## Supplementary material for "Detection and characterization of the SARS-CoV-2 lineage B.1.526 in New York": Supp. Table 3 GISAID Acknowledgment part 2: gisaid_hcov-19_acknowledgement_table_2021_02_13_00-9.pdf

We gratefully acknowledge the following Authors from the Originating laboratories responsible for obtaining the specimens, as well as the Submitting laboratories where the genome data were generated and shared via GISAID, on which this research is based.

All Submitters of data may be contacted directly via www.gisaid.org

Authors are sorted alphabetically.

| Accession ID | Originating Laboratory | Submitting Laboratory | Authors |
|---|---|---|---|
| EPI_ISL_645133, EPI_ISL_645134, EPI_ISL_649108, EPI_ISL_649110 | Respiratory Virus Unit, Microbiology Services Colindale, Public Health England | COVID-19 Genomics UK (COG-UK) Consortium | PHE Covid Sequencing Team |
| EPI_ISL_649125 | Wellington SCL (WN) | Institute of Environmental Science and Research (ESR) | Xiaoyun Ren, Matt Storey, Nikki Freed, Muhammad Faisal, Jing Wang, Hermes Perez, Anja Werno, Antje van der Linden, Arlo Upton, Chris Mansell, David Hammer, Dragana Drinkovic, Gary McAuliffe, Hana Sofia Andersson, James Ussher, Jill Sherwood, Josh Freeman, Julia Howard, Juliet Elvy, Mary DeAlmeida, Matt Blakiston, Matthew Rogers, Max Bloomfield, Michael Addidle, Michelle Balm, Sally Roberts, Sarah Jefferies, Sharmini Muttaiyah, Susan Morpeth, Susan Taylor, Timothy Blackmore, Vani Sathyendran, Veronica Playle, Virginia Hope, Erasmus Smit, Lauren Jelly, Olin Silander, Joep de Ligt |
| EPI_ISL_649936, EPI_ISL_649937 | Respiratory Virus Unit, Microbiology Services Colindale, Public Health England | COVID-19 Genomics UK (COG-UK) Consortium | PHE Covid Sequencing Team |
| EPI_ISL_650178, EPI_ISL_650187, EPI_ISL_650266, EPI_ISL_650276, EPI_ISL_650314, EPI_ISL_650443 | Queens Medical Centre, Clinical Microbiology Department / DeepSeq Nottingham | COVID-19 Genomics UK (COG-UK) Consortium | Gemma Clark, Wendy Smith, Manjinder Khakh, Vicki M Fleming, Michelle M Lister, Hannah Howson-Wells, Jonathan Ball, Patrick McClure, Joseph Chappell, Theocharis Tsoleridis, Nadine Holmes, Matthew Carlisle, Christopher Moore, Fei Sang, Johnny Debebe, Victoria Wright, Matthew Loose |
| EPI_ISL_650459 | Quadram Institute Bioscience | COVID-19 Genomics UK (COG-UK) Consortium | Dave J. Baker, Gemma L. Kay, Alp Aydin, Thanh Le-Viet, Steven Rudder, Ana P. Tedim, Anastasia Kolyva, Maria Diaz, Leonardo de Oliveira Martins, Nabil-Fareed Alikhan, Lizzie Meadows, Rachael Stanley, Ngozi Elumogo, Muhammed Yasir, Nicholas M. Thomson, Alexander J Trotter, Rachel Gilroy, Samuel Bloomfield, Claire Stuart, Andrew Bell, Reenesh Prakash, Samir Dervisevic, Alison E. Mather, John Wain, Mark Webber, Andrew J. Page, Justin O'Grady |
| EPI_ISL_650786, EPI_ISL_650830, EPI_ISL_650835, EPI_ISL_650979, EPI_ISL_650987, EPI_ISL_651065, EPI_ISL_651107, EPI_ISL_651189, EPI_ISL_651286, EPI_ISL_651401, EPI_ISL_651408, EPI_ISL_652401, EPI_ISL_652402, EPI_ISL_652403, EPI_ISL_652404, EPI_ISL_652405, EPI_ISL_652406, EPI_ISL_652407, EPI_ISL_652408, EPI_ISL_652409, EPI_ISL_652410, EPI_ISL_652411, EPI_ISL_652412, EPI_ISL_652413 | | | |
| see above | Queens Medical Centre, Clinical Microbiology Department / DeepSeq Nottingham | COVID-19 Genomics UK (COG-UK) Consortium | Gemma Clark, Wendy Smith, Manjinder Khakh, Vicki M Fleming, Michelle M Lister, Hannah Howson-Wells, Jonathan Ball, Patrick McClure, Joseph Chappell, Theocharis Tsoleridis, Nadine Holmes, Matthew Carlisle, Christopher Moore, Fei Sang, Johnny Debebe, Victoria Wright, Matthew Loose |
| EPI_ISL_654799, EPI_ISL_654800, EPI_ISL_654801, EPI_ISL_654802, EPI_ISL_654803, EPI_ISL_654804, EPI_ISL_654805, EPI_ISL_654806, EPI_ISL_654807, EPI_ISL_654808, EPI_ISL_654809 | | | |
| see above | SA Pathology | SA Pathology | Lex Leong, Julien Soubrier, Chuan Kok Lim, Song Gao, Mark Turra, Karin Kassahn, Ivan Bastian, Geoff Higgins |
| EPI_ISL_660228, EPI_ISL_660229, EPI_ISL_660230, EPI_ISL_660231, EPI_ISL_660232, EPI_ISL_660256, EPI_ISL_660257, EPI_ISL_660258 | KRISP, KZN Research Innovation and Sequencing Platform | KRISP, KZN Research Innovation and Sequencing Platform | Giandhari J, Pillay S, Lessells R, Mdlalose K, York D, Khan S, Tegally H, Wilkinson E, de Oliveira T |
| EPI_ISL_661173 | Gundersen Molecular Diagnostics Laboratory | Kabara Cancer Research Institute | Craig S. Richmond, Paraic A. Kenny |
| EPI_ISL_661201, EPI_ISL_661204, EPI_ISL_661205, EPI_ISL_661206, EPI_ISL_661207, EPI_ISL_661208, EPI_ISL_661209, EPI_ISL_661210, EPI_ISL_661211, EPI_ISL_661212, EPI_ISL_661213, EPI_ISL_661214, EPI_ISL_661218, EPI_ISL_661219, EPI_ISL_661220, EPI_ISL_661221, EPI_ISL_661222, EPI_ISL_661223, EPI_ISL_661224, EPI_ISL_661225, EPI_ISL_661226, EPI_ISL_661227, EPI_ISL_661228, EPI_ISL_661229, EPI_ISL_661230, EPI_ISL_661231, EPI_ISL_661232, EPI_ISL_661233, EPI_ISL_661234, EPI_ISL_661235, EPI_ISL_661236, EPI_ISL_661237, EPI_ISL_661238, EPI_ISL_661239, EPI_ISL_661240, EPI_ISL_661241, EPI_ISL_661242, EPI_ISL_661243, EPI_ISL_661244, EPI_ISL_661245 | | | |
| see above | Department of Clinical Microbiology | GIGA Medical Genomics | Keith Durkin, Maria Artesi, Sébastien Bontems, Raphaël Boreux, Bouchra Boujemla, Cécile Meex, Pierrette Melin, Marie-Pierre Hayette, Vincent Bours |
| EPI_ISL_661257, EPI_ISL_661258, EPI_ISL_661259, EPI_ISL_661262, EPI_ISL_661263 | LabPLUS | Institute of Environmental Science and Research (ESR) | Xiaoyun Ren, Matt Storey, Nikki Freed, Muhammad Faisal, Jing Wang, Hermes Perez, Anja Werno, Antje van der Linden, Arlo Upton, Chris Mansell, David Hammer, Dragana Drinkovic, Gary McAuliffe, Hana Sofia Andersson, James Ussher, Jill Sherwood, Josh Freeman, Julia Howard, Juliet Elvy, Mary DeAlmeida, Matt Blakiston, Matthew Rogers, Max Bloomfield, Michael Addidle, Michelle Balm, Sally Roberts, Sarah Jefferies, Sharmini Muttaiyah, Susan Morpeth, Susan Taylor, Timothy Blackmore, Vani Sathyendran, Veronica Playle, Virginia Hope, Erasmus Smit, Lauren Jelly, Olin Silander, Joep de Ligt |
| EPI_ISL_664120, EPI_ISL_664139, EPI_ISL_664199, EPI_ISL_664200, EPI_ISL_664234, EPI_ISL_664291, EPI_ISL_664304, EPI_ISL_664316, EPI_ISL_664317, EPI_ISL_664380, EPI_ISL_664382, EPI_ISL_664383, EPI_ISL_664384, EPI_ISL_664446, EPI_ISL_664455, EPI_ISL_664470, EPI_ISL_664516, EPI_ISL_664527, EPI_ISL_664529, EPI_ISL_664535, EPI_ISL_664536, EPI_ISL_664537, EPI_ISL_664538, EPI_ISL_664588, EPI_ISL_664589, EPI_ISL_664601, EPI_ISL_664616, EPI_ISL_664635, EPI_ISL_664656, EPI_ISL_664731, EPI_ISL_664749, EPI_ISL_664995, EPI_ISL_665005, EPI_ISL_665006, EPI_ISL_665007, EPI_ISL_665008, EPI_ISL_665009, EPI_ISL_665010, EPI_ISL_665011, EPI_ISL_665012, EPI_ISL_665020, EPI_ISL_665021, EPI_ISL_665022 | | | |
| see above | Department of Pathology, University of Cambridge | COVID-19 Genomics UK (COG-UK) Consortium | Aminu S. Jahun, Yasmin Chaudhry, Grant Hall, Iliana Georgana, Myra Hosmillo, Martin D. Curran, Malte Pinckert, Surendra Parmar, Ian Goodfellow |
| EPI_ISL_665268 | Quadram Institute Bioscience | COVID-19 Genomics UK (COG-UK) Consortium | Dave J. Baker, Gemma L. Kay, Alp Aydin, Thanh Le-Viet, Steven Rudder, Ana P. Tedim, Anastasia Kolyva, Maria Diaz, Leonardo de Oliveira Martins, Nabil-Fareed Alikhan, Lizzie Meadows, Rachael Stanley, Ngozi Elumogo, Muhammed Yasir, Nicholas M. Thomson, Alexander J Trotter, Rachel Gilroy, Samuel Bloomfield, Claire Stuart, Andrew Bell, Reenesh Prakash, Samir Dervisevic, Alison E. Mather, John Wain, Mark Webber, Andrew J. Page, Justin O'Grady |
| EPI_ISL_665291 | Queens Medical Centre, Clinical Microbiology Department / DeepSeq Nottingham | COVID-19 Genomics UK (COG-UK) Consortium | Gemma Clark, Wendy Smith, Manjinder Khakh, Vicki M Fleming, Michelle M Lister, Hannah Howson-Wells, Jonathan Ball, Patrick McClure, Joseph Chappell, Theocharis Tsoleridis, Nadine Holmes, Matthew Carlisle, Christopher Moore, Fei Sang, Johnny Debebe, Victoria Wright, Matthew Loose |
| EPI_ISL_665316, EPI_ISL_665336 | Quadram Institute Bioscience | COVID-19 Genomics UK (COG-UK) Consortium | Dave J. Baker, Gemma L. Kay, Alp Aydin, Thanh Le-Viet, Steven Rudder, Ana P. Tedim, Anastasia Kolyva, Maria Diaz, Leonardo de Oliveira Martins, Nabil-Fareed Alikhan, Lizzie Meadows, Rachael Stanley, Ngozi Elumogo, Muhammed Yasir, Nicholas M. Thomson, Alexander J Trotter, Rachel Gilroy, Samuel Bloomfield, Claire Stuart, Andrew Bell, Reenesh Prakash, Samir Dervisevic, Alison E. Mather, John Wain, Mark Webber, Andrew J. Page, Justin O'Grady |
| EPI_ISL_665345 | Queens Medical Centre, Clinical Microbiology Department / DeepSeq Nottingham | COVID-19 Genomics UK (COG-UK) Consortium | Gemma Clark, Wendy Smith, Manjinder Khakh, Vicki M Fleming, Michelle M Lister, Hannah Howson-Wells, Jonathan Ball, Patrick McClure, Joseph Chappell, Theocharis Tsoleridis, Nadine Holmes, Matthew Carlisle, Christopher Moore, Fei Sang, Johnny Debebe, Victoria Wright, Matthew Loose |
| EPI_ISL_665362, EPI_ISL_665369, EPI_ISL_665370 | Quadram Institute Bioscience | COVID-19 Genomics UK (COG-UK) Consortium | Dave J. Baker, Gemma L. Kay, Alp Aydin, Thanh Le-Viet, Steven Rudder, Ana P. Tedim, Anastasia Kolyva, Maria Diaz, Leonardo de Oliveira Martins, Nabil-Fareed Alikhan, Lizzie Meadows, Rachael Stanley, Ngozi Elumogo, Muhammed Yasir, Nicholas M. Thomson, Alexander J Trotter, Rachel Gilroy, Samuel Bloomfield, Claire Stuart, Andrew Bell, Reenesh Prakash, Samir Dervisevic, Alison E. Mather, John Wain, Mark Webber, Andrew J. Page, Justin O'Grady |
| EPI_ISL_665420, EPI_ISL_665421, EPI_ISL_665424, EPI_ISL_665426 | Queens Medical Centre, Clinical Microbiology Department / DeepSeq Nottingham | COVID-19 Genomics UK (COG-UK) Consortium | Gemma Clark, Wendy Smith, Manjinder Khakh, Vicki M Fleming, Michelle M Lister, Hannah Howson-Wells, Jonathan Ball, Patrick McClure, Joseph Chappell, Theocharis Tsoleridis, Nadine Holmes, Matthew Carlisle, Christopher Moore, Fei Sang, Johnny Debebe, Victoria Wright, Matthew Loose |
| EPI_ISL_665427, EPI_ISL_665429 | Quadram Institute Bioscience | COVID-19 Genomics UK (COG-UK) Consortium | Dave J. Baker, Gemma L. Kay, Alp Aydin, Thanh Le-Viet, Steven Rudder, Ana P. Tedim, Anastasia Kolyva, Maria Diaz, Leonardo de Oliveira Martins, Nabil-Fareed Alikhan, Lizzie Meadows, Rachael Stanley, Ngozi Elumogo, Muhammed Yasir, Nicholas M. Thomson, Alexander J Trotter, Rachel Gilroy, Samuel Bloomfield, Claire Stuart, Andrew Bell, Reenesh Prakash, Samir Dervisevic, Alison E. Mather, John Wain, Mark Webber, Andrew J. Page, Justin O'Grady |
| EPI_ISL_665430 | Queens Medical Centre, Clinical Microbiology Department / DeepSeq Nottingham | COVID-19 Genomics UK (COG-UK) Consortium | Gemma Clark, Wendy Smith, Manjinder Khakh, Vicki M Fleming, Michelle M Lister, Hannah Howson-Wells, Jonathan Ball, Patrick McClure, Joseph Chappell, Theocharis Tsoleridis, Nadine Holmes, Matthew Carlisle, Christopher Moore, Fei Sang, Johnny Debebe, Victoria Wright, Matthew Loose |
| EPI_ISL_665436 | West of Scotland Specialist Virology Centre, NHSGGC / | COVID-19 Genomics UK (COG-UK) Consortium | Ana da Silva Filipe, Natasha Johnson, Kathy Smollett, Daniel Mair, Stephen Carmichael, Alice Broos, Lily Tong, Jenna Nichols, Kyriaki Nomikou; Sarah |

| | | | |
|---|---|---|---|
| | MRC-University of Glasgow Centre for Virus Research | | McDonald; Richard Orton, Joseph Hughes, Sreenu Vattipally, David L Robertson; Alasdair MacLean, Rory Gunson; Sharif Shaaban, Matthew Holden; Rachel Blacow, Guy Mollett, Kathy Li, James Shepherd, Antonia Ho, Emma Thomson |
| EPI_ISL_665438 | Quadram Institute Bioscience | COVID-19 Genomics UK (COG-UK) Consortium | Dave J. Baker, Gemma L. Kay, Alp Aydin, Thanh Le-Viet, Steven Rudder, Ana P. Tedim, Anastasia Kolyva, Maria Diaz, Leonardo de Oliveira Martins, Nabil-Fareed Alikhan, Lizzie Meadows, Rachael Stanley, Ngozi Elumogo, Muhammed Yasir, Nicholas M. Thomson, Alexander J Trotter, Rachel Gilroy, Samuel Bloomfield, Claire Stuart, Andrew Bell, Reenesh Prakash, Samir Dervisevic, Alison E. Mather, John Wain, Mark Webber, Andrew J. Page, Justin O'Grady |
| EPI_ISL_665443, EPI_ISL_665452 | Queens Medical Centre, Clinical Microbiology Department / DeepSeq Nottingham | COVID-19 Genomics UK (COG-UK) Consortium | Gemma Clark, Wendy Smith, Manjinder Khakh, Vicki M Fleming, Michelle M Lister, Hannah Howson-Wells, Jonathan Ball, Patrick McClure, Joseph Chappell, Theocharis Tsoleridis, Nadine Holmes, Matthew Carlisle, Christopher Moore, Fei Sang, Johnny Debebe, Victoria Wright, Matthew Loose |
| EPI_ISL_665453, EPI_ISL_665463, EPI_ISL_665465, EPI_ISL_665475, EPI_ISL_665519 | Quadram Institute Bioscience | COVID-19 Genomics UK (COG-UK) Consortium | Dave J. Baker, Gemma L. Kay, Alp Aydin, Thanh Le-Viet, Steven Rudder, Ana P. Tedim, Anastasia Kolyva, Maria Diaz, Leonardo de Oliveira Martins, Nabil-Fareed Alikhan, Lizzie Meadows, Rachael Stanley, Ngozi Elumogo, Muhammed Yasir, Nicholas M. Thomson, Alexander J Trotter, Rachel Gilroy, Samuel Bloomfield, Claire Stuart, Andrew Bell, Reenesh Prakash, Samir Dervisevic, Alison E. Mather, John Wain, Mark Webber, Andrew J. Page, Justin O'Grady |
| EPI_ISL_665521 | West of Scotland Specialist Virology Centre, NHSGGC / MRC-University of Glasgow Centre for Virus Research | COVID-19 Genomics UK (COG-UK) Consortium | Ana da Silva Filipe, Natasha Johnson, Kathy Smollett, Daniel Mair, Stephen Carmichael, Alice Broos, Lily Tong, Jenna Nichols, Kyriaki Nomikou; Sarah McDonald; Richard Orton, Joseph Hughes, Sreenu Vattipally, David L Robertson; Alasdair MacLean, Rory Gunson; Sharif Shaaban, Matthew Holden; Rachel Blacow, Guy Mollett, Kathy Li, James Shepherd, Antonia Ho, Emma Thomson |
| EPI_ISL_665534 | Quadram Institute Bioscience | COVID-19 Genomics UK (COG-UK) Consortium | Dave J. Baker, Gemma L. Kay, Alp Aydin, Thanh Le-Viet, Steven Rudder, Ana P. Tedim, Anastasia Kolyva, Maria Diaz, Leonardo de Oliveira Martins, Nabil-Fareed Alikhan, Lizzie Meadows, Rachael Stanley, Ngozi Elumogo, Muhammed Yasir, Nicholas M. Thomson, Alexander J Trotter, Rachel Gilroy, Samuel Bloomfield, Claire Stuart, Andrew Bell, Reenesh Prakash, Samir Dervisevic, Alison E. Mather, John Wain, Mark Webber, Andrew J. Page, Justin O'Grady |
| EPI_ISL_665538 | West of Scotland Specialist Virology Centre, NHSGGC / MRC-University of Glasgow Centre for Virus Research | COVID-19 Genomics UK (COG-UK) Consortium | Ana da Silva Filipe, Natasha Johnson, Kathy Smollett, Daniel Mair, Stephen Carmichael, Alice Broos, Lily Tong, Jenna Nichols, Kyriaki Nomikou; Sarah McDonald; Richard Orton, Joseph Hughes, Sreenu Vattipally, David L Robertson; Alasdair MacLean, Rory Gunson; Sharif Shaaban, Matthew Holden; Rachel Blacow, Guy Mollett, Kathy Li, James Shepherd, Antonia Ho, Emma Thomson |
| EPI_ISL_665543, EPI_ISL_665544, EPI_ISL_665550, EPI_ISL_665551 | Queens Medical Centre, Clinical Microbiology Department / DeepSeq Nottingham | COVID-19 Genomics UK (COG-UK) Consortium | Gemma Clark, Wendy Smith, Manjinder Khakh, Vicki M Fleming, Michelle M Lister, Hannah Howson-Wells, Jonathan Ball, Patrick McClure, Joseph Chappell, Theocharis Tsoleridis, Nadine Holmes, Matthew Carlisle, Christopher Moore, Fei Sang, Johnny Debebe, Victoria Wright, Matthew Loose |
| EPI_ISL_665560 | Quadram Institute Bioscience | COVID-19 Genomics UK (COG-UK) Consortium | Dave J. Baker, Gemma L. Kay, Alp Aydin, Thanh Le-Viet, Steven Rudder, Ana P. Tedim, Anastasia Kolyva, Maria Diaz, Leonardo de Oliveira Martins, Nabil-Fareed Alikhan, Lizzie Meadows, Rachael Stanley, Ngozi Elumogo, Muhammed Yasir, Nicholas M. Thomson, Alexander J Trotter, Rachel Gilroy, Samuel Bloomfield, Claire Stuart, Andrew Bell, Reenesh Prakash, Samir Dervisevic, Alison E. Mather, John Wain, Mark Webber, Andrew J. Page, Justin O'Grady |
| EPI_ISL_665561, EPI_ISL_665562 | Queens Medical Centre, Clinical Microbiology Department / DeepSeq Nottingham | COVID-19 Genomics UK (COG-UK) Consortium | Gemma Clark, Wendy Smith, Manjinder Khakh, Vicki M Fleming, Michelle M Lister, Hannah Howson-Wells, Jonathan Ball, Patrick McClure, Joseph Chappell, Theocharis Tsoleridis, Nadine Holmes, Matthew Carlisle, Christopher Moore, Fei Sang, Johnny Debebe, Victoria Wright, Matthew Loose |
| EPI_ISL_665599 | Quadram Institute Bioscience | COVID-19 Genomics UK (COG-UK) Consortium | Dave J. Baker, Gemma L. Kay, Alp Aydin, Thanh Le-Viet, Steven Rudder, Ana P. Tedim, Anastasia Kolyva, Maria Diaz, Leonardo de Oliveira Martins, Nabil-Fareed Alikhan, Lizzie Meadows, Rachael Stanley, Ngozi Elumogo, Muhammed Yasir, Nicholas M. Thomson, Alexander J Trotter, Rachel Gilroy, Samuel Bloomfield, Claire Stuart, Andrew Bell, Reenesh Prakash, Samir Dervisevic, Alison E. Mather, John Wain, Mark Webber, Andrew J. Page, Justin O'Grady |
| EPI_ISL_665601 | Queens Medical Centre, Clinical Microbiology Department / DeepSeq Nottingham | COVID-19 Genomics UK (COG-UK) Consortium | Gemma Clark, Wendy Smith, Manjinder Khakh, Vicki M Fleming, Michelle M Lister, Hannah Howson-Wells, Jonathan Ball, Patrick McClure, Joseph Chappell, Theocharis Tsoleridis, Nadine Holmes, Matthew Carlisle, Christopher Moore, Fei Sang, Johnny Debebe, Victoria Wright, Matthew Loose |
| EPI_ISL_665609 | West of Scotland Specialist Virology Centre, NHSGGC / MRC-University of Glasgow Centre for Virus Research | COVID-19 Genomics UK (COG-UK) Consortium | Ana da Silva Filipe, Natasha Johnson, Kathy Smollett, Daniel Mair, Stephen Carmichael, Alice Broos, Lily Tong, Jenna Nichols, Kyriaki Nomikou; Sarah McDonald; Richard Orton, Joseph Hughes, Sreenu Vattipally, David L Robertson; Alasdair MacLean, Rory Gunson; Sharif Shaaban, Matthew Holden; Rachel Blacow, Guy Mollett, Kathy Li, James Shepherd, Antonia Ho, Emma Thomson |
| EPI_ISL_665612 | Queens Medical Centre, Clinical Microbiology Department / DeepSeq Nottingham | COVID-19 Genomics UK (COG-UK) Consortium | Gemma Clark, Wendy Smith, Manjinder Khakh, Vicki M Fleming, Michelle M Lister, Hannah Howson-Wells, Jonathan Ball, Patrick McClure, Joseph Chappell, Theocharis Tsoleridis, Nadine Holmes, Matthew Carlisle, Christopher Moore, Fei Sang, Johnny Debebe, Victoria Wright, Matthew Loose |
| EPI_ISL_665616 | Centre for Enzyme Innovation, University of Portsmouth / Translational Research Laboratory, Portsmouth Hospitals NHS Trust | COVID-19 Genomics UK (COG-UK) Consortium | Angela Beckett,Yann Bourgeois,Garry Scarlett,Sharon Glaysher,Scott Elliott,Kelly Bicknell,Robert Impey,Allyson Lloyd,Sarah Wyllie,Ethan Butcher,Anoop Chauhan,Samuel Robson |
| EPI_ISL_665622 | Queens Medical Centre, Clinical Microbiology Department / DeepSeq Nottingham | COVID-19 Genomics UK (COG-UK) Consortium | Gemma Clark, Wendy Smith, Manjinder Khakh, Vicki M Fleming, Michelle M Lister, Hannah Howson-Wells, Jonathan Ball, Patrick McClure, Joseph Chappell, Theocharis Tsoleridis, Nadine Holmes, Matthew Carlisle, Christopher Moore, Fei Sang, Johnny Debebe, Victoria Wright, Matthew Loose |
| EPI_ISL_665625 | Centre for Enzyme Innovation, University of Portsmouth / Translational Research Laboratory, Portsmouth Hospitals NHS Trust | COVID-19 Genomics UK (COG-UK) Consortium | Angela Beckett,Yann Bourgeois,Garry Scarlett,Sharon Glaysher,Scott Elliott,Kelly Bicknell,Robert Impey,Allyson Lloyd,Sarah Wyllie,Ethan Butcher,Anoop Chauhan,Samuel Robson |
| EPI_ISL_665636 | University College London Hospital | COVID-19 Genomics UK (COG-UK) Consortium | Judith Heaney, Matthew Byott, Catherine Houlihan, Dan Frampton, Stuart Kirk, Moira Spyer and Eleni Nastouli |
| EPI_ISL_665638 | Quadram Institute Bioscience | COVID-19 Genomics UK (COG-UK) Consortium | Dave J. Baker, Gemma L. Kay, Alp Aydin, Thanh Le-Viet, Steven Rudder, Ana P. Tedim, Anastasia Kolyva, Maria Diaz, Leonardo de Oliveira Martins, Nabil-Fareed Alikhan, Lizzie Meadows, Rachael Stanley, Ngozi Elumogo, Muhammed Yasir, Nicholas M. Thomson, Alexander J Trotter, Rachel Gilroy, Samuel Bloomfield, Claire Stuart, Andrew Bell, Reenesh Prakash, Samir Dervisevic, Alison E. Mather, John Wain, Mark Webber, Andrew J. Page, Justin O'Grady |
| EPI_ISL_665641 | West of Scotland Specialist Virology Centre, NHSGGC / MRC-University of Glasgow Centre for Virus Research | COVID-19 Genomics UK (COG-UK) Consortium | Ana da Silva Filipe, Natasha Johnson, Kathy Smollett, Daniel Mair, Stephen Carmichael, Alice Broos, Lily Tong, Jenna Nichols, Kyriaki Nomikou; Sarah McDonald; Richard Orton, Joseph Hughes, Sreenu Vattipally, David L Robertson; Alasdair MacLean, Rory Gunson; Sharif Shaaban, Matthew Holden; Rachel Blacow, Guy Mollett, Kathy Li, James Shepherd, Antonia Ho, Emma Thomson |
| EPI_ISL_665643 | University College London Hospital | COVID-19 Genomics UK (COG-UK) Consortium | Judith Heaney, Matthew Byott, Catherine Houlihan, Dan Frampton, Stuart Kirk, Moira Spyer and Eleni Nastouli |
| EPI_ISL_665668, EPI_ISL_665678, EPI_ISL_665692, EPI_ISL_665693, EPI_ISL_665697, EPI_ISL_665714, EPI_ISL_665730 | Quadram Institute Bioscience | COVID-19 Genomics UK (COG-UK) Consortium | Dave J. Baker, Gemma L. Kay, Alp Aydin, Thanh Le-Viet, Steven Rudder, Ana P. Tedim, Anastasia Kolyva, Maria Diaz, Leonardo de Oliveira Martins, Nabil-Fareed Alikhan, Lizzie Meadows, Rachael Stanley, Ngozi Elumogo, Muhammed Yasir, Nicholas M. Thomson, Alexander J Trotter, Rachel Gilroy, Samuel Bloomfield, Claire Stuart, Andrew Bell, Reenesh Prakash, Samir Dervisevic, Alison E. Mather, John Wain, Mark Webber, Andrew J. Page, Justin O'Grady |
| EPI_ISL_665740 | Queens Medical Centre, Clinical Microbiology Department / DeepSeq Nottingham | COVID-19 Genomics UK (COG-UK) Consortium | Gemma Clark, Wendy Smith, Manjinder Khakh, Vicki M Fleming, Michelle M Lister, Hannah Howson-Wells, Jonathan Ball, Patrick McClure, Joseph Chappell, Theocharis Tsoleridis, Nadine Holmes, Matthew Carlisle, Christopher Moore, Fei Sang, Johnny Debebe, Victoria Wright, Matthew Loose |
| EPI_ISL_665746 | Quadram Institute Bioscience | COVID-19 Genomics UK (COG-UK) Consortium | Dave J. Baker, Gemma L. Kay, Alp Aydin, Thanh Le-Viet, Steven Rudder, Ana P. Tedim, Anastasia Kolyva, Maria Diaz, Leonardo de Oliveira Martins, Nabil-Fareed Alikhan, Lizzie Meadows, Rachael Stanley, Ngozi Elumogo, Muhammed Yasir, Nicholas M. Thomson, Alexander J Trotter, Rachel Gilroy, Samuel Bloomfield, Claire Stuart, Andrew Bell, Reenesh Prakash, Samir Dervisevic, Alison E. Mather, John Wain, Mark Webber, Andrew J. Page, Justin O'Grady |
| EPI_ISL_665789 | Queens Medical Centre, Clinical Microbiology Department / DeepSeq Nottingham | COVID-19 Genomics UK (COG-UK) Consortium | Gemma Clark, Wendy Smith, Manjinder Khakh, Vicki M Fleming, Michelle M Lister, Hannah Howson-Wells, Jonathan Ball, Patrick McClure, Joseph Chappell, Theocharis Tsoleridis, Nadine Holmes, Matthew Carlisle, Christopher Moore, Fei Sang, Johnny Debebe, Victoria Wright, Matthew Loose |
| EPI_ISL_665825, EPI_ISL_665839, EPI_ISL_665840, EPI_ISL_665841, EPI_ISL_665847, EPI_ISL_665848 | University College London Hospital | COVID-19 Genomics UK (COG-UK) Consortium | Judith Heaney, Matthew Byott, Catherine Houlihan, Dan Frampton, Stuart Kirk, Moira Spyer and Eleni Nastouli |
| EPI_ISL_665876 | Quadram Institute Bioscience | COVID-19 Genomics UK (COG-UK) Consortium | Dave J. Baker, Gemma L. Kay, Alp Aydin, Thanh Le-Viet, Steven Rudder, Ana P. Tedim, Anastasia Kolyva, Maria Diaz, Leonardo de Oliveira Martins, |

| | | | |
|---|---|---|---|
| | | | Nabil-Fareed Alikhan, Lizzie Meadows, Rachael Stanley, Ngozi Elumogo, Muhammed Yasir, Nicholas M. Thomson, Alexander J Trotter, Rachel Gilroy, Samuel Bloomfield, Claire Stuart, Andrew Bell, Reenesh Prakash, Samir Dervisevic, Alison E. Mather, John Wain, Mark Webber, Andrew J. Page, Justin O'Grady |
| EPI_ISL_665901, EPI_ISL_665902, EPI_ISL_665903, EPI_ISL_665904, EPI_ISL_665905, EPI_ISL_665906, EPI_ISL_665907, EPI_ISL_665908, EPI_ISL_665942 | Queens Medical Centre, Clinical Microbiology Department / DeepSeq Nottingham | COVID-19 Genomics UK (COG-UK) Consortium | Gemma Clark, Wendy Smith, Manjinder Khakh, Vicki M Fleming, Michelle M Lister, Hannah Howson-Wells, Jonathan Ball, Patrick McClure, Joseph Chappell, Theocharis Tsoleridis, Nadine Holmes, Matthew Carlisle, Christopher Moore, Fei Sang, Johnny Debebe, Victoria Wright, Matthew Loose |
| EPI_ISL_666041, EPI_ISL_666042, EPI_ISL_666043, EPI_ISL_666044, EPI_ISL_666045, EPI_ISL_666046, EPI_ISL_666047, EPI_ISL_666048, EPI_ISL_666049, EPI_ISL_666050, EPI_ISL_666051, EPI_ISL_666052, EPI_ISL_666053, EPI_ISL_666054, EPI_ISL_666055, EPI_ISL_666056, EPI_ISL_666057, EPI_ISL_666058, EPI_ISL_666059 | | | |
| see above | West of Scotland Specialist Virology Centre, NHSGGC / MRC-University of Glasgow Centre for Virus Research | COVID-19 Genomics UK (COG-UK) Consortium | Ana da Silva Filipe, Natasha Johnson, Kathy Smollett, Daniel Mair, Stephen Carmichael, Alice Broos, Lily Tong, Jenna Nichols, Kyriaki Nomikou; Sarah McDonald; Richard Orton, Joseph Hughes, Sreenu Vattipally, David L Robertson; Alasdair MacLean, Rory Gunson; Sharif Shaaban, Matthew Holden; Rachel Blacow, Guy Mollett, Kathy Li, James Shepherd, Antonia Ho, Emma Thomson |
| EPI_ISL_667549 | OHSU Lab Services Molecular Microbiology Lab | Oregon SARS-CoV-2 Genome Sequencing Center | Brendan L. O'Connell, Ruth V. Nichols, Sally Grindstaff, Alec J. Hirsch, Donna Hansel, Guang Fan, Daniel N. Streblow, William B. Messer, Andrew C. Adey, Benjamin N. Bimber, Brian J. O'Roak |
| EPI_ISL_668465, EPI_ISL_668467, EPI_ISL_668468, EPI_ISL_668501, EPI_ISL_668502, EPI_ISL_668510, EPI_ISL_668517, EPI_ISL_668518, EPI_ISL_668519, EPI_ISL_668521, EPI_ISL_668522, EPI_ISL_668523, EPI_ISL_668524, EPI_ISL_668525, EPI_ISL_668526, EPI_ISL_668527, EPI_ISL_668530, EPI_ISL_668534, EPI_ISL_668535, EPI_ISL_668536, EPI_ISL_668537, EPI_ISL_668544, EPI_ISL_668546, EPI_ISL_668558, EPI_ISL_668559, EPI_ISL_668561, EPI_ISL_668562, EPI_ISL_668563, EPI_ISL_668572, EPI_ISL_668579, EPI_ISL_668580, EPI_ISL_668581, EPI_ISL_668592, EPI_ISL_668595, EPI_ISL_668596, EPI_ISL_668597, EPI_ISL_668601, EPI_ISL_668606, EPI_ISL_668609, EPI_ISL_668610, EPI_ISL_668612, EPI_ISL_668615, EPI_ISL_668633, EPI_ISL_668635, EPI_ISL_668636, EPI_ISL_668643, EPI_ISL_668652, EPI_ISL_668653, EPI_ISL_668663, EPI_ISL_668664, EPI_ISL_668665, EPI_ISL_668666, EPI_ISL_668667, EPI_ISL_668668, EPI_ISL_668669, EPI_ISL_668670, EPI_ISL_668671, EPI_ISL_668672, EPI_ISL_668673, EPI_ISL_668674, EPI_ISL_668718, EPI_ISL_668719, EPI_ISL_668727, EPI_ISL_668728, EPI_ISL_668730, EPI_ISL_668735, EPI_ISL_668738, EPI_ISL_668743, EPI_ISL_668744, EPI_ISL_668757, EPI_ISL_668758, EPI_ISL_668759, EPI_ISL_668760, EPI_ISL_668761, EPI_ISL_668762, EPI_ISL_668763, EPI_ISL_668764, EPI_ISL_668765, EPI_ISL_668766, EPI_ISL_668767, EPI_ISL_668768, EPI_ISL_668769, EPI_ISL_668770, EPI_ISL_668771, EPI_ISL_668772, EPI_ISL_668773, EPI_ISL_668774, EPI_ISL_668829, EPI_ISL_668830, EPI_ISL_668839, EPI_ISL_668840, EPI_ISL_668850, EPI_ISL_668853, EPI_ISL_668854, EPI_ISL_668855, EPI_ISL_668856, EPI_ISL_668857, EPI_ISL_668858, EPI_ISL_668859, EPI_ISL_668860, EPI_ISL_668861, EPI_ISL_668862, EPI_ISL_668863, EPI_ISL_668864, EPI_ISL_668865, EPI_ISL_668866, EPI_ISL_668867, EPI_ISL_668868, EPI_ISL_668869, EPI_ISL_668870, EPI_ISL_668871, EPI_ISL_668872, EPI_ISL_668873, EPI_ISL_668874, EPI_ISL_668875, EPI_ISL_668876, EPI_ISL_668877, EPI_ISL_668878, EPI_ISL_668879, EPI_ISL_668880, EPI_ISL_668881, EPI_ISL_668882, EPI_ISL_668883, EPI_ISL_668884, EPI_ISL_668885, EPI_ISL_668886, EPI_ISL_668887, EPI_ISL_668888, EPI_ISL_668889, EPI_ISL_668890, EPI_ISL_668891, EPI_ISL_668892, EPI_ISL_668893, EPI_ISL_668894, EPI_ISL_668895, EPI_ISL_668896, EPI_ISL_668897, EPI_ISL_668898, EPI_ISL_668899, EPI_ISL_668900, EPI_ISL_668901, EPI_ISL_668902, EPI_ISL_668903, EPI_ISL_668904, EPI_ISL_668905, EPI_ISL_668906, EPI_ISL_668907, EPI_ISL_668908, EPI_ISL_668909, EPI_ISL_668910, EPI_ISL_668911, EPI_ISL_668912, EPI_ISL_668913, EPI_ISL_668914, EPI_ISL_668915, EPI_ISL_668916, EPI_ISL_668917, EPI_ISL_668918, EPI_ISL_668919, EPI_ISL_668920, EPI_ISL_668921, EPI_ISL_668922, EPI_ISL_668923, EPI_ISL_668924, EPI_ISL_668925, EPI_ISL_668926, EPI_ISL_668927, EPI_ISL_668928, EPI_ISL_668929, EPI_ISL_668930, EPI_ISL_668931, EPI_ISL_668932, EPI_ISL_668933, EPI_ISL_668934, EPI_ISL_668935, EPI_ISL_668936, EPI_ISL_668937, EPI_ISL_668938, EPI_ISL_668939, EPI_ISL_668940, EPI_ISL_668941, EPI_ISL_668942, EPI_ISL_668943, EPI_ISL_668944, EPI_ISL_668945, EPI_ISL_668946, EPI_ISL_668947, EPI_ISL_668948, EPI_ISL_668949, EPI_ISL_668950, EPI_ISL_668951, EPI_ISL_668952, EPI_ISL_668953, EPI_ISL_668954, EPI_ISL_668955, EPI_ISL_668956 | | | |
| see above | Department of Virus and Microbiological Special Diagnostics, Statens Serum Institut, Copenhagen, Denmark | Albertsen Lab, Department of Chemistry and Bioscience, Aalborg University, Denmark | Danish Covid-19 Genome Consortium |
| EPI_ISL_671883 | National Virus Reference Laboratory | National Virus Reference Laboratory | Michael Carr, Gabriel Gonzalez, Jonathan Dean, Daniel Hare, Cillian F De Gascun |
| EPI_ISL_672655, EPI_ISL_672656, EPI_ISL_672657, EPI_ISL_672658, EPI_ISL_672659 | South Eastern Area Laboratory Services (SEALS) | NSW Health Pathology - Institute of Clinical Pathology and Medical Research; Westmead Hospital; University of Sydney | CIDM-PH et al. |
| EPI_ISL_672752, EPI_ISL_672753, EPI_ISL_672754, EPI_ISL_672755, EPI_ISL_672756, EPI_ISL_672757, EPI_ISL_672758, EPI_ISL_672759, EPI_ISL_672760, EPI_ISL_672761, EPI_ISL_672762, EPI_ISL_672763, EPI_ISL_672764, EPI_ISL_672765, EPI_ISL_672766, EPI_ISL_672767, EPI_ISL_672768, EPI_ISL_672769, EPI_ISL_672770, EPI_ISL_672771, EPI_ISL_672772, EPI_ISL_672773, EPI_ISL_672774, EPI_ISL_672775, EPI_ISL_672776, EPI_ISL_672777, EPI_ISL_672778, EPI_ISL_672779, EPI_ISL_672780, EPI_ISL_672781, EPI_ISL_672782, EPI_ISL_672783, EPI_ISL_672784, EPI_ISL_672785, EPI_ISL_672786, EPI_ISL_672787, EPI_ISL_672788, EPI_ISL_672789, EPI_ISL_672790, EPI_ISL_672791, EPI_ISL_672792, EPI_ISL_672793, EPI_ISL_672794, EPI_ISL_672795, EPI_ISL_672796, EPI_ISL_672797, EPI_ISL_672798, EPI_ISL_672799, EPI_ISL_672800, EPI_ISL_672801, EPI_ISL_672802, EPI_ISL_672803, EPI_ISL_672804, EPI_ISL_672805, EPI_ISL_672806, EPI_ISL_672807, EPI_ISL_672808, EPI_ISL_672809, EPI_ISL_672810, EPI_ISL_672811, EPI_ISL_672812, EPI_ISL_672813, EPI_ISL_672814, EPI_ISL_672815, EPI_ISL_672816, EPI_ISL_672817, EPI_ISL_672818, EPI_ISL_672819, EPI_ISL_672820, EPI_ISL_672821, EPI_ISL_672822, EPI_ISL_672823, EPI_ISL_672824, EPI_ISL_672825, EPI_ISL_672826, EPI_ISL_672827, EPI_ISL_672828, EPI_ISL_672829, EPI_ISL_672830, EPI_ISL_672831, EPI_ISL_672832, EPI_ISL_672833, EPI_ISL_672834, EPI_ISL_672835, EPI_ISL_672836, EPI_ISL_672837, EPI_ISL_672838, EPI_ISL_672839, EPI_ISL_672840, EPI_ISL_672841, EPI_ISL_672842, EPI_ISL_672843, EPI_ISL_672844, EPI_ISL_672845, EPI_ISL_672846, EPI_ISL_672847, EPI_ISL_672848, EPI_ISL_672849, EPI_ISL_672850, EPI_ISL_672851, EPI_ISL_672852, EPI_ISL_672853, EPI_ISL_672854, EPI_ISL_672855, EPI_ISL_672856, EPI_ISL_672857, EPI_ISL_672858, EPI_ISL_672859, EPI_ISL_672860, EPI_ISL_672861, EPI_ISL_672862, EPI_ISL_672863, EPI_ISL_672864, EPI_ISL_672865, EPI_ISL_672866, EPI_ISL_672867, EPI_ISL_672868, EPI_ISL_672869, EPI_ISL_672870, EPI_ISL_672871, EPI_ISL_672872, EPI_ISL_672873, EPI_ISL_672874, EPI_ISL_672875, EPI_ISL_672876, EPI_ISL_672877, EPI_ISL_672878, EPI_ISL_672879, EPI_ISL_672880, EPI_ISL_672881, EPI_ISL_672882, EPI_ISL_672883, EPI_ISL_672884, EPI_ISL_672885, EPI_ISL_672886, EPI_ISL_672887, EPI_ISL_672888, EPI_ISL_672889, EPI_ISL_672890, EPI_ISL_672891, EPI_ISL_672892, EPI_ISL_672893, EPI_ISL_672894, EPI_ISL_672895, EPI_ISL_672896, EPI_ISL_672897, EPI_ISL_672898, EPI_ISL_672899, EPI_ISL_672900, EPI_ISL_672901, EPI_ISL_672902, EPI_ISL_672903, EPI_ISL_672904, EPI_ISL_672905, EPI_ISL_672906, EPI_ISL_672907, EPI_ISL_672908, EPI_ISL_672909, EPI_ISL_672910, EPI_ISL_672911 | | | |
| see above | Lighthouse Lab in Glasgow | Wellcome Sanger Institute for the COVID-19 Genomics UK (COG-UK) Consortium | Harper VanSteenhouse, Yumi Kasai, David Gray, Carol Clugston, Anna Dominiczak and Alex Alderton, Roberto Amato, Sonia Goncalves, Ewan Harrison, David K. Jackson, Ian Johnston, Dominic Kwiatkowski, Cordelia Langford, John Sillitoe on behalf of the Wellcome Sanger Institute COVID-19 Surveillance Team |
| EPI_ISL_673284, EPI_ISL_673288, EPI_ISL_673443, EPI_ISL_673483, EPI_ISL_673597, EPI_ISL_673629, EPI_ISL_673705, EPI_ISL_673771 | Lighthouse Lab in Cambridge | Wellcome Sanger Institute for the COVID-19 Genomics UK (COG-UK) Consortium | Rob Howes, The Lighthouse Lab in Cambridge and Alex Alderton, Roberto Amato, Sonia Goncalves, Ewan Harrison, David K. Jackson, Ian Johnston, Dominic Kwiatkowski, Cordelia Langford, John Sillitoe on behalf of the Wellcome Sanger Institute COVID-19 Surveillance Team |
| EPI_ISL_676511 | Respiratory Virus Unit, Microbiology Services Colindale, Public Health England | COVID-19 Genomics UK (COG-UK) Consortium | PHE Covid Sequencing Team |
| EPI_ISL_678383, EPI_ISL_678384, EPI_ISL_678385 | Area of Virology, Serology and Virology Division (SAViD), New South Wales Health Pathology Randwick | Virology Research Laboratory; Area of Virology, Serology and Virology Division (SAViD), New South Wales Health Pathology Randwick | Foster, C.; Au, J.; Ruiz Silva, M.; Deveson, I.; Bull, R.; Van Hal, S.; Rawlinson, W. |
| EPI_ISL_678548, EPI_ISL_678549, EPI_ISL_678550, EPI_ISL_678551, EPI_ISL_678552, EPI_ISL_678553, EPI_ISL_678554, EPI_ISL_678555, EPI_ISL_678556, EPI_ISL_678557, EPI_ISL_678558, EPI_ISL_678559, EPI_ISL_678560, EPI_ISL_678561, EPI_ISL_678562, EPI_ISL_678563, EPI_ISL_678564, EPI_ISL_678565, EPI_ISL_678567, EPI_ISL_678568, EPI_ISL_678571, EPI_ISL_678572, EPI_ISL_678573, EPI_ISL_678576, EPI_ISL_678577, EPI_ISL_678578, EPI_ISL_678579, EPI_ISL_678580, EPI_ISL_678581, EPI_ISL_678582, EPI_ISL_678583, EPI_ISL_678584, EPI_ISL_678585, EPI_ISL_678586, EPI_ISL_678587, EPI_ISL_678588, EPI_ISL_678589, EPI_ISL_678590, EPI_ISL_678591, EPI_ISL_678592, EPI_ISL_678593, EPI_ISL_678594, EPI_ISL_678595, EPI_ISL_678596, EPI_ISL_678597 | | | |
| see above | Netcare | KRISP, KZN Research Innovation and Sequencing Platform | Giandhari J, Pillay S, Lessells R, ChimukangaraB, Mdlalose K, York D, Khan S, Tegally H, Wilkinson E, de Oliveira T |
| EPI_ISL_678605, EPI_ISL_678607, EPI_ISL_678613, EPI_ISL_678614, EPI_ISL_678616, EPI_ISL_678617, EPI_ISL_678618 | NHLS-IALCH | KRISP, KZN Research Innovation and Sequencing Platform | Giandhari J, Pillay S, Lessells R, ChimukangaraB, Mdlalose K, York D, Khan S, Tegally H, Wilkinson E, de Oliveira T |
| EPI_ISL_678706, EPI_ISL_678709, EPI_ISL_678710, EPI_ISL_678711, EPI_ISL_678713, EPI_ISL_678715, EPI_ISL_678716, EPI_ISL_678717, EPI_ISL_678718, EPI_ISL_678719, EPI_ISL_678720, EPI_ISL_678721, EPI_ISL_678722, EPI_ISL_678723, EPI_ISL_678724, EPI_ISL_678725, EPI_ISL_678726, EPI_ISL_678727, EPI_ISL_678728, EPI_ISL_678729, EPI_ISL_678730, EPI_ISL_678731, EPI_ISL_678732, EPI_ISL_678733, EPI_ISL_678734, EPI_ISL_678735, EPI_ISL_678736, EPI_ISL_678737, EPI_ISL_678738, EPI_ISL_678739, EPI_ISL_678740, EPI_ISL_678741, EPI_ISL_678742, EPI_ISL_678743, EPI_ISL_678744, EPI_ISL_678745, EPI_ISL_678746, EPI_ISL_678749, EPI_ISL_678750, EPI_ISL_678752, EPI_ISL_678753, EPI_ISL_678755, EPI_ISL_678760, EPI_ISL_678761 | | | |
| see above | Respiratory Virus Unit, Microbiology Services Colindale, Public Health England | COVID-19 Genomics UK (COG-UK) Consortium | PHE Covid Sequencing Team |
| EPI_ISL_679162, EPI_ISL_679163, EPI_ISL_679164, EPI_ISL_679165, EPI_ISL_679166, EPI_ISL_679167, EPI_ISL_679168, EPI_ISL_679169, EPI_ISL_679170, EPI_ISL_679171, EPI_ISL_679172, EPI_ISL_679173, EPI_ISL_679174, EPI_ISL_679175, EPI_ISL_679176, EPI_ISL_679179, EPI_ISL_679181, EPI_ISL_679183, EPI_ISL_679185, EPI_ISL_679187, EPI_ISL_679189, EPI_ISL_679191, EPI_ISL_679196, EPI_ISL_679197, EPI_ISL_679198, EPI_ISL_679199, EPI_ISL_679200, EPI_ISL_679201, EPI_ISL_679202, EPI_ISL_679203, EPI_ISL_679204, EPI_ISL_679206, EPI_ISL_679208, EPI_ISL_679210, EPI_ISL_679212, EPI_ISL_679214, EPI_ISL_679216, EPI_ISL_679218 | | | |
| see above | Department of Pathology, University of Cambridge | COVID-19 Genomics UK (COG-UK) Consortium | Aminu S. Jahun, Yasmin Chaudhry, Grant Hall, Iliana Georgana, Myra Hosmillo, Martin D. Curran, Malte Pinckert, Surendra Parmar, Ian Goodfellow |
| EPI_ISL_679292, EPI_ISL_679293, EPI_ISL_679294, EPI_ISL_679296, EPI_ISL_679308, EPI_ISL_679309, EPI_ISL_679310, EPI_ISL_679311 | Queens Medical Centre, Clinical Microbiology Department / DeepSeq Nottingham | COVID-19 Genomics UK (COG-UK) Consortium | Gemma Clark, Wendy Smith, Manjinder Khakh, Vicki M Fleming, Michelle M Lister, Hannah Howson-Wells, Jonathan Ball, Patrick McClure, Joseph Chappell, Theocharis Tsoleridis, Nadine Holmes, Matthew Carlisle, Christopher Moore, Fei Sang, Johnny Debebe, Victoria Wright, Matthew Loose |
| EPI_ISL_679416, EPI_ISL_679417, EPI_ISL_679418, EPI_ISL_679419, EPI_ISL_679420, EPI_ISL_679421, EPI_ISL_679430, EPI_ISL_679431, EPI_ISL_679434, EPI_ISL_679435, EPI_ISL_679436, EPI_ISL_679437, EPI_ISL_679438, EPI_ISL_679439 | | | |
| see above | University College London Hospital | COVID-19 Genomics UK (COG-UK) Consortium | Judith Heaney, Matthew Byott, Catherine Houlihan, Dan Frampton, Stuart Kirk, Moira Spyer and Eleni Nastouli |

| | | | |
|---|---|---|---|
| EPI_ISL_679545, EPI_ISL_679546, EPI_ISL_679547, EPI_ISL_679548, EPI_ISL_679549, EPI_ISL_679575, EPI_ISL_679576, EPI_ISL_679577, EPI_ISL_679578, EPI_ISL_679579, EPI_ISL_679580, EPI_ISL_679581, EPI_ISL_679582, EPI_ISL_679583, EPI_ISL_679585, EPI_ISL_679586, EPI_ISL_679588, EPI_ISL_679589, EPI_ISL_679590, EPI_ISL_679591, EPI_ISL_679592, EPI_ISL_679596 | | | |
| see above | University College London, Great Ormond Street Hospital for Children NHS Foundation Trust, Imperial College Healthcare NHS Trust | COVID-19 Genomics UK (COG-UK) Consortium | Sergi Castellano, Rachel Williams, Mark Kristiansen, Paola Resende Silva, Sunando Roy, Tony Brooks, Helena Tutill, Paola Niola, Patricia Dyal, Charlotte Williams, Leysa Forrest, Yasmin Panchbhaya, Jacqueline Findlay, Samuel Weeks, Julianne Brown, Kathryn Harris, Paul Randell, James Price, Alison Holmes, Judith Breuer |
| EPI_ISL_679933 | Queens Medical Centre, Clinical Microbiology Department / DeepSeq Nottingham | COVID-19 Genomics UK (COG-UK) Consortium | Gemma Clark, Wendy Smith, Manjinder Khakh, Vicki M Fleming, Michelle M Lister, Hannah Howson-Wells, Jonathan Ball, Patrick McClure, Joseph Chappell, Theocharis Tsoleridis, Nadine Holmes, Matthew Carlisle, Christopher Moore, Fei Sang, Johnny Debebe, Victoria Wright, Matthew Loose |
| EPI_ISL_680102, EPI_ISL_680103, EPI_ISL_680104, EPI_ISL_680105, EPI_ISL_680106, EPI_ISL_680107, EPI_ISL_680108, EPI_ISL_680109, EPI_ISL_680113, EPI_ISL_680114, EPI_ISL_680119, EPI_ISL_680128, EPI_ISL_680137, EPI_ISL_680187, EPI_ISL_680192 | | | |
| see above | Virology Department, Sheffield Teaching Hospitals NHS Foundation Trust/Department of Infection, Immunity and Cardiovascular Disease, The Medical School, University of Sheffield | COVID-19 Genomics UK (COG-UK) Consortium | Thushan de Silva, Matthew Parker, Nikki Smith, Adri Angyal, Rebecca Brown, Luke Green, Rachel Tucker, Paul Parsons, Danielle Groves, Katie Johnson, Laura Carrilero, Alex Keeley, Dave Partridge, Matthew Wyles, Benjamin Lindsey, Mehmet Yavuz, Mohammad Raza, Cariad Evans |
| EPI_ISL_680449, EPI_ISL_680450, EPI_ISL_680451, EPI_ISL_680452 | West of Scotland Specialist Virology Centre, NHSGGC / MRC-University of Glasgow Centre for Virus Research | COVID-19 Genomics UK (COG-UK) Consortium | Ana da Silva Filipe, Natasha Johnson, Kathy Smollett, Daniel Mair, Stephen Carmichael, Alice Broos, Lily Tong, Jenna Nichols, Kyriaki Nomikou; Sarah McDonald; Richard Orton, Joseph Hughes, Sreenu Vattipally, David L Robertson; Alasdair MacLean, Rory Gunson; Sharif Shaaban, Matthew Holden; Rachel Blacow, Guy Mollett, Kathy Li, James Shepherd, Antonia Ho, Emma Thomson |
| EPI_ISL_680461, EPI_ISL_680462, EPI_ISL_680463, EPI_ISL_680541, EPI_ISL_680542, EPI_ISL_680543, EPI_ISL_680548, EPI_ISL_680549, EPI_ISL_680550, EPI_ISL_680551, EPI_ISL_680552, EPI_ISL_680553, EPI_ISL_680554, EPI_ISL_680555 | | | |
| see above | Virology Department, Royal Infirmary of Edinburgh, NHS Lothian / School of Biological Sciences, University of Edinburgh / Institute of Genetics and Molecular Medicine, University of Edinburgh | COVID-19 Genomics UK (COG-UK) Consortium | McHugh M, Dewar R, Rooke S, Gallagher M, Balcaza C, O'Toole Á, Scher E, Hill V, McCrone JT, Colquhoun R, Yu X, Jackson B, Rambaut A, Williams TC, Templeton K |
| EPI_ISL_680591, EPI_ISL_680610, EPI_ISL_680654, EPI_ISL_680662, EPI_ISL_680663, EPI_ISL_680664, EPI_ISL_680665, EPI_ISL_680666, EPI_ISL_680667, EPI_ISL_680668, EPI_ISL_680669, EPI_ISL_680670 | | | |
| see above | Wales Specialist Virology Centre Sequencing lab: Pathogen Genomics Unit | COVID-19 Genomics UK (COG-UK) Consortium | Catherine Moore, Johnathan Evans, Laura Gifford, Malorie Perry, Simon Cottrell, Angela Marchbank, Alec Birchley, Alexander Adams, Amy Gaskin, Bree Gatica-Wilcox, Jason Coombes, Joel Southgate, Lauren Gilbert, Lee Graham, Nicole Pacchiarini, Sara Kumziene-Summerhayes, Sarah Taylor, Sophie Jones, Sara Rey, Matthew Bull, Joanne Watkins, Sally Corden, Tom Connor |
| EPI_ISL_681313, EPI_ISL_681314 | Communicable Disease Laboratory, Public Health Directorate | Communicable Disease Laboratory, Public Health Directorate | Alwasti,H., Altaif,Z., AlHujairi,Z., AlAbbas,Z. |
| EPI_ISL_681325, EPI_ISL_681326, EPI_ISL_681327, EPI_ISL_681328, EPI_ISL_681329, EPI_ISL_681330, EPI_ISL_681331, EPI_ISL_681332, EPI_ISL_681333, EPI_ISL_681334, EPI_ISL_681335, EPI_ISL_681336, EPI_ISL_681337, EPI_ISL_681338, EPI_ISL_681339, EPI_ISL_681340, EPI_ISL_681341, EPI_ISL_681342, EPI_ISL_681343, EPI_ISL_681344, EPI_ISL_681345, EPI_ISL_681346, EPI_ISL_681347, EPI_ISL_681348, EPI_ISL_681349, EPI_ISL_681350, EPI_ISL_681351, EPI_ISL_681352, EPI_ISL_681353, EPI_ISL_681354, EPI_ISL_681355, EPI_ISL_681356, EPI_ISL_681357, EPI_ISL_681358, EPI_ISL_681359, EPI_ISL_681360, EPI_ISL_681361, EPI_ISL_681362, EPI_ISL_681363, EPI_ISL_681364, EPI_ISL_681365, EPI_ISL_681366, EPI_ISL_681367, EPI_ISL_681368, EPI_ISL_681369, EPI_ISL_681370, EPI_ISL_681371, EPI_ISL_681372, EPI_ISL_681373, EPI_ISL_681374, EPI_ISL_681375, EPI_ISL_681376, EPI_ISL_681377, EPI_ISL_681378, EPI_ISL_681379, EPI_ISL_681380, EPI_ISL_681381, EPI_ISL_681382, EPI_ISL_681383, EPI_ISL_681384, EPI_ISL_681385, EPI_ISL_681386, EPI_ISL_681387, EPI_ISL_681388, EPI_ISL_681389, EPI_ISL_681390, EPI_ISL_681391, EPI_ISL_681392, EPI_ISL_681393, EPI_ISL_681394, EPI_ISL_681395, EPI_ISL_681396, EPI_ISL_681397, EPI_ISL_681398, EPI_ISL_681399, EPI_ISL_681400, EPI_ISL_681401, EPI_ISL_681402, EPI_ISL_681403, EPI_ISL_681404, EPI_ISL_681405, EPI_ISL_681406, EPI_ISL_681407, EPI_ISL_681408, EPI_ISL_681409, EPI_ISL_681410, EPI_ISL_681411, EPI_ISL_681412, EPI_ISL_681413, EPI_ISL_681414, EPI_ISL_681415, EPI_ISL_681416, EPI_ISL_681417, EPI_ISL_681418, EPI_ISL_681419, EPI_ISL_681420, EPI_ISL_681421, EPI_ISL_681422, EPI_ISL_681423, EPI_ISL_681424, EPI_ISL_681425, EPI_ISL_681426, EPI_ISL_681427, EPI_ISL_681428, EPI_ISL_681429, EPI_ISL_681430, EPI_ISL_681431, EPI_ISL_681432, EPI_ISL_681433, EPI_ISL_681434, EPI_ISL_681435, EPI_ISL_681436, EPI_ISL_681437, EPI_ISL_681438, EPI_ISL_681439, EPI_ISL_681440, EPI_ISL_681441, EPI_ISL_681442, EPI_ISL_681443, EPI_ISL_681444, EPI_ISL_681445, EPI_ISL_681446, EPI_ISL_681447, EPI_ISL_681448, EPI_ISL_681449, EPI_ISL_681450, EPI_ISL_681451, EPI_ISL_681452, EPI_ISL_681453, EPI_ISL_681454, EPI_ISL_681455, EPI_ISL_681456, EPI_ISL_681457, EPI_ISL_681458, EPI_ISL_681459, EPI_ISL_681460, EPI_ISL_681461, EPI_ISL_681462, EPI_ISL_681463, EPI_ISL_681464, EPI_ISL_681465, EPI_ISL_681467, EPI_ISL_681468, EPI_ISL_681469, EPI_ISL_681470, EPI_ISL_681471, EPI_ISL_681472, EPI_ISL_681473, EPI_ISL_681474, EPI_ISL_681475, EPI_ISL_681476, EPI_ISL_681477, EPI_ISL_681478, EPI_ISL_681479, EPI_ISL_681480, EPI_ISL_681481, EPI_ISL_681482, EPI_ISL_681483, EPI_ISL_681484, EPI_ISL_681485, EPI_ISL_681486, EPI_ISL_681487, EPI_ISL_681488, EPI_ISL_681489, EPI_ISL_681490, EPI_ISL_681491, EPI_ISL_681492, EPI_ISL_681493, EPI_ISL_681494, EPI_ISL_681495, EPI_ISL_681496, EPI_ISL_681497, EPI_ISL_681498, EPI_ISL_681499, EPI_ISL_681500, EPI_ISL_681501, EPI_ISL_681502, EPI_ISL_681503, EPI_ISL_681504, EPI_ISL_681505, EPI_ISL_681506, EPI_ISL_681507, EPI_ISL_681508, EPI_ISL_681509, EPI_ISL_681510, EPI_ISL_681511, EPI_ISL_681512, EPI_ISL_681513, EPI_ISL_681514, EPI_ISL_681515, EPI_ISL_681516, EPI_ISL_681517, EPI_ISL_681518, EPI_ISL_681519, EPI_ISL_681520, EPI_ISL_681521, EPI_ISL_681522, EPI_ISL_681523, EPI_ISL_681524, EPI_ISL_681525, EPI_ISL_681526, EPI_ISL_681527, EPI_ISL_681528, EPI_ISL_681529, EPI_ISL_681530, EPI_ISL_681531, EPI_ISL_681532, EPI_ISL_681533, EPI_ISL_681534, EPI_ISL_681535, EPI_ISL_681536, EPI_ISL_681537, EPI_ISL_681538, EPI_ISL_681539, EPI_ISL_681540, EPI_ISL_681541, EPI_ISL_681542, EPI_ISL_681543, EPI_ISL_681544, EPI_ISL_681545, EPI_ISL_681546, EPI_ISL_681547, EPI_ISL_681548, EPI_ISL_681549, EPI_ISL_681550, EPI_ISL_681551, EPI_ISL_681552, EPI_ISL_681553, EPI_ISL_681554, EPI_ISL_681555, EPI_ISL_681556, EPI_ISL_681557, EPI_ISL_681558, EPI_ISL_681559, EPI_ISL_681560, EPI_ISL_681561, EPI_ISL_681562, EPI_ISL_681563, EPI_ISL_681564, EPI_ISL_681565, EPI_ISL_681566, EPI_ISL_681567, EPI_ISL_681568, EPI_ISL_681569, EPI_ISL_681570, EPI_ISL_681571, EPI_ISL_681572, EPI_ISL_681573, EPI_ISL_681574, EPI_ISL_681575, EPI_ISL_681576, EPI_ISL_681577, EPI_ISL_681578, EPI_ISL_681579, EPI_ISL_681580, EPI_ISL_681581, EPI_ISL_681582, EPI_ISL_681583, EPI_ISL_681584, EPI_ISL_681585, EPI_ISL_681586, EPI_ISL_681587, EPI_ISL_681588, EPI_ISL_681589, EPI_ISL_681590, EPI_ISL_681591, EPI_ISL_681592, EPI_ISL_681593, EPI_ISL_681594, EPI_ISL_681595, EPI_ISL_681596, EPI_ISL_681597, EPI_ISL_681598, EPI_ISL_681599, EPI_ISL_681600, EPI_ISL_681601, EPI_ISL_681602, EPI_ISL_681603, EPI_ISL_681604, EPI_ISL_681605, EPI_ISL_681606, EPI_ISL_681607, EPI_ISL_681608, EPI_ISL_681609, EPI_ISL_681610, EPI_ISL_681611, EPI_ISL_681612, EPI_ISL_681613, EPI_ISL_681614, EPI_ISL_681615, EPI_ISL_681616, EPI_ISL_681617, EPI_ISL_681618, EPI_ISL_681619, EPI_ISL_681620, EPI_ISL_681621, EPI_ISL_681622, EPI_ISL_681623, EPI_ISL_681624, EPI_ISL_681625, EPI_ISL_681626, EPI_ISL_681627, EPI_ISL_681628, EPI_ISL_681629, EPI_ISL_681630, EPI_ISL_681631, EPI_ISL_681632, EPI_ISL_681633, EPI_ISL_681634, EPI_ISL_681635, EPI_ISL_681636, EPI_ISL_681637, EPI_ISL_681638, EPI_ISL_681639, EPI_ISL_681640, EPI_ISL_681641, EPI_ISL_681642, EPI_ISL_681643, EPI_ISL_681644, EPI_ISL_681645, EPI_ISL_681646, EPI_ISL_681647, EPI_ISL_681648, EPI_ISL_681649, EPI_ISL_681650, EPI_ISL_681651, EPI_ISL_681652, EPI_ISL_681653, EPI_ISL_681654, EPI_ISL_681655 | | | |
| see above | Lighthouse Lab in Milton Keynes | Wellcome Sanger Institute for the COVID-19 Genomics UK (COG-UK) Consortium | The Lighthouse Lab in Milton Keynes and Alex Alderton, Roberto Amato, Sonia Goncalves, Ewan Harrison, David K. Jackson, Ian Johnston, Dominic Kwiatkowski, Cordelia Langford, John Sillitoe on behalf of the Wellcome Sanger Institute COVID-19 Surveillance Team |
| EPI_ISL_682279 | Middlemore Hospital | Institute of Environmental Science and Research (ESR) | Xiaoyun Ren, Matt Storey, Nikki Freed, Muhammad Faisal, Jing Wang, Hermes Perez, Anja Werno, Antje van der Linden, Arlo Upton, Chris Mansell, David Hammer, Dragana Drinkovic, Gary McAuliffe, Hana Sofia Andersson, James Ussher, Jill Sherwood, Josh Freeman, Julia Howard, Juliet Elvy, Mary DeAlmeida, Matt Blakiston, Matthew Rogers, Max Bloomfield, Michael Addidle, Michelle Balm, Sally Roberts, Sarah Jefferies, Sharmini Muttaiyah, Susan Morpeth, Susan Taylor, Timothy Blackmore, Vani Sathyendran, Veronica Playle, Virginia Hope, Erasmus Smit, Lauren Jelly, Olin Silander, Joep de Ligt |
| EPI_ISL_682315, EPI_ISL_682316, EPI_ISL_682319, EPI_ISL_682320, EPI_ISL_682321 | Communicable Disease Laboratory, Public Health Directorate | Communicable Disease Laboratory, Public Health Directorate | Alwasti,H., Altaif,Z., AlHujairi,Z., AlAbbas,Z. |
| EPI_ISL_682363, EPI_ISL_682364, EPI_ISL_682365, EPI_ISL_682366, EPI_ISL_682367, EPI_ISL_682368, EPI_ISL_682369, EPI_ISL_682370, EPI_ISL_682371, EPI_ISL_682372, EPI_ISL_682373, EPI_ISL_682374, EPI_ISL_682375, EPI_ISL_682376, EPI_ISL_682377, EPI_ISL_682378, EPI_ISL_682379, EPI_ISL_682380, EPI_ISL_682381, EPI_ISL_682382, EPI_ISL_682383, EPI_ISL_682384, EPI_ISL_682385, EPI_ISL_682386, EPI_ISL_682387, EPI_ISL_682388, EPI_ISL_682389, EPI_ISL_682390, EPI_ISL_682391, EPI_ISL_682392, EPI_ISL_682393, EPI_ISL_682394, EPI_ISL_682395, EPI_ISL_682396, EPI_ISL_682397, EPI_ISL_682398, EPI_ISL_682399, EPI_ISL_682400, EPI_ISL_682401, EPI_ISL_682402, EPI_ISL_682403, EPI_ISL_682404, EPI_ISL_682405, EPI_ISL_682406, EPI_ISL_682407, EPI_ISL_682408, EPI_ISL_682409, EPI_ISL_682410, EPI_ISL_682411, EPI_ISL_682412, EPI_ISL_682413, EPI_ISL_682414, EPI_ISL_682415, EPI_ISL_682416, EPI_ISL_682417, EPI_ISL_682418, EPI_ISL_682419, EPI_ISL_682420, EPI_ISL_682421, EPI_ISL_682422, EPI_ISL_682423, EPI_ISL_682424, EPI_ISL_682425, EPI_ISL_682426, EPI_ISL_682427, EPI_ISL_682428, EPI_ISL_682429, EPI_ISL_682430, EPI_ISL_682431, EPI_ISL_682432, EPI_ISL_682433, EPI_ISL_682434, EPI_ISL_682435, EPI_ISL_682436, EPI_ISL_682437, EPI_ISL_682438, EPI_ISL_682439, EPI_ISL_682440, EPI_ISL_682441, EPI_ISL_682442, EPI_ISL_682443, EPI_ISL_682444, EPI_ISL_682445, EPI_ISL_682446, EPI_ISL_682447, EPI_ISL_682448, EPI_ISL_682449, EPI_ISL_682450, EPI_ISL_682451, EPI_ISL_682452, EPI_ISL_682453, EPI_ISL_682454, EPI_ISL_682455, EPI_ISL_682456, EPI_ISL_682457, EPI_ISL_682458, EPI_ISL_682459, EPI_ISL_682460, EPI_ISL_682461, EPI_ISL_682462, EPI_ISL_682463, EPI_ISL_682464, EPI_ISL_682465, EPI_ISL_682466, EPI_ISL_682467, EPI_ISL_682468, EPI_ISL_682469, EPI_ISL_682470, EPI_ISL_682471, EPI_ISL_682472, EPI_ISL_682473, EPI_ISL_682474, EPI_ISL_682475, EPI_ISL_682476, EPI_ISL_682477, EPI_ISL_682478, EPI_ISL_682479, EPI_ISL_682480, EPI_ISL_682481, EPI_ISL_682482, EPI_ISL_682483, EPI_ISL_682484, EPI_ISL_682485, EPI_ISL_682486, EPI_ISL_682487, EPI_ISL_682488, EPI_ISL_682489, EPI_ISL_682490, EPI_ISL_682491, EPI_ISL_682492, EPI_ISL_682493, EPI_ISL_682494, EPI_ISL_682495, EPI_ISL_682496, EPI_ISL_682497, EPI_ISL_682498, EPI_ISL_682499, EPI_ISL_682500, EPI_ISL_682501, EPI_ISL_682502, EPI_ISL_682503, EPI_ISL_682504, EPI_ISL_682505, EPI_ISL_682506, EPI_ISL_682507, EPI_ISL_682508, EPI_ISL_682509, EPI_ISL_682510, EPI_ISL_682511, EPI_ISL_682512, EPI_ISL_682513, EPI_ISL_682514, EPI_ISL_682515, EPI_ISL_682516, EPI_ISL_682517, EPI_ISL_682518, EPI_ISL_682519, EPI_ISL_682520, EPI_ISL_682521, EPI_ISL_682522, EPI_ISL_682523, EPI_ISL_682524, EPI_ISL_682525, EPI_ISL_682526, EPI_ISL_682527, EPI_ISL_682528, EPI_ISL_682529, EPI_ISL_682530, EPI_ISL_682531, EPI_ISL_682532, EPI_ISL_682533, EPI_ISL_682534, EPI_ISL_682535, EPI_ISL_682536, EPI_ISL_682537, EPI_ISL_682538, EPI_ISL_682539, EPI_ISL_682540, EPI_ISL_682541, EPI_ISL_682542, EPI_ISL_682543, EPI_ISL_682544, EPI_ISL_682545, EPI_ISL_682546, EPI_ISL_682547, EPI_ISL_682548, EPI_ISL_682549, EPI_ISL_682550, EPI_ISL_682551, EPI_ISL_682552, EPI_ISL_682553, EPI_ISL_682554, EPI_ISL_682555, EPI_ISL_682556, EPI_ISL_682557, EPI_ISL_682558, EPI_ISL_682559, EPI_ISL_682560, EPI_ISL_682561, EPI_ISL_682562, EPI_ISL_682563, EPI_ISL_682564, EPI_ISL_682565, EPI_ISL_682566, EPI_ISL_682567, EPI_ISL_682568, EPI_ISL_682569, EPI_ISL_682570, EPI_ISL_682571, EPI_ISL_682572, EPI_ISL_682573, EPI_ISL_682574, EPI_ISL_682575, EPI_ISL_682576, EPI_ISL_682577, EPI_ISL_682578, EPI_ISL_682579, EPI_ISL_682580, EPI_ISL_682581, EPI_ISL_682582, EPI_ISL_682583, EPI_ISL_682584, EPI_ISL_682585, EPI_ISL_682586, EPI_ISL_682587, EPI_ISL_682588, EPI_ISL_682589, EPI_ISL_682590, EPI_ISL_682591, EPI_ISL_682592, EPI_ISL_682593, EPI_ISL_682594, EPI_ISL_682595, EPI_ISL_682596, EPI_ISL_682597, EPI_ISL_682598, EPI_ISL_682599, EPI_ISL_682600, EPI_ISL_682601, EPI_ISL_682602, EPI_ISL_682603, EPI_ISL_682604, EPI_ISL_682605, EPI_ISL_682606, EPI_ISL_682607, EPI_ISL_682608, EPI_ISL_682609, EPI_ISL_682610, EPI_ISL_682611, EPI_ISL_682612, EPI_ISL_682613, EPI_ISL_682614, EPI_ISL_682615, EPI_ISL_682616, EPI_ISL_682617, EPI_ISL_682618, EPI_ISL_682619, EPI_ISL_682620, EPI_ISL_682621, EPI_ISL_682622, EPI_ISL_682623, EPI_ISL_682624, EPI_ISL_682625, EPI_ISL_682626, EPI_ISL_682627, EPI_ISL_682628, EPI_ISL_682629, EPI_ISL_682630, EPI_ISL_682631, EPI_ISL_682632, EPI_ISL_682633, EPI_ISL_682634, EPI_ISL_682635, EPI_ISL_682636, EPI_ISL_682637, EPI_ISL_682638, EPI_ISL_682639, EPI_ISL_682640, EPI_ISL_682641, EPI_ISL_682642, EPI_ISL_682643, EPI_ISL_682644, EPI_ISL_682645, EPI_ISL_682646, EPI_ISL_682647, EPI_ISL_682648, EPI_ISL_682649, EPI_ISL_682650, EPI_ISL_682651, EPI_ISL_682652, EPI_ISL_682653, EPI_ISL_682654, EPI_ISL_682655, EPI_ISL_682656, EPI_ISL_682657, EPI_ISL_682658, EPI_ISL_682659, EPI_ISL_682660, EPI_ISL_682661, EPI_ISL_682662, EPI_ISL_682663, EPI_ISL_682664, EPI_ISL_682665, EPI_ISL_682666, EPI_ISL_682667, EPI_ISL_682668, EPI_ISL_682669, EPI_ISL_682670, EPI_ISL_682671, EPI_ISL_682672, EPI_ISL_682673, EPI_ISL_682674, EPI_ISL_682675, EPI_ISL_682676, EPI_ISL_682677, EPI_ISL_682678, EPI_ISL_682679, EPI_ISL_682680, EPI_ISL_682681, EPI_ISL_682682, EPI_ISL_682683, EPI_ISL_682684, EPI_ISL_682685, EPI_ISL_682686, EPI_ISL_682687, EPI_ISL_682688, EPI_ISL_682689, EPI_ISL_682690, EPI_ISL_682691, EPI_ISL_682692, EPI_ISL_682693, EPI_ISL_682694, EPI_ISL_682695, EPI_ISL_682696, EPI_ISL_682697, EPI_ISL_682698, EPI_ISL_682699, EPI_ISL_682700, EPI_ISL_682701, EPI_ISL_682702, EPI_ISL_682703, EPI_ISL_682704, EPI_ISL_682705, EPI_ISL_682706, EPI_ISL_682707, EPI_ISL_682708, EPI_ISL_682709, EPI_ISL_682710, EPI_ISL_682711, EPI_ISL_682712, EPI_ISL_682713, EPI_ISL_682714, EPI_ISL_682715, EPI_ISL_682716, EPI_ISL_682717, EPI_ISL_682718, EPI_ISL_682719, EPI_ISL_682720, EPI_ISL_682721, EPI_ISL_682722, EPI_ISL_682723, EPI_ISL_682724, EPI_ISL_682725, EPI_ISL_682726, EPI_ISL_682727, EPI_ISL_682728, EPI_ISL_682729, EPI_ISL_682730, EPI_ISL_682731, EPI_ISL_682732, EPI_ISL_682733, EPI_ISL_682734, EPI_ISL_682735, EPI_ISL_682736, EPI_ISL_682737, EPI_ISL_682738, EPI_ISL_682739, EPI_ISL_682740, EPI_ISL_682741, EPI_ISL_682742, EPI_ISL_682743, EPI_ISL_682744, EPI_ISL_682745, EPI_ISL_682746, EPI_ISL_682747, EPI_ISL_682748, EPI_ISL_682749, EPI_ISL_682750, EPI_ISL_682751, EPI_ISL_682752, EPI_ISL_682753, EPI_ISL_682754, EPI_ISL_682755, EPI_ISL_682756, EPI_ISL_682757, EPI_ISL_682758, EPI_ISL_682759, EPI_ISL_682760, EPI_ISL_682761, EPI_ISL_682762, EPI_ISL_682763, EPI_ISL_682764, EPI_ISL_682765, EPI_ISL_682766, EPI_ISL_682767, EPI_ISL_682768, EPI_ISL_682769, EPI_ISL_682770, EPI_ISL_682771, EPI_ISL_682772, EPI_ISL_682773, EPI_ISL_682774, EPI_ISL_682775, EPI_ISL_682776, EPI_ISL_682777, EPI_ISL_682778, EPI_ISL_682779, EPI_ISL_682780, EPI_ISL_682781, EPI_ISL_682782, EPI_ISL_682783, EPI_ISL_682784, EPI_ISL_682785, EPI_ISL_682786, EPI_ISL_682787, EPI_ISL_682788, EPI_ISL_682789, EPI_ISL_682790, EPI_ISL_682791, EPI_ISL_682792, EPI_ISL_682793, EPI_ISL_682794, EPI_ISL_682795, EPI_ISL_682796, EPI_ISL_682797, EPI_ISL_682798, EPI_ISL_682799, EPI_ISL_682800, EPI_ISL_682801, EPI_ISL_682802, EPI_ISL_682803, EPI_ISL_682804, EPI_ISL_682805, EPI_ISL_682806, EPI_ISL_682807, EPI_ISL_682808, EPI_ISL_682809, EPI_ISL_682810, EPI_ISL_682811, EPI_ISL_682812, EPI_ISL_682813, EPI_ISL_682814, EPI_ISL_682815, EPI_ISL_682816, EPI_ISL_682817, EPI_ISL_682818, EPI_ISL_682819, EPI_ISL_682820, EPI_ISL_682821, EPI_ISL_682822, EPI_ISL_682823, EPI_ISL_682824, EPI_ISL_682825, EPI_ISL_682826, EPI_ISL_682827, EPI_ISL_682828, EPI_ISL_682829, EPI_ISL_682830, EPI_ISL_682831, EPI_ISL_682832, EPI_ISL_682833, EPI_ISL_682834, EPI_ISL_682835, EPI_ISL_682836, EPI_ISL_682837, EPI_ISL_682838, EPI_ISL_682839, EPI_ISL_682840, EPI_ISL_682841, EPI_ISL_682842, EPI_ISL_682843, EPI_ISL_682844, EPI_ISL_682845, EPI_ISL_682846, EPI_ISL_682847, EPI_ISL_682848, | | | |

| | | | |
|---|---|---|---|
| EPI_ISL_682849, EPI_ISL_682850, EPI_ISL_682851, EPI_ISL_682852, EPI_ISL_682853, EPI_ISL_682854, EPI_ISL_682855, EPI_ISL_682856, EPI_ISL_682857, EPI_ISL_682858, EPI_ISL_682859, EPI_ISL_682860, EPI_ISL_682861, EPI_ISL_682862, EPI_ISL_682863, EPI_ISL_682864, EPI_ISL_682865, EPI_ISL_682866, EPI_ISL_682867, EPI_ISL_682868, EPI_ISL_682869, EPI_ISL_682870, EPI_ISL_682871, EPI_ISL_682872, EPI_ISL_682873, EPI_ISL_682874, EPI_ISL_682875, EPI_ISL_682876, EPI_ISL_682877, EPI_ISL_682878, EPI_ISL_682879, EPI_ISL_682880, EPI_ISL_682881, EPI_ISL_682882, EPI_ISL_682883, EPI_ISL_682884, EPI_ISL_682885, EPI_ISL_682886, EPI_ISL_682887, EPI_ISL_682888, EPI_ISL_682889, EPI_ISL_682890, EPI_ISL_682891, EPI_ISL_682892, EPI_ISL_682893, EPI_ISL_682894, EPI_ISL_682895, EPI_ISL_682896, EPI_ISL_682897, EPI_ISL_682898, EPI_ISL_682899, EPI_ISL_682900, EPI_ISL_682901, EPI_ISL_682902, EPI_ISL_682903, EPI_ISL_682904, EPI_ISL_682905, EPI_ISL_682906, EPI_ISL_682907, EPI_ISL_682908, EPI_ISL_682909, EPI_ISL_682910, EPI_ISL_682911, EPI_ISL_682912, EPI_ISL_682913, EPI_ISL_682914, EPI_ISL_682915, EPI_ISL_682916, EPI_ISL_682917, EPI_ISL_682918, EPI_ISL_682919, EPI_ISL_682920, EPI_ISL_682921, EPI_ISL_682922, EPI_ISL_682923, EPI_ISL_682924, EPI_ISL_682925, EPI_ISL_682926, EPI_ISL_682927, EPI_ISL_682928, EPI_ISL_682929, EPI_ISL_683161, EPI_ISL_683162, EPI_ISL_683163, EPI_ISL_683164, EPI_ISL_683165, EPI_ISL_683166, EPI_ISL_683255, EPI_ISL_683256, EPI_ISL_683257, EPI_ISL_683258, EPI_ISL_683259, EPI_ISL_683260 | | | |
| see above | Department of Virus and Microbiological Special Diagnostics, Statens Serum Institut, Copenhagen, Denmark | Albertsen Lab, Department of Chemistry and Bioscience, Aalborg University, Denmark | Danish Covid-19 Genome Consortium |
| EPI_ISL_683457, EPI_ISL_683475, EPI_ISL_683476, EPI_ISL_683478, EPI_ISL_683484, EPI_ISL_683489, EPI_ISL_683490, EPI_ISL_683534, EPI_ISL_683535, EPI_ISL_683536, EPI_ISL_683537, EPI_ISL_683538, EPI_ISL_683539, EPI_ISL_683540, EPI_ISL_683541, EPI_ISL_683542, EPI_ISL_683543, EPI_ISL_683544, EPI_ISL_683545, EPI_ISL_683546, EPI_ISL_683547, EPI_ISL_683548, EPI_ISL_683549, EPI_ISL_683550, EPI_ISL_683551, EPI_ISL_683552, EPI_ISL_683553, EPI_ISL_683554, EPI_ISL_683555, EPI_ISL_683556, EPI_ISL_683557, EPI_ISL_683558, EPI_ISL_683559, EPI_ISL_683560, EPI_ISL_683561, EPI_ISL_683562, EPI_ISL_683563, EPI_ISL_683564, EPI_ISL_683565 | | | |
| see above | Respiratory Virus Unit, Microbiology Services Colindale, Public Health England | COVID-19 Genomics UK (COG-UK) Consortium | PHE Covid Sequencing Team |
| EPI_ISL_683608, EPI_ISL_683609, EPI_ISL_683623, EPI_ISL_683636 | Servicio de Microbiología, Laboratori Clínic Metropolitana Nord. Hospital Universitari Germans Trias i Pujol. Institut d'Investigació en Ciències de la Salut Germans Trias i Pujol (IGTP) | SeqCOVID-SPAIN consortium/IBV(CSIC) | Elisa Martró, Antoni E. Bordoy, Anna Not, Adrián Antuori, Anabel Fernández, Nona Romaní, Verónica Saludes, Cristina Casañ and SeqCOVID-SPAIN consortium |
| EPI_ISL_683689 | Hennepin County Medical Center | Minnesota Department of Health, Public Health Laboratory | Alexandra Lorentz, Jacob Garfin, Matt Plumb, and Xiong Wang |
| EPI_ISL_683729 | Minnesota Department of Health, Public Health Laboratory | Minnesota Department of Health, Public Health Laboratory | Alexandra Lorentz, Jacob Garfin, Matt Plumb, and Xiong Wang |
| EPI_ISL_683753 | DOHMH Corona | New York City Public Health Laboratory | Jade Wang, et al. |
| EPI_ISL_683755 | DOHMH PHL | New York City Public Health Laboratory | Jade Wang, et al. |
| EPI_ISL_683756 | DOHMH Central Harlem | New York City Public Health Laboratory | Jade Wang, et al. |
| EPI_ISL_683757, EPI_ISL_683758 | DOHMH Corona | New York City Public Health Laboratory | Jade Wang, et al. |
| EPI_ISL_683759 | DOHMH PHL | New York City Public Health Laboratory | Jade Wang, et al. |
| EPI_ISL_683764, EPI_ISL_683779 | DOHMH Corona | New York City Public Health Laboratory | Jade Wang, et al. |
| EPI_ISL_683780 | DOHMH Chelsea | New York City Public Health Laboratory | Jade Wang, et al. |
| EPI_ISL_683781, EPI_ISL_683782 | DOHMH Jamaica | New York City Public Health Laboratory | Jade Wang, et al. |
| EPI_ISL_683783 | DOHMH PHL | New York City Public Health Laboratory | Jade Wang, et al. |
| EPI_ISL_683784 | DOHMH Riverside | New York City Public Health Laboratory | Jade Wang, et al. |
| EPI_ISL_683785 | DOHMH Corona | New York City Public Health Laboratory | Jade Wang, et al. |
| EPI_ISL_683786 | DOHMH Jamaica | New York City Public Health Laboratory | Jade Wang, et al. |
| EPI_ISL_683787 | DOHMH PHL | New York City Public Health Laboratory | Jade Wang, et al. |
| EPI_ISL_683788 | DOHMH Chelsea | New York City Public Health Laboratory | Jade Wang, et al. |
| EPI_ISL_683789 | DOHMH PHL | New York City Public Health Laboratory | Jade Wang, et al. |
| EPI_ISL_683790 | DOHMH Morrisania | New York City Public Health Laboratory | Jade Wang, et al. |
| EPI_ISL_683791 | DOHMH Jamaica | New York City Public Health Laboratory | Jade Wang, et al. |
| EPI_ISL_683792 | DOHMH Chelsea | New York City Public Health Laboratory | Jade Wang, et al. |
| EPI_ISL_683837 | DOHMH Jamaica | New York City Public Health Laboratory | Jade Wang, et al. |
| EPI_ISL_683842 | DOHMH Central Harlem | New York City Public Health Laboratory | Jade Wang, et al. |
| EPI_ISL_683847 | DOHMH Corona | New York City Public Health Laboratory | Jade Wang, et al. |
| EPI_ISL_683869 | DOHMH Morrisania | New York City Public Health Laboratory | Jade Wang, et al. |
| EPI_ISL_683871, EPI_ISL_683875 | DOHMH PHL | New York City Public Health Laboratory | Jade Wang, et al. |
| EPI_ISL_683881 | DOHMH Corona | New York City Public Health Laboratory | Jade Wang, et al. |
| EPI_ISL_683927, EPI_ISL_683928 | DOHMH Central Harlem | New York City Public Health Laboratory | Jade Wang, et al. |
| EPI_ISL_683929 | DOHMH Corona | New York City Public Health Laboratory | Jade Wang, et al. |
| EPI_ISL_683930 | DOHMH Jamaica | New York City Public Health Laboratory | Jade Wang, et al. |
| EPI_ISL_683931 | DOHMH Chelsea | New York City Public Health Laboratory | Jade Wang, et al. |
| EPI_ISL_683932 | DOHMH Corona | New York City Public Health Laboratory | Jade Wang, et al. |
| EPI_ISL_683933 | DOHMH Central Harlem | New York City Public Health Laboratory | Jade Wang, et al. |
| EPI_ISL_683934 | DOHMH Corona | New York City Public Health Laboratory | Jade Wang, et al. |
| EPI_ISL_683935 | DOHMH Crown Heights | New York City Public Health Laboratory | Jade Wang, et al. |
| EPI_ISL_683936 | DOHMH Chelsea | New York City Public Health Laboratory | Jade Wang, et al. |
| EPI_ISL_683978, EPI_ISL_683979 | DOHMH Corona | New York City Public Health Laboratory | Jade Wang, et al. |
| EPI_ISL_683980, EPI_ISL_683981 | DOHMH Chelsea | New York City Public Health Laboratory | Jade Wang, et al. |
| EPI_ISL_683982, EPI_ISL_683983, EPI_ISL_683984, EPI_ISL_683985 | DOHMH Central Harlem | New York City Public Health Laboratory | Jade Wang, et al. |
| EPI_ISL_683986 | DOHMH Jamaica | New York City Public Health Laboratory | Jade Wang, et al. |
| EPI_ISL_683987, EPI_ISL_683988 | DOHMH PHL | New York City Public Health Laboratory | Jade Wang, et al. |
| EPI_ISL_683989 | DOHMH Chelsea | New York City Public Health Laboratory | Jade Wang, et al. |
| EPI_ISL_683990 | DOHMH Jamaica | New York City Public Health Laboratory | Jade Wang, et al. |
| EPI_ISL_683991 | DOHMH Central Harlem | New York City Public Health Laboratory | Jade Wang, et al. |
| EPI_ISL_683992 | DOHMH Morrisania | New York City Public Health Laboratory | Jade Wang, et al. |
| EPI_ISL_683993 | DOHMH PHL | New York City Public Health Laboratory | Jade Wang, et al. |

| | | | |
|---|---|---|---|
| EPI_ISL_683994 | DOHMH Riverside | New York City Public Health Laboratory | Jade Wang, et al. |
| EPI_ISL_683995 | DOHMH PHL | New York City Public Health Laboratory | Jade Wang, et al. |
| EPI_ISL_683996 | DOHMH Fort Greene | New York City Public Health Laboratory | Jade Wang, et al. |
| EPI_ISL_684028, EPI_ISL_684029, EPI_ISL_684036 | Communicable Disease Laboratory, Public Health Directorate | Communicable Disease Laboratory, Public Health Directorate | Alwasti,H., Altaif,Z., AlHujairi,Z., AlAbbas,Z. |
| EPI_ISL_686601, EPI_ISL_686602, EPI_ISL_686603, EPI_ISL_686604, EPI_ISL_686605, EPI_ISL_686606, EPI_ISL_686607, EPI_ISL_686608, EPI_ISL_686609, EPI_ISL_686610 | Respiratory Virus Unit, Microbiology Services Colindale, Public Health England | COVID-19 Genomics UK (COG-UK) Consortium | PHE Covid Sequencing Team |
| EPI_ISL_692774, EPI_ISL_692776, EPI_ISL_692777, EPI_ISL_692778, EPI_ISL_692779, EPI_ISL_692780, EPI_ISL_692782, EPI_ISL_692785 | Massachusetts State Public Health Laboratory | Massachusetts State Public Health Laboratory | Andrew Lang, Timelia Fink, Glen Gallagher, Sandra Smole |
| EPI_ISL_693303 | Department of Laboratory Medicine, National Taiwan University Hospital | Microbial Genomics Core Lab, National Taiwan University Centers of Genomic and Precision Medicine | Shiou-Hwei Yeh, You-Yu Lin, Ya-Yun Lai, Chiao-Ling Li, Shan-Chwen Chang, Pei-Jer Chen, Sui-Yuan Chang |
| EPI_ISL_693307, EPI_ISL_693308, EPI_ISL_693309, EPI_ISL_693310, EPI_ISL_693311, EPI_ISL_693312, EPI_ISL_693314, EPI_ISL_693317 | National Public Health Laboratory, National Centre for Infectious Diseases | National Public Health Laboratory, National Centre for Infectious Diseases | Tze Minn Mak, Sophie Octavia, Zhenyang Zhou, Lin Cui, Raymond Tzer Pin Lin |
| EPI_ISL_693332, EPI_ISL_693333 | Lighthouse Lab in Alderley Park | Wellcome Sanger Institute for the COVID-19 Genomics UK (COG-UK) Consortium | Jacquelyn Wynn, Mairead Hyland, The Lighthouse Lab in Alderley Park and Alex Alderton, Roberto Amato, Sonia Goncalves, Ewan Harrison, David K. Jackson, Ian Johnston, Dominic Kwiatkowski, Cordelia Langford, John Sillitoe on behalf of the Wellcome Sanger Institute COVID-19 Surveillance Team |
| EPI_ISL_693397, EPI_ISL_693404, EPI_ISL_693406, EPI_ISL_693407, EPI_ISL_693461, EPI_ISL_693462, EPI_ISL_693463, EPI_ISL_693464, EPI_ISL_693465, EPI_ISL_693466, EPI_ISL_693468 | | | |
| see above | Respiratory Virus Unit, Microbiology Services Colindale, Public Health England | COVID-19 Genomics UK (COG-UK) Consortium | PHE Covid Sequencing Team |
| EPI_ISL_693770 | General practitioner | National Reference Center for Viruses of Respiratory Infections, Institut Pasteur, Paris | Marion Barbet, Sylvie Behillil, Méline Bizard, Angela Brisebarre, Camille Capel, Etienne Simon-Lorière, Vincent Enouf, Maud Vanpeene, Sylvie van der Werf |
| EPI_ISL_693948, EPI_ISL_693949, EPI_ISL_693950, EPI_ISL_693951, EPI_ISL_693952, EPI_ISL_693953, EPI_ISL_693954, EPI_ISL_693955, EPI_ISL_693956, EPI_ISL_693957, EPI_ISL_693958, EPI_ISL_693959, EPI_ISL_693960, EPI_ISL_693961, EPI_ISL_693962, EPI_ISL_693963, EPI_ISL_693964, EPI_ISL_693965, EPI_ISL_693966, EPI_ISL_693967, EPI_ISL_693968, EPI_ISL_693969, EPI_ISL_693970, EPI_ISL_693971, EPI_ISL_693972, EPI_ISL_693973, EPI_ISL_693974, EPI_ISL_693975, EPI_ISL_693976, EPI_ISL_693977, EPI_ISL_693978, EPI_ISL_693979, EPI_ISL_693980, EPI_ISL_693981, EPI_ISL_693982, EPI_ISL_693983, EPI_ISL_693984, EPI_ISL_693985, EPI_ISL_693986, EPI_ISL_693987, EPI_ISL_693988, EPI_ISL_693989, EPI_ISL_693990, EPI_ISL_693991, EPI_ISL_693992, EPI_ISL_693993, EPI_ISL_693994, EPI_ISL_693995, EPI_ISL_693996, EPI_ISL_693997, EPI_ISL_693998, EPI_ISL_693999, EPI_ISL_694000, EPI_ISL_694001, EPI_ISL_694002, EPI_ISL_694003, EPI_ISL_694004, EPI_ISL_694005, EPI_ISL_694006, EPI_ISL_694007, EPI_ISL_694008 | | | |
| see above | Viollier AG | Department of Biosystems Science and Engineering, ETH Zürich | Christian Beisel, Sarah Nadeau, Chaoran Chen, Ivan Topolsky, Pedro Ferreira, Philipp Jablonski, Susana Posada-Céspedes, Tobias Schär, Ina Nissen, Natascha Santacroce, Elodie Burcklen, Christiane Beckmann, Maurice Redondo, Olivier Kobel, Christoph Noppen, Sophie Seidel, Noemie Santamaria de Souza, Niko Beerenwinkel, Tanja Stadler |
| EPI_ISL_696452, EPI_ISL_696455 | Thembalethu CDC wc THC & NHLS/UCT | KRISP, KZN Research Innovation and Sequencing Platform | Arash Iranzadeh, Deelan Doolabh, Lynn Tyers, Bruna Galvao, Innocent Mudau, Marvin Hsiao, Kruger Marais, Jennifer Giandhari, Sureshnee Pillay, Houriiyah Tegally, Emanuel James San, Tulio de Oliveira, Diana Hardie, Stephen Korsman, Carolyn Williamson |
| EPI_ISL_696457 | Hornlee Clinic wc HLC & NHLS/UCT | KRISP, KZN Research Innovation and Sequencing Platform | Arash Iranzadeh, Deelan Doolabh, Lynn Tyers, Bruna Galvao, Innocent Mudau, Marvin Hsiao, Kruger Marais, Jennifer Giandhari, Sureshnee Pillay, Houriiyah Tegally, Emanuel James San, Tulio de Oliveira, Diana Hardie, Stephen Korsman, Carolyn Williamson |
| EPI_ISL_696461 | Thembalethu CDC wc THC & NHLS/UCT | KRISP, KZN Research Innovation and Sequencing Platform | Arash Iranzadeh, Deelan Doolabh, Lynn Tyers, Bruna Galvao, Innocent Mudau, Marvin Hsiao, Kruger Marais, Jennifer Giandhari, Sureshnee Pillay, Houriiyah Tegally, Emanuel James San, Tulio de Oliveira, Diana Hardie, Stephen Korsman, Carolyn Williamson |
| EPI_ISL_696462 | Sedgefield Clinic wc SGE & NHLS/UCT | KRISP, KZN Research Innovation and Sequencing Platform | Arash Iranzadeh, Deelan Doolabh, Lynn Tyers, Bruna Galvao, Innocent Mudau, Marvin Hsiao, Kruger Marais, Jennifer Giandhari, Sureshnee Pillay, Houriiyah Tegally, Emanuel James San, Tulio de Oliveira, Diana Hardie, Stephen Korsman, Carolyn Williamson |
| EPI_ISL_696463 | Knysna Hospital wc KNY & NHLS/UCT | KRISP, KZN Research Innovation and Sequencing Platform | Arash Iranzadeh, Deelan Doolabh, Lynn Tyers, Bruna Galvao, Innocent Mudau, Marvin Hsiao, Kruger Marais, Jennifer Giandhari, Sureshnee Pillay, Houriiyah Tegally, Emanuel James San, Tulio de Oliveira, Diana Hardie, Stephen Korsman, Carolyn Williamson |
| EPI_ISL_696464 | Groote Schuur Hospital wc GSH & NHLS/UCT | KRISP, KZN Research Innovation and Sequencing Platform | Arash Iranzadeh, Deelan Doolabh, Lynn Tyers, Bruna Galvao, Innocent Mudau, Marvin Hsiao, Kruger Marais, Jennifer Giandhari, Sureshnee Pillay, Houriiyah Tegally, Emanuel James San, Tulio de Oliveira, Diana Hardie, Stephen Korsman, Carolyn Williamson |
| EPI_ISL_696465 | Kwanokuthula CDC wc KWA & NHLS/UCT | KRISP, KZN Research Innovation and Sequencing Platform | Arash Iranzadeh, Deelan Doolabh, Lynn Tyers, Bruna Galvao, Innocent Mudau, Marvin Hsiao, Kruger Marais, Jennifer Giandhari, Sureshnee Pillay, Houriiyah Tegally, Emanuel James San, Tulio de Oliveira, Diana Hardie, Stephen Korsman, Carolyn Williamson |
| EPI_ISL_696466 | Great Brak River Clinic wc GBC & NHLS/UCT | KRISP, KZN Research Innovation and Sequencing Platform | Arash Iranzadeh, Deelan Doolabh, Lynn Tyers, Bruna Galvao, Innocent Mudau, Marvin Hsiao, Kruger Marais, Jennifer Giandhari, Sureshnee Pillay, Houriiyah Tegally, Emanuel James San, Tulio de Oliveira, Diana Hardie, Stephen Korsman, Carolyn Williamson |
| EPI_ISL_696467 | Knysna CDC wc WLC & NHLS/UCT | KRISP, KZN Research Innovation and Sequencing Platform | Arash Iranzadeh, Deelan Doolabh, Lynn Tyers, Bruna Galvao, Innocent Mudau, Marvin Hsiao, Kruger Marais, Jennifer Giandhari, Sureshnee Pillay, Houriiyah Tegally, Emanuel James San, Tulio de Oliveira, Diana Hardie, Stephen Korsman, Carolyn Williamson |
| EPI_ISL_696468 | New Horizon Clinic wc NZC & NHLS/UCT | KRISP, KZN Research Innovation and Sequencing Platform | Arash Iranzadeh, Deelan Doolabh, Lynn Tyers, Bruna Galvao, Innocent Mudau, Marvin Hsiao, Kruger Marais, Jennifer Giandhari, Sureshnee Pillay, Houriiyah Tegally, Emanuel James San, Tulio de Oliveira, Diana Hardie, Stephen Korsman, Carolyn Williamson |
| EPI_ISL_696469 | Knysna Hospital wc KNY & NHLS/UCT | KRISP, KZN Research Innovation and Sequencing Platform | Arash Iranzadeh, Deelan Doolabh, Lynn Tyers, Bruna Galvao, Innocent Mudau, Marvin Hsiao, Kruger Marais, Jennifer Giandhari, Sureshnee Pillay, Houriiyah Tegally, Emanuel James San, Tulio de Oliveira, Diana Hardie, Stephen Korsman, Carolyn Williamson |
| EPI_ISL_696470 | Pacaltsdorp Clinic wc PAC & NHLS/UCT | KRISP, KZN Research Innovation and Sequencing Platform | Arash Iranzadeh, Deelan Doolabh, Lynn Tyers, Bruna Galvao, Innocent Mudau, Marvin Hsiao, Kruger Marais, Jennifer Giandhari, Sureshnee Pillay, Houriiyah Tegally, Emanuel James San, Tulio de Oliveira, Diana Hardie, Stephen Korsman, Carolyn Williamson |
| EPI_ISL_696471 | George Hospital wc GRH & NHLS/UCT | KRISP, KZN Research Innovation and Sequencing Platform | Arash Iranzadeh, Deelan Doolabh, Lynn Tyers, Bruna Galvao, Innocent Mudau, Marvin Hsiao, Kruger Marais, Jennifer Giandhari, Sureshnee Pillay, Houriiyah Tegally, Emanuel James San, Tulio de Oliveira, Diana Hardie, Stephen Korsman, Carolyn Williamson |
| EPI_ISL_696472 | Conville CDC wc CVC & NHLS/UCT | KRISP, KZN Research Innovation and Sequencing Platform | Arash Iranzadeh, Deelan Doolabh, Lynn Tyers, Bruna Galvao, Innocent Mudau, Marvin Hsiao, Kruger Marais, Jennifer Giandhari, Sureshnee Pillay, Houriiyah Tegally, Emanuel James San, Tulio de Oliveira, Diana Hardie, Stephen Korsman, Carolyn Williamson |
| EPI_ISL_696473 | George Hospital wc GRH & NHLS/UCT | KRISP, KZN Research Innovation and Sequencing Platform | Arash Iranzadeh, Deelan Doolabh, Lynn Tyers, Bruna Galvao, Innocent Mudau, Marvin Hsiao, Kruger Marais, Jennifer Giandhari, Sureshnee Pillay, Houriiyah Tegally, Emanuel James San, Tulio de Oliveira, Diana Hardie, Stephen Korsman, Carolyn Williamson |
| EPI_ISL_696474 | Pacaltsdorp Clinic wc PAC & NHLS/UCT | KRISP, KZN Research Innovation and Sequencing Platform | Arash Iranzadeh, Deelan Doolabh, Lynn Tyers, Bruna Galvao, Innocent Mudau, Marvin Hsiao, Kruger Marais, Jennifer Giandhari, Sureshnee Pillay, Houriiyah Tegally, Emanuel James San, Tulio de Oliveira, Diana Hardie, Stephen Korsman, Carolyn Williamson |
| EPI_ISL_696475 | Sedgefield Clinic wc SGE & NHLS/UCT | KRISP, KZN Research Innovation and Sequencing Platform | Arash Iranzadeh, Deelan Doolabh, Lynn Tyers, Bruna Galvao, Innocent Mudau, Marvin Hsiao, Kruger Marais, Jennifer Giandhari, Sureshnee Pillay, Houriiyah Tegally, Emanuel James San, Tulio de Oliveira, Diana Hardie, Stephen Korsman, Carolyn Williamson |
| EPI_ISL_696476 | Plettenberg Bay Clinic wc PLC & NHLS/UCT | KRISP, KZN Research Innovation and Sequencing Platform | Arash Iranzadeh, Deelan Doolabh, Lynn Tyers, Bruna Galvao, Innocent Mudau, Marvin Hsiao, Kruger Marais, Jennifer Giandhari, Sureshnee Pillay, Houriiyah Tegally, Emanuel James San, Tulio de Oliveira, Diana Hardie, Stephen Korsman, Carolyn Williamson |
| EPI_ISL_696477, EPI_ISL_696478 | Knysna Hospital wc KNY & NHLS/UCT | KRISP, KZN Research Innovation and Sequencing Platform | Arash Iranzadeh, Deelan Doolabh, Lynn Tyers, Bruna Galvao, Innocent Mudau, Marvin Hsiao, Kruger Marais, Jennifer Giandhari, Sureshnee Pillay, Houriiyah Tegally, Emanuel James San, Tulio de Oliveira, Diana Hardie, Stephen Korsman, Carolyn Williamson |

| | | | |
|---|---|---|---|
| EPI_ISL_696480 | Crags Clinic wc CRG & NHLS/UCT | KRISP, KZN Research Innovation and Sequencing Platform | Arash Iranzadeh, Deelan Doolabh, Lynn Tyers, Bruna Galvao, Innocent Mudau, Marvin Hsiao, Kruger Marais, Jennifer Giandhari, Sureshnee Pillay, Houriiyah Tegally, Emanuel James San, Tulio de Oliveira, Diana Hardie, Stephen Korsman, Carolyn Williamson |
| EPI_ISL_696481 | George Hospital wc GRH & NHLS/UCT | KRISP, KZN Research Innovation and Sequencing Platform | Arash Iranzadeh, Deelan Doolabh, Lynn Tyers, Bruna Galvao, Innocent Mudau, Marvin Hsiao, Kruger Marais, Jennifer Giandhari, Sureshnee Pillay, Houriiyah Tegally, Emanuel James San, Tulio de Oliveira, Diana Hardie, Stephen Korsman, Carolyn Williamson |
| EPI_ISL_696485 | Touwsranten Clinic wc TST & NHLS/UCT | KRISP, KZN Research Innovation and Sequencing Platform | Arash Iranzadeh, Deelan Doolabh, Lynn Tyers, Bruna Galvao, Innocent Mudau, Marvin Hsiao, Kruger Marais, Jennifer Giandhari, Sureshnee Pillay, Houriiyah Tegally, Emanuel James San, Tulio de Oliveira, Diana Hardie, Stephen Korsman, Carolyn Williamson |
| EPI_ISL_696491 | Knysna Hospital wc KNY & NHLS/UCT | KRISP, KZN Research Innovation and Sequencing Platform | Arash Iranzadeh, Deelan Doolabh, Lynn Tyers, Bruna Galvao, Innocent Mudau, Marvin Hsiao, Kruger Marais, Jennifer Giandhari, Sureshnee Pillay, Houriiyah Tegally, Emanuel James San, Tulio de Oliveira, Diana Hardie, Stephen Korsman, Carolyn Williamson |
| EPI_ISL_701675, EPI_ISL_701681, EPI_ISL_701683, EPI_ISL_701697, EPI_ISL_701698, EPI_ISL_701701, EPI_ISL_701713, EPI_ISL_701714, EPI_ISL_701722, EPI_ISL_701726, EPI_ISL_701729, EPI_ISL_701737, EPI_ISL_701739, EPI_ISL_701743, EPI_ISL_701746, EPI_ISL_701749, EPI_ISL_701754, EPI_ISL_701759, EPI_ISL_701762, EPI_ISL_701774, EPI_ISL_701775, EPI_ISL_701781, EPI_ISL_701788, EPI_ISL_701792, EPI_ISL_701795, EPI_ISL_701800, EPI_ISL_701803, EPI_ISL_701819, EPI_ISL_701829, EPI_ISL_701837, EPI_ISL_701838, EPI_ISL_701839, EPI_ISL_701841, EPI_ISL_701843, EPI_ISL_701847, EPI_ISL_701848, EPI_ISL_701850, EPI_ISL_701853, EPI_ISL_701870, EPI_ISL_701871, EPI_ISL_701875, EPI_ISL_701876, EPI_ISL_701878, EPI_ISL_701881, EPI_ISL_701913, EPI_ISL_701914, EPI_ISL_701916, EPI_ISL_701920, EPI_ISL_701923, EPI_ISL_701927, EPI_ISL_701930, EPI_ISL_701932, EPI_ISL_701943, EPI_ISL_701947, EPI_ISL_701963, EPI_ISL_701975, EPI_ISL_701976, EPI_ISL_701980, EPI_ISL_701987, EPI_ISL_701993, EPI_ISL_701994, EPI_ISL_701999, EPI_ISL_702001, EPI_ISL_702002, EPI_ISL_702014, EPI_ISL_702015, EPI_ISL_702016, EPI_ISL_702376, EPI_ISL_702381, EPI_ISL_702384, EPI_ISL_702413, EPI_ISL_702423, EPI_ISL_702424, EPI_ISL_702427, EPI_ISL_702443, EPI_ISL_702447, EPI_ISL_702453, EPI_ISL_702486 | | | |
| see above | Lighthouse Lab in Alderley Park | Wellcome Sanger Institute for the COVID-19 Genomics UK (COG-UK) Consortium | Jacquelyn Wynn, Mairead Hyland, The Lighthouse Lab in Alderley Park and Alex Alderton, Roberto Amato, Sonia Goncalves, Ewan Harrison, David K. Jackson, Ian Johnston, Dominic Kwiatkowski, Cordelia Langford, John Sillitoe on behalf of the Wellcome Sanger Institute COVID-19 Surveillance Team |
| EPI_ISL_702495 | Northumbria University / South Tees Hospitals NHS Foundation Trust / North Cumbria Integrated Care NHS Foundation Trust / North Tees and Hartlepool NHS Foundation Trust / Newcastle Hospitals NHS Foundation Trust | COVID-19 Genomics UK (COG-UK) Consortium | Darren L Smith,Andrew Nelson,Matthew Bashton,Greg R Young,Joshua Loh,John Allan,Mohammad A Tariq,Giles S Holt,Gary Black,Wen C Yew,Lynn Dover,Paul Baker,Steve Liggett,Sarah Essex,Jane Greenaway,Debra Padgett,Clive Graham,Garren Scott,Edward Barton,Emma Swindells,Brendan Payne,Jennifer Collins,Yusri Taha,Gary Eltringham |
| EPI_ISL_702519, EPI_ISL_702521, EPI_ISL_702537 | Lighthouse Lab in Alderley Park | Wellcome Sanger Institute for the COVID-19 Genomics UK (COG-UK) Consortium | Jacquelyn Wynn, Mairead Hyland, The Lighthouse Lab in Alderley Park and Alex Alderton, Roberto Amato, Sonia Goncalves, Ewan Harrison, David K. Jackson, Ian Johnston, Dominic Kwiatkowski, Cordelia Langford, John Sillitoe on behalf of the Wellcome Sanger Institute COVID-19 Surveillance Team |
| EPI_ISL_702539 | Northumbria University / South Tees Hospitals NHS Foundation Trust / North Cumbria Integrated Care NHS Foundation Trust / North Tees and Hartlepool NHS Foundation Trust / Newcastle Hospitals NHS Foundation Trust | COVID-19 Genomics UK (COG-UK) Consortium | Darren L Smith,Andrew Nelson,Matthew Bashton,Greg R Young,Joshua Loh,John Allan,Mohammad A Tariq,Giles S Holt,Gary Black,Wen C Yew,Lynn Dover,Paul Baker,Steve Liggett,Sarah Essex,Jane Greenaway,Debra Padgett,Clive Graham,Garren Scott,Edward Barton,Emma Swindells,Brendan Payne,Jennifer Collins,Yusri Taha,Gary Eltringham |
| EPI_ISL_702543 | Lighthouse Lab in Alderley Park | Wellcome Sanger Institute for the COVID-19 Genomics UK (COG-UK) Consortium | Jacquelyn Wynn, Mairead Hyland, The Lighthouse Lab in Alderley Park and Alex Alderton, Roberto Amato, Sonia Goncalves, Ewan Harrison, David K. Jackson, Ian Johnston, Dominic Kwiatkowski, Cordelia Langford, John Sillitoe on behalf of the Wellcome Sanger Institute COVID-19 Surveillance Team |
| EPI_ISL_702544 | University College London, Great Ormond Street Hospital for Children NHS Foundation Trust, Imperial College Healthcare NHS Trust | COVID-19 Genomics UK (COG-UK) Consortium | Sergi Castellano, Rachel Williams, Mark Kristiansen, Paola Resende Silva, Sunando Roy, Tony Brooks, Helena Tutill, Paola Niola, Patricia Dyal, Charlotte Williams, Leysa Forrest, Yasmin Panchbhaya, Jacqueline Findlay, Samuel Weeks, Julianne Brown, Kathryn Harris, Paul Randell, James Price, Alison Holmes, Judith Breuer |
| EPI_ISL_702547 | Northumbria University / South Tees Hospitals NHS Foundation Trust / North Cumbria Integrated Care NHS Foundation Trust / North Tees and Hartlepool NHS Foundation Trust / Newcastle Hospitals NHS Foundation Trust | COVID-19 Genomics UK (COG-UK) Consortium | Darren L Smith,Andrew Nelson,Matthew Bashton,Greg R Young,Joshua Loh,John Allan,Mohammad A Tariq,Giles S Holt,Gary Black,Wen C Yew,Lynn Dover,Paul Baker,Steve Liggett,Sarah Essex,Jane Greenaway,Debra Padgett,Clive Graham,Garren Scott,Edward Barton,Emma Swindells,Brendan Payne,Jennifer Collins,Yusri Taha,Gary Eltringham |
| EPI_ISL_702553 | Wales Specialist Virology Centre Sequencing lab: Pathogen Genomics Unit | COVID-19 Genomics UK (COG-UK) Consortium | Catherine Moore, Johnathan Evans, Laura Gifford, Malorie Perry, Simon Cottrell, Angela Marchbank, Alec Birchley, Alexander Adams, Amy Gaskin, Bree Gatica-Wilcox, Jason Coombes, Joel Southgate, Lauren Gilbert, Lee Graham, Nicole Pacchiarini, Sara Kumziene-Summerhayes, Sarah Taylor, Sophie Jones, Sara Rey, Matthew Bull, Joanne Watkins, Sally Corden, Tom Connor |
| EPI_ISL_702571, EPI_ISL_702591 | Liverpool Clinical Laboratories | COVID-19 Genomics UK (COG-UK) Consortium | Sam Haldenby, Anita Lucaci, Steve Paterson, Julian Hiscox, Alistair Darby, M Almsaud, A Alrezaihi, Muhannad Alruwaili, Stuart D Armstrong, Jones Benjamin, Eleanor G Bentley, Anu Chawla, Jordan J Clark, Angela Cowell, Richard Eccles, Isabel García-Dorival, Matthew Gemmell, Alessandro Gerada, PKF Gilmore, Richard Gregory, Ximeng Han, Catherine Hartley, Margaret Hughes, Miren Iturriza-Gomara, James Johnson, L Luu, Jenifer Manson, Charlotte Nelson, Elaine O'Toole, Cassie Olateju, Rebekah Penrice-Randal , Lucille Rainbow, N.P Randle, Trevor Ian Robinson, Parul Sharma, Ghada T Shawli, James P Stewart, Neil Swainston, Ecaterina Vamos, Joanne Watts, Mark Whitehead |
| EPI_ISL_702610 | University College London, Great Ormond Street Hospital for Children NHS Foundation Trust, Imperial College Healthcare NHS Trust | COVID-19 Genomics UK (COG-UK) Consortium | Sergi Castellano, Rachel Williams, Mark Kristiansen, Paola Resende Silva, Sunando Roy, Tony Brooks, Helena Tutill, Paola Niola, Patricia Dyal, Charlotte Williams, Leysa Forrest, Yasmin Panchbhaya, Jacqueline Findlay, Samuel Weeks, Julianne Brown, Kathryn Harris, Paul Randell, James Price, Alison Holmes, Judith Breuer |
| EPI_ISL_702615 | Liverpool Clinical Laboratories | COVID-19 Genomics UK (COG-UK) Consortium | Sam Haldenby, Anita Lucaci, Steve Paterson, Julian Hiscox, Alistair Darby, M Almsaud, A Alrezaihi, Muhannad Alruwaili, Stuart D Armstrong, Jones Benjamin, Eleanor G Bentley, Anu Chawla, Jordan J Clark, Angela Cowell, Richard Eccles, Isabel García-Dorival, Matthew Gemmell, Alessandro Gerada, PKF Gilmore, Richard Gregory, Ximeng Han, Catherine Hartley, Margaret Hughes, Miren Iturriza-Gomara, James Johnson, L Luu, Jenifer Manson, Charlotte Nelson, Elaine O'Toole, Cassie Olateju, Rebekah Penrice-Randal , Lucille Rainbow, N.P Randle, Trevor Ian Robinson, Parul Sharma, Ghada T Shawli, James P Stewart, Neil Swainston, Ecaterina Vamos, Joanne Watts, Mark Whitehead |
| EPI_ISL_702619, EPI_ISL_702622 | Lighthouse Lab in Alderley Park | Wellcome Sanger Institute for the COVID-19 Genomics UK (COG-UK) Consortium | Jacquelyn Wynn, Mairead Hyland, The Lighthouse Lab in Alderley Park and Alex Alderton, Roberto Amato, Sonia Goncalves, Ewan Harrison, David K. Jackson, Ian Johnston, Dominic Kwiatkowski, Cordelia Langford, John Sillitoe on behalf of the Wellcome Sanger Institute COVID-19 Surveillance Team |
| EPI_ISL_702623, EPI_ISL_702628, EPI_ISL_702631 | Liverpool Clinical Laboratories | COVID-19 Genomics UK (COG-UK) Consortium | Sam Haldenby, Anita Lucaci, Steve Paterson, Julian Hiscox, Alistair Darby, M Almsaud, A Alrezaihi, Muhannad Alruwaili, Stuart D Armstrong, Jones Benjamin, Eleanor G Bentley, Anu Chawla, Jordan J Clark, Angela Cowell, Richard Eccles, Isabel García-Dorival, Matthew Gemmell, Alessandro Gerada, PKF Gilmore, Richard Gregory, Ximeng Han, Catherine Hartley, Margaret Hughes, Miren Iturriza-Gomara, James Johnson, L Luu, Jenifer Manson, Charlotte Nelson, Elaine O'Toole, Cassie Olateju, Rebekah Penrice-Randal , Lucille Rainbow, N.P Randle, Trevor Ian Robinson, Parul Sharma, Ghada T Shawli, James P Stewart, Neil Swainston, Ecaterina Vamos, Joanne Watts, Mark Whitehead |
| EPI_ISL_702634, EPI_ISL_702635, EPI_ISL_702637, EPI_ISL_702638, EPI_ISL_702640, EPI_ISL_702658 | Lighthouse Lab in Alderley Park | Wellcome Sanger Institute for the COVID-19 Genomics UK (COG-UK) Consortium | Jacquelyn Wynn, Mairead Hyland, The Lighthouse Lab in Alderley Park and Alex Alderton, Roberto Amato, Sonia Goncalves, Ewan Harrison, David K. Jackson, Ian Johnston, Dominic Kwiatkowski, Cordelia Langford, John Sillitoe on behalf of the Wellcome Sanger Institute COVID-19 Surveillance Team |
| EPI_ISL_702659 | West of Scotland Specialist Virology Centre, NHSGGC / MRC-University of Glasgow Centre for Virus Research | COVID-19 Genomics UK (COG-UK) Consortium | Ana da Silva Filipe, Natasha Johnson, Kathy Smollett, Daniel Mair, Stephen Carmichael, Alice Broos, Lily Tong, Jenna Nichols, Kyriaki Nomikou; Sarah McDonald; Richard Orton, Joseph Hughes, Sreenu Vattipally, David L Robertson; Alasdair MacLean, Rory Gunson; Sharif Shaaban, Matthew Holden; Rachel Blacow, Guy Mollett, Kathy Li, James Shepherd, Antonia Ho, Emma Thomson |
| EPI_ISL_702663, EPI_ISL_702668, EPI_ISL_702678, EPI_ISL_702681, EPI_ISL_702683, EPI_ISL_702701, EPI_ISL_702704, EPI_ISL_702707, EPI_ISL_702711, EPI_ISL_702712 | Lighthouse Lab in Alderley Park | Wellcome Sanger Institute for the COVID-19 Genomics UK (COG-UK) Consortium | Jacquelyn Wynn, Mairead Hyland, The Lighthouse Lab in Alderley Park and Alex Alderton, Roberto Amato, Sonia Goncalves, Ewan Harrison, David K. Jackson, Ian Johnston, Dominic Kwiatkowski, Cordelia Langford, John Sillitoe on behalf of the Wellcome Sanger Institute COVID-19 Surveillance Team |
| EPI_ISL_702713 | Wales Specialist Virology Centre Sequencing lab: Pathogen Genomics Unit | COVID-19 Genomics UK (COG-UK) Consortium | Catherine Moore, Johnathan Evans, Laura Gifford, Malorie Perry, Simon Cottrell, Angela Marchbank, Alec Birchley, Alexander Adams, Amy Gaskin, Bree Gatica-Wilcox, Jason Coombes, Joel Southgate, Lauren Gilbert, Lee Graham, Nicole Pacchiarini, Sara Kumziene-Summerhayes, Sarah Taylor, Sophie Jones, Sara Rey, Matthew Bull, Joanne Watkins, Sally Corden, Tom Connor |
| EPI_ISL_702719, EPI_ISL_702720, EPI_ISL_702725 | Lighthouse Lab in Alderley Park | Wellcome Sanger Institute for the COVID-19 Genomics UK (COG-UK) Consortium | Jacquelyn Wynn, Mairead Hyland, The Lighthouse Lab in Alderley Park and Alex Alderton, Roberto Amato, Sonia Goncalves, Ewan Harrison, David K. Jackson, Ian Johnston, Dominic Kwiatkowski, Cordelia Langford, John Sillitoe on behalf of the Wellcome Sanger Institute COVID-19 Surveillance Team |
| EPI_ISL_702729 | Liverpool Clinical Laboratories | COVID-19 Genomics UK (COG-UK) Consortium | Sam Haldenby, Anita Lucaci, Steve Paterson, Julian Hiscox, Alistair Darby, M Almsaud, A Alrezaihi, Muhannad Alruwaili, Stuart D Armstrong, Jones Benjamin, Eleanor G Bentley, Anu Chawla, Jordan J Clark, Angela Cowell, Richard Eccles, Isabel García-Dorival, Matthew Gemmell, Alessandro Gerada, |

| | | | |
|---|---|---|---|
| | | | PKF Gilmore, Richard Gregory, Ximeng Han, Catherine Hartley, Margaret Hughes, Miren Iturriza-Gomara, James Johnson, L Luu, Jenifer Manson, Charlotte Nelson, Elaine O'Toole, Cassie Olateju, Rebekah Penrice-Randal , Lucille Rainbow, N.P Randle, Trevor Ian Robinson, Parul Sharma, Ghada T Shawli, James P Stewart, Neil Swainston, Ecaterina Vamos, Joanne Watts, Mark Whitehead |
| EPI_ISL_702731 | University College London, Great Ormond Street Hospital for Children NHS Foundation Trust, Imperial College Healthcare NHS Trust | COVID-19 Genomics UK (COG-UK) Consortium | Sergi Castellano, Rachel Williams, Mark Kristiansen, Paola Resende Silva, Sunando Roy, Tony Brooks, Helena Tutill, Paola Niola, Patricia Dyal, Charlotte Williams, Leysa Forrest, Yasmin Panchbhaya, Jacqueline Findlay, Samuel Weeks, Julianne Brown, Kathryn Harris, Paul Randell, James Price, Alison Holmes, Judith Breuer |
| EPI_ISL_702732, EPI_ISL_702733, EPI_ISL_702741 | Lighthouse Lab in Alderley Park | Wellcome Sanger Institute for the COVID-19 Genomics UK (COG-UK) Consortium | Jacquelyn Wynn, Mairead Hyland, The Lighthouse Lab in Alderley Park and Alex Alderton, Roberto Amato, Sonia Goncalves, Ewan Harrison, David K. Jackson, Ian Johnston, Dominic Kwiatkowski, Cordelia Langford, John Sillitoe on behalf of the Wellcome Sanger Institute COVID-19 Surveillance Team |
| EPI_ISL_702742 | Wales Specialist Virology Centre Sequencing lab: Pathogen Genomics Unit | COVID-19 Genomics UK (COG-UK) Consortium | Catherine Moore, Johnathan Evans, Laura Gifford, Malorie Perry, Simon Cottrell, Angela Marchbank, Alec Birchley, Alexander Adams, Amy Gaskin, Bree Gatica-Wilcox, Jason Coombes, Joel Southgate, Lauren Gilbert, Lee Graham, Nicole Pacchiarini, Sara Kumziene-Summerhayes, Sarah Taylor, Sophie Jones, Sara Rey, Matthew Bull, Joanne Watkins, Sally Corden, Tom Connor |
| EPI_ISL_702745, EPI_ISL_702752, EPI_ISL_702754, EPI_ISL_702755 | Lighthouse Lab in Alderley Park | Wellcome Sanger Institute for the COVID-19 Genomics UK (COG-UK) Consortium | Jacquelyn Wynn, Mairead Hyland, The Lighthouse Lab in Alderley Park and Alex Alderton, Roberto Amato, Sonia Goncalves, Ewan Harrison, David K. Jackson, Ian Johnston, Dominic Kwiatkowski, Cordelia Langford, John Sillitoe on behalf of the Wellcome Sanger Institute COVID-19 Surveillance Team |
| EPI_ISL_702769 | Liverpool Clinical Laboratories | COVID-19 Genomics UK (COG-UK) Consortium | Sam Haldenby, Anita Lucaci, Steve Paterson, Julian Hiscox, Alistair Darby, M Almsaud, A Alrezaihi, Muhannad Alruwaili, Stuart D Armstrong, Jones Benjamin, Eleanor G Bentley, Anu Chawla, Jordan J Clark, Angela Cowell, Richard Eccles, Isabel García-Dorival, Matthew Gemmell, Alessandro Gerada, PKF Gilmore, Richard Gregory, Ximeng Han, Catherine Hartley, Margaret Hughes, Miren Iturriza-Gomara, James Johnson, L Luu, Jenifer Manson, Charlotte Nelson, Elaine O'Toole, Cassie Olateju, Rebekah Penrice-Randal , Lucille Rainbow, N.P Randle, Trevor Ian Robinson, Parul Sharma, Ghada T Shawli, James P Stewart, Neil Swainston, Ecaterina Vamos, Joanne Watts, Mark Whitehead |
| EPI_ISL_702773, EPI_ISL_702785, EPI_ISL_702791, EPI_ISL_702796, EPI_ISL_702797, EPI_ISL_702800, EPI_ISL_702804 | Lighthouse Lab in Alderley Park | Wellcome Sanger Institute for the COVID-19 Genomics UK (COG-UK) Consortium | Jacquelyn Wynn, Mairead Hyland, The Lighthouse Lab in Alderley Park and Alex Alderton, Roberto Amato, Sonia Goncalves, Ewan Harrison, David K. Jackson, Ian Johnston, Dominic Kwiatkowski, Cordelia Langford, John Sillitoe on behalf of the Wellcome Sanger Institute COVID-19 Surveillance Team |
| EPI_ISL_702812, EPI_ISL_702815 | Northumbria University / South Tees Hospitals NHS Foundation Trust / North Cumbria Integrated Care NHS Foundation Trust / North Tees and Hartlepool NHS Foundation Trust / Newcastle Hospitals NHS Foundation Trust | COVID-19 Genomics UK (COG-UK) Consortium | Darren L Smith,Andrew Nelson,Matthew Bashton,Greg R Young,Joshua Loh,John Allan,Mohammad A Tariq,Giles S Holt,Gary Black,Wen C Yew,Lynn Dover,Paul Baker,Steve Liggett,Sarah Essex,Jane Greenaway,Debra Padgett,Clive Graham,Garren Scott,Edward Barton,Emma Swindells,Brendan Payne,Jennifer Collins,Yusri Taha,Gary Eltringham |
| EPI_ISL_702816 | Lighthouse Lab in Alderley Park | Wellcome Sanger Institute for the COVID-19 Genomics UK (COG-UK) Consortium | Jacquelyn Wynn, Mairead Hyland, The Lighthouse Lab in Alderley Park and Alex Alderton, Roberto Amato, Sonia Goncalves, Ewan Harrison, David K. Jackson, Ian Johnston, Dominic Kwiatkowski, Cordelia Langford, John Sillitoe on behalf of the Wellcome Sanger Institute COVID-19 Surveillance Team |
| EPI_ISL_702817 | Wales Specialist Virology Centre Sequencing lab: Pathogen Genomics Unit | COVID-19 Genomics UK (COG-UK) Consortium | Catherine Moore, Johnathan Evans, Laura Gifford, Malorie Perry, Simon Cottrell, Angela Marchbank, Alec Birchley, Alexander Adams, Amy Gaskin, Bree Gatica-Wilcox, Jason Coombes, Joel Southgate, Lauren Gilbert, Lee Graham, Nicole Pacchiarini, Sara Kumziene-Summerhayes, Sarah Taylor, Sophie Jones, Sara Rey, Matthew Bull, Joanne Watkins, Sally Corden, Tom Connor |
| EPI_ISL_702821 | Lighthouse Lab in Alderley Park | Wellcome Sanger Institute for the COVID-19 Genomics UK (COG-UK) Consortium | Jacquelyn Wynn, Mairead Hyland, The Lighthouse Lab in Alderley Park and Alex Alderton, Roberto Amato, Sonia Goncalves, Ewan Harrison, David K. Jackson, Ian Johnston, Dominic Kwiatkowski, Cordelia Langford, John Sillitoe on behalf of the Wellcome Sanger Institute COVID-19 Surveillance Team |
| EPI_ISL_702823, EPI_ISL_702827 | Northumbria University / South Tees Hospitals NHS Foundation Trust / North Cumbria Integrated Care NHS Foundation Trust / North Tees and Hartlepool NHS Foundation Trust / Newcastle Hospitals NHS Foundation Trust | COVID-19 Genomics UK (COG-UK) Consortium | Darren L Smith,Andrew Nelson,Matthew Bashton,Greg R Young,Joshua Loh,John Allan,Mohammad A Tariq,Giles S Holt,Gary Black,Wen C Yew,Lynn Dover,Paul Baker,Steve Liggett,Sarah Essex,Jane Greenaway,Debra Padgett,Clive Graham,Garren Scott,Edward Barton,Emma Swindells,Brendan Payne,Jennifer Collins,Yusri Taha,Gary Eltringham |
| EPI_ISL_702838 | Liverpool Clinical Laboratories | COVID-19 Genomics UK (COG-UK) Consortium | Sam Haldenby, Anita Lucaci, Steve Paterson, Julian Hiscox, Alistair Darby, M Almsaud, A Alrezaihi, Muhannad Alruwaili, Stuart D Armstrong, Jones Benjamin, Eleanor G Bentley, Anu Chawla, Jordan J Clark, Angela Cowell, Richard Eccles, Isabel García-Dorival, Matthew Gemmell, Alessandro Gerada, PKF Gilmore, Richard Gregory, Ximeng Han, Catherine Hartley, Margaret Hughes, Miren Iturriza-Gomara, James Johnson, L Luu, Jenifer Manson, Charlotte Nelson, Elaine O'Toole, Cassie Olateju, Rebekah Penrice-Randal , Lucille Rainbow, N.P Randle, Trevor Ian Robinson, Parul Sharma, Ghada T Shawli, James P Stewart, Neil Swainston, Ecaterina Vamos, Joanne Watts, Mark Whitehead |
| EPI_ISL_702857, EPI_ISL_702859 | Wales Specialist Virology Centre Sequencing lab: Pathogen Genomics Unit | COVID-19 Genomics UK (COG-UK) Consortium | Catherine Moore, Johnathan Evans, Laura Gifford, Malorie Perry, Simon Cottrell, Angela Marchbank, Alec Birchley, Alexander Adams, Amy Gaskin, Bree Gatica-Wilcox, Jason Coombes, Joel Southgate, Lauren Gilbert, Lee Graham, Nicole Pacchiarini, Sara Kumziene-Summerhayes, Sarah Taylor, Sophie Jones, Sara Rey, Matthew Bull, Joanne Watkins, Sally Corden, Tom Connor |
| EPI_ISL_702867 | West of Scotland Specialist Virology Centre, NHSGGC / MRC-University of Glasgow Centre for Virus Research | COVID-19 Genomics UK (COG-UK) Consortium | Ana da Silva Filipe, Natasha Johnson, Kathy Smollett, Daniel Mair, Stephen Carmichael, Alice Broos, Lily Tong, Jenna Nichols, Kyriaki Nomikou; Sarah McDonald; Richard Orton, Joseph Hughes, Sreenu Vattipally, David L Robertson; Alasdair MacLean, Rory Gunson; Sharif Shaaban, Matthew Holden; Rachel Blacow, Guy Mollett, Kathy Li, James Shepherd, Antonia Ho, Emma Thomson |
| EPI_ISL_702895, EPI_ISL_702905 | Wales Specialist Virology Centre Sequencing lab: Pathogen Genomics Unit | COVID-19 Genomics UK (COG-UK) Consortium | Catherine Moore, Johnathan Evans, Laura Gifford, Malorie Perry, Simon Cottrell, Angela Marchbank, Alec Birchley, Alexander Adams, Amy Gaskin, Bree Gatica-Wilcox, Jason Coombes, Joel Southgate, Lauren Gilbert, Lee Graham, Nicole Pacchiarini, Sara Kumziene-Summerhayes, Sarah Taylor, Sophie Jones, Sara Rey, Matthew Bull, Joanne Watkins, Sally Corden, Tom Connor |
| EPI_ISL_702906, EPI_ISL_702908, EPI_ISL_702916, EPI_ISL_702918, EPI_ISL_702923, EPI_ISL_702927, EPI_ISL_702929 | Lighthouse Lab in Alderley Park | Wellcome Sanger Institute for the COVID-19 Genomics UK (COG-UK) Consortium | Jacquelyn Wynn, Mairead Hyland, The Lighthouse Lab in Alderley Park and Alex Alderton, Roberto Amato, Sonia Goncalves, Ewan Harrison, David K. Jackson, Ian Johnston, Dominic Kwiatkowski, Cordelia Langford, John Sillitoe on behalf of the Wellcome Sanger Institute COVID-19 Surveillance Team |
| EPI_ISL_702930 | University College London, Great Ormond Street Hospital for Children NHS Foundation Trust, Imperial College Healthcare NHS Trust | COVID-19 Genomics UK (COG-UK) Consortium | Sergi Castellano, Rachel Williams, Mark Kristiansen, Paola Resende Silva, Sunando Roy, Tony Brooks, Helena Tutill, Paola Niola, Patricia Dyal, Charlotte Williams, Leysa Forrest, Yasmin Panchbhaya, Jacqueline Findlay, Samuel Weeks, Julianne Brown, Kathryn Harris, Paul Randell, James Price, Alison Holmes, Judith Breuer |
| EPI_ISL_702940, EPI_ISL_702942 | Lighthouse Lab in Alderley Park | Wellcome Sanger Institute for the COVID-19 Genomics UK (COG-UK) Consortium | Jacquelyn Wynn, Mairead Hyland, The Lighthouse Lab in Alderley Park and Alex Alderton, Roberto Amato, Sonia Goncalves, Ewan Harrison, David K. Jackson, Ian Johnston, Dominic Kwiatkowski, Cordelia Langford, John Sillitoe on behalf of the Wellcome Sanger Institute COVID-19 Surveillance Team |
| EPI_ISL_702947 | Liverpool Clinical Laboratories | COVID-19 Genomics UK (COG-UK) Consortium | Sam Haldenby, Anita Lucaci, Steve Paterson, Julian Hiscox, Alistair Darby, M Almsaud, A Alrezaihi, Muhannad Alruwaili, Stuart D Armstrong, Jones Benjamin, Eleanor G Bentley, Anu Chawla, Jordan J Clark, Angela Cowell, Richard Eccles, Isabel García-Dorival, Matthew Gemmell, Alessandro Gerada, PKF Gilmore, Richard Gregory, Ximeng Han, Catherine Hartley, Margaret Hughes, Miren Iturriza-Gomara, James Johnson, L Luu, Jenifer Manson, Charlotte Nelson, Elaine O'Toole, Cassie Olateju, Rebekah Penrice-Randal , Lucille Rainbow, N.P Randle, Trevor Ian Robinson, Parul Sharma, Ghada T Shawli, James P Stewart, Neil Swainston, Ecaterina Vamos, Joanne Watts, Mark Whitehead |
| EPI_ISL_702952, EPI_ISL_702954 | Lighthouse Lab in Alderley Park | Wellcome Sanger Institute for the COVID-19 Genomics UK (COG-UK) Consortium | Jacquelyn Wynn, Mairead Hyland, The Lighthouse Lab in Alderley Park and Alex Alderton, Roberto Amato, Sonia Goncalves, Ewan Harrison, David K. Jackson, Ian Johnston, Dominic Kwiatkowski, Cordelia Langford, John Sillitoe on behalf of the Wellcome Sanger Institute COVID-19 Surveillance Team |
| EPI_ISL_702959 | Northumbria University / South Tees Hospitals NHS Foundation Trust / North Cumbria Integrated Care NHS Foundation Trust / North Tees and Hartlepool NHS Foundation Trust / Newcastle Hospitals NHS Foundation Trust | COVID-19 Genomics UK (COG-UK) Consortium | Darren L Smith,Andrew Nelson,Matthew Bashton,Greg R Young,Joshua Loh,John Allan,Mohammad A Tariq,Giles S Holt,Gary Black,Wen C Yew,Lynn Dover,Paul Baker,Steve Liggett,Sarah Essex,Jane Greenaway,Debra Padgett,Clive Graham,Garren Scott,Edward Barton,Emma Swindells,Brendan Payne,Jennifer Collins,Yusri Taha,Gary Eltringham |
| EPI_ISL_702963, EPI_ISL_702965, EPI_ISL_702967, EPI_ISL_702968, EPI_ISL_702970, EPI_ISL_702972, EPI_ISL_702976, EPI_ISL_702979 | Lighthouse Lab in Alderley Park | Wellcome Sanger Institute for the COVID-19 Genomics UK (COG-UK) Consortium | Jacquelyn Wynn, Mairead Hyland, The Lighthouse Lab in Alderley Park and Alex Alderton, Roberto Amato, Sonia Goncalves, Ewan Harrison, David K. Jackson, Ian Johnston, Dominic Kwiatkowski, Cordelia Langford, John Sillitoe on behalf of the Wellcome Sanger Institute COVID-19 Surveillance Team |
| EPI_ISL_702980 | Liverpool Clinical Laboratories | COVID-19 Genomics UK (COG-UK) Consortium | Sam Haldenby, Anita Lucaci, Steve Paterson, Julian Hiscox, Alistair Darby, M Almsaud, A Alrezaihi, Muhannad Alruwaili, Stuart D Armstrong, Jones |

| | | | |
|---|---|---|---|
| | | | Benjamin, Eleanor G Bentley, Anu Chawla, Jordan J Clark, Angela Cowell, Richard Eccles, Isabel García-Dorival, Matthew Gemmell, Alessandro Gerada, PKF Gilmore, Richard Gregory, Ximeng Han, Catherine Hartley, Margaret Hughes, Miren Iturriza-Gomara, James Johnson, L Luu, Jenifer Manson, Charlotte Nelson, Elaine O'Toole, Cassie Olateju, Rebekah Penrice-Randal , Lucille Rainbow, N.P Randle, Trevor Ian Robinson, Parul Sharma, Ghada T Shawli, James P Stewart, Neil Swainston, Ecaterina Vamos, Joanne Watts, Mark Whitehead |
| EPI_ISL_702981, EPI_ISL_702983, EPI_ISL_702986, EPI_ISL_702988, EPI_ISL_702991, EPI_ISL_702993 | Lighthouse Lab in Alderley Park | Wellcome Sanger Institute for the COVID-19 Genomics UK (COG-UK) Consortium | Jacquelyn Wynn, Mairead Hyland, The Lighthouse Lab in Alderley Park and Alex Alderton, Roberto Amato, Sonia Goncalves, Ewan Harrison, David K. Jackson, Ian Johnston, Dominic Kwiatkowski, Cordelia Langford, John Sillitoe on behalf of the Wellcome Sanger Institute COVID-19 Surveillance Team |
| EPI_ISL_702996 | Liverpool Clinical Laboratories | COVID-19 Genomics UK (COG-UK) Consortium | Sam Haldenby, Anita Lucaci, Steve Paterson, Julian Hiscox, Alistair Darby, M Almsaud, A Alrezaihi, Muhannad Alruwaili, Stuart D Armstrong, Jones Benjamin, Eleanor G Bentley, Anu Chawla, Jordan J Clark, Angela Cowell, Richard Eccles, Isabel García-Dorival, Matthew Gemmell, Alessandro Gerada, PKF Gilmore, Richard Gregory, Ximeng Han, Catherine Hartley, Margaret Hughes, Miren Iturriza-Gomara, James Johnson, L Luu, Jenifer Manson, Charlotte Nelson, Elaine O'Toole, Cassie Olateju, Rebekah Penrice-Randal , Lucille Rainbow, N.P Randle, Trevor Ian Robinson, Parul Sharma, Ghada T Shawli, James P Stewart, Neil Swainston, Ecaterina Vamos, Joanne Watts, Mark Whitehead |
| EPI_ISL_702997 | Lighthouse Lab in Alderley Park | Wellcome Sanger Institute for the COVID-19 Genomics UK (COG-UK) Consortium | Jacquelyn Wynn, Mairead Hyland, The Lighthouse Lab in Alderley Park and Alex Alderton, Roberto Amato, Sonia Goncalves, Ewan Harrison, David K. Jackson, Ian Johnston, Dominic Kwiatkowski, Cordelia Langford, John Sillitoe on behalf of the Wellcome Sanger Institute COVID-19 Surveillance Team |
| EPI_ISL_702999 | University College London, Great Ormond Street Hospital for Children NHS Foundation Trust, Imperial College Healthcare NHS Trust | COVID-19 Genomics UK (COG-UK) Consortium | Sergi Castellano, Rachel Williams, Mark Kristiansen, Paola Resende Silva, Sunando Roy, Tony Brooks, Helena Tutill, Paola Niola, Patricia Dyal, Charlotte Williams, Leysa Forrest, Yasmin Panchbhaya, Jacqueline Findlay, Samuel Weeks, Julianne Brown, Kathryn Harris, Paul Randell, James Price, Alison Holmes, Judith Breuer |
| EPI_ISL_703010 | Northumbria University / South Tees Hospitals NHS Foundation Trust / North Cumbria Integrated Care NHS Foundation Trust / North Tees and Hartlepool NHS Foundation Trust / Newcastle Hospitals NHS Foundation Trust | COVID-19 Genomics UK (COG-UK) Consortium | Darren L Smith,Andrew Nelson,Matthew Bashton,Greg R Young,Joshua Loh,John Allan,Mohammad A Tariq,Giles S Holt,Gary Black,Wen C Yew,Lynn Dover,Paul Baker,Steve Liggett,Sarah Essex,Jane Greenaway,Debra Padgett,Clive Graham,Garren Scott,Edward Barton,Emma Swindells,Brendan Payne,Jennifer Collins,Yusri Taha,Gary Eltringham |
| EPI_ISL_703012 | Lighthouse Lab in Alderley Park | Wellcome Sanger Institute for the COVID-19 Genomics UK (COG-UK) Consortium | Jacquelyn Wynn, Mairead Hyland, The Lighthouse Lab in Alderley Park and Alex Alderton, Roberto Amato, Sonia Goncalves, Ewan Harrison, David K. Jackson, Ian Johnston, Dominic Kwiatkowski, Cordelia Langford, John Sillitoe on behalf of the Wellcome Sanger Institute COVID-19 Surveillance Team |
| EPI_ISL_703013 | Northumbria University / South Tees Hospitals NHS Foundation Trust / North Cumbria Integrated Care NHS Foundation Trust / North Tees and Hartlepool NHS Foundation Trust / Newcastle Hospitals NHS Foundation Trust | COVID-19 Genomics UK (COG-UK) Consortium | Darren L Smith,Andrew Nelson,Matthew Bashton,Greg R Young,Joshua Loh,John Allan,Mohammad A Tariq,Giles S Holt,Gary Black,Wen C Yew,Lynn Dover,Paul Baker,Steve Liggett,Sarah Essex,Jane Greenaway,Debra Padgett,Clive Graham,Garren Scott,Edward Barton,Emma Swindells,Brendan Payne,Jennifer Collins,Yusri Taha,Gary Eltringham |
| EPI_ISL_703021, EPI_ISL_703022, EPI_ISL_703027, EPI_ISL_703030, EPI_ISL_703033 | Lighthouse Lab in Alderley Park | Wellcome Sanger Institute for the COVID-19 Genomics UK (COG-UK) Consortium | Jacquelyn Wynn, Mairead Hyland, The Lighthouse Lab in Alderley Park and Alex Alderton, Roberto Amato, Sonia Goncalves, Ewan Harrison, David K. Jackson, Ian Johnston, Dominic Kwiatkowski, Cordelia Langford, John Sillitoe on behalf of the Wellcome Sanger Institute COVID-19 Surveillance Team |
| EPI_ISL_703034 | Northumbria University / South Tees Hospitals NHS Foundation Trust / North Cumbria Integrated Care NHS Foundation Trust / North Tees and Hartlepool NHS Foundation Trust / Newcastle Hospitals NHS Foundation Trust | COVID-19 Genomics UK (COG-UK) Consortium | Darren L Smith,Andrew Nelson,Matthew Bashton,Greg R Young,Joshua Loh,John Allan,Mohammad A Tariq,Giles S Holt,Gary Black,Wen C Yew,Lynn Dover,Paul Baker,Steve Liggett,Sarah Essex,Jane Greenaway,Debra Padgett,Clive Graham,Garren Scott,Edward Barton,Emma Swindells,Brendan Payne,Jennifer Collins,Yusri Taha,Gary Eltringham |
| EPI_ISL_703035, EPI_ISL_703039, EPI_ISL_703044, EPI_ISL_703049 | Lighthouse Lab in Alderley Park | Wellcome Sanger Institute for the COVID-19 Genomics UK (COG-UK) Consortium | Jacquelyn Wynn, Mairead Hyland, The Lighthouse Lab in Alderley Park and Alex Alderton, Roberto Amato, Sonia Goncalves, Ewan Harrison, David K. Jackson, Ian Johnston, Dominic Kwiatkowski, Cordelia Langford, John Sillitoe on behalf of the Wellcome Sanger Institute COVID-19 Surveillance Team |
| EPI_ISL_703051 | Liverpool Clinical Laboratories | COVID-19 Genomics UK (COG-UK) Consortium | Sam Haldenby, Anita Lucaci, Steve Paterson, Julian Hiscox, Alistair Darby, M Almsaud, A Alrezaihi, Muhannad Alruwaili, Stuart D Armstrong, Jones Benjamin, Eleanor G Bentley, Anu Chawla, Jordan J Clark, Angela Cowell, Richard Eccles, Isabel García-Dorival, Matthew Gemmell, Alessandro Gerada, PKF Gilmore, Richard Gregory, Ximeng Han, Catherine Hartley, Margaret Hughes, Miren Iturriza-Gomara, James Johnson, L Luu, Jenifer Manson, Charlotte Nelson, Elaine O'Toole, Cassie Olateju, Rebekah Penrice-Randal , Lucille Rainbow, N.P Randle, Trevor Ian Robinson, Parul Sharma, Ghada T Shawli, James P Stewart, Neil Swainston, Ecaterina Vamos, Joanne Watts, Mark Whitehead |
| EPI_ISL_703054, EPI_ISL_703055, EPI_ISL_703057, EPI_ISL_703058 | Lighthouse Lab in Alderley Park | Wellcome Sanger Institute for the COVID-19 Genomics UK (COG-UK) Consortium | Jacquelyn Wynn, Mairead Hyland, The Lighthouse Lab in Alderley Park and Alex Alderton, Roberto Amato, Sonia Goncalves, Ewan Harrison, David K. Jackson, Ian Johnston, Dominic Kwiatkowski, Cordelia Langford, John Sillitoe on behalf of the Wellcome Sanger Institute COVID-19 Surveillance Team |
| EPI_ISL_703064 | Northumbria University / South Tees Hospitals NHS Foundation Trust / North Cumbria Integrated Care NHS Foundation Trust / North Tees and Hartlepool NHS Foundation Trust / Newcastle Hospitals NHS Foundation Trust | COVID-19 Genomics UK (COG-UK) Consortium | Darren L Smith,Andrew Nelson,Matthew Bashton,Greg R Young,Joshua Loh,John Allan,Mohammad A Tariq,Giles S Holt,Gary Black,Wen C Yew,Lynn Dover,Paul Baker,Steve Liggett,Sarah Essex,Jane Greenaway,Debra Padgett,Clive Graham,Garren Scott,Edward Barton,Emma Swindells,Brendan Payne,Jennifer Collins,Yusri Taha,Gary Eltringham |
| EPI_ISL_703066, EPI_ISL_703074 | Wales Specialist Virology Centre Sequencing lab: Pathogen Genomics Unit | COVID-19 Genomics UK (COG-UK) Consortium | Catherine Moore, Johnathan Evans, Laura Gifford, Malorie Perry, Simon Cottrell, Angela Marchbank, Alec Birchley, Alexander Adams, Amy Gaskin, Bree Gatica-Wilcox, Jason Coombes, Joel Southgate, Lauren Gilbert, Lee Graham, Nicole Pacchiarini, Sara Kumziene-Summerhayes, Sarah Taylor, Sophie Jones, Sara Rey, Matthew Bull, Joanne Watkins, Sally Corden, Tom Connor |
| EPI_ISL_703075, EPI_ISL_703081, EPI_ISL_703087, EPI_ISL_703089, EPI_ISL_703094, EPI_ISL_703101 | Lighthouse Lab in Alderley Park | Wellcome Sanger Institute for the COVID-19 Genomics UK (COG-UK) Consortium | Jacquelyn Wynn, Mairead Hyland, The Lighthouse Lab in Alderley Park and Alex Alderton, Roberto Amato, Sonia Goncalves, Ewan Harrison, David K. Jackson, Ian Johnston, Dominic Kwiatkowski, Cordelia Langford, John Sillitoe on behalf of the Wellcome Sanger Institute COVID-19 Surveillance Team |
| EPI_ISL_703105 | Northumbria University / South Tees Hospitals NHS Foundation Trust / North Cumbria Integrated Care NHS Foundation Trust / North Tees and Hartlepool NHS Foundation Trust / Newcastle Hospitals NHS Foundation Trust | COVID-19 Genomics UK (COG-UK) Consortium | Darren L Smith,Andrew Nelson,Matthew Bashton,Greg R Young,Joshua Loh,John Allan,Mohammad A Tariq,Giles S Holt,Gary Black,Wen C Yew,Lynn Dover,Paul Baker,Steve Liggett,Sarah Essex,Jane Greenaway,Debra Padgett,Clive Graham,Garren Scott,Edward Barton,Emma Swindells,Brendan Payne,Jennifer Collins,Yusri Taha,Gary Eltringham |
| EPI_ISL_703109, EPI_ISL_703111, EPI_ISL_703112, EPI_ISL_703114, EPI_ISL_703116, EPI_ISL_703117, EPI_ISL_703119, EPI_ISL_703125, EPI_ISL_703134, EPI_ISL_703137, EPI_ISL_703140, EPI_ISL_703145, EPI_ISL_703148, EPI_ISL_703151, EPI_ISL_703157, EPI_ISL_703164, EPI_ISL_703166, EPI_ISL_703168, EPI_ISL_703171, EPI_ISL_703175 | | | |
| see above | Lighthouse Lab in Alderley Park | Wellcome Sanger Institute for the COVID-19 Genomics UK (COG-UK) Consortium | Jacquelyn Wynn, Mairead Hyland, The Lighthouse Lab in Alderley Park and Alex Alderton, Roberto Amato, Sonia Goncalves, Ewan Harrison, David K. Jackson, Ian Johnston, Dominic Kwiatkowski, Cordelia Langford, John Sillitoe on behalf of the Wellcome Sanger Institute COVID-19 Surveillance Team |
| EPI_ISL_703176 | Liverpool Clinical Laboratories | COVID-19 Genomics UK (COG-UK) Consortium | Sam Haldenby, Anita Lucaci, Steve Paterson, Julian Hiscox, Alistair Darby, M Almsaud, A Alrezaihi, Muhannad Alruwaili, Stuart D Armstrong, Jones Benjamin, Eleanor G Bentley, Anu Chawla, Jordan J Clark, Angela Cowell, Richard Eccles, Isabel García-Dorival, Matthew Gemmell, Alessandro Gerada, PKF Gilmore, Richard Gregory, Ximeng Han, Catherine Hartley, Margaret Hughes, Miren Iturriza-Gomara, James Johnson, L Luu, Jenifer Manson, Charlotte Nelson, Elaine O'Toole, Cassie Olateju, Rebekah Penrice-Randal , Lucille Rainbow, N.P Randle, Trevor Ian Robinson, Parul Sharma, Ghada T Shawli, James P Stewart, Neil Swainston, Ecaterina Vamos, Joanne Watts, Mark Whitehead |
| EPI_ISL_703177, EPI_ISL_703178, EPI_ISL_703180, EPI_ISL_703181 | Lighthouse Lab in Alderley Park | Wellcome Sanger Institute for the COVID-19 Genomics UK (COG-UK) Consortium | Jacquelyn Wynn, Mairead Hyland, The Lighthouse Lab in Alderley Park and Alex Alderton, Roberto Amato, Sonia Goncalves, Ewan Harrison, David K. Jackson, Ian Johnston, Dominic Kwiatkowski, Cordelia Langford, John Sillitoe on behalf of the Wellcome Sanger Institute COVID-19 Surveillance Team |
| EPI_ISL_703182 | Liverpool Clinical Laboratories | COVID-19 Genomics UK (COG-UK) Consortium | Sam Haldenby, Anita Lucaci, Steve Paterson, Julian Hiscox, Alistair Darby, M Almsaud, A Alrezaihi, Muhannad Alruwaili, Stuart D Armstrong, Jones Benjamin, Eleanor G Bentley, Anu Chawla, Jordan J Clark, Angela Cowell, Richard Eccles, Isabel García-Dorival, Matthew Gemmell, Alessandro Gerada, PKF Gilmore, Richard Gregory, Ximeng Han, Catherine Hartley, Margaret Hughes, Miren Iturriza-Gomara, James Johnson, L Luu, Jenifer Manson, Charlotte Nelson, Elaine O'Toole, Cassie Olateju, Rebekah Penrice-Randal , Lucille Rainbow, N.P Randle, Trevor Ian Robinson, Parul Sharma, Ghada T Shawli, James P Stewart, Neil Swainston, Ecaterina Vamos, Joanne Watts, Mark Whitehead |
| EPI_ISL_703185, EPI_ISL_703189, EPI_ISL_703191, EPI_ISL_703194, EPI_ISL_703196, EPI_ISL_703199, | Lighthouse Lab in Alderley Park | Wellcome Sanger Institute for the COVID-19 Genomics UK (COG-UK) Consortium | Jacquelyn Wynn, Mairead Hyland, The Lighthouse Lab in Alderley Park and Alex Alderton, Roberto Amato, Sonia Goncalves, Ewan Harrison, David K. Jackson, Ian Johnston, Dominic Kwiatkowski, Cordelia Langford, John Sillitoe on behalf of the Wellcome Sanger Institute COVID-19 Surveillance Team |

| | | | |
|---|---|---|---|
| EPI_ISL_703202, EPI_ISL_703205, EPI_ISL_703214, EPI_ISL_703218 | | | |
| EPI_ISL_703219 | Liverpool Clinical Laboratories | COVID-19 Genomics UK (COG-UK) Consortium | Sam Haldenby, Anita Lucaci, Steve Paterson, Julian Hiscox, Alistair Darby, M Almsaud, A Alrezaihi, Muhannad Alruwaili, Stuart D Armstrong, Jones Benjamin, Eleanor G Bentley, Anu Chawla, Jordan J Clark, Angela Cowell, Richard Eccles, Isabel García-Dorival, Matthew Gemmell, Alessandro Gerada, PKF Gilmore, Richard Gregory, Ximeng Han, Catherine Hartley, Margaret Hughes, Miren Iturriza-Gomara, James Johnson, L Luu, Jenifer Manson, Charlotte Nelson, Elaine O'Toole, Cassie Olateju, Rebekah Penrice-Randal , Lucille Rainbow, N.P Randle, Trevor Ian Robinson, Parul Sharma, Ghada T Shawli, James P Stewart, Neil Swainston, Ecaterina Vamos, Joanne Watts, Mark Whitehead |
| EPI_ISL_703223, EPI_ISL_703229, EPI_ISL_703234 | Lighthouse Lab in Alderley Park | Wellcome Sanger Institute for the COVID-19 Genomics UK (COG-UK) Consortium | Jacquelyn Wynn, Mairead Hyland, The Lighthouse Lab in Alderley Park and Alex Alderton, Roberto Amato, Sonia Goncalves, Ewan Harrison, David K. Jackson, Ian Johnston, Dominic Kwiatkowski, Cordelia Langford, John Sillitoe on behalf of the Wellcome Sanger Institute COVID-19 Surveillance Team |
| EPI_ISL_703235 | Liverpool Clinical Laboratories | COVID-19 Genomics UK (COG-UK) Consortium | Sam Haldenby, Anita Lucaci, Steve Paterson, Julian Hiscox, Alistair Darby, M Almsaud, A Alrezaihi, Muhannad Alruwaili, Stuart D Armstrong, Jones Benjamin, Eleanor G Bentley, Anu Chawla, Jordan J Clark, Angela Cowell, Richard Eccles, Isabel García-Dorival, Matthew Gemmell, Alessandro Gerada, PKF Gilmore, Richard Gregory, Ximeng Han, Catherine Hartley, Margaret Hughes, Miren Iturriza-Gomara, James Johnson, L Luu, Jenifer Manson, Charlotte Nelson, Elaine O'Toole, Cassie Olateju, Rebekah Penrice-Randal , Lucille Rainbow, N.P Randle, Trevor Ian Robinson, Parul Sharma, Ghada T Shawli, James P Stewart, Neil Swainston, Ecaterina Vamos, Joanne Watts, Mark Whitehead |
| EPI_ISL_703236, EPI_ISL_703239, EPI_ISL_703244 | Lighthouse Lab in Alderley Park | Wellcome Sanger Institute for the COVID-19 Genomics UK (COG-UK) Consortium | Jacquelyn Wynn, Mairead Hyland, The Lighthouse Lab in Alderley Park and Alex Alderton, Roberto Amato, Sonia Goncalves, Ewan Harrison, David K. Jackson, Ian Johnston, Dominic Kwiatkowski, Cordelia Langford, John Sillitoe on behalf of the Wellcome Sanger Institute COVID-19 Surveillance Team |
| EPI_ISL_703246 | West of Scotland Specialist Virology Centre, NHSGGC / MRC-University of Glasgow Centre for Virus Research | COVID-19 Genomics UK (COG-UK) Consortium | Ana da Silva Filipe, Natasha Johnson, Kathy Smollett, Daniel Mair, Stephen Carmichael, Alice Broos, Lily Tong, Jenna Nichols, Kyriaki Nomikou; Sarah McDonald; Richard Orton, Joseph Hughes, Sreenu Vattipally, David L Robertson; Alasdair MacLean, Rory Gunson; Sharif Shaaban, Matthew Holden; Rachel Blacow, Guy Mollett, Kathy Li, James Shepherd, Antonia Ho, Emma Thomson |
| EPI_ISL_703251 | Wales Specialist Virology Centre Sequencing lab: Pathogen Genomics Unit | COVID-19 Genomics UK (COG-UK) Consortium | Catherine Moore, Johnathan Evans, Laura Gifford, Malorie Perry, Simon Cottrell, Angela Marchbank, Alec Birchley, Alexander Adams, Amy Gaskin, Bree Gatica-Wilcox, Jason Coombes, Joel Southgate, Lauren Gilbert, Lee Graham, Nicole Pacchiarini, Sara Kumziene-Summerhayes, Sarah Taylor, Sophie Jones, Sara Rey, Matthew Bull, Joanne Watkins, Sally Corden, Tom Connor |
| EPI_ISL_703269 | West of Scotland Specialist Virology Centre, NHSGGC / MRC-University of Glasgow Centre for Virus Research | COVID-19 Genomics UK (COG-UK) Consortium | Ana da Silva Filipe, Natasha Johnson, Kathy Smollett, Daniel Mair, Stephen Carmichael, Alice Broos, Lily Tong, Jenna Nichols, Kyriaki Nomikou; Sarah McDonald; Richard Orton, Joseph Hughes, Sreenu Vattipally, David L Robertson; Alasdair MacLean, Rory Gunson; Sharif Shaaban, Matthew Holden; Rachel Blacow, Guy Mollett, Kathy Li, James Shepherd, Antonia Ho, Emma Thomson |
| EPI_ISL_703272 | Liverpool Clinical Laboratories | COVID-19 Genomics UK (COG-UK) Consortium | Sam Haldenby, Anita Lucaci, Steve Paterson, Julian Hiscox, Alistair Darby, M Almsaud, A Alrezaihi, Muhannad Alruwaili, Stuart D Armstrong, Jones Benjamin, Eleanor G Bentley, Anu Chawla, Jordan J Clark, Angela Cowell, Richard Eccles, Isabel García-Dorival, Matthew Gemmell, Alessandro Gerada, PKF Gilmore, Richard Gregory, Ximeng Han, Catherine Hartley, Margaret Hughes, Miren Iturriza-Gomara, James Johnson, L Luu, Jenifer Manson, Charlotte Nelson, Elaine O'Toole, Cassie Olateju, Rebekah Penrice-Randal , Lucille Rainbow, N.P Randle, Trevor Ian Robinson, Parul Sharma, Ghada T Shawli, James P Stewart, Neil Swainston, Ecaterina Vamos, Joanne Watts, Mark Whitehead |
| EPI_ISL_703278 | Wales Specialist Virology Centre Sequencing lab: Pathogen Genomics Unit | COVID-19 Genomics UK (COG-UK) Consortium | Catherine Moore, Johnathan Evans, Laura Gifford, Malorie Perry, Simon Cottrell, Angela Marchbank, Alec Birchley, Alexander Adams, Amy Gaskin, Bree Gatica-Wilcox, Jason Coombes, Joel Southgate, Lauren Gilbert, Lee Graham, Nicole Pacchiarini, Sara Kumziene-Summerhayes, Sarah Taylor, Sophie Jones, Sara Rey, Matthew Bull, Joanne Watkins, Sally Corden, Tom Connor |
| EPI_ISL_703319 | University College London, Great Ormond Street Hospital for Children NHS Foundation Trust, Imperial College Healthcare NHS Trust | COVID-19 Genomics UK (COG-UK) Consortium | Sergi Castellano, Rachel Williams, Mark Kristiansen, Paola Resende Silva, Sunando Roy, Tony Brooks, Helena Tutill, Paola Niola, Patricia Dyal, Charlotte Williams, Leysa Forrest, Yasmin Panchbhaya, Jacqueline Findlay, Samuel Weeks, Julianne Brown, Kathryn Harris, Paul Randell, James Price, Alison Holmes, Judith Breuer |
| EPI_ISL_703321 | Wales Specialist Virology Centre Sequencing lab: Pathogen Genomics Unit | COVID-19 Genomics UK (COG-UK) Consortium | Catherine Moore, Johnathan Evans, Laura Gifford, Malorie Perry, Simon Cottrell, Angela Marchbank, Alec Birchley, Alexander Adams, Amy Gaskin, Bree Gatica-Wilcox, Jason Coombes, Joel Southgate, Lauren Gilbert, Lee Graham, Nicole Pacchiarini, Sara Kumziene-Summerhayes, Sarah Taylor, Sophie Jones, Sara Rey, Matthew Bull, Joanne Watkins, Sally Corden, Tom Connor |
| EPI_ISL_703324 | Liverpool Clinical Laboratories | COVID-19 Genomics UK (COG-UK) Consortium | Sam Haldenby, Anita Lucaci, Steve Paterson, Julian Hiscox, Alistair Darby, M Almsaud, A Alrezaihi, Muhannad Alruwaili, Stuart D Armstrong, Jones Benjamin, Eleanor G Bentley, Anu Chawla, Jordan J Clark, Angela Cowell, Richard Eccles, Isabel García-Dorival, Matthew Gemmell, Alessandro Gerada, PKF Gilmore, Richard Gregory, Ximeng Han, Catherine Hartley, Margaret Hughes, Miren Iturriza-Gomara, James Johnson, L Luu, Jenifer Manson, Charlotte Nelson, Elaine O'Toole, Cassie Olateju, Rebekah Penrice-Randal , Lucille Rainbow, N.P Randle, Trevor Ian Robinson, Parul Sharma, Ghada T Shawli, James P Stewart, Neil Swainston, Ecaterina Vamos, Joanne Watts, Mark Whitehead |
| EPI_ISL_703330 | Wales Specialist Virology Centre Sequencing lab: Pathogen Genomics Unit | COVID-19 Genomics UK (COG-UK) Consortium | Catherine Moore, Johnathan Evans, Laura Gifford, Malorie Perry, Simon Cottrell, Angela Marchbank, Alec Birchley, Alexander Adams, Amy Gaskin, Bree Gatica-Wilcox, Jason Coombes, Joel Southgate, Lauren Gilbert, Lee Graham, Nicole Pacchiarini, Sara Kumziene-Summerhayes, Sarah Taylor, Sophie Jones, Sara Rey, Matthew Bull, Joanne Watkins, Sally Corden, Tom Connor |
| EPI_ISL_703335 | University College London, Great Ormond Street Hospital for Children NHS Foundation Trust, Imperial College Healthcare NHS Trust | COVID-19 Genomics UK (COG-UK) Consortium | Sergi Castellano, Rachel Williams, Mark Kristiansen, Paola Resende Silva, Sunando Roy, Tony Brooks, Helena Tutill, Paola Niola, Patricia Dyal, Charlotte Williams, Leysa Forrest, Yasmin Panchbhaya, Jacqueline Findlay, Samuel Weeks, Julianne Brown, Kathryn Harris, Paul Randell, James Price, Alison Holmes, Judith Breuer |
| EPI_ISL_703343, EPI_ISL_703359 | Wales Specialist Virology Centre Sequencing lab: Pathogen Genomics Unit | COVID-19 Genomics UK (COG-UK) Consortium | Catherine Moore, Johnathan Evans, Laura Gifford, Malorie Perry, Simon Cottrell, Angela Marchbank, Alec Birchley, Alexander Adams, Amy Gaskin, Bree Gatica-Wilcox, Jason Coombes, Joel Southgate, Lauren Gilbert, Lee Graham, Nicole Pacchiarini, Sara Kumziene-Summerhayes, Sarah Taylor, Sophie Jones, Sara Rey, Matthew Bull, Joanne Watkins, Sally Corden, Tom Connor |
| EPI_ISL_703364, EPI_ISL_703375 | Liverpool Clinical Laboratories | COVID-19 Genomics UK (COG-UK) Consortium | Sam Haldenby, Anita Lucaci, Steve Paterson, Julian Hiscox, Alistair Darby, M Almsaud, A Alrezaihi, Muhannad Alruwaili, Stuart D Armstrong, Jones Benjamin, Eleanor G Bentley, Anu Chawla, Jordan J Clark, Angela Cowell, Richard Eccles, Isabel García-Dorival, Matthew Gemmell, Alessandro Gerada, PKF Gilmore, Richard Gregory, Ximeng Han, Catherine Hartley, Margaret Hughes, Miren Iturriza-Gomara, James Johnson, L Luu, Jenifer Manson, Charlotte Nelson, Elaine O'Toole, Cassie Olateju, Rebekah Penrice-Randal , Lucille Rainbow, N.P Randle, Trevor Ian Robinson, Parul Sharma, Ghada T Shawli, James P Stewart, Neil Swainston, Ecaterina Vamos, Joanne Watts, Mark Whitehead |
| EPI_ISL_703377 | Northumbria University / South Tees Hospitals NHS Foundation Trust / North Cumbria Integrated Care NHS Foundation Trust / North Tees and Hartlepool NHS Foundation Trust / Newcastle Hospitals NHS Foundation Trust | COVID-19 Genomics UK (COG-UK) Consortium | Darren L Smith,Andrew Nelson,Matthew Bashton,Greg R Young,Joshua Loh,John Allan,Mohammad A Tariq,Giles S Holt,Gary Black,Wen C Yew,Lynn Dover,Paul Baker,Steve Liggett,Sarah Essex,Jane Greenaway,Debra Padgett,Clive Graham,Garren Scott,Edward Barton,Emma Swindells,Brendan Payne,Jennifer Collins,Yusri Taha,Gary Eltringham |
| EPI_ISL_703383, EPI_ISL_703385 | University College London, Great Ormond Street Hospital for Children NHS Foundation Trust, Imperial College Healthcare NHS Trust | COVID-19 Genomics UK (COG-UK) Consortium | Sergi Castellano, Rachel Williams, Mark Kristiansen, Paola Resende Silva, Sunando Roy, Tony Brooks, Helena Tutill, Paola Niola, Patricia Dyal, Charlotte Williams, Leysa Forrest, Yasmin Panchbhaya, Jacqueline Findlay, Samuel Weeks, Julianne Brown, Kathryn Harris, Paul Randell, James Price, Alison Holmes, Judith Breuer |
| EPI_ISL_703387 | Liverpool Clinical Laboratories | COVID-19 Genomics UK (COG-UK) Consortium | Sam Haldenby, Anita Lucaci, Steve Paterson, Julian Hiscox, Alistair Darby, M Almsaud, A Alrezaihi, Muhannad Alruwaili, Stuart D Armstrong, Jones Benjamin, Eleanor G Bentley, Anu Chawla, Jordan J Clark, Angela Cowell, Richard Eccles, Isabel García-Dorival, Matthew Gemmell, Alessandro Gerada, PKF Gilmore, Richard Gregory, Ximeng Han, Catherine Hartley, Margaret Hughes, Miren Iturriza-Gomara, James Johnson, L Luu, Jenifer Manson, Charlotte Nelson, Elaine O'Toole, Cassie Olateju, Rebekah Penrice-Randal , Lucille Rainbow, N.P Randle, Trevor Ian Robinson, Parul Sharma, Ghada T Shawli, James P Stewart, Neil Swainston, Ecaterina Vamos, Joanne Watts, Mark Whitehead |
| EPI_ISL_703390 | Northumbria University / South Tees Hospitals NHS Foundation Trust / North Cumbria Integrated Care NHS Foundation Trust / North Tees and Hartlepool NHS Foundation Trust / Newcastle Hospitals NHS Foundation Trust | COVID-19 Genomics UK (COG-UK) Consortium | Darren L Smith,Andrew Nelson,Matthew Bashton,Greg R Young,Joshua Loh,John Allan,Mohammad A Tariq,Giles S Holt,Gary Black,Wen C Yew,Lynn Dover,Paul Baker,Steve Liggett,Sarah Essex,Jane Greenaway,Debra Padgett,Clive Graham,Garren Scott,Edward Barton,Emma Swindells,Brendan Payne,Jennifer Collins,Yusri Taha,Gary Eltringham |
| EPI_ISL_703395, EPI_ISL_703398 | Wales Specialist Virology Centre Sequencing lab: Pathogen Genomics Unit | COVID-19 Genomics UK (COG-UK) Consortium | Catherine Moore, Johnathan Evans, Laura Gifford, Malorie Perry, Simon Cottrell, Angela Marchbank, Alec Birchley, Alexander Adams, Amy Gaskin, Bree Gatica-Wilcox, Jason Coombes, Joel Southgate, Lauren Gilbert, Lee Graham, Nicole Pacchiarini, Sara Kumziene-Summerhayes, Sarah Taylor, Sophie Jones, Sara Rey, Matthew Bull, Joanne Watkins, Sally Corden, Tom Connor |

| | | | |
|---|---|---|---|
| EPI_ISL_703484, EPI_ISL_703542, EPI_ISL_703545 | Northumbria University / South Tees Hospitals NHS Foundation Trust / North Cumbria Integrated Care NHS Foundation Trust / North Tees and Hartlepool NHS Foundation Trust / Newcastle Hospitals NHS Foundation Trust | COVID-19 Genomics UK (COG-UK) Consortium | Darren L Smith,Andrew Nelson,Matthew Bashton,Greg R Young,Joshua Loh,John Allan,Mohammad A Tariq,Giles S Holt,Gary Black,Wen C Yew,Lynn Dover,Paul Baker,Steve Liggett,Sarah Essex,Jane Greenaway,Debra Padgett,Clive Graham,Garren Scott,Edward Barton,Emma Swindells,Brendan Payne,Jennifer Collins,Yusri Taha,Gary Eltringham |
| EPI_ISL_703577 | Liverpool Clinical Laboratories | COVID-19 Genomics UK (COG-UK) Consortium | Sam Haldenby, Anita Lucaci, Steve Paterson, Julian Hiscox, Alistair Darby, M Almsaud, A Alrezaihi, Muhannad Alruwaili, Stuart D Armstrong, Jones Benjamin, Eleanor G Bentley, Anu Chawla, Jordan J Clark, Angela Cowell, Richard Eccles, Isabel García-Dorival, Matthew Gemmell, Alessandro Gerada, PKF Gilmore, Richard Gregory, Ximeng Han, Catherine Hartley, Margaret Hughes, Miren Iturriza-Gomara, James Johnson, L Luu, Jenifer Manson, Charlotte Nelson, Elaine O'Toole, Cassie Olateju, Rebekah Penrice-Randal , Lucille Rainbow, N.P Randle, Trevor Ian Robinson, Parul Sharma, Ghada T Shawli, James P Stewart, Neil Swainston, Ecaterina Vamos, Joanne Watts, Mark Whitehead |
| EPI_ISL_703614 | Wales Specialist Virology Centre Sequencing lab: Pathogen Genomics Unit | COVID-19 Genomics UK (COG-UK) Consortium | Catherine Moore, Johnathan Evans, Laura Gifford, Malorie Perry, Simon Cottrell, Angela Marchbank, Alec Birchley, Alexander Adams, Amy Gaskin, Bree Gatica-Wilcox, Jason Coombes, Joel Southgate, Lauren Gilbert, Lee Graham, Nicole Pacchiarini, Sara Kumziene-Summerhayes, Sarah Taylor, Sophie Jones, Sara Rey, Matthew Bull, Joanne Watkins, Sally Corden, Tom Connor |
| EPI_ISL_703624, EPI_ISL_703698 | Liverpool Clinical Laboratories | COVID-19 Genomics UK (COG-UK) Consortium | Sam Haldenby, Anita Lucaci, Steve Paterson, Julian Hiscox, Alistair Darby, M Almsaud, A Alrezaihi, Muhannad Alruwaili, Stuart D Armstrong, Jones Benjamin, Eleanor G Bentley, Anu Chawla, Jordan J Clark, Angela Cowell, Richard Eccles, Isabel García-Dorival, Matthew Gemmell, Alessandro Gerada, PKF Gilmore, Richard Gregory, Ximeng Han, Catherine Hartley, Margaret Hughes, Miren Iturriza-Gomara, James Johnson, L Luu, Jenifer Manson, Charlotte Nelson, Elaine O'Toole, Cassie Olateju, Rebekah Penrice-Randal , Lucille Rainbow, N.P Randle, Trevor Ian Robinson, Parul Sharma, Ghada T Shawli, James P Stewart, Neil Swainston, Ecaterina Vamos, Joanne Watts, Mark Whitehead |
| EPI_ISL_703704 | Northumbria University / South Tees Hospitals NHS Foundation Trust / North Cumbria Integrated Care NHS Foundation Trust / North Tees and Hartlepool NHS Foundation Trust / Newcastle Hospitals NHS Foundation Trust | COVID-19 Genomics UK (COG-UK) Consortium | Darren L Smith,Andrew Nelson,Matthew Bashton,Greg R Young,Joshua Loh,John Allan,Mohammad A Tariq,Giles S Holt,Gary Black,Wen C Yew,Lynn Dover,Paul Baker,Steve Liggett,Sarah Essex,Jane Greenaway,Debra Padgett,Clive Graham,Garren Scott,Edward Barton,Emma Swindells,Brendan Payne,Jennifer Collins,Yusri Taha,Gary Eltringham |
| EPI_ISL_703706 | University College London, Great Ormond Street Hospital for Children NHS Foundation Trust, Imperial College Healthcare NHS Trust | COVID-19 Genomics UK (COG-UK) Consortium | Sergi Castellano, Rachel Williams, Mark Kristiansen, Paola Resende Silva, Sunando Roy, Tony Brooks, Helena Tutill, Paola Niola, Patricia Dyal, Charlotte Williams, Leysa Forrest, Yasmin Panchbhaya, Jacqueline Findlay, Samuel Weeks, Julianne Brown, Kathryn Harris, Paul Randell, James Price, Alison Holmes, Judith Breuer |
| EPI_ISL_703717 | West of Scotland Specialist Virology Centre, NHSGGC / MRC-University of Glasgow Centre for Virus Research | COVID-19 Genomics UK (COG-UK) Consortium | Ana da Silva Filipe, Natasha Johnson, Kathy Smollett, Daniel Mair, Stephen Carmichael, Alice Broos, Lily Tong, Jenna Nichols, Kyriaki Nomikou; Sarah McDonald; Richard Orton, Joseph Hughes, Sreenu Vattipally, David L Robertson; Alasdair MacLean, Rory Gunson; Sharif Shaaban, Matthew Holden; Rachel Blacow, Guy Mollett, Kathy Li, James Shepherd, Antonia Ho, Emma Thomson |
| EPI_ISL_703728 | Northumbria University / South Tees Hospitals NHS Foundation Trust / North Cumbria Integrated Care NHS Foundation Trust / North Tees and Hartlepool NHS Foundation Trust / Newcastle Hospitals NHS Foundation Trust | COVID-19 Genomics UK (COG-UK) Consortium | Darren L Smith,Andrew Nelson,Matthew Bashton,Greg R Young,Joshua Loh,John Allan,Mohammad A Tariq,Giles S Holt,Gary Black,Wen C Yew,Lynn Dover,Paul Baker,Steve Liggett,Sarah Essex,Jane Greenaway,Debra Padgett,Clive Graham,Garren Scott,Edward Barton,Emma Swindells,Brendan Payne,Jennifer Collins,Yusri Taha,Gary Eltringham |
| EPI_ISL_703732 | Liverpool Clinical Laboratories | COVID-19 Genomics UK (COG-UK) Consortium | Sam Haldenby, Anita Lucaci, Steve Paterson, Julian Hiscox, Alistair Darby, M Almsaud, A Alrezaihi, Muhannad Alruwaili, Stuart D Armstrong, Jones Benjamin, Eleanor G Bentley, Anu Chawla, Jordan J Clark, Angela Cowell, Richard Eccles, Isabel García-Dorival, Matthew Gemmell, Alessandro Gerada, PKF Gilmore, Richard Gregory, Ximeng Han, Catherine Hartley, Margaret Hughes, Miren Iturriza-Gomara, James Johnson, L Luu, Jenifer Manson, Charlotte Nelson, Elaine O'Toole, Cassie Olateju, Rebekah Penrice-Randal , Lucille Rainbow, N.P Randle, Trevor Ian Robinson, Parul Sharma, Ghada T Shawli, James P Stewart, Neil Swainston, Ecaterina Vamos, Joanne Watts, Mark Whitehead |
| EPI_ISL_703739 | University College London, Great Ormond Street Hospital for Children NHS Foundation Trust, Imperial College Healthcare NHS Trust | COVID-19 Genomics UK (COG-UK) Consortium | Sergi Castellano, Rachel Williams, Mark Kristiansen, Paola Resende Silva, Sunando Roy, Tony Brooks, Helena Tutill, Paola Niola, Patricia Dyal, Charlotte Williams, Leysa Forrest, Yasmin Panchbhaya, Jacqueline Findlay, Samuel Weeks, Julianne Brown, Kathryn Harris, Paul Randell, James Price, Alison Holmes, Judith Breuer |
| EPI_ISL_703746 | Wales Specialist Virology Centre Sequencing lab: Pathogen Genomics Unit | COVID-19 Genomics UK (COG-UK) Consortium | Catherine Moore, Johnathan Evans, Laura Gifford, Malorie Perry, Simon Cottrell, Angela Marchbank, Alec Birchley, Alexander Adams, Amy Gaskin, Bree Gatica-Wilcox, Jason Coombes, Joel Southgate, Lauren Gilbert, Lee Graham, Nicole Pacchiarini, Sara Kumziene-Summerhayes, Sarah Taylor, Sophie Jones, Sara Rey, Matthew Bull, Joanne Watkins, Sally Corden, Tom Connor |
| EPI_ISL_703748, EPI_ISL_703749, EPI_ISL_703756 | West of Scotland Specialist Virology Centre, NHSGGC / MRC-University of Glasgow Centre for Virus Research | COVID-19 Genomics UK (COG-UK) Consortium | Ana da Silva Filipe, Natasha Johnson, Kathy Smollett, Daniel Mair, Stephen Carmichael, Alice Broos, Lily Tong, Jenna Nichols, Kyriaki Nomikou; Sarah McDonald; Richard Orton, Joseph Hughes, Sreenu Vattipally, David L Robertson; Alasdair MacLean, Rory Gunson; Sharif Shaaban, Matthew Holden; Rachel Blacow, Guy Mollett, Kathy Li, James Shepherd, Antonia Ho, Emma Thomson |
| EPI_ISL_703760, EPI_ISL_703761 | Wales Specialist Virology Centre Sequencing lab: Pathogen Genomics Unit | COVID-19 Genomics UK (COG-UK) Consortium | Catherine Moore, Johnathan Evans, Laura Gifford, Malorie Perry, Simon Cottrell, Angela Marchbank, Alec Birchley, Alexander Adams, Amy Gaskin, Bree Gatica-Wilcox, Jason Coombes, Joel Southgate, Lauren Gilbert, Lee Graham, Nicole Pacchiarini, Sara Kumziene-Summerhayes, Sarah Taylor, Sophie Jones, Sara Rey, Matthew Bull, Joanne Watkins, Sally Corden, Tom Connor |
| EPI_ISL_703767 | University College London, Great Ormond Street Hospital for Children NHS Foundation Trust, Imperial College Healthcare NHS Trust | COVID-19 Genomics UK (COG-UK) Consortium | Sergi Castellano, Rachel Williams, Mark Kristiansen, Paola Resende Silva, Sunando Roy, Tony Brooks, Helena Tutill, Paola Niola, Patricia Dyal, Charlotte Williams, Leysa Forrest, Yasmin Panchbhaya, Jacqueline Findlay, Samuel Weeks, Julianne Brown, Kathryn Harris, Paul Randell, James Price, Alison Holmes, Judith Breuer |
| EPI_ISL_703768 | Wales Specialist Virology Centre Sequencing lab: Pathogen Genomics Unit | COVID-19 Genomics UK (COG-UK) Consortium | Catherine Moore, Johnathan Evans, Laura Gifford, Malorie Perry, Simon Cottrell, Angela Marchbank, Alec Birchley, Alexander Adams, Amy Gaskin, Bree Gatica-Wilcox, Jason Coombes, Joel Southgate, Lauren Gilbert, Lee Graham, Nicole Pacchiarini, Sara Kumziene-Summerhayes, Sarah Taylor, Sophie Jones, Sara Rey, Matthew Bull, Joanne Watkins, Sally Corden, Tom Connor |
| EPI_ISL_703770 | Liverpool Clinical Laboratories | COVID-19 Genomics UK (COG-UK) Consortium | Sam Haldenby, Anita Lucaci, Steve Paterson, Julian Hiscox, Alistair Darby, M Almsaud, A Alrezaihi, Muhannad Alruwaili, Stuart D Armstrong, Jones Benjamin, Eleanor G Bentley, Anu Chawla, Jordan J Clark, Angela Cowell, Richard Eccles, Isabel García-Dorival, Matthew Gemmell, Alessandro Gerada, PKF Gilmore, Richard Gregory, Ximeng Han, Catherine Hartley, Margaret Hughes, Miren Iturriza-Gomara, James Johnson, L Luu, Jenifer Manson, Charlotte Nelson, Elaine O'Toole, Cassie Olateju, Rebekah Penrice-Randal , Lucille Rainbow, N.P Randle, Trevor Ian Robinson, Parul Sharma, Ghada T Shawli, James P Stewart, Neil Swainston, Ecaterina Vamos, Joanne Watts, Mark Whitehead |
| EPI_ISL_703810 | Virology Department, Royal Infirmary of Edinburgh, NHS Lothian / School of Biological Sciences, University of Edinburgh / Institute of Genetics and Molecular Medicine, University of Edinburgh | COVID-19 Genomics UK (COG-UK) Consortium | McHugh M, Dewar R, Rooke S, Gallagher M, Balcaza C, O'Toole Á, Scher E, Hill V, McCrone JT, Colquhoun R, Yu X, Jackson B, Rambaut A, Williams TC, Templeton K |
| EPI_ISL_703858, EPI_ISL_703860 | Wales Specialist Virology Centre Sequencing lab: Pathogen Genomics Unit | COVID-19 Genomics UK (COG-UK) Consortium | Catherine Moore, Johnathan Evans, Laura Gifford, Malorie Perry, Simon Cottrell, Angela Marchbank, Alec Birchley, Alexander Adams, Amy Gaskin, Bree Gatica-Wilcox, Jason Coombes, Joel Southgate, Lauren Gilbert, Lee Graham, Nicole Pacchiarini, Sara Kumziene-Summerhayes, Sarah Taylor, Sophie Jones, Sara Rey, Matthew Bull, Joanne Watkins, Sally Corden, Tom Connor |
| EPI_ISL_703865 | University College London, Great Ormond Street Hospital for Children NHS Foundation Trust, Imperial College Healthcare NHS Trust | COVID-19 Genomics UK (COG-UK) Consortium | Sergi Castellano, Rachel Williams, Mark Kristiansen, Paola Resende Silva, Sunando Roy, Tony Brooks, Helena Tutill, Paola Niola, Patricia Dyal, Charlotte Williams, Leysa Forrest, Yasmin Panchbhaya, Jacqueline Findlay, Samuel Weeks, Julianne Brown, Kathryn Harris, Paul Randell, James Price, Alison Holmes, Judith Breuer |
| EPI_ISL_703870, EPI_ISL_703873 | Wales Specialist Virology Centre Sequencing lab: Pathogen Genomics Unit | COVID-19 Genomics UK (COG-UK) Consortium | Catherine Moore, Johnathan Evans, Laura Gifford, Malorie Perry, Simon Cottrell, Angela Marchbank, Alec Birchley, Alexander Adams, Amy Gaskin, Bree Gatica-Wilcox, Jason Coombes, Joel Southgate, Lauren Gilbert, Lee Graham, Nicole Pacchiarini, Sara Kumziene-Summerhayes, Sarah Taylor, Sophie Jones, Sara Rey, Matthew Bull, Joanne Watkins, Sally Corden, Tom Connor |
| EPI_ISL_703879 | University College London, Great Ormond Street Hospital for Children NHS Foundation Trust, Imperial College Healthcare NHS Trust | COVID-19 Genomics UK (COG-UK) Consortium | Sergi Castellano, Rachel Williams, Mark Kristiansen, Paola Resende Silva, Sunando Roy, Tony Brooks, Helena Tutill, Paola Niola, Patricia Dyal, Charlotte Williams, Leysa Forrest, Yasmin Panchbhaya, Jacqueline Findlay, Samuel Weeks, Julianne Brown, Kathryn Harris, Paul Randell, James Price, Alison Holmes, Judith Breuer |
| EPI_ISL_703884, EPI_ISL_703891, EPI_ISL_703897 | Liverpool Clinical Laboratories | COVID-19 Genomics UK (COG-UK) Consortium | Sam Haldenby, Anita Lucaci, Steve Paterson, Julian Hiscox, Alistair Darby, M Almsaud, A Alrezaihi, Muhannad Alruwaili, Stuart D Armstrong, Jones Benjamin, Eleanor G Bentley, Anu Chawla, Jordan J Clark, Angela Cowell, Richard Eccles, Isabel García-Dorival, Matthew Gemmell, Alessandro Gerada, |

| | | | |
|---|---|---|---|
| | | | PKF Gilmore, Richard Gregory, Ximeng Han, Catherine Hartley, Margaret Hughes, Miren Iturriza-Gomara, James Johnson, L Luu, Jenifer Manson, Charlotte Nelson, Elaine O'Toole, Cassie Olateju, Rebekah Penrice-Randal , Lucille Rainbow, N.P Randle, Trevor Ian Robinson, Parul Sharma, Ghada T Shawli, James P Stewart, Neil Swainston, Ecaterina Vamos, Joanne Watts, Mark Whitehead |
| EPI_ISL_703911 | West of Scotland Specialist Virology Centre, NHSGGC / MRC-University of Glasgow Centre for Virus Research | COVID-19 Genomics UK (COG-UK) Consortium | Ana da Silva Filipe, Natasha Johnson, Kathy Smollett, Daniel Mair, Stephen Carmichael, Alice Broos, Lily Tong, Jenna Nichols, Kyriaki Nomikou; Sarah McDonald; Richard Orton, Joseph Hughes, Sreenu Vattipally, David L Robertson; Alasdair MacLean, Rory Gunson; Sharif Shaaban, Matthew Holden; Rachel Blacow, Guy Mollett, Kathy Li, James Shepherd, Antonia Ho, Emma Thomson |
| EPI_ISL_703937 | Liverpool Clinical Laboratories | COVID-19 Genomics UK (COG-UK) Consortium | Sam Haldenby, Anita Lucaci, Steve Paterson, Julian Hiscox, Alistair Darby, M Almsaud, A Alrezaihi, Muhannad Alruwaili, Stuart D Armstrong, Jones Benjamin, Eleanor G Bentley, Anu Chawla, Jordan J Clark, Angela Cowell, Richard Eccles, Isabel García-Dorival, Matthew Gemmell, Alessandro Gerada, PKF Gilmore, Richard Gregory, Ximeng Han, Catherine Hartley, Margaret Hughes, Miren Iturriza-Gomara, James Johnson, L Luu, Jenifer Manson, Charlotte Nelson, Elaine O'Toole, Cassie Olateju, Rebekah Penrice-Randal , Lucille Rainbow, N.P Randle, Trevor Ian Robinson, Parul Sharma, Ghada T Shawli, James P Stewart, Neil Swainston, Ecaterina Vamos, Joanne Watts, Mark Whitehead |
| EPI_ISL_703955 | Wales Specialist Virology Centre Sequencing lab: Pathogen Genomics Unit | COVID-19 Genomics UK (COG-UK) Consortium | Catherine Moore, Johnathan Evans, Laura Gifford, Malorie Perry, Simon Cottrell, Angela Marchbank, Alec Birchley, Alexander Adams, Amy Gaskin, Bree Gatica-Wilcox, Jason Coombes, Joel Southgate, Lauren Gilbert, Lee Graham, Nicole Pacchiarini, Sara Kumziene-Summerhayes, Sarah Taylor, Sophie Jones, Sara Rey, Matthew Bull, Joanne Watkins, Sally Corden, Tom Connor |
| EPI_ISL_703977 | Liverpool Clinical Laboratories | COVID-19 Genomics UK (COG-UK) Consortium | Sam Haldenby, Anita Lucaci, Steve Paterson, Julian Hiscox, Alistair Darby, M Almsaud, A Alrezaihi, Muhannad Alruwaili, Stuart D Armstrong, Jones Benjamin, Eleanor G Bentley, Anu Chawla, Jordan J Clark, Angela Cowell, Richard Eccles, Isabel García-Dorival, Matthew Gemmell, Alessandro Gerada, PKF Gilmore, Richard Gregory, Ximeng Han, Catherine Hartley, Margaret Hughes, Miren Iturriza-Gomara, James Johnson, L Luu, Jenifer Manson, Charlotte Nelson, Elaine O'Toole, Cassie Olateju, Rebekah Penrice-Randal , Lucille Rainbow, N.P Randle, Trevor Ian Robinson, Parul Sharma, Ghada T Shawli, James P Stewart, Neil Swainston, Ecaterina Vamos, Joanne Watts, Mark Whitehead |
| EPI_ISL_704001, EPI_ISL_704014 | Wales Specialist Virology Centre Sequencing lab: Pathogen Genomics Unit | COVID-19 Genomics UK (COG-UK) Consortium | Catherine Moore, Johnathan Evans, Laura Gifford, Malorie Perry, Simon Cottrell, Angela Marchbank, Alec Birchley, Alexander Adams, Amy Gaskin, Bree Gatica-Wilcox, Jason Coombes, Joel Southgate, Lauren Gilbert, Lee Graham, Nicole Pacchiarini, Sara Kumziene-Summerhayes, Sarah Taylor, Sophie Jones, Sara Rey, Matthew Bull, Joanne Watkins, Sally Corden, Tom Connor |
| EPI_ISL_704025 | University College London, Great Ormond Street Hospital for Children NHS Foundation Trust, Imperial College Healthcare NHS Trust | COVID-19 Genomics UK (COG-UK) Consortium | Sergi Castellano, Rachel Williams, Mark Kristiansen, Paola Resende Silva, Sunando Roy, Tony Brooks, Helena Tutill, Paola Niola, Patricia Dyal, Charlotte Williams, Leysa Forrest, Yasmin Panchbhaya, Jacqueline Findlay, Samuel Weeks, Julianne Brown, Kathryn Harris, Paul Randell, James Price, Alison Holmes, Judith Breuer |
| EPI_ISL_704030 | Wales Specialist Virology Centre Sequencing lab: Pathogen Genomics Unit | COVID-19 Genomics UK (COG-UK) Consortium | Catherine Moore, Johnathan Evans, Laura Gifford, Malorie Perry, Simon Cottrell, Angela Marchbank, Alec Birchley, Alexander Adams, Amy Gaskin, Bree Gatica-Wilcox, Jason Coombes, Joel Southgate, Lauren Gilbert, Lee Graham, Nicole Pacchiarini, Sara Kumziene-Summerhayes, Sarah Taylor, Sophie Jones, Sara Rey, Matthew Bull, Joanne Watkins, Sally Corden, Tom Connor |
| EPI_ISL_704036 | University College London, Great Ormond Street Hospital for Children NHS Foundation Trust, Imperial College Healthcare NHS Trust | COVID-19 Genomics UK (COG-UK) Consortium | Sergi Castellano, Rachel Williams, Mark Kristiansen, Paola Resende Silva, Sunando Roy, Tony Brooks, Helena Tutill, Paola Niola, Patricia Dyal, Charlotte Williams, Leysa Forrest, Yasmin Panchbhaya, Jacqueline Findlay, Samuel Weeks, Julianne Brown, Kathryn Harris, Paul Randell, James Price, Alison Holmes, Judith Breuer |
| EPI_ISL_704038, EPI_ISL_704044 | Wales Specialist Virology Centre Sequencing lab: Pathogen Genomics Unit | COVID-19 Genomics UK (COG-UK) Consortium | Catherine Moore, Johnathan Evans, Laura Gifford, Malorie Perry, Simon Cottrell, Angela Marchbank, Alec Birchley, Alexander Adams, Amy Gaskin, Bree Gatica-Wilcox, Jason Coombes, Joel Southgate, Lauren Gilbert, Lee Graham, Nicole Pacchiarini, Sara Kumziene-Summerhayes, Sarah Taylor, Sophie Jones, Sara Rey, Matthew Bull, Joanne Watkins, Sally Corden, Tom Connor |
| EPI_ISL_704049 | Liverpool Clinical Laboratories | COVID-19 Genomics UK (COG-UK) Consortium | Sam Haldenby, Anita Lucaci, Steve Paterson, Julian Hiscox, Alistair Darby, M Almsaud, A Alrezaihi, Muhannad Alruwaili, Stuart D Armstrong, Jones Benjamin, Eleanor G Bentley, Anu Chawla, Jordan J Clark, Angela Cowell, Richard Eccles, Isabel García-Dorival, Matthew Gemmell, Alessandro Gerada, PKF Gilmore, Richard Gregory, Ximeng Han, Catherine Hartley, Margaret Hughes, Miren Iturriza-Gomara, James Johnson, L Luu, Jenifer Manson, Charlotte Nelson, Elaine O'Toole, Cassie Olateju, Rebekah Penrice-Randal , Lucille Rainbow, N.P Randle, Trevor Ian Robinson, Parul Sharma, Ghada T Shawli, James P Stewart, Neil Swainston, Ecaterina Vamos, Joanne Watts, Mark Whitehead |
| EPI_ISL_704086 | University College London, Great Ormond Street Hospital for Children NHS Foundation Trust, Imperial College Healthcare NHS Trust | COVID-19 Genomics UK (COG-UK) Consortium | Sergi Castellano, Rachel Williams, Mark Kristiansen, Paola Resende Silva, Sunando Roy, Tony Brooks, Helena Tutill, Paola Niola, Patricia Dyal, Charlotte Williams, Leysa Forrest, Yasmin Panchbhaya, Jacqueline Findlay, Samuel Weeks, Julianne Brown, Kathryn Harris, Paul Randell, James Price, Alison Holmes, Judith Breuer |
| EPI_ISL_704111, EPI_ISL_704161 | Wales Specialist Virology Centre Sequencing lab: Pathogen Genomics Unit | COVID-19 Genomics UK (COG-UK) Consortium | Catherine Moore, Johnathan Evans, Laura Gifford, Malorie Perry, Simon Cottrell, Angela Marchbank, Alec Birchley, Alexander Adams, Amy Gaskin, Bree Gatica-Wilcox, Jason Coombes, Joel Southgate, Lauren Gilbert, Lee Graham, Nicole Pacchiarini, Sara Kumziene-Summerhayes, Sarah Taylor, Sophie Jones, Sara Rey, Matthew Bull, Joanne Watkins, Sally Corden, Tom Connor |
| EPI_ISL_704179 | University College London, Great Ormond Street Hospital for Children NHS Foundation Trust, Imperial College Healthcare NHS Trust | COVID-19 Genomics UK (COG-UK) Consortium | Sergi Castellano, Rachel Williams, Mark Kristiansen, Paola Resende Silva, Sunando Roy, Tony Brooks, Helena Tutill, Paola Niola, Patricia Dyal, Charlotte Williams, Leysa Forrest, Yasmin Panchbhaya, Jacqueline Findlay, Samuel Weeks, Julianne Brown, Kathryn Harris, Paul Randell, James Price, Alison Holmes, Judith Breuer |
| EPI_ISL_704209 | Liverpool Clinical Laboratories | COVID-19 Genomics UK (COG-UK) Consortium | Sam Haldenby, Anita Lucaci, Steve Paterson, Julian Hiscox, Alistair Darby, M Almsaud, A Alrezaihi, Muhannad Alruwaili, Stuart D Armstrong, Jones Benjamin, Eleanor G Bentley, Anu Chawla, Jordan J Clark, Angela Cowell, Richard Eccles, Isabel García-Dorival, Matthew Gemmell, Alessandro Gerada, PKF Gilmore, Richard Gregory, Ximeng Han, Catherine Hartley, Margaret Hughes, Miren Iturriza-Gomara, James Johnson, L Luu, Jenifer Manson, Charlotte Nelson, Elaine O'Toole, Cassie Olateju, Rebekah Penrice-Randal , Lucille Rainbow, N.P Randle, Trevor Ian Robinson, Parul Sharma, Ghada T Shawli, James P Stewart, Neil Swainston, Ecaterina Vamos, Joanne Watts, Mark Whitehead |
| EPI_ISL_704312, EPI_ISL_704372 | University College London, Great Ormond Street Hospital for Children NHS Foundation Trust, Imperial College Healthcare NHS Trust | COVID-19 Genomics UK (COG-UK) Consortium | Sergi Castellano, Rachel Williams, Mark Kristiansen, Paola Resende Silva, Sunando Roy, Tony Brooks, Helena Tutill, Paola Niola, Patricia Dyal, Charlotte Williams, Leysa Forrest, Yasmin Panchbhaya, Jacqueline Findlay, Samuel Weeks, Julianne Brown, Kathryn Harris, Paul Randell, James Price, Alison Holmes, Judith Breuer |
| EPI_ISL_704382, EPI_ISL_704502, EPI_ISL_704505, EPI_ISL_704507 | Department of Pathology, University of Cambridge | COVID-19 Genomics UK (COG-UK) Consortium | Aminu S. Jahun, Yasmin Chaudhry, Grant Hall, Iliana Georgana, Myra Hosmillo, Martin D. Curran, Malte Pinckert, Surendra Parmar, Ian Goodfellow |
| EPI_ISL_704572, EPI_ISL_704573, EPI_ISL_704574, EPI_ISL_704578 | Northumbria University / South Tees Hospitals NHS Foundation Trust / North Cumbria Integrated Care NHS Foundation Trust / North Tees and Hartlepool NHS Foundation Trust / Newcastle Hospitals NHS Foundation Trust | COVID-19 Genomics UK (COG-UK) Consortium | Darren L Smith,Andrew Nelson,Matthew Bashton,Greg R Young,Joshua Loh,John Allan,Mohammad A Tariq,Giles S Holt,Gary Black,Wen C Yew,Lynn Dover,Paul Baker,Steve Liggett,Sarah Essex,Jane Greenaway,Debra Padgett,Clive Graham,Garren Scott,Edward Barton,Emma Swindells,Brendan Payne,Jennifer Collins,Yusri Taha,Gary Eltringham |
| EPI_ISL_704584 | Wales Specialist Virology Centre Sequencing lab: Pathogen Genomics Unit | COVID-19 Genomics UK (COG-UK) Consortium | Catherine Moore, Johnathan Evans, Laura Gifford, Malorie Perry, Simon Cottrell, Angela Marchbank, Alec Birchley, Alexander Adams, Amy Gaskin, Bree Gatica-Wilcox, Jason Coombes, Joel Southgate, Lauren Gilbert, Lee Graham, Nicole Pacchiarini, Sara Kumziene-Summerhayes, Sarah Taylor, Sophie Jones, Sara Rey, Matthew Bull, Joanne Watkins, Sally Corden, Tom Connor |
| EPI_ISL_704595 | West of Scotland Specialist Virology Centre, NHSGGC / MRC-University of Glasgow Centre for Virus Research | COVID-19 Genomics UK (COG-UK) Consortium | Ana da Silva Filipe, Natasha Johnson, Kathy Smollett, Daniel Mair, Stephen Carmichael, Alice Broos, Lily Tong, Jenna Nichols, Kyriaki Nomikou; Sarah McDonald; Richard Orton, Joseph Hughes, Sreenu Vattipally, David L Robertson; Alasdair MacLean, Rory Gunson; Sharif Shaaban, Matthew Holden; Rachel Blacow, Guy Mollett, Kathy Li, James Shepherd, Antonia Ho, Emma Thomson |
| EPI_ISL_704605 | Virology Department, Sheffield Teaching Hospitals NHS Foundation Trust/Department of Infection, Immunity and Cardiovascular Disease, The Medical School, University of Sheffield | COVID-19 Genomics UK (COG-UK) Consortium | Thushan de Silva, Matthew Parker, Nikki Smith, Adri Angyal, Rebecca Brown, Luke Green, Rachel Tucker, Paul Parsons, Danielle Groves, Katie Johnson, Laura Carrilero, Alex Keeley, Dave Partridge, Matthew Wyles, Benjamin Lindsey, Mehmet Yavuz, Mohammad Raza, Cariad Evans |
| EPI_ISL_704628 | Quadram Institute Bioscience | COVID-19 Genomics UK (COG-UK) Consortium | Dave J. Baker, Gemma L. Kay, Alp Aydin, Thanh Le-Viet, Steven Rudder, Ana P. Tedim, Anastasia Kolyva, Maria Diaz, Leonardo de Oliveira Martins, Nabil-Fareed Alikhan, Lizzie Meadows, Rachael Stanley, Ngozi Elumogo, Muhammed Yasir, Nicholas M. Thomson, Alexander J Trotter, Rachel Gilroy, Samuel Bloomfield, Claire Stuart, Andrew Bell, Reenesh Prakash, Samir Dervisevic, Alison E. Mather, John Wain, Mark Webber, Andrew J. Page, Justin O'Grady |

| | | | |
|---|---|---|---|
| EPI_ISL_704750, EPI_ISL_704867 | Lighthouse Lab in Cambridge | Wellcome Sanger Institute for the COVID-19 Genomics UK (COG-UK) Consortium | Rob Howes, The Lighthouse Lab in Cambridge and Alex Alderton, Roberto Amato, Sonia Goncalves, Ewan Harrison, David K. Jackson, Ian Johnston, Dominic Kwiatkowski, Cordelia Langford, John Sillitoe on behalf of the Wellcome Sanger Institute COVID-19 Surveillance Team |
| EPI_ISL_704905 | Liverpool Clinical Laboratories | COVID-19 Genomics UK (COG-UK) Consortium | Sam Haldenby, Anita Lucaci, Steve Paterson, Julian Hiscox, Alistair Darby, M Almsaud, A Alrezaihi, Muhannad Alruwaili, Stuart D Armstrong, Jones Benjamin, Eleanor G Bentley, Anu Chawla, Jordan J Clark, Angela Cowell, Richard Eccles, Isabel García-Dorival, Matthew Gemmell, Alessandro Gerada, PKF Gilmore, Richard Gregory, Ximeng Han, Catherine Hartley, Margaret Hughes, Miren Iturriza-Gomara, James Johnson, L Luu, Jenifer Manson, Charlotte Nelson, Elaine O'Toole, Cassie Olateju, Rebekah Penrice-Randal , Lucille Rainbow, N.P Randle, Trevor Ian Robinson, Parul Sharma, Ghada T Shawli, James P Stewart, Neil Swainston, Ecaterina Vamos, Joanne Watts, Mark Whitehead |
| EPI_ISL_704931 | Wales Specialist Virology Centre Sequencing lab: Pathogen Genomics Unit | COVID-19 Genomics UK (COG-UK) Consortium | Catherine Moore, Johnathan Evans, Laura Gifford, Malorie Perry, Simon Cottrell, Angela Marchbank, Alec Birchley, Alexander Adams, Amy Gaskin, Bree Gatica-Wilcox, Jason Coombes, Joel Southgate, Lauren Gilbert, Lee Graham, Nicole Pacchiarini, Sara Kumziene-Summerhayes, Sarah Taylor, Sophie Jones, Sara Rey, Matthew Bull, Joanne Watkins, Sally Corden, Tom Connor |
| EPI_ISL_704932 | Lighthouse Lab in Cambridge | Wellcome Sanger Institute for the COVID-19 Genomics UK (COG-UK) Consortium | Rob Howes, The Lighthouse Lab in Cambridge and Alex Alderton, Roberto Amato, Sonia Goncalves, Ewan Harrison, David K. Jackson, Ian Johnston, Dominic Kwiatkowski, Cordelia Langford, John Sillitoe on behalf of the Wellcome Sanger Institute COVID-19 Surveillance Team |
| EPI_ISL_704950 | Liverpool Clinical Laboratories | COVID-19 Genomics UK (COG-UK) Consortium | Sam Haldenby, Anita Lucaci, Steve Paterson, Julian Hiscox, Alistair Darby, M Almsaud, A Alrezaihi, Muhannad Alruwaili, Stuart D Armstrong, Jones Benjamin, Eleanor G Bentley, Anu Chawla, Jordan J Clark, Angela Cowell, Richard Eccles, Isabel García-Dorival, Matthew Gemmell, Alessandro Gerada, PKF Gilmore, Richard Gregory, Ximeng Han, Catherine Hartley, Margaret Hughes, Miren Iturriza-Gomara, James Johnson, L Luu, Jenifer Manson, Charlotte Nelson, Elaine O'Toole, Cassie Olateju, Rebekah Penrice-Randal , Lucille Rainbow, N.P Randle, Trevor Ian Robinson, Parul Sharma, Ghada T Shawli, James P Stewart, Neil Swainston, Ecaterina Vamos, Joanne Watts, Mark Whitehead |
| EPI_ISL_704955, EPI_ISL_704965 | University College London, Great Ormond Street Hospital for Children NHS Foundation Trust, Imperial College Healthcare NHS Trust | COVID-19 Genomics UK (COG-UK) Consortium | Sergi Castellano, Rachel Williams, Mark Kristiansen, Paola Resende Silva, Sunando Roy, Tony Brooks, Helena Tutill, Paola Niola, Patricia Dyal, Charlotte Williams, Leysa Forrest, Yasmin Panchbhaya, Jacqueline Findlay, Samuel Weeks, Julianne Brown, Kathryn Harris, Paul Randell, James Price, Alison Holmes, Judith Breuer |
| EPI_ISL_704968, EPI_ISL_704973 | Liverpool Clinical Laboratories | COVID-19 Genomics UK (COG-UK) Consortium | Sam Haldenby, Anita Lucaci, Steve Paterson, Julian Hiscox, Alistair Darby, M Almsaud, A Alrezaihi, Muhannad Alruwaili, Stuart D Armstrong, Jones Benjamin, Eleanor G Bentley, Anu Chawla, Jordan J Clark, Angela Cowell, Richard Eccles, Isabel García-Dorival, Matthew Gemmell, Alessandro Gerada, PKF Gilmore, Richard Gregory, Ximeng Han, Catherine Hartley, Margaret Hughes, Miren Iturriza-Gomara, James Johnson, L Luu, Jenifer Manson, Charlotte Nelson, Elaine O'Toole, Cassie Olateju, Rebekah Penrice-Randal , Lucille Rainbow, N.P Randle, Trevor Ian Robinson, Parul Sharma, Ghada T Shawli, James P Stewart, Neil Swainston, Ecaterina Vamos, Joanne Watts, Mark Whitehead |
| EPI_ISL_705010 | Northumbria University / South Tees Hospitals NHS Foundation Trust / North Cumbria Integrated Care NHS Foundation Trust / North Tees and Hartlepool NHS Foundation Trust / Newcastle Hospitals NHS Foundation Trust | COVID-19 Genomics UK (COG-UK) Consortium | Darren L Smith,Andrew Nelson,Matthew Bashton,Greg R Young,Joshua Loh,John Allan,Mohammad A Tariq,Giles S Holt,Gary Black,Wen C Yew,Lynn Dover,Paul Baker,Steve Liggett,Sarah Essex,Jane Greenaway,Debra Padgett,Clive Graham,Garren Scott,Edward Barton,Emma Swindells,Brendan Payne,Jennifer Collins,Yusri Taha,Gary Eltringham |
| EPI_ISL_705054 | Liverpool Clinical Laboratories | COVID-19 Genomics UK (COG-UK) Consortium | Sam Haldenby, Anita Lucaci, Steve Paterson, Julian Hiscox, Alistair Darby, M Almsaud, A Alrezaihi, Muhannad Alruwaili, Stuart D Armstrong, Jones Benjamin, Eleanor G Bentley, Anu Chawla, Jordan J Clark, Angela Cowell, Richard Eccles, Isabel García-Dorival, Matthew Gemmell, Alessandro Gerada, PKF Gilmore, Richard Gregory, Ximeng Han, Catherine Hartley, Margaret Hughes, Miren Iturriza-Gomara, James Johnson, L Luu, Jenifer Manson, Charlotte Nelson, Elaine O'Toole, Cassie Olateju, Rebekah Penrice-Randal , Lucille Rainbow, N.P Randle, Trevor Ian Robinson, Parul Sharma, Ghada T Shawli, James P Stewart, Neil Swainston, Ecaterina Vamos, Joanne Watts, Mark Whitehead |
| EPI_ISL_705061 | Lighthouse Lab in Cambridge | Wellcome Sanger Institute for the COVID-19 Genomics UK (COG-UK) Consortium | Rob Howes, The Lighthouse Lab in Cambridge and Alex Alderton, Roberto Amato, Sonia Goncalves, Ewan Harrison, David K. Jackson, Ian Johnston, Dominic Kwiatkowski, Cordelia Langford, John Sillitoe on behalf of the Wellcome Sanger Institute COVID-19 Surveillance Team |
| EPI_ISL_705086 | Liverpool Clinical Laboratories | COVID-19 Genomics UK (COG-UK) Consortium | Sam Haldenby, Anita Lucaci, Steve Paterson, Julian Hiscox, Alistair Darby, M Almsaud, A Alrezaihi, Muhannad Alruwaili, Stuart D Armstrong, Jones Benjamin, Eleanor G Bentley, Anu Chawla, Jordan J Clark, Angela Cowell, Richard Eccles, Isabel García-Dorival, Matthew Gemmell, Alessandro Gerada, PKF Gilmore, Richard Gregory, Ximeng Han, Catherine Hartley, Margaret Hughes, Miren Iturriza-Gomara, James Johnson, L Luu, Jenifer Manson, Charlotte Nelson, Elaine O'Toole, Cassie Olateju, Rebekah Penrice-Randal , Lucille Rainbow, N.P Randle, Trevor Ian Robinson, Parul Sharma, Ghada T Shawli, James P Stewart, Neil Swainston, Ecaterina Vamos, Joanne Watts, Mark Whitehead |
| EPI_ISL_705104 | Lighthouse Lab in Cambridge | Wellcome Sanger Institute for the COVID-19 Genomics UK (COG-UK) Consortium | Rob Howes, The Lighthouse Lab in Cambridge and Alex Alderton, Roberto Amato, Sonia Goncalves, Ewan Harrison, David K. Jackson, Ian Johnston, Dominic Kwiatkowski, Cordelia Langford, John Sillitoe on behalf of the Wellcome Sanger Institute COVID-19 Surveillance Team |
| EPI_ISL_705137 | Northumbria University / South Tees Hospitals NHS Foundation Trust / North Cumbria Integrated Care NHS Foundation Trust / North Tees and Hartlepool NHS Foundation Trust / Newcastle Hospitals NHS Foundation Trust | COVID-19 Genomics UK (COG-UK) Consortium | Darren L Smith,Andrew Nelson,Matthew Bashton,Greg R Young,Joshua Loh,John Allan,Mohammad A Tariq,Giles S Holt,Gary Black,Wen C Yew,Lynn Dover,Paul Baker,Steve Liggett,Sarah Essex,Jane Greenaway,Debra Padgett,Clive Graham,Garren Scott,Edward Barton,Emma Swindells,Brendan Payne,Jennifer Collins,Yusri Taha,Gary Eltringham |
| EPI_ISL_705143 | University College London, Great Ormond Street Hospital for Children NHS Foundation Trust, Imperial College Healthcare NHS Trust | COVID-19 Genomics UK (COG-UK) Consortium | Sergi Castellano, Rachel Williams, Mark Kristiansen, Paola Resende Silva, Sunando Roy, Tony Brooks, Helena Tutill, Paola Niola, Patricia Dyal, Charlotte Williams, Leysa Forrest, Yasmin Panchbhaya, Jacqueline Findlay, Samuel Weeks, Julianne Brown, Kathryn Harris, Paul Randell, James Price, Alison Holmes, Judith Breuer |
| EPI_ISL_705148 | Northumbria University / South Tees Hospitals NHS Foundation Trust / North Cumbria Integrated Care NHS Foundation Trust / North Tees and Hartlepool NHS Foundation Trust / Newcastle Hospitals NHS Foundation Trust | COVID-19 Genomics UK (COG-UK) Consortium | Darren L Smith,Andrew Nelson,Matthew Bashton,Greg R Young,Joshua Loh,John Allan,Mohammad A Tariq,Giles S Holt,Gary Black,Wen C Yew,Lynn Dover,Paul Baker,Steve Liggett,Sarah Essex,Jane Greenaway,Debra Padgett,Clive Graham,Garren Scott,Edward Barton,Emma Swindells,Brendan Payne,Jennifer Collins,Yusri Taha,Gary Eltringham |
| EPI_ISL_705168, EPI_ISL_705176 | Liverpool Clinical Laboratories | COVID-19 Genomics UK (COG-UK) Consortium | Sam Haldenby, Anita Lucaci, Steve Paterson, Julian Hiscox, Alistair Darby, M Almsaud, A Alrezaihi, Muhannad Alruwaili, Stuart D Armstrong, Jones Benjamin, Eleanor G Bentley, Anu Chawla, Jordan J Clark, Angela Cowell, Richard Eccles, Isabel García-Dorival, Matthew Gemmell, Alessandro Gerada, PKF Gilmore, Richard Gregory, Ximeng Han, Catherine Hartley, Margaret Hughes, Miren Iturriza-Gomara, James Johnson, L Luu, Jenifer Manson, Charlotte Nelson, Elaine O'Toole, Cassie Olateju, Rebekah Penrice-Randal , Lucille Rainbow, N.P Randle, Trevor Ian Robinson, Parul Sharma, Ghada T Shawli, James P Stewart, Neil Swainston, Ecaterina Vamos, Joanne Watts, Mark Whitehead |
| EPI_ISL_705191, EPI_ISL_705192 | Lighthouse Lab in Milton Keynes | Wellcome Sanger Institute for the COVID-19 Genomics UK (COG-UK) Consortium | The Lighthouse Lab in Milton Keynes and Alex Alderton, Roberto Amato, Sonia Goncalves, Ewan Harrison, David K. Jackson, Ian Johnston, Dominic Kwiatkowski, Cordelia Langford, John Sillitoe on behalf of the Wellcome Sanger Institute COVID-19 Surveillance Team |
| EPI_ISL_705193 | Liverpool Clinical Laboratories | COVID-19 Genomics UK (COG-UK) Consortium | Sam Haldenby, Anita Lucaci, Steve Paterson, Julian Hiscox, Alistair Darby, M Almsaud, A Alrezaihi, Muhannad Alruwaili, Stuart D Armstrong, Jones Benjamin, Eleanor G Bentley, Anu Chawla, Jordan J Clark, Angela Cowell, Richard Eccles, Isabel García-Dorival, Matthew Gemmell, Alessandro Gerada, PKF Gilmore, Richard Gregory, Ximeng Han, Catherine Hartley, Margaret Hughes, Miren Iturriza-Gomara, James Johnson, L Luu, Jenifer Manson, Charlotte Nelson, Elaine O'Toole, Cassie Olateju, Rebekah Penrice-Randal , Lucille Rainbow, N.P Randle, Trevor Ian Robinson, Parul Sharma, Ghada T Shawli, James P Stewart, Neil Swainston, Ecaterina Vamos, Joanne Watts, Mark Whitehead |
| EPI_ISL_705194, EPI_ISL_705196, EPI_ISL_705197, EPI_ISL_705199, EPI_ISL_705200, EPI_ISL_705202 | Lighthouse Lab in Milton Keynes | Wellcome Sanger Institute for the COVID-19 Genomics UK (COG-UK) Consortium | The Lighthouse Lab in Milton Keynes and Alex Alderton, Roberto Amato, Sonia Goncalves, Ewan Harrison, David K. Jackson, Ian Johnston, Dominic Kwiatkowski, Cordelia Langford, John Sillitoe on behalf of the Wellcome Sanger Institute COVID-19 Surveillance Team |
| EPI_ISL_705203, EPI_ISL_705208 | Liverpool Clinical Laboratories | COVID-19 Genomics UK (COG-UK) Consortium | Sam Haldenby, Anita Lucaci, Steve Paterson, Julian Hiscox, Alistair Darby, M Almsaud, A Alrezaihi, Muhannad Alruwaili, Stuart D Armstrong, Jones Benjamin, Eleanor G Bentley, Anu Chawla, Jordan J Clark, Angela Cowell, Richard Eccles, Isabel García-Dorival, Matthew Gemmell, Alessandro Gerada, PKF Gilmore, Richard Gregory, Ximeng Han, Catherine Hartley, Margaret Hughes, Miren Iturriza-Gomara, James Johnson, L Luu, Jenifer Manson, Charlotte Nelson, Elaine O'Toole, Cassie Olateju, Rebekah Penrice-Randal , Lucille Rainbow, N.P Randle, Trevor Ian Robinson, Parul Sharma, Ghada T Shawli, James P Stewart, Neil Swainston, Ecaterina Vamos, Joanne Watts, Mark Whitehead |
| EPI_ISL_705209 | University College London, Great Ormond Street Hospital for Children NHS Foundation Trust, Imperial College Healthcare NHS Trust | COVID-19 Genomics UK (COG-UK) Consortium | Sergi Castellano, Rachel Williams, Mark Kristiansen, Paola Resende Silva, Sunando Roy, Tony Brooks, Helena Tutill, Paola Niola, Patricia Dyal, Charlotte Williams, Leysa Forrest, Yasmin Panchbhaya, Jacqueline Findlay, Samuel Weeks, Julianne Brown, Kathryn Harris, Paul Randell, James Price, Alison Holmes, Judith Breuer |

| | | | |
|---|---|---|---|
| EPI_ISL_705214 | Northumbria University / South Tees Hospitals NHS Foundation Trust / North Cumbria Integrated Care NHS Foundation Trust / North Tees and Hartlepool NHS Foundation Trust / Newcastle Hospitals NHS Foundation Trust | COVID-19 Genomics UK (COG-UK) Consortium | Darren L Smith,Andrew Nelson,Matthew Bashton,Greg R Young,Joshua Loh,John Allan,Mohammad A Tariq,Giles S Holt,Gary Black,Wen C Yew,Lynn Dover,Paul Baker,Steve Liggett,Sarah Essex,Jane Greenaway,Debra Padgett,Clive Graham,Garren Scott,Edward Barton,Emma Swindells,Brendan Payne,Jennifer Collins,Yusri Taha,Gary Eltringham |
| EPI_ISL_705231 | Liverpool Clinical Laboratories | COVID-19 Genomics UK (COG-UK) Consortium | Sam Haldenby, Anita Lucaci, Steve Paterson, Julian Hiscox, Alistair Darby, M Almsaud, A Alrezaihi, Muhannad Alruwaili, Stuart D Armstrong, Jones Benjamin, Eleanor G Bentley, Anu Chawla, Jordan J Clark, Angela Cowell, Richard Eccles, Isabel García-Dorival, Matthew Gemmell, Alessandro Gerada, PKF Gilmore, Richard Gregory, Ximeng Han, Catherine Hartley, Margaret Hughes, Miren Iturriza-Gomara, James Johnson, L Luu, Jenifer Manson, Charlotte Nelson, Elaine O'Toole, Cassie Olateju, Rebekah Penrice-Randal , Lucille Rainbow, N.P Randle, Trevor Ian Robinson, Parul Sharma, Ghada T Shawli, James P Stewart, Neil Swainston, Ecaterina Vamos, Joanne Watts, Mark Whitehead |
| EPI_ISL_705239, EPI_ISL_705244 | Wales Specialist Virology Centre Sequencing lab: Pathogen Genomics Unit | COVID-19 Genomics UK (COG-UK) Consortium | Catherine Moore, Johnathan Evans, Laura Gifford, Malorie Perry, Simon Cottrell, Angela Marchbank, Alec Birchley, Alexander Adams, Amy Gaskin, Bree Gatica-Wilcox, Jason Coombes, Joel Southgate, Lauren Gilbert, Lee Graham, Nicole Pacchiarini, Sara Kumziene-Summerhayes, Sarah Taylor, Sophie Jones, Sara Rey, Matthew Bull, Joanne Watkins, Sally Corden, Tom Connor |
| EPI_ISL_705266, EPI_ISL_705282 | Northumbria University / South Tees Hospitals NHS Foundation Trust / North Cumbria Integrated Care NHS Foundation Trust / North Tees and Hartlepool NHS Foundation Trust / Newcastle Hospitals NHS Foundation Trust | COVID-19 Genomics UK (COG-UK) Consortium | Darren L Smith,Andrew Nelson,Matthew Bashton,Greg R Young,Joshua Loh,John Allan,Mohammad A Tariq,Giles S Holt,Gary Black,Wen C Yew,Lynn Dover,Paul Baker,Steve Liggett,Sarah Essex,Jane Greenaway,Debra Padgett,Clive Graham,Garren Scott,Edward Barton,Emma Swindells,Brendan Payne,Jennifer Collins,Yusri Taha,Gary Eltringham |
| EPI_ISL_705287 | Liverpool Clinical Laboratories | COVID-19 Genomics UK (COG-UK) Consortium | Sam Haldenby, Anita Lucaci, Steve Paterson, Julian Hiscox, Alistair Darby, M Almsaud, A Alrezaihi, Muhannad Alruwaili, Stuart D Armstrong, Jones Benjamin, Eleanor G Bentley, Anu Chawla, Jordan J Clark, Angela Cowell, Richard Eccles, Isabel García-Dorival, Matthew Gemmell, Alessandro Gerada, PKF Gilmore, Richard Gregory, Ximeng Han, Catherine Hartley, Margaret Hughes, Miren Iturriza-Gomara, James Johnson, L Luu, Jenifer Manson, Charlotte Nelson, Elaine O'Toole, Cassie Olateju, Rebekah Penrice-Randal , Lucille Rainbow, N.P Randle, Trevor Ian Robinson, Parul Sharma, Ghada T Shawli, James P Stewart, Neil Swainston, Ecaterina Vamos, Joanne Watts, Mark Whitehead |
| EPI_ISL_705295 | Northumbria University / South Tees Hospitals NHS Foundation Trust / North Cumbria Integrated Care NHS Foundation Trust / North Tees and Hartlepool NHS Foundation Trust / Newcastle Hospitals NHS Foundation Trust | COVID-19 Genomics UK (COG-UK) Consortium | Darren L Smith,Andrew Nelson,Matthew Bashton,Greg R Young,Joshua Loh,John Allan,Mohammad A Tariq,Giles S Holt,Gary Black,Wen C Yew,Lynn Dover,Paul Baker,Steve Liggett,Sarah Essex,Jane Greenaway,Debra Padgett,Clive Graham,Garren Scott,Edward Barton,Emma Swindells,Brendan Payne,Jennifer Collins,Yusri Taha,Gary Eltringham |
| EPI_ISL_705310 | University College London, Great Ormond Street Hospital for Children NHS Foundation Trust, Imperial College Healthcare NHS Trust | COVID-19 Genomics UK (COG-UK) Consortium | Sergi Castellano, Rachel Williams, Mark Kristiansen, Paola Resende Silva, Sunando Roy, Tony Brooks, Helena Tutill, Paola Niola, Patricia Dyal, Charlotte Williams, Leysa Forrest, Yasmin Panchbhaya, Jacqueline Findlay, Samuel Weeks, Julianne Brown, Kathryn Harris, Paul Randell, James Price, Alison Holmes, Judith Breuer |
| EPI_ISL_705329 | Liverpool Clinical Laboratories | COVID-19 Genomics UK (COG-UK) Consortium | Sam Haldenby, Anita Lucaci, Steve Paterson, Julian Hiscox, Alistair Darby, M Almsaud, A Alrezaihi, Muhannad Alruwaili, Stuart D Armstrong, Jones Benjamin, Eleanor G Bentley, Anu Chawla, Jordan J Clark, Angela Cowell, Richard Eccles, Isabel García-Dorival, Matthew Gemmell, Alessandro Gerada, PKF Gilmore, Richard Gregory, Ximeng Han, Catherine Hartley, Margaret Hughes, Miren Iturriza-Gomara, James Johnson, L Luu, Jenifer Manson, Charlotte Nelson, Elaine O'Toole, Cassie Olateju, Rebekah Penrice-Randal , Lucille Rainbow, N.P Randle, Trevor Ian Robinson, Parul Sharma, Ghada T Shawli, James P Stewart, Neil Swainston, Ecaterina Vamos, Joanne Watts, Mark Whitehead |
| EPI_ISL_705342 | University College London, Great Ormond Street Hospital for Children NHS Foundation Trust, Imperial College Healthcare NHS Trust | COVID-19 Genomics UK (COG-UK) Consortium | Sergi Castellano, Rachel Williams, Mark Kristiansen, Paola Resende Silva, Sunando Roy, Tony Brooks, Helena Tutill, Paola Niola, Patricia Dyal, Charlotte Williams, Leysa Forrest, Yasmin Panchbhaya, Jacqueline Findlay, Samuel Weeks, Julianne Brown, Kathryn Harris, Paul Randell, James Price, Alison Holmes, Judith Breuer |
| EPI_ISL_705343, EPI_ISL_705353 | Northumbria University / South Tees Hospitals NHS Foundation Trust / North Cumbria Integrated Care NHS Foundation Trust / North Tees and Hartlepool NHS Foundation Trust / Newcastle Hospitals NHS Foundation Trust | COVID-19 Genomics UK (COG-UK) Consortium | Darren L Smith,Andrew Nelson,Matthew Bashton,Greg R Young,Joshua Loh,John Allan,Mohammad A Tariq,Giles S Holt,Gary Black,Wen C Yew,Lynn Dover,Paul Baker,Steve Liggett,Sarah Essex,Jane Greenaway,Debra Padgett,Clive Graham,Garren Scott,Edward Barton,Emma Swindells,Brendan Payne,Jennifer Collins,Yusri Taha,Gary Eltringham |
| EPI_ISL_705358 | Liverpool Clinical Laboratories | COVID-19 Genomics UK (COG-UK) Consortium | Sam Haldenby, Anita Lucaci, Steve Paterson, Julian Hiscox, Alistair Darby, M Almsaud, A Alrezaihi, Muhannad Alruwaili, Stuart D Armstrong, Jones Benjamin, Eleanor G Bentley, Anu Chawla, Jordan J Clark, Angela Cowell, Richard Eccles, Isabel García-Dorival, Matthew Gemmell, Alessandro Gerada, PKF Gilmore, Richard Gregory, Ximeng Han, Catherine Hartley, Margaret Hughes, Miren Iturriza-Gomara, James Johnson, L Luu, Jenifer Manson, Charlotte Nelson, Elaine O'Toole, Cassie Olateju, Rebekah Penrice-Randal , Lucille Rainbow, N.P Randle, Trevor Ian Robinson, Parul Sharma, Ghada T Shawli, James P Stewart, Neil Swainston, Ecaterina Vamos, Joanne Watts, Mark Whitehead |
| EPI_ISL_705364, EPI_ISL_705367 | Northumbria University / South Tees Hospitals NHS Foundation Trust / North Cumbria Integrated Care NHS Foundation Trust / North Tees and Hartlepool NHS Foundation Trust / Newcastle Hospitals NHS Foundation Trust | COVID-19 Genomics UK (COG-UK) Consortium | Darren L Smith,Andrew Nelson,Matthew Bashton,Greg R Young,Joshua Loh,John Allan,Mohammad A Tariq,Giles S Holt,Gary Black,Wen C Yew,Lynn Dover,Paul Baker,Steve Liggett,Sarah Essex,Jane Greenaway,Debra Padgett,Clive Graham,Garren Scott,Edward Barton,Emma Swindells,Brendan Payne,Jennifer Collins,Yusri Taha,Gary Eltringham |
| EPI_ISL_705384 | Wales Specialist Virology Centre Sequencing lab: Pathogen Genomics Unit | COVID-19 Genomics UK (COG-UK) Consortium | Catherine Moore, Johnathan Evans, Laura Gifford, Malorie Perry, Simon Cottrell, Angela Marchbank, Alec Birchley, Alexander Adams, Amy Gaskin, Bree Gatica-Wilcox, Jason Coombes, Joel Southgate, Lauren Gilbert, Lee Graham, Nicole Pacchiarini, Sara Kumziene-Summerhayes, Sarah Taylor, Sophie Jones, Sara Rey, Matthew Bull, Joanne Watkins, Sally Corden, Tom Connor |
| EPI_ISL_705388 | Liverpool Clinical Laboratories | COVID-19 Genomics UK (COG-UK) Consortium | Sam Haldenby, Anita Lucaci, Steve Paterson, Julian Hiscox, Alistair Darby, M Almsaud, A Alrezaihi, Muhannad Alruwaili, Stuart D Armstrong, Jones Benjamin, Eleanor G Bentley, Anu Chawla, Jordan J Clark, Angela Cowell, Richard Eccles, Isabel García-Dorival, Matthew Gemmell, Alessandro Gerada, PKF Gilmore, Richard Gregory, Ximeng Han, Catherine Hartley, Margaret Hughes, Miren Iturriza-Gomara, James Johnson, L Luu, Jenifer Manson, Charlotte Nelson, Elaine O'Toole, Cassie Olateju, Rebekah Penrice-Randal , Lucille Rainbow, N.P Randle, Trevor Ian Robinson, Parul Sharma, Ghada T Shawli, James P Stewart, Neil Swainston, Ecaterina Vamos, Joanne Watts, Mark Whitehead |
| EPI_ISL_705397 | University College London, Great Ormond Street Hospital for Children NHS Foundation Trust, Imperial College Healthcare NHS Trust | COVID-19 Genomics UK (COG-UK) Consortium | Sergi Castellano, Rachel Williams, Mark Kristiansen, Paola Resende Silva, Sunando Roy, Tony Brooks, Helena Tutill, Paola Niola, Patricia Dyal, Charlotte Williams, Leysa Forrest, Yasmin Panchbhaya, Jacqueline Findlay, Samuel Weeks, Julianne Brown, Kathryn Harris, Paul Randell, James Price, Alison Holmes, Judith Breuer |
| EPI_ISL_705412 | Wales Specialist Virology Centre Sequencing lab: Pathogen Genomics Unit | COVID-19 Genomics UK (COG-UK) Consortium | Catherine Moore, Johnathan Evans, Laura Gifford, Malorie Perry, Simon Cottrell, Angela Marchbank, Alec Birchley, Alexander Adams, Amy Gaskin, Bree Gatica-Wilcox, Jason Coombes, Joel Southgate, Lauren Gilbert, Lee Graham, Nicole Pacchiarini, Sara Kumziene-Summerhayes, Sarah Taylor, Sophie Jones, Sara Rey, Matthew Bull, Joanne Watkins, Sally Corden, Tom Connor |
| EPI_ISL_705422 | Quadram Institute Bioscience | COVID-19 Genomics UK (COG-UK) Consortium | Dave J. Baker, Gemma L. Kay, Alp Aydin, Thanh Le-Viet, Steven Rudder, Ana P. Tedim, Anastasia Kolyva, Maria Diaz, Leonardo de Oliveira Martins, Nabil-Fareed Alikhan, Lizzie Meadows, Rachael Stanley, Ngozi Elumogo, Muhammed Yasir, Nicholas M. Thomson, Alexander J Trotter, Rachel Gilroy, Samuel Bloomfield, Claire Stuart, Andrew Bell, Reenesh Prakash, Samir Dervisevic, Alison E. Mather, John Wain, Mark Webber, Andrew J. Page, Justin O'Grady |
| EPI_ISL_705435, EPI_ISL_705440, EPI_ISL_705441, EPI_ISL_705442 | Wales Specialist Virology Centre Sequencing lab: Pathogen Genomics Unit | COVID-19 Genomics UK (COG-UK) Consortium | Catherine Moore, Johnathan Evans, Laura Gifford, Malorie Perry, Simon Cottrell, Angela Marchbank, Alec Birchley, Alexander Adams, Amy Gaskin, Bree Gatica-Wilcox, Jason Coombes, Joel Southgate, Lauren Gilbert, Lee Graham, Nicole Pacchiarini, Sara Kumziene-Summerhayes, Sarah Taylor, Sophie Jones, Sara Rey, Matthew Bull, Joanne Watkins, Sally Corden, Tom Connor |
| EPI_ISL_705490 | West of Scotland Specialist Virology Centre, NHSGGC / MRC-University of Glasgow Centre for Virus Research | COVID-19 Genomics UK (COG-UK) Consortium | Ana da Silva Filipe, Natasha Johnson, Kathy Smollett, Daniel Mair, Stephen Carmichael, Alice Broos, Lily Tong, Jenna Nichols, Kyriaki Nomikou; Sarah McDonald; Richard Orton, Joseph Hughes, Sreenu Vattipally, David L Robertson; Alasdair MacLean, Rory Gunson; Sharif Shaaban, Matthew Holden; Rachel Blacow, Guy Mollett, Kathy Li, James Shepherd, Antonia Ho, Emma Thomson |
| EPI_ISL_705496 | Northumbria University / South Tees Hospitals NHS Foundation Trust / North Cumbria Integrated Care NHS Foundation Trust / North Tees and Hartlepool NHS Foundation Trust / Newcastle Hospitals NHS Foundation Trust | COVID-19 Genomics UK (COG-UK) Consortium | Darren L Smith,Andrew Nelson,Matthew Bashton,Greg R Young,Joshua Loh,John Allan,Mohammad A Tariq,Giles S Holt,Gary Black,Wen C Yew,Lynn Dover,Paul Baker,Steve Liggett,Sarah Essex,Jane Greenaway,Debra Padgett,Clive Graham,Garren Scott,Edward Barton,Emma Swindells,Brendan Payne,Jennifer Collins,Yusri Taha,Gary Eltringham |

| | | | |
|---|---|---|---|
| EPI_ISL_705560 | Department of Pathology, University of Cambridge | COVID-19 Genomics UK (COG-UK) Consortium | Aminu S. Jahun, Yasmin Chaudhry, Grant Hall, Iliana Georgana, Myra Hosmillo, Martin D. Curran, Malte Pinckert, Surendra Parmar, Ian Goodfellow |
| EPI_ISL_705572, EPI_ISL_705593, EPI_ISL_705594 | Quadram Institute Bioscience | COVID-19 Genomics UK (COG-UK) Consortium | Dave J. Baker, Gemma L. Kay, Alp Aydin, Thanh Le-Viet, Steven Rudder, Ana P. Tedim, Anastasia Kolyva, Maria Diaz, Leonardo de Oliveira Martins, Nabil-Fareed Alikhan, Lizzie Meadows, Rachael Stanley, Ngozi Elumogo, Muhammed Yasir, Nicholas M. Thomson, Alexander J Trotter, Rachel Gilroy, Samuel Bloomfield, Claire Stuart, Andrew Bell, Reenesh Prakash, Samir Dervisevic, Alison E. Mather, John Wain, Mark Webber, Andrew J. Page, Justin O'Grady |
| EPI_ISL_705626, EPI_ISL_705627, EPI_ISL_705628 | Virology Department, Sheffield Teaching Hospitals NHS Foundation Trust/Department of Infection, Immunity and Cardiovascular Disease, The Medical School, University of Sheffield | COVID-19 Genomics UK (COG-UK) Consortium | Thushan de Silva, Matthew Parker, Nikki Smith, Adri Angyal, Rebecca Brown, Luke Green, Rachel Tucker, Paul Parsons, Danielle Groves, Katie Johnson, Laura Carrilero, Alex Keeley, Dave Partridge, Matthew Wyles, Benjamin Lindsey, Mehmet Yavuz, Mohammad Raza, Cariad Evans |
| EPI_ISL_705641, EPI_ISL_705642, EPI_ISL_705643, EPI_ISL_705644, EPI_ISL_705645, EPI_ISL_705646, EPI_ISL_705647, EPI_ISL_705648, EPI_ISL_705649, EPI_ISL_705650, EPI_ISL_705651, EPI_ISL_705652, EPI_ISL_705653, EPI_ISL_705654, EPI_ISL_705655, EPI_ISL_705656, EPI_ISL_705657, EPI_ISL_705658, EPI_ISL_705659, EPI_ISL_705660, EPI_ISL_705661, EPI_ISL_705662, EPI_ISL_705663, EPI_ISL_705664, EPI_ISL_705665, EPI_ISL_705666, EPI_ISL_705708, EPI_ISL_705710, EPI_ISL_705730 | | | |
| see above | West of Scotland Specialist Virology Centre, NHSGGC / MRC-University of Glasgow Centre for Virus Research | COVID-19 Genomics UK (COG-UK) Consortium | Ana da Silva Filipe, Natasha Johnson, Kathy Smollett, Daniel Mair, Stephen Carmichael, Alice Broos, Lily Tong, Jenna Nichols, Kyriaki Nomikou; Sarah McDonald; Richard Orton, Joseph Hughes, Sreenu Vattipally, David L Robertson; Alasdair MacLean, Rory Gunson; Sharif Shaaban, Matthew Holden; Rachel Blacow, Guy Mollett, Kathy Li, James Shepherd, Antonia Ho, Emma Thomson |
| EPI_ISL_705804 | Virology Department, Royal Infirmary of Edinburgh, NHS Lothian / School of Biological Sciences, University of Edinburgh / Institute of Genetics and Molecular Medicine, University of Edinburgh | COVID-19 Genomics UK (COG-UK) Consortium | McHugh M, Dewar R, Rooke S, Gallagher M, Balcaza C, O'Toole Á, Scher E, Hill V, McCrone JT, Colquhoun R, Yu X, Jackson B, Rambaut A, Williams TC, Templeton K |
| EPI_ISL_705810, EPI_ISL_705812, EPI_ISL_705818, EPI_ISL_705929, EPI_ISL_705930, EPI_ISL_705931, EPI_ISL_705932, EPI_ISL_705933, EPI_ISL_705934, EPI_ISL_705935, EPI_ISL_705936, EPI_ISL_705937, EPI_ISL_705938, EPI_ISL_705939, EPI_ISL_705940 | | | |
| see above | Liverpool Clinical Laboratories | COVID-19 Genomics UK (COG-UK) Consortium | Sam Haldenby, Anita Lucaci, Steve Paterson, Julian Hiscox, Alistair Darby, M Almsaud, A Alrezaihi, Muhannad Alruwaili, Stuart D Armstrong, Jones Benjamin, Eleanor G Bentley, Anu Chawla, Jordan J Clark, Angela Cowell, Richard Eccles, Isabel García-Dorival, Matthew Gemmell, Alessandro Gerada, PKF Gilmore, Richard Gregory, Ximeng Han, Catherine Hartley, Margaret Hughes, Miren Iturriza-Gomara, James Johnson, L Luu, Jenifer Manson, Charlotte Nelson, Elaine O'Toole, Cassie Olateju, Rebekah Penrice-Randal , Lucille Rainbow, N.P Randle, Trevor Ian Robinson, Parul Sharma, Ghada T Shawli, James P Stewart, Neil Swainston, Ecaterina Vamos, Joanne Watts, Mark Whitehead |
| EPI_ISL_705949 | University College London Hospital | COVID-19 Genomics UK (COG-UK) Consortium | Judith Heaney, Matthew Byott, Catherine Houlihan, Dan Frampton, Stuart Kirk, Moira Spyer and Eleni Nastouli |
| EPI_ISL_705961, EPI_ISL_705962, EPI_ISL_705963, EPI_ISL_705964, EPI_ISL_705965, EPI_ISL_705966, EPI_ISL_705967, EPI_ISL_705968, EPI_ISL_705969, EPI_ISL_705970, EPI_ISL_705971 | | | |
| see above | University College London, Great Ormond Street Hospital for Children NHS Foundation Trust, Imperial College Healthcare NHS Trust | COVID-19 Genomics UK (COG-UK) Consortium | Sergi Castellano, Rachel Williams, Mark Kristiansen, Paola Resende Silva, Sunando Roy, Tony Brooks, Helena Tutill, Paola Niola, Patricia Dyal, Charlotte Williams, Leysa Forrest, Yasmin Panchbhaya, Jacqueline Findlay, Samuel Weeks, Julianne Brown, Kathryn Harris, Paul Randell, James Price, Alison Holmes, Judith Breuer |
| EPI_ISL_706312, EPI_ISL_706313, EPI_ISL_706314, EPI_ISL_706316, EPI_ISL_706317, EPI_ISL_706318, EPI_ISL_706319, EPI_ISL_706322, EPI_ISL_706334, EPI_ISL_706335, EPI_ISL_706336, EPI_ISL_706337, EPI_ISL_706338, EPI_ISL_706339, EPI_ISL_706341, EPI_ISL_706342, EPI_ISL_706343, EPI_ISL_706344, EPI_ISL_706345, EPI_ISL_706346, EPI_ISL_706347, EPI_ISL_706348, EPI_ISL_706349, EPI_ISL_706350, EPI_ISL_706351, EPI_ISL_706352 | | | |
| see above | Northumbria University / South Tees Hospitals NHS Foundation Trust / North Cumbria Integrated Care NHS Foundation Trust / North Tees and Hartlepool NHS Foundation Trust / Newcastle Hospitals NHS Foundation Trust | COVID-19 Genomics UK (COG-UK) Consortium | Darren L Smith,Andrew Nelson,Matthew Bashton,Greg R Young,Joshua Loh,John Allan,Mohammad A Tariq,Giles S Holt,Gary Black,Wen C Yew,Lynn Dover,Paul Baker,Steve Liggett,Sarah Essex,Jane Greenaway,Debra Padgett,Clive Graham,Garren Scott,Edward Barton,Emma Swindells,Brendan Payne,Jennifer Collins,Yusri Taha,Gary Eltringham |
| EPI_ISL_706445, EPI_ISL_706449, EPI_ISL_706453, EPI_ISL_706454, EPI_ISL_706455, EPI_ISL_706458, EPI_ISL_706459, EPI_ISL_706460, EPI_ISL_706461, EPI_ISL_706462, EPI_ISL_706527, EPI_ISL_706529, EPI_ISL_706535, EPI_ISL_706538, EPI_ISL_706542, EPI_ISL_706554, EPI_ISL_706555, EPI_ISL_706556, EPI_ISL_706557, EPI_ISL_706558, EPI_ISL_706559, EPI_ISL_706560, EPI_ISL_706561, EPI_ISL_706562, EPI_ISL_706563, EPI_ISL_706564, EPI_ISL_706565, EPI_ISL_706566, EPI_ISL_706567, EPI_ISL_706568, EPI_ISL_706569, EPI_ISL_706570, EPI_ISL_706571, EPI_ISL_706572, EPI_ISL_706573, EPI_ISL_706574, EPI_ISL_706575, EPI_ISL_706576, EPI_ISL_706577, EPI_ISL_706578, EPI_ISL_706579, EPI_ISL_706580, EPI_ISL_706581, EPI_ISL_706582, EPI_ISL_706583, EPI_ISL_706584, EPI_ISL_706585, EPI_ISL_706586, EPI_ISL_706587, EPI_ISL_706588, EPI_ISL_706596, EPI_ISL_706597, EPI_ISL_706598, EPI_ISL_706599, EPI_ISL_706600, EPI_ISL_706601, EPI_ISL_706602, EPI_ISL_706608, EPI_ISL_706609, EPI_ISL_706610, EPI_ISL_706611, EPI_ISL_706612, EPI_ISL_706613, EPI_ISL_706620, EPI_ISL_706621, EPI_ISL_706622, EPI_ISL_706623, EPI_ISL_706624, EPI_ISL_706625, EPI_ISL_706626, EPI_ISL_706630, EPI_ISL_706631, EPI_ISL_706632, EPI_ISL_706633, EPI_ISL_706634, EPI_ISL_706635, EPI_ISL_706639, EPI_ISL_706640, EPI_ISL_706641, EPI_ISL_706642, EPI_ISL_706643, EPI_ISL_706644, EPI_ISL_706645, EPI_ISL_706646, EPI_ISL_706647, EPI_ISL_706648, EPI_ISL_706649, EPI_ISL_706650, EPI_ISL_706651, EPI_ISL_706652, EPI_ISL_706653, EPI_ISL_706654, EPI_ISL_706655, EPI_ISL_706656, EPI_ISL_706657, EPI_ISL_706662, EPI_ISL_706776, EPI_ISL_706783, EPI_ISL_706784, EPI_ISL_706785, EPI_ISL_706786, EPI_ISL_706787, EPI_ISL_706788, EPI_ISL_706789, EPI_ISL_706791, EPI_ISL_706792, EPI_ISL_706793, EPI_ISL_706794, EPI_ISL_706796, EPI_ISL_706797, EPI_ISL_706800, EPI_ISL_706801, EPI_ISL_706802, EPI_ISL_706803, EPI_ISL_706804, EPI_ISL_706805, EPI_ISL_706806, EPI_ISL_706807, EPI_ISL_706808, EPI_ISL_706809, EPI_ISL_706810, EPI_ISL_706811, EPI_ISL_706812, EPI_ISL_706813, EPI_ISL_706814, EPI_ISL_706815, EPI_ISL_706816, EPI_ISL_706817, EPI_ISL_706818, EPI_ISL_706819, EPI_ISL_706820, EPI_ISL_706821, EPI_ISL_706822, EPI_ISL_706823, EPI_ISL_706824, EPI_ISL_706825, EPI_ISL_706826, EPI_ISL_706827, EPI_ISL_706828, EPI_ISL_706829, EPI_ISL_706830, EPI_ISL_706831, EPI_ISL_706832, EPI_ISL_706833, EPI_ISL_706834, EPI_ISL_706835, EPI_ISL_706836, EPI_ISL_706837, EPI_ISL_706838, EPI_ISL_706839, EPI_ISL_706840, EPI_ISL_706841, EPI_ISL_706842, EPI_ISL_706843, EPI_ISL_706844, EPI_ISL_706845, EPI_ISL_706846, EPI_ISL_706847, EPI_ISL_706848, EPI_ISL_706849, EPI_ISL_706850, EPI_ISL_706851, EPI_ISL_706852, EPI_ISL_706853, EPI_ISL_706854, EPI_ISL_706855, EPI_ISL_706856, EPI_ISL_706857, EPI_ISL_706858, EPI_ISL_706859, EPI_ISL_706860, EPI_ISL_706861, EPI_ISL_706862, EPI_ISL_706863 | | | |
| see above | Wales Specialist Virology Centre Sequencing lab: Pathogen Genomics Unit | COVID-19 Genomics UK (COG-UK) Consortium | Catherine Moore, Johnathan Evans, Laura Gifford, Malorie Perry, Simon Cottrell, Angela Marchbank, Alec Birchley, Alexander Adams, Amy Gaskin, Bree Gatica-Wilcox, Jason Coombes, Joel Southgate, Lauren Gilbert, Lee Graham, Nicole Pacchiarini, Sara Kumziene-Summerhayes, Sarah Taylor, Sophie Jones, Sara Rey, Matthew Bull, Joanne Watkins, Sally Corden, Tom Connor |
| EPI_ISL_706988 | Virology Department, Sheffield Teaching Hospitals NHS Foundation Trust/Department of Infection, Immunity and Cardiovascular Disease, The Medical School, University of Sheffield | COVID-19 Genomics UK (COG-UK) Consortium | Thushan de Silva, Matthew Parker, Nikki Smith, Adri Angyal, Rebecca Brown, Luke Green, Rachel Tucker, Paul Parsons, Danielle Groves, Katie Johnson, Laura Carrilero, Alex Keeley, Dave Partridge, Matthew Wyles, Benjamin Lindsey, Mehmet Yavuz, Mohammad Raza, Cariad Evans |
| EPI_ISL_708136, EPI_ISL_708137, EPI_ISL_708138, EPI_ISL_708139 | Medical Microbiology Unit, Department for Laboratory Medicine, Drammen Hospital, Vestre Viken Health Trust, | Norwegian Institute of Public Health, Department of Virology | Kathrine Stene-Johansen, Kamilla Heddeland Instefjord, Hilde Elshaug, Marie Paulsen Madsen, Rasmus Riis Kopperud, Hilde Vollan, Karoline Bragstad, Olav Hungnes |
| EPI_ISL_708140, EPI_ISL_708169, EPI_ISL_708170 | Norwegian Institute of Public Health, Department of Virology | Norwegian Institute of Public Health, Department of Virology | Kathrine Stene-Johansen, Kamilla Heddeland Instefjord, Hilde Elshaug, Marie Paulsen Madsen, Rasmus Riis Kopperud, Hilde Vollan, Karoline Bragstad, Olav Hungnes |
| EPI_ISL_708172, EPI_ISL_708173, EPI_ISL_708174, EPI_ISL_708175 | Oslo University Hospital, Department of Medical Microbiology | Norwegian Institute of Public Health, Department of Virology | Kathrine Stene-Johansen, Kamilla Heddeland Instefjord, Hilde Elshaug, Marie Paulsen Madsen, Rasmus Riis Kopperud, Hilde Vollan, Karoline Bragstad, Olav Hungnes |
| EPI_ISL_708180, EPI_ISL_708181 | Norwegian Institute of Public Health, Department of Virology | Norwegian Institute of Public Health, Department of Virology | Kathrine Stene-Johansen, Kamilla Heddeland Instefjord, Hilde Elshaug, Marie Paulsen Madsen, Rasmus Riis Kopperud, Hilde Vollan, Karoline Bragstad, Olav Hungnes |
| EPI_ISL_708200, EPI_ISL_708201, EPI_ISL_708202, EPI_ISL_708203, EPI_ISL_708204, EPI_ISL_708205, EPI_ISL_708206, EPI_ISL_708207, EPI_ISL_708208, EPI_ISL_708209, EPI_ISL_708210, EPI_ISL_708211, EPI_ISL_708212, EPI_ISL_708213, EPI_ISL_708214, EPI_ISL_708215, EPI_ISL_708216, EPI_ISL_708217, EPI_ISL_708218, EPI_ISL_708219, EPI_ISL_708220, EPI_ISL_708221, EPI_ISL_708222, EPI_ISL_708223, EPI_ISL_708224, EPI_ISL_708225, EPI_ISL_708226, EPI_ISL_708227, EPI_ISL_708228, EPI_ISL_708229, EPI_ISL_708230, EPI_ISL_708231, EPI_ISL_708232, EPI_ISL_708233, EPI_ISL_708235, EPI_ISL_708236, EPI_ISL_708237, EPI_ISL_708238, EPI_ISL_708239, EPI_ISL_708240, EPI_ISL_708241, EPI_ISL_708242, EPI_ISL_708244, EPI_ISL_708247, EPI_ISL_708249, EPI_ISL_708251, EPI_ISL_708252, EPI_ISL_708253, EPI_ISL_708261, EPI_ISL_708262, EPI_ISL_708263, EPI_ISL_708264, EPI_ISL_708266, EPI_ISL_708267, EPI_ISL_708268, EPI_ISL_708269, EPI_ISL_708270, EPI_ISL_708272, EPI_ISL_708273, EPI_ISL_708318, EPI_ISL_708320, EPI_ISL_708332, EPI_ISL_708333, EPI_ISL_708345, EPI_ISL_708358, EPI_ISL_708359, EPI_ISL_708362, EPI_ISL_708366, EPI_ISL_708368, EPI_ISL_708369 | | | |
| see above | Michigan Department of Health and Human Services, Bureau of Laboratories | Michigan Department of Health and Human Services, Bureau of Laboratories | Blankenship HM, Riner D, Soehnlen MK |
| EPI_ISL_708456, EPI_ISL_708457, EPI_ISL_708458, EPI_ISL_708459 | Department of Clinical Microbiology | GIGA Medical Genomics | Keith Durkin, Maria Artesi, Justine Defêche, Gilles Darcis, Michel Moutschen, Sébastien Bontems, Raphaël Boreux, Bouchra Boujemla, Cécile Meex, Pierrette Melin, Marie-Pierre Hayette, Vincent Bours |
| EPI_ISL_709882 | Lighthouse Lab in Milton Keynes | Wellcome Sanger Institute for the COVID-19 Genomics UK (COG-UK) Consortium | The Lighthouse Lab in Milton Keynes and Alex Alderton, Roberto Amato, Sonia Goncalves, Ewan Harrison, David K. Jackson, Ian Johnston, Dominic Kwiatkowski, Cordelia Langford, John Sillitoe on behalf of the Wellcome Sanger Institute COVID-19 Surveillance Team |
| EPI_ISL_710123 | South Eastern Area Laboratory Services (SEALS) | CIDM-PH et al. | CIDM-PH et al. |
| EPI_ISL_710589, EPI_ISL_710590 | Narhalsan Fjallbacka VC | The Public Health Agency of Sweden | Department of Microbiology, The Public Health Agency of Sweden |
| EPI_ISL_710607 | Halsomedicinskt Center | The Public Health Agency of Sweden | Department of Microbiology, The Public Health Agency of Sweden |
| EPI_ISL_710830, EPI_ISL_710831, | Lighthouse Lab in Glasgow | Wellcome Sanger Institute for the COVID-19 Genomics UK | Harper VanSteenhouse, Yumi Kasai, David Gray, Carol Clugston, Anna Dominiczak and Alex Alderton, Roberto Amato, Sonia Goncalves, Ewan Harrison, |

| | | | |
|---|---|---|---|
| EPI_ISL_710832, EPI_ISL_710833, EPI_ISL_710834 | | (COG-UK) Consortium | David K. Jackson, Ian Johnston, Dominic Kwiatkowski, Cordelia Langford, John Sillitoe on behalf of the Wellcome Sanger Institute COVID-19 Surveillance Team |
| EPI_ISL_714098, EPI_ISL_714100, EPI_ISL_714101, EPI_ISL_714102, EPI_ISL_714103, EPI_ISL_714104, EPI_ISL_714105, EPI_ISL_714106, EPI_ISL_714108, EPI_ISL_714109, EPI_ISL_714110, EPI_ISL_714111, EPI_ISL_714112, EPI_ISL_714113, EPI_ISL_714114, EPI_ISL_714124, EPI_ISL_714125, EPI_ISL_714126, EPI_ISL_714127, EPI_ISL_714128, EPI_ISL_714129, EPI_ISL_714130, EPI_ISL_714131, EPI_ISL_714132, EPI_ISL_714133, EPI_ISL_714134, EPI_ISL_714135, EPI_ISL_714136, EPI_ISL_714137, EPI_ISL_714138, EPI_ISL_714139, EPI_ISL_714140, EPI_ISL_714141, EPI_ISL_714142, EPI_ISL_714143, EPI_ISL_714144, EPI_ISL_714145, EPI_ISL_714146, EPI_ISL_714147, EPI_ISL_714148, EPI_ISL_714149, EPI_ISL_714150, EPI_ISL_714151, EPI_ISL_714152, EPI_ISL_714153, EPI_ISL_714154, EPI_ISL_714155, EPI_ISL_714156, EPI_ISL_714157, EPI_ISL_714158, EPI_ISL_714159, EPI_ISL_714160, EPI_ISL_714161, EPI_ISL_714162, EPI_ISL_714163, EPI_ISL_714164, EPI_ISL_714165, EPI_ISL_714166, EPI_ISL_714167, EPI_ISL_714168, EPI_ISL_714169, EPI_ISL_714170, EPI_ISL_714171, EPI_ISL_714172, EPI_ISL_714173, EPI_ISL_714175, EPI_ISL_714176, EPI_ISL_714177, EPI_ISL_714178, EPI_ISL_714179, EPI_ISL_714180, EPI_ISL_714181, EPI_ISL_714183, EPI_ISL_714184, EPI_ISL_714185, EPI_ISL_714186, EPI_ISL_714187, EPI_ISL_714188, EPI_ISL_714189, EPI_ISL_714190, EPI_ISL_714192, EPI_ISL_714193, EPI_ISL_714194, EPI_ISL_714195, EPI_ISL_714199, EPI_ISL_714200, EPI_ISL_714201, EPI_ISL_714202, EPI_ISL_714203, EPI_ISL_714204, EPI_ISL_714205, EPI_ISL_714206, EPI_ISL_714207, EPI_ISL_714208, EPI_ISL_714209, EPI_ISL_714211, EPI_ISL_714212, EPI_ISL_714213, EPI_ISL_714214, EPI_ISL_714215, EPI_ISL_714216, EPI_ISL_714221, EPI_ISL_714223, EPI_ISL_714224, EPI_ISL_714225, EPI_ISL_714226, EPI_ISL_714228, EPI_ISL_714229, EPI_ISL_714230, EPI_ISL_714231, EPI_ISL_714232, EPI_ISL_714233, EPI_ISL_714234, EPI_ISL_714237, EPI_ISL_714238, EPI_ISL_714239, EPI_ISL_714242, EPI_ISL_714243, EPI_ISL_714244, EPI_ISL_714245, EPI_ISL_714246, EPI_ISL_714247, EPI_ISL_714248, EPI_ISL_714249, EPI_ISL_714250, EPI_ISL_714251, EPI_ISL_714255, EPI_ISL_714256, EPI_ISL_714257, EPI_ISL_714261, EPI_ISL_714262, EPI_ISL_714263, EPI_ISL_714265, EPI_ISL_714266, EPI_ISL_714267, EPI_ISL_714268, EPI_ISL_714270, EPI_ISL_714271, EPI_ISL_714273, EPI_ISL_714274, EPI_ISL_714277, EPI_ISL_714278, EPI_ISL_714280, EPI_ISL_714281, EPI_ISL_714283, EPI_ISL_714284, EPI_ISL_714285, EPI_ISL_714286, EPI_ISL_714287, EPI_ISL_714288, EPI_ISL_714290, EPI_ISL_714291, EPI_ISL_714292, EPI_ISL_714293, EPI_ISL_714294, EPI_ISL_714295, EPI_ISL_714296, EPI_ISL_714297, EPI_ISL_714298, EPI_ISL_714299, EPI_ISL_714320, EPI_ISL_714321, EPI_ISL_714322, EPI_ISL_714323, EPI_ISL_714324, EPI_ISL_714325, EPI_ISL_714326, EPI_ISL_714327, EPI_ISL_714328, EPI_ISL_714329, EPI_ISL_714330, EPI_ISL_714331, EPI_ISL_714332, EPI_ISL_714333, EPI_ISL_714334, EPI_ISL_714335, EPI_ISL_714336, EPI_ISL_714337, EPI_ISL_714338, EPI_ISL_714339, EPI_ISL_714340, EPI_ISL_714341, EPI_ISL_714342, EPI_ISL_714343, EPI_ISL_714344, EPI_ISL_714345, EPI_ISL_714346, EPI_ISL_714347, EPI_ISL_714348, EPI_ISL_714349, EPI_ISL_714350, EPI_ISL_714351, EPI_ISL_714352, EPI_ISL_714353, EPI_ISL_714354, EPI_ISL_714355, EPI_ISL_714356, EPI_ISL_714357, EPI_ISL_714358, EPI_ISL_714359, EPI_ISL_714360, EPI_ISL_714361, EPI_ISL_714362, EPI_ISL_714365, EPI_ISL_714367, EPI_ISL_714368, EPI_ISL_714370, EPI_ISL_714371, EPI_ISL_714372, EPI_ISL_714373, EPI_ISL_714374, EPI_ISL_714375, EPI_ISL_714376, EPI_ISL_714378, EPI_ISL_714379, EPI_ISL_714380, EPI_ISL_714381, EPI_ISL_714382, EPI_ISL_714383, EPI_ISL_714384, EPI_ISL_714392, EPI_ISL_714393, EPI_ISL_714394, EPI_ISL_714395, EPI_ISL_714396, EPI_ISL_714397, EPI_ISL_714398, EPI_ISL_714399, EPI_ISL_714400, EPI_ISL_714401, EPI_ISL_714402, EPI_ISL_714403, EPI_ISL_714404, EPI_ISL_714405, EPI_ISL_714406, EPI_ISL_714407, EPI_ISL_714408, EPI_ISL_714409, EPI_ISL_714410, EPI_ISL_714413, EPI_ISL_714414, EPI_ISL_714415, EPI_ISL_714417, EPI_ISL_714418, EPI_ISL_714444, EPI_ISL_714445, EPI_ISL_714446, EPI_ISL_714447, EPI_ISL_714448, EPI_ISL_714449, EPI_ISL_714451, EPI_ISL_714452, EPI_ISL_714453, EPI_ISL_714454, EPI_ISL_714455, EPI_ISL_714456, EPI_ISL_714457, EPI_ISL_714458, EPI_ISL_714459, EPI_ISL_714460, EPI_ISL_714461, EPI_ISL_714465, EPI_ISL_714466, EPI_ISL_714467, EPI_ISL_714468, EPI_ISL_714469, EPI_ISL_714470, EPI_ISL_714471, EPI_ISL_714472, EPI_ISL_714473, EPI_ISL_714474, EPI_ISL_714477, EPI_ISL_714566, EPI_ISL_714567, EPI_ISL_714568, EPI_ISL_714569, EPI_ISL_714570, EPI_ISL_714571, EPI_ISL_714572, EPI_ISL_714573, EPI_ISL_714574, EPI_ISL_714575, EPI_ISL_714576, EPI_ISL_714577, EPI_ISL_714578, EPI_ISL_714579, EPI_ISL_714580, EPI_ISL_714581, EPI_ISL_714582, EPI_ISL_714583, EPI_ISL_714584, EPI_ISL_714585, EPI_ISL_714586, EPI_ISL_714587, EPI_ISL_714588, EPI_ISL_714589, EPI_ISL_714590, EPI_ISL_714591, EPI_ISL_714592, EPI_ISL_714593, EPI_ISL_714594, EPI_ISL_714595, EPI_ISL_714596, EPI_ISL_714597, EPI_ISL_714598, EPI_ISL_714599, EPI_ISL_714600, EPI_ISL_714601, EPI_ISL_714602, EPI_ISL_714603, EPI_ISL_714604, EPI_ISL_714605, EPI_ISL_714606, EPI_ISL_714607, EPI_ISL_714608, EPI_ISL_714609, EPI_ISL_714610, EPI_ISL_714611, EPI_ISL_714612, EPI_ISL_714613, EPI_ISL_714614, EPI_ISL_714615, EPI_ISL_714616, EPI_ISL_714617, EPI_ISL_714618, EPI_ISL_714619, EPI_ISL_714620, EPI_ISL_714621, EPI_ISL_714622, EPI_ISL_714623, EPI_ISL_714624, EPI_ISL_714625, EPI_ISL_714626, EPI_ISL_714627, EPI_ISL_714628, EPI_ISL_714629, EPI_ISL_714630, EPI_ISL_714631, EPI_ISL_714632, EPI_ISL_714633, EPI_ISL_714634, EPI_ISL_714635, EPI_ISL_714636, EPI_ISL_714637, EPI_ISL_714638, EPI_ISL_714639, EPI_ISL_714640, EPI_ISL_714641, EPI_ISL_714642, EPI_ISL_714643, EPI_ISL_714644, EPI_ISL_714645, EPI_ISL_714646, EPI_ISL_714647, EPI_ISL_714648, EPI_ISL_714649, EPI_ISL_714650, EPI_ISL_714651, EPI_ISL_714652, EPI_ISL_714653, EPI_ISL_714654, EPI_ISL_714655, EPI_ISL_714656, EPI_ISL_714657, EPI_ISL_714658, EPI_ISL_714659, EPI_ISL_714660, EPI_ISL_714661, EPI_ISL_714662, EPI_ISL_714663, EPI_ISL_714664, EPI_ISL_714665, EPI_ISL_714666, EPI_ISL_714667, EPI_ISL_714668, EPI_ISL_714669, EPI_ISL_714670, EPI_ISL_714671, EPI_ISL_714672, EPI_ISL_714673, EPI_ISL_714674, EPI_ISL_714675, EPI_ISL_714676, EPI_ISL_714677, EPI_ISL_714678, EPI_ISL_714679, EPI_ISL_714680, EPI_ISL_714681, EPI_ISL_714682, EPI_ISL_714683, EPI_ISL_714684, EPI_ISL_714685, EPI_ISL_714686, EPI_ISL_714687, EPI_ISL_714688, EPI_ISL_714689, EPI_ISL_714690, EPI_ISL_714691, EPI_ISL_714692, EPI_ISL_714693, EPI_ISL_714694, EPI_ISL_714695, EPI_ISL_714696, EPI_ISL_714697, EPI_ISL_714698, EPI_ISL_714699, EPI_ISL_714700, EPI_ISL_714701, EPI_ISL_714702, EPI_ISL_714703, EPI_ISL_714704, EPI_ISL_714705, EPI_ISL_714706, EPI_ISL_714707, EPI_ISL_714708, EPI_ISL_714709, EPI_ISL_714710, EPI_ISL_714711, EPI_ISL_714712, EPI_ISL_714713, EPI_ISL_714714, EPI_ISL_714715, EPI_ISL_714716, EPI_ISL_714717, EPI_ISL_714718, EPI_ISL_714719, EPI_ISL_714720, EPI_ISL_714721, EPI_ISL_714722, EPI_ISL_714723, EPI_ISL_714724, EPI_ISL_714725, EPI_ISL_714726, EPI_ISL_714727, EPI_ISL_714728, EPI_ISL_714729, EPI_ISL_714730, EPI_ISL_714731, EPI_ISL_714732, EPI_ISL_714733, EPI_ISL_714734, EPI_ISL_714735, EPI_ISL_714736, EPI_ISL_714737, EPI_ISL_714738, EPI_ISL_714739, EPI_ISL_714740, EPI_ISL_714741, EPI_ISL_714742, EPI_ISL_714743, EPI_ISL_714744, EPI_ISL_714745, EPI_ISL_714746, EPI_ISL_714747, EPI_ISL_714748, EPI_ISL_714749, EPI_ISL_714750, EPI_ISL_714751, EPI_ISL_714752, EPI_ISL_714753, EPI_ISL_714754, EPI_ISL_714755, EPI_ISL_714756, EPI_ISL_714757, EPI_ISL_714758, EPI_ISL_714759, EPI_ISL_714760, EPI_ISL_714761, EPI_ISL_714762, EPI_ISL_714763, EPI_ISL_714764, EPI_ISL_714765, EPI_ISL_714766, EPI_ISL_714767, EPI_ISL_714768, EPI_ISL_714769, EPI_ISL_714770, EPI_ISL_714771, EPI_ISL_714772, EPI_ISL_714773, EPI_ISL_714774, EPI_ISL_714775, EPI_ISL_714776, EPI_ISL_714777, EPI_ISL_714778, EPI_ISL_714779, EPI_ISL_714780, EPI_ISL_714781, EPI_ISL_714782, EPI_ISL_714783, EPI_ISL_714784, EPI_ISL_714785, EPI_ISL_714786, EPI_ISL_714787, EPI_ISL_714788, EPI_ISL_714789, EPI_ISL_714790, EPI_ISL_714791, EPI_ISL_714792, EPI_ISL_714793, EPI_ISL_714794, EPI_ISL_714795, EPI_ISL_714796, EPI_ISL_714797, EPI_ISL_714798, EPI_ISL_714799, EPI_ISL_714800, EPI_ISL_714801, EPI_ISL_714802, EPI_ISL_714803, EPI_ISL_714804, EPI_ISL_714805, EPI_ISL_714806, EPI_ISL_714807, EPI_ISL_714808, EPI_ISL_714809, EPI_ISL_714810, EPI_ISL_714811, EPI_ISL_714812, EPI_ISL_714813, EPI_ISL_714814, EPI_ISL_714815, EPI_ISL_714816, EPI_ISL_714817, EPI_ISL_714818, EPI_ISL_714819, EPI_ISL_714820, EPI_ISL_714821, EPI_ISL_714822, EPI_ISL_714823, EPI_ISL_714824, EPI_ISL_714825, EPI_ISL_714826, EPI_ISL_714827, EPI_ISL_714828, EPI_ISL_714829, EPI_ISL_714830, EPI_ISL_714831, EPI_ISL_714832, EPI_ISL_714833, EPI_ISL_714834, EPI_ISL_714835, EPI_ISL_714836, EPI_ISL_714837, EPI_ISL_714838, EPI_ISL_714839, EPI_ISL_714840, EPI_ISL_714841, EPI_ISL_714842, EPI_ISL_714843, EPI_ISL_714844, EPI_ISL_714845, EPI_ISL_714846, EPI_ISL_714847, EPI_ISL_714848, EPI_ISL_714849, EPI_ISL_714850, EPI_ISL_714851, EPI_ISL_714852, EPI_ISL_714853, EPI_ISL_714854, EPI_ISL_714855, EPI_ISL_714856, EPI_ISL_714857, EPI_ISL_714858, EPI_ISL_714859, EPI_ISL_714860, EPI_ISL_714861, EPI_ISL_714862, EPI_ISL_714863, EPI_ISL_714864, EPI_ISL_714865, EPI_ISL_714866, EPI_ISL_714867, EPI_ISL_714868, EPI_ISL_714869, EPI_ISL_714870, EPI_ISL_714871, EPI_ISL_714872, EPI_ISL_714873, EPI_ISL_714874, EPI_ISL_714875, EPI_ISL_714876, EPI_ISL_714877, EPI_ISL_714878, EPI_ISL_714879, EPI_ISL_714880, EPI_ISL_714881, EPI_ISL_714882, EPI_ISL_714883, EPI_ISL_714884, EPI_ISL_714885, EPI_ISL_714886, EPI_ISL_714887, EPI_ISL_714888, EPI_ISL_714889, EPI_ISL_714890, EPI_ISL_714891, EPI_ISL_714892, EPI_ISL_714893, EPI_ISL_714894, EPI_ISL_714895, EPI_ISL_714896, EPI_ISL_714897, EPI_ISL_714898, EPI_ISL_714899, EPI_ISL_714900, EPI_ISL_714901, EPI_ISL_714902, EPI_ISL_714903 | | | |
| see above | Department of Virus and Microbiological Special Diagnostics, Statens Serum Institut, Copenhagen, Denmark | Albertsen Lab, Department of Chemistry and Bioscience, Aalborg University, Denmark | Danish Covid-19 Genome Consortium |
| EPI_ISL_717620, EPI_ISL_717621 | Lab voor klinische biologie | Onderzoeksgroep Virologie | Nick Vereecke, Laurens Lambrechts, Marthe Pauwels, Bruno Verhasselt, Linos Vandekerckhove, Hans Nauwynck, Sebastiaan Theuns |
| EPI_ISL_717985 | Lab voor klinische biologie | Onderzoeksgroep Virologie | Laurens Lambrechts, Nick Vereecke, Marthe Pauwels, Bruno Verhasselt, Linos Vandekerckhove, Hans Nauwynck, Sebastiaan Theuns |
| EPI_ISL_717987 | Lab voor klinische biologie | Onderzoeksgroep Virologie | Nick Vereecke, Laurens Lambrechts, Marthe Pauwels, Bruno Verhasselt, Linos Vandekerckhove, Hans Nauwynck, Sebastiaan Theuns |
| EPI_ISL_717989 | Lab voor klinische biologie | Onderzoeksgroep Virologie | Laurens Lambrechts, Nick Vereecke, Marthe Pauwels, Bruno Verhasselt, Linos Vandekerckhove, Hans Nauwynck, Sebastiaan Theuns |
| EPI_ISL_718008 | Lab voor klinische biologie | Onderzoeksgroep Virologie | Nick Vereecke, Laurens Lambrechts, Marthe Pauwels, Bruno Verhasselt, Linos Vandekerckhove, Hans Nauwynck, Sebastiaan Theuns |
| EPI_ISL_718010, EPI_ISL_718012 | Lab voor klinische biologie | Onderzoeksgroep Virologie | Laurens Lambrechts, Nick Vereecke, Marthe Pauwels, Bruno Verhasselt, Linos Vandekerckhove, Hans Nauwynck, Sebastiaan Theuns |
| EPI_ISL_718016, EPI_ISL_718017, EPI_ISL_718018 | Lab voor klinische biologie | Onderzoeksgroep Virologie | Nick Vereecke, Laurens Lambrechts, Marthe Pauwels, Bruno Verhasselt, Linos Vandekerckhove, Hans Nauwynck, Sebastiaan Theuns |
| EPI_ISL_718172, EPI_ISL_718200 | Ministry of Health Hospitals | Institute of Health and Community Medicine | David Perera, Ooi Mong How, Chua Hock Hin, Tonnii Sia Loong Loong, Wong Jyn Shan, Wong Kiing Aik, Chan Chia Jui |
| EPI_ISL_718257 | Institute of Virology, Biomedical Research Center of the Slovak Academy of Sciences, Bratislava | Faculty of Natural Sciences, Comenius University, Bratislava | Broa Brejová, Viktória Hodorová, Kristína Boršová, Viktória abanová, Sabina Fumaová Havlíková, Juraj Kopáek, Martina Liková, ubomíra Lukáiková, Martina Neboháová, Monika Sláviková, Tomáš Vina, Jozef Nosek, Boris Klempa |
| EPI_ISL_718258 | Institute of Virology, Biomedical Research Center of the Slovak Academy of Sciences, Bratislava | Faculty of Natural Sciences, Comenius University, Bratislava | Viktória Hodorová, Kristína Boršová, Broa Brejová, Viktória abanová, Sabina Fumaová Havlíková, Juraj Kopáek, Martina Liková, ubomíra Lukáiková, Martina Neboháová, Monika Sláviková, Tomáš Vina, Jozef Nosek, Boris Klempa |
| EPI_ISL_719605, EPI_ISL_719606, EPI_ISL_719607, EPI_ISL_719608, EPI_ISL_719609, EPI_ISL_719610, EPI_ISL_719611, EPI_ISL_719612, EPI_ISL_719613, EPI_ISL_719614, EPI_ISL_719615, EPI_ISL_719616, EPI_ISL_719617, EPI_ISL_719618, EPI_ISL_719619, EPI_ISL_719620, EPI_ISL_719621, EPI_ISL_719622, EPI_ISL_719623, EPI_ISL_719624, EPI_ISL_719625, EPI_ISL_719626, EPI_ISL_719627, EPI_ISL_719628, EPI_ISL_719629, EPI_ISL_719630, EPI_ISL_719631, EPI_ISL_719632, EPI_ISL_719633, EPI_ISL_719634, EPI_ISL_719635, EPI_ISL_719636, EPI_ISL_719637, EPI_ISL_719638, EPI_ISL_719639, EPI_ISL_719640, EPI_ISL_719641, EPI_ISL_719642, EPI_ISL_719643, EPI_ISL_719644, EPI_ISL_719645, EPI_ISL_719646, EPI_ISL_719647, EPI_ISL_719648, EPI_ISL_719649, EPI_ISL_719650, EPI_ISL_719651, EPI_ISL_719652, EPI_ISL_719653, EPI_ISL_719654, EPI_ISL_719655, EPI_ISL_719656, EPI_ISL_719657, EPI_ISL_719658, EPI_ISL_719659, EPI_ISL_719660, EPI_ISL_719661, EPI_ISL_719662, EPI_ISL_719663, EPI_ISL_719664, EPI_ISL_719665, EPI_ISL_719666, EPI_ISL_719667, EPI_ISL_719668, EPI_ISL_719669, EPI_ISL_719670, EPI_ISL_719671, EPI_ISL_719672, EPI_ISL_719673, EPI_ISL_719674, EPI_ISL_719675, EPI_ISL_719676, EPI_ISL_719677, EPI_ISL_719678, EPI_ISL_719679, EPI_ISL_719680, EPI_ISL_719681, EPI_ISL_719682, EPI_ISL_719683, EPI_ISL_719684, EPI_ISL_719685, EPI_ISL_719686, EPI_ISL_719687, EPI_ISL_719688, EPI_ISL_719689, EPI_ISL_719690, EPI_ISL_719691, EPI_ISL_719692, EPI_ISL_719693, EPI_ISL_719694, EPI_ISL_719695, EPI_ISL_719696, EPI_ISL_719697, EPI_ISL_719698, EPI_ISL_719699, EPI_ISL_719700, EPI_ISL_719701, EPI_ISL_719702, EPI_ISL_719703, EPI_ISL_719704, EPI_ISL_719705, EPI_ISL_719706, EPI_ISL_719707, EPI_ISL_719708, EPI_ISL_719709, EPI_ISL_719710, EPI_ISL_719711, EPI_ISL_719712, EPI_ISL_719713, EPI_ISL_719714, EPI_ISL_719715, EPI_ISL_719716, EPI_ISL_719717, EPI_ISL_719718, EPI_ISL_719719, EPI_ISL_719720, EPI_ISL_719721, EPI_ISL_719722, EPI_ISL_719723, EPI_ISL_719724, EPI_ISL_719725, EPI_ISL_719726, EPI_ISL_719727, EPI_ISL_719728, EPI_ISL_719729, EPI_ISL_719730, EPI_ISL_719731, EPI_ISL_719732, EPI_ISL_719733, EPI_ISL_719734, EPI_ISL_719735, EPI_ISL_719736, EPI_ISL_719737, EPI_ISL_719738, EPI_ISL_719739, EPI_ISL_719740, EPI_ISL_719741, EPI_ISL_719742, EPI_ISL_719743, EPI_ISL_719744, EPI_ISL_719745, EPI_ISL_719746, EPI_ISL_719747, EPI_ISL_719748, EPI_ISL_719749, EPI_ISL_719750, EPI_ISL_719751, EPI_ISL_719752, EPI_ISL_719753, EPI_ISL_719754, EPI_ISL_719755, EPI_ISL_719756, EPI_ISL_719757, EPI_ISL_719758, EPI_ISL_719759, EPI_ISL_719760, EPI_ISL_719761, EPI_ISL_719762, EPI_ISL_719763, EPI_ISL_719764, EPI_ISL_719765, EPI_ISL_719766, EPI_ISL_719767, EPI_ISL_719768, EPI_ISL_719769, EPI_ISL_719770, EPI_ISL_719771, EPI_ISL_719772, EPI_ISL_719773, EPI_ISL_719774, EPI_ISL_719775, EPI_ISL_719776, EPI_ISL_719777, EPI_ISL_719778, EPI_ISL_719779, EPI_ISL_719780, EPI_ISL_719781, EPI_ISL_719782, EPI_ISL_719783, EPI_ISL_719784, EPI_ISL_719785, EPI_ISL_719786, EPI_ISL_719787, EPI_ISL_719788, EPI_ISL_719789, EPI_ISL_719790, EPI_ISL_719791, EPI_ISL_719792, EPI_ISL_719793, EPI_ISL_719794, EPI_ISL_719795, EPI_ISL_719796, EPI_ISL_719797, EPI_ISL_719798, EPI_ISL_719799, EPI_ISL_719800, EPI_ISL_719801, EPI_ISL_719802, EPI_ISL_719803, EPI_ISL_719804, EPI_ISL_719805, EPI_ISL_719806, EPI_ISL_719807, EPI_ISL_719808, EPI_ISL_719809, EPI_ISL_719810, EPI_ISL_719811, EPI_ISL_719812, EPI_ISL_719813, EPI_ISL_719814, EPI_ISL_719815, EPI_ISL_719816, EPI_ISL_719817, EPI_ISL_719818, EPI_ISL_719819, EPI_ISL_719820, EPI_ISL_719821, EPI_ISL_719822, EPI_ISL_719823, EPI_ISL_719824, EPI_ISL_719825, EPI_ISL_719826, EPI_ISL_719827, EPI_ISL_719828, EPI_ISL_719829, EPI_ISL_719830, EPI_ISL_719831, EPI_ISL_719832, EPI_ISL_719833, EPI_ISL_719834, EPI_ISL_719835, EPI_ISL_719836, EPI_ISL_719837, EPI_ISL_719838, EPI_ISL_719839, EPI_ISL_719840, EPI_ISL_719841, EPI_ISL_719842, EPI_ISL_719843, EPI_ISL_719844, EPI_ISL_719845, EPI_ISL_719846, EPI_ISL_719847, EPI_ISL_719848, EPI_ISL_719849, EPI_ISL_719850, EPI_ISL_719851, EPI_ISL_719852, EPI_ISL_719853, EPI_ISL_719854, EPI_ISL_719855, EPI_ISL_719856, EPI_ISL_719857, EPI_ISL_719858, EPI_ISL_719859, EPI_ISL_719860, EPI_ISL_719861, EPI_ISL_719862, EPI_ISL_719863, EPI_ISL_719864, EPI_ISL_719865, EPI_ISL_719866, EPI_ISL_719867, EPI_ISL_719868, EPI_ISL_719869, EPI_ISL_719870, EPI_ISL_719871, EPI_ISL_719872, EPI_ISL_719873, EPI_ISL_719874, EPI_ISL_719875, EPI_ISL_719876, EPI_ISL_719877, EPI_ISL_719878, EPI_ISL_719879, EPI_ISL_719880, EPI_ISL_719881, EPI_ISL_719882, EPI_ISL_719883, EPI_ISL_719884, EPI_ISL_719885, EPI_ISL_719886, EPI_ISL_719887, EPI_ISL_719888, EPI_ISL_719889, EPI_ISL_719890, EPI_ISL_719891, EPI_ISL_719892, EPI_ISL_719893, EPI_ISL_719894, EPI_ISL_719895, EPI_ISL_719896, EPI_ISL_719897, EPI_ISL_719898, EPI_ISL_719899, EPI_ISL_719900, EPI_ISL_719901, EPI_ISL_719902, EPI_ISL_719903, EPI_ISL_719904, EPI_ISL_719905, EPI_ISL_719906, EPI_ISL_719907, EPI_ISL_719908, EPI_ISL_719909, EPI_ISL_719910, EPI_ISL_719911, EPI_ISL_719912, EPI_ISL_719913, EPI_ISL_719914, EPI_ISL_719915, EPI_ISL_719916, EPI_ISL_719917, EPI_ISL_719918, EPI_ISL_719919, EPI_ISL_719920, EPI_ISL_719921, EPI_ISL_719922, EPI_ISL_719923, EPI_ISL_719924, EPI_ISL_719925, EPI_ISL_719926, EPI_ISL_719927, EPI_ISL_719928, EPI_ISL_719929, EPI_ISL_719930, EPI_ISL_719931, EPI_ISL_719932, EPI_ISL_719933, EPI_ISL_719934, EPI_ISL_719935, EPI_ISL_719936, EPI_ISL_719937, EPI_ISL_719938, EPI_ISL_719939, EPI_ISL_719940, EPI_ISL_719941, EPI_ISL_719942, EPI_ISL_719943, EPI_ISL_719944, EPI_ISL_719945, EPI_ISL_719946, EPI_ISL_719947, EPI_ISL_719948, EPI_ISL_719949, EPI_ISL_719950, EPI_ISL_719951, EPI_ISL_719952, EPI_ISL_719953, EPI_ISL_719954, EPI_ISL_719955, EPI_ISL_719956, EPI_ISL_719957, EPI_ISL_719958, EPI_ISL_719959, EPI_ISL_719960, EPI_ISL_719961, EPI_ISL_719962, EPI_ISL_719963, EPI_ISL_719964, EPI_ISL_719965, EPI_ISL_719966, EPI_ISL_719967, EPI_ISL_719968, EPI_ISL_719969, EPI_ISL_719970, EPI_ISL_719971, EPI_ISL_719972, EPI_ISL_719973, EPI_ISL_719974, EPI_ISL_719975, EPI_ISL_719976, EPI_ISL_719977, EPI_ISL_719978, EPI_ISL_719979, EPI_ISL_719980, EPI_ISL_719981, EPI_ISL_719982, EPI_ISL_719983, EPI_ISL_719984, EPI_ISL_719985, EPI_ISL_719986 | | | |
| see above | Lighthouse Lab in Glasgow | Wellcome Sanger Institute for the COVID-19 Genomics UK (COG-UK) Consortium | Harper VanSteenhouse, Yumi Kasai, David Gray, Carol Clugston, Anna Dominiczak and Alex Alderton, Roberto Amato, Sonia Goncalves, Ewan Harrison, David K. Jackson, Ian Johnston, Dominic Kwiatkowski, Cordelia Langford, John Sillitoe on behalf of the Wellcome Sanger Institute COVID-19 Surveillance Team |
| EPI_ISL_719987, EPI_ISL_719988, EPI_ISL_719989, EPI_ISL_719990, EPI_ISL_719991, EPI_ISL_719992, EPI_ISL_719993, EPI_ISL_719994, EPI_ISL_719995, EPI_ISL_719996, EPI_ISL_719997, EPI_ISL_719998, EPI_ISL_719999, EPI_ISL_720000, EPI_ISL_720001, EPI_ISL_720002, EPI_ISL_720003, EPI_ISL_720004, EPI_ISL_720005, EPI_ISL_720006, EPI_ISL_720007, EPI_ISL_720008, EPI_ISL_720009, EPI_ISL_720010, EPI_ISL_720011, EPI_ISL_720012, EPI_ISL_720013, EPI_ISL_720014, EPI_ISL_720015, EPI_ISL_720016, EPI_ISL_720017, EPI_ISL_720018, EPI_ISL_720019, EPI_ISL_720020, EPI_ISL_720021, EPI_ISL_720022, EPI_ISL_720023, EPI_ISL_720024, EPI_ISL_720025, EPI_ISL_720026, EPI_ISL_720027, EPI_ISL_720028, EPI_ISL_720029, EPI_ISL_720030, EPI_ISL_720031, EPI_ISL_720032, EPI_ISL_720033, EPI_ISL_720034, EPI_ISL_720035, EPI_ISL_720036, EPI_ISL_720037, EPI_ISL_720038, EPI_ISL_720039, EPI_ISL_720040, EPI_ISL_720041, EPI_ISL_720042, EPI_ISL_720043, EPI_ISL_720044, EPI_ISL_720045, EPI_ISL_720046, EPI_ISL_720047, EPI_ISL_720048, EPI_ISL_720049, EPI_ISL_720050, EPI_ISL_720051, EPI_ISL_720052, EPI_ISL_720053, EPI_ISL_720054, EPI_ISL_720055, EPI_ISL_720056, EPI_ISL_720057, EPI_ISL_720058, | | | |

| | | | |
|---|---|---|---|
| EPI_ISL_720059, EPI_ISL_720060, EPI_ISL_720061, EPI_ISL_720062, EPI_ISL_720063, EPI_ISL_720064, EPI_ISL_720065, EPI_ISL_720066, EPI_ISL_720067, EPI_ISL_720068, EPI_ISL_720069, EPI_ISL_720070, EPI_ISL_720071, EPI_ISL_720072, EPI_ISL_720073, EPI_ISL_720074, EPI_ISL_720075, EPI_ISL_720076, EPI_ISL_720077, EPI_ISL_720078, EPI_ISL_720079, EPI_ISL_720080, EPI_ISL_720081, EPI_ISL_720082, EPI_ISL_720083, EPI_ISL_720084, EPI_ISL_720085, EPI_ISL_720086, EPI_ISL_720087, EPI_ISL_720088, EPI_ISL_720089, EPI_ISL_720090, EPI_ISL_720091, EPI_ISL_720092, EPI_ISL_720093, EPI_ISL_720094, EPI_ISL_720095, EPI_ISL_720096, EPI_ISL_720097, EPI_ISL_720098, EPI_ISL_720099, EPI_ISL_720100, EPI_ISL_720101, EPI_ISL_720102, EPI_ISL_720103, EPI_ISL_720104, EPI_ISL_720105, EPI_ISL_720106, EPI_ISL_720107, EPI_ISL_720108, EPI_ISL_720109, EPI_ISL_720110, EPI_ISL_720111, EPI_ISL_720112, EPI_ISL_720113, EPI_ISL_720114, EPI_ISL_720115, EPI_ISL_720116, EPI_ISL_720117, EPI_ISL_720118, EPI_ISL_720119, EPI_ISL_720120, EPI_ISL_720121, EPI_ISL_720122, EPI_ISL_720123, EPI_ISL_720124, EPI_ISL_720125, EPI_ISL_720126, EPI_ISL_720127, EPI_ISL_720128, EPI_ISL_720129, EPI_ISL_720130, EPI_ISL_720131, EPI_ISL_720132, EPI_ISL_720133, EPI_ISL_720134, EPI_ISL_720135, EPI_ISL_720136, EPI_ISL_720137, EPI_ISL_720138, EPI_ISL_720139, EPI_ISL_720140, EPI_ISL_720141, EPI_ISL_720142, EPI_ISL_720143, EPI_ISL_720144, EPI_ISL_720145, EPI_ISL_720147, EPI_ISL_720148, EPI_ISL_720149, EPI_ISL_720150, EPI_ISL_720151, EPI_ISL_720152, EPI_ISL_720153, EPI_ISL_720154, EPI_ISL_720155, EPI_ISL_720156, EPI_ISL_720157, EPI_ISL_720158, EPI_ISL_720159, EPI_ISL_720160, EPI_ISL_720161, EPI_ISL_720162, EPI_ISL_720163, EPI_ISL_720164, EPI_ISL_720165, EPI_ISL_720166, EPI_ISL_720167, EPI_ISL_720168, EPI_ISL_720169, EPI_ISL_720170, EPI_ISL_720171, EPI_ISL_720172, EPI_ISL_720173, EPI_ISL_720174, EPI_ISL_720175, EPI_ISL_720176, EPI_ISL_720177, EPI_ISL_720178, EPI_ISL_720179, EPI_ISL_720180, EPI_ISL_720181, EPI_ISL_720182, EPI_ISL_720183, EPI_ISL_720184, EPI_ISL_720185, EPI_ISL_720186, EPI_ISL_720187, EPI_ISL_720188, EPI_ISL_720189, EPI_ISL_720190, EPI_ISL_720191, EPI_ISL_720192, EPI_ISL_720193, EPI_ISL_720194, EPI_ISL_720195, EPI_ISL_720196, EPI_ISL_720197, EPI_ISL_720198, EPI_ISL_720199, EPI_ISL_720200, EPI_ISL_720201, EPI_ISL_720202, EPI_ISL_720203, EPI_ISL_720204, EPI_ISL_720205, EPI_ISL_720206, EPI_ISL_720207, EPI_ISL_720208, EPI_ISL_720209, EPI_ISL_720210, EPI_ISL_720211, EPI_ISL_720212, EPI_ISL_720213, EPI_ISL_720214, EPI_ISL_720215, EPI_ISL_720216, EPI_ISL_720217, EPI_ISL_720218, EPI_ISL_720219, EPI_ISL_720220, EPI_ISL_720221, EPI_ISL_720222, EPI_ISL_720223, EPI_ISL_720224, EPI_ISL_720225, EPI_ISL_720226, EPI_ISL_720227, EPI_ISL_720228, EPI_ISL_720229, EPI_ISL_720230, EPI_ISL_720231, EPI_ISL_720232, EPI_ISL_720233, EPI_ISL_720234, EPI_ISL_720235, EPI_ISL_720236, EPI_ISL_720237, EPI_ISL_720238, EPI_ISL_720239, EPI_ISL_720240, EPI_ISL_720241, EPI_ISL_720242, EPI_ISL_720243, EPI_ISL_720244, EPI_ISL_720245, EPI_ISL_720246, EPI_ISL_720247, EPI_ISL_720248, EPI_ISL_720249, EPI_ISL_720250, EPI_ISL_720251, EPI_ISL_720252, EPI_ISL_720253, EPI_ISL_720254, EPI_ISL_720255, EPI_ISL_720256, EPI_ISL_720257, EPI_ISL_720258, EPI_ISL_720259, EPI_ISL_720260, EPI_ISL_720261, EPI_ISL_720262, EPI_ISL_720263, EPI_ISL_720264, EPI_ISL_720265, EPI_ISL_720266, EPI_ISL_720267, EPI_ISL_720268, EPI_ISL_720269, EPI_ISL_720270, EPI_ISL_720271, EPI_ISL_720272, EPI_ISL_720273, EPI_ISL_720274, EPI_ISL_720275, EPI_ISL_720276, EPI_ISL_720277, EPI_ISL_720278, EPI_ISL_720279, EPI_ISL_720280, EPI_ISL_720281, EPI_ISL_720282, EPI_ISL_720283, EPI_ISL_720284, EPI_ISL_720285, EPI_ISL_720286, EPI_ISL_720287, EPI_ISL_720288, EPI_ISL_720289, EPI_ISL_720290, EPI_ISL_720291, EPI_ISL_720292, EPI_ISL_720293, EPI_ISL_720294, EPI_ISL_720295, EPI_ISL_720296, EPI_ISL_720297, EPI_ISL_720298, EPI_ISL_720299, EPI_ISL_720300, EPI_ISL_720301, EPI_ISL_720302, EPI_ISL_720303, EPI_ISL_720304, EPI_ISL_720305, EPI_ISL_720306, EPI_ISL_720307, EPI_ISL_720308, EPI_ISL_720309, EPI_ISL_720310, EPI_ISL_720311, EPI_ISL_720312, EPI_ISL_720313, EPI_ISL_720314, EPI_ISL_720315, EPI_ISL_720316, EPI_ISL_720317, EPI_ISL_720318, EPI_ISL_720319, EPI_ISL_720320, EPI_ISL_720321, EPI_ISL_720322, EPI_ISL_720323, EPI_ISL_720324, EPI_ISL_720325, EPI_ISL_720326, EPI_ISL_720327, EPI_ISL_720328, EPI_ISL_720329, EPI_ISL_720330, EPI_ISL_720331, EPI_ISL_720332, EPI_ISL_720333, EPI_ISL_720334, EPI_ISL_720335, EPI_ISL_720336, EPI_ISL_720337, EPI_ISL_720338, EPI_ISL_720339, EPI_ISL_720340, EPI_ISL_720341, EPI_ISL_720342, EPI_ISL_720343, EPI_ISL_720344, EPI_ISL_720345, EPI_ISL_720346, EPI_ISL_720347, EPI_ISL_720348, EPI_ISL_720349, EPI_ISL_720350, EPI_ISL_720351, EPI_ISL_720352, EPI_ISL_720353, EPI_ISL_720354, EPI_ISL_720355, EPI_ISL_720356, EPI_ISL_720357, EPI_ISL_720358, EPI_ISL_720359, EPI_ISL_720360, EPI_ISL_720361, EPI_ISL_720362, EPI_ISL_720363, EPI_ISL_720364, EPI_ISL_720365, EPI_ISL_720366, EPI_ISL_720367, EPI_ISL_720368, EPI_ISL_720369, EPI_ISL_720370, EPI_ISL_720371, EPI_ISL_720372, EPI_ISL_720373, EPI_ISL_720374, EPI_ISL_720375, EPI_ISL_720376, EPI_ISL_720377, EPI_ISL_720378, EPI_ISL_720379, EPI_ISL_720380, EPI_ISL_720381, EPI_ISL_720382, EPI_ISL_720383, EPI_ISL_720384, EPI_ISL_720385, EPI_ISL_720386, EPI_ISL_720387, EPI_ISL_720388, EPI_ISL_720389, EPI_ISL_720390, EPI_ISL_720391, EPI_ISL_720392, EPI_ISL_720393, EPI_ISL_720394, EPI_ISL_720395, EPI_ISL_720396, EPI_ISL_720397, EPI_ISL_720398, EPI_ISL_720399, EPI_ISL_720400, EPI_ISL_720401, EPI_ISL_720402, EPI_ISL_720403, EPI_ISL_720404, EPI_ISL_720405, EPI_ISL_720406, EPI_ISL_720407, EPI_ISL_720408, EPI_ISL_720409, EPI_ISL_720410, EPI_ISL_720411, EPI_ISL_720412, EPI_ISL_720413, EPI_ISL_720414, EPI_ISL_720415, EPI_ISL_720416, EPI_ISL_720417, EPI_ISL_720418, EPI_ISL_720419, EPI_ISL_720420, EPI_ISL_720421, EPI_ISL_720422, EPI_ISL_720423, EPI_ISL_720424, EPI_ISL_720425, EPI_ISL_720426, EPI_ISL_720427, EPI_ISL_720428, EPI_ISL_720429, EPI_ISL_720430, EPI_ISL_720431, EPI_ISL_720432, EPI_ISL_720433, EPI_ISL_720434, EPI_ISL_720435, EPI_ISL_720436, EPI_ISL_720437, EPI_ISL_720438, EPI_ISL_720439, EPI_ISL_720440, EPI_ISL_720441, EPI_ISL_720442, EPI_ISL_720443, EPI_ISL_720444, EPI_ISL_720445, EPI_ISL_720446, EPI_ISL_720447, EPI_ISL_720448, EPI_ISL_720449, EPI_ISL_720450, EPI_ISL_720451, EPI_ISL_720452, EPI_ISL_720453, EPI_ISL_720454, EPI_ISL_720455, EPI_ISL_720456, EPI_ISL_720457, EPI_ISL_720458, EPI_ISL_720459, EPI_ISL_720460, EPI_ISL_720461, EPI_ISL_720462, EPI_ISL_720463, EPI_ISL_720464, EPI_ISL_720465, EPI_ISL_720466, EPI_ISL_720467, EPI_ISL_720468, EPI_ISL_720469, EPI_ISL_720470, EPI_ISL_720471, EPI_ISL_720472, EPI_ISL_720473, EPI_ISL_720474, EPI_ISL_720475, EPI_ISL_720476, EPI_ISL_720477, EPI_ISL_720478, EPI_ISL_720479, EPI_ISL_720480, EPI_ISL_720481, EPI_ISL_720482, EPI_ISL_720483, EPI_ISL_720484, EPI_ISL_720485, EPI_ISL_720486, EPI_ISL_720487, EPI_ISL_720488, EPI_ISL_720489, EPI_ISL_720490, EPI_ISL_720491, EPI_ISL_720492, EPI_ISL_720493, EPI_ISL_720494, EPI_ISL_720495, EPI_ISL_720496, EPI_ISL_720497, EPI_ISL_720498, EPI_ISL_720499, EPI_ISL_720500, EPI_ISL_720501, EPI_ISL_720502, EPI_ISL_720503, EPI_ISL_720504, EPI_ISL_720505, EPI_ISL_720506, EPI_ISL_720507, EPI_ISL_720508, EPI_ISL_720509, EPI_ISL_720510, EPI_ISL_720511, EPI_ISL_720512, EPI_ISL_720513, EPI_ISL_720514, EPI_ISL_720515, EPI_ISL_720516, EPI_ISL_720517, EPI_ISL_720518, EPI_ISL_720519, EPI_ISL_720520, EPI_ISL_720521, EPI_ISL_720522, EPI_ISL_720523, EPI_ISL_720524, EPI_ISL_720525, EPI_ISL_720526, EPI_ISL_720527, EPI_ISL_720528, EPI_ISL_720529, EPI_ISL_720530, EPI_ISL_720531, EPI_ISL_720532, EPI_ISL_720533, EPI_ISL_720534, EPI_ISL_720535, EPI_ISL_720536, EPI_ISL_720537, EPI_ISL_720538, EPI_ISL_720539, EPI_ISL_720540, EPI_ISL_720541, EPI_ISL_720542, EPI_ISL_720543, EPI_ISL_720544, EPI_ISL_720545, EPI_ISL_720546, EPI_ISL_720547, EPI_ISL_720548, EPI_ISL_720549, EPI_ISL_720550, EPI_ISL_720551, EPI_ISL_720552, EPI_ISL_720553, EPI_ISL_720554, EPI_ISL_720555, EPI_ISL_720556, EPI_ISL_720557, EPI_ISL_720558, EPI_ISL_720559, EPI_ISL_720560, EPI_ISL_720561, EPI_ISL_720562, EPI_ISL_720563, EPI_ISL_720564, EPI_ISL_720565, EPI_ISL_720566, EPI_ISL_720567, EPI_ISL_720568, EPI_ISL_720569, EPI_ISL_720570, EPI_ISL_720571, EPI_ISL_720572, EPI_ISL_720573, EPI_ISL_720574, EPI_ISL_720575, EPI_ISL_720576, EPI_ISL_720577, EPI_ISL_720578, EPI_ISL_720579, EPI_ISL_720580, EPI_ISL_720581, EPI_ISL_720582, EPI_ISL_720583, EPI_ISL_720584, EPI_ISL_720585, EPI_ISL_720586, EPI_ISL_720587, EPI_ISL_720588, EPI_ISL_720589, EPI_ISL_720590, EPI_ISL_720591, EPI_ISL_720592, EPI_ISL_720593, EPI_ISL_720594, EPI_ISL_720595, EPI_ISL_720596, EPI_ISL_720597, EPI_ISL_720598, EPI_ISL_720599, EPI_ISL_720600, EPI_ISL_720601, EPI_ISL_720602, EPI_ISL_720603, EPI_ISL_720604, EPI_ISL_720605, EPI_ISL_720606, EPI_ISL_720607, EPI_ISL_720608, EPI_ISL_720609, EPI_ISL_720610, EPI_ISL_720611, EPI_ISL_720612, EPI_ISL_720613, EPI_ISL_720614, EPI_ISL_720615, EPI_ISL_720616, EPI_ISL_720617, EPI_ISL_720618, EPI_ISL_720619, EPI_ISL_720620, EPI_ISL_720621, EPI_ISL_720622, EPI_ISL_720623, EPI_ISL_720624, EPI_ISL_720625, EPI_ISL_720626, EPI_ISL_720627, EPI_ISL_720628, EPI_ISL_720629, EPI_ISL_720630, EPI_ISL_720631, EPI_ISL_720632, EPI_ISL_720633, EPI_ISL_720634, EPI_ISL_720635, EPI_ISL_720636, EPI_ISL_720637, EPI_ISL_720638, EPI_ISL_720639, EPI_ISL_720640, EPI_ISL_720641, EPI_ISL_720642, EPI_ISL_720643, EPI_ISL_720644, EPI_ISL_720645, EPI_ISL_720646, EPI_ISL_720647, EPI_ISL_720648, EPI_ISL_720649, EPI_ISL_720650, EPI_ISL_720651, EPI_ISL_720652, EPI_ISL_720653, EPI_ISL_720654, EPI_ISL_720655, EPI_ISL_720656, EPI_ISL_720657, EPI_ISL_720658, EPI_ISL_720659, EPI_ISL_720660, EPI_ISL_720661, EPI_ISL_720662, EPI_ISL_720663, EPI_ISL_720664, EPI_ISL_720665, EPI_ISL_720666, EPI_ISL_720667 | | | |
| see above | Lighthouse Lab in Milton Keynes | Wellcome Sanger Institute for the COVID-19 Genomics UK (COG-UK) Consortium | The Lighthouse Lab in Milton Keynes and Alex Alderton, Roberto Amato, Sonia Goncalves, Ewan Harrison, David K. Jackson, Ian Johnston, Dominic Kwiatkowski, Cordelia Langford, John Sillitoe on behalf of the Wellcome Sanger Institute COVID-19 Surveillance Team |
| EPI_ISL_720909, EPI_ISL_720910, EPI_ISL_720911, EPI_ISL_720912, EPI_ISL_720913, EPI_ISL_720914, EPI_ISL_720915, EPI_ISL_720916, EPI_ISL_720917, EPI_ISL_720918, EPI_ISL_720919, EPI_ISL_720920, EPI_ISL_720921, EPI_ISL_720922, EPI_ISL_720923, EPI_ISL_720924, EPI_ISL_720925, EPI_ISL_720926, EPI_ISL_720927, EPI_ISL_720928, EPI_ISL_720929, EPI_ISL_720930, EPI_ISL_720931, EPI_ISL_720932, EPI_ISL_720933, EPI_ISL_720934, EPI_ISL_720935, EPI_ISL_720936, EPI_ISL_720937, EPI_ISL_720938, EPI_ISL_720939, EPI_ISL_720940, EPI_ISL_720941, EPI_ISL_720942, EPI_ISL_720943, EPI_ISL_720944, EPI_ISL_720945, EPI_ISL_720946, EPI_ISL_720947, EPI_ISL_720948, EPI_ISL_720949, EPI_ISL_720950, EPI_ISL_720951, EPI_ISL_720952, EPI_ISL_720953, EPI_ISL_720954, EPI_ISL_720955, EPI_ISL_720956, EPI_ISL_720957, EPI_ISL_720958, EPI_ISL_720959, EPI_ISL_720960, EPI_ISL_720961, EPI_ISL_720962, EPI_ISL_720963, EPI_ISL_720964, EPI_ISL_720965, EPI_ISL_720966, EPI_ISL_720967, EPI_ISL_720968, EPI_ISL_720969, EPI_ISL_720970, EPI_ISL_720971, EPI_ISL_720972, EPI_ISL_720973, EPI_ISL_720974, EPI_ISL_720975, EPI_ISL_720976, EPI_ISL_720977, EPI_ISL_720978, EPI_ISL_720979, EPI_ISL_720980, EPI_ISL_720981, EPI_ISL_720982, EPI_ISL_720983, EPI_ISL_720984, EPI_ISL_720985, EPI_ISL_720986, EPI_ISL_720987, EPI_ISL_720988, EPI_ISL_720989, EPI_ISL_720990, EPI_ISL_720991, EPI_ISL_720992, EPI_ISL_720993, EPI_ISL_720994, EPI_ISL_720995, EPI_ISL_720996, EPI_ISL_720997, EPI_ISL_720998, EPI_ISL_720999, EPI_ISL_721000, EPI_ISL_721001, EPI_ISL_721002, EPI_ISL_721003, EPI_ISL_721004, EPI_ISL_721005, EPI_ISL_721006, EPI_ISL_721007, EPI_ISL_721008, EPI_ISL_721009, EPI_ISL_721010, EPI_ISL_721011, EPI_ISL_721012, EPI_ISL_721013, EPI_ISL_721014, EPI_ISL_721015, EPI_ISL_721016, EPI_ISL_721017, EPI_ISL_721018, EPI_ISL_721019, EPI_ISL_721020, EPI_ISL_721021, EPI_ISL_721022, EPI_ISL_721023, EPI_ISL_721024, EPI_ISL_721025, EPI_ISL_721026, EPI_ISL_721027, EPI_ISL_721028, EPI_ISL_721029, EPI_ISL_721030, EPI_ISL_721031, EPI_ISL_721032, EPI_ISL_721033, EPI_ISL_721034, EPI_ISL_721035, EPI_ISL_721036, EPI_ISL_721037, EPI_ISL_721038, EPI_ISL_721039, EPI_ISL_721040, EPI_ISL_721041, EPI_ISL_721042, EPI_ISL_721043, EPI_ISL_721044, EPI_ISL_721045, EPI_ISL_721046, EPI_ISL_721047, EPI_ISL_721048, EPI_ISL_721049, EPI_ISL_721050, EPI_ISL_721051, EPI_ISL_721052, EPI_ISL_721053, EPI_ISL_721054, EPI_ISL_721055, EPI_ISL_721056, EPI_ISL_721057, EPI_ISL_721058, EPI_ISL_721059, EPI_ISL_721060, EPI_ISL_721061, EPI_ISL_721062, EPI_ISL_721063, EPI_ISL_721064, EPI_ISL_721065, EPI_ISL_721066, EPI_ISL_721067, EPI_ISL_721068, EPI_ISL_721069, EPI_ISL_721070, EPI_ISL_721071, EPI_ISL_721072, EPI_ISL_721073, EPI_ISL_721074, EPI_ISL_721075, EPI_ISL_721076, EPI_ISL_721077, EPI_ISL_721078, EPI_ISL_721079, EPI_ISL_721080, EPI_ISL_721081, EPI_ISL_721082, EPI_ISL_721083, EPI_ISL_721084, EPI_ISL_721085, EPI_ISL_721086, EPI_ISL_721087, EPI_ISL_721088, EPI_ISL_721089, EPI_ISL_721090, EPI_ISL_721091, EPI_ISL_721092, EPI_ISL_721093, EPI_ISL_721094, EPI_ISL_721095, EPI_ISL_721096, EPI_ISL_721097, EPI_ISL_721098, EPI_ISL_721099, EPI_ISL_721100, EPI_ISL_721101, EPI_ISL_721102, EPI_ISL_721103, EPI_ISL_721104, EPI_ISL_721105, EPI_ISL_721106, EPI_ISL_721107, EPI_ISL_721108, EPI_ISL_721109, EPI_ISL_721110, EPI_ISL_721111, EPI_ISL_721112, EPI_ISL_721113, EPI_ISL_721114, EPI_ISL_721115, EPI_ISL_721116, EPI_ISL_721117, EPI_ISL_721118, EPI_ISL_721119, EPI_ISL_721120, EPI_ISL_721121, EPI_ISL_721122, EPI_ISL_721123, EPI_ISL_721124, EPI_ISL_721125, EPI_ISL_721126, EPI_ISL_721127, EPI_ISL_721128, EPI_ISL_721129, EPI_ISL_721130, EPI_ISL_721131, EPI_ISL_721132, EPI_ISL_721133, EPI_ISL_721134, EPI_ISL_721135, EPI_ISL_721136, EPI_ISL_721137, EPI_ISL_721138, EPI_ISL_721139, EPI_ISL_721140, EPI_ISL_721141, EPI_ISL_721142, EPI_ISL_721143, EPI_ISL_721144, EPI_ISL_721145, EPI_ISL_721146, EPI_ISL_721147, EPI_ISL_721148, EPI_ISL_721149, EPI_ISL_721150, EPI_ISL_721151, EPI_ISL_721152, EPI_ISL_721153, EPI_ISL_721154, EPI_ISL_721155, EPI_ISL_721156, EPI_ISL_721157, EPI_ISL_721158, EPI_ISL_721159, EPI_ISL_721160, EPI_ISL_721161, EPI_ISL_721162, EPI_ISL_721163, EPI_ISL_721164, EPI_ISL_721165, EPI_ISL_721166, EPI_ISL_721167, EPI_ISL_721168, EPI_ISL_721169, EPI_ISL_721170, EPI_ISL_721171, EPI_ISL_721172, EPI_ISL_721173, EPI_ISL_721174, EPI_ISL_721175, EPI_ISL_721176, EPI_ISL_721177, EPI_ISL_721178, EPI_ISL_721179, EPI_ISL_721180, EPI_ISL_721181, EPI_ISL_721182, EPI_ISL_721183, EPI_ISL_721184 | | | |
| see above | Lighthouse Lab in Glasgow | Wellcome Sanger Institute for the COVID-19 Genomics UK (COG-UK) Consortium | Harper VanSteenhouse, Yumi Kasai, David Gray, Carol Clugston, Anna Dominiczak and Alex Alderton, Roberto Amato, Sonia Goncalves, Ewan Harrison, David K. Jackson, Ian Johnston, Dominic Kwiatkowski, Cordelia Langford, John Sillitoe on behalf of the Wellcome Sanger Institute COVID-19 Surveillance Team |
| EPI_ISL_721306, EPI_ISL_721370, EPI_ISL_721456, EPI_ISL_721460 | Lighthouse Lab in Alderley Park | Wellcome Sanger Institute for the COVID-19 Genomics UK (COG-UK) Consortium | Jacquelyn Wynn, Mairead Hyland, The Lighthouse Lab in Alderley Park and Alex Alderton, Roberto Amato, Sonia Goncalves, Ewan Harrison, David K. Jackson, Ian Johnston, Dominic Kwiatkowski, Cordelia Langford, John Sillitoe on behalf of the Wellcome Sanger Institute COVID-19 Surveillance Team |
| EPI_ISL_721624 | National Institute for Infectious Diseases, INMI, "L. Spallanzani" IRCCS | National Institute for Infectious Diseases, INMI, "L. Spallanzani" IRCCS | C.E.M Gruber, B Bartolini, M Rueca, F Messina, E Giombini, A Di Caro, MR Capobianchi |
| EPI_ISL_721625 | National Institute for Infectious Diseases, INMI, "L. Spallanzani" IRCCS | National Institute for Infectious Diseases, INMI, "L. Spallanzani" IRCCS | B Bartolini, C.E.M Gruber, M Rueca, F Messina, E Giombini, MR Capobianchi, A Di Caro |
| EPI_ISL_721628 | National Institute for Infectious Diseases, INMI, "L. Spallanzani" IRCCS | National Institute for Infectious Diseases, INMI, "L. Spallanzani" IRCCS | E Giombini, M Rueca, B Bartolini, C.E.M Gruber, F Messina, A Di Caro, MR Capobianchi |
| EPI_ISL_722276, EPI_ISL_722277, EPI_ISL_722279, EPI_ISL_722284, EPI_ISL_722288, EPI_ISL_722302, EPI_ISL_722306, EPI_ISL_722321, EPI_ISL_722326, EPI_ISL_722327, EPI_ISL_722331, EPI_ISL_722340, EPI_ISL_722345, EPI_ISL_722346, EPI_ISL_722350, EPI_ISL_722364, EPI_ISL_722376, EPI_ISL_722391, EPI_ISL_722393, EPI_ISL_722395, EPI_ISL_722408, EPI_ISL_722419, EPI_ISL_722432, EPI_ISL_722648, EPI_ISL_722649, EPI_ISL_722650, EPI_ISL_722651, EPI_ISL_722652, EPI_ISL_722653, EPI_ISL_722654, EPI_ISL_722655, EPI_ISL_722656, EPI_ISL_722657, EPI_ISL_722658, EPI_ISL_722659, EPI_ISL_722660, EPI_ISL_722661, EPI_ISL_722662, EPI_ISL_722663, EPI_ISL_722664, EPI_ISL_722665, EPI_ISL_722666, EPI_ISL_722667, EPI_ISL_722668, EPI_ISL_722669, EPI_ISL_722670, EPI_ISL_722671, EPI_ISL_722672, EPI_ISL_722673, EPI_ISL_722674, EPI_ISL_722675, EPI_ISL_722676, EPI_ISL_722677, EPI_ISL_722678 | | | |
| see above | Dutch COVID-19 response team | Erasmus Medical Center | Bas Oude Munnink, Reina Sikkema, David Nieuwenhuijse, Irina Chestakova, Anne van der Linden, Marjan Boter, Emmanuelle Munger, Corine GeurtsvanKessel, Annemiek van der Eijk, Richard Molenkamp, Marion Koopmans, on behalf of the Dutch national COVID-19 response team. |
| EPI_ISL_723123 | Minnesota Department of Health, Public Health Laboratory | Minnesota Department of Health, Public Health Laboratory | Alexandra Lorentz, Jacob Garfin, Matt Plumb, and Xiong Wang |
| EPI_ISL_723145 | Mayo Clinic & Mayo Clinic Laboratories | Minnesota Department of Health, Public Health Laboratory | Alexandra Lorentz, Jacob Garfin, Matt Plumb, and Xiong Wang |
| EPI_ISL_723157, EPI_ISL_723179, EPI_ISL_723225, EPI_ISL_723251, EPI_ISL_723272, EPI_ISL_723273, EPI_ISL_723274, EPI_ISL_723292, EPI_ISL_723293, EPI_ISL_723294, EPI_ISL_723295, EPI_ISL_723296, EPI_ISL_723297, EPI_ISL_723324, EPI_ISL_723325, EPI_ISL_723406, EPI_ISL_723407, EPI_ISL_723408, EPI_ISL_723409, EPI_ISL_723410, EPI_ISL_723411, EPI_ISL_723424, EPI_ISL_723428, EPI_ISL_723430, EPI_ISL_723431, EPI_ISL_723432, EPI_ISL_723433, EPI_ISL_723434, EPI_ISL_723435, EPI_ISL_723436, EPI_ISL_723440, EPI_ISL_723444, EPI_ISL_723459 | | | |
| see above | Dutch COVID-19 response team | National Institute for Public Health and the Environment (RIVM) | Adam Meijer, Harry Vennema, Jeroen Cremer, Sharon van den Brink, Bas van der Veer, AnneMarie van den Brandt, Florian Zwagemaker, Dennis Schmitz, Chantal Reusken, on behalf of the national COVID-19 response team |
| EPI_ISL_723493, EPI_ISL_723512, EPI_ISL_723513, EPI_ISL_723514 | Virginia Division of Consolidated Laboratory Services (DCLS) | Virginia Division of Consolidated Laboratory Services (DCLS) | Virginia DCLS |
| EPI_ISL_724063 | Northumbria University / South Tees Hospitals NHS Foundation Trust / North Cumbria Integrated Care NHS Foundation Trust / North Tees and Hartlepool NHS Foundation Trust / Newcastle Hospitals NHS Foundation Trust | COVID-19 Genomics UK (COG-UK) Consortium | Darren L Smith,Andrew Nelson,Matthew Bashton,Greg R Young,Joshua Loh,John Allan,Mohammad A Tariq,Giles S Holt,Gary Black,Wen C Yew,Lynn Dover,Paul Baker,Steve Liggett,Sarah Essex,Jane Greenaway,Debra Padgett,Clive Graham,Garren Scott,Edward Barton,Emma Swindells,Brendan Payne,Jennifer Collins,Yusri Taha,Gary Eltringham |

| | | | |
|---|---|---|---|
| EPI_ISL_724125, EPI_ISL_724126, EPI_ISL_724127, EPI_ISL_724128, EPI_ISL_724129, EPI_ISL_724148, EPI_ISL_724149, EPI_ISL_724150, EPI_ISL_724188, EPI_ISL_724200, EPI_ISL_724201, EPI_ISL_724202, EPI_ISL_724203, EPI_ISL_724204, EPI_ISL_724205, EPI_ISL_724206, EPI_ISL_724207, EPI_ISL_724208, EPI_ISL_724209 | | | |
| see above | Quadram Institute Bioscience | COVID-19 Genomics UK (COG-UK) Consortium | Dave J. Baker, Gemma L. Kay, Alp Aydin, Thanh Le-Viet, Steven Rudder, Ana P. Tedim, Anastasia Kolyva, Maria Diaz, Leonardo de Oliveira Martins, Nabil-Fareed Alikhan, Lizzie Meadows, Rachael Stanley, Ngozi Elumogo, Muhammed Yasir, Nicholas M. Thomson, Alexander J Trotter, Rachel Gilroy, Samuel Bloomfield, Claire Stuart, Andrew Bell, Reenesh Prakash, Samir Dervisevic, Alison E. Mather, John Wain, Mark Webber, Andrew J. Page, Justin O'Grady |
| EPI_ISL_724212 | West of Scotland Specialist Virology Centre, NHSGGC / MRC-University of Glasgow Centre for Virus Research | COVID-19 Genomics UK (COG-UK) Consortium | Ana da Silva Filipe, Natasha Johnson, Kathy Smollett, Daniel Mair, Stephen Carmichael, Alice Broos, Lily Tong, Jenna Nichols, Kyriaki Nomikou; Sarah McDonald; Richard Orton, Joseph Hughes, Sreenu Vattipally, David L Robertson; Alasdair MacLean, Rory Gunson; Sharif Shaaban, Matthew Holden; Rachel Blacow, Guy Mollett, Kathy Li, James Shepherd, Antonia Ho, Emma Thomson |
| EPI_ISL_724541, EPI_ISL_724549, EPI_ISL_724612, EPI_ISL_724616, EPI_ISL_724619, EPI_ISL_724641, EPI_ISL_724653, EPI_ISL_724654, EPI_ISL_724663, EPI_ISL_724668, EPI_ISL_724669, EPI_ISL_724670, EPI_ISL_724688, EPI_ISL_724711, EPI_ISL_724714, EPI_ISL_724721, EPI_ISL_724768, EPI_ISL_724769, EPI_ISL_724770, EPI_ISL_724772, EPI_ISL_724774, EPI_ISL_724775 | | | |
| see above | University College London, Great Ormond Street Hospital for Children NHS Foundation Trust, Imperial College Healthcare NHS Trust | COVID-19 Genomics UK (COG-UK) Consortium | Sergi Castellano, Rachel Williams, Mark Kristiansen, Paola Resende Silva, Sunando Roy, Tony Brooks, Helena Tutill, Paola Niola, Patricia Dyal, Charlotte Williams, Leysa Forrest, Yasmin Panchbhaya, Jacqueline Findlay, Samuel Weeks, Julianne Brown, Kathryn Harris, Paul Randell, James Price, Alison Holmes, Judith Breuer |
| EPI_ISL_724924, EPI_ISL_724925, EPI_ISL_724926, EPI_ISL_724928, EPI_ISL_724929, EPI_ISL_724931, EPI_ISL_725007, EPI_ISL_725008, EPI_ISL_725009, EPI_ISL_725010, EPI_ISL_725011 | | | |
| see above | Northumbria University / South Tees Hospitals NHS Foundation Trust / North Cumbria Integrated Care NHS Foundation Trust / North Tees and Hartlepool NHS Foundation Trust / Newcastle Hospitals NHS Foundation Trust | COVID-19 Genomics UK (COG-UK) Consortium | Darren L Smith,Andrew Nelson,Matthew Bashton,Greg R Young,Joshua Loh,John Allan,Mohammad A Tariq,Giles S Holt,Gary Black,Wen C Yew,Lynn Dover,Paul Baker,Steve Liggett,Sarah Essex,Jane Greenaway,Debra Padgett,Clive Graham,Garren Scott,Edward Barton,Emma Swindells,Brendan Payne,Jennifer Collins,Yusri Taha,Gary Eltringham |
| EPI_ISL_725160, EPI_ISL_725174, EPI_ISL_725177, EPI_ISL_725178, EPI_ISL_725179, EPI_ISL_725183, EPI_ISL_725184, EPI_ISL_725186, EPI_ISL_725188, EPI_ISL_725189, EPI_ISL_725194, EPI_ISL_725195, EPI_ISL_725196, EPI_ISL_725197, EPI_ISL_725199, EPI_ISL_725200, EPI_ISL_725202, EPI_ISL_725203, EPI_ISL_725207, EPI_ISL_725208, EPI_ISL_725209, EPI_ISL_725211, EPI_ISL_725213, EPI_ISL_725214, EPI_ISL_725215, EPI_ISL_725216, EPI_ISL_725217, EPI_ISL_725218, EPI_ISL_725219, EPI_ISL_725221, EPI_ISL_725222, EPI_ISL_725228, EPI_ISL_725233, EPI_ISL_725237, EPI_ISL_725238, EPI_ISL_725239, EPI_ISL_725240, EPI_ISL_725241, EPI_ISL_725242, EPI_ISL_725248, EPI_ISL_725249, EPI_ISL_725251, EPI_ISL_725252, EPI_ISL_725254, EPI_ISL_725258, EPI_ISL_725260, EPI_ISL_725262, EPI_ISL_725281, EPI_ISL_725283 | | | |
| see above | Quadram Institute Bioscience | COVID-19 Genomics UK (COG-UK) Consortium | Dave J. Baker, Gemma L. Kay, Alp Aydin, Thanh Le-Viet, Steven Rudder, Ana P. Tedim, Anastasia Kolyva, Maria Diaz, Leonardo de Oliveira Martins, Nabil-Fareed Alikhan, Lizzie Meadows, Rachael Stanley, Ngozi Elumogo, Muhammed Yasir, Nicholas M. Thomson, Alexander J Trotter, Rachel Gilroy, Samuel Bloomfield, Claire Stuart, Andrew Bell, Reenesh Prakash, Samir Dervisevic, Alison E. Mather, John Wain, Mark Webber, Andrew J. Page, Justin O'Grady |
| EPI_ISL_725679, EPI_ISL_725718, EPI_ISL_725720, EPI_ISL_725721, EPI_ISL_725722, EPI_ISL_725723, EPI_ISL_725724, EPI_ISL_725982, EPI_ISL_725984, EPI_ISL_725994, EPI_ISL_725995, EPI_ISL_725996, EPI_ISL_725997, EPI_ISL_725998, EPI_ISL_726002, EPI_ISL_726003, EPI_ISL_726004, EPI_ISL_726005, EPI_ISL_726006, EPI_ISL_726010, EPI_ISL_726011, EPI_ISL_726012, EPI_ISL_726013, EPI_ISL_726014, EPI_ISL_726015, EPI_ISL_726016, EPI_ISL_726025, EPI_ISL_726032, EPI_ISL_726053, EPI_ISL_726063, EPI_ISL_726064, EPI_ISL_726065, EPI_ISL_726066, EPI_ISL_726067, EPI_ISL_726068, EPI_ISL_726069, EPI_ISL_726070, EPI_ISL_726071, EPI_ISL_726072, EPI_ISL_726073, EPI_ISL_726074, EPI_ISL_726075, EPI_ISL_726076, EPI_ISL_726077, EPI_ISL_726078, EPI_ISL_726079, EPI_ISL_726080, EPI_ISL_726081, EPI_ISL_726082, EPI_ISL_726083, EPI_ISL_726084, EPI_ISL_726085, EPI_ISL_726086, EPI_ISL_726087, EPI_ISL_726088, EPI_ISL_726089, EPI_ISL_726090, EPI_ISL_726091, EPI_ISL_726092, EPI_ISL_726093, EPI_ISL_726094, EPI_ISL_726095, EPI_ISL_726096, EPI_ISL_726097, EPI_ISL_726098, EPI_ISL_726099, EPI_ISL_726100, EPI_ISL_726101, EPI_ISL_726102, EPI_ISL_726103, EPI_ISL_726104, EPI_ISL_726105, EPI_ISL_726106, EPI_ISL_726108, EPI_ISL_726109, EPI_ISL_726110, EPI_ISL_726112, EPI_ISL_726113, EPI_ISL_726114, EPI_ISL_726115, EPI_ISL_726116, EPI_ISL_726117, EPI_ISL_726118, EPI_ISL_726119, EPI_ISL_726121, EPI_ISL_726122, EPI_ISL_726124, EPI_ISL_726125, EPI_ISL_726126, EPI_ISL_726127, EPI_ISL_726128, EPI_ISL_726130, EPI_ISL_726208, EPI_ISL_726213, EPI_ISL_726253, EPI_ISL_726254, EPI_ISL_726255, EPI_ISL_726256, EPI_ISL_726280, EPI_ISL_726306, EPI_ISL_726308, EPI_ISL_726309, EPI_ISL_726310, EPI_ISL_726522, EPI_ISL_726537, EPI_ISL_726570, EPI_ISL_726571, EPI_ISL_726605, EPI_ISL_726606, EPI_ISL_726607, EPI_ISL_726609, EPI_ISL_726610, EPI_ISL_726611, EPI_ISL_726613, EPI_ISL_726615, EPI_ISL_726616, EPI_ISL_726618, EPI_ISL_726619 | | | |
| see above | Wales Specialist Virology Centre Sequencing lab: Pathogen Genomics Unit | COVID-19 Genomics UK (COG-UK) Consortium | Catherine Moore, Johnathan Evans, Laura Gifford, Malorie Perry, Simon Cottrell, Angela Marchbank, Alec Birchley, Alexander Adams, Amy Gaskin, Bree Gatica-Wilcox, Jason Coombes, Joel Southgate, Lauren Gilbert, Lee Graham, Nicole Pacchiarini, Sara Kumziene-Summerhayes, Sarah Taylor, Sophie Jones, Sara Rey, Matthew Bull, Joanne Watkins, Sally Corden, Tom Connor |
| EPI_ISL_727734 | Centre for Enzyme Innovation, University of Portsmouth / Translational Research Laboratory, Portsmouth Hospitals NHS Trust | COVID-19 Genomics UK (COG-UK) Consortium | Angela Beckett,Yann Bourgeois,Garry Scarlett,Sharon Glaysher,Scott Elliott,Kelly Bicknell,Robert Impey,Allyson Lloyd,Sarah Wyllie,Ethan Butcher,Anoop Chauhan,Samuel Robson |
| EPI_ISL_727991 | Virology Department, Sheffield Teaching Hospitals NHS Foundation Trust/Department of Infection, Immunity and Cardiovascular Disease, The Medical School, University of Sheffield | COVID-19 Genomics UK (COG-UK) Consortium | Thushan de Silva, Matthew Parker, Nikki Smith, Adri Angyal, Rebecca Brown, Luke Green, Rachel Tucker, Paul Parsons, Danielle Groves, Katie Johnson, Laura Carrilero, Alex Keeley, Dave Partridge, Matthew Wyles, Benjamin Lindsey, Mehmet Yavuz, Mohammad Raza, Cariad Evans |
| EPI_ISL_728006, EPI_ISL_728008, EPI_ISL_728038, EPI_ISL_728039, EPI_ISL_728040, EPI_ISL_728047, EPI_ISL_728048, EPI_ISL_728049, EPI_ISL_728094, EPI_ISL_728098, EPI_ISL_728099, EPI_ISL_728100, EPI_ISL_728101, EPI_ISL_728102, EPI_ISL_728108, EPI_ISL_728109, EPI_ISL_728112, EPI_ISL_728113, EPI_ISL_728116, EPI_ISL_728117, EPI_ISL_728118, EPI_ISL_728120, EPI_ISL_728121, EPI_ISL_728122, EPI_ISL_728124, EPI_ISL_728125, EPI_ISL_728126, EPI_ISL_728127, EPI_ISL_728129, EPI_ISL_728133 | | | |
| see above | University of Wisconsin-Madison AIDS Vaccine Research Laboratories | University of Wisconsin-Madison AIDS Vaccine Research Laboratories | Gage Moreno, Katarina Braun, et al. AIDS Vaccine Research Laboratories |
| EPI_ISL_728248, EPI_ISL_728249, EPI_ISL_728250, EPI_ISL_728251, EPI_ISL_728252 | Institute for Medical Research, Infectious Disease Research Centre, National Institutes of Health, Ministry of Health Malaysia | Institute for Medical Research, Infectious Disease Research Centre, National Institutes of Health, Ministry of Health Malaysia | Suppiah J, Kamel K, Mohd-Zawawi Z, Thayan R |
| EPI_ISL_728300 | Lighthouse Lab in Glasgow | Wellcome Sanger Institute for the COVID-19 Genomics UK (COG-UK) Consortium | Harper VanSteenhouse, Yumi Kasai, David Gray, Carol Clugston, Anna Dominiczak and Alex Alderton, Roberto Amato, Sonia Goncalves, Ewan Harrison, David K. Jackson, Ian Johnston, Dominic Kwiatkowski, Cordelia Langford, John Sillitoe on behalf of the Wellcome Sanger Institute COVID-19 Surveillance Team |
| EPI_ISL_728311, EPI_ISL_728312, EPI_ISL_728313, EPI_ISL_728314, EPI_ISL_728315, EPI_ISL_728316, EPI_ISL_728317, EPI_ISL_728318, EPI_ISL_728319, EPI_ISL_728320, EPI_ISL_728516, EPI_ISL_728517, EPI_ISL_728518, EPI_ISL_728519, EPI_ISL_728520, EPI_ISL_728521, EPI_ISL_728522, EPI_ISL_728523, EPI_ISL_728524, EPI_ISL_728525, EPI_ISL_728526, EPI_ISL_728527, EPI_ISL_728528, EPI_ISL_728529, EPI_ISL_728530, EPI_ISL_728531, EPI_ISL_728532, EPI_ISL_728533 | | | |
| see above | CNR Virus des Infections Respiratoires - France SUD | CNR Virus des Infections Respiratoires - France SUD | Antonin Bal, Gregory Destras, Claudia Gonzalez, Gwendolyne Burfin, Quentin Semanas, Martine Valette, Bruno Lina, Laurence Josset |
| EPI_ISL_728553 | University of Wisconsin-Madison AIDS Vaccine Research Laboratories | University of Wisconsin-Madison AIDS Vaccine Research Laboratories | Gage Moreno, Katarina Braun, et al. AIDS Vaccine Research Laboratories |
| EPI_ISL_728676, EPI_ISL_728677, EPI_ISL_728678, EPI_ISL_728679, EPI_ISL_728680, EPI_ISL_728681, EPI_ISL_728684, EPI_ISL_728713, EPI_ISL_728714, EPI_ISL_728715, EPI_ISL_728716, EPI_ISL_728717, EPI_ISL_728718, EPI_ISL_728719, EPI_ISL_728720, EPI_ISL_728723, EPI_ISL_728724, EPI_ISL_728753, EPI_ISL_728754, EPI_ISL_728755 | | | |
| see above | Dutch COVID-19 response team | National Institute for Public Health and the Environment (RIVM) | Adam Meijer, Harry Vennema, Jeroen Cremer, Sharon van den Brink, Bas van der Veer, AnneMarie van den Brandt, Florian Zwagemaker, Dennis Schmitz, Chantal Reusken, on behalf of the national COVID-19 response team |
| EPI_ISL_728968, EPI_ISL_729008 | Viollier AG | Department of Biosystems Science and Engineering, ETH Zürich | Chaoran Chen, Sarah Nadeau, Catharine Aquino, Ivan Topolsky, Pedro Ferreira, Philipp Jablonski, Susana Posada-Céspedes, Andreia Cabral de Gouvea, Maria Domenica Moccia, Simon Grüter, Timothy Sykes, Lennart Opitz, Ralph Schlapbach, Christiane Beckmann, Maurice Redondo, Olivier Kobel, Christoph Noppen, Sophie Seidel, Noemie Santamaria de Souza, Niko Beerenwinkel, Tanja Stadler |
| EPI_ISL_729338, EPI_ISL_729371, EPI_ISL_729443, EPI_ISL_729445, EPI_ISL_729446, EPI_ISL_729467, EPI_ISL_729645, EPI_ISL_729646, EPI_ISL_729647, EPI_ISL_729648, EPI_ISL_729649, EPI_ISL_729650, EPI_ISL_729651, EPI_ISL_729652, EPI_ISL_729671, EPI_ISL_729672, EPI_ISL_729673, EPI_ISL_729674, EPI_ISL_729675, EPI_ISL_729676, EPI_ISL_729677, EPI_ISL_729678, EPI_ISL_729679, EPI_ISL_729680, EPI_ISL_729681, EPI_ISL_729682 | | | |
| see above | A. Krumbholz, Labor Dr. Krause und Kollegen MVZ GmbH, Kiel | Charité Universitätsmedizin Berlin, Institut für Virologie | Victor M Corman, Barbara Mühlemann, Jörn Beheim-Schwarzbach, Talitha Veith, Julia Schneider, Terry Jones, Christian Drosten |
| EPI_ISL_731251, EPI_ISL_731252, EPI_ISL_731253 | Lighthouse Lab in Milton Keynes | Wellcome Sanger Institute for the COVID-19 Genomics UK (COG-UK) Consortium | The Lighthouse Lab in Milton Keynes and Alex Alderton, Roberto Amato, Sonia Goncalves, Ewan Harrison, David K. Jackson, Ian Johnston, Dominic Kwiatkowski, Cordelia Langford, John Sillitoe on behalf of the Wellcome Sanger Institute COVID-19 Surveillance Team |
| EPI_ISL_731894 | Lighthouse Lab in Glasgow | Wellcome Sanger Institute for the COVID-19 Genomics UK (COG-UK) Consortium | Harper VanSteenhouse, Yumi Kasai, David Gray, Carol Clugston, Anna Dominiczak and Alex Alderton, Roberto Amato, Sonia Goncalves, Ewan Harrison, David K. Jackson, Ian Johnston, Dominic Kwiatkowski, Cordelia Langford, John Sillitoe on behalf of the Wellcome Sanger Institute COVID-19 Surveillance Team |

| | | | |
|---|---|---|---|
| EPI_ISL_731897 | Instituto Nacional de Saude (INSA) and Instituto Gulbenkian de Ciencia (IGC) | Instituto Nacional de Saude (INSA) and Instituto Gulbenkian de Ciencia (IGC) | Borges et al |
| EPI_ISL_731920, EPI_ISL_731921 | Instituto Nacional de Saude (INSA) | Instituto Nacional de Saude (INSA) | Borges et al |
| EPI_ISL_731925, EPI_ISL_731927, EPI_ISL_731928 | Instituto Nacional de Saude (INSA) and Instituto Gulbenkian de Ciencia (IGC) | Instituto Nacional de Saude (INSA) and Instituto Gulbenkian de Ciencia (IGC) | Borges et al |
| EPI_ISL_731951, EPI_ISL_731998, EPI_ISL_731999, EPI_ISL_732008, EPI_ISL_732009, EPI_ISL_732010, EPI_ISL_732011 | Instituto Nacional de Saude (INSA) | Instituto Nacional de Saude (INSA) | Borges et al |
| EPI_ISL_732218, EPI_ISL_732219, EPI_ISL_732220, EPI_ISL_732232, EPI_ISL_732233, EPI_ISL_732234, EPI_ISL_732235, EPI_ISL_732236, EPI_ISL_732237, EPI_ISL_732238, EPI_ISL_732239, EPI_ISL_732240 | | | |
| see above | Instituto Nacional de Saude (INSA) and Instituto Gulbenkian de Ciencia (IGC) | Instituto Nacional de Saude (INSA) and Instituto Gulbenkian de Ciencia (IGC) | Borges et al |
| EPI_ISL_732382, EPI_ISL_732383, EPI_ISL_732384, EPI_ISL_732394, EPI_ISL_732403, EPI_ISL_732404, EPI_ISL_732408, EPI_ISL_732409, EPI_ISL_732423, EPI_ISL_732427, EPI_ISL_732445, EPI_ISL_732448, EPI_ISL_732449, EPI_ISL_732451, EPI_ISL_732452, EPI_ISL_732453, EPI_ISL_732493, EPI_ISL_732503, EPI_ISL_732504, EPI_ISL_732505, EPI_ISL_732506, EPI_ISL_732507, EPI_ISL_732508, EPI_ISL_732509 | | | |
| see above | National Virus Reference Laboratory | National Virus Reference Laboratory | Michael Carr, Gabriel Gonzalez, Jonathan Dean, Daniel Hare, Cillian F De Gascun |
| EPI_ISL_732686, EPI_ISL_732687, EPI_ISL_732688 | CNR Virus des Infections Respiratoires - France SUD | CNR Virus des Infections Respiratoires - France SUD | Antonin Bal, Gregory Destras, Claudia Gonzalez, Gwendolyne Burfin, Quentin Semanas, Martine Valette, Bruno Lina, Laurence Josset |
| EPI_ISL_732948, EPI_ISL_732949, EPI_ISL_732950, EPI_ISL_732951, EPI_ISL_732952, EPI_ISL_732953, EPI_ISL_732954, EPI_ISL_732955, EPI_ISL_732956, EPI_ISL_732957, EPI_ISL_732958, EPI_ISL_732959 | | | |
| see above | Department of Tropical Parasitology | Laboratory of Recombinant Vaccines | Lukasz Rabalski, Maciej Kosinski, Teemu Smura, Kirsi Aaltonen, Ravi Kant, Tarja Sironen, Boguslaw Szewczyk, Maciej Grzybek |
| EPI_ISL_732974, EPI_ISL_732975, EPI_ISL_732976, EPI_ISL_732977, EPI_ISL_732978, EPI_ISL_732979, EPI_ISL_732980, EPI_ISL_732981, EPI_ISL_732982, EPI_ISL_732983, EPI_ISL_732984, EPI_ISL_732985, EPI_ISL_732986, EPI_ISL_732987, EPI_ISL_732988, EPI_ISL_733459, EPI_ISL_733460, EPI_ISL_733461, EPI_ISL_733464, EPI_ISL_733476, EPI_ISL_733477 | | | |
| see above | WHO National Influenza Centre Russian Federation | WHO National Influenza Centre Russian Federation | Andrey Komissarov, Artem Fadeev, Anna Ivanova, Kseniya Komissarova, Dmitry Bazhenov, Daria Danilenko, Ksenia Safina, Elena Nabieva, Georgii Bazykin, Dmitry Lioznov |
| EPI_ISL_733492, EPI_ISL_733493 | Omsk Research Institute of Natural Focal Infections | WHO National Influenza Centre Russian Federation | Andrey Komissarov, Artem Fadeev, Anna Ivanova, Kseniya Komissarova, Dmitry Bazhenov, Daria Danilenko, Ekaterina Gradoboeva, Elena Poleshchuk, Aleksei Vasilenko, Valery Yakimenko, Ksenia Safina, Elena Nabieva, Georgii Bazykin, Dmitry Lioznov |
| EPI_ISL_733494, EPI_ISL_733495 | WHO National Influenza Centre Russian Federation | WHO National Influenza Centre Russian Federation | Andrey Komissarov, Artem Fadeev, Anna Ivanova, Kseniya Komissarova, Dmitry Bazhenov, Daria Danilenko, Ksenia Safina, Elena Nabieva, Georgii Bazykin, Dmitry Lioznov |
| EPI_ISL_734170, EPI_ISL_734171, EPI_ISL_734172, EPI_ISL_734173, EPI_ISL_734174, EPI_ISL_734175, EPI_ISL_734176, EPI_ISL_734180, EPI_ISL_734181, EPI_ISL_734182, EPI_ISL_734183, EPI_ISL_734184, EPI_ISL_734185, EPI_ISL_734186, EPI_ISL_734187, EPI_ISL_734188, EPI_ISL_734189, EPI_ISL_734190, EPI_ISL_734191, EPI_ISL_734192, EPI_ISL_734199, EPI_ISL_734200, EPI_ISL_734201, EPI_ISL_734202, EPI_ISL_734203, EPI_ISL_734204, EPI_ISL_734205, EPI_ISL_734206, EPI_ISL_734224, EPI_ISL_734225, EPI_ISL_734254, EPI_ISL_734255, EPI_ISL_734256, EPI_ISL_734269, EPI_ISL_734280, EPI_ISL_734281, EPI_ISL_734282, EPI_ISL_734283, EPI_ISL_734284, EPI_ISL_734285, EPI_ISL_734286 | | | |
| see above | Virginia Division of Consolidated Laboratory Services (DCLS) | Virginia Division of Consolidated Laboratory Services (DCLS) | Virginia DCLS |
| EPI_ISL_734481 | Masonic Medical Research Institute | Wadsworth Center, New York State Department.of Health | Kirsten St. George, Nathan Tucker, Ryan D. Pfeiffer, Daryl M. Lamson, Alexis Russel, Jonathan Plitnick, Navjot Singh, John Kelly, Sara Griesemer, Erasmus Schneider, Erica Lasek-Nesselquist |
| EPI_ISL_736936, EPI_ISL_736937 | Netcare | KRISP, KZN Research Innovation and Sequencing Platform | Giandhari J, Pillay S, Lessells R, ChimukangaraB, Mdlalose K, York D, Khan S, Tegally H, Wilkinson E, de Oliveira T |
| EPI_ISL_736960 | NHLS-IALCH | KRISP, KZN Research Innovation and Sequencing Platform | Giandhari J, Pillay S, Lessells R, ChimukangaraB, Mdlalose K, York D, Khan S, Tegally H, Wilkinson E, de Oliveira T |
| EPI_ISL_736961, EPI_ISL_736962, EPI_ISL_736963, EPI_ISL_736964, EPI_ISL_736965 | Netcare | KRISP, KZN Research Innovation and Sequencing Platform | Giandhari J, Pillay S, Lessells R, ChimukangaraB, Mdlalose K, York D, Khan S, Tegally H, Wilkinson E, de Oliveira T |
| EPI_ISL_737154, EPI_ISL_737160, EPI_ISL_737163, EPI_ISL_737164, EPI_ISL_737169, EPI_ISL_737178, EPI_ISL_737179, EPI_ISL_737180, EPI_ISL_737182, EPI_ISL_737193, EPI_ISL_737264, EPI_ISL_737265, EPI_ISL_737268, EPI_ISL_737269, EPI_ISL_737281, EPI_ISL_737284, EPI_ISL_737287, EPI_ISL_737290, EPI_ISL_737299, EPI_ISL_737306, EPI_ISL_737307 | | | |
| see above | Michigan Department of Health and Human Services, Bureau of Laboratories | Michigan Department of Health and Human Services, Bureau of Laboratories | Blankenship HM, Riner D, Soehnlen MK |
| EPI_ISL_738503 | Orange County Public Health Lab | Chan-Zuckerberg Biohub | CZB Cliahub Consortium |
| EPI_ISL_738504 | Humboldt County Public Health Laboratory | Chan-Zuckerberg Biohub | CZB Cliahub Consortium |
| EPI_ISL_738516, EPI_ISL_738532 | Santa Clara County Public Health Laboratory | Chan-Zuckerberg Biohub | CZB Cliahub Consortium |
| EPI_ISL_738539 | Humboldt County Public Health Laboratory | Chan-Zuckerberg Biohub | CZB Cliahub Consortium |
| EPI_ISL_738556 | Santa Clara County Public Health Laboratory | Chan-Zuckerberg Biohub | CZB Cliahub Consortium |
| EPI_ISL_738559 | Orange County Public Health Lab | Chan-Zuckerberg Biohub | CZB Cliahub Consortium |
| EPI_ISL_738563 | Humboldt County Public Health Laboratory | Chan-Zuckerberg Biohub | CZB Cliahub Consortium |
| EPI_ISL_738568, EPI_ISL_738581, EPI_ISL_738586, EPI_ISL_738590 | Santa Clara County Public Health Laboratory | Chan-Zuckerberg Biohub | CZB Cliahub Consortium |
| EPI_ISL_738594, EPI_ISL_738599 | Orange County Public Health Lab | Chan-Zuckerberg Biohub | CZB Cliahub Consortium |
| EPI_ISL_738601 | Santa Clara County Public Health Laboratory | Chan-Zuckerberg Biohub | CZB Cliahub Consortium |
| EPI_ISL_738603 | Orange County Public Health Lab | Chan-Zuckerberg Biohub | CZB Cliahub Consortium |
| EPI_ISL_738612, EPI_ISL_738618 | Santa Clara County Public Health Laboratory | Chan-Zuckerberg Biohub | CZB Cliahub Consortium |
| EPI_ISL_738621, EPI_ISL_738632 | Orange County Public Health Lab | Chan-Zuckerberg Biohub | CZB Cliahub Consortium |
| EPI_ISL_738645, EPI_ISL_738670, EPI_ISL_738681, EPI_ISL_738682, EPI_ISL_738685, EPI_ISL_738691 | Santa Clara County Public Health Laboratory | Chan-Zuckerberg Biohub | CZB Cliahub Consortium |
| EPI_ISL_738693, EPI_ISL_738694 | Orange County Public Health Lab | Chan-Zuckerberg Biohub | CZB Cliahub Consortium |
| EPI_ISL_738698, EPI_ISL_738701 | Santa Clara County Public Health Laboratory | Chan-Zuckerberg Biohub | CZB Cliahub Consortium |
| EPI_ISL_738709 | Orange County Public Health Lab | Chan-Zuckerberg Biohub | CZB Cliahub Consortium |
| EPI_ISL_738710 | Santa Clara County Public Health Laboratory | Chan-Zuckerberg Biohub | CZB Cliahub Consortium |
| EPI_ISL_738722, EPI_ISL_738729, EPI_ISL_738755, EPI_ISL_738761 | Orange County Public Health Lab | Chan-Zuckerberg Biohub | CZB Cliahub Consortium |
| EPI_ISL_738769, EPI_ISL_738771, EPI_ISL_738780, EPI_ISL_738798 | Santa Clara County Public Health Laboratory | Chan-Zuckerberg Biohub | CZB Cliahub Consortium |

| | | | |
|---|---|---|---|
| EPI_ISL_738799 | Orange County Public Health Lab | Chan-Zuckerberg Biohub | CZB Cliahub Consortium |
| EPI_ISL_738801, EPI_ISL_738806, EPI_ISL_738807 | Humboldt County Public Health Laboratory | Chan-Zuckerberg Biohub | CZB Cliahub Consortium |
| EPI_ISL_738811, EPI_ISL_738818, EPI_ISL_738824, EPI_ISL_738830 | Orange County Public Health Lab | Chan-Zuckerberg Biohub | CZB Cliahub Consortium |
| EPI_ISL_738842 | Santa Clara County Public Health Laboratory | Chan-Zuckerberg Biohub | CZB Cliahub Consortium |
| EPI_ISL_738845 | Orange County Public Health Lab | Chan-Zuckerberg Biohub | CZB Cliahub Consortium |
| EPI_ISL_738849 | Humboldt County Public Health Laboratory | Chan-Zuckerberg Biohub | CZB Cliahub Consortium |
| EPI_ISL_738860, EPI_ISL_738862 | Orange County Public Health Lab | Chan-Zuckerberg Biohub | CZB Cliahub Consortium |
| EPI_ISL_738864 | Santa Clara County Public Health Laboratory | Chan-Zuckerberg Biohub | CZB Cliahub Consortium |
| EPI_ISL_738865 | Humboldt County Public Health Laboratory | Chan-Zuckerberg Biohub | CZB Cliahub Consortium |
| EPI_ISL_738867 | Santa Clara County Public Health Laboratory | Chan-Zuckerberg Biohub | CZB Cliahub Consortium |
| EPI_ISL_738869, EPI_ISL_738873 | Orange County Public Health Lab | Chan-Zuckerberg Biohub | CZB Cliahub Consortium |
| EPI_ISL_738875 | Humboldt County Public Health Laboratory | Chan-Zuckerberg Biohub | CZB Cliahub Consortium |
| EPI_ISL_738884 | Orange County Public Health Lab | Chan-Zuckerberg Biohub | CZB Cliahub Consortium |
| EPI_ISL_738887 | Santa Clara County Public Health Laboratory | Chan-Zuckerberg Biohub | CZB Cliahub Consortium |
| EPI_ISL_738902 | Humboldt County Public Health Laboratory | Chan-Zuckerberg Biohub | CZB Cliahub Consortium |
| EPI_ISL_738903 | Santa Clara County Public Health Laboratory | Chan-Zuckerberg Biohub | CZB Cliahub Consortium |
| EPI_ISL_738905 | Orange County Public Health Lab | Chan-Zuckerberg Biohub | CZB Cliahub Consortium |
| EPI_ISL_738917 | Humboldt County Public Health Laboratory | Chan-Zuckerberg Biohub | CZB Cliahub Consortium |
| EPI_ISL_738921 | Orange County Public Health Lab | Chan-Zuckerberg Biohub | CZB Cliahub Consortium |
| EPI_ISL_738924 | Santa Clara County Public Health Laboratory | Chan-Zuckerberg Biohub | CZB Cliahub Consortium |
| EPI_ISL_738935 | Orange County Public Health Lab | Chan-Zuckerberg Biohub | CZB Cliahub Consortium |
| EPI_ISL_738936 | Santa Clara County Public Health Laboratory | Chan-Zuckerberg Biohub | CZB Cliahub Consortium |
| EPI_ISL_738938 | Humboldt County Public Health Laboratory | Chan-Zuckerberg Biohub | CZB Cliahub Consortium |
| EPI_ISL_738944 | Orange County Public Health Lab | Chan-Zuckerberg Biohub | CZB Cliahub Consortium |
| EPI_ISL_738960, EPI_ISL_738965 | Santa Clara County Public Health Laboratory | Chan-Zuckerberg Biohub | CZB Cliahub Consortium |
| EPI_ISL_738966, EPI_ISL_738968 | Orange County Public Health Lab | Chan-Zuckerberg Biohub | CZB Cliahub Consortium |
| EPI_ISL_738969 | Humboldt County Public Health Laboratory | Chan-Zuckerberg Biohub | CZB Cliahub Consortium |
| EPI_ISL_738970, EPI_ISL_738973, EPI_ISL_738975 | Orange County Public Health Lab | Chan-Zuckerberg Biohub | CZB Cliahub Consortium |
| EPI_ISL_738980, EPI_ISL_739000, EPI_ISL_739001, EPI_ISL_739007 | Humboldt County Public Health Laboratory | Chan-Zuckerberg Biohub | CZB Cliahub Consortium |
| EPI_ISL_739015, EPI_ISL_739026 | Santa Clara County Public Health Laboratory | Chan-Zuckerberg Biohub | CZB Cliahub Consortium |
| EPI_ISL_739028 | Orange County Public Health Lab | Chan-Zuckerberg Biohub | CZB Cliahub Consortium |
| EPI_ISL_739031 | Santa Clara County Public Health Laboratory | Chan-Zuckerberg Biohub | CZB Cliahub Consortium |
| EPI_ISL_739043 | Humboldt County Public Health Laboratory | Chan-Zuckerberg Biohub | CZB Cliahub Consortium |
| EPI_ISL_739048, EPI_ISL_739055 | Orange County Public Health Lab | Chan-Zuckerberg Biohub | CZB Cliahub Consortium |
| EPI_ISL_739060 | Santa Clara County Public Health Laboratory | Chan-Zuckerberg Biohub | CZB Cliahub Consortium |
| EPI_ISL_739063, EPI_ISL_739071 | Orange County Public Health Lab | Chan-Zuckerberg Biohub | CZB Cliahub Consortium |
| EPI_ISL_739074 | Santa Clara County Public Health Laboratory | Chan-Zuckerberg Biohub | CZB Cliahub Consortium |
| EPI_ISL_739078 | Orange County Public Health Lab | Chan-Zuckerberg Biohub | CZB Cliahub Consortium |
| EPI_ISL_739091 | Santa Clara County Public Health Laboratory | Chan-Zuckerberg Biohub | CZB Cliahub Consortium |
| EPI_ISL_739092, EPI_ISL_739099 | Orange County Public Health Lab | Chan-Zuckerberg Biohub | CZB Cliahub Consortium |
| EPI_ISL_739103 | Santa Clara County Public Health Laboratory | Chan-Zuckerberg Biohub | CZB Cliahub Consortium |
| EPI_ISL_739112, EPI_ISL_739114, EPI_ISL_739123 | Humboldt County Public Health Laboratory | Chan-Zuckerberg Biohub | CZB Cliahub Consortium |
| EPI_ISL_739147 | Santa Clara County Public Health Laboratory | Chan-Zuckerberg Biohub | CZB Cliahub Consortium |
| EPI_ISL_739162 | Orange County Public Health Lab | Chan-Zuckerberg Biohub | CZB Cliahub Consortium |
| EPI_ISL_739170 | Humboldt County Public Health Laboratory | Chan-Zuckerberg Biohub | CZB Cliahub Consortium |
| EPI_ISL_739195, EPI_ISL_739204 | Santa Clara County Public Health Laboratory | Chan-Zuckerberg Biohub | CZB Cliahub Consortium |
| EPI_ISL_739205 | Orange County Public Health Lab | Chan-Zuckerberg Biohub | CZB Cliahub Consortium |
| EPI_ISL_739210 | Humboldt County Public Health Laboratory | Chan-Zuckerberg Biohub | CZB Cliahub Consortium |
| EPI_ISL_739227 | Orange County Public Health Lab | Chan-Zuckerberg Biohub | CZB Cliahub Consortium |
| EPI_ISL_739228 | Humboldt County Public Health Laboratory | Chan-Zuckerberg Biohub | CZB Cliahub Consortium |
| EPI_ISL_739230 | Orange County Public Health Lab | Chan-Zuckerberg Biohub | CZB Cliahub Consortium |
| EPI_ISL_739232 | Santa Clara County Public Health Laboratory | Chan-Zuckerberg Biohub | CZB Cliahub Consortium |
| EPI_ISL_739238 | Orange County Public Health Lab | Chan-Zuckerberg Biohub | CZB Cliahub Consortium |
| EPI_ISL_739240, EPI_ISL_739242 | Humboldt County Public Health Laboratory | Chan-Zuckerberg Biohub | CZB Cliahub Consortium |
| EPI_ISL_739246, EPI_ISL_739253 | Santa Clara County Public Health Laboratory | Chan-Zuckerberg Biohub | CZB Cliahub Consortium |

| | | | |
|---|---|---|---|
| EPI_ISL_739255, EPI_ISL_739271 | Orange County Public Health Lab | Chan-Zuckerberg Biohub | CZB Cliahub Consortium |
| EPI_ISL_739274 | Humboldt County Public Health Laboratory | Chan-Zuckerberg Biohub | CZB Cliahub Consortium |
| EPI_ISL_739275 | Santa Clara County Public Health Laboratory | Chan-Zuckerberg Biohub | CZB Cliahub Consortium |
| EPI_ISL_739276 | Orange County Public Health Lab | Chan-Zuckerberg Biohub | CZB Cliahub Consortium |
| EPI_ISL_739284 | Santa Clara County Public Health Laboratory | Chan-Zuckerberg Biohub | CZB Cliahub Consortium |
| EPI_ISL_739287 | Humboldt County Public Health Laboratory | Chan-Zuckerberg Biohub | CZB Cliahub Consortium |
| EPI_ISL_739289 | Orange County Public Health Lab | Chan-Zuckerberg Biohub | CZB Cliahub Consortium |
| EPI_ISL_739293 | Santa Clara County Public Health Laboratory | Chan-Zuckerberg Biohub | CZB Cliahub Consortium |
| EPI_ISL_739296 | Orange County Public Health Lab | Chan-Zuckerberg Biohub | CZB Cliahub Consortium |
| EPI_ISL_739305 | Santa Clara County Public Health Laboratory | Chan-Zuckerberg Biohub | CZB Cliahub Consortium |
| EPI_ISL_739313 | Humboldt County Public Health Laboratory | Chan-Zuckerberg Biohub | CZB Cliahub Consortium |
| EPI_ISL_739327 | Santa Clara County Public Health Laboratory | Chan-Zuckerberg Biohub | CZB Cliahub Consortium |
| EPI_ISL_739331 | Orange County Public Health Lab | Chan-Zuckerberg Biohub | CZB Cliahub Consortium |
| EPI_ISL_739332 | Santa Clara County Public Health Laboratory | Chan-Zuckerberg Biohub | CZB Cliahub Consortium |
| EPI_ISL_739334 | Orange County Public Health Lab | Chan-Zuckerberg Biohub | CZB Cliahub Consortium |
| EPI_ISL_739335, EPI_ISL_739343 | Santa Clara County Public Health Laboratory | Chan-Zuckerberg Biohub | CZB Cliahub Consortium |
| EPI_ISL_739345, EPI_ISL_739350 | Orange County Public Health Lab | Chan-Zuckerberg Biohub | CZB Cliahub Consortium |
| EPI_ISL_739352 | Santa Clara County Public Health Laboratory | Chan-Zuckerberg Biohub | CZB Cliahub Consortium |
| EPI_ISL_739353, EPI_ISL_739362 | Orange County Public Health Lab | Chan-Zuckerberg Biohub | CZB Cliahub Consortium |
| EPI_ISL_739365 | Santa Clara County Public Health Laboratory | Chan-Zuckerberg Biohub | CZB Cliahub Consortium |
| EPI_ISL_739373 | Orange County Public Health Lab | Chan-Zuckerberg Biohub | CZB Cliahub Consortium |
| EPI_ISL_739376 | Santa Clara County Public Health Laboratory | Chan-Zuckerberg Biohub | CZB Cliahub Consortium |
| EPI_ISL_739379 | Orange County Public Health Lab | Chan-Zuckerberg Biohub | CZB Cliahub Consortium |
| EPI_ISL_739380 | Santa Clara County Public Health Laboratory | Chan-Zuckerberg Biohub | CZB Cliahub Consortium |
| EPI_ISL_739389 | Humboldt County Public Health Laboratory | Chan-Zuckerberg Biohub | CZB Cliahub Consortium |
| EPI_ISL_739397 | Orange County Public Health Lab | Chan-Zuckerberg Biohub | CZB Cliahub Consortium |
| EPI_ISL_739403 | Humboldt County Public Health Laboratory | Chan-Zuckerberg Biohub | CZB Cliahub Consortium |
| EPI_ISL_739404 | Santa Clara County Public Health Laboratory | Chan-Zuckerberg Biohub | CZB Cliahub Consortium |
| EPI_ISL_739405 | Humboldt County Public Health Laboratory | Chan-Zuckerberg Biohub | CZB Cliahub Consortium |
| EPI_ISL_739409 | Santa Clara County Public Health Laboratory | Chan-Zuckerberg Biohub | CZB Cliahub Consortium |
| EPI_ISL_739410, EPI_ISL_739412 | Orange County Public Health Lab | Chan-Zuckerberg Biohub | CZB Cliahub Consortium |
| EPI_ISL_739417, EPI_ISL_739425 | Humboldt County Public Health Laboratory | Chan-Zuckerberg Biohub | CZB Cliahub Consortium |
| EPI_ISL_739428 | Orange County Public Health Lab | Chan-Zuckerberg Biohub | CZB Cliahub Consortium |
| EPI_ISL_739436 | Humboldt County Public Health Laboratory | Chan-Zuckerberg Biohub | CZB Cliahub Consortium |
| EPI_ISL_739438 | Orange County Public Health Lab | Chan-Zuckerberg Biohub | CZB Cliahub Consortium |
| EPI_ISL_739453 | Santa Clara County Public Health Laboratory | Chan-Zuckerberg Biohub | CZB Cliahub Consortium |
| EPI_ISL_739461 | Humboldt County Public Health Laboratory | Chan-Zuckerberg Biohub | CZB Cliahub Consortium |
| EPI_ISL_739465 | Orange County Public Health Lab | Chan-Zuckerberg Biohub | CZB Cliahub Consortium |
| EPI_ISL_739475 | Humboldt County Public Health Laboratory | Chan-Zuckerberg Biohub | CZB Cliahub Consortium |
| EPI_ISL_739476, EPI_ISL_739479, EPI_ISL_739481 | Santa Clara County Public Health Laboratory | Chan-Zuckerberg Biohub | CZB Cliahub Consortium |
| EPI_ISL_739482, EPI_ISL_739493 | Orange County Public Health Lab | Chan-Zuckerberg Biohub | CZB Cliahub Consortium |
| EPI_ISL_739502, EPI_ISL_739510, EPI_ISL_739521 | Santa Clara County Public Health Laboratory | Chan-Zuckerberg Biohub | CZB Cliahub Consortium |
| EPI_ISL_739531 | Humboldt County Public Health Laboratory | Chan-Zuckerberg Biohub | CZB Cliahub Consortium |
| EPI_ISL_739532, EPI_ISL_739534 | Orange County Public Health Lab | Chan-Zuckerberg Biohub | CZB Cliahub Consortium |
| EPI_ISL_739542 | Humboldt County Public Health Laboratory | Chan-Zuckerberg Biohub | CZB Cliahub Consortium |
| EPI_ISL_739543 | Santa Clara County Public Health Laboratory | Chan-Zuckerberg Biohub | CZB Cliahub Consortium |
| EPI_ISL_739547 | Orange County Public Health Lab | Chan-Zuckerberg Biohub | CZB Cliahub Consortium |
| EPI_ISL_739560 | Santa Clara County Public Health Laboratory | Chan-Zuckerberg Biohub | CZB Cliahub Consortium |
| EPI_ISL_739568, EPI_ISL_739570, EPI_ISL_739573 | Orange County Public Health Lab | Chan-Zuckerberg Biohub | CZB Cliahub Consortium |
| EPI_ISL_739574 | Santa Clara County Public Health Laboratory | Chan-Zuckerberg Biohub | CZB Cliahub Consortium |
| EPI_ISL_739582, EPI_ISL_739591 | Orange County Public Health Lab | Chan-Zuckerberg Biohub | CZB Cliahub Consortium |
| EPI_ISL_739596 | Santa Clara County Public Health Laboratory | Chan-Zuckerberg Biohub | CZB Cliahub Consortium |
| EPI_ISL_739611 | Humboldt County Public Health Laboratory | Chan-Zuckerberg Biohub | CZB Cliahub Consortium |
| EPI_ISL_739612 | Santa Clara County Public Health Laboratory | Chan-Zuckerberg Biohub | CZB Cliahub Consortium |
| EPI_ISL_739614 | Orange County Public Health Lab | Chan-Zuckerberg Biohub | CZB Cliahub Consortium |
| EPI_ISL_739624 | Santa Clara County Public Health Laboratory | Chan-Zuckerberg Biohub | CZB Cliahub Consortium |
| EPI_ISL_739625, EPI_ISL_739627, | Orange County Public Health Lab | Chan-Zuckerberg Biohub | CZB Cliahub Consortium |

| | | | |
|---|---|---|---|
| EPI_ISL_739636 | | | |
| EPI_ISL_739651 | Humboldt County Public Health Laboratory | Chan-Zuckerberg Biohub | CZB Cliahub Consortium |
| EPI_ISL_739706, EPI_ISL_739707, EPI_ISL_739708, EPI_ISL_739710, EPI_ISL_739719, EPI_ISL_739741, EPI_ISL_739752, EPI_ISL_739772, EPI_ISL_739775, EPI_ISL_739805, EPI_ISL_739811, EPI_ISL_739842, EPI_ISL_739876, EPI_ISL_739879, EPI_ISL_739887, EPI_ISL_739888, EPI_ISL_739893, EPI_ISL_739899, EPI_ISL_739932, EPI_ISL_739938, EPI_ISL_739942, EPI_ISL_739959, EPI_ISL_739979, EPI_ISL_739989, EPI_ISL_740009, EPI_ISL_740035, EPI_ISL_740065, EPI_ISL_740075, EPI_ISL_740088, EPI_ISL_740089, EPI_ISL_740102, EPI_ISL_740112, EPI_ISL_740128, EPI_ISL_740143, EPI_ISL_740151, EPI_ISL_740152, EPI_ISL_740157, EPI_ISL_740164, EPI_ISL_740183, EPI_ISL_740216, EPI_ISL_740235, EPI_ISL_740253, EPI_ISL_740255, EPI_ISL_740285, EPI_ISL_740297, EPI_ISL_740303, EPI_ISL_740308, EPI_ISL_740311, EPI_ISL_740312, EPI_ISL_740315, EPI_ISL_740350, EPI_ISL_740367, EPI_ISL_740369, EPI_ISL_740379, EPI_ISL_740422, EPI_ISL_740426, EPI_ISL_740459, EPI_ISL_740465, EPI_ISL_740476, EPI_ISL_740483, EPI_ISL_740484, EPI_ISL_740517, EPI_ISL_740524 | | | |
| see above | Laboratoire national de santé, Microbiology, Virology | Laboratoire national de santé, Microbiology, Microbial Genomics Platform | Anke Wienecke-Baldacchino, Catherine Ragimbeau,Jessica Tapp, Fatu Djabi, Lise Pignon, Raoul Salmon, Tamir Abdelrahman |
| EPI_ISL_741516 | Quadram Institute Bioscience | COVID-19 Genomics UK (COG-UK) Consortium | Dave J. Baker, Gemma L. Kay, Alp Aydin, Thanh Le-Viet, Steven Rudder, Ana P. Tedim, Anastasia Kolyva, Maria Diaz, Leonardo de Oliveira Martins, Nabil-Fareed Alikhan, Lizzie Meadows, Rachael Stanley, Ngozi Elumogo, Muhammed Yasir, Nicholas M. Thomson, Alexander J Trotter, Rachel Gilroy, Samuel Bloomfield, Claire Stuart, Andrew Bell, Reenesh Prakash, Samir Dervisevic, Alison E. Mather, John Wain, Mark Webber, Andrew J. Page, Justin O'Grady |
| EPI_ISL_741716, EPI_ISL_741717, EPI_ISL_741718, EPI_ISL_741719, EPI_ISL_741740, EPI_ISL_741741 | Centre for Enzyme Innovation, University of Portsmouth / Translational Research Laboratory, Portsmouth Hospitals NHS Trust | COVID-19 Genomics UK (COG-UK) Consortium | Angela Beckett,Yann Bourgeois,Garry Scarlett,Sharon Glaysher,Scott Elliott,Kelly Bicknell,Robert Impey,Allyson Lloyd,Sarah Wyllie,Ethan Butcher,Anoop Chauhan,Samuel Robson |
| EPI_ISL_742541, EPI_ISL_742542, EPI_ISL_742543, EPI_ISL_742544, EPI_ISL_742547, EPI_ISL_742548, EPI_ISL_742549, EPI_ISL_742559, EPI_ISL_742561, EPI_ISL_742563, EPI_ISL_742564, EPI_ISL_742585, EPI_ISL_742586, EPI_ISL_742588, EPI_ISL_742589, EPI_ISL_742590, EPI_ISL_742591, EPI_ISL_742592, EPI_ISL_742593, EPI_ISL_742594, EPI_ISL_742595, EPI_ISL_742596, EPI_ISL_742597, EPI_ISL_742598, EPI_ISL_742599, EPI_ISL_742600, EPI_ISL_742601, EPI_ISL_742602, EPI_ISL_742603, EPI_ISL_742604, EPI_ISL_742605, EPI_ISL_742606, EPI_ISL_742607, EPI_ISL_742608, EPI_ISL_742609, EPI_ISL_742610, EPI_ISL_742611, EPI_ISL_742612, EPI_ISL_742613, EPI_ISL_742614, EPI_ISL_742615, EPI_ISL_742616, EPI_ISL_742617, EPI_ISL_742618, EPI_ISL_742619, EPI_ISL_742620, EPI_ISL_742621, EPI_ISL_742622, EPI_ISL_742623, EPI_ISL_742624, EPI_ISL_742625, EPI_ISL_742626, EPI_ISL_742627, EPI_ISL_742628, EPI_ISL_742629, EPI_ISL_742630, EPI_ISL_742631, EPI_ISL_742632, EPI_ISL_742633, EPI_ISL_742634, EPI_ISL_742635, EPI_ISL_742636 | | | |
| see above | Wales Specialist Virology Centre Sequencing lab: Pathogen Genomics Unit | COVID-19 Genomics UK (COG-UK) Consortium | Catherine Moore, Johnathan Evans, Laura Gifford, Malorie Perry, Simon Cottrell, Angela Marchbank, Alec Birchley, Alexander Adams, Amy Gaskin, Bree Gatica-Wilcox, Jason Coombes, Joel Southgate, Lauren Gilbert, Lee Graham, Nicole Pacchiarini, Sara Kumziene-Summerhayes, Sarah Taylor, Sophie Jones, Sara Rey, Matthew Bull, Joanne Watkins, Sally Corden, Tom Connor |
| EPI_ISL_744137, EPI_ISL_744148, EPI_ISL_744177, EPI_ISL_744179, EPI_ISL_744186, EPI_ISL_744244, EPI_ISL_744248, EPI_ISL_744250, EPI_ISL_744251, EPI_ISL_744254, EPI_ISL_744256, EPI_ISL_744263, EPI_ISL_744268, EPI_ISL_744271, EPI_ISL_744277, EPI_ISL_744310, EPI_ISL_744329, EPI_ISL_744336, EPI_ISL_744349, EPI_ISL_744353, EPI_ISL_744407, EPI_ISL_744415, EPI_ISL_744422, EPI_ISL_744424, EPI_ISL_744426, EPI_ISL_744453, EPI_ISL_744462, EPI_ISL_744476, EPI_ISL_744494, EPI_ISL_744499, EPI_ISL_744526, EPI_ISL_744545, EPI_ISL_744562, EPI_ISL_744603, EPI_ISL_744614, EPI_ISL_744629, EPI_ISL_744648, EPI_ISL_744656, EPI_ISL_744657, EPI_ISL_744667, EPI_ISL_744692, EPI_ISL_744704, EPI_ISL_744708, EPI_ISL_744740, EPI_ISL_744776, EPI_ISL_744781, EPI_ISL_744795, EPI_ISL_744797, EPI_ISL_744802, EPI_ISL_744834, EPI_ISL_744844, EPI_ISL_744853, EPI_ISL_744880, EPI_ISL_744923, EPI_ISL_744933, EPI_ISL_744953, EPI_ISL_744957, EPI_ISL_744982, EPI_ISL_744983, EPI_ISL_745000, EPI_ISL_745003, EPI_ISL_745021, EPI_ISL_745026 | | | |
| see above | Laboratoire national de santé, Microbiology, Virology | Laboratoire national de santé, Microbiology, Microbial Genomics Platform | Anke Wienecke-Baldacchino, Catherine Ragimbeau,Jessica Tapp, Fatu Djabi, Lise Pignon, Raoul Salmon, Tamir Abdelrahman |
| EPI_ISL_745399 | DOHMH Morrisania | New York City Public Health Laboratory | Jade Wang, et al. |
| EPI_ISL_745404 | DOHMH Central Harlem | New York City Public Health Laboratory | Jade Wang, et al. |
| EPI_ISL_746878, EPI_ISL_746965, EPI_ISL_746969, EPI_ISL_746982, EPI_ISL_746996 | Utah Public Health Laboratory | Utah Public Health Laboratory | Erin Young, Kelly Oakeson, Tara Gallagher |
| EPI_ISL_747155, EPI_ISL_747156, EPI_ISL_747157, EPI_ISL_747158, EPI_ISL_747159, EPI_ISL_747160, EPI_ISL_747161, EPI_ISL_747162, EPI_ISL_747163, EPI_ISL_747164, EPI_ISL_747165, EPI_ISL_747166, EPI_ISL_747167, EPI_ISL_747169, EPI_ISL_747179, EPI_ISL_747180, EPI_ISL_747191, EPI_ISL_747193 | | | |
| see above | Respiratory Viruses Branch, Centers for Disease Control and Prevention | Respiratory Viruses Branch, Centers for Disease Control and Prevention | Queen,K., Li,Y., Tao,Y., Uehara,A., Montmayeur,A., Paden,C.R., Cook,P.W., Marine,R., Sheth,M., Wang,H., Lee,J., Tong,S. |
| EPI_ISL_747384, EPI_ISL_747386, EPI_ISL_747392, EPI_ISL_747393, EPI_ISL_747397, EPI_ISL_747400, EPI_ISL_747401, EPI_ISL_747402, EPI_ISL_747439, EPI_ISL_747446, EPI_ISL_747447, EPI_ISL_747448, EPI_ISL_747451, EPI_ISL_747452 | | | |
| see above | Division of Emerging Infectious Diseases, Bureau of Infectious Diseases Diagnosis Control, Korea Disease Control and Prevention Agency | Division of Emerging Infectious Diseases, Bureau of Infectious Diseases Diagnosis Control, Korea Disease Control and Prevention Agency | Ae Kyung Park, Il-Hwan Kim, Heui Man Kim, Jeong-Min Kim, Namjoo Lee, Chaeyoung Lee, Sang Hee Woo, Eun-Jin Kim |
| EPI_ISL_747588, EPI_ISL_747604, EPI_ISL_747605, EPI_ISL_747649, EPI_ISL_747653, EPI_ISL_747720, EPI_ISL_747744, EPI_ISL_747751, EPI_ISL_748113, EPI_ISL_748114, EPI_ISL_748115, EPI_ISL_748116, EPI_ISL_748117, EPI_ISL_748118 | | | |
| see above | Department of Virus and Microbiological Special Diagnostics, Statens Serum Institut, Copenhagen, Denmark | Albertsen Lab, Department of Chemistry and Bioscience, Aalborg University, Denmark | Danish Covid-19 Genome Consortium |
| EPI_ISL_751454, EPI_ISL_751455, EPI_ISL_751456, EPI_ISL_751457, EPI_ISL_751463, EPI_ISL_751465, EPI_ISL_751474, EPI_ISL_751475, EPI_ISL_751476 | CHU Purpan - Laboratoire de Virologie - Institut Fédératif de Biologie | CHU Purpan - Laboratoire de Virologie - Institut Fédératif de Biologie | Latour J., Ranger N., Dubois M., Carcenac R., Harter A., Boyer P., Tremeaux P., Izopet J. |
| EPI_ISL_751546 | VA-Division of Consolidated Laboratory Services | Genomics and Discovery, Respiratory Viruses Branch, Division of Viral Diseases, Centers for Disease Control and Prevention | Krista Queen, Yan Li, Ying Tao, Jing Zhang, Anna Uehara, Anna Montmayeur, Clinton R. Paden, Peter W. Cook,Rachel Marine, Mili Sheth, Haibin Wang, Justin Lee, Suxiang Tong |
| EPI_ISL_751558 | NH Dept. of Health and Human Services Public Health Labs | Genomics and Discovery, Respiratory Viruses Branch, Division of Viral Diseases, Centers for Disease Control and Prevention | Krista Queen, Yan Li, Ying Tao, Jing Zhang, Anna Uehara, Anna Montmayeur, Clinton R. Paden, Peter W. Cook,Rachel Marine, Mili Sheth, Haibin Wang, Justin Lee, Suxiang Tong |
| EPI_ISL_751560 | NYSDOH Wadsworth Center, Virology Lab | Genomics and Discovery, Respiratory Viruses Branch, Division of Viral Diseases, Centers for Disease Control and Prevention | Krista Queen, Yan Li, Ying Tao, Jing Zhang, Anna Uehara, Anna Montmayeur, Clinton R. Paden, Peter W. Cook,Rachel Marine, Mili Sheth, Haibin Wang, Justin Lee, Suxiang Tong |
| EPI_ISL_751564 | VA-Division of Consolidated Laboratory Services | Genomics and Discovery, Respiratory Viruses Branch, Division of Viral Diseases, Centers for Disease Control and Prevention | Krista Queen, Yan Li, Ying Tao, Jing Zhang, Anna Uehara, Anna Montmayeur, Clinton R. Paden, Peter W. Cook,Rachel Marine, Mili Sheth, Haibin Wang, Justin Lee, Suxiang Tong |
| EPI_ISL_751576 | MN PHL Division, Minnesota Department of Health | Genomics and Discovery, Respiratory Viruses Branch, Division of Viral Diseases, Centers for Disease Control and Prevention | Krista Queen, Yan Li, Ying Tao, Jing Zhang, Anna Uehara, Anna Montmayeur, Clinton R. Paden, Peter W. Cook,Rachel Marine, Mili Sheth, Haibin Wang, Justin Lee, Suxiang Tong |
| EPI_ISL_751588 | VA-Division of Consolidated Laboratory Services | Genomics and Discovery, Respiratory Viruses Branch, Division of Viral Diseases, Centers for Disease Control and Prevention | Krista Queen, Yan Li, Ying Tao, Jing Zhang, Anna Uehara, Anna Montmayeur, Clinton R. Paden, Peter W. Cook,Rachel Marine, Mili Sheth, Haibin Wang, Justin Lee, Suxiang Tong |
| EPI_ISL_751590 | ID Bureau of Laboratories | Genomics and Discovery, Respiratory Viruses Branch, Division of Viral Diseases, Centers for Disease Control and Prevention | Krista Queen, Yan Li, Ying Tao, Jing Zhang, Anna Uehara, Anna Montmayeur, Clinton R. Paden, Peter W. Cook,Rachel Marine, Mili Sheth, Haibin Wang, Justin Lee, Suxiang Tong |
| EPI_ISL_751592, EPI_ISL_751598 | AK State Public Health Lab, State Health Department | Genomics and Discovery, Respiratory Viruses Branch, Division of Viral Diseases, Centers for Disease Control and Prevention | Krista Queen, Yan Li, Ying Tao, Jing Zhang, Anna Uehara, Anna Montmayeur, Clinton R. Paden, Peter W. Cook,Rachel Marine, Mili Sheth, Haibin Wang, Justin Lee, Suxiang Tong |
| EPI_ISL_751602 | DE Public Health Laboratory | Genomics and Discovery, Respiratory Viruses Branch, | Krista Queen, Yan Li, Ying Tao, Jing Zhang, Anna Uehara, Anna Montmayeur, Clinton R. Paden, Peter W. Cook,Rachel Marine, Mili Sheth, Haibin Wang, |

| | | | |
|---|---|---|---|
| | | Division of Viral Diseases, Centers for Disease Control and Prevention | Justin Lee, Suxiang Tong |
| EPI_ISL_751610 | OH Department of Health Laboratory | Genomics and Discovery, Respiratory Viruses Branch, Division of Viral Diseases, Centers for Disease Control and Prevention | Krista Queen, Yan Li, Ying Tao, Jing Zhang, Anna Uehara, Anna Montmayeur, Clinton R. Paden, Peter W. Cook,Rachel Marine, Mili Sheth, Haibin Wang, Justin Lee, Suxiang Tong |
| EPI_ISL_751612 | ID Bureau of Laboratories | Genomics and Discovery, Respiratory Viruses Branch, Division of Viral Diseases, Centers for Disease Control and Prevention | Krista Queen, Yan Li, Ying Tao, Jing Zhang, Anna Uehara, Anna Montmayeur, Clinton R. Paden, Peter W. Cook,Rachel Marine, Mili Sheth, Haibin Wang, Justin Lee, Suxiang Tong |
| EPI_ISL_751613 | OH Department of Health Laboratory | Genomics and Discovery, Respiratory Viruses Branch, Division of Viral Diseases, Centers for Disease Control and Prevention | Krista Queen, Yan Li, Ying Tao, Jing Zhang, Anna Uehara, Anna Montmayeur, Clinton R. Paden, Peter W. Cook,Rachel Marine, Mili Sheth, Haibin Wang, Justin Lee, Suxiang Tong |
| EPI_ISL_751614 | ID Bureau of Laboratories | Genomics and Discovery, Respiratory Viruses Branch, Division of Viral Diseases, Centers for Disease Control and Prevention | Krista Queen, Yan Li, Ying Tao, Jing Zhang, Anna Uehara, Anna Montmayeur, Clinton R. Paden, Peter W. Cook,Rachel Marine, Mili Sheth, Haibin Wang, Justin Lee, Suxiang Tong |
| EPI_ISL_751616 | OH Department of Health Laboratory | Genomics and Discovery, Respiratory Viruses Branch, Division of Viral Diseases, Centers for Disease Control and Prevention | Krista Queen, Yan Li, Ying Tao, Jing Zhang, Anna Uehara, Anna Montmayeur, Clinton R. Paden, Peter W. Cook,Rachel Marine, Mili Sheth, Haibin Wang, Justin Lee, Suxiang Tong |
| EPI_ISL_751618 | ID Bureau of Laboratories | Genomics and Discovery, Respiratory Viruses Branch, Division of Viral Diseases, Centers for Disease Control and Prevention | Krista Queen, Yan Li, Ying Tao, Jing Zhang, Anna Uehara, Anna Montmayeur, Clinton R. Paden, Peter W. Cook,Rachel Marine, Mili Sheth, Haibin Wang, Justin Lee, Suxiang Tong |
| EPI_ISL_751641 | MN PHL Division, Minnesota Department of Health | Genomics and Discovery, Respiratory Viruses Branch, Division of Viral Diseases, Centers for Disease Control and Prevention | Krista Queen, Yan Li, Ying Tao, Jing Zhang, Anna Uehara, Anna Montmayeur, Clinton R. Paden, Peter W. Cook,Rachel Marine, Mili Sheth, Haibin Wang, Justin Lee, Suxiang Tong |
| EPI_ISL_751656 | MO State Public Health Laboratory | Genomics and Discovery, Respiratory Viruses Branch, Division of Viral Diseases, Centers for Disease Control and Prevention | Krista Queen, Yan Li, Ying Tao, Jing Zhang, Anna Uehara, Anna Montmayeur, Clinton R. Paden, Peter W. Cook,Rachel Marine, Mili Sheth, Haibin Wang, Justin Lee, Suxiang Tong |
| EPI_ISL_751659, EPI_ISL_751662 | NJ Public Health and Environmental Laboratories | Genomics and Discovery, Respiratory Viruses Branch, Division of Viral Diseases, Centers for Disease Control and Prevention | Krista Queen, Yan Li, Ying Tao, Jing Zhang, Anna Uehara, Anna Montmayeur, Clinton R. Paden, Peter W. Cook,Rachel Marine, Mili Sheth, Haibin Wang, Justin Lee, Suxiang Tong |
| EPI_ISL_751694 | AK State Public Health Lab, State Health Department | Genomics and Discovery, Respiratory Viruses Branch, Division of Viral Diseases, Centers for Disease Control and Prevention | Krista Queen, Yan Li, Ying Tao, Jing Zhang, Anna Uehara, Anna Montmayeur, Clinton R. Paden, Peter W. Cook,Rachel Marine, Mili Sheth, Haibin Wang, Justin Lee, Suxiang Tong |
| EPI_ISL_751716 | MN PHL Division, Minnesota Department of Health | Genomics and Discovery, Respiratory Viruses Branch, Division of Viral Diseases, Centers for Disease Control and Prevention | Krista Queen, Yan Li, Ying Tao, Jing Zhang, Anna Uehara, Anna Montmayeur, Clinton R. Paden, Peter W. Cook,Rachel Marine, Mili Sheth, Haibin Wang, Justin Lee, Suxiang Tong |
| EPI_ISL_751720 | TX DSHS, Lab Services Section MC 1947 | Genomics and Discovery, Respiratory Viruses Branch, Division of Viral Diseases, Centers for Disease Control and Prevention | Krista Queen, Yan Li, Ying Tao, Jing Zhang, Anna Uehara, Anna Montmayeur, Clinton R. Paden, Peter W. Cook,Rachel Marine, Mili Sheth, Haibin Wang, Justin Lee, Suxiang Tong |
| EPI_ISL_751734 | NE Public Health Laboratory | Genomics and Discovery, Respiratory Viruses Branch, Division of Viral Diseases, Centers for Disease Control and Prevention | Krista Queen, Yan Li, Ying Tao, Jing Zhang, Anna Uehara, Anna Montmayeur, Clinton R. Paden, Peter W. Cook,Rachel Marine, Mili Sheth, Haibin Wang, Justin Lee, Suxiang Tong |
| EPI_ISL_751741 | OH Department of Health Laboratory | Genomics and Discovery, Respiratory Viruses Branch, Division of Viral Diseases, Centers for Disease Control and Prevention | Krista Queen, Yan Li, Ying Tao, Jing Zhang, Anna Uehara, Anna Montmayeur, Clinton R. Paden, Peter W. Cook,Rachel Marine, Mili Sheth, Haibin Wang, Justin Lee, Suxiang Tong |
| EPI_ISL_751748, EPI_ISL_751782 | ID Bureau of Laboratories | Genomics and Discovery, Respiratory Viruses Branch, Division of Viral Diseases, Centers for Disease Control and Prevention | Krista Queen, Yan Li, Ying Tao, Jing Zhang, Anna Uehara, Anna Montmayeur, Clinton R. Paden, Peter W. Cook,Rachel Marine, Mili Sheth, Haibin Wang, Justin Lee, Suxiang Tong |
| EPI_ISL_751784 | VA-Division of Consolidated Laboratory Services | Genomics and Discovery, Respiratory Viruses Branch, Division of Viral Diseases, Centers for Disease Control and Prevention | Krista Queen, Yan Li, Ying Tao, Jing Zhang, Anna Uehara, Anna Montmayeur, Clinton R. Paden, Peter W. Cook,Rachel Marine, Mili Sheth, Haibin Wang, Justin Lee, Suxiang Tong |
| EPI_ISL_751789 | ID Bureau of Laboratories | Genomics and Discovery, Respiratory Viruses Branch, Division of Viral Diseases, Centers for Disease Control and Prevention | Krista Queen, Yan Li, Ying Tao, Jing Zhang, Anna Uehara, Anna Montmayeur, Clinton R. Paden, Peter W. Cook,Rachel Marine, Mili Sheth, Haibin Wang, Justin Lee, Suxiang Tong |
| EPI_ISL_751793 | AK State Public Health Lab, State Health Department | Genomics and Discovery, Respiratory Viruses Branch, Division of Viral Diseases, Centers for Disease Control and Prevention | Krista Queen, Yan Li, Ying Tao, Jing Zhang, Anna Uehara, Anna Montmayeur, Clinton R. Paden, Peter W. Cook,Rachel Marine, Mili Sheth, Haibin Wang, Justin Lee, Suxiang Tong |
| EPI_ISL_753114 | State Laboratories Division, Hawaii State Department of Health | State Laboratories Division, Hawaii State Department of Health | Pamela O'Brien, Sabrina Diemert, Drew Kuwazaki, Razvan Sultana, Edward Desmond |
| EPI_ISL_753345, EPI_ISL_753346, EPI_ISL_753347, EPI_ISL_753348, EPI_ISL_753349, EPI_ISL_753350, EPI_ISL_753351, EPI_ISL_753352, EPI_ISL_753353, EPI_ISL_753354, EPI_ISL_753355, EPI_ISL_753356, EPI_ISL_753357, EPI_ISL_753358, EPI_ISL_753359, EPI_ISL_753360, EPI_ISL_753361, EPI_ISL_753362, EPI_ISL_753363, EPI_ISL_753364, EPI_ISL_753365, EPI_ISL_753366, EPI_ISL_753367, EPI_ISL_753368, EPI_ISL_753369, EPI_ISL_753370, EPI_ISL_753371, EPI_ISL_753372, EPI_ISL_753373, EPI_ISL_753374, EPI_ISL_753375, EPI_ISL_753376, EPI_ISL_753377, EPI_ISL_753378, EPI_ISL_753379, EPI_ISL_753380, EPI_ISL_753381, EPI_ISL_753382, EPI_ISL_753383, EPI_ISL_753384, EPI_ISL_753385, EPI_ISL_753386, EPI_ISL_753387, EPI_ISL_753389, EPI_ISL_753390, EPI_ISL_753391, EPI_ISL_753392, EPI_ISL_753403, EPI_ISL_753404, EPI_ISL_753405, EPI_ISL_753406, EPI_ISL_753407, EPI_ISL_753408 | | | |
| see above | Clinical virology Laboratory, Children's Hospital Los Angeles | Center for Personalized Medicine, Children's Hospital Los Angeles | Gai et al |
| EPI_ISL_753714, EPI_ISL_753769, EPI_ISL_753827, EPI_ISL_753831, EPI_ISL_753833 | Charité Universitätsmedizin Berlin, Institut für Virologie/Labor Berlin | Charité Universitätsmedizin Berlin, Institut für Virologie | Victor M Corman, Jörn Beheim-Schwarzbach, Barbara Mühlemann, Julia Schneider, Talitha Veith, Terry Jones, Christian Drosten |
| EPI_ISL_754141, EPI_ISL_754161 | Laboratoire de Microbiologie | National Reference Center for Viruses of Respiratory Infections, Institut Pasteur, Paris | Marion Barbet, Sylvie Behillil, Méline Bizard, Angela Brisebarre, Camille Capel, Etienne Simon-Lorière, Vincent Enouf, Maud Vanpeene, Sylvie van der Werf, Marie-Sarah Fangous |
| EPI_ISL_754546, EPI_ISL_754579, EPI_ISL_754580, EPI_ISL_754581, EPI_ISL_754582, EPI_ISL_754583, EPI_ISL_754584, EPI_ISL_754585, EPI_ISL_754586 | Wadsworth Center, New York State Department.of Health | Wadsworth Center, New York State Department.of Health | Kirsten St. George, Daryl M. Lamson, Alexis Russel, Matthew Shudt, Melissa A Leisner, Jonathan Plitnick, Navjot Singh, John Kelly, Sara Griesemer, Erasmus Schneider, Erica Lasek-Nesselquist |
| EPI_ISL_754771, EPI_ISL_754773, EPI_ISL_754774 | Emory Molecular Diagnostics Laboratory, Emory Healthcare | Piantadosi Lab, Emory Department of Pathology | Ahmed Babiker, Anne Piantadosi |

| | | | |
|---|---|---|---|
| EPI_ISL_755117, EPI_ISL_755119, EPI_ISL_755130, EPI_ISL_755134, EPI_ISL_755136, EPI_ISL_755270, EPI_ISL_755271, EPI_ISL_755274, EPI_ISL_755275, EPI_ISL_755280, EPI_ISL_755290, EPI_ISL_755295 | | | |
| see above | San Diego County Public Health Laboratory | Andersen lab at Scripps Research | SEARCH Alliance San Diego with Tracy Basler, Jovan Shephard, Brett Austin |
| EPI_ISL_755805, EPI_ISL_755806, EPI_ISL_755807 | Toronto Invasive Bacterial Diseases Network | McMaster University | Allison McGeer, Patryk Aftanas, Hooman Derakhshani, Angel Li, Kuganya Nirmalarajah, Emily Panousis, Ahmed Draia, Jalees Nasir, Michael Surette, Samira Mubareka, Andrew G. McArthur |
| EPI_ISL_756311 | The Caribbean Public Health Agency | Carrington Lab, Department of PreClinical Sciences, Faculty of Medical Sciences, The University of the West Indies | Nikita S. D. Sahadeo, Arianne Brown-Jordan, Sarah Hill, Vernie Ramkissoon, Roshan Parasram, Naresh Nandram, Avery Hinds, Jerome Foster, Stanley Giddings, Karla Georges, Marsha Ivey, Rahul Naidu, Risha Singh, SueMin Nathaniel, Rajini Haraksingh, Jaya Jayaraman, Chinna Chinnadurai, Adesh Ramsubhag, Nuno Faria, Oliver Pybus, Christopher Oura, Gabriel Escobar, Christine V. F. Carrington |
| EPI_ISL_756360 | Caribbean Regional Public Health Authority | Carrington Lab, Department of PreClinical Sciences, Faculty of Medical Sciences, The University of the West Indies | Nikita S. D. Sahadeo, Gabriel Escobar, Sarah Hill, Vernie Ramkissoon, Risha Singh, SueMin Nathaniel, Simone Keizer-Beache, Arianne Brown-Jordan, Naresh Nandram, Avery Hinds, Jerome Foster, Stanley Giddings, Karla Georges, Marsha Ivey, Rahul Naidu, , Rajini Haraksingh, Jaya Jayaraman, Chinna Chinnadurai, Adesh Ramsubhag, Nuno Faria, Oliver Pybus, Christopher Oura, Christine V. F. Carrington |
| EPI_ISL_757187, EPI_ISL_757188, EPI_ISL_757189, EPI_ISL_757190, EPI_ISL_757191, EPI_ISL_757192, EPI_ISL_757193, EPI_ISL_757194, EPI_ISL_757195, EPI_ISL_757196, EPI_ISL_757197, EPI_ISL_757198, EPI_ISL_757199, EPI_ISL_757200, EPI_ISL_757201, EPI_ISL_757202, EPI_ISL_757203, EPI_ISL_757204, EPI_ISL_757205, EPI_ISL_757206, EPI_ISL_757207, EPI_ISL_757208, EPI_ISL_757209, EPI_ISL_757210, EPI_ISL_757211, EPI_ISL_757212, EPI_ISL_757213, EPI_ISL_757214, EPI_ISL_757215, EPI_ISL_757216, EPI_ISL_757217, EPI_ISL_757218, EPI_ISL_757219, EPI_ISL_757220, EPI_ISL_757221, EPI_ISL_757222, EPI_ISL_757223, EPI_ISL_757224, EPI_ISL_757273, EPI_ISL_757274, EPI_ISL_757275, EPI_ISL_757276, EPI_ISL_757277, EPI_ISL_757278, EPI_ISL_757279, EPI_ISL_757280, EPI_ISL_757281, EPI_ISL_757282, EPI_ISL_757283, EPI_ISL_757284 | | | |
| see above | Lighthouse Lab in Glasgow | Wellcome Sanger Institute for the COVID-19 Genomics UK (COG-UK) Consortium | Harper VanSteenhouse, Yumi Kasai, David Gray, Carol Clugston, Anna Dominiczak and Alex Alderton, Roberto Amato, Sonia Goncalves, Ewan Harrison, David K. Jackson, Ian Johnston, Dominic Kwiatkowski, Cordelia Langford, John Sillitoe on behalf of the Wellcome Sanger Institute COVID-19 Surveillance Team |
| EPI_ISL_757296, EPI_ISL_757297, EPI_ISL_757298, EPI_ISL_757299, EPI_ISL_757300, EPI_ISL_757301, EPI_ISL_757302, EPI_ISL_757303, EPI_ISL_757304, EPI_ISL_757305, EPI_ISL_757306, EPI_ISL_757307, EPI_ISL_757308, EPI_ISL_757309, EPI_ISL_757310, EPI_ISL_757311, EPI_ISL_757312, EPI_ISL_757313, EPI_ISL_757314, EPI_ISL_757315, EPI_ISL_757316, EPI_ISL_757317, EPI_ISL_757318, EPI_ISL_757319, EPI_ISL_757320, EPI_ISL_757321, EPI_ISL_757322, EPI_ISL_757323, EPI_ISL_757324, EPI_ISL_757325, EPI_ISL_757326, EPI_ISL_757327, EPI_ISL_757328, EPI_ISL_757329, EPI_ISL_757330, EPI_ISL_757331, EPI_ISL_757332, EPI_ISL_757333, EPI_ISL_757334, EPI_ISL_757335, EPI_ISL_757336, EPI_ISL_757337, EPI_ISL_757338, EPI_ISL_757339, EPI_ISL_757340, EPI_ISL_757341 | | | |
| see above | Department of Virology and Immunology, University of Helsinki and Helsinki University Hospital, Huslab Finland | Department of Virology, Faculty of Medicine, University of Helsinki, Helsinki, Finland | Teemu Smura, Ravi Kant, Phuoc Truong, Hussein Alburkat, Hannimari Kallio-Kokko, Jenni Virtanen, Maija Suvanto, Sari Hannula, Harri Kangas, Pekka Ellonen, Olli Vapalahti |
| EPI_ISL_759722 | Instituto Nacional de Saude (INSA) | Instituto Nacional de Saude (INSA) | Borges et al |
| EPI_ISL_759973, EPI_ISL_759974 | Furst Medical Laboratory | Norwegian Institute of Public Health, Department of Virology | Kathrine Stene-Johansen, Kamilla Heddeland Instefjord, Hilde Elshaug, Marie Paulsen Madsen, Rasmus Riis Kopperud, Hilde Vollan, Karoline Bragstad, Olav Hungnes |
| EPI_ISL_759975 | Haukeland University Hospital, Dept. of Microbiology | Norwegian Institute of Public Health, Department of Virology | Kathrine Stene-Johansen, Kamilla Heddeland Instefjord, Hilde Elshaug, Marie Paulsen Madsen, Rasmus Riis Kopperud, Hilde Vollan, Karoline Bragstad, Olav Hungnes |
| EPI_ISL_759976 | Hospital of Southern Norway - Kristiansand, Department of Medical Microbiology | Norwegian Institute of Public Health, Department of Virology | Kathrine Stene-Johansen, Kamilla Heddeland Instefjord, Hilde Elshaug, Marie Paulsen Madsen, Rasmus Riis Kopperud, Hilde Vollan, Karoline Bragstad, Olav Hungnes |
| EPI_ISL_759977 | Medical Microbiology Unit, Department for Laboratory Medicine, Drammen Hospital, Vestre Viken Health Trust, | Norwegian Institute of Public Health, Department of Virology | Kathrine Stene-Johansen, Kamilla Heddeland Instefjord, Hilde Elshaug, Marie Paulsen Madsen, Rasmus Riis Kopperud, Hilde Vollan, Karoline Bragstad, Olav Hungnes |
| EPI_ISL_760230, EPI_ISL_760231, EPI_ISL_760232, EPI_ISL_760233, EPI_ISL_760234, EPI_ISL_760235, EPI_ISL_760236, EPI_ISL_760237, EPI_ISL_760238, EPI_ISL_760239, EPI_ISL_760240, EPI_ISL_760241, EPI_ISL_760242, EPI_ISL_760243, EPI_ISL_760244, EPI_ISL_760245 | | | |
| see above | Division of Emerging Infectious Diseases, Bureau of Infectious Diseases Diagnosis Control, Korea Disease Control and Prevention Agency | Division of Emerging Infectious Diseases, Bureau of Infectious Diseases Diagnosis Control, Korea Disease Control and Prevention Agency | Ae Kyung Park, Il-Hwan Kim, Heui Man Kim, Jeong-Min Kim, Namjoo Lee, Chaeyoung Lee, Sang Hee Woo, Eun-Jin Kim |
| EPI_ISL_763111, EPI_ISL_763163, EPI_ISL_763172, EPI_ISL_763235, EPI_ISL_763305 | Dutch COVID-19 response team | Erasmus Medical Center | Bas Oude Munnink, Reina Sikkema, David Nieuwenhuijse, Irina Chestakova, Anne van der Linden, Marjan Boter, Emmanuelle Munger, Corine GeurtsvanKessel, Annemiek van der Eijk, Richard Molenkamp, Marion Koopmans, on behalf of the Dutch national COVID-19 response team. |
| EPI_ISL_763331, EPI_ISL_763337, EPI_ISL_763338, EPI_ISL_763339, EPI_ISL_763340, EPI_ISL_763341, EPI_ISL_763342 | Florida Bureau of Public Health Laboratories | Florida Bureau of Public Health Laboratories | Sarah Schmedes, Jason Blanton |
| EPI_ISL_765748, EPI_ISL_765749, EPI_ISL_765844, EPI_ISL_765845, EPI_ISL_765846, EPI_ISL_765847, EPI_ISL_765848, EPI_ISL_765849, EPI_ISL_765850, EPI_ISL_765851, EPI_ISL_765852, EPI_ISL_765853 | | | |
| see above | Massachusetts General Hospital | Infectious Disease Program, Broad Institute of Harvard and MIT | Lemieux,J.E., Siddle,K.J., Shaw,B., Adams,G., Pierce,V., Turbett,S., Anahtar,M., Branda,J., Slater,D., Harris,J., Lin,A.E., Gladden-Young,A., Lagerborg,K., Rudy,M., DeRuff,K., Carter,A., Normandin,E., Bauer,M., Reilly,S., Tomkins-Tinch,C., Loreth,C., Chaluvadi,S., Neumann,A., Cusick,C., Chapman,S.B., Gnirke,A., Flowers,K., Cerrato,F., Birren,B.W., Gallagher,G., Smole,S., Park,D.J., MacInnis,B.L., Ryan,E., LaRocque,R., Rosenberg,E. and Sabeti,P.C. |
| EPI_ISL_766071, EPI_ISL_766077, EPI_ISL_766094, EPI_ISL_766098, EPI_ISL_766271, EPI_ISL_766273, EPI_ISL_766274, EPI_ISL_766275 | Respiratory Virus Unit, National Infection Service, Public Health England | COVID-19 Genomics UK (COG-UK) Consortium | PHE Covid Sequencing Team |
| EPI_ISL_766621 | Klinisk mikrobiologi | The Public Health Agency of Sweden | Department of Microbiology, The Public Health Agency of Sweden |
| EPI_ISL_766742 | Texas Department of State Health Services | Texas Department of State Health Services | Rashmi Tuladhar, Bonnie Oh, Jenny Zhang, Maliha Rahman, Anita Pokharel, Myong Koag, Chung Wang, Rachel Lee, Grace Kubin, Mayela Pedrueza, James Daniel Bonser |
| EPI_ISL_767001, EPI_ISL_767004 | Delaware Public Health Laboratory | Delaware Public Health Laboratory | Gregory Hovan |
| EPI_ISL_767239 | Lighthouse Lab in Milton Keynes | Wellcome Sanger Institute for the COVID-19 Genomics UK (COG-UK) Consortium | The Lighthouse Lab in Milton Keynes and Alex Alderton, Roberto Amato, Sonia Goncalves, Ewan Harrison, David K. Jackson, Ian Johnston, Dominic Kwiatkowski, Cordelia Langford, John Sillitoe on behalf of the Wellcome Sanger Institute COVID-19 Surveillance Team |
| EPI_ISL_767351, EPI_ISL_767353, EPI_ISL_767356 | Michigan Department of Health and Human Services, Bureau of Laboratories | Michigan Department of Health and Human Services, Bureau of Laboratories | Blankenship HM, Riner D, Soehnlen MK |
| EPI_ISL_767861, EPI_ISL_767862 | Sydney South West Pathology Service (SSWPS) - Royal Prince Alfred Hospital - NSW Health Pathology | NSW Health Pathology - Institute of Clinical Pathology and Medical Research; Westmead Hospital; University of Sydney | CIDM-PH et al. |
| EPI_ISL_768522 | Synphaet Hospital | National Institute of Health, Department of Medical Sciences, Ministry of Public Health, Thailand | Pilailuk Okada; Siripaporn Phuygun; Sittiporn Parnmen; Ratana Tacharoenmuang; Pakorn Piromtong; Natchaya Khiadsang; Thanutsapa Thanadachakul; Warawan Wongboot; sirikanda wimol; Sunthareeya Waicharoen; |
| EPI_ISL_768727 | Jena University Hospital, Institute for Infectious Diseases and Infection Control | Institute of infectious medicine & hospital hygiene, CaSe-Group | Spott, Riccardo; Marquet, Mike; Pletz, Matthias W.; Brandt, Christian |
| EPI_ISL_769865 | Respiratory Virus Unit, National Infection Service, Public Health England | COVID-19 Genomics UK (COG-UK) Consortium | PHE Covid Sequencing Team |
| EPI_ISL_769986 | Area De Salud Catedral Noreste | Inciensa, Instituto Costarricense de Investigación y Enseñanza en Nutrición y Salud | Francisco Duarte, Hebleen Porras, Claudio Soto-Garita, Estela Cordero, Adriana Godínez, Melany Calderón & Mariel López |
| EPI_ISL_769988 | Area De Salud San Francisco-San Antonio (Coopesana) | Inciensa, Instituto Costarricense de Investigación y Enseñanza en Nutrición y Salud | Francisco Duarte, Hebleen Porras, Claudio Soto-Garita, Estela Cordero, Adriana Godínez, Melany Calderón & Mariel López |
| EPI_ISL_770021 | Area De Salud Catedral Noreste | Inciensa, Instituto Costarricense de Investigación y Enseñanza en Nutrición y Salud | Francisco Duarte, Hebleen Porras, Claudio Soto-Garita, Estela Cordero, Adriana Godínez, Melany Calderón & Mariel López |

| | | | |
|---|---|---|---|
| EPI_ISL_770024 | Hle - Asociacion Hogar De Ancianos Santiago Crespo Calvo | Inciensa, Instituto Costarricense de Investigación y Enseñanza en Nutrición y Salud | Francisco Duarte, Hebleen Porras, Claudio Soto-Garita, Estela Cordero, Adriana Godínez, Melany Calderón & Mariel López |
| EPI_ISL_770026 | Area De Salud Catedral Noreste | Inciensa, Instituto Costarricense de Investigación y Enseñanza en Nutrición y Salud | Francisco Duarte, Hebleen Porras, Claudio Soto-Garita, Estela Cordero, Adriana Godínez, Melany Calderón & Mariel López |
| EPI_ISL_770635 | WHO National Influenza Centre Russian Federation | WHO National Influenza Centre Russian Federation | Andrey Komissarov, Artem Fadeev, Anna Ivanova, Kseniya Komissarova, Dmitry Bazhenov, Daria Danilenko, Ksenia Safina, Elena Nabieva, Georgii Bazykin, Dmitry Lioznov |
| EPI_ISL_770780, EPI_ISL_770781, EPI_ISL_770782 | Minnesota Department of Health, Public Health Laboratory | Minnesota Department of Health, Public Health Laboratory | Alexandra Lorentz, Jacob Garfin, Matt Plumb, and Xiong Wang |
| EPI_ISL_770808, EPI_ISL_770809, EPI_ISL_770812 | Essentia Health-St. Mary's Medical Center | Minnesota Department of Health, Public Health Laboratory | Alexandra Lorentz, Jacob Garfin, Matt Plumb, and Xiong Wang |
| EPI_ISL_771152, EPI_ISL_771154, EPI_ISL_771155, EPI_ISL_771158, EPI_ISL_771159, EPI_ISL_771161, EPI_ISL_771163, EPI_ISL_771164, EPI_ISL_771171, EPI_ISL_771172, EPI_ISL_771181, EPI_ISL_771184, EPI_ISL_771189, EPI_ISL_771200, EPI_ISL_771203, EPI_ISL_771206, EPI_ISL_771207, EPI_ISL_771209, EPI_ISL_771210, EPI_ISL_771211, EPI_ISL_771213, EPI_ISL_771216, EPI_ISL_771220, EPI_ISL_771247, EPI_ISL_771248, EPI_ISL_771249, EPI_ISL_771250, EPI_ISL_771251, EPI_ISL_771252, EPI_ISL_771253, EPI_ISL_771254, EPI_ISL_771255, EPI_ISL_771256, EPI_ISL_771257, EPI_ISL_771258, EPI_ISL_771259, EPI_ISL_771260, EPI_ISL_771261, EPI_ISL_771262, EPI_ISL_771263, EPI_ISL_771264, EPI_ISL_771265, EPI_ISL_771266, EPI_ISL_771267, EPI_ISL_771268, EPI_ISL_771269, EPI_ISL_771270, EPI_ISL_771271, EPI_ISL_771272, EPI_ISL_771273, EPI_ISL_771274, EPI_ISL_771275, EPI_ISL_771276, EPI_ISL_771277, EPI_ISL_771278, EPI_ISL_771279, EPI_ISL_771280, EPI_ISL_771281, EPI_ISL_771282, EPI_ISL_771283, EPI_ISL_771284, EPI_ISL_771285, EPI_ISL_771286, EPI_ISL_771287, EPI_ISL_771288, EPI_ISL_771289, EPI_ISL_771290, EPI_ISL_771292, EPI_ISL_771293, EPI_ISL_771294, EPI_ISL_771295, EPI_ISL_771296, EPI_ISL_771297, EPI_ISL_771298, EPI_ISL_771299, EPI_ISL_771302, EPI_ISL_771307, EPI_ISL_771331, EPI_ISL_771332, EPI_ISL_771350 | | | |
| see above | Colorado Department of Public Health and Environment | Colorado Department of Puplic Health and Environment | Laura Bankers, Molly C. Hetherington-Rauth, Diana Ir, Shannon Ely, Shannon R. Matzinger, Sarah Elizabeth Totten, Emily A. Travanty |
| EPI_ISL_775271, EPI_ISL_775294 | Furst Medical Laboratory | Norwegian Institute of Public Health, Department of Virology | Kathrine Stene-Johansen, Kamilla Heddeland Instefjord, Hilde Elshaug, Atiya R Ali,Marie Paulsen Madsen, Rasmus Riis Kopperud, Hilde Vollan, Karoline Bragstad, Olav Hungnes |
| EPI_ISL_775297 | Nordland Hospital - Bodo, Laboratory Department, Molecular Biology Unit | Norwegian Institute of Public Health, Department of Virology | Kathrine Stene-Johansen, Kamilla Heddeland Instefjord, Hilde Elshaug, Atiya R Ali,Marie Paulsen Madsen, Rasmus Riis Kopperud, Hilde Vollan, Karoline Bragstad, Olav Hungnes |
| EPI_ISL_775335, EPI_ISL_775336 | Haukeland University Hospital, Dept. of Microbiology | Norwegian Institute of Public Health, Department of Virology | Kathrine Stene-Johansen, Kamilla Heddeland Instefjord, Hilde Elshaug, Atiya R Ali,Marie Paulsen Madsen, Rasmus Riis Kopperud, Hilde Vollan, Karoline Bragstad, Olav Hungnes |
| EPI_ISL_775349, EPI_ISL_775350, EPI_ISL_775351, EPI_ISL_775352 | Unilabs Laboratory Medicine | Norwegian Institute of Public Health, Department of Virology | Kathrine Stene-Johansen, Kamilla Heddeland Instefjord, Hilde Elshaug, Atiya R Ali,Marie Paulsen Madsen, Rasmus Riis Kopperud, Hilde Vollan, Karoline Bragstad, Olav Hungnes |
| EPI_ISL_775354 | Dept. of Medical Microbiology, Stavanger University Hospital, Helse Stavanger HF | Norwegian Institute of Public Health, Department of Virology | Kathrine Stene-Johansen, Kamilla Heddeland Instefjord, Hilde Elshaug, Atiya R Ali,Marie Paulsen Madsen, Rasmus Riis Kopperud, Hilde Vollan, Karoline Bragstad, Olav Hungnes |
| EPI_ISL_775363, EPI_ISL_775364, EPI_ISL_775365, EPI_ISL_775366, EPI_ISL_775367 | Foerde Hospital, Department of Microbiology | Norwegian Institute of Public Health, Department of Virology | Kathrine Stene-Johansen, Kamilla Heddeland Instefjord, Hilde Elshaug, Atiya R Ali,Marie Paulsen Madsen, Rasmus Riis Kopperud, Hilde Vollan, Karoline Bragstad, Olav Hungnes |
| EPI_ISL_775388 | Akershus University Hospital, Department for Microbiology and Infectious Disease Control | Norwegian Institute of Public Health, Department of Virology | Kathrine Stene-Johansen, Kamilla Heddeland Instefjord, Hilde Elshaug, Atiya R Ali,Marie Paulsen Madsen, Rasmus Riis Kopperud, Hilde Vollan, Karoline Bragstad, Olav Hungnes |
| EPI_ISL_775457 | Oslo University Hospital, Department of Medical Microbiology | Norwegian Institute of Public Health, Department of Virology | Kathrine Stene-Johansen, Kamilla Heddeland Instefjord, Hilde Elshaug, Atiya R Ali,Marie Paulsen Madsen, Rasmus Riis Kopperud, Hilde Vollan, Karoline Bragstad, Olav Hungnes |
| EPI_ISL_775484, EPI_ISL_775489, EPI_ISL_775490 | Furst Medical Laboratory | Norwegian Institute of Public Health, Department of Virology | Kathrine Stene-Johansen, Kamilla Heddeland Instefjord, Hilde Elshaug, Atiya R Ali,Marie Paulsen Madsen, Rasmus Riis Kopperud, Hilde Vollan, Karoline Bragstad, Olav Hungnes |
| EPI_ISL_775498, EPI_ISL_775499 | Vestfold Hospital, Toensberg Department of Microbiology | Norwegian Institute of Public Health, Department of Virology | Kathrine Stene-Johansen, Kamilla Heddeland Instefjord, Hilde Elshaug, Atiya R Ali,Marie Paulsen Madsen, Rasmus Riis Kopperud, Hilde Vollan, Karoline Bragstad, Olav Hungnes |
| EPI_ISL_775514 | Norwegian Institute of Public Health, Department of Virology | Norwegian Institute of Public Health, Department of Virology | Kathrine Stene-Johansen, Kamilla Heddeland Instefjord, Hilde Elshaug, Atiya R Ali,Marie Paulsen Madsen, Rasmus Riis Kopperud, Hilde Vollan, Karoline Bragstad, Olav Hungnes |
| EPI_ISL_778834 | AIID | Irish Coronavirus Sequencing Consortium-Teagasc Grange | Matthew McCabe, Aljandro Abner Garcia Leon, Fiona Crispie, Calum Walsh, Michael Carr, John Kenny, Paul Cotter, Patrick Mallon, Gabriel Gonzalez |
| EPI_ISL_779185, EPI_ISL_779186, EPI_ISL_779188, EPI_ISL_779192, EPI_ISL_779196 | Laboratorio Estatal de Salud Pública de Nuevo León | Laboratorio de Infectología Molecular, Departamento de Bioquímica y Medicina Molecular,Facultad de Medicina - Universidad Autónoma de Nuevo León | Kame A. Galán-Huerta, María F. Herrera-Saldivar, Natalia Martínez-Acuña, Sonia A. Lozano-Sepúlveda, Daniel Arellanos-Soto, Ana M. Rivas-Estilla, Samuel Buentello-Wong, Else del Carmen García-García, Gloria A. Jasso-de-la-Peña, Roberto Montes-de-Oca, Consuelo Treviño-Garza, Manuel E. de-la-O-Cavazos |
| EPI_ISL_779781, EPI_ISL_779782, EPI_ISL_779821, EPI_ISL_779822, EPI_ISL_779823, EPI_ISL_779824, EPI_ISL_779825, EPI_ISL_779826, EPI_ISL_779827, EPI_ISL_779828, EPI_ISL_779829, EPI_ISL_779830, EPI_ISL_779831, EPI_ISL_779832, EPI_ISL_779833, EPI_ISL_779834, EPI_ISL_779835 | | | |
| see above | Laboratoire de virologie, CHU de Grenoble | CNR Virus des Infections Respiratoires - France SUD | Antonin Bal, Gregory Destras, Gwendolyne Burfin, Hadrien Règue, Quentin Semanas, Martine Valette, Bruno Lina, Sylvie Larrat, Laurence Josset |
| EPI_ISL_779955, EPI_ISL_779959, EPI_ISL_779963, EPI_ISL_779969, EPI_ISL_779971, EPI_ISL_779972, EPI_ISL_779973, EPI_ISL_779975, EPI_ISL_779976, EPI_ISL_780012, EPI_ISL_780013, EPI_ISL_780014, EPI_ISL_780015, EPI_ISL_780016, EPI_ISL_780017, EPI_ISL_780019, EPI_ISL_780020, EPI_ISL_780029, EPI_ISL_780032, EPI_ISL_780034, EPI_ISL_780039, EPI_ISL_780042, EPI_ISL_780045, EPI_ISL_780050, EPI_ISL_780055, EPI_ISL_780056, EPI_ISL_780060, EPI_ISL_780063, EPI_ISL_780065, EPI_ISL_780067, EPI_ISL_780075, EPI_ISL_780076, EPI_ISL_780078, EPI_ISL_780079, EPI_ISL_780080, EPI_ISL_780081, EPI_ISL_780082, EPI_ISL_780083, EPI_ISL_780084, EPI_ISL_780085, EPI_ISL_780086, EPI_ISL_780087, EPI_ISL_780088, EPI_ISL_780089 | | | |
| see above | Hospital General Universitario Gregorio Marañón | SeqCOVID-SPAIN consortium/IBV(CSIC) | Darío García de Viedma, Laura Pérez-Lago, Marta Herranz, Jon Sicilia, Julia Suárez, Pilar Catalán, Patricia Muñoz and SeqCOVID-SPAIN consortium |
| EPI_ISL_780397, EPI_ISL_780398, EPI_ISL_780399, EPI_ISL_780400, EPI_ISL_780401 | Bermuda Government Molecular Diagnostics Laboratory (MDL) | Respiratory Virus Unit, National Infection Service, Public Health England | PHE Covid Sequencing Team, Dr Carika Weldon (Bermuda), Dr Ayoola Oyinloye (Bermuda) |
| EPI_ISL_783069, EPI_ISL_783070, EPI_ISL_783071, EPI_ISL_783072, EPI_ISL_783073, EPI_ISL_783074, EPI_ISL_783075, EPI_ISL_783076, EPI_ISL_783077, EPI_ISL_783078, EPI_ISL_783079, EPI_ISL_783080, EPI_ISL_783081, EPI_ISL_783082, EPI_ISL_783083, EPI_ISL_783084, EPI_ISL_783085, EPI_ISL_783086, EPI_ISL_783087, EPI_ISL_783088, EPI_ISL_783089, EPI_ISL_783090, EPI_ISL_783091, EPI_ISL_783092, EPI_ISL_783093, EPI_ISL_783094, EPI_ISL_783095, EPI_ISL_783096, EPI_ISL_783097, EPI_ISL_783098, EPI_ISL_783099, EPI_ISL_783100, EPI_ISL_783101, EPI_ISL_783102, EPI_ISL_783103, EPI_ISL_783104, EPI_ISL_783105, EPI_ISL_783106, EPI_ISL_783107, EPI_ISL_783108, EPI_ISL_783109, EPI_ISL_783110, EPI_ISL_783111, EPI_ISL_783112, EPI_ISL_783113, EPI_ISL_783114, EPI_ISL_783115, EPI_ISL_783116, EPI_ISL_783117, EPI_ISL_783118, EPI_ISL_783119, EPI_ISL_783120, EPI_ISL_783121, EPI_ISL_783122, EPI_ISL_783123, EPI_ISL_783124, EPI_ISL_783125, EPI_ISL_783126, EPI_ISL_783127, EPI_ISL_783128, EPI_ISL_783129, EPI_ISL_783130, EPI_ISL_783131, EPI_ISL_783132, EPI_ISL_783133, EPI_ISL_783134, EPI_ISL_783135, EPI_ISL_783136, EPI_ISL_783137, EPI_ISL_783138, EPI_ISL_783139, EPI_ISL_783140, EPI_ISL_783141, EPI_ISL_783142, EPI_ISL_783143, EPI_ISL_783144, EPI_ISL_783145, EPI_ISL_783146, EPI_ISL_783147, EPI_ISL_783148, EPI_ISL_783149, EPI_ISL_783150, EPI_ISL_783151, EPI_ISL_783152, EPI_ISL_783153, EPI_ISL_783154, EPI_ISL_783155, EPI_ISL_783156, EPI_ISL_783157, EPI_ISL_783158, EPI_ISL_783159, EPI_ISL_783160, EPI_ISL_783161, EPI_ISL_783162, EPI_ISL_783163, EPI_ISL_783164, EPI_ISL_783165, EPI_ISL_783166, EPI_ISL_783167, EPI_ISL_783168, EPI_ISL_783169, EPI_ISL_783170, EPI_ISL_783171, EPI_ISL_783172, EPI_ISL_783173, EPI_ISL_783174, EPI_ISL_783175, EPI_ISL_783176, EPI_ISL_783177, EPI_ISL_783178, EPI_ISL_783179, EPI_ISL_783180, EPI_ISL_783181, EPI_ISL_783182, EPI_ISL_783183, EPI_ISL_783184, EPI_ISL_783185, EPI_ISL_783186, EPI_ISL_783187, EPI_ISL_783188, EPI_ISL_783189, EPI_ISL_783190, EPI_ISL_783191, EPI_ISL_783192, EPI_ISL_783193, EPI_ISL_783194, EPI_ISL_783195, EPI_ISL_783196, EPI_ISL_783197, EPI_ISL_783198, EPI_ISL_783199, EPI_ISL_783200, EPI_ISL_783201, EPI_ISL_783202, EPI_ISL_783203, EPI_ISL_783204, EPI_ISL_783205, EPI_ISL_783206, EPI_ISL_783207, EPI_ISL_783208, EPI_ISL_783209, EPI_ISL_783210, EPI_ISL_783211, EPI_ISL_783212, EPI_ISL_783213, EPI_ISL_783214, EPI_ISL_783215, EPI_ISL_783216, EPI_ISL_783217, EPI_ISL_783218, EPI_ISL_783219, EPI_ISL_783220, EPI_ISL_783221, EPI_ISL_783222, EPI_ISL_783223, EPI_ISL_783224, EPI_ISL_783225, EPI_ISL_783226, EPI_ISL_783227, EPI_ISL_783228, EPI_ISL_783229, EPI_ISL_783230, EPI_ISL_783231, EPI_ISL_783232, EPI_ISL_783233, EPI_ISL_783234, EPI_ISL_783235, EPI_ISL_783236, EPI_ISL_783237, EPI_ISL_783238, EPI_ISL_783239, EPI_ISL_783240, EPI_ISL_783241, EPI_ISL_783242, EPI_ISL_783243, EPI_ISL_783244, EPI_ISL_783245, EPI_ISL_783246, EPI_ISL_783247, EPI_ISL_783248, EPI_ISL_783249, EPI_ISL_783250, EPI_ISL_783251, EPI_ISL_783252, EPI_ISL_783253, EPI_ISL_783254, EPI_ISL_783255, EPI_ISL_783256, EPI_ISL_783257, EPI_ISL_783258, EPI_ISL_783259, EPI_ISL_783260, EPI_ISL_783261, EPI_ISL_783262, EPI_ISL_783263, EPI_ISL_783264, EPI_ISL_783265, EPI_ISL_783266, EPI_ISL_783267, EPI_ISL_783268, EPI_ISL_783269, EPI_ISL_783270, EPI_ISL_783271, EPI_ISL_783272, EPI_ISL_783273, EPI_ISL_783274, EPI_ISL_783275, EPI_ISL_783276, EPI_ISL_783277, EPI_ISL_783278, EPI_ISL_783279, EPI_ISL_783280, EPI_ISL_783281, EPI_ISL_783282, EPI_ISL_783283, EPI_ISL_783284, EPI_ISL_783285, EPI_ISL_783286, EPI_ISL_783287, EPI_ISL_783288, EPI_ISL_783289, EPI_ISL_783290, EPI_ISL_783291, EPI_ISL_783292, EPI_ISL_783293, EPI_ISL_783294, EPI_ISL_783295, EPI_ISL_783296, EPI_ISL_783297, EPI_ISL_783298, EPI_ISL_783299, EPI_ISL_783300, EPI_ISL_783301, EPI_ISL_783302, EPI_ISL_783303, EPI_ISL_783304, EPI_ISL_783305, EPI_ISL_783306, EPI_ISL_783307, EPI_ISL_783308, EPI_ISL_783309, EPI_ISL_783310, EPI_ISL_783311, EPI_ISL_783312, EPI_ISL_783313, EPI_ISL_783314, EPI_ISL_783315, EPI_ISL_783316, EPI_ISL_783317, EPI_ISL_783318, EPI_ISL_783319, EPI_ISL_783320, EPI_ISL_783321, EPI_ISL_783322, EPI_ISL_783323, EPI_ISL_783324, EPI_ISL_783325, EPI_ISL_783326, EPI_ISL_783327, EPI_ISL_783328, EPI_ISL_783329, EPI_ISL_783330, EPI_ISL_783331, EPI_ISL_783332, EPI_ISL_783333, EPI_ISL_783334, EPI_ISL_783335, EPI_ISL_783336, EPI_ISL_783337, EPI_ISL_783338, EPI_ISL_783339, EPI_ISL_783340, EPI_ISL_783341, EPI_ISL_783342, EPI_ISL_783343, EPI_ISL_783344, EPI_ISL_783345, EPI_ISL_783346, EPI_ISL_783347, EPI_ISL_783348, EPI_ISL_783349, EPI_ISL_783350, EPI_ISL_783351, EPI_ISL_783352, EPI_ISL_783353, EPI_ISL_783354, EPI_ISL_783355, EPI_ISL_783356, EPI_ISL_783357, EPI_ISL_783358, EPI_ISL_783359, EPI_ISL_783360, EPI_ISL_783361, EPI_ISL_783362, EPI_ISL_783363, EPI_ISL_783364, EPI_ISL_783365, EPI_ISL_783366 | | | |
| see above | Lighthouse Lab in Milton Keynes | Wellcome Sanger Institute for the COVID-19 Genomics UK (COG-UK) Consortium | The Lighthouse Lab in Milton Keynes and Alex Alderton, Roberto Amato, Sonia Goncalves, Ewan Harrison, David K. Jackson, Ian Johnston, Dominic Kwiatkowski, Cordelia Langford, John Sillitoe on behalf of the Wellcome Sanger Institute COVID-19 Surveillance Team |
| EPI_ISL_783537, EPI_ISL_783540, EPI_ISL_783545, EPI_ISL_783547, EPI_ISL_783549, EPI_ISL_783551, | Houston Methodist Hospital | Houston Methodist Hospital | S. Wesley Long, Randall J. Olsen, Paul A. Christensen, David W. Bernard, James J. Davis, Maulik Shukla, Marcus Nguyen, Matthew Ojeda Saavedra, Prasanti Yerramilli, Layne Pruitt, Sishir Subedi, Heather Hendrickson, and James M. Musser |

| | | | |
|---|---|---|---|
| EPI_ISL_783553, EPI_ISL_783569 | | | |
| EPI_ISL_783850 | Bermuda Government Molecular Diagnostics Laboratory (MDL) | Respiratory Virus Unit, National Infection Service, Public Health England | PHE Covid Sequencing Team, Dr Carika Weldon (Bermuda), Dr Ayoola Oyinloye (Bermuda) |
| EPI_ISL_783869, EPI_ISL_783870, EPI_ISL_783871, EPI_ISL_783872, EPI_ISL_783873, EPI_ISL_783874, EPI_ISL_783879, EPI_ISL_783880, EPI_ISL_783881, EPI_ISL_783882, EPI_ISL_783883, EPI_ISL_783889, EPI_ISL_783890, EPI_ISL_783891, EPI_ISL_783892, EPI_ISL_783893, EPI_ISL_783898, EPI_ISL_783899, EPI_ISL_783900, EPI_ISL_783901, EPI_ISL_783902, EPI_ISL_783903, EPI_ISL_783908, EPI_ISL_783909, EPI_ISL_783910, EPI_ISL_783911, EPI_ISL_783912, EPI_ISL_783913, EPI_ISL_783914, EPI_ISL_783919, EPI_ISL_783920, EPI_ISL_783921, EPI_ISL_783922, EPI_ISL_783923, EPI_ISL_783924, EPI_ISL_783928, EPI_ISL_783929, EPI_ISL_783930, EPI_ISL_783931, EPI_ISL_783932, EPI_ISL_783933, EPI_ISL_783940, EPI_ISL_783941, EPI_ISL_783942, EPI_ISL_783943, EPI_ISL_783944, EPI_ISL_783945, EPI_ISL_783946, EPI_ISL_783950, EPI_ISL_783953, EPI_ISL_783957, EPI_ISL_783963, EPI_ISL_783964, EPI_ISL_783967, EPI_ISL_783971, EPI_ISL_783977, EPI_ISL_783979, EPI_ISL_783981, EPI_ISL_783982, EPI_ISL_783983, EPI_ISL_783985, EPI_ISL_783991, EPI_ISL_783992, EPI_ISL_784001, EPI_ISL_784002, EPI_ISL_784004, EPI_ISL_784008, EPI_ISL_784013, EPI_ISL_784019, EPI_ISL_784022, EPI_ISL_784024, EPI_ISL_784025, EPI_ISL_784026, EPI_ISL_784027, EPI_ISL_784035, EPI_ISL_784040, EPI_ISL_784045, EPI_ISL_784052, EPI_ISL_784059, EPI_ISL_784060, EPI_ISL_784066, EPI_ISL_784069, EPI_ISL_784073, EPI_ISL_784076, EPI_ISL_784077, EPI_ISL_784078, EPI_ISL_784081, EPI_ISL_784085, EPI_ISL_784087, EPI_ISL_784092, EPI_ISL_784099, EPI_ISL_784106, EPI_ISL_784112, EPI_ISL_784113, EPI_ISL_784123, EPI_ISL_784134, EPI_ISL_784146, EPI_ISL_784150, EPI_ISL_784154, EPI_ISL_784162, EPI_ISL_784170, EPI_ISL_784171, EPI_ISL_784399, EPI_ISL_784400, EPI_ISL_784401, EPI_ISL_784402, EPI_ISL_784403, EPI_ISL_784410, EPI_ISL_784411, EPI_ISL_784412, EPI_ISL_784413, EPI_ISL_784414, EPI_ISL_784415, EPI_ISL_784423, EPI_ISL_784424, EPI_ISL_784425, EPI_ISL_784426, EPI_ISL_784431, EPI_ISL_784435, EPI_ISL_784436, EPI_ISL_784437, EPI_ISL_784438, EPI_ISL_784446, EPI_ISL_784447, EPI_ISL_784448, EPI_ISL_784449, EPI_ISL_784450, EPI_ISL_784454, EPI_ISL_784456, EPI_ISL_784458, EPI_ISL_784459, EPI_ISL_784460, EPI_ISL_784461, EPI_ISL_784462, EPI_ISL_784469, EPI_ISL_784470, EPI_ISL_784471, EPI_ISL_784472, EPI_ISL_784473, EPI_ISL_784474, EPI_ISL_784481, EPI_ISL_784482, EPI_ISL_784492, EPI_ISL_784509, EPI_ISL_784510, EPI_ISL_784514, EPI_ISL_784524, EPI_ISL_784568, EPI_ISL_784570, EPI_ISL_784572, EPI_ISL_784575, EPI_ISL_784577, EPI_ISL_784578, EPI_ISL_784580, EPI_ISL_784582, EPI_ISL_784583, EPI_ISL_784584, EPI_ISL_784586, EPI_ISL_784587, EPI_ISL_784588, EPI_ISL_784589, EPI_ISL_784590, EPI_ISL_784593, EPI_ISL_784594, EPI_ISL_784595, EPI_ISL_784596, EPI_ISL_784597, EPI_ISL_784600, EPI_ISL_784601, EPI_ISL_784602, EPI_ISL_784603, EPI_ISL_784605, EPI_ISL_784606, EPI_ISL_784607, EPI_ISL_784608, EPI_ISL_784609, EPI_ISL_784610, EPI_ISL_784611, EPI_ISL_784612, EPI_ISL_784613, EPI_ISL_784614, EPI_ISL_784615, EPI_ISL_784618, EPI_ISL_784619, EPI_ISL_784620, EPI_ISL_784621, EPI_ISL_784622, EPI_ISL_784623, EPI_ISL_784624, EPI_ISL_784625, EPI_ISL_784626, EPI_ISL_784627, EPI_ISL_784628, EPI_ISL_784629, EPI_ISL_784630, EPI_ISL_784631, EPI_ISL_784632, EPI_ISL_784633, EPI_ISL_784634, EPI_ISL_784635, EPI_ISL_784636, EPI_ISL_784637, EPI_ISL_784638, EPI_ISL_784641, EPI_ISL_784642, EPI_ISL_784643, EPI_ISL_784644, EPI_ISL_784645, EPI_ISL_784646, EPI_ISL_784647, EPI_ISL_784648, EPI_ISL_784649, EPI_ISL_784650, EPI_ISL_784653, EPI_ISL_784654, EPI_ISL_784655, EPI_ISL_784656, EPI_ISL_784657, EPI_ISL_784658, EPI_ISL_784659, EPI_ISL_784660, EPI_ISL_784661, EPI_ISL_784665, EPI_ISL_784666, EPI_ISL_784667, EPI_ISL_784668, EPI_ISL_784669, EPI_ISL_784670, EPI_ISL_784671, EPI_ISL_784672, EPI_ISL_784673, EPI_ISL_784674, EPI_ISL_784675, EPI_ISL_784676, EPI_ISL_784677, EPI_ISL_784678, EPI_ISL_784679, EPI_ISL_784680, EPI_ISL_784681, EPI_ISL_784682, EPI_ISL_784683, EPI_ISL_784684, EPI_ISL_784685, EPI_ISL_784686, EPI_ISL_784687, EPI_ISL_784688, EPI_ISL_784689, EPI_ISL_784690, EPI_ISL_784691, EPI_ISL_784692, EPI_ISL_784693, EPI_ISL_784694, EPI_ISL_784695, EPI_ISL_784696, EPI_ISL_784697, EPI_ISL_784698, EPI_ISL_784699, EPI_ISL_784700, EPI_ISL_784701, EPI_ISL_784702, EPI_ISL_784703, EPI_ISL_784704, EPI_ISL_784705, EPI_ISL_784706, EPI_ISL_784708, EPI_ISL_784709, EPI_ISL_784710, EPI_ISL_784711, EPI_ISL_784712, EPI_ISL_784713, EPI_ISL_784714, EPI_ISL_784715, EPI_ISL_784716, EPI_ISL_784717, EPI_ISL_784718, EPI_ISL_784719, EPI_ISL_784720, EPI_ISL_784721, EPI_ISL_784722, EPI_ISL_784723, EPI_ISL_784724, EPI_ISL_784725, EPI_ISL_784726, EPI_ISL_784727, EPI_ISL_784728, EPI_ISL_784729, EPI_ISL_784730, EPI_ISL_784731, EPI_ISL_784732, EPI_ISL_784733, EPI_ISL_784734, EPI_ISL_784735, EPI_ISL_784736, EPI_ISL_784737, EPI_ISL_784738, EPI_ISL_784739, EPI_ISL_784740, EPI_ISL_784741, EPI_ISL_784742, EPI_ISL_784743, EPI_ISL_784744, EPI_ISL_784745, EPI_ISL_784746, EPI_ISL_784747, EPI_ISL_784748, EPI_ISL_784749, EPI_ISL_784750, EPI_ISL_784751, EPI_ISL_784752, EPI_ISL_784753, EPI_ISL_784754, EPI_ISL_784755, EPI_ISL_784756, EPI_ISL_784757, EPI_ISL_784758, EPI_ISL_784759, EPI_ISL_784760, EPI_ISL_784761, EPI_ISL_784762, EPI_ISL_784763, EPI_ISL_784764, EPI_ISL_784765, EPI_ISL_784766, EPI_ISL_784767, EPI_ISL_784768, EPI_ISL_784769, EPI_ISL_784770, EPI_ISL_784771, EPI_ISL_784772, EPI_ISL_784774, EPI_ISL_784776, EPI_ISL_784778, EPI_ISL_784780, EPI_ISL_784783, EPI_ISL_784784, EPI_ISL_784785, EPI_ISL_784787, EPI_ISL_784788, EPI_ISL_784789, EPI_ISL_784791, EPI_ISL_784792, EPI_ISL_784793, EPI_ISL_784795, EPI_ISL_784796, EPI_ISL_784797, EPI_ISL_784798, EPI_ISL_784799, EPI_ISL_784800, EPI_ISL_784801, EPI_ISL_784803, EPI_ISL_784804, EPI_ISL_784805, EPI_ISL_784806, EPI_ISL_784807, EPI_ISL_784808, EPI_ISL_784809, EPI_ISL_784810, EPI_ISL_784811, EPI_ISL_784812, EPI_ISL_784816, EPI_ISL_784819, EPI_ISL_784820, EPI_ISL_784821, EPI_ISL_784822, EPI_ISL_784823, EPI_ISL_784825, EPI_ISL_784827, EPI_ISL_784829, EPI_ISL_784831, EPI_ISL_784833, EPI_ISL_784834, EPI_ISL_784835, EPI_ISL_784836, EPI_ISL_784837, EPI_ISL_784838, EPI_ISL_784840, EPI_ISL_784843, EPI_ISL_784844, EPI_ISL_784845, EPI_ISL_784846, EPI_ISL_784847, EPI_ISL_784848, EPI_ISL_784850, EPI_ISL_784851, EPI_ISL_784853, EPI_ISL_784855, EPI_ISL_784856, EPI_ISL_784858, EPI_ISL_784859, EPI_ISL_784863, EPI_ISL_784864, EPI_ISL_784867, EPI_ISL_784868, EPI_ISL_784869, EPI_ISL_784870, EPI_ISL_784872, EPI_ISL_784873, EPI_ISL_784874, EPI_ISL_784875, EPI_ISL_784878, EPI_ISL_784879, EPI_ISL_784881, EPI_ISL_784883, EPI_ISL_784885, EPI_ISL_784887, EPI_ISL_784892, EPI_ISL_784895, EPI_ISL_784896, EPI_ISL_784897, EPI_ISL_784899, EPI_ISL_784901, EPI_ISL_784902, EPI_ISL_784903, EPI_ISL_784904, EPI_ISL_784905, EPI_ISL_784908, EPI_ISL_784911, EPI_ISL_784912, EPI_ISL_784913, EPI_ISL_784915, EPI_ISL_784916, EPI_ISL_784919, EPI_ISL_784921, EPI_ISL_784923, EPI_ISL_784925, EPI_ISL_784926, EPI_ISL_784927, EPI_ISL_784928, EPI_ISL_784933, EPI_ISL_784936, EPI_ISL_784937, EPI_ISL_784938, EPI_ISL_784940, EPI_ISL_784941, EPI_ISL_784942, EPI_ISL_784943, EPI_ISL_784944, EPI_ISL_784946, EPI_ISL_784947, EPI_ISL_784948, EPI_ISL_784949, EPI_ISL_784950, EPI_ISL_784953, EPI_ISL_784956, EPI_ISL_784958, EPI_ISL_784960, EPI_ISL_784962, EPI_ISL_784964, EPI_ISL_784967, EPI_ISL_784975, EPI_ISL_784977, EPI_ISL_784987, EPI_ISL_784992, EPI_ISL_784998, EPI_ISL_785000, EPI_ISL_785007, EPI_ISL_785012, EPI_ISL_785013, EPI_ISL_785016, EPI_ISL_785023, EPI_ISL_785025, EPI_ISL_785026, EPI_ISL_785027, EPI_ISL_785028, EPI_ISL_785034, EPI_ISL_785038, EPI_ISL_785048, EPI_ISL_785049, EPI_ISL_785050, EPI_ISL_785051, EPI_ISL_785052, EPI_ISL_785074, EPI_ISL_785106, EPI_ISL_785122, EPI_ISL_786091, EPI_ISL_786092, EPI_ISL_786093, EPI_ISL_786094, EPI_ISL_786095, EPI_ISL_786096, EPI_ISL_786097, EPI_ISL_786098, EPI_ISL_786099, EPI_ISL_786100, EPI_ISL_786101, EPI_ISL_786102, EPI_ISL_786103, EPI_ISL_786104, EPI_ISL_786105, EPI_ISL_786106, EPI_ISL_786107, EPI_ISL_786108, EPI_ISL_786109, EPI_ISL_786110, EPI_ISL_786111, EPI_ISL_786112, EPI_ISL_786113, EPI_ISL_786114, EPI_ISL_786115, EPI_ISL_786116, EPI_ISL_786117, EPI_ISL_786118, EPI_ISL_786119, EPI_ISL_786120, EPI_ISL_786121, EPI_ISL_786122, EPI_ISL_786123, EPI_ISL_786124, EPI_ISL_786125, EPI_ISL_786126, EPI_ISL_786127, EPI_ISL_786128, EPI_ISL_786129, EPI_ISL_786130, EPI_ISL_786131, EPI_ISL_786132, EPI_ISL_786133, EPI_ISL_786134, EPI_ISL_786135, EPI_ISL_786136, EPI_ISL_786137, EPI_ISL_786138, EPI_ISL_786139, EPI_ISL_786140, EPI_ISL_786141, EPI_ISL_786142, EPI_ISL_786143, EPI_ISL_786144, EPI_ISL_786145, EPI_ISL_786146, EPI_ISL_786147, EPI_ISL_786148, EPI_ISL_786149, EPI_ISL_786150, EPI_ISL_786151, EPI_ISL_786152, EPI_ISL_786153, EPI_ISL_786154, EPI_ISL_786155, EPI_ISL_786156, EPI_ISL_786157, EPI_ISL_786158, EPI_ISL_786159, EPI_ISL_786160, EPI_ISL_786161, EPI_ISL_786162, EPI_ISL_786163, EPI_ISL_786164, EPI_ISL_786165, EPI_ISL_786166, EPI_ISL_786167, EPI_ISL_786168, EPI_ISL_786169, EPI_ISL_786170, EPI_ISL_786171, EPI_ISL_786172, EPI_ISL_786173, EPI_ISL_786174, EPI_ISL_786175, EPI_ISL_786176, EPI_ISL_786177, EPI_ISL_786178, EPI_ISL_786179, EPI_ISL_786180, EPI_ISL_786181, EPI_ISL_786182, EPI_ISL_786183, EPI_ISL_786184, EPI_ISL_786185, EPI_ISL_786186, EPI_ISL_786187, EPI_ISL_786188, EPI_ISL_786189, EPI_ISL_786190, EPI_ISL_786191 | | | |
| see above | Houston Methodist Hospital | Houston Methodist Hospital | S. Wesley Long, Randall J. Olsen, Paul A. Christensen, David W. Bernard, James J. Davis, Maulik Shukla, Marcus Nguyen, Matthew Ojeda Saavedra, Prasanti Yerramilli, Layne Pruitt, Sishir Subedi, Heather Hendrickson, and James M. Musser |
| EPI_ISL_790631, EPI_ISL_790674 | Dutch COVID-19 response team | National Institute for Public Health and the Environment (RIVM) | Adam Meijer, Harry Vennema, Jeroen Cremer, Sharon van den Brink, Bas van der Veer, AnneMarie van den Brandt, Florian Zwagemaker, Dennis Schmitz, Chantal Reusken, on behalf of the national COVID-19 response team |
| EPI_ISL_791298 | National Virus Reference Laboratory | Irish Coronavirus Sequencing Consortium - Teagasc Moorepark | Alejandro Abner Garcia Leon, Paul Cotter, Fiona Crispie, John Kenny, Paddy Mallon, Calum Walsh |
| EPI_ISL_791366, EPI_ISL_791369, EPI_ISL_791380, EPI_ISL_791407, EPI_ISL_791409, EPI_ISL_791410 | Johns Hopkins Hospital Department of Pathology | Johns Hopkins Hospital Department of Pathology | C. Paul Morris, Chun Huai Luo, Heba H. Mostafa |
| EPI_ISL_792558 | Grubaugh Lab - Yale School of Public Health | Grubaugh Lab - Yale School of Public Health | Joseph Fauver, Tara Alpert, Anderson Brito, Nathan Grubaugh |
| EPI_ISL_792649 | LACEN-PR | Laboratory of Respiratory Viruses and Measles, Oswaldo Cruz Institute, FIOCRUZ | Paola Resende, Luciana Appolinario, Fernando Motta, Anna Carolina Paixao, Ana Carolina Mendonca, Maria do Carmo Debur, Irina Nastassja Riediger, Marilda Siqueira |
| EPI_ISL_792686, EPI_ISL_792690, EPI_ISL_792693, EPI_ISL_792694, EPI_ISL_792705, EPI_ISL_792706 | The National Institute of Public Health | State Veterinary Institute Prague | Nagy,A;Jirincova,H;Trnka,D;Vecerova,J |
| EPI_ISL_792828, EPI_ISL_793012 | Department of Virus and Microbiological Special Diagnostics, Statens Serum Institut, Copenhagen, Denmark | Albertsen Lab, Department of Chemistry and Bioscience, Aalborg University, Denmark | Danish Covid-19 Genome Consortium |
| EPI_ISL_796777 | Instituto Nacional de Saude (INSA) | Instituto Nacional de Saude (INSA) | Borges et al |
| EPI_ISL_800897, EPI_ISL_800920, EPI_ISL_800924, EPI_ISL_800967, EPI_ISL_800971, EPI_ISL_800995, EPI_ISL_801002, EPI_ISL_801055, EPI_ISL_801063, EPI_ISL_801110, EPI_ISL_801125, EPI_ISL_801148 | | | |
| see above | Lighthouse Lab in Cambridge | Wellcome Sanger Institute for the COVID-19 Genomics UK (COG-UK) Consortium | Rob Howes, The Lighthouse Lab in Cambridge and Alex Alderton, Roberto Amato, Sonia Goncalves, Ewan Harrison, David K. Jackson, Ian Johnston, Dominic Kwiatkowski, Cordelia Langford, John Sillitoe on behalf of the Wellcome Sanger Institute COVID-19 Surveillance Team |
| EPI_ISL_801182, EPI_ISL_801183, EPI_ISL_801184, EPI_ISL_801185, EPI_ISL_801186, EPI_ISL_801187, EPI_ISL_801188, EPI_ISL_801189, EPI_ISL_801190, EPI_ISL_801191, EPI_ISL_801192, EPI_ISL_801193, EPI_ISL_801194, EPI_ISL_801195, EPI_ISL_801196, EPI_ISL_801197, EPI_ISL_801198, EPI_ISL_801199, EPI_ISL_801200, EPI_ISL_801201, EPI_ISL_801202, EPI_ISL_801203, EPI_ISL_801204, EPI_ISL_801205, EPI_ISL_801206, EPI_ISL_801207, EPI_ISL_801208, EPI_ISL_801209, EPI_ISL_801210, EPI_ISL_801211, EPI_ISL_801212, EPI_ISL_801213, EPI_ISL_801214, EPI_ISL_801215, EPI_ISL_801216, EPI_ISL_801217, EPI_ISL_801218, EPI_ISL_801219, EPI_ISL_801220, EPI_ISL_801221, EPI_ISL_801222, EPI_ISL_801223, EPI_ISL_801224, EPI_ISL_801225, EPI_ISL_801226, EPI_ISL_801227, EPI_ISL_801228, EPI_ISL_801229, EPI_ISL_801230 | | | |
| see above | Lighthouse Lab in Milton Keynes | Wellcome Sanger Institute for the COVID-19 Genomics UK (COG-UK) Consortium | The Lighthouse Lab in Milton Keynes and Alex Alderton, Roberto Amato, Sonia Goncalves, Ewan Harrison, David K. Jackson, Ian Johnston, Dominic Kwiatkowski, Cordelia Langford, John Sillitoe on behalf of the Wellcome Sanger Institute COVID-19 Surveillance Team |
| EPI_ISL_802855 | Vilnius University Hospital Santaros Klinikos, Vilnius University | Institute of Biotechnology, Life Sciences Center, Vilnius University | Emilija Vasiliunaite, Milda Norkiene, Albertas Timinskas, Alma Gedvilaite, Aurelija Zvirbliene, Daniel Naumovas, Laimonas Griskevicius |
| EPI_ISL_803101, EPI_ISL_803104 | Respiratory Viruses Branch, Centers for Disease Control and Prevention | Respiratory Viruses Branch, Centers for Disease Control and Prevention | Queen,K., Li,Y., Tao,Y., Uehara,A., Montmayeur,A., Paden,C.R., Cook,P.W., Marine,R., Sheth,M., Wang,H., Lee,J., Tong,S. |
| EPI_ISL_804582 | MEPHI, Aix Marseille University | MEPHI, Aix Marseille University | Anthony LEVASSEUR |
| EPI_ISL_804602, EPI_ISL_804608, EPI_ISL_804614, EPI_ISL_804615, EPI_ISL_804624, EPI_ISL_804631, EPI_ISL_804632, EPI_ISL_804634, EPI_ISL_804635, EPI_ISL_804641, EPI_ISL_804644, EPI_ISL_804645, EPI_ISL_804650, EPI_ISL_804651, EPI_ISL_804655, EPI_ISL_804657, EPI_ISL_804658, EPI_ISL_804661, EPI_ISL_804662, EPI_ISL_804666, EPI_ISL_804671, EPI_ISL_804673, EPI_ISL_804674, EPI_ISL_804675, EPI_ISL_804678, EPI_ISL_804682, EPI_ISL_804684, EPI_ISL_804688, EPI_ISL_804690, EPI_ISL_804692, EPI_ISL_804693, EPI_ISL_804698, EPI_ISL_804699, EPI_ISL_804702, EPI_ISL_804704, EPI_ISL_804710, EPI_ISL_804743, EPI_ISL_804754, EPI_ISL_804756, EPI_ISL_804757, EPI_ISL_804758, EPI_ISL_804759, EPI_ISL_804779, EPI_ISL_804781, EPI_ISL_804786, EPI_ISL_804790, EPI_ISL_804792, EPI_ISL_804806, EPI_ISL_804808 | | | |
| see above | Michigan Department of Health and Human Services, Bureau of Laboratories | Michigan Department of Health and Human Services, Bureau of Laboratories | Blankenship HM, Riner D, Soehnlen MK |
| EPI_ISL_806819, EPI_ISL_806820, EPI_ISL_806821 | Alaska State Virology Laboratory | Alaska State Virology Laboratory | Stephanie DeRonde, Lisa Smith, Ph.D., Devin M. Drown, Ph.D., Jack Chen, Ph.D. |
| EPI_ISL_807015, EPI_ISL_807021, EPI_ISL_807055, EPI_ISL_807056, EPI_ISL_807057, EPI_ISL_807058, EPI_ISL_807059, EPI_ISL_807060, EPI_ISL_807061, EPI_ISL_807062, EPI_ISL_807063, EPI_ISL_807064, EPI_ISL_807065, EPI_ISL_807066, EPI_ISL_807067, EPI_ISL_807068, EPI_ISL_807069, EPI_ISL_807070, EPI_ISL_807071, EPI_ISL_807072, EPI_ISL_807073, EPI_ISL_807074, EPI_ISL_807075, EPI_ISL_807076, EPI_ISL_807077, EPI_ISL_807078, EPI_ISL_807079, EPI_ISL_807080, EPI_ISL_807081, EPI_ISL_807082, EPI_ISL_807083, EPI_ISL_807084, EPI_ISL_807085, EPI_ISL_807086, EPI_ISL_807087, EPI_ISL_807089, EPI_ISL_807090, EPI_ISL_807091, EPI_ISL_807092, EPI_ISL_807093, EPI_ISL_807095, EPI_ISL_807096, EPI_ISL_807097, EPI_ISL_807098, EPI_ISL_807099, EPI_ISL_807100, EPI_ISL_807101, EPI_ISL_807102, EPI_ISL_807103, EPI_ISL_807104, EPI_ISL_807105, EPI_ISL_807106, EPI_ISL_807107, EPI_ISL_807109, | | | |

| | | | |
|---|---|---|---|
| EPI_ISL_807110, EPI_ISL_807111, EPI_ISL_807112, EPI_ISL_807115 | | | |
| see above | Washington State Department of Health | Seattle Flu Study | Deborah A. Nickerson, Chris D. Frazar, Jover Lee, Benjamin Pelle, Matthew Richardson, Amanda Adler, Elisabeth Brandstetter, Peter D. Han, Kairsten Fay, Misja Ilcisin, Kirsten Lacombe, Thomas R. Sibley, Melissa Truong, Caitlin R. Wolf, Romesh Gautom, Geoff Melly, Brian Hiatt, Philip Dykema, Scott Lindquist, Michael Boeckh, Janet A. Englund, Michael Famulare, Barry R. Lutz, Mark J. Rieder, Lea M. Starita, Matthew Thompson, Helen Y. Chu, Jay Shendure, Trevor Bedford |
| EPI_ISL_812132, EPI_ISL_812139 | TN Division of Laboratory Services | Pathogen Discovery, Respiratory Viruses Branch, Division of Viral Diseases, Centers for Disease Control and Prevention | Yan Li, Ying Tao, Anna Montmayeur, Jing Zhang, Brian Lynch, Krista Queen, Anna Uehara, Rachel Marine, Peter Cook, Clinton R. Paden, Haibin Wang, Suxiang Tong |
| EPI_ISL_812151, EPI_ISL_812152, EPI_ISL_812153, EPI_ISL_812154, EPI_ISL_812155, EPI_ISL_812197 | GA Department of Public Health Laboratory | Pathogen Discovery, Respiratory Viruses Branch, Division of Viral Diseases, Centers for Disease Control and Prevention | Yan Li, Ying Tao, Anna Montmayeur, Jing Zhang, Brian Lynch, Krista Queen, Anna Uehara, Rachel Marine, Peter Cook, Clinton R. Paden, Haibin Wang, Suxiang Tong |
| EPI_ISL_812204, EPI_ISL_812205, EPI_ISL_812206, EPI_ISL_812207, EPI_ISL_812208, EPI_ISL_812209, EPI_ISL_812210, EPI_ISL_812211, EPI_ISL_812212, EPI_ISL_812213, EPI_ISL_812214, EPI_ISL_812215, EPI_ISL_812216, EPI_ISL_812217, EPI_ISL_812218, EPI_ISL_812220, EPI_ISL_812221, EPI_ISL_812222, EPI_ISL_812223, EPI_ISL_812224, EPI_ISL_812225, EPI_ISL_812226, EPI_ISL_812227, EPI_ISL_812228, EPI_ISL_812229, EPI_ISL_812230, EPI_ISL_812231 | | | |
| see above | TN Division of Laboratory Services | Pathogen Discovery, Respiratory Viruses Branch, Division of Viral Diseases, Centers for Disease Control and Prevention | Yan Li, Ying Tao, Anna Montmayeur, Jing Zhang, Brian Lynch, Krista Queen, Anna Uehara, Rachel Marine, Peter Cook, Clinton R. Paden, Haibin Wang, Suxiang Tong |
| EPI_ISL_813751 | Liverpool Clinical Laboratories | COVID-19 Genomics UK (COG-UK) Consortium | Sam Haldenby, Anita Lucaci, Steve Paterson, Julian Hiscox, Alistair Darby, M Almsaud, A Alrezaihi, Muhannad Alruwaili, Stuart D Armstrong, Jones Benjamin, Eleanor G Bentley, Anu Chawla, Jordan J Clark, Angela Cowell, Richard Eccles, Isabel García-Dorival, Matthew Gemmell, Alessandro Gerada, PKF Gilmore, Richard Gregory, Ximeng Han, Catherine Hartley, Margaret Hughes, Miren Iturriza-Gomara, James Johnson, L Luu, Jenifer Manson, Charlotte Nelson, Elaine O'Toole, Cassie Olateju, Rebekah Penrice-Randal , Lucille Rainbow, N.P Randle, Trevor Ian Robinson, Parul Sharma, Ghada T Shawli, James P Stewart, Neil Swainston, Ecaterina Vamos, Joanne Watts, Mark Whitehead |
| EPI_ISL_814534, EPI_ISL_814571 | Virology Department, Royal Infirmary of Edinburgh, NHS Lothian / School of Biological Sciences, University of Edinburgh / Institute of Genetics and Molecular Medicine, University of Edinburgh | COVID-19 Genomics UK (COG-UK) Consortium | McHugh M, Dewar R, Rooke S, Gallagher M, Balcaza C, O'Toole Á, Scher E, Hill V, McCrone JT, Colquhoun R, Yu X, Jackson B, Rambaut A, Williams TC, Templeton K |
| EPI_ISL_814796, EPI_ISL_814797 | Wales Specialist Virology Centre Sequencing lab: Pathogen Genomics Unit | COVID-19 Genomics UK (COG-UK) Consortium | Catherine Moore, Johnathan Evans, Laura Gifford, Malorie Perry, Simon Cottrell, Angela Marchbank, Alec Birchley, Alexander Adams, Amy Gaskin, Bree Gatica-Wilcox, Jason Coombes, Joel Southgate, Lauren Gilbert, Lee Graham, Nicole Pacchiarini, Sara Kumziene-Summerhayes, Sarah Taylor, Sophie Jones, Sara Rey, Matthew Bull, Joanne Watkins, Sally Corden, Tom Connor |
| EPI_ISL_815463, EPI_ISL_815466, EPI_ISL_815484, EPI_ISL_815558, EPI_ISL_815600, EPI_ISL_816095, EPI_ISL_816096, EPI_ISL_816097, EPI_ISL_816098, EPI_ISL_816099, EPI_ISL_816100, EPI_ISL_816101, EPI_ISL_816102, EPI_ISL_816103, EPI_ISL_816104, EPI_ISL_816105, EPI_ISL_816106, EPI_ISL_816107, EPI_ISL_816108, EPI_ISL_816109, EPI_ISL_816110 | | | |
| see above | Department of Virus and Microbiological Special Diagnostics, Statens Serum Institut, Copenhagen, Denmark | Albertsen Lab, Department of Chemistry and Bioscience, Aalborg University, Denmark | Danish Covid-19 Genome Consortium |
| EPI_ISL_816324, EPI_ISL_816453 | Virology Department, Sheffield Teaching Hospitals NHS Foundation Trust/Department of Infection, Immunity and Cardiovascular Disease, The Medical School, University of Sheffield | COVID-19 Genomics UK (COG-UK) Consortium | Thushan de Silva, Matthew Parker, Nikki Smith, Adri Angyal, Rebecca Brown, Luke Green, Rachel Tucker, Paul Parsons, Danielle Groves, Katie Johnson, Laura Carrilero, Alex Keeley, Dave Partridge, Matthew Wyles, Benjamin Lindsey, Mehmet Yavuz, Mohammad Raza, Cariad Evans |
| EPI_ISL_823970 | Department of Homeless Services | New York City Public Health Laboratory | Jade Wang, et al. |
| EPI_ISL_823971 | OCME Office Of Chief Medical Examiner | New York City Public Health Laboratory | Jade Wang, et al. |
| EPI_ISL_825035 | B.J. Medical College and Civil hospital, Ahmedabad | Gujarat Biotechnology Research Centre | Nilima Shah, Sanjay Kapadia, Nitin Savaliya, Dinesh Kumar, Zuber Saiyed, Labdhi Pandya, Afzal Ansari, Nikha Trivedi, Apurvasinh Puvar, Ramesh Pandit, Janvi Raval, Zarna Patel, Kamlesh J Upadhyay, Pranay Shah, Dipa Kinariwala, Chaitanya Joshi, Madhvi Joshi |
| EPI_ISL_825092, EPI_ISL_825096, EPI_ISL_825097, EPI_ISL_825098, EPI_ISL_825099, EPI_ISL_825101, EPI_ISL_825104, EPI_ISL_825106, EPI_ISL_825107, EPI_ISL_825108 | NHLS-IALCH | KRISP, KZN Research Innovation and Sequencing Platform | Giandhari J, Pillay S, Lessells R, Mdlalose K, York D, Khan S, Tegally H, Wilkinson E, de Oliveira T |
| EPI_ISL_825330, EPI_ISL_825331, EPI_ISL_825358, EPI_ISL_825359, EPI_ISL_825360 | Hospital Universitari Vall d'Hebron - Vall d'Hebron Institut de Rercerca | Hospital Universitari Vall d'Hebron | Cristina Andrés, Maria Piñana, Josep F Abril, Damir Garcia-Cehic, Ariadna Rando, Juliana Esperalba, Maria Gema Codina, Carla Castillo, Maria Carmen Martin, Tomàs Pumarola, Josep Quer, Andrés Antón |
| EPI_ISL_825597, EPI_ISL_825598, EPI_ISL_825599, EPI_ISL_825600, EPI_ISL_825601, EPI_ISL_825602, EPI_ISL_825603, EPI_ISL_825604 | Respiratory Virus Unit, National Infection Service, Public Health England | COVID-19 Genomics UK (COG-UK) Consortium | PHE Covid Sequencing Team |
| EPI_ISL_826569 | Texas Department of State Health Services | Texas Department of State Health Services | Rashmi Tuladhar, Bonnie Oh, Jenny Zhang, Maliha Rahman, Anita Pokharel, Myong Koag, Chung Wang, Rachel Lee, Grace Kubin, Mayela Pedrueza, James Daniel Bonser |
| EPI_ISL_826723, EPI_ISL_826995, EPI_ISL_826998, EPI_ISL_827267 | deCODE genetics | deCODE genetics | Daniel F Gudbjartsson; Agnar Helgason; Hakon Jonsson; Olafur T Magnusson; Pall Melsted; Gudmundur L Norddahl; Jona Saemundsdottir; Asgeir Sigurdsson; Patrick Sulem; Arna B Agustsdottir; Hannes Eggertsson; Berglind Eiriksdottir; Run Fridriksdottir; Elisabet E Gardarsdottir; Gudmundur Georgsson; Olafia S Gretarsdottir; Kjartan R Gudmundsson; Thora R Gunnarsdottir; Arnaldur Gylfason; Hilma Holm; Brynjar O Jensson; Aslaug Jonasdottir; Kamilla S Josefsdottir; Thordur Kristjansson; Droplaug N Magnusdottir; Solvi Rognvaldsson; Louise le Roux; Gudrun Sigmundsdottir; Gardar Sveinbjornsson; Kristin E Sveinsdottir; Maney Sveinsdottir; Emil A Thorarensen; Bjarni Thorbjornsson; Gisli Masson; Ingileif Jonsdottir; Alma Moller; Thorolfur Gudnason; Karl G Kristinsson; Unnur Thorsteinsdottir; Kari Stefansson |
| EPI_ISL_827395 | The National University Hospital of Iceland | deCODE genetics | Daniel F Gudbjartsson; Agnar Helgason; Hakon Jonsson; Olafur T Magnusson; Pall Melsted; Gudmundur L Norddahl; Jona Saemundsdottir; Asgeir Sigurdsson; Patrick Sulem; Arna B Agustsdottir; Hannes Eggertsson; Berglind Eiriksdottir; Run Fridriksdottir; Elisabet E Gardarsdottir; Gudmundur Georgsson; Olafia S Gretarsdottir; Kjartan R Gudmundsson; Thora R Gunnarsdottir; Arnaldur Gylfason; Hilma Holm; Brynjar O Jensson; Aslaug Jonasdottir; Kamilla S Josefsdottir; Thordur Kristjansson; Droplaug N Magnusdottir; Solvi Rognvaldsson; Louise le Roux; Gudrun Sigmundsdottir; Gardar Sveinbjornsson; Kristin E Sveinsdottir; Maney Sveinsdottir; Emil A Thorarensen; Bjarni Thorbjornsson; Gisli Masson; Ingileif Jonsdottir; Alma Moller; Thorolfur Gudnason; Karl G Kristinsson; Unnur Thorsteinsdottir; Kari Stefansson |
| EPI_ISL_827397, EPI_ISL_827398, EPI_ISL_827399, EPI_ISL_827400, EPI_ISL_827402, EPI_ISL_827442, EPI_ISL_827458, EPI_ISL_827830, EPI_ISL_827852, EPI_ISL_827861, EPI_ISL_828110 | | | |
| see above | deCODE genetics | deCODE genetics | Daniel F Gudbjartsson; Agnar Helgason; Hakon Jonsson; Olafur T Magnusson; Pall Melsted; Gudmundur L Norddahl; Jona Saemundsdottir; Asgeir Sigurdsson; Patrick Sulem; Arna B Agustsdottir; Hannes Eggertsson; Berglind Eiriksdottir; Run Fridriksdottir; Elisabet E Gardarsdottir; Gudmundur Georgsson; Olafia S Gretarsdottir; Kjartan R Gudmundsson; Thora R Gunnarsdottir; Arnaldur Gylfason; Hilma Holm; Brynjar O Jensson; Aslaug Jonasdottir; Kamilla S Josefsdottir; Thordur Kristjansson; Droplaug N Magnusdottir; Solvi Rognvaldsson; Louise le Roux; Gudrun Sigmundsdottir; Gardar Sveinbjornsson; Kristin E Sveinsdottir; Maney Sveinsdottir; Emil A Thorarensen; Bjarni Thorbjornsson; Gisli Masson; Ingileif Jonsdottir; Alma Moller; Thorolfur Gudnason; Karl G Kristinsson; Unnur Thorsteinsdottir; Kari Stefansson |
| EPI_ISL_828802, EPI_ISL_828804 | The National University Hospital of Iceland | deCODE genetics | Daniel F Gudbjartsson; Agnar Helgason; Hakon Jonsson; Olafur T Magnusson; Pall Melsted; Gudmundur L Norddahl; Jona Saemundsdottir; Asgeir Sigurdsson; Patrick Sulem; Arna B Agustsdottir; Hannes Eggertsson; Berglind Eiriksdottir; Run Fridriksdottir; Elisabet E Gardarsdottir; Gudmundur Georgsson; Olafia S Gretarsdottir; Kjartan R Gudmundsson; Thora R Gunnarsdottir; Arnaldur Gylfason; Hilma Holm; Brynjar O Jensson; Aslaug |

| | | | |
|---|---|---|---|
| | | | Jonasdottir; Kamilla S Josefsdottir; Thordur Kristjansson; Droplaug N Magnusdottir; Solvi Rognvaldsson; Louise le Roux; Gudrun Sigmundsdottir; Gardar Sveinbjornsson; Kristin E Sveinsdottir; Maney Sveinsdottir; Emil A Thorarensen; Bjarni Thorbjornsson; Gisli Masson; Ingileif Jonsdottir; Alma Moller; Thorolfur Gudnason; Karl G Kristinsson; Unnur Thorsteinsdottir; Kari Stefansson |
| EPI_ISL_828964, EPI_ISL_829041 | deCODE genetics | deCODE genetics | Daniel F Gudbjartsson; Agnar Helgason; Hakon Jonsson; Olafur T Magnusson; Pall Melsted; Gudmundur L Norddahl; Jona Saemundsdottir; Asgeir Sigurdsson; Patrick Sulem; Arna B Agustsdottir; Hannes Eggertsson; Berglind Eiriksdottir; Run Fridriksdottir; Elisabet E Gardarsdottir; Gudmundur Georgsson; Olafia S Gretarsdottir; Kjartan R Gudmundsson; Thora R Gunnarsdottir; Arnaldur Gylfason; Hilma Holm; Brynjar O Jensson; Aslaug Jonasdottir; Kamilla S Josefsdottir; Thordur Kristjansson; Droplaug N Magnusdottir; Solvi Rognvaldsson; Louise le Roux; Gudrun Sigmundsdottir; Gardar Sveinbjornsson; Kristin E Sveinsdottir; Maney Sveinsdottir; Emil A Thorarensen; Bjarni Thorbjornsson; Gisli Masson; Ingileif Jonsdottir; Alma Moller; Thorolfur Gudnason; Karl G Kristinsson; Unnur Thorsteinsdottir; Kari Stefansson |
| EPI_ISL_829105 | The National University Hospital of Iceland | deCODE genetics | Daniel F Gudbjartsson; Agnar Helgason; Hakon Jonsson; Olafur T Magnusson; Pall Melsted; Gudmundur L Norddahl; Jona Saemundsdottir; Asgeir Sigurdsson; Patrick Sulem; Arna B Agustsdottir; Hannes Eggertsson; Berglind Eiriksdottir; Run Fridriksdottir; Elisabet E Gardarsdottir; Gudmundur Georgsson; Olafia S Gretarsdottir; Kjartan R Gudmundsson; Thora R Gunnarsdottir; Arnaldur Gylfason; Hilma Holm; Brynjar O Jensson; Aslaug Jonasdottir; Kamilla S Josefsdottir; Thordur Kristjansson; Droplaug N Magnusdottir; Solvi Rognvaldsson; Louise le Roux; Gudrun Sigmundsdottir; Gardar Sveinbjornsson; Kristin E Sveinsdottir; Maney Sveinsdottir; Emil A Thorarensen; Bjarni Thorbjornsson; Gisli Masson; Ingileif Jonsdottir; Alma Moller; Thorolfur Gudnason; Karl G Kristinsson; Unnur Thorsteinsdottir; Kari Stefansson |
| EPI_ISL_829198, EPI_ISL_829235 | deCODE genetics | deCODE genetics | Daniel F Gudbjartsson; Agnar Helgason; Hakon Jonsson; Olafur T Magnusson; Pall Melsted; Gudmundur L Norddahl; Jona Saemundsdottir; Asgeir Sigurdsson; Patrick Sulem; Arna B Agustsdottir; Hannes Eggertsson; Berglind Eiriksdottir; Run Fridriksdottir; Elisabet E Gardarsdottir; Gudmundur Georgsson; Olafia S Gretarsdottir; Kjartan R Gudmundsson; Thora R Gunnarsdottir; Arnaldur Gylfason; Hilma Holm; Brynjar O Jensson; Aslaug Jonasdottir; Kamilla S Josefsdottir; Thordur Kristjansson; Droplaug N Magnusdottir; Solvi Rognvaldsson; Louise le Roux; Gudrun Sigmundsdottir; Gardar Sveinbjornsson; Kristin E Sveinsdottir; Maney Sveinsdottir; Emil A Thorarensen; Bjarni Thorbjornsson; Gisli Masson; Ingileif Jonsdottir; Alma Moller; Thorolfur Gudnason; Karl G Kristinsson; Unnur Thorsteinsdottir; Kari Stefansson |
| EPI_ISL_829617 | The National University Hospital of Iceland | deCODE genetics | Daniel F Gudbjartsson; Agnar Helgason; Hakon Jonsson; Olafur T Magnusson; Pall Melsted; Gudmundur L Norddahl; Jona Saemundsdottir; Asgeir Sigurdsson; Patrick Sulem; Arna B Agustsdottir; Hannes Eggertsson; Berglind Eiriksdottir; Run Fridriksdottir; Elisabet E Gardarsdottir; Gudmundur Georgsson; Olafia S Gretarsdottir; Kjartan R Gudmundsson; Thora R Gunnarsdottir; Arnaldur Gylfason; Hilma Holm; Brynjar O Jensson; Aslaug Jonasdottir; Kamilla S Josefsdottir; Thordur Kristjansson; Droplaug N Magnusdottir; Solvi Rognvaldsson; Louise le Roux; Gudrun Sigmundsdottir; Gardar Sveinbjornsson; Kristin E Sveinsdottir; Maney Sveinsdottir; Emil A Thorarensen; Bjarni Thorbjornsson; Gisli Masson; Ingileif Jonsdottir; Alma Moller; Thorolfur Gudnason; Karl G Kristinsson; Unnur Thorsteinsdottir; Kari Stefansson |
| EPI_ISL_829726, EPI_ISL_829727, EPI_ISL_829728, EPI_ISL_829729, EPI_ISL_829730, EPI_ISL_829731, EPI_ISL_829976, EPI_ISL_829980, EPI_ISL_829983, EPI_ISL_829985, EPI_ISL_830157, EPI_ISL_830158, EPI_ISL_830159, EPI_ISL_830160, EPI_ISL_830161 | | | |
| see above | deCODE genetics | deCODE genetics | Daniel F Gudbjartsson; Agnar Helgason; Hakon Jonsson; Olafur T Magnusson; Pall Melsted; Gudmundur L Norddahl; Jona Saemundsdottir; Asgeir Sigurdsson; Patrick Sulem; Arna B Agustsdottir; Hannes Eggertsson; Berglind Eiriksdottir; Run Fridriksdottir; Elisabet E Gardarsdottir; Gudmundur Georgsson; Olafia S Gretarsdottir; Kjartan R Gudmundsson; Thora R Gunnarsdottir; Arnaldur Gylfason; Hilma Holm; Brynjar O Jensson; Aslaug Jonasdottir; Kamilla S Josefsdottir; Thordur Kristjansson; Droplaug N Magnusdottir; Solvi Rognvaldsson; Louise le Roux; Gudrun Sigmundsdottir; Gardar Sveinbjornsson; Kristin E Sveinsdottir; Maney Sveinsdottir; Emil A Thorarensen; Bjarni Thorbjornsson; Gisli Masson; Ingileif Jonsdottir; Alma Moller; Thorolfur Gudnason; Karl G Kristinsson; Unnur Thorsteinsdottir; Kari Stefansson |
| EPI_ISL_830235, EPI_ISL_830237, EPI_ISL_830240, EPI_ISL_830242, EPI_ISL_830245 | The National University Hospital of Iceland | deCODE genetics | Daniel F Gudbjartsson; Agnar Helgason; Hakon Jonsson; Olafur T Magnusson; Pall Melsted; Gudmundur L Norddahl; Jona Saemundsdottir; Asgeir Sigurdsson; Patrick Sulem; Arna B Agustsdottir; Hannes Eggertsson; Berglind Eiriksdottir; Run Fridriksdottir; Elisabet E Gardarsdottir; Gudmundur Georgsson; Olafia S Gretarsdottir; Kjartan R Gudmundsson; Thora R Gunnarsdottir; Arnaldur Gylfason; Hilma Holm; Brynjar O Jensson; Aslaug Jonasdottir; Kamilla S Josefsdottir; Thordur Kristjansson; Droplaug N Magnusdottir; Solvi Rognvaldsson; Louise le Roux; Gudrun Sigmundsdottir; Gardar Sveinbjornsson; Kristin E Sveinsdottir; Maney Sveinsdottir; Emil A Thorarensen; Bjarni Thorbjornsson; Gisli Masson; Ingileif Jonsdottir; Alma Moller; Thorolfur Gudnason; Karl G Kristinsson; Unnur Thorsteinsdottir; Kari Stefansson |
| EPI_ISL_831032, EPI_ISL_831202, EPI_ISL_831205, EPI_ISL_831206 | Hospital Universitario La Paz (Madrid) | SeqCOVID-SPAIN consortium/IBV(CSIC) | María Rodríguez-Tejedor, Elias Dahdouh, Fernando Lázaro-Perona, Jesús Mingorance and SeqCOVID-SPAIN consortium |
| EPI_ISL_831707, EPI_ISL_831717, EPI_ISL_831720, EPI_ISL_831721, EPI_ISL_831722, EPI_ISL_831724, EPI_ISL_831725, EPI_ISL_831726, EPI_ISL_831727, EPI_ISL_831728, EPI_ISL_831729, EPI_ISL_831744, EPI_ISL_831753, EPI_ISL_831755, EPI_ISL_831756, EPI_ISL_831803, EPI_ISL_831804, EPI_ISL_831805, EPI_ISL_831806, EPI_ISL_831807, EPI_ISL_831808, EPI_ISL_831809, EPI_ISL_831810, EPI_ISL_831811, EPI_ISL_831812, EPI_ISL_831822, EPI_ISL_831823, EPI_ISL_831824 | | | |
| see above | United States Air Force School of Aerospace Medicine | United States Air Force School of Aerospace Medicine | Anthony Fries, Jennifer Meyer, William Gruner, Amanda Javorina, Sarah Purves, Clarise Starr, Elizabeth Macias |
| EPI_ISL_832758, EPI_ISL_832759, EPI_ISL_832760, EPI_ISL_832776, EPI_ISL_832777, EPI_ISL_832778, EPI_ISL_832779, EPI_ISL_832780, EPI_ISL_832783, EPI_ISL_832784, EPI_ISL_832785, EPI_ISL_832786, EPI_ISL_832787, EPI_ISL_832788, EPI_ISL_832789, EPI_ISL_832790, EPI_ISL_832791, EPI_ISL_832792, EPI_ISL_832793 | | | |
| see above | OHSU Lab Services Molecular Microbiology Lab | Oregon SARS-CoV-2 Genome Sequencing Center | Brendan L. O'Connell, Ruth V. Nichols, Sally Grindstaff, Alex J. Hirsch, Donna Hansel, Guang Fan, Daniel N. Streblow, William B. Messer, Andrew C. Adey, Benjamin N. Bimber, Brian J. O'Roak |
| EPI_ISL_833157 | Instituto Adolfo Lutz - Regional de Santo Andre | Instituto Adolfo Lutz, Interdiciplinary Procedures Center, Strategic Laboratory | Claudio Tavares Sacchi, Claudia Regina Gonçalves, Erica Valessa Ramos Gomes, Karoline Rodrigues Campos |
| EPI_ISL_833163 | Instituto Adolfo Lutz - Regional de Marilia | Instituto Adolfo Lutz, Interdiciplinary Procedures Center, Strategic Laboratory | Claudio Tavares Sacchi, Claudia Regina Gonçalves, Erica Valessa Ramos Gomes, Karoline Rodrigues Campos |
| EPI_ISL_833463, EPI_ISL_833471 | CHU Purpan - Laboratoire de Virologie - Institut Fédératif de Biologie | CHU Purpan - Laboratoire de Virologie - Institut Fédératif de Biologie | Latour J., Ranger N., Dubois M., Carcenac R., Harter A., Boyer P., Tremeaux P., Izopet J. |
| EPI_ISL_838349 | Queens Medical Centre, Clinical Microbiology Department / DeepSeq Nottingham | COVID-19 Genomics UK (COG-UK) Consortium | Gemma Clark, Wendy Smith, Manjinder Khakh, Vicki M Fleming, Michelle M Lister, Hannah Howson-Wells, Jonathan Ball, Patrick McClure, Joseph Chappell, Theocharis Tsoleridis, Nadine Holmes, Matthew Carlisle, Christopher Moore, Fei Sang, Johnny Debebe, Victoria Wright, Matthew Loose |
| EPI_ISL_840186, EPI_ISL_840187, EPI_ISL_840190, EPI_ISL_840191, EPI_ISL_840192, EPI_ISL_840193, EPI_ISL_840194, EPI_ISL_840195, EPI_ISL_840196, EPI_ISL_840197, EPI_ISL_840198, EPI_ISL_840199, EPI_ISL_840207 | | | |
| see above | Oxford Viromics, NDM, University of Oxford; Oxford University Hospitals; Basingstoke and North Hampshire Hospital | COVID-19 Genomics UK (COG-UK) Consortium | Tanya Golubchik, David Bonsall, George Macintyre, Amy Trebes, Mariateresa de Cesare, Catrin Moore, Alex Mobbs, Anita Justice, Robert Shaw, Monique Andersson, Timothy Peto, Emma Wise, Nathan Moore, Jessica Lynch, Nick Cortes, Matilde Mori, Stephen Kidd, David Buck, John Todd, Christophe Fraser |
| EPI_ISL_842204 | Virology Department, Sheffield Teaching Hospitals NHS Foundation Trust/Department of Infection, Immunity and Cardiovascular Disease, The Medical School, University of Sheffield | COVID-19 Genomics UK (COG-UK) Consortium | Thushan de Silva, Matthew Parker, Nikki Smith, Adri Angyal, Rebecca Brown, Luke Green, Rachel Tucker, Paul Parsons, Danielle Groves, Katie Johnson, Laura Carrilero, Alex Keeley, Dave Partridge, Matthew Wyles, Benjamin Lindsey, Mehmet Yavuz, Mohammad Raza, Cariad Evans |
| EPI_ISL_842910, EPI_ISL_842911, EPI_ISL_842912, EPI_ISL_842913, EPI_ISL_842914, EPI_ISL_842915 | Barts Health NHS Trust | COVID-19 Genomics UK (COG-UK) Consortium | CUTINO-MOGUEL, Maria-Teresa; HARRINGTON, David; OWOYEMI, Dola; SHYLINI, Raghavendran; BROAD, Claire; KELE, Beatrix |
| EPI_ISL_844234, EPI_ISL_844299, EPI_ISL_844300, EPI_ISL_844539, EPI_ISL_844540 | Department of Virus and Microbiological Special Diagnostics, Statens Serum Institut, Copenhagen, Denmark | Albertsen Lab, Department of Chemistry and Bioscience, Aalborg University, Denmark | Danish Covid-19 Genome Consortium |
| EPI_ISL_845557, EPI_ISL_845558, EPI_ISL_845559, EPI_ISL_845560, EPI_ISL_845561, EPI_ISL_845562 | National Public Health Laboratory, Cameroon | African Centre of Excellence for Genomics of Infectious Diseases (ACEGID), Redeemer's University | Oluniyi P.E. et al |
| EPI_ISL_846656, EPI_ISL_846659, EPI_ISL_846664, EPI_ISL_846665, EPI_ISL_846666, EPI_ISL_846667, EPI_ISL_846669, EPI_ISL_846670, EPI_ISL_846671, EPI_ISL_846676, EPI_ISL_846677, EPI_ISL_846682, EPI_ISL_846684, EPI_ISL_846685, EPI_ISL_846686, EPI_ISL_846690, EPI_ISL_846691, EPI_ISL_846693, EPI_ISL_846695, EPI_ISL_846696, EPI_ISL_846698, EPI_ISL_846702, EPI_ISL_846703, EPI_ISL_846736, EPI_ISL_846737, EPI_ISL_846738, EPI_ISL_846739, EPI_ISL_846740, EPI_ISL_846741, EPI_ISL_846742, EPI_ISL_846743, EPI_ISL_846744, EPI_ISL_846745, EPI_ISL_846747, EPI_ISL_846767, EPI_ISL_846768, EPI_ISL_846769, EPI_ISL_846770, EPI_ISL_846771, EPI_ISL_846772, EPI_ISL_846773 | | | |

| | | | |
|---|---|---|---|
| see above | University of Michigan Clinical Microbiology Laboratory | Lauring Lab, University of Michigan, Department of Microbiology and Immunology | Valesano |
| EPI_ISL_847524, EPI_ISL_847539, EPI_ISL_847576, EPI_ISL_847582, EPI_ISL_847586, EPI_ISL_847695, EPI_ISL_847696, EPI_ISL_847697, EPI_ISL_847698, EPI_ISL_847699, EPI_ISL_847700, EPI_ISL_847701, EPI_ISL_847702, EPI_ISL_847727, EPI_ISL_847732, EPI_ISL_847754, EPI_ISL_847816, EPI_ISL_847817, EPI_ISL_847818 | | | |
| see above | California Department of Public Health | Chiu Laboratory, University of California, San Francisco | Charles Chiu, Xianding (Wayne) Deng, Candace Wang, Brian Bushnell, Scot Federman, Jill Hacker, Debra Wadford |
| EPI_ISL_848211, EPI_ISL_848213, EPI_ISL_848214, EPI_ISL_848217, EPI_ISL_848221, EPI_ISL_848225, EPI_ISL_848233, EPI_ISL_848240, EPI_ISL_848241, EPI_ISL_848242, EPI_ISL_848250, EPI_ISL_848256, EPI_ISL_848257, EPI_ISL_848258, EPI_ISL_848264, EPI_ISL_848273, EPI_ISL_848385, EPI_ISL_848386, EPI_ISL_848387, EPI_ISL_848388, EPI_ISL_848389, EPI_ISL_848390, EPI_ISL_848391, EPI_ISL_848392, EPI_ISL_848393, EPI_ISL_848394, EPI_ISL_848395, EPI_ISL_848396, EPI_ISL_848442, EPI_ISL_848444, EPI_ISL_848454, EPI_ISL_848465, EPI_ISL_848524, EPI_ISL_848525, EPI_ISL_848526 | | | |
| see above | Illinois Department of Public Health | Gagnon Lab, Southern Illinois University | Keith Gagnon |
| EPI_ISL_848758, EPI_ISL_848759, EPI_ISL_848760, EPI_ISL_848761, EPI_ISL_848762, EPI_ISL_848763, EPI_ISL_848764, EPI_ISL_848765, EPI_ISL_848766, EPI_ISL_848767, EPI_ISL_848768, EPI_ISL_848769, EPI_ISL_848770, EPI_ISL_848771, EPI_ISL_848772, EPI_ISL_848773, EPI_ISL_848774, EPI_ISL_848775, EPI_ISL_848776, EPI_ISL_848777, EPI_ISL_848778, EPI_ISL_848779, EPI_ISL_848780, EPI_ISL_848781, EPI_ISL_848782, EPI_ISL_848783, EPI_ISL_848784, EPI_ISL_848785, EPI_ISL_848786, EPI_ISL_848787, EPI_ISL_848788, EPI_ISL_848789, EPI_ISL_848790, EPI_ISL_848791, EPI_ISL_848792, EPI_ISL_848793, EPI_ISL_848794, EPI_ISL_848795, EPI_ISL_848796, EPI_ISL_848797, EPI_ISL_848798, EPI_ISL_848799, EPI_ISL_848800, EPI_ISL_848801, EPI_ISL_848802, EPI_ISL_848803, EPI_ISL_848804, EPI_ISL_848805, EPI_ISL_848806, EPI_ISL_848807, EPI_ISL_848808, EPI_ISL_848809 | | | |
| see above | Florida Bureau of Public Health Laboratories | Florida Bureau of Public Health Laboratories | Sarah Schmedes, Jason Blanton |
| EPI_ISL_849516, EPI_ISL_849517 | Seattle Flu Study | Seattle Flu Study | Deborah A. Nickerson, Chris D. Frazar, Jover Lee, Benjamin Pelle, Matthew Richardson, Amanda Adler, Elisabeth Brandstetter, Peter D. Han, Kairsten Fay, Misja Ilcisin, Kirsten Lacombe, Thomas R. Sibley, Melissa Truong, Caitlin R. Wolf, Michael Boeckh, Janet A. Englund, Michael Famulare, Barry R. Lutz, Mark J. Rieder, Lea M. Starita, Matthew Thompson, Jay Shendure, Trevor Bedford, Helen Y. Chu |
| EPI_ISL_849686 | unknown | PHV-FSS | Son Nguyen et al. |
| EPI_ISL_849948, EPI_ISL_849951 | UC Davis- Department of Pathology and Laboratory Medicine | Chan-Zuckerberg Biohub | CZB Cliahub Consortium |
| EPI_ISL_853329, EPI_ISL_853332 | UPMC Clinical Microbiology Laboratory | Microbial Genome Sequencing Center; Microbial Genomic Epidemiology Laboratory | Mustapha M. Mustapha, Jane W. Marsh, Dan Snyder, Marissa P. Griffith, Stephanie L. Mitchell, Vatsala R. Srinivasa, Kady D. Waggle, Chinelo Ezeonwuku, Vaughn S. Cooper, Lee H. Harrison |
| EPI_ISL_853767, EPI_ISL_853792, EPI_ISL_853933, EPI_ISL_853942, EPI_ISL_853945 | Department of Microbiology, University Innsbruck | Bergthaler laboratory, CeMM Research Center for Molecular Medicine of the Austrian Academy of Sciences | Lukas Endler, Alexandra Popa, Benedikt Agerer, Jakob-Wendelin Genger, Alexander Lercher, Anna Schedl, Thomas Penz, Michael Schuster, Jan Laine, Martin Senekowitsch, Christoph Bock, Andreas Bergthaler |
| EPI_ISL_855389, EPI_ISL_855390, EPI_ISL_855391 | California Department of Public Health | Chiu Laboratory, University of California, San Francisco | Charles Chiu, Xianding (Wayne) Deng, Candace Wang, Brian Bushnell, Scot Federman, Jill Hacker, Debra Wadford |
| EPI_ISL_855499, EPI_ISL_855516, EPI_ISL_855521, EPI_ISL_855528, EPI_ISL_855529 | KEMRI-Wellcome Trust Research Programme/KEMRI-CGMR-C Kilifi | KEMRI-Wellcome Trust Research Programme/KEMRI-CGMR-C Kilifi | Githinji et al |
| EPI_ISL_856760 | Servicio Virosis Respiratorias-Departamento Virología-INEI | Instituto Nacional Enfermedades Infecciosas C.G.Malbran | Baumeister E., Avaro M., Benedetti E., Russo M., Dattero ME, Pontoriero A., Cisterna D., Molina V., Perandones C., Tuduri E., Lorenzo F., Poklepovich T., Campos J. |
| EPI_ISL_857049 | Colorado Department of Public Health and Environment | Colorado Department of Puplic Health and Environment | Laura Bankers, Molly C. Hetherington-Rauth, Diana Ir, Shannon Ely, Shannon R. Matzinger, Sarah Elizabeth Totten, Emily A. Travanty |
| EPI_ISL_857496, EPI_ISL_857497, EPI_ISL_857513 | Swiss National Reference Centre for Influenza | Swiss National Reference Centre for Influenza | Ana Rita Goncalves,Samuel Cordey, Laurent Kaiser, Lorenzo Cerutti, Henri Pegeot, Melyssa Elies, Keith Harshman, Ioannis Xenarios, Emmanouil Dermitzakis |
| EPI_ISL_857520, EPI_ISL_857540 | Swiss National Reference Centre for Influenza | Swiss National Reference Centre for Influenza | Tim Roloff, Ana Rita Gonçalves, Madlen Stange, Helena MB Seth-Smith, Alfredo Mari, Karoline Leuzinger, Julia Bielicki, Manuel Battegay, Hans Hirsch, Laurent Kaiser, Adrian Egli |
| EPI_ISL_859555, EPI_ISL_859560 | Lighthouse Lab in Cambridge | Wellcome Sanger Institute for the COVID-19 Genomics UK (COG-UK) Consortium | Rob Howes, The Lighthouse Lab in Cambridge and Alex Alderton, Roberto Amato, Sonia Goncalves, Ewan Harrison, David K. Jackson, Ian Johnston, Dominic Kwiatkowski, Cordelia Langford, John Sillitoe on behalf of the Wellcome Sanger Institute COVID-19 Surveillance Team |
| EPI_ISL_859846, EPI_ISL_859847, EPI_ISL_859848, EPI_ISL_859849, EPI_ISL_859850, EPI_ISL_859851, EPI_ISL_859852, EPI_ISL_859853, EPI_ISL_859865, EPI_ISL_859866, EPI_ISL_859867, EPI_ISL_859868, EPI_ISL_859869, EPI_ISL_859870, EPI_ISL_859871, EPI_ISL_859872, EPI_ISL_859873, EPI_ISL_859874, EPI_ISL_859875, EPI_ISL_859887, EPI_ISL_859902, EPI_ISL_859903, EPI_ISL_859924, EPI_ISL_859925, EPI_ISL_859926, EPI_ISL_859927, EPI_ISL_859928 | | | |
| see above | BTC, Khalifa University | BTC, Khalifa University | Al Safar et al |
| EPI_ISL_860123, EPI_ISL_860134, EPI_ISL_860135, EPI_ISL_860149, EPI_ISL_860152, EPI_ISL_860153, EPI_ISL_860156, EPI_ISL_860158, EPI_ISL_860159 | Keio University School of Medicine | Keio University School of Medicine | Kenjiro Kosaki, Yuka Iwasaki, Hirotsugu Ishizu, Haruhiko Siomi, Kodai Abe |
| EPI_ISL_861887, EPI_ISL_861888 | LATE - Laboratório de Técnicas Especiais - Hospital Israelita Albert Einstein | LATE - Laboratório de Técnicas Especiais - Hospital Israelita Albert Einstein | Deyvid Amgarten, Fernanda de Mello Malta, Raquel Riyuzo, Ana Paula Moreira Salles, Pedro Henrique Sebe Rodrigues, João Renato Rebello Pinho |
| EPI_ISL_864573 | Institute of Medical Microbiology and Hospital Hygiene | Institute of Medical Microbiology and Hospital Hygiene | Prof. Dr. Achim Kaasch, Aljoscha Tersteegen |
| EPI_ISL_865173 | Liverpool Clinical Laboratories | COVID-19 Genomics UK (COG-UK) Consortium | Sam Haldenby, Anita Lucaci, Steve Paterson, Julian Hiscox, Alistair Darby, M Almsaud, A Alrezaihi, Muhannad Alruwaili, Stuart D Armstrong, Jones Benjamin, Eleanor G Bentley, Anu Chawla, Jordan J Clark, Angela Cowell, Richard Eccles, Isabel García-Dorival, Matthew Gemmell, Alessandro Gerada, PKF Gilmore, Richard Gregory, Ximeng Han, Catherine Hartley, Margaret Hughes, Miren Iturriza-Gomara, James Johnson, L Luu, Jenifer Manson, Charlotte Nelson, Elaine O'Toole, Cassie Olateju, Rebekah Penrice-Randal , Lucille Rainbow, N.P Randle, Trevor Ian Robinson, Parul Sharma, Ghada T Shawli, James P Stewart, Neil Swainston, Ecaterina Vamos, Joanne Watts, Mark Whitehead |
| EPI_ISL_865683, EPI_ISL_865697, EPI_ISL_865698, EPI_ISL_865699, EPI_ISL_865700, EPI_ISL_865701 | University College London, Great Ormond Street Hospital for Children NHS Foundation Trust, Imperial College Healthcare NHS Trust | COVID-19 Genomics UK (COG-UK) Consortium | Sergi Castellano, Rachel Williams, Mark Kristiansen, Paola Resende Silva, Sunando Roy, Tony Brooks, Helena Tutill, Paola Niola, Patricia Dyal, Charlotte Williams, Leysa Forrest, Yasmin Panchbhaya, Jacqueline Findlay, Samuel Weeks, Julianne Brown, Kathryn Harris, Paul Randell, James Price, Alison Holmes, Judith Breuer |
| EPI_ISL_866054 | University College London Hospital | COVID-19 Genomics UK (COG-UK) Consortium | Judith Heaney, Matthew Byott, Catherine Houlihan, Dan Frampton, Stuart Kirk, Moira Spyer and Eleni Nastouli |
| EPI_ISL_872607 | Nigeria Centre for Disease Control (NCDC) | African Centre of Excellence for Genomics of Infectious Diseases (ACEGID), Redeemer's University | Oluniyi P.E. et al |
| EPI_ISL_872692, EPI_ISL_872693, EPI_ISL_872694, EPI_ISL_872695, EPI_ISL_872696 | Rhode Island Department of Health | Infectious Disease Program, Broad Institute of Harvard and MIT | Lemieux,J.E., Siddle,K.J., Huard,R., King,E., Azevedo,K., Miller,A., Adams,G., Gladden-Young,A., Lagerborg,K., Rudy,M., DeRuff,K., Carter,A., Normandin,E., Bauer,M., Reilly,S., Tomkins-Tinch,C., Loreth,C., Chaluvadi,S., Birren,B.W., Gallagher,G., Smole,S., Park,D.J., MacInnis,B.L., and Sabeti,P.C. |
| EPI_ISL_873132, EPI_ISL_873135, EPI_ISL_873138, EPI_ISL_873140, EPI_ISL_873143, EPI_ISL_873144, EPI_ISL_873146, EPI_ISL_873154 | University of Michigan Clinical Microbiology Laboratory | Lauring Lab, University of Michigan, Department of Microbiology and Immunology | Valesano |
| EPI_ISL_876189, EPI_ISL_876325, EPI_ISL_876326, EPI_ISL_876327 | Massachusetts State Public Health Laboratory | Massachusetts State Public Health Laboratory | Andrew Lang, Timelia Fink, Glen Gallagher, Sandra Smole |
| EPI_ISL_876567 | Florida Bureau of Public Health Laboratories | Florida Bureau of Public Health Laboratories | Sarah Schmedes, Jason Blanton |
| EPI_ISL_877331, EPI_ISL_877332, EPI_ISL_877333, EPI_ISL_877334, EPI_ISL_877335, EPI_ISL_877336, EPI_ISL_877337, EPI_ISL_877338, EPI_ISL_877339, EPI_ISL_877340, EPI_ISL_877341, EPI_ISL_877342, EPI_ISL_877343, EPI_ISL_877344, EPI_ISL_877345, EPI_ISL_877346, EPI_ISL_877347, EPI_ISL_877348, EPI_ISL_877349, EPI_ISL_877350, EPI_ISL_877351, EPI_ISL_877352, EPI_ISL_877353, EPI_ISL_877354, EPI_ISL_877355, EPI_ISL_877356, EPI_ISL_877357, EPI_ISL_877358, EPI_ISL_877359, EPI_ISL_877360, EPI_ISL_877361, EPI_ISL_877362, EPI_ISL_877363, EPI_ISL_877364, EPI_ISL_877365, EPI_ISL_877366, EPI_ISL_877367, EPI_ISL_877368, EPI_ISL_877369, EPI_ISL_877370, EPI_ISL_877371, EPI_ISL_877372, EPI_ISL_877373, EPI_ISL_877374, EPI_ISL_877375, EPI_ISL_877376, EPI_ISL_877377, EPI_ISL_877378, EPI_ISL_877379, EPI_ISL_877380, EPI_ISL_877381, EPI_ISL_877382, EPI_ISL_877383, EPI_ISL_877384, | | | |

| | | | |
|---|---|---|---|
| EPI_ISL_877385, EPI_ISL_877386, EPI_ISL_877387, EPI_ISL_877388, EPI_ISL_877389, EPI_ISL_877390, EPI_ISL_877391, EPI_ISL_877392, EPI_ISL_877393, EPI_ISL_877394, EPI_ISL_877395, EPI_ISL_877396, EPI_ISL_877397, EPI_ISL_877398, EPI_ISL_877399, EPI_ISL_877400, EPI_ISL_877401, EPI_ISL_877402, EPI_ISL_877403, EPI_ISL_877404, EPI_ISL_877405, EPI_ISL_877406, EPI_ISL_877407, EPI_ISL_877408, EPI_ISL_877409, EPI_ISL_877410 | | | |
| see above | Eurofins Diatherix | Hudsonalpha Genome Sequencing Center | Jane Grimwood, Melissa Williams, Lori H. Handley, Joshua Stough, Leslie Malone, Stefan Brzezinski, Ada Stewart, Teresa Jones, Jenell Webber, John Lovell, Jennifer Cart, and Jeremy Schmutz |
| EPI_ISL_878399, EPI_ISL_878403, EPI_ISL_878405, EPI_ISL_878407, EPI_ISL_878431 | San Diego County Public Health Laboratory | Andersen lab at Scripps Research | SEARCH Alliance San Diego with Tracy Basler, Jovan Shephard, Brett Austin |
| EPI_ISL_878547 | Robert Garry lab | Andersen lab at Scripps Research | Allison Smither, Gilberto Sabino-Santos, Patricia Snarski, Lilia Melnik, Antoinette Bell, Kaylynn Genemaras, Arnaud Drouin, Dahlene Fusco, Robert Garry with SEARCH Alliance San Diego |
| EPI_ISL_878682, EPI_ISL_878684, EPI_ISL_878726 | Rady's Childrens Hospital | Andersen lab at Scripps Research | SEARCH Alliance San Diego with Nanda Radamchar, David Dimmock, Linda Luo, Christina Clarke, Kathryn Bouic, Teresa Mueller, Denise Malicki |
| EPI_ISL_882688, EPI_ISL_882689, EPI_ISL_882697, EPI_ISL_882698, EPI_ISL_882699, EPI_ISL_882700, EPI_ISL_882701, EPI_ISL_882704, EPI_ISL_882705, EPI_ISL_882706, EPI_ISL_882707, EPI_ISL_882708 | | | |
| see above | 1.AO Universitaria 'S. Giovanni di Dio e Ruggi D'Aragona, Scuola Medica Salernitana' Hospital / 2.UOC di Virologia e Microbiologia, Università della Campania 'L. Vanvitelli' / 3.AO Universitaria 'Federico II' Napoli Hospital / 4.AORN 'San Giuseppe Moscati' Avellino Hospital / 5.AO 'San Pio - presidio G. Rummo' Benevento Hospital / 6.AO 'Sant'Anna e San Sebastiano' Caserta Hospital / 7.PO 'Maria Santissima Addolorata' Eboli Hospital / 8.Biogem Istituto di Ricerche Genetiche | 1. Genome Research Center for Health (CRGS) / 2. Laboratory of Molecular Medicine and Genomics(LMMGe) / 3. Center for Research in Pure and Applied Mathematics (CRMPA) | Giorgio Giurato, Francesca Rizzo, Alessandro Weisz, Gianluigi Franci, Giovanni Nassa, Pasquale Pagliano, Roberta Tarallo, Elena Alexandrova, Ylenia D'Agostino, Carlo Ferravante, Jessica Lamberti, Viola Melone, Domenico Memoli, Valeria Mirici Cappa, Domenico Palumbo, Giovanni Pecoraro, Assunta Sellitto, Oriana Strianese, Ilaria Terenzi, Giuseppe Fenza, Aniello Gentile, Antonello Saccomanno, Sonia Amabile, Teresa Rocco, Annamaria Salvati, Emilia Vaccaro, Massimiliano Galdiero, Michele Cennamo, Giuseppe Portella, Maria Grazia Foti, Mariarosaria Ingino, Maria Landi, Maurizio Fumi, Vincenzo Rocco, Rita Greco, Vittoria Letizia, Arnolfo Petruzziello, Maddalena Schioppa, Gregorio Goffredi, Francesca Marciano, Michele Caraglia, Alessia Cossu, Marianna Scrima |
| EPI_ISL_882957 | Specjalistyczny Szpital im. Dra A. Sokoowskiego | National Institute of Public Health - National Institute of Hygiene | Wokowicz Tomasz, Zacharczuk Katarzyna, Gawor Jan |
| EPI_ISL_885130 | California Institute of Technology | Chan-Zuckerberg Biohub | CZB Cliahub Consortium |
| EPI_ISL_888836 | Michigan Department of Health and Human Services, Bureau of Laboratories | Michigan Department of Health and Human Services, Bureau of Laboratories | Blankenship HM, Riner D, Soehnlen MK |
| EPI_ISL_888988 | RS Qadr | Eijkman Institute for Molecular Biology, Ministry of Research and Technology/National Agency for Research and Innovation | Frilasita A Yudhaputri, Hidayat Trimarsanto, Iskandar Adnan, Lydia V. Panggalo, Sukma Oktavianthi, Willy Agustine, Edison Johar, Safarina G Malik, Khin Saw Myint, Amin Soebandrio |
| EPI_ISL_888989 | RS Pondok Indah Bintaro Jaya | Eijkman Institute for Molecular Biology, Ministry of Research and Technology/National Agency for Research and Innovation | Frilasita A Yudhaputri, Hidayat Trimarsanto, Iskandar Adnan, Lydia V. Panggalo, Sukma Oktavianthi, Willy Agustine, Edison Johar, Safarina G Malik, Khin Saw Myint, Amin Soebandrio |
| EPI_ISL_889010 | RSU Medika Dramaga | Eijkman Institute for Molecular Biology, Ministry of Research and Technology/National Agency for Research and Innovation | Frilasita A Yudhaputri, Hidayat Trimarsanto, Iskandar Adnan, Lydia V. Panggalo, Sukma Oktavianthi, Willy Agustine, Edison Johar, Safarina G Malik, Khin Saw Myint, Amin Soebandrio |
| EPI_ISL_891195 | DPH, Massachusetts State Public Health Lab | DPH, Massachusetts State Public Health Lab | Lang,A.S., Fink,T., Gallagher,G.R., Smole,S.C. |
| EPI_ISL_902996, EPI_ISL_902997, EPI_ISL_902998, EPI_ISL_902999, EPI_ISL_903000, EPI_ISL_903001, EPI_ISL_903003 | Seattle Flu Study | Seattle Flu Study | Deborah A. Nickerson, Chris D. Frazar, Jover Lee, Benjamin Pelle, Erica Ryke, Matthew Richardson, Amanda Adler, Elisabeth Brandstetter, Peter D. Han, Kairsten Fay, Misja Ilcisin, Kirsten Lacombe, Thomas R. Sibley, Melissa Truong, Caitlin R. Wolf, Michael Boeckh, Janet A. Englund, Michael Famulare, Barry R. Lutz, Mark J. Rieder, Lea M. Starita, Matthew Thompson, Jay Shendure, Trevor Bedford, Helen Y. Chu |
| EPI_ISL_903633 | WA State Department of Health | Genomics and Discovery, Respiratory Viruses Branch, Division of Viral Diseases, Centers for Disease Control and Prevention | Krista Queen, Yan Li, Ying Tao, Jing Zhang, Anna Uehara, Anna Montmayeur, Clinton R. Paden, Peter W. Cook, Rachel Marine, Mili Sheth, Jasmine Padilla, Sarah Nobles, Mark Burroughs, Lori Rowe, Haibin Wang, Ben L. Rambo-Martin, Dhwani Batra, Justin Lee, Suxiang Tong |
| EPI_ISL_903748, EPI_ISL_903785, EPI_ISL_903820, EPI_ISL_903854 | PA Department of Health, Bureau of Laboratories | Genomics and Discovery, Respiratory Viruses Branch, Division of Viral Diseases, Centers for Disease Control and Prevention | Krista Queen, Yan Li, Ying Tao, Jing Zhang, Anna Uehara, Anna Montmayeur, Clinton R. Paden, Peter W. Cook, Rachel Marine, Mili Sheth, Jasmine Padilla, Sarah Nobles, Mark Burroughs, Lori Rowe, Haibin Wang, Ben L. Rambo-Martin, Dhwani Batra, Justin Lee, Suxiang Tong |
| EPI_ISL_903949, EPI_ISL_903952 | WA State Department of Health | Genomics and Discovery, Respiratory Viruses Branch, Division of Viral Diseases, Centers for Disease Control and Prevention | Krista Queen, Yan Li, Ying Tao, Jing Zhang, Anna Uehara, Anna Montmayeur, Clinton R. Paden, Peter W. Cook, Rachel Marine, Mili Sheth, Jasmine Padilla, Sarah Nobles, Mark Burroughs, Lori Rowe, Haibin Wang, Ben L. Rambo-Martin, Dhwani Batra, Justin Lee, Suxiang Tong |
| EPI_ISL_904735 | Dutch COVID-19 response team | National Institute for Public Health and the Environment (RIVM) | Adam Meijer, Harry Vennema, Dirk Eggink, Jeroen Cremer, Sharon van den Brink, Bas van der Veer, AnneMarie van den Brandt, Florian Zwagemaker, Dennis Schmitz, Chantal Reusken, on behalf of the national COVID-19 response team |
| EPI_ISL_904947, EPI_ISL_904948 | Vilnius University Hospital Santaros Klinikos, Vilnius University | Institute of Biotechnology, Life Sciences Center, Vilnius University | Emilija Vasiliunaite, Milda Norkiene, Albertas Timinskas, Alma Gedvilaite, Aurelija Zvirbliene, Daniel Naumovas, Laimonas Griskevicius |
| EPI_ISL_906105 | Child Health Research Foundation | Child Health Research Foundation | Senjuti Saha, Sharmistha Goswami, Afroza Akter Tanni, Syed Muktadir Al Sium, Arif Mohammad Tanmoy, Roly Malaker, Md Hafizur Rahman, Samir K Saha |
| EPI_ISL_906130 | Institute of Microbiology and Immunology, Faculty of Medicine, University of Ljubljana | Institute of Microbiology and Immunology, Faculty of Medicine, University of Ljubljana | Samo Zakotnik, Tomaž Mark Zorec, Matic Brvar, Miša Korva, Mario Poljak, Tatjana Avši - Županc |
| EPI_ISL_906263 | University of Wisconsin-Madison AIDS Vaccine Research Laboratories | University of Wisconsin-Madison AIDS Vaccine Research Laboratories | Gage Moreno, Katarina Braun, et al. AIDS Vaccine Research Laboratories |
| EPI_ISL_906852 | Respiratory Viruses Branch, Centers for Disease Control and Prevention | Respiratory Viruses Branch, Centers for Disease Control and Prevention | Tao,Y., Li,Y., Zhang,J., Queen,K., Uehara,A., Cook,P., Paden,C.R., Wang,H., Tong,S. |
| EPI_ISL_907077, EPI_ISL_907078, EPI_ISL_907079, EPI_ISL_907080, EPI_ISL_907081, EPI_ISL_907082, EPI_ISL_907083, EPI_ISL_907084, EPI_ISL_907085 | Biology, MCL | Biology, MCL | Seadawy,M.G., Shamel,M.D., Gad,A.F., Elhoseiny,M.F., Zekri,A.N. |
| EPI_ISL_911239, EPI_ISL_911240 | Laboratoire national de sante, Microbiology, Virology | Laboratoire national de sante, Microbiology, Microbial Genomics Platform | Anke Wienecke-Baldacchino, Catherine Ragimbeau,Jessica Tapp, Fatu Djabi, Lise Pignon, Raoul Salmon, Tamir Abdelrahman |
| EPI_ISL_911687 | Alaska State Virology Laboratory | Alaska State Virology Laboratory | Stephanie DeRonde, Lisa Smith, Ph.D., Jack Chen, Ph.D. |
| EPI_ISL_912466, EPI_ISL_912472, EPI_ISL_912475, EPI_ISL_912479, EPI_ISL_912481, EPI_ISL_912487, EPI_ISL_912536 | NHLS Universitas Academic | UFS Virology | PA Bester, MM Nyaga, P Nthiga, MT Mogotsi, D Goedhals, T de Oliveira |
| EPI_ISL_912925 | Hôpital Pitié-Salpêtrière | Department of Virology, Henri Mondor University Hospital, Assistance Publique Hôpitaux de Paris, Université Paris-Est Créteil, INSERM U955 | Christophe Rodriguez, Slim Fourati, Vanessa Demontant, Guillaume Gricourt, Melissa N'Debi, Alexandre Soulier, Elisabeth Trawinski, Jean-Michel Pawlotsky |

| | | | |
|---|---|---|---|
| EPI_ISL_913088 | Center for Virology | Center for Virology | Jeremy V. Camp, Irene Goerzer, Monika Redlberger-Fritz, Stephan W. Aberle |
| EPI_ISL_913117, EPI_ISL_913122, EPI_ISL_913123, EPI_ISL_913124, EPI_ISL_913132, EPI_ISL_913140, EPI_ISL_913145, EPI_ISL_913147, EPI_ISL_913148, EPI_ISL_913151, EPI_ISL_913152, EPI_ISL_913153, EPI_ISL_913154, EPI_ISL_913157, EPI_ISL_913160, EPI_ISL_913201, EPI_ISL_913202, EPI_ISL_913203, EPI_ISL_913204, EPI_ISL_913205, EPI_ISL_913206, EPI_ISL_913207, EPI_ISL_913208, EPI_ISL_913209, EPI_ISL_913210, EPI_ISL_913211, EPI_ISL_913212, EPI_ISL_913213, EPI_ISL_913214, EPI_ISL_913215, EPI_ISL_913216, EPI_ISL_913217, EPI_ISL_913218, EPI_ISL_913219, EPI_ISL_913220, EPI_ISL_913221, EPI_ISL_913222, EPI_ISL_913223, EPI_ISL_913224, EPI_ISL_913225, EPI_ISL_913226, EPI_ISL_913227, EPI_ISL_913228, EPI_ISL_913229, EPI_ISL_913230, EPI_ISL_913231, EPI_ISL_913232, EPI_ISL_913233, EPI_ISL_913234, EPI_ISL_913235, EPI_ISL_913236, EPI_ISL_913237, EPI_ISL_913238, EPI_ISL_913239, EPI_ISL_913240, EPI_ISL_913241, EPI_ISL_913242, EPI_ISL_913243, EPI_ISL_913244, EPI_ISL_913245, EPI_ISL_913246, EPI_ISL_913247, EPI_ISL_913248, EPI_ISL_913249, EPI_ISL_913250, EPI_ISL_913251, EPI_ISL_913252, EPI_ISL_913253, EPI_ISL_913254, EPI_ISL_913255 | | | |
| see above | University of Michigan Clinical Microbiology Laboratory | Lauring Lab, University of Michigan, Department of Microbiology and Immunology | Valesano |
| EPI_ISL_913480 | Klinisk mikrobiologi | The Public Health Agency of Sweden | Anna-Malin Linde, Maria Lind Karlberg, Carlo Berg, Oskar Karlsson Lindsjo, Sofia Stamouli, Reza Advani, Mattias Haukland, Petra Holmstrom, Noura Walai, Petra Edquist, Mia Brytting, Anna Risberg, Karin Tegmark-Wisell |
| EPI_ISL_913517, EPI_ISL_913518 | University of Michigan Clinical Microbiology Laboratory | Lauring Lab, University of Michigan, Department of Microbiology and Immunology | Valesano |
| EPI_ISL_913933, EPI_ISL_913958 | Instituto de Diagnostico y Referencia Epidemiologicos INDRE_RNLSP | Instituto de Diagnostico y Referencia Epidemiologicos (INDRE) | Claudia Wong-Arambula, Abril Rodriguez-Maldonado, Fabiola Garces-Ayala, Adnan Araiza-Rodriguez, David Fragoso-Fonseca, Sergio Rangel-Guerrero, Mayra Jimenez-Morales, Nancy Munoz-Hernandez, Natividad Cruz-Ortiz, Tatiana Nunez-Garcia, Gisela Barrera-Badillo, Lucia Hernandez-Rivas, Irma Lopez-Martinez, Ernesto Ramirez-Gonzalez. |
| EPI_ISL_915365, EPI_ISL_915366, EPI_ISL_915371, EPI_ISL_915378, EPI_ISL_915412 | Keio University School of Medicine | Keio University School of Medicine | Kenjiro Kosaki, Yuka Iwasaki, Hirotsugu Ishizu, Haruhiko Siomi, Kodai Abe |
| EPI_ISL_915656, EPI_ISL_916068, EPI_ISL_916071, EPI_ISL_916075, EPI_ISL_916076, EPI_ISL_916079, EPI_ISL_916080, EPI_ISL_916082, EPI_ISL_916086, EPI_ISL_916087, EPI_ISL_916088, EPI_ISL_916091, EPI_ISL_916092, EPI_ISL_916094, EPI_ISL_916098, EPI_ISL_916099, EPI_ISL_916100, EPI_ISL_916105, EPI_ISL_916106, EPI_ISL_916107, EPI_ISL_916108, EPI_ISL_916109, EPI_ISL_916110, EPI_ISL_916111, EPI_ISL_916113, EPI_ISL_916121, EPI_ISL_916124, EPI_ISL_916127, EPI_ISL_916129, EPI_ISL_916131, EPI_ISL_916132, EPI_ISL_916134, EPI_ISL_916137, EPI_ISL_916144, EPI_ISL_916145, EPI_ISL_916147, EPI_ISL_916151, EPI_ISL_916152, EPI_ISL_916154, EPI_ISL_916159, EPI_ISL_916161, EPI_ISL_916162, EPI_ISL_916164, EPI_ISL_916165, EPI_ISL_916166, EPI_ISL_916167, EPI_ISL_916169, EPI_ISL_916171, EPI_ISL_916174 | | | |
| see above | Lighthouse Lab in Alderley Park | Wellcome Sanger Institute for the COVID-19 Genomics UK (COG-UK) Consortium | Jacquelyn Wynn, Mairead Hyland, The Lighthouse Lab in Alderley Park and Alex Alderton, Roberto Amato, Sonia Goncalves, Ewan Harrison, David K. Jackson, Ian Johnston, Dominic Kwiatkowski, Cordelia Langford, John Sillitoe on behalf of the Wellcome Sanger Institute COVID-19 Surveillance Team |
| EPI_ISL_918184, EPI_ISL_918202, EPI_ISL_918205, EPI_ISL_918206, EPI_ISL_918207, EPI_ISL_918209, EPI_ISL_918211, EPI_ISL_918214, EPI_ISL_918216 | Innovative Genomics Institute, UC Berkeley | Innovative Genomics Institute, UC Berkeley | Stacia Wyman, Haridha Shivram, Phil Frankino, Liana Lareau, Shana McDevitt, Justin Choi |
| EPI_ISL_918515 | LACEN - Laboratório Central de Saúde Pública do Para | Evandro Chagas Institute | Santos, M.C.; Silva, A.M.; Junior, W.D.C.; Barbagelata, L.S.; Ferreira, J.A.; Sousa, E.M.A.; da Silva, P.S.; Pinheiro, K.C.; L.C.; Sousa Junior, E.C. |
| EPI_ISL_918564, EPI_ISL_918565, EPI_ISL_918566, EPI_ISL_918567, EPI_ISL_918568, EPI_ISL_918585, EPI_ISL_918586, EPI_ISL_918587, EPI_ISL_918588, EPI_ISL_918590, EPI_ISL_918665 | | | |
| see above | University of Birmingham | COVID-19 Genomics UK (COG-UK) Consortium | Institute of Microbiology, University of Birmingham: Claire McMurray, Joanne Stockton, Samuel Nicholls, Radoslaw Poplawski, Will Rowe, Josh Quick, Nicholas Loman. University of Birmingham Testing Laboratory: Celina M Whalley, Andrew Bosworth, Charlotte Poxon, Kasun Wanigasooriya, Oliver Pickles, Mike Kidd, Alex Richter, Andrew D Beggs PHE Heartlands Lab: Husam Osman, Andrew Bosworth. Queen Elizabeth Hospital: Anna Casey |
| EPI_ISL_920268, EPI_ISL_920278, EPI_ISL_920355, EPI_ISL_920364, EPI_ISL_920376, EPI_ISL_920517 | University College London Hospital | COVID-19 Genomics UK (COG-UK) Consortium | Judith Heaney, Matthew Byott, Catherine Houlihan, Dan Frampton, Stuart Kirk, Moira Spyer and Eleni Nastouli |
| EPI_ISL_925181, EPI_ISL_925182, EPI_ISL_925183, EPI_ISL_925184, EPI_ISL_925185, EPI_ISL_925186, EPI_ISL_925187, EPI_ISL_925188, EPI_ISL_925189, EPI_ISL_925190, EPI_ISL_925191, EPI_ISL_925192, EPI_ISL_925193, EPI_ISL_925194, EPI_ISL_925195, EPI_ISL_925196, EPI_ISL_925197, EPI_ISL_925201, EPI_ISL_925202, EPI_ISL_925203 | | | |
| see above | Virginia DCLS | Virginia DCLS | Virginia DCLS |
| EPI_ISL_925376, EPI_ISL_925377, EPI_ISL_925378, EPI_ISL_925466, EPI_ISL_925495 | Department of Clinical Microbiology | GIGA Medical Genomics | Keith Durkin, Maria Artesi, Sébastien Bontems, Raphaël Boreux, Bouchra Boujemla, Cécile Meex, Pierrette Melin, Marie-Pierre Hayette, Vincent Bours |
| EPI_ISL_925992, EPI_ISL_926388, EPI_ISL_926432, EPI_ISL_926494, EPI_ISL_926504, EPI_ISL_926768, EPI_ISL_926787, EPI_ISL_926811, EPI_ISL_927079, EPI_ISL_927118, EPI_ISL_927211, EPI_ISL_927252, EPI_ISL_927295, EPI_ISL_927345, EPI_ISL_927431, EPI_ISL_927603, EPI_ISL_928118, EPI_ISL_928179, EPI_ISL_928282, EPI_ISL_928285, EPI_ISL_928297, EPI_ISL_928380, EPI_ISL_928422, EPI_ISL_928513, EPI_ISL_928606, EPI_ISL_928628, EPI_ISL_928630, EPI_ISL_928655, EPI_ISL_928670, EPI_ISL_928722, EPI_ISL_928884, EPI_ISL_928954, EPI_ISL_929022, EPI_ISL_929213, EPI_ISL_929286, EPI_ISL_929299, EPI_ISL_929391, EPI_ISL_929450, EPI_ISL_929476, EPI_ISL_929657, EPI_ISL_929678, EPI_ISL_929797, EPI_ISL_929900, EPI_ISL_929916, EPI_ISL_930088, EPI_ISL_930112, EPI_ISL_930187, EPI_ISL_930202, EPI_ISL_930220, EPI_ISL_930376, EPI_ISL_930403, EPI_ISL_930562 | | | |
| see above | Department of Virus and Microbiological Special Diagnostics, Statens Serum Institut, Copenhagen, Denmark | Aalborg University | Danish Covid-19 Genome Consortium |
| EPI_ISL_931343, EPI_ISL_931344, EPI_ISL_931345, EPI_ISL_931346, EPI_ISL_931347, EPI_ISL_931348, EPI_ISL_931349, EPI_ISL_931350, EPI_ISL_931351, EPI_ISL_931352, EPI_ISL_931353, EPI_ISL_931354, EPI_ISL_931355, EPI_ISL_931356, EPI_ISL_931357, EPI_ISL_931358, EPI_ISL_931359, EPI_ISL_931360, EPI_ISL_931361, EPI_ISL_931362, EPI_ISL_931370, EPI_ISL_931371, EPI_ISL_931372, EPI_ISL_931373, EPI_ISL_931374, EPI_ISL_931375 | | | |
| see above | University Hospital Basel, Clinical Virology | University Hospital Basel, Clinical Bacteriology | Tim Roloff, Madlen Stange, Helena MB Seth-Smith, Alfredo Mari, Karoline Leuzinger, Julia Bielicki, Manuel Battegay, Hans Hirsch, Adrian Egli |
| EPI_ISL_933495 | Lighthouse Lab in Milton Keynes | Wellcome Sanger Institute for the COVID-19 Genomics UK (COG-UK) Consortium | The Lighthouse Lab in Milton Keynes and Alex Alderton, Roberto Amato, Sonia Goncalves, Ewan Harrison, David K. Jackson, Ian Johnston, Dominic Kwiatkowski, Cordelia Langford, John Sillitoe on behalf of the Wellcome Sanger Institute COVID-19 Surveillance Team |
| EPI_ISL_933540, EPI_ISL_933542, EPI_ISL_933543, EPI_ISL_933544, EPI_ISL_933545, EPI_ISL_933546 | Arizona State Public Health Laboratory | Arizona State Public Health Laboratory | Trung Huynh, Jessica Escobar, Katherine Fullerton, Nobuko Fukushima, Stacy White, Linda Getsinger, Victor Waddell |
| EPI_ISL_933707 | Instituto de Diagnostico y Referencia Epidemiologicos INDRE_RNLSP | Instituto de Diagnostico y Referencia Epidemiologicos (INDRE) | Claudia Wong-Arambula, Abril Rodriguez-Maldonado, Fabiola Garces-Ayala, Adnan Araiza-Rodriguez, David Fragoso-Fonseca, Sergio Rangel-Guerrero, Mayra Jimenez-Morales, Nancy Munoz-Hernandez, Natividad Cruz-Ortiz, Tatiana Nunez-Garcia, Gisela Barrera-Badillo, Lucia Hernandez-Rivas, Irma Lopez-Martinez, Ernesto Ramirez-Gonzalez. |
| EPI_ISL_934542, EPI_ISL_934544 | Institut für Virologie am Department für Hygiene, Mikrobiologie und Public Health | Bergthaler laboratory, CeMM Research Center for Molecular Medicine of the Austrian Academy of Sciences | Lukas Endler, Anna Schedl, Thomas Penz, Benedikt Agerer, Maelle Le Moing, Michael Schuster, Bekir Erguner, Jan Laine, Martin Senekowitsch, Christoph Bock, Andreas Bergthaler |
| EPI_ISL_935107, EPI_ISL_935108, EPI_ISL_935119, EPI_ISL_935120, EPI_ISL_935121, EPI_ISL_935125, EPI_ISL_935128 | 1.AO Universitaria 'S. Giovanni di Dio e Ruggi D'Aragona, Scuola Medica Salernitana' Hospital / 2.UOC di Virologia e Microbiologia, Università della Campania 'L. Vanvitelli' / 3.AO Universitaria 'Federico II' Napoli Hospital / 4.AORN 'San Giuseppe Moscati' Avellino Hospital / 5.AO 'San Pio - presidio G. Rummo' Benevento Hospital / 6.AO 'Sant'Anna e San Sebastiano' Caserta Hospital / 7.PO 'Maria Santissima Addolorata' Eboli Hospital / 8.Biogem Istituto di Ricerche Genetiche | 1. Genome Research Center for Health (CRGS) / 2. Laboratory of Molecular Medicine and Genomics(LMMGe) / 3. Center for Research in Pure and Applied Mathematics (CRMPA) | Giorgio Giurato (Corresponding Author), Francesca Rizzo (Corresponding Author), Alessandro Weisz (Corresponding Author), Gianluigi Franci, Giovanni Nassa, Pasquale Pagliano, Roberta Tarallo, Elena Alexandrova, Ylenia D'Agostino, Carlo Ferravante, Jessica Lamberti, Viola Melone, Domenico Memoli, Valeria Mirici Cappa, Domenico Palumbo, Giovanni Pecoraro, Assunta Sellitto, Oriana Strianese, Ilaria Terenzi, Giuseppe Fenza, Aniello Gentile, Antonello Saccomanno, Sonia Amabile, Teresa Rocco, Annamaria Salvati, Emilia Vaccaro, Massimiliano Galdiero, Michele Cennamo, Giuseppe Portella, Maria Grazia Foti, Mariarosaria Ingino, Maria Landi, Maurizio Fumi, Vincenzo Rocco, Rita Greco, Vittoria Letizia, Arnolfo Petruzziello, Maddalena Schioppa, Gregorio Goffredi, Francesca Marciano, Michele Caraglia, Alessia Cossu, Marianna Scrima |
| EPI_ISL_935415, EPI_ISL_935416, EPI_ISL_935417, EPI_ISL_935418, EPI_ISL_935419, EPI_ISL_935420, EPI_ISL_935421, EPI_ISL_935422, EPI_ISL_935423, EPI_ISL_935424, EPI_ISL_935425, EPI_ISL_935426, EPI_ISL_935427, EPI_ISL_935428, EPI_ISL_935429, EPI_ISL_935430, EPI_ISL_935431, EPI_ISL_935432, EPI_ISL_935433, EPI_ISL_935434, EPI_ISL_935435, EPI_ISL_935436, EPI_ISL_935437, EPI_ISL_935438, EPI_ISL_935439, EPI_ISL_935440, EPI_ISL_935441, EPI_ISL_935442, EPI_ISL_935443, EPI_ISL_935444, EPI_ISL_935445, EPI_ISL_935446, EPI_ISL_935447, EPI_ISL_935448, EPI_ISL_935449, EPI_ISL_935450, EPI_ISL_935451, EPI_ISL_935452, EPI_ISL_935453, EPI_ISL_935454, EPI_ISL_935455, EPI_ISL_935456, EPI_ISL_935457, EPI_ISL_935458, EPI_ISL_935459, EPI_ISL_935460, EPI_ISL_935461, EPI_ISL_935462, EPI_ISL_935463 | | | |
| see above | Florida Bureau of Public Health Laboratories | Florida Bureau of Public Health Laboratories | Sarah Schmedes, Jason Blanton |
| EPI_ISL_935881, EPI_ISL_935900, EPI_ISL_935904, EPI_ISL_935908, EPI_ISL_935919, EPI_ISL_935920, EPI_ISL_935922, EPI_ISL_935925, EPI_ISL_935926, EPI_ISL_935927, EPI_ISL_935928, EPI_ISL_935929, EPI_ISL_935931, EPI_ISL_935932, EPI_ISL_935934, EPI_ISL_935935, EPI_ISL_935936, EPI_ISL_935937, | | | |

| | | | |
|---|---|---|---|
| EPI_ISL_935938, EPI_ISL_935939, EPI_ISL_935940, EPI_ISL_935941, EPI_ISL_935942, EPI_ISL_935943, EPI_ISL_935944, EPI_ISL_935945, EPI_ISL_935946, EPI_ISL_935947, EPI_ISL_935948, EPI_ISL_935949, EPI_ISL_935950, EPI_ISL_935951, EPI_ISL_935952 | | | |
| see above | Cadham Provincial laboratory | National Microbiology Laboratory (NML) | Anna Majer, Shari Tyson, Grace Seo, Philip Mabon, Elsie Grudeski, Rhiannon Huzarewich, Russell Mandes, Anneliese Landgraff, Jennifer Tanner, Natalie Knox, Morag Graham, Gary Van Domselaar, Paul Van Caeseele, Jared Bullard, David Alexander, Kerry Dust, Nathalie Bastien, Yan Li, Timothy Booth, Darian Hole, Madison Chapel, Kirsten Biggar, CanCOGeN's metadata curation team, Public Health Agency of Canada CanCOGeN team |
| EPI_ISL_936613, EPI_ISL_936615, EPI_ISL_936616, EPI_ISL_936617, EPI_ISL_936618 | Northwestern Memorial Hospital | Ozer Lab | Ramon Lorenzo-Redondo, Lacy M. Simons, Chad J. Achenbach, Lawrence J. Jennings, Michael G. Ison, Judd F. Hultquist, Egon A. Ozer |
| EPI_ISL_940427, EPI_ISL_940451 | Hôpital Bichat Claude Bernard, Laboratoire de Virologie | IAME UMR1137 Inserm, Université de Paris, Hôpital Bichat | Antoine Bridier-Nahmias, Amélie Recoing, Quentin Le Hingrat, Lena Daniel, Siham Hamri, Gilles Collin, Alexandre Storto, Mélanie Bertine, Charlotte Charpentier, Nadhira Houhou-Fidouh, Diane Descamps, Benoit Visseaux |
| EPI_ISL_940855 | Vaccines and Infectious Diseases Analytics Research Unit (VIDA) | KRISP, KZN Research Innovation and Sequencing Platform | Baillie Vicky, du Plessis Jeanine, Giandhari Jennifer, Pillay Sureshnee, Naidoo Yeshnee, Tegally Houriiyah, de Oliveira Tulio, Madhi Shabir |
| EPI_ISL_941206 | Servicio de Microbiología. Hospital Clínico Universitario de Valencia | SeqCOVID-SPAIN consortium/IBV(CSIC) | David Navarro Ortega, Eliseo Albert Vicent, Ignacio Torres and SeqCOVID-SPAIN consortium |
| EPI_ISL_941936, EPI_ISL_941937 | Florida Bureau of Public Health Laboratories | Florida Bureau of Public Health Laboratories | Sarah Schmedes, Jason Blanton |
| EPI_ISL_942454, EPI_ISL_942455, EPI_ISL_942456, EPI_ISL_942457, EPI_ISL_942459, EPI_ISL_942460, EPI_ISL_942461, EPI_ISL_942462, EPI_ISL_942463, EPI_ISL_942464, EPI_ISL_942465, EPI_ISL_942466, EPI_ISL_942467, EPI_ISL_942468, EPI_ISL_942469, EPI_ISL_942470, EPI_ISL_942471, EPI_ISL_942472, EPI_ISL_942473, EPI_ISL_942474, EPI_ISL_942475, EPI_ISL_942476, EPI_ISL_942477, EPI_ISL_942478, EPI_ISL_942479 | | | |
| see above | Gundersen Molecular Diagnostics Laboratory | Kabara Cancer Research Institute | Craig S. Richmond, Paraic A. Kenny |
| EPI_ISL_943602, EPI_ISL_943603, EPI_ISL_943605, EPI_ISL_943606, EPI_ISL_943608, EPI_ISL_943610, EPI_ISL_943612 | Lacen_RS | State Center for Health Surveillance. Rio Grande do Sul State Secretary of Health | Aline Campos, Amanda da Silva, Anelise Schaurich, Claudia Dornelles, Cynthia Molina, Fernanda Godinho, Lara Crescente, Leticia Garay, Regina Barcellos, Richard Salvato, Tatiana Gregianini, Vagner Fonseca |
| EPI_ISL_947265 | RS Sari Asih Ciledug | Eijkman Institute for Molecular Biology, Ministry of Research and Technology/National Agency for Research and Innovation | Frilasita A Yudhaputri, Hidayat Trimarsanto, Iskandar Adnan, Lydia V. Panggalo, Sukma Oktavianthi, Willy Agustine, Edison Johar, Safarina G Malik, Khin Saw Myint, Amin Soebandrio |
| EPI_ISL_947268 | RSU Anggrek Mas | Eijkman Institute for Molecular Biology, Ministry of Research and Technology/National Agency for Research and Innovation | Frilasita A Yudhaputri, Hidayat Trimarsanto, Iskandar Adnan, Lydia V. Panggalo, Sukma Oktavianthi, Willy Agustine, Edison Johar, Safarina G Malik, Khin Saw Myint, Amin Soebandrio |
| EPI_ISL_947270 | RS Mitra Keluarga Gading Serpong | Eijkman Institute for Molecular Biology, Ministry of Research and Technology/National Agency for Research and Innovation | Frilasita A Yudhaputri, Hidayat Trimarsanto, Iskandar Adnan, Lydia V. Panggalo, Sukma Oktavianthi, Willy Agustine, Edison Johar, Safarina G Malik, Khin Saw Myint, Amin Soebandrio |
| EPI_ISL_949347, EPI_ISL_949349, EPI_ISL_949350, EPI_ISL_949351, EPI_ISL_949352, EPI_ISL_949353, EPI_ISL_949354, EPI_ISL_949355, EPI_ISL_949356, EPI_ISL_949358, EPI_ISL_949359, EPI_ISL_949360, EPI_ISL_949361, EPI_ISL_949362, EPI_ISL_949363, EPI_ISL_949364, EPI_ISL_949365, EPI_ISL_949366, EPI_ISL_949367, EPI_ISL_949368, EPI_ISL_949369, EPI_ISL_949370, EPI_ISL_949371, EPI_ISL_949372, EPI_ISL_949373, EPI_ISL_949374, EPI_ISL_949375, EPI_ISL_949376, EPI_ISL_949377, EPI_ISL_949378, EPI_ISL_949379, EPI_ISL_949380, EPI_ISL_949381, EPI_ISL_949382, EPI_ISL_949383, EPI_ISL_949384, EPI_ISL_949385, EPI_ISL_949386, EPI_ISL_949387, EPI_ISL_949388, EPI_ISL_949389, EPI_ISL_949390 | | | |
| see above | University of Birmingham | COVID-19 Genomics UK (COG-UK) Consortium | Institute of Microbiology, University of Birmingham: Claire McMurray, Joanne Stockton, Samuel Nicholls, Radoslaw Poplawski, Will Rowe, Josh Quick, Nicholas Loman. University of Birmingham Testing Laboratory: Celina M Whalley, Andrew Bosworth, Charlotte Poxon, Kasun Wanigasooriya, Oliver Pickles, Mike Kidd, Alex Richter, Andrew D Beggs PHE Heartlands Lab: Husam Osman, Andrew Bosworth. Queen Elizabeth Hospital: Anna Casey |
| EPI_ISL_952431 | Centre for Enzyme Innovation, University of Portsmouth / Translational Research Laboratory, Portsmouth Hospitals NHS Trust | COVID-19 Genomics UK (COG-UK) Consortium | Angela Beckett,Salman Goudarzi,Christopher Fearn,Kate Cook,Katie Loveson,Sharon Glaysher,Scott Elliott,Samuel Robson |
| EPI_ISL_955165, EPI_ISL_955166 | Institute for Biocides and Medical Ecology | Institute of microbiology and Immunology, Faculty of Medicine, University of Belgrade | Knezevic,A., Jankovic,M., Vidanovic,D., Milicevic,O., Tesovic,B., Sekler,M., Jovanovic,T. |
| EPI_ISL_955189, EPI_ISL_955190, EPI_ISL_955191, EPI_ISL_955194, EPI_ISL_955195, EPI_ISL_955197, EPI_ISL_955198, EPI_ISL_955199, EPI_ISL_955200, EPI_ISL_955201, EPI_ISL_955202, EPI_ISL_955203, EPI_ISL_955204, EPI_ISL_955205, EPI_ISL_955206, EPI_ISL_955207, EPI_ISL_955208, EPI_ISL_955209 | | | |
| see above | Instituto Nacional de Medicina Genomica | Instituto Nacional de Medicina Genomica | Hidalgo-Miranda A, Mendoza-Vargas A, Reyes-Grajeda JP, Cisneros-Villanueva M, Cedro-Tanda A,Peñaloza-Figueroa F, Herrera-Montalvo LA |
| EPI_ISL_955939, EPI_ISL_955946, EPI_ISL_955947, EPI_ISL_955948, EPI_ISL_955950, EPI_ISL_955954, EPI_ISL_955955, EPI_ISL_955960, EPI_ISL_955961, EPI_ISL_955962, EPI_ISL_955963, EPI_ISL_955964, EPI_ISL_955965, EPI_ISL_955966, EPI_ISL_955967, EPI_ISL_955968, EPI_ISL_955969, EPI_ISL_955970, EPI_ISL_955971, EPI_ISL_955976, EPI_ISL_955977, EPI_ISL_955978 | | | |
| see above | Division of Emerging Infectious Diseases, Bureau of Infectious Diseases Diagnosis Control, Korea Disease Control and Prevention Agency | Division of Emerging Infectious Diseases, Bureau of Infectious Diseases Diagnosis Control, Korea Disease Control and Prevention Agency | Ae Kyung Park, Il-Hwan Kim, Heui Man Kim, Jeong-Min Kim, Namjoo Lee, Chae Young Lee, Sang Hee Woo, Eun-Jin Kim |
| EPI_ISL_959302, EPI_ISL_959303 | Centre de Recerca en Salut Animal (IRTA/CRESA) | IrsiCaixa - Can Ruti CovidSeq | Marc Noguera-Julian, Mariona Parera, Maria Casadellà, Pilar Armengol, Francesc Catala-Moll, Roger Paredes, Bonaventura Clotet J. Segalés, M. Puig, J. Rodon, C. Avila-Nieto, J. Carrillo, G. Cantero, M.T. Terrón, S. Cruz, N. Izquierdo-Useros, E. Vidal, J. Blanco, B. Clotet, J. Vergara-Alert |
| EPI_ISL_959479, EPI_ISL_959480, EPI_ISL_959481, EPI_ISL_959482, EPI_ISL_959501, EPI_ISL_959506 | Division of Emerging Infectious Diseases, Bureau of Infectious Diseases Diagnosis Control, Korea Disease Control and Prevention Agency | Division of Emerging Infectious Diseases, Bureau of Infectious Diseases Diagnosis Control, Korea Disease Control and Prevention Agency | Ae Kyung Park, Il-Hwan Kim, Heui Man Kim, Jeong-Min Kim, Namjoo Lee, Chae Young Lee, Sang Hee Woo, Eun-Jin Kim |
| EPI_ISL_960343, EPI_ISL_960352, EPI_ISL_960375, EPI_ISL_960381, EPI_ISL_960386 | University of Wisconsin-Madison AIDS Vaccine Research Laboratories | University of Wisconsin-Madison AIDS Vaccine Research Laboratories | Gage Moreno, Katarina Braun, et al. AIDS Vaccine Research Laboratories |
| EPI_ISL_960445, EPI_ISL_960450, EPI_ISL_960483, EPI_ISL_960484, EPI_ISL_960485, EPI_ISL_960502, EPI_ISL_960557, EPI_ISL_960568, EPI_ISL_960569, EPI_ISL_960578, EPI_ISL_960579, EPI_ISL_960580, EPI_ISL_960646, EPI_ISL_960660, EPI_ISL_960661 | | | |
| see above | Istituto Zooprofilattico Sperimentale del Mezzogiorno | TIGEM | Patrizia Annunziata, Andrea Ballabio, Valentina Bouche, Davide Cacchiarelli, Pellegrino Cerino, Chiara Colantuono, Maria Concetta Cuomo, Denise Di Concilio, Lucio Di Filippo, Antonio Grimaldi, Antonio Limone, Anna Manfredi, Francesco Panariello, Biancamaria Pierri, Marcello Salvi |
| EPI_ISL_961667 | Hôpital Georges L. Dumont | National Microbiology Laboratory (NML) | Anna Majer, Shari Tyson, Grace Seo, Philip Mabon, Elsie Grudeski, Rhiannon Huzarewich, Russell Mandes, Anneliese Landgraff, Jennifer Tanner, Natalie Knox, Morag Graham, Gary Van Domselaar, Richard Garceau, Guillaume Desnoyers, Nathalie Bastien, Yan Li, Timothy Booth, Darian Hole, Madison Chapel, Kirsten Biggar, CanCOGeN's metadata curation team, Public Health Agency of Canada CanCOGeN team |
| EPI_ISL_962208, EPI_ISL_962209, EPI_ISL_962210, EPI_ISL_962211, EPI_ISL_962212, EPI_ISL_962213, EPI_ISL_962214, EPI_ISL_962215, EPI_ISL_962216, EPI_ISL_962217, EPI_ISL_962218, EPI_ISL_962219, EPI_ISL_962220, EPI_ISL_962221, EPI_ISL_962222, EPI_ISL_962223 | | | |
| see above | Seattle Flu Study | Seattle Flu Study | Deborah A. Nickerson, Chris D. Frazar, Jover Lee, Benjamin Pelle, Erica Ryke, Matthew Richardson, Amanda Adler, Elisabeth Brandstetter, Peter D. Han, Kairsten Fay, Misja Ilcisin, Kirsten Lacombe, Thomas R. Sibley, Melissa Truong, Caitlin R. Wolf, Karen Cowgill, Stephanie Schrag, Jeff Duchin, Michael Boeckh, Janet A. Englund, Michael Famulare, Barry R. Lutz, Mark J. Rieder, Lea M. Starita, Matthew Thompson, Helen Y. Chu, Trevor Bedford, Jay Shendure |
| EPI_ISL_962771, EPI_ISL_962775, EPI_ISL_962781, EPI_ISL_962782, EPI_ISL_962789, EPI_ISL_962794, EPI_ISL_962798, EPI_ISL_962804 | San Diego County Public Health Laboratory | Andersen lab at Scripps Research | SEARCH Alliance San Diego with Tracy Basler, Jovan Shephard, Brett Austin |
| EPI_ISL_965341, EPI_ISL_965344, EPI_ISL_965347, EPI_ISL_965348, EPI_ISL_965349, EPI_ISL_965353, EPI_ISL_965354, EPI_ISL_965360, EPI_ISL_965363, EPI_ISL_965366, EPI_ISL_965367, EPI_ISL_965368, EPI_ISL_965369, EPI_ISL_965370, EPI_ISL_965372, EPI_ISL_965373, EPI_ISL_965374, EPI_ISL_965385, EPI_ISL_965391, EPI_ISL_965461, EPI_ISL_965464 | | | |
| see above | University of Liège COVID-19 testing center | GIGA Medical Genomics | Keith Durkin, Maria Artesi, Bouchra Boujemla, Emmanuel André, Marc Van Ranst, Fabrice Bureau, Laurent Gillet, Wouter Coppieters, Vincent Bours |
| EPI_ISL_965532, EPI_ISL_965543, | Dutch COVID-19 response team | Medical Microbiology, Maastricht University Medical Centre | Jozef Dingemans*, Brian van der Veer*, Erik Beuken, Carmen Reumkens, Lieke van Alphen, Christian Hoebe, Paul Savelkoul |

| | | | |
|---|---|---|---|
| EPI_ISL_965663 | | | |
| EPI_ISL_968199, EPI_ISL_968200, EPI_ISL_968201, EPI_ISL_968202, EPI_ISL_968203, EPI_ISL_968204 | Arizona State Public Health Laboratory | Arizona State Public Health Laboratory | Trung Huynh, Jessica Escobar, Katherine Fullerton, Nobuko Fukushima, Stacy White, Linda Getsinger, Victor Waddell |
| EPI_ISL_968808, EPI_ISL_968810, EPI_ISL_968925, EPI_ISL_968926, EPI_ISL_968927, EPI_ISL_968993 | KEMRI-Wellcome Trust Research Programme/KEMRI-CGMR-C Kilifi | KEMRI-Wellcome Trust Research Programme/KEMRI-CGMR-C Kilifi | Githinji et al |
| EPI_ISL_975495, EPI_ISL_975496, EPI_ISL_975497, EPI_ISL_975498, EPI_ISL_975499, EPI_ISL_975500, EPI_ISL_975501, EPI_ISL_975502, EPI_ISL_975503, EPI_ISL_975504, EPI_ISL_975505, EPI_ISL_975506, EPI_ISL_975507, EPI_ISL_975508, EPI_ISL_975509, EPI_ISL_975510, EPI_ISL_975511, EPI_ISL_975512, EPI_ISL_975513, EPI_ISL_975514, EPI_ISL_975515, EPI_ISL_975516, EPI_ISL_975517, EPI_ISL_975518, EPI_ISL_975519, EPI_ISL_975520, EPI_ISL_975521, EPI_ISL_975522, EPI_ISL_975523, EPI_ISL_975524, EPI_ISL_975525, EPI_ISL_975526, EPI_ISL_975527, EPI_ISL_975528, EPI_ISL_975529, EPI_ISL_975530, EPI_ISL_975531, EPI_ISL_975532, EPI_ISL_975533, EPI_ISL_975534, EPI_ISL_975535, EPI_ISL_975536, EPI_ISL_975537, EPI_ISL_975538, EPI_ISL_975539, EPI_ISL_975540, EPI_ISL_975541, EPI_ISL_975542, EPI_ISL_975543, EPI_ISL_975544, EPI_ISL_975545, EPI_ISL_975546, EPI_ISL_975547, EPI_ISL_975548, EPI_ISL_975549, EPI_ISL_975550, EPI_ISL_975551, EPI_ISL_975552, EPI_ISL_975553, EPI_ISL_975554, EPI_ISL_975555, EPI_ISL_975556, EPI_ISL_975557, EPI_ISL_975558, EPI_ISL_975559, EPI_ISL_975560, EPI_ISL_975561, EPI_ISL_975562, EPI_ISL_975563, EPI_ISL_975564, EPI_ISL_975565, EPI_ISL_975566, EPI_ISL_975567, EPI_ISL_975568, EPI_ISL_975569, EPI_ISL_975570, EPI_ISL_975571, EPI_ISL_975572, EPI_ISL_975573, EPI_ISL_975574, EPI_ISL_975575, EPI_ISL_975576, EPI_ISL_975577, EPI_ISL_975578, EPI_ISL_975579, EPI_ISL_975580, EPI_ISL_975581, EPI_ISL_975582, EPI_ISL_975583, EPI_ISL_975584, EPI_ISL_975585, EPI_ISL_975586, EPI_ISL_975587, EPI_ISL_975588, EPI_ISL_975589, EPI_ISL_975590, EPI_ISL_975591, EPI_ISL_975592, EPI_ISL_975593, EPI_ISL_975594, EPI_ISL_975595, EPI_ISL_975596, EPI_ISL_975597, EPI_ISL_975598, EPI_ISL_975599, EPI_ISL_975600, EPI_ISL_975601, EPI_ISL_975602, EPI_ISL_975603, EPI_ISL_975604, EPI_ISL_975605, EPI_ISL_975606, EPI_ISL_975607, EPI_ISL_975608, EPI_ISL_975609, EPI_ISL_975610, EPI_ISL_975611, EPI_ISL_975612, EPI_ISL_975613, EPI_ISL_975614, EPI_ISL_975615, EPI_ISL_975616, EPI_ISL_975617, EPI_ISL_975618, EPI_ISL_975619, EPI_ISL_975620, EPI_ISL_975621, EPI_ISL_975622, EPI_ISL_975623, EPI_ISL_975624, EPI_ISL_975625, EPI_ISL_975626, EPI_ISL_975627, EPI_ISL_975628, EPI_ISL_975629, EPI_ISL_975630, EPI_ISL_975631, EPI_ISL_975632, EPI_ISL_975633, EPI_ISL_975634, EPI_ISL_975635, EPI_ISL_975636, EPI_ISL_975637, EPI_ISL_975638, EPI_ISL_975639, EPI_ISL_975640, EPI_ISL_975641, EPI_ISL_975642, EPI_ISL_975643, EPI_ISL_975644, EPI_ISL_975645, EPI_ISL_975646, EPI_ISL_975647, EPI_ISL_975648, EPI_ISL_975649, EPI_ISL_975650, EPI_ISL_975651, EPI_ISL_975652, EPI_ISL_975653, EPI_ISL_975654, EPI_ISL_975655, EPI_ISL_975656, EPI_ISL_975657, EPI_ISL_975658, EPI_ISL_975659, EPI_ISL_975660 | | | |
| see above | BCCDC Public Health Laboratory | BCCDC Public Health Laboratory | Prystajecky Natalie, Linda Hoang, Dan Fornika, John Tyson, Shannon Russell, Kim Macdonald, Kimia Kamelian, Ana Pacagnella, Corrinne Ng, Loretta Janz, Robert Azana Terry Snutch, Mel Krajden |
| EPI_ISL_981176 | Johns Hopkins Hospital Department of Pathology | Johns Hopkins Hospital Department of Pathology | C. Paul Morris, Chun Huai Luo, Adannaya Amadi, Matthew Schwartz, Nicholas Gallagher, Heba H. Mostafa |
| EPI_ISL_982771, EPI_ISL_982772, EPI_ISL_982773, EPI_ISL_982774, EPI_ISL_982775 | Kentucky State Public Health Lab | Kentucky State Public Health Lab | Stephanie Lunn, Karim George, Joshua Tobias, William Grooms, Vaneet Arora, Matthew Johnson, Rachel Zinner, Rhonda Lucas |
| EPI_ISL_983699, EPI_ISL_983700 | Essentia Health-St. Mary's Medical Center | Minnesota Department of Health, Public Health Laboratory | Alexandra Lorentz, Jacob Garfin, Matt Plumb, and Xiong Wang |
