## Supplementary material for "Detection and characterization of the SARS-CoV-2 lineage B.1.526 in New York": Supp. Table 3 GISAID Acknowledgment part 2: gisaid_hcov-19_acknowledgement_table_2021_02_13_00-10.pdf

We gratefully acknowledge the following Authors from the Originating laboratories responsible for obtaining the specimens, as well as the Submitting laboratories where the genome data were generated and shared via GISAID, on which this research is based.

All Submitters of data may be contacted directly via www.gisaid.org

Authors are sorted alphabetically.

| Accession ID | Originating Laboratory | Submitting Laboratory | Authors |
|---|---|---|---|
| EPI_ISL_660794, EPI_ISL_661175, EPI_ISL_661176 | Respiratory Virus Unit, Microbiology Services Colindale, Public Health England | COVID-19 Genomics UK (COG-UK) Consortium | PHE Covid Sequencing Team |
| EPI_ISL_661260 | Canterbury Health Laboratories | Institute of Environmental Science and Research (ESR) | Xiaoyun Ren, Matt Storey, Nikki Freed, Muhammad Faisal, Jing Wang, Hermes Perez, Anja Werno, Antje van der Linden, Arlo Upton, Chris Mansell, David Hammer, Dragana Drinkovic, Gary McAuliffe, Hana Sofia Andersson, James Ussher, Jill Sherwood, Josh Freeman, Julia Howard, Juliet Elvy, Mary DeAlmeida, Matt Blakiston, Matthew Rogers, Max Bloomfield, Michael Addidle, Michelle Balm, Sally Roberts, Sarah Jefferies, Sharmini Muttaiyah, Susan Morpeth, Susan Taylor, Timothy Blackmore, Vani Sathyendran, Veronica Playle, Virginia Hope, Erasmus Smit, Lauren Jelly, Olin Silander, Joep de Ligt |
| EPI_ISL_661261, EPI_ISL_661264 | LabPLUS | Institute of Environmental Science and Research (ESR) | Xiaoyun Ren, Matt Storey, Nikki Freed, Muhammad Faisal, Jing Wang, Hermes Perez, Anja Werno, Antje van der Linden, Arlo Upton, Chris Mansell, David Hammer, Dragana Drinkovic, Gary McAuliffe, Hana Sofia Andersson, James Ussher, Jill Sherwood, Josh Freeman, Julia Howard, Juliet Elvy, Mary DeAlmeida, Matt Blakiston, Matthew Rogers, Max Bloomfield, Michael Addidle, Michelle Balm, Sally Roberts, Sarah Jefferies, Sharmini Muttaiyah, Susan Morpeth, Susan Taylor, Timothy Blackmore, Vani Sathyendran, Veronica Playle, Virginia Hope, Erasmus Smit, Lauren Jelly, Olin Silander, Joep de Ligt |
| EPI_ISL_664149, EPI_ISL_664150, EPI_ISL_664326, EPI_ISL_664381, EPI_ISL_664546, EPI_ISL_664591, EPI_ISL_664610, EPI_ISL_665013, EPI_ISL_665014, EPI_ISL_665015, EPI_ISL_665016, EPI_ISL_665017, EPI_ISL_665018, EPI_ISL_665019 | | | |
| see above | Department of Pathology, University of Cambridge | COVID-19 Genomics UK (COG-UK) Consortium | Aminu S. Jahun, Yasmin Chaudhry, Grant Hall, Iliana Georgana, Myra Hosmillo, Martin D. Curran, Malte Pinckert, Surendra Parmar, Ian Goodfellow |
| EPI_ISL_665290 | Queens Medical Centre, Clinical Microbiology Department / DeepSeq Nottingham | COVID-19 Genomics UK (COG-UK) Consortium | Gemma Clark, Wendy Smith, Manjinder Khakh, Vicki M Fleming, Michelle M Lister, Hannah Howson-Wells, Jonathan Ball, Patrick McClure, Joseph Chappell, Theocharis Tsoleridis, Nadine Holmes, Matthew Carlisle, Christopher Moore, Fei Sang, Johnny Debebe, Victoria Wright, Matthew Loose |
| EPI_ISL_665295 | Quadram Institute Bioscience | COVID-19 Genomics UK (COG-UK) Consortium | Dave J. Baker, Gemma L. Kay, Alp Aydin, Thanh Le-Viet, Steven Rudder, Ana P. Tedim, Anastasia Kolyva, Maria Diaz, Leonardo de Oliveira Martins, Nabil-Fareed Alikhan, Lizzie Meadows, Rachael Stanley, Ngozi Elumogo, Muhammed Yasir, Nicholas M. Thomson, Alexander J Trotter, Rachel Gilroy, Samuel Bloomfield, Claire Stuart, Andrew Bell, Reenesh Prakash, Samir Dervisevic, Alison E. Mather, John Wain, Mark Webber, Andrew J. Page, Justin O'Grady |
| EPI_ISL_665344, EPI_ISL_665366 | Queens Medical Centre, Clinical Microbiology Department / DeepSeq Nottingham | COVID-19 Genomics UK (COG-UK) Consortium | Gemma Clark, Wendy Smith, Manjinder Khakh, Vicki M Fleming, Michelle M Lister, Hannah Howson-Wells, Jonathan Ball, Patrick McClure, Joseph Chappell, Theocharis Tsoleridis, Nadine Holmes, Matthew Carlisle, Christopher Moore, Fei Sang, Johnny Debebe, Victoria Wright, Matthew Loose |
| EPI_ISL_665377, EPI_ISL_665380, EPI_ISL_665381 | Quadram Institute Bioscience | COVID-19 Genomics UK (COG-UK) Consortium | Dave J. Baker, Gemma L. Kay, Alp Aydin, Thanh Le-Viet, Steven Rudder, Ana P. Tedim, Anastasia Kolyva, Maria Diaz, Leonardo de Oliveira Martins, Nabil-Fareed Alikhan, Lizzie Meadows, Rachael Stanley, Ngozi Elumogo, Muhammed Yasir, Nicholas M. Thomson, Alexander J Trotter, Rachel Gilroy, Samuel Bloomfield, Claire Stuart, Andrew Bell, Reenesh Prakash, Samir Dervisevic, Alison E. Mather, John Wain, Mark Webber, Andrew J. Page, Justin O'Grady |
| EPI_ISL_665415 | Queens Medical Centre, Clinical Microbiology Department / DeepSeq Nottingham | COVID-19 Genomics UK (COG-UK) Consortium | Gemma Clark, Wendy Smith, Manjinder Khakh, Vicki M Fleming, Michelle M Lister, Hannah Howson-Wells, Jonathan Ball, Patrick McClure, Joseph Chappell, Theocharis Tsoleridis, Nadine Holmes, Matthew Carlisle, Christopher Moore, Fei Sang, Johnny Debebe, Victoria Wright, Matthew Loose |
| EPI_ISL_665423, EPI_ISL_665440, EPI_ISL_665450, EPI_ISL_665508 | Quadram Institute Bioscience | COVID-19 Genomics UK (COG-UK) Consortium | Dave J. Baker, Gemma L. Kay, Alp Aydin, Thanh Le-Viet, Steven Rudder, Ana P. Tedim, Anastasia Kolyva, Maria Diaz, Leonardo de Oliveira Martins, Nabil-Fareed Alikhan, Lizzie Meadows, Rachael Stanley, Ngozi Elumogo, Muhammed Yasir, Nicholas M. Thomson, Alexander J Trotter, Rachel Gilroy, Samuel Bloomfield, Claire Stuart, Andrew Bell, Reenesh Prakash, Samir Dervisevic, Alison E. Mather, John Wain, Mark Webber, Andrew J. Page, Justin O'Grady |
| EPI_ISL_665512, EPI_ISL_665513 | Queens Medical Centre, Clinical Microbiology Department / DeepSeq Nottingham | COVID-19 Genomics UK (COG-UK) Consortium | Gemma Clark, Wendy Smith, Manjinder Khakh, Vicki M Fleming, Michelle M Lister, Hannah Howson-Wells, Jonathan Ball, Patrick McClure, Joseph Chappell, Theocharis Tsoleridis, Nadine Holmes, Matthew Carlisle, Christopher Moore, Fei Sang, Johnny Debebe, Victoria Wright, Matthew Loose |
| EPI_ISL_665518 | Quadram Institute Bioscience | COVID-19 Genomics UK (COG-UK) Consortium | Dave J. Baker, Gemma L. Kay, Alp Aydin, Thanh Le-Viet, Steven Rudder, Ana P. Tedim, Anastasia Kolyva, Maria Diaz, Leonardo de Oliveira Martins, Nabil-Fareed Alikhan, Lizzie Meadows, Rachael Stanley, Ngozi Elumogo, Muhammed Yasir, Nicholas M. Thomson, Alexander J Trotter, Rachel Gilroy, Samuel Bloomfield, Claire Stuart, Andrew Bell, Reenesh Prakash, Samir Dervisevic, Alison E. Mather, John Wain, Mark Webber, Andrew J. Page, Justin O'Grady |
| EPI_ISL_665528, EPI_ISL_665545 | Queens Medical Centre, Clinical Microbiology Department / DeepSeq Nottingham | COVID-19 Genomics UK (COG-UK) Consortium | Gemma Clark, Wendy Smith, Manjinder Khakh, Vicki M Fleming, Michelle M Lister, Hannah Howson-Wells, Jonathan Ball, Patrick McClure, Joseph Chappell, Theocharis Tsoleridis, Nadine Holmes, Matthew Carlisle, Christopher Moore, Fei Sang, Johnny Debebe, Victoria Wright, Matthew Loose |
| EPI_ISL_665574, EPI_ISL_665594, EPI_ISL_665596, EPI_ISL_665603 | Quadram Institute Bioscience | COVID-19 Genomics UK (COG-UK) Consortium | Dave J. Baker, Gemma L. Kay, Alp Aydin, Thanh Le-Viet, Steven Rudder, Ana P. Tedim, Anastasia Kolyva, Maria Diaz, Leonardo de Oliveira Martins, Nabil-Fareed Alikhan, Lizzie Meadows, Rachael Stanley, Ngozi Elumogo, Muhammed Yasir, Nicholas M. Thomson, Alexander J Trotter, Rachel Gilroy, Samuel Bloomfield, Claire Stuart, Andrew Bell, Reenesh Prakash, Samir Dervisevic, Alison E. Mather, John Wain, Mark Webber, Andrew J. Page, Justin O'Grady |
| EPI_ISL_665610, EPI_ISL_665623 | Queens Medical Centre, Clinical Microbiology Department / DeepSeq Nottingham | COVID-19 Genomics UK (COG-UK) Consortium | Gemma Clark, Wendy Smith, Manjinder Khakh, Vicki M Fleming, Michelle M Lister, Hannah Howson-Wells, Jonathan Ball, Patrick McClure, Joseph Chappell, Theocharis Tsoleridis, Nadine Holmes, Matthew Carlisle, Christopher Moore, Fei Sang, Johnny Debebe, Victoria Wright, Matthew Loose |
| EPI_ISL_665639, EPI_ISL_665665, EPI_ISL_665689, EPI_ISL_665712, EPI_ISL_665732, EPI_ISL_665737, EPI_ISL_665749, EPI_ISL_665897 | Quadram Institute Bioscience | COVID-19 Genomics UK (COG-UK) Consortium | Dave J. Baker, Gemma L. Kay, Alp Aydin, Thanh Le-Viet, Steven Rudder, Ana P. Tedim, Anastasia Kolyva, Maria Diaz, Leonardo de Oliveira Martins, Nabil-Fareed Alikhan, Lizzie Meadows, Rachael Stanley, Ngozi Elumogo, Muhammed Yasir, Nicholas M. Thomson, Alexander J Trotter, Rachel Gilroy, Samuel Bloomfield, Claire Stuart, Andrew Bell, Reenesh Prakash, Samir Dervisevic, Alison E. Mather, John Wain, Mark Webber, Andrew J. Page, Justin O'Grady |
| EPI_ISL_665909, EPI_ISL_665910, EPI_ISL_665911, EPI_ISL_665912, EPI_ISL_665913, EPI_ISL_665914, EPI_ISL_665915, EPI_ISL_665916, EPI_ISL_665917, EPI_ISL_665918, EPI_ISL_665919, EPI_ISL_665920, EPI_ISL_665921, EPI_ISL_665922, EPI_ISL_665923, EPI_ISL_665924, EPI_ISL_665925, EPI_ISL_665926, EPI_ISL_665927, EPI_ISL_665928, EPI_ISL_665929, EPI_ISL_665930, EPI_ISL_665931, EPI_ISL_665932, EPI_ISL_665933, EPI_ISL_665934, EPI_ISL_665935, EPI_ISL_665936, EPI_ISL_665937, EPI_ISL_665938, EPI_ISL_665939 | | | |
| see above | Queens Medical Centre, Clinical Microbiology Department / DeepSeq Nottingham | COVID-19 Genomics UK (COG-UK) Consortium | Gemma Clark, Wendy Smith, Manjinder Khakh, Vicki M Fleming, Michelle M Lister, Hannah Howson-Wells, Jonathan Ball, Patrick McClure, Joseph Chappell, Theocharis Tsoleridis, Nadine Holmes, Matthew Carlisle, Christopher Moore, Fei Sang, Johnny Debebe, Victoria Wright, Matthew Loose |
| EPI_ISL_674863 | Lighthouse Lab in Milton Keynes | Wellcome Sanger Institute for the COVID-19 Genomics UK (COG-UK) Consortium | The Lighthouse Lab in Milton Keynes and Alex Alderton, Roberto Amato, Sonia Goncalves, Ewan Harrison, David K. Jackson, Ian Johnston, Dominic Kwiatkowski, Cordelia Langford, John Sillitoe on behalf of the Wellcome Sanger Institute COVID-19 Surveillance Team |
| EPI_ISL_676491, EPI_ISL_676502, EPI_ISL_676503, EPI_ISL_676504 | Klinisk mikrobiologi | The Public Health Agency of Sweden | Department of Microbiology, The Public Health Agency of Sweden |
| EPI_ISL_678261, EPI_ISL_678262, EPI_ISL_678263, EPI_ISL_678264, EPI_ISL_678265, EPI_ISL_678266, EPI_ISL_678267, EPI_ISL_678268 | Communicable Disease Laboratory, Public Health Directorate | Communicable Disease Laboratory, Public Health Directorate | Alwasti,H., Altaif,Z., AlHujairi,Z., AlAbbas,Z. |
| EPI_ISL_678598 | NHLS-IALCH | KRISP, KZN Research Innovation and Sequencing Platform | Giandhari J, Pillay S, Lessells R, ChimukangaraB, Mdlalose K, York D, Khan S, Tegally H, Wilkinson E, de Oliveira T |
| EPI_ISL_678747, EPI_ISL_678748, EPI_ISL_678754, EPI_ISL_678756, EPI_ISL_678757, EPI_ISL_678758, EPI_ISL_678759, EPI_ISL_678762, EPI_ISL_678763, EPI_ISL_678764, EPI_ISL_678765, EPI_ISL_678766, EPI_ISL_678767, EPI_ISL_678768, EPI_ISL_678769, EPI_ISL_678770, EPI_ISL_678771, EPI_ISL_678772, EPI_ISL_678774, EPI_ISL_678775, EPI_ISL_678776 | | | |

| | | | |
|---|---|---|---|
| see above | Respiratory Virus Unit, Microbiology Services Colindale, Public Health England | COVID-19 Genomics UK (COG-UK) Consortium | PHE Covid Sequencing Team |
| EPI_ISL_679160, EPI_ISL_679161, EPI_ISL_679193, EPI_ISL_679195 | Department of Pathology, University of Cambridge | COVID-19 Genomics UK (COG-UK) Consortium | Aminu S. Jahun, Yasmin Chaudhry, Grant Hall, Iliana Georgana, Myra Hosmillo, Martin D. Curran, Malte Pinckert, Surendra Parmar, Ian Goodfellow |
| EPI_ISL_679249, EPI_ISL_679250, EPI_ISL_679251, EPI_ISL_679252, EPI_ISL_679253, EPI_ISL_679254, EPI_ISL_679255, EPI_ISL_679256, EPI_ISL_679257, EPI_ISL_679258, EPI_ISL_679259, EPI_ISL_679260, EPI_ISL_679261, EPI_ISL_679262, EPI_ISL_679263, EPI_ISL_679264, EPI_ISL_679265, EPI_ISL_679266, EPI_ISL_679267, EPI_ISL_679268, EPI_ISL_679269, EPI_ISL_679270, EPI_ISL_679271, EPI_ISL_679272, EPI_ISL_679273, EPI_ISL_679274, EPI_ISL_679275, EPI_ISL_679276, EPI_ISL_679277, EPI_ISL_679278, EPI_ISL_679279, EPI_ISL_679280, EPI_ISL_679281, EPI_ISL_679282, EPI_ISL_679283, EPI_ISL_679284, EPI_ISL_679285, EPI_ISL_679286, EPI_ISL_679287, EPI_ISL_679288, EPI_ISL_679289, EPI_ISL_679290, EPI_ISL_679291, EPI_ISL_679295, EPI_ISL_679297, EPI_ISL_679298, EPI_ISL_679299, EPI_ISL_679300, EPI_ISL_679301, EPI_ISL_679302, EPI_ISL_679303, EPI_ISL_679304, EPI_ISL_679305, EPI_ISL_679306, EPI_ISL_679307, EPI_ISL_679312, EPI_ISL_679313, EPI_ISL_679314 | | | |
| see above | Queens Medical Centre, Clinical Microbiology Department / DeepSeq Nottingham | COVID-19 Genomics UK (COG-UK) Consortium | Gemma Clark, Wendy Smith, Manjinder Khakh, Vicki M Fleming, Michelle M Lister, Hannah Howson-Wells, Jonathan Ball, Patrick McClure, Joseph Chappell, Theocharis Tsoleridis, Nadine Holmes, Matthew Carlisle, Christopher Moore, Fei Sang, Johnny Debebe, Victoria Wright, Matthew Loose |
| EPI_ISL_679393, EPI_ISL_679394, EPI_ISL_679395, EPI_ISL_679396, EPI_ISL_679397, EPI_ISL_679398, EPI_ISL_679399, EPI_ISL_679400, EPI_ISL_679401, EPI_ISL_679402, EPI_ISL_679403, EPI_ISL_679404, EPI_ISL_679405 | | | |
| see above | Liverpool Clinical Laboratories | COVID-19 Genomics UK (COG-UK) Consortium | Sam Haldenby, Anita Lucaci, Steve Paterson, Julian Hiscox, Alistair Darby, M Almsaud, A Alrezaihi, Muhannad Alruwaili, Stuart D Armstrong, Jones Benjamin, Eleanor G Bentley, Anu Chawla, Jordan J Clark, Angela Cowell, Richard Eccles, Isabel García-Dorival, Matthew Gemmell, Alessandro Gerada, PKF Gilmore, Richard Gregory, Ximeng Han, Catherine Hartley, Margaret Hughes, Miren Iturriza-Gomara, James Johnson, L Luu, Jenifer Manson, Charlotte Nelson, Elaine O'Toole, Cassie Olateju, Rebekah Penrice-Randal , Lucille Rainbow, N.P Randle, Trevor Ian Robinson, Parul Sharma, Ghada T Shawli, James P Stewart, Neil Swainston, Ecaterina Vamos, Joanne Watts, Mark Whitehead |
| EPI_ISL_679406, EPI_ISL_679407, EPI_ISL_679422, EPI_ISL_679423, EPI_ISL_679424, EPI_ISL_679425, EPI_ISL_679426, EPI_ISL_679427, EPI_ISL_679428, EPI_ISL_679429, EPI_ISL_679432, EPI_ISL_679433, EPI_ISL_679440, EPI_ISL_679441, EPI_ISL_679442, EPI_ISL_679443, EPI_ISL_679444, EPI_ISL_679445, EPI_ISL_679446 | | | |
| see above | University College London Hospital | COVID-19 Genomics UK (COG-UK) Consortium | Judith Heaney, Matthew Byott, Catherine Houlihan, Dan Frampton, Stuart Kirk, Moira Spyer and Eleni Nastouli |
| EPI_ISL_679593, EPI_ISL_679594, EPI_ISL_679595, EPI_ISL_679597, EPI_ISL_679598, EPI_ISL_679599, EPI_ISL_679600, EPI_ISL_679602, EPI_ISL_679603 | University College London, Great Ormond Street Hospital for Children NHS Foundation Trust, Imperial College Healthcare NHS Trust | COVID-19 Genomics UK (COG-UK) Consortium | Sergi Castellano, Rachel Williams, Mark Kristiansen, Paola Resende Silva, Sunando Roy, Tony Brooks, Helena Tutill, Paola Niola, Patricia Dyal, Charlotte Williams, Leysa Forrest, Yasmin Panchbhaya, Jacqueline Findlay, Samuel Weeks, Julianne Brown, Kathryn Harris, Paul Randell, James Price, Alison Holmes, Judith Breuer |
| EPI_ISL_679934 | Queens Medical Centre, Clinical Microbiology Department / DeepSeq Nottingham | COVID-19 Genomics UK (COG-UK) Consortium | Gemma Clark, Wendy Smith, Manjinder Khakh, Vicki M Fleming, Michelle M Lister, Hannah Howson-Wells, Jonathan Ball, Patrick McClure, Joseph Chappell, Theocharis Tsoleridis, Nadine Holmes, Matthew Carlisle, Christopher Moore, Fei Sang, Johnny Debebe, Victoria Wright, Matthew Loose |
| EPI_ISL_679946, EPI_ISL_679947, EPI_ISL_679948, EPI_ISL_679949, EPI_ISL_679950, EPI_ISL_679951, EPI_ISL_679952, EPI_ISL_679953, EPI_ISL_679954, EPI_ISL_679955 | Lincolnshire Hospitals and DeepSeq Nottingham | COVID-19 Genomics UK (COG-UK) Consortium | Nichola Duckworth, Tim Sloan, Sarah Walsh, Jonathan Ball, Patrick McClure, Joeseph Chappell, Nadine Holmes, Matthew Carlisle, Christopher Moore, Fei Sang, Johnny Debebe, Victoria Wright, Matthew Loose |
| EPI_ISL_679971 | Centre for Enzyme Innovation, University of Portsmouth / Translational Research Laboratory, Portsmouth Hospitals NHS Trust | COVID-19 Genomics UK (COG-UK) Consortium | Angela Beckett,Yann Bourgeois,Garry Scarlett,Sharon Glaysher,Scott Elliott,Kelly Bicknell,Robert Impey,Allyson Lloyd,Sarah Wyllie,Ethan Butcher,Anoop Chauhan,Samuel Robson |
| EPI_ISL_680110, EPI_ISL_680111, EPI_ISL_680112, EPI_ISL_680121, EPI_ISL_680135, EPI_ISL_680136, EPI_ISL_680141, EPI_ISL_680144, EPI_ISL_680147, EPI_ISL_680148, EPI_ISL_680150, EPI_ISL_680154, EPI_ISL_680155, EPI_ISL_680158, EPI_ISL_680168, EPI_ISL_680170, EPI_ISL_680171, EPI_ISL_680178, EPI_ISL_680183, EPI_ISL_680194 | | | |
| see above | Virology Department, Sheffield Teaching Hospitals NHS Foundation Trust/Department of Infection, Immunity and Cardiovascular Disease, The Medical School, University of Sheffield | COVID-19 Genomics UK (COG-UK) Consortium | Thushan de Silva, Matthew Parker, Nikki Smith, Adri Angyal, Rebecca Brown, Luke Green, Rachel Tucker, Paul Parsons, Danielle Groves, Katie Johnson, Laura Carrilero, Alex Keeley, Dave Partridge, Matthew Wyles, Benjamin Lindsey, Mehmet Yavuz, Mohammad Raza, Cariad Evans |
| EPI_ISL_680464, EPI_ISL_680465, EPI_ISL_680466, EPI_ISL_680467, EPI_ISL_680468, EPI_ISL_680486, EPI_ISL_680487, EPI_ISL_680488, EPI_ISL_680489, EPI_ISL_680490, EPI_ISL_680491, EPI_ISL_680492, EPI_ISL_680493, EPI_ISL_680494 | | | |
| see above | Virology Department, Royal Infirmary of Edinburgh, NHS Lothian / School of Biological Sciences, University of Edinburgh / Institute of Genetics and Molecular Medicine, University of Edinburgh | COVID-19 Genomics UK (COG-UK) Consortium | McHugh M, Dewar R, Rooke S, Gallagher M, Balcaza C, O'Toole Á, Scher E, Hill V, McCrone JT, Colquhoun R, Yu X, Jackson B, Rambaut A, Williams TC, Templeton K |
| EPI_ISL_681466 | Lighthouse Lab in Milton Keynes | Wellcome Sanger Institute for the COVID-19 Genomics UK (COG-UK) Consortium | The Lighthouse Lab in Milton Keynes and Alex Alderton, Roberto Amato, Sonia Goncalves, Ewan Harrison, David K. Jackson, Ian Johnston, Dominic Kwiatkowski, Cordelia Langford, John Sillitoe on behalf of the Wellcome Sanger Institute COVID-19 Surveillance Team |
| EPI_ISL_682280, EPI_ISL_682281, EPI_ISL_682282 | Middlemore Hospital | Institute of Environmental Science and Research (ESR) | Xiaoyun Ren, Matt Storey, Nikki Freed, Muhammad Faisal, Jing Wang, Hermes Perez, Anja Werno, Antje van der Linden, Arlo Upton, Chris Mansell, David Hammer, Dragana Drinkovic, Gary McAuliffe, Hana Sofia Andersson, James Ussher, Jill Sherwood, Josh Freeman, Julia Howard, Juliet Elvy, Mary DeAlmeida, Matt Blakiston, Matthew Rogers, Max Bloomfield, Michael Addidle, Michelle Balm, Sally Roberts, Sarah Jefferies, Sharmini Muttaiyah, Susan Morpeth, Susan Taylor, Timothy Blackmore, Vani Sathyendran, Veronica Playle, Virginia Hope, Erasmus Smit, Lauren Jelly, Olin Silander, Joep de Ligt |
| EPI_ISL_682322 | Communicable Disease Laboratory, Public Health Directorate | Communicable Disease Laboratory, Public Health Directorate | Alwasti,H., Altaif,Z., AlHujairi,Z., AlAbbas,Z. |
| EPI_ISL_682353, EPI_ISL_682354, EPI_ISL_682355, EPI_ISL_682356, EPI_ISL_682357, EPI_ISL_682358, EPI_ISL_682359, EPI_ISL_682360, EPI_ISL_682361, EPI_ISL_682362 | Department of Virus and Microbiological Special Diagnostics, Statens Serum Institut, Copenhagen, Denmark | Albertsen Lab, Department of Chemistry and Bioscience, Aalborg University, Denmark | Danish Covid-19 Genome Consortium |
| EPI_ISL_683482, EPI_ISL_683566, EPI_ISL_683567, EPI_ISL_683568, EPI_ISL_683569, EPI_ISL_683570, EPI_ISL_683571, EPI_ISL_683572, EPI_ISL_683576, EPI_ISL_683577, EPI_ISL_683585 | | | |
| see above | Respiratory Virus Unit, Microbiology Services Colindale, Public Health England | COVID-19 Genomics UK (COG-UK) Consortium | PHE Covid Sequencing Team |
| EPI_ISL_683754 | DOHMH Morrisania | New York City Public Health Laboratory | Jade Wang, et al. |
| EPI_ISL_683761 | DOHMH Corona | New York City Public Health Laboratory | Jade Wang, et al. |
| EPI_ISL_683762 | DOHMH Morrisania | New York City Public Health Laboratory | Jade Wang, et al. |
| EPI_ISL_683765 | DOHMH Chelsea | New York City Public Health Laboratory | Jade Wang, et al. |
| EPI_ISL_683793, EPI_ISL_683794 | DOHMH Corona | New York City Public Health Laboratory | Jade Wang, et al. |
| EPI_ISL_683795 | DOHMH Central Harlem | New York City Public Health Laboratory | Jade Wang, et al. |
| EPI_ISL_683796 | DOHMH Corona | New York City Public Health Laboratory | Jade Wang, et al. |
| EPI_ISL_683797 | DOHMH PHL | New York City Public Health Laboratory | Jade Wang, et al. |
| EPI_ISL_683798, EPI_ISL_683799, EPI_ISL_683800 | DOHMH Corona | New York City Public Health Laboratory | Jade Wang, et al. |
| EPI_ISL_683801 | DOHMH Morrisania | New York City Public Health Laboratory | Jade Wang, et al. |
| EPI_ISL_683802 | DOHMH Corona | New York City Public Health Laboratory | Jade Wang, et al. |

| | | | |
|---|---|---|---|
| EPI_ISL_683803, EPI_ISL_683804, EPI_ISL_683805, EPI_ISL_683806 | DOHMH Morrisania | New York City Public Health Laboratory | Jade Wang, et al. |
| EPI_ISL_683807, EPI_ISL_683808 | DOHMH Chelsea | New York City Public Health Laboratory | Jade Wang, et al. |
| EPI_ISL_686611 | Respiratory Virus Unit, Microbiology Services Colindale, Public Health England | COVID-19 Genomics UK (COG-UK) Consortium | PHE Covid Sequencing Team |
| EPI_ISL_692769, EPI_ISL_692772, EPI_ISL_692773 | Massachusetts State Public Health Laboratory | Massachusetts State Public Health Laboratory | Andrew Lang, Timelia Fink, Glen Gallagher, Sandra Smole |
| EPI_ISL_693304, EPI_ISL_693305, EPI_ISL_693306 | Department of Laboratory Medicine, National Taiwan University Hospital | Microbial Genomics Core Lab, National Taiwan University Centers of Genomic and Precision Medicine | Shiou-Hwei Yeh, You-Yu Lin, Ya-Yun Lai, Chiao-Ling Li, Shan-Chwen Chang, Pei-Jer Chen, Sui-Yuan Chang |
| EPI_ISL_693313, EPI_ISL_693315, EPI_ISL_693316, EPI_ISL_693320, EPI_ISL_693321, EPI_ISL_693324, EPI_ISL_693326, EPI_ISL_693327 | National Public Health Laboratory, National Centre for Infectious Diseases | National Public Health Laboratory, National Centre for Infectious Diseases | Tze Minn Mak, Sophie Octavia, Zhenyang Zhou, Lin Cui, Raymond Tzer Pin Lin |
| EPI_ISL_696453 | Pacaltsdorp Clinic wc PAC & NHLS/UCT | KRISP, KZN Research Innovation and Sequencing Platform | Arash Iranzadeh, Deelan Doolabh, Lynn Tyers, Bruna Galvao, Innocent Mudau, Marvin Hsiao, Kruger Marais, Jennifer Giandhari, Sureshnee Pillay, Houriiyah Tegally, Emanuel James San, Tulio de Oliveira, Diana Hardie, Stephen Korsman, Carolyn Williamson |
| EPI_ISL_696454 | Plettenberg Bay Clinic wc PLC & NHLS/UCT | KRISP, KZN Research Innovation and Sequencing Platform | Arash Iranzadeh, Deelan Doolabh, Lynn Tyers, Bruna Galvao, Innocent Mudau, Marvin Hsiao, Kruger Marais, Jennifer Giandhari, Sureshnee Pillay, Houriiyah Tegally, Emanuel James San, Tulio de Oliveira, Diana Hardie, Stephen Korsman, Carolyn Williamson |
| EPI_ISL_696456 | Pacaltsdorp Clinic wc PAC & NHLS/UCT | KRISP, KZN Research Innovation and Sequencing Platform | Arash Iranzadeh, Deelan Doolabh, Lynn Tyers, Bruna Galvao, Innocent Mudau, Marvin Hsiao, Kruger Marais, Jennifer Giandhari, Sureshnee Pillay, Houriiyah Tegally, Emanuel James San, Tulio de Oliveira, Diana Hardie, Stephen Korsman, Carolyn Williamson |
| EPI_ISL_696458 | George Hospital wc GRH & NHLS/UCT | KRISP, KZN Research Innovation and Sequencing Platform | Arash Iranzadeh, Deelan Doolabh, Lynn Tyers, Bruna Galvao, Innocent Mudau, Marvin Hsiao, Kruger Marais, Jennifer Giandhari, Sureshnee Pillay, Houriiyah Tegally, Emanuel James San, Tulio de Oliveira, Diana Hardie, Stephen Korsman, Carolyn Williamson |
| EPI_ISL_696459 | Pacaltsdorp Clinic wc PAC & NHLS/UCT | KRISP, KZN Research Innovation and Sequencing Platform | Arash Iranzadeh, Deelan Doolabh, Lynn Tyers, Bruna Galvao, Innocent Mudau, Marvin Hsiao, Kruger Marais, Jennifer Giandhari, Sureshnee Pillay, Houriiyah Tegally, Emanuel James San, Tulio de Oliveira, Diana Hardie, Stephen Korsman, Carolyn Williamson |
| EPI_ISL_696460 | Kranshoek Clinic wc KSH & NHLS/UCT | KRISP, KZN Research Innovation and Sequencing Platform | Arash Iranzadeh, Deelan Doolabh, Lynn Tyers, Bruna Galvao, Innocent Mudau, Marvin Hsiao, Kruger Marais, Jennifer Giandhari, Sureshnee Pillay, Houriiyah Tegally, Emanuel James San, Tulio de Oliveira, Diana Hardie, Stephen Korsman, Carolyn Williamson |
| EPI_ISL_696479 | Hornlee Clinic wc HLC & NHLS/UCT | KRISP, KZN Research Innovation and Sequencing Platform | Arash Iranzadeh, Deelan Doolabh, Lynn Tyers, Bruna Galvao, Innocent Mudau, Marvin Hsiao, Kruger Marais, Jennifer Giandhari, Sureshnee Pillay, Houriiyah Tegally, Emanuel James San, Tulio de Oliveira, Diana Hardie, Stephen Korsman, Carolyn Williamson |
| EPI_ISL_696482 | Knysna Hospital wc KNY & NHLS/UCT | KRISP, KZN Research Innovation and Sequencing Platform | Arash Iranzadeh, Deelan Doolabh, Lynn Tyers, Bruna Galvao, Innocent Mudau, Marvin Hsiao, Kruger Marais, Jennifer Giandhari, Sureshnee Pillay, Houriiyah Tegally, Emanuel James San, Tulio de Oliveira, Diana Hardie, Stephen Korsman, Carolyn Williamson |
| EPI_ISL_696483 | Hornlee Clinic wc HLC & NHLS/UCT | KRISP, KZN Research Innovation and Sequencing Platform | Arash Iranzadeh, Deelan Doolabh, Lynn Tyers, Bruna Galvao, Innocent Mudau, Marvin Hsiao, Kruger Marais, Jennifer Giandhari, Sureshnee Pillay, Houriiyah Tegally, Emanuel James San, Tulio de Oliveira, Diana Hardie, Stephen Korsman, Carolyn Williamson |
| EPI_ISL_696484 | Khayelethu Clinic wc KLC & NHLS/UCT | KRISP, KZN Research Innovation and Sequencing Platform | Arash Iranzadeh, Deelan Doolabh, Lynn Tyers, Bruna Galvao, Innocent Mudau, Marvin Hsiao, Kruger Marais, Jennifer Giandhari, Sureshnee Pillay, Houriiyah Tegally, Emanuel James San, Tulio de Oliveira, Diana Hardie, Stephen Korsman, Carolyn Williamson |
| EPI_ISL_696486 | Hornlee Clinic wc HLC & NHLS/UCT | KRISP, KZN Research Innovation and Sequencing Platform | Arash Iranzadeh, Deelan Doolabh, Lynn Tyers, Bruna Galvao, Innocent Mudau, Marvin Hsiao, Kruger Marais, Jennifer Giandhari, Sureshnee Pillay, Houriiyah Tegally, Emanuel James San, Tulio de Oliveira, Diana Hardie, Stephen Korsman, Carolyn Williamson |
| EPI_ISL_696487, EPI_ISL_696488 | Sedgefield Clinic wc SGE & NHLS/UCT | KRISP, KZN Research Innovation and Sequencing Platform | Arash Iranzadeh, Deelan Doolabh, Lynn Tyers, Bruna Galvao, Innocent Mudau, Marvin Hsiao, Kruger Marais, Jennifer Giandhari, Sureshnee Pillay, Houriiyah Tegally, Emanuel James San, Tulio de Oliveira, Diana Hardie, Stephen Korsman, Carolyn Williamson |
| EPI_ISL_696489 | New Horizon Clinic wc NZC & NHLS/UCT | KRISP, KZN Research Innovation and Sequencing Platform | Arash Iranzadeh, Deelan Doolabh, Lynn Tyers, Bruna Galvao, Innocent Mudau, Marvin Hsiao, Kruger Marais, Jennifer Giandhari, Sureshnee Pillay, Houriiyah Tegally, Emanuel James San, Tulio de Oliveira, Diana Hardie, Stephen Korsman, Carolyn Williamson |
| EPI_ISL_696490 | Touwsranten Clinic wc TST & NHLS/UCT | KRISP, KZN Research Innovation and Sequencing Platform | Arash Iranzadeh, Deelan Doolabh, Lynn Tyers, Bruna Galvao, Innocent Mudau, Marvin Hsiao, Kruger Marais, Jennifer Giandhari, Sureshnee Pillay, Houriiyah Tegally, Emanuel James San, Tulio de Oliveira, Diana Hardie, Stephen Korsman, Carolyn Williamson |
| EPI_ISL_699655 | 1-Laboratory of Microbiology, National Reference Lab, Charles Nicolle Hospital; 2-University of Tunis ElManar, Faculty of Medicine of Tunis, LR99ES09, Tunis, Tunisia | 1-Clinical and Experimental Pharmacology Lab, LR16SP02, National Center of Pharmacovigilance, University of Tunis El Manar, Tunis, Tunisia. 2-Neurodegenerative diseases and psychiatric troubles, LR18SP03, Razi Hospital, University of Tunis El Manar, Tunis, Tunisia. 3- Ministry of Health, National Observatory of New and Emerging Diseases, 1006, Tunis, Tunisia | Ilhem Boutiba-Ben Boubaker, Sameh Trabelsi, Nissaf Ben Alaya, Maher Kharrat, Alia Ben Kahla, Jalila Ben Khelil, Salma Abid, Sana Ferjani, Asma Ferjani, Mouna Ben Sassi, Mouna Safer, Guedi Berrabeh,Salwa Mrabet, Hanen ElJebari, Gaies Emna, Riadh Daghfous, Riadh Gouider. |
| EPI_ISL_701673, EPI_ISL_701674, EPI_ISL_701676, EPI_ISL_701677, EPI_ISL_701678, EPI_ISL_701679, EPI_ISL_701680, EPI_ISL_701682, EPI_ISL_701684, EPI_ISL_701685, EPI_ISL_701686, EPI_ISL_701687, EPI_ISL_701688, EPI_ISL_701689, EPI_ISL_701690, EPI_ISL_701691, EPI_ISL_701692, EPI_ISL_701693, EPI_ISL_701694, EPI_ISL_701695, EPI_ISL_701696, EPI_ISL_701699, EPI_ISL_701700, EPI_ISL_701702, EPI_ISL_701703, EPI_ISL_701704, EPI_ISL_701705, EPI_ISL_701706, EPI_ISL_701707, EPI_ISL_701708, EPI_ISL_701709, EPI_ISL_701710, EPI_ISL_701711, EPI_ISL_701712, EPI_ISL_701715, EPI_ISL_701716, EPI_ISL_701717, EPI_ISL_701718, EPI_ISL_701719, EPI_ISL_701720, EPI_ISL_701721, EPI_ISL_701723, EPI_ISL_701725, EPI_ISL_701727, EPI_ISL_701728, EPI_ISL_701730, EPI_ISL_701731, EPI_ISL_701732, EPI_ISL_701733, EPI_ISL_701734, EPI_ISL_701735, EPI_ISL_701736, EPI_ISL_701738, EPI_ISL_701740, EPI_ISL_701741, EPI_ISL_701742, EPI_ISL_701744, EPI_ISL_701745, EPI_ISL_701747, EPI_ISL_701748, EPI_ISL_701750, EPI_ISL_701751, EPI_ISL_701752, EPI_ISL_701753, EPI_ISL_701755, EPI_ISL_701756, EPI_ISL_701757, EPI_ISL_701758, EPI_ISL_701760, EPI_ISL_701761, EPI_ISL_701763, EPI_ISL_701764, EPI_ISL_701765, EPI_ISL_701766, EPI_ISL_701767, EPI_ISL_701768, EPI_ISL_701769, EPI_ISL_701770, EPI_ISL_701771, EPI_ISL_701772, EPI_ISL_701773, EPI_ISL_701776, EPI_ISL_701777, EPI_ISL_701778, EPI_ISL_701779, EPI_ISL_701780, EPI_ISL_701782, EPI_ISL_701783, EPI_ISL_701784, EPI_ISL_701785, EPI_ISL_701786, EPI_ISL_701787, EPI_ISL_701789, EPI_ISL_701790, EPI_ISL_701791, EPI_ISL_701793, EPI_ISL_701794, EPI_ISL_701796, EPI_ISL_701797, EPI_ISL_701798, EPI_ISL_701799, EPI_ISL_701801, EPI_ISL_701802, EPI_ISL_701804, EPI_ISL_701805, EPI_ISL_701806, EPI_ISL_701807, EPI_ISL_701808, EPI_ISL_701809, EPI_ISL_701810, EPI_ISL_701811, EPI_ISL_701812, EPI_ISL_701813, EPI_ISL_701814, EPI_ISL_701815, EPI_ISL_701816, EPI_ISL_701817, EPI_ISL_701818, EPI_ISL_701820, EPI_ISL_701821, EPI_ISL_701822, EPI_ISL_701823, EPI_ISL_701824, EPI_ISL_701825, EPI_ISL_701826, EPI_ISL_701827, EPI_ISL_701828, EPI_ISL_701830, EPI_ISL_701831, EPI_ISL_701832, EPI_ISL_701833, EPI_ISL_701834, EPI_ISL_701835, EPI_ISL_701836, EPI_ISL_701840, EPI_ISL_701842, EPI_ISL_701844, EPI_ISL_701845, EPI_ISL_701846, EPI_ISL_701849, EPI_ISL_701851, EPI_ISL_701852, EPI_ISL_701854, EPI_ISL_701855, EPI_ISL_701856, EPI_ISL_701857, EPI_ISL_701858, EPI_ISL_701859, EPI_ISL_701860, EPI_ISL_701861, EPI_ISL_701862, EPI_ISL_701863, EPI_ISL_701864, EPI_ISL_701865, EPI_ISL_701866, EPI_ISL_701867, EPI_ISL_701868, EPI_ISL_701869, EPI_ISL_701872, EPI_ISL_701873, EPI_ISL_701874, EPI_ISL_701877, EPI_ISL_701879, EPI_ISL_701880, EPI_ISL_701882, EPI_ISL_701883, EPI_ISL_701884, EPI_ISL_701885, EPI_ISL_701886, EPI_ISL_701887, EPI_ISL_701888, EPI_ISL_701889, EPI_ISL_701890, EPI_ISL_701891, EPI_ISL_701892, EPI_ISL_701893, EPI_ISL_701894, EPI_ISL_701895, EPI_ISL_701896, EPI_ISL_701897, EPI_ISL_701898, EPI_ISL_701899, EPI_ISL_701900, EPI_ISL_701901, EPI_ISL_701902, EPI_ISL_701903, EPI_ISL_701904, EPI_ISL_701905, EPI_ISL_701906, EPI_ISL_701907, EPI_ISL_701908, EPI_ISL_701909, EPI_ISL_701910, EPI_ISL_701911, EPI_ISL_701912, EPI_ISL_701915, EPI_ISL_701917, EPI_ISL_701918, EPI_ISL_701919, EPI_ISL_701921, EPI_ISL_701922, EPI_ISL_701924, EPI_ISL_701925, EPI_ISL_701926, EPI_ISL_701928, EPI_ISL_701929, EPI_ISL_701931, EPI_ISL_701933, EPI_ISL_701934, EPI_ISL_701935, EPI_ISL_701936, EPI_ISL_701937, EPI_ISL_701938, EPI_ISL_701939, EPI_ISL_701940, EPI_ISL_701941, EPI_ISL_701942, EPI_ISL_701944, EPI_ISL_701945, EPI_ISL_701946, EPI_ISL_701948, EPI_ISL_701949, EPI_ISL_701950, EPI_ISL_701951, EPI_ISL_701952, EPI_ISL_701953, EPI_ISL_701954, EPI_ISL_701955, EPI_ISL_701956, EPI_ISL_701957, EPI_ISL_701958, EPI_ISL_701959, EPI_ISL_701960, EPI_ISL_701962, EPI_ISL_701964, EPI_ISL_701965, EPI_ISL_701966, EPI_ISL_701967, EPI_ISL_701968, EPI_ISL_701969, EPI_ISL_701970, EPI_ISL_701971, EPI_ISL_701972, EPI_ISL_701973, EPI_ISL_701974, EPI_ISL_701977, EPI_ISL_701978, EPI_ISL_701979, EPI_ISL_701981, EPI_ISL_701982, EPI_ISL_701983, EPI_ISL_701984, EPI_ISL_701985, EPI_ISL_701986, EPI_ISL_701988, EPI_ISL_701989, EPI_ISL_701990, EPI_ISL_701991, EPI_ISL_701992, EPI_ISL_701995, EPI_ISL_701996, EPI_ISL_701997, EPI_ISL_701998, EPI_ISL_702000, EPI_ISL_702003, EPI_ISL_702004, EPI_ISL_702005, EPI_ISL_702006, EPI_ISL_702007, EPI_ISL_702008, EPI_ISL_702009, EPI_ISL_702010, EPI_ISL_702011, EPI_ISL_702012, EPI_ISL_702013, EPI_ISL_702017, EPI_ISL_702018, EPI_ISL_702019, EPI_ISL_702020, EPI_ISL_702021, EPI_ISL_702022, EPI_ISL_702370, EPI_ISL_702371, EPI_ISL_702372, EPI_ISL_702373, EPI_ISL_702374, EPI_ISL_702375, EPI_ISL_702377, EPI_ISL_702378, EPI_ISL_702379, EPI_ISL_702380, EPI_ISL_702382, EPI_ISL_702383, EPI_ISL_702386, EPI_ISL_702387, EPI_ISL_702388, EPI_ISL_702389, EPI_ISL_702390, EPI_ISL_702391, EPI_ISL_702392, EPI_ISL_702393, EPI_ISL_702394, EPI_ISL_702395, EPI_ISL_702396, EPI_ISL_702397, EPI_ISL_702398, EPI_ISL_702399, EPI_ISL_702400, EPI_ISL_702401, EPI_ISL_702402, EPI_ISL_702403, EPI_ISL_702404, EPI_ISL_702405, EPI_ISL_702406, EPI_ISL_702407, EPI_ISL_702408, EPI_ISL_702409, EPI_ISL_702410, EPI_ISL_702412, EPI_ISL_702414, EPI_ISL_702415, EPI_ISL_702416, EPI_ISL_702417, EPI_ISL_702418, EPI_ISL_702419, EPI_ISL_702420, EPI_ISL_702421, EPI_ISL_702422, EPI_ISL_702425, EPI_ISL_702426, EPI_ISL_702428, EPI_ISL_702429, EPI_ISL_702430, EPI_ISL_702431, EPI_ISL_702432, EPI_ISL_702433, EPI_ISL_702434, EPI_ISL_702435, EPI_ISL_702436, EPI_ISL_702437, EPI_ISL_702439, EPI_ISL_702440, EPI_ISL_702441, EPI_ISL_702442, EPI_ISL_702444, EPI_ISL_702445, EPI_ISL_702446, EPI_ISL_702448, EPI_ISL_702449, EPI_ISL_702450, EPI_ISL_702451, EPI_ISL_702452, EPI_ISL_702454, EPI_ISL_702455, EPI_ISL_702456, EPI_ISL_702457, EPI_ISL_702458, EPI_ISL_702459, EPI_ISL_702460, EPI_ISL_702461, EPI_ISL_702462, EPI_ISL_702463, EPI_ISL_702465, EPI_ISL_702466, EPI_ISL_702467, EPI_ISL_702468, EPI_ISL_702470 | | | |
| see above | Lighthouse Lab in Alderley Park | Wellcome Sanger Institute for the COVID-19 Genomics UK (COG-UK) Consortium | Jacquelyn Wynn, Mairead Hyland, The Lighthouse Lab in Alderley Park and Alex Alderton, Roberto Amato, Sonia Goncalves, Ewan Harrison, David K. Jackson, Ian Johnston, Dominic Kwiatkowski, Cordelia Langford, John Sillitoe on behalf of the Wellcome Sanger Institute COVID-19 Surveillance Team |
| EPI_ISL_702471 | Liverpool Clinical Laboratories | COVID-19 Genomics UK (COG-UK) Consortium | Sam Haldenby, Anita Lucaci, Steve Paterson, Julian Hiscox, Alistair Darby, M Almsaud, A Alrezaihi, Muhannad Alruwaili, Stuart D Armstrong, Jones Benjamin, Eleanor G Bentley, Anu Chawla, Jordan J Clark, Angela Cowell, Richard Eccles, Isabel García-Dorival, Matthew Gemmell, Alessandro Gerada, |

| | | | |
|---|---|---|---|
| | | | PKF Gilmore, Richard Gregory, Ximeng Han, Catherine Hartley, Margaret Hughes, Miren Iturriza-Gomara, James Johnson, L Luu, Jenifer Manson, Charlotte Nelson, Elaine O'Toole, Cassie Olateju, Rebekah Penrice-Randal , Lucille Rainbow, N.P Randle, Trevor Ian Robinson, Parul Sharma, Ghada T Shawli, James P Stewart, Neil Swainston, Ecaterina Vamos, Joanne Watts, Mark Whitehead |
| EPI_ISL_702472, EPI_ISL_702473, EPI_ISL_702475 | Lighthouse Lab in Alderley Park | Wellcome Sanger Institute for the COVID-19 Genomics UK (COG-UK) Consortium | Jacquelyn Wynn, Mairead Hyland, The Lighthouse Lab in Alderley Park and Alex Alderton, Roberto Amato, Sonia Goncalves, Ewan Harrison, David K. Jackson, Ian Johnston, Dominic Kwiatkowski, Cordelia Langford, John Sillitoe on behalf of the Wellcome Sanger Institute COVID-19 Surveillance Team |
| EPI_ISL_702476 | Virology Department, Sheffield Teaching Hospitals NHS Foundation Trust/Department of Infection, Immunity and Cardiovascular Disease, The Medical School, University of Sheffield | COVID-19 Genomics UK (COG-UK) Consortium | Thushan de Silva, Matthew Parker, Nikki Smith, Adri Angyal, Rebecca Brown, Luke Green, Rachel Tucker, Paul Parsons, Danielle Groves, Katie Johnson, Laura Carrilero, Alex Keeley, Dave Partridge, Matthew Wyles, Benjamin Lindsey, Mehmet Yavuz, Mohammad Raza, Cariad Evans |
| EPI_ISL_702477, EPI_ISL_702479, EPI_ISL_702480, EPI_ISL_702482, EPI_ISL_702483, EPI_ISL_702485, EPI_ISL_702488 | Lighthouse Lab in Alderley Park | Wellcome Sanger Institute for the COVID-19 Genomics UK (COG-UK) Consortium | Jacquelyn Wynn, Mairead Hyland, The Lighthouse Lab in Alderley Park and Alex Alderton, Roberto Amato, Sonia Goncalves, Ewan Harrison, David K. Jackson, Ian Johnston, Dominic Kwiatkowski, Cordelia Langford, John Sillitoe on behalf of the Wellcome Sanger Institute COVID-19 Surveillance Team |
| EPI_ISL_702489 | Department of Pathology, University of Cambridge | COVID-19 Genomics UK (COG-UK) Consortium | Aminu S. Jahun, Yasmin Chaudhry, Grant Hall, Iliana Georgana, Myra Hosmillo, Martin D. Curran, Malte Pinckert, Surendra Parmar, Ian Goodfellow |
| EPI_ISL_702490, EPI_ISL_702491 | Lighthouse Lab in Alderley Park | Wellcome Sanger Institute for the COVID-19 Genomics UK (COG-UK) Consortium | Jacquelyn Wynn, Mairead Hyland, The Lighthouse Lab in Alderley Park and Alex Alderton, Roberto Amato, Sonia Goncalves, Ewan Harrison, David K. Jackson, Ian Johnston, Dominic Kwiatkowski, Cordelia Langford, John Sillitoe on behalf of the Wellcome Sanger Institute COVID-19 Surveillance Team |
| EPI_ISL_702492 | Liverpool Clinical Laboratories | COVID-19 Genomics UK (COG-UK) Consortium | Sam Haldenby, Anita Lucaci, Steve Paterson, Julian Hiscox, Alistair Darby, M Almsaud, A Alrezaihi, Muhannad Alruwaili, Stuart D Armstrong, Jones Benjamin, Eleanor G Bentley, Anu Chawla, Jordan J Clark, Angela Cowell, Richard Eccles, Isabel García-Dorival, Matthew Gemmell, Alessandro Gerada, PKF Gilmore, Richard Gregory, Ximeng Han, Catherine Hartley, Margaret Hughes, Miren Iturriza-Gomara, James Johnson, L Luu, Jenifer Manson, Charlotte Nelson, Elaine O'Toole, Cassie Olateju, Rebekah Penrice-Randal , Lucille Rainbow, N.P Randle, Trevor Ian Robinson, Parul Sharma, Ghada T Shawli, James P Stewart, Neil Swainston, Ecaterina Vamos, Joanne Watts, Mark Whitehead |
| EPI_ISL_702493, EPI_ISL_702494, EPI_ISL_702496, EPI_ISL_702497, EPI_ISL_702499 | Lighthouse Lab in Alderley Park | Wellcome Sanger Institute for the COVID-19 Genomics UK (COG-UK) Consortium | Jacquelyn Wynn, Mairead Hyland, The Lighthouse Lab in Alderley Park and Alex Alderton, Roberto Amato, Sonia Goncalves, Ewan Harrison, David K. Jackson, Ian Johnston, Dominic Kwiatkowski, Cordelia Langford, John Sillitoe on behalf of the Wellcome Sanger Institute COVID-19 Surveillance Team |
| EPI_ISL_702500 | Department of Pathology, University of Cambridge | COVID-19 Genomics UK (COG-UK) Consortium | Aminu S. Jahun, Yasmin Chaudhry, Grant Hall, Iliana Georgana, Myra Hosmillo, Martin D. Curran, Malte Pinckert, Surendra Parmar, Ian Goodfellow |
| EPI_ISL_702501, EPI_ISL_702502, EPI_ISL_702503, EPI_ISL_702506, EPI_ISL_702508, EPI_ISL_702509 | Lighthouse Lab in Alderley Park | Wellcome Sanger Institute for the COVID-19 Genomics UK (COG-UK) Consortium | Jacquelyn Wynn, Mairead Hyland, The Lighthouse Lab in Alderley Park and Alex Alderton, Roberto Amato, Sonia Goncalves, Ewan Harrison, David K. Jackson, Ian Johnston, Dominic Kwiatkowski, Cordelia Langford, John Sillitoe on behalf of the Wellcome Sanger Institute COVID-19 Surveillance Team |
| EPI_ISL_702510 | Liverpool Clinical Laboratories | COVID-19 Genomics UK (COG-UK) Consortium | Sam Haldenby, Anita Lucaci, Steve Paterson, Julian Hiscox, Alistair Darby, M Almsaud, A Alrezaihi, Muhannad Alruwaili, Stuart D Armstrong, Jones Benjamin, Eleanor G Bentley, Anu Chawla, Jordan J Clark, Angela Cowell, Richard Eccles, Isabel García-Dorival, Matthew Gemmell, Alessandro Gerada, PKF Gilmore, Richard Gregory, Ximeng Han, Catherine Hartley, Margaret Hughes, Miren Iturriza-Gomara, James Johnson, L Luu, Jenifer Manson, Charlotte Nelson, Elaine O'Toole, Cassie Olateju, Rebekah Penrice-Randal , Lucille Rainbow, N.P Randle, Trevor Ian Robinson, Parul Sharma, Ghada T Shawli, James P Stewart, Neil Swainston, Ecaterina Vamos, Joanne Watts, Mark Whitehead |
| EPI_ISL_702511, EPI_ISL_702513, EPI_ISL_702514, EPI_ISL_702516, EPI_ISL_702517 | Lighthouse Lab in Alderley Park | Wellcome Sanger Institute for the COVID-19 Genomics UK (COG-UK) Consortium | Jacquelyn Wynn, Mairead Hyland, The Lighthouse Lab in Alderley Park and Alex Alderton, Roberto Amato, Sonia Goncalves, Ewan Harrison, David K. Jackson, Ian Johnston, Dominic Kwiatkowski, Cordelia Langford, John Sillitoe on behalf of the Wellcome Sanger Institute COVID-19 Surveillance Team |
| EPI_ISL_702518 | Virology Department, Sheffield Teaching Hospitals NHS Foundation Trust/Department of Infection, Immunity and Cardiovascular Disease, The Medical School, University of Sheffield | COVID-19 Genomics UK (COG-UK) Consortium | Thushan de Silva, Matthew Parker, Nikki Smith, Adri Angyal, Rebecca Brown, Luke Green, Rachel Tucker, Paul Parsons, Danielle Groves, Katie Johnson, Laura Carrilero, Alex Keeley, Dave Partridge, Matthew Wyles, Benjamin Lindsey, Mehmet Yavuz, Mohammad Raza, Cariad Evans |
| EPI_ISL_702522, EPI_ISL_702524, EPI_ISL_702526, EPI_ISL_702527, EPI_ISL_702529, EPI_ISL_702530, EPI_ISL_702532, EPI_ISL_702534, EPI_ISL_702535, EPI_ISL_702538, EPI_ISL_702540, EPI_ISL_702542, EPI_ISL_702545, EPI_ISL_702546, EPI_ISL_702548, EPI_ISL_702551, EPI_ISL_702552, EPI_ISL_702554, EPI_ISL_702555, EPI_ISL_702557, EPI_ISL_702559 | | | |
| see above | Lighthouse Lab in Alderley Park | Wellcome Sanger Institute for the COVID-19 Genomics UK (COG-UK) Consortium | Jacquelyn Wynn, Mairead Hyland, The Lighthouse Lab in Alderley Park and Alex Alderton, Roberto Amato, Sonia Goncalves, Ewan Harrison, David K. Jackson, Ian Johnston, Dominic Kwiatkowski, Cordelia Langford, John Sillitoe on behalf of the Wellcome Sanger Institute COVID-19 Surveillance Team |
| EPI_ISL_702561 | Department of Pathology, University of Cambridge | COVID-19 Genomics UK (COG-UK) Consortium | Aminu S. Jahun, Yasmin Chaudhry, Grant Hall, Iliana Georgana, Myra Hosmillo, Martin D. Curran, Malte Pinckert, Surendra Parmar, Ian Goodfellow |
| EPI_ISL_702562 | University of Exeter | COVID-19 Genomics UK (COG-UK) Consortium | Ben Temperton,Aaron Jeffries,Michelle Michelsen,Joanna Warwick-Dugdale,Audrey Farbos,Robyn Manley,Stephen Michell,Jane Masoli |
| EPI_ISL_702563, EPI_ISL_702565, EPI_ISL_702567, EPI_ISL_702568, EPI_ISL_702570, EPI_ISL_702572, EPI_ISL_702573 | Lighthouse Lab in Alderley Park | Wellcome Sanger Institute for the COVID-19 Genomics UK (COG-UK) Consortium | Jacquelyn Wynn, Mairead Hyland, The Lighthouse Lab in Alderley Park and Alex Alderton, Roberto Amato, Sonia Goncalves, Ewan Harrison, David K. Jackson, Ian Johnston, Dominic Kwiatkowski, Cordelia Langford, John Sillitoe on behalf of the Wellcome Sanger Institute COVID-19 Surveillance Team |
| EPI_ISL_702574 | Liverpool Clinical Laboratories | COVID-19 Genomics UK (COG-UK) Consortium | Sam Haldenby, Anita Lucaci, Steve Paterson, Julian Hiscox, Alistair Darby, M Almsaud, A Alrezaihi, Muhannad Alruwaili, Stuart D Armstrong, Jones Benjamin, Eleanor G Bentley, Anu Chawla, Jordan J Clark, Angela Cowell, Richard Eccles, Isabel García-Dorival, Matthew Gemmell, Alessandro Gerada, PKF Gilmore, Richard Gregory, Ximeng Han, Catherine Hartley, Margaret Hughes, Miren Iturriza-Gomara, James Johnson, L Luu, Jenifer Manson, Charlotte Nelson, Elaine O'Toole, Cassie Olateju, Rebekah Penrice-Randal , Lucille Rainbow, N.P Randle, Trevor Ian Robinson, Parul Sharma, Ghada T Shawli, James P Stewart, Neil Swainston, Ecaterina Vamos, Joanne Watts, Mark Whitehead |
| EPI_ISL_702575, EPI_ISL_702576 | Lighthouse Lab in Alderley Park | Wellcome Sanger Institute for the COVID-19 Genomics UK (COG-UK) Consortium | Jacquelyn Wynn, Mairead Hyland, The Lighthouse Lab in Alderley Park and Alex Alderton, Roberto Amato, Sonia Goncalves, Ewan Harrison, David K. Jackson, Ian Johnston, Dominic Kwiatkowski, Cordelia Langford, John Sillitoe on behalf of the Wellcome Sanger Institute COVID-19 Surveillance Team |
| EPI_ISL_702577 | Liverpool Clinical Laboratories | COVID-19 Genomics UK (COG-UK) Consortium | Sam Haldenby, Anita Lucaci, Steve Paterson, Julian Hiscox, Alistair Darby, M Almsaud, A Alrezaihi, Muhannad Alruwaili, Stuart D Armstrong, Jones Benjamin, Eleanor G Bentley, Anu Chawla, Jordan J Clark, Angela Cowell, Richard Eccles, Isabel García-Dorival, Matthew Gemmell, Alessandro Gerada, PKF Gilmore, Richard Gregory, Ximeng Han, Catherine Hartley, Margaret Hughes, Miren Iturriza-Gomara, James Johnson, L Luu, Jenifer Manson, Charlotte Nelson, Elaine O'Toole, Cassie Olateju, Rebekah Penrice-Randal , Lucille Rainbow, N.P Randle, Trevor Ian Robinson, Parul Sharma, Ghada T Shawli, James P Stewart, Neil Swainston, Ecaterina Vamos, Joanne Watts, Mark Whitehead |
| EPI_ISL_702578, EPI_ISL_702580, EPI_ISL_702581, EPI_ISL_702583, EPI_ISL_702584, EPI_ISL_702586, EPI_ISL_702588, EPI_ISL_702589, EPI_ISL_702592, EPI_ISL_702593, EPI_ISL_702595, EPI_ISL_702596, EPI_ISL_702598, EPI_ISL_702599, EPI_ISL_702601, EPI_ISL_702603, EPI_ISL_702604, EPI_ISL_702606, EPI_ISL_702608, EPI_ISL_702609, EPI_ISL_702611, EPI_ISL_702613, EPI_ISL_702614, EPI_ISL_702616, EPI_ISL_702617, EPI_ISL_702621, EPI_ISL_702624, EPI_ISL_702625, EPI_ISL_702627, EPI_ISL_702629, EPI_ISL_702630, EPI_ISL_702632 | | | |
| see above | Lighthouse Lab in Alderley Park | Wellcome Sanger Institute for the COVID-19 Genomics UK (COG-UK) Consortium | Jacquelyn Wynn, Mairead Hyland, The Lighthouse Lab in Alderley Park and Alex Alderton, Roberto Amato, Sonia Goncalves, Ewan Harrison, David K. Jackson, Ian Johnston, Dominic Kwiatkowski, Cordelia Langford, John Sillitoe on behalf of the Wellcome Sanger Institute COVID-19 Surveillance Team |
| EPI_ISL_702639 | Liverpool Clinical Laboratories | COVID-19 Genomics UK (COG-UK) Consortium | Sam Haldenby, Anita Lucaci, Steve Paterson, Julian Hiscox, Alistair Darby, M Almsaud, A Alrezaihi, Muhannad Alruwaili, Stuart D Armstrong, Jones Benjamin, Eleanor G Bentley, Anu Chawla, Jordan J Clark, Angela Cowell, Richard Eccles, Isabel García-Dorival, Matthew Gemmell, Alessandro Gerada, PKF Gilmore, Richard Gregory, Ximeng Han, Catherine Hartley, Margaret Hughes, Miren Iturriza-Gomara, James Johnson, L Luu, Jenifer Manson, Charlotte Nelson, Elaine O'Toole, Cassie Olateju, Rebekah Penrice-Randal , Lucille Rainbow, N.P Randle, Trevor Ian Robinson, Parul Sharma, Ghada T Shawli, James P Stewart, Neil Swainston, Ecaterina Vamos, Joanne Watts, Mark Whitehead |
| EPI_ISL_702641, EPI_ISL_702643, EPI_ISL_702645, EPI_ISL_702646, EPI_ISL_702648 | Lighthouse Lab in Alderley Park | Wellcome Sanger Institute for the COVID-19 Genomics UK (COG-UK) Consortium | Jacquelyn Wynn, Mairead Hyland, The Lighthouse Lab in Alderley Park and Alex Alderton, Roberto Amato, Sonia Goncalves, Ewan Harrison, David K. Jackson, Ian Johnston, Dominic Kwiatkowski, Cordelia Langford, John Sillitoe on behalf of the Wellcome Sanger Institute COVID-19 Surveillance Team |
| EPI_ISL_702649 | Liverpool Clinical Laboratories | COVID-19 Genomics UK (COG-UK) Consortium | Sam Haldenby, Anita Lucaci, Steve Paterson, Julian Hiscox, Alistair Darby, M Almsaud, A Alrezaihi, Muhannad Alruwaili, Stuart D Armstrong, Jones |

| | | | |
|---|---|---|---|
| | | | Benjamin, Eleanor G Bentley, Anu Chawla, Jordan J Clark, Angela Cowell, Richard Eccles, Isabel García-Dorival, Matthew Gemmell, Alessandro Gerada, PKF Gilmore, Richard Gregory, Ximeng Han, Catherine Hartley, Margaret Hughes, Miren Iturriza-Gomara, James Johnson, L Luu, Jenifer Manson, Charlotte Nelson, Elaine O'Toole, Cassie Olateju, Rebekah Penrice-Randal , Lucille Rainbow, N.P Randle, Trevor Ian Robinson, Parul Sharma, Ghada T Shawli, James P Stewart, Neil Swainston, Ecaterina Vamos, Joanne Watts, Mark Whitehead |
| EPI_ISL_702650, EPI_ISL_702651, EPI_ISL_702653, EPI_ISL_702654, EPI_ISL_702657, EPI_ISL_702660, EPI_ISL_702662, EPI_ISL_702665, EPI_ISL_702666 | Lighthouse Lab in Alderley Park | Wellcome Sanger Institute for the COVID-19 Genomics UK (COG-UK) Consortium | Jacquelyn Wynn, Mairead Hyland, The Lighthouse Lab in Alderley Park and Alex Alderton, Roberto Amato, Sonia Goncalves, Ewan Harrison, David K. Jackson, Ian Johnston, Dominic Kwiatkowski, Cordelia Langford, John Sillitoe on behalf of the Wellcome Sanger Institute COVID-19 Surveillance Team |
| EPI_ISL_702669 | University College London Hospital | COVID-19 Genomics UK (COG-UK) Consortium | Judith Heaney, Matthew Byott, Catherine Houlihan, Dan Frampton, Stuart Kirk, Moira Spyer and Eleni Nastouli |
| EPI_ISL_702670, EPI_ISL_702671, EPI_ISL_702673, EPI_ISL_702675, EPI_ISL_702676, EPI_ISL_702680, EPI_ISL_702685, EPI_ISL_702686, EPI_ISL_702688, EPI_ISL_702689, EPI_ISL_702691, EPI_ISL_702693, EPI_ISL_702695, EPI_ISL_702696, EPI_ISL_702699, EPI_ISL_702702 | | | |
| see above | Lighthouse Lab in Alderley Park | Wellcome Sanger Institute for the COVID-19 Genomics UK (COG-UK) Consortium | Jacquelyn Wynn, Mairead Hyland, The Lighthouse Lab in Alderley Park and Alex Alderton, Roberto Amato, Sonia Goncalves, Ewan Harrison, David K. Jackson, Ian Johnston, Dominic Kwiatkowski, Cordelia Langford, John Sillitoe on behalf of the Wellcome Sanger Institute COVID-19 Surveillance Team |
| EPI_ISL_702703 | Department of Pathology, University of Cambridge | COVID-19 Genomics UK (COG-UK) Consortium | Aminu S. Jahun, Yasmin Chaudhry, Grant Hall, Iliana Georgana, Myra Hosmillo, Martin D. Curran, Malte Pinckert, Surendra Parmar, Ian Goodfellow |
| EPI_ISL_702706, EPI_ISL_702709, EPI_ISL_702714, EPI_ISL_702715, EPI_ISL_702717 | Lighthouse Lab in Alderley Park | Wellcome Sanger Institute for the COVID-19 Genomics UK (COG-UK) Consortium | Jacquelyn Wynn, Mairead Hyland, The Lighthouse Lab in Alderley Park and Alex Alderton, Roberto Amato, Sonia Goncalves, Ewan Harrison, David K. Jackson, Ian Johnston, Dominic Kwiatkowski, Cordelia Langford, John Sillitoe on behalf of the Wellcome Sanger Institute COVID-19 Surveillance Team |
| EPI_ISL_702718 | Liverpool Clinical Laboratories | COVID-19 Genomics UK (COG-UK) Consortium | Sam Haldenby, Anita Lucaci, Steve Paterson, Julian Hiscox, Alistair Darby, M Almsaud, A Alrezaihi, Muhannad Alruwaili, Stuart D Armstrong, Jones Benjamin, Eleanor G Bentley, Anu Chawla, Jordan J Clark, Angela Cowell, Richard Eccles, Isabel García-Dorival, Matthew Gemmell, Alessandro Gerada, PKF Gilmore, Richard Gregory, Ximeng Han, Catherine Hartley, Margaret Hughes, Miren Iturriza-Gomara, James Johnson, L Luu, Jenifer Manson, Charlotte Nelson, Elaine O'Toole, Cassie Olateju, Rebekah Penrice-Randal , Lucille Rainbow, N.P Randle, Trevor Ian Robinson, Parul Sharma, Ghada T Shawli, James P Stewart, Neil Swainston, Ecaterina Vamos, Joanne Watts, Mark Whitehead |
| EPI_ISL_702722, EPI_ISL_702724, EPI_ISL_702727, EPI_ISL_702728, EPI_ISL_702730 | Lighthouse Lab in Alderley Park | Wellcome Sanger Institute for the COVID-19 Genomics UK (COG-UK) Consortium | Jacquelyn Wynn, Mairead Hyland, The Lighthouse Lab in Alderley Park and Alex Alderton, Roberto Amato, Sonia Goncalves, Ewan Harrison, David K. Jackson, Ian Johnston, Dominic Kwiatkowski, Cordelia Langford, John Sillitoe on behalf of the Wellcome Sanger Institute COVID-19 Surveillance Team |
| EPI_ISL_702734 | Department of Pathology, University of Cambridge | COVID-19 Genomics UK (COG-UK) Consortium | Aminu S. Jahun, Yasmin Chaudhry, Grant Hall, Iliana Georgana, Myra Hosmillo, Martin D. Curran, Malte Pinckert, Surendra Parmar, Ian Goodfellow |
| EPI_ISL_702735, EPI_ISL_702737, EPI_ISL_702738, EPI_ISL_702740, EPI_ISL_702743, EPI_ISL_702746 | Lighthouse Lab in Alderley Park | Wellcome Sanger Institute for the COVID-19 Genomics UK (COG-UK) Consortium | Jacquelyn Wynn, Mairead Hyland, The Lighthouse Lab in Alderley Park and Alex Alderton, Roberto Amato, Sonia Goncalves, Ewan Harrison, David K. Jackson, Ian Johnston, Dominic Kwiatkowski, Cordelia Langford, John Sillitoe on behalf of the Wellcome Sanger Institute COVID-19 Surveillance Team |
| EPI_ISL_702747 | Northumbria University / South Tees Hospitals NHS Foundation Trust / North Cumbria Integrated Care NHS Foundation Trust / North Tees and Hartlepool NHS Foundation Trust / Newcastle Hospitals NHS Foundation Trust | COVID-19 Genomics UK (COG-UK) Consortium | Darren L Smith,Andrew Nelson,Matthew Bashton,Greg R Young,Joshua Loh,John Allan,Mohammad A Tariq,Giles S Holt,Gary Black,Wen C Yew,Lynn Dover,Paul Baker,Steve Liggett,Sarah Essex,Jane Greenaway,Debra Padgett,Clive Graham,Garren Scott,Edward Barton,Emma Swindells,Brendan Payne,Jennifer Collins,Yusri Taha,Gary Eltringham |
| EPI_ISL_702748, EPI_ISL_702750, EPI_ISL_702757, EPI_ISL_702759, EPI_ISL_702760 | Lighthouse Lab in Alderley Park | Wellcome Sanger Institute for the COVID-19 Genomics UK (COG-UK) Consortium | Jacquelyn Wynn, Mairead Hyland, The Lighthouse Lab in Alderley Park and Alex Alderton, Roberto Amato, Sonia Goncalves, Ewan Harrison, David K. Jackson, Ian Johnston, Dominic Kwiatkowski, Cordelia Langford, John Sillitoe on behalf of the Wellcome Sanger Institute COVID-19 Surveillance Team |
| EPI_ISL_702761 | Virology Department, Sheffield Teaching Hospitals NHS Foundation Trust/Department of Infection, Immunity and Cardiovascular Disease, The Medical School, University of Sheffield | COVID-19 Genomics UK (COG-UK) Consortium | Thushan de Silva, Matthew Parker, Nikki Smith, Adri Angyal, Rebecca Brown, Luke Green, Rachel Tucker, Paul Parsons, Danielle Groves, Katie Johnson, Laura Carrilero, Alex Keeley, Dave Partridge, Matthew Wyles, Benjamin Lindsey, Mehmet Yavuz, Mohammad Raza, Cariad Evans |
| EPI_ISL_702762, EPI_ISL_702764, EPI_ISL_702765, EPI_ISL_702767, EPI_ISL_702768, EPI_ISL_702770 | Lighthouse Lab in Alderley Park | Wellcome Sanger Institute for the COVID-19 Genomics UK (COG-UK) Consortium | Jacquelyn Wynn, Mairead Hyland, The Lighthouse Lab in Alderley Park and Alex Alderton, Roberto Amato, Sonia Goncalves, Ewan Harrison, David K. Jackson, Ian Johnston, Dominic Kwiatkowski, Cordelia Langford, John Sillitoe on behalf of the Wellcome Sanger Institute COVID-19 Surveillance Team |
| EPI_ISL_702771 | West of Scotland Specialist Virology Centre, NHSGGC / MRC-University of Glasgow Centre for Virus Research | COVID-19 Genomics UK (COG-UK) Consortium | Ana da Silva Filipe, Natasha Johnson, Kathy Smollett, Daniel Mair, Stephen Carmichael, Alice Broos, Lily Tong, Jenna Nichols, Kyriaki Nomikou; Sarah McDonald; Richard Orton, Joseph Hughes, Sreenu Vattipally, David L Robertson; Alasdair MacLean, Rory Gunson; Sharif Shaaban, Matthew Holden; Rachel Blacow, Guy Mollett, Kathy Li, James Shepherd, Antonia Ho, Emma Thomson |
| EPI_ISL_702772, EPI_ISL_702775, EPI_ISL_702777, EPI_ISL_702778, EPI_ISL_702780, EPI_ISL_702781, EPI_ISL_702783, EPI_ISL_702786, EPI_ISL_702788, EPI_ISL_702789 | Lighthouse Lab in Alderley Park | Wellcome Sanger Institute for the COVID-19 Genomics UK (COG-UK) Consortium | Jacquelyn Wynn, Mairead Hyland, The Lighthouse Lab in Alderley Park and Alex Alderton, Roberto Amato, Sonia Goncalves, Ewan Harrison, David K. Jackson, Ian Johnston, Dominic Kwiatkowski, Cordelia Langford, John Sillitoe on behalf of the Wellcome Sanger Institute COVID-19 Surveillance Team |
| EPI_ISL_702790 | University of Exeter | COVID-19 Genomics UK (COG-UK) Consortium | Ben Temperton,Aaron Jeffries,Michelle Michelsen,Joanna Warwick-Dugdale,Audrey Farbos,Robyn Manley,Stephen Michell,Jane Masoli |
| EPI_ISL_702793 | Lighthouse Lab in Alderley Park | Wellcome Sanger Institute for the COVID-19 Genomics UK (COG-UK) Consortium | Jacquelyn Wynn, Mairead Hyland, The Lighthouse Lab in Alderley Park and Alex Alderton, Roberto Amato, Sonia Goncalves, Ewan Harrison, David K. Jackson, Ian Johnston, Dominic Kwiatkowski, Cordelia Langford, John Sillitoe on behalf of the Wellcome Sanger Institute COVID-19 Surveillance Team |
| EPI_ISL_702798 | Virology Department, Sheffield Teaching Hospitals NHS Foundation Trust/Department of Infection, Immunity and Cardiovascular Disease, The Medical School, University of Sheffield | COVID-19 Genomics UK (COG-UK) Consortium | Thushan de Silva, Matthew Parker, Nikki Smith, Adri Angyal, Rebecca Brown, Luke Green, Rachel Tucker, Paul Parsons, Danielle Groves, Katie Johnson, Laura Carrilero, Alex Keeley, Dave Partridge, Matthew Wyles, Benjamin Lindsey, Mehmet Yavuz, Mohammad Raza, Cariad Evans |
| EPI_ISL_702799 | Lighthouse Lab in Alderley Park | Wellcome Sanger Institute for the COVID-19 Genomics UK (COG-UK) Consortium | Jacquelyn Wynn, Mairead Hyland, The Lighthouse Lab in Alderley Park and Alex Alderton, Roberto Amato, Sonia Goncalves, Ewan Harrison, David K. Jackson, Ian Johnston, Dominic Kwiatkowski, Cordelia Langford, John Sillitoe on behalf of the Wellcome Sanger Institute COVID-19 Surveillance Team |
| EPI_ISL_702801 | Liverpool Clinical Laboratories | COVID-19 Genomics UK (COG-UK) Consortium | Sam Haldenby, Anita Lucaci, Steve Paterson, Julian Hiscox, Alistair Darby, M Almsaud, A Alrezaihi, Muhannad Alruwaili, Stuart D Armstrong, Jones Benjamin, Eleanor G Bentley, Anu Chawla, Jordan J Clark, Angela Cowell, Richard Eccles, Isabel García-Dorival, Matthew Gemmell, Alessandro Gerada, PKF Gilmore, Richard Gregory, Ximeng Han, Catherine Hartley, Margaret Hughes, Miren Iturriza-Gomara, James Johnson, L Luu, Jenifer Manson, Charlotte Nelson, Elaine O'Toole, Cassie Olateju, Rebekah Penrice-Randal , Lucille Rainbow, N.P Randle, Trevor Ian Robinson, Parul Sharma, Ghada T Shawli, James P Stewart, Neil Swainston, Ecaterina Vamos, Joanne Watts, Mark Whitehead |
| EPI_ISL_702802 | Lighthouse Lab in Alderley Park | Wellcome Sanger Institute for the COVID-19 Genomics UK (COG-UK) Consortium | Jacquelyn Wynn, Mairead Hyland, The Lighthouse Lab in Alderley Park and Alex Alderton, Roberto Amato, Sonia Goncalves, Ewan Harrison, David K. Jackson, Ian Johnston, Dominic Kwiatkowski, Cordelia Langford, John Sillitoe on behalf of the Wellcome Sanger Institute COVID-19 Surveillance Team |
| EPI_ISL_702803 | Northumbria University / South Tees Hospitals NHS Foundation Trust / North Cumbria Integrated Care NHS Foundation Trust / North Tees and Hartlepool NHS Foundation Trust / Newcastle Hospitals NHS Foundation Trust | COVID-19 Genomics UK (COG-UK) Consortium | Darren L Smith,Andrew Nelson,Matthew Bashton,Greg R Young,Joshua Loh,John Allan,Mohammad A Tariq,Giles S Holt,Gary Black,Wen C Yew,Lynn Dover,Paul Baker,Steve Liggett,Sarah Essex,Jane Greenaway,Debra Padgett,Clive Graham,Garren Scott,Edward Barton,Emma Swindells,Brendan Payne,Jennifer Collins,Yusri Taha,Gary Eltringham |
| EPI_ISL_702805, EPI_ISL_702807, EPI_ISL_702808, EPI_ISL_702810, | Lighthouse Lab in Alderley Park | Wellcome Sanger Institute for the COVID-19 Genomics UK (COG-UK) Consortium | Jacquelyn Wynn, Mairead Hyland, The Lighthouse Lab in Alderley Park and Alex Alderton, Roberto Amato, Sonia Goncalves, Ewan Harrison, David K. Jackson, Ian Johnston, Dominic Kwiatkowski, Cordelia Langford, John Sillitoe on behalf of the Wellcome Sanger Institute COVID-19 Surveillance Team |

| | | | |
|---|---|---|---|
| EPI_ISL_702811, EPI_ISL_702813, EPI_ISL_702814, EPI_ISL_702818, EPI_ISL_702819, EPI_ISL_702822 | | | |
| EPI_ISL_702826 | Liverpool Clinical Laboratories | COVID-19 Genomics UK (COG-UK) Consortium | Sam Haldenby, Anita Lucaci, Steve Paterson, Julian Hiscox, Alistair Darby, M Almsaud, A Alrezaihi, Muhannad Alruwaili, Stuart D Armstrong, Jones Benjamin, Eleanor G Bentley, Anu Chawla, Jordan J Clark, Angela Cowell, Richard Eccles, Isabel García-Dorival, Matthew Gemmell, Alessandro Gerada, PKF Gilmore, Richard Gregory, Ximeng Han, Catherine Hartley, Margaret Hughes, Miren Iturriza-Gomara, James Johnson, L Luu, Jenifer Manson, Charlotte Nelson, Elaine O'Toole, Cassie Olateju, Rebekah Penrice-Randal , Lucille Rainbow, N.P Randle, Trevor Ian Robinson, Parul Sharma, Ghada T Shawli, James P Stewart, Neil Swainston, Ecaterina Vamos, Joanne Watts, Mark Whitehead |
| EPI_ISL_702829, EPI_ISL_702830, EPI_ISL_702833 | Virology Department, Sheffield Teaching Hospitals NHS Foundation Trust/Department of Infection, Immunity and Cardiovascular Disease, The Medical School, University of Sheffield | COVID-19 Genomics UK (COG-UK) Consortium | Thushan de Silva, Matthew Parker, Nikki Smith, Adri Angyal, Rebecca Brown, Luke Green, Rachel Tucker, Paul Parsons, Danielle Groves, Katie Johnson, Laura Carrilero, Alex Keeley, Dave Partridge, Matthew Wyles, Benjamin Lindsey, Mehmet Yavuz, Mohammad Raza, Cariad Evans |
| EPI_ISL_702844 | University of Exeter | COVID-19 Genomics UK (COG-UK) Consortium | Ben Temperton,Aaron Jeffries,Michelle Michelsen,Joanna Warwick-Dugdale,Audrey Farbos,Robyn Manley,Stephen Michell,Jane Masoli |
| EPI_ISL_702845 | Liverpool Clinical Laboratories | COVID-19 Genomics UK (COG-UK) Consortium | Sam Haldenby, Anita Lucaci, Steve Paterson, Julian Hiscox, Alistair Darby, M Almsaud, A Alrezaihi, Muhannad Alruwaili, Stuart D Armstrong, Jones Benjamin, Eleanor G Bentley, Anu Chawla, Jordan J Clark, Angela Cowell, Richard Eccles, Isabel García-Dorival, Matthew Gemmell, Alessandro Gerada, PKF Gilmore, Richard Gregory, Ximeng Han, Catherine Hartley, Margaret Hughes, Miren Iturriza-Gomara, James Johnson, L Luu, Jenifer Manson, Charlotte Nelson, Elaine O'Toole, Cassie Olateju, Rebekah Penrice-Randal , Lucille Rainbow, N.P Randle, Trevor Ian Robinson, Parul Sharma, Ghada T Shawli, James P Stewart, Neil Swainston, Ecaterina Vamos, Joanne Watts, Mark Whitehead |
| EPI_ISL_702848, EPI_ISL_702876 | Department of Pathology, University of Cambridge | COVID-19 Genomics UK (COG-UK) Consortium | Aminu S. Jahun, Yasmin Chaudhry, Grant Hall, Iliana Georgana, Myra Hosmillo, Martin D. Curran, Malte Pinckert, Surendra Parmar, Ian Goodfellow |
| EPI_ISL_702880 | Virology Department, Sheffield Teaching Hospitals NHS Foundation Trust/Department of Infection, Immunity and Cardiovascular Disease, The Medical School, University of Sheffield | COVID-19 Genomics UK (COG-UK) Consortium | Thushan de Silva, Matthew Parker, Nikki Smith, Adri Angyal, Rebecca Brown, Luke Green, Rachel Tucker, Paul Parsons, Danielle Groves, Katie Johnson, Laura Carrilero, Alex Keeley, Dave Partridge, Matthew Wyles, Benjamin Lindsey, Mehmet Yavuz, Mohammad Raza, Cariad Evans |
| EPI_ISL_702882 | Quadram Institute Bioscience | COVID-19 Genomics UK (COG-UK) Consortium | Dave J. Baker, Gemma L. Kay, Alp Aydin, Thanh Le-Viet, Steven Rudder, Ana P. Tedim, Anastasia Kolyva, Maria Diaz, Leonardo de Oliveira Martins, Nabil-Fareed Alikhan, Lizzie Meadows, Rachael Stanley, Ngozi Elumogo, Muhammed Yasir, Nicholas M. Thomson, Alexander J Trotter, Rachel Gilroy, Samuel Bloomfield, Claire Stuart, Andrew Bell, Reenesh Prakash, Samir Dervisevic, Alison E. Mather, John Wain, Mark Webber, Andrew J. Page, Justin O'Grady |
| EPI_ISL_702883, EPI_ISL_702886, EPI_ISL_702891 | Department of Pathology, University of Cambridge | COVID-19 Genomics UK (COG-UK) Consortium | Aminu S. Jahun, Yasmin Chaudhry, Grant Hall, Iliana Georgana, Myra Hosmillo, Martin D. Curran, Malte Pinckert, Surendra Parmar, Ian Goodfellow |
| EPI_ISL_702893 | West of Scotland Specialist Virology Centre, NHSGGC / MRC-University of Glasgow Centre for Virus Research | COVID-19 Genomics UK (COG-UK) Consortium | Ana da Silva Filipe, Natasha Johnson, Kathy Smollett, Daniel Mair, Stephen Carmichael, Alice Broos, Lily Tong, Jenna Nichols, Kyriaki Nomikou; Sarah McDonald; Richard Orton, Joseph Hughes, Sreenu Vattipally, David L Robertson; Alasdair MacLean, Rory Gunson; Sharif Shaaban, Matthew Holden; Rachel Blacow, Guy Mollett, Kathy Li, James Shepherd, Antonia Ho, Emma Thomson |
| EPI_ISL_702898, EPI_ISL_702899, EPI_ISL_702901, EPI_ISL_702903, EPI_ISL_702904 | Lighthouse Lab in Alderley Park | Wellcome Sanger Institute for the COVID-19 Genomics UK (COG-UK) Consortium | Jacquelyn Wynn, Mairead Hyland, The Lighthouse Lab in Alderley Park and Alex Alderton, Roberto Amato, Sonia Goncalves, Ewan Harrison, David K. Jackson, Ian Johnston, Dominic Kwiatkowski, Cordelia Langford, John Sillitoe on behalf of the Wellcome Sanger Institute COVID-19 Surveillance Team |
| EPI_ISL_702907 | Liverpool Clinical Laboratories | COVID-19 Genomics UK (COG-UK) Consortium | Sam Haldenby, Anita Lucaci, Steve Paterson, Julian Hiscox, Alistair Darby, M Almsaud, A Alrezaihi, Muhannad Alruwaili, Stuart D Armstrong, Jones Benjamin, Eleanor G Bentley, Anu Chawla, Jordan J Clark, Angela Cowell, Richard Eccles, Isabel García-Dorival, Matthew Gemmell, Alessandro Gerada, PKF Gilmore, Richard Gregory, Ximeng Han, Catherine Hartley, Margaret Hughes, Miren Iturriza-Gomara, James Johnson, L Luu, Jenifer Manson, Charlotte Nelson, Elaine O'Toole, Cassie Olateju, Rebekah Penrice-Randal , Lucille Rainbow, N.P Randle, Trevor Ian Robinson, Parul Sharma, Ghada T Shawli, James P Stewart, Neil Swainston, Ecaterina Vamos, Joanne Watts, Mark Whitehead |
| EPI_ISL_702909, EPI_ISL_702912, EPI_ISL_702914, EPI_ISL_702919, EPI_ISL_702921, EPI_ISL_702924, EPI_ISL_702926, EPI_ISL_702931, EPI_ISL_702933, EPI_ISL_702934, EPI_ISL_702936, EPI_ISL_702938, EPI_ISL_702943, EPI_ISL_702945, EPI_ISL_702946, EPI_ISL_702948, EPI_ISL_702951, EPI_ISL_702955, EPI_ISL_702957, EPI_ISL_702958, EPI_ISL_702960, EPI_ISL_702962 | | | |
| see above | Lighthouse Lab in Alderley Park | Wellcome Sanger Institute for the COVID-19 Genomics UK (COG-UK) Consortium | Jacquelyn Wynn, Mairead Hyland, The Lighthouse Lab in Alderley Park and Alex Alderton, Roberto Amato, Sonia Goncalves, Ewan Harrison, David K. Jackson, Ian Johnston, Dominic Kwiatkowski, Cordelia Langford, John Sillitoe on behalf of the Wellcome Sanger Institute COVID-19 Surveillance Team |
| EPI_ISL_702964 | Virology Department, Sheffield Teaching Hospitals NHS Foundation Trust/Department of Infection, Immunity and Cardiovascular Disease, The Medical School, University of Sheffield | COVID-19 Genomics UK (COG-UK) Consortium | Thushan de Silva, Matthew Parker, Nikki Smith, Adri Angyal, Rebecca Brown, Luke Green, Rachel Tucker, Paul Parsons, Danielle Groves, Katie Johnson, Laura Carrilero, Alex Keeley, Dave Partridge, Matthew Wyles, Benjamin Lindsey, Mehmet Yavuz, Mohammad Raza, Cariad Evans |
| EPI_ISL_702973, EPI_ISL_702975, EPI_ISL_702978, EPI_ISL_702984, EPI_ISL_702989 | Lighthouse Lab in Alderley Park | Wellcome Sanger Institute for the COVID-19 Genomics UK (COG-UK) Consortium | Jacquelyn Wynn, Mairead Hyland, The Lighthouse Lab in Alderley Park and Alex Alderton, Roberto Amato, Sonia Goncalves, Ewan Harrison, David K. Jackson, Ian Johnston, Dominic Kwiatkowski, Cordelia Langford, John Sillitoe on behalf of the Wellcome Sanger Institute COVID-19 Surveillance Team |
| EPI_ISL_702994 | University of Exeter | COVID-19 Genomics UK (COG-UK) Consortium | Ben Temperton,Aaron Jeffries,Michelle Michelsen,Joanna Warwick-Dugdale,Audrey Farbos,Robyn Manley,Stephen Michell,Jane Masoli |
| EPI_ISL_702995, EPI_ISL_702998, EPI_ISL_703000, EPI_ISL_703001, EPI_ISL_703003, EPI_ISL_703004, EPI_ISL_703006, EPI_ISL_703008, EPI_ISL_703009, EPI_ISL_703011, EPI_ISL_703014, EPI_ISL_703016, EPI_ISL_703017 | | | |
| see above | Lighthouse Lab in Alderley Park | Wellcome Sanger Institute for the COVID-19 Genomics UK (COG-UK) Consortium | Jacquelyn Wynn, Mairead Hyland, The Lighthouse Lab in Alderley Park and Alex Alderton, Roberto Amato, Sonia Goncalves, Ewan Harrison, David K. Jackson, Ian Johnston, Dominic Kwiatkowski, Cordelia Langford, John Sillitoe on behalf of the Wellcome Sanger Institute COVID-19 Surveillance Team |
| EPI_ISL_703018 | Virology Department, Sheffield Teaching Hospitals NHS Foundation Trust/Department of Infection, Immunity and Cardiovascular Disease, The Medical School, University of Sheffield | COVID-19 Genomics UK (COG-UK) Consortium | Thushan de Silva, Matthew Parker, Nikki Smith, Adri Angyal, Rebecca Brown, Luke Green, Rachel Tucker, Paul Parsons, Danielle Groves, Katie Johnson, Laura Carrilero, Alex Keeley, Dave Partridge, Matthew Wyles, Benjamin Lindsey, Mehmet Yavuz, Mohammad Raza, Cariad Evans |
| EPI_ISL_703019 | Lighthouse Lab in Alderley Park | Wellcome Sanger Institute for the COVID-19 Genomics UK (COG-UK) Consortium | Jacquelyn Wynn, Mairead Hyland, The Lighthouse Lab in Alderley Park and Alex Alderton, Roberto Amato, Sonia Goncalves, Ewan Harrison, David K. Jackson, Ian Johnston, Dominic Kwiatkowski, Cordelia Langford, John Sillitoe on behalf of the Wellcome Sanger Institute COVID-19 Surveillance Team |
| EPI_ISL_703020 | Virology Department, Sheffield Teaching Hospitals NHS Foundation Trust/Department of Infection, Immunity and Cardiovascular Disease, The Medical School, University of Sheffield | COVID-19 Genomics UK (COG-UK) Consortium | Thushan de Silva, Matthew Parker, Nikki Smith, Adri Angyal, Rebecca Brown, Luke Green, Rachel Tucker, Paul Parsons, Danielle Groves, Katie Johnson, Laura Carrilero, Alex Keeley, Dave Partridge, Matthew Wyles, Benjamin Lindsey, Mehmet Yavuz, Mohammad Raza, Cariad Evans |
| EPI_ISL_703024, EPI_ISL_703025 | Lighthouse Lab in Alderley Park | Wellcome Sanger Institute for the COVID-19 Genomics UK (COG-UK) Consortium | Jacquelyn Wynn, Mairead Hyland, The Lighthouse Lab in Alderley Park and Alex Alderton, Roberto Amato, Sonia Goncalves, Ewan Harrison, David K. Jackson, Ian Johnston, Dominic Kwiatkowski, Cordelia Langford, John Sillitoe on behalf of the Wellcome Sanger Institute COVID-19 Surveillance Team |
| EPI_ISL_703026 | Virology Department, Sheffield Teaching Hospitals NHS Foundation Trust/Department of Infection, Immunity and Cardiovascular Disease, The Medical School, University of Sheffield | COVID-19 Genomics UK (COG-UK) Consortium | Thushan de Silva, Matthew Parker, Nikki Smith, Adri Angyal, Rebecca Brown, Luke Green, Rachel Tucker, Paul Parsons, Danielle Groves, Katie Johnson, Laura Carrilero, Alex Keeley, Dave Partridge, Matthew Wyles, Benjamin Lindsey, Mehmet Yavuz, Mohammad Raza, Cariad Evans |
| EPI_ISL_703028 | Lighthouse Lab in Alderley Park | Wellcome Sanger Institute for the COVID-19 Genomics UK (COG-UK) Consortium | Jacquelyn Wynn, Mairead Hyland, The Lighthouse Lab in Alderley Park and Alex Alderton, Roberto Amato, Sonia Goncalves, Ewan Harrison, David K. Jackson, Ian Johnston, Dominic Kwiatkowski, Cordelia Langford, John Sillitoe on behalf of the Wellcome Sanger Institute COVID-19 Surveillance Team |
| EPI_ISL_703029 | Virology Department, Sheffield Teaching Hospitals NHS Foundation Trust/Department of Infection, Immunity and | COVID-19 Genomics UK (COG-UK) Consortium | Thushan de Silva, Matthew Parker, Nikki Smith, Adri Angyal, Rebecca Brown, Luke Green, Rachel Tucker, Paul Parsons, Danielle Groves, Katie Johnson, Laura Carrilero, Alex Keeley, Dave Partridge, Matthew Wyles, Benjamin Lindsey, Mehmet Yavuz, Mohammad Raza, Cariad Evans |

| | | | |
|---|---|---|---|
| | Cardiovascular Disease, The Medical School, University of Sheffield | | |
| EPI_ISL_703031 | Lighthouse Lab in Alderley Park | Wellcome Sanger Institute for the COVID-19 Genomics UK (COG-UK) Consortium | Jacquelyn Wynn, Mairead Hyland, The Lighthouse Lab in Alderley Park and Alex Alderton, Roberto Amato, Sonia Goncalves, Ewan Harrison, David K. Jackson, Ian Johnston, Dominic Kwiatkowski, Cordelia Langford, John Sillitoe on behalf of the Wellcome Sanger Institute COVID-19 Surveillance Team |
| EPI_ISL_703032 | Department of Pathology, University of Cambridge | COVID-19 Genomics UK (COG-UK) Consortium | Aminu S. Jahun, Yasmin Chaudhry, Grant Hall, Iliana Georgana, Myra Hosmillo, Martin D. Curran, Malte Pinckert, Surendra Parmar, Ian Goodfellow |
| EPI_ISL_703037, EPI_ISL_703041, EPI_ISL_703042, EPI_ISL_703045, EPI_ISL_703047, EPI_ISL_703050, EPI_ISL_703052, EPI_ISL_703060, EPI_ISL_703062, EPI_ISL_703063, EPI_ISL_703065, EPI_ISL_703067, EPI_ISL_703068, EPI_ISL_703070, EPI_ISL_703072, EPI_ISL_703073 | | | |
| see above | Lighthouse Lab in Alderley Park | Wellcome Sanger Institute for the COVID-19 Genomics UK (COG-UK) Consortium | Jacquelyn Wynn, Mairead Hyland, The Lighthouse Lab in Alderley Park and Alex Alderton, Roberto Amato, Sonia Goncalves, Ewan Harrison, David K. Jackson, Ian Johnston, Dominic Kwiatkowski, Cordelia Langford, John Sillitoe on behalf of the Wellcome Sanger Institute COVID-19 Surveillance Team |
| EPI_ISL_703076 | Department of Pathology, University of Cambridge | COVID-19 Genomics UK (COG-UK) Consortium | Aminu S. Jahun, Yasmin Chaudhry, Grant Hall, Iliana Georgana, Myra Hosmillo, Martin D. Curran, Malte Pinckert, Surendra Parmar, Ian Goodfellow |
| EPI_ISL_703078, EPI_ISL_703079, EPI_ISL_703083, EPI_ISL_703084, EPI_ISL_703086, EPI_ISL_703091, EPI_ISL_703092, EPI_ISL_703095, EPI_ISL_703097, EPI_ISL_703098, EPI_ISL_703100, EPI_ISL_703103, EPI_ISL_703104, EPI_ISL_703106, EPI_ISL_703108 | | | |
| see above | Lighthouse Lab in Alderley Park | Wellcome Sanger Institute for the COVID-19 Genomics UK (COG-UK) Consortium | Jacquelyn Wynn, Mairead Hyland, The Lighthouse Lab in Alderley Park and Alex Alderton, Roberto Amato, Sonia Goncalves, Ewan Harrison, David K. Jackson, Ian Johnston, Dominic Kwiatkowski, Cordelia Langford, John Sillitoe on behalf of the Wellcome Sanger Institute COVID-19 Surveillance Team |
| EPI_ISL_703113 | Quadram Institute Bioscience | COVID-19 Genomics UK (COG-UK) Consortium | Dave J. Baker, Gemma L. Kay, Alp Aydin, Thanh Le-Viet, Steven Rudder, Ana P. Tedim, Anastasia Kolyva, Maria Diaz, Leonardo de Oliveira Martins, Nabil-Fareed Alikhan, Lizzie Meadows, Rachael Stanley, Ngozi Elumogo, Muhammed Yasir, Nicholas M. Thomson, Alexander J Trotter, Rachel Gilroy, Samuel Bloomfield, Claire Stuart, Andrew Bell, Reenesh Prakash, Samir Dervisevic, Alison E. Mather, John Wain, Mark Webber, Andrew J. Page, Justin O'Grady |
| EPI_ISL_703118 | Department of Pathology, University of Cambridge | COVID-19 Genomics UK (COG-UK) Consortium | Aminu S. Jahun, Yasmin Chaudhry, Grant Hall, Iliana Georgana, Myra Hosmillo, Martin D. Curran, Malte Pinckert, Surendra Parmar, Ian Goodfellow |
| EPI_ISL_703120, EPI_ISL_703122, EPI_ISL_703123, EPI_ISL_703126 | Lighthouse Lab in Alderley Park | Wellcome Sanger Institute for the COVID-19 Genomics UK (COG-UK) Consortium | Jacquelyn Wynn, Mairead Hyland, The Lighthouse Lab in Alderley Park and Alex Alderton, Roberto Amato, Sonia Goncalves, Ewan Harrison, David K. Jackson, Ian Johnston, Dominic Kwiatkowski, Cordelia Langford, John Sillitoe on behalf of the Wellcome Sanger Institute COVID-19 Surveillance Team |
| EPI_ISL_703127 | Department of Pathology, University of Cambridge | COVID-19 Genomics UK (COG-UK) Consortium | Aminu S. Jahun, Yasmin Chaudhry, Grant Hall, Iliana Georgana, Myra Hosmillo, Martin D. Curran, Malte Pinckert, Surendra Parmar, Ian Goodfellow |
| EPI_ISL_703128, EPI_ISL_703130, EPI_ISL_703131, EPI_ISL_703135, EPI_ISL_703138 | Lighthouse Lab in Alderley Park | Wellcome Sanger Institute for the COVID-19 Genomics UK (COG-UK) Consortium | Jacquelyn Wynn, Mairead Hyland, The Lighthouse Lab in Alderley Park and Alex Alderton, Roberto Amato, Sonia Goncalves, Ewan Harrison, David K. Jackson, Ian Johnston, Dominic Kwiatkowski, Cordelia Langford, John Sillitoe on behalf of the Wellcome Sanger Institute COVID-19 Surveillance Team |
| EPI_ISL_703141 | West of Scotland Specialist Virology Centre, NHSGGC / MRC-University of Glasgow Centre for Virus Research | COVID-19 Genomics UK (COG-UK) Consortium | Ana da Silva Filipe, Natasha Johnson, Kathy Smollett, Daniel Mair, Stephen Carmichael, Alice Broos, Lily Tong, Jenna Nichols, Kyriaki Nomikou; Sarah McDonald; Richard Orton, Joseph Hughes, Sreenu Vattipally, David L Robertson; Alasdair MacLean, Rory Gunson; Sharif Shaaban, Matthew Holden; Rachel Blacow, Guy Mollett, Kathy Li, James Shepherd, Antonia Ho, Emma Thomson |
| EPI_ISL_703142, EPI_ISL_703143, EPI_ISL_703146 | Lighthouse Lab in Alderley Park | Wellcome Sanger Institute for the COVID-19 Genomics UK (COG-UK) Consortium | Jacquelyn Wynn, Mairead Hyland, The Lighthouse Lab in Alderley Park and Alex Alderton, Roberto Amato, Sonia Goncalves, Ewan Harrison, David K. Jackson, Ian Johnston, Dominic Kwiatkowski, Cordelia Langford, John Sillitoe on behalf of the Wellcome Sanger Institute COVID-19 Surveillance Team |
| EPI_ISL_703147 | Liverpool Clinical Laboratories | COVID-19 Genomics UK (COG-UK) Consortium | Sam Haldenby, Anita Lucaci, Steve Paterson, Julian Hiscox, Alistair Darby, M Almsaud, A Alrezaihi, Muhannad Alruwaili, Stuart D Armstrong, Jones Benjamin, Eleanor G Bentley, Anu Chawla, Jordan J Clark, Angela Cowell, Richard Eccles, Isabel García-Dorival, Matthew Gemmell, Alessandro Gerada, PKF Gilmore, Richard Gregory, Ximeng Han, Catherine Hartley, Margaret Hughes, Miren Iturriza-Gomara, James Johnson, L Luu, Jenifer Manson, Charlotte Nelson, Elaine O'Toole, Cassie Olateju, Rebekah Penrice-Randal , Lucille Rainbow, N.P Randle, Trevor Ian Robinson, Parul Sharma, Ghada T Shawli, James P Stewart, Neil Swainston, Ecaterina Vamos, Joanne Watts, Mark Whitehead |
| EPI_ISL_703149, EPI_ISL_703153, EPI_ISL_703154, EPI_ISL_703156, EPI_ISL_703159, EPI_ISL_703160, EPI_ISL_703161 | Lighthouse Lab in Alderley Park | Wellcome Sanger Institute for the COVID-19 Genomics UK (COG-UK) Consortium | Jacquelyn Wynn, Mairead Hyland, The Lighthouse Lab in Alderley Park and Alex Alderton, Roberto Amato, Sonia Goncalves, Ewan Harrison, David K. Jackson, Ian Johnston, Dominic Kwiatkowski, Cordelia Langford, John Sillitoe on behalf of the Wellcome Sanger Institute COVID-19 Surveillance Team |
| EPI_ISL_703162 | Liverpool Clinical Laboratories | COVID-19 Genomics UK (COG-UK) Consortium | Sam Haldenby, Anita Lucaci, Steve Paterson, Julian Hiscox, Alistair Darby, M Almsaud, A Alrezaihi, Muhannad Alruwaili, Stuart D Armstrong, Jones Benjamin, Eleanor G Bentley, Anu Chawla, Jordan J Clark, Angela Cowell, Richard Eccles, Isabel García-Dorival, Matthew Gemmell, Alessandro Gerada, PKF Gilmore, Richard Gregory, Ximeng Han, Catherine Hartley, Margaret Hughes, Miren Iturriza-Gomara, James Johnson, L Luu, Jenifer Manson, Charlotte Nelson, Elaine O'Toole, Cassie Olateju, Rebekah Penrice-Randal , Lucille Rainbow, N.P Randle, Trevor Ian Robinson, Parul Sharma, Ghada T Shawli, James P Stewart, Neil Swainston, Ecaterina Vamos, Joanne Watts, Mark Whitehead |
| EPI_ISL_703163, EPI_ISL_703169, EPI_ISL_703173, EPI_ISL_703183 | Lighthouse Lab in Alderley Park | Wellcome Sanger Institute for the COVID-19 Genomics UK (COG-UK) Consortium | Jacquelyn Wynn, Mairead Hyland, The Lighthouse Lab in Alderley Park and Alex Alderton, Roberto Amato, Sonia Goncalves, Ewan Harrison, David K. Jackson, Ian Johnston, Dominic Kwiatkowski, Cordelia Langford, John Sillitoe on behalf of the Wellcome Sanger Institute COVID-19 Surveillance Team |
| EPI_ISL_703184 | Liverpool Clinical Laboratories | COVID-19 Genomics UK (COG-UK) Consortium | Sam Haldenby, Anita Lucaci, Steve Paterson, Julian Hiscox, Alistair Darby, M Almsaud, A Alrezaihi, Muhannad Alruwaili, Stuart D Armstrong, Jones Benjamin, Eleanor G Bentley, Anu Chawla, Jordan J Clark, Angela Cowell, Richard Eccles, Isabel García-Dorival, Matthew Gemmell, Alessandro Gerada, PKF Gilmore, Richard Gregory, Ximeng Han, Catherine Hartley, Margaret Hughes, Miren Iturriza-Gomara, James Johnson, L Luu, Jenifer Manson, Charlotte Nelson, Elaine O'Toole, Cassie Olateju, Rebekah Penrice-Randal , Lucille Rainbow, N.P Randle, Trevor Ian Robinson, Parul Sharma, Ghada T Shawli, James P Stewart, Neil Swainston, Ecaterina Vamos, Joanne Watts, Mark Whitehead |
| EPI_ISL_703186 | Lighthouse Lab in Alderley Park | Wellcome Sanger Institute for the COVID-19 Genomics UK (COG-UK) Consortium | Jacquelyn Wynn, Mairead Hyland, The Lighthouse Lab in Alderley Park and Alex Alderton, Roberto Amato, Sonia Goncalves, Ewan Harrison, David K. Jackson, Ian Johnston, Dominic Kwiatkowski, Cordelia Langford, John Sillitoe on behalf of the Wellcome Sanger Institute COVID-19 Surveillance Team |
| EPI_ISL_703187 | Quadram Institute Bioscience | COVID-19 Genomics UK (COG-UK) Consortium | Dave J. Baker, Gemma L. Kay, Alp Aydin, Thanh Le-Viet, Steven Rudder, Ana P. Tedim, Anastasia Kolyva, Maria Diaz, Leonardo de Oliveira Martins, Nabil-Fareed Alikhan, Lizzie Meadows, Rachael Stanley, Ngozi Elumogo, Muhammed Yasir, Nicholas M. Thomson, Alexander J Trotter, Rachel Gilroy, Samuel Bloomfield, Claire Stuart, Andrew Bell, Reenesh Prakash, Samir Dervisevic, Alison E. Mather, John Wain, Mark Webber, Andrew J. Page, Justin O'Grady |
| EPI_ISL_703188, EPI_ISL_703193, EPI_ISL_703197, EPI_ISL_703201, EPI_ISL_703204, EPI_ISL_703207 | Lighthouse Lab in Alderley Park | Wellcome Sanger Institute for the COVID-19 Genomics UK (COG-UK) Consortium | Jacquelyn Wynn, Mairead Hyland, The Lighthouse Lab in Alderley Park and Alex Alderton, Roberto Amato, Sonia Goncalves, Ewan Harrison, David K. Jackson, Ian Johnston, Dominic Kwiatkowski, Cordelia Langford, John Sillitoe on behalf of the Wellcome Sanger Institute COVID-19 Surveillance Team |
| EPI_ISL_703208 | Liverpool Clinical Laboratories | COVID-19 Genomics UK (COG-UK) Consortium | Sam Haldenby, Anita Lucaci, Steve Paterson, Julian Hiscox, Alistair Darby, M Almsaud, A Alrezaihi, Muhannad Alruwaili, Stuart D Armstrong, Jones Benjamin, Eleanor G Bentley, Anu Chawla, Jordan J Clark, Angela Cowell, Richard Eccles, Isabel García-Dorival, Matthew Gemmell, Alessandro Gerada, PKF Gilmore, Richard Gregory, Ximeng Han, Catherine Hartley, Margaret Hughes, Miren Iturriza-Gomara, James Johnson, L Luu, Jenifer Manson, Charlotte Nelson, Elaine O'Toole, Cassie Olateju, Rebekah Penrice-Randal , Lucille Rainbow, N.P Randle, Trevor Ian Robinson, Parul Sharma, Ghada T Shawli, James P Stewart, Neil Swainston, Ecaterina Vamos, Joanne Watts, Mark Whitehead |
| EPI_ISL_703209, EPI_ISL_703210, EPI_ISL_703212 | Lighthouse Lab in Alderley Park | Wellcome Sanger Institute for the COVID-19 Genomics UK (COG-UK) Consortium | Jacquelyn Wynn, Mairead Hyland, The Lighthouse Lab in Alderley Park and Alex Alderton, Roberto Amato, Sonia Goncalves, Ewan Harrison, David K. Jackson, Ian Johnston, Dominic Kwiatkowski, Cordelia Langford, John Sillitoe on behalf of the Wellcome Sanger Institute COVID-19 Surveillance Team |
| EPI_ISL_703213 | Department of Pathology, University of Cambridge | COVID-19 Genomics UK (COG-UK) Consortium | Aminu S. Jahun, Yasmin Chaudhry, Grant Hall, Iliana Georgana, Myra Hosmillo, Martin D. Curran, Malte Pinckert, Surendra Parmar, Ian Goodfellow |
| EPI_ISL_703215, EPI_ISL_703217, EPI_ISL_703220, EPI_ISL_703221 | Lighthouse Lab in Alderley Park | Wellcome Sanger Institute for the COVID-19 Genomics UK (COG-UK) Consortium | Jacquelyn Wynn, Mairead Hyland, The Lighthouse Lab in Alderley Park and Alex Alderton, Roberto Amato, Sonia Goncalves, Ewan Harrison, David K. Jackson, Ian Johnston, Dominic Kwiatkowski, Cordelia Langford, John Sillitoe on behalf of the Wellcome Sanger Institute COVID-19 Surveillance Team |
| EPI_ISL_703222 | University College London Hospital | COVID-19 Genomics UK (COG-UK) Consortium | Judith Heaney, Matthew Byott, Catherine Houlihan, Dan Frampton, Stuart Kirk, Moira Spyer and Eleni Nastouli |
| EPI_ISL_703224, EPI_ISL_703226, EPI_ISL_703228, EPI_ISL_703231, EPI_ISL_703232, EPI_ISL_703237 | Lighthouse Lab in Alderley Park | Wellcome Sanger Institute for the COVID-19 Genomics UK (COG-UK) Consortium | Jacquelyn Wynn, Mairead Hyland, The Lighthouse Lab in Alderley Park and Alex Alderton, Roberto Amato, Sonia Goncalves, Ewan Harrison, David K. Jackson, Ian Johnston, Dominic Kwiatkowski, Cordelia Langford, John Sillitoe on behalf of the Wellcome Sanger Institute COVID-19 Surveillance Team |
| EPI_ISL_703238 | Department of Pathology, University of Cambridge | COVID-19 Genomics UK (COG-UK) Consortium | Aminu S. Jahun, Yasmin Chaudhry, Grant Hall, Iliana Georgana, Myra Hosmillo, Martin D. Curran, Malte Pinckert, Surendra Parmar, Ian Goodfellow |

| | | | |
|---|---|---|---|
| EPI_ISL_703240 | West of Scotland Specialist Virology Centre, NHSGGC / MRC-University of Glasgow Centre for Virus Research | COVID-19 Genomics UK (COG-UK) Consortium | Ana da Silva Filipe, Natasha Johnson, Kathy Smollett, Daniel Mair, Stephen Carmichael, Alice Broos, Lily Tong, Jenna Nichols, Kyriaki Nomikou; Sarah McDonald; Richard Orton, Joseph Hughes, Sreenu Vattipally, David L Robertson; Alasdair MacLean, Rory Gunson; Sharif Shaaban, Matthew Holden; Rachel Blacow, Guy Mollett, Kathy Li, James Shepherd, Antonia Ho, Emma Thomson |
| EPI_ISL_703241, EPI_ISL_703242 | Lighthouse Lab in Alderley Park | Wellcome Sanger Institute for the COVID-19 Genomics UK (COG-UK) Consortium | Jacquelyn Wynn, Mairead Hyland, The Lighthouse Lab in Alderley Park and Alex Alderton, Roberto Amato, Sonia Goncalves, Ewan Harrison, David K. Jackson, Ian Johnston, Dominic Kwiatkowski, Cordelia Langford, John Sillitoe on behalf of the Wellcome Sanger Institute COVID-19 Surveillance Team |
| EPI_ISL_703243 | West of Scotland Specialist Virology Centre, NHSGGC / MRC-University of Glasgow Centre for Virus Research | COVID-19 Genomics UK (COG-UK) Consortium | Ana da Silva Filipe, Natasha Johnson, Kathy Smollett, Daniel Mair, Stephen Carmichael, Alice Broos, Lily Tong, Jenna Nichols, Kyriaki Nomikou; Sarah McDonald; Richard Orton, Joseph Hughes, Sreenu Vattipally, David L Robertson; Alasdair MacLean, Rory Gunson; Sharif Shaaban, Matthew Holden; Rachel Blacow, Guy Mollett, Kathy Li, James Shepherd, Antonia Ho, Emma Thomson |
| EPI_ISL_703245, EPI_ISL_703247, EPI_ISL_703248, EPI_ISL_703250, EPI_ISL_703252, EPI_ISL_703253, EPI_ISL_703255, EPI_ISL_703257, EPI_ISL_703258, EPI_ISL_703260 | Lighthouse Lab in Alderley Park | Wellcome Sanger Institute for the COVID-19 Genomics UK (COG-UK) Consortium | Jacquelyn Wynn, Mairead Hyland, The Lighthouse Lab in Alderley Park and Alex Alderton, Roberto Amato, Sonia Goncalves, Ewan Harrison, David K. Jackson, Ian Johnston, Dominic Kwiatkowski, Cordelia Langford, John Sillitoe on behalf of the Wellcome Sanger Institute COVID-19 Surveillance Team |
| EPI_ISL_703261 | Liverpool Clinical Laboratories | COVID-19 Genomics UK (COG-UK) Consortium | Sam Haldenby, Anita Lucaci, Steve Paterson, Julian Hiscox, Alistair Darby, M Almsaud, A Alrezaihi, Muhannad Alruwaili, Stuart D Armstrong, Jones Benjamin, Eleanor G Bentley, Anu Chawla, Jordan J Clark, Angela Cowell, Richard Eccles, Isabel García-Dorival, Matthew Gemmell, Alessandro Gerada, PKF Gilmore, Richard Gregory, Ximeng Han, Catherine Hartley, Margaret Hughes, Miren Iturriza-Gomara, James Johnson, L Luu, Jenifer Manson, Charlotte Nelson, Elaine O'Toole, Cassie Olateju, Rebekah Penrice-Randal , Lucille Rainbow, N.P Randle, Trevor Ian Robinson, Parul Sharma, Ghada T Shawli, James P Stewart, Neil Swainston, Ecaterina Vamos, Joanne Watts, Mark Whitehead |
| EPI_ISL_703265, EPI_ISL_703266, EPI_ISL_703268, EPI_ISL_703271, EPI_ISL_703274 | Lighthouse Lab in Alderley Park | Wellcome Sanger Institute for the COVID-19 Genomics UK (COG-UK) Consortium | Jacquelyn Wynn, Mairead Hyland, The Lighthouse Lab in Alderley Park and Alex Alderton, Roberto Amato, Sonia Goncalves, Ewan Harrison, David K. Jackson, Ian Johnston, Dominic Kwiatkowski, Cordelia Langford, John Sillitoe on behalf of the Wellcome Sanger Institute COVID-19 Surveillance Team |
| EPI_ISL_703275 | Liverpool Clinical Laboratories | COVID-19 Genomics UK (COG-UK) Consortium | Sam Haldenby, Anita Lucaci, Steve Paterson, Julian Hiscox, Alistair Darby, M Almsaud, A Alrezaihi, Muhannad Alruwaili, Stuart D Armstrong, Jones Benjamin, Eleanor G Bentley, Anu Chawla, Jordan J Clark, Angela Cowell, Richard Eccles, Isabel García-Dorival, Matthew Gemmell, Alessandro Gerada, PKF Gilmore, Richard Gregory, Ximeng Han, Catherine Hartley, Margaret Hughes, Miren Iturriza-Gomara, James Johnson, L Luu, Jenifer Manson, Charlotte Nelson, Elaine O'Toole, Cassie Olateju, Rebekah Penrice-Randal , Lucille Rainbow, N.P Randle, Trevor Ian Robinson, Parul Sharma, Ghada T Shawli, James P Stewart, Neil Swainston, Ecaterina Vamos, Joanne Watts, Mark Whitehead |
| EPI_ISL_703276, EPI_ISL_703277, EPI_ISL_703282 | Lighthouse Lab in Alderley Park | Wellcome Sanger Institute for the COVID-19 Genomics UK (COG-UK) Consortium | Jacquelyn Wynn, Mairead Hyland, The Lighthouse Lab in Alderley Park and Alex Alderton, Roberto Amato, Sonia Goncalves, Ewan Harrison, David K. Jackson, Ian Johnston, Dominic Kwiatkowski, Cordelia Langford, John Sillitoe on behalf of the Wellcome Sanger Institute COVID-19 Surveillance Team |
| EPI_ISL_703286 | Lighthouse Lab in Glasgow | Wellcome Sanger Institute for the COVID-19 Genomics UK (COG-UK) Consortium | Harper VanSteenhouse, Yumi Kasai, David Gray, Carol Clugston, Anna Dominiczak and Alex Alderton, Roberto Amato, Sonia Goncalves, Ewan Harrison, David K. Jackson, Ian Johnston, Dominic Kwiatkowski, Cordelia Langford, John Sillitoe on behalf of the Wellcome Sanger Institute COVID-19 Surveillance Team |
| EPI_ISL_703289 | Liverpool Clinical Laboratories | COVID-19 Genomics UK (COG-UK) Consortium | Sam Haldenby, Anita Lucaci, Steve Paterson, Julian Hiscox, Alistair Darby, M Almsaud, A Alrezaihi, Muhannad Alruwaili, Stuart D Armstrong, Jones Benjamin, Eleanor G Bentley, Anu Chawla, Jordan J Clark, Angela Cowell, Richard Eccles, Isabel García-Dorival, Matthew Gemmell, Alessandro Gerada, PKF Gilmore, Richard Gregory, Ximeng Han, Catherine Hartley, Margaret Hughes, Miren Iturriza-Gomara, James Johnson, L Luu, Jenifer Manson, Charlotte Nelson, Elaine O'Toole, Cassie Olateju, Rebekah Penrice-Randal , Lucille Rainbow, N.P Randle, Trevor Ian Robinson, Parul Sharma, Ghada T Shawli, James P Stewart, Neil Swainston, Ecaterina Vamos, Joanne Watts, Mark Whitehead |
| EPI_ISL_703293, EPI_ISL_703294, EPI_ISL_703298, EPI_ISL_703301, EPI_ISL_703306, EPI_ISL_703307, EPI_ISL_703311, EPI_ISL_703312, EPI_ISL_703314, EPI_ISL_703317, EPI_ISL_703318, EPI_ISL_703322, EPI_ISL_703326, EPI_ISL_703328, EPI_ISL_703329, EPI_ISL_703333, EPI_ISL_703336, EPI_ISL_703339, EPI_ISL_703341, EPI_ISL_703342, EPI_ISL_703344 | | | |
| see above | Lighthouse Lab in Alderley Park | Wellcome Sanger Institute for the COVID-19 Genomics UK (COG-UK) Consortium | Jacquelyn Wynn, Mairead Hyland, The Lighthouse Lab in Alderley Park and Alex Alderton, Roberto Amato, Sonia Goncalves, Ewan Harrison, David K. Jackson, Ian Johnston, Dominic Kwiatkowski, Cordelia Langford, John Sillitoe on behalf of the Wellcome Sanger Institute COVID-19 Surveillance Team |
| EPI_ISL_703345 | Lighthouse Lab in Glasgow | Wellcome Sanger Institute for the COVID-19 Genomics UK (COG-UK) Consortium | Harper VanSteenhouse, Yumi Kasai, David Gray, Carol Clugston, Anna Dominiczak and Alex Alderton, Roberto Amato, Sonia Goncalves, Ewan Harrison, David K. Jackson, Ian Johnston, Dominic Kwiatkowski, Cordelia Langford, John Sillitoe on behalf of the Wellcome Sanger Institute COVID-19 Surveillance Team |
| EPI_ISL_703347 | Lighthouse Lab in Alderley Park | Wellcome Sanger Institute for the COVID-19 Genomics UK (COG-UK) Consortium | Jacquelyn Wynn, Mairead Hyland, The Lighthouse Lab in Alderley Park and Alex Alderton, Roberto Amato, Sonia Goncalves, Ewan Harrison, David K. Jackson, Ian Johnston, Dominic Kwiatkowski, Cordelia Langford, John Sillitoe on behalf of the Wellcome Sanger Institute COVID-19 Surveillance Team |
| EPI_ISL_703353 | Liverpool Clinical Laboratories | COVID-19 Genomics UK (COG-UK) Consortium | Sam Haldenby, Anita Lucaci, Steve Paterson, Julian Hiscox, Alistair Darby, M Almsaud, A Alrezaihi, Muhannad Alruwaili, Stuart D Armstrong, Jones Benjamin, Eleanor G Bentley, Anu Chawla, Jordan J Clark, Angela Cowell, Richard Eccles, Isabel García-Dorival, Matthew Gemmell, Alessandro Gerada, PKF Gilmore, Richard Gregory, Ximeng Han, Catherine Hartley, Margaret Hughes, Miren Iturriza-Gomara, James Johnson, L Luu, Jenifer Manson, Charlotte Nelson, Elaine O'Toole, Cassie Olateju, Rebekah Penrice-Randal , Lucille Rainbow, N.P Randle, Trevor Ian Robinson, Parul Sharma, Ghada T Shawli, James P Stewart, Neil Swainston, Ecaterina Vamos, Joanne Watts, Mark Whitehead |
| EPI_ISL_703357, EPI_ISL_703363, EPI_ISL_703365 | Lighthouse Lab in Alderley Park | Wellcome Sanger Institute for the COVID-19 Genomics UK (COG-UK) Consortium | Jacquelyn Wynn, Mairead Hyland, The Lighthouse Lab in Alderley Park and Alex Alderton, Roberto Amato, Sonia Goncalves, Ewan Harrison, David K. Jackson, Ian Johnston, Dominic Kwiatkowski, Cordelia Langford, John Sillitoe on behalf of the Wellcome Sanger Institute COVID-19 Surveillance Team |
| EPI_ISL_703366 | Lighthouse Lab in Glasgow | Wellcome Sanger Institute for the COVID-19 Genomics UK (COG-UK) Consortium | Harper VanSteenhouse, Yumi Kasai, David Gray, Carol Clugston, Anna Dominiczak and Alex Alderton, Roberto Amato, Sonia Goncalves, Ewan Harrison, David K. Jackson, Ian Johnston, Dominic Kwiatkowski, Cordelia Langford, John Sillitoe on behalf of the Wellcome Sanger Institute COVID-19 Surveillance Team |
| EPI_ISL_703368, EPI_ISL_703373, EPI_ISL_703374, EPI_ISL_703376 | Lighthouse Lab in Alderley Park | Wellcome Sanger Institute for the COVID-19 Genomics UK (COG-UK) Consortium | Jacquelyn Wynn, Mairead Hyland, The Lighthouse Lab in Alderley Park and Alex Alderton, Roberto Amato, Sonia Goncalves, Ewan Harrison, David K. Jackson, Ian Johnston, Dominic Kwiatkowski, Cordelia Langford, John Sillitoe on behalf of the Wellcome Sanger Institute COVID-19 Surveillance Team |
| EPI_ISL_703380 | Liverpool Clinical Laboratories | COVID-19 Genomics UK (COG-UK) Consortium | Sam Haldenby, Anita Lucaci, Steve Paterson, Julian Hiscox, Alistair Darby, M Almsaud, A Alrezaihi, Muhannad Alruwaili, Stuart D Armstrong, Jones Benjamin, Eleanor G Bentley, Anu Chawla, Jordan J Clark, Angela Cowell, Richard Eccles, Isabel García-Dorival, Matthew Gemmell, Alessandro Gerada, PKF Gilmore, Richard Gregory, Ximeng Han, Catherine Hartley, Margaret Hughes, Miren Iturriza-Gomara, James Johnson, L Luu, Jenifer Manson, Charlotte Nelson, Elaine O'Toole, Cassie Olateju, Rebekah Penrice-Randal , Lucille Rainbow, N.P Randle, Trevor Ian Robinson, Parul Sharma, Ghada T Shawli, James P Stewart, Neil Swainston, Ecaterina Vamos, Joanne Watts, Mark Whitehead |
| EPI_ISL_703386, EPI_ISL_703388, EPI_ISL_703394, EPI_ISL_703396, EPI_ISL_703397 | Lighthouse Lab in Alderley Park | Wellcome Sanger Institute for the COVID-19 Genomics UK (COG-UK) Consortium | Jacquelyn Wynn, Mairead Hyland, The Lighthouse Lab in Alderley Park and Alex Alderton, Roberto Amato, Sonia Goncalves, Ewan Harrison, David K. Jackson, Ian Johnston, Dominic Kwiatkowski, Cordelia Langford, John Sillitoe on behalf of the Wellcome Sanger Institute COVID-19 Surveillance Team |
| EPI_ISL_703399, EPI_ISL_703401 | Lighthouse Lab in Glasgow | Wellcome Sanger Institute for the COVID-19 Genomics UK (COG-UK) Consortium | Harper VanSteenhouse, Yumi Kasai, David Gray, Carol Clugston, Anna Dominiczak and Alex Alderton, Roberto Amato, Sonia Goncalves, Ewan Harrison, David K. Jackson, Ian Johnston, Dominic Kwiatkowski, Cordelia Langford, John Sillitoe on behalf of the Wellcome Sanger Institute COVID-19 Surveillance Team |
| EPI_ISL_703402, EPI_ISL_703404, EPI_ISL_703405, EPI_ISL_703413, EPI_ISL_703416, EPI_ISL_703418, EPI_ISL_703420 | Lighthouse Lab in Alderley Park | Wellcome Sanger Institute for the COVID-19 Genomics UK (COG-UK) Consortium | Jacquelyn Wynn, Mairead Hyland, The Lighthouse Lab in Alderley Park and Alex Alderton, Roberto Amato, Sonia Goncalves, Ewan Harrison, David K. Jackson, Ian Johnston, Dominic Kwiatkowski, Cordelia Langford, John Sillitoe on behalf of the Wellcome Sanger Institute COVID-19 Surveillance Team |
| EPI_ISL_703424 | Quadram Institute Bioscience | COVID-19 Genomics UK (COG-UK) Consortium | Dave J. Baker, Gemma L. Kay, Alp Aydin, Thanh Le-Viet, Steven Rudder, Ana P. Tedim, Anastasia Kolyva, Maria Diaz, Leonardo de Oliveira Martins, Nabil-Fareed Alikhan, Lizzie Meadows, Rachael Stanley, Ngozi Elumogo, Muhammed Yasir, Nicholas M. Thomson, Alexander J Trotter, Rachel Gilroy, Samuel Bloomfield, Claire Stuart, Andrew Bell, Reenesh Prakash, Samir Dervisevic, Alison E. Mather, John Wain, Mark Webber, Andrew J. Page, Justin O'Grady |

| | | | |
|---|---|---|---|
| EPI_ISL_703425, EPI_ISL_703433, EPI_ISL_703434, EPI_ISL_703436, EPI_ISL_703437, EPI_ISL_703441 | Lighthouse Lab in Alderley Park | Wellcome Sanger Institute for the COVID-19 Genomics UK (COG-UK) Consortium | Jacquelyn Wynn, Mairead Hyland, The Lighthouse Lab in Alderley Park and Alex Alderton, Roberto Amato, Sonia Goncalves, Ewan Harrison, David K. Jackson, Ian Johnston, Dominic Kwiatkowski, Cordelia Langford, John Sillitoe on behalf of the Wellcome Sanger Institute COVID-19 Surveillance Team |
| EPI_ISL_703445 | Lighthouse Lab in Glasgow | Wellcome Sanger Institute for the COVID-19 Genomics UK (COG-UK) Consortium | Harper VanSteenhouse, Yumi Kasai, David Gray, Carol Clugston, Anna Dominiczak and Alex Alderton, Roberto Amato, Sonia Goncalves, Ewan Harrison, David K. Jackson, Ian Johnston, Dominic Kwiatkowski, Cordelia Langford, John Sillitoe on behalf of the Wellcome Sanger Institute COVID-19 Surveillance Team |
| EPI_ISL_703448, EPI_ISL_703450 | Lighthouse Lab in Alderley Park | Wellcome Sanger Institute for the COVID-19 Genomics UK (COG-UK) Consortium | Jacquelyn Wynn, Mairead Hyland, The Lighthouse Lab in Alderley Park and Alex Alderton, Roberto Amato, Sonia Goncalves, Ewan Harrison, David K. Jackson, Ian Johnston, Dominic Kwiatkowski, Cordelia Langford, John Sillitoe on behalf of the Wellcome Sanger Institute COVID-19 Surveillance Team |
| EPI_ISL_703451 | Department of Pathology, University of Cambridge | COVID-19 Genomics UK (COG-UK) Consortium | Aminu S. Jahun, Yasmin Chaudhry, Grant Hall, Iliana Georgana, Myra Hosmillo, Martin D. Curran, Malte Pinckert, Surendra Parmar, Ian Goodfellow |
| EPI_ISL_703452, EPI_ISL_703455 | Lighthouse Lab in Alderley Park | Wellcome Sanger Institute for the COVID-19 Genomics UK (COG-UK) Consortium | Jacquelyn Wynn, Mairead Hyland, The Lighthouse Lab in Alderley Park and Alex Alderton, Roberto Amato, Sonia Goncalves, Ewan Harrison, David K. Jackson, Ian Johnston, Dominic Kwiatkowski, Cordelia Langford, John Sillitoe on behalf of the Wellcome Sanger Institute COVID-19 Surveillance Team |
| EPI_ISL_703457 | Department of Pathology, University of Cambridge | COVID-19 Genomics UK (COG-UK) Consortium | Aminu S. Jahun, Yasmin Chaudhry, Grant Hall, Iliana Georgana, Myra Hosmillo, Martin D. Curran, Malte Pinckert, Surendra Parmar, Ian Goodfellow |
| EPI_ISL_703460, EPI_ISL_703461, EPI_ISL_703464 | Lighthouse Lab in Alderley Park | Wellcome Sanger Institute for the COVID-19 Genomics UK (COG-UK) Consortium | Jacquelyn Wynn, Mairead Hyland, The Lighthouse Lab in Alderley Park and Alex Alderton, Roberto Amato, Sonia Goncalves, Ewan Harrison, David K. Jackson, Ian Johnston, Dominic Kwiatkowski, Cordelia Langford, John Sillitoe on behalf of the Wellcome Sanger Institute COVID-19 Surveillance Team |
| EPI_ISL_703466 | Lighthouse Lab in Glasgow | Wellcome Sanger Institute for the COVID-19 Genomics UK (COG-UK) Consortium | Harper VanSteenhouse, Yumi Kasai, David Gray, Carol Clugston, Anna Dominiczak and Alex Alderton, Roberto Amato, Sonia Goncalves, Ewan Harrison, David K. Jackson, Ian Johnston, Dominic Kwiatkowski, Cordelia Langford, John Sillitoe on behalf of the Wellcome Sanger Institute COVID-19 Surveillance Team |
| EPI_ISL_703467, EPI_ISL_703470, EPI_ISL_703475 | Lighthouse Lab in Alderley Park | Wellcome Sanger Institute for the COVID-19 Genomics UK (COG-UK) Consortium | Jacquelyn Wynn, Mairead Hyland, The Lighthouse Lab in Alderley Park and Alex Alderton, Roberto Amato, Sonia Goncalves, Ewan Harrison, David K. Jackson, Ian Johnston, Dominic Kwiatkowski, Cordelia Langford, John Sillitoe on behalf of the Wellcome Sanger Institute COVID-19 Surveillance Team |
| EPI_ISL_703476 | Department of Pathology, University of Cambridge | COVID-19 Genomics UK (COG-UK) Consortium | Aminu S. Jahun, Yasmin Chaudhry, Grant Hall, Iliana Georgana, Myra Hosmillo, Martin D. Curran, Malte Pinckert, Surendra Parmar, Ian Goodfellow |
| EPI_ISL_703477, EPI_ISL_703478 | Lighthouse Lab in Alderley Park | Wellcome Sanger Institute for the COVID-19 Genomics UK (COG-UK) Consortium | Jacquelyn Wynn, Mairead Hyland, The Lighthouse Lab in Alderley Park and Alex Alderton, Roberto Amato, Sonia Goncalves, Ewan Harrison, David K. Jackson, Ian Johnston, Dominic Kwiatkowski, Cordelia Langford, John Sillitoe on behalf of the Wellcome Sanger Institute COVID-19 Surveillance Team |
| EPI_ISL_703479 | Department of Pathology, University of Cambridge | COVID-19 Genomics UK (COG-UK) Consortium | Aminu S. Jahun, Yasmin Chaudhry, Grant Hall, Iliana Georgana, Myra Hosmillo, Martin D. Curran, Malte Pinckert, Surendra Parmar, Ian Goodfellow |
| EPI_ISL_703480, EPI_ISL_703483, EPI_ISL_703485 | Lighthouse Lab in Alderley Park | Wellcome Sanger Institute for the COVID-19 Genomics UK (COG-UK) Consortium | Jacquelyn Wynn, Mairead Hyland, The Lighthouse Lab in Alderley Park and Alex Alderton, Roberto Amato, Sonia Goncalves, Ewan Harrison, David K. Jackson, Ian Johnston, Dominic Kwiatkowski, Cordelia Langford, John Sillitoe on behalf of the Wellcome Sanger Institute COVID-19 Surveillance Team |
| EPI_ISL_703487 | Northumbria University / South Tees Hospitals NHS Foundation Trust / North Cumbria Integrated Care NHS Foundation Trust / North Tees and Hartlepool NHS Foundation Trust / Newcastle Hospitals NHS Foundation Trust | COVID-19 Genomics UK (COG-UK) Consortium | Darren L Smith,Andrew Nelson,Matthew Bashton,Greg R Young,Joshua Loh,John Allan,Mohammad A Tariq,Giles S Holt,Gary Black,Wen C Yew,Lynn Dover,Paul Baker,Steve Liggett,Sarah Essex,Jane Greenaway,Debra Padgett,Clive Graham,Garren Scott,Edward Barton,Emma Swindells,Brendan Payne,Jennifer Collins,Yusri Taha,Gary Eltringham |
| EPI_ISL_703488, EPI_ISL_703490, EPI_ISL_703493, EPI_ISL_703494, EPI_ISL_703498 | Lighthouse Lab in Alderley Park | Wellcome Sanger Institute for the COVID-19 Genomics UK (COG-UK) Consortium | Jacquelyn Wynn, Mairead Hyland, The Lighthouse Lab in Alderley Park and Alex Alderton, Roberto Amato, Sonia Goncalves, Ewan Harrison, David K. Jackson, Ian Johnston, Dominic Kwiatkowski, Cordelia Langford, John Sillitoe on behalf of the Wellcome Sanger Institute COVID-19 Surveillance Team |
| EPI_ISL_703501 | Lighthouse Lab in Glasgow | Wellcome Sanger Institute for the COVID-19 Genomics UK (COG-UK) Consortium | Harper VanSteenhouse, Yumi Kasai, David Gray, Carol Clugston, Anna Dominiczak and Alex Alderton, Roberto Amato, Sonia Goncalves, Ewan Harrison, David K. Jackson, Ian Johnston, Dominic Kwiatkowski, Cordelia Langford, John Sillitoe on behalf of the Wellcome Sanger Institute COVID-19 Surveillance Team |
| EPI_ISL_703503, EPI_ISL_703504 | Lighthouse Lab in Alderley Park | Wellcome Sanger Institute for the COVID-19 Genomics UK (COG-UK) Consortium | Jacquelyn Wynn, Mairead Hyland, The Lighthouse Lab in Alderley Park and Alex Alderton, Roberto Amato, Sonia Goncalves, Ewan Harrison, David K. Jackson, Ian Johnston, Dominic Kwiatkowski, Cordelia Langford, John Sillitoe on behalf of the Wellcome Sanger Institute COVID-19 Surveillance Team |
| EPI_ISL_703505 | Virology Department, Sheffield Teaching Hospitals NHS Foundation Trust/Department of Infection, Immunity and Cardiovascular Disease, The Medical School, University of Sheffield | COVID-19 Genomics UK (COG-UK) Consortium | Thushan de Silva, Matthew Parker, Nikki Smith, Adri Angyal, Rebecca Brown, Luke Green, Rachel Tucker, Paul Parsons, Danielle Groves, Katie Johnson, Laura Carrilero, Alex Keeley, Dave Partridge, Matthew Wyles, Benjamin Lindsey, Mehmet Yavuz, Mohammad Raza, Cariad Evans |
| EPI_ISL_703507 | Lighthouse Lab in Alderley Park | Wellcome Sanger Institute for the COVID-19 Genomics UK (COG-UK) Consortium | Jacquelyn Wynn, Mairead Hyland, The Lighthouse Lab in Alderley Park and Alex Alderton, Roberto Amato, Sonia Goncalves, Ewan Harrison, David K. Jackson, Ian Johnston, Dominic Kwiatkowski, Cordelia Langford, John Sillitoe on behalf of the Wellcome Sanger Institute COVID-19 Surveillance Team |
| EPI_ISL_703508 | Department of Pathology, University of Cambridge | COVID-19 Genomics UK (COG-UK) Consortium | Aminu S. Jahun, Yasmin Chaudhry, Grant Hall, Iliana Georgana, Myra Hosmillo, Martin D. Curran, Malte Pinckert, Surendra Parmar, Ian Goodfellow |
| EPI_ISL_703509, EPI_ISL_703518, EPI_ISL_703520, EPI_ISL_703523 | Lighthouse Lab in Alderley Park | Wellcome Sanger Institute for the COVID-19 Genomics UK (COG-UK) Consortium | Jacquelyn Wynn, Mairead Hyland, The Lighthouse Lab in Alderley Park and Alex Alderton, Roberto Amato, Sonia Goncalves, Ewan Harrison, David K. Jackson, Ian Johnston, Dominic Kwiatkowski, Cordelia Langford, John Sillitoe on behalf of the Wellcome Sanger Institute COVID-19 Surveillance Team |
| EPI_ISL_703524 | Department of Pathology, University of Cambridge | COVID-19 Genomics UK (COG-UK) Consortium | Aminu S. Jahun, Yasmin Chaudhry, Grant Hall, Iliana Georgana, Myra Hosmillo, Martin D. Curran, Malte Pinckert, Surendra Parmar, Ian Goodfellow |
| EPI_ISL_703526 | Lighthouse Lab in Alderley Park | Wellcome Sanger Institute for the COVID-19 Genomics UK (COG-UK) Consortium | Jacquelyn Wynn, Mairead Hyland, The Lighthouse Lab in Alderley Park and Alex Alderton, Roberto Amato, Sonia Goncalves, Ewan Harrison, David K. Jackson, Ian Johnston, Dominic Kwiatkowski, Cordelia Langford, John Sillitoe on behalf of the Wellcome Sanger Institute COVID-19 Surveillance Team |
| EPI_ISL_703527 | Liverpool Clinical Laboratories | COVID-19 Genomics UK (COG-UK) Consortium | Sam Haldenby, Anita Lucaci, Steve Paterson, Julian Hiscox, Alistair Darby, M Almsaud, A Alrezaihi, Muhannad Alruwaili, Stuart D Armstrong, Jones Benjamin, Eleanor G Bentley, Anu Chawla, Jordan J Clark, Angela Cowell, Richard Eccles, Isabel García-Dorival, Matthew Gemmell, Alessandro Gerada, PKF Gilmore, Richard Gregory, Ximeng Han, Catherine Hartley, Margaret Hughes, Miren Iturriza-Gomara, James Johnson, L Luu, Jenifer Manson, Charlotte Nelson, Elaine O'Toole, Cassie Olateju, Rebekah Penrice-Randal , Lucille Rainbow, N.P Randle, Trevor Ian Robinson, Parul Sharma, Ghada T Shawli, James P Stewart, Neil Swainston, Ecaterina Vamos, Joanne Watts, Mark Whitehead |
| EPI_ISL_703528, EPI_ISL_703531, EPI_ISL_703532, EPI_ISL_703536, EPI_ISL_703539, EPI_ISL_703543, EPI_ISL_703544, EPI_ISL_703547, EPI_ISL_703549 | Lighthouse Lab in Alderley Park | Wellcome Sanger Institute for the COVID-19 Genomics UK (COG-UK) Consortium | Jacquelyn Wynn, Mairead Hyland, The Lighthouse Lab in Alderley Park and Alex Alderton, Roberto Amato, Sonia Goncalves, Ewan Harrison, David K. Jackson, Ian Johnston, Dominic Kwiatkowski, Cordelia Langford, John Sillitoe on behalf of the Wellcome Sanger Institute COVID-19 Surveillance Team |
| EPI_ISL_703550 | Lighthouse Lab in Glasgow | Wellcome Sanger Institute for the COVID-19 Genomics UK (COG-UK) Consortium | Harper VanSteenhouse, Yumi Kasai, David Gray, Carol Clugston, Anna Dominiczak and Alex Alderton, Roberto Amato, Sonia Goncalves, Ewan Harrison, David K. Jackson, Ian Johnston, Dominic Kwiatkowski, Cordelia Langford, John Sillitoe on behalf of the Wellcome Sanger Institute COVID-19 Surveillance Team |
| EPI_ISL_703552, EPI_ISL_703555, EPI_ISL_703558 | Lighthouse Lab in Alderley Park | Wellcome Sanger Institute for the COVID-19 Genomics UK (COG-UK) Consortium | Jacquelyn Wynn, Mairead Hyland, The Lighthouse Lab in Alderley Park and Alex Alderton, Roberto Amato, Sonia Goncalves, Ewan Harrison, David K. Jackson, Ian Johnston, Dominic Kwiatkowski, Cordelia Langford, John Sillitoe on behalf of the Wellcome Sanger Institute COVID-19 Surveillance Team |
| EPI_ISL_703560 | Lighthouse Lab in Glasgow | Wellcome Sanger Institute for the COVID-19 Genomics UK (COG-UK) Consortium | Harper VanSteenhouse, Yumi Kasai, David Gray, Carol Clugston, Anna Dominiczak and Alex Alderton, Roberto Amato, Sonia Goncalves, Ewan Harrison, David K. Jackson, Ian Johnston, Dominic Kwiatkowski, Cordelia Langford, John Sillitoe on behalf of the Wellcome Sanger Institute COVID-19 Surveillance Team |
| EPI_ISL_703563, EPI_ISL_703565, EPI_ISL_703566, EPI_ISL_703568 | Lighthouse Lab in Alderley Park | Wellcome Sanger Institute for the COVID-19 Genomics UK (COG-UK) Consortium | Jacquelyn Wynn, Mairead Hyland, The Lighthouse Lab in Alderley Park and Alex Alderton, Roberto Amato, Sonia Goncalves, Ewan Harrison, David K. Jackson, Ian Johnston, Dominic Kwiatkowski, Cordelia Langford, John Sillitoe on behalf of the Wellcome Sanger Institute COVID-19 Surveillance Team |
| EPI_ISL_703569 | Virology Department, Sheffield Teaching Hospitals NHS Foundation Trust/Department of Infection, Immunity and Cardiovascular Disease, The Medical School, University of Sheffield | COVID-19 Genomics UK (COG-UK) Consortium | Thushan de Silva, Matthew Parker, Nikki Smith, Adri Angyal, Rebecca Brown, Luke Green, Rachel Tucker, Paul Parsons, Danielle Groves, Katie Johnson, Laura Carrilero, Alex Keeley, Dave Partridge, Matthew Wyles, Benjamin Lindsey, Mehmet Yavuz, Mohammad Raza, Cariad Evans |

| | | | |
|---|---|---|---|
| EPI_ISL_703570, EPI_ISL_703571, EPI_ISL_703573, EPI_ISL_703574 | Lighthouse Lab in Alderley Park | Wellcome Sanger Institute for the COVID-19 Genomics UK (COG-UK) Consortium | Jacquelyn Wynn, Mairead Hyland, The Lighthouse Lab in Alderley Park and Alex Alderton, Roberto Amato, Sonia Goncalves, Ewan Harrison, David K. Jackson, Ian Johnston, Dominic Kwiatkowski, Cordelia Langford, John Sillitoe on behalf of the Wellcome Sanger Institute COVID-19 Surveillance Team |
| EPI_ISL_703575 | Virology Department, Sheffield Teaching Hospitals NHS Foundation Trust/Department of Infection, Immunity and Cardiovascular Disease, The Medical School, University of Sheffield | COVID-19 Genomics UK (COG-UK) Consortium | Thushan de Silva, Matthew Parker, Nikki Smith, Adri Angyal, Rebecca Brown, Luke Green, Rachel Tucker, Paul Parsons, Danielle Groves, Katie Johnson, Laura Carrilero, Alex Keeley, Dave Partridge, Matthew Wyles, Benjamin Lindsey, Mehmet Yavuz, Mohammad Raza, Cariad Evans |
| EPI_ISL_703576, EPI_ISL_703580, EPI_ISL_703581 | Lighthouse Lab in Alderley Park | Wellcome Sanger Institute for the COVID-19 Genomics UK (COG-UK) Consortium | Jacquelyn Wynn, Mairead Hyland, The Lighthouse Lab in Alderley Park and Alex Alderton, Roberto Amato, Sonia Goncalves, Ewan Harrison, David K. Jackson, Ian Johnston, Dominic Kwiatkowski, Cordelia Langford, John Sillitoe on behalf of the Wellcome Sanger Institute COVID-19 Surveillance Team |
| EPI_ISL_703582 | Virology Department, Sheffield Teaching Hospitals NHS Foundation Trust/Department of Infection, Immunity and Cardiovascular Disease, The Medical School, University of Sheffield | COVID-19 Genomics UK (COG-UK) Consortium | Thushan de Silva, Matthew Parker, Nikki Smith, Adri Angyal, Rebecca Brown, Luke Green, Rachel Tucker, Paul Parsons, Danielle Groves, Katie Johnson, Laura Carrilero, Alex Keeley, Dave Partridge, Matthew Wyles, Benjamin Lindsey, Mehmet Yavuz, Mohammad Raza, Cariad Evans |
| EPI_ISL_703584 | Lighthouse Lab in Glasgow | Wellcome Sanger Institute for the COVID-19 Genomics UK (COG-UK) Consortium | Harper VanSteenhouse, Yumi Kasai, David Gray, Carol Clugston, Anna Dominiczak and Alex Alderton, Roberto Amato, Sonia Goncalves, Ewan Harrison, David K. Jackson, Ian Johnston, Dominic Kwiatkowski, Cordelia Langford, John Sillitoe on behalf of the Wellcome Sanger Institute COVID-19 Surveillance Team |
| EPI_ISL_703585 | Virology Department, Sheffield Teaching Hospitals NHS Foundation Trust/Department of Infection, Immunity and Cardiovascular Disease, The Medical School, University of Sheffield | COVID-19 Genomics UK (COG-UK) Consortium | Thushan de Silva, Matthew Parker, Nikki Smith, Adri Angyal, Rebecca Brown, Luke Green, Rachel Tucker, Paul Parsons, Danielle Groves, Katie Johnson, Laura Carrilero, Alex Keeley, Dave Partridge, Matthew Wyles, Benjamin Lindsey, Mehmet Yavuz, Mohammad Raza, Cariad Evans |
| EPI_ISL_703586 | Lighthouse Lab in Alderley Park | Wellcome Sanger Institute for the COVID-19 Genomics UK (COG-UK) Consortium | Jacquelyn Wynn, Mairead Hyland, The Lighthouse Lab in Alderley Park and Alex Alderton, Roberto Amato, Sonia Goncalves, Ewan Harrison, David K. Jackson, Ian Johnston, Dominic Kwiatkowski, Cordelia Langford, John Sillitoe on behalf of the Wellcome Sanger Institute COVID-19 Surveillance Team |
| EPI_ISL_703587 | Virology Department, Sheffield Teaching Hospitals NHS Foundation Trust/Department of Infection, Immunity and Cardiovascular Disease, The Medical School, University of Sheffield | COVID-19 Genomics UK (COG-UK) Consortium | Thushan de Silva, Matthew Parker, Nikki Smith, Adri Angyal, Rebecca Brown, Luke Green, Rachel Tucker, Paul Parsons, Danielle Groves, Katie Johnson, Laura Carrilero, Alex Keeley, Dave Partridge, Matthew Wyles, Benjamin Lindsey, Mehmet Yavuz, Mohammad Raza, Cariad Evans |
| EPI_ISL_703588, EPI_ISL_703589 | Lighthouse Lab in Alderley Park | Wellcome Sanger Institute for the COVID-19 Genomics UK (COG-UK) Consortium | Jacquelyn Wynn, Mairead Hyland, The Lighthouse Lab in Alderley Park and Alex Alderton, Roberto Amato, Sonia Goncalves, Ewan Harrison, David K. Jackson, Ian Johnston, Dominic Kwiatkowski, Cordelia Langford, John Sillitoe on behalf of the Wellcome Sanger Institute COVID-19 Surveillance Team |
| EPI_ISL_703590 | Quadram Institute Bioscience | COVID-19 Genomics UK (COG-UK) Consortium | Dave J. Baker, Gemma L. Kay, Alp Aydin, Thanh Le-Viet, Steven Rudder, Ana P. Tedim, Anastasia Kolyva, Maria Diaz, Leonardo de Oliveira Martins, Nabil-Fareed Alikhan, Lizzie Meadows, Rachael Stanley, Ngozi Elumogo, Muhammed Yasir, Nicholas M. Thomson, Alexander J Trotter, Rachel Gilroy, Samuel Bloomfield, Claire Stuart, Andrew Bell, Reenesh Prakash, Samir Dervisevic, Alison E. Mather, John Wain, Mark Webber, Andrew J. Page, Justin O'Grady |
| EPI_ISL_703591, EPI_ISL_703592 | Lighthouse Lab in Alderley Park | Wellcome Sanger Institute for the COVID-19 Genomics UK (COG-UK) Consortium | Jacquelyn Wynn, Mairead Hyland, The Lighthouse Lab in Alderley Park and Alex Alderton, Roberto Amato, Sonia Goncalves, Ewan Harrison, David K. Jackson, Ian Johnston, Dominic Kwiatkowski, Cordelia Langford, John Sillitoe on behalf of the Wellcome Sanger Institute COVID-19 Surveillance Team |
| EPI_ISL_703593 | Virology Department, Sheffield Teaching Hospitals NHS Foundation Trust/Department of Infection, Immunity and Cardiovascular Disease, The Medical School, University of Sheffield | COVID-19 Genomics UK (COG-UK) Consortium | Thushan de Silva, Matthew Parker, Nikki Smith, Adri Angyal, Rebecca Brown, Luke Green, Rachel Tucker, Paul Parsons, Danielle Groves, Katie Johnson, Laura Carrilero, Alex Keeley, Dave Partridge, Matthew Wyles, Benjamin Lindsey, Mehmet Yavuz, Mohammad Raza, Cariad Evans |
| EPI_ISL_703595, EPI_ISL_703597, EPI_ISL_703602, EPI_ISL_703603, EPI_ISL_703605, EPI_ISL_703613, EPI_ISL_703615, EPI_ISL_703618 | Lighthouse Lab in Alderley Park | Wellcome Sanger Institute for the COVID-19 Genomics UK (COG-UK) Consortium | Jacquelyn Wynn, Mairead Hyland, The Lighthouse Lab in Alderley Park and Alex Alderton, Roberto Amato, Sonia Goncalves, Ewan Harrison, David K. Jackson, Ian Johnston, Dominic Kwiatkowski, Cordelia Langford, John Sillitoe on behalf of the Wellcome Sanger Institute COVID-19 Surveillance Team |
| EPI_ISL_703619 | Department of Pathology, University of Cambridge | COVID-19 Genomics UK (COG-UK) Consortium | Aminu S. Jahun, Yasmin Chaudhry, Grant Hall, Iliana Georgana, Myra Hosmillo, Martin D. Curran, Malte Pinckert, Surendra Parmar, Ian Goodfellow |
| EPI_ISL_703620, EPI_ISL_703623, EPI_ISL_703625, EPI_ISL_703628 | Lighthouse Lab in Alderley Park | Wellcome Sanger Institute for the COVID-19 Genomics UK (COG-UK) Consortium | Jacquelyn Wynn, Mairead Hyland, The Lighthouse Lab in Alderley Park and Alex Alderton, Roberto Amato, Sonia Goncalves, Ewan Harrison, David K. Jackson, Ian Johnston, Dominic Kwiatkowski, Cordelia Langford, John Sillitoe on behalf of the Wellcome Sanger Institute COVID-19 Surveillance Team |
| EPI_ISL_703629 | Department of Pathology, University of Cambridge | COVID-19 Genomics UK (COG-UK) Consortium | Aminu S. Jahun, Yasmin Chaudhry, Grant Hall, Iliana Georgana, Myra Hosmillo, Martin D. Curran, Malte Pinckert, Surendra Parmar, Ian Goodfellow |
| EPI_ISL_703630 | Lighthouse Lab in Alderley Park | Wellcome Sanger Institute for the COVID-19 Genomics UK (COG-UK) Consortium | Jacquelyn Wynn, Mairead Hyland, The Lighthouse Lab in Alderley Park and Alex Alderton, Roberto Amato, Sonia Goncalves, Ewan Harrison, David K. Jackson, Ian Johnston, Dominic Kwiatkowski, Cordelia Langford, John Sillitoe on behalf of the Wellcome Sanger Institute COVID-19 Surveillance Team |
| EPI_ISL_703632, EPI_ISL_703635 | Department of Pathology, University of Cambridge | COVID-19 Genomics UK (COG-UK) Consortium | Aminu S. Jahun, Yasmin Chaudhry, Grant Hall, Iliana Georgana, Myra Hosmillo, Martin D. Curran, Malte Pinckert, Surendra Parmar, Ian Goodfellow |
| EPI_ISL_703636 | Virology Department, Sheffield Teaching Hospitals NHS Foundation Trust/Department of Infection, Immunity and Cardiovascular Disease, The Medical School, University of Sheffield | COVID-19 Genomics UK (COG-UK) Consortium | Thushan de Silva, Matthew Parker, Nikki Smith, Adri Angyal, Rebecca Brown, Luke Green, Rachel Tucker, Paul Parsons, Danielle Groves, Katie Johnson, Laura Carrilero, Alex Keeley, Dave Partridge, Matthew Wyles, Benjamin Lindsey, Mehmet Yavuz, Mohammad Raza, Cariad Evans |
| EPI_ISL_703637, EPI_ISL_703638 | Lighthouse Lab in Alderley Park | Wellcome Sanger Institute for the COVID-19 Genomics UK (COG-UK) Consortium | Jacquelyn Wynn, Mairead Hyland, The Lighthouse Lab in Alderley Park and Alex Alderton, Roberto Amato, Sonia Goncalves, Ewan Harrison, David K. Jackson, Ian Johnston, Dominic Kwiatkowski, Cordelia Langford, John Sillitoe on behalf of the Wellcome Sanger Institute COVID-19 Surveillance Team |
| EPI_ISL_703639 | Virology Department, Sheffield Teaching Hospitals NHS Foundation Trust/Department of Infection, Immunity and Cardiovascular Disease, The Medical School, University of Sheffield | COVID-19 Genomics UK (COG-UK) Consortium | Thushan de Silva, Matthew Parker, Nikki Smith, Adri Angyal, Rebecca Brown, Luke Green, Rachel Tucker, Paul Parsons, Danielle Groves, Katie Johnson, Laura Carrilero, Alex Keeley, Dave Partridge, Matthew Wyles, Benjamin Lindsey, Mehmet Yavuz, Mohammad Raza, Cariad Evans |
| EPI_ISL_703646, EPI_ISL_703648 | Lighthouse Lab in Alderley Park | Wellcome Sanger Institute for the COVID-19 Genomics UK (COG-UK) Consortium | Jacquelyn Wynn, Mairead Hyland, The Lighthouse Lab in Alderley Park and Alex Alderton, Roberto Amato, Sonia Goncalves, Ewan Harrison, David K. Jackson, Ian Johnston, Dominic Kwiatkowski, Cordelia Langford, John Sillitoe on behalf of the Wellcome Sanger Institute COVID-19 Surveillance Team |
| EPI_ISL_703650, EPI_ISL_703659 | Lighthouse Lab in Glasgow | Wellcome Sanger Institute for the COVID-19 Genomics UK (COG-UK) Consortium | Harper VanSteenhouse, Yumi Kasai, David Gray, Carol Clugston, Anna Dominiczak and Alex Alderton, Roberto Amato, Sonia Goncalves, Ewan Harrison, David K. Jackson, Ian Johnston, Dominic Kwiatkowski, Cordelia Langford, John Sillitoe on behalf of the Wellcome Sanger Institute COVID-19 Surveillance Team |
| EPI_ISL_703662, EPI_ISL_703664, EPI_ISL_703665, EPI_ISL_703667, EPI_ISL_703670, EPI_ISL_703675, EPI_ISL_703678 | Lighthouse Lab in Alderley Park | Wellcome Sanger Institute for the COVID-19 Genomics UK (COG-UK) Consortium | Jacquelyn Wynn, Mairead Hyland, The Lighthouse Lab in Alderley Park and Alex Alderton, Roberto Amato, Sonia Goncalves, Ewan Harrison, David K. Jackson, Ian Johnston, Dominic Kwiatkowski, Cordelia Langford, John Sillitoe on behalf of the Wellcome Sanger Institute COVID-19 Surveillance Team |
| EPI_ISL_703679 | Department of Pathology, University of Cambridge | COVID-19 Genomics UK (COG-UK) Consortium | Aminu S. Jahun, Yasmin Chaudhry, Grant Hall, Iliana Georgana, Myra Hosmillo, Martin D. Curran, Malte Pinckert, Surendra Parmar, Ian Goodfellow |
| EPI_ISL_703680, EPI_ISL_703681, EPI_ISL_703683, EPI_ISL_703686, EPI_ISL_703688, EPI_ISL_703689, EPI_ISL_703691, EPI_ISL_703692 | Lighthouse Lab in Alderley Park | Wellcome Sanger Institute for the COVID-19 Genomics UK (COG-UK) Consortium | Jacquelyn Wynn, Mairead Hyland, The Lighthouse Lab in Alderley Park and Alex Alderton, Roberto Amato, Sonia Goncalves, Ewan Harrison, David K. Jackson, Ian Johnston, Dominic Kwiatkowski, Cordelia Langford, John Sillitoe on behalf of the Wellcome Sanger Institute COVID-19 Surveillance Team |
| EPI_ISL_703693 | Department of Pathology, University of Cambridge | COVID-19 Genomics UK (COG-UK) Consortium | Aminu S. Jahun, Yasmin Chaudhry, Grant Hall, Iliana Georgana, Myra Hosmillo, Martin D. Curran, Malte Pinckert, Surendra Parmar, Ian Goodfellow |
| EPI_ISL_703696 | Lighthouse Lab in Alderley Park | Wellcome Sanger Institute for the COVID-19 Genomics UK | Jacquelyn Wynn, Mairead Hyland, The Lighthouse Lab in Alderley Park and Alex Alderton, Roberto Amato, Sonia Goncalves, Ewan Harrison, David K. |

| | | | |
|---|---|---|---|
| | | (COG-UK) Consortium | Jackson, Ian Johnston, Dominic Kwiatkowski, Cordelia Langford, John Sillitoe on behalf of the Wellcome Sanger Institute COVID-19 Surveillance Team |
| EPI_ISL_703697 | Lighthouse Lab in Glasgow | Wellcome Sanger Institute for the COVID-19 Genomics UK (COG-UK) Consortium | Harper VanSteenhouse, Yumi Kasai, David Gray, Carol Clugston, Anna Dominiczak and Alex Alderton, Roberto Amato, Sonia Goncalves, Ewan Harrison, David K. Jackson, Ian Johnston, Dominic Kwiatkowski, Cordelia Langford, John Sillitoe on behalf of the Wellcome Sanger Institute COVID-19 Surveillance Team |
| EPI_ISL_703699, EPI_ISL_703702, EPI_ISL_703705, EPI_ISL_703707 | Lighthouse Lab in Alderley Park | Wellcome Sanger Institute for the COVID-19 Genomics UK (COG-UK) Consortium | Jacquelyn Wynn, Mairead Hyland, The Lighthouse Lab in Alderley Park and Alex Alderton, Roberto Amato, Sonia Goncalves, Ewan Harrison, David K. Jackson, Ian Johnston, Dominic Kwiatkowski, Cordelia Langford, John Sillitoe on behalf of the Wellcome Sanger Institute COVID-19 Surveillance Team |
| EPI_ISL_703708, EPI_ISL_703709, EPI_ISL_703710, EPI_ISL_703713, EPI_ISL_703716 | Department of Pathology, University of Cambridge | COVID-19 Genomics UK (COG-UK) Consortium | Aminu S. Jahun, Yasmin Chaudhry, Grant Hall, Iliana Georgana, Myra Hosmillo, Martin D. Curran, Malte Pinckert, Surendra Parmar, Ian Goodfellow |
| EPI_ISL_703723 | Virology Department, Sheffield Teaching Hospitals NHS Foundation Trust/Department of Infection, Immunity and Cardiovascular Disease, The Medical School, University of Sheffield | COVID-19 Genomics UK (COG-UK) Consortium | Thushan de Silva, Matthew Parker, Nikki Smith, Adri Angyal, Rebecca Brown, Luke Green, Rachel Tucker, Paul Parsons, Danielle Groves, Katie Johnson, Laura Carrilero, Alex Keeley, Dave Partridge, Matthew Wyles, Benjamin Lindsey, Mehmet Yavuz, Mohammad Raza, Cariad Evans |
| EPI_ISL_703724 | Department of Pathology, University of Cambridge | COVID-19 Genomics UK (COG-UK) Consortium | Aminu S. Jahun, Yasmin Chaudhry, Grant Hall, Iliana Georgana, Myra Hosmillo, Martin D. Curran, Malte Pinckert, Surendra Parmar, Ian Goodfellow |
| EPI_ISL_703738 | Liverpool Clinical Laboratories | COVID-19 Genomics UK (COG-UK) Consortium | Sam Haldenby, Anita Lucaci, Steve Paterson, Julian Hiscox, Alistair Darby, M Almsaud, A Alrezaihi, Muhannad Alruwaili, Stuart D Armstrong, Jones Benjamin, Eleanor G Bentley, Anu Chawla, Jordan J Clark, Angela Cowell, Richard Eccles, Isabel García-Dorival, Matthew Gemmell, Alessandro Gerada, PKF Gilmore, Richard Gregory, Ximeng Han, Catherine Hartley, Margaret Hughes, Miren Iturriza-Gomara, James Johnson, L Luu, Jenifer Manson, Charlotte Nelson, Elaine O'Toole, Cassie Olateju, Rebekah Penrice-Randal , Lucille Rainbow, N.P Randle, Trevor Ian Robinson, Parul Sharma, Ghada T Shawli, James P Stewart, Neil Swainston, Ecaterina Vamos, Joanne Watts, Mark Whitehead |
| EPI_ISL_703741, EPI_ISL_703742 | Department of Pathology, University of Cambridge | COVID-19 Genomics UK (COG-UK) Consortium | Aminu S. Jahun, Yasmin Chaudhry, Grant Hall, Iliana Georgana, Myra Hosmillo, Martin D. Curran, Malte Pinckert, Surendra Parmar, Ian Goodfellow |
| EPI_ISL_703743 | West of Scotland Specialist Virology Centre, NHSGGC / MRC-University of Glasgow Centre for Virus Research | COVID-19 Genomics UK (COG-UK) Consortium | Ana da Silva Filipe, Natasha Johnson, Kathy Smollett, Daniel Mair, Stephen Carmichael, Alice Broos, Lily Tong, Jenna Nichols, Kyriaki Nomikou; Sarah McDonald; Richard Orton, Joseph Hughes, Sreenu Vattipally, David L Robertson; Alasdair MacLean, Rory Gunson; Sharif Shaaban, Matthew Holden; Rachel Blacow, Guy Mollett, Kathy Li, James Shepherd, Antonia Ho, Emma Thomson |
| EPI_ISL_703747 | Department of Pathology, University of Cambridge | COVID-19 Genomics UK (COG-UK) Consortium | Aminu S. Jahun, Yasmin Chaudhry, Grant Hall, Iliana Georgana, Myra Hosmillo, Martin D. Curran, Malte Pinckert, Surendra Parmar, Ian Goodfellow |
| EPI_ISL_703750, EPI_ISL_703751, EPI_ISL_703752 | West of Scotland Specialist Virology Centre, NHSGGC / MRC-University of Glasgow Centre for Virus Research | COVID-19 Genomics UK (COG-UK) Consortium | Ana da Silva Filipe, Natasha Johnson, Kathy Smollett, Daniel Mair, Stephen Carmichael, Alice Broos, Lily Tong, Jenna Nichols, Kyriaki Nomikou; Sarah McDonald; Richard Orton, Joseph Hughes, Sreenu Vattipally, David L Robertson; Alasdair MacLean, Rory Gunson; Sharif Shaaban, Matthew Holden; Rachel Blacow, Guy Mollett, Kathy Li, James Shepherd, Antonia Ho, Emma Thomson |
| EPI_ISL_703759, EPI_ISL_703762 | Liverpool Clinical Laboratories | COVID-19 Genomics UK (COG-UK) Consortium | Sam Haldenby, Anita Lucaci, Steve Paterson, Julian Hiscox, Alistair Darby, M Almsaud, A Alrezaihi, Muhannad Alruwaili, Stuart D Armstrong, Jones Benjamin, Eleanor G Bentley, Anu Chawla, Jordan J Clark, Angela Cowell, Richard Eccles, Isabel García-Dorival, Matthew Gemmell, Alessandro Gerada, PKF Gilmore, Richard Gregory, Ximeng Han, Catherine Hartley, Margaret Hughes, Miren Iturriza-Gomara, James Johnson, L Luu, Jenifer Manson, Charlotte Nelson, Elaine O'Toole, Cassie Olateju, Rebekah Penrice-Randal , Lucille Rainbow, N.P Randle, Trevor Ian Robinson, Parul Sharma, Ghada T Shawli, James P Stewart, Neil Swainston, Ecaterina Vamos, Joanne Watts, Mark Whitehead |
| EPI_ISL_703769 | Quadram Institute Bioscience | COVID-19 Genomics UK (COG-UK) Consortium | Dave J. Baker, Gemma L. Kay, Alp Aydin, Thanh Le-Viet, Steven Rudder, Ana P. Tedim, Anastasia Kolyva, Maria Diaz, Leonardo de Oliveira Martins, Nabil-Fareed Alikhan, Lizzie Meadows, Rachael Stanley, Ngozi Elumogo, Muhammed Yasir, Nicholas M. Thomson, Alexander J Trotter, Rachel Gilroy, Samuel Bloomfield, Claire Stuart, Andrew Bell, Reenesh Prakash, Samir Dervisevic, Alison E. Mather, John Wain, Mark Webber, Andrew J. Page, Justin O'Grady |
| EPI_ISL_703773 | West of Scotland Specialist Virology Centre, NHSGGC / MRC-University of Glasgow Centre for Virus Research | COVID-19 Genomics UK (COG-UK) Consortium | Ana da Silva Filipe, Natasha Johnson, Kathy Smollett, Daniel Mair, Stephen Carmichael, Alice Broos, Lily Tong, Jenna Nichols, Kyriaki Nomikou; Sarah McDonald; Richard Orton, Joseph Hughes, Sreenu Vattipally, David L Robertson; Alasdair MacLean, Rory Gunson; Sharif Shaaban, Matthew Holden; Rachel Blacow, Guy Mollett, Kathy Li, James Shepherd, Antonia Ho, Emma Thomson |
| EPI_ISL_703777, EPI_ISL_703778, EPI_ISL_703780, EPI_ISL_703781, EPI_ISL_703783, EPI_ISL_703785, EPI_ISL_703788, EPI_ISL_703791, EPI_ISL_703793, EPI_ISL_703795, EPI_ISL_703796, EPI_ISL_703798, EPI_ISL_703803, EPI_ISL_703804, EPI_ISL_703809, EPI_ISL_703812 | | | |
| see above | Lighthouse Lab in Alderley Park | Wellcome Sanger Institute for the COVID-19 Genomics UK (COG-UK) Consortium | Jacquelyn Wynn, Mairead Hyland, The Lighthouse Lab in Alderley Park and Alex Alderton, Roberto Amato, Sonia Goncalves, Ewan Harrison, David K. Jackson, Ian Johnston, Dominic Kwiatkowski, Cordelia Langford, John Sillitoe on behalf of the Wellcome Sanger Institute COVID-19 Surveillance Team |
| EPI_ISL_703813 | Virology Department, Royal Infirmary of Edinburgh, NHS Lothian / School of Biological Sciences, University of Edinburgh / Institute of Genetics and Molecular Medicine, University of Edinburgh | COVID-19 Genomics UK (COG-UK) Consortium | McHugh M, Dewar R, Rooke S, Gallagher M, Balcaza C, O'Toole Á, Scher E, Hill V, McCrone JT, Colquhoun R, Yu X, Jackson B, Rambaut A, Williams TC, Templeton K |
| EPI_ISL_703814, EPI_ISL_703817 | Lighthouse Lab in Alderley Park | Wellcome Sanger Institute for the COVID-19 Genomics UK (COG-UK) Consortium | Jacquelyn Wynn, Mairead Hyland, The Lighthouse Lab in Alderley Park and Alex Alderton, Roberto Amato, Sonia Goncalves, Ewan Harrison, David K. Jackson, Ian Johnston, Dominic Kwiatkowski, Cordelia Langford, John Sillitoe on behalf of the Wellcome Sanger Institute COVID-19 Surveillance Team |
| EPI_ISL_703821, EPI_ISL_703823 | Liverpool Clinical Laboratories | COVID-19 Genomics UK (COG-UK) Consortium | Sam Haldenby, Anita Lucaci, Steve Paterson, Julian Hiscox, Alistair Darby, M Almsaud, A Alrezaihi, Muhannad Alruwaili, Stuart D Armstrong, Jones Benjamin, Eleanor G Bentley, Anu Chawla, Jordan J Clark, Angela Cowell, Richard Eccles, Isabel García-Dorival, Matthew Gemmell, Alessandro Gerada, PKF Gilmore, Richard Gregory, Ximeng Han, Catherine Hartley, Margaret Hughes, Miren Iturriza-Gomara, James Johnson, L Luu, Jenifer Manson, Charlotte Nelson, Elaine O'Toole, Cassie Olateju, Rebekah Penrice-Randal , Lucille Rainbow, N.P Randle, Trevor Ian Robinson, Parul Sharma, Ghada T Shawli, James P Stewart, Neil Swainston, Ecaterina Vamos, Joanne Watts, Mark Whitehead |
| EPI_ISL_703842 | West of Scotland Specialist Virology Centre, NHSGGC / MRC-University of Glasgow Centre for Virus Research | COVID-19 Genomics UK (COG-UK) Consortium | Ana da Silva Filipe, Natasha Johnson, Kathy Smollett, Daniel Mair, Stephen Carmichael, Alice Broos, Lily Tong, Jenna Nichols, Kyriaki Nomikou; Sarah McDonald; Richard Orton, Joseph Hughes, Sreenu Vattipally, David L Robertson; Alasdair MacLean, Rory Gunson; Sharif Shaaban, Matthew Holden; Rachel Blacow, Guy Mollett, Kathy Li, James Shepherd, Antonia Ho, Emma Thomson |
| EPI_ISL_703876, EPI_ISL_703913, EPI_ISL_704006 | Liverpool Clinical Laboratories | COVID-19 Genomics UK (COG-UK) Consortium | Sam Haldenby, Anita Lucaci, Steve Paterson, Julian Hiscox, Alistair Darby, M Almsaud, A Alrezaihi, Muhannad Alruwaili, Stuart D Armstrong, Jones Benjamin, Eleanor G Bentley, Anu Chawla, Jordan J Clark, Angela Cowell, Richard Eccles, Isabel García-Dorival, Matthew Gemmell, Alessandro Gerada, PKF Gilmore, Richard Gregory, Ximeng Han, Catherine Hartley, Margaret Hughes, Miren Iturriza-Gomara, James Johnson, L Luu, Jenifer Manson, Charlotte Nelson, Elaine O'Toole, Cassie Olateju, Rebekah Penrice-Randal , Lucille Rainbow, N.P Randle, Trevor Ian Robinson, Parul Sharma, Ghada T Shawli, James P Stewart, Neil Swainston, Ecaterina Vamos, Joanne Watts, Mark Whitehead |
| EPI_ISL_704060 | Department of Pathology, University of Cambridge | COVID-19 Genomics UK (COG-UK) Consortium | Aminu S. Jahun, Yasmin Chaudhry, Grant Hall, Iliana Georgana, Myra Hosmillo, Martin D. Curran, Malte Pinckert, Surendra Parmar, Ian Goodfellow |
| EPI_ISL_704100 | Quadram Institute Bioscience | COVID-19 Genomics UK (COG-UK) Consortium | Dave J. Baker, Gemma L. Kay, Alp Aydin, Thanh Le-Viet, Steven Rudder, Ana P. Tedim, Anastasia Kolyva, Maria Diaz, Leonardo de Oliveira Martins, Nabil-Fareed Alikhan, Lizzie Meadows, Rachael Stanley, Ngozi Elumogo, Muhammed Yasir, Nicholas M. Thomson, Alexander J Trotter, Rachel Gilroy, Samuel Bloomfield, Claire Stuart, Andrew Bell, Reenesh Prakash, Samir Dervisevic, Alison E. Mather, John Wain, Mark Webber, Andrew J. Page, Justin O'Grady |
| EPI_ISL_704102 | University College London Hospital | COVID-19 Genomics UK (COG-UK) Consortium | Judith Heaney, Matthew Byott, Catherine Houlihan, Dan Frampton, Stuart Kirk, Moira Spyer and Eleni Nastouli |
| EPI_ISL_704115 | Department of Pathology, University of Cambridge | COVID-19 Genomics UK (COG-UK) Consortium | Aminu S. Jahun, Yasmin Chaudhry, Grant Hall, Iliana Georgana, Myra Hosmillo, Martin D. Curran, Malte Pinckert, Surendra Parmar, Ian Goodfellow |
| EPI_ISL_704131, EPI_ISL_704137, EPI_ISL_704153, EPI_ISL_704156 | Liverpool Clinical Laboratories | COVID-19 Genomics UK (COG-UK) Consortium | Sam Haldenby, Anita Lucaci, Steve Paterson, Julian Hiscox, Alistair Darby, M Almsaud, A Alrezaihi, Muhannad Alruwaili, Stuart D Armstrong, Jones Benjamin, Eleanor G Bentley, Anu Chawla, Jordan J Clark, Angela Cowell, Richard Eccles, Isabel García-Dorival, Matthew Gemmell, Alessandro Gerada, PKF Gilmore, Richard Gregory, Ximeng Han, Catherine Hartley, Margaret Hughes, Miren Iturriza-Gomara, James Johnson, L Luu, Jenifer Manson, Charlotte Nelson, Elaine O'Toole, Cassie Olateju, Rebekah Penrice-Randal , Lucille Rainbow, N.P Randle, Trevor Ian Robinson, Parul Sharma, Ghada T Shawli, James P Stewart, Neil Swainston, Ecaterina Vamos, Joanne Watts, Mark Whitehead |
| EPI_ISL_704190 | Virology Department, Sheffield Teaching Hospitals NHS | COVID-19 Genomics UK (COG-UK) Consortium | Thushan de Silva, Matthew Parker, Nikki Smith, Adri Angyal, Rebecca Brown, Luke Green, Rachel Tucker, Paul Parsons, Danielle Groves, Katie Johnson, |

| | | | |
|---|---|---|---|
| | Foundation Trust/Department of Infection, Immunity and Cardiovascular Disease, The Medical School, University of Sheffield | | Laura Carrilero, Alex Keeley, Dave Partridge, Matthew Wyles, Benjamin Lindsey, Mehmet Yavuz, Mohammad Raza, Cariad Evans |
| EPI_ISL_704198 | Northumbria University / South Tees Hospitals NHS Foundation Trust / North Cumbria Integrated Care NHS Foundation Trust / North Tees and Hartlepool NHS Foundation Trust / Newcastle Hospitals NHS Foundation Trust | COVID-19 Genomics UK (COG-UK) Consortium | Darren L Smith,Andrew Nelson,Matthew Bashton,Greg R Young,Joshua Loh,John Allan,Mohammad A Tariq,Giles S Holt,Gary Black,Wen C Yew,Lynn Dover,Paul Baker,Steve Liggett,Sarah Essex,Jane Greenaway,Debra Padgett,Clive Graham,Garren Scott,Edward Barton,Emma Swindells,Brendan Payne,Jennifer Collins,Yusri Taha,Gary Eltringham |
| EPI_ISL_704203 | Department of Pathology, University of Cambridge | COVID-19 Genomics UK (COG-UK) Consortium | Aminu S. Jahun, Yasmin Chaudhry, Grant Hall, Iliana Georgana, Myra Hosmillo, Martin D. Curran, Malte Pinckert, Surendra Parmar, Ian Goodfellow |
| EPI_ISL_704228 | West of Scotland Specialist Virology Centre, NHSGGC / MRC-University of Glasgow Centre for Virus Research | COVID-19 Genomics UK (COG-UK) Consortium | Ana da Silva Filipe, Natasha Johnson, Kathy Smollett, Daniel Mair, Stephen Carmichael, Alice Broos, Lily Tong, Jenna Nichols, Kyriaki Nomikou; Sarah McDonald; Richard Orton, Joseph Hughes, Sreenu Vattipally, David L Robertson; Alasdair MacLean, Rory Gunson; Sharif Shaaban, Matthew Holden; Rachel Blacow, Guy Mollett, Kathy Li, James Shepherd, Antonia Ho, Emma Thomson |
| EPI_ISL_704247, EPI_ISL_704250, EPI_ISL_704252, EPI_ISL_704255, EPI_ISL_704257, EPI_ISL_704269, EPI_ISL_704289, EPI_ISL_704291, EPI_ISL_704294, EPI_ISL_704297, EPI_ISL_704299, EPI_ISL_704302, EPI_ISL_704305, EPI_ISL_704307, EPI_ISL_704309, EPI_ISL_704328, EPI_ISL_704331, EPI_ISL_704333, EPI_ISL_704336, EPI_ISL_704339, EPI_ISL_704341, EPI_ISL_704344, EPI_ISL_704347, EPI_ISL_704350, EPI_ISL_704353, EPI_ISL_704356, EPI_ISL_704359, EPI_ISL_704361, EPI_ISL_704364, EPI_ISL_704366, EPI_ISL_704369, EPI_ISL_704374, EPI_ISL_704377, EPI_ISL_704379, EPI_ISL_704384 | | | |
| see above | Department of Pathology, University of Cambridge | COVID-19 Genomics UK (COG-UK) Consortium | Aminu S. Jahun, Yasmin Chaudhry, Grant Hall, Iliana Georgana, Myra Hosmillo, Martin D. Curran, Malte Pinckert, Surendra Parmar, Ian Goodfellow |
| EPI_ISL_704422, EPI_ISL_704425 | Northumbria University / South Tees Hospitals NHS Foundation Trust / North Cumbria Integrated Care NHS Foundation Trust / North Tees and Hartlepool NHS Foundation Trust / Newcastle Hospitals NHS Foundation Trust | COVID-19 Genomics UK (COG-UK) Consortium | Darren L Smith,Andrew Nelson,Matthew Bashton,Greg R Young,Joshua Loh,John Allan,Mohammad A Tariq,Giles S Holt,Gary Black,Wen C Yew,Lynn Dover,Paul Baker,Steve Liggett,Sarah Essex,Jane Greenaway,Debra Padgett,Clive Graham,Garren Scott,Edward Barton,Emma Swindells,Brendan Payne,Jennifer Collins,Yusri Taha,Gary Eltringham |
| EPI_ISL_704478, EPI_ISL_704481, EPI_ISL_704484, EPI_ISL_704486, EPI_ISL_704489, EPI_ISL_704492, EPI_ISL_704494, EPI_ISL_704497, EPI_ISL_704499, EPI_ISL_704510, EPI_ISL_704513, EPI_ISL_704516, EPI_ISL_704518, EPI_ISL_704521, EPI_ISL_704524, EPI_ISL_704526, EPI_ISL_704529, EPI_ISL_704532, EPI_ISL_704535, EPI_ISL_704537, EPI_ISL_704540, EPI_ISL_704543, EPI_ISL_704545, EPI_ISL_704548, EPI_ISL_704551, EPI_ISL_704554, EPI_ISL_704557 | | | |
| see above | Department of Pathology, University of Cambridge | COVID-19 Genomics UK (COG-UK) Consortium | Aminu S. Jahun, Yasmin Chaudhry, Grant Hall, Iliana Georgana, Myra Hosmillo, Martin D. Curran, Malte Pinckert, Surendra Parmar, Ian Goodfellow |
| EPI_ISL_704562 | Liverpool Clinical Laboratories | COVID-19 Genomics UK (COG-UK) Consortium | Sam Haldenby, Anita Lucaci, Steve Paterson, Julian Hiscox, Alistair Darby, M Almsaud, A Alrezaihi, Muhannad Alruwaili, Stuart D Armstrong, Jones Benjamin, Eleanor G Bentley, Anu Chawla, Jordan J Clark, Angela Cowell, Richard Eccles, Isabel García-Dorival, Matthew Gemmell, Alessandro Gerada, PKF Gilmore, Richard Gregory, Ximeng Han, Catherine Hartley, Margaret Hughes, Miren Iturriza-Gomara, James Johnson, L Luu, Jenifer Manson, Charlotte Nelson, Elaine O'Toole, Cassie Olateju, Rebekah Penrice-Randal , Lucille Rainbow, N.P Randle, Trevor Ian Robinson, Parul Sharma, Ghada T Shawli, James P Stewart, Neil Swainston, Ecaterina Vamos, Joanne Watts, Mark Whitehead |
| EPI_ISL_704575, EPI_ISL_704576 | Northumbria University / South Tees Hospitals NHS Foundation Trust / North Cumbria Integrated Care NHS Foundation Trust / North Tees and Hartlepool NHS Foundation Trust / Newcastle Hospitals NHS Foundation Trust | COVID-19 Genomics UK (COG-UK) Consortium | Darren L Smith,Andrew Nelson,Matthew Bashton,Greg R Young,Joshua Loh,John Allan,Mohammad A Tariq,Giles S Holt,Gary Black,Wen C Yew,Lynn Dover,Paul Baker,Steve Liggett,Sarah Essex,Jane Greenaway,Debra Padgett,Clive Graham,Garren Scott,Edward Barton,Emma Swindells,Brendan Payne,Jennifer Collins,Yusri Taha,Gary Eltringham |
| EPI_ISL_704580, EPI_ISL_704598, EPI_ISL_704599 | Quadram Institute Bioscience | COVID-19 Genomics UK (COG-UK) Consortium | Dave J. Baker, Gemma L. Kay, Alp Aydin, Thanh Le-Viet, Steven Rudder, Ana P. Tedim, Anastasia Kolyva, Maria Diaz, Leonardo de Oliveira Martins, Nabil-Fareed Alikhan, Lizzie Meadows, Rachael Stanley, Ngozi Elumogo, Muhammed Yasir, Nicholas M. Thomson, Alexander J Trotter, Rachel Gilroy, Samuel Bloomfield, Claire Stuart, Andrew Bell, Reenesh Prakash, Samir Dervisevic, Alison E. Mather, John Wain, Mark Webber, Andrew J. Page, Justin O'Grady |
| EPI_ISL_704600, EPI_ISL_704601, EPI_ISL_704602, EPI_ISL_704603, EPI_ISL_704604 | Virology Department, Sheffield Teaching Hospitals NHS Foundation Trust/Department of Infection, Immunity and Cardiovascular Disease, The Medical School, University of Sheffield | COVID-19 Genomics UK (COG-UK) Consortium | Thushan de Silva, Matthew Parker, Nikki Smith, Adri Angyal, Rebecca Brown, Luke Green, Rachel Tucker, Paul Parsons, Danielle Groves, Katie Johnson, Laura Carrilero, Alex Keeley, Dave Partridge, Matthew Wyles, Benjamin Lindsey, Mehmet Yavuz, Mohammad Raza, Cariad Evans |
| EPI_ISL_704606, EPI_ISL_704617, EPI_ISL_704618 | Department of Pathology, University of Cambridge | COVID-19 Genomics UK (COG-UK) Consortium | Aminu S. Jahun, Yasmin Chaudhry, Grant Hall, Iliana Georgana, Myra Hosmillo, Martin D. Curran, Malte Pinckert, Surendra Parmar, Ian Goodfellow |
| EPI_ISL_704619, EPI_ISL_704620, EPI_ISL_704621, EPI_ISL_704622, EPI_ISL_704623, EPI_ISL_704624, EPI_ISL_704625, EPI_ISL_704626, EPI_ISL_704627, EPI_ISL_704629, EPI_ISL_704630, EPI_ISL_704631, EPI_ISL_704632, EPI_ISL_704633, EPI_ISL_704634, EPI_ISL_704635 | | | |
| see above | Quadram Institute Bioscience | COVID-19 Genomics UK (COG-UK) Consortium | Dave J. Baker, Gemma L. Kay, Alp Aydin, Thanh Le-Viet, Steven Rudder, Ana P. Tedim, Anastasia Kolyva, Maria Diaz, Leonardo de Oliveira Martins, Nabil-Fareed Alikhan, Lizzie Meadows, Rachael Stanley, Ngozi Elumogo, Muhammed Yasir, Nicholas M. Thomson, Alexander J Trotter, Rachel Gilroy, Samuel Bloomfield, Claire Stuart, Andrew Bell, Reenesh Prakash, Samir Dervisevic, Alison E. Mather, John Wain, Mark Webber, Andrew J. Page, Justin O'Grady |
| EPI_ISL_704636, EPI_ISL_704637, EPI_ISL_704638, EPI_ISL_704639 | Virology Department, Sheffield Teaching Hospitals NHS Foundation Trust/Department of Infection, Immunity and Cardiovascular Disease, The Medical School, University of Sheffield | COVID-19 Genomics UK (COG-UK) Consortium | Thushan de Silva, Matthew Parker, Nikki Smith, Adri Angyal, Rebecca Brown, Luke Green, Rachel Tucker, Paul Parsons, Danielle Groves, Katie Johnson, Laura Carrilero, Alex Keeley, Dave Partridge, Matthew Wyles, Benjamin Lindsey, Mehmet Yavuz, Mohammad Raza, Cariad Evans |
| EPI_ISL_704640, EPI_ISL_704641, EPI_ISL_704643, EPI_ISL_704646, EPI_ISL_704648, EPI_ISL_704651, EPI_ISL_704654, EPI_ISL_704656, EPI_ISL_704659, EPI_ISL_704662, EPI_ISL_704664 | | | |
| see above | Department of Pathology, University of Cambridge | COVID-19 Genomics UK (COG-UK) Consortium | Aminu S. Jahun, Yasmin Chaudhry, Grant Hall, Iliana Georgana, Myra Hosmillo, Martin D. Curran, Malte Pinckert, Surendra Parmar, Ian Goodfellow |
| EPI_ISL_704688, EPI_ISL_704689, EPI_ISL_704691, EPI_ISL_704693, EPI_ISL_704694, EPI_ISL_704696, EPI_ISL_704697, EPI_ISL_704699, EPI_ISL_704700, EPI_ISL_704702, EPI_ISL_704704, EPI_ISL_704705, EPI_ISL_704707, EPI_ISL_704708, EPI_ISL_704710, EPI_ISL_704712, EPI_ISL_704713, EPI_ISL_704715, EPI_ISL_704717, EPI_ISL_704718, EPI_ISL_704720, EPI_ISL_704721, EPI_ISL_704723, EPI_ISL_704724, EPI_ISL_704726, EPI_ISL_704727, EPI_ISL_704729, EPI_ISL_704730, EPI_ISL_704732, EPI_ISL_704733, EPI_ISL_704735, EPI_ISL_704737, EPI_ISL_704738, EPI_ISL_704740, EPI_ISL_704742, EPI_ISL_704743, EPI_ISL_704745, EPI_ISL_704747, EPI_ISL_704748, EPI_ISL_704751, EPI_ISL_704753, EPI_ISL_704755, EPI_ISL_704756, EPI_ISL_704758, EPI_ISL_704760, EPI_ISL_704761 | | | |
| see above | Lighthouse Lab in Cambridge | Wellcome Sanger Institute for the COVID-19 Genomics UK (COG-UK) Consortium | Rob Howes, The Lighthouse Lab in Cambridge and Alex Alderton, Roberto Amato, Sonia Goncalves, Ewan Harrison, David K. Jackson, Ian Johnston, Dominic Kwiatkowski, Cordelia Langford, John Sillitoe on behalf of the Wellcome Sanger Institute COVID-19 Surveillance Team |
| EPI_ISL_704762 | Quadram Institute Bioscience | COVID-19 Genomics UK (COG-UK) Consortium | Dave J. Baker, Gemma L. Kay, Alp Aydin, Thanh Le-Viet, Steven Rudder, Ana P. Tedim, Anastasia Kolyva, Maria Diaz, Leonardo de Oliveira Martins, Nabil-Fareed Alikhan, Lizzie Meadows, Rachael Stanley, Ngozi Elumogo, Muhammed Yasir, Nicholas M. Thomson, Alexander J Trotter, Rachel Gilroy, Samuel Bloomfield, Claire Stuart, Andrew Bell, Reenesh Prakash, Samir Dervisevic, Alison E. Mather, John Wain, Mark Webber, Andrew J. Page, Justin O'Grady |
| EPI_ISL_704763, EPI_ISL_704764 | Lighthouse Lab in Cambridge | Wellcome Sanger Institute for the COVID-19 Genomics UK (COG-UK) Consortium | Rob Howes, The Lighthouse Lab in Cambridge and Alex Alderton, Roberto Amato, Sonia Goncalves, Ewan Harrison, David K. Jackson, Ian Johnston, Dominic Kwiatkowski, Cordelia Langford, John Sillitoe on behalf of the Wellcome Sanger Institute COVID-19 Surveillance Team |
| EPI_ISL_704765 | Quadram Institute Bioscience | COVID-19 Genomics UK (COG-UK) Consortium | Dave J. Baker, Gemma L. Kay, Alp Aydin, Thanh Le-Viet, Steven Rudder, Ana P. Tedim, Anastasia Kolyva, Maria Diaz, Leonardo de Oliveira Martins, Nabil-Fareed Alikhan, Lizzie Meadows, Rachael Stanley, Ngozi Elumogo, Muhammed Yasir, Nicholas M. Thomson, Alexander J Trotter, Rachel Gilroy, Samuel Bloomfield, Claire Stuart, Andrew Bell, Reenesh Prakash, Samir Dervisevic, Alison E. Mather, John Wain, Mark Webber, Andrew J. Page, Justin O'Grady |
| EPI_ISL_704766, EPI_ISL_704767 | Lighthouse Lab in Cambridge | Wellcome Sanger Institute for the COVID-19 Genomics UK (COG-UK) Consortium | Rob Howes, The Lighthouse Lab in Cambridge and Alex Alderton, Roberto Amato, Sonia Goncalves, Ewan Harrison, David K. Jackson, Ian Johnston, Dominic Kwiatkowski, Cordelia Langford, John Sillitoe on behalf of the Wellcome Sanger Institute COVID-19 Surveillance Team |
| EPI_ISL_704768 | Quadram Institute Bioscience | COVID-19 Genomics UK (COG-UK) Consortium | Dave J. Baker, Gemma L. Kay, Alp Aydin, Thanh Le-Viet, Steven Rudder, Ana P. Tedim, Anastasia Kolyva, Maria Diaz, Leonardo de Oliveira Martins, Nabil-Fareed Alikhan, Lizzie Meadows, Rachael Stanley, Ngozi Elumogo, Muhammed Yasir, Nicholas M. Thomson, Alexander J Trotter, Rachel Gilroy, Samuel Bloomfield, Claire Stuart, Andrew Bell, Reenesh Prakash, Samir Dervisevic, Alison E. Mather, John Wain, Mark Webber, Andrew J. Page, Justin O'Grady |
| EPI_ISL_704769 | Lighthouse Lab in Cambridge | Wellcome Sanger Institute for the COVID-19 Genomics UK (COG-UK) Consortium | Rob Howes, The Lighthouse Lab in Cambridge and Alex Alderton, Roberto Amato, Sonia Goncalves, Ewan Harrison, David K. Jackson, Ian Johnston, Dominic Kwiatkowski, Cordelia Langford, John Sillitoe on behalf of the Wellcome Sanger Institute COVID-19 Surveillance Team |

| | | | |
|---|---|---|---|
| EPI_ISL_704770 | Quadram Institute Bioscience | COVID-19 Genomics UK (COG-UK) Consortium | Dave J. Baker, Gemma L. Kay, Alp Aydin, Thanh Le-Viet, Steven Rudder, Ana P. Tedim, Anastasia Kolyva, Maria Diaz, Leonardo de Oliveira Martins, Nabil-Fareed Alikhan, Lizzie Meadows, Rachael Stanley, Ngozi Elumogo, Muhammed Yasir, Nicholas M. Thomson, Alexander J Trotter, Rachel Gilroy, Samuel Bloomfield, Claire Stuart, Andrew Bell, Reenesh Prakash, Samir Dervisevic, Alison E. Mather, John Wain, Mark Webber, Andrew J. Page, Justin O'Grady |
| EPI_ISL_704771, EPI_ISL_704773, EPI_ISL_704774, EPI_ISL_704776, EPI_ISL_704778, EPI_ISL_704779, EPI_ISL_704781, EPI_ISL_704782, EPI_ISL_704784, EPI_ISL_704785, EPI_ISL_704787, EPI_ISL_704789, EPI_ISL_704790, EPI_ISL_704792, EPI_ISL_704793, EPI_ISL_704795, EPI_ISL_704796, EPI_ISL_704798, EPI_ISL_704800, EPI_ISL_704801, EPI_ISL_704803, EPI_ISL_704805, EPI_ISL_704806, EPI_ISL_704808, EPI_ISL_704810, EPI_ISL_704811, EPI_ISL_704813, EPI_ISL_704814, EPI_ISL_704816, EPI_ISL_704818, EPI_ISL_704819 | | | |
| see above | Lighthouse Lab in Cambridge | Wellcome Sanger Institute for the COVID-19 Genomics UK (COG-UK) Consortium | Rob Howes, The Lighthouse Lab in Cambridge and Alex Alderton, Roberto Amato, Sonia Goncalves, Ewan Harrison, David K. Jackson, Ian Johnston, Dominic Kwiatkowski, Cordelia Langford, John Sillitoe on behalf of the Wellcome Sanger Institute COVID-19 Surveillance Team |
| EPI_ISL_704820 | Quadram Institute Bioscience | COVID-19 Genomics UK (COG-UK) Consortium | Dave J. Baker, Gemma L. Kay, Alp Aydin, Thanh Le-Viet, Steven Rudder, Ana P. Tedim, Anastasia Kolyva, Maria Diaz, Leonardo de Oliveira Martins, Nabil-Fareed Alikhan, Lizzie Meadows, Rachael Stanley, Ngozi Elumogo, Muhammed Yasir, Nicholas M. Thomson, Alexander J Trotter, Rachel Gilroy, Samuel Bloomfield, Claire Stuart, Andrew Bell, Reenesh Prakash, Samir Dervisevic, Alison E. Mather, John Wain, Mark Webber, Andrew J. Page, Justin O'Grady |
| EPI_ISL_704821, EPI_ISL_704822 | Lighthouse Lab in Cambridge | Wellcome Sanger Institute for the COVID-19 Genomics UK (COG-UK) Consortium | Rob Howes, The Lighthouse Lab in Cambridge and Alex Alderton, Roberto Amato, Sonia Goncalves, Ewan Harrison, David K. Jackson, Ian Johnston, Dominic Kwiatkowski, Cordelia Langford, John Sillitoe on behalf of the Wellcome Sanger Institute COVID-19 Surveillance Team |
| EPI_ISL_704823 | Quadram Institute Bioscience | COVID-19 Genomics UK (COG-UK) Consortium | Dave J. Baker, Gemma L. Kay, Alp Aydin, Thanh Le-Viet, Steven Rudder, Ana P. Tedim, Anastasia Kolyva, Maria Diaz, Leonardo de Oliveira Martins, Nabil-Fareed Alikhan, Lizzie Meadows, Rachael Stanley, Ngozi Elumogo, Muhammed Yasir, Nicholas M. Thomson, Alexander J Trotter, Rachel Gilroy, Samuel Bloomfield, Claire Stuart, Andrew Bell, Reenesh Prakash, Samir Dervisevic, Alison E. Mather, John Wain, Mark Webber, Andrew J. Page, Justin O'Grady |
| EPI_ISL_704824, EPI_ISL_704825 | Lighthouse Lab in Cambridge | Wellcome Sanger Institute for the COVID-19 Genomics UK (COG-UK) Consortium | Rob Howes, The Lighthouse Lab in Cambridge and Alex Alderton, Roberto Amato, Sonia Goncalves, Ewan Harrison, David K. Jackson, Ian Johnston, Dominic Kwiatkowski, Cordelia Langford, John Sillitoe on behalf of the Wellcome Sanger Institute COVID-19 Surveillance Team |
| EPI_ISL_704826 | Quadram Institute Bioscience | COVID-19 Genomics UK (COG-UK) Consortium | Dave J. Baker, Gemma L. Kay, Alp Aydin, Thanh Le-Viet, Steven Rudder, Ana P. Tedim, Anastasia Kolyva, Maria Diaz, Leonardo de Oliveira Martins, Nabil-Fareed Alikhan, Lizzie Meadows, Rachael Stanley, Ngozi Elumogo, Muhammed Yasir, Nicholas M. Thomson, Alexander J Trotter, Rachel Gilroy, Samuel Bloomfield, Claire Stuart, Andrew Bell, Reenesh Prakash, Samir Dervisevic, Alison E. Mather, John Wain, Mark Webber, Andrew J. Page, Justin O'Grady |
| EPI_ISL_704827 | Lighthouse Lab in Cambridge | Wellcome Sanger Institute for the COVID-19 Genomics UK (COG-UK) Consortium | Rob Howes, The Lighthouse Lab in Cambridge and Alex Alderton, Roberto Amato, Sonia Goncalves, Ewan Harrison, David K. Jackson, Ian Johnston, Dominic Kwiatkowski, Cordelia Langford, John Sillitoe on behalf of the Wellcome Sanger Institute COVID-19 Surveillance Team |
| EPI_ISL_704828 | Virology Department, Sheffield Teaching Hospitals NHS Foundation Trust/Department of Infection, Immunity and Cardiovascular Disease, The Medical School, University of Sheffield | COVID-19 Genomics UK (COG-UK) Consortium | Thushan de Silva, Matthew Parker, Nikki Smith, Adri Angyal, Rebecca Brown, Luke Green, Rachel Tucker, Paul Parsons, Danielle Groves, Katie Johnson, Laura Carrilero, Alex Keeley, Dave Partridge, Matthew Wyles, Benjamin Lindsey, Mehmet Yavuz, Mohammad Raza, Cariad Evans |
| EPI_ISL_704829, EPI_ISL_704830 | Lighthouse Lab in Cambridge | Wellcome Sanger Institute for the COVID-19 Genomics UK (COG-UK) Consortium | Rob Howes, The Lighthouse Lab in Cambridge and Alex Alderton, Roberto Amato, Sonia Goncalves, Ewan Harrison, David K. Jackson, Ian Johnston, Dominic Kwiatkowski, Cordelia Langford, John Sillitoe on behalf of the Wellcome Sanger Institute COVID-19 Surveillance Team |
| EPI_ISL_704831 | Quadram Institute Bioscience | COVID-19 Genomics UK (COG-UK) Consortium | Dave J. Baker, Gemma L. Kay, Alp Aydin, Thanh Le-Viet, Steven Rudder, Ana P. Tedim, Anastasia Kolyva, Maria Diaz, Leonardo de Oliveira Martins, Nabil-Fareed Alikhan, Lizzie Meadows, Rachael Stanley, Ngozi Elumogo, Muhammed Yasir, Nicholas M. Thomson, Alexander J Trotter, Rachel Gilroy, Samuel Bloomfield, Claire Stuart, Andrew Bell, Reenesh Prakash, Samir Dervisevic, Alison E. Mather, John Wain, Mark Webber, Andrew J. Page, Justin O'Grady |
| EPI_ISL_704832, EPI_ISL_704833 | Lighthouse Lab in Cambridge | Wellcome Sanger Institute for the COVID-19 Genomics UK (COG-UK) Consortium | Rob Howes, The Lighthouse Lab in Cambridge and Alex Alderton, Roberto Amato, Sonia Goncalves, Ewan Harrison, David K. Jackson, Ian Johnston, Dominic Kwiatkowski, Cordelia Langford, John Sillitoe on behalf of the Wellcome Sanger Institute COVID-19 Surveillance Team |
| EPI_ISL_704834 | Department of Pathology, University of Cambridge | COVID-19 Genomics UK (COG-UK) Consortium | Aminu S. Jahun, Yasmin Chaudhry, Grant Hall, Iliana Georgana, Myra Hosmillo, Martin D. Curran, Malte Pinckert, Surendra Parmar, Ian Goodfellow |
| EPI_ISL_704835, EPI_ISL_704837, EPI_ISL_704838, EPI_ISL_704840, EPI_ISL_704841 | Lighthouse Lab in Cambridge | Wellcome Sanger Institute for the COVID-19 Genomics UK (COG-UK) Consortium | Rob Howes, The Lighthouse Lab in Cambridge and Alex Alderton, Roberto Amato, Sonia Goncalves, Ewan Harrison, David K. Jackson, Ian Johnston, Dominic Kwiatkowski, Cordelia Langford, John Sillitoe on behalf of the Wellcome Sanger Institute COVID-19 Surveillance Team |
| EPI_ISL_704842 | Quadram Institute Bioscience | COVID-19 Genomics UK (COG-UK) Consortium | Dave J. Baker, Gemma L. Kay, Alp Aydin, Thanh Le-Viet, Steven Rudder, Ana P. Tedim, Anastasia Kolyva, Maria Diaz, Leonardo de Oliveira Martins, Nabil-Fareed Alikhan, Lizzie Meadows, Rachael Stanley, Ngozi Elumogo, Muhammed Yasir, Nicholas M. Thomson, Alexander J Trotter, Rachel Gilroy, Samuel Bloomfield, Claire Stuart, Andrew Bell, Reenesh Prakash, Samir Dervisevic, Alison E. Mather, John Wain, Mark Webber, Andrew J. Page, Justin O'Grady |
| EPI_ISL_704843, EPI_ISL_704844, EPI_ISL_704846, EPI_ISL_704848, EPI_ISL_704849, EPI_ISL_704851, EPI_ISL_704852, EPI_ISL_704854, EPI_ISL_704856, EPI_ISL_704857, EPI_ISL_704859, EPI_ISL_704860, EPI_ISL_704862, EPI_ISL_704863, EPI_ISL_704865, EPI_ISL_704868, EPI_ISL_704870, EPI_ISL_704871, EPI_ISL_704873, EPI_ISL_704875, EPI_ISL_704876, EPI_ISL_704878, EPI_ISL_704880, EPI_ISL_704881, EPI_ISL_704883, EPI_ISL_704884, EPI_ISL_704886, EPI_ISL_704888, EPI_ISL_704889, EPI_ISL_704891, EPI_ISL_704893, EPI_ISL_704894, EPI_ISL_704896, EPI_ISL_704898, EPI_ISL_704899, EPI_ISL_704901, EPI_ISL_704902 | | | |
| see above | Lighthouse Lab in Cambridge | Wellcome Sanger Institute for the COVID-19 Genomics UK (COG-UK) Consortium | Rob Howes, The Lighthouse Lab in Cambridge and Alex Alderton, Roberto Amato, Sonia Goncalves, Ewan Harrison, David K. Jackson, Ian Johnston, Dominic Kwiatkowski, Cordelia Langford, John Sillitoe on behalf of the Wellcome Sanger Institute COVID-19 Surveillance Team |
| EPI_ISL_704903 | Liverpool Clinical Laboratories | COVID-19 Genomics UK (COG-UK) Consortium | Sam Haldenby, Anita Lucaci, Steve Paterson, Julian Hiscox, Alistair Darby, M Almsaud, A Alrezaihi, Muhannad Alruwaili, Stuart D Armstrong, Jones Benjamin, Eleanor G Bentley, Anu Chawla, Jordan J Clark, Angela Cowell, Richard Eccles, Isabel García-Dorival, Matthew Gemmell, Alessandro Gerada, PKF Gilmore, Richard Gregory, Ximeng Han, Catherine Hartley, Margaret Hughes, Miren Iturriza-Gomara, James Johnson, L Luu, Jenifer Manson, Charlotte Nelson, Elaine O'Toole, Cassie Olateju, Rebekah Penrice-Randal , Lucille Rainbow, N.P Randle, Trevor Ian Robinson, Parul Sharma, Ghada T Shawli, James P Stewart, Neil Swainston, Ecaterina Vamos, Joanne Watts, Mark Whitehead |
| EPI_ISL_704904, EPI_ISL_704906, EPI_ISL_704907, EPI_ISL_704909, EPI_ISL_704910, EPI_ISL_704912, EPI_ISL_704913 | Lighthouse Lab in Cambridge | Wellcome Sanger Institute for the COVID-19 Genomics UK (COG-UK) Consortium | Rob Howes, The Lighthouse Lab in Cambridge and Alex Alderton, Roberto Amato, Sonia Goncalves, Ewan Harrison, David K. Jackson, Ian Johnston, Dominic Kwiatkowski, Cordelia Langford, John Sillitoe on behalf of the Wellcome Sanger Institute COVID-19 Surveillance Team |
| EPI_ISL_704914 | Quadram Institute Bioscience | COVID-19 Genomics UK (COG-UK) Consortium | Dave J. Baker, Gemma L. Kay, Alp Aydin, Thanh Le-Viet, Steven Rudder, Ana P. Tedim, Anastasia Kolyva, Maria Diaz, Leonardo de Oliveira Martins, Nabil-Fareed Alikhan, Lizzie Meadows, Rachael Stanley, Ngozi Elumogo, Muhammed Yasir, Nicholas M. Thomson, Alexander J Trotter, Rachel Gilroy, Samuel Bloomfield, Claire Stuart, Andrew Bell, Reenesh Prakash, Samir Dervisevic, Alison E. Mather, John Wain, Mark Webber, Andrew J. Page, Justin O'Grady |
| EPI_ISL_704915, EPI_ISL_704917, EPI_ISL_704919, EPI_ISL_704920, EPI_ISL_704922, EPI_ISL_704924, EPI_ISL_704925, EPI_ISL_704927, EPI_ISL_704928, EPI_ISL_704930, EPI_ISL_704933, EPI_ISL_704935, EPI_ISL_704936, EPI_ISL_704938, EPI_ISL_704940, EPI_ISL_704941, EPI_ISL_704943, EPI_ISL_704944, EPI_ISL_704946, EPI_ISL_704948, EPI_ISL_704949, EPI_ISL_704951, EPI_ISL_704953, EPI_ISL_704954, EPI_ISL_704956, EPI_ISL_704958, EPI_ISL_704959, EPI_ISL_704961, EPI_ISL_704962, EPI_ISL_704964, EPI_ISL_704966, EPI_ISL_704967, EPI_ISL_704969, EPI_ISL_704971, EPI_ISL_704972, EPI_ISL_704974, EPI_ISL_704975 | | | |
| see above | Lighthouse Lab in Cambridge | Wellcome Sanger Institute for the COVID-19 Genomics UK (COG-UK) Consortium | Rob Howes, The Lighthouse Lab in Cambridge and Alex Alderton, Roberto Amato, Sonia Goncalves, Ewan Harrison, David K. Jackson, Ian Johnston, Dominic Kwiatkowski, Cordelia Langford, John Sillitoe on behalf of the Wellcome Sanger Institute COVID-19 Surveillance Team |
| EPI_ISL_704976 | Northumbria University / South Tees Hospitals NHS Foundation Trust / North Cumbria Integrated Care NHS Foundation Trust / North Tees and Hartlepool NHS Foundation Trust / Newcastle Hospitals NHS Foundation Trust | COVID-19 Genomics UK (COG-UK) Consortium | Darren L Smith,Andrew Nelson,Matthew Bashton,Greg R Young,Joshua Loh,John Allan,Mohammad A Tariq,Giles S Holt,Gary Black,Wen C Yew,Lynn Dover,Paul Baker,Steve Liggett,Sarah Essex,Jane Greenaway,Debra Padgett,Clive Graham,Garren Scott,Edward Barton,Emma Swindells,Brendan Payne,Jennifer Collins,Yusri Taha,Gary Eltringham |
| EPI_ISL_704977, EPI_ISL_704979, EPI_ISL_704980, EPI_ISL_704982 | Lighthouse Lab in Cambridge | Wellcome Sanger Institute for the COVID-19 Genomics UK (COG-UK) Consortium | Rob Howes, The Lighthouse Lab in Cambridge and Alex Alderton, Roberto Amato, Sonia Goncalves, Ewan Harrison, David K. Jackson, Ian Johnston, Dominic Kwiatkowski, Cordelia Langford, John Sillitoe on behalf of the Wellcome Sanger Institute COVID-19 Surveillance Team |

| | | | |
|---|---|---|---|
| EPI_ISL_704983 | Liverpool Clinical Laboratories | COVID-19 Genomics UK (COG-UK) Consortium | Sam Haldenby, Anita Lucaci, Steve Paterson, Julian Hiscox, Alistair Darby, M Almsaud, A Alrezaihi, Muhannad Alruwaili, Stuart D Armstrong, Jones Benjamin, Eleanor G Bentley, Anu Chawla, Jordan J Clark, Angela Cowell, Richard Eccles, Isabel García-Dorival, Matthew Gemmell, Alessandro Gerada, PKF Gilmore, Richard Gregory, Ximeng Han, Catherine Hartley, Margaret Hughes, Miren Iturriza-Gomara, James Johnson, L Luu, Jenifer Manson, Charlotte Nelson, Elaine O'Toole, Cassie Olateju, Rebekah Penrice-Randal , Lucille Rainbow, N.P Randle, Trevor Ian Robinson, Parul Sharma, Ghada T Shawli, James P Stewart, Neil Swainston, Ecaterina Vamos, Joanne Watts, Mark Whitehead |
| EPI_ISL_704984, EPI_ISL_704985, EPI_ISL_704987, EPI_ISL_704988 | Lighthouse Lab in Cambridge | Wellcome Sanger Institute for the COVID-19 Genomics UK (COG-UK) Consortium | Rob Howes, The Lighthouse Lab in Cambridge and Alex Alderton, Roberto Amato, Sonia Goncalves, Ewan Harrison, David K. Jackson, Ian Johnston, Dominic Kwiatkowski, Cordelia Langford, John Sillitoe on behalf of the Wellcome Sanger Institute COVID-19 Surveillance Team |
| EPI_ISL_704989 | Liverpool Clinical Laboratories | COVID-19 Genomics UK (COG-UK) Consortium | Sam Haldenby, Anita Lucaci, Steve Paterson, Julian Hiscox, Alistair Darby, M Almsaud, A Alrezaihi, Muhannad Alruwaili, Stuart D Armstrong, Jones Benjamin, Eleanor G Bentley, Anu Chawla, Jordan J Clark, Angela Cowell, Richard Eccles, Isabel García-Dorival, Matthew Gemmell, Alessandro Gerada, PKF Gilmore, Richard Gregory, Ximeng Han, Catherine Hartley, Margaret Hughes, Miren Iturriza-Gomara, James Johnson, L Luu, Jenifer Manson, Charlotte Nelson, Elaine O'Toole, Cassie Olateju, Rebekah Penrice-Randal , Lucille Rainbow, N.P Randle, Trevor Ian Robinson, Parul Sharma, Ghada T Shawli, James P Stewart, Neil Swainston, Ecaterina Vamos, Joanne Watts, Mark Whitehead |
| EPI_ISL_704990, EPI_ISL_704992, EPI_ISL_704993, EPI_ISL_704995, EPI_ISL_704996 | Lighthouse Lab in Cambridge | Wellcome Sanger Institute for the COVID-19 Genomics UK (COG-UK) Consortium | Rob Howes, The Lighthouse Lab in Cambridge and Alex Alderton, Roberto Amato, Sonia Goncalves, Ewan Harrison, David K. Jackson, Ian Johnston, Dominic Kwiatkowski, Cordelia Langford, John Sillitoe on behalf of the Wellcome Sanger Institute COVID-19 Surveillance Team |
| EPI_ISL_704997 | University College London, Great Ormond Street Hospital for Children NHS Foundation Trust, Imperial College Healthcare NHS Trust | COVID-19 Genomics UK (COG-UK) Consortium | Sergi Castellano, Rachel Williams, Mark Kristiansen, Paola Resende Silva, Sunando Roy, Tony Brooks, Helena Tutill, Paola Niola, Patricia Dyal, Charlotte Williams, Leysa Forrest, Yasmin Panchbhaya, Jacqueline Findlay, Samuel Weeks, Julianne Brown, Kathryn Harris, Paul Randell, James Price, Alison Holmes, Judith Breuer |
| EPI_ISL_704998 | Lighthouse Lab in Cambridge | Wellcome Sanger Institute for the COVID-19 Genomics UK (COG-UK) Consortium | Rob Howes, The Lighthouse Lab in Cambridge and Alex Alderton, Roberto Amato, Sonia Goncalves, Ewan Harrison, David K. Jackson, Ian Johnston, Dominic Kwiatkowski, Cordelia Langford, John Sillitoe on behalf of the Wellcome Sanger Institute COVID-19 Surveillance Team |
| EPI_ISL_704999 | University College London, Great Ormond Street Hospital for Children NHS Foundation Trust, Imperial College Healthcare NHS Trust | COVID-19 Genomics UK (COG-UK) Consortium | Sergi Castellano, Rachel Williams, Mark Kristiansen, Paola Resende Silva, Sunando Roy, Tony Brooks, Helena Tutill, Paola Niola, Patricia Dyal, Charlotte Williams, Leysa Forrest, Yasmin Panchbhaya, Jacqueline Findlay, Samuel Weeks, Julianne Brown, Kathryn Harris, Paul Randell, James Price, Alison Holmes, Judith Breuer |
| EPI_ISL_705000, EPI_ISL_705001, EPI_ISL_705003, EPI_ISL_705004 | Lighthouse Lab in Cambridge | Wellcome Sanger Institute for the COVID-19 Genomics UK (COG-UK) Consortium | Rob Howes, The Lighthouse Lab in Cambridge and Alex Alderton, Roberto Amato, Sonia Goncalves, Ewan Harrison, David K. Jackson, Ian Johnston, Dominic Kwiatkowski, Cordelia Langford, John Sillitoe on behalf of the Wellcome Sanger Institute COVID-19 Surveillance Team |
| EPI_ISL_705005 | Liverpool Clinical Laboratories | COVID-19 Genomics UK (COG-UK) Consortium | Sam Haldenby, Anita Lucaci, Steve Paterson, Julian Hiscox, Alistair Darby, M Almsaud, A Alrezaihi, Muhannad Alruwaili, Stuart D Armstrong, Jones Benjamin, Eleanor G Bentley, Anu Chawla, Jordan J Clark, Angela Cowell, Richard Eccles, Isabel García-Dorival, Matthew Gemmell, Alessandro Gerada, PKF Gilmore, Richard Gregory, Ximeng Han, Catherine Hartley, Margaret Hughes, Miren Iturriza-Gomara, James Johnson, L Luu, Jenifer Manson, Charlotte Nelson, Elaine O'Toole, Cassie Olateju, Rebekah Penrice-Randal , Lucille Rainbow, N.P Randle, Trevor Ian Robinson, Parul Sharma, Ghada T Shawli, James P Stewart, Neil Swainston, Ecaterina Vamos, Joanne Watts, Mark Whitehead |
| EPI_ISL_705006, EPI_ISL_705008, EPI_ISL_705009, EPI_ISL_705011, EPI_ISL_705012, EPI_ISL_705014, EPI_ISL_705016, EPI_ISL_705017, EPI_ISL_705019, EPI_ISL_705020, EPI_ISL_705022, EPI_ISL_705023, EPI_ISL_705025, EPI_ISL_705027, EPI_ISL_705028, EPI_ISL_705029, EPI_ISL_705031, EPI_ISL_705033, EPI_ISL_705034, EPI_ISL_705036, EPI_ISL_705037, EPI_ISL_705039, EPI_ISL_705041, EPI_ISL_705042, EPI_ISL_705044, EPI_ISL_705045, EPI_ISL_705047, EPI_ISL_705049, EPI_ISL_705050, EPI_ISL_705052, EPI_ISL_705053, EPI_ISL_705055, EPI_ISL_705057, EPI_ISL_705058 | | | |
| see above | Lighthouse Lab in Cambridge | Wellcome Sanger Institute for the COVID-19 Genomics UK (COG-UK) Consortium | Rob Howes, The Lighthouse Lab in Cambridge and Alex Alderton, Roberto Amato, Sonia Goncalves, Ewan Harrison, David K. Jackson, Ian Johnston, Dominic Kwiatkowski, Cordelia Langford, John Sillitoe on behalf of the Wellcome Sanger Institute COVID-19 Surveillance Team |
| EPI_ISL_705059 | Liverpool Clinical Laboratories | COVID-19 Genomics UK (COG-UK) Consortium | Sam Haldenby, Anita Lucaci, Steve Paterson, Julian Hiscox, Alistair Darby, M Almsaud, A Alrezaihi, Muhannad Alruwaili, Stuart D Armstrong, Jones Benjamin, Eleanor G Bentley, Anu Chawla, Jordan J Clark, Angela Cowell, Richard Eccles, Isabel García-Dorival, Matthew Gemmell, Alessandro Gerada, PKF Gilmore, Richard Gregory, Ximeng Han, Catherine Hartley, Margaret Hughes, Miren Iturriza-Gomara, James Johnson, L Luu, Jenifer Manson, Charlotte Nelson, Elaine O'Toole, Cassie Olateju, Rebekah Penrice-Randal , Lucille Rainbow, N.P Randle, Trevor Ian Robinson, Parul Sharma, Ghada T Shawli, James P Stewart, Neil Swainston, Ecaterina Vamos, Joanne Watts, Mark Whitehead |
| EPI_ISL_705060, EPI_ISL_705063, EPI_ISL_705065, EPI_ISL_705066 | Lighthouse Lab in Cambridge | Wellcome Sanger Institute for the COVID-19 Genomics UK (COG-UK) Consortium | Rob Howes, The Lighthouse Lab in Cambridge and Alex Alderton, Roberto Amato, Sonia Goncalves, Ewan Harrison, David K. Jackson, Ian Johnston, Dominic Kwiatkowski, Cordelia Langford, John Sillitoe on behalf of the Wellcome Sanger Institute COVID-19 Surveillance Team |
| EPI_ISL_705067 | Northumbria University / South Tees Hospitals NHS Foundation Trust / North Cumbria Integrated Care NHS Foundation Trust / North Tees and Hartlepool NHS Foundation Trust / Newcastle Hospitals NHS Foundation Trust | COVID-19 Genomics UK (COG-UK) Consortium | Darren L Smith,Andrew Nelson,Matthew Bashton,Greg R Young,Joshua Loh,John Allan,Mohammad A Tariq,Giles S Holt,Gary Black,Wen C Yew,Lynn Dover,Paul Baker,Steve Liggett,Sarah Essex,Jane Greenaway,Debra Padgett,Clive Graham,Garren Scott,Edward Barton,Emma Swindells,Brendan Payne,Jennifer Collins,Yusri Taha,Gary Eltringham |
| EPI_ISL_705068, EPI_ISL_705069, EPI_ISL_705071 | Lighthouse Lab in Cambridge | Wellcome Sanger Institute for the COVID-19 Genomics UK (COG-UK) Consortium | Rob Howes, The Lighthouse Lab in Cambridge and Alex Alderton, Roberto Amato, Sonia Goncalves, Ewan Harrison, David K. Jackson, Ian Johnston, Dominic Kwiatkowski, Cordelia Langford, John Sillitoe on behalf of the Wellcome Sanger Institute COVID-19 Surveillance Team |
| EPI_ISL_705072 | University College London, Great Ormond Street Hospital for Children NHS Foundation Trust, Imperial College Healthcare NHS Trust | COVID-19 Genomics UK (COG-UK) Consortium | Sergi Castellano, Rachel Williams, Mark Kristiansen, Paola Resende Silva, Sunando Roy, Tony Brooks, Helena Tutill, Paola Niola, Patricia Dyal, Charlotte Williams, Leysa Forrest, Yasmin Panchbhaya, Jacqueline Findlay, Samuel Weeks, Julianne Brown, Kathryn Harris, Paul Randell, James Price, Alison Holmes, Judith Breuer |
| EPI_ISL_705073, EPI_ISL_705074, EPI_ISL_705076, EPI_ISL_705077, EPI_ISL_705079, EPI_ISL_705080 | Lighthouse Lab in Cambridge | Wellcome Sanger Institute for the COVID-19 Genomics UK (COG-UK) Consortium | Rob Howes, The Lighthouse Lab in Cambridge and Alex Alderton, Roberto Amato, Sonia Goncalves, Ewan Harrison, David K. Jackson, Ian Johnston, Dominic Kwiatkowski, Cordelia Langford, John Sillitoe on behalf of the Wellcome Sanger Institute COVID-19 Surveillance Team |
| EPI_ISL_705081 | Department of Pathology, University of Cambridge | COVID-19 Genomics UK (COG-UK) Consortium | Aminu S. Jahun, Yasmin Chaudhry, Grant Hall, Iliana Georgana, Myra Hosmillo, Martin D. Curran, Malte Pinckert, Surendra Parmar, Ian Goodfellow |
| EPI_ISL_705082, EPI_ISL_705084, EPI_ISL_705085, EPI_ISL_705087, EPI_ISL_705089, EPI_ISL_705090, EPI_ISL_705092, EPI_ISL_705093, EPI_ISL_705095, EPI_ISL_705097, EPI_ISL_705098, EPI_ISL_705100, EPI_ISL_705101, EPI_ISL_705103, EPI_ISL_705106, EPI_ISL_705107, EPI_ISL_705109 | | | |
| see above | Lighthouse Lab in Cambridge | Wellcome Sanger Institute for the COVID-19 Genomics UK (COG-UK) Consortium | Rob Howes, The Lighthouse Lab in Cambridge and Alex Alderton, Roberto Amato, Sonia Goncalves, Ewan Harrison, David K. Jackson, Ian Johnston, Dominic Kwiatkowski, Cordelia Langford, John Sillitoe on behalf of the Wellcome Sanger Institute COVID-19 Surveillance Team |
| EPI_ISL_705110 | Department of Pathology, University of Cambridge | COVID-19 Genomics UK (COG-UK) Consortium | Aminu S. Jahun, Yasmin Chaudhry, Grant Hall, Iliana Georgana, Myra Hosmillo, Martin D. Curran, Malte Pinckert, Surendra Parmar, Ian Goodfellow |
| EPI_ISL_705111, EPI_ISL_705112, EPI_ISL_705114, EPI_ISL_705116, EPI_ISL_705117, EPI_ISL_705119, EPI_ISL_705120, EPI_ISL_705122, EPI_ISL_705123, EPI_ISL_705125, EPI_ISL_705127, EPI_ISL_705128, EPI_ISL_705130, EPI_ISL_705131, EPI_ISL_705133 | | | |
| see above | Lighthouse Lab in Cambridge | Wellcome Sanger Institute for the COVID-19 Genomics UK (COG-UK) Consortium | Rob Howes, The Lighthouse Lab in Cambridge and Alex Alderton, Roberto Amato, Sonia Goncalves, Ewan Harrison, David K. Jackson, Ian Johnston, Dominic Kwiatkowski, Cordelia Langford, John Sillitoe on behalf of the Wellcome Sanger Institute COVID-19 Surveillance Team |
| EPI_ISL_705134 | University of Exeter | COVID-19 Genomics UK (COG-UK) Consortium | Ben Temperton,Aaron Jeffries,Michelle Michelsen,Joanna Warwick-Dugdale,Audrey Farbos,Robyn Manley,Stephen Michell,Jane Masoli |
| EPI_ISL_705135, EPI_ISL_705136, EPI_ISL_705138, EPI_ISL_705139, EPI_ISL_705141, EPI_ISL_705142, EPI_ISL_705144, EPI_ISL_705146, EPI_ISL_705147, EPI_ISL_705149 | Lighthouse Lab in Cambridge | Wellcome Sanger Institute for the COVID-19 Genomics UK (COG-UK) Consortium | Rob Howes, The Lighthouse Lab in Cambridge and Alex Alderton, Roberto Amato, Sonia Goncalves, Ewan Harrison, David K. Jackson, Ian Johnston, Dominic Kwiatkowski, Cordelia Langford, John Sillitoe on behalf of the Wellcome Sanger Institute COVID-19 Surveillance Team |
| EPI_ISL_705150 | Liverpool Clinical Laboratories | COVID-19 Genomics UK (COG-UK) Consortium | Sam Haldenby, Anita Lucaci, Steve Paterson, Julian Hiscox, Alistair Darby, M Almsaud, A Alrezaihi, Muhannad Alruwaili, Stuart D Armstrong, Jones Benjamin, Eleanor G Bentley, Anu Chawla, Jordan J Clark, Angela Cowell, Richard Eccles, Isabel García-Dorival, Matthew Gemmell, Alessandro Gerada, PKF Gilmore, Richard Gregory, Ximeng Han, Catherine Hartley, Margaret Hughes, Miren Iturriza-Gomara, James Johnson, L Luu, Jenifer Manson, Charlotte Nelson, Elaine O'Toole, Cassie Olateju, Rebekah Penrice-Randal , Lucille Rainbow, N.P Randle, Trevor Ian Robinson, Parul Sharma, Ghada T Shawli, James P Stewart, Neil Swainston, Ecaterina Vamos, Joanne Watts, Mark Whitehead |
| EPI_ISL_705151, EPI_ISL_705152, EPI_ISL_705154, EPI_ISL_705155, | Lighthouse Lab in Cambridge | Wellcome Sanger Institute for the COVID-19 Genomics UK (COG-UK) Consortium | Rob Howes, The Lighthouse Lab in Cambridge and Alex Alderton, Roberto Amato, Sonia Goncalves, Ewan Harrison, David K. Jackson, Ian Johnston, Dominic Kwiatkowski, Cordelia Langford, John Sillitoe on behalf of the Wellcome Sanger Institute COVID-19 Surveillance Team |

| | | | |
|---|---|---|---|
| EPI_ISL_705157, EPI_ISL_705159, EPI_ISL_705160, EPI_ISL_705162 | | | |
| EPI_ISL_705163 | Virology Department, Sheffield Teaching Hospitals NHS Foundation Trust/Department of Infection, Immunity and Cardiovascular Disease, The Medical School, University of Sheffield | COVID-19 Genomics UK (COG-UK) Consortium | Thushan de Silva, Matthew Parker, Nikki Smith, Adri Angyal, Rebecca Brown, Luke Green, Rachel Tucker, Paul Parsons, Danielle Groves, Katie Johnson, Laura Carrilero, Alex Keeley, Dave Partridge, Matthew Wyles, Benjamin Lindsey, Mehmet Yavuz, Mohammad Raza, Cariad Evans |
| EPI_ISL_705164, EPI_ISL_705165, EPI_ISL_705167, EPI_ISL_705169, EPI_ISL_705170, EPI_ISL_705172, EPI_ISL_705173, EPI_ISL_705175, EPI_ISL_705177, EPI_ISL_705178, EPI_ISL_705180, EPI_ISL_705181, EPI_ISL_705183, EPI_ISL_705185, EPI_ISL_705186, EPI_ISL_705188, EPI_ISL_705189 | | | |
| see above | Lighthouse Lab in Cambridge | Wellcome Sanger Institute for the COVID-19 Genomics UK (COG-UK) Consortium | Rob Howes, The Lighthouse Lab in Cambridge and Alex Alderton, Roberto Amato, Sonia Goncalves, Ewan Harrison, David K. Jackson, Ian Johnston, Dominic Kwiatkowski, Cordelia Langford, John Sillitoe on behalf of the Wellcome Sanger Institute COVID-19 Surveillance Team |
| EPI_ISL_705195 | Virology Department, Sheffield Teaching Hospitals NHS Foundation Trust/Department of Infection, Immunity and Cardiovascular Disease, The Medical School, University of Sheffield | COVID-19 Genomics UK (COG-UK) Consortium | Thushan de Silva, Matthew Parker, Nikki Smith, Adri Angyal, Rebecca Brown, Luke Green, Rachel Tucker, Paul Parsons, Danielle Groves, Katie Johnson, Laura Carrilero, Alex Keeley, Dave Partridge, Matthew Wyles, Benjamin Lindsey, Mehmet Yavuz, Mohammad Raza, Cariad Evans |
| EPI_ISL_705201 | Northumbria University / South Tees Hospitals NHS Foundation Trust / North Cumbria Integrated Care NHS Foundation Trust / North Tees and Hartlepool NHS Foundation Trust / Newcastle Hospitals NHS Foundation Trust | COVID-19 Genomics UK (COG-UK) Consortium | Darren L Smith,Andrew Nelson,Matthew Bashton,Greg R Young,Joshua Loh,John Allan,Mohammad A Tariq,Giles S Holt,Gary Black,Wen C Yew,Lynn Dover,Paul Baker,Steve Liggett,Sarah Essex,Jane Greenaway,Debra Padgett,Clive Graham,Garren Scott,Edward Barton,Emma Swindells,Brendan Payne,Jennifer Collins,Yusri Taha,Gary Eltringham |
| EPI_ISL_705211 | Quadram Institute Bioscience | COVID-19 Genomics UK (COG-UK) Consortium | Dave J. Baker, Gemma L. Kay, Alp Aydin, Thanh Le-Viet, Steven Rudder, Ana P. Tedim, Anastasia Kolyva, Maria Diaz, Leonardo de Oliveira Martins, Nabil-Fareed Alikhan, Lizzie Meadows, Rachael Stanley, Ngozi Elumogo, Muhammed Yasir, Nicholas M. Thomson, Alexander J Trotter, Rachel Gilroy, Samuel Bloomfield, Claire Stuart, Andrew Bell, Reenesh Prakash, Samir Dervisevic, Alison E. Mather, John Wain, Mark Webber, Andrew J. Page, Justin O'Grady |
| EPI_ISL_705215, EPI_ISL_705216 | University College London, Great Ormond Street Hospital for Children NHS Foundation Trust, Imperial College Healthcare NHS Trust | COVID-19 Genomics UK (COG-UK) Consortium | Sergi Castellano, Rachel Williams, Mark Kristiansen, Paola Resende Silva, Sunando Roy, Tony Brooks, Helena Tutill, Paola Niola, Patricia Dyal, Charlotte Williams, Leysa Forrest, Yasmin Panchbhaya, Jacqueline Findlay, Samuel Weeks, Julianne Brown, Kathryn Harris, Paul Randell, James Price, Alison Holmes, Judith Breuer |
| EPI_ISL_705217 | Department of Pathology, University of Cambridge | COVID-19 Genomics UK (COG-UK) Consortium | Aminu S. Jahun, Yasmin Chaudhry, Grant Hall, Iliana Georgana, Myra Hosmillo, Martin D. Curran, Malte Pinckert, Surendra Parmar, Ian Goodfellow |
| EPI_ISL_705228 | Quadram Institute Bioscience | COVID-19 Genomics UK (COG-UK) Consortium | Dave J. Baker, Gemma L. Kay, Alp Aydin, Thanh Le-Viet, Steven Rudder, Ana P. Tedim, Anastasia Kolyva, Maria Diaz, Leonardo de Oliveira Martins, Nabil-Fareed Alikhan, Lizzie Meadows, Rachael Stanley, Ngozi Elumogo, Muhammed Yasir, Nicholas M. Thomson, Alexander J Trotter, Rachel Gilroy, Samuel Bloomfield, Claire Stuart, Andrew Bell, Reenesh Prakash, Samir Dervisevic, Alison E. Mather, John Wain, Mark Webber, Andrew J. Page, Justin O'Grady |
| EPI_ISL_705237 | University of Exeter | COVID-19 Genomics UK (COG-UK) Consortium | Ben Temperton,Aaron Jeffries,Michelle Michelsen,Joanna Warwick-Dugdale,Audrey Farbos,Robyn Manley,Stephen Michell,Jane Masoli |
| EPI_ISL_705241, EPI_ISL_705248 | Liverpool Clinical Laboratories | COVID-19 Genomics UK (COG-UK) Consortium | Sam Haldenby, Anita Lucaci, Steve Paterson, Julian Hiscox, Alistair Darby, M Almsaud, A Alrezaihi, Muhannad Alruwaili, Stuart D Armstrong, Jones Benjamin, Eleanor G Bentley, Anu Chawla, Jordan J Clark, Angela Cowell, Richard Eccles, Isabel García-Dorival, Matthew Gemmell, Alessandro Gerada, PKF Gilmore, Richard Gregory, Ximeng Han, Catherine Hartley, Margaret Hughes, Miren Iturriza-Gomara, James Johnson, L Luu, Jenifer Manson, Charlotte Nelson, Elaine O'Toole, Cassie Olateju, Rebekah Penrice-Randal , Lucille Rainbow, N.P Randle, Trevor Ian Robinson, Parul Sharma, Ghada T Shawli, James P Stewart, Neil Swainston, Ecaterina Vamos, Joanne Watts, Mark Whitehead |
| EPI_ISL_705254 | University of Exeter | COVID-19 Genomics UK (COG-UK) Consortium | Ben Temperton,Aaron Jeffries,Michelle Michelsen,Joanna Warwick-Dugdale,Audrey Farbos,Robyn Manley,Stephen Michell,Jane Masoli |
| EPI_ISL_705262 | Quadram Institute Bioscience | COVID-19 Genomics UK (COG-UK) Consortium | Dave J. Baker, Gemma L. Kay, Alp Aydin, Thanh Le-Viet, Steven Rudder, Ana P. Tedim, Anastasia Kolyva, Maria Diaz, Leonardo de Oliveira Martins, Nabil-Fareed Alikhan, Lizzie Meadows, Rachael Stanley, Ngozi Elumogo, Muhammed Yasir, Nicholas M. Thomson, Alexander J Trotter, Rachel Gilroy, Samuel Bloomfield, Claire Stuart, Andrew Bell, Reenesh Prakash, Samir Dervisevic, Alison E. Mather, John Wain, Mark Webber, Andrew J. Page, Justin O'Grady |
| EPI_ISL_705269 | Liverpool Clinical Laboratories | COVID-19 Genomics UK (COG-UK) Consortium | Sam Haldenby, Anita Lucaci, Steve Paterson, Julian Hiscox, Alistair Darby, M Almsaud, A Alrezaihi, Muhannad Alruwaili, Stuart D Armstrong, Jones Benjamin, Eleanor G Bentley, Anu Chawla, Jordan J Clark, Angela Cowell, Richard Eccles, Isabel García-Dorival, Matthew Gemmell, Alessandro Gerada, PKF Gilmore, Richard Gregory, Ximeng Han, Catherine Hartley, Margaret Hughes, Miren Iturriza-Gomara, James Johnson, L Luu, Jenifer Manson, Charlotte Nelson, Elaine O'Toole, Cassie Olateju, Rebekah Penrice-Randal , Lucille Rainbow, N.P Randle, Trevor Ian Robinson, Parul Sharma, Ghada T Shawli, James P Stewart, Neil Swainston, Ecaterina Vamos, Joanne Watts, Mark Whitehead |
| EPI_ISL_705270 | Virology Department, Sheffield Teaching Hospitals NHS Foundation Trust/Department of Infection, Immunity and Cardiovascular Disease, The Medical School, University of Sheffield | COVID-19 Genomics UK (COG-UK) Consortium | Thushan de Silva, Matthew Parker, Nikki Smith, Adri Angyal, Rebecca Brown, Luke Green, Rachel Tucker, Paul Parsons, Danielle Groves, Katie Johnson, Laura Carrilero, Alex Keeley, Dave Partridge, Matthew Wyles, Benjamin Lindsey, Mehmet Yavuz, Mohammad Raza, Cariad Evans |
| EPI_ISL_705273 | Northumbria University / South Tees Hospitals NHS Foundation Trust / North Cumbria Integrated Care NHS Foundation Trust / North Tees and Hartlepool NHS Foundation Trust / Newcastle Hospitals NHS Foundation Trust | COVID-19 Genomics UK (COG-UK) Consortium | Darren L Smith,Andrew Nelson,Matthew Bashton,Greg R Young,Joshua Loh,John Allan,Mohammad A Tariq,Giles S Holt,Gary Black,Wen C Yew,Lynn Dover,Paul Baker,Steve Liggett,Sarah Essex,Jane Greenaway,Debra Padgett,Clive Graham,Garren Scott,Edward Barton,Emma Swindells,Brendan Payne,Jennifer Collins,Yusri Taha,Gary Eltringham |
| EPI_ISL_705274 | Liverpool Clinical Laboratories | COVID-19 Genomics UK (COG-UK) Consortium | Sam Haldenby, Anita Lucaci, Steve Paterson, Julian Hiscox, Alistair Darby, M Almsaud, A Alrezaihi, Muhannad Alruwaili, Stuart D Armstrong, Jones Benjamin, Eleanor G Bentley, Anu Chawla, Jordan J Clark, Angela Cowell, Richard Eccles, Isabel García-Dorival, Matthew Gemmell, Alessandro Gerada, PKF Gilmore, Richard Gregory, Ximeng Han, Catherine Hartley, Margaret Hughes, Miren Iturriza-Gomara, James Johnson, L Luu, Jenifer Manson, Charlotte Nelson, Elaine O'Toole, Cassie Olateju, Rebekah Penrice-Randal , Lucille Rainbow, N.P Randle, Trevor Ian Robinson, Parul Sharma, Ghada T Shawli, James P Stewart, Neil Swainston, Ecaterina Vamos, Joanne Watts, Mark Whitehead |
| EPI_ISL_705278 | University College London, Great Ormond Street Hospital for Children NHS Foundation Trust, Imperial College Healthcare NHS Trust | COVID-19 Genomics UK (COG-UK) Consortium | Sergi Castellano, Rachel Williams, Mark Kristiansen, Paola Resende Silva, Sunando Roy, Tony Brooks, Helena Tutill, Paola Niola, Patricia Dyal, Charlotte Williams, Leysa Forrest, Yasmin Panchbhaya, Jacqueline Findlay, Samuel Weeks, Julianne Brown, Kathryn Harris, Paul Randell, James Price, Alison Holmes, Judith Breuer |
| EPI_ISL_705286 | Liverpool Clinical Laboratories | COVID-19 Genomics UK (COG-UK) Consortium | Sam Haldenby, Anita Lucaci, Steve Paterson, Julian Hiscox, Alistair Darby, M Almsaud, A Alrezaihi, Muhannad Alruwaili, Stuart D Armstrong, Jones Benjamin, Eleanor G Bentley, Anu Chawla, Jordan J Clark, Angela Cowell, Richard Eccles, Isabel García-Dorival, Matthew Gemmell, Alessandro Gerada, PKF Gilmore, Richard Gregory, Ximeng Han, Catherine Hartley, Margaret Hughes, Miren Iturriza-Gomara, James Johnson, L Luu, Jenifer Manson, Charlotte Nelson, Elaine O'Toole, Cassie Olateju, Rebekah Penrice-Randal , Lucille Rainbow, N.P Randle, Trevor Ian Robinson, Parul Sharma, Ghada T Shawli, James P Stewart, Neil Swainston, Ecaterina Vamos, Joanne Watts, Mark Whitehead |
| EPI_ISL_705289 | Department of Pathology, University of Cambridge | COVID-19 Genomics UK (COG-UK) Consortium | Aminu S. Jahun, Yasmin Chaudhry, Grant Hall, Iliana Georgana, Myra Hosmillo, Martin D. Curran, Malte Pinckert, Surendra Parmar, Ian Goodfellow |
| EPI_ISL_705299 | Liverpool Clinical Laboratories | COVID-19 Genomics UK (COG-UK) Consortium | Sam Haldenby, Anita Lucaci, Steve Paterson, Julian Hiscox, Alistair Darby, M Almsaud, A Alrezaihi, Muhannad Alruwaili, Stuart D Armstrong, Jones Benjamin, Eleanor G Bentley, Anu Chawla, Jordan J Clark, Angela Cowell, Richard Eccles, Isabel García-Dorival, Matthew Gemmell, Alessandro Gerada, PKF Gilmore, Richard Gregory, Ximeng Han, Catherine Hartley, Margaret Hughes, Miren Iturriza-Gomara, James Johnson, L Luu, Jenifer Manson, Charlotte Nelson, Elaine O'Toole, Cassie Olateju, Rebekah Penrice-Randal , Lucille Rainbow, N.P Randle, Trevor Ian Robinson, Parul Sharma, Ghada T Shawli, James P Stewart, Neil Swainston, Ecaterina Vamos, Joanne Watts, Mark Whitehead |
| EPI_ISL_705300 | West of Scotland Specialist Virology Centre, NHSGGC / MRC-University of Glasgow Centre for Virus Research | COVID-19 Genomics UK (COG-UK) Consortium | Ana da Silva Filipe, Natasha Johnson, Kathy Smollett, Daniel Mair, Stephen Carmichael, Alice Broos, Lily Tong, Jenna Nichols, Kyriaki Nomikou; Sarah McDonald; Richard Orton, Joseph Hughes, Sreenu Vattipally, David L Robertson; Alasdair MacLean, Rory Gunson; Sharif Shaaban, Matthew Holden; Rachel Blacow, Guy Mollett, Kathy Li, James Shepherd, Antonia Ho, Emma Thomson |
| EPI_ISL_705313 | Liverpool Clinical Laboratories | COVID-19 Genomics UK (COG-UK) Consortium | Sam Haldenby, Anita Lucaci, Steve Paterson, Julian Hiscox, Alistair Darby, M Almsaud, A Alrezaihi, Muhannad Alruwaili, Stuart D Armstrong, Jones |

| | | | |
|---|---|---|---|
| | | | Benjamin, Eleanor G Bentley, Anu Chawla, Jordan J Clark, Angela Cowell, Richard Eccles, Isabel García-Dorival, Matthew Gemmell, Alessandro Gerada, PKF Gilmore, Richard Gregory, Ximeng Han, Catherine Hartley, Margaret Hughes, Miren Iturriza-Gomara, James Johnson, L Luu, Jenifer Manson, Charlotte Nelson, Elaine O'Toole, Cassie Olateju, Rebekah Penrice-Randal , Lucille Rainbow, N.P Randle, Trevor Ian Robinson, Parul Sharma, Ghada T Shawli, James P Stewart, Neil Swainston, Ecaterina Vamos, Joanne Watts, Mark Whitehead |
| EPI_ISL_705317, EPI_ISL_705319 | Virology Department, Sheffield Teaching Hospitals NHS Foundation Trust/Department of Infection, Immunity and Cardiovascular Disease, The Medical School, University of Sheffield | COVID-19 Genomics UK (COG-UK) Consortium | Thushan de Silva, Matthew Parker, Nikki Smith, Adri Angyal, Rebecca Brown, Luke Green, Rachel Tucker, Paul Parsons, Danielle Groves, Katie Johnson, Laura Carrilero, Alex Keeley, Dave Partridge, Matthew Wyles, Benjamin Lindsey, Mehmet Yavuz, Mohammad Raza, Cariad Evans |
| EPI_ISL_705334 | Liverpool Clinical Laboratories | COVID-19 Genomics UK (COG-UK) Consortium | Sam Haldenby, Anita Lucaci, Steve Paterson, Julian Hiscox, Alistair Darby, M Almsaud, A Alrezaihi, Muhannad Alruwaili, Stuart D Armstrong, Jones Benjamin, Eleanor G Bentley, Anu Chawla, Jordan J Clark, Angela Cowell, Richard Eccles, Isabel García-Dorival, Matthew Gemmell, Alessandro Gerada, PKF Gilmore, Richard Gregory, Ximeng Han, Catherine Hartley, Margaret Hughes, Miren Iturriza-Gomara, James Johnson, L Luu, Jenifer Manson, Charlotte Nelson, Elaine O'Toole, Cassie Olateju, Rebekah Penrice-Randal , Lucille Rainbow, N.P Randle, Trevor Ian Robinson, Parul Sharma, Ghada T Shawli, James P Stewart, Neil Swainston, Ecaterina Vamos, Joanne Watts, Mark Whitehead |
| EPI_ISL_705335 | Virology Department, Sheffield Teaching Hospitals NHS Foundation Trust/Department of Infection, Immunity and Cardiovascular Disease, The Medical School, University of Sheffield | COVID-19 Genomics UK (COG-UK) Consortium | Thushan de Silva, Matthew Parker, Nikki Smith, Adri Angyal, Rebecca Brown, Luke Green, Rachel Tucker, Paul Parsons, Danielle Groves, Katie Johnson, Laura Carrilero, Alex Keeley, Dave Partridge, Matthew Wyles, Benjamin Lindsey, Mehmet Yavuz, Mohammad Raza, Cariad Evans |
| EPI_ISL_705339 | Department of Pathology, University of Cambridge | COVID-19 Genomics UK (COG-UK) Consortium | Aminu S. Jahun, Yasmin Chaudhry, Grant Hall, Iliana Georgana, Myra Hosmillo, Martin D. Curran, Malte Pinckert, Surendra Parmar, Ian Goodfellow |
| EPI_ISL_705356 | Northumbria University / South Tees Hospitals NHS Foundation Trust / North Cumbria Integrated Care NHS Foundation Trust / North Tees and Hartlepool NHS Foundation Trust / Newcastle Hospitals NHS Foundation Trust | COVID-19 Genomics UK (COG-UK) Consortium | Darren L Smith,Andrew Nelson,Matthew Bashton,Greg R Young,Joshua Loh,John Allan,Mohammad A Tariq,Giles S Holt,Gary Black,Wen C Yew,Lynn Dover,Paul Baker,Steve Liggett,Sarah Essex,Jane Greenaway,Debra Padgett,Clive Graham,Garren Scott,Edward Barton,Emma Swindells,Brendan Payne,Jennifer Collins,Yusri Taha,Gary Eltringham |
| EPI_ISL_705371 | Virology Department, Sheffield Teaching Hospitals NHS Foundation Trust/Department of Infection, Immunity and Cardiovascular Disease, The Medical School, University of Sheffield | COVID-19 Genomics UK (COG-UK) Consortium | Thushan de Silva, Matthew Parker, Nikki Smith, Adri Angyal, Rebecca Brown, Luke Green, Rachel Tucker, Paul Parsons, Danielle Groves, Katie Johnson, Laura Carrilero, Alex Keeley, Dave Partridge, Matthew Wyles, Benjamin Lindsey, Mehmet Yavuz, Mohammad Raza, Cariad Evans |
| EPI_ISL_705373 | University of Exeter | COVID-19 Genomics UK (COG-UK) Consortium | Ben Temperton,Aaron Jeffries,Michelle Michelsen,Joanna Warwick-Dugdale,Audrey Farbos,Robyn Manley,Stephen Michell,Jane Masoli |
| EPI_ISL_705375 | Quadram Institute Bioscience | COVID-19 Genomics UK (COG-UK) Consortium | Dave J. Baker, Gemma L. Kay, Alp Aydin, Thanh Le-Viet, Steven Rudder, Ana P. Tedim, Anastasia Kolyva, Maria Diaz, Leonardo de Oliveira Martins, Nabil-Fareed Alikhan, Lizzie Meadows, Rachael Stanley, Ngozi Elumogo, Muhammed Yasir, Nicholas M. Thomson, Alexander J Trotter, Rachel Gilroy, Samuel Bloomfield, Claire Stuart, Andrew Bell, Reenesh Prakash, Samir Dervisevic, Alison E. Mather, John Wain, Mark Webber, Andrew J. Page, Justin O'Grady |
| EPI_ISL_705376 | Department of Pathology, University of Cambridge | COVID-19 Genomics UK (COG-UK) Consortium | Aminu S. Jahun, Yasmin Chaudhry, Grant Hall, Iliana Georgana, Myra Hosmillo, Martin D. Curran, Malte Pinckert, Surendra Parmar, Ian Goodfellow |
| EPI_ISL_705383 | University of Exeter | COVID-19 Genomics UK (COG-UK) Consortium | Ben Temperton,Aaron Jeffries,Michelle Michelsen,Joanna Warwick-Dugdale,Audrey Farbos,Robyn Manley,Stephen Michell,Jane Masoli |
| EPI_ISL_705398 | Liverpool Clinical Laboratories | COVID-19 Genomics UK (COG-UK) Consortium | Sam Haldenby, Anita Lucaci, Steve Paterson, Julian Hiscox, Alistair Darby, M Almsaud, A Alrezaihi, Muhannad Alruwaili, Stuart D Armstrong, Jones Benjamin, Eleanor G Bentley, Anu Chawla, Jordan J Clark, Angela Cowell, Richard Eccles, Isabel García-Dorival, Matthew Gemmell, Alessandro Gerada, PKF Gilmore, Richard Gregory, Ximeng Han, Catherine Hartley, Margaret Hughes, Miren Iturriza-Gomara, James Johnson, L Luu, Jenifer Manson, Charlotte Nelson, Elaine O'Toole, Cassie Olateju, Rebekah Penrice-Randal , Lucille Rainbow, N.P Randle, Trevor Ian Robinson, Parul Sharma, Ghada T Shawli, James P Stewart, Neil Swainston, Ecaterina Vamos, Joanne Watts, Mark Whitehead |
| EPI_ISL_705405 | Virology Department, Sheffield Teaching Hospitals NHS Foundation Trust/Department of Infection, Immunity and Cardiovascular Disease, The Medical School, University of Sheffield | COVID-19 Genomics UK (COG-UK) Consortium | Thushan de Silva, Matthew Parker, Nikki Smith, Adri Angyal, Rebecca Brown, Luke Green, Rachel Tucker, Paul Parsons, Danielle Groves, Katie Johnson, Laura Carrilero, Alex Keeley, Dave Partridge, Matthew Wyles, Benjamin Lindsey, Mehmet Yavuz, Mohammad Raza, Cariad Evans |
| EPI_ISL_705410 | Department of Pathology, University of Cambridge | COVID-19 Genomics UK (COG-UK) Consortium | Aminu S. Jahun, Yasmin Chaudhry, Grant Hall, Iliana Georgana, Myra Hosmillo, Martin D. Curran, Malte Pinckert, Surendra Parmar, Ian Goodfellow |
| EPI_ISL_705471 | Liverpool Clinical Laboratories | COVID-19 Genomics UK (COG-UK) Consortium | Sam Haldenby, Anita Lucaci, Steve Paterson, Julian Hiscox, Alistair Darby, M Almsaud, A Alrezaihi, Muhannad Alruwaili, Stuart D Armstrong, Jones Benjamin, Eleanor G Bentley, Anu Chawla, Jordan J Clark, Angela Cowell, Richard Eccles, Isabel García-Dorival, Matthew Gemmell, Alessandro Gerada, PKF Gilmore, Richard Gregory, Ximeng Han, Catherine Hartley, Margaret Hughes, Miren Iturriza-Gomara, James Johnson, L Luu, Jenifer Manson, Charlotte Nelson, Elaine O'Toole, Cassie Olateju, Rebekah Penrice-Randal , Lucille Rainbow, N.P Randle, Trevor Ian Robinson, Parul Sharma, Ghada T Shawli, James P Stewart, Neil Swainston, Ecaterina Vamos, Joanne Watts, Mark Whitehead |
| EPI_ISL_705472 | University College London Hospital | COVID-19 Genomics UK (COG-UK) Consortium | Judith Heaney, Matthew Byott, Catherine Houlihan, Dan Frampton, Stuart Kirk, Moira Spyer and Eleni Nastouli |
| EPI_ISL_705486 | Northumbria University / South Tees Hospitals NHS Foundation Trust / North Cumbria Integrated Care NHS Foundation Trust / North Tees and Hartlepool NHS Foundation Trust / Newcastle Hospitals NHS Foundation Trust | COVID-19 Genomics UK (COG-UK) Consortium | Darren L Smith,Andrew Nelson,Matthew Bashton,Greg R Young,Joshua Loh,John Allan,Mohammad A Tariq,Giles S Holt,Gary Black,Wen C Yew,Lynn Dover,Paul Baker,Steve Liggett,Sarah Essex,Jane Greenaway,Debra Padgett,Clive Graham,Garren Scott,Edward Barton,Emma Swindells,Brendan Payne,Jennifer Collins,Yusri Taha,Gary Eltringham |
| EPI_ISL_705487 | Queens Medical Centre, Clinical Microbiology Department / DeepSeq Nottingham | COVID-19 Genomics UK (COG-UK) Consortium | Gemma Clark, Wendy Smith, Manjinder Khakh, Vicki M Fleming, Michelle M Lister, Hannah Howson-Wells, Jonathan Ball, Patrick McClure, Joseph Chappell, Theocharis Tsoleridis, Nadine Holmes, Matthew Carlisle, Christopher Moore, Fei Sang, Johnny Debebe, Victoria Wright, Matthew Loose |
| EPI_ISL_705491 | Quadram Institute Bioscience | COVID-19 Genomics UK (COG-UK) Consortium | Dave J. Baker, Gemma L. Kay, Alp Aydin, Thanh Le-Viet, Steven Rudder, Ana P. Tedim, Anastasia Kolyva, Maria Diaz, Leonardo de Oliveira Martins, Nabil-Fareed Alikhan, Lizzie Meadows, Rachael Stanley, Ngozi Elumogo, Muhammed Yasir, Nicholas M. Thomson, Alexander J Trotter, Rachel Gilroy, Samuel Bloomfield, Claire Stuart, Andrew Bell, Reenesh Prakash, Samir Dervisevic, Alison E. Mather, John Wain, Mark Webber, Andrew J. Page, Justin O'Grady |
| EPI_ISL_705494, EPI_ISL_705495 | Northumbria University / South Tees Hospitals NHS Foundation Trust / North Cumbria Integrated Care NHS Foundation Trust / North Tees and Hartlepool NHS Foundation Trust / Newcastle Hospitals NHS Foundation Trust | COVID-19 Genomics UK (COG-UK) Consortium | Darren L Smith,Andrew Nelson,Matthew Bashton,Greg R Young,Joshua Loh,John Allan,Mohammad A Tariq,Giles S Holt,Gary Black,Wen C Yew,Lynn Dover,Paul Baker,Steve Liggett,Sarah Essex,Jane Greenaway,Debra Padgett,Clive Graham,Garren Scott,Edward Barton,Emma Swindells,Brendan Payne,Jennifer Collins,Yusri Taha,Gary Eltringham |
| EPI_ISL_705551, EPI_ISL_705557, EPI_ISL_705559, EPI_ISL_705561, EPI_ISL_705562, EPI_ISL_705563, EPI_ISL_705564, EPI_ISL_705565, EPI_ISL_705566 | Department of Pathology, University of Cambridge | COVID-19 Genomics UK (COG-UK) Consortium | Aminu S. Jahun, Yasmin Chaudhry, Grant Hall, Iliana Georgana, Myra Hosmillo, Martin D. Curran, Malte Pinckert, Surendra Parmar, Ian Goodfellow |
| EPI_ISL_705567, EPI_ISL_705568, EPI_ISL_705569, EPI_ISL_705570, EPI_ISL_705571, EPI_ISL_705573, EPI_ISL_705574, EPI_ISL_705575, EPI_ISL_705576, EPI_ISL_705577, EPI_ISL_705578, EPI_ISL_705579, EPI_ISL_705580, EPI_ISL_705581, EPI_ISL_705582, EPI_ISL_705583, EPI_ISL_705584, EPI_ISL_705585, EPI_ISL_705586, EPI_ISL_705587, EPI_ISL_705588, EPI_ISL_705589, EPI_ISL_705590, EPI_ISL_705591, EPI_ISL_705592, EPI_ISL_705595, EPI_ISL_705596, EPI_ISL_705597, EPI_ISL_705598, EPI_ISL_705599, EPI_ISL_705600, EPI_ISL_705601, EPI_ISL_705602, EPI_ISL_705603, EPI_ISL_705604, EPI_ISL_705605, EPI_ISL_705606, EPI_ISL_705607, EPI_ISL_705608, EPI_ISL_705609, EPI_ISL_705610, EPI_ISL_705611, EPI_ISL_705612 | | | |
| see above | Quadram Institute Bioscience | COVID-19 Genomics UK (COG-UK) Consortium | Dave J. Baker, Gemma L. Kay, Alp Aydin, Thanh Le-Viet, Steven Rudder, Ana P. Tedim, Anastasia Kolyva, Maria Diaz, Leonardo de Oliveira Martins, Nabil-Fareed Alikhan, Lizzie Meadows, Rachael Stanley, Ngozi Elumogo, Muhammed Yasir, Nicholas M. Thomson, Alexander J Trotter, Rachel Gilroy, Samuel Bloomfield, Claire Stuart, Andrew Bell, Reenesh Prakash, Samir Dervisevic, Alison E. Mather, John Wain, Mark Webber, Andrew J. Page, Justin O'Grady |
| EPI_ISL_705613, EPI_ISL_705614 | Queens Medical Centre, Clinical Microbiology Department / | COVID-19 Genomics UK (COG-UK) Consortium | Gemma Clark, Wendy Smith, Manjinder Khakh, Vicki M Fleming, Michelle M Lister, Hannah Howson-Wells, Jonathan Ball, Patrick McClure, Joseph |

| | | | |
|---|---|---|---|
| | DeepSeq Nottingham | | Chappell, Theocharis Tsoleridis, Nadine Holmes, Matthew Carlisle, Christopher Moore, Fei Sang, Johnny Debebe, Victoria Wright, Matthew Loose |
| EPI_ISL_705615, EPI_ISL_705616, EPI_ISL_705617, EPI_ISL_705618, EPI_ISL_705619, EPI_ISL_705620, EPI_ISL_705621, EPI_ISL_705622, EPI_ISL_705623, EPI_ISL_705624, EPI_ISL_705625 | | | |
| see above | Virology Department, Sheffield Teaching Hospitals NHS Foundation Trust/Department of Infection, Immunity and Cardiovascular Disease, The Medical School, University of Sheffield | COVID-19 Genomics UK (COG-UK) Consortium | Thushan de Silva, Matthew Parker, Nikki Smith, Adri Angyal, Rebecca Brown, Luke Green, Rachel Tucker, Paul Parsons, Danielle Groves, Katie Johnson, Laura Carrilero, Alex Keeley, Dave Partridge, Matthew Wyles, Benjamin Lindsey, Mehmet Yavuz, Mohammad Raza, Cariad Evans |
| EPI_ISL_705629, EPI_ISL_705630, EPI_ISL_705631, EPI_ISL_705632, EPI_ISL_705633, EPI_ISL_705634, EPI_ISL_705635 | Department of Pathology, University of Cambridge | COVID-19 Genomics UK (COG-UK) Consortium | Aminu S. Jahun, Yasmin Chaudhry, Grant Hall, Iliana Georgana, Myra Hosmillo, Martin D. Curran, Malte Pinckert, Surendra Parmar, Ian Goodfellow |
| EPI_ISL_705667, EPI_ISL_705668, EPI_ISL_705669, EPI_ISL_705670, EPI_ISL_705671, EPI_ISL_705672, EPI_ISL_705673, EPI_ISL_705674, EPI_ISL_705675, EPI_ISL_705676, EPI_ISL_705677, EPI_ISL_705678, EPI_ISL_705679, EPI_ISL_705680, EPI_ISL_705681, EPI_ISL_705682, EPI_ISL_705683, EPI_ISL_705684, EPI_ISL_705685, EPI_ISL_705686, EPI_ISL_705687, EPI_ISL_705688, EPI_ISL_705689, EPI_ISL_705690, EPI_ISL_705691, EPI_ISL_705692, EPI_ISL_705693, EPI_ISL_705694, EPI_ISL_705695, EPI_ISL_705696, EPI_ISL_705697, EPI_ISL_705701, EPI_ISL_705709, EPI_ISL_705729 | | | |
| see above | West of Scotland Specialist Virology Centre, NHSGGC / MRC-University of Glasgow Centre for Virus Research | COVID-19 Genomics UK (COG-UK) Consortium | Ana da Silva Filipe, Natasha Johnson, Kathy Smollett, Daniel Mair, Stephen Carmichael, Alice Broos, Lily Tong, Jenna Nichols, Kyriaki Nomikou; Sarah McDonald; Richard Orton, Joseph Hughes, Sreenu Vattipally, David L Robertson; Alasdair MacLean, Rory Gunson; Sharif Shaaban, Matthew Holden; Rachel Blacow, Guy Mollett, Kathy Li, James Shepherd, Antonia Ho, Emma Thomson |
| EPI_ISL_705737, EPI_ISL_705738 | Virology Department, Royal Infirmary of Edinburgh, NHS Lothian / School of Biological Sciences, University of Edinburgh / Institute of Genetics and Molecular Medicine, University of Edinburgh | COVID-19 Genomics UK (COG-UK) Consortium | McHugh M, Dewar R, Rooke S, Gallagher M, Balcaza C, O'Toole Á, Scher E, Hill V, McCrone JT, Colquhoun R, Yu X, Jackson B, Rambaut A, Williams TC, Templeton K |
| EPI_ISL_705809, EPI_ISL_705811, EPI_ISL_705813, EPI_ISL_705814, EPI_ISL_705815, EPI_ISL_705816, EPI_ISL_705817, EPI_ISL_705819, EPI_ISL_705820, EPI_ISL_705821, EPI_ISL_705822, EPI_ISL_705823, EPI_ISL_705826, EPI_ISL_705829, EPI_ISL_705831, EPI_ISL_705836, EPI_ISL_705838, EPI_ISL_705859, EPI_ISL_705860, EPI_ISL_705861, EPI_ISL_705862, EPI_ISL_705863, EPI_ISL_705864, EPI_ISL_705925 | | | |
| see above | Liverpool Clinical Laboratories | COVID-19 Genomics UK (COG-UK) Consortium | Sam Haldenby, Anita Lucaci, Steve Paterson, Julian Hiscox, Alistair Darby, M Almsaud, A Alrezaihi, Muhannad Alruwaili, Stuart D Armstrong, Jones Benjamin, Eleanor G Bentley, Anu Chawla, Jordan J Clark, Angela Cowell, Richard Eccles, Isabel García-Dorival, Matthew Gemmell, Alessandro Gerada, PKF Gilmore, Richard Gregory, Ximeng Han, Catherine Hartley, Margaret Hughes, Miren Iturriza-Gomara, James Johnson, L Luu, Jenifer Manson, Charlotte Nelson, Elaine O'Toole, Cassie Olateju, Rebekah Penrice-Randal , Lucille Rainbow, N.P Randle, Trevor Ian Robinson, Parul Sharma, Ghada T Shawli, James P Stewart, Neil Swainston, Ecaterina Vamos, Joanne Watts, Mark Whitehead |
| EPI_ISL_705942, EPI_ISL_705943, EPI_ISL_705944 | University College London Hospital | COVID-19 Genomics UK (COG-UK) Consortium | Judith Heaney, Matthew Byott, Catherine Houlihan, Dan Frampton, Stuart Kirk, Moira Spyer and Eleni Nastouli |
| EPI_ISL_706320, EPI_ISL_706321, EPI_ISL_706323, EPI_ISL_706324, EPI_ISL_706325, EPI_ISL_706326, EPI_ISL_706327, EPI_ISL_706328, EPI_ISL_706329, EPI_ISL_706330 | Northumbria University / South Tees Hospitals NHS Foundation Trust / North Cumbria Integrated Care NHS Foundation Trust / North Tees and Hartlepool NHS Foundation Trust / Newcastle Hospitals NHS Foundation Trust | COVID-19 Genomics UK (COG-UK) Consortium | Darren L Smith,Andrew Nelson,Matthew Bashton,Greg R Young,Joshua Loh,John Allan,Mohammad A Tariq,Giles S Holt,Gary Black,Wen C Yew,Lynn Dover,Paul Baker,Steve Liggett,Sarah Essex,Jane Greenaway,Debra Padgett,Clive Graham,Garren Scott,Edward Barton,Emma Swindells,Brendan Payne,Jennifer Collins,Yusri Taha,Gary Eltringham |
| EPI_ISL_706361, EPI_ISL_706362, EPI_ISL_706408 | Quadram Institute Bioscience | COVID-19 Genomics UK (COG-UK) Consortium | Dave J. Baker, Gemma L. Kay, Alp Aydin, Thanh Le-Viet, Steven Rudder, Ana P. Tedim, Anastasia Kolyva, Maria Diaz, Leonardo de Oliveira Martins, Nabil-Fareed Alikhan, Lizzie Meadows, Rachael Stanley, Ngozi Elumogo, Muhammed Yasir, Nicholas M. Thomson, Alexander J Trotter, Rachel Gilroy, Samuel Bloomfield, Claire Stuart, Andrew Bell, Reenesh Prakash, Samir Dervisevic, Alison E. Mather, John Wain, Mark Webber, Andrew J. Page, Justin O'Grady |
| EPI_ISL_706420, EPI_ISL_706421 | Queens Medical Centre, Clinical Microbiology Department / DeepSeq Nottingham | COVID-19 Genomics UK (COG-UK) Consortium | Gemma Clark, Wendy Smith, Manjinder Khakh, Vicki M Fleming, Michelle M Lister, Hannah Howson-Wells, Jonathan Ball, Patrick McClure, Joseph Chappell, Theocharis Tsoleridis, Nadine Holmes, Matthew Carlisle, Christopher Moore, Fei Sang, Johnny Debebe, Victoria Wright, Matthew Loose |
| EPI_ISL_706463 | Wales Specialist Virology Centre Sequencing lab: Pathogen Genomics Unit | COVID-19 Genomics UK (COG-UK) Consortium | Catherine Moore, Johnathan Evans, Laura Gifford, Malorie Perry, Simon Cottrell, Angela Marchbank, Alec Birchley, Alexander Adams, Amy Gaskin, Bree Gatica-Wilcox, Jason Coombes, Joel Southgate, Lauren Gilbert, Lee Graham, Nicole Pacchiarini, Sara Kumziene-Summerhayes, Sarah Taylor, Sophie Jones, Sara Rey, Matthew Bull, Joanne Watkins, Sally Corden, Tom Connor |
| EPI_ISL_706983 | Virology Department, Sheffield Teaching Hospitals NHS Foundation Trust/Department of Infection, Immunity and Cardiovascular Disease, The Medical School, University of Sheffield | COVID-19 Genomics UK (COG-UK) Consortium | Thushan de Silva, Matthew Parker, Nikki Smith, Adri Angyal, Rebecca Brown, Luke Green, Rachel Tucker, Paul Parsons, Danielle Groves, Katie Johnson, Laura Carrilero, Alex Keeley, Dave Partridge, Matthew Wyles, Benjamin Lindsey, Mehmet Yavuz, Mohammad Raza, Cariad Evans |
| EPI_ISL_707697, EPI_ISL_707698, EPI_ISL_707699, EPI_ISL_707700 | 1-Laboratory of Microbiology, National Reference Lab, Charles Nicolle Hospital; 2-University of Tunis ElManar, Faculty of Medicine of Tunis, LR99ES09, Tunis, Tunisia | 1-Clinical and Experimental Pharmacology Lab, LR16SP02, National Center of Pharmacovigilance, University of Tunis El Manar, Tunis, Tunisia. 2-Neurodegenerative diseases and psychiatric troubles, LR18SP03, Razi Hospital, University of Tunis El Manar, Tunis, Tunisia. 3- Ministry of Health, National Observatory of New and Emerging Diseases, 1006, Tunis, Tunisia | Ilhem Boutiba-Ben Boubaker, Sameh Trabelsi, Nissaf Ben Alaya, Maher Kharrat, Alia Ben Kahla, Jalila Ben Khelil, Salma Abid, Sana Ferjani, Mouna Ben Sassi, Mouna Safer, Zaineb Hamzaoui, Habiba Ben Romdhane, Souissi Amira, Rouaa Ben Othman, Hanen El Jebari, Asma Ferjani, Gaies Emna, Riadh Daghfous, Riadh Guider. |
| EPI_ISL_708485, EPI_ISL_708486, EPI_ISL_708488 | Minnesota Department of Health, Public Health Laboratory | Minnesota Department of Health, Public Health Laboratory | Alexandra Lorentz, Jacob Garfin, Matt Plumb, and Xiong Wang |
| EPI_ISL_708530 | Secretaria Municipal de Saude de Fernandópolis | Instituto Adolfo Lutz, Interdiciplinary Procedures Center, Strategic Laboratory | Claudio Tavares Sacchi, Claudia Regina Gonçalves, Erica Valessa Ramos Gomes, Carlos Henrique Camargo, Karoline Rodrigues Campos, Fernanda Modesto Tolentino Binhardi, Maricelia Navarro Pinheiro Flores, Marcia Maria Costa Nunes Soares, Janaina Other Martins Montanha |
| EPI_ISL_710424, EPI_ISL_710427, EPI_ISL_710428 | Los Angeles County PHL | Los Angeles County PHL | P. Hemarajata et al. |
| EPI_ISL_710577 | Vardcentralen Brinken | The Public Health Agency of Sweden | Department of Microbiology, The Public Health Agency of Sweden |
| EPI_ISL_710612 | Capio Bro VC | The Public Health Agency of Sweden | Department of Microbiology, The Public Health Agency of Sweden |
| EPI_ISL_710699, EPI_ISL_710700, EPI_ISL_710701, EPI_ISL_710702, EPI_ISL_710703, EPI_ISL_710704, EPI_ISL_710705, EPI_ISL_710706, EPI_ISL_710707, EPI_ISL_710708, EPI_ISL_710709, EPI_ISL_710710, EPI_ISL_710711, EPI_ISL_710712, EPI_ISL_710713, EPI_ISL_710714, EPI_ISL_710715, EPI_ISL_710716, EPI_ISL_710717, EPI_ISL_710718, EPI_ISL_710719, EPI_ISL_710720, EPI_ISL_710721, EPI_ISL_710722, EPI_ISL_710723, EPI_ISL_710724, EPI_ISL_710725, EPI_ISL_710726, EPI_ISL_710727, EPI_ISL_710728, EPI_ISL_710729, EPI_ISL_710730, EPI_ISL_710731, EPI_ISL_710732, EPI_ISL_710733, EPI_ISL_710734, EPI_ISL_710735, EPI_ISL_710736, EPI_ISL_710737, EPI_ISL_710738, EPI_ISL_710739, EPI_ISL_710740, EPI_ISL_710741, EPI_ISL_710742, EPI_ISL_710743, EPI_ISL_710744, EPI_ISL_710745, EPI_ISL_710746, EPI_ISL_710747, EPI_ISL_710749, EPI_ISL_710750, EPI_ISL_710751, EPI_ISL_710752 | | | |
| see above | Lighthouse Lab in Alderley Park | Wellcome Sanger Institute for the COVID-19 Genomics UK (COG-UK) Consortium | Jacquelyn Wynn, Mairead Hyland, The Lighthouse Lab in Alderley Park and Alex Alderton, Roberto Amato, Sonia Goncalves, Ewan Harrison, David K. Jackson, Ian Johnston, Dominic Kwiatkowski, Cordelia Langford, John Sillitoe on behalf of the Wellcome Sanger Institute COVID-19 Surveillance Team |
| EPI_ISL_710821, EPI_ISL_710822, EPI_ISL_710823, EPI_ISL_710824, EPI_ISL_710825, EPI_ISL_710826, EPI_ISL_710827, EPI_ISL_710828, EPI_ISL_710829 | Lighthouse Lab in Cambridge | Wellcome Sanger Institute for the COVID-19 Genomics UK (COG-UK) Consortium | Rob Howes, The Lighthouse Lab in Cambridge and Alex Alderton, Roberto Amato, Sonia Goncalves, Ewan Harrison, David K. Jackson, Ian Johnston, Dominic Kwiatkowski, Cordelia Langford, John Sillitoe on behalf of the Wellcome Sanger Institute COVID-19 Surveillance Team |
| EPI_ISL_711067, EPI_ISL_711074, EPI_ISL_711076, EPI_ISL_711077, EPI_ISL_711078, EPI_ISL_711149, EPI_ISL_711150, EPI_ISL_711151, EPI_ISL_711152, EPI_ISL_711153, EPI_ISL_711154, EPI_ISL_711155, EPI_ISL_711156, EPI_ISL_711157, EPI_ISL_711158, EPI_ISL_711159, EPI_ISL_711160, EPI_ISL_711161, EPI_ISL_711162, EPI_ISL_711163, EPI_ISL_711164, EPI_ISL_711165, EPI_ISL_711166, EPI_ISL_711167, EPI_ISL_711168, EPI_ISL_711169, EPI_ISL_711170, EPI_ISL_711171, EPI_ISL_711172, EPI_ISL_711173, EPI_ISL_711174, EPI_ISL_711175, EPI_ISL_711177, EPI_ISL_711178, EPI_ISL_711179, EPI_ISL_711180, EPI_ISL_711181, EPI_ISL_711185, EPI_ISL_711190, EPI_ISL_711191, EPI_ISL_711192, EPI_ISL_711194, EPI_ISL_711195, EPI_ISL_711198, EPI_ISL_711199, EPI_ISL_711201, EPI_ISL_711207, EPI_ISL_711208, EPI_ISL_711209, EPI_ISL_711210, EPI_ISL_711211, EPI_ISL_711212, EPI_ISL_711213, EPI_ISL_711216, EPI_ISL_711217, EPI_ISL_711218, EPI_ISL_711220, EPI_ISL_711228, EPI_ISL_711232, EPI_ISL_711256, EPI_ISL_711257, EPI_ISL_711258, EPI_ISL_711259, EPI_ISL_711260, EPI_ISL_711261, EPI_ISL_711262, EPI_ISL_711263, EPI_ISL_711264, EPI_ISL_711265, EPI_ISL_711268, EPI_ISL_711271, EPI_ISL_711272, | | | |

EPI_ISL_711273, EPI_ISL_711274, EPI_ISL_711276, EPI_ISL_711277, EPI_ISL_711281, EPI_ISL_711295, EPI_ISL_711296, EPI_ISL_711297, EPI_ISL_711298, EPI_ISL_711361, EPI_ISL_711362, EPI_ISL_711363, EPI_ISL_711364, EPI_ISL_711365, EPI_ISL_711366, EPI_ISL_711367, EPI_ISL_711368, EPI_ISL_711369,
EPI_ISL_711370, EPI_ISL_711371, EPI_ISL_711372, EPI_ISL_711373, EPI_ISL_711374, EPI_ISL_711375, EPI_ISL_711376, EPI_ISL_711377, EPI_ISL_711378, EPI_ISL_711379, EPI_ISL_711380, EPI_ISL_711381, EPI_ISL_711382, EPI_ISL_711383, EPI_ISL_711390, EPI_ISL_711391, EPI_ISL_711392, EPI_ISL_711395,
EPI_ISL_711396, EPI_ISL_711397, EPI_ISL_711407, EPI_ISL_711408, EPI_ISL_711411, EPI_ISL_711429, EPI_ISL_711430, EPI_ISL_711438, EPI_ISL_711439, EPI_ISL_711440, EPI_ISL_711441, EPI_ISL_711442, EPI_ISL_711443, EPI_ISL_711444, EPI_ISL_711850, EPI_ISL_711851, EPI_ISL_711852, EPI_ISL_711853,
EPI_ISL_711854, EPI_ISL_711855, EPI_ISL_711856, EPI_ISL_711857, EPI_ISL_711858, EPI_ISL_711859, EPI_ISL_711860, EPI_ISL_711861, EPI_ISL_711862, EPI_ISL_711863, EPI_ISL_711864, EPI_ISL_711865, EPI_ISL_711866, EPI_ISL_711867, EPI_ISL_711868, EPI_ISL_711869, EPI_ISL_711870, EPI_ISL_711871,
EPI_ISL_711872, EPI_ISL_711873, EPI_ISL_711874, EPI_ISL_711875, EPI_ISL_711876, EPI_ISL_711877, EPI_ISL_711878, EPI_ISL_711879, EPI_ISL_711880, EPI_ISL_711881, EPI_ISL_711882, EPI_ISL_711883, EPI_ISL_711884, EPI_ISL_711885, EPI_ISL_711886, EPI_ISL_711887, EPI_ISL_711888, EPI_ISL_711889,
EPI_ISL_711890, EPI_ISL_711891, EPI_ISL_711892, EPI_ISL_711893, EPI_ISL_711894, EPI_ISL_711895, EPI_ISL_711896, EPI_ISL_711897, EPI_ISL_711898, EPI_ISL_711899, EPI_ISL_711900, EPI_ISL_711901, EPI_ISL_711902, EPI_ISL_711903, EPI_ISL_711904, EPI_ISL_711905, EPI_ISL_711906, EPI_ISL_711907,
EPI_ISL_711908, EPI_ISL_711909, EPI_ISL_711910, EPI_ISL_711911, EPI_ISL_711912, EPI_ISL_711913, EPI_ISL_711914, EPI_ISL_711915, EPI_ISL_711916, EPI_ISL_711917, EPI_ISL_711918, EPI_ISL_711919, EPI_ISL_711920, EPI_ISL_711921, EPI_ISL_711922, EPI_ISL_711923, EPI_ISL_711924, EPI_ISL_711925,
EPI_ISL_711926, EPI_ISL_711927, EPI_ISL_711928, EPI_ISL_711929, EPI_ISL_711930, EPI_ISL_711931, EPI_ISL_711932, EPI_ISL_711933, EPI_ISL_711934, EPI_ISL_711935, EPI_ISL_711936, EPI_ISL_711937, EPI_ISL_711938, EPI_ISL_711939, EPI_ISL_711940, EPI_ISL_711941, EPI_ISL_711942, EPI_ISL_711943,
EPI_ISL_711944, EPI_ISL_711945, EPI_ISL_711946, EPI_ISL_711947, EPI_ISL_711948, EPI_ISL_711949, EPI_ISL_711950, EPI_ISL_711951, EPI_ISL_711952, EPI_ISL_711953, EPI_ISL_711954, EPI_ISL_711955, EPI_ISL_711956, EPI_ISL_711957, EPI_ISL_711958, EPI_ISL_711959, EPI_ISL_711960, EPI_ISL_711961,
EPI_ISL_711962, EPI_ISL_711963, EPI_ISL_711964, EPI_ISL_711965, EPI_ISL_711966, EPI_ISL_711967, EPI_ISL_711968, EPI_ISL_711969, EPI_ISL_711970, EPI_ISL_711971, EPI_ISL_711972, EPI_ISL_711973, EPI_ISL_711974, EPI_ISL_711975, EPI_ISL_711976, EPI_ISL_711977, EPI_ISL_711978, EPI_ISL_711979,
EPI_ISL_711980, EPI_ISL_711981, EPI_ISL_711982, EPI_ISL_711983, EPI_ISL_711984, EPI_ISL_711985, EPI_ISL_711986, EPI_ISL_711987, EPI_ISL_711988, EPI_ISL_711989, EPI_ISL_711990, EPI_ISL_711991, EPI_ISL_711992, EPI_ISL_711993, EPI_ISL_711994, EPI_ISL_711995, EPI_ISL_711996, EPI_ISL_711997,
EPI_ISL_711998, EPI_ISL_711999, EPI_ISL_712000, EPI_ISL_712001, EPI_ISL_712002, EPI_ISL_712003, EPI_ISL_712004, EPI_ISL_712005, EPI_ISL_712006, EPI_ISL_712007, EPI_ISL_712008, EPI_ISL_712009, EPI_ISL_712010, EPI_ISL_712011, EPI_ISL_712012, EPI_ISL_712013, EPI_ISL_712014, EPI_ISL_712015,
EPI_ISL_712016, EPI_ISL_712017, EPI_ISL_712018, EPI_ISL_712019, EPI_ISL_712020, EPI_ISL_712021, EPI_ISL_712022, EPI_ISL_712023, EPI_ISL_712024, EPI_ISL_712025, EPI_ISL_712026, EPI_ISL_712027, EPI_ISL_712028, EPI_ISL_712029, EPI_ISL_712030, EPI_ISL_712031, EPI_ISL_712032, EPI_ISL_712033,
EPI_ISL_712034, EPI_ISL_712035, EPI_ISL_712036, EPI_ISL_712037, EPI_ISL_712038, EPI_ISL_712039, EPI_ISL_712040, EPI_ISL_712041, EPI_ISL_712042, EPI_ISL_712043, EPI_ISL_712044, EPI_ISL_712045, EPI_ISL_712046, EPI_ISL_712047, EPI_ISL_712048, EPI_ISL_712049, EPI_ISL_712050, EPI_ISL_712051,
EPI_ISL_712052, EPI_ISL_712053, EPI_ISL_712054, EPI_ISL_712055, EPI_ISL_712056, EPI_ISL_712057, EPI_ISL_712058, EPI_ISL_712059, EPI_ISL_713098, EPI_ISL_713099, EPI_ISL_713100, EPI_ISL_713101, EPI_ISL_713102, EPI_ISL_713103, EPI_ISL_713104, EPI_ISL_713105, EPI_ISL_713106, EPI_ISL_713107,
EPI_ISL_713108, EPI_ISL_713109, EPI_ISL_713110, EPI_ISL_713111, EPI_ISL_713112, EPI_ISL_713113, EPI_ISL_713114, EPI_ISL_713115, EPI_ISL_713116, EPI_ISL_713117, EPI_ISL_713118, EPI_ISL_713119, EPI_ISL_713120, EPI_ISL_713121, EPI_ISL_713122, EPI_ISL_713123, EPI_ISL_713124, EPI_ISL_713125,
EPI_ISL_713126, EPI_ISL_713127, EPI_ISL_713128, EPI_ISL_713129, EPI_ISL_713130, EPI_ISL_713131, EPI_ISL_713132, EPI_ISL_713133, EPI_ISL_713134, EPI_ISL_713135, EPI_ISL_713136, EPI_ISL_713137, EPI_ISL_713138, EPI_ISL_713139, EPI_ISL_713140, EPI_ISL_713141, EPI_ISL_713142, EPI_ISL_713143,
EPI_ISL_713144, EPI_ISL_713145, EPI_ISL_713146, EPI_ISL_713147, EPI_ISL_713148, EPI_ISL_713149, EPI_ISL_713150, EPI_ISL_713151, EPI_ISL_713152, EPI_ISL_713153, EPI_ISL_713154, EPI_ISL_713155, EPI_ISL_713156, EPI_ISL_713157, EPI_ISL_713158, EPI_ISL_713159, EPI_ISL_713160, EPI_ISL_713161,
EPI_ISL_713162, EPI_ISL_713163, EPI_ISL_713164, EPI_ISL_713165, EPI_ISL_713166, EPI_ISL_713167, EPI_ISL_713168, EPI_ISL_713169, EPI_ISL_713170, EPI_ISL_713171, EPI_ISL_713172, EPI_ISL_713173, EPI_ISL_713174, EPI_ISL_713175, EPI_ISL_713176, EPI_ISL_713177, EPI_ISL_713178, EPI_ISL_713179,
EPI_ISL_713180, EPI_ISL_713181, EPI_ISL_713182, EPI_ISL_713183, EPI_ISL_713184, EPI_ISL_713185, EPI_ISL_713186, EPI_ISL_713187, EPI_ISL_713188, EPI_ISL_713189, EPI_ISL_713190, EPI_ISL_713191, EPI_ISL_713192, EPI_ISL_713193, EPI_ISL_713194, EPI_ISL_713195, EPI_ISL_713196, EPI_ISL_713197,
EPI_ISL_713198, EPI_ISL_713199, EPI_ISL_713200, EPI_ISL_713201, EPI_ISL_713202, EPI_ISL_713203, EPI_ISL_713204, EPI_ISL_713205, EPI_ISL_713206, EPI_ISL_713207, EPI_ISL_713208, EPI_ISL_713209, EPI_ISL_713210, EPI_ISL_713211, EPI_ISL_713212, EPI_ISL_713213, EPI_ISL_713214, EPI_ISL_713215,
EPI_ISL_713216, EPI_ISL_713217, EPI_ISL_713218, EPI_ISL_713219, EPI_ISL_713220, EPI_ISL_713221, EPI_ISL_713222, EPI_ISL_713223, EPI_ISL_713224, EPI_ISL_713225, EPI_ISL_713226, EPI_ISL_713227, EPI_ISL_713228, EPI_ISL_713229, EPI_ISL_713230, EPI_ISL_713231, EPI_ISL_713232, EPI_ISL_713233,
EPI_ISL_713234, EPI_ISL_713235, EPI_ISL_713236, EPI_ISL_713237, EPI_ISL_713238, EPI_ISL_713239, EPI_ISL_713240, EPI_ISL_713241, EPI_ISL_713242, EPI_ISL_713243, EPI_ISL_713244, EPI_ISL_713245, EPI_ISL_713246, EPI_ISL_713247, EPI_ISL_713248, EPI_ISL_713249, EPI_ISL_713250, EPI_ISL_713251,
EPI_ISL_713252, EPI_ISL_713253, EPI_ISL_713254, EPI_ISL_713255, EPI_ISL_713256, EPI_ISL_713257, EPI_ISL_713258, EPI_ISL_713259, EPI_ISL_713260, EPI_ISL_713261, EPI_ISL_713262, EPI_ISL_713263, EPI_ISL_713264, EPI_ISL_713265, EPI_ISL_713266, EPI_ISL_713267, EPI_ISL_713268, EPI_ISL_713269,
EPI_ISL_713270, EPI_ISL_713271, EPI_ISL_713272, EPI_ISL_713273, EPI_ISL_713274, EPI_ISL_713275, EPI_ISL_713276, EPI_ISL_713277, EPI_ISL_713278, EPI_ISL_713279, EPI_ISL_713280, EPI_ISL_713281, EPI_ISL_713282, EPI_ISL_713283, EPI_ISL_713284, EPI_ISL_713285, EPI_ISL_713286, EPI_ISL_713287,
EPI_ISL_713288, EPI_ISL_713289, EPI_ISL_713290, EPI_ISL_713291, EPI_ISL_713292, EPI_ISL_713293, EPI_ISL_713294, EPI_ISL_713295, EPI_ISL_713296, EPI_ISL_713297, EPI_ISL_713298, EPI_ISL_713299, EPI_ISL_713300, EPI_ISL_713301, EPI_ISL_713302, EPI_ISL_713303, EPI_ISL_713304, EPI_ISL_713305,
EPI_ISL_713306, EPI_ISL_713307, EPI_ISL_713308, EPI_ISL_713309, EPI_ISL_713310, EPI_ISL_713311, EPI_ISL_713312, EPI_ISL_713313, EPI_ISL_713314, EPI_ISL_713315, EPI_ISL_713316, EPI_ISL_713317, EPI_ISL_713318, EPI_ISL_713319, EPI_ISL_713320, EPI_ISL_713321, EPI_ISL_713322, EPI_ISL_713323,
EPI_ISL_713324, EPI_ISL_713325, EPI_ISL_713326, EPI_ISL_713327, EPI_ISL_713328, EPI_ISL_713329, EPI_ISL_713330, EPI_ISL_713331, EPI_ISL_713332, EPI_ISL_713333, EPI_ISL_713334, EPI_ISL_713335, EPI_ISL_713336, EPI_ISL_713337, EPI_ISL_713338, EPI_ISL_713339, EPI_ISL_713340, EPI_ISL_713341,
EPI_ISL_713342, EPI_ISL_713343, EPI_ISL_713344, EPI_ISL_713345, EPI_ISL_713346, EPI_ISL_713347, EPI_ISL_713348, EPI_ISL_713349, EPI_ISL_713350, EPI_ISL_713351, EPI_ISL_713352, EPI_ISL_713353, EPI_ISL_713354, EPI_ISL_713355, EPI_ISL_713356, EPI_ISL_713357, EPI_ISL_713358, EPI_ISL_713359,
EPI_ISL_713360, EPI_ISL_713361, EPI_ISL_713362, EPI_ISL_713363, EPI_ISL_713364, EPI_ISL_713365, EPI_ISL_713366, EPI_ISL_713367, EPI_ISL_713368, EPI_ISL_713369, EPI_ISL_713370, EPI_ISL_713371, EPI_ISL_713372, EPI_ISL_713373, EPI_ISL_713374, EPI_ISL_713375, EPI_ISL_713376, EPI_ISL_713377,
EPI_ISL_713378, EPI_ISL_713379, EPI_ISL_713380, EPI_ISL_713381, EPI_ISL_713382, EPI_ISL_713383, EPI_ISL_713384, EPI_ISL_713385, EPI_ISL_713386, EPI_ISL_713387, EPI_ISL_713388, EPI_ISL_713389, EPI_ISL_713390, EPI_ISL_713391, EPI_ISL_713392, EPI_ISL_713393, EPI_ISL_713394, EPI_ISL_713395,
EPI_ISL_713396, EPI_ISL_713397, EPI_ISL_713398, EPI_ISL_713399, EPI_ISL_713400, EPI_ISL_713401, EPI_ISL_713402, EPI_ISL_713403, EPI_ISL_713404, EPI_ISL_713405, EPI_ISL_713406, EPI_ISL_713407, EPI_ISL_713408, EPI_ISL_713409, EPI_ISL_713410, EPI_ISL_713411, EPI_ISL_713412, EPI_ISL_713413,
EPI_ISL_713414, EPI_ISL_713415, EPI_ISL_713416, EPI_ISL_713417, EPI_ISL_713418, EPI_ISL_713419, EPI_ISL_713420, EPI_ISL_713421, EPI_ISL_713422, EPI_ISL_713423, EPI_ISL_713424, EPI_ISL_713425, EPI_ISL_713426, EPI_ISL_713427, EPI_ISL_713428, EPI_ISL_713429, EPI_ISL_713430, EPI_ISL_713431,
EPI_ISL_713432, EPI_ISL_713433, EPI_ISL_713434, EPI_ISL_713435, EPI_ISL_713436, EPI_ISL_713437, EPI_ISL_713438, EPI_ISL_713439, EPI_ISL_713440, EPI_ISL_713441, EPI_ISL_713442, EPI_ISL_713443, EPI_ISL_713444, EPI_ISL_713445, EPI_ISL_713446, EPI_ISL_713447, EPI_ISL_713448, EPI_ISL_713449,
EPI_ISL_713450, EPI_ISL_713451, EPI_ISL_713452, EPI_ISL_713453, EPI_ISL_713454, EPI_ISL_713455, EPI_ISL_713456, EPI_ISL_713457, EPI_ISL_713458, EPI_ISL_713459, EPI_ISL_713460, EPI_ISL_713461, EPI_ISL_713462, EPI_ISL_713463, EPI_ISL_713464, EPI_ISL_713465, EPI_ISL_713466, EPI_ISL_713467,
EPI_ISL_713468, EPI_ISL_713469, EPI_ISL_713470, EPI_ISL_713471, EPI_ISL_713472, EPI_ISL_713473, EPI_ISL_713474, EPI_ISL_713475, EPI_ISL_713476, EPI_ISL_713477, EPI_ISL_713478, EPI_ISL_713479, EPI_ISL_713480, EPI_ISL_713481, EPI_ISL_713482, EPI_ISL_713483, EPI_ISL_713484, EPI_ISL_713485,
EPI_ISL_713486, EPI_ISL_713487, EPI_ISL_713488, EPI_ISL_713489, EPI_ISL_713490, EPI_ISL_713491, EPI_ISL_713492, EPI_ISL_713493, EPI_ISL_713494, EPI_ISL_713495, EPI_ISL_713496, EPI_ISL_713497, EPI_ISL_713498, EPI_ISL_713499, EPI_ISL_713500, EPI_ISL_713501, EPI_ISL_713502, EPI_ISL_713503,
EPI_ISL_713504, EPI_ISL_713505, EPI_ISL_713506, EPI_ISL_713507, EPI_ISL_713508, EPI_ISL_713509, EPI_ISL_713510, EPI_ISL_713511, EPI_ISL_713512, EPI_ISL_713513, EPI_ISL_713514, EPI_ISL_713515, EPI_ISL_713516, EPI_ISL_713517, EPI_ISL_713518, EPI_ISL_713519, EPI_ISL_713520, EPI_ISL_713521,
EPI_ISL_713522, EPI_ISL_713523, EPI_ISL_713524, EPI_ISL_713525, EPI_ISL_713526, EPI_ISL_713527, EPI_ISL_713528, EPI_ISL_713529, EPI_ISL_713530, EPI_ISL_713531, EPI_ISL_713532, EPI_ISL_713533, EPI_ISL_713534, EPI_ISL_713535, EPI_ISL_713536, EPI_ISL_713537, EPI_ISL_713538, EPI_ISL_713539,
EPI_ISL_713540, EPI_ISL_713541, EPI_ISL_713542, EPI_ISL_713543, EPI_ISL_713544, EPI_ISL_713545, EPI_ISL_713546, EPI_ISL_713547, EPI_ISL_713548, EPI_ISL_713549, EPI_ISL_713550, EPI_ISL_713551, EPI_ISL_713552, EPI_ISL_713553, EPI_ISL_713554, EPI_ISL_713555, EPI_ISL_713556, EPI_ISL_713557,
EPI_ISL_713558, EPI_ISL_713559, EPI_ISL_713560, EPI_ISL_713561, EPI_ISL_713562, EPI_ISL_713563, EPI_ISL_713564, EPI_ISL_713565, EPI_ISL_713566, EPI_ISL_713567, EPI_ISL_713568, EPI_ISL_713569, EPI_ISL_713570, EPI_ISL_713571, EPI_ISL_713572, EPI_ISL_713573, EPI_ISL_713574, EPI_ISL_713575,
EPI_ISL_713576, EPI_ISL_713577, EPI_ISL_713578, EPI_ISL_713579, EPI_ISL_713580, EPI_ISL_713581, EPI_ISL_713582, EPI_ISL_713583, EPI_ISL_713584, EPI_ISL_713585, EPI_ISL_713586, EPI_ISL_713587, EPI_ISL_713588, EPI_ISL_713589, EPI_ISL_713590, EPI_ISL_713591, EPI_ISL_713592, EPI_ISL_713593,
EPI_ISL_713594, EPI_ISL_713595, EPI_ISL_713596, EPI_ISL_713597, EPI_ISL_713598, EPI_ISL_713599, EPI_ISL_713600, EPI_ISL_713601, EPI_ISL_713602, EPI_ISL_713603, EPI_ISL_713604, EPI_ISL_713605, EPI_ISL_713606, EPI_ISL_713607, EPI_ISL_713608, EPI_ISL_713609, EPI_ISL_713610, EPI_ISL_713611,
EPI_ISL_713612, EPI_ISL_713613, EPI_ISL_713614, EPI_ISL_713615, EPI_ISL_713616, EPI_ISL_713617, EPI_ISL_713618, EPI_ISL_713619, EPI_ISL_713620, EPI_ISL_713621, EPI_ISL_713622, EPI_ISL_713623, EPI_ISL_713624, EPI_ISL_713625, EPI_ISL_713626, EPI_ISL_713627, EPI_ISL_713628, EPI_ISL_713629,
EPI_ISL_713630, EPI_ISL_713631, EPI_ISL_713632, EPI_ISL_713633, EPI_ISL_713634, EPI_ISL_713635, EPI_ISL_713636, EPI_ISL_713637, EPI_ISL_713638, EPI_ISL_713639, EPI_ISL_713640, EPI_ISL_713641, EPI_ISL_713642, EPI_ISL_713643, EPI_ISL_713644, EPI_ISL_713645, EPI_ISL_713646, EPI_ISL_713647,
EPI_ISL_713648, EPI_ISL_713649, EPI_ISL_713650, EPI_ISL_713651, EPI_ISL_713652, EPI_ISL_713653, EPI_ISL_713654, EPI_ISL_713655, EPI_ISL_713656, EPI_ISL_713657, EPI_ISL_713658, EPI_ISL_713659, EPI_ISL_713660, EPI_ISL_713661, EPI_ISL_713662, EPI_ISL_713663, EPI_ISL_713664, EPI_ISL_713665,
EPI_ISL_713666, EPI_ISL_713667, EPI_ISL_713668, EPI_ISL_713669, EPI_ISL_713670, EPI_ISL_713671, EPI_ISL_713672, EPI_ISL_713673, EPI_ISL_713674, EPI_ISL_713675, EPI_ISL_713676, EPI_ISL_713677, EPI_ISL_713678, EPI_ISL_713679, EPI_ISL_713680, EPI_ISL_713681, EPI_ISL_713682, EPI_ISL_713683,
EPI_ISL_713684, EPI_ISL_713685, EPI_ISL_713686, EPI_ISL_713687, EPI_ISL_713688, EPI_ISL_713689, EPI_ISL_713690, EPI_ISL_713691, EPI_ISL_713692, EPI_ISL_713693, EPI_ISL_713694, EPI_ISL_713695, EPI_ISL_713696, EPI_ISL_713697, EPI_ISL_713698, EPI_ISL_713699, EPI_ISL_713700, EPI_ISL_713701,
EPI_ISL_713702, EPI_ISL_713703, EPI_ISL_713704, EPI_ISL_713705, EPI_ISL_713706, EPI_ISL_713707, EPI_ISL_713708, EPI_ISL_713709, EPI_ISL_713710, EPI_ISL_713711, EPI_ISL_713712, EPI_ISL_713713, EPI_ISL_713714, EPI_ISL_713715, EPI_ISL_713716, EPI_ISL_713717, EPI_ISL_713718, EPI_ISL_713719,
EPI_ISL_713720, EPI_ISL_713721, EPI_ISL_713722, EPI_ISL_713723, EPI_ISL_713724, EPI_ISL_713725, EPI_ISL_713726, EPI_ISL_713727, EPI_ISL_713728, EPI_ISL_713729, EPI_ISL_713730, EPI_ISL_713731, EPI_ISL_713732, EPI_ISL_713733, EPI_ISL_713734, EPI_ISL_713735, EPI_ISL_713736, EPI_ISL_713737,
EPI_ISL_713738, EPI_ISL_713739, EPI_ISL_713740, EPI_ISL_713741, EPI_ISL_713742, EPI_ISL_713743, EPI_ISL_713744, EPI_ISL_713745, EPI_ISL_713746, EPI_ISL_713747, EPI_ISL_713748, EPI_ISL_713749, EPI_ISL_713750, EPI_ISL_713751, EPI_ISL_713752, EPI_ISL_713753, EPI_ISL_713754, EPI_ISL_713755,
EPI_ISL_713756, EPI_ISL_713757, EPI_ISL_713758, EPI_ISL_713759, EPI_ISL_713760, EPI_ISL_713761, EPI_ISL_713762, EPI_ISL_713763, EPI_ISL_713764, EPI_ISL_713765, EPI_ISL_713766, EPI_ISL_713767, EPI_ISL_713768, EPI_ISL_713769, EPI_ISL_713770, EPI_ISL_713771, EPI_ISL_713772, EPI_ISL_713773,
EPI_ISL_713774, EPI_ISL_713775, EPI_ISL_713776, EPI_ISL_713777, EPI_ISL_713778, EPI_ISL_713779, EPI_ISL_713780, EPI_ISL_713781, EPI_ISL_713782, EPI_ISL_713783, EPI_ISL_713784, EPI_ISL_713785, EPI_ISL_713786, EPI_ISL_713787, EPI_ISL_713788, EPI_ISL_713789, EPI_ISL_713790, EPI_ISL_713791,
EPI_ISL_713792, EPI_ISL_713793, EPI_ISL_713794, EPI_ISL_713795, EPI_ISL_713796, EPI_ISL_713797, EPI_ISL_713798, EPI_ISL_713799, EPI_ISL_713800, EPI_ISL_713801, EPI_ISL_713802, EPI_ISL_713803, EPI_ISL_713804, EPI_ISL_713805, EPI_ISL_713806, EPI_ISL_713807, EPI_ISL_713808, EPI_ISL_713809,
EPI_ISL_713810, EPI_ISL_713811, EPI_ISL_713812, EPI_ISL_713813, EPI_ISL_713814, EPI_ISL_713815, EPI_ISL_713816, EPI_ISL_713817, EPI_ISL_713818, EPI_ISL_713819, EPI_ISL_713820, EPI_ISL_713821, EPI_ISL_713822, EPI_ISL_713823, EPI_ISL_713824, EPI_ISL_713825, EPI_ISL_713826, EPI_ISL_713827,
EPI_ISL_713828, EPI_ISL_713829, EPI_ISL_713830, EPI_ISL_713831, EPI_ISL_713832, EPI_ISL_713833, EPI_ISL_713834, EPI_ISL_713835, EPI_ISL_713836, EPI_ISL_713837, EPI_ISL_713838, EPI_ISL_713839, EPI_ISL_713840, EPI_ISL_713841, EPI_ISL_713842, EPI_ISL_713843, EPI_ISL_713844, EPI_ISL_713845,
EPI_ISL_713846, EPI_ISL_713847, EPI_ISL_713848, EPI_ISL_713849, EPI_ISL_713850, EPI_ISL_713851, EPI_ISL_713852, EPI_ISL_713853, EPI_ISL_713854, EPI_ISL_713855, EPI_ISL_713856, EPI_ISL_713857, EPI_ISL_713858, EPI_ISL_713859, EPI_ISL_713860, EPI_ISL_713861, EPI_ISL_713862, EPI_ISL_713863,
EPI_ISL_713864, EPI_ISL_713865, EPI_ISL_713866, EPI_ISL_713867, EPI_ISL_713868, EPI_ISL_713869, EPI_ISL_713870, EPI_ISL_713871, EPI_ISL_713872, EPI_ISL_713873, EPI_ISL_713874, EPI_ISL_713875, EPI_ISL_713876, EPI_ISL_713877, EPI_ISL_713878, EPI_ISL_713879, EPI_ISL_713880, EPI_ISL_713881,
EPI_ISL_713882, EPI_ISL_713883, EPI_ISL_713884, EPI_ISL_713885, EPI_ISL_713886, EPI_ISL_713887, EPI_ISL_713888, EPI_ISL_713889, EPI_ISL_713890, EPI_ISL_713891, EPI_ISL_713892, EPI_ISL_713893, EPI_ISL_713894, EPI_ISL_713895, EPI_ISL_713896, EPI_ISL_713897, EPI_ISL_713898, EPI_ISL_713899,
EPI_ISL_713900, EPI_ISL_713901, EPI_ISL_713902, EPI_ISL_713903, EPI_ISL_713904, EPI_ISL_713905, EPI_ISL_713906, EPI_ISL_713907, EPI_ISL_713908, EPI_ISL_713909, EPI_ISL_713910, EPI_ISL_713911, EPI_ISL_713912, EPI_ISL_713913, EPI_ISL_713914, EPI_ISL_713915, EPI_ISL_713916, EPI_ISL_713917,
EPI_ISL_713918, EPI_ISL_713919, EPI_ISL_713920, EPI_ISL_713921, EPI_ISL_713922, EPI_ISL_713923, EPI_ISL_713924, EPI_ISL_713925, EPI_ISL_713926, EPI_ISL_713927, EPI_ISL_713928, EPI_ISL_713929, EPI_ISL_713930, EPI_ISL_713931, EPI_ISL_713932, EPI_ISL_713933, EPI_ISL_713934, EPI_ISL_713935,
EPI_ISL_713936, EPI_ISL_713937, EPI_ISL_713938, EPI_ISL_713939, EPI_ISL_713940, EPI_ISL_713941, EPI_ISL_713942, EPI_ISL_713943, EPI_ISL_713944, EPI_ISL_713945, EPI_ISL_713946, EPI_ISL_713947, EPI_ISL_713948, EPI_ISL_713949, EPI_ISL_713950, EPI_ISL_713951, EPI_ISL_713952, EPI_ISL_713953,
EPI_ISL_713954, EPI_ISL_713955, EPI_ISL_713956, EPI_ISL_713957, EPI_ISL_713958, EPI_ISL_713959, EPI_ISL_713960, EPI_ISL_713961, EPI_ISL_713962, EPI_ISL_713963, EPI_ISL_713964, EPI_ISL_713965, EPI_ISL_713966, EPI_ISL_713967, EPI_ISL_713968, EPI_ISL_713969, EPI_ISL_713970, EPI_ISL_713971,
EPI_ISL_713972, EPI_ISL_713973, EPI_ISL_713974, EPI_ISL_713975, EPI_ISL_713976, EPI_ISL_713977, EPI_ISL_713978, EPI_ISL_713979, EPI_ISL_713980, EPI_ISL_713981, EPI_ISL_713982, EPI_ISL_713983, EPI_ISL_713984, EPI_ISL_713985, EPI_ISL_713986, EPI_ISL_713987, EPI_ISL_713988, EPI_ISL_713989,
EPI_ISL_713990, EPI_ISL_713991, EPI_ISL_713992, EPI_ISL_713993, EPI_ISL_713994, EPI_ISL_713995, EPI_ISL_713996, EPI_ISL_713997, EPI_ISL_713998, EPI_ISL_713999, EPI_ISL_714000, EPI_ISL_714001, EPI_ISL_714002, EPI_ISL_714003, EPI_ISL_714004, EPI_ISL_714005, EPI_ISL_714006, EPI_ISL_714007,
EPI_ISL_714008, EPI_ISL_714009, EPI_ISL_714010, EPI_ISL_714011, EPI_ISL_714012, EPI_ISL_714013, EPI_ISL_714014, EPI_ISL_714015, EPI_ISL_714016, EPI_ISL_714017, EPI_ISL_714018, EPI_ISL_714019, EPI_ISL_714020, EPI_ISL_714021, EPI_ISL_714022, EPI_ISL_714023, EPI_ISL_714024, EPI_ISL_714025,
EPI_ISL_714026, EPI_ISL_714027, EPI_ISL_714028, EPI_ISL_714029, EPI_ISL_714030, EPI_ISL_714031, EPI_ISL_714032, EPI_ISL_714033, EPI_ISL_714034, EPI_ISL_714035, EPI_ISL_714036, EPI_ISL_714037, EPI_ISL_714038, EPI_ISL_714039, EPI_ISL_714040, EPI_ISL_714041, EPI_ISL_714042, EPI_ISL_714043,
EPI_ISL_714044, EPI_ISL_714045, EPI_ISL_714046, EPI_ISL_714047, EPI_ISL_714048, EPI_ISL_714049, EPI_ISL_714050, EPI_ISL_714051, EPI_ISL_714052, EPI_ISL_714053, EPI_ISL_714054, EPI_ISL_714055, EPI_ISL_714056, EPI_ISL_714057, EPI_ISL_714058, EPI_ISL_714059, EPI_ISL_714060, EPI_ISL_714061,
EPI_ISL_714062, EPI_ISL_714063, EPI_ISL_714064, EPI_ISL_714065, EPI_ISL_714066, EPI_ISL_714067, EPI_ISL_714068, EPI_ISL_714069, EPI_ISL_714070, EPI_ISL_714071, EPI_ISL_714072, EPI_ISL_714073, EPI_ISL_714074, EPI_ISL_714075, EPI_ISL_714076, EPI_ISL_714077, EPI_ISL_714078, EPI_ISL_714079,
EPI_ISL_714080, EPI_ISL_714081, EPI_ISL_714082, EPI_ISL_714083, EPI_ISL_714084, EPI_ISL_714085, EPI_ISL_714086, EPI_ISL_714087, EPI_ISL_714088, EPI_ISL_714089, EPI_ISL_714090, EPI_ISL_714091, EPI_ISL_714092, EPI_ISL_714093, EPI_ISL_714094, EPI_ISL_714095, EPI_ISL_714096, EPI_ISL_714097,
EPI_ISL_714099, EPI_ISL_714115, EPI_ISL_714116, EPI_ISL_714117, EPI_ISL_714118, EPI_ISL_714119, EPI_ISL_714120, EPI_ISL_714121, EPI_ISL_714122, EPI_ISL_714123, EPI_ISL_714182, EPI_ISL_714196, EPI_ISL_714197, EPI_ISL_714198, EPI_ISL_714222, EPI_ISL_714235, EPI_ISL_714236, EPI_ISL_714240,
EPI_ISL_714241, EPI_ISL_714254, EPI_ISL_714259, EPI_ISL_714260, EPI_ISL_714264, EPI_ISL_714272, EPI_ISL_714276, EPI_ISL_714279, EPI_ISL_714289, EPI_ISL_714300, EPI_ISL_714301, EPI_ISL_714302, EPI_ISL_714303, EPI_ISL_714304, EPI_ISL_714305, EPI_ISL_714306, EPI_ISL_714307, EPI_ISL_714308,
EPI_ISL_714309, EPI_ISL_714310, EPI_ISL_714311, EPI_ISL_714312, EPI_ISL_714313, EPI_ISL_714314, EPI_ISL_714315, EPI_ISL_714316, EPI_ISL_714317, EPI_ISL_714318, EPI_ISL_714319, EPI_ISL_714366, EPI_ISL_714369, EPI_ISL_714385, EPI_ISL_714386, EPI_ISL_714387, EPI_ISL_714388, EPI_ISL_714389,
EPI_ISL_714390, EPI_ISL_714391, EPI_ISL_714416, EPI_ISL_714450, EPI_ISL_714462, EPI_ISL_714463, EPI_ISL_714464, EPI_ISL_714478, EPI_ISL_714479, EPI_ISL_714480, EPI_ISL_714481, EPI_ISL_714482, EPI_ISL_714483, EPI_ISL_714484, EPI_ISL_714485, EPI_ISL_714486, EPI_ISL_714487, EPI_ISL_714488,
EPI_ISL_714489, EPI_ISL_714490, EPI_ISL_714491, EPI_ISL_714492, EPI_ISL_714493, EPI_ISL_714494, EPI_ISL_714495, EPI_ISL_714496, EPI_ISL_714497, EPI_ISL_714498, EPI_ISL_714499, EPI_ISL_714500, EPI_ISL_714501, EPI_ISL_714502, EPI_ISL_714503, EPI_ISL_714504, EPI_ISL_714505, EPI_ISL_714506,
EPI_ISL_714507, EPI_ISL_714508, EPI_ISL_714509, EPI_ISL_714510, EPI_ISL_714511, EPI_ISL_714512, EPI_ISL_714513, EPI_ISL_714514, EPI_ISL_714515, EPI_ISL_714516, EPI_ISL_714517, EPI_ISL_714518, EPI_ISL_714519, EPI_ISL_714520, EPI_ISL_714521, EPI_ISL_714522, EPI_ISL_714523, EPI_ISL_714524,
EPI_ISL_714525, EPI_ISL_714526, EPI_ISL_714527, EPI_ISL_714528, EPI_ISL_714529, EPI_ISL_714530, EPI_ISL_714531, EPI_ISL_714532, EPI_ISL_714533, EPI_ISL_714534, EPI_ISL_714535, EPI_ISL_714536, EPI_ISL_714537, EPI_ISL_714538, EPI_ISL_714539, EPI_ISL_714540, EPI_ISL_714541, EPI_ISL_714542,
EPI_ISL_714543, EPI_ISL_714544, EPI_ISL_714545, EPI_ISL_714546, EPI_ISL_714547, EPI_ISL_714548, EPI_ISL_714549, EPI_ISL_714550, EPI_ISL_714551, EPI_ISL_714552, EPI_ISL_714553, EPI_ISL_714554, EPI_ISL_714555, EPI_ISL_714556, EPI_ISL_714557, EPI_ISL_714558, EPI_ISL_714559, EPI_ISL_714560,
EPI_ISL_714561, EPI_ISL_714562, EPI_ISL_714563, EPI_ISL_714564, EPI_ISL_714565

| | | | |
|---|---|---|---|
| see above | Department of Virus and Microbiological Special Diagnostics, Statens Serum Institut, Copenhagen, Denmark | Albertsen Lab, Department of Chemistry and Bioscience, Aalborg University, Denmark | Danish Covid-19 Genome Consortium |
| EPI_ISL_717979 | Laboratory of Microbiology and Infectious Diseases, Faculty of Veterinary Medicine, Aristotle University of Thessaloniki, University Campus, 541 24, Thessaloniki, Greece. | Laboratory of Biology, Department of Medicine, Democritus University of Thrace, Alexandroupolis, Greece | Dovrolis N., Chaintoutis S., Dimitriou M., Bampali M., Dovas C., Karakasiliotis I. |
| EPI_ISL_717980, EPI_ISL_718009, EPI_ISL_718011, EPI_ISL_718013 | Lab voor klinische biologie | Onderzoeksgroep Virologie | Laurens Lambrechts, Nick Vereecke, Marthe Pauwels, Bruno Verhasselt, Linos Vandekerckhove, Hans Nauwynck, Sebastiaan Theuns |

| | | | |
|---|---|---|---|
| EPI_ISL_718050, EPI_ISL_718059 | ZOTZ KLIMAS MVZ Düsseldorf-Centrum GbR ÜBAG für Labormedizin, Genetik, Zytologie, Pathologie | Center of Medical Microbiology, Virology, and Hospital Hygiene, University of Duesseldorf | Maximilian Damagnez, Alexander Dilthey, Ashley-Jane Duplessis, Patrick Finzer, Katrin Hoffmann, Torsten Houwaart, Lisanna Hülse, Malte Kohns Vasconcelos, Marek Korencak, Nadine Lübke, Jessica Nicolai, Klaus Pfeffer, Daniel Strelow, Jörg Timm, Andreas Walker, Tobias Wienemann, Rainer Zotz |
| EPI_ISL_718202 | Ministry of Health Hospitals | Institute of Health and Community Medicine | David Perera, Ooi Mong How, Chua Hock Hin, Tonnii Sia Loong Loong, Wong Jyn Shan, Wong Kiing Aik, Chan Chia Jui |
| EPI_ISL_718250 | Institute of Virology, Biomedical Research Center of the Slovak Academy of Sciences, Bratislava | Faculty of Natural Sciences, Comenius University, Bratislava | Kristína Boršová, Viktória Hodorová, Broa Brejová, Viktória abanová, Sabina Fumaová Havlíková, Juraj Kopáek, Martina Liková, ubomíra Lukáiková, Martina Neboháová, Monika Sláviková, Tomáš Vina, Boris Klempa, Jozef Nosek |
| EPI_ISL_721185, EPI_ISL_721186, EPI_ISL_721187, EPI_ISL_721188, EPI_ISL_721189, EPI_ISL_721190, EPI_ISL_721191, EPI_ISL_721192, EPI_ISL_721193, EPI_ISL_721194, EPI_ISL_721195, EPI_ISL_721196, EPI_ISL_721197, EPI_ISL_721198, EPI_ISL_721199, EPI_ISL_721200, EPI_ISL_721201, EPI_ISL_721202, EPI_ISL_721203, EPI_ISL_721204, EPI_ISL_721205, EPI_ISL_721206, EPI_ISL_721207, EPI_ISL_721208, EPI_ISL_721209, EPI_ISL_721210, EPI_ISL_721211, EPI_ISL_721212, EPI_ISL_721213, EPI_ISL_721214, EPI_ISL_721215, EPI_ISL_721216, EPI_ISL_721217, EPI_ISL_721218, EPI_ISL_721219, EPI_ISL_721220, EPI_ISL_721221, EPI_ISL_721222, EPI_ISL_721223, EPI_ISL_721224, EPI_ISL_721225, EPI_ISL_721226, EPI_ISL_721227, EPI_ISL_721228, EPI_ISL_721229, EPI_ISL_721230, EPI_ISL_721231, EPI_ISL_721232, EPI_ISL_721233, EPI_ISL_721234, EPI_ISL_721235, EPI_ISL_721236, EPI_ISL_721237, EPI_ISL_721238, EPI_ISL_721239, EPI_ISL_721240, EPI_ISL_721241, EPI_ISL_721242, EPI_ISL_721243, EPI_ISL_721244, EPI_ISL_721245, EPI_ISL_721246, EPI_ISL_721247, EPI_ISL_721248, EPI_ISL_721249, EPI_ISL_721250, EPI_ISL_721251, EPI_ISL_721252, EPI_ISL_721253, EPI_ISL_721254, EPI_ISL_721255, EPI_ISL_721256, EPI_ISL_721257, EPI_ISL_721258, EPI_ISL_721259, EPI_ISL_721260, EPI_ISL_721261, EPI_ISL_721262, EPI_ISL_721263, EPI_ISL_721264, EPI_ISL_721265, EPI_ISL_721266, EPI_ISL_721267, EPI_ISL_721268, EPI_ISL_721269, EPI_ISL_721270, EPI_ISL_721271, EPI_ISL_721272, EPI_ISL_721273, EPI_ISL_721274, EPI_ISL_721275, EPI_ISL_721276, EPI_ISL_721277, EPI_ISL_721278, EPI_ISL_721279, EPI_ISL_721280, EPI_ISL_721281, EPI_ISL_721282, EPI_ISL_721283, EPI_ISL_721284, EPI_ISL_721285, EPI_ISL_721286, EPI_ISL_721287, EPI_ISL_721288, EPI_ISL_721289, EPI_ISL_721290, EPI_ISL_721291, EPI_ISL_721292, EPI_ISL_721293, EPI_ISL_721294, EPI_ISL_721295, EPI_ISL_721296, EPI_ISL_721297, EPI_ISL_721298, EPI_ISL_721299, EPI_ISL_721300, EPI_ISL_721301, EPI_ISL_721302, EPI_ISL_721303, EPI_ISL_721304, EPI_ISL_721305, EPI_ISL_721307, EPI_ISL_721308, EPI_ISL_721309, EPI_ISL_721310, EPI_ISL_721311, EPI_ISL_721312, EPI_ISL_721313, EPI_ISL_721314, EPI_ISL_721315, EPI_ISL_721316, EPI_ISL_721317, EPI_ISL_721318, EPI_ISL_721319, EPI_ISL_721320, EPI_ISL_721321, EPI_ISL_721322, EPI_ISL_721323, EPI_ISL_721324, EPI_ISL_721325, EPI_ISL_721326, EPI_ISL_721327, EPI_ISL_721328, EPI_ISL_721329, EPI_ISL_721330, EPI_ISL_721331, EPI_ISL_721332, EPI_ISL_721333, EPI_ISL_721334, EPI_ISL_721335, EPI_ISL_721336, EPI_ISL_721337, EPI_ISL_721338, EPI_ISL_721339, EPI_ISL_721340, EPI_ISL_721341, EPI_ISL_721342, EPI_ISL_721343, EPI_ISL_721344, EPI_ISL_721345, EPI_ISL_721346, EPI_ISL_721347, EPI_ISL_721348, EPI_ISL_721349, EPI_ISL_721350, EPI_ISL_721351, EPI_ISL_721352, EPI_ISL_721353, EPI_ISL_721354, EPI_ISL_721355, EPI_ISL_721356, EPI_ISL_721357, EPI_ISL_721358, EPI_ISL_721359, EPI_ISL_721360, EPI_ISL_721361, EPI_ISL_721362, EPI_ISL_721363, EPI_ISL_721364, EPI_ISL_721365, EPI_ISL_721366, EPI_ISL_721367, EPI_ISL_721368, EPI_ISL_721369, EPI_ISL_721371, EPI_ISL_721372, EPI_ISL_721373, EPI_ISL_721374, EPI_ISL_721375, EPI_ISL_721376, EPI_ISL_721377, EPI_ISL_721378, EPI_ISL_721379, EPI_ISL_721380, EPI_ISL_721381, EPI_ISL_721382, EPI_ISL_721383, EPI_ISL_721384, EPI_ISL_721385, EPI_ISL_721386, EPI_ISL_721387, EPI_ISL_721388, EPI_ISL_721389, EPI_ISL_721390, EPI_ISL_721391, EPI_ISL_721392, EPI_ISL_721393, EPI_ISL_721394, EPI_ISL_721395, EPI_ISL_721396, EPI_ISL_721397, EPI_ISL_721398, EPI_ISL_721399, EPI_ISL_721400, EPI_ISL_721401, EPI_ISL_721402, EPI_ISL_721403, EPI_ISL_721404, EPI_ISL_721405, EPI_ISL_721406, EPI_ISL_721407, EPI_ISL_721408, EPI_ISL_721409, EPI_ISL_721410, EPI_ISL_721411, EPI_ISL_721412, EPI_ISL_721413, EPI_ISL_721414, EPI_ISL_721415, EPI_ISL_721416, EPI_ISL_721417, EPI_ISL_721418, EPI_ISL_721419, EPI_ISL_721420, EPI_ISL_721421, EPI_ISL_721422, EPI_ISL_721423, EPI_ISL_721424, EPI_ISL_721425, EPI_ISL_721426, EPI_ISL_721427, EPI_ISL_721428, EPI_ISL_721429, EPI_ISL_721430, EPI_ISL_721431, EPI_ISL_721432, EPI_ISL_721433, EPI_ISL_721434, EPI_ISL_721435, EPI_ISL_721436, EPI_ISL_721437, EPI_ISL_721438, EPI_ISL_721439, EPI_ISL_721440, EPI_ISL_721441, EPI_ISL_721442, EPI_ISL_721443, EPI_ISL_721444, EPI_ISL_721445, EPI_ISL_721446, EPI_ISL_721447, EPI_ISL_721448, EPI_ISL_721449, EPI_ISL_721450, EPI_ISL_721451, EPI_ISL_721452, EPI_ISL_721453, EPI_ISL_721454, EPI_ISL_721455, EPI_ISL_721457, EPI_ISL_721458, EPI_ISL_721459, EPI_ISL_721461, EPI_ISL_721462, EPI_ISL_721463, EPI_ISL_721464, EPI_ISL_721465, EPI_ISL_721466, EPI_ISL_721467, EPI_ISL_721468, EPI_ISL_721469, EPI_ISL_721470, EPI_ISL_721471, EPI_ISL_721472, EPI_ISL_721473, EPI_ISL_721474, EPI_ISL_721475, EPI_ISL_721476, EPI_ISL_721477, EPI_ISL_721478, EPI_ISL_721479, EPI_ISL_721480, EPI_ISL_721481, EPI_ISL_721482, EPI_ISL_721483, EPI_ISL_721484, EPI_ISL_721485, EPI_ISL_721486, EPI_ISL_721487, EPI_ISL_721488, EPI_ISL_721489, EPI_ISL_721490, EPI_ISL_721491, EPI_ISL_721492, EPI_ISL_721493, EPI_ISL_721494, EPI_ISL_721495, EPI_ISL_721496, EPI_ISL_721497, EPI_ISL_721498, EPI_ISL_721499, EPI_ISL_721500, EPI_ISL_721501, EPI_ISL_721502, EPI_ISL_721503, EPI_ISL_721504, EPI_ISL_721505, EPI_ISL_721506, EPI_ISL_721507, EPI_ISL_721508, EPI_ISL_721509, EPI_ISL_721510, EPI_ISL_721511, EPI_ISL_721512, EPI_ISL_721513, EPI_ISL_721514, EPI_ISL_721515, EPI_ISL_721516 | | | |
| see above | Lighthouse Lab in Alderley Park | Wellcome Sanger Institute for the COVID-19 Genomics UK (COG-UK) Consortium | Jacquelyn Wynn, Mairead Hyland, The Lighthouse Lab in Alderley Park and Alex Alderton, Roberto Amato, Sonia Goncalves, Ewan Harrison, David K. Jackson, Ian Johnston, Dominic Kwiatkowski, Cordelia Langford, John Sillitoe on behalf of the Wellcome Sanger Institute COVID-19 Surveillance Team |
| EPI_ISL_721626 | National Institute for Infectious Diseases, INMI, "L. Spallanzani" IRCCS | National Institute for Infectious Diseases, INMI, "L. Spallanzani" IRCCS | M Rueca, B Bartolini, C.E.M Gruber, F Messina, E Giombini, A Di Caro, MR Capobianchi |
| EPI_ISL_721627 | National Institute for Infectious Diseases, INMI, "L. Spallanzani" IRCCS | National Institute for Infectious Diseases, INMI, "L. Spallanzani" IRCCS | F Messina, E Giombini, M Rueca, B Bartolini, C.E.M Gruber, MR Capobianchi, A Di Caro |
| EPI_ISL_721671, EPI_ISL_721677, EPI_ISL_721679, EPI_ISL_721682, EPI_ISL_721684, EPI_ISL_721691, EPI_ISL_721693, EPI_ISL_721696, EPI_ISL_721701, EPI_ISL_721711, EPI_ISL_721712, EPI_ISL_721713, EPI_ISL_721714, EPI_ISL_721715, EPI_ISL_721757, EPI_ISL_721758, EPI_ISL_721759, EPI_ISL_721760, EPI_ISL_721767, EPI_ISL_721768, EPI_ISL_721769, EPI_ISL_721770, EPI_ISL_721778, EPI_ISL_721779, EPI_ISL_721780, EPI_ISL_721781, EPI_ISL_721782, EPI_ISL_721783, EPI_ISL_721784, EPI_ISL_721785, EPI_ISL_721786, EPI_ISL_721788, EPI_ISL_721789, EPI_ISL_721790, EPI_ISL_721791, EPI_ISL_721792, EPI_ISL_721793, EPI_ISL_721794, EPI_ISL_721795, EPI_ISL_721796, EPI_ISL_721797, EPI_ISL_721798, EPI_ISL_721799, EPI_ISL_721800, EPI_ISL_721801, EPI_ISL_721802, EPI_ISL_721803, EPI_ISL_721804, EPI_ISL_721805, EPI_ISL_721813, EPI_ISL_721814, EPI_ISL_721826, EPI_ISL_721844, EPI_ISL_721845, EPI_ISL_721846, EPI_ISL_721847, EPI_ISL_721858, EPI_ISL_721859, EPI_ISL_721860, EPI_ISL_721861, EPI_ISL_721862, EPI_ISL_721863, EPI_ISL_721864, EPI_ISL_721869, EPI_ISL_721870, EPI_ISL_721871, EPI_ISL_721872, EPI_ISL_721873, EPI_ISL_721874, EPI_ISL_721885, EPI_ISL_721886, EPI_ISL_721887, EPI_ISL_721915, EPI_ISL_721916, EPI_ISL_721940, EPI_ISL_721942, EPI_ISL_721943, EPI_ISL_721944, EPI_ISL_721945, EPI_ISL_721946, EPI_ISL_721947, EPI_ISL_721948, EPI_ISL_721949, EPI_ISL_721950, EPI_ISL_721951, EPI_ISL_721952, EPI_ISL_721953, EPI_ISL_721954, EPI_ISL_721955, EPI_ISL_721956, EPI_ISL_721957, EPI_ISL_721958 | | | |
| see above | Viollier AG | Department of Biosystems Science and Engineering, ETH Zürich | Christian Beisel |
| EPI_ISL_722201 | Vitalis Mostar | Alea Genetic Center | Salihefendic L., Pecar D., Konjhodzic R. |
| EPI_ISL_722209 | Dom Zdravlja Sarajevo | Alea Genetic Center | Salihefendic L., Pecar D., Konjhodzic R. |
| EPI_ISL_722290, EPI_ISL_722328, EPI_ISL_722334, EPI_ISL_722351, EPI_ISL_722363, EPI_ISL_722388, EPI_ISL_722406, EPI_ISL_722682, EPI_ISL_722683, EPI_ISL_722684, EPI_ISL_722685, EPI_ISL_722686 | | | |
| see above | Dutch COVID-19 response team | Erasmus Medical Center | Bas Oude Munnink, Reina Sikkema, David Nieuwenhuijse, Irina Chestakova, Anne van der Linden, Marjan Boter, Emmanuelle Munger, Corine GeurtsvanKessel, Annemiek van der Eijk, Richard Molenkamp, Marion Koopmans, on behalf of the Dutch national COVID-19 response team. |
| EPI_ISL_722910 | Istituto Zooprofilattico Sperimentale della Puglia e della Basilicata | Istituto Zooprofilattico Sperimentale della Puglia e della Basilicata | Parisi A., Bianco A., Capozzi L., Del Sambro L., Manzulli V, Rondinone V., Pace L., Cipolletta D., Galante D. |
| EPI_ISL_723124, EPI_ISL_723125, EPI_ISL_723126 | Minnesota Department of Health, Public Health Laboratory | Minnesota Department of Health, Public Health Laboratory | Alexandra Lorentz, Jacob Garfin, Matt Plumb, and Xiong Wang |
| EPI_ISL_723150 | Vault Health | Minnesota Department of Health, Public Health Laboratory | Alexandra Lorentz, Jacob Garfin, Matt Plumb, and Xiong Wang |
| EPI_ISL_723156, EPI_ISL_723200, EPI_ISL_723202, EPI_ISL_723203, EPI_ISL_723204, EPI_ISL_723205, EPI_ISL_723206, EPI_ISL_723207, EPI_ISL_723208, EPI_ISL_723209, EPI_ISL_723210, EPI_ISL_723211, EPI_ISL_723213, EPI_ISL_723214, EPI_ISL_723215, EPI_ISL_723216, EPI_ISL_723217, EPI_ISL_723220, EPI_ISL_723226, EPI_ISL_723227, EPI_ISL_723229, EPI_ISL_723230, EPI_ISL_723231, EPI_ISL_723266, EPI_ISL_723267, EPI_ISL_723268, EPI_ISL_723289, EPI_ISL_723316, EPI_ISL_723317, EPI_ISL_723323, EPI_ISL_723356, EPI_ISL_723358, EPI_ISL_723359, EPI_ISL_723383, EPI_ISL_723394, EPI_ISL_723395, EPI_ISL_723396, EPI_ISL_723397, EPI_ISL_723412, EPI_ISL_723413, EPI_ISL_723423, EPI_ISL_723429, EPI_ISL_723441, EPI_ISL_723442, EPI_ISL_723443, EPI_ISL_723445, EPI_ISL_723449, EPI_ISL_723450, EPI_ISL_723451 | | | |
| see above | Dutch COVID-19 response team | National Institute for Public Health and the Environment (RIVM) | Adam Meijer, Harry Vennema, Jeroen Cremer, Sharon van den Brink, Bas van der Veer, AnneMarie van den Brandt, Florian Zwagemaker, Dennis Schmitz, Chantal Reusken, on behalf of the national COVID-19 response team |
| EPI_ISL_723480, EPI_ISL_723481, EPI_ISL_723482, EPI_ISL_723483, EPI_ISL_723484, EPI_ISL_723485, EPI_ISL_723486, EPI_ISL_723487, EPI_ISL_723488, EPI_ISL_723489, EPI_ISL_723490, EPI_ISL_723491 | | | |
| see above | Virginia DCLS | Virginia DCLS | Virginia DCLS |
| EPI_ISL_723852, EPI_ISL_723854, EPI_ISL_723855, EPI_ISL_723856 | Department of Pathology, University of Cambridge | COVID-19 Genomics UK (COG-UK) Consortium | Aminu S. Jahun, Yasmin Chaudhry, Grant Hall, Iliana Georgana, Myra Hosmillo, Martin D. Curran, Malte Pinckert, Surendra Parmar, Ian Goodfellow |
| EPI_ISL_724064, EPI_ISL_724065, EPI_ISL_724066, EPI_ISL_724067, EPI_ISL_724068, EPI_ISL_724069, EPI_ISL_724070, EPI_ISL_724071, EPI_ISL_724072, EPI_ISL_724073, EPI_ISL_724074, EPI_ISL_724075, EPI_ISL_724076, EPI_ISL_724077, EPI_ISL_724078, EPI_ISL_724079, EPI_ISL_724080, EPI_ISL_724081, EPI_ISL_724082, EPI_ISL_724083, EPI_ISL_724084, EPI_ISL_724085, EPI_ISL_724086, EPI_ISL_724087, EPI_ISL_724088, EPI_ISL_724089, EPI_ISL_724090, EPI_ISL_724091, EPI_ISL_724092, EPI_ISL_724093, EPI_ISL_724094, EPI_ISL_724095, EPI_ISL_724096, EPI_ISL_724097, EPI_ISL_724098, EPI_ISL_724099, EPI_ISL_724100, EPI_ISL_724101, EPI_ISL_724102, EPI_ISL_724103, EPI_ISL_724104, EPI_ISL_724105, EPI_ISL_724106, EPI_ISL_724107, EPI_ISL_724108, EPI_ISL_724109, EPI_ISL_724110, EPI_ISL_724111, EPI_ISL_724112, EPI_ISL_724113, EPI_ISL_724114, EPI_ISL_724115, EPI_ISL_724116, EPI_ISL_724117, EPI_ISL_724118, EPI_ISL_724119, EPI_ISL_724120, EPI_ISL_724121, EPI_ISL_724122, EPI_ISL_724123 | | | |
| see above | Northumbria University / South Tees Hospitals NHS Foundation Trust / North Cumbria Integrated Care NHS Foundation Trust / North Tees and Hartlepool NHS Foundation Trust / Newcastle Hospitals NHS Foundation Trust | COVID-19 Genomics UK (COG-UK) Consortium | Darren L Smith,Andrew Nelson,Matthew Bashton,Greg R Young,Joshua Loh,John Allan,Mohammad A Tariq,Giles S Holt,Gary Black,Wen C Yew,Lynn Dover,Paul Baker,Steve Liggett,Sarah Essex,Jane Greenaway,Debra Padgett,Clive Graham,Garren Scott,Edward Barton,Emma Swindells,Brendan Payne,Jennifer Collins,Yusri Taha,Gary Eltringham |
| EPI_ISL_724124, EPI_ISL_724130, EPI_ISL_724131, EPI_ISL_724132, EPI_ISL_724133, EPI_ISL_724134, EPI_ISL_724135, EPI_ISL_724136, EPI_ISL_724137, EPI_ISL_724138, EPI_ISL_724139, EPI_ISL_724140, EPI_ISL_724141, EPI_ISL_724142, EPI_ISL_724143, EPI_ISL_724144, EPI_ISL_724145, EPI_ISL_724146, EPI_ISL_724147, EPI_ISL_724151, EPI_ISL_724152, EPI_ISL_724153, EPI_ISL_724154, EPI_ISL_724155, EPI_ISL_724156, EPI_ISL_724157, EPI_ISL_724158, EPI_ISL_724159, EPI_ISL_724160, EPI_ISL_724161, EPI_ISL_724162, EPI_ISL_724163, EPI_ISL_724164, EPI_ISL_724165, EPI_ISL_724166, EPI_ISL_724167, EPI_ISL_724168, EPI_ISL_724169, EPI_ISL_724170, EPI_ISL_724171, EPI_ISL_724172, EPI_ISL_724173, EPI_ISL_724174, EPI_ISL_724175, EPI_ISL_724176, EPI_ISL_724177, EPI_ISL_724178, EPI_ISL_724179, EPI_ISL_724180, EPI_ISL_724181, EPI_ISL_724182, EPI_ISL_724183, EPI_ISL_724184, EPI_ISL_724185, EPI_ISL_724186, EPI_ISL_724187, EPI_ISL_724189, EPI_ISL_724190, EPI_ISL_724191, EPI_ISL_724192, EPI_ISL_724193, EPI_ISL_724194, EPI_ISL_724195, EPI_ISL_724196, EPI_ISL_724197, EPI_ISL_724198, EPI_ISL_724199 | | | |
| see above | Quadram Institute Bioscience | COVID-19 Genomics UK (COG-UK) Consortium | Dave J. Baker, Gemma L. Kay, Alp Aydin, Thanh Le-Viet, Steven Rudder, Ana P. Tedim, Anastasia Kolyva, Maria Diaz, Leonardo de Oliveira Martins, Nabil-Fareed Alikhan, Lizzie Meadows, Rachael Stanley, Ngozi Elumogo, Muhammed Yasir, Nicholas M. Thomson, Alexander J Trotter, Rachel Gilroy, Samuel Bloomfield, Claire Stuart, Andrew Bell, Reenesh Prakash, Samir Dervisevic, Alison E. Mather, John Wain, Mark Webber, Andrew J. Page, Justin O'Grady |
| EPI_ISL_724222 | Virology Department, Royal Infirmary of Edinburgh, NHS Lothian / School of Biological Sciences, University of | COVID-19 Genomics UK (COG-UK) Consortium | McHugh M, Dewar R, Rooke S, Gallagher M, Balcaza C, O'Toole Á, Scher E, Hill V, McCrone JT, Colquhoun R, Yu X, Jackson B, Rambaut A, Williams TC, Templeton K |

| | | | |
|---|---|---|---|
| | Edinburgh / Institute of Genetics and Molecular Medicine, University of Edinburgh | | |
| EPI_ISL_724397, EPI_ISL_724398, EPI_ISL_724400, EPI_ISL_724402, EPI_ISL_724403, EPI_ISL_724404, EPI_ISL_724405, EPI_ISL_724407, EPI_ISL_724418, EPI_ISL_724424, EPI_ISL_724427, EPI_ISL_724430, EPI_ISL_724433, EPI_ISL_724435, EPI_ISL_724438, EPI_ISL_724467, EPI_ISL_724468, EPI_ISL_724469, EPI_ISL_724470, EPI_ISL_724471, EPI_ISL_724472, EPI_ISL_724473, EPI_ISL_724474, EPI_ISL_724475, EPI_ISL_724476, EPI_ISL_724478, EPI_ISL_724480, EPI_ISL_724481, EPI_ISL_724482, EPI_ISL_724484, EPI_ISL_724486, EPI_ISL_724487, EPI_ISL_724488, EPI_ISL_724489, EPI_ISL_724490, EPI_ISL_724491, EPI_ISL_724492, EPI_ISL_724494, EPI_ISL_724495 | | | |
| see above | Liverpool Clinical Laboratories | COVID-19 Genomics UK (COG-UK) Consortium | Sam Haldenby, Anita Lucaci, Steve Paterson, Julian Hiscox, Alistair Darby, M Almsaud, A Alrezaihi, Muhannad Alruwaili, Stuart D Armstrong, Jones Benjamin, Eleanor G Bentley, Anu Chawla, Jordan J Clark, Angela Cowell, Richard Eccles, Isabel García-Dorival, Matthew Gemmell, Alessandro Gerada, PKF Gilmore, Richard Gregory, Ximeng Han, Catherine Hartley, Margaret Hughes, Miren Iturriza-Gomara, James Johnson, L Luu, Jenifer Manson, Charlotte Nelson, Elaine O'Toole, Cassie Olateju, Rebekah Penrice-Randal , Lucille Rainbow, N.P Randle, Trevor Ian Robinson, Parul Sharma, Ghada T Shawli, James P Stewart, Neil Swainston, Ecaterina Vamos, Joanne Watts, Mark Whitehead |
| EPI_ISL_724542, EPI_ISL_724543, EPI_ISL_724544, EPI_ISL_724545, EPI_ISL_724546, EPI_ISL_724547, EPI_ISL_724548, EPI_ISL_724550, EPI_ISL_724551, EPI_ISL_724552, EPI_ISL_724553, EPI_ISL_724554, EPI_ISL_724555, EPI_ISL_724556, EPI_ISL_724557, EPI_ISL_724558, EPI_ISL_724559, EPI_ISL_724560, EPI_ISL_724561, EPI_ISL_724562, EPI_ISL_724563, EPI_ISL_724564, EPI_ISL_724565, EPI_ISL_724566, EPI_ISL_724567, EPI_ISL_724568, EPI_ISL_724569, EPI_ISL_724570, EPI_ISL_724571, EPI_ISL_724572, EPI_ISL_724573, EPI_ISL_724574, EPI_ISL_724575, EPI_ISL_724580, EPI_ISL_724581, EPI_ISL_724582 | | | |
| see above | University College London, Great Ormond Street Hospital for Children NHS Foundation Trust, Imperial College Healthcare NHS Trust | COVID-19 Genomics UK (COG-UK) Consortium | Sergi Castellano, Rachel Williams, Mark Kristiansen, Paola Resende Silva, Sunando Roy, Tony Brooks, Helena Tutill, Paola Niola, Patricia Dyal, Charlotte Williams, Leysa Forrest, Yasmin Panchbhaya, Jacqueline Findlay, Samuel Weeks, Julianne Brown, Kathryn Harris, Paul Randell, James Price, Alison Holmes, Judith Breuer |
| EPI_ISL_724927, EPI_ISL_724930, EPI_ISL_724932, EPI_ISL_724933, EPI_ISL_724934, EPI_ISL_724935, EPI_ISL_724936, EPI_ISL_724937, EPI_ISL_724938, EPI_ISL_724939, EPI_ISL_724940, EPI_ISL_724941, EPI_ISL_724942, EPI_ISL_724943, EPI_ISL_724944, EPI_ISL_724945, EPI_ISL_724946, EPI_ISL_724947, EPI_ISL_724948, EPI_ISL_724949, EPI_ISL_724950, EPI_ISL_724951, EPI_ISL_724952, EPI_ISL_724970, EPI_ISL_725012, EPI_ISL_725013, EPI_ISL_725014, EPI_ISL_725015, EPI_ISL_725016, EPI_ISL_725017 | | | |
| see above | Northumbria University / South Tees Hospitals NHS Foundation Trust / North Cumbria Integrated Care NHS Foundation Trust / North Tees and Hartlepool NHS Foundation Trust / Newcastle Hospitals NHS Foundation Trust | COVID-19 Genomics UK (COG-UK) Consortium | Darren L Smith,Andrew Nelson,Matthew Bashton,Greg R Young,Joshua Loh,John Allan,Mohammad A Tariq,Giles S Holt,Gary Black,Wen C Yew,Lynn Dover,Paul Baker,Steve Liggett,Sarah Essex,Jane Greenaway,Debra Padgett,Clive Graham,Garren Scott,Edward Barton,Emma Swindells,Brendan Payne,Jennifer Collins,Yusri Taha,Gary Eltringham |
| EPI_ISL_725192, EPI_ISL_725201, EPI_ISL_725205, EPI_ISL_725206, EPI_ISL_725210, EPI_ISL_725220, EPI_ISL_725223, EPI_ISL_725224, EPI_ISL_725225, EPI_ISL_725226, EPI_ISL_725229, EPI_ISL_725232, EPI_ISL_725234, EPI_ISL_725235, EPI_ISL_725236, EPI_ISL_725243, EPI_ISL_725244, EPI_ISL_725253, EPI_ISL_725259, EPI_ISL_725284, EPI_ISL_725288 | | | |
| see above | Quadram Institute Bioscience | COVID-19 Genomics UK (COG-UK) Consortium | Dave J. Baker, Gemma L. Kay, Alp Aydin, Thanh Le-Viet, Steven Rudder, Ana P. Tedim, Anastasia Kolyva, Maria Diaz, Leonardo de Oliveira Martins, Nabil-Fareed Alikhan, Lizzie Meadows, Rachael Stanley, Ngozi Elumogo, Muhammed Yasir, Nicholas M. Thomson, Alexander J Trotter, Rachel Gilroy, Samuel Bloomfield, Claire Stuart, Andrew Bell, Reenesh Prakash, Samir Dervisevic, Alison E. Mather, John Wain, Mark Webber, Andrew J. Page, Justin O'Grady |
| EPI_ISL_725519 | Queens Medical Centre, Clinical Microbiology Department / DeepSeq Nottingham | COVID-19 Genomics UK (COG-UK) Consortium | Gemma Clark, Wendy Smith, Manjinder Khakh, Vicki M Fleming, Michelle M Lister, Hannah Howson-Wells, Jonathan Ball, Patrick McClure, Joseph Chappell, Theocharis Tsoleridis, Nadine Holmes, Matthew Carlisle, Christopher Moore, Fei Sang, Johnny Debebe, Victoria Wright, Matthew Loose |
| EPI_ISL_725677, EPI_ISL_725678, EPI_ISL_725680, EPI_ISL_725681, EPI_ISL_725682, EPI_ISL_725683, EPI_ISL_725684, EPI_ISL_725685, EPI_ISL_725716, EPI_ISL_725719, EPI_ISL_725725, EPI_ISL_725727, EPI_ISL_725728, EPI_ISL_725729, EPI_ISL_725730, EPI_ISL_725731, EPI_ISL_725732, EPI_ISL_725733, EPI_ISL_725734, EPI_ISL_725735, EPI_ISL_725736, EPI_ISL_725737, EPI_ISL_725738, EPI_ISL_725983, EPI_ISL_726000, EPI_ISL_726001, EPI_ISL_726007, EPI_ISL_726008, EPI_ISL_726009, EPI_ISL_726022, EPI_ISL_726023, EPI_ISL_726024, EPI_ISL_726026, EPI_ISL_726027, EPI_ISL_726028, EPI_ISL_726029, EPI_ISL_726030, EPI_ISL_726031, EPI_ISL_726107, EPI_ISL_726111, EPI_ISL_726120, EPI_ISL_726123, EPI_ISL_726129, EPI_ISL_726131, EPI_ISL_726132, EPI_ISL_726133, EPI_ISL_726134, EPI_ISL_726135, EPI_ISL_726136, EPI_ISL_726137, EPI_ISL_726138, EPI_ISL_726139, EPI_ISL_726140, EPI_ISL_726141, EPI_ISL_726142, EPI_ISL_726143, EPI_ISL_726144, EPI_ISL_726145, EPI_ISL_726146, EPI_ISL_726147, EPI_ISL_726148, EPI_ISL_726149, EPI_ISL_726150, EPI_ISL_726151, EPI_ISL_726152, EPI_ISL_726153, EPI_ISL_726154, EPI_ISL_726155, EPI_ISL_726156, EPI_ISL_726157, EPI_ISL_726158, EPI_ISL_726159, EPI_ISL_726160, EPI_ISL_726161, EPI_ISL_726162, EPI_ISL_726163, EPI_ISL_726164, EPI_ISL_726165, EPI_ISL_726166, EPI_ISL_726167, EPI_ISL_726168, EPI_ISL_726169, EPI_ISL_726170, EPI_ISL_726171, EPI_ISL_726172, EPI_ISL_726173, EPI_ISL_726174, EPI_ISL_726175, EPI_ISL_726176, EPI_ISL_726177, EPI_ISL_726178, EPI_ISL_726179, EPI_ISL_726180, EPI_ISL_726181, EPI_ISL_726182, EPI_ISL_726183, EPI_ISL_726184, EPI_ISL_726185, EPI_ISL_726186, EPI_ISL_726187, EPI_ISL_726188, EPI_ISL_726189, EPI_ISL_726190, EPI_ISL_726191, EPI_ISL_726192, EPI_ISL_726193, EPI_ISL_726194, EPI_ISL_726195, EPI_ISL_726196, EPI_ISL_726197, EPI_ISL_726198, EPI_ISL_726199, EPI_ISL_726200, EPI_ISL_726201, EPI_ISL_726202, EPI_ISL_726203, EPI_ISL_726204, EPI_ISL_726205, EPI_ISL_726206, EPI_ISL_726207, EPI_ISL_726209, EPI_ISL_726210, EPI_ISL_726211, EPI_ISL_726214, EPI_ISL_726215, EPI_ISL_726216, EPI_ISL_726218, EPI_ISL_726219, EPI_ISL_726222, EPI_ISL_726223, EPI_ISL_726224, EPI_ISL_726225, EPI_ISL_726226, EPI_ISL_726227, EPI_ISL_726228, EPI_ISL_726229, EPI_ISL_726230, EPI_ISL_726231, EPI_ISL_726232, EPI_ISL_726233, EPI_ISL_726234, EPI_ISL_726235, EPI_ISL_726242, EPI_ISL_726243, EPI_ISL_726244, EPI_ISL_726245, EPI_ISL_726246, EPI_ISL_726247, EPI_ISL_726248, EPI_ISL_726249, EPI_ISL_726250, EPI_ISL_726252, EPI_ISL_726257, EPI_ISL_726258, EPI_ISL_726259, EPI_ISL_726260, EPI_ISL_726261, EPI_ISL_726262, EPI_ISL_726263, EPI_ISL_726264, EPI_ISL_726265, EPI_ISL_726268, EPI_ISL_726269, EPI_ISL_726270, EPI_ISL_726271, EPI_ISL_726276, EPI_ISL_726277, EPI_ISL_726278, EPI_ISL_726279, EPI_ISL_726305, EPI_ISL_726307, EPI_ISL_726311, EPI_ISL_726312, EPI_ISL_726313, EPI_ISL_726314, EPI_ISL_726315, EPI_ISL_726316, EPI_ISL_726317, EPI_ISL_726318, EPI_ISL_726319, EPI_ISL_726320, EPI_ISL_726321, EPI_ISL_726322, EPI_ISL_726323, EPI_ISL_726324, EPI_ISL_726325, EPI_ISL_726326, EPI_ISL_726327, EPI_ISL_726328, EPI_ISL_726329, EPI_ISL_726330, EPI_ISL_726331, EPI_ISL_726332, EPI_ISL_726333, EPI_ISL_726334, EPI_ISL_726335, EPI_ISL_726336, EPI_ISL_726337, EPI_ISL_726338, EPI_ISL_726339, EPI_ISL_726340, EPI_ISL_726341, EPI_ISL_726342, EPI_ISL_726343, EPI_ISL_726344, EPI_ISL_726345, EPI_ISL_726346, EPI_ISL_726347, EPI_ISL_726348, EPI_ISL_726349, EPI_ISL_726350, EPI_ISL_726351, EPI_ISL_726352, EPI_ISL_726353, EPI_ISL_726354, EPI_ISL_726355, EPI_ISL_726356, EPI_ISL_726358, EPI_ISL_726359, EPI_ISL_726360, EPI_ISL_726361, EPI_ISL_726362, EPI_ISL_726363, EPI_ISL_726364, EPI_ISL_726365, EPI_ISL_726366, EPI_ISL_726367, EPI_ISL_726368, EPI_ISL_726369, EPI_ISL_726370, EPI_ISL_726371, EPI_ISL_726372, EPI_ISL_726373, EPI_ISL_726374, EPI_ISL_726375, EPI_ISL_726376, EPI_ISL_726377, EPI_ISL_726378, EPI_ISL_726379, EPI_ISL_726380, EPI_ISL_726381, EPI_ISL_726382, EPI_ISL_726383, EPI_ISL_726384, EPI_ISL_726385, EPI_ISL_726386, EPI_ISL_726387, EPI_ISL_726388, EPI_ISL_726389, EPI_ISL_726390, EPI_ISL_726391, EPI_ISL_726392, EPI_ISL_726393, EPI_ISL_726394, EPI_ISL_726395, EPI_ISL_726396, EPI_ISL_726397, EPI_ISL_726398, EPI_ISL_726399, EPI_ISL_726401, EPI_ISL_726499, EPI_ISL_726500, EPI_ISL_726501, EPI_ISL_726502, EPI_ISL_726503, EPI_ISL_726504, EPI_ISL_726505, EPI_ISL_726506, EPI_ISL_726507, EPI_ISL_726508, EPI_ISL_726520, EPI_ISL_726521, EPI_ISL_726523, EPI_ISL_726524, EPI_ISL_726525, EPI_ISL_726526, EPI_ISL_726538, EPI_ISL_726539, EPI_ISL_726540, EPI_ISL_726541, EPI_ISL_726542, EPI_ISL_726546, EPI_ISL_726547, EPI_ISL_726548, EPI_ISL_726555, EPI_ISL_726556, EPI_ISL_726557, EPI_ISL_726558, EPI_ISL_726559, EPI_ISL_726567, EPI_ISL_726568, EPI_ISL_726572, EPI_ISL_726573, EPI_ISL_726574, EPI_ISL_726575, EPI_ISL_726576, EPI_ISL_726577, EPI_ISL_726578, EPI_ISL_726579, EPI_ISL_726580, EPI_ISL_726581, EPI_ISL_726582, EPI_ISL_726583, EPI_ISL_726584, EPI_ISL_726585, EPI_ISL_726587, EPI_ISL_726589, EPI_ISL_726591, EPI_ISL_726594, EPI_ISL_726599, EPI_ISL_726601, EPI_ISL_726603, EPI_ISL_726608, EPI_ISL_726612, EPI_ISL_726614, EPI_ISL_726617, EPI_ISL_726620, EPI_ISL_726621, EPI_ISL_726622, EPI_ISL_726623, EPI_ISL_726624, EPI_ISL_726625, EPI_ISL_726626, EPI_ISL_726627, EPI_ISL_726628, EPI_ISL_726629, EPI_ISL_726630, EPI_ISL_726631, EPI_ISL_726632, EPI_ISL_726633, EPI_ISL_726634, EPI_ISL_726635, EPI_ISL_726636, EPI_ISL_726637, EPI_ISL_726638, EPI_ISL_726639, EPI_ISL_726640, EPI_ISL_726641, EPI_ISL_726642, EPI_ISL_726643, EPI_ISL_726644, EPI_ISL_726645, EPI_ISL_726646, EPI_ISL_726647, EPI_ISL_726648, EPI_ISL_726649, EPI_ISL_726650, EPI_ISL_726651, EPI_ISL_726652, EPI_ISL_726653, EPI_ISL_726654, EPI_ISL_726655, EPI_ISL_726656, EPI_ISL_726657, EPI_ISL_726658, EPI_ISL_726659, EPI_ISL_726660, EPI_ISL_726661, EPI_ISL_726662, EPI_ISL_726663, EPI_ISL_726664, EPI_ISL_726665, EPI_ISL_726666, EPI_ISL_726667, EPI_ISL_726668, EPI_ISL_726670, EPI_ISL_726671, EPI_ISL_726672, EPI_ISL_726673, EPI_ISL_726674, EPI_ISL_726675, EPI_ISL_726676, EPI_ISL_726677, EPI_ISL_726678, EPI_ISL_726679, EPI_ISL_726680, EPI_ISL_726681, EPI_ISL_726682, EPI_ISL_726683, EPI_ISL_726684, EPI_ISL_726685, EPI_ISL_726686, EPI_ISL_726687, EPI_ISL_726688, EPI_ISL_726689, EPI_ISL_726691, EPI_ISL_726692, EPI_ISL_726693, EPI_ISL_726694, EPI_ISL_726695, EPI_ISL_726697, EPI_ISL_726698, EPI_ISL_726699, EPI_ISL_726700, EPI_ISL_726701, EPI_ISL_726702, EPI_ISL_726706, EPI_ISL_726707, EPI_ISL_726708, EPI_ISL_726709, EPI_ISL_726710, EPI_ISL_726711, EPI_ISL_726713, EPI_ISL_726714, EPI_ISL_726865, EPI_ISL_726866, EPI_ISL_726867, EPI_ISL_726868, EPI_ISL_726869, EPI_ISL_726870, EPI_ISL_726871, EPI_ISL_726872, EPI_ISL_726873, EPI_ISL_726874, EPI_ISL_726875, EPI_ISL_726876, EPI_ISL_726877, EPI_ISL_726878, EPI_ISL_726879, EPI_ISL_726880, EPI_ISL_726881, EPI_ISL_726882, EPI_ISL_726883, EPI_ISL_726884, EPI_ISL_726885, EPI_ISL_726886, EPI_ISL_726887, EPI_ISL_726888, EPI_ISL_726889, EPI_ISL_726890, EPI_ISL_726891, EPI_ISL_726892, EPI_ISL_726893, EPI_ISL_726894, EPI_ISL_726895, EPI_ISL_726896, EPI_ISL_726897, EPI_ISL_726898, EPI_ISL_726899, EPI_ISL_726900, EPI_ISL_726901, EPI_ISL_726902, EPI_ISL_726903, EPI_ISL_726904, EPI_ISL_726905, EPI_ISL_726906, EPI_ISL_726907, EPI_ISL_726908, EPI_ISL_726909, EPI_ISL_726910, EPI_ISL_726911, EPI_ISL_726912, EPI_ISL_726913, EPI_ISL_726914, EPI_ISL_726915, EPI_ISL_726916, EPI_ISL_726917, EPI_ISL_726918, EPI_ISL_726919, EPI_ISL_726920, EPI_ISL_726921, EPI_ISL_726922, EPI_ISL_726923, EPI_ISL_726924, EPI_ISL_726925, EPI_ISL_726926, EPI_ISL_726927, EPI_ISL_726928, EPI_ISL_726929, EPI_ISL_726930, EPI_ISL_726931, EPI_ISL_726932, EPI_ISL_726933, EPI_ISL_726934, EPI_ISL_726935, EPI_ISL_726936, EPI_ISL_726937, EPI_ISL_726938, EPI_ISL_726939, EPI_ISL_726940, EPI_ISL_726941, EPI_ISL_726942, EPI_ISL_727103, EPI_ISL_727104, EPI_ISL_727105, EPI_ISL_727106, EPI_ISL_727107, EPI_ISL_727109, EPI_ISL_727113, EPI_ISL_727114, EPI_ISL_727181, EPI_ISL_727182, EPI_ISL_727183, EPI_ISL_727185 | | | |
| see above | Wales Specialist Virology Centre Sequencing lab: Pathogen Genomics Unit | COVID-19 Genomics UK (COG-UK) Consortium | Catherine Moore, Johnathan Evans, Laura Gifford, Malorie Perry, Simon Cottrell, Angela Marchbank, Alec Birchley, Alexander Adams, Amy Gaskin, Bree Gatica-Wilcox, Jason Coombes, Joel Southgate, Lauren Gilbert, Lee Graham, Nicole Pacchiarini, Sara Kumziene-Summerhayes, Sarah Taylor, Sophie Jones, Sara Rey, Matthew Bull, Joanne Watkins, Sally Corden, Tom Connor |
| EPI_ISL_727908, EPI_ISL_727917, EPI_ISL_727978, EPI_ISL_727981, EPI_ISL_727996 | Virology Department, Sheffield Teaching Hospitals NHS Foundation Trust/Department of Infection, Immunity and Cardiovascular Disease, The Medical School, University of Sheffield | COVID-19 Genomics UK (COG-UK) Consortium | Thushan de Silva, Matthew Parker, Nikki Smith, Adri Angyal, Rebecca Brown, Luke Green, Rachel Tucker, Paul Parsons, Danielle Groves, Katie Johnson, Laura Carrilero, Alex Keeley, Dave Partridge, Matthew Wyles, Benjamin Lindsey, Mehmet Yavuz, Mohammad Raza, Cariad Evans |
| EPI_ISL_728058, EPI_ISL_728061, EPI_ISL_728062, EPI_ISL_728063, EPI_ISL_728090 | University of Wisconsin-Madison AIDS Vaccine Research Laboratories | University of Wisconsin-Madison AIDS Vaccine Research Laboratories | Gage Moreno, Katarina Braun, et al. AIDS Vaccine Research Laboratories |
| EPI_ISL_728534, EPI_ISL_728535, EPI_ISL_728536, EPI_ISL_728537, EPI_ISL_728538, EPI_ISL_728539, EPI_ISL_728540, EPI_ISL_728541, EPI_ISL_728542, EPI_ISL_728543, EPI_ISL_728544, EPI_ISL_728545 | | | |
| see above | CNR Virus des Infections Respiratoires - France SUD | CNR Virus des Infections Respiratoires - France SUD | Antonin Bal, Gregory Destras, Claudia Gonzalez, Gwendolyne Burfin, Quentin Semanas, Martine Valette, Bruno Lina, Laurence Josset |
| EPI_ISL_728674, EPI_ISL_728675, EPI_ISL_728685, EPI_ISL_728709, EPI_ISL_728710, EPI_ISL_728711, EPI_ISL_728712, EPI_ISL_728727, EPI_ISL_728734, EPI_ISL_728735, EPI_ISL_728748, EPI_ISL_728749, EPI_ISL_728750, EPI_ISL_728751, EPI_ISL_728752, EPI_ISL_728757 | | | |
| see above | Dutch COVID-19 response team | National Institute for Public Health and the Environment (RIVM) | Adam Meijer, Harry Vennema, Jeroen Cremer, Sharon van den Brink, Bas van der Veer, AnneMarie van den Brandt, Florian Zwagemaker, Dennis Schmitz, Chantal Reusken, on behalf of the national COVID-19 response team |
| EPI_ISL_729385, EPI_ISL_729684 | A. Krumbholz, Labor Dr. Krause und Kollegen MVZ GmbH, Kiel | Charité Universitätsmedizin Berlin, Institut für Virologie | Victor M Corman, Barbara Mühlemann, Jörn Beheim-Schwarzbach, Talitha Veith, Julia Schneider, Terry Jones, Christian Drosten |
| EPI_ISL_731900, EPI_ISL_731929, EPI_ISL_731930 | Instituto Nacional de Saude (INSA) and Instituto Gulbenkian de Ciencia (IGC) | Instituto Nacional de Saude (INSA) and Instituto Gulbenkian de Ciencia (IGC) | Borges et al |
| EPI_ISL_732111, EPI_ISL_732112, | Instituto Nacional de Saude (INSA) | Instituto Nacional de Saude (INSA) | Borges et al |

| | | | |
|---|---|---|---|
| EPI_ISL_732113 | | | |
| EPI_ISL_732186, EPI_ISL_732187, EPI_ISL_732188, EPI_ISL_732241, EPI_ISL_732242, EPI_ISL_732243, EPI_ISL_732244, EPI_ISL_732245 | Instituto Nacional de Saude (INSA) and Instituto Gulbenkian de Ciencia (IGC) | Instituto Nacional de Saude (INSA) and Instituto Gulbenkian de Ciencia (IGC) | Borges et al |
| EPI_ISL_732376, EPI_ISL_732377, EPI_ISL_732387, EPI_ISL_732392, EPI_ISL_732400, EPI_ISL_732428, EPI_ISL_732435, EPI_ISL_732443, EPI_ISL_732447, EPI_ISL_732510, EPI_ISL_732511, EPI_ISL_732512, EPI_ISL_732513 | | | |
| see above | National Virus Reference Laboratory | National Virus Reference Laboratory | Michael Carr, Gabriel Gonzalez, Jonathan Dean, Daniel Hare, Cillian F De Gascun |
| EPI_ISL_732689, EPI_ISL_732690 | CNR Virus des Infections Respiratoires - France SUD | CNR Virus des Infections Respiratoires - France SUD | Antonin Bal, Gregory Destras, Claudia Gonzalez, Gwendolyne Burfin, Quentin Semanas, Martine Valette, Bruno Lina, Laurence Josset |
| EPI_ISL_733462, EPI_ISL_733463, EPI_ISL_733465, EPI_ISL_733468, EPI_ISL_733470, EPI_ISL_733471, EPI_ISL_733472, EPI_ISL_733474 | WHO National Influenza Centre Russian Federation | WHO National Influenza Centre Russian Federation | Andrey Komissarov, Artem Fadeev, Anna Ivanova, Kseniya Komissarova, Dmitry Bazhenov, Daria Danilenko, Ksenia Safina, Elena Nabieva, Georgii Bazykin, Dmitry Lioznov |
| EPI_ISL_733496 | University Hospital Zürich | Institute of Medical Virology, University of Zurich | Stefan Schmutz, Verena Kufner, Maryam Zaheri, Gabriela Ziltener, Thomas Scheier, Jürg Böni, Michael Huber, Alexandra Trkola |
| EPI_ISL_734177, EPI_ISL_734178, EPI_ISL_734179, EPI_ISL_734193, EPI_ISL_734194, EPI_ISL_734195, EPI_ISL_734196, EPI_ISL_734197, EPI_ISL_734198, EPI_ISL_734226, EPI_ISL_734227, EPI_ISL_734228, EPI_ISL_734229, EPI_ISL_734231 | | | |
| see above | Virginia Division of Consolidated Laboratory Services (DCLS) | Virginia Division of Consolidated Laboratory Services (DCLS) | Virginia DCLS |
| EPI_ISL_735493 | Cox's Bazar Medicila College | Central Biological Research Laboratory and Department of Biochemistry and Molecular Biology Central Biological Research Laboratory and Department of Biochemistry and Molecular Biology | Robiul Hasan Bhuiyan, Md. Imranul Hoq, Md. Khondakar Raziur Rahman, Imam Hossen, Sajib Rudra, Md. Arif Hossain, Shanta Paul, Md. Omer Faruq, H. M. Abdullah Al Masud, Mohammad Omar Faruque |
| EPI_ISL_735496 | Chattogram Veterinary and Animal Sciences University | Central Biological Research Laboratory and Department of Biochemistry and Molecular Biology | Md. Imranul Hoq, Robiul Hasan Bhuiyan, Md. Khondakar Raziur Rahman, Imam Hossen, Sajib Rudra, Md. Arif Hossain, Shanta Paul, Md. Omer Faruq, Mohammad Omar Faruque, H. M. Abdullah Al Masud |
| EPI_ISL_735497 | Chattogram Veterinary and Animal Sciences University | Central Biological Research Laboratory and Department of Biochemistry and Molecular Biology | Mohammad Omar Faruque, H. M. Abdullah Al Masud, Imam Hossen, Md. Khondakar Raziur Rahman, Sajib Rudra, Md. Arif Hossain, Shanta Paul, Md. Omer Faruq, Robiul Hasan Bhuiyan, Md. Imranul Hoq |
| EPI_ISL_736957, EPI_ISL_736958, EPI_ISL_736959 | NHLS-IALCH | KRISP, KZN Research Innovation and Sequencing Platform | Giandhari J, Pillay S, Lessells R, ChimukangaraB, Mdlalose K, York D, Khan S, Tegally H, Wilkinson E, de Oliveira T |
| EPI_ISL_737103, EPI_ISL_737115, EPI_ISL_737125, EPI_ISL_737131, EPI_ISL_737134, EPI_ISL_737135, EPI_ISL_737141, EPI_ISL_737142, EPI_ISL_737143, EPI_ISL_737144, EPI_ISL_737146, EPI_ISL_737147, EPI_ISL_737148, EPI_ISL_737151, EPI_ISL_737156, EPI_ISL_737157, EPI_ISL_737167, EPI_ISL_737175, EPI_ISL_737183, EPI_ISL_737194, EPI_ISL_737196, EPI_ISL_737286, EPI_ISL_737288, EPI_ISL_737292, EPI_ISL_737294, EPI_ISL_737303, EPI_ISL_737304 | | | |
| see above | Michigan Department of Health and Human Services, Bureau of Laboratories | Michigan Department of Health and Human Services, Bureau of Laboratories | Blankenship HM, Riner D, Soehnlen MK |
| EPI_ISL_738505 | Humboldt County Public Health Laboratory | Chan-Zuckerberg Biohub | CZB Cliahub Consortium |
| EPI_ISL_738508, EPI_ISL_738525, EPI_ISL_738533, EPI_ISL_738534, EPI_ISL_738543, EPI_ISL_738549, EPI_ISL_738551, EPI_ISL_738552, EPI_ISL_738558, EPI_ISL_738582, EPI_ISL_738583, EPI_ISL_738589, EPI_ISL_738608, EPI_ISL_738611, EPI_ISL_738631, EPI_ISL_738643 | | | |
| see above | Santa Clara County Public Health Laboratory | Chan-Zuckerberg Biohub | CZB Cliahub Consortium |
| EPI_ISL_738647, EPI_ISL_738649 | Humboldt County Public Health Laboratory | Chan-Zuckerberg Biohub | CZB Cliahub Consortium |
| EPI_ISL_738650 | Santa Clara County Public Health Laboratory | Chan-Zuckerberg Biohub | CZB Cliahub Consortium |
| EPI_ISL_738651 | Humboldt County Public Health Laboratory | Chan-Zuckerberg Biohub | CZB Cliahub Consortium |
| EPI_ISL_738652, EPI_ISL_738653, EPI_ISL_738657, EPI_ISL_738661, EPI_ISL_738663, EPI_ISL_738689, EPI_ISL_738697 | Santa Clara County Public Health Laboratory | Chan-Zuckerberg Biohub | CZB Cliahub Consortium |
| EPI_ISL_738723 | Renegade | Chan-Zuckerberg Biohub | CZB Cliahub Consortium |
| EPI_ISL_738746, EPI_ISL_738763, EPI_ISL_738772, EPI_ISL_738789 | Santa Clara County Public Health Laboratory | Chan-Zuckerberg Biohub | CZB Cliahub Consortium |
| EPI_ISL_738790 | Orange County Public Health Lab | Chan-Zuckerberg Biohub | CZB Cliahub Consortium |
| EPI_ISL_738791, EPI_ISL_738794, EPI_ISL_738803 | Santa Clara County Public Health Laboratory | Chan-Zuckerberg Biohub | CZB Cliahub Consortium |
| EPI_ISL_738809 | Humboldt County Public Health Laboratory | Chan-Zuckerberg Biohub | CZB Cliahub Consortium |
| EPI_ISL_738812 | Santa Clara County Public Health Laboratory | Chan-Zuckerberg Biohub | CZB Cliahub Consortium |
| EPI_ISL_738854 | Humboldt County Public Health Laboratory | Chan-Zuckerberg Biohub | CZB Cliahub Consortium |
| EPI_ISL_738883, EPI_ISL_738889, EPI_ISL_738906, EPI_ISL_738909 | Santa Clara County Public Health Laboratory | Chan-Zuckerberg Biohub | CZB Cliahub Consortium |
| EPI_ISL_738929 | Humboldt County Public Health Laboratory | Chan-Zuckerberg Biohub | CZB Cliahub Consortium |
| EPI_ISL_738931, EPI_ISL_738933, EPI_ISL_738941, EPI_ISL_738942, EPI_ISL_738948 | Santa Clara County Public Health Laboratory | Chan-Zuckerberg Biohub | CZB Cliahub Consortium |
| EPI_ISL_738958 | Humboldt County Public Health Laboratory | Chan-Zuckerberg Biohub | CZB Cliahub Consortium |
| EPI_ISL_738971, EPI_ISL_738984, EPI_ISL_738987 | Santa Clara County Public Health Laboratory | Chan-Zuckerberg Biohub | CZB Cliahub Consortium |
| EPI_ISL_738991 | Orange County Public Health Lab | Chan-Zuckerberg Biohub | CZB Cliahub Consortium |
| EPI_ISL_738999 | Humboldt County Public Health Laboratory | Chan-Zuckerberg Biohub | CZB Cliahub Consortium |
| EPI_ISL_739008, EPI_ISL_739012, EPI_ISL_739021, EPI_ISL_739041, EPI_ISL_739065, EPI_ISL_739070 | Santa Clara County Public Health Laboratory | Chan-Zuckerberg Biohub | CZB Cliahub Consortium |
| EPI_ISL_739072 | Orange County Public Health Lab | Chan-Zuckerberg Biohub | CZB Cliahub Consortium |
| EPI_ISL_739085 | Santa Clara County Public Health Laboratory | Chan-Zuckerberg Biohub | CZB Cliahub Consortium |
| EPI_ISL_739106 | Humboldt County Public Health Laboratory | Chan-Zuckerberg Biohub | CZB Cliahub Consortium |
| EPI_ISL_739117, EPI_ISL_739121 | Santa Clara County Public Health Laboratory | Chan-Zuckerberg Biohub | CZB Cliahub Consortium |
| EPI_ISL_739126 | Humboldt County Public Health Laboratory | Chan-Zuckerberg Biohub | CZB Cliahub Consortium |

| | | | |
|---|---|---|---|
| EPI_ISL_739128, EPI_ISL_739129 | Santa Clara County Public Health Laboratory | Chan-Zuckerberg Biohub | CZB Cliahub Consortium |
| EPI_ISL_739131 | Humboldt County Public Health Laboratory | Chan-Zuckerberg Biohub | CZB Cliahub Consortium |
| EPI_ISL_739138 | Santa Clara County Public Health Laboratory | Chan-Zuckerberg Biohub | CZB Cliahub Consortium |
| EPI_ISL_739155, EPI_ISL_739159 | Humboldt County Public Health Laboratory | Chan-Zuckerberg Biohub | CZB Cliahub Consortium |
| EPI_ISL_739180 | Santa Clara County Public Health Laboratory | Chan-Zuckerberg Biohub | CZB Cliahub Consortium |
| EPI_ISL_739183 | Humboldt County Public Health Laboratory | Chan-Zuckerberg Biohub | CZB Cliahub Consortium |
| EPI_ISL_739187 | Santa Clara County Public Health Laboratory | Chan-Zuckerberg Biohub | CZB Cliahub Consortium |
| EPI_ISL_739198 | Humboldt County Public Health Laboratory | Chan-Zuckerberg Biohub | CZB Cliahub Consortium |
| EPI_ISL_739219, EPI_ISL_739234 | Santa Clara County Public Health Laboratory | Chan-Zuckerberg Biohub | CZB Cliahub Consortium |
| EPI_ISL_739235 | Humboldt County Public Health Laboratory | Chan-Zuckerberg Biohub | CZB Cliahub Consortium |
| EPI_ISL_739237 | Santa Clara County Public Health Laboratory | Chan-Zuckerberg Biohub | CZB Cliahub Consortium |
| EPI_ISL_739250 | Humboldt County Public Health Laboratory | Chan-Zuckerberg Biohub | CZB Cliahub Consortium |
| EPI_ISL_739252, EPI_ISL_739254 | Santa Clara County Public Health Laboratory | Chan-Zuckerberg Biohub | CZB Cliahub Consortium |
| EPI_ISL_739269 | Humboldt County Public Health Laboratory | Chan-Zuckerberg Biohub | CZB Cliahub Consortium |
| EPI_ISL_739273, EPI_ISL_739278 | Santa Clara County Public Health Laboratory | Chan-Zuckerberg Biohub | CZB Cliahub Consortium |
| EPI_ISL_739299 | Humboldt County Public Health Laboratory | Chan-Zuckerberg Biohub | CZB Cliahub Consortium |
| EPI_ISL_739310, EPI_ISL_739360, EPI_ISL_739377, EPI_ISL_739382 | Santa Clara County Public Health Laboratory | Chan-Zuckerberg Biohub | CZB Cliahub Consortium |
| EPI_ISL_739394 | Renegade | Chan-Zuckerberg Biohub | CZB Cliahub Consortium |
| EPI_ISL_739413, EPI_ISL_739416 | Santa Clara County Public Health Laboratory | Chan-Zuckerberg Biohub | CZB Cliahub Consortium |
| EPI_ISL_739418 | Humboldt County Public Health Laboratory | Chan-Zuckerberg Biohub | CZB Cliahub Consortium |
| EPI_ISL_739431 | Santa Clara County Public Health Laboratory | Chan-Zuckerberg Biohub | CZB Cliahub Consortium |
| EPI_ISL_739441 | Orange County Public Health Lab | Chan-Zuckerberg Biohub | CZB Cliahub Consortium |
| EPI_ISL_739443 | Santa Clara County Public Health Laboratory | Chan-Zuckerberg Biohub | CZB Cliahub Consortium |
| EPI_ISL_739445 | Orange County Public Health Lab | Chan-Zuckerberg Biohub | CZB Cliahub Consortium |
| EPI_ISL_739452 | Humboldt County Public Health Laboratory | Chan-Zuckerberg Biohub | CZB Cliahub Consortium |
| EPI_ISL_739458, EPI_ISL_739491, EPI_ISL_739498 | Santa Clara County Public Health Laboratory | Chan-Zuckerberg Biohub | CZB Cliahub Consortium |
| EPI_ISL_739499 | Humboldt County Public Health Laboratory | Chan-Zuckerberg Biohub | CZB Cliahub Consortium |
| EPI_ISL_739501, EPI_ISL_739508, EPI_ISL_739509, EPI_ISL_739520, EPI_ISL_739537, EPI_ISL_739549 | Santa Clara County Public Health Laboratory | Chan-Zuckerberg Biohub | CZB Cliahub Consortium |
| EPI_ISL_739559, EPI_ISL_739581 | Orange County Public Health Lab | Chan-Zuckerberg Biohub | CZB Cliahub Consortium |
| EPI_ISL_739585 | Humboldt County Public Health Laboratory | Chan-Zuckerberg Biohub | CZB Cliahub Consortium |
| EPI_ISL_739594, EPI_ISL_739597, EPI_ISL_739602 | Santa Clara County Public Health Laboratory | Chan-Zuckerberg Biohub | CZB Cliahub Consortium |
| EPI_ISL_739607 | Orange County Public Health Lab | Chan-Zuckerberg Biohub | CZB Cliahub Consortium |
| EPI_ISL_739608 | Humboldt County Public Health Laboratory | Chan-Zuckerberg Biohub | CZB Cliahub Consortium |
| EPI_ISL_739626, EPI_ISL_739629, EPI_ISL_739647 | Santa Clara County Public Health Laboratory | Chan-Zuckerberg Biohub | CZB Cliahub Consortium |
| EPI_ISL_739672 | Instituto Nacional de Salud, Bogotá, Colombia | Instituto Nacional de Salud, Bogotá, Colombia | Katherine Laiton-Donato, Diego A. Álvarez-Díaz, Carlos Franco-Muñoz, Mauricio Pacheco-Montealegre, Jonathan Reales, Diego Andrés Prada, Sheryl Corchuelo, Magdalena Weisner, Martha Lucia Ospina Martinez, Marcela Mercado-Reyes |
| EPI_ISL_739694, EPI_ISL_739697, EPI_ISL_739721, EPI_ISL_739724, EPI_ISL_739730, EPI_ISL_739740, EPI_ISL_739764, EPI_ISL_739771, EPI_ISL_739846, EPI_ISL_739857, EPI_ISL_739858, EPI_ISL_739900, EPI_ISL_739901, EPI_ISL_739921, EPI_ISL_739966, EPI_ISL_739972, EPI_ISL_739976, EPI_ISL_739984, EPI_ISL_739991, EPI_ISL_740012, EPI_ISL_740072, EPI_ISL_740094, EPI_ISL_740119, EPI_ISL_740158, EPI_ISL_740167, EPI_ISL_740212, EPI_ISL_740213, EPI_ISL_740236, EPI_ISL_740284, EPI_ISL_740299, EPI_ISL_740309, EPI_ISL_740313, EPI_ISL_740323, EPI_ISL_740335, EPI_ISL_740339, EPI_ISL_740341, EPI_ISL_740351, EPI_ISL_740365, EPI_ISL_740368, EPI_ISL_740376, EPI_ISL_740378, EPI_ISL_740380, EPI_ISL_740389, EPI_ISL_740392, EPI_ISL_740393, EPI_ISL_740395, EPI_ISL_740416, EPI_ISL_740420, EPI_ISL_740421, EPI_ISL_740428, EPI_ISL_740437, EPI_ISL_740439, EPI_ISL_740463, EPI_ISL_740466, EPI_ISL_740472, EPI_ISL_740478, EPI_ISL_740494, EPI_ISL_740505, EPI_ISL_740523, EPI_ISL_740529, EPI_ISL_740541 | | | |
| see above | Laboratoire national de santé, Microbiology, Virology | Laboratoire national de santé, Microbiology, Microbial Genomics Platform | Anke Wienecke-Baldacchino, Catherine Ragimbeau,Jessica Tapp, Fatu Djabi, Lise Pignon, Raoul Salmon, Tamir Abdelrahman |
| EPI_ISL_741759 | Centre for Enzyme Innovation, University of Portsmouth / Translational Research Laboratory, Portsmouth Hospitals NHS Trust | COVID-19 Genomics UK (COG-UK) Consortium | Angela Beckett,Yann Bourgeois,Garry Scarlett,Sharon Glaysher,Scott Elliott,Kelly Bicknell,Robert Impey,Allyson Lloyd,Sarah Wyllie,Ethan Butcher,Anoop Chauhan,Samuel Robson |
| EPI_ISL_742174 | Regional Virus Laboratory, Belfast Health and Social Care Trust | COVID-19 Genomics UK (COG-UK) Consortium | Conall McCaughey, James McKenna, Tanya Curran, Susan Feeney, Alison Watt, Ciara Cox, Mairead Connor, Zoltan Molnar, David Simpson, Derek Fairley |
| EPI_ISL_743611, EPI_ISL_743612, EPI_ISL_743613 | Wales Specialist Virology Centre Sequencing lab: Pathogen Genomics Unit | COVID-19 Genomics UK (COG-UK) Consortium | Catherine Moore, Johnathan Evans, Laura Gifford, Malorie Perry, Simon Cottrell, Angela Marchbank, Alec Birchley, Alexander Adams, Amy Gaskin, Bree Gatica-Wilcox, Jason Coombes, Joel Southgate, Lauren Gilbert, Lee Graham, Nicole Pacchiarini, Sara Kumziene-Summerhayes, Sarah Taylor, Sophie Jones, Sara Rey, Matthew Bull, Joanne Watkins, Sally Corden, Tom Connor |
| EPI_ISL_744151, EPI_ISL_744176, EPI_ISL_744183, EPI_ISL_744210, EPI_ISL_744230, EPI_ISL_744273, EPI_ISL_744328, EPI_ISL_744330, EPI_ISL_744333, EPI_ISL_744335, EPI_ISL_744338, EPI_ISL_744342, EPI_ISL_744360, EPI_ISL_744377, EPI_ISL_744468, EPI_ISL_744479, EPI_ISL_744485, EPI_ISL_744504, EPI_ISL_744508, EPI_ISL_744518, EPI_ISL_744523, EPI_ISL_744542, EPI_ISL_744554, EPI_ISL_744558, EPI_ISL_744564, EPI_ISL_744576, EPI_ISL_744580, EPI_ISL_744606, EPI_ISL_744608, EPI_ISL_744689, EPI_ISL_744717, EPI_ISL_744728, EPI_ISL_744730, EPI_ISL_744743, EPI_ISL_744747, EPI_ISL_744786, EPI_ISL_744787, EPI_ISL_744817, EPI_ISL_744838, EPI_ISL_744839, EPI_ISL_744840, EPI_ISL_744850, EPI_ISL_744856, EPI_ISL_744871, EPI_ISL_744884, EPI_ISL_744905, EPI_ISL_744915, EPI_ISL_744927, EPI_ISL_744930, EPI_ISL_744931, EPI_ISL_744944, EPI_ISL_744959, EPI_ISL_744965, EPI_ISL_744966, EPI_ISL_744980, EPI_ISL_744991, EPI_ISL_745008, EPI_ISL_745009 | | | |
| see above | Laboratoire national de santé, Microbiology, Virology | Laboratoire national de santé, Microbiology, Microbial Genomics Platform | Anke Wienecke-Baldacchino, Catherine Ragimbeau,Jessica Tapp, Fatu Djabi, Lise Pignon, Raoul Salmon, Tamir Abdelrahman |
| EPI_ISL_745335 | CHU Clermont-Ferrand | CNR Virus des Infections Respiratoires - France SUD | Antonin Bal, Gregory Destras, Gwendolyne Burfin, Hadrien Règue, Quentin Semanas, Martine Valette, Bruno Lina, Christine Archimbaud, Amélie Brebion, Hélène Chabrolles, Martine Chambon, Audrey Mirand, Christel Regagnon, Maxime Bisseux, Patricia Combes, Cécile Henquell, Laurence Josset |
| EPI_ISL_746839, EPI_ISL_746853, EPI_ISL_746899, EPI_ISL_746914, | Utah Public Health Laboratory | Utah Public Health Laboratory | Erin Young, Kelly Oakeson, Tara Gallagher |

| | | | |
|---|---|---|---|
| EPI_ISL_746963, EPI_ISL_746974, EPI_ISL_746979 | | | |
| EPI_ISL_747168, EPI_ISL_747170, EPI_ISL_747171, EPI_ISL_747181, EPI_ISL_747182 | Respiratory Viruses Branch, Centers for Disease Control and Prevention | Respiratory Viruses Branch, Centers for Disease Control and Prevention | Queen,K., Li,Y., Tao,Y., Uehara,A., Montmayeur,A., Paden,C.R., Cook,P.W., Marine,R., Sheth,M., Wang,H., Lee,J., Tong,S. |
| EPI_ISL_747391, EPI_ISL_747398, EPI_ISL_747399, EPI_ISL_747408, EPI_ISL_747425, EPI_ISL_747426, EPI_ISL_747427, EPI_ISL_747428, EPI_ISL_747429, EPI_ISL_747438, EPI_ISL_747458 | | | |
| see above | Division of Emerging Infectious Diseases, Bureau of Infectious Diseases Diagnosis Control, Korea Disease Control and Prevention Agency | Division of Emerging Infectious Diseases, Bureau of Infectious Diseases Diagnosis Control, Korea Disease Control and Prevention Agency | Ae Kyung Park, Il-Hwan Kim, Heui Man Kim, Jeong-Min Kim, Namjoo Lee, Chaeyoung Lee, Sang Hee Woo, Eun-Jin Kim |
| EPI_ISL_747471, EPI_ISL_747472, EPI_ISL_747473, EPI_ISL_747474, EPI_ISL_747475 | ULSS9 Distretto di Bussolengo | Istituto Zooprofilattico Sperimentale delle Venezie | Adelaide Milani, Alessia Schivo, Annalisa Salviato, Erika Giorgia Quaranta, Ambra Pastori, Bianca Zecchin, Alice Fusaro, Isabella Monne, Calogero Terregino, Antonia Ricci |
| EPI_ISL_747531, EPI_ISL_747585, EPI_ISL_747586, EPI_ISL_747587, EPI_ISL_747598, EPI_ISL_747609, EPI_ISL_747629, EPI_ISL_747630, EPI_ISL_747631, EPI_ISL_747680, EPI_ISL_748099, EPI_ISL_748100, EPI_ISL_748101, EPI_ISL_748102, EPI_ISL_748103, EPI_ISL_748104, EPI_ISL_748105, EPI_ISL_748106, EPI_ISL_748107, EPI_ISL_748108, EPI_ISL_748109, EPI_ISL_748110, EPI_ISL_748111, EPI_ISL_748112 | | | |
| see above | Department of Virus and Microbiological Special Diagnostics, Statens Serum Institut, Copenhagen, Denmark | Albertsen Lab, Department of Chemistry and Bioscience, Aalborg University, Denmark | Danish Covid-19 Genome Consortium |
| EPI_ISL_751458, EPI_ISL_751459, EPI_ISL_751460, EPI_ISL_751464, EPI_ISL_751473 | CHU Purpan - Laboratoire de Virologie - Institut Fédératif de Biologie | CHU Purpan - Laboratoire de Virologie - Institut Fédératif de Biologie | Latour J., Ranger N., Dubois M., Carcenac R., Harter A., Boyer P., Tremeaux P., Izopet J. |
| EPI_ISL_751562 | NYSDOH Wadsworth Center, Virology Lab | Genomics and Discovery, Respiratory Viruses Branch, Division of Viral Diseases, Centers for Disease Control and Prevention | Krista Queen, Yan Li, Ying Tao, Jing Zhang, Anna Uehara, Anna Montmayeur, Clinton R. Paden, Peter W. Cook,Rachel Marine, Mili Sheth, Haibin Wang, Justin Lee, Suxiang Tong |
| EPI_ISL_751577 | MD DOH Laboratories Administration | Genomics and Discovery, Respiratory Viruses Branch, Division of Viral Diseases, Centers for Disease Control and Prevention | Krista Queen, Yan Li, Ying Tao, Jing Zhang, Anna Uehara, Anna Montmayeur, Clinton R. Paden, Peter W. Cook,Rachel Marine, Mili Sheth, Haibin Wang, Justin Lee, Suxiang Tong |
| EPI_ISL_751581 | NYSDOH Wadsworth Center, Virology Lab | Genomics and Discovery, Respiratory Viruses Branch, Division of Viral Diseases, Centers for Disease Control and Prevention | Krista Queen, Yan Li, Ying Tao, Jing Zhang, Anna Uehara, Anna Montmayeur, Clinton R. Paden, Peter W. Cook,Rachel Marine, Mili Sheth, Haibin Wang, Justin Lee, Suxiang Tong |
| EPI_ISL_751606, EPI_ISL_751607 | MD DOH Laboratories Administration | Genomics and Discovery, Respiratory Viruses Branch, Division of Viral Diseases, Centers for Disease Control and Prevention | Krista Queen, Yan Li, Ying Tao, Jing Zhang, Anna Uehara, Anna Montmayeur, Clinton R. Paden, Peter W. Cook,Rachel Marine, Mili Sheth, Haibin Wang, Justin Lee, Suxiang Tong |
| EPI_ISL_751638 | NE Public Health Laboratory | Genomics and Discovery, Respiratory Viruses Branch, Division of Viral Diseases, Centers for Disease Control and Prevention | Krista Queen, Yan Li, Ying Tao, Jing Zhang, Anna Uehara, Anna Montmayeur, Clinton R. Paden, Peter W. Cook,Rachel Marine, Mili Sheth, Haibin Wang, Justin Lee, Suxiang Tong |
| EPI_ISL_751649, EPI_ISL_751650 | CT-Dr. Katherine A. Kelley State Public Health Lab | Genomics and Discovery, Respiratory Viruses Branch, Division of Viral Diseases, Centers for Disease Control and Prevention | Krista Queen, Yan Li, Ying Tao, Jing Zhang, Anna Uehara, Anna Montmayeur, Clinton R. Paden, Peter W. Cook,Rachel Marine, Mili Sheth, Haibin Wang, Justin Lee, Suxiang Tong |
| EPI_ISL_751673 | HI Dept. of Health, State Laboratories Division | Genomics and Discovery, Respiratory Viruses Branch, Division of Viral Diseases, Centers for Disease Control and Prevention | Krista Queen, Yan Li, Ying Tao, Jing Zhang, Anna Uehara, Anna Montmayeur, Clinton R. Paden, Peter W. Cook,Rachel Marine, Mili Sheth, Haibin Wang, Justin Lee, Suxiang Tong |
| EPI_ISL_751702 | MD DOH Laboratories Administration | Genomics and Discovery, Respiratory Viruses Branch, Division of Viral Diseases, Centers for Disease Control and Prevention | Krista Queen, Yan Li, Ying Tao, Jing Zhang, Anna Uehara, Anna Montmayeur, Clinton R. Paden, Peter W. Cook,Rachel Marine, Mili Sheth, Haibin Wang, Justin Lee, Suxiang Tong |
| EPI_ISL_751709 | NJ Public Health and Environmental Laboratories | Genomics and Discovery, Respiratory Viruses Branch, Division of Viral Diseases, Centers for Disease Control and Prevention | Krista Queen, Yan Li, Ying Tao, Jing Zhang, Anna Uehara, Anna Montmayeur, Clinton R. Paden, Peter W. Cook,Rachel Marine, Mili Sheth, Haibin Wang, Justin Lee, Suxiang Tong |
| EPI_ISL_751725 | MD DOH Laboratories Administration | Genomics and Discovery, Respiratory Viruses Branch, Division of Viral Diseases, Centers for Disease Control and Prevention | Krista Queen, Yan Li, Ying Tao, Jing Zhang, Anna Uehara, Anna Montmayeur, Clinton R. Paden, Peter W. Cook,Rachel Marine, Mili Sheth, Haibin Wang, Justin Lee, Suxiang Tong |
| EPI_ISL_751767 | HI Dept. of Health, State Laboratories Division | Genomics and Discovery, Respiratory Viruses Branch, Division of Viral Diseases, Centers for Disease Control and Prevention | Krista Queen, Yan Li, Ying Tao, Jing Zhang, Anna Uehara, Anna Montmayeur, Clinton R. Paden, Peter W. Cook,Rachel Marine, Mili Sheth, Haibin Wang, Justin Lee, Suxiang Tong |
| EPI_ISL_751787 | TX DSHS, Lab Services Section MC 1947 | Genomics and Discovery, Respiratory Viruses Branch, Division of Viral Diseases, Centers for Disease Control and Prevention | Krista Queen, Yan Li, Ying Tao, Jing Zhang, Anna Uehara, Anna Montmayeur, Clinton R. Paden, Peter W. Cook,Rachel Marine, Mili Sheth, Haibin Wang, Justin Lee, Suxiang Tong |
| EPI_ISL_751788 | OH Department of Health Laboratory | Genomics and Discovery, Respiratory Viruses Branch, Division of Viral Diseases, Centers for Disease Control and Prevention | Krista Queen, Yan Li, Ying Tao, Jing Zhang, Anna Uehara, Anna Montmayeur, Clinton R. Paden, Peter W. Cook,Rachel Marine, Mili Sheth, Haibin Wang, Justin Lee, Suxiang Tong |
| EPI_ISL_752531, EPI_ISL_752532, EPI_ISL_752533, EPI_ISL_752534, EPI_ISL_752535, EPI_ISL_752536, EPI_ISL_752537, EPI_ISL_752538, EPI_ISL_752539, EPI_ISL_752540, EPI_ISL_752541, EPI_ISL_752542, EPI_ISL_752543, EPI_ISL_752544, EPI_ISL_752545, EPI_ISL_752546, EPI_ISL_752547, EPI_ISL_752548, EPI_ISL_752549, EPI_ISL_752550, EPI_ISL_752551, EPI_ISL_752552, EPI_ISL_752553, EPI_ISL_752554, EPI_ISL_752555, EPI_ISL_752556, EPI_ISL_752557, EPI_ISL_752558, EPI_ISL_752559, EPI_ISL_752560, EPI_ISL_752561, EPI_ISL_752562, EPI_ISL_752563, EPI_ISL_752564, EPI_ISL_752565, EPI_ISL_752566, EPI_ISL_752567, EPI_ISL_752568, EPI_ISL_752569, EPI_ISL_752570, EPI_ISL_752571, EPI_ISL_752572, EPI_ISL_752573, EPI_ISL_752574, EPI_ISL_752575, EPI_ISL_752576, EPI_ISL_752577, EPI_ISL_752578, EPI_ISL_752579, EPI_ISL_752580, EPI_ISL_752581, EPI_ISL_752582, EPI_ISL_752593 | | | |
| see above | National Virus Reference Laboratory | National Virus Reference Laboratory | Michael Carr, Gabriel Gonzalez, Jonathan Dean, Daniel Hare, Cillian F De Gascun |
| EPI_ISL_753116, EPI_ISL_753117, EPI_ISL_753118, EPI_ISL_753119, EPI_ISL_753120, EPI_ISL_753129, EPI_ISL_753130, EPI_ISL_753131, EPI_ISL_753132, EPI_ISL_753133, EPI_ISL_753134, EPI_ISL_753135, EPI_ISL_753136, EPI_ISL_753137, EPI_ISL_753138, EPI_ISL_753139, EPI_ISL_753140, EPI_ISL_753141, EPI_ISL_753142, EPI_ISL_753143, EPI_ISL_753144, EPI_ISL_753145, EPI_ISL_753146, EPI_ISL_753154, EPI_ISL_753155, EPI_ISL_753156, EPI_ISL_753157, EPI_ISL_753158, EPI_ISL_753159, EPI_ISL_753160, EPI_ISL_753161, EPI_ISL_753162, EPI_ISL_753163, EPI_ISL_753164, EPI_ISL_753165, EPI_ISL_753166, EPI_ISL_753167, EPI_ISL_753168, EPI_ISL_753169, EPI_ISL_753170, EPI_ISL_753171, EPI_ISL_753172, EPI_ISL_753173 | | | |
| see above | State Laboratories Division, Hawaii State Department of Health | State Laboratories Division, Hawaii State Department of Health | Pamela O'Brien, Sabrina Diemert, Drew Kuwazaki, Razvan Sultana, Edward Desmond |
| EPI_ISL_753393, EPI_ISL_753395, EPI_ISL_753396, EPI_ISL_753397, EPI_ISL_753398, EPI_ISL_753399, EPI_ISL_753400, EPI_ISL_753401, EPI_ISL_753402, EPI_ISL_753409, EPI_ISL_753410, EPI_ISL_753411, EPI_ISL_753412 | | | |
| see above | Clinical virology Laboratory, Children's Hospital Los Angeles | Center for Personalized Medicine, Children's Hospital Los Angeles | Gai et al |
| EPI_ISL_753741, EPI_ISL_753828, EPI_ISL_753829, EPI_ISL_753830 | Charité Universitätsmedizin Berlin, Institut für Virologie/Labor Berlin | Charité Universitätsmedizin Berlin, Institut für Virologie | Victor M Corman, Jörn Beheim-Schwarzbach, Barbara Mühlemann, Julia Schneider, Talitha Veith, Terry Jones, Christian Drosten |
| EPI_ISL_754143, EPI_ISL_754144, | Laboratoire de Microbiologie | National Reference Center for Viruses of Respiratory | Marion Barbet, Sylvie Behillil, Méline Bizard, Angela Brisebarre, Camille Capel, Etienne Simon-Lorière, Vincent Enouf, Maud Vanpeene, Sylvie van der |

| | | | |
|---|---|---|---|
| EPI_ISL_754152, EPI_ISL_754153, EPI_ISL_754154 | | Infections, Institut Pasteur, Paris | Werf, Marie-Sarah Fangous |
| EPI_ISL_754233 | The Republican Research and Practical Center for Epidemiology and Microbiology (RRPCEM) | WHO National Influenza Centre Russian Federation | Elena Gasich, Kirill Bulda, Anatoly Krasko, Andrey Komissarov, Artem Fadeev, Anna Ivanova, Kseniya Komissarova, Dmitry Bazhenov, Daria Danilenko, Ksenia Safina, Elena Nabieva, Georgii Bazykin, Dmitry Lioznov |
| EPI_ISL_754405, EPI_ISL_754406, EPI_ISL_754477, EPI_ISL_754478, EPI_ISL_754479, EPI_ISL_754480, EPI_ISL_754481, EPI_ISL_754488, EPI_ISL_754587, EPI_ISL_754588, EPI_ISL_754589, EPI_ISL_754590, EPI_ISL_754591, EPI_ISL_754592, EPI_ISL_754593, EPI_ISL_754594, EPI_ISL_754595, EPI_ISL_754596, EPI_ISL_754597, EPI_ISL_754598, EPI_ISL_754599, EPI_ISL_754600, EPI_ISL_754601, EPI_ISL_754602, EPI_ISL_754603, EPI_ISL_754604, EPI_ISL_754605, EPI_ISL_754606, EPI_ISL_754607, EPI_ISL_754608, EPI_ISL_754609, EPI_ISL_754610, EPI_ISL_754611, EPI_ISL_754612, EPI_ISL_754613, EPI_ISL_754614, EPI_ISL_754615, EPI_ISL_754616, EPI_ISL_754617, EPI_ISL_754618 | | | |
| see above | Wadsworth Center, New York State Department.of Health | Wadsworth Center, New York State Department.of Health | Kirsten St. George, Daryl M. Lamson, Alexis Russel, Matthew Shudt, Melissa A Leisner, Jonathan Plitnick, Navjot Singh, John Kelly, Sara Griesemer, Erasmus Schneider, Erica Lasek-Nesselquist |
| EPI_ISL_755135, EPI_ISL_755272, EPI_ISL_755281, EPI_ISL_755284, EPI_ISL_755299 | San Diego County Public Health Laboratory | Andersen lab at Scripps Research | SEARCH Alliance San Diego with Tracy Basler, Jovan Shephard, Brett Austin |
| EPI_ISL_755474, EPI_ISL_755475, EPI_ISL_755476 | Maine Health and Environmental Testing Laboratory | Tewhey Lab, The Jackson Laboratory | Matluk,N., Dewey,H., Iosue,F., Barter,M., Lynch,R., Munger,H. and Tewhey,R. |
| EPI_ISL_756279 | State Laboratories Division, Hawaii State Department of Health | State Laboratories Division, Hawaii State Department of Health | Pamela O'Brien, Sabrina Diemert, Drew Kuwazaki, Razvan Sultana, Edward Desmond |
| EPI_ISL_759978, EPI_ISL_759979, EPI_ISL_759980 | Medical Microbiology Unit, Department for Laboratory Medicine, Drammen Hospital, Vestre Viken Health Trust, | Norwegian Institute of Public Health, Department of Virology | Kathrine Stene-Johansen, Kamilla Heddeland Instefjord, Hilde Elshaug, Marie Paulsen Madsen, Rasmus Riis Kopperud, Hilde Vollan, Karoline Bragstad, Olav Hungnes |
| EPI_ISL_759981 | Norwegian Institute of Public Health, Department of Virology | Norwegian Institute of Public Health, Department of Virology | Kathrine Stene-Johansen, Kamilla Heddeland Instefjord, Hilde Elshaug, Marie Paulsen Madsen, Rasmus Riis Kopperud, Hilde Vollan, Karoline Bragstad, Olav Hungnes |
| EPI_ISL_760490 | Lighthouse Lab in Glasgow | Wellcome Sanger Institute for the COVID-19 Genomics UK (COG-UK) Consortium | Harper VanSteenhouse, Yumi Kasai, David Gray, Carol Clugston, Anna Dominiczak and Alex Alderton, Roberto Amato, Sonia Goncalves, Ewan Harrison, David K. Jackson, Ian Johnston, Dominic Kwiatkowski, Cordelia Langford, John Sillitoe on behalf of the Wellcome Sanger Institute COVID-19 Surveillance Team |
| EPI_ISL_763026, EPI_ISL_763030, EPI_ISL_763051 | Unit 17: Influenza & Other Respiratory Viruses, German National Influenza Center | Project group Epidemiology of Highly Pathogenic Microorganisms, Robert Koch-Institute | Ariane Düx, Andreas Sachse, Grit Schubert, Sébastien Calvignac-Spencer, Fabian Leendertz, Thorsten Wolff, Ralf Dürrwald, Djin-Ye Oh, Marianne Wedde |
| EPI_ISL_763100, EPI_ISL_763101, EPI_ISL_763102, EPI_ISL_763103, EPI_ISL_763112, EPI_ISL_763113, EPI_ISL_763114, EPI_ISL_763115, EPI_ISL_763116, EPI_ISL_763117, EPI_ISL_763118, EPI_ISL_763164, EPI_ISL_763165, EPI_ISL_763166, EPI_ISL_763167, EPI_ISL_763171, EPI_ISL_763236, EPI_ISL_763275, EPI_ISL_763276, EPI_ISL_763277, EPI_ISL_763313 | | | |
| see above | Dutch COVID-19 response team | Erasmus Medical Center | Bas Oude Munnink, Reina Sikkema, David Nieuwenhuijse, Irina Chestakova, Anne van der Linden, Marjan Boter, Emmanuelle Munger, Corine GeurtsvanKessel, Annemiek van der Eijk, Richard Molenkamp, Marion Koopmans, on behalf of the Dutch national COVID-19 response team. |
| EPI_ISL_763332, EPI_ISL_763343 | Florida Bureau of Public Health Laboratories | Florida Bureau of Public Health Laboratories | Sarah Schmedes, Jason Blanton |
| EPI_ISL_763359 | Dutch COVID-19 response team | Erasmus Medical Center | Bas Oude Munnink, Reina Sikkema, David Nieuwenhuijse, Irina Chestakova, Anne van der Linden, Marjan Boter, Emmanuelle Munger, Corine GeurtsvanKessel, Annemiek van der Eijk, Richard Molenkamp, Marion Koopmans, on behalf of the Dutch national COVID-19 response team. |
| EPI_ISL_764360 | Virology Department, Royal Infirmary of Edinburgh, NHS Lothian / School of Biological Sciences, University of Edinburgh / Institute of Genetics and Molecular Medicine, University of Edinburgh | COVID-19 Genomics UK (COG-UK) Consortium | McHugh M, Dewar R, Rooke S, Gallagher M, Balcaza C, O'Toole Á, Scher E, Hill V, McCrone JT, Colquhoun R, Yu X, Jackson B, Rambaut A, Williams TC, Templeton K |
| EPI_ISL_765217 | Instituto Nacional de Saude (INSA) and Instituto Gulbenkian de Ciencia (IGC) | Instituto Nacional de Saude (INSA) and Instituto Gulbenkian de Ciencia (IGC) | Borges et al |
| EPI_ISL_765779, EPI_ISL_765780, EPI_ISL_765854, EPI_ISL_765855, EPI_ISL_765856, EPI_ISL_765857, EPI_ISL_765858, EPI_ISL_765859, EPI_ISL_765860, EPI_ISL_765861, EPI_ISL_765862 | | | |
| see above | Massachusetts General Hospital | Infectious Disease Program, Broad Institute of Harvard and MIT | Lemieux,J.E., Siddle,K.J., Shaw,B., Adams,G., Pierce,V., Turbett,S., Anahtar,M., Branda,J., Slater,D., Harris,J., Lin,A.E., Gladden-Young,A., Lagerborg,K., Rudy,M., DeRuff,K., Carter,A., Normandin,E., Bauer,M., Reilly,S., Tomkins-Tinch,C., Loreth,C., Chaluvadi,S., Neumann,A., Cusick,C., Chapman,S.B., Gnirke,A., Flowers,K., Cerrato,F., Birren,B.W., Gallagher,G., Smole,S., Park,D.J., MacInnis,B.L., Ryan,E., LaRocque,R., Rosenberg,E. and Sabeti,P.C. |
| EPI_ISL_766276, EPI_ISL_766277 | Respiratory Virus Unit, National Infection Service, Public Health England | COVID-19 Genomics UK (COG-UK) Consortium | PHE Covid Sequencing Team |
| EPI_ISL_766617, EPI_ISL_766618, EPI_ISL_766620 | Klinisk mikrobiologi | The Public Health Agency of Sweden | Department of Microbiology, The Public Health Agency of Sweden |
| EPI_ISL_766642 | Vardcentralen Brinken | The Public Health Agency of Sweden | Department of Microbiology, The Public Health Agency of Sweden |
| EPI_ISL_766659, EPI_ISL_766660, EPI_ISL_766680 | Texas Department of State Health Services | Texas Department of State Health Services | Rashmi Tuladhar, Bonnie Oh, Jenny Zhang, Maliha Rahman, Anita Pokharel, Myong Koag, Chung Wang, Rachel Lee, Grace Kubin, Mayela Pedrueza, James Daniel Bonser |
| EPI_ISL_766976, EPI_ISL_766993, EPI_ISL_766999 | Delaware Public Health Laboratory | Delaware Public Health Laboratory | Gregory Hovan |
| EPI_ISL_767319, EPI_ISL_767320, EPI_ISL_767321, EPI_ISL_767322, EPI_ISL_767323, EPI_ISL_767324, EPI_ISL_767325, EPI_ISL_767326, EPI_ISL_767327, EPI_ISL_767328, EPI_ISL_767329, EPI_ISL_767330, EPI_ISL_767331, EPI_ISL_767332, EPI_ISL_767333 | | | |
| see above | Lighthouse Lab in Alderley Park | Wellcome Sanger Institute for the COVID-19 Genomics UK (COG-UK) Consortium | Jacquelyn Wynn, Mairead Hyland, The Lighthouse Lab in Alderley Park and Alex Alderton, Roberto Amato, Sonia Goncalves, Ewan Harrison, David K. Jackson, Ian Johnston, Dominic Kwiatkowski, Cordelia Langford, John Sillitoe on behalf of the Wellcome Sanger Institute COVID-19 Surveillance Team |
| EPI_ISL_767582, EPI_ISL_767590, EPI_ISL_767626, EPI_ISL_767627 | WHITE PLAINS HOSPITAL CENTER LABORATORY | Wadsworth Center, New York State Department.of Health | Kirsten St. George, Daryl M. Lamson, Alexis Russel, Matthew Shudt, Melissa A Leisner, Jonathan Plitnick, Navjot Singh, John Kelly, Sara Griesemer, Erasmus Schneider, Erica Lasek-Nesselquist |
| EPI_ISL_767851, EPI_ISL_767852 | South Eastern Area Laboratory Services (SEALS) | NSW Health Pathology - Institute of Clinical Pathology and Medical Research; Westmead Hospital; University of Sydney | CIDM-PH et al. |
| EPI_ISL_767858, EPI_ISL_767859, EPI_ISL_767860 | Sydney South West Pathology Service (SSWPS) - Royal Prince Alfred Hospital - NSW Health Pathology | NSW Health Pathology - Institute of Clinical Pathology and Medical Research; Westmead Hospital; University of Sydney | CIDM-PH et al. |
| EPI_ISL_768353, EPI_ISL_768354, EPI_ISL_768355, EPI_ISL_768356 | LSUHS Emerging Viral Threat Laboratory | Microbial Genome Sequencing Center | Jeremy P. Kamil, Jennifer L. Carroll, Camille F. Abshire, Maarten Van Diest, Andrew D. Yurochko, Martin J. Sapp, Rona S. Scott, Christopher G. Kevil, Daniel J. Snyder, Vaughn S. Cooper, John A. Vanchiere |
| EPI_ISL_770022 | Area De Salud Perez Zeledon | Inciensa, Instituto Costarricense de Investigación y Enseñanza en Nutrición y Salud | Francisco Duarte, Hebleen Porras, Claudio Soto-Garita, Estela Cordero, Adriana Godínez, Melany Calderón & Mariel López |
| EPI_ISL_770067, EPI_ISL_770068, EPI_ISL_770069, EPI_ISL_770086, EPI_ISL_770087, EPI_ISL_770088, EPI_ISL_770089, EPI_ISL_770090, EPI_ISL_770091, EPI_ISL_770092, EPI_ISL_770093, EPI_ISL_770108, EPI_ISL_770128 | | | |
| see above | Wyoming Public Health Laboratory | Wyoming Public Health Laboratory | Noah Hull, Taylor Fearing, Lynette Gumbleton, Channing Weber, Ashley Norberg, Bailey Bowcutt, and Wanda Manley |
| EPI_ISL_770555, EPI_ISL_770556, EPI_ISL_770573, EPI_ISL_770574, EPI_ISL_770575, EPI_ISL_770576 | Laboratório de Microbiologia Molecular - Universidade FEEVALE | Bioinformatics Laboratory / LNCC | Felipe Benites, Fernando Rosado Spilki, Alana Witt Hansen, Juliane Deise Fleck, Juliana Schons, Meriane Demoliner, Ana Karolina Eisen Antunes, Fagner Henrique Heldt, Larissa Mallmann, Bruna Hermann, Ana Luiza Ziulkoski, Vyctoria Goes, Karoline Schallenberger, Matheus Nunes Weber, Paula Rodrigues de Almeida, Alessandra Pavan Lamarca da Silva, Ronaldo da Silva F Jr , Luiz G P de Almeida, Alexandra L Gerber , Ana Paula de C Guimarães,Ana Tereza R de Vasconcelos |
| EPI_ISL_770658, EPI_ISL_770673 | WHO National Influenza Centre Russian Federation | WHO National Influenza Centre Russian Federation | Andrey Komissarov, Artem Fadeev, Anna Ivanova, Kseniya Komissarova, Dmitry Bazhenov, Daria Danilenko, Ksenia Safina, Elena Nabieva, Georgii |

| | | | |
|---|---|---|---|
| | | | Bazykin, Dmitry Lioznov |
| EPI_ISL_771157, EPI_ISL_771160, EPI_ISL_771162, EPI_ISL_771165, EPI_ISL_771167, EPI_ISL_771168, EPI_ISL_771169, EPI_ISL_771170, EPI_ISL_771173, EPI_ISL_771176, EPI_ISL_771179, EPI_ISL_771180, EPI_ISL_771182, EPI_ISL_771183, EPI_ISL_771191, EPI_ISL_771192, EPI_ISL_771193, EPI_ISL_771198, EPI_ISL_771199, EPI_ISL_771201, EPI_ISL_771208, EPI_ISL_771212, EPI_ISL_771222, EPI_ISL_771291, EPI_ISL_771300, EPI_ISL_771301, EPI_ISL_771303, EPI_ISL_771304, EPI_ISL_771305, EPI_ISL_771306, EPI_ISL_771308, EPI_ISL_771309, EPI_ISL_771310, EPI_ISL_771311, EPI_ISL_771312, EPI_ISL_771313, EPI_ISL_771314, EPI_ISL_771315, EPI_ISL_771316, EPI_ISL_771317, EPI_ISL_771318, EPI_ISL_771319, EPI_ISL_771320, EPI_ISL_771321, EPI_ISL_771322, EPI_ISL_771323, EPI_ISL_771324, EPI_ISL_771325, EPI_ISL_771326, EPI_ISL_771327, EPI_ISL_771328, EPI_ISL_771330, EPI_ISL_771333, EPI_ISL_771334, EPI_ISL_771351, EPI_ISL_771352 | | | |
| see above | Colorado Department of Public Health and Environment | Colorado Department of Puplic Health and Environment | Laura Bankers, Molly C. Hetherington-Rauth, Diana Ir, Shannon Ely, Shannon R. Matzinger, Sarah Elizabeth Totten, Emily A. Travanty |
| EPI_ISL_775286 | Nordland Hospital - Bodo, Laboratory Department, Molecular Biology Unit | Norwegian Institute of Public Health, Department of Virology | Kathrine Stene-Johansen, Kamilla Heddeland Instefjord, Hilde Elshaug, Atiya R Ali,Marie Paulsen Madsen, Rasmus Riis Kopperud, Hilde Vollan, Karoline Bragstad, Olav Hungnes |
| EPI_ISL_775306 | Department of Medical Microbiology, St. Olavs hospital | Norwegian Institute of Public Health, Department of Virology | Kathrine Stene-Johansen, Kamilla Heddeland Instefjord, Hilde Elshaug, Atiya R Ali,Marie Paulsen Madsen, Rasmus Riis Kopperud, Hilde Vollan, Karoline Bragstad, Olav Hungnes |
| EPI_ISL_775348 | Medical Microbiology Unit, Department for Laboratory Medicine, Drammen Hospital, Vestre Viken Health Trust, | Norwegian Institute of Public Health, Department of Virology | Kathrine Stene-Johansen, Kamilla Heddeland Instefjord, Hilde Elshaug, Atiya R Ali,Marie Paulsen Madsen, Rasmus Riis Kopperud, Hilde Vollan, Karoline Bragstad, Olav Hungnes |
| EPI_ISL_775358, EPI_ISL_775359, EPI_ISL_775360, EPI_ISL_775361 | Unilabs Laboratory Medicine | Norwegian Institute of Public Health, Department of Virology | Kathrine Stene-Johansen, Kamilla Heddeland Instefjord, Hilde Elshaug, Atiya R Ali,Marie Paulsen Madsen, Rasmus Riis Kopperud, Hilde Vollan, Karoline Bragstad, Olav Hungnes |
| EPI_ISL_775374, EPI_ISL_775376, EPI_ISL_775398 | Nordland Hospital - Bodo, Laboratory Department, Molecular Biology Unit | Norwegian Institute of Public Health, Department of Virology | Kathrine Stene-Johansen, Kamilla Heddeland Instefjord, Hilde Elshaug, Atiya R Ali,Marie Paulsen Madsen, Rasmus Riis Kopperud, Hilde Vollan, Karoline Bragstad, Olav Hungnes |
| EPI_ISL_775488 | Furst Medical Laboratory | Norwegian Institute of Public Health, Department of Virology | Kathrine Stene-Johansen, Kamilla Heddeland Instefjord, Hilde Elshaug, Atiya R Ali,Marie Paulsen Madsen, Rasmus Riis Kopperud, Hilde Vollan, Karoline Bragstad, Olav Hungnes |
| EPI_ISL_775493, EPI_ISL_775494 | Department of Medical Microbiology, St. Olavs hospital | Norwegian Institute of Public Health, Department of Virology | Kathrine Stene-Johansen, Kamilla Heddeland Instefjord, Hilde Elshaug, Atiya R Ali,Marie Paulsen Madsen, Rasmus Riis Kopperud, Hilde Vollan, Karoline Bragstad, Olav Hungnes |
| EPI_ISL_778843 | Servicio Virosis Respiratorias-Departamento Virología-INEI | Instituto Nacional Enfermedades Infecciosas C.G.Malbran | Baumeister E., Avaro M., Benedetti E., Russo M., Dattero ME, Pontoriero A., Cisterna D., Molina V., Perandones C., Tuduri E., Lorenzo F., Poklepovich T., Campos J. |
| EPI_ISL_779155, EPI_ISL_779156 | Laboratório de Microbiologia Molecular - Universidade FEEVALE | Bioinformatics Laboratory / LNCC | Felipe Benites, Fernando Rosado Spilki, Alana Witt Hansen, Juliane Deise Fleck, Juliana Schons, Meriane Demoliner, Ana Karolina Eisen Antunes, Fagner Henrique Heldt, Larissa Mallmann, Bruna Hermann, Ana Luiza Ziulkoski, Vyctoria Goes, Karoline Schallenberger, Matheus Nunes Weber, Paula Rodrigues de Almeida, Alessandra Pavan Lamarca da Silva, Ronaldo da Silva F Jr , Luiz G P de Almeida, Alexandra L Gerber , Ana Paula de C Guimarães,Ana Tereza R de Vasconcelos |
| EPI_ISL_779198 | Laboratorio Estatal de Salud Pública de Nuevo León | Laboratorio de Infectología Molecular, Departamento de Bioquímica y Medicina Molecular,Facultad de Medicina - Universidad Autónoma de Nuevo León | Kame A. Galán-Huerta, María F. Herrera-Saldivar, Natalia Martínez-Acuña, Sonia A. Lozano-Sepúlveda, Daniel Arellanos-Soto, Ana M. Rivas-Estilla, Samuel Buentello-Wong, Else del Carmen García-García, Gloria A. Jasso-de-la-Peña, Roberto Montes-de-Oca, Consuelo Treviño-Garza, Manuel E. de-la-O-Cavazos |
| EPI_ISL_779403 | Institute of Virology, Biomedical Research Center of the Slovak Academy of Sciences, Bratislava | Faculty of Natural Sciences, Comenius University, Bratislava | Viktória Hodorová, Kristína Boršová, Broa Brejová, Viktória abanová, Sabina Fumaová Havlíková, Juraj Kopáek, Martina Liková, ubomíra Lukáiková, Martina Neboháová, Monika Sláviková, Tomáš Vina, Jozef Nosek, Boris Klempa |
| EPI_ISL_779956, EPI_ISL_779957, EPI_ISL_779958, EPI_ISL_779960, EPI_ISL_779962, EPI_ISL_779964, EPI_ISL_779965, EPI_ISL_779966, EPI_ISL_779967, EPI_ISL_779968, EPI_ISL_779974, EPI_ISL_779977, EPI_ISL_779978, EPI_ISL_779979, EPI_ISL_779980, EPI_ISL_779981, EPI_ISL_779982, EPI_ISL_779983, EPI_ISL_779984, EPI_ISL_779985, EPI_ISL_779986, EPI_ISL_779987, EPI_ISL_779988, EPI_ISL_779989, EPI_ISL_779990, EPI_ISL_779991, EPI_ISL_779992, EPI_ISL_779993, EPI_ISL_779994, EPI_ISL_779995, EPI_ISL_779996, EPI_ISL_779997, EPI_ISL_779998, EPI_ISL_779999, EPI_ISL_780000, EPI_ISL_780001, EPI_ISL_780002, EPI_ISL_780003, EPI_ISL_780004, EPI_ISL_780005, EPI_ISL_780006, EPI_ISL_780007, EPI_ISL_780008, EPI_ISL_780009, EPI_ISL_780010, EPI_ISL_780011, EPI_ISL_780033, EPI_ISL_780049, EPI_ISL_780061, EPI_ISL_780069 | | | |
| see above | Hospital General Universitario Gregorio Marañón | SeqCOVID-SPAIN consortium/IBV(CSIC) | Darío García de Viedma, Laura Pérez-Lago, Marta Herranz, Jon Sicilia, Julia Suárez, Pilar Catalán, Patricia Muñoz and SeqCOVID-SPAIN consortium |
| EPI_ISL_783538, EPI_ISL_783544, EPI_ISL_783550, EPI_ISL_783552, EPI_ISL_783560, EPI_ISL_783566, EPI_ISL_783571, EPI_ISL_783594, EPI_ISL_783595, EPI_ISL_783597, EPI_ISL_783603, EPI_ISL_783604, EPI_ISL_783605, EPI_ISL_783607, EPI_ISL_783613, EPI_ISL_783621, EPI_ISL_783622, EPI_ISL_783623, EPI_ISL_783624, EPI_ISL_783626, EPI_ISL_783629, EPI_ISL_783631, EPI_ISL_783632, EPI_ISL_783633, EPI_ISL_783634, EPI_ISL_783636, EPI_ISL_783637, EPI_ISL_783638, EPI_ISL_783639, EPI_ISL_783640, EPI_ISL_783642, EPI_ISL_783643, EPI_ISL_783645, EPI_ISL_783646, EPI_ISL_783648, EPI_ISL_783650, EPI_ISL_783654, EPI_ISL_783655, EPI_ISL_783656, EPI_ISL_783657, EPI_ISL_783658, EPI_ISL_783659, EPI_ISL_783660, EPI_ISL_783661, EPI_ISL_783662, EPI_ISL_783663, EPI_ISL_783664, EPI_ISL_783665, EPI_ISL_783666, EPI_ISL_783668, EPI_ISL_783670, EPI_ISL_783672, EPI_ISL_783673, EPI_ISL_783675, EPI_ISL_783676, EPI_ISL_783678, EPI_ISL_783681, EPI_ISL_783688, EPI_ISL_783689, EPI_ISL_783697, EPI_ISL_783704, EPI_ISL_783711, EPI_ISL_783712, EPI_ISL_783713, EPI_ISL_783720, EPI_ISL_783723, EPI_ISL_783724, EPI_ISL_783725, EPI_ISL_783734, EPI_ISL_783735, EPI_ISL_783954, EPI_ISL_783956, EPI_ISL_783959, EPI_ISL_783960, EPI_ISL_783962, EPI_ISL_783965, EPI_ISL_783966, EPI_ISL_783968, EPI_ISL_783974, EPI_ISL_783987, EPI_ISL_783988, EPI_ISL_783995, EPI_ISL_784007, EPI_ISL_784015, EPI_ISL_784018, EPI_ISL_784020, EPI_ISL_784023, EPI_ISL_784028, EPI_ISL_784029, EPI_ISL_784033, EPI_ISL_784034, EPI_ISL_784038, EPI_ISL_784041, EPI_ISL_784050, EPI_ISL_784051, EPI_ISL_784053, EPI_ISL_784055, EPI_ISL_784058, EPI_ISL_784061, EPI_ISL_784063, EPI_ISL_784065, EPI_ISL_784067, EPI_ISL_784068, EPI_ISL_784075, EPI_ISL_784079, EPI_ISL_784082, EPI_ISL_784083, EPI_ISL_784084, EPI_ISL_784091, EPI_ISL_784093, EPI_ISL_784094, EPI_ISL_784096, EPI_ISL_784097, EPI_ISL_784101, EPI_ISL_784103, EPI_ISL_784105, EPI_ISL_784107, EPI_ISL_784108, EPI_ISL_784116, EPI_ISL_784121, EPI_ISL_784122, EPI_ISL_784124, EPI_ISL_784126, EPI_ISL_784129, EPI_ISL_784130, EPI_ISL_784131, EPI_ISL_784132, EPI_ISL_784137, EPI_ISL_784138, EPI_ISL_784141, EPI_ISL_784142, EPI_ISL_784143, EPI_ISL_784144, EPI_ISL_784145, EPI_ISL_784148, EPI_ISL_784157, EPI_ISL_784160, EPI_ISL_784161, EPI_ISL_784166, EPI_ISL_784167, EPI_ISL_784168, EPI_ISL_784169, EPI_ISL_784172, EPI_ISL_784173, EPI_ISL_784176, EPI_ISL_784179, EPI_ISL_784181, EPI_ISL_784183, EPI_ISL_784707, EPI_ISL_784773, EPI_ISL_784775, EPI_ISL_784777, EPI_ISL_784779, EPI_ISL_784781, EPI_ISL_784782, EPI_ISL_784786, EPI_ISL_784790, EPI_ISL_784794, EPI_ISL_784802, EPI_ISL_784813, EPI_ISL_784814, EPI_ISL_784815, EPI_ISL_784817, EPI_ISL_784818, EPI_ISL_784824, EPI_ISL_784826, EPI_ISL_784828, EPI_ISL_784830, EPI_ISL_784832, EPI_ISL_784839, EPI_ISL_784841, EPI_ISL_784842, EPI_ISL_784849, EPI_ISL_784852, EPI_ISL_784854, EPI_ISL_784857, EPI_ISL_784860, EPI_ISL_784861, EPI_ISL_784862, EPI_ISL_784865, EPI_ISL_784866, EPI_ISL_784871, EPI_ISL_784876, EPI_ISL_784877, EPI_ISL_784880, EPI_ISL_784882, EPI_ISL_784884, EPI_ISL_784886, EPI_ISL_784888, EPI_ISL_784889, EPI_ISL_784890, EPI_ISL_784891, EPI_ISL_784893, EPI_ISL_784894, EPI_ISL_784898, EPI_ISL_784900, EPI_ISL_784906, EPI_ISL_784907, EPI_ISL_784909, EPI_ISL_784910, EPI_ISL_784914, EPI_ISL_784917, EPI_ISL_784918, EPI_ISL_784920, EPI_ISL_784922, EPI_ISL_784924, EPI_ISL_784929, EPI_ISL_784930, EPI_ISL_784931, EPI_ISL_784932, EPI_ISL_784934, EPI_ISL_784935, EPI_ISL_784939, EPI_ISL_784945, EPI_ISL_784951, EPI_ISL_784952, EPI_ISL_784954, EPI_ISL_784955, EPI_ISL_784957, EPI_ISL_784959, EPI_ISL_784961, EPI_ISL_784963, EPI_ISL_784965, EPI_ISL_784966, EPI_ISL_784968, EPI_ISL_784969, EPI_ISL_784970, EPI_ISL_784971, EPI_ISL_784972, EPI_ISL_784973, EPI_ISL_784974, EPI_ISL_784976, EPI_ISL_784978, EPI_ISL_784979, EPI_ISL_784980, EPI_ISL_784981, EPI_ISL_784982, EPI_ISL_784983, EPI_ISL_784984, EPI_ISL_784985, EPI_ISL_784986, EPI_ISL_784988, EPI_ISL_784989, EPI_ISL_784990, EPI_ISL_784991, EPI_ISL_784993, EPI_ISL_784994, EPI_ISL_784995, EPI_ISL_784996, EPI_ISL_784997, EPI_ISL_784999, EPI_ISL_785001, EPI_ISL_785002, EPI_ISL_785003, EPI_ISL_785004, EPI_ISL_785005, EPI_ISL_785006, EPI_ISL_785008, EPI_ISL_785009, EPI_ISL_785010, EPI_ISL_785011, EPI_ISL_785014, EPI_ISL_785015, EPI_ISL_785017, EPI_ISL_785018, EPI_ISL_785019, EPI_ISL_785020, EPI_ISL_785021, EPI_ISL_785022, EPI_ISL_785024, EPI_ISL_785029, EPI_ISL_785030, EPI_ISL_785031, EPI_ISL_785032, EPI_ISL_785033, EPI_ISL_785035, EPI_ISL_785036, EPI_ISL_785037, EPI_ISL_785039, EPI_ISL_785040, EPI_ISL_785041, EPI_ISL_785042, EPI_ISL_785043, EPI_ISL_785044, EPI_ISL_785045, EPI_ISL_785046, EPI_ISL_785047, EPI_ISL_785053, EPI_ISL_785054, EPI_ISL_785055, EPI_ISL_785056, EPI_ISL_785057, EPI_ISL_785058, EPI_ISL_785059, EPI_ISL_785060, EPI_ISL_785061, EPI_ISL_785062, EPI_ISL_785063, EPI_ISL_785064, EPI_ISL_785065, EPI_ISL_785066, EPI_ISL_785067, EPI_ISL_785068, EPI_ISL_785069, EPI_ISL_785070, EPI_ISL_785071, EPI_ISL_785072, EPI_ISL_785073, EPI_ISL_785075, EPI_ISL_785078, EPI_ISL_785079, EPI_ISL_785080, EPI_ISL_785081, EPI_ISL_785082, EPI_ISL_785083, EPI_ISL_785084, EPI_ISL_785086, EPI_ISL_785087, EPI_ISL_785088, EPI_ISL_785089, EPI_ISL_785090, EPI_ISL_785091, EPI_ISL_785092, EPI_ISL_785093, EPI_ISL_785094, EPI_ISL_785095, EPI_ISL_785096, EPI_ISL_785097, EPI_ISL_785098, EPI_ISL_785099, EPI_ISL_785100, EPI_ISL_785103, EPI_ISL_785104, EPI_ISL_785105, EPI_ISL_785107, EPI_ISL_785108, EPI_ISL_785109, EPI_ISL_785110, EPI_ISL_785112, EPI_ISL_785114, EPI_ISL_785115, EPI_ISL_785116, EPI_ISL_785117, EPI_ISL_785118, EPI_ISL_785119, EPI_ISL_785120, EPI_ISL_785123, EPI_ISL_785124, EPI_ISL_785125, EPI_ISL_785126, EPI_ISL_785127, EPI_ISL_785128, EPI_ISL_785129, EPI_ISL_785132, EPI_ISL_785133, EPI_ISL_785134, EPI_ISL_785135, EPI_ISL_785136, EPI_ISL_785137, EPI_ISL_785138, EPI_ISL_785144, EPI_ISL_785155, EPI_ISL_785162, EPI_ISL_785172, EPI_ISL_785184, EPI_ISL_785185, EPI_ISL_785192, EPI_ISL_785194, EPI_ISL_785200, EPI_ISL_785201, EPI_ISL_785280, EPI_ISL_785288 | | | |
| see above | Houston Methodist Hospital | Houston Methodist Hospital | S. Wesley Long, Randall J. Olsen, Paul A. Christensen, David W. Bernard, James J. Davis, Maulik Shukla, Marcus Nguyen, Matthew Ojeda Saavedra, Prasanti Yerramilli, Layne Pruitt, Sishir Subedi, Heather Hendrickson, and James M. Musser |
| EPI_ISL_788985 | Institute of Virology, Biomedical Research Center of the Slovak Academy of Sciences, Bratislava | Faculty of Natural Sciences, Comenius University, Bratislava | Broa Brejová, Viktória abanová, Kristína Boršová, Viktória Hodorová, Sabina Fumaová Havlíková, Juraj Kopáek, Martina Liková, ubomíra Lukáiková, Martina Neboháová, Monika Sláviková, Alena Košálová, Peter Sabaka, Tomáš Vina, Jozef Nosek, Boris Klempa |
| EPI_ISL_790695, EPI_ISL_790697 | Dutch COVID-19 response team | National Institute for Public Health and the Environment (RIVM) | Adam Meijer, Harry Vennema, Jeroen Cremer, Sharon van den Brink, Bas van der Veer, AnneMarie van den Brandt, Florian Zwagemaker, Dennis Schmitz, Chantal Reusken, on behalf of the national COVID-19 response team |
| EPI_ISL_791299 | National Virus Reference Laboratory | Irish Coronavirus Sequencing Consortium - Teagasc Moorepark | Alejandro Abner Garcia Leon, Paul Cotter, Fiona Crispie, John Kenny, Paddy Mallon, Calum Walsh |
| EPI_ISL_791343, EPI_ISL_791350, EPI_ISL_791392, EPI_ISL_791393, EPI_ISL_791395, EPI_ISL_791404, EPI_ISL_791405, EPI_ISL_791408, EPI_ISL_791412, EPI_ISL_791414 | Johns Hopkins Hospital Department of Pathology | Johns Hopkins Hospital Department of Pathology | C. Paul Morris, Chun Huai Luo, Heba H. Mostafa |
| EPI_ISL_791990 | Institute of Virology, Biomedical Research Center of the Slovak Academy of Sciences, Bratislava | Faculty of Natural Sciences, Comenius University, Bratislava | Viktória abanová, Kristína Boršová, Broa Brejová, Viktória Hodorová, Sabina Fumaová Havlíková, Juraj Kopáek, Martina Liková, ubomíra Lukáiková, Martina Neboháová, Monika Sláviková, Alena Košálová, Peter Sabaka, Tomáš Vina, Jozef Nosek, Boris Klempa |
| EPI_ISL_791992 | CHU - Hôpital Cavale Blanche | National Reference Center for Viruses of Respiratory Infections, Institut Pasteur, Paris | Marion Barbet, Sylvie Behillil, Méline Bizard, Angela Brisebarre, Camille Capel, Etienne Simon-Lorière, Vincent Enouf, Maud Vanpeene, Sylvie van der Werf, Léa Pilorgé |
| EPI_ISL_792639, EPI_ISL_792641 | LACEN-AL | Laboratory of Respiratory Viruses and Measles, Oswaldo Cruz Institute, FIOCRUZ | Paola Resende, Luciana Appolinario, Fernando Motta, Anna Carolina Paixao, Ana Carolina Mendonca, Anderson Brandao Leite, Marilda Siqueira |

| | | | |
|---|---|---|---|
| EPI_ISL_792650, EPI_ISL_792651, EPI_ISL_792654 | LACEN-PR | Laboratory of Respiratory Viruses and Measles, Oswaldo Cruz Institute, FIOCRUZ | Paola Resende, Luciana Appolinario, Fernando Motta, Anna Carolina Paixao, Ana Carolina Mendonca, Maria do Carmo Debur, Irina Nastassja Riediger, Marilda Siqueira |
| EPI_ISL_792688, EPI_ISL_792695 | The National Institute of Public Health | State Veterinary Institute Prague | Nagy,A;Jirincova,H;Trnka,D;Vecerova,J |
| EPI_ISL_793011, EPI_ISL_794589, EPI_ISL_794590, EPI_ISL_794591, EPI_ISL_795010 | Department of Virus and Microbiological Special Diagnostics, Statens Serum Institut, Copenhagen, Denmark | Albertsen Lab, Department of Chemistry and Bioscience, Aalborg University, Denmark | Danish Covid-19 Genome Consortium |
| EPI_ISL_800885, EPI_ISL_800886, EPI_ISL_800887, EPI_ISL_800888, EPI_ISL_800889, EPI_ISL_800890, EPI_ISL_800891, EPI_ISL_800892, EPI_ISL_800893, EPI_ISL_800894, EPI_ISL_800895, EPI_ISL_800896, EPI_ISL_800898, EPI_ISL_800899, EPI_ISL_800900, EPI_ISL_800901, EPI_ISL_800902, EPI_ISL_800903, EPI_ISL_800904, EPI_ISL_800905, EPI_ISL_800906, EPI_ISL_800907, EPI_ISL_800908, EPI_ISL_800909, EPI_ISL_800910, EPI_ISL_800911, EPI_ISL_800912, EPI_ISL_800913, EPI_ISL_800914, EPI_ISL_800915, EPI_ISL_800916, EPI_ISL_800917, EPI_ISL_800918, EPI_ISL_800919, EPI_ISL_800921, EPI_ISL_800922, EPI_ISL_800923, EPI_ISL_800925, EPI_ISL_800926, EPI_ISL_800927, EPI_ISL_800928, EPI_ISL_800929, EPI_ISL_800930, EPI_ISL_800931, EPI_ISL_800932, EPI_ISL_800933, EPI_ISL_800934, EPI_ISL_800935, EPI_ISL_800936, EPI_ISL_800937, EPI_ISL_800938, EPI_ISL_800939, EPI_ISL_800940, EPI_ISL_800941, EPI_ISL_800942, EPI_ISL_800943, EPI_ISL_800944, EPI_ISL_800945, EPI_ISL_800946, EPI_ISL_800947, EPI_ISL_800948, EPI_ISL_800949, EPI_ISL_800950, EPI_ISL_800951, EPI_ISL_800952, EPI_ISL_800953, EPI_ISL_800954, EPI_ISL_800955, EPI_ISL_800956, EPI_ISL_800957, EPI_ISL_800958, EPI_ISL_800959, EPI_ISL_800960, EPI_ISL_800961, EPI_ISL_800962, EPI_ISL_800963, EPI_ISL_800964, EPI_ISL_800965, EPI_ISL_800966, EPI_ISL_800968, EPI_ISL_800969, EPI_ISL_800970, EPI_ISL_800972, EPI_ISL_800973, EPI_ISL_800974, EPI_ISL_800975, EPI_ISL_800976, EPI_ISL_800977, EPI_ISL_800978, EPI_ISL_800979, EPI_ISL_800980, EPI_ISL_800981, EPI_ISL_800982, EPI_ISL_800983, EPI_ISL_800984, EPI_ISL_800985, EPI_ISL_800986, EPI_ISL_800987, EPI_ISL_800988, EPI_ISL_800989, EPI_ISL_800990, EPI_ISL_800991, EPI_ISL_800992, EPI_ISL_800993, EPI_ISL_800994, EPI_ISL_800996, EPI_ISL_800997, EPI_ISL_800998, EPI_ISL_800999, EPI_ISL_801000, EPI_ISL_801001, EPI_ISL_801003, EPI_ISL_801004, EPI_ISL_801005, EPI_ISL_801006, EPI_ISL_801007, EPI_ISL_801008, EPI_ISL_801009, EPI_ISL_801010, EPI_ISL_801011, EPI_ISL_801012, EPI_ISL_801013, EPI_ISL_801014, EPI_ISL_801015, EPI_ISL_801016, EPI_ISL_801017, EPI_ISL_801018, EPI_ISL_801019, EPI_ISL_801020, EPI_ISL_801021, EPI_ISL_801022, EPI_ISL_801023, EPI_ISL_801024, EPI_ISL_801025, EPI_ISL_801026, EPI_ISL_801027, EPI_ISL_801028, EPI_ISL_801029, EPI_ISL_801030, EPI_ISL_801031, EPI_ISL_801032, EPI_ISL_801033, EPI_ISL_801034, EPI_ISL_801035, EPI_ISL_801036, EPI_ISL_801037, EPI_ISL_801038, EPI_ISL_801039, EPI_ISL_801040, EPI_ISL_801041, EPI_ISL_801042, EPI_ISL_801043, EPI_ISL_801044, EPI_ISL_801045, EPI_ISL_801046, EPI_ISL_801047, EPI_ISL_801048, EPI_ISL_801049, EPI_ISL_801050, EPI_ISL_801051, EPI_ISL_801052, EPI_ISL_801053, EPI_ISL_801054, EPI_ISL_801056, EPI_ISL_801057, EPI_ISL_801058, EPI_ISL_801059, EPI_ISL_801060, EPI_ISL_801061, EPI_ISL_801062, EPI_ISL_801064, EPI_ISL_801065, EPI_ISL_801066, EPI_ISL_801067, EPI_ISL_801068, EPI_ISL_801069, EPI_ISL_801070, EPI_ISL_801071, EPI_ISL_801072, EPI_ISL_801073, EPI_ISL_801074, EPI_ISL_801075, EPI_ISL_801076, EPI_ISL_801077, EPI_ISL_801078, EPI_ISL_801079, EPI_ISL_801080, EPI_ISL_801081, EPI_ISL_801082, EPI_ISL_801083, EPI_ISL_801084, EPI_ISL_801085, EPI_ISL_801086, EPI_ISL_801087, EPI_ISL_801088, EPI_ISL_801089, EPI_ISL_801090, EPI_ISL_801091, EPI_ISL_801092, EPI_ISL_801093, EPI_ISL_801094, EPI_ISL_801095, EPI_ISL_801096, EPI_ISL_801097, EPI_ISL_801098, EPI_ISL_801099, EPI_ISL_801100, EPI_ISL_801101, EPI_ISL_801102, EPI_ISL_801103, EPI_ISL_801104, EPI_ISL_801105, EPI_ISL_801106, EPI_ISL_801107, EPI_ISL_801108, EPI_ISL_801109, EPI_ISL_801111, EPI_ISL_801112, EPI_ISL_801113, EPI_ISL_801114, EPI_ISL_801115, EPI_ISL_801116, EPI_ISL_801117, EPI_ISL_801118, EPI_ISL_801119, EPI_ISL_801120, EPI_ISL_801121, EPI_ISL_801122, EPI_ISL_801123, EPI_ISL_801124, EPI_ISL_801126, EPI_ISL_801127, EPI_ISL_801128, EPI_ISL_801129, EPI_ISL_801130, EPI_ISL_801131, EPI_ISL_801132, EPI_ISL_801133, EPI_ISL_801134, EPI_ISL_801135, EPI_ISL_801136, EPI_ISL_801137, EPI_ISL_801138, EPI_ISL_801139, EPI_ISL_801140, EPI_ISL_801141, EPI_ISL_801142, EPI_ISL_801143, EPI_ISL_801144, EPI_ISL_801145, EPI_ISL_801146, EPI_ISL_801147, EPI_ISL_801149, EPI_ISL_801150, EPI_ISL_801151, EPI_ISL_801152, EPI_ISL_801153, EPI_ISL_801154, EPI_ISL_801155, EPI_ISL_801156, EPI_ISL_801157, EPI_ISL_801158, EPI_ISL_801159, EPI_ISL_801160, EPI_ISL_801161, EPI_ISL_801162, EPI_ISL_801163, EPI_ISL_801164, EPI_ISL_801165, EPI_ISL_801166, EPI_ISL_801167, EPI_ISL_801168, EPI_ISL_801169, EPI_ISL_801170, EPI_ISL_801171, EPI_ISL_801172, EPI_ISL_801173, EPI_ISL_801174, EPI_ISL_801175, EPI_ISL_801176, EPI_ISL_801177, EPI_ISL_801178 | | | |
| see above | Lighthouse Lab in Cambridge | Wellcome Sanger Institute for the COVID-19 Genomics UK (COG-UK) Consortium | Rob Howes, The Lighthouse Lab in Cambridge and Alex Alderton, Roberto Amato, Sonia Goncalves, Ewan Harrison, David K. Jackson, Ian Johnston, Dominic Kwiatkowski, Cordelia Langford, John Sillitoe on behalf of the Wellcome Sanger Institute COVID-19 Surveillance Team |
| EPI_ISL_802567 | Essentia Health-St. Mary's Medical Center | Minnesota Department of Health, Public Health Laboratory | Alexandra Lorentz, Jacob Garfin, Matt Plumb, and Xiong Wang |
| EPI_ISL_802853, EPI_ISL_802854 | Vilnius University Hospital Santaros Klinikos, Vilnius University | Institute of Biotechnology, Life Sciences Center, Vilnius University | Emilija Vasiliunaite, Milda Norkiene, Albertas Timinskas, Alma Gedvilaite, Aurelija Zvirbliene, Daniel Naumovas, Laimonas Griskevicius |
| EPI_ISL_804600, EPI_ISL_804605, EPI_ISL_804616, EPI_ISL_804628, EPI_ISL_804630, EPI_ISL_804633, EPI_ISL_804638, EPI_ISL_804643, EPI_ISL_804652, EPI_ISL_804659, EPI_ISL_804665, EPI_ISL_804670, EPI_ISL_804676, EPI_ISL_804681, EPI_ISL_804683, EPI_ISL_804686, EPI_ISL_804691, EPI_ISL_804695, EPI_ISL_804696, EPI_ISL_804697, EPI_ISL_804700, EPI_ISL_804711, EPI_ISL_804761, EPI_ISL_804762, EPI_ISL_804763, EPI_ISL_804764, EPI_ISL_804765, EPI_ISL_804766, EPI_ISL_804768, EPI_ISL_804769, EPI_ISL_804773, EPI_ISL_804774, EPI_ISL_804775, EPI_ISL_804776, EPI_ISL_804777, EPI_ISL_804778, EPI_ISL_804780, EPI_ISL_804782, EPI_ISL_804783, EPI_ISL_804787, EPI_ISL_804788, EPI_ISL_804789, EPI_ISL_804791, EPI_ISL_804794, EPI_ISL_804795, EPI_ISL_804796, EPI_ISL_804797, EPI_ISL_804801, EPI_ISL_804802, EPI_ISL_804803, EPI_ISL_804804, EPI_ISL_804805, EPI_ISL_804807, EPI_ISL_804809, EPI_ISL_804810, EPI_ISL_804811 | | | |
| see above | Michigan Department of Health and Human Services, Bureau of Laboratories | Michigan Department of Health and Human Services, Bureau of Laboratories | Blankenship HM, Riner D, Soehnlen MK |
| EPI_ISL_806816, EPI_ISL_806817, EPI_ISL_806818 | Alaska State Virology Laboratory | Alaska State Virology Laboratory | Stephanie DeRonde, Lisa Smith, Ph.D., Devin M. Drown, Ph.D., Jack Chen, Ph.D. |
| EPI_ISL_807094, EPI_ISL_807108, EPI_ISL_807113, EPI_ISL_807114, EPI_ISL_807116, EPI_ISL_807117, EPI_ISL_807118, EPI_ISL_807119, EPI_ISL_807120, EPI_ISL_807121, EPI_ISL_807122, EPI_ISL_807123, EPI_ISL_807124, EPI_ISL_807125, EPI_ISL_807126, EPI_ISL_807127, EPI_ISL_807128, EPI_ISL_807129, EPI_ISL_807130, EPI_ISL_807131, EPI_ISL_807132, EPI_ISL_807133, EPI_ISL_807134, EPI_ISL_807135, EPI_ISL_807136, EPI_ISL_807137, EPI_ISL_807138, EPI_ISL_807139, EPI_ISL_807140, EPI_ISL_807141, EPI_ISL_807142, EPI_ISL_807143, EPI_ISL_807144, EPI_ISL_807145, EPI_ISL_807146, EPI_ISL_807147 | | | |
| see above | Washington State Department of Health | Seattle Flu Study | Deborah A. Nickerson, Chris D. Frazar, Jover Lee, Benjamin Pelle, Matthew Richardson, Amanda Adler, Elisabeth Brandstetter, Peter D. Han, Kairsten Fay, Misja Ilcisin, Kirsten Lacombe, Thomas R. Sibley, Melissa Truong, Caitlin R. Wolf, Romesh Gautom, Geoff Melly, Brian Hiatt, Philip Dykema, Scott Lindquist, Michael Boeckh, Janet A. Englund, Michael Famulare, Barry R. Lutz, Mark J. Rieder, Lea M. Starita, Matthew Thompson, Helen Y. Chu, Jay Shendure, Trevor Bedford |
| EPI_ISL_814321, EPI_ISL_814343, EPI_ISL_814695, EPI_ISL_814764, EPI_ISL_814765, EPI_ISL_814766, EPI_ISL_814767, EPI_ISL_814768, EPI_ISL_814769, EPI_ISL_814770, EPI_ISL_814771, EPI_ISL_814772, EPI_ISL_814773, EPI_ISL_814774, EPI_ISL_814775, EPI_ISL_814776, EPI_ISL_814777 | | | |
| see above | Wales Specialist Virology Centre Sequencing lab: Pathogen Genomics Unit | COVID-19 Genomics UK (COG-UK) Consortium | Catherine Moore, Johnathan Evans, Laura Gifford, Malorie Perry, Simon Cottrell, Angela Marchbank, Alec Birchley, Alexander Adams, Amy Gaskin, Bree Gatica-Wilcox, Jason Coombes, Joel Southgate, Lauren Gilbert, Lee Graham, Nicole Pacchiarini, Sara Kumziene-Summerhayes, Sarah Taylor, Sophie Jones, Sara Rey, Matthew Bull, Joanne Watkins, Sally Corden, Tom Connor |
| EPI_ISL_815525, EPI_ISL_815530, EPI_ISL_815599, EPI_ISL_815640, EPI_ISL_815654, EPI_ISL_815660, EPI_ISL_816073, EPI_ISL_816074, EPI_ISL_816075, EPI_ISL_816076, EPI_ISL_816077, EPI_ISL_816078, EPI_ISL_816079, EPI_ISL_816080, EPI_ISL_816081, EPI_ISL_816082, EPI_ISL_816083, EPI_ISL_816084, EPI_ISL_816085, EPI_ISL_816086, EPI_ISL_816087, EPI_ISL_816088, EPI_ISL_816089, EPI_ISL_816090, EPI_ISL_816091, EPI_ISL_816092, EPI_ISL_816093, EPI_ISL_816094 | | | |
| see above | Department of Virus and Microbiological Special Diagnostics, Statens Serum Institut, Copenhagen, Denmark | Albertsen Lab, Department of Chemistry and Bioscience, Aalborg University, Denmark | Danish Covid-19 Genome Consortium |
| EPI_ISL_819462, EPI_ISL_819463 | University College London, Great Ormond Street Hospital for Children NHS Foundation Trust, Imperial College Healthcare NHS Trust | COVID-19 Genomics UK (COG-UK) Consortium | Sergi Castellano, Rachel Williams, Mark Kristiansen, Paola Resende Silva, Sunando Roy, Tony Brooks, Helena Tutill, Paola Niola, Patricia Dyal, Charlotte Williams, Leysa Forrest, Yasmin Panchbhaya, Jacqueline Findlay, Samuel Weeks, Julianne Brown, Kathryn Harris, Paul Randell, James Price, Alison Holmes, Judith Breuer |
| EPI_ISL_823973 | Department of Homeless Services | New York City Public Health Laboratory | Jade Wang, et al. |
| EPI_ISL_824000 | Dutch COVID-19 response team | National Institute for Public Health and the Environment (RIVM) | Adam Meijer, Harry Vennema, Jeroen Cremer, Sharon van den Brink, Bas van der Veer, AnneMarie van den Brandt, Florian Zwagemaker, Dennis Schmitz, Chantal Reusken, on behalf of the national COVID-19 response team |
| EPI_ISL_824334 | Michigan Department of Health and Human Services, Bureau of Laboratories | Michigan Department of Health and Human Services, Bureau of Laboratories | Blankenship HM, Riner D, Soehnlen MK |
| EPI_ISL_824395, EPI_ISL_824399 | California Department of Public Health | California Department of Public Health | CDPH IDLB COVIDNet |
| EPI_ISL_824517, EPI_ISL_824518 | New Mexico Department of Health Scientific Laboratory | New Mexico Department of Health Scientific Laboratory | Ellie Johnson, Anastacia Griego-Fisher, D'eldra Malone |
| EPI_ISL_824659, EPI_ISL_824670, EPI_ISL_824682, EPI_ISL_824694, EPI_ISL_824705, EPI_ISL_824706, EPI_ISL_824717, EPI_ISL_824718, EPI_ISL_824729, EPI_ISL_824740, EPI_ISL_824741 | | | |
| see above | Cedars-Sinai Medical Center, Department of Pathology & Laboratory Medicine, Molecular Pathology Laboratory | Cedars-Sinai Medical Center, Molecular Pathology Laboratory of Department of Pathology & Laboratory Medicine and Genomic Core | Wenjuan Zhang, Brian Davis, Stephanie Chen, Jorge Mario Sincuir Martinez, Jasmine T Plummer, Eric Vail |
| EPI_ISL_825034 | B.J. Medical College and Civil hospital, Ahmedabad | Gujarat Biotechnology Research Centre | Kamlesh J Upadhyay, Zarna Patel, Nitin Savaliya, Dinesh Kumar, Zuber Saiyed, Labdhi Pandya, Afzal Ansari, Nikha Trivedi, Apurvasinh Puvar, Ramesh Pandit, Janvi Raval, Pranay Shah, Nilima Shah, Sanjay Kapadia, Dipa Kinariwala, Chaitanya Joshi, Madhvi Joshi |
| EPI_ISL_825090, EPI_ISL_825091, EPI_ISL_825093, EPI_ISL_825094, EPI_ISL_825095, EPI_ISL_825100, EPI_ISL_825102, EPI_ISL_825103, EPI_ISL_825105, EPI_ISL_825121, EPI_ISL_825122, EPI_ISL_825123 | | | |
| see above | NHLS-IALCH | KRISP, KZN Research Innovation and Sequencing Platform | Giandhari J, Pillay S, Lessells R, Mdlalose K, York D, Khan S, Tegally H, Wilkinson E, de Oliveira T |
| EPI_ISL_825334, EPI_ISL_825361, EPI_ISL_825362, EPI_ISL_825363 | Hospital Universitari Vall d'Hebron - Vall d'Hebron Institut de Rercerca | Hospital Universitari Vall d'Hebron | Cristina Andrés, Maria Piñana, Josep F Abril, Damir Garcia-Cehic, Ariadna Rando, Juliana Esperalba, Maria Gema Codina, Carla Castillo, Maria Carmen Martin, Tomàs Pumarola, Josep Quer, Andrés Antón |
| EPI_ISL_825552, EPI_ISL_826281 | Respiratory Virus Unit, National Infection Service, Public Health England | COVID-19 Genomics UK (COG-UK) Consortium | PHE Covid Sequencing Team |
| EPI_ISL_826879, EPI_ISL_826882, EPI_ISL_827068, EPI_ISL_827662, | deCODE genetics | deCODE genetics | Daniel F Gudbjartsson; Agnar Helgason; Hakon Jonsson; Olafur T Magnusson; Pall Melsted; Gudmundur L Norddahl; Jona Saemundsdottir; Asgeir Sigurdsson; Patrick Sulem; Arna B Agustsdottir; Hannes Eggertsson; Berglind Eiriksdottir; Run Fridriksdottir; Elisabet E Gardarsdottir; Gudmundur |

| | | | |
|---|---|---|---|
| EPI_ISL_827826, EPI_ISL_828173, EPI_ISL_828179, EPI_ISL_828233, EPI_ISL_828249, EPI_ISL_828321 | | | Georgsson; Olafia S Gretarsdottir; Kjartan R Gudmundsson; Thora R Gunnarsdottir; Arnaldur Gylfason; Hilma Holm; Brynjar O Jensson; Aslaug Jonasdottir; Kamilla S Josefsdottir; Thordur Kristjansson; Droplaug N Magnusdottir; Solvi Rognvaldsson; Louise le Roux; Gudrun Sigmundsdottir; Gardar Sveinbjornsson; Kristin E Sveinsdottir; Maney Sveinsdottir; Emil A Thorarensen; Bjarni Thorbjornsson; Gisli Masson; Ingileif Jonsdottir; Alma Moller; Thorolfur Gudnason; Karl G Kristinsson; Unnur Thorsteinsdottir; Kari Stefansson |
| EPI_ISL_828483 | The National University Hospital of Iceland | deCODE genetics | Daniel F Gudbjartsson; Agnar Helgason; Hakon Jonsson; Olafur T Magnusson; Pall Melsted; Gudmundur L Norddahl; Jona Saemundsdottir; Asgeir Sigurdsson; Patrick Sulem; Arna B Agustsdottir; Hannes Eggertsson; Berglind Eiriksdottir; Run Fridriksdottir; Elisabet E Gardarsdottir; Gudmundur Georgsson; Olafia S Gretarsdottir; Kjartan R Gudmundsson; Thora R Gunnarsdottir; Arnaldur Gylfason; Hilma Holm; Brynjar O Jensson; Aslaug Jonasdottir; Kamilla S Josefsdottir; Thordur Kristjansson; Droplaug N Magnusdottir; Solvi Rognvaldsson; Louise le Roux; Gudrun Sigmundsdottir; Gardar Sveinbjornsson; Kristin E Sveinsdottir; Maney Sveinsdottir; Emil A Thorarensen; Bjarni Thorbjornsson; Gisli Masson; Ingileif Jonsdottir; Alma Moller; Thorolfur Gudnason; Karl G Kristinsson; Unnur Thorsteinsdottir; Kari Stefansson |
| EPI_ISL_828920, EPI_ISL_829051, EPI_ISL_829052, EPI_ISL_829137, EPI_ISL_829138, EPI_ISL_829145, EPI_ISL_829148, EPI_ISL_829150, EPI_ISL_829151, EPI_ISL_829176, EPI_ISL_829177, EPI_ISL_829178, EPI_ISL_829182, EPI_ISL_829184, EPI_ISL_829185, EPI_ISL_829684, EPI_ISL_829904, EPI_ISL_829905, EPI_ISL_829906, EPI_ISL_829908, EPI_ISL_829911, EPI_ISL_829912, EPI_ISL_829913, EPI_ISL_830174, EPI_ISL_830175 | | | |
| see above | deCODE genetics | deCODE genetics | Daniel F Gudbjartsson; Agnar Helgason; Hakon Jonsson; Olafur T Magnusson; Pall Melsted; Gudmundur L Norddahl; Jona Saemundsdottir; Asgeir Sigurdsson; Patrick Sulem; Arna B Agustsdottir; Hannes Eggertsson; Berglind Eiriksdottir; Run Fridriksdottir; Elisabet E Gardarsdottir; Gudmundur Georgsson; Olafia S Gretarsdottir; Kjartan R Gudmundsson; Thora R Gunnarsdottir; Arnaldur Gylfason; Hilma Holm; Brynjar O Jensson; Aslaug Jonasdottir; Kamilla S Josefsdottir; Thordur Kristjansson; Droplaug N Magnusdottir; Solvi Rognvaldsson; Louise le Roux; Gudrun Sigmundsdottir; Gardar Sveinbjornsson; Kristin E Sveinsdottir; Maney Sveinsdottir; Emil A Thorarensen; Bjarni Thorbjornsson; Gisli Masson; Ingileif Jonsdottir; Alma Moller; Thorolfur Gudnason; Karl G Kristinsson; Unnur Thorsteinsdottir; Kari Stefansson |
| EPI_ISL_831701, EPI_ISL_831708, EPI_ISL_831709, EPI_ISL_831713, EPI_ISL_831743, EPI_ISL_831825, EPI_ISL_831826, EPI_ISL_831827, EPI_ISL_831828, EPI_ISL_831830, EPI_ISL_831831 | | | |
| see above | United States Air Force School of Aerospace Medicine | United States Air Force School of Aerospace Medicine | Anthony Fries, Jennifer Meyer, William Gruner, Amanda Javorina, Sarah Purves, Clarise Starr, Elizabeth Macias |
| EPI_ISL_832032, EPI_ISL_832033, EPI_ISL_832034 | Wyoming Public Health Laboratory | Wyoming Public Health Laboratory | Noah Hull, Taylor Fearing, Lynette Gumbleton, Channing Weber, Ashley Norberg, Bailey Bowcutt, and Wanda Manley |
| EPI_ISL_832146, EPI_ISL_832147, EPI_ISL_832148 | Hospital | National Reference Center for Viruses of Respiratory Infections, Institut Pasteur, Paris | Marion Barbet, Sylvie Behillil, Méline Bizard, Angela Brisebarre, Camille Capel, Etienne Simon-Lorière, Vincent Enouf, Maud Vanpeene, Sylvie van der Werf, Léa Pilorge |
| EPI_ISL_832781, EPI_ISL_832782, EPI_ISL_832794, EPI_ISL_832795, EPI_ISL_832796, EPI_ISL_832797, EPI_ISL_832798, EPI_ISL_832799, EPI_ISL_832800, EPI_ISL_832811, EPI_ISL_832812 | | | |
| see above | OHSU Lab Services Molecular Microbiology Lab | Oregon SARS-CoV-2 Genome Sequencing Center | Brendan L. O'Connell, Ruth V. Nichols, Sally Grindstaff, Alec J. Hirsch, Donna Hansel, Guang Fan, Daniel N. Streblow, William B. Messer, Andrew C. Adey, Benjamin N. Bimber, Brian J. O'Roak |
| EPI_ISL_833455 | SC (UCO) Igiene e Sanità Pubblica (funzione integrata con SC Microbiologia e Virologia) e Laboratory of Molecular Virology of the International Centre for Genetic Engineering and Biotechnology (ICGEB) | ARGO Laboratorio Genomica ed Epigenomica | Licastro D, Dal Monego S, Degasperi M, Marcello A, D'Agaro P |
| EPI_ISL_833458, EPI_ISL_833475, EPI_ISL_833476, EPI_ISL_833479 | CHU Purpan - Laboratoire de Virologie - Institut Fédératif de Biologie | CHU Purpan - Laboratoire de Virologie - Institut Fédératif de Biologie | Latour J., Ranger N., Dubois M., Carcenac R., Harter A., Boyer P., Tremeaux P., Izopet J. |
| EPI_ISL_838281 | Virology Department, Royal Infirmary of Edinburgh, NHS Lothian / School of Biological Sciences, University of Edinburgh / Institute of Genetics and Molecular Medicine, University of Edinburgh | COVID-19 Genomics UK (COG-UK) Consortium | McHugh M, Dewar R, Rooke S, Gallagher M, Balcaza C, O'Toole Á, Scher E, Hill V, McCrone JT, Colquhoun R, Yu X, Jackson B, Rambaut A, Williams TC, Templeton K |
| EPI_ISL_838344, EPI_ISL_838345, EPI_ISL_838346 | Queens Medical Centre, Clinical Microbiology Department / DeepSeq Nottingham | COVID-19 Genomics UK (COG-UK) Consortium | Gemma Clark, Wendy Smith, Manjinder Khakh, Vicki M Fleming, Michelle M Lister, Hannah Howson-Wells, Jonathan Ball, Patrick McClure, Joseph Chappell, Theocharis Tsoleridis, Nadine Holmes, Matthew Carlisle, Christopher Moore, Fei Sang, Johnny Debebe, Victoria Wright, Matthew Loose |
| EPI_ISL_840188, EPI_ISL_840189, EPI_ISL_840204, EPI_ISL_840205, EPI_ISL_840206, EPI_ISL_840216, EPI_ISL_840217 | Oxford Viromics, NDM, University of Oxford; Oxford University Hospitals; Basingstoke and North Hampshire Hospital | COVID-19 Genomics UK (COG-UK) Consortium | Tanya Golubchik, David Bonsall, George Macintyre, Amy Trebes, Mariateresa de Cesare, Catrin Moore, Alex Mobbs, Anita Justice, Robert Shaw, Monique Andersson, Timothy Peto, Emma Wise, Nathan Moore, Jessica Lynch, Nick Cortes, Matilde Mori, Stephen Kidd, David Buck, John Todd, Christophe Fraser |
| EPI_ISL_842916, EPI_ISL_842917, EPI_ISL_842918, EPI_ISL_842925, EPI_ISL_842926, EPI_ISL_842927, EPI_ISL_842928, EPI_ISL_842929, EPI_ISL_842930, EPI_ISL_842931, EPI_ISL_842933, EPI_ISL_842934, EPI_ISL_842935, EPI_ISL_842936, EPI_ISL_842943, EPI_ISL_842944, EPI_ISL_842946, EPI_ISL_842947, EPI_ISL_842948, EPI_ISL_842949, EPI_ISL_842953, EPI_ISL_842956, EPI_ISL_842963, EPI_ISL_842964, EPI_ISL_842965, EPI_ISL_842967, EPI_ISL_842968, EPI_ISL_842984, EPI_ISL_842985, EPI_ISL_842986, EPI_ISL_842987, EPI_ISL_842988, EPI_ISL_842989, EPI_ISL_842990, EPI_ISL_842991, EPI_ISL_842992, EPI_ISL_842993, EPI_ISL_842995, EPI_ISL_842996, EPI_ISL_842997, EPI_ISL_842998, EPI_ISL_842999, EPI_ISL_843001, EPI_ISL_843002, EPI_ISL_843003, EPI_ISL_843004 | | | |
| see above | Barts Health NHS Trust | COVID-19 Genomics UK (COG-UK) Consortium | CUTINO-MOGUEL, Maria-Teresa; HARRINGTON, David; OWOYEMI, Dola; SHYLINI, Raghavendran; BROAD, Claire; KELE, Beatrix |
| EPI_ISL_844204, EPI_ISL_844223, EPI_ISL_844229, EPI_ISL_844239, EPI_ISL_844292, EPI_ISL_844293, EPI_ISL_844294, EPI_ISL_844296, EPI_ISL_844528, EPI_ISL_844529, EPI_ISL_844530, EPI_ISL_844531, EPI_ISL_844532, EPI_ISL_844533, EPI_ISL_844534, EPI_ISL_844535, EPI_ISL_844536, EPI_ISL_844537, EPI_ISL_844538 | | | |
| see above | Department of Virus and Microbiological Special Diagnostics, Statens Serum Institut, Copenhagen, Denmark | Albertsen Lab, Department of Chemistry and Bioscience, Aalborg University, Denmark | Danish Covid-19 Genome Consortium |
| EPI_ISL_847516, EPI_ISL_847521, EPI_ISL_847523, EPI_ISL_847552, EPI_ISL_847554, EPI_ISL_847561, EPI_ISL_847573, EPI_ISL_847588, EPI_ISL_847674, EPI_ISL_847675, EPI_ISL_847676, EPI_ISL_847677, EPI_ISL_847678, EPI_ISL_847679, EPI_ISL_847680, EPI_ISL_847681, EPI_ISL_847682, EPI_ISL_847683, EPI_ISL_847684, EPI_ISL_847685, EPI_ISL_847686, EPI_ISL_847687, EPI_ISL_847688, EPI_ISL_847689, EPI_ISL_847690, EPI_ISL_847691, EPI_ISL_847692, EPI_ISL_847693, EPI_ISL_847694, EPI_ISL_847706, EPI_ISL_847709, EPI_ISL_847748, EPI_ISL_847804, EPI_ISL_847805, EPI_ISL_847806, EPI_ISL_847807, EPI_ISL_847808, EPI_ISL_847809, EPI_ISL_847810, EPI_ISL_847811, EPI_ISL_847812, EPI_ISL_847813, EPI_ISL_847814, EPI_ISL_847815 | | | |
| see above | California Department of Public Health | Chiu Laboratory, University of California, San Francisco | Charles Chiu, Xianding (Wayne) Deng, Candace Wang, Brian Bushnell, Scot Federman, Jill Hacker, Debra Wadford |
| EPI_ISL_848231 | Illinois Department of Public Health | Gagnon Lab, Southern Illinois University | Keith Gagnon |
| EPI_ISL_848734, EPI_ISL_848735, EPI_ISL_848736, EPI_ISL_848737, EPI_ISL_848738, EPI_ISL_848739, EPI_ISL_848740, EPI_ISL_848741, EPI_ISL_848742, EPI_ISL_848743, EPI_ISL_848744, EPI_ISL_848745, EPI_ISL_848746, EPI_ISL_848747, EPI_ISL_848748, EPI_ISL_848749, EPI_ISL_848750, EPI_ISL_848751, EPI_ISL_848752, EPI_ISL_848753, EPI_ISL_848754, EPI_ISL_848755, EPI_ISL_848756, EPI_ISL_848757 | | | |
| see above | Florida Bureau of Public Health Laboratories | Florida Bureau of Public Health Laboratories | Sarah Schmedes, Jason Blanton |
| EPI_ISL_849421, EPI_ISL_849448, EPI_ISL_849518, EPI_ISL_849519, EPI_ISL_849520, EPI_ISL_849575, EPI_ISL_849618 | Washington State Department of Health | Seattle Flu Study | Deborah A. Nickerson, Chris D. Frazar, Jover Lee, Benjamin Pelle, Matthew Richardson, Amanda Adler, Elisabeth Brandstetter, Peter D. Han, Kairsten Fay, Misja Ilcisin, Kirsten Lacombe, Thomas R. Sibley, Melissa Truong, Caitlin R. Wolf, Romesh Gautom, Geoff Melly, Brian Hiatt, Philip Dykema, Scott Lindquist, Michael Boeckh, Janet A. Englund, Michael Famulare, Barry R. Lutz, Mark J. Rieder, Lea M. Starita, Matthew Thompson, Helen Y. Chu, Jay Shendure, Trevor Bedford |
| EPI_ISL_850019, EPI_ISL_850020, EPI_ISL_850021, EPI_ISL_850022, EPI_ISL_850023, EPI_ISL_850024, EPI_ISL_850025, EPI_ISL_850026, EPI_ISL_850027, EPI_ISL_850028, EPI_ISL_850029, EPI_ISL_850030, EPI_ISL_850031, EPI_ISL_850032, EPI_ISL_850033, EPI_ISL_850034, EPI_ISL_850035, EPI_ISL_850036, EPI_ISL_850037, EPI_ISL_850038 | | | |
| see above | Renegade | Chan-Zuckerberg Biohub | CZB Cliahub Consortium |
| EPI_ISL_850663 | The National Institute of Public Health | State Veterinary Institute Prague | Nagy,A;Jirincova,H;Trnka,D;Vecerova,J;Trinklova,M |
| EPI_ISL_853336 | UPMC Clinical Microbiology Laboratory | Microbial Genome Sequencing Center; Microbial Genomic Epidemiology Laboratory | Mustapha M. Mustapha, Jane W. Marsh, Dan Snyder, Marissa P. Griffith, Stephanie L. Mitchell, Vatsala R. Srinivasa, Kady D. Waggle, Chinelo Ezeonwuku, Vaughn S. Cooper, Lee H. Harrison |
| EPI_ISL_853733, EPI_ISL_853779, EPI_ISL_853780 | Austrian Agency for Health and Food Safety (AGES) | Bergthaler laboratory, CeMM Research Center for Molecular Medicine of the Austrian Academy of Sciences | Lukas Endler, Alexandra Popa, Benedikt Agerer, Jakob-Wendelin Genger, Alexander Lercher, Anna Schedl, Thomas Penz, Michael Schuster, Jan Laine, Martin Senekowitsch, Christoph Bock, Andreas Bergthaler |
| EPI_ISL_853948 | Department of Microbiology, University Innsbruck | Bergthaler laboratory, CeMM Research Center for Molecular Medicine of the Austrian Academy of Sciences | Lukas Endler, Alexandra Popa, Benedikt Agerer, Jakob-Wendelin Genger, Alexander Lercher, Anna Schedl, Thomas Penz, Michael Schuster, Jan Laine, Martin Senekowitsch, Christoph Bock, Andreas Bergthaler |

| | | | |
|---|---|---|---|
| EPI_ISL_853976, EPI_ISL_854113, EPI_ISL_854114, EPI_ISL_854115, EPI_ISL_854116, EPI_ISL_854117, EPI_ISL_854118, EPI_ISL_854119, EPI_ISL_854120, EPI_ISL_854121, EPI_ISL_854243, EPI_ISL_854285, EPI_ISL_854297 | | | |
| see above | Austrian Agency for Health and Food Safety (AGES) | Bergthaler laboratory, CeMM Research Center for Molecular Medicine of the Austrian Academy of Sciences | Lukas Endler, Alexandra Popa, Benedikt Agerer, Jakob-Wendelin Genger, Alexander Lercher, Anna Schedl, Thomas Penz, Michael Schuster, Jan Laine, Martin Senekowitsch, Christoph Bock, Andreas Bergthaler |
| EPI_ISL_856761, EPI_ISL_856762, EPI_ISL_856763 | Servicio Virosis Respiratorias-Departamento Virología-INEI | Instituto Nacional Enfermedades Infecciosas C.G.Malbran | Baumeister E., Avaro M., Benedetti E., Russo M., Dattero ME, Pontoriero A., Cisterna D., Molina V., Perandones C., Tuduri E., Lorenzo F., Poklepovich T., Campos J. |
| EPI_ISL_857521, EPI_ISL_857522, EPI_ISL_857523 | Swiss National Reference Centre for Influenza | Swiss National Reference Centre for Influenza | Tim Roloff, Ana Rita Gonçalves, Madlen Stange, Helena MB Seth-Smith, Alfredo Mari, Karoline Leuzinger, Julia Bielicki, Manuel Battegay, Hans Hirsch, Laurent Kaiser, Adrian Egli |
| EPI_ISL_859554, EPI_ISL_859556, EPI_ISL_859557, EPI_ISL_859558, EPI_ISL_859559 | Lighthouse Lab in Cambridge | Wellcome Sanger Institute for the COVID-19 Genomics UK (COG-UK) Consortium | Rob Howes, The Lighthouse Lab in Cambridge and Alex Alderton, Roberto Amato, Sonia Goncalves, Ewan Harrison, David K. Jackson, Ian Johnston, Dominic Kwiatkowski, Cordelia Langford, John Sillitoe on behalf of the Wellcome Sanger Institute COVID-19 Surveillance Team |
| EPI_ISL_859854, EPI_ISL_859855, EPI_ISL_859856, EPI_ISL_859857, EPI_ISL_859858, EPI_ISL_859859, EPI_ISL_859860, EPI_ISL_859861, EPI_ISL_859862, EPI_ISL_859863, EPI_ISL_859864, EPI_ISL_859876, EPI_ISL_859877, EPI_ISL_859878, EPI_ISL_859879, EPI_ISL_859880, EPI_ISL_859881, EPI_ISL_859904, EPI_ISL_859905, EPI_ISL_859906, EPI_ISL_859907, EPI_ISL_859908, EPI_ISL_859909, EPI_ISL_859910, EPI_ISL_859911, EPI_ISL_859912, EPI_ISL_859913, EPI_ISL_859920, EPI_ISL_859921, EPI_ISL_859922, EPI_ISL_859923, EPI_ISL_859934, EPI_ISL_859935 | | | |
| see above | BTC, Khalifa University | BTC, Khalifa University | Al Safar et al |
| EPI_ISL_860161, EPI_ISL_860164 | Keio University School of Medicine | Keio University School of Medicine | Kenjiro Kosaki, Yuka Iwasaki, Hirotsugu Ishizu, Haruhiko Siomi, Kodai Abe |
| EPI_ISL_861460 | UHAS COVID-19 Lab | UHAS COVID-19 Lab | Kwabena O. Duedu, Jones Gyamfi, Reuben Ayivor-Djanie, John O. Gyapong and the UHAS COVID-19 Lab Team |
| EPI_ISL_861868, EPI_ISL_861881, EPI_ISL_861883, EPI_ISL_861894, EPI_ISL_861903, EPI_ISL_861904, EPI_ISL_861908, EPI_ISL_861909 | LATE - Laboratório de Técnicas Especiais - Hospital Israelita Albert Einstein | LATE - Laboratório de Técnicas Especiais - Hospital Israelita Albert Einstein | Deyvid Amgarten, Fernanda de Mello Malta, Raquel Riyuzo, Ana Paula Moreira Salles, Pedro Henrique Sebe Rodrigues, João Renato Rebello Pinho |
| EPI_ISL_861916, EPI_ISL_861917, EPI_ISL_861918, EPI_ISL_861919, EPI_ISL_861920, EPI_ISL_861921, EPI_ISL_861922, EPI_ISL_861923, EPI_ISL_861924 | OHSU Lab Services Molecular Microbiology Lab | Oregon SARS-CoV-2 Genome Sequencing Center | Brendan L. O'Connell, Sally Grindstaff, Kayla Carter, Ruth V. Nichols, Alec J. Hirsch, Donna Hansel, Guang Fan, Xuan Qin, Daniel N. Streblow, William B. Messer, Andrew C. Adey, Benjamin N. Bimber, Brian J. O'Roak |
| EPI_ISL_864570, EPI_ISL_864571 | Institute of Medical Microbiology and Hospital Hygiene | Institute of Medical Microbiology and Hospital Hygiene | Prof. Dr. Achim Kaasch, Aljoscha Tersteegen |
| EPI_ISL_868760, EPI_ISL_868763, EPI_ISL_868766, EPI_ISL_868772, EPI_ISL_868773, EPI_ISL_868776, EPI_ISL_868780, EPI_ISL_868781, EPI_ISL_868782, EPI_ISL_868783, EPI_ISL_868784, EPI_ISL_868788 | | | |
| see above | Bioinformatics and Biostatistics Lab, Advanced Sequencing Facility | COVID-19 Genomics UK (COG-UK) Consortium | Aengus Stewart,Jerome Nicod,Chelsea Sawyer,Laura Cubitt,Harshil Patel,Margaret Crawford |
| EPI_ISL_869250, EPI_ISL_869262, EPI_ISL_869263, EPI_ISL_869264 | Krasnoyarsk Regional Clinical Hospital | WHO National Influenza Centre Russian Federation | Andrey Komissarov, Artem Fadeev, Anna Ivanova, Kseniya Komissarova, Dmitry Bazhenov, Mikhail Bakaev, Daria Danilenko, Ksenia Safina, Elena Nabieva, Georgii Bazykin, Dmitry Lioznov |
| EPI_ISL_871783, EPI_ISL_871784, EPI_ISL_871785 | Department of Virus and Microbiological Special Diagnostics, Statens Serum Institut, Copenhagen, Denmark | Aalborg University | Danish Covid-19 Genome Consortium |
| EPI_ISL_872570 | Krasnoyarsk Regional Clinical Hospital | WHO National Influenza Centre Russian Federation | Andrey Komissarov, Artem Fadeev, Anna Ivanova, Kseniya Komissarova, Dmitry Bazhenov, Mikhail Bakaev, Daria Danilenko, Ksenia Safina, Elena Nabieva, Georgii Bazykin, Dmitry Lioznov |
| EPI_ISL_872697, EPI_ISL_872698 | Rhode Island Department of Health | Infectious Disease Program, Broad Institute of Harvard and MIT | Lemieux,J.E., Siddle,K.J., Huard,R., King,E., Azevedo,K., Miller,A., Adams,G., Gladden-Young,A., Lagerborg,K., Rudy,M., DeRuff,K., Carter,A., Normandin,E., Bauer,M., Reilly,S., Tomkins-Tinch,C., Loreth,C., Chaluvadi,S., Birren,B.W., Gallagher,G., Smole,S., Park,D.J., MacInnis,B.L., and Sabeti,P.C. |
| EPI_ISL_873126, EPI_ISL_873127, EPI_ISL_873128, EPI_ISL_873131, EPI_ISL_873136, EPI_ISL_873137, EPI_ISL_873141, EPI_ISL_873142, EPI_ISL_873145 | University of Michigan Clinical Microbiology Laboratory | Lauring Lab, University of Michigan, Department of Microbiology and Immunology | Valesano |
| EPI_ISL_875553 | ULSS 2 Treviso | Istituto Zooprofilattico Sperimentale delle Venezie | Adelaide Milani, Alessia Schivo, Annalisa Salviato, Erika Giorgia Quaranta, Ambra Pastori, Bianca Zecchin, Alice Fusaro, Isabella Monne, Calogero Terregino, Antonia Ricci |
| EPI_ISL_876074, EPI_ISL_876104, EPI_ISL_876113, EPI_ISL_876119, EPI_ISL_876135, EPI_ISL_876138, EPI_ISL_876187, EPI_ISL_876322, EPI_ISL_876323, EPI_ISL_876324 | Massachusetts State Public Health Laboratory | Massachusetts State Public Health Laboratory | Andrew Lang, Timelia Fink, Glen Gallagher, Sandra Smole |
| EPI_ISL_876523, EPI_ISL_876527, EPI_ISL_876529, EPI_ISL_876530, EPI_ISL_876555, EPI_ISL_876556, EPI_ISL_876557, EPI_ISL_876558, EPI_ISL_876559, EPI_ISL_876560, EPI_ISL_876561, EPI_ISL_876562, EPI_ISL_876563, EPI_ISL_876564, EPI_ISL_876565, EPI_ISL_876566 | | | |
| see above | Florida Bureau of Public Health Laboratories | Florida Bureau of Public Health Laboratories | Sarah Schmedes, Jason Blanton |
| EPI_ISL_878276, EPI_ISL_878279, EPI_ISL_878300, EPI_ISL_878303, EPI_ISL_878308, EPI_ISL_878319, EPI_ISL_878322, EPI_ISL_878330, EPI_ISL_878336, EPI_ISL_878345, EPI_ISL_878346, EPI_ISL_878351, EPI_ISL_878362, EPI_ISL_878365, EPI_ISL_878382, EPI_ISL_878400, EPI_ISL_878401, EPI_ISL_878402, EPI_ISL_878404, EPI_ISL_878406, EPI_ISL_878408, EPI_ISL_878416, EPI_ISL_878418, EPI_ISL_878423, EPI_ISL_878426, EPI_ISL_878434, EPI_ISL_878437 | | | |
| see above | San Diego County Public Health Laboratory | Andersen lab at Scripps Research | SEARCH Alliance San Diego with Tracy Basler, Jovan Shephard, Brett Austin |
| EPI_ISL_878544 | Robert Garry lab | Andersen lab at Scripps Research | Allison Smither, Gilberto Sabino-Santos, Patricia Snarski, Lilia Melnik, Antoinette Bell, Kaylynn Genemaras, Arnaud Drouin, Dahlene Fusco, Robert Garry with SEARCH Alliance San Diego |
| EPI_ISL_878648, EPI_ISL_878661, EPI_ISL_878705 | Rady's Childrens Hospital | Andersen lab at Scripps Research | SEARCH Alliance San Diego with Nanda Radamchar, David Dimmock, Linda Luo, Christina Clarke, Kathryn Bouic, Teresa Mueller, Denise Malicki |
| EPI_ISL_882658 | Secretaria Municipal de Saude | Instituto Adolfo Lutz, Interdiciplinary Procedures Center, Strategic Laboratory | Claudio Tavares Sacchi, Claudia Regina Gonçalves, Erica Valessa Ramos Gomes, Karoline Rodrigues Campos |
| EPI_ISL_887164 | Massachusetts General Hospital | Infectious Disease Program, Broad Institute of Harvard and MIT | Lemieux,J.E., Siddle,K.J., Shaw,B., Adams,G., Pierce,V., Turbett,S., Anahtar,M., Branda,J., Slater,D., Harris,J., Lin,A.E., Gladden-Young,A., Lagerborg,K., Rudy,M., DeRuff,K., Carter,A., Normandin,E., Bauer,M., Reilly,S., Tomkins-Tinch,C., Loreth,C., Chaluvadi,S., Neumann,A., Cusick,C., Chapman,S.B., Gnirke,A., Flowers,K., Cerrato,F., Birren,B.W., Gallagher,G., Smole,S., Park,D.J., MacInnis,B.L., Ryan,E., LaRocque,R., Rosenberg,E. and Sabeti,P.C. |
| EPI_ISL_887425, EPI_ISL_887461, EPI_ISL_887462, EPI_ISL_887463, EPI_ISL_887465 | Instituto Nacional de Saude (INS), Mozambique | KRISP, KZN Research Innovation and Sequencing Platform | Nalia Ismael, Nadia Sitoe, Paulo Arnaldo, Nedio Mabunda, Giandhari J, Pillay S, Tegally H, Wilkinson E, de Oliveira T |
| EPI_ISL_887589 | SC Dept of Health and Env. Control-Bureau of Laboratories | Centers for Disease Control and Prevention Division of Viral Diseases, Pathogen Discovery | Yan Li, Anna Montmayeur, Ying Tao, Jing Zhang, Krista Queen, Anna Uehara, Rachel Marine, Peter Cook, Clinton R. Paden, Haibin Wang, Suxiang Tong |
| EPI_ISL_888990 | RS Pondok Indah Bintaro Jaya | Eijkman Institute for Molecular Biology, Ministry of Research and Technology/National Agency for Research and Innovation | Frilasita A Yudhaputri, Hidayat Trimarsanto, Iskandar Adnan, Lydia V. Panggalo, Sukma Oktavianthi, Willy Agustine, Edison Johar, Safarina G Malik, Khin Saw Myint, Amin Soebandrio |
| EPI_ISL_890297 | KU Leuven, Rega Institute, Clinical and Epidemiological | KU Leuven, Rega Institute, Clinical and Epidemiological | Tony Wawina-Bokalanga, Bert Vanmechelen, Joan Marti-Carerras, Piet Maes |

| | | | |
|---|---|---|---|
| | Virology | Virology | |
| EPI_ISL_891221 | University of Medicine and Pharmacy of Craiova | "Stefan cel Mare" University Metagenomics Lab | Lobiuc Andrei, Gheorghita Roxana |
| EPI_ISL_891229 | Suceava County Emergency Hospital | "Stefan cel Mare" University Metagenomics Lab | Lobiuc Andrei, Gheorghita Roxana |
| EPI_ISL_893721, EPI_ISL_893722, EPI_ISL_893723 | Oregon State Public Health Laboratory | Pathogen Discovery, Respiratory Viruses Branch, Division of Viral Diseases, Centers for Disease Control and Prevention | Ying Tao, Yan Li, Jing Zhang, Krista Queen, Anna Uehara, Peter Cook, Clinton R. Paden, Haibin Wang, Suxiang Tong |
| EPI_ISL_896206 | MEPHI, Aix Marseille University | MEPHI, Aix Marseille University | Anthony LEVASSEUR |
| EPI_ISL_896376 | University of Medicine and Pharmacy of Craiova | "Stefan cel Mare" University Metagenomics Lab | Lobiuc Andrei, Gheorghita Roxana |
| EPI_ISL_896378, EPI_ISL_896379 | Suceava County Emergency Hospital | "Stefan cel Mare" University Metagenomics Lab | Lobiuc Andrei, Gheorghita Roxana |
| EPI_ISL_903002, EPI_ISL_903004, EPI_ISL_903005, EPI_ISL_903006 | Seattle Flu Study | Seattle Flu Study | Deborah A. Nickerson, Chris D. Frazar, Jover Lee, Benjamin Pelle, Erica Ryke, Matthew Richardson, Amanda Adler, Elisabeth Brandstetter, Peter D. Han, Kairsten Fay, Misja Ilcisin, Kirsten Lacombe, Thomas R. Sibley, Melissa Truong, Caitlin R. Wolf, Michael Boeckh, Janet A. Englund, Michael Famulare, Barry R. Lutz, Mark J. Rieder, Lea M. Starita, Matthew Thompson, Jay Shendure, Trevor Bedford, Helen Y. Chu |
| EPI_ISL_903613, EPI_ISL_903627 | LA Office of Public Health Laboratories | Genomics and Discovery, Respiratory Viruses Branch, Division of Viral Diseases, Centers for Disease Control and Prevention | Krista Queen, Yan Li, Ying Tao, Jing Zhang, Anna Uehara, Anna Montmayeur, Clinton R. Paden, Peter W. Cook, Rachel Marine, Mili Sheth, Jasmine Padilla, Sarah Nobles, Mark Burroughs, Lori Rowe, Haibin Wang, Ben L. Rambo-Martin, Dhwani Batra, Justin Lee, Suxiang Tong |
| EPI_ISL_903632 | NC State Laboratory of Public Health | Genomics and Discovery, Respiratory Viruses Branch, Division of Viral Diseases, Centers for Disease Control and Prevention | Krista Queen, Yan Li, Ying Tao, Jing Zhang, Anna Uehara, Anna Montmayeur, Clinton R. Paden, Peter W. Cook, Rachel Marine, Mili Sheth, Jasmine Padilla, Sarah Nobles, Mark Burroughs, Lori Rowe, Haibin Wang, Ben L. Rambo-Martin, Dhwani Batra, Justin Lee, Suxiang Tong |
| EPI_ISL_903720, EPI_ISL_903765, EPI_ISL_903906 | LA Office of Public Health Laboratories | Genomics and Discovery, Respiratory Viruses Branch, Division of Viral Diseases, Centers for Disease Control and Prevention | Krista Queen, Yan Li, Ying Tao, Jing Zhang, Anna Uehara, Anna Montmayeur, Clinton R. Paden, Peter W. Cook, Rachel Marine, Mili Sheth, Jasmine Padilla, Sarah Nobles, Mark Burroughs, Lori Rowe, Haibin Wang, Ben L. Rambo-Martin, Dhwani Batra, Justin Lee, Suxiang Tong |
| EPI_ISL_903954, EPI_ISL_903960, EPI_ISL_903972, EPI_ISL_903975, EPI_ISL_903978 | WA State Department of Health | Genomics and Discovery, Respiratory Viruses Branch, Division of Viral Diseases, Centers for Disease Control and Prevention | Krista Queen, Yan Li, Ying Tao, Jing Zhang, Anna Uehara, Anna Montmayeur, Clinton R. Paden, Peter W. Cook, Rachel Marine, Mili Sheth, Jasmine Padilla, Sarah Nobles, Mark Burroughs, Lori Rowe, Haibin Wang, Ben L. Rambo-Martin, Dhwani Batra, Justin Lee, Suxiang Tong |
| EPI_ISL_904736 | Dutch COVID-19 response team | National Institute for Public Health and the Environment (RIVM) | Adam Meijer, Harry Vennema, Dirk Eggink, Jeroen Cremer, Sharon van den Brink, Bas van der Veer, AnneMarie van den Brandt, Florian Zwagemaker, Dennis Schmitz, Chantal Reusken, on behalf of the national COVID-19 response team |
| EPI_ISL_904941, EPI_ISL_904949, EPI_ISL_904950, EPI_ISL_904954 | Vilnius University Hospital Santaros Klinikos, Vilnius University | Institute of Biotechnology, Life Sciences Center, Vilnius University | Emilija Vasiliunaite, Milda Norkiene, Albertas Timinskas, Alma Gedvilaite, Aurelija Zvirbliene, Daniel Naumovas, Laimonas Griskevicius |
| EPI_ISL_906096 | Child Health Research Foundation | Child Health Research Foundation | Senjuti Saha, Afroza Akter Tanni, Sharmistha Goswami, Syed Muktadir Al Sium, Arif Mohammad Tanmoy, Roly Malaker, Md Hafizur Rahman, Samir K Saha |
| EPI_ISL_909958, EPI_ISL_909962 | Apollo Hospitals | CSIR-Centre for Cellular and Molecular Biology | Onkar Kulkarni, Suneetha Narreddy, Lamuk Zaveri, Irawathy Goud, Sofia Banu, Payel Mukherjee, Karthik Bharadwaj Tallapaka, Divya Tej Sowpati |
| EPI_ISL_911679 | Alaska State Virology Laboratory | Alaska State Virology Laboratory | Stephanie DeRonde, Lisa Smith, Ph.D., Jack Chen, Ph.D. |
| EPI_ISL_912465, EPI_ISL_912505, EPI_ISL_912511, EPI_ISL_912527 | NHLS Universitas Academic | UFS Virology | PA Bester, MM Nyaga, P Nthiga, MT Mogotsi, D Goedhals, T de Oliveira |
| EPI_ISL_913934, EPI_ISL_913935, EPI_ISL_913970 | Instituto de Diagnostico y Referencia Epidemiologicos INDRE_RNLSP | Instituto de Diagnostico y Referencia Epidemiologicos (INDRE) | Claudia Wong-Arambula, Abril Rodriguez-Maldonado, Fabiola Garces-Ayala, Adnan Araiza-Rodriguez, David Fragoso-Fonseca, Sergio Rangel-Guerrero, Mayra Jimenez-Morales, Nancy Munoz-Hernandez, Natividad Cruz-Ortiz, Tatiana Nunez-Garcia, Gisela Barrera-Badillo, Lucia Hernandez-Rivas, Irma Lopez-Martinez, Ernesto Ramirez-Gonzalez. |
| EPI_ISL_914882 | Vilnius university hospital Santaros Klinikos, Center of Laboratory Medicine | Vilnius University Hospital Santaros Klinikos | Ingrida Olendraite, Daniel Naumovas, Rimvydas Norvilas, Dovil Ežerskyt, Justinas Šlikas |
| EPI_ISL_915367, EPI_ISL_915370, EPI_ISL_915413, EPI_ISL_915414 | Keio University School of Medicine | Keio University School of Medicine | Kenjiro Kosaki, Yuka Iwasaki, Hirotsugu Ishizu, Haruhiko Siomi, Kodai Abe |

EPI_ISL_915440, EPI_ISL_915441, EPI_ISL_915442, EPI_ISL_915443, EPI_ISL_915444, EPI_ISL_915445, EPI_ISL_915446, EPI_ISL_915447, EPI_ISL_915448, EPI_ISL_915449, EPI_ISL_915450, EPI_ISL_915451, EPI_ISL_915452, EPI_ISL_915453, EPI_ISL_915454, EPI_ISL_915455, EPI_ISL_915456, EPI_ISL_915457,
EPI_ISL_915458, EPI_ISL_915459, EPI_ISL_915460, EPI_ISL_915461, EPI_ISL_915462, EPI_ISL_915463, EPI_ISL_915464, EPI_ISL_915465, EPI_ISL_915466, EPI_ISL_915467, EPI_ISL_915468, EPI_ISL_915469, EPI_ISL_915470, EPI_ISL_915471, EPI_ISL_915472, EPI_ISL_915473, EPI_ISL_915474, EPI_ISL_915475,
EPI_ISL_915476, EPI_ISL_915477, EPI_ISL_915478, EPI_ISL_915479, EPI_ISL_915480, EPI_ISL_915481, EPI_ISL_915482, EPI_ISL_915483, EPI_ISL_915484, EPI_ISL_915485, EPI_ISL_915486, EPI_ISL_915487, EPI_ISL_915488, EPI_ISL_915489, EPI_ISL_915490, EPI_ISL_915491, EPI_ISL_915492, EPI_ISL_915493,
EPI_ISL_915494, EPI_ISL_915495, EPI_ISL_915496, EPI_ISL_915497, EPI_ISL_915498, EPI_ISL_915499, EPI_ISL_915500, EPI_ISL_915501, EPI_ISL_915502, EPI_ISL_915503, EPI_ISL_915504, EPI_ISL_915505, EPI_ISL_915506, EPI_ISL_915507, EPI_ISL_915508, EPI_ISL_915509, EPI_ISL_915510, EPI_ISL_915511,
EPI_ISL_915512, EPI_ISL_915513, EPI_ISL_915514, EPI_ISL_915515, EPI_ISL_915516, EPI_ISL_915517, EPI_ISL_915518, EPI_ISL_915519, EPI_ISL_915520, EPI_ISL_915521, EPI_ISL_915522, EPI_ISL_915523, EPI_ISL_915524, EPI_ISL_915525, EPI_ISL_915526, EPI_ISL_915527, EPI_ISL_915528, EPI_ISL_915529,
EPI_ISL_915530, EPI_ISL_915531, EPI_ISL_915532, EPI_ISL_915533, EPI_ISL_915534, EPI_ISL_915535, EPI_ISL_915536, EPI_ISL_915537, EPI_ISL_915538, EPI_ISL_915539, EPI_ISL_915540, EPI_ISL_915541, EPI_ISL_915542, EPI_ISL_915543, EPI_ISL_915544, EPI_ISL_915545, EPI_ISL_915546, EPI_ISL_915547,
EPI_ISL_915548, EPI_ISL_915549, EPI_ISL_915550, EPI_ISL_915551, EPI_ISL_915552, EPI_ISL_915553, EPI_ISL_915554, EPI_ISL_915555, EPI_ISL_915556, EPI_ISL_915557, EPI_ISL_915558, EPI_ISL_915559, EPI_ISL_915560, EPI_ISL_915561, EPI_ISL_915562, EPI_ISL_915563, EPI_ISL_915564, EPI_ISL_915565,
EPI_ISL_915566, EPI_ISL_915567, EPI_ISL_915568, EPI_ISL_915569, EPI_ISL_915570, EPI_ISL_915571, EPI_ISL_915572, EPI_ISL_915573, EPI_ISL_915574, EPI_ISL_915575, EPI_ISL_915576, EPI_ISL_915577, EPI_ISL_915578, EPI_ISL_915579, EPI_ISL_915580, EPI_ISL_915581, EPI_ISL_915582, EPI_ISL_915583,
EPI_ISL_915584, EPI_ISL_915585, EPI_ISL_915586, EPI_ISL_915587, EPI_ISL_915588, EPI_ISL_915589, EPI_ISL_915590, EPI_ISL_915591, EPI_ISL_915592, EPI_ISL_915593, EPI_ISL_915594, EPI_ISL_915595, EPI_ISL_915596, EPI_ISL_915597, EPI_ISL_915598, EPI_ISL_915599, EPI_ISL_915600, EPI_ISL_915601,
EPI_ISL_915602, EPI_ISL_915603, EPI_ISL_915604, EPI_ISL_915605, EPI_ISL_915606, EPI_ISL_915607, EPI_ISL_915608, EPI_ISL_915609, EPI_ISL_915610, EPI_ISL_915611, EPI_ISL_915612, EPI_ISL_915613, EPI_ISL_915614, EPI_ISL_915615, EPI_ISL_915616, EPI_ISL_915617, EPI_ISL_915618, EPI_ISL_915619,
EPI_ISL_915620, EPI_ISL_915621, EPI_ISL_915622, EPI_ISL_915623, EPI_ISL_915624, EPI_ISL_915625, EPI_ISL_915626, EPI_ISL_915627, EPI_ISL_915628, EPI_ISL_915629, EPI_ISL_915630, EPI_ISL_915631, EPI_ISL_915632, EPI_ISL_915633, EPI_ISL_915634, EPI_ISL_915635, EPI_ISL_915636, EPI_ISL_915637,
EPI_ISL_915638, EPI_ISL_915639, EPI_ISL_915640, EPI_ISL_915641, EPI_ISL_915642, EPI_ISL_915643, EPI_ISL_915644, EPI_ISL_915645, EPI_ISL_915646, EPI_ISL_915647, EPI_ISL_915648, EPI_ISL_915649, EPI_ISL_915650, EPI_ISL_915651, EPI_ISL_915652, EPI_ISL_915653, EPI_ISL_915654, EPI_ISL_915655,
EPI_ISL_915657, EPI_ISL_915658, EPI_ISL_915659, EPI_ISL_915660, EPI_ISL_915661, EPI_ISL_915662, EPI_ISL_915663, EPI_ISL_915664, EPI_ISL_915665, EPI_ISL_915666, EPI_ISL_915667, EPI_ISL_915668, EPI_ISL_915669, EPI_ISL_915670, EPI_ISL_915671, EPI_ISL_915672, EPI_ISL_915673, EPI_ISL_915674,
EPI_ISL_915675, EPI_ISL_915676, EPI_ISL_915677, EPI_ISL_915678, EPI_ISL_915679, EPI_ISL_915680, EPI_ISL_915681, EPI_ISL_915682, EPI_ISL_915683, EPI_ISL_915684, EPI_ISL_915685, EPI_ISL_915686, EPI_ISL_915687, EPI_ISL_915688, EPI_ISL_915689, EPI_ISL_915690, EPI_ISL_915691, EPI_ISL_915692,
EPI_ISL_915693, EPI_ISL_915694, EPI_ISL_915695, EPI_ISL_915696, EPI_ISL_915697, EPI_ISL_915698, EPI_ISL_915699, EPI_ISL_915700, EPI_ISL_915701, EPI_ISL_915702, EPI_ISL_915703, EPI_ISL_915704, EPI_ISL_915705, EPI_ISL_915706, EPI_ISL_915707, EPI_ISL_915708, EPI_ISL_915709, EPI_ISL_915710,
EPI_ISL_915711, EPI_ISL_915712, EPI_ISL_915713, EPI_ISL_915714, EPI_ISL_915715, EPI_ISL_915716, EPI_ISL_915717, EPI_ISL_915718, EPI_ISL_915719, EPI_ISL_915720, EPI_ISL_915721, EPI_ISL_915722, EPI_ISL_915723, EPI_ISL_915724, EPI_ISL_915725, EPI_ISL_915726, EPI_ISL_915727, EPI_ISL_915728,
EPI_ISL_915729, EPI_ISL_915730, EPI_ISL_915731, EPI_ISL_915732, EPI_ISL_915733, EPI_ISL_915734, EPI_ISL_915735, EPI_ISL_915736, EPI_ISL_915737, EPI_ISL_915738, EPI_ISL_915739, EPI_ISL_915740, EPI_ISL_915741, EPI_ISL_915742, EPI_ISL_915743, EPI_ISL_915744, EPI_ISL_915745, EPI_ISL_915746,
EPI_ISL_915747, EPI_ISL_915748, EPI_ISL_915749, EPI_ISL_915750, EPI_ISL_915751, EPI_ISL_915752, EPI_ISL_915753, EPI_ISL_915754, EPI_ISL_915755, EPI_ISL_915756, EPI_ISL_915757, EPI_ISL_915758, EPI_ISL_915759, EPI_ISL_915760, EPI_ISL_915761, EPI_ISL_915762, EPI_ISL_915763, EPI_ISL_915764,
EPI_ISL_915765, EPI_ISL_915766, EPI_ISL_915767, EPI_ISL_915768, EPI_ISL_915769, EPI_ISL_915770, EPI_ISL_915771, EPI_ISL_915772, EPI_ISL_915773, EPI_ISL_915774, EPI_ISL_915775, EPI_ISL_915776, EPI_ISL_915777, EPI_ISL_915778, EPI_ISL_915779, EPI_ISL_915780, EPI_ISL_915781, EPI_ISL_915782,
EPI_ISL_915783, EPI_ISL_915784, EPI_ISL_915785, EPI_ISL_915786, EPI_ISL_915787, EPI_ISL_915788, EPI_ISL_915789, EPI_ISL_915790, EPI_ISL_915791, EPI_ISL_915792, EPI_ISL_915793, EPI_ISL_915794, EPI_ISL_915795, EPI_ISL_915796, EPI_ISL_915797, EPI_ISL_915798, EPI_ISL_915799, EPI_ISL_915800,
EPI_ISL_915801, EPI_ISL_915802, EPI_ISL_915803, EPI_ISL_915804, EPI_ISL_915805, EPI_ISL_915806, EPI_ISL_915807, EPI_ISL_915808, EPI_ISL_915809, EPI_ISL_915810, EPI_ISL_915811, EPI_ISL_915812, EPI_ISL_915813, EPI_ISL_915814, EPI_ISL_915815, EPI_ISL_915816, EPI_ISL_915817, EPI_ISL_915818,
EPI_ISL_915819, EPI_ISL_915820, EPI_ISL_915821, EPI_ISL_915822, EPI_ISL_915823, EPI_ISL_915824, EPI_ISL_915825, EPI_ISL_915826, EPI_ISL_915827, EPI_ISL_915828, EPI_ISL_915829, EPI_ISL_915830, EPI_ISL_915831, EPI_ISL_915832, EPI_ISL_915833, EPI_ISL_915834, EPI_ISL_915835, EPI_ISL_915836,
EPI_ISL_915837, EPI_ISL_915838, EPI_ISL_915839, EPI_ISL_915840, EPI_ISL_915841, EPI_ISL_915842, EPI_ISL_915843, EPI_ISL_915844, EPI_ISL_915845, EPI_ISL_915846, EPI_ISL_915847, EPI_ISL_915848, EPI_ISL_915849, EPI_ISL_915850, EPI_ISL_915851, EPI_ISL_915852, EPI_ISL_915853, EPI_ISL_915854,
EPI_ISL_915855, EPI_ISL_915856, EPI_ISL_915857, EPI_ISL_915858, EPI_ISL_915859, EPI_ISL_915860, EPI_ISL_915861, EPI_ISL_915862, EPI_ISL_915863, EPI_ISL_915864, EPI_ISL_915865, EPI_ISL_915866, EPI_ISL_915867, EPI_ISL_915868, EPI_ISL_915869, EPI_ISL_915870, EPI_ISL_915871, EPI_ISL_915872,
EPI_ISL_915873, EPI_ISL_915874, EPI_ISL_915875, EPI_ISL_915876, EPI_ISL_915877, EPI_ISL_915878, EPI_ISL_915879, EPI_ISL_915880, EPI_ISL_915881, EPI_ISL_915882, EPI_ISL_915883, EPI_ISL_915884, EPI_ISL_915885, EPI_ISL_915886, EPI_ISL_915887, EPI_ISL_915888, EPI_ISL_915889, EPI_ISL_915890,
EPI_ISL_915891, EPI_ISL_915892, EPI_ISL_915893, EPI_ISL_915894, EPI_ISL_915895, EPI_ISL_915896, EPI_ISL_915897, EPI_ISL_915898, EPI_ISL_915899, EPI_ISL_915900, EPI_ISL_915901, EPI_ISL_915902, EPI_ISL_915903, EPI_ISL_915904, EPI_ISL_915905, EPI_ISL_915906, EPI_ISL_915907, EPI_ISL_915908,
EPI_ISL_915909, EPI_ISL_915910, EPI_ISL_915912, EPI_ISL_915913, EPI_ISL_915914, EPI_ISL_915915, EPI_ISL_915916, EPI_ISL_915917, EPI_ISL_915918, EPI_ISL_915919, EPI_ISL_915920, EPI_ISL_915921, EPI_ISL_915922, EPI_ISL_915923, EPI_ISL_915924, EPI_ISL_915925, EPI_ISL_915926, EPI_ISL_915927,
EPI_ISL_915928, EPI_ISL_915929, EPI_ISL_915930, EPI_ISL_915931, EPI_ISL_915932, EPI_ISL_915933, EPI_ISL_915934, EPI_ISL_915935, EPI_ISL_915936, EPI_ISL_915937, EPI_ISL_915938, EPI_ISL_915939, EPI_ISL_915940, EPI_ISL_915941, EPI_ISL_915942, EPI_ISL_915943, EPI_ISL_915944, EPI_ISL_915945,
EPI_ISL_915946, EPI_ISL_915947, EPI_ISL_915948, EPI_ISL_915949, EPI_ISL_915950, EPI_ISL_915951, EPI_ISL_915952, EPI_ISL_915953, EPI_ISL_915954, EPI_ISL_915955, EPI_ISL_915956, EPI_ISL_915957, EPI_ISL_915958, EPI_ISL_915959, EPI_ISL_915960, EPI_ISL_915961, EPI_ISL_915962, EPI_ISL_915963,
EPI_ISL_915964, EPI_ISL_915965, EPI_ISL_915966, EPI_ISL_915967, EPI_ISL_915968, EPI_ISL_915969, EPI_ISL_915970, EPI_ISL_915971, EPI_ISL_915972, EPI_ISL_915973, EPI_ISL_915974, EPI_ISL_915975, EPI_ISL_915976, EPI_ISL_915977, EPI_ISL_915978, EPI_ISL_915979, EPI_ISL_915980, EPI_ISL_915981,
EPI_ISL_915982, EPI_ISL_915983, EPI_ISL_915984, EPI_ISL_915985, EPI_ISL_915986, EPI_ISL_915987, EPI_ISL_915988, EPI_ISL_915989, EPI_ISL_915990, EPI_ISL_915991, EPI_ISL_915992, EPI_ISL_915993, EPI_ISL_915994, EPI_ISL_915995, EPI_ISL_915996, EPI_ISL_915997, EPI_ISL_915998, EPI_ISL_915999,
EPI_ISL_916000, EPI_ISL_916001, EPI_ISL_916002, EPI_ISL_916003, EPI_ISL_916004, EPI_ISL_916005, EPI_ISL_916006, EPI_ISL_916007, EPI_ISL_916008, EPI_ISL_916009, EPI_ISL_916010, EPI_ISL_916011, EPI_ISL_916012, EPI_ISL_916013, EPI_ISL_916014, EPI_ISL_916015, EPI_ISL_916016, EPI_ISL_916017,
EPI_ISL_916018, EPI_ISL_916019, EPI_ISL_916020, EPI_ISL_916021, EPI_ISL_916022, EPI_ISL_916023, EPI_ISL_916024, EPI_ISL_916025, EPI_ISL_916026, EPI_ISL_916027, EPI_ISL_916028, EPI_ISL_916029, EPI_ISL_916030, EPI_ISL_916031, EPI_ISL_916032, EPI_ISL_916033, EPI_ISL_916034, EPI_ISL_916035,
EPI_ISL_916036, EPI_ISL_916037, EPI_ISL_916038, EPI_ISL_916039, EPI_ISL_916040, EPI_ISL_916041, EPI_ISL_916042, EPI_ISL_916043, EPI_ISL_916044, EPI_ISL_916045, EPI_ISL_916046, EPI_ISL_916047, EPI_ISL_916048, EPI_ISL_916049, EPI_ISL_916050, EPI_ISL_916051, EPI_ISL_916052, EPI_ISL_916053,
EPI_ISL_916054, EPI_ISL_916055, EPI_ISL_916056, EPI_ISL_916057, EPI_ISL_916058, EPI_ISL_916059, EPI_ISL_916060, EPI_ISL_916061, EPI_ISL_916062, EPI_ISL_916063, EPI_ISL_916064, EPI_ISL_916065, EPI_ISL_916066, EPI_ISL_916067, EPI_ISL_916069, EPI_ISL_916070, EPI_ISL_916072, EPI_ISL_916073,
EPI_ISL_916074, EPI_ISL_916077, EPI_ISL_916078, EPI_ISL_916081, EPI_ISL_916083, EPI_ISL_916084, EPI_ISL_916085, EPI_ISL_916089, EPI_ISL_916090, EPI_ISL_916093, EPI_ISL_916095, EPI_ISL_916096, EPI_ISL_916097, EPI_ISL_916101, EPI_ISL_916102, EPI_ISL_916103, EPI_ISL_916104, EPI_ISL_916112,
EPI_ISL_916114, EPI_ISL_916115, EPI_ISL_916116, EPI_ISL_916117, EPI_ISL_916118, EPI_ISL_916119, EPI_ISL_916120, EPI_ISL_916122, EPI_ISL_916123, EPI_ISL_916125, EPI_ISL_916126, EPI_ISL_916128, EPI_ISL_916130, EPI_ISL_916133, EPI_ISL_916135, EPI_ISL_916136, EPI_ISL_916138, EPI_ISL_916139,

| | | | |
|---|---|---|---|
| EPI_ISL_916140, EPI_ISL_916141, EPI_ISL_916142, EPI_ISL_916143, EPI_ISL_916146, EPI_ISL_916148, EPI_ISL_916149, EPI_ISL_916150, EPI_ISL_916153, EPI_ISL_916155, EPI_ISL_916156, EPI_ISL_916157, EPI_ISL_916158, EPI_ISL_916160, EPI_ISL_916163, EPI_ISL_916168, EPI_ISL_916170, EPI_ISL_916172, EPI_ISL_916173, EPI_ISL_916175, EPI_ISL_916176, EPI_ISL_916177, EPI_ISL_916178, EPI_ISL_916179 | | | |
| see above | Lighthouse Lab in Alderley Park | Wellcome Sanger Institute for the COVID-19 Genomics UK (COG-UK) Consortium | Jacquelyn Wynn, Mairead Hyland, The Lighthouse Lab in Alderley Park and Alex Alderton, Roberto Amato, Sonia Goncalves, Ewan Harrison, David K. Jackson, Ian Johnston, Dominic Kwiatkowski, Cordelia Langford, John Sillitoe on behalf of the Wellcome Sanger Institute COVID-19 Surveillance Team |
| EPI_ISL_918516 | LACEN - Laboratório Central de Saúde Pública do Para | Evandro Chagas Institute | Santos, M.C.; Silva, A.M.; Junior, W.D.C.; Barbagelata, L.S.; Ferreira, J.A.; Sousa, E.M.A.; da Silva, P.S.; Pinheiro, K.C.; L.C.; Sousa Junior, E.C. |
| EPI_ISL_918569, EPI_ISL_918570, EPI_ISL_918571, EPI_ISL_918572, EPI_ISL_918573, EPI_ISL_918589, EPI_ISL_918591, EPI_ISL_918592, EPI_ISL_918593, EPI_ISL_918594, EPI_ISL_918595 | | | |
| see above | University of Birmingham | COVID-19 Genomics UK (COG-UK) Consortium | Institute of Microbiology, University of Birmingham: Claire McMurray, Joanne Stockton, Samuel Nicholls, Radoslaw Poplawski, Will Rowe, Josh Quick, Nicholas Loman. University of Birmingham Testing Laboratory: Celina M Whalley, Andrew Bosworth, Charlotte Poxon, Kasun Wanigasooriya, Oliver Pickles, Mike Kidd, Alex Richter, Andrew D Beggs PHE Heartlands Lab: Husam Osman, Andrew Bosworth. Queen Elizabeth Hospital: Anna Casey |
| EPI_ISL_919272 | West of Scotland Specialist Virology Centre, NHSGGC / MRC-University of Glasgow Centre for Virus Research | COVID-19 Genomics UK (COG-UK) Consortium | Ana da Silva Filipe, Natasha Johnson, Kathy Smollett, Daniel Mair, Stephen Carmichael, Alice Broos, Lily Tong, Jenna Nichols, Kyriaki Nomikou; Sarah McDonald; Richard Orton, Joseph Hughes, Sreenu Vattipally, David L Robertson; Alasdair MacLean, Rory Gunson; Sharif Shaaban, Matthew Holden; Rachel Blacow, Guy Mollett, Kathy Li, James Shepherd, Antonia Ho, Emma Thomson |
| EPI_ISL_920482 | University College London Hospital | COVID-19 Genomics UK (COG-UK) Consortium | Judith Heaney, Matthew Byott, Catherine Houlihan, Dan Frampton, Stuart Kirk, Moira Spyer and Eleni Nastouli |
| EPI_ISL_924783, EPI_ISL_924784, EPI_ISL_924785, EPI_ISL_924786, EPI_ISL_924789, EPI_ISL_924790, EPI_ISL_924791, EPI_ISL_924792, EPI_ISL_924793 | Bioinformatics and Biostatistics Lab, Advanced Sequencing Facility | COVID-19 Genomics UK (COG-UK) Consortium | Aengus Stewart,Jerome Nicod,Chelsea Sawyer,Laura Cubitt,Harshil Patel,Margaret Crawford |
| EPI_ISL_925496 | Department of Clinical Microbiology | GIGA Medical Genomics | Keith Durkin, Maria Artesi, Sébastien Bontems, Raphaël Boreux, Bouchra Boujemla, Cécile Meex, Pierrette Melin, Marie-Pierre Hayette, Vincent Bours |
| EPI_ISL_926137, EPI_ISL_926166, EPI_ISL_926339, EPI_ISL_926776, EPI_ISL_926882, EPI_ISL_927025, EPI_ISL_927325, EPI_ISL_927352, EPI_ISL_927461, EPI_ISL_927491, EPI_ISL_927502, EPI_ISL_927662, EPI_ISL_927843, EPI_ISL_927872, EPI_ISL_927895, EPI_ISL_927932, EPI_ISL_928073, EPI_ISL_928106, EPI_ISL_928112, EPI_ISL_928148, EPI_ISL_928208, EPI_ISL_928240, EPI_ISL_928261, EPI_ISL_928316, EPI_ISL_928322, EPI_ISL_928335, EPI_ISL_928696, EPI_ISL_928718, EPI_ISL_928939, EPI_ISL_929027, EPI_ISL_929116, EPI_ISL_929357, EPI_ISL_929467, EPI_ISL_929695, EPI_ISL_929937, EPI_ISL_929995, EPI_ISL_930105, EPI_ISL_930111, EPI_ISL_930265, EPI_ISL_930431, EPI_ISL_930434 | | | |
| see above | Department of Virus and Microbiological Special Diagnostics, Statens Serum Institut, Copenhagen, Denmark | Aalborg University | Danish Covid-19 Genome Consortium |
| EPI_ISL_931363, EPI_ISL_931364, EPI_ISL_931389 | University Hospital Basel, Clinical Virology | University Hospital Basel, Clinical Bacteriology | Tim Roloff, Madlen Stange, Helena MB Seth-Smith, Alfredo Mari, Karoline Leuzinger, Julia Bielicki, Manuel Battegay, Hans Hirsch, Adrian Egli |
| EPI_ISL_933516, EPI_ISL_933518 | Instituto Nacional de Medicina Genomica | Instituto Nacional de Medicina Genomica | Hidalgo-Miranda A, Mendoza-Vargas A, Reyes-Grajeda JP, Cisneros-Villanueva M, Cedro-Tanda A,Peñaloza-Figueroa F, Herrera-Montalvo LA |
| EPI_ISL_933547, EPI_ISL_933548, EPI_ISL_933549, EPI_ISL_933550, EPI_ISL_933551, EPI_ISL_933552, EPI_ISL_933553, EPI_ISL_933558 | Arizona State Public Health Laboratory | Arizona State Public Health Laboratory | Trung Huynh, Jessica Escobar, Katherine Fullerton, Nobuko Fukushima, Stacy White, Linda Getsinger, Victor Waddell |
| EPI_ISL_933668 | Instituto de Diagnostico y Referencia Epidemiologicos INDRE_RNLSP | Instituto de Diagnostico y Referencia Epidemiologicos (INDRE) | Claudia Wong-Arambula, Abril Rodriguez-Maldonado, Fabiola Garces-Ayala, Adnan Araiza-Rodriguez, David Fragoso-Fonseca, Sergio Rangel-Guerrero, Mayra Jimenez-Morales, Nancy Munoz-Hernandez, Natividad Cruz-Ortiz, Tatiana Nunez-Garcia, Gisela Barrera-Badillo, Lucia Hernandez-Rivas, Irma Lopez-Martinez, Ernesto Ramirez-Gonzalez. |
| EPI_ISL_935464, EPI_ISL_935465 | Florida Bureau of Public Health Laboratories | Florida Bureau of Public Health Laboratories | Sarah Schmedes, Jason Blanton |
| EPI_ISL_936489 | Hospital Raja Perempuan Zainab II | Institute for Medical Research, Infectious Disease Research Centre, National Institutes of Health, Ministry of Health Malaysia | Suppiah J, Kamel K, Azizan MA, Thayan R |
| EPI_ISL_936619, EPI_ISL_936620, EPI_ISL_936621, EPI_ISL_936623, EPI_ISL_936624, EPI_ISL_937015 | Northwestern Memorial Hospital | Ozer Lab | Ramon Lorenzo-Redondo, Lacy M. Simons, Chad J. Achenbach, Lawrence J. Jennings, Michael G. Ison, Judd F. Hultquist, Egon A. Ozer |
| EPI_ISL_940187, EPI_ISL_940501 | Hôpital Bichat Claude Bernard, Laboratoire de Virologie | IAME UMR1137 Inserm, Université de Paris, Hôpital Bichat | Antoine Bridier-Nahmias, Amélie Recoing, Quentin Le Hingrat, Lena Daniel, Siham Hamri, Gilles Collin, Alexandre Storto, Mélanie Bertine, Charlotte Charpentier, Nadhira Houhou-Fidouh, Diane Descamps, Benoit Visseaux |
| EPI_ISL_941276, EPI_ISL_941278 | Nigeria Centre for Disease Control (NCDC) | African Centre of Excellence for Genomics of Infectious Diseases (ACEGID), Redeemer's University | Oluniyi P.E. et al |
| EPI_ISL_941909, EPI_ISL_941912 | Virginia DCLS | Virginia DCLS | Virginia DCLS |
| EPI_ISL_941930, EPI_ISL_941934, EPI_ISL_941939 | Florida Bureau of Public Health Laboratories | Florida Bureau of Public Health Laboratories | Sarah Schmedes, Jason Blanton |
| EPI_ISL_942403 | Virginia DCLS | Virginia DCLS | Virginia DCLS |
| EPI_ISL_942480, EPI_ISL_942481 | Gundersen Molecular Diagnostics Laboratory | Kabara Cancer Research Institute | Craig S. Richmond, Paraic A. Kenny |
| EPI_ISL_942482, EPI_ISL_942483, EPI_ISL_942484, EPI_ISL_942485, EPI_ISL_942486, EPI_ISL_942487, EPI_ISL_942488, EPI_ISL_942489, EPI_ISL_942490, EPI_ISL_942491, EPI_ISL_942492, EPI_ISL_942493, EPI_ISL_942494 | | | |
| see above | Gundersen Clinical Microbiology Laboratory | Kabara Cancer Research Institute | Craig S. Richmond, Paraic A. Kenny |
| EPI_ISL_942495, EPI_ISL_942496, EPI_ISL_942497, EPI_ISL_942498, EPI_ISL_942499, EPI_ISL_942500, EPI_ISL_942501, EPI_ISL_942503 | Gundersen Molecular Diagnostics Laboratory | Kabara Cancer Research Institute | Craig S. Richmond, Paraic A. Kenny |
| EPI_ISL_942941 | Gundersen Clinical Microbiology Laboratory | Kabara Cancer Research Institute | Craig S. Richmond, Paraic A. Kenny |
| EPI_ISL_943604, EPI_ISL_943607, EPI_ISL_943609, EPI_ISL_943611, EPI_ISL_943613 | Lacen_RS | State Center for Health Surveillance. Rio Grande do Sul State Secretary of Health | Aline Campos, Amanda da Silva, Anelise Schaurich, Claudia Dornelles, Cynthia Molina, Fernanda Godinho, Lara Crescente, Leticia Garay, Regina Barcellos, Richard Salvato, Tatiana Gregianini, Vagner Fonseca |
| EPI_ISL_951788 | Originating lab: Wales Specialist Virology Centre Sequencing lab: Pathogen Genomics Unit | Public Health Wales Microbiology Cardiff Wales Specialist Virology Centre | Catherine Moore, Johnathan Evans, Laura Gifford, Malorie Perry, Simon Cottrell, Angela Marchbank, Alec Birchley, Alexander Adams, Amy Gaskin, Bree Gatica-Wilcox, Jason Coombes, Joel Southgate, Lauren Gilbert, Lee Graham, Nicole Pacchiarini, Sara Kumziene-Summerhayes, Sarah Taylor, Sophie Jones, Sara Rey, Matthew Bull, Joanne Watkins, Sally Corden, Tom Connor |
| EPI_ISL_952404 | Centre for Enzyme Innovation, University of Portsmouth / Translational Research Laboratory, Portsmouth Hospitals NHS Trust | COVID-19 Genomics UK (COG-UK) Consortium | Angela Beckett,Salman Goudarzi,Christopher Fearn,Kate Cook,Katie Loveson,Sharon Glaysher,Scott Elliott,Samuel Robson |
| EPI_ISL_955167, EPI_ISL_955168 | Institute for Biocides and Medical Ecology | Institute of microbiology and Immunology, Faculty of Medicine, University of Belgrade | Knezevic,A., Jankovic,M., Vidanovic,D., Milicevic,O., Tesovic,B., Sekler,M., Jovanovic,T. |
| EPI_ISL_955192, EPI_ISL_955193, EPI_ISL_955196 | Instituto Nacional de Medicina Genomica | Instituto Nacional de Medicina Genomica | Hidalgo-Miranda A, Mendoza-Vargas A, Reyes-Grajeda JP, Cisneros-Villanueva M, Cedro-Tanda A,Peñaloza-Figueroa F, Herrera-Montalvo LA |
| EPI_ISL_955337, EPI_ISL_955338, | UC Davis- Department of Pathology and Laboratory Medicine | Chan-Zuckerberg Biohub | CZB Cliahub Consortium |

| | | | |
|---|---|---|---|
| EPI_ISL_955339, EPI_ISL_955340 | | | |
| EPI_ISL_955945, EPI_ISL_955956, EPI_ISL_955957, EPI_ISL_955958, EPI_ISL_955959, EPI_ISL_955979, EPI_ISL_955980, EPI_ISL_959483, EPI_ISL_959484, EPI_ISL_959507, EPI_ISL_959508, EPI_ISL_959509, EPI_ISL_959510, EPI_ISL_959511, EPI_ISL_959512, EPI_ISL_959513 | | | |
| see above | Division of Emerging Infectious Diseases, Bureau of Infectious Diseases Diagnosis Control, Korea Disease Control and Prevention Agency | Division of Emerging Infectious Diseases, Bureau of Infectious Diseases Diagnosis Control, Korea Disease Control and Prevention Agency | Ae Kyung Park, Il-Hwan Kim, Heui Man Kim, Jeong-Min Kim, Namjoo Lee, Chae Young Lee, Sang Hee Woo, Eun-Jin Kim |
| EPI_ISL_960340, EPI_ISL_960358, EPI_ISL_960359, EPI_ISL_960376 | University of Wisconsin-Madison AIDS Vaccine Research Laboratories | University of Wisconsin-Madison AIDS Vaccine Research Laboratories | Gage Moreno, Katarina Braun, et al. AIDS Vaccine Research Laboratories |
| EPI_ISL_960556, EPI_ISL_960581, EPI_ISL_960647 | Istituto Zooprofilattico Sperimentale del Mezzogiorno | TIGEM | Patrizia Annunziata, Andrea Ballabio, Valentina Bouche, Davide Cacchiarelli, Pellegrino Cerino, Chiara Colantuono, Maria Concetta Cuomo, Denise Di Concilio, Lucio Di Filippo, Antonio Grimaldi, Antonio Limone, Anna Manfredi, Francesco Panariello, Biancamaria Pierri, Marcello Salvi |
| EPI_ISL_962224, EPI_ISL_962225, EPI_ISL_962226, EPI_ISL_962227, EPI_ISL_962228, EPI_ISL_962229, EPI_ISL_962230, EPI_ISL_962231, EPI_ISL_962232, EPI_ISL_962233, EPI_ISL_962234, EPI_ISL_962235, EPI_ISL_962236, EPI_ISL_962237, EPI_ISL_962238, EPI_ISL_962239, EPI_ISL_962240, EPI_ISL_962241, EPI_ISL_962242, EPI_ISL_962243, EPI_ISL_962244, EPI_ISL_962260, EPI_ISL_962261 | | | |
| see above | Seattle Flu Study | Seattle Flu Study | Deborah A. Nickerson, Chris D. Frazar, Jover Lee, Benjamin Pelle, Erica Ryke, Matthew Richardson, Amanda Adler, Elisabeth Brandstetter, Peter D. Han, Kairsten Fay, Misja Ilcisin, Kirsten Lacombe, Thomas R. Sibley, Melissa Truong, Caitlin R. Wolf, Karen Cowgill, Stephanie Schrag, Jeff Duchin, Michael Boeckh, Janet A. Englund, Michael Famulare, Barry R. Lutz, Mark J. Rieder, Lea M. Starita, Matthew Thompson, Helen Y. Chu, Trevor Bedford, Jay Shendure |
| EPI_ISL_965345, EPI_ISL_965389, EPI_ISL_965390 | University of Liège COVID-19 testing center | GIGA Medical Genomics | Keith Durkin, Maria Artesi, Bouchra Boujemla, Emmanuel André, Marc Van Ranst, Fabrice Bureau, Laurent Gillet, Wouter Coppieters, Vincent Bours |
| EPI_ISL_965535, EPI_ISL_965554, EPI_ISL_965611, EPI_ISL_965702, EPI_ISL_965703, EPI_ISL_965790 | Dutch COVID-19 response team | Medical Microbiology, Maastricht University Medical Centre | Jozef Dingemans*, Brian van der Veer*, Erik Beuken, Carmen Reumkens, Lieke van Alphen, Christian Hoebe, Paul Savelkoul |
| EPI_ISL_968205, EPI_ISL_968206, EPI_ISL_968207, EPI_ISL_968208, EPI_ISL_968209, EPI_ISL_968210, EPI_ISL_968211 | Arizona State Public Health Laboratory | Arizona State Public Health Laboratory | Trung Huynh, Jessica Escobar, Katherine Fullerton, Nobuko Fukushima, Stacy White, Linda Getsinger, Victor Waddell |
| EPI_ISL_968842, EPI_ISL_968930 | KEMRI-Wellcome Trust Research Programme/KEMRI-CGMR-C Kilifi | KEMRI-Wellcome Trust Research Programme/KEMRI-CGMR-C Kilifi | Githinji et al |
| EPI_ISL_975661, EPI_ISL_975662, EPI_ISL_975663, EPI_ISL_975664, EPI_ISL_975665, EPI_ISL_975666, EPI_ISL_975667, EPI_ISL_975668, EPI_ISL_975669, EPI_ISL_975670, EPI_ISL_975671, EPI_ISL_975672, EPI_ISL_975673, EPI_ISL_975674, EPI_ISL_975675, EPI_ISL_975676, EPI_ISL_975677, EPI_ISL_975678, EPI_ISL_975679, EPI_ISL_975680, EPI_ISL_975681, EPI_ISL_975682, EPI_ISL_975683, EPI_ISL_975684, EPI_ISL_975685, EPI_ISL_975686, EPI_ISL_975687, EPI_ISL_975688, EPI_ISL_975689, EPI_ISL_975690, EPI_ISL_975691, EPI_ISL_975692, EPI_ISL_975693, EPI_ISL_975694, EPI_ISL_975695, EPI_ISL_975696, EPI_ISL_975697, EPI_ISL_975698, EPI_ISL_975699, EPI_ISL_975700, EPI_ISL_975701, EPI_ISL_975702, EPI_ISL_975703, EPI_ISL_975704, EPI_ISL_975705, EPI_ISL_975706, EPI_ISL_975707, EPI_ISL_975708, EPI_ISL_975709, EPI_ISL_975710, EPI_ISL_975711, EPI_ISL_975712, EPI_ISL_975713, EPI_ISL_975714, EPI_ISL_975715, EPI_ISL_975716, EPI_ISL_975717, EPI_ISL_975718, EPI_ISL_975719, EPI_ISL_975720, EPI_ISL_975721, EPI_ISL_975722, EPI_ISL_975723, EPI_ISL_975724, EPI_ISL_975725, EPI_ISL_975726, EPI_ISL_975727, EPI_ISL_975728, EPI_ISL_975729, EPI_ISL_975730, EPI_ISL_975731, EPI_ISL_975732, EPI_ISL_975733, EPI_ISL_975734, EPI_ISL_975735, EPI_ISL_975736, EPI_ISL_975737, EPI_ISL_975738, EPI_ISL_975739, EPI_ISL_975740, EPI_ISL_975741, EPI_ISL_975742, EPI_ISL_975743, EPI_ISL_975744, EPI_ISL_975745, EPI_ISL_975746, EPI_ISL_975747, EPI_ISL_975748, EPI_ISL_975749, EPI_ISL_975750, EPI_ISL_975751, EPI_ISL_975752, EPI_ISL_975753, EPI_ISL_975754, EPI_ISL_975755, EPI_ISL_975756, EPI_ISL_975757, EPI_ISL_975758, EPI_ISL_975759, EPI_ISL_975760, EPI_ISL_975761, EPI_ISL_975762, EPI_ISL_975763, EPI_ISL_975764, EPI_ISL_975765, EPI_ISL_975766, EPI_ISL_975767, EPI_ISL_975768, EPI_ISL_975769, EPI_ISL_975770, EPI_ISL_975771, EPI_ISL_975772, EPI_ISL_975773, EPI_ISL_975774, EPI_ISL_975776, EPI_ISL_975777, EPI_ISL_975778, EPI_ISL_975779, EPI_ISL_975780, EPI_ISL_975781, EPI_ISL_975782, EPI_ISL_975783, EPI_ISL_975784, EPI_ISL_975785, EPI_ISL_975786, EPI_ISL_975787, EPI_ISL_975788, EPI_ISL_975789, EPI_ISL_975790, EPI_ISL_975791, EPI_ISL_975792, EPI_ISL_975793, EPI_ISL_975794 | | | |
| see above | BCCDC Public Health Laboratory | BCCDC Public Health Laboratory | Prystajecky Natalie, Linda Hoang, Dan Fornika, John Tyson, Shannon Russell, Kim Macdonald, Kimia Kamelian, Ana Pacagnella, Corrinne Ng, Loretta Janz, Robert Azana Terry Snutch, Mel Krajden |
| EPI_ISL_977154 | ULSS 2 Treviso | Istituto Zooprofilattico Sperimentale delle Venezie | Adelaide Milani, Alessia Schivo, Annalisa Salviato, Erika Giorgia Quaranta, Ambra Pastori, Bianca Zecchin, Alice Fusaro, Isabella Monne, Calogero Terregino, Antonia Ricci |
| EPI_ISL_977177, EPI_ISL_977178, EPI_ISL_977179 | Microbiologia e Virologia | Istituto Zooprofilattico Sperimentale delle Venezie | Adelaide Milani, Alessia Schivo, Annalisa Salviato, Erika Giorgia Quaranta, Ambra Pastori, Bianca Zecchin, Alice Fusaro, Isabella Monne, Calogero Terregino, Antonia Ricci |
| EPI_ISL_977188, EPI_ISL_977189, EPI_ISL_977192, EPI_ISL_977193 | ULSS 2 Treviso | Istituto Zooprofilattico Sperimentale delle Venezie | Adelaide Milani, Alessia Schivo, Annalisa Salviato, Erika Giorgia Quaranta, Ambra Pastori, Bianca Zecchin, Alice Fusaro, Isabella Monne, Calogero Terregino, Antonia Ricci |
| EPI_ISL_977225, EPI_ISL_977226 | ULSS 1 Dolomiti | Istituto Zooprofilattico Sperimentale delle Venezie | Adelaide Milani, Alessia Schivo, Annalisa Salviato, Erika Giorgia Quaranta, Ambra Pastori, Bianca Zecchin, Alice Fusaro, Isabella Monne, Calogero Terregino, Antonia Ricci |
