## Supplementary material for "Detection and characterization of the SARS-CoV-2 lineage B.1.526 in New York": Supp. Table 3 GISAID Acknowledgment part 2: gisaid_hcov-19_acknowledgement_table_2021_02_13_00-11.pdf

We gratefully acknowledge the following Authors from the Originating laboratories responsible for obtaining the specimens, as well as the Submitting laboratories where the genome data were generated and shared via GISAID, on which this research is based.

All Submitters of data may be contacted directly via www.gisaid.org

Authors are sorted alphabetically.

| Accession ID | Originating Laboratory | Submitting Laboratory | Authors |
|---|---|---|---|
| EPI_ISL_660070 | Zurita & Zurita Laboratorios | Zurita & Zurita Laboratorios | Gabriela Sevillano Camilo Zurita-Salinas Karen Loaiza David Ortega-Paredes Jeannete Zurita |
| EPI_ISL_661265 | Middlemore Hospital | Institute of Environmental Science and Research (ESR) | Xiaoyun Ren, Matt Storey, Nikki Freed, Muhammad Faisal, Jing Wang, Hermes Perez, Anja Werno, Antje van der Linden, Arlo Upton, Chris Mansell, David Hammer, Dragana Drinkovic, Gary McAuliffe, Hana Sofia Andersson, James Ussher, Jill Sherwood, Josh Freeman, Julia Howard, Juliet Elvy, Mary DeAlmeida, Matt Blakiston, Matthew Rogers, Max Bloomfield, Michael Addidle, Michelle Balm, Sally Roberts, Sarah Jefferies, Sharmini Muttaiyah, Susan Morpeth, Susan Taylor, Timothy Blackmore, Vani Sathyendran, Veronica Playle, Virginia Hope, Erasmus Smit, Lauren Jelly, Olin Silander, Joep de Ligt |
| EPI_ISL_665365, EPI_ISL_665367, EPI_ISL_665510, EPI_ISL_665724, EPI_ISL_665943, EPI_ISL_665944, EPI_ISL_665945 | Queens Medical Centre, Clinical Microbiology Department / DeepSeq Nottingham | COVID-19 Genomics UK (COG-UK) Consortium | Gemma Clark, Wendy Smith, Manjinder Khakh, Vicki M Fleming, Michelle M Lister, Hannah Howson-Wells, Jonathan Ball, Patrick McClure, Joseph Chappell, Theocharis Tsoleridis, Nadine Holmes, Matthew Carlisle, Christopher Moore, Fei Sang, Johnny Debebe, Victoria Wright, Matthew Loose |
| EPI_ISL_678456, EPI_ISL_678457, EPI_ISL_678458, EPI_ISL_678459, EPI_ISL_678460, EPI_ISL_678461, EPI_ISL_678462, EPI_ISL_678463, EPI_ISL_678464, EPI_ISL_678465, EPI_ISL_678466, EPI_ISL_678467, EPI_ISL_678468, EPI_ISL_678469, EPI_ISL_678470, EPI_ISL_678471, EPI_ISL_678472, EPI_ISL_678473, EPI_ISL_678474, EPI_ISL_678475, EPI_ISL_678476, EPI_ISL_678477, EPI_ISL_678478, EPI_ISL_678479, EPI_ISL_678480, EPI_ISL_678481, EPI_ISL_678482, EPI_ISL_678483, EPI_ISL_678484, EPI_ISL_678485 | | | |
| see above | Department of Clinical Microbiology | GIGA Medical Genomics | Keith Durkin, Maria Artesi, Sébastien Bontems, Raphaël Boreux, Bouchra Boujemla, Cécile Meex, Pierrette Melin, Marie-Pierre Hayette, Vincent Bours |
| EPI_ISL_678773, EPI_ISL_678777, EPI_ISL_678778, EPI_ISL_678779, EPI_ISL_678780, EPI_ISL_678781, EPI_ISL_678782, EPI_ISL_678783, EPI_ISL_678784, EPI_ISL_678785, EPI_ISL_678786, EPI_ISL_678828 | | | |
| see above | Respiratory Virus Unit, Microbiology Services Colindale, Public Health England | COVID-19 Genomics UK (COG-UK) Consortium | PHE Covid Sequencing Team |
| EPI_ISL_679601, EPI_ISL_679604, EPI_ISL_679605, EPI_ISL_679606, EPI_ISL_679607, EPI_ISL_679608, EPI_ISL_679609, EPI_ISL_679610, EPI_ISL_679611, EPI_ISL_679612, EPI_ISL_679613, EPI_ISL_679614, EPI_ISL_679615, EPI_ISL_679616, EPI_ISL_679617, EPI_ISL_679618, EPI_ISL_679619, EPI_ISL_679620 | | | |
| see above | University College London, Great Ormond Street Hospital for Children NHS Foundation Trust, Imperial College Healthcare NHS Trust | COVID-19 Genomics UK (COG-UK) Consortium | Sergi Castellano, Rachel Williams, Mark Kristiansen, Paola Resende Silva, Sunando Roy, Tony Brooks, Helena Tutill, Paola Niola, Patricia Dyal, Charlotte Williams, Leysa Forrest, Yasmin Panchbhaya, Jacqueline Findlay, Samuel Weeks, Julianne Brown, Kathryn Harris, Paul Randell, James Price, Alison Holmes, Judith Breuer |
| EPI_ISL_679935, EPI_ISL_679936, EPI_ISL_679937, EPI_ISL_679938, EPI_ISL_679939, EPI_ISL_679940, EPI_ISL_679941, EPI_ISL_679942, EPI_ISL_679943, EPI_ISL_679944 | Queens Medical Centre, Clinical Microbiology Department / DeepSeq Nottingham | COVID-19 Genomics UK (COG-UK) Consortium | Gemma Clark, Wendy Smith, Manjinder Khakh, Vicki M Fleming, Michelle M Lister, Hannah Howson-Wells, Jonathan Ball, Patrick McClure, Joseph Chappell, Theocharis Tsoleridis, Nadine Holmes, Matthew Carlisle, Christopher Moore, Fei Sang, Johnny Debebe, Victoria Wright, Matthew Loose |
| EPI_ISL_679956, EPI_ISL_679957, EPI_ISL_679958, EPI_ISL_679959, EPI_ISL_679960, EPI_ISL_679961, EPI_ISL_679962, EPI_ISL_679963, EPI_ISL_679964, EPI_ISL_679965, EPI_ISL_679966 | | | |
| see above | Lincolnshire Hospitals and DeepSeq Nottingham | COVID-19 Genomics UK (COG-UK) Consortium | Nichola Duckworth, Tim Sloan, Sarah Walsh, Jonathan Ball, Patrick McClure, Joeseph Chappell, Nadine Holmes, Matthew Carlisle, Christopher Moore, Fei Sang, Johnny Debebe, Victoria Wright, Matthew Loose |
| EPI_ISL_680115, EPI_ISL_680118, EPI_ISL_680134, EPI_ISL_680142, EPI_ISL_680146, EPI_ISL_680181, EPI_ISL_680186, EPI_ISL_680190 | Virology Department, Sheffield Teaching Hospitals NHS Foundation Trust/Department of Infection, Immunity and Cardiovascular Disease, The Medical School, University of Sheffield | COVID-19 Genomics UK (COG-UK) Consortium | Thushan de Silva, Matthew Parker, Nikki Smith, Adri Angyal, Rebecca Brown, Luke Green, Rachel Tucker, Paul Parsons, Danielle Groves, Katie Johnson, Laura Carrilero, Alex Keeley, Dave Partridge, Matthew Wyles, Benjamin Lindsey, Mehmet Yavuz, Mohammad Raza, Cariad Evans |
| EPI_ISL_680440 | Regional Virus Laboratory, Belfast Health and Social Care Trust | COVID-19 Genomics UK (COG-UK) Consortium | Conall McCaughey, James McKenna, Tanya Curran, Susan Feeney, Alison Watt, Ciara Cox, Mairead Connor, Zoltan Molnar, David Simpson, Derek Fairley |
| EPI_ISL_680453, EPI_ISL_680454, EPI_ISL_680455, EPI_ISL_680456, EPI_ISL_680457, EPI_ISL_680458 | West of Scotland Specialist Virology Centre, NHSGGC / MRC-University of Glasgow Centre for Virus Research | COVID-19 Genomics UK (COG-UK) Consortium | Ana da Silva Filipe, Natasha Johnson, Kathy Smollett, Daniel Mair, Stephen Carmichael, Alice Broos, Lily Tong, Jenna Nichols, Kyriaki Nomikou; Sarah McDonald; Richard Orton, Joseph Hughes, Sreenu Vattipally, David L Robertson; Alasdair MacLean, Rory Gunson; Sharif Shaaban, Matthew Holden; Rachel Blacow, Guy Mollett, Kathy Li, James Shepherd, Antonia Ho, Emma Thomson |
| EPI_ISL_681270 | Department of Clinical Microbiology | GIGA Medical Genomics | Keith Durkin, Maria Artesi, Sébastien Bontems, Raphaël Boreux, Bouchra Boujemla, Cécile Meex, Pierrette Melin, Marie-Pierre Hayette, Vincent Bours |
| EPI_ISL_682283, EPI_ISL_682284, EPI_ISL_682285, EPI_ISL_682286 | Canterbury Health Laboratories | Institute of Environmental Science and Research (ESR) | Xiaoyun Ren, Matt Storey, Nikki Freed, Muhammad Faisal, Jing Wang, Hermes Perez, Anja Werno, Antje van der Linden, Arlo Upton, Chris Mansell, David Hammer, Dragana Drinkovic, Gary McAuliffe, Hana Sofia Andersson, James Ussher, Jill Sherwood, Josh Freeman, Julia Howard, Juliet Elvy, Mary DeAlmeida, Matt Blakiston, Matthew Rogers, Max Bloomfield, Michael Addidle, Michelle Balm, Sally Roberts, Sarah Jefferies, Sharmini Muttaiyah, Susan Morpeth, Susan Taylor, Timothy Blackmore, Vani Sathyendran, Veronica Playle, Virginia Hope, Erasmus Smit, Lauren Jelly, Olin Silander, Joep de Ligt |
| EPI_ISL_682287, EPI_ISL_682288, EPI_ISL_682289, EPI_ISL_682290 | Middlemore Hospital | Institute of Environmental Science and Research (ESR) | Xiaoyun Ren, Matt Storey, Nikki Freed, Muhammad Faisal, Jing Wang, Hermes Perez, Anja Werno, Antje van der Linden, Arlo Upton, Chris Mansell, David Hammer, Dragana Drinkovic, Gary McAuliffe, Hana Sofia Andersson, James Ussher, Jill Sherwood, Josh Freeman, Julia Howard, Juliet Elvy, Mary DeAlmeida, Matt Blakiston, Matthew Rogers, Max Bloomfield, Michael Addidle, Michelle Balm, Sally Roberts, Sarah Jefferies, Sharmini Muttaiyah, Susan Morpeth, Susan Taylor, Timothy Blackmore, Vani Sathyendran, Veronica Playle, Virginia Hope, Erasmus Smit, Lauren Jelly, Olin Silander, Joep de Ligt |
| EPI_ISL_682291, EPI_ISL_682292, EPI_ISL_682293 | LabPLUS | Institute of Environmental Science and Research (ESR) | Xiaoyun Ren, Matt Storey, Nikki Freed, Muhammad Faisal, Jing Wang, Hermes Perez, Anja Werno, Antje van der Linden, Arlo Upton, Chris Mansell, David Hammer, Dragana Drinkovic, Gary McAuliffe, Hana Sofia Andersson, James Ussher, Jill Sherwood, Josh Freeman, Julia Howard, Juliet Elvy, Mary DeAlmeida, Matt Blakiston, Matthew Rogers, Max Bloomfield, Michael Addidle, Michelle Balm, Sally Roberts, Sarah Jefferies, Sharmini Muttaiyah, Susan Morpeth, Susan Taylor, Timothy Blackmore, Vani Sathyendran, Veronica Playle, Virginia Hope, Erasmus Smit, Lauren Jelly, Olin Silander, Joep de Ligt |
| EPI_ISL_682295 | Canterbury Health Laboratories | Institute of Environmental Science and Research (ESR) | Xiaoyun Ren, Matt Storey, Nikki Freed, Muhammad Faisal, Jing Wang, Hermes Perez, Anja Werno, Antje van der Linden, Arlo Upton, Chris Mansell, David Hammer, Dragana Drinkovic, Gary McAuliffe, Hana Sofia Andersson, James Ussher, Jill Sherwood, Josh Freeman, Julia Howard, Juliet Elvy, Mary DeAlmeida, Matt Blakiston, Matthew Rogers, Max Bloomfield, Michael Addidle, Michelle Balm, Sally Roberts, Sarah Jefferies, Sharmini Muttaiyah, Susan Morpeth, Susan Taylor, Timothy Blackmore, Vani Sathyendran, Veronica Playle, Virginia Hope, Erasmus Smit, Lauren Jelly, Olin Silander, Joep de Ligt |
| EPI_ISL_683453, EPI_ISL_683459, EPI_ISL_683460, EPI_ISL_683462, EPI_ISL_683463, EPI_ISL_683464, EPI_ISL_683465, EPI_ISL_683466, EPI_ISL_683467, EPI_ISL_683468, EPI_ISL_683469, EPI_ISL_683470, EPI_ISL_683471, EPI_ISL_683472, EPI_ISL_683473, EPI_ISL_683474, EPI_ISL_683483, EPI_ISL_683573, EPI_ISL_683574, EPI_ISL_683575, EPI_ISL_683578, EPI_ISL_683579, EPI_ISL_683580, EPI_ISL_683581, EPI_ISL_683582, EPI_ISL_683583, EPI_ISL_683584, EPI_ISL_683586, EPI_ISL_683587, EPI_ISL_683588, EPI_ISL_683589, EPI_ISL_683590, EPI_ISL_683591 | | | |
| see above | Respiratory Virus Unit, Microbiology Services Colindale, Public Health England | COVID-19 Genomics UK (COG-UK) Consortium | PHE Covid Sequencing Team |
| EPI_ISL_683760, EPI_ISL_683763 | DOHMH Morrisania | New York City Public Health Laboratory | Jade Wang, et al. |
| EPI_ISL_683766 | DOHMH Corona | New York City Public Health Laboratory | Jade Wang, et al. |
| EPI_ISL_683767, EPI_ISL_683768, EPI_ISL_683769 | DOHMH Riverside | New York City Public Health Laboratory | Jade Wang, et al. |
| EPI_ISL_683770, EPI_ISL_683811, EPI_ISL_683812 | DOHMH Corona | New York City Public Health Laboratory | Jade Wang, et al. |

| | | | |
|---|---|---|---|
| EPI_ISL_683813 | DOHMH Central Harlem | New York City Public Health Laboratory | Jade Wang, et al. |
| EPI_ISL_683814 | DOHMH Morrisania | New York City Public Health Laboratory | Jade Wang, et al. |
| EPI_ISL_683815 | DOHMH Corona | New York City Public Health Laboratory | Jade Wang, et al. |
| EPI_ISL_683816 | DOHMH Fort Greene | New York City Public Health Laboratory | Jade Wang, et al. |
| EPI_ISL_683817, EPI_ISL_683818 | DOHMH Corona | New York City Public Health Laboratory | Jade Wang, et al. |
| EPI_ISL_683819, EPI_ISL_683820 | DOHMH Riverside | New York City Public Health Laboratory | Jade Wang, et al. |
| EPI_ISL_683821 | DOHMH Crown Heights | New York City Public Health Laboratory | Jade Wang, et al. |
| EPI_ISL_683822 | DOHMH Jamaica | New York City Public Health Laboratory | Jade Wang, et al. |
| EPI_ISL_683823 | DOHMH Morrisania | New York City Public Health Laboratory | Jade Wang, et al. |
| EPI_ISL_683824 | DOHMH Corona | New York City Public Health Laboratory | Jade Wang, et al. |
| EPI_ISL_683825, EPI_ISL_683826 | DOHMH Morrisania | New York City Public Health Laboratory | Jade Wang, et al. |
| EPI_ISL_683827 | DOHMH Crown Heights | New York City Public Health Laboratory | Jade Wang, et al. |
| EPI_ISL_683828 | DOHMH Chelsea | New York City Public Health Laboratory | Jade Wang, et al. |
| EPI_ISL_683829 | DOHMH Central Harlem | New York City Public Health Laboratory | Jade Wang, et al. |
| EPI_ISL_683830 | DOHMH Chelsea | New York City Public Health Laboratory | Jade Wang, et al. |
| EPI_ISL_683831 | DOHMH Corona | New York City Public Health Laboratory | Jade Wang, et al. |
| EPI_ISL_683832 | DOHMH Fort Greene | New York City Public Health Laboratory | Jade Wang, et al. |
| EPI_ISL_683833 | DOHMH Corona | New York City Public Health Laboratory | Jade Wang, et al. |
| EPI_ISL_683834 | DOHMH Central Harlem | New York City Public Health Laboratory | Jade Wang, et al. |
| EPI_ISL_692775 | Massachusetts State Public Health Laboratory | Massachusetts State Public Health Laboratory | Andrew Lang, Timelia Fink, Glen Gallagher, Sandra Smole |
| EPI_ISL_693318, EPI_ISL_693319, EPI_ISL_693323, EPI_ISL_693325, EPI_ISL_693328, EPI_ISL_693329, EPI_ISL_693331 | National Public Health Laboratory, National Centre for Infectious Diseases | National Public Health Laboratory, National Centre for Infectious Diseases | Tze Minn Mak, Sophie Octavia, Zhenyang Zhou, Lin Cui, Raymond Tzer Pin Lin |
| EPI_ISL_693766, EPI_ISL_693774 | General practitioner | National Reference Center for Viruses of Respiratory Infections, Institut Pasteur, Paris | Marion Barbet, Sylvie Behillil, Méline Bizard, Angela Brisebarre, Camille Capel, Etienne Simon-Lorière, Vincent Enouf, Maud Vanpeene, Sylvie van der Werf |
| EPI_ISL_696492 | Conville CDC wc CVC & NHLS/UCT | KRISP, KZN Research Innovation and Sequencing Platform | Arash Iranzadeh, Deelan Doolabh, Lynn Tyers, Bruna Galvao, Innocent Mudau, Marvin Hsiao, Kruger Marais, Jennifer Giandhari, Sureshnee Pillay, Houriiyah Tegally, Emanuel James San, Tulio de Oliveira, Diana Hardie, Stephen Korsman, Carolyn Williamson |
| EPI_ISL_696493, EPI_ISL_696494 | Thembalethu CDC wc THC & NHLS/UCT | KRISP, KZN Research Innovation and Sequencing Platform | Arash Iranzadeh, Deelan Doolabh, Lynn Tyers, Bruna Galvao, Innocent Mudau, Marvin Hsiao, Kruger Marais, Jennifer Giandhari, Sureshnee Pillay, Houriiyah Tegally, Emanuel James San, Tulio de Oliveira, Diana Hardie, Stephen Korsman, Carolyn Williamson |
| EPI_ISL_696495 | Knysna CDC wc WLC & NHLS/UCT | KRISP, KZN Research Innovation and Sequencing Platform | Arash Iranzadeh, Deelan Doolabh, Lynn Tyers, Bruna Galvao, Innocent Mudau, Marvin Hsiao, Kruger Marais, Jennifer Giandhari, Sureshnee Pillay, Houriiyah Tegally, Emanuel James San, Tulio de Oliveira, Diana Hardie, Stephen Korsman, Carolyn Williamson |
| EPI_ISL_696496 | Thembalethu CDC wc THC & NHLS/UCT | KRISP, KZN Research Innovation and Sequencing Platform | Arash Iranzadeh, Deelan Doolabh, Lynn Tyers, Bruna Galvao, Innocent Mudau, Marvin Hsiao, Kruger Marais, Jennifer Giandhari, Sureshnee Pillay, Houriiyah Tegally, Emanuel James San, Tulio de Oliveira, Diana Hardie, Stephen Korsman, Carolyn Williamson |
| EPI_ISL_696497 | Great Brak River Clinic wc GBC & NHLS/UCT | KRISP, KZN Research Innovation and Sequencing Platform | Arash Iranzadeh, Deelan Doolabh, Lynn Tyers, Bruna Galvao, Innocent Mudau, Marvin Hsiao, Kruger Marais, Jennifer Giandhari, Sureshnee Pillay, Houriiyah Tegally, Emanuel James San, Tulio de Oliveira, Diana Hardie, Stephen Korsman, Carolyn Williamson |
| EPI_ISL_696498 | Kranshoek Clinic wc KSH & NHLS/UCT | KRISP, KZN Research Innovation and Sequencing Platform | Arash Iranzadeh, Deelan Doolabh, Lynn Tyers, Bruna Galvao, Innocent Mudau, Marvin Hsiao, Kruger Marais, Jennifer Giandhari, Sureshnee Pillay, Houriiyah Tegally, Emanuel James San, Tulio de Oliveira, Diana Hardie, Stephen Korsman, Carolyn Williamson |
| EPI_ISL_696499, EPI_ISL_696500 | Thembalethu CDC wc THC & NHLS/UCT | KRISP, KZN Research Innovation and Sequencing Platform | Arash Iranzadeh, Deelan Doolabh, Lynn Tyers, Bruna Galvao, Innocent Mudau, Marvin Hsiao, Kruger Marais, Jennifer Giandhari, Sureshnee Pillay, Houriiyah Tegally, Emanuel James San, Tulio de Oliveira, Diana Hardie, Stephen Korsman, Carolyn Williamson |
| EPI_ISL_696501 | Great Brak River Clinic wc GBC & NHLS/UCT | KRISP, KZN Research Innovation and Sequencing Platform | Arash Iranzadeh, Deelan Doolabh, Lynn Tyers, Bruna Galvao, Innocent Mudau, Marvin Hsiao, Kruger Marais, Jennifer Giandhari, Sureshnee Pillay, Houriiyah Tegally, Emanuel James San, Tulio de Oliveira, Diana Hardie, Stephen Korsman, Carolyn Williamson |
| EPI_ISL_696502 | Hornlee Clinic wc HLC & NHLS/UCT | KRISP, KZN Research Innovation and Sequencing Platform | Arash Iranzadeh, Deelan Doolabh, Lynn Tyers, Bruna Galvao, Innocent Mudau, Marvin Hsiao, Kruger Marais, Jennifer Giandhari, Sureshnee Pillay, Houriiyah Tegally, Emanuel James San, Tulio de Oliveira, Diana Hardie, Stephen Korsman, Carolyn Williamson |
| EPI_ISL_696503 | George Hospital wc GRH & NHLS/UCT | KRISP, KZN Research Innovation and Sequencing Platform | Arash Iranzadeh, Deelan Doolabh, Lynn Tyers, Bruna Galvao, Innocent Mudau, Marvin Hsiao, Kruger Marais, Jennifer Giandhari, Sureshnee Pillay, Houriiyah Tegally, Emanuel James San, Tulio de Oliveira, Diana Hardie, Stephen Korsman, Carolyn Williamson |
| EPI_ISL_696504 | Pacaltsdorp Clinic wc PAC & NHLS/UCT | KRISP, KZN Research Innovation and Sequencing Platform | Arash Iranzadeh, Deelan Doolabh, Lynn Tyers, Bruna Galvao, Innocent Mudau, Marvin Hsiao, Kruger Marais, Jennifer Giandhari, Sureshnee Pillay, Houriiyah Tegally, Emanuel James San, Tulio de Oliveira, Diana Hardie, Stephen Korsman, Carolyn Williamson |
| EPI_ISL_696505 | Khayelethu Clinic wc KLC & NHLS/UCT | KRISP, KZN Research Innovation and Sequencing Platform | Arash Iranzadeh, Deelan Doolabh, Lynn Tyers, Bruna Galvao, Innocent Mudau, Marvin Hsiao, Kruger Marais, Jennifer Giandhari, Sureshnee Pillay, Houriiyah Tegally, Emanuel James San, Tulio de Oliveira, Diana Hardie, Stephen Korsman, Carolyn Williamson |
| EPI_ISL_696506 | Knysna CDC wc WLC & NHLS/UCT | KRISP, KZN Research Innovation and Sequencing Platform | Arash Iranzadeh, Deelan Doolabh, Lynn Tyers, Bruna Galvao, Innocent Mudau, Marvin Hsiao, Kruger Marais, Jennifer Giandhari, Sureshnee Pillay, Houriiyah Tegally, Emanuel James San, Tulio de Oliveira, Diana Hardie, Stephen Korsman, Carolyn Williamson |
| EPI_ISL_696507 | Great Brak River Clinic wc GBC & NHLS/UCT | KRISP, KZN Research Innovation and Sequencing Platform | Arash Iranzadeh, Deelan Doolabh, Lynn Tyers, Bruna Galvao, Innocent Mudau, Marvin Hsiao, Kruger Marais, Jennifer Giandhari, Sureshnee Pillay, Houriiyah Tegally, Emanuel James San, Tulio de Oliveira, Diana Hardie, Stephen Korsman, Carolyn Williamson |
| EPI_ISL_696508, EPI_ISL_696509 | Conville CDC wc CVC & NHLS/UCT | KRISP, KZN Research Innovation and Sequencing Platform | Arash Iranzadeh, Deelan Doolabh, Lynn Tyers, Bruna Galvao, Innocent Mudau, Marvin Hsiao, Kruger Marais, Jennifer Giandhari, Sureshnee Pillay, Houriiyah Tegally, Emanuel James San, Tulio de Oliveira, Diana Hardie, Stephen Korsman, Carolyn Williamson |
| EPI_ISL_696510 | Sedgefield Clinic wc SGE & NHLS/UCT | KRISP, KZN Research Innovation and Sequencing Platform | Arash Iranzadeh, Deelan Doolabh, Lynn Tyers, Bruna Galvao, Innocent Mudau, Marvin Hsiao, Kruger Marais, Jennifer Giandhari, Sureshnee Pillay, Houriiyah Tegally, Emanuel James San, Tulio de Oliveira, Diana Hardie, Stephen Korsman, Carolyn Williamson |
| EPI_ISL_696511 | Keurhoek Clinic wc KTT & NHLS/UCT | KRISP, KZN Research Innovation and Sequencing Platform | Arash Iranzadeh, Deelan Doolabh, Lynn Tyers, Bruna Galvao, Innocent Mudau, Marvin Hsiao, Kruger Marais, Jennifer Giandhari, Sureshnee Pillay, Houriiyah Tegally, Emanuel James San, Tulio de Oliveira, Diana Hardie, Stephen Korsman, Carolyn Williamson |
| EPI_ISL_696512 | Conville CDC wc CVC & NHLS/UCT | KRISP, KZN Research Innovation and Sequencing Platform | Arash Iranzadeh, Deelan Doolabh, Lynn Tyers, Bruna Galvao, Innocent Mudau, Marvin Hsiao, Kruger Marais, Jennifer Giandhari, Sureshnee Pillay, Houriiyah Tegally, Emanuel James San, Tulio de Oliveira, Diana Hardie, Stephen Korsman, Carolyn Williamson |
| EPI_ISL_696513 | Pacaltsdorp Clinic wc PAC & NHLS/UCT | KRISP, KZN Research Innovation and Sequencing Platform | Arash Iranzadeh, Deelan Doolabh, Lynn Tyers, Bruna Galvao, Innocent Mudau, Marvin Hsiao, Kruger Marais, Jennifer Giandhari, Sureshnee Pillay, Houriiyah Tegally, Emanuel James San, Tulio de Oliveira, Diana Hardie, Stephen Korsman, Carolyn Williamson |
| EPI_ISL_696514, EPI_ISL_696515, EPI_ISL_696516, EPI_ISL_696517, EPI_ISL_696518 | Sedgefield Clinic wc SGE & NHLS/UCT | KRISP, KZN Research Innovation and Sequencing Platform | Arash Iranzadeh, Deelan Doolabh, Lynn Tyers, Bruna Galvao, Innocent Mudau, Marvin Hsiao, Kruger Marais, Jennifer Giandhari, Sureshnee Pillay, Houriiyah Tegally, Emanuel James San, Tulio de Oliveira, Diana Hardie, Stephen Korsman, Carolyn Williamson |

| | | | |
|---|---|---|---|
| EPI_ISL_696519, EPI_ISL_696520 | Thembalethu CDC wc THC & NHLS/UCT | KRISP, KZN Research Innovation and Sequencing Platform | Arash Iranzadeh, Deelan Doolabh, Lynn Tyers, Bruna Galvao, Innocent Mudau, Marvin Hsiao, Kruger Marais, Jennifer Giandhari, Sureshnee Pillay, Houriiyah Tegally, Emanuel James San, Tulio de Oliveira, Diana Hardie, Stephen Korsman, Carolyn Williamson |
| EPI_ISL_696521 | Thembalethu CDC wc THC & NHLS/UC | KRISP, KZN Research Innovation and Sequencing Platform | Arash Iranzadeh, Deelan Doolabh, Lynn Tyers, Bruna Galvao, Innocent Mudau, Marvin Hsiao, Kruger Marais, Jennifer Giandhari, Sureshnee Pillay, Houriiyah Tegally, Emanuel James San, Tulio de Oliveira, Diana Hardie, Stephen Korsman, Carolyn Williamson |
| EPI_ISL_698079, EPI_ISL_698080, EPI_ISL_698081, EPI_ISL_698082, EPI_ISL_698083, EPI_ISL_698084, EPI_ISL_698085, EPI_ISL_698086, EPI_ISL_698087, EPI_ISL_698088, EPI_ISL_698089, EPI_ISL_698090, EPI_ISL_698091, EPI_ISL_698092, EPI_ISL_698093, EPI_ISL_698094, EPI_ISL_698095, EPI_ISL_698096, EPI_ISL_698097, EPI_ISL_698098, EPI_ISL_698099, EPI_ISL_698100, EPI_ISL_698101, EPI_ISL_698102, EPI_ISL_698103, EPI_ISL_698104 | | | |
| see above | Unity Health Toronto | Ontario Institute for Cancer Research | Ramzi Fattouh, Larissa M. Matukas, Yan Chen,Mark Downing, Trina Otterman, Karel Boissinot, Le Luu, Samira Mubareka, TIBDN, Ilinca Lungu, Bernard Lam, Jeremy Johns, Paul Krzyzanowski, Richard de Borja, Felicia Vincelli, Philip Zuzarte, Jared T. Simpson |
| EPI_ISL_702023, EPI_ISL_702024, EPI_ISL_702025, EPI_ISL_702026, EPI_ISL_702027, EPI_ISL_702028, EPI_ISL_702029, EPI_ISL_702030, EPI_ISL_702031, EPI_ISL_702032, EPI_ISL_702033, EPI_ISL_702034, EPI_ISL_702035, EPI_ISL_702036, EPI_ISL_702037, EPI_ISL_702038, EPI_ISL_702039, EPI_ISL_702040, EPI_ISL_702041, EPI_ISL_702042, EPI_ISL_702043, EPI_ISL_702044, EPI_ISL_702045, EPI_ISL_702046, EPI_ISL_702047, EPI_ISL_702048, EPI_ISL_702049, EPI_ISL_702050, EPI_ISL_702051, EPI_ISL_702052, EPI_ISL_702053, EPI_ISL_702054, EPI_ISL_702055, EPI_ISL_702056, EPI_ISL_702057, EPI_ISL_702058, EPI_ISL_702059, EPI_ISL_702060, EPI_ISL_702061, EPI_ISL_702062, EPI_ISL_702063, EPI_ISL_702064, EPI_ISL_702065, EPI_ISL_702066, EPI_ISL_702067, EPI_ISL_702068, EPI_ISL_702069, EPI_ISL_702070, EPI_ISL_702071, EPI_ISL_702072, EPI_ISL_702073, EPI_ISL_702074, EPI_ISL_702075, EPI_ISL_702076, EPI_ISL_702077, EPI_ISL_702078, EPI_ISL_702079, EPI_ISL_702080, EPI_ISL_702081, EPI_ISL_702082, EPI_ISL_702083, EPI_ISL_702084, EPI_ISL_702085, EPI_ISL_702086, EPI_ISL_702087, EPI_ISL_702088, EPI_ISL_702089, EPI_ISL_702090, EPI_ISL_702091, EPI_ISL_702092, EPI_ISL_702093, EPI_ISL_702094, EPI_ISL_702095, EPI_ISL_702096, EPI_ISL_702098, EPI_ISL_702099, EPI_ISL_702100, EPI_ISL_702101, EPI_ISL_702102, EPI_ISL_702103, EPI_ISL_702104, EPI_ISL_702105, EPI_ISL_702106, EPI_ISL_702107, EPI_ISL_702108, EPI_ISL_702109, EPI_ISL_702110, EPI_ISL_702111, EPI_ISL_702112, EPI_ISL_702113, EPI_ISL_702114, EPI_ISL_702115, EPI_ISL_702116, EPI_ISL_702117, EPI_ISL_702118, EPI_ISL_702119, EPI_ISL_702120, EPI_ISL_702121, EPI_ISL_702122, EPI_ISL_702123, EPI_ISL_702124, EPI_ISL_702125, EPI_ISL_702126, EPI_ISL_702127, EPI_ISL_702128, EPI_ISL_702129, EPI_ISL_702130, EPI_ISL_702131, EPI_ISL_702132, EPI_ISL_702133, EPI_ISL_702134, EPI_ISL_702135, EPI_ISL_702136, EPI_ISL_702137, EPI_ISL_702138, EPI_ISL_702139, EPI_ISL_702140, EPI_ISL_702141, EPI_ISL_702142, EPI_ISL_702143, EPI_ISL_702144, EPI_ISL_702145, EPI_ISL_702146, EPI_ISL_702147, EPI_ISL_702148, EPI_ISL_702149, EPI_ISL_702150, EPI_ISL_702151, EPI_ISL_702152, EPI_ISL_702153, EPI_ISL_702154, EPI_ISL_702155, EPI_ISL_702156, EPI_ISL_702157, EPI_ISL_702158, EPI_ISL_702159, EPI_ISL_702160, EPI_ISL_702161, EPI_ISL_702162, EPI_ISL_702163, EPI_ISL_702164, EPI_ISL_702165, EPI_ISL_702166, EPI_ISL_702167, EPI_ISL_702168, EPI_ISL_702169, EPI_ISL_702170, EPI_ISL_702171, EPI_ISL_702172, EPI_ISL_702173, EPI_ISL_702174, EPI_ISL_702175, EPI_ISL_702177, EPI_ISL_702178, EPI_ISL_702179, EPI_ISL_702180, EPI_ISL_702181, EPI_ISL_702182, EPI_ISL_702183, EPI_ISL_702184, EPI_ISL_702185, EPI_ISL_702186, EPI_ISL_702187, EPI_ISL_702188, EPI_ISL_702189, EPI_ISL_702190, EPI_ISL_702191, EPI_ISL_702192, EPI_ISL_702193, EPI_ISL_702194, EPI_ISL_702195, EPI_ISL_702196, EPI_ISL_702197, EPI_ISL_702198, EPI_ISL_702199, EPI_ISL_702200, EPI_ISL_702201, EPI_ISL_702202, EPI_ISL_702203, EPI_ISL_702204, EPI_ISL_702205, EPI_ISL_702206, EPI_ISL_702207, EPI_ISL_702208, EPI_ISL_702209, EPI_ISL_702210, EPI_ISL_702211, EPI_ISL_702212, EPI_ISL_702213, EPI_ISL_702214, EPI_ISL_702215, EPI_ISL_702216, EPI_ISL_702217, EPI_ISL_702218, EPI_ISL_702219, EPI_ISL_702220, EPI_ISL_702221, EPI_ISL_702222, EPI_ISL_702223, EPI_ISL_702224, EPI_ISL_702225, EPI_ISL_702226, EPI_ISL_702227, EPI_ISL_702228, EPI_ISL_702229, EPI_ISL_702230, EPI_ISL_702231, EPI_ISL_702232, EPI_ISL_702233, EPI_ISL_702234, EPI_ISL_702235, EPI_ISL_702236, EPI_ISL_702237, EPI_ISL_702238, EPI_ISL_702239, EPI_ISL_702240, EPI_ISL_702241, EPI_ISL_702242, EPI_ISL_702243, EPI_ISL_702245, EPI_ISL_702246, EPI_ISL_702247, EPI_ISL_702248, EPI_ISL_702249, EPI_ISL_702250, EPI_ISL_702251, EPI_ISL_702252, EPI_ISL_702253, EPI_ISL_702254, EPI_ISL_702255, EPI_ISL_702256, EPI_ISL_702257, EPI_ISL_702258, EPI_ISL_702259, EPI_ISL_702260, EPI_ISL_702261, EPI_ISL_702262, EPI_ISL_702263, EPI_ISL_702264, EPI_ISL_702265, EPI_ISL_702266, EPI_ISL_702267, EPI_ISL_702268, EPI_ISL_702269, EPI_ISL_702270, EPI_ISL_702271, EPI_ISL_702272, EPI_ISL_702273, EPI_ISL_702274, EPI_ISL_702275, EPI_ISL_702276, EPI_ISL_702277, EPI_ISL_702278, EPI_ISL_702279, EPI_ISL_702280, EPI_ISL_702281, EPI_ISL_702282, EPI_ISL_702283, EPI_ISL_702284, EPI_ISL_702285, EPI_ISL_702286, EPI_ISL_702287, EPI_ISL_702288, EPI_ISL_702289, EPI_ISL_702290, EPI_ISL_702291, EPI_ISL_702292, EPI_ISL_702293, EPI_ISL_702294, EPI_ISL_702295, EPI_ISL_702296, EPI_ISL_702297, EPI_ISL_702298, EPI_ISL_702299, EPI_ISL_702300, EPI_ISL_702301, EPI_ISL_702302, EPI_ISL_702303, EPI_ISL_702304, EPI_ISL_702305, EPI_ISL_702306, EPI_ISL_702307, EPI_ISL_702308, EPI_ISL_702309, EPI_ISL_702310, EPI_ISL_702311, EPI_ISL_702312, EPI_ISL_702313, EPI_ISL_702314, EPI_ISL_702315, EPI_ISL_702316, EPI_ISL_702317, EPI_ISL_702318, EPI_ISL_702319, EPI_ISL_702320, EPI_ISL_702321, EPI_ISL_702322, EPI_ISL_702323, EPI_ISL_702324, EPI_ISL_702325, EPI_ISL_702326, EPI_ISL_702327, EPI_ISL_702328, EPI_ISL_702329, EPI_ISL_702330, EPI_ISL_702331, EPI_ISL_702332, EPI_ISL_702333, EPI_ISL_702334, EPI_ISL_702335, EPI_ISL_702336, EPI_ISL_702337, EPI_ISL_702338, EPI_ISL_702339, EPI_ISL_702340, EPI_ISL_702341, EPI_ISL_702342, EPI_ISL_702343, EPI_ISL_702344, EPI_ISL_702345, EPI_ISL_702346, EPI_ISL_702347, EPI_ISL_702348, EPI_ISL_702349, EPI_ISL_702350, EPI_ISL_702351, EPI_ISL_702352, EPI_ISL_702353, EPI_ISL_702354, EPI_ISL_702355, EPI_ISL_702356, EPI_ISL_702357, EPI_ISL_702358, EPI_ISL_702359, EPI_ISL_702360, EPI_ISL_702361, EPI_ISL_702362, EPI_ISL_702363, EPI_ISL_702364, EPI_ISL_702365, EPI_ISL_702366, EPI_ISL_702367, EPI_ISL_702368, EPI_ISL_702369 | | | |
| see above | Lighthouse Lab in Cambridge | Wellcome Sanger Institute for the COVID-19 Genomics UK (COG-UK) Consortium | Rob Howes, The Lighthouse Lab in Cambridge and Alex Alderton, Roberto Amato, Sonia Goncalves, Ewan Harrison, David K. Jackson, Ian Johnston, Dominic Kwiatkowski, Cordelia Langford, John Sillitoe on behalf of the Wellcome Sanger Institute COVID-19 Surveillance Team |
| EPI_ISL_702469, EPI_ISL_702481 | Liverpool Clinical Laboratories | COVID-19 Genomics UK (COG-UK) Consortium | Sam Haldenby, Anita Lucaci, Steve Paterson, Julian Hiscox, Alistair Darby, M Almsaud, A Alrezaihi, Muhannad Alruwaili, Stuart D Armstrong, Jones Benjamin, Eleanor G Bentley, Anu Chawla, Jordan J Clark, Angela Cowell, Richard Eccles, Isabel García-Dorival, Matthew Gemmell, Alessandro Gerada, PKF Gilmore, Richard Gregory, Ximeng Han, Catherine Hartley, Margaret Hughes, Miren Iturriza-Gomara, James Johnson, L Luu, Jenifer Manson, Charlotte Nelson, Elaine O'Toole, Cassie Olateju, Rebekah Penrice-Randal , Lucille Rainbow, N.P Randle, Trevor Ian Robinson, Parul Sharma, Ghada T Shawli, James P Stewart, Neil Swainston, Ecaterina Vamos, Joanne Watts, Mark Whitehead |
| EPI_ISL_702507 | University College London Hospital | COVID-19 Genomics UK (COG-UK) Consortium | Judith Heaney, Matthew Byott, Catherine Houlihan, Dan Frampton, Stuart Kirk, Moira Spyer and Eleni Nastouli |
| EPI_ISL_702520 | Northumbria University / South Tees Hospitals NHS Foundation Trust / North Cumbria Integrated Care NHS Foundation Trust / North Tees and Hartlepool NHS Foundation Trust / Newcastle Hospitals NHS Foundation Trust | COVID-19 Genomics UK (COG-UK) Consortium | Darren L Smith,Andrew Nelson,Matthew Bashton,Greg R Young,Joshua Loh,John Allan,Mohammad A Tariq,Giles S Holt,Gary Black,Wen C Yew,Lynn Dover,Paul Baker,Steve Liggett,Sarah Essex,Jane Greenaway,Debra Padgett,Clive Graham,Garren Scott,Edward Barton,Emma Swindells,Brendan Payne,Jennifer Collins,Yusri Taha,Gary Eltringham |
| EPI_ISL_702523 | Liverpool Clinical Laboratories | COVID-19 Genomics UK (COG-UK) Consortium | Sam Haldenby, Anita Lucaci, Steve Paterson, Julian Hiscox, Alistair Darby, M Almsaud, A Alrezaihi, Muhannad Alruwaili, Stuart D Armstrong, Jones Benjamin, Eleanor G Bentley, Anu Chawla, Jordan J Clark, Angela Cowell, Richard Eccles, Isabel García-Dorival, Matthew Gemmell, Alessandro Gerada, PKF Gilmore, Richard Gregory, Ximeng Han, Catherine Hartley, Margaret Hughes, Miren Iturriza-Gomara, James Johnson, L Luu, Jenifer Manson, Charlotte Nelson, Elaine O'Toole, Cassie Olateju, Rebekah Penrice-Randal , Lucille Rainbow, N.P Randle, Trevor Ian Robinson, Parul Sharma, Ghada T Shawli, James P Stewart, Neil Swainston, Ecaterina Vamos, Joanne Watts, Mark Whitehead |
| EPI_ISL_702531, EPI_ISL_702550, EPI_ISL_702564 | Virology Department, Sheffield Teaching Hospitals NHS Foundation Trust/Department of Infection, Immunity and Cardiovascular Disease, The Medical School, University of Sheffield | COVID-19 Genomics UK (COG-UK) Consortium | Thushan de Silva, Matthew Parker, Nikki Smith, Adri Angyal, Rebecca Brown, Luke Green, Rachel Tucker, Paul Parsons, Danielle Groves, Katie Johnson, Laura Carrilero, Alex Keeley, Dave Partridge, Matthew Wyles, Benjamin Lindsey, Mehmet Yavuz, Mohammad Raza, Cariad Evans |
| EPI_ISL_702602 | University College London Hospital | COVID-19 Genomics UK (COG-UK) Consortium | Judith Heaney, Matthew Byott, Catherine Houlihan, Dan Frampton, Stuart Kirk, Moira Spyer and Eleni Nastouli |
| EPI_ISL_702605, EPI_ISL_702626, EPI_ISL_702644, EPI_ISL_702674, EPI_ISL_702677 | Liverpool Clinical Laboratories | COVID-19 Genomics UK (COG-UK) Consortium | Sam Haldenby, Anita Lucaci, Steve Paterson, Julian Hiscox, Alistair Darby, M Almsaud, A Alrezaihi, Muhannad Alruwaili, Stuart D Armstrong, Jones Benjamin, Eleanor G Bentley, Anu Chawla, Jordan J Clark, Angela Cowell, Richard Eccles, Isabel García-Dorival, Matthew Gemmell, Alessandro Gerada, PKF Gilmore, Richard Gregory, Ximeng Han, Catherine Hartley, Margaret Hughes, Miren Iturriza-Gomara, James Johnson, L Luu, Jenifer Manson, Charlotte Nelson, Elaine O'Toole, Cassie Olateju, Rebekah Penrice-Randal , Lucille Rainbow, N.P Randle, Trevor Ian Robinson, Parul Sharma, Ghada T Shawli, James P Stewart, Neil Swainston, Ecaterina Vamos, Joanne Watts, Mark Whitehead |
| EPI_ISL_702700 | Quadram Institute Bioscience | COVID-19 Genomics UK (COG-UK) Consortium | Dave J. Baker, Gemma L. Kay, Alp Aydin, Thanh Le-Viet, Steven Rudder, Ana P. Tedim, Anastasia Kolyva, Maria Diaz, Leonardo de Oliveira Martins, Nabil-Fareed Alikhan, Lizzie Meadows, Rachael Stanley, Ngozi Elumogo, Muhammed Yasir, Nicholas M. Thomson, Alexander J Trotter, Rachel Gilroy, Samuel Bloomfield, Claire Stuart, Andrew Bell, Reenesh Prakash, Samir Dervisevic, Alison E. Mather, John Wain, Mark Webber, Andrew J. Page, Justin O'Grady |
| EPI_ISL_702705 | Virology Department, Sheffield Teaching Hospitals NHS Foundation Trust/Department of Infection, Immunity and Cardiovascular Disease, The Medical School, University of Sheffield | COVID-19 Genomics UK (COG-UK) Consortium | Thushan de Silva, Matthew Parker, Nikki Smith, Adri Angyal, Rebecca Brown, Luke Green, Rachel Tucker, Paul Parsons, Danielle Groves, Katie Johnson, Laura Carrilero, Alex Keeley, Dave Partridge, Matthew Wyles, Benjamin Lindsey, Mehmet Yavuz, Mohammad Raza, Cariad Evans |
| EPI_ISL_702716 | Quadram Institute Bioscience | COVID-19 Genomics UK (COG-UK) Consortium | Dave J. Baker, Gemma L. Kay, Alp Aydin, Thanh Le-Viet, Steven Rudder, Ana P. Tedim, Anastasia Kolyva, Maria Diaz, Leonardo de Oliveira Martins, Nabil-Fareed Alikhan, Lizzie Meadows, Rachael Stanley, Ngozi Elumogo, Muhammed Yasir, Nicholas M. Thomson, Alexander J Trotter, Rachel Gilroy, Samuel Bloomfield, Claire Stuart, Andrew Bell, Reenesh Prakash, Samir Dervisevic, Alison E. Mather, John Wain, Mark Webber, Andrew J. Page, Justin O'Grady |
| EPI_ISL_702776 | Virology Department, Sheffield Teaching Hospitals NHS Foundation Trust/Department of Infection, Immunity and Cardiovascular Disease, The Medical School, University of Sheffield | COVID-19 Genomics UK (COG-UK) Consortium | Thushan de Silva, Matthew Parker, Nikki Smith, Adri Angyal, Rebecca Brown, Luke Green, Rachel Tucker, Paul Parsons, Danielle Groves, Katie Johnson, Laura Carrilero, Alex Keeley, Dave Partridge, Matthew Wyles, Benjamin Lindsey, Mehmet Yavuz, Mohammad Raza, Cariad Evans |
| EPI_ISL_702806 | Liverpool Clinical Laboratories | COVID-19 Genomics UK (COG-UK) Consortium | Sam Haldenby, Anita Lucaci, Steve Paterson, Julian Hiscox, Alistair Darby, M Almsaud, A Alrezaihi, Muhannad Alruwaili, Stuart D Armstrong, Jones Benjamin, Eleanor G Bentley, Anu Chawla, Jordan J Clark, Angela Cowell, Richard Eccles, Isabel García-Dorival, Matthew Gemmell, Alessandro Gerada, PKF Gilmore, Richard Gregory, Ximeng Han, Catherine Hartley, Margaret Hughes, Miren Iturriza-Gomara, James Johnson, L Luu, Jenifer Manson, Charlotte Nelson, Elaine O'Toole, Cassie Olateju, Rebekah Penrice-Randal , Lucille Rainbow, N.P Randle, Trevor Ian Robinson, Parul Sharma, Ghada T Shawli, James P Stewart, Neil Swainston, Ecaterina Vamos, Joanne Watts, Mark Whitehead |
| EPI_ISL_702842, EPI_ISL_702843 | Virology Department, Sheffield Teaching Hospitals NHS Foundation Trust/Department of Infection, Immunity and | COVID-19 Genomics UK (COG-UK) Consortium | Thushan de Silva, Matthew Parker, Nikki Smith, Adri Angyal, Rebecca Brown, Luke Green, Rachel Tucker, Paul Parsons, Danielle Groves, Katie Johnson, Laura Carrilero, Alex Keeley, Dave Partridge, Matthew Wyles, Benjamin Lindsey, Mehmet Yavuz, Mohammad Raza, Cariad Evans |

| | | | |
|---|---|---|---|
| | Cardiovascular Disease, The Medical School, University of Sheffield | | |
| EPI_ISL_702846 | Queens Medical Centre, Clinical Microbiology Department / DeepSeq Nottingham | COVID-19 Genomics UK (COG-UK) Consortium | Gemma Clark, Wendy Smith, Manjinder Khakh, Vicki M Fleming, Michelle M Lister, Hannah Howson-Wells, Jonathan Ball, Patrick McClure, Joseph Chappell, Theocharis Tsoleridis, Nadine Holmes, Matthew Carlisle, Christopher Moore, Fei Sang, Johnny Debebe, Victoria Wright, Matthew Loose |
| EPI_ISL_702856 | Liverpool Clinical Laboratories | COVID-19 Genomics UK (COG-UK) Consortium | Sam Haldenby, Anita Lucaci, Steve Paterson, Julian Hiscox, Alistair Darby, M Almsaud, A Alrezaihi, Muhannad Alruwaili, Stuart D Armstrong, Jones Benjamin, Eleanor G Bentley, Anu Chawla, Jordan J Clark, Angela Cowell, Richard Eccles, Isabel García-Dorival, Matthew Gemmell, Alessandro Gerada, PKF Gilmore, Richard Gregory, Ximeng Han, Catherine Hartley, Margaret Hughes, Miren Iturriza-Gomara, James Johnson, L Luu, Jenifer Manson, Charlotte Nelson, Elaine O'Toole, Cassie Olateju, Rebekah Penrice-Randal , Lucille Rainbow, N.P Randle, Trevor Ian Robinson, Parul Sharma, Ghada T Shawli, James P Stewart, Neil Swainston, Ecaterina Vamos, Joanne Watts, Mark Whitehead |
| EPI_ISL_702861 | Quadram Institute Bioscience | COVID-19 Genomics UK (COG-UK) Consortium | Dave J. Baker, Gemma L. Kay, Alp Aydin, Thanh Le-Viet, Steven Rudder, Ana P. Tedim, Anastasia Kolyva, Maria Diaz, Leonardo de Oliveira Martins, Nabil-Fareed Alikhan, Lizzie Meadows, Rachael Stanley, Ngozi Elumogo, Muhammed Yasir, Nicholas M. Thomson, Alexander J Trotter, Rachel Gilroy, Samuel Bloomfield, Claire Stuart, Andrew Bell, Reenesh Prakash, Samir Dervisevic, Alison E. Mather, John Wain, Mark Webber, Andrew J. Page, Justin O'Grady |
| EPI_ISL_702866, EPI_ISL_702871 | Liverpool Clinical Laboratories | COVID-19 Genomics UK (COG-UK) Consortium | Sam Haldenby, Anita Lucaci, Steve Paterson, Julian Hiscox, Alistair Darby, M Almsaud, A Alrezaihi, Muhannad Alruwaili, Stuart D Armstrong, Jones Benjamin, Eleanor G Bentley, Anu Chawla, Jordan J Clark, Angela Cowell, Richard Eccles, Isabel García-Dorival, Matthew Gemmell, Alessandro Gerada, PKF Gilmore, Richard Gregory, Ximeng Han, Catherine Hartley, Margaret Hughes, Miren Iturriza-Gomara, James Johnson, L Luu, Jenifer Manson, Charlotte Nelson, Elaine O'Toole, Cassie Olateju, Rebekah Penrice-Randal , Lucille Rainbow, N.P Randle, Trevor Ian Robinson, Parul Sharma, Ghada T Shawli, James P Stewart, Neil Swainston, Ecaterina Vamos, Joanne Watts, Mark Whitehead |
| EPI_ISL_702875 | Quadram Institute Bioscience | COVID-19 Genomics UK (COG-UK) Consortium | Dave J. Baker, Gemma L. Kay, Alp Aydin, Thanh Le-Viet, Steven Rudder, Ana P. Tedim, Anastasia Kolyva, Maria Diaz, Leonardo de Oliveira Martins, Nabil-Fareed Alikhan, Lizzie Meadows, Rachael Stanley, Ngozi Elumogo, Muhammed Yasir, Nicholas M. Thomson, Alexander J Trotter, Rachel Gilroy, Samuel Bloomfield, Claire Stuart, Andrew Bell, Reenesh Prakash, Samir Dervisevic, Alison E. Mather, John Wain, Mark Webber, Andrew J. Page, Justin O'Grady |
| EPI_ISL_702878 | Liverpool Clinical Laboratories | COVID-19 Genomics UK (COG-UK) Consortium | Sam Haldenby, Anita Lucaci, Steve Paterson, Julian Hiscox, Alistair Darby, M Almsaud, A Alrezaihi, Muhannad Alruwaili, Stuart D Armstrong, Jones Benjamin, Eleanor G Bentley, Anu Chawla, Jordan J Clark, Angela Cowell, Richard Eccles, Isabel García-Dorival, Matthew Gemmell, Alessandro Gerada, PKF Gilmore, Richard Gregory, Ximeng Han, Catherine Hartley, Margaret Hughes, Miren Iturriza-Gomara, James Johnson, L Luu, Jenifer Manson, Charlotte Nelson, Elaine O'Toole, Cassie Olateju, Rebekah Penrice-Randal , Lucille Rainbow, N.P Randle, Trevor Ian Robinson, Parul Sharma, Ghada T Shawli, James P Stewart, Neil Swainston, Ecaterina Vamos, Joanne Watts, Mark Whitehead |
| EPI_ISL_702879 | Queens Medical Centre, Clinical Microbiology Department / DeepSeq Nottingham | COVID-19 Genomics UK (COG-UK) Consortium | Gemma Clark, Wendy Smith, Manjinder Khakh, Vicki M Fleming, Michelle M Lister, Hannah Howson-Wells, Jonathan Ball, Patrick McClure, Joseph Chappell, Theocharis Tsoleridis, Nadine Holmes, Matthew Carlisle, Christopher Moore, Fei Sang, Johnny Debebe, Victoria Wright, Matthew Loose |
| EPI_ISL_702881 | Virology Department, Sheffield Teaching Hospitals NHS Foundation Trust/Department of Infection, Immunity and Cardiovascular Disease, The Medical School, University of Sheffield | COVID-19 Genomics UK (COG-UK) Consortium | Thushan de Silva, Matthew Parker, Nikki Smith, Adri Angyal, Rebecca Brown, Luke Green, Rachel Tucker, Paul Parsons, Danielle Groves, Katie Johnson, Laura Carrilero, Alex Keeley, Dave Partridge, Matthew Wyles, Benjamin Lindsey, Mehmet Yavuz, Mohammad Raza, Cariad Evans |
| EPI_ISL_702884 | Liverpool Clinical Laboratories | COVID-19 Genomics UK (COG-UK) Consortium | Sam Haldenby, Anita Lucaci, Steve Paterson, Julian Hiscox, Alistair Darby, M Almsaud, A Alrezaihi, Muhannad Alruwaili, Stuart D Armstrong, Jones Benjamin, Eleanor G Bentley, Anu Chawla, Jordan J Clark, Angela Cowell, Richard Eccles, Isabel García-Dorival, Matthew Gemmell, Alessandro Gerada, PKF Gilmore, Richard Gregory, Ximeng Han, Catherine Hartley, Margaret Hughes, Miren Iturriza-Gomara, James Johnson, L Luu, Jenifer Manson, Charlotte Nelson, Elaine O'Toole, Cassie Olateju, Rebekah Penrice-Randal , Lucille Rainbow, N.P Randle, Trevor Ian Robinson, Parul Sharma, Ghada T Shawli, James P Stewart, Neil Swainston, Ecaterina Vamos, Joanne Watts, Mark Whitehead |
| EPI_ISL_702889 | Queens Medical Centre, Clinical Microbiology Department / DeepSeq Nottingham | COVID-19 Genomics UK (COG-UK) Consortium | Gemma Clark, Wendy Smith, Manjinder Khakh, Vicki M Fleming, Michelle M Lister, Hannah Howson-Wells, Jonathan Ball, Patrick McClure, Joseph Chappell, Theocharis Tsoleridis, Nadine Holmes, Matthew Carlisle, Christopher Moore, Fei Sang, Johnny Debebe, Victoria Wright, Matthew Loose |
| EPI_ISL_702894 | Liverpool Clinical Laboratories | COVID-19 Genomics UK (COG-UK) Consortium | Sam Haldenby, Anita Lucaci, Steve Paterson, Julian Hiscox, Alistair Darby, M Almsaud, A Alrezaihi, Muhannad Alruwaili, Stuart D Armstrong, Jones Benjamin, Eleanor G Bentley, Anu Chawla, Jordan J Clark, Angela Cowell, Richard Eccles, Isabel García-Dorival, Matthew Gemmell, Alessandro Gerada, PKF Gilmore, Richard Gregory, Ximeng Han, Catherine Hartley, Margaret Hughes, Miren Iturriza-Gomara, James Johnson, L Luu, Jenifer Manson, Charlotte Nelson, Elaine O'Toole, Cassie Olateju, Rebekah Penrice-Randal , Lucille Rainbow, N.P Randle, Trevor Ian Robinson, Parul Sharma, Ghada T Shawli, James P Stewart, Neil Swainston, Ecaterina Vamos, Joanne Watts, Mark Whitehead |
| EPI_ISL_702902, EPI_ISL_702925 | Quadram Institute Bioscience | COVID-19 Genomics UK (COG-UK) Consortium | Dave J. Baker, Gemma L. Kay, Alp Aydin, Thanh Le-Viet, Steven Rudder, Ana P. Tedim, Anastasia Kolyva, Maria Diaz, Leonardo de Oliveira Martins, Nabil-Fareed Alikhan, Lizzie Meadows, Rachael Stanley, Ngozi Elumogo, Muhammed Yasir, Nicholas M. Thomson, Alexander J Trotter, Rachel Gilroy, Samuel Bloomfield, Claire Stuart, Andrew Bell, Reenesh Prakash, Samir Dervisevic, Alison E. Mather, John Wain, Mark Webber, Andrew J. Page, Justin O'Grady |
| EPI_ISL_702932 | Liverpool Clinical Laboratories | COVID-19 Genomics UK (COG-UK) Consortium | Sam Haldenby, Anita Lucaci, Steve Paterson, Julian Hiscox, Alistair Darby, M Almsaud, A Alrezaihi, Muhannad Alruwaili, Stuart D Armstrong, Jones Benjamin, Eleanor G Bentley, Anu Chawla, Jordan J Clark, Angela Cowell, Richard Eccles, Isabel García-Dorival, Matthew Gemmell, Alessandro Gerada, PKF Gilmore, Richard Gregory, Ximeng Han, Catherine Hartley, Margaret Hughes, Miren Iturriza-Gomara, James Johnson, L Luu, Jenifer Manson, Charlotte Nelson, Elaine O'Toole, Cassie Olateju, Rebekah Penrice-Randal , Lucille Rainbow, N.P Randle, Trevor Ian Robinson, Parul Sharma, Ghada T Shawli, James P Stewart, Neil Swainston, Ecaterina Vamos, Joanne Watts, Mark Whitehead |
| EPI_ISL_702941, EPI_ISL_702944 | Quadram Institute Bioscience | COVID-19 Genomics UK (COG-UK) Consortium | Dave J. Baker, Gemma L. Kay, Alp Aydin, Thanh Le-Viet, Steven Rudder, Ana P. Tedim, Anastasia Kolyva, Maria Diaz, Leonardo de Oliveira Martins, Nabil-Fareed Alikhan, Lizzie Meadows, Rachael Stanley, Ngozi Elumogo, Muhammed Yasir, Nicholas M. Thomson, Alexander J Trotter, Rachel Gilroy, Samuel Bloomfield, Claire Stuart, Andrew Bell, Reenesh Prakash, Samir Dervisevic, Alison E. Mather, John Wain, Mark Webber, Andrew J. Page, Justin O'Grady |
| EPI_ISL_702950, EPI_ISL_702956, EPI_ISL_702969, EPI_ISL_702974, EPI_ISL_703023 | Liverpool Clinical Laboratories | COVID-19 Genomics UK (COG-UK) Consortium | Sam Haldenby, Anita Lucaci, Steve Paterson, Julian Hiscox, Alistair Darby, M Almsaud, A Alrezaihi, Muhannad Alruwaili, Stuart D Armstrong, Jones Benjamin, Eleanor G Bentley, Anu Chawla, Jordan J Clark, Angela Cowell, Richard Eccles, Isabel García-Dorival, Matthew Gemmell, Alessandro Gerada, PKF Gilmore, Richard Gregory, Ximeng Han, Catherine Hartley, Margaret Hughes, Miren Iturriza-Gomara, James Johnson, L Luu, Jenifer Manson, Charlotte Nelson, Elaine O'Toole, Cassie Olateju, Rebekah Penrice-Randal , Lucille Rainbow, N.P Randle, Trevor Ian Robinson, Parul Sharma, Ghada T Shawli, James P Stewart, Neil Swainston, Ecaterina Vamos, Joanne Watts, Mark Whitehead |
| EPI_ISL_703040 | Virology Department, Sheffield Teaching Hospitals NHS Foundation Trust/Department of Infection, Immunity and Cardiovascular Disease, The Medical School, University of Sheffield | COVID-19 Genomics UK (COG-UK) Consortium | Thushan de Silva, Matthew Parker, Nikki Smith, Adri Angyal, Rebecca Brown, Luke Green, Rachel Tucker, Paul Parsons, Danielle Groves, Katie Johnson, Laura Carrilero, Alex Keeley, Dave Partridge, Matthew Wyles, Benjamin Lindsey, Mehmet Yavuz, Mohammad Raza, Cariad Evans |
| EPI_ISL_703043 | Liverpool Clinical Laboratories | COVID-19 Genomics UK (COG-UK) Consortium | Sam Haldenby, Anita Lucaci, Steve Paterson, Julian Hiscox, Alistair Darby, M Almsaud, A Alrezaihi, Muhannad Alruwaili, Stuart D Armstrong, Jones Benjamin, Eleanor G Bentley, Anu Chawla, Jordan J Clark, Angela Cowell, Richard Eccles, Isabel García-Dorival, Matthew Gemmell, Alessandro Gerada, PKF Gilmore, Richard Gregory, Ximeng Han, Catherine Hartley, Margaret Hughes, Miren Iturriza-Gomara, James Johnson, L Luu, Jenifer Manson, Charlotte Nelson, Elaine O'Toole, Cassie Olateju, Rebekah Penrice-Randal , Lucille Rainbow, N.P Randle, Trevor Ian Robinson, Parul Sharma, Ghada T Shawli, James P Stewart, Neil Swainston, Ecaterina Vamos, Joanne Watts, Mark Whitehead |
| EPI_ISL_703056 | Virology Department, Sheffield Teaching Hospitals NHS Foundation Trust/Department of Infection, Immunity and Cardiovascular Disease, The Medical School, University of Sheffield | COVID-19 Genomics UK (COG-UK) Consortium | Thushan de Silva, Matthew Parker, Nikki Smith, Adri Angyal, Rebecca Brown, Luke Green, Rachel Tucker, Paul Parsons, Danielle Groves, Katie Johnson, Laura Carrilero, Alex Keeley, Dave Partridge, Matthew Wyles, Benjamin Lindsey, Mehmet Yavuz, Mohammad Raza, Cariad Evans |
| EPI_ISL_703088, EPI_ISL_703090 | Liverpool Clinical Laboratories | COVID-19 Genomics UK (COG-UK) Consortium | Sam Haldenby, Anita Lucaci, Steve Paterson, Julian Hiscox, Alistair Darby, M Almsaud, A Alrezaihi, Muhannad Alruwaili, Stuart D Armstrong, Jones Benjamin, Eleanor G Bentley, Anu Chawla, Jordan J Clark, Angela Cowell, Richard Eccles, Isabel García-Dorival, Matthew Gemmell, Alessandro Gerada, |

| | | | |
|---|---|---|---|
| | | | PKF Gilmore, Richard Gregory, Ximeng Han, Catherine Hartley, Margaret Hughes, Miren Iturriza-Gomara, James Johnson, L Luu, Jenifer Manson, Charlotte Nelson, Elaine O'Toole, Cassie Olateju, Rebekah Penrice-Randal , Lucille Rainbow, N.P Randle, Trevor Ian Robinson, Parul Sharma, Ghada T Shawli, James P Stewart, Neil Swainston, Ecaterina Vamos, Joanne Watts, Mark Whitehead |
| EPI_ISL_703096 | Northumbria University / South Tees Hospitals NHS Foundation Trust / North Cumbria Integrated Care NHS Foundation Trust / North Tees and Hartlepool NHS Foundation Trust / Newcastle Hospitals NHS Foundation Trust | COVID-19 Genomics UK (COG-UK) Consortium | Darren L Smith,Andrew Nelson,Matthew Bashton,Greg R Young,Joshua Loh,John Allan,Mohammad A Tariq,Giles S Holt,Gary Black,Wen C Yew,Lynn Dover,Paul Baker,Steve Liggett,Sarah Essex,Jane Greenaway,Debra Padgett,Clive Graham,Garren Scott,Edward Barton,Emma Swindells,Brendan Payne,Jennifer Collins,Yusri Taha,Gary Eltringham |
| EPI_ISL_703099 | Quadram Institute Bioscience | COVID-19 Genomics UK (COG-UK) Consortium | Dave J. Baker, Gemma L. Kay, Alp Aydin, Thanh Le-Viet, Steven Rudder, Ana P. Tedim, Anastasia Kolyva, Maria Diaz, Leonardo de Oliveira Martins, Nabil-Fareed Alikhan, Lizzie Meadows, Rachael Stanley, Ngozi Elumogo, Muhammed Yasir, Nicholas M. Thomson, Alexander J Trotter, Rachel Gilroy, Samuel Bloomfield, Claire Stuart, Andrew Bell, Reenesh Prakash, Samir Dervisevic, Alison E. Mather, John Wain, Mark Webber, Andrew J. Page, Justin O'Grady |
| EPI_ISL_703102, EPI_ISL_703110 | Liverpool Clinical Laboratories | COVID-19 Genomics UK (COG-UK) Consortium | Sam Haldenby, Anita Lucaci, Steve Paterson, Julian Hiscox, Alistair Darby, M Almsaud, A Alrezaihi, Muhannad Alruwaili, Stuart D Armstrong, Jones Benjamin, Eleanor G Bentley, Anu Chawla, Jordan J Clark, Angela Cowell, Richard Eccles, Isabel García-Dorival, Matthew Gemmell, Alessandro Gerada, PKF Gilmore, Richard Gregory, Ximeng Han, Catherine Hartley, Margaret Hughes, Miren Iturriza-Gomara, James Johnson, L Luu, Jenifer Manson, Charlotte Nelson, Elaine O'Toole, Cassie Olateju, Rebekah Penrice-Randal , Lucille Rainbow, N.P Randle, Trevor Ian Robinson, Parul Sharma, Ghada T Shawli, James P Stewart, Neil Swainston, Ecaterina Vamos, Joanne Watts, Mark Whitehead |
| EPI_ISL_703124 | Queens Medical Centre, Clinical Microbiology Department / DeepSeq Nottingham | COVID-19 Genomics UK (COG-UK) Consortium | Gemma Clark, Wendy Smith, Manjinder Khakh, Vicki M Fleming, Michelle M Lister, Hannah Howson-Wells, Jonathan Ball, Patrick McClure, Joseph Chappell, Theocharis Tsoleridis, Nadine Holmes, Matthew Carlisle, Christopher Moore, Fei Sang, Johnny Debebe, Victoria Wright, Matthew Loose |
| EPI_ISL_703155 | West of Scotland Specialist Virology Centre, NHSGGC / MRC-University of Glasgow Centre for Virus Research | COVID-19 Genomics UK (COG-UK) Consortium | Ana da Silva Filipe, Natasha Johnson, Kathy Smollett, Daniel Mair, Stephen Carmichael, Alice Broos, Lily Tong, Jenna Nichols, Kyriaki Nomikou; Sarah McDonald; Richard Orton, Joseph Hughes, Sreenu Vattipally, David L Robertson; Alasdair MacLean, Rory Gunson; Sharif Shaaban, Matthew Holden; Rachel Blacow, Guy Mollett, Kathy Li, James Shepherd, Antonia Ho, Emma Thomson |
| EPI_ISL_703165, EPI_ISL_703167, EPI_ISL_703170 | Liverpool Clinical Laboratories | COVID-19 Genomics UK (COG-UK) Consortium | Sam Haldenby, Anita Lucaci, Steve Paterson, Julian Hiscox, Alistair Darby, M Almsaud, A Alrezaihi, Muhannad Alruwaili, Stuart D Armstrong, Jones Benjamin, Eleanor G Bentley, Anu Chawla, Jordan J Clark, Angela Cowell, Richard Eccles, Isabel García-Dorival, Matthew Gemmell, Alessandro Gerada, PKF Gilmore, Richard Gregory, Ximeng Han, Catherine Hartley, Margaret Hughes, Miren Iturriza-Gomara, James Johnson, L Luu, Jenifer Manson, Charlotte Nelson, Elaine O'Toole, Cassie Olateju, Rebekah Penrice-Randal , Lucille Rainbow, N.P Randle, Trevor Ian Robinson, Parul Sharma, Ghada T Shawli, James P Stewart, Neil Swainston, Ecaterina Vamos, Joanne Watts, Mark Whitehead |
| EPI_ISL_703200 | Quadram Institute Bioscience | COVID-19 Genomics UK (COG-UK) Consortium | Dave J. Baker, Gemma L. Kay, Alp Aydin, Thanh Le-Viet, Steven Rudder, Ana P. Tedim, Anastasia Kolyva, Maria Diaz, Leonardo de Oliveira Martins, Nabil-Fareed Alikhan, Lizzie Meadows, Rachael Stanley, Ngozi Elumogo, Muhammed Yasir, Nicholas M. Thomson, Alexander J Trotter, Rachel Gilroy, Samuel Bloomfield, Claire Stuart, Andrew Bell, Reenesh Prakash, Samir Dervisevic, Alison E. Mather, John Wain, Mark Webber, Andrew J. Page, Justin O'Grady |
| EPI_ISL_703206 | Liverpool Clinical Laboratories | COVID-19 Genomics UK (COG-UK) Consortium | Sam Haldenby, Anita Lucaci, Steve Paterson, Julian Hiscox, Alistair Darby, M Almsaud, A Alrezaihi, Muhannad Alruwaili, Stuart D Armstrong, Jones Benjamin, Eleanor G Bentley, Anu Chawla, Jordan J Clark, Angela Cowell, Richard Eccles, Isabel García-Dorival, Matthew Gemmell, Alessandro Gerada, PKF Gilmore, Richard Gregory, Ximeng Han, Catherine Hartley, Margaret Hughes, Miren Iturriza-Gomara, James Johnson, L Luu, Jenifer Manson, Charlotte Nelson, Elaine O'Toole, Cassie Olateju, Rebekah Penrice-Randal , Lucille Rainbow, N.P Randle, Trevor Ian Robinson, Parul Sharma, Ghada T Shawli, James P Stewart, Neil Swainston, Ecaterina Vamos, Joanne Watts, Mark Whitehead |
| EPI_ISL_703211 | Quadram Institute Bioscience | COVID-19 Genomics UK (COG-UK) Consortium | Dave J. Baker, Gemma L. Kay, Alp Aydin, Thanh Le-Viet, Steven Rudder, Ana P. Tedim, Anastasia Kolyva, Maria Diaz, Leonardo de Oliveira Martins, Nabil-Fareed Alikhan, Lizzie Meadows, Rachael Stanley, Ngozi Elumogo, Muhammed Yasir, Nicholas M. Thomson, Alexander J Trotter, Rachel Gilroy, Samuel Bloomfield, Claire Stuart, Andrew Bell, Reenesh Prakash, Samir Dervisevic, Alison E. Mather, John Wain, Mark Webber, Andrew J. Page, Justin O'Grady |
| EPI_ISL_703216 | Liverpool Clinical Laboratories | COVID-19 Genomics UK (COG-UK) Consortium | Sam Haldenby, Anita Lucaci, Steve Paterson, Julian Hiscox, Alistair Darby, M Almsaud, A Alrezaihi, Muhannad Alruwaili, Stuart D Armstrong, Jones Benjamin, Eleanor G Bentley, Anu Chawla, Jordan J Clark, Angela Cowell, Richard Eccles, Isabel García-Dorival, Matthew Gemmell, Alessandro Gerada, PKF Gilmore, Richard Gregory, Ximeng Han, Catherine Hartley, Margaret Hughes, Miren Iturriza-Gomara, James Johnson, L Luu, Jenifer Manson, Charlotte Nelson, Elaine O'Toole, Cassie Olateju, Rebekah Penrice-Randal , Lucille Rainbow, N.P Randle, Trevor Ian Robinson, Parul Sharma, Ghada T Shawli, James P Stewart, Neil Swainston, Ecaterina Vamos, Joanne Watts, Mark Whitehead |
| EPI_ISL_703262, EPI_ISL_703263 | Lighthouse Lab in Glasgow | Wellcome Sanger Institute for the COVID-19 Genomics UK (COG-UK) Consortium | Harper VanSteenhouse, Yumi Kasai, David Gray, Carol Clugston, Anna Dominiczak and Alex Alderton, Roberto Amato, Sonia Goncalves, Ewan Harrison, David K. Jackson, Ian Johnston, Dominic Kwiatkowski, Cordelia Langford, John Sillitoe on behalf of the Wellcome Sanger Institute COVID-19 Surveillance Team |
| EPI_ISL_703264 | Quadram Institute Bioscience | COVID-19 Genomics UK (COG-UK) Consortium | Dave J. Baker, Gemma L. Kay, Alp Aydin, Thanh Le-Viet, Steven Rudder, Ana P. Tedim, Anastasia Kolyva, Maria Diaz, Leonardo de Oliveira Martins, Nabil-Fareed Alikhan, Lizzie Meadows, Rachael Stanley, Ngozi Elumogo, Muhammed Yasir, Nicholas M. Thomson, Alexander J Trotter, Rachel Gilroy, Samuel Bloomfield, Claire Stuart, Andrew Bell, Reenesh Prakash, Samir Dervisevic, Alison E. Mather, John Wain, Mark Webber, Andrew J. Page, Justin O'Grady |
| EPI_ISL_703270, EPI_ISL_703273, EPI_ISL_703280, EPI_ISL_703283, EPI_ISL_703285, EPI_ISL_703288, EPI_ISL_703290, EPI_ISL_703291 | Lighthouse Lab in Glasgow | Wellcome Sanger Institute for the COVID-19 Genomics UK (COG-UK) Consortium | Harper VanSteenhouse, Yumi Kasai, David Gray, Carol Clugston, Anna Dominiczak and Alex Alderton, Roberto Amato, Sonia Goncalves, Ewan Harrison, David K. Jackson, Ian Johnston, Dominic Kwiatkowski, Cordelia Langford, John Sillitoe on behalf of the Wellcome Sanger Institute COVID-19 Surveillance Team |
| EPI_ISL_703292 | Liverpool Clinical Laboratories | COVID-19 Genomics UK (COG-UK) Consortium | Sam Haldenby, Anita Lucaci, Steve Paterson, Julian Hiscox, Alistair Darby, M Almsaud, A Alrezaihi, Muhannad Alruwaili, Stuart D Armstrong, Jones Benjamin, Eleanor G Bentley, Anu Chawla, Jordan J Clark, Angela Cowell, Richard Eccles, Isabel García-Dorival, Matthew Gemmell, Alessandro Gerada, PKF Gilmore, Richard Gregory, Ximeng Han, Catherine Hartley, Margaret Hughes, Miren Iturriza-Gomara, James Johnson, L Luu, Jenifer Manson, Charlotte Nelson, Elaine O'Toole, Cassie Olateju, Rebekah Penrice-Randal , Lucille Rainbow, N.P Randle, Trevor Ian Robinson, Parul Sharma, Ghada T Shawli, James P Stewart, Neil Swainston, Ecaterina Vamos, Joanne Watts, Mark Whitehead |
| EPI_ISL_703296, EPI_ISL_703300, EPI_ISL_703303, EPI_ISL_703304 | Lighthouse Lab in Glasgow | Wellcome Sanger Institute for the COVID-19 Genomics UK (COG-UK) Consortium | Harper VanSteenhouse, Yumi Kasai, David Gray, Carol Clugston, Anna Dominiczak and Alex Alderton, Roberto Amato, Sonia Goncalves, Ewan Harrison, David K. Jackson, Ian Johnston, Dominic Kwiatkowski, Cordelia Langford, John Sillitoe on behalf of the Wellcome Sanger Institute COVID-19 Surveillance Team |
| EPI_ISL_703305 | Liverpool Clinical Laboratories | COVID-19 Genomics UK (COG-UK) Consortium | Sam Haldenby, Anita Lucaci, Steve Paterson, Julian Hiscox, Alistair Darby, M Almsaud, A Alrezaihi, Muhannad Alruwaili, Stuart D Armstrong, Jones Benjamin, Eleanor G Bentley, Anu Chawla, Jordan J Clark, Angela Cowell, Richard Eccles, Isabel García-Dorival, Matthew Gemmell, Alessandro Gerada, PKF Gilmore, Richard Gregory, Ximeng Han, Catherine Hartley, Margaret Hughes, Miren Iturriza-Gomara, James Johnson, L Luu, Jenifer Manson, Charlotte Nelson, Elaine O'Toole, Cassie Olateju, Rebekah Penrice-Randal , Lucille Rainbow, N.P Randle, Trevor Ian Robinson, Parul Sharma, Ghada T Shawli, James P Stewart, Neil Swainston, Ecaterina Vamos, Joanne Watts, Mark Whitehead |
| EPI_ISL_703309 | Lighthouse Lab in Glasgow | Wellcome Sanger Institute for the COVID-19 Genomics UK (COG-UK) Consortium | Harper VanSteenhouse, Yumi Kasai, David Gray, Carol Clugston, Anna Dominiczak and Alex Alderton, Roberto Amato, Sonia Goncalves, Ewan Harrison, David K. Jackson, Ian Johnston, Dominic Kwiatkowski, Cordelia Langford, John Sillitoe on behalf of the Wellcome Sanger Institute COVID-19 Surveillance Team |
| EPI_ISL_703313 | Liverpool Clinical Laboratories | COVID-19 Genomics UK (COG-UK) Consortium | Sam Haldenby, Anita Lucaci, Steve Paterson, Julian Hiscox, Alistair Darby, M Almsaud, A Alrezaihi, Muhannad Alruwaili, Stuart D Armstrong, Jones Benjamin, Eleanor G Bentley, Anu Chawla, Jordan J Clark, Angela Cowell, Richard Eccles, Isabel García-Dorival, Matthew Gemmell, Alessandro Gerada, PKF Gilmore, Richard Gregory, Ximeng Han, Catherine Hartley, Margaret Hughes, Miren Iturriza-Gomara, James Johnson, L Luu, Jenifer Manson, Charlotte Nelson, Elaine O'Toole, Cassie Olateju, Rebekah Penrice-Randal , Lucille Rainbow, N.P Randle, Trevor Ian Robinson, Parul Sharma, Ghada T Shawli, James P Stewart, Neil Swainston, Ecaterina Vamos, Joanne Watts, Mark Whitehead |
| EPI_ISL_703315, EPI_ISL_703320, EPI_ISL_703323, EPI_ISL_703325, | Lighthouse Lab in Glasgow | Wellcome Sanger Institute for the COVID-19 Genomics UK (COG-UK) Consortium | Harper VanSteenhouse, Yumi Kasai, David Gray, Carol Clugston, Anna Dominiczak and Alex Alderton, Roberto Amato, Sonia Goncalves, Ewan Harrison, David K. Jackson, Ian Johnston, Dominic Kwiatkowski, Cordelia Langford, John Sillitoe on behalf of the Wellcome Sanger Institute COVID-19 Surveillance Team |

| | | | |
|---|---|---|---|
| EPI_ISL_703331, EPI_ISL_703334, EPI_ISL_703337 | | | Team |
| EPI_ISL_703348 | Liverpool Clinical Laboratories | COVID-19 Genomics UK (COG-UK) Consortium | Sam Haldenby, Anita Lucaci, Steve Paterson, Julian Hiscox, Alistair Darby, M Almsaud, A Alrezaihi, Muhannad Alruwaili, Stuart D Armstrong, Jones Benjamin, Eleanor G Bentley, Anu Chawla, Jordan J Clark, Angela Cowell, Richard Eccles, Isabel García-Dorival, Matthew Gemmell, Alessandro Gerada, PKF Gilmore, Richard Gregory, Ximeng Han, Catherine Hartley, Margaret Hughes, Miren Iturriza-Gomara, James Johnson, L Luu, Jenifer Manson, Charlotte Nelson, Elaine O'Toole, Cassie Olateju, Rebekah Penrice-Randal , Lucille Rainbow, N.P Randle, Trevor Ian Robinson, Parul Sharma, Ghada T Shawli, James P Stewart, Neil Swainston, Ecaterina Vamos, Joanne Watts, Mark Whitehead |
| EPI_ISL_703349, EPI_ISL_703350, EPI_ISL_703352, EPI_ISL_703354, EPI_ISL_703355 | Lighthouse Lab in Glasgow | Wellcome Sanger Institute for the COVID-19 Genomics UK (COG-UK) Consortium | Harper VanSteenhouse, Yumi Kasai, David Gray, Carol Clugston, Anna Dominiczak and Alex Alderton, Roberto Amato, Sonia Goncalves, Ewan Harrison, David K. Jackson, Ian Johnston, Dominic Kwiatkowski, Cordelia Langford, John Sillitoe on behalf of the Wellcome Sanger Institute COVID-19 Surveillance Team |
| EPI_ISL_703356 | Liverpool Clinical Laboratories | COVID-19 Genomics UK (COG-UK) Consortium | Sam Haldenby, Anita Lucaci, Steve Paterson, Julian Hiscox, Alistair Darby, M Almsaud, A Alrezaihi, Muhannad Alruwaili, Stuart D Armstrong, Jones Benjamin, Eleanor G Bentley, Anu Chawla, Jordan J Clark, Angela Cowell, Richard Eccles, Isabel García-Dorival, Matthew Gemmell, Alessandro Gerada, PKF Gilmore, Richard Gregory, Ximeng Han, Catherine Hartley, Margaret Hughes, Miren Iturriza-Gomara, James Johnson, L Luu, Jenifer Manson, Charlotte Nelson, Elaine O'Toole, Cassie Olateju, Rebekah Penrice-Randal , Lucille Rainbow, N.P Randle, Trevor Ian Robinson, Parul Sharma, Ghada T Shawli, James P Stewart, Neil Swainston, Ecaterina Vamos, Joanne Watts, Mark Whitehead |
| EPI_ISL_703358, EPI_ISL_703360 | Lighthouse Lab in Glasgow | Wellcome Sanger Institute for the COVID-19 Genomics UK (COG-UK) Consortium | Harper VanSteenhouse, Yumi Kasai, David Gray, Carol Clugston, Anna Dominiczak and Alex Alderton, Roberto Amato, Sonia Goncalves, Ewan Harrison, David K. Jackson, Ian Johnston, Dominic Kwiatkowski, Cordelia Langford, John Sillitoe on behalf of the Wellcome Sanger Institute COVID-19 Surveillance Team |
| EPI_ISL_703361 | Liverpool Clinical Laboratories | COVID-19 Genomics UK (COG-UK) Consortium | Sam Haldenby, Anita Lucaci, Steve Paterson, Julian Hiscox, Alistair Darby, M Almsaud, A Alrezaihi, Muhannad Alruwaili, Stuart D Armstrong, Jones Benjamin, Eleanor G Bentley, Anu Chawla, Jordan J Clark, Angela Cowell, Richard Eccles, Isabel García-Dorival, Matthew Gemmell, Alessandro Gerada, PKF Gilmore, Richard Gregory, Ximeng Han, Catherine Hartley, Margaret Hughes, Miren Iturriza-Gomara, James Johnson, L Luu, Jenifer Manson, Charlotte Nelson, Elaine O'Toole, Cassie Olateju, Rebekah Penrice-Randal , Lucille Rainbow, N.P Randle, Trevor Ian Robinson, Parul Sharma, Ghada T Shawli, James P Stewart, Neil Swainston, Ecaterina Vamos, Joanne Watts, Mark Whitehead |
| EPI_ISL_703362, EPI_ISL_703370, EPI_ISL_703371, EPI_ISL_703378, EPI_ISL_703379, EPI_ISL_703381, EPI_ISL_703382, EPI_ISL_703384, EPI_ISL_703389, EPI_ISL_703391, EPI_ISL_703393 | | | |
| see above | Lighthouse Lab in Glasgow | Wellcome Sanger Institute for the COVID-19 Genomics UK (COG-UK) Consortium | Harper VanSteenhouse, Yumi Kasai, David Gray, Carol Clugston, Anna Dominiczak and Alex Alderton, Roberto Amato, Sonia Goncalves, Ewan Harrison, David K. Jackson, Ian Johnston, Dominic Kwiatkowski, Cordelia Langford, John Sillitoe on behalf of the Wellcome Sanger Institute COVID-19 Surveillance Team |
| EPI_ISL_703400 | Virology Department, Sheffield Teaching Hospitals NHS Foundation Trust/Department of Infection, Immunity and Cardiovascular Disease, The Medical School, University of Sheffield | COVID-19 Genomics UK (COG-UK) Consortium | Thushan de Silva, Matthew Parker, Nikki Smith, Adri Angyal, Rebecca Brown, Luke Green, Rachel Tucker, Paul Parsons, Danielle Groves, Katie Johnson, Laura Carrilero, Alex Keeley, Dave Partridge, Matthew Wyles, Benjamin Lindsey, Mehmet Yavuz, Mohammad Raza, Cariad Evans |
| EPI_ISL_703407, EPI_ISL_703408, EPI_ISL_703410, EPI_ISL_703411, EPI_ISL_703415 | Lighthouse Lab in Glasgow | Wellcome Sanger Institute for the COVID-19 Genomics UK (COG-UK) Consortium | Harper VanSteenhouse, Yumi Kasai, David Gray, Carol Clugston, Anna Dominiczak and Alex Alderton, Roberto Amato, Sonia Goncalves, Ewan Harrison, David K. Jackson, Ian Johnston, Dominic Kwiatkowski, Cordelia Langford, John Sillitoe on behalf of the Wellcome Sanger Institute COVID-19 Surveillance Team |
| EPI_ISL_703421 | Liverpool Clinical Laboratories | COVID-19 Genomics UK (COG-UK) Consortium | Sam Haldenby, Anita Lucaci, Steve Paterson, Julian Hiscox, Alistair Darby, M Almsaud, A Alrezaihi, Muhannad Alruwaili, Stuart D Armstrong, Jones Benjamin, Eleanor G Bentley, Anu Chawla, Jordan J Clark, Angela Cowell, Richard Eccles, Isabel García-Dorival, Matthew Gemmell, Alessandro Gerada, PKF Gilmore, Richard Gregory, Ximeng Han, Catherine Hartley, Margaret Hughes, Miren Iturriza-Gomara, James Johnson, L Luu, Jenifer Manson, Charlotte Nelson, Elaine O'Toole, Cassie Olateju, Rebekah Penrice-Randal , Lucille Rainbow, N.P Randle, Trevor Ian Robinson, Parul Sharma, Ghada T Shawli, James P Stewart, Neil Swainston, Ecaterina Vamos, Joanne Watts, Mark Whitehead |
| EPI_ISL_703422, EPI_ISL_703423, EPI_ISL_703426 | Lighthouse Lab in Glasgow | Wellcome Sanger Institute for the COVID-19 Genomics UK (COG-UK) Consortium | Harper VanSteenhouse, Yumi Kasai, David Gray, Carol Clugston, Anna Dominiczak and Alex Alderton, Roberto Amato, Sonia Goncalves, Ewan Harrison, David K. Jackson, Ian Johnston, Dominic Kwiatkowski, Cordelia Langford, John Sillitoe on behalf of the Wellcome Sanger Institute COVID-19 Surveillance Team |
| EPI_ISL_703427 | Liverpool Clinical Laboratories | COVID-19 Genomics UK (COG-UK) Consortium | Sam Haldenby, Anita Lucaci, Steve Paterson, Julian Hiscox, Alistair Darby, M Almsaud, A Alrezaihi, Muhannad Alruwaili, Stuart D Armstrong, Jones Benjamin, Eleanor G Bentley, Anu Chawla, Jordan J Clark, Angela Cowell, Richard Eccles, Isabel García-Dorival, Matthew Gemmell, Alessandro Gerada, PKF Gilmore, Richard Gregory, Ximeng Han, Catherine Hartley, Margaret Hughes, Miren Iturriza-Gomara, James Johnson, L Luu, Jenifer Manson, Charlotte Nelson, Elaine O'Toole, Cassie Olateju, Rebekah Penrice-Randal , Lucille Rainbow, N.P Randle, Trevor Ian Robinson, Parul Sharma, Ghada T Shawli, James P Stewart, Neil Swainston, Ecaterina Vamos, Joanne Watts, Mark Whitehead |
| EPI_ISL_703428, EPI_ISL_703429, EPI_ISL_703431, EPI_ISL_703439, EPI_ISL_703442, EPI_ISL_703444, EPI_ISL_703447, EPI_ISL_703453, EPI_ISL_703456, EPI_ISL_703458, EPI_ISL_703463, EPI_ISL_703469, EPI_ISL_703472, EPI_ISL_703474 | | | |
| see above | Lighthouse Lab in Glasgow | Wellcome Sanger Institute for the COVID-19 Genomics UK (COG-UK) Consortium | Harper VanSteenhouse, Yumi Kasai, David Gray, Carol Clugston, Anna Dominiczak and Alex Alderton, Roberto Amato, Sonia Goncalves, Ewan Harrison, David K. Jackson, Ian Johnston, Dominic Kwiatkowski, Cordelia Langford, John Sillitoe on behalf of the Wellcome Sanger Institute COVID-19 Surveillance Team |
| EPI_ISL_703481 | Liverpool Clinical Laboratories | COVID-19 Genomics UK (COG-UK) Consortium | Sam Haldenby, Anita Lucaci, Steve Paterson, Julian Hiscox, Alistair Darby, M Almsaud, A Alrezaihi, Muhannad Alruwaili, Stuart D Armstrong, Jones Benjamin, Eleanor G Bentley, Anu Chawla, Jordan J Clark, Angela Cowell, Richard Eccles, Isabel García-Dorival, Matthew Gemmell, Alessandro Gerada, PKF Gilmore, Richard Gregory, Ximeng Han, Catherine Hartley, Margaret Hughes, Miren Iturriza-Gomara, James Johnson, L Luu, Jenifer Manson, Charlotte Nelson, Elaine O'Toole, Cassie Olateju, Rebekah Penrice-Randal , Lucille Rainbow, N.P Randle, Trevor Ian Robinson, Parul Sharma, Ghada T Shawli, James P Stewart, Neil Swainston, Ecaterina Vamos, Joanne Watts, Mark Whitehead |
| EPI_ISL_703482, EPI_ISL_703491, EPI_ISL_703496, EPI_ISL_703499, EPI_ISL_703506, EPI_ISL_703510, EPI_ISL_703512, EPI_ISL_703514, EPI_ISL_703515, EPI_ISL_703517, EPI_ISL_703522, EPI_ISL_703525, EPI_ISL_703529, EPI_ISL_703534, EPI_ISL_703538, EPI_ISL_703541, EPI_ISL_703546, EPI_ISL_703553, EPI_ISL_703557, EPI_ISL_703561 | | | |
| see above | Lighthouse Lab in Glasgow | Wellcome Sanger Institute for the COVID-19 Genomics UK (COG-UK) Consortium | Harper VanSteenhouse, Yumi Kasai, David Gray, Carol Clugston, Anna Dominiczak and Alex Alderton, Roberto Amato, Sonia Goncalves, Ewan Harrison, David K. Jackson, Ian Johnston, Dominic Kwiatkowski, Cordelia Langford, John Sillitoe on behalf of the Wellcome Sanger Institute COVID-19 Surveillance Team |
| EPI_ISL_703572 | Virology Department, Royal Infirmary of Edinburgh, NHS Lothian / School of Biological Sciences, University of Edinburgh / Institute of Genetics and Molecular Medicine, University of Edinburgh | COVID-19 Genomics UK (COG-UK) Consortium | McHugh M, Dewar R, Rooke S, Gallagher M, Balcaza C, O'Toole Á, Scher E, Hill V, McCrone JT, Colquhoun R, Yu X, Jackson B, Rambaut A, Williams TC, Templeton K |
| EPI_ISL_703578 | Lighthouse Lab in Glasgow | Wellcome Sanger Institute for the COVID-19 Genomics UK (COG-UK) Consortium | Harper VanSteenhouse, Yumi Kasai, David Gray, Carol Clugston, Anna Dominiczak and Alex Alderton, Roberto Amato, Sonia Goncalves, Ewan Harrison, David K. Jackson, Ian Johnston, Dominic Kwiatkowski, Cordelia Langford, John Sillitoe on behalf of the Wellcome Sanger Institute COVID-19 Surveillance Team |
| EPI_ISL_703579 | Virology Department, Sheffield Teaching Hospitals NHS Foundation Trust/Department of Infection, Immunity and Cardiovascular Disease, The Medical School, University of Sheffield | COVID-19 Genomics UK (COG-UK) Consortium | Thushan de Silva, Matthew Parker, Nikki Smith, Adri Angyal, Rebecca Brown, Luke Green, Rachel Tucker, Paul Parsons, Danielle Groves, Katie Johnson, Laura Carrilero, Alex Keeley, Dave Partridge, Matthew Wyles, Benjamin Lindsey, Mehmet Yavuz, Mohammad Raza, Cariad Evans |
| EPI_ISL_703583 | Lighthouse Lab in Glasgow | Wellcome Sanger Institute for the COVID-19 Genomics UK (COG-UK) Consortium | Harper VanSteenhouse, Yumi Kasai, David Gray, Carol Clugston, Anna Dominiczak and Alex Alderton, Roberto Amato, Sonia Goncalves, Ewan Harrison, David K. Jackson, Ian Johnston, Dominic Kwiatkowski, Cordelia Langford, John Sillitoe on behalf of the Wellcome Sanger Institute COVID-19 Surveillance Team |
| EPI_ISL_703594 | Lighthouse Lab in Alderley Park | Wellcome Sanger Institute for the COVID-19 Genomics UK | Jacquelyn Wynn, Mairead Hyland, The Lighthouse Lab in Alderley Park and Alex Alderton, Roberto Amato, Sonia Goncalves, Ewan Harrison, David K. |

| | | | |
|---|---|---|---|
| | | (COG-UK) Consortium | Jackson, Ian Johnston, Dominic Kwiatkowski, Cordelia Langford, John Sillitoe on behalf of the Wellcome Sanger Institute COVID-19 Surveillance Team |
| EPI_ISL_703598, EPI_ISL_703600 | Lighthouse Lab in Glasgow | Wellcome Sanger Institute for the COVID-19 Genomics UK (COG-UK) Consortium | Harper VanSteenhouse, Yumi Kasai, David Gray, Carol Clugston, Anna Dominiczak and Alex Alderton, Roberto Amato, Sonia Goncalves, Ewan Harrison, David K. Jackson, Ian Johnston, Dominic Kwiatkowski, Cordelia Langford, John Sillitoe on behalf of the Wellcome Sanger Institute COVID-19 Surveillance Team |
| EPI_ISL_703601, EPI_ISL_703604, EPI_ISL_703606 | Virology Department, Sheffield Teaching Hospitals NHS Foundation Trust/Department of Infection, Immunity and Cardiovascular Disease, The Medical School, University of Sheffield | COVID-19 Genomics UK (COG-UK) Consortium | Thushan de Silva, Matthew Parker, Nikki Smith, Adri Angyal, Rebecca Brown, Luke Green, Rachel Tucker, Paul Parsons, Danielle Groves, Katie Johnson, Laura Carrilero, Alex Keeley, Dave Partridge, Matthew Wyles, Benjamin Lindsey, Mehmet Yavuz, Mohammad Raza, Cariad Evans |
| EPI_ISL_703607, EPI_ISL_703608, EPI_ISL_703610, EPI_ISL_703612 | Lighthouse Lab in Glasgow | Wellcome Sanger Institute for the COVID-19 Genomics UK (COG-UK) Consortium | Harper VanSteenhouse, Yumi Kasai, David Gray, Carol Clugston, Anna Dominiczak and Alex Alderton, Roberto Amato, Sonia Goncalves, Ewan Harrison, David K. Jackson, Ian Johnston, Dominic Kwiatkowski, Cordelia Langford, John Sillitoe on behalf of the Wellcome Sanger Institute COVID-19 Surveillance Team |
| EPI_ISL_703616 | Quadram Institute Bioscience | COVID-19 Genomics UK (COG-UK) Consortium | Dave J. Baker, Gemma L. Kay, Alp Aydin, Thanh Le-Viet, Steven Rudder, Ana P. Tedim, Anastasia Kolyva, Maria Diaz, Leonardo de Oliveira Martins, Nabil-Fareed Alikhan, Lizzie Meadows, Rachael Stanley, Ngozi Elumogo, Muhammed Yasir, Nicholas M. Thomson, Alexander J Trotter, Rachel Gilroy, Samuel Bloomfield, Claire Stuart, Andrew Bell, Reenesh Prakash, Samir Dervisevic, Alison E. Mather, John Wain, Mark Webber, Andrew J. Page, Justin O'Grady |
| EPI_ISL_703617, EPI_ISL_703621, EPI_ISL_703626, EPI_ISL_703631, EPI_ISL_703633, EPI_ISL_703634, EPI_ISL_703640, EPI_ISL_703641, EPI_ISL_703643 | Lighthouse Lab in Glasgow | Wellcome Sanger Institute for the COVID-19 Genomics UK (COG-UK) Consortium | Harper VanSteenhouse, Yumi Kasai, David Gray, Carol Clugston, Anna Dominiczak and Alex Alderton, Roberto Amato, Sonia Goncalves, Ewan Harrison, David K. Jackson, Ian Johnston, Dominic Kwiatkowski, Cordelia Langford, John Sillitoe on behalf of the Wellcome Sanger Institute COVID-19 Surveillance Team |
| EPI_ISL_703644 | Virology Department, Sheffield Teaching Hospitals NHS Foundation Trust/Department of Infection, Immunity and Cardiovascular Disease, The Medical School, University of Sheffield | COVID-19 Genomics UK (COG-UK) Consortium | Thushan de Silva, Matthew Parker, Nikki Smith, Adri Angyal, Rebecca Brown, Luke Green, Rachel Tucker, Paul Parsons, Danielle Groves, Katie Johnson, Laura Carrilero, Alex Keeley, Dave Partridge, Matthew Wyles, Benjamin Lindsey, Mehmet Yavuz, Mohammad Raza, Cariad Evans |
| EPI_ISL_703645 | Lighthouse Lab in Glasgow | Wellcome Sanger Institute for the COVID-19 Genomics UK (COG-UK) Consortium | Harper VanSteenhouse, Yumi Kasai, David Gray, Carol Clugston, Anna Dominiczak and Alex Alderton, Roberto Amato, Sonia Goncalves, Ewan Harrison, David K. Jackson, Ian Johnston, Dominic Kwiatkowski, Cordelia Langford, John Sillitoe on behalf of the Wellcome Sanger Institute COVID-19 Surveillance Team |
| EPI_ISL_703647 | Virology Department, Sheffield Teaching Hospitals NHS Foundation Trust/Department of Infection, Immunity and Cardiovascular Disease, The Medical School, University of Sheffield | COVID-19 Genomics UK (COG-UK) Consortium | Thushan de Silva, Matthew Parker, Nikki Smith, Adri Angyal, Rebecca Brown, Luke Green, Rachel Tucker, Paul Parsons, Danielle Groves, Katie Johnson, Laura Carrilero, Alex Keeley, Dave Partridge, Matthew Wyles, Benjamin Lindsey, Mehmet Yavuz, Mohammad Raza, Cariad Evans |
| EPI_ISL_703651, EPI_ISL_703653, EPI_ISL_703654 | Lighthouse Lab in Glasgow | Wellcome Sanger Institute for the COVID-19 Genomics UK (COG-UK) Consortium | Harper VanSteenhouse, Yumi Kasai, David Gray, Carol Clugston, Anna Dominiczak and Alex Alderton, Roberto Amato, Sonia Goncalves, Ewan Harrison, David K. Jackson, Ian Johnston, Dominic Kwiatkowski, Cordelia Langford, John Sillitoe on behalf of the Wellcome Sanger Institute COVID-19 Surveillance Team |
| EPI_ISL_703655 | Virology Department, Sheffield Teaching Hospitals NHS Foundation Trust/Department of Infection, Immunity and Cardiovascular Disease, The Medical School, University of Sheffield | COVID-19 Genomics UK (COG-UK) Consortium | Thushan de Silva, Matthew Parker, Nikki Smith, Adri Angyal, Rebecca Brown, Luke Green, Rachel Tucker, Paul Parsons, Danielle Groves, Katie Johnson, Laura Carrilero, Alex Keeley, Dave Partridge, Matthew Wyles, Benjamin Lindsey, Mehmet Yavuz, Mohammad Raza, Cariad Evans |
| EPI_ISL_703656, EPI_ISL_703657 | Lighthouse Lab in Glasgow | Wellcome Sanger Institute for the COVID-19 Genomics UK (COG-UK) Consortium | Harper VanSteenhouse, Yumi Kasai, David Gray, Carol Clugston, Anna Dominiczak and Alex Alderton, Roberto Amato, Sonia Goncalves, Ewan Harrison, David K. Jackson, Ian Johnston, Dominic Kwiatkowski, Cordelia Langford, John Sillitoe on behalf of the Wellcome Sanger Institute COVID-19 Surveillance Team |
| EPI_ISL_703658, EPI_ISL_703660 | Virology Department, Sheffield Teaching Hospitals NHS Foundation Trust/Department of Infection, Immunity and Cardiovascular Disease, The Medical School, University of Sheffield | COVID-19 Genomics UK (COG-UK) Consortium | Thushan de Silva, Matthew Parker, Nikki Smith, Adri Angyal, Rebecca Brown, Luke Green, Rachel Tucker, Paul Parsons, Danielle Groves, Katie Johnson, Laura Carrilero, Alex Keeley, Dave Partridge, Matthew Wyles, Benjamin Lindsey, Mehmet Yavuz, Mohammad Raza, Cariad Evans |
| EPI_ISL_703661, EPI_ISL_703669, EPI_ISL_703672, EPI_ISL_703673 | Lighthouse Lab in Glasgow | Wellcome Sanger Institute for the COVID-19 Genomics UK (COG-UK) Consortium | Harper VanSteenhouse, Yumi Kasai, David Gray, Carol Clugston, Anna Dominiczak and Alex Alderton, Roberto Amato, Sonia Goncalves, Ewan Harrison, David K. Jackson, Ian Johnston, Dominic Kwiatkowski, Cordelia Langford, John Sillitoe on behalf of the Wellcome Sanger Institute COVID-19 Surveillance Team |
| EPI_ISL_703676 | Virology Department, Sheffield Teaching Hospitals NHS Foundation Trust/Department of Infection, Immunity and Cardiovascular Disease, The Medical School, University of Sheffield | COVID-19 Genomics UK (COG-UK) Consortium | Thushan de Silva, Matthew Parker, Nikki Smith, Adri Angyal, Rebecca Brown, Luke Green, Rachel Tucker, Paul Parsons, Danielle Groves, Katie Johnson, Laura Carrilero, Alex Keeley, Dave Partridge, Matthew Wyles, Benjamin Lindsey, Mehmet Yavuz, Mohammad Raza, Cariad Evans |
| EPI_ISL_703677 | Lighthouse Lab in Glasgow | Wellcome Sanger Institute for the COVID-19 Genomics UK (COG-UK) Consortium | Harper VanSteenhouse, Yumi Kasai, David Gray, Carol Clugston, Anna Dominiczak and Alex Alderton, Roberto Amato, Sonia Goncalves, Ewan Harrison, David K. Jackson, Ian Johnston, Dominic Kwiatkowski, Cordelia Langford, John Sillitoe on behalf of the Wellcome Sanger Institute COVID-19 Surveillance Team |
| EPI_ISL_703684 | Virology Department, Sheffield Teaching Hospitals NHS Foundation Trust/Department of Infection, Immunity and Cardiovascular Disease, The Medical School, University of Sheffield | COVID-19 Genomics UK (COG-UK) Consortium | Thushan de Silva, Matthew Parker, Nikki Smith, Adri Angyal, Rebecca Brown, Luke Green, Rachel Tucker, Paul Parsons, Danielle Groves, Katie Johnson, Laura Carrilero, Alex Keeley, Dave Partridge, Matthew Wyles, Benjamin Lindsey, Mehmet Yavuz, Mohammad Raza, Cariad Evans |
| EPI_ISL_703685 | Lighthouse Lab in Glasgow | Wellcome Sanger Institute for the COVID-19 Genomics UK (COG-UK) Consortium | Harper VanSteenhouse, Yumi Kasai, David Gray, Carol Clugston, Anna Dominiczak and Alex Alderton, Roberto Amato, Sonia Goncalves, Ewan Harrison, David K. Jackson, Ian Johnston, Dominic Kwiatkowski, Cordelia Langford, John Sillitoe on behalf of the Wellcome Sanger Institute COVID-19 Surveillance Team |
| EPI_ISL_703690 | Liverpool Clinical Laboratories | COVID-19 Genomics UK (COG-UK) Consortium | Sam Haldenby, Anita Lucaci, Steve Paterson, Julian Hiscox, Alistair Darby, M Almsaud, A Alrezaihi, Muhannad Alruwaili, Stuart D Armstrong, Jones Benjamin, Eleanor G Bentley, Anu Chawla, Jordan J Clark, Angela Cowell, Richard Eccles, Isabel García-Dorival, Matthew Gemmell, Alessandro Gerada, PKF Gilmore, Richard Gregory, Ximeng Han, Catherine Hartley, Margaret Hughes, Miren Iturriza-Gomara, James Johnson, L Luu, Jenifer Manson, Charlotte Nelson, Elaine O'Toole, Cassie Olateju, Rebekah Penrice-Randal , Lucille Rainbow, N.P Randle, Trevor Ian Robinson, Parul Sharma, Ghada T Shawli, James P Stewart, Neil Swainston, Ecaterina Vamos, Joanne Watts, Mark Whitehead |
| EPI_ISL_703694 | Lighthouse Lab in Glasgow | Wellcome Sanger Institute for the COVID-19 Genomics UK (COG-UK) Consortium | Harper VanSteenhouse, Yumi Kasai, David Gray, Carol Clugston, Anna Dominiczak and Alex Alderton, Roberto Amato, Sonia Goncalves, Ewan Harrison, David K. Jackson, Ian Johnston, Dominic Kwiatkowski, Cordelia Langford, John Sillitoe on behalf of the Wellcome Sanger Institute COVID-19 Surveillance Team |
| EPI_ISL_703695 | Liverpool Clinical Laboratories | COVID-19 Genomics UK (COG-UK) Consortium | Sam Haldenby, Anita Lucaci, Steve Paterson, Julian Hiscox, Alistair Darby, M Almsaud, A Alrezaihi, Muhannad Alruwaili, Stuart D Armstrong, Jones Benjamin, Eleanor G Bentley, Anu Chawla, Jordan J Clark, Angela Cowell, Richard Eccles, Isabel García-Dorival, Matthew Gemmell, Alessandro Gerada, PKF Gilmore, Richard Gregory, Ximeng Han, Catherine Hartley, Margaret Hughes, Miren Iturriza-Gomara, James Johnson, L Luu, Jenifer Manson, Charlotte Nelson, Elaine O'Toole, Cassie Olateju, Rebekah Penrice-Randal , Lucille Rainbow, N.P Randle, Trevor Ian Robinson, Parul Sharma, Ghada T Shawli, James P Stewart, Neil Swainston, Ecaterina Vamos, Joanne Watts, Mark Whitehead |
| EPI_ISL_703700, EPI_ISL_703703 | Lighthouse Lab in Glasgow | Wellcome Sanger Institute for the COVID-19 Genomics UK (COG-UK) Consortium | Harper VanSteenhouse, Yumi Kasai, David Gray, Carol Clugston, Anna Dominiczak and Alex Alderton, Roberto Amato, Sonia Goncalves, Ewan Harrison, David K. Jackson, Ian Johnston, Dominic Kwiatkowski, Cordelia Langford, John Sillitoe on behalf of the Wellcome Sanger Institute COVID-19 Surveillance |

| | | | |
|---|---|---|---|
| | | | Team |
| EPI_ISL_703714 | Quadram Institute Bioscience | COVID-19 Genomics UK (COG-UK) Consortium | Dave J. Baker, Gemma L. Kay, Alp Aydin, Thanh Le-Viet, Steven Rudder, Ana P. Tedim, Anastasia Kolyva, Maria Diaz, Leonardo de Oliveira Martins, Nabil-Fareed Alikhan, Lizzie Meadows, Rachael Stanley, Ngozi Elumogo, Muhammed Yasir, Nicholas M. Thomson, Alexander J Trotter, Rachel Gilroy, Samuel Bloomfield, Claire Stuart, Andrew Bell, Reenesh Prakash, Samir Dervisevic, Alison E. Mather, John Wain, Mark Webber, Andrew J. Page, Justin O'Grady |
| EPI_ISL_703719 | West of Scotland Specialist Virology Centre, NHSGGC / MRC-University of Glasgow Centre for Virus Research | COVID-19 Genomics UK (COG-UK) Consortium | Ana da Silva Filipe, Natasha Johnson, Kathy Smollett, Daniel Mair, Stephen Carmichael, Alice Broos, Lily Tong, Jenna Nichols, Kyriaki Nomikou; Sarah McDonald; Richard Orton, Joseph Hughes, Sreenu Vattipally, David L Robertson; Alasdair MacLean, Rory Gunson; Sharif Shaaban, Matthew Holden; Rachel Blacow, Guy Mollett, Kathy Li, James Shepherd, Antonia Ho, Emma Thomson |
| EPI_ISL_703720 | Queens Medical Centre, Clinical Microbiology Department / DeepSeq Nottingham | COVID-19 Genomics UK (COG-UK) Consortium | Gemma Clark, Wendy Smith, Manjinder Khakh, Vicki M Fleming, Michelle M Lister, Hannah Howson-Wells, Jonathan Ball, Patrick McClure, Joseph Chappell, Theocharis Tsoleridis, Nadine Holmes, Matthew Carlisle, Christopher Moore, Fei Sang, Johnny Debebe, Victoria Wright, Matthew Loose |
| EPI_ISL_703727 | Northumbria University / South Tees Hospitals NHS Foundation Trust / North Cumbria Integrated Care NHS Foundation Trust / North Tees and Hartlepool NHS Foundation Trust / Newcastle Hospitals NHS Foundation Trust | COVID-19 Genomics UK (COG-UK) Consortium | Darren L Smith,Andrew Nelson,Matthew Bashton,Greg R Young,Joshua Loh,John Allan,Mohammad A Tariq,Giles S Holt,Gary Black,Wen C Yew,Lynn Dover,Paul Baker,Steve Liggett,Sarah Essex,Jane Greenaway,Debra Padgett,Clive Graham,Garren Scott,Edward Barton,Emma Swindells,Brendan Payne,Jennifer Collins,Yusri Taha,Gary Eltringham |
| EPI_ISL_703733 | Quadram Institute Bioscience | COVID-19 Genomics UK (COG-UK) Consortium | Dave J. Baker, Gemma L. Kay, Alp Aydin, Thanh Le-Viet, Steven Rudder, Ana P. Tedim, Anastasia Kolyva, Maria Diaz, Leonardo de Oliveira Martins, Nabil-Fareed Alikhan, Lizzie Meadows, Rachael Stanley, Ngozi Elumogo, Muhammed Yasir, Nicholas M. Thomson, Alexander J Trotter, Rachel Gilroy, Samuel Bloomfield, Claire Stuart, Andrew Bell, Reenesh Prakash, Samir Dervisevic, Alison E. Mather, John Wain, Mark Webber, Andrew J. Page, Justin O'Grady |
| EPI_ISL_703734, EPI_ISL_703735 | Liverpool Clinical Laboratories | COVID-19 Genomics UK (COG-UK) Consortium | Sam Haldenby, Anita Lucaci, Steve Paterson, Julian Hiscox, Alistair Darby, M Almsaud, A Alrezaihi, Muhannad Alruwaili, Stuart D Armstrong, Jones Benjamin, Eleanor G Bentley, Anu Chawla, Jordan J Clark, Angela Cowell, Richard Eccles, Isabel García-Dorival, Matthew Gemmell, Alessandro Gerada, PKF Gilmore, Richard Gregory, Ximeng Han, Catherine Hartley, Margaret Hughes, Miren Iturriza-Gomara, James Johnson, L Luu, Jenifer Manson, Charlotte Nelson, Elaine O'Toole, Cassie Olateju, Rebekah Penrice-Randal , Lucille Rainbow, N.P Randle, Trevor Ian Robinson, Parul Sharma, Ghada T Shawli, James P Stewart, Neil Swainston, Ecaterina Vamos, Joanne Watts, Mark Whitehead |
| EPI_ISL_703736 | Quadram Institute Bioscience | COVID-19 Genomics UK (COG-UK) Consortium | Dave J. Baker, Gemma L. Kay, Alp Aydin, Thanh Le-Viet, Steven Rudder, Ana P. Tedim, Anastasia Kolyva, Maria Diaz, Leonardo de Oliveira Martins, Nabil-Fareed Alikhan, Lizzie Meadows, Rachael Stanley, Ngozi Elumogo, Muhammed Yasir, Nicholas M. Thomson, Alexander J Trotter, Rachel Gilroy, Samuel Bloomfield, Claire Stuart, Andrew Bell, Reenesh Prakash, Samir Dervisevic, Alison E. Mather, John Wain, Mark Webber, Andrew J. Page, Justin O'Grady |
| EPI_ISL_703744 | Liverpool Clinical Laboratories | COVID-19 Genomics UK (COG-UK) Consortium | Sam Haldenby, Anita Lucaci, Steve Paterson, Julian Hiscox, Alistair Darby, M Almsaud, A Alrezaihi, Muhannad Alruwaili, Stuart D Armstrong, Jones Benjamin, Eleanor G Bentley, Anu Chawla, Jordan J Clark, Angela Cowell, Richard Eccles, Isabel García-Dorival, Matthew Gemmell, Alessandro Gerada, PKF Gilmore, Richard Gregory, Ximeng Han, Catherine Hartley, Margaret Hughes, Miren Iturriza-Gomara, James Johnson, L Luu, Jenifer Manson, Charlotte Nelson, Elaine O'Toole, Cassie Olateju, Rebekah Penrice-Randal , Lucille Rainbow, N.P Randle, Trevor Ian Robinson, Parul Sharma, Ghada T Shawli, James P Stewart, Neil Swainston, Ecaterina Vamos, Joanne Watts, Mark Whitehead |
| EPI_ISL_703753, EPI_ISL_703754, EPI_ISL_703757 | West of Scotland Specialist Virology Centre, NHSGGC / MRC-University of Glasgow Centre for Virus Research | COVID-19 Genomics UK (COG-UK) Consortium | Ana da Silva Filipe, Natasha Johnson, Kathy Smollett, Daniel Mair, Stephen Carmichael, Alice Broos, Lily Tong, Jenna Nichols, Kyriaki Nomikou; Sarah McDonald; Richard Orton, Joseph Hughes, Sreenu Vattipally, David L Robertson; Alasdair MacLean, Rory Gunson; Sharif Shaaban, Matthew Holden; Rachel Blacow, Guy Mollett, Kathy Li, James Shepherd, Antonia Ho, Emma Thomson |
| EPI_ISL_703763 | Liverpool Clinical Laboratories | COVID-19 Genomics UK (COG-UK) Consortium | Sam Haldenby, Anita Lucaci, Steve Paterson, Julian Hiscox, Alistair Darby, M Almsaud, A Alrezaihi, Muhannad Alruwaili, Stuart D Armstrong, Jones Benjamin, Eleanor G Bentley, Anu Chawla, Jordan J Clark, Angela Cowell, Richard Eccles, Isabel García-Dorival, Matthew Gemmell, Alessandro Gerada, PKF Gilmore, Richard Gregory, Ximeng Han, Catherine Hartley, Margaret Hughes, Miren Iturriza-Gomara, James Johnson, L Luu, Jenifer Manson, Charlotte Nelson, Elaine O'Toole, Cassie Olateju, Rebekah Penrice-Randal , Lucille Rainbow, N.P Randle, Trevor Ian Robinson, Parul Sharma, Ghada T Shawli, James P Stewart, Neil Swainston, Ecaterina Vamos, Joanne Watts, Mark Whitehead |
| EPI_ISL_703764 | Quadram Institute Bioscience | COVID-19 Genomics UK (COG-UK) Consortium | Dave J. Baker, Gemma L. Kay, Alp Aydin, Thanh Le-Viet, Steven Rudder, Ana P. Tedim, Anastasia Kolyva, Maria Diaz, Leonardo de Oliveira Martins, Nabil-Fareed Alikhan, Lizzie Meadows, Rachael Stanley, Ngozi Elumogo, Muhammed Yasir, Nicholas M. Thomson, Alexander J Trotter, Rachel Gilroy, Samuel Bloomfield, Claire Stuart, Andrew Bell, Reenesh Prakash, Samir Dervisevic, Alison E. Mather, John Wain, Mark Webber, Andrew J. Page, Justin O'Grady |
| EPI_ISL_703766 | Virology Department, Sheffield Teaching Hospitals NHS Foundation Trust/Department of Infection, Immunity and Cardiovascular Disease, The Medical School, University of Sheffield | COVID-19 Genomics UK (COG-UK) Consortium | Thushan de Silva, Matthew Parker, Nikki Smith, Adri Angyal, Rebecca Brown, Luke Green, Rachel Tucker, Paul Parsons, Danielle Groves, Katie Johnson, Laura Carrilero, Alex Keeley, Dave Partridge, Matthew Wyles, Benjamin Lindsey, Mehmet Yavuz, Mohammad Raza, Cariad Evans |
| EPI_ISL_703775, EPI_ISL_703776 | Queens Medical Centre, Clinical Microbiology Department / DeepSeq Nottingham | COVID-19 Genomics UK (COG-UK) Consortium | Gemma Clark, Wendy Smith, Manjinder Khakh, Vicki M Fleming, Michelle M Lister, Hannah Howson-Wells, Jonathan Ball, Patrick McClure, Joseph Chappell, Theocharis Tsoleridis, Nadine Holmes, Matthew Carlisle, Christopher Moore, Fei Sang, Johnny Debebe, Victoria Wright, Matthew Loose |
| EPI_ISL_703786 | Lighthouse Lab in Glasgow | Wellcome Sanger Institute for the COVID-19 Genomics UK (COG-UK) Consortium | Harper VanSteenhouse, Yumi Kasai, David Gray, Carol Clugston, Anna Dominiczak and Alex Alderton, Roberto Amato, Sonia Goncalves, Ewan Harrison, David K. Jackson, Ian Johnston, Dominic Kwiatkowski, Cordelia Langford, John Sillitoe on behalf of the Wellcome Sanger Institute COVID-19 Surveillance Team |
| EPI_ISL_703789 | Queens Medical Centre, Clinical Microbiology Department / DeepSeq Nottingham | COVID-19 Genomics UK (COG-UK) Consortium | Gemma Clark, Wendy Smith, Manjinder Khakh, Vicki M Fleming, Michelle M Lister, Hannah Howson-Wells, Jonathan Ball, Patrick McClure, Joseph Chappell, Theocharis Tsoleridis, Nadine Holmes, Matthew Carlisle, Christopher Moore, Fei Sang, Johnny Debebe, Victoria Wright, Matthew Loose |
| EPI_ISL_703790 | Lighthouse Lab in Glasgow | Wellcome Sanger Institute for the COVID-19 Genomics UK (COG-UK) Consortium | Harper VanSteenhouse, Yumi Kasai, David Gray, Carol Clugston, Anna Dominiczak and Alex Alderton, Roberto Amato, Sonia Goncalves, Ewan Harrison, David K. Jackson, Ian Johnston, Dominic Kwiatkowski, Cordelia Langford, John Sillitoe on behalf of the Wellcome Sanger Institute COVID-19 Surveillance Team |
| EPI_ISL_703792, EPI_ISL_703794 | Queens Medical Centre, Clinical Microbiology Department / DeepSeq Nottingham | COVID-19 Genomics UK (COG-UK) Consortium | Gemma Clark, Wendy Smith, Manjinder Khakh, Vicki M Fleming, Michelle M Lister, Hannah Howson-Wells, Jonathan Ball, Patrick McClure, Joseph Chappell, Theocharis Tsoleridis, Nadine Holmes, Matthew Carlisle, Christopher Moore, Fei Sang, Johnny Debebe, Victoria Wright, Matthew Loose |
| EPI_ISL_703799 | Lighthouse Lab in Glasgow | Wellcome Sanger Institute for the COVID-19 Genomics UK (COG-UK) Consortium | Harper VanSteenhouse, Yumi Kasai, David Gray, Carol Clugston, Anna Dominiczak and Alex Alderton, Roberto Amato, Sonia Goncalves, Ewan Harrison, David K. Jackson, Ian Johnston, Dominic Kwiatkowski, Cordelia Langford, John Sillitoe on behalf of the Wellcome Sanger Institute COVID-19 Surveillance Team |
| EPI_ISL_703800 | Quadram Institute Bioscience | COVID-19 Genomics UK (COG-UK) Consortium | Dave J. Baker, Gemma L. Kay, Alp Aydin, Thanh Le-Viet, Steven Rudder, Ana P. Tedim, Anastasia Kolyva, Maria Diaz, Leonardo de Oliveira Martins, Nabil-Fareed Alikhan, Lizzie Meadows, Rachael Stanley, Ngozi Elumogo, Muhammed Yasir, Nicholas M. Thomson, Alexander J Trotter, Rachel Gilroy, Samuel Bloomfield, Claire Stuart, Andrew Bell, Reenesh Prakash, Samir Dervisevic, Alison E. Mather, John Wain, Mark Webber, Andrew J. Page, Justin O'Grady |
| EPI_ISL_703801 | Lighthouse Lab in Glasgow | Wellcome Sanger Institute for the COVID-19 Genomics UK (COG-UK) Consortium | Harper VanSteenhouse, Yumi Kasai, David Gray, Carol Clugston, Anna Dominiczak and Alex Alderton, Roberto Amato, Sonia Goncalves, Ewan Harrison, David K. Jackson, Ian Johnston, Dominic Kwiatkowski, Cordelia Langford, John Sillitoe on behalf of the Wellcome Sanger Institute COVID-19 Surveillance Team |
| EPI_ISL_703802, EPI_ISL_703805 | Quadram Institute Bioscience | COVID-19 Genomics UK (COG-UK) Consortium | Dave J. Baker, Gemma L. Kay, Alp Aydin, Thanh Le-Viet, Steven Rudder, Ana P. Tedim, Anastasia Kolyva, Maria Diaz, Leonardo de Oliveira Martins, Nabil-Fareed Alikhan, Lizzie Meadows, Rachael Stanley, Ngozi Elumogo, Muhammed Yasir, Nicholas M. Thomson, Alexander J Trotter, Rachel Gilroy, Samuel Bloomfield, Claire Stuart, Andrew Bell, Reenesh Prakash, Samir Dervisevic, Alison E. Mather, John Wain, Mark Webber, Andrew J. Page, Justin O'Grady |
| EPI_ISL_703806 | Lighthouse Lab in Glasgow | Wellcome Sanger Institute for the COVID-19 Genomics UK (COG-UK) Consortium | Harper VanSteenhouse, Yumi Kasai, David Gray, Carol Clugston, Anna Dominiczak and Alex Alderton, Roberto Amato, Sonia Goncalves, Ewan Harrison, David K. Jackson, Ian Johnston, Dominic Kwiatkowski, Cordelia Langford, John Sillitoe on behalf of the Wellcome Sanger Institute COVID-19 Surveillance |

| | | | |
|---|---|---|---|
| | | | Team |
| EPI_ISL_703807 | Liverpool Clinical Laboratories | COVID-19 Genomics UK (COG-UK) Consortium | Sam Haldenby, Anita Lucaci, Steve Paterson, Julian Hiscox, Alistair Darby, M Almsaud, A Alrezaihi, Muhannad Alruwaili, Stuart D Armstrong, Jones Benjamin, Eleanor G Bentley, Anu Chawla, Jordan J Clark, Angela Cowell, Richard Eccles, Isabel García-Dorival, Matthew Gemmell, Alessandro Gerada, PKF Gilmore, Richard Gregory, Ximeng Han, Catherine Hartley, Margaret Hughes, Miren Iturriza-Gomara, James Johnson, L Luu, Jenifer Manson, Charlotte Nelson, Elaine O'Toole, Cassie Olateju, Rebekah Penrice-Randal , Lucille Rainbow, N.P Randle, Trevor Ian Robinson, Parul Sharma, Ghada T Shawli, James P Stewart, Neil Swainston, Ecaterina Vamos, Joanne Watts, Mark Whitehead |
| EPI_ISL_703808, EPI_ISL_703811 | Lighthouse Lab in Glasgow | Wellcome Sanger Institute for the COVID-19 Genomics UK (COG-UK) Consortium | Harper VanSteenhouse, Yumi Kasai, David Gray, Carol Clugston, Anna Dominiczak and Alex Alderton, Roberto Amato, Sonia Goncalves, Ewan Harrison, David K. Jackson, Ian Johnston, Dominic Kwiatkowski, Cordelia Langford, John Sillitoe on behalf of the Wellcome Sanger Institute COVID-19 Surveillance Team |
| EPI_ISL_703815 | Liverpool Clinical Laboratories | COVID-19 Genomics UK (COG-UK) Consortium | Sam Haldenby, Anita Lucaci, Steve Paterson, Julian Hiscox, Alistair Darby, M Almsaud, A Alrezaihi, Muhannad Alruwaili, Stuart D Armstrong, Jones Benjamin, Eleanor G Bentley, Anu Chawla, Jordan J Clark, Angela Cowell, Richard Eccles, Isabel García-Dorival, Matthew Gemmell, Alessandro Gerada, PKF Gilmore, Richard Gregory, Ximeng Han, Catherine Hartley, Margaret Hughes, Miren Iturriza-Gomara, James Johnson, L Luu, Jenifer Manson, Charlotte Nelson, Elaine O'Toole, Cassie Olateju, Rebekah Penrice-Randal , Lucille Rainbow, N.P Randle, Trevor Ian Robinson, Parul Sharma, Ghada T Shawli, James P Stewart, Neil Swainston, Ecaterina Vamos, Joanne Watts, Mark Whitehead |
| EPI_ISL_703816 | Lighthouse Lab in Glasgow | Wellcome Sanger Institute for the COVID-19 Genomics UK (COG-UK) Consortium | Harper VanSteenhouse, Yumi Kasai, David Gray, Carol Clugston, Anna Dominiczak and Alex Alderton, Roberto Amato, Sonia Goncalves, Ewan Harrison, David K. Jackson, Ian Johnston, Dominic Kwiatkowski, Cordelia Langford, John Sillitoe on behalf of the Wellcome Sanger Institute COVID-19 Surveillance Team |
| EPI_ISL_703818 | Liverpool Clinical Laboratories | COVID-19 Genomics UK (COG-UK) Consortium | Sam Haldenby, Anita Lucaci, Steve Paterson, Julian Hiscox, Alistair Darby, M Almsaud, A Alrezaihi, Muhannad Alruwaili, Stuart D Armstrong, Jones Benjamin, Eleanor G Bentley, Anu Chawla, Jordan J Clark, Angela Cowell, Richard Eccles, Isabel García-Dorival, Matthew Gemmell, Alessandro Gerada, PKF Gilmore, Richard Gregory, Ximeng Han, Catherine Hartley, Margaret Hughes, Miren Iturriza-Gomara, James Johnson, L Luu, Jenifer Manson, Charlotte Nelson, Elaine O'Toole, Cassie Olateju, Rebekah Penrice-Randal , Lucille Rainbow, N.P Randle, Trevor Ian Robinson, Parul Sharma, Ghada T Shawli, James P Stewart, Neil Swainston, Ecaterina Vamos, Joanne Watts, Mark Whitehead |
| EPI_ISL_703819, EPI_ISL_703820, EPI_ISL_703822, EPI_ISL_703824, EPI_ISL_703825, EPI_ISL_703827, EPI_ISL_703828, EPI_ISL_703830, EPI_ISL_703832, EPI_ISL_703833, EPI_ISL_703835, EPI_ISL_703836, EPI_ISL_703838, EPI_ISL_703840, EPI_ISL_703841, EPI_ISL_703843, EPI_ISL_703844 | | | |
| see above | Lighthouse Lab in Glasgow | Wellcome Sanger Institute for the COVID-19 Genomics UK (COG-UK) Consortium | Harper VanSteenhouse, Yumi Kasai, David Gray, Carol Clugston, Anna Dominiczak and Alex Alderton, Roberto Amato, Sonia Goncalves, Ewan Harrison, David K. Jackson, Ian Johnston, Dominic Kwiatkowski, Cordelia Langford, John Sillitoe on behalf of the Wellcome Sanger Institute COVID-19 Surveillance Team |
| EPI_ISL_703845 | West of Scotland Specialist Virology Centre, NHSGGC / MRC-University of Glasgow Centre for Virus Research | COVID-19 Genomics UK (COG-UK) Consortium | Ana da Silva Filipe, Natasha Johnson, Kathy Smollett, Daniel Mair, Stephen Carmichael, Alice Broos, Lily Tong, Jenna Nichols, Kyriaki Nomikou; Sarah McDonald; Richard Orton, Joseph Hughes, Sreenu Vattipally, David L Robertson; Alasdair MacLean, Rory Gunson; Sharif Shaaban, Matthew Holden; Rachel Blacow, Guy Mollett, Kathy Li, James Shepherd, Antonia Ho, Emma Thomson |
| EPI_ISL_703846 | Lighthouse Lab in Glasgow | Wellcome Sanger Institute for the COVID-19 Genomics UK (COG-UK) Consortium | Harper VanSteenhouse, Yumi Kasai, David Gray, Carol Clugston, Anna Dominiczak and Alex Alderton, Roberto Amato, Sonia Goncalves, Ewan Harrison, David K. Jackson, Ian Johnston, Dominic Kwiatkowski, Cordelia Langford, John Sillitoe on behalf of the Wellcome Sanger Institute COVID-19 Surveillance Team |
| EPI_ISL_703847 | Liverpool Clinical Laboratories | COVID-19 Genomics UK (COG-UK) Consortium | Sam Haldenby, Anita Lucaci, Steve Paterson, Julian Hiscox, Alistair Darby, M Almsaud, A Alrezaihi, Muhannad Alruwaili, Stuart D Armstrong, Jones Benjamin, Eleanor G Bentley, Anu Chawla, Jordan J Clark, Angela Cowell, Richard Eccles, Isabel García-Dorival, Matthew Gemmell, Alessandro Gerada, PKF Gilmore, Richard Gregory, Ximeng Han, Catherine Hartley, Margaret Hughes, Miren Iturriza-Gomara, James Johnson, L Luu, Jenifer Manson, Charlotte Nelson, Elaine O'Toole, Cassie Olateju, Rebekah Penrice-Randal , Lucille Rainbow, N.P Randle, Trevor Ian Robinson, Parul Sharma, Ghada T Shawli, James P Stewart, Neil Swainston, Ecaterina Vamos, Joanne Watts, Mark Whitehead |
| EPI_ISL_703848, EPI_ISL_703849 | Lighthouse Lab in Glasgow | Wellcome Sanger Institute for the COVID-19 Genomics UK (COG-UK) Consortium | Harper VanSteenhouse, Yumi Kasai, David Gray, Carol Clugston, Anna Dominiczak and Alex Alderton, Roberto Amato, Sonia Goncalves, Ewan Harrison, David K. Jackson, Ian Johnston, Dominic Kwiatkowski, Cordelia Langford, John Sillitoe on behalf of the Wellcome Sanger Institute COVID-19 Surveillance Team |
| EPI_ISL_703850 | Liverpool Clinical Laboratories | COVID-19 Genomics UK (COG-UK) Consortium | Sam Haldenby, Anita Lucaci, Steve Paterson, Julian Hiscox, Alistair Darby, M Almsaud, A Alrezaihi, Muhannad Alruwaili, Stuart D Armstrong, Jones Benjamin, Eleanor G Bentley, Anu Chawla, Jordan J Clark, Angela Cowell, Richard Eccles, Isabel García-Dorival, Matthew Gemmell, Alessandro Gerada, PKF Gilmore, Richard Gregory, Ximeng Han, Catherine Hartley, Margaret Hughes, Miren Iturriza-Gomara, James Johnson, L Luu, Jenifer Manson, Charlotte Nelson, Elaine O'Toole, Cassie Olateju, Rebekah Penrice-Randal , Lucille Rainbow, N.P Randle, Trevor Ian Robinson, Parul Sharma, Ghada T Shawli, James P Stewart, Neil Swainston, Ecaterina Vamos, Joanne Watts, Mark Whitehead |
| EPI_ISL_703851, EPI_ISL_703852 | Lighthouse Lab in Glasgow | Wellcome Sanger Institute for the COVID-19 Genomics UK (COG-UK) Consortium | Harper VanSteenhouse, Yumi Kasai, David Gray, Carol Clugston, Anna Dominiczak and Alex Alderton, Roberto Amato, Sonia Goncalves, Ewan Harrison, David K. Jackson, Ian Johnston, Dominic Kwiatkowski, Cordelia Langford, John Sillitoe on behalf of the Wellcome Sanger Institute COVID-19 Surveillance Team |
| EPI_ISL_703853 | Queens Medical Centre, Clinical Microbiology Department / DeepSeq Nottingham | COVID-19 Genomics UK (COG-UK) Consortium | Gemma Clark, Wendy Smith, Manjinder Khakh, Vicki M Fleming, Michelle M Lister, Hannah Howson-Wells, Jonathan Ball, Patrick McClure, Joseph Chappell, Theocharis Tsoleridis, Nadine Holmes, Matthew Carlisle, Christopher Moore, Fei Sang, Johnny Debebe, Victoria Wright, Matthew Loose |
| EPI_ISL_703854, EPI_ISL_703856, EPI_ISL_703857, EPI_ISL_703859, EPI_ISL_703861, EPI_ISL_703862, EPI_ISL_703864, EPI_ISL_703866, EPI_ISL_703867, EPI_ISL_703869, EPI_ISL_703871, EPI_ISL_703872, EPI_ISL_703874, EPI_ISL_703875, EPI_ISL_703877, EPI_ISL_703878, EPI_ISL_703880, EPI_ISL_703882, EPI_ISL_703883, EPI_ISL_703885, EPI_ISL_703887, EPI_ISL_703889, EPI_ISL_703890, EPI_ISL_703892, EPI_ISL_703893 | | | |
| see above | Lighthouse Lab in Glasgow | Wellcome Sanger Institute for the COVID-19 Genomics UK (COG-UK) Consortium | Harper VanSteenhouse, Yumi Kasai, David Gray, Carol Clugston, Anna Dominiczak and Alex Alderton, Roberto Amato, Sonia Goncalves, Ewan Harrison, David K. Jackson, Ian Johnston, Dominic Kwiatkowski, Cordelia Langford, John Sillitoe on behalf of the Wellcome Sanger Institute COVID-19 Surveillance Team |
| EPI_ISL_703894 | Virology Department, Sheffield Teaching Hospitals NHS Foundation Trust/Department of Infection, Immunity and Cardiovascular Disease, The Medical School, University of Sheffield | COVID-19 Genomics UK (COG-UK) Consortium | Thushan de Silva, Matthew Parker, Nikki Smith, Adri Angyal, Rebecca Brown, Luke Green, Rachel Tucker, Paul Parsons, Danielle Groves, Katie Johnson, Laura Carrilero, Alex Keeley, Dave Partridge, Matthew Wyles, Benjamin Lindsey, Mehmet Yavuz, Mohammad Raza, Cariad Evans |
| EPI_ISL_703895, EPI_ISL_703896, EPI_ISL_703898, EPI_ISL_703900, EPI_ISL_703901, EPI_ISL_703903, EPI_ISL_703904 | Lighthouse Lab in Glasgow | Wellcome Sanger Institute for the COVID-19 Genomics UK (COG-UK) Consortium | Harper VanSteenhouse, Yumi Kasai, David Gray, Carol Clugston, Anna Dominiczak and Alex Alderton, Roberto Amato, Sonia Goncalves, Ewan Harrison, David K. Jackson, Ian Johnston, Dominic Kwiatkowski, Cordelia Langford, John Sillitoe on behalf of the Wellcome Sanger Institute COVID-19 Surveillance Team |
| EPI_ISL_703905 | Quadram Institute Bioscience | COVID-19 Genomics UK (COG-UK) Consortium | Dave J. Baker, Gemma L. Kay, Alp Aydin, Thanh Le-Viet, Steven Rudder, Ana P. Tedim, Anastasia Kolyva, Maria Diaz, Leonardo de Oliveira Martins, Nabil-Fareed Alikhan, Lizzie Meadows, Rachael Stanley, Ngozi Elumogo, Muhammed Yasir, Nicholas M. Thomson, Alexander J Trotter, Rachel Gilroy, Samuel Bloomfield, Claire Stuart, Andrew Bell, Reenesh Prakash, Samir Dervisevic, Alison E. Mather, John Wain, Mark Webber, Andrew J. Page, Justin O'Grady |
| EPI_ISL_703906, EPI_ISL_703907, EPI_ISL_703909, EPI_ISL_703910, EPI_ISL_703912, EPI_ISL_703914, EPI_ISL_703915 | Lighthouse Lab in Glasgow | Wellcome Sanger Institute for the COVID-19 Genomics UK (COG-UK) Consortium | Harper VanSteenhouse, Yumi Kasai, David Gray, Carol Clugston, Anna Dominiczak and Alex Alderton, Roberto Amato, Sonia Goncalves, Ewan Harrison, David K. Jackson, Ian Johnston, Dominic Kwiatkowski, Cordelia Langford, John Sillitoe on behalf of the Wellcome Sanger Institute COVID-19 Surveillance Team |
| EPI_ISL_703916 | Liverpool Clinical Laboratories | COVID-19 Genomics UK (COG-UK) Consortium | Sam Haldenby, Anita Lucaci, Steve Paterson, Julian Hiscox, Alistair Darby, M Almsaud, A Alrezaihi, Muhannad Alruwaili, Stuart D Armstrong, Jones Benjamin, Eleanor G Bentley, Anu Chawla, Jordan J Clark, Angela Cowell, Richard Eccles, Isabel García-Dorival, Matthew Gemmell, Alessandro Gerada, PKF Gilmore, Richard Gregory, Ximeng Han, Catherine Hartley, Margaret Hughes, Miren Iturriza-Gomara, James Johnson, L Luu, Jenifer Manson, Charlotte Nelson, Elaine O'Toole, Cassie Olateju, Rebekah Penrice-Randal , Lucille Rainbow, N.P Randle, Trevor Ian Robinson, Parul Sharma, Ghada T Shawli, James P Stewart, Neil Swainston, Ecaterina Vamos, Joanne Watts, Mark Whitehead |

| | | | |
|---|---|---|---|
| EPI_ISL_703917, EPI_ISL_703919, EPI_ISL_703920 | Lighthouse Lab in Glasgow | Wellcome Sanger Institute for the COVID-19 Genomics UK (COG-UK) Consortium | Harper VanSteenhouse, Yumi Kasai, David Gray, Carol Clugston, Anna Dominiczak and Alex Alderton, Roberto Amato, Sonia Goncalves, Ewan Harrison, David K. Jackson, Ian Johnston, Dominic Kwiatkowski, Cordelia Langford, John Sillitoe on behalf of the Wellcome Sanger Institute COVID-19 Surveillance Team |
| EPI_ISL_703921 | West of Scotland Specialist Virology Centre, NHSGGC / MRC-University of Glasgow Centre for Virus Research | COVID-19 Genomics UK (COG-UK) Consortium | Ana da Silva Filipe, Natasha Johnson, Kathy Smollett, Daniel Mair, Stephen Carmichael, Alice Broos, Lily Tong, Jenna Nichols, Kyriaki Nomikou; Sarah McDonald; Richard Orton, Joseph Hughes, Sreenu Vattipally, David L Robertson; Alasdair MacLean, Rory Gunson; Sharif Shaaban, Matthew Holden; Rachel Blacow, Guy Mollett, Kathy Li, James Shepherd, Antonia Ho, Emma Thomson |
| EPI_ISL_703922, EPI_ISL_703923 | Lighthouse Lab in Glasgow | Wellcome Sanger Institute for the COVID-19 Genomics UK (COG-UK) Consortium | Harper VanSteenhouse, Yumi Kasai, David Gray, Carol Clugston, Anna Dominiczak and Alex Alderton, Roberto Amato, Sonia Goncalves, Ewan Harrison, David K. Jackson, Ian Johnston, Dominic Kwiatkowski, Cordelia Langford, John Sillitoe on behalf of the Wellcome Sanger Institute COVID-19 Surveillance Team |
| EPI_ISL_703924 | West of Scotland Specialist Virology Centre, NHSGGC / MRC-University of Glasgow Centre for Virus Research | COVID-19 Genomics UK (COG-UK) Consortium | Ana da Silva Filipe, Natasha Johnson, Kathy Smollett, Daniel Mair, Stephen Carmichael, Alice Broos, Lily Tong, Jenna Nichols, Kyriaki Nomikou; Sarah McDonald; Richard Orton, Joseph Hughes, Sreenu Vattipally, David L Robertson; Alasdair MacLean, Rory Gunson; Sharif Shaaban, Matthew Holden; Rachel Blacow, Guy Mollett, Kathy Li, James Shepherd, Antonia Ho, Emma Thomson |
| EPI_ISL_703925, EPI_ISL_703926, EPI_ISL_703928, EPI_ISL_703930, EPI_ISL_703931 | Lighthouse Lab in Glasgow | Wellcome Sanger Institute for the COVID-19 Genomics UK (COG-UK) Consortium | Harper VanSteenhouse, Yumi Kasai, David Gray, Carol Clugston, Anna Dominiczak and Alex Alderton, Roberto Amato, Sonia Goncalves, Ewan Harrison, David K. Jackson, Ian Johnston, Dominic Kwiatkowski, Cordelia Langford, John Sillitoe on behalf of the Wellcome Sanger Institute COVID-19 Surveillance Team |
| EPI_ISL_703932 | Quadram Institute Bioscience | COVID-19 Genomics UK (COG-UK) Consortium | Dave J. Baker, Gemma L. Kay, Alp Aydin, Thanh Le-Viet, Steven Rudder, Ana P. Tedim, Anastasia Kolyva, Maria Diaz, Leonardo de Oliveira Martins, Nabil-Fareed Alikhan, Lizzie Meadows, Rachael Stanley, Ngozi Elumogo, Muhammed Yasir, Nicholas M. Thomson, Alexander J Trotter, Rachel Gilroy, Samuel Bloomfield, Claire Stuart, Andrew Bell, Reenesh Prakash, Samir Dervisevic, Alison E. Mather, John Wain, Mark Webber, Andrew J. Page, Justin O'Grady |
| EPI_ISL_703933, EPI_ISL_703934, EPI_ISL_703936, EPI_ISL_703938, EPI_ISL_703939, EPI_ISL_703941, EPI_ISL_703943, EPI_ISL_703944 | Lighthouse Lab in Glasgow | Wellcome Sanger Institute for the COVID-19 Genomics UK (COG-UK) Consortium | Harper VanSteenhouse, Yumi Kasai, David Gray, Carol Clugston, Anna Dominiczak and Alex Alderton, Roberto Amato, Sonia Goncalves, Ewan Harrison, David K. Jackson, Ian Johnston, Dominic Kwiatkowski, Cordelia Langford, John Sillitoe on behalf of the Wellcome Sanger Institute COVID-19 Surveillance Team |
| EPI_ISL_703945 | Quadram Institute Bioscience | COVID-19 Genomics UK (COG-UK) Consortium | Dave J. Baker, Gemma L. Kay, Alp Aydin, Thanh Le-Viet, Steven Rudder, Ana P. Tedim, Anastasia Kolyva, Maria Diaz, Leonardo de Oliveira Martins, Nabil-Fareed Alikhan, Lizzie Meadows, Rachael Stanley, Ngozi Elumogo, Muhammed Yasir, Nicholas M. Thomson, Alexander J Trotter, Rachel Gilroy, Samuel Bloomfield, Claire Stuart, Andrew Bell, Reenesh Prakash, Samir Dervisevic, Alison E. Mather, John Wain, Mark Webber, Andrew J. Page, Justin O'Grady |
| EPI_ISL_703946 | Lighthouse Lab in Glasgow | Wellcome Sanger Institute for the COVID-19 Genomics UK (COG-UK) Consortium | Harper VanSteenhouse, Yumi Kasai, David Gray, Carol Clugston, Anna Dominiczak and Alex Alderton, Roberto Amato, Sonia Goncalves, Ewan Harrison, David K. Jackson, Ian Johnston, Dominic Kwiatkowski, Cordelia Langford, John Sillitoe on behalf of the Wellcome Sanger Institute COVID-19 Surveillance Team |
| EPI_ISL_703947 | Quadram Institute Bioscience | COVID-19 Genomics UK (COG-UK) Consortium | Dave J. Baker, Gemma L. Kay, Alp Aydin, Thanh Le-Viet, Steven Rudder, Ana P. Tedim, Anastasia Kolyva, Maria Diaz, Leonardo de Oliveira Martins, Nabil-Fareed Alikhan, Lizzie Meadows, Rachael Stanley, Ngozi Elumogo, Muhammed Yasir, Nicholas M. Thomson, Alexander J Trotter, Rachel Gilroy, Samuel Bloomfield, Claire Stuart, Andrew Bell, Reenesh Prakash, Samir Dervisevic, Alison E. Mather, John Wain, Mark Webber, Andrew J. Page, Justin O'Grady |
| EPI_ISL_703948, EPI_ISL_703949, EPI_ISL_703951, EPI_ISL_703953, EPI_ISL_703954, EPI_ISL_703956, EPI_ISL_703957, EPI_ISL_703959, EPI_ISL_703960, EPI_ISL_703962 | Lighthouse Lab in Glasgow | Wellcome Sanger Institute for the COVID-19 Genomics UK (COG-UK) Consortium | Harper VanSteenhouse, Yumi Kasai, David Gray, Carol Clugston, Anna Dominiczak and Alex Alderton, Roberto Amato, Sonia Goncalves, Ewan Harrison, David K. Jackson, Ian Johnston, Dominic Kwiatkowski, Cordelia Langford, John Sillitoe on behalf of the Wellcome Sanger Institute COVID-19 Surveillance Team |
| EPI_ISL_703963 | Quadram Institute Bioscience | COVID-19 Genomics UK (COG-UK) Consortium | Dave J. Baker, Gemma L. Kay, Alp Aydin, Thanh Le-Viet, Steven Rudder, Ana P. Tedim, Anastasia Kolyva, Maria Diaz, Leonardo de Oliveira Martins, Nabil-Fareed Alikhan, Lizzie Meadows, Rachael Stanley, Ngozi Elumogo, Muhammed Yasir, Nicholas M. Thomson, Alexander J Trotter, Rachel Gilroy, Samuel Bloomfield, Claire Stuart, Andrew Bell, Reenesh Prakash, Samir Dervisevic, Alison E. Mather, John Wain, Mark Webber, Andrew J. Page, Justin O'Grady |
| EPI_ISL_703964, EPI_ISL_703965 | Lighthouse Lab in Glasgow | Wellcome Sanger Institute for the COVID-19 Genomics UK (COG-UK) Consortium | Harper VanSteenhouse, Yumi Kasai, David Gray, Carol Clugston, Anna Dominiczak and Alex Alderton, Roberto Amato, Sonia Goncalves, Ewan Harrison, David K. Jackson, Ian Johnston, Dominic Kwiatkowski, Cordelia Langford, John Sillitoe on behalf of the Wellcome Sanger Institute COVID-19 Surveillance Team |
| EPI_ISL_703966 | University College London Hospital | COVID-19 Genomics UK (COG-UK) Consortium | Judith Heaney, Matthew Byott, Catherine Houlihan, Dan Frampton, Stuart Kirk, Moira Spyer and Eleni Nastouli |
| EPI_ISL_703967 | Lighthouse Lab in Glasgow | Wellcome Sanger Institute for the COVID-19 Genomics UK (COG-UK) Consortium | Harper VanSteenhouse, Yumi Kasai, David Gray, Carol Clugston, Anna Dominiczak and Alex Alderton, Roberto Amato, Sonia Goncalves, Ewan Harrison, David K. Jackson, Ian Johnston, Dominic Kwiatkowski, Cordelia Langford, John Sillitoe on behalf of the Wellcome Sanger Institute COVID-19 Surveillance Team |
| EPI_ISL_703968 | West of Scotland Specialist Virology Centre, NHSGGC / MRC-University of Glasgow Centre for Virus Research | COVID-19 Genomics UK (COG-UK) Consortium | Ana da Silva Filipe, Natasha Johnson, Kathy Smollett, Daniel Mair, Stephen Carmichael, Alice Broos, Lily Tong, Jenna Nichols, Kyriaki Nomikou; Sarah McDonald; Richard Orton, Joseph Hughes, Sreenu Vattipally, David L Robertson; Alasdair MacLean, Rory Gunson; Sharif Shaaban, Matthew Holden; Rachel Blacow, Guy Mollett, Kathy Li, James Shepherd, Antonia Ho, Emma Thomson |
| EPI_ISL_703969, EPI_ISL_703970 | Lighthouse Lab in Glasgow | Wellcome Sanger Institute for the COVID-19 Genomics UK (COG-UK) Consortium | Harper VanSteenhouse, Yumi Kasai, David Gray, Carol Clugston, Anna Dominiczak and Alex Alderton, Roberto Amato, Sonia Goncalves, Ewan Harrison, David K. Jackson, Ian Johnston, Dominic Kwiatkowski, Cordelia Langford, John Sillitoe on behalf of the Wellcome Sanger Institute COVID-19 Surveillance Team |
| EPI_ISL_703971 | Liverpool Clinical Laboratories | COVID-19 Genomics UK (COG-UK) Consortium | Sam Haldenby, Anita Lucaci, Steve Paterson, Julian Hiscox, Alistair Darby, M Almsaud, A Alrezaihi, Muhannad Alruwaili, Stuart D Armstrong, Jones Benjamin, Eleanor G Bentley, Anu Chawla, Jordan J Clark, Angela Cowell, Richard Eccles, Isabel García-Dorival, Matthew Gemmell, Alessandro Gerada, PKF Gilmore, Richard Gregory, Ximeng Han, Catherine Hartley, Margaret Hughes, Miren Iturriza-Gomara, James Johnson, L Luu, Jenifer Manson, Charlotte Nelson, Elaine O'Toole, Cassie Olateju, Rebekah Penrice-Randal , Lucille Rainbow, N.P Randle, Trevor Ian Robinson, Parul Sharma, Ghada T Shawli, James P Stewart, Neil Swainston, Ecaterina Vamos, Joanne Watts, Mark Whitehead |
| EPI_ISL_703972, EPI_ISL_703973 | Lighthouse Lab in Glasgow | Wellcome Sanger Institute for the COVID-19 Genomics UK (COG-UK) Consortium | Harper VanSteenhouse, Yumi Kasai, David Gray, Carol Clugston, Anna Dominiczak and Alex Alderton, Roberto Amato, Sonia Goncalves, Ewan Harrison, David K. Jackson, Ian Johnston, Dominic Kwiatkowski, Cordelia Langford, John Sillitoe on behalf of the Wellcome Sanger Institute COVID-19 Surveillance Team |
| EPI_ISL_703974 | Quadram Institute Bioscience | COVID-19 Genomics UK (COG-UK) Consortium | Dave J. Baker, Gemma L. Kay, Alp Aydin, Thanh Le-Viet, Steven Rudder, Ana P. Tedim, Anastasia Kolyva, Maria Diaz, Leonardo de Oliveira Martins, Nabil-Fareed Alikhan, Lizzie Meadows, Rachael Stanley, Ngozi Elumogo, Muhammed Yasir, Nicholas M. Thomson, Alexander J Trotter, Rachel Gilroy, Samuel Bloomfield, Claire Stuart, Andrew Bell, Reenesh Prakash, Samir Dervisevic, Alison E. Mather, John Wain, Mark Webber, Andrew J. Page, Justin O'Grady |
| EPI_ISL_703975, EPI_ISL_703976, EPI_ISL_703978 | Lighthouse Lab in Glasgow | Wellcome Sanger Institute for the COVID-19 Genomics UK (COG-UK) Consortium | Harper VanSteenhouse, Yumi Kasai, David Gray, Carol Clugston, Anna Dominiczak and Alex Alderton, Roberto Amato, Sonia Goncalves, Ewan Harrison, David K. Jackson, Ian Johnston, Dominic Kwiatkowski, Cordelia Langford, John Sillitoe on behalf of the Wellcome Sanger Institute COVID-19 Surveillance Team |
| EPI_ISL_703979 | Quadram Institute Bioscience | COVID-19 Genomics UK (COG-UK) Consortium | Dave J. Baker, Gemma L. Kay, Alp Aydin, Thanh Le-Viet, Steven Rudder, Ana P. Tedim, Anastasia Kolyva, Maria Diaz, Leonardo de Oliveira Martins, Nabil-Fareed Alikhan, Lizzie Meadows, Rachael Stanley, Ngozi Elumogo, Muhammed Yasir, Nicholas M. Thomson, Alexander J Trotter, Rachel Gilroy, Samuel Bloomfield, Claire Stuart, Andrew Bell, Reenesh Prakash, Samir Dervisevic, Alison E. Mather, John Wain, Mark Webber, Andrew J. Page, Justin O'Grady |

EPI_ISL_703980, EPI_ISL_703981, EPI_ISL_703983, EPI_ISL_703984, EPI_ISL_703986, EPI_ISL_703988, EPI_ISL_703989, EPI_ISL_703991, EPI_ISL_703993, EPI_ISL_703994, EPI_ISL_703996, EPI_ISL_703997

| | | | |
|---|---|---|---|
| see above | Lighthouse Lab in Glasgow | Wellcome Sanger Institute for the COVID-19 Genomics UK (COG-UK) Consortium | Harper VanSteenhouse, Yumi Kasai, David Gray, Carol Clugston, Anna Dominiczak and Alex Alderton, Roberto Amato, Sonia Goncalves, Ewan Harrison, David K. Jackson, Ian Johnston, Dominic Kwiatkowski, Cordelia Langford, John Sillitoe on behalf of the Wellcome Sanger Institute COVID-19 Surveillance Team |
| EPI_ISL_703998 | Quadram Institute Bioscience | COVID-19 Genomics UK (COG-UK) Consortium | Dave J. Baker, Gemma L. Kay, Alp Aydin, Thanh Le-Viet, Steven Rudder, Ana P. Tedim, Anastasia Kolyva, Maria Diaz, Leonardo de Oliveira Martins, Nabil-Fareed Alikhan, Lizzie Meadows, Rachael Stanley, Ngozi Elumogo, Muhammed Yasir, Nicholas M. Thomson, Alexander J Trotter, Rachel Gilroy, Samuel Bloomfield, Claire Stuart, Andrew Bell, Reenesh Prakash, Samir Dervisevic, Alison E. Mather, John Wain, Mark Webber, Andrew J. Page, Justin O'Grady |
| EPI_ISL_703999, EPI_ISL_704000, EPI_ISL_704002 | Lighthouse Lab in Glasgow | Wellcome Sanger Institute for the COVID-19 Genomics UK (COG-UK) Consortium | Harper VanSteenhouse, Yumi Kasai, David Gray, Carol Clugston, Anna Dominiczak and Alex Alderton, Roberto Amato, Sonia Goncalves, Ewan Harrison, David K. Jackson, Ian Johnston, Dominic Kwiatkowski, Cordelia Langford, John Sillitoe on behalf of the Wellcome Sanger Institute COVID-19 Surveillance Team |
| EPI_ISL_704003 | Liverpool Clinical Laboratories | COVID-19 Genomics UK (COG-UK) Consortium | Sam Haldenby, Anita Lucaci, Steve Paterson, Julian Hiscox, Alistair Darby, M Almsaud, A Alrezaihi, Muhannad Alruwaili, Stuart D Armstrong, Jones Benjamin, Eleanor G Bentley, Anu Chawla, Jordan J Clark, Angela Cowell, Richard Eccles, Isabel García-Dorival, Matthew Gemmell, Alessandro Gerada, PKF Gilmore, Richard Gregory, Ximeng Han, Catherine Hartley, Margaret Hughes, Miren Iturriza-Gomara, James Johnson, L Luu, Jenifer Manson, Charlotte Nelson, Elaine O'Toole, Cassie Olateju, Rebekah Penrice-Randal , Lucille Rainbow, N.P Randle, Trevor Ian Robinson, Parul Sharma, Ghada T Shawli, James P Stewart, Neil Swainston, Ecaterina Vamos, Joanne Watts, Mark Whitehead |
| EPI_ISL_704004, EPI_ISL_704005, EPI_ISL_704007, EPI_ISL_704008 | Lighthouse Lab in Glasgow | Wellcome Sanger Institute for the COVID-19 Genomics UK (COG-UK) Consortium | Harper VanSteenhouse, Yumi Kasai, David Gray, Carol Clugston, Anna Dominiczak and Alex Alderton, Roberto Amato, Sonia Goncalves, Ewan Harrison, David K. Jackson, Ian Johnston, Dominic Kwiatkowski, Cordelia Langford, John Sillitoe on behalf of the Wellcome Sanger Institute COVID-19 Surveillance Team |
| EPI_ISL_704009 | Liverpool Clinical Laboratories | COVID-19 Genomics UK (COG-UK) Consortium | Sam Haldenby, Anita Lucaci, Steve Paterson, Julian Hiscox, Alistair Darby, M Almsaud, A Alrezaihi, Muhannad Alruwaili, Stuart D Armstrong, Jones Benjamin, Eleanor G Bentley, Anu Chawla, Jordan J Clark, Angela Cowell, Richard Eccles, Isabel García-Dorival, Matthew Gemmell, Alessandro Gerada, PKF Gilmore, Richard Gregory, Ximeng Han, Catherine Hartley, Margaret Hughes, Miren Iturriza-Gomara, James Johnson, L Luu, Jenifer Manson, Charlotte Nelson, Elaine O'Toole, Cassie Olateju, Rebekah Penrice-Randal , Lucille Rainbow, N.P Randle, Trevor Ian Robinson, Parul Sharma, Ghada T Shawli, James P Stewart, Neil Swainston, Ecaterina Vamos, Joanne Watts, Mark Whitehead |
| EPI_ISL_704010, EPI_ISL_704011, EPI_ISL_704013, EPI_ISL_704015, EPI_ISL_704016, EPI_ISL_704018, EPI_ISL_704019, EPI_ISL_704021, EPI_ISL_704023, EPI_ISL_704024, EPI_ISL_704026, EPI_ISL_704027, EPI_ISL_704029, EPI_ISL_704031, EPI_ISL_704032, EPI_ISL_704034, EPI_ISL_704035, EPI_ISL_704037, EPI_ISL_704039, EPI_ISL_704040, EPI_ISL_704042, EPI_ISL_704043, EPI_ISL_704045, EPI_ISL_704047, EPI_ISL_704048, EPI_ISL_704050, EPI_ISL_704051 | | | |
| see above | Lighthouse Lab in Glasgow | Wellcome Sanger Institute for the COVID-19 Genomics UK (COG-UK) Consortium | Harper VanSteenhouse, Yumi Kasai, David Gray, Carol Clugston, Anna Dominiczak and Alex Alderton, Roberto Amato, Sonia Goncalves, Ewan Harrison, David K. Jackson, Ian Johnston, Dominic Kwiatkowski, Cordelia Langford, John Sillitoe on behalf of the Wellcome Sanger Institute COVID-19 Surveillance Team |
| EPI_ISL_704052 | University College London Hospital | COVID-19 Genomics UK (COG-UK) Consortium | Judith Heaney, Matthew Byott, Catherine Houlihan, Dan Frampton, Stuart Kirk, Moira Spyer and Eleni Nastouli |
| EPI_ISL_704053 | Lighthouse Lab in Glasgow | Wellcome Sanger Institute for the COVID-19 Genomics UK (COG-UK) Consortium | Harper VanSteenhouse, Yumi Kasai, David Gray, Carol Clugston, Anna Dominiczak and Alex Alderton, Roberto Amato, Sonia Goncalves, Ewan Harrison, David K. Jackson, Ian Johnston, Dominic Kwiatkowski, Cordelia Langford, John Sillitoe on behalf of the Wellcome Sanger Institute COVID-19 Surveillance Team |
| EPI_ISL_704054 | Quadram Institute Bioscience | COVID-19 Genomics UK (COG-UK) Consortium | Dave J. Baker, Gemma L. Kay, Alp Aydin, Thanh Le-Viet, Steven Rudder, Ana P. Tedim, Anastasia Kolyva, Maria Diaz, Leonardo de Oliveira Martins, Nabil-Fareed Alikhan, Lizzie Meadows, Rachael Stanley, Ngozi Elumogo, Muhammed Yasir, Nicholas M. Thomson, Alexander J Trotter, Rachel Gilroy, Samuel Bloomfield, Claire Stuart, Andrew Bell, Reenesh Prakash, Samir Dervisevic, Alison E. Mather, John Wain, Mark Webber, Andrew J. Page, Justin O'Grady |
| EPI_ISL_704055, EPI_ISL_704056, EPI_ISL_704058, EPI_ISL_704059, EPI_ISL_704061, EPI_ISL_704063, EPI_ISL_704064, EPI_ISL_704066, EPI_ISL_704067, EPI_ISL_704069, EPI_ISL_704071, EPI_ISL_704072 | | | |
| see above | Lighthouse Lab in Glasgow | Wellcome Sanger Institute for the COVID-19 Genomics UK (COG-UK) Consortium | Harper VanSteenhouse, Yumi Kasai, David Gray, Carol Clugston, Anna Dominiczak and Alex Alderton, Roberto Amato, Sonia Goncalves, Ewan Harrison, David K. Jackson, Ian Johnston, Dominic Kwiatkowski, Cordelia Langford, John Sillitoe on behalf of the Wellcome Sanger Institute COVID-19 Surveillance Team |
| EPI_ISL_704073 | Quadram Institute Bioscience | COVID-19 Genomics UK (COG-UK) Consortium | Dave J. Baker, Gemma L. Kay, Alp Aydin, Thanh Le-Viet, Steven Rudder, Ana P. Tedim, Anastasia Kolyva, Maria Diaz, Leonardo de Oliveira Martins, Nabil-Fareed Alikhan, Lizzie Meadows, Rachael Stanley, Ngozi Elumogo, Muhammed Yasir, Nicholas M. Thomson, Alexander J Trotter, Rachel Gilroy, Samuel Bloomfield, Claire Stuart, Andrew Bell, Reenesh Prakash, Samir Dervisevic, Alison E. Mather, John Wain, Mark Webber, Andrew J. Page, Justin O'Grady |
| EPI_ISL_704074, EPI_ISL_704075, EPI_ISL_704077, EPI_ISL_704078, EPI_ISL_704080, EPI_ISL_704082, EPI_ISL_704083, EPI_ISL_704085, EPI_ISL_704087, EPI_ISL_704088, EPI_ISL_704090, EPI_ISL_704091, EPI_ISL_704093, EPI_ISL_704094, EPI_ISL_704096, EPI_ISL_704098, EPI_ISL_704099, EPI_ISL_704101, EPI_ISL_704103, EPI_ISL_704104, EPI_ISL_704106, EPI_ISL_704107, EPI_ISL_704109, EPI_ISL_704110, EPI_ISL_704113, EPI_ISL_704114, EPI_ISL_704116, EPI_ISL_704117 | | | |
| see above | Lighthouse Lab in Glasgow | Wellcome Sanger Institute for the COVID-19 Genomics UK (COG-UK) Consortium | Harper VanSteenhouse, Yumi Kasai, David Gray, Carol Clugston, Anna Dominiczak and Alex Alderton, Roberto Amato, Sonia Goncalves, Ewan Harrison, David K. Jackson, Ian Johnston, Dominic Kwiatkowski, Cordelia Langford, John Sillitoe on behalf of the Wellcome Sanger Institute COVID-19 Surveillance Team |
| EPI_ISL_704118 | West of Scotland Specialist Virology Centre, NHSGGC / MRC-University of Glasgow Centre for Virus Research | COVID-19 Genomics UK (COG-UK) Consortium | Ana da Silva Filipe, Natasha Johnson, Kathy Smollett, Daniel Mair, Stephen Carmichael, Alice Broos, Lily Tong, Jenna Nichols, Kyriaki Nomikou; Sarah McDonald; Richard Orton, Joseph Hughes, Sreenu Vattipally, David L Robertson; Alasdair MacLean, Rory Gunson; Sharif Shaaban, Matthew Holden; Rachel Blacow, Guy Mollett, Kathy Li, James Shepherd, Antonia Ho, Emma Thomson |
| EPI_ISL_704119, EPI_ISL_704121, EPI_ISL_704122 | Lighthouse Lab in Glasgow | Wellcome Sanger Institute for the COVID-19 Genomics UK (COG-UK) Consortium | Harper VanSteenhouse, Yumi Kasai, David Gray, Carol Clugston, Anna Dominiczak and Alex Alderton, Roberto Amato, Sonia Goncalves, Ewan Harrison, David K. Jackson, Ian Johnston, Dominic Kwiatkowski, Cordelia Langford, John Sillitoe on behalf of the Wellcome Sanger Institute COVID-19 Surveillance Team |
| EPI_ISL_704123 | Quadram Institute Bioscience | COVID-19 Genomics UK (COG-UK) Consortium | Dave J. Baker, Gemma L. Kay, Alp Aydin, Thanh Le-Viet, Steven Rudder, Ana P. Tedim, Anastasia Kolyva, Maria Diaz, Leonardo de Oliveira Martins, Nabil-Fareed Alikhan, Lizzie Meadows, Rachael Stanley, Ngozi Elumogo, Muhammed Yasir, Nicholas M. Thomson, Alexander J Trotter, Rachel Gilroy, Samuel Bloomfield, Claire Stuart, Andrew Bell, Reenesh Prakash, Samir Dervisevic, Alison E. Mather, John Wain, Mark Webber, Andrew J. Page, Justin O'Grady |
| EPI_ISL_704124, EPI_ISL_704125, EPI_ISL_704127, EPI_ISL_704128, EPI_ISL_704130, EPI_ISL_704132, EPI_ISL_704133 | Lighthouse Lab in Glasgow | Wellcome Sanger Institute for the COVID-19 Genomics UK (COG-UK) Consortium | Harper VanSteenhouse, Yumi Kasai, David Gray, Carol Clugston, Anna Dominiczak and Alex Alderton, Roberto Amato, Sonia Goncalves, Ewan Harrison, David K. Jackson, Ian Johnston, Dominic Kwiatkowski, Cordelia Langford, John Sillitoe on behalf of the Wellcome Sanger Institute COVID-19 Surveillance Team |
| EPI_ISL_704134 | Liverpool Clinical Laboratories | COVID-19 Genomics UK (COG-UK) Consortium | Sam Haldenby, Anita Lucaci, Steve Paterson, Julian Hiscox, Alistair Darby, M Almsaud, A Alrezaihi, Muhannad Alruwaili, Stuart D Armstrong, Jones Benjamin, Eleanor G Bentley, Anu Chawla, Jordan J Clark, Angela Cowell, Richard Eccles, Isabel García-Dorival, Matthew Gemmell, Alessandro Gerada, PKF Gilmore, Richard Gregory, Ximeng Han, Catherine Hartley, Margaret Hughes, Miren Iturriza-Gomara, James Johnson, L Luu, Jenifer Manson, Charlotte Nelson, Elaine O'Toole, Cassie Olateju, Rebekah Penrice-Randal , Lucille Rainbow, N.P Randle, Trevor Ian Robinson, Parul Sharma, Ghada T Shawli, James P Stewart, Neil Swainston, Ecaterina Vamos, Joanne Watts, Mark Whitehead |
| EPI_ISL_704135, EPI_ISL_704136, EPI_ISL_704138 | Lighthouse Lab in Glasgow | Wellcome Sanger Institute for the COVID-19 Genomics UK (COG-UK) Consortium | Harper VanSteenhouse, Yumi Kasai, David Gray, Carol Clugston, Anna Dominiczak and Alex Alderton, Roberto Amato, Sonia Goncalves, Ewan Harrison, David K. Jackson, Ian Johnston, Dominic Kwiatkowski, Cordelia Langford, John Sillitoe on behalf of the Wellcome Sanger Institute COVID-19 Surveillance Team |
| EPI_ISL_704139 | Quadram Institute Bioscience | COVID-19 Genomics UK (COG-UK) Consortium | Dave J. Baker, Gemma L. Kay, Alp Aydin, Thanh Le-Viet, Steven Rudder, Ana P. Tedim, Anastasia Kolyva, Maria Diaz, Leonardo de Oliveira Martins, Nabil-Fareed Alikhan, Lizzie Meadows, Rachael Stanley, Ngozi Elumogo, Muhammed Yasir, Nicholas M. Thomson, Alexander J Trotter, Rachel Gilroy, Samuel Bloomfield, Claire Stuart, Andrew Bell, Reenesh Prakash, Samir Dervisevic, Alison E. Mather, John Wain, Mark Webber, Andrew J. Page, Justin O'Grady |
| EPI_ISL_704140, EPI_ISL_704141, | Lighthouse Lab in Glasgow | Wellcome Sanger Institute for the COVID-19 Genomics UK | Harper VanSteenhouse, Yumi Kasai, David Gray, Carol Clugston, Anna Dominiczak and Alex Alderton, Roberto Amato, Sonia Goncalves, Ewan Harrison, |

| | | | |
|---|---|---|---|
| EPI_ISL_704143, EPI_ISL_704144, EPI_ISL_704146, EPI_ISL_704147, EPI_ISL_704149 | | (COG-UK) Consortium | David K. Jackson, Ian Johnston, Dominic Kwiatkowski, Cordelia Langford, John Sillitoe on behalf of the Wellcome Sanger Institute COVID-19 Surveillance Team |
| EPI_ISL_704150 | Quadram Institute Bioscience | COVID-19 Genomics UK (COG-UK) Consortium | Dave J. Baker, Gemma L. Kay, Alp Aydin, Thanh Le-Viet, Steven Rudder, Ana P. Tedim, Anastasia Kolyva, Maria Diaz, Leonardo de Oliveira Martins, Nabil-Fareed Alikhan, Lizzie Meadows, Rachael Stanley, Ngozi Elumogo, Muhammed Yasir, Nicholas M. Thomson, Alexander J Trotter, Rachel Gilroy, Samuel Bloomfield, Claire Stuart, Andrew Bell, Reenesh Prakash, Samir Dervisevic, Alison E. Mather, John Wain, Mark Webber, Andrew J. Page, Justin O'Grady |
| EPI_ISL_704151, EPI_ISL_704152, EPI_ISL_704154, EPI_ISL_704155, EPI_ISL_704157 | Lighthouse Lab in Glasgow | Wellcome Sanger Institute for the COVID-19 Genomics UK (COG-UK) Consortium | Harper VanSteenhouse, Yumi Kasai, David Gray, Carol Clugston, Anna Dominiczak and Alex Alderton, Roberto Amato, Sonia Goncalves, Ewan Harrison, David K. Jackson, Ian Johnston, Dominic Kwiatkowski, Cordelia Langford, John Sillitoe on behalf of the Wellcome Sanger Institute COVID-19 Surveillance Team |
| EPI_ISL_704158 | Quadram Institute Bioscience | COVID-19 Genomics UK (COG-UK) Consortium | Dave J. Baker, Gemma L. Kay, Alp Aydin, Thanh Le-Viet, Steven Rudder, Ana P. Tedim, Anastasia Kolyva, Maria Diaz, Leonardo de Oliveira Martins, Nabil-Fareed Alikhan, Lizzie Meadows, Rachael Stanley, Ngozi Elumogo, Muhammed Yasir, Nicholas M. Thomson, Alexander J Trotter, Rachel Gilroy, Samuel Bloomfield, Claire Stuart, Andrew Bell, Reenesh Prakash, Samir Dervisevic, Alison E. Mather, John Wain, Mark Webber, Andrew J. Page, Justin O'Grady |
| EPI_ISL_704159, EPI_ISL_704160, EPI_ISL_704162, EPI_ISL_704164, EPI_ISL_704165, EPI_ISL_704167, EPI_ISL_704169, EPI_ISL_704170, EPI_ISL_704172, EPI_ISL_704173 | Lighthouse Lab in Glasgow | Wellcome Sanger Institute for the COVID-19 Genomics UK (COG-UK) Consortium | Harper VanSteenhouse, Yumi Kasai, David Gray, Carol Clugston, Anna Dominiczak and Alex Alderton, Roberto Amato, Sonia Goncalves, Ewan Harrison, David K. Jackson, Ian Johnston, Dominic Kwiatkowski, Cordelia Langford, John Sillitoe on behalf of the Wellcome Sanger Institute COVID-19 Surveillance Team |
| EPI_ISL_704174 | Quadram Institute Bioscience | COVID-19 Genomics UK (COG-UK) Consortium | Dave J. Baker, Gemma L. Kay, Alp Aydin, Thanh Le-Viet, Steven Rudder, Ana P. Tedim, Anastasia Kolyva, Maria Diaz, Leonardo de Oliveira Martins, Nabil-Fareed Alikhan, Lizzie Meadows, Rachael Stanley, Ngozi Elumogo, Muhammed Yasir, Nicholas M. Thomson, Alexander J Trotter, Rachel Gilroy, Samuel Bloomfield, Claire Stuart, Andrew Bell, Reenesh Prakash, Samir Dervisevic, Alison E. Mather, John Wain, Mark Webber, Andrew J. Page, Justin O'Grady |
| EPI_ISL_704175, EPI_ISL_704177, EPI_ISL_704178, EPI_ISL_704180 | Lighthouse Lab in Glasgow | Wellcome Sanger Institute for the COVID-19 Genomics UK (COG-UK) Consortium | Harper VanSteenhouse, Yumi Kasai, David Gray, Carol Clugston, Anna Dominiczak and Alex Alderton, Roberto Amato, Sonia Goncalves, Ewan Harrison, David K. Jackson, Ian Johnston, Dominic Kwiatkowski, Cordelia Langford, John Sillitoe on behalf of the Wellcome Sanger Institute COVID-19 Surveillance Team |
| EPI_ISL_704181 | Quadram Institute Bioscience | COVID-19 Genomics UK (COG-UK) Consortium | Dave J. Baker, Gemma L. Kay, Alp Aydin, Thanh Le-Viet, Steven Rudder, Ana P. Tedim, Anastasia Kolyva, Maria Diaz, Leonardo de Oliveira Martins, Nabil-Fareed Alikhan, Lizzie Meadows, Rachael Stanley, Ngozi Elumogo, Muhammed Yasir, Nicholas M. Thomson, Alexander J Trotter, Rachel Gilroy, Samuel Bloomfield, Claire Stuart, Andrew Bell, Reenesh Prakash, Samir Dervisevic, Alison E. Mather, John Wain, Mark Webber, Andrew J. Page, Justin O'Grady |
| EPI_ISL_704182, EPI_ISL_704183 | Lighthouse Lab in Glasgow | Wellcome Sanger Institute for the COVID-19 Genomics UK (COG-UK) Consortium | Harper VanSteenhouse, Yumi Kasai, David Gray, Carol Clugston, Anna Dominiczak and Alex Alderton, Roberto Amato, Sonia Goncalves, Ewan Harrison, David K. Jackson, Ian Johnston, Dominic Kwiatkowski, Cordelia Langford, John Sillitoe on behalf of the Wellcome Sanger Institute COVID-19 Surveillance Team |
| EPI_ISL_704184 | West of Scotland Specialist Virology Centre, NHSGGC / MRC-University of Glasgow Centre for Virus Research | COVID-19 Genomics UK (COG-UK) Consortium | Ana da Silva Filipe, Natasha Johnson, Kathy Smollett, Daniel Mair, Stephen Carmichael, Alice Broos, Lily Tong, Jenna Nichols, Kyriaki Nomikou; Sarah McDonald; Richard Orton, Joseph Hughes, Sreenu Vattipally, David L Robertson; Alasdair MacLean, Rory Gunson; Sharif Shaaban, Matthew Holden; Rachel Blacow, Guy Mollett, Kathy Li, James Shepherd, Antonia Ho, Emma Thomson |
| EPI_ISL_704185, EPI_ISL_704186, EPI_ISL_704188, EPI_ISL_704189, EPI_ISL_704191, EPI_ISL_704192, EPI_ISL_704194 | Lighthouse Lab in Glasgow | Wellcome Sanger Institute for the COVID-19 Genomics UK (COG-UK) Consortium | Harper VanSteenhouse, Yumi Kasai, David Gray, Carol Clugston, Anna Dominiczak and Alex Alderton, Roberto Amato, Sonia Goncalves, Ewan Harrison, David K. Jackson, Ian Johnston, Dominic Kwiatkowski, Cordelia Langford, John Sillitoe on behalf of the Wellcome Sanger Institute COVID-19 Surveillance Team |
| EPI_ISL_704195 | Liverpool Clinical Laboratories | COVID-19 Genomics UK (COG-UK) Consortium | Sam Haldenby, Anita Lucaci, Steve Paterson, Julian Hiscox, Alistair Darby, M Almsaud, A Alrezaihi, Muhannad Alruwaili, Stuart D Armstrong, Jones Benjamin, Eleanor G Bentley, Anu Chawla, Jordan J Clark, Angela Cowell, Richard Eccles, Isabel García-Dorival, Matthew Gemmell, Alessandro Gerada, PKF Gilmore, Richard Gregory, Ximeng Han, Catherine Hartley, Margaret Hughes, Miren Iturriza-Gomara, James Johnson, L Luu, Jenifer Manson, Charlotte Nelson, Elaine O'Toole, Cassie Olateju, Rebekah Penrice-Randal , Lucille Rainbow, N.P Randle, Trevor Ian Robinson, Parul Sharma, Ghada T Shawli, James P Stewart, Neil Swainston, Ecaterina Vamos, Joanne Watts, Mark Whitehead |
| EPI_ISL_704196, EPI_ISL_704197, EPI_ISL_704199, EPI_ISL_704200, EPI_ISL_704202, EPI_ISL_704204, EPI_ISL_704205, EPI_ISL_704207, EPI_ISL_704208, EPI_ISL_704210, EPI_ISL_704211, EPI_ISL_704212, EPI_ISL_704213, EPI_ISL_704214, EPI_ISL_704215, EPI_ISL_704216, EPI_ISL_704218, EPI_ISL_704220, EPI_ISL_704221, EPI_ISL_704223, EPI_ISL_704224, EPI_ISL_704226, EPI_ISL_704227, EPI_ISL_704229, EPI_ISL_704230, EPI_ISL_704232, EPI_ISL_704234, EPI_ISL_704235 | | | |
| see above | Lighthouse Lab in Glasgow | Wellcome Sanger Institute for the COVID-19 Genomics UK (COG-UK) Consortium | Harper VanSteenhouse, Yumi Kasai, David Gray, Carol Clugston, Anna Dominiczak and Alex Alderton, Roberto Amato, Sonia Goncalves, Ewan Harrison, David K. Jackson, Ian Johnston, Dominic Kwiatkowski, Cordelia Langford, John Sillitoe on behalf of the Wellcome Sanger Institute COVID-19 Surveillance Team |
| EPI_ISL_704236 | Quadram Institute Bioscience | COVID-19 Genomics UK (COG-UK) Consortium | Dave J. Baker, Gemma L. Kay, Alp Aydin, Thanh Le-Viet, Steven Rudder, Ana P. Tedim, Anastasia Kolyva, Maria Diaz, Leonardo de Oliveira Martins, Nabil-Fareed Alikhan, Lizzie Meadows, Rachael Stanley, Ngozi Elumogo, Muhammed Yasir, Nicholas M. Thomson, Alexander J Trotter, Rachel Gilroy, Samuel Bloomfield, Claire Stuart, Andrew Bell, Reenesh Prakash, Samir Dervisevic, Alison E. Mather, John Wain, Mark Webber, Andrew J. Page, Justin O'Grady |
| EPI_ISL_704237, EPI_ISL_704239, EPI_ISL_704240 | Lighthouse Lab in Glasgow | Wellcome Sanger Institute for the COVID-19 Genomics UK (COG-UK) Consortium | Harper VanSteenhouse, Yumi Kasai, David Gray, Carol Clugston, Anna Dominiczak and Alex Alderton, Roberto Amato, Sonia Goncalves, Ewan Harrison, David K. Jackson, Ian Johnston, Dominic Kwiatkowski, Cordelia Langford, John Sillitoe on behalf of the Wellcome Sanger Institute COVID-19 Surveillance Team |
| EPI_ISL_704241 | Quadram Institute Bioscience | COVID-19 Genomics UK (COG-UK) Consortium | Dave J. Baker, Gemma L. Kay, Alp Aydin, Thanh Le-Viet, Steven Rudder, Ana P. Tedim, Anastasia Kolyva, Maria Diaz, Leonardo de Oliveira Martins, Nabil-Fareed Alikhan, Lizzie Meadows, Rachael Stanley, Ngozi Elumogo, Muhammed Yasir, Nicholas M. Thomson, Alexander J Trotter, Rachel Gilroy, Samuel Bloomfield, Claire Stuart, Andrew Bell, Reenesh Prakash, Samir Dervisevic, Alison E. Mather, John Wain, Mark Webber, Andrew J. Page, Justin O'Grady |
| EPI_ISL_704242, EPI_ISL_704244, EPI_ISL_704245, EPI_ISL_704246, EPI_ISL_704248, EPI_ISL_704249, EPI_ISL_704251, EPI_ISL_704253, EPI_ISL_704254, EPI_ISL_704256, EPI_ISL_704258, EPI_ISL_704259 | | | |
| see above | Lighthouse Lab in Glasgow | Wellcome Sanger Institute for the COVID-19 Genomics UK (COG-UK) Consortium | Harper VanSteenhouse, Yumi Kasai, David Gray, Carol Clugston, Anna Dominiczak and Alex Alderton, Roberto Amato, Sonia Goncalves, Ewan Harrison, David K. Jackson, Ian Johnston, Dominic Kwiatkowski, Cordelia Langford, John Sillitoe on behalf of the Wellcome Sanger Institute COVID-19 Surveillance Team |
| EPI_ISL_704260 | Liverpool Clinical Laboratories | COVID-19 Genomics UK (COG-UK) Consortium | Sam Haldenby, Anita Lucaci, Steve Paterson, Julian Hiscox, Alistair Darby, M Almsaud, A Alrezaihi, Muhannad Alruwaili, Stuart D Armstrong, Jones Benjamin, Eleanor G Bentley, Anu Chawla, Jordan J Clark, Angela Cowell, Richard Eccles, Isabel García-Dorival, Matthew Gemmell, Alessandro Gerada, PKF Gilmore, Richard Gregory, Ximeng Han, Catherine Hartley, Margaret Hughes, Miren Iturriza-Gomara, James Johnson, L Luu, Jenifer Manson, Charlotte Nelson, Elaine O'Toole, Cassie Olateju, Rebekah Penrice-Randal , Lucille Rainbow, N.P Randle, Trevor Ian Robinson, Parul Sharma, Ghada T Shawli, James P Stewart, Neil Swainston, Ecaterina Vamos, Joanne Watts, Mark Whitehead |
| EPI_ISL_704261 | Lighthouse Lab in Glasgow | Wellcome Sanger Institute for the COVID-19 Genomics UK (COG-UK) Consortium | Harper VanSteenhouse, Yumi Kasai, David Gray, Carol Clugston, Anna Dominiczak and Alex Alderton, Roberto Amato, Sonia Goncalves, Ewan Harrison, David K. Jackson, Ian Johnston, Dominic Kwiatkowski, Cordelia Langford, John Sillitoe on behalf of the Wellcome Sanger Institute COVID-19 Surveillance Team |
| EPI_ISL_704262 | Northumbria University / South Tees Hospitals NHS Foundation Trust / North Cumbria Integrated Care NHS Foundation Trust / North Tees and Hartlepool NHS Foundation Trust / Newcastle Hospitals NHS Foundation Trust | COVID-19 Genomics UK (COG-UK) Consortium | Darren L Smith,Andrew Nelson,Matthew Bashton,Greg R Young,Joshua Loh,John Allan,Mohammad A Tariq,Giles S Holt,Gary Black,Wen C Yew,Lynn Dover,Paul Baker,Steve Liggett,Sarah Essex,Jane Greenaway,Debra Padgett,Clive Graham,Garren Scott,Edward Barton,Emma Swindells,Brendan Payne,Jennifer Collins,Yusri Taha,Gary Eltringham |

| | | | |
|---|---|---|---|
| EPI_ISL_704263, EPI_ISL_704264, EPI_ISL_704265, EPI_ISL_704267, EPI_ISL_704268, EPI_ISL_704270, EPI_ISL_704272, EPI_ISL_704274, EPI_ISL_704275, EPI_ISL_704277, EPI_ISL_704278, EPI_ISL_704279, EPI_ISL_704281, EPI_ISL_704282, EPI_ISL_704284, EPI_ISL_704285, EPI_ISL_704287, EPI_ISL_704288, EPI_ISL_704290, EPI_ISL_704292, EPI_ISL_704293, EPI_ISL_704295, EPI_ISL_704296, EPI_ISL_704298, EPI_ISL_704300, EPI_ISL_704301, EPI_ISL_704303, EPI_ISL_704304, EPI_ISL_704306, EPI_ISL_704308, EPI_ISL_704310, EPI_ISL_704311, EPI_ISL_704313, EPI_ISL_704314, EPI_ISL_704316, EPI_ISL_704318, EPI_ISL_704319, EPI_ISL_704321, EPI_ISL_704323, EPI_ISL_704324, EPI_ISL_704326, EPI_ISL_704327, EPI_ISL_704329, EPI_ISL_704330, EPI_ISL_704332, EPI_ISL_704334, EPI_ISL_704335, EPI_ISL_704337, EPI_ISL_704338, EPI_ISL_704340, EPI_ISL_704342, EPI_ISL_704343, EPI_ISL_704345, EPI_ISL_704346, EPI_ISL_704348, EPI_ISL_704349, EPI_ISL_704351, EPI_ISL_704352, EPI_ISL_704354, EPI_ISL_704355, EPI_ISL_704357, EPI_ISL_704358, EPI_ISL_704360, EPI_ISL_704362, EPI_ISL_704363, EPI_ISL_704365, EPI_ISL_704367, EPI_ISL_704368, EPI_ISL_704370, EPI_ISL_704371, EPI_ISL_704373, EPI_ISL_704375, EPI_ISL_704376, EPI_ISL_704378, EPI_ISL_704380, EPI_ISL_704381, EPI_ISL_704383, EPI_ISL_704385, EPI_ISL_704386, EPI_ISL_704388, EPI_ISL_704390, EPI_ISL_704391, EPI_ISL_704393, EPI_ISL_704394, EPI_ISL_704396, EPI_ISL_704397, EPI_ISL_704399, EPI_ISL_704400, EPI_ISL_704402, EPI_ISL_704404, EPI_ISL_704405, EPI_ISL_704407, EPI_ISL_704409, EPI_ISL_704410, EPI_ISL_704412, EPI_ISL_704414, EPI_ISL_704415, EPI_ISL_704417, EPI_ISL_704418, EPI_ISL_704420, EPI_ISL_704421, EPI_ISL_704423, EPI_ISL_704424, EPI_ISL_704426, EPI_ISL_704428, EPI_ISL_704429, EPI_ISL_704431, EPI_ISL_704432, EPI_ISL_704434, EPI_ISL_704436, EPI_ISL_704437, EPI_ISL_704439, EPI_ISL_704440, EPI_ISL_704442, EPI_ISL_704444, EPI_ISL_704445, EPI_ISL_704447, EPI_ISL_704448, EPI_ISL_704450, EPI_ISL_704451, EPI_ISL_704453, EPI_ISL_704455, EPI_ISL_704456, EPI_ISL_704458, EPI_ISL_704459, EPI_ISL_704461, EPI_ISL_704463, EPI_ISL_704464, EPI_ISL_704466, EPI_ISL_704468, EPI_ISL_704469, EPI_ISL_704471, EPI_ISL_704472, EPI_ISL_704474, EPI_ISL_704476, EPI_ISL_704477, EPI_ISL_704479, EPI_ISL_704480, EPI_ISL_704482, EPI_ISL_704483, EPI_ISL_704485, EPI_ISL_704487, EPI_ISL_704488, EPI_ISL_704490, EPI_ISL_704491, EPI_ISL_704493, EPI_ISL_704495, EPI_ISL_704496, EPI_ISL_704498, EPI_ISL_704500, EPI_ISL_704501, EPI_ISL_704503, EPI_ISL_704504, EPI_ISL_704506, EPI_ISL_704508, EPI_ISL_704509, EPI_ISL_704511, EPI_ISL_704512, EPI_ISL_704514, EPI_ISL_704515, EPI_ISL_704517, EPI_ISL_704519, EPI_ISL_704520, EPI_ISL_704522, EPI_ISL_704523, EPI_ISL_704525, EPI_ISL_704527, EPI_ISL_704528, EPI_ISL_704530, EPI_ISL_704531, EPI_ISL_704533, EPI_ISL_704534, EPI_ISL_704536, EPI_ISL_704538, EPI_ISL_704539, EPI_ISL_704541, EPI_ISL_704542, EPI_ISL_704544, EPI_ISL_704546, EPI_ISL_704547, EPI_ISL_704549, EPI_ISL_704550, EPI_ISL_704552, EPI_ISL_704553, EPI_ISL_704555, EPI_ISL_704556, EPI_ISL_704558 | | | |
| see above | Lighthouse Lab in Glasgow | Wellcome Sanger Institute for the COVID-19 Genomics UK (COG-UK) Consortium | Harper VanSteenhouse, Yumi Kasai, David Gray, Carol Clugston, Anna Dominiczak and Alex Alderton, Roberto Amato, Sonia Goncalves, Ewan Harrison, David K. Jackson, Ian Johnston, Dominic Kwiatkowski, Cordelia Langford, John Sillitoe on behalf of the Wellcome Sanger Institute COVID-19 Surveillance Team |
| EPI_ISL_704559 | Virology Department, Royal Infirmary of Edinburgh, NHS Lothian / School of Biological Sciences, University of Edinburgh / Institute of Genetics and Molecular Medicine, University of Edinburgh | COVID-19 Genomics UK (COG-UK) Consortium | McHugh M, Dewar R, Rooke S, Gallagher M, Balcaza C, O'Toole Á, Scher E, Hill V, McCrone JT, Colquhoun R, Yu X, Jackson B, Rambaut A, Williams TC, Templeton K |
| EPI_ISL_704560, EPI_ISL_704561, EPI_ISL_704563 | Lighthouse Lab in Glasgow | Wellcome Sanger Institute for the COVID-19 Genomics UK (COG-UK) Consortium | Harper VanSteenhouse, Yumi Kasai, David Gray, Carol Clugston, Anna Dominiczak and Alex Alderton, Roberto Amato, Sonia Goncalves, Ewan Harrison, David K. Jackson, Ian Johnston, Dominic Kwiatkowski, Cordelia Langford, John Sillitoe on behalf of the Wellcome Sanger Institute COVID-19 Surveillance Team |
| EPI_ISL_704564 | Northumbria University / South Tees Hospitals NHS Foundation Trust / North Cumbria Integrated Care NHS Foundation Trust / North Tees and Hartlepool NHS Foundation Trust / Newcastle Hospitals NHS Foundation Trust | COVID-19 Genomics UK (COG-UK) Consortium | Darren L Smith,Andrew Nelson,Matthew Bashton,Greg R Young,Joshua Loh,John Allan,Mohammad A Tariq,Giles S Holt,Gary Black,Wen C Yew,Lynn Dover,Paul Baker,Steve Liggett,Sarah Essex,Jane Greenaway,Debra Padgett,Clive Graham,Garren Scott,Edward Barton,Emma Swindells,Brendan Payne,Jennifer Collins,Yusri Taha,Gary Eltringham |
| EPI_ISL_704565, EPI_ISL_704566, EPI_ISL_704568, EPI_ISL_704569, EPI_ISL_704571 | Lighthouse Lab in Glasgow | Wellcome Sanger Institute for the COVID-19 Genomics UK (COG-UK) Consortium | Harper VanSteenhouse, Yumi Kasai, David Gray, Carol Clugston, Anna Dominiczak and Alex Alderton, Roberto Amato, Sonia Goncalves, Ewan Harrison, David K. Jackson, Ian Johnston, Dominic Kwiatkowski, Cordelia Langford, John Sillitoe on behalf of the Wellcome Sanger Institute COVID-19 Surveillance Team |
| EPI_ISL_704582 | Queens Medical Centre, Clinical Microbiology Department / DeepSeq Nottingham | COVID-19 Genomics UK (COG-UK) Consortium | Gemma Clark, Wendy Smith, Manjinder Khakh, Vicki M Fleming, Michelle M Lister, Hannah Howson-Wells, Jonathan Ball, Patrick McClure, Joseph Chappell, Theocharis Tsoleridis, Nadine Holmes, Matthew Carlisle, Christopher Moore, Fei Sang, Johnny Debebe, Victoria Wright, Matthew Loose |
| EPI_ISL_704607 | University College London Hospital | COVID-19 Genomics UK (COG-UK) Consortium | Judith Heaney, Matthew Byott, Catherine Houlihan, Dan Frampton, Stuart Kirk, Moira Spyer and Eleni Nastouli |
| EPI_ISL_704610, EPI_ISL_704611, EPI_ISL_704612, EPI_ISL_704613, EPI_ISL_704616 | Department of Pathology, University of Cambridge | COVID-19 Genomics UK (COG-UK) Consortium | Aminu S. Jahun, Yasmin Chaudhry, Grant Hall, Iliana Georgana, Myra Hosmillo, Martin D. Curran, Malte Pinckert, Surendra Parmar, Ian Goodfellow |
| EPI_ISL_704642, EPI_ISL_704644, EPI_ISL_704645, EPI_ISL_704647, EPI_ISL_704649, EPI_ISL_704650, EPI_ISL_704652, EPI_ISL_704653, EPI_ISL_704655, EPI_ISL_704657, EPI_ISL_704658, EPI_ISL_704660, EPI_ISL_704661, EPI_ISL_704663, EPI_ISL_704665 | | | |
| see above | Lighthouse Lab in Glasgow | Wellcome Sanger Institute for the COVID-19 Genomics UK (COG-UK) Consortium | Harper VanSteenhouse, Yumi Kasai, David Gray, Carol Clugston, Anna Dominiczak and Alex Alderton, Roberto Amato, Sonia Goncalves, Ewan Harrison, David K. Jackson, Ian Johnston, Dominic Kwiatkowski, Cordelia Langford, John Sillitoe on behalf of the Wellcome Sanger Institute COVID-19 Surveillance Team |
| EPI_ISL_704666 | University College London Hospital | COVID-19 Genomics UK (COG-UK) Consortium | Judith Heaney, Matthew Byott, Catherine Houlihan, Dan Frampton, Stuart Kirk, Moira Spyer and Eleni Nastouli |
| EPI_ISL_704667, EPI_ISL_704668, EPI_ISL_704670, EPI_ISL_704672, EPI_ISL_704673, EPI_ISL_704675, EPI_ISL_704677, EPI_ISL_704678, EPI_ISL_704680, EPI_ISL_704681, EPI_ISL_704683, EPI_ISL_704684, EPI_ISL_704686 | | | |
| see above | Lighthouse Lab in Glasgow | Wellcome Sanger Institute for the COVID-19 Genomics UK (COG-UK) Consortium | Harper VanSteenhouse, Yumi Kasai, David Gray, Carol Clugston, Anna Dominiczak and Alex Alderton, Roberto Amato, Sonia Goncalves, Ewan Harrison, David K. Jackson, Ian Johnston, Dominic Kwiatkowski, Cordelia Langford, John Sillitoe on behalf of the Wellcome Sanger Institute COVID-19 Surveillance Team |
| EPI_ISL_704719, EPI_ISL_704722, EPI_ISL_704725, EPI_ISL_704749, EPI_ISL_704752, EPI_ISL_704754, EPI_ISL_704757, EPI_ISL_704759, EPI_ISL_704772, EPI_ISL_704775, EPI_ISL_704777, EPI_ISL_704786, EPI_ISL_704797, EPI_ISL_704799, EPI_ISL_704802 | | | |
| see above | Quadram Institute Bioscience | COVID-19 Genomics UK (COG-UK) Consortium | Dave J. Baker, Gemma L. Kay, Alp Aydin, Thanh Le-Viet, Steven Rudder, Ana P. Tedim, Anastasia Kolyva, Maria Diaz, Leonardo de Oliveira Martins, Nabil-Fareed Alikhan, Lizzie Meadows, Rachael Stanley, Ngozi Elumogo, Muhammed Yasir, Nicholas M. Thomson, Alexander J Trotter, Rachel Gilroy, Samuel Bloomfield, Claire Stuart, Andrew Bell, Reenesh Prakash, Samir Dervisevic, Alison E. Mather, John Wain, Mark Webber, Andrew J. Page, Justin O'Grady |
| EPI_ISL_704809 | West of Scotland Specialist Virology Centre, NHSGGC / MRC-University of Glasgow Centre for Virus Research | COVID-19 Genomics UK (COG-UK) Consortium | Ana da Silva Filipe, Natasha Johnson, Kathy Smollett, Daniel Mair, Stephen Carmichael, Alice Broos, Lily Tong, Jenna Nichols, Kyriaki Nomikou; Sarah McDonald; Richard Orton, Joseph Hughes, Sreenu Vattipally, David L Robertson; Alasdair MacLean, Rory Gunson; Sharif Shaaban, Matthew Holden; Rachel Blacow, Guy Mollett, Kathy Li, James Shepherd, Antonia Ho, Emma Thomson |
| EPI_ISL_704847, EPI_ISL_704855, EPI_ISL_704858, EPI_ISL_704892 | Quadram Institute Bioscience | COVID-19 Genomics UK (COG-UK) Consortium | Dave J. Baker, Gemma L. Kay, Alp Aydin, Thanh Le-Viet, Steven Rudder, Ana P. Tedim, Anastasia Kolyva, Maria Diaz, Leonardo de Oliveira Martins, Nabil-Fareed Alikhan, Lizzie Meadows, Rachael Stanley, Ngozi Elumogo, Muhammed Yasir, Nicholas M. Thomson, Alexander J Trotter, Rachel Gilroy, Samuel Bloomfield, Claire Stuart, Andrew Bell, Reenesh Prakash, Samir Dervisevic, Alison E. Mather, John Wain, Mark Webber, Andrew J. Page, Justin O'Grady |
| EPI_ISL_704895, EPI_ISL_704897, EPI_ISL_704900 | Liverpool Clinical Laboratories | COVID-19 Genomics UK (COG-UK) Consortium | Sam Haldenby, Anita Lucaci, Steve Paterson, Julian Hiscox, Alistair Darby, M Almsaud, A Alrezaihi, Muhannad Alruwaili, Stuart D Armstrong, Jones Benjamin, Eleanor G Bentley, Anu Chawla, Jordan J Clark, Angela Cowell, Richard Eccles, Isabel García-Dorival, Matthew Gemmell, Alessandro Gerada, PKF Gilmore, Richard Gregory, Ximeng Han, Catherine Hartley, Margaret Hughes, Miren Iturriza-Gomara, James Johnson, L Luu, Jenifer Manson, Charlotte Nelson, Elaine O'Toole, Cassie Olateju, Rebekah Penrice-Randal , Lucille Rainbow, N.P Randle, Trevor Ian Robinson, Parul Sharma, Ghada T Shawli, James P Stewart, Neil Swainston, Ecaterina Vamos, Joanne Watts, Mark Whitehead |
| EPI_ISL_704908, EPI_ISL_704911, EPI_ISL_704916 | Quadram Institute Bioscience | COVID-19 Genomics UK (COG-UK) Consortium | Dave J. Baker, Gemma L. Kay, Alp Aydin, Thanh Le-Viet, Steven Rudder, Ana P. Tedim, Anastasia Kolyva, Maria Diaz, Leonardo de Oliveira Martins, Nabil-Fareed Alikhan, Lizzie Meadows, Rachael Stanley, Ngozi Elumogo, Muhammed Yasir, Nicholas M. Thomson, Alexander J Trotter, Rachel Gilroy, Samuel Bloomfield, Claire Stuart, Andrew Bell, Reenesh Prakash, Samir Dervisevic, Alison E. Mather, John Wain, Mark Webber, Andrew J. Page, Justin O'Grady |
| EPI_ISL_704945 | Liverpool Clinical Laboratories | COVID-19 Genomics UK (COG-UK) Consortium | Sam Haldenby, Anita Lucaci, Steve Paterson, Julian Hiscox, Alistair Darby, M Almsaud, A Alrezaihi, Muhannad Alruwaili, Stuart D Armstrong, Jones Benjamin, Eleanor G Bentley, Anu Chawla, Jordan J Clark, Angela Cowell, Richard Eccles, Isabel García-Dorival, Matthew Gemmell, Alessandro Gerada, PKF Gilmore, Richard Gregory, Ximeng Han, Catherine Hartley, Margaret Hughes, Miren Iturriza-Gomara, James Johnson, L Luu, Jenifer Manson, Charlotte Nelson, Elaine O'Toole, Cassie Olateju, Rebekah Penrice-Randal , Lucille Rainbow, N.P Randle, Trevor Ian Robinson, Parul Sharma, Ghada T Shawli, James P Stewart, Neil Swainston, Ecaterina Vamos, Joanne Watts, Mark Whitehead |
| EPI_ISL_704960 | Virology Department, Sheffield Teaching Hospitals NHS Foundation Trust/Department of Infection, Immunity and Cardiovascular Disease, The Medical School, University of Sheffield | COVID-19 Genomics UK (COG-UK) Consortium | Thushan de Silva, Matthew Parker, Nikki Smith, Adri Angyal, Rebecca Brown, Luke Green, Rachel Tucker, Paul Parsons, Danielle Groves, Katie Johnson, Laura Carrilero, Alex Keeley, Dave Partridge, Matthew Wyles, Benjamin Lindsey, Mehmet Yavuz, Mohammad Raza, Cariad Evans |
| EPI_ISL_705024, EPI_ISL_705030, EPI_ISL_705038, EPI_ISL_705043, EPI_ISL_705046, EPI_ISL_705056, | Liverpool Clinical Laboratories | COVID-19 Genomics UK (COG-UK) Consortium | Sam Haldenby, Anita Lucaci, Steve Paterson, Julian Hiscox, Alistair Darby, M Almsaud, A Alrezaihi, Muhannad Alruwaili, Stuart D Armstrong, Jones Benjamin, Eleanor G Bentley, Anu Chawla, Jordan J Clark, Angela Cowell, Richard Eccles, Isabel García-Dorival, Matthew Gemmell, Alessandro Gerada, PKF Gilmore, Richard Gregory, Ximeng Han, Catherine Hartley, Margaret Hughes, Miren Iturriza-Gomara, James Johnson, L Luu, Jenifer Manson, |

| | | | |
|---|---|---|---|
| EPI_ISL_705083 | | | Charlotte Nelson, Elaine O'Toole, Cassie Olateju, Rebekah Penrice-Randal , Lucille Rainbow, N.P Randle, Trevor Ian Robinson, Parul Sharma, Ghada T Shawli, James P Stewart, Neil Swainston, Ecaterina Vamos, Joanne Watts, Mark Whitehead |
| EPI_ISL_705088 | Virology Department, Sheffield Teaching Hospitals NHS Foundation Trust/Department of Infection, Immunity and Cardiovascular Disease, The Medical School, University of Sheffield | COVID-19 Genomics UK (COG-UK) Consortium | Thushan de Silva, Matthew Parker, Nikki Smith, Adri Angyal, Rebecca Brown, Luke Green, Rachel Tucker, Paul Parsons, Danielle Groves, Katie Johnson, Laura Carrilero, Alex Keeley, Dave Partridge, Matthew Wyles, Benjamin Lindsey, Mehmet Yavuz, Mohammad Raza, Cariad Evans |
| EPI_ISL_705094 | Queens Medical Centre, Clinical Microbiology Department / DeepSeq Nottingham | COVID-19 Genomics UK (COG-UK) Consortium | Gemma Clark, Wendy Smith, Manjinder Khakh, Vicki M Fleming, Michelle M Lister, Hannah Howson-Wells, Jonathan Ball, Patrick McClure, Joseph Chappell, Theocharis Tsoleridis, Nadine Holmes, Matthew Carlisle, Christopher Moore, Fei Sang, Johnny Debebe, Victoria Wright, Matthew Loose |
| EPI_ISL_705096 | Virology Department, Sheffield Teaching Hospitals NHS Foundation Trust/Department of Infection, Immunity and Cardiovascular Disease, The Medical School, University of Sheffield | COVID-19 Genomics UK (COG-UK) Consortium | Thushan de Silva, Matthew Parker, Nikki Smith, Adri Angyal, Rebecca Brown, Luke Green, Rachel Tucker, Paul Parsons, Danielle Groves, Katie Johnson, Laura Carrilero, Alex Keeley, Dave Partridge, Matthew Wyles, Benjamin Lindsey, Mehmet Yavuz, Mohammad Raza, Cariad Evans |
| EPI_ISL_705099 | University of Exeter | COVID-19 Genomics UK (COG-UK) Consortium | Ben Temperton,Aaron Jeffries,Michelle Michelsen,Joanna Warwick-Dugdale,Audrey Farbos,Robyn Manley,Stephen Michell,Jane Masoli |
| EPI_ISL_705108 | Liverpool Clinical Laboratories | COVID-19 Genomics UK (COG-UK) Consortium | Sam Haldenby, Anita Lucaci, Steve Paterson, Julian Hiscox, Alistair Darby, M Almsaud, A Alrezaihi, Muhannad Alruwaili, Stuart D Armstrong, Jones Benjamin, Eleanor G Bentley, Anu Chawla, Jordan J Clark, Angela Cowell, Richard Eccles, Isabel García-Dorival, Matthew Gemmell, Alessandro Gerada, PKF Gilmore, Richard Gregory, Ximeng Han, Catherine Hartley, Margaret Hughes, Miren Iturriza-Gomara, James Johnson, L Luu, Jenifer Manson, Charlotte Nelson, Elaine O'Toole, Cassie Olateju, Rebekah Penrice-Randal , Lucille Rainbow, N.P Randle, Trevor Ian Robinson, Parul Sharma, Ghada T Shawli, James P Stewart, Neil Swainston, Ecaterina Vamos, Joanne Watts, Mark Whitehead |
| EPI_ISL_705118 | Virology Department, Royal Infirmary of Edinburgh, NHS Lothian / School of Biological Sciences, University of Edinburgh / Institute of Genetics and Molecular Medicine, University of Edinburgh | COVID-19 Genomics UK (COG-UK) Consortium | McHugh M, Dewar R, Rooke S, Gallagher M, Balcaza C, O'Toole Á, Scher E, Hill V, McCrone JT, Colquhoun R, Yu X, Jackson B, Rambaut A, Williams TC, Templeton K |
| EPI_ISL_705121 | Liverpool Clinical Laboratories | COVID-19 Genomics UK (COG-UK) Consortium | Sam Haldenby, Anita Lucaci, Steve Paterson, Julian Hiscox, Alistair Darby, M Almsaud, A Alrezaihi, Muhannad Alruwaili, Stuart D Armstrong, Jones Benjamin, Eleanor G Bentley, Anu Chawla, Jordan J Clark, Angela Cowell, Richard Eccles, Isabel García-Dorival, Matthew Gemmell, Alessandro Gerada, PKF Gilmore, Richard Gregory, Ximeng Han, Catherine Hartley, Margaret Hughes, Miren Iturriza-Gomara, James Johnson, L Luu, Jenifer Manson, Charlotte Nelson, Elaine O'Toole, Cassie Olateju, Rebekah Penrice-Randal , Lucille Rainbow, N.P Randle, Trevor Ian Robinson, Parul Sharma, Ghada T Shawli, James P Stewart, Neil Swainston, Ecaterina Vamos, Joanne Watts, Mark Whitehead |
| EPI_ISL_705145 | Northumbria University / South Tees Hospitals NHS Foundation Trust / North Cumbria Integrated Care NHS Foundation Trust / North Tees and Hartlepool NHS Foundation Trust / Newcastle Hospitals NHS Foundation Trust | COVID-19 Genomics UK (COG-UK) Consortium | Darren L Smith,Andrew Nelson,Matthew Bashton,Greg R Young,Joshua Loh,John Allan,Mohammad A Tariq,Giles S Holt,Gary Black,Wen C Yew,Lynn Dover,Paul Baker,Steve Liggett,Sarah Essex,Jane Greenaway,Debra Padgett,Clive Graham,Garren Scott,Edward Barton,Emma Swindells,Brendan Payne,Jennifer Collins,Yusri Taha,Gary Eltringham |
| EPI_ISL_705206 | Liverpool Clinical Laboratories | COVID-19 Genomics UK (COG-UK) Consortium | Sam Haldenby, Anita Lucaci, Steve Paterson, Julian Hiscox, Alistair Darby, M Almsaud, A Alrezaihi, Muhannad Alruwaili, Stuart D Armstrong, Jones Benjamin, Eleanor G Bentley, Anu Chawla, Jordan J Clark, Angela Cowell, Richard Eccles, Isabel García-Dorival, Matthew Gemmell, Alessandro Gerada, PKF Gilmore, Richard Gregory, Ximeng Han, Catherine Hartley, Margaret Hughes, Miren Iturriza-Gomara, James Johnson, L Luu, Jenifer Manson, Charlotte Nelson, Elaine O'Toole, Cassie Olateju, Rebekah Penrice-Randal , Lucille Rainbow, N.P Randle, Trevor Ian Robinson, Parul Sharma, Ghada T Shawli, James P Stewart, Neil Swainston, Ecaterina Vamos, Joanne Watts, Mark Whitehead |
| EPI_ISL_705207 | Northumbria University / South Tees Hospitals NHS Foundation Trust / North Cumbria Integrated Care NHS Foundation Trust / North Tees and Hartlepool NHS Foundation Trust / Newcastle Hospitals NHS Foundation Trust | COVID-19 Genomics UK (COG-UK) Consortium | Darren L Smith,Andrew Nelson,Matthew Bashton,Greg R Young,Joshua Loh,John Allan,Mohammad A Tariq,Giles S Holt,Gary Black,Wen C Yew,Lynn Dover,Paul Baker,Steve Liggett,Sarah Essex,Jane Greenaway,Debra Padgett,Clive Graham,Garren Scott,Edward Barton,Emma Swindells,Brendan Payne,Jennifer Collins,Yusri Taha,Gary Eltringham |
| EPI_ISL_705210 | Virology Department, Sheffield Teaching Hospitals NHS Foundation Trust/Department of Infection, Immunity and Cardiovascular Disease, The Medical School, University of Sheffield | COVID-19 Genomics UK (COG-UK) Consortium | Thushan de Silva, Matthew Parker, Nikki Smith, Adri Angyal, Rebecca Brown, Luke Green, Rachel Tucker, Paul Parsons, Danielle Groves, Katie Johnson, Laura Carrilero, Alex Keeley, Dave Partridge, Matthew Wyles, Benjamin Lindsey, Mehmet Yavuz, Mohammad Raza, Cariad Evans |
| EPI_ISL_705213, EPI_ISL_705223 | Queens Medical Centre, Clinical Microbiology Department / DeepSeq Nottingham | COVID-19 Genomics UK (COG-UK) Consortium | Gemma Clark, Wendy Smith, Manjinder Khakh, Vicki M Fleming, Michelle M Lister, Hannah Howson-Wells, Jonathan Ball, Patrick McClure, Joseph Chappell, Theocharis Tsoleridis, Nadine Holmes, Matthew Carlisle, Christopher Moore, Fei Sang, Johnny Debebe, Victoria Wright, Matthew Loose |
| EPI_ISL_705242, EPI_ISL_705246 | Liverpool Clinical Laboratories | COVID-19 Genomics UK (COG-UK) Consortium | Sam Haldenby, Anita Lucaci, Steve Paterson, Julian Hiscox, Alistair Darby, M Almsaud, A Alrezaihi, Muhannad Alruwaili, Stuart D Armstrong, Jones Benjamin, Eleanor G Bentley, Anu Chawla, Jordan J Clark, Angela Cowell, Richard Eccles, Isabel García-Dorival, Matthew Gemmell, Alessandro Gerada, PKF Gilmore, Richard Gregory, Ximeng Han, Catherine Hartley, Margaret Hughes, Miren Iturriza-Gomara, James Johnson, L Luu, Jenifer Manson, Charlotte Nelson, Elaine O'Toole, Cassie Olateju, Rebekah Penrice-Randal , Lucille Rainbow, N.P Randle, Trevor Ian Robinson, Parul Sharma, Ghada T Shawli, James P Stewart, Neil Swainston, Ecaterina Vamos, Joanne Watts, Mark Whitehead |
| EPI_ISL_705247 | University College London Hospital | COVID-19 Genomics UK (COG-UK) Consortium | Judith Heaney, Matthew Byott, Catherine Houlihan, Dan Frampton, Stuart Kirk, Moira Spyer and Eleni Nastouli |
| EPI_ISL_705251 | University of Exeter | COVID-19 Genomics UK (COG-UK) Consortium | Ben Temperton,Aaron Jeffries,Michelle Michelsen,Joanna Warwick-Dugdale,Audrey Farbos,Robyn Manley,Stephen Michell,Jane Masoli |
| EPI_ISL_705272, EPI_ISL_705288 | Virology Department, Sheffield Teaching Hospitals NHS Foundation Trust/Department of Infection, Immunity and Cardiovascular Disease, The Medical School, University of Sheffield | COVID-19 Genomics UK (COG-UK) Consortium | Thushan de Silva, Matthew Parker, Nikki Smith, Adri Angyal, Rebecca Brown, Luke Green, Rachel Tucker, Paul Parsons, Danielle Groves, Katie Johnson, Laura Carrilero, Alex Keeley, Dave Partridge, Matthew Wyles, Benjamin Lindsey, Mehmet Yavuz, Mohammad Raza, Cariad Evans |
| EPI_ISL_705291, EPI_ISL_705292, EPI_ISL_705303, EPI_ISL_705315, EPI_ISL_705321, EPI_ISL_705324 | Liverpool Clinical Laboratories | COVID-19 Genomics UK (COG-UK) Consortium | Sam Haldenby, Anita Lucaci, Steve Paterson, Julian Hiscox, Alistair Darby, M Almsaud, A Alrezaihi, Muhannad Alruwaili, Stuart D Armstrong, Jones Benjamin, Eleanor G Bentley, Anu Chawla, Jordan J Clark, Angela Cowell, Richard Eccles, Isabel García-Dorival, Matthew Gemmell, Alessandro Gerada, PKF Gilmore, Richard Gregory, Ximeng Han, Catherine Hartley, Margaret Hughes, Miren Iturriza-Gomara, James Johnson, L Luu, Jenifer Manson, Charlotte Nelson, Elaine O'Toole, Cassie Olateju, Rebekah Penrice-Randal , Lucille Rainbow, N.P Randle, Trevor Ian Robinson, Parul Sharma, Ghada T Shawli, James P Stewart, Neil Swainston, Ecaterina Vamos, Joanne Watts, Mark Whitehead |
| EPI_ISL_705332 | Quadram Institute Bioscience | COVID-19 Genomics UK (COG-UK) Consortium | Dave J. Baker, Gemma L. Kay, Alp Aydin, Thanh Le-Viet, Steven Rudder, Ana P. Tedim, Anastasia Kolyva, Maria Diaz, Leonardo de Oliveira Martins, Nabil-Fareed Alikhan, Lizzie Meadows, Rachael Stanley, Ngozi Elumogo, Muhammed Yasir, Nicholas M. Thomson, Alexander J Trotter, Rachel Gilroy, Samuel Bloomfield, Claire Stuart, Andrew Bell, Reenesh Prakash, Samir Dervisevic, Alison E. Mather, John Wain, Mark Webber, Andrew J. Page, Justin O'Grady |
| EPI_ISL_705336 | Liverpool Clinical Laboratories | COVID-19 Genomics UK (COG-UK) Consortium | Sam Haldenby, Anita Lucaci, Steve Paterson, Julian Hiscox, Alistair Darby, M Almsaud, A Alrezaihi, Muhannad Alruwaili, Stuart D Armstrong, Jones Benjamin, Eleanor G Bentley, Anu Chawla, Jordan J Clark, Angela Cowell, Richard Eccles, Isabel García-Dorival, Matthew Gemmell, Alessandro Gerada, PKF Gilmore, Richard Gregory, Ximeng Han, Catherine Hartley, Margaret Hughes, Miren Iturriza-Gomara, James Johnson, L Luu, Jenifer Manson, Charlotte Nelson, Elaine O'Toole, Cassie Olateju, Rebekah Penrice-Randal , Lucille Rainbow, N.P Randle, Trevor Ian Robinson, Parul Sharma, Ghada T Shawli, James P Stewart, Neil Swainston, Ecaterina Vamos, Joanne Watts, Mark Whitehead |
| EPI_ISL_705337 | University College London, Great Ormond Street Hospital for Children NHS Foundation Trust, Imperial College Healthcare NHS Trust | COVID-19 Genomics UK (COG-UK) Consortium | Sergi Castellano, Rachel Williams, Mark Kristiansen, Paola Resende Silva, Sunando Roy, Tony Brooks, Helena Tutill, Paola Niola, Patricia Dyal, Charlotte Williams, Leysa Forrest, Yasmin Panchbhaya, Jacqueline Findlay, Samuel Weeks, Julianne Brown, Kathryn Harris, Paul Randell, James Price, Alison Holmes, Judith Breuer |
| EPI_ISL_705349 | Liverpool Clinical Laboratories | COVID-19 Genomics UK (COG-UK) Consortium | Sam Haldenby, Anita Lucaci, Steve Paterson, Julian Hiscox, Alistair Darby, M Almsaud, A Alrezaihi, Muhannad Alruwaili, Stuart D Armstrong, Jones Benjamin, Eleanor G Bentley, Anu Chawla, Jordan J Clark, Angela Cowell, Richard Eccles, Isabel García-Dorival, Matthew Gemmell, Alessandro Gerada, PKF Gilmore, Richard Gregory, Ximeng Han, Catherine Hartley, Margaret Hughes, Miren Iturriza-Gomara, James Johnson, L Luu, Jenifer Manson, |

| | | | |
|---|---|---|---|
| | | | Charlotte Nelson, Elaine O'Toole, Cassie Olateju, Rebekah Penrice-Randal , Lucille Rainbow, N.P Randle, Trevor Ian Robinson, Parul Sharma, Ghada T Shawli, James P Stewart, Neil Swainston, Ecaterina Vamos, Joanne Watts, Mark Whitehead |
| EPI_ISL_705351 | Virology Department, Sheffield Teaching Hospitals NHS Foundation Trust/Department of Infection, Immunity and Cardiovascular Disease, The Medical School, University of Sheffield | COVID-19 Genomics UK (COG-UK) Consortium | Thushan de Silva, Matthew Parker, Nikki Smith, Adri Angyal, Rebecca Brown, Luke Green, Rachel Tucker, Paul Parsons, Danielle Groves, Katie Johnson, Laura Carrilero, Alex Keeley, Dave Partridge, Matthew Wyles, Benjamin Lindsey, Mehmet Yavuz, Mohammad Raza, Cariad Evans |
| EPI_ISL_705357 | Northumbria University / South Tees Hospitals NHS Foundation Trust / North Cumbria Integrated Care NHS Foundation Trust / North Tees and Hartlepool NHS Foundation Trust / Newcastle Hospitals NHS Foundation Trust | COVID-19 Genomics UK (COG-UK) Consortium | Darren L Smith,Andrew Nelson,Matthew Bashton,Greg R Young,Joshua Loh,John Allan,Mohammad A Tariq,Giles S Holt,Gary Black,Wen C Yew,Lynn Dover,Paul Baker,Steve Liggett,Sarah Essex,Jane Greenaway,Debra Padgett,Clive Graham,Garren Scott,Edward Barton,Emma Swindells,Brendan Payne,Jennifer Collins,Yusri Taha,Gary Eltringham |
| EPI_ISL_705372, EPI_ISL_705377 | Liverpool Clinical Laboratories | COVID-19 Genomics UK (COG-UK) Consortium | Sam Haldenby, Anita Lucaci, Steve Paterson, Julian Hiscox, Alistair Darby, M Almsaud, A Alrezaihi, Muhannad Alruwaili, Stuart D Armstrong, Jones Benjamin, Eleanor G Bentley, Anu Chawla, Jordan J Clark, Angela Cowell, Richard Eccles, Isabel García-Dorival, Matthew Gemmell, Alessandro Gerada, PKF Gilmore, Richard Gregory, Ximeng Han, Catherine Hartley, Margaret Hughes, Miren Iturriza-Gomara, James Johnson, L Luu, Jenifer Manson, Charlotte Nelson, Elaine O'Toole, Cassie Olateju, Rebekah Penrice-Randal , Lucille Rainbow, N.P Randle, Trevor Ian Robinson, Parul Sharma, Ghada T Shawli, James P Stewart, Neil Swainston, Ecaterina Vamos, Joanne Watts, Mark Whitehead |
| EPI_ISL_705380 | Queens Medical Centre, Clinical Microbiology Department / DeepSeq Nottingham | COVID-19 Genomics UK (COG-UK) Consortium | Gemma Clark, Wendy Smith, Manjinder Khakh, Vicki M Fleming, Michelle M Lister, Hannah Howson-Wells, Jonathan Ball, Patrick McClure, Joseph Chappell, Theocharis Tsoleridis, Nadine Holmes, Matthew Carlisle, Christopher Moore, Fei Sang, Johnny Debebe, Victoria Wright, Matthew Loose |
| EPI_ISL_705385, EPI_ISL_705386, EPI_ISL_705387 | Liverpool Clinical Laboratories | COVID-19 Genomics UK (COG-UK) Consortium | Sam Haldenby, Anita Lucaci, Steve Paterson, Julian Hiscox, Alistair Darby, M Almsaud, A Alrezaihi, Muhannad Alruwaili, Stuart D Armstrong, Jones Benjamin, Eleanor G Bentley, Anu Chawla, Jordan J Clark, Angela Cowell, Richard Eccles, Isabel García-Dorival, Matthew Gemmell, Alessandro Gerada, PKF Gilmore, Richard Gregory, Ximeng Han, Catherine Hartley, Margaret Hughes, Miren Iturriza-Gomara, James Johnson, L Luu, Jenifer Manson, Charlotte Nelson, Elaine O'Toole, Cassie Olateju, Rebekah Penrice-Randal , Lucille Rainbow, N.P Randle, Trevor Ian Robinson, Parul Sharma, Ghada T Shawli, James P Stewart, Neil Swainston, Ecaterina Vamos, Joanne Watts, Mark Whitehead |
| EPI_ISL_705399 | West of Scotland Specialist Virology Centre, NHSGGC / MRC-University of Glasgow Centre for Virus Research | COVID-19 Genomics UK (COG-UK) Consortium | Ana da Silva Filipe, Natasha Johnson, Kathy Smollett, Daniel Mair, Stephen Carmichael, Alice Broos, Lily Tong, Jenna Nichols, Kyriaki Nomikou; Sarah McDonald; Richard Orton, Joseph Hughes, Sreenu Vattipally, David L Robertson; Alasdair MacLean, Rory Gunson; Sharif Shaaban, Matthew Holden; Rachel Blacow, Guy Mollett, Kathy Li, James Shepherd, Antonia Ho, Emma Thomson |
| EPI_ISL_705400 | Liverpool Clinical Laboratories | COVID-19 Genomics UK (COG-UK) Consortium | Sam Haldenby, Anita Lucaci, Steve Paterson, Julian Hiscox, Alistair Darby, M Almsaud, A Alrezaihi, Muhannad Alruwaili, Stuart D Armstrong, Jones Benjamin, Eleanor G Bentley, Anu Chawla, Jordan J Clark, Angela Cowell, Richard Eccles, Isabel García-Dorival, Matthew Gemmell, Alessandro Gerada, PKF Gilmore, Richard Gregory, Ximeng Han, Catherine Hartley, Margaret Hughes, Miren Iturriza-Gomara, James Johnson, L Luu, Jenifer Manson, Charlotte Nelson, Elaine O'Toole, Cassie Olateju, Rebekah Penrice-Randal , Lucille Rainbow, N.P Randle, Trevor Ian Robinson, Parul Sharma, Ghada T Shawli, James P Stewart, Neil Swainston, Ecaterina Vamos, Joanne Watts, Mark Whitehead |
| EPI_ISL_705401 | Quadram Institute Bioscience | COVID-19 Genomics UK (COG-UK) Consortium | Dave J. Baker, Gemma L. Kay, Alp Aydin, Thanh Le-Viet, Steven Rudder, Ana P. Tedim, Anastasia Kolyva, Maria Diaz, Leonardo de Oliveira Martins, Nabil-Fareed Alikhan, Lizzie Meadows, Rachael Stanley, Ngozi Elumogo, Muhammed Yasir, Nicholas M. Thomson, Alexander J Trotter, Rachel Gilroy, Samuel Bloomfield, Claire Stuart, Andrew Bell, Reenesh Prakash, Samir Dervisevic, Alison E. Mather, John Wain, Mark Webber, Andrew J. Page, Justin O'Grady |
| EPI_ISL_705403 | Liverpool Clinical Laboratories | COVID-19 Genomics UK (COG-UK) Consortium | Sam Haldenby, Anita Lucaci, Steve Paterson, Julian Hiscox, Alistair Darby, M Almsaud, A Alrezaihi, Muhannad Alruwaili, Stuart D Armstrong, Jones Benjamin, Eleanor G Bentley, Anu Chawla, Jordan J Clark, Angela Cowell, Richard Eccles, Isabel García-Dorival, Matthew Gemmell, Alessandro Gerada, PKF Gilmore, Richard Gregory, Ximeng Han, Catherine Hartley, Margaret Hughes, Miren Iturriza-Gomara, James Johnson, L Luu, Jenifer Manson, Charlotte Nelson, Elaine O'Toole, Cassie Olateju, Rebekah Penrice-Randal , Lucille Rainbow, N.P Randle, Trevor Ian Robinson, Parul Sharma, Ghada T Shawli, James P Stewart, Neil Swainston, Ecaterina Vamos, Joanne Watts, Mark Whitehead |
| EPI_ISL_705432 | Quadram Institute Bioscience | COVID-19 Genomics UK (COG-UK) Consortium | Dave J. Baker, Gemma L. Kay, Alp Aydin, Thanh Le-Viet, Steven Rudder, Ana P. Tedim, Anastasia Kolyva, Maria Diaz, Leonardo de Oliveira Martins, Nabil-Fareed Alikhan, Lizzie Meadows, Rachael Stanley, Ngozi Elumogo, Muhammed Yasir, Nicholas M. Thomson, Alexander J Trotter, Rachel Gilroy, Samuel Bloomfield, Claire Stuart, Andrew Bell, Reenesh Prakash, Samir Dervisevic, Alison E. Mather, John Wain, Mark Webber, Andrew J. Page, Justin O'Grady |
| EPI_ISL_705457 | Northumbria University / South Tees Hospitals NHS Foundation Trust / North Cumbria Integrated Care NHS Foundation Trust / North Tees and Hartlepool NHS Foundation Trust / Newcastle Hospitals NHS Foundation Trust | COVID-19 Genomics UK (COG-UK) Consortium | Darren L Smith,Andrew Nelson,Matthew Bashton,Greg R Young,Joshua Loh,John Allan,Mohammad A Tariq,Giles S Holt,Gary Black,Wen C Yew,Lynn Dover,Paul Baker,Steve Liggett,Sarah Essex,Jane Greenaway,Debra Padgett,Clive Graham,Garren Scott,Edward Barton,Emma Swindells,Brendan Payne,Jennifer Collins,Yusri Taha,Gary Eltringham |
| EPI_ISL_705536, EPI_ISL_705555, EPI_ISL_705556 | Department of Pathology, University of Cambridge | COVID-19 Genomics UK (COG-UK) Consortium | Aminu S. Jahun, Yasmin Chaudhry, Grant Hall, Iliana Georgana, Myra Hosmillo, Martin D. Curran, Malte Pinckert, Surendra Parmar, Ian Goodfellow |
| EPI_ISL_705698, EPI_ISL_705699, EPI_ISL_705700, EPI_ISL_705702, EPI_ISL_705703, EPI_ISL_705704, EPI_ISL_705705, EPI_ISL_705706, EPI_ISL_705707, EPI_ISL_705712, EPI_ISL_705713, EPI_ISL_705714, EPI_ISL_705716, EPI_ISL_705718, EPI_ISL_705720, EPI_ISL_705722, EPI_ISL_705726, EPI_ISL_705727, EPI_ISL_705731 | | | |
| see above | West of Scotland Specialist Virology Centre, NHSGGC / MRC-University of Glasgow Centre for Virus Research | COVID-19 Genomics UK (COG-UK) Consortium | Ana da Silva Filipe, Natasha Johnson, Kathy Smollett, Daniel Mair, Stephen Carmichael, Alice Broos, Lily Tong, Jenna Nichols, Kyriaki Nomikou; Sarah McDonald; Richard Orton, Joseph Hughes, Sreenu Vattipally, David L Robertson; Alasdair MacLean, Rory Gunson; Sharif Shaaban, Matthew Holden; Rachel Blacow, Guy Mollett, Kathy Li, James Shepherd, Antonia Ho, Emma Thomson |
| EPI_ISL_705734, EPI_ISL_705735, EPI_ISL_705736, EPI_ISL_705739, EPI_ISL_705740, EPI_ISL_705742, EPI_ISL_705743, EPI_ISL_705744, EPI_ISL_705745, EPI_ISL_705746, EPI_ISL_705747, EPI_ISL_705748, EPI_ISL_705749, EPI_ISL_705750, EPI_ISL_705774, EPI_ISL_705775, EPI_ISL_705776 | | | |
| see above | Virology Department, Royal Infirmary of Edinburgh, NHS Lothian / School of Biological Sciences, University of Edinburgh / Institute of Genetics and Molecular Medicine, University of Edinburgh | COVID-19 Genomics UK (COG-UK) Consortium | McHugh M, Dewar R, Rooke S, Gallagher M, Balcaza C, O'Toole Á, Scher E, Hill V, McCrone JT, Colquhoun R, Yu X, Jackson B, Rambaut A, Williams TC, Templeton K |
| EPI_ISL_705824, EPI_ISL_705825, EPI_ISL_705827, EPI_ISL_705828, EPI_ISL_705830, EPI_ISL_705832, EPI_ISL_705833, EPI_ISL_705834, EPI_ISL_705835, EPI_ISL_705837, EPI_ISL_705839, EPI_ISL_705840, EPI_ISL_705841, EPI_ISL_705842, EPI_ISL_705843, EPI_ISL_705844, EPI_ISL_705845, EPI_ISL_705846, EPI_ISL_705847, EPI_ISL_705848, EPI_ISL_705849, EPI_ISL_705850, EPI_ISL_705851, EPI_ISL_705852, EPI_ISL_705853, EPI_ISL_705854, EPI_ISL_705855, EPI_ISL_705856, EPI_ISL_705857, EPI_ISL_705858, EPI_ISL_705921, EPI_ISL_705922, EPI_ISL_705923, EPI_ISL_705924, EPI_ISL_705926, EPI_ISL_705927 | | | |
| see above | Liverpool Clinical Laboratories | COVID-19 Genomics UK (COG-UK) Consortium | Sam Haldenby, Anita Lucaci, Steve Paterson, Julian Hiscox, Alistair Darby, M Almsaud, A Alrezaihi, Muhannad Alruwaili, Stuart D Armstrong, Jones Benjamin, Eleanor G Bentley, Anu Chawla, Jordan J Clark, Angela Cowell, Richard Eccles, Isabel García-Dorival, Matthew Gemmell, Alessandro Gerada, PKF Gilmore, Richard Gregory, Ximeng Han, Catherine Hartley, Margaret Hughes, Miren Iturriza-Gomara, James Johnson, L Luu, Jenifer Manson, Charlotte Nelson, Elaine O'Toole, Cassie Olateju, Rebekah Penrice-Randal , Lucille Rainbow, N.P Randle, Trevor Ian Robinson, Parul Sharma, Ghada T Shawli, James P Stewart, Neil Swainston, Ecaterina Vamos, Joanne Watts, Mark Whitehead |
| EPI_ISL_705945, EPI_ISL_705946, EPI_ISL_705947, EPI_ISL_705948, EPI_ISL_705950, EPI_ISL_705951, EPI_ISL_705952, EPI_ISL_705953, EPI_ISL_705954, EPI_ISL_705955, EPI_ISL_705956 | | | |
| see above | University College London Hospital | COVID-19 Genomics UK (COG-UK) Consortium | Judith Heaney, Matthew Byott, Catherine Houlihan, Dan Frampton, Stuart Kirk, Moira Spyer and Eleni Nastouli |
| EPI_ISL_706280, EPI_ISL_706281, EPI_ISL_706282, EPI_ISL_706284, EPI_ISL_706285, EPI_ISL_706331 | Northumbria University / South Tees Hospitals NHS Foundation Trust / North Cumbria Integrated Care NHS Foundation Trust / North Tees and Hartlepool NHS Foundation Trust / Newcastle Hospitals NHS Foundation Trust | COVID-19 Genomics UK (COG-UK) Consortium | Darren L Smith,Andrew Nelson,Matthew Bashton,Greg R Young,Joshua Loh,John Allan,Mohammad A Tariq,Giles S Holt,Gary Black,Wen C Yew,Lynn Dover,Paul Baker,Steve Liggett,Sarah Essex,Jane Greenaway,Debra Padgett,Clive Graham,Garren Scott,Edward Barton,Emma Swindells,Brendan Payne,Jennifer Collins,Yusri Taha,Gary Eltringham |
| EPI_ISL_706356, EPI_ISL_706359, EPI_ISL_706363, EPI_ISL_706364, EPI_ISL_706365, EPI_ISL_706366, EPI_ISL_706384, EPI_ISL_706385, EPI_ISL_706386, EPI_ISL_706387, EPI_ISL_706388, EPI_ISL_706389, EPI_ISL_706390, EPI_ISL_706391, EPI_ISL_706392, EPI_ISL_706393, EPI_ISL_706394, EPI_ISL_706395, EPI_ISL_706396, EPI_ISL_706400, EPI_ISL_706402, EPI_ISL_706414, EPI_ISL_706417 | | | |
| see above | Quadram Institute Bioscience | COVID-19 Genomics UK (COG-UK) Consortium | Dave J. Baker, Gemma L. Kay, Alp Aydin, Thanh Le-Viet, Steven Rudder, Ana P. Tedim, Anastasia Kolyva, Maria Diaz, Leonardo de Oliveira Martins, Nabil-Fareed Alikhan, Lizzie Meadows, Rachael Stanley, Ngozi Elumogo, Muhammed Yasir, Nicholas M. Thomson, Alexander J Trotter, Rachel Gilroy, |

| | | | |
|---|---|---|---|
| | | | Samuel Bloomfield, Claire Stuart, Andrew Bell, Reenesh Prakash, Samir Dervisevic, Alison E. Mather, John Wain, Mark Webber, Andrew J. Page, Justin O'Grady |
| EPI_ISL_706422, EPI_ISL_706423, EPI_ISL_706424, EPI_ISL_706425, EPI_ISL_706426, EPI_ISL_706427, EPI_ISL_706428, EPI_ISL_706431, EPI_ISL_706432, EPI_ISL_706433, EPI_ISL_706434 | | | |
| see above | Queens Medical Centre, Clinical Microbiology Department / DeepSeq Nottingham | COVID-19 Genomics UK (COG-UK) Consortium | Gemma Clark, Wendy Smith, Manjinder Khakh, Vicki M Fleming, Michelle M Lister, Hannah Howson-Wells, Jonathan Ball, Patrick McClure, Joseph Chappell, Theocharis Tsoleridis, Nadine Holmes, Matthew Carlisle, Christopher Moore, Fei Sang, Johnny Debebe, Victoria Wright, Matthew Loose |
| EPI_ISL_706987, EPI_ISL_706989, EPI_ISL_706992, EPI_ISL_706993, EPI_ISL_706995, EPI_ISL_706996, EPI_ISL_706997, EPI_ISL_706998, EPI_ISL_706999, EPI_ISL_707000, EPI_ISL_707001, EPI_ISL_707002, EPI_ISL_707003, EPI_ISL_707005, EPI_ISL_707006 | | | |
| see above | Virology Department, Sheffield Teaching Hospitals NHS Foundation Trust/Department of Infection, Immunity and Cardiovascular Disease, The Medical School, University of Sheffield | COVID-19 Genomics UK (COG-UK) Consortium | Thushan de Silva, Matthew Parker, Nikki Smith, Adri Angyal, Rebecca Brown, Luke Green, Rachel Tucker, Paul Parsons, Danielle Groves, Katie Johnson, Laura Carrilero, Alex Keeley, Dave Partridge, Matthew Wyles, Benjamin Lindsey, Mehmet Yavuz, Mohammad Raza, Cariad Evans |
| EPI_ISL_707707 | Institute for Urban Disease Control and Prevention | COVID-19 Network Investigations (CONI) Alliance | Kamolthip Atsawawaranunt, Elizabeth Batty, Wasun Chantratita, Thanat Chookajorn, Stefan Fernandez, Angkana Huang, Anthony R. Jones, Khajohn Joonsalak, Prayuth Kaewmalang, Amornmas Kongklieng, Chonticha Klungtong, Theerarat Kochakarn, Namfon Kotanan, Krittikorn Kumpornsin, Wudtichai Manasatienkij, Anek Mungaomklang, Bhakbhoom Panthan, Ekawat Pasomsub, Pukkaporn Parnwijitkul, Kingkan Rakmanee, Insee Sensorn, Janjira Thaipadungpanit, Arporn Wangwiwatsin,Treewat Watthanachockchai |
| EPI_ISL_707794 | Waikato Hospital | Institute of Environmental Science and Research (ESR) | Xiaoyun Ren, Matt Storey, Nikki Freed, Muhammad Faisal, Jing Wang, Hermes Perez, Anja Werno, Antje van der Linden, Arlo Upton, Chris Mansell, David Hammer, Dragana Drinkovic, Gary McAuliffe, Hana Sofia Andersson, James Ussher, Jill Sherwood, Josh Freeman, Julia Howard, Juliet Elvy, Mary DeAlmeida, Matt Blakiston, Matthew Rogers, Max Bloomfield, Michael Addidle, Michelle Balm, Sally Roberts, Sarah Jefferies, Sharmini Muttaiyah, Susan Morpeth, Susan Taylor, Timothy Blackmore, Vani Sathyendran, Veronica Playle, Virginia Hope, Erasmus Smit, Lauren Jelly, Olin Silander, Joep de Ligt |
| EPI_ISL_708487, EPI_ISL_708489, EPI_ISL_708490, EPI_ISL_708491, EPI_ISL_708492 | Minnesota Department of Health, Public Health Laboratory | Minnesota Department of Health, Public Health Laboratory | Alexandra Lorentz, Jacob Garfin, Matt Plumb, and Xiong Wang |
| EPI_ISL_708788, EPI_ISL_708789, EPI_ISL_708790, EPI_ISL_708791, EPI_ISL_708792, EPI_ISL_708794 | PathWest Laboratory Medicine WA | PathWest Laboratory Medicine WA Microbial Surveillance Unit | PathWest Laboratory Medicine WA Microbial Surveillance Unit |
| EPI_ISL_708863 | Lighthouse Lab in Alderley Park | Wellcome Sanger Institute for the COVID-19 Genomics UK (COG-UK) Consortium | Jacquelyn Wynn, Mairead Hyland, The Lighthouse Lab in Alderley Park and Alex Alderton, Roberto Amato, Sonia Goncalves, Ewan Harrison, David K. Jackson, Ian Johnston, Dominic Kwiatkowski, Cordelia Langford, John Sillitoe on behalf of the Wellcome Sanger Institute COVID-19 Surveillance Team |
| EPI_ISL_708998, EPI_ISL_709048 | Lighthouse Lab in Glasgow | Wellcome Sanger Institute for the COVID-19 Genomics UK (COG-UK) Consortium | Harper VanSteenhouse, Yumi Kasai, David Gray, Carol Clugston, Anna Dominiczak and Alex Alderton, Roberto Amato, Sonia Goncalves, Ewan Harrison, David K. Jackson, Ian Johnston, Dominic Kwiatkowski, Cordelia Langford, John Sillitoe on behalf of the Wellcome Sanger Institute COVID-19 Surveillance Team |
| EPI_ISL_709567, EPI_ISL_709568, EPI_ISL_709569, EPI_ISL_709570, EPI_ISL_709571, EPI_ISL_709572, EPI_ISL_709573, EPI_ISL_709574, EPI_ISL_709575, EPI_ISL_709576, EPI_ISL_709577, EPI_ISL_709578, EPI_ISL_709579, EPI_ISL_709580, EPI_ISL_709581, EPI_ISL_709582, EPI_ISL_709583, EPI_ISL_709584, EPI_ISL_709585, EPI_ISL_709586, EPI_ISL_709587, EPI_ISL_709588, EPI_ISL_709589, EPI_ISL_709590, EPI_ISL_709591, EPI_ISL_709592, EPI_ISL_709593, EPI_ISL_709594, EPI_ISL_709595, EPI_ISL_709596, EPI_ISL_709597, EPI_ISL_709598, EPI_ISL_709599, EPI_ISL_709600, EPI_ISL_709601, EPI_ISL_709602, EPI_ISL_709603, EPI_ISL_709604, EPI_ISL_709605, EPI_ISL_709606, EPI_ISL_709607, EPI_ISL_709608, EPI_ISL_709609, EPI_ISL_709610, EPI_ISL_709611, EPI_ISL_709612, EPI_ISL_709613, EPI_ISL_709614, EPI_ISL_709615, EPI_ISL_709616, EPI_ISL_709617, EPI_ISL_709618, EPI_ISL_709619, EPI_ISL_709620, EPI_ISL_709621, EPI_ISL_709622, EPI_ISL_709623, EPI_ISL_709624, EPI_ISL_709625, EPI_ISL_709626, EPI_ISL_709627, EPI_ISL_709628, EPI_ISL_709629, EPI_ISL_709630, EPI_ISL_709631, EPI_ISL_709632, EPI_ISL_709633, EPI_ISL_709634, EPI_ISL_709635, EPI_ISL_709636, EPI_ISL_709637, EPI_ISL_709638, EPI_ISL_709639, EPI_ISL_709640, EPI_ISL_709641, EPI_ISL_709642, EPI_ISL_709643, EPI_ISL_709644, EPI_ISL_709645, EPI_ISL_709646, EPI_ISL_709647, EPI_ISL_709648, EPI_ISL_709649, EPI_ISL_709650, EPI_ISL_709651, EPI_ISL_709652, EPI_ISL_709653, EPI_ISL_709654, EPI_ISL_709655, EPI_ISL_709656, EPI_ISL_709657, EPI_ISL_709658, EPI_ISL_709659, EPI_ISL_709660, EPI_ISL_709661, EPI_ISL_709662, EPI_ISL_709663, EPI_ISL_709664, EPI_ISL_709665, EPI_ISL_709666, EPI_ISL_709667, EPI_ISL_709668, EPI_ISL_709669, EPI_ISL_709670, EPI_ISL_709671, EPI_ISL_709672, EPI_ISL_709673, EPI_ISL_709674, EPI_ISL_709675, EPI_ISL_709676, EPI_ISL_709677, EPI_ISL_709678, EPI_ISL_709679, EPI_ISL_709680, EPI_ISL_709681, EPI_ISL_709682, EPI_ISL_709683, EPI_ISL_709684, EPI_ISL_709685, EPI_ISL_709686, EPI_ISL_709687, EPI_ISL_709688, EPI_ISL_709689, EPI_ISL_709690, EPI_ISL_709691, EPI_ISL_709692, EPI_ISL_709693, EPI_ISL_709694, EPI_ISL_709695, EPI_ISL_709696, EPI_ISL_709697, EPI_ISL_709698, EPI_ISL_709699, EPI_ISL_709700, EPI_ISL_709701, EPI_ISL_709702, EPI_ISL_709703, EPI_ISL_709704, EPI_ISL_709705, EPI_ISL_709706, EPI_ISL_709707, EPI_ISL_709708, EPI_ISL_709709, EPI_ISL_709710, EPI_ISL_709711, EPI_ISL_709712, EPI_ISL_709713, EPI_ISL_709714, EPI_ISL_709715, EPI_ISL_709716, EPI_ISL_709717, EPI_ISL_709718, EPI_ISL_709719, EPI_ISL_709720, EPI_ISL_709721, EPI_ISL_709722, EPI_ISL_709723, EPI_ISL_709724, EPI_ISL_709725, EPI_ISL_709726, EPI_ISL_709727, EPI_ISL_709728, EPI_ISL_709729, EPI_ISL_709730, EPI_ISL_709731, EPI_ISL_709732, EPI_ISL_709733, EPI_ISL_709734, EPI_ISL_709735, EPI_ISL_709736, EPI_ISL_709737, EPI_ISL_709738, EPI_ISL_709739, EPI_ISL_709740, EPI_ISL_709741, EPI_ISL_709742, EPI_ISL_709743, EPI_ISL_709744, EPI_ISL_709745, EPI_ISL_709746, EPI_ISL_709747, EPI_ISL_709748, EPI_ISL_709749, EPI_ISL_709750, EPI_ISL_709751, EPI_ISL_709752, EPI_ISL_709753, EPI_ISL_709754, EPI_ISL_709755, EPI_ISL_709756, EPI_ISL_709757, EPI_ISL_709758, EPI_ISL_709759, EPI_ISL_709760, EPI_ISL_709761, EPI_ISL_709762, EPI_ISL_709763, EPI_ISL_709764, EPI_ISL_709765, EPI_ISL_709766, EPI_ISL_709767, EPI_ISL_709768, EPI_ISL_709769, EPI_ISL_709770, EPI_ISL_709771, EPI_ISL_709772, EPI_ISL_709773, EPI_ISL_709774, EPI_ISL_709775, EPI_ISL_709776, EPI_ISL_709777, EPI_ISL_709778, EPI_ISL_709779, EPI_ISL_709780, EPI_ISL_709781, EPI_ISL_709782, EPI_ISL_709783, EPI_ISL_709784, EPI_ISL_709785, EPI_ISL_709786, EPI_ISL_709787, EPI_ISL_709788, EPI_ISL_709789, EPI_ISL_709790, EPI_ISL_709791, EPI_ISL_709792, EPI_ISL_709793, EPI_ISL_709794, EPI_ISL_709795, EPI_ISL_709796, EPI_ISL_709797, EPI_ISL_709798, EPI_ISL_709799, EPI_ISL_709801, EPI_ISL_709802, EPI_ISL_709803 | | | |
| see above | Lighthouse Lab in Milton Keynes | Wellcome Sanger Institute for the COVID-19 Genomics UK (COG-UK) Consortium | The Lighthouse Lab in Milton Keynes and Alex Alderton, Roberto Amato, Sonia Goncalves, Ewan Harrison, David K. Jackson, Ian Johnston, Dominic Kwiatkowski, Cordelia Langford, John Sillitoe on behalf of the Wellcome Sanger Institute COVID-19 Surveillance Team |
| EPI_ISL_709805, EPI_ISL_709807, EPI_ISL_709808, EPI_ISL_709810, EPI_ISL_709813, EPI_ISL_709814 | Lighthouse Lab in Cambridge | Wellcome Sanger Institute for the COVID-19 Genomics UK (COG-UK) Consortium | Rob Howes, The Lighthouse Lab in Cambridge and Alex Alderton, Roberto Amato, Sonia Goncalves, Ewan Harrison, David K. Jackson, Ian Johnston, Dominic Kwiatkowski, Cordelia Langford, John Sillitoe on behalf of the Wellcome Sanger Institute COVID-19 Surveillance Team |
| EPI_ISL_709815 | Lighthouse Lab in Milton Keynes | Wellcome Sanger Institute for the COVID-19 Genomics UK (COG-UK) Consortium | The Lighthouse Lab in Milton Keynes and Alex Alderton, Roberto Amato, Sonia Goncalves, Ewan Harrison, David K. Jackson, Ian Johnston, Dominic Kwiatkowski, Cordelia Langford, John Sillitoe on behalf of the Wellcome Sanger Institute COVID-19 Surveillance Team |
| EPI_ISL_709817 | Lighthouse Lab in Cambridge | Wellcome Sanger Institute for the COVID-19 Genomics UK (COG-UK) Consortium | Rob Howes, The Lighthouse Lab in Cambridge and Alex Alderton, Roberto Amato, Sonia Goncalves, Ewan Harrison, David K. Jackson, Ian Johnston, Dominic Kwiatkowski, Cordelia Langford, John Sillitoe on behalf of the Wellcome Sanger Institute COVID-19 Surveillance Team |
| EPI_ISL_709819 | Lighthouse Lab in Milton Keynes | Wellcome Sanger Institute for the COVID-19 Genomics UK (COG-UK) Consortium | The Lighthouse Lab in Milton Keynes and Alex Alderton, Roberto Amato, Sonia Goncalves, Ewan Harrison, David K. Jackson, Ian Johnston, Dominic Kwiatkowski, Cordelia Langford, John Sillitoe on behalf of the Wellcome Sanger Institute COVID-19 Surveillance Team |
| EPI_ISL_709822 | Lighthouse Lab in Cambridge | Wellcome Sanger Institute for the COVID-19 Genomics UK (COG-UK) Consortium | Rob Howes, The Lighthouse Lab in Cambridge and Alex Alderton, Roberto Amato, Sonia Goncalves, Ewan Harrison, David K. Jackson, Ian Johnston, Dominic Kwiatkowski, Cordelia Langford, John Sillitoe on behalf of the Wellcome Sanger Institute COVID-19 Surveillance Team |
| EPI_ISL_709823, EPI_ISL_709824 | Lighthouse Lab in Milton Keynes | Wellcome Sanger Institute for the COVID-19 Genomics UK (COG-UK) Consortium | The Lighthouse Lab in Milton Keynes and Alex Alderton, Roberto Amato, Sonia Goncalves, Ewan Harrison, David K. Jackson, Ian Johnston, Dominic Kwiatkowski, Cordelia Langford, John Sillitoe on behalf of the Wellcome Sanger Institute COVID-19 Surveillance Team |
| EPI_ISL_709825 | Lighthouse Lab in Cambridge | Wellcome Sanger Institute for the COVID-19 Genomics UK (COG-UK) Consortium | Rob Howes, The Lighthouse Lab in Cambridge and Alex Alderton, Roberto Amato, Sonia Goncalves, Ewan Harrison, David K. Jackson, Ian Johnston, Dominic Kwiatkowski, Cordelia Langford, John Sillitoe on behalf of the Wellcome Sanger Institute COVID-19 Surveillance Team |
| EPI_ISL_709828 | Lighthouse Lab in Milton Keynes | Wellcome Sanger Institute for the COVID-19 Genomics UK (COG-UK) Consortium | The Lighthouse Lab in Milton Keynes and Alex Alderton, Roberto Amato, Sonia Goncalves, Ewan Harrison, David K. Jackson, Ian Johnston, Dominic Kwiatkowski, Cordelia Langford, John Sillitoe on behalf of the Wellcome Sanger Institute COVID-19 Surveillance Team |
| EPI_ISL_709829 | Lighthouse Lab in Cambridge | Wellcome Sanger Institute for the COVID-19 Genomics UK (COG-UK) Consortium | Rob Howes, The Lighthouse Lab in Cambridge and Alex Alderton, Roberto Amato, Sonia Goncalves, Ewan Harrison, David K. Jackson, Ian Johnston, Dominic Kwiatkowski, Cordelia Langford, John Sillitoe on behalf of the Wellcome Sanger Institute COVID-19 Surveillance Team |
| EPI_ISL_709830, EPI_ISL_709833 | Lighthouse Lab in Milton Keynes | Wellcome Sanger Institute for the COVID-19 Genomics UK (COG-UK) Consortium | The Lighthouse Lab in Milton Keynes and Alex Alderton, Roberto Amato, Sonia Goncalves, Ewan Harrison, David K. Jackson, Ian Johnston, Dominic Kwiatkowski, Cordelia Langford, John Sillitoe on behalf of the Wellcome Sanger Institute COVID-19 Surveillance Team |
| EPI_ISL_709837 | Lighthouse Lab in Cambridge | Wellcome Sanger Institute for the COVID-19 Genomics UK (COG-UK) Consortium | Rob Howes, The Lighthouse Lab in Cambridge and Alex Alderton, Roberto Amato, Sonia Goncalves, Ewan Harrison, David K. Jackson, Ian Johnston, Dominic Kwiatkowski, Cordelia Langford, John Sillitoe on behalf of the Wellcome Sanger Institute COVID-19 Surveillance Team |
| EPI_ISL_709838, EPI_ISL_709839, EPI_ISL_709841 | Lighthouse Lab in Milton Keynes | Wellcome Sanger Institute for the COVID-19 Genomics UK (COG-UK) Consortium | The Lighthouse Lab in Milton Keynes and Alex Alderton, Roberto Amato, Sonia Goncalves, Ewan Harrison, David K. Jackson, Ian Johnston, Dominic Kwiatkowski, Cordelia Langford, John Sillitoe on behalf of the Wellcome Sanger Institute COVID-19 Surveillance Team |
| EPI_ISL_709842, EPI_ISL_709843, EPI_ISL_709844, EPI_ISL_709846 | Lighthouse Lab in Cambridge | Wellcome Sanger Institute for the COVID-19 Genomics UK (COG-UK) Consortium | Rob Howes, The Lighthouse Lab in Cambridge and Alex Alderton, Roberto Amato, Sonia Goncalves, Ewan Harrison, David K. Jackson, Ian Johnston, Dominic Kwiatkowski, Cordelia Langford, John Sillitoe on behalf of the Wellcome Sanger Institute COVID-19 Surveillance Team |
| EPI_ISL_709847 | Lighthouse Lab in Milton Keynes | Wellcome Sanger Institute for the COVID-19 Genomics UK (COG-UK) Consortium | The Lighthouse Lab in Milton Keynes and Alex Alderton, Roberto Amato, Sonia Goncalves, Ewan Harrison, David K. Jackson, Ian Johnston, Dominic Kwiatkowski, Cordelia Langford, John Sillitoe on behalf of the Wellcome Sanger Institute COVID-19 Surveillance Team |

| | | | |
|---|---|---|---|
| EPI_ISL_709850 | Lighthouse Lab in Cambridge | Wellcome Sanger Institute for the COVID-19 Genomics UK (COG-UK) Consortium | Rob Howes, The Lighthouse Lab in Cambridge and Alex Alderton, Roberto Amato, Sonia Goncalves, Ewan Harrison, David K. Jackson, Ian Johnston, Dominic Kwiatkowski, Cordelia Langford, John Sillitoe on behalf of the Wellcome Sanger Institute COVID-19 Surveillance Team |
| EPI_ISL_709852, EPI_ISL_709853, EPI_ISL_709856, EPI_ISL_709860, EPI_ISL_709863 | Lighthouse Lab in Milton Keynes | Wellcome Sanger Institute for the COVID-19 Genomics UK (COG-UK) Consortium | The Lighthouse Lab in Milton Keynes and Alex Alderton, Roberto Amato, Sonia Goncalves, Ewan Harrison, David K. Jackson, Ian Johnston, Dominic Kwiatkowski, Cordelia Langford, John Sillitoe on behalf of the Wellcome Sanger Institute COVID-19 Surveillance Team |
| EPI_ISL_709864, EPI_ISL_709865 | Lighthouse Lab in Cambridge | Wellcome Sanger Institute for the COVID-19 Genomics UK (COG-UK) Consortium | Rob Howes, The Lighthouse Lab in Cambridge and Alex Alderton, Roberto Amato, Sonia Goncalves, Ewan Harrison, David K. Jackson, Ian Johnston, Dominic Kwiatkowski, Cordelia Langford, John Sillitoe on behalf of the Wellcome Sanger Institute COVID-19 Surveillance Team |
| EPI_ISL_709866 | Lighthouse Lab in Milton Keynes | Wellcome Sanger Institute for the COVID-19 Genomics UK (COG-UK) Consortium | The Lighthouse Lab in Milton Keynes and Alex Alderton, Roberto Amato, Sonia Goncalves, Ewan Harrison, David K. Jackson, Ian Johnston, Dominic Kwiatkowski, Cordelia Langford, John Sillitoe on behalf of the Wellcome Sanger Institute COVID-19 Surveillance Team |
| EPI_ISL_709867 | Lighthouse Lab in Cambridge | Wellcome Sanger Institute for the COVID-19 Genomics UK (COG-UK) Consortium | Rob Howes, The Lighthouse Lab in Cambridge and Alex Alderton, Roberto Amato, Sonia Goncalves, Ewan Harrison, David K. Jackson, Ian Johnston, Dominic Kwiatkowski, Cordelia Langford, John Sillitoe on behalf of the Wellcome Sanger Institute COVID-19 Surveillance Team |
| EPI_ISL_709868 | Lighthouse Lab in Milton Keynes | Wellcome Sanger Institute for the COVID-19 Genomics UK (COG-UK) Consortium | The Lighthouse Lab in Milton Keynes and Alex Alderton, Roberto Amato, Sonia Goncalves, Ewan Harrison, David K. Jackson, Ian Johnston, Dominic Kwiatkowski, Cordelia Langford, John Sillitoe on behalf of the Wellcome Sanger Institute COVID-19 Surveillance Team |
| EPI_ISL_709869, EPI_ISL_709870 | Lighthouse Lab in Cambridge | Wellcome Sanger Institute for the COVID-19 Genomics UK (COG-UK) Consortium | Rob Howes, The Lighthouse Lab in Cambridge and Alex Alderton, Roberto Amato, Sonia Goncalves, Ewan Harrison, David K. Jackson, Ian Johnston, Dominic Kwiatkowski, Cordelia Langford, John Sillitoe on behalf of the Wellcome Sanger Institute COVID-19 Surveillance Team |
| EPI_ISL_709871, EPI_ISL_709872 | Lighthouse Lab in Milton Keynes | Wellcome Sanger Institute for the COVID-19 Genomics UK (COG-UK) Consortium | The Lighthouse Lab in Milton Keynes and Alex Alderton, Roberto Amato, Sonia Goncalves, Ewan Harrison, David K. Jackson, Ian Johnston, Dominic Kwiatkowski, Cordelia Langford, John Sillitoe on behalf of the Wellcome Sanger Institute COVID-19 Surveillance Team |
| EPI_ISL_709873 | Lighthouse Lab in Cambridge | Wellcome Sanger Institute for the COVID-19 Genomics UK (COG-UK) Consortium | Rob Howes, The Lighthouse Lab in Cambridge and Alex Alderton, Roberto Amato, Sonia Goncalves, Ewan Harrison, David K. Jackson, Ian Johnston, Dominic Kwiatkowski, Cordelia Langford, John Sillitoe on behalf of the Wellcome Sanger Institute COVID-19 Surveillance Team |
| EPI_ISL_709874 | Lighthouse Lab in Milton Keynes | Wellcome Sanger Institute for the COVID-19 Genomics UK (COG-UK) Consortium | The Lighthouse Lab in Milton Keynes and Alex Alderton, Roberto Amato, Sonia Goncalves, Ewan Harrison, David K. Jackson, Ian Johnston, Dominic Kwiatkowski, Cordelia Langford, John Sillitoe on behalf of the Wellcome Sanger Institute COVID-19 Surveillance Team |
| EPI_ISL_709876, EPI_ISL_709877, EPI_ISL_709878 | Lighthouse Lab in Cambridge | Wellcome Sanger Institute for the COVID-19 Genomics UK (COG-UK) Consortium | Rob Howes, The Lighthouse Lab in Cambridge and Alex Alderton, Roberto Amato, Sonia Goncalves, Ewan Harrison, David K. Jackson, Ian Johnston, Dominic Kwiatkowski, Cordelia Langford, John Sillitoe on behalf of the Wellcome Sanger Institute COVID-19 Surveillance Team |
| EPI_ISL_709879 | Lighthouse Lab in Milton Keynes | Wellcome Sanger Institute for the COVID-19 Genomics UK (COG-UK) Consortium | The Lighthouse Lab in Milton Keynes and Alex Alderton, Roberto Amato, Sonia Goncalves, Ewan Harrison, David K. Jackson, Ian Johnston, Dominic Kwiatkowski, Cordelia Langford, John Sillitoe on behalf of the Wellcome Sanger Institute COVID-19 Surveillance Team |
| EPI_ISL_709880 | Lighthouse Lab in Cambridge | Wellcome Sanger Institute for the COVID-19 Genomics UK (COG-UK) Consortium | Rob Howes, The Lighthouse Lab in Cambridge and Alex Alderton, Roberto Amato, Sonia Goncalves, Ewan Harrison, David K. Jackson, Ian Johnston, Dominic Kwiatkowski, Cordelia Langford, John Sillitoe on behalf of the Wellcome Sanger Institute COVID-19 Surveillance Team |
| EPI_ISL_709881 | Lighthouse Lab in Milton Keynes | Wellcome Sanger Institute for the COVID-19 Genomics UK (COG-UK) Consortium | The Lighthouse Lab in Milton Keynes and Alex Alderton, Roberto Amato, Sonia Goncalves, Ewan Harrison, David K. Jackson, Ian Johnston, Dominic Kwiatkowski, Cordelia Langford, John Sillitoe on behalf of the Wellcome Sanger Institute COVID-19 Surveillance Team |
| EPI_ISL_710124 | South Eastern Area Laboratory Services (SEALS) | CIDM-PH et al. | CIDM-PH et al. |
| EPI_ISL_710483, EPI_ISL_710484 | Department of Medical Laboratory Sciences, Arab American University | Department of Medical Laboratory Sciences, Arab American University | Al-Jawabreh,A., Nasereddin,A., Dumaidi,K., Al-Jawabreh,H., Ereqat,S. |
| EPI_ISL_710576 | VC Sorgenfrimottagningen | The Public Health Agency of Sweden | Department of Microbiology, The Public Health Agency of Sweden |
| EPI_ISL_710578 | Vardcentralen Brinken | The Public Health Agency of Sweden | Department of Microbiology, The Public Health Agency of Sweden |
| EPI_ISL_710581, EPI_ISL_710582, EPI_ISL_710583 | The Public Health Agency of Sweden | The Public Health Agency of Sweden | Department of Microbiology, The Public Health Agency of Sweden |
| EPI_ISL_710585, EPI_ISL_710586 | Orsa VC | The Public Health Agency of Sweden | Department of Microbiology, The Public Health Agency of Sweden |
| EPI_ISL_710588 | Omtanken Grimmered | The Public Health Agency of Sweden | Department of Microbiology, The Public Health Agency of Sweden |
| EPI_ISL_710604 | Klinisk mikrobiologi | The Public Health Agency of Sweden | Department of Microbiology, The Public Health Agency of Sweden |
| EPI_ISL_710748, EPI_ISL_710753, EPI_ISL_710754, EPI_ISL_710755, EPI_ISL_710756, EPI_ISL_710757, EPI_ISL_710758, EPI_ISL_710759, EPI_ISL_710760, EPI_ISL_710761, EPI_ISL_710762, EPI_ISL_710763, EPI_ISL_710764, EPI_ISL_710765, EPI_ISL_710766, EPI_ISL_710767, EPI_ISL_710768, EPI_ISL_710769, EPI_ISL_710770, EPI_ISL_710771, EPI_ISL_710772, EPI_ISL_710773, EPI_ISL_710774, EPI_ISL_710775, EPI_ISL_710776, EPI_ISL_710777, EPI_ISL_710778, EPI_ISL_710779, EPI_ISL_710780, EPI_ISL_710781, EPI_ISL_710782, EPI_ISL_710783, EPI_ISL_710784, EPI_ISL_710785, EPI_ISL_710786, EPI_ISL_710787, EPI_ISL_710788, EPI_ISL_710789, EPI_ISL_710790, EPI_ISL_710791, EPI_ISL_710792, EPI_ISL_710793, EPI_ISL_710794, EPI_ISL_710795, EPI_ISL_710796, EPI_ISL_710797, EPI_ISL_710798, EPI_ISL_710799, EPI_ISL_710800, EPI_ISL_710801, EPI_ISL_710802, EPI_ISL_710803, EPI_ISL_710804, EPI_ISL_710805, EPI_ISL_710806, EPI_ISL_710807, EPI_ISL_710808, EPI_ISL_710809, EPI_ISL_710810, EPI_ISL_710811, EPI_ISL_710812, EPI_ISL_710813, EPI_ISL_710814, EPI_ISL_710815, EPI_ISL_710816, EPI_ISL_710817, EPI_ISL_710818, EPI_ISL_710819, EPI_ISL_710820 | | | |
| see above | Lighthouse Lab in Glasgow | Wellcome Sanger Institute for the COVID-19 Genomics UK (COG-UK) Consortium | Harper VanSteenhouse, Yumi Kasai, David Gray, Carol Clugston, Anna Dominiczak and Alex Alderton, Roberto Amato, Sonia Goncalves, Ewan Harrison, David K. Jackson, Ian Johnston, Dominic Kwiatkowski, Cordelia Langford, John Sillitoe on behalf of the Wellcome Sanger Institute COVID-19 Surveillance Team |
| EPI_ISL_717734, EPI_ISL_717737, EPI_ISL_717741, EPI_ISL_717752 | Kingston Health Sciences Centre and Queen's University | Ontario Institute for Cancer Research | Prameet M. Sheth,Calvin Sjaarda,Robert Colautti,Katya Douchant,Ilinca Lungu,Bernard Lam,Paul Krzyzanowski,Michael Laszloffy,Lawrence E. Heisler,Richard de Borja,Jared T. Simpson |
| EPI_ISL_717981, EPI_ISL_717990, EPI_ISL_718014, EPI_ISL_718015 | Lab voor klinische biologie | Onderzoeksgroep Virologie | Laurens Lambrechts, Nick Vereecke, Marthe Pauwels, Bruno Verhasselt, Linos Vandekerckhove, Hans Nauwynck, Sebastiaan Theuns |
| EPI_ISL_718019, EPI_ISL_718020, EPI_ISL_718021, EPI_ISL_718022 | Lab voor klinische biologie | Onderzoeksgroep Virologie | Nick Vereecke, Laurens Lambrechts, Marthe Pauwels, Bruno Verhasselt, Linos Vandekerckhove, Hans Nauwynck, Sebastiaan Theuns |
| EPI_ISL_718023 | Lab voor klinische biologie | Onderzoeksgroep Virologie | Laurens Lambrechts, Nick Vereecke, Marthe Pauwels, Bruno Verhasselt, Linos Vandekerckhove, Hans Nauwynck, Sebastiaan Theuns |
| EPI_ISL_718025 | Lab voor klinische biologie | Onderzoeksgroep Virologie | Nick Vereecke, Laurens Lambrechts, Marthe Pauwels, Bruno Verhasselt, Linos Vandekerckhove, Hans Nauwynck, Sebastiaan Theuns |
| EPI_ISL_718040, EPI_ISL_718041, EPI_ISL_718042, EPI_ISL_718043, EPI_ISL_718044, EPI_ISL_718045, EPI_ISL_718046, EPI_ISL_718047, EPI_ISL_718048, EPI_ISL_718049, EPI_ISL_718051, EPI_ISL_718052, EPI_ISL_718053, EPI_ISL_718054, EPI_ISL_718055, EPI_ISL_718056, EPI_ISL_718057, EPI_ISL_718058 | | | |
| see above | ZOTZ KLIMAS MVZ Düsseldorf-Centrum GbR ÜBAG für Labormedizin, Genetik, Zytologie, Pathologie | Center of Medical Microbiology, Virology, and Hospital Hygiene, University of Duesseldorf | Maximilian Damagnez, Alexander Dilthey, Ashley-Jane Duplessis, Patrick Finzer, Katrin Hoffmann, Torsten Houwaart, Lisanna Hülse, Malte Kohns Vasconcelos, Marek Korencak, Nadine Lübke, Jessica Nicolai, Klaus Pfeffer, Daniel Strelow, Jörg Timm, Andreas Walker, Tobias Wienemann, Rainer Zotz |
| EPI_ISL_718251 | Institute of Virology, Biomedical Research Center of the Slovak Academy of Sciences, Bratislava | Faculty of Natural Sciences, Comenius University, Bratislava | Viktória Hodorová, Kristína Boršová, Broa Brejová, Viktória abanová, Sabina Fumaová Havlíková, Juraj Kopáek, Martina Liková, ubomíra Lukáiková, Martina Neboháová, Monika Sláviková, Tomáš Vina, Jozef Nosek, Boris Klempa |
| EPI_ISL_720673 | Lighthouse Lab in Milton Keynes | Wellcome Sanger Institute for the COVID-19 Genomics UK (COG-UK) Consortium | The Lighthouse Lab in Milton Keynes and Alex Alderton, Roberto Amato, Sonia Goncalves, Ewan Harrison, David K. Jackson, Ian Johnston, Dominic Kwiatkowski, Cordelia Langford, John Sillitoe on behalf of the Wellcome Sanger Institute COVID-19 Surveillance Team |
| EPI_ISL_720732 | Lighthouse Lab in Glasgow | Wellcome Sanger Institute for the COVID-19 Genomics UK (COG-UK) Consortium | Harper VanSteenhouse, Yumi Kasai, David Gray, Carol Clugston, Anna Dominiczak and Alex Alderton, Roberto Amato, Sonia Goncalves, Ewan Harrison, David K. Jackson, Ian Johnston, Dominic Kwiatkowski, Cordelia Langford, John Sillitoe on behalf of the Wellcome Sanger Institute COVID-19 Surveillance Team |
| EPI_ISL_721528, EPI_ISL_721529, EPI_ISL_721530, EPI_ISL_721531, EPI_ISL_721532 | Lighthouse Lab in Milton Keynes | Wellcome Sanger Institute for the COVID-19 Genomics UK (COG-UK) Consortium | The Lighthouse Lab in Milton Keynes and Alex Alderton, Roberto Amato, Sonia Goncalves, Ewan Harrison, David K. Jackson, Ian Johnston, Dominic Kwiatkowski, Cordelia Langford, John Sillitoe on behalf of the Wellcome Sanger Institute COVID-19 Surveillance Team |
| EPI_ISL_721533 | Lighthouse Lab in Glasgow | Wellcome Sanger Institute for the COVID-19 Genomics UK (COG-UK) Consortium | Harper VanSteenhouse, Yumi Kasai, David Gray, Carol Clugston, Anna Dominiczak and Alex Alderton, Roberto Amato, Sonia Goncalves, Ewan Harrison, David K. Jackson, Ian Johnston, Dominic Kwiatkowski, Cordelia Langford, John Sillitoe on behalf of the Wellcome Sanger Institute COVID-19 Surveillance Team |

| | | | |
|---|---|---|---|
| EPI_ISL_721673, EPI_ISL_721674, EPI_ISL_721681, EPI_ISL_721686, EPI_ISL_721689, EPI_ISL_721690, EPI_ISL_721692, EPI_ISL_721694, EPI_ISL_721716, EPI_ISL_721717, EPI_ISL_721718, EPI_ISL_721719, EPI_ISL_721720, EPI_ISL_721721, EPI_ISL_721722, EPI_ISL_721723, EPI_ISL_721724, EPI_ISL_721725, EPI_ISL_721726, EPI_ISL_721727, EPI_ISL_721728, EPI_ISL_721729, EPI_ISL_721730, EPI_ISL_721731, EPI_ISL_721732, EPI_ISL_721733, EPI_ISL_721734, EPI_ISL_721735, EPI_ISL_721736, EPI_ISL_721740, EPI_ISL_721761, EPI_ISL_721762, EPI_ISL_721763, EPI_ISL_721764, EPI_ISL_721765, EPI_ISL_721766, EPI_ISL_721771, EPI_ISL_721787, EPI_ISL_721827, EPI_ISL_721828, EPI_ISL_721829, EPI_ISL_721830, EPI_ISL_721831, EPI_ISL_721832, EPI_ISL_721833, EPI_ISL_721834, EPI_ISL_721835, EPI_ISL_721836, EPI_ISL_721837, EPI_ISL_721838, EPI_ISL_721839, EPI_ISL_721843, EPI_ISL_721848, EPI_ISL_721849, EPI_ISL_721850, EPI_ISL_721865, EPI_ISL_721866, EPI_ISL_721875, EPI_ISL_721876, EPI_ISL_721877, EPI_ISL_721878, EPI_ISL_721879, EPI_ISL_721880, EPI_ISL_721881, EPI_ISL_721882, EPI_ISL_721883, EPI_ISL_721884, EPI_ISL_721888, EPI_ISL_721905, EPI_ISL_721910, EPI_ISL_721911, EPI_ISL_721912, EPI_ISL_721917, EPI_ISL_721922, EPI_ISL_721923, EPI_ISL_721924, EPI_ISL_721925, EPI_ISL_721926, EPI_ISL_721927, EPI_ISL_721941, EPI_ISL_721959, EPI_ISL_721960, EPI_ISL_721961, EPI_ISL_721962, EPI_ISL_721963, EPI_ISL_721964, EPI_ISL_721965 | | | |
| see above | Viollier AG | Department of Biosystems Science and Engineering, ETH Zürich | Christian Beisel |
| EPI_ISL_722281, EPI_ISL_722311, EPI_ISL_722317, EPI_ISL_722329, EPI_ISL_722339, EPI_ISL_722352, EPI_ISL_722394, EPI_ISL_722687, EPI_ISL_722688, EPI_ISL_722689, EPI_ISL_722690, EPI_ISL_722691, EPI_ISL_722692, EPI_ISL_722693, EPI_ISL_722694, EPI_ISL_722695, EPI_ISL_722696, EPI_ISL_722697 | | | |
| see above | Dutch COVID-19 response team | Erasmus Medical Center | Bas Oude Munnink, Reina Sikkema, David Nieuwenhuijse, Irina Chestakova, Anne van der Linden, Marjan Boter, Emmanuelle Munger, Corine GeurtsvanKessel, Annemiek van der Eijk, Richard Molenkamp, Marion Koopmans, on behalf of the Dutch national COVID-19 response team. |
| EPI_ISL_722934, EPI_ISL_722936, EPI_ISL_722937, EPI_ISL_722938, EPI_ISL_722939, EPI_ISL_722940, EPI_ISL_722941, EPI_ISL_722942, EPI_ISL_722943 | Department of Clinical Microbiology | GIGA Medical Genomics | Keith Durkin, Maria Artesi, Sébastien Bontems, Raphaël Boreux, Bouchra Boujemla, Cécile Meex, Pierrette Melin, Marie-Pierre Hayette, Vincent Bours |
| EPI_ISL_722971, EPI_ISL_722972, EPI_ISL_722973, EPI_ISL_722975, EPI_ISL_723016, EPI_ISL_723017, EPI_ISL_723018, EPI_ISL_723019, EPI_ISL_723020, EPI_ISL_723021 | Respiratory Virus Unit, National Infection Service, Public Health England | COVID-19 Genomics UK (COG-UK) Consortium | PHE Covid Sequencing Team |
| EPI_ISL_723127, EPI_ISL_723128, EPI_ISL_723129, EPI_ISL_723130, EPI_ISL_723131, EPI_ISL_723132, EPI_ISL_723133, EPI_ISL_723134, EPI_ISL_723135, EPI_ISL_723136 | Minnesota Department of Health, Public Health Laboratory | Minnesota Department of Health, Public Health Laboratory | Alexandra Lorentz, Jacob Garfin, Matt Plumb, and Xiong Wang |
| EPI_ISL_723147, EPI_ISL_723148, EPI_ISL_723149, EPI_ISL_723151, EPI_ISL_723152, EPI_ISL_723153, EPI_ISL_723154 | Vault Health | Minnesota Department of Health, Public Health Laboratory | Alexandra Lorentz, Jacob Garfin, Matt Plumb, and Xiong Wang |
| EPI_ISL_723160, EPI_ISL_723181, EPI_ISL_723196, EPI_ISL_723197, EPI_ISL_723198, EPI_ISL_723218, EPI_ISL_723221, EPI_ISL_723222, EPI_ISL_723228, EPI_ISL_723242, EPI_ISL_723243, EPI_ISL_723244, EPI_ISL_723245, EPI_ISL_723246, EPI_ISL_723247, EPI_ISL_723248, EPI_ISL_723249, EPI_ISL_723250, EPI_ISL_723263, EPI_ISL_723264, EPI_ISL_723265, EPI_ISL_723269, EPI_ISL_723270, EPI_ISL_723288, EPI_ISL_723290, EPI_ISL_723291, EPI_ISL_723306, EPI_ISL_723307, EPI_ISL_723308, EPI_ISL_723309, EPI_ISL_723310, EPI_ISL_723311, EPI_ISL_723313, EPI_ISL_723314, EPI_ISL_723315, EPI_ISL_723318, EPI_ISL_723320, EPI_ISL_723321, EPI_ISL_723322, EPI_ISL_723327, EPI_ISL_723329, EPI_ISL_723330, EPI_ISL_723331, EPI_ISL_723333, EPI_ISL_723341, EPI_ISL_723342, EPI_ISL_723343, EPI_ISL_723344, EPI_ISL_723345, EPI_ISL_723347, EPI_ISL_723348, EPI_ISL_723349, EPI_ISL_723350, EPI_ISL_723351, EPI_ISL_723352, EPI_ISL_723353, EPI_ISL_723357, EPI_ISL_723360, EPI_ISL_723361, EPI_ISL_723362, EPI_ISL_723363, EPI_ISL_723364, EPI_ISL_723365, EPI_ISL_723398, EPI_ISL_723399, EPI_ISL_723400, EPI_ISL_723401, EPI_ISL_723402, EPI_ISL_723403, EPI_ISL_723404, EPI_ISL_723405, EPI_ISL_723414, EPI_ISL_723415, EPI_ISL_723416, EPI_ISL_723417, EPI_ISL_723418, EPI_ISL_723419, EPI_ISL_723420, EPI_ISL_723421, EPI_ISL_723422 | | | |
| see above | Dutch COVID-19 response team | National Institute for Public Health and the Environment (RIVM) | Adam Meijer, Harry Vennema, Jeroen Cremer, Sharon van den Brink, Bas van der Veer, AnneMarie van den Brandt, Florian Zwagemaker, Dennis Schmitz, Chantal Reusken, on behalf of the national COVID-19 response team |
| EPI_ISL_723473, EPI_ISL_723474, EPI_ISL_723475, EPI_ISL_723476, EPI_ISL_723477, EPI_ISL_723478, EPI_ISL_723479 | Virginia DCLS | Virginia DCLS | Virginia DCLS |
| EPI_ISL_723853, EPI_ISL_723857, EPI_ISL_723858, EPI_ISL_723859, EPI_ISL_723860, EPI_ISL_723861, EPI_ISL_723862, EPI_ISL_723863, EPI_ISL_723864, EPI_ISL_723865, EPI_ISL_723866, EPI_ISL_723867, EPI_ISL_723869, EPI_ISL_723870, EPI_ISL_723871, EPI_ISL_723872, EPI_ISL_723873, EPI_ISL_723874, EPI_ISL_723875, EPI_ISL_723876, EPI_ISL_723877, EPI_ISL_723878, EPI_ISL_723879, EPI_ISL_723880, EPI_ISL_723881, EPI_ISL_723882, EPI_ISL_723883, EPI_ISL_723885, EPI_ISL_723889, EPI_ISL_723890, EPI_ISL_723891, EPI_ISL_723892, EPI_ISL_723893, EPI_ISL_723894, EPI_ISL_723895, EPI_ISL_723896 | | | |
| see above | Department of Pathology, University of Cambridge | COVID-19 Genomics UK (COG-UK) Consortium | Aminu S. Jahun, Yasmin Chaudhry, Grant Hall, Iliana Georgana, Myra Hosmillo, Martin D. Curran, Malte Pinckert, Surendra Parmar, Ian Goodfellow |
| EPI_ISL_724213 | West of Scotland Specialist Virology Centre, NHSGGC / MRC-University of Glasgow Centre for Virus Research | COVID-19 Genomics UK (COG-UK) Consortium | Ana da Silva Filipe, Natasha Johnson, Kathy Smollett, Daniel Mair, Stephen Carmichael, Alice Broos, Lily Tong, Jenna Nichols, Kyriaki Nomikou; Sarah McDonald; Richard Orton, Joseph Hughes, Sreenu Vattipally, David L Robertson; Alasdair MacLean, Rory Gunson; Sharif Shaaban, Matthew Holden; Rachel Blacow, Guy Mollett, Kathy Li, James Shepherd, Antonia Ho, Emma Thomson |
| EPI_ISL_724223, EPI_ISL_724224, EPI_ISL_724225, EPI_ISL_724226 | Virology Department, Royal Infirmary of Edinburgh, NHS Lothian / School of Biological Sciences, University of Edinburgh / Institute of Genetics and Molecular Medicine, University of Edinburgh | COVID-19 Genomics UK (COG-UK) Consortium | McHugh M, Dewar R, Rooke S, Gallagher M, Balcaza C, O'Toole Á, Scher E, Hill V, McCrone JT, Colquhoun R, Yu X, Jackson B, Rambaut A, Williams TC, Templeton K |
| EPI_ISL_724262, EPI_ISL_724263, EPI_ISL_724264, EPI_ISL_724265, EPI_ISL_724266, EPI_ISL_724267 | University of Exeter | COVID-19 Genomics UK (COG-UK) Consortium | Ben Temperton,Aaron Jeffries,Michelle Michelsen,Joanna Warwick-Dugdale,Audrey Farbos,Robyn Manley,Stephen Michell,Jane Masoli |
| EPI_ISL_724396, EPI_ISL_724399, EPI_ISL_724401, EPI_ISL_724406, EPI_ISL_724408, EPI_ISL_724409, EPI_ISL_724410, EPI_ISL_724411, EPI_ISL_724412, EPI_ISL_724413, EPI_ISL_724414, EPI_ISL_724415, EPI_ISL_724416, EPI_ISL_724417, EPI_ISL_724419, EPI_ISL_724420, EPI_ISL_724421, EPI_ISL_724422, EPI_ISL_724423, EPI_ISL_724425, EPI_ISL_724426, EPI_ISL_724428, EPI_ISL_724429, EPI_ISL_724431, EPI_ISL_724432, EPI_ISL_724434, EPI_ISL_724436, EPI_ISL_724437, EPI_ISL_724439, EPI_ISL_724440, EPI_ISL_724441, EPI_ISL_724442, EPI_ISL_724443, EPI_ISL_724444, EPI_ISL_724445, EPI_ISL_724446, EPI_ISL_724447, EPI_ISL_724448, EPI_ISL_724449, EPI_ISL_724450, EPI_ISL_724451, EPI_ISL_724452, EPI_ISL_724453, EPI_ISL_724454, EPI_ISL_724455, EPI_ISL_724456, EPI_ISL_724457, EPI_ISL_724458, EPI_ISL_724459, EPI_ISL_724460, EPI_ISL_724461, EPI_ISL_724462, EPI_ISL_724463, EPI_ISL_724464, EPI_ISL_724465, EPI_ISL_724466, EPI_ISL_724477, EPI_ISL_724479, EPI_ISL_724483, EPI_ISL_724485, EPI_ISL_724493, EPI_ISL_724496, EPI_ISL_724497, EPI_ISL_724498, EPI_ISL_724499, EPI_ISL_724500, EPI_ISL_724501, EPI_ISL_724502, EPI_ISL_724503, EPI_ISL_724504, EPI_ISL_724505, EPI_ISL_724506, EPI_ISL_724507, EPI_ISL_724508, EPI_ISL_724509, EPI_ISL_724510, EPI_ISL_724511, EPI_ISL_724512, EPI_ISL_724513, EPI_ISL_724514, EPI_ISL_724515, EPI_ISL_724516, EPI_ISL_724517, EPI_ISL_724518, EPI_ISL_724519, EPI_ISL_724520, EPI_ISL_724521, EPI_ISL_724522, EPI_ISL_724523, EPI_ISL_724524 | | | |
| see above | Liverpool Clinical Laboratories | COVID-19 Genomics UK (COG-UK) Consortium | Sam Haldenby, Anita Lucaci, Steve Paterson, Julian Hiscox, Alistair Darby, M Almsaud, A Alrezaihi, Muhannad Alruwaili, Stuart D Armstrong, Jones Benjamin, Eleanor G Bentley, Anu Chawla, Jordan J Clark, Angela Cowell, Richard Eccles, Isabel García-Dorival, Matthew Gemmell, Alessandro Gerada, PKF Gilmore, Richard Gregory, Ximeng Han, Catherine Hartley, Margaret Hughes, Miren Iturriza-Gomara, James Johnson, L Luu, Jenifer Manson, Charlotte Nelson, Elaine O'Toole, Cassie Olateju, Rebekah Penrice-Randal , Lucille Rainbow, N.P Randle, Trevor Ian Robinson, Parul Sharma, Ghada T Shawli, James P Stewart, Neil Swainston, Ecaterina Vamos, Joanne Watts, Mark Whitehead |
| EPI_ISL_724576, EPI_ISL_724577, EPI_ISL_724578, EPI_ISL_724579, EPI_ISL_724583, EPI_ISL_724584, EPI_ISL_724585, EPI_ISL_724586, EPI_ISL_724587, EPI_ISL_724588, EPI_ISL_724589, EPI_ISL_724590, EPI_ISL_724591, EPI_ISL_724592, EPI_ISL_724595, EPI_ISL_724596 | | | |
| see above | University College London, Great Ormond Street Hospital for Children NHS Foundation Trust, Imperial College Healthcare NHS Trust | COVID-19 Genomics UK (COG-UK) Consortium | Sergi Castellano, Rachel Williams, Mark Kristiansen, Paola Resende Silva, Sunando Roy, Tony Brooks, Helena Tutill, Paola Niola, Patricia Dyal, Charlotte Williams, Leysa Forrest, Yasmin Panchbhaya, Jacqueline Findlay, Samuel Weeks, Julianne Brown, Kathryn Harris, Paul Randell, James Price, Alison Holmes, Judith Breuer |
| EPI_ISL_724855, EPI_ISL_724856, EPI_ISL_724857, EPI_ISL_724858, EPI_ISL_724859, EPI_ISL_724860, EPI_ISL_724861, EPI_ISL_724862, EPI_ISL_724863, EPI_ISL_724864, EPI_ISL_724865, EPI_ISL_724866, EPI_ISL_724867, EPI_ISL_724868, EPI_ISL_724869, EPI_ISL_724870, EPI_ISL_724871, EPI_ISL_724872, EPI_ISL_724873, EPI_ISL_724874, EPI_ISL_724875, EPI_ISL_724876, EPI_ISL_724877, EPI_ISL_724878, EPI_ISL_724879, EPI_ISL_724880, EPI_ISL_724881, EPI_ISL_724885, EPI_ISL_724886, EPI_ISL_724953, EPI_ISL_724954, EPI_ISL_724955, EPI_ISL_724956, EPI_ISL_724957, EPI_ISL_724958, EPI_ISL_724959, EPI_ISL_724960, EPI_ISL_724961, EPI_ISL_724962, EPI_ISL_725018, EPI_ISL_725019 | | | |
| see above | Northumbria University / South Tees Hospitals NHS Foundation Trust / North Cumbria Integrated Care NHS Foundation Trust / North Tees and Hartlepool NHS Foundation Trust / Newcastle Hospitals NHS Foundation Trust | COVID-19 Genomics UK (COG-UK) Consortium | Darren L Smith,Andrew Nelson,Matthew Bashton,Greg R Young,Joshua Loh,John Allan,Mohammad A Tariq,Giles S Holt,Gary Black,Wen C Yew,Lynn Dover,Paul Baker,Steve Liggett,Sarah Essex,Jane Greenaway,Debra Padgett,Clive Graham,Garren Scott,Edward Barton,Emma Swindells,Brendan Payne,Jennifer Collins,Yusri Taha,Gary Eltringham |
| EPI_ISL_725370, EPI_ISL_725371, EPI_ISL_725372, EPI_ISL_725373, EPI_ISL_725374, EPI_ISL_725375, EPI_ISL_725376, EPI_ISL_725377, EPI_ISL_725378, EPI_ISL_725379, EPI_ISL_725425, EPI_ISL_725513 | | | |
| see above | Quadram Institute Bioscience | COVID-19 Genomics UK (COG-UK) Consortium | Dave J. Baker, Gemma L. Kay, Alp Aydin, Thanh Le-Viet, Steven Rudder, Ana P. Tedim, Anastasia Kolyva, Maria Diaz, Leonardo de Oliveira Martins, |

| | | | |
|---|---|---|---|
| | | | Nabil-Fareed Alikhan, Lizzie Meadows, Rachael Stanley, Ngozi Elumogo, Muhammed Yasir, Nicholas M. Thomson, Alexander J Trotter, Rachel Gilroy, Samuel Bloomfield, Claire Stuart, Andrew Bell, Reenesh Prakash, Samir Dervisevic, Alison E. Mather, John Wain, Mark Webber, Andrew J. Page, Justin O'Grady |

EPI_ISL_725520, EPI_ISL_725521, EPI_ISL_725522, EPI_ISL_725523, EPI_ISL_725524, EPI_ISL_725525, EPI_ISL_725526, EPI_ISL_725527, EPI_ISL_725528, EPI_ISL_725529, EPI_ISL_725530, EPI_ISL_725531, EPI_ISL_725532, EPI_ISL_725533, EPI_ISL_725534, EPI_ISL_725535, EPI_ISL_725536, EPI_ISL_725537

| | | | |
|---|---|---|---|
| see above | Queens Medical Centre, Clinical Microbiology Department / DeepSeq Nottingham | COVID-19 Genomics UK (COG-UK) Consortium | Gemma Clark, Wendy Smith, Manjinder Khakh, Vicki M Fleming, Michelle M Lister, Hannah Howson-Wells, Jonathan Ball, Patrick McClure, Joseph Chappell, Theocharis Tsoleridis, Nadine Holmes, Matthew Carlisle, Christopher Moore, Fei Sang, Johnny Debebe, Victoria Wright, Matthew Loose |

EPI_ISL_725540, EPI_ISL_725541, EPI_ISL_725542, EPI_ISL_725543, EPI_ISL_725544, EPI_ISL_725545, EPI_ISL_725546, EPI_ISL_725547, EPI_ISL_725548, EPI_ISL_725549, EPI_ISL_725550, EPI_ISL_725551, EPI_ISL_725552, EPI_ISL_725553, EPI_ISL_725554, EPI_ISL_725555, EPI_ISL_725556, EPI_ISL_725557, EPI_ISL_725558, EPI_ISL_725559, EPI_ISL_725560, EPI_ISL_725561, EPI_ISL_725562, EPI_ISL_725563, EPI_ISL_725564, EPI_ISL_725565, EPI_ISL_725566, EPI_ISL_725567, EPI_ISL_725568, EPI_ISL_725569, EPI_ISL_725570, EPI_ISL_725571, EPI_ISL_725572, EPI_ISL_725573, EPI_ISL_725574, EPI_ISL_725575, EPI_ISL_725576, EPI_ISL_725577, EPI_ISL_725578, EPI_ISL_725579, EPI_ISL_725580, EPI_ISL_725581, EPI_ISL_725582, EPI_ISL_725583, EPI_ISL_725584, EPI_ISL_725585, EPI_ISL_725586, EPI_ISL_725587, EPI_ISL_725588, EPI_ISL_725589, EPI_ISL_725590, EPI_ISL_725591, EPI_ISL_725592, EPI_ISL_725593, EPI_ISL_725594, EPI_ISL_725595, EPI_ISL_725596, EPI_ISL_725597, EPI_ISL_725598, EPI_ISL_725599, EPI_ISL_725600, EPI_ISL_725601, EPI_ISL_725602, EPI_ISL_725603, EPI_ISL_725604

| | | | |
|---|---|---|---|
| see above | Lincolnshire Hospitals and DeepSeq Nottingham | COVID-19 Genomics UK (COG-UK) Consortium | Nichola Duckworth, Tim Sloan, Sarah Walsh, Jonathan Ball, Patrick McClure, Joeseph Chappell, Nadine Holmes, Matthew Carlisle, Christopher Moore, Fei Sang, Johnny Debebe, Victoria Wright, Matthew Loose |

EPI_ISL_725672, EPI_ISL_725673, EPI_ISL_725674, EPI_ISL_725675, EPI_ISL_725676, EPI_ISL_725686, EPI_ISL_725687, EPI_ISL_725688, EPI_ISL_725689, EPI_ISL_725690, EPI_ISL_725691, EPI_ISL_725692, EPI_ISL_725693, EPI_ISL_725694, EPI_ISL_725695, EPI_ISL_725696, EPI_ISL_725697, EPI_ISL_725698, EPI_ISL_725699, EPI_ISL_725700, EPI_ISL_725702, EPI_ISL_725703, EPI_ISL_725704, EPI_ISL_725705, EPI_ISL_725706, EPI_ISL_725707, EPI_ISL_725708, EPI_ISL_725711, EPI_ISL_725712, EPI_ISL_725713, EPI_ISL_725714, EPI_ISL_725715, EPI_ISL_725717, EPI_ISL_725726, EPI_ISL_725739, EPI_ISL_725740, EPI_ISL_725741, EPI_ISL_725742, EPI_ISL_725743, EPI_ISL_725744, EPI_ISL_725745, EPI_ISL_725746, EPI_ISL_725747, EPI_ISL_725748, EPI_ISL_725749, EPI_ISL_725750, EPI_ISL_725751, EPI_ISL_725752, EPI_ISL_725753, EPI_ISL_725754, EPI_ISL_725755, EPI_ISL_725756, EPI_ISL_725757, EPI_ISL_725758, EPI_ISL_725759, EPI_ISL_725760, EPI_ISL_725761, EPI_ISL_725762, EPI_ISL_725763, EPI_ISL_725764, EPI_ISL_725765, EPI_ISL_725766, EPI_ISL_725767, EPI_ISL_725768, EPI_ISL_725769, EPI_ISL_725770, EPI_ISL_725771, EPI_ISL_725772, EPI_ISL_725773, EPI_ISL_725774, EPI_ISL_725775, EPI_ISL_725776, EPI_ISL_725777, EPI_ISL_725778, EPI_ISL_725779, EPI_ISL_725780, EPI_ISL_725781, EPI_ISL_725782, EPI_ISL_725783, EPI_ISL_725784, EPI_ISL_725785, EPI_ISL_725786, EPI_ISL_725787, EPI_ISL_725788, EPI_ISL_725789, EPI_ISL_725790, EPI_ISL_725791, EPI_ISL_725792, EPI_ISL_725793, EPI_ISL_725794, EPI_ISL_725795, EPI_ISL_725796, EPI_ISL_725797, EPI_ISL_725798, EPI_ISL_725799, EPI_ISL_725800, EPI_ISL_725801, EPI_ISL_725802, EPI_ISL_725803, EPI_ISL_725804, EPI_ISL_725805, EPI_ISL_725806, EPI_ISL_725807, EPI_ISL_725808, EPI_ISL_725809, EPI_ISL_725810, EPI_ISL_725811, EPI_ISL_725812, EPI_ISL_725813, EPI_ISL_725814, EPI_ISL_725815, EPI_ISL_725816, EPI_ISL_725817, EPI_ISL_725818, EPI_ISL_725819, EPI_ISL_725820, EPI_ISL_725821, EPI_ISL_725822, EPI_ISL_725823, EPI_ISL_725824, EPI_ISL_725825, EPI_ISL_725826, EPI_ISL_725827, EPI_ISL_725828, EPI_ISL_725829, EPI_ISL_725830, EPI_ISL_725831, EPI_ISL_725832, EPI_ISL_725833, EPI_ISL_725834, EPI_ISL_725835, EPI_ISL_725836, EPI_ISL_725837, EPI_ISL_725838, EPI_ISL_725839, EPI_ISL_725840, EPI_ISL_725841, EPI_ISL_725842, EPI_ISL_725843, EPI_ISL_725844, EPI_ISL_725845, EPI_ISL_725846, EPI_ISL_725847, EPI_ISL_725848, EPI_ISL_725849, EPI_ISL_725850, EPI_ISL_725851, EPI_ISL_725852, EPI_ISL_725853, EPI_ISL_725854, EPI_ISL_725855, EPI_ISL_725856, EPI_ISL_725857, EPI_ISL_725858, EPI_ISL_725859, EPI_ISL_725860, EPI_ISL_725861, EPI_ISL_725862, EPI_ISL_725863, EPI_ISL_725864, EPI_ISL_725865, EPI_ISL_725866, EPI_ISL_725867, EPI_ISL_725868, EPI_ISL_725869, EPI_ISL_725870, EPI_ISL_725871, EPI_ISL_725872, EPI_ISL_725873, EPI_ISL_725874, EPI_ISL_725875, EPI_ISL_725876, EPI_ISL_725877, EPI_ISL_725878, EPI_ISL_725879, EPI_ISL_725880, EPI_ISL_725881, EPI_ISL_725882, EPI_ISL_725883, EPI_ISL_725884, EPI_ISL_725885, EPI_ISL_725886, EPI_ISL_725887, EPI_ISL_725888, EPI_ISL_725889, EPI_ISL_725890, EPI_ISL_725891, EPI_ISL_725892, EPI_ISL_725894, EPI_ISL_725896, EPI_ISL_725897, EPI_ISL_725898, EPI_ISL_725899, EPI_ISL_725900, EPI_ISL_725901, EPI_ISL_725902, EPI_ISL_725903, EPI_ISL_725904, EPI_ISL_725905, EPI_ISL_725906, EPI_ISL_725907, EPI_ISL_725908, EPI_ISL_725909, EPI_ISL_725910, EPI_ISL_725911, EPI_ISL_725912, EPI_ISL_725913, EPI_ISL_725914, EPI_ISL_725915, EPI_ISL_725916, EPI_ISL_725917, EPI_ISL_725918, EPI_ISL_725920, EPI_ISL_725921, EPI_ISL_725922, EPI_ISL_725923, EPI_ISL_725924, EPI_ISL_725925, EPI_ISL_725926, EPI_ISL_725927, EPI_ISL_725928, EPI_ISL_725929, EPI_ISL_725930, EPI_ISL_725931, EPI_ISL_725932, EPI_ISL_725933, EPI_ISL_725934, EPI_ISL_725935, EPI_ISL_725936, EPI_ISL_725937, EPI_ISL_725938, EPI_ISL_725939, EPI_ISL_725940, EPI_ISL_725941, EPI_ISL_725942, EPI_ISL_725943, EPI_ISL_725944, EPI_ISL_725945, EPI_ISL_725946, EPI_ISL_725947, EPI_ISL_725948, EPI_ISL_725949, EPI_ISL_725950, EPI_ISL_725951, EPI_ISL_725952, EPI_ISL_725953, EPI_ISL_725954, EPI_ISL_726212, EPI_ISL_726217, EPI_ISL_726220, EPI_ISL_726221, EPI_ISL_726236, EPI_ISL_726237, EPI_ISL_726238, EPI_ISL_726239, EPI_ISL_726240, EPI_ISL_726241, EPI_ISL_726251, EPI_ISL_726266, EPI_ISL_726267, EPI_ISL_726272, EPI_ISL_726273, EPI_ISL_726274, EPI_ISL_726275, EPI_ISL_726281, EPI_ISL_726282, EPI_ISL_726283, EPI_ISL_726284, EPI_ISL_726285, EPI_ISL_726286, EPI_ISL_726287, EPI_ISL_726288, EPI_ISL_726289, EPI_ISL_726290, EPI_ISL_726291, EPI_ISL_726292, EPI_ISL_726293, EPI_ISL_726294, EPI_ISL_726295, EPI_ISL_726296, EPI_ISL_726297, EPI_ISL_726298, EPI_ISL_726299, EPI_ISL_726300, EPI_ISL_726301, EPI_ISL_726302, EPI_ISL_726303, EPI_ISL_726304, EPI_ISL_726357, EPI_ISL_726400, EPI_ISL_726402, EPI_ISL_726403, EPI_ISL_726404, EPI_ISL_726405, EPI_ISL_726406, EPI_ISL_726407, EPI_ISL_726408, EPI_ISL_726409, EPI_ISL_726410, EPI_ISL_726411, EPI_ISL_726412, EPI_ISL_726413, EPI_ISL_726414, EPI_ISL_726415, EPI_ISL_726416, EPI_ISL_726417, EPI_ISL_726418, EPI_ISL_726419, EPI_ISL_726420, EPI_ISL_726421, EPI_ISL_726422, EPI_ISL_726423, EPI_ISL_726424, EPI_ISL_726425, EPI_ISL_726426, EPI_ISL_726427, EPI_ISL_726428, EPI_ISL_726429, EPI_ISL_726430, EPI_ISL_726431, EPI_ISL_726432, EPI_ISL_726433, EPI_ISL_726434, EPI_ISL_726435, EPI_ISL_726436, EPI_ISL_726437, EPI_ISL_726438, EPI_ISL_726439, EPI_ISL_726440, EPI_ISL_726441, EPI_ISL_726442, EPI_ISL_726443, EPI_ISL_726444, EPI_ISL_726445, EPI_ISL_726446, EPI_ISL_726447, EPI_ISL_726448, EPI_ISL_726449, EPI_ISL_726450, EPI_ISL_726451, EPI_ISL_726452, EPI_ISL_726453, EPI_ISL_726454, EPI_ISL_726455, EPI_ISL_726456, EPI_ISL_726457, EPI_ISL_726458, EPI_ISL_726459, EPI_ISL_726460, EPI_ISL_726461, EPI_ISL_726462, EPI_ISL_726463, EPI_ISL_726464, EPI_ISL_726465, EPI_ISL_726466, EPI_ISL_726467, EPI_ISL_726468, EPI_ISL_726469, EPI_ISL_726470, EPI_ISL_726471, EPI_ISL_726472, EPI_ISL_726473, EPI_ISL_726474, EPI_ISL_726475, EPI_ISL_726476, EPI_ISL_726477, EPI_ISL_726478, EPI_ISL_726479, EPI_ISL_726480, EPI_ISL_726481, EPI_ISL_726482, EPI_ISL_726483, EPI_ISL_726484, EPI_ISL_726485, EPI_ISL_726486, EPI_ISL_726487, EPI_ISL_726488, EPI_ISL_726489, EPI_ISL_726490, EPI_ISL_726491, EPI_ISL_726492, EPI_ISL_726493, EPI_ISL_726494, EPI_ISL_726495, EPI_ISL_726496, EPI_ISL_726497, EPI_ISL_726498, EPI_ISL_726509, EPI_ISL_726510, EPI_ISL_726511, EPI_ISL_726512, EPI_ISL_726513, EPI_ISL_726514, EPI_ISL_726515, EPI_ISL_726516, EPI_ISL_726517, EPI_ISL_726518, EPI_ISL_726519, EPI_ISL_726527, EPI_ISL_726528, EPI_ISL_726529, EPI_ISL_726530, EPI_ISL_726531, EPI_ISL_726532, EPI_ISL_726533, EPI_ISL_726534, EPI_ISL_726535, EPI_ISL_726536, EPI_ISL_726543, EPI_ISL_726544, EPI_ISL_726545, EPI_ISL_726549, EPI_ISL_726550, EPI_ISL_726551, EPI_ISL_726552, EPI_ISL_726553, EPI_ISL_726554, EPI_ISL_726560, EPI_ISL_726561, EPI_ISL_726562, EPI_ISL_726563, EPI_ISL_726564, EPI_ISL_726565, EPI_ISL_726566, EPI_ISL_726569, EPI_ISL_726586, EPI_ISL_726588, EPI_ISL_726590, EPI_ISL_726592, EPI_ISL_726593, EPI_ISL_726595, EPI_ISL_726596, EPI_ISL_726597, EPI_ISL_726598, EPI_ISL_726600, EPI_ISL_726602, EPI_ISL_726604, EPI_ISL_726690, EPI_ISL_726696, EPI_ISL_726703, EPI_ISL_726704, EPI_ISL_726705, EPI_ISL_726712, EPI_ISL_726715, EPI_ISL_726716, EPI_ISL_726717, EPI_ISL_726718, EPI_ISL_726719, EPI_ISL_726720, EPI_ISL_726721, EPI_ISL_726723, EPI_ISL_726724, EPI_ISL_726725, EPI_ISL_726726, EPI_ISL_726727, EPI_ISL_726728, EPI_ISL_726729, EPI_ISL_726730, EPI_ISL_726731, EPI_ISL_726732, EPI_ISL_726733, EPI_ISL_726734, EPI_ISL_726735, EPI_ISL_726736, EPI_ISL_726737, EPI_ISL_726738, EPI_ISL_726739, EPI_ISL_726740, EPI_ISL_726741, EPI_ISL_726742, EPI_ISL_726743, EPI_ISL_726744, EPI_ISL_726745, EPI_ISL_726746, EPI_ISL_726747, EPI_ISL_726748, EPI_ISL_726749, EPI_ISL_726750, EPI_ISL_726751, EPI_ISL_726752, EPI_ISL_726753, EPI_ISL_726754, EPI_ISL_726755, EPI_ISL_726756, EPI_ISL_726757, EPI_ISL_726758, EPI_ISL_726759, EPI_ISL_726760, EPI_ISL_726761, EPI_ISL_726762, EPI_ISL_726763, EPI_ISL_726764, EPI_ISL_726765, EPI_ISL_726766, EPI_ISL_726767, EPI_ISL_726768, EPI_ISL_726769, EPI_ISL_726770, EPI_ISL_726771, EPI_ISL_726772, EPI_ISL_726773, EPI_ISL_726774, EPI_ISL_726775, EPI_ISL_726776, EPI_ISL_726777, EPI_ISL_726778, EPI_ISL_726779, EPI_ISL_726780, EPI_ISL_726781, EPI_ISL_726782, EPI_ISL_726783, EPI_ISL_726784, EPI_ISL_726785, EPI_ISL_726786, EPI_ISL_726787, EPI_ISL_726788, EPI_ISL_726789, EPI_ISL_726790, EPI_ISL_726791, EPI_ISL_726792, EPI_ISL_726793, EPI_ISL_726794, EPI_ISL_726795, EPI_ISL_726796, EPI_ISL_726797, EPI_ISL_726798, EPI_ISL_726799, EPI_ISL_726800, EPI_ISL_726801, EPI_ISL_726802, EPI_ISL_726803, EPI_ISL_726804, EPI_ISL_726805, EPI_ISL_726806, EPI_ISL_726807, EPI_ISL_726808, EPI_ISL_726809, EPI_ISL_726810, EPI_ISL_726811, EPI_ISL_726812, EPI_ISL_726813, EPI_ISL_726814, EPI_ISL_726815, EPI_ISL_726816, EPI_ISL_726817, EPI_ISL_726818, EPI_ISL_726819, EPI_ISL_726820, EPI_ISL_726821, EPI_ISL_726822, EPI_ISL_726823, EPI_ISL_726824, EPI_ISL_726825, EPI_ISL_726826, EPI_ISL_726827, EPI_ISL_726828, EPI_ISL_726829, EPI_ISL_726830, EPI_ISL_726831, EPI_ISL_726832, EPI_ISL_726833, EPI_ISL_726834, EPI_ISL_726835, EPI_ISL_726836, EPI_ISL_726837, EPI_ISL_726839, EPI_ISL_726943, EPI_ISL_726944, EPI_ISL_726945, EPI_ISL_726946, EPI_ISL_726947, EPI_ISL_726948, EPI_ISL_726949, EPI_ISL_726950, EPI_ISL_726951, EPI_ISL_726952, EPI_ISL_726953, EPI_ISL_726954, EPI_ISL_726955, EPI_ISL_726956, EPI_ISL_726957, EPI_ISL_726958, EPI_ISL_726959, EPI_ISL_726960, EPI_ISL_726962, EPI_ISL_727108, EPI_ISL_727110, EPI_ISL_727111, EPI_ISL_727112, EPI_ISL_727115, EPI_ISL_727116, EPI_ISL_727117, EPI_ISL_727124, EPI_ISL_727125, EPI_ISL_727126, EPI_ISL_727127, EPI_ISL_727128, EPI_ISL_727129, EPI_ISL_727130, EPI_ISL_727131, EPI_ISL_727132, EPI_ISL_727133, EPI_ISL_727134, EPI_ISL_727135, EPI_ISL_727136, EPI_ISL_727137, EPI_ISL_727138, EPI_ISL_727139, EPI_ISL_727140, EPI_ISL_727141, EPI_ISL_727142, EPI_ISL_727184, EPI_ISL_727186, EPI_ISL_727188, EPI_ISL_727189, EPI_ISL_727190, EPI_ISL_727191, EPI_ISL_727192, EPI_ISL_727193, EPI_ISL_727194, EPI_ISL_727195, EPI_ISL_727196, EPI_ISL_727197, EPI_ISL_727198, EPI_ISL_727199, EPI_ISL_727200, EPI_ISL_727201, EPI_ISL_727202, EPI_ISL_727203, EPI_ISL_727204, EPI_ISL_727206, EPI_ISL_727207, EPI_ISL_727208, EPI_ISL_727209, EPI_ISL_727210, EPI_ISL_727211, EPI_ISL_727212, EPI_ISL_727213, EPI_ISL_727214, EPI_ISL_727215, EPI_ISL_727216, EPI_ISL_727217, EPI_ISL_727218, EPI_ISL_727219, EPI_ISL_727220, EPI_ISL_727221, EPI_ISL_727222, EPI_ISL_727409, EPI_ISL_727410, EPI_ISL_727411, EPI_ISL_727412, EPI_ISL_727413, EPI_ISL_727414, EPI_ISL_727415, EPI_ISL_727416, EPI_ISL_727432, EPI_ISL_727433, EPI_ISL_727434, EPI_ISL_727435, EPI_ISL_727436, EPI_ISL_727437, EPI_ISL_727438, EPI_ISL_727439, EPI_ISL_727440, EPI_ISL_727441, EPI_ISL_727442, EPI_ISL_727443, EPI_ISL_727444, EPI_ISL_727445, EPI_ISL_727446, EPI_ISL_727447, EPI_ISL_727448, EPI_ISL_727449, EPI_ISL_727450, EPI_ISL_727451, EPI_ISL_727452, EPI_ISL_727453, EPI_ISL_727454, EPI_ISL_727455, EPI_ISL_727456, EPI_ISL_727457, EPI_ISL_727458, EPI_ISL_727459, EPI_ISL_727460, EPI_ISL_727461, EPI_ISL_727462, EPI_ISL_727463, EPI_ISL_727464, EPI_ISL_727465, EPI_ISL_727466, EPI_ISL_727467, EPI_ISL_727468, EPI_ISL_727469, EPI_ISL_727470, EPI_ISL_727471, EPI_ISL_727472, EPI_ISL_727473, EPI_ISL_727474, EPI_ISL_727475, EPI_ISL_727476, EPI_ISL_727477, EPI_ISL_727478, EPI_ISL_727479, EPI_ISL_727480, EPI_ISL_727481, EPI_ISL_727482, EPI_ISL_727483, EPI_ISL_727484

| | | | |
|---|---|---|---|
| see above | Wales Specialist Virology Centre Sequencing lab: Pathogen Genomics Unit | COVID-19 Genomics UK (COG-UK) Consortium | Catherine Moore, Johnathan Evans, Laura Gifford, Malorie Perry, Simon Cottrell, Angela Marchbank, Alec Birchley, Alexander Adams, Amy Gaskin, Bree Gatica-Wilcox, Jason Coombes, Joel Southgate, Lauren Gilbert, Lee Graham, Nicole Pacchiarini, Sara Kumziene-Summerhayes, Sarah Taylor, Sophie Jones, Sara Rey, Matthew Bull, Joanne Watkins, Sally Corden, Tom Connor |

EPI_ISL_727868, EPI_ISL_727883, EPI_ISL_727895, EPI_ISL_727931, EPI_ISL_727934, EPI_ISL_727940, EPI_ISL_727943, EPI_ISL_727967, EPI_ISL_727968, EPI_ISL_727969, EPI_ISL_727976

| | | | |
|---|---|---|---|
| see above | Virology Department, Sheffield Teaching Hospitals NHS Foundation Trust/Department of Infection, Immunity and Cardiovascular Disease, The Medical School, University of Sheffield | COVID-19 Genomics UK (COG-UK) Consortium | Thushan de Silva, Matthew Parker, Nikki Smith, Adri Angyal, Rebecca Brown, Luke Green, Rachel Tucker, Paul Parsons, Danielle Groves, Katie Johnson, Laura Carrilero, Alex Keeley, Dave Partridge, Matthew Wyles, Benjamin Lindsey, Mehmet Yavuz, Mohammad Raza, Cariad Evans |

EPI_ISL_728059, EPI_ISL_728064, EPI_ISL_728065, EPI_ISL_728066, EPI_ISL_728067, EPI_ISL_728068, EPI_ISL_728069, EPI_ISL_728070, EPI_ISL_728071, EPI_ISL_728072, EPI_ISL_728073, EPI_ISL_728074, EPI_ISL_728095

| | | | |
|---|---|---|---|
| see above | University of Wisconsin-Madison AIDS Vaccine Research Laboratories | University of Wisconsin-Madison AIDS Vaccine Research Laboratories | Gage Moreno, Katarina Braun, et al. AIDS Vaccine Research Laboratories |
| EPI_ISL_728202 | Institute of Microbiology, Universidad San Francisco de Quito | Institute of Microbiology, Universidad San Francisco de Quito | Sully Márquez, Belén Prado-Vivar, Juan José Guadalupe, Monica Becerra-Wong, Bernardo Gutiérrez, Tania Guayasamin, Patricio Reyes, Verónica Barragán, Patricio Rojas-Silva, Gabriel Trueba, Michelle Grunauer, Paúl Cárdenas |
| EPI_ISL_728286 | National Institute for Infectious Diseases, INMI, "L. Spallanzani" IRCCS | National Institute for Infectious Diseases, INMI, "L. Spallanzani" IRCCS | E Giombini, M Rueca, B Bartolini, C.E.M Gruber, F Messina, A Di Caro, MR Capobianchi |
| EPI_ISL_728546, EPI_ISL_728547, EPI_ISL_728548, EPI_ISL_728549, EPI_ISL_728550, EPI_ISL_728551 | CNR Virus des Infections Respiratoires - France SUD | CNR Virus des Infections Respiratoires - France SUD | Antonin Bal, Gregory Destras, Claudia Gonzalez, Gwendolyne Burfin, Quentin Semanas, Martine Valette, Bruno Lina, Laurence Josset |

EPI_ISL_728564, EPI_ISL_728638, EPI_ISL_728639, EPI_ISL_728687, EPI_ISL_728688, EPI_ISL_728708, EPI_ISL_728733, EPI_ISL_728736, EPI_ISL_728737, EPI_ISL_728758, EPI_ISL_728759, EPI_ISL_728760, EPI_ISL_728761, EPI_ISL_728762

| | | | |
|---|---|---|---|
| see above | Dutch COVID-19 response team | National Institute for Public Health and the Environment (RIVM) | Adam Meijer, Harry Vennema, Jeroen Cremer, Sharon van den Brink, Bas van der Veer, AnneMarie van den Brandt, Florian Zwagemaker, Dennis Schmitz, Chantal Reusken, on behalf of the national COVID-19 response team |

EPI_ISL_728766, EPI_ISL_728767, EPI_ISL_728795, EPI_ISL_728806, EPI_ISL_728897, EPI_ISL_728906, EPI_ISL_728908, EPI_ISL_728909, EPI_ISL_728911, EPI_ISL_728921, EPI_ISL_728923, EPI_ISL_728929, EPI_ISL_728930, EPI_ISL_728944, EPI_ISL_728972, EPI_ISL_728973, EPI_ISL_728974, EPI_ISL_728975, EPI_ISL_728977, EPI_ISL_728979, EPI_ISL_728986, EPI_ISL_728991, EPI_ISL_728992, EPI_ISL_728994, EPI_ISL_728996, EPI_ISL_728998, EPI_ISL_729006, EPI_ISL_729015, EPI_ISL_729018, EPI_ISL_729021, EPI_ISL_729029, EPI_ISL_729037, EPI_ISL_729038, EPI_ISL_729039, EPI_ISL_729045, EPI_ISL_729046

| | | | |
|---|---|---|---|
| see above | Viollier AG | Department of Biosystems Science and Engineering, ETH Zürich | Chaoran Chen, Sarah Nadeau, Catharine Aquino, Ivan Topolsky, Pedro Ferreira, Philipp Jablonski, Susana Posada-Céspedes, Andreia Cabral de Gouvea, Maria Domenica Moccia, Simon Grüter, Timothy Sykes, Lennart Opitz, Ralph Schlapbach, Christiane Beckmann, Maurice Redondo, Olivier Kobel, Christoph Noppen, Sophie Seidel, Noemie Santamaria de Souza, Niko Beerenwinkel, Tanja Stadler |

| | | | |
|---|---|---|---|
| EPI_ISL_729259, EPI_ISL_729260, EPI_ISL_729261, EPI_ISL_729262, EPI_ISL_729263, EPI_ISL_729264, EPI_ISL_729265, EPI_ISL_729266, EPI_ISL_729267, EPI_ISL_729268, EPI_ISL_729269, EPI_ISL_729270, EPI_ISL_729271, EPI_ISL_729272, EPI_ISL_729273, EPI_ISL_729274, EPI_ISL_729275, EPI_ISL_729276, EPI_ISL_729277, EPI_ISL_729278, EPI_ISL_729279, EPI_ISL_729280, EPI_ISL_729281, EPI_ISL_729282, EPI_ISL_729283, EPI_ISL_729284, EPI_ISL_729285, EPI_ISL_729286, EPI_ISL_729287, EPI_ISL_729288, EPI_ISL_729289, EPI_ISL_729290, EPI_ISL_729291, EPI_ISL_729292, EPI_ISL_729293, EPI_ISL_729294, EPI_ISL_729295, EPI_ISL_729296, EPI_ISL_729297, EPI_ISL_729298, EPI_ISL_729299, EPI_ISL_729300, EPI_ISL_729301, EPI_ISL_729302, EPI_ISL_729303, EPI_ISL_729304, EPI_ISL_729305, EPI_ISL_729306, EPI_ISL_729307, EPI_ISL_729308, EPI_ISL_729309, EPI_ISL_729310, EPI_ISL_729311, EPI_ISL_729312, EPI_ISL_729313, EPI_ISL_729314, EPI_ISL_729315, EPI_ISL_729316, EPI_ISL_729317, EPI_ISL_729318, EPI_ISL_729319, EPI_ISL_729320, EPI_ISL_729321, EPI_ISL_729322, EPI_ISL_729323, EPI_ISL_729324, EPI_ISL_729325, EPI_ISL_729326, EPI_ISL_729327, EPI_ISL_729328, EPI_ISL_729329, EPI_ISL_729330, EPI_ISL_729331, EPI_ISL_729332, EPI_ISL_729333, EPI_ISL_729334, EPI_ISL_729335 | | | |
| see above | Toronto Invasive Bacterial Diseases Network | McMaster University | Allison McGeer, Patryk Aftanas, Hooman Derakhshani, Angel Li, Kuganya Nirmalarajah, Emily Panousis, Ahmed Draia, Jalees Nasir, Michael Surette, Samira Mubareka, Andrew G. McArthur |
| EPI_ISL_729420, EPI_ISL_729447, EPI_ISL_729448, EPI_ISL_729464, EPI_ISL_729685, EPI_ISL_729686, EPI_ISL_729687, EPI_ISL_729688, EPI_ISL_729689, EPI_ISL_729690, EPI_ISL_729691, EPI_ISL_729692, EPI_ISL_729693, EPI_ISL_729694, EPI_ISL_729695, EPI_ISL_729696, EPI_ISL_729704, EPI_ISL_729720, EPI_ISL_729723, EPI_ISL_729734 | | | |
| see above | A. Krumbholz, Labor Dr. Krause und Kollegen MVZ GmbH, Kiel | Charité Universitätsmedizin Berlin, Institut für Virologie | Victor M Corman, Barbara Mühlemann, Jörn Beheim-Schwarzbach, Talitha Veith, Julia Schneider, Terry Jones, Christian Drosten |
| EPI_ISL_730320, EPI_ISL_730322, EPI_ISL_730324, EPI_ISL_730328, EPI_ISL_730336 | San Diego County Public Health Laboratory | Andersen lab at Scripps Research | SEARCH Alliance San Diego with Tracy Basler, Jovan Shephard, Brett Austin |
| EPI_ISL_732189, EPI_ISL_732190, EPI_ISL_732191, EPI_ISL_732192, EPI_ISL_732193, EPI_ISL_732194, EPI_ISL_732195 | Instituto Nacional de Saude (INSA) and Instituto Gulbenkian de Ciencia (IGC) | Instituto Nacional de Saude (INSA) and Instituto Gulbenkian de Ciencia (IGC) | Borges et al |
| EPI_ISL_732379, EPI_ISL_732386, EPI_ISL_732388, EPI_ISL_732390, EPI_ISL_732391, EPI_ISL_732405, EPI_ISL_732407, EPI_ISL_732440, EPI_ISL_732442, EPI_ISL_732444, EPI_ISL_732446, EPI_ISL_732455, EPI_ISL_732457, EPI_ISL_732472, EPI_ISL_732484, EPI_ISL_732495, EPI_ISL_732497, EPI_ISL_732498, EPI_ISL_732499, EPI_ISL_732500, EPI_ISL_732501, EPI_ISL_732514, EPI_ISL_732515, EPI_ISL_732522 | | | |
| see above | National Virus Reference Laboratory | National Virus Reference Laboratory | Michael Carr, Gabriel Gonzalez, Jonathan Dean, Daniel Hare, Cillian F De Gascun |
| EPI_ISL_732654 | Klinisk mikrobiologi | The Public Health Agency of Sweden | Department of Microbiology, The Public Health Agency of Sweden |
| EPI_ISL_732655 | Narhalsan Fjallbacka VC | The Public Health Agency of Sweden | Department of Microbiology, The Public Health Agency of Sweden |
| EPI_ISL_732691, EPI_ISL_732692 | CNR Virus des Infections Respiratoires - France SUD | CNR Virus des Infections Respiratoires - France SUD | Antonin Bal, Gregory Destras, Claudia Gonzalez, Gwendolyne Burfin, Quentin Semanas, Martine Valette, Bruno Lina, Laurence Josset |
| EPI_ISL_733466, EPI_ISL_733467, EPI_ISL_733469, EPI_ISL_733473, EPI_ISL_733475 | WHO National Influenza Centre Russian Federation | WHO National Influenza Centre Russian Federation | Andrey Komissarov, Artem Fadeev, Anna Ivanova, Kseniya Komissarova, Dmitry Bazhenov, Daria Danilenko, Ksenia Safina, Elena Nabieva, Georgii Bazykin, Dmitry Lioznov |
| EPI_ISL_733478, EPI_ISL_733479, EPI_ISL_733480, EPI_ISL_733481 | SOGAZ PROFMEDICINA LLC | WHO National Influenza Centre Russian Federation | Andrey Komissarov, Artem Fadeev, Anna Ivanova, Kseniya Komissarova, Dmitry Bazhenov, Daria Danilenko, Ksenia Safina, Elena Nabieva, Georgii Bazykin, Dmitry Lioznov |
| EPI_ISL_733569 | CHAI WAN FAMILIES CLINIC | Hong Kong Department of Health | Alan K.L. Tsang, Peter C.W. Yip, Edman T.K. Lam, Rickjason C.W. Chan, Dominic N.C. Tsang |
| EPI_ISL_733591 | Respiratory Virus Unit, National Infection Service, Public Health England | COVID-19 Genomics UK (COG-UK) Consortium | PHE Covid Sequencing Team |
| EPI_ISL_735221, EPI_ISL_735222, EPI_ISL_735223, EPI_ISL_735224, EPI_ISL_735225, EPI_ISL_735226, EPI_ISL_735227, EPI_ISL_735228, EPI_ISL_735229, EPI_ISL_735230, EPI_ISL_735231, EPI_ISL_735232, EPI_ISL_735233, EPI_ISL_735234, EPI_ISL_735235 | | | |
| see above | UZ Leuven, National Reference Laboratory for Coronaviruses, Laboratory Medicine, Leuven, Belgium | KU Leuven, Rega Institute, Clinical and Epidemiological Virology | Tony Wawina-Bokalanga, Joan Marti-Carerras, Bert Vanmechelen, Piet Maes |
| EPI_ISL_735490 | Rangamati General Hospital RT-PCR lab, | Central Biological Research Laboratory and Department of Biochemistry and Molecular Biology | Robiul Hasan Bhuiyan, Md. Imranul Hoq, Md. Khondakar Raziur Rahman, Imam Hossen, Sajib Rudra, Md. Arif Hossain, Shanta Paul, Md. Omer Faruq, H. M. Abdullah Al Masud, Mohammad Omar Faruque |
| EPI_ISL_735494 | Cox's Bazar Medical College | Central Biological Research Laboratory and Department of Biochemistry and Molecular Biology | Md. Imranul Hoq, Robiul Hasan Bhuiyan, Md. Khondakar Raziur Rahman, Imam Hossen, Sajib Rudra, Md. Arif Hossain, Shanta Paul, Md. Omer Faruq, Mohammad Omar Faruque, H. M. Abdullah Al Masud |
| EPI_ISL_736925, EPI_ISL_736926 | NHLS-IALCH | KRISP, KZN Research Innovation and Sequencing Platform | Giandhari J, Pillay S, Lessells R, ChimukangaraB, Mdlalose K, York D, Khan S, Tegally H, Wilkinson E, de Oliveira T |
| EPI_ISL_736928, EPI_ISL_736929 | NHLS-UCT | KRISP, KZN Research Innovation and Sequencing Platform | Arash Iranzadeh, Deelan Doolabh, Lynn Tyers, Bruna Galvao, Innocent Mudau, Marvin Hsiao, Kruger Marais, Jennifer Giandhari, Sureshnee Pillay, Houriiyah Tegally, Emanuel James San, Tulio de Oliveira, Diana Hardie, Stephen Korsman, Carolyn Williamson |
| EPI_ISL_736931, EPI_ISL_736932, EPI_ISL_736933, EPI_ISL_736934, EPI_ISL_736935 | NHLS-IALCH | KRISP, KZN Research Innovation and Sequencing Platform | Giandhari J, Pillay S, Lessells R, ChimukangaraB, Mdlalose K, York D, Khan S, Tegally H, Wilkinson E, de Oliveira T |
| EPI_ISL_736939 | NHLS-UCT | KRISP, KZN Research Innovation and Sequencing Platform | Arash Iranzadeh, Deelan Doolabh, Lynn Tyers, Bruna Galvao, Innocent Mudau, Marvin Hsiao, Kruger Marais, Jennifer Giandhari, Sureshnee Pillay, Houriiyah Tegally, Emanuel James San, Tulio de Oliveira, Diana Hardie, Stephen Korsman, Carolyn Williamson |
| EPI_ISL_736940, EPI_ISL_736941, EPI_ISL_736942, EPI_ISL_736943, EPI_ISL_736944, EPI_ISL_736945, EPI_ISL_736946, EPI_ISL_736947, EPI_ISL_736948, EPI_ISL_736949, EPI_ISL_736950, EPI_ISL_736951, EPI_ISL_736952, EPI_ISL_736953, EPI_ISL_736954, EPI_ISL_736955, EPI_ISL_736956 | | | |
| see above | NHLS-IALCH | KRISP, KZN Research Innovation and Sequencing Platform | Giandhari J, Pillay S, Lessells R, ChimukangaraB, Mdlalose K, York D, Khan S, Tegally H, Wilkinson E, de Oliveira T |
| EPI_ISL_736981, EPI_ISL_736982, EPI_ISL_736983 | NHLS-UCT | KRISP, KZN Research Innovation and Sequencing Platform | Arash Iranzadeh, Deelan Doolabh, Lynn Tyers, Bruna Galvao, Innocent Mudau, Marvin Hsiao, Kruger Marais, Jennifer Giandhari, Sureshnee Pillay, Houriiyah Tegally, Emanuel James San, Tulio de Oliveira, Diana Hardie, Stephen Korsman, Carolyn Williamson |
| EPI_ISL_736984, EPI_ISL_736985, EPI_ISL_736986, EPI_ISL_736987, EPI_ISL_736988, EPI_ISL_736989, EPI_ISL_736990, EPI_ISL_736991, EPI_ISL_736992, EPI_ISL_736993, EPI_ISL_736994 | | | |
| see above | NHLS-IALCH | KRISP, KZN Research Innovation and Sequencing Platform | Giandhari J, Pillay S, Lessells R, ChimukangaraB, Mdlalose K, York D, Khan S, Tegally H, Wilkinson E, de Oliveira T |
| EPI_ISL_737109, EPI_ISL_737124, EPI_ISL_737149, EPI_ISL_737153, EPI_ISL_737155, EPI_ISL_737159, EPI_ISL_737161, EPI_ISL_737166, EPI_ISL_737170, EPI_ISL_737174, EPI_ISL_737176, EPI_ISL_737177, EPI_ISL_737185, EPI_ISL_737188, EPI_ISL_737197 | | | |
| see above | Michigan Department of Health and Human Services, Bureau of Laboratories | Michigan Department of Health and Human Services, Bureau of Laboratories | Blankenship HM, Riner D, Soehnlen MK |
| EPI_ISL_737236 | Los Angeles County PHL | Los Angeles County PHL | P. Hemarajata et al. |
| EPI_ISL_737283, EPI_ISL_737285, EPI_ISL_737289, EPI_ISL_737291, EPI_ISL_737293, EPI_ISL_737295, EPI_ISL_737297, EPI_ISL_737300, EPI_ISL_737302, EPI_ISL_737305, EPI_ISL_737308 | | | |
| see above | Michigan Department of Health and Human Services, Bureau of Laboratories | Michigan Department of Health and Human Services, Bureau of Laboratories | Blankenship HM, Riner D, Soehnlen MK |
| EPI_ISL_737553, EPI_ISL_737563, EPI_ISL_737598, EPI_ISL_737599, EPI_ISL_737609, EPI_ISL_737618, EPI_ISL_737629, EPI_ISL_737636, EPI_ISL_737648, EPI_ISL_737660, EPI_ISL_737679, EPI_ISL_737680, EPI_ISL_737681, EPI_ISL_737728, EPI_ISL_737729, EPI_ISL_737730, EPI_ISL_737732, EPI_ISL_737733, EPI_ISL_737734, EPI_ISL_737735, EPI_ISL_737736, EPI_ISL_737737, EPI_ISL_737738, EPI_ISL_737761, EPI_ISL_737772, EPI_ISL_737773, EPI_ISL_737774, EPI_ISL_737775, EPI_ISL_737776, EPI_ISL_737777, EPI_ISL_737778, EPI_ISL_737779, EPI_ISL_737811, EPI_ISL_737812, EPI_ISL_737813, EPI_ISL_737815, EPI_ISL_737848, EPI_ISL_737849, EPI_ISL_737850, EPI_ISL_737851, EPI_ISL_737852, EPI_ISL_737853, EPI_ISL_737854, EPI_ISL_737855, EPI_ISL_737856, EPI_ISL_737903, EPI_ISL_737904, EPI_ISL_737905, EPI_ISL_737906, EPI_ISL_737919, EPI_ISL_737920, EPI_ISL_737921, EPI_ISL_737922, EPI_ISL_737923, EPI_ISL_737924 | | | |
| see above | Viollier AG | Department of Biosystems Science and Engineering, ETH Zürich | Chaoran Chen, Sarah Nadeau, Catharine Aquino, Ivan Topolsky, Philipp Jablonski, Lara Fuhrmann, David Dreifuss, Katharina Jahn, Andreia Cabral de Gouvea, Maria Domenica Moccia, Simon Grüter, Timothy Sykes, Lennart Opitz, Griffin White, Laura Neff, Doris Popovic, Andrea Patrignani, Jay Tracy, Ralph Schlapbach, Christiane Beckmann, Maurice Redondo, Olivier Kobel, Christoph Noppen, Sophie Seidel, Noemie Santamaria de Souza, Niko Beerenwinkel, Tanja Stadler |
| EPI_ISL_738064, EPI_ISL_738065 | Department of Laboratory Medicine, National Taiwan University Hospital | Microbial Genomics Core Lab, National Taiwan University Centers of Genomic and Precision Medicine | Shiou-Hwei Yeh, You-Yu Lin, Ya-Yun Lai, Chiao-Ling Li, Shan-Chwen Chang, Pei-Jer Chen, Sui-Yuan Chang |

| | | | |
|---|---|---|---|
| EPI_ISL_738317, EPI_ISL_738318, EPI_ISL_738319, EPI_ISL_738320, EPI_ISL_738321, EPI_ISL_738322, EPI_ISL_738323, EPI_ISL_738324, EPI_ISL_738325, EPI_ISL_738326, EPI_ISL_738327, EPI_ISL_738328, EPI_ISL_738329, EPI_ISL_738330, EPI_ISL_738331, EPI_ISL_738332, EPI_ISL_738333, EPI_ISL_738334, EPI_ISL_738335, EPI_ISL_738336, EPI_ISL_738337, EPI_ISL_738338, EPI_ISL_738339, EPI_ISL_738340, EPI_ISL_738341, EPI_ISL_738342, EPI_ISL_738343, EPI_ISL_738344, EPI_ISL_738345, EPI_ISL_738346, EPI_ISL_738347, EPI_ISL_738348, EPI_ISL_738349 | | | |
| see above | Landstuhl Regional Medical Center | United States Air Force School of Aerospace Medicine | Anthony Fries, Jennifer Meyer, Amanda Javorina, Sarah Purves, William Gruner, Clarise Starr, Elizabeth Macias, Fritz Castillo, Cole Anderson |
| EPI_ISL_738501, EPI_ISL_738502, EPI_ISL_738511, EPI_ISL_738520, EPI_ISL_738546 | Santa Clara County Public Health Laboratory | Chan-Zuckerberg Biohub | CZB Cliahub Consortium |
| EPI_ISL_738575, EPI_ISL_738576 | Renegade | Chan-Zuckerberg Biohub | CZB Cliahub Consortium |
| EPI_ISL_738593, EPI_ISL_738595, EPI_ISL_738596, EPI_ISL_738598 | Santa Clara County Public Health Laboratory | Chan-Zuckerberg Biohub | CZB Cliahub Consortium |
| EPI_ISL_738602 | Orange County Public Health Lab | Chan-Zuckerberg Biohub | CZB Cliahub Consortium |
| EPI_ISL_738610 | Santa Clara County Public Health Laboratory | Chan-Zuckerberg Biohub | CZB Cliahub Consortium |
| EPI_ISL_738655 | Orange County Public Health Lab | Chan-Zuckerberg Biohub | CZB Cliahub Consortium |
| EPI_ISL_738660 | Renegade | Chan-Zuckerberg Biohub | CZB Cliahub Consortium |
| EPI_ISL_738664 | Orange County Public Health Lab | Chan-Zuckerberg Biohub | CZB Cliahub Consortium |
| EPI_ISL_738667 | Santa Clara County Public Health Laboratory | Chan-Zuckerberg Biohub | CZB Cliahub Consortium |
| EPI_ISL_738669 | Orange County Public Health Lab | Chan-Zuckerberg Biohub | CZB Cliahub Consortium |
| EPI_ISL_738673, EPI_ISL_738683, EPI_ISL_738684, EPI_ISL_738690, EPI_ISL_738695, EPI_ISL_738700, EPI_ISL_738702 | Santa Clara County Public Health Laboratory | Chan-Zuckerberg Biohub | CZB Cliahub Consortium |
| EPI_ISL_738703 | Orange County Public Health Lab | Chan-Zuckerberg Biohub | CZB Cliahub Consortium |
| EPI_ISL_738705 | Santa Clara County Public Health Laboratory | Chan-Zuckerberg Biohub | CZB Cliahub Consortium |
| EPI_ISL_738713, EPI_ISL_738714, EPI_ISL_738727 | Renegade | Chan-Zuckerberg Biohub | CZB Cliahub Consortium |
| EPI_ISL_738732, EPI_ISL_738734, EPI_ISL_738735, EPI_ISL_738736, EPI_ISL_738737, EPI_ISL_738738, EPI_ISL_738739, EPI_ISL_738740, EPI_ISL_738741 | Orange County Public Health Lab | Chan-Zuckerberg Biohub | CZB Cliahub Consortium |
| EPI_ISL_738764 | Santa Clara County Public Health Laboratory | Chan-Zuckerberg Biohub | CZB Cliahub Consortium |
| EPI_ISL_738782, EPI_ISL_738823, EPI_ISL_738826 | Orange County Public Health Lab | Chan-Zuckerberg Biohub | CZB Cliahub Consortium |
| EPI_ISL_738839 | Santa Clara County Public Health Laboratory | Chan-Zuckerberg Biohub | CZB Cliahub Consortium |
| EPI_ISL_738840, EPI_ISL_738841 | Orange County Public Health Lab | Chan-Zuckerberg Biohub | CZB Cliahub Consortium |
| EPI_ISL_738859 | Santa Clara County Public Health Laboratory | Chan-Zuckerberg Biohub | CZB Cliahub Consortium |
| EPI_ISL_738863 | Orange County Public Health Lab | Chan-Zuckerberg Biohub | CZB Cliahub Consortium |
| EPI_ISL_738877 | Renegade | Chan-Zuckerberg Biohub | CZB Cliahub Consortium |
| EPI_ISL_738893 | Orange County Public Health Lab | Chan-Zuckerberg Biohub | CZB Cliahub Consortium |
| EPI_ISL_738894 | Santa Clara County Public Health Laboratory | Chan-Zuckerberg Biohub | CZB Cliahub Consortium |
| EPI_ISL_738899 | Orange County Public Health Lab | Chan-Zuckerberg Biohub | CZB Cliahub Consortium |
| EPI_ISL_738901 | Renegade | Chan-Zuckerberg Biohub | CZB Cliahub Consortium |
| EPI_ISL_738907, EPI_ISL_738908 | Orange County Public Health Lab | Chan-Zuckerberg Biohub | CZB Cliahub Consortium |
| EPI_ISL_738915, EPI_ISL_738927 | Santa Clara County Public Health Laboratory | Chan-Zuckerberg Biohub | CZB Cliahub Consortium |
| EPI_ISL_738928 | Orange County Public Health Lab | Chan-Zuckerberg Biohub | CZB Cliahub Consortium |
| EPI_ISL_738979, EPI_ISL_738994 | Santa Clara County Public Health Laboratory | Chan-Zuckerberg Biohub | CZB Cliahub Consortium |
| EPI_ISL_738995 | Orange County Public Health Lab | Chan-Zuckerberg Biohub | CZB Cliahub Consortium |
| EPI_ISL_738996, EPI_ISL_738997, EPI_ISL_739005, EPI_ISL_739009 | Santa Clara County Public Health Laboratory | Chan-Zuckerberg Biohub | CZB Cliahub Consortium |
| EPI_ISL_739013 | Orange County Public Health Lab | Chan-Zuckerberg Biohub | CZB Cliahub Consortium |
| EPI_ISL_739020 | Santa Clara County Public Health Laboratory | Chan-Zuckerberg Biohub | CZB Cliahub Consortium |
| EPI_ISL_739022 | Renegade | Chan-Zuckerberg Biohub | CZB Cliahub Consortium |
| EPI_ISL_739025, EPI_ISL_739030 | Santa Clara County Public Health Laboratory | Chan-Zuckerberg Biohub | CZB Cliahub Consortium |
| EPI_ISL_739034, EPI_ISL_739044 | Orange County Public Health Lab | Chan-Zuckerberg Biohub | CZB Cliahub Consortium |
| EPI_ISL_739045, EPI_ISL_739066, EPI_ISL_739067 | Santa Clara County Public Health Laboratory | Chan-Zuckerberg Biohub | CZB Cliahub Consortium |
| EPI_ISL_739069 | Orange County Public Health Lab | Chan-Zuckerberg Biohub | CZB Cliahub Consortium |
| EPI_ISL_739076 | Santa Clara County Public Health Laboratory | Chan-Zuckerberg Biohub | CZB Cliahub Consortium |
| EPI_ISL_739089, EPI_ISL_739094, EPI_ISL_739097 | Orange County Public Health Lab | Chan-Zuckerberg Biohub | CZB Cliahub Consortium |
| EPI_ISL_739098 | Santa Clara County Public Health Laboratory | Chan-Zuckerberg Biohub | CZB Cliahub Consortium |
| EPI_ISL_739125, EPI_ISL_739132, EPI_ISL_739142 | Orange County Public Health Lab | Chan-Zuckerberg Biohub | CZB Cliahub Consortium |
| EPI_ISL_739146, EPI_ISL_739152, EPI_ISL_739153 | Santa Clara County Public Health Laboratory | Chan-Zuckerberg Biohub | CZB Cliahub Consortium |
| EPI_ISL_739154, EPI_ISL_739191, | Orange County Public Health Lab | Chan-Zuckerberg Biohub | CZB Cliahub Consortium |

| | | | |
|---|---|---|---|
| EPI_ISL_739211 | | | |
| EPI_ISL_739214 | Santa Clara County Public Health Laboratory | Chan-Zuckerberg Biohub | CZB Cliahub Consortium |
| EPI_ISL_739216 | Orange County Public Health Lab | Chan-Zuckerberg Biohub | CZB Cliahub Consortium |
| EPI_ISL_739221 | Santa Clara County Public Health Laboratory | Chan-Zuckerberg Biohub | CZB Cliahub Consortium |
| EPI_ISL_739222 | Orange County Public Health Lab | Chan-Zuckerberg Biohub | CZB Cliahub Consortium |
| EPI_ISL_739247, EPI_ISL_739248 | Santa Clara County Public Health Laboratory | Chan-Zuckerberg Biohub | CZB Cliahub Consortium |
| EPI_ISL_739266, EPI_ISL_739267 | Orange County Public Health Lab | Chan-Zuckerberg Biohub | CZB Cliahub Consortium |
| EPI_ISL_739277 | Santa Clara County Public Health Laboratory | Chan-Zuckerberg Biohub | CZB Cliahub Consortium |
| EPI_ISL_739279 | Orange County Public Health Lab | Chan-Zuckerberg Biohub | CZB Cliahub Consortium |
| EPI_ISL_739282 | Santa Clara County Public Health Laboratory | Chan-Zuckerberg Biohub | CZB Cliahub Consortium |
| EPI_ISL_739316 | Renegade | Chan-Zuckerberg Biohub | CZB Cliahub Consortium |
| EPI_ISL_739317, EPI_ISL_739320 | Santa Clara County Public Health Laboratory | Chan-Zuckerberg Biohub | CZB Cliahub Consortium |
| EPI_ISL_739322, EPI_ISL_739347 | Orange County Public Health Lab | Chan-Zuckerberg Biohub | CZB Cliahub Consortium |
| EPI_ISL_739381, EPI_ISL_739392, EPI_ISL_739415 | Santa Clara County Public Health Laboratory | Chan-Zuckerberg Biohub | CZB Cliahub Consortium |
| EPI_ISL_739437 | Orange County Public Health Lab | Chan-Zuckerberg Biohub | CZB Cliahub Consortium |
| EPI_ISL_739440 | Santa Clara County Public Health Laboratory | Chan-Zuckerberg Biohub | CZB Cliahub Consortium |
| EPI_ISL_739450, EPI_ISL_739455 | Orange County Public Health Lab | Chan-Zuckerberg Biohub | CZB Cliahub Consortium |
| EPI_ISL_739456, EPI_ISL_739457 | Santa Clara County Public Health Laboratory | Chan-Zuckerberg Biohub | CZB Cliahub Consortium |
| EPI_ISL_739464 | Orange County Public Health Lab | Chan-Zuckerberg Biohub | CZB Cliahub Consortium |
| EPI_ISL_739466 | Renegade | Chan-Zuckerberg Biohub | CZB Cliahub Consortium |
| EPI_ISL_739497 | Orange County Public Health Lab | Chan-Zuckerberg Biohub | CZB Cliahub Consortium |
| EPI_ISL_739512 | Santa Clara County Public Health Laboratory | Chan-Zuckerberg Biohub | CZB Cliahub Consortium |
| EPI_ISL_739513, EPI_ISL_739517, EPI_ISL_739529 | Orange County Public Health Lab | Chan-Zuckerberg Biohub | CZB Cliahub Consortium |
| EPI_ISL_739550 | Santa Clara County Public Health Laboratory | Chan-Zuckerberg Biohub | CZB Cliahub Consortium |
| EPI_ISL_739556 | Orange County Public Health Lab | Chan-Zuckerberg Biohub | CZB Cliahub Consortium |
| EPI_ISL_739562 | Santa Clara County Public Health Laboratory | Chan-Zuckerberg Biohub | CZB Cliahub Consortium |
| EPI_ISL_739564 | Renegade | Chan-Zuckerberg Biohub | CZB Cliahub Consortium |
| EPI_ISL_739566, EPI_ISL_739577 | Orange County Public Health Lab | Chan-Zuckerberg Biohub | CZB Cliahub Consortium |
| EPI_ISL_739579, EPI_ISL_739584, EPI_ISL_739589 | Santa Clara County Public Health Laboratory | Chan-Zuckerberg Biohub | CZB Cliahub Consortium |
| EPI_ISL_739630 | Renegade | Chan-Zuckerberg Biohub | CZB Cliahub Consortium |
| EPI_ISL_739642 | Santa Clara County Public Health Laboratory | Chan-Zuckerberg Biohub | CZB Cliahub Consortium |
| EPI_ISL_739646 | Orange County Public Health Lab | Chan-Zuckerberg Biohub | CZB Cliahub Consortium |
| EPI_ISL_739649 | Santa Clara County Public Health Laboratory | Chan-Zuckerberg Biohub | CZB Cliahub Consortium |
| EPI_ISL_739655 | Renegade | Chan-Zuckerberg Biohub | CZB Cliahub Consortium |
| EPI_ISL_739659, EPI_ISL_739660, EPI_ISL_739661 | Al-Quds Nutrition and Health Research Institute, Al-Quds University | Al-Quds Nutrition and Health Research Institute, Al-Quds University | Nasereddin,A., Ereqat,S., Al-Jawabreh,A., Rishmawi,C. |
| EPI_ISL_739723, EPI_ISL_739742, EPI_ISL_739759, EPI_ISL_739763, EPI_ISL_739776, EPI_ISL_739793, EPI_ISL_739797, EPI_ISL_739801, EPI_ISL_739821, EPI_ISL_739823, EPI_ISL_739847, EPI_ISL_739849, EPI_ISL_739865, EPI_ISL_739872, EPI_ISL_739885, EPI_ISL_739890, EPI_ISL_739895, EPI_ISL_739907, EPI_ISL_739928, EPI_ISL_739935, EPI_ISL_739950, EPI_ISL_739951, EPI_ISL_739960, EPI_ISL_739961, EPI_ISL_739986, EPI_ISL_739994, EPI_ISL_740011, EPI_ISL_740029, EPI_ISL_740059, EPI_ISL_740083, EPI_ISL_740096, EPI_ISL_740140, EPI_ISL_740148, EPI_ISL_740163, EPI_ISL_740174, EPI_ISL_740177, EPI_ISL_740211, EPI_ISL_740223, EPI_ISL_740228, EPI_ISL_740241, EPI_ISL_740252, EPI_ISL_740257, EPI_ISL_740287, EPI_ISL_740294, EPI_ISL_740301, EPI_ISL_740325, EPI_ISL_740363, EPI_ISL_740371, EPI_ISL_740374, EPI_ISL_740377, EPI_ISL_740385, EPI_ISL_740400, EPI_ISL_740411, EPI_ISL_740413, EPI_ISL_740446, EPI_ISL_740452, EPI_ISL_740454, EPI_ISL_740458, EPI_ISL_740493 | | | |
| see above | Laboratoire national de santé, Microbiology, Virology | Laboratoire national de santé, Microbiology, Microbial Genomics Platform | Anke Wienecke-Baldacchino, Catherine Ragimbeau,Jessica Tapp, Fatu Djabi, Lise Pignon, Raoul Salmon, Tamir Abdelrahman |
| EPI_ISL_740547 | Department of Laboratory Medicine, National Taiwan University Hospital | Microbial Genomics Core Lab, National Taiwan University Centers of Genomic and Precision Medicine | Shiou-Hwei Yeh, You-Yu Lin, Ya-Yun Lai, Chiao-Ling Li, Shan-Chwen Chang, Pei-Jer Chen, Sui-Yuan Chang |
| EPI_ISL_740864 | South Eastern Area Laboratory Services (SEALS) | NSW Health Pathology - Institute of Clinical Pathology and Medical Research; Westmead Hospital; University of Sydney | CIDM-PH et al. |
| EPI_ISL_740953, EPI_ISL_740954 | Department of Pathology, University of Cambridge | COVID-19 Genomics UK (COG-UK) Consortium | Aminu S. Jahun, Yasmin Chaudhry, Grant Hall, Iliana Georgana, Myra Hosmillo, Martin D. Curran, Malte Pinckert, Surendra Parmar, Ian Goodfellow |
| EPI_ISL_741304 | University College London, Great Ormond Street Hospital for Children NHS Foundation Trust, Imperial College Healthcare NHS Trust | COVID-19 Genomics UK (COG-UK) Consortium | Sergi Castellano, Rachel Williams, Mark Kristiansen, Paola Resende Silva, Sunando Roy, Tony Brooks, Helena Tutill, Paola Niola, Patricia Dyal, Charlotte Williams, Leysa Forrest, Yasmin Panchbhaya, Jacqueline Findlay, Samuel Weeks, Julianne Brown, Kathryn Harris, Paul Randell, James Price, Alison Holmes, Judith Breuer |
| EPI_ISL_741436, EPI_ISL_741437, EPI_ISL_741438, EPI_ISL_741439, EPI_ISL_741440, EPI_ISL_741441, EPI_ISL_741442, EPI_ISL_741443, EPI_ISL_741444, EPI_ISL_741445, EPI_ISL_741446, EPI_ISL_741447, EPI_ISL_741448, EPI_ISL_741449, EPI_ISL_741450, EPI_ISL_741451, EPI_ISL_741452 | | | |
| see above | Northumbria University / South Tees Hospitals NHS Foundation Trust / North Cumbria Integrated Care NHS Foundation Trust / North Tees and Hartlepool NHS Foundation Trust / Newcastle Hospitals NHS Foundation Trust | COVID-19 Genomics UK (COG-UK) Consortium | Darren L Smith,Andrew Nelson,Matthew Bashton,Greg R Young,Joshua Loh,John Allan,Mohammad A Tariq,Giles S Holt,Gary Black,Wen C Yew,Lynn Dover,Paul Baker,Steve Liggett,Sarah Essex,Jane Greenaway,Debra Padgett,Clive Graham,Garren Scott,Edward Barton,Emma Swindells,Brendan Payne,Jennifer Collins,Yusri Taha,Gary Eltringham |
| EPI_ISL_741544 | Quadram Institute Bioscience | COVID-19 Genomics UK (COG-UK) Consortium | Dave J. Baker, Gemma L. Kay, Alp Aydin, Thanh Le-Viet, Steven Rudder, Ana P. Tedim, Anastasia Kolyva, Maria Diaz, Leonardo de Oliveira Martins, Nabil-Fareed Alikhan, Lizzie Meadows, Rachael Stanley, Ngozi Elumogo, Muhammed Yasir, Nicholas M. Thomson, Alexander J Trotter, Rachel Gilroy, Samuel Bloomfield, Claire Stuart, Andrew Bell, Reenesh Prakash, Samir Dervisevic, Alison E. Mather, John Wain, Mark Webber, Andrew J. Page, Justin O'Grady |
| EPI_ISL_741610 | Queens Medical Centre, Clinical Microbiology Department / DeepSeq Nottingham | COVID-19 Genomics UK (COG-UK) Consortium | Gemma Clark, Wendy Smith, Manjinder Khakh, Vicki M Fleming, Michelle M Lister, Hannah Howson-Wells, Jonathan Ball, Patrick McClure, Joseph Chappell, Theocharis Tsoleridis, Nadine Holmes, Matthew Carlisle, Christopher Moore, Fei Sang, Johnny Debebe, Victoria Wright, Matthew Loose |
| EPI_ISL_741738, EPI_ISL_741739, | Centre for Enzyme Innovation, University of Portsmouth / | COVID-19 Genomics UK (COG-UK) Consortium | Angela Beckett,Yann Bourgeois,Garry Scarlett,Sharon Glaysher,Scott Elliott,Kelly Bicknell,Robert Impey,Allyson Lloyd,Sarah Wyllie,Ethan Butcher,Anoop |

| | | | |
|---|---|---|---|
| EPI_ISL_741756, EPI_ISL_741757, EPI_ISL_741758 | Translational Research Laboratory, Portsmouth Hospitals NHS Trust | | Chauhan,Samuel Robson |
| EPI_ISL_742840, EPI_ISL_742880, EPI_ISL_742881, EPI_ISL_742882, EPI_ISL_742883, EPI_ISL_742884, EPI_ISL_742885, EPI_ISL_742886, EPI_ISL_742887, EPI_ISL_742888, EPI_ISL_742889, EPI_ISL_743117, EPI_ISL_743126, EPI_ISL_743129, EPI_ISL_743130, EPI_ISL_743131, EPI_ISL_743161, EPI_ISL_743162, EPI_ISL_743380, EPI_ISL_743381, EPI_ISL_743592, EPI_ISL_743593, EPI_ISL_743594, EPI_ISL_743595, EPI_ISL_743606, EPI_ISL_743607, EPI_ISL_743608, EPI_ISL_743609, EPI_ISL_743610, EPI_ISL_743614, EPI_ISL_743615, EPI_ISL_743616, EPI_ISL_743617, EPI_ISL_743618, EPI_ISL_743619, EPI_ISL_743620, EPI_ISL_743621, EPI_ISL_743622, EPI_ISL_743623, EPI_ISL_743624, EPI_ISL_743625, EPI_ISL_743626, EPI_ISL_743627, EPI_ISL_743628, EPI_ISL_743629, EPI_ISL_743630, EPI_ISL_743631, EPI_ISL_743632, EPI_ISL_743633, EPI_ISL_743634, EPI_ISL_743635, EPI_ISL_743636, EPI_ISL_743637, EPI_ISL_743638, EPI_ISL_743639 | | | |
| see above | Wales Specialist Virology Centre Sequencing lab: Pathogen Genomics Unit | COVID-19 Genomics UK (COG-UK) Consortium | Catherine Moore, Johnathan Evans, Laura Gifford, Malorie Perry, Simon Cottrell, Angela Marchbank, Alec Birchley, Alexander Adams, Amy Gaskin, Bree Gatica-Wilcox, Jason Coombes, Joel Southgate, Lauren Gilbert, Lee Graham, Nicole Pacchiarini, Sara Kumziene-Summerhayes, Sarah Taylor, Sophie Jones, Sara Rey, Matthew Bull, Joanne Watkins, Sally Corden, Tom Connor |
| EPI_ISL_744143, EPI_ISL_744162, EPI_ISL_744173, EPI_ISL_744189, EPI_ISL_744199, EPI_ISL_744225, EPI_ISL_744239, EPI_ISL_744247, EPI_ISL_744255, EPI_ISL_744267, EPI_ISL_744276, EPI_ISL_744279, EPI_ISL_744286, EPI_ISL_744293, EPI_ISL_744299, EPI_ISL_744314, EPI_ISL_744318, EPI_ISL_744319, EPI_ISL_744322, EPI_ISL_744331, EPI_ISL_744345, EPI_ISL_744346, EPI_ISL_744348, EPI_ISL_744355, EPI_ISL_744356, EPI_ISL_744366, EPI_ISL_744368, EPI_ISL_744390, EPI_ISL_744403, EPI_ISL_744404, EPI_ISL_744421, EPI_ISL_744427, EPI_ISL_744431, EPI_ISL_744432, EPI_ISL_744449, EPI_ISL_744450, EPI_ISL_744469, EPI_ISL_744477, EPI_ISL_744525, EPI_ISL_744540, EPI_ISL_744544, EPI_ISL_744560, EPI_ISL_744573, EPI_ISL_744592, EPI_ISL_744593, EPI_ISL_744634, EPI_ISL_744635, EPI_ISL_744665, EPI_ISL_744677, EPI_ISL_744691, EPI_ISL_744698, EPI_ISL_744707, EPI_ISL_744714, EPI_ISL_744715, EPI_ISL_744737, EPI_ISL_744765, EPI_ISL_744768, EPI_ISL_744771, EPI_ISL_744798, EPI_ISL_744803, EPI_ISL_744807, EPI_ISL_744827, EPI_ISL_744845, EPI_ISL_744867, EPI_ISL_744878, EPI_ISL_744889, EPI_ISL_744898, EPI_ISL_744900, EPI_ISL_744910, EPI_ISL_744921, EPI_ISL_744940, EPI_ISL_744950, EPI_ISL_744970, EPI_ISL_744972, EPI_ISL_744975, EPI_ISL_744976, EPI_ISL_744977, EPI_ISL_744986, EPI_ISL_744996 | | | |
| see above | Laboratoire national de santé, Microbiology, Virology | Laboratoire national de santé, Microbiology, Microbial Genomics Platform | Anke Wienecke-Baldacchino, Catherine Ragimbeau,Jessica Tapp, Fatu Djabi, Lise Pignon, Raoul Salmon, Tamir Abdelrahman |
| EPI_ISL_745128, EPI_ISL_745153 | CoVid EC Nelson Mandela Bay Metro | National Health Laboratory Service (NHLS), Tygerberg | Susan Engelbrecht, Kayla Delaney, Bronwyn Kleinhans, Houriiyah Tegally, Eduan Wilkindon, Gert van Zyl, Wolfgang Preiser, Tulio de Oliveira |
| EPI_ISL_745307, EPI_ISL_745308, EPI_ISL_745316, EPI_ISL_745320, EPI_ISL_745321, EPI_ISL_745337, EPI_ISL_745338, EPI_ISL_745339, EPI_ISL_745340, EPI_ISL_745341, EPI_ISL_745342, EPI_ISL_745343, EPI_ISL_745344, EPI_ISL_745345, EPI_ISL_745396 | | | |
| see above | CNR Virus des Infections Respiratoires - France SUD | CNR Virus des Infections Respiratoires - France SUD | Antonin Bal, Gregory Destras, Claudia Gonzalez, Gwendolyne Burfin, Quentin Semanas, Martine Valette, Bruno Lina, Laurence Josset |
| EPI_ISL_745398, EPI_ISL_745407 | DOHMH Jamaica | New York City Public Health Laboratory | Jade Wang, et al. |
| EPI_ISL_745413 | DOHMH Central Harlem | New York City Public Health Laboratory | Jade Wang, et al. |
| EPI_ISL_745414, EPI_ISL_745415, EPI_ISL_745416, EPI_ISL_745417, EPI_ISL_745418 | DOHMH Jamaica | New York City Public Health Laboratory | Jade Wang, et al. |
| EPI_ISL_745419 | DOHMH Crown Heights | New York City Public Health Laboratory | Jade Wang, et al. |
| EPI_ISL_746408, EPI_ISL_746467, EPI_ISL_746836, EPI_ISL_746840, EPI_ISL_746844, EPI_ISL_746845, EPI_ISL_746851, EPI_ISL_746881, EPI_ISL_746886, EPI_ISL_746888, EPI_ISL_746894, EPI_ISL_746898, EPI_ISL_746904, EPI_ISL_746905, EPI_ISL_746908, EPI_ISL_746911, EPI_ISL_746919, EPI_ISL_746922, EPI_ISL_746923, EPI_ISL_746939, EPI_ISL_746940, EPI_ISL_746943, EPI_ISL_746947, EPI_ISL_746951, EPI_ISL_746952, EPI_ISL_746953, EPI_ISL_746954, EPI_ISL_746955, EPI_ISL_746970, EPI_ISL_746998, EPI_ISL_746999, EPI_ISL_747000, EPI_ISL_747001, EPI_ISL_747002, EPI_ISL_747003, EPI_ISL_747004, EPI_ISL_747005, EPI_ISL_747006, EPI_ISL_747007, EPI_ISL_747008, EPI_ISL_747009, EPI_ISL_747010, EPI_ISL_747011, EPI_ISL_747012, EPI_ISL_747013, EPI_ISL_747014 | | | |
| see above | Utah Public Health Laboratory | Utah Public Health Laboratory | Erin Young, Kelly Oakeson, Tara Gallagher |
| EPI_ISL_747172, EPI_ISL_747183, EPI_ISL_747184, EPI_ISL_747185, EPI_ISL_747186, EPI_ISL_747187, EPI_ISL_747188 | Respiratory Viruses Branch, Centers for Disease Control and Prevention | Respiratory Viruses Branch, Centers for Disease Control and Prevention | Queen,K., Li,Y., Tao,Y., Uehara,A., Montmayeur,A., Paden,C.R., Cook,P.W., Marine,R., Sheth,M., Wang,H., Lee,J., Tong,S. |
| EPI_ISL_747431, EPI_ISL_747432, EPI_ISL_747433, EPI_ISL_747434, EPI_ISL_747435, EPI_ISL_747436, EPI_ISL_747437, EPI_ISL_747440, EPI_ISL_747441, EPI_ISL_747442, EPI_ISL_747443, EPI_ISL_747444, EPI_ISL_747445, EPI_ISL_747453, EPI_ISL_747454, EPI_ISL_747457 | | | |
| see above | Division of Emerging Infectious Diseases, Bureau of Infectious Diseases Diagnosis Control, Korea Disease Control and Prevention Agency | Division of Emerging Infectious Diseases, Bureau of Infectious Diseases Diagnosis Control, Korea Disease Control and Prevention Agency | Ae Kyung Park, Il-Hwan Kim, Heui Man Kim, Jeong-Min Kim, Namjoo Lee, Chaeyoung Lee, Sang Hee Woo, Eun-Jin Kim |
| EPI_ISL_747459, EPI_ISL_747460, EPI_ISL_747461, EPI_ISL_747462, EPI_ISL_747463 | Ospedale San Bonifacio | Istituto Zooprofilattico Sperimentale delle Venezie | Adelaide Milani, Alessia Schivo, Annalisa Salviato, Erika Giorgia Quaranta, Ambra Pastori, Bianca Zecchin, Alice Fusaro, Isabella Monne, Calogero Terregino, Antonia Ricci |
| EPI_ISL_747476 | ULSS9 Distretto di Bussolengo | Istituto Zooprofilattico Sperimentale delle Venezie | Adelaide Milani, Alessia Schivo, Annalisa Salviato, Erika Giorgia Quaranta, Ambra Pastori, Bianca Zecchin, Alice Fusaro, Isabella Monne, Calogero Terregino, Antonia Ricci |
| EPI_ISL_747478, EPI_ISL_747479, EPI_ISL_747480 | ULSS 5 Polesana | Istituto Zooprofilattico Sperimentale delle Venezie | Adelaide Milani, Alessia Schivo, Annalisa Salviato, Erika Giorgia Quaranta, Ambra Pastori, Bianca Zecchin, Alice Fusaro, Isabella Monne, Calogero Terregino, Antonia Ricci |
| EPI_ISL_747482 | Ospedale San Bonifacio | Istituto Zooprofilattico Sperimentale delle Venezie | Adelaide Milani, Alessia Schivo, Annalisa Salviato, Erika Giorgia Quaranta, Ambra Pastori, Bianca Zecchin, Alice Fusaro, Isabella Monne, Calogero Terregino, Antonia Ricci |
| EPI_ISL_747486, EPI_ISL_747487, EPI_ISL_747488, EPI_ISL_747489 | ULSS 5 Polesana | Istituto Zooprofilattico Sperimentale delle Venezie | Adelaide Milani, Alessia Schivo, Annalisa Salviato, Erika Giorgia Quaranta, Ambra Pastori, Bianca Zecchin, Alice Fusaro, Isabella Monne, Calogero Terregino, Antonia Ricci |
| EPI_ISL_747492 | Respiratory Virus Unit, National Infection Service, Public Health England | COVID-19 Genomics UK (COG-UK) Consortium | PHE Covid Sequencing Team |
| EPI_ISL_747522 | Dutch COVID-19 response team | National Institute for Public Health and the Environment (RIVM) | Adam Meijer, Harry Vennema, Jeroen Cremer, Sharon van den Brink, Bas van der Veer, AnneMarie van den Brandt, Florian Zwagemaker, Dennis Schmitz, Chantal Reusken, on behalf of the national COVID-19 response team |
| EPI_ISL_751472 | CHU Purpan - Laboratoire de Virologie - Institut Fédératif de Biologie | CHU Purpan - Laboratoire de Virologie - Institut Fédératif de Biologie | Latour J., Ranger N., Dubois M., Carcenac R., Harter A., Boyer P., Tremeaux P., Izopet J. |
| EPI_ISL_751551 | MS Public Health Laboratory | Genomics and Discovery, Respiratory Viruses Branch, Division of Viral Diseases, Centers for Disease Control and Prevention | Krista Queen, Yan Li, Ying Tao, Jing Zhang, Anna Uehara, Anna Montmayeur, Clinton R. Paden, Peter W. Cook,Rachel Marine, Mili Sheth, Haibin Wang, Justin Lee, Suxiang Tong |
| EPI_ISL_751553, EPI_ISL_751565 | IA State Hygienic Laboratory | Genomics and Discovery, Respiratory Viruses Branch, Division of Viral Diseases, Centers for Disease Control and Prevention | Krista Queen, Yan Li, Ying Tao, Jing Zhang, Anna Uehara, Anna Montmayeur, Clinton R. Paden, Peter W. Cook,Rachel Marine, Mili Sheth, Haibin Wang, Justin Lee, Suxiang Tong |
| EPI_ISL_751578 | HI Dept. of Health, State Laboratories Division | Genomics and Discovery, Respiratory Viruses Branch, Division of Viral Diseases, Centers for Disease Control and Prevention | Krista Queen, Yan Li, Ying Tao, Jing Zhang, Anna Uehara, Anna Montmayeur, Clinton R. Paden, Peter W. Cook,Rachel Marine, Mili Sheth, Haibin Wang, Justin Lee, Suxiang Tong |
| EPI_ISL_751580 | NYSDOH Wadsworth Center, Virology Lab | Genomics and Discovery, Respiratory Viruses Branch, Division of Viral Diseases, Centers for Disease Control and Prevention | Krista Queen, Yan Li, Ying Tao, Jing Zhang, Anna Uehara, Anna Montmayeur, Clinton R. Paden, Peter W. Cook,Rachel Marine, Mili Sheth, Haibin Wang, Justin Lee, Suxiang Tong |
| EPI_ISL_751585 | NE Public Health Laboratory | Genomics and Discovery, Respiratory Viruses Branch, Division of Viral Diseases, Centers for Disease Control and Prevention | Krista Queen, Yan Li, Ying Tao, Jing Zhang, Anna Uehara, Anna Montmayeur, Clinton R. Paden, Peter W. Cook,Rachel Marine, Mili Sheth, Haibin Wang, Justin Lee, Suxiang Tong |
| EPI_ISL_751594 | KY State Public Health Lab | Genomics and Discovery, Respiratory Viruses Branch, Division of Viral Diseases, Centers for Disease Control and Prevention | Krista Queen, Yan Li, Ying Tao, Jing Zhang, Anna Uehara, Anna Montmayeur, Clinton R. Paden, Peter W. Cook,Rachel Marine, Mili Sheth, Haibin Wang, Justin Lee, Suxiang Tong |

| | | | |
|---|---|---|---|
| EPI_ISL_751603 | HI Dept. of Health, State Laboratories Division | Genomics and Discovery, Respiratory Viruses Branch, Division of Viral Diseases, Centers for Disease Control and Prevention | Krista Queen, Yan Li, Ying Tao, Jing Zhang, Anna Uehara, Anna Montmayeur, Clinton R. Paden, Peter W. Cook,Rachel Marine, Mili Sheth, Haibin Wang, Justin Lee, Suxiang Tong |
| EPI_ISL_751629 | MD DOH Laboratories Administration | Genomics and Discovery, Respiratory Viruses Branch, Division of Viral Diseases, Centers for Disease Control and Prevention | Krista Queen, Yan Li, Ying Tao, Jing Zhang, Anna Uehara, Anna Montmayeur, Clinton R. Paden, Peter W. Cook,Rachel Marine, Mili Sheth, Haibin Wang, Justin Lee, Suxiang Tong |
| EPI_ISL_751651 | HI Dept. of Health, State Laboratories Division | Genomics and Discovery, Respiratory Viruses Branch, Division of Viral Diseases, Centers for Disease Control and Prevention | Krista Queen, Yan Li, Ying Tao, Jing Zhang, Anna Uehara, Anna Montmayeur, Clinton R. Paden, Peter W. Cook,Rachel Marine, Mili Sheth, Haibin Wang, Justin Lee, Suxiang Tong |
| EPI_ISL_751666 | IA State Hygienic Laboratory | Genomics and Discovery, Respiratory Viruses Branch, Division of Viral Diseases, Centers for Disease Control and Prevention | Krista Queen, Yan Li, Ying Tao, Jing Zhang, Anna Uehara, Anna Montmayeur, Clinton R. Paden, Peter W. Cook,Rachel Marine, Mili Sheth, Haibin Wang, Justin Lee, Suxiang Tong |
| EPI_ISL_751683 | MS Public Health Laboratory | Genomics and Discovery, Respiratory Viruses Branch, Division of Viral Diseases, Centers for Disease Control and Prevention | Krista Queen, Yan Li, Ying Tao, Jing Zhang, Anna Uehara, Anna Montmayeur, Clinton R. Paden, Peter W. Cook,Rachel Marine, Mili Sheth, Haibin Wang, Justin Lee, Suxiang Tong |
| EPI_ISL_751687 | IA State Hygienic Laboratory | Genomics and Discovery, Respiratory Viruses Branch, Division of Viral Diseases, Centers for Disease Control and Prevention | Krista Queen, Yan Li, Ying Tao, Jing Zhang, Anna Uehara, Anna Montmayeur, Clinton R. Paden, Peter W. Cook,Rachel Marine, Mili Sheth, Haibin Wang, Justin Lee, Suxiang Tong |
| EPI_ISL_751689 | OR State PHL-Virology/Immunology Section | Genomics and Discovery, Respiratory Viruses Branch, Division of Viral Diseases, Centers for Disease Control and Prevention | Krista Queen, Yan Li, Ying Tao, Jing Zhang, Anna Uehara, Anna Montmayeur, Clinton R. Paden, Peter W. Cook,Rachel Marine, Mili Sheth, Haibin Wang, Justin Lee, Suxiang Tong |
| EPI_ISL_751692 | IA State Hygienic Laboratory | Genomics and Discovery, Respiratory Viruses Branch, Division of Viral Diseases, Centers for Disease Control and Prevention | Krista Queen, Yan Li, Ying Tao, Jing Zhang, Anna Uehara, Anna Montmayeur, Clinton R. Paden, Peter W. Cook,Rachel Marine, Mili Sheth, Haibin Wang, Justin Lee, Suxiang Tong |
| EPI_ISL_751715 | MO State Public Health Laboratory | Genomics and Discovery, Respiratory Viruses Branch, Division of Viral Diseases, Centers for Disease Control and Prevention | Krista Queen, Yan Li, Ying Tao, Jing Zhang, Anna Uehara, Anna Montmayeur, Clinton R. Paden, Peter W. Cook,Rachel Marine, Mili Sheth, Haibin Wang, Justin Lee, Suxiang Tong |
| EPI_ISL_751721 | IA State Hygienic Laboratory | Genomics and Discovery, Respiratory Viruses Branch, Division of Viral Diseases, Centers for Disease Control and Prevention | Krista Queen, Yan Li, Ying Tao, Jing Zhang, Anna Uehara, Anna Montmayeur, Clinton R. Paden, Peter W. Cook,Rachel Marine, Mili Sheth, Haibin Wang, Justin Lee, Suxiang Tong |
| EPI_ISL_751729 | CT-Dr. Katherine A. Kelley State Public Health Lab | Genomics and Discovery, Respiratory Viruses Branch, Division of Viral Diseases, Centers for Disease Control and Prevention | Krista Queen, Yan Li, Ying Tao, Jing Zhang, Anna Uehara, Anna Montmayeur, Clinton R. Paden, Peter W. Cook,Rachel Marine, Mili Sheth, Haibin Wang, Justin Lee, Suxiang Tong |
| EPI_ISL_751733, EPI_ISL_751739 | MN PHL Division, Minnesota Department of Health | Genomics and Discovery, Respiratory Viruses Branch, Division of Viral Diseases, Centers for Disease Control and Prevention | Krista Queen, Yan Li, Ying Tao, Jing Zhang, Anna Uehara, Anna Montmayeur, Clinton R. Paden, Peter W. Cook,Rachel Marine, Mili Sheth, Haibin Wang, Justin Lee, Suxiang Tong |
| EPI_ISL_751742 | IA State Hygienic Laboratory | Genomics and Discovery, Respiratory Viruses Branch, Division of Viral Diseases, Centers for Disease Control and Prevention | Krista Queen, Yan Li, Ying Tao, Jing Zhang, Anna Uehara, Anna Montmayeur, Clinton R. Paden, Peter W. Cook,Rachel Marine, Mili Sheth, Haibin Wang, Justin Lee, Suxiang Tong |
| EPI_ISL_751758 | HI Dept. of Health, State Laboratories Division | Genomics and Discovery, Respiratory Viruses Branch, Division of Viral Diseases, Centers for Disease Control and Prevention | Krista Queen, Yan Li, Ying Tao, Jing Zhang, Anna Uehara, Anna Montmayeur, Clinton R. Paden, Peter W. Cook,Rachel Marine, Mili Sheth, Haibin Wang, Justin Lee, Suxiang Tong |
| EPI_ISL_751779, EPI_ISL_751783 | IA State Hygienic Laboratory | Genomics and Discovery, Respiratory Viruses Branch, Division of Viral Diseases, Centers for Disease Control and Prevention | Krista Queen, Yan Li, Ying Tao, Jing Zhang, Anna Uehara, Anna Montmayeur, Clinton R. Paden, Peter W. Cook,Rachel Marine, Mili Sheth, Haibin Wang, Justin Lee, Suxiang Tong |
| EPI_ISL_752583, EPI_ISL_752584, EPI_ISL_752585, EPI_ISL_752586, EPI_ISL_752587, EPI_ISL_752588, EPI_ISL_752589, EPI_ISL_752590, EPI_ISL_752591, EPI_ISL_752592, EPI_ISL_752594, EPI_ISL_752595, EPI_ISL_752596, EPI_ISL_752597 | | | |
| see above | National Virus Reference Laboratory | National Virus Reference Laboratory | Michael Carr, Gabriel Gonzalez, Jonathan Dean, Daniel Hare, Cillian F De Gascun |
| EPI_ISL_752605 | Faculty of Medicine, Al-Quds University | Faculty of Medicine, Al-Quds University | Ereqat,S., Nasereddin,A. and Al-Jawabreh,A. |
| EPI_ISL_753174, EPI_ISL_753175, EPI_ISL_753196, EPI_ISL_753197, EPI_ISL_753198, EPI_ISL_753199, EPI_ISL_753214, EPI_ISL_753221, EPI_ISL_753222 | State Laboratories Division, Hawaii State Department of Health | State Laboratories Division, Hawaii State Department of Health | Pamela O'Brien, Sabrina Diemert, Drew Kuwazaki, Razvan Sultana, Edward Desmond |
| EPI_ISL_753527, EPI_ISL_753528, EPI_ISL_753529, EPI_ISL_753530, EPI_ISL_753531, EPI_ISL_753532, EPI_ISL_753533, EPI_ISL_753643, EPI_ISL_753644, EPI_ISL_753645, EPI_ISL_753647, EPI_ISL_753648, EPI_ISL_753649, EPI_ISL_753650, EPI_ISL_753651, EPI_ISL_753652 | | | |
| see above | Clinical virology Laboratory, Children's Hospital Los Angeles | Center for Personalized Medicine, Children's Hospital Los Angeles | Gai et al |
| EPI_ISL_754145, EPI_ISL_754155, EPI_ISL_754162 | Laboratoire de Microbiologie | National Reference Center for Viruses of Respiratory Infections, Institut Pasteur, Paris | Marion Barbet, Sylvie Behillil, Méline Bizard, Angela Brisebarre, Camille Capel, Etienne Simon-Lorière, Vincent Enouf, Maud Vanpeene, Sylvie van der Werf, Marie-Sarah Fangous |
| EPI_ISL_754173 | Specialized Lab for COVID-19 Detection, Department of Genetic Engineering and Biotechnology | Specialized Lab for COVID-19 Detection, Department of Genetic Engineering and Biotechnology | Md. Nazmul Hasan, Shamsul H. Prodhan, Hammadul Hoque , Nurnabi Azad Jewel, Mumtahin Ahammed, Mahedi Hasan Ridoy, Avishek Sarkar |
| EPI_ISL_754177 | Specialized Lab for COVID-19 Detection, Department of Genetic Engineering and Biotechnology | Specialized Lab for COVID-19 Detection, Department of Genetic Engineering and Biotechnology | Nurnabi Azad Jewel, Md. Nazmul Hasan, Shamsul H. Prodhan, Hammadul Hoque, Asim Debnath, Sudipto Sakib Duranto |
| EPI_ISL_754232 | The Republican Research and Practical Center for Epidemiology and Microbiology (RRPCEM) | WHO National Influenza Centre Russian Federation | Elena Gasich, Kirill Bulda, Anatoly Krasko, Andrey Komissarov, Artem Fadeev, Anna Ivanova, Kseniya Komissarova, Dmitry Bazhenov, Daria Danilenko, Ksenia Safina, Elena Nabieva, Georgii Bazykin, Dmitry Lioznov |
| EPI_ISL_754247, EPI_ISL_754257, EPI_ISL_754287, EPI_ISL_754294, EPI_ISL_754295, EPI_ISL_754296, EPI_ISL_754297, EPI_ISL_754303, EPI_ISL_754332, EPI_ISL_754333, EPI_ISL_754334, EPI_ISL_754335, EPI_ISL_754336, EPI_ISL_754337, EPI_ISL_754338, EPI_ISL_754343 | | | |
| see above | Respiratory Virus Unit, National Infection Service, Public Health England | COVID-19 Genomics UK (COG-UK) Consortium | PHE Covid Sequencing Team |
| EPI_ISL_754402, EPI_ISL_754408, EPI_ISL_754482, EPI_ISL_754483, EPI_ISL_754484, EPI_ISL_754485, EPI_ISL_754487, EPI_ISL_754489, EPI_ISL_754490, EPI_ISL_754491, EPI_ISL_754492, EPI_ISL_754493, EPI_ISL_754494, EPI_ISL_754495, EPI_ISL_754496, EPI_ISL_754497, EPI_ISL_754498, EPI_ISL_754499, EPI_ISL_754500, EPI_ISL_754501, EPI_ISL_754502, EPI_ISL_754503, EPI_ISL_754504 | | | |
| see above | Wadsworth Center, New York State Department.of Health | Wadsworth Center, New York State Department.of Health | Kirsten St. George, Daryl M. Lamson, Alexis Russel, Matthew Shudt, Melissa A Leisner, Jonathan Plitnick, Navjot Singh, John Kelly, Sara Griesemer, Erasmus Schneider, Erica Lasek-Nesselquist |
| EPI_ISL_754914, EPI_ISL_754942, EPI_ISL_754973, EPI_ISL_755002, EPI_ISL_755009, EPI_ISL_755024, EPI_ISL_755026, EPI_ISL_755092, EPI_ISL_755093, EPI_ISL_755109, EPI_ISL_755110, EPI_ISL_755111 | | | |
| see above | California Department of Public Health | California Department of Public Health | CDPH IDLB COVIDNet |

| | | | |
|---|---|---|---|
| EPI_ISL_755115, EPI_ISL_755116, EPI_ISL_755139 | San Diego County Public Health Laboratory | Andersen lab at Scripps Research | SEARCH Alliance San Diego with Tracy Basler, Jovan Shephard, Brett Austin |
| EPI_ISL_755222, EPI_ISL_755223, EPI_ISL_755224, EPI_ISL_755225, EPI_ISL_755229 | UCSD EXCITE lab | Andersen lab at Scripps Research | SEARCH Alliance San Diego |
| EPI_ISL_755269, EPI_ISL_755273, EPI_ISL_755276, EPI_ISL_755277, EPI_ISL_755278, EPI_ISL_755282, EPI_ISL_755287, EPI_ISL_755288, EPI_ISL_755291, EPI_ISL_755292, EPI_ISL_755293, EPI_ISL_755296, EPI_ISL_755297, EPI_ISL_755298, EPI_ISL_755300 | | | |
| see above | San Diego County Public Health Laboratory | Andersen lab at Scripps Research | SEARCH Alliance San Diego with Tracy Basler, Jovan Shephard, Brett Austin |
| EPI_ISL_755456, EPI_ISL_755457, EPI_ISL_755458, EPI_ISL_755459, EPI_ISL_755460, EPI_ISL_755461, EPI_ISL_755462, EPI_ISL_755463, EPI_ISL_755464, EPI_ISL_755465, EPI_ISL_755466, EPI_ISL_755467, EPI_ISL_755468, EPI_ISL_755469, EPI_ISL_755470, EPI_ISL_755471, EPI_ISL_755472, EPI_ISL_755473, EPI_ISL_755477, EPI_ISL_755478, EPI_ISL_755479, EPI_ISL_755480, EPI_ISL_755481 | | | |
| see above | Maine Health and Environmental Testing Laboratory | Tewhey Lab, The Jackson Laboratory | Matluk,N., Dewey,H., Iosue,F., Barter,M., Lynch,R., Munger,H. and Tewhey,R. |
| EPI_ISL_755656, EPI_ISL_755657, EPI_ISL_755658, EPI_ISL_755659, EPI_ISL_755660, EPI_ISL_755661, EPI_ISL_755662, EPI_ISL_755663, EPI_ISL_755664, EPI_ISL_755665, EPI_ISL_755666, EPI_ISL_755667, EPI_ISL_755668, EPI_ISL_755669, EPI_ISL_755670, EPI_ISL_755671, EPI_ISL_755672, EPI_ISL_755673, EPI_ISL_755674, EPI_ISL_755675, EPI_ISL_755676, EPI_ISL_755677, EPI_ISL_755678, EPI_ISL_755679, EPI_ISL_755680, EPI_ISL_755681, EPI_ISL_755682, EPI_ISL_755683, EPI_ISL_755684, EPI_ISL_755685, EPI_ISL_755686, EPI_ISL_755687, EPI_ISL_755688, EPI_ISL_755689, EPI_ISL_755690, EPI_ISL_755691, EPI_ISL_755692, EPI_ISL_755693, EPI_ISL_755694, EPI_ISL_755695, EPI_ISL_755696, EPI_ISL_755697, EPI_ISL_755698, EPI_ISL_755699, EPI_ISL_755700, EPI_ISL_755701, EPI_ISL_755702, EPI_ISL_755703, EPI_ISL_755704, EPI_ISL_755705, EPI_ISL_755706, EPI_ISL_755707, EPI_ISL_755708, EPI_ISL_755709, EPI_ISL_755710, EPI_ISL_755711 | | | |
| see above | Toronto Invasive Bacterial Diseases Network | McMaster University | Allison McGeer, Patryk Aftanas, Hooman Derakhshani, Angel Li, Kuganya Nirmalarajah, Emily Panousis, Ahmed Draia, Jalees Nasir, Michael Surette, Samira Mubareka, Andrew G. McArthur |
| EPI_ISL_755712, EPI_ISL_755713, EPI_ISL_755714, EPI_ISL_755715, EPI_ISL_755716, EPI_ISL_755717, EPI_ISL_755718, EPI_ISL_755719, EPI_ISL_755720, EPI_ISL_755721, EPI_ISL_755722, EPI_ISL_755723, EPI_ISL_755724, EPI_ISL_755725, EPI_ISL_755726, EPI_ISL_755727, EPI_ISL_755728, EPI_ISL_755729, EPI_ISL_755730, EPI_ISL_755731, EPI_ISL_755732, EPI_ISL_755733, EPI_ISL_755734, EPI_ISL_755735, EPI_ISL_755736 | | | |
| see above | Toronto Invasive Bacterial Diseases Network | McMaster University | David Richardson, Allison McGeer, Patryk Aftanas, Hooman Derakhshani, Angel Li, Kuganya Nirmalarajah, Emily Panousis, Ahmed Draia, Jalees Nasir, Michael Surette, Samira Mubareka, Andrew G. McArthur |
| EPI_ISL_755737, EPI_ISL_755738, EPI_ISL_755739, EPI_ISL_755802 | Toronto Invasive Bacterial Diseases Network | McMaster University | Allison McGeer, Patryk Aftanas, Hooman Derakhshani, Angel Li, Kuganya Nirmalarajah, Emily Panousis, Ahmed Draia, Jalees Nasir, Michael Surette, Samira Mubareka, Andrew G. McArthur |
| EPI_ISL_756280 | State Laboratories Division, Hawaii State Department of Health | State Laboratories Division, Hawaii State Department of Health | Pamela O'Brien, Sabrina Diemert, Drew Kuwazaki, Razvan Sultana, Edward Desmond |
| EPI_ISL_756357 | The Caribbean Public Health Agency | Carrington Lab, Department of PreClinical Sciences, Faculty of Medical Sciences, The University of the West Indies | Nikita S. D. Sahadeo, Arianne Brown-Jordan, Sarah Hill, Vernie Ramkissoon, Roshan Parasram, Naresh Nandram, Avery Hinds, Jerome Foster, Stanley Giddings, Karla Georges, Marsha Ivey, Rahul Naidu, Risha Singh, SueMin Nathaniel, Rajini Haraksingh, Jaya Jayaraman, Chinna Chinnadurai, Adesh Ramsubhag, Nuno Faria, Oliver Pybus, Christopher Oura, Gabriel Escobar, Christine V. F. Carrington |
| EPI_ISL_759982 | Oslo University Hospital, Department of Medical Microbiology | Norwegian Institute of Public Health, Department of Virology | Kathrine Stene-Johansen, Kamilla Heddeland Instefjord, Hilde Elshaug, Marie Paulsen Madsen, Rasmus Riis Kopperud, Hilde Vollan, Karoline Bragstad, Olav Hungnes |
| EPI_ISL_759983 | Medical Microbiology Unit, Department for Laboratory Medicine, Drammen Hospital, Vestre Viken Health Trust, | Norwegian Institute of Public Health, Department of Virology | Kathrine Stene-Johansen, Kamilla Heddeland Instefjord, Hilde Elshaug, Marie Paulsen Madsen, Rasmus Riis Kopperud, Hilde Vollan, Karoline Bragstad, Olav Hungnes |
| EPI_ISL_763029 | Unit 17: Influenza & Other Respiratory Viruses, German National Influenza Center | Project group Epidemiology of Highly Pathogenic Microorganisms, Robert Koch-Institute | Ariane Düx, Andreas Sachse, Grit Schubert, Sébastien Calvignac-Spencer, Fabian Leendertz, Thorsten Wolff, Ralf Dürrwald, Djin-Ye Oh, Marianne Wedde |
| EPI_ISL_763099, EPI_ISL_763119, EPI_ISL_763120, EPI_ISL_763121, EPI_ISL_763122, EPI_ISL_763123, EPI_ISL_763124, EPI_ISL_763125, EPI_ISL_763168, EPI_ISL_763169, EPI_ISL_763170, EPI_ISL_763237 | | | |
| see above | Dutch COVID-19 response team | Erasmus Medical Center | Bas Oude Munnink, Reina Sikkema, David Nieuwenhuijse, Irina Chestakova, Anne van der Linden, Marjan Boter, Emmanuelle Munger, Corine GeurtsvanKessel, Annemiek van der Eijk, Richard Molenkamp, Marion Koopmans, on behalf of the Dutch national COVID-19 response team. |
| EPI_ISL_763333, EPI_ISL_763334, EPI_ISL_763335, EPI_ISL_763336 | Florida Bureau of Public Health Laboratories | Florida Bureau of Public Health Laboratories | Sarah Schmedes, Jason Blanton |
| EPI_ISL_763790, EPI_ISL_763873, EPI_ISL_764302, EPI_ISL_764327 | Department of Pathology, University of Cambridge | COVID-19 Genomics UK (COG-UK) Consortium | Aminu S. Jahun, Yasmin Chaudhry, Grant Hall, Iliana Georgana, Myra Hosmillo, Martin D. Curran, Malte Pinckert, Surendra Parmar, Ian Goodfellow |
| EPI_ISL_765863, EPI_ISL_765864, EPI_ISL_765865, EPI_ISL_765866, EPI_ISL_765867, EPI_ISL_765868, EPI_ISL_765869, EPI_ISL_765870, EPI_ISL_765871, EPI_ISL_765872, EPI_ISL_765873, EPI_ISL_765874, EPI_ISL_765875, EPI_ISL_765876, EPI_ISL_765877, EPI_ISL_765878, EPI_ISL_765879, EPI_ISL_765880, EPI_ISL_765881 | | | |
| see above | Massachusetts General Hospital | Infectious Disease Program, Broad Institute of Harvard and MIT | Lemieux,J.E., Siddle,K.J., Shaw,B., Adams,G., Pierce,V., Turbett,S., Anahtar,M., Branda,J., Slater,D., Harris,J., Lin,A.E., Gladden-Young,A., Lagerborg,K., Rudy,M., DeRuff,K., Carter,A., Normandin,E., Bauer,M., Reilly,S., Tomkins-Tinch,C., Loreth,C., Chaluvadi,S., Neumann,A., Cusick,C., Chapman,S.B., Gnirke,A., Flowers,K., Cerrato,F., Birren,B.W., Gallagher,G., Smole,S., Park,D.J., MacInnis,B.L., Ryan,E., LaRocque,R., Rosenberg,E. and Sabeti,P.C. |
| EPI_ISL_766087, EPI_ISL_766115, EPI_ISL_766116, EPI_ISL_766233, EPI_ISL_766235, EPI_ISL_766278 | Respiratory Virus Unit, National Infection Service, Public Health England | COVID-19 Genomics UK (COG-UK) Consortium | PHE Covid Sequencing Team |
| EPI_ISL_766571, EPI_ISL_766572, EPI_ISL_766573 | ULSS 7 Pedemontana - Distretto 1 | Istituto Zooprofilattico Sperimentale delle Venezie | Adelaide Milani, Alessia Schivo, Annalisa Salviato, Erika Giorgia Quaranta, Ambra Pastori, Bianca Zecchin, Alice Fusaro, Isabella Monne, Calogero Terregino, Antonia Ricci |
| EPI_ISL_766614 | unknown | The Public Health Agency of Sweden | Department of Microbiology, The Public Health Agency of Sweden |
| EPI_ISL_766643 | Vardcentralen Brinken | The Public Health Agency of Sweden | Department of Microbiology, The Public Health Agency of Sweden |
| EPI_ISL_766645 | Barnakuten | The Public Health Agency of Sweden | Department of Microbiology, The Public Health Agency of Sweden |
| EPI_ISL_766879, EPI_ISL_766880, EPI_ISL_766881 | Wyoming Public Health Laboratory | Wyoming Public Health Laboratory | Noah Hull, Taylor Fearing, Lynette Gumbleton, Channing Weber, Ashley Norberg, Bailey Bowcutt, and Wanda Manley |
| EPI_ISL_766975, EPI_ISL_766989, EPI_ISL_767003, EPI_ISL_767006 | Delaware Public Health Laboratory | Delaware Public Health Laboratory | Gregory Hovan |
| EPI_ISL_767857 | Sydney South West Pathology Service (SSWPS) - Royal Prince Alfred Hospital - NSW Health Pathology | NSW Health Pathology - Institute of Clinical Pathology and Medical Research; Westmead Hospital; University of Sydney | CIDM-PH et al. |
| EPI_ISL_768325, EPI_ISL_768326, EPI_ISL_768327, EPI_ISL_768328, EPI_ISL_768329, EPI_ISL_768330, EPI_ISL_768331, EPI_ISL_768332, EPI_ISL_768333, EPI_ISL_768334, EPI_ISL_768335, EPI_ISL_768336, EPI_ISL_768337, EPI_ISL_768338, EPI_ISL_768339, EPI_ISL_768340, EPI_ISL_768341, EPI_ISL_768342, EPI_ISL_768343, EPI_ISL_768344, EPI_ISL_768345, EPI_ISL_768346, EPI_ISL_768347, EPI_ISL_768348, EPI_ISL_768349, EPI_ISL_768350, EPI_ISL_768351, EPI_ISL_768352 | | | |
| see above | LSUHS Emerging Viral Threat Laboratory | Microbial Genome Sequencing Center | Jeremy P. Kamil, Jennifer L. Carroll, Camille F. Abshire, Maarten Van Diest, Andrew D. Yurochko, Martin J. Sapp, Rona S. Scott, Christopher G. Kevil, Daniel J. Snyder, Vaughn S. Cooper, John A. Vanchiere |
| EPI_ISL_768524 | Regional Medical Sciences Center 1/1 Chiang Rai | National Institute of Health, Department of Medical Sciences, Ministry of Public Health, Thailand | Pilailuk Okada; Siripaporn Phuygun; Sittiporn Parnmen; Ratana Tacharoenmuang; Pakorn Piromtong; Natchaya Khiadsang; Thanutsapa Thanadachakul; Warawan Wongboot; sirikanda wimol; Sunthareeya Waicharoen; |
| EPI_ISL_768610, EPI_ISL_768611 | Regional medical sciences center 6 Chonburi | National Institute of Health, Department of Medical Sciences, Ministry of Public Health, Thailand | Pilailuk Okada; Siripaporn Phuygun; Sittiporn Parnmen; Ratana Tacharoenmuang; Pakorn Piromtong; Natchaya Khiadsang; Thanutsapa Thanadachakul; Warawan Wongboot; sirikanda wimol; Sunthareeya Waicharoen; |
| EPI_ISL_768616 | Rajavithi Hospital | National Institute of Health, Department of Medical Sciences, Ministry of Public Health, Thailand | Pilailuk Okada; Siripaporn Phuygun; Sittiporn Parnmen; Ratana Tacharoenmuang; Pakorn Piromtong; Natchaya Khiadsang; Thanutsapa Thanadachakul; Warawan Wongboot; sirikanda wimol; Sunthareeya Waicharoen; |
| EPI_ISL_769866 | Respiratory Virus Unit, National Infection Service, Public | COVID-19 Genomics UK (COG-UK) Consortium | PHE Covid Sequencing Team |

| | | | |
|---|---|---|---|
| | Health England | | |
| EPI_ISL_769999 | Area De Salud Paraiso-Cervantes | Inciensa, Instituto Costarricense de Investigación y Enseñanza en Nutrición y Salud | Francisco Duarte, Hebleen Porras, Claudio Soto-Garita, Estela Cordero, Adriana Godínez, Melany Calderón & Mariel López |
| EPI_ISL_770005 | Area De Salud Moravia | Inciensa, Instituto Costarricense de Investigación y Enseñanza en Nutrición y Salud | Francisco Duarte, Hebleen Porras, Claudio Soto-Garita, Estela Cordero, Adriana Godínez, Melany Calderón & Mariel López |
| EPI_ISL_770023 | Area De Salud San Francisco-San Antonio (Coopesana) | Inciensa, Instituto Costarricense de Investigación y Enseñanza en Nutrición y Salud | Francisco Duarte, Hebleen Porras, Claudio Soto-Garita, Estela Cordero, Adriana Godínez, Melany Calderón & Mariel López |
| EPI_ISL_770063, EPI_ISL_770064, EPI_ISL_770066, EPI_ISL_770071, EPI_ISL_770094, EPI_ISL_770095, EPI_ISL_770096, EPI_ISL_770097, EPI_ISL_770098, EPI_ISL_770099, EPI_ISL_770100, EPI_ISL_770101, EPI_ISL_770102, EPI_ISL_770103, EPI_ISL_770104, EPI_ISL_770105, EPI_ISL_770106, EPI_ISL_770107, EPI_ISL_770109, EPI_ISL_770110, EPI_ISL_770111, EPI_ISL_770112 | | | |
| see above | Wyoming Public Health Laboratory | Wyoming Public Health Laboratory | Noah Hull, Taylor Fearing, Lynette Gumbleton, Channing Weber, Ashley Norberg, Bailey Bowcutt, and Wanda Manley |
| EPI_ISL_770551, EPI_ISL_770552, EPI_ISL_770553, EPI_ISL_770554, EPI_ISL_770557, EPI_ISL_770558, EPI_ISL_770559, EPI_ISL_770560, EPI_ISL_770561, EPI_ISL_770562, EPI_ISL_770563, EPI_ISL_770564, EPI_ISL_770565, EPI_ISL_770566, EPI_ISL_770567, EPI_ISL_770568, EPI_ISL_770569, EPI_ISL_770570, EPI_ISL_770571, EPI_ISL_770572, EPI_ISL_770577, EPI_ISL_770578, EPI_ISL_770579, EPI_ISL_770580, EPI_ISL_770581, EPI_ISL_770582, EPI_ISL_770583, EPI_ISL_770584, EPI_ISL_770585, EPI_ISL_770586, EPI_ISL_770587, EPI_ISL_770588, EPI_ISL_770589, EPI_ISL_770590, EPI_ISL_770591, EPI_ISL_770592, EPI_ISL_770593, EPI_ISL_770594, EPI_ISL_770595, EPI_ISL_770596, EPI_ISL_770597, EPI_ISL_770598, EPI_ISL_770599, EPI_ISL_770600, EPI_ISL_770601, EPI_ISL_770602, EPI_ISL_770603, EPI_ISL_770604, EPI_ISL_770605, EPI_ISL_770606, EPI_ISL_770607, EPI_ISL_770608, EPI_ISL_770609, EPI_ISL_770610, EPI_ISL_770611, EPI_ISL_770612, EPI_ISL_770613, EPI_ISL_770614, EPI_ISL_770615, EPI_ISL_770616, EPI_ISL_770617, EPI_ISL_770618, EPI_ISL_770619, EPI_ISL_770620, EPI_ISL_770621, EPI_ISL_770622, EPI_ISL_770623, EPI_ISL_770624, EPI_ISL_770625, EPI_ISL_770626, EPI_ISL_770627, EPI_ISL_770628, EPI_ISL_770629 | | | |
| see above | Laboratório de Microbiologia Molecular - Universidade FEEVALE | Bioinformatics Laboratory / LNCC | Felipe Benites, Fernando Rosado Spilki, Alana Witt Hansen, Juliane Deise Fleck, Juliana Schons, Meriane Demoliner, Ana Karolina Eisen Antunes, Fagner Henrique Heldt, Larissa Mallmann, Bruna Hermann, Ana Luiza Ziulkoski, Vyctoria Goes, Karoline Schallenberger, Matheus Nunes Weber, Paula Rodrigues de Almeida, Alessandra Pavan Lamarca da Silva, Ronaldo da Silva F Jr , Luiz G P de Almeida, Alexandra L Gerber , Ana Paula de C Guimarães,Ana Tereza R de Vasconcelos |
| EPI_ISL_770633, EPI_ISL_770661, EPI_ISL_770674 | WHO National Influenza Centre Russian Federation | WHO National Influenza Centre Russian Federation | Andrey Komissarov, Artem Fadeev, Anna Ivanova, Kseniya Komissarova, Dmitry Bazhenov, Daria Danilenko, Ksenia Safina, Elena Nabieva, Georgii Bazykin, Dmitry Lioznov |
| EPI_ISL_770890, EPI_ISL_770891, EPI_ISL_770897, EPI_ISL_770915, EPI_ISL_770916, EPI_ISL_770917, EPI_ISL_770918, EPI_ISL_770919, EPI_ISL_770920, EPI_ISL_770921, EPI_ISL_770922, EPI_ISL_770923, EPI_ISL_770924, EPI_ISL_770925, EPI_ISL_770926, EPI_ISL_770927, EPI_ISL_770928, EPI_ISL_770929 | | | |
| see above | Laboratoire national de santé, Microbiology, Virology | Laboratoire national de santé, Microbiology, Microbial Genomics Platform | Anke Wienecke-Baldacchino, Catherine Ragimbeau,Jessica Tapp, Fatu Djabi, Lise Pignon, Raoul Salmon, Tamir Abdelrahman |
| EPI_ISL_771153, EPI_ISL_771166, EPI_ISL_771174, EPI_ISL_771175, EPI_ISL_771177, EPI_ISL_771186, EPI_ISL_771187, EPI_ISL_771196, EPI_ISL_771214, EPI_ISL_771215, EPI_ISL_771217, EPI_ISL_771223, EPI_ISL_771329, EPI_ISL_771335, EPI_ISL_771336, EPI_ISL_771337, EPI_ISL_771338, EPI_ISL_771339, EPI_ISL_771340, EPI_ISL_771341, EPI_ISL_771342, EPI_ISL_771343, EPI_ISL_771344, EPI_ISL_771345, EPI_ISL_771346, EPI_ISL_771347, EPI_ISL_771348, EPI_ISL_771349, EPI_ISL_771353, EPI_ISL_771354, EPI_ISL_771355, EPI_ISL_771356, EPI_ISL_771357, EPI_ISL_771358, EPI_ISL_771359, EPI_ISL_771360 | | | |
| see above | Colorado Department of Public Health and Environment | Colorado Department of Puplic Health and Environment | Laura Bankers, Molly C. Hetherington-Rauth, Diana Ir, Shannon Ely, Shannon R. Matzinger, Sarah Elizabeth Totten, Emily A. Travanty |
| EPI_ISL_775276 | Haukeland University Hospital, Dept. of Microbiology | Norwegian Institute of Public Health, Department of Virology | Kathrine Stene-Johansen, Kamilla Heddeland Instefjord, Hilde Elshaug, Atiya R Ali,Marie Paulsen Madsen, Rasmus Riis Kopperud, Hilde Vollan, Karoline Bragstad, Olav Hungnes |
| EPI_ISL_775284 | Oslo University Hospital, Department of Medical Microbiology | Norwegian Institute of Public Health, Department of Virology | Kathrine Stene-Johansen, Kamilla Heddeland Instefjord, Hilde Elshaug, Atiya R Ali,Marie Paulsen Madsen, Rasmus Riis Kopperud, Hilde Vollan, Karoline Bragstad, Olav Hungnes |
| EPI_ISL_775287, EPI_ISL_775288, EPI_ISL_775289, EPI_ISL_775290 | Nordland Hospital - Bodo, Laboratory Department, Molecular Biology Unit | Norwegian Institute of Public Health, Department of Virology | Kathrine Stene-Johansen, Kamilla Heddeland Instefjord, Hilde Elshaug, Atiya R Ali,Marie Paulsen Madsen, Rasmus Riis Kopperud, Hilde Vollan, Karoline Bragstad, Olav Hungnes |
| EPI_ISL_775356 | Norwegian Institute of Public Health, Department of Virology | Norwegian Institute of Public Health, Department of Virology | Kathrine Stene-Johansen, Kamilla Heddeland Instefjord, Hilde Elshaug, Atiya R Ali,Marie Paulsen Madsen, Rasmus Riis Kopperud, Hilde Vollan, Karoline Bragstad, Olav Hungnes |
| EPI_ISL_775357 | Haukeland University Hospital, Dept. of Microbiology | Norwegian Institute of Public Health, Department of Virology | Kathrine Stene-Johansen, Kamilla Heddeland Instefjord, Hilde Elshaug, Atiya R Ali,Marie Paulsen Madsen, Rasmus Riis Kopperud, Hilde Vollan, Karoline Bragstad, Olav Hungnes |
| EPI_ISL_775362, EPI_ISL_775368, EPI_ISL_775369, EPI_ISL_775370 | Foerde Hospital, Department of Microbiology | Norwegian Institute of Public Health, Department of Virology | Kathrine Stene-Johansen, Kamilla Heddeland Instefjord, Hilde Elshaug, Atiya R Ali,Marie Paulsen Madsen, Rasmus Riis Kopperud, Hilde Vollan, Karoline Bragstad, Olav Hungnes |
| EPI_ISL_775373, EPI_ISL_775375, EPI_ISL_775377, EPI_ISL_775383 | Nordland Hospital - Bodo, Laboratory Department, Molecular Biology Unit | Norwegian Institute of Public Health, Department of Virology | Kathrine Stene-Johansen, Kamilla Heddeland Instefjord, Hilde Elshaug, Atiya R Ali,Marie Paulsen Madsen, Rasmus Riis Kopperud, Hilde Vollan, Karoline Bragstad, Olav Hungnes |
| EPI_ISL_775384 | Unilabs Laboratory Medicine | Norwegian Institute of Public Health, Department of Virology | Kathrine Stene-Johansen, Kamilla Heddeland Instefjord, Hilde Elshaug, Atiya R Ali,Marie Paulsen Madsen, Rasmus Riis Kopperud, Hilde Vollan, Karoline Bragstad, Olav Hungnes |
| EPI_ISL_775387, EPI_ISL_775389, EPI_ISL_775390 | Akershus University Hospital, Department for Microbiology and Infectious Disease Control | Norwegian Institute of Public Health, Department of Virology | Kathrine Stene-Johansen, Kamilla Heddeland Instefjord, Hilde Elshaug, Atiya R Ali,Marie Paulsen Madsen, Rasmus Riis Kopperud, Hilde Vollan, Karoline Bragstad, Olav Hungnes |
| EPI_ISL_775415 | Oslo University Hospital, Department of Medical Microbiology | Norwegian Institute of Public Health, Department of Virology | Kathrine Stene-Johansen, Kamilla Heddeland Instefjord, Hilde Elshaug, Atiya R Ali,Marie Paulsen Madsen, Rasmus Riis Kopperud, Hilde Vollan, Karoline Bragstad, Olav Hungnes |
| EPI_ISL_775462 | Norwegian Institute of Public Health, Department of Virology | Norwegian Institute of Public Health, Department of Virology | Kathrine Stene-Johansen, Kamilla Heddeland Instefjord, Hilde Elshaug, Atiya R Ali,Marie Paulsen Madsen, Rasmus Riis Kopperud, Hilde Vollan, Karoline Bragstad, Olav Hungnes |
| EPI_ISL_775491 | Department of Medical Microbiology, St. Olavs hospital | Norwegian Institute of Public Health, Department of Virology | Kathrine Stene-Johansen, Kamilla Heddeland Instefjord, Hilde Elshaug, Atiya R Ali,Marie Paulsen Madsen, Rasmus Riis Kopperud, Hilde Vollan, Karoline Bragstad, Olav Hungnes |
| EPI_ISL_775495 | Unilabs Laboratory Medicine | Norwegian Institute of Public Health, Department of Virology | Kathrine Stene-Johansen, Kamilla Heddeland Instefjord, Hilde Elshaug, Atiya R Ali,Marie Paulsen Madsen, Rasmus Riis Kopperud, Hilde Vollan, Karoline Bragstad, Olav Hungnes |
| EPI_ISL_775511, EPI_ISL_775513 | Norwegian Institute of Public Health, Department of Virology | Norwegian Institute of Public Health, Department of Virology | Kathrine Stene-Johansen, Kamilla Heddeland Instefjord, Hilde Elshaug, Atiya R Ali,Marie Paulsen Madsen, Rasmus Riis Kopperud, Hilde Vollan, Karoline Bragstad, Olav Hungnes |
| EPI_ISL_776766 | Instituto Adolfo Lutz - Regional de Santo Andre | Instituto Adolfo Lutz, Interdiciplinary Procedures Center, Strategic Laboratory | Claudio Tavares Sacchi, Claudia Regina Gonçalves, Erica Valessa Ramos Gomes, Karoline Rodrigues Campos |
| EPI_ISL_776767 | Instituto Adolfo Lutz - Regional de Marilia | Instituto Adolfo Lutz, Interdiciplinary Procedures Center, Strategic Laboratory | Claudio Tavares Sacchi, Claudia Regina Gonçalves, Erica Valessa Ramos Gomes, Karoline Rodrigues Campos |
| EPI_ISL_778820, EPI_ISL_778835 | AIID | Irish Coronavirus Sequencing Consortium-Teagasc Grange | Matthew McCabe, Aljandro Abner Garcia Leon, Fiona Crispie, Calum Walsh, Michael Carr, John Kenny, Paul Cotter, Patrick Mallon, Gabriel Gonzalez |
| EPI_ISL_778892 | Maryland Public Health Laboratory | Maryland Public Health Laboratory | Maryland Department of Health Laboratories Administration |
| EPI_ISL_779131, EPI_ISL_779132, EPI_ISL_779133, EPI_ISL_779134, EPI_ISL_779136 | Wyoming Public Health Laboratory | Wyoming Public Health Laboratory | Noah Hull, Taylor Fearing, Lynette Gumbleton, Channing Weber, Ashley Norberg, Bailey Bowcutt, and Wanda Manley |
| EPI_ISL_779157, EPI_ISL_779158, EPI_ISL_779159, EPI_ISL_779160, EPI_ISL_779161, EPI_ISL_779162, EPI_ISL_779163, EPI_ISL_779164, EPI_ISL_779165, EPI_ISL_779166, EPI_ISL_779167, EPI_ISL_779168 | | | |
| see above | Laboratório de Microbiologia Molecular - Universidade FEEVALE | Bioinformatics Laboratory / LNCC | Felipe Benites, Fernando Rosado Spilki, Alana Witt Hansen, Juliane Deise Fleck, Juliana Schons, Meriane Demoliner, Ana Karolina Eisen Antunes, Fagner Henrique Heldt, Larissa Mallmann, Bruna Hermann, Ana Luiza Ziulkoski, Vyctoria Goes, Karoline Schallenberger, Matheus Nunes Weber, Paula Rodrigues de Almeida, Alessandra Pavan Lamarca da Silva, Ronaldo da Silva F Jr , Luiz G P de Almeida, Alexandra L Gerber , Ana Paula de C Guimarães,Ana Tereza R de Vasconcelos |
| EPI_ISL_779189 | Laboratorio Estatal de Salud Pública de Nuevo León | Laboratorio de Infectología Molecular, Departamento de Bioquímica y Medicina Molecular,Facultad de Medicina - | Kame A. Galán-Huerta, María F. Herrera-Saldivar, Natalia Martínez-Acuña, Sonia A. Lozano-Sepúlveda, Daniel Arellanos-Soto, Ana M. Rivas-Estilla, Samuel Buentello-Wong, Else del Carmen García-García, Gloria A. Jasso-de-la-Peña, Roberto Montes-de-Oca, Consuelo Treviño-Garza, Manuel E. |

| | | | |
|---|---|---|---|
| | | Universidad Autónoma de Nuevo León | de-la-O-Cavazos |
| EPI_ISL_779302, EPI_ISL_779311, EPI_ISL_779318 | Utah Public Health Laboratory, Utah Public Health Laboratory Infectious Disease submission group | Utah Public Health Laboratory, Utah Public Health Laboratory Infectious Disease submission group | Gallagher,T., Young,E.L., Oakeson,K.F. |
| EPI_ISL_779407 | Royal Darwin Hospital Pathology | MDU-PHL | Meumann, E., Caly L., Seemann T., Sait, M.L., Druce J., Sherry, N.L. |
| EPI_ISL_779838 | CHU Bordeaux | CNR Virus des Infections Respiratoires - France SUD | Antonin Bal, Gregory Destras, Gwendolyne Burfin, Hadrien Règue, Quentin Semanas, Martine Valette, Bruno Lina, Pantxika Bellecave, Camille Ciccone, Isabelle Garrigue, Marie-Edith Lafon, Pascale Trimoulet, Laurence Josset |
| EPI_ISL_780030, EPI_ISL_780031, EPI_ISL_780035, EPI_ISL_780037, EPI_ISL_780038, EPI_ISL_780040, EPI_ISL_780044, EPI_ISL_780046, EPI_ISL_780052, EPI_ISL_780053, EPI_ISL_780054, EPI_ISL_780066, EPI_ISL_780070, EPI_ISL_780071, EPI_ISL_780072, EPI_ISL_780077, EPI_ISL_780094, EPI_ISL_780095 | | | |
| see above | Hospital General Universitario Gregorio Marañón | SeqCOVID-SPAIN consortium/IBV(CSIC) | Darío García de Viedma, Laura Pérez-Lago, Marta Herranz, Jon Sicilia, Julia Suárez, Pilar Catalán, Patricia Muñoz and SeqCOVID-SPAIN consortium |
| EPI_ISL_783546, EPI_ISL_783548, EPI_ISL_783562, EPI_ISL_783563, EPI_ISL_783564, EPI_ISL_783565, EPI_ISL_783568, EPI_ISL_783580, EPI_ISL_783596, EPI_ISL_783598, EPI_ISL_783599, EPI_ISL_783600, EPI_ISL_783601, EPI_ISL_783602, EPI_ISL_783606, EPI_ISL_783608, EPI_ISL_783609, EPI_ISL_783610, EPI_ISL_783611, EPI_ISL_783612, EPI_ISL_783614, EPI_ISL_783615, EPI_ISL_783616, EPI_ISL_783617, EPI_ISL_783618, EPI_ISL_783619, EPI_ISL_783620, EPI_ISL_783625, EPI_ISL_783627, EPI_ISL_783628, EPI_ISL_783630, EPI_ISL_783635, EPI_ISL_783641, EPI_ISL_783644, EPI_ISL_783647, EPI_ISL_783649, EPI_ISL_783651, EPI_ISL_783652, EPI_ISL_783653, EPI_ISL_783667, EPI_ISL_783669, EPI_ISL_783671, EPI_ISL_783674, EPI_ISL_783677, EPI_ISL_783679, EPI_ISL_783680, EPI_ISL_783682, EPI_ISL_783683, EPI_ISL_783684, EPI_ISL_783685, EPI_ISL_783686, EPI_ISL_783687, EPI_ISL_783690, EPI_ISL_783691, EPI_ISL_783692, EPI_ISL_783693, EPI_ISL_783694, EPI_ISL_783695, EPI_ISL_783696, EPI_ISL_783698, EPI_ISL_783699, EPI_ISL_783700, EPI_ISL_783701, EPI_ISL_783702, EPI_ISL_783703, EPI_ISL_783705, EPI_ISL_783706, EPI_ISL_783707, EPI_ISL_783708, EPI_ISL_783709, EPI_ISL_783710, EPI_ISL_783714, EPI_ISL_783715, EPI_ISL_783716, EPI_ISL_783717, EPI_ISL_783718, EPI_ISL_783719, EPI_ISL_783721, EPI_ISL_783722, EPI_ISL_783726, EPI_ISL_783727, EPI_ISL_783728, EPI_ISL_783729, EPI_ISL_783730, EPI_ISL_783731, EPI_ISL_783732, EPI_ISL_783733, EPI_ISL_783736, EPI_ISL_783737, EPI_ISL_783738, EPI_ISL_783739, EPI_ISL_783740, EPI_ISL_783741, EPI_ISL_783742, EPI_ISL_783743, EPI_ISL_783744, EPI_ISL_783745, EPI_ISL_783746, EPI_ISL_783747, EPI_ISL_783748, EPI_ISL_783749, EPI_ISL_783750, EPI_ISL_783751, EPI_ISL_783752, EPI_ISL_783753, EPI_ISL_783754, EPI_ISL_783755, EPI_ISL_783756, EPI_ISL_783757, EPI_ISL_783758, EPI_ISL_783759, EPI_ISL_783760, EPI_ISL_783761, EPI_ISL_783762, EPI_ISL_783763, EPI_ISL_783764, EPI_ISL_783765, EPI_ISL_783766, EPI_ISL_783767, EPI_ISL_783768, EPI_ISL_783769, EPI_ISL_783770, EPI_ISL_783771, EPI_ISL_783772, EPI_ISL_783773, EPI_ISL_783774, EPI_ISL_783776, EPI_ISL_783777, EPI_ISL_783778, EPI_ISL_783779, EPI_ISL_783780, EPI_ISL_783781, EPI_ISL_783782, EPI_ISL_783783, EPI_ISL_783784, EPI_ISL_783785, EPI_ISL_783786, EPI_ISL_783787, EPI_ISL_783790, EPI_ISL_783791, EPI_ISL_783792, EPI_ISL_783793, EPI_ISL_783794, EPI_ISL_783795, EPI_ISL_783796, EPI_ISL_783797, EPI_ISL_783799, EPI_ISL_783800, EPI_ISL_783801, EPI_ISL_783802, EPI_ISL_783803, EPI_ISL_783804, EPI_ISL_783805, EPI_ISL_783806, EPI_ISL_783807, EPI_ISL_783814, EPI_ISL_783820, EPI_ISL_783832, EPI_ISL_783833, EPI_ISL_783834, EPI_ISL_783840, EPI_ISL_783845, EPI_ISL_783853, EPI_ISL_783857, EPI_ISL_783862, EPI_ISL_783863, EPI_ISL_783894, EPI_ISL_783915, EPI_ISL_783934, EPI_ISL_783947, EPI_ISL_783948, EPI_ISL_783949, EPI_ISL_783951, EPI_ISL_783952, EPI_ISL_783955, EPI_ISL_783958, EPI_ISL_783961, EPI_ISL_783969, EPI_ISL_783970, EPI_ISL_783972, EPI_ISL_783973, EPI_ISL_783975, EPI_ISL_783976, EPI_ISL_783978, EPI_ISL_783980, EPI_ISL_783984, EPI_ISL_783986, EPI_ISL_783989, EPI_ISL_783990, EPI_ISL_783993, EPI_ISL_783994, EPI_ISL_783996, EPI_ISL_783997, EPI_ISL_783998, EPI_ISL_783999, EPI_ISL_784000, EPI_ISL_784003, EPI_ISL_784005, EPI_ISL_784006, EPI_ISL_784009, EPI_ISL_784010, EPI_ISL_784011, EPI_ISL_784012, EPI_ISL_784014, EPI_ISL_784016, EPI_ISL_784017, EPI_ISL_784021, EPI_ISL_784030, EPI_ISL_784036, EPI_ISL_784037, EPI_ISL_784039, EPI_ISL_784042, EPI_ISL_784043, EPI_ISL_784044, EPI_ISL_784046, EPI_ISL_784048, EPI_ISL_784049, EPI_ISL_784054, EPI_ISL_784056, EPI_ISL_784057, EPI_ISL_784062, EPI_ISL_784064, EPI_ISL_784070, EPI_ISL_784071, EPI_ISL_784072, EPI_ISL_784080, EPI_ISL_784086, EPI_ISL_784088, EPI_ISL_784089, EPI_ISL_784090, EPI_ISL_784095, EPI_ISL_784098, EPI_ISL_784100, EPI_ISL_784102, EPI_ISL_784104, EPI_ISL_784109, EPI_ISL_784110, EPI_ISL_784111, EPI_ISL_784114, EPI_ISL_784115, EPI_ISL_784117, EPI_ISL_784118, EPI_ISL_784119, EPI_ISL_784120, EPI_ISL_784125, EPI_ISL_784127, EPI_ISL_784128, EPI_ISL_784133, EPI_ISL_784135, EPI_ISL_784136, EPI_ISL_784140, EPI_ISL_784149, EPI_ISL_784151, EPI_ISL_784152, EPI_ISL_784153, EPI_ISL_784155, EPI_ISL_784156, EPI_ISL_784158, EPI_ISL_784159, EPI_ISL_784163, EPI_ISL_784164, EPI_ISL_784165, EPI_ISL_784174, EPI_ISL_784175, EPI_ISL_784177, EPI_ISL_784178, EPI_ISL_784180, EPI_ISL_784182, EPI_ISL_785076, EPI_ISL_785077, EPI_ISL_785085, EPI_ISL_785101, EPI_ISL_785102, EPI_ISL_785111, EPI_ISL_785113, EPI_ISL_785121, EPI_ISL_785130, EPI_ISL_785131, EPI_ISL_785139, EPI_ISL_785140, EPI_ISL_785141, EPI_ISL_785142, EPI_ISL_785143, EPI_ISL_785145, EPI_ISL_785146, EPI_ISL_785147, EPI_ISL_785148, EPI_ISL_785149, EPI_ISL_785150, EPI_ISL_785151, EPI_ISL_785152, EPI_ISL_785153, EPI_ISL_785154, EPI_ISL_785156, EPI_ISL_785157, EPI_ISL_785158, EPI_ISL_785159, EPI_ISL_785160, EPI_ISL_785161, EPI_ISL_785163, EPI_ISL_785164, EPI_ISL_785165, EPI_ISL_785166, EPI_ISL_785167, EPI_ISL_785168, EPI_ISL_785169, EPI_ISL_785170, EPI_ISL_785171, EPI_ISL_785173, EPI_ISL_785174, EPI_ISL_785175, EPI_ISL_785176, EPI_ISL_785177, EPI_ISL_785178, EPI_ISL_785179, EPI_ISL_785180, EPI_ISL_785181, EPI_ISL_785182, EPI_ISL_785183, EPI_ISL_785186, EPI_ISL_785187, EPI_ISL_785188, EPI_ISL_785189, EPI_ISL_785190, EPI_ISL_785191, EPI_ISL_785193, EPI_ISL_785195, EPI_ISL_785196, EPI_ISL_785197, EPI_ISL_785198, EPI_ISL_785199, EPI_ISL_785202, EPI_ISL_785203, EPI_ISL_785204, EPI_ISL_785205, EPI_ISL_785206, EPI_ISL_785207, EPI_ISL_785208, EPI_ISL_785209, EPI_ISL_785210, EPI_ISL_785211, EPI_ISL_785212, EPI_ISL_785213, EPI_ISL_785214, EPI_ISL_785215, EPI_ISL_785216, EPI_ISL_785217, EPI_ISL_785218, EPI_ISL_785219, EPI_ISL_785220, EPI_ISL_785221, EPI_ISL_785222, EPI_ISL_785223, EPI_ISL_785224, EPI_ISL_785225, EPI_ISL_785226, EPI_ISL_785227, EPI_ISL_785228, EPI_ISL_785229, EPI_ISL_785230, EPI_ISL_785231, EPI_ISL_785232, EPI_ISL_785233, EPI_ISL_785234, EPI_ISL_785235, EPI_ISL_785236, EPI_ISL_785237, EPI_ISL_785238, EPI_ISL_785239, EPI_ISL_785240, EPI_ISL_785241, EPI_ISL_785242, EPI_ISL_785243, EPI_ISL_785244, EPI_ISL_785245, EPI_ISL_785246, EPI_ISL_785247, EPI_ISL_785248, EPI_ISL_785249, EPI_ISL_785250, EPI_ISL_785251, EPI_ISL_785252, EPI_ISL_785253, EPI_ISL_785254, EPI_ISL_785255, EPI_ISL_785256, EPI_ISL_785257, EPI_ISL_785258, EPI_ISL_785259, EPI_ISL_785260, EPI_ISL_785261, EPI_ISL_785262, EPI_ISL_785263, EPI_ISL_785264, EPI_ISL_785265, EPI_ISL_785266, EPI_ISL_785267, EPI_ISL_785268, EPI_ISL_785269, EPI_ISL_785270, EPI_ISL_785271, EPI_ISL_785272, EPI_ISL_785273, EPI_ISL_785274, EPI_ISL_785275, EPI_ISL_785276, EPI_ISL_785277, EPI_ISL_785278, EPI_ISL_785279, EPI_ISL_785281, EPI_ISL_785282, EPI_ISL_785283, EPI_ISL_785284, EPI_ISL_785285, EPI_ISL_785286, EPI_ISL_785287, EPI_ISL_785289, EPI_ISL_785290, EPI_ISL_785291, EPI_ISL_785292, EPI_ISL_785293, EPI_ISL_785294, EPI_ISL_785295, EPI_ISL_785296, EPI_ISL_785297 | | | |
| see above | Houston Methodist Hospital | Houston Methodist Hospital | S. Wesley Long, Randall J. Olsen, Paul A. Christensen, David W. Bernard, James J. Davis, Maulik Shukla, Marcus Nguyen, Matthew Ojeda Saavedra, Prasanti Yerramilli, Layne Pruitt, Sishir Subedi, Heather Hendrickson, and James M. Musser |
| EPI_ISL_788983 | Institute of Virology, Biomedical Research Center of the Slovak Academy of Sciences, Bratislava | Faculty of Natural Sciences, Comenius University, Bratislava | Kristína Boršová, Viktória abanová, Broa Brejová, Viktória Hodorová, Sabina Fumaová Havlíková, Juraj Kopáek, Martina Liková, ubomíra Lukáiková, Martina Neboháová, Monika Sláviková, Alena Košálová, Peter Sabaka, Tomáš Vina, Boris Klempa, Jozef Nosek |
| EPI_ISL_790595 | Dutch COVID-19 response team | National Institute for Public Health and the Environment (RIVM) | Adam Meijer, Harry Vennema, Jeroen Cremer, Sharon van den Brink, Bas van der Veer, AnneMarie van den Brandt, Florian Zwagemaker, Dennis Schmitz, Chantal Reusken, on behalf of the national COVID-19 response team |
| EPI_ISL_791317, EPI_ISL_791318 | National Virus Reference Laboratory | Irish Coronavirus Sequencing Consortium - Teagasc Moorepark | Alejandro Abner Garcia Leon, Paul Cotter, Fiona Crispie, John Kenny, Paddy Mallon, Calum Walsh |
| EPI_ISL_791349, EPI_ISL_791385, EPI_ISL_791386, EPI_ISL_791387, EPI_ISL_791389, EPI_ISL_791394, EPI_ISL_791397, EPI_ISL_791399, EPI_ISL_791400, EPI_ISL_791402, EPI_ISL_791406 | | | |
| see above | Johns Hopkins Hospital Department of Pathology | Johns Hopkins Hospital Department of Pathology | C. Paul Morris, Chun Huai Luo, Heba H. Mostafa |
| EPI_ISL_791488, EPI_ISL_791489, EPI_ISL_791490, EPI_ISL_791491, EPI_ISL_791492, EPI_ISL_791493, EPI_ISL_791494, EPI_ISL_791495, EPI_ISL_791496, EPI_ISL_791497, EPI_ISL_791498, EPI_ISL_791499, EPI_ISL_791500, EPI_ISL_791501, EPI_ISL_791502, EPI_ISL_791503, EPI_ISL_791504, EPI_ISL_791505, EPI_ISL_791506, EPI_ISL_791507, EPI_ISL_791508, EPI_ISL_791509, EPI_ISL_791510, EPI_ISL_791511, EPI_ISL_791512, EPI_ISL_791513 | | | |
| see above | The Hosptial for Sick Children | The Hospital for Sick Children | Aaron Campigotto, Christian Marshall, Lynette Lau |
| EPI_ISL_792059, EPI_ISL_792060, EPI_ISL_792061, EPI_ISL_792062, EPI_ISL_792063, EPI_ISL_792064, EPI_ISL_792065, EPI_ISL_792066, EPI_ISL_792067, EPI_ISL_792068, EPI_ISL_792069, EPI_ISL_792070, EPI_ISL_792071, EPI_ISL_792072, EPI_ISL_792073, EPI_ISL_792074, EPI_ISL_792075, EPI_ISL_792076, EPI_ISL_792077, EPI_ISL_792078, EPI_ISL_792079, EPI_ISL_792080, EPI_ISL_792081, EPI_ISL_792082, EPI_ISL_792083 | | | |
| see above | Toronto Invasive Bacterial Diseases Network | McMaster University | David Richardson, Allison McGeer, Patryk Aftanas, Hooman Derakhshani, Angel Li, Kuganya Nirmalarajah, Emily Panousis, Ahmed Draia, Jalees Nasir, Michael Surette, Samira Mubareka, Andrew G. McArthur |
| EPI_ISL_792547, EPI_ISL_792548, EPI_ISL_792552, EPI_ISL_792553, EPI_ISL_792554 | Centre for Dengue Research and AICBU, Department of Immunology and Molecular Medicine | Centre for Dengue Research and AICBU, Department of Immunology and Molecular Medicine | Chandima Jeewandara, Deshni Jayathilaka, Dinuka Ariyaratne, Diyanath Ranasinghe, Laksiri Gomes, Gathsaurie Neelika Malavige |
| EPI_ISL_792603 | LACEN-PB | Laboratory of Respiratory Viruses and Measles, Oswaldo Cruz Institute, FIOCRUZ | Paola Resende, Luciana Appolinario, Fernando Motta, Anna Carolina Paixao, Ana Carolina Mendonca, João Felipe Bezerra, Romero Henrique Teixeira de Vasconcelos, Dalane Loudal Florentino Teixeira, Thiago Franco de Oliveira Carneiro, Marilda Siqueira |
| EPI_ISL_792652 | LACEN-PR | Laboratory of Respiratory Viruses and Measles, Oswaldo Cruz Institute, FIOCRUZ | Paola Resende, Luciana Appolinario, Fernando Motta, Anna Carolina Paixao, Ana Carolina Mendonca, Maria do Carmo Debur, Irina Nastassja Riediger, Marilda Siqueira |
| EPI_ISL_792696, EPI_ISL_792707 | The National Institute of Public Health | State Veterinary Institute Prague | Nagy,A;Jirincova,H;Trnka,D;Vecerova,J |
| EPI_ISL_794718 | PathWest Laboratory Medicine WA | PathWest Laboratory Medicine WA Microbial Surveillance Unit | PathWest Laboratory Medicine WA Microbial Surveillance Unit |
| EPI_ISL_794749, EPI_ISL_794755 | Istituto Zooprofilattico Sperimentale della Puglia e della Basilicata | Istituto Zooprofilattico Sperimentale della Puglia e della Basilicata | Parisi A., Bianco A., Capozzi L., Del Sambro L., Manzulli V, Rondinone V., Pace L., Cipolletta D., Galante D. |
| EPI_ISL_796778 | Instituto Nacional de Saude (INSA) | Instituto Nacional de Saude (INSA) | Borges et al |
| EPI_ISL_800220 | Lighthouse Lab in Cambridge | Wellcome Sanger Institute for the COVID-19 Genomics UK (COG-UK) Consortium | Rob Howes, The Lighthouse Lab in Cambridge and Alex Alderton, Roberto Amato, Sonia Goncalves, Ewan Harrison, David K. Jackson, Ian Johnston, Dominic Kwiatkowski, Cordelia Langford, John Sillitoe on behalf of the Wellcome Sanger Institute COVID-19 Surveillance Team |
| EPI_ISL_801416, EPI_ISL_801417, EPI_ISL_801418, EPI_ISL_801450, EPI_ISL_801460, EPI_ISL_801461, EPI_ISL_801462, EPI_ISL_801463, EPI_ISL_801464, EPI_ISL_801465, EPI_ISL_801466, EPI_ISL_801467, EPI_ISL_801468, EPI_ISL_801469, EPI_ISL_801470, EPI_ISL_801477, EPI_ISL_801490 | | | |
| see above | Dutch COVID-19 response team | Erasmus Medical Center | Bas Oude Munnink, Reina Sikkema, David Nieuwenhuijse, Irina Chestakova, Anne van der Linden, Marjan Boter, Emmanuelle Munger, Corine GeurtsvanKessel, Annemiek van der Eijk, Richard Molenkamp, Marion Koopmans, on behalf of the Dutch national COVID-19 response team. |
| EPI_ISL_802856, EPI_ISL_802857, EPI_ISL_802858, EPI_ISL_802859 | Vilnius University Hospital Santaros Klinikos, Vilnius University | Institute of Biotechnology, Life Sciences Center, Vilnius University | Emilija Vasiliunaite, Milda Norkiene, Albertas Timinskas, Alma Gedvilaite, Aurelija Zvirbliene, Daniel Naumovas, Laimonas Griskevicius |
| EPI_ISL_803100 | Respiratory Viruses Branch, Centers for Disease Control and Prevention | Respiratory Viruses Branch, Centers for Disease Control and Prevention | Queen,K., Li,Y., Tao,Y., Uehara,A., Montmayeur,A., Paden,C.R., Cook,P.W., Marine,R., Sheth,M., Wang,H., Lee,J., Tong,S. |
| EPI_ISL_803871 | Institute of Medical Microbiology and Hospital Hygiene | Institute of Medical Microbiology and Hospital Hygiene | Prof. Dr. Achim Kaasch, Aljoscha Tersteegen |
| EPI_ISL_804606, EPI_ISL_804607, EPI_ISL_804612, EPI_ISL_804613, EPI_ISL_804619, EPI_ISL_804623, EPI_ISL_804627, EPI_ISL_804629, EPI_ISL_804646, EPI_ISL_804647, EPI_ISL_804653, EPI_ISL_804660, EPI_ISL_804668, EPI_ISL_804672, EPI_ISL_804679, EPI_ISL_804703, EPI_ISL_804744, EPI_ISL_804745, EPI_ISL_804746, EPI_ISL_804747, EPI_ISL_804748, EPI_ISL_804749, EPI_ISL_804750, EPI_ISL_804751, EPI_ISL_804752, EPI_ISL_804755, EPI_ISL_804767, EPI_ISL_804770, EPI_ISL_804771, EPI_ISL_804772, EPI_ISL_804784, EPI_ISL_804785, EPI_ISL_804793, EPI_ISL_804798, EPI_ISL_804799, EPI_ISL_804800 | | | |

| | | | |
|---|---|---|---|
| see above | Michigan Department of Health and Human Services, Bureau of Laboratories | Michigan Department of Health and Human Services, Bureau of Laboratories | Blankenship HM, Riner D, Soehnlen MK |
| EPI_ISL_804855 | DC Public Health Lab/ Dept. of Forensic Sciences | DC Public Health Lab/ Dept. of Forensic Sciences | Scott Nguyen, Elizabeth Zelaya, Connie Maza, Monica Mann, Brittany Hamilton, David Payne, Jocelyn Hauser |
| EPI_ISL_806810, EPI_ISL_806811, EPI_ISL_806823, EPI_ISL_806824, EPI_ISL_806825, EPI_ISL_806826, EPI_ISL_806827, EPI_ISL_806828, EPI_ISL_806829, EPI_ISL_806832, EPI_ISL_806833 | | | |
| see above | Alaska State Virology Laboratory | Alaska State Virology Laboratory | Stephanie DeRonde, Lisa Smith, Ph.D., Devin M. Drown, Ph.D., Jack Chen, Ph.D. |
| EPI_ISL_806991 | Washington State Department of Health | Seattle Flu Study | Deborah A. Nickerson, Chris D. Frazar, Jover Lee, Benjamin Pelle, Matthew Richardson, Amanda Adler, Elisabeth Brandstetter, Peter D. Han, Kairsten Fay, Misja Ilcisin, Kirsten Lacombe, Thomas R. Sibley, Melissa Truong, Caitlin R. Wolf, Romesh Gautom, Geoff Melly, Brian Hiatt, Philip Dykema, Scott Lindquist, Michael Boeckh, Janet A. Englund, Michael Famulare, Barry R. Lutz, Mark J. Rieder, Lea M. Starita, Matthew Thompson, Helen Y. Chu, Jay Shendure, Trevor Bedford |
| EPI_ISL_813045, EPI_ISL_813046, EPI_ISL_813047, EPI_ISL_813048, EPI_ISL_813049, EPI_ISL_813050, EPI_ISL_813051, EPI_ISL_813053, EPI_ISL_813054, EPI_ISL_813055, EPI_ISL_813056, EPI_ISL_813057, EPI_ISL_813058, EPI_ISL_813059 | | | |
| see above | University of Birmingham | COVID-19 Genomics UK (COG-UK) Consortium | Institute of Microbiology, University of Birmingham: Claire McMurray, Joanne Stockton, Samuel Nicholls, Radoslaw Poplawski, Will Rowe, Josh Quick, Nicholas Loman. University of Birmingham Testing Laboratory: Celina M Whalley, Andrew Bosworth, Charlotte Poxon, Kasun Wanigasooriya, Oliver Pickles, Mike Kidd, Alex Richter, Andrew D Beggs PHE Heartlands Lab: Husam Osman, Andrew Bosworth. Queen Elizabeth Hospital: Anna Casey |
| EPI_ISL_813753 | Liverpool Clinical Laboratories | COVID-19 Genomics UK (COG-UK) Consortium | Sam Haldenby, Anita Lucaci, Steve Paterson, Julian Hiscox, Alistair Darby, M Almsaud, A Alrezaihi, Muhannad Alruwaili, Stuart D Armstrong, Jones Benjamin, Eleanor G Bentley, Anu Chawla, Jordan J Clark, Angela Cowell, Richard Eccles, Isabel García-Dorival, Matthew Gemmell, Alessandro Gerada, PKF Gilmore, Richard Gregory, Ximeng Han, Catherine Hartley, Margaret Hughes, Miren Iturriza-Gomara, James Johnson, L Luu, Jenifer Manson, Charlotte Nelson, Elaine O'Toole, Cassie Olateju, Rebekah Penrice-Randal , Lucille Rainbow, N.P Randle, Trevor Ian Robinson, Parul Sharma, Ghada T Shawli, James P Stewart, Neil Swainston, Ecaterina Vamos, Joanne Watts, Mark Whitehead |
| EPI_ISL_814276, EPI_ISL_814416 | Wales Specialist Virology Centre Sequencing lab: Pathogen Genomics Unit | COVID-19 Genomics UK (COG-UK) Consortium | Catherine Moore, Johnathan Evans, Laura Gifford, Malorie Perry, Simon Cottrell, Angela Marchbank, Alec Birchley, Alexander Adams, Amy Gaskin, Bree Gatica-Wilcox, Jason Coombes, Joel Southgate, Lauren Gilbert, Lee Graham, Nicole Pacchiarini, Sara Kumziene-Summerhayes, Sarah Taylor, Sophie Jones, Sara Rey, Matthew Bull, Joanne Watkins, Sally Corden, Tom Connor |
| EPI_ISL_814680 | Virology Department, Royal Infirmary of Edinburgh, NHS Lothian / School of Biological Sciences, University of Edinburgh / Institute of Genetics and Molecular Medicine, University of Edinburgh | COVID-19 Genomics UK (COG-UK) Consortium | McHugh M, Dewar R, Rooke S, Gallagher M, Balcaza C, O'Toole Á, Scher E, Hill V, McCrone JT, Colquhoun R, Yu X, Jackson B, Rambaut A, Williams TC, Templeton K |
| EPI_ISL_814778, EPI_ISL_814779, EPI_ISL_814780, EPI_ISL_814781, EPI_ISL_814782, EPI_ISL_814783, EPI_ISL_814784, EPI_ISL_814785, EPI_ISL_814786, EPI_ISL_814787, EPI_ISL_814788, EPI_ISL_814789, EPI_ISL_814790, EPI_ISL_814791, EPI_ISL_814792, EPI_ISL_814793 | | | |
| see above | Wales Specialist Virology Centre Sequencing lab: Pathogen Genomics Unit | COVID-19 Genomics UK (COG-UK) Consortium | Catherine Moore, Johnathan Evans, Laura Gifford, Malorie Perry, Simon Cottrell, Angela Marchbank, Alec Birchley, Alexander Adams, Amy Gaskin, Bree Gatica-Wilcox, Jason Coombes, Joel Southgate, Lauren Gilbert, Lee Graham, Nicole Pacchiarini, Sara Kumziene-Summerhayes, Sarah Taylor, Sophie Jones, Sara Rey, Matthew Bull, Joanne Watkins, Sally Corden, Tom Connor |
| EPI_ISL_816468 | Virology Department, Sheffield Teaching Hospitals NHS Foundation Trust/Department of Infection, Immunity and Cardiovascular Disease, The Medical School, University of Sheffield | COVID-19 Genomics UK (COG-UK) Consortium | Thushan de Silva, Matthew Parker, Nikki Smith, Adri Angyal, Rebecca Brown, Luke Green, Rachel Tucker, Paul Parsons, Danielle Groves, Katie Johnson, Laura Carrilero, Alex Keeley, Dave Partridge, Matthew Wyles, Benjamin Lindsey, Mehmet Yavuz, Mohammad Raza, Cariad Evans |
| EPI_ISL_819464 | Queens Medical Centre, Clinical Microbiology Department / DeepSeq Nottingham | COVID-19 Genomics UK (COG-UK) Consortium | Gemma Clark, Wendy Smith, Manjinder Khakh, Vicki M Fleming, Michelle M Lister, Hannah Howson-Wells, Jonathan Ball, Patrick McClure, Joseph Chappell, Theocharis Tsoleridis, Nadine Holmes, Matthew Carlisle, Christopher Moore, Fei Sang, Johnny Debebe, Victoria Wright, Matthew Loose |
| EPI_ISL_819465, EPI_ISL_819466, EPI_ISL_819467, EPI_ISL_819468, EPI_ISL_819469, EPI_ISL_819470, EPI_ISL_819471, EPI_ISL_819472, EPI_ISL_819473, EPI_ISL_819474, EPI_ISL_819475 | | | |
| see above | University College London, Great Ormond Street Hospital for Children NHS Foundation Trust, Imperial College Healthcare NHS Trust | COVID-19 Genomics UK (COG-UK) Consortium | Sergi Castellano, Rachel Williams, Mark Kristiansen, Paola Resende Silva, Sunando Roy, Tony Brooks, Helena Tutill, Paola Niola, Patricia Dyal, Charlotte Williams, Leysa Forrest, Yasmin Panchbhaya, Jacqueline Findlay, Samuel Weeks, Julianne Brown, Kathryn Harris, Paul Randell, James Price, Alison Holmes, Judith Breuer |
| EPI_ISL_819476 | Lincolnshire Hospitals and DeepSeq Nottingham | COVID-19 Genomics UK (COG-UK) Consortium | Nichola Duckworth, Tim Sloan, Sarah Walsh, Jonathan Ball, Patrick McClure, Joeseph Chappell, Nadine Holmes, Matthew Carlisle, Christopher Moore, Fei Sang, Johnny Debebe, Victoria Wright, Matthew Loose |
| EPI_ISL_819477 | University College London, Great Ormond Street Hospital for Children NHS Foundation Trust, Imperial College Healthcare NHS Trust | COVID-19 Genomics UK (COG-UK) Consortium | Sergi Castellano, Rachel Williams, Mark Kristiansen, Paola Resende Silva, Sunando Roy, Tony Brooks, Helena Tutill, Paola Niola, Patricia Dyal, Charlotte Williams, Leysa Forrest, Yasmin Panchbhaya, Jacqueline Findlay, Samuel Weeks, Julianne Brown, Kathryn Harris, Paul Randell, James Price, Alison Holmes, Judith Breuer |
| EPI_ISL_819478, EPI_ISL_819479, EPI_ISL_819480, EPI_ISL_819481, EPI_ISL_819482 | Lincolnshire Hospitals and DeepSeq Nottingham | COVID-19 Genomics UK (COG-UK) Consortium | Nichola Duckworth, Tim Sloan, Sarah Walsh, Jonathan Ball, Patrick McClure, Joeseph Chappell, Nadine Holmes, Matthew Carlisle, Christopher Moore, Fei Sang, Johnny Debebe, Victoria Wright, Matthew Loose |
| EPI_ISL_819483, EPI_ISL_819484, EPI_ISL_819485, EPI_ISL_819486 | University College London, Great Ormond Street Hospital for Children NHS Foundation Trust, Imperial College Healthcare NHS Trust | COVID-19 Genomics UK (COG-UK) Consortium | Sergi Castellano, Rachel Williams, Mark Kristiansen, Paola Resende Silva, Sunando Roy, Tony Brooks, Helena Tutill, Paola Niola, Patricia Dyal, Charlotte Williams, Leysa Forrest, Yasmin Panchbhaya, Jacqueline Findlay, Samuel Weeks, Julianne Brown, Kathryn Harris, Paul Randell, James Price, Alison Holmes, Judith Breuer |
| EPI_ISL_819487, EPI_ISL_819488, EPI_ISL_819489 | Lincolnshire Hospitals and DeepSeq Nottingham | COVID-19 Genomics UK (COG-UK) Consortium | Nichola Duckworth, Tim Sloan, Sarah Walsh, Jonathan Ball, Patrick McClure, Joeseph Chappell, Nadine Holmes, Matthew Carlisle, Christopher Moore, Fei Sang, Johnny Debebe, Victoria Wright, Matthew Loose |
| EPI_ISL_819490, EPI_ISL_819491, EPI_ISL_819492 | University College London, Great Ormond Street Hospital for Children NHS Foundation Trust, Imperial College Healthcare NHS Trust | COVID-19 Genomics UK (COG-UK) Consortium | Sergi Castellano, Rachel Williams, Mark Kristiansen, Paola Resende Silva, Sunando Roy, Tony Brooks, Helena Tutill, Paola Niola, Patricia Dyal, Charlotte Williams, Leysa Forrest, Yasmin Panchbhaya, Jacqueline Findlay, Samuel Weeks, Julianne Brown, Kathryn Harris, Paul Randell, James Price, Alison Holmes, Judith Breuer |
| EPI_ISL_819493 | Lincolnshire Hospitals and DeepSeq Nottingham | COVID-19 Genomics UK (COG-UK) Consortium | Nichola Duckworth, Tim Sloan, Sarah Walsh, Jonathan Ball, Patrick McClure, Joeseph Chappell, Nadine Holmes, Matthew Carlisle, Christopher Moore, Fei Sang, Johnny Debebe, Victoria Wright, Matthew Loose |
| EPI_ISL_819494, EPI_ISL_819495, EPI_ISL_819496, EPI_ISL_819497, EPI_ISL_819498 | University College London, Great Ormond Street Hospital for Children NHS Foundation Trust, Imperial College Healthcare NHS Trust | COVID-19 Genomics UK (COG-UK) Consortium | Sergi Castellano, Rachel Williams, Mark Kristiansen, Paola Resende Silva, Sunando Roy, Tony Brooks, Helena Tutill, Paola Niola, Patricia Dyal, Charlotte Williams, Leysa Forrest, Yasmin Panchbhaya, Jacqueline Findlay, Samuel Weeks, Julianne Brown, Kathryn Harris, Paul Randell, James Price, Alison Holmes, Judith Breuer |
| EPI_ISL_819499 | Lincolnshire Hospitals and DeepSeq Nottingham | COVID-19 Genomics UK (COG-UK) Consortium | Nichola Duckworth, Tim Sloan, Sarah Walsh, Jonathan Ball, Patrick McClure, Joeseph Chappell, Nadine Holmes, Matthew Carlisle, Christopher Moore, Fei Sang, Johnny Debebe, Victoria Wright, Matthew Loose |
| EPI_ISL_819500, EPI_ISL_819501, EPI_ISL_819502 | University College London, Great Ormond Street Hospital for Children NHS Foundation Trust, Imperial College Healthcare NHS Trust | COVID-19 Genomics UK (COG-UK) Consortium | Sergi Castellano, Rachel Williams, Mark Kristiansen, Paola Resende Silva, Sunando Roy, Tony Brooks, Helena Tutill, Paola Niola, Patricia Dyal, Charlotte Williams, Leysa Forrest, Yasmin Panchbhaya, Jacqueline Findlay, Samuel Weeks, Julianne Brown, Kathryn Harris, Paul Randell, James Price, Alison Holmes, Judith Breuer |
| EPI_ISL_819503, EPI_ISL_819504 | Lincolnshire Hospitals and DeepSeq Nottingham | COVID-19 Genomics UK (COG-UK) Consortium | Nichola Duckworth, Tim Sloan, Sarah Walsh, Jonathan Ball, Patrick McClure, Joeseph Chappell, Nadine Holmes, Matthew Carlisle, Christopher Moore, Fei Sang, Johnny Debebe, Victoria Wright, Matthew Loose |
| EPI_ISL_824001, EPI_ISL_824008, EPI_ISL_824017, EPI_ISL_824050 | Dutch COVID-19 response team | National Institute for Public Health and the Environment (RIVM) | Adam Meijer, Harry Vennema, Jeroen Cremer, Sharon van den Brink, Bas van der Veer, AnneMarie van den Brandt, Florian Zwagemaker, Dennis Schmitz, Chantal Reusken, on behalf of the national COVID-19 response team |
| EPI_ISL_824296 | DOHMH Jamaica | New York City Public Health Laboratory | Jade Wang, et al. |
| EPI_ISL_824298 | DOHMH Central Harlem | New York City Public Health Laboratory | Jade Wang, et al. |
| EPI_ISL_824392, EPI_ISL_824393, | California Department of Public Health | California Department of Public Health | CDPH IDLB COVIDNet |

| | | | |
|---|---|---|---|
| EPI_ISL_824394, EPI_ISL_824398 | | | |
| EPI_ISL_825042 | B.J. Medical College and Civil hospital, Ahmedabad | Gujarat Biotechnology Research Centre | Apurvasinh Puvar, Ramesh Pandit, Janvi Raval, Zarna Patel, Nitin Savaliya, Dinesh Kumar, Zuber Saiyed, Labdhi Pandya, Afzal Ansari, Nikha Trivedi, Pranay Shah, Kamlesh J Upadhyay, Sanjay Kapadia, Dipa Kinariwala, Chaitanya Joshi, Madhvi Joshi |
| EPI_ISL_825109, EPI_ISL_825110, EPI_ISL_825111, EPI_ISL_825112, EPI_ISL_825113, EPI_ISL_825114, EPI_ISL_825115, EPI_ISL_825116, EPI_ISL_825117, EPI_ISL_825118, EPI_ISL_825119, EPI_ISL_825120, EPI_ISL_825124, EPI_ISL_825125, EPI_ISL_825126, EPI_ISL_825127, EPI_ISL_825128, EPI_ISL_825129, EPI_ISL_825130 | | | |
| see above | NHLS-IALCH | KRISP, KZN Research Innovation and Sequencing Platform | Giandhari J, Pillay S, Lessells R, Mdlalose K, York D, Khan S, Tegally H, Wilkinson E, de Oliveira T |
| EPI_ISL_825327, EPI_ISL_825364, EPI_ISL_825365, EPI_ISL_825384 | Hospital Universitari Vall d'Hebron - Vall d'Hebron Institut de Rercerca | Hospital Universitari Vall d'Hebron | Cristina Andrés, Maria Piñana, Josep F Abril, Damir Garcia-Cehic, Ariadna Rando, Juliana Esperalba, Maria Gema Codina, Carla Castillo, Maria Carmen Martin, Tomàs Pumarola, Josep Quer, Andrés Antón |
| EPI_ISL_825714 | Nigeria Centre For Disease Control | National reference Laboratory, NCDC, Gaduwa, Abuja | Dr Ndodo Nnaemeka, Olusola Anuoluwapo Akanbi, Chimaobi Chukwu, Dr Omoare Adesuyi, Esebanmen Grace, Anthony Ahumibe, Naidoo Dhamari, Nwando Mba, Dr Chikwe Ihekweazu |
| EPI_ISL_826553 | Texas Department of State Health Services | Texas Department of State Health Services | Rashmi Tuladhar, Bonnie Oh, Jenny Zhang, Maliha Rahman, Anita Pokharel, Myong Koag, Chung Wang, Rachel Lee, Grace Kubin, Mayela Pedrueza, James Daniel Bonser |
| EPI_ISL_826673, EPI_ISL_826883 | deCODE genetics | deCODE genetics | Daniel F Gudbjartsson; Agnar Helgason; Hakon Jonsson; Olafur T Magnusson; Pall Melsted; Gudmundur L Norddahl; Jona Saemundsdottir; Asgeir Sigurdsson; Patrick Sulem; Arna B Agustsdottir; Hannes Eggertsson; Berglind Eiriksdottir; Run Fridriksdottir; Elisabet E Gardarsdottir; Gudmundur Georgsson; Olafia S Gretarsdottir; Kjartan R Gudmundsson; Thora R Gunnarsdottir; Arnaldur Gylfason; Hilma Holm; Brynjar O Jensson; Aslaug Jonasdottir; Kamilla S Josefsdottir; Thordur Kristjansson; Droplaug N Magnusdottir; Solvi Rognvaldsson; Louise le Roux; Gudrun Sigmundsdottir; Gardar Sveinbjornsson; Kristin E Sveinsdottir; Maney Sveinsdottir; Emil A Thorarensen; Bjarni Thorbjornsson; Gisli Masson; Ingileif Jonsdottir; Alma Moller; Thorolfur Gudnason; Karl G Kristinsson; Unnur Thorsteinsdottir; Kari Stefansson |
| EPI_ISL_826884 | The National University Hospital of Iceland | deCODE genetics | Daniel F Gudbjartsson; Agnar Helgason; Hakon Jonsson; Olafur T Magnusson; Pall Melsted; Gudmundur L Norddahl; Jona Saemundsdottir; Asgeir Sigurdsson; Patrick Sulem; Arna B Agustsdottir; Hannes Eggertsson; Berglind Eiriksdottir; Run Fridriksdottir; Elisabet E Gardarsdottir; Gudmundur Georgsson; Olafia S Gretarsdottir; Kjartan R Gudmundsson; Thora R Gunnarsdottir; Arnaldur Gylfason; Hilma Holm; Brynjar O Jensson; Aslaug Jonasdottir; Kamilla S Josefsdottir; Thordur Kristjansson; Droplaug N Magnusdottir; Solvi Rognvaldsson; Louise le Roux; Gudrun Sigmundsdottir; Gardar Sveinbjornsson; Kristin E Sveinsdottir; Maney Sveinsdottir; Emil A Thorarensen; Bjarni Thorbjornsson; Gisli Masson; Ingileif Jonsdottir; Alma Moller; Thorolfur Gudnason; Karl G Kristinsson; Unnur Thorsteinsdottir; Kari Stefansson |
| EPI_ISL_826913, EPI_ISL_826914, EPI_ISL_826932, EPI_ISL_827127, EPI_ISL_827128, EPI_ISL_827891, EPI_ISL_827894, EPI_ISL_827906, EPI_ISL_827964, EPI_ISL_827965, EPI_ISL_827969, EPI_ISL_828026 | | | |
| see above | deCODE genetics | deCODE genetics | Daniel F Gudbjartsson; Agnar Helgason; Hakon Jonsson; Olafur T Magnusson; Pall Melsted; Gudmundur L Norddahl; Jona Saemundsdottir; Asgeir Sigurdsson; Patrick Sulem; Arna B Agustsdottir; Hannes Eggertsson; Berglind Eiriksdottir; Run Fridriksdottir; Elisabet E Gardarsdottir; Gudmundur Georgsson; Olafia S Gretarsdottir; Kjartan R Gudmundsson; Thora R Gunnarsdottir; Arnaldur Gylfason; Hilma Holm; Brynjar O Jensson; Aslaug Jonasdottir; Kamilla S Josefsdottir; Thordur Kristjansson; Droplaug N Magnusdottir; Solvi Rognvaldsson; Louise le Roux; Gudrun Sigmundsdottir; Gardar Sveinbjornsson; Kristin E Sveinsdottir; Maney Sveinsdottir; Emil A Thorarensen; Bjarni Thorbjornsson; Gisli Masson; Ingileif Jonsdottir; Alma Moller; Thorolfur Gudnason; Karl G Kristinsson; Unnur Thorsteinsdottir; Kari Stefansson |
| EPI_ISL_828066 | The National University Hospital of Iceland | deCODE genetics | Daniel F Gudbjartsson; Agnar Helgason; Hakon Jonsson; Olafur T Magnusson; Pall Melsted; Gudmundur L Norddahl; Jona Saemundsdottir; Asgeir Sigurdsson; Patrick Sulem; Arna B Agustsdottir; Hannes Eggertsson; Berglind Eiriksdottir; Run Fridriksdottir; Elisabet E Gardarsdottir; Gudmundur Georgsson; Olafia S Gretarsdottir; Kjartan R Gudmundsson; Thora R Gunnarsdottir; Arnaldur Gylfason; Hilma Holm; Brynjar O Jensson; Aslaug Jonasdottir; Kamilla S Josefsdottir; Thordur Kristjansson; Droplaug N Magnusdottir; Solvi Rognvaldsson; Louise le Roux; Gudrun Sigmundsdottir; Gardar Sveinbjornsson; Kristin E Sveinsdottir; Maney Sveinsdottir; Emil A Thorarensen; Bjarni Thorbjornsson; Gisli Masson; Ingileif Jonsdottir; Alma Moller; Thorolfur Gudnason; Karl G Kristinsson; Unnur Thorsteinsdottir; Kari Stefansson |
| EPI_ISL_828090, EPI_ISL_828091, EPI_ISL_828092, EPI_ISL_828093, EPI_ISL_828094, EPI_ISL_828096, EPI_ISL_828097, EPI_ISL_828123, EPI_ISL_828138, EPI_ISL_828139, EPI_ISL_828141, EPI_ISL_828146, EPI_ISL_828148, EPI_ISL_828151, EPI_ISL_828152 | | | |
| see above | deCODE genetics | deCODE genetics | Daniel F Gudbjartsson; Agnar Helgason; Hakon Jonsson; Olafur T Magnusson; Pall Melsted; Gudmundur L Norddahl; Jona Saemundsdottir; Asgeir Sigurdsson; Patrick Sulem; Arna B Agustsdottir; Hannes Eggertsson; Berglind Eiriksdottir; Run Fridriksdottir; Elisabet E Gardarsdottir; Gudmundur Georgsson; Olafia S Gretarsdottir; Kjartan R Gudmundsson; Thora R Gunnarsdottir; Arnaldur Gylfason; Hilma Holm; Brynjar O Jensson; Aslaug Jonasdottir; Kamilla S Josefsdottir; Thordur Kristjansson; Droplaug N Magnusdottir; Solvi Rognvaldsson; Louise le Roux; Gudrun Sigmundsdottir; Gardar Sveinbjornsson; Kristin E Sveinsdottir; Maney Sveinsdottir; Emil A Thorarensen; Bjarni Thorbjornsson; Gisli Masson; Ingileif Jonsdottir; Alma Moller; Thorolfur Gudnason; Karl G Kristinsson; Unnur Thorsteinsdottir; Kari Stefansson |
| EPI_ISL_828175, EPI_ISL_828188, EPI_ISL_828244 | The National University Hospital of Iceland | deCODE genetics | Daniel F Gudbjartsson; Agnar Helgason; Hakon Jonsson; Olafur T Magnusson; Pall Melsted; Gudmundur L Norddahl; Jona Saemundsdottir; Asgeir Sigurdsson; Patrick Sulem; Arna B Agustsdottir; Hannes Eggertsson; Berglind Eiriksdottir; Run Fridriksdottir; Elisabet E Gardarsdottir; Gudmundur Georgsson; Olafia S Gretarsdottir; Kjartan R Gudmundsson; Thora R Gunnarsdottir; Arnaldur Gylfason; Hilma Holm; Brynjar O Jensson; Aslaug Jonasdottir; Kamilla S Josefsdottir; Thordur Kristjansson; Droplaug N Magnusdottir; Solvi Rognvaldsson; Louise le Roux; Gudrun Sigmundsdottir; Gardar Sveinbjornsson; Kristin E Sveinsdottir; Maney Sveinsdottir; Emil A Thorarensen; Bjarni Thorbjornsson; Gisli Masson; Ingileif Jonsdottir; Alma Moller; Thorolfur Gudnason; Karl G Kristinsson; Unnur Thorsteinsdottir; Kari Stefansson |
| EPI_ISL_828409, EPI_ISL_828411, EPI_ISL_828712 | deCODE genetics | deCODE genetics | Daniel F Gudbjartsson; Agnar Helgason; Hakon Jonsson; Olafur T Magnusson; Pall Melsted; Gudmundur L Norddahl; Jona Saemundsdottir; Asgeir Sigurdsson; Patrick Sulem; Arna B Agustsdottir; Hannes Eggertsson; Berglind Eiriksdottir; Run Fridriksdottir; Elisabet E Gardarsdottir; Gudmundur Georgsson; Olafia S Gretarsdottir; Kjartan R Gudmundsson; Thora R Gunnarsdottir; Arnaldur Gylfason; Hilma Holm; Brynjar O Jensson; Aslaug Jonasdottir; Kamilla S Josefsdottir; Thordur Kristjansson; Droplaug N Magnusdottir; Solvi Rognvaldsson; Louise le Roux; Gudrun Sigmundsdottir; Gardar Sveinbjornsson; Kristin E Sveinsdottir; Maney Sveinsdottir; Emil A Thorarensen; Bjarni Thorbjornsson; Gisli Masson; Ingileif Jonsdottir; Alma Moller; Thorolfur Gudnason; Karl G Kristinsson; Unnur Thorsteinsdottir; Kari Stefansson |
| EPI_ISL_828826 | The National University Hospital of Iceland | deCODE genetics | Daniel F Gudbjartsson; Agnar Helgason; Hakon Jonsson; Olafur T Magnusson; Pall Melsted; Gudmundur L Norddahl; Jona Saemundsdottir; Asgeir Sigurdsson; Patrick Sulem; Arna B Agustsdottir; Hannes Eggertsson; Berglind Eiriksdottir; Run Fridriksdottir; Elisabet E Gardarsdottir; Gudmundur Georgsson; Olafia S Gretarsdottir; Kjartan R Gudmundsson; Thora R Gunnarsdottir; Arnaldur Gylfason; Hilma Holm; Brynjar O Jensson; Aslaug Jonasdottir; Kamilla S Josefsdottir; Thordur Kristjansson; Droplaug N Magnusdottir; Solvi Rognvaldsson; Louise le Roux; Gudrun Sigmundsdottir; Gardar Sveinbjornsson; Kristin E Sveinsdottir; Maney Sveinsdottir; Emil A Thorarensen; Bjarni Thorbjornsson; Gisli Masson; Ingileif Jonsdottir; Alma Moller; Thorolfur Gudnason; Karl G Kristinsson; Unnur Thorsteinsdottir; Kari Stefansson |
| EPI_ISL_828947, EPI_ISL_828949, EPI_ISL_828950, EPI_ISL_828959, EPI_ISL_828963, EPI_ISL_828967, EPI_ISL_829057 | deCODE genetics | deCODE genetics | Daniel F Gudbjartsson; Agnar Helgason; Hakon Jonsson; Olafur T Magnusson; Pall Melsted; Gudmundur L Norddahl; Jona Saemundsdottir; Asgeir Sigurdsson; Patrick Sulem; Arna B Agustsdottir; Hannes Eggertsson; Berglind Eiriksdottir; Run Fridriksdottir; Elisabet E Gardarsdottir; Gudmundur Georgsson; Olafia S Gretarsdottir; Kjartan R Gudmundsson; Thora R Gunnarsdottir; Arnaldur Gylfason; Hilma Holm; Brynjar O Jensson; Aslaug Jonasdottir; Kamilla S Josefsdottir; Thordur Kristjansson; Droplaug N Magnusdottir; Solvi Rognvaldsson; Louise le Roux; Gudrun Sigmundsdottir; Gardar Sveinbjornsson; Kristin E Sveinsdottir; Maney Sveinsdottir; Emil A Thorarensen; Bjarni Thorbjornsson; Gisli Masson; Ingileif Jonsdottir; Alma Moller; Thorolfur Gudnason; Karl G Kristinsson; Unnur Thorsteinsdottir; Kari Stefansson |
| EPI_ISL_829220 | The National University Hospital of Iceland | deCODE genetics | Daniel F Gudbjartsson; Agnar Helgason; Hakon Jonsson; Olafur T Magnusson; Pall Melsted; Gudmundur L Norddahl; Jona Saemundsdottir; Asgeir Sigurdsson; Patrick Sulem; Arna B Agustsdottir; Hannes Eggertsson; Berglind Eiriksdottir; Run Fridriksdottir; Elisabet E Gardarsdottir; Gudmundur Georgsson; Olafia S Gretarsdottir; Kjartan R Gudmundsson; Thora R Gunnarsdottir; Arnaldur Gylfason; Hilma Holm; Brynjar O Jensson; Aslaug Jonasdottir; Kamilla S Josefsdottir; Thordur Kristjansson; Droplaug N Magnusdottir; Solvi Rognvaldsson; Louise le Roux; Gudrun Sigmundsdottir; Gardar Sveinbjornsson; Kristin E Sveinsdottir; Maney Sveinsdottir; Emil A Thorarensen; Bjarni Thorbjornsson; Gisli Masson; Ingileif Jonsdottir; Alma Moller; Thorolfur Gudnason; Karl G Kristinsson; Unnur Thorsteinsdottir; Kari Stefansson |
| EPI_ISL_829340, EPI_ISL_829341, EPI_ISL_829342, EPI_ISL_829343, EPI_ISL_829344, EPI_ISL_829345, EPI_ISL_829346, EPI_ISL_829347, EPI_ISL_829368, EPI_ISL_829380, EPI_ISL_829390, EPI_ISL_829421, EPI_ISL_829437, EPI_ISL_829462, EPI_ISL_829584, EPI_ISL_829586, EPI_ISL_829587, EPI_ISL_829588, EPI_ISL_829589, EPI_ISL_829591 | | | |
| see above | deCODE genetics | deCODE genetics | Daniel F Gudbjartsson; Agnar Helgason; Hakon Jonsson; Olafur T Magnusson; Pall Melsted; Gudmundur L Norddahl; Jona Saemundsdottir; Asgeir |

| | | | |
|---|---|---|---|
| | | | Sigurdsson; Patrick Sulem; Arna B Agustsdottir; Hannes Eggertsson; Berglind Eiriksdottir; Run Fridriksdottir; Elisabet E Gardarsdottir; Gudmundur Georgsson; Olafia S Gretarsdottir; Kjartan R Gudmundsson; Thora R Gunnarsdottir; Arnaldur Gylfason; Hilma Holm; Brynjar O Jensson; Aslaug Jonasdottir; Kamilla S Josefsdottir; Thordur Kristjansson; Droplaug N Magnusdottir; Solvi Rognvaldsson; Louise le Roux; Gudrun Sigmundsdottir; Gardar Sveinbjornsson; Kristin E Sveinsdottir; Maney Sveinsdottir; Emil A Thorarensen; Bjarni Thorbjornsson; Gisli Masson; Ingileif Jonsdottir; Alma Moller; Thorolfur Gudnason; Karl G Kristinsson; Unnur Thorsteinsdottir; Kari Stefansson |
| EPI_ISL_829851 | The National University Hospital of Iceland | deCODE genetics | Daniel F Gudbjartsson; Agnar Helgason; Hakon Jonsson; Olafur T Magnusson; Pall Melsted; Gudmundur L Norddahl; Jona Saemundsdottir; Asgeir Sigurdsson; Patrick Sulem; Arna B Agustsdottir; Hannes Eggertsson; Berglind Eiriksdottir; Run Fridriksdottir; Elisabet E Gardarsdottir; Gudmundur Georgsson; Olafia S Gretarsdottir; Kjartan R Gudmundsson; Thora R Gunnarsdottir; Arnaldur Gylfason; Hilma Holm; Brynjar O Jensson; Aslaug Jonasdottir; Kamilla S Josefsdottir; Thordur Kristjansson; Droplaug N Magnusdottir; Solvi Rognvaldsson; Louise le Roux; Gudrun Sigmundsdottir; Gardar Sveinbjornsson; Kristin E Sveinsdottir; Maney Sveinsdottir; Emil A Thorarensen; Bjarni Thorbjornsson; Gisli Masson; Ingileif Jonsdottir; Alma Moller; Thorolfur Gudnason; Karl G Kristinsson; Unnur Thorsteinsdottir; Kari Stefansson |
| EPI_ISL_829852, EPI_ISL_829853, EPI_ISL_829854, EPI_ISL_829855, EPI_ISL_829856, EPI_ISL_829857, EPI_ISL_829858 | deCODE genetics | deCODE genetics | Daniel F Gudbjartsson; Agnar Helgason; Hakon Jonsson; Olafur T Magnusson; Pall Melsted; Gudmundur L Norddahl; Jona Saemundsdottir; Asgeir Sigurdsson; Patrick Sulem; Arna B Agustsdottir; Hannes Eggertsson; Berglind Eiriksdottir; Run Fridriksdottir; Elisabet E Gardarsdottir; Gudmundur Georgsson; Olafia S Gretarsdottir; Kjartan R Gudmundsson; Thora R Gunnarsdottir; Arnaldur Gylfason; Hilma Holm; Brynjar O Jensson; Aslaug Jonasdottir; Kamilla S Josefsdottir; Thordur Kristjansson; Droplaug N Magnusdottir; Solvi Rognvaldsson; Louise le Roux; Gudrun Sigmundsdottir; Gardar Sveinbjornsson; Kristin E Sveinsdottir; Maney Sveinsdottir; Emil A Thorarensen; Bjarni Thorbjornsson; Gisli Masson; Ingileif Jonsdottir; Alma Moller; Thorolfur Gudnason; Karl G Kristinsson; Unnur Thorsteinsdottir; Kari Stefansson |
| EPI_ISL_829859, EPI_ISL_829860 | The National University Hospital of Iceland | deCODE genetics | Daniel F Gudbjartsson; Agnar Helgason; Hakon Jonsson; Olafur T Magnusson; Pall Melsted; Gudmundur L Norddahl; Jona Saemundsdottir; Asgeir Sigurdsson; Patrick Sulem; Arna B Agustsdottir; Hannes Eggertsson; Berglind Eiriksdottir; Run Fridriksdottir; Elisabet E Gardarsdottir; Gudmundur Georgsson; Olafia S Gretarsdottir; Kjartan R Gudmundsson; Thora R Gunnarsdottir; Arnaldur Gylfason; Hilma Holm; Brynjar O Jensson; Aslaug Jonasdottir; Kamilla S Josefsdottir; Thordur Kristjansson; Droplaug N Magnusdottir; Solvi Rognvaldsson; Louise le Roux; Gudrun Sigmundsdottir; Gardar Sveinbjornsson; Kristin E Sveinsdottir; Maney Sveinsdottir; Emil A Thorarensen; Bjarni Thorbjornsson; Gisli Masson; Ingileif Jonsdottir; Alma Moller; Thorolfur Gudnason; Karl G Kristinsson; Unnur Thorsteinsdottir; Kari Stefansson |
| EPI_ISL_829951, EPI_ISL_829963 | deCODE genetics | deCODE genetics | Daniel F Gudbjartsson; Agnar Helgason; Hakon Jonsson; Olafur T Magnusson; Pall Melsted; Gudmundur L Norddahl; Jona Saemundsdottir; Asgeir Sigurdsson; Patrick Sulem; Arna B Agustsdottir; Hannes Eggertsson; Berglind Eiriksdottir; Run Fridriksdottir; Elisabet E Gardarsdottir; Gudmundur Georgsson; Olafia S Gretarsdottir; Kjartan R Gudmundsson; Thora R Gunnarsdottir; Arnaldur Gylfason; Hilma Holm; Brynjar O Jensson; Aslaug Jonasdottir; Kamilla S Josefsdottir; Thordur Kristjansson; Droplaug N Magnusdottir; Solvi Rognvaldsson; Louise le Roux; Gudrun Sigmundsdottir; Gardar Sveinbjornsson; Kristin E Sveinsdottir; Maney Sveinsdottir; Emil A Thorarensen; Bjarni Thorbjornsson; Gisli Masson; Ingileif Jonsdottir; Alma Moller; Thorolfur Gudnason; Karl G Kristinsson; Unnur Thorsteinsdottir; Kari Stefansson |
| EPI_ISL_830102, EPI_ISL_830103, EPI_ISL_830104 | The National University Hospital of Iceland | deCODE genetics | Daniel F Gudbjartsson; Agnar Helgason; Hakon Jonsson; Olafur T Magnusson; Pall Melsted; Gudmundur L Norddahl; Jona Saemundsdottir; Asgeir Sigurdsson; Patrick Sulem; Arna B Agustsdottir; Hannes Eggertsson; Berglind Eiriksdottir; Run Fridriksdottir; Elisabet E Gardarsdottir; Gudmundur Georgsson; Olafia S Gretarsdottir; Kjartan R Gudmundsson; Thora R Gunnarsdottir; Arnaldur Gylfason; Hilma Holm; Brynjar O Jensson; Aslaug Jonasdottir; Kamilla S Josefsdottir; Thordur Kristjansson; Droplaug N Magnusdottir; Solvi Rognvaldsson; Louise le Roux; Gudrun Sigmundsdottir; Gardar Sveinbjornsson; Kristin E Sveinsdottir; Maney Sveinsdottir; Emil A Thorarensen; Bjarni Thorbjornsson; Gisli Masson; Ingileif Jonsdottir; Alma Moller; Thorolfur Gudnason; Karl G Kristinsson; Unnur Thorsteinsdottir; Kari Stefansson |
| EPI_ISL_830163, EPI_ISL_830197 | deCODE genetics | deCODE genetics | Daniel F Gudbjartsson; Agnar Helgason; Hakon Jonsson; Olafur T Magnusson; Pall Melsted; Gudmundur L Norddahl; Jona Saemundsdottir; Asgeir Sigurdsson; Patrick Sulem; Arna B Agustsdottir; Hannes Eggertsson; Berglind Eiriksdottir; Run Fridriksdottir; Elisabet E Gardarsdottir; Gudmundur Georgsson; Olafia S Gretarsdottir; Kjartan R Gudmundsson; Thora R Gunnarsdottir; Arnaldur Gylfason; Hilma Holm; Brynjar O Jensson; Aslaug Jonasdottir; Kamilla S Josefsdottir; Thordur Kristjansson; Droplaug N Magnusdottir; Solvi Rognvaldsson; Louise le Roux; Gudrun Sigmundsdottir; Gardar Sveinbjornsson; Kristin E Sveinsdottir; Maney Sveinsdottir; Emil A Thorarensen; Bjarni Thorbjornsson; Gisli Masson; Ingileif Jonsdottir; Alma Moller; Thorolfur Gudnason; Karl G Kristinsson; Unnur Thorsteinsdottir; Kari Stefansson |
| EPI_ISL_830415 | The National University Hospital of Iceland | deCODE genetics | Daniel F Gudbjartsson; Agnar Helgason; Hakon Jonsson; Olafur T Magnusson; Pall Melsted; Gudmundur L Norddahl; Jona Saemundsdottir; Asgeir Sigurdsson; Patrick Sulem; Arna B Agustsdottir; Hannes Eggertsson; Berglind Eiriksdottir; Run Fridriksdottir; Elisabet E Gardarsdottir; Gudmundur Georgsson; Olafia S Gretarsdottir; Kjartan R Gudmundsson; Thora R Gunnarsdottir; Arnaldur Gylfason; Hilma Holm; Brynjar O Jensson; Aslaug Jonasdottir; Kamilla S Josefsdottir; Thordur Kristjansson; Droplaug N Magnusdottir; Solvi Rognvaldsson; Louise le Roux; Gudrun Sigmundsdottir; Gardar Sveinbjornsson; Kristin E Sveinsdottir; Maney Sveinsdottir; Emil A Thorarensen; Bjarni Thorbjornsson; Gisli Masson; Ingileif Jonsdottir; Alma Moller; Thorolfur Gudnason; Karl G Kristinsson; Unnur Thorsteinsdottir; Kari Stefansson |
| EPI_ISL_830561 | deCODE genetics | deCODE genetics | Daniel F Gudbjartsson; Agnar Helgason; Hakon Jonsson; Olafur T Magnusson; Pall Melsted; Gudmundur L Norddahl; Jona Saemundsdottir; Asgeir Sigurdsson; Patrick Sulem; Arna B Agustsdottir; Hannes Eggertsson; Berglind Eiriksdottir; Run Fridriksdottir; Elisabet E Gardarsdottir; Gudmundur Georgsson; Olafia S Gretarsdottir; Kjartan R Gudmundsson; Thora R Gunnarsdottir; Arnaldur Gylfason; Hilma Holm; Brynjar O Jensson; Aslaug Jonasdottir; Kamilla S Josefsdottir; Thordur Kristjansson; Droplaug N Magnusdottir; Solvi Rognvaldsson; Louise le Roux; Gudrun Sigmundsdottir; Gardar Sveinbjornsson; Kristin E Sveinsdottir; Maney Sveinsdottir; Emil A Thorarensen; Bjarni Thorbjornsson; Gisli Masson; Ingileif Jonsdottir; Alma Moller; Thorolfur Gudnason; Karl G Kristinsson; Unnur Thorsteinsdottir; Kari Stefansson |
| EPI_ISL_831029, EPI_ISL_831203, EPI_ISL_831204, EPI_ISL_831207, EPI_ISL_831208, EPI_ISL_831209, EPI_ISL_831210, EPI_ISL_831211, EPI_ISL_831212, EPI_ISL_831245, EPI_ISL_831246 | | | |
| see above | Hospital Universitario La Paz (Madrid) | SeqCOVID-SPAIN consortium/IBV(CSIC) | María Rodríguez-Tejedor, Elias Dahdouh, Fernando Lázaro-Perona, Jesús Mingorance and SeqCOVID-SPAIN consortium |
| EPI_ISL_831735, EPI_ISL_831740, EPI_ISL_831829, EPI_ISL_831832, EPI_ISL_831833 | United States Air Force School of Aerospace Medicine | United States Air Force School of Aerospace Medicine | Anthony Fries, Jennifer Meyer, William Gruner, Amanda Javorina, Sarah Purves, Clarise Starr, Elizabeth Macias |
| EPI_ISL_832035, EPI_ISL_832036, EPI_ISL_832037, EPI_ISL_832038, EPI_ISL_832039, EPI_ISL_832040, EPI_ISL_832041, EPI_ISL_832042 | Wyoming Public Health Laboratory | Wyoming Public Health Laboratory | Noah Hull, Taylor Fearing, Lynette Gumbleton, Channing Weber, Ashley Norberg, Bailey Bowcutt, and Wanda Manley |
| EPI_ISL_832149, EPI_ISL_832150, EPI_ISL_832151, EPI_ISL_832152, EPI_ISL_832153, EPI_ISL_832154, EPI_ISL_832155, EPI_ISL_832157, EPI_ISL_832158, EPI_ISL_832159 | Hospital | National Reference Center for Viruses of Respiratory Infections, Institut Pasteur, Paris | Marion Barbet, Sylvie Behillil, Méline Bizard, Angela Brisebarre, Camille Capel, Etienne Simon-Lorière, Vincent Enouf, Maud Vanpeene, Sylvie van der Werf, Léa Pilorge |
| EPI_ISL_832801, EPI_ISL_832802, EPI_ISL_832803, EPI_ISL_832805, EPI_ISL_832806, EPI_ISL_832807, EPI_ISL_832809, EPI_ISL_832810, EPI_ISL_832813 | OHSU Lab Services Molecular Microbiology Lab | Oregon SARS-CoV-2 Genome Sequencing Center | Brendan L. O'Connell, Ruth V. Nichols, Sally Grindstaff, Alec J. Hirsch, Donna Hansel, Guang Fan, Daniel N. Streblow, William B. Messer, Andrew C. Adey, Benjamin N. Bimber, Brian J. O'Roak |
| EPI_ISL_833161 | Instituto Adolfo Lutz - Central | Instituto Adolfo Lutz, Interdiciplinary Procedures Center, Strategic Laboratory | Claudio Tavares Sacchi, Claudia Regina Gonçalves, Erica Valessa Ramos Gomes, Karoline Rodrigues Campos |
| EPI_ISL_833162 | Lab LOC - Itapecerica da Serra | Instituto Adolfo Lutz, Interdiciplinary Procedures Center, Strategic Laboratory | Claudio Tavares Sacchi, Claudia Regina Gonçalves, Erica Valessa Ramos Gomes, Karoline Rodrigues Campos |
| EPI_ISL_833448, EPI_ISL_833449, EPI_ISL_833450, EPI_ISL_833451, EPI_ISL_833452, EPI_ISL_833453, | SC (UCO) Igiene e Sanità Pubblica (funzione integrata con SC Microbiologia e Virologia) e Laboratory of Molecular Virology of the International Centre for Genetic Engineering and | ARGO Laboratorio Genomica ed Epigenomica | Licastro D, Dal Monego S, Degasperi M, Marcello A, D'Agaro P |

| | | | |
|---|---|---|---|
| EPI_ISL_833454 | Biotechnology (ICGEB) | | |
| EPI_ISL_837249, EPI_ISL_837251, EPI_ISL_837256, EPI_ISL_837257, EPI_ISL_837258, EPI_ISL_837259, EPI_ISL_837260, EPI_ISL_837261, EPI_ISL_837262, EPI_ISL_837263, EPI_ISL_837264, EPI_ISL_837265, EPI_ISL_837266, EPI_ISL_837267, EPI_ISL_837268, EPI_ISL_837434, EPI_ISL_837436, EPI_ISL_837437, EPI_ISL_837441, EPI_ISL_837443, EPI_ISL_837445, EPI_ISL_837447, EPI_ISL_837457, EPI_ISL_837458, EPI_ISL_837459, EPI_ISL_837460, EPI_ISL_837461, EPI_ISL_837462, EPI_ISL_837463, EPI_ISL_837464, EPI_ISL_837465, EPI_ISL_837466, EPI_ISL_837467, EPI_ISL_837468, EPI_ISL_837469, EPI_ISL_837470, EPI_ISL_837471, EPI_ISL_837472, EPI_ISL_837479, EPI_ISL_837480, EPI_ISL_837486, EPI_ISL_837487, EPI_ISL_837488 | | | |
| see above | Istituto Zooprofilattico Sperimentale del Mezzogiorno | TIGEM | Patrizia Annunziata, Andrea Ballabio, Valentina Bouche, Davide Cacchiarelli (CorrespAuthor), Pellegrino Cerino, Chiara Colantuono, Lucio Di Filippo, Antonio Grimaldi, Antonio Limone, Gabriella Loconte, Anna Manfredi, Francesco Panariello, Biancamaria Pierri, Marcello Salvi, Lucia Vassallo |
| EPI_ISL_838055, EPI_ISL_838056, EPI_ISL_838057, EPI_ISL_838058, EPI_ISL_838059, EPI_ISL_838060, EPI_ISL_838061, EPI_ISL_838062, EPI_ISL_838063, EPI_ISL_838064, EPI_ISL_838066, EPI_ISL_838067, EPI_ISL_838068, EPI_ISL_838069, EPI_ISL_838070, EPI_ISL_838071 | | | |
| see above | West of Scotland Specialist Virology Centre, NHSGGC / MRC-University of Glasgow Centre for Virus Research | COVID-19 Genomics UK (COG-UK) Consortium | Ana da Silva Filipe, Natasha Johnson, Kathy Smollett, Daniel Mair, Stephen Carmichael, Alice Broos, Lily Tong, Jenna Nichols, Kyriaki Nomikou; Sarah McDonald; Richard Orton, Joseph Hughes, Sreenu Vattipally, David L Robertson; Alasdair MacLean, Rory Gunson; Sharif Shaaban, Matthew Holden; Rachel Blacow, Guy Mollett, Kathy Li, James Shepherd, Antonia Ho, Emma Thomson |
| EPI_ISL_838214, EPI_ISL_838282 | Virology Department, Royal Infirmary of Edinburgh, NHS Lothian / School of Biological Sciences, University of Edinburgh / Institute of Genetics and Molecular Medicine, University of Edinburgh | COVID-19 Genomics UK (COG-UK) Consortium | McHugh M, Dewar R, Rooke S, Gallagher M, Balcaza C, O'Toole Á, Scher E, Hill V, McCrone JT, Colquhoun R, Yu X, Jackson B, Rambaut A, Williams TC, Templeton K |
| EPI_ISL_838347, EPI_ISL_838348, EPI_ISL_838350 | Queens Medical Centre, Clinical Microbiology Department / DeepSeq Nottingham | COVID-19 Genomics UK (COG-UK) Consortium | Gemma Clark, Wendy Smith, Manjinder Khakh, Vicki M Fleming, Michelle M Lister, Hannah Howson-Wells, Jonathan Ball, Patrick McClure, Joseph Chappell, Theocharis Tsoleridis, Nadine Holmes, Matthew Carlisle, Christopher Moore, Fei Sang, Johnny Debebe, Victoria Wright, Matthew Loose |
| EPI_ISL_838351, EPI_ISL_838360, EPI_ISL_838371, EPI_ISL_838374, EPI_ISL_838402, EPI_ISL_838430, EPI_ISL_838431, EPI_ISL_838434, EPI_ISL_838435, EPI_ISL_838436, EPI_ISL_838437, EPI_ISL_838441, EPI_ISL_838442, EPI_ISL_838443, EPI_ISL_838444, EPI_ISL_838447, EPI_ISL_838448, EPI_ISL_838449, EPI_ISL_838450 | | | |
| see above | Liverpool Clinical Laboratories | COVID-19 Genomics UK (COG-UK) Consortium | Sam Haldenby, Anita Lucaci, Steve Paterson, Julian Hiscox, Alistair Darby, M Almsaud, A Alrezaihi, Muhannad Alruwaili, Stuart D Armstrong, Jones Benjamin, Eleanor G Bentley, Anu Chawla, Jordan J Clark, Angela Cowell, Richard Eccles, Isabel García-Dorival, Matthew Gemmell, Alessandro Gerada, PKF Gilmore, Richard Gregory, Ximeng Han, Catherine Hartley, Margaret Hughes, Miren Iturriza-Gomara, James Johnson, L Luu, Jenifer Manson, Charlotte Nelson, Elaine O'Toole, Cassie Olateju, Rebekah Penrice-Randal , Lucille Rainbow, N.P Randle, Trevor Ian Robinson, Parul Sharma, Ghada T Shawli, James P Stewart, Neil Swainston, Ecaterina Vamos, Joanne Watts, Mark Whitehead |
| EPI_ISL_840017, EPI_ISL_840018 | Lincolnshire Hospitals and DeepSeq Nottingham | COVID-19 Genomics UK (COG-UK) Consortium | Nichola Duckworth, Tim Sloan, Sarah Walsh, Jonathan Ball, Patrick McClure, Joeseph Chappell, Nadine Holmes, Matthew Carlisle, Christopher Moore, Fei Sang, Johnny Debebe, Victoria Wright, Matthew Loose |
| EPI_ISL_840209, EPI_ISL_840210, EPI_ISL_840211, EPI_ISL_840212, EPI_ISL_840213, EPI_ISL_840214, EPI_ISL_840215, EPI_ISL_840220, EPI_ISL_840221 | Oxford Viromics, NDM, University of Oxford; Oxford University Hospitals; Basingstoke and North Hampshire Hospital | COVID-19 Genomics UK (COG-UK) Consortium | Tanya Golubchik, David Bonsall, George Macintyre, Amy Trebes, Mariateresa de Cesare, Catrin Moore, Alex Mobbs, Anita Justice, Robert Shaw, Monique Andersson, Timothy Peto, Emma Wise, Nathan Moore, Jessica Lynch, Nick Cortes, Matilde Mori, Stephen Kidd, David Buck, John Todd, Christophe Fraser |
| EPI_ISL_841531, EPI_ISL_841559, EPI_ISL_841560, EPI_ISL_841561, EPI_ISL_841563, EPI_ISL_841564, EPI_ISL_841565, EPI_ISL_841566, EPI_ISL_841568, EPI_ISL_841569, EPI_ISL_841570, EPI_ISL_841571, EPI_ISL_841572 | | | |
| see above | Originating lab: Wales Specialist Virology Centre Sequencing lab: Pathogen Genomics Unit | Public Health Wales Microbiology Cardiff Wales Specialist Virology Centre | Catherine Moore, Johnathan Evans, Laura Gifford, Malorie Perry, Simon Cottrell, Angela Marchbank, Alec Birchley, Alexander Adams, Amy Gaskin, Bree Gatica-Wilcox, Jason Coombes, Joel Southgate, Lauren Gilbert, Lee Graham, Nicole Pacchiarini, Sara Kumziene-Summerhayes, Sarah Taylor, Sophie Jones, Sara Rey, Matthew Bull, Joanne Watkins, Sally Corden, Tom Connor |
| EPI_ISL_842937, EPI_ISL_842938, EPI_ISL_842940, EPI_ISL_842941, EPI_ISL_842942, EPI_ISL_842950, EPI_ISL_842951, EPI_ISL_842952, EPI_ISL_842954, EPI_ISL_842955, EPI_ISL_842957, EPI_ISL_842958, EPI_ISL_842959, EPI_ISL_842960, EPI_ISL_842961, EPI_ISL_842962, EPI_ISL_842966, EPI_ISL_842969, EPI_ISL_842970, EPI_ISL_842971, EPI_ISL_842972, EPI_ISL_842973, EPI_ISL_842974, EPI_ISL_842975, EPI_ISL_842976, EPI_ISL_842977, EPI_ISL_842978, EPI_ISL_842979, EPI_ISL_842980, EPI_ISL_842981, EPI_ISL_842982, EPI_ISL_842983, EPI_ISL_843000, EPI_ISL_843005, EPI_ISL_843007, EPI_ISL_843013, EPI_ISL_843014, EPI_ISL_843015, EPI_ISL_843016, EPI_ISL_843017, EPI_ISL_843018, EPI_ISL_843019, EPI_ISL_843020, EPI_ISL_843021, EPI_ISL_843022, EPI_ISL_843023, EPI_ISL_843024, EPI_ISL_843041 | | | |
| see above | Barts Health NHS Trust | COVID-19 Genomics UK (COG-UK) Consortium | CUTINO-MOGUEL, Maria-Teresa; HARRINGTON, David; OWOYEMI, Dola; SHYLINI, Raghavendran; BROAD, Claire; KELE, Beatrix |
| EPI_ISL_845786, EPI_ISL_845787, EPI_ISL_845788, EPI_ISL_845789, EPI_ISL_845790, EPI_ISL_845791, EPI_ISL_845792, EPI_ISL_845793, EPI_ISL_845795 | South Eastern Area Laboratory Services (SEALS) | NSW Health Pathology - Institute of Clinical Pathology and Medical Research; Westmead Hospital; University of Sydney | CIDM-PH et al. |
| EPI_ISL_847519, EPI_ISL_847535, EPI_ISL_847551, EPI_ISL_847584, EPI_ISL_847672, EPI_ISL_847673, EPI_ISL_847707, EPI_ISL_847708, EPI_ISL_847803 | California Department of Public Health | Chiu Laboratory, University of California, San Francisco | Charles Chiu, Xianding (Wayne) Deng, Candace Wang, Brian Bushnell, Scot Federman, Jill Hacker, Debra Wadford |
| EPI_ISL_847978, EPI_ISL_847994, EPI_ISL_847995, EPI_ISL_847997, EPI_ISL_847999, EPI_ISL_848001, EPI_ISL_848004, EPI_ISL_848010, EPI_ISL_848011, EPI_ISL_848020, EPI_ISL_848037, EPI_ISL_848038, EPI_ISL_848039, EPI_ISL_848040, EPI_ISL_848041, EPI_ISL_848042, EPI_ISL_848043, EPI_ISL_848046, EPI_ISL_848052, EPI_ISL_848062 | | | |
| see above | Michigan Department of Health and Human Services, Bureau of Laboratories | Michigan Department of Health and Human Services, Bureau of Laboratories | Blankenship HM, Riner D, Soehnlen MK |
| EPI_ISL_848203, EPI_ISL_848209, EPI_ISL_848229, EPI_ISL_848271, EPI_ISL_848397, EPI_ISL_848398, EPI_ISL_848399, EPI_ISL_848400, EPI_ISL_848401, EPI_ISL_848402, EPI_ISL_848403, EPI_ISL_848404, EPI_ISL_848405, EPI_ISL_848406, EPI_ISL_848407, EPI_ISL_848408, EPI_ISL_848409, EPI_ISL_848410, EPI_ISL_848411, EPI_ISL_848412, EPI_ISL_848413, EPI_ISL_848449, EPI_ISL_848456, EPI_ISL_848461, EPI_ISL_848464, EPI_ISL_848527, EPI_ISL_848528, EPI_ISL_848529 | | | |
| see above | Illinois Department of Public Health | Gagnon Lab, Southern Illinois University | Keith Gagnon |
| EPI_ISL_848717, EPI_ISL_848718, EPI_ISL_848719, EPI_ISL_848720, EPI_ISL_848721, EPI_ISL_848722, EPI_ISL_848723, EPI_ISL_848724, EPI_ISL_848725, EPI_ISL_848726, EPI_ISL_848727, EPI_ISL_848728, EPI_ISL_848729, EPI_ISL_848730, EPI_ISL_848731, EPI_ISL_848732, EPI_ISL_848733 | | | |
| see above | Florida Bureau of Public Health Laboratories | Florida Bureau of Public Health Laboratories | Sarah Schmedes, Jason Blanton |
| EPI_ISL_849383, EPI_ISL_849386, EPI_ISL_849389, EPI_ISL_849393, EPI_ISL_849395, EPI_ISL_849406, EPI_ISL_849407, EPI_ISL_849409, EPI_ISL_849411, EPI_ISL_849416, EPI_ISL_849417, EPI_ISL_849420, EPI_ISL_849422, EPI_ISL_849427, EPI_ISL_849433, EPI_ISL_849435, EPI_ISL_849437, EPI_ISL_849438, EPI_ISL_849443, EPI_ISL_849445, EPI_ISL_849446, EPI_ISL_849450, EPI_ISL_849451, EPI_ISL_849452, EPI_ISL_849454, EPI_ISL_849456, EPI_ISL_849458, EPI_ISL_849459 | | | |
| see above | Washington State Department of Health | Seattle Flu Study | Deborah A. Nickerson, Chris D. Frazar, Jover Lee, Benjamin Pelle, Matthew Richardson, Amanda Adler, Elisabeth Brandstetter, Peter D. Han, Kairsten Fay, Misja Ilcisin, Kirsten Lacombe, Thomas R. Sibley, Melissa Truong, Caitlin R. Wolf, Romesh Gautom, Geoff Melly, Brian Hiatt, Philip Dykema, Scott Lindquist, Michael Boeckh, Janet A. Englund, Michael Famulare, Barry R. Lutz, Mark J. Rieder, Lea M. Starita, Matthew Thompson, Helen Y. Chu, Jay Shendure, Trevor Bedford |
| EPI_ISL_849462 | Seattle Flu Study | Seattle Flu Study | Deborah A. Nickerson, Chris D. Frazar, Jover Lee, Benjamin Pelle, Matthew Richardson, Amanda Adler, Elisabeth Brandstetter, Peter D. Han, Kairsten Fay, Misja Ilcisin, Kirsten Lacombe, Thomas R. Sibley, Melissa Truong, Caitlin R. Wolf, Michael Boeckh, Janet A. Englund, Michael Famulare, Barry R. Lutz, Mark J. Rieder, Lea M. Starita, Matthew Thompson, Jay Shendure, Trevor Bedford, Helen Y. Chu |
| EPI_ISL_849464, EPI_ISL_849466, EPI_ISL_849522, EPI_ISL_849523, EPI_ISL_849525, EPI_ISL_849556, EPI_ISL_849558, EPI_ISL_849567, EPI_ISL_849568, EPI_ISL_849570, EPI_ISL_849577, EPI_ISL_849582, EPI_ISL_849620, EPI_ISL_849621, EPI_ISL_849622 | | | |
| see above | Washington State Department of Health | Seattle Flu Study | Deborah A. Nickerson, Chris D. Frazar, Jover Lee, Benjamin Pelle, Matthew Richardson, Amanda Adler, Elisabeth Brandstetter, Peter D. Han, Kairsten Fay, Misja Ilcisin, Kirsten Lacombe, Thomas R. Sibley, Melissa Truong, Caitlin R. Wolf, Romesh Gautom, Geoff Melly, Brian Hiatt, Philip Dykema, Scott Lindquist, Michael Boeckh, Janet A. Englund, Michael Famulare, Barry R. Lutz, Mark J. Rieder, Lea M. Starita, Matthew Thompson, Helen Y. Chu, Jay Shendure, Trevor Bedford |
| EPI_ISL_849952 | UC Davis- Department of Pathology and Laboratory Medicine | Chan-Zuckerberg Biohub | CZB Cliahub Consortium |
| EPI_ISL_850039, EPI_ISL_850040, EPI_ISL_850041, EPI_ISL_850042, EPI_ISL_850043, EPI_ISL_850044, EPI_ISL_850045, EPI_ISL_850046, EPI_ISL_850047, EPI_ISL_850048, EPI_ISL_850049, EPI_ISL_850050, EPI_ISL_850051, EPI_ISL_850052, EPI_ISL_850053, EPI_ISL_850054, EPI_ISL_850055, EPI_ISL_850056, | | | |

| | | | |
|---|---|---|---|
| EPI_ISL_850057, EPI_ISL_850058, EPI_ISL_850059, EPI_ISL_850060, EPI_ISL_850061, EPI_ISL_850062, EPI_ISL_850063, EPI_ISL_850064, EPI_ISL_850065, EPI_ISL_850066, EPI_ISL_850067, EPI_ISL_850068, EPI_ISL_850069, EPI_ISL_850070, EPI_ISL_850071, EPI_ISL_850072, EPI_ISL_850073, EPI_ISL_850074, EPI_ISL_850075, EPI_ISL_850076, EPI_ISL_850077, EPI_ISL_850078, EPI_ISL_850079, EPI_ISL_850080, EPI_ISL_850081, EPI_ISL_850082, EPI_ISL_850083, EPI_ISL_850084, EPI_ISL_850085, EPI_ISL_850086, EPI_ISL_850087, EPI_ISL_850088, EPI_ISL_850089, EPI_ISL_850090, EPI_ISL_850091, EPI_ISL_850092, EPI_ISL_850093, EPI_ISL_850094, EPI_ISL_850095, EPI_ISL_850096, EPI_ISL_850097, EPI_ISL_850098, EPI_ISL_850099, EPI_ISL_850100, EPI_ISL_850101, EPI_ISL_850102, EPI_ISL_850103, EPI_ISL_850104, EPI_ISL_850105, EPI_ISL_850106, EPI_ISL_850107, EPI_ISL_850108, EPI_ISL_850109, EPI_ISL_850110, EPI_ISL_850111, EPI_ISL_850112, EPI_ISL_850113, EPI_ISL_850176, EPI_ISL_850177, EPI_ISL_850178 | | | |
| see above | Renegade | Chan-Zuckerberg Biohub | CZB Cliahub Consortium |
| EPI_ISL_852598, EPI_ISL_852601 | Max von Pettenkofer Institute, Virology, National Reference Center for Retroviruses, LMU München | Laboratory for Functional Genome Analysis, Dept. Genomics, Gene Center of the LMU Munich | Max Muenchhoff, Stefan Krebs, Alexander Graf, Oliver Keppler, Helmut Blum |
| EPI_ISL_852633, EPI_ISL_852634, EPI_ISL_852635, EPI_ISL_852636, EPI_ISL_852637, EPI_ISL_852638, EPI_ISL_852639, EPI_ISL_852640, EPI_ISL_852641, EPI_ISL_852642, EPI_ISL_852643, EPI_ISL_852644, EPI_ISL_852645, EPI_ISL_852646, EPI_ISL_852647, EPI_ISL_852648, EPI_ISL_852649 | | | |
| see above | Institute of Virology, Medical Center, University of Freiburg, Freiburg, Germany | Institute of Virology, Clinial Virus Genomics, Medical Center, University of Freiburg, Freiburg, Germany | Jonas Fuchs, Lisa Kern, Sandra Reuter, Hajo Grundmann, Marcus Panning |
| EPI_ISL_852843, EPI_ISL_852844, EPI_ISL_852845, EPI_ISL_852846, EPI_ISL_852847, EPI_ISL_852848, EPI_ISL_852849, EPI_ISL_852850, EPI_ISL_852851, EPI_ISL_852852, EPI_ISL_852853, EPI_ISL_852854, EPI_ISL_852855, EPI_ISL_852856, EPI_ISL_852857, EPI_ISL_852862, EPI_ISL_852863, EPI_ISL_852864, EPI_ISL_852865, EPI_ISL_852866, EPI_ISL_852867, EPI_ISL_852868, EPI_ISL_852869, EPI_ISL_852870, EPI_ISL_852871, EPI_ISL_852872, EPI_ISL_852873, EPI_ISL_852874, EPI_ISL_852875, EPI_ISL_852876, EPI_ISL_852877, EPI_ISL_852878, EPI_ISL_852879, EPI_ISL_852880, EPI_ISL_852881, EPI_ISL_852882, EPI_ISL_852883, EPI_ISL_852884, EPI_ISL_852885, EPI_ISL_852886, EPI_ISL_852887, EPI_ISL_852888, EPI_ISL_852889, EPI_ISL_852890, EPI_ISL_852891, EPI_ISL_852892, EPI_ISL_852893, EPI_ISL_852894, EPI_ISL_852895, EPI_ISL_852896, EPI_ISL_852897, EPI_ISL_852898, EPI_ISL_852899, EPI_ISL_852900, EPI_ISL_852901, EPI_ISL_852902, EPI_ISL_852903, EPI_ISL_852904, EPI_ISL_852905, EPI_ISL_852906, EPI_ISL_852907, EPI_ISL_852908, EPI_ISL_852909, EPI_ISL_852910, EPI_ISL_852911, EPI_ISL_852912, EPI_ISL_852913, EPI_ISL_852914, EPI_ISL_852915, EPI_ISL_852916, EPI_ISL_852917, EPI_ISL_852918, EPI_ISL_852919, EPI_ISL_852920, EPI_ISL_852921, EPI_ISL_852922, EPI_ISL_852923, EPI_ISL_852924, EPI_ISL_852925, EPI_ISL_852926, EPI_ISL_852927, EPI_ISL_852928, EPI_ISL_852929, EPI_ISL_852930, EPI_ISL_852931, EPI_ISL_852932, EPI_ISL_852933, EPI_ISL_852934, EPI_ISL_852935, EPI_ISL_852936, EPI_ISL_852937, EPI_ISL_852938, EPI_ISL_852939, EPI_ISL_852940, EPI_ISL_852941, EPI_ISL_852942, EPI_ISL_852943, EPI_ISL_852944, EPI_ISL_852945, EPI_ISL_852946, EPI_ISL_853033, EPI_ISL_853034, EPI_ISL_853035, EPI_ISL_853036, EPI_ISL_853037, EPI_ISL_853038, EPI_ISL_853039, EPI_ISL_853040, EPI_ISL_853041, EPI_ISL_853042, EPI_ISL_853043, EPI_ISL_853044, EPI_ISL_853045, EPI_ISL_853046, EPI_ISL_853047, EPI_ISL_853048, EPI_ISL_853049, EPI_ISL_853050, EPI_ISL_853051, EPI_ISL_853052, EPI_ISL_853053, EPI_ISL_853054, EPI_ISL_853055, EPI_ISL_853056, EPI_ISL_853057, EPI_ISL_853058, EPI_ISL_853059, EPI_ISL_853060, EPI_ISL_853061, EPI_ISL_853062, EPI_ISL_853063, EPI_ISL_853064, EPI_ISL_853065, EPI_ISL_853066, EPI_ISL_853067, EPI_ISL_853068, EPI_ISL_853069, EPI_ISL_853070, EPI_ISL_853071, EPI_ISL_853072, EPI_ISL_853073, EPI_ISL_853074, EPI_ISL_853075, EPI_ISL_853076, EPI_ISL_853077, EPI_ISL_853078, EPI_ISL_853079, EPI_ISL_853080, EPI_ISL_853081, EPI_ISL_853082, EPI_ISL_853083, EPI_ISL_853084, EPI_ISL_853085, EPI_ISL_853086, EPI_ISL_853087, EPI_ISL_853088, EPI_ISL_853089, EPI_ISL_853090, EPI_ISL_853091, EPI_ISL_853092, EPI_ISL_853093, EPI_ISL_853094, EPI_ISL_853095, EPI_ISL_853096, EPI_ISL_853097, EPI_ISL_853098, EPI_ISL_853099, EPI_ISL_853100, EPI_ISL_853101, EPI_ISL_853102, EPI_ISL_853103, EPI_ISL_853104, EPI_ISL_853105, EPI_ISL_853106, EPI_ISL_853107, EPI_ISL_853108, EPI_ISL_853109, EPI_ISL_853110, EPI_ISL_853111, EPI_ISL_853112, EPI_ISL_853113, EPI_ISL_853114, EPI_ISL_853115, EPI_ISL_853116, EPI_ISL_853117, EPI_ISL_853118, EPI_ISL_853119, EPI_ISL_853120, EPI_ISL_853121, EPI_ISL_853122, EPI_ISL_853123, EPI_ISL_853124, EPI_ISL_853125, EPI_ISL_853126, EPI_ISL_853127, EPI_ISL_853128, EPI_ISL_853129, EPI_ISL_853130, EPI_ISL_853131, EPI_ISL_853132, EPI_ISL_853133, EPI_ISL_853134, EPI_ISL_853135, EPI_ISL_853136, EPI_ISL_853137, EPI_ISL_853138, EPI_ISL_853139, EPI_ISL_853140, EPI_ISL_853141, EPI_ISL_853142, EPI_ISL_853143, EPI_ISL_853144, EPI_ISL_853145, EPI_ISL_853146, EPI_ISL_853147, EPI_ISL_853148, EPI_ISL_853149, EPI_ISL_853150, EPI_ISL_853151, EPI_ISL_853152, EPI_ISL_853153, EPI_ISL_853154, EPI_ISL_853155, EPI_ISL_853156, EPI_ISL_853157, EPI_ISL_853158, EPI_ISL_853159, EPI_ISL_853160, EPI_ISL_853161, EPI_ISL_853162, EPI_ISL_853163, EPI_ISL_853164, EPI_ISL_853165, EPI_ISL_853166, EPI_ISL_853167, EPI_ISL_853168, EPI_ISL_853169, EPI_ISL_853170, EPI_ISL_853171, EPI_ISL_853172, EPI_ISL_853173, EPI_ISL_853174, EPI_ISL_853175, EPI_ISL_853176, EPI_ISL_853177, EPI_ISL_853178, EPI_ISL_853179, EPI_ISL_853180, EPI_ISL_853181, EPI_ISL_853182, EPI_ISL_853183, EPI_ISL_853184, EPI_ISL_853185, EPI_ISL_853186, EPI_ISL_853187, EPI_ISL_853188, EPI_ISL_853189, EPI_ISL_853190, EPI_ISL_853191, EPI_ISL_853192, EPI_ISL_853193, EPI_ISL_853194, EPI_ISL_853195, EPI_ISL_853196, EPI_ISL_853197, EPI_ISL_853198, EPI_ISL_853199, EPI_ISL_853200, EPI_ISL_853201, EPI_ISL_853202, EPI_ISL_853203, EPI_ISL_853204, EPI_ISL_853205, EPI_ISL_853206, EPI_ISL_853207, EPI_ISL_853208, EPI_ISL_853209, EPI_ISL_853210, EPI_ISL_853211, EPI_ISL_853212, EPI_ISL_853213, EPI_ISL_853214, EPI_ISL_853215, EPI_ISL_853216, EPI_ISL_853217, EPI_ISL_853218, EPI_ISL_853219, EPI_ISL_853220, EPI_ISL_853221, EPI_ISL_853222, EPI_ISL_853223, EPI_ISL_853224, EPI_ISL_853225, EPI_ISL_853226, EPI_ISL_853227, EPI_ISL_853228, EPI_ISL_853229, EPI_ISL_853230, EPI_ISL_853231, EPI_ISL_853232, EPI_ISL_853233, EPI_ISL_853234, EPI_ISL_853235, EPI_ISL_853236, EPI_ISL_853237, EPI_ISL_853238, EPI_ISL_853239, EPI_ISL_853240, EPI_ISL_853241, EPI_ISL_853242, EPI_ISL_853243, EPI_ISL_853244, EPI_ISL_853245, EPI_ISL_853246, EPI_ISL_853247, EPI_ISL_853248, EPI_ISL_853249, EPI_ISL_853250, EPI_ISL_853251, EPI_ISL_853252, EPI_ISL_853253, EPI_ISL_853254, EPI_ISL_853255, EPI_ISL_853256, EPI_ISL_853257, EPI_ISL_853258, EPI_ISL_853259, EPI_ISL_853260, EPI_ISL_853261, EPI_ISL_853262, EPI_ISL_853263, EPI_ISL_853264, EPI_ISL_853265, EPI_ISL_853266, EPI_ISL_853267, EPI_ISL_853268, EPI_ISL_853269, EPI_ISL_853270, EPI_ISL_853271, EPI_ISL_853272, EPI_ISL_853273 | | | |
| see above | Toronto Invasive Bacterial Diseases Network | Ontario Institute for Cancer Research | Allison McGeer, Patryk Aftanas, Angel Li, Kuganya Nirmalarajah, Samira Mubareka, Ilinca Lungu, Lubaina Kothari, Bernard Lam, Paul Krzyzanowski, Michael Laszloffy, Lawrence E. Heisler, Richard de Borja, Jared T. Simpson |
| EPI_ISL_853327, EPI_ISL_853328, EPI_ISL_853335 | UPMC Clinical Microbiology Laboratory | Microbial Genome Sequencing Center; Microbial Genomic Epidemiology Laboratory | Mustapha M. Mustapha, Jane W. Marsh, Dan Snyder, Marissa P. Griffith, Stephanie L. Mitchell, Vatsala R. Srinivasa, Kady D. Waggle, Chinelo Ezeonwuku, Vaughn S. Cooper, Lee H. Harrison |
| EPI_ISL_853566, EPI_ISL_853567, EPI_ISL_853568, EPI_ISL_853569, EPI_ISL_853570, EPI_ISL_853571, EPI_ISL_853572, EPI_ISL_853573, EPI_ISL_853574, EPI_ISL_853575, EPI_ISL_853576, EPI_ISL_853577, EPI_ISL_853578, EPI_ISL_853579, EPI_ISL_853580, EPI_ISL_853581, EPI_ISL_853582, EPI_ISL_853583, EPI_ISL_853584, EPI_ISL_853585, EPI_ISL_853586, EPI_ISL_853587, EPI_ISL_853588, EPI_ISL_853589, EPI_ISL_853590, EPI_ISL_853591, EPI_ISL_853592, EPI_ISL_853593, EPI_ISL_853594, EPI_ISL_853595, EPI_ISL_853596, EPI_ISL_853597, EPI_ISL_853598, EPI_ISL_853599, EPI_ISL_853600, EPI_ISL_853601, EPI_ISL_853602, EPI_ISL_853603, EPI_ISL_853604, EPI_ISL_853605, EPI_ISL_853606, EPI_ISL_853607, EPI_ISL_853608, EPI_ISL_853609, EPI_ISL_853610, EPI_ISL_853611, EPI_ISL_853612, EPI_ISL_853613, EPI_ISL_853614, EPI_ISL_853615, EPI_ISL_853616, EPI_ISL_853617, EPI_ISL_853618, EPI_ISL_853619, EPI_ISL_853620, EPI_ISL_853621, EPI_ISL_853622, EPI_ISL_853623, EPI_ISL_853624, EPI_ISL_853625, EPI_ISL_853626, EPI_ISL_853627, EPI_ISL_853628, EPI_ISL_853629, EPI_ISL_853630, EPI_ISL_853631, EPI_ISL_853632, EPI_ISL_853633, EPI_ISL_853634, EPI_ISL_853635, EPI_ISL_853636, EPI_ISL_853637, EPI_ISL_853638, EPI_ISL_853639, EPI_ISL_853640, EPI_ISL_853641, EPI_ISL_853642, EPI_ISL_853643, EPI_ISL_853644, EPI_ISL_853645, EPI_ISL_853646, EPI_ISL_853647 | | | |
| see above | Toronto Invasive Bacterial Diseases Network | Ontario Institute for Cancer Research | Allison McGeer, Patryk Aftanas, Angel Li, Kuganya Nirmalarajah, Samira Mubareka, Ilinca Lungu, Lubaina Kothari, Bernard Lam, Paul Krzyzanowski, Michael Laszloffy, Lawrence E. Heisler, Richard de Borja, Jared T. Simpson |
| EPI_ISL_853723 | Institute for Water Quality and Resource Management, Technical University Vienna | Bergthaler laboratory, CeMM Research Center for Molecular Medicine of the Austrian Academy of Sciences | Lukas Endler, Alexandra Popa, Benedikt Agerer, Jakob-Wendelin Genger, Alexander Lercher, Anna Schedl, Thomas Penz, Michael Schuster, Jan Laine, Martin Senekowitsch, Christoph Bock, Andreas Bergthaler |
| EPI_ISL_853738, EPI_ISL_853749, EPI_ISL_853791, EPI_ISL_853793, EPI_ISL_853794, EPI_ISL_853797, EPI_ISL_853815 | Austrian Agency for Health and Food Safety (AGES) | Bergthaler laboratory, CeMM Research Center for Molecular Medicine of the Austrian Academy of Sciences | Lukas Endler, Alexandra Popa, Benedikt Agerer, Jakob-Wendelin Genger, Alexander Lercher, Anna Schedl, Thomas Penz, Michael Schuster, Jan Laine, Martin Senekowitsch, Christoph Bock, Andreas Bergthaler |
| EPI_ISL_853953 | Department of Microbiology, University Innsbruck | Bergthaler laboratory, CeMM Research Center for Molecular Medicine of the Austrian Academy of Sciences | Lukas Endler, Alexandra Popa, Benedikt Agerer, Jakob-Wendelin Genger, Alexander Lercher, Anna Schedl, Thomas Penz, Michael Schuster, Jan Laine, Martin Senekowitsch, Christoph Bock, Andreas Bergthaler |
| EPI_ISL_853968, EPI_ISL_853969, EPI_ISL_854122, EPI_ISL_854123, EPI_ISL_854124, EPI_ISL_854125, EPI_ISL_854126, EPI_ISL_854127, EPI_ISL_854128, EPI_ISL_854129, EPI_ISL_854130, EPI_ISL_854131, EPI_ISL_854132, EPI_ISL_854133, EPI_ISL_854134, EPI_ISL_854135, EPI_ISL_854136, EPI_ISL_854137, EPI_ISL_854138, EPI_ISL_854139, EPI_ISL_854140, EPI_ISL_854141, EPI_ISL_854142, EPI_ISL_854143, EPI_ISL_854144, EPI_ISL_854145, EPI_ISL_854146, EPI_ISL_854147, EPI_ISL_854209, EPI_ISL_854215, EPI_ISL_854244, EPI_ISL_854245, EPI_ISL_854246, EPI_ISL_854247, EPI_ISL_854248, EPI_ISL_854263, EPI_ISL_854264, EPI_ISL_854286, EPI_ISL_854287, EPI_ISL_854288, EPI_ISL_854289, EPI_ISL_854290, EPI_ISL_854291, EPI_ISL_854292, EPI_ISL_854304 | | | |
| see above | Austrian Agency for Health and Food Safety (AGES) | Bergthaler laboratory, CeMM Research Center for Molecular Medicine of the Austrian Academy of Sciences | Lukas Endler, Alexandra Popa, Benedikt Agerer, Jakob-Wendelin Genger, Alexander Lercher, Anna Schedl, Thomas Penz, Michael Schuster, Jan Laine, Martin Senekowitsch, Christoph Bock, Andreas Bergthaler |
| EPI_ISL_854441, EPI_ISL_854442 | SARATOGA HOSPITAL LABORATORY | Wadsworth Center, New York State Department of Health | Kirsten St. George, Daryl M. Lamson, Alexis Russel, Matthew Shudt, Melissa A Leisner, Jonathan Plitnick, Navjot Singh, John Kelly, Erasmus Schneider, Erica Lasek-Nesselquist |
| EPI_ISL_854763, EPI_ISL_854764, EPI_ISL_854765, EPI_ISL_854766, EPI_ISL_854767, EPI_ISL_854768, EPI_ISL_854769, EPI_ISL_854770, EPI_ISL_854771, EPI_ISL_854772, EPI_ISL_854773, EPI_ISL_854774, EPI_ISL_854775, EPI_ISL_854776, EPI_ISL_854777, EPI_ISL_854778, EPI_ISL_854779, EPI_ISL_854780, EPI_ISL_854781, EPI_ISL_854782, EPI_ISL_854783, EPI_ISL_854784, EPI_ISL_854785, EPI_ISL_854786, EPI_ISL_854787, EPI_ISL_854788, EPI_ISL_854789, EPI_ISL_854790, EPI_ISL_854791, EPI_ISL_854792, EPI_ISL_854793, EPI_ISL_854794, EPI_ISL_854795, EPI_ISL_854796, EPI_ISL_854797, EPI_ISL_854798, EPI_ISL_854799, EPI_ISL_854800, EPI_ISL_854801, EPI_ISL_854802, EPI_ISL_854803, EPI_ISL_854804, EPI_ISL_854805, EPI_ISL_854806, EPI_ISL_854807, EPI_ISL_854808, EPI_ISL_854809 | | | |
| see above | Ontario's COVID-19 Genomics Rapid Response Coalition | McMaster University | Ana Cabrera, Allison McGeer, Patryk Aftanas, Hooman Derakhshani, Angel Li, Kuganya Nirmalarajah, Emily Panousis, Ahmed Draia, Jalees Nasir, Michael Surette, Samira Mubareka, Andrew G. McArthur |
| EPI_ISL_855494 | KEMRI-Wellcome Trust Research Programme/KEMRI-CGMR-C Kilifi | KEMRI-Wellcome Trust Research Programme/KEMRI-CGMR-C Kilifi | Githinji et al |
| EPI_ISL_856764, EPI_ISL_856765, EPI_ISL_856766, EPI_ISL_856767, EPI_ISL_856768, EPI_ISL_856769, EPI_ISL_856770, EPI_ISL_856771 | Servicio Virosis Respiratorias-Departamento Virología-INEI | Instituto Nacional Enfermedades Infecciosas C.G.Malbran | Baumeister E., Avaro M., Benedetti E., Russo M., Dattero ME, Pontoriero A., Cisterna D., Molina V., Perandones C., Tuduri E., Lorenzo F., Poklepovich T., Campos J. |
| EPI_ISL_857048 | Colorado Department of Public Health and Environment | Colorado Department of Puplic Health and Environment | Laura Bankers, Molly C. Hetherington-Rauth, Diana Ir, Shannon Ely, Shannon R. Matzinger, Sarah Elizabeth Totten, Emily A. Travanty |
| EPI_ISL_857498, EPI_ISL_857499 | Swiss National Reference Centre for Influenza | Swiss National Reference Centre for Influenza | Ana Rita Goncalves,Samuel Cordey, Laurent Kaiser, Lorenzo Cerutti, Henri Pegeot, Melyssa Elies, Keith Harshman, Ioannis Xenarios, Emmanouil Dermitzakis |
| EPI_ISL_857524, EPI_ISL_857525, EPI_ISL_857526 | Swiss National Reference Centre for Influenza | Swiss National Reference Centre for Influenza | Tim Roloff, Ana Rita Gonçalves, Madlen Stange, Helena MB Seth-Smith, Alfredo Mari, Karoline Leuzinger, Julia Bielicki, Manuel Battegay, Hans Hirsch, Laurent Kaiser, Adrian Egli |
| EPI_ISL_859882, EPI_ISL_859883, EPI_ISL_859884, EPI_ISL_859885, EPI_ISL_859886, EPI_ISL_859895, EPI_ISL_859896, EPI_ISL_859897, EPI_ISL_859898, EPI_ISL_859899, EPI_ISL_859900, EPI_ISL_859901, EPI_ISL_859916, EPI_ISL_859917, EPI_ISL_859918, EPI_ISL_859919, EPI_ISL_859929, EPI_ISL_859930 | | | |
| see above | BTC, Khalifa University | BTC, Khalifa University | Al Safar et al |
| EPI_ISL_860117, EPI_ISL_860118, EPI_ISL_860154, EPI_ISL_860162, EPI_ISL_860163, EPI_ISL_860165, EPI_ISL_860166, EPI_ISL_860167, EPI_ISL_860168, EPI_ISL_860169, EPI_ISL_860170, EPI_ISL_860171, EPI_ISL_860172, EPI_ISL_860173 | | | |

| | | | |
|---|---|---|---|
| see above | Keio University School of Medicine | Keio University School of Medicine | Kenjiro Kosaki, Yuka Iwasaki, Hirotsugu Ishizu, Haruhiko Siomi, Kodai Abe |
| EPI_ISL_861462, EPI_ISL_861464, EPI_ISL_861467 | UHAS COVID-19 Lab | UHAS COVID-19 Lab | Kwabena O. Duedu, Jones Gyamfi, Reuben Ayivor-Djanie, John O. Gyapong and the UHAS COVID-19 Lab Team |
| EPI_ISL_861668 | Centro de Triagem Covid19 | Instituto Adolfo Lutz, Interdiciplinary Procedures Center, Strategic Laboratory | Claudio Tavares Sacchi, Claudia Regina Gonçalves, Erica Valessa Ramos Gomes, Karoline Rodrigues Campos |
| EPI_ISL_861882, EPI_ISL_861886 | LATE - Laboratório de Técnicas Especiais - Hospital Israelita Albert Einstein | LATE - Laboratório de Técnicas Especiais - Hospital Israelita Albert Einstein | Deyvid Amgarten, Fernanda de Mello Malta, Raquel Riyuzo, Ana Paula Moreira Salles, Pedro Henrique Sebe Rodrigues, João Renato Rebello Pinho |
| EPI_ISL_861925, EPI_ISL_861926, EPI_ISL_861927, EPI_ISL_861928, EPI_ISL_861929, EPI_ISL_861930, EPI_ISL_861931, EPI_ISL_861932, EPI_ISL_861933, EPI_ISL_861934, EPI_ISL_861935, EPI_ISL_861936, EPI_ISL_861937, EPI_ISL_861938, EPI_ISL_861939, EPI_ISL_861940, EPI_ISL_861941, EPI_ISL_861942, EPI_ISL_861943, EPI_ISL_861944, EPI_ISL_861945, EPI_ISL_861946, EPI_ISL_861947, EPI_ISL_861948, EPI_ISL_861949, EPI_ISL_861950, EPI_ISL_861951, EPI_ISL_861952, EPI_ISL_861953, EPI_ISL_861954 | | | |
| see above | OHSU Lab Services Molecular Microbiology Lab | Oregon SARS-CoV-2 Genome Sequencing Center | Brendan L. O'Connell, Sally Grindstaff, Kayla Carter, Ruth V. Nichols, Alec J. Hirsch, Donna Hansel, Guang Fan, Xuan Qin, Daniel N. Streblow, William B. Messer, Andrew C. Adey, Benjamin N. Bimber, Brian J. O'Roak |
| EPI_ISL_862076 | National Influenza Center, Virology Department | National Influenza Center | NZ Shafiei Jandaghi, V Salimi, A Nejati, K Sadeghi, J Yavarian, N Ghavvami,F Ajaminejad and T Mokhtari Azad |
| EPI_ISL_862080 | National Influenza Center, Virology Department | National Influenza Center | J Yavarian, NZ Shafiei Jandaghi, V Salimi, A Nejati, K Sadeghi, F Ajaminejad,N Ghavvami and T Mokhtari Azad |
| EPI_ISL_862081 | National Influenza Center, Virology Department | National Influenza Center | NZ Shafiei Jandaghi, V Salimi, A Nejati, K Sadeghi, J Yavarian, F Ajaminejad,N Ghavvami and T Mokhtari Azad |
| EPI_ISL_864563 | Institute of Medical Microbiology and Hospital Hygiene | Institute of Medical Microbiology and Hospital Hygiene | Prof. Dr. Achim Kaasch, Aljoscha Tersteegen |
| EPI_ISL_865174 | Liverpool Clinical Laboratories | COVID-19 Genomics UK (COG-UK) Consortium | Sam Haldenby, Anita Lucaci, Steve Paterson, Julian Hiscox, Alistair Darby, M Almsaud, A Alrezaihi, Muhannad Alruwaili, Stuart D Armstrong, Jones Benjamin, Eleanor G Bentley, Anu Chawla, Jordan J Clark, Angela Cowell, Richard Eccles, Isabel García-Dorival, Matthew Gemmell, Alessandro Gerada, PKF Gilmore, Richard Gregory, Ximeng Han, Catherine Hartley, Margaret Hughes, Miren Iturriza-Gomara, James Johnson, L Luu, Jenifer Manson, Charlotte Nelson, Elaine O'Toole, Cassie Olateju, Rebekah Penrice-Randal , Lucille Rainbow, N.P Randle, Trevor Ian Robinson, Parul Sharma, Ghada T Shawli, James P Stewart, Neil Swainston, Ecaterina Vamos, Joanne Watts, Mark Whitehead |
| EPI_ISL_865702, EPI_ISL_865703, EPI_ISL_865704, EPI_ISL_865705, EPI_ISL_865706, EPI_ISL_865707 | University College London, Great Ormond Street Hospital for Children NHS Foundation Trust, Imperial College Healthcare NHS Trust | COVID-19 Genomics UK (COG-UK) Consortium | Sergi Castellano, Rachel Williams, Mark Kristiansen, Paola Resende Silva, Sunando Roy, Tony Brooks, Helena Tutill, Paola Niola, Patricia Dyal, Charlotte Williams, Leysa Forrest, Yasmin Panchbhaya, Jacqueline Findlay, Samuel Weeks, Julianne Brown, Kathryn Harris, Paul Randell, James Price, Alison Holmes, Judith Breuer |
| EPI_ISL_866055, EPI_ISL_866056, EPI_ISL_866057 | University College London Hospital | COVID-19 Genomics UK (COG-UK) Consortium | Judith Heaney, Matthew Byott, Catherine Houlihan, Dan Frampton, Stuart Kirk, Moira Spyer and Eleni Nastouli |
| EPI_ISL_866822 | Quadram Institute Bioscience | COVID-19 Genomics UK (COG-UK) Consortium | Dave J. Baker, Gemma L. Kay, Alp Aydin, Thanh Le-Viet, Steven Rudder, Ana P. Tedim, Anastasia Kolyva, Maria Diaz, Leonardo de Oliveira Martins, Nabil-Fareed Alikhan, Lizzie Meadows, Rachael Stanley, Ngozi Elumogo, Muhammed Yasir, Nicholas M. Thomson, Alexander J Trotter, Rachel Gilroy, Samuel Bloomfield, Claire Stuart, Andrew Bell, Reenesh Prakash, Samir Dervisevic, Alison E. Mather, John Wain, Mark Webber, Andrew J. Page, Justin O'Grady |
| EPI_ISL_867271 | Originating lab: Wales Specialist Virology Centre Sequencing lab: Pathogen Genomics Unit | Public Health Wales Microbiology Cardiff Wales Specialist Virology Centre | Catherine Moore, Johnathan Evans, Laura Gifford, Malorie Perry, Simon Cottrell, Angela Marchbank, Alec Birchley, Alexander Adams, Amy Gaskin, Bree Gatica-Wilcox, Jason Coombes, Joel Southgate, Lauren Gilbert, Lee Graham, Nicole Pacchiarini, Sara Kumziene-Summerhayes, Sarah Taylor, Sophie Jones, Sara Rey, Matthew Bull, Joanne Watkins, Sally Corden, Tom Connor |
| EPI_ISL_867950, EPI_ISL_867951 | Centre for Enzyme Innovation, University of Portsmouth / Translational Research Laboratory, Portsmouth Hospitals NHS Trust | COVID-19 Genomics UK (COG-UK) Consortium | Angela Beckett,Yann Bourgeois,Garry Scarlett,Sharon Glaysher,Scott Elliott,Kelly Bicknell,Robert Impey,Allyson Lloyd,Sarah Wyllie,Ethan Butcher,Anoop Chauhan,Samuel Robson |
| EPI_ISL_868878, EPI_ISL_868879, EPI_ISL_868880, EPI_ISL_868881, EPI_ISL_868882, EPI_ISL_868883, EPI_ISL_868884, EPI_ISL_868885, EPI_ISL_868886, EPI_ISL_868887, EPI_ISL_868888, EPI_ISL_868890, EPI_ISL_868891, EPI_ISL_868892, EPI_ISL_868893, EPI_ISL_868894, EPI_ISL_868895, EPI_ISL_868896, EPI_ISL_868897, EPI_ISL_868898, EPI_ISL_868899, EPI_ISL_868900, EPI_ISL_868981, EPI_ISL_868982, EPI_ISL_868983, EPI_ISL_868984, EPI_ISL_868985, EPI_ISL_868986, EPI_ISL_868987, EPI_ISL_868988, EPI_ISL_868989, EPI_ISL_868990, EPI_ISL_868991, EPI_ISL_868992, EPI_ISL_868993, EPI_ISL_868994, EPI_ISL_868995, EPI_ISL_868996, EPI_ISL_868997, EPI_ISL_868998, EPI_ISL_868999, EPI_ISL_869000, EPI_ISL_869001, EPI_ISL_869002, EPI_ISL_869003, EPI_ISL_869004, EPI_ISL_869005, EPI_ISL_869006, EPI_ISL_869007, EPI_ISL_869008, EPI_ISL_869009, EPI_ISL_869010, EPI_ISL_869011, EPI_ISL_869012, EPI_ISL_869013 | | | |
| see above | Bioinformatics and Biostatistics Lab, Advanced Sequencing Facility | COVID-19 Genomics UK (COG-UK) Consortium | Aengus Stewart,Jerome Nicod,Chelsea Sawyer,Laura Cubitt,Harshil Patel,Margaret Crawford |
| EPI_ISL_869247 | Krasnoyarsk Regional Clinical Hospital | WHO National Influenza Centre Russian Federation | Andrey Komissarov, Artem Fadeev, Anna Ivanova, Kseniya Komissarova, Dmitry Bazhenov, Mikhail Bakaev, Daria Danilenko, Ksenia Safina, Elena Nabieva, Georgii Bazykin, Dmitry Lioznov |
| EPI_ISL_872608, EPI_ISL_872609 | Nigeria Centre for Disease Control (NCDC) | African Centre of Excellence for Genomics of Infectious Diseases (ACEGID), Redeemer's University | Oluniyi P.E. et al |
| EPI_ISL_872699, EPI_ISL_872700, EPI_ISL_872701, EPI_ISL_872702, EPI_ISL_872703 | Rhode Island Department of Health | Infectious Disease Program, Broad Institute of Harvard and MIT | Lemieux,J.E., Siddle,K.J., Huard,R., King,E., Azevedo,K., Miller,A., Adams,G., Gladden-Young,A., Lagerborg,K., Rudy,M., DeRuff,K., Carter,A., Normandin,E., Bauer,M., Reilly,S., Tomkins-Tinch,C., Loreth,C., Chaluvadi,S., Birren,B.W., Gallagher,G., Smole,S., Park,D.J., MacInnis,B.L., and Sabeti,P.C. |
| EPI_ISL_875554 | ULSS 2 Treviso | Istituto Zooprofilattico Sperimentale delle Venezie | Adelaide Milani, Alessia Schivo, Annalisa Salviato, Erika Giorgia Quaranta, Ambra Pastori, Bianca Zecchin, Alice Fusaro, Isabella Monne, Calogero Terregino, Antonia Ricci |
| EPI_ISL_876049, EPI_ISL_876051, EPI_ISL_876055, EPI_ISL_876079, EPI_ISL_876086, EPI_ISL_876092, EPI_ISL_876098, EPI_ISL_876102, EPI_ISL_876107, EPI_ISL_876120, EPI_ISL_876121, EPI_ISL_876123, EPI_ISL_876188, EPI_ISL_876195, EPI_ISL_876318, EPI_ISL_876319, EPI_ISL_876320, EPI_ISL_876321 | | | |
| see above | Massachusetts State Public Health Laboratory | Massachusetts State Public Health Laboratory | Andrew Lang, Timelia Fink, Glen Gallagher, Sandra Smole |
| EPI_ISL_876521, EPI_ISL_876522, EPI_ISL_876526, EPI_ISL_876532, EPI_ISL_876552, EPI_ISL_876553, EPI_ISL_876554 | Florida Bureau of Public Health Laboratories | Florida Bureau of Public Health Laboratories | Sarah Schmedes, Jason Blanton |
| EPI_ISL_877461, EPI_ISL_877462, EPI_ISL_877463, EPI_ISL_877464, EPI_ISL_877465, EPI_ISL_877466, EPI_ISL_877467, EPI_ISL_877468, EPI_ISL_877469, EPI_ISL_877470, EPI_ISL_877471, EPI_ISL_877472, EPI_ISL_877473, EPI_ISL_877474, EPI_ISL_877475, EPI_ISL_877476, EPI_ISL_877477, EPI_ISL_877478, EPI_ISL_877479, EPI_ISL_877480, EPI_ISL_877481, EPI_ISL_877482, EPI_ISL_877483, EPI_ISL_877484, EPI_ISL_877485, EPI_ISL_877486, EPI_ISL_877487, EPI_ISL_877488, EPI_ISL_877489, EPI_ISL_877490, EPI_ISL_877491, EPI_ISL_877492, EPI_ISL_877493, EPI_ISL_877494, EPI_ISL_877495, EPI_ISL_877496, EPI_ISL_877497, EPI_ISL_877498, EPI_ISL_877499, EPI_ISL_877500, EPI_ISL_877501, EPI_ISL_877502, EPI_ISL_877503, EPI_ISL_877504, EPI_ISL_877505, EPI_ISL_877506, EPI_ISL_877507, EPI_ISL_877508, EPI_ISL_877509, EPI_ISL_877510, EPI_ISL_877511, EPI_ISL_877512, EPI_ISL_877513, EPI_ISL_877514, EPI_ISL_877515, EPI_ISL_877516, EPI_ISL_877517, EPI_ISL_877518, EPI_ISL_877519, EPI_ISL_877520, EPI_ISL_877521, EPI_ISL_877522, EPI_ISL_877523, EPI_ISL_877524, EPI_ISL_877525, EPI_ISL_877526, EPI_ISL_877527, EPI_ISL_877528, EPI_ISL_877529, EPI_ISL_877530, EPI_ISL_877531, EPI_ISL_877532, EPI_ISL_877533, EPI_ISL_877534, EPI_ISL_877535, EPI_ISL_877536, EPI_ISL_877537, EPI_ISL_877539, EPI_ISL_877540, EPI_ISL_877541, EPI_ISL_877542, EPI_ISL_877543, EPI_ISL_877544, EPI_ISL_877545, EPI_ISL_877546, EPI_ISL_877547, EPI_ISL_877548, EPI_ISL_877549, EPI_ISL_877550 | | | |
| see above | Eurofins Diatherix | Hudsonalpha Genome Sequencing Center | Jane Grimwood, Melissa Williams, Lori H. Handley, Joshua Stough, Leslie Malone, Stefan Brzezinski, Ada Stewart, Teresa Jones, Jenell Webber, John Lovell, Jennifer Cart, and Jeremy Schmutz |
| EPI_ISL_878281, EPI_ISL_878284, EPI_ISL_878287, EPI_ISL_878290, EPI_ISL_878292, EPI_ISL_878295, EPI_ISL_878298, EPI_ISL_878305, EPI_ISL_878311, EPI_ISL_878314, EPI_ISL_878316, EPI_ISL_878325, EPI_ISL_878327, EPI_ISL_878333, EPI_ISL_878338, EPI_ISL_878341, EPI_ISL_878342, EPI_ISL_878343, EPI_ISL_878344, EPI_ISL_878347, EPI_ISL_878348, EPI_ISL_878349, EPI_ISL_878350, EPI_ISL_878352, EPI_ISL_878353, EPI_ISL_878354, EPI_ISL_878356, EPI_ISL_878357, EPI_ISL_878358, EPI_ISL_878359, EPI_ISL_878363, EPI_ISL_878364, EPI_ISL_878366, EPI_ISL_878368, EPI_ISL_878369, EPI_ISL_878370, EPI_ISL_878372, EPI_ISL_878373, EPI_ISL_878375, EPI_ISL_878376, EPI_ISL_878377, EPI_ISL_878378, EPI_ISL_878379, EPI_ISL_878380, EPI_ISL_878383, EPI_ISL_878385, EPI_ISL_878387, EPI_ISL_878388, EPI_ISL_878390, EPI_ISL_878391, EPI_ISL_878394, EPI_ISL_878395, EPI_ISL_878396, EPI_ISL_878397, EPI_ISL_878411, EPI_ISL_878413, EPI_ISL_878421, EPI_ISL_878429 | | | |
| see above | San Diego County Public Health Laboratory | Andersen lab at Scripps Research | SEARCH Alliance San Diego with Tracy Basler, Jovan Shephard, Brett Austin |
| EPI_ISL_878666, EPI_ISL_878713, EPI_ISL_878716, EPI_ISL_878719, EPI_ISL_878729 | Rady's Childrens Hospital | Andersen lab at Scripps Research | SEARCH Alliance San Diego with Nanda Radamchar, David Dimmock, Linda Luo, Christina Clarke, Kathryn Bouic, Teresa Mueller, Denise Malicki |
| EPI_ISL_879924, EPI_ISL_879937, EPI_ISL_879956, EPI_ISL_879961, EPI_ISL_879964, EPI_ISL_879967, EPI_ISL_879972, EPI_ISL_879980, EPI_ISL_879982, EPI_ISL_879985, EPI_ISL_879987, EPI_ISL_879993, EPI_ISL_879998, EPI_ISL_880000, EPI_ISL_880009, EPI_ISL_880011, EPI_ISL_880014, EPI_ISL_880019, EPI_ISL_880027, EPI_ISL_880029, EPI_ISL_880032, EPI_ISL_880034 | | | |

| | | | |
|---|---|---|---|
| see above | San Diego County Public Health Laboratory | Andersen lab at Scripps Research | SEARCH Alliance San Diego with Tracy Basler, Jovan Shephard, Brett Austin |
| EPI_ISL_882659 | Centro de Triagem Covid19 | Instituto Adolfo Lutz, Interdiciplinary Procedures Center, Strategic Laboratory | Claudio Tavares Sacchi, Claudia Regina Gonçalves, Erica Valessa Ramos Gomes, Karoline Rodrigues Campos |
| EPI_ISL_882660 | Hospital Municipal Prefeito Waldemar Costa Filho | Instituto Adolfo Lutz, Interdiciplinary Procedures Center, Strategic Laboratory | Claudio Tavares Sacchi, Claudia Regina Gonçalves, Erica Valessa Ramos Gomes, Karoline Rodrigues Campos |
| EPI_ISL_883066, EPI_ISL_883067, EPI_ISL_883068, EPI_ISL_883069, EPI_ISL_883070, EPI_ISL_883071, EPI_ISL_883072, EPI_ISL_883073, EPI_ISL_883074, EPI_ISL_883075, EPI_ISL_883076, EPI_ISL_883077, EPI_ISL_883078, EPI_ISL_883079, EPI_ISL_883080, EPI_ISL_883081, EPI_ISL_883082, EPI_ISL_883083, EPI_ISL_883084, EPI_ISL_883085, EPI_ISL_883086, EPI_ISL_883087, EPI_ISL_883088, EPI_ISL_883089, EPI_ISL_883090, EPI_ISL_883091, EPI_ISL_883092, EPI_ISL_883093, EPI_ISL_883094, EPI_ISL_883095, EPI_ISL_883096, EPI_ISL_883097, EPI_ISL_883098, EPI_ISL_883099, EPI_ISL_883100, EPI_ISL_883101, EPI_ISL_883102, EPI_ISL_883103, EPI_ISL_883104, EPI_ISL_883105, EPI_ISL_883106, EPI_ISL_883107, EPI_ISL_883108, EPI_ISL_883109, EPI_ISL_883110, EPI_ISL_883111, EPI_ISL_883112, EPI_ISL_883113, EPI_ISL_883114, EPI_ISL_883115, EPI_ISL_883116, EPI_ISL_883117, EPI_ISL_883118, EPI_ISL_883119, EPI_ISL_883120, EPI_ISL_883121, EPI_ISL_883122, EPI_ISL_883123, EPI_ISL_883124, EPI_ISL_883125, EPI_ISL_883126, EPI_ISL_883127, EPI_ISL_883128, EPI_ISL_883129, EPI_ISL_883130, EPI_ISL_883131, EPI_ISL_883132, EPI_ISL_883133, EPI_ISL_883134, EPI_ISL_883135, EPI_ISL_883136, EPI_ISL_883137, EPI_ISL_883138, EPI_ISL_883139, EPI_ISL_883140, EPI_ISL_883141, EPI_ISL_883142, EPI_ISL_883143, EPI_ISL_883144, EPI_ISL_883145, EPI_ISL_883146, EPI_ISL_883147, EPI_ISL_883148, EPI_ISL_883149, EPI_ISL_883150, EPI_ISL_883151, EPI_ISL_883152, EPI_ISL_883153, EPI_ISL_883154, EPI_ISL_883195, EPI_ISL_883197, EPI_ISL_883226, EPI_ISL_883268 | | | |
| see above | Eurofins Diatherix | Hudsonalpha Genome Sequencing Center | Jane Grimwood, Melissa Williams, Lori H. Handley, Joshua Stough, Leslie Malone, Stefan Brzezinski, Ada Stewart, Teresa Jones, Jenell Webber, John Lovell, Jennifer Cart, and Jeremy Schmutz |
| EPI_ISL_884899 | Santa Clara County Public Health Laboratory | Chan-Zuckerberg Biohub | CZB Cliahub Consortium |
| EPI_ISL_887165 | Massachusetts General Hospital | Infectious Disease Program, Broad Institute of Harvard and MIT | Lemieux,J.E., Siddle,K.J., Shaw,B., Adams,G., Pierce,V., Turbett,S., Anahtar,M., Branda,J., Slater,D., Harris,J., Lin,A.E., Gladden-Young,A., Lagerborg,K., Rudy,M., DeRuff,K., Carter,A., Normandin,E., Bauer,M., Reilly,S., Tomkins-Tinch,C., Loreth,C., Chaluvadi,S., Neumann,A., Cusick,C., Chapman,S.B., Gnirke,A., Flowers,K., Cerrato,F., Birren,B.W., Gallagher,G., Smole,S., Park,D.J., MacInnis,B.L., Ryan,E., LaRocque,R., Rosenberg,E. and Sabeti,P.C. |
| EPI_ISL_887435, EPI_ISL_887436, EPI_ISL_887438, EPI_ISL_887455, EPI_ISL_887456, EPI_ISL_887457, EPI_ISL_887458, EPI_ISL_887459, EPI_ISL_887460, EPI_ISL_887464 | Instituto Nacional de Saude (INS), Mozambique | KRISP, KZN Research Innovation and Sequencing Platform | Nalia Ismael, Nadia Sitoe, Paulo Arnaldo, Nedio Mabunda, Giandhari J, Pillay S, Tegally H, Wilkinson E, de Oliveira T |
| EPI_ISL_888734, EPI_ISL_888757, EPI_ISL_888758 | KU Leuven, Rega Institute, Clinical and Epidemiological Virology | KU Leuven, Rega Institute, Clinical and Epidemiological Virology | Tony Wawina-Bokalanga, Bert Vanmechelen, Joan Marti-Carerras, Piet Maes |
| EPI_ISL_888837, EPI_ISL_888838, EPI_ISL_888839, EPI_ISL_888840, EPI_ISL_888841, EPI_ISL_888877, EPI_ISL_888878 | Michigan Department of Health and Human Services, Bureau of Laboratories | Michigan Department of Health and Human Services, Bureau of Laboratories | Blankenship HM, Riner D, Soehnlen MK |
| EPI_ISL_888991 | RS Mitra Keluarga Gading Serpong | Eijkman Institute for Molecular Biology, Ministry of Research and Technology/National Agency for Research and Innovation | Frilasita A Yudhaputri, Hidayat Trimarsanto, Iskandar Adnan, Lydia V. Panggalo, Sukma Oktavianthi, Willy Agustine, Edison Johar, Safarina G Malik, Khin Saw Myint, Amin Soebandrio |
| EPI_ISL_889011 | RSU Permata Cibubur | Eijkman Institute for Molecular Biology, Ministry of Research and Technology/National Agency for Research and Innovation | Hidayat Trimarsanto, Iskandar Adnan, Lydia V. Panggalo, Sukma Oktavianthi, Willy Agustine, Edison Johar, Frilasita A Yudhaputri, Safarina G Malik, Khin Saw Myint, Amin Soebandrio |
| EPI_ISL_889443 | Ohio Department of Health Laboratory | Ohio Department of Health Laboratory | Holmes, Jennifer; Eric Brandt, Keoni Omura, Glen McGillivary, Caitlin McDonnell, Jade Mowery, Stephanie Mccracken, Tyler Payne, Kirtana Ramadugu, Erica Leasure, Brent Lee, Kelsey Florek, Heather Blankenship, Quanta Brown, and Tammy Bannerman |
| EPI_ISL_890303 | KU Leuven, Rega Institute, Clinical and Epidemiological Virology | KU Leuven, Rega Institute, Clinical and Epidemiological Virology | Tony Wawina-Bokalanga, Bert Vanmechelen, Joan Marti-Carerras, Piet Maes |
| EPI_ISL_890896, EPI_ISL_890897, EPI_ISL_890898, EPI_ISL_890899, EPI_ISL_890900, EPI_ISL_890901, EPI_ISL_890902, EPI_ISL_890903, EPI_ISL_890904, EPI_ISL_890905, EPI_ISL_890906, EPI_ISL_890910, EPI_ISL_890911, EPI_ISL_890912, EPI_ISL_890925 | | | |
| see above | Seattle Flu Study | Seattle Flu Study | Deborah A. Nickerson, Chris D. Frazar, Jover Lee, Benjamin Pelle, Erica Ryke, Matthew Richardson, Amanda Adler, Elisabeth Brandstetter, Peter D. Han, Kairsten Fay, Misja Ilcisin, Kirsten Lacombe, Thomas R. Sibley, Melissa Truong, Caitlin R. Wolf, Karen Cowgill, Stephanie Schrag, Jeff Duchin, Michael Boeckh, Janet A. Englund, Michael Famulare, Barry R. Lutz, Mark J. Rieder, Lea M. Starita, Matthew Thompson, Helen Y. Chu, Trevor Bedford, Jay Shendure |
| EPI_ISL_891196 | DPH, Massachusetts State Public Health Lab | DPH, Massachusetts State Public Health Lab | Lang,A.S., Fink,T., Gallagher,G.R., Smole,S.C. |
| EPI_ISL_891228 | University of Medicine and Pharmacy of Craiova | "Stefan cel Mare" University Metagenomics Lab | Lobiuc Andrei, Gheorghita Roxana |
| EPI_ISL_896107 | Viollier AG | University Hospital Basel, Clinical Bacteriology | Tim Roloff, Madlen Stange, Helena MB Seth-Smith, Alfredo Mari, Karoline Leuzinger, Julia Bielicki, Christiane Beckmann, Manuel Battegay, Hans Hirsch, Adrian Egli |
| EPI_ISL_896186 | MEPHI, Aix Marseille University | MEPHI, Aix Marseille University | Anthony LEVASSEUR |
| EPI_ISL_899515 | Viollier AG | Department of Biosystems Science and Engineering, ETH Zürich | Christian Beisel, Sarah Nadeau, Chaoran Chen, Ivan Topolsky, Philipp Jablonski, Lara Fuhrmann, David Dreifuss, Katharina Jahn, Tobias Schär, Ina Nissen, Natascha Santacroce, Elodie Burcklen, Christiane Beckmann, Maurice Redondo, Olivier Kobel, Christoph Noppen, Sophie Seidel, Noemie Santamaria de Souza, Niko Beerenwinkel, Tanja Stadler |
| EPI_ISL_902768, EPI_ISL_902769, EPI_ISL_902770, EPI_ISL_902771, EPI_ISL_902772, EPI_ISL_902773, EPI_ISL_902774, EPI_ISL_902775, EPI_ISL_902776, EPI_ISL_902777, EPI_ISL_902778, EPI_ISL_902779, EPI_ISL_902780, EPI_ISL_902781, EPI_ISL_902782, EPI_ISL_902783, EPI_ISL_902784, EPI_ISL_902785, EPI_ISL_902786, EPI_ISL_902787, EPI_ISL_902788, EPI_ISL_902789, EPI_ISL_902790, EPI_ISL_902791, EPI_ISL_902792, EPI_ISL_902793, EPI_ISL_902794, EPI_ISL_902795, EPI_ISL_902796, EPI_ISL_902797, EPI_ISL_902798, EPI_ISL_902799, EPI_ISL_902800, EPI_ISL_902801, EPI_ISL_902802, EPI_ISL_902803, EPI_ISL_902804, EPI_ISL_902805, EPI_ISL_902806, EPI_ISL_902807, EPI_ISL_902808, EPI_ISL_902809, EPI_ISL_902810, EPI_ISL_902811, EPI_ISL_902812, EPI_ISL_902813, EPI_ISL_902814, EPI_ISL_902815, EPI_ISL_902816, EPI_ISL_902817, EPI_ISL_902818, EPI_ISL_902819, EPI_ISL_902820, EPI_ISL_902821, EPI_ISL_902822, EPI_ISL_902823, EPI_ISL_902824, EPI_ISL_902825, EPI_ISL_902826, EPI_ISL_902827, EPI_ISL_902828, EPI_ISL_902829, EPI_ISL_902830, EPI_ISL_902831, EPI_ISL_902832, EPI_ISL_902833, EPI_ISL_902834, EPI_ISL_902835, EPI_ISL_902836, EPI_ISL_902837, EPI_ISL_902838, EPI_ISL_902839, EPI_ISL_902840, EPI_ISL_902841, EPI_ISL_902842, EPI_ISL_902843, EPI_ISL_902844, EPI_ISL_902845, EPI_ISL_902846, EPI_ISL_902847, EPI_ISL_902848, EPI_ISL_902849, EPI_ISL_902850, EPI_ISL_902851, EPI_ISL_902852, EPI_ISL_902853, EPI_ISL_902854, EPI_ISL_902855 | | | |
| see above | Department of Virology and Immunology, University of Helsinki and Helsinki University Hospital, Huslab Finland | Department of Virology, Faculty of Medicine, University of Helsinki, Helsinki, Finland | Teemu Smura, Ravi Kant, Phuoc Truong, Hussein Alburkat, Hannimari Kallio-Kokko, Jenni Virtanen, Maija Suvanto, Essi Korhonen, Sari Hannula, Harri Kangas, Hanna Liimatainen, Satu Kurkela, Hanna Jarva, Maija Lappalainen, Pekka Ellonen, Olli Vapalahti |
| EPI_ISL_903007, EPI_ISL_903008, EPI_ISL_903009, EPI_ISL_903010, EPI_ISL_903011, EPI_ISL_903012 | Seattle Flu Study | Seattle Flu Study | Deborah A. Nickerson, Chris D. Frazar, Jover Lee, Benjamin Pelle, Erica Ryke, Matthew Richardson, Amanda Adler, Elisabeth Brandstetter, Peter D. Han, Kairsten Fay, Misja Ilcisin, Kirsten Lacombe, Thomas R. Sibley, Melissa Truong, Caitlin R. Wolf, Michael Boeckh, Janet A. Englund, Michael Famulare, Barry R. Lutz, Mark J. Rieder, Lea M. Starita, Matthew Thompson, Jay Shendure, Trevor Bedford, Helen Y. Chu |
| EPI_ISL_903608 | MN PHL Division, Minnesota Department of Health | Genomics and Discovery, Respiratory Viruses Branch, Division of Viral Diseases, Centers for Disease Control and Prevention | Krista Queen, Yan Li, Ying Tao, Jing Zhang, Anna Uehara, Anna Montmayeur, Clinton R. Paden, Peter W. Cook, Rachel Marine, Mili Sheth, Jasmine Padilla, Sarah Nobles, Mark Burroughs, Lori Rowe, Haibin Wang, Ben L. Rambo-Martin, Dhwani Batra, Justin Lee, Suxiang Tong |
| EPI_ISL_903650, EPI_ISL_903718 | VT Dept. of Health Laboratory | Genomics and Discovery, Respiratory Viruses Branch, Division of Viral Diseases, Centers for Disease Control and Prevention | Krista Queen, Yan Li, Ying Tao, Jing Zhang, Anna Uehara, Anna Montmayeur, Clinton R. Paden, Peter W. Cook, Rachel Marine, Mili Sheth, Jasmine Padilla, Sarah Nobles, Mark Burroughs, Lori Rowe, Haibin Wang, Ben L. Rambo-Martin, Dhwani Batra, Justin Lee, Suxiang Tong |
| EPI_ISL_903851 | CT-Dr. Katherine A. Kelley State Public Health Lab | Genomics and Discovery, Respiratory Viruses Branch, Division of Viral Diseases, Centers for Disease Control and Prevention | Krista Queen, Yan Li, Ying Tao, Jing Zhang, Anna Uehara, Anna Montmayeur, Clinton R. Paden, Peter W. Cook, Rachel Marine, Mili Sheth, Jasmine Padilla, Sarah Nobles, Mark Burroughs, Lori Rowe, Haibin Wang, Ben L. Rambo-Martin, Dhwani Batra, Justin Lee, Suxiang Tong |
| EPI_ISL_903879 | UT-Unified State Labs: Public Health Utah DOH | Genomics and Discovery, Respiratory Viruses Branch, Division of Viral Diseases, Centers for Disease Control and Prevention | Krista Queen, Yan Li, Ying Tao, Jing Zhang, Anna Uehara, Anna Montmayeur, Clinton R. Paden, Peter W. Cook, Rachel Marine, Mili Sheth, Jasmine Padilla, Sarah Nobles, Mark Burroughs, Lori Rowe, Haibin Wang, Ben L. Rambo-Martin, Dhwani Batra, Justin Lee, Suxiang Tong |
| EPI_ISL_904005, EPI_ISL_904006 | Scientific Veterinary Institute Novi Sad | Veterinary Specialized Institute "Kraljevo", Serbia | Vidanovic,D., Tesovic,B., Knezevic,A., Jovanovic,T., Jankovic,M., Sekler,M., Banovic Djeri,B., Petrovic,T., Volkening,J., Afonso,C. |
| EPI_ISL_904017 | NCSLPH | NCSLPH | Kitty Chase, Michael C Miller, Shermalyn Greene, William Glover |

| | | | |
|---|---|---|---|
| EPI_ISL_904035 | New Mexico Department of Health Scientific Laboratory | New Mexico Department of Health Scientific Laboratory | Ellie Johnson, Anastacia Griego-Fisher, D'eldra Malone |
| EPI_ISL_904167, EPI_ISL_904469, EPI_ISL_904470, EPI_ISL_904471, EPI_ISL_904472, EPI_ISL_904473, EPI_ISL_904474 | Dutch COVID-19 response team | Erasmus Medical Center | Bas Oude Munnink, Reina Sikkema, David Nieuwenhuijse, Irina Chestakova, Anne van der Linden, Marjan Boter, Emmanuelle Munger, Corine GeurtsvanKessel, Annemiek van der Eijk, Richard Molenkamp, Marion Koopmans, on behalf of the Dutch national COVID-19 response team. |
| EPI_ISL_904737, EPI_ISL_904738, EPI_ISL_904797, EPI_ISL_904798 | Dutch COVID-19 response team | National Institute for Public Health and the Environment (RIVM) | Adam Meijer, Harry Vennema, Dirk Eggink, Jeroen Cremer, Sharon van den Brink, Bas van der Veer, AnneMarie van den Brandt, Florian Zwagemaker, Dennis Schmitz, Chantal Reusken, on behalf of the national COVID-19 response team |
| EPI_ISL_904935, EPI_ISL_904940, EPI_ISL_904951, EPI_ISL_904952, EPI_ISL_904953, EPI_ISL_904955, EPI_ISL_904956, EPI_ISL_904957 | Vilnius University Hospital Santaros Klinikos, Vilnius University | Institute of Biotechnology, Life Sciences Center, Vilnius University | Emilija Vasiliunaite, Milda Norkiene, Albertas Timinskas, Alma Gedvilaite, Aurelija Zvirbliene, Daniel Naumovas, Laimonas Griskevicius |
| EPI_ISL_911683, EPI_ISL_911697 | Alaska State Virology Laboratory | Alaska State Virology Laboratory | Stephanie DeRonde, Lisa Smith, Ph.D., Jack Chen, Ph.D. |
| EPI_ISL_911708 | Tanjungpura University Hospital | Tanjungpura University Hospital | Puji Astuti; Andriani; Mahyarudin; Delima Fajar Liana; Virhan Novianry ; Sofi Siti Shofiyah |
| EPI_ISL_911750 | Tanjungpura University Hospital | Tanjungpura University Hospital | Virhan Novianry; Delima Fajar Liana; Mahyarudin; Andriani; Sofi Siti Shofiyah; Puji Astuti; Muhammad Kahtan; Ambar Rialita; Eka Ardiani Putri; Wiwik Windarti; Helmi Sastriawan, Rini Andriani |
| EPI_ISL_912460, EPI_ISL_912476, EPI_ISL_912480, EPI_ISL_912494, EPI_ISL_912496, EPI_ISL_912506, EPI_ISL_912529, EPI_ISL_912538 | NHLS Universitas Academic | UFS Virology | PA Bester, MM Nyaga, P Nthiga, MT Mogotsi, D Goedhals, T de Oliveira |
| EPI_ISL_913278, EPI_ISL_913279, EPI_ISL_913285, EPI_ISL_913313 | Klinisk mikrobiologi | The Public Health Agency of Sweden | Anna-Malin Linde, Maria Lind Karlberg, Carlo Berg, Oskar Karlsson Lindsjo, Sofia Stamouli, Reza Advani, Mattias Haukland, Petra Holmstrom, Noura Walai, Petra Edquist, Mia Brytting, Anna Risberg, Karin Tegmark-Wisell |
| EPI_ISL_913343 | Synlab Medilab, Mikrobiologi | The Public Health Agency of Sweden | Anna-Malin Linde, Maria Lind Karlberg, Carlo Berg, Oskar Karlsson Lindsjo, Sofia Stamouli, Reza Advani, Mattias Haukland, Petra Holmstrom, Noura Walai, Petra Edquist, Mia Brytting, Anna Risberg, Karin Tegmark-Wisell |
| EPI_ISL_913344, EPI_ISL_913345, EPI_ISL_913346, EPI_ISL_913347, EPI_ISL_913354, EPI_ISL_913355, EPI_ISL_913356, EPI_ISL_913461 | Klinisk mikrobiologi | The Public Health Agency of Sweden | Anna-Malin Linde, Maria Lind Karlberg, Carlo Berg, Oskar Karlsson Lindsjo, Sofia Stamouli, Reza Advani, Mattias Haukland, Petra Holmstrom, Noura Walai, Petra Edquist, Mia Brytting, Anna Risberg, Karin Tegmark-Wisell |
| EPI_ISL_913936, EPI_ISL_913937, EPI_ISL_913938, EPI_ISL_913939, EPI_ISL_913959, EPI_ISL_913971 | Instituto de Diagnostico y Referencia Epidemiologicos INDRE_RNLSP | Instituto de Diagnostico y Referencia Epidemiologicos (INDRE) | Claudia Wong-Arambula, Abril Rodriguez-Maldonado, Fabiola Garces-Ayala, Adnan Araiza-Rodriguez, David Fragoso-Fonseca, Sergio Rangel-Guerrero, Mayra Jimenez-Morales, Nancy Munoz-Hernandez, Natividad Cruz-Ortiz, Tatiana Nunez-Garcia, Gisela Barrera-Badillo, Lucia Hernandez-Rivas, Irma Lopez-Martinez, Ernesto Ramirez-Gonzalez. |
| EPI_ISL_915372, EPI_ISL_915373, EPI_ISL_915375, EPI_ISL_915376 | Keio University School of Medicine | Keio University School of Medicine | Kenjiro Kosaki, Yuka Iwasaki, Hirotsugu Ishizu, Haruhiko Siomi, Kodai Abe |
| EPI_ISL_918366, EPI_ISL_918367, EPI_ISL_918368, EPI_ISL_918369, EPI_ISL_918370, EPI_ISL_918371, EPI_ISL_918372 | Virology Unit, Institut Pasteur du Cambodge | Virology Unit, Institut Pasteur du Cambodge | Sokhoun Yann, Ly Sovann, Kraing Sidonn, Yi Sengdoeurn, Chin Savuth, Chau Darapheak, Etienne Simon-Loriere, Veasna Duong, Erik A Karlsson |
| EPI_ISL_918523 | LACEN - Laboratório Central de Saúde Pública do Para | Evandro Chagas Institute | Santos, M.C.; Silva, A.M.; Junior, W.D.C.; Barbagelata, L.S.; Ferreira, J.A.; Sousa, E.M.A.; da Silva, P.S.; Pinheiro, K.C.; L.C.; Sousa Junior, E.C. |
| EPI_ISL_918574, EPI_ISL_918575, EPI_ISL_918576, EPI_ISL_918577, EPI_ISL_918596, EPI_ISL_918597, EPI_ISL_918598, EPI_ISL_918656, EPI_ISL_918657, EPI_ISL_918666, EPI_ISL_918667, EPI_ISL_918668, EPI_ISL_918669, EPI_ISL_918670, EPI_ISL_918671, EPI_ISL_918672, EPI_ISL_918673, EPI_ISL_918674, EPI_ISL_918675, EPI_ISL_918676, EPI_ISL_918677, EPI_ISL_918678, EPI_ISL_918679, EPI_ISL_918680, EPI_ISL_918681, EPI_ISL_918682, EPI_ISL_918683, EPI_ISL_918684 | | | |
| see above | University of Birmingham | COVID-19 Genomics UK (COG-UK) Consortium | Institute of Microbiology, University of Birmingham: Claire McMurray, Joanne Stockton, Samuel Nicholls, Radoslaw Poplawski, Will Rowe, Josh Quick, Nicholas Loman. University of Birmingham Testing Laboratory: Celina M Whalley, Andrew Bosworth, Charlotte Poxon, Kasun Wanigasooriya, Oliver Pickles, Mike Kidd, Alex Richter, Andrew D Beggs PHE Heartlands Lab: Husam Osman, Andrew Bosworth. Queen Elizabeth Hospital: Anna Casey |
| EPI_ISL_919239, EPI_ISL_919270, EPI_ISL_919273, EPI_ISL_919274, EPI_ISL_919275 | West of Scotland Specialist Virology Centre, NHSGGC / MRC-University of Glasgow Centre for Virus Research | COVID-19 Genomics UK (COG-UK) Consortium | Ana da Silva Filipe, Natasha Johnson, Kathy Smollett, Daniel Mair, Stephen Carmichael, Alice Broos, Lily Tong, Jenna Nichols, Kyriaki Nomikou; Sarah McDonald; Richard Orton, Joseph Hughes, Sreenu Vattipally, David L Robertson; Alasdair MacLean, Rory Gunson; Sharif Shaaban, Matthew Holden; Rachel Blacow, Guy Mollett, Kathy Li, James Shepherd, Antonia Ho, Emma Thomson |
| EPI_ISL_919347 | Virology Department, Royal Infirmary of Edinburgh, NHS Lothian / School of Biological Sciences, University of Edinburgh / Institute of Genetics and Molecular Medicine, University of Edinburgh | COVID-19 Genomics UK (COG-UK) Consortium | McHugh M, Dewar R, Rooke S, Gallagher M, Balcaza C, O'Toole Á, Scher E, Hill V, McCrone JT, Colquhoun R, Yu X, Jackson B, Rambaut A, Williams TC, Templeton K |
| EPI_ISL_920491, EPI_ISL_920509 | University College London Hospital | COVID-19 Genomics UK (COG-UK) Consortium | Judith Heaney, Matthew Byott, Catherine Houlihan, Dan Frampton, Stuart Kirk, Moira Spyer and Eleni Nastouli |
| EPI_ISL_923219 | Centre for Enzyme Innovation, University of Portsmouth / Translational Research Laboratory, Portsmouth Hospitals NHS Trust | COVID-19 Genomics UK (COG-UK) Consortium | Angela Beckett,Salman Goudarzi,Christopher Fearn,Kate Cook,Katie Loveson,Sharon Glaysher,Scott Elliott,Samuel Robson |
| EPI_ISL_924788 | Bioinformatics and Biostatistics Lab, Advanced Sequencing Facility | COVID-19 Genomics UK (COG-UK) Consortium | Aengus Stewart,Jerome Nicod,Chelsea Sawyer,Laura Cubitt,Harshil Patel,Margaret Crawford |
| EPI_ISL_925112, EPI_ISL_925134 | 1.AO Universitaria 'S. Giovanni di Dio e Ruggi D'Aragona, Scuola Medica Salernitana' Hospital / 2.UOC di Virologia e Microbiologia, Università della Campania 'L. Vanvitelli' / 3.AO Universitaria 'Federico II' Napoli Hospital / 4.AORN 'San Giuseppe Moscati' Avellino Hospital / 5.AO 'San Pio - presidio G. Rummo' Benevento Hospital / 6.AO 'Sant'Anna e San Sebastiano' Caserta Hospital / 7.PO 'Maria Santissima Addolorata' Eboli Hospital / 8.Biogem Istituto di Ricerche Genetiche | 1. Genome Research Center for Health (CRGS) / 2. Laboratory of Molecular Medicine and Genomics(LMMGe) / 3. Center for Research in Pure and Applied Mathematics (CRMPA) | Giorgio Giurato, Francesca Rizzo, Alessandro Weisz, Gianluigi Franci, Giovanni Nassa, Pasquale Pagliano, Roberta Tarallo, Elena Alexandrova, Ylenia D'Agostino, Carlo Ferravante, Jessica Lamberti, Viola Melone, Domenico Memoli, Valeria Mirici Cappa, Domenico Palumbo, Giovanni Pecoraro, Assunta Sellitto, Oriana Strianese, Ilaria Terenzi, Giuseppe Fenza, Aniello Gentile, Antonello Saccomanno, Sonia Amabile, Teresa Rocco, Annamaria Salvati, Emilia Vaccaro, Massimiliano Galdiero, Michele Cennamo, Giuseppe Portella, Maria Grazia Foti, Mariarosaria Ingino, Maria Landi, Maurizio Fumi, Vincenzo Rocco, Rita Greco, Vittoria Letizia, Arnolfo Petruzziello, Maddalena Schioppa, Gregorio Goffredi, Francesca Marciano, Michele Caraglia, Alessia Cossu, Marianna Scrima |
| EPI_ISL_925361 | NCSLPH | NCSLPH | Chase K, Miller MC, Greene S, Glover W |
| EPI_ISL_925380, EPI_ISL_925386 | Department of Clinical Microbiology | GIGA Medical Genomics | Keith Durkin, Maria Artesi, Sébastien Bontems, Raphaël Boreux, Bouchra Boujemla, Cécile Meex, Pierrette Melin, Marie-Pierre Hayette, Vincent Bours |
| EPI_ISL_925900 | Nucleic Acid Testing, National Reference Laboratory | GIGA Medical Genomics | Yvan Butera, Keith Durkin, Maria Artesi, Bouchra Boujemla, Robert Rutayisire, Patrick Tuyisenge, Esperence Umumararungu, Sébastien Bontems, Marie-Pierre Hayette, Nathalie Renotte, Swaibu Gatare, Jacob Souopgui, Sabin Nsanzimana, Vincent Bours, Léon Mutesa |
| EPI_ISL_929763 | NCSLPH | NCSLPH | Chase K, Miller MC, Greene S, Glover W |
| EPI_ISL_930783 | Instituto Nacional de Medicina Genomica | Instituto Nacional de Medicina Genomica | Hidalgo-Miranda A, Mendoza-Vargas A, Reyes-Grajeda JP, Cisneros-Villanueva M, Cedro-Tanda A,Peñaloza-Figueroa F, Herrera-Montalvo LA |
| EPI_ISL_931239, EPI_ISL_931240, EPI_ISL_931241, EPI_ISL_931242, EPI_ISL_931365, EPI_ISL_931366, | University Hospital Basel, Clinical Virology | University Hospital Basel, Clinical Bacteriology | Tim Roloff, Madlen Stange, Helena MB Seth-Smith, Alfredo Mari, Karoline Leuzinger, Julia Bielicki, Manuel Battegay, Hans Hirsch, Adrian Egli |

| | | | |
|---|---|---|---|
| EPI_ISL_931390 | | | |
| EPI_ISL_933512, EPI_ISL_933513, EPI_ISL_933514, EPI_ISL_933515, EPI_ISL_933517 | Instituto Nacional de Medicina Genomica | Instituto Nacional de Medicina Genomica | Hidalgo-Miranda A, Mendoza-Vargas A, Reyes-Grajeda JP, Cisneros-Villanueva M, Cedro-Tanda A,Peñaloza-Figueroa F, Herrera-Montalvo LA |
| EPI_ISL_933559, EPI_ISL_933560, EPI_ISL_933561, EPI_ISL_933562, EPI_ISL_933563, EPI_ISL_933598, EPI_ISL_933599, EPI_ISL_933600, EPI_ISL_933601 | Arizona State Public Health Laboratory | Arizona State Public Health Laboratory | Trung Huynh, Jessica Escobar, Katherine Fullerton, Nobuko Fukushima, Stacy White, Linda Getsinger, Victor Waddell |
| EPI_ISL_934337 | Klinisk mikrobiologi | The Public Health Agency of Sweden | Anna-Malin Linde, Maria Lind Karlberg, Carlo Berg, Oskar Karlsson Lindsjo, Sofia Stamouli, Reza Advani, Mattias Haukland, Petra Holmstrom, Noura Walai, Petra Edquist, Mia Brytting, Anna Risberg, Karin Tegmark-Wisell |
| EPI_ISL_935877, EPI_ISL_935878, EPI_ISL_935879, EPI_ISL_935880, EPI_ISL_935882, EPI_ISL_935883, EPI_ISL_935884, EPI_ISL_935885, EPI_ISL_935886, EPI_ISL_935888, EPI_ISL_935889, EPI_ISL_935893, EPI_ISL_935894, EPI_ISL_935895, EPI_ISL_935898, EPI_ISL_935899, EPI_ISL_935901, EPI_ISL_935902, EPI_ISL_935907, EPI_ISL_935910, EPI_ISL_935911, EPI_ISL_935912, EPI_ISL_935913, EPI_ISL_935915, EPI_ISL_935916, EPI_ISL_935917, EPI_ISL_935921, EPI_ISL_935924, EPI_ISL_935930, EPI_ISL_935933 | | | |
| see above | Cadham Provincial laboratory | National Microbiology Laboratory (NML) | Anna Majer, Shari Tyson, Grace Seo, Philip Mabon, Elsie Grudeski, Rhiannon Huzarewich, Russell Mandes, Anneliese Landgraff, Jennifer Tanner, Natalie Knox, Morag Graham, Gary Van Domselaar, Paul Van Caeseele, Jared Bullard, David Alexander, Kerry Dust, Nathalie Bastien, Yan Li, Timothy Booth, Darian Hole, Madison Chapel, Kirsten Biggar, CanCOGeN's metadata curation team, Public Health Agency of Canada CanCOGeN team |
| EPI_ISL_936490 | Makmal Kesihatan Awam Kota Bharu | Institute for Medical Research, Infectious Disease Research Centre, National Institutes of Health, Ministry of Health Malaysia | Suppiah J, Kamel K, Azizan MA, Thayan R |
| EPI_ISL_936622, EPI_ISL_936625, EPI_ISL_936626, EPI_ISL_936627, EPI_ISL_936628, EPI_ISL_936629, EPI_ISL_936630, EPI_ISL_936631 | Northwestern Memorial Hospital | Ozer Lab | Ramon Lorenzo-Redondo, Lacy M. Simons, Chad J. Achenbach, Lawrence J. Jennings, Michael G. Ison, Judd F. Hultquist, Egon A. Ozer |
| EPI_ISL_937326 | Utah Public Health Laboratory | Utah Public Health Laboratory | Erin L. Young, Kelly F. Oakeson, Tara Gallagher |
| EPI_ISL_940119 | Charlotte Maxeke Johannesburg Academic Hospital, National Health Laboratory Services, Gauteng, South Africa | National Institute for Communicable Diseases of the National Health Laboratory Service | Amoako DG, Mohale T, Ntuli N, Mahlangu B, Allam M, Ismail A, Bhiman JN |
| EPI_ISL_940141, EPI_ISL_940142, EPI_ISL_940143 | National Institute for Communicable Diseases,National Health Laboratory Services, Gauteng, South Africa | National Institute for Communicable Diseases of the National Health Laboratory Service | Amoako DG, Mohale T, Ntuli N, Mahlangu B, Allam M, Ismail A, Bhiman JN |
| EPI_ISL_940159, EPI_ISL_940161, EPI_ISL_940208, EPI_ISL_940234, EPI_ISL_940244, EPI_ISL_940258 | Hôpital Bichat Claude Bernard, Laboratoire de Virologie | IAME UMR1137 Inserm, Université de Paris, Hôpital Bichat | Antoine Bridier-Nahmias, Amélie Recoing, Quentin Le Hingrat, Lena Daniel, Siham Hamri, Gilles Collin, Alexandre Storto, Mélanie Bertine, Charlotte Charpentier, Nadhira Houhou-Fidouh, Diane Descamps, Benoit Visseaux |
| EPI_ISL_941207 | Servicio de Microbiología. Hospital Clínico Universitario de Valencia | SeqCOVID-SPAIN consortium/IBV(CSIC) | David Navarro Ortega, Eliseo Albert Vicent, Ignacio Torres and SeqCOVID-SPAIN consortium |
| EPI_ISL_941275, EPI_ISL_941277, EPI_ISL_941279, EPI_ISL_941280, EPI_ISL_941281 | Nigeria Centre for Disease Control (NCDC) | African Centre of Excellence for Genomics of Infectious Diseases (ACEGID), Redeemer's University | Oluniyi P.E. et al |
| EPI_ISL_941340 | Instituto Nacional de Saude (INSA) | Instituto Nacional de Saude (INSA) | Borges et al |
| EPI_ISL_941903, EPI_ISL_941904, EPI_ISL_941905, EPI_ISL_941906, EPI_ISL_941907, EPI_ISL_941908, EPI_ISL_941910, EPI_ISL_941911, EPI_ISL_941921, EPI_ISL_942401, EPI_ISL_942402, EPI_ISL_942404, EPI_ISL_942405 | | | |
| see above | Virginia DCLS | Virginia DCLS | Virginia DCLS |
| EPI_ISL_942502, EPI_ISL_942504 | Gundersen Molecular Diagnostics Laboratory | Kabara Cancer Research Institute | Craig S. Richmond, Paraic A. Kenny |
| EPI_ISL_942505, EPI_ISL_942506 | Gundersen Clinical Microbiology Laboratory | Kabara Cancer Research Institute | Craig S. Richmond, Paraic A. Kenny |
| EPI_ISL_942507, EPI_ISL_942508, EPI_ISL_942509, EPI_ISL_942510, EPI_ISL_942511, EPI_ISL_942512, EPI_ISL_942513, EPI_ISL_942514, EPI_ISL_942516, EPI_ISL_942517, EPI_ISL_942518, EPI_ISL_942519, EPI_ISL_942520, EPI_ISL_942521, EPI_ISL_942522, EPI_ISL_942523, EPI_ISL_942524, EPI_ISL_942525, EPI_ISL_942526, EPI_ISL_942527, EPI_ISL_942528, EPI_ISL_942529, EPI_ISL_942530, EPI_ISL_942531, EPI_ISL_942532, EPI_ISL_942533, EPI_ISL_942534, EPI_ISL_942535, EPI_ISL_942536, EPI_ISL_942537, EPI_ISL_942538, EPI_ISL_942539, EPI_ISL_942540, EPI_ISL_942541, EPI_ISL_942542, EPI_ISL_942543 | | | |
| see above | Gundersen Molecular Diagnostics Laboratory | Kabara Cancer Research Institute | Craig S. Richmond, Paraic A. Kenny |
| EPI_ISL_947273 | RSIA Sayyidah | Eijkman Institute for Molecular Biology, Ministry of Research and Technology/National Agency for Research and Innovation | Frilasita A Yudhaputri, Hidayat Trimarsanto, Iskandar Adnan, Lydia V. Panggalo, Sukma Oktavianthi, Willy Agustine, Edison Johar, Safarina G Malik, Khin Saw Myint, Amin Soebandrio |
| EPI_ISL_949193, EPI_ISL_949199, EPI_ISL_949200, EPI_ISL_949201, EPI_ISL_949210, EPI_ISL_949214, EPI_ISL_949220 | Departamento de Microbiología, CDB, Hospital Clínic, Barcelona | SeqCOVID-SPAIN consortium/IBV(CSIC) | Andrea Vergara, Mikel Martínez, Elisa Rubio, Jéssica Navero, Aida Peiró and SeqCOVID-SPAIN consortium |
| EPI_ISL_950603, EPI_ISL_950613, EPI_ISL_950614, EPI_ISL_950617, EPI_ISL_950618, EPI_ISL_950619, EPI_ISL_950620, EPI_ISL_950621, EPI_ISL_950622, EPI_ISL_950623, EPI_ISL_950624, EPI_ISL_950625, EPI_ISL_950626, EPI_ISL_950627, EPI_ISL_950628, EPI_ISL_950629, EPI_ISL_950630, EPI_ISL_950631, EPI_ISL_950632, EPI_ISL_950634, EPI_ISL_950635 | | | |
| see above | Queens Medical Centre, Clinical Microbiology Department / DeepSeq Nottingham | COVID-19 Genomics UK (COG-UK) Consortium | Gemma Clark, Wendy Smith, Manjinder Khakh, Vicki M Fleming, Michelle M Lister, Hannah Howson-Wells, Jonathan Ball, Patrick McClure, Joseph Chappell, Theocharis Tsoleridis, Nadine Holmes, Matthew Carlisle, Christopher Moore, Fei Sang, Johnny Debebe, Victoria Wright, Matthew Loose |
| EPI_ISL_952390, EPI_ISL_952399 | Centre for Enzyme Innovation, University of Portsmouth / Translational Research Laboratory, Portsmouth Hospitals NHS Trust | COVID-19 Genomics UK (COG-UK) Consortium | Angela Beckett,Salman Goudarzi,Christopher Fearn,Kate Cook,Katie Loveson,Sharon Glaysher,Scott Elliott,Samuel Robson |
| EPI_ISL_955156 | University of Sarajevo, Veterinary Faculty, Laboratory for Molecular Diagnostic and Research Laboratory | University of Sarajevo, Veterinary Faculty, Laboratory for Molecular Diagnostic and Research Laboratory | Goleti T., Goleti Š., Softi A., Ali-Šeho A., Jaži A., Šabi E., Nicevi M., Hodži A., Terzi I. |
| EPI_ISL_955341, EPI_ISL_955342, EPI_ISL_955343, EPI_ISL_955344, EPI_ISL_955345, EPI_ISL_955346, EPI_ISL_955347, EPI_ISL_955348, EPI_ISL_955349, EPI_ISL_955350, EPI_ISL_955351, EPI_ISL_955352 | | | |
| see above | UC Davis- Department of Pathology and Laboratory Medicine | Chan-Zuckerberg Biohub | CZB Cliahub Consortium |
| EPI_ISL_955496, EPI_ISL_955497, EPI_ISL_955498, EPI_ISL_955499, EPI_ISL_955500, EPI_ISL_955501, EPI_ISL_955502, EPI_ISL_955503, EPI_ISL_955504, EPI_ISL_955505, EPI_ISL_955506, EPI_ISL_955507, EPI_ISL_955508, EPI_ISL_955509, EPI_ISL_955510, EPI_ISL_955512, EPI_ISL_955513, EPI_ISL_955514, EPI_ISL_955515, EPI_ISL_955516, EPI_ISL_955517, EPI_ISL_955518, EPI_ISL_955519, EPI_ISL_955520, EPI_ISL_955521, EPI_ISL_955522, EPI_ISL_955533 | | | |
| see above | Humboldt County Public Health Laboratory | Chan-Zuckerberg Biohub | CZB Cliahub Consortium |
| EPI_ISL_955561, EPI_ISL_955562, EPI_ISL_955563, EPI_ISL_955564, EPI_ISL_955565, EPI_ISL_955566, EPI_ISL_955567 | Santa Clara County Public Health Laboratory | Chan-Zuckerberg Biohub | CZB Cliahub Consortium |
| EPI_ISL_955628, EPI_ISL_955629, EPI_ISL_955630, EPI_ISL_955631 | Orange County Public Health Lab | Chan-Zuckerberg Biohub | CZB Cliahub Consortium |
| EPI_ISL_959476, EPI_ISL_959477, EPI_ISL_959478, EPI_ISL_959485, EPI_ISL_959486, EPI_ISL_959487, EPI_ISL_959488, EPI_ISL_959489, EPI_ISL_959490, EPI_ISL_959491, EPI_ISL_959492, EPI_ISL_959493, EPI_ISL_959500, EPI_ISL_959502, EPI_ISL_959514, EPI_ISL_959515, EPI_ISL_959516, EPI_ISL_959517, | | | |

| | | | |
|---|---|---|---|
| EPI_ISL_959518, EPI_ISL_959519, EPI_ISL_959520, EPI_ISL_959521, EPI_ISL_959522, EPI_ISL_959523, EPI_ISL_959524, EPI_ISL_959525, EPI_ISL_959526, EPI_ISL_959527, EPI_ISL_959528 | | | |
| see above | Division of Emerging Infectious Diseases, Bureau of Infectious Diseases Diagnosis Control, Korea Disease Control and Prevention Agency | Division of Emerging Infectious Diseases, Bureau of Infectious Diseases Diagnosis Control, Korea Disease Control and Prevention Agency | Ae Kyung Park, Il-Hwan Kim, Heui Man Kim, Jeong-Min Kim, Namjoo Lee, Chae Young Lee, Sang Hee Woo, Eun-Jin Kim |
| EPI_ISL_960112 | False Bay Hospital wc FBH | National Health Laboratory Service/UCT | Arash Iranzadeh, Deelan Doolabh, Lynn Tyers, Bruna Galvao, Innocent Mudau, Marvin Hsiao, Kruger Marais, Diana Hardie, Stephen Korsman, Carolyn Williamson |
| EPI_ISL_960113 | Victoria Hospital wc VHW | National Health Laboratory Service/UCT | Arash Iranzadeh, Deelan Doolabh, Lynn Tyers, Bruna Galvao, Innocent Mudau, Marvin Hsiao, Kruger Marais, Diana Hardie, Stephen Korsman, Carolyn Williamson |
| EPI_ISL_960348, EPI_ISL_960360, EPI_ISL_960393 | University of Wisconsin-Madison AIDS Vaccine Research Laboratories | University of Wisconsin-Madison AIDS Vaccine Research Laboratories | Gage Moreno, Katarina Braun, et al. AIDS Vaccine Research Laboratories |
| EPI_ISL_960451, EPI_ISL_960455, EPI_ISL_960458, EPI_ISL_960486, EPI_ISL_960487, EPI_ISL_960488, EPI_ISL_960489, EPI_ISL_960490, EPI_ISL_960491, EPI_ISL_960493, EPI_ISL_960494, EPI_ISL_960495, EPI_ISL_960501, EPI_ISL_960582, EPI_ISL_960583, EPI_ISL_960584, EPI_ISL_960585, EPI_ISL_960586, EPI_ISL_960587, EPI_ISL_960588 | | | |
| see above | Istituto Zooprofilattico Sperimentale del Mezzogiorno | TIGEM | Patrizia Annunziata, Andrea Ballabio, Valentina Bouche, Davide Cacchiarelli, Pellegrino Cerino, Chiara Colantuono, Maria Concetta Cuomo, Denise Di Concilio, Lucio Di Filippo, Antonio Grimaldi, Antonio Limone, Anna Manfredi, Francesco Panariello, Biancamaria Pierri, Marcello Salvi |
| EPI_ISL_962106, EPI_ISL_962107 | Illinois Department of Public Health | Gagnon Lab, Southern Illinois University | Keith Gagnon |
| EPI_ISL_962245, EPI_ISL_962246, EPI_ISL_962247, EPI_ISL_962248, EPI_ISL_962249, EPI_ISL_962250, EPI_ISL_962251, EPI_ISL_962252, EPI_ISL_962253, EPI_ISL_962254, EPI_ISL_962255, EPI_ISL_962256, EPI_ISL_962257, EPI_ISL_962258, EPI_ISL_962259, EPI_ISL_962262 | | | |
| see above | Seattle Flu Study | Seattle Flu Study | Deborah A. Nickerson, Chris D. Frazar, Jover Lee, Benjamin Pelle, Erica Ryke, Matthew Richardson, Amanda Adler, Elisabeth Brandstetter, Peter D. Han, Kairsten Fay, Misja Ilcisin, Kirsten Lacombe, Thomas R. Sibley, Melissa Truong, Caitlin R. Wolf, Karen Cowgill, Stephanie Schrag, Jeff Duchin, Michael Boeckh, Janet A. Englund, Michael Famulare, Barry R. Lutz, Mark J. Rieder, Lea M. Starita, Matthew Thompson, Helen Y. Chu, Trevor Bedford, Jay Shendure |
| EPI_ISL_962892 | ULSS 03 Venezia | Istituto Zooprofilattico Sperimentale delle Venezie | Natale Alda, Mazzotta Elisa, Mason Nicoletta, Ceglie Letizia, Lucchese L, Stefani Annalisa, Bonfante Francesco, Bortolami Alessio, Fincato Alice, Monne Isabella, Bellinati Laura, Mion Monica, Guadagno Carmine, Quaranta Erika, Pastori Ambra, Terregino Calogero |
| EPI_ISL_965540, EPI_ISL_965541, EPI_ISL_965545, EPI_ISL_965557, EPI_ISL_965571, EPI_ISL_965598, EPI_ISL_965600, EPI_ISL_965609, EPI_ISL_965613, EPI_ISL_965617, EPI_ISL_965621, EPI_ISL_965628, EPI_ISL_965630, EPI_ISL_965644, EPI_ISL_965646, EPI_ISL_965648, EPI_ISL_965650, EPI_ISL_965658, EPI_ISL_965665, EPI_ISL_965670, EPI_ISL_965677, EPI_ISL_965682, EPI_ISL_965685, EPI_ISL_965691, EPI_ISL_965704, EPI_ISL_965706, EPI_ISL_965709, EPI_ISL_965712, EPI_ISL_965722, EPI_ISL_965723, EPI_ISL_965727, EPI_ISL_965728, EPI_ISL_965732, EPI_ISL_965733, EPI_ISL_965737, EPI_ISL_965767, EPI_ISL_965771, EPI_ISL_965775, EPI_ISL_965787, EPI_ISL_965788, EPI_ISL_965791, EPI_ISL_965794 | | | |
| see above | Dutch COVID-19 response team | Medical Microbiology, Maastricht University Medical Centre | Jozef Dingemans*, Brian van der Veer*, Erik Beuken, Carmen Reumkens, Lieke van Alphen, Christian Hoebe, Paul Savelkoul |
| EPI_ISL_968815, EPI_ISL_968843, EPI_ISL_968941, EPI_ISL_968942, EPI_ISL_968943, EPI_ISL_968955, EPI_ISL_968996, EPI_ISL_968997 | KEMRI-Wellcome Trust Research Programme/KEMRI-CGMR-C Kilifi | KEMRI-Wellcome Trust Research Programme/KEMRI-CGMR-C Kilifi | Githinji et al |
| EPI_ISL_975795, EPI_ISL_975796, EPI_ISL_975797, EPI_ISL_975798, EPI_ISL_975799, EPI_ISL_975800, EPI_ISL_975801, EPI_ISL_975802, EPI_ISL_975803, EPI_ISL_975804, EPI_ISL_975805, EPI_ISL_975806, EPI_ISL_975807, EPI_ISL_975808, EPI_ISL_975809, EPI_ISL_975810, EPI_ISL_975811, EPI_ISL_975812, EPI_ISL_975813, EPI_ISL_975814, EPI_ISL_975815, EPI_ISL_975816, EPI_ISL_975817, EPI_ISL_975818, EPI_ISL_975819, EPI_ISL_975820, EPI_ISL_975821, EPI_ISL_975822, EPI_ISL_975823, EPI_ISL_975824, EPI_ISL_975825, EPI_ISL_975826, EPI_ISL_975827, EPI_ISL_975828, EPI_ISL_975829, EPI_ISL_975830, EPI_ISL_975831, EPI_ISL_975832, EPI_ISL_975833, EPI_ISL_975834, EPI_ISL_975835, EPI_ISL_975836, EPI_ISL_975837, EPI_ISL_975838, EPI_ISL_975839, EPI_ISL_975840, EPI_ISL_975841, EPI_ISL_975842, EPI_ISL_975843, EPI_ISL_975844, EPI_ISL_975845, EPI_ISL_975846, EPI_ISL_975847, EPI_ISL_975848, EPI_ISL_975849, EPI_ISL_975850, EPI_ISL_975851, EPI_ISL_975852, EPI_ISL_975853, EPI_ISL_975854, EPI_ISL_975855, EPI_ISL_975856, EPI_ISL_975857, EPI_ISL_975858, EPI_ISL_975859, EPI_ISL_975860, EPI_ISL_975861, EPI_ISL_975862, EPI_ISL_975863, EPI_ISL_975864, EPI_ISL_975865, EPI_ISL_975866, EPI_ISL_975867, EPI_ISL_975868, EPI_ISL_975869, EPI_ISL_975870, EPI_ISL_975871, EPI_ISL_975872, EPI_ISL_975873, EPI_ISL_975874, EPI_ISL_975875, EPI_ISL_975876, EPI_ISL_975877, EPI_ISL_975878, EPI_ISL_975879, EPI_ISL_975880, EPI_ISL_975881, EPI_ISL_975882, EPI_ISL_975883, EPI_ISL_975884, EPI_ISL_975885, EPI_ISL_975886, EPI_ISL_975887, EPI_ISL_975888, EPI_ISL_975889, EPI_ISL_975890, EPI_ISL_975891, EPI_ISL_975892, EPI_ISL_975893, EPI_ISL_975894, EPI_ISL_975895, EPI_ISL_975896, EPI_ISL_975897, EPI_ISL_975898, EPI_ISL_975899, EPI_ISL_975900, EPI_ISL_975901, EPI_ISL_975902, EPI_ISL_975903, EPI_ISL_975904, EPI_ISL_975905, EPI_ISL_975906, EPI_ISL_975907, EPI_ISL_975908, EPI_ISL_975909, EPI_ISL_975910, EPI_ISL_975911, EPI_ISL_975912, EPI_ISL_975913, EPI_ISL_975914, EPI_ISL_975915, EPI_ISL_975916, EPI_ISL_975917, EPI_ISL_975918, EPI_ISL_975919, EPI_ISL_975920, EPI_ISL_975921, EPI_ISL_975922, EPI_ISL_975923, EPI_ISL_975924, EPI_ISL_975925, EPI_ISL_975926, EPI_ISL_975927, EPI_ISL_975928, EPI_ISL_975929, EPI_ISL_975930, EPI_ISL_975931, EPI_ISL_975932, EPI_ISL_975933, EPI_ISL_975934, EPI_ISL_975935, EPI_ISL_975936, EPI_ISL_975937, EPI_ISL_975938, EPI_ISL_975939, EPI_ISL_975940, EPI_ISL_975941, EPI_ISL_975942, EPI_ISL_975943, EPI_ISL_975944, EPI_ISL_975945, EPI_ISL_975946, EPI_ISL_975947, EPI_ISL_975948, EPI_ISL_975949, EPI_ISL_975950, EPI_ISL_975951, EPI_ISL_975952, EPI_ISL_975953, EPI_ISL_975954, EPI_ISL_975955, EPI_ISL_975956, EPI_ISL_975957, EPI_ISL_975958, EPI_ISL_975959, EPI_ISL_975960, EPI_ISL_975961, EPI_ISL_975962, EPI_ISL_975963, EPI_ISL_975964, EPI_ISL_975965, EPI_ISL_975966, EPI_ISL_975967, EPI_ISL_975968, EPI_ISL_975969, EPI_ISL_975970, EPI_ISL_975971, EPI_ISL_975972 | | | |
| see above | BCCDC Public Health Laboratory | BCCDC Public Health Laboratory | Prystajecky Natalie, Linda Hoang, Dan Fornika, John Tyson, Shannon Russell, Kim Macdonald, Kimia Kamelian, Ana Pacagnella, Corrinne Ng, Loretta Janz, Robert Azana Terry Snutch, Mel Krajden |
| EPI_ISL_977190, EPI_ISL_977191 | ULSS 2 Treviso | Istituto Zooprofilattico Sperimentale delle Venezie | Adelaide Milani, Alessia Schivo, Annalisa Salviato, Erika Giorgia Quaranta, Ambra Pastori, Bianca Zecchin, Alice Fusaro, Isabella Monne, Calogero Terregino, Antonia Ricci |
| EPI_ISL_977514, EPI_ISL_977515, EPI_ISL_977536 | NCSLPH | NCSLPH | Chase K, Miller MC, Greene S, Glover W |
| EPI_ISL_977885, EPI_ISL_977886, EPI_ISL_977887, EPI_ISL_977888, EPI_ISL_977889, EPI_ISL_977890, EPI_ISL_977891, EPI_ISL_978006, EPI_ISL_978007, EPI_ISL_978008, EPI_ISL_978160, EPI_ISL_978161, EPI_ISL_978162, EPI_ISL_978163, EPI_ISL_978164, EPI_ISL_978165, EPI_ISL_978166, EPI_ISL_978167, EPI_ISL_978168, EPI_ISL_978967, EPI_ISL_978968, EPI_ISL_978969 | | | |
| see above | Chiu Laboratory, University of California, San Francisco | Chiu Laboratory, University of California, San Francisco | Charles Chiu, Xianding (Wayne) Deng, Candace Wang, Venice Servellita, Jill Hacker, Debra Wadford |
| EPI_ISL_982903 | MEPHI Aix Marseille University (AMU) | MEPHI Aix Marseille University (AMU) | Anthony LEVASSEUR |
| EPI_ISL_983648 | North Carolina State Laboratory of Public Health | North Carolina State Laboratory of Public Health | Chase K, Miller MC, Greene S, Glover W |
