## Supplementary material for "Detection and characterization of the SARS-CoV-2 lineage B.1.526 in New York": Supp. Table 3 GISAID Acknowledgment part 2: gisaid_hcov-19_acknowledgement_table_2021_02_13_00-12.pdf

We gratefully acknowledge the following Authors from the Originating laboratories responsible for obtaining the specimens, as well as the Submitting laboratories where the genome data were generated and shared via GISAID, on which this research is based.

All Submitters of data may be contacted directly via www.gisaid.org

Authors are sorted alphabetically.

| Accession ID | Originating Laboratory | Submitting Laboratory | Authors |
|---|---|---|---|
| EPI_ISL_672574 | Histopath | NSW Health Pathology - Institute of Clinical Pathology and Medical Research; Westmead Hospital; University of Sydney | CIDM-PH et al. |
| EPI_ISL_678386 | Area of Virology, Serology and Virology Division (SAViD), New South Wales Health Pathology Randwick | Virology Research Laboratory; Area of Virology, Serology and Virology Division (SAViD), New South Wales Health Pathology Randwick | Foster, C.; Au, J.; Ruiz Silva, M.; Deveson, I.; Bull, R.; Van Hal, S.; Rawlinson, W. |
| EPI_ISL_678830 | Respiratory Virus Unit, Microbiology Services Colindale, Public Health England | COVID-19 Genomics UK (COG-UK) Consortium | PHE Covid Sequencing Team |
| EPI_ISL_679945 | Queens Medical Centre, Clinical Microbiology Department / DeepSeq Nottingham | COVID-19 Genomics UK (COG-UK) Consortium | Gemma Clark, Wendy Smith, Manjinder Khakh, Vicki M Fleming, Michelle M Lister, Hannah Howson-Wells, Jonathan Ball, Patrick McClure, Joseph Chappell, Theocharis Tsoleridis, Nadine Holmes, Matthew Carlisle, Christopher Moore, Fei Sang, Johnny Debebe, Victoria Wright, Matthew Loose |
| EPI_ISL_680459, EPI_ISL_680460 | West of Scotland Specialist Virology Centre, NHSGGC / MRC-University of Glasgow Centre for Virus Research | COVID-19 Genomics UK (COG-UK) Consortium | Ana da Silva Filipe, Natasha Johnson, Kathy Smollett, Daniel Mair, Stephen Carmichael, Alice Broos, Lily Tong, Jenna Nichols, Kyriaki Nomikou; Sarah McDonald; Richard Orton, Joseph Hughes, Sreenu Vattipally, David L Robertson; Alasdair MacLean, Rory Gunson; Sharif Shaaban, Matthew Holden; Rachel Blacow, Guy Mollett, Kathy Li, James Shepherd, Antonia Ho, Emma Thomson |
| EPI_ISL_682296 | Canterbury Health Laboratories | Institute of Environmental Science and Research (ESR) | Xiaoyun Ren, Matt Storey, Nikki Freed, Muhammad Faisal, Jing Wang, Hermes Perez, Anja Werno, Antje van der Linden, Arlo Upton, Chris Mansell, David Hammer, Dragana Drinkovic, Gary McAuliffe, Hana Sofia Andersson, James Ussher, Jill Sherwood, Josh Freeman, Julia Howard, Juliet Elvy, Mary DeAlmeida, Matt Blakiston, Matthew Rogers, Max Bloomfield, Michael Addidle, Michelle Balm, Sally Roberts, Sarah Jefferies, Sharmini Muttaiyah, Susan Morpeth, Susan Taylor, Timothy Blackmore, Vani Sathyendran, Veronica Playle, Virginia Hope, Erasmus Smit, Lauren Jelly, Olin Silander, Joep de Ligt |
| EPI_ISL_684006, EPI_ISL_684007, EPI_ISL_684008, EPI_ISL_684009, EPI_ISL_684010, EPI_ISL_684011, EPI_ISL_684012, EPI_ISL_684013, EPI_ISL_684015, EPI_ISL_684016, EPI_ISL_684017, EPI_ISL_684018, EPI_ISL_684019, EPI_ISL_684020, EPI_ISL_684021, EPI_ISL_684022, EPI_ISL_684023, EPI_ISL_684024, EPI_ISL_684025, EPI_ISL_684026, EPI_ISL_684027 | | | |
| see above | Utah Public Health Laboratory | Utah Public Health Laboratory | Erin Young, Kelly Oakeson |
| EPI_ISL_692770, EPI_ISL_692771 | Massachusetts State Public Health Laboratory | Massachusetts State Public Health Laboratory | Andrew Lang, Timelia Fink, Glen Gallagher, Sandra Smole |
| EPI_ISL_693322, EPI_ISL_693330 | National Public Health Laboratory, National Centre for Infectious Diseases | National Public Health Laboratory, National Centre for Infectious Diseases | Tze Minn Mak, Sophie Octavia, Zhenyang Zhou, Lin Cui, Raymond Tzer Pin Lin |
| EPI_ISL_697798, EPI_ISL_697799 | Institute of Microbiology, Universidad San Francisco de Quito | Institute of Microbiology, Universidad San Francisco de Quito | Belén Prado-Vivar, Sully Márquez, Juan José Guadalupe, Monica Becerra-Wong, Diana Zambrano, Fredy Loor, Juan Zuñiga, Edison Chavez, Bernardo Gutiérrez, Verónica Barragán, Patricio Rojas-Silva, Gabriel Trueba, Michelle Grunauer, Paúl Cárdenas |
| EPI_ISL_697800 | Institute of Microbiology, Universidad San Francisco de Quito | Institute of Microbiology, Universidad San Francisco de Quito | Belén Prado-Vivar, Sully Márquez, Juan José Guadalupe, Monica Becerra-Wong, Maureen Mosquera, Bernardo Gutiérrez, Verónica Barragán, Patricio Rojas-Silva, Gabriel Trueba, Michelle Grunauer, Paúl Cárdenas |
| EPI_ISL_702478 | Quadram Institute Bioscience | COVID-19 Genomics UK (COG-UK) Consortium | Dave J. Baker, Gemma L. Kay, Alp Aydin, Thanh Le-Viet, Steven Rudder, Ana P. Tedim, Anastasia Kolyva, Maria Diaz, Leonardo de Oliveira Martins, Nabil-Fareed Alikhan, Lizzie Meadows, Rachael Stanley, Ngozi Elumogo, Muhammed Yasir, Nicholas M. Thomson, Alexander J Trotter, Rachel Gilroy, Samuel Bloomfield, Claire Stuart, Andrew Bell, Reenesh Prakash, Samir Dervisevic, Alison E. Mather, John Wain, Mark Webber, Andrew J. Page, Justin O'Grady |
| EPI_ISL_702515, EPI_ISL_702558 | Liverpool Clinical Laboratories | COVID-19 Genomics UK (COG-UK) Consortium | Sam Haldenby, Anita Lucaci, Steve Paterson, Julian Hiscox, Alistair Darby, M Almsaud, A Alrezaihi, Muhannad Alruwaili, Stuart D Armstrong, Jones Benjamin, Eleanor G Bentley, Anu Chawla, Jordan J Clark, Angela Cowell, Richard Eccles, Isabel García-Dorival, Matthew Gemmell, Alessandro Gerada, PKF Gilmore, Richard Gregory, Ximeng Han, Catherine Hartley, Margaret Hughes, Miren Iturriza-Gomara, James Johnson, L Luu, Jenifer Manson, Charlotte Nelson, Elaine O'Toole, Cassie Olateju, Rebekah Penrice-Randal , Lucille Rainbow, N.P Randle, Trevor Ian Robinson, Parul Sharma, Ghada T Shawli, James P Stewart, Neil Swainston, Ecaterina Vamos, Joanne Watts, Mark Whitehead |
| EPI_ISL_702607 | Virology Department, Sheffield Teaching Hospitals NHS Foundation Trust/Department of Infection, Immunity and Cardiovascular Disease, The Medical School, University of Sheffield | COVID-19 Genomics UK (COG-UK) Consortium | Thushan de Silva, Matthew Parker, Nikki Smith, Adri Angyal, Rebecca Brown, Luke Green, Rachel Tucker, Paul Parsons, Danielle Groves, Katie Johnson, Laura Carrilero, Alex Keeley, Dave Partridge, Matthew Wyles, Benjamin Lindsey, Mehmet Yavuz, Mohammad Raza, Cariad Evans |
| EPI_ISL_702652 | Quadram Institute Bioscience | COVID-19 Genomics UK (COG-UK) Consortium | Dave J. Baker, Gemma L. Kay, Alp Aydin, Thanh Le-Viet, Steven Rudder, Ana P. Tedim, Anastasia Kolyva, Maria Diaz, Leonardo de Oliveira Martins, Nabil-Fareed Alikhan, Lizzie Meadows, Rachael Stanley, Ngozi Elumogo, Muhammed Yasir, Nicholas M. Thomson, Alexander J Trotter, Rachel Gilroy, Samuel Bloomfield, Claire Stuart, Andrew Bell, Reenesh Prakash, Samir Dervisevic, Alison E. Mather, John Wain, Mark Webber, Andrew J. Page, Justin O'Grady |
| EPI_ISL_702661 | University College London, Great Ormond Street Hospital for Children NHS Foundation Trust, Imperial College Healthcare NHS Trust | COVID-19 Genomics UK (COG-UK) Consortium | Sergi Castellano, Rachel Williams, Mark Kristiansen, Paola Resende Silva, Sunando Roy, Tony Brooks, Helena Tutill, Paola Niola, Patricia Dyal, Charlotte Williams, Leysa Forrest, Yasmin Panchbhaya, Jacqueline Findlay, Samuel Weeks, Julianne Brown, Kathryn Harris, Paul Randell, James Price, Alison Holmes, Judith Breuer |
| EPI_ISL_702694, EPI_ISL_702749, EPI_ISL_702825, EPI_ISL_702840 | Quadram Institute Bioscience | COVID-19 Genomics UK (COG-UK) Consortium | Dave J. Baker, Gemma L. Kay, Alp Aydin, Thanh Le-Viet, Steven Rudder, Ana P. Tedim, Anastasia Kolyva, Maria Diaz, Leonardo de Oliveira Martins, Nabil-Fareed Alikhan, Lizzie Meadows, Rachael Stanley, Ngozi Elumogo, Muhammed Yasir, Nicholas M. Thomson, Alexander J Trotter, Rachel Gilroy, Samuel Bloomfield, Claire Stuart, Andrew Bell, Reenesh Prakash, Samir Dervisevic, Alison E. Mather, John Wain, Mark Webber, Andrew J. Page, Justin O'Grady |
| EPI_ISL_702852 | Virology Department, Sheffield Teaching Hospitals NHS Foundation Trust/Department of Infection, Immunity and Cardiovascular Disease, The Medical School, University of Sheffield | COVID-19 Genomics UK (COG-UK) Consortium | Thushan de Silva, Matthew Parker, Nikki Smith, Adri Angyal, Rebecca Brown, Luke Green, Rachel Tucker, Paul Parsons, Danielle Groves, Katie Johnson, Laura Carrilero, Alex Keeley, Dave Partridge, Matthew Wyles, Benjamin Lindsey, Mehmet Yavuz, Mohammad Raza, Cariad Evans |
| EPI_ISL_702873 | Quadram Institute Bioscience | COVID-19 Genomics UK (COG-UK) Consortium | Dave J. Baker, Gemma L. Kay, Alp Aydin, Thanh Le-Viet, Steven Rudder, Ana P. Tedim, Anastasia Kolyva, Maria Diaz, Leonardo de Oliveira Martins, Nabil-Fareed Alikhan, Lizzie Meadows, Rachael Stanley, Ngozi Elumogo, Muhammed Yasir, Nicholas M. Thomson, Alexander J Trotter, Rachel Gilroy, Samuel Bloomfield, Claire Stuart, Andrew Bell, Reenesh Prakash, Samir Dervisevic, Alison E. Mather, John Wain, Mark Webber, Andrew J. Page, Justin O'Grady |
| EPI_ISL_703069, EPI_ISL_703107 | University College London, Great Ormond Street Hospital for Children NHS Foundation Trust, Imperial College Healthcare NHS Trust | COVID-19 Genomics UK (COG-UK) Consortium | Sergi Castellano, Rachel Williams, Mark Kristiansen, Paola Resende Silva, Sunando Roy, Tony Brooks, Helena Tutill, Paola Niola, Patricia Dyal, Charlotte Williams, Leysa Forrest, Yasmin Panchbhaya, Jacqueline Findlay, Samuel Weeks, Julianne Brown, Kathryn Harris, Paul Randell, James Price, Alison Holmes, Judith Breuer |
| EPI_ISL_703133 | Virology Department, Sheffield Teaching Hospitals NHS Foundation Trust/Department of Infection, Immunity and Cardiovascular Disease, The Medical School, University of Sheffield | COVID-19 Genomics UK (COG-UK) Consortium | Thushan de Silva, Matthew Parker, Nikki Smith, Adri Angyal, Rebecca Brown, Luke Green, Rachel Tucker, Paul Parsons, Danielle Groves, Katie Johnson, Laura Carrilero, Alex Keeley, Dave Partridge, Matthew Wyles, Benjamin Lindsey, Mehmet Yavuz, Mohammad Raza, Cariad Evans |

| | | | |
|---|---|---|---|
| EPI_ISL_703136 | West of Scotland Specialist Virology Centre, NHSGGC / MRC-University of Glasgow Centre for Virus Research | COVID-19 Genomics UK (COG-UK) Consortium | Ana da Silva Filipe, Natasha Johnson, Kathy Smollett, Daniel Mair, Stephen Carmichael, Alice Broos, Lily Tong, Jenna Nichols, Kyriaki Nomikou; Sarah McDonald; Richard Orton, Joseph Hughes, Sreenu Vattipally, David L Robertson; Alasdair MacLean, Rory Gunson; Sharif Shaaban, Matthew Holden; Rachel Blacow, Guy Mollett, Kathy Li, James Shepherd, Antonia Ho, Emma Thomson |
| EPI_ISL_703295 | Quadram Institute Bioscience | COVID-19 Genomics UK (COG-UK) Consortium | Dave J. Baker, Gemma L. Kay, Alp Aydin, Thanh Le-Viet, Steven Rudder, Ana P. Tedim, Anastasia Kolyva, Maria Diaz, Leonardo de Oliveira Martins, Nabil-Fareed Alikhan, Lizzie Meadows, Rachael Stanley, Ngozi Elumogo, Muhammed Yasir, Nicholas M. Thomson, Alexander J Trotter, Rachel Gilroy, Samuel Bloomfield, Claire Stuart, Andrew Bell, Reenesh Prakash, Samir Dervisevic, Alison E. Mather, John Wain, Mark Webber, Andrew J. Page, Justin O'Grady |
| EPI_ISL_703310 | Liverpool Clinical Laboratories | COVID-19 Genomics UK (COG-UK) Consortium | Sam Haldenby, Anita Lucaci, Steve Paterson, Julian Hiscox, Alistair Darby, M Almsaud, A Alrezaihi, Muhannad Alruwaili, Stuart D Armstrong, Jones Benjamin, Eleanor G Bentley, Anu Chawla, Jordan J Clark, Angela Cowell, Richard Eccles, Isabel García-Dorival, Matthew Gemmell, Alessandro Gerada, PKF Gilmore, Richard Gregory, Ximeng Han, Catherine Hartley, Margaret Hughes, Miren Iturriza-Gomara, James Johnson, L Luu, Jenifer Manson, Charlotte Nelson, Elaine O'Toole, Cassie Olateju, Rebekah Penrice-Randal , Lucille Rainbow, N.P Randle, Trevor Ian Robinson, Parul Sharma, Ghada T Shawli, James P Stewart, Neil Swainston, Ecaterina Vamos, Joanne Watts, Mark Whitehead |
| EPI_ISL_703369 | University College London Hospital | COVID-19 Genomics UK (COG-UK) Consortium | Judith Heaney, Matthew Byott, Catherine Houlihan, Dan Frampton, Stuart Kirk, Moira Spyer and Eleni Nastouli |
| EPI_ISL_703409 | Liverpool Clinical Laboratories | COVID-19 Genomics UK (COG-UK) Consortium | Sam Haldenby, Anita Lucaci, Steve Paterson, Julian Hiscox, Alistair Darby, M Almsaud, A Alrezaihi, Muhannad Alruwaili, Stuart D Armstrong, Jones Benjamin, Eleanor G Bentley, Anu Chawla, Jordan J Clark, Angela Cowell, Richard Eccles, Isabel García-Dorival, Matthew Gemmell, Alessandro Gerada, PKF Gilmore, Richard Gregory, Ximeng Han, Catherine Hartley, Margaret Hughes, Miren Iturriza-Gomara, James Johnson, L Luu, Jenifer Manson, Charlotte Nelson, Elaine O'Toole, Cassie Olateju, Rebekah Penrice-Randal , Lucille Rainbow, N.P Randle, Trevor Ian Robinson, Parul Sharma, Ghada T Shawli, James P Stewart, Neil Swainston, Ecaterina Vamos, Joanne Watts, Mark Whitehead |
| EPI_ISL_703414 | Virology Department, Royal Infirmary of Edinburgh, NHS Lothian / School of Biological Sciences, University of Edinburgh / Institute of Genetics and Molecular Medicine, University of Edinburgh | COVID-19 Genomics UK (COG-UK) Consortium | McHugh M, Dewar R, Rooke S, Gallagher M, Balcaza C, O'Toole Á, Scher E, Hill V, McCrone JT, Colquhoun R, Yu X, Jackson B, Rambaut A, Williams TC, Templeton K |
| EPI_ISL_703611 | Liverpool Clinical Laboratories | COVID-19 Genomics UK (COG-UK) Consortium | Sam Haldenby, Anita Lucaci, Steve Paterson, Julian Hiscox, Alistair Darby, M Almsaud, A Alrezaihi, Muhannad Alruwaili, Stuart D Armstrong, Jones Benjamin, Eleanor G Bentley, Anu Chawla, Jordan J Clark, Angela Cowell, Richard Eccles, Isabel García-Dorival, Matthew Gemmell, Alessandro Gerada, PKF Gilmore, Richard Gregory, Ximeng Han, Catherine Hartley, Margaret Hughes, Miren Iturriza-Gomara, James Johnson, L Luu, Jenifer Manson, Charlotte Nelson, Elaine O'Toole, Cassie Olateju, Rebekah Penrice-Randal , Lucille Rainbow, N.P Randle, Trevor Ian Robinson, Parul Sharma, Ghada T Shawli, James P Stewart, Neil Swainston, Ecaterina Vamos, Joanne Watts, Mark Whitehead |
| EPI_ISL_703649 | Virology Department, Sheffield Teaching Hospitals NHS Foundation Trust/Department of Infection, Immunity and Cardiovascular Disease, The Medical School, University of Sheffield | COVID-19 Genomics UK (COG-UK) Consortium | Thushan de Silva, Matthew Parker, Nikki Smith, Adri Angyal, Rebecca Brown, Luke Green, Rachel Tucker, Paul Parsons, Danielle Groves, Katie Johnson, Laura Carrilero, Alex Keeley, Dave Partridge, Matthew Wyles, Benjamin Lindsey, Mehmet Yavuz, Mohammad Raza, Cariad Evans |
| EPI_ISL_703758 | West of Scotland Specialist Virology Centre, NHSGGC / MRC-University of Glasgow Centre for Virus Research | COVID-19 Genomics UK (COG-UK) Consortium | Ana da Silva Filipe, Natasha Johnson, Kathy Smollett, Daniel Mair, Stephen Carmichael, Alice Broos, Lily Tong, Jenna Nichols, Kyriaki Nomikou; Sarah McDonald; Richard Orton, Joseph Hughes, Sreenu Vattipally, David L Robertson; Alasdair MacLean, Rory Gunson; Sharif Shaaban, Matthew Holden; Rachel Blacow, Guy Mollett, Kathy Li, James Shepherd, Antonia Ho, Emma Thomson |
| EPI_ISL_703765 | Virology Department, Sheffield Teaching Hospitals NHS Foundation Trust/Department of Infection, Immunity and Cardiovascular Disease, The Medical School, University of Sheffield | COVID-19 Genomics UK (COG-UK) Consortium | Thushan de Silva, Matthew Parker, Nikki Smith, Adri Angyal, Rebecca Brown, Luke Green, Rachel Tucker, Paul Parsons, Danielle Groves, Katie Johnson, Laura Carrilero, Alex Keeley, Dave Partridge, Matthew Wyles, Benjamin Lindsey, Mehmet Yavuz, Mohammad Raza, Cariad Evans |
| EPI_ISL_703886 | University College London Hospital | COVID-19 Genomics UK (COG-UK) Consortium | Judith Heaney, Matthew Byott, Catherine Houlihan, Dan Frampton, Stuart Kirk, Moira Spyer and Eleni Nastouli |
| EPI_ISL_703927 | West of Scotland Specialist Virology Centre, NHSGGC / MRC-University of Glasgow Centre for Virus Research | COVID-19 Genomics UK (COG-UK) Consortium | Ana da Silva Filipe, Natasha Johnson, Kathy Smollett, Daniel Mair, Stephen Carmichael, Alice Broos, Lily Tong, Jenna Nichols, Kyriaki Nomikou; Sarah McDonald; Richard Orton, Joseph Hughes, Sreenu Vattipally, David L Robertson; Alasdair MacLean, Rory Gunson; Sharif Shaaban, Matthew Holden; Rachel Blacow, Guy Mollett, Kathy Li, James Shepherd, Antonia Ho, Emma Thomson |
| EPI_ISL_703982, EPI_ISL_704070 | Quadram Institute Bioscience | COVID-19 Genomics UK (COG-UK) Consortium | Dave J. Baker, Gemma L. Kay, Alp Aydin, Thanh Le-Viet, Steven Rudder, Ana P. Tedim, Anastasia Kolyva, Maria Diaz, Leonardo de Oliveira Martins, Nabil-Fareed Alikhan, Lizzie Meadows, Rachael Stanley, Ngozi Elumogo, Muhammed Yasir, Nicholas M. Thomson, Alexander J Trotter, Rachel Gilroy, Samuel Bloomfield, Claire Stuart, Andrew Bell, Reenesh Prakash, Samir Dervisevic, Alison E. Mather, John Wain, Mark Webber, Andrew J. Page, Justin O'Grady |
| EPI_ISL_704105 | University College London Hospital | COVID-19 Genomics UK (COG-UK) Consortium | Judith Heaney, Matthew Byott, Catherine Houlihan, Dan Frampton, Stuart Kirk, Moira Spyer and Eleni Nastouli |
| EPI_ISL_704142, EPI_ISL_704145, EPI_ISL_704148, EPI_ISL_704176, EPI_ISL_704217, EPI_ISL_704233, EPI_ISL_704238 | Quadram Institute Bioscience | COVID-19 Genomics UK (COG-UK) Consortium | Dave J. Baker, Gemma L. Kay, Alp Aydin, Thanh Le-Viet, Steven Rudder, Ana P. Tedim, Anastasia Kolyva, Maria Diaz, Leonardo de Oliveira Martins, Nabil-Fareed Alikhan, Lizzie Meadows, Rachael Stanley, Ngozi Elumogo, Muhammed Yasir, Nicholas M. Thomson, Alexander J Trotter, Rachel Gilroy, Samuel Bloomfield, Claire Stuart, Andrew Bell, Reenesh Prakash, Samir Dervisevic, Alison E. Mather, John Wain, Mark Webber, Andrew J. Page, Justin O'Grady |
| EPI_ISL_704596, EPI_ISL_704597 | West of Scotland Specialist Virology Centre, NHSGGC / MRC-University of Glasgow Centre for Virus Research | COVID-19 Genomics UK (COG-UK) Consortium | Ana da Silva Filipe, Natasha Johnson, Kathy Smollett, Daniel Mair, Stephen Carmichael, Alice Broos, Lily Tong, Jenna Nichols, Kyriaki Nomikou; Sarah McDonald; Richard Orton, Joseph Hughes, Sreenu Vattipally, David L Robertson; Alasdair MacLean, Rory Gunson; Sharif Shaaban, Matthew Holden; Rachel Blacow, Guy Mollett, Kathy Li, James Shepherd, Antonia Ho, Emma Thomson |
| EPI_ISL_704608 | University College London Hospital | COVID-19 Genomics UK (COG-UK) Consortium | Judith Heaney, Matthew Byott, Catherine Houlihan, Dan Frampton, Stuart Kirk, Moira Spyer and Eleni Nastouli |
| EPI_ISL_704609 | Quadram Institute Bioscience | COVID-19 Genomics UK (COG-UK) Consortium | Dave J. Baker, Gemma L. Kay, Alp Aydin, Thanh Le-Viet, Steven Rudder, Ana P. Tedim, Anastasia Kolyva, Maria Diaz, Leonardo de Oliveira Martins, Nabil-Fareed Alikhan, Lizzie Meadows, Rachael Stanley, Ngozi Elumogo, Muhammed Yasir, Nicholas M. Thomson, Alexander J Trotter, Rachel Gilroy, Samuel Bloomfield, Claire Stuart, Andrew Bell, Reenesh Prakash, Samir Dervisevic, Alison E. Mather, John Wain, Mark Webber, Andrew J. Page, Justin O'Grady |
| EPI_ISL_704669, EPI_ISL_704674 | University College London Hospital | COVID-19 Genomics UK (COG-UK) Consortium | Judith Heaney, Matthew Byott, Catherine Houlihan, Dan Frampton, Stuart Kirk, Moira Spyer and Eleni Nastouli |
| EPI_ISL_704709, EPI_ISL_704711, EPI_ISL_704714, EPI_ISL_704716, EPI_ISL_704728, EPI_ISL_704731, EPI_ISL_704734, EPI_ISL_704736, EPI_ISL_704739, EPI_ISL_704746, EPI_ISL_704788, EPI_ISL_704791, EPI_ISL_704794 | | | |
| see above | Quadram Institute Bioscience | COVID-19 Genomics UK (COG-UK) Consortium | Dave J. Baker, Gemma L. Kay, Alp Aydin, Thanh Le-Viet, Steven Rudder, Ana P. Tedim, Anastasia Kolyva, Maria Diaz, Leonardo de Oliveira Martins, Nabil-Fareed Alikhan, Lizzie Meadows, Rachael Stanley, Ngozi Elumogo, Muhammed Yasir, Nicholas M. Thomson, Alexander J Trotter, Rachel Gilroy, Samuel Bloomfield, Claire Stuart, Andrew Bell, Reenesh Prakash, Samir Dervisevic, Alison E. Mather, John Wain, Mark Webber, Andrew J. Page, Justin O'Grady |
| EPI_ISL_704804, EPI_ISL_704807 | West of Scotland Specialist Virology Centre, NHSGGC / MRC-University of Glasgow Centre for Virus Research | COVID-19 Genomics UK (COG-UK) Consortium | Ana da Silva Filipe, Natasha Johnson, Kathy Smollett, Daniel Mair, Stephen Carmichael, Alice Broos, Lily Tong, Jenna Nichols, Kyriaki Nomikou; Sarah McDonald; Richard Orton, Joseph Hughes, Sreenu Vattipally, David L Robertson; Alasdair MacLean, Rory Gunson; Sharif Shaaban, Matthew Holden; Rachel Blacow, Guy Mollett, Kathy Li, James Shepherd, Antonia Ho, Emma Thomson |
| EPI_ISL_704850, EPI_ISL_704853, EPI_ISL_704887, EPI_ISL_704890, EPI_ISL_704923, EPI_ISL_704926, EPI_ISL_704929 | Quadram Institute Bioscience | COVID-19 Genomics UK (COG-UK) Consortium | Dave J. Baker, Gemma L. Kay, Alp Aydin, Thanh Le-Viet, Steven Rudder, Ana P. Tedim, Anastasia Kolyva, Maria Diaz, Leonardo de Oliveira Martins, Nabil-Fareed Alikhan, Lizzie Meadows, Rachael Stanley, Ngozi Elumogo, Muhammed Yasir, Nicholas M. Thomson, Alexander J Trotter, Rachel Gilroy, Samuel Bloomfield, Claire Stuart, Andrew Bell, Reenesh Prakash, Samir Dervisevic, Alison E. Mather, John Wain, Mark Webber, Andrew J. Page, Justin O'Grady |
| EPI_ISL_704963 | University College London, Great Ormond Street Hospital for Children NHS Foundation Trust, Imperial College Healthcare NHS Trust | COVID-19 Genomics UK (COG-UK) Consortium | Sergi Castellano, Rachel Williams, Mark Kristiansen, Paola Resende Silva, Sunando Roy, Tony Brooks, Helena Tutill, Paola Niola, Patricia Dyal, Charlotte Williams, Leysa Forrest, Yasmin Panchbhaya, Jacqueline Findlay, Samuel Weeks, Julianne Brown, Kathryn Harris, Paul Randell, James Price, Alison Holmes, Judith Breuer |

| | | | |
|---|---|---|---|
| EPI_ISL_705002 | West of Scotland Specialist Virology Centre, NHSGGC / MRC-University of Glasgow Centre for Virus Research | COVID-19 Genomics UK (COG-UK) Consortium | Ana da Silva Filipe, Natasha Johnson, Kathy Smollett, Daniel Mair, Stephen Carmichael, Alice Broos, Lily Tong, Jenna Nichols, Kyriaki Nomikou; Sarah McDonald; Richard Orton, Joseph Hughes, Sreenu Vattipally, David L Robertson; Alasdair MacLean, Rory Gunson; Sharif Shaaban, Matthew Holden; Rachel Blacow, Guy Mollett, Kathy Li, James Shepherd, Antonia Ho, Emma Thomson |
| EPI_ISL_705091 | Quadram Institute Bioscience | COVID-19 Genomics UK (COG-UK) Consortium | Dave J. Baker, Gemma L. Kay, Alp Aydin, Thanh Le-Viet, Steven Rudder, Ana P. Tedim, Anastasia Kolyva, Maria Diaz, Leonardo de Oliveira Martins, Nabil-Fareed Alikhan, Lizzie Meadows, Rachael Stanley, Ngozi Elumogo, Muhammed Yasir, Nicholas M. Thomson, Alexander J Trotter, Rachel Gilroy, Samuel Bloomfield, Claire Stuart, Andrew Bell, Reenesh Prakash, Samir Dervisevic, Alison E. Mather, John Wain, Mark Webber, Andrew J. Page, Justin O'Grady |
| EPI_ISL_705129, EPI_ISL_705220 | Virology Department, Sheffield Teaching Hospitals NHS Foundation Trust/Department of Infection, Immunity and Cardiovascular Disease, The Medical School, University of Sheffield | COVID-19 Genomics UK (COG-UK) Consortium | Thushan de Silva, Matthew Parker, Nikki Smith, Adri Angyal, Rebecca Brown, Luke Green, Rachel Tucker, Paul Parsons, Danielle Groves, Katie Johnson, Laura Carrilero, Alex Keeley, Dave Partridge, Matthew Wyles, Benjamin Lindsey, Mehmet Yavuz, Mohammad Raza, Cariad Evans |
| EPI_ISL_705232 | University College London, Great Ormond Street Hospital for Children NHS Foundation Trust, Imperial College Healthcare NHS Trust | COVID-19 Genomics UK (COG-UK) Consortium | Sergi Castellano, Rachel Williams, Mark Kristiansen, Paola Resende Silva, Sunando Roy, Tony Brooks, Helena Tutill, Paola Niola, Patricia Dyal, Charlotte Williams, Leysa Forrest, Yasmin Panchbhaya, Jacqueline Findlay, Samuel Weeks, Julianne Brown, Kathryn Harris, Paul Randell, James Price, Alison Holmes, Judith Breuer |
| EPI_ISL_705245 | Liverpool Clinical Laboratories | COVID-19 Genomics UK (COG-UK) Consortium | Sam Haldenby, Anita Lucaci, Steve Paterson, Julian Hiscox, Alistair Darby, M Almsaud, A Alrezaihi, Muhannad Alruwaili, Stuart D Armstrong, Jones Benjamin, Eleanor G Bentley, Anu Chawla, Jordan J Clark, Angela Cowell, Richard Eccles, Isabel García-Dorival, Matthew Gemmell, Alessandro Gerada, PKF Gilmore, Richard Gregory, Ximeng Han, Catherine Hartley, Margaret Hughes, Miren Iturriza-Gomara, James Johnson, L Luu, Jenifer Manson, Charlotte Nelson, Elaine O'Toole, Cassie Olateju, Rebekah Penrice-Randal , Lucille Rainbow, N.P Randle, Trevor Ian Robinson, Parul Sharma, Ghada T Shawli, James P Stewart, Neil Swainston, Ecaterina Vamos, Joanne Watts, Mark Whitehead |
| EPI_ISL_705255, EPI_ISL_705256, EPI_ISL_705261 | Quadram Institute Bioscience | COVID-19 Genomics UK (COG-UK) Consortium | Dave J. Baker, Gemma L. Kay, Alp Aydin, Thanh Le-Viet, Steven Rudder, Ana P. Tedim, Anastasia Kolyva, Maria Diaz, Leonardo de Oliveira Martins, Nabil-Fareed Alikhan, Lizzie Meadows, Rachael Stanley, Ngozi Elumogo, Muhammed Yasir, Nicholas M. Thomson, Alexander J Trotter, Rachel Gilroy, Samuel Bloomfield, Claire Stuart, Andrew Bell, Reenesh Prakash, Samir Dervisevic, Alison E. Mather, John Wain, Mark Webber, Andrew J. Page, Justin O'Grady |
| EPI_ISL_705285, EPI_ISL_705297 | University College London, Great Ormond Street Hospital for Children NHS Foundation Trust, Imperial College Healthcare NHS Trust | COVID-19 Genomics UK (COG-UK) Consortium | Sergi Castellano, Rachel Williams, Mark Kristiansen, Paola Resende Silva, Sunando Roy, Tony Brooks, Helena Tutill, Paola Niola, Patricia Dyal, Charlotte Williams, Leysa Forrest, Yasmin Panchbhaya, Jacqueline Findlay, Samuel Weeks, Julianne Brown, Kathryn Harris, Paul Randell, James Price, Alison Holmes, Judith Breuer |
| EPI_ISL_705345, EPI_ISL_705394 | Liverpool Clinical Laboratories | COVID-19 Genomics UK (COG-UK) Consortium | Sam Haldenby, Anita Lucaci, Steve Paterson, Julian Hiscox, Alistair Darby, M Almsaud, A Alrezaihi, Muhannad Alruwaili, Stuart D Armstrong, Jones Benjamin, Eleanor G Bentley, Anu Chawla, Jordan J Clark, Angela Cowell, Richard Eccles, Isabel García-Dorival, Matthew Gemmell, Alessandro Gerada, PKF Gilmore, Richard Gregory, Ximeng Han, Catherine Hartley, Margaret Hughes, Miren Iturriza-Gomara, James Johnson, L Luu, Jenifer Manson, Charlotte Nelson, Elaine O'Toole, Cassie Olateju, Rebekah Penrice-Randal , Lucille Rainbow, N.P Randle, Trevor Ian Robinson, Parul Sharma, Ghada T Shawli, James P Stewart, Neil Swainston, Ecaterina Vamos, Joanne Watts, Mark Whitehead |
| EPI_ISL_705431 | Quadram Institute Bioscience | COVID-19 Genomics UK (COG-UK) Consortium | Dave J. Baker, Gemma L. Kay, Alp Aydin, Thanh Le-Viet, Steven Rudder, Ana P. Tedim, Anastasia Kolyva, Maria Diaz, Leonardo de Oliveira Martins, Nabil-Fareed Alikhan, Lizzie Meadows, Rachael Stanley, Ngozi Elumogo, Muhammed Yasir, Nicholas M. Thomson, Alexander J Trotter, Rachel Gilroy, Samuel Bloomfield, Claire Stuart, Andrew Bell, Reenesh Prakash, Samir Dervisevic, Alison E. Mather, John Wain, Mark Webber, Andrew J. Page, Justin O'Grady |
| EPI_ISL_705492 | Virology Department, Royal Infirmary of Edinburgh, NHS Lothian / School of Biological Sciences, University of Edinburgh / Institute of Genetics and Molecular Medicine, University of Edinburgh | COVID-19 Genomics UK (COG-UK) Consortium | McHugh M, Dewar R, Rooke S, Gallagher M, Balcaza C, O'Toole Á, Scher E, Hill V, McCrone JT, Colquhoun R, Yu X, Jackson B, Rambaut A, Williams TC, Templeton K |
| EPI_ISL_705715, EPI_ISL_705717, EPI_ISL_705721, EPI_ISL_705723, EPI_ISL_705724, EPI_ISL_705725, EPI_ISL_705728 | West of Scotland Specialist Virology Centre, NHSGGC / MRC-University of Glasgow Centre for Virus Research | COVID-19 Genomics UK (COG-UK) Consortium | Ana da Silva Filipe, Natasha Johnson, Kathy Smollett, Daniel Mair, Stephen Carmichael, Alice Broos, Lily Tong, Jenna Nichols, Kyriaki Nomikou; Sarah McDonald; Richard Orton, Joseph Hughes, Sreenu Vattipally, David L Robertson; Alasdair MacLean, Rory Gunson; Sharif Shaaban, Matthew Holden; Rachel Blacow, Guy Mollett, Kathy Li, James Shepherd, Antonia Ho, Emma Thomson |
| EPI_ISL_705777, EPI_ISL_705780, EPI_ISL_705781, EPI_ISL_705782, EPI_ISL_705783, EPI_ISL_705784, EPI_ISL_705785, EPI_ISL_705786, EPI_ISL_705787, EPI_ISL_705788, EPI_ISL_705789, EPI_ISL_705790, EPI_ISL_705791, EPI_ISL_705792 | | | |
| see above | Virology Department, Royal Infirmary of Edinburgh, NHS Lothian / School of Biological Sciences, University of Edinburgh / Institute of Genetics and Molecular Medicine, University of Edinburgh | COVID-19 Genomics UK (COG-UK) Consortium | McHugh M, Dewar R, Rooke S, Gallagher M, Balcaza C, O'Toole Á, Scher E, Hill V, McCrone JT, Colquhoun R, Yu X, Jackson B, Rambaut A, Williams TC, Templeton K |
| EPI_ISL_705928 | Liverpool Clinical Laboratories | COVID-19 Genomics UK (COG-UK) Consortium | Sam Haldenby, Anita Lucaci, Steve Paterson, Julian Hiscox, Alistair Darby, M Almsaud, A Alrezaihi, Muhannad Alruwaili, Stuart D Armstrong, Jones Benjamin, Eleanor G Bentley, Anu Chawla, Jordan J Clark, Angela Cowell, Richard Eccles, Isabel García-Dorival, Matthew Gemmell, Alessandro Gerada, PKF Gilmore, Richard Gregory, Ximeng Han, Catherine Hartley, Margaret Hughes, Miren Iturriza-Gomara, James Johnson, L Luu, Jenifer Manson, Charlotte Nelson, Elaine O'Toole, Cassie Olateju, Rebekah Penrice-Randal , Lucille Rainbow, N.P Randle, Trevor Ian Robinson, Parul Sharma, Ghada T Shawli, James P Stewart, Neil Swainston, Ecaterina Vamos, Joanne Watts, Mark Whitehead |
| EPI_ISL_705957, EPI_ISL_705959 | University College London Hospital | COVID-19 Genomics UK (COG-UK) Consortium | Judith Heaney, Matthew Byott, Catherine Houlihan, Dan Frampton, Stuart Kirk, Moira Spyer and Eleni Nastouli |
| EPI_ISL_706357, EPI_ISL_706358, EPI_ISL_706360, EPI_ISL_706367, EPI_ISL_706368, EPI_ISL_706369, EPI_ISL_706370, EPI_ISL_706371, EPI_ISL_706372, EPI_ISL_706373, EPI_ISL_706374, EPI_ISL_706375, EPI_ISL_706376, EPI_ISL_706377, EPI_ISL_706378, EPI_ISL_706379, EPI_ISL_706381, EPI_ISL_706382, EPI_ISL_706397, EPI_ISL_706398, EPI_ISL_706399, EPI_ISL_706401, EPI_ISL_706403, EPI_ISL_706404, EPI_ISL_706405, EPI_ISL_706406, EPI_ISL_706407, EPI_ISL_706409, EPI_ISL_706410, EPI_ISL_706411, EPI_ISL_706413, EPI_ISL_706415, EPI_ISL_706416, EPI_ISL_706418, EPI_ISL_706419 | | | |
| see above | Quadram Institute Bioscience | COVID-19 Genomics UK (COG-UK) Consortium | Dave J. Baker, Gemma L. Kay, Alp Aydin, Thanh Le-Viet, Steven Rudder, Ana P. Tedim, Anastasia Kolyva, Maria Diaz, Leonardo de Oliveira Martins, Nabil-Fareed Alikhan, Lizzie Meadows, Rachael Stanley, Ngozi Elumogo, Muhammed Yasir, Nicholas M. Thomson, Alexander J Trotter, Rachel Gilroy, Samuel Bloomfield, Claire Stuart, Andrew Bell, Reenesh Prakash, Samir Dervisevic, Alison E. Mather, John Wain, Mark Webber, Andrew J. Page, Justin O'Grady |
| EPI_ISL_706429, EPI_ISL_706430 | Queens Medical Centre, Clinical Microbiology Department / DeepSeq Nottingham | COVID-19 Genomics UK (COG-UK) Consortium | Gemma Clark, Wendy Smith, Manjinder Khakh, Vicki M Fleming, Michelle M Lister, Hannah Howson-Wells, Jonathan Ball, Patrick McClure, Joseph Chappell, Theocharis Tsoleridis, Nadine Holmes, Matthew Carlisle, Christopher Moore, Fei Sang, Johnny Debebe, Victoria Wright, Matthew Loose |
| EPI_ISL_706975, EPI_ISL_706978, EPI_ISL_706990, EPI_ISL_707004 | Virology Department, Sheffield Teaching Hospitals NHS Foundation Trust/Department of Infection, Immunity and Cardiovascular Disease, The Medical School, University of Sheffield | COVID-19 Genomics UK (COG-UK) Consortium | Thushan de Silva, Matthew Parker, Nikki Smith, Adri Angyal, Rebecca Brown, Luke Green, Rachel Tucker, Paul Parsons, Danielle Groves, Katie Johnson, Laura Carrilero, Alex Keeley, Dave Partridge, Matthew Wyles, Benjamin Lindsey, Mehmet Yavuz, Mohammad Raza, Cariad Evans |
| EPI_ISL_707708 | Institute for Urban Disease Control and Prevention | COVID-19 Network Investigations (CONI) Alliance | Kamolthip Atsawawaranunt, Elizabeth Batty, Wasun Chantratita, Thanat Chookajorn, Stefan Fernandez, Angkana Huang, Anthony R. Jones, Khajohn Joonsalak, Prayuth Kaewmalang, Amornmas Kongklieng, Chonticha Klungtong, Theerarat Kochakarn, Namfon Kotanan, Krittikorn Kumpornsin, Wudtichai Manasatienkij, Anek Mungaomklang, Bhakbhoom Panthan, Ekawat Pasomsub, Pukkaporn Parnwijitkul, Kingkan Rakmanee, Insee Sensorn, Janjira Thaipadungpanit, Arporn Wangwiwatsin,Treewat Watthanachockchai |
| EPI_ISL_707795, EPI_ISL_707796, EPI_ISL_707797, EPI_ISL_707798 | LabPLUS | Institute of Environmental Science and Research (ESR) | Xiaoyun Ren, Matt Storey, Nikki Freed, Muhammad Faisal, Jing Wang, Hermes Perez, Anja Werno, Antje van der Linden, Arlo Upton, Chris Mansell, David Hammer, Dragana Drinkovic, Gary McAuliffe, Hana Sofia Andersson, James Ussher, Jill Sherwood, Josh Freeman, Julia Howard, Juliet Elvy, Mary DeAlmeida, Matt Blakiston, Matthew Rogers, Max Bloomfield, Michael Addidle, Michelle Balm, Sally Roberts, Sarah Jefferies, Sharmini Muttaiyah, Susan Morpeth, Susan Taylor, Timothy Blackmore, Vani Sathyendran, Veronica Playle, Virginia Hope, Erasmus Smit, Lauren Jelly, Olin Silander, Joep de Ligt |
| EPI_ISL_708493, EPI_ISL_708494, EPI_ISL_708495, EPI_ISL_708496, EPI_ISL_708497, EPI_ISL_708498, | Minnesota Department of Health, Public Health Laboratory | Minnesota Department of Health, Public Health Laboratory | Alexandra Lorentz, Jacob Garfin, Matt Plumb, and Xiong Wang |

| | | | |
|---|---|---|---|
| EPI_ISL_708499, EPI_ISL_708503 | | | |
| EPI_ISL_708793, EPI_ISL_708795 | PathWest Laboratory Medicine WA | PathWest Laboratory Medicine WA Microbial Surveillance Unit | PathWest Laboratory Medicine WA Microbial Surveillance Unit |
| EPI_ISL_708844, EPI_ISL_708847, EPI_ISL_708848, EPI_ISL_708849, EPI_ISL_708850 | Lighthouse Lab in Milton Keynes | Wellcome Sanger Institute for the COVID-19 Genomics UK (COG-UK) Consortium | The Lighthouse Lab in Milton Keynes and Alex Alderton, Roberto Amato, Sonia Goncalves, Ewan Harrison, David K. Jackson, Ian Johnston, Dominic Kwiatkowski, Cordelia Langford, John Sillitoe on behalf of the Wellcome Sanger Institute COVID-19 Surveillance Team |
| EPI_ISL_708851 | Lighthouse Lab in Alderley Park | Wellcome Sanger Institute for the COVID-19 Genomics UK (COG-UK) Consortium | Jacquelyn Wynn, Mairead Hyland, The Lighthouse Lab in Alderley Park and Alex Alderton, Roberto Amato, Sonia Goncalves, Ewan Harrison, David K. Jackson, Ian Johnston, Dominic Kwiatkowski, Cordelia Langford, John Sillitoe on behalf of the Wellcome Sanger Institute COVID-19 Surveillance Team |
| EPI_ISL_708856, EPI_ISL_708857, EPI_ISL_708858, EPI_ISL_708862, EPI_ISL_708869, EPI_ISL_708871 | Lighthouse Lab in Milton Keynes | Wellcome Sanger Institute for the COVID-19 Genomics UK (COG-UK) Consortium | The Lighthouse Lab in Milton Keynes and Alex Alderton, Roberto Amato, Sonia Goncalves, Ewan Harrison, David K. Jackson, Ian Johnston, Dominic Kwiatkowski, Cordelia Langford, John Sillitoe on behalf of the Wellcome Sanger Institute COVID-19 Surveillance Team |
| EPI_ISL_708873, EPI_ISL_708875 | Lighthouse Lab in Alderley Park | Wellcome Sanger Institute for the COVID-19 Genomics UK (COG-UK) Consortium | Jacquelyn Wynn, Mairead Hyland, The Lighthouse Lab in Alderley Park and Alex Alderton, Roberto Amato, Sonia Goncalves, Ewan Harrison, David K. Jackson, Ian Johnston, Dominic Kwiatkowski, Cordelia Langford, John Sillitoe on behalf of the Wellcome Sanger Institute COVID-19 Surveillance Team |
| EPI_ISL_708879 | Lighthouse Lab in Milton Keynes | Wellcome Sanger Institute for the COVID-19 Genomics UK (COG-UK) Consortium | The Lighthouse Lab in Milton Keynes and Alex Alderton, Roberto Amato, Sonia Goncalves, Ewan Harrison, David K. Jackson, Ian Johnston, Dominic Kwiatkowski, Cordelia Langford, John Sillitoe on behalf of the Wellcome Sanger Institute COVID-19 Surveillance Team |
| EPI_ISL_708880 | Lighthouse Lab in Alderley Park | Wellcome Sanger Institute for the COVID-19 Genomics UK (COG-UK) Consortium | Jacquelyn Wynn, Mairead Hyland, The Lighthouse Lab in Alderley Park and Alex Alderton, Roberto Amato, Sonia Goncalves, Ewan Harrison, David K. Jackson, Ian Johnston, Dominic Kwiatkowski, Cordelia Langford, John Sillitoe on behalf of the Wellcome Sanger Institute COVID-19 Surveillance Team |
| EPI_ISL_708881, EPI_ISL_708883 | Lighthouse Lab in Milton Keynes | Wellcome Sanger Institute for the COVID-19 Genomics UK (COG-UK) Consortium | The Lighthouse Lab in Milton Keynes and Alex Alderton, Roberto Amato, Sonia Goncalves, Ewan Harrison, David K. Jackson, Ian Johnston, Dominic Kwiatkowski, Cordelia Langford, John Sillitoe on behalf of the Wellcome Sanger Institute COVID-19 Surveillance Team |
| EPI_ISL_708885, EPI_ISL_708886 | Lighthouse Lab in Alderley Park | Wellcome Sanger Institute for the COVID-19 Genomics UK (COG-UK) Consortium | Jacquelyn Wynn, Mairead Hyland, The Lighthouse Lab in Alderley Park and Alex Alderton, Roberto Amato, Sonia Goncalves, Ewan Harrison, David K. Jackson, Ian Johnston, Dominic Kwiatkowski, Cordelia Langford, John Sillitoe on behalf of the Wellcome Sanger Institute COVID-19 Surveillance Team |
| EPI_ISL_708889 | Lighthouse Lab in Milton Keynes | Wellcome Sanger Institute for the COVID-19 Genomics UK (COG-UK) Consortium | The Lighthouse Lab in Milton Keynes and Alex Alderton, Roberto Amato, Sonia Goncalves, Ewan Harrison, David K. Jackson, Ian Johnston, Dominic Kwiatkowski, Cordelia Langford, John Sillitoe on behalf of the Wellcome Sanger Institute COVID-19 Surveillance Team |
| EPI_ISL_708893 | Lighthouse Lab in Glasgow | Wellcome Sanger Institute for the COVID-19 Genomics UK (COG-UK) Consortium | Harper VanSteenhouse, Yumi Kasai, David Gray, Carol Clugston, Anna Dominiczak and Alex Alderton, Roberto Amato, Sonia Goncalves, Ewan Harrison, David K. Jackson, Ian Johnston, Dominic Kwiatkowski, Cordelia Langford, John Sillitoe on behalf of the Wellcome Sanger Institute COVID-19 Surveillance Team |
| EPI_ISL_708894 | Lighthouse Lab in Alderley Park | Wellcome Sanger Institute for the COVID-19 Genomics UK (COG-UK) Consortium | Jacquelyn Wynn, Mairead Hyland, The Lighthouse Lab in Alderley Park and Alex Alderton, Roberto Amato, Sonia Goncalves, Ewan Harrison, David K. Jackson, Ian Johnston, Dominic Kwiatkowski, Cordelia Langford, John Sillitoe on behalf of the Wellcome Sanger Institute COVID-19 Surveillance Team |
| EPI_ISL_708896, EPI_ISL_708897, EPI_ISL_708898 | Lighthouse Lab in Milton Keynes | Wellcome Sanger Institute for the COVID-19 Genomics UK (COG-UK) Consortium | The Lighthouse Lab in Milton Keynes and Alex Alderton, Roberto Amato, Sonia Goncalves, Ewan Harrison, David K. Jackson, Ian Johnston, Dominic Kwiatkowski, Cordelia Langford, John Sillitoe on behalf of the Wellcome Sanger Institute COVID-19 Surveillance Team |
| EPI_ISL_708899 | Lighthouse Lab in Alderley Park | Wellcome Sanger Institute for the COVID-19 Genomics UK (COG-UK) Consortium | Jacquelyn Wynn, Mairead Hyland, The Lighthouse Lab in Alderley Park and Alex Alderton, Roberto Amato, Sonia Goncalves, Ewan Harrison, David K. Jackson, Ian Johnston, Dominic Kwiatkowski, Cordelia Langford, John Sillitoe on behalf of the Wellcome Sanger Institute COVID-19 Surveillance Team |
| EPI_ISL_708900, EPI_ISL_708901, EPI_ISL_708908, EPI_ISL_708912, EPI_ISL_708913 | Lighthouse Lab in Milton Keynes | Wellcome Sanger Institute for the COVID-19 Genomics UK (COG-UK) Consortium | The Lighthouse Lab in Milton Keynes and Alex Alderton, Roberto Amato, Sonia Goncalves, Ewan Harrison, David K. Jackson, Ian Johnston, Dominic Kwiatkowski, Cordelia Langford, John Sillitoe on behalf of the Wellcome Sanger Institute COVID-19 Surveillance Team |
| EPI_ISL_708915 | Lighthouse Lab in Alderley Park | Wellcome Sanger Institute for the COVID-19 Genomics UK (COG-UK) Consortium | Jacquelyn Wynn, Mairead Hyland, The Lighthouse Lab in Alderley Park and Alex Alderton, Roberto Amato, Sonia Goncalves, Ewan Harrison, David K. Jackson, Ian Johnston, Dominic Kwiatkowski, Cordelia Langford, John Sillitoe on behalf of the Wellcome Sanger Institute COVID-19 Surveillance Team |
| EPI_ISL_708916 | Lighthouse Lab in Milton Keynes | Wellcome Sanger Institute for the COVID-19 Genomics UK (COG-UK) Consortium | The Lighthouse Lab in Milton Keynes and Alex Alderton, Roberto Amato, Sonia Goncalves, Ewan Harrison, David K. Jackson, Ian Johnston, Dominic Kwiatkowski, Cordelia Langford, John Sillitoe on behalf of the Wellcome Sanger Institute COVID-19 Surveillance Team |
| EPI_ISL_708918 | Lighthouse Lab in Alderley Park | Wellcome Sanger Institute for the COVID-19 Genomics UK (COG-UK) Consortium | Jacquelyn Wynn, Mairead Hyland, The Lighthouse Lab in Alderley Park and Alex Alderton, Roberto Amato, Sonia Goncalves, Ewan Harrison, David K. Jackson, Ian Johnston, Dominic Kwiatkowski, Cordelia Langford, John Sillitoe on behalf of the Wellcome Sanger Institute COVID-19 Surveillance Team |
| EPI_ISL_708920 | Lighthouse Lab in Milton Keynes | Wellcome Sanger Institute for the COVID-19 Genomics UK (COG-UK) Consortium | The Lighthouse Lab in Milton Keynes and Alex Alderton, Roberto Amato, Sonia Goncalves, Ewan Harrison, David K. Jackson, Ian Johnston, Dominic Kwiatkowski, Cordelia Langford, John Sillitoe on behalf of the Wellcome Sanger Institute COVID-19 Surveillance Team |
| EPI_ISL_708921, EPI_ISL_708925 | Lighthouse Lab in Glasgow | Wellcome Sanger Institute for the COVID-19 Genomics UK (COG-UK) Consortium | Harper VanSteenhouse, Yumi Kasai, David Gray, Carol Clugston, Anna Dominiczak and Alex Alderton, Roberto Amato, Sonia Goncalves, Ewan Harrison, David K. Jackson, Ian Johnston, Dominic Kwiatkowski, Cordelia Langford, John Sillitoe on behalf of the Wellcome Sanger Institute COVID-19 Surveillance Team |
| EPI_ISL_708926, EPI_ISL_708927 | Lighthouse Lab in Alderley Park | Wellcome Sanger Institute for the COVID-19 Genomics UK (COG-UK) Consortium | Jacquelyn Wynn, Mairead Hyland, The Lighthouse Lab in Alderley Park and Alex Alderton, Roberto Amato, Sonia Goncalves, Ewan Harrison, David K. Jackson, Ian Johnston, Dominic Kwiatkowski, Cordelia Langford, John Sillitoe on behalf of the Wellcome Sanger Institute COVID-19 Surveillance Team |
| EPI_ISL_708928, EPI_ISL_708930 | Lighthouse Lab in Milton Keynes | Wellcome Sanger Institute for the COVID-19 Genomics UK (COG-UK) Consortium | The Lighthouse Lab in Milton Keynes and Alex Alderton, Roberto Amato, Sonia Goncalves, Ewan Harrison, David K. Jackson, Ian Johnston, Dominic Kwiatkowski, Cordelia Langford, John Sillitoe on behalf of the Wellcome Sanger Institute COVID-19 Surveillance Team |
| EPI_ISL_708934, EPI_ISL_708936, EPI_ISL_708938, EPI_ISL_708939 | Lighthouse Lab in Alderley Park | Wellcome Sanger Institute for the COVID-19 Genomics UK (COG-UK) Consortium | Jacquelyn Wynn, Mairead Hyland, The Lighthouse Lab in Alderley Park and Alex Alderton, Roberto Amato, Sonia Goncalves, Ewan Harrison, David K. Jackson, Ian Johnston, Dominic Kwiatkowski, Cordelia Langford, John Sillitoe on behalf of the Wellcome Sanger Institute COVID-19 Surveillance Team |
| EPI_ISL_708941, EPI_ISL_708945, EPI_ISL_708948, EPI_ISL_708951 | Lighthouse Lab in Milton Keynes | Wellcome Sanger Institute for the COVID-19 Genomics UK (COG-UK) Consortium | The Lighthouse Lab in Milton Keynes and Alex Alderton, Roberto Amato, Sonia Goncalves, Ewan Harrison, David K. Jackson, Ian Johnston, Dominic Kwiatkowski, Cordelia Langford, John Sillitoe on behalf of the Wellcome Sanger Institute COVID-19 Surveillance Team |
| EPI_ISL_708952 | Lighthouse Lab in Alderley Park | Wellcome Sanger Institute for the COVID-19 Genomics UK (COG-UK) Consortium | Jacquelyn Wynn, Mairead Hyland, The Lighthouse Lab in Alderley Park and Alex Alderton, Roberto Amato, Sonia Goncalves, Ewan Harrison, David K. Jackson, Ian Johnston, Dominic Kwiatkowski, Cordelia Langford, John Sillitoe on behalf of the Wellcome Sanger Institute COVID-19 Surveillance Team |
| EPI_ISL_708958, EPI_ISL_708963, EPI_ISL_708964 | Lighthouse Lab in Milton Keynes | Wellcome Sanger Institute for the COVID-19 Genomics UK (COG-UK) Consortium | The Lighthouse Lab in Milton Keynes and Alex Alderton, Roberto Amato, Sonia Goncalves, Ewan Harrison, David K. Jackson, Ian Johnston, Dominic Kwiatkowski, Cordelia Langford, John Sillitoe on behalf of the Wellcome Sanger Institute COVID-19 Surveillance Team |
| EPI_ISL_708967 | Lighthouse Lab in Alderley Park | Wellcome Sanger Institute for the COVID-19 Genomics UK (COG-UK) Consortium | Jacquelyn Wynn, Mairead Hyland, The Lighthouse Lab in Alderley Park and Alex Alderton, Roberto Amato, Sonia Goncalves, Ewan Harrison, David K. Jackson, Ian Johnston, Dominic Kwiatkowski, Cordelia Langford, John Sillitoe on behalf of the Wellcome Sanger Institute COVID-19 Surveillance Team |
| EPI_ISL_708969 | Lighthouse Lab in Milton Keynes | Wellcome Sanger Institute for the COVID-19 Genomics UK (COG-UK) Consortium | The Lighthouse Lab in Milton Keynes and Alex Alderton, Roberto Amato, Sonia Goncalves, Ewan Harrison, David K. Jackson, Ian Johnston, Dominic Kwiatkowski, Cordelia Langford, John Sillitoe on behalf of the Wellcome Sanger Institute COVID-19 Surveillance Team |
| EPI_ISL_708970, EPI_ISL_708971 | Lighthouse Lab in Alderley Park | Wellcome Sanger Institute for the COVID-19 Genomics UK (COG-UK) Consortium | Jacquelyn Wynn, Mairead Hyland, The Lighthouse Lab in Alderley Park and Alex Alderton, Roberto Amato, Sonia Goncalves, Ewan Harrison, David K. Jackson, Ian Johnston, Dominic Kwiatkowski, Cordelia Langford, John Sillitoe on behalf of the Wellcome Sanger Institute COVID-19 Surveillance Team |
| EPI_ISL_708975, EPI_ISL_708979 | Lighthouse Lab in Milton Keynes | Wellcome Sanger Institute for the COVID-19 Genomics UK (COG-UK) Consortium | The Lighthouse Lab in Milton Keynes and Alex Alderton, Roberto Amato, Sonia Goncalves, Ewan Harrison, David K. Jackson, Ian Johnston, Dominic Kwiatkowski, Cordelia Langford, John Sillitoe on behalf of the Wellcome Sanger Institute COVID-19 Surveillance Team |
| EPI_ISL_708981 | Lighthouse Lab in Alderley Park | Wellcome Sanger Institute for the COVID-19 Genomics UK (COG-UK) Consortium | Jacquelyn Wynn, Mairead Hyland, The Lighthouse Lab in Alderley Park and Alex Alderton, Roberto Amato, Sonia Goncalves, Ewan Harrison, David K. Jackson, Ian Johnston, Dominic Kwiatkowski, Cordelia Langford, John Sillitoe on behalf of the Wellcome Sanger Institute COVID-19 Surveillance Team |
| EPI_ISL_708983, EPI_ISL_708984 | Lighthouse Lab in Milton Keynes | Wellcome Sanger Institute for the COVID-19 Genomics UK (COG-UK) Consortium | The Lighthouse Lab in Milton Keynes and Alex Alderton, Roberto Amato, Sonia Goncalves, Ewan Harrison, David K. Jackson, Ian Johnston, Dominic Kwiatkowski, Cordelia Langford, John Sillitoe on behalf of the Wellcome Sanger Institute COVID-19 Surveillance Team |
| EPI_ISL_708985 | Lighthouse Lab in Glasgow | Wellcome Sanger Institute for the COVID-19 Genomics UK (COG-UK) Consortium | Harper VanSteenhouse, Yumi Kasai, David Gray, Carol Clugston, Anna Dominiczak and Alex Alderton, Roberto Amato, Sonia Goncalves, Ewan Harrison, David K. Jackson, Ian Johnston, Dominic Kwiatkowski, Cordelia Langford, John Sillitoe on behalf of the Wellcome Sanger Institute COVID-19 Surveillance |

| | | | |
|---|---|---|---|
| | | | Team |
| EPI_ISL_708986 | Lighthouse Lab in Milton Keynes | Wellcome Sanger Institute for the COVID-19 Genomics UK (COG-UK) Consortium | The Lighthouse Lab in Milton Keynes and Alex Alderton, Roberto Amato, Sonia Goncalves, Ewan Harrison, David K. Jackson, Ian Johnston, Dominic Kwiatkowski, Cordelia Langford, John Sillitoe on behalf of the Wellcome Sanger Institute COVID-19 Surveillance Team |
| EPI_ISL_709480 | Lighthouse Lab in Alderley Park | Wellcome Sanger Institute for the COVID-19 Genomics UK (COG-UK) Consortium | Jacquelyn Wynn, Mairead Hyland, The Lighthouse Lab in Alderley Park and Alex Alderton, Roberto Amato, Sonia Goncalves, Ewan Harrison, David K. Jackson, Ian Johnston, Dominic Kwiatkowski, Cordelia Langford, John Sillitoe on behalf of the Wellcome Sanger Institute COVID-19 Surveillance Team |
| EPI_ISL_709800 | Lighthouse Lab in Glasgow | Wellcome Sanger Institute for the COVID-19 Genomics UK (COG-UK) Consortium | Harper VanSteenhouse, Yumi Kasai, David Gray, Carol Clugston, Anna Dominiczak and Alex Alderton, Roberto Amato, Sonia Goncalves, Ewan Harrison, David K. Jackson, Ian Johnston, Dominic Kwiatkowski, Cordelia Langford, John Sillitoe on behalf of the Wellcome Sanger Institute COVID-19 Surveillance Team |
| EPI_ISL_709804 | Lighthouse Lab in Cambridge | Wellcome Sanger Institute for the COVID-19 Genomics UK (COG-UK) Consortium | Rob Howes, The Lighthouse Lab in Cambridge and Alex Alderton, Roberto Amato, Sonia Goncalves, Ewan Harrison, David K. Jackson, Ian Johnston, Dominic Kwiatkowski, Cordelia Langford, John Sillitoe on behalf of the Wellcome Sanger Institute COVID-19 Surveillance Team |
| EPI_ISL_709806, EPI_ISL_709809, EPI_ISL_709811, EPI_ISL_709812, EPI_ISL_709816, EPI_ISL_709818, EPI_ISL_709820 | Lighthouse Lab in Glasgow | Wellcome Sanger Institute for the COVID-19 Genomics UK (COG-UK) Consortium | Harper VanSteenhouse, Yumi Kasai, David Gray, Carol Clugston, Anna Dominiczak and Alex Alderton, Roberto Amato, Sonia Goncalves, Ewan Harrison, David K. Jackson, Ian Johnston, Dominic Kwiatkowski, Cordelia Langford, John Sillitoe on behalf of the Wellcome Sanger Institute COVID-19 Surveillance Team |
| EPI_ISL_709821 | Lighthouse Lab in Cambridge | Wellcome Sanger Institute for the COVID-19 Genomics UK (COG-UK) Consortium | Rob Howes, The Lighthouse Lab in Cambridge and Alex Alderton, Roberto Amato, Sonia Goncalves, Ewan Harrison, David K. Jackson, Ian Johnston, Dominic Kwiatkowski, Cordelia Langford, John Sillitoe on behalf of the Wellcome Sanger Institute COVID-19 Surveillance Team |
| EPI_ISL_709826 | Lighthouse Lab in Glasgow | Wellcome Sanger Institute for the COVID-19 Genomics UK (COG-UK) Consortium | Harper VanSteenhouse, Yumi Kasai, David Gray, Carol Clugston, Anna Dominiczak and Alex Alderton, Roberto Amato, Sonia Goncalves, Ewan Harrison, David K. Jackson, Ian Johnston, Dominic Kwiatkowski, Cordelia Langford, John Sillitoe on behalf of the Wellcome Sanger Institute COVID-19 Surveillance Team |
| EPI_ISL_709827 | Lighthouse Lab in Cambridge | Wellcome Sanger Institute for the COVID-19 Genomics UK (COG-UK) Consortium | Rob Howes, The Lighthouse Lab in Cambridge and Alex Alderton, Roberto Amato, Sonia Goncalves, Ewan Harrison, David K. Jackson, Ian Johnston, Dominic Kwiatkowski, Cordelia Langford, John Sillitoe on behalf of the Wellcome Sanger Institute COVID-19 Surveillance Team |
| EPI_ISL_709831, EPI_ISL_709832, EPI_ISL_709834, EPI_ISL_709835 | Lighthouse Lab in Glasgow | Wellcome Sanger Institute for the COVID-19 Genomics UK (COG-UK) Consortium | Harper VanSteenhouse, Yumi Kasai, David Gray, Carol Clugston, Anna Dominiczak and Alex Alderton, Roberto Amato, Sonia Goncalves, Ewan Harrison, David K. Jackson, Ian Johnston, Dominic Kwiatkowski, Cordelia Langford, John Sillitoe on behalf of the Wellcome Sanger Institute COVID-19 Surveillance Team |
| EPI_ISL_709836 | Lighthouse Lab in Cambridge | Wellcome Sanger Institute for the COVID-19 Genomics UK (COG-UK) Consortium | Rob Howes, The Lighthouse Lab in Cambridge and Alex Alderton, Roberto Amato, Sonia Goncalves, Ewan Harrison, David K. Jackson, Ian Johnston, Dominic Kwiatkowski, Cordelia Langford, John Sillitoe on behalf of the Wellcome Sanger Institute COVID-19 Surveillance Team |
| EPI_ISL_709840, EPI_ISL_709845, EPI_ISL_709848, EPI_ISL_709849, EPI_ISL_709851 | Lighthouse Lab in Glasgow | Wellcome Sanger Institute for the COVID-19 Genomics UK (COG-UK) Consortium | Harper VanSteenhouse, Yumi Kasai, David Gray, Carol Clugston, Anna Dominiczak and Alex Alderton, Roberto Amato, Sonia Goncalves, Ewan Harrison, David K. Jackson, Ian Johnston, Dominic Kwiatkowski, Cordelia Langford, John Sillitoe on behalf of the Wellcome Sanger Institute COVID-19 Surveillance Team |
| EPI_ISL_709854 | Lighthouse Lab in Cambridge | Wellcome Sanger Institute for the COVID-19 Genomics UK (COG-UK) Consortium | Rob Howes, The Lighthouse Lab in Cambridge and Alex Alderton, Roberto Amato, Sonia Goncalves, Ewan Harrison, David K. Jackson, Ian Johnston, Dominic Kwiatkowski, Cordelia Langford, John Sillitoe on behalf of the Wellcome Sanger Institute COVID-19 Surveillance Team |
| EPI_ISL_709855, EPI_ISL_709857, EPI_ISL_709858, EPI_ISL_709859, EPI_ISL_709861, EPI_ISL_709862, EPI_ISL_709875 | Lighthouse Lab in Glasgow | Wellcome Sanger Institute for the COVID-19 Genomics UK (COG-UK) Consortium | Harper VanSteenhouse, Yumi Kasai, David Gray, Carol Clugston, Anna Dominiczak and Alex Alderton, Roberto Amato, Sonia Goncalves, Ewan Harrison, David K. Jackson, Ian Johnston, Dominic Kwiatkowski, Cordelia Langford, John Sillitoe on behalf of the Wellcome Sanger Institute COVID-19 Surveillance Team |
| EPI_ISL_709948 | Microbiology, Department of Pathology, St. Bernard's Hospital, Gibraltar Health Authority | Respiratory Virus Unit, National Infection Service, Public Health England | PHE Covid Sequencing Team, Dr Nicholas Cortes (Gibraltar), Charlotte Gillborn-Jones (Gibraltar) |
| EPI_ISL_709949 | Respiratory Virus Unit, National Infection Service, Public Health England | COVID-19 Genomics UK (COG-UK) Consortium | PHE Covid Sequencing Team |
| EPI_ISL_709950, EPI_ISL_709951, EPI_ISL_709952, EPI_ISL_709953, EPI_ISL_709954, EPI_ISL_709955, EPI_ISL_709956, EPI_ISL_709957 | Microbiology, Department of Pathology, St. Bernard's Hospital, Gibraltar Health Authority | Respiratory Virus Unit, National Infection Service, Public Health England | PHE Covid Sequencing Team, Dr Nicholas Cortes (Gibraltar), Charlotte Gillborn-Jones (Gibraltar) |
| EPI_ISL_709958 | Respiratory Virus Unit, National Infection Service, Public Health England | COVID-19 Genomics UK (COG-UK) Consortium | PHE Covid Sequencing Team |
| EPI_ISL_709959, EPI_ISL_709960, EPI_ISL_709961, EPI_ISL_709962, EPI_ISL_709963, EPI_ISL_709964, EPI_ISL_709965, EPI_ISL_709966, EPI_ISL_709967, EPI_ISL_709968, EPI_ISL_709969, EPI_ISL_709970, EPI_ISL_709971, EPI_ISL_709972, EPI_ISL_709973, EPI_ISL_709974, EPI_ISL_709975, EPI_ISL_709976, EPI_ISL_709977, EPI_ISL_709978, EPI_ISL_709979, EPI_ISL_709980, EPI_ISL_709981, EPI_ISL_709982, EPI_ISL_709983, EPI_ISL_709984, EPI_ISL_709985, EPI_ISL_709986, EPI_ISL_709987, EPI_ISL_709988, EPI_ISL_709989, EPI_ISL_709990, EPI_ISL_709991, EPI_ISL_709992, EPI_ISL_709993, EPI_ISL_709994, EPI_ISL_709995, EPI_ISL_709996, EPI_ISL_709997, EPI_ISL_709998, EPI_ISL_709999, EPI_ISL_710000, EPI_ISL_710001, EPI_ISL_710002, EPI_ISL_710003, EPI_ISL_710004, EPI_ISL_710005, EPI_ISL_710006, EPI_ISL_710007, EPI_ISL_710008, EPI_ISL_710009, EPI_ISL_710010, EPI_ISL_710011, EPI_ISL_710012, EPI_ISL_710013, EPI_ISL_710014, EPI_ISL_710015, EPI_ISL_710016, EPI_ISL_710017, EPI_ISL_710018, EPI_ISL_710019, EPI_ISL_710020, EPI_ISL_710021, EPI_ISL_710022, EPI_ISL_710023, EPI_ISL_710024, EPI_ISL_710025, EPI_ISL_710026, EPI_ISL_710027, EPI_ISL_710028, EPI_ISL_710029, EPI_ISL_710030, EPI_ISL_710031, EPI_ISL_710032, EPI_ISL_710033 | | | |
| see above | Microbiology, Department of Pathology, St. Bernard's Hospital, Gibraltar Health Authority | Respiratory Virus Unit, National Infection Service, Public Health England | PHE Covid Sequencing Team, Dr Nicholas Cortes (Gibraltar), Charlotte Gillborn-Jones (Gibraltar) |
| EPI_ISL_710125, EPI_ISL_710127, EPI_ISL_710128 | South Eastern Area Laboratory Services (SEALS) | CIDM-PH et al. | CIDM-PH et al. |
| EPI_ISL_710485, EPI_ISL_710486 | Utah Public Health Laboratory, Utah Public Health Laboratory Infectious Disease submission group | Utah Public Health Laboratory, Utah Public Health Laboratory Infectious Disease submission group | Young,E.L., Oakeson,K.F. |
| EPI_ISL_710535, EPI_ISL_710536 | Microbiology, Department of Pathology, St. Bernard's Hospital, Gibraltar Health Authority | Respiratory Virus Unit, National Infection Service, Public Health England | PHE Covid Sequencing Team, Dr Nicholas Cortes (Gibraltar), Charlotte Gillborn-Jones (Gibraltar) |
| EPI_ISL_710616, EPI_ISL_710617, EPI_ISL_710625, EPI_ISL_710628, EPI_ISL_710636, EPI_ISL_710645, EPI_ISL_710648 | Lighthouse Lab in Alderley Park | Wellcome Sanger Institute for the COVID-19 Genomics UK (COG-UK) Consortium | Jacquelyn Wynn, Mairead Hyland, The Lighthouse Lab in Alderley Park and Alex Alderton, Roberto Amato, Sonia Goncalves, Ewan Harrison, David K. Jackson, Ian Johnston, Dominic Kwiatkowski, Cordelia Langford, John Sillitoe on behalf of the Wellcome Sanger Institute COVID-19 Surveillance Team |
| EPI_ISL_711060, EPI_ISL_711061, EPI_ISL_711062, EPI_ISL_711063, EPI_ISL_711064, EPI_ISL_711065, EPI_ISL_711066, EPI_ISL_711068, EPI_ISL_711069, EPI_ISL_711070, EPI_ISL_711071, EPI_ISL_711072, EPI_ISL_711073, EPI_ISL_711075, EPI_ISL_711079, EPI_ISL_711080, EPI_ISL_711081, EPI_ISL_711082, EPI_ISL_711083, EPI_ISL_711084, EPI_ISL_711085, EPI_ISL_711086, EPI_ISL_711087, EPI_ISL_711088, EPI_ISL_711089, EPI_ISL_711090, EPI_ISL_711091, EPI_ISL_711092, EPI_ISL_711093, EPI_ISL_711094, EPI_ISL_711095, EPI_ISL_711096, EPI_ISL_711097, EPI_ISL_711098, EPI_ISL_711099, EPI_ISL_711100, EPI_ISL_711101, EPI_ISL_711102, EPI_ISL_711103, EPI_ISL_711104, EPI_ISL_711105, EPI_ISL_711106, EPI_ISL_711107, EPI_ISL_711108, EPI_ISL_711109, EPI_ISL_711110, EPI_ISL_711111, EPI_ISL_711112, EPI_ISL_711113, EPI_ISL_711114, EPI_ISL_711115, EPI_ISL_711116, EPI_ISL_711117, EPI_ISL_711118, EPI_ISL_711119, EPI_ISL_711120, EPI_ISL_711121, EPI_ISL_711122, EPI_ISL_711123, EPI_ISL_711124, EPI_ISL_711125, EPI_ISL_711126, EPI_ISL_711127, EPI_ISL_711128, EPI_ISL_711129, EPI_ISL_711130, EPI_ISL_711131, EPI_ISL_711132, EPI_ISL_711133, EPI_ISL_711134, EPI_ISL_711135, EPI_ISL_711136, EPI_ISL_711137, EPI_ISL_711138, EPI_ISL_711139, EPI_ISL_711140, EPI_ISL_711141, EPI_ISL_711142, EPI_ISL_711143, EPI_ISL_711144, EPI_ISL_711145, EPI_ISL_711146, EPI_ISL_711147, EPI_ISL_711148, EPI_ISL_711176, EPI_ISL_711182, EPI_ISL_711183, EPI_ISL_711184, EPI_ISL_711186, EPI_ISL_711187, EPI_ISL_711188, EPI_ISL_711189, EPI_ISL_711193, EPI_ISL_711196, EPI_ISL_711197, EPI_ISL_711200, EPI_ISL_711202, EPI_ISL_711203, EPI_ISL_711204, EPI_ISL_711205, EPI_ISL_711206, EPI_ISL_711214, EPI_ISL_711215, EPI_ISL_711219, EPI_ISL_711221, EPI_ISL_711222, EPI_ISL_711223, EPI_ISL_711224, EPI_ISL_711225, EPI_ISL_711226, EPI_ISL_711227, EPI_ISL_711229, EPI_ISL_711230, EPI_ISL_711231, EPI_ISL_711233, EPI_ISL_711234, EPI_ISL_711235, EPI_ISL_711236, EPI_ISL_711237, EPI_ISL_711238, EPI_ISL_711239, EPI_ISL_711240, EPI_ISL_711241, EPI_ISL_711242, EPI_ISL_711243, EPI_ISL_711244, EPI_ISL_711245, EPI_ISL_711246, EPI_ISL_711247, EPI_ISL_711248, EPI_ISL_711249, EPI_ISL_711250, EPI_ISL_711251, EPI_ISL_711252, EPI_ISL_711253, EPI_ISL_711254, EPI_ISL_711255, EPI_ISL_711266, EPI_ISL_711267, EPI_ISL_711269, EPI_ISL_711270, EPI_ISL_711275, EPI_ISL_711278, EPI_ISL_711279, EPI_ISL_711280, EPI_ISL_711282, EPI_ISL_711283, EPI_ISL_711284, EPI_ISL_711285, EPI_ISL_711286, EPI_ISL_711287, EPI_ISL_711288, EPI_ISL_711289, EPI_ISL_711290, EPI_ISL_711291, EPI_ISL_711292, EPI_ISL_711293, EPI_ISL_711294, EPI_ISL_711299, EPI_ISL_711300, EPI_ISL_711301, EPI_ISL_711302, EPI_ISL_711303, EPI_ISL_711304, EPI_ISL_711305, EPI_ISL_711306, EPI_ISL_711307, EPI_ISL_711308, EPI_ISL_711309, EPI_ISL_711310, EPI_ISL_711311, EPI_ISL_711312, EPI_ISL_711313, EPI_ISL_711314, EPI_ISL_711315, EPI_ISL_711316, EPI_ISL_711317, EPI_ISL_711318, EPI_ISL_711319, EPI_ISL_711320, EPI_ISL_711321, EPI_ISL_711322, EPI_ISL_711323, EPI_ISL_711324, EPI_ISL_711325, EPI_ISL_711326, EPI_ISL_711327, EPI_ISL_711328, EPI_ISL_711329, EPI_ISL_711330, EPI_ISL_711331, EPI_ISL_711332, EPI_ISL_711333, EPI_ISL_711334, EPI_ISL_711335, EPI_ISL_711336, EPI_ISL_711337, EPI_ISL_711338, EPI_ISL_711339, EPI_ISL_711340, EPI_ISL_711341, EPI_ISL_711342, EPI_ISL_711343, EPI_ISL_711344, EPI_ISL_711345, EPI_ISL_711346, EPI_ISL_711347, EPI_ISL_711348, EPI_ISL_711349, EPI_ISL_711350, EPI_ISL_711351, EPI_ISL_711352, EPI_ISL_711353, EPI_ISL_711354, EPI_ISL_711355, EPI_ISL_711356, EPI_ISL_711357, EPI_ISL_711358, EPI_ISL_711359, EPI_ISL_711360, EPI_ISL_711384, EPI_ISL_711385, EPI_ISL_711386, EPI_ISL_711387, EPI_ISL_711388, EPI_ISL_711389, EPI_ISL_711393, EPI_ISL_711394, EPI_ISL_711398, EPI_ISL_711399, EPI_ISL_711400, EPI_ISL_711401, EPI_ISL_711402, EPI_ISL_711403, | | | |

| | | | |
|---|---|---|---|
| EPI_ISL_711404, EPI_ISL_711405, EPI_ISL_711406, EPI_ISL_711409, EPI_ISL_711410, EPI_ISL_711412, EPI_ISL_711413, EPI_ISL_711414, EPI_ISL_711415, EPI_ISL_711416, EPI_ISL_711417, EPI_ISL_711418, EPI_ISL_711419, EPI_ISL_711420, EPI_ISL_711421, EPI_ISL_711422, EPI_ISL_711423, EPI_ISL_711424, EPI_ISL_711425, EPI_ISL_711426, EPI_ISL_711427, EPI_ISL_711428, EPI_ISL_711431, EPI_ISL_711432, EPI_ISL_711433, EPI_ISL_711434, EPI_ISL_711435, EPI_ISL_711436, EPI_ISL_711437, EPI_ISL_711445, EPI_ISL_711446, EPI_ISL_711447, EPI_ISL_711448, EPI_ISL_711449, EPI_ISL_711450, EPI_ISL_711451, EPI_ISL_711452, EPI_ISL_711453, EPI_ISL_711454, EPI_ISL_711455, EPI_ISL_711456, EPI_ISL_711457, EPI_ISL_711458, EPI_ISL_711459, EPI_ISL_711460, EPI_ISL_711461, EPI_ISL_711462, EPI_ISL_711463, EPI_ISL_711464, EPI_ISL_711465, EPI_ISL_711466, EPI_ISL_711467, EPI_ISL_711468, EPI_ISL_711469, EPI_ISL_711470, EPI_ISL_711471, EPI_ISL_711472, EPI_ISL_711473, EPI_ISL_711474, EPI_ISL_711475, EPI_ISL_711476, EPI_ISL_711477, EPI_ISL_711478, EPI_ISL_711479, EPI_ISL_711480, EPI_ISL_711481, EPI_ISL_711482, EPI_ISL_711483, EPI_ISL_711484, EPI_ISL_711485, EPI_ISL_711486, EPI_ISL_711487, EPI_ISL_711488, EPI_ISL_711489, EPI_ISL_711490, EPI_ISL_711491, EPI_ISL_711492, EPI_ISL_711493, EPI_ISL_711494, EPI_ISL_711495, EPI_ISL_711496, EPI_ISL_711497, EPI_ISL_711498, EPI_ISL_711499, EPI_ISL_711500, EPI_ISL_711501, EPI_ISL_711502, EPI_ISL_711503, EPI_ISL_711504, EPI_ISL_711505, EPI_ISL_711506, EPI_ISL_711507, EPI_ISL_711508, EPI_ISL_711509, EPI_ISL_711510, EPI_ISL_711511, EPI_ISL_711512, EPI_ISL_711513, EPI_ISL_711514, EPI_ISL_711515, EPI_ISL_711516, EPI_ISL_711517, EPI_ISL_711518, EPI_ISL_711519, EPI_ISL_711520, EPI_ISL_711521, EPI_ISL_711522, EPI_ISL_711523, EPI_ISL_711524, EPI_ISL_711525, EPI_ISL_711526, EPI_ISL_711527, EPI_ISL_711528, EPI_ISL_711529, EPI_ISL_711530, EPI_ISL_711531, EPI_ISL_711532, EPI_ISL_711533, EPI_ISL_711534, EPI_ISL_711535, EPI_ISL_711536, EPI_ISL_711537, EPI_ISL_711538, EPI_ISL_711539, EPI_ISL_711540, EPI_ISL_711541, EPI_ISL_711542, EPI_ISL_711543, EPI_ISL_711544, EPI_ISL_711545, EPI_ISL_711546, EPI_ISL_711547, EPI_ISL_711548, EPI_ISL_711549, EPI_ISL_711550, EPI_ISL_711551, EPI_ISL_711552, EPI_ISL_711553, EPI_ISL_711554, EPI_ISL_711555, EPI_ISL_711556, EPI_ISL_711557, EPI_ISL_711558, EPI_ISL_711559, EPI_ISL_711560, EPI_ISL_711561, EPI_ISL_711562, EPI_ISL_711563, EPI_ISL_711564, EPI_ISL_711565, EPI_ISL_711566, EPI_ISL_711567, EPI_ISL_711568, EPI_ISL_711569, EPI_ISL_711570, EPI_ISL_711571, EPI_ISL_711572, EPI_ISL_711573, EPI_ISL_711574, EPI_ISL_711575, EPI_ISL_711576, EPI_ISL_711577, EPI_ISL_711578, EPI_ISL_711579, EPI_ISL_711580, EPI_ISL_711581, EPI_ISL_711582, EPI_ISL_711583, EPI_ISL_711584, EPI_ISL_711585, EPI_ISL_711586, EPI_ISL_711587, EPI_ISL_711588, EPI_ISL_711589, EPI_ISL_711590, EPI_ISL_711591, EPI_ISL_711592, EPI_ISL_711593, EPI_ISL_711594, EPI_ISL_711595, EPI_ISL_711596, EPI_ISL_711597, EPI_ISL_711598, EPI_ISL_711599, EPI_ISL_711600, EPI_ISL_711601, EPI_ISL_711602, EPI_ISL_711603, EPI_ISL_711604, EPI_ISL_711605, EPI_ISL_711606, EPI_ISL_711607, EPI_ISL_711608, EPI_ISL_711609, EPI_ISL_711610, EPI_ISL_711611, EPI_ISL_711612, EPI_ISL_711613, EPI_ISL_711614, EPI_ISL_711615, EPI_ISL_711616, EPI_ISL_711617, EPI_ISL_711618, EPI_ISL_711619, EPI_ISL_711620, EPI_ISL_711621, EPI_ISL_711622, EPI_ISL_711623, EPI_ISL_711624, EPI_ISL_711625, EPI_ISL_711626, EPI_ISL_711627, EPI_ISL_711628, EPI_ISL_711629, EPI_ISL_711630, EPI_ISL_711631, EPI_ISL_711632, EPI_ISL_711633, EPI_ISL_711634, EPI_ISL_711635, EPI_ISL_711636, EPI_ISL_711637, EPI_ISL_711638, EPI_ISL_711639, EPI_ISL_711640, EPI_ISL_711641, EPI_ISL_711642, EPI_ISL_711643, EPI_ISL_711644, EPI_ISL_711645, EPI_ISL_711646, EPI_ISL_711647, EPI_ISL_711648, EPI_ISL_711649, EPI_ISL_711650, EPI_ISL_711651, EPI_ISL_711652, EPI_ISL_711653, EPI_ISL_711654, EPI_ISL_711655, EPI_ISL_711656, EPI_ISL_711657, EPI_ISL_711658, EPI_ISL_711659, EPI_ISL_711660, EPI_ISL_711661, EPI_ISL_711662, EPI_ISL_711663, EPI_ISL_711664, EPI_ISL_711665, EPI_ISL_711666, EPI_ISL_711667, EPI_ISL_711668, EPI_ISL_711669, EPI_ISL_711670, EPI_ISL_711671, EPI_ISL_711672, EPI_ISL_711673, EPI_ISL_711674, EPI_ISL_711675, EPI_ISL_711676, EPI_ISL_711677, EPI_ISL_711678, EPI_ISL_711679, EPI_ISL_711680, EPI_ISL_711681, EPI_ISL_711682, EPI_ISL_711683, EPI_ISL_711684, EPI_ISL_711685, EPI_ISL_711686, EPI_ISL_711687, EPI_ISL_711688, EPI_ISL_711689, EPI_ISL_711690, EPI_ISL_711691, EPI_ISL_711692, EPI_ISL_711693, EPI_ISL_711694, EPI_ISL_711695, EPI_ISL_711696, EPI_ISL_711697, EPI_ISL_711698, EPI_ISL_711699, EPI_ISL_711700, EPI_ISL_711701, EPI_ISL_711702, EPI_ISL_711703, EPI_ISL_711704, EPI_ISL_711705, EPI_ISL_711706, EPI_ISL_711707, EPI_ISL_711708, EPI_ISL_711709, EPI_ISL_711710, EPI_ISL_711711, EPI_ISL_711712, EPI_ISL_711713, EPI_ISL_711714, EPI_ISL_711715, EPI_ISL_711716, EPI_ISL_711717, EPI_ISL_711718, EPI_ISL_711719, EPI_ISL_711720, EPI_ISL_711721, EPI_ISL_711722, EPI_ISL_711723, EPI_ISL_711724, EPI_ISL_711725, EPI_ISL_711726, EPI_ISL_711727, EPI_ISL_711728, EPI_ISL_711729, EPI_ISL_711730, EPI_ISL_711731, EPI_ISL_711732, EPI_ISL_711733, EPI_ISL_711734, EPI_ISL_711735, EPI_ISL_711736, EPI_ISL_711737, EPI_ISL_711738, EPI_ISL_711739, EPI_ISL_711740, EPI_ISL_711741, EPI_ISL_711742, EPI_ISL_711743, EPI_ISL_711744, EPI_ISL_711745, EPI_ISL_711746, EPI_ISL_711747, EPI_ISL_711748, EPI_ISL_711749, EPI_ISL_711750, EPI_ISL_711751, EPI_ISL_711752, EPI_ISL_711753, EPI_ISL_711754, EPI_ISL_711755, EPI_ISL_711756, EPI_ISL_711757, EPI_ISL_711758, EPI_ISL_711759, EPI_ISL_711760, EPI_ISL_711761, EPI_ISL_711762, EPI_ISL_711763, EPI_ISL_711764, EPI_ISL_711765, EPI_ISL_711766, EPI_ISL_711767, EPI_ISL_711768, EPI_ISL_711769, EPI_ISL_711770, EPI_ISL_711771, EPI_ISL_711772, EPI_ISL_711773, EPI_ISL_711774, EPI_ISL_711775, EPI_ISL_711776, EPI_ISL_711777, EPI_ISL_711778, EPI_ISL_711779, EPI_ISL_711780, EPI_ISL_711781, EPI_ISL_711782, EPI_ISL_711783, EPI_ISL_711784, EPI_ISL_711785, EPI_ISL_711786, EPI_ISL_711787, EPI_ISL_711788, EPI_ISL_711789, EPI_ISL_711790, EPI_ISL_711791, EPI_ISL_711792, EPI_ISL_711793, EPI_ISL_711794, EPI_ISL_711795, EPI_ISL_711796, EPI_ISL_711797, EPI_ISL_711798, EPI_ISL_711799, EPI_ISL_711800, EPI_ISL_711801, EPI_ISL_711802, EPI_ISL_711803, EPI_ISL_711804, EPI_ISL_711805, EPI_ISL_711806, EPI_ISL_711807, EPI_ISL_711808, EPI_ISL_711809, EPI_ISL_711810, EPI_ISL_711811, EPI_ISL_711812, EPI_ISL_711813, EPI_ISL_711814, EPI_ISL_711815, EPI_ISL_711816, EPI_ISL_711817, EPI_ISL_711818, EPI_ISL_711819, EPI_ISL_711820, EPI_ISL_711821, EPI_ISL_711822, EPI_ISL_711823, EPI_ISL_711824, EPI_ISL_711825, EPI_ISL_711826, EPI_ISL_711827, EPI_ISL_711828, EPI_ISL_711829, EPI_ISL_711830, EPI_ISL_711831, EPI_ISL_711832, EPI_ISL_711833, EPI_ISL_711834, EPI_ISL_711835, EPI_ISL_711836, EPI_ISL_711837, EPI_ISL_711838, EPI_ISL_711839, EPI_ISL_711840, EPI_ISL_711841, EPI_ISL_711842, EPI_ISL_711843, EPI_ISL_711844, EPI_ISL_711845, EPI_ISL_711846, EPI_ISL_711847, EPI_ISL_711848, EPI_ISL_711849, EPI_ISL_712097, EPI_ISL_712098, EPI_ISL_712101, EPI_ISL_712102, EPI_ISL_712103, EPI_ISL_712104, EPI_ISL_712105, EPI_ISL_712106, EPI_ISL_712107, EPI_ISL_712108, EPI_ISL_712109, EPI_ISL_712110, EPI_ISL_712111, EPI_ISL_712112, EPI_ISL_712113, EPI_ISL_712114, EPI_ISL_712115, EPI_ISL_712116, EPI_ISL_712117, EPI_ISL_712118, EPI_ISL_712129, EPI_ISL_712130, EPI_ISL_712131, EPI_ISL_712132, EPI_ISL_712133, EPI_ISL_712134, EPI_ISL_712135, EPI_ISL_712136, EPI_ISL_712137, EPI_ISL_712138, EPI_ISL_712139, EPI_ISL_712140, EPI_ISL_712141, EPI_ISL_712142, EPI_ISL_712143, EPI_ISL_712144, EPI_ISL_712145, EPI_ISL_712146, EPI_ISL_712147, EPI_ISL_712148, EPI_ISL_712149, EPI_ISL_712150, EPI_ISL_712151, EPI_ISL_712152, EPI_ISL_712153, EPI_ISL_712154, EPI_ISL_712155, EPI_ISL_712156, EPI_ISL_712157, EPI_ISL_712158, EPI_ISL_712159, EPI_ISL_712160, EPI_ISL_712161, EPI_ISL_712162, EPI_ISL_712163, EPI_ISL_712164, EPI_ISL_712165, EPI_ISL_712166, EPI_ISL_712167, EPI_ISL_712168, EPI_ISL_712169, EPI_ISL_712170, EPI_ISL_712171, EPI_ISL_712172, EPI_ISL_712173, EPI_ISL_712174, EPI_ISL_712175, EPI_ISL_712176, EPI_ISL_712177, EPI_ISL_712178, EPI_ISL_712179, EPI_ISL_712180, EPI_ISL_712181, EPI_ISL_712182, EPI_ISL_712183, EPI_ISL_712184, EPI_ISL_712185, EPI_ISL_712186, EPI_ISL_712187, EPI_ISL_712188, EPI_ISL_712189, EPI_ISL_712190, EPI_ISL_712191, EPI_ISL_712192, EPI_ISL_712193, EPI_ISL_712194, EPI_ISL_712195, EPI_ISL_712196, EPI_ISL_712197, EPI_ISL_712198, EPI_ISL_712199, EPI_ISL_712200, EPI_ISL_712201, EPI_ISL_712202, EPI_ISL_712203, EPI_ISL_712204, EPI_ISL_712205, EPI_ISL_712206, EPI_ISL_712207, EPI_ISL_712208, EPI_ISL_712209, EPI_ISL_712210, EPI_ISL_712211, EPI_ISL_712212, EPI_ISL_712213, EPI_ISL_712214, EPI_ISL_712215, EPI_ISL_712216, EPI_ISL_712217, EPI_ISL_712218, EPI_ISL_712219, EPI_ISL_712220, EPI_ISL_712221, EPI_ISL_712222, EPI_ISL_712223, EPI_ISL_712224, EPI_ISL_712225, EPI_ISL_712226, EPI_ISL_712227, EPI_ISL_712228, EPI_ISL_712229, EPI_ISL_712230, EPI_ISL_712231, EPI_ISL_712232, EPI_ISL_712233, EPI_ISL_712234, EPI_ISL_712235, EPI_ISL_712236, EPI_ISL_712237, EPI_ISL_712238, EPI_ISL_712239, EPI_ISL_712240, EPI_ISL_712241, EPI_ISL_712242, EPI_ISL_712243, EPI_ISL_712244, EPI_ISL_712245, EPI_ISL_712246, EPI_ISL_712247, EPI_ISL_712248, EPI_ISL_712249, EPI_ISL_712254, EPI_ISL_712255, EPI_ISL_712256, EPI_ISL_712257, EPI_ISL_712258, EPI_ISL_712259, EPI_ISL_712260, EPI_ISL_712261, EPI_ISL_712262, EPI_ISL_712263, EPI_ISL_712264, EPI_ISL_712265, EPI_ISL_712266, EPI_ISL_712267, EPI_ISL_712268, EPI_ISL_712269, EPI_ISL_712270, EPI_ISL_712271, EPI_ISL_712272, EPI_ISL_712273, EPI_ISL_712274, EPI_ISL_712275, EPI_ISL_712276, EPI_ISL_712277, EPI_ISL_712278, EPI_ISL_712279, EPI_ISL_712280, EPI_ISL_712281, EPI_ISL_712282, EPI_ISL_712283, EPI_ISL_712284, EPI_ISL_712285, EPI_ISL_712286, EPI_ISL_712287, EPI_ISL_712288, EPI_ISL_712289, EPI_ISL_712290, EPI_ISL_712291, EPI_ISL_712292, EPI_ISL_712293, EPI_ISL_712294, EPI_ISL_712296, EPI_ISL_712297, EPI_ISL_712298, EPI_ISL_712299, EPI_ISL_712300, EPI_ISL_712301, EPI_ISL_712302, EPI_ISL_712303, EPI_ISL_712304, EPI_ISL_712305, EPI_ISL_712306, EPI_ISL_712307, EPI_ISL_712308, EPI_ISL_712309, EPI_ISL_712310, EPI_ISL_712311, EPI_ISL_712312, EPI_ISL_712313, EPI_ISL_712314, EPI_ISL_712315, EPI_ISL_712316, EPI_ISL_712317, EPI_ISL_712318, EPI_ISL_712319, EPI_ISL_712320, EPI_ISL_712322, EPI_ISL_712323, EPI_ISL_712324, EPI_ISL_712325, EPI_ISL_712326, EPI_ISL_712327, EPI_ISL_712328, EPI_ISL_712329, EPI_ISL_712330, EPI_ISL_712331, EPI_ISL_712332, EPI_ISL_712333, EPI_ISL_712334, EPI_ISL_712335, EPI_ISL_712336, EPI_ISL_712337, EPI_ISL_712338, EPI_ISL_712339, EPI_ISL_712347, EPI_ISL_712348, EPI_ISL_712349, EPI_ISL_712350, EPI_ISL_712351, EPI_ISL_712352, EPI_ISL_712353, EPI_ISL_712354, EPI_ISL_712355, EPI_ISL_712356, EPI_ISL_712357, EPI_ISL_712358, EPI_ISL_712359, EPI_ISL_712360, EPI_ISL_712361, EPI_ISL_712362, EPI_ISL_712363, EPI_ISL_712364, EPI_ISL_712365, EPI_ISL_712366, EPI_ISL_712367, EPI_ISL_712368, EPI_ISL_712369, EPI_ISL_712370, EPI_ISL_712371, EPI_ISL_712372, EPI_ISL_712373, EPI_ISL_712374, EPI_ISL_712375, EPI_ISL_712376, EPI_ISL_712377, EPI_ISL_712378, EPI_ISL_712379, EPI_ISL_712380, EPI_ISL_712381, EPI_ISL_712382, EPI_ISL_712383, EPI_ISL_712384, EPI_ISL_712385, EPI_ISL_712386, EPI_ISL_712387, EPI_ISL_712388, EPI_ISL_712389, EPI_ISL_712390, EPI_ISL_712391, EPI_ISL_712392, EPI_ISL_712393, EPI_ISL_712394, EPI_ISL_712395, EPI_ISL_712396, EPI_ISL_712397, EPI_ISL_712398, EPI_ISL_712399, EPI_ISL_712400, EPI_ISL_712401, EPI_ISL_712402, EPI_ISL_712403, EPI_ISL_712404, EPI_ISL_712405, EPI_ISL_712406, EPI_ISL_712407, EPI_ISL_712408, EPI_ISL_712409, EPI_ISL_712410, EPI_ISL_712411, EPI_ISL_712412, EPI_ISL_712413, EPI_ISL_712414, EPI_ISL_712415, EPI_ISL_712416, EPI_ISL_712418, EPI_ISL_712419, EPI_ISL_712420, EPI_ISL_712421, EPI_ISL_712422, EPI_ISL_712423, EPI_ISL_712424, EPI_ISL_712425, EPI_ISL_712426, EPI_ISL_712427, EPI_ISL_712428, EPI_ISL_712429, EPI_ISL_712430, EPI_ISL_712432, EPI_ISL_712433, EPI_ISL_712434, EPI_ISL_712435, EPI_ISL_712436, EPI_ISL_712437, EPI_ISL_712438, EPI_ISL_712440, EPI_ISL_712441, EPI_ISL_712442, EPI_ISL_712443, EPI_ISL_712444, EPI_ISL_712445, EPI_ISL_712449, EPI_ISL_712450, EPI_ISL_712451, EPI_ISL_712452, EPI_ISL_712453, EPI_ISL_712454, EPI_ISL_712531, EPI_ISL_712532, EPI_ISL_712533, EPI_ISL_712534, EPI_ISL_712535, EPI_ISL_712536, EPI_ISL_712537, EPI_ISL_712538, EPI_ISL_712539, EPI_ISL_712540, EPI_ISL_712541, EPI_ISL_712542, EPI_ISL_712543, EPI_ISL_712544, EPI_ISL_712545, EPI_ISL_712546, EPI_ISL_712547, EPI_ISL_712548, EPI_ISL_712549, EPI_ISL_712550, EPI_ISL_712551, EPI_ISL_712552, EPI_ISL_712553, EPI_ISL_712554, EPI_ISL_712555, EPI_ISL_712556, EPI_ISL_712557, EPI_ISL_712558, EPI_ISL_712559, EPI_ISL_712560, EPI_ISL_712561, EPI_ISL_712562, EPI_ISL_712563, EPI_ISL_712564, EPI_ISL_712565, EPI_ISL_712566, EPI_ISL_712567, EPI_ISL_712569, EPI_ISL_712570, EPI_ISL_712571, EPI_ISL_712572, EPI_ISL_712573, EPI_ISL_712574, EPI_ISL_712575, EPI_ISL_712576, EPI_ISL_712577, EPI_ISL_712578, EPI_ISL_712579, EPI_ISL_712580, EPI_ISL_712581, EPI_ISL_712582, EPI_ISL_712583, EPI_ISL_712584, EPI_ISL_712585, EPI_ISL_712586, EPI_ISL_712587, EPI_ISL_712588, EPI_ISL_712589, EPI_ISL_712590, EPI_ISL_712591, EPI_ISL_712592, EPI_ISL_712593, EPI_ISL_712594, EPI_ISL_712595, EPI_ISL_712596, EPI_ISL_712597, EPI_ISL_712598, EPI_ISL_712599, EPI_ISL_712600, EPI_ISL_712601, EPI_ISL_712602, EPI_ISL_712603, EPI_ISL_712604, EPI_ISL_712605, EPI_ISL_712606, EPI_ISL_712607, EPI_ISL_712608, EPI_ISL_712609, EPI_ISL_712610, EPI_ISL_712611, EPI_ISL_712612, EPI_ISL_712613, EPI_ISL_712614, EPI_ISL_712615, EPI_ISL_712616, EPI_ISL_712617, EPI_ISL_712618, EPI_ISL_712619, EPI_ISL_712620, EPI_ISL_712621, EPI_ISL_712622, EPI_ISL_712623, EPI_ISL_712624, EPI_ISL_712625, EPI_ISL_712626, EPI_ISL_712627, EPI_ISL_712628, EPI_ISL_712629, EPI_ISL_712630, EPI_ISL_712631, EPI_ISL_712632, EPI_ISL_712633, EPI_ISL_712634, EPI_ISL_712635, EPI_ISL_712636, EPI_ISL_712637, EPI_ISL_712638, EPI_ISL_712639, EPI_ISL_712640, EPI_ISL_712641, EPI_ISL_712642, EPI_ISL_712643, EPI_ISL_712644, EPI_ISL_712645, EPI_ISL_712646, EPI_ISL_712647, EPI_ISL_712648, EPI_ISL_712649, EPI_ISL_712650, EPI_ISL_712651, EPI_ISL_712652, EPI_ISL_712653, EPI_ISL_712654, EPI_ISL_712655, EPI_ISL_712656, EPI_ISL_712657, EPI_ISL_712658, EPI_ISL_712659, EPI_ISL_712660, EPI_ISL_712661, EPI_ISL_712662, EPI_ISL_712663, EPI_ISL_712664, EPI_ISL_712665, EPI_ISL_712666, EPI_ISL_712667, EPI_ISL_712668, EPI_ISL_712669, EPI_ISL_712670, EPI_ISL_712671, EPI_ISL_712672, EPI_ISL_712673, EPI_ISL_712674, EPI_ISL_712675, EPI_ISL_712676, EPI_ISL_712677, EPI_ISL_712678, EPI_ISL_712679, EPI_ISL_712680, EPI_ISL_712681, EPI_ISL_712682, EPI_ISL_712683, EPI_ISL_712684, EPI_ISL_712685, EPI_ISL_712686, EPI_ISL_712687, EPI_ISL_712688, EPI_ISL_712689, EPI_ISL_712690, EPI_ISL_712691, EPI_ISL_712692, EPI_ISL_712693, EPI_ISL_712694, EPI_ISL_712695, EPI_ISL_712696, EPI_ISL_712697, EPI_ISL_712698, EPI_ISL_712699, EPI_ISL_712700, EPI_ISL_712701, EPI_ISL_712702, EPI_ISL_712703, EPI_ISL_712704, EPI_ISL_712705, EPI_ISL_712706, EPI_ISL_712707, EPI_ISL_712708, EPI_ISL_712709, EPI_ISL_712710, EPI_ISL_712711, EPI_ISL_712712, EPI_ISL_712713, EPI_ISL_712714, EPI_ISL_712715, EPI_ISL_712716, EPI_ISL_712717, EPI_ISL_712718, EPI_ISL_712719, EPI_ISL_712720, EPI_ISL_712721, EPI_ISL_712722, EPI_ISL_712723, EPI_ISL_712724, EPI_ISL_712725, EPI_ISL_712726, EPI_ISL_712727, EPI_ISL_712728, EPI_ISL_712729, EPI_ISL_712730, EPI_ISL_712731, EPI_ISL_712732, EPI_ISL_712733, EPI_ISL_712734, EPI_ISL_712735, EPI_ISL_712736, EPI_ISL_712737, EPI_ISL_712738, EPI_ISL_712739, EPI_ISL_712740, EPI_ISL_712741, EPI_ISL_712742, EPI_ISL_712743, EPI_ISL_712744, EPI_ISL_712745, EPI_ISL_712746, EPI_ISL_712747, EPI_ISL_712748, EPI_ISL_712749, EPI_ISL_712750, EPI_ISL_712751, EPI_ISL_712752, EPI_ISL_712753, EPI_ISL_712754, EPI_ISL_712755, EPI_ISL_712756, EPI_ISL_712757, EPI_ISL_712758, EPI_ISL_712759, EPI_ISL_712760, EPI_ISL_712761, EPI_ISL_712762, EPI_ISL_712763, EPI_ISL_712764, EPI_ISL_712765, EPI_ISL_712766, EPI_ISL_712767, EPI_ISL_712768, EPI_ISL_712769, EPI_ISL_712770, EPI_ISL_712771, EPI_ISL_712772, EPI_ISL_712773, EPI_ISL_712774, EPI_ISL_712775, EPI_ISL_712776, EPI_ISL_712777, EPI_ISL_712778, EPI_ISL_712779, EPI_ISL_712780, EPI_ISL_712781, EPI_ISL_712782, EPI_ISL_712783, EPI_ISL_712784, EPI_ISL_712785, EPI_ISL_712786, EPI_ISL_712787, EPI_ISL_712788, EPI_ISL_712789, EPI_ISL_712790, EPI_ISL_712791, EPI_ISL_712792, EPI_ISL_712793, EPI_ISL_712794, EPI_ISL_712795, EPI_ISL_712796, EPI_ISL_712797, EPI_ISL_712798, EPI_ISL_712799, EPI_ISL_712800, EPI_ISL_712801, EPI_ISL_712802, EPI_ISL_712803, EPI_ISL_712804, EPI_ISL_712805, EPI_ISL_712806, EPI_ISL_712807, EPI_ISL_712808, EPI_ISL_712809, EPI_ISL_712810, EPI_ISL_712811, EPI_ISL_712812, EPI_ISL_712813, EPI_ISL_712814, EPI_ISL_712815, EPI_ISL_712816, EPI_ISL_712817, EPI_ISL_712818, EPI_ISL_712819, EPI_ISL_712820, EPI_ISL_712821, EPI_ISL_712822, EPI_ISL_712823, EPI_ISL_712824, EPI_ISL_712825, EPI_ISL_712826, EPI_ISL_712827, EPI_ISL_712828, EPI_ISL_712829, EPI_ISL_712830, EPI_ISL_712831, EPI_ISL_712832, EPI_ISL_712833, EPI_ISL_712834, EPI_ISL_712835, EPI_ISL_712836, EPI_ISL_712837, EPI_ISL_712838, EPI_ISL_712839, EPI_ISL_712840, EPI_ISL_712841, EPI_ISL_712842, EPI_ISL_712843, EPI_ISL_712844, EPI_ISL_712845, EPI_ISL_712846, EPI_ISL_712847, EPI_ISL_712848, EPI_ISL_712849, EPI_ISL_712850, EPI_ISL_712851, EPI_ISL_712852, EPI_ISL_712853, EPI_ISL_712854, EPI_ISL_712855, EPI_ISL_712856, EPI_ISL_712857, EPI_ISL_712858, EPI_ISL_712859, EPI_ISL_712860, EPI_ISL_712861, EPI_ISL_712862, EPI_ISL_712863, EPI_ISL_712864, EPI_ISL_712865, EPI_ISL_712866, EPI_ISL_712867, EPI_ISL_712868, EPI_ISL_712869, EPI_ISL_712870, EPI_ISL_712871, EPI_ISL_712872, EPI_ISL_712873, EPI_ISL_712874, EPI_ISL_712875, EPI_ISL_712876, EPI_ISL_712877, EPI_ISL_712878, EPI_ISL_712879, EPI_ISL_712880, EPI_ISL_712881, EPI_ISL_712882, EPI_ISL_712883, EPI_ISL_712884, EPI_ISL_712885, EPI_ISL_712886, EPI_ISL_712887, EPI_ISL_712888, EPI_ISL_712889, EPI_ISL_712890, EPI_ISL_712891, EPI_ISL_712892, EPI_ISL_712893, EPI_ISL_712894, EPI_ISL_712895, EPI_ISL_712896, EPI_ISL_712897, EPI_ISL_712898, EPI_ISL_712899, EPI_ISL_712900, EPI_ISL_712901, EPI_ISL_712902, EPI_ISL_712903, EPI_ISL_712904, EPI_ISL_712905, EPI_ISL_712906, EPI_ISL_712907, EPI_ISL_712908, EPI_ISL_712909, EPI_ISL_712910, EPI_ISL_712911, EPI_ISL_712912, EPI_ISL_712913, EPI_ISL_712914, EPI_ISL_712915, EPI_ISL_712916, EPI_ISL_712917, EPI_ISL_712918, EPI_ISL_712919, EPI_ISL_712920, EPI_ISL_712921, EPI_ISL_712922, EPI_ISL_712923, EPI_ISL_712924, EPI_ISL_712925, EPI_ISL_712926, EPI_ISL_712927, EPI_ISL_712928, EPI_ISL_712929, EPI_ISL_712930, EPI_ISL_712931, EPI_ISL_712932, EPI_ISL_712933, EPI_ISL_712934, EPI_ISL_712935, EPI_ISL_712936, EPI_ISL_712937, EPI_ISL_712938, EPI_ISL_712939, EPI_ISL_712940, EPI_ISL_712941, EPI_ISL_712942, EPI_ISL_712943, EPI_ISL_712944, EPI_ISL_712945, EPI_ISL_712946, EPI_ISL_712947, EPI_ISL_712948, EPI_ISL_712949, EPI_ISL_712950, EPI_ISL_712951, EPI_ISL_712952, EPI_ISL_712953, EPI_ISL_712954, EPI_ISL_712955, EPI_ISL_712956, EPI_ISL_712957, EPI_ISL_712958, EPI_ISL_712959, EPI_ISL_712960, EPI_ISL_712961, EPI_ISL_712962, EPI_ISL_712963, EPI_ISL_712964, EPI_ISL_712965, EPI_ISL_712966, EPI_ISL_712967, EPI_ISL_712968, EPI_ISL_712969, EPI_ISL_712970, EPI_ISL_712971, EPI_ISL_712972, EPI_ISL_712973, EPI_ISL_712974, EPI_ISL_712975, EPI_ISL_712976, EPI_ISL_712977, EPI_ISL_712978, EPI_ISL_712979, EPI_ISL_712980, EPI_ISL_712981, EPI_ISL_712982, EPI_ISL_712983, EPI_ISL_712984, EPI_ISL_712985, EPI_ISL_712986, EPI_ISL_712987, EPI_ISL_712988, EPI_ISL_712989, EPI_ISL_712990, EPI_ISL_712991, EPI_ISL_712992, EPI_ISL_712993, EPI_ISL_712994, EPI_ISL_712995, EPI_ISL_712996, EPI_ISL_712997, EPI_ISL_712998, EPI_ISL_712999, EPI_ISL_713000, EPI_ISL_713001, EPI_ISL_713002, EPI_ISL_713003, EPI_ISL_713004, EPI_ISL_713005, EPI_ISL_713006, EPI_ISL_713007, EPI_ISL_713008, EPI_ISL_713009, EPI_ISL_713010, EPI_ISL_713011, EPI_ISL_713012, EPI_ISL_713013, EPI_ISL_713014, EPI_ISL_713015, EPI_ISL_713016, EPI_ISL_713017, EPI_ISL_713018, EPI_ISL_713019, EPI_ISL_713020, EPI_ISL_713021, EPI_ISL_713022, EPI_ISL_713023, EPI_ISL_713024, EPI_ISL_713025, EPI_ISL_713026, EPI_ISL_713027, EPI_ISL_713028, EPI_ISL_713029, EPI_ISL_713030, EPI_ISL_713031, EPI_ISL_713032, EPI_ISL_713033, EPI_ISL_713034, EPI_ISL_713035, EPI_ISL_713036, EPI_ISL_713037, EPI_ISL_713038, EPI_ISL_713039, EPI_ISL_713040, EPI_ISL_713041, EPI_ISL_713042, EPI_ISL_713043, EPI_ISL_713044, EPI_ISL_713045, EPI_ISL_713046, EPI_ISL_713047, EPI_ISL_713048, EPI_ISL_713049, EPI_ISL_713050, EPI_ISL_713051, EPI_ISL_713052, EPI_ISL_713053, EPI_ISL_713054, EPI_ISL_713055, EPI_ISL_713056, EPI_ISL_713057, EPI_ISL_713058, EPI_ISL_713059, EPI_ISL_713060, EPI_ISL_713061, EPI_ISL_713062, EPI_ISL_713063, EPI_ISL_713064, EPI_ISL_713065, EPI_ISL_713066, EPI_ISL_713067, EPI_ISL_713068, EPI_ISL_713069, EPI_ISL_713070, EPI_ISL_713071, EPI_ISL_713072, EPI_ISL_713073, EPI_ISL_713074, EPI_ISL_713075, EPI_ISL_713076, EPI_ISL_713077, EPI_ISL_713078, EPI_ISL_713079, EPI_ISL_713080, EPI_ISL_713081, EPI_ISL_713082, EPI_ISL_713083, EPI_ISL_713084, EPI_ISL_713085, EPI_ISL_713086, EPI_ISL_713087, EPI_ISL_713088, EPI_ISL_713089, EPI_ISL_713090, EPI_ISL_713091, EPI_ISL_713092, EPI_ISL_713093, EPI_ISL_713094, EPI_ISL_713095, EPI_ISL_713096, EPI_ISL_713097 | | | |
| see above | Department of Virus and Microbiological Special Diagnostics, Statens Serum Institut, Copenhagen, Denmark | Albertsen Lab, Department of Chemistry and Bioscience, Aalborg University, Denmark | Danish Covid-19 Genome Consortium |
| EPI_ISL_717982, EPI_ISL_717986 | Lab voor klinische biologie | Onderzoeksgroep Virologie | Nick Vereecke, Laurens Lambrechts, Marthe Pauwels, Bruno Verhasselt, Linos Vandekerckhove, Hans Nauwynck, Sebastiaan Theuns |
| EPI_ISL_718024 | Lab voor klinische biologie | Onderzoeksgroep Virologie | Laurens Lambrechts, Nick Vereecke, Marthe Pauwels, Bruno Verhasselt, Linos Vandekerckhove, Hans Nauwynck, Sebastiaan Theuns |
| EPI_ISL_718026 | Lab voor klinische biologie | Onderzoeksgroep Virologie | Nick Vereecke, Laurens Lambrechts, Marthe Pauwels, Bruno Verhasselt, Linos Vandekerckhove, Hans Nauwynck, Sebastiaan Theuns |
| EPI_ISL_718252 | Institute of Virology, Biomedical Research Center of the Slovak Academy of Sciences, Bratislava | Faculty of Natural Sciences, Comenius University, Bratislava | Kristína Boršová, Viktória Hodorová, Broa Brejová, Viktória abanová, Sabina Fumaová Havlíková, Juraj Kopáek, Martina Liková, ubomíra Lukáiková, Martina Neboháová, Monika Sláviková, Tomáš Vina, Boris Klempa, Jozef Nosek |
| EPI_ISL_718253 | Institute of Virology, Biomedical Research Center of the Slovak Academy of Sciences, Bratislava | Faculty of Natural Sciences, Comenius University, Bratislava | Viktória Hodorová, Kristína Boršová, Broa Brejová, Viktória abanová, Sabina Fumaová Havlíková, Juraj Kopáek, Martina Liková, ubomíra Lukáiková, Martina Neboháová, Monika Sláviková, Tomáš Vina, Jozef Nosek, Boris Klempa |
| EPI_ISL_718254 | Institute of Virology, Biomedical Research Center of the | Faculty of Natural Sciences, Comenius University, Bratislava | Kristína Boršová, Viktória Hodorová, Broa Brejová, Viktória abanová, Sabina Fumaová Havlíková, Juraj Kopáek, Martina Liková, ubomíra Lukáiková, |

| | | | |
|---|---|---|---|
| | Slovak Academy of Sciences, Bratislava | | Martina Neboháová, Monika Sláviková, Tomáš Vina, Boris Klempa, Jozef Nosek |
| EPI_ISL_718255, EPI_ISL_718256 | Institute of Virology, Biomedical Research Center of the Slovak Academy of Sciences, Bratislava | Faculty of Natural Sciences, Comenius University, Bratislava | Broa Brejová, Viktória Hodorová, Kristína Boršová, Viktória abanová, Sabina Fumaová Havlíková, Juraj Kopáek, Martina Liková, ubomíra Lukáiková, Martina Neboháová, Monika Sláviková, Tomáš Vina, Jozef Nosek, Boris Klempa |
| EPI_ISL_720669, EPI_ISL_720670, EPI_ISL_720675, EPI_ISL_720676, EPI_ISL_720677, EPI_ISL_720679, EPI_ISL_720680, EPI_ISL_720681, EPI_ISL_720682, EPI_ISL_720687, EPI_ISL_720692, EPI_ISL_720696, EPI_ISL_720698, EPI_ISL_720699, EPI_ISL_720701, EPI_ISL_720703, EPI_ISL_720705, EPI_ISL_720706, EPI_ISL_720708, EPI_ISL_720710, EPI_ISL_720712, EPI_ISL_720714, EPI_ISL_720716, EPI_ISL_720717, EPI_ISL_720718, EPI_ISL_720720, EPI_ISL_720721, EPI_ISL_720723, EPI_ISL_720724, EPI_ISL_720725, EPI_ISL_720727, EPI_ISL_720729, EPI_ISL_720730, EPI_ISL_720731, EPI_ISL_720735, EPI_ISL_720736, EPI_ISL_720739, EPI_ISL_720742, EPI_ISL_720743, EPI_ISL_720744, EPI_ISL_720746, EPI_ISL_720747, EPI_ISL_720748, EPI_ISL_720751, EPI_ISL_720753, EPI_ISL_720754, EPI_ISL_720758, EPI_ISL_720759, EPI_ISL_720760, EPI_ISL_720763, EPI_ISL_720768, EPI_ISL_720772, EPI_ISL_720774, EPI_ISL_720775, EPI_ISL_720778, EPI_ISL_720781, EPI_ISL_720782, EPI_ISL_720785, EPI_ISL_720789, EPI_ISL_720791, EPI_ISL_720794, EPI_ISL_720797, EPI_ISL_720798, EPI_ISL_720799, EPI_ISL_720800, EPI_ISL_720801, EPI_ISL_720804, EPI_ISL_720812, EPI_ISL_720813, EPI_ISL_720815, EPI_ISL_720817, EPI_ISL_720818, EPI_ISL_720820, EPI_ISL_720822, EPI_ISL_720824, EPI_ISL_720826, EPI_ISL_720829, EPI_ISL_720830, EPI_ISL_720832, EPI_ISL_720835, EPI_ISL_720838, EPI_ISL_720840, EPI_ISL_720843, EPI_ISL_720844, EPI_ISL_720847, EPI_ISL_720852, EPI_ISL_720854, EPI_ISL_720855, EPI_ISL_720859, EPI_ISL_720861, EPI_ISL_720863 | | | |
| see above | Lighthouse Lab in Milton Keynes | Wellcome Sanger Institute for the COVID-19 Genomics UK (COG-UK) Consortium | The Lighthouse Lab in Milton Keynes and Alex Alderton, Roberto Amato, Sonia Goncalves, Ewan Harrison, David K. Jackson, Ian Johnston, Dominic Kwiatkowski, Cordelia Langford, John Sillitoe on behalf of the Wellcome Sanger Institute COVID-19 Surveillance Team |
| EPI_ISL_720864 | Lighthouse Lab in Alderley Park | Wellcome Sanger Institute for the COVID-19 Genomics UK (COG-UK) Consortium | Jacquelyn Wynn, Mairead Hyland, The Lighthouse Lab in Alderley Park and Alex Alderton, Roberto Amato, Sonia Goncalves, Ewan Harrison, David K. Jackson, Ian Johnston, Dominic Kwiatkowski, Cordelia Langford, John Sillitoe on behalf of the Wellcome Sanger Institute COVID-19 Surveillance Team |
| EPI_ISL_720865, EPI_ISL_720866, EPI_ISL_720868, EPI_ISL_720873, EPI_ISL_720876, EPI_ISL_720878, EPI_ISL_720879, EPI_ISL_720881, EPI_ISL_720882, EPI_ISL_720883, EPI_ISL_720884, EPI_ISL_720887, EPI_ISL_720890, EPI_ISL_720892, EPI_ISL_720895, EPI_ISL_720899, EPI_ISL_720901, EPI_ISL_720905, EPI_ISL_720906, EPI_ISL_720908, EPI_ISL_721526, EPI_ISL_721527 | | | |
| see above | Lighthouse Lab in Milton Keynes | Wellcome Sanger Institute for the COVID-19 Genomics UK (COG-UK) Consortium | The Lighthouse Lab in Milton Keynes and Alex Alderton, Roberto Amato, Sonia Goncalves, Ewan Harrison, David K. Jackson, Ian Johnston, Dominic Kwiatkowski, Cordelia Langford, John Sillitoe on behalf of the Wellcome Sanger Institute COVID-19 Surveillance Team |
| EPI_ISL_721568, EPI_ISL_721573, EPI_ISL_721575, EPI_ISL_721598, EPI_ISL_721599, EPI_ISL_721600, EPI_ISL_721611, EPI_ISL_721612, EPI_ISL_721613, EPI_ISL_721614, EPI_ISL_721615, EPI_ISL_721616, EPI_ISL_721617, EPI_ISL_721622, EPI_ISL_721623 | | | |
| see above | Pathogen Genomics Center, National Institute of Infectious Diseases | Pathogen Genomics Center, National Institute of Infectious Diseases | Tsuyoshi Sekizuka, Kentaro Itokawa, Rina Tanaka, Masanori Hashino, Makoto Kuroda |
| EPI_ISL_721670, EPI_ISL_721675, EPI_ISL_721695, EPI_ISL_721702, EPI_ISL_721741, EPI_ISL_721772, EPI_ISL_721818, EPI_ISL_721819, EPI_ISL_721820, EPI_ISL_721821, EPI_ISL_721822, EPI_ISL_721840, EPI_ISL_721841, EPI_ISL_721842, EPI_ISL_721851, EPI_ISL_721852, EPI_ISL_721853, EPI_ISL_721889, EPI_ISL_721890, EPI_ISL_721891, EPI_ISL_721892, EPI_ISL_721893, EPI_ISL_721894, EPI_ISL_721895, EPI_ISL_721896, EPI_ISL_721901, EPI_ISL_721902, EPI_ISL_721906, EPI_ISL_721907, EPI_ISL_721908, EPI_ISL_721928, EPI_ISL_721966, EPI_ISL_721967, EPI_ISL_721968, EPI_ISL_721969, EPI_ISL_721970 | | | |
| see above | Viollier AG | Department of Biosystems Science and Engineering, ETH Zürich | Christian Beisel |
| EPI_ISL_722338, EPI_ISL_722372, EPI_ISL_722384, EPI_ISL_722454, EPI_ISL_722711, EPI_ISL_722712, EPI_ISL_722713, EPI_ISL_722758, EPI_ISL_722759, EPI_ISL_722760, EPI_ISL_722761, EPI_ISL_722762, EPI_ISL_722784, EPI_ISL_722785, EPI_ISL_722786 | | | |
| see above | Dutch COVID-19 response team | Erasmus Medical Center | Bas Oude Munnink, Reina Sikkema, David Nieuwenhuijse, Irina Chestakova, Anne van der Linden, Marjan Boter, Emmanuelle Munger, Corine GeurtsvanKessel, Annemiek van der Eijk, Richard Molenkamp, Marion Koopmans, on behalf of the Dutch national COVID-19 response team. |
| EPI_ISL_722926, EPI_ISL_722927, EPI_ISL_722928, EPI_ISL_722929, EPI_ISL_722930, EPI_ISL_722931, EPI_ISL_722932, EPI_ISL_722933, EPI_ISL_722935, EPI_ISL_722944, EPI_ISL_722945, EPI_ISL_722946, EPI_ISL_722947, EPI_ISL_722948, EPI_ISL_722949, EPI_ISL_722950, EPI_ISL_722951, EPI_ISL_722952, EPI_ISL_722953, EPI_ISL_722954, EPI_ISL_722955, EPI_ISL_722956, EPI_ISL_722957, EPI_ISL_722958, EPI_ISL_722959, EPI_ISL_722960, EPI_ISL_722961, EPI_ISL_722962, EPI_ISL_722963, EPI_ISL_722964, EPI_ISL_722965, EPI_ISL_722966 | | | |
| see above | Department of Clinical Microbiology | GIGA Medical Genomics | Keith Durkin, Maria Artesi, Sébastien Bontems, Raphaël Boreux, Bouchra Boujemla, Cécile Meex, Pierrette Melin, Marie-Pierre Hayette, Vincent Bours |
| EPI_ISL_722967, EPI_ISL_722968, EPI_ISL_722969, EPI_ISL_722970, EPI_ISL_722976, EPI_ISL_722977, EPI_ISL_722978, EPI_ISL_723008, EPI_ISL_723009, EPI_ISL_723010, EPI_ISL_723011, EPI_ISL_723012, EPI_ISL_723013, EPI_ISL_723014, EPI_ISL_723022, EPI_ISL_723023, EPI_ISL_723024 | | | |
| see above | Respiratory Virus Unit, National Infection Service, Public Health England | COVID-19 Genomics UK (COG-UK) Consortium | PHE Covid Sequencing Team |
| EPI_ISL_723137 | Minnesota Department of Health, Public Health Laboratory | Minnesota Department of Health, Public Health Laboratory | Alexandra Lorentz, Jacob Garfin, Matt Plumb, and Xiong Wang |
| EPI_ISL_723155, EPI_ISL_723158, EPI_ISL_723159, EPI_ISL_723161, EPI_ISL_723162, EPI_ISL_723163, EPI_ISL_723165, EPI_ISL_723169, EPI_ISL_723175, EPI_ISL_723176, EPI_ISL_723177, EPI_ISL_723201, EPI_ISL_723252, EPI_ISL_723277, EPI_ISL_723278, EPI_ISL_723279, EPI_ISL_723312, EPI_ISL_723319, EPI_ISL_723328, EPI_ISL_723332, EPI_ISL_723334, EPI_ISL_723335, EPI_ISL_723336, EPI_ISL_723337, EPI_ISL_723338, EPI_ISL_723339, EPI_ISL_723346, EPI_ISL_723354, EPI_ISL_723355, EPI_ISL_723366, EPI_ISL_723367, EPI_ISL_723368, EPI_ISL_723369, EPI_ISL_723381, EPI_ISL_723382, EPI_ISL_723384, EPI_ISL_723386, EPI_ISL_723387, EPI_ISL_723388, EPI_ISL_723389, EPI_ISL_723390, EPI_ISL_723391, EPI_ISL_723392, EPI_ISL_723393, EPI_ISL_723446, EPI_ISL_723447, EPI_ISL_723454, EPI_ISL_723466, EPI_ISL_723467 | | | |
| see above | Dutch COVID-19 response team | National Institute for Public Health and the Environment (RIVM) | Adam Meijer, Harry Vennema, Jeroen Cremer, Sharon van den Brink, Bas van der Veer, AnneMarie van den Brandt, Florian Zwagemaker, Dennis Schmitz, Chantal Reusken, on behalf of the national COVID-19 response team |
| EPI_ISL_723886, EPI_ISL_723887, EPI_ISL_723888, EPI_ISL_723916, EPI_ISL_723920 | Department of Pathology, University of Cambridge | COVID-19 Genomics UK (COG-UK) Consortium | Aminu S. Jahun, Yasmin Chaudhry, Grant Hall, Iliana Georgana, Myra Hosmillo, Martin D. Curran, Malte Pinckert, Surendra Parmar, Ian Goodfellow |
| EPI_ISL_724210, EPI_ISL_724211 | West of Scotland Specialist Virology Centre, NHSGGC / MRC-University of Glasgow Centre for Virus Research | COVID-19 Genomics UK (COG-UK) Consortium | Ana da Silva Filipe, Natasha Johnson, Kathy Smollett, Daniel Mair, Stephen Carmichael, Alice Broos, Lily Tong, Jenna Nichols, Kyriaki Nomikou; Sarah McDonald; Richard Orton, Joseph Hughes, Sreenu Vattipally, David L Robertson; Alasdair MacLean, Rory Gunson; Sharif Shaaban, Matthew Holden; Rachel Blacow, Guy Mollett, Kathy Li, James Shepherd, Antonia Ho, Emma Thomson |
| EPI_ISL_724268, EPI_ISL_724276, EPI_ISL_724277, EPI_ISL_724278, EPI_ISL_724279, EPI_ISL_724280, EPI_ISL_724281, EPI_ISL_724282, EPI_ISL_724283, EPI_ISL_724284, EPI_ISL_724285, EPI_ISL_724286, EPI_ISL_724287, EPI_ISL_724288, EPI_ISL_724290, EPI_ISL_724292, EPI_ISL_724293 | | | |
| see above | University of Exeter | COVID-19 Genomics UK (COG-UK) Consortium | Ben Temperton,Aaron Jeffries,Michelle Michelsen,Joanna Warwick-Dugdale,Audrey Farbos,Robyn Manley,Stephen Michell,Jane Masoli |
| EPI_ISL_724349, EPI_ISL_724350, EPI_ISL_724352, EPI_ISL_724353 | Liverpool Clinical Laboratories | COVID-19 Genomics UK (COG-UK) Consortium | Sam Haldenby, Anita Lucaci, Steve Paterson, Julian Hiscox, Alistair Darby, M Almsaud, A Alrezaihi, Muhannad Alruwaili, Stuart D Armstrong, Jones Benjamin, Eleanor G Bentley, Anu Chawla, Jordan J Clark, Angela Cowell, Richard Eccles, Isabel García-Dorival, Matthew Gemmell, Alessandro Gerada, PKF Gilmore, Richard Gregory, Ximeng Han, Catherine Hartley, Margaret Hughes, Miren Iturriza-Gomara, James Johnson, L Luu, Jenifer Manson, Charlotte Nelson, Elaine O'Toole, Cassie Olateju, Rebekah Penrice-Randal , Lucille Rainbow, N.P Randle, Trevor Ian Robinson, Parul Sharma, Ghada T Shawli, James P Stewart, Neil Swainston, Ecaterina Vamos, Joanne Watts, Mark Whitehead |
| EPI_ISL_724597 | University College London, Great Ormond Street Hospital for Children NHS Foundation Trust, Imperial College Healthcare NHS Trust | COVID-19 Genomics UK (COG-UK) Consortium | Sergi Castellano, Rachel Williams, Mark Kristiansen, Paola Resende Silva, Sunando Roy, Tony Brooks, Helena Tutill, Paola Niola, Patricia Dyal, Charlotte Williams, Leysa Forrest, Yasmin Panchbhaya, Jacqueline Findlay, Samuel Weeks, Julianne Brown, Kathryn Harris, Paul Randell, James Price, Alison Holmes, Judith Breuer |
| EPI_ISL_724837, EPI_ISL_724841, EPI_ISL_724842, EPI_ISL_724848, EPI_ISL_724849, EPI_ISL_724850, EPI_ISL_724882, EPI_ISL_724883, EPI_ISL_724884, EPI_ISL_724887, EPI_ISL_724888, EPI_ISL_724889, EPI_ISL_724890, EPI_ISL_724891, EPI_ISL_724892, EPI_ISL_724893, EPI_ISL_724894 | | | |
| see above | Northumbria University / South Tees Hospitals NHS Foundation Trust / North Cumbria Integrated Care NHS Foundation Trust / North Tees and Hartlepool NHS Foundation Trust / Newcastle Hospitals NHS Foundation Trust | COVID-19 Genomics UK (COG-UK) Consortium | Darren L Smith,Andrew Nelson,Matthew Bashton,Greg R Young,Joshua Loh,John Allan,Mohammad A Tariq,Giles S Holt,Gary Black,Wen C Yew,Lynn Dover,Paul Baker,Steve Liggett,Sarah Essex,Jane Greenaway,Debra Padgett,Clive Graham,Garren Scott,Edward Barton,Emma Swindells,Brendan Payne,Jennifer Collins,Yusri Taha,Gary Eltringham |
| EPI_ISL_725380, EPI_ISL_725381, EPI_ISL_725382, EPI_ISL_725383, EPI_ISL_725384, EPI_ISL_725385, EPI_ISL_725386, EPI_ISL_725387, EPI_ISL_725388, EPI_ISL_725389, EPI_ISL_725390, EPI_ISL_725391, EPI_ISL_725392, EPI_ISL_725393, EPI_ISL_725394, EPI_ISL_725395, EPI_ISL_725396, EPI_ISL_725397, EPI_ISL_725398, EPI_ISL_725399, EPI_ISL_725400, EPI_ISL_725401, EPI_ISL_725402, EPI_ISL_725403, EPI_ISL_725404 | | | |
| see above | Quadram Institute Bioscience | COVID-19 Genomics UK (COG-UK) Consortium | Dave J. Baker, Gemma L. Kay, Alp Aydin, Thanh Le-Viet, Steven Rudder, Ana P. Tedim, Anastasia Kolyva, Maria Diaz, Leonardo de Oliveira Martins, Nabil-Fareed Alikhan, Lizzie Meadows, Rachael Stanley, Ngozi Elumogo, Muhammed Yasir, Nicholas M. Thomson, Alexander J Trotter, Rachel Gilroy, Samuel Bloomfield, Claire Stuart, Andrew Bell, Reenesh Prakash, Samir Dervisevic, Alison E. Mather, John Wain, Mark Webber, Andrew J. Page, Justin O'Grady |
| EPI_ISL_725538, EPI_ISL_725539 | Queens Medical Centre, Clinical Microbiology Department / DeepSeq Nottingham | COVID-19 Genomics UK (COG-UK) Consortium | Gemma Clark, Wendy Smith, Manjinder Khakh, Vicki M Fleming, Michelle M Lister, Hannah Howson-Wells, Jonathan Ball, Patrick McClure, Joseph Chappell, Theocharis Tsoleridis, Nadine Holmes, Matthew Carlisle, Christopher Moore, Fei Sang, Johnny Debebe, Victoria Wright, Matthew Loose |
| EPI_ISL_725671, EPI_ISL_725701, EPI_ISL_725709, EPI_ISL_725710, EPI_ISL_725893, EPI_ISL_725895, EPI_ISL_725919, EPI_ISL_725955, EPI_ISL_725956, EPI_ISL_725957, EPI_ISL_725958, EPI_ISL_725959, EPI_ISL_725960, EPI_ISL_725961, EPI_ISL_725962, EPI_ISL_725963, EPI_ISL_725964, EPI_ISL_725965, EPI_ISL_725966, EPI_ISL_725967, EPI_ISL_725968, EPI_ISL_725969, EPI_ISL_725970, EPI_ISL_725971, EPI_ISL_725972, EPI_ISL_725973, EPI_ISL_725974, EPI_ISL_725975, EPI_ISL_725976, EPI_ISL_725977, EPI_ISL_725978, EPI_ISL_725979, EPI_ISL_725980, EPI_ISL_725981, EPI_ISL_726838, EPI_ISL_726840, EPI_ISL_726841, EPI_ISL_726842, EPI_ISL_726843, EPI_ISL_726844, EPI_ISL_726845, EPI_ISL_726846, EPI_ISL_726847, EPI_ISL_726848, EPI_ISL_726849, EPI_ISL_726850, EPI_ISL_726851, EPI_ISL_726852, EPI_ISL_726853, EPI_ISL_726854, EPI_ISL_726855, EPI_ISL_726856, EPI_ISL_726857, EPI_ISL_726858, EPI_ISL_726859, EPI_ISL_726860, EPI_ISL_726861, EPI_ISL_726862, EPI_ISL_726864, EPI_ISL_726961, EPI_ISL_726963, EPI_ISL_726964, EPI_ISL_726965, EPI_ISL_726966, EPI_ISL_726967, EPI_ISL_726968, EPI_ISL_726969, EPI_ISL_726970, EPI_ISL_726971, EPI_ISL_726972, EPI_ISL_726973, EPI_ISL_726974, | | | |

| | | | |
|---|---|---|---|
| EPI_ISL_726975, EPI_ISL_726976, EPI_ISL_726977, EPI_ISL_726978, EPI_ISL_726979, EPI_ISL_726980, EPI_ISL_726981, EPI_ISL_726982, EPI_ISL_726983, EPI_ISL_726984, EPI_ISL_726985, EPI_ISL_726986, EPI_ISL_726987, EPI_ISL_726988, EPI_ISL_726989, EPI_ISL_726990, EPI_ISL_726991, EPI_ISL_726992, EPI_ISL_726993, EPI_ISL_726994, EPI_ISL_726995, EPI_ISL_726996, EPI_ISL_726997, EPI_ISL_726998, EPI_ISL_726999, EPI_ISL_727000, EPI_ISL_727001, EPI_ISL_727002, EPI_ISL_727003, EPI_ISL_727004, EPI_ISL_727005, EPI_ISL_727006, EPI_ISL_727007, EPI_ISL_727008, EPI_ISL_727009, EPI_ISL_727010, EPI_ISL_727011, EPI_ISL_727012, EPI_ISL_727013, EPI_ISL_727014, EPI_ISL_727015, EPI_ISL_727016, EPI_ISL_727017, EPI_ISL_727018, EPI_ISL_727019, EPI_ISL_727020, EPI_ISL_727021, EPI_ISL_727022, EPI_ISL_727023, EPI_ISL_727024, EPI_ISL_727025, EPI_ISL_727026, EPI_ISL_727027, EPI_ISL_727028, EPI_ISL_727029, EPI_ISL_727030, EPI_ISL_727031, EPI_ISL_727032, EPI_ISL_727033, EPI_ISL_727034, EPI_ISL_727035, EPI_ISL_727036, EPI_ISL_727037, EPI_ISL_727038, EPI_ISL_727039, EPI_ISL_727040, EPI_ISL_727041, EPI_ISL_727042, EPI_ISL_727043, EPI_ISL_727044, EPI_ISL_727045, EPI_ISL_727046, EPI_ISL_727047, EPI_ISL_727048, EPI_ISL_727049, EPI_ISL_727050, EPI_ISL_727051, EPI_ISL_727052, EPI_ISL_727053, EPI_ISL_727054, EPI_ISL_727055, EPI_ISL_727056, EPI_ISL_727057, EPI_ISL_727058, EPI_ISL_727059, EPI_ISL_727060, EPI_ISL_727061, EPI_ISL_727062, EPI_ISL_727063, EPI_ISL_727064, EPI_ISL_727065, EPI_ISL_727066, EPI_ISL_727067, EPI_ISL_727068, EPI_ISL_727069, EPI_ISL_727070, EPI_ISL_727071, EPI_ISL_727072, EPI_ISL_727073, EPI_ISL_727074, EPI_ISL_727075, EPI_ISL_727076, EPI_ISL_727077, EPI_ISL_727078, EPI_ISL_727079, EPI_ISL_727080, EPI_ISL_727081, EPI_ISL_727082, EPI_ISL_727083, EPI_ISL_727084, EPI_ISL_727085, EPI_ISL_727086, EPI_ISL_727087, EPI_ISL_727088, EPI_ISL_727089, EPI_ISL_727090, EPI_ISL_727091, EPI_ISL_727092, EPI_ISL_727093, EPI_ISL_727094, EPI_ISL_727095, EPI_ISL_727096, EPI_ISL_727097, EPI_ISL_727101, EPI_ISL_727118, EPI_ISL_727119, EPI_ISL_727120, EPI_ISL_727121, EPI_ISL_727147, EPI_ISL_727148, EPI_ISL_727149, EPI_ISL_727178, EPI_ISL_727179, EPI_ISL_727187, EPI_ISL_727223, EPI_ISL_727224, EPI_ISL_727225, EPI_ISL_727226, EPI_ISL_727227, EPI_ISL_727228, EPI_ISL_727229, EPI_ISL_727230, EPI_ISL_727231, EPI_ISL_727232, EPI_ISL_727233, EPI_ISL_727234, EPI_ISL_727235, EPI_ISL_727236, EPI_ISL_727238, EPI_ISL_727239, EPI_ISL_727240, EPI_ISL_727241, EPI_ISL_727242, EPI_ISL_727243, EPI_ISL_727245, EPI_ISL_727246, EPI_ISL_727247, EPI_ISL_727248, EPI_ISL_727249, EPI_ISL_727250, EPI_ISL_727251, EPI_ISL_727252, EPI_ISL_727253, EPI_ISL_727254, EPI_ISL_727257, EPI_ISL_727258, EPI_ISL_727259, EPI_ISL_727260, EPI_ISL_727261, EPI_ISL_727262, EPI_ISL_727263, EPI_ISL_727264, EPI_ISL_727265, EPI_ISL_727266, EPI_ISL_727267, EPI_ISL_727268, EPI_ISL_727269, EPI_ISL_727270, EPI_ISL_727271, EPI_ISL_727272, EPI_ISL_727273, EPI_ISL_727274, EPI_ISL_727275, EPI_ISL_727277, EPI_ISL_727278, EPI_ISL_727280, EPI_ISL_727417, EPI_ISL_727419, EPI_ISL_727421, EPI_ISL_727422, EPI_ISL_727423, EPI_ISL_727485, EPI_ISL_727486, EPI_ISL_727487, EPI_ISL_727488, EPI_ISL_727489, EPI_ISL_727490, EPI_ISL_727491, EPI_ISL_727492, EPI_ISL_727493, EPI_ISL_727494, EPI_ISL_727495, EPI_ISL_727496, EPI_ISL_727497, EPI_ISL_727498, EPI_ISL_727499, EPI_ISL_727500, EPI_ISL_727501, EPI_ISL_727502, EPI_ISL_727503, EPI_ISL_727504, EPI_ISL_727505, EPI_ISL_727506, EPI_ISL_727507, EPI_ISL_727508, EPI_ISL_727509, EPI_ISL_727510, EPI_ISL_727511, EPI_ISL_727512, EPI_ISL_727513, EPI_ISL_727514, EPI_ISL_727515, EPI_ISL_727516, EPI_ISL_727517, EPI_ISL_727518, EPI_ISL_727519, EPI_ISL_727520, EPI_ISL_727521, EPI_ISL_727522, EPI_ISL_727523, EPI_ISL_727524, EPI_ISL_727525, EPI_ISL_727526, EPI_ISL_727527, EPI_ISL_727528, EPI_ISL_727529, EPI_ISL_727530, EPI_ISL_727531, EPI_ISL_727532, EPI_ISL_727533, EPI_ISL_727534, EPI_ISL_727535, EPI_ISL_727536, EPI_ISL_727537, EPI_ISL_727538, EPI_ISL_727539, EPI_ISL_727540, EPI_ISL_727541, EPI_ISL_727542, EPI_ISL_727543, EPI_ISL_727544, EPI_ISL_727545, EPI_ISL_727546, EPI_ISL_727547, EPI_ISL_727548, EPI_ISL_727549, EPI_ISL_727550, EPI_ISL_727551, EPI_ISL_727553, EPI_ISL_727554, EPI_ISL_727555, EPI_ISL_727556, EPI_ISL_727557, EPI_ISL_727558, EPI_ISL_727559, EPI_ISL_727560, EPI_ISL_727561, EPI_ISL_727562, EPI_ISL_727563, EPI_ISL_727564, EPI_ISL_727565, EPI_ISL_727566, EPI_ISL_727567, EPI_ISL_727568, EPI_ISL_727569, EPI_ISL_727570, EPI_ISL_727571, EPI_ISL_727572, EPI_ISL_727573, EPI_ISL_727574, EPI_ISL_727575, EPI_ISL_727576, EPI_ISL_727577, EPI_ISL_727578, EPI_ISL_727579, EPI_ISL_727580, EPI_ISL_727581, EPI_ISL_727582, EPI_ISL_727583, EPI_ISL_727584, EPI_ISL_727585, EPI_ISL_727586, EPI_ISL_727587, EPI_ISL_727588, EPI_ISL_727589, EPI_ISL_727590, EPI_ISL_727591, EPI_ISL_727592, EPI_ISL_727593, EPI_ISL_727594, EPI_ISL_727595, EPI_ISL_727596, EPI_ISL_727597, EPI_ISL_727598, EPI_ISL_727599, EPI_ISL_727600, EPI_ISL_727601, EPI_ISL_727602, EPI_ISL_727603, EPI_ISL_727604, EPI_ISL_727605, EPI_ISL_727606, EPI_ISL_727607, EPI_ISL_727608, EPI_ISL_727609, EPI_ISL_727610, EPI_ISL_727611, EPI_ISL_727612, EPI_ISL_727613, EPI_ISL_727615, EPI_ISL_727616, EPI_ISL_727617, EPI_ISL_727618, EPI_ISL_727619, EPI_ISL_727620, EPI_ISL_727621 | | | |
| see above | Wales Specialist Virology Centre Sequencing lab: Pathogen Genomics Unit | COVID-19 Genomics UK (COG-UK) Consortium | Catherine Moore, Johnathan Evans, Laura Gifford, Malorie Perry, Simon Cottrell, Angela Marchbank, Alec Birchley, Alexander Adams, Amy Gaskin, Bree Gatica-Wilcox, Jason Coombes, Joel Southgate, Lauren Gilbert, Lee Graham, Nicole Pacchiarini, Sara Kumziene-Summerhayes, Sarah Taylor, Sophie Jones, Sara Rey, Matthew Bull, Joanne Watkins, Sally Corden, Tom Connor |
| EPI_ISL_727863, EPI_ISL_727869, EPI_ISL_727875, EPI_ISL_727888, EPI_ISL_727892, EPI_ISL_727903, EPI_ISL_727918, EPI_ISL_727929, EPI_ISL_727971, EPI_ISL_727983, EPI_ISL_727985, EPI_ISL_727986 | | | |
| see above | Virology Department, Sheffield Teaching Hospitals NHS Foundation Trust/Department of Infection, Immunity and Cardiovascular Disease, The Medical School, University of Sheffield | COVID-19 Genomics UK (COG-UK) Consortium | Thushan de Silva, Matthew Parker, Nikki Smith, Adri Angyal, Rebecca Brown, Luke Green, Rachel Tucker, Paul Parsons, Danielle Groves, Katie Johnson, Laura Carrilero, Alex Keeley, Dave Partridge, Matthew Wyles, Benjamin Lindsey, Mehmet Yavuz, Mohammad Raza, Cariad Evans |
| EPI_ISL_728188 | National Public Health Laboratory, National Centre for Infectious Diseases | National Public Health Laboratory, National Centre for Infectious Diseases | Tze Minn Mak, Sophie Octavia, Zhenyang Zhou, Lin Cui, Raymond Tzer Pin Lin |
| EPI_ISL_728204 | Institute of Microbiology, Universidad San Francisco de Quito | Institute of Microbiology, Universidad San Francisco de Quito | Sully Márquez, Belén Prado-Vivar, Juan José Guadalupe, Monica Becerra-Wong, Bernardo Gutiérrez, Tania Guayasamin, Patricio Reyes, Verónica Barragán, Patricio Rojas-Silva, Gabriel Trueba, Michelle Grunauer, Paúl Cárdenas |
| EPI_ISL_728279 | National Institute for Infectious Diseases, INMI, "L. Spallanzani" IRCCS | National Institute for Infectious Diseases, INMI, "L. Spallanzani" IRCCS | E Giombini, C.E.M Gruber, M Rueca, B Bartolini, F Messina, A Di Caro, MR Capobianchi |
| EPI_ISL_728280 | National Institute for Infectious Diseases, INMI, "L. Spallanzani" IRCCS | National Institute for Infectious Diseases, INMI, "L. Spallanzani" IRCCS | F Messina, M Rueca, B Bartolini, C.E.M Gruber, E Giombini, MR Capobianchi, A Di Caro |
| EPI_ISL_728281 | National Institute for Infectious Diseases, INMI, "L. Spallanzani" IRCCS | National Institute for Infectious Diseases, INMI, "L. Spallanzani" IRCCS | B Bartolini, M Rueca, C.E.M Gruber, F Messina, E Giombini, A Di Caro, MR Capobianchi |
| EPI_ISL_728282 | National Institute for Infectious Diseases, INMI, "L. Spallanzani" IRCCS | National Institute for Infectious Diseases, INMI, "L. Spallanzani" IRCCS | M. Rueca, C.E.M Gruber, B Bartolini, F Messina, E Giombini, A Di Caro, MR Capobianchi |
| EPI_ISL_728283 | National Institute for Infectious Diseases, INMI, "L. Spallanzani" IRCCS | National Institute for Infectious Diseases, INMI, "L. Spallanzani" IRCCS | B Bartolini, C.E.M Gruber, M Rueca, F Messina, E Giombini, MR Capobianchi, A Di Caro |
| EPI_ISL_728284 | National Institute for Infectious Diseases, INMI, "L. Spallanzani" IRCCS | National Institute for Infectious Diseases, INMI, "L. Spallanzani" IRCCS | M Rueca, B Bartolini, C.E.M Gruber, F Messina, E Giombini, A Di Caro, MR Capobianchi |
| EPI_ISL_728285 | National Institute for Infectious Diseases, INMI, "L. Spallanzani" IRCCS | National Institute for Infectious Diseases, INMI, "L. Spallanzani" IRCCS | F Messina, E Giombini, M Rueca, B Bartolini, C.E.M Gruber, MR Capobianchi, A Di Caro |
| EPI_ISL_728287 | National Institute for Infectious Diseases, INMI, "L. Spallanzani" IRCCS | National Institute for Infectious Diseases, INMI, "L. Spallanzani" IRCCS | C.E.M Gruber, F Messina, M Rueca, B Bartolini, E Giombini, MR Capobianchi, A Di Caro |
| EPI_ISL_728324, EPI_ISL_728331, EPI_ISL_728333 | Jena University Hospital, Institute for Infectious Diseases and Infection Control | Institute of infectious medicine & hospital hygiene, CaSe-Group | Spott, Riccardo; Marquet, Mike; Pletz, Matthias W.; Brandt, Christian |
| EPI_ISL_728334, EPI_ISL_728336, EPI_ISL_728339 | Laboratoire Biolife | Laboratoire de Biotechnologie | Mouna Ouadghiri, Tarik Aanniz, Mohammed Walid Chemao Elfihri, Mohamed Chenaoui, Hanae Dakka, Afaf Alaoui, Otmane Touzani, Bouchra Belfquih, Lahcen belyamani, Saaid Amzazi and Azeddine Ibrahimi |
| EPI_ISL_728340 | Laboratoire Biolife | Laboratoire de Biotechnologie | Mouna Ouadghiri, Tarik Aanniz, Mohammed Walid Chemao Elfihri, Mohamed Chenaoui, Hanae Dakka, Afaf Alaoui, Otmane Touzani, Bouchra Belfquih, Lahcen belyamani, Saaid Amzazi and Azeddine Ibrahim |
| EPI_ISL_728342, EPI_ISL_728344, EPI_ISL_728347, EPI_ISL_728349, EPI_ISL_728352, EPI_ISL_728353, EPI_ISL_728355, EPI_ISL_728359, EPI_ISL_728360, EPI_ISL_728363, EPI_ISL_728366, EPI_ISL_728367 | | | |
| see above | Laboratoire Biolife | Laboratoire de Biotechnologie | Mouna Ouadghiri, Tarik Aanniz, Mohammed Walid Chemao Elfihri, Mohamed Chenaoui, Hanae Dakka, Afaf Alaoui, Otmane Touzani, Bouchra Belfquih, Lahcen belyamani, Saaid Amzazi and Azeddine Ibrahimi |
| EPI_ISL_728635, EPI_ISL_728636, EPI_ISL_728637, EPI_ISL_728640, EPI_ISL_728641, EPI_ISL_728643, EPI_ISL_728645, EPI_ISL_728646, EPI_ISL_728647, EPI_ISL_728648, EPI_ISL_728696, EPI_ISL_728745 | | | |
| see above | Dutch COVID-19 response team | National Institute for Public Health and the Environment (RIVM) | Adam Meijer, Harry Vennema, Jeroen Cremer, Sharon van den Brink, Bas van der Veer, AnneMarie van den Brandt, Florian Zwagemaker, Dennis Schmitz, Chantal Reusken, on behalf of the national COVID-19 response team |
| EPI_ISL_728765, EPI_ISL_728768, EPI_ISL_728770, EPI_ISL_728783, EPI_ISL_728813, EPI_ISL_728885, EPI_ISL_728892, EPI_ISL_728896, EPI_ISL_728900, EPI_ISL_728907, EPI_ISL_728914, EPI_ISL_728916, EPI_ISL_728919, EPI_ISL_728922, EPI_ISL_728924, EPI_ISL_728928, EPI_ISL_728932, EPI_ISL_728939, EPI_ISL_728942, EPI_ISL_728945, EPI_ISL_728947, EPI_ISL_728948, EPI_ISL_728955, EPI_ISL_728956, EPI_ISL_728957, EPI_ISL_728963, EPI_ISL_728978, EPI_ISL_728995, EPI_ISL_729002, EPI_ISL_729011, EPI_ISL_729019, EPI_ISL_729032, EPI_ISL_729035, EPI_ISL_729036, EPI_ISL_729040, EPI_ISL_729041, EPI_ISL_729068 | | | |
| see above | Viollier AG | Department of Biosystems Science and Engineering, ETH Zürich | Chaoran Chen, Sarah Nadeau, Catharine Aquino, Ivan Topolsky, Pedro Ferreira, Philipp Jablonski, Susana Posada-Céspedes, Andreia Cabral de Gouvea, Maria Domenica Moccia, Simon Grüter, Timothy Sykes, Lennart Opitz, Ralph Schlapbach, Christiane Beckmann, Maurice Redondo, Olivier Kobel, Christoph Noppen, Sophie Seidel, Noemie Santamaria de Souza, Niko Beerenwinkel, Tanja Stadler |
| EPI_ISL_729347, EPI_ISL_729392, EPI_ISL_729416, EPI_ISL_729449, EPI_ISL_729450, EPI_ISL_729451, EPI_ISL_729452, EPI_ISL_729453, EPI_ISL_729454, EPI_ISL_729455, EPI_ISL_729456, EPI_ISL_729459, EPI_ISL_729460, EPI_ISL_729463, EPI_ISL_729697, EPI_ISL_729698, EPI_ISL_729699, EPI_ISL_729700, EPI_ISL_729701, EPI_ISL_729702, EPI_ISL_729703, EPI_ISL_729705, EPI_ISL_729706, EPI_ISL_729707, EPI_ISL_729708, EPI_ISL_729709, EPI_ISL_729710, EPI_ISL_729711, EPI_ISL_729712, EPI_ISL_729713, EPI_ISL_729714, EPI_ISL_729715, EPI_ISL_729716, EPI_ISL_729717, EPI_ISL_729719, EPI_ISL_729721, EPI_ISL_729722, EPI_ISL_729728, EPI_ISL_729729, EPI_ISL_729730, EPI_ISL_729731, EPI_ISL_729732, EPI_ISL_729733 | | | |
| see above | A. Krumbholz, Labor Dr. Krause und Kollegen MVZ GmbH, Kiel | Charité Universitätsmedizin Berlin, Institut für Virologie | Victor M Corman, Barbara Mühlemann, Jörn Beheim-Schwarzbach, Talitha Veith, Julia Schneider, Terry Jones, Christian Drosten |
| EPI_ISL_730303, EPI_ISL_730312, EPI_ISL_730319, EPI_ISL_730325, EPI_ISL_730332, EPI_ISL_730340, EPI_ISL_730349 | San Diego County Public Health Laboratory | Andersen lab at Scripps Research | SEARCH Alliance San Diego with Tracy Basler, Jovan Shephard, Brett Austin |

| | | | |
|---|---|---|---|
| EPI_ISL_731891, EPI_ISL_731892, EPI_ISL_731893 | Lighthouse Lab in Milton Keynes | Wellcome Sanger Institute for the COVID-19 Genomics UK (COG-UK) Consortium | The Lighthouse Lab in Milton Keynes and Alex Alderton, Roberto Amato, Sonia Goncalves, Ewan Harrison, David K. Jackson, Ian Johnston, Dominic Kwiatkowski, Cordelia Langford, John Sillitoe on behalf of the Wellcome Sanger Institute COVID-19 Surveillance Team |
| EPI_ISL_732114, EPI_ISL_732115, EPI_ISL_732116, EPI_ISL_732117 | Instituto Nacional de Saude (INSA) | Instituto Nacional de Saude (INSA) | Borges et al |
| EPI_ISL_732426, EPI_ISL_732433 | National Virus Reference Laboratory | National Virus Reference Laboratory | Michael Carr, Gabriel Gonzalez, Jonathan Dean, Daniel Hare, Cillian F De Gascun |
| EPI_ISL_732567, EPI_ISL_732570, EPI_ISL_732572, EPI_ISL_732576, EPI_ISL_732577, EPI_ISL_732578, EPI_ISL_732579, EPI_ISL_732580, EPI_ISL_732581, EPI_ISL_732582, EPI_ISL_732583, EPI_ISL_732584, EPI_ISL_732597, EPI_ISL_732598, EPI_ISL_732600, EPI_ISL_732612, EPI_ISL_732613, EPI_ISL_732614, EPI_ISL_732615 | | | |
| see above | Department of Virology and Immunology, University of Helsinki and Helsinki University Hospital, Huslab Finland | Department of Virology, Faculty of Medicine, University of Helsinki, Helsinki, Finland | Teemu Smura, Ravi Kant, Phuoc Truong, Hussein Alburkat, Hannimari Kallio-Kokko, Jenni Virtanen, Maija Suvanto, Sari Hannula, Harri Kangas, Pekka Ellonen, Olli Vapalahti |
| EPI_ISL_733499 | 1-Laboratory of Microbiology, National Reference Lab, Charles Nicolle Hospital; 2-University of Tunis ElManar, Faculty of Medicine of Tunis, LR99ES09, Tunis, Tunisia | 1-Clinical and Experimental Pharmacology Lab, LR16SP02, National Center of Pharmacovigilance, University of Tunis El Manar, Tunis, Tunisia. 2-Neurodegenerative diseases and psychiatric troubles, LR18SP03, Razi Hospital, University of Tunis El Manar, Tunis, Tunisia. 3- Ministry of Health, National Observatory of New and Emerging Diseases, 1006, Tunis, Tunisia | Ilhem Boutiba-Ben Boubaker, Sameh Trabelsi, Nissaf Ben Alaya, Maher Kharrat, Alia Ben Kahla, Jalila Ben Khelil, Salma Abid, Sana Ferjani, Mouna Ben Sassi, Mouna Safer, Guedi Ali Barreh, Habiba Ben Romdhane, Souissi Amira, Sarra Chamman, Hanen El Jebari, Asma Ferjani, Gaies Emna, Riadh Daghfous, Riadh Gouider. |
| EPI_ISL_733576 | Temporary Specimen Collection Centre | Hong Kong Department of Health | Alan K.L. Tsang, Peter C.W. Yip, Edman T.K. Lam, Rickjason C.W. Chan, Dominic N.C. Tsang |
| EPI_ISL_733606, EPI_ISL_733753, EPI_ISL_733883, EPI_ISL_733899, EPI_ISL_733906, EPI_ISL_733907 | Lighthouse Lab in Cambridge | Wellcome Sanger Institute for the COVID-19 Genomics UK (COG-UK) Consortium | Rob Howes, The Lighthouse Lab in Cambridge and Alex Alderton, Roberto Amato, Sonia Goncalves, Ewan Harrison, David K. Jackson, Ian Johnston, Dominic Kwiatkowski, Cordelia Langford, John Sillitoe on behalf of the Wellcome Sanger Institute COVID-19 Surveillance Team |
| EPI_ISL_735236, EPI_ISL_735237, EPI_ISL_735238, EPI_ISL_735239, EPI_ISL_735240 | UZ Leuven, National Reference Laboratory for Coronaviruses, Laboratory Medicine, Leuven, Belgium | KU Leuven, Rega Institute, Clinical and Epidemiological Virology | Tony Wawina-Bokalanga, Joan Marti-Carerras, Bert Vanmechelen, Piet Maes |
| EPI_ISL_735439, EPI_ISL_735440, EPI_ISL_735441, EPI_ISL_735442, EPI_ISL_735443 | Pathogen Genomics Center, National Institute of Infectious Diseases | Pathogen Genomics Center, National Institute of Infectious Diseases | Tsuyoshi Sekizuka, Kentaro Itokawa, Rina Tanaka, Masanori Hashino, Makoto Kuroda |
| EPI_ISL_735500 | University of Chittagong | Central Biological Research Laboratory and Department of Biochemistry and Molecular Biology | H. M. Abdullah Al Masud, Mohammad Omar Faruque, Sajib Rudra, Md. Khondakar Raziur Rahman, Imam Hossen, Md. Arif Hossain, Shanta Paul, Md. Omer Faruq, Md. Imranul Hoq, Robiul Hasan Bhuiyan |
| EPI_ISL_736891, EPI_ISL_736892, EPI_ISL_736893, EPI_ISL_736894, EPI_ISL_736895, EPI_ISL_736896, EPI_ISL_736897, EPI_ISL_736898, EPI_ISL_736899, EPI_ISL_736900, EPI_ISL_736901, EPI_ISL_736902, EPI_ISL_736903, EPI_ISL_736904, EPI_ISL_736905, EPI_ISL_736906, EPI_ISL_736907, EPI_ISL_736908, EPI_ISL_736909, EPI_ISL_736910, EPI_ISL_736911, EPI_ISL_736912, EPI_ISL_736913, EPI_ISL_736914, EPI_ISL_736915, EPI_ISL_736916, EPI_ISL_736917, EPI_ISL_736918, EPI_ISL_736919, EPI_ISL_736920, EPI_ISL_736921, EPI_ISL_736922, EPI_ISL_736923, EPI_ISL_736924 | | | |
| see above | Pathogen Genomics Center, National Institute of Infectious Diseases | Pathogen Genomics Center, National Institute of Infectious Diseases | Tsuyoshi Sekizuka, Kentaro Itokawa, Rina Tanaka, Masanori Hashino, Makoto Kuroda |
| EPI_ISL_736969 | NHLS-IALCH | KRISP, KZN Research Innovation and Sequencing Platform | Giandhari J, Pillay S, Lessells R, ChimukangaraB, Mdlalose K, York D, Khan S, Tegally H, Wilkinson E, de Oliveira T |
| EPI_ISL_737100, EPI_ISL_737101, EPI_ISL_737102, EPI_ISL_737107, EPI_ISL_737111, EPI_ISL_737112, EPI_ISL_737113, EPI_ISL_737114, EPI_ISL_737117, EPI_ISL_737118, EPI_ISL_737120, EPI_ISL_737121, EPI_ISL_737123, EPI_ISL_737127, EPI_ISL_737128, EPI_ISL_737129, EPI_ISL_737132, EPI_ISL_737145, EPI_ISL_737150, EPI_ISL_737152, EPI_ISL_737162, EPI_ISL_737165, EPI_ISL_737168, EPI_ISL_737181, EPI_ISL_737198 | | | |
| see above | Michigan Department of Health and Human Services, Bureau of Laboratories | Michigan Department of Health and Human Services, Bureau of Laboratories | Blankenship HM, Riner D, Soehnlen MK |
| EPI_ISL_737208 | Pathogen Genomics Center, National Institute of Infectious Diseases | Pathogen Genomics Center, National Institute of Infectious Diseases | Tsuyoshi Sekizuka, Kentaro Itokawa, Rina Tanaka, Masanori Hashino, Makoto Kuroda |
| EPI_ISL_737243, EPI_ISL_737244 | Los Angeles County PHL | Los Angeles County PHL | P. Hemarajata et al. |
| EPI_ISL_737605, EPI_ISL_737637, EPI_ISL_737638, EPI_ISL_737661, EPI_ISL_737747, EPI_ISL_737748, EPI_ISL_737749, EPI_ISL_737750, EPI_ISL_737758, EPI_ISL_737762, EPI_ISL_737816, EPI_ISL_737871, EPI_ISL_737872, EPI_ISL_737873, EPI_ISL_737874, EPI_ISL_737907, EPI_ISL_737925, EPI_ISL_737926, EPI_ISL_737927, EPI_ISL_737928, EPI_ISL_737929 | | | |
| see above | Viollier AG | Department of Biosystems Science and Engineering, ETH Zürich | Chaoran Chen, Sarah Nadeau, Catharine Aquino, Ivan Topolsky, Philipp Jablonski, Lara Fuhrmann, David Dreifuss, Katharina Jahn, Andreia Cabral de Gouvea, Maria Domenica Moccia, Simon Grüter, Timothy Sykes, Lennart Opitz, Griffin White, Laura Neff, Doris Popovic, Andrea Patrignani, Jay Tracy, Ralph Schlapbach, Christiane Beckmann, Maurice Redondo, Olivier Kobel, Christoph Noppen, Sophie Seidel, Noemie Santamaria de Souza, Niko Beerenwinkel, Tanja Stadler |
| EPI_ISL_738507 | Orange County Public Health Lab | Chan-Zuckerberg Biohub | CZB Cliahub Consortium |
| EPI_ISL_738521, EPI_ISL_738542, EPI_ISL_738550, EPI_ISL_738555, EPI_ISL_738557 | Santa Clara County Public Health Laboratory | Chan-Zuckerberg Biohub | CZB Cliahub Consortium |
| EPI_ISL_738567 | Renegade | Chan-Zuckerberg Biohub | CZB Cliahub Consortium |
| EPI_ISL_738569, EPI_ISL_738570 | Orange County Public Health Lab | Chan-Zuckerberg Biohub | CZB Cliahub Consortium |
| EPI_ISL_738574 | Renegade | Chan-Zuckerberg Biohub | CZB Cliahub Consortium |
| EPI_ISL_738579 | Santa Clara County Public Health Laboratory | Chan-Zuckerberg Biohub | CZB Cliahub Consortium |
| EPI_ISL_738588 | Orange County Public Health Lab | Chan-Zuckerberg Biohub | CZB Cliahub Consortium |
| EPI_ISL_738591 | Renegade | Chan-Zuckerberg Biohub | CZB Cliahub Consortium |
| EPI_ISL_738620, EPI_ISL_738624, EPI_ISL_738630, EPI_ISL_738634, EPI_ISL_738635, EPI_ISL_738636, EPI_ISL_738637, EPI_ISL_738638, EPI_ISL_738639, EPI_ISL_738640, EPI_ISL_738641, EPI_ISL_738646 | | | |
| see above | Orange County Public Health Lab | Chan-Zuckerberg Biohub | CZB Cliahub Consortium |
| EPI_ISL_738662 | Renegade | Chan-Zuckerberg Biohub | CZB Cliahub Consortium |
| EPI_ISL_738696 | Santa Clara County Public Health Laboratory | Chan-Zuckerberg Biohub | CZB Cliahub Consortium |
| EPI_ISL_738712, EPI_ISL_738725 | Renegade | Chan-Zuckerberg Biohub | CZB Cliahub Consortium |
| EPI_ISL_738747, EPI_ISL_738750 | Santa Clara County Public Health Laboratory | Chan-Zuckerberg Biohub | CZB Cliahub Consortium |
| EPI_ISL_738758 | Orange County Public Health Lab | Chan-Zuckerberg Biohub | CZB Cliahub Consortium |
| EPI_ISL_738770 | Renegade | Chan-Zuckerberg Biohub | CZB Cliahub Consortium |
| EPI_ISL_738784 | Orange County Public Health Lab | Chan-Zuckerberg Biohub | CZB Cliahub Consortium |
| EPI_ISL_738805 | Santa Clara County Public Health Laboratory | Chan-Zuckerberg Biohub | CZB Cliahub Consortium |
| EPI_ISL_738816 | Orange County Public Health Lab | Chan-Zuckerberg Biohub | CZB Cliahub Consortium |
| EPI_ISL_738834 | Santa Clara County Public Health Laboratory | Chan-Zuckerberg Biohub | CZB Cliahub Consortium |

| | | | |
|---|---|---|---|
| EPI_ISL_738851 | Renegade | Chan-Zuckerberg Biohub | CZB Cliahub Consortium |
| EPI_ISL_738855 | Santa Clara County Public Health Laboratory | Chan-Zuckerberg Biohub | CZB Cliahub Consortium |
| EPI_ISL_738874, EPI_ISL_738878, EPI_ISL_738949 | Orange County Public Health Lab | Chan-Zuckerberg Biohub | CZB Cliahub Consortium |
| EPI_ISL_738955 | Santa Clara County Public Health Laboratory | Chan-Zuckerberg Biohub | CZB Cliahub Consortium |
| EPI_ISL_738977 | Renegade | Chan-Zuckerberg Biohub | CZB Cliahub Consortium |
| EPI_ISL_738978, EPI_ISL_738981, EPI_ISL_739002 | Orange County Public Health Lab | Chan-Zuckerberg Biohub | CZB Cliahub Consortium |
| EPI_ISL_739014 | Renegade | Chan-Zuckerberg Biohub | CZB Cliahub Consortium |
| EPI_ISL_739027, EPI_ISL_739037, EPI_ISL_739040, EPI_ISL_739050, EPI_ISL_739053, EPI_ISL_739058, EPI_ISL_739068, EPI_ISL_739087, EPI_ISL_739104 | Santa Clara County Public Health Laboratory | Chan-Zuckerberg Biohub | CZB Cliahub Consortium |
| EPI_ISL_739116 | Orange County Public Health Lab | Chan-Zuckerberg Biohub | CZB Cliahub Consortium |
| EPI_ISL_739141, EPI_ISL_739166, EPI_ISL_739186 | Santa Clara County Public Health Laboratory | Chan-Zuckerberg Biohub | CZB Cliahub Consortium |
| EPI_ISL_739259 | Orange County Public Health Lab | Chan-Zuckerberg Biohub | CZB Cliahub Consortium |
| EPI_ISL_739263, EPI_ISL_739298, EPI_ISL_739319 | Santa Clara County Public Health Laboratory | Chan-Zuckerberg Biohub | CZB Cliahub Consortium |
| EPI_ISL_739341 | Orange County Public Health Lab | Chan-Zuckerberg Biohub | CZB Cliahub Consortium |
| EPI_ISL_739344 | Santa Clara County Public Health Laboratory | Chan-Zuckerberg Biohub | CZB Cliahub Consortium |
| EPI_ISL_739348 | Renegade | Chan-Zuckerberg Biohub | CZB Cliahub Consortium |
| EPI_ISL_739354 | Santa Clara County Public Health Laboratory | Chan-Zuckerberg Biohub | CZB Cliahub Consortium |
| EPI_ISL_739355 | Renegade | Chan-Zuckerberg Biohub | CZB Cliahub Consortium |
| EPI_ISL_739370, EPI_ISL_739371, EPI_ISL_739398, EPI_ISL_739419 | Santa Clara County Public Health Laboratory | Chan-Zuckerberg Biohub | CZB Cliahub Consortium |
| EPI_ISL_739462 | Orange County Public Health Lab | Chan-Zuckerberg Biohub | CZB Cliahub Consortium |
| EPI_ISL_739467, EPI_ISL_739478 | Renegade | Chan-Zuckerberg Biohub | CZB Cliahub Consortium |
| EPI_ISL_739485, EPI_ISL_739487 | Santa Clara County Public Health Laboratory | Chan-Zuckerberg Biohub | CZB Cliahub Consortium |
| EPI_ISL_739496 | Orange County Public Health Lab | Chan-Zuckerberg Biohub | CZB Cliahub Consortium |
| EPI_ISL_739505 | Santa Clara County Public Health Laboratory | Chan-Zuckerberg Biohub | CZB Cliahub Consortium |
| EPI_ISL_739524 | Orange County Public Health Lab | Chan-Zuckerberg Biohub | CZB Cliahub Consortium |
| EPI_ISL_739548 | Santa Clara County Public Health Laboratory | Chan-Zuckerberg Biohub | CZB Cliahub Consortium |
| EPI_ISL_739609 | Orange County Public Health Lab | Chan-Zuckerberg Biohub | CZB Cliahub Consortium |
| EPI_ISL_739617, EPI_ISL_739650, EPI_ISL_739653 | Santa Clara County Public Health Laboratory | Chan-Zuckerberg Biohub | CZB Cliahub Consortium |
| EPI_ISL_739662 | BCCDC Public Health Laboratory | BCCDC Public Health Laboratory | Prystajecky Natalie, Linda Hoang, Dan Fornika, Shannon Russell, Kim Macdonald, Kimia Kamelian, John Tyson, Inna Sekirov, Mel Krajden |
| EPI_ISL_740863, EPI_ISL_740865 | South Eastern Area Laboratory Services (SEALS) | NSW Health Pathology - Institute of Clinical Pathology and Medical Research; Westmead Hospital; University of Sydney | CIDM-PH et al. |
| EPI_ISL_740955 | Department of Pathology, University of Cambridge | COVID-19 Genomics UK (COG-UK) Consortium | Aminu S. Jahun, Yasmin Chaudhry, Grant Hall, Iliana Georgana, Myra Hosmillo, Martin D. Curran, Malte Pinckert, Surendra Parmar, Ian Goodfellow |
| EPI_ISL_741256, EPI_ISL_741257, EPI_ISL_741258 | University College London, Great Ormond Street Hospital for Children NHS Foundation Trust, Imperial College Healthcare NHS Trust | COVID-19 Genomics UK (COG-UK) Consortium | Sergi Castellano, Rachel Williams, Mark Kristiansen, Paola Resende Silva, Sunando Roy, Tony Brooks, Helena Tutill, Paola Niola, Patricia Dyal, Charlotte Williams, Leysa Forrest, Yasmin Panchbhaya, Jacqueline Findlay, Samuel Weeks, Julianne Brown, Kathryn Harris, Paul Randell, James Price, Alison Holmes, Judith Breuer |
| EPI_ISL_741425, EPI_ISL_741429, EPI_ISL_741430, EPI_ISL_741431, EPI_ISL_741432, EPI_ISL_741433, EPI_ISL_741434, EPI_ISL_741435 | Northumbria University / South Tees Hospitals NHS Foundation Trust / North Cumbria Integrated Care NHS Foundation Trust / North Tees and Hartlepool NHS Foundation Trust / Newcastle Hospitals NHS Foundation Trust | COVID-19 Genomics UK (COG-UK) Consortium | Darren L Smith,Andrew Nelson,Matthew Bashton,Greg R Young,Joshua Loh,John Allan,Mohammad A Tariq,Giles S Holt,Gary Black,Wen C Yew,Lynn Dover,Paul Baker,Steve Liggett,Sarah Essex,Jane Greenaway,Debra Padgett,Clive Graham,Garren Scott,Edward Barton,Emma Swindells,Brendan Payne,Jennifer Collins,Yusri Taha,Gary Eltringham |
| EPI_ISL_741611, EPI_ISL_741612 | Queens Medical Centre, Clinical Microbiology Department / DeepSeq Nottingham | COVID-19 Genomics UK (COG-UK) Consortium | Gemma Clark, Wendy Smith, Manjinder Khakh, Vicki M Fleming, Michelle M Lister, Hannah Howson-Wells, Jonathan Ball, Patrick McClure, Joseph Chappell, Theocharis Tsoleridis, Nadine Holmes, Matthew Carlisle, Christopher Moore, Fei Sang, Johnny Debebe, Victoria Wright, Matthew Loose |
| EPI_ISL_741737, EPI_ISL_741754 | Centre for Enzyme Innovation, University of Portsmouth / Translational Research Laboratory, Portsmouth Hospitals NHS Trust | COVID-19 Genomics UK (COG-UK) Consortium | Angela Beckett,Yann Bourgeois,Garry Scarlett,Sharon Glaysher,Scott Elliott,Kelly Bicknell,Robert Impey,Allyson Lloyd,Sarah Wyllie,Ethan Butcher,Anoop Chauhan,Samuel Robson |
| EPI_ISL_741944, EPI_ISL_741957, EPI_ISL_741963, EPI_ISL_741980, EPI_ISL_741997, EPI_ISL_741998, EPI_ISL_742026, EPI_ISL_742032, EPI_ISL_742037, EPI_ISL_742073, EPI_ISL_742087, EPI_ISL_742102 | | | |
| see above | Virology Department, Sheffield Teaching Hospitals NHS Foundation Trust/Department of Infection, Immunity and Cardiovascular Disease, The Medical School, University of Sheffield | COVID-19 Genomics UK (COG-UK) Consortium | Thushan de Silva, Matthew Parker, Nikki Smith, Adri Angyal, Rebecca Brown, Luke Green, Rachel Tucker, Paul Parsons, Danielle Groves, Katie Johnson, Laura Carrilero, Alex Keeley, Dave Partridge, Matthew Wyles, Benjamin Lindsey, Mehmet Yavuz, Mohammad Raza, Cariad Evans |
| EPI_ISL_742175 | Virology Department, Royal Infirmary of Edinburgh, NHS Lothian / School of Biological Sciences, University of Edinburgh / Institute of Genetics and Molecular Medicine, University of Edinburgh | COVID-19 Genomics UK (COG-UK) Consortium | McHugh M, Dewar R, Rooke S, Gallagher M, Balcaza C, O'Toole Á, Scher E, Hill V, McCrone JT, Colquhoun R, Yu X, Jackson B, Rambaut A, Williams TC, Templeton K |
| EPI_ISL_742267, EPI_ISL_742268, EPI_ISL_742351, EPI_ISL_742353, EPI_ISL_742354, EPI_ISL_742355, EPI_ISL_742841, EPI_ISL_742842, EPI_ISL_742843, EPI_ISL_742844, EPI_ISL_742845, EPI_ISL_742846, EPI_ISL_742890, EPI_ISL_742891, EPI_ISL_742892, EPI_ISL_742893, EPI_ISL_742894, EPI_ISL_742895, EPI_ISL_742896, EPI_ISL_742897, EPI_ISL_742898, EPI_ISL_742899, EPI_ISL_742900, EPI_ISL_742901, EPI_ISL_742902, EPI_ISL_742903, EPI_ISL_742904, EPI_ISL_742905, EPI_ISL_742906, EPI_ISL_743118, EPI_ISL_743124, EPI_ISL_743125, EPI_ISL_743132, EPI_ISL_743133, EPI_ISL_743134, EPI_ISL_743135, EPI_ISL_743166, EPI_ISL_743167, EPI_ISL_743168, EPI_ISL_743169, EPI_ISL_743170, EPI_ISL_743171, EPI_ISL_743172, EPI_ISL_743173, EPI_ISL_743342, EPI_ISL_743343, EPI_ISL_743353, EPI_ISL_743354, EPI_ISL_743358, EPI_ISL_743359, EPI_ISL_743360, EPI_ISL_743361, EPI_ISL_743382, EPI_ISL_743383, EPI_ISL_743384, EPI_ISL_743385, EPI_ISL_743386, EPI_ISL_743387, EPI_ISL_743388, EPI_ISL_743389, EPI_ISL_743390, EPI_ISL_743391, EPI_ISL_743392, EPI_ISL_743393, EPI_ISL_743394, EPI_ISL_743395, EPI_ISL_743396, EPI_ISL_743398, EPI_ISL_743399, EPI_ISL_743400, EPI_ISL_743401, EPI_ISL_743402, EPI_ISL_743403, EPI_ISL_743404, EPI_ISL_743405 | | | |
| see above | Wales Specialist Virology Centre Sequencing lab: Pathogen Genomics Unit | COVID-19 Genomics UK (COG-UK) Consortium | Catherine Moore, Johnathan Evans, Laura Gifford, Malorie Perry, Simon Cottrell, Angela Marchbank, Alec Birchley, Alexander Adams, Amy Gaskin, Bree Gatica-Wilcox, Jason Coombes, Joel Southgate, Lauren Gilbert, Lee Graham, Nicole Pacchiarini, Sara Kumziene-Summerhayes, Sarah Taylor, Sophie |

| | | | |
|---|---|---|---|
| | | | Jones, Sara Rey, Matthew Bull, Joanne Watkins, Sally Corden, Tom Connor |
| EPI_ISL_745034, EPI_ISL_745035, EPI_ISL_745036, EPI_ISL_745037, EPI_ISL_745038 | Israel Central Virology laboratory | Israel Central Virology laboratory | Neta Zuckerman, Efrat Dahan Bucris, Oran Erster, Michal Mandelboim, Orna Mor, Ella Mendelson |
| EPI_ISL_745113 | Dr Ivan Toms Clinic wc IVT | National Health Laboratory Service (NHLS), Tygerberg | Susan Engelbrecht, Kayla Delaney, Bronwyn Kleinhans, Houriiyah Tegally, Eduan Wilkindon, Gert van Zyl, Wolfgang Preiser, Tulio de Oliveira |
| EPI_ISL_745121 | Calvinia Hospital | National Health Laboratory Service (NHLS), Tygerberg | Susan Engelbrecht, Kayla Delaney, Bronwyn Kleinhans, Houriiyah Tegally, Eduan Wilkindon, Gert van Zyl, Wolfgang Preiser, Tulio de Oliveira |
| EPI_ISL_745124 | Laetitia Bam CHC | National Health Laboratory Service (NHLS), Tygerberg | Susan Engelbrecht, Kayla Delaney, Bronwyn Kleinhans, Houriiyah Tegally, Eduan Wilkindon, Gert van Zyl, Wolfgang Preiser, Tulio de Oliveira |
| EPI_ISL_745126 | Okiep CHC | National Health Laboratory Service (NHLS), Tygerberg | Susan Engelbrecht, Kayla Delaney, Bronwyn Kleinhans, Houriiyah Tegally, Eduan Wilkindon, Gert van Zyl, Wolfgang Preiser, Tulio de Oliveira |
| EPI_ISL_745129 | Port Nolloth Hospital | National Health Laboratory Service (NHLS), Tygerberg | Susan Engelbrecht, Kayla Delaney, Bronwyn Kleinhans, Houriiyah Tegally, Eduan Wilkindon, Gert van Zyl, Wolfgang Preiser, Tulio de Oliveira |
| EPI_ISL_745137 | Edameni Clinic | National Health Laboratory Service (NHLS), Tygerberg | Susan Engelbrecht, Kayla Delaney, Bronwyn Kleinhans, Houriiyah Tegally, Eduan Wilkindon, Gert van Zyl, Wolfgang Preiser, Tulio de Oliveira |
| EPI_ISL_745138 | Dr Van Niekerk Hospital | National Health Laboratory Service (NHLS), Tygerberg | Susan Engelbrecht, Kayla Delaney, Bronwyn Kleinhans, Houriiyah Tegally, Eduan Wilkindon, Gert van Zyl, Wolfgang Preiser, Tulio de Oliveira |
| EPI_ISL_745142, EPI_ISL_745143 | Nomangesi Jayiya Clinic | National Health Laboratory Service (NHLS), Tygerberg | Susan Engelbrecht, Kayla Delaney, Bronwyn Kleinhans, Houriiyah Tegally, Eduan Wilkindon, Gert van Zyl, Wolfgang Preiser, Tulio de Oliveira |
| EPI_ISL_745151 | Edameni Clinic | National Health Laboratory Service (NHLS), Tygerberg | Susan Engelbrecht, Kayla Delaney, Bronwyn Kleinhans, Houriiyah Tegally, Eduan Wilkindon, Gert van Zyl, Wolfgang Preiser, Tulio de Oliveira |
| EPI_ISL_745152 | Laetitia Bam CHC | National Health Laboratory Service (NHLS), Tygerberg | Susan Engelbrecht, Kayla Delaney, Bronwyn Kleinhans, Houriiyah Tegally, Eduan Wilkindon, Gert van Zyl, Wolfgang Preiser, Tulio de Oliveira |
| EPI_ISL_745154 | Dr Ivan Toms Clinic wc IVT | National Health Laboratory Service (NHLS), Tygerberg | Susan Engelbrecht, Kayla Delaney, Bronwyn Kleinhans, Houriiyah Tegally, Eduan Wilkindon, Gert van Zyl, Wolfgang Preiser, Tulio de Oliveira |
| EPI_ISL_745155 | Langebaan Clinic wc LBC | National Health Laboratory Service (NHLS), Tygerberg | Susan Engelbrecht, Kayla Delaney, Bronwyn Kleinhans, Houriiyah Tegally, Eduan Wilkindon, Gert van Zyl, Wolfgang Preiser, Tulio de Oliveira |
| EPI_ISL_745156 | SAS Saldanha | National Health Laboratory Service (NHLS), Tygerberg | Susan Engelbrecht, Kayla Delaney, Bronwyn Kleinhans, Houriiyah Tegally, Eduan Wilkindon, Gert van Zyl, Wolfgang Preiser, Tulio de Oliveira |
| EPI_ISL_745202 | Lab voor klinische biologie | Onderzoeksgroep Virologie | Laurens Lambrechts, Nick Vereecke, Marthe Pauwels, Bruno Verhasselt, Linos Vandekerckhove, Hans Nauwynck, Sebastiaan Theuns |
| EPI_ISL_745317, EPI_ISL_745326, EPI_ISL_745327, EPI_ISL_745328, EPI_ISL_745329 | CNR Virus des Infections Respiratoires - France SUD | CNR Virus des Infections Respiratoires - France SUD | Antonin Bal, Gregory Destras, Claudia Gonzalez, Gwendolyne Burfin, Quentin Semanas, Martine Valette, Bruno Lina, Laurence Josset |
| EPI_ISL_745336 | CHU Clermont-Ferrand | CNR Virus des Infections Respiratoires - France SUD | Antonin Bal, Gregory Destras, Gwendolyne Burfin, Hadrien Règue, Quentin Semanas, Martine Valette, Bruno Lina, Christine Archimbaud, Amélie Brebion, Hélène Chabrolles, Martine Chambon, Audrey Mirand, Christel Regagnon, Maxime Bisseux, Patricia Combes, Cécile Henquell, Laurence Josset |
| EPI_ISL_745346, EPI_ISL_745347, EPI_ISL_745348, EPI_ISL_745349, EPI_ISL_745350, EPI_ISL_745351, EPI_ISL_745352, EPI_ISL_745353, EPI_ISL_745354, EPI_ISL_745355, EPI_ISL_745356 | | | |
| see above | CNR Virus des Infections Respiratoires - France SUD | CNR Virus des Infections Respiratoires - France SUD | Antonin Bal, Gregory Destras, Claudia Gonzalez, Gwendolyne Burfin, Quentin Semanas, Martine Valette, Bruno Lina, Laurence Josset |
| EPI_ISL_745400 | DOHMH Central Harlem | New York City Public Health Laboratory | Jade Wang, et al. |
| EPI_ISL_745401 | DOHMH Corona | New York City Public Health Laboratory | Jade Wang, et al. |
| EPI_ISL_745402 | DOHMH Jamaica | New York City Public Health Laboratory | Jade Wang, et al. |
| EPI_ISL_745403 | DOHMH Corona | New York City Public Health Laboratory | Jade Wang, et al. |
| EPI_ISL_745405 | DOHMH Jamaica | New York City Public Health Laboratory | Jade Wang, et al. |
| EPI_ISL_745406 | DOHMH Morrisania | New York City Public Health Laboratory | Jade Wang, et al. |
| EPI_ISL_745408 | DOHMH Riverside | New York City Public Health Laboratory | Jade Wang, et al. |
| EPI_ISL_745409 | DOHMH Corona | New York City Public Health Laboratory | Jade Wang, et al. |
| EPI_ISL_745410 | DOHMH Chelsea | New York City Public Health Laboratory | Jade Wang, et al. |
| EPI_ISL_745411 | DOHMH Corona | New York City Public Health Laboratory | Jade Wang, et al. |
| EPI_ISL_745412 | DOHMH Morrisania | New York City Public Health Laboratory | Jade Wang, et al. |
| EPI_ISL_745420 | DOHMH Crown Heights | New York City Public Health Laboratory | Jade Wang, et al. |
| EPI_ISL_745421, EPI_ISL_745422 | DOHMH Chelsea | New York City Public Health Laboratory | Jade Wang, et al. |
| EPI_ISL_745423, EPI_ISL_745424 | DOHMH Corona | New York City Public Health Laboratory | Jade Wang, et al. |
| EPI_ISL_745425 | DOHMH Central Harlem | New York City Public Health Laboratory | Jade Wang, et al. |
| EPI_ISL_745426, EPI_ISL_745427 | DOHMH Corona | New York City Public Health Laboratory | Jade Wang, et al. |
| EPI_ISL_745428, EPI_ISL_745429 | DOHMH Riverside | New York City Public Health Laboratory | Jade Wang, et al. |
| EPI_ISL_745430, EPI_ISL_745431, EPI_ISL_745432, EPI_ISL_745433 | DOHMH Morrisania | New York City Public Health Laboratory | Jade Wang, et al. |
| EPI_ISL_745434, EPI_ISL_745435 | DOHMH Corona | New York City Public Health Laboratory | Jade Wang, et al. |
| EPI_ISL_745436 | DOHMH Chelsea | New York City Public Health Laboratory | Jade Wang, et al. |
| EPI_ISL_745437, EPI_ISL_745438 | DOHMH PHL | New York City Public Health Laboratory | Jade Wang, et al. |
| EPI_ISL_745439 | DOHMH Riverside | New York City Public Health Laboratory | Jade Wang, et al. |
| EPI_ISL_745440 | DOHMH Jamaica | New York City Public Health Laboratory | Jade Wang, et al. |
| EPI_ISL_745445 | DOHMH Crown Heights | New York City Public Health Laboratory | Jade Wang, et al. |
| EPI_ISL_745446, EPI_ISL_745447 | DOHMH Corona | New York City Public Health Laboratory | Jade Wang, et al. |
| EPI_ISL_745448 | DOHMH Crown Heights | New York City Public Health Laboratory | Jade Wang, et al. |
| EPI_ISL_745449, EPI_ISL_745450 | DOHMH Morrisania | New York City Public Health Laboratory | Jade Wang, et al. |
| EPI_ISL_745451 | DOHMH Central Harlem | New York City Public Health Laboratory | Jade Wang, et al. |
| EPI_ISL_745452 | DOHMH Crown Heights | New York City Public Health Laboratory | Jade Wang, et al. |
| EPI_ISL_745453 | DOHMH Chelsea | New York City Public Health Laboratory | Jade Wang, et al. |
| EPI_ISL_745454 | DOHMH Crown Heights | New York City Public Health Laboratory | Jade Wang, et al. |
| EPI_ISL_745455 | DOHMH Corona | New York City Public Health Laboratory | Jade Wang, et al. |
| EPI_ISL_745456 | DOHMH PHL | New York City Public Health Laboratory | Jade Wang, et al. |
| EPI_ISL_745457 | DOHMH Chelsea | New York City Public Health Laboratory | Jade Wang, et al. |
| EPI_ISL_745458, EPI_ISL_745459, | DOHMH Corona | New York City Public Health Laboratory | Jade Wang, et al. |

| | | | |
|---|---|---|---|
| EPI_ISL_745460, EPI_ISL_745461, EPI_ISL_745462 | | | |
| EPI_ISL_745463 | DOHMH Fort Greene | New York City Public Health Laboratory | Jade Wang, et al. |
| EPI_ISL_745464 | DOHMH Corona | New York City Public Health Laboratory | Jade Wang, et al. |
| EPI_ISL_745465 | DOHMH PHL | New York City Public Health Laboratory | Jade Wang, et al. |
| EPI_ISL_745466, EPI_ISL_745467 | DOHMH Corona | New York City Public Health Laboratory | Jade Wang, et al. |
| EPI_ISL_745468 | DOHMH Riverside | New York City Public Health Laboratory | Jade Wang, et al. |
| EPI_ISL_745469 | DOHMH Central Harlem | New York City Public Health Laboratory | Jade Wang, et al. |
| EPI_ISL_746390, EPI_ISL_746398, EPI_ISL_746402, EPI_ISL_746403, EPI_ISL_746406, EPI_ISL_746413, EPI_ISL_746416, EPI_ISL_746422, EPI_ISL_746426, EPI_ISL_746428, EPI_ISL_746430, EPI_ISL_746431, EPI_ISL_746432, EPI_ISL_746434, EPI_ISL_746440, EPI_ISL_746446, EPI_ISL_746454, EPI_ISL_746455, EPI_ISL_746460, EPI_ISL_746461, EPI_ISL_746463, EPI_ISL_746466, EPI_ISL_746469, EPI_ISL_746837, EPI_ISL_746838, EPI_ISL_746841, EPI_ISL_746842, EPI_ISL_746843, EPI_ISL_746846, EPI_ISL_746847, EPI_ISL_746848, EPI_ISL_746849, EPI_ISL_746852, EPI_ISL_746854, EPI_ISL_746858, EPI_ISL_746860, EPI_ISL_746861, EPI_ISL_746864, EPI_ISL_746865, EPI_ISL_746868, EPI_ISL_746870, EPI_ISL_746873, EPI_ISL_746875, EPI_ISL_746876, EPI_ISL_746877, EPI_ISL_746884, EPI_ISL_746889, EPI_ISL_746891, EPI_ISL_746892, EPI_ISL_746893, EPI_ISL_746896, EPI_ISL_746897, EPI_ISL_746900, EPI_ISL_746901, EPI_ISL_746902, EPI_ISL_746903, EPI_ISL_746906, EPI_ISL_746912, EPI_ISL_746915, EPI_ISL_746916, EPI_ISL_746917, EPI_ISL_746918, EPI_ISL_746924, EPI_ISL_746926, EPI_ISL_746927, EPI_ISL_746928, EPI_ISL_746929, EPI_ISL_746931, EPI_ISL_746932, EPI_ISL_746934, EPI_ISL_746941, EPI_ISL_746942, EPI_ISL_746945, EPI_ISL_746949, EPI_ISL_746956, EPI_ISL_746957, EPI_ISL_746959, EPI_ISL_746960, EPI_ISL_746961, EPI_ISL_746962, EPI_ISL_746967, EPI_ISL_746968, EPI_ISL_746971, EPI_ISL_746973, EPI_ISL_746975, EPI_ISL_746977, EPI_ISL_746978, EPI_ISL_746980, EPI_ISL_746981, EPI_ISL_746985, EPI_ISL_746987, EPI_ISL_746988, EPI_ISL_746989, EPI_ISL_746990, EPI_ISL_746991, EPI_ISL_746993, EPI_ISL_746994, EPI_ISL_746995, EPI_ISL_747015, EPI_ISL_747016, EPI_ISL_747017, EPI_ISL_747018, EPI_ISL_747019, EPI_ISL_747024, EPI_ISL_747025, EPI_ISL_747026, EPI_ISL_747027, EPI_ISL_747028 | | | |
| see above | Utah Public Health Laboratory | Utah Public Health Laboratory | Erin Young, Kelly Oakeson, Tara Gallagher |
| EPI_ISL_747464, EPI_ISL_747483 | Ospedale San Bonifacio | Istituto Zooprofilattico Sperimentale delle Venezie | Adelaide Milani, Alessia Schivo, Annalisa Salviato, Erika Giorgia Quaranta, Ambra Pastori, Bianca Zecchin, Alice Fusaro, Isabella Monne, Calogero Terregino, Antonia Ricci |
| EPI_ISL_747490, EPI_ISL_747491, EPI_ISL_747493, EPI_ISL_747494, EPI_ISL_747495, EPI_ISL_747496, EPI_ISL_747497 | Respiratory Virus Unit, National Infection Service, Public Health England | COVID-19 Genomics UK (COG-UK) Consortium | PHE Covid Sequencing Team |
| EPI_ISL_747530, EPI_ISL_747532, EPI_ISL_747534, EPI_ISL_747537, EPI_ISL_747538, EPI_ISL_747539, EPI_ISL_747577, EPI_ISL_747578, EPI_ISL_747579, EPI_ISL_747580, EPI_ISL_747581, EPI_ISL_747582, EPI_ISL_747583, EPI_ISL_747584, EPI_ISL_747596, EPI_ISL_747603, EPI_ISL_747606, EPI_ISL_747624, EPI_ISL_747625, EPI_ISL_747626, EPI_ISL_747627, EPI_ISL_747628, EPI_ISL_747633, EPI_ISL_747634, EPI_ISL_747635, EPI_ISL_747641, EPI_ISL_747646, EPI_ISL_747650, EPI_ISL_747654, EPI_ISL_747657, EPI_ISL_747675, EPI_ISL_747676, EPI_ISL_747677, EPI_ISL_747678, EPI_ISL_747679, EPI_ISL_747709, EPI_ISL_747710, EPI_ISL_747711, EPI_ISL_747712, EPI_ISL_747713, EPI_ISL_747714, EPI_ISL_747715, EPI_ISL_747716, EPI_ISL_747717, EPI_ISL_747718, EPI_ISL_747719, EPI_ISL_747724, EPI_ISL_747725, EPI_ISL_747726, EPI_ISL_747727, EPI_ISL_747730, EPI_ISL_747735, EPI_ISL_747736, EPI_ISL_747737, EPI_ISL_747738, EPI_ISL_747755, EPI_ISL_748021, EPI_ISL_748022, EPI_ISL_748023, EPI_ISL_748024, EPI_ISL_748025, EPI_ISL_748026, EPI_ISL_748027, EPI_ISL_748028, EPI_ISL_748029, EPI_ISL_748030, EPI_ISL_748031, EPI_ISL_748032, EPI_ISL_748033, EPI_ISL_748034, EPI_ISL_748035, EPI_ISL_748036, EPI_ISL_748037, EPI_ISL_748038, EPI_ISL_748039, EPI_ISL_748040, EPI_ISL_748041, EPI_ISL_748042, EPI_ISL_748043, EPI_ISL_748044, EPI_ISL_748045, EPI_ISL_748046, EPI_ISL_748047, EPI_ISL_748048, EPI_ISL_748049, EPI_ISL_748050, EPI_ISL_748051, EPI_ISL_748052, EPI_ISL_748053, EPI_ISL_748054, EPI_ISL_748055, EPI_ISL_748056, EPI_ISL_748057, EPI_ISL_748058, EPI_ISL_748059, EPI_ISL_748060, EPI_ISL_748061, EPI_ISL_748062, EPI_ISL_748063, EPI_ISL_748064, EPI_ISL_748065, EPI_ISL_748066, EPI_ISL_748067, EPI_ISL_748068, EPI_ISL_748069, EPI_ISL_748070, EPI_ISL_748071, EPI_ISL_748072, EPI_ISL_748073, EPI_ISL_748074, EPI_ISL_748075, EPI_ISL_748076, EPI_ISL_748077, EPI_ISL_748078, EPI_ISL_748079, EPI_ISL_748080, EPI_ISL_748081, EPI_ISL_748082, EPI_ISL_748083, EPI_ISL_748084, EPI_ISL_748085, EPI_ISL_748086, EPI_ISL_748087, EPI_ISL_748088, EPI_ISL_748089, EPI_ISL_748090, EPI_ISL_748091, EPI_ISL_748092, EPI_ISL_748093, EPI_ISL_748094, EPI_ISL_748095, EPI_ISL_748096, EPI_ISL_748097, EPI_ISL_748098 | | | |
| see above | Department of Virus and Microbiological Special Diagnostics, Statens Serum Institut, Copenhagen, Denmark | Albertsen Lab, Department of Chemistry and Bioscience, Aalborg University, Denmark | Danish Covid-19 Genome Consortium |
| EPI_ISL_751270 | Devki Devi Foundation, a unit of Max Healthcare | CSIR-IGIB/Max | Rajesh Pandey#, Samreen Siddiqui, Janani Srinivasa Vasudevan, Akshay Kanakan, Ranjeet Maurya, Uzma Shamim, Bansidhar Tarai, Akansha Tyagi, Mitali Mukerji, Poonam Das, Sujeet Jha, Mohammed Faruq, Anurag Agrawal |
| EPI_ISL_751542 | SC Dept of Health and Env. Control-Bureau of Laboratories | Genomics and Discovery, Respiratory Viruses Branch, Division of Viral Diseases, Centers for Disease Control and Prevention | Krista Queen, Yan Li, Ying Tao, Jing Zhang, Anna Uehara, Anna Montmayeur, Clinton R. Paden, Peter W. Cook,Rachel Marine, Mili Sheth, Haibin Wang, Justin Lee, Suxiang Tong |
| EPI_ISL_751549 | MS Public Health Laboratory | Genomics and Discovery, Respiratory Viruses Branch, Division of Viral Diseases, Centers for Disease Control and Prevention | Krista Queen, Yan Li, Ying Tao, Jing Zhang, Anna Uehara, Anna Montmayeur, Clinton R. Paden, Peter W. Cook,Rachel Marine, Mili Sheth, Haibin Wang, Justin Lee, Suxiang Tong |
| EPI_ISL_751550 | SC Dept of Health and Env. Control-Bureau of Laboratories | Genomics and Discovery, Respiratory Viruses Branch, Division of Viral Diseases, Centers for Disease Control and Prevention | Krista Queen, Yan Li, Ying Tao, Jing Zhang, Anna Uehara, Anna Montmayeur, Clinton R. Paden, Peter W. Cook,Rachel Marine, Mili Sheth, Haibin Wang, Justin Lee, Suxiang Tong |
| EPI_ISL_751559 | MD DOH Laboratories Administration | Genomics and Discovery, Respiratory Viruses Branch, Division of Viral Diseases, Centers for Disease Control and Prevention | Krista Queen, Yan Li, Ying Tao, Jing Zhang, Anna Uehara, Anna Montmayeur, Clinton R. Paden, Peter W. Cook,Rachel Marine, Mili Sheth, Haibin Wang, Justin Lee, Suxiang Tong |
| EPI_ISL_751566, EPI_ISL_751589 | WVDHHR - Office of Laboratory Services | Genomics and Discovery, Respiratory Viruses Branch, Division of Viral Diseases, Centers for Disease Control and Prevention | Krista Queen, Yan Li, Ying Tao, Jing Zhang, Anna Uehara, Anna Montmayeur, Clinton R. Paden, Peter W. Cook,Rachel Marine, Mili Sheth, Haibin Wang, Justin Lee, Suxiang Tong |
| EPI_ISL_751611 | SC Dept of Health and Env. Control-Bureau of Laboratories | Genomics and Discovery, Respiratory Viruses Branch, Division of Viral Diseases, Centers for Disease Control and Prevention | Krista Queen, Yan Li, Ying Tao, Jing Zhang, Anna Uehara, Anna Montmayeur, Clinton R. Paden, Peter W. Cook,Rachel Marine, Mili Sheth, Haibin Wang, Justin Lee, Suxiang Tong |
| EPI_ISL_751615 | OR State PHL-Virology/Immunology Section | Genomics and Discovery, Respiratory Viruses Branch, Division of Viral Diseases, Centers for Disease Control and Prevention | Krista Queen, Yan Li, Ying Tao, Jing Zhang, Anna Uehara, Anna Montmayeur, Clinton R. Paden, Peter W. Cook,Rachel Marine, Mili Sheth, Haibin Wang, Justin Lee, Suxiang Tong |
| EPI_ISL_751617 | TX DSHS, Lab Services Section MC 1947 | Genomics and Discovery, Respiratory Viruses Branch, Division of Viral Diseases, Centers for Disease Control and Prevention | Krista Queen, Yan Li, Ying Tao, Jing Zhang, Anna Uehara, Anna Montmayeur, Clinton R. Paden, Peter W. Cook,Rachel Marine, Mili Sheth, Haibin Wang, Justin Lee, Suxiang Tong |
| EPI_ISL_751620 | LA Office of Public Health Laboratories | Genomics and Discovery, Respiratory Viruses Branch, Division of Viral Diseases, Centers for Disease Control and Prevention | Krista Queen, Yan Li, Ying Tao, Jing Zhang, Anna Uehara, Anna Montmayeur, Clinton R. Paden, Peter W. Cook,Rachel Marine, Mili Sheth, Haibin Wang, Justin Lee, Suxiang Tong |
| EPI_ISL_751633 | MT Public Health Laboratory | Genomics and Discovery, Respiratory Viruses Branch, Division of Viral Diseases, Centers for Disease Control and Prevention | Krista Queen, Yan Li, Ying Tao, Jing Zhang, Anna Uehara, Anna Montmayeur, Clinton R. Paden, Peter W. Cook,Rachel Marine, Mili Sheth, Haibin Wang, Justin Lee, Suxiang Tong |
| EPI_ISL_751634 | MO State Public Health Laboratory | Genomics and Discovery, Respiratory Viruses Branch, Division of Viral Diseases, Centers for Disease Control and Prevention | Krista Queen, Yan Li, Ying Tao, Jing Zhang, Anna Uehara, Anna Montmayeur, Clinton R. Paden, Peter W. Cook,Rachel Marine, Mili Sheth, Haibin Wang, Justin Lee, Suxiang Tong |
| EPI_ISL_751640 | MT Public Health Laboratory | Genomics and Discovery, Respiratory Viruses Branch, Division of Viral Diseases, Centers for Disease Control and Prevention | Krista Queen, Yan Li, Ying Tao, Jing Zhang, Anna Uehara, Anna Montmayeur, Clinton R. Paden, Peter W. Cook,Rachel Marine, Mili Sheth, Haibin Wang, Justin Lee, Suxiang Tong |
| EPI_ISL_751645 | TX DSHS, Lab Services Section MC 1947 | Genomics and Discovery, Respiratory Viruses Branch, Division of Viral Diseases, Centers for Disease Control and Prevention | Krista Queen, Yan Li, Ying Tao, Jing Zhang, Anna Uehara, Anna Montmayeur, Clinton R. Paden, Peter W. Cook,Rachel Marine, Mili Sheth, Haibin Wang, Justin Lee, Suxiang Tong |
| EPI_ISL_751655 | MN PHL Division, Minnesota Department of Health | Genomics and Discovery, Respiratory Viruses Branch, Division of Viral Diseases, Centers for Disease Control and | Krista Queen, Yan Li, Ying Tao, Jing Zhang, Anna Uehara, Anna Montmayeur, Clinton R. Paden, Peter W. Cook,Rachel Marine, Mili Sheth, Haibin Wang, Justin Lee, Suxiang Tong |

| | | | |
|---|---|---|---|
| | | Prevention | |
| EPI_ISL_751661 | MS Public Health Laboratory | Genomics and Discovery, Respiratory Viruses Branch, Division of Viral Diseases, Centers for Disease Control and Prevention | Krista Queen, Yan Li, Ying Tao, Jing Zhang, Anna Uehara, Anna Montmayeur, Clinton R. Paden, Peter W. Cook,Rachel Marine, Mili Sheth, Haibin Wang, Justin Lee, Suxiang Tong |
| EPI_ISL_751664, EPI_ISL_751665 | SC Dept of Health and Env. Control-Bureau of Laboratories | Genomics and Discovery, Respiratory Viruses Branch, Division of Viral Diseases, Centers for Disease Control and Prevention | Krista Queen, Yan Li, Ying Tao, Jing Zhang, Anna Uehara, Anna Montmayeur, Clinton R. Paden, Peter W. Cook,Rachel Marine, Mili Sheth, Haibin Wang, Justin Lee, Suxiang Tong |
| EPI_ISL_751693 | OR State PHL-Virology/Immunology Section | Genomics and Discovery, Respiratory Viruses Branch, Division of Viral Diseases, Centers for Disease Control and Prevention | Krista Queen, Yan Li, Ying Tao, Jing Zhang, Anna Uehara, Anna Montmayeur, Clinton R. Paden, Peter W. Cook,Rachel Marine, Mili Sheth, Haibin Wang, Justin Lee, Suxiang Tong |
| EPI_ISL_751696 | LA Office of Public Health Laboratories | Genomics and Discovery, Respiratory Viruses Branch, Division of Viral Diseases, Centers for Disease Control and Prevention | Krista Queen, Yan Li, Ying Tao, Jing Zhang, Anna Uehara, Anna Montmayeur, Clinton R. Paden, Peter W. Cook,Rachel Marine, Mili Sheth, Haibin Wang, Justin Lee, Suxiang Tong |
| EPI_ISL_751703 | MD DOH Laboratories Administration | Genomics and Discovery, Respiratory Viruses Branch, Division of Viral Diseases, Centers for Disease Control and Prevention | Krista Queen, Yan Li, Ying Tao, Jing Zhang, Anna Uehara, Anna Montmayeur, Clinton R. Paden, Peter W. Cook,Rachel Marine, Mili Sheth, Haibin Wang, Justin Lee, Suxiang Tong |
| EPI_ISL_751713 | MT Public Health Laboratory | Genomics and Discovery, Respiratory Viruses Branch, Division of Viral Diseases, Centers for Disease Control and Prevention | Krista Queen, Yan Li, Ying Tao, Jing Zhang, Anna Uehara, Anna Montmayeur, Clinton R. Paden, Peter W. Cook,Rachel Marine, Mili Sheth, Haibin Wang, Justin Lee, Suxiang Tong |
| EPI_ISL_751718 | SC Dept of Health and Env. Control-Bureau of Laboratories | Genomics and Discovery, Respiratory Viruses Branch, Division of Viral Diseases, Centers for Disease Control and Prevention | Krista Queen, Yan Li, Ying Tao, Jing Zhang, Anna Uehara, Anna Montmayeur, Clinton R. Paden, Peter W. Cook,Rachel Marine, Mili Sheth, Haibin Wang, Justin Lee, Suxiang Tong |
| EPI_ISL_751728 | CO Dept. of Public Health and Environment, Lab Services Division | Genomics and Discovery, Respiratory Viruses Branch, Division of Viral Diseases, Centers for Disease Control and Prevention | Krista Queen, Yan Li, Ying Tao, Jing Zhang, Anna Uehara, Anna Montmayeur, Clinton R. Paden, Peter W. Cook,Rachel Marine, Mili Sheth, Haibin Wang, Justin Lee, Suxiang Tong |
| EPI_ISL_751740 | MO State Public Health Laboratory | Genomics and Discovery, Respiratory Viruses Branch, Division of Viral Diseases, Centers for Disease Control and Prevention | Krista Queen, Yan Li, Ying Tao, Jing Zhang, Anna Uehara, Anna Montmayeur, Clinton R. Paden, Peter W. Cook,Rachel Marine, Mili Sheth, Haibin Wang, Justin Lee, Suxiang Tong |
| EPI_ISL_751744 | SC Dept of Health and Env. Control-Bureau of Laboratories | Genomics and Discovery, Respiratory Viruses Branch, Division of Viral Diseases, Centers for Disease Control and Prevention | Krista Queen, Yan Li, Ying Tao, Jing Zhang, Anna Uehara, Anna Montmayeur, Clinton R. Paden, Peter W. Cook,Rachel Marine, Mili Sheth, Haibin Wang, Justin Lee, Suxiang Tong |
| EPI_ISL_751746 | WVDHHR - Office of Laboratory Services | Genomics and Discovery, Respiratory Viruses Branch, Division of Viral Diseases, Centers for Disease Control and Prevention | Krista Queen, Yan Li, Ying Tao, Jing Zhang, Anna Uehara, Anna Montmayeur, Clinton R. Paden, Peter W. Cook,Rachel Marine, Mili Sheth, Haibin Wang, Justin Lee, Suxiang Tong |
| EPI_ISL_751749 | TX DSHS, Lab Services Section MC 1947 | Genomics and Discovery, Respiratory Viruses Branch, Division of Viral Diseases, Centers for Disease Control and Prevention | Krista Queen, Yan Li, Ying Tao, Jing Zhang, Anna Uehara, Anna Montmayeur, Clinton R. Paden, Peter W. Cook,Rachel Marine, Mili Sheth, Haibin Wang, Justin Lee, Suxiang Tong |
| EPI_ISL_751752 | LA Office of Public Health Laboratories | Genomics and Discovery, Respiratory Viruses Branch, Division of Viral Diseases, Centers for Disease Control and Prevention | Krista Queen, Yan Li, Ying Tao, Jing Zhang, Anna Uehara, Anna Montmayeur, Clinton R. Paden, Peter W. Cook,Rachel Marine, Mili Sheth, Haibin Wang, Justin Lee, Suxiang Tong |
| EPI_ISL_751763 | CO Dept. of Public Health and Environment, Lab Services Division | Genomics and Discovery, Respiratory Viruses Branch, Division of Viral Diseases, Centers for Disease Control and Prevention | Krista Queen, Yan Li, Ying Tao, Jing Zhang, Anna Uehara, Anna Montmayeur, Clinton R. Paden, Peter W. Cook,Rachel Marine, Mili Sheth, Haibin Wang, Justin Lee, Suxiang Tong |
| EPI_ISL_751771 | MD DOH Laboratories Administration | Genomics and Discovery, Respiratory Viruses Branch, Division of Viral Diseases, Centers for Disease Control and Prevention | Krista Queen, Yan Li, Ying Tao, Jing Zhang, Anna Uehara, Anna Montmayeur, Clinton R. Paden, Peter W. Cook,Rachel Marine, Mili Sheth, Haibin Wang, Justin Lee, Suxiang Tong |
| EPI_ISL_751772 | HI Dept. of Health, State Laboratories Division | Genomics and Discovery, Respiratory Viruses Branch, Division of Viral Diseases, Centers for Disease Control and Prevention | Krista Queen, Yan Li, Ying Tao, Jing Zhang, Anna Uehara, Anna Montmayeur, Clinton R. Paden, Peter W. Cook,Rachel Marine, Mili Sheth, Haibin Wang, Justin Lee, Suxiang Tong |
| EPI_ISL_751775 | MT Public Health Laboratory | Genomics and Discovery, Respiratory Viruses Branch, Division of Viral Diseases, Centers for Disease Control and Prevention | Krista Queen, Yan Li, Ying Tao, Jing Zhang, Anna Uehara, Anna Montmayeur, Clinton R. Paden, Peter W. Cook,Rachel Marine, Mili Sheth, Haibin Wang, Justin Lee, Suxiang Tong |
| EPI_ISL_751780 | ID Bureau of Laboratories | Genomics and Discovery, Respiratory Viruses Branch, Division of Viral Diseases, Centers for Disease Control and Prevention | Krista Queen, Yan Li, Ying Tao, Jing Zhang, Anna Uehara, Anna Montmayeur, Clinton R. Paden, Peter W. Cook,Rachel Marine, Mili Sheth, Haibin Wang, Justin Lee, Suxiang Tong |
| EPI_ISL_751785, EPI_ISL_751786 | WVDHHR - Office of Laboratory Services | Genomics and Discovery, Respiratory Viruses Branch, Division of Viral Diseases, Centers for Disease Control and Prevention | Krista Queen, Yan Li, Ying Tao, Jing Zhang, Anna Uehara, Anna Montmayeur, Clinton R. Paden, Peter W. Cook,Rachel Marine, Mili Sheth, Haibin Wang, Justin Lee, Suxiang Tong |
| EPI_ISL_751790 | RI State Health Laboratories | Genomics and Discovery, Respiratory Viruses Branch, Division of Viral Diseases, Centers for Disease Control and Prevention | Krista Queen, Yan Li, Ying Tao, Jing Zhang, Anna Uehara, Anna Montmayeur, Clinton R. Paden, Peter W. Cook,Rachel Marine, Mili Sheth, Haibin Wang, Justin Lee, Suxiang Tong |
| EPI_ISL_751794 | KY State Public Health Lab | Genomics and Discovery, Respiratory Viruses Branch, Division of Viral Diseases, Centers for Disease Control and Prevention | Krista Queen, Yan Li, Ying Tao, Jing Zhang, Anna Uehara, Anna Montmayeur, Clinton R. Paden, Peter W. Cook,Rachel Marine, Mili Sheth, Haibin Wang, Justin Lee, Suxiang Tong |
| EPI_ISL_751796, EPI_ISL_751797, EPI_ISL_751798 | Public Health Ontario Laboratory | Public Health Ontario Laboratory | Vanessa G Allen, Philip Banh, Richard de Borja, Yao Chen, Alireza Eshaghi, Nahuel Fittipaldi, Christine Frantz, Jonathan B Gubbay, Jennifer L Guthrie, Lawrence Heisler, Esha Joshi, Michael Laszloffy, Aimin Li, Michael CY Li, Dean Maxwell, Sandeep Nagra, Samir N Patel, Heather Rilkoff, Jared Simpson, Karthikeyan Sivaraman, Yogi Sundaravadanam, Sarah Teatero, Andre Villegas, Sandra Zittermann |
| EPI_ISL_753176, EPI_ISL_753177, EPI_ISL_753178, EPI_ISL_753179, EPI_ISL_753180, EPI_ISL_753181, EPI_ISL_753182, EPI_ISL_753183, EPI_ISL_753184, EPI_ISL_753185, EPI_ISL_753186, EPI_ISL_753187, EPI_ISL_753188, EPI_ISL_753189, EPI_ISL_753190, EPI_ISL_753191, EPI_ISL_753200, EPI_ISL_753201, EPI_ISL_753202, EPI_ISL_753203, EPI_ISL_753204, EPI_ISL_753205, EPI_ISL_753206, EPI_ISL_753207, EPI_ISL_753208, EPI_ISL_753209, EPI_ISL_753210, EPI_ISL_753211, EPI_ISL_753212, EPI_ISL_753213, EPI_ISL_753216, EPI_ISL_753217, EPI_ISL_753218, EPI_ISL_753219, EPI_ISL_753220, EPI_ISL_753240 | | | |
| see above | State Laboratories Division, Hawaii State Department of Health | State Laboratories Division, Hawaii State Department of Health | Pamela O'Brien, Sabrina Diemert, Drew Kuwazaki, Razvan Sultana, Edward Desmond |
| EPI_ISL_753413, EPI_ISL_753414, EPI_ISL_753415, EPI_ISL_753416, EPI_ISL_753417, EPI_ISL_753418, EPI_ISL_753419, EPI_ISL_753420, EPI_ISL_753421, EPI_ISL_753422, EPI_ISL_753423, EPI_ISL_753424, EPI_ISL_753425, EPI_ISL_753426, EPI_ISL_753427, EPI_ISL_753428, EPI_ISL_753429, EPI_ISL_753430, EPI_ISL_753431, EPI_ISL_753432, EPI_ISL_753433, EPI_ISL_753434, EPI_ISL_753435, EPI_ISL_753436, EPI_ISL_753437, EPI_ISL_753438, EPI_ISL_753439, EPI_ISL_753440, EPI_ISL_753441, EPI_ISL_753442, EPI_ISL_753443, EPI_ISL_753444, EPI_ISL_753445, EPI_ISL_753446, EPI_ISL_753447, EPI_ISL_753448, EPI_ISL_753449, EPI_ISL_753450, EPI_ISL_753451, EPI_ISL_753452, EPI_ISL_753453, EPI_ISL_753454, EPI_ISL_753455, EPI_ISL_753456, EPI_ISL_753457, EPI_ISL_753458, EPI_ISL_753459, EPI_ISL_753460, EPI_ISL_753461, EPI_ISL_753462, EPI_ISL_753463, EPI_ISL_753464, EPI_ISL_753465, EPI_ISL_753466, EPI_ISL_753467, EPI_ISL_753468, EPI_ISL_753469, EPI_ISL_753470, EPI_ISL_753471, EPI_ISL_753472, EPI_ISL_753473, EPI_ISL_753474, EPI_ISL_753475, EPI_ISL_753476, EPI_ISL_753477, EPI_ISL_753478, EPI_ISL_753479, EPI_ISL_753480, EPI_ISL_753481, EPI_ISL_753482, EPI_ISL_753483, EPI_ISL_753484, | | | |

| | | | |
|---|---|---|---|
| EPI_ISL_753485, EPI_ISL_753486, EPI_ISL_753487, EPI_ISL_753488, EPI_ISL_753489, EPI_ISL_753490, EPI_ISL_753491, EPI_ISL_753492, EPI_ISL_753493, EPI_ISL_753494, EPI_ISL_753495, EPI_ISL_753496, EPI_ISL_753497, EPI_ISL_753498, EPI_ISL_753499, EPI_ISL_753500, EPI_ISL_753501, EPI_ISL_753502, EPI_ISL_753503, EPI_ISL_753504, EPI_ISL_753505, EPI_ISL_753506, EPI_ISL_753507, EPI_ISL_753508, EPI_ISL_753509, EPI_ISL_753510, EPI_ISL_753511, EPI_ISL_753512, EPI_ISL_753513, EPI_ISL_753514, EPI_ISL_753515, EPI_ISL_753516, EPI_ISL_753517, EPI_ISL_753518, EPI_ISL_753519, EPI_ISL_753520, EPI_ISL_753521, EPI_ISL_753522, EPI_ISL_753523, EPI_ISL_753524, EPI_ISL_753525, EPI_ISL_753526, EPI_ISL_753534, EPI_ISL_753535, EPI_ISL_753536, EPI_ISL_753537, EPI_ISL_753538, EPI_ISL_753539, EPI_ISL_753540, EPI_ISL_753541, EPI_ISL_753542, EPI_ISL_753543, EPI_ISL_753544, EPI_ISL_753545, EPI_ISL_753546, EPI_ISL_753547, EPI_ISL_753548, EPI_ISL_753549, EPI_ISL_753550, EPI_ISL_753551, EPI_ISL_753552, EPI_ISL_753553, EPI_ISL_753554, EPI_ISL_753555, EPI_ISL_753556, EPI_ISL_753557, EPI_ISL_753558, EPI_ISL_753559, EPI_ISL_753560, EPI_ISL_753561, EPI_ISL_753562, EPI_ISL_753563, EPI_ISL_753564, EPI_ISL_753565, EPI_ISL_753566, EPI_ISL_753567, EPI_ISL_753568, EPI_ISL_753569, EPI_ISL_753570, EPI_ISL_753571, EPI_ISL_753572, EPI_ISL_753573, EPI_ISL_753574, EPI_ISL_753575, EPI_ISL_753576, EPI_ISL_753577, EPI_ISL_753578, EPI_ISL_753579, EPI_ISL_753580, EPI_ISL_753581, EPI_ISL_753582, EPI_ISL_753583, EPI_ISL_753584, EPI_ISL_753585, EPI_ISL_753586, EPI_ISL_753587, EPI_ISL_753588, EPI_ISL_753589, EPI_ISL_753590, EPI_ISL_753591, EPI_ISL_753592, EPI_ISL_753593, EPI_ISL_753594, EPI_ISL_753624, EPI_ISL_753625, EPI_ISL_753627, EPI_ISL_753628, EPI_ISL_753630, EPI_ISL_753633, EPI_ISL_753634, EPI_ISL_753646, EPI_ISL_753653 | | | |
| see above | Clinical virology Laboratory, Children's Hospital Los Angeles | Center for Personalized Medicine, Children's Hospital Los Angeles | Gai et al |
| EPI_ISL_753697 | Specialized Lab for COVID-19 Detection, Department of Genetic Engineering and Biotechnology | Specialized Lab for COVID-19 Detection, Department of Genetic Engineering and Biotechnology | Shamsul H. Prodhan, Md. Asraful Jahan, Hammadul Hoque, Nurnabi Azad Jewel, Md. Nazmul Hasan, Hafiz Al Ashad, Rahatul Islam, Salman Ahmed |
| EPI_ISL_753700 | Specialized Lab for COVID-19 Detection, Department of Genetic Engineering and Biotechnology | Specialized Lab for COVID-19 Detection, Department of Genetic Engineering and Biotechnology | Hammadul Hoque, Ajit Ghosh, Nurnabi Azad Jewel, Md. Nazmul Hasan, Shamsul H. Prodhan, Amit Kumar Mondal, Fahmid H. Bhuiyan, Md. Rakib Wazed Nayon |
| EPI_ISL_753701 | Specialized Lab for COVID-19 Detection, Department of Genetic Engineering and Biotechnology | Specialized Lab for COVID-19 Detection, Department of Genetic Engineering and Biotechnology | Nurnabi Azad Jewel, Ziaul Faruque Joy , Md. Nazmul Hasan, Shamsul H. Prodhan, Hammadul Hoque , Md. Mobarok Karim, Shahrear Arefin, Md. Tahsin Khan |
| EPI_ISL_754059 | Specialized Lab for COVID-19 Detection, Department of Genetic Engineering and Biotechnology | Specialized Lab for COVID-19 Detection, Department of Genetic Engineering and Biotechnology | Md. Nazmul Hasan, Md. Akkas Ali, Shamsul H. Prodhan, Hammadul Hoque , Nurnabi Azad Jewel, Md Mohsin Bapary, Diptha Chakraborthy |
| EPI_ISL_754109 | Specialized Lab for COVID-19 Detection, Department of Genetic Engineering and Biotechnology | Specialized Lab for COVID-19 Detection, Department of Genetic Engineering and Biotechnology | Hammadul Hoque , Nurnabi Azad Jewel, Md. Nazmul Hasan, Shamsul H. Prodhan, Md. Mosarof Hossen, Md. Mashiur Rahaman |
| EPI_ISL_754148, EPI_ISL_754156, EPI_ISL_754157, EPI_ISL_754158, EPI_ISL_754160, EPI_ISL_754164 | Laboratoire de Microbiologie | National Reference Center for Viruses of Respiratory Infections, Institut Pasteur, Paris | Marion Barbet, Sylvie Behillil, Méline Bizard, Angela Brisebarre, Camille Capel, Etienne Simon-Lorière, Vincent Enouf, Maud Vanpeene, Sylvie van der Werf, Marie-Sarah Fangous |
| EPI_ISL_754176 | Specialized Lab for COVID-19 Detection, Department of Genetic Engineering and Biotechnology | Specialized Lab for COVID-19 Detection, Department of Genetic Engineering and Biotechnology | Shamsul H. Prodhan, Hammadul Hoque, Nurnabi Azad Jewel, Md. Nazmul Hasan, Md. Mainul Hossain Bakul, Md. Rabiul Awal |
| EPI_ISL_754242, EPI_ISL_754244, EPI_ISL_754248, EPI_ISL_754249, EPI_ISL_754250, EPI_ISL_754258, EPI_ISL_754260, EPI_ISL_754290, EPI_ISL_754298, EPI_ISL_754299, EPI_ISL_754300, EPI_ISL_754301, EPI_ISL_754304, EPI_ISL_754339, EPI_ISL_754340, EPI_ISL_754341, EPI_ISL_754342, EPI_ISL_754344, EPI_ISL_754345, EPI_ISL_754346, EPI_ISL_754347, EPI_ISL_754351, EPI_ISL_754354, EPI_ISL_754355 | | | |
| see above | Respiratory Virus Unit, National Infection Service, Public Health England | COVID-19 Genomics UK (COG-UK) Consortium | PHE Covid Sequencing Team |
| EPI_ISL_754388, EPI_ISL_754389 | Wyoming Public Health Laboratory | Wyoming Public Health Laboratory | Noah Hull, Taylor Fearing, Channing Weber, Ashley Norberg, Bailey Bowcutt, and Wanda Manley |
| EPI_ISL_754395, EPI_ISL_754396, EPI_ISL_754397, EPI_ISL_754398, EPI_ISL_754403, EPI_ISL_754407, EPI_ISL_754409, EPI_ISL_754410, EPI_ISL_754411, EPI_ISL_754412, EPI_ISL_754413, EPI_ISL_754414, EPI_ISL_754415, EPI_ISL_754416, EPI_ISL_754417, EPI_ISL_754418, EPI_ISL_754419, EPI_ISL_754420, EPI_ISL_754421, EPI_ISL_754422, EPI_ISL_754423, EPI_ISL_754424, EPI_ISL_754425, EPI_ISL_754426, EPI_ISL_754427, EPI_ISL_754428, EPI_ISL_754429, EPI_ISL_754430, EPI_ISL_754431, EPI_ISL_754432, EPI_ISL_754433, EPI_ISL_754434, EPI_ISL_754435, EPI_ISL_754436, EPI_ISL_754437, EPI_ISL_754438, EPI_ISL_754505, EPI_ISL_754506, EPI_ISL_754507, EPI_ISL_754508, EPI_ISL_754509 | | | |
| see above | Wadsworth Center, New York State Department.of Health | Wadsworth Center, New York State Department.of Health | Kirsten St. George, Daryl M. Lamson, Alexis Russel, Matthew Shudt, Melissa A Leisner, Jonathan Plitnick, Navjot Singh, John Kelly, Sara Griesemer, Erasmus Schneider, Erica Lasek-Nesselquist |
| EPI_ISL_754657, EPI_ISL_754658 | University of Wisconsin-Madison AIDS Vaccine Research Laboratories | University of Wisconsin-Madison AIDS Vaccine Research Laboratories | Gage Moreno, Katarina Braun, et al. AIDS Vaccine Research Laboratories |
| EPI_ISL_754772, EPI_ISL_754775 | Emory Molecular Diagnostics Laboratory, Emory Healthcare | Piantadosi Lab, Emory Department of Pathology | Ahmed Babiker, Anne Piantadosi |
| EPI_ISL_755132, EPI_ISL_755138 | San Diego County Public Health Laboratory | Andersen lab at Scripps Research | SEARCH Alliance San Diego with Tracy Basler, Jovan Shephard, Brett Austin |
| EPI_ISL_755198, EPI_ISL_755199, EPI_ISL_755200, EPI_ISL_755201, EPI_ISL_755202, EPI_ISL_755203, EPI_ISL_755204, EPI_ISL_755205, EPI_ISL_755209, EPI_ISL_755216, EPI_ISL_755217, EPI_ISL_755218, EPI_ISL_755219, EPI_ISL_755220, EPI_ISL_755226, EPI_ISL_755227, EPI_ISL_755228, EPI_ISL_755230, EPI_ISL_755231 | | | |
| see above | UCSD EXCITE lab | Andersen lab at Scripps Research | SEARCH Alliance San Diego |
| EPI_ISL_755279, EPI_ISL_755283, EPI_ISL_755285, EPI_ISL_755286, EPI_ISL_755289, EPI_ISL_755294 | San Diego County Public Health Laboratory | Andersen lab at Scripps Research | SEARCH Alliance San Diego with Tracy Basler, Jovan Shephard, Brett Austin |
| EPI_ISL_755434, EPI_ISL_755435, EPI_ISL_755436, EPI_ISL_755437, EPI_ISL_755438, EPI_ISL_755439, EPI_ISL_755440, EPI_ISL_755441, EPI_ISL_755442, EPI_ISL_755443, EPI_ISL_755444, EPI_ISL_755445, EPI_ISL_755446, EPI_ISL_755447, EPI_ISL_755448, EPI_ISL_755449, EPI_ISL_755450, EPI_ISL_755451, EPI_ISL_755452, EPI_ISL_755453, EPI_ISL_755454, EPI_ISL_755455, EPI_ISL_755482, EPI_ISL_755483, EPI_ISL_755484, EPI_ISL_755485, EPI_ISL_755486, EPI_ISL_755487, EPI_ISL_755488 | | | |
| see above | Maine Health and Environmental Testing Laboratory | Tewhey Lab, The Jackson Laboratory | Matluk,N., Dewey,H., Iosue,F., Barter,M., Lynch,R., Munger,H. and Tewhey,R. |
| EPI_ISL_755579 | Utah Public Health Laboratory, Utah Public Health Laboratory Infectious Disease submission group | Utah Public Health Laboratory, Utah Public Health Laboratory Infectious Disease submission group | Gallagher,T., Young,E.L., Oakeson,K.F. |
| EPI_ISL_755641 | Instituto Adolfo Lutz - Regional de Santo Andre | Instituto Adolfo Lutz, Interdiciplinary Procedures Center, Strategic Laboratory | Claudio Tavares Sacchi, Claudia Regina Gonçalves, Erica Valessa Ramos Gomes, Karoline Rodrigues Campos |
| EPI_ISL_755803 | Toronto Invasive Bacterial Diseases Network | McMaster University | Allison McGeer, Patryk Aftanas, Hooman Derakhshani, Angel Li, Kuganya Nirmalarajah, Emily Panousis, Ahmed Draia, Jalees Nasir, Michael Surette, Samira Mubareka, Andrew G. McArthur |
| EPI_ISL_756126, EPI_ISL_756127, EPI_ISL_756128, EPI_ISL_756129, EPI_ISL_756130, EPI_ISL_756131, EPI_ISL_756132, EPI_ISL_756133, EPI_ISL_756134, EPI_ISL_756164, EPI_ISL_756184, EPI_ISL_756185, EPI_ISL_756186 | | | |
| see above | Wyoming Public Health Laboratory | Wyoming Public Health Laboratory | Noah Hull, Taylor Fearing, Channing Weber, Ashley Norberg, Bailey Bowcutt, and Wanda Manley |
| EPI_ISL_756281 | State Laboratories Division, Hawaii State Department of Health | State Laboratories Division, Hawaii State Department of Health | Pamela O'Brien, Sabrina Diemert, Drew Kuwazaki, Razvan Sultana, Edward Desmond |
| EPI_ISL_756765, EPI_ISL_756860, EPI_ISL_756865, EPI_ISL_756876, EPI_ISL_756925, EPI_ISL_757014 | Lighthouse Lab in Milton Keynes | Wellcome Sanger Institute for the COVID-19 Genomics UK (COG-UK) Consortium | The Lighthouse Lab in Milton Keynes and Alex Alderton, Roberto Amato, Sonia Goncalves, Ewan Harrison, David K. Jackson, Ian Johnston, Dominic Kwiatkowski, Cordelia Langford, John Sillitoe on behalf of the Wellcome Sanger Institute COVID-19 Surveillance Team |
| EPI_ISL_757699 | Department of Virus and Microbiological Special Diagnostics, Statens Serum Institut, Copenhagen, Denmark | Albertsen Lab, Department of Chemistry and Bioscience, Aalborg University, Denmark | Danish Covid-19 Genome Consortium |
| EPI_ISL_759758 | University of Wisconsin-Madison AIDS Vaccine Research Laboratories | University of Wisconsin-Madison AIDS Vaccine Research Laboratories | Gage Moreno, Katarina Braun, et al. AIDS Vaccine Research Laboratories |
| EPI_ISL_759985 | Furst Medical Laboratory | Norwegian Institute of Public Health, Department of Virology | Kathrine Stene-Johansen, Kamilla Heddeland Instefjord, Hilde Elshaug, Marie Paulsen Madsen, Rasmus Riis Kopperud, Hilde Vollan, Karoline Bragstad, Olav Hungnes |
| EPI_ISL_759986, EPI_ISL_759987 | University Hospital of Northern Norway, Department for Microbiology and Infectious Disease Control | Norwegian Institute of Public Health, Department of Virology | Kathrine Stene-Johansen, Kamilla Heddeland Instefjord, Hilde Elshaug, Marie Paulsen Madsen, Rasmus Riis Kopperud, Hilde Vollan, Karoline Bragstad, Olav Hungnes |
| EPI_ISL_759988 | Dept. of Medical Microbiology, Stavanger University Hospital, Helse Stavanger HF | Norwegian Institute of Public Health, Department of Virology | Kathrine Stene-Johansen, Kamilla Heddeland Instefjord, Hilde Elshaug, Marie Paulsen Madsen, Rasmus Riis Kopperud, Hilde Vollan, Karoline Bragstad, Olav Hungnes |
| EPI_ISL_760247 | Pasteur Institute in Ho Chi Minh city | Department of Microbiology and Immunology - Pasteur | Lan Trong Phan, Manh Huy Dao, Hang Thi Thu Pham, Nhung Pham Hong Vu, Hieu Minh Le, Thang Minh Cao, Loan Thi Kim Huynh, Long Thanh Nguyen, |

| | | | |
|---|---|---|---|
| | | Institute in Ho Chi Minh city | Anh Hoang Nguyen, Hieu Trung Nguyen, Thao Thi Ngoc Nguyen, Quang Duy Pham, Quang Chan Luong, Thuong Vu Nguyen |
| EPI_ISL_760813, EPI_ISL_760833, EPI_ISL_760858, EPI_ISL_760859 | Lighthouse Lab in Cambridge | Wellcome Sanger Institute for the COVID-19 Genomics UK (COG-UK) Consortium | Rob Howes, The Lighthouse Lab in Cambridge and Alex Alderton, Roberto Amato, Sonia Goncalves, Ewan Harrison, David K. Jackson, Ian Johnston, Dominic Kwiatkowski, Cordelia Langford, John Sillitoe on behalf of the Wellcome Sanger Institute COVID-19 Surveillance Team |
| EPI_ISL_760876 | Lighthouse Lab in Glasgow | Wellcome Sanger Institute for the COVID-19 Genomics UK (COG-UK) Consortium | Harper VanSteenhouse, Yumi Kasai, David Gray, Carol Clugston, Anna Dominiczak and Alex Alderton, Roberto Amato, Sonia Goncalves, Ewan Harrison, David K. Jackson, Ian Johnston, Dominic Kwiatkowski, Cordelia Langford, John Sillitoe on behalf of the Wellcome Sanger Institute COVID-19 Surveillance Team |
| EPI_ISL_760881 | Lighthouse Lab in Milton Keynes | Wellcome Sanger Institute for the COVID-19 Genomics UK (COG-UK) Consortium | The Lighthouse Lab in Milton Keynes and Alex Alderton, Roberto Amato, Sonia Goncalves, Ewan Harrison, David K. Jackson, Ian Johnston, Dominic Kwiatkowski, Cordelia Langford, John Sillitoe on behalf of the Wellcome Sanger Institute COVID-19 Surveillance Team |
| EPI_ISL_760888 | Lighthouse Lab in Alderley Park | Wellcome Sanger Institute for the COVID-19 Genomics UK (COG-UK) Consortium | Jacquelyn Wynn, Mairead Hyland, The Lighthouse Lab in Alderley Park and Alex Alderton, Roberto Amato, Sonia Goncalves, Ewan Harrison, David K. Jackson, Ian Johnston, Dominic Kwiatkowski, Cordelia Langford, John Sillitoe on behalf of the Wellcome Sanger Institute COVID-19 Surveillance Team |
| EPI_ISL_760943, EPI_ISL_760975 | Lighthouse Lab in Milton Keynes | Wellcome Sanger Institute for the COVID-19 Genomics UK (COG-UK) Consortium | The Lighthouse Lab in Milton Keynes and Alex Alderton, Roberto Amato, Sonia Goncalves, Ewan Harrison, David K. Jackson, Ian Johnston, Dominic Kwiatkowski, Cordelia Langford, John Sillitoe on behalf of the Wellcome Sanger Institute COVID-19 Surveillance Team |
| EPI_ISL_760976 | Lighthouse Lab in Cambridge | Wellcome Sanger Institute for the COVID-19 Genomics UK (COG-UK) Consortium | Rob Howes, The Lighthouse Lab in Cambridge and Alex Alderton, Roberto Amato, Sonia Goncalves, Ewan Harrison, David K. Jackson, Ian Johnston, Dominic Kwiatkowski, Cordelia Langford, John Sillitoe on behalf of the Wellcome Sanger Institute COVID-19 Surveillance Team |
| EPI_ISL_761023, EPI_ISL_761039 | Lighthouse Lab in Milton Keynes | Wellcome Sanger Institute for the COVID-19 Genomics UK (COG-UK) Consortium | The Lighthouse Lab in Milton Keynes and Alex Alderton, Roberto Amato, Sonia Goncalves, Ewan Harrison, David K. Jackson, Ian Johnston, Dominic Kwiatkowski, Cordelia Langford, John Sillitoe on behalf of the Wellcome Sanger Institute COVID-19 Surveillance Team |
| EPI_ISL_761042 | Lighthouse Lab in Alderley Park | Wellcome Sanger Institute for the COVID-19 Genomics UK (COG-UK) Consortium | Jacquelyn Wynn, Mairead Hyland, The Lighthouse Lab in Alderley Park and Alex Alderton, Roberto Amato, Sonia Goncalves, Ewan Harrison, David K. Jackson, Ian Johnston, Dominic Kwiatkowski, Cordelia Langford, John Sillitoe on behalf of the Wellcome Sanger Institute COVID-19 Surveillance Team |
| EPI_ISL_761043, EPI_ISL_761067 | Lighthouse Lab in Glasgow | Wellcome Sanger Institute for the COVID-19 Genomics UK (COG-UK) Consortium | Harper VanSteenhouse, Yumi Kasai, David Gray, Carol Clugston, Anna Dominiczak and Alex Alderton, Roberto Amato, Sonia Goncalves, Ewan Harrison, David K. Jackson, Ian Johnston, Dominic Kwiatkowski, Cordelia Langford, John Sillitoe on behalf of the Wellcome Sanger Institute COVID-19 Surveillance Team |
| EPI_ISL_763001, EPI_ISL_763007, EPI_ISL_763020, EPI_ISL_763025, EPI_ISL_763027, EPI_ISL_763028, EPI_ISL_763033, EPI_ISL_763034, EPI_ISL_763035, EPI_ISL_763047, EPI_ISL_763048, EPI_ISL_763049, EPI_ISL_763050 | | | |
| see above | Unit 17: Influenza & Other Respiratory Viruses, German National Influenza Center | Project group Epidemiology of Highly Pathogenic Microorganisms, Robert Koch-Institute | Ariane Düx, Andreas Sachse, Grit Schubert, Sébastien Calvignac-Spencer, Fabian Leendertz, Thorsten Wolff, Ralf Dürrwald, Djin-Ye Oh, Marianne Wedde |
| EPI_ISL_763344, EPI_ISL_763345, EPI_ISL_763346 | Florida Bureau of Public Health Laboratories | Florida Bureau of Public Health Laboratories | Sarah Schmedes, Jason Blanton |
| EPI_ISL_763375, EPI_ISL_763438, EPI_ISL_763569, EPI_ISL_763593, EPI_ISL_763641, EPI_ISL_763792, EPI_ISL_763793, EPI_ISL_763794, EPI_ISL_763815, EPI_ISL_763872, EPI_ISL_764159, EPI_ISL_764171, EPI_ISL_764174, EPI_ISL_764176, EPI_ISL_764177, EPI_ISL_764191, EPI_ISL_764209, EPI_ISL_764281, EPI_ISL_764284, EPI_ISL_764285, EPI_ISL_764286, EPI_ISL_764287, EPI_ISL_764288, EPI_ISL_764289, EPI_ISL_764290, EPI_ISL_764291, EPI_ISL_764292, EPI_ISL_764293, EPI_ISL_764294, EPI_ISL_764295, EPI_ISL_764296, EPI_ISL_764297, EPI_ISL_764298, EPI_ISL_764299, EPI_ISL_764300, EPI_ISL_764301, EPI_ISL_764305, EPI_ISL_764307, EPI_ISL_764309, EPI_ISL_764312, EPI_ISL_764313, EPI_ISL_764314, EPI_ISL_764322, EPI_ISL_764323, EPI_ISL_764324, EPI_ISL_764325, EPI_ISL_764326 | | | |
| see above | Department of Pathology, University of Cambridge | COVID-19 Genomics UK (COG-UK) Consortium | Aminu S. Jahun, Yasmin Chaudhry, Grant Hall, Iliana Georgana, Myra Hosmillo, Martin D. Curran, Malte Pinckert, Surendra Parmar, Ian Goodfellow |
| EPI_ISL_764766 | Wales Specialist Virology Centre Sequencing lab: Pathogen Genomics Unit | COVID-19 Genomics UK (COG-UK) Consortium | Catherine Moore, Johnathan Evans, Laura Gifford, Malorie Perry, Simon Cottrell, Angela Marchbank, Alec Birchley, Alexander Adams, Amy Gaskin, Bree Gatica-Wilcox, Jason Coombes, Joel Southgate, Lauren Gilbert, Lee Graham, Nicole Pacchiarini, Sara Kumziene-Summerhayes, Sarah Taylor, Sophie Jones, Sara Rey, Matthew Bull, Joanne Watkins, Sally Corden, Tom Connor |
| EPI_ISL_765882, EPI_ISL_765883, EPI_ISL_765884, EPI_ISL_765885, EPI_ISL_765886, EPI_ISL_765887, EPI_ISL_765888, EPI_ISL_765889, EPI_ISL_765890, EPI_ISL_765891, EPI_ISL_765892 | | | |
| see above | Massachusetts General Hospital | Infectious Disease Program, Broad Institute of Harvard and MIT | Lemieux,J.E., Siddle,K.J., Shaw,B., Adams,G., Pierce,V., Turbett,S., Anahtar,M., Branda,J., Slater,D., Harris,J., Lin,A.E., Gladden-Young,A., Lagerborg,K., Rudy,M., DeRuff,K., Carter,A., Normandin,E., Bauer,M., Reilly,S., Tomkins-Tinch,C., Loreth,C., Chaluvadi,S., Neumann,A., Cusick,C., Chapman,S.B., Gnirke,A., Flowers,K., Cerrato,F., Birren,B.W., Gallagher,G., Smole,S., Park,D.J., MacInnis,B.L., Ryan,E., LaRocque,R., Rosenberg,E. and Sabeti,P.C. |
| EPI_ISL_766028 | Hannover Medical School, Institute of Virology | Hannover Medical School, Institute of Virology | Lars Steinbrück |
| EPI_ISL_766055 | Department of Virology | Department of Virology | Massab Umair, Muhammad Salman, Sana Tamim, Nazish Badar, Zaira Rehman, Adnan Khurshid, Samee Ullah, Abdul Ahad, Hamza Ahmed, Aamer Ikram |
| EPI_ISL_766064, EPI_ISL_766239, EPI_ISL_766279, EPI_ISL_766280 | Respiratory Virus Unit, National Infection Service, Public Health England | COVID-19 Genomics UK (COG-UK) Consortium | PHE Covid Sequencing Team |
| EPI_ISL_766576 | ULSS 8 Berica | Istituto Zooprofilattico Sperimentale delle Venezie | Adelaide Milani, Alessia Schivo, Annalisa Salviato, Erika Giorgia Quaranta, Ambra Pastori, Bianca Zecchin, Alice Fusaro, Isabella Monne, Calogero Terregino, Antonia Ricci |
| EPI_ISL_766591 | Halsomedicinskt Center | The Public Health Agency of Sweden | Department of Microbiology, The Public Health Agency of Sweden |
| EPI_ISL_766661, EPI_ISL_766666, EPI_ISL_766674, EPI_ISL_766734, EPI_ISL_766751, EPI_ISL_766759 | Texas Department of State Health Services | Texas Department of State Health Services | Rashmi Tuladhar, Bonnie Oh, Jenny Zhang, Maliha Rahman, Anita Pokharel, Myong Koag, Chung Wang, Rachel Lee, Grace Kubin, Mayela Pedrueza, James Daniel Bonser |
| EPI_ISL_766972, EPI_ISL_766973, EPI_ISL_766974, EPI_ISL_766978, EPI_ISL_766979, EPI_ISL_766981, EPI_ISL_766982, EPI_ISL_766983, EPI_ISL_766985, EPI_ISL_766986, EPI_ISL_766987, EPI_ISL_766990, EPI_ISL_766991, EPI_ISL_766992, EPI_ISL_766994, EPI_ISL_766997, EPI_ISL_767002, EPI_ISL_767007, EPI_ISL_767009, EPI_ISL_767010, EPI_ISL_767011, EPI_ISL_767012 | | | |
| see above | Delaware Public Health Laboratory | Delaware Public Health Laboratory | Gregory Hovan |
| EPI_ISL_767291, EPI_ISL_767299, EPI_ISL_767318 | Lighthouse Lab in Alderley Park | Wellcome Sanger Institute for the COVID-19 Genomics UK (COG-UK) Consortium | Jacquelyn Wynn, Mairead Hyland, The Lighthouse Lab in Alderley Park and Alex Alderton, Roberto Amato, Sonia Goncalves, Ewan Harrison, David K. Jackson, Ian Johnston, Dominic Kwiatkowski, Cordelia Langford, John Sillitoe on behalf of the Wellcome Sanger Institute COVID-19 Surveillance Team |
| EPI_ISL_767583, EPI_ISL_767588, EPI_ISL_767591, EPI_ISL_767592, EPI_ISL_767593, EPI_ISL_767596, EPI_ISL_767597, EPI_ISL_767629, EPI_ISL_767630, EPI_ISL_767631, EPI_ISL_767632, EPI_ISL_767633, EPI_ISL_767634, EPI_ISL_767635, EPI_ISL_767636, EPI_ISL_767637, EPI_ISL_767638, EPI_ISL_767639, EPI_ISL_767640, EPI_ISL_767641, EPI_ISL_767642, EPI_ISL_767643, EPI_ISL_767644, EPI_ISL_767659, EPI_ISL_767660, EPI_ISL_767661, EPI_ISL_767662, EPI_ISL_767663, EPI_ISL_767664 | | | |
| see above | MEMORIAL SLOAN KETTERING CANCER CENTER | Wadsworth Center, New York State Department.of Health | Kirsten St. George, Daryl M. Lamson, Alexis Russel, Matthew Shudt, Melissa A Leisner, Jonathan Plitnick, Navjot Singh, John Kelly, Sara Griesemer, Erasmus Schneider, Erica Lasek-Nesselquist |
| EPI_ISL_767854, EPI_ISL_767855, EPI_ISL_767856 | Sydney South West Pathology Service (SSWPS) - Royal Prince Alfred Hospital - NSW Health Pathology | NSW Health Pathology - Institute of Clinical Pathology and Medical Research; Westmead Hospital; University of Sydney | CIDM-PH et al. |
| EPI_ISL_768357, EPI_ISL_768358, EPI_ISL_768359, EPI_ISL_768360, EPI_ISL_768361, EPI_ISL_768362, EPI_ISL_768363, EPI_ISL_768364, EPI_ISL_768365, EPI_ISL_768366, EPI_ISL_768367, EPI_ISL_768368, EPI_ISL_768369, EPI_ISL_768370, EPI_ISL_768371, EPI_ISL_768372 | | | |
| see above | LSUHS Emerging Viral Threat Laboratory | Microbial Genome Sequencing Center | Jeremy P. Kamil, Jennifer L. Carroll, Camille F. Abshire, Maarten Van Diest, Andrew D. Yurochko, Martin J. Sapp, Rona S. Scott, Christopher G. Kevil, Daniel J. Snyder, Vaughn S. Cooper, John A. Vanchiere |
| EPI_ISL_768521 | Regional medical sciences center 6 Chonburi | National Institute of Health, Department of Medical Sciences, Ministry of Public Health, Thailand | Pilailuk Okada; Siripaporn Phuygun; Sittiporn Parnmen; Ratana Tacharoenmuang; Pakorn Piromtong; Natchaya Khiadsang; Thanutsapa Thanadachakul; Warawan Wongboot; sirikanda wimol; Sunthareeya Waicharoen; |
| EPI_ISL_768526, EPI_ISL_768527 | Department of Virology I, National Institute of Infectious Diseases | Pathogen Genomics Center, National Institute of Infectious Diseases | Tsuyoshi Sekizuka, Shuetsu Fukushi, Souichi Yamada, Kentaro Itokawa, Rina Tanaka, Masanori Hashino, Makoto Kuroda |
| EPI_ISL_768528, EPI_ISL_768614 | Regional Medical Sciences Center 1/1 Chiang Rai | National Institute of Health, Department of Medical Sciences, Ministry of Public Health, Thailand | Pilailuk Okada; Siripaporn Phuygun; Sittiporn Parnmen; Ratana Tacharoenmuang; Pakorn Piromtong; Natchaya Khiadsang; Thanutsapa Thanadachakul; Warawan Wongboot; sirikanda wimol; Sunthareeya Waicharoen; |
| EPI_ISL_768640, EPI_ISL_768641, EPI_ISL_768642, EPI_ISL_768655, EPI_ISL_768656, EPI_ISL_768657, EPI_ISL_768658, EPI_ISL_768661, EPI_ISL_768662, EPI_ISL_768663, EPI_ISL_768664, EPI_ISL_768665, EPI_ISL_768666, EPI_ISL_768668, EPI_ISL_768669, EPI_ISL_768670, EPI_ISL_768671, EPI_ISL_768672, EPI_ISL_768673, EPI_ISL_768674, EPI_ISL_768675, EPI_ISL_768676, EPI_ISL_768677, EPI_ISL_768678, EPI_ISL_768679, EPI_ISL_768680, EPI_ISL_768681, EPI_ISL_768682, EPI_ISL_768683, EPI_ISL_768684, EPI_ISL_768685, EPI_ISL_768686, EPI_ISL_768687, EPI_ISL_768688, EPI_ISL_768689, EPI_ISL_768690, EPI_ISL_768691, EPI_ISL_768692, EPI_ISL_768693, EPI_ISL_768694, EPI_ISL_768695, EPI_ISL_768696, EPI_ISL_768697, EPI_ISL_768698, EPI_ISL_768699, EPI_ISL_768700, EPI_ISL_768701, EPI_ISL_768703, EPI_ISL_768704, EPI_ISL_768705, EPI_ISL_768706, EPI_ISL_768707, EPI_ISL_768708, EPI_ISL_768709, | | | |

| | | | |
|---|---|---|---|
| EPI_ISL_768710, EPI_ISL_768711, EPI_ISL_768712, EPI_ISL_768713, EPI_ISL_768714, EPI_ISL_768715, EPI_ISL_768716, EPI_ISL_768717, EPI_ISL_768718, EPI_ISL_768719, EPI_ISL_768720, EPI_ISL_768721, EPI_ISL_768722, EPI_ISL_768723, EPI_ISL_768724 | | | |
| see above | Pathogen Genomics Center, National Institute of Infectious Diseases | Pathogen Genomics Center, National Institute of Infectious Diseases | Tsuyoshi Sekizuka, Kentaro Itokawa, Rina Tanaka, Masanori Hashino, Makoto Kuroda |
| EPI_ISL_768725 | Jena University Hospital, Institute for Infectious Diseases and Infection Control | Institute of infectious medicine & hospital hygiene, CaSe-Group | Spott, Riccardo; Marquet, Mike; Pletz, Matthias W.; Brandt, Christian |
| EPI_ISL_768831, EPI_ISL_768832, EPI_ISL_768834, EPI_ISL_768835, EPI_ISL_768836, EPI_ISL_768837 | Laboratoire Biolife | Laboratoire de Biotechnologie | Mouna Ouadghiri, Tarik Aanniz, Mohammed Walid Chemao Elfihri, Mohamed Chenaoui, Hanae Dakka, Afaf Alaoui, Otmane Touzani, Amina Benouda, Bouchra Belfquih, Lahcen belyamani, Saaid Amzazi and Azeddine Ibrahimi |
| EPI_ISL_770070, EPI_ISL_770113, EPI_ISL_770114, EPI_ISL_770115, EPI_ISL_770116, EPI_ISL_770117, EPI_ISL_770118, EPI_ISL_770119, EPI_ISL_770120 | Wyoming Public Health Laboratory | Wyoming Public Health Laboratory | Noah Hull, Taylor Fearing, Lynette Gumbleton, Channing Weber, Ashley Norberg, Bailey Bowcutt, and Wanda Manley |
| EPI_ISL_770542, EPI_ISL_770543, EPI_ISL_770544, EPI_ISL_770545, EPI_ISL_770546, EPI_ISL_770547, EPI_ISL_770548, EPI_ISL_770549 | Private lab | Lithuanian University of Health Sciences, Molecular cardiology lab. | Lukas Zemaitis, Ingrida Olendrait, Arnoldas Pautienius, Kamile Tamusauskaite, Dovydas Gecys, Laura Pareckaite, Vaiva Lesauskaite, Astra Vitkauskiene |
| EPI_ISL_770630 | Laboratório de Microbiologia Molecular - Universidade FEEVALE | Bioinformatics Laboratory / LNCC | Felipe Benites, Fernando Rosado Spilki, Alana Witt Hansen, Juliane Deise Fleck, Juliana Schons, Meriane Demoliner, Ana Karolina Eisen Antunes, Fagner Henrique Heldt, Larissa Mallmann, Bruna Hermann, Ana Luiza Ziulkoski, Vyctoria Goes, Karoline Schallenberger, Matheus Nunes Weber, Paula Rodrigues de Almeida, Alessandra Pavan Lamarca da Silva, Ronaldo da Silva F Jr , Luiz G P de Almeida, Alexandra L Gerber , Ana Paula de C Guimarães,Ana Tereza R de Vasconcelos |
| EPI_ISL_770634, EPI_ISL_770636, EPI_ISL_770637, EPI_ISL_770639, EPI_ISL_770640, EPI_ISL_770641, EPI_ISL_770643, EPI_ISL_770644, EPI_ISL_770645, EPI_ISL_770646, EPI_ISL_770653, EPI_ISL_770660, EPI_ISL_770666, EPI_ISL_770669, EPI_ISL_770675, EPI_ISL_770676, EPI_ISL_770677, EPI_ISL_770678, EPI_ISL_770679, EPI_ISL_770680, EPI_ISL_770681, EPI_ISL_770682, EPI_ISL_770683, EPI_ISL_770684, EPI_ISL_770685, EPI_ISL_770686, EPI_ISL_770687 | | | |
| see above | WHO National Influenza Centre Russian Federation | WHO National Influenza Centre Russian Federation | Andrey Komissarov, Artem Fadeev, Anna Ivanova, Kseniya Komissarova, Dmitry Bazhenov, Daria Danilenko, Ksenia Safina, Elena Nabieva, Georgii Bazykin, Dmitry Lioznov |
| EPI_ISL_770715, EPI_ISL_770717, EPI_ISL_770718, EPI_ISL_770719 | ZOTZ KLIMAS MVZ Düsseldorf-Centrum GbR ÜBAG für Labormedizin, Genetik, Zytologie, Pathologie | Center of Medical Microbiology, Virology, and Hospital Hygiene, University of Duesseldorf | Maximilian Damagnez, Alexander Dilthey, Ashley-Jane Duplessis, Patrick Finzer, Katrin Hoffmann, Torsten Houwaart, Lisanna Hülse, Malte Kohns Vasconcelos, Marek Korencak, Nadine Lübke, Jessica Nicolai, Klaus Pfeffer, Daniel Strelow, Jörg Timm, Andreas Walker, Tobias Wienemann, Rainer Zotz |
| EPI_ISL_770783, EPI_ISL_770784, EPI_ISL_770785, EPI_ISL_770786, EPI_ISL_770787, EPI_ISL_770788, EPI_ISL_770789 | Minnesota Department of Health, Public Health Laboratory | Minnesota Department of Health, Public Health Laboratory | Alexandra Lorentz, Jacob Garfin, Matt Plumb, and Xiong Wang |
| EPI_ISL_770804, EPI_ISL_770805, EPI_ISL_770806, EPI_ISL_770807, EPI_ISL_770814 | Vault Health | Minnesota Department of Health, Public Health Laboratory | Alexandra Lorentz, Jacob Garfin, Matt Plumb, and Xiong Wang |
| EPI_ISL_770848, EPI_ISL_770854, EPI_ISL_770862, EPI_ISL_770866, EPI_ISL_770867, EPI_ISL_770869, EPI_ISL_770874, EPI_ISL_770875, EPI_ISL_770878, EPI_ISL_770879, EPI_ISL_770892, EPI_ISL_770893, EPI_ISL_770894, EPI_ISL_770896, EPI_ISL_770898, EPI_ISL_770899, EPI_ISL_770900, EPI_ISL_770906, EPI_ISL_770907, EPI_ISL_770908, EPI_ISL_770909, EPI_ISL_770910, EPI_ISL_770911, EPI_ISL_770912, EPI_ISL_770913, EPI_ISL_770914, EPI_ISL_770930, EPI_ISL_770931, EPI_ISL_770932, EPI_ISL_770933, EPI_ISL_770934, EPI_ISL_770935, EPI_ISL_770936, EPI_ISL_770937, EPI_ISL_770938, EPI_ISL_770939, EPI_ISL_770940, EPI_ISL_770941, EPI_ISL_770942, EPI_ISL_770943, EPI_ISL_770944, EPI_ISL_770945, EPI_ISL_770946, EPI_ISL_770947, EPI_ISL_770948, EPI_ISL_770949, EPI_ISL_770950, EPI_ISL_770951, EPI_ISL_770952, EPI_ISL_770953, EPI_ISL_770954, EPI_ISL_770955, EPI_ISL_770956, EPI_ISL_770957, EPI_ISL_770958, EPI_ISL_770959, EPI_ISL_770960, EPI_ISL_770961, EPI_ISL_770962, EPI_ISL_770963, EPI_ISL_770964, EPI_ISL_770965, EPI_ISL_770966, EPI_ISL_770967, EPI_ISL_770968, EPI_ISL_770969, EPI_ISL_770970, EPI_ISL_770971, EPI_ISL_770972, EPI_ISL_770973, EPI_ISL_770974, EPI_ISL_770975, EPI_ISL_770976, EPI_ISL_770977, EPI_ISL_770978, EPI_ISL_770979, EPI_ISL_770985, EPI_ISL_770986, EPI_ISL_770987, EPI_ISL_770988, EPI_ISL_770989, EPI_ISL_770991, EPI_ISL_770992, EPI_ISL_770993, EPI_ISL_770994, EPI_ISL_770995, EPI_ISL_770996, EPI_ISL_770997, EPI_ISL_770998, EPI_ISL_770999, EPI_ISL_771000, EPI_ISL_771001, EPI_ISL_771002, EPI_ISL_771003, EPI_ISL_771004, EPI_ISL_771005, EPI_ISL_771006, EPI_ISL_771007, EPI_ISL_771008, EPI_ISL_771009, EPI_ISL_771010, EPI_ISL_771011, EPI_ISL_771012, EPI_ISL_771013, EPI_ISL_771014, EPI_ISL_771015, EPI_ISL_771016, EPI_ISL_771017, EPI_ISL_771018, EPI_ISL_771019, EPI_ISL_771020, EPI_ISL_771022, EPI_ISL_771024, EPI_ISL_771025, EPI_ISL_771030, EPI_ISL_771031, EPI_ISL_771032, EPI_ISL_771033, EPI_ISL_771034, EPI_ISL_771035, EPI_ISL_771036, EPI_ISL_771037, EPI_ISL_771038 | | | |
| see above | Laboratoire national de santé, Microbiology, Virology | Laboratoire national de santé, Microbiology, Microbial Genomics Platform | Anke Wienecke-Baldacchino, Catherine Ragimbeau,Jessica Tapp, Fatu Djabi, Lise Pignon, Raoul Salmon, Tamir Abdelrahman |
| EPI_ISL_775223, EPI_ISL_775224, EPI_ISL_775226, EPI_ISL_775247, EPI_ISL_775249, EPI_ISL_775254 | Laboratoire Biolife | Laboratoire de Biotechnologie | Mouna Ouadghiri, Tarik Aanniz, Mohammed Walid Chemao Elfihri, Mohamed Chenaoui, Hanae Dakka, Afaf Alaoui, Otmane Touzani, Amina Benouda, Bouchra Belfquih, Lahcen belyamani, Saaid Amzazi and Azeddine Ibrahimi |
| EPI_ISL_775285, EPI_ISL_775371 | Hospital of Southern Norway - Kristiansand, Department of Medical Microbiology | Norwegian Institute of Public Health, Department of Virology | Kathrine Stene-Johansen, Kamilla Heddeland Instefjord, Hilde Elshaug, Atiya R Ali,Marie Paulsen Madsen, Rasmus Riis Kopperud, Hilde Vollan, Karoline Bragstad, Olav Hungnes |
| EPI_ISL_775385 | Unilabs Laboratory Medicine | Norwegian Institute of Public Health, Department of Virology | Kathrine Stene-Johansen, Kamilla Heddeland Instefjord, Hilde Elshaug, Atiya R Ali,Marie Paulsen Madsen, Rasmus Riis Kopperud, Hilde Vollan, Karoline Bragstad, Olav Hungnes |
| EPI_ISL_775400, EPI_ISL_775402 | Nordland Hospital - Bodo, Laboratory Department, Molecular Biology Unit | Norwegian Institute of Public Health, Department of Virology | Kathrine Stene-Johansen, Kamilla Heddeland Instefjord, Hilde Elshaug, Atiya R Ali,Marie Paulsen Madsen, Rasmus Riis Kopperud, Hilde Vollan, Karoline Bragstad, Olav Hungnes |
| EPI_ISL_775417, EPI_ISL_775418, EPI_ISL_775419, EPI_ISL_775420, EPI_ISL_775421, EPI_ISL_775422, EPI_ISL_775423 | Oslo University Hospital, Department of Medical Microbiology | Norwegian Institute of Public Health, Department of Virology | Kathrine Stene-Johansen, Kamilla Heddeland Instefjord, Hilde Elshaug, Atiya R Ali,Marie Paulsen Madsen, Rasmus Riis Kopperud, Hilde Vollan, Karoline Bragstad, Olav Hungnes |
| EPI_ISL_775492 | Department of Medical Microbiology, St. Olavs hospital | Norwegian Institute of Public Health, Department of Virology | Kathrine Stene-Johansen, Kamilla Heddeland Instefjord, Hilde Elshaug, Atiya R Ali,Marie Paulsen Madsen, Rasmus Riis Kopperud, Hilde Vollan, Karoline Bragstad, Olav Hungnes |
| EPI_ISL_776768 | Instituto Adolfo Lutz - Regional de Aracatuba | Instituto Adolfo Lutz, Interdiciplinary Procedures Center, Strategic Laboratory | Claudio Tavares Sacchi, Claudia Regina Gonçalves, Erica Valessa Ramos Gomes, Karoline Rodrigues Campos |
| EPI_ISL_776769 | Instituto Adolfo Lutz - Regional de Santo Andre | Instituto Adolfo Lutz, Interdiciplinary Procedures Center, Strategic Laboratory | Claudio Tavares Sacchi, Claudia Regina Gonçalves, Erica Valessa Ramos Gomes, Karoline Rodrigues Campos |
| EPI_ISL_777056 | Lighthouse Lab in Milton Keynes | Wellcome Sanger Institute for the COVID-19 Genomics UK (COG-UK) Consortium | The Lighthouse Lab in Milton Keynes and Alex Alderton, Roberto Amato, Sonia Goncalves, Ewan Harrison, David K. Jackson, Ian Johnston, Dominic Kwiatkowski, Cordelia Langford, John Sillitoe on behalf of the Wellcome Sanger Institute COVID-19 Surveillance Team |
| EPI_ISL_778825, EPI_ISL_778826, EPI_ISL_778827, EPI_ISL_778836 | AIID | Irish Coronavirus Sequencing Consortium-Teagasc Grange | Matthew McCabe, Aljandro Abner Garcia Leon, Fiona Crispie, Calum Walsh, Michael Carr, John Kenny, Paul Cotter, Patrick Mallon, Gabriel Gonzalez |
| EPI_ISL_778871, EPI_ISL_778872, EPI_ISL_778873, EPI_ISL_778874, EPI_ISL_778875, EPI_ISL_778876, EPI_ISL_778877, EPI_ISL_778891, EPI_ISL_778893, EPI_ISL_778894, EPI_ISL_778895, EPI_ISL_778896, EPI_ISL_778897, EPI_ISL_778898, EPI_ISL_778899, EPI_ISL_778900, EPI_ISL_778901, EPI_ISL_778902, EPI_ISL_778903, EPI_ISL_778904, EPI_ISL_778905, EPI_ISL_778906, EPI_ISL_778907, EPI_ISL_778908, EPI_ISL_778909, EPI_ISL_778910, EPI_ISL_778911, EPI_ISL_778912, EPI_ISL_778913, EPI_ISL_778914, EPI_ISL_778915, EPI_ISL_778916, EPI_ISL_778917 | | | |
| see above | Maryland Public Health Laboratory | Maryland Public Health Laboratory | Maryland Department of Health Laboratories Administration |
| EPI_ISL_778918 | LSUHS Emerging Viral Threat Laboratory | Microbial Genome Sequencing Center | Jeremy P. Kamil, Jennifer L. Carroll, Camille F. Abshire, Maarten Van Diest, Andrew D. Yurochko, Martin J. Sapp, Rona S. Scott, Christopher G. Kevil, Daniel J. Snyder, Vaughn S. Cooper, John A. Vanchiere |
| EPI_ISL_779135 | Wyoming Public Health Laboratory | Wyoming Public Health Laboratory | Noah Hull, Taylor Fearing, Lynette Gumbleton, Channing Weber, Ashley Norberg, Bailey Bowcutt, and Wanda Manley |
| EPI_ISL_779205, EPI_ISL_779206, EPI_ISL_779207, EPI_ISL_779208, EPI_ISL_779209, EPI_ISL_779210, EPI_ISL_779211, EPI_ISL_779212, EPI_ISL_779213, EPI_ISL_779214, EPI_ISL_779215, EPI_ISL_779216, EPI_ISL_779217, EPI_ISL_779232, EPI_ISL_779233, EPI_ISL_779234, EPI_ISL_779235, EPI_ISL_779236, | | | |

| | | | |
|---|---|---|---|
| EPI_ISL_779237, EPI_ISL_779238 | | | |
| see above | Pathogen Genomics Center, National Institute of Infectious Diseases | Pathogen Genomics Center, National Institute of Infectious Diseases | Tsuyoshi Sekizuka, Kentaro Itokawa, Rina Tanaka, Masanori Hashino, Makoto Kuroda |
| EPI_ISL_779291, EPI_ISL_779292, EPI_ISL_779293, EPI_ISL_779294, EPI_ISL_779295, EPI_ISL_779297, EPI_ISL_779298, EPI_ISL_779299, EPI_ISL_779300, EPI_ISL_779301, EPI_ISL_779303, EPI_ISL_779304, EPI_ISL_779305, EPI_ISL_779306, EPI_ISL_779307, EPI_ISL_779308, EPI_ISL_779309, EPI_ISL_779312, EPI_ISL_779313, EPI_ISL_779314, EPI_ISL_779317 | | | |
| see above | Utah Public Health Laboratory, Utah Public Health Laboratory Infectious Disease submission group | Utah Public Health Laboratory, Utah Public Health Laboratory Infectious Disease submission group | Gallagher,T., Young,E.L., Oakeson,K.F. |
| EPI_ISL_779697, EPI_ISL_779698 | Pathogen Genomics Center, National Institute of Infectious Diseases | Pathogen Genomics Center, National Institute of Infectious Diseases | Tsuyoshi Sekizuka, Kentaro Itokawa, Rina Tanaka, Masanori Hashino, Makoto Kuroda |
| EPI_ISL_779871, EPI_ISL_779875, EPI_ISL_779881, EPI_ISL_779886, EPI_ISL_779893, EPI_ISL_779899, EPI_ISL_779916 | Servicio de Microbiología, Hospital Universitario Son Espases | SeqCOVID-SPAIN consortium/IBV(CSIC) | Carla López-Causapé, Jordi Reina, Antonio Oliver and SeqCOVID-SPAIN consortium |
| EPI_ISL_783543, EPI_ISL_783561, EPI_ISL_783798, EPI_ISL_783808, EPI_ISL_783809, EPI_ISL_783810, EPI_ISL_783811, EPI_ISL_783812, EPI_ISL_783813, EPI_ISL_783815, EPI_ISL_783816, EPI_ISL_783817, EPI_ISL_783818, EPI_ISL_783819, EPI_ISL_783821, EPI_ISL_783822, EPI_ISL_783823, EPI_ISL_783824, EPI_ISL_783825, EPI_ISL_783826, EPI_ISL_783827, EPI_ISL_783828, EPI_ISL_783829, EPI_ISL_783830, EPI_ISL_783831, EPI_ISL_783835, EPI_ISL_783836, EPI_ISL_783837, EPI_ISL_783838, EPI_ISL_783839, EPI_ISL_783841, EPI_ISL_783842, EPI_ISL_783843, EPI_ISL_783844, EPI_ISL_783846, EPI_ISL_783847, EPI_ISL_783848, EPI_ISL_783849, EPI_ISL_783851, EPI_ISL_783852, EPI_ISL_783854, EPI_ISL_783855, EPI_ISL_783856, EPI_ISL_783858, EPI_ISL_783859, EPI_ISL_783860, EPI_ISL_783861, EPI_ISL_783864, EPI_ISL_783865, EPI_ISL_783866, EPI_ISL_783867, EPI_ISL_783868, EPI_ISL_783875, EPI_ISL_783876, EPI_ISL_783877, EPI_ISL_783878, EPI_ISL_783884, EPI_ISL_783885, EPI_ISL_783886, EPI_ISL_783887, EPI_ISL_783888, EPI_ISL_783895, EPI_ISL_783896, EPI_ISL_783897, EPI_ISL_783904, EPI_ISL_783905, EPI_ISL_783906, EPI_ISL_783907, EPI_ISL_783916, EPI_ISL_783917, EPI_ISL_783918, EPI_ISL_783925, EPI_ISL_783926, EPI_ISL_783927, EPI_ISL_783935, EPI_ISL_783936, EPI_ISL_783937, EPI_ISL_783938, EPI_ISL_783939, EPI_ISL_784031, EPI_ISL_784032, EPI_ISL_784047, EPI_ISL_784074, EPI_ISL_784139, EPI_ISL_784147 | | | |
| see above | Houston Methodist Hospital | Houston Methodist Hospital | S. Wesley Long, Randall J. Olsen, Paul A. Christensen, David W. Bernard, James J. Davis, Maulik Shukla, Marcus Nguyen, Matthew Ojeda Saavedra, Prasanti Yerramilli, Layne Pruitt, Sishir Subedi, Heather Hendrickson, and James M. Musser |
| EPI_ISL_788984, EPI_ISL_788986 | Institute of Virology, Biomedical Research Center of the Slovak Academy of Sciences, Bratislava | Faculty of Natural Sciences, Comenius University, Bratislava | Broa Brejová, Viktória abanová, Kristína Boršová, Viktória Hodorová, Sabina Fumaová Havlíková, Juraj Kopáek, Martina Liková, ubomíra Lukáiková, Martina Neboháová, Monika Sláviková, Alena Košálová, Peter Sabaka, Tomáš Vina, Jozef Nosek, Boris Klempa |
| EPI_ISL_788987 | Institute of Virology, Biomedical Research Center of the Slovak Academy of Sciences, Bratislava | Faculty of Natural Sciences, Comenius University, Bratislava | Viktória abanová, Kristína Boršová, Broa Brejová, Viktória Hodorová, Sabina Fumaová Havlíková, Juraj Kopáek, Martina Liková, ubomíra Lukáiková, Martina Neboháová, Monika Sláviková, Alena Košálová, Peter Sabaka, Tomáš Vina, Jozef Nosek, Boris Klempa |
| EPI_ISL_788988 | Institute of Virology, Biomedical Research Center of the Slovak Academy of Sciences, Bratislava | Faculty of Natural Sciences, Comenius University, Bratislava | Kristína Boršová, Viktória abanová, Broa Brejová, Viktória Hodorová, Sabina Fumaová Havlíková, Juraj Kopáek, Martina Liková, ubomíra Lukáiková, Martina Neboháová, Monika Sláviková, Tomáš Vina, Boris Klempa, Jozef Nosek |
| EPI_ISL_791067 | Dutch COVID-19 response team | National Institute for Public Health and the Environment (RIVM) | Adam Meijer, Harry Vennema, Jeroen Cremer, Sharon van den Brink, Bas van der Veer, AnneMarie van den Brandt, Florian Zwagemaker, Dennis Schmitz, Chantal Reusken, on behalf of the national COVID-19 response team |
| EPI_ISL_791281, EPI_ISL_791288, EPI_ISL_791289, EPI_ISL_791303, EPI_ISL_791304, EPI_ISL_791305, EPI_ISL_791319, EPI_ISL_791320 | National Virus Reference Laboratory | Irish Coronavirus Sequencing Consortium - Teagasc Moorepark | Alejandro Abner Garcia Leon, Paul Cotter, Fiona Crispie, John Kenny, Paddy Mallon, Calum Walsh |
| EPI_ISL_791351, EPI_ISL_791391, EPI_ISL_791401 | Johns Hopkins Hospital Department of Pathology | Johns Hopkins Hospital Department of Pathology | C. Paul Morris, Chun Huai Luo, Heba H. Mostafa |
| EPI_ISL_792549 | Centre for Dengue Research and AICBU, Department of Immunology and Molecular Medicine | Centre for Dengue Research and AICBU, Department of Immunology and Molecular Medicine | Chandima Jeewandara, Deshni Jayathilaka, Dinuka Ariyaratne, Diyanath Ranasinghe, Laksiri Gomes, Gathsaurie Neelika Malavige |
| EPI_ISL_792697, EPI_ISL_792702 | The National Institute of Public Health | State Veterinary Institute Prague | Nagy,A;Jirincova,H;Trnka,D;Vecerova,J |
| EPI_ISL_792838, EPI_ISL_793010 | Department of Virus and Microbiological Special Diagnostics, Statens Serum Institut, Copenhagen, Denmark | Albertsen Lab, Department of Chemistry and Bioscience, Aalborg University, Denmark | Danish Covid-19 Genome Consortium |
| EPI_ISL_794321, EPI_ISL_794322, EPI_ISL_794323 | UZ Leuven, National Reference Laboratory for Coronaviruses, Laboratory Medicine, Leuven, Belgium | KU Leuven, Rega Institute, Clinical and Epidemiological Virology | Tony Wawina-Bokalanga, Joan Marti-Carerras, Bert Vanmechelen, Piet Maes |
| EPI_ISL_794337, EPI_ISL_794338, EPI_ISL_794342, EPI_ISL_794352, EPI_ISL_794370, EPI_ISL_794372, EPI_ISL_794373, EPI_ISL_794374, EPI_ISL_794378, EPI_ISL_794412, EPI_ISL_794413, EPI_ISL_794414, EPI_ISL_794415, EPI_ISL_794416, EPI_ISL_794417, EPI_ISL_794418, EPI_ISL_794419, EPI_ISL_794424, EPI_ISL_794431, EPI_ISL_794435, EPI_ISL_794437, EPI_ISL_794440, EPI_ISL_794441, EPI_ISL_794555, EPI_ISL_794556, EPI_ISL_794557, EPI_ISL_794558, EPI_ISL_794559, EPI_ISL_794560, EPI_ISL_794561, EPI_ISL_794562, EPI_ISL_794563, EPI_ISL_794564, EPI_ISL_794565, EPI_ISL_794566, EPI_ISL_794567, EPI_ISL_794568, EPI_ISL_794569, EPI_ISL_794570, EPI_ISL_794571, EPI_ISL_794572, EPI_ISL_794573, EPI_ISL_794574, EPI_ISL_794575, EPI_ISL_794576, EPI_ISL_794577, EPI_ISL_794578, EPI_ISL_794579, EPI_ISL_794580, EPI_ISL_794581, EPI_ISL_794582, EPI_ISL_794583, EPI_ISL_794584, EPI_ISL_794585, EPI_ISL_794586, EPI_ISL_794587, EPI_ISL_794588 | | | |
| see above | Department of Virus and Microbiological Special Diagnostics, Statens Serum Institut, Copenhagen, Denmark | Albertsen Lab, Department of Chemistry and Bioscience, Aalborg University, Denmark | Danish Covid-19 Genome Consortium |
| EPI_ISL_794634 | Biology, MCL | Biology, MCL | Seadawy,M.G., Shamel,M.D., EL-hosieny,M.F., Gad,A.F., EL-harty,B.S. and EL-Safty,A.S. |
| EPI_ISL_794721 | PathWest Laboratory Medicine WA | PathWest Laboratory Medicine WA Microbial Surveillance Unit | PathWest Laboratory Medicine WA Microbial Surveillance Unit |
| EPI_ISL_794829, EPI_ISL_794858, EPI_ISL_794859, EPI_ISL_794870, EPI_ISL_794902, EPI_ISL_795008, EPI_ISL_795009 | Department of Virus and Microbiological Special Diagnostics, Statens Serum Institut, Copenhagen, Denmark | Albertsen Lab, Department of Chemistry and Bioscience, Aalborg University, Denmark | Danish Covid-19 Genome Consortium |
| EPI_ISL_796673 | Unilabs Laboratory Medicine | Norwegian Institute of Public Health, Department of Virology | Kathrine Stene-Johansen, Kamilla Heddeland Instefjord, Hilde Elshaug, Atiya R Ali,Marie Paulsen Madsen, Rasmus Riis Kopperud, Hilde Vollan, Karoline Bragstad, Olav Hungnes |
| EPI_ISL_796779 | Instituto Nacional de Saude (INSA) | Instituto Nacional de Saude (INSA) | Borges et al |
| EPI_ISL_801414, EPI_ISL_801427, EPI_ISL_801428, EPI_ISL_801451, EPI_ISL_801478, EPI_ISL_801480 | Dutch COVID-19 response team | Erasmus Medical Center | Bas Oude Munnink, Reina Sikkema, David Nieuwenhuijse, Irina Chestakova, Anne van der Linden, Marjan Boter, Emmanuelle Munger, Corine GeurtsvanKessel, Annemiek van der Eijk, Richard Molenkamp, Marion Koopmans, on behalf of the Dutch national COVID-19 response team. |
| EPI_ISL_802438 | Wadsworth Center, New York State Department.of Health | Wadsworth Center, New York State Department.of Health | Kirsten St. George, Daryl M. Lamson, Alexis Russel, Matthew Shudt, Melissa A Leisner, Jonathan Plitnick, Navjot Singh, John Kelly, Sara Griesemer, Erasmus Schneider, Erica Lasek-Nesselquist |
| EPI_ISL_802571 | Essentia Health-St. Mary's Medical Center | Minnesota Department of Health, Public Health Laboratory | Alexandra Lorentz, Jacob Garfin, Matt Plumb, and Xiong Wang |
| EPI_ISL_803699 | Wisconsin State Laboratory of Hygiene Communicable Disease Division | Wisconsin State Laboratory of Hygiene Communicable Disease Division | Kelsey R. Florek, Abigail C. Shockey |
| EPI_ISL_803920, EPI_ISL_803940, EPI_ISL_803941, EPI_ISL_803942, EPI_ISL_803943, EPI_ISL_803944, EPI_ISL_803945, EPI_ISL_803946, EPI_ISL_803947, EPI_ISL_803948, EPI_ISL_803949 | | | |
| see above | Pathogen Genomics Center, National Institute of Infectious Diseases | Pathogen Genomics Center, National Institute of Infectious Diseases | Tsuyoshi Sekizuka, Kentaro Itokawa, Rina Tanaka, Masanori Hashino, Makoto Kuroda |
| EPI_ISL_804007, EPI_ISL_804008 | Department of Virology I, National Institute of Infectious Diseases | Pathogen Genomics Center, National Institute of Infectious Diseases | Tsuyoshi Sekizuka, Shuetsu Fukushi, Souichi Yamada, Kentaro Itokawa, Rina Tanaka, Masanori Hashino, Makoto Kuroda |
| EPI_ISL_804226, EPI_ISL_804265 | Respiratory Virus Unit, National Infection Service, Public Health England | COVID-19 Genomics UK (COG-UK) Consortium | PHE Covid Sequencing Team |

| | | | |
|---|---|---|---|
| EPI_ISL_804374 | CHU Purpan - Laboratoire de Virologie - Institut Fédératif de Biologie | CHU Purpan - Laboratoire de Virologie - Institut Fédératif de Biologie | Latour J., Ranger N., Dubois M., Carcenac R., Harter A., Boyer P., Tremeaux P., Izopet J. |
| EPI_ISL_806830, EPI_ISL_806831, EPI_ISL_806834, EPI_ISL_806835, EPI_ISL_806836, EPI_ISL_806840, EPI_ISL_806847 | Alaska State Virology Laboratory | Alaska State Virology Laboratory | Stephanie DeRonde, Lisa Smith, Ph.D., Devin M. Drown, Ph.D., Jack Chen, Ph.D. |
| EPI_ISL_806998, EPI_ISL_807006, EPI_ISL_807018, EPI_ISL_807022 | Washington State Department of Health | Seattle Flu Study | Deborah A. Nickerson, Chris D. Frazar, Jover Lee, Benjamin Pelle, Matthew Richardson, Amanda Adler, Elisabeth Brandstetter, Peter D. Han, Kairsten Fay, Misja Ilcisin, Kirsten Lacombe, Thomas R. Sibley, Melissa Truong, Caitlin R. Wolf, Romesh Gautom, Geoff Melly, Brian Hiatt, Philip Dykema, Scott Lindquist, Michael Boeckh, Janet A. Englund, Michael Famulare, Barry R. Lutz, Mark J. Rieder, Lea M. Starita, Matthew Thompson, Helen Y. Chu, Jay Shendure, Trevor Bedford |
| EPI_ISL_812350, EPI_ISL_812351 | Santa Clara County Public Health Laboratory | Santa Clara County Public Health Laboratory | Santa Clara County Public Health Department |
| EPI_ISL_812404 | Utah Public Health Laboratory | Utah Public Health Laboratory | Erin L. Young, Kelly F. Oakeson, Tara Gallagher |
| EPI_ISL_812488 | Laboratorio de Referencia Nacional de Virus Respiratorios, Instituto Nacional de Salud Peru | Laboratorio de Genómica Microbiana, Universidad Peruana Cayetano Heredia | Pablo Tsukayama, Alejandra Dávila-Barclay, Guillermo Salvatierra, Luis González, Pedro E. Romero, Brenda Ayzanoa, Janet Huancachoque, Pool Marcos, Camila Castillo-Vilcahuamán, Oscar Escalante, Priscila Lope, Nancy Rojas |
| EPI_ISL_813706, EPI_ISL_813710, EPI_ISL_813712, EPI_ISL_813715, EPI_ISL_813716, EPI_ISL_813717, EPI_ISL_813718, EPI_ISL_813719, EPI_ISL_813720, EPI_ISL_813721, EPI_ISL_813722, EPI_ISL_813752, EPI_ISL_813754, EPI_ISL_813763 | | | |
| see above | Liverpool Clinical Laboratories | COVID-19 Genomics UK (COG-UK) Consortium | Sam Haldenby, Anita Lucaci, Steve Paterson, Julian Hiscox, Alistair Darby, M Almsaud, A Alrezaihi, Muhannad Alruwaili, Stuart D Armstrong, Jones Benjamin, Eleanor G Bentley, Anu Chawla, Jordan J Clark, Angela Cowell, Richard Eccles, Isabel García-Dorival, Matthew Gemmell, Alessandro Gerada, PKF Gilmore, Richard Gregory, Ximeng Han, Catherine Hartley, Margaret Hughes, Miren Iturriza-Gomara, James Johnson, L Luu, Jenifer Manson, Charlotte Nelson, Elaine O'Toole, Cassie Olateju, Rebekah Penrice-Randal , Lucille Rainbow, N.P Randle, Trevor Ian Robinson, Parul Sharma, Ghada T Shawli, James P Stewart, Neil Swainston, Ecaterina Vamos, Joanne Watts, Mark Whitehead |
| EPI_ISL_814297, EPI_ISL_814324, EPI_ISL_814346, EPI_ISL_814390, EPI_ISL_814399, EPI_ISL_814472, EPI_ISL_814514, EPI_ISL_814516 | Wales Specialist Virology Centre Sequencing lab: Pathogen Genomics Unit | COVID-19 Genomics UK (COG-UK) Consortium | Catherine Moore, Johnathan Evans, Laura Gifford, Malorie Perry, Simon Cottrell, Angela Marchbank, Alec Birchley, Alexander Adams, Amy Gaskin, Bree Gatica-Wilcox, Jason Coombes, Joel Southgate, Lauren Gilbert, Lee Graham, Nicole Pacchiarini, Sara Kumziene-Summerhayes, Sarah Taylor, Sophie Jones, Sara Rey, Matthew Bull, Joanne Watkins, Sally Corden, Tom Connor |
| EPI_ISL_814533 | Virology Department, Royal Infirmary of Edinburgh, NHS Lothian / School of Biological Sciences, University of Edinburgh / Institute of Genetics and Molecular Medicine, University of Edinburgh | COVID-19 Genomics UK (COG-UK) Consortium | McHugh M, Dewar R, Rooke S, Gallagher M, Balcaza C, O'Toole Á, Scher E, Hill V, McCrone JT, Colquhoun R, Yu X, Jackson B, Rambaut A, Williams TC, Templeton K |
| EPI_ISL_814562, EPI_ISL_814563, EPI_ISL_814564, EPI_ISL_814565, EPI_ISL_814576, EPI_ISL_814577, EPI_ISL_814603 | Wales Specialist Virology Centre Sequencing lab: Pathogen Genomics Unit | COVID-19 Genomics UK (COG-UK) Consortium | Catherine Moore, Johnathan Evans, Laura Gifford, Malorie Perry, Simon Cottrell, Angela Marchbank, Alec Birchley, Alexander Adams, Amy Gaskin, Bree Gatica-Wilcox, Jason Coombes, Joel Southgate, Lauren Gilbert, Lee Graham, Nicole Pacchiarini, Sara Kumziene-Summerhayes, Sarah Taylor, Sophie Jones, Sara Rey, Matthew Bull, Joanne Watkins, Sally Corden, Tom Connor |
| EPI_ISL_814665, EPI_ISL_814666, EPI_ISL_814667, EPI_ISL_814668, EPI_ISL_814669, EPI_ISL_814670 | Virology Department, Royal Infirmary of Edinburgh, NHS Lothian / School of Biological Sciences, University of Edinburgh / Institute of Genetics and Molecular Medicine, University of Edinburgh | COVID-19 Genomics UK (COG-UK) Consortium | McHugh M, Dewar R, Rooke S, Gallagher M, Balcaza C, O'Toole Á, Scher E, Hill V, McCrone JT, Colquhoun R, Yu X, Jackson B, Rambaut A, Williams TC, Templeton K |
| EPI_ISL_814754, EPI_ISL_814755, EPI_ISL_814794, EPI_ISL_814795, EPI_ISL_814800, EPI_ISL_814801, EPI_ISL_814802, EPI_ISL_814803, EPI_ISL_814804, EPI_ISL_814805, EPI_ISL_814806, EPI_ISL_814807, EPI_ISL_814808, EPI_ISL_814809, EPI_ISL_814810, EPI_ISL_814811, EPI_ISL_814812, EPI_ISL_814813, EPI_ISL_814814, EPI_ISL_814815, EPI_ISL_814816, EPI_ISL_814817, EPI_ISL_814818, EPI_ISL_814819, EPI_ISL_814820, EPI_ISL_814821, EPI_ISL_814822, EPI_ISL_814823, EPI_ISL_814824, EPI_ISL_814825, EPI_ISL_814826, EPI_ISL_814827, EPI_ISL_814828, EPI_ISL_814829, EPI_ISL_814830, EPI_ISL_814831, EPI_ISL_814832, EPI_ISL_814833, EPI_ISL_814834, EPI_ISL_814835, EPI_ISL_814837, EPI_ISL_814838, EPI_ISL_814839, EPI_ISL_814840, EPI_ISL_814841, EPI_ISL_814842, EPI_ISL_814843, EPI_ISL_814844, EPI_ISL_814845, EPI_ISL_814846, EPI_ISL_814847, EPI_ISL_814848, EPI_ISL_814849, EPI_ISL_814850, EPI_ISL_814851, EPI_ISL_814852, EPI_ISL_814853, EPI_ISL_814854, EPI_ISL_814855, EPI_ISL_814856, EPI_ISL_814857, EPI_ISL_814858, EPI_ISL_814859 | | | |
| see above | Wales Specialist Virology Centre Sequencing lab: Pathogen Genomics Unit | COVID-19 Genomics UK (COG-UK) Consortium | Catherine Moore, Johnathan Evans, Laura Gifford, Malorie Perry, Simon Cottrell, Angela Marchbank, Alec Birchley, Alexander Adams, Amy Gaskin, Bree Gatica-Wilcox, Jason Coombes, Joel Southgate, Lauren Gilbert, Lee Graham, Nicole Pacchiarini, Sara Kumziene-Summerhayes, Sarah Taylor, Sophie Jones, Sara Rey, Matthew Bull, Joanne Watkins, Sally Corden, Tom Connor |
| EPI_ISL_815597, EPI_ISL_815598, EPI_ISL_816067, EPI_ISL_816068, EPI_ISL_816069, EPI_ISL_816070, EPI_ISL_816071, EPI_ISL_816072 | Department of Virus and Microbiological Special Diagnostics, Statens Serum Institut, Copenhagen, Denmark | Albertsen Lab, Department of Chemistry and Bioscience, Aalborg University, Denmark | Danish Covid-19 Genome Consortium |
| EPI_ISL_816241, EPI_ISL_816295, EPI_ISL_816370, EPI_ISL_816428, EPI_ISL_816443, EPI_ISL_816512, EPI_ISL_816522, EPI_ISL_816551 | Virology Department, Sheffield Teaching Hospitals NHS Foundation Trust/Department of Infection, Immunity and Cardiovascular Disease, The Medical School, University of Sheffield | COVID-19 Genomics UK (COG-UK) Consortium | Thushan de Silva, Matthew Parker, Nikki Smith, Adri Angyal, Rebecca Brown, Luke Green, Rachel Tucker, Paul Parsons, Danielle Groves, Katie Johnson, Laura Carrilero, Alex Keeley, Dave Partridge, Matthew Wyles, Benjamin Lindsey, Mehmet Yavuz, Mohammad Raza, Cariad Evans |
| EPI_ISL_819505 | University College London, Great Ormond Street Hospital for Children NHS Foundation Trust, Imperial College Healthcare NHS Trust | COVID-19 Genomics UK (COG-UK) Consortium | Sergi Castellano, Rachel Williams, Mark Kristiansen, Paola Resende Silva, Sunando Roy, Tony Brooks, Helena Tutill, Paola Niola, Patricia Dyal, Charlotte Williams, Leysa Forrest, Yasmin Panchbhaya, Jacqueline Findlay, Samuel Weeks, Julianne Brown, Kathryn Harris, Paul Randell, James Price, Alison Holmes, Judith Breuer |
| EPI_ISL_819506, EPI_ISL_819507 | Lincolnshire Hospitals and DeepSeq Nottingham | COVID-19 Genomics UK (COG-UK) Consortium | Nichola Duckworth, Tim Sloan, Sarah Walsh, Jonathan Ball, Patrick McClure, Joeseph Chappell, Nadine Holmes, Matthew Carlisle, Christopher Moore, Fei Sang, Johnny Debebe, Victoria Wright, Matthew Loose |
| EPI_ISL_819508, EPI_ISL_819509, EPI_ISL_819510, EPI_ISL_819511, EPI_ISL_819512, EPI_ISL_819513, EPI_ISL_819514, EPI_ISL_819515, EPI_ISL_819516, EPI_ISL_819517, EPI_ISL_819518, EPI_ISL_819519 | | | |
| see above | University College London, Great Ormond Street Hospital for Children NHS Foundation Trust, Imperial College Healthcare NHS Trust | COVID-19 Genomics UK (COG-UK) Consortium | Sergi Castellano, Rachel Williams, Mark Kristiansen, Paola Resende Silva, Sunando Roy, Tony Brooks, Helena Tutill, Paola Niola, Patricia Dyal, Charlotte Williams, Leysa Forrest, Yasmin Panchbhaya, Jacqueline Findlay, Samuel Weeks, Julianne Brown, Kathryn Harris, Paul Randell, James Price, Alison Holmes, Judith Breuer |
| EPI_ISL_819520, EPI_ISL_819521, EPI_ISL_819522, EPI_ISL_819523 | Lincolnshire Hospitals and DeepSeq Nottingham | COVID-19 Genomics UK (COG-UK) Consortium | Nichola Duckworth, Tim Sloan, Sarah Walsh, Jonathan Ball, Patrick McClure, Joeseph Chappell, Nadine Holmes, Matthew Carlisle, Christopher Moore, Fei Sang, Johnny Debebe, Victoria Wright, Matthew Loose |
| EPI_ISL_819524, EPI_ISL_819525, EPI_ISL_819526, EPI_ISL_819527, EPI_ISL_819528 | University College London, Great Ormond Street Hospital for Children NHS Foundation Trust, Imperial College Healthcare NHS Trust | COVID-19 Genomics UK (COG-UK) Consortium | Sergi Castellano, Rachel Williams, Mark Kristiansen, Paola Resende Silva, Sunando Roy, Tony Brooks, Helena Tutill, Paola Niola, Patricia Dyal, Charlotte Williams, Leysa Forrest, Yasmin Panchbhaya, Jacqueline Findlay, Samuel Weeks, Julianne Brown, Kathryn Harris, Paul Randell, James Price, Alison Holmes, Judith Breuer |
| EPI_ISL_819529 | Lincolnshire Hospitals and DeepSeq Nottingham | COVID-19 Genomics UK (COG-UK) Consortium | Nichola Duckworth, Tim Sloan, Sarah Walsh, Jonathan Ball, Patrick McClure, Joeseph Chappell, Nadine Holmes, Matthew Carlisle, Christopher Moore, Fei Sang, Johnny Debebe, Victoria Wright, Matthew Loose |
| EPI_ISL_819530, EPI_ISL_819531 | University College London, Great Ormond Street Hospital for Children NHS Foundation Trust, Imperial College Healthcare NHS Trust | COVID-19 Genomics UK (COG-UK) Consortium | Sergi Castellano, Rachel Williams, Mark Kristiansen, Paola Resende Silva, Sunando Roy, Tony Brooks, Helena Tutill, Paola Niola, Patricia Dyal, Charlotte Williams, Leysa Forrest, Yasmin Panchbhaya, Jacqueline Findlay, Samuel Weeks, Julianne Brown, Kathryn Harris, Paul Randell, James Price, Alison Holmes, Judith Breuer |
| EPI_ISL_819532 | Lincolnshire Hospitals and DeepSeq Nottingham | COVID-19 Genomics UK (COG-UK) Consortium | Nichola Duckworth, Tim Sloan, Sarah Walsh, Jonathan Ball, Patrick McClure, Joeseph Chappell, Nadine Holmes, Matthew Carlisle, Christopher Moore, Fei Sang, Johnny Debebe, Victoria Wright, Matthew Loose |
| EPI_ISL_819533, EPI_ISL_819534, EPI_ISL_819535, EPI_ISL_819536, EPI_ISL_819537, EPI_ISL_819538, EPI_ISL_819539, EPI_ISL_819540, EPI_ISL_819541, EPI_ISL_819542, EPI_ISL_819543, EPI_ISL_819544, EPI_ISL_819545, EPI_ISL_819546, EPI_ISL_819547, EPI_ISL_819548, EPI_ISL_819549 | | | |
| see above | University College London, Great Ormond Street Hospital for | COVID-19 Genomics UK (COG-UK) Consortium | Sergi Castellano, Rachel Williams, Mark Kristiansen, Paola Resende Silva, Sunando Roy, Tony Brooks, Helena Tutill, Paola Niola, Patricia Dyal, Charlotte |

| | | | |
|---|---|---|---|
| | Children NHS Foundation Trust, Imperial College Healthcare NHS Trust | | Williams, Leysa Forrest, Yasmin Panchbhaya, Jacqueline Findlay, Samuel Weeks, Julianne Brown, Kathryn Harris, Paul Randell, James Price, Alison Holmes, Judith Breuer |
| EPI_ISL_822093 | Lighthouse Lab in Alderley Park | Wellcome Sanger Institute for the COVID-19 Genomics UK (COG-UK) Consortium | Jacquelyn Wynn, Mairead Hyland, The Lighthouse Lab in Alderley Park and Alex Alderton, Roberto Amato, Sonia Goncalves, Ewan Harrison, David K. Jackson, Ian Johnston, Dominic Kwiatkowski, Cordelia Langford, John Sillitoe on behalf of the Wellcome Sanger Institute COVID-19 Surveillance Team |
| EPI_ISL_823967 | DOHMH Corona | New York City Public Health Laboratory | Jade Wang, et al. |
| EPI_ISL_823968, EPI_ISL_823969 | DOHMH Jamaica | New York City Public Health Laboratory | Jade Wang, et al. |
| EPI_ISL_824004, EPI_ISL_824005, EPI_ISL_824007, EPI_ISL_824009, EPI_ISL_824011, EPI_ISL_824012, EPI_ISL_824014, EPI_ISL_824018, EPI_ISL_824126, EPI_ISL_824127, EPI_ISL_824131 | | | |
| see above | Dutch COVID-19 response team | National Institute for Public Health and the Environment (RIVM) | Adam Meijer, Harry Vennema, Jeroen Cremer, Sharon van den Brink, Bas van der Veer, AnneMarie van den Brandt, Florian Zwagemaker, Dennis Schmitz, Chantal Reusken, on behalf of the national COVID-19 response team |
| EPI_ISL_824335, EPI_ISL_824344, EPI_ISL_824351, EPI_ISL_824352, EPI_ISL_824355, EPI_ISL_824356, EPI_ISL_824357, EPI_ISL_824366, EPI_ISL_824371, EPI_ISL_824374, EPI_ISL_824375, EPI_ISL_824376, EPI_ISL_824379, EPI_ISL_824382 | | | |
| see above | Michigan Department of Health and Human Services, Bureau of Laboratories | Michigan Department of Health and Human Services, Bureau of Laboratories | Blankenship HM, Riner D, Soehnlen MK |
| EPI_ISL_824396, EPI_ISL_824397 | California Department of Public Health | California Department of Public Health | CDPH IDLB COVIDNet |
| EPI_ISL_825084, EPI_ISL_825086, EPI_ISL_825087, EPI_ISL_825088, EPI_ISL_825089 | Indian Council of Medical Research-National Institute of Virology,Maximum Containment Laboratory | Indian Council of Medical Research-National Institute of Virology,Maximum Containment Laboratory | Pragya D Yadav, Dimpal A Nyayanit, Rima R Sahay, Prasad Sarkale, Jayshri Pethani, Savita Patil, Shrikant Baradkar, Varsha Potdar, Deepak Y Patil |
| EPI_ISL_825333, EPI_ISL_825335, EPI_ISL_825366, EPI_ISL_825367, EPI_ISL_825368, EPI_ISL_825369, EPI_ISL_825370, EPI_ISL_825371 | Hospital Universitari Vall d'Hebron - Vall d'Hebron Institut de Rercerca | Hospital Universitari Vall d'Hebron | Cristina Andrés, Maria Piñana, Josep F Abril, Damir Garcia-Cehic, Ariadna Rando, Juliana Esperalba, Maria Gema Codina, Carla Castillo, Maria Carmen Martin, Tomàs Pumarola, Josep Quer, Andrés Antón |
| EPI_ISL_826554 | Texas Department of State Health Services | Texas Department of State Health Services | Rashmi Tuladhar, Bonnie Oh, Jenny Zhang, Maliha Rahman, Anita Pokharel, Myong Koag, Chung Wang, Rachel Lee, Grace Kubin, Mayela Pedrueza, James Daniel Bonser |
| EPI_ISL_826881, EPI_ISL_826892, EPI_ISL_827078, EPI_ISL_827152, EPI_ISL_827170, EPI_ISL_827434 | deCODE genetics | deCODE genetics | Daniel F Gudbjartsson; Agnar Helgason; Hakon Jonsson; Olafur T Magnusson; Pall Melsted; Gudmundur L Norddahl; Jona Saemundsdottir; Asgeir Sigurdsson; Patrick Sulem; Arna B Agustsdottir; Hannes Eggertsson; Berglind Eiriksdottir; Run Fridriksdottir; Elisabet E Gardarsdottir; Gudmundur Georgsson; Olafia S Gretarsdottir; Kjartan R Gudmundsson; Thora R Gunnarsdottir; Arnaldur Gylfason; Hilma Holm; Brynjar O Jensson; Aslaug Jonasdottir; Kamilla S Josefsdottir; Thordur Kristjansson; Droplaug N Magnusdottir; Solvi Rognvaldsson; Louise le Roux; Gudrun Sigmundsdottir; Gardar Sveinbjornsson; Kristin E Sveinsdottir; Maney Sveinsdottir; Emil A Thorarensen; Bjarni Thorbjornsson; Gisli Masson; Ingileif Jonsdottir; Alma Moller; Thorolfur Gudnason; Karl G Kristinsson; Unnur Thorsteinsdottir; Kari Stefansson |
| EPI_ISL_827692 | The National University Hospital of Iceland | deCODE genetics | Daniel F Gudbjartsson; Agnar Helgason; Hakon Jonsson; Olafur T Magnusson; Pall Melsted; Gudmundur L Norddahl; Jona Saemundsdottir; Asgeir Sigurdsson; Patrick Sulem; Arna B Agustsdottir; Hannes Eggertsson; Berglind Eiriksdottir; Run Fridriksdottir; Elisabet E Gardarsdottir; Gudmundur Georgsson; Olafia S Gretarsdottir; Kjartan R Gudmundsson; Thora R Gunnarsdottir; Arnaldur Gylfason; Hilma Holm; Brynjar O Jensson; Aslaug Jonasdottir; Kamilla S Josefsdottir; Thordur Kristjansson; Droplaug N Magnusdottir; Solvi Rognvaldsson; Louise le Roux; Gudrun Sigmundsdottir; Gardar Sveinbjornsson; Kristin E Sveinsdottir; Maney Sveinsdottir; Emil A Thorarensen; Bjarni Thorbjornsson; Gisli Masson; Ingileif Jonsdottir; Alma Moller; Thorolfur Gudnason; Karl G Kristinsson; Unnur Thorsteinsdottir; Kari Stefansson |
| EPI_ISL_827858, EPI_ISL_827912, EPI_ISL_827920, EPI_ISL_827926, EPI_ISL_828050, EPI_ISL_828208, EPI_ISL_828520, EPI_ISL_828521, EPI_ISL_828524, EPI_ISL_828535, EPI_ISL_828536, EPI_ISL_828632, EPI_ISL_828860, EPI_ISL_829056, EPI_ISL_829209, EPI_ISL_829210, EPI_ISL_829211, EPI_ISL_829212, EPI_ISL_829213, EPI_ISL_829214, EPI_ISL_829215, EPI_ISL_829216, EPI_ISL_829329, EPI_ISL_829330, EPI_ISL_829331, EPI_ISL_829332, EPI_ISL_829333, EPI_ISL_829334 | | | |
| see above | deCODE genetics | deCODE genetics | Daniel F Gudbjartsson; Agnar Helgason; Hakon Jonsson; Olafur T Magnusson; Pall Melsted; Gudmundur L Norddahl; Jona Saemundsdottir; Asgeir Sigurdsson; Patrick Sulem; Arna B Agustsdottir; Hannes Eggertsson; Berglind Eiriksdottir; Run Fridriksdottir; Elisabet E Gardarsdottir; Gudmundur Georgsson; Olafia S Gretarsdottir; Kjartan R Gudmundsson; Thora R Gunnarsdottir; Arnaldur Gylfason; Hilma Holm; Brynjar O Jensson; Aslaug Jonasdottir; Kamilla S Josefsdottir; Thordur Kristjansson; Droplaug N Magnusdottir; Solvi Rognvaldsson; Louise le Roux; Gudrun Sigmundsdottir; Gardar Sveinbjornsson; Kristin E Sveinsdottir; Maney Sveinsdottir; Emil A Thorarensen; Bjarni Thorbjornsson; Gisli Masson; Ingileif Jonsdottir; Alma Moller; Thorolfur Gudnason; Karl G Kristinsson; Unnur Thorsteinsdottir; Kari Stefansson |
| EPI_ISL_829607 | The National University Hospital of Iceland | deCODE genetics | Daniel F Gudbjartsson; Agnar Helgason; Hakon Jonsson; Olafur T Magnusson; Pall Melsted; Gudmundur L Norddahl; Jona Saemundsdottir; Asgeir Sigurdsson; Patrick Sulem; Arna B Agustsdottir; Hannes Eggertsson; Berglind Eiriksdottir; Run Fridriksdottir; Elisabet E Gardarsdottir; Gudmundur Georgsson; Olafia S Gretarsdottir; Kjartan R Gudmundsson; Thora R Gunnarsdottir; Arnaldur Gylfason; Hilma Holm; Brynjar O Jensson; Aslaug Jonasdottir; Kamilla S Josefsdottir; Thordur Kristjansson; Droplaug N Magnusdottir; Solvi Rognvaldsson; Louise le Roux; Gudrun Sigmundsdottir; Gardar Sveinbjornsson; Kristin E Sveinsdottir; Maney Sveinsdottir; Emil A Thorarensen; Bjarni Thorbjornsson; Gisli Masson; Ingileif Jonsdottir; Alma Moller; Thorolfur Gudnason; Karl G Kristinsson; Unnur Thorsteinsdottir; Kari Stefansson |
| EPI_ISL_829742, EPI_ISL_829966, EPI_ISL_829968, EPI_ISL_829969, EPI_ISL_829974, EPI_ISL_830251 | deCODE genetics | deCODE genetics | Daniel F Gudbjartsson; Agnar Helgason; Hakon Jonsson; Olafur T Magnusson; Pall Melsted; Gudmundur L Norddahl; Jona Saemundsdottir; Asgeir Sigurdsson; Patrick Sulem; Arna B Agustsdottir; Hannes Eggertsson; Berglind Eiriksdottir; Run Fridriksdottir; Elisabet E Gardarsdottir; Gudmundur Georgsson; Olafia S Gretarsdottir; Kjartan R Gudmundsson; Thora R Gunnarsdottir; Arnaldur Gylfason; Hilma Holm; Brynjar O Jensson; Aslaug Jonasdottir; Kamilla S Josefsdottir; Thordur Kristjansson; Droplaug N Magnusdottir; Solvi Rognvaldsson; Louise le Roux; Gudrun Sigmundsdottir; Gardar Sveinbjornsson; Kristin E Sveinsdottir; Maney Sveinsdottir; Emil A Thorarensen; Bjarni Thorbjornsson; Gisli Masson; Ingileif Jonsdottir; Alma Moller; Thorolfur Gudnason; Karl G Kristinsson; Unnur Thorsteinsdottir; Kari Stefansson |
| EPI_ISL_830254 | The National University Hospital of Iceland | deCODE genetics | Daniel F Gudbjartsson; Agnar Helgason; Hakon Jonsson; Olafur T Magnusson; Pall Melsted; Gudmundur L Norddahl; Jona Saemundsdottir; Asgeir Sigurdsson; Patrick Sulem; Arna B Agustsdottir; Hannes Eggertsson; Berglind Eiriksdottir; Run Fridriksdottir; Elisabet E Gardarsdottir; Gudmundur Georgsson; Olafia S Gretarsdottir; Kjartan R Gudmundsson; Thora R Gunnarsdottir; Arnaldur Gylfason; Hilma Holm; Brynjar O Jensson; Aslaug Jonasdottir; Kamilla S Josefsdottir; Thordur Kristjansson; Droplaug N Magnusdottir; Solvi Rognvaldsson; Louise le Roux; Gudrun Sigmundsdottir; Gardar Sveinbjornsson; Kristin E Sveinsdottir; Maney Sveinsdottir; Emil A Thorarensen; Bjarni Thorbjornsson; Gisli Masson; Ingileif Jonsdottir; Alma Moller; Thorolfur Gudnason; Karl G Kristinsson; Unnur Thorsteinsdottir; Kari Stefansson |
| EPI_ISL_830256, EPI_ISL_830260, EPI_ISL_830262 | deCODE genetics | deCODE genetics | Daniel F Gudbjartsson; Agnar Helgason; Hakon Jonsson; Olafur T Magnusson; Pall Melsted; Gudmundur L Norddahl; Jona Saemundsdottir; Asgeir Sigurdsson; Patrick Sulem; Arna B Agustsdottir; Hannes Eggertsson; Berglind Eiriksdottir; Run Fridriksdottir; Elisabet E Gardarsdottir; Gudmundur Georgsson; Olafia S Gretarsdottir; Kjartan R Gudmundsson; Thora R Gunnarsdottir; Arnaldur Gylfason; Hilma Holm; Brynjar O Jensson; Aslaug Jonasdottir; Kamilla S Josefsdottir; Thordur Kristjansson; Droplaug N Magnusdottir; Solvi Rognvaldsson; Louise le Roux; Gudrun Sigmundsdottir; Gardar Sveinbjornsson; Kristin E Sveinsdottir; Maney Sveinsdottir; Emil A Thorarensen; Bjarni Thorbjornsson; Gisli Masson; Ingileif Jonsdottir; Alma Moller; Thorolfur Gudnason; Karl G Kristinsson; Unnur Thorsteinsdottir; Kari Stefansson |
| EPI_ISL_830268 | The National University Hospital of Iceland | deCODE genetics | Daniel F Gudbjartsson; Agnar Helgason; Hakon Jonsson; Olafur T Magnusson; Pall Melsted; Gudmundur L Norddahl; Jona Saemundsdottir; Asgeir Sigurdsson; Patrick Sulem; Arna B Agustsdottir; Hannes Eggertsson; Berglind Eiriksdottir; Run Fridriksdottir; Elisabet E Gardarsdottir; Gudmundur Georgsson; Olafia S Gretarsdottir; Kjartan R Gudmundsson; Thora R Gunnarsdottir; Arnaldur Gylfason; Hilma Holm; Brynjar O Jensson; Aslaug Jonasdottir; Kamilla S Josefsdottir; Thordur Kristjansson; Droplaug N Magnusdottir; Solvi Rognvaldsson; Louise le Roux; Gudrun Sigmundsdottir; Gardar Sveinbjornsson; Kristin E Sveinsdottir; Maney Sveinsdottir; Emil A Thorarensen; Bjarni Thorbjornsson; Gisli Masson; Ingileif Jonsdottir; Alma Moller; Thorolfur Gudnason; Karl G Kristinsson; Unnur Thorsteinsdottir; Kari Stefansson |
| EPI_ISL_831420, EPI_ISL_831421, EPI_ISL_831422 | Labor Dr. Wisplinghoff - Köln | Robert Koch Institute, Influenza and respiratory viruses FG17 & Bioinformatics MF1, Berlin, Germany | Dr. R. Grosser, Stephan Fuchs, Stefan Kroeger, Marianne Wedde, Oliver Drechsel, Aleksandar Radonic, Rene Kmiecinski, Ralf Duerrwald, Thorsten Wolff |
| EPI_ISL_831423 | Labor Dr. Krause & Kollegen MVZ GmbH | Robert Koch Institute, Influenza and respiratory viruses FG17 & Bioinformatics MF1, Berlin, Germany | Dr. Lorentz, Stephan Fuchs, Stefan Kroeger, Marianne Wedde, Oliver Drechsel, Aleksandar Radonic, Rene Kmiecinski, Ralf Duerrwald, Thorsten Wolff |

| | | | |
|---|---|---|---|
| EPI_ISL_831424, EPI_ISL_831425 | Labor Dr. Wisplinghoff - Köln | Robert Koch Institute, Influenza and respiratory viruses FG17 & Bioinformatics MF1, Berlin, Germany | Dr. R. Grosser, Stephan Fuchs, Stefan Kroeger, Marianne Wedde, Oliver Drechsel, Aleksandar Radonic, Rene Kmiecinski, Ralf Duerrwald, Thorsten Wolff |
| EPI_ISL_831426 | Labor Dr. Krause & Kollegen MVZ GmbH | Robert Koch Institute, Influenza and respiratory viruses FG17 & Bioinformatics MF1, Berlin, Germany | Dr. Lorentz, Stephan Fuchs, Stefan Kroeger, Marianne Wedde, Oliver Drechsel, Aleksandar Radonic, Rene Kmiecinski, Ralf Duerrwald, Thorsten Wolff |
| EPI_ISL_831662, EPI_ISL_831684, EPI_ISL_831686 | Santa Clara County Public Health Laboratory | Santa Clara County Public Health Laboratory | Santa Clara County Public Health Department |
| EPI_ISL_831697, EPI_ISL_831706, EPI_ISL_831715, EPI_ISL_831730, EPI_ISL_831731, EPI_ISL_831736, EPI_ISL_831741, EPI_ISL_831834, EPI_ISL_831835, EPI_ISL_831836, EPI_ISL_831837, EPI_ISL_831838, EPI_ISL_831839, EPI_ISL_831840, EPI_ISL_831843, EPI_ISL_831844, EPI_ISL_831845, EPI_ISL_831855, EPI_ISL_831856, EPI_ISL_831857, EPI_ISL_831858, EPI_ISL_831859, EPI_ISL_831860, EPI_ISL_831864 | | | |
| see above | United States Air Force School of Aerospace Medicine | United States Air Force School of Aerospace Medicine | Anthony Fries, Jennifer Meyer, William Gruner, Amanda Javorina, Sarah Purves, Clarise Starr, Elizabeth Macias |
| EPI_ISL_832156, EPI_ISL_832160, EPI_ISL_832161, EPI_ISL_832162 | Hospital | National Reference Center for Viruses of Respiratory Infections, Institut Pasteur, Paris | Marion Barbet, Sylvie Behillil, Méline Bizard, Angela Brisebarre, Camille Capel, Etienne Simon-Lorière, Vincent Enouf, Maud Vanpeene, Sylvie van der Werf, Léa Pilorge |
| EPI_ISL_832814 | OHSU Lab Services Molecular Microbiology Lab | Oregon SARS-CoV-2 Genome Sequencing Center | Brendan L. O'Connell, Ruth V. Nichols, Sally Grindstaff, Alec J. Hirsch, Donna Hansel, Guang Fan, Daniel N. Streblow, William B. Messer, Andrew C. Adey, Benjamin N. Bimber, Brian J. O'Roak |
| EPI_ISL_833131 | Laboratorio de Ecologia de Doencas Transmissiveis na Amazonia, Instituto Leonidas e Maria Deane - Fiocruz Amazonia | Laboratorio de Ecologia de Doencas Transmissiveis na Amazonia, Instituto Leonidas e Maria Deane - Fiocruz Amazonia | Valdinete Nascimento, Victor Souza, André Corado, Fernanda Nascimento, George Silva, Ágatha Costa, Debora Duarte, Karina Pessoa, Matilde Mejía, Luciana Gonçalves, Maria Júlia Brandão, Michele Jesus, Felipe Naveca |
| EPI_ISL_833158 | Instituto Adolfo Lutz - Regional de Santo Andre | Instituto Adolfo Lutz, Interdiciplinary Procedures Center, Strategic Laboratory | Claudio Tavares Sacchi, Claudia Regina Gonçalves, Erica Valessa Ramos Gomes, Karoline Rodrigues Campos |
| EPI_ISL_833343 | Buenavista Rural Health Unit | Research Institute for Tropical Medicine | Hannah Leah Morito, Othoniel Jan Onza, John Leonard Chan, Ma Angelica Tujan, Francisco Gerardo Polotan, Inez Andrea Medado, Kirstyn Brunker, Edelwisa Mercado, Daria Manalo, Catalino Demetria |
| EPI_ISL_833356, EPI_ISL_833357, EPI_ISL_833358, EPI_ISL_833359, EPI_ISL_833360, EPI_ISL_833361, EPI_ISL_833362, EPI_ISL_833363 | Microbiology Division, South Carolina Department of Health and Environmental Control (SC DHEC) | Microbiology Division, South Carolina Department of Health and Environmental Control (SC DHEC) | Flores,H., Freeman,J. |
| EPI_ISL_835474 | Lighthouse Lab in Alderley Park | Wellcome Sanger Institute for the COVID-19 Genomics UK (COG-UK) Consortium | Jacquelyn Wynn, Mairead Hyland, The Lighthouse Lab in Alderley Park and Alex Alderton, Roberto Amato, Sonia Goncalves, Ewan Harrison, David K. Jackson, Ian Johnston, Dominic Kwiatkowski, Cordelia Langford, John Sillitoe on behalf of the Wellcome Sanger Institute COVID-19 Surveillance Team |
| EPI_ISL_837107, EPI_ISL_837108, EPI_ISL_837109, EPI_ISL_837110, EPI_ISL_837111, EPI_ISL_837198 | Respiratory Virus Unit, National Infection Service, Public Health England | COVID-19 Genomics UK (COG-UK) Consortium | PHE Covid Sequencing Team |
| EPI_ISL_837450, EPI_ISL_837451, EPI_ISL_837452, EPI_ISL_837453, EPI_ISL_837454, EPI_ISL_837455, EPI_ISL_837456, EPI_ISL_837481, EPI_ISL_837482 | Istituto Zooprofilattico Sperimentale del Mezzogiorno | TIGEM | Patrizia Annunziata, Andrea Ballabio, Valentina Bouche, Davide Cacchiarelli (CorrespAuthor), Pellegrino Cerino, Chiara Colantuono, Lucio Di Filippo, Antonio Grimaldi, Antonio Limone, Gabriella Loconte, Anna Manfredi, Francesco Panariello, Biancamaria Pierri, Marcello Salvi, Lucia Vassallo |
| EPI_ISL_838065, EPI_ISL_838073, EPI_ISL_838074, EPI_ISL_838075, EPI_ISL_838076, EPI_ISL_838077, EPI_ISL_838078, EPI_ISL_838079 | West of Scotland Specialist Virology Centre, NHSGGC / MRC-University of Glasgow Centre for Virus Research | COVID-19 Genomics UK (COG-UK) Consortium | Ana da Silva Filipe, Natasha Johnson, Kathy Smollett, Daniel Mair, Stephen Carmichael, Alice Broos, Lily Tong, Jenna Nichols, Kyriaki Nomikou; Sarah McDonald; Richard Orton, Joseph Hughes, Sreenu Vattipally, David L Robertson; Alasdair MacLean, Rory Gunson; Sharif Shaaban, Matthew Holden; Rachel Blacow, Guy Mollett, Kathy Li, James Shepherd, Antonia Ho, Emma Thomson |
| EPI_ISL_838215 | Virology Department, Royal Infirmary of Edinburgh, NHS Lothian / School of Biological Sciences, University of Edinburgh / Institute of Genetics and Molecular Medicine, University of Edinburgh | COVID-19 Genomics UK (COG-UK) Consortium | McHugh M, Dewar R, Rooke S, Gallagher M, Balcaza C, O'Toole Á, Scher E, Hill V, McCrone JT, Colquhoun R, Yu X, Jackson B, Rambaut A, Williams TC, Templeton K |
| EPI_ISL_838353, EPI_ISL_838354, EPI_ISL_838355, EPI_ISL_838356, EPI_ISL_838357, EPI_ISL_838358, EPI_ISL_838359, EPI_ISL_838361, EPI_ISL_838362, EPI_ISL_838363, EPI_ISL_838364, EPI_ISL_838366, EPI_ISL_838367, EPI_ISL_838368, EPI_ISL_838369, EPI_ISL_838372, EPI_ISL_838376, EPI_ISL_838377, EPI_ISL_838378, EPI_ISL_838379, EPI_ISL_838380, EPI_ISL_838382, EPI_ISL_838383, EPI_ISL_838384, EPI_ISL_838385, EPI_ISL_838386, EPI_ISL_838388, EPI_ISL_838393, EPI_ISL_838394, EPI_ISL_838395, EPI_ISL_838398, EPI_ISL_838399, EPI_ISL_838401, EPI_ISL_838405, EPI_ISL_838422, EPI_ISL_838424, EPI_ISL_838427, EPI_ISL_838440, EPI_ISL_838446, EPI_ISL_838453, EPI_ISL_838454 | | | |
| see above | Liverpool Clinical Laboratories | COVID-19 Genomics UK (COG-UK) Consortium | Sam Haldenby, Anita Lucaci, Steve Paterson, Julian Hiscox, Alistair Darby, M Almsaud, A Alrezaihi, Muhannad Alruwaili, Stuart D Armstrong, Jones Benjamin, Eleanor G Bentley, Anu Chawla, Jordan J Clark, Angela Cowell, Richard Eccles, Isabel García-Dorival, Matthew Gemmell, Alessandro Gerada, PKF Gilmore, Richard Gregory, Ximeng Han, Catherine Hartley, Margaret Hughes, Miren Iturriza-Gomara, James Johnson, L Luu, Jenifer Manson, Charlotte Nelson, Elaine O'Toole, Cassie Olateju, Rebekah Penrice-Randal , Lucille Rainbow, N.P Randle, Trevor Ian Robinson, Parul Sharma, Ghada T Shawli, James P Stewart, Neil Swainston, Ecaterina Vamos, Joanne Watts, Mark Whitehead |
| EPI_ISL_839338, EPI_ISL_839341 | University College London, Great Ormond Street Hospital for Children NHS Foundation Trust, Imperial College Healthcare NHS Trust | COVID-19 Genomics UK (COG-UK) Consortium | Sergi Castellano, Rachel Williams, Mark Kristiansen, Paola Resende Silva, Sunando Roy, Tony Brooks, Helena Tutill, Paola Niola, Patricia Dyal, Charlotte Williams, Leysa Forrest, Yasmin Panchbhaya, Jacqueline Findlay, Samuel Weeks, Julianne Brown, Kathryn Harris, Paul Randell, James Price, Alison Holmes, Judith Breuer |
| EPI_ISL_839987 | Queens Medical Centre, Clinical Microbiology Department / DeepSeq Nottingham | COVID-19 Genomics UK (COG-UK) Consortium | Gemma Clark, Wendy Smith, Manjinder Khakh, Vicki M Fleming, Michelle M Lister, Hannah Howson-Wells, Jonathan Ball, Patrick McClure, Joseph Chappell, Theocharis Tsoleridis, Nadine Holmes, Matthew Carlisle, Christopher Moore, Fei Sang, Johnny Debebe, Victoria Wright, Matthew Loose |
| EPI_ISL_840019, EPI_ISL_840020, EPI_ISL_840021, EPI_ISL_840025 | Lincolnshire Hospitals and DeepSeq Nottingham | COVID-19 Genomics UK (COG-UK) Consortium | Nichola Duckworth, Tim Sloan, Sarah Walsh, Jonathan Ball, Patrick McClure, Joeseph Chappell, Nadine Holmes, Matthew Carlisle, Christopher Moore, Fei Sang, Johnny Debebe, Victoria Wright, Matthew Loose |
| EPI_ISL_840222, EPI_ISL_840223, EPI_ISL_840224 | Oxford Viromics, NDM, University of Oxford; Oxford University Hospitals; Basingstoke and North Hampshire Hospital | COVID-19 Genomics UK (COG-UK) Consortium | Tanya Golubchik, David Bonsall, George Macintyre, Amy Trebes, Mariateresa de Cesare, Catrin Moore, Alex Mobbs, Anita Justice, Robert Shaw, Monique Andersson, Timothy Peto, Emma Wise, Nathan Moore, Jessica Lynch, Nick Cortes, Matilde Mori, Stephen Kidd, David Buck, John Todd, Christophe Fraser |
| EPI_ISL_841503, EPI_ISL_841529, EPI_ISL_841562, EPI_ISL_841573, EPI_ISL_841574, EPI_ISL_841575, EPI_ISL_841576, EPI_ISL_841577, EPI_ISL_841578 | Originating lab: Wales Specialist Virology Centre Sequencing lab: Pathogen Genomics Unit | Public Health Wales Microbiology Cardiff Wales Specialist Virology Centre | Catherine Moore, Johnathan Evans, Laura Gifford, Malorie Perry, Simon Cottrell, Angela Marchbank, Alec Birchley, Alexander Adams, Amy Gaskin, Bree Gatica-Wilcox, Jason Coombes, Joel Southgate, Lauren Gilbert, Lee Graham, Nicole Pacchiarini, Sara Kumziene-Summerhayes, Sarah Taylor, Sophie Jones, Sara Rey, Matthew Bull, Joanne Watkins, Sally Corden, Tom Connor |
| EPI_ISL_842662, EPI_ISL_842696 | University College London Hospital | COVID-19 Genomics UK (COG-UK) Consortium | Judith Heaney, Matthew Byott, Catherine Houlihan, Dan Frampton, Stuart Kirk, Moira Spyer and Eleni Nastouli |
| EPI_ISL_843008, EPI_ISL_843009, EPI_ISL_843012, EPI_ISL_843025, EPI_ISL_843026, EPI_ISL_843027, EPI_ISL_843028, EPI_ISL_843029, EPI_ISL_843030, EPI_ISL_843031, EPI_ISL_843038, EPI_ISL_843040, EPI_ISL_843042, EPI_ISL_843043, EPI_ISL_843045, EPI_ISL_843047, EPI_ISL_843067, EPI_ISL_843068, EPI_ISL_843070 | | | |
| see above | Barts Health NHS Trust | COVID-19 Genomics UK (COG-UK) Consortium | CUTINO-MOGUEL, Maria-Teresa; HARRINGTON, David; OWOYEMI, Dola; SHYLINI, Raghavendran; BROAD, Claire; KELE, Beatrix |
| EPI_ISL_844220, EPI_ISL_844221, EPI_ISL_844222, EPI_ISL_844228, EPI_ISL_844232, EPI_ISL_844233, EPI_ISL_844236, EPI_ISL_844238, EPI_ISL_844251, EPI_ISL_844252, EPI_ISL_844263, EPI_ISL_844264, EPI_ISL_844266, EPI_ISL_844269, EPI_ISL_844287, EPI_ISL_844288, EPI_ISL_844289, EPI_ISL_844290, EPI_ISL_844291, EPI_ISL_844297, EPI_ISL_844302, EPI_ISL_844303, EPI_ISL_844304, EPI_ISL_844305, EPI_ISL_844462, EPI_ISL_844463, EPI_ISL_844464, EPI_ISL_844465, EPI_ISL_844466, EPI_ISL_844467, EPI_ISL_844468, EPI_ISL_844469, EPI_ISL_844470, EPI_ISL_844471, EPI_ISL_844472, EPI_ISL_844473, EPI_ISL_844474, EPI_ISL_844475, EPI_ISL_844476, EPI_ISL_844477, EPI_ISL_844478, EPI_ISL_844479, EPI_ISL_844480, EPI_ISL_844481, EPI_ISL_844482, EPI_ISL_844483, EPI_ISL_844484, EPI_ISL_844485, EPI_ISL_844486, EPI_ISL_844487, EPI_ISL_844488, EPI_ISL_844489, EPI_ISL_844490, EPI_ISL_844491, EPI_ISL_844492, EPI_ISL_844493, EPI_ISL_844494, EPI_ISL_844495, EPI_ISL_844496, EPI_ISL_844497, EPI_ISL_844498, EPI_ISL_844499, EPI_ISL_844500, EPI_ISL_844501, EPI_ISL_844502, EPI_ISL_844503, EPI_ISL_844504, EPI_ISL_844505, EPI_ISL_844506, EPI_ISL_844507, EPI_ISL_844508, EPI_ISL_844509, EPI_ISL_844510, EPI_ISL_844511, EPI_ISL_844512, EPI_ISL_844513, EPI_ISL_844514, EPI_ISL_844515, EPI_ISL_844516, EPI_ISL_844517, EPI_ISL_844518, EPI_ISL_844519, EPI_ISL_844520, EPI_ISL_844521, EPI_ISL_844522, EPI_ISL_844523, EPI_ISL_844524, EPI_ISL_844525, EPI_ISL_844526, EPI_ISL_844527 | | | |
| see above | Department of Virus and Microbiological Special Diagnostics, | Albertsen Lab, Department of Chemistry and Bioscience, | Danish Covid-19 Genome Consortium |

| | | | |
|---|---|---|---|
| | Statens Serum Institut, Copenhagen, Denmark | Aalborg University, Denmark | |
| EPI_ISL_845563 | National Public Health Laboratory, Cameroon | African Centre of Excellence for Genomics of Infectious Diseases (ACEGID), Redeemer's University | Oluniyi P.E. et al |
| EPI_ISL_845680, EPI_ISL_845681, EPI_ISL_845682, EPI_ISL_845683, EPI_ISL_845684, EPI_ISL_845685, EPI_ISL_845686, EPI_ISL_845687, EPI_ISL_845688, EPI_ISL_845689, EPI_ISL_845690, EPI_ISL_845691, EPI_ISL_845692, EPI_ISL_845693, EPI_ISL_845694, EPI_ISL_845695, EPI_ISL_845696, EPI_ISL_845697, EPI_ISL_845698, EPI_ISL_845699, EPI_ISL_845700, EPI_ISL_845701, EPI_ISL_845702, EPI_ISL_845703, EPI_ISL_845704, EPI_ISL_845705, EPI_ISL_845706, EPI_ISL_845707, EPI_ISL_845708, EPI_ISL_845709, EPI_ISL_845710, EPI_ISL_845711, EPI_ISL_845712, EPI_ISL_845713, EPI_ISL_845714 | | | |
| see above | Ontario's COVID-19 Genomics Rapid Response Coalition | McMaster University | Allison McGeer, Patryk Aftanas, Hooman Derakhshani, Angel Li, Kuganya Nirmalarajah, Emily Panousis, Ahmed Draia, Jalees Nasir, Michael Surette, Samira Mubareka, Andrew G. McArthur |
| EPI_ISL_845794 | South Eastern Area Laboratory Services (SEALS) | NSW Health Pathology - Institute of Clinical Pathology and Medical Research; Westmead Hospital; University of Sydney | CIDM-PH et al. |
| EPI_ISL_846588 | Respiratory Virus Unit, National Infection Service, Public Health England | COVID-19 Genomics UK (COG-UK) Consortium | PHE Covid Sequencing Team |
| EPI_ISL_847179 | Department of Virus and Microbiological Special Diagnostics, Statens Serum Institut, Copenhagen, Denmark | Albertsen Lab, Department of Chemistry and Bioscience, Aalborg University, Denmark | Danish Covid-19 Genome Consortium |
| EPI_ISL_847558, EPI_ISL_847560, EPI_ISL_847566, EPI_ISL_847571, EPI_ISL_847583, EPI_ISL_847664, EPI_ISL_847665, EPI_ISL_847666, EPI_ISL_847667, EPI_ISL_847668, EPI_ISL_847669, EPI_ISL_847670, EPI_ISL_847671, EPI_ISL_847715, EPI_ISL_847749, EPI_ISL_847797, EPI_ISL_847798, EPI_ISL_847799, EPI_ISL_847800, EPI_ISL_847801, EPI_ISL_847802 | | | |
| see above | California Department of Public Health | Chiu Laboratory, University of California, San Francisco | Charles Chiu, Xianding (Wayne) Deng, Candace Wang, Brian Bushnell, Scot Federman, Jill Hacker, Debra Wadford |
| EPI_ISL_847980, EPI_ISL_847981, EPI_ISL_847985, EPI_ISL_847986, EPI_ISL_848005, EPI_ISL_848007, EPI_ISL_848008, EPI_ISL_848015, EPI_ISL_848017, EPI_ISL_848018, EPI_ISL_848019, EPI_ISL_848024, EPI_ISL_848025, EPI_ISL_848026, EPI_ISL_848027, EPI_ISL_848028, EPI_ISL_848029, EPI_ISL_848030, EPI_ISL_848031, EPI_ISL_848032, EPI_ISL_848033, EPI_ISL_848034, EPI_ISL_848035, EPI_ISL_848036, EPI_ISL_848045, EPI_ISL_848047, EPI_ISL_848054, EPI_ISL_848059, EPI_ISL_848060, EPI_ISL_848061 | | | |
| see above | Michigan Department of Health and Human Services, Bureau of Laboratories | Michigan Department of Health and Human Services, Bureau of Laboratories | Blankenship HM, Riner D, Soehnlen MK |
| EPI_ISL_848223, EPI_ISL_848227, EPI_ISL_848228, EPI_ISL_848236, EPI_ISL_848414, EPI_ISL_848415, EPI_ISL_848416, EPI_ISL_848417, EPI_ISL_848418, EPI_ISL_848419, EPI_ISL_848420, EPI_ISL_848421, EPI_ISL_848422, EPI_ISL_848423, EPI_ISL_848424, EPI_ISL_848439, EPI_ISL_848447, EPI_ISL_848462, EPI_ISL_848530, EPI_ISL_848531, EPI_ISL_848532, EPI_ISL_848533, EPI_ISL_848534 | | | |
| see above | Illinois Department of Public Health | Gagnon Lab, Southern Illinois University | Keith Gagnon |
| EPI_ISL_848674, EPI_ISL_848675, EPI_ISL_848676, EPI_ISL_848677, EPI_ISL_848678, EPI_ISL_848679, EPI_ISL_848680, EPI_ISL_848681, EPI_ISL_848682, EPI_ISL_848683, EPI_ISL_848684, EPI_ISL_848685, EPI_ISL_848686, EPI_ISL_848687, EPI_ISL_848688, EPI_ISL_848689, EPI_ISL_848690, EPI_ISL_848691, EPI_ISL_848692, EPI_ISL_848693, EPI_ISL_848694, EPI_ISL_848695, EPI_ISL_848696, EPI_ISL_848697, EPI_ISL_848698, EPI_ISL_848699, EPI_ISL_848700, EPI_ISL_848701, EPI_ISL_848702, EPI_ISL_848703, EPI_ISL_848704, EPI_ISL_848705, EPI_ISL_848706, EPI_ISL_848707, EPI_ISL_848708, EPI_ISL_848709, EPI_ISL_848710, EPI_ISL_848711, EPI_ISL_848712, EPI_ISL_848713, EPI_ISL_848714, EPI_ISL_848715, EPI_ISL_848716 | | | |
| see above | Florida Bureau of Public Health Laboratories | Florida Bureau of Public Health Laboratories | Sarah Schmedes, Jason Blanton |
| EPI_ISL_849387, EPI_ISL_849390, EPI_ISL_849392, EPI_ISL_849396, EPI_ISL_849397, EPI_ISL_849399, EPI_ISL_849400, EPI_ISL_849401, EPI_ISL_849402, EPI_ISL_849405, EPI_ISL_849414, EPI_ISL_849419, EPI_ISL_849423, EPI_ISL_849432, EPI_ISL_849439, EPI_ISL_849461, EPI_ISL_849463, EPI_ISL_849526, EPI_ISL_849527, EPI_ISL_849528, EPI_ISL_849529, EPI_ISL_849530, EPI_ISL_849531, EPI_ISL_849563, EPI_ISL_849573, EPI_ISL_849574, EPI_ISL_849587, EPI_ISL_849624, EPI_ISL_849625, EPI_ISL_849626 | | | |
| see above | Washington State Department of Health | Seattle Flu Study | Deborah A. Nickerson, Chris D. Frazar, Jover Lee, Benjamin Pelle, Matthew Richardson, Amanda Adler, Elisabeth Brandstetter, Peter D. Han, Kairsten Fay, Misja Ilcisin, Kirsten Lacombe, Thomas R. Sibley, Melissa Truong, Caitlin R. Wolf, Romesh Gautom, Geoff Melly, Brian Hiatt, Philip Dykema, Scott Lindquist, Michael Boeckh, Janet A. Englund, Michael Famulare, Barry R. Lutz, Mark J. Rieder, Lea M. Starita, Matthew Thompson, Helen Y. Chu, Jay Shendure, Trevor Bedford |
| EPI_ISL_849953, EPI_ISL_849954, EPI_ISL_850012, EPI_ISL_850013, EPI_ISL_850014, EPI_ISL_850015, EPI_ISL_850016, EPI_ISL_850017, EPI_ISL_850018 | Santa Clara County Public Health Laboratory | Chan-Zuckerberg Biohub | CZB Cliahub Consortium |
| EPI_ISL_850114, EPI_ISL_850115, EPI_ISL_850116, EPI_ISL_850117, EPI_ISL_850118, EPI_ISL_850119, EPI_ISL_850120, EPI_ISL_850121, EPI_ISL_850122, EPI_ISL_850123, EPI_ISL_850124, EPI_ISL_850125, EPI_ISL_850126, EPI_ISL_850127, EPI_ISL_850128, EPI_ISL_850129, EPI_ISL_850130, EPI_ISL_850131, EPI_ISL_850132, EPI_ISL_850133, EPI_ISL_850134, EPI_ISL_850135, EPI_ISL_850136, EPI_ISL_850137, EPI_ISL_850138, EPI_ISL_850139, EPI_ISL_850140, EPI_ISL_850141, EPI_ISL_850142, EPI_ISL_850143, EPI_ISL_850144, EPI_ISL_850145, EPI_ISL_850146, EPI_ISL_850147, EPI_ISL_850148, EPI_ISL_850149, EPI_ISL_850150, EPI_ISL_850151, EPI_ISL_850152, EPI_ISL_850153, EPI_ISL_850154, EPI_ISL_850155, EPI_ISL_850156, EPI_ISL_850157, EPI_ISL_850158, EPI_ISL_850159, EPI_ISL_850160, EPI_ISL_850161, EPI_ISL_850162, EPI_ISL_850163, EPI_ISL_850164, EPI_ISL_850165, EPI_ISL_850166, EPI_ISL_850167, EPI_ISL_850168, EPI_ISL_850169, EPI_ISL_850170, EPI_ISL_850171, EPI_ISL_850172, EPI_ISL_850173, EPI_ISL_850174, EPI_ISL_850175, EPI_ISL_850179 | | | |
| see above | Renegade | Chan-Zuckerberg Biohub | CZB Cliahub Consortium |
| EPI_ISL_852602, EPI_ISL_852603 | Max von Pettenkofer Institute, Virology, National Reference Center for Retroviruses, LMU München | Laboratory for Functional Genome Analysis, Dept. Genomics, Gene Center of the LMU Munich | Max Muenchhoff, Stefan Krebs, Alexander Graf, Oliver Keppler, Helmut Blum |
| EPI_ISL_852953 | Hospital General Universitario Gregorio Marañón | SeqCOVID-SPAIN consortium/IBV(CSIC) | Darío García de Viedma, Laura Pérez-Lago, Pedro J Sola-Campoy, Sergio Buenestado-Serrano, Marta Herranz, Victor Manuel de la Cueva, Julia Suárez, Pilar Catalán, Patricia Muñoz and SeqCOVID-SPAIN consortium |
| EPI_ISL_853330, EPI_ISL_853333, EPI_ISL_853334, EPI_ISL_853337, EPI_ISL_853339, EPI_ISL_853340 | UPMC Clinical Microbiology Laboratory | Microbial Genome Sequencing Center; Microbial Genomic Epidemiology Laboratory | Mustapha M. Mustapha, Jane W. Marsh, Dan Snyder, Marissa P. Griffith, Stephanie L. Mitchell, Vatsala R. Srinivasa, Kady D. Waggle, Chinelo Ezeonwuku, Vaughn S. Cooper, Lee H. Harrison |
| EPI_ISL_853417, EPI_ISL_853418, EPI_ISL_853419, EPI_ISL_853420, EPI_ISL_853421, EPI_ISL_853422, EPI_ISL_853423, EPI_ISL_853424, EPI_ISL_853425, EPI_ISL_853428, EPI_ISL_853429, EPI_ISL_853430, EPI_ISL_853431, EPI_ISL_853432, EPI_ISL_853433, EPI_ISL_853434, EPI_ISL_853435, EPI_ISL_853436, EPI_ISL_853437, EPI_ISL_853438, EPI_ISL_853439, EPI_ISL_853440, EPI_ISL_853441, EPI_ISL_853442, EPI_ISL_853443, EPI_ISL_853444, EPI_ISL_853445, EPI_ISL_853446, EPI_ISL_853447, EPI_ISL_853448, EPI_ISL_853449, EPI_ISL_853450, EPI_ISL_853451, EPI_ISL_853452, EPI_ISL_853453, EPI_ISL_853454, EPI_ISL_853455, EPI_ISL_853456, EPI_ISL_853457, EPI_ISL_853458, EPI_ISL_853459, EPI_ISL_853460, EPI_ISL_853461, EPI_ISL_853462, EPI_ISL_853463, EPI_ISL_853464, EPI_ISL_853465, EPI_ISL_853466, EPI_ISL_853467, EPI_ISL_853468, EPI_ISL_853469, EPI_ISL_853470, EPI_ISL_853471, EPI_ISL_853472, EPI_ISL_853473, EPI_ISL_853474, EPI_ISL_853475, EPI_ISL_853476, EPI_ISL_853477, EPI_ISL_853478, EPI_ISL_853479, EPI_ISL_853480, EPI_ISL_853481, EPI_ISL_853482, EPI_ISL_853483, EPI_ISL_853484, EPI_ISL_853485, EPI_ISL_853486, EPI_ISL_853487, EPI_ISL_853488, EPI_ISL_853489, EPI_ISL_853490, EPI_ISL_853491, EPI_ISL_853492, EPI_ISL_853493, EPI_ISL_853494, EPI_ISL_853495, EPI_ISL_853496, EPI_ISL_853497, EPI_ISL_853498, EPI_ISL_853499, EPI_ISL_853500, EPI_ISL_853501, EPI_ISL_853502, EPI_ISL_853503, EPI_ISL_853504, EPI_ISL_853505, EPI_ISL_853506, EPI_ISL_853507, EPI_ISL_853508, EPI_ISL_853509, EPI_ISL_853510, EPI_ISL_853511, EPI_ISL_853512, EPI_ISL_853513, EPI_ISL_853514, EPI_ISL_853515, EPI_ISL_853516, EPI_ISL_853517, EPI_ISL_853518, EPI_ISL_853519, EPI_ISL_853520, EPI_ISL_853521, EPI_ISL_853522, EPI_ISL_853523, EPI_ISL_853524, EPI_ISL_853525, EPI_ISL_853526, EPI_ISL_853527, EPI_ISL_853528, EPI_ISL_853529, EPI_ISL_853530, EPI_ISL_853531, EPI_ISL_853532, EPI_ISL_853534, EPI_ISL_853535, EPI_ISL_853536, EPI_ISL_853537, EPI_ISL_853538, EPI_ISL_853539, EPI_ISL_853540, EPI_ISL_853541, EPI_ISL_853542, EPI_ISL_853543, EPI_ISL_853544, EPI_ISL_853545, EPI_ISL_853560, EPI_ISL_853561, EPI_ISL_853562, EPI_ISL_853563, EPI_ISL_853564, EPI_ISL_853565 | | | |
| see above | Santa Clara County Public Health Laboratory | Chan-Zuckerberg Biohub | CZB Cliahub Consortium |
| EPI_ISL_853814, EPI_ISL_853930, EPI_ISL_853934, EPI_ISL_853940 | Department of Microbiology, University Innsbruck | Bergthaler laboratory, CeMM Research Center for Molecular Medicine of the Austrian Academy of Sciences | Lukas Endler, Alexandra Popa, Benedikt Agerer, Jakob-Wendelin Genger, Alexander Lercher, Anna Schedl, Thomas Penz, Michael Schuster, Jan Laine, Martin Senekowitsch, Christoph Bock, Andreas Bergthaler |
| EPI_ISL_854440 | SARATOGA HOSPITAL LABORATORY | Wadsworth Center, New York State Department of Health | Kirsten St. George, Daryl M. Lamson, Alexis Russel, Matthew Shudt, Melissa A Leisner, Jonathan Plitnick, Navjot Singh, John Kelly, Erasmus Schneider, Erica Lasek-Nesselquist |
| EPI_ISL_854625, EPI_ISL_854626, EPI_ISL_854627, EPI_ISL_854628, EPI_ISL_854629, EPI_ISL_854630, EPI_ISL_854631, EPI_ISL_854632, EPI_ISL_854633, EPI_ISL_854634, EPI_ISL_854635, EPI_ISL_854636, EPI_ISL_854637, EPI_ISL_854638, EPI_ISL_854639, EPI_ISL_854640, EPI_ISL_854641, EPI_ISL_854642, EPI_ISL_854643, EPI_ISL_854644, EPI_ISL_854645, EPI_ISL_854646, EPI_ISL_854647, EPI_ISL_854648, EPI_ISL_854649, EPI_ISL_854650, EPI_ISL_854651, EPI_ISL_854652, EPI_ISL_854653, EPI_ISL_854654, EPI_ISL_854655, EPI_ISL_854656, EPI_ISL_854657, EPI_ISL_854658, EPI_ISL_854659, EPI_ISL_854660, EPI_ISL_854661, EPI_ISL_854662, EPI_ISL_854663, EPI_ISL_854664, EPI_ISL_854665, EPI_ISL_854666, EPI_ISL_854667, EPI_ISL_854668, EPI_ISL_854669, EPI_ISL_854670, EPI_ISL_854671, EPI_ISL_854672, EPI_ISL_854673, EPI_ISL_854674, EPI_ISL_854675, EPI_ISL_854676, EPI_ISL_854677, EPI_ISL_854678, EPI_ISL_854679, EPI_ISL_854680, EPI_ISL_854681, EPI_ISL_854682, EPI_ISL_854683, EPI_ISL_854684, EPI_ISL_854685, EPI_ISL_854686, EPI_ISL_854687, EPI_ISL_854688, EPI_ISL_854689, EPI_ISL_854690, EPI_ISL_854691, EPI_ISL_854692, EPI_ISL_854693, EPI_ISL_854694, EPI_ISL_854695, EPI_ISL_854696, EPI_ISL_854697, EPI_ISL_854698, EPI_ISL_854699, EPI_ISL_854700, EPI_ISL_854701, EPI_ISL_854702, EPI_ISL_854703, EPI_ISL_854704, EPI_ISL_854705, EPI_ISL_854706, EPI_ISL_854707, EPI_ISL_854708, EPI_ISL_854709, EPI_ISL_854710, EPI_ISL_854711, EPI_ISL_854712, EPI_ISL_854713, EPI_ISL_854716, EPI_ISL_854717, EPI_ISL_854718, EPI_ISL_854719, EPI_ISL_854720, EPI_ISL_854721, EPI_ISL_854722, EPI_ISL_854723, EPI_ISL_854724, EPI_ISL_854725, EPI_ISL_854726, EPI_ISL_854727, EPI_ISL_854728, EPI_ISL_854729, EPI_ISL_854730, EPI_ISL_854731, EPI_ISL_854732, EPI_ISL_854733, EPI_ISL_854734, EPI_ISL_854735, EPI_ISL_854737, EPI_ISL_854738, EPI_ISL_854739, EPI_ISL_854740, EPI_ISL_854741, EPI_ISL_854742, EPI_ISL_854743, EPI_ISL_854744 | | | |
| see above | Public Health Ontario Laboratory | Public Health Ontario Laboratory | Vanessa G Allen, Philip Banh, Yao Chen, Richard de Borja, Alireza Eshaghi, Nahuel Fittipaldi, Christine Frantz, Jonathan B Gubbay, Jennifer L Guthrie, Lawrence Heisler, Esha Joshi, Michael Laszloffy, Aimin Li, Michael CY Li, Dean Maxwell, Sandeep Nagra, Samir N Patel, Jared Simpson, Karthikeyan Sivaraman, Ashleigh Sullivan, Yogi Sundaravadanam, Sarah Teatero, Matthew Watson, Andre Villegas, Sandra Zittermann |

| | | | |
|---|---|---|---|
| EPI_ISL_856772, EPI_ISL_856773 | Servicio Virosis Respiratorias-Departamento Virología-INEI | Instituto Nacional Enfermedades Infecciosas C.G.Malbran | Baumeister E., Avaro M., Benedetti E., Russo M., Dattero ME, Pontoriero A., Cisterna D., Molina V., Perandones C., Tuduri E., Lorenzo F., Poklepovich T., Campos J. |
| EPI_ISL_857500, EPI_ISL_857501, EPI_ISL_857502, EPI_ISL_857503, EPI_ISL_857504 | Swiss National Reference Centre for Influenza | Swiss National Reference Centre for Influenza | Ana Rita Goncalves,Samuel Cordey, Laurent Kaiser, Lorenzo Cerutti, Henri Pegeot, Melyssa Elies, Keith Harshman, Ioannis Xenarios, Emmanouil Dermitzakis |
| EPI_ISL_857527 | Swiss National Reference Centre for Influenza | Swiss National Reference Centre for Influenza | Tim Roloff, Ana Rita Gonçalves, Madlen Stange, Helena MB Seth-Smith, Alfredo Mari, Karoline Leuzinger, Julia Bielicki, Manuel Battegay, Hans Hirsch, Laurent Kaiser, Adrian Egli |
| EPI_ISL_859931, EPI_ISL_859932, EPI_ISL_859933, EPI_ISL_860090, EPI_ISL_860091 | BTC, Khalifa University | BTC, Khalifa University | Al Safar et al |
| EPI_ISL_860804 | WHO/Minsk | Charité Universitätsmedizin Berlin, Institut für Virologie | Victor M Corman, Barbara Mühlemann, Jörn Beheim-Schwarzbach, Talitha Veith, Julia Tesch, Tobias Bleicker, Julia Schneider, Shmialiova Natallia, Sivets Natallia, Terry Jones, Christian Drosten |
| EPI_ISL_861780 | Hospital General Universitario Gregorio Marañón | SeqCOVID-SPAIN consortium/IBV(CSIC) | Darío García de Viedma, Laura Pérez-Lago, Pedro J Sola-Campoy, Sergio Buenestado-Serrano, Marta Herranz, Victor Manuel de la Cueva, Julia Suárez, Pilar Catalán, Patricia Muñoz and SeqCOVID-SPAIN consortium |
| EPI_ISL_862588 | Hospital Clínic | Instituto de Salud Carlos III | Iglesias-Caballero, M.Camarero, S. Molinero Calamita, M. González-Esguevillas, M. Pozo, F. Casas, I. Jiménez, P. Jiménez, M. Zaballos, A. Monzón, S. Varona, S. Juliá, M. Cuesta, I. Marcos, M.A. |
| EPI_ISL_864577 | Institute of Medical Microbiology and Hospital Hygiene | Institute of Medical Microbiology and Hospital Hygiene | Prof. Dr. Achim Kaasch, Aljoscha Tersteegen |
| EPI_ISL_865708, EPI_ISL_865709, EPI_ISL_865710, EPI_ISL_865711, EPI_ISL_865712 | University College London, Great Ormond Street Hospital for Children NHS Foundation Trust, Imperial College Healthcare NHS Trust | COVID-19 Genomics UK (COG-UK) Consortium | Sergi Castellano, Rachel Williams, Mark Kristiansen, Paola Resende Silva, Sunando Roy, Tony Brooks, Helena Tutill, Paola Niola, Patricia Dyal, Charlotte Williams, Leysa Forrest, Yasmin Panchbhaya, Jacqueline Findlay, Samuel Weeks, Julianne Brown, Kathryn Harris, Paul Randell, James Price, Alison Holmes, Judith Breuer |
| EPI_ISL_866453 | Northumbria University / South Tees Hospitals NHS Foundation Trust / North Cumbria Integrated Care NHS Foundation Trust / North Tees and Hartlepool NHS Foundation Trust / Newcastle Hospitals NHS Foundation Trust | COVID-19 Genomics UK (COG-UK) Consortium | Darren L Smith,Andrew Nelson,Matthew Bashton,Greg R Young,Joshua Loh,John Allan,Mohammad A Tariq,Giles S Holt,Gary Black,Wen C Yew,Lynn Dover,Paul Baker,Steve Liggett,Sarah Essex,Jane Greenaway,Debra Padgett,Clive Graham,Garren Scott,Edward Barton,Emma Swindells,Brendan Payne,Jennifer Collins,Yusri Taha,Gary Eltringham |
| EPI_ISL_866804, EPI_ISL_866813, EPI_ISL_866827 | Quadram Institute Bioscience | COVID-19 Genomics UK (COG-UK) Consortium | Dave J. Baker, Gemma L. Kay, Alp Aydin, Thanh Le-Viet, Steven Rudder, Ana P. Tedim, Anastasia Kolyva, Maria Diaz, Leonardo de Oliveira Martins, Nabil-Fareed Alikhan, Lizzie Meadows, Rachael Stanley, Ngozi Elumogo, Muhammed Yasir, Nicholas M. Thomson, Alexander J Trotter, Rachel Gilroy, Samuel Bloomfield, Claire Stuart, Andrew Bell, Reenesh Prakash, Samir Dervisevic, Alison E. Mather, John Wain, Mark Webber, Andrew J. Page, Justin O'Grady |
| EPI_ISL_867272, EPI_ISL_867273, EPI_ISL_867274, EPI_ISL_867275, EPI_ISL_867276, EPI_ISL_867277, EPI_ISL_867427 | Originating lab: Wales Specialist Virology Centre Sequencing lab: Pathogen Genomics Unit | Public Health Wales Microbiology Cardiff Wales Specialist Virology Centre | Catherine Moore, Johnathan Evans, Laura Gifford, Malorie Perry, Simon Cottrell, Angela Marchbank, Alec Birchley, Alexander Adams, Amy Gaskin, Bree Gatica-Wilcox, Jason Coombes, Joel Southgate, Lauren Gilbert, Lee Graham, Nicole Pacchiarini, Sara Kumziene-Summerhayes, Sarah Taylor, Sophie Jones, Sara Rey, Matthew Bull, Joanne Watkins, Sally Corden, Tom Connor |
| EPI_ISL_871781, EPI_ISL_871782 | Department of Virus and Microbiological Special Diagnostics, Statens Serum Institut, Copenhagen, Denmark | Aalborg University | Danish Covid-19 Genome Consortium |
| EPI_ISL_872571 | Kamchatka Regional Children's Infectious Diseases Hospital | WHO National Influenza Centre Russian Federation | Andrey Komissarov, Artem Fadeev, Anna Ivanova, Kseniya Komissarova, Dmitry Bazhenov, Mikhail Bakaev, Daria Danilenko, Ksenia Safina, Elena Nabieva, Georgii Bazykin, Dmitry Lioznov |
| EPI_ISL_872611, EPI_ISL_872612 | Nigeria Centre for Disease Control (NCDC) | African Centre of Excellence for Genomics of Infectious Diseases (ACEGID), Redeemer's University | Oluniyi P.E. et al |
| EPI_ISL_872644, EPI_ISL_872648, EPI_ISL_872654, EPI_ISL_872667, EPI_ISL_872673, EPI_ISL_872677 | University of Massachusetts Medical School | Infectious Disease Program, Broad Institute of Harvard and MIT | Lemieux,J.E., Siddle,K.J., Ward,D., Ellison,R., Adams,G., Gladden-Young,A., Lagerborg,K., Rudy,M., DeRuff,K., Carter,A., Normandin,E., Bauer,M., Reilly,S., Tomkins-Tinch,C., Loreth,C., Chaluvadi,S., Birren,B.W., Gallagher,G., Smole,S., Park,D.J., MacInnis,B.L., and Sabeti,P.C. |
| EPI_ISL_872704, EPI_ISL_872705, EPI_ISL_872706, EPI_ISL_872707, EPI_ISL_872708, EPI_ISL_872709, EPI_ISL_872710, EPI_ISL_872711, EPI_ISL_872712, EPI_ISL_872713, EPI_ISL_872714, EPI_ISL_872715, EPI_ISL_872716, EPI_ISL_872718, EPI_ISL_872719, EPI_ISL_872721, EPI_ISL_872723 | | | |
| see above | Rhode Island Department of Health | Infectious Disease Program, Broad Institute of Harvard and MIT | Lemieux,J.E., Siddle,K.J., Huard,R., King,E., Azevedo,K., Miller,A., Adams,G., Gladden-Young,A., Lagerborg,K., Rudy,M., DeRuff,K., Carter,A., Normandin,E., Bauer,M., Reilly,S., Tomkins-Tinch,C., Loreth,C., Chaluvadi,S., Birren,B.W., Gallagher,G., Smole,S., Park,D.J., MacInnis,B.L., and Sabeti,P.C. |
| EPI_ISL_872918 | WHO National Influenza Centre Russian Federation | WHO National Influenza Centre Russian Federation | Andrey Komissarov, Artem Fadeev, Anna Ivanova, Kseniya Komissarova, Dmitry Bazhenov, Mikhail Bakaev, Daria Danilenko, Ksenia Safina, Elena Nabieva, Georgii Bazykin, Dmitry Lioznov |
| EPI_ISL_873124, EPI_ISL_873129, EPI_ISL_873130, EPI_ISL_873133, EPI_ISL_873134, EPI_ISL_873139, EPI_ISL_873147 | University of Michigan Clinical Microbiology Laboratory | Lauring Lab, University of Michigan, Department of Microbiology and Immunology | Valesano |
| EPI_ISL_873211, EPI_ISL_873212 | M Health Fairview | Minnesota Department of Health, Public Health Laboratory | Alexandra Lorentz, Jacob Garfin, Matt Plumb, and Xiong Wang |
| EPI_ISL_875690, EPI_ISL_875691, EPI_ISL_875692, EPI_ISL_875693, EPI_ISL_875694, EPI_ISL_875695, EPI_ISL_875696, EPI_ISL_875697, EPI_ISL_875698, EPI_ISL_875699, EPI_ISL_875700, EPI_ISL_875701, EPI_ISL_875702, EPI_ISL_875703, EPI_ISL_875704, EPI_ISL_875705, EPI_ISL_875706, EPI_ISL_875707, EPI_ISL_875708, EPI_ISL_875709, EPI_ISL_875710, EPI_ISL_875711, EPI_ISL_875712, EPI_ISL_875713, EPI_ISL_875714, EPI_ISL_875715, EPI_ISL_875716, EPI_ISL_875717, EPI_ISL_875718, EPI_ISL_875719, EPI_ISL_875720, EPI_ISL_875721, EPI_ISL_875722, EPI_ISL_875723, EPI_ISL_875724, EPI_ISL_875725, EPI_ISL_875726, EPI_ISL_875727, EPI_ISL_875728, EPI_ISL_875729, EPI_ISL_875730, EPI_ISL_875731, EPI_ISL_875732, EPI_ISL_875733, EPI_ISL_875734, EPI_ISL_875735, EPI_ISL_875736, EPI_ISL_875737, EPI_ISL_875738, EPI_ISL_875739, EPI_ISL_875740, EPI_ISL_875741, EPI_ISL_875742, EPI_ISL_875743, EPI_ISL_875744, EPI_ISL_875745, EPI_ISL_875746, EPI_ISL_875747, EPI_ISL_875748, EPI_ISL_875749, EPI_ISL_875750, EPI_ISL_875751, EPI_ISL_875752, EPI_ISL_875753, EPI_ISL_875754, EPI_ISL_875755, EPI_ISL_875756, EPI_ISL_875757, EPI_ISL_875758, EPI_ISL_875759, EPI_ISL_875760, EPI_ISL_875761, EPI_ISL_875762, EPI_ISL_875763, EPI_ISL_875764, EPI_ISL_875765, EPI_ISL_875766, EPI_ISL_875767, EPI_ISL_875768, EPI_ISL_875769, EPI_ISL_875770, EPI_ISL_875771, EPI_ISL_875772, EPI_ISL_875773, EPI_ISL_875774, EPI_ISL_875775, EPI_ISL_875776, EPI_ISL_875777, EPI_ISL_875778, EPI_ISL_875779, EPI_ISL_875780, EPI_ISL_875781, EPI_ISL_875782, EPI_ISL_875783, EPI_ISL_875784, EPI_ISL_875785, EPI_ISL_875786, EPI_ISL_875787, EPI_ISL_875788, EPI_ISL_875789, EPI_ISL_875790, EPI_ISL_875791, EPI_ISL_875792, EPI_ISL_875793, EPI_ISL_875794, EPI_ISL_875795, EPI_ISL_875796, EPI_ISL_875797, EPI_ISL_875798, EPI_ISL_875799, EPI_ISL_875800, EPI_ISL_875801, EPI_ISL_875802, EPI_ISL_875803, EPI_ISL_875804, EPI_ISL_875805, EPI_ISL_875806, EPI_ISL_875807, EPI_ISL_875808, EPI_ISL_875809, EPI_ISL_875810, EPI_ISL_875811, EPI_ISL_875812, EPI_ISL_875813, EPI_ISL_875816, EPI_ISL_875817, EPI_ISL_875818, EPI_ISL_875819, EPI_ISL_875820, EPI_ISL_875821, EPI_ISL_875822, EPI_ISL_875823, EPI_ISL_875824, EPI_ISL_875825, EPI_ISL_875826, EPI_ISL_875827, EPI_ISL_875828, EPI_ISL_875829, EPI_ISL_875830, EPI_ISL_875831, EPI_ISL_875832, EPI_ISL_875833, EPI_ISL_875834, EPI_ISL_875835, EPI_ISL_875836, EPI_ISL_875837, EPI_ISL_875838, EPI_ISL_875839, EPI_ISL_875840, EPI_ISL_875841, EPI_ISL_875842, EPI_ISL_875843, EPI_ISL_875844, EPI_ISL_875845, EPI_ISL_875846, EPI_ISL_875847, EPI_ISL_875848, EPI_ISL_875849, EPI_ISL_875850, EPI_ISL_875851, EPI_ISL_875852, EPI_ISL_875853, EPI_ISL_875854, EPI_ISL_875855, EPI_ISL_875856, EPI_ISL_875857, EPI_ISL_875858, EPI_ISL_875859, EPI_ISL_875860, EPI_ISL_875861, EPI_ISL_875862, EPI_ISL_875863, EPI_ISL_875864, EPI_ISL_875865, EPI_ISL_875866, EPI_ISL_875867, EPI_ISL_875868, EPI_ISL_875869, EPI_ISL_875870 | | | |
| see above | Public Health Ontario Laboratory | Public Health Ontario Laboratory | Vanessa G Allen, Philip Banh, Yao Chen, Richard de Borja, Alireza Eshaghi, Nahuel Fittipaldi, Christine Frantz, Jonathan B Gubbay, Jennifer L Guthrie, Lawrence Heisler, Esha Joshi, Michael Laszloffy, Aimin Li, Michael CY Li, Dean Maxwell, Sandeep Nagra, Samir N Patel, Jared Simpson, Karthikeyan Sivaraman, Ashleigh Sullivan, Yogi Sundaravadanam, Sarah Teatero, Matthew Watson, Andre Villegas, Sandra Zittermann |
| EPI_ISL_876177, EPI_ISL_876192, EPI_ISL_876317 | Massachusetts State Public Health Laboratory | Massachusetts State Public Health Laboratory | Andrew Lang, Timelia Fink, Glen Gallagher, Sandra Smole |
| EPI_ISL_876549, EPI_ISL_876550, EPI_ISL_876551 | Florida Bureau of Public Health Laboratories | Florida Bureau of Public Health Laboratories | Sarah Schmedes, Jason Blanton |
| EPI_ISL_878157, EPI_ISL_878167, EPI_ISL_878170, EPI_ISL_878180, EPI_ISL_878183, EPI_ISL_878191, EPI_ISL_878202, EPI_ISL_878205, EPI_ISL_878212, EPI_ISL_878215, EPI_ISL_878226, EPI_ISL_878229, EPI_ISL_878232, EPI_ISL_878239, EPI_ISL_878248, EPI_ISL_878250, EPI_ISL_878253, EPI_ISL_878258, EPI_ISL_878260, EPI_ISL_878263, EPI_ISL_878274, EPI_ISL_878355, EPI_ISL_878360, EPI_ISL_878361, EPI_ISL_878367, EPI_ISL_878371, EPI_ISL_878374, EPI_ISL_878381, EPI_ISL_878386, EPI_ISL_878392, EPI_ISL_878393, EPI_ISL_878398 | | | |
| see above | San Diego County Public Health Laboratory | Andersen lab at Scripps Research | SEARCH Alliance San Diego with Tracy Basler, Jovan Shephard, Brett Austin |
| EPI_ISL_878557 | Robert Garry lab | Andersen lab at Scripps Research | Allison Smither, Gilberto Sabino-Santos, Patricia Snarski, Lilia Melnik, Antoinette Bell, Kaylynn Genemaras, Arnaud Drouin, Dahlene Fusco, Robert Garry with SEARCH Alliance San Diego |

| | | | |
|---|---|---|---|
| EPI_ISL_878674, EPI_ISL_878742 | Rady's Childrens Hospital | Andersen lab at Scripps Research | SEARCH Alliance San Diego with Nanda Radamchar, David Dimmock, Linda Luo, Christina Clarke, Kathryn Bouic, Teresa Mueller, Denise Malicki |
| EPI_ISL_879926, EPI_ISL_879932, EPI_ISL_879934, EPI_ISL_879942, EPI_ISL_879947, EPI_ISL_879950, EPI_ISL_879953, EPI_ISL_879959, EPI_ISL_879969, EPI_ISL_879975, EPI_ISL_879977, EPI_ISL_879990, EPI_ISL_879995, EPI_ISL_880003, EPI_ISL_880017, EPI_ISL_880022 | | | |
| see above | San Diego County Public Health Laboratory | Andersen lab at Scripps Research | SEARCH Alliance San Diego with Tracy Basler, Jovan Shephard, Brett Austin |
| EPI_ISL_882333 | Lighthouse Lab in Alderley Park | Wellcome Sanger Institute for the COVID-19 Genomics UK (COG-UK) Consortium | Jacquelyn Wynn, Mairead Hyland, The Lighthouse Lab in Alderley Park and Alex Alderton, Roberto Amato, Sonia Goncalves, Ewan Harrison, David K. Jackson, Ian Johnston, Dominic Kwiatkowski, Cordelia Langford, John Sillitoe on behalf of the Wellcome Sanger Institute COVID-19 Surveillance Team |
| EPI_ISL_882648 | Swiss National Reference Centre for Influenza Virology laboratory, CNRI | Swiss National Reference Centre for Influenza Virology laboratory, CNRI | Tim Roloff, Ana Rita Gonçalves, Madlen Stange, Helena MB Seth-Smith, Alfredo Mari, Karoline Leuzinger, Julia Bielicki, Manuel Battegay, Hans Hirsch, Laurent Kaiser, Adrian Egli |
| EPI_ISL_882764 | 1.AO Universitaria 'S. Giovanni di Dio e Ruggi D'Aragona, Scuola Medica Salernitana' Hospital / 2.UOC di Virologia e Microbiologia, Università della Campania 'L. Vanvitelli' / 3.AO Universitaria 'Federico II' Napoli Hospital / 4.AORN 'San Giuseppe Moscati' Avellino Hospital / 5.AO 'San Pio - presidio G. Rummo' Benevento Hospital / 6.AO 'Sant'Anna e San Sebastiano' Caserta Hospital / 7.PO 'Maria Santissima Addolorata' Eboli Hospital / 8.Biogem Istituto di Ricerche Genetiche | 1. Genome Research Center for Health (CRGS) / 2. Laboratory of Molecular Medicine and Genomics(LMMGe) / 3. Center for Research in Pure and Applied Mathematics (CRMPA) | Giorgio Giurato, Francesca Rizzo, Alessandro Weisz, Gianluigi Franci, Giovanni Nassa, Pasquale Pagliano, Roberta Tarallo, Elena Alexandrova, Ylenia D'Agostino, Carlo Ferravante, Jessica Lamberti, Viola Melone, Domenico Memoli, Valeria Mirici Cappa, Domenico Palumbo, Giovanni Pecoraro, Assunta Sellitto, Oriana Strianese, Ilaria Terenzi, Giuseppe Fenza, Aniello Gentile, Antonello Saccomanno, Sonia Amabile, Teresa Rocco, Annamaria Salvati, Emilia Vaccaro, Massimiliano Galdiero, Michele Cennamo, Giuseppe Portella, Maria Grazia Foti, Mariarosaria Ingino, Maria Landi, Maurizio Fumi, Vincenzo Rocco, Rita Greco, Vittoria Letizia, Arnolfo Petruzziello, Maddalena Schioppa, Gregorio Goffredi, Francesca Marciano, Michele Caraglia, Alessia Cossu, Marianna Scrima |
| EPI_ISL_883196, EPI_ISL_883198, EPI_ISL_883199, EPI_ISL_883200, EPI_ISL_883201, EPI_ISL_883202, EPI_ISL_883203, EPI_ISL_883204, EPI_ISL_883205, EPI_ISL_883206, EPI_ISL_883207, EPI_ISL_883208, EPI_ISL_883209, EPI_ISL_883210, EPI_ISL_883211, EPI_ISL_883212, EPI_ISL_883213, EPI_ISL_883214, EPI_ISL_883215, EPI_ISL_883216, EPI_ISL_883217, EPI_ISL_883218, EPI_ISL_883219, EPI_ISL_883220, EPI_ISL_883221, EPI_ISL_883222, EPI_ISL_883223, EPI_ISL_883224, EPI_ISL_883225, EPI_ISL_883227, EPI_ISL_883228, EPI_ISL_883229, EPI_ISL_883230, EPI_ISL_883231, EPI_ISL_883232, EPI_ISL_883233, EPI_ISL_883234, EPI_ISL_883235, EPI_ISL_883236, EPI_ISL_883237, EPI_ISL_883238, EPI_ISL_883239, EPI_ISL_883240, EPI_ISL_883241, EPI_ISL_883242, EPI_ISL_883243, EPI_ISL_883244, EPI_ISL_883245, EPI_ISL_883246, EPI_ISL_883247, EPI_ISL_883248, EPI_ISL_883249, EPI_ISL_883250, EPI_ISL_883251, EPI_ISL_883252, EPI_ISL_883253, EPI_ISL_883254, EPI_ISL_883255, EPI_ISL_883256, EPI_ISL_883257, EPI_ISL_883258, EPI_ISL_883259, EPI_ISL_883260, EPI_ISL_883261, EPI_ISL_883262, EPI_ISL_883263, EPI_ISL_883264, EPI_ISL_883265, EPI_ISL_883266, EPI_ISL_883267, EPI_ISL_883269, EPI_ISL_883270, EPI_ISL_883271, EPI_ISL_883272, EPI_ISL_883273, EPI_ISL_883274, EPI_ISL_883275, EPI_ISL_883276, EPI_ISL_883277, EPI_ISL_883278, EPI_ISL_883279, EPI_ISL_883280, EPI_ISL_883281, EPI_ISL_883282, EPI_ISL_883283, EPI_ISL_883284, EPI_ISL_883501, EPI_ISL_883502, EPI_ISL_883503, EPI_ISL_883504, EPI_ISL_883505, EPI_ISL_883506, EPI_ISL_883507, EPI_ISL_883508, EPI_ISL_883509, EPI_ISL_883510, EPI_ISL_883511, EPI_ISL_883512, EPI_ISL_883513, EPI_ISL_883514, EPI_ISL_883515, EPI_ISL_883516, EPI_ISL_883517, EPI_ISL_883519, EPI_ISL_883520, EPI_ISL_883521, EPI_ISL_883522, EPI_ISL_883523, EPI_ISL_883524, EPI_ISL_883525, EPI_ISL_883526, EPI_ISL_883527, EPI_ISL_883528, EPI_ISL_883529, EPI_ISL_883530, EPI_ISL_883531, EPI_ISL_883532, EPI_ISL_883533, EPI_ISL_883534, EPI_ISL_883535, EPI_ISL_883536, EPI_ISL_883537, EPI_ISL_883538, EPI_ISL_883539, EPI_ISL_883540, EPI_ISL_883541, EPI_ISL_883542, EPI_ISL_883543, EPI_ISL_883544, EPI_ISL_883545, EPI_ISL_883546, EPI_ISL_883547, EPI_ISL_883548, EPI_ISL_883549, EPI_ISL_883550, EPI_ISL_883551, EPI_ISL_883552, EPI_ISL_883553, EPI_ISL_883554, EPI_ISL_883555, EPI_ISL_883556, EPI_ISL_883557, EPI_ISL_883558, EPI_ISL_883559, EPI_ISL_883560, EPI_ISL_883561, EPI_ISL_883562, EPI_ISL_883563, EPI_ISL_883564, EPI_ISL_883565, EPI_ISL_883566, EPI_ISL_883567, EPI_ISL_883568, EPI_ISL_883569, EPI_ISL_883570, EPI_ISL_883571, EPI_ISL_883572, EPI_ISL_883573, EPI_ISL_883574, EPI_ISL_883575, EPI_ISL_883576, EPI_ISL_883577, EPI_ISL_883578, EPI_ISL_883579, EPI_ISL_883580, EPI_ISL_883581, EPI_ISL_883582, EPI_ISL_883583, EPI_ISL_883584, EPI_ISL_883585, EPI_ISL_883586, EPI_ISL_883587, EPI_ISL_883588, EPI_ISL_883589, EPI_ISL_883590, EPI_ISL_883591, EPI_ISL_883592, EPI_ISL_883593, EPI_ISL_883594, EPI_ISL_883595, EPI_ISL_883596, EPI_ISL_883597, EPI_ISL_883598, EPI_ISL_883599, EPI_ISL_883600, EPI_ISL_883601, EPI_ISL_883602, EPI_ISL_883603, EPI_ISL_883604, EPI_ISL_883605, EPI_ISL_883606, EPI_ISL_883607, EPI_ISL_883608, EPI_ISL_883609, EPI_ISL_883610, EPI_ISL_883611, EPI_ISL_883612, EPI_ISL_883613, EPI_ISL_883614, EPI_ISL_883615, EPI_ISL_883616, EPI_ISL_883617, EPI_ISL_883618, EPI_ISL_883619, EPI_ISL_883620, EPI_ISL_883621, EPI_ISL_883622, EPI_ISL_883623, EPI_ISL_883624, EPI_ISL_883625, EPI_ISL_883626, EPI_ISL_883627, EPI_ISL_883628, EPI_ISL_883629, EPI_ISL_883630, EPI_ISL_883631, EPI_ISL_883632, EPI_ISL_883633, EPI_ISL_883634, EPI_ISL_883635, EPI_ISL_883636, EPI_ISL_883637, EPI_ISL_883638, EPI_ISL_883639, EPI_ISL_883640, EPI_ISL_883641, EPI_ISL_883642, EPI_ISL_883643, EPI_ISL_883644, EPI_ISL_883645, EPI_ISL_883646, EPI_ISL_883647, EPI_ISL_883648, EPI_ISL_883649, EPI_ISL_883650, EPI_ISL_883656, EPI_ISL_883657, EPI_ISL_883658, EPI_ISL_883659, EPI_ISL_883660, EPI_ISL_883661, EPI_ISL_883662, EPI_ISL_883663, EPI_ISL_883664, EPI_ISL_883665, EPI_ISL_883666, EPI_ISL_883667, EPI_ISL_883677, EPI_ISL_883678, EPI_ISL_883679, EPI_ISL_883680, EPI_ISL_883681, EPI_ISL_883686, EPI_ISL_883687, EPI_ISL_883688, EPI_ISL_883689, EPI_ISL_883690, EPI_ISL_883691, EPI_ISL_883692, EPI_ISL_883693, EPI_ISL_883695, EPI_ISL_883723, EPI_ISL_883724, EPI_ISL_883725, EPI_ISL_883741, EPI_ISL_883742, EPI_ISL_883743, EPI_ISL_883755, EPI_ISL_883757 | | | |
| see above | Eurofins Diatherix | Hudsonalpha Genome Sequencing Center | Jane Grimwood, Melissa Williams, Lori H. Handley, Joshua Stough, Leslie Malone, Stefan Brzezinski, Ada Stewart, Teresa Jones, Jenell Webber, John Lovell, Jennifer Cart, and Jeremy Schmutz |
| EPI_ISL_884088 | Wazed Mia Science Research Center, Jahangirnagar University in collaboration with GRMDRC, Dhaka and Invent Technologies Ltd, Dhaka, Bangladesh | COVID Research Cell (CRC), Wazed Miah Science Research Center | Md. Abdul Matin, Bishajit Sarkar, Md. Asad Ullah, Yusha Araf, Nihad Adnan, Abdullah Al Nahid, Rashedul Islam, Mohammad Shahedur Rahman |
| EPI_ISL_884216, EPI_ISL_884219, EPI_ISL_884220 | Alaska State Virology Laboratory (Alaska DHHS) | Alaska State Virology Laboratory (Alaska DHHS) | Stephanie DeRonde, Lisa Smith, PhD. Jack Chen, PhD. |
| EPI_ISL_884295 | Microbiology Division, South Carolina Department of Health and Environmental Control Public Health Laboratory (SC DHEC PHL) | Microbiology Division, South Carolina Department of Health and Environmental Control Public Health Laboratory (SC DHEC PHL) | Haley Flores, Jessica Freeman |
| EPI_ISL_884884, EPI_ISL_884885, EPI_ISL_884887, EPI_ISL_884888, EPI_ISL_884898 | Santa Clara County Public Health Laboratory | Chan-Zuckerberg Biohub | CZB Cliahub Consortium |
| EPI_ISL_885135, EPI_ISL_885136 | California Institute of Technology | Chan-Zuckerberg Biohub | CZB Cliahub Consortium |
| EPI_ISL_888685, EPI_ISL_888755, EPI_ISL_888756 | KU Leuven, Rega Institute, Clinical and Epidemiological Virology | KU Leuven, Rega Institute, Clinical and Epidemiological Virology | Tony Wawina-Bokalanga, Bert Vanmechelen, Joan Marti-Carerras, Piet Maes |
| EPI_ISL_888842, EPI_ISL_888876 | Michigan Department of Health and Human Services, Bureau of Laboratories | Michigan Department of Health and Human Services, Bureau of Laboratories | Blankenship HM, Riner D, Soehnlen MK |
| EPI_ISL_888992 | RS Mitra Keluarga Gading Serpong | Eijkman Institute for Molecular Biology, Ministry of Research and Technology/National Agency for Research and Innovation | Hidayat Trimarsanto, Iskandar Adnan, Lydia V. Panggalo, Sukma Oktavianthi, Willy Agustine, Edison Johar, Frilasita A Yudhaputri, Safarina G Malik, Khin Saw Myint, Amin Soebandrio |
| EPI_ISL_889021 | RSIA Kenari Graha Medika | Eijkman Institute for Molecular Biology, Ministry of Research and Technology/National Agency for Research and Innovation | Hidayat Trimarsanto, Iskandar Adnan, Lydia V. Panggalo, Sukma Oktavianthi, Willy Agustine, Edison Johar, Frilasita A Yudhaputri, Safarina G Malik, Khin Saw Myint, Amin Soebandrio |
| EPI_ISL_889022 | RS Anna | Eijkman Institute for Molecular Biology, Ministry of Research and Technology/National Agency for Research and Innovation | Hidayat Trimarsanto, Iskandar Adnan, Lydia V. Panggalo, Sukma Oktavianthi, Willy Agustine, Edison Johar, Frilasita A Yudhaputri, Safarina G Malik, Khin Saw Myint, Amin Soebandrio |
| EPI_ISL_889542, EPI_ISL_890174, EPI_ISL_890175, EPI_ISL_890176, EPI_ISL_890178, EPI_ISL_890179, EPI_ISL_890180 | Ohio Department of Health Laboratory | Ohio Department of Health Laboratory | Holmes, Jennifer; Eric Brandt, Keoni Omura, Glen McGillivary, Caitlin McDonnell, Jade Mowery, Stephanie Mccracken, Tyler Payne, Kirtana Ramadugu, Erica Leasure, Brent Lee, Kelsey Florek, Heather Blankenship, Quanta Brown, and Tammy Bannerman |
| EPI_ISL_890185 | Academic Hospital of Gadjah Mada University (RSA UGM) | Genetics Working Group (Pokja Genetik) Faculty of Medicine, Public Health and Nursing Universitas Gadjah Mada (FK-KMK UGM); Disease Investigation Center Wates Ministry of Agriculture Indonesia; Department of Microbiology FK-KMK UGM; Laboratorium Diagnostik Yayasan Tahija World Mosquito Program (WMP) Yogyakarta Center for Tropical Medicine FK-KMK UGM; Integrated Research center FK-KMK UGM; Department of Computer Science and Electronics FMIPA UGM | Gunadi, Hendra Wibawa, Marcellus, Mohamad S. Hakim, Edwin W. Daniwijaya, Ludhang P. Rizki, Endah Supriyati, Eggi Arguni, Titik Nuryastuti, Tri Wibawa, Dwi AA Nugrahaningsih, Afiahayati, Siswanto, Kristy Iskandar, Nungki Anggorowati, Aditya Rifqi Fauzi, Fadil Fahri |
| EPI_ISL_890301, EPI_ISL_890302 | KU Leuven, Rega Institute, Clinical and Epidemiological Virology | KU Leuven, Rega Institute, Clinical and Epidemiological Virology | Tony Wawina-Bokalanga, Bert Vanmechelen, Joan Marti-Carerras, Piet Maes |
| EPI_ISL_890907, EPI_ISL_890908, EPI_ISL_890909, EPI_ISL_890913, EPI_ISL_890914, EPI_ISL_890915, EPI_ISL_890916, EPI_ISL_890917, EPI_ISL_890918 | Seattle Flu Study | Seattle Flu Study | Deborah A. Nickerson, Chris D. Frazar, Jover Lee, Benjamin Pelle, Erica Ryke, Matthew Richardson, Amanda Adler, Elisabeth Brandstetter, Peter D. Han, Kairsten Fay, Misja Ilcisin, Kirsten Lacombe, Thomas R. Sibley, Melissa Truong, Caitlin R. Wolf, Karen Cowgill, Stephanie Schrag, Jeff Duchin, Michael Boeckh, Janet A. Englund, Michael Famulare, Barry R. Lutz, Mark J. Rieder, Lea M. Starita, Matthew Thompson, Helen Y. Chu, Trevor Bedford, Jay Shendure |
| EPI_ISL_890919, EPI_ISL_890920 | Seattle Flu Study | Seattle Flu Study | Deborah A. Nickerson, Chris D. Frazar, Jover Lee, Benjamin Pelle, Erica Ryke, Matthew Richardson, Amanda Adler, Elisabeth Brandstetter, Peter D. Han, |

| | | | |
|---|---|---|---|
| | | | Kairsten Fay, Misja Ilcisin, Kirsten Lacombe, Thomas R. Sibley, Melissa Truong, Caitlin R. Wolf, Michael Boeckh, Janet A. Englund, Michael Famulare, Barry R. Lutz, Mark J. Rieder, Lea M. Starita, Matthew Thompson, Jay Shendure, Trevor Bedford, Helen Y. Chu |
| EPI_ISL_890921, EPI_ISL_890922, EPI_ISL_890923, EPI_ISL_890924, EPI_ISL_890926 | Seattle Flu Study | Seattle Flu Study | Deborah A. Nickerson, Chris D. Frazar, Jover Lee, Benjamin Pelle, Erica Ryke, Matthew Richardson, Amanda Adler, Elisabeth Brandstetter, Peter D. Han, Kairsten Fay, Misja Ilcisin, Kirsten Lacombe, Thomas R. Sibley, Melissa Truong, Caitlin R. Wolf, Karen Cowgill, Stephanie Schrag, Jeff Duchin, Michael Boeckh, Janet A. Englund, Michael Famulare, Barry R. Lutz, Mark J. Rieder, Lea M. Starita, Matthew Thompson, Helen Y. Chu, Trevor Bedford, Jay Shendure |
| EPI_ISL_890956 | Seattle Flu Study | Seattle Flu Study | Deborah A. Nickerson, Chris D. Frazar, Jover Lee, Benjamin Pelle, Erica Ryke, Matthew Richardson, Amanda Adler, Elisabeth Brandstetter, Peter D. Han, Kairsten Fay, Misja Ilcisin, Kirsten Lacombe, Thomas R. Sibley, Melissa Truong, Caitlin R. Wolf, Michael Boeckh, Janet A. Englund, Michael Famulare, Barry R. Lutz, Mark J. Rieder, Lea M. Starita, Matthew Thompson, Jay Shendure, Trevor Bedford, Helen Y. Chu |
| EPI_ISL_892280, EPI_ISL_892281, EPI_ISL_892282, EPI_ISL_892283, EPI_ISL_892284, EPI_ISL_892285, EPI_ISL_892286, EPI_ISL_892287 | Servicio de Microbiología. Hospital Universitario Donostia. OSI Donostialdea. Área de Enfermedades Infecciosas, Grupo de Infección Respiratoria y Resistencia Antimicrobiana. Instituto de Investigación Sanitaria Biodonostia | SeqCOVID-SPAIN consortium/IBV(CSIC) | Gustavo Cilla Eguiluz, Milagrosa Montes Ros, Luis Piñeiro Vázquez, Ane Sorrarain, Jose Maria Marimón and SeqCOVID-SPAIN consortium |
| EPI_ISL_892325, EPI_ISL_892326, EPI_ISL_892327, EPI_ISL_892328, EPI_ISL_892329, EPI_ISL_892330, EPI_ISL_892331, EPI_ISL_892332 | Servicio de Microbiología Clínica (Complejo Hospitalario de Navarra, Pamplona), Instituto de Investigación Sanitaria de Navarra (IdiSNA) | SeqCOVID-SPAIN consortium/IBV(CSIC) | Carmen Ezpeleta Baquedano, Ana Navascués, Ana Miqueleiz and SeqCOVID-SPAIN consortium |
| EPI_ISL_892369, EPI_ISL_892370, EPI_ISL_892371, EPI_ISL_892372, EPI_ISL_892378, EPI_ISL_892379, EPI_ISL_892380, EPI_ISL_892381, EPI_ISL_892382, EPI_ISL_892383, EPI_ISL_892384, EPI_ISL_892385, EPI_ISL_892386, EPI_ISL_892387, EPI_ISL_892388, EPI_ISL_892389, EPI_ISL_892390, EPI_ISL_892391, EPI_ISL_892392, EPI_ISL_892393, EPI_ISL_892394, EPI_ISL_892395, EPI_ISL_892396, EPI_ISL_892397, EPI_ISL_892398, EPI_ISL_892399, EPI_ISL_892400, EPI_ISL_892401, EPI_ISL_892402, EPI_ISL_892403, EPI_ISL_892404, EPI_ISL_892405, EPI_ISL_892406, EPI_ISL_892415, EPI_ISL_892416, EPI_ISL_892417, EPI_ISL_892418, EPI_ISL_892419, EPI_ISL_892420, EPI_ISL_892421, EPI_ISL_892422, EPI_ISL_892423, EPI_ISL_892424, EPI_ISL_892425, EPI_ISL_892426, EPI_ISL_892427, EPI_ISL_892428, EPI_ISL_892429, EPI_ISL_892430, EPI_ISL_892431, EPI_ISL_892432, EPI_ISL_892433, EPI_ISL_892434, EPI_ISL_892435, EPI_ISL_892436, EPI_ISL_892437, EPI_ISL_892438, EPI_ISL_892439, EPI_ISL_892440, EPI_ISL_892441, EPI_ISL_892442, EPI_ISL_892443, EPI_ISL_892444, EPI_ISL_892445, EPI_ISL_892446, EPI_ISL_892447, EPI_ISL_892448, EPI_ISL_892449, EPI_ISL_892450, EPI_ISL_892451, EPI_ISL_892452, EPI_ISL_892453, EPI_ISL_892454, EPI_ISL_892455, EPI_ISL_892456, EPI_ISL_892457, EPI_ISL_892458, EPI_ISL_892459, EPI_ISL_892460, EPI_ISL_892461, EPI_ISL_892462, EPI_ISL_892463, EPI_ISL_892464, EPI_ISL_892465, EPI_ISL_892466, EPI_ISL_892467, EPI_ISL_892468, EPI_ISL_892469, EPI_ISL_892470, EPI_ISL_892471, EPI_ISL_892472, EPI_ISL_892473, EPI_ISL_892474, EPI_ISL_892475, EPI_ISL_892476, EPI_ISL_892477, EPI_ISL_892478, EPI_ISL_892479, EPI_ISL_892480, EPI_ISL_892481, EPI_ISL_892482, EPI_ISL_892483, EPI_ISL_892484, EPI_ISL_892485, EPI_ISL_892486, EPI_ISL_892487, EPI_ISL_892488, EPI_ISL_892489, EPI_ISL_892490, EPI_ISL_892491, EPI_ISL_892492, EPI_ISL_892493, EPI_ISL_892494, EPI_ISL_892495, EPI_ISL_892496, EPI_ISL_892497, EPI_ISL_892498, EPI_ISL_892499, EPI_ISL_892500, EPI_ISL_892501, EPI_ISL_892502, EPI_ISL_892503, EPI_ISL_892504, EPI_ISL_892505, EPI_ISL_892506, EPI_ISL_892507, EPI_ISL_892508, EPI_ISL_892509, EPI_ISL_892510, EPI_ISL_892511, EPI_ISL_892512, EPI_ISL_892513, EPI_ISL_892514, EPI_ISL_892534, EPI_ISL_892535, EPI_ISL_892536, EPI_ISL_892537, EPI_ISL_892538, EPI_ISL_892539, EPI_ISL_892540, EPI_ISL_892541, EPI_ISL_892542, EPI_ISL_892543, EPI_ISL_892544, EPI_ISL_892545, EPI_ISL_892546, EPI_ISL_892547, EPI_ISL_892548, EPI_ISL_892549, EPI_ISL_892550, EPI_ISL_892551, EPI_ISL_892552, EPI_ISL_892553, EPI_ISL_892554, EPI_ISL_892555, EPI_ISL_892556, EPI_ISL_892557, EPI_ISL_892558, EPI_ISL_892559, EPI_ISL_892560 | | | |
| see above | Pathogen Genomics Center, National Institute of Infectious Diseases | Pathogen Genomics Center, National Institute of Infectious Diseases | Tsuyoshi Sekizuka, Kentaro Itokawa, Rina Tanaka, Masanori Hashino, Makoto Kuroda |
| EPI_ISL_892561, EPI_ISL_892562, EPI_ISL_892563, EPI_ISL_892564, EPI_ISL_892565 | Osaka Institute of Public Health, Morinomiya Center | Pathogen Genomics Center, National Institute of Infectious Diseases | Tsuyoshi Sekizuka, Kentaro Itokawa, Rina Tanaka, Masanori Hashino, Makoto Kuroda |
| EPI_ISL_892585, EPI_ISL_892586, EPI_ISL_892587, EPI_ISL_892588, EPI_ISL_892611, EPI_ISL_892612, EPI_ISL_892613, EPI_ISL_892614, EPI_ISL_892615, EPI_ISL_892616, EPI_ISL_892617, EPI_ISL_892618, EPI_ISL_892619, EPI_ISL_892620, EPI_ISL_892621, EPI_ISL_892622, EPI_ISL_892623, EPI_ISL_892624, EPI_ISL_892625, EPI_ISL_892626, EPI_ISL_892627, EPI_ISL_892628, EPI_ISL_892629, EPI_ISL_892630, EPI_ISL_892631, EPI_ISL_892632, EPI_ISL_892633, EPI_ISL_892634, EPI_ISL_892635, EPI_ISL_892636, EPI_ISL_892637, EPI_ISL_892638, EPI_ISL_892639, EPI_ISL_892640, EPI_ISL_892641, EPI_ISL_892642, EPI_ISL_892643, EPI_ISL_892644, EPI_ISL_892645, EPI_ISL_892646, EPI_ISL_892647, EPI_ISL_892648, EPI_ISL_892649, EPI_ISL_892650, EPI_ISL_892651, EPI_ISL_892652, EPI_ISL_892653, EPI_ISL_892654, EPI_ISL_892655, EPI_ISL_892656, EPI_ISL_892657, EPI_ISL_892658, EPI_ISL_892659, EPI_ISL_892660, EPI_ISL_892661, EPI_ISL_892662, EPI_ISL_892663, EPI_ISL_892664, EPI_ISL_892665, EPI_ISL_892666, EPI_ISL_892667, EPI_ISL_892668, EPI_ISL_892669, EPI_ISL_892670, EPI_ISL_892671, EPI_ISL_892672, EPI_ISL_892673, EPI_ISL_892674, EPI_ISL_892675, EPI_ISL_892676, EPI_ISL_892677, EPI_ISL_892678, EPI_ISL_892679, EPI_ISL_892680, EPI_ISL_892681, EPI_ISL_892682, EPI_ISL_892683, EPI_ISL_892684, EPI_ISL_892685, EPI_ISL_892686, EPI_ISL_892687, EPI_ISL_892688, EPI_ISL_892689, EPI_ISL_892690, EPI_ISL_892691, EPI_ISL_892692, EPI_ISL_892693, EPI_ISL_892694, EPI_ISL_892695, EPI_ISL_892696, EPI_ISL_892697, EPI_ISL_892698, EPI_ISL_892699, EPI_ISL_892701, EPI_ISL_892702, EPI_ISL_892703, EPI_ISL_892704, EPI_ISL_893242, EPI_ISL_893243, EPI_ISL_893244, EPI_ISL_893245, EPI_ISL_893246, EPI_ISL_893247, EPI_ISL_893248, EPI_ISL_893249, EPI_ISL_893250, EPI_ISL_893251, EPI_ISL_893255, EPI_ISL_893256, EPI_ISL_893257, EPI_ISL_893258, EPI_ISL_893259, EPI_ISL_893260, EPI_ISL_893261, EPI_ISL_893262, EPI_ISL_893263, EPI_ISL_893264, EPI_ISL_893265, EPI_ISL_893266, EPI_ISL_893281, EPI_ISL_893282, EPI_ISL_893283, EPI_ISL_893284, EPI_ISL_893285, EPI_ISL_893286, EPI_ISL_893287, EPI_ISL_893288, EPI_ISL_893289, EPI_ISL_893290, EPI_ISL_893291, EPI_ISL_893292, EPI_ISL_893293, EPI_ISL_893294, EPI_ISL_893295, EPI_ISL_893296, EPI_ISL_893297, EPI_ISL_893298, EPI_ISL_893299, EPI_ISL_893300, EPI_ISL_893301, EPI_ISL_893302, EPI_ISL_893303, EPI_ISL_893304, EPI_ISL_893305, EPI_ISL_893306, EPI_ISL_893307, EPI_ISL_893308, EPI_ISL_893309, EPI_ISL_893310, EPI_ISL_893311, EPI_ISL_893312, EPI_ISL_893313, EPI_ISL_893314, EPI_ISL_893315, EPI_ISL_893316, EPI_ISL_893317, EPI_ISL_893318, EPI_ISL_893319, EPI_ISL_893320, EPI_ISL_893321, EPI_ISL_893322, EPI_ISL_893323, EPI_ISL_893324, EPI_ISL_893325, EPI_ISL_893326, EPI_ISL_893327, EPI_ISL_893328, EPI_ISL_893329, EPI_ISL_893330, EPI_ISL_893331, EPI_ISL_893332, EPI_ISL_893333, EPI_ISL_893334, EPI_ISL_893335, EPI_ISL_893336, EPI_ISL_893337, EPI_ISL_893338, EPI_ISL_893339, EPI_ISL_893340, EPI_ISL_893341, EPI_ISL_893342, EPI_ISL_893343, EPI_ISL_893344, EPI_ISL_893345, EPI_ISL_893346, EPI_ISL_893347, EPI_ISL_893348, EPI_ISL_893349, EPI_ISL_893350, EPI_ISL_893351, EPI_ISL_893352, EPI_ISL_893353, EPI_ISL_893354, EPI_ISL_893355, EPI_ISL_893356, EPI_ISL_893357, EPI_ISL_893358, EPI_ISL_893359, EPI_ISL_893360, EPI_ISL_893361, EPI_ISL_893362, EPI_ISL_893363, EPI_ISL_893364, EPI_ISL_893365, EPI_ISL_893366 | | | |
| see above | Pathogen Genomics Center, National Institute of Infectious Diseases | Pathogen Genomics Center, National Institute of Infectious Diseases | Tsuyoshi Sekizuka, Kentaro Itokawa, Rina Tanaka, Masanori Hashino, Makoto Kuroda |
| EPI_ISL_893371, EPI_ISL_893608, EPI_ISL_893609, EPI_ISL_893610, EPI_ISL_893611, EPI_ISL_893612, EPI_ISL_893613, EPI_ISL_893614, EPI_ISL_893615, EPI_ISL_893616, EPI_ISL_893617, EPI_ISL_893618, EPI_ISL_893619, EPI_ISL_893620, EPI_ISL_893621, EPI_ISL_893622, EPI_ISL_893623, EPI_ISL_893624, EPI_ISL_893625, EPI_ISL_893626, EPI_ISL_893627, EPI_ISL_893628, EPI_ISL_893629, EPI_ISL_893630, EPI_ISL_893631, EPI_ISL_893632, EPI_ISL_893633, EPI_ISL_893634, EPI_ISL_893635, EPI_ISL_893636, EPI_ISL_893637, EPI_ISL_893638, EPI_ISL_893639, EPI_ISL_893640, EPI_ISL_893641, EPI_ISL_893642, EPI_ISL_893643, EPI_ISL_893644, EPI_ISL_893645, EPI_ISL_893646, EPI_ISL_893647, EPI_ISL_893648, EPI_ISL_893649, EPI_ISL_893650, EPI_ISL_893651, EPI_ISL_893652, EPI_ISL_893653, EPI_ISL_893654, EPI_ISL_893655, EPI_ISL_893656, EPI_ISL_893657, EPI_ISL_893658, EPI_ISL_893659, EPI_ISL_893660, EPI_ISL_893661, EPI_ISL_893662, EPI_ISL_893663, EPI_ISL_893664, EPI_ISL_893665, EPI_ISL_893666, EPI_ISL_893667, EPI_ISL_893668, EPI_ISL_893669, EPI_ISL_893670, EPI_ISL_893671, EPI_ISL_893672, EPI_ISL_893677, EPI_ISL_893678, EPI_ISL_893679, EPI_ISL_893680, EPI_ISL_893681, EPI_ISL_893682, EPI_ISL_893683, EPI_ISL_893684, EPI_ISL_893685, EPI_ISL_893686, EPI_ISL_893687, EPI_ISL_893688, EPI_ISL_893689, EPI_ISL_893690, EPI_ISL_893691, EPI_ISL_893692, EPI_ISL_893693, EPI_ISL_893694, EPI_ISL_893695, EPI_ISL_893696, EPI_ISL_893697, EPI_ISL_893698, EPI_ISL_893699, EPI_ISL_893700, EPI_ISL_893701, EPI_ISL_893702, EPI_ISL_893703, EPI_ISL_893704, EPI_ISL_893705, EPI_ISL_893706, EPI_ISL_893707, EPI_ISL_893708, EPI_ISL_893709, EPI_ISL_893710, EPI_ISL_893711, EPI_ISL_893712, EPI_ISL_893713, EPI_ISL_893714, EPI_ISL_893715, EPI_ISL_893716, EPI_ISL_893717, EPI_ISL_893718, EPI_ISL_893719 | | | |
| see above | Department of Infectious Diseases, Kobe Institute of Health | Department of Infectious Diseases, Kobe Institute of Health | Ryohei Nomoto, Tomotada Iwamoto, Tsuyoshi Sekizuka, Kentaro Itokawa, Rina Tanaka, Masanori Hashino, Makoto Kuroda |
| EPI_ISL_894278, EPI_ISL_894279 | Fukuoka Institute of Health and Environmental Sciences | Pathogen Genomics Center, National Institute of Infectious Diseases | Tsuyoshi Sekizuka, Kentaro Itokawa, Rina Tanaka, Masanori Hashino, Makoto Kuroda |
| EPI_ISL_894281, EPI_ISL_894282, EPI_ISL_894283, EPI_ISL_894284, EPI_ISL_894285, EPI_ISL_894286, EPI_ISL_894289, EPI_ISL_894292, EPI_ISL_894293, EPI_ISL_894294, EPI_ISL_894295, EPI_ISL_894298, EPI_ISL_894299, EPI_ISL_894305, EPI_ISL_894306, EPI_ISL_894307, EPI_ISL_894308, EPI_ISL_894309, EPI_ISL_894310, EPI_ISL_894311, EPI_ISL_894312, EPI_ISL_894313, EPI_ISL_894314, EPI_ISL_894315, EPI_ISL_894316, EPI_ISL_894323, EPI_ISL_894324, EPI_ISL_894325, EPI_ISL_894326, EPI_ISL_894327, EPI_ISL_894328, EPI_ISL_894329, EPI_ISL_894333, EPI_ISL_894334, EPI_ISL_894335, EPI_ISL_894336, EPI_ISL_894337, EPI_ISL_894338, EPI_ISL_894339, EPI_ISL_894340, EPI_ISL_894341, EPI_ISL_894342, EPI_ISL_894343, EPI_ISL_894344, EPI_ISL_894345, EPI_ISL_894346, EPI_ISL_894347, EPI_ISL_894348, EPI_ISL_894349, EPI_ISL_894350, EPI_ISL_894351, EPI_ISL_894352, EPI_ISL_894353 | | | |
| see above | Pathogen Genomics Center, National Institute of Infectious Diseases | Pathogen Genomics Center, National Institute of Infectious Diseases | Tsuyoshi Sekizuka, Kentaro Itokawa, Rina Tanaka, Masanori Hashino, Makoto Kuroda |
| EPI_ISL_894354, EPI_ISL_894355, EPI_ISL_894356, EPI_ISL_894357, EPI_ISL_894358, EPI_ISL_894359, EPI_ISL_894360, EPI_ISL_894361, EPI_ISL_894362, EPI_ISL_894363 | Department of Infectious Diseases, Kobe Institute of Health | Pathogen Genomics Center, National Institute of Infectious Diseases | Tsuyoshi Sekizuka, Kentaro Itokawa, Rina Tanaka, Masanori Hashino, Makoto Kuroda |
| EPI_ISL_894624, EPI_ISL_894625, EPI_ISL_894626, EPI_ISL_894627, EPI_ISL_894628, EPI_ISL_894629, EPI_ISL_894630, EPI_ISL_894631, EPI_ISL_894632, EPI_ISL_894633, EPI_ISL_894634, EPI_ISL_894635, EPI_ISL_894636, EPI_ISL_894637, EPI_ISL_894638, EPI_ISL_894639, EPI_ISL_894640, EPI_ISL_894641, EPI_ISL_894642, EPI_ISL_894643, EPI_ISL_894644, EPI_ISL_894645, EPI_ISL_894646, EPI_ISL_894647, EPI_ISL_894648, EPI_ISL_894649, EPI_ISL_894650, EPI_ISL_894651, EPI_ISL_894652, EPI_ISL_894653, EPI_ISL_894654, EPI_ISL_894655, EPI_ISL_894656, EPI_ISL_894657, EPI_ISL_894658, EPI_ISL_894659, EPI_ISL_894660, EPI_ISL_894661, EPI_ISL_894662, EPI_ISL_894663, EPI_ISL_894664, EPI_ISL_894665, EPI_ISL_894666, EPI_ISL_894667, EPI_ISL_894668, EPI_ISL_894669, EPI_ISL_894670, EPI_ISL_894671 | | | |
| see above | Pathogen Genomics Center, National Institute of Infectious Diseases | Pathogen Genomics Center, National Institute of Infectious Diseases | Tsuyoshi Sekizuka, Kentaro Itokawa, Rina Tanaka, Masanori Hashino, Makoto Kuroda |
| EPI_ISL_894695, EPI_ISL_894696, EPI_ISL_894697, EPI_ISL_894698, EPI_ISL_894699, EPI_ISL_894700, EPI_ISL_894701, EPI_ISL_894702, EPI_ISL_894703, EPI_ISL_894704, EPI_ISL_894705, EPI_ISL_894706, EPI_ISL_894707, EPI_ISL_894708, EPI_ISL_894709, EPI_ISL_894710, EPI_ISL_894711, EPI_ISL_894712, EPI_ISL_894713, EPI_ISL_894714, EPI_ISL_894715, EPI_ISL_894716, EPI_ISL_894717, EPI_ISL_894718, EPI_ISL_894719, EPI_ISL_894720, EPI_ISL_894721, EPI_ISL_894722, EPI_ISL_894723, EPI_ISL_894724, EPI_ISL_894725, EPI_ISL_894726, EPI_ISL_894727, EPI_ISL_894728, EPI_ISL_894729, EPI_ISL_894730, EPI_ISL_894731, EPI_ISL_894732, EPI_ISL_894733, EPI_ISL_894734, EPI_ISL_894735, EPI_ISL_894736, EPI_ISL_894737, EPI_ISL_894738, EPI_ISL_894739, EPI_ISL_894740, EPI_ISL_894741, EPI_ISL_894742, EPI_ISL_894743, EPI_ISL_894744, EPI_ISL_894745, EPI_ISL_894746, EPI_ISL_894747, EPI_ISL_894748, EPI_ISL_894749, EPI_ISL_894750, EPI_ISL_894751, EPI_ISL_894752, EPI_ISL_894753, EPI_ISL_894754, EPI_ISL_894755, EPI_ISL_894756, EPI_ISL_894757, EPI_ISL_894758, EPI_ISL_894759, EPI_ISL_894760, EPI_ISL_894761, EPI_ISL_894762, EPI_ISL_894763, EPI_ISL_894764, EPI_ISL_894765, EPI_ISL_894766, EPI_ISL_894767, EPI_ISL_894768, EPI_ISL_894769, EPI_ISL_894770, EPI_ISL_894771, EPI_ISL_894772, EPI_ISL_894773, EPI_ISL_894774, EPI_ISL_894775, EPI_ISL_894776, EPI_ISL_894777, EPI_ISL_894778, EPI_ISL_894779, EPI_ISL_894780, EPI_ISL_894781, EPI_ISL_894782, EPI_ISL_894783, EPI_ISL_894816 | | | |
| see above | Fukuoka Institute of Health and Environmental Sciences | Pathogen Genomics Center, National Institute of Infectious Diseases | Tsuyoshi Sekizuka, Kentaro Itokawa, Rina Tanaka, Masanori Hashino, Makoto Kuroda |

| | | | |
|---|---|---|---|
| EPI_ISL_894820, EPI_ISL_894821, EPI_ISL_894822, EPI_ISL_894823, EPI_ISL_894824, EPI_ISL_894825, EPI_ISL_894826, EPI_ISL_894827, EPI_ISL_894828, EPI_ISL_894829, EPI_ISL_894830, EPI_ISL_894831, EPI_ISL_894832, EPI_ISL_894833, EPI_ISL_894834, EPI_ISL_894835, EPI_ISL_894836, EPI_ISL_894837, EPI_ISL_894838, EPI_ISL_894839, EPI_ISL_894840, EPI_ISL_894841, EPI_ISL_894842, EPI_ISL_894843, EPI_ISL_894844, EPI_ISL_894845, EPI_ISL_894846, EPI_ISL_894847, EPI_ISL_894848, EPI_ISL_894849, EPI_ISL_894850, EPI_ISL_894851, EPI_ISL_894852, EPI_ISL_894853, EPI_ISL_894854, EPI_ISL_894855, EPI_ISL_894856, EPI_ISL_894857, EPI_ISL_894858, EPI_ISL_894859, EPI_ISL_894860, EPI_ISL_894861, EPI_ISL_894862, EPI_ISL_894863, EPI_ISL_894864, EPI_ISL_894865, EPI_ISL_894866, EPI_ISL_894867, EPI_ISL_894868, EPI_ISL_894869, EPI_ISL_894870, EPI_ISL_894871, EPI_ISL_894872, EPI_ISL_894873, EPI_ISL_894874, EPI_ISL_894875, EPI_ISL_894876, EPI_ISL_894877, EPI_ISL_894878, EPI_ISL_894879, EPI_ISL_894880, EPI_ISL_894881, EPI_ISL_894882, EPI_ISL_894883, EPI_ISL_894884, EPI_ISL_894885, EPI_ISL_894886, EPI_ISL_894887, EPI_ISL_894888, EPI_ISL_894913, EPI_ISL_894914, EPI_ISL_894915, EPI_ISL_894916, EPI_ISL_894917, EPI_ISL_894918, EPI_ISL_894919, EPI_ISL_894920, EPI_ISL_894921, EPI_ISL_894922, EPI_ISL_894923, EPI_ISL_894924, EPI_ISL_894925, EPI_ISL_894926, EPI_ISL_894927, EPI_ISL_894928, EPI_ISL_894929, EPI_ISL_894930, EPI_ISL_894931, EPI_ISL_894933, EPI_ISL_894934, EPI_ISL_894935, EPI_ISL_894936, EPI_ISL_894937, EPI_ISL_894938, EPI_ISL_894939, EPI_ISL_894940, EPI_ISL_894941, EPI_ISL_894942, EPI_ISL_894959, EPI_ISL_894960, EPI_ISL_894961, EPI_ISL_894962, EPI_ISL_894963 | | | |
| see above | Pathogen Genomics Center, National Institute of Infectious Diseases | Pathogen Genomics Center, National Institute of Infectious Diseases | Tsuyoshi Sekizuka, Kentaro Itokawa, Rina Tanaka, Masanori Hashino, Makoto Kuroda |
| EPI_ISL_895013 | Niigata Prefectural Institute of Public Health and Environmental Sciences | Pathogen Genomics Center, National Institute of Infectious Diseases | Tsuyoshi Sekizuka, Kentaro Itokawa, Rina Tanaka, Masanori Hashino, Makoto Kuroda |
| EPI_ISL_895023, EPI_ISL_895024, EPI_ISL_895025, EPI_ISL_895026, EPI_ISL_895027, EPI_ISL_895028, EPI_ISL_895029, EPI_ISL_895030, EPI_ISL_895031, EPI_ISL_895032, EPI_ISL_895033, EPI_ISL_895034, EPI_ISL_895035, EPI_ISL_895036, EPI_ISL_895037, EPI_ISL_895038, EPI_ISL_895039, EPI_ISL_895040, EPI_ISL_895041, EPI_ISL_895042, EPI_ISL_895043, EPI_ISL_895044, EPI_ISL_895045, EPI_ISL_895046, EPI_ISL_895047, EPI_ISL_895048, EPI_ISL_895049, EPI_ISL_895050, EPI_ISL_895051, EPI_ISL_895052, EPI_ISL_895053, EPI_ISL_895054, EPI_ISL_895055, EPI_ISL_895056, EPI_ISL_895057, EPI_ISL_895058, EPI_ISL_895059, EPI_ISL_895060, EPI_ISL_895061, EPI_ISL_895062, EPI_ISL_895063, EPI_ISL_895064, EPI_ISL_895065, EPI_ISL_895066, EPI_ISL_895067, EPI_ISL_895068, EPI_ISL_895069, EPI_ISL_895089, EPI_ISL_895090, EPI_ISL_895091, EPI_ISL_895092, EPI_ISL_895094 | | | |
| see above | Department of Infectious Diseases, Kobe Institute of Health | Pathogen Genomics Center, National Institute of Infectious Diseases | Tsuyoshi Sekizuka, Kentaro Itokawa, Rina Tanaka, Masanori Hashino, Makoto Kuroda |
| EPI_ISL_895151, EPI_ISL_895152, EPI_ISL_895153, EPI_ISL_895154, EPI_ISL_895155, EPI_ISL_895156, EPI_ISL_895157, EPI_ISL_895158, EPI_ISL_895159, EPI_ISL_895266, EPI_ISL_895267, EPI_ISL_895268, EPI_ISL_895269, EPI_ISL_895270, EPI_ISL_895271, EPI_ISL_895272, EPI_ISL_895273, EPI_ISL_895274, EPI_ISL_895275, EPI_ISL_895276, EPI_ISL_895277, EPI_ISL_895278, EPI_ISL_895279, EPI_ISL_895280, EPI_ISL_895281, EPI_ISL_895282, EPI_ISL_895283, EPI_ISL_895284, EPI_ISL_895285, EPI_ISL_895286, EPI_ISL_895287, EPI_ISL_895288, EPI_ISL_895290, EPI_ISL_895291, EPI_ISL_895292, EPI_ISL_895293, EPI_ISL_895294, EPI_ISL_895295, EPI_ISL_895296, EPI_ISL_895297, EPI_ISL_895298, EPI_ISL_895299, EPI_ISL_895300, EPI_ISL_895301, EPI_ISL_895302, EPI_ISL_895303, EPI_ISL_895304, EPI_ISL_895305, EPI_ISL_895306, EPI_ISL_895307, EPI_ISL_895308, EPI_ISL_895309, EPI_ISL_895310, EPI_ISL_895311, EPI_ISL_895312, EPI_ISL_895313, EPI_ISL_895314, EPI_ISL_895315, EPI_ISL_895316, EPI_ISL_895317, EPI_ISL_895318, EPI_ISL_895319, EPI_ISL_895320, EPI_ISL_895321, EPI_ISL_895322, EPI_ISL_895323, EPI_ISL_895324, EPI_ISL_895325, EPI_ISL_895326, EPI_ISL_895327, EPI_ISL_895328, EPI_ISL_895329, EPI_ISL_895330, EPI_ISL_895331, EPI_ISL_895332, EPI_ISL_895333, EPI_ISL_895334, EPI_ISL_895335, EPI_ISL_895336, EPI_ISL_895337, EPI_ISL_895338, EPI_ISL_895339, EPI_ISL_895340, EPI_ISL_895341, EPI_ISL_895342, EPI_ISL_895343, EPI_ISL_895344, EPI_ISL_895345, EPI_ISL_895346, EPI_ISL_895347, EPI_ISL_895348, EPI_ISL_895349, EPI_ISL_895350, EPI_ISL_895351, EPI_ISL_895352, EPI_ISL_895353, EPI_ISL_895354, EPI_ISL_895355, EPI_ISL_895356, EPI_ISL_895357, EPI_ISL_895358, EPI_ISL_895359, EPI_ISL_895360, EPI_ISL_895361, EPI_ISL_895362, EPI_ISL_895363, EPI_ISL_895364, EPI_ISL_895365, EPI_ISL_895366, EPI_ISL_895367, EPI_ISL_895368, EPI_ISL_895369, EPI_ISL_895370, EPI_ISL_895371, EPI_ISL_895372, EPI_ISL_895373, EPI_ISL_895374, EPI_ISL_895375, EPI_ISL_895376, EPI_ISL_895377, EPI_ISL_895378, EPI_ISL_895379, EPI_ISL_895380, EPI_ISL_895381, EPI_ISL_895382, EPI_ISL_895383, EPI_ISL_895384, EPI_ISL_895385, EPI_ISL_895386, EPI_ISL_895387, EPI_ISL_895388, EPI_ISL_895389, EPI_ISL_895390, EPI_ISL_895391, EPI_ISL_895392, EPI_ISL_895393, EPI_ISL_895394, EPI_ISL_895395, EPI_ISL_895467, EPI_ISL_895468, EPI_ISL_895469, EPI_ISL_895470, EPI_ISL_895471, EPI_ISL_895472, EPI_ISL_895473, EPI_ISL_895474, EPI_ISL_895475, EPI_ISL_895476, EPI_ISL_895477, EPI_ISL_895478, EPI_ISL_895479, EPI_ISL_895480, EPI_ISL_895481, EPI_ISL_895482, EPI_ISL_895483, EPI_ISL_895484, EPI_ISL_895485, EPI_ISL_895486, EPI_ISL_895488, EPI_ISL_895489, EPI_ISL_895508, EPI_ISL_895509, EPI_ISL_895510, EPI_ISL_895511, EPI_ISL_895512, EPI_ISL_895513, EPI_ISL_895514, EPI_ISL_895515, EPI_ISL_895516, EPI_ISL_895517, EPI_ISL_895533, EPI_ISL_895534, EPI_ISL_895535, EPI_ISL_895567, EPI_ISL_895568, EPI_ISL_895569, EPI_ISL_895570, EPI_ISL_895571, EPI_ISL_895572, EPI_ISL_895573, EPI_ISL_895574, EPI_ISL_895575, EPI_ISL_895576, EPI_ISL_895577, EPI_ISL_895578, EPI_ISL_895579, EPI_ISL_895580, EPI_ISL_895581, EPI_ISL_895582, EPI_ISL_895583, EPI_ISL_895584, EPI_ISL_895585, EPI_ISL_895586, EPI_ISL_895587, EPI_ISL_895588, EPI_ISL_895589, EPI_ISL_895590, EPI_ISL_895591, EPI_ISL_895592, EPI_ISL_895593, EPI_ISL_895594, EPI_ISL_895595, EPI_ISL_895596, EPI_ISL_895597, EPI_ISL_895598, EPI_ISL_895599, EPI_ISL_895600, EPI_ISL_895601, EPI_ISL_895602, EPI_ISL_895603, EPI_ISL_895604, EPI_ISL_895605, EPI_ISL_895606, EPI_ISL_895607, EPI_ISL_895658, EPI_ISL_895659, EPI_ISL_895660, EPI_ISL_895661, EPI_ISL_895662, EPI_ISL_895663, EPI_ISL_895664, EPI_ISL_895665, EPI_ISL_895666, EPI_ISL_895667, EPI_ISL_895668, EPI_ISL_895669, EPI_ISL_895670, EPI_ISL_895671, EPI_ISL_895672, EPI_ISL_895673, EPI_ISL_895674, EPI_ISL_895675, EPI_ISL_895676, EPI_ISL_895677, EPI_ISL_895678, EPI_ISL_895679, EPI_ISL_895680, EPI_ISL_895681, EPI_ISL_895682, EPI_ISL_895683, EPI_ISL_895684, EPI_ISL_895685, EPI_ISL_895686, EPI_ISL_895687, EPI_ISL_895688, EPI_ISL_895689, EPI_ISL_895690, EPI_ISL_895691, EPI_ISL_895692, EPI_ISL_895693, EPI_ISL_895694, EPI_ISL_895695, EPI_ISL_895696, EPI_ISL_895697, EPI_ISL_895698, EPI_ISL_895699, EPI_ISL_895700, EPI_ISL_895701, EPI_ISL_895702, EPI_ISL_895703, EPI_ISL_895704, EPI_ISL_895705, EPI_ISL_895706, EPI_ISL_895707, EPI_ISL_895708, EPI_ISL_895709, EPI_ISL_895710, EPI_ISL_895711, EPI_ISL_895712, EPI_ISL_895713, EPI_ISL_895714, EPI_ISL_895715, EPI_ISL_895716, EPI_ISL_895717, EPI_ISL_895718, EPI_ISL_895719, EPI_ISL_895720, EPI_ISL_895721, EPI_ISL_895722, EPI_ISL_895723, EPI_ISL_895724, EPI_ISL_895725, EPI_ISL_895726, EPI_ISL_895849, EPI_ISL_895850, EPI_ISL_895851, EPI_ISL_895852, EPI_ISL_895853, EPI_ISL_895854, EPI_ISL_895855, EPI_ISL_895856, EPI_ISL_895857, EPI_ISL_895858, EPI_ISL_895859, EPI_ISL_895860, EPI_ISL_895861, EPI_ISL_895862, EPI_ISL_895863, EPI_ISL_895864, EPI_ISL_895865, EPI_ISL_895866, EPI_ISL_895867, EPI_ISL_895868, EPI_ISL_895869, EPI_ISL_895870, EPI_ISL_895871, EPI_ISL_895872, EPI_ISL_895873, EPI_ISL_895874, EPI_ISL_895875, EPI_ISL_895876, EPI_ISL_895877, EPI_ISL_895878, EPI_ISL_895879, EPI_ISL_895880, EPI_ISL_895881, EPI_ISL_895882, EPI_ISL_895883, EPI_ISL_895884, EPI_ISL_895885, EPI_ISL_895886, EPI_ISL_895887, EPI_ISL_895888, EPI_ISL_895889, EPI_ISL_895890, EPI_ISL_895891, EPI_ISL_895892, EPI_ISL_895893, EPI_ISL_895894, EPI_ISL_895895, EPI_ISL_895896, EPI_ISL_895897, EPI_ISL_895898, EPI_ISL_895899, EPI_ISL_895900, EPI_ISL_895901, EPI_ISL_895902, EPI_ISL_895903, EPI_ISL_895904, EPI_ISL_895905, EPI_ISL_895906, EPI_ISL_895907, EPI_ISL_895908, EPI_ISL_895909, EPI_ISL_895910, EPI_ISL_895911, EPI_ISL_895912, EPI_ISL_895913, EPI_ISL_895914, EPI_ISL_895915, EPI_ISL_895916, EPI_ISL_895917, EPI_ISL_895918, EPI_ISL_895919, EPI_ISL_895920, EPI_ISL_895921, EPI_ISL_895922, EPI_ISL_895923, EPI_ISL_895924, EPI_ISL_895925, EPI_ISL_895926, EPI_ISL_895927, EPI_ISL_895928, EPI_ISL_895929, EPI_ISL_895930, EPI_ISL_895931, EPI_ISL_895932, EPI_ISL_895933, EPI_ISL_895934, EPI_ISL_895935, EPI_ISL_895936, EPI_ISL_895937, EPI_ISL_895938, EPI_ISL_895939, EPI_ISL_895940, EPI_ISL_895941, EPI_ISL_895942, EPI_ISL_895943, EPI_ISL_895944, EPI_ISL_895945, EPI_ISL_895946, EPI_ISL_895947, EPI_ISL_895948, EPI_ISL_895949, EPI_ISL_895950, EPI_ISL_895951, EPI_ISL_895952, EPI_ISL_895953, EPI_ISL_895954, EPI_ISL_895955, EPI_ISL_895956, EPI_ISL_895957, EPI_ISL_895958, EPI_ISL_895959, EPI_ISL_895960, EPI_ISL_895961, EPI_ISL_895962, EPI_ISL_895963, EPI_ISL_895964, EPI_ISL_895965, EPI_ISL_895966, EPI_ISL_895967, EPI_ISL_895968, EPI_ISL_895969, EPI_ISL_895970, EPI_ISL_895971, EPI_ISL_895972, EPI_ISL_895973, EPI_ISL_895974, EPI_ISL_895975, EPI_ISL_895976, EPI_ISL_895977, EPI_ISL_895978 | | | |
| see above | Pathogen Genomics Center, National Institute of Infectious Diseases | Pathogen Genomics Center, National Institute of Infectious Diseases | Tsuyoshi Sekizuka, Kentaro Itokawa, Rina Tanaka, Masanori Hashino, Makoto Kuroda |
| EPI_ISL_895989, EPI_ISL_895990, EPI_ISL_895991, EPI_ISL_895992, EPI_ISL_895993, EPI_ISL_895994, EPI_ISL_895995, EPI_ISL_895996, EPI_ISL_895997, EPI_ISL_895998, EPI_ISL_895999, EPI_ISL_896000, EPI_ISL_896001, EPI_ISL_896002, EPI_ISL_896003, EPI_ISL_896004, EPI_ISL_896005, EPI_ISL_896006, EPI_ISL_896007, EPI_ISL_896008, EPI_ISL_896009, EPI_ISL_896010, EPI_ISL_896011, EPI_ISL_896012 | | | |
| see above | Tokyo Metropolitan Institute of Public Health | Pathogen Genomics Center, National Institute of Infectious Diseases | Tsuyoshi Sekizuka, Kentaro Itokawa, Rina Tanaka, Masanori Hashino, Makoto Kuroda |
| EPI_ISL_896064, EPI_ISL_896065, EPI_ISL_896066, EPI_ISL_896067, EPI_ISL_896068 | Pathogen Genomics Center, National Institute of Infectious Diseases | Pathogen Genomics Center, National Institute of Infectious Diseases | Tsuyoshi Sekizuka, Kentaro Itokawa, Rina Tanaka, Masanori Hashino, Makoto Kuroda |
| EPI_ISL_896093 | Viollier AG | University Hospital Basel, Clinical Bacteriology | Tim Roloff, Madlen Stange, Helena MB Seth-Smith, Alfredo Mari, Karoline Leuzinger, Julia Bielicki, Christiane Beckmann, Manuel Battegay, Hans Hirsch, Adrian Egli |
| EPI_ISL_896184, EPI_ISL_896207, EPI_ISL_896208 | MEPHI, Aix Marseille University | MEPHI, Aix Marseille University | Anthony LEVASSEUR |
| EPI_ISL_897097, EPI_ISL_897135, EPI_ISL_897136, EPI_ISL_897137, EPI_ISL_897138, EPI_ISL_897248, EPI_ISL_897249, EPI_ISL_897250, EPI_ISL_897252, EPI_ISL_897253, EPI_ISL_897254, EPI_ISL_897255, EPI_ISL_897256, EPI_ISL_897257, EPI_ISL_897258, EPI_ISL_897259, EPI_ISL_897260, EPI_ISL_897261, EPI_ISL_897262, EPI_ISL_897263, EPI_ISL_897264, EPI_ISL_897265, EPI_ISL_897266, EPI_ISL_897267, EPI_ISL_897268, EPI_ISL_897269, EPI_ISL_897270, EPI_ISL_897271, EPI_ISL_897272, EPI_ISL_897313, EPI_ISL_897314, EPI_ISL_897315, EPI_ISL_897434, EPI_ISL_897435, EPI_ISL_897436, EPI_ISL_897437, EPI_ISL_897438, EPI_ISL_897439, EPI_ISL_897440, EPI_ISL_897441, EPI_ISL_897442, EPI_ISL_897443, EPI_ISL_897444 | | | |
| see above | Pathogen Genomics Center, National Institute of Infectious Diseases | Pathogen Genomics Center, National Institute of Infectious Diseases | Tsuyoshi Sekizuka, Kentaro Itokawa, Rina Tanaka, Masanori Hashino, Makoto Kuroda |
| EPI_ISL_900173, EPI_ISL_900229, EPI_ISL_900273, EPI_ISL_900477 | MEPHI, Aix Marseille University | MEPHI, Aix Marseille University | Anthony LEVASSEUR |
| EPI_ISL_902933, EPI_ISL_902934, EPI_ISL_902949 | Maryland Public Health Laboratory | Maryland Public Health Laboratory | Maryland Department of Health Laboratories Administration |
| EPI_ISL_903013, EPI_ISL_903014, EPI_ISL_903015, EPI_ISL_903016, EPI_ISL_903017, EPI_ISL_903018, EPI_ISL_903019 | Seattle Flu Study | Seattle Flu Study | Deborah A. Nickerson, Chris D. Frazar, Jover Lee, Benjamin Pelle, Erica Ryke, Matthew Richardson, Amanda Adler, Elisabeth Brandstetter, Peter D. Han, Kairsten Fay, Misja Ilcisin, Kirsten Lacombe, Thomas R. Sibley, Melissa Truong, Caitlin R. Wolf, Michael Boeckh, Janet A. Englund, Michael Famulare, Barry R. Lutz, Mark J. Rieder, Lea M. Starita, Matthew Thompson, Jay Shendure, Trevor Bedford, Helen Y. Chu |
| EPI_ISL_903568, EPI_ISL_903569 | NC State Laboratory of Public Health | Genomics and Discovery, Respiratory Viruses Branch, Division of Viral Diseases, Centers for Disease Control and Prevention | Krista Queen, Yan Li, Ying Tao, Jing Zhang, Anna Uehara, Anna Montmayeur, Clinton R. Paden, Peter W. Cook, Rachel Marine, Mili Sheth, Jasmine Padilla, Sarah Nobles, Mark Burroughs, Lori Rowe, Haibin Wang, Ben L. Rambo-Martin, Dhwani Batra, Justin Lee, Suxiang Tong |
| EPI_ISL_903598, EPI_ISL_903617 | UT-Unified State Labs: Public Health Utah DOH | Genomics and Discovery, Respiratory Viruses Branch, Division of Viral Diseases, Centers for Disease Control and Prevention | Krista Queen, Yan Li, Ying Tao, Jing Zhang, Anna Uehara, Anna Montmayeur, Clinton R. Paden, Peter W. Cook, Rachel Marine, Mili Sheth, Jasmine Padilla, Sarah Nobles, Mark Burroughs, Lori Rowe, Haibin Wang, Ben L. Rambo-Martin, Dhwani Batra, Justin Lee, Suxiang Tong |
| EPI_ISL_903684 | WI State Laboratory of Hygiene | Genomics and Discovery, Respiratory Viruses Branch, Division of Viral Diseases, Centers for Disease Control and Prevention | Krista Queen, Yan Li, Ying Tao, Jing Zhang, Anna Uehara, Anna Montmayeur, Clinton R. Paden, Peter W. Cook, Rachel Marine, Mili Sheth, Jasmine Padilla, Sarah Nobles, Mark Burroughs, Lori Rowe, Haibin Wang, Ben L. Rambo-Martin, Dhwani Batra, Justin Lee, Suxiang Tong |
| EPI_ISL_903722 | OK Public Health Laboratory, Oklahoma State DOH | Genomics and Discovery, Respiratory Viruses Branch, Division of Viral Diseases, Centers for Disease Control and Prevention | Krista Queen, Yan Li, Ying Tao, Jing Zhang, Anna Uehara, Anna Montmayeur, Clinton R. Paden, Peter W. Cook, Rachel Marine, Mili Sheth, Jasmine Padilla, Sarah Nobles, Mark Burroughs, Lori Rowe, Haibin Wang, Ben L. Rambo-Martin, Dhwani Batra, Justin Lee, Suxiang Tong |
| EPI_ISL_903741 | VA-Division of Consolidated Laboratory Services | Genomics and Discovery, Respiratory Viruses Branch, | Krista Queen, Yan Li, Ying Tao, Jing Zhang, Anna Uehara, Anna Montmayeur, Clinton R. Paden, Peter W. Cook, Rachel Marine, Mili Sheth, Jasmine |

| | | | |
|---|---|---|---|
| | | Division of Viral Diseases, Centers for Disease Control and Prevention | Padilla, Sarah Nobles, Mark Burroughs, Lori Rowe, Haibin Wang, Ben L. Rambo-Martin, Dhwani Batra, Justin Lee, Suxiang Tong |
| EPI_ISL_903756, EPI_ISL_903758 | OK Public Health Laboratory, Oklahoma State DOH | Genomics and Discovery, Respiratory Viruses Branch, Division of Viral Diseases, Centers for Disease Control and Prevention | Krista Queen, Yan Li, Ying Tao, Jing Zhang, Anna Uehara, Anna Montmayeur, Clinton R. Paden, Peter W. Cook, Rachel Marine, Mili Sheth, Jasmine Padilla, Sarah Nobles, Mark Burroughs, Lori Rowe, Haibin Wang, Ben L. Rambo-Martin, Dhwani Batra, Justin Lee, Suxiang Tong |
| EPI_ISL_903766 | UT-Unified State Labs: Public Health Utah DOH | Genomics and Discovery, Respiratory Viruses Branch, Division of Viral Diseases, Centers for Disease Control and Prevention | Krista Queen, Yan Li, Ying Tao, Jing Zhang, Anna Uehara, Anna Montmayeur, Clinton R. Paden, Peter W. Cook, Rachel Marine, Mili Sheth, Jasmine Padilla, Sarah Nobles, Mark Burroughs, Lori Rowe, Haibin Wang, Ben L. Rambo-Martin, Dhwani Batra, Justin Lee, Suxiang Tong |
| EPI_ISL_903770 | OK Public Health Laboratory, Oklahoma State DOH | Genomics and Discovery, Respiratory Viruses Branch, Division of Viral Diseases, Centers for Disease Control and Prevention | Krista Queen, Yan Li, Ying Tao, Jing Zhang, Anna Uehara, Anna Montmayeur, Clinton R. Paden, Peter W. Cook, Rachel Marine, Mili Sheth, Jasmine Padilla, Sarah Nobles, Mark Burroughs, Lori Rowe, Haibin Wang, Ben L. Rambo-Martin, Dhwani Batra, Justin Lee, Suxiang Tong |
| EPI_ISL_903771 | VA-Division of Consolidated Laboratory Services | Genomics and Discovery, Respiratory Viruses Branch, Division of Viral Diseases, Centers for Disease Control and Prevention | Krista Queen, Yan Li, Ying Tao, Jing Zhang, Anna Uehara, Anna Montmayeur, Clinton R. Paden, Peter W. Cook, Rachel Marine, Mili Sheth, Jasmine Padilla, Sarah Nobles, Mark Burroughs, Lori Rowe, Haibin Wang, Ben L. Rambo-Martin, Dhwani Batra, Justin Lee, Suxiang Tong |
| EPI_ISL_903783 | OK Public Health Laboratory, Oklahoma State DOH | Genomics and Discovery, Respiratory Viruses Branch, Division of Viral Diseases, Centers for Disease Control and Prevention | Krista Queen, Yan Li, Ying Tao, Jing Zhang, Anna Uehara, Anna Montmayeur, Clinton R. Paden, Peter W. Cook, Rachel Marine, Mili Sheth, Jasmine Padilla, Sarah Nobles, Mark Burroughs, Lori Rowe, Haibin Wang, Ben L. Rambo-Martin, Dhwani Batra, Justin Lee, Suxiang Tong |
| EPI_ISL_903792 | VA-Division of Consolidated Laboratory Services | Genomics and Discovery, Respiratory Viruses Branch, Division of Viral Diseases, Centers for Disease Control and Prevention | Krista Queen, Yan Li, Ying Tao, Jing Zhang, Anna Uehara, Anna Montmayeur, Clinton R. Paden, Peter W. Cook, Rachel Marine, Mili Sheth, Jasmine Padilla, Sarah Nobles, Mark Burroughs, Lori Rowe, Haibin Wang, Ben L. Rambo-Martin, Dhwani Batra, Justin Lee, Suxiang Tong |
| EPI_ISL_903801 | WI State Laboratory of Hygiene | Genomics and Discovery, Respiratory Viruses Branch, Division of Viral Diseases, Centers for Disease Control and Prevention | Krista Queen, Yan Li, Ying Tao, Jing Zhang, Anna Uehara, Anna Montmayeur, Clinton R. Paden, Peter W. Cook, Rachel Marine, Mili Sheth, Jasmine Padilla, Sarah Nobles, Mark Burroughs, Lori Rowe, Haibin Wang, Ben L. Rambo-Martin, Dhwani Batra, Justin Lee, Suxiang Tong |
| EPI_ISL_903830 | VA-Division of Consolidated Laboratory Services | Genomics and Discovery, Respiratory Viruses Branch, Division of Viral Diseases, Centers for Disease Control and Prevention | Krista Queen, Yan Li, Ying Tao, Jing Zhang, Anna Uehara, Anna Montmayeur, Clinton R. Paden, Peter W. Cook, Rachel Marine, Mili Sheth, Jasmine Padilla, Sarah Nobles, Mark Burroughs, Lori Rowe, Haibin Wang, Ben L. Rambo-Martin, Dhwani Batra, Justin Lee, Suxiang Tong |
| EPI_ISL_903841 | UT-Unified State Labs: Public Health Utah DOH | Genomics and Discovery, Respiratory Viruses Branch, Division of Viral Diseases, Centers for Disease Control and Prevention | Krista Queen, Yan Li, Ying Tao, Jing Zhang, Anna Uehara, Anna Montmayeur, Clinton R. Paden, Peter W. Cook, Rachel Marine, Mili Sheth, Jasmine Padilla, Sarah Nobles, Mark Burroughs, Lori Rowe, Haibin Wang, Ben L. Rambo-Martin, Dhwani Batra, Justin Lee, Suxiang Tong |
| EPI_ISL_903846 | WI State Laboratory of Hygiene | Genomics and Discovery, Respiratory Viruses Branch, Division of Viral Diseases, Centers for Disease Control and Prevention | Krista Queen, Yan Li, Ying Tao, Jing Zhang, Anna Uehara, Anna Montmayeur, Clinton R. Paden, Peter W. Cook, Rachel Marine, Mili Sheth, Jasmine Padilla, Sarah Nobles, Mark Burroughs, Lori Rowe, Haibin Wang, Ben L. Rambo-Martin, Dhwani Batra, Justin Lee, Suxiang Tong |
| EPI_ISL_903891 | UT-Unified State Labs: Public Health Utah DOH | Genomics and Discovery, Respiratory Viruses Branch, Division of Viral Diseases, Centers for Disease Control and Prevention | Krista Queen, Yan Li, Ying Tao, Jing Zhang, Anna Uehara, Anna Montmayeur, Clinton R. Paden, Peter W. Cook, Rachel Marine, Mili Sheth, Jasmine Padilla, Sarah Nobles, Mark Burroughs, Lori Rowe, Haibin Wang, Ben L. Rambo-Martin, Dhwani Batra, Justin Lee, Suxiang Tong |
| EPI_ISL_903903 | WI State Laboratory of Hygiene | Genomics and Discovery, Respiratory Viruses Branch, Division of Viral Diseases, Centers for Disease Control and Prevention | Krista Queen, Yan Li, Ying Tao, Jing Zhang, Anna Uehara, Anna Montmayeur, Clinton R. Paden, Peter W. Cook, Rachel Marine, Mili Sheth, Jasmine Padilla, Sarah Nobles, Mark Burroughs, Lori Rowe, Haibin Wang, Ben L. Rambo-Martin, Dhwani Batra, Justin Lee, Suxiang Tong |
| EPI_ISL_903910, EPI_ISL_903938 | UT-Unified State Labs: Public Health Utah DOH | Genomics and Discovery, Respiratory Viruses Branch, Division of Viral Diseases, Centers for Disease Control and Prevention | Krista Queen, Yan Li, Ying Tao, Jing Zhang, Anna Uehara, Anna Montmayeur, Clinton R. Paden, Peter W. Cook, Rachel Marine, Mili Sheth, Jasmine Padilla, Sarah Nobles, Mark Burroughs, Lori Rowe, Haibin Wang, Ben L. Rambo-Martin, Dhwani Batra, Justin Lee, Suxiang Tong |
| EPI_ISL_903940 | WI State Laboratory of Hygiene | Genomics and Discovery, Respiratory Viruses Branch, Division of Viral Diseases, Centers for Disease Control and Prevention | Krista Queen, Yan Li, Ying Tao, Jing Zhang, Anna Uehara, Anna Montmayeur, Clinton R. Paden, Peter W. Cook, Rachel Marine, Mili Sheth, Jasmine Padilla, Sarah Nobles, Mark Burroughs, Lori Rowe, Haibin Wang, Ben L. Rambo-Martin, Dhwani Batra, Justin Lee, Suxiang Tong |
| EPI_ISL_903951, EPI_ISL_903953 | MN PHL Division, Minnesota Department of Health | Genomics and Discovery, Respiratory Viruses Branch, Division of Viral Diseases, Centers for Disease Control and Prevention | Krista Queen, Yan Li, Ying Tao, Jing Zhang, Anna Uehara, Anna Montmayeur, Clinton R. Paden, Peter W. Cook, Rachel Marine, Mili Sheth, Jasmine Padilla, Sarah Nobles, Mark Burroughs, Lori Rowe, Haibin Wang, Ben L. Rambo-Martin, Dhwani Batra, Justin Lee, Suxiang Tong |
| EPI_ISL_904140, EPI_ISL_904176, EPI_ISL_904475, EPI_ISL_904476, EPI_ISL_904477, EPI_ISL_904478 | Dutch COVID-19 response team | Erasmus Medical Center | Bas Oude Munnink, Reina Sikkema, David Nieuwenhuijse, Irina Chestakova, Anne van der Linden, Marjan Boter, Emmanuelle Munger, Corine GeurtsvanKessel, Annemiek van der Eijk, Richard Molenkamp, Marion Koopmans, on behalf of the Dutch national COVID-19 response team. |
| EPI_ISL_904861, EPI_ISL_904864, EPI_ISL_905080, EPI_ISL_905083 | Dutch COVID-19 response team | National Institute for Public Health and the Environment (RIVM) | Adam Meijer, Harry Vennema, Dirk Eggink, Jeroen Cremer, Sharon van den Brink, Bas van der Veer, AnneMarie van den Brandt, Florian Zwagemaker, Dennis Schmitz, Chantal Reusken, on behalf of the national COVID-19 response team |
| EPI_ISL_906082 | Child Health Research Foundation | Child Health Research Foundation | Senjuti Saha, Arif Mohammad Tanmoy, Sharmistha Goswami, Afroza Akter Tanni, Syed Muktadir Al Sium, Roly Malaker, Md Hafizur Rahman, Samir K Saha |
| EPI_ISL_906085, EPI_ISL_906089 | Child Health Research Foundation | Child Health Research Foundation | Senjuti Saha, Sharmistha Goswami, Afroza Akter Tanni, Syed Muktadir Al Sium, Arif Mohammad Tanmoy, Roly Malaker, Md Hafizur Rahman, Samir K Saha |
| EPI_ISL_906090 | Child Health Research Foundation | Child Health Research Foundation | Senjuti Saha, Afroza Akter Tanni, Sharmistha Goswami, Syed Muktadir Al Sium, Arif Mohammad Tanmoy, Roly Malaker, Md Hafizur Rahman, Samir K Saha |
| EPI_ISL_906094, EPI_ISL_906102 | Child Health Research Foundation | Child Health Research Foundation | Senjuti Saha, Syed Muktadir Al Sium, Sharmistha Goswami, Afroza Akter Tanni, Arif Mohammad Tanmoy, Roly Malaker, Md Hafizur Rahman, Samir K Saha |
| EPI_ISL_906104 | Child Health Research Foundation | Child Health Research Foundation | Senjuti Saha, Arif Mohammad Tanmoy, Sharmistha Goswami, Afroza Akter Tanni, Syed Muktadir Al Sium, Roly Malaker, Md Hafizur Rahman, Samir K Saha |
| EPI_ISL_906107 | Child Health Research Foundation | Child Health Research Foundation | Senjuti Saha, Afroza Akter Tanni, Sharmistha Goswami, Syed Muktadir Al Sium, Arif Mohammad Tanmoy, Roly Malaker, Md Hafizur Rahman, Samir K Saha |
| EPI_ISL_906259, EPI_ISL_906260, EPI_ISL_906261 | University of Wisconsin-Madison AIDS Vaccine Research Laboratories | University of Wisconsin-Madison AIDS Vaccine Research Laboratories | Gage Moreno, Katarina Braun, et al. AIDS Vaccine Research Laboratories |
| EPI_ISL_906285 | Nigeria Centre for Disease Control (NCDC) | African Centre of Excellence for Genomics of Infectious Diseases (ACEGID), Redeemer's University | Oluniyi P.E. et al |
| EPI_ISL_906523 | Department of Infectious Diseases, Kobe Institute of Health | Pathogen Genomics Center, National Institute of Infectious Diseases | Tsuyoshi Sekizuka, Kentaro Itokawa, Rina Tanaka, Masanori Hashino, Makoto Kuroda |
| EPI_ISL_906526 | Pathogen Genomics Center, National Institute of Infectious Diseases | Pathogen Genomics Center, National Institute of Infectious Diseases | Tsuyoshi Sekizuka, Kentaro Itokawa, Rina Tanaka, Masanori Hashino, Makoto Kuroda |
| EPI_ISL_906533, EPI_ISL_906542 | Laboratorio de Virologia-Instituto Nacional de Salud | Instituto Nacional de Salud- Dirección de Investigación en | Katherine Laiton-Donato, Diego A. Álvarez-Díaz, Carlos Franco-Muñoz, Mauricio Pacheco-Montealegre, Héctor Alejandro Ruiz-Moreno, Maria T. |

| | | | |
|---|---|---|---|
| | | Salud Pública, Universidad de los Andes- Applied genomics research group, Vicerrectoria de Investigación y Creación, Universidad de los Andes- Systems and Computing Engineering Department | Herrera-Sepúlveda, Diego Andrés Prada, Jhonnatan Reales-González, Sheryll Corchuelo, Julian Naizaque, Gerardo Santamaría Jorge Duitama, Laura Natalia Gonzalez, Jorge Ivan Diaz, Silvia Restrepo-Restrepo, Magdalena Wiesner, Martha Lucia Ospina Martinez, Marcela Mercado-Reyes |
| EPI_ISL_910360, EPI_ISL_910386, EPI_ISL_910387, EPI_ISL_910388, EPI_ISL_910389, EPI_ISL_910457, EPI_ISL_910458, EPI_ISL_910459, EPI_ISL_910460, EPI_ISL_910461, EPI_ISL_910462, EPI_ISL_910647, EPI_ISL_910700, EPI_ISL_910701, EPI_ISL_910702, EPI_ISL_910703, EPI_ISL_910705, EPI_ISL_910714, EPI_ISL_910715, EPI_ISL_910716, EPI_ISL_910717, EPI_ISL_910718, EPI_ISL_910719, EPI_ISL_910720, EPI_ISL_910723, EPI_ISL_910774, EPI_ISL_910775, EPI_ISL_910776, EPI_ISL_911235 | | | |
| see above | Laboratoire national de sante, Microbiology, Virology | Laboratoire national de sante, Microbiology, Microbial Genomics Platform | Anke Wienecke-Baldacchino, Catherine Ragimbeau,Jessica Tapp, Fatu Djabi, Lise Pignon, Raoul Salmon, Tamir Abdelrahman |
| EPI_ISL_911532, EPI_ISL_911533, EPI_ISL_911534, EPI_ISL_911535, EPI_ISL_911536, EPI_ISL_911537, EPI_ISL_911538, EPI_ISL_911539, EPI_ISL_911540, EPI_ISL_911541, EPI_ISL_911542, EPI_ISL_911543, EPI_ISL_911544, EPI_ISL_911546, EPI_ISL_911547 | | | |
| see above | CSIR-IGIB | CSIR-IGIB | Rajesh Pandey, Akshay Kanakan, Janani SV, Ranjeet Maurya, Aparna S Murali, Shweta Sahni, Azka Khan, Partha Chattopadhyay, Priti Devi, Priyanka Mehta, Anil Kumar, Nisha Rawat |
| EPI_ISL_911677, EPI_ISL_911680, EPI_ISL_911682, EPI_ISL_911700, EPI_ISL_911703 | Alaska State Virology Laboratory | Alaska State Virology Laboratory | Stephanie DeRonde, Lisa Smith, Ph.D., Jack Chen, Ph.D. |
| EPI_ISL_912467, EPI_ISL_912489, EPI_ISL_912515, EPI_ISL_912523, EPI_ISL_912530 | NHLS Universitas Academic | UFS Virology | PA Bester, MM Nyaga, P Nthiga, MT Mogotsi, D Goedhals, T de Oliveira |
| EPI_ISL_913306, EPI_ISL_913462 | Klinisk mikrobiologi | The Public Health Agency of Sweden | Anna-Malin Linde, Maria Lind Karlberg, Carlo Berg, Oskar Karlsson Lindsjo, Sofia Stamouli, Reza Advani, Mattias Haukland, Petra Holmstrom, Noura Walai, Petra Edquist, Mia Brytting, Anna Risberg, Karin Tegmark-Wisell |
| EPI_ISL_914794 | AREA DE SALUD BUENOS AIRES | Inciensa, Instituto Costarricense de Investigación y Enseñanza en Nutrición y Salud | Francisco Duarte, Hebleen Porras, Claudio Soto-Garita, Estela Cordero, Adriana Godínez, Melany Calderón & Mariel López |
| EPI_ISL_914795 | HOSPITAL DR. FERNANDO ESCALANTE PRADILLA | Inciensa, Instituto Costarricense de Investigación y Enseñanza en Nutrición y Salud | Francisco Duarte, Hebleen Porras, Claudio Soto-Garita, Estela Cordero, Adriana Godínez, Melany Calderón & Mariel López |
| EPI_ISL_915374, EPI_ISL_915377, EPI_ISL_915415, EPI_ISL_915416 | Keio University School of Medicine | Keio University School of Medicine | Kenjiro Kosaki, Yuka Iwasaki, Hirotsugu Ishizu, Haruhiko Siomi, Kodai Abe |
| EPI_ISL_918167 | Department of Infectious Diseases and Immunology, National Hospital Organization Nagoya Medical Center | Clinical Research Center, National Hospital Organization Nagoya Medical Center | Yoshihiro Nakata, Hirotaka Ode, Mai Kubota, Masakazu Matsuda, Kazuhiro Matsuoka, Miho Nakasuji, Mikiko Mori, Mayumi Imahashi, Yoshiyuki Yokomaku, Yasumasa Iwatani |
| EPI_ISL_918359, EPI_ISL_918365 | Virology Unit, Institut Pasteur du Cambodge | Virology Unit, Institut Pasteur du Cambodge | Sokhoun Yann, Ly Sovann, Kraing Sidonn, Yi Sengdoeurn, Chin Savuth, Chau Darapheak, Etienne Simon-Loriere, Veasna Duong, Erik A Karlsson |
| EPI_ISL_918578, EPI_ISL_918636, EPI_ISL_918637, EPI_ISL_918638, EPI_ISL_918639, EPI_ISL_918640, EPI_ISL_918641, EPI_ISL_918642, EPI_ISL_918643, EPI_ISL_918644, EPI_ISL_918645, EPI_ISL_918646, EPI_ISL_918648, EPI_ISL_918649, EPI_ISL_918650, EPI_ISL_918651, EPI_ISL_918652, EPI_ISL_918653, EPI_ISL_918654, EPI_ISL_918655, EPI_ISL_918658, EPI_ISL_918659, EPI_ISL_918660, EPI_ISL_918661, EPI_ISL_918662, EPI_ISL_918663, EPI_ISL_918664, EPI_ISL_918685 | | | |
| see above | University of Birmingham | COVID-19 Genomics UK (COG-UK) Consortium | Institute of Microbiology, University of Birmingham: Claire McMurray, Joanne Stockton, Samuel Nicholls, Radoslaw Poplawski, Will Rowe, Josh Quick, Nicholas Loman. University of Birmingham Testing Laboratory: Celina M Whalley, Andrew Bosworth, Charlotte Poxon, Kasun Wanigasooriya, Oliver Pickles, Mike Kidd, Alex Richter, Andrew D Beggs PHE Heartlands Lab: Husam Osman, Andrew Bosworth. Queen Elizabeth Hospital: Anna Casey |
| EPI_ISL_919271 | West of Scotland Specialist Virology Centre, NHSGGC / MRC-University of Glasgow Centre for Virus Research | COVID-19 Genomics UK (COG-UK) Consortium | Ana da Silva Filipe, Natasha Johnson, Kathy Smollett, Daniel Mair, Stephen Carmichael, Alice Broos, Lily Tong, Jenna Nichols, Kyriaki Nomikou; Sarah McDonald; Richard Orton, Joseph Hughes, Sreenu Vattipally, David L Robertson; Alasdair MacLean, Rory Gunson; Sharif Shaaban, Matthew Holden; Rachel Blacow, Guy Mollett, Kathy Li, James Shepherd, Antonia Ho, Emma Thomson |
| EPI_ISL_919348 | Virology Department, Royal Infirmary of Edinburgh, NHS Lothian / School of Biological Sciences, University of Edinburgh / Institute of Genetics and Molecular Medicine, University of Edinburgh | COVID-19 Genomics UK (COG-UK) Consortium | McHugh M, Dewar R, Rooke S, Gallagher M, Balcaza C, O'Toole Á, Scher E, Hill V, McCrone JT, Colquhoun R, Yu X, Jackson B, Rambaut A, Williams TC, Templeton K |
| EPI_ISL_919445 | Liverpool Clinical Laboratories | COVID-19 Genomics UK (COG-UK) Consortium | Sam Haldenby, Anita Lucaci, Steve Paterson, Julian Hiscox, Alistair Darby, M Almsaud, A Alrezaihi, Muhannad Alruwaili, Stuart D Armstrong, Jones Benjamin, Eleanor G Bentley, Anu Chawla, Jordan J Clark, Angela Cowell, Richard Eccles, Isabel García-Dorival, Matthew Gemmell, Alessandro Gerada, PKF Gilmore, Richard Gregory, Ximeng Han, Catherine Hartley, Margaret Hughes, Miren Iturriza-Gomara, James Johnson, L Luu, Jenifer Manson, Charlotte Nelson, Elaine O'Toole, Cassie Olateju, Rebekah Penrice-Randal , Lucille Rainbow, N.P Randle, Trevor Ian Robinson, Parul Sharma, Ghada T Shawli, James P Stewart, Neil Swainston, Ecaterina Vamos, Joanne Watts, Mark Whitehead |
| EPI_ISL_920223 | University College London Hospital | COVID-19 Genomics UK (COG-UK) Consortium | Judith Heaney, Matthew Byott, Catherine Houlihan, Dan Frampton, Stuart Kirk, Moira Spyer and Eleni Nastouli |
| EPI_ISL_923221, EPI_ISL_923222 | Centre for Enzyme Innovation, University of Portsmouth / Translational Research Laboratory, Portsmouth Hospitals NHS Trust | COVID-19 Genomics UK (COG-UK) Consortium | Angela Beckett,Salman Goudarzi,Christopher Fearn,Kate Cook,Katie Loveson,Sharon Glaysher,Scott Elliott,Samuel Robson |
| EPI_ISL_924141 | Virology Department, Sheffield Teaching Hospitals NHS Foundation Trust/Department of Infection, Immunity and Cardiovascular Disease, The Medical School, University of Sheffield | COVID-19 Genomics UK (COG-UK) Consortium | Thushan de Silva, Matthew Parker, Nikki Smith, Adri Angyal, Rebecca Brown, Luke Green, Rachel Tucker, Paul Parsons, Danielle Groves, Katie Johnson, Laura Carrilero, Alex Keeley, Dave Partridge, Matthew Wyles, Benjamin Lindsey, Mehmet Yavuz, Mohammad Raza, Cariad Evans |
| EPI_ISL_925151, EPI_ISL_925152, EPI_ISL_925153, EPI_ISL_925154, EPI_ISL_925155, EPI_ISL_925156, EPI_ISL_925157, EPI_ISL_925158, EPI_ISL_925159, EPI_ISL_925164, EPI_ISL_925165, EPI_ISL_925166 | | | |
| see above | Virginia DCLS | Virginia DCLS | Virginia DCLS |
| EPI_ISL_925539, EPI_ISL_925597, EPI_ISL_925598, EPI_ISL_925599, EPI_ISL_925600, EPI_ISL_925601, EPI_ISL_925602, EPI_ISL_925603, EPI_ISL_925604, EPI_ISL_925605, EPI_ISL_925606, EPI_ISL_925607, EPI_ISL_925608, EPI_ISL_925609, EPI_ISL_925610, EPI_ISL_925611, EPI_ISL_925612, EPI_ISL_925613, EPI_ISL_925614, EPI_ISL_925615, EPI_ISL_925616, EPI_ISL_925617, EPI_ISL_925618, EPI_ISL_925619, EPI_ISL_925620, EPI_ISL_925621, EPI_ISL_925622, EPI_ISL_925623, EPI_ISL_925624, EPI_ISL_925625, EPI_ISL_925626, EPI_ISL_925627, EPI_ISL_925628, EPI_ISL_925629, EPI_ISL_925630, EPI_ISL_925631, EPI_ISL_925632, EPI_ISL_925633, EPI_ISL_925634, EPI_ISL_925635, EPI_ISL_925636, EPI_ISL_925637, EPI_ISL_925638, EPI_ISL_925639, EPI_ISL_925640, EPI_ISL_925641, EPI_ISL_925642, EPI_ISL_925643, EPI_ISL_925644, EPI_ISL_925645, EPI_ISL_925646, EPI_ISL_925647, EPI_ISL_925648, EPI_ISL_925649, EPI_ISL_925650, EPI_ISL_925651, EPI_ISL_925652, EPI_ISL_925653, EPI_ISL_925654, EPI_ISL_925655, EPI_ISL_925656, EPI_ISL_925657, EPI_ISL_925658, EPI_ISL_925659, EPI_ISL_925660, EPI_ISL_925661, EPI_ISL_925662, EPI_ISL_925663, EPI_ISL_925664, EPI_ISL_925665, EPI_ISL_925666, EPI_ISL_925667, EPI_ISL_925668, EPI_ISL_925669, EPI_ISL_925670, EPI_ISL_925671, EPI_ISL_925672, EPI_ISL_925673, EPI_ISL_925674, EPI_ISL_925675, EPI_ISL_925676, EPI_ISL_925677, EPI_ISL_925701, EPI_ISL_925702, EPI_ISL_925703, EPI_ISL_925704, EPI_ISL_925705, EPI_ISL_925706, EPI_ISL_925707, EPI_ISL_925708, EPI_ISL_925709, EPI_ISL_925710, EPI_ISL_925711, EPI_ISL_925712, EPI_ISL_925713, EPI_ISL_925714, EPI_ISL_925715, EPI_ISL_925716, EPI_ISL_925717, EPI_ISL_925718, EPI_ISL_925719, EPI_ISL_925720, EPI_ISL_925721, EPI_ISL_925722, EPI_ISL_925723, EPI_ISL_925724, EPI_ISL_925725, EPI_ISL_925730, EPI_ISL_925763 | | | |
| see above | Public Health Ontario Laboratory | Public Health Ontario Laboratory | Vanessa G Allen, Philip Banh, Yao Chen, Richard de Borja, Alireza Eshaghi, Nahuel Fittipaldi, Christine Frantz, Jonathan B Gubbay, Jennifer L Guthrie, Lawrence Heisler, Esha Joshi, Michael Laszloffy, Aimin Li, Michael CY Li, Dean Maxwell, Sandeep Nagra, Samir N Patel, Jared Simpson, Karthikeyan Sivaraman, Ashleigh Sullivan, Yogi Sundaravadanam, Sarah Teatero, Matthew Watson, Andre Villegas, Sandra Zittermann |
| EPI_ISL_925979, EPI_ISL_926193, EPI_ISL_926252, EPI_ISL_926292, EPI_ISL_926323, EPI_ISL_926334, EPI_ISL_926515, EPI_ISL_926536, EPI_ISL_926697, EPI_ISL_926878, EPI_ISL_926995, EPI_ISL_927223, EPI_ISL_927239, EPI_ISL_927808, EPI_ISL_927824, EPI_ISL_928081, EPI_ISL_928230, EPI_ISL_928687, EPI_ISL_928888, EPI_ISL_928988, EPI_ISL_929006, EPI_ISL_929069, EPI_ISL_929355, EPI_ISL_929513, EPI_ISL_929658, EPI_ISL_929662, EPI_ISL_929738, EPI_ISL_929781, EPI_ISL_929790, EPI_ISL_929830, EPI_ISL_930065, EPI_ISL_930158, EPI_ISL_930178, EPI_ISL_930180, EPI_ISL_930212, EPI_ISL_930324, EPI_ISL_930347, EPI_ISL_930458, EPI_ISL_930497, EPI_ISL_930518, EPI_ISL_930523 | | | |
| see above | Department of Virus and Microbiological Special Diagnostics, Statens Serum Institut, Copenhagen, Denmark | Aalborg University | Danish Covid-19 Genome Consortium |
| EPI_ISL_931367, EPI_ISL_931368, EPI_ISL_931369 | University Hospital Basel, Clinical Virology | University Hospital Basel, Clinical Bacteriology | Tim Roloff, Madlen Stange, Helena MB Seth-Smith, Alfredo Mari, Karoline Leuzinger, Julia Bielicki, Manuel Battegay, Hans Hirsch, Adrian Egli |
| EPI_ISL_933496, EPI_ISL_933497, EPI_ISL_933498, EPI_ISL_933499, EPI_ISL_933500, EPI_ISL_933501, EPI_ISL_933502, EPI_ISL_933503, EPI_ISL_933504, EPI_ISL_933505, EPI_ISL_933506, EPI_ISL_933507, EPI_ISL_933508, EPI_ISL_933509 | | | |

| | | | |
|---|---|---|---|
| see above | Lighthouse Lab in Milton Keynes | Wellcome Sanger Institute for the COVID-19 Genomics UK (COG-UK) Consortium | The Lighthouse Lab in Milton Keynes and Alex Alderton, Roberto Amato, Sonia Goncalves, Ewan Harrison, David K. Jackson, Ian Johnston, Dominic Kwiatkowski, Cordelia Langford, John Sillitoe on behalf of the Wellcome Sanger Institute COVID-19 Surveillance Team |
| EPI_ISL_933519, EPI_ISL_933520, EPI_ISL_933521, EPI_ISL_933522, EPI_ISL_933523, EPI_ISL_933524, EPI_ISL_933525, EPI_ISL_933526, EPI_ISL_933527, EPI_ISL_933528, EPI_ISL_933529, EPI_ISL_933530, EPI_ISL_933531 | | | |
| see above | Instituto Nacional de Medicina Genomica | Instituto Nacional de Medicina Genomica | Hidalgo-Miranda A, Mendoza-Vargas A, Reyes-Grajeda JP, Cisneros-Villanueva M, Cedro-Tanda A,Peñaloza-Figueroa F, Herrera-Montalvo LA |
| EPI_ISL_933554, EPI_ISL_933555, EPI_ISL_933556, EPI_ISL_933557, EPI_ISL_933580, EPI_ISL_933581, EPI_ISL_933582, EPI_ISL_933583, EPI_ISL_933584, EPI_ISL_933585, EPI_ISL_933586, EPI_ISL_933587, EPI_ISL_933588, EPI_ISL_933589, EPI_ISL_933590, EPI_ISL_933591, EPI_ISL_933592, EPI_ISL_933593, EPI_ISL_933595 | | | |
| see above | Arizona State Public Health Laboratory | Arizona State Public Health Laboratory | Trung Huynh, Jessica Escobar, Katherine Fullerton, Nobuko Fukushima, Stacy White, Linda Getsinger, Victor Waddell |
| EPI_ISL_933805, EPI_ISL_933806, EPI_ISL_933807, EPI_ISL_933808, EPI_ISL_933809, EPI_ISL_933810, EPI_ISL_933811, EPI_ISL_933812, EPI_ISL_933813, EPI_ISL_933814, EPI_ISL_933815, EPI_ISL_933816, EPI_ISL_933817, EPI_ISL_933818, EPI_ISL_933819, EPI_ISL_933820, EPI_ISL_933821, EPI_ISL_933822, EPI_ISL_933823, EPI_ISL_933824, EPI_ISL_933825, EPI_ISL_933826, EPI_ISL_933827, EPI_ISL_933828, EPI_ISL_933829, EPI_ISL_933830, EPI_ISL_933831, EPI_ISL_933832, EPI_ISL_933833, EPI_ISL_933834, EPI_ISL_933835, EPI_ISL_933836, EPI_ISL_933837, EPI_ISL_933838, EPI_ISL_933839, EPI_ISL_933840, EPI_ISL_933841, EPI_ISL_933842, EPI_ISL_933843, EPI_ISL_933844, EPI_ISL_933845, EPI_ISL_933846, EPI_ISL_933847, EPI_ISL_933848, EPI_ISL_933849, EPI_ISL_933850, EPI_ISL_933851, EPI_ISL_933852, EPI_ISL_933853, EPI_ISL_933854, EPI_ISL_933855, EPI_ISL_933856, EPI_ISL_933857, EPI_ISL_933858, EPI_ISL_933859, EPI_ISL_933860, EPI_ISL_933861, EPI_ISL_933862, EPI_ISL_933863, EPI_ISL_933864, EPI_ISL_933865, EPI_ISL_933866, EPI_ISL_933867, EPI_ISL_933868, EPI_ISL_933869, EPI_ISL_933870, EPI_ISL_933871, EPI_ISL_933872, EPI_ISL_933873, EPI_ISL_933874, EPI_ISL_933875, EPI_ISL_933876, EPI_ISL_933877, EPI_ISL_933941, EPI_ISL_933942, EPI_ISL_933943, EPI_ISL_933947 | | | |
| see above | Vilnius university hospital Santaros Klinikos, Center of Laboratory Medicine | Vilnius university hospital Santaros Klinikos, Center of Laboratory Medicine | Ingrida Olendraite, Daniel Naumovas, Rimvydas Norvilas, Dovile Ezerskyte, Justinas Slikas, Gytis Dudas |
| EPI_ISL_935215 | KU Leuven, Rega Institute, Clinical and Epidemiological Virology | KU Leuven, Rega Institute, Clinical and Epidemiological Virology | Tony Wawina-Bokalanga, Bert Vanmechelen, Joan Marti-Carerras, Piet Maes |
| EPI_ISL_935466, EPI_ISL_935467, EPI_ISL_935468, EPI_ISL_935469, EPI_ISL_935470, EPI_ISL_935471, EPI_ISL_935472, EPI_ISL_935473, EPI_ISL_935474, EPI_ISL_935475, EPI_ISL_935476, EPI_ISL_935477, EPI_ISL_935478, EPI_ISL_935479, EPI_ISL_935480, EPI_ISL_935481, EPI_ISL_935482, EPI_ISL_935483, EPI_ISL_935484, EPI_ISL_935485, EPI_ISL_935486, EPI_ISL_935487, EPI_ISL_935488, EPI_ISL_935489, EPI_ISL_935490, EPI_ISL_935491, EPI_ISL_935492, EPI_ISL_935493, EPI_ISL_935494 | | | |
| see above | Florida Bureau of Public Health Laboratories | Florida Bureau of Public Health Laboratories | Sarah Schmedes, Jason Blanton |
| EPI_ISL_936632 | Northwestern Memorial Hospital | Ozer Lab | Ramon Lorenzo-Redondo, Lacy M. Simons, Chad J. Achenbach, Lawrence J. Jennings, Michael G. Ison, Judd F. Hultquist, Egon A. Ozer |
| EPI_ISL_940153, EPI_ISL_940154 | NHLS Universitas Academic | UFS Virology | PA Bester, MM Nyaga, P Nthiga, MT Mogotsi, D Goedhals, T de Oliveira |
| EPI_ISL_940259, EPI_ISL_940262, EPI_ISL_940263, EPI_ISL_940275 | Hôpital Bichat Claude Bernard, Laboratoire de Virologie | IAME UMR1137 Inserm, Université de Paris, Hôpital Bichat | Antoine Bridier-Nahmias, Amélie Recoing, Quentin Le Hingrat, Lena Daniel, Siham Hamri, Gilles Collin, Alexandre Storto, Mélanie Bertine, Charlotte Charpentier, Nadhira Houhou-Fidouh, Diane Descamps, Benoit Visseaux |
| EPI_ISL_940694, EPI_ISL_940699, EPI_ISL_940702, EPI_ISL_940707 | National Laboratory for Health, Environment and Food | National Laboratory for Health, Environment and Food | Aleksander Mahnic, Sandra Janezic, Maja Rupnik |
| EPI_ISL_940830, EPI_ISL_940831, EPI_ISL_941918, EPI_ISL_941919, EPI_ISL_941920 | Virginia DCLS | Virginia DCLS | Virginia DCLS |
| EPI_ISL_942097, EPI_ISL_942098, EPI_ISL_942099, EPI_ISL_942100, EPI_ISL_942101, EPI_ISL_942102, EPI_ISL_942103, EPI_ISL_942104, EPI_ISL_942105, EPI_ISL_942106, EPI_ISL_942107, EPI_ISL_942108, EPI_ISL_942120, EPI_ISL_942127, EPI_ISL_942143, EPI_ISL_942144, EPI_ISL_942145, EPI_ISL_942146 | | | |
| see above | Wisconsin State Laboratory of Hygiene Communicable Disease Division | Wisconsin State Laboratory of Hygiene Communicable Disease Division | Kelsey R. Florek, Abigail C. Shockey |
| EPI_ISL_942544, EPI_ISL_942545, EPI_ISL_942546, EPI_ISL_942547, EPI_ISL_942548, EPI_ISL_942549, EPI_ISL_942550, EPI_ISL_942551, EPI_ISL_942552, EPI_ISL_942553, EPI_ISL_942554, EPI_ISL_942555, EPI_ISL_942556, EPI_ISL_942557, EPI_ISL_942558, EPI_ISL_942559, EPI_ISL_942560, EPI_ISL_942561, EPI_ISL_942562, EPI_ISL_942563, EPI_ISL_942564, EPI_ISL_942565, EPI_ISL_942566, EPI_ISL_942567, EPI_ISL_942568, EPI_ISL_942569, EPI_ISL_942570, EPI_ISL_942571, EPI_ISL_942572, EPI_ISL_942573, EPI_ISL_942574, EPI_ISL_942575, EPI_ISL_942576, EPI_ISL_942577, EPI_ISL_942578, EPI_ISL_942579, EPI_ISL_942580, EPI_ISL_942581, EPI_ISL_942582 | | | |
| see above | Gundersen Molecular Diagnostics Laboratory | Kabara Cancer Research Institute | Craig S. Richmond, Paraic A. Kenny |
| EPI_ISL_944596 | Connecticut Department of Public Health | Grubaugh Lab - Yale School of Public Health | Joseph Fauver, Tara Alpert, Anderson Brito, Mallery Breban, Anne Wyllie, Chantal Vogels, Mary Petrone, Annie Watkins, Chaney Kalinich, Isabel Ott, Nathan Grubaugh |
| EPI_ISL_947275 | RS Umum Mulia Pajajaran | Eijkman Institute for Molecular Biology, Ministry of Research and Technology/National Agency for Research and Innovation | Frilasita A Yudhaputri, Hidayat Trimarsanto, Iskandar Adnan, Lydia V. Panggalo, Sukma Oktavianthi, Willy Agustine, Edison Johar, Safarina G Malik, Khin Saw Myint, Amin Soebandrio |
| EPI_ISL_947278 | RS Umum Ciputra Hospital Citragarden City | Eijkman Institute for Molecular Biology, Ministry of Research and Technology/National Agency for Research and Innovation | Hidayat Trimarsanto, Iskandar Adnan, Lydia V. Panggalo, Sukma Oktavianthi, Willy Agustine, Edison Johar, Frilasita A Yudhaputri, Safarina G Malik, Khin Saw Myint, Amin Soebandrio |
| EPI_ISL_947281 | RSU Medika Dramaga | Eijkman Institute for Molecular Biology, Ministry of Research and Technology/National Agency for Research and Innovation | Hidayat Trimarsanto, Iskandar Adnan, Lydia V. Panggalo, Sukma Oktavianthi, Willy Agustine, Edison Johar, Frilasita A Yudhaputri, Safarina G Malik, Khin Saw Myint, Amin Soebandrio |
| EPI_ISL_947287 | RSIA PKU Muhammadiyah | Eijkman Institute for Molecular Biology, Ministry of Research and Technology/National Agency for Research and Innovation | Hidayat Trimarsanto, Iskandar Adnan, Lydia V. Panggalo, Sukma Oktavianthi, Willy Agustine, Edison Johar, Frilasita A Yudhaputri, Safarina G Malik, Khin Saw Myint, Amin Soebandrio |
| EPI_ISL_947290 | RS Hermina Bitung | Eijkman Institute for Molecular Biology, Ministry of Research and Technology/National Agency for Research and Innovation | Hidayat Trimarsanto, Iskandar Adnan, Lydia V. Panggalo, Sukma Oktavianthi, Willy Agustine, Edison Johar, Frilasita A Yudhaputri, Safarina G Malik, Khin Saw Myint, Amin Soebandrio |
| EPI_ISL_949391 | University of Birmingham | COVID-19 Genomics UK (COG-UK) Consortium | Institute of Microbiology, University of Birmingham: Claire McMurray, Joanne Stockton, Samuel Nicholls, Radoslaw Poplawski, Will Rowe, Josh Quick, Nicholas Loman. University of Birmingham Testing Laboratory: Celina M Whalley, Andrew Bosworth, Charlotte Poxon, Kasun Wanigasooriya, Oliver Pickles, Mike Kidd, Alex Richter, Andrew D Beggs PHE Heartlands Lab: Husam Osman, Andrew Bosworth. Queen Elizabeth Hospital: Anna Casey |
| EPI_ISL_949631, EPI_ISL_949633, EPI_ISL_949635, EPI_ISL_949636 | West of Scotland Specialist Virology Centre, NHSGGC / MRC-University of Glasgow Centre for Virus Research | COVID-19 Genomics UK (COG-UK) Consortium | Ana da Silva Filipe, Natasha Johnson, Kathy Smollett, Daniel Mair, Stephen Carmichael, Alice Broos, Lily Tong, Jenna Nichols, Kyriaki Nomikou; Sarah McDonald; Richard Orton, Joseph Hughes, Sreenu Vattipally, David L Robertson; Alasdair MacLean, Rory Gunson; Sharif Shaaban, Matthew Holden; Rachel Blacow, Guy Mollett, Kathy Li, James Shepherd, Antonia Ho, Emma Thomson |
| EPI_ISL_955511, EPI_ISL_955523, EPI_ISL_955524, EPI_ISL_955525, EPI_ISL_955526, EPI_ISL_955527, EPI_ISL_955528, EPI_ISL_955529, EPI_ISL_955530, EPI_ISL_955531, EPI_ISL_955532, EPI_ISL_955534, EPI_ISL_955535, EPI_ISL_955536, EPI_ISL_955537, EPI_ISL_955538, EPI_ISL_955539, EPI_ISL_955540, EPI_ISL_955541, EPI_ISL_955542, EPI_ISL_955543, EPI_ISL_955544, EPI_ISL_955545, EPI_ISL_955546, EPI_ISL_955547, EPI_ISL_955548, EPI_ISL_955549, EPI_ISL_955550, EPI_ISL_955551, EPI_ISL_955552, EPI_ISL_955553, EPI_ISL_955554, EPI_ISL_955555, EPI_ISL_955556, EPI_ISL_955557, EPI_ISL_955558, EPI_ISL_955559, EPI_ISL_955560 | | | |
| see above | Humboldt County Public Health Laboratory | Chan-Zuckerberg Biohub | CZB Cliahub Consortium |
| EPI_ISL_955568, EPI_ISL_955569, EPI_ISL_955570, EPI_ISL_955571 | Santa Clara County Public Health Laboratory | Chan-Zuckerberg Biohub | CZB Cliahub Consortium |
| EPI_ISL_955632 | Orange County Public Health Lab | Chan-Zuckerberg Biohub | CZB Cliahub Consortium |
| EPI_ISL_956323 | Laboratory Medicine | Department of Laboratory Medicine, Lin-Kou Chang Gung Memorial Hospital, Taoyuan, Taiwan | Kuo-Chien Tsao, Yu-Nong Gong, Shu-Li Yang, Yi-Chun Liu, Chung-Guei Huang, Mei-Jen Hsiao, Po-Wei Huang, Cheng-Ta Yang, Cheng-Hsun Chiu, Peng-Nien Huang, Kuo-Ming Lee, Guang-Wu Chen, Shin-Ru Shih |
| EPI_ISL_959494, EPI_ISL_959495, EPI_ISL_959496, EPI_ISL_959529, EPI_ISL_959530, EPI_ISL_959531, EPI_ISL_959532, EPI_ISL_959533, EPI_ISL_959534, EPI_ISL_959535 | Division of Emerging Infectious Diseases, Bureau of Infectious Diseases Diagnosis Control, Korea Disease Control and Prevention Agency | Division of Emerging Infectious Diseases, Bureau of Infectious Diseases Diagnosis Control, Korea Disease Control and Prevention Agency | Ae Kyung Park, Il-Hwan Kim, Heui Man Kim, Jeong-Min Kim, Namjoo Lee, Chae Young Lee, Sang Hee Woo, Eun-Jin Kim |
| EPI_ISL_960097, EPI_ISL_960098 | Mitchells Plain Hospital wc MPH | National Health Laboratory Service/UCT | Arash Iranzadeh, Deelan Doolabh, Lynn Tyers, Bruna Galvao, Innocent Mudau, Marvin Hsiao, Kruger Marais, Diana Hardie, Stephen Korsman, Carolyn Williamson |
| EPI_ISL_960099 | Victoria Hospital wc VHW | National Health Laboratory Service/UCT | Arash Iranzadeh, Deelan Doolabh, Lynn Tyers, Bruna Galvao, Innocent Mudau, Marvin Hsiao, Kruger Marais, Diana Hardie, Stephen Korsman, Carolyn Williamson |

| | | | |
|---|---|---|---|
| EPI_ISL_960100 | Mitchells Plain Hospital wc MPH | National Health Laboratory Service/UCT | Arash Iranzadeh, Deelan Doolabh, Lynn Tyers, Bruna Galvao, Innocent Mudau, Marvin Hsiao, Kruger Marais, Diana Hardie, Stephen Korsman, Carolyn Williamson |
| EPI_ISL_960101 | Victoria Hospital wc VHW | National Health Laboratory Service/UCT | Arash Iranzadeh, Deelan Doolabh, Lynn Tyers, Bruna Galvao, Innocent Mudau, Marvin Hsiao, Kruger Marais, Diana Hardie, Stephen Korsman, Carolyn Williamson |
| EPI_ISL_960102, EPI_ISL_960103 | Groote Schuur Hospital wc GSH | National Health Laboratory Service/UCT | Arash Iranzadeh, Deelan Doolabh, Lynn Tyers, Bruna Galvao, Innocent Mudau, Marvin Hsiao, Kruger Marais, Diana Hardie, Stephen Korsman, Carolyn Williamson |
| EPI_ISL_960104 | Mowbray Maternity Hospital wc MMH | National Health Laboratory Service/UCT | Arash Iranzadeh, Deelan Doolabh, Lynn Tyers, Bruna Galvao, Innocent Mudau, Marvin Hsiao, Kruger Marais, Diana Hardie, Stephen Korsman, Carolyn Williamson |
| EPI_ISL_960105 | False Bay Hospital wc FBH | National Health Laboratory Service/UCT | Arash Iranzadeh, Deelan Doolabh, Lynn Tyers, Bruna Galvao, Innocent Mudau, Marvin Hsiao, Kruger Marais, Diana Hardie, Stephen Korsman, Carolyn Williamson |
| EPI_ISL_960106 | Red Cross Children's Hospital wc RXH | National Health Laboratory Service/UCT | Arash Iranzadeh, Deelan Doolabh, Lynn Tyers, Bruna Galvao, Innocent Mudau, Marvin Hsiao, Kruger Marais, Diana Hardie, Stephen Korsman, Carolyn Williamson |
| EPI_ISL_960107 | Groote Schuur Hospital wc GSH | National Health Laboratory Service/UCT | Arash Iranzadeh, Deelan Doolabh, Lynn Tyers, Bruna Galvao, Innocent Mudau, Marvin Hsiao, Kruger Marais, Diana Hardie, Stephen Korsman, Carolyn Williamson |
| EPI_ISL_960108 | Victoria Hospital wc VHW | National Health Laboratory Service/UCT | Arash Iranzadeh, Deelan Doolabh, Lynn Tyers, Bruna Galvao, Innocent Mudau, Marvin Hsiao, Kruger Marais, Diana Hardie, Stephen Korsman, Carolyn Williamson |
| EPI_ISL_960109 | Groote Schuur Hospital wc GSH | National Health Laboratory Service/UCT | Arash Iranzadeh, Deelan Doolabh, Lynn Tyers, Bruna Galvao, Innocent Mudau, Marvin Hsiao, Kruger Marais, Diana Hardie, Stephen Korsman, Carolyn Williamson |
| EPI_ISL_960110 | False Bay Hospital wc FBH | National Health Laboratory Service/UCT | Arash Iranzadeh, Deelan Doolabh, Lynn Tyers, Bruna Galvao, Innocent Mudau, Marvin Hsiao, Kruger Marais, Diana Hardie, Stephen Korsman, Carolyn Williamson |
| EPI_ISL_960111 | Pelican Park CDC wc PAX | National Health Laboratory Service/UCT | Arash Iranzadeh, Deelan Doolabh, Lynn Tyers, Bruna Galvao, Innocent Mudau, Marvin Hsiao, Kruger Marais, Diana Hardie, Stephen Korsman, Carolyn Williamson |
| EPI_ISL_961364 | Toronto Invasive Bacterial Diseases Network | McMaster University | Allison McGeer, Patryk Aftanas, Hooman Derakhshani, Angel Li, Kuganya Nirmalarajah, Emily Panousis, Ahmed Draia, Jalees Nasir, Michael Surette, Samira Mubareka, Andrew G. McArthur |
| EPI_ISL_962263 | Seattle Flu Study | Seattle Flu Study | Deborah A. Nickerson, Chris D. Frazar, Jover Lee, Benjamin Pelle, Erica Ryke, Matthew Richardson, Amanda Adler, Elisabeth Brandstetter, Peter D. Han, Kairsten Fay, Misja Ilcisin, Kirsten Lacombe, Thomas R. Sibley, Melissa Truong, Caitlin R. Wolf, Karen Cowgill, Stephanie Schrag, Jeff Duchin, Michael Boeckh, Janet A. Englund, Michael Famulare, Barry R. Lutz, Mark J. Rieder, Lea M. Starita, Matthew Thompson, Helen Y. Chu, Trevor Bedford, Jay Shendure |
| EPI_ISL_962522 | Tokyo Metropolitan Institute of Public Health | Tokyo Metropolitan Institute of Public Health | Masaki Hayashi, Takako Yamazaki, Mami Nagashima,Takushi Fujiwara, Takashi Chiba, Kenji Sadamasu |
| EPI_ISL_962609, EPI_ISL_962614, EPI_ISL_962616, EPI_ISL_962617 | San Diego County Public Health Laboratory | Andersen lab at Scripps Research | SEARCH Alliance San Diego with Tracy Basler, Jovan Shephard, Brett Austin |
| EPI_ISL_965553, EPI_ISL_965577, EPI_ISL_965580, EPI_ISL_965590, EPI_ISL_965639, EPI_ISL_965688, EPI_ISL_965689, EPI_ISL_965705, EPI_ISL_965714, EPI_ISL_965769, EPI_ISL_965799 | | | |
| see above | Dutch COVID-19 response team | Medical Microbiology, Maastricht University Medical Centre | Jozef Dingemans*, Brian van der Veer*, Erik Beuken, Carmen Reumkens, Lieke van Alphen, Christian Hoebe, Paul Savelkoul |
| EPI_ISL_971260, EPI_ISL_972244, EPI_ISL_973148 | Department of Virus and Microbiological Special Diagnostics, Statens Serum Institut, Copenhagen, Denmark | Aalborg University | Danish Covid-19 Genome Consortium |
| EPI_ISL_975973, EPI_ISL_975974, EPI_ISL_975975, EPI_ISL_975976, EPI_ISL_975977, EPI_ISL_975978, EPI_ISL_975979, EPI_ISL_975980, EPI_ISL_975981, EPI_ISL_975982, EPI_ISL_975983, EPI_ISL_975984, EPI_ISL_975985, EPI_ISL_975986, EPI_ISL_975987, EPI_ISL_975988, EPI_ISL_975989, EPI_ISL_975990, EPI_ISL_975991, EPI_ISL_975992, EPI_ISL_975993, EPI_ISL_975994, EPI_ISL_975995, EPI_ISL_975996, EPI_ISL_975997, EPI_ISL_975998, EPI_ISL_975999, EPI_ISL_976000, EPI_ISL_976001, EPI_ISL_976002, EPI_ISL_976003, EPI_ISL_976004, EPI_ISL_976005, EPI_ISL_976006, EPI_ISL_976007, EPI_ISL_976008, EPI_ISL_976009, EPI_ISL_976010, EPI_ISL_976011, EPI_ISL_976012, EPI_ISL_976013, EPI_ISL_976014, EPI_ISL_976015, EPI_ISL_976016, EPI_ISL_976017, EPI_ISL_976018, EPI_ISL_976019, EPI_ISL_976020, EPI_ISL_976021, EPI_ISL_976022, EPI_ISL_976023, EPI_ISL_976024, EPI_ISL_976025, EPI_ISL_976026, EPI_ISL_976027, EPI_ISL_976028, EPI_ISL_976029, EPI_ISL_976030, EPI_ISL_976031, EPI_ISL_976032, EPI_ISL_976033, EPI_ISL_976034, EPI_ISL_976035, EPI_ISL_976036, EPI_ISL_976037, EPI_ISL_976038, EPI_ISL_976039, EPI_ISL_976040, EPI_ISL_976041, EPI_ISL_976042, EPI_ISL_976043, EPI_ISL_976044, EPI_ISL_976045, EPI_ISL_976046, EPI_ISL_976047, EPI_ISL_976048, EPI_ISL_976049, EPI_ISL_976050, EPI_ISL_976051, EPI_ISL_976052, EPI_ISL_976053, EPI_ISL_976054, EPI_ISL_976055, EPI_ISL_976056, EPI_ISL_976057, EPI_ISL_976058, EPI_ISL_976059, EPI_ISL_976060, EPI_ISL_976061, EPI_ISL_976062, EPI_ISL_976063 | | | |
| see above | BCCDC Public Health Laboratory | BCCDC Public Health Laboratory | Prystajecky Natalie, Linda Hoang, Dan Fornika, John Tyson, Shannon Russell, Kim Macdonald, Kimia Kamelian, Ana Pacagnella, Corrinne Ng, Loretta Janz, Robert Azana Terry Snutch, Mel Krajden |
| EPI_ISL_977204 | ULSS 7 Pedemontana - Distretto 1 | Istituto Zooprofilattico Sperimentale delle Venezie | Adelaide Milani, Alessia Schivo, Annalisa Salviato, Erika Giorgia Quaranta, Ambra Pastori, Bianca Zecchin, Alice Fusaro, Isabella Monne, Calogero Terregino, Antonia Ricci |
| EPI_ISL_977352, EPI_ISL_977353, EPI_ISL_977354 | University of Zambia, School of Veterinary Medicine | UNZAVET and PATH | Mulenga Mwenda-Chimfwembe, Ngonda Saasa, Daniel Bridges |
| EPI_ISL_977999, EPI_ISL_978000, EPI_ISL_978002, EPI_ISL_978004 | Chiu Laboratory, University of California, San Francisco | Chiu Laboratory, University of California, San Francisco | Charles Chiu, Xianding (Wayne) Deng, Candace Wang, Venice Servellita, Jill Hacker, Debra Wadford |
| EPI_ISL_979247 | Institute of Microbiology and Immunology, Faculty of Medicine, University of Ljubljana | Institute of Microbiology and Immunology, Faculty of Medicine, University of Ljubljana | Samo Zakotnik, Tomaž Mark Zorec, Matic Brvar, Doroteja Vlaj, Patricija Pozvek,Špela Pleh, Miša Korva, Mario Poljak, Tatjana Avši - Županc |
| EPI_ISL_979716, EPI_ISL_979717 | Humboldt County Public Health Laboratory | Chan-Zuckerberg Biohub | CZB Cliahub Consortium |
| EPI_ISL_981850, EPI_ISL_981851, EPI_ISL_981852, EPI_ISL_981853, EPI_ISL_981854, EPI_ISL_981855, EPI_ISL_981856, EPI_ISL_981857, EPI_ISL_981858, EPI_ISL_981859, EPI_ISL_981860, EPI_ISL_981861, EPI_ISL_981862, EPI_ISL_981863 | | | |
| see above | Microbiology Service, Hospital Universitario Clinico San Cecilio, Granada | Microbiology Service, Hospital Universitario Clinico San Cecilio, Granada | Adolfo de Salazar, Natalia Chueca, Laura Viñuela, Ana Fuentes, Federico García |
| EPI_ISL_982116, EPI_ISL_982117, EPI_ISL_982119, EPI_ISL_982122, EPI_ISL_982124, EPI_ISL_982126, EPI_ISL_982129, EPI_ISL_982150, EPI_ISL_982151, EPI_ISL_982152, EPI_ISL_982157, EPI_ISL_982160, EPI_ISL_982163, EPI_ISL_982167, EPI_ISL_982168, EPI_ISL_982172, EPI_ISL_982173, EPI_ISL_982184, EPI_ISL_982185, EPI_ISL_982186, EPI_ISL_982187, EPI_ISL_982188, EPI_ISL_982189, EPI_ISL_982190, EPI_ISL_982191, EPI_ISL_982192, EPI_ISL_982193, EPI_ISL_982194, EPI_ISL_982195, EPI_ISL_982196, EPI_ISL_982197, EPI_ISL_982198, EPI_ISL_982199 | | | |
| see above | Hôpital Henri Mondor | Department of Virology, Henri Mondor University Hospital, Assistance Publique Hôpitaux de Paris, Université Paris-Est Créteil, INSERM U955 | Christophe Rodriguez, Slim Fourati, Vanessa Demontant, Guillaume Gricourt, Melissa N'Debi, Alexandre Soulier, Elisabeth Trawinski, Jean-Michel Pawlotsky |
| EPI_ISL_982776, EPI_ISL_982777 | Kentucky State Public Health Lab | Kentucky State Public Health Lab | Stephanie Lunn, Karim George, Joshua Tobias, William Grooms, Vaneet Arora, Matthew Johnson, Rachel Zinner, Rhonda Lucas |
| EPI_ISL_983861, EPI_ISL_983862 | Colorado Department of Public Health and Environment | Colorado Department of Puplic Health and Environment | Laura Bankers, Molly C. Hetherington-Rauth, Diana Ir, Shannon Ely, Shannon R. Matzinger, Sarah Elizabeth Totten, Emily A. Travanty |
