## Supplementary material for "Detection and characterization of the SARS-CoV-2 lineage B.1.526 in New York": Supp. Table 3 GISAID Acknowledgment part 2: gisaid_hcov-19_acknowledgement_table_2021_02_13_00-14.pdf

We gratefully acknowledge the following Authors from the Originating laboratories responsible for obtaining the specimens, as well as the Submitting laboratories where the genome data were generated and shared via GISAID, on which this research is based.

All Submitters of data may be contacted directly via www.gisaid.org

Authors are sorted alphabetically.

| Accession ID | Originating Laboratory | Submitting Laboratory | Authors |
|---|---|---|---|
| EPI_ISL_702633 | Virology Department, Sheffield Teaching Hospitals NHS Foundation Trust/Department of Infection, Immunity and Cardiovascular Disease, The Medical School, University of Sheffield | COVID-19 Genomics UK (COG-UK) Consortium | Thushan de Silva, Matthew Parker, Nikki Smith, Adri Angyal, Rebecca Brown, Luke Green, Rachel Tucker, Paul Parsons, Danielle Groves, Katie Johnson, Laura Carrilero, Alex Keeley, Dave Partridge, Matthew Wyles, Benjamin Lindsey, Mehmet Yavuz, Mohammad Raza, Cariad Evans |
| EPI_ISL_704583, EPI_ISL_704780 | Queens Medical Centre, Clinical Microbiology Department / DeepSeq Nottingham | COVID-19 Genomics UK (COG-UK) Consortium | Gemma Clark, Wendy Smith, Manjinder Khakh, Vicki M Fleming, Michelle M Lister, Hannah Howson-Wells, Jonathan Ball, Patrick McClure, Joseph Chappell, Theocharis Tsoleridis, Nadine Holmes, Matthew Carlisle, Christopher Moore, Fei Sang, Johnny Debebe, Victoria Wright, Matthew Loose |
| EPI_ISL_705204, EPI_ISL_705421 | Virology Department, Sheffield Teaching Hospitals NHS Foundation Trust/Department of Infection, Immunity and Cardiovascular Disease, The Medical School, University of Sheffield | COVID-19 Genomics UK (COG-UK) Consortium | Thushan de Silva, Matthew Parker, Nikki Smith, Adri Angyal, Rebecca Brown, Luke Green, Rachel Tucker, Paul Parsons, Danielle Groves, Katie Johnson, Laura Carrilero, Alex Keeley, Dave Partridge, Matthew Wyles, Benjamin Lindsey, Mehmet Yavuz, Mohammad Raza, Cariad Evans |
| EPI_ISL_706435, EPI_ISL_706436, EPI_ISL_706437 | Queens Medical Centre, Clinical Microbiology Department / DeepSeq Nottingham | COVID-19 Genomics UK (COG-UK) Consortium | Gemma Clark, Wendy Smith, Manjinder Khakh, Vicki M Fleming, Michelle M Lister, Hannah Howson-Wells, Jonathan Ball, Patrick McClure, Joseph Chappell, Theocharis Tsoleridis, Nadine Holmes, Matthew Carlisle, Christopher Moore, Fei Sang, Johnny Debebe, Victoria Wright, Matthew Loose |
| EPI_ISL_706976, EPI_ISL_706984, EPI_ISL_706985, EPI_ISL_706986 | Virology Department, Sheffield Teaching Hospitals NHS Foundation Trust/Department of Infection, Immunity and Cardiovascular Disease, The Medical School, University of Sheffield | COVID-19 Genomics UK (COG-UK) Consortium | Thushan de Silva, Matthew Parker, Nikki Smith, Adri Angyal, Rebecca Brown, Luke Green, Rachel Tucker, Paul Parsons, Danielle Groves, Katie Johnson, Laura Carrilero, Alex Keeley, Dave Partridge, Matthew Wyles, Benjamin Lindsey, Mehmet Yavuz, Mohammad Raza, Cariad Evans |
| EPI_ISL_712099, EPI_ISL_712100, EPI_ISL_712119, EPI_ISL_712120, EPI_ISL_712121, EPI_ISL_712122, EPI_ISL_712123, EPI_ISL_712124, EPI_ISL_712125, EPI_ISL_712126, EPI_ISL_712127, EPI_ISL_712128, EPI_ISL_712250, EPI_ISL_712251, EPI_ISL_712252, EPI_ISL_712253, EPI_ISL_712295, EPI_ISL_712321, EPI_ISL_712340, EPI_ISL_712341, EPI_ISL_712342, EPI_ISL_712343, EPI_ISL_712344, EPI_ISL_712345, EPI_ISL_712346, EPI_ISL_712417, EPI_ISL_712431, EPI_ISL_712439, EPI_ISL_712446, EPI_ISL_712447, EPI_ISL_712448, EPI_ISL_712455, EPI_ISL_712456, EPI_ISL_712457, EPI_ISL_712458, EPI_ISL_712459, EPI_ISL_712460, EPI_ISL_712461, EPI_ISL_712462, EPI_ISL_712463, EPI_ISL_712464, EPI_ISL_712465, EPI_ISL_712466, EPI_ISL_712467, EPI_ISL_712468, EPI_ISL_712469, EPI_ISL_712470, EPI_ISL_712471, EPI_ISL_712472, EPI_ISL_712473, EPI_ISL_712474, EPI_ISL_712475, EPI_ISL_712476, EPI_ISL_712477, EPI_ISL_712478, EPI_ISL_712479, EPI_ISL_712480, EPI_ISL_712481, EPI_ISL_712482, EPI_ISL_712483, EPI_ISL_712484, EPI_ISL_712485, EPI_ISL_712486, EPI_ISL_712487, EPI_ISL_712488, EPI_ISL_712489, EPI_ISL_712490, EPI_ISL_712491, EPI_ISL_712492, EPI_ISL_712493, EPI_ISL_712494, EPI_ISL_712495, EPI_ISL_712496, EPI_ISL_712497, EPI_ISL_712498, EPI_ISL_712499, EPI_ISL_712500, EPI_ISL_712501, EPI_ISL_712502, EPI_ISL_712503, EPI_ISL_712504, EPI_ISL_712505, EPI_ISL_712506, EPI_ISL_712507, EPI_ISL_712508, EPI_ISL_712509, EPI_ISL_712510, EPI_ISL_712511, EPI_ISL_712512, EPI_ISL_712513, EPI_ISL_712514, EPI_ISL_712515, EPI_ISL_712516, EPI_ISL_712517, EPI_ISL_712518, EPI_ISL_712519, EPI_ISL_712520, EPI_ISL_712521, EPI_ISL_712522, EPI_ISL_712523, EPI_ISL_712524, EPI_ISL_712525, EPI_ISL_712526, EPI_ISL_712527, EPI_ISL_712528, EPI_ISL_712529, EPI_ISL_712530 | | | |
| see above | Department of Virus and Microbiological Special Diagnostics, Statens Serum Institut, Copenhagen, Denmark | Albertsen Lab, Department of Chemistry and Bioscience, Aalborg University, Denmark | Danish Covid-19 Genome Consortium |
| EPI_ISL_717703, EPI_ISL_717708, EPI_ISL_717709, EPI_ISL_717710, EPI_ISL_717711, EPI_ISL_717712, EPI_ISL_717713 | Area of Virology, Serology and Virology Division (SAViD), New South Wales Health Pathology Randwick | Virology Research Laboratory; Area of Virology, Serology and Virology Division (SAViD), New South Wales Health Pathology Randwick | Foster, C.; Au, J.; Ruiz Silva, M.; Deveson, I.; Bull, R.; Van Hal, S.; Rawlinson, W. |
| EPI_ISL_722299, EPI_ISL_722315, EPI_ISL_722402, EPI_ISL_722461, EPI_ISL_722462, EPI_ISL_722796, EPI_ISL_722797, EPI_ISL_722798, EPI_ISL_722799 | Dutch COVID-19 response team | Erasmus Medical Center | Bas Oude Munnink, Reina Sikkema, David Nieuwenhuijse, Irina Chestakova, Anne van der Linden, Marjan Boter, Emmanuelle Munger, Corine GeurtsvanKessel, Annemiek van der Eijk, Richard Molenkamp, Marion Koopmans, on behalf of the Dutch national COVID-19 response team. |
| EPI_ISL_722974, EPI_ISL_722996, EPI_ISL_722997, EPI_ISL_722998, EPI_ISL_723001, EPI_ISL_723002, EPI_ISL_723003, EPI_ISL_723004, EPI_ISL_723005, EPI_ISL_723038, EPI_ISL_723039, EPI_ISL_723040, EPI_ISL_723041, EPI_ISL_723042, EPI_ISL_723043 | | | |
| see above | Respiratory Virus Unit, National Infection Service, Public Health England | COVID-19 Genomics UK (COG-UK) Consortium | PHE Covid Sequencing Team |
| EPI_ISL_723139, EPI_ISL_723140, EPI_ISL_723146 | Minnesota Department of Health, Public Health Laboratory | Minnesota Department of Health, Public Health Laboratory | Alexandra Lorentz, Jacob Garfin, Matt Plumb, and Xiong Wang |
| EPI_ISL_723933, EPI_ISL_723935, EPI_ISL_723936, EPI_ISL_723947, EPI_ISL_723948 | Department of Pathology, University of Cambridge | COVID-19 Genomics UK (COG-UK) Consortium | Aminu S. Jahun, Yasmin Chaudhry, Grant Hall, Iliana Georgana, Myra Hosmillo, Martin D. Curran, Malte Pinckert, Surendra Parmar, Ian Goodfellow |
| EPI_ISL_724217, EPI_ISL_724219, EPI_ISL_724220 | West of Scotland Specialist Virology Centre, NHSGGC / MRC-University of Glasgow Centre for Virus Research | COVID-19 Genomics UK (COG-UK) Consortium | Ana da Silva Filipe, Natasha Johnson, Kathy Smollett, Daniel Mair, Stephen Carmichael, Alice Broos, Lily Tong, Jenna Nichols, Kyriaki Nomikou; Sarah McDonald; Richard Orton, Joseph Hughes, Sreenu Vattipally, David L Robertson; Alasdair MacLean, Rory Gunson; Sharif Shaaban, Matthew Holden; Rachel Blacow, Guy Mollett, Kathy Li, James Shepherd, Antonia Ho, Emma Thomson |
| EPI_ISL_724228, EPI_ISL_724229, EPI_ISL_724231, EPI_ISL_724232, EPI_ISL_724233, EPI_ISL_724234, EPI_ISL_724235, EPI_ISL_724236, EPI_ISL_724237, EPI_ISL_724238, EPI_ISL_724239, EPI_ISL_724240, EPI_ISL_724241, EPI_ISL_724242, EPI_ISL_724243, EPI_ISL_724244, EPI_ISL_724246, EPI_ISL_724247 | | | |
| see above | Virology Department, Royal Infirmary of Edinburgh, NHS Lothian / School of Biological Sciences, University of Edinburgh / Institute of Genetics and Molecular Medicine, University of Edinburgh | COVID-19 Genomics UK (COG-UK) Consortium | McHugh M, Dewar R, Rooke S, Gallagher M, Balcaza C, O'Toole Á, Scher E, Hill V, McCrone JT, Colquhoun R, Yu X, Jackson B, Rambaut A, Williams TC, Templeton K |
| EPI_ISL_724357, EPI_ISL_724358, EPI_ISL_724361, EPI_ISL_724363, EPI_ISL_724367, EPI_ISL_724368, EPI_ISL_724373, EPI_ISL_724374, EPI_ISL_724376, EPI_ISL_724379, EPI_ISL_724381, EPI_ISL_724385, EPI_ISL_724386, EPI_ISL_724387, EPI_ISL_724388, EPI_ISL_724389, EPI_ISL_724390, EPI_ISL_724391, EPI_ISL_724392, EPI_ISL_724393, EPI_ISL_724394, EPI_ISL_724395 | | | |
| see above | Liverpool Clinical Laboratories | COVID-19 Genomics UK (COG-UK) Consortium | Sam Haldenby, Anita Lucaci, Steve Paterson, Julian Hiscox, Alistair Darby, M Almsaud, A Alrezaihi, Muhannad Alruwaili, Stuart D Armstrong, Jones Benjamin, Eleanor G Bentley, Anu Chawla, Jordan J Clark, Angela Cowell, Richard Eccles, Isabel García-Dorival, Matthew Gemmell, Alessandro Gerada, PKF Gilmore, Richard Gregory, Ximeng Han, Catherine Hartley, Margaret Hughes, Miren Iturriza-Gomara, James Johnson, L Luu, Jenifer Manson, Charlotte Nelson, Elaine O'Toole, Cassie Olateju, Rebekah Penrice-Randal , Lucille Rainbow, N.P Randle, Trevor Ian Robinson, Parul Sharma, Ghada T Shawli, James P Stewart, Neil Swainston, Ecaterina Vamos, Joanne Watts, Mark Whitehead |
| EPI_ISL_724598, EPI_ISL_724599, EPI_ISL_724600, EPI_ISL_724603, EPI_ISL_724604, EPI_ISL_724605 | University College London, Great Ormond Street Hospital for Children NHS Foundation Trust, Imperial College Healthcare NHS Trust | COVID-19 Genomics UK (COG-UK) Consortium | Sergi Castellano, Rachel Williams, Mark Kristiansen, Paola Resende Silva, Sunando Roy, Tony Brooks, Helena Tutill, Paola Niola, Patricia Dyal, Charlotte Williams, Leysa Forrest, Yasmin Panchbhaya, Jacqueline Findlay, Samuel Weeks, Julianne Brown, Kathryn Harris, Paul Randell, James Price, Alison Holmes, Judith Breuer |
| EPI_ISL_724908, EPI_ISL_724912, EPI_ISL_724914, EPI_ISL_724917, EPI_ISL_724918, EPI_ISL_724919, EPI_ISL_724920, EPI_ISL_724921, EPI_ISL_724922, EPI_ISL_724923 | Northumbria University / South Tees Hospitals NHS Foundation Trust / North Cumbria Integrated Care NHS Foundation Trust / North Tees and Hartlepool NHS Foundation Trust / Newcastle Hospitals NHS Foundation Trust | COVID-19 Genomics UK (COG-UK) Consortium | Darren L Smith,Andrew Nelson,Matthew Bashton,Greg R Young,Joshua Loh,John Allan,Mohammad A Tariq,Giles S Holt,Gary Black,Wen C Yew,Lynn Dover,Paul Baker,Steve Liggett,Sarah Essex,Jane Greenaway,Debra Padgett,Clive Graham,Garren Scott,Edward Barton,Emma Swindells,Brendan Payne,Jennifer Collins,Yusri Taha,Gary Eltringham |
| EPI_ISL_725029, EPI_ISL_725030, EPI_ISL_725031, EPI_ISL_725032, EPI_ISL_725036, EPI_ISL_725037, EPI_ISL_725039, EPI_ISL_725040, EPI_ISL_725042, EPI_ISL_725043, EPI_ISL_725044, EPI_ISL_725045, EPI_ISL_725046, EPI_ISL_725047, EPI_ISL_725048, EPI_ISL_725049, EPI_ISL_725050, EPI_ISL_725051, | | | |

| | | | |
|---|---|---|---|
| EPI_ISL_725052, EPI_ISL_725053, EPI_ISL_725054, EPI_ISL_725056, EPI_ISL_725057, EPI_ISL_725058, EPI_ISL_725059, EPI_ISL_725060, EPI_ISL_725061, EPI_ISL_725139, EPI_ISL_725140, EPI_ISL_725142, EPI_ISL_725145, EPI_ISL_725154, EPI_ISL_725158, EPI_ISL_725302, EPI_ISL_725303, EPI_ISL_725307, EPI_ISL_725309, EPI_ISL_725310, EPI_ISL_725311, EPI_ISL_725312, EPI_ISL_725313, EPI_ISL_725314, EPI_ISL_725321, EPI_ISL_725324, EPI_ISL_725325, EPI_ISL_725327, EPI_ISL_725328, EPI_ISL_725331, EPI_ISL_725365, EPI_ISL_725415, EPI_ISL_725416, EPI_ISL_725418, EPI_ISL_725421, EPI_ISL_725426, EPI_ISL_725427, EPI_ISL_725429, EPI_ISL_725431, EPI_ISL_725432, EPI_ISL_725434, EPI_ISL_725435, EPI_ISL_725436, EPI_ISL_725437, EPI_ISL_725439, EPI_ISL_725440, EPI_ISL_725441, EPI_ISL_725442, EPI_ISL_725443, EPI_ISL_725444, EPI_ISL_725445, EPI_ISL_725446, EPI_ISL_725447, EPI_ISL_725448, EPI_ISL_725449, EPI_ISL_725450, EPI_ISL_725451, EPI_ISL_725452, EPI_ISL_725453, EPI_ISL_725454, EPI_ISL_725455, EPI_ISL_725456, EPI_ISL_725457, EPI_ISL_725460, EPI_ISL_725461, EPI_ISL_725462, EPI_ISL_725465, EPI_ISL_725470, EPI_ISL_725473, EPI_ISL_725474, EPI_ISL_725477, EPI_ISL_725478, EPI_ISL_725479, EPI_ISL_725482, EPI_ISL_725483, EPI_ISL_725484, EPI_ISL_725485, EPI_ISL_725486, EPI_ISL_725488, EPI_ISL_725489, EPI_ISL_725490, EPI_ISL_725491, EPI_ISL_725511, EPI_ISL_725514 | | | |
| see above | Quadram Institute Bioscience | COVID-19 Genomics UK (COG-UK) Consortium | Dave J. Baker, Gemma L. Kay, Alp Aydin, Thanh Le-Viet, Steven Rudder, Ana P. Tedim, Anastasia Kolyva, Maria Diaz, Leonardo de Oliveira Martins, Nabil-Fareed Alikhan, Lizzie Meadows, Rachael Stanley, Ngozi Elumogo, Muhammed Yasir, Nicholas M. Thomson, Alexander J Trotter, Rachel Gilroy, Samuel Bloomfield, Claire Stuart, Andrew Bell, Reenesh Prakash, Samir Dervisevic, Alison E. Mather, John Wain, Mark Webber, Andrew J. Page, Justin O'Grady |
| EPI_ISL_725606, EPI_ISL_725607, EPI_ISL_725608, EPI_ISL_725609, EPI_ISL_725610, EPI_ISL_725611, EPI_ISL_725612, EPI_ISL_725613, EPI_ISL_725614, EPI_ISL_725615, EPI_ISL_725616, EPI_ISL_725617, EPI_ISL_725618, EPI_ISL_725619, EPI_ISL_725620, EPI_ISL_725621, EPI_ISL_725622, EPI_ISL_725623, EPI_ISL_725624, EPI_ISL_725625, EPI_ISL_725626, EPI_ISL_725627, EPI_ISL_725628, EPI_ISL_725629, EPI_ISL_725630, EPI_ISL_725631, EPI_ISL_725632, EPI_ISL_725633, EPI_ISL_725634, EPI_ISL_725635, EPI_ISL_725636, EPI_ISL_725637, EPI_ISL_725638, EPI_ISL_725639, EPI_ISL_725640, EPI_ISL_725641, EPI_ISL_725642 | | | |
| see above | Queens Medical Centre, Clinical Microbiology Department / DeepSeq Nottingham | COVID-19 Genomics UK (COG-UK) Consortium | Gemma Clark, Wendy Smith, Manjinder Khakh, Vicki M Fleming, Michelle M Lister, Hannah Howson-Wells, Jonathan Ball, Patrick McClure, Joseph Chappell, Theocharis Tsoleridis, Nadine Holmes, Matthew Carlisle, Christopher Moore, Fei Sang, Johnny Debebe, Victoria Wright, Matthew Loose |
| EPI_ISL_727871, EPI_ISL_727885, EPI_ISL_727905, EPI_ISL_727906, EPI_ISL_727920, EPI_ISL_727922, EPI_ISL_727923, EPI_ISL_727964, EPI_ISL_727966, EPI_ISL_727975, EPI_ISL_727987 | | | |
| see above | Virology Department, Sheffield Teaching Hospitals NHS Foundation Trust/Department of Infection, Immunity and Cardiovascular Disease, The Medical School, University of Sheffield | COVID-19 Genomics UK (COG-UK) Consortium | Thushan de Silva, Matthew Parker, Nikki Smith, Adri Angyal, Rebecca Brown, Luke Green, Rachel Tucker, Paul Parsons, Danielle Groves, Katie Johnson, Laura Carrilero, Alex Keeley, Dave Partridge, Matthew Wyles, Benjamin Lindsey, Mehmet Yavuz, Mohammad Raza, Cariad Evans |
| EPI_ISL_728180, EPI_ISL_728183, EPI_ISL_728184, EPI_ISL_728189, EPI_ISL_728191, EPI_ISL_728201 | National Public Health Laboratory, National Centre for Infectious Diseases | National Public Health Laboratory, National Centre for Infectious Diseases | Tze Minn Mak, Sophie Octavia, Zhenyang Zhou, Lin Cui, Raymond Tzer Pin Lin |
| EPI_ISL_728205 | Institute of Microbiology, Universidad San Francisco de Quito | Institute of Microbiology, Universidad San Francisco de Quito | Belén Prado-Vivar, Sully Márquez, Juan José Guadalupe, Monica Becerra-Wong, Diana Zambrano, Fredy Loor, Juan Zuñiga, Edison Chavez, Bernardo Gutiérrez, Verónica Barragán, Patricio Rojas-Silva, Gabriel Trueba, Michelle Grunauer, Paúl Cárdenas |
| EPI_ISL_728557, EPI_ISL_728558, EPI_ISL_728559, EPI_ISL_728560, EPI_ISL_728561, EPI_ISL_728562, EPI_ISL_728563 | Institute for Urban Disease Control and Prevention | COVID-19 Network Investigations (CONI) Alliance | Kamolthip Atsawawaranunt, Elizabeth Batty, Wasun Chantratita, Thanat Chookajorn, Stefan Fernandez, Angkana Huang, Anthony R. Jones, Khajohn Joonsalak, Chonticha Klungtong, Theerarat Kochakarn, Prayuth Kaewmalang, Amornmas Kongklieng, Namfon Kotanan, Krittikorn Kumpornsin, Wudtichai Manasatienkij, Anek Mungaomklang, Bhakbhoom Panthan, Pukkaporn Parnwijitkul, Ekawat Pasomsub, Kingkan Rakmanee, Insee Sensorn, Janjira Thaipadungpanit, Arporn Wangwiwatsin, Treewat Watthanachockchai |
| EPI_ISL_728570, EPI_ISL_728571, EPI_ISL_728572, EPI_ISL_728573, EPI_ISL_728574, EPI_ISL_728575, EPI_ISL_728576, EPI_ISL_728578, EPI_ISL_728579, EPI_ISL_728581, EPI_ISL_728582, EPI_ISL_728583, EPI_ISL_728584, EPI_ISL_728585, EPI_ISL_728588, EPI_ISL_728589, EPI_ISL_728590, EPI_ISL_728591, EPI_ISL_728595, EPI_ISL_728596, EPI_ISL_728600, EPI_ISL_728601, EPI_ISL_728602, EPI_ISL_728603, EPI_ISL_728606, EPI_ISL_728607, EPI_ISL_728608, EPI_ISL_728609, EPI_ISL_728610, EPI_ISL_728611, EPI_ISL_728612, EPI_ISL_728613, EPI_ISL_728618, EPI_ISL_728619, EPI_ISL_728622, EPI_ISL_728623, EPI_ISL_728624, EPI_ISL_728625, EPI_ISL_728626, EPI_ISL_728627, EPI_ISL_728628, EPI_ISL_728629, EPI_ISL_728630, EPI_ISL_728631, EPI_ISL_728649, EPI_ISL_728650, EPI_ISL_728652, EPI_ISL_728653, EPI_ISL_728654, EPI_ISL_728655, EPI_ISL_728656, EPI_ISL_728657, EPI_ISL_728658, EPI_ISL_728659, EPI_ISL_728662, EPI_ISL_728663, EPI_ISL_728664, EPI_ISL_728665, EPI_ISL_728666, EPI_ISL_728667, EPI_ISL_728695, EPI_ISL_728698, EPI_ISL_728699, EPI_ISL_728700, EPI_ISL_728702, EPI_ISL_728703, EPI_ISL_728721, EPI_ISL_728722, EPI_ISL_728746 | | | |
| see above | Dutch COVID-19 response team | National Institute for Public Health and the Environment (RIVM) | Adam Meijer, Harry Vennema, Jeroen Cremer, Sharon van den Brink, Bas van der Veer, AnneMarie van den Brandt, Florian Zwagemaker, Dennis Schmitz, Chantal Reusken, on behalf of the national COVID-19 response team |
| EPI_ISL_728764, EPI_ISL_728769, EPI_ISL_728771, EPI_ISL_728772 | Viollier AG | Department of Biosystems Science and Engineering, ETH Zürich | Chaoran Chen, Sarah Nadeau, Catharine Aquino, Ivan Topolsky, Pedro Ferreira, Philipp Jablonski, Susana Posada-Céspedes, Andreia Cabral de Gouvea, Maria Domenica Moccia, Simon Grüter, Timothy Sykes, Lennart Opitz, Ralph Schlapbach, Christiane Beckmann, Maurice Redondo, Olivier Kobel, Christoph Noppen, Sophie Seidel, Noemie Santamaria de Souza, Niko Beerenwinkel, Tanja Stadler |
| EPI_ISL_728776 | Viollier AG | Department of Biosystems Science and Engineering, ETH Zürich | Christian Beisel, Sarah Nadeau, Chaoran Chen, Ivan Topolsky, Pedro Ferreira, Philipp Jablonski, Susana Posada-Céspedes, Tobias Schär, Ina Nissen, Natascha Santacroce, Elodie Burcklen, Christiane Beckmann, Maurice Redondo, Olivier Kobel, Christoph Noppen, Sophie Seidel, Noemie Santamaria de Souza, Niko Beerenwinkel, Tanja Stadler |
| EPI_ISL_728779 | Viollier AG | Department of Biosystems Science and Engineering, ETH Zürich | Chaoran Chen, Sarah Nadeau, Catharine Aquino, Ivan Topolsky, Pedro Ferreira, Philipp Jablonski, Susana Posada-Céspedes, Andreia Cabral de Gouvea, Maria Domenica Moccia, Simon Grüter, Timothy Sykes, Lennart Opitz, Ralph Schlapbach, Christiane Beckmann, Maurice Redondo, Olivier Kobel, Christoph Noppen, Sophie Seidel, Noemie Santamaria de Souza, Niko Beerenwinkel, Tanja Stadler |
| EPI_ISL_728808 | Viollier AG | Department of Biosystems Science and Engineering, ETH Zürich | Christian Beisel, Sarah Nadeau, Chaoran Chen, Ivan Topolsky, Pedro Ferreira, Philipp Jablonski, Susana Posada-Céspedes, Tobias Schär, Ina Nissen, Natascha Santacroce, Elodie Burcklen, Christiane Beckmann, Maurice Redondo, Olivier Kobel, Christoph Noppen, Sophie Seidel, Noemie Santamaria de Souza, Niko Beerenwinkel, Tanja Stadler |
| EPI_ISL_728809 | Viollier AG | Department of Biosystems Science and Engineering, ETH Zürich | Chaoran Chen, Sarah Nadeau, Catharine Aquino, Ivan Topolsky, Pedro Ferreira, Philipp Jablonski, Susana Posada-Céspedes, Andreia Cabral de Gouvea, Maria Domenica Moccia, Simon Grüter, Timothy Sykes, Lennart Opitz, Ralph Schlapbach, Christiane Beckmann, Maurice Redondo, Olivier Kobel, Christoph Noppen, Sophie Seidel, Noemie Santamaria de Souza, Niko Beerenwinkel, Tanja Stadler |
| EPI_ISL_728828, EPI_ISL_728882 | Viollier AG | Department of Biosystems Science and Engineering, ETH Zürich | Christian Beisel, Sarah Nadeau, Chaoran Chen, Ivan Topolsky, Pedro Ferreira, Philipp Jablonski, Susana Posada-Céspedes, Tobias Schär, Ina Nissen, Natascha Santacroce, Elodie Burcklen, Christiane Beckmann, Maurice Redondo, Olivier Kobel, Christoph Noppen, Sophie Seidel, Noemie Santamaria de Souza, Niko Beerenwinkel, Tanja Stadler |
| EPI_ISL_728886, EPI_ISL_728887, EPI_ISL_728888, EPI_ISL_728889, EPI_ISL_728890, EPI_ISL_728891, EPI_ISL_728894, EPI_ISL_728895, EPI_ISL_728898, EPI_ISL_728899, EPI_ISL_728901, EPI_ISL_728902, EPI_ISL_728903, EPI_ISL_728904, EPI_ISL_728905, EPI_ISL_728910, EPI_ISL_728917, EPI_ISL_728920, EPI_ISL_728925, EPI_ISL_728926, EPI_ISL_728927, EPI_ISL_728933, EPI_ISL_728936, EPI_ISL_728937, EPI_ISL_728949, EPI_ISL_728952, EPI_ISL_728953, EPI_ISL_728954, EPI_ISL_728958, EPI_ISL_728962, EPI_ISL_728964, EPI_ISL_728969, EPI_ISL_728980, EPI_ISL_728988, EPI_ISL_728990, EPI_ISL_728997, EPI_ISL_728999, EPI_ISL_729001, EPI_ISL_729003, EPI_ISL_729005, EPI_ISL_729009, EPI_ISL_729010, EPI_ISL_729012, EPI_ISL_729013, EPI_ISL_729014, EPI_ISL_729016, EPI_ISL_729017, EPI_ISL_729020, EPI_ISL_729023, EPI_ISL_729024, EPI_ISL_729026, EPI_ISL_729027, EPI_ISL_729028, EPI_ISL_729031, EPI_ISL_729033, EPI_ISL_729043, EPI_ISL_729044 | | | |
| see above | Viollier AG | Department of Biosystems Science and Engineering, ETH Zürich | Chaoran Chen, Sarah Nadeau, Catharine Aquino, Ivan Topolsky, Pedro Ferreira, Philipp Jablonski, Susana Posada-Céspedes, Andreia Cabral de Gouvea, Maria Domenica Moccia, Simon Grüter, Timothy Sykes, Lennart Opitz, Ralph Schlapbach, Christiane Beckmann, Maurice Redondo, Olivier Kobel, Christoph Noppen, Sophie Seidel, Noemie Santamaria de Souza, Niko Beerenwinkel, Tanja Stadler |
| EPI_ISL_729050, EPI_ISL_729057, EPI_ISL_729059, EPI_ISL_729060, EPI_ISL_729061, EPI_ISL_729062, EPI_ISL_729063, EPI_ISL_729064, EPI_ISL_729065, EPI_ISL_729066 | Viollier AG | Department of Biosystems Science and Engineering, ETH Zürich | Christian Beisel, Sarah Nadeau, Chaoran Chen, Ivan Topolsky, Pedro Ferreira, Philipp Jablonski, Susana Posada-Céspedes, Tobias Schär, Ina Nissen, Natascha Santacroce, Elodie Burcklen, Christiane Beckmann, Maurice Redondo, Olivier Kobel, Christoph Noppen, Sophie Seidel, Noemie Santamaria de Souza, Niko Beerenwinkel, Tanja Stadler |
| EPI_ISL_729139, EPI_ISL_729140, EPI_ISL_729141, EPI_ISL_729142, EPI_ISL_729143, EPI_ISL_729144, EPI_ISL_729145, EPI_ISL_729146, EPI_ISL_729147, EPI_ISL_729148, EPI_ISL_729149, EPI_ISL_729150, EPI_ISL_729151 | | | |
| see above | Viollier AG | Department of Biosystems Science and Engineering, ETH Zürich | Chaoran Chen, Sarah Nadeau, Catharine Aquino, Ivan Topolsky, Pedro Ferreira, Philipp Jablonski, Susana Posada-Céspedes, Andreia Cabral de Gouvea, Maria Domenica Moccia, Simon Grüter, Timothy Sykes, Lennart Opitz, Ralph Schlapbach, Christiane Beckmann, Maurice Redondo, Olivier Kobel, Christoph Noppen, Sophie Seidel, Noemie Santamaria de Souza, Niko Beerenwinkel, Tanja Stadler |
| EPI_ISL_729156, EPI_ISL_729157, EPI_ISL_729158, EPI_ISL_729159, EPI_ISL_729160, EPI_ISL_729161, EPI_ISL_729162, EPI_ISL_729163, EPI_ISL_729164, EPI_ISL_729165, EPI_ISL_729166, EPI_ISL_729167, EPI_ISL_729168, EPI_ISL_729169, EPI_ISL_729170, EPI_ISL_729171, EPI_ISL_729172, EPI_ISL_729173, EPI_ISL_729174, EPI_ISL_729175, EPI_ISL_729176, EPI_ISL_729177, EPI_ISL_729178, EPI_ISL_729179, EPI_ISL_729180, EPI_ISL_729181, EPI_ISL_729182, EPI_ISL_729183, EPI_ISL_729184, EPI_ISL_729185, EPI_ISL_729186 | | | |
| see above | Viollier AG | Department of Biosystems Science and Engineering, ETH Zürich | Christian Beisel, Sarah Nadeau, Chaoran Chen, Ivan Topolsky, Pedro Ferreira, Philipp Jablonski, Susana Posada-Céspedes, Tobias Schär, Ina Nissen, Natascha Santacroce, Elodie Burcklen, Christiane Beckmann, Maurice Redondo, Olivier Kobel, Christoph Noppen, Sophie Seidel, Noemie Santamaria de Souza, Niko Beerenwinkel, Tanja Stadler |
| EPI_ISL_730654, EPI_ISL_730660, EPI_ISL_730661, EPI_ISL_730662, EPI_ISL_730663, EPI_ISL_730665, EPI_ISL_730666, EPI_ISL_730669, EPI_ISL_730670, EPI_ISL_730671, EPI_ISL_730672, EPI_ISL_730674, EPI_ISL_730677, EPI_ISL_730679, EPI_ISL_730681, EPI_ISL_730684, EPI_ISL_730687, EPI_ISL_730688, EPI_ISL_730689, EPI_ISL_730690, EPI_ISL_730691, EPI_ISL_730693, EPI_ISL_730694, EPI_ISL_730698, EPI_ISL_730700, EPI_ISL_730701, EPI_ISL_730703, EPI_ISL_730706, EPI_ISL_730707, EPI_ISL_730708, EPI_ISL_730713, EPI_ISL_730714, EPI_ISL_730715, EPI_ISL_730716, EPI_ISL_730717, EPI_ISL_730719, | | | |

| | | | |
|---|---|---|---|
| EPI_ISL_730720, EPI_ISL_730721 | | | |
| see above | Lighthouse Lab in Alderley Park | Wellcome Sanger Institute for the COVID-19 Genomics UK (COG-UK) Consortium | Jacquelyn Wynn, Mairead Hyland, The Lighthouse Lab in Alderley Park and Alex Alderton, Roberto Amato, Sonia Goncalves, Ewan Harrison, David K. Jackson, Ian Johnston, Dominic Kwiatkowski, Cordelia Langford, John Sillitoe on behalf of the Wellcome Sanger Institute COVID-19 Surveillance Team |
| EPI_ISL_730723 | Lighthouse Lab in Glasgow | Wellcome Sanger Institute for the COVID-19 Genomics UK (COG-UK) Consortium | Harper VanSteenhouse, Yumi Kasai, David Gray, Carol Clugston, Anna Dominiczak and Alex Alderton, Roberto Amato, Sonia Goncalves, Ewan Harrison, David K. Jackson, Ian Johnston, Dominic Kwiatkowski, Cordelia Langford, John Sillitoe on behalf of the Wellcome Sanger Institute COVID-19 Surveillance Team |
| EPI_ISL_730724, EPI_ISL_730725, EPI_ISL_730726, EPI_ISL_730727, EPI_ISL_730728, EPI_ISL_730730, EPI_ISL_730735, EPI_ISL_730736, EPI_ISL_730738, EPI_ISL_730739, EPI_ISL_730740, EPI_ISL_730741, EPI_ISL_730744, EPI_ISL_730745, EPI_ISL_730746, EPI_ISL_730747, EPI_ISL_730748, EPI_ISL_730749, EPI_ISL_730755, EPI_ISL_730756, EPI_ISL_730757, EPI_ISL_730760, EPI_ISL_730761, EPI_ISL_730762, EPI_ISL_730763, EPI_ISL_730764, EPI_ISL_730766, EPI_ISL_730767, EPI_ISL_730768, EPI_ISL_730769, EPI_ISL_730770, EPI_ISL_730772, EPI_ISL_730775, EPI_ISL_730776, EPI_ISL_730779, EPI_ISL_730781, EPI_ISL_730782, EPI_ISL_730783, EPI_ISL_730787, EPI_ISL_730790, EPI_ISL_730791, EPI_ISL_730792, EPI_ISL_730797, EPI_ISL_730798, EPI_ISL_730799, EPI_ISL_730800, EPI_ISL_730807 | | | |
| see above | Lighthouse Lab in Alderley Park | Wellcome Sanger Institute for the COVID-19 Genomics UK (COG-UK) Consortium | Jacquelyn Wynn, Mairead Hyland, The Lighthouse Lab in Alderley Park and Alex Alderton, Roberto Amato, Sonia Goncalves, Ewan Harrison, David K. Jackson, Ian Johnston, Dominic Kwiatkowski, Cordelia Langford, John Sillitoe on behalf of the Wellcome Sanger Institute COVID-19 Surveillance Team |
| EPI_ISL_730808 | Lighthouse Lab in Glasgow | Wellcome Sanger Institute for the COVID-19 Genomics UK (COG-UK) Consortium | Harper VanSteenhouse, Yumi Kasai, David Gray, Carol Clugston, Anna Dominiczak and Alex Alderton, Roberto Amato, Sonia Goncalves, Ewan Harrison, David K. Jackson, Ian Johnston, Dominic Kwiatkowski, Cordelia Langford, John Sillitoe on behalf of the Wellcome Sanger Institute COVID-19 Surveillance Team |
| EPI_ISL_730810, EPI_ISL_730811, EPI_ISL_730812, EPI_ISL_730816, EPI_ISL_730817, EPI_ISL_730820, EPI_ISL_730821, EPI_ISL_730822, EPI_ISL_730827, EPI_ISL_730835, EPI_ISL_730836, EPI_ISL_730837, EPI_ISL_730838, EPI_ISL_730841, EPI_ISL_730842, EPI_ISL_730843, EPI_ISL_730851, EPI_ISL_730853, EPI_ISL_730856, EPI_ISL_730857, EPI_ISL_730863, EPI_ISL_730866, EPI_ISL_730868, EPI_ISL_730871, EPI_ISL_730872, EPI_ISL_730874, EPI_ISL_730879, EPI_ISL_730880, EPI_ISL_730882, EPI_ISL_730883, EPI_ISL_730884, EPI_ISL_730885, EPI_ISL_730888, EPI_ISL_730891, EPI_ISL_730892, EPI_ISL_730895, EPI_ISL_730896, EPI_ISL_730897, EPI_ISL_730899, EPI_ISL_730901, EPI_ISL_730903, EPI_ISL_730905, EPI_ISL_730906, EPI_ISL_730907, EPI_ISL_730909, EPI_ISL_730912, EPI_ISL_730915, EPI_ISL_730916, EPI_ISL_730921, EPI_ISL_730924, EPI_ISL_730929, EPI_ISL_730935, EPI_ISL_730939, EPI_ISL_730948, EPI_ISL_730959, EPI_ISL_730966, EPI_ISL_730967, EPI_ISL_730969, EPI_ISL_730970, EPI_ISL_730971, EPI_ISL_730973, EPI_ISL_730980, EPI_ISL_730987, EPI_ISL_730989, EPI_ISL_730994, EPI_ISL_731000, EPI_ISL_731006, EPI_ISL_731017, EPI_ISL_731019, EPI_ISL_731020, EPI_ISL_731024, EPI_ISL_731026, EPI_ISL_731027, EPI_ISL_731033, EPI_ISL_731035, EPI_ISL_731040, EPI_ISL_731045, EPI_ISL_731050, EPI_ISL_731052, EPI_ISL_731054, EPI_ISL_731055, EPI_ISL_731057, EPI_ISL_731059, EPI_ISL_731065, EPI_ISL_731066, EPI_ISL_731074, EPI_ISL_731076, EPI_ISL_731079, EPI_ISL_731080, EPI_ISL_731082, EPI_ISL_731083, EPI_ISL_731086, EPI_ISL_731087, EPI_ISL_731088, EPI_ISL_731094, EPI_ISL_731096, EPI_ISL_731104, EPI_ISL_731106, EPI_ISL_731107, EPI_ISL_731110, EPI_ISL_731116, EPI_ISL_731119, EPI_ISL_731120, EPI_ISL_731123, EPI_ISL_731126, EPI_ISL_731129, EPI_ISL_731130, EPI_ISL_731131, EPI_ISL_731133, EPI_ISL_731135, EPI_ISL_731137, EPI_ISL_731141, EPI_ISL_731143, EPI_ISL_731147, EPI_ISL_731153, EPI_ISL_731154, EPI_ISL_731157, EPI_ISL_731158, EPI_ISL_731160, EPI_ISL_731161, EPI_ISL_731169, EPI_ISL_731171, EPI_ISL_731181, EPI_ISL_731183, EPI_ISL_731188, EPI_ISL_731196, EPI_ISL_731199, EPI_ISL_731200, EPI_ISL_731204, EPI_ISL_731206, EPI_ISL_731214, EPI_ISL_731215, EPI_ISL_731233, EPI_ISL_731235, EPI_ISL_731240, EPI_ISL_731247, EPI_ISL_731248, EPI_ISL_731254, EPI_ISL_731260 | | | |
| see above | Lighthouse Lab in Alderley Park | Wellcome Sanger Institute for the COVID-19 Genomics UK (COG-UK) Consortium | Jacquelyn Wynn, Mairead Hyland, The Lighthouse Lab in Alderley Park and Alex Alderton, Roberto Amato, Sonia Goncalves, Ewan Harrison, David K. Jackson, Ian Johnston, Dominic Kwiatkowski, Cordelia Langford, John Sillitoe on behalf of the Wellcome Sanger Institute COVID-19 Surveillance Team |
| EPI_ISL_731261, EPI_ISL_731262 | Lighthouse Lab in Glasgow | Wellcome Sanger Institute for the COVID-19 Genomics UK (COG-UK) Consortium | Harper VanSteenhouse, Yumi Kasai, David Gray, Carol Clugston, Anna Dominiczak and Alex Alderton, Roberto Amato, Sonia Goncalves, Ewan Harrison, David K. Jackson, Ian Johnston, Dominic Kwiatkowski, Cordelia Langford, John Sillitoe on behalf of the Wellcome Sanger Institute COVID-19 Surveillance Team |
| EPI_ISL_731263 | Lighthouse Lab in Alderley Park | Wellcome Sanger Institute for the COVID-19 Genomics UK (COG-UK) Consortium | Jacquelyn Wynn, Mairead Hyland, The Lighthouse Lab in Alderley Park and Alex Alderton, Roberto Amato, Sonia Goncalves, Ewan Harrison, David K. Jackson, Ian Johnston, Dominic Kwiatkowski, Cordelia Langford, John Sillitoe on behalf of the Wellcome Sanger Institute COVID-19 Surveillance Team |
| EPI_ISL_731264 | Lighthouse Lab in Glasgow | Wellcome Sanger Institute for the COVID-19 Genomics UK (COG-UK) Consortium | Harper VanSteenhouse, Yumi Kasai, David Gray, Carol Clugston, Anna Dominiczak and Alex Alderton, Roberto Amato, Sonia Goncalves, Ewan Harrison, David K. Jackson, Ian Johnston, Dominic Kwiatkowski, Cordelia Langford, John Sillitoe on behalf of the Wellcome Sanger Institute COVID-19 Surveillance Team |
| EPI_ISL_731265, EPI_ISL_731266, EPI_ISL_731268, EPI_ISL_731269, EPI_ISL_731270, EPI_ISL_731272, EPI_ISL_731274, EPI_ISL_731275, EPI_ISL_731276, EPI_ISL_731277, EPI_ISL_731278, EPI_ISL_731279 | | | |
| see above | Lighthouse Lab in Alderley Park | Wellcome Sanger Institute for the COVID-19 Genomics UK (COG-UK) Consortium | Jacquelyn Wynn, Mairead Hyland, The Lighthouse Lab in Alderley Park and Alex Alderton, Roberto Amato, Sonia Goncalves, Ewan Harrison, David K. Jackson, Ian Johnston, Dominic Kwiatkowski, Cordelia Langford, John Sillitoe on behalf of the Wellcome Sanger Institute COVID-19 Surveillance Team |
| EPI_ISL_731283, EPI_ISL_731284 | Lighthouse Lab in Glasgow | Wellcome Sanger Institute for the COVID-19 Genomics UK (COG-UK) Consortium | Harper VanSteenhouse, Yumi Kasai, David Gray, Carol Clugston, Anna Dominiczak and Alex Alderton, Roberto Amato, Sonia Goncalves, Ewan Harrison, David K. Jackson, Ian Johnston, Dominic Kwiatkowski, Cordelia Langford, John Sillitoe on behalf of the Wellcome Sanger Institute COVID-19 Surveillance Team |
| EPI_ISL_731285, EPI_ISL_731288, EPI_ISL_731289 | Lighthouse Lab in Alderley Park | Wellcome Sanger Institute for the COVID-19 Genomics UK (COG-UK) Consortium | Jacquelyn Wynn, Mairead Hyland, The Lighthouse Lab in Alderley Park and Alex Alderton, Roberto Amato, Sonia Goncalves, Ewan Harrison, David K. Jackson, Ian Johnston, Dominic Kwiatkowski, Cordelia Langford, John Sillitoe on behalf of the Wellcome Sanger Institute COVID-19 Surveillance Team |
| EPI_ISL_731290 | Lighthouse Lab in Glasgow | Wellcome Sanger Institute for the COVID-19 Genomics UK (COG-UK) Consortium | Harper VanSteenhouse, Yumi Kasai, David Gray, Carol Clugston, Anna Dominiczak and Alex Alderton, Roberto Amato, Sonia Goncalves, Ewan Harrison, David K. Jackson, Ian Johnston, Dominic Kwiatkowski, Cordelia Langford, John Sillitoe on behalf of the Wellcome Sanger Institute COVID-19 Surveillance Team |
| EPI_ISL_731292 | Lighthouse Lab in Alderley Park | Wellcome Sanger Institute for the COVID-19 Genomics UK (COG-UK) Consortium | Jacquelyn Wynn, Mairead Hyland, The Lighthouse Lab in Alderley Park and Alex Alderton, Roberto Amato, Sonia Goncalves, Ewan Harrison, David K. Jackson, Ian Johnston, Dominic Kwiatkowski, Cordelia Langford, John Sillitoe on behalf of the Wellcome Sanger Institute COVID-19 Surveillance Team |
| EPI_ISL_731293 | Lighthouse Lab in Glasgow | Wellcome Sanger Institute for the COVID-19 Genomics UK (COG-UK) Consortium | Harper VanSteenhouse, Yumi Kasai, David Gray, Carol Clugston, Anna Dominiczak and Alex Alderton, Roberto Amato, Sonia Goncalves, Ewan Harrison, David K. Jackson, Ian Johnston, Dominic Kwiatkowski, Cordelia Langford, John Sillitoe on behalf of the Wellcome Sanger Institute COVID-19 Surveillance Team |
| EPI_ISL_731294, EPI_ISL_731296, EPI_ISL_731298 | Lighthouse Lab in Alderley Park | Wellcome Sanger Institute for the COVID-19 Genomics UK (COG-UK) Consortium | Jacquelyn Wynn, Mairead Hyland, The Lighthouse Lab in Alderley Park and Alex Alderton, Roberto Amato, Sonia Goncalves, Ewan Harrison, David K. Jackson, Ian Johnston, Dominic Kwiatkowski, Cordelia Langford, John Sillitoe on behalf of the Wellcome Sanger Institute COVID-19 Surveillance Team |
| EPI_ISL_731299 | Lighthouse Lab in Glasgow | Wellcome Sanger Institute for the COVID-19 Genomics UK (COG-UK) Consortium | Harper VanSteenhouse, Yumi Kasai, David Gray, Carol Clugston, Anna Dominiczak and Alex Alderton, Roberto Amato, Sonia Goncalves, Ewan Harrison, David K. Jackson, Ian Johnston, Dominic Kwiatkowski, Cordelia Langford, John Sillitoe on behalf of the Wellcome Sanger Institute COVID-19 Surveillance Team |
| EPI_ISL_731300, EPI_ISL_731301, EPI_ISL_731302, EPI_ISL_731303, EPI_ISL_731304 | Lighthouse Lab in Alderley Park | Wellcome Sanger Institute for the COVID-19 Genomics UK (COG-UK) Consortium | Jacquelyn Wynn, Mairead Hyland, The Lighthouse Lab in Alderley Park and Alex Alderton, Roberto Amato, Sonia Goncalves, Ewan Harrison, David K. Jackson, Ian Johnston, Dominic Kwiatkowski, Cordelia Langford, John Sillitoe on behalf of the Wellcome Sanger Institute COVID-19 Surveillance Team |
| EPI_ISL_731306 | Lighthouse Lab in Glasgow | Wellcome Sanger Institute for the COVID-19 Genomics UK (COG-UK) Consortium | Harper VanSteenhouse, Yumi Kasai, David Gray, Carol Clugston, Anna Dominiczak and Alex Alderton, Roberto Amato, Sonia Goncalves, Ewan Harrison, David K. Jackson, Ian Johnston, Dominic Kwiatkowski, Cordelia Langford, John Sillitoe on behalf of the Wellcome Sanger Institute COVID-19 Surveillance Team |
| EPI_ISL_731307, EPI_ISL_731311, EPI_ISL_731312, EPI_ISL_731313, EPI_ISL_731315, EPI_ISL_731316, EPI_ISL_731317, EPI_ISL_731318, EPI_ISL_731323, EPI_ISL_731324, EPI_ISL_731325, EPI_ISL_731328, EPI_ISL_731329, EPI_ISL_731331 | | | |
| see above | Lighthouse Lab in Alderley Park | Wellcome Sanger Institute for the COVID-19 Genomics UK (COG-UK) Consortium | Jacquelyn Wynn, Mairead Hyland, The Lighthouse Lab in Alderley Park and Alex Alderton, Roberto Amato, Sonia Goncalves, Ewan Harrison, David K. Jackson, Ian Johnston, Dominic Kwiatkowski, Cordelia Langford, John Sillitoe on behalf of the Wellcome Sanger Institute COVID-19 Surveillance Team |
| EPI_ISL_731333, EPI_ISL_731334 | Lighthouse Lab in Glasgow | Wellcome Sanger Institute for the COVID-19 Genomics UK (COG-UK) Consortium | Harper VanSteenhouse, Yumi Kasai, David Gray, Carol Clugston, Anna Dominiczak and Alex Alderton, Roberto Amato, Sonia Goncalves, Ewan Harrison, David K. Jackson, Ian Johnston, Dominic Kwiatkowski, Cordelia Langford, John Sillitoe on behalf of the Wellcome Sanger Institute COVID-19 Surveillance Team |
| EPI_ISL_731335, EPI_ISL_731336, EPI_ISL_731339, EPI_ISL_731342, EPI_ISL_731343, EPI_ISL_731344, EPI_ISL_731348 | Lighthouse Lab in Alderley Park | Wellcome Sanger Institute for the COVID-19 Genomics UK (COG-UK) Consortium | Jacquelyn Wynn, Mairead Hyland, The Lighthouse Lab in Alderley Park and Alex Alderton, Roberto Amato, Sonia Goncalves, Ewan Harrison, David K. Jackson, Ian Johnston, Dominic Kwiatkowski, Cordelia Langford, John Sillitoe on behalf of the Wellcome Sanger Institute COVID-19 Surveillance Team |
| EPI_ISL_731349 | Lighthouse Lab in Glasgow | Wellcome Sanger Institute for the COVID-19 Genomics UK (COG-UK) Consortium | Harper VanSteenhouse, Yumi Kasai, David Gray, Carol Clugston, Anna Dominiczak and Alex Alderton, Roberto Amato, Sonia Goncalves, Ewan Harrison, David K. Jackson, Ian Johnston, Dominic Kwiatkowski, Cordelia Langford, John Sillitoe on behalf of the Wellcome Sanger Institute COVID-19 Surveillance Team |
| EPI_ISL_731350, EPI_ISL_731351, EPI_ISL_731352 | Lighthouse Lab in Alderley Park | Wellcome Sanger Institute for the COVID-19 Genomics UK (COG-UK) Consortium | Jacquelyn Wynn, Mairead Hyland, The Lighthouse Lab in Alderley Park and Alex Alderton, Roberto Amato, Sonia Goncalves, Ewan Harrison, David K. Jackson, Ian Johnston, Dominic Kwiatkowski, Cordelia Langford, John Sillitoe on behalf of the Wellcome Sanger Institute COVID-19 Surveillance Team |

| | | | |
|---|---|---|---|
| EPI_ISL_731353 | Lighthouse Lab in Glasgow | Wellcome Sanger Institute for the COVID-19 Genomics UK (COG-UK) Consortium | Harper VanSteenhouse, Yumi Kasai, David Gray, Carol Clugston, Anna Dominiczak and Alex Alderton, Roberto Amato, Sonia Goncalves, Ewan Harrison, David K. Jackson, Ian Johnston, Dominic Kwiatkowski, Cordelia Langford, John Sillitoe on behalf of the Wellcome Sanger Institute COVID-19 Surveillance Team |
| EPI_ISL_731356, EPI_ISL_731358, EPI_ISL_731359, EPI_ISL_731360, EPI_ISL_731362, EPI_ISL_731363, EPI_ISL_731364, EPI_ISL_731365, EPI_ISL_731366, EPI_ISL_731369, EPI_ISL_731370, EPI_ISL_731371, EPI_ISL_731374, EPI_ISL_731375, EPI_ISL_731377 | | | |
| see above | Lighthouse Lab in Alderley Park | Wellcome Sanger Institute for the COVID-19 Genomics UK (COG-UK) Consortium | Jacquelyn Wynn, Mairead Hyland, The Lighthouse Lab in Alderley Park and Alex Alderton, Roberto Amato, Sonia Goncalves, Ewan Harrison, David K. Jackson, Ian Johnston, Dominic Kwiatkowski, Cordelia Langford, John Sillitoe on behalf of the Wellcome Sanger Institute COVID-19 Surveillance Team |
| EPI_ISL_731378 | Lighthouse Lab in Glasgow | Wellcome Sanger Institute for the COVID-19 Genomics UK (COG-UK) Consortium | Harper VanSteenhouse, Yumi Kasai, David Gray, Carol Clugston, Anna Dominiczak and Alex Alderton, Roberto Amato, Sonia Goncalves, Ewan Harrison, David K. Jackson, Ian Johnston, Dominic Kwiatkowski, Cordelia Langford, John Sillitoe on behalf of the Wellcome Sanger Institute COVID-19 Surveillance Team |
| EPI_ISL_731381, EPI_ISL_731384, EPI_ISL_731385, EPI_ISL_731387, EPI_ISL_731388, EPI_ISL_731389, EPI_ISL_731391 | Lighthouse Lab in Alderley Park | Wellcome Sanger Institute for the COVID-19 Genomics UK (COG-UK) Consortium | Jacquelyn Wynn, Mairead Hyland, The Lighthouse Lab in Alderley Park and Alex Alderton, Roberto Amato, Sonia Goncalves, Ewan Harrison, David K. Jackson, Ian Johnston, Dominic Kwiatkowski, Cordelia Langford, John Sillitoe on behalf of the Wellcome Sanger Institute COVID-19 Surveillance Team |
| EPI_ISL_731393 | Lighthouse Lab in Glasgow | Wellcome Sanger Institute for the COVID-19 Genomics UK (COG-UK) Consortium | Harper VanSteenhouse, Yumi Kasai, David Gray, Carol Clugston, Anna Dominiczak and Alex Alderton, Roberto Amato, Sonia Goncalves, Ewan Harrison, David K. Jackson, Ian Johnston, Dominic Kwiatkowski, Cordelia Langford, John Sillitoe on behalf of the Wellcome Sanger Institute COVID-19 Surveillance Team |
| EPI_ISL_731394, EPI_ISL_731395 | Lighthouse Lab in Alderley Park | Wellcome Sanger Institute for the COVID-19 Genomics UK (COG-UK) Consortium | Jacquelyn Wynn, Mairead Hyland, The Lighthouse Lab in Alderley Park and Alex Alderton, Roberto Amato, Sonia Goncalves, Ewan Harrison, David K. Jackson, Ian Johnston, Dominic Kwiatkowski, Cordelia Langford, John Sillitoe on behalf of the Wellcome Sanger Institute COVID-19 Surveillance Team |
| EPI_ISL_731396, EPI_ISL_731398 | Lighthouse Lab in Glasgow | Wellcome Sanger Institute for the COVID-19 Genomics UK (COG-UK) Consortium | Harper VanSteenhouse, Yumi Kasai, David Gray, Carol Clugston, Anna Dominiczak and Alex Alderton, Roberto Amato, Sonia Goncalves, Ewan Harrison, David K. Jackson, Ian Johnston, Dominic Kwiatkowski, Cordelia Langford, John Sillitoe on behalf of the Wellcome Sanger Institute COVID-19 Surveillance Team |
| EPI_ISL_731399 | Lighthouse Lab in Alderley Park | Wellcome Sanger Institute for the COVID-19 Genomics UK (COG-UK) Consortium | Jacquelyn Wynn, Mairead Hyland, The Lighthouse Lab in Alderley Park and Alex Alderton, Roberto Amato, Sonia Goncalves, Ewan Harrison, David K. Jackson, Ian Johnston, Dominic Kwiatkowski, Cordelia Langford, John Sillitoe on behalf of the Wellcome Sanger Institute COVID-19 Surveillance Team |
| EPI_ISL_731400, EPI_ISL_731405 | Lighthouse Lab in Glasgow | Wellcome Sanger Institute for the COVID-19 Genomics UK (COG-UK) Consortium | Harper VanSteenhouse, Yumi Kasai, David Gray, Carol Clugston, Anna Dominiczak and Alex Alderton, Roberto Amato, Sonia Goncalves, Ewan Harrison, David K. Jackson, Ian Johnston, Dominic Kwiatkowski, Cordelia Langford, John Sillitoe on behalf of the Wellcome Sanger Institute COVID-19 Surveillance Team |
| EPI_ISL_731409, EPI_ISL_731410, EPI_ISL_731412, EPI_ISL_731413, EPI_ISL_731415, EPI_ISL_731418, EPI_ISL_731419 | Lighthouse Lab in Alderley Park | Wellcome Sanger Institute for the COVID-19 Genomics UK (COG-UK) Consortium | Jacquelyn Wynn, Mairead Hyland, The Lighthouse Lab in Alderley Park and Alex Alderton, Roberto Amato, Sonia Goncalves, Ewan Harrison, David K. Jackson, Ian Johnston, Dominic Kwiatkowski, Cordelia Langford, John Sillitoe on behalf of the Wellcome Sanger Institute COVID-19 Surveillance Team |
| EPI_ISL_731421 | Lighthouse Lab in Glasgow | Wellcome Sanger Institute for the COVID-19 Genomics UK (COG-UK) Consortium | Harper VanSteenhouse, Yumi Kasai, David Gray, Carol Clugston, Anna Dominiczak and Alex Alderton, Roberto Amato, Sonia Goncalves, Ewan Harrison, David K. Jackson, Ian Johnston, Dominic Kwiatkowski, Cordelia Langford, John Sillitoe on behalf of the Wellcome Sanger Institute COVID-19 Surveillance Team |
| EPI_ISL_731422, EPI_ISL_731423, EPI_ISL_731424, EPI_ISL_731427, EPI_ISL_731428, EPI_ISL_731430 | Lighthouse Lab in Alderley Park | Wellcome Sanger Institute for the COVID-19 Genomics UK (COG-UK) Consortium | Jacquelyn Wynn, Mairead Hyland, The Lighthouse Lab in Alderley Park and Alex Alderton, Roberto Amato, Sonia Goncalves, Ewan Harrison, David K. Jackson, Ian Johnston, Dominic Kwiatkowski, Cordelia Langford, John Sillitoe on behalf of the Wellcome Sanger Institute COVID-19 Surveillance Team |
| EPI_ISL_731433 | Lighthouse Lab in Glasgow | Wellcome Sanger Institute for the COVID-19 Genomics UK (COG-UK) Consortium | Harper VanSteenhouse, Yumi Kasai, David Gray, Carol Clugston, Anna Dominiczak and Alex Alderton, Roberto Amato, Sonia Goncalves, Ewan Harrison, David K. Jackson, Ian Johnston, Dominic Kwiatkowski, Cordelia Langford, John Sillitoe on behalf of the Wellcome Sanger Institute COVID-19 Surveillance Team |
| EPI_ISL_731434, EPI_ISL_731436 | Lighthouse Lab in Alderley Park | Wellcome Sanger Institute for the COVID-19 Genomics UK (COG-UK) Consortium | Jacquelyn Wynn, Mairead Hyland, The Lighthouse Lab in Alderley Park and Alex Alderton, Roberto Amato, Sonia Goncalves, Ewan Harrison, David K. Jackson, Ian Johnston, Dominic Kwiatkowski, Cordelia Langford, John Sillitoe on behalf of the Wellcome Sanger Institute COVID-19 Surveillance Team |
| EPI_ISL_731438 | Lighthouse Lab in Glasgow | Wellcome Sanger Institute for the COVID-19 Genomics UK (COG-UK) Consortium | Harper VanSteenhouse, Yumi Kasai, David Gray, Carol Clugston, Anna Dominiczak and Alex Alderton, Roberto Amato, Sonia Goncalves, Ewan Harrison, David K. Jackson, Ian Johnston, Dominic Kwiatkowski, Cordelia Langford, John Sillitoe on behalf of the Wellcome Sanger Institute COVID-19 Surveillance Team |
| EPI_ISL_731439, EPI_ISL_731440, EPI_ISL_731441, EPI_ISL_731442, EPI_ISL_731443, EPI_ISL_731445, EPI_ISL_731446, EPI_ISL_731448, EPI_ISL_731451, EPI_ISL_731455 | Lighthouse Lab in Alderley Park | Wellcome Sanger Institute for the COVID-19 Genomics UK (COG-UK) Consortium | Jacquelyn Wynn, Mairead Hyland, The Lighthouse Lab in Alderley Park and Alex Alderton, Roberto Amato, Sonia Goncalves, Ewan Harrison, David K. Jackson, Ian Johnston, Dominic Kwiatkowski, Cordelia Langford, John Sillitoe on behalf of the Wellcome Sanger Institute COVID-19 Surveillance Team |
| EPI_ISL_731456 | Lighthouse Lab in Glasgow | Wellcome Sanger Institute for the COVID-19 Genomics UK (COG-UK) Consortium | Harper VanSteenhouse, Yumi Kasai, David Gray, Carol Clugston, Anna Dominiczak and Alex Alderton, Roberto Amato, Sonia Goncalves, Ewan Harrison, David K. Jackson, Ian Johnston, Dominic Kwiatkowski, Cordelia Langford, John Sillitoe on behalf of the Wellcome Sanger Institute COVID-19 Surveillance Team |
| EPI_ISL_731457 | Lighthouse Lab in Alderley Park | Wellcome Sanger Institute for the COVID-19 Genomics UK (COG-UK) Consortium | Jacquelyn Wynn, Mairead Hyland, The Lighthouse Lab in Alderley Park and Alex Alderton, Roberto Amato, Sonia Goncalves, Ewan Harrison, David K. Jackson, Ian Johnston, Dominic Kwiatkowski, Cordelia Langford, John Sillitoe on behalf of the Wellcome Sanger Institute COVID-19 Surveillance Team |
| EPI_ISL_731459 | Lighthouse Lab in Glasgow | Wellcome Sanger Institute for the COVID-19 Genomics UK (COG-UK) Consortium | Harper VanSteenhouse, Yumi Kasai, David Gray, Carol Clugston, Anna Dominiczak and Alex Alderton, Roberto Amato, Sonia Goncalves, Ewan Harrison, David K. Jackson, Ian Johnston, Dominic Kwiatkowski, Cordelia Langford, John Sillitoe on behalf of the Wellcome Sanger Institute COVID-19 Surveillance Team |
| EPI_ISL_731460, EPI_ISL_731461, EPI_ISL_731462, EPI_ISL_731463, EPI_ISL_731464, EPI_ISL_731465 | Lighthouse Lab in Alderley Park | Wellcome Sanger Institute for the COVID-19 Genomics UK (COG-UK) Consortium | Jacquelyn Wynn, Mairead Hyland, The Lighthouse Lab in Alderley Park and Alex Alderton, Roberto Amato, Sonia Goncalves, Ewan Harrison, David K. Jackson, Ian Johnston, Dominic Kwiatkowski, Cordelia Langford, John Sillitoe on behalf of the Wellcome Sanger Institute COVID-19 Surveillance Team |
| EPI_ISL_731466 | Lighthouse Lab in Glasgow | Wellcome Sanger Institute for the COVID-19 Genomics UK (COG-UK) Consortium | Harper VanSteenhouse, Yumi Kasai, David Gray, Carol Clugston, Anna Dominiczak and Alex Alderton, Roberto Amato, Sonia Goncalves, Ewan Harrison, David K. Jackson, Ian Johnston, Dominic Kwiatkowski, Cordelia Langford, John Sillitoe on behalf of the Wellcome Sanger Institute COVID-19 Surveillance Team |
| EPI_ISL_731467, EPI_ISL_731468, EPI_ISL_731470, EPI_ISL_731473, EPI_ISL_731474, EPI_ISL_731475, EPI_ISL_731478 | Lighthouse Lab in Alderley Park | Wellcome Sanger Institute for the COVID-19 Genomics UK (COG-UK) Consortium | Jacquelyn Wynn, Mairead Hyland, The Lighthouse Lab in Alderley Park and Alex Alderton, Roberto Amato, Sonia Goncalves, Ewan Harrison, David K. Jackson, Ian Johnston, Dominic Kwiatkowski, Cordelia Langford, John Sillitoe on behalf of the Wellcome Sanger Institute COVID-19 Surveillance Team |
| EPI_ISL_731479 | Lighthouse Lab in Glasgow | Wellcome Sanger Institute for the COVID-19 Genomics UK (COG-UK) Consortium | Harper VanSteenhouse, Yumi Kasai, David Gray, Carol Clugston, Anna Dominiczak and Alex Alderton, Roberto Amato, Sonia Goncalves, Ewan Harrison, David K. Jackson, Ian Johnston, Dominic Kwiatkowski, Cordelia Langford, John Sillitoe on behalf of the Wellcome Sanger Institute COVID-19 Surveillance Team |
| EPI_ISL_731480, EPI_ISL_731481, EPI_ISL_731483, EPI_ISL_731487, EPI_ISL_731489, EPI_ISL_731490, EPI_ISL_731492, EPI_ISL_731495, EPI_ISL_731498, EPI_ISL_731500, EPI_ISL_731502, EPI_ISL_731503, EPI_ISL_731504, EPI_ISL_731506, EPI_ISL_731509, EPI_ISL_731510, EPI_ISL_731511, EPI_ISL_731512, EPI_ISL_731514, EPI_ISL_731515, EPI_ISL_731516, EPI_ISL_731518, EPI_ISL_731520, EPI_ISL_731521 | | | |
| see above | Lighthouse Lab in Alderley Park | Wellcome Sanger Institute for the COVID-19 Genomics UK (COG-UK) Consortium | Jacquelyn Wynn, Mairead Hyland, The Lighthouse Lab in Alderley Park and Alex Alderton, Roberto Amato, Sonia Goncalves, Ewan Harrison, David K. Jackson, Ian Johnston, Dominic Kwiatkowski, Cordelia Langford, John Sillitoe on behalf of the Wellcome Sanger Institute COVID-19 Surveillance Team |

| | | | |
|---|---|---|---|
| EPI_ISL_731522 | Lighthouse Lab in Glasgow | Wellcome Sanger Institute for the COVID-19 Genomics UK (COG-UK) Consortium | Harper VanSteenhouse, Yumi Kasai, David Gray, Carol Clugston, Anna Dominiczak and Alex Alderton, Roberto Amato, Sonia Goncalves, Ewan Harrison, David K. Jackson, Ian Johnston, Dominic Kwiatkowski, Cordelia Langford, John Sillitoe on behalf of the Wellcome Sanger Institute COVID-19 Surveillance Team |
| EPI_ISL_731524, EPI_ISL_731525, EPI_ISL_731526, EPI_ISL_731527, EPI_ISL_731528, EPI_ISL_731529, EPI_ISL_731530, EPI_ISL_731531, EPI_ISL_731532, EPI_ISL_731533, EPI_ISL_731534 | | | |
| see above | Lighthouse Lab in Alderley Park | Wellcome Sanger Institute for the COVID-19 Genomics UK (COG-UK) Consortium | Jacquelyn Wynn, Mairead Hyland, The Lighthouse Lab in Alderley Park and Alex Alderton, Roberto Amato, Sonia Goncalves, Ewan Harrison, David K. Jackson, Ian Johnston, Dominic Kwiatkowski, Cordelia Langford, John Sillitoe on behalf of the Wellcome Sanger Institute COVID-19 Surveillance Team |
| EPI_ISL_731535 | Lighthouse Lab in Glasgow | Wellcome Sanger Institute for the COVID-19 Genomics UK (COG-UK) Consortium | Harper VanSteenhouse, Yumi Kasai, David Gray, Carol Clugston, Anna Dominiczak and Alex Alderton, Roberto Amato, Sonia Goncalves, Ewan Harrison, David K. Jackson, Ian Johnston, Dominic Kwiatkowski, Cordelia Langford, John Sillitoe on behalf of the Wellcome Sanger Institute COVID-19 Surveillance Team |
| EPI_ISL_731538, EPI_ISL_731539, EPI_ISL_731540 | Lighthouse Lab in Alderley Park | Wellcome Sanger Institute for the COVID-19 Genomics UK (COG-UK) Consortium | Jacquelyn Wynn, Mairead Hyland, The Lighthouse Lab in Alderley Park and Alex Alderton, Roberto Amato, Sonia Goncalves, Ewan Harrison, David K. Jackson, Ian Johnston, Dominic Kwiatkowski, Cordelia Langford, John Sillitoe on behalf of the Wellcome Sanger Institute COVID-19 Surveillance Team |
| EPI_ISL_731541 | Lighthouse Lab in Glasgow | Wellcome Sanger Institute for the COVID-19 Genomics UK (COG-UK) Consortium | Harper VanSteenhouse, Yumi Kasai, David Gray, Carol Clugston, Anna Dominiczak and Alex Alderton, Roberto Amato, Sonia Goncalves, Ewan Harrison, David K. Jackson, Ian Johnston, Dominic Kwiatkowski, Cordelia Langford, John Sillitoe on behalf of the Wellcome Sanger Institute COVID-19 Surveillance Team |
| EPI_ISL_731542, EPI_ISL_731543, EPI_ISL_731545, EPI_ISL_731546, EPI_ISL_731547, EPI_ISL_731549, EPI_ISL_731555, EPI_ISL_731565, EPI_ISL_731570, EPI_ISL_731571, EPI_ISL_731576, EPI_ISL_731580, EPI_ISL_731584, EPI_ISL_731585, EPI_ISL_731588, EPI_ISL_731592, EPI_ISL_731599, EPI_ISL_731611, EPI_ISL_731624, EPI_ISL_731626, EPI_ISL_731629, EPI_ISL_731632, EPI_ISL_731635, EPI_ISL_731639, EPI_ISL_731640, EPI_ISL_731641, EPI_ISL_731644, EPI_ISL_731647, EPI_ISL_731648, EPI_ISL_731649, EPI_ISL_731655, EPI_ISL_731656, EPI_ISL_731664, EPI_ISL_731665, EPI_ISL_731667, EPI_ISL_731671, EPI_ISL_731680, EPI_ISL_731684, EPI_ISL_731686, EPI_ISL_731688, EPI_ISL_731690, EPI_ISL_731691, EPI_ISL_731692, EPI_ISL_731693, EPI_ISL_731697, EPI_ISL_731701, EPI_ISL_731705, EPI_ISL_731707, EPI_ISL_731709, EPI_ISL_731711, EPI_ISL_731714, EPI_ISL_731719, EPI_ISL_731721, EPI_ISL_731722, EPI_ISL_731724, EPI_ISL_731731, EPI_ISL_731736, EPI_ISL_731737, EPI_ISL_731763, EPI_ISL_731767, EPI_ISL_731770, EPI_ISL_731775, EPI_ISL_731785, EPI_ISL_731787, EPI_ISL_731788, EPI_ISL_731792, EPI_ISL_731795, EPI_ISL_731796, EPI_ISL_731799, EPI_ISL_731800, EPI_ISL_731806, EPI_ISL_731808, EPI_ISL_731810, EPI_ISL_731819, EPI_ISL_731824, EPI_ISL_731827, EPI_ISL_731836, EPI_ISL_731838, EPI_ISL_731840, EPI_ISL_731841, EPI_ISL_731842, EPI_ISL_731846, EPI_ISL_731850, EPI_ISL_731854, EPI_ISL_731858, EPI_ISL_731863, EPI_ISL_731868, EPI_ISL_731874, EPI_ISL_731875, EPI_ISL_731877, EPI_ISL_731881, EPI_ISL_731884, EPI_ISL_731885, EPI_ISL_731887, EPI_ISL_731888, EPI_ISL_731889 | | | |
| see above | Lighthouse Lab in Alderley Park | Wellcome Sanger Institute for the COVID-19 Genomics UK (COG-UK) Consortium | Jacquelyn Wynn, Mairead Hyland, The Lighthouse Lab in Alderley Park and Alex Alderton, Roberto Amato, Sonia Goncalves, Ewan Harrison, David K. Jackson, Ian Johnston, Dominic Kwiatkowski, Cordelia Langford, John Sillitoe on behalf of the Wellcome Sanger Institute COVID-19 Surveillance Team |
| EPI_ISL_732657 | Barnakuten | The Public Health Agency of Sweden | Department of Microbiology, The Public Health Agency of Sweden |
| EPI_ISL_732962 | LabPLUS | Institute of Environmental Science and Research (ESR) | Xiaoyun Ren, Matt Storey, Nikki Freed, Muhammad Faisal, Jing Wang, Hermes Perez, Anja Werno, Antje van der Linden, Arlo Upton, Chris Mansell, David Hammer, Dragana Drinkovic, Gary McAuliffe, Hana Sofia Andersson, James Ussher, Jill Sherwood, Josh Freeman, Julia Howard, Juliet Elvy, Mary DeAlmeida, Matt Blakiston, Matthew Rogers, Max Bloomfield, Michael Addidle, Michelle Balm, Sally Roberts, Sarah Jefferies, Sharmini Muttaiyah, Susan Morpeth, Susan Taylor, Timothy Blackmore, Vani Sathyendran, Veronica Playle, Virginia Hope, Erasmus Smit, Lauren Jelly, Olin Silander, Joep de Ligt |
| EPI_ISL_732968, EPI_ISL_732969, EPI_ISL_732970 | Middlemore Hospital | Institute of Environmental Science and Research (ESR) | Xiaoyun Ren, Matt Storey, Nikki Freed, Muhammad Faisal, Jing Wang, Hermes Perez, Anja Werno, Antje van der Linden, Arlo Upton, Chris Mansell, David Hammer, Dragana Drinkovic, Gary McAuliffe, Hana Sofia Andersson, James Ussher, Jill Sherwood, Josh Freeman, Julia Howard, Juliet Elvy, Mary DeAlmeida, Matt Blakiston, Matthew Rogers, Max Bloomfield, Michael Addidle, Michelle Balm, Sally Roberts, Sarah Jefferies, Sharmini Muttaiyah, Susan Morpeth, Susan Taylor, Timothy Blackmore, Vani Sathyendran, Veronica Playle, Virginia Hope, Erasmus Smit, Lauren Jelly, Olin Silander, Joep de Ligt |
| EPI_ISL_733573, EPI_ISL_733578, EPI_ISL_733579 | Temporary Specimen Collection Centre | Hong Kong Department of Health | Alan K.L. Tsang, Peter C.W. Yip, Edman T.K. Lam, Rickjason C.W. Chan, Dominic N.C. Tsang |
| EPI_ISL_733589, EPI_ISL_733592 | Respiratory Virus Unit, National Infection Service, Public Health England | COVID-19 Genomics UK (COG-UK) Consortium | PHE Covid Sequencing Team |
| EPI_ISL_733596, EPI_ISL_733597, EPI_ISL_733599 | Lighthouse Lab in Cambridge | Wellcome Sanger Institute for the COVID-19 Genomics UK (COG-UK) Consortium | Rob Howes, The Lighthouse Lab in Cambridge and Alex Alderton, Roberto Amato, Sonia Goncalves, Ewan Harrison, David K. Jackson, Ian Johnston, Dominic Kwiatkowski, Cordelia Langford, John Sillitoe on behalf of the Wellcome Sanger Institute COVID-19 Surveillance Team |
| EPI_ISL_733600 | Lighthouse Lab in Milton Keynes | Wellcome Sanger Institute for the COVID-19 Genomics UK (COG-UK) Consortium | The Lighthouse Lab in Milton Keynes and Alex Alderton, Roberto Amato, Sonia Goncalves, Ewan Harrison, David K. Jackson, Ian Johnston, Dominic Kwiatkowski, Cordelia Langford, John Sillitoe on behalf of the Wellcome Sanger Institute COVID-19 Surveillance Team |
| EPI_ISL_733601, EPI_ISL_733602, EPI_ISL_733604 | Lighthouse Lab in Cambridge | Wellcome Sanger Institute for the COVID-19 Genomics UK (COG-UK) Consortium | Rob Howes, The Lighthouse Lab in Cambridge and Alex Alderton, Roberto Amato, Sonia Goncalves, Ewan Harrison, David K. Jackson, Ian Johnston, Dominic Kwiatkowski, Cordelia Langford, John Sillitoe on behalf of the Wellcome Sanger Institute COVID-19 Surveillance Team |
| EPI_ISL_733607 | Lighthouse Lab in Milton Keynes | Wellcome Sanger Institute for the COVID-19 Genomics UK (COG-UK) Consortium | The Lighthouse Lab in Milton Keynes and Alex Alderton, Roberto Amato, Sonia Goncalves, Ewan Harrison, David K. Jackson, Ian Johnston, Dominic Kwiatkowski, Cordelia Langford, John Sillitoe on behalf of the Wellcome Sanger Institute COVID-19 Surveillance Team |
| EPI_ISL_733610, EPI_ISL_733612, EPI_ISL_733614, EPI_ISL_733615 | Lighthouse Lab in Cambridge | Wellcome Sanger Institute for the COVID-19 Genomics UK (COG-UK) Consortium | Rob Howes, The Lighthouse Lab in Cambridge and Alex Alderton, Roberto Amato, Sonia Goncalves, Ewan Harrison, David K. Jackson, Ian Johnston, Dominic Kwiatkowski, Cordelia Langford, John Sillitoe on behalf of the Wellcome Sanger Institute COVID-19 Surveillance Team |
| EPI_ISL_733616, EPI_ISL_733617 | Lighthouse Lab in Milton Keynes | Wellcome Sanger Institute for the COVID-19 Genomics UK (COG-UK) Consortium | The Lighthouse Lab in Milton Keynes and Alex Alderton, Roberto Amato, Sonia Goncalves, Ewan Harrison, David K. Jackson, Ian Johnston, Dominic Kwiatkowski, Cordelia Langford, John Sillitoe on behalf of the Wellcome Sanger Institute COVID-19 Surveillance Team |
| EPI_ISL_733619, EPI_ISL_733620, EPI_ISL_733622, EPI_ISL_733624, EPI_ISL_733626, EPI_ISL_733627 | Lighthouse Lab in Cambridge | Wellcome Sanger Institute for the COVID-19 Genomics UK (COG-UK) Consortium | Rob Howes, The Lighthouse Lab in Cambridge and Alex Alderton, Roberto Amato, Sonia Goncalves, Ewan Harrison, David K. Jackson, Ian Johnston, Dominic Kwiatkowski, Cordelia Langford, John Sillitoe on behalf of the Wellcome Sanger Institute COVID-19 Surveillance Team |
| EPI_ISL_733628 | Lighthouse Lab in Milton Keynes | Wellcome Sanger Institute for the COVID-19 Genomics UK (COG-UK) Consortium | The Lighthouse Lab in Milton Keynes and Alex Alderton, Roberto Amato, Sonia Goncalves, Ewan Harrison, David K. Jackson, Ian Johnston, Dominic Kwiatkowski, Cordelia Langford, John Sillitoe on behalf of the Wellcome Sanger Institute COVID-19 Surveillance Team |
| EPI_ISL_733629 | Lighthouse Lab in Cambridge | Wellcome Sanger Institute for the COVID-19 Genomics UK (COG-UK) Consortium | Rob Howes, The Lighthouse Lab in Cambridge and Alex Alderton, Roberto Amato, Sonia Goncalves, Ewan Harrison, David K. Jackson, Ian Johnston, Dominic Kwiatkowski, Cordelia Langford, John Sillitoe on behalf of the Wellcome Sanger Institute COVID-19 Surveillance Team |
| EPI_ISL_733630 | Lighthouse Lab in Milton Keynes | Wellcome Sanger Institute for the COVID-19 Genomics UK (COG-UK) Consortium | The Lighthouse Lab in Milton Keynes and Alex Alderton, Roberto Amato, Sonia Goncalves, Ewan Harrison, David K. Jackson, Ian Johnston, Dominic Kwiatkowski, Cordelia Langford, John Sillitoe on behalf of the Wellcome Sanger Institute COVID-19 Surveillance Team |
| EPI_ISL_733631, EPI_ISL_733632, EPI_ISL_733633, EPI_ISL_733634 | Lighthouse Lab in Cambridge | Wellcome Sanger Institute for the COVID-19 Genomics UK (COG-UK) Consortium | Rob Howes, The Lighthouse Lab in Cambridge and Alex Alderton, Roberto Amato, Sonia Goncalves, Ewan Harrison, David K. Jackson, Ian Johnston, Dominic Kwiatkowski, Cordelia Langford, John Sillitoe on behalf of the Wellcome Sanger Institute COVID-19 Surveillance Team |
| EPI_ISL_733636 | Lighthouse Lab in Milton Keynes | Wellcome Sanger Institute for the COVID-19 Genomics UK (COG-UK) Consortium | The Lighthouse Lab in Milton Keynes and Alex Alderton, Roberto Amato, Sonia Goncalves, Ewan Harrison, David K. Jackson, Ian Johnston, Dominic Kwiatkowski, Cordelia Langford, John Sillitoe on behalf of the Wellcome Sanger Institute COVID-19 Surveillance Team |
| EPI_ISL_733637, EPI_ISL_733640 | Lighthouse Lab in Cambridge | Wellcome Sanger Institute for the COVID-19 Genomics UK (COG-UK) Consortium | Rob Howes, The Lighthouse Lab in Cambridge and Alex Alderton, Roberto Amato, Sonia Goncalves, Ewan Harrison, David K. Jackson, Ian Johnston, Dominic Kwiatkowski, Cordelia Langford, John Sillitoe on behalf of the Wellcome Sanger Institute COVID-19 Surveillance Team |
| EPI_ISL_733644, EPI_ISL_733645 | Lighthouse Lab in Milton Keynes | Wellcome Sanger Institute for the COVID-19 Genomics UK (COG-UK) Consortium | The Lighthouse Lab in Milton Keynes and Alex Alderton, Roberto Amato, Sonia Goncalves, Ewan Harrison, David K. Jackson, Ian Johnston, Dominic Kwiatkowski, Cordelia Langford, John Sillitoe on behalf of the Wellcome Sanger Institute COVID-19 Surveillance Team |
| EPI_ISL_733646 | Lighthouse Lab in Cambridge | Wellcome Sanger Institute for the COVID-19 Genomics UK (COG-UK) Consortium | Rob Howes, The Lighthouse Lab in Cambridge and Alex Alderton, Roberto Amato, Sonia Goncalves, Ewan Harrison, David K. Jackson, Ian Johnston, Dominic Kwiatkowski, Cordelia Langford, John Sillitoe on behalf of the Wellcome Sanger Institute COVID-19 Surveillance Team |
| EPI_ISL_733648 | Lighthouse Lab in Milton Keynes | Wellcome Sanger Institute for the COVID-19 Genomics UK (COG-UK) Consortium | The Lighthouse Lab in Milton Keynes and Alex Alderton, Roberto Amato, Sonia Goncalves, Ewan Harrison, David K. Jackson, Ian Johnston, Dominic Kwiatkowski, Cordelia Langford, John Sillitoe on behalf of the Wellcome Sanger Institute COVID-19 Surveillance Team |
| EPI_ISL_733649, EPI_ISL_733650, EPI_ISL_733651 | Lighthouse Lab in Cambridge | Wellcome Sanger Institute for the COVID-19 Genomics UK (COG-UK) Consortium | Rob Howes, The Lighthouse Lab in Cambridge and Alex Alderton, Roberto Amato, Sonia Goncalves, Ewan Harrison, David K. Jackson, Ian Johnston, Dominic Kwiatkowski, Cordelia Langford, John Sillitoe on behalf of the Wellcome Sanger Institute COVID-19 Surveillance Team |
| EPI_ISL_733652 | Lighthouse Lab in Milton Keynes | Wellcome Sanger Institute for the COVID-19 Genomics UK (COG-UK) Consortium | The Lighthouse Lab in Milton Keynes and Alex Alderton, Roberto Amato, Sonia Goncalves, Ewan Harrison, David K. Jackson, Ian Johnston, Dominic Kwiatkowski, Cordelia Langford, John Sillitoe on behalf of the Wellcome Sanger Institute COVID-19 Surveillance Team |

| | | | |
|---|---|---|---|
| EPI_ISL_733655, EPI_ISL_733656 | Lighthouse Lab in Cambridge | Wellcome Sanger Institute for the COVID-19 Genomics UK (COG-UK) Consortium | Rob Howes, The Lighthouse Lab in Cambridge and Alex Alderton, Roberto Amato, Sonia Goncalves, Ewan Harrison, David K. Jackson, Ian Johnston, Dominic Kwiatkowski, Cordelia Langford, John Sillitoe on behalf of the Wellcome Sanger Institute COVID-19 Surveillance Team |
| EPI_ISL_733657, EPI_ISL_733658 | Lighthouse Lab in Milton Keynes | Wellcome Sanger Institute for the COVID-19 Genomics UK (COG-UK) Consortium | The Lighthouse Lab in Milton Keynes and Alex Alderton, Roberto Amato, Sonia Goncalves, Ewan Harrison, David K. Jackson, Ian Johnston, Dominic Kwiatkowski, Cordelia Langford, John Sillitoe on behalf of the Wellcome Sanger Institute COVID-19 Surveillance Team |
| EPI_ISL_733660, EPI_ISL_733661, EPI_ISL_733662, EPI_ISL_733663 | Lighthouse Lab in Cambridge | Wellcome Sanger Institute for the COVID-19 Genomics UK (COG-UK) Consortium | Rob Howes, The Lighthouse Lab in Cambridge and Alex Alderton, Roberto Amato, Sonia Goncalves, Ewan Harrison, David K. Jackson, Ian Johnston, Dominic Kwiatkowski, Cordelia Langford, John Sillitoe on behalf of the Wellcome Sanger Institute COVID-19 Surveillance Team |
| EPI_ISL_733664 | Lighthouse Lab in Milton Keynes | Wellcome Sanger Institute for the COVID-19 Genomics UK (COG-UK) Consortium | The Lighthouse Lab in Milton Keynes and Alex Alderton, Roberto Amato, Sonia Goncalves, Ewan Harrison, David K. Jackson, Ian Johnston, Dominic Kwiatkowski, Cordelia Langford, John Sillitoe on behalf of the Wellcome Sanger Institute COVID-19 Surveillance Team |
| EPI_ISL_733665 | Lighthouse Lab in Cambridge | Wellcome Sanger Institute for the COVID-19 Genomics UK (COG-UK) Consortium | Rob Howes, The Lighthouse Lab in Cambridge and Alex Alderton, Roberto Amato, Sonia Goncalves, Ewan Harrison, David K. Jackson, Ian Johnston, Dominic Kwiatkowski, Cordelia Langford, John Sillitoe on behalf of the Wellcome Sanger Institute COVID-19 Surveillance Team |
| EPI_ISL_733666 | Lighthouse Lab in Milton Keynes | Wellcome Sanger Institute for the COVID-19 Genomics UK (COG-UK) Consortium | The Lighthouse Lab in Milton Keynes and Alex Alderton, Roberto Amato, Sonia Goncalves, Ewan Harrison, David K. Jackson, Ian Johnston, Dominic Kwiatkowski, Cordelia Langford, John Sillitoe on behalf of the Wellcome Sanger Institute COVID-19 Surveillance Team |
| EPI_ISL_733667, EPI_ISL_733668 | Lighthouse Lab in Cambridge | Wellcome Sanger Institute for the COVID-19 Genomics UK (COG-UK) Consortium | Rob Howes, The Lighthouse Lab in Cambridge and Alex Alderton, Roberto Amato, Sonia Goncalves, Ewan Harrison, David K. Jackson, Ian Johnston, Dominic Kwiatkowski, Cordelia Langford, John Sillitoe on behalf of the Wellcome Sanger Institute COVID-19 Surveillance Team |
| EPI_ISL_733669 | Lighthouse Lab in Milton Keynes | Wellcome Sanger Institute for the COVID-19 Genomics UK (COG-UK) Consortium | The Lighthouse Lab in Milton Keynes and Alex Alderton, Roberto Amato, Sonia Goncalves, Ewan Harrison, David K. Jackson, Ian Johnston, Dominic Kwiatkowski, Cordelia Langford, John Sillitoe on behalf of the Wellcome Sanger Institute COVID-19 Surveillance Team |
| EPI_ISL_733670, EPI_ISL_733671, EPI_ISL_733672 | Lighthouse Lab in Cambridge | Wellcome Sanger Institute for the COVID-19 Genomics UK (COG-UK) Consortium | Rob Howes, The Lighthouse Lab in Cambridge and Alex Alderton, Roberto Amato, Sonia Goncalves, Ewan Harrison, David K. Jackson, Ian Johnston, Dominic Kwiatkowski, Cordelia Langford, John Sillitoe on behalf of the Wellcome Sanger Institute COVID-19 Surveillance Team |
| EPI_ISL_733673 | Lighthouse Lab in Milton Keynes | Wellcome Sanger Institute for the COVID-19 Genomics UK (COG-UK) Consortium | The Lighthouse Lab in Milton Keynes and Alex Alderton, Roberto Amato, Sonia Goncalves, Ewan Harrison, David K. Jackson, Ian Johnston, Dominic Kwiatkowski, Cordelia Langford, John Sillitoe on behalf of the Wellcome Sanger Institute COVID-19 Surveillance Team |
| EPI_ISL_733674, EPI_ISL_733675, EPI_ISL_733676, EPI_ISL_733677 | Lighthouse Lab in Cambridge | Wellcome Sanger Institute for the COVID-19 Genomics UK (COG-UK) Consortium | Rob Howes, The Lighthouse Lab in Cambridge and Alex Alderton, Roberto Amato, Sonia Goncalves, Ewan Harrison, David K. Jackson, Ian Johnston, Dominic Kwiatkowski, Cordelia Langford, John Sillitoe on behalf of the Wellcome Sanger Institute COVID-19 Surveillance Team |
| EPI_ISL_733678, EPI_ISL_733679 | Lighthouse Lab in Milton Keynes | Wellcome Sanger Institute for the COVID-19 Genomics UK (COG-UK) Consortium | The Lighthouse Lab in Milton Keynes and Alex Alderton, Roberto Amato, Sonia Goncalves, Ewan Harrison, David K. Jackson, Ian Johnston, Dominic Kwiatkowski, Cordelia Langford, John Sillitoe on behalf of the Wellcome Sanger Institute COVID-19 Surveillance Team |
| EPI_ISL_733680, EPI_ISL_733681, EPI_ISL_733684 | Lighthouse Lab in Cambridge | Wellcome Sanger Institute for the COVID-19 Genomics UK (COG-UK) Consortium | Rob Howes, The Lighthouse Lab in Cambridge and Alex Alderton, Roberto Amato, Sonia Goncalves, Ewan Harrison, David K. Jackson, Ian Johnston, Dominic Kwiatkowski, Cordelia Langford, John Sillitoe on behalf of the Wellcome Sanger Institute COVID-19 Surveillance Team |
| EPI_ISL_733686, EPI_ISL_733687 | Lighthouse Lab in Milton Keynes | Wellcome Sanger Institute for the COVID-19 Genomics UK (COG-UK) Consortium | The Lighthouse Lab in Milton Keynes and Alex Alderton, Roberto Amato, Sonia Goncalves, Ewan Harrison, David K. Jackson, Ian Johnston, Dominic Kwiatkowski, Cordelia Langford, John Sillitoe on behalf of the Wellcome Sanger Institute COVID-19 Surveillance Team |
| EPI_ISL_733688, EPI_ISL_733689, EPI_ISL_733690, EPI_ISL_733691 | Lighthouse Lab in Cambridge | Wellcome Sanger Institute for the COVID-19 Genomics UK (COG-UK) Consortium | Rob Howes, The Lighthouse Lab in Cambridge and Alex Alderton, Roberto Amato, Sonia Goncalves, Ewan Harrison, David K. Jackson, Ian Johnston, Dominic Kwiatkowski, Cordelia Langford, John Sillitoe on behalf of the Wellcome Sanger Institute COVID-19 Surveillance Team |
| EPI_ISL_733692 | Lighthouse Lab in Milton Keynes | Wellcome Sanger Institute for the COVID-19 Genomics UK (COG-UK) Consortium | The Lighthouse Lab in Milton Keynes and Alex Alderton, Roberto Amato, Sonia Goncalves, Ewan Harrison, David K. Jackson, Ian Johnston, Dominic Kwiatkowski, Cordelia Langford, John Sillitoe on behalf of the Wellcome Sanger Institute COVID-19 Surveillance Team |
| EPI_ISL_733694, EPI_ISL_733695, EPI_ISL_733696, EPI_ISL_733698, EPI_ISL_733700, EPI_ISL_733702, EPI_ISL_733704 | Lighthouse Lab in Cambridge | Wellcome Sanger Institute for the COVID-19 Genomics UK (COG-UK) Consortium | Rob Howes, The Lighthouse Lab in Cambridge and Alex Alderton, Roberto Amato, Sonia Goncalves, Ewan Harrison, David K. Jackson, Ian Johnston, Dominic Kwiatkowski, Cordelia Langford, John Sillitoe on behalf of the Wellcome Sanger Institute COVID-19 Surveillance Team |
| EPI_ISL_733707 | Lighthouse Lab in Milton Keynes | Wellcome Sanger Institute for the COVID-19 Genomics UK (COG-UK) Consortium | The Lighthouse Lab in Milton Keynes and Alex Alderton, Roberto Amato, Sonia Goncalves, Ewan Harrison, David K. Jackson, Ian Johnston, Dominic Kwiatkowski, Cordelia Langford, John Sillitoe on behalf of the Wellcome Sanger Institute COVID-19 Surveillance Team |
| EPI_ISL_733708, EPI_ISL_733709 | Lighthouse Lab in Cambridge | Wellcome Sanger Institute for the COVID-19 Genomics UK (COG-UK) Consortium | Rob Howes, The Lighthouse Lab in Cambridge and Alex Alderton, Roberto Amato, Sonia Goncalves, Ewan Harrison, David K. Jackson, Ian Johnston, Dominic Kwiatkowski, Cordelia Langford, John Sillitoe on behalf of the Wellcome Sanger Institute COVID-19 Surveillance Team |
| EPI_ISL_733711, EPI_ISL_733713 | Lighthouse Lab in Milton Keynes | Wellcome Sanger Institute for the COVID-19 Genomics UK (COG-UK) Consortium | The Lighthouse Lab in Milton Keynes and Alex Alderton, Roberto Amato, Sonia Goncalves, Ewan Harrison, David K. Jackson, Ian Johnston, Dominic Kwiatkowski, Cordelia Langford, John Sillitoe on behalf of the Wellcome Sanger Institute COVID-19 Surveillance Team |
| EPI_ISL_733714, EPI_ISL_733715, EPI_ISL_733716, EPI_ISL_733717, EPI_ISL_733719, EPI_ISL_733720, EPI_ISL_733721, EPI_ISL_733722, EPI_ISL_733723, EPI_ISL_733724, EPI_ISL_733726, EPI_ISL_733727, EPI_ISL_733728, EPI_ISL_733729, EPI_ISL_733730, EPI_ISL_733732, EPI_ISL_733733 | | | |
| see above | Lighthouse Lab in Cambridge | Wellcome Sanger Institute for the COVID-19 Genomics UK (COG-UK) Consortium | Rob Howes, The Lighthouse Lab in Cambridge and Alex Alderton, Roberto Amato, Sonia Goncalves, Ewan Harrison, David K. Jackson, Ian Johnston, Dominic Kwiatkowski, Cordelia Langford, John Sillitoe on behalf of the Wellcome Sanger Institute COVID-19 Surveillance Team |
| EPI_ISL_733735 | Lighthouse Lab in Milton Keynes | Wellcome Sanger Institute for the COVID-19 Genomics UK (COG-UK) Consortium | The Lighthouse Lab in Milton Keynes and Alex Alderton, Roberto Amato, Sonia Goncalves, Ewan Harrison, David K. Jackson, Ian Johnston, Dominic Kwiatkowski, Cordelia Langford, John Sillitoe on behalf of the Wellcome Sanger Institute COVID-19 Surveillance Team |
| EPI_ISL_733736, EPI_ISL_733738 | Lighthouse Lab in Cambridge | Wellcome Sanger Institute for the COVID-19 Genomics UK (COG-UK) Consortium | Rob Howes, The Lighthouse Lab in Cambridge and Alex Alderton, Roberto Amato, Sonia Goncalves, Ewan Harrison, David K. Jackson, Ian Johnston, Dominic Kwiatkowski, Cordelia Langford, John Sillitoe on behalf of the Wellcome Sanger Institute COVID-19 Surveillance Team |
| EPI_ISL_733739 | Lighthouse Lab in Milton Keynes | Wellcome Sanger Institute for the COVID-19 Genomics UK (COG-UK) Consortium | The Lighthouse Lab in Milton Keynes and Alex Alderton, Roberto Amato, Sonia Goncalves, Ewan Harrison, David K. Jackson, Ian Johnston, Dominic Kwiatkowski, Cordelia Langford, John Sillitoe on behalf of the Wellcome Sanger Institute COVID-19 Surveillance Team |
| EPI_ISL_733740, EPI_ISL_733741 | Lighthouse Lab in Cambridge | Wellcome Sanger Institute for the COVID-19 Genomics UK (COG-UK) Consortium | Rob Howes, The Lighthouse Lab in Cambridge and Alex Alderton, Roberto Amato, Sonia Goncalves, Ewan Harrison, David K. Jackson, Ian Johnston, Dominic Kwiatkowski, Cordelia Langford, John Sillitoe on behalf of the Wellcome Sanger Institute COVID-19 Surveillance Team |
| EPI_ISL_733743 | Lighthouse Lab in Milton Keynes | Wellcome Sanger Institute for the COVID-19 Genomics UK (COG-UK) Consortium | The Lighthouse Lab in Milton Keynes and Alex Alderton, Roberto Amato, Sonia Goncalves, Ewan Harrison, David K. Jackson, Ian Johnston, Dominic Kwiatkowski, Cordelia Langford, John Sillitoe on behalf of the Wellcome Sanger Institute COVID-19 Surveillance Team |
| EPI_ISL_733744, EPI_ISL_733745 | Lighthouse Lab in Cambridge | Wellcome Sanger Institute for the COVID-19 Genomics UK (COG-UK) Consortium | Rob Howes, The Lighthouse Lab in Cambridge and Alex Alderton, Roberto Amato, Sonia Goncalves, Ewan Harrison, David K. Jackson, Ian Johnston, Dominic Kwiatkowski, Cordelia Langford, John Sillitoe on behalf of the Wellcome Sanger Institute COVID-19 Surveillance Team |
| EPI_ISL_733746, EPI_ISL_733747, EPI_ISL_733748, EPI_ISL_733749 | Lighthouse Lab in Milton Keynes | Wellcome Sanger Institute for the COVID-19 Genomics UK (COG-UK) Consortium | The Lighthouse Lab in Milton Keynes and Alex Alderton, Roberto Amato, Sonia Goncalves, Ewan Harrison, David K. Jackson, Ian Johnston, Dominic Kwiatkowski, Cordelia Langford, John Sillitoe on behalf of the Wellcome Sanger Institute COVID-19 Surveillance Team |
| EPI_ISL_733750, EPI_ISL_733751, EPI_ISL_733752 | Lighthouse Lab in Cambridge | Wellcome Sanger Institute for the COVID-19 Genomics UK (COG-UK) Consortium | Rob Howes, The Lighthouse Lab in Cambridge and Alex Alderton, Roberto Amato, Sonia Goncalves, Ewan Harrison, David K. Jackson, Ian Johnston, Dominic Kwiatkowski, Cordelia Langford, John Sillitoe on behalf of the Wellcome Sanger Institute COVID-19 Surveillance Team |
| EPI_ISL_733756, EPI_ISL_733757 | Lighthouse Lab in Milton Keynes | Wellcome Sanger Institute for the COVID-19 Genomics UK (COG-UK) Consortium | The Lighthouse Lab in Milton Keynes and Alex Alderton, Roberto Amato, Sonia Goncalves, Ewan Harrison, David K. Jackson, Ian Johnston, Dominic Kwiatkowski, Cordelia Langford, John Sillitoe on behalf of the Wellcome Sanger Institute COVID-19 Surveillance Team |
| EPI_ISL_733758 | Lighthouse Lab in Cambridge | Wellcome Sanger Institute for the COVID-19 Genomics UK (COG-UK) Consortium | Rob Howes, The Lighthouse Lab in Cambridge and Alex Alderton, Roberto Amato, Sonia Goncalves, Ewan Harrison, David K. Jackson, Ian Johnston, Dominic Kwiatkowski, Cordelia Langford, John Sillitoe on behalf of the Wellcome Sanger Institute COVID-19 Surveillance Team |
| EPI_ISL_733760, EPI_ISL_733761 | Lighthouse Lab in Milton Keynes | Wellcome Sanger Institute for the COVID-19 Genomics UK (COG-UK) Consortium | The Lighthouse Lab in Milton Keynes and Alex Alderton, Roberto Amato, Sonia Goncalves, Ewan Harrison, David K. Jackson, Ian Johnston, Dominic Kwiatkowski, Cordelia Langford, John Sillitoe on behalf of the Wellcome Sanger Institute COVID-19 Surveillance Team |
| EPI_ISL_733763 | Lighthouse Lab in Cambridge | Wellcome Sanger Institute for the COVID-19 Genomics UK (COG-UK) Consortium | Rob Howes, The Lighthouse Lab in Cambridge and Alex Alderton, Roberto Amato, Sonia Goncalves, Ewan Harrison, David K. Jackson, Ian Johnston, Dominic Kwiatkowski, Cordelia Langford, John Sillitoe on behalf of the Wellcome Sanger Institute COVID-19 Surveillance Team |
| EPI_ISL_733764 | Lighthouse Lab in Milton Keynes | Wellcome Sanger Institute for the COVID-19 Genomics UK (COG-UK) Consortium | The Lighthouse Lab in Milton Keynes and Alex Alderton, Roberto Amato, Sonia Goncalves, Ewan Harrison, David K. Jackson, Ian Johnston, Dominic Kwiatkowski, Cordelia Langford, John Sillitoe on behalf of the Wellcome Sanger Institute COVID-19 Surveillance Team |

| | | | |
|---|---|---|---|
| EPI_ISL_733765, EPI_ISL_733767 | Lighthouse Lab in Cambridge | Wellcome Sanger Institute for the COVID-19 Genomics UK (COG-UK) Consortium | Rob Howes, The Lighthouse Lab in Cambridge and Alex Alderton, Roberto Amato, Sonia Goncalves, Ewan Harrison, David K. Jackson, Ian Johnston, Dominic Kwiatkowski, Cordelia Langford, John Sillitoe on behalf of the Wellcome Sanger Institute COVID-19 Surveillance Team |
| EPI_ISL_733768 | Lighthouse Lab in Milton Keynes | Wellcome Sanger Institute for the COVID-19 Genomics UK (COG-UK) Consortium | The Lighthouse Lab in Milton Keynes and Alex Alderton, Roberto Amato, Sonia Goncalves, Ewan Harrison, David K. Jackson, Ian Johnston, Dominic Kwiatkowski, Cordelia Langford, John Sillitoe on behalf of the Wellcome Sanger Institute COVID-19 Surveillance Team |
| EPI_ISL_733770 | Lighthouse Lab in Cambridge | Wellcome Sanger Institute for the COVID-19 Genomics UK (COG-UK) Consortium | Rob Howes, The Lighthouse Lab in Cambridge and Alex Alderton, Roberto Amato, Sonia Goncalves, Ewan Harrison, David K. Jackson, Ian Johnston, Dominic Kwiatkowski, Cordelia Langford, John Sillitoe on behalf of the Wellcome Sanger Institute COVID-19 Surveillance Team |
| EPI_ISL_733771 | Lighthouse Lab in Milton Keynes | Wellcome Sanger Institute for the COVID-19 Genomics UK (COG-UK) Consortium | The Lighthouse Lab in Milton Keynes and Alex Alderton, Roberto Amato, Sonia Goncalves, Ewan Harrison, David K. Jackson, Ian Johnston, Dominic Kwiatkowski, Cordelia Langford, John Sillitoe on behalf of the Wellcome Sanger Institute COVID-19 Surveillance Team |
| EPI_ISL_733772, EPI_ISL_733773, EPI_ISL_733774, EPI_ISL_733775 | Lighthouse Lab in Cambridge | Wellcome Sanger Institute for the COVID-19 Genomics UK (COG-UK) Consortium | Rob Howes, The Lighthouse Lab in Cambridge and Alex Alderton, Roberto Amato, Sonia Goncalves, Ewan Harrison, David K. Jackson, Ian Johnston, Dominic Kwiatkowski, Cordelia Langford, John Sillitoe on behalf of the Wellcome Sanger Institute COVID-19 Surveillance Team |
| EPI_ISL_733776 | Lighthouse Lab in Milton Keynes | Wellcome Sanger Institute for the COVID-19 Genomics UK (COG-UK) Consortium | The Lighthouse Lab in Milton Keynes and Alex Alderton, Roberto Amato, Sonia Goncalves, Ewan Harrison, David K. Jackson, Ian Johnston, Dominic Kwiatkowski, Cordelia Langford, John Sillitoe on behalf of the Wellcome Sanger Institute COVID-19 Surveillance Team |
| EPI_ISL_733778, EPI_ISL_733779 | Lighthouse Lab in Cambridge | Wellcome Sanger Institute for the COVID-19 Genomics UK (COG-UK) Consortium | Rob Howes, The Lighthouse Lab in Cambridge and Alex Alderton, Roberto Amato, Sonia Goncalves, Ewan Harrison, David K. Jackson, Ian Johnston, Dominic Kwiatkowski, Cordelia Langford, John Sillitoe on behalf of the Wellcome Sanger Institute COVID-19 Surveillance Team |
| EPI_ISL_733780, EPI_ISL_733781 | Lighthouse Lab in Milton Keynes | Wellcome Sanger Institute for the COVID-19 Genomics UK (COG-UK) Consortium | The Lighthouse Lab in Milton Keynes and Alex Alderton, Roberto Amato, Sonia Goncalves, Ewan Harrison, David K. Jackson, Ian Johnston, Dominic Kwiatkowski, Cordelia Langford, John Sillitoe on behalf of the Wellcome Sanger Institute COVID-19 Surveillance Team |
| EPI_ISL_733782, EPI_ISL_733784, EPI_ISL_733786, EPI_ISL_733787, EPI_ISL_733788 | Lighthouse Lab in Cambridge | Wellcome Sanger Institute for the COVID-19 Genomics UK (COG-UK) Consortium | Rob Howes, The Lighthouse Lab in Cambridge and Alex Alderton, Roberto Amato, Sonia Goncalves, Ewan Harrison, David K. Jackson, Ian Johnston, Dominic Kwiatkowski, Cordelia Langford, John Sillitoe on behalf of the Wellcome Sanger Institute COVID-19 Surveillance Team |
| EPI_ISL_733789 | Lighthouse Lab in Milton Keynes | Wellcome Sanger Institute for the COVID-19 Genomics UK (COG-UK) Consortium | The Lighthouse Lab in Milton Keynes and Alex Alderton, Roberto Amato, Sonia Goncalves, Ewan Harrison, David K. Jackson, Ian Johnston, Dominic Kwiatkowski, Cordelia Langford, John Sillitoe on behalf of the Wellcome Sanger Institute COVID-19 Surveillance Team |
| EPI_ISL_733790, EPI_ISL_733791, EPI_ISL_733792, EPI_ISL_733793 | Lighthouse Lab in Cambridge | Wellcome Sanger Institute for the COVID-19 Genomics UK (COG-UK) Consortium | Rob Howes, The Lighthouse Lab in Cambridge and Alex Alderton, Roberto Amato, Sonia Goncalves, Ewan Harrison, David K. Jackson, Ian Johnston, Dominic Kwiatkowski, Cordelia Langford, John Sillitoe on behalf of the Wellcome Sanger Institute COVID-19 Surveillance Team |
| EPI_ISL_733796, EPI_ISL_733798 | Lighthouse Lab in Milton Keynes | Wellcome Sanger Institute for the COVID-19 Genomics UK (COG-UK) Consortium | The Lighthouse Lab in Milton Keynes and Alex Alderton, Roberto Amato, Sonia Goncalves, Ewan Harrison, David K. Jackson, Ian Johnston, Dominic Kwiatkowski, Cordelia Langford, John Sillitoe on behalf of the Wellcome Sanger Institute COVID-19 Surveillance Team |
| EPI_ISL_733799, EPI_ISL_733800 | Lighthouse Lab in Cambridge | Wellcome Sanger Institute for the COVID-19 Genomics UK (COG-UK) Consortium | Rob Howes, The Lighthouse Lab in Cambridge and Alex Alderton, Roberto Amato, Sonia Goncalves, Ewan Harrison, David K. Jackson, Ian Johnston, Dominic Kwiatkowski, Cordelia Langford, John Sillitoe on behalf of the Wellcome Sanger Institute COVID-19 Surveillance Team |
| EPI_ISL_733801 | Lighthouse Lab in Milton Keynes | Wellcome Sanger Institute for the COVID-19 Genomics UK (COG-UK) Consortium | The Lighthouse Lab in Milton Keynes and Alex Alderton, Roberto Amato, Sonia Goncalves, Ewan Harrison, David K. Jackson, Ian Johnston, Dominic Kwiatkowski, Cordelia Langford, John Sillitoe on behalf of the Wellcome Sanger Institute COVID-19 Surveillance Team |
| EPI_ISL_733802, EPI_ISL_733804, EPI_ISL_733805, EPI_ISL_733809, EPI_ISL_733810 | Lighthouse Lab in Cambridge | Wellcome Sanger Institute for the COVID-19 Genomics UK (COG-UK) Consortium | Rob Howes, The Lighthouse Lab in Cambridge and Alex Alderton, Roberto Amato, Sonia Goncalves, Ewan Harrison, David K. Jackson, Ian Johnston, Dominic Kwiatkowski, Cordelia Langford, John Sillitoe on behalf of the Wellcome Sanger Institute COVID-19 Surveillance Team |
| EPI_ISL_733811 | Lighthouse Lab in Milton Keynes | Wellcome Sanger Institute for the COVID-19 Genomics UK (COG-UK) Consortium | The Lighthouse Lab in Milton Keynes and Alex Alderton, Roberto Amato, Sonia Goncalves, Ewan Harrison, David K. Jackson, Ian Johnston, Dominic Kwiatkowski, Cordelia Langford, John Sillitoe on behalf of the Wellcome Sanger Institute COVID-19 Surveillance Team |
| EPI_ISL_733812 | Lighthouse Lab in Cambridge | Wellcome Sanger Institute for the COVID-19 Genomics UK (COG-UK) Consortium | Rob Howes, The Lighthouse Lab in Cambridge and Alex Alderton, Roberto Amato, Sonia Goncalves, Ewan Harrison, David K. Jackson, Ian Johnston, Dominic Kwiatkowski, Cordelia Langford, John Sillitoe on behalf of the Wellcome Sanger Institute COVID-19 Surveillance Team |
| EPI_ISL_733813 | Lighthouse Lab in Milton Keynes | Wellcome Sanger Institute for the COVID-19 Genomics UK (COG-UK) Consortium | The Lighthouse Lab in Milton Keynes and Alex Alderton, Roberto Amato, Sonia Goncalves, Ewan Harrison, David K. Jackson, Ian Johnston, Dominic Kwiatkowski, Cordelia Langford, John Sillitoe on behalf of the Wellcome Sanger Institute COVID-19 Surveillance Team |
| EPI_ISL_733814, EPI_ISL_733816, EPI_ISL_733817 | Lighthouse Lab in Cambridge | Wellcome Sanger Institute for the COVID-19 Genomics UK (COG-UK) Consortium | Rob Howes, The Lighthouse Lab in Cambridge and Alex Alderton, Roberto Amato, Sonia Goncalves, Ewan Harrison, David K. Jackson, Ian Johnston, Dominic Kwiatkowski, Cordelia Langford, John Sillitoe on behalf of the Wellcome Sanger Institute COVID-19 Surveillance Team |
| EPI_ISL_733820 | Lighthouse Lab in Milton Keynes | Wellcome Sanger Institute for the COVID-19 Genomics UK (COG-UK) Consortium | The Lighthouse Lab in Milton Keynes and Alex Alderton, Roberto Amato, Sonia Goncalves, Ewan Harrison, David K. Jackson, Ian Johnston, Dominic Kwiatkowski, Cordelia Langford, John Sillitoe on behalf of the Wellcome Sanger Institute COVID-19 Surveillance Team |
| EPI_ISL_733821, EPI_ISL_733822, EPI_ISL_733823, EPI_ISL_733824 | Lighthouse Lab in Cambridge | Wellcome Sanger Institute for the COVID-19 Genomics UK (COG-UK) Consortium | Rob Howes, The Lighthouse Lab in Cambridge and Alex Alderton, Roberto Amato, Sonia Goncalves, Ewan Harrison, David K. Jackson, Ian Johnston, Dominic Kwiatkowski, Cordelia Langford, John Sillitoe on behalf of the Wellcome Sanger Institute COVID-19 Surveillance Team |
| EPI_ISL_733825, EPI_ISL_733826 | Lighthouse Lab in Milton Keynes | Wellcome Sanger Institute for the COVID-19 Genomics UK (COG-UK) Consortium | The Lighthouse Lab in Milton Keynes and Alex Alderton, Roberto Amato, Sonia Goncalves, Ewan Harrison, David K. Jackson, Ian Johnston, Dominic Kwiatkowski, Cordelia Langford, John Sillitoe on behalf of the Wellcome Sanger Institute COVID-19 Surveillance Team |
| EPI_ISL_733827 | Lighthouse Lab in Cambridge | Wellcome Sanger Institute for the COVID-19 Genomics UK (COG-UK) Consortium | Rob Howes, The Lighthouse Lab in Cambridge and Alex Alderton, Roberto Amato, Sonia Goncalves, Ewan Harrison, David K. Jackson, Ian Johnston, Dominic Kwiatkowski, Cordelia Langford, John Sillitoe on behalf of the Wellcome Sanger Institute COVID-19 Surveillance Team |
| EPI_ISL_733828, EPI_ISL_733829 | Lighthouse Lab in Milton Keynes | Wellcome Sanger Institute for the COVID-19 Genomics UK (COG-UK) Consortium | The Lighthouse Lab in Milton Keynes and Alex Alderton, Roberto Amato, Sonia Goncalves, Ewan Harrison, David K. Jackson, Ian Johnston, Dominic Kwiatkowski, Cordelia Langford, John Sillitoe on behalf of the Wellcome Sanger Institute COVID-19 Surveillance Team |
| EPI_ISL_733831 | Lighthouse Lab in Cambridge | Wellcome Sanger Institute for the COVID-19 Genomics UK (COG-UK) Consortium | Rob Howes, The Lighthouse Lab in Cambridge and Alex Alderton, Roberto Amato, Sonia Goncalves, Ewan Harrison, David K. Jackson, Ian Johnston, Dominic Kwiatkowski, Cordelia Langford, John Sillitoe on behalf of the Wellcome Sanger Institute COVID-19 Surveillance Team |
| EPI_ISL_733832 | Lighthouse Lab in Milton Keynes | Wellcome Sanger Institute for the COVID-19 Genomics UK (COG-UK) Consortium | The Lighthouse Lab in Milton Keynes and Alex Alderton, Roberto Amato, Sonia Goncalves, Ewan Harrison, David K. Jackson, Ian Johnston, Dominic Kwiatkowski, Cordelia Langford, John Sillitoe on behalf of the Wellcome Sanger Institute COVID-19 Surveillance Team |
| EPI_ISL_733834, EPI_ISL_733835 | Lighthouse Lab in Cambridge | Wellcome Sanger Institute for the COVID-19 Genomics UK (COG-UK) Consortium | Rob Howes, The Lighthouse Lab in Cambridge and Alex Alderton, Roberto Amato, Sonia Goncalves, Ewan Harrison, David K. Jackson, Ian Johnston, Dominic Kwiatkowski, Cordelia Langford, John Sillitoe on behalf of the Wellcome Sanger Institute COVID-19 Surveillance Team |
| EPI_ISL_733836, EPI_ISL_733838 | Lighthouse Lab in Milton Keynes | Wellcome Sanger Institute for the COVID-19 Genomics UK (COG-UK) Consortium | The Lighthouse Lab in Milton Keynes and Alex Alderton, Roberto Amato, Sonia Goncalves, Ewan Harrison, David K. Jackson, Ian Johnston, Dominic Kwiatkowski, Cordelia Langford, John Sillitoe on behalf of the Wellcome Sanger Institute COVID-19 Surveillance Team |
| EPI_ISL_733839, EPI_ISL_733840, EPI_ISL_733841 | Lighthouse Lab in Cambridge | Wellcome Sanger Institute for the COVID-19 Genomics UK (COG-UK) Consortium | Rob Howes, The Lighthouse Lab in Cambridge and Alex Alderton, Roberto Amato, Sonia Goncalves, Ewan Harrison, David K. Jackson, Ian Johnston, Dominic Kwiatkowski, Cordelia Langford, John Sillitoe on behalf of the Wellcome Sanger Institute COVID-19 Surveillance Team |
| EPI_ISL_733842, EPI_ISL_733845 | Lighthouse Lab in Milton Keynes | Wellcome Sanger Institute for the COVID-19 Genomics UK (COG-UK) Consortium | The Lighthouse Lab in Milton Keynes and Alex Alderton, Roberto Amato, Sonia Goncalves, Ewan Harrison, David K. Jackson, Ian Johnston, Dominic Kwiatkowski, Cordelia Langford, John Sillitoe on behalf of the Wellcome Sanger Institute COVID-19 Surveillance Team |
| EPI_ISL_733846, EPI_ISL_733847, EPI_ISL_733849 | Lighthouse Lab in Cambridge | Wellcome Sanger Institute for the COVID-19 Genomics UK (COG-UK) Consortium | Rob Howes, The Lighthouse Lab in Cambridge and Alex Alderton, Roberto Amato, Sonia Goncalves, Ewan Harrison, David K. Jackson, Ian Johnston, Dominic Kwiatkowski, Cordelia Langford, John Sillitoe on behalf of the Wellcome Sanger Institute COVID-19 Surveillance Team |
| EPI_ISL_733851 | Lighthouse Lab in Milton Keynes | Wellcome Sanger Institute for the COVID-19 Genomics UK (COG-UK) Consortium | The Lighthouse Lab in Milton Keynes and Alex Alderton, Roberto Amato, Sonia Goncalves, Ewan Harrison, David K. Jackson, Ian Johnston, Dominic Kwiatkowski, Cordelia Langford, John Sillitoe on behalf of the Wellcome Sanger Institute COVID-19 Surveillance Team |
| EPI_ISL_733852, EPI_ISL_733853, EPI_ISL_733854 | Lighthouse Lab in Cambridge | Wellcome Sanger Institute for the COVID-19 Genomics UK (COG-UK) Consortium | Rob Howes, The Lighthouse Lab in Cambridge and Alex Alderton, Roberto Amato, Sonia Goncalves, Ewan Harrison, David K. Jackson, Ian Johnston, Dominic Kwiatkowski, Cordelia Langford, John Sillitoe on behalf of the Wellcome Sanger Institute COVID-19 Surveillance Team |
| EPI_ISL_733855 | Lighthouse Lab in Milton Keynes | Wellcome Sanger Institute for the COVID-19 Genomics UK (COG-UK) Consortium | The Lighthouse Lab in Milton Keynes and Alex Alderton, Roberto Amato, Sonia Goncalves, Ewan Harrison, David K. Jackson, Ian Johnston, Dominic Kwiatkowski, Cordelia Langford, John Sillitoe on behalf of the Wellcome Sanger Institute COVID-19 Surveillance Team |

| | | | |
|---|---|---|---|
| EPI_ISL_733856, EPI_ISL_733857 | Lighthouse Lab in Cambridge | Wellcome Sanger Institute for the COVID-19 Genomics UK (COG-UK) Consortium | Rob Howes, The Lighthouse Lab in Cambridge and Alex Alderton, Roberto Amato, Sonia Goncalves, Ewan Harrison, David K. Jackson, Ian Johnston, Dominic Kwiatkowski, Cordelia Langford, John Sillitoe on behalf of the Wellcome Sanger Institute COVID-19 Surveillance Team |
| EPI_ISL_733858 | Lighthouse Lab in Milton Keynes | Wellcome Sanger Institute for the COVID-19 Genomics UK (COG-UK) Consortium | The Lighthouse Lab in Milton Keynes and Alex Alderton, Roberto Amato, Sonia Goncalves, Ewan Harrison, David K. Jackson, Ian Johnston, Dominic Kwiatkowski, Cordelia Langford, John Sillitoe on behalf of the Wellcome Sanger Institute COVID-19 Surveillance Team |
| EPI_ISL_733860 | Lighthouse Lab in Cambridge | Wellcome Sanger Institute for the COVID-19 Genomics UK (COG-UK) Consortium | Rob Howes, The Lighthouse Lab in Cambridge and Alex Alderton, Roberto Amato, Sonia Goncalves, Ewan Harrison, David K. Jackson, Ian Johnston, Dominic Kwiatkowski, Cordelia Langford, John Sillitoe on behalf of the Wellcome Sanger Institute COVID-19 Surveillance Team |
| EPI_ISL_733861 | Lighthouse Lab in Milton Keynes | Wellcome Sanger Institute for the COVID-19 Genomics UK (COG-UK) Consortium | The Lighthouse Lab in Milton Keynes and Alex Alderton, Roberto Amato, Sonia Goncalves, Ewan Harrison, David K. Jackson, Ian Johnston, Dominic Kwiatkowski, Cordelia Langford, John Sillitoe on behalf of the Wellcome Sanger Institute COVID-19 Surveillance Team |
| EPI_ISL_733862 | Lighthouse Lab in Cambridge | Wellcome Sanger Institute for the COVID-19 Genomics UK (COG-UK) Consortium | Rob Howes, The Lighthouse Lab in Cambridge and Alex Alderton, Roberto Amato, Sonia Goncalves, Ewan Harrison, David K. Jackson, Ian Johnston, Dominic Kwiatkowski, Cordelia Langford, John Sillitoe on behalf of the Wellcome Sanger Institute COVID-19 Surveillance Team |
| EPI_ISL_733863, EPI_ISL_733864, EPI_ISL_733865 | Lighthouse Lab in Milton Keynes | Wellcome Sanger Institute for the COVID-19 Genomics UK (COG-UK) Consortium | The Lighthouse Lab in Milton Keynes and Alex Alderton, Roberto Amato, Sonia Goncalves, Ewan Harrison, David K. Jackson, Ian Johnston, Dominic Kwiatkowski, Cordelia Langford, John Sillitoe on behalf of the Wellcome Sanger Institute COVID-19 Surveillance Team |
| EPI_ISL_733866, EPI_ISL_733868, EPI_ISL_733869, EPI_ISL_733870, EPI_ISL_733871 | Lighthouse Lab in Cambridge | Wellcome Sanger Institute for the COVID-19 Genomics UK (COG-UK) Consortium | Rob Howes, The Lighthouse Lab in Cambridge and Alex Alderton, Roberto Amato, Sonia Goncalves, Ewan Harrison, David K. Jackson, Ian Johnston, Dominic Kwiatkowski, Cordelia Langford, John Sillitoe on behalf of the Wellcome Sanger Institute COVID-19 Surveillance Team |
| EPI_ISL_733872, EPI_ISL_733873 | Lighthouse Lab in Milton Keynes | Wellcome Sanger Institute for the COVID-19 Genomics UK (COG-UK) Consortium | The Lighthouse Lab in Milton Keynes and Alex Alderton, Roberto Amato, Sonia Goncalves, Ewan Harrison, David K. Jackson, Ian Johnston, Dominic Kwiatkowski, Cordelia Langford, John Sillitoe on behalf of the Wellcome Sanger Institute COVID-19 Surveillance Team |
| EPI_ISL_733874 | Lighthouse Lab in Cambridge | Wellcome Sanger Institute for the COVID-19 Genomics UK (COG-UK) Consortium | Rob Howes, The Lighthouse Lab in Cambridge and Alex Alderton, Roberto Amato, Sonia Goncalves, Ewan Harrison, David K. Jackson, Ian Johnston, Dominic Kwiatkowski, Cordelia Langford, John Sillitoe on behalf of the Wellcome Sanger Institute COVID-19 Surveillance Team |
| EPI_ISL_733875 | Lighthouse Lab in Milton Keynes | Wellcome Sanger Institute for the COVID-19 Genomics UK (COG-UK) Consortium | The Lighthouse Lab in Milton Keynes and Alex Alderton, Roberto Amato, Sonia Goncalves, Ewan Harrison, David K. Jackson, Ian Johnston, Dominic Kwiatkowski, Cordelia Langford, John Sillitoe on behalf of the Wellcome Sanger Institute COVID-19 Surveillance Team |
| EPI_ISL_733876, EPI_ISL_733878, EPI_ISL_733881 | Lighthouse Lab in Cambridge | Wellcome Sanger Institute for the COVID-19 Genomics UK (COG-UK) Consortium | Rob Howes, The Lighthouse Lab in Cambridge and Alex Alderton, Roberto Amato, Sonia Goncalves, Ewan Harrison, David K. Jackson, Ian Johnston, Dominic Kwiatkowski, Cordelia Langford, John Sillitoe on behalf of the Wellcome Sanger Institute COVID-19 Surveillance Team |
| EPI_ISL_733884 | Lighthouse Lab in Milton Keynes | Wellcome Sanger Institute for the COVID-19 Genomics UK (COG-UK) Consortium | The Lighthouse Lab in Milton Keynes and Alex Alderton, Roberto Amato, Sonia Goncalves, Ewan Harrison, David K. Jackson, Ian Johnston, Dominic Kwiatkowski, Cordelia Langford, John Sillitoe on behalf of the Wellcome Sanger Institute COVID-19 Surveillance Team |
| EPI_ISL_733886 | Lighthouse Lab in Cambridge | Wellcome Sanger Institute for the COVID-19 Genomics UK (COG-UK) Consortium | Rob Howes, The Lighthouse Lab in Cambridge and Alex Alderton, Roberto Amato, Sonia Goncalves, Ewan Harrison, David K. Jackson, Ian Johnston, Dominic Kwiatkowski, Cordelia Langford, John Sillitoe on behalf of the Wellcome Sanger Institute COVID-19 Surveillance Team |
| EPI_ISL_733887 | Lighthouse Lab in Milton Keynes | Wellcome Sanger Institute for the COVID-19 Genomics UK (COG-UK) Consortium | The Lighthouse Lab in Milton Keynes and Alex Alderton, Roberto Amato, Sonia Goncalves, Ewan Harrison, David K. Jackson, Ian Johnston, Dominic Kwiatkowski, Cordelia Langford, John Sillitoe on behalf of the Wellcome Sanger Institute COVID-19 Surveillance Team |
| EPI_ISL_733888 | Lighthouse Lab in Cambridge | Wellcome Sanger Institute for the COVID-19 Genomics UK (COG-UK) Consortium | Rob Howes, The Lighthouse Lab in Cambridge and Alex Alderton, Roberto Amato, Sonia Goncalves, Ewan Harrison, David K. Jackson, Ian Johnston, Dominic Kwiatkowski, Cordelia Langford, John Sillitoe on behalf of the Wellcome Sanger Institute COVID-19 Surveillance Team |
| EPI_ISL_733889, EPI_ISL_733890, EPI_ISL_733891 | Lighthouse Lab in Milton Keynes | Wellcome Sanger Institute for the COVID-19 Genomics UK (COG-UK) Consortium | The Lighthouse Lab in Milton Keynes and Alex Alderton, Roberto Amato, Sonia Goncalves, Ewan Harrison, David K. Jackson, Ian Johnston, Dominic Kwiatkowski, Cordelia Langford, John Sillitoe on behalf of the Wellcome Sanger Institute COVID-19 Surveillance Team |
| EPI_ISL_733892 | Lighthouse Lab in Cambridge | Wellcome Sanger Institute for the COVID-19 Genomics UK (COG-UK) Consortium | Rob Howes, The Lighthouse Lab in Cambridge and Alex Alderton, Roberto Amato, Sonia Goncalves, Ewan Harrison, David K. Jackson, Ian Johnston, Dominic Kwiatkowski, Cordelia Langford, John Sillitoe on behalf of the Wellcome Sanger Institute COVID-19 Surveillance Team |
| EPI_ISL_733893, EPI_ISL_733894 | Lighthouse Lab in Milton Keynes | Wellcome Sanger Institute for the COVID-19 Genomics UK (COG-UK) Consortium | The Lighthouse Lab in Milton Keynes and Alex Alderton, Roberto Amato, Sonia Goncalves, Ewan Harrison, David K. Jackson, Ian Johnston, Dominic Kwiatkowski, Cordelia Langford, John Sillitoe on behalf of the Wellcome Sanger Institute COVID-19 Surveillance Team |
| EPI_ISL_733895, EPI_ISL_733896, EPI_ISL_733898 | Lighthouse Lab in Cambridge | Wellcome Sanger Institute for the COVID-19 Genomics UK (COG-UK) Consortium | Rob Howes, The Lighthouse Lab in Cambridge and Alex Alderton, Roberto Amato, Sonia Goncalves, Ewan Harrison, David K. Jackson, Ian Johnston, Dominic Kwiatkowski, Cordelia Langford, John Sillitoe on behalf of the Wellcome Sanger Institute COVID-19 Surveillance Team |
| EPI_ISL_733900, EPI_ISL_733902, EPI_ISL_733903, EPI_ISL_733904, EPI_ISL_733905 | Lighthouse Lab in Milton Keynes | Wellcome Sanger Institute for the COVID-19 Genomics UK (COG-UK) Consortium | The Lighthouse Lab in Milton Keynes and Alex Alderton, Roberto Amato, Sonia Goncalves, Ewan Harrison, David K. Jackson, Ian Johnston, Dominic Kwiatkowski, Cordelia Langford, John Sillitoe on behalf of the Wellcome Sanger Institute COVID-19 Surveillance Team |
| EPI_ISL_733908, EPI_ISL_733909, EPI_ISL_733911 | Lighthouse Lab in Cambridge | Wellcome Sanger Institute for the COVID-19 Genomics UK (COG-UK) Consortium | Rob Howes, The Lighthouse Lab in Cambridge and Alex Alderton, Roberto Amato, Sonia Goncalves, Ewan Harrison, David K. Jackson, Ian Johnston, Dominic Kwiatkowski, Cordelia Langford, John Sillitoe on behalf of the Wellcome Sanger Institute COVID-19 Surveillance Team |
| EPI_ISL_733914 | Lighthouse Lab in Milton Keynes | Wellcome Sanger Institute for the COVID-19 Genomics UK (COG-UK) Consortium | The Lighthouse Lab in Milton Keynes and Alex Alderton, Roberto Amato, Sonia Goncalves, Ewan Harrison, David K. Jackson, Ian Johnston, Dominic Kwiatkowski, Cordelia Langford, John Sillitoe on behalf of the Wellcome Sanger Institute COVID-19 Surveillance Team |
| EPI_ISL_733915 | Lighthouse Lab in Cambridge | Wellcome Sanger Institute for the COVID-19 Genomics UK (COG-UK) Consortium | Rob Howes, The Lighthouse Lab in Cambridge and Alex Alderton, Roberto Amato, Sonia Goncalves, Ewan Harrison, David K. Jackson, Ian Johnston, Dominic Kwiatkowski, Cordelia Langford, John Sillitoe on behalf of the Wellcome Sanger Institute COVID-19 Surveillance Team |
| EPI_ISL_733928, EPI_ISL_733950, EPI_ISL_734041, EPI_ISL_734044, EPI_ISL_734093, EPI_ISL_734117, EPI_ISL_734131 | Lighthouse Lab in Glasgow | Wellcome Sanger Institute for the COVID-19 Genomics UK (COG-UK) Consortium | Harper VanSteenhouse, Yumi Kasai, David Gray, Carol Clugston, Anna Dominiczak and Alex Alderton, Roberto Amato, Sonia Goncalves, Ewan Harrison, David K. Jackson, Ian Johnston, Dominic Kwiatkowski, Cordelia Langford, John Sillitoe on behalf of the Wellcome Sanger Institute COVID-19 Surveillance Team |
| EPI_ISL_736202, EPI_ISL_736240, EPI_ISL_736252, EPI_ISL_736264, EPI_ISL_736274, EPI_ISL_736289, EPI_ISL_736292, EPI_ISL_736293, EPI_ISL_736303, EPI_ISL_736349, EPI_ISL_736369, EPI_ISL_736380, EPI_ISL_736392, EPI_ISL_736408, EPI_ISL_736449 | | | |
| see above | Lighthouse Lab in Milton Keynes | Wellcome Sanger Institute for the COVID-19 Genomics UK (COG-UK) Consortium | The Lighthouse Lab in Milton Keynes and Alex Alderton, Roberto Amato, Sonia Goncalves, Ewan Harrison, David K. Jackson, Ian Johnston, Dominic Kwiatkowski, Cordelia Langford, John Sillitoe on behalf of the Wellcome Sanger Institute COVID-19 Surveillance Team |
| EPI_ISL_736458 | Lighthouse Lab in Glasgow | Wellcome Sanger Institute for the COVID-19 Genomics UK (COG-UK) Consortium | Harper VanSteenhouse, Yumi Kasai, David Gray, Carol Clugston, Anna Dominiczak and Alex Alderton, Roberto Amato, Sonia Goncalves, Ewan Harrison, David K. Jackson, Ian Johnston, Dominic Kwiatkowski, Cordelia Langford, John Sillitoe on behalf of the Wellcome Sanger Institute COVID-19 Surveillance Team |
| EPI_ISL_736469 | Lighthouse Lab in Milton Keynes | Wellcome Sanger Institute for the COVID-19 Genomics UK (COG-UK) Consortium | The Lighthouse Lab in Milton Keynes and Alex Alderton, Roberto Amato, Sonia Goncalves, Ewan Harrison, David K. Jackson, Ian Johnston, Dominic Kwiatkowski, Cordelia Langford, John Sillitoe on behalf of the Wellcome Sanger Institute COVID-19 Surveillance Team |
| EPI_ISL_736477 | Lighthouse Lab in Glasgow | Wellcome Sanger Institute for the COVID-19 Genomics UK (COG-UK) Consortium | Harper VanSteenhouse, Yumi Kasai, David Gray, Carol Clugston, Anna Dominiczak and Alex Alderton, Roberto Amato, Sonia Goncalves, Ewan Harrison, David K. Jackson, Ian Johnston, Dominic Kwiatkowski, Cordelia Langford, John Sillitoe on behalf of the Wellcome Sanger Institute COVID-19 Surveillance Team |
| EPI_ISL_736547, EPI_ISL_736686, EPI_ISL_736694, EPI_ISL_736749 | Lighthouse Lab in Milton Keynes | Wellcome Sanger Institute for the COVID-19 Genomics UK (COG-UK) Consortium | The Lighthouse Lab in Milton Keynes and Alex Alderton, Roberto Amato, Sonia Goncalves, Ewan Harrison, David K. Jackson, Ian Johnston, Dominic Kwiatkowski, Cordelia Langford, John Sillitoe on behalf of the Wellcome Sanger Institute COVID-19 Surveillance Team |
| EPI_ISL_736758 | Lighthouse Lab in Alderley Park | Wellcome Sanger Institute for the COVID-19 Genomics UK (COG-UK) Consortium | Jacquelyn Wynn, Mairead Hyland, The Lighthouse Lab in Alderley Park and Alex Alderton, Roberto Amato, Sonia Goncalves, Ewan Harrison, David K. Jackson, Ian Johnston, Dominic Kwiatkowski, Cordelia Langford, John Sillitoe on behalf of the Wellcome Sanger Institute COVID-19 Surveillance Team |
| EPI_ISL_736927 | MDS | KRISP, KZN Research Innovation and Sequencing Platform | Giandhari J, Pillay S, Lessells R, ChimukangaraB, Mdlalose K, York D, Khan S, Tegally H, Wilkinson E, de Oliveira T |
| EPI_ISL_736938, EPI_ISL_736970, EPI_ISL_736971, EPI_ISL_736973, EPI_ISL_736974, EPI_ISL_736975, EPI_ISL_736976, EPI_ISL_736977, EPI_ISL_736978, EPI_ISL_736979, EPI_ISL_736980 | | | |
| see above | NHLS-IALCH | KRISP, KZN Research Innovation and Sequencing Platform | Giandhari J, Pillay S, Lessells R, ChimukangaraB, Mdlalose K, York D, Khan S, Tegally H, Wilkinson E, de Oliveira T |
| EPI_ISL_736995 | MDS | KRISP, KZN Research Innovation and Sequencing Platform | Giandhari J, Pillay S, Lessells R, ChimukangaraB, Mdlalose K, York D, Khan S, Tegally H, Wilkinson E, de Oliveira T |

| | | | |
|---|---|---|---|
| EPI_ISL_737105, EPI_ISL_737282, EPI_ISL_737301 | Michigan Department of Health and Human Services, Bureau of Laboratories | Michigan Department of Health and Human Services, Bureau of Laboratories | Blankenship HM, Riner D, Soehnlen MK |
| EPI_ISL_737630, EPI_ISL_737857, EPI_ISL_737859, EPI_ISL_737860, EPI_ISL_737861, EPI_ISL_737862 | Viollier AG | Department of Biosystems Science and Engineering, ETH Zürich | Chaoran Chen, Sarah Nadeau, Catharine Aquino, Ivan Topolsky, Philipp Jablonski, Lara Fuhrmann, David Dreifuss, Katharina Jahn, Andreia Cabral de Gouvea, Maria Domenica Moccia, Simon Grüter, Timothy Sykes, Lennart Opitz, Griffin White, Laura Neff, Doris Popovic, Andrea Patrignani, Jay Tracy, Ralph Schlapbach, Christiane Beckmann, Maurice Redondo, Olivier Kobel, Christoph Noppen, Sophie Seidel, Noemie Santamaria de Souza, Niko Beerenwinkel, Tanja Stadler |
| EPI_ISL_738108, EPI_ISL_738112 | Instituto Nacional de Saude (INSA) | Instituto Nacional de Saude (INSA) | Borges et al |
| EPI_ISL_738361, EPI_ISL_738371, EPI_ISL_738389, EPI_ISL_738414, EPI_ISL_738431, EPI_ISL_738437 | UZ Leuven, National Reference Laboratory for Coronaviruses, Laboratory Medicine, Leuven, Belgium | KU Leuven, Rega Institute, Clinical and Epidemiological Virology | Tony Wawina-Bokalanga, Joan Marti-Carerras, Bert Vanmechelen, Piet Maes |
| EPI_ISL_738658 | Monterey County Public Health Lab | Chan-Zuckerberg Biohub | CZB Cliahub Consortium |
| EPI_ISL_740871, EPI_ISL_740872, EPI_ISL_740873, EPI_ISL_740874 | South Eastern Area Laboratory Services (SEALS) | NSW Health Pathology - Institute of Clinical Pathology and Medical Research; Westmead Hospital; University of Sydney | CIDM-PH et al. |
| EPI_ISL_740958, EPI_ISL_740963, EPI_ISL_740964, EPI_ISL_740965, EPI_ISL_740966, EPI_ISL_740967, EPI_ISL_740968, EPI_ISL_740969, EPI_ISL_740970, EPI_ISL_740972, EPI_ISL_740973, EPI_ISL_740974, EPI_ISL_740975, EPI_ISL_740976, EPI_ISL_740977, EPI_ISL_740978, EPI_ISL_740979, EPI_ISL_740980, EPI_ISL_740981, EPI_ISL_740982, EPI_ISL_740983, EPI_ISL_740984, EPI_ISL_740985, EPI_ISL_740990, EPI_ISL_740991, EPI_ISL_740993 | | | |
| see above | Department of Pathology, University of Cambridge | COVID-19 Genomics UK (COG-UK) Consortium | Aminu S. Jahun, Yasmin Chaudhry, Grant Hall, Iliana Georgana, Myra Hosmillo, Martin D. Curran, Malte Pinckert, Surendra Parmar, Ian Goodfellow |
| EPI_ISL_741187, EPI_ISL_741207, EPI_ISL_741255, EPI_ISL_741264, EPI_ISL_741265, EPI_ISL_741266, EPI_ISL_741267, EPI_ISL_741268, EPI_ISL_741269, EPI_ISL_741271, EPI_ISL_741272, EPI_ISL_741273, EPI_ISL_741274, EPI_ISL_741276, EPI_ISL_741278, EPI_ISL_741279, EPI_ISL_741280, EPI_ISL_741281, EPI_ISL_741282, EPI_ISL_741283, EPI_ISL_741284, EPI_ISL_741285, EPI_ISL_741286, EPI_ISL_741288, EPI_ISL_741289, EPI_ISL_741290, EPI_ISL_741291, EPI_ISL_741292, EPI_ISL_741293, EPI_ISL_741295, EPI_ISL_741300, EPI_ISL_741303, EPI_ISL_741308 | | | |
| see above | University College London, Great Ormond Street Hospital for Children NHS Foundation Trust, Imperial College Healthcare NHS Trust | COVID-19 Genomics UK (COG-UK) Consortium | Sergi Castellano, Rachel Williams, Mark Kristiansen, Paola Resende Silva, Sunando Roy, Tony Brooks, Helena Tutill, Paola Niola, Patricia Dyal, Charlotte Williams, Leysa Forrest, Yasmin Panchbhaya, Jacqueline Findlay, Samuel Weeks, Julianne Brown, Kathryn Harris, Paul Randell, James Price, Alison Holmes, Judith Breuer |
| EPI_ISL_741613 | Queens Medical Centre, Clinical Microbiology Department / DeepSeq Nottingham | COVID-19 Genomics UK (COG-UK) Consortium | Gemma Clark, Wendy Smith, Manjinder Khakh, Vicki M Fleming, Michelle M Lister, Hannah Howson-Wells, Jonathan Ball, Patrick McClure, Joseph Chappell, Theocharis Tsoleridis, Nadine Holmes, Matthew Carlisle, Christopher Moore, Fei Sang, Johnny Debebe, Victoria Wright, Matthew Loose |
| EPI_ISL_741734, EPI_ISL_741750, EPI_ISL_741751, EPI_ISL_741752 | Centre for Enzyme Innovation, University of Portsmouth / Translational Research Laboratory, Portsmouth Hospitals NHS Trust | COVID-19 Genomics UK (COG-UK) Consortium | Angela Beckett,Yann Bourgeois,Garry Scarlett,Sharon Glaysher,Scott Elliott,Kelly Bicknell,Robert Impey,Allyson Lloyd,Sarah Wyllie,Ethan Butcher,Anoop Chauhan,Samuel Robson |
| EPI_ISL_741939, EPI_ISL_741940, EPI_ISL_741941, EPI_ISL_741949, EPI_ISL_741955, EPI_ISL_741959, EPI_ISL_741973, EPI_ISL_741979, EPI_ISL_741987, EPI_ISL_741988, EPI_ISL_741993, EPI_ISL_742001, EPI_ISL_742006, EPI_ISL_742008, EPI_ISL_742011, EPI_ISL_742021, EPI_ISL_742027, EPI_ISL_742029, EPI_ISL_742036, EPI_ISL_742038, EPI_ISL_742042, EPI_ISL_742043, EPI_ISL_742045, EPI_ISL_742053, EPI_ISL_742057, EPI_ISL_742058, EPI_ISL_742085, EPI_ISL_742088, EPI_ISL_742091, EPI_ISL_742093, EPI_ISL_742095, EPI_ISL_742110 | | | |
| see above | Virology Department, Sheffield Teaching Hospitals NHS Foundation Trust/Department of Infection, Immunity and Cardiovascular Disease, The Medical School, University of Sheffield | COVID-19 Genomics UK (COG-UK) Consortium | Thushan de Silva, Matthew Parker, Nikki Smith, Adri Angyal, Rebecca Brown, Luke Green, Rachel Tucker, Paul Parsons, Danielle Groves, Katie Johnson, Laura Carrilero, Alex Keeley, Dave Partridge, Matthew Wyles, Benjamin Lindsey, Mehmet Yavuz, Mohammad Raza, Cariad Evans |
| EPI_ISL_742180 | Virology Department, Royal Infirmary of Edinburgh, NHS Lothian / School of Biological Sciences, University of Edinburgh / Institute of Genetics and Molecular Medicine, University of Edinburgh | COVID-19 Genomics UK (COG-UK) Consortium | McHugh M, Dewar R, Rooke S, Gallagher M, Balcaza C, O'Toole Á, Scher E, Hill V, McCrone JT, Colquhoun R, Yu X, Jackson B, Rambaut A, Williams TC, Templeton K |
| EPI_ISL_742263, EPI_ISL_742266, EPI_ISL_742288, EPI_ISL_742289, EPI_ISL_742290, EPI_ISL_742291, EPI_ISL_742292, EPI_ISL_742293, EPI_ISL_742294, EPI_ISL_742295, EPI_ISL_742296, EPI_ISL_742297, EPI_ISL_742356, EPI_ISL_742359, EPI_ISL_742360, EPI_ISL_742361, EPI_ISL_742362, EPI_ISL_742363, EPI_ISL_742488, EPI_ISL_742489, EPI_ISL_742490, EPI_ISL_742491, EPI_ISL_742492, EPI_ISL_742493, EPI_ISL_742494, EPI_ISL_742495, EPI_ISL_742496, EPI_ISL_742497, EPI_ISL_742498, EPI_ISL_742499, EPI_ISL_742500, EPI_ISL_742501, EPI_ISL_742502, EPI_ISL_742503, EPI_ISL_742504, EPI_ISL_742505, EPI_ISL_742506, EPI_ISL_742507, EPI_ISL_742508, EPI_ISL_742509, EPI_ISL_742510, EPI_ISL_742511, EPI_ISL_742512, EPI_ISL_742513, EPI_ISL_742514, EPI_ISL_742515, EPI_ISL_742516, EPI_ISL_742517, EPI_ISL_742518, EPI_ISL_742519, EPI_ISL_742520, EPI_ISL_742521, EPI_ISL_742522, EPI_ISL_742523, EPI_ISL_742524, EPI_ISL_742525, EPI_ISL_742526, EPI_ISL_742527, EPI_ISL_742528, EPI_ISL_742529, EPI_ISL_742531, EPI_ISL_742532, EPI_ISL_742533, EPI_ISL_742534, EPI_ISL_742535, EPI_ISL_742536, EPI_ISL_742537, EPI_ISL_742538, EPI_ISL_742539, EPI_ISL_742540, EPI_ISL_742545, EPI_ISL_742550, EPI_ISL_742551, EPI_ISL_742552, EPI_ISL_742553, EPI_ISL_742554, EPI_ISL_742565, EPI_ISL_742637, EPI_ISL_742638, EPI_ISL_742639, EPI_ISL_742640, EPI_ISL_742646, EPI_ISL_742649, EPI_ISL_742651, EPI_ISL_742653, EPI_ISL_742654, EPI_ISL_742655, EPI_ISL_742656, EPI_ISL_742657, EPI_ISL_742658, EPI_ISL_742659, EPI_ISL_742660, EPI_ISL_742661, EPI_ISL_742662, EPI_ISL_742663, EPI_ISL_742664, EPI_ISL_742665, EPI_ISL_742666, EPI_ISL_742667, EPI_ISL_742668, EPI_ISL_742669, EPI_ISL_742670, EPI_ISL_742671, EPI_ISL_742672, EPI_ISL_742673, EPI_ISL_742674, EPI_ISL_742675, EPI_ISL_742676, EPI_ISL_742677, EPI_ISL_742678, EPI_ISL_742679, EPI_ISL_742683, EPI_ISL_742684, EPI_ISL_742685, EPI_ISL_742686, EPI_ISL_742687, EPI_ISL_742688, EPI_ISL_742689, EPI_ISL_742690, EPI_ISL_742691, EPI_ISL_742694, EPI_ISL_742695, EPI_ISL_742696, EPI_ISL_742697, EPI_ISL_742700, EPI_ISL_742701, EPI_ISL_742702, EPI_ISL_742704, EPI_ISL_742705, EPI_ISL_742707, EPI_ISL_742709, EPI_ISL_742710, EPI_ISL_742713, EPI_ISL_742714, EPI_ISL_742716, EPI_ISL_742717, EPI_ISL_742718, EPI_ISL_742719, EPI_ISL_742720, EPI_ISL_742849, EPI_ISL_742850, EPI_ISL_742851, EPI_ISL_742852, EPI_ISL_742853, EPI_ISL_742854, EPI_ISL_742855, EPI_ISL_742858, EPI_ISL_742859, EPI_ISL_742860, EPI_ISL_742861, EPI_ISL_742862, EPI_ISL_742863, EPI_ISL_742864, EPI_ISL_742865, EPI_ISL_742866, EPI_ISL_742867, EPI_ISL_742868, EPI_ISL_742869, EPI_ISL_742870, EPI_ISL_742871, EPI_ISL_742872, EPI_ISL_742873, EPI_ISL_742874, EPI_ISL_742875, EPI_ISL_742876, EPI_ISL_742877, EPI_ISL_742878, EPI_ISL_742879, EPI_ISL_742911, EPI_ISL_742912, EPI_ISL_742915, EPI_ISL_742924, EPI_ISL_742925, EPI_ISL_742939, EPI_ISL_742946, EPI_ISL_742947, EPI_ISL_742948, EPI_ISL_742949, EPI_ISL_742950, EPI_ISL_742951, EPI_ISL_742952, EPI_ISL_742953, EPI_ISL_742954, EPI_ISL_742955, EPI_ISL_742956, EPI_ISL_742957, EPI_ISL_742958, EPI_ISL_742959, EPI_ISL_742960, EPI_ISL_742961, EPI_ISL_742962, EPI_ISL_742963, EPI_ISL_742964, EPI_ISL_742965, EPI_ISL_742966, EPI_ISL_742967, EPI_ISL_742968, EPI_ISL_742969, EPI_ISL_742970, EPI_ISL_742971, EPI_ISL_742972, EPI_ISL_742973, EPI_ISL_742974, EPI_ISL_742975, EPI_ISL_742976, EPI_ISL_742977, EPI_ISL_742978, EPI_ISL_742979, EPI_ISL_742980, EPI_ISL_742981, EPI_ISL_742982, EPI_ISL_742983, EPI_ISL_742984, EPI_ISL_742985, EPI_ISL_742986, EPI_ISL_742987, EPI_ISL_742988, EPI_ISL_742989, EPI_ISL_742990, EPI_ISL_742991, EPI_ISL_742992, EPI_ISL_742993, EPI_ISL_742994, EPI_ISL_742995, EPI_ISL_742996, EPI_ISL_742997, EPI_ISL_742998, EPI_ISL_742999, EPI_ISL_743000, EPI_ISL_743001, EPI_ISL_743002, EPI_ISL_743003, EPI_ISL_743004, EPI_ISL_743005, EPI_ISL_743006, EPI_ISL_743007, EPI_ISL_743008, EPI_ISL_743009, EPI_ISL_743010, EPI_ISL_743011, EPI_ISL_743012, EPI_ISL_743013, EPI_ISL_743014, EPI_ISL_743015, EPI_ISL_743016, EPI_ISL_743017, EPI_ISL_743018, EPI_ISL_743019, EPI_ISL_743020, EPI_ISL_743021, EPI_ISL_743022, EPI_ISL_743023, EPI_ISL_743024, EPI_ISL_743025, EPI_ISL_743026, EPI_ISL_743027, EPI_ISL_743028, EPI_ISL_743029, EPI_ISL_743030, EPI_ISL_743031, EPI_ISL_743032, EPI_ISL_743033, EPI_ISL_743034, EPI_ISL_743035, EPI_ISL_743036, EPI_ISL_743037, EPI_ISL_743038, EPI_ISL_743039, EPI_ISL_743040, EPI_ISL_743041, EPI_ISL_743042, EPI_ISL_743043, EPI_ISL_743044, EPI_ISL_743045, EPI_ISL_743046, EPI_ISL_743047, EPI_ISL_743048, EPI_ISL_743049, EPI_ISL_743050, EPI_ISL_743051, EPI_ISL_743052, EPI_ISL_743053, EPI_ISL_743054, EPI_ISL_743055, EPI_ISL_743056, EPI_ISL_743057, EPI_ISL_743058, EPI_ISL_743059, EPI_ISL_743060, EPI_ISL_743061, EPI_ISL_743062, EPI_ISL_743063, EPI_ISL_743064, EPI_ISL_743065, EPI_ISL_743066, EPI_ISL_743067, EPI_ISL_743068, EPI_ISL_743069, EPI_ISL_743070, EPI_ISL_743071, EPI_ISL_743072, EPI_ISL_743073, EPI_ISL_743074, EPI_ISL_743075, EPI_ISL_743076, EPI_ISL_743077, EPI_ISL_743078, EPI_ISL_743079, EPI_ISL_743080, EPI_ISL_743081, EPI_ISL_743082, EPI_ISL_743083, EPI_ISL_743084, EPI_ISL_743085, EPI_ISL_743086, EPI_ISL_743087, EPI_ISL_743088, EPI_ISL_743089, EPI_ISL_743090, EPI_ISL_743091, EPI_ISL_743092, EPI_ISL_743093, EPI_ISL_743094, EPI_ISL_743095, EPI_ISL_743096, EPI_ISL_743097, EPI_ISL_743098, EPI_ISL_743099, EPI_ISL_743100, EPI_ISL_743101, EPI_ISL_743102, EPI_ISL_743103, EPI_ISL_743104, EPI_ISL_743105, EPI_ISL_743106, EPI_ISL_743107, EPI_ISL_743108, EPI_ISL_743109, EPI_ISL_743110, EPI_ISL_743111, EPI_ISL_743112, EPI_ISL_743113, EPI_ISL_743114, EPI_ISL_743115, EPI_ISL_743116, EPI_ISL_743121, EPI_ISL_743122, EPI_ISL_743123, EPI_ISL_743128, EPI_ISL_743137, EPI_ISL_743138, EPI_ISL_743139, EPI_ISL_743140, EPI_ISL_743141, EPI_ISL_743142, EPI_ISL_743143, EPI_ISL_743144, EPI_ISL_743145, EPI_ISL_743159, EPI_ISL_743160, EPI_ISL_743163, EPI_ISL_743164, EPI_ISL_743165, EPI_ISL_743175, EPI_ISL_743176, EPI_ISL_743177, EPI_ISL_743178, EPI_ISL_743179, EPI_ISL_743180, EPI_ISL_743181, EPI_ISL_743182, EPI_ISL_743190, EPI_ISL_743192, EPI_ISL_743193, EPI_ISL_743194, EPI_ISL_743195, EPI_ISL_743196, EPI_ISL_743197, EPI_ISL_743198, EPI_ISL_743199, EPI_ISL_743200, EPI_ISL_743201, EPI_ISL_743202, EPI_ISL_743203, EPI_ISL_743204, EPI_ISL_743205, EPI_ISL_743206, EPI_ISL_743207, EPI_ISL_743208, EPI_ISL_743209, EPI_ISL_743210, EPI_ISL_743211, EPI_ISL_743212, EPI_ISL_743213, EPI_ISL_743214, EPI_ISL_743215, EPI_ISL_743216, EPI_ISL_743217, EPI_ISL_743218, EPI_ISL_743219, EPI_ISL_743220, EPI_ISL_743221, EPI_ISL_743222, EPI_ISL_743223, EPI_ISL_743224, EPI_ISL_743225, EPI_ISL_743226, EPI_ISL_743227, EPI_ISL_743228, EPI_ISL_743229, EPI_ISL_743230, EPI_ISL_743231, EPI_ISL_743232, EPI_ISL_743233, EPI_ISL_743234, EPI_ISL_743235, EPI_ISL_743236, EPI_ISL_743237, EPI_ISL_743238, EPI_ISL_743239, EPI_ISL_743240, EPI_ISL_743241, EPI_ISL_743242, EPI_ISL_743243, EPI_ISL_743244, EPI_ISL_743245, EPI_ISL_743246, EPI_ISL_743247, EPI_ISL_743248, EPI_ISL_743249, EPI_ISL_743250, EPI_ISL_743251, EPI_ISL_743252, EPI_ISL_743253, EPI_ISL_743254, EPI_ISL_743255, EPI_ISL_743256, EPI_ISL_743257, EPI_ISL_743258, EPI_ISL_743259, EPI_ISL_743260, EPI_ISL_743261, EPI_ISL_743262, EPI_ISL_743263, EPI_ISL_743264, EPI_ISL_743265, EPI_ISL_743266, EPI_ISL_743267, EPI_ISL_743268, EPI_ISL_743269, EPI_ISL_743270, EPI_ISL_743271, EPI_ISL_743272, EPI_ISL_743273, EPI_ISL_743274, EPI_ISL_743275, EPI_ISL_743276, EPI_ISL_743277, EPI_ISL_743278, EPI_ISL_743279, EPI_ISL_743280, EPI_ISL_743281, EPI_ISL_743282, EPI_ISL_743283, EPI_ISL_743284, EPI_ISL_743285, EPI_ISL_743286, EPI_ISL_743287, EPI_ISL_743288, EPI_ISL_743289, EPI_ISL_743290, EPI_ISL_743291, EPI_ISL_743292, EPI_ISL_743293, EPI_ISL_743294, EPI_ISL_743295, EPI_ISL_743296, EPI_ISL_743297, EPI_ISL_743298, EPI_ISL_743299, EPI_ISL_743300, EPI_ISL_743301, EPI_ISL_743302, EPI_ISL_743303, EPI_ISL_743304, EPI_ISL_743305, EPI_ISL_743306, EPI_ISL_743307, EPI_ISL_743308, EPI_ISL_743309, EPI_ISL_743310, EPI_ISL_743311, EPI_ISL_743312, EPI_ISL_743313, EPI_ISL_743314, EPI_ISL_743315, EPI_ISL_743316, EPI_ISL_743317, EPI_ISL_743318, EPI_ISL_743319, EPI_ISL_743320, EPI_ISL_743323, EPI_ISL_743324, EPI_ISL_743325, EPI_ISL_743327, EPI_ISL_743328, EPI_ISL_743329, EPI_ISL_743330, EPI_ISL_743331, EPI_ISL_743332, EPI_ISL_743333, EPI_ISL_743334, EPI_ISL_743335, EPI_ISL_743336, EPI_ISL_743337, EPI_ISL_743338, EPI_ISL_743339, EPI_ISL_743340, EPI_ISL_743346, EPI_ISL_743347, EPI_ISL_743348, EPI_ISL_743349, EPI_ISL_743350, EPI_ISL_743351, EPI_ISL_743356, EPI_ISL_743357, EPI_ISL_743411, EPI_ISL_743454, EPI_ISL_743455, EPI_ISL_743456, EPI_ISL_743457, EPI_ISL_743458, EPI_ISL_743459, EPI_ISL_743460, EPI_ISL_743461, EPI_ISL_743462, EPI_ISL_743463, EPI_ISL_743464, EPI_ISL_743465, EPI_ISL_743466, EPI_ISL_743467, EPI_ISL_743468, EPI_ISL_743469, EPI_ISL_743470, EPI_ISL_743471, EPI_ISL_743472, EPI_ISL_743473, EPI_ISL_743474, EPI_ISL_743475, EPI_ISL_743476, EPI_ISL_743477, EPI_ISL_743478, EPI_ISL_743479, EPI_ISL_743480, EPI_ISL_743481, EPI_ISL_743482, EPI_ISL_743483, EPI_ISL_743484, EPI_ISL_743485, EPI_ISL_743486, EPI_ISL_743487, EPI_ISL_743488, EPI_ISL_743489, EPI_ISL_743490, EPI_ISL_743491, EPI_ISL_743492, EPI_ISL_743493, EPI_ISL_743494, EPI_ISL_743495, EPI_ISL_743496, EPI_ISL_743497, EPI_ISL_743498, EPI_ISL_743499, EPI_ISL_743500, EPI_ISL_743501, EPI_ISL_743502, EPI_ISL_743503, EPI_ISL_743504, EPI_ISL_743505, EPI_ISL_743506, EPI_ISL_743507, EPI_ISL_743508, EPI_ISL_743510, EPI_ISL_743568, EPI_ISL_743580, EPI_ISL_743640, EPI_ISL_743642, EPI_ISL_743643, EPI_ISL_743644, EPI_ISL_743645, EPI_ISL_743646, EPI_ISL_743647, EPI_ISL_743648, EPI_ISL_743649, EPI_ISL_743650, EPI_ISL_743651, EPI_ISL_743652, EPI_ISL_743653, EPI_ISL_743654, EPI_ISL_743655, EPI_ISL_743656, EPI_ISL_743657, EPI_ISL_743658, EPI_ISL_743659, EPI_ISL_743660, EPI_ISL_743661, EPI_ISL_743665, EPI_ISL_743666, EPI_ISL_743667, EPI_ISL_743668, EPI_ISL_743669, EPI_ISL_743670, EPI_ISL_743671, EPI_ISL_743672, EPI_ISL_743673, EPI_ISL_743674, EPI_ISL_743676, EPI_ISL_743677, EPI_ISL_743679, EPI_ISL_743901, EPI_ISL_743943, EPI_ISL_743945, EPI_ISL_743946, EPI_ISL_743947, EPI_ISL_743948, EPI_ISL_743949, EPI_ISL_743950, EPI_ISL_743951, EPI_ISL_743953, EPI_ISL_743954, EPI_ISL_743955, EPI_ISL_743956, EPI_ISL_743957 | | | |
| see above | Wales Specialist Virology Centre Sequencing lab: Pathogen Genomics Unit | COVID-19 Genomics UK (COG-UK) Consortium | Catherine Moore, Johnathan Evans, Laura Gifford, Malorie Perry, Simon Cottrell, Angela Marchbank, Alec Birchley, Alexander Adams, Amy Gaskin, Bree Gatica-Wilcox, Jason Coombes, Joel Southgate, Lauren Gilbert, Lee Graham, Nicole Pacchiarini, Sara Kumziene-Summerhayes, Sarah Taylor, Sophie Jones, Sara Rey, Matthew Bull, Joanne Watkins, Sally Corden, Tom Connor |
| EPI_ISL_745043, EPI_ISL_745044, EPI_ISL_745045, EPI_ISL_745046, EPI_ISL_745047, EPI_ISL_745048, EPI_ISL_745049, EPI_ISL_745050, EPI_ISL_745051, EPI_ISL_745052, EPI_ISL_745053, EPI_ISL_745054, EPI_ISL_745055, EPI_ISL_745057 | | | |
| see above | Israel Central Virology laboratory | Israel Central Virology laboratory | Neta Zuckerman, Efrat Dahan Bucris, Oran Erster, Michal Mandelboim, Orna Mor, Ella Mendelson |

| | | | |
|---|---|---|---|
| EPI_ISL_745127 | Khayelitsha (Site B) CHC wc KHC | National Health Laboratory Service (NHLS), Tygerberg | Susan Engelbrecht, Kayla Delaney, Bronwyn Kleinhans, Houriiyah Tegally, Eduan Wilkindon, Gert van Zyl, Wolfgang Preiser, Tulio de Oliveira |
| EPI_ISL_745131, EPI_ISL_745133, EPI_ISL_745136 | Kraaifontein CHC wc KFP | National Health Laboratory Service (NHLS), Tygerberg | Susan Engelbrecht, Kayla Delaney, Bronwyn Kleinhans, Houriiyah Tegally, Eduan Wilkindon, Gert van Zyl, Wolfgang Preiser, Tulio de Oliveira |
| EPI_ISL_745140, EPI_ISL_745141, EPI_ISL_745147, EPI_ISL_745148, EPI_ISL_745149, EPI_ISL_745150 | Tygerberg Hospital wc TBH | National Health Laboratory Service (NHLS), Tygerberg | Susan Engelbrecht, Kayla Delaney, Bronwyn Kleinhans, Houriiyah Tegally, Eduan Wilkindon, Gert van Zyl, Wolfgang Preiser, Tulio de Oliveira |
| EPI_ISL_745168, EPI_ISL_745169 | Bishop Lavis CDC wc BLP | National Health Laboratory Service (NHLS), Tygerberg | Susan Engelbrecht, Kayla Delaney, Bronwyn Kleinhans, Houriiyah Tegally, Eduan Wilkindon, Gert van Zyl, Wolfgang Preiser, Tulio de Oliveira |
| EPI_ISL_745170 | Brackengate Field Hospital COVID-19 wc BRG | National Health Laboratory Service (NHLS), Tygerberg | Susan Engelbrecht, Kayla Delaney, Bronwyn Kleinhans, Houriiyah Tegally, Eduan Wilkindon, Gert van Zyl, Wolfgang Preiser, Tulio de Oliveira |
| EPI_ISL_745171, EPI_ISL_745172 | Delft CHC wc DFP | National Health Laboratory Service (NHLS), Tygerberg | Susan Engelbrecht, Kayla Delaney, Bronwyn Kleinhans, Houriiyah Tegally, Eduan Wilkindon, Gert van Zyl, Wolfgang Preiser, Tulio de Oliveira |
| EPI_ISL_745173, EPI_ISL_745174, EPI_ISL_745175, EPI_ISL_745176, EPI_ISL_745177, EPI_ISL_745178 | Eerste River Hospital wc ERH | National Health Laboratory Service (NHLS), Tygerberg | Susan Engelbrecht, Kayla Delaney, Bronwyn Kleinhans, Houriiyah Tegally, Eduan Wilkindon, Gert van Zyl, Wolfgang Preiser, Tulio de Oliveira |
| EPI_ISL_745179 | Kraaifontein CHC wc KFP | National Health Laboratory Service (NHLS), Tygerberg | Susan Engelbrecht, Kayla Delaney, Bronwyn Kleinhans, Houriiyah Tegally, Eduan Wilkindon, Gert van Zyl, Wolfgang Preiser, Tulio de Oliveira |
| EPI_ISL_745180, EPI_ISL_745181, EPI_ISL_745182, EPI_ISL_745183, EPI_ISL_745184, EPI_ISL_745185 | Tygerberg Hospital wc TBH | National Health Laboratory Service (NHLS), Tygerberg | Susan Engelbrecht, Kayla Delaney, Bronwyn Kleinhans, Houriiyah Tegally, Eduan Wilkindon, Gert van Zyl, Wolfgang Preiser, Tulio de Oliveira |
| EPI_ISL_745186 | Wallacedene Clinic wc WAL | National Health Laboratory Service (NHLS), Tygerberg | Susan Engelbrecht, Kayla Delaney, Bronwyn Kleinhans, Houriiyah Tegally, Eduan Wilkindon, Gert van Zyl, Wolfgang Preiser, Tulio de Oliveira |
| EPI_ISL_745190 | Brackengate Field Hospital COVID-19 wc BRG | National Health Laboratory Service (NHLS), Tygerberg | Susan Engelbrecht, Kayla Delaney, Bronwyn Kleinhans, Houriiyah Tegally, Eduan Wilkindon, Gert van Zyl, Wolfgang Preiser, Tulio de Oliveira |
| EPI_ISL_745208, EPI_ISL_745209, EPI_ISL_745210 | Lab voor klinische biologie | Onderzoeksgroep Virologie | Laurens Lambrechts, Nick Vereecke, Marthe Pauwels, Bruno Verhasselt, Linos Vandekerckhove, Hans Nauwynck, Sebastiaan Theuns |
| EPI_ISL_745309, EPI_ISL_745330, EPI_ISL_745368, EPI_ISL_745369, EPI_ISL_745370, EPI_ISL_745371, EPI_ISL_745372, EPI_ISL_745373, EPI_ISL_745374, EPI_ISL_745375, EPI_ISL_745376, EPI_ISL_745377, EPI_ISL_745397 | | | |
| see above | CNR Virus des Infections Respiratoires - France SUD | CNR Virus des Infections Respiratoires - France SUD | Antonin Bal, Gregory Destras, Claudia Gonzalez, Gwendolyne Burfin, Quentin Semanas, Martine Valette, Bruno Lina, Laurence Josset |
| EPI_ISL_746966 | Utah Public Health Laboratory | Utah Public Health Laboratory | Erin Young, Kelly Oakeson, Tara Gallagher |
| EPI_ISL_747195, EPI_ISL_747196, EPI_ISL_747197, EPI_ISL_747198, EPI_ISL_747199, EPI_ISL_747200, EPI_ISL_747201, EPI_ISL_747202, EPI_ISL_747203, EPI_ISL_747204, EPI_ISL_747205, EPI_ISL_747206, EPI_ISL_747207, EPI_ISL_747208, EPI_ISL_747209, EPI_ISL_747210, EPI_ISL_747211, EPI_ISL_747212, EPI_ISL_747213, EPI_ISL_747214, EPI_ISL_747215, EPI_ISL_747216, EPI_ISL_747217, EPI_ISL_747218, EPI_ISL_747219, EPI_ISL_747220, EPI_ISL_747221, EPI_ISL_747222, EPI_ISL_747223, EPI_ISL_747224, EPI_ISL_747225, EPI_ISL_747226, EPI_ISL_747227, EPI_ISL_747228, EPI_ISL_747229, EPI_ISL_747230, EPI_ISL_747231, EPI_ISL_747232 | | | |
| see above | UW Virology Lab | UW Virology Lab | Pavitra Roychoudhury, Hong Xie, Lasata Shrestha, Michelle Lin, Meei-Li Huang, Keith R Jerome, Alexander Greninger |
| EPI_ISL_747527, EPI_ISL_747528, EPI_ISL_747529, EPI_ISL_747533, EPI_ISL_747535, EPI_ISL_747536, EPI_ISL_747540, EPI_ISL_747541, EPI_ISL_747542, EPI_ISL_747543, EPI_ISL_747544, EPI_ISL_747545, EPI_ISL_747546, EPI_ISL_747547, EPI_ISL_747548, EPI_ISL_747549, EPI_ISL_747550, EPI_ISL_747551, EPI_ISL_747552, EPI_ISL_747553, EPI_ISL_747554, EPI_ISL_747555, EPI_ISL_747556, EPI_ISL_747557, EPI_ISL_747558, EPI_ISL_747559, EPI_ISL_747560, EPI_ISL_747561, EPI_ISL_747562, EPI_ISL_747563, EPI_ISL_747564, EPI_ISL_747565, EPI_ISL_747566, EPI_ISL_747567, EPI_ISL_747568, EPI_ISL_747569, EPI_ISL_747570, EPI_ISL_747571, EPI_ISL_747572, EPI_ISL_747573, EPI_ISL_747574, EPI_ISL_747575, EPI_ISL_747576, EPI_ISL_747589, EPI_ISL_747590, EPI_ISL_747591, EPI_ISL_747592, EPI_ISL_747593, EPI_ISL_747594, EPI_ISL_747595, EPI_ISL_747599, EPI_ISL_747601, EPI_ISL_747602, EPI_ISL_747607, EPI_ISL_747608, EPI_ISL_747610, EPI_ISL_747611, EPI_ISL_747612, EPI_ISL_747613, EPI_ISL_747614, EPI_ISL_747616, EPI_ISL_747617, EPI_ISL_747618, EPI_ISL_747619, EPI_ISL_747620, EPI_ISL_747621, EPI_ISL_747622, EPI_ISL_747623, EPI_ISL_747632, EPI_ISL_747636, EPI_ISL_747637, EPI_ISL_747638, EPI_ISL_747639, EPI_ISL_747640, EPI_ISL_747642, EPI_ISL_747643, EPI_ISL_747644, EPI_ISL_747645, EPI_ISL_747648, EPI_ISL_747652, EPI_ISL_747655, EPI_ISL_747656, EPI_ISL_747658, EPI_ISL_747659, EPI_ISL_747660, EPI_ISL_747661, EPI_ISL_747662, EPI_ISL_747663, EPI_ISL_747664, EPI_ISL_747665, EPI_ISL_747666, EPI_ISL_747667, EPI_ISL_747668, EPI_ISL_747669, EPI_ISL_747670, EPI_ISL_747671, EPI_ISL_747672, EPI_ISL_747673, EPI_ISL_747674, EPI_ISL_747681, EPI_ISL_747682, EPI_ISL_747683, EPI_ISL_747684, EPI_ISL_747685, EPI_ISL_747686, EPI_ISL_747687, EPI_ISL_747688, EPI_ISL_747689, EPI_ISL_747690, EPI_ISL_747691, EPI_ISL_747692, EPI_ISL_747693, EPI_ISL_747694, EPI_ISL_747695, EPI_ISL_747696, EPI_ISL_747697, EPI_ISL_747698, EPI_ISL_747699, EPI_ISL_747700, EPI_ISL_747701, EPI_ISL_747702, EPI_ISL_747703, EPI_ISL_747704, EPI_ISL_747705, EPI_ISL_747706, EPI_ISL_747707, EPI_ISL_747708, EPI_ISL_747721, EPI_ISL_747722, EPI_ISL_747723, EPI_ISL_747728, EPI_ISL_747729, EPI_ISL_747731, EPI_ISL_747732, EPI_ISL_747733, EPI_ISL_747734, EPI_ISL_747739, EPI_ISL_747740, EPI_ISL_747741, EPI_ISL_747742, EPI_ISL_747743, EPI_ISL_747745, EPI_ISL_747746, EPI_ISL_747748, EPI_ISL_747750, EPI_ISL_747752, EPI_ISL_747753, EPI_ISL_747754, EPI_ISL_747756, EPI_ISL_747757, EPI_ISL_747758, EPI_ISL_747759, EPI_ISL_747760, EPI_ISL_747761, EPI_ISL_747762, EPI_ISL_747763, EPI_ISL_747764, EPI_ISL_747765, EPI_ISL_747766, EPI_ISL_747767, EPI_ISL_747768, EPI_ISL_747769, EPI_ISL_747770, EPI_ISL_747771, EPI_ISL_747772, EPI_ISL_747773, EPI_ISL_747774, EPI_ISL_747775, EPI_ISL_747776, EPI_ISL_747777, EPI_ISL_747778, EPI_ISL_747779, EPI_ISL_747780, EPI_ISL_747781, EPI_ISL_747782, EPI_ISL_747783, EPI_ISL_747784, EPI_ISL_747785, EPI_ISL_747786, EPI_ISL_747787, EPI_ISL_747788, EPI_ISL_747789, EPI_ISL_747790, EPI_ISL_747791, EPI_ISL_747792, EPI_ISL_747793, EPI_ISL_747794, EPI_ISL_747795, EPI_ISL_747796, EPI_ISL_747797, EPI_ISL_747798, EPI_ISL_747799, EPI_ISL_747800, EPI_ISL_747801, EPI_ISL_747802, EPI_ISL_747803, EPI_ISL_747804, EPI_ISL_747805, EPI_ISL_747806, EPI_ISL_747807, EPI_ISL_747808, EPI_ISL_747809, EPI_ISL_747810, EPI_ISL_747811, EPI_ISL_747812, EPI_ISL_747813, EPI_ISL_747814, EPI_ISL_747815, EPI_ISL_747816, EPI_ISL_747817, EPI_ISL_747818, EPI_ISL_747819, EPI_ISL_747820, EPI_ISL_747821, EPI_ISL_747822, EPI_ISL_747823, EPI_ISL_747824, EPI_ISL_747825, EPI_ISL_747826, EPI_ISL_747827, EPI_ISL_747828, EPI_ISL_747829, EPI_ISL_747830, EPI_ISL_747831, EPI_ISL_747832, EPI_ISL_747833, EPI_ISL_747834, EPI_ISL_747835, EPI_ISL_747836, EPI_ISL_747837, EPI_ISL_747838, EPI_ISL_747839, EPI_ISL_747840, EPI_ISL_747841, EPI_ISL_747842, EPI_ISL_747843, EPI_ISL_747844, EPI_ISL_747845, EPI_ISL_747846, EPI_ISL_747847, EPI_ISL_747848, EPI_ISL_747849, EPI_ISL_747850, EPI_ISL_747851, EPI_ISL_747852, EPI_ISL_747853, EPI_ISL_747854, EPI_ISL_747855, EPI_ISL_747856, EPI_ISL_747857, EPI_ISL_747858, EPI_ISL_747859, EPI_ISL_747860, EPI_ISL_747861, EPI_ISL_747862, EPI_ISL_747863, EPI_ISL_747864, EPI_ISL_747865, EPI_ISL_747866, EPI_ISL_747867, EPI_ISL_747868, EPI_ISL_747869, EPI_ISL_747870, EPI_ISL_747871, EPI_ISL_747872, EPI_ISL_747873, EPI_ISL_747874, EPI_ISL_747875, EPI_ISL_747876, EPI_ISL_747877, EPI_ISL_747878, EPI_ISL_747879, EPI_ISL_747880, EPI_ISL_747881, EPI_ISL_747882, EPI_ISL_747883, EPI_ISL_747884, EPI_ISL_747885, EPI_ISL_747886, EPI_ISL_747887, EPI_ISL_747888, EPI_ISL_747889, EPI_ISL_747890, EPI_ISL_747891, EPI_ISL_747892, EPI_ISL_747893, EPI_ISL_747894, EPI_ISL_747895, EPI_ISL_747896, EPI_ISL_747897, EPI_ISL_747898, EPI_ISL_747899, EPI_ISL_747900, EPI_ISL_747901, EPI_ISL_747902, EPI_ISL_747903, EPI_ISL_747904, EPI_ISL_747905, EPI_ISL_747906, EPI_ISL_747907, EPI_ISL_747908, EPI_ISL_747909, EPI_ISL_747910, EPI_ISL_747911, EPI_ISL_747912, EPI_ISL_747913, EPI_ISL_747914, EPI_ISL_747915, EPI_ISL_747916, EPI_ISL_747917, EPI_ISL_747918, EPI_ISL_747919, EPI_ISL_747920, EPI_ISL_747921, EPI_ISL_747922, EPI_ISL_747923, EPI_ISL_747924, EPI_ISL_747925, EPI_ISL_747926, EPI_ISL_747927, EPI_ISL_747928, EPI_ISL_747929, EPI_ISL_747930, EPI_ISL_747931, EPI_ISL_747932, EPI_ISL_747933, EPI_ISL_747934, EPI_ISL_747935, EPI_ISL_747936, EPI_ISL_747937, EPI_ISL_747938, EPI_ISL_747939, EPI_ISL_747940, EPI_ISL_747941, EPI_ISL_747942, EPI_ISL_747943, EPI_ISL_747944, EPI_ISL_747945, EPI_ISL_747946, EPI_ISL_747947, EPI_ISL_747948, EPI_ISL_747949, EPI_ISL_747950, EPI_ISL_747951, EPI_ISL_747952, EPI_ISL_747953, EPI_ISL_747954, EPI_ISL_747955, EPI_ISL_747956, EPI_ISL_747957, EPI_ISL_747958, EPI_ISL_747959, EPI_ISL_747960, EPI_ISL_747961, EPI_ISL_747962, EPI_ISL_747963, EPI_ISL_747964, EPI_ISL_747965, EPI_ISL_747966, EPI_ISL_747967, EPI_ISL_747968, EPI_ISL_747969, EPI_ISL_747970, EPI_ISL_747971, EPI_ISL_747972, EPI_ISL_747973, EPI_ISL_747974, EPI_ISL_747975, EPI_ISL_747976, EPI_ISL_747977, EPI_ISL_747978, EPI_ISL_747979, EPI_ISL_747980, EPI_ISL_747981, EPI_ISL_747982, EPI_ISL_747983, EPI_ISL_747984, EPI_ISL_747985, EPI_ISL_747986, EPI_ISL_747987, EPI_ISL_747988, EPI_ISL_747989, EPI_ISL_747990, EPI_ISL_747991, EPI_ISL_747992, EPI_ISL_747993, EPI_ISL_747994, EPI_ISL_747995, EPI_ISL_747996, EPI_ISL_747997, EPI_ISL_747998, EPI_ISL_747999, EPI_ISL_748000, EPI_ISL_748001, EPI_ISL_748002, EPI_ISL_748003, EPI_ISL_748004, EPI_ISL_748005, EPI_ISL_748006, EPI_ISL_748007, EPI_ISL_748008, EPI_ISL_748009, EPI_ISL_748010, EPI_ISL_748011, EPI_ISL_748012, EPI_ISL_748013, EPI_ISL_748014, EPI_ISL_748015, EPI_ISL_748016, EPI_ISL_748017, EPI_ISL_748018, EPI_ISL_748019, EPI_ISL_748020, EPI_ISL_748146, EPI_ISL_748147, EPI_ISL_748148, EPI_ISL_748149, EPI_ISL_748150, EPI_ISL_748151, EPI_ISL_748152, EPI_ISL_748153, EPI_ISL_748154, EPI_ISL_748155, EPI_ISL_748156, EPI_ISL_748157, EPI_ISL_748158, EPI_ISL_748159, EPI_ISL_748160, EPI_ISL_748161, EPI_ISL_748162, EPI_ISL_748163, EPI_ISL_748164, EPI_ISL_748165, EPI_ISL_748166, EPI_ISL_748167, EPI_ISL_748168, EPI_ISL_748169, EPI_ISL_748170, EPI_ISL_748171, EPI_ISL_748172, EPI_ISL_748173, EPI_ISL_748174, EPI_ISL_748175, EPI_ISL_748176, EPI_ISL_748177, EPI_ISL_748178, EPI_ISL_748179, EPI_ISL_748180, EPI_ISL_748181, EPI_ISL_748182, EPI_ISL_748183, EPI_ISL_748184, EPI_ISL_748185, EPI_ISL_748186, EPI_ISL_748187, EPI_ISL_748188, EPI_ISL_748189, EPI_ISL_748190, EPI_ISL_748191, EPI_ISL_748192, EPI_ISL_748193, EPI_ISL_748194, EPI_ISL_748195, EPI_ISL_748196, EPI_ISL_748197, EPI_ISL_748198, EPI_ISL_748199, EPI_ISL_748200, EPI_ISL_748201, EPI_ISL_748202, EPI_ISL_748203, EPI_ISL_748204, EPI_ISL_748205, EPI_ISL_748206, EPI_ISL_748207, EPI_ISL_748208, EPI_ISL_748209, EPI_ISL_748210, EPI_ISL_748211, EPI_ISL_748212, EPI_ISL_748213, EPI_ISL_748214, EPI_ISL_748215, EPI_ISL_748216, EPI_ISL_748217, EPI_ISL_748218, EPI_ISL_748219, EPI_ISL_748220, EPI_ISL_748221, EPI_ISL_748222, EPI_ISL_748223, EPI_ISL_748224, EPI_ISL_748225, EPI_ISL_748226, EPI_ISL_748227, EPI_ISL_748228, EPI_ISL_748229, EPI_ISL_748230, EPI_ISL_748231, EPI_ISL_748232, EPI_ISL_748233, EPI_ISL_748234, EPI_ISL_748235, EPI_ISL_748236, EPI_ISL_748237, EPI_ISL_748238, EPI_ISL_748239, EPI_ISL_748240, EPI_ISL_748241, EPI_ISL_748242, EPI_ISL_748243, EPI_ISL_748244, EPI_ISL_748245, EPI_ISL_748246, EPI_ISL_748247, EPI_ISL_748248, EPI_ISL_748249, EPI_ISL_748250, EPI_ISL_748251, EPI_ISL_748252, EPI_ISL_748253, EPI_ISL_748254, EPI_ISL_748255, EPI_ISL_748256, EPI_ISL_748257, EPI_ISL_748258, EPI_ISL_748259, EPI_ISL_748260, EPI_ISL_748261, EPI_ISL_748262, EPI_ISL_748263, EPI_ISL_748264, EPI_ISL_748265, EPI_ISL_748266, EPI_ISL_748267, EPI_ISL_748268, EPI_ISL_748269, EPI_ISL_748270, EPI_ISL_748271, EPI_ISL_748272, EPI_ISL_748273, EPI_ISL_748274, EPI_ISL_748275, EPI_ISL_748276, EPI_ISL_748277, EPI_ISL_748278, EPI_ISL_748279, EPI_ISL_748280, EPI_ISL_748281, EPI_ISL_748282, EPI_ISL_748283, EPI_ISL_748284, EPI_ISL_748285, EPI_ISL_748286, EPI_ISL_748287, EPI_ISL_748288, EPI_ISL_748289, EPI_ISL_748290, EPI_ISL_748291, EPI_ISL_748292, EPI_ISL_748293, EPI_ISL_748294, EPI_ISL_748295, EPI_ISL_748296, EPI_ISL_748297, EPI_ISL_748298, EPI_ISL_748299, EPI_ISL_748300, EPI_ISL_748301, EPI_ISL_748302, EPI_ISL_748303, EPI_ISL_748304, EPI_ISL_748305, EPI_ISL_748306, EPI_ISL_748307, EPI_ISL_748308, EPI_ISL_748309, EPI_ISL_748310, EPI_ISL_748311, EPI_ISL_748312, EPI_ISL_748313, EPI_ISL_748314, EPI_ISL_748315, EPI_ISL_748316, EPI_ISL_748317, EPI_ISL_748318, EPI_ISL_748319, EPI_ISL_748320, EPI_ISL_748321, EPI_ISL_748322, EPI_ISL_748323, EPI_ISL_748324, EPI_ISL_748325, EPI_ISL_748326, EPI_ISL_748327, EPI_ISL_748328, EPI_ISL_748329, EPI_ISL_748330, EPI_ISL_748331, EPI_ISL_748332, EPI_ISL_748333, EPI_ISL_748334, EPI_ISL_748335, EPI_ISL_748336, EPI_ISL_748337, EPI_ISL_748338, EPI_ISL_748339, EPI_ISL_748340, EPI_ISL_748341, EPI_ISL_748342, EPI_ISL_748343, EPI_ISL_748344, EPI_ISL_748345, EPI_ISL_748346, EPI_ISL_748347, EPI_ISL_748348, EPI_ISL_748349, EPI_ISL_748350, EPI_ISL_748351, EPI_ISL_748352, EPI_ISL_748353, EPI_ISL_748354, EPI_ISL_748355, EPI_ISL_748356, EPI_ISL_748357, EPI_ISL_748358, EPI_ISL_748359, EPI_ISL_748360, EPI_ISL_748361, EPI_ISL_748362, EPI_ISL_748363, EPI_ISL_748364, EPI_ISL_748365, EPI_ISL_748366, EPI_ISL_748367, EPI_ISL_748368, EPI_ISL_748369, EPI_ISL_748370, EPI_ISL_748371, EPI_ISL_748372, EPI_ISL_748373, EPI_ISL_748374, EPI_ISL_748375, EPI_ISL_748376, EPI_ISL_748377, EPI_ISL_748378, EPI_ISL_748379, EPI_ISL_748380, EPI_ISL_748381, EPI_ISL_748382, EPI_ISL_748383, EPI_ISL_748384, EPI_ISL_748385, EPI_ISL_748386, EPI_ISL_748387, EPI_ISL_748388, EPI_ISL_748389, EPI_ISL_748390, EPI_ISL_748391, EPI_ISL_748392, EPI_ISL_748393, EPI_ISL_748394, EPI_ISL_748395, EPI_ISL_748396, EPI_ISL_748397, EPI_ISL_748398, EPI_ISL_748399, EPI_ISL_748400, EPI_ISL_748401, EPI_ISL_748402, EPI_ISL_748403, EPI_ISL_748404, EPI_ISL_748405, EPI_ISL_748406, EPI_ISL_748407, EPI_ISL_748408, EPI_ISL_748409, EPI_ISL_748410, EPI_ISL_748411, EPI_ISL_748412, EPI_ISL_748413, EPI_ISL_748414, EPI_ISL_748415, EPI_ISL_748416, EPI_ISL_748417, EPI_ISL_748418, EPI_ISL_748419, EPI_ISL_748420, EPI_ISL_748421, EPI_ISL_748422, EPI_ISL_748423, EPI_ISL_748424, EPI_ISL_748425, EPI_ISL_748426, EPI_ISL_748427, EPI_ISL_748428, EPI_ISL_748429, EPI_ISL_748430, EPI_ISL_748431, EPI_ISL_748432, EPI_ISL_748433, EPI_ISL_748434, EPI_ISL_748435, EPI_ISL_748436, EPI_ISL_748437, EPI_ISL_748438, EPI_ISL_748439, EPI_ISL_748440, EPI_ISL_748441, EPI_ISL_748442, EPI_ISL_748443, EPI_ISL_748444, EPI_ISL_748445, EPI_ISL_748446, EPI_ISL_748447, EPI_ISL_748448, EPI_ISL_748449, EPI_ISL_748450, EPI_ISL_748451, EPI_ISL_748452, EPI_ISL_748453, EPI_ISL_748454, EPI_ISL_748455, EPI_ISL_748456, EPI_ISL_748457, EPI_ISL_748458, EPI_ISL_748459, EPI_ISL_748460, EPI_ISL_748461, EPI_ISL_748462, EPI_ISL_748463, EPI_ISL_748464, EPI_ISL_748465, EPI_ISL_748466, EPI_ISL_748467, EPI_ISL_748468, EPI_ISL_748469, EPI_ISL_748470, EPI_ISL_748471, EPI_ISL_748472, EPI_ISL_748473, EPI_ISL_748474, EPI_ISL_748475, EPI_ISL_748476, EPI_ISL_748477, EPI_ISL_748478, EPI_ISL_748479, EPI_ISL_748480, EPI_ISL_748481, EPI_ISL_748482, EPI_ISL_748483, EPI_ISL_748484, EPI_ISL_748485, EPI_ISL_748486, EPI_ISL_748487, EPI_ISL_748488, EPI_ISL_748489, EPI_ISL_748490, EPI_ISL_748491, EPI_ISL_748492, EPI_ISL_748493, EPI_ISL_748494, EPI_ISL_748495, EPI_ISL_748496, EPI_ISL_748497, EPI_ISL_748498, EPI_ISL_748499, EPI_ISL_748500, EPI_ISL_748501, EPI_ISL_748502, EPI_ISL_748503, EPI_ISL_748504, EPI_ISL_748505, EPI_ISL_748506, EPI_ISL_748507, EPI_ISL_748508, EPI_ISL_748509, EPI_ISL_748510, EPI_ISL_748511, EPI_ISL_748512, EPI_ISL_748513, EPI_ISL_748514, EPI_ISL_748515, EPI_ISL_748516, EPI_ISL_748517, EPI_ISL_748518, EPI_ISL_748519, EPI_ISL_748520, EPI_ISL_748521, EPI_ISL_748522, EPI_ISL_748523, EPI_ISL_748524, EPI_ISL_748525, EPI_ISL_748526, EPI_ISL_748527, EPI_ISL_748528, EPI_ISL_748529, EPI_ISL_748530, EPI_ISL_748531, EPI_ISL_748532, EPI_ISL_748533, EPI_ISL_748534, EPI_ISL_748535, EPI_ISL_748536, EPI_ISL_748537, EPI_ISL_748538, EPI_ISL_748539, EPI_ISL_748540, EPI_ISL_748541, EPI_ISL_748542, EPI_ISL_748543, EPI_ISL_748544, EPI_ISL_748545, EPI_ISL_748546, EPI_ISL_748547, EPI_ISL_748548, EPI_ISL_748549, EPI_ISL_748550, EPI_ISL_748551, EPI_ISL_748552, EPI_ISL_748553, EPI_ISL_748554, EPI_ISL_748555, EPI_ISL_748556, EPI_ISL_748557, EPI_ISL_748558, EPI_ISL_748559, EPI_ISL_748560, EPI_ISL_748561, EPI_ISL_748562, EPI_ISL_748563, EPI_ISL_748564, EPI_ISL_748565, EPI_ISL_748566, EPI_ISL_748567, EPI_ISL_748568, EPI_ISL_748569, EPI_ISL_748570, EPI_ISL_748571, EPI_ISL_748572, EPI_ISL_748573, EPI_ISL_748574, EPI_ISL_748575, EPI_ISL_748576, EPI_ISL_748577, EPI_ISL_748578, EPI_ISL_748579, EPI_ISL_748580, EPI_ISL_748581, EPI_ISL_748582, EPI_ISL_748583, EPI_ISL_748584, EPI_ISL_748585, EPI_ISL_748586, EPI_ISL_748587, EPI_ISL_748588, EPI_ISL_748589, EPI_ISL_748590, EPI_ISL_748591, EPI_ISL_748592, EPI_ISL_748593, EPI_ISL_748594, EPI_ISL_748595, EPI_ISL_748596, EPI_ISL_748597, EPI_ISL_748598, EPI_ISL_748599, EPI_ISL_748600, EPI_ISL_748601, EPI_ISL_748602, EPI_ISL_748603, EPI_ISL_748604, EPI_ISL_748605, EPI_ISL_748606, EPI_ISL_748607, EPI_ISL_748608, EPI_ISL_748609, EPI_ISL_748610, EPI_ISL_748611, EPI_ISL_748612, EPI_ISL_748613, EPI_ISL_748614, EPI_ISL_748615, EPI_ISL_748616, EPI_ISL_748617, EPI_ISL_748618, EPI_ISL_748619, EPI_ISL_748620, EPI_ISL_748621, EPI_ISL_748622, EPI_ISL_748623, EPI_ISL_748624, EPI_ISL_748625, EPI_ISL_748626, EPI_ISL_748627, EPI_ISL_748628, EPI_ISL_748629, EPI_ISL_748630, EPI_ISL_748631, EPI_ISL_748632, EPI_ISL_748633, EPI_ISL_748634, EPI_ISL_748635, EPI_ISL_748636, EPI_ISL_748637, EPI_ISL_748638, EPI_ISL_748639, EPI_ISL_748640, EPI_ISL_748641, EPI_ISL_748642, EPI_ISL_748643, EPI_ISL_748644, EPI_ISL_748645, EPI_ISL_748646, EPI_ISL_748647, EPI_ISL_748648, EPI_ISL_748649, EPI_ISL_748650, EPI_ISL_748651, EPI_ISL_748652, EPI_ISL_748653, EPI_ISL_748654, EPI_ISL_748655, EPI_ISL_748656, EPI_ISL_748657, EPI_ISL_748658, EPI_ISL_748659, EPI_ISL_748660, EPI_ISL_748661, EPI_ISL_748662, EPI_ISL_748663, EPI_ISL_748664, EPI_ISL_748665, EPI_ISL_748666, EPI_ISL_748668, EPI_ISL_748669, | | | |

| | | | |
|---|---|---|---|
| EPI_ISL_748670, EPI_ISL_748671, EPI_ISL_748672, EPI_ISL_748673, EPI_ISL_748674, EPI_ISL_748675, EPI_ISL_748676, EPI_ISL_748677, EPI_ISL_748678, EPI_ISL_748679, EPI_ISL_748680, EPI_ISL_748681, EPI_ISL_748682, EPI_ISL_748683, EPI_ISL_748684, EPI_ISL_748685, EPI_ISL_748686, EPI_ISL_748687, EPI_ISL_748688, EPI_ISL_748689, EPI_ISL_748690, EPI_ISL_748691, EPI_ISL_748692, EPI_ISL_748693, EPI_ISL_748694, EPI_ISL_748695, EPI_ISL_748696, EPI_ISL_748697, EPI_ISL_748698, EPI_ISL_748699, EPI_ISL_748700, EPI_ISL_748701, EPI_ISL_748702, EPI_ISL_748703, EPI_ISL_748704, EPI_ISL_748705, EPI_ISL_748706, EPI_ISL_748707, EPI_ISL_748708, EPI_ISL_748709, EPI_ISL_748710, EPI_ISL_748711, EPI_ISL_748712, EPI_ISL_748713, EPI_ISL_748714, EPI_ISL_748715, EPI_ISL_748716, EPI_ISL_748717, EPI_ISL_748718, EPI_ISL_748719, EPI_ISL_748720, EPI_ISL_748721, EPI_ISL_748722, EPI_ISL_748723, EPI_ISL_748724, EPI_ISL_748725, EPI_ISL_748726, EPI_ISL_748727, EPI_ISL_748728, EPI_ISL_748729, EPI_ISL_748730, EPI_ISL_748731, EPI_ISL_748732, EPI_ISL_748733, EPI_ISL_748734, EPI_ISL_748735, EPI_ISL_748736, EPI_ISL_748737, EPI_ISL_748738, EPI_ISL_748739, EPI_ISL_748740, EPI_ISL_748741, EPI_ISL_748742, EPI_ISL_748743, EPI_ISL_748744, EPI_ISL_748745, EPI_ISL_748746, EPI_ISL_748747, EPI_ISL_748748, EPI_ISL_748749, EPI_ISL_748750, EPI_ISL_748751, EPI_ISL_748752, EPI_ISL_748753, EPI_ISL_748754, EPI_ISL_748755, EPI_ISL_748756, EPI_ISL_748757, EPI_ISL_748758, EPI_ISL_748759, EPI_ISL_748760, EPI_ISL_748761, EPI_ISL_748762, EPI_ISL_748763, EPI_ISL_748764, EPI_ISL_748765, EPI_ISL_748766, EPI_ISL_748767, EPI_ISL_748768, EPI_ISL_748769, EPI_ISL_748770, EPI_ISL_748771, EPI_ISL_748772, EPI_ISL_748773, EPI_ISL_748774, EPI_ISL_748775, EPI_ISL_748776, EPI_ISL_748777, EPI_ISL_748778, EPI_ISL_748779, EPI_ISL_748780, EPI_ISL_748781, EPI_ISL_748782, EPI_ISL_748783, EPI_ISL_748784, EPI_ISL_748785, EPI_ISL_748786, EPI_ISL_748787, EPI_ISL_748788, EPI_ISL_748789, EPI_ISL_748790, EPI_ISL_748791, EPI_ISL_748792, EPI_ISL_748793, EPI_ISL_748794, EPI_ISL_748795, EPI_ISL_748796, EPI_ISL_748797, EPI_ISL_748798, EPI_ISL_748799, EPI_ISL_748800, EPI_ISL_748801, EPI_ISL_748802, EPI_ISL_748803, EPI_ISL_748804, EPI_ISL_748805, EPI_ISL_748806, EPI_ISL_748807, EPI_ISL_748808, EPI_ISL_748809, EPI_ISL_748810, EPI_ISL_748811, EPI_ISL_748812, EPI_ISL_748813, EPI_ISL_748814, EPI_ISL_748815, EPI_ISL_748816, EPI_ISL_748817, EPI_ISL_748818, EPI_ISL_748819, EPI_ISL_748820, EPI_ISL_748821, EPI_ISL_748822, EPI_ISL_748823, EPI_ISL_748824, EPI_ISL_748825, EPI_ISL_748826, EPI_ISL_748827, EPI_ISL_748828, EPI_ISL_748829, EPI_ISL_748830, EPI_ISL_748831, EPI_ISL_748832, EPI_ISL_748833, EPI_ISL_748834, EPI_ISL_748835, EPI_ISL_748836, EPI_ISL_748837, EPI_ISL_748838, EPI_ISL_748839, EPI_ISL_748840, EPI_ISL_748841, EPI_ISL_748842, EPI_ISL_748843, EPI_ISL_748844, EPI_ISL_748845, EPI_ISL_748846, EPI_ISL_748847, EPI_ISL_748848, EPI_ISL_748849, EPI_ISL_748850, EPI_ISL_748851, EPI_ISL_748852, EPI_ISL_748853, EPI_ISL_748854, EPI_ISL_748855, EPI_ISL_748856, EPI_ISL_748857, EPI_ISL_748858, EPI_ISL_748859, EPI_ISL_748860, EPI_ISL_748861, EPI_ISL_748862, EPI_ISL_748863, EPI_ISL_748864, EPI_ISL_748865, EPI_ISL_748866, EPI_ISL_748867, EPI_ISL_748868, EPI_ISL_748869, EPI_ISL_748870, EPI_ISL_748871, EPI_ISL_748872, EPI_ISL_748873, EPI_ISL_748874, EPI_ISL_748875, EPI_ISL_748876, EPI_ISL_748877, EPI_ISL_748878, EPI_ISL_748879, EPI_ISL_748880, EPI_ISL_748881, EPI_ISL_748882, EPI_ISL_748883, EPI_ISL_748884, EPI_ISL_748885, EPI_ISL_748886, EPI_ISL_748887, EPI_ISL_748888, EPI_ISL_748889, EPI_ISL_748890, EPI_ISL_748891, EPI_ISL_748892, EPI_ISL_748893, EPI_ISL_748894, EPI_ISL_748895, EPI_ISL_748896, EPI_ISL_748897, EPI_ISL_748898, EPI_ISL_748899, EPI_ISL_748900, EPI_ISL_748901, EPI_ISL_748902, EPI_ISL_748903, EPI_ISL_748904, EPI_ISL_748905, EPI_ISL_748906, EPI_ISL_748907, EPI_ISL_748908, EPI_ISL_748909, EPI_ISL_748910, EPI_ISL_748911, EPI_ISL_748912, EPI_ISL_748913, EPI_ISL_748914, EPI_ISL_748915, EPI_ISL_748916, EPI_ISL_748917, EPI_ISL_748918, EPI_ISL_748919, EPI_ISL_748920, EPI_ISL_748921, EPI_ISL_748922, EPI_ISL_748923, EPI_ISL_748924, EPI_ISL_748925, EPI_ISL_748926, EPI_ISL_748927, EPI_ISL_748928, EPI_ISL_748929, EPI_ISL_748930, EPI_ISL_748931, EPI_ISL_748932, EPI_ISL_748933, EPI_ISL_748934, EPI_ISL_748935, EPI_ISL_748936, EPI_ISL_748937, EPI_ISL_748938, EPI_ISL_748939, EPI_ISL_748940, EPI_ISL_748941, EPI_ISL_748942, EPI_ISL_748943, EPI_ISL_748944, EPI_ISL_748945, EPI_ISL_748946, EPI_ISL_748947, EPI_ISL_748948, EPI_ISL_748949, EPI_ISL_748950, EPI_ISL_748951, EPI_ISL_748952, EPI_ISL_748953, EPI_ISL_748954, EPI_ISL_748955, EPI_ISL_748956, EPI_ISL_748957, EPI_ISL_748958, EPI_ISL_748959, EPI_ISL_748960, EPI_ISL_748961, EPI_ISL_748962, EPI_ISL_748963, EPI_ISL_748964, EPI_ISL_748965, EPI_ISL_748966, EPI_ISL_748967, EPI_ISL_748968, EPI_ISL_748969, EPI_ISL_748970, EPI_ISL_748971, EPI_ISL_748972, EPI_ISL_748973, EPI_ISL_748974, EPI_ISL_748975, EPI_ISL_748976, EPI_ISL_748977, EPI_ISL_748978, EPI_ISL_748979, EPI_ISL_748980, EPI_ISL_748981, EPI_ISL_748982, EPI_ISL_748983, EPI_ISL_748984, EPI_ISL_748985, EPI_ISL_748986, EPI_ISL_748987, EPI_ISL_748988, EPI_ISL_748989, EPI_ISL_748990, EPI_ISL_748991, EPI_ISL_748992, EPI_ISL_748993, EPI_ISL_748994, EPI_ISL_748995, EPI_ISL_748996, EPI_ISL_748997, EPI_ISL_748998, EPI_ISL_748999, EPI_ISL_749000, EPI_ISL_749001, EPI_ISL_749002, EPI_ISL_749003, EPI_ISL_749004, EPI_ISL_749005, EPI_ISL_749006, EPI_ISL_749007, EPI_ISL_749008, EPI_ISL_749009, EPI_ISL_749010, EPI_ISL_749011, EPI_ISL_749012, EPI_ISL_749013, EPI_ISL_749014, EPI_ISL_749015, EPI_ISL_749016, EPI_ISL_749017, EPI_ISL_749018, EPI_ISL_749019, EPI_ISL_749020, EPI_ISL_749021, EPI_ISL_749022, EPI_ISL_749023, EPI_ISL_749024, EPI_ISL_749025, EPI_ISL_749026, EPI_ISL_749027, EPI_ISL_749028, EPI_ISL_749029, EPI_ISL_749030, EPI_ISL_749031, EPI_ISL_749032, EPI_ISL_749033, EPI_ISL_749034, EPI_ISL_749035, EPI_ISL_749037, EPI_ISL_749038, EPI_ISL_749039, EPI_ISL_749040, EPI_ISL_749041, EPI_ISL_749042, EPI_ISL_749043, EPI_ISL_749044, EPI_ISL_749045, EPI_ISL_749046, EPI_ISL_749047, EPI_ISL_749048, EPI_ISL_749049, EPI_ISL_749050, EPI_ISL_749051, EPI_ISL_749052, EPI_ISL_749053, EPI_ISL_749054, EPI_ISL_749055, EPI_ISL_749056, EPI_ISL_749057, EPI_ISL_749058, EPI_ISL_749059, EPI_ISL_749060, EPI_ISL_749061, EPI_ISL_749062, EPI_ISL_749063, EPI_ISL_749064, EPI_ISL_749065, EPI_ISL_749066, EPI_ISL_749067, EPI_ISL_749068, EPI_ISL_749069, EPI_ISL_749070, EPI_ISL_749071, EPI_ISL_749072, EPI_ISL_749073, EPI_ISL_749074, EPI_ISL_749075, EPI_ISL_749076, EPI_ISL_749077, EPI_ISL_749078, EPI_ISL_749079, EPI_ISL_749080, EPI_ISL_749081, EPI_ISL_749082, EPI_ISL_749083, EPI_ISL_749084, EPI_ISL_749085, EPI_ISL_749086, EPI_ISL_749087, EPI_ISL_749088, EPI_ISL_749089, EPI_ISL_749090, EPI_ISL_749091, EPI_ISL_749092, EPI_ISL_749093, EPI_ISL_749094, EPI_ISL_749095, EPI_ISL_749096, EPI_ISL_749097, EPI_ISL_749098, EPI_ISL_749099, EPI_ISL_749100, EPI_ISL_749101, EPI_ISL_749102, EPI_ISL_749103, EPI_ISL_749104, EPI_ISL_749105, EPI_ISL_749106, EPI_ISL_749107, EPI_ISL_749108, EPI_ISL_749109, EPI_ISL_749110, EPI_ISL_749111, EPI_ISL_749112, EPI_ISL_749113, EPI_ISL_749114, EPI_ISL_749115, EPI_ISL_749116, EPI_ISL_749117, EPI_ISL_749118, EPI_ISL_749119, EPI_ISL_749120, EPI_ISL_749121, EPI_ISL_749122, EPI_ISL_749123, EPI_ISL_749124, EPI_ISL_749125, EPI_ISL_749126, EPI_ISL_749127, EPI_ISL_749128, EPI_ISL_749129, EPI_ISL_749130, EPI_ISL_749131, EPI_ISL_749132, EPI_ISL_749133, EPI_ISL_749134, EPI_ISL_749135, EPI_ISL_749136, EPI_ISL_749137, EPI_ISL_749138, EPI_ISL_749139, EPI_ISL_749140, EPI_ISL_749141, EPI_ISL_749142, EPI_ISL_749143, EPI_ISL_749144, EPI_ISL_749145, EPI_ISL_749146, EPI_ISL_749147, EPI_ISL_749158, EPI_ISL_749167, EPI_ISL_749169, EPI_ISL_749170, EPI_ISL_749171, EPI_ISL_749176, EPI_ISL_749177, EPI_ISL_749181, EPI_ISL_749223, EPI_ISL_749224, EPI_ISL_749225, EPI_ISL_749226, EPI_ISL_749227, EPI_ISL_749228, EPI_ISL_749229, EPI_ISL_749230, EPI_ISL_749231, EPI_ISL_749232, EPI_ISL_749233, EPI_ISL_749234, EPI_ISL_749235, EPI_ISL_749236, EPI_ISL_749237, EPI_ISL_749239, EPI_ISL_749240, EPI_ISL_749241, EPI_ISL_749242, EPI_ISL_749243, EPI_ISL_749244, EPI_ISL_749245, EPI_ISL_749246, EPI_ISL_749247, EPI_ISL_749248, EPI_ISL_749249, EPI_ISL_749250, EPI_ISL_749251, EPI_ISL_749252, EPI_ISL_749253, EPI_ISL_749254, EPI_ISL_749255, EPI_ISL_749256, EPI_ISL_749257, EPI_ISL_749258, EPI_ISL_749259, EPI_ISL_749260, EPI_ISL_749261, EPI_ISL_749262, EPI_ISL_749267, EPI_ISL_749268, EPI_ISL_749271, EPI_ISL_749273, EPI_ISL_749274, EPI_ISL_749276, EPI_ISL_749279, EPI_ISL_749287, EPI_ISL_749290, EPI_ISL_749291, EPI_ISL_749292, EPI_ISL_749293, EPI_ISL_749294, EPI_ISL_749295, EPI_ISL_749321, EPI_ISL_749322, EPI_ISL_749323, EPI_ISL_749324, EPI_ISL_749325, EPI_ISL_749326, EPI_ISL_749327, EPI_ISL_749328, EPI_ISL_749329, EPI_ISL_749330, EPI_ISL_749331, EPI_ISL_749332, EPI_ISL_749333, EPI_ISL_749334, EPI_ISL_749335, EPI_ISL_749336, EPI_ISL_749337, EPI_ISL_749338, EPI_ISL_749339, EPI_ISL_749340, EPI_ISL_749341, EPI_ISL_749342, EPI_ISL_749343, EPI_ISL_749344, EPI_ISL_749345, EPI_ISL_749346, EPI_ISL_749347, EPI_ISL_749348, EPI_ISL_749349, EPI_ISL_749350, EPI_ISL_749351, EPI_ISL_749352, EPI_ISL_749353, EPI_ISL_749354, EPI_ISL_749357, EPI_ISL_749361, EPI_ISL_749369, EPI_ISL_749374, EPI_ISL_749375, EPI_ISL_749390, EPI_ISL_749391, EPI_ISL_749392, EPI_ISL_749393, EPI_ISL_749394, EPI_ISL_749395, EPI_ISL_749396, EPI_ISL_749397, EPI_ISL_749398, EPI_ISL_749399, EPI_ISL_749400, EPI_ISL_749401, EPI_ISL_749402, EPI_ISL_749403, EPI_ISL_749404, EPI_ISL_749405, EPI_ISL_749406, EPI_ISL_749452, EPI_ISL_749453, EPI_ISL_749454, EPI_ISL_749455, EPI_ISL_749456, EPI_ISL_749457, EPI_ISL_749458, EPI_ISL_749459, EPI_ISL_749460, EPI_ISL_749461, EPI_ISL_749462, EPI_ISL_749463, EPI_ISL_749464, EPI_ISL_749465, EPI_ISL_749466, EPI_ISL_749467, EPI_ISL_749468, EPI_ISL_749469, EPI_ISL_749470, EPI_ISL_749471, EPI_ISL_749472, EPI_ISL_749473, EPI_ISL_749475, EPI_ISL_749476, EPI_ISL_749477, EPI_ISL_749478, EPI_ISL_749479, EPI_ISL_749480, EPI_ISL_749481, EPI_ISL_749482, EPI_ISL_749483, EPI_ISL_749491, EPI_ISL_749492, EPI_ISL_749494, EPI_ISL_749495, EPI_ISL_749496, EPI_ISL_749497, EPI_ISL_749499, EPI_ISL_749506, EPI_ISL_749508, EPI_ISL_749509, EPI_ISL_749510, EPI_ISL_749513, EPI_ISL_749514, EPI_ISL_749520, EPI_ISL_749521, EPI_ISL_749522, EPI_ISL_749525, EPI_ISL_749526, EPI_ISL_749527, EPI_ISL_749540, EPI_ISL_749541, EPI_ISL_749859, EPI_ISL_749860, EPI_ISL_749861, EPI_ISL_749862, EPI_ISL_749863, EPI_ISL_749864, EPI_ISL_749865, EPI_ISL_749866, EPI_ISL_749867, EPI_ISL_749868, EPI_ISL_749869, EPI_ISL_749870, EPI_ISL_749871, EPI_ISL_749872, EPI_ISL_749873, EPI_ISL_749874, EPI_ISL_749875, EPI_ISL_749876, EPI_ISL_749877, EPI_ISL_749878, EPI_ISL_749879, EPI_ISL_749880, EPI_ISL_749881, EPI_ISL_749882, EPI_ISL_749883, EPI_ISL_749884, EPI_ISL_749885, EPI_ISL_749886, EPI_ISL_749887, EPI_ISL_749888, EPI_ISL_749889, EPI_ISL_749890, EPI_ISL_749891, EPI_ISL_749892, EPI_ISL_749893, EPI_ISL_749894, EPI_ISL_749895, EPI_ISL_749896, EPI_ISL_749897, EPI_ISL_749898, EPI_ISL_749899, EPI_ISL_749900, EPI_ISL_749901, EPI_ISL_749902, EPI_ISL_749903, EPI_ISL_749904, EPI_ISL_749905, EPI_ISL_749907, EPI_ISL_749908, EPI_ISL_749909, EPI_ISL_749910, EPI_ISL_749911, EPI_ISL_749912, EPI_ISL_749913, EPI_ISL_749914, EPI_ISL_749915, EPI_ISL_749916, EPI_ISL_749917, EPI_ISL_749918, EPI_ISL_749919, EPI_ISL_749920, EPI_ISL_749921, EPI_ISL_749922, EPI_ISL_749923, EPI_ISL_749924, EPI_ISL_749925, EPI_ISL_749926, EPI_ISL_749927, EPI_ISL_749928, EPI_ISL_749929, EPI_ISL_749930, EPI_ISL_749931, EPI_ISL_749932, EPI_ISL_749933, EPI_ISL_749934, EPI_ISL_749935, EPI_ISL_749936, EPI_ISL_749937, EPI_ISL_749938, EPI_ISL_749939, EPI_ISL_749940, EPI_ISL_749941, EPI_ISL_749942, EPI_ISL_749943, EPI_ISL_749944, EPI_ISL_749945, EPI_ISL_749946, EPI_ISL_749947, EPI_ISL_749948, EPI_ISL_749949, EPI_ISL_749950, EPI_ISL_749951, EPI_ISL_749952, EPI_ISL_749953, EPI_ISL_749954, EPI_ISL_749955, EPI_ISL_749956, EPI_ISL_749957, EPI_ISL_749958, EPI_ISL_749959, EPI_ISL_749960, EPI_ISL_749961, EPI_ISL_749962, EPI_ISL_749963, EPI_ISL_749964, EPI_ISL_749965, EPI_ISL_749966, EPI_ISL_749967, EPI_ISL_749968, EPI_ISL_749969, EPI_ISL_749970, EPI_ISL_749971, EPI_ISL_749972, EPI_ISL_749973, EPI_ISL_749974, EPI_ISL_749975, EPI_ISL_749976, EPI_ISL_749977, EPI_ISL_749978, EPI_ISL_749979, EPI_ISL_749980, EPI_ISL_749981, EPI_ISL_749982, EPI_ISL_749983, EPI_ISL_749984, EPI_ISL_749985, EPI_ISL_749986, EPI_ISL_749987, EPI_ISL_749988, EPI_ISL_749989, EPI_ISL_749990, EPI_ISL_749991, EPI_ISL_749992, EPI_ISL_749993, EPI_ISL_749994, EPI_ISL_749995, EPI_ISL_749996, EPI_ISL_749997, EPI_ISL_749998, EPI_ISL_749999, EPI_ISL_750000, EPI_ISL_750001, EPI_ISL_750002, EPI_ISL_750003, EPI_ISL_750004, EPI_ISL_750005, EPI_ISL_750006, EPI_ISL_750007, EPI_ISL_750008, EPI_ISL_750009, EPI_ISL_750010, EPI_ISL_750011, EPI_ISL_750012, EPI_ISL_750013, EPI_ISL_750014, EPI_ISL_750015, EPI_ISL_750016, EPI_ISL_750017, EPI_ISL_750018, EPI_ISL_750019, EPI_ISL_750020, EPI_ISL_750021, EPI_ISL_750022, EPI_ISL_750023, EPI_ISL_750024, EPI_ISL_750025, EPI_ISL_750026, EPI_ISL_750027, EPI_ISL_750028, EPI_ISL_750029, EPI_ISL_750030, EPI_ISL_750031, EPI_ISL_750032, EPI_ISL_750033, EPI_ISL_750034, EPI_ISL_750035, EPI_ISL_750036, EPI_ISL_750037, EPI_ISL_750038, EPI_ISL_750039, EPI_ISL_750040, EPI_ISL_750041, EPI_ISL_750042, EPI_ISL_750043, EPI_ISL_750044, EPI_ISL_750045, EPI_ISL_750046, EPI_ISL_750047, EPI_ISL_750048, EPI_ISL_750049, EPI_ISL_750050, EPI_ISL_750051, EPI_ISL_750052, EPI_ISL_750053, EPI_ISL_750054, EPI_ISL_750055, EPI_ISL_750056, EPI_ISL_750057, EPI_ISL_750058, EPI_ISL_750059, EPI_ISL_750060, EPI_ISL_750061, EPI_ISL_750062, EPI_ISL_750063, EPI_ISL_750064, EPI_ISL_750065, EPI_ISL_750066, EPI_ISL_750067, EPI_ISL_750068, EPI_ISL_750069, EPI_ISL_750070, EPI_ISL_750071, EPI_ISL_750072, EPI_ISL_750073, EPI_ISL_750074, EPI_ISL_750075, EPI_ISL_750076, EPI_ISL_750077, EPI_ISL_750078, EPI_ISL_750079, EPI_ISL_750080, EPI_ISL_750081, EPI_ISL_750082, EPI_ISL_750083, EPI_ISL_750084, EPI_ISL_750085, EPI_ISL_750086, EPI_ISL_750087, EPI_ISL_750088, EPI_ISL_750089, EPI_ISL_750090, EPI_ISL_750091, EPI_ISL_750092, EPI_ISL_750093, EPI_ISL_750094, EPI_ISL_750095, EPI_ISL_750096, EPI_ISL_750097, EPI_ISL_750098, EPI_ISL_750099, EPI_ISL_750100, EPI_ISL_750101, EPI_ISL_750102, EPI_ISL_750103, EPI_ISL_750104, EPI_ISL_750105, EPI_ISL_750106, EPI_ISL_750107, EPI_ISL_750109, EPI_ISL_750110, EPI_ISL_750111, EPI_ISL_750112, EPI_ISL_750113, EPI_ISL_750114, EPI_ISL_750115, EPI_ISL_750116, EPI_ISL_750117, EPI_ISL_750118, EPI_ISL_750119, EPI_ISL_750120, EPI_ISL_750121, EPI_ISL_750122, EPI_ISL_750123, EPI_ISL_750124, EPI_ISL_750125, EPI_ISL_750126, EPI_ISL_750127, EPI_ISL_750128, EPI_ISL_750129, EPI_ISL_750130, EPI_ISL_750131, EPI_ISL_750132, EPI_ISL_750133, EPI_ISL_750134, EPI_ISL_750135, EPI_ISL_750136, EPI_ISL_750137, EPI_ISL_750138, EPI_ISL_750139, EPI_ISL_750140, EPI_ISL_750141, EPI_ISL_750142, EPI_ISL_750143, EPI_ISL_750144, EPI_ISL_750145, EPI_ISL_750146, EPI_ISL_750147, EPI_ISL_750148, EPI_ISL_750149, EPI_ISL_750150, EPI_ISL_750151, EPI_ISL_750152, EPI_ISL_750153, EPI_ISL_750154, EPI_ISL_750155, EPI_ISL_750156, EPI_ISL_750157, EPI_ISL_750158, EPI_ISL_750159, EPI_ISL_750160 | | | |
| see above | Department of Virus and Microbiological Special Diagnostics, Statens Serum Institut, Copenhagen, Denmark | Albertsen Lab, Department of Chemistry and Bioscience, Aalborg University, Denmark | Danish Covid-19 Genome Consortium |
| EPI_ISL_751191 | Limmattal Hospital | Institute of Medical Virology, University of Zurich | Stefan Schmutz, Kevin Steiner, Verena Kufner, Maryam Zaheri, Gabriela Ziltener, Jürg Böni, Michael Huber, Alexandra Trkola, Roberto Buonomano |
| EPI_ISL_751799 | Labor Dr. Wisplinghoff - Köln | Robert Koch Institute, Bioinformatics MF1, Berlin, Germany | Dr. R. Grosser, Stephan Fuchs, Stefan Kroeger, Marianne Wedde, Oliver Drechsel, Aleksandar Radonic, Rene Kmiecinski, Ralf Duerrwald, Thorsten Wolff |
| EPI_ISL_753225, EPI_ISL_753226, EPI_ISL_753227, EPI_ISL_753228, EPI_ISL_753229, EPI_ISL_753230, EPI_ISL_753231, EPI_ISL_753232, EPI_ISL_753233, EPI_ISL_753234, EPI_ISL_753235, EPI_ISL_753236, EPI_ISL_753237, EPI_ISL_753241, EPI_ISL_753242, EPI_ISL_753243, EPI_ISL_753244, EPI_ISL_753245, EPI_ISL_753246, EPI_ISL_753247, EPI_ISL_753248, EPI_ISL_753249, EPI_ISL_753250, EPI_ISL_753251, EPI_ISL_753252, EPI_ISL_753253, EPI_ISL_753254, EPI_ISL_753255, EPI_ISL_753256, EPI_ISL_753257, EPI_ISL_753258, EPI_ISL_753259, EPI_ISL_753260, EPI_ISL_753261 | | | |
| see above | State Laboratories Division, Hawaii State Department of Health | State Laboratories Division, Hawaii State Department of Health | Pamela O'Brien, Sabrina Diemert, Drew Kuwazaki, Razvan Sultana, Edward Desmond |
| EPI_ISL_753661, EPI_ISL_753662, EPI_ISL_753664, EPI_ISL_753665, EPI_ISL_753666, EPI_ISL_753667, EPI_ISL_753668, EPI_ISL_753670 | Clinical virology Laboratory, Children's Hospital Los Angeles | Center for Personalized Medicine, Children's Hospital Los Angeles | Gai et al |
| EPI_ISL_754149, EPI_ISL_754150, EPI_ISL_754163, EPI_ISL_754165, EPI_ISL_754166 | Laboratoire de Microbiologie | National Reference Center for Viruses of Respiratory Infections, Institut Pasteur, Paris | Marion Barbet, Sylvie Behillil, Méline Bizard, Angela Brisebarre, Camille Capel, Etienne Simon-Lorière, Vincent Enouf, Maud Vanpeene, Sylvie van der Werf, Marie-Sarah Fangous |
| EPI_ISL_754275 | Respiratory Virus Unit, National Infection Service, Public Health England | COVID-19 Genomics UK (COG-UK) Consortium | PHE Covid Sequencing Team |
| EPI_ISL_754394, EPI_ISL_754399, EPI_ISL_754448, EPI_ISL_754449, EPI_ISL_754450, EPI_ISL_754451, EPI_ISL_754452, EPI_ISL_754453, EPI_ISL_754454, EPI_ISL_754455, EPI_ISL_754456, EPI_ISL_754457, EPI_ISL_754458, EPI_ISL_754459, EPI_ISL_754460, EPI_ISL_754461, EPI_ISL_754462, EPI_ISL_754463, EPI_ISL_754464, EPI_ISL_754465, EPI_ISL_754466, EPI_ISL_754467, EPI_ISL_754468, EPI_ISL_754469, EPI_ISL_754470, EPI_ISL_754471, EPI_ISL_754472, EPI_ISL_754473, EPI_ISL_754474, EPI_ISL_754475, EPI_ISL_754476, EPI_ISL_754547, EPI_ISL_754548, EPI_ISL_754549, EPI_ISL_754572 | | | |
| see above | Wadsworth Center, New York State Department.of Health | Wadsworth Center, New York State Department.of Health | Kirsten St. George, Daryl M. Lamson, Alexis Russel, Matthew Shudt, Melissa A Leisner, Jonathan Plitnick, Navjot Singh, John Kelly, Sara Griesemer, Erasmus Schneider, Erica Lasek-Nesselquist |
| EPI_ISL_754624, EPI_ISL_754673, EPI_ISL_754674, EPI_ISL_754675 | University of Wisconsin-Madison AIDS Vaccine Research Laboratories | University of Wisconsin-Madison AIDS Vaccine Research Laboratories | Gage Moreno, Katarina Braun, et al. AIDS Vaccine Research Laboratories |
| EPI_ISL_755133, EPI_ISL_755140, EPI_ISL_755147, EPI_ISL_755148, EPI_ISL_755149, EPI_ISL_755150, EPI_ISL_755151, EPI_ISL_755152, EPI_ISL_755153, EPI_ISL_755154, EPI_ISL_755155, EPI_ISL_755156, EPI_ISL_755157, EPI_ISL_755158, EPI_ISL_755159, EPI_ISL_755160, EPI_ISL_755161, EPI_ISL_755162 | | | |
| see above | UCSD EXCITE lab | Andersen lab at Scripps Research | SEARCH Alliance San Diego |
| EPI_ISL_755489, EPI_ISL_755490, EPI_ISL_755491, EPI_ISL_755492, | Maine Health and Environmental Testing Laboratory | Tewhey Lab, The Jackson Laboratory | Matluk,N., Dewey,H., Iosue,F., Barter,M., Lynch,R., Munger,H. and Tewhey,R. |

| | | | |
|---|---|---|---|
| EPI_ISL_755493, EPI_ISL_755496 | | | |
| EPI_ISL_755643 | Instituto Adolfo Lutz - Central | Instituto Adolfo Lutz, Interdiciplinary Procedures Center, Strategic Laboratory | Claudio Tavares Sacchi, Claudia Regina Gonçalves, Erica Valessa Ramos Gomes, Karoline Rodrigues Campos |
| EPI_ISL_755644, EPI_ISL_755645 | Lab LOC - Itapecerica da Serra | Instituto Adolfo Lutz, Interdiciplinary Procedures Center, Strategic Laboratory | Claudio Tavares Sacchi, Claudia Regina Gonçalves, Erica Valessa Ramos Gomes, Karoline Rodrigues Campos |
| EPI_ISL_755646, EPI_ISL_755647 | Instituto Adolfo Lutz - Regional de Santo Andre | Instituto Adolfo Lutz, Interdiciplinary Procedures Center, Strategic Laboratory | Claudio Tavares Sacchi, Claudia Regina Gonçalves, Erica Valessa Ramos Gomes, Karoline Rodrigues Campos |
| EPI_ISL_755648, EPI_ISL_755650 | Instituto Adolfo Lutz - Regional de Taubate | Instituto Adolfo Lutz, Interdiciplinary Procedures Center, Strategic Laboratory | Claudio Tavares Sacchi, Claudia Regina Gonçalves, Erica Valessa Ramos Gomes, Karoline Rodrigues Campos |
| EPI_ISL_755652 | Lab LOC - Itapecerica da Serra | Instituto Adolfo Lutz, Interdiciplinary Procedures Center, Strategic Laboratory | Claudio Tavares Sacchi, Claudia Regina Gonçalves, Erica Valessa Ramos Gomes, Karoline Rodrigues Campos |
| EPI_ISL_756202, EPI_ISL_756203, EPI_ISL_756204, EPI_ISL_756205, EPI_ISL_756207, EPI_ISL_756298 | UW Virology Lab | UW Virology Lab | Pavitra Roychoudhury, Hong Xie, Lasata Shrestha, Meei-Li Huang, Keith R Jerome, Alexander Greninger |
| EPI_ISL_757226, EPI_ISL_757227, EPI_ISL_757231 | Lighthouse Lab in Glasgow | Wellcome Sanger Institute for the COVID-19 Genomics UK (COG-UK) Consortium | Harper VanSteenhouse, Yumi Kasai, David Gray, Carol Clugston, Anna Dominiczak and Alex Alderton, Roberto Amato, Sonia Goncalves, Ewan Harrison, David K. Jackson, Ian Johnston, Dominic Kwiatkowski, Cordelia Langford, John Sillitoe on behalf of the Wellcome Sanger Institute COVID-19 Surveillance Team |
| EPI_ISL_757443, EPI_ISL_757448, EPI_ISL_757491, EPI_ISL_757688, EPI_ISL_757689, EPI_ISL_757690, EPI_ISL_757691, EPI_ISL_757692, EPI_ISL_757693, EPI_ISL_757694, EPI_ISL_757695, EPI_ISL_757696, EPI_ISL_757697, EPI_ISL_757698 | | | |
| see above | Department of Virus and Microbiological Special Diagnostics, Statens Serum Institut, Copenhagen, Denmark | Albertsen Lab, Department of Chemistry and Bioscience, Aalborg University, Denmark | Danish Covid-19 Genome Consortium |
| EPI_ISL_759989 | Norwegian Institute of Public Health, Department of Virology | Norwegian Institute of Public Health, Department of Virology | Kathrine Stene-Johansen, Kamilla Heddeland Instefjord, Hilde Elshaug, Marie Paulsen Madsen, Rasmus Riis Kopperud, Hilde Vollan, Karoline Bragstad, Olav Hungnes |
| EPI_ISL_760822 | Lighthouse Lab in Cambridge | Wellcome Sanger Institute for the COVID-19 Genomics UK (COG-UK) Consortium | Rob Howes, The Lighthouse Lab in Cambridge and Alex Alderton, Roberto Amato, Sonia Goncalves, Ewan Harrison, David K. Jackson, Ian Johnston, Dominic Kwiatkowski, Cordelia Langford, John Sillitoe on behalf of the Wellcome Sanger Institute COVID-19 Surveillance Team |
| EPI_ISL_760844, EPI_ISL_760871 | Lighthouse Lab in Milton Keynes | Wellcome Sanger Institute for the COVID-19 Genomics UK (COG-UK) Consortium | The Lighthouse Lab in Milton Keynes and Alex Alderton, Roberto Amato, Sonia Goncalves, Ewan Harrison, David K. Jackson, Ian Johnston, Dominic Kwiatkowski, Cordelia Langford, John Sillitoe on behalf of the Wellcome Sanger Institute COVID-19 Surveillance Team |
| EPI_ISL_760891, EPI_ISL_760929 | Lighthouse Lab in Alderley Park | Wellcome Sanger Institute for the COVID-19 Genomics UK (COG-UK) Consortium | Jacquelyn Wynn, Mairead Hyland, The Lighthouse Lab in Alderley Park and Alex Alderton, Roberto Amato, Sonia Goncalves, Ewan Harrison, David K. Jackson, Ian Johnston, Dominic Kwiatkowski, Cordelia Langford, John Sillitoe on behalf of the Wellcome Sanger Institute COVID-19 Surveillance Team |
| EPI_ISL_760939 | Lighthouse Lab in Cambridge | Wellcome Sanger Institute for the COVID-19 Genomics UK (COG-UK) Consortium | Rob Howes, The Lighthouse Lab in Cambridge and Alex Alderton, Roberto Amato, Sonia Goncalves, Ewan Harrison, David K. Jackson, Ian Johnston, Dominic Kwiatkowski, Cordelia Langford, John Sillitoe on behalf of the Wellcome Sanger Institute COVID-19 Surveillance Team |
| EPI_ISL_760982 | Lighthouse Lab in Alderley Park | Wellcome Sanger Institute for the COVID-19 Genomics UK (COG-UK) Consortium | Jacquelyn Wynn, Mairead Hyland, The Lighthouse Lab in Alderley Park and Alex Alderton, Roberto Amato, Sonia Goncalves, Ewan Harrison, David K. Jackson, Ian Johnston, Dominic Kwiatkowski, Cordelia Langford, John Sillitoe on behalf of the Wellcome Sanger Institute COVID-19 Surveillance Team |
| EPI_ISL_760986 | Lighthouse Lab in Glasgow | Wellcome Sanger Institute for the COVID-19 Genomics UK (COG-UK) Consortium | Harper VanSteenhouse, Yumi Kasai, David Gray, Carol Clugston, Anna Dominiczak and Alex Alderton, Roberto Amato, Sonia Goncalves, Ewan Harrison, David K. Jackson, Ian Johnston, Dominic Kwiatkowski, Cordelia Langford, John Sillitoe on behalf of the Wellcome Sanger Institute COVID-19 Surveillance Team |
| EPI_ISL_760988 | Lighthouse Lab in Cambridge | Wellcome Sanger Institute for the COVID-19 Genomics UK (COG-UK) Consortium | Rob Howes, The Lighthouse Lab in Cambridge and Alex Alderton, Roberto Amato, Sonia Goncalves, Ewan Harrison, David K. Jackson, Ian Johnston, Dominic Kwiatkowski, Cordelia Langford, John Sillitoe on behalf of the Wellcome Sanger Institute COVID-19 Surveillance Team |
| EPI_ISL_761013, EPI_ISL_761046 | Lighthouse Lab in Alderley Park | Wellcome Sanger Institute for the COVID-19 Genomics UK (COG-UK) Consortium | Jacquelyn Wynn, Mairead Hyland, The Lighthouse Lab in Alderley Park and Alex Alderton, Roberto Amato, Sonia Goncalves, Ewan Harrison, David K. Jackson, Ian Johnston, Dominic Kwiatkowski, Cordelia Langford, John Sillitoe on behalf of the Wellcome Sanger Institute COVID-19 Surveillance Team |
| EPI_ISL_761047 | Lighthouse Lab in Glasgow | Wellcome Sanger Institute for the COVID-19 Genomics UK (COG-UK) Consortium | Harper VanSteenhouse, Yumi Kasai, David Gray, Carol Clugston, Anna Dominiczak and Alex Alderton, Roberto Amato, Sonia Goncalves, Ewan Harrison, David K. Jackson, Ian Johnston, Dominic Kwiatkowski, Cordelia Langford, John Sillitoe on behalf of the Wellcome Sanger Institute COVID-19 Surveillance Team |
| EPI_ISL_761050 | Lighthouse Lab in Milton Keynes | Wellcome Sanger Institute for the COVID-19 Genomics UK (COG-UK) Consortium | The Lighthouse Lab in Milton Keynes and Alex Alderton, Roberto Amato, Sonia Goncalves, Ewan Harrison, David K. Jackson, Ian Johnston, Dominic Kwiatkowski, Cordelia Langford, John Sillitoe on behalf of the Wellcome Sanger Institute COVID-19 Surveillance Team |
| EPI_ISL_763004, EPI_ISL_763005, EPI_ISL_763006, EPI_ISL_763014, EPI_ISL_763015, EPI_ISL_763016, EPI_ISL_763017, EPI_ISL_763018, EPI_ISL_763019, EPI_ISL_763031, EPI_ISL_763032, EPI_ISL_763043, EPI_ISL_763044, EPI_ISL_763046, EPI_ISL_763060, EPI_ISL_763061 | | | |
| see above | Unit 17: Influenza & Other Respiratory Viruses, German National Influenza Center | Project group Epidemiology of Highly Pathogenic Microorganisms, Robert Koch-Institute | Ariane Düx, Andreas Sachse, Grit Schubert, Sébastien Calvignac-Spencer, Fabian Leendertz, Thorsten Wolff, Ralf Dürrwald, Djin-Ye Oh, Marianne Wedde |
| EPI_ISL_763126, EPI_ISL_763174, EPI_ISL_763176, EPI_ISL_763191, EPI_ISL_763192, EPI_ISL_763193, EPI_ISL_763200 | Dutch COVID-19 response team | Erasmus Medical Center | Bas Oude Munnink, Reina Sikkema, David Nieuwenhuijse, Irina Chestakova, Anne van der Linden, Marjan Boter, Emmanuelle Munger, Corine GeurtsvanKessel, Annemiek van der Eijk, Richard Molenkamp, Marion Koopmans, on behalf of the Dutch national COVID-19 response team. |
| EPI_ISL_763370 | Department of Pathology, University of Cambridge | COVID-19 Genomics UK (COG-UK) Consortium | Aminu S. Jahun, Yasmin Chaudhry, Grant Hall, Iliana Georgana, Myra Hosmillo, Martin D. Curran, Malte Pinckert, Surendra Parmar, Ian Goodfellow |
| EPI_ISL_763392, EPI_ISL_763409, EPI_ISL_763416, EPI_ISL_763420, EPI_ISL_763422, EPI_ISL_763423, EPI_ISL_763454, EPI_ISL_763462, EPI_ISL_763467, EPI_ISL_763470, EPI_ISL_763479, EPI_ISL_763480, EPI_ISL_763481, EPI_ISL_763489, EPI_ISL_763492, EPI_ISL_763565, EPI_ISL_763572, EPI_ISL_763590, EPI_ISL_763658, EPI_ISL_763721, EPI_ISL_763725, EPI_ISL_763750, EPI_ISL_763760, EPI_ISL_763782, EPI_ISL_763821 | | | |
| see above | Wales Specialist Virology Centre Sequencing lab: Pathogen Genomics Unit | COVID-19 Genomics UK (COG-UK) Consortium | Catherine Moore, Johnathan Evans, Laura Gifford, Malorie Perry, Simon Cottrell, Angela Marchbank, Alec Birchley, Alexander Adams, Amy Gaskin, Bree Gatica-Wilcox, Jason Coombes, Joel Southgate, Lauren Gilbert, Lee Graham, Nicole Pacchiarini, Sara Kumziene-Summerhayes, Sarah Taylor, Sophie Jones, Sara Rey, Matthew Bull, Joanne Watkins, Sally Corden, Tom Connor |
| EPI_ISL_763903 | Virology Department, Royal Infirmary of Edinburgh, NHS Lothian / School of Biological Sciences, University of Edinburgh / Institute of Genetics and Molecular Medicine, University of Edinburgh | COVID-19 Genomics UK (COG-UK) Consortium | McHugh M, Dewar R, Rooke S, Gallagher M, Balcaza C, O'Toole Á, Scher E, Hill V, McCrone JT, Colquhoun R, Yu X, Jackson B, Rambaut A, Williams TC, Templeton K |
| EPI_ISL_763934, EPI_ISL_763935, EPI_ISL_763940, EPI_ISL_763941, EPI_ISL_763945 | Oxford Viromics, NDM, University of Oxford; Oxford University Hospitals; Basingstoke and North Hampshire Hospital | COVID-19 Genomics UK (COG-UK) Consortium | Tanya Golubchik, David Bonsall, George Macintyre, Amy Trebes, Mariateresa de Cesare, Catrin Moore, Alex Mobbs, Anita Justice, Robert Shaw, Monique Andersson, Timothy Peto, Emma Wise, Nathan Moore, Jessica Lynch, Nick Cortes, Matilde Mori, Stephen Kidd, David Buck, John Todd, Christophe Fraser |
| EPI_ISL_764064 | Wales Specialist Virology Centre Sequencing lab: Pathogen Genomics Unit | COVID-19 Genomics UK (COG-UK) Consortium | Catherine Moore, Johnathan Evans, Laura Gifford, Malorie Perry, Simon Cottrell, Angela Marchbank, Alec Birchley, Alexander Adams, Amy Gaskin, Bree Gatica-Wilcox, Jason Coombes, Joel Southgate, Lauren Gilbert, Lee Graham, Nicole Pacchiarini, Sara Kumziene-Summerhayes, Sarah Taylor, Sophie Jones, Sara Rey, Matthew Bull, Joanne Watkins, Sally Corden, Tom Connor |
| EPI_ISL_764066, EPI_ISL_764087 | Oxford Viromics, NDM, University of Oxford; Oxford University Hospitals; Basingstoke and North Hampshire Hospital | COVID-19 Genomics UK (COG-UK) Consortium | Tanya Golubchik, David Bonsall, George Macintyre, Amy Trebes, Mariateresa de Cesare, Catrin Moore, Alex Mobbs, Anita Justice, Robert Shaw, Monique Andersson, Timothy Peto, Emma Wise, Nathan Moore, Jessica Lynch, Nick Cortes, Matilde Mori, Stephen Kidd, David Buck, John Todd, Christophe Fraser |
| EPI_ISL_764133, EPI_ISL_764138, EPI_ISL_764155, EPI_ISL_764157, EPI_ISL_764192, EPI_ISL_764200, | Wales Specialist Virology Centre Sequencing lab: Pathogen Genomics Unit | COVID-19 Genomics UK (COG-UK) Consortium | Catherine Moore, Johnathan Evans, Laura Gifford, Malorie Perry, Simon Cottrell, Angela Marchbank, Alec Birchley, Alexander Adams, Amy Gaskin, Bree Gatica-Wilcox, Jason Coombes, Joel Southgate, Lauren Gilbert, Lee Graham, Nicole Pacchiarini, Sara Kumziene-Summerhayes, Sarah Taylor, Sophie Jones, Sara Rey, Matthew Bull, Joanne Watkins, Sally Corden, Tom Connor |

| | | | |
|---|---|---|---|
| EPI_ISL_764201, EPI_ISL_764235 | | | |
| EPI_ISL_764238 | University College London, Great Ormond Street Hospital for Children NHS Foundation Trust, Imperial College Healthcare NHS Trust | COVID-19 Genomics UK (COG-UK) Consortium | Sergi Castellano, Rachel Williams, Mark Kristiansen, Paola Resende Silva, Sunando Roy, Tony Brooks, Helena Tutill, Paola Niola, Patricia Dyal, Charlotte Williams, Leysa Forrest, Yasmin Panchbhaya, Jacqueline Findlay, Samuel Weeks, Julianne Brown, Kathryn Harris, Paul Randell, James Price, Alison Holmes, Judith Breuer |
| EPI_ISL_764255 | Wales Specialist Virology Centre Sequencing lab: Pathogen Genomics Unit | COVID-19 Genomics UK (COG-UK) Consortium | Catherine Moore, Johnathan Evans, Laura Gifford, Malorie Perry, Simon Cottrell, Angela Marchbank, Alec Birchley, Alexander Adams, Amy Gaskin, Bree Gatica-Wilcox, Jason Coombes, Joel Southgate, Lauren Gilbert, Lee Graham, Nicole Pacchiarini, Sara Kumziene-Summerhayes, Sarah Taylor, Sophie Jones, Sara Rey, Matthew Bull, Joanne Watkins, Sally Corden, Tom Connor |
| EPI_ISL_764303, EPI_ISL_764304 | Department of Pathology, University of Cambridge | COVID-19 Genomics UK (COG-UK) Consortium | Aminu S. Jahun, Yasmin Chaudhry, Grant Hall, Iliana Georgana, Myra Hosmillo, Martin D. Curran, Malte Pinckert, Surendra Parmar, Ian Goodfellow |
| EPI_ISL_764532, EPI_ISL_764533, EPI_ISL_764538, EPI_ISL_764539 | Oxford Viromics, NDM, University of Oxford; Oxford University Hospitals; Basingstoke and North Hampshire Hospital | COVID-19 Genomics UK (COG-UK) Consortium | Tanya Golubchik, David Bonsall, George Macintyre, Amy Trebes, Mariateresa de Cesare, Catrin Moore, Alex Mobbs, Anita Justice, Robert Shaw, Monique Andersson, Timothy Peto, Emma Wise, Nathan Moore, Jessica Lynch, Nick Cortes, Matilde Mori, Stephen Kidd, David Buck, John Todd, Christophe Fraser |
| EPI_ISL_764582, EPI_ISL_764746, EPI_ISL_764747, EPI_ISL_764753, EPI_ISL_764754, EPI_ISL_764755, EPI_ISL_764769, EPI_ISL_764770, EPI_ISL_764771, EPI_ISL_764772, EPI_ISL_764773, EPI_ISL_764774, EPI_ISL_764775, EPI_ISL_764776, EPI_ISL_764777, EPI_ISL_764778, EPI_ISL_764779, EPI_ISL_764780, EPI_ISL_764781, EPI_ISL_764782, EPI_ISL_764783, EPI_ISL_764784, EPI_ISL_764785, EPI_ISL_764786, EPI_ISL_764787, EPI_ISL_764788, EPI_ISL_764789, EPI_ISL_764790, EPI_ISL_764791, EPI_ISL_764792, EPI_ISL_764793, EPI_ISL_764794, EPI_ISL_764795, EPI_ISL_764796, EPI_ISL_764797, EPI_ISL_764798, EPI_ISL_764799, EPI_ISL_764800, EPI_ISL_764801, EPI_ISL_764802, EPI_ISL_764803, EPI_ISL_764804 | | | |
| see above | Wales Specialist Virology Centre Sequencing lab: Pathogen Genomics Unit | COVID-19 Genomics UK (COG-UK) Consortium | Catherine Moore, Johnathan Evans, Laura Gifford, Malorie Perry, Simon Cottrell, Angela Marchbank, Alec Birchley, Alexander Adams, Amy Gaskin, Bree Gatica-Wilcox, Jason Coombes, Joel Southgate, Lauren Gilbert, Lee Graham, Nicole Pacchiarini, Sara Kumziene-Summerhayes, Sarah Taylor, Sophie Jones, Sara Rey, Matthew Bull, Joanne Watkins, Sally Corden, Tom Connor |
| EPI_ISL_765818, EPI_ISL_765819, EPI_ISL_765820, EPI_ISL_765821, EPI_ISL_765822, EPI_ISL_765823 | North Shore Medical Center | Infectious Disease Program, Broad Institute of Harvard and MIT | Lemieux,J.E., Siddle,K.J., Shaw,B., Adams,G., Pierce,V., Turbett,S., Anahtar,M., Branda,J., Slater,D., Harris,J., Lin,A.E., Gladden-Young,A., Lagerborg,K., Rudy,M., DeRuff,K., Carter,A., Normandin,E., Bauer,M., Reilly,S., Tomkins-Tinch,C., Loreth,C., Chaluvadi,S., Neumann,A., Cusick,C., Chapman,S.B., Gnirke,A., Flowers,K., Cerrato,F., Birren,B.W., Gallagher,G., Smole,S., Park,D.J., MacInnis,B.L., Ryan,E., LaRocque,R., Rosenberg,E. and Sabeti,P.C. |
| EPI_ISL_765893, EPI_ISL_765894 | Unit 17: Influenza & Other Respiratory Viruses, German National Influenza Center | Project group Epidemiology of Highly Pathogenic Microorganisms, Robert Koch-Institute | Ariane Düx, Andreas Sachse, Grit Schubert, Sébastien Calvignac-Spencer, Fabian Leendertz, Thorsten Wolff, Ralf Dürrwald, Djin-Ye Oh, Marianne Wedde |
| EPI_ISL_765960, EPI_ISL_765961, EPI_ISL_765962, EPI_ISL_765963, EPI_ISL_765964, EPI_ISL_765965, EPI_ISL_765966, EPI_ISL_765967, EPI_ISL_765968, EPI_ISL_765969, EPI_ISL_765970, EPI_ISL_765971, EPI_ISL_765972, EPI_ISL_765973, EPI_ISL_765974, EPI_ISL_765975, EPI_ISL_765976, EPI_ISL_765977, EPI_ISL_765978 | | | |
| see above | Wyoming Public Health Laboratory | Wyoming Public Health Laboratory | Noah Hull, Taylor Fearing, Channing Weber, Ashley Norberg, Bailey Bowcutt, and Wanda Manley |
| EPI_ISL_766096, EPI_ISL_766125, EPI_ISL_766129, EPI_ISL_766130, EPI_ISL_766213, EPI_ISL_766246, EPI_ISL_766247, EPI_ISL_766281, EPI_ISL_766282, EPI_ISL_766283, EPI_ISL_766285, EPI_ISL_766289, EPI_ISL_766291, EPI_ISL_766292, EPI_ISL_766506, EPI_ISL_766507, EPI_ISL_766510, EPI_ISL_766511, EPI_ISL_766512, EPI_ISL_766513, EPI_ISL_766514, EPI_ISL_766515, EPI_ISL_766516, EPI_ISL_766558, EPI_ISL_766559, EPI_ISL_766561 | | | |
| see above | Respiratory Virus Unit, National Infection Service, Public Health England | COVID-19 Genomics UK (COG-UK) Consortium | PHE Covid Sequencing Team |
| EPI_ISL_766630 | Klinisk mikrobiologi, Viruslab | The Public Health Agency of Sweden | Department of Microbiology, The Public Health Agency of Sweden |
| EPI_ISL_766651, EPI_ISL_766658 | Texas Department of State Health Services | Texas Department of State Health Services | Rashmi Tuladhar, Bonnie Oh, Jenny Zhang, Maliha Rahman, Anita Pokharel, Myong Koag, Chung Wang, Rachel Lee, Grace Kubin, Mayela Pedrueza, James Daniel Bonser |
| EPI_ISL_766853 | Respiratory Virus Unit, National Infection Service, Public Health England | COVID-19 Genomics UK (COG-UK) Consortium | PHE Covid Sequencing Team |
| EPI_ISL_767134 | Lighthouse Lab in Milton Keynes | Wellcome Sanger Institute for the COVID-19 Genomics UK (COG-UK) Consortium | The Lighthouse Lab in Milton Keynes and Alex Alderton, Roberto Amato, Sonia Goncalves, Ewan Harrison, David K. Jackson, Ian Johnston, Dominic Kwiatkowski, Cordelia Langford, John Sillitoe on behalf of the Wellcome Sanger Institute COVID-19 Surveillance Team |
| EPI_ISL_767151, EPI_ISL_767152, EPI_ISL_767153, EPI_ISL_767154, EPI_ISL_767155, EPI_ISL_767156, EPI_ISL_767157, EPI_ISL_767158, EPI_ISL_767159, EPI_ISL_767160, EPI_ISL_767161, EPI_ISL_767162, EPI_ISL_767163, EPI_ISL_767164, EPI_ISL_767165, EPI_ISL_767166, EPI_ISL_767167, EPI_ISL_767168, EPI_ISL_767169, EPI_ISL_767170, EPI_ISL_767171, EPI_ISL_767172, EPI_ISL_767173, EPI_ISL_767174, EPI_ISL_767175, EPI_ISL_767176, EPI_ISL_767177, EPI_ISL_767179, EPI_ISL_767180, EPI_ISL_767181, EPI_ISL_767182, EPI_ISL_767183, EPI_ISL_767184, EPI_ISL_767185, EPI_ISL_767186, EPI_ISL_767187, EPI_ISL_767188, EPI_ISL_767189, EPI_ISL_767190, EPI_ISL_767191, EPI_ISL_767192, EPI_ISL_767194, EPI_ISL_767195, EPI_ISL_767196, EPI_ISL_767197, EPI_ISL_767198, EPI_ISL_767199, EPI_ISL_767200, EPI_ISL_767201, EPI_ISL_767202, EPI_ISL_767203, EPI_ISL_767204, EPI_ISL_767206, EPI_ISL_767207, EPI_ISL_767208, EPI_ISL_767209, EPI_ISL_767210, EPI_ISL_767211, EPI_ISL_767212, EPI_ISL_767213, EPI_ISL_767214, EPI_ISL_767215, EPI_ISL_767221, EPI_ISL_767230, EPI_ISL_767251, EPI_ISL_767252, EPI_ISL_767253, EPI_ISL_767254, EPI_ISL_767255, EPI_ISL_767256, EPI_ISL_767257, EPI_ISL_767258, EPI_ISL_767259, EPI_ISL_767260, EPI_ISL_767261, EPI_ISL_767262, EPI_ISL_767263, EPI_ISL_767264, EPI_ISL_767265, EPI_ISL_767266, EPI_ISL_767267, EPI_ISL_767268, EPI_ISL_767269, EPI_ISL_767270, EPI_ISL_767271, EPI_ISL_767272, EPI_ISL_767273, EPI_ISL_767274, EPI_ISL_767275, EPI_ISL_767276, EPI_ISL_767278, EPI_ISL_767284 | | | |
| see above | Lighthouse Lab in Alderley Park | Wellcome Sanger Institute for the COVID-19 Genomics UK (COG-UK) Consortium | Jacquelyn Wynn, Mairead Hyland, The Lighthouse Lab in Alderley Park and Alex Alderton, Roberto Amato, Sonia Goncalves, Ewan Harrison, David K. Jackson, Ian Johnston, Dominic Kwiatkowski, Cordelia Langford, John Sillitoe on behalf of the Wellcome Sanger Institute COVID-19 Surveillance Team |
| EPI_ISL_767344, EPI_ISL_767345, EPI_ISL_767347, EPI_ISL_767349, EPI_ISL_767350, EPI_ISL_767354, EPI_ISL_767358, EPI_ISL_767359, EPI_ISL_767361 | Michigan Department of Health and Human Services, Bureau of Laboratories | Michigan Department of Health and Human Services, Bureau of Laboratories | Blankenship HM, Riner D, Soehnlen MK |
| EPI_ISL_767655, EPI_ISL_767658, EPI_ISL_767690, EPI_ISL_767691, EPI_ISL_767692, EPI_ISL_767693, EPI_ISL_767694, EPI_ISL_767695, EPI_ISL_767696, EPI_ISL_767697, EPI_ISL_767698, EPI_ISL_767699, EPI_ISL_767700, EPI_ISL_767701, EPI_ISL_767702, EPI_ISL_767703, EPI_ISL_767704, EPI_ISL_767705, EPI_ISL_767706, EPI_ISL_767707, EPI_ISL_767708, EPI_ISL_767709 | | | |
| see above | MEMORIAL SLOAN KETTERING CANCER CENTER | Wadsworth Center, New York State Department.of Health | Kirsten St. George, Daryl M. Lamson, Alexis Russel, Matthew Shudt, Melissa A Leisner, Jonathan Plitnick, Navjot Singh, John Kelly, Sara Griesemer, Erasmus Schneider, Erica Lasek-Nesselquist |
| EPI_ISL_768398, EPI_ISL_768399, EPI_ISL_768400, EPI_ISL_768401, EPI_ISL_768402, EPI_ISL_768403, EPI_ISL_768404, EPI_ISL_768405, EPI_ISL_768406, EPI_ISL_768407, EPI_ISL_768408, EPI_ISL_768409, EPI_ISL_768410, EPI_ISL_768411, EPI_ISL_768412, EPI_ISL_768413, EPI_ISL_768414, EPI_ISL_768415, EPI_ISL_768416, EPI_ISL_768417, EPI_ISL_768418 | | | |
| see above | LSUHS Emerging Viral Threat Laboratory | Microbial Genome Sequencing Center | Jeremy P. Kamil, Jennifer L. Carroll, Camille F. Abshire, Maarten Van Diest, Andrew D. Yurochko, Martin J. Sapp, Rona S. Scott, Christopher G. Kevil, Daniel J. Snyder, Vaughn S. Cooper, John A. Vanchiere |
| EPI_ISL_768633 | Jena University Hospital, Institute for Infectious Diseases and Infection Control | Institute of infectious medicine & hospital hygiene, CaSe-Group | Spott, Riccardo; Marquet, Mike; Pletz, Matthias W.; Brandt, Christian |
| EPI_ISL_768801, EPI_ISL_768802, EPI_ISL_768803 | Respiratory Virus Unit, National Infection Service, Public Health England | COVID-19 Genomics UK (COG-UK) Consortium | PHE Covid Sequencing Team |
| EPI_ISL_769545, EPI_ISL_769584, EPI_ISL_769590, EPI_ISL_769597, EPI_ISL_769646, EPI_ISL_769715, EPI_ISL_769744, EPI_ISL_769764, EPI_ISL_769776, EPI_ISL_769845, EPI_ISL_769852, EPI_ISL_769857 | | | |
| see above | Lighthouse Lab in Milton Keynes | Wellcome Sanger Institute for the COVID-19 Genomics UK (COG-UK) Consortium | The Lighthouse Lab in Milton Keynes and Alex Alderton, Roberto Amato, Sonia Goncalves, Ewan Harrison, David K. Jackson, Ian Johnston, Dominic Kwiatkowski, Cordelia Langford, John Sillitoe on behalf of the Wellcome Sanger Institute COVID-19 Surveillance Team |
| EPI_ISL_769868, EPI_ISL_769870, EPI_ISL_769878 | Respiratory Virus Unit, National Infection Service, Public Health England | COVID-19 Genomics UK (COG-UK) Consortium | PHE Covid Sequencing Team |
| EPI_ISL_770791, EPI_ISL_770792 | Minnesota Department of Health, Public Health Laboratory | Minnesota Department of Health, Public Health Laboratory | Alexandra Lorentz, Jacob Garfin, Matt Plumb, and Xiong Wang |
| EPI_ISL_775020 | Gonoshasthya-RNA Molecular Research Center | Gonoshasthya-RNA Molecular Research Center | Mohd. Raeed Jamiruddin, Nihad Adnan, Md. Ahsanul Haq, Mohib Ullah Khondoker, Nafisa Azmuda, Firoz Ahmed, Shahana Sharmin, Salma Akter, Taslin Jahan Mou, Mahfuza Marzan, Sayeda Moriam Liza, Nowshin Jahan, Tamanna Ali, Shahad Saif Khandker, Maha Jamiruddin, Mousumi Chaity, Mumtarin Jannat Oishee |
| EPI_ISL_775213 | Gonoshasthya-RNA Molecular Research Center | Gonoshasthya-RNA Molecular Research Center | Nihad Adnan, Mohd. Raeed Jamiruddin, Md. Ahsanul Haq, Mohib Ullah Khondoker, Nafisa Azmuda, Firoz Ahmed, Shahana Sharmin, Salma Akter, Taslin Jahan Mou, Mahfuza Marzan, Sayeda Moriam Liza, Nowshin Jahan, Tamanna Ali, Maha Jamiruddin, Mousumi Chaity, Shahad Saif Khandker, Mumtarin Jannat Oishee |

| | | | |
|---|---|---|---|
| EPI_ISL_775293 | Oslo University Hospital, Department of Medical Microbiology | Norwegian Institute of Public Health, Department of Virology | Kathrine Stene-Johansen, Kamilla Heddeland Instefjord, Hilde Elshaug, Atiya R Ali,Marie Paulsen Madsen, Rasmus Riis Kopperud, Hilde Vollan, Karoline Bragstad, Olav Hungnes |
| EPI_ISL_775403 | Nordland Hospital - Bodo, Laboratory Department, Molecular Biology Unit | Norwegian Institute of Public Health, Department of Virology | Kathrine Stene-Johansen, Kamilla Heddeland Instefjord, Hilde Elshaug, Atiya R Ali,Marie Paulsen Madsen, Rasmus Riis Kopperud, Hilde Vollan, Karoline Bragstad, Olav Hungnes |
| EPI_ISL_775406, EPI_ISL_775463 | Unilabs Laboratory Medicine | Norwegian Institute of Public Health, Department of Virology | Kathrine Stene-Johansen, Kamilla Heddeland Instefjord, Hilde Elshaug, Atiya R Ali,Marie Paulsen Madsen, Rasmus Riis Kopperud, Hilde Vollan, Karoline Bragstad, Olav Hungnes |
| EPI_ISL_775505, EPI_ISL_775506, EPI_ISL_775507, EPI_ISL_775508 | Department of Medical Microbiology, St. Olavs hospital | Norwegian Institute of Public Health, Department of Virology | Kathrine Stene-Johansen, Kamilla Heddeland Instefjord, Hilde Elshaug, Atiya R Ali,Marie Paulsen Madsen, Rasmus Riis Kopperud, Hilde Vollan, Karoline Bragstad, Olav Hungnes |
| EPI_ISL_775516, EPI_ISL_775517 | Unilabs Laboratory Medicine | Norwegian Institute of Public Health, Department of Virology | Kathrine Stene-Johansen, Kamilla Heddeland Instefjord, Hilde Elshaug, Atiya R Ali,Marie Paulsen Madsen, Rasmus Riis Kopperud, Hilde Vollan, Karoline Bragstad, Olav Hungnes |
| EPI_ISL_775520 | Hospital of Southern Norway - Kristiansand, Department of Medical Microbiology | Norwegian Institute of Public Health, Department of Virology | Kathrine Stene-Johansen, Kamilla Heddeland Instefjord, Hilde Elshaug, Atiya R Ali,Marie Paulsen Madsen, Rasmus Riis Kopperud, Hilde Vollan, Karoline Bragstad, Olav Hungnes |
| EPI_ISL_775531, EPI_ISL_775544, EPI_ISL_775545 | Klinisk mikrobiologi, Viruslab | The Public Health Agency of Sweden | Department of Microbiology, The Public Health Agency of Sweden |
| EPI_ISL_775549, EPI_ISL_775592 | Klinisk mikrobiologi | The Public Health Agency of Sweden | Department of Microbiology, The Public Health Agency of Sweden |
| EPI_ISL_776696, EPI_ISL_776697, EPI_ISL_776698, EPI_ISL_776699, EPI_ISL_776700, EPI_ISL_776701, EPI_ISL_776702, EPI_ISL_776703, EPI_ISL_776704, EPI_ISL_776705, EPI_ISL_776730, EPI_ISL_776746 | | | |
| see above | UW Virology Lab | UW Virology Lab | Pavitra Roychoudhury, Hong Xie, Lasata Shrestha, Meei-Li Huang, Keith R Jerome, Alexander Greninger |
| EPI_ISL_777013 | Lighthouse Lab in Glasgow | Wellcome Sanger Institute for the COVID-19 Genomics UK (COG-UK) Consortium | Harper VanSteenhouse, Yumi Kasai, David Gray, Carol Clugston, Anna Dominiczak and Alex Alderton, Roberto Amato, Sonia Goncalves, Ewan Harrison, David K. Jackson, Ian Johnston, Dominic Kwiatkowski, Cordelia Langford, John Sillitoe on behalf of the Wellcome Sanger Institute COVID-19 Surveillance Team |
| EPI_ISL_777039 | Lighthouse Lab in Milton Keynes | Wellcome Sanger Institute for the COVID-19 Genomics UK (COG-UK) Consortium | The Lighthouse Lab in Milton Keynes and Alex Alderton, Roberto Amato, Sonia Goncalves, Ewan Harrison, David K. Jackson, Ian Johnston, Dominic Kwiatkowski, Cordelia Langford, John Sillitoe on behalf of the Wellcome Sanger Institute COVID-19 Surveillance Team |
| EPI_ISL_777081 | Lighthouse Lab in Glasgow | Wellcome Sanger Institute for the COVID-19 Genomics UK (COG-UK) Consortium | Harper VanSteenhouse, Yumi Kasai, David Gray, Carol Clugston, Anna Dominiczak and Alex Alderton, Roberto Amato, Sonia Goncalves, Ewan Harrison, David K. Jackson, Ian Johnston, Dominic Kwiatkowski, Cordelia Langford, John Sillitoe on behalf of the Wellcome Sanger Institute COVID-19 Surveillance Team |
| EPI_ISL_777085 | Lighthouse Lab in Milton Keynes | Wellcome Sanger Institute for the COVID-19 Genomics UK (COG-UK) Consortium | The Lighthouse Lab in Milton Keynes and Alex Alderton, Roberto Amato, Sonia Goncalves, Ewan Harrison, David K. Jackson, Ian Johnston, Dominic Kwiatkowski, Cordelia Langford, John Sillitoe on behalf of the Wellcome Sanger Institute COVID-19 Surveillance Team |
| EPI_ISL_777101 | Lighthouse Lab in Glasgow | Wellcome Sanger Institute for the COVID-19 Genomics UK (COG-UK) Consortium | Harper VanSteenhouse, Yumi Kasai, David Gray, Carol Clugston, Anna Dominiczak and Alex Alderton, Roberto Amato, Sonia Goncalves, Ewan Harrison, David K. Jackson, Ian Johnston, Dominic Kwiatkowski, Cordelia Langford, John Sillitoe on behalf of the Wellcome Sanger Institute COVID-19 Surveillance Team |
| EPI_ISL_777102, EPI_ISL_777108 | Lighthouse Lab in Milton Keynes | Wellcome Sanger Institute for the COVID-19 Genomics UK (COG-UK) Consortium | The Lighthouse Lab in Milton Keynes and Alex Alderton, Roberto Amato, Sonia Goncalves, Ewan Harrison, David K. Jackson, Ian Johnston, Dominic Kwiatkowski, Cordelia Langford, John Sillitoe on behalf of the Wellcome Sanger Institute COVID-19 Surveillance Team |
| EPI_ISL_777142 | Lighthouse Lab in Glasgow | Wellcome Sanger Institute for the COVID-19 Genomics UK (COG-UK) Consortium | Harper VanSteenhouse, Yumi Kasai, David Gray, Carol Clugston, Anna Dominiczak and Alex Alderton, Roberto Amato, Sonia Goncalves, Ewan Harrison, David K. Jackson, Ian Johnston, Dominic Kwiatkowski, Cordelia Langford, John Sillitoe on behalf of the Wellcome Sanger Institute COVID-19 Surveillance Team |
| EPI_ISL_777145 | Lighthouse Lab in Milton Keynes | Wellcome Sanger Institute for the COVID-19 Genomics UK (COG-UK) Consortium | The Lighthouse Lab in Milton Keynes and Alex Alderton, Roberto Amato, Sonia Goncalves, Ewan Harrison, David K. Jackson, Ian Johnston, Dominic Kwiatkowski, Cordelia Langford, John Sillitoe on behalf of the Wellcome Sanger Institute COVID-19 Surveillance Team |
| EPI_ISL_777169 | Lighthouse Lab in Alderley Park | Wellcome Sanger Institute for the COVID-19 Genomics UK (COG-UK) Consortium | Jacquelyn Wynn, Mairead Hyland, The Lighthouse Lab in Alderley Park and Alex Alderton, Roberto Amato, Sonia Goncalves, Ewan Harrison, David K. Jackson, Ian Johnston, Dominic Kwiatkowski, Cordelia Langford, John Sillitoe on behalf of the Wellcome Sanger Institute COVID-19 Surveillance Team |
| EPI_ISL_777192 | Lighthouse Lab in Glasgow | Wellcome Sanger Institute for the COVID-19 Genomics UK (COG-UK) Consortium | Harper VanSteenhouse, Yumi Kasai, David Gray, Carol Clugston, Anna Dominiczak and Alex Alderton, Roberto Amato, Sonia Goncalves, Ewan Harrison, David K. Jackson, Ian Johnston, Dominic Kwiatkowski, Cordelia Langford, John Sillitoe on behalf of the Wellcome Sanger Institute COVID-19 Surveillance Team |
| EPI_ISL_777201 | Lighthouse Lab in Cambridge | Wellcome Sanger Institute for the COVID-19 Genomics UK (COG-UK) Consortium | Rob Howes, The Lighthouse Lab in Cambridge and Alex Alderton, Roberto Amato, Sonia Goncalves, Ewan Harrison, David K. Jackson, Ian Johnston, Dominic Kwiatkowski, Cordelia Langford, John Sillitoe on behalf of the Wellcome Sanger Institute COVID-19 Surveillance Team |
| EPI_ISL_777295, EPI_ISL_777313 | Lighthouse Lab in Milton Keynes | Wellcome Sanger Institute for the COVID-19 Genomics UK (COG-UK) Consortium | The Lighthouse Lab in Milton Keynes and Alex Alderton, Roberto Amato, Sonia Goncalves, Ewan Harrison, David K. Jackson, Ian Johnston, Dominic Kwiatkowski, Cordelia Langford, John Sillitoe on behalf of the Wellcome Sanger Institute COVID-19 Surveillance Team |
| EPI_ISL_777323 | Lighthouse Lab in Cambridge | Wellcome Sanger Institute for the COVID-19 Genomics UK (COG-UK) Consortium | Rob Howes, The Lighthouse Lab in Cambridge and Alex Alderton, Roberto Amato, Sonia Goncalves, Ewan Harrison, David K. Jackson, Ian Johnston, Dominic Kwiatkowski, Cordelia Langford, John Sillitoe on behalf of the Wellcome Sanger Institute COVID-19 Surveillance Team |
| EPI_ISL_778837, EPI_ISL_778838 | AIID | Irish Coronavirus Sequencing Consortium-Teagasc Grange | Matthew McCabe, Aljandro Abner Garcia Leon, Fiona Crispie, Calum Walsh, Michael Carr, John Kenny, Paul Cotter, Patrick Mallon, Gabriel Gonzalez |
| EPI_ISL_779813 | CNR Virus des Infections Respiratoires - France SUD | CNR Virus des Infections Respiratoires - France SUD | Antonin Bal, Gregory Destras, Gwendolyne Burfin, Hadrien Règue, Quentin Semanas, Martine Valette, Bruno Lina, Laurence Josset |
| EPI_ISL_779837 | CHU Bordeaux | CNR Virus des Infections Respiratoires - France SUD | Antonin Bal, Gregory Destras, Gwendolyne Burfin, Hadrien Règue, Quentin Semanas, Martine Valette, Bruno Lina, Pantxika Bellecave, Camille Ciccone, Isabelle Garrigue, Marie-Edith Lafon, Pascale Trimoulet, Laurence Josset |
| EPI_ISL_779845 | CNR Virus des Infections Respiratoires - France SUD | CNR Virus des Infections Respiratoires - France SUD | Antonin Bal, Gregory Destras, Gwendolyne Burfin, Hadrien Règue, Quentin Semanas, Martine Valette, Bruno Lina, Laurence Josset |
| EPI_ISL_779862, EPI_ISL_779867, EPI_ISL_779870, EPI_ISL_779873, EPI_ISL_779876, EPI_ISL_779877, EPI_ISL_779887, EPI_ISL_779900, EPI_ISL_779903, EPI_ISL_779904, EPI_ISL_779920 | | | |
| see above | Servicio de Microbiología, Hospital Universitario Son Espases | SeqCOVID-SPAIN consortium/IBV(CSIC) | Carla López-Causapé, Jordi Reina, Antonio Oliver and SeqCOVID-SPAIN consortium |
| EPI_ISL_782486, EPI_ISL_782494, EPI_ISL_782500, EPI_ISL_782504, EPI_ISL_782512, EPI_ISL_782536, EPI_ISL_782538, EPI_ISL_782568, EPI_ISL_782580, EPI_ISL_782584, EPI_ISL_782596, EPI_ISL_782605, EPI_ISL_782618, EPI_ISL_782624, EPI_ISL_782631, EPI_ISL_782633, EPI_ISL_782674, EPI_ISL_782679, EPI_ISL_782689, EPI_ISL_782710 | | | |
| see above | Lighthouse Lab in Glasgow | Wellcome Sanger Institute for the COVID-19 Genomics UK (COG-UK) Consortium | Harper VanSteenhouse, Yumi Kasai, David Gray, Carol Clugston, Anna Dominiczak and Alex Alderton, Roberto Amato, Sonia Goncalves, Ewan Harrison, David K. Jackson, Ian Johnston, Dominic Kwiatkowski, Cordelia Langford, John Sillitoe on behalf of the Wellcome Sanger Institute COVID-19 Surveillance Team |
| EPI_ISL_782734, EPI_ISL_782735, EPI_ISL_782736, EPI_ISL_782738 | Lighthouse Lab in Milton Keynes | Wellcome Sanger Institute for the COVID-19 Genomics UK (COG-UK) Consortium | The Lighthouse Lab in Milton Keynes and Alex Alderton, Roberto Amato, Sonia Goncalves, Ewan Harrison, David K. Jackson, Ian Johnston, Dominic Kwiatkowski, Cordelia Langford, John Sillitoe on behalf of the Wellcome Sanger Institute COVID-19 Surveillance Team |
| EPI_ISL_782739 | Lighthouse Lab in Cambridge | Wellcome Sanger Institute for the COVID-19 Genomics UK (COG-UK) Consortium | Rob Howes, The Lighthouse Lab in Cambridge and Alex Alderton, Roberto Amato, Sonia Goncalves, Ewan Harrison, David K. Jackson, Ian Johnston, Dominic Kwiatkowski, Cordelia Langford, John Sillitoe on behalf of the Wellcome Sanger Institute COVID-19 Surveillance Team |
| EPI_ISL_782740, EPI_ISL_782741, EPI_ISL_782742, EPI_ISL_782743, EPI_ISL_782744 | Lighthouse Lab in Milton Keynes | Wellcome Sanger Institute for the COVID-19 Genomics UK (COG-UK) Consortium | The Lighthouse Lab in Milton Keynes and Alex Alderton, Roberto Amato, Sonia Goncalves, Ewan Harrison, David K. Jackson, Ian Johnston, Dominic Kwiatkowski, Cordelia Langford, John Sillitoe on behalf of the Wellcome Sanger Institute COVID-19 Surveillance Team |
| EPI_ISL_782746 | Lighthouse Lab in Cambridge | Wellcome Sanger Institute for the COVID-19 Genomics UK (COG-UK) Consortium | Rob Howes, The Lighthouse Lab in Cambridge and Alex Alderton, Roberto Amato, Sonia Goncalves, Ewan Harrison, David K. Jackson, Ian Johnston, Dominic Kwiatkowski, Cordelia Langford, John Sillitoe on behalf of the Wellcome Sanger Institute COVID-19 Surveillance Team |
| EPI_ISL_782749, EPI_ISL_782750, | Lighthouse Lab in Milton Keynes | Wellcome Sanger Institute for the COVID-19 Genomics UK | The Lighthouse Lab in Milton Keynes and Alex Alderton, Roberto Amato, Sonia Goncalves, Ewan Harrison, David K. Jackson, Ian Johnston, Dominic |

| | | | |
|---|---|---|---|
| EPI_ISL_782751, EPI_ISL_782752, EPI_ISL_782753, EPI_ISL_782754, EPI_ISL_782755, EPI_ISL_782756 | | (COG-UK) Consortium | Kwiatkowski, Cordelia Langford, John Sillitoe on behalf of the Wellcome Sanger Institute COVID-19 Surveillance Team |
| EPI_ISL_782757, EPI_ISL_782759 | Lighthouse Lab in Cambridge | Wellcome Sanger Institute for the COVID-19 Genomics UK (COG-UK) Consortium | Rob Howes, The Lighthouse Lab in Cambridge and Alex Alderton, Roberto Amato, Sonia Goncalves, Ewan Harrison, David K. Jackson, Ian Johnston, Dominic Kwiatkowski, Cordelia Langford, John Sillitoe on behalf of the Wellcome Sanger Institute COVID-19 Surveillance Team |
| EPI_ISL_782760 | Lighthouse Lab in Milton Keynes | Wellcome Sanger Institute for the COVID-19 Genomics UK (COG-UK) Consortium | The Lighthouse Lab in Milton Keynes and Alex Alderton, Roberto Amato, Sonia Goncalves, Ewan Harrison, David K. Jackson, Ian Johnston, Dominic Kwiatkowski, Cordelia Langford, John Sillitoe on behalf of the Wellcome Sanger Institute COVID-19 Surveillance Team |
| EPI_ISL_782761 | Lighthouse Lab in Cambridge | Wellcome Sanger Institute for the COVID-19 Genomics UK (COG-UK) Consortium | Rob Howes, The Lighthouse Lab in Cambridge and Alex Alderton, Roberto Amato, Sonia Goncalves, Ewan Harrison, David K. Jackson, Ian Johnston, Dominic Kwiatkowski, Cordelia Langford, John Sillitoe on behalf of the Wellcome Sanger Institute COVID-19 Surveillance Team |
| EPI_ISL_782762 | Lighthouse Lab in Milton Keynes | Wellcome Sanger Institute for the COVID-19 Genomics UK (COG-UK) Consortium | The Lighthouse Lab in Milton Keynes and Alex Alderton, Roberto Amato, Sonia Goncalves, Ewan Harrison, David K. Jackson, Ian Johnston, Dominic Kwiatkowski, Cordelia Langford, John Sillitoe on behalf of the Wellcome Sanger Institute COVID-19 Surveillance Team |
| EPI_ISL_782763, EPI_ISL_782764 | Lighthouse Lab in Cambridge | Wellcome Sanger Institute for the COVID-19 Genomics UK (COG-UK) Consortium | Rob Howes, The Lighthouse Lab in Cambridge and Alex Alderton, Roberto Amato, Sonia Goncalves, Ewan Harrison, David K. Jackson, Ian Johnston, Dominic Kwiatkowski, Cordelia Langford, John Sillitoe on behalf of the Wellcome Sanger Institute COVID-19 Surveillance Team |
| EPI_ISL_782765, EPI_ISL_782766, EPI_ISL_782767 | Lighthouse Lab in Milton Keynes | Wellcome Sanger Institute for the COVID-19 Genomics UK (COG-UK) Consortium | The Lighthouse Lab in Milton Keynes and Alex Alderton, Roberto Amato, Sonia Goncalves, Ewan Harrison, David K. Jackson, Ian Johnston, Dominic Kwiatkowski, Cordelia Langford, John Sillitoe on behalf of the Wellcome Sanger Institute COVID-19 Surveillance Team |
| EPI_ISL_782769 | Lighthouse Lab in Cambridge | Wellcome Sanger Institute for the COVID-19 Genomics UK (COG-UK) Consortium | Rob Howes, The Lighthouse Lab in Cambridge and Alex Alderton, Roberto Amato, Sonia Goncalves, Ewan Harrison, David K. Jackson, Ian Johnston, Dominic Kwiatkowski, Cordelia Langford, John Sillitoe on behalf of the Wellcome Sanger Institute COVID-19 Surveillance Team |
| EPI_ISL_782770, EPI_ISL_782772, EPI_ISL_782773, EPI_ISL_782775 | Lighthouse Lab in Milton Keynes | Wellcome Sanger Institute for the COVID-19 Genomics UK (COG-UK) Consortium | The Lighthouse Lab in Milton Keynes and Alex Alderton, Roberto Amato, Sonia Goncalves, Ewan Harrison, David K. Jackson, Ian Johnston, Dominic Kwiatkowski, Cordelia Langford, John Sillitoe on behalf of the Wellcome Sanger Institute COVID-19 Surveillance Team |
| EPI_ISL_782776, EPI_ISL_782777 | Lighthouse Lab in Cambridge | Wellcome Sanger Institute for the COVID-19 Genomics UK (COG-UK) Consortium | Rob Howes, The Lighthouse Lab in Cambridge and Alex Alderton, Roberto Amato, Sonia Goncalves, Ewan Harrison, David K. Jackson, Ian Johnston, Dominic Kwiatkowski, Cordelia Langford, John Sillitoe on behalf of the Wellcome Sanger Institute COVID-19 Surveillance Team |
| EPI_ISL_782778, EPI_ISL_782779 | Lighthouse Lab in Milton Keynes | Wellcome Sanger Institute for the COVID-19 Genomics UK (COG-UK) Consortium | The Lighthouse Lab in Milton Keynes and Alex Alderton, Roberto Amato, Sonia Goncalves, Ewan Harrison, David K. Jackson, Ian Johnston, Dominic Kwiatkowski, Cordelia Langford, John Sillitoe on behalf of the Wellcome Sanger Institute COVID-19 Surveillance Team |
| EPI_ISL_782780 | Lighthouse Lab in Cambridge | Wellcome Sanger Institute for the COVID-19 Genomics UK (COG-UK) Consortium | Rob Howes, The Lighthouse Lab in Cambridge and Alex Alderton, Roberto Amato, Sonia Goncalves, Ewan Harrison, David K. Jackson, Ian Johnston, Dominic Kwiatkowski, Cordelia Langford, John Sillitoe on behalf of the Wellcome Sanger Institute COVID-19 Surveillance Team |
| EPI_ISL_782781, EPI_ISL_782782, EPI_ISL_782785, EPI_ISL_782786 | Lighthouse Lab in Milton Keynes | Wellcome Sanger Institute for the COVID-19 Genomics UK (COG-UK) Consortium | The Lighthouse Lab in Milton Keynes and Alex Alderton, Roberto Amato, Sonia Goncalves, Ewan Harrison, David K. Jackson, Ian Johnston, Dominic Kwiatkowski, Cordelia Langford, John Sillitoe on behalf of the Wellcome Sanger Institute COVID-19 Surveillance Team |
| EPI_ISL_782788 | Lighthouse Lab in Cambridge | Wellcome Sanger Institute for the COVID-19 Genomics UK (COG-UK) Consortium | Rob Howes, The Lighthouse Lab in Cambridge and Alex Alderton, Roberto Amato, Sonia Goncalves, Ewan Harrison, David K. Jackson, Ian Johnston, Dominic Kwiatkowski, Cordelia Langford, John Sillitoe on behalf of the Wellcome Sanger Institute COVID-19 Surveillance Team |
| EPI_ISL_782789, EPI_ISL_782790, EPI_ISL_782793, EPI_ISL_782794 | Lighthouse Lab in Milton Keynes | Wellcome Sanger Institute for the COVID-19 Genomics UK (COG-UK) Consortium | The Lighthouse Lab in Milton Keynes and Alex Alderton, Roberto Amato, Sonia Goncalves, Ewan Harrison, David K. Jackson, Ian Johnston, Dominic Kwiatkowski, Cordelia Langford, John Sillitoe on behalf of the Wellcome Sanger Institute COVID-19 Surveillance Team |
| EPI_ISL_782797 | Lighthouse Lab in Cambridge | Wellcome Sanger Institute for the COVID-19 Genomics UK (COG-UK) Consortium | Rob Howes, The Lighthouse Lab in Cambridge and Alex Alderton, Roberto Amato, Sonia Goncalves, Ewan Harrison, David K. Jackson, Ian Johnston, Dominic Kwiatkowski, Cordelia Langford, John Sillitoe on behalf of the Wellcome Sanger Institute COVID-19 Surveillance Team |
| EPI_ISL_782798, EPI_ISL_782799 | Lighthouse Lab in Milton Keynes | Wellcome Sanger Institute for the COVID-19 Genomics UK (COG-UK) Consortium | The Lighthouse Lab in Milton Keynes and Alex Alderton, Roberto Amato, Sonia Goncalves, Ewan Harrison, David K. Jackson, Ian Johnston, Dominic Kwiatkowski, Cordelia Langford, John Sillitoe on behalf of the Wellcome Sanger Institute COVID-19 Surveillance Team |
| EPI_ISL_782800 | Lighthouse Lab in Cambridge | Wellcome Sanger Institute for the COVID-19 Genomics UK (COG-UK) Consortium | Rob Howes, The Lighthouse Lab in Cambridge and Alex Alderton, Roberto Amato, Sonia Goncalves, Ewan Harrison, David K. Jackson, Ian Johnston, Dominic Kwiatkowski, Cordelia Langford, John Sillitoe on behalf of the Wellcome Sanger Institute COVID-19 Surveillance Team |
| EPI_ISL_782802 | Lighthouse Lab in Milton Keynes | Wellcome Sanger Institute for the COVID-19 Genomics UK (COG-UK) Consortium | The Lighthouse Lab in Milton Keynes and Alex Alderton, Roberto Amato, Sonia Goncalves, Ewan Harrison, David K. Jackson, Ian Johnston, Dominic Kwiatkowski, Cordelia Langford, John Sillitoe on behalf of the Wellcome Sanger Institute COVID-19 Surveillance Team |
| EPI_ISL_782803 | Lighthouse Lab in Cambridge | Wellcome Sanger Institute for the COVID-19 Genomics UK (COG-UK) Consortium | Rob Howes, The Lighthouse Lab in Cambridge and Alex Alderton, Roberto Amato, Sonia Goncalves, Ewan Harrison, David K. Jackson, Ian Johnston, Dominic Kwiatkowski, Cordelia Langford, John Sillitoe on behalf of the Wellcome Sanger Institute COVID-19 Surveillance Team |
| EPI_ISL_782804, EPI_ISL_782807 | Lighthouse Lab in Milton Keynes | Wellcome Sanger Institute for the COVID-19 Genomics UK (COG-UK) Consortium | The Lighthouse Lab in Milton Keynes and Alex Alderton, Roberto Amato, Sonia Goncalves, Ewan Harrison, David K. Jackson, Ian Johnston, Dominic Kwiatkowski, Cordelia Langford, John Sillitoe on behalf of the Wellcome Sanger Institute COVID-19 Surveillance Team |
| EPI_ISL_782809, EPI_ISL_782810 | Lighthouse Lab in Cambridge | Wellcome Sanger Institute for the COVID-19 Genomics UK (COG-UK) Consortium | Rob Howes, The Lighthouse Lab in Cambridge and Alex Alderton, Roberto Amato, Sonia Goncalves, Ewan Harrison, David K. Jackson, Ian Johnston, Dominic Kwiatkowski, Cordelia Langford, John Sillitoe on behalf of the Wellcome Sanger Institute COVID-19 Surveillance Team |
| EPI_ISL_782811, EPI_ISL_782812 | Lighthouse Lab in Milton Keynes | Wellcome Sanger Institute for the COVID-19 Genomics UK (COG-UK) Consortium | The Lighthouse Lab in Milton Keynes and Alex Alderton, Roberto Amato, Sonia Goncalves, Ewan Harrison, David K. Jackson, Ian Johnston, Dominic Kwiatkowski, Cordelia Langford, John Sillitoe on behalf of the Wellcome Sanger Institute COVID-19 Surveillance Team |
| EPI_ISL_782814 | Lighthouse Lab in Cambridge | Wellcome Sanger Institute for the COVID-19 Genomics UK (COG-UK) Consortium | Rob Howes, The Lighthouse Lab in Cambridge and Alex Alderton, Roberto Amato, Sonia Goncalves, Ewan Harrison, David K. Jackson, Ian Johnston, Dominic Kwiatkowski, Cordelia Langford, John Sillitoe on behalf of the Wellcome Sanger Institute COVID-19 Surveillance Team |
| EPI_ISL_782815 | Lighthouse Lab in Milton Keynes | Wellcome Sanger Institute for the COVID-19 Genomics UK (COG-UK) Consortium | The Lighthouse Lab in Milton Keynes and Alex Alderton, Roberto Amato, Sonia Goncalves, Ewan Harrison, David K. Jackson, Ian Johnston, Dominic Kwiatkowski, Cordelia Langford, John Sillitoe on behalf of the Wellcome Sanger Institute COVID-19 Surveillance Team |
| EPI_ISL_782816, EPI_ISL_782817, EPI_ISL_782818 | Lighthouse Lab in Cambridge | Wellcome Sanger Institute for the COVID-19 Genomics UK (COG-UK) Consortium | Rob Howes, The Lighthouse Lab in Cambridge and Alex Alderton, Roberto Amato, Sonia Goncalves, Ewan Harrison, David K. Jackson, Ian Johnston, Dominic Kwiatkowski, Cordelia Langford, John Sillitoe on behalf of the Wellcome Sanger Institute COVID-19 Surveillance Team |
| EPI_ISL_782819, EPI_ISL_782821 | Lighthouse Lab in Milton Keynes | Wellcome Sanger Institute for the COVID-19 Genomics UK (COG-UK) Consortium | The Lighthouse Lab in Milton Keynes and Alex Alderton, Roberto Amato, Sonia Goncalves, Ewan Harrison, David K. Jackson, Ian Johnston, Dominic Kwiatkowski, Cordelia Langford, John Sillitoe on behalf of the Wellcome Sanger Institute COVID-19 Surveillance Team |
| EPI_ISL_782822 | Lighthouse Lab in Cambridge | Wellcome Sanger Institute for the COVID-19 Genomics UK (COG-UK) Consortium | Rob Howes, The Lighthouse Lab in Cambridge and Alex Alderton, Roberto Amato, Sonia Goncalves, Ewan Harrison, David K. Jackson, Ian Johnston, Dominic Kwiatkowski, Cordelia Langford, John Sillitoe on behalf of the Wellcome Sanger Institute COVID-19 Surveillance Team |
| EPI_ISL_782823, EPI_ISL_782824, EPI_ISL_782826, EPI_ISL_782828, EPI_ISL_782829 | Lighthouse Lab in Milton Keynes | Wellcome Sanger Institute for the COVID-19 Genomics UK (COG-UK) Consortium | The Lighthouse Lab in Milton Keynes and Alex Alderton, Roberto Amato, Sonia Goncalves, Ewan Harrison, David K. Jackson, Ian Johnston, Dominic Kwiatkowski, Cordelia Langford, John Sillitoe on behalf of the Wellcome Sanger Institute COVID-19 Surveillance Team |
| EPI_ISL_782830 | Lighthouse Lab in Cambridge | Wellcome Sanger Institute for the COVID-19 Genomics UK (COG-UK) Consortium | Rob Howes, The Lighthouse Lab in Cambridge and Alex Alderton, Roberto Amato, Sonia Goncalves, Ewan Harrison, David K. Jackson, Ian Johnston, Dominic Kwiatkowski, Cordelia Langford, John Sillitoe on behalf of the Wellcome Sanger Institute COVID-19 Surveillance Team |
| EPI_ISL_782831, EPI_ISL_782833 | Lighthouse Lab in Milton Keynes | Wellcome Sanger Institute for the COVID-19 Genomics UK (COG-UK) Consortium | The Lighthouse Lab in Milton Keynes and Alex Alderton, Roberto Amato, Sonia Goncalves, Ewan Harrison, David K. Jackson, Ian Johnston, Dominic Kwiatkowski, Cordelia Langford, John Sillitoe on behalf of the Wellcome Sanger Institute COVID-19 Surveillance Team |
| EPI_ISL_782834 | Lighthouse Lab in Cambridge | Wellcome Sanger Institute for the COVID-19 Genomics UK (COG-UK) Consortium | Rob Howes, The Lighthouse Lab in Cambridge and Alex Alderton, Roberto Amato, Sonia Goncalves, Ewan Harrison, David K. Jackson, Ian Johnston, Dominic Kwiatkowski, Cordelia Langford, John Sillitoe on behalf of the Wellcome Sanger Institute COVID-19 Surveillance Team |
| EPI_ISL_782835, EPI_ISL_782836, EPI_ISL_782837, EPI_ISL_782839 | Lighthouse Lab in Milton Keynes | Wellcome Sanger Institute for the COVID-19 Genomics UK (COG-UK) Consortium | The Lighthouse Lab in Milton Keynes and Alex Alderton, Roberto Amato, Sonia Goncalves, Ewan Harrison, David K. Jackson, Ian Johnston, Dominic Kwiatkowski, Cordelia Langford, John Sillitoe on behalf of the Wellcome Sanger Institute COVID-19 Surveillance Team |
| EPI_ISL_782840, EPI_ISL_782842 | Lighthouse Lab in Cambridge | Wellcome Sanger Institute for the COVID-19 Genomics UK (COG-UK) Consortium | Rob Howes, The Lighthouse Lab in Cambridge and Alex Alderton, Roberto Amato, Sonia Goncalves, Ewan Harrison, David K. Jackson, Ian Johnston, Dominic Kwiatkowski, Cordelia Langford, John Sillitoe on behalf of the Wellcome Sanger Institute COVID-19 Surveillance Team |
| EPI_ISL_782845 | Lighthouse Lab in Milton Keynes | Wellcome Sanger Institute for the COVID-19 Genomics UK | The Lighthouse Lab in Milton Keynes and Alex Alderton, Roberto Amato, Sonia Goncalves, Ewan Harrison, David K. Jackson, Ian Johnston, Dominic |

| | | | |
|---|---|---|---|
| | | (COG-UK) Consortium | Kwiatkowski, Cordelia Langford, John Sillitoe on behalf of the Wellcome Sanger Institute COVID-19 Surveillance Team |
| EPI_ISL_782846 | Lighthouse Lab in Cambridge | Wellcome Sanger Institute for the COVID-19 Genomics UK (COG-UK) Consortium | Rob Howes, The Lighthouse Lab in Cambridge and Alex Alderton, Roberto Amato, Sonia Goncalves, Ewan Harrison, David K. Jackson, Ian Johnston, Dominic Kwiatkowski, Cordelia Langford, John Sillitoe on behalf of the Wellcome Sanger Institute COVID-19 Surveillance Team |
| EPI_ISL_782847, EPI_ISL_782848 | Lighthouse Lab in Milton Keynes | Wellcome Sanger Institute for the COVID-19 Genomics UK (COG-UK) Consortium | The Lighthouse Lab in Milton Keynes and Alex Alderton, Roberto Amato, Sonia Goncalves, Ewan Harrison, David K. Jackson, Ian Johnston, Dominic Kwiatkowski, Cordelia Langford, John Sillitoe on behalf of the Wellcome Sanger Institute COVID-19 Surveillance Team |
| EPI_ISL_782850 | Lighthouse Lab in Cambridge | Wellcome Sanger Institute for the COVID-19 Genomics UK (COG-UK) Consortium | Rob Howes, The Lighthouse Lab in Cambridge and Alex Alderton, Roberto Amato, Sonia Goncalves, Ewan Harrison, David K. Jackson, Ian Johnston, Dominic Kwiatkowski, Cordelia Langford, John Sillitoe on behalf of the Wellcome Sanger Institute COVID-19 Surveillance Team |
| EPI_ISL_782851, EPI_ISL_782852 | Lighthouse Lab in Milton Keynes | Wellcome Sanger Institute for the COVID-19 Genomics UK (COG-UK) Consortium | The Lighthouse Lab in Milton Keynes and Alex Alderton, Roberto Amato, Sonia Goncalves, Ewan Harrison, David K. Jackson, Ian Johnston, Dominic Kwiatkowski, Cordelia Langford, John Sillitoe on behalf of the Wellcome Sanger Institute COVID-19 Surveillance Team |
| EPI_ISL_782853, EPI_ISL_782854, EPI_ISL_782857 | Lighthouse Lab in Cambridge | Wellcome Sanger Institute for the COVID-19 Genomics UK (COG-UK) Consortium | Rob Howes, The Lighthouse Lab in Cambridge and Alex Alderton, Roberto Amato, Sonia Goncalves, Ewan Harrison, David K. Jackson, Ian Johnston, Dominic Kwiatkowski, Cordelia Langford, John Sillitoe on behalf of the Wellcome Sanger Institute COVID-19 Surveillance Team |
| EPI_ISL_782858, EPI_ISL_782859 | Lighthouse Lab in Milton Keynes | Wellcome Sanger Institute for the COVID-19 Genomics UK (COG-UK) Consortium | The Lighthouse Lab in Milton Keynes and Alex Alderton, Roberto Amato, Sonia Goncalves, Ewan Harrison, David K. Jackson, Ian Johnston, Dominic Kwiatkowski, Cordelia Langford, John Sillitoe on behalf of the Wellcome Sanger Institute COVID-19 Surveillance Team |
| EPI_ISL_782860 | Lighthouse Lab in Cambridge | Wellcome Sanger Institute for the COVID-19 Genomics UK (COG-UK) Consortium | Rob Howes, The Lighthouse Lab in Cambridge and Alex Alderton, Roberto Amato, Sonia Goncalves, Ewan Harrison, David K. Jackson, Ian Johnston, Dominic Kwiatkowski, Cordelia Langford, John Sillitoe on behalf of the Wellcome Sanger Institute COVID-19 Surveillance Team |
| EPI_ISL_782861 | Lighthouse Lab in Milton Keynes | Wellcome Sanger Institute for the COVID-19 Genomics UK (COG-UK) Consortium | The Lighthouse Lab in Milton Keynes and Alex Alderton, Roberto Amato, Sonia Goncalves, Ewan Harrison, David K. Jackson, Ian Johnston, Dominic Kwiatkowski, Cordelia Langford, John Sillitoe on behalf of the Wellcome Sanger Institute COVID-19 Surveillance Team |
| EPI_ISL_782862 | Lighthouse Lab in Cambridge | Wellcome Sanger Institute for the COVID-19 Genomics UK (COG-UK) Consortium | Rob Howes, The Lighthouse Lab in Cambridge and Alex Alderton, Roberto Amato, Sonia Goncalves, Ewan Harrison, David K. Jackson, Ian Johnston, Dominic Kwiatkowski, Cordelia Langford, John Sillitoe on behalf of the Wellcome Sanger Institute COVID-19 Surveillance Team |
| EPI_ISL_782863, EPI_ISL_782864, EPI_ISL_782865 | Lighthouse Lab in Milton Keynes | Wellcome Sanger Institute for the COVID-19 Genomics UK (COG-UK) Consortium | The Lighthouse Lab in Milton Keynes and Alex Alderton, Roberto Amato, Sonia Goncalves, Ewan Harrison, David K. Jackson, Ian Johnston, Dominic Kwiatkowski, Cordelia Langford, John Sillitoe on behalf of the Wellcome Sanger Institute COVID-19 Surveillance Team |
| EPI_ISL_782866, EPI_ISL_782867, EPI_ISL_782868 | Lighthouse Lab in Cambridge | Wellcome Sanger Institute for the COVID-19 Genomics UK (COG-UK) Consortium | Rob Howes, The Lighthouse Lab in Cambridge and Alex Alderton, Roberto Amato, Sonia Goncalves, Ewan Harrison, David K. Jackson, Ian Johnston, Dominic Kwiatkowski, Cordelia Langford, John Sillitoe on behalf of the Wellcome Sanger Institute COVID-19 Surveillance Team |
| EPI_ISL_782870, EPI_ISL_782871, EPI_ISL_782874, EPI_ISL_782875, EPI_ISL_782876, EPI_ISL_782879 | Lighthouse Lab in Milton Keynes | Wellcome Sanger Institute for the COVID-19 Genomics UK (COG-UK) Consortium | The Lighthouse Lab in Milton Keynes and Alex Alderton, Roberto Amato, Sonia Goncalves, Ewan Harrison, David K. Jackson, Ian Johnston, Dominic Kwiatkowski, Cordelia Langford, John Sillitoe on behalf of the Wellcome Sanger Institute COVID-19 Surveillance Team |
| EPI_ISL_782880 | Lighthouse Lab in Cambridge | Wellcome Sanger Institute for the COVID-19 Genomics UK (COG-UK) Consortium | Rob Howes, The Lighthouse Lab in Cambridge and Alex Alderton, Roberto Amato, Sonia Goncalves, Ewan Harrison, David K. Jackson, Ian Johnston, Dominic Kwiatkowski, Cordelia Langford, John Sillitoe on behalf of the Wellcome Sanger Institute COVID-19 Surveillance Team |
| EPI_ISL_782881 | Lighthouse Lab in Milton Keynes | Wellcome Sanger Institute for the COVID-19 Genomics UK (COG-UK) Consortium | The Lighthouse Lab in Milton Keynes and Alex Alderton, Roberto Amato, Sonia Goncalves, Ewan Harrison, David K. Jackson, Ian Johnston, Dominic Kwiatkowski, Cordelia Langford, John Sillitoe on behalf of the Wellcome Sanger Institute COVID-19 Surveillance Team |
| EPI_ISL_782882, EPI_ISL_782883 | Lighthouse Lab in Cambridge | Wellcome Sanger Institute for the COVID-19 Genomics UK (COG-UK) Consortium | Rob Howes, The Lighthouse Lab in Cambridge and Alex Alderton, Roberto Amato, Sonia Goncalves, Ewan Harrison, David K. Jackson, Ian Johnston, Dominic Kwiatkowski, Cordelia Langford, John Sillitoe on behalf of the Wellcome Sanger Institute COVID-19 Surveillance Team |
| EPI_ISL_782884 | Lighthouse Lab in Milton Keynes | Wellcome Sanger Institute for the COVID-19 Genomics UK (COG-UK) Consortium | The Lighthouse Lab in Milton Keynes and Alex Alderton, Roberto Amato, Sonia Goncalves, Ewan Harrison, David K. Jackson, Ian Johnston, Dominic Kwiatkowski, Cordelia Langford, John Sillitoe on behalf of the Wellcome Sanger Institute COVID-19 Surveillance Team |
| EPI_ISL_782885, EPI_ISL_782886, EPI_ISL_782887 | Lighthouse Lab in Cambridge | Wellcome Sanger Institute for the COVID-19 Genomics UK (COG-UK) Consortium | Rob Howes, The Lighthouse Lab in Cambridge and Alex Alderton, Roberto Amato, Sonia Goncalves, Ewan Harrison, David K. Jackson, Ian Johnston, Dominic Kwiatkowski, Cordelia Langford, John Sillitoe on behalf of the Wellcome Sanger Institute COVID-19 Surveillance Team |
| EPI_ISL_782888 | Lighthouse Lab in Milton Keynes | Wellcome Sanger Institute for the COVID-19 Genomics UK (COG-UK) Consortium | The Lighthouse Lab in Milton Keynes and Alex Alderton, Roberto Amato, Sonia Goncalves, Ewan Harrison, David K. Jackson, Ian Johnston, Dominic Kwiatkowski, Cordelia Langford, John Sillitoe on behalf of the Wellcome Sanger Institute COVID-19 Surveillance Team |
| EPI_ISL_782889, EPI_ISL_782890 | Lighthouse Lab in Cambridge | Wellcome Sanger Institute for the COVID-19 Genomics UK (COG-UK) Consortium | Rob Howes, The Lighthouse Lab in Cambridge and Alex Alderton, Roberto Amato, Sonia Goncalves, Ewan Harrison, David K. Jackson, Ian Johnston, Dominic Kwiatkowski, Cordelia Langford, John Sillitoe on behalf of the Wellcome Sanger Institute COVID-19 Surveillance Team |
| EPI_ISL_782891, EPI_ISL_782895, EPI_ISL_782896, EPI_ISL_782898, EPI_ISL_782900, EPI_ISL_782901, EPI_ISL_782906 | Lighthouse Lab in Milton Keynes | Wellcome Sanger Institute for the COVID-19 Genomics UK (COG-UK) Consortium | The Lighthouse Lab in Milton Keynes and Alex Alderton, Roberto Amato, Sonia Goncalves, Ewan Harrison, David K. Jackson, Ian Johnston, Dominic Kwiatkowski, Cordelia Langford, John Sillitoe on behalf of the Wellcome Sanger Institute COVID-19 Surveillance Team |
| EPI_ISL_782907, EPI_ISL_782908, EPI_ISL_782909, EPI_ISL_782910, EPI_ISL_782912, EPI_ISL_782913 | Lighthouse Lab in Cambridge | Wellcome Sanger Institute for the COVID-19 Genomics UK (COG-UK) Consortium | Rob Howes, The Lighthouse Lab in Cambridge and Alex Alderton, Roberto Amato, Sonia Goncalves, Ewan Harrison, David K. Jackson, Ian Johnston, Dominic Kwiatkowski, Cordelia Langford, John Sillitoe on behalf of the Wellcome Sanger Institute COVID-19 Surveillance Team |
| EPI_ISL_782914, EPI_ISL_782915, EPI_ISL_782916, EPI_ISL_782917 | Lighthouse Lab in Milton Keynes | Wellcome Sanger Institute for the COVID-19 Genomics UK (COG-UK) Consortium | The Lighthouse Lab in Milton Keynes and Alex Alderton, Roberto Amato, Sonia Goncalves, Ewan Harrison, David K. Jackson, Ian Johnston, Dominic Kwiatkowski, Cordelia Langford, John Sillitoe on behalf of the Wellcome Sanger Institute COVID-19 Surveillance Team |
| EPI_ISL_782920 | Lighthouse Lab in Cambridge | Wellcome Sanger Institute for the COVID-19 Genomics UK (COG-UK) Consortium | Rob Howes, The Lighthouse Lab in Cambridge and Alex Alderton, Roberto Amato, Sonia Goncalves, Ewan Harrison, David K. Jackson, Ian Johnston, Dominic Kwiatkowski, Cordelia Langford, John Sillitoe on behalf of the Wellcome Sanger Institute COVID-19 Surveillance Team |
| EPI_ISL_782921, EPI_ISL_782922 | Lighthouse Lab in Milton Keynes | Wellcome Sanger Institute for the COVID-19 Genomics UK (COG-UK) Consortium | The Lighthouse Lab in Milton Keynes and Alex Alderton, Roberto Amato, Sonia Goncalves, Ewan Harrison, David K. Jackson, Ian Johnston, Dominic Kwiatkowski, Cordelia Langford, John Sillitoe on behalf of the Wellcome Sanger Institute COVID-19 Surveillance Team |
| EPI_ISL_782923 | Lighthouse Lab in Cambridge | Wellcome Sanger Institute for the COVID-19 Genomics UK (COG-UK) Consortium | Rob Howes, The Lighthouse Lab in Cambridge and Alex Alderton, Roberto Amato, Sonia Goncalves, Ewan Harrison, David K. Jackson, Ian Johnston, Dominic Kwiatkowski, Cordelia Langford, John Sillitoe on behalf of the Wellcome Sanger Institute COVID-19 Surveillance Team |
| EPI_ISL_782925, EPI_ISL_782927 | Lighthouse Lab in Milton Keynes | Wellcome Sanger Institute for the COVID-19 Genomics UK (COG-UK) Consortium | The Lighthouse Lab in Milton Keynes and Alex Alderton, Roberto Amato, Sonia Goncalves, Ewan Harrison, David K. Jackson, Ian Johnston, Dominic Kwiatkowski, Cordelia Langford, John Sillitoe on behalf of the Wellcome Sanger Institute COVID-19 Surveillance Team |
| EPI_ISL_782928, EPI_ISL_782930, EPI_ISL_782932 | Lighthouse Lab in Cambridge | Wellcome Sanger Institute for the COVID-19 Genomics UK (COG-UK) Consortium | Rob Howes, The Lighthouse Lab in Cambridge and Alex Alderton, Roberto Amato, Sonia Goncalves, Ewan Harrison, David K. Jackson, Ian Johnston, Dominic Kwiatkowski, Cordelia Langford, John Sillitoe on behalf of the Wellcome Sanger Institute COVID-19 Surveillance Team |
| EPI_ISL_782933, EPI_ISL_782934, EPI_ISL_782935 | Lighthouse Lab in Milton Keynes | Wellcome Sanger Institute for the COVID-19 Genomics UK (COG-UK) Consortium | The Lighthouse Lab in Milton Keynes and Alex Alderton, Roberto Amato, Sonia Goncalves, Ewan Harrison, David K. Jackson, Ian Johnston, Dominic Kwiatkowski, Cordelia Langford, John Sillitoe on behalf of the Wellcome Sanger Institute COVID-19 Surveillance Team |
| EPI_ISL_782937 | Lighthouse Lab in Cambridge | Wellcome Sanger Institute for the COVID-19 Genomics UK (COG-UK) Consortium | Rob Howes, The Lighthouse Lab in Cambridge and Alex Alderton, Roberto Amato, Sonia Goncalves, Ewan Harrison, David K. Jackson, Ian Johnston, Dominic Kwiatkowski, Cordelia Langford, John Sillitoe on behalf of the Wellcome Sanger Institute COVID-19 Surveillance Team |
| EPI_ISL_782938, EPI_ISL_782939, EPI_ISL_782940 | Lighthouse Lab in Milton Keynes | Wellcome Sanger Institute for the COVID-19 Genomics UK (COG-UK) Consortium | The Lighthouse Lab in Milton Keynes and Alex Alderton, Roberto Amato, Sonia Goncalves, Ewan Harrison, David K. Jackson, Ian Johnston, Dominic Kwiatkowski, Cordelia Langford, John Sillitoe on behalf of the Wellcome Sanger Institute COVID-19 Surveillance Team |
| EPI_ISL_782941, EPI_ISL_782942 | Lighthouse Lab in Cambridge | Wellcome Sanger Institute for the COVID-19 Genomics UK (COG-UK) Consortium | Rob Howes, The Lighthouse Lab in Cambridge and Alex Alderton, Roberto Amato, Sonia Goncalves, Ewan Harrison, David K. Jackson, Ian Johnston, Dominic Kwiatkowski, Cordelia Langford, John Sillitoe on behalf of the Wellcome Sanger Institute COVID-19 Surveillance Team |
| EPI_ISL_782943, EPI_ISL_782944 | Lighthouse Lab in Milton Keynes | Wellcome Sanger Institute for the COVID-19 Genomics UK (COG-UK) Consortium | The Lighthouse Lab in Milton Keynes and Alex Alderton, Roberto Amato, Sonia Goncalves, Ewan Harrison, David K. Jackson, Ian Johnston, Dominic Kwiatkowski, Cordelia Langford, John Sillitoe on behalf of the Wellcome Sanger Institute COVID-19 Surveillance Team |
| EPI_ISL_782947 | Lighthouse Lab in Cambridge | Wellcome Sanger Institute for the COVID-19 Genomics UK (COG-UK) Consortium | Rob Howes, The Lighthouse Lab in Cambridge and Alex Alderton, Roberto Amato, Sonia Goncalves, Ewan Harrison, David K. Jackson, Ian Johnston, Dominic Kwiatkowski, Cordelia Langford, John Sillitoe on behalf of the Wellcome Sanger Institute COVID-19 Surveillance Team |

| | | | |
|---|---|---|---|
| EPI_ISL_782948, EPI_ISL_782949, EPI_ISL_782950, EPI_ISL_782951, EPI_ISL_782952, EPI_ISL_782953 | Lighthouse Lab in Milton Keynes | Wellcome Sanger Institute for the COVID-19 Genomics UK (COG-UK) Consortium | The Lighthouse Lab in Milton Keynes and Alex Alderton, Roberto Amato, Sonia Goncalves, Ewan Harrison, David K. Jackson, Ian Johnston, Dominic Kwiatkowski, Cordelia Langford, John Sillitoe on behalf of the Wellcome Sanger Institute COVID-19 Surveillance Team |
| EPI_ISL_782954 | Lighthouse Lab in Cambridge | Wellcome Sanger Institute for the COVID-19 Genomics UK (COG-UK) Consortium | Rob Howes, The Lighthouse Lab in Cambridge and Alex Alderton, Roberto Amato, Sonia Goncalves, Ewan Harrison, David K. Jackson, Ian Johnston, Dominic Kwiatkowski, Cordelia Langford, John Sillitoe on behalf of the Wellcome Sanger Institute COVID-19 Surveillance Team |
| EPI_ISL_782955, EPI_ISL_782957, EPI_ISL_782958 | Lighthouse Lab in Milton Keynes | Wellcome Sanger Institute for the COVID-19 Genomics UK (COG-UK) Consortium | The Lighthouse Lab in Milton Keynes and Alex Alderton, Roberto Amato, Sonia Goncalves, Ewan Harrison, David K. Jackson, Ian Johnston, Dominic Kwiatkowski, Cordelia Langford, John Sillitoe on behalf of the Wellcome Sanger Institute COVID-19 Surveillance Team |
| EPI_ISL_782959, EPI_ISL_782960 | Lighthouse Lab in Cambridge | Wellcome Sanger Institute for the COVID-19 Genomics UK (COG-UK) Consortium | Rob Howes, The Lighthouse Lab in Cambridge and Alex Alderton, Roberto Amato, Sonia Goncalves, Ewan Harrison, David K. Jackson, Ian Johnston, Dominic Kwiatkowski, Cordelia Langford, John Sillitoe on behalf of the Wellcome Sanger Institute COVID-19 Surveillance Team |
| EPI_ISL_782961, EPI_ISL_782962, EPI_ISL_782965 | Lighthouse Lab in Milton Keynes | Wellcome Sanger Institute for the COVID-19 Genomics UK (COG-UK) Consortium | The Lighthouse Lab in Milton Keynes and Alex Alderton, Roberto Amato, Sonia Goncalves, Ewan Harrison, David K. Jackson, Ian Johnston, Dominic Kwiatkowski, Cordelia Langford, John Sillitoe on behalf of the Wellcome Sanger Institute COVID-19 Surveillance Team |
| EPI_ISL_782966 | Lighthouse Lab in Cambridge | Wellcome Sanger Institute for the COVID-19 Genomics UK (COG-UK) Consortium | Rob Howes, The Lighthouse Lab in Cambridge and Alex Alderton, Roberto Amato, Sonia Goncalves, Ewan Harrison, David K. Jackson, Ian Johnston, Dominic Kwiatkowski, Cordelia Langford, John Sillitoe on behalf of the Wellcome Sanger Institute COVID-19 Surveillance Team |
| EPI_ISL_782968 | Lighthouse Lab in Milton Keynes | Wellcome Sanger Institute for the COVID-19 Genomics UK (COG-UK) Consortium | The Lighthouse Lab in Milton Keynes and Alex Alderton, Roberto Amato, Sonia Goncalves, Ewan Harrison, David K. Jackson, Ian Johnston, Dominic Kwiatkowski, Cordelia Langford, John Sillitoe on behalf of the Wellcome Sanger Institute COVID-19 Surveillance Team |
| EPI_ISL_782971 | Lighthouse Lab in Cambridge | Wellcome Sanger Institute for the COVID-19 Genomics UK (COG-UK) Consortium | Rob Howes, The Lighthouse Lab in Cambridge and Alex Alderton, Roberto Amato, Sonia Goncalves, Ewan Harrison, David K. Jackson, Ian Johnston, Dominic Kwiatkowski, Cordelia Langford, John Sillitoe on behalf of the Wellcome Sanger Institute COVID-19 Surveillance Team |
| EPI_ISL_782972, EPI_ISL_782973 | Lighthouse Lab in Milton Keynes | Wellcome Sanger Institute for the COVID-19 Genomics UK (COG-UK) Consortium | The Lighthouse Lab in Milton Keynes and Alex Alderton, Roberto Amato, Sonia Goncalves, Ewan Harrison, David K. Jackson, Ian Johnston, Dominic Kwiatkowski, Cordelia Langford, John Sillitoe on behalf of the Wellcome Sanger Institute COVID-19 Surveillance Team |
| EPI_ISL_782974 | Lighthouse Lab in Cambridge | Wellcome Sanger Institute for the COVID-19 Genomics UK (COG-UK) Consortium | Rob Howes, The Lighthouse Lab in Cambridge and Alex Alderton, Roberto Amato, Sonia Goncalves, Ewan Harrison, David K. Jackson, Ian Johnston, Dominic Kwiatkowski, Cordelia Langford, John Sillitoe on behalf of the Wellcome Sanger Institute COVID-19 Surveillance Team |
| EPI_ISL_782975, EPI_ISL_782977, EPI_ISL_782978 | Lighthouse Lab in Milton Keynes | Wellcome Sanger Institute for the COVID-19 Genomics UK (COG-UK) Consortium | The Lighthouse Lab in Milton Keynes and Alex Alderton, Roberto Amato, Sonia Goncalves, Ewan Harrison, David K. Jackson, Ian Johnston, Dominic Kwiatkowski, Cordelia Langford, John Sillitoe on behalf of the Wellcome Sanger Institute COVID-19 Surveillance Team |
| EPI_ISL_782979 | Lighthouse Lab in Cambridge | Wellcome Sanger Institute for the COVID-19 Genomics UK (COG-UK) Consortium | Rob Howes, The Lighthouse Lab in Cambridge and Alex Alderton, Roberto Amato, Sonia Goncalves, Ewan Harrison, David K. Jackson, Ian Johnston, Dominic Kwiatkowski, Cordelia Langford, John Sillitoe on behalf of the Wellcome Sanger Institute COVID-19 Surveillance Team |
| EPI_ISL_782980, EPI_ISL_782981, EPI_ISL_782982 | Lighthouse Lab in Milton Keynes | Wellcome Sanger Institute for the COVID-19 Genomics UK (COG-UK) Consortium | The Lighthouse Lab in Milton Keynes and Alex Alderton, Roberto Amato, Sonia Goncalves, Ewan Harrison, David K. Jackson, Ian Johnston, Dominic Kwiatkowski, Cordelia Langford, John Sillitoe on behalf of the Wellcome Sanger Institute COVID-19 Surveillance Team |
| EPI_ISL_782983, EPI_ISL_782984, EPI_ISL_782985 | Lighthouse Lab in Cambridge | Wellcome Sanger Institute for the COVID-19 Genomics UK (COG-UK) Consortium | Rob Howes, The Lighthouse Lab in Cambridge and Alex Alderton, Roberto Amato, Sonia Goncalves, Ewan Harrison, David K. Jackson, Ian Johnston, Dominic Kwiatkowski, Cordelia Langford, John Sillitoe on behalf of the Wellcome Sanger Institute COVID-19 Surveillance Team |
| EPI_ISL_782986 | Lighthouse Lab in Milton Keynes | Wellcome Sanger Institute for the COVID-19 Genomics UK (COG-UK) Consortium | The Lighthouse Lab in Milton Keynes and Alex Alderton, Roberto Amato, Sonia Goncalves, Ewan Harrison, David K. Jackson, Ian Johnston, Dominic Kwiatkowski, Cordelia Langford, John Sillitoe on behalf of the Wellcome Sanger Institute COVID-19 Surveillance Team |
| EPI_ISL_782987 | Lighthouse Lab in Cambridge | Wellcome Sanger Institute for the COVID-19 Genomics UK (COG-UK) Consortium | Rob Howes, The Lighthouse Lab in Cambridge and Alex Alderton, Roberto Amato, Sonia Goncalves, Ewan Harrison, David K. Jackson, Ian Johnston, Dominic Kwiatkowski, Cordelia Langford, John Sillitoe on behalf of the Wellcome Sanger Institute COVID-19 Surveillance Team |
| EPI_ISL_782989, EPI_ISL_782990 | Lighthouse Lab in Milton Keynes | Wellcome Sanger Institute for the COVID-19 Genomics UK (COG-UK) Consortium | The Lighthouse Lab in Milton Keynes and Alex Alderton, Roberto Amato, Sonia Goncalves, Ewan Harrison, David K. Jackson, Ian Johnston, Dominic Kwiatkowski, Cordelia Langford, John Sillitoe on behalf of the Wellcome Sanger Institute COVID-19 Surveillance Team |
| EPI_ISL_782991 | Lighthouse Lab in Cambridge | Wellcome Sanger Institute for the COVID-19 Genomics UK (COG-UK) Consortium | Rob Howes, The Lighthouse Lab in Cambridge and Alex Alderton, Roberto Amato, Sonia Goncalves, Ewan Harrison, David K. Jackson, Ian Johnston, Dominic Kwiatkowski, Cordelia Langford, John Sillitoe on behalf of the Wellcome Sanger Institute COVID-19 Surveillance Team |
| EPI_ISL_782992, EPI_ISL_782995 | Lighthouse Lab in Milton Keynes | Wellcome Sanger Institute for the COVID-19 Genomics UK (COG-UK) Consortium | The Lighthouse Lab in Milton Keynes and Alex Alderton, Roberto Amato, Sonia Goncalves, Ewan Harrison, David K. Jackson, Ian Johnston, Dominic Kwiatkowski, Cordelia Langford, John Sillitoe on behalf of the Wellcome Sanger Institute COVID-19 Surveillance Team |
| EPI_ISL_782996, EPI_ISL_782998 | Lighthouse Lab in Cambridge | Wellcome Sanger Institute for the COVID-19 Genomics UK (COG-UK) Consortium | Rob Howes, The Lighthouse Lab in Cambridge and Alex Alderton, Roberto Amato, Sonia Goncalves, Ewan Harrison, David K. Jackson, Ian Johnston, Dominic Kwiatkowski, Cordelia Langford, John Sillitoe on behalf of the Wellcome Sanger Institute COVID-19 Surveillance Team |
| EPI_ISL_782999, EPI_ISL_783001, EPI_ISL_783003, EPI_ISL_783004, EPI_ISL_783005, EPI_ISL_783006, EPI_ISL_783009, EPI_ISL_783010, EPI_ISL_783011 | Lighthouse Lab in Milton Keynes | Wellcome Sanger Institute for the COVID-19 Genomics UK (COG-UK) Consortium | The Lighthouse Lab in Milton Keynes and Alex Alderton, Roberto Amato, Sonia Goncalves, Ewan Harrison, David K. Jackson, Ian Johnston, Dominic Kwiatkowski, Cordelia Langford, John Sillitoe on behalf of the Wellcome Sanger Institute COVID-19 Surveillance Team |
| EPI_ISL_783012 | Lighthouse Lab in Cambridge | Wellcome Sanger Institute for the COVID-19 Genomics UK (COG-UK) Consortium | Rob Howes, The Lighthouse Lab in Cambridge and Alex Alderton, Roberto Amato, Sonia Goncalves, Ewan Harrison, David K. Jackson, Ian Johnston, Dominic Kwiatkowski, Cordelia Langford, John Sillitoe on behalf of the Wellcome Sanger Institute COVID-19 Surveillance Team |
| EPI_ISL_783013, EPI_ISL_783016, EPI_ISL_783017, EPI_ISL_783018 | Lighthouse Lab in Milton Keynes | Wellcome Sanger Institute for the COVID-19 Genomics UK (COG-UK) Consortium | The Lighthouse Lab in Milton Keynes and Alex Alderton, Roberto Amato, Sonia Goncalves, Ewan Harrison, David K. Jackson, Ian Johnston, Dominic Kwiatkowski, Cordelia Langford, John Sillitoe on behalf of the Wellcome Sanger Institute COVID-19 Surveillance Team |
| EPI_ISL_783019 | Lighthouse Lab in Cambridge | Wellcome Sanger Institute for the COVID-19 Genomics UK (COG-UK) Consortium | Rob Howes, The Lighthouse Lab in Cambridge and Alex Alderton, Roberto Amato, Sonia Goncalves, Ewan Harrison, David K. Jackson, Ian Johnston, Dominic Kwiatkowski, Cordelia Langford, John Sillitoe on behalf of the Wellcome Sanger Institute COVID-19 Surveillance Team |
| EPI_ISL_783020, EPI_ISL_783021, EPI_ISL_783022 | Lighthouse Lab in Milton Keynes | Wellcome Sanger Institute for the COVID-19 Genomics UK (COG-UK) Consortium | The Lighthouse Lab in Milton Keynes and Alex Alderton, Roberto Amato, Sonia Goncalves, Ewan Harrison, David K. Jackson, Ian Johnston, Dominic Kwiatkowski, Cordelia Langford, John Sillitoe on behalf of the Wellcome Sanger Institute COVID-19 Surveillance Team |
| EPI_ISL_783023 | Lighthouse Lab in Cambridge | Wellcome Sanger Institute for the COVID-19 Genomics UK (COG-UK) Consortium | Rob Howes, The Lighthouse Lab in Cambridge and Alex Alderton, Roberto Amato, Sonia Goncalves, Ewan Harrison, David K. Jackson, Ian Johnston, Dominic Kwiatkowski, Cordelia Langford, John Sillitoe on behalf of the Wellcome Sanger Institute COVID-19 Surveillance Team |
| EPI_ISL_783024, EPI_ISL_783025, EPI_ISL_783027 | Lighthouse Lab in Milton Keynes | Wellcome Sanger Institute for the COVID-19 Genomics UK (COG-UK) Consortium | The Lighthouse Lab in Milton Keynes and Alex Alderton, Roberto Amato, Sonia Goncalves, Ewan Harrison, David K. Jackson, Ian Johnston, Dominic Kwiatkowski, Cordelia Langford, John Sillitoe on behalf of the Wellcome Sanger Institute COVID-19 Surveillance Team |
| EPI_ISL_783028 | Lighthouse Lab in Cambridge | Wellcome Sanger Institute for the COVID-19 Genomics UK (COG-UK) Consortium | Rob Howes, The Lighthouse Lab in Cambridge and Alex Alderton, Roberto Amato, Sonia Goncalves, Ewan Harrison, David K. Jackson, Ian Johnston, Dominic Kwiatkowski, Cordelia Langford, John Sillitoe on behalf of the Wellcome Sanger Institute COVID-19 Surveillance Team |
| EPI_ISL_783029 | Lighthouse Lab in Milton Keynes | Wellcome Sanger Institute for the COVID-19 Genomics UK (COG-UK) Consortium | The Lighthouse Lab in Milton Keynes and Alex Alderton, Roberto Amato, Sonia Goncalves, Ewan Harrison, David K. Jackson, Ian Johnston, Dominic Kwiatkowski, Cordelia Langford, John Sillitoe on behalf of the Wellcome Sanger Institute COVID-19 Surveillance Team |
| EPI_ISL_783031 | Lighthouse Lab in Cambridge | Wellcome Sanger Institute for the COVID-19 Genomics UK (COG-UK) Consortium | Rob Howes, The Lighthouse Lab in Cambridge and Alex Alderton, Roberto Amato, Sonia Goncalves, Ewan Harrison, David K. Jackson, Ian Johnston, Dominic Kwiatkowski, Cordelia Langford, John Sillitoe on behalf of the Wellcome Sanger Institute COVID-19 Surveillance Team |
| EPI_ISL_783035 | Lighthouse Lab in Milton Keynes | Wellcome Sanger Institute for the COVID-19 Genomics UK (COG-UK) Consortium | The Lighthouse Lab in Milton Keynes and Alex Alderton, Roberto Amato, Sonia Goncalves, Ewan Harrison, David K. Jackson, Ian Johnston, Dominic Kwiatkowski, Cordelia Langford, John Sillitoe on behalf of the Wellcome Sanger Institute COVID-19 Surveillance Team |
| EPI_ISL_783036, EPI_ISL_783037, EPI_ISL_783038 | Lighthouse Lab in Cambridge | Wellcome Sanger Institute for the COVID-19 Genomics UK (COG-UK) Consortium | Rob Howes, The Lighthouse Lab in Cambridge and Alex Alderton, Roberto Amato, Sonia Goncalves, Ewan Harrison, David K. Jackson, Ian Johnston, Dominic Kwiatkowski, Cordelia Langford, John Sillitoe on behalf of the Wellcome Sanger Institute COVID-19 Surveillance Team |
| EPI_ISL_783039, EPI_ISL_783040, EPI_ISL_783041 | Lighthouse Lab in Milton Keynes | Wellcome Sanger Institute for the COVID-19 Genomics UK (COG-UK) Consortium | The Lighthouse Lab in Milton Keynes and Alex Alderton, Roberto Amato, Sonia Goncalves, Ewan Harrison, David K. Jackson, Ian Johnston, Dominic Kwiatkowski, Cordelia Langford, John Sillitoe on behalf of the Wellcome Sanger Institute COVID-19 Surveillance Team |
| EPI_ISL_783042, EPI_ISL_783044 | Lighthouse Lab in Cambridge | Wellcome Sanger Institute for the COVID-19 Genomics UK | Rob Howes, The Lighthouse Lab in Cambridge and Alex Alderton, Roberto Amato, Sonia Goncalves, Ewan Harrison, David K. Jackson, Ian Johnston, |

| | | | |
|---|---|---|---|
| | | (COG-UK) Consortium | Dominic Kwiatkowski, Cordelia Langford, John Sillitoe on behalf of the Wellcome Sanger Institute COVID-19 Surveillance Team |
| EPI_ISL_783048, EPI_ISL_783049, EPI_ISL_783050 | Lighthouse Lab in Milton Keynes | Wellcome Sanger Institute for the COVID-19 Genomics UK (COG-UK) Consortium | The Lighthouse Lab in Milton Keynes and Alex Alderton, Roberto Amato, Sonia Goncalves, Ewan Harrison, David K. Jackson, Ian Johnston, Dominic Kwiatkowski, Cordelia Langford, John Sillitoe on behalf of the Wellcome Sanger Institute COVID-19 Surveillance Team |
| EPI_ISL_783051 | Lighthouse Lab in Cambridge | Wellcome Sanger Institute for the COVID-19 Genomics UK (COG-UK) Consortium | Rob Howes, The Lighthouse Lab in Cambridge and Alex Alderton, Roberto Amato, Sonia Goncalves, Ewan Harrison, David K. Jackson, Ian Johnston, Dominic Kwiatkowski, Cordelia Langford, John Sillitoe on behalf of the Wellcome Sanger Institute COVID-19 Surveillance Team |
| EPI_ISL_783053, EPI_ISL_783054 | Lighthouse Lab in Milton Keynes | Wellcome Sanger Institute for the COVID-19 Genomics UK (COG-UK) Consortium | The Lighthouse Lab in Milton Keynes and Alex Alderton, Roberto Amato, Sonia Goncalves, Ewan Harrison, David K. Jackson, Ian Johnston, Dominic Kwiatkowski, Cordelia Langford, John Sillitoe on behalf of the Wellcome Sanger Institute COVID-19 Surveillance Team |
| EPI_ISL_783055, EPI_ISL_783057 | Lighthouse Lab in Cambridge | Wellcome Sanger Institute for the COVID-19 Genomics UK (COG-UK) Consortium | Rob Howes, The Lighthouse Lab in Cambridge and Alex Alderton, Roberto Amato, Sonia Goncalves, Ewan Harrison, David K. Jackson, Ian Johnston, Dominic Kwiatkowski, Cordelia Langford, John Sillitoe on behalf of the Wellcome Sanger Institute COVID-19 Surveillance Team |
| EPI_ISL_783058, EPI_ISL_783059, EPI_ISL_783060, EPI_ISL_783061, EPI_ISL_783062, EPI_ISL_783063, EPI_ISL_783064, EPI_ISL_783065 | Lighthouse Lab in Milton Keynes | Wellcome Sanger Institute for the COVID-19 Genomics UK (COG-UK) Consortium | The Lighthouse Lab in Milton Keynes and Alex Alderton, Roberto Amato, Sonia Goncalves, Ewan Harrison, David K. Jackson, Ian Johnston, Dominic Kwiatkowski, Cordelia Langford, John Sillitoe on behalf of the Wellcome Sanger Institute COVID-19 Surveillance Team |
| EPI_ISL_783066 | Lighthouse Lab in Cambridge | Wellcome Sanger Institute for the COVID-19 Genomics UK (COG-UK) Consortium | Rob Howes, The Lighthouse Lab in Cambridge and Alex Alderton, Roberto Amato, Sonia Goncalves, Ewan Harrison, David K. Jackson, Ian Johnston, Dominic Kwiatkowski, Cordelia Langford, John Sillitoe on behalf of the Wellcome Sanger Institute COVID-19 Surveillance Team |
| EPI_ISL_783067 | Lighthouse Lab in Milton Keynes | Wellcome Sanger Institute for the COVID-19 Genomics UK (COG-UK) Consortium | The Lighthouse Lab in Milton Keynes and Alex Alderton, Roberto Amato, Sonia Goncalves, Ewan Harrison, David K. Jackson, Ian Johnston, Dominic Kwiatkowski, Cordelia Langford, John Sillitoe on behalf of the Wellcome Sanger Institute COVID-19 Surveillance Team |
| EPI_ISL_789037, EPI_ISL_789039 | Klinisk mikrobiologi | The Public Health Agency of Sweden | Department of Microbiology, The Public Health Agency of Sweden |
| EPI_ISL_790626, EPI_ISL_790647, EPI_ISL_790651, EPI_ISL_790652, EPI_ISL_790653, EPI_ISL_790665, EPI_ISL_790675, EPI_ISL_790676, EPI_ISL_790677, EPI_ISL_790678, EPI_ISL_790686, EPI_ISL_790688, EPI_ISL_790689, EPI_ISL_790693, EPI_ISL_790745, EPI_ISL_790746, EPI_ISL_790747, EPI_ISL_790814, EPI_ISL_790823, EPI_ISL_791034 | | | |
| see above | Dutch COVID-19 response team | National Institute for Public Health and the Environment (RIVM) | Adam Meijer, Harry Vennema, Jeroen Cremer, Sharon van den Brink, Bas van der Veer, AnneMarie van den Brandt, Florian Zwagemaker, Dennis Schmitz, Chantal Reusken, on behalf of the national COVID-19 response team |
| EPI_ISL_791093, EPI_ISL_791094, EPI_ISL_791095 | Instituto Nacional de Salud - Unidad de Secuenciación y Análisis Genómico | Instituto Nacional de Salud - Dirección de Investigación en Salud Pública | Katherine Laiton-Donato, Diego A. Álvarez-Díaz, Carlos Franco-Muñoz, Mauricio Pacheco-Montealegre, Jonathan Reales, Sheryl Corchuelo, Maria T. Herrera, Julian Naizaque, Gerardo Santamaría, Paola Muñoz-Laiton, Diego Andrés Prada, Magdalena Wiesner, Martha Lucia Ospina Martinez, Marcela Mercado-Reyes |
| EPI_ISL_791314, EPI_ISL_791322 | National Virus Reference Laboratory | Irish Coronavirus Sequencing Consortium - Teagasc Moorepark | Alejandro Abner Garcia Leon, Paul Cotter, Fiona Crispie, John Kenny, Paddy Mallon, Calum Walsh |
| EPI_ISL_791994, EPI_ISL_791995 | CHU - Hôpital Cavale Blanche | National Reference Center for Viruses of Respiratory Infections, Institut Pasteur, Paris | Marion Barbet, Sylvie Behillil, Méline Bizard, Angela Brisebarre, Camille Capel, Etienne Simon-Lorière, Vincent Enouf, Maud Vanpeene, Sylvie van der Werf, Léa Pilorgé |
| EPI_ISL_792556 | Centre for Dengue Research and AICBU, Department of Immunology and Molecular Medicine | Centre for Dengue Research and AICBU, Department of Immunology and Molecular Medicine | Chandima Jeewandara, Deshni Jayathilaka, Dinuka Ariyaratne, Diyanath Ranasinghe, Laksiri Gomes, Gathsaurie Neelika Malavige |
| EPI_ISL_792699, EPI_ISL_792704 | The National Institute of Public Health | State Veterinary Institute Prague | Nagy,A;Jirincova,H;Trnka,D;Vecerova,J |
| EPI_ISL_792710, EPI_ISL_792713, EPI_ISL_792740, EPI_ISL_792741, EPI_ISL_792742, EPI_ISL_792743, EPI_ISL_792744, EPI_ISL_792745, EPI_ISL_792746, EPI_ISL_792747, EPI_ISL_792748, EPI_ISL_792754, EPI_ISL_792759, EPI_ISL_792780, EPI_ISL_792783, EPI_ISL_792784, EPI_ISL_792793, EPI_ISL_792825, EPI_ISL_792826, EPI_ISL_792827, EPI_ISL_792831, EPI_ISL_792832, EPI_ISL_792833, EPI_ISL_792836, EPI_ISL_792837, EPI_ISL_792980, EPI_ISL_792981, EPI_ISL_792982, EPI_ISL_792983, EPI_ISL_792984, EPI_ISL_792985, EPI_ISL_792986, EPI_ISL_792987, EPI_ISL_792988, EPI_ISL_792989, EPI_ISL_792990, EPI_ISL_792991, EPI_ISL_792992, EPI_ISL_792993, EPI_ISL_792994, EPI_ISL_792995, EPI_ISL_792996, EPI_ISL_792997, EPI_ISL_792998, EPI_ISL_792999, EPI_ISL_793000, EPI_ISL_793001, EPI_ISL_793002, EPI_ISL_793003, EPI_ISL_793004, EPI_ISL_793005, EPI_ISL_793006, EPI_ISL_793007, EPI_ISL_793008, EPI_ISL_793009, EPI_ISL_794324, EPI_ISL_794329, EPI_ISL_794330, EPI_ISL_794331, EPI_ISL_794332, EPI_ISL_794333, EPI_ISL_794334, EPI_ISL_794335, EPI_ISL_794336, EPI_ISL_794341, EPI_ISL_794343, EPI_ISL_794344, EPI_ISL_794345, EPI_ISL_794346, EPI_ISL_794347, EPI_ISL_794349, EPI_ISL_794350, EPI_ISL_794351, EPI_ISL_794361, EPI_ISL_794362, EPI_ISL_794363, EPI_ISL_794364, EPI_ISL_794365, EPI_ISL_794366, EPI_ISL_794367, EPI_ISL_794369, EPI_ISL_794371, EPI_ISL_794375, EPI_ISL_794376, EPI_ISL_794377, EPI_ISL_794383, EPI_ISL_794384, EPI_ISL_794385, EPI_ISL_794386, EPI_ISL_794387, EPI_ISL_794388, EPI_ISL_794398, EPI_ISL_794399, EPI_ISL_794400, EPI_ISL_794401, EPI_ISL_794402, EPI_ISL_794403, EPI_ISL_794404, EPI_ISL_794405, EPI_ISL_794406, EPI_ISL_794407, EPI_ISL_794408, EPI_ISL_794409, EPI_ISL_794410, EPI_ISL_794411, EPI_ISL_794420, EPI_ISL_794421, EPI_ISL_794423, EPI_ISL_794425, EPI_ISL_794426, EPI_ISL_794430, EPI_ISL_794434, EPI_ISL_794496, EPI_ISL_794497, EPI_ISL_794498, EPI_ISL_794499, EPI_ISL_794500, EPI_ISL_794501, EPI_ISL_794502, EPI_ISL_794503, EPI_ISL_794504, EPI_ISL_794505, EPI_ISL_794506, EPI_ISL_794507, EPI_ISL_794508, EPI_ISL_794509, EPI_ISL_794510, EPI_ISL_794511, EPI_ISL_794512, EPI_ISL_794513, EPI_ISL_794514, EPI_ISL_794515, EPI_ISL_794516, EPI_ISL_794517, EPI_ISL_794518, EPI_ISL_794519, EPI_ISL_794520, EPI_ISL_794521, EPI_ISL_794522, EPI_ISL_794523, EPI_ISL_794524, EPI_ISL_794525, EPI_ISL_794526, EPI_ISL_794527, EPI_ISL_794528, EPI_ISL_794529, EPI_ISL_794530, EPI_ISL_794531, EPI_ISL_794532, EPI_ISL_794533, EPI_ISL_794534, EPI_ISL_794535, EPI_ISL_794536, EPI_ISL_794537, EPI_ISL_794538, EPI_ISL_794539, EPI_ISL_794540, EPI_ISL_794541, EPI_ISL_794542, EPI_ISL_794543, EPI_ISL_794544, EPI_ISL_794545, EPI_ISL_794546, EPI_ISL_794547, EPI_ISL_794548, EPI_ISL_794549, EPI_ISL_794550, EPI_ISL_794551, EPI_ISL_794552, EPI_ISL_794553, EPI_ISL_794554 | | | |
| see above | Department of Virus and Microbiological Special Diagnostics, Statens Serum Institut, Copenhagen, Denmark | Albertsen Lab, Department of Chemistry and Bioscience, Aalborg University, Denmark | Danish Covid-19 Genome Consortium |
| EPI_ISL_794716, EPI_ISL_794730 | PathWest Laboratory Medicine WA | PathWest Laboratory Medicine WA Microbial Surveillance Unit | PathWest Laboratory Medicine WA Microbial Surveillance Unit |
| EPI_ISL_794850, EPI_ISL_794851, EPI_ISL_794852, EPI_ISL_794853, EPI_ISL_794854, EPI_ISL_794855, EPI_ISL_794856, EPI_ISL_794857, EPI_ISL_794860, EPI_ISL_794862, EPI_ISL_794864, EPI_ISL_794871, EPI_ISL_794876, EPI_ISL_794883, EPI_ISL_794895, EPI_ISL_794896, EPI_ISL_794897, EPI_ISL_794898, EPI_ISL_794899, EPI_ISL_794901, EPI_ISL_794905, EPI_ISL_794983, EPI_ISL_794984, EPI_ISL_794985, EPI_ISL_794986, EPI_ISL_794987, EPI_ISL_794988, EPI_ISL_794989, EPI_ISL_794990, EPI_ISL_794991, EPI_ISL_794992, EPI_ISL_794993, EPI_ISL_794994, EPI_ISL_794995, EPI_ISL_794996, EPI_ISL_794997, EPI_ISL_794998, EPI_ISL_794999, EPI_ISL_795000, EPI_ISL_795001, EPI_ISL_795002, EPI_ISL_795003, EPI_ISL_795004, EPI_ISL_795005, EPI_ISL_795006, EPI_ISL_795007 | | | |
| see above | Department of Virus and Microbiological Special Diagnostics, Statens Serum Institut, Copenhagen, Denmark | Albertsen Lab, Department of Chemistry and Bioscience, Aalborg University, Denmark | Danish Covid-19 Genome Consortium |
| EPI_ISL_796650 | Nordland Hospital - Bodo, Laboratory Department, Molecular Biology Unit | Norwegian Institute of Public Health, Department of Virology | Kathrine Stene-Johansen, Kamilla Heddeland Instefjord, Hilde Elshaug, Atiya R Ali,Marie Paulsen Madsen, Rasmus Riis Kopperud, Hilde Vollan, Karoline Bragstad, Olav Hungnes |
| EPI_ISL_796663 | Dept. of Medical Microbiology, Stavanger University Hospital, Helse Stavanger HF | Norwegian Institute of Public Health, Department of Virology | Kathrine Stene-Johansen, Kamilla Heddeland Instefjord, Hilde Elshaug, Atiya R Ali,Marie Paulsen Madsen, Rasmus Riis Kopperud, Hilde Vollan, Karoline Bragstad, Olav Hungnes |
| EPI_ISL_796667, EPI_ISL_796668, EPI_ISL_796669, EPI_ISL_796672, EPI_ISL_796691, EPI_ISL_796692, EPI_ISL_796693, EPI_ISL_796694 | Nordland Hospital - Bodo, Laboratory Department, Molecular Biology Unit | Norwegian Institute of Public Health, Department of Virology | Kathrine Stene-Johansen, Kamilla Heddeland Instefjord, Hilde Elshaug, Atiya R Ali,Marie Paulsen Madsen, Rasmus Riis Kopperud, Hilde Vollan, Karoline Bragstad, Olav Hungnes |
| EPI_ISL_796695, EPI_ISL_796696, EPI_ISL_796697 | Ostfold Hospital Trust - Kalnes, Centre for Laboratory Medicine, Section for gene technology and infection serology | Norwegian Institute of Public Health, Department of Virology | Kathrine Stene-Johansen, Kamilla Heddeland Instefjord, Hilde Elshaug, Atiya R Ali,Marie Paulsen Madsen, Rasmus Riis Kopperud, Hilde Vollan, Karoline Bragstad, Olav Hungnes |
| EPI_ISL_796698 | Dept. of Medical Microbiology, Stavanger University Hospital, Helse Stavanger HF | Norwegian Institute of Public Health, Department of Virology | Kathrine Stene-Johansen, Kamilla Heddeland Instefjord, Hilde Elshaug, Atiya R Ali,Marie Paulsen Madsen, Rasmus Riis Kopperud, Hilde Vollan, Karoline Bragstad, Olav Hungnes |
| EPI_ISL_796702 | Unilabs Laboratory Medicine | Norwegian Institute of Public Health, Department of Virology | Kathrine Stene-Johansen, Kamilla Heddeland Instefjord, Hilde Elshaug, Atiya R Ali,Marie Paulsen Madsen, Rasmus Riis Kopperud, Hilde Vollan, Karoline Bragstad, Olav Hungnes |
| EPI_ISL_796730, EPI_ISL_796731, EPI_ISL_796732 | Furst Medical Laboratory | Norwegian Institute of Public Health, Department of Virology | Kathrine Stene-Johansen, Kamilla Heddeland Instefjord, Hilde Elshaug, Atiya R Ali,Marie Paulsen Madsen, Rasmus Riis Kopperud, Hilde Vollan, Karoline Bragstad, Olav Hungnes |
| EPI_ISL_800089, EPI_ISL_800157, EPI_ISL_800203, EPI_ISL_800586, EPI_ISL_800587, EPI_ISL_800589, EPI_ISL_800590 | Lighthouse Lab in Cambridge | Wellcome Sanger Institute for the COVID-19 Genomics UK (COG-UK) Consortium | Rob Howes, The Lighthouse Lab in Cambridge and Alex Alderton, Roberto Amato, Sonia Goncalves, Ewan Harrison, David K. Jackson, Ian Johnston, Dominic Kwiatkowski, Cordelia Langford, John Sillitoe on behalf of the Wellcome Sanger Institute COVID-19 Surveillance Team |
| EPI_ISL_800591 | Lighthouse Lab in Glasgow | Wellcome Sanger Institute for the COVID-19 Genomics UK (COG-UK) Consortium | Harper VanSteenhouse, Yumi Kasai, David Gray, Carol Clugston, Anna Dominiczak and Alex Alderton, Roberto Amato, Sonia Goncalves, Ewan Harrison, David K. Jackson, Ian Johnston, Dominic Kwiatkowski, Cordelia Langford, John Sillitoe on behalf of the Wellcome Sanger Institute COVID-19 Surveillance Team |

| | | | |
|---|---|---|---|
| EPI_ISL_800592 | Lighthouse Lab in Cambridge | Wellcome Sanger Institute for the COVID-19 Genomics UK (COG-UK) Consortium | Rob Howes, The Lighthouse Lab in Cambridge and Alex Alderton, Roberto Amato, Sonia Goncalves, Ewan Harrison, David K. Jackson, Ian Johnston, Dominic Kwiatkowski, Cordelia Langford, John Sillitoe on behalf of the Wellcome Sanger Institute COVID-19 Surveillance Team |
| EPI_ISL_800594, EPI_ISL_800596 | Lighthouse Lab in Glasgow | Wellcome Sanger Institute for the COVID-19 Genomics UK (COG-UK) Consortium | Harper VanSteenhouse, Yumi Kasai, David Gray, Carol Clugston, Anna Dominiczak and Alex Alderton, Roberto Amato, Sonia Goncalves, Ewan Harrison, David K. Jackson, Ian Johnston, Dominic Kwiatkowski, Cordelia Langford, John Sillitoe on behalf of the Wellcome Sanger Institute COVID-19 Surveillance Team |
| EPI_ISL_800597, EPI_ISL_800600 | Lighthouse Lab in Cambridge | Wellcome Sanger Institute for the COVID-19 Genomics UK (COG-UK) Consortium | Rob Howes, The Lighthouse Lab in Cambridge and Alex Alderton, Roberto Amato, Sonia Goncalves, Ewan Harrison, David K. Jackson, Ian Johnston, Dominic Kwiatkowski, Cordelia Langford, John Sillitoe on behalf of the Wellcome Sanger Institute COVID-19 Surveillance Team |
| EPI_ISL_800601, EPI_ISL_800602, EPI_ISL_800603, EPI_ISL_800604, EPI_ISL_800605, EPI_ISL_800607, EPI_ISL_800608 | Lighthouse Lab in Glasgow | Wellcome Sanger Institute for the COVID-19 Genomics UK (COG-UK) Consortium | Harper VanSteenhouse, Yumi Kasai, David Gray, Carol Clugston, Anna Dominiczak and Alex Alderton, Roberto Amato, Sonia Goncalves, Ewan Harrison, David K. Jackson, Ian Johnston, Dominic Kwiatkowski, Cordelia Langford, John Sillitoe on behalf of the Wellcome Sanger Institute COVID-19 Surveillance Team |
| EPI_ISL_800609, EPI_ISL_800610 | Lighthouse Lab in Cambridge | Wellcome Sanger Institute for the COVID-19 Genomics UK (COG-UK) Consortium | Rob Howes, The Lighthouse Lab in Cambridge and Alex Alderton, Roberto Amato, Sonia Goncalves, Ewan Harrison, David K. Jackson, Ian Johnston, Dominic Kwiatkowski, Cordelia Langford, John Sillitoe on behalf of the Wellcome Sanger Institute COVID-19 Surveillance Team |
| EPI_ISL_800611, EPI_ISL_800612, EPI_ISL_800614 | Lighthouse Lab in Glasgow | Wellcome Sanger Institute for the COVID-19 Genomics UK (COG-UK) Consortium | Harper VanSteenhouse, Yumi Kasai, David Gray, Carol Clugston, Anna Dominiczak and Alex Alderton, Roberto Amato, Sonia Goncalves, Ewan Harrison, David K. Jackson, Ian Johnston, Dominic Kwiatkowski, Cordelia Langford, John Sillitoe on behalf of the Wellcome Sanger Institute COVID-19 Surveillance Team |
| EPI_ISL_800615 | Lighthouse Lab in Cambridge | Wellcome Sanger Institute for the COVID-19 Genomics UK (COG-UK) Consortium | Rob Howes, The Lighthouse Lab in Cambridge and Alex Alderton, Roberto Amato, Sonia Goncalves, Ewan Harrison, David K. Jackson, Ian Johnston, Dominic Kwiatkowski, Cordelia Langford, John Sillitoe on behalf of the Wellcome Sanger Institute COVID-19 Surveillance Team |
| EPI_ISL_800616, EPI_ISL_800618, EPI_ISL_800620, EPI_ISL_800621 | Lighthouse Lab in Glasgow | Wellcome Sanger Institute for the COVID-19 Genomics UK (COG-UK) Consortium | Harper VanSteenhouse, Yumi Kasai, David Gray, Carol Clugston, Anna Dominiczak and Alex Alderton, Roberto Amato, Sonia Goncalves, Ewan Harrison, David K. Jackson, Ian Johnston, Dominic Kwiatkowski, Cordelia Langford, John Sillitoe on behalf of the Wellcome Sanger Institute COVID-19 Surveillance Team |
| EPI_ISL_800622 | Lighthouse Lab in Cambridge | Wellcome Sanger Institute for the COVID-19 Genomics UK (COG-UK) Consortium | Rob Howes, The Lighthouse Lab in Cambridge and Alex Alderton, Roberto Amato, Sonia Goncalves, Ewan Harrison, David K. Jackson, Ian Johnston, Dominic Kwiatkowski, Cordelia Langford, John Sillitoe on behalf of the Wellcome Sanger Institute COVID-19 Surveillance Team |
| EPI_ISL_800625 | Lighthouse Lab in Glasgow | Wellcome Sanger Institute for the COVID-19 Genomics UK (COG-UK) Consortium | Harper VanSteenhouse, Yumi Kasai, David Gray, Carol Clugston, Anna Dominiczak and Alex Alderton, Roberto Amato, Sonia Goncalves, Ewan Harrison, David K. Jackson, Ian Johnston, Dominic Kwiatkowski, Cordelia Langford, John Sillitoe on behalf of the Wellcome Sanger Institute COVID-19 Surveillance Team |
| EPI_ISL_800626 | Lighthouse Lab in Cambridge | Wellcome Sanger Institute for the COVID-19 Genomics UK (COG-UK) Consortium | Rob Howes, The Lighthouse Lab in Cambridge and Alex Alderton, Roberto Amato, Sonia Goncalves, Ewan Harrison, David K. Jackson, Ian Johnston, Dominic Kwiatkowski, Cordelia Langford, John Sillitoe on behalf of the Wellcome Sanger Institute COVID-19 Surveillance Team |
| EPI_ISL_800627, EPI_ISL_800628, EPI_ISL_800629, EPI_ISL_800630, EPI_ISL_800631, EPI_ISL_800633, EPI_ISL_800634, EPI_ISL_800635 | Lighthouse Lab in Glasgow | Wellcome Sanger Institute for the COVID-19 Genomics UK (COG-UK) Consortium | Harper VanSteenhouse, Yumi Kasai, David Gray, Carol Clugston, Anna Dominiczak and Alex Alderton, Roberto Amato, Sonia Goncalves, Ewan Harrison, David K. Jackson, Ian Johnston, Dominic Kwiatkowski, Cordelia Langford, John Sillitoe on behalf of the Wellcome Sanger Institute COVID-19 Surveillance Team |
| EPI_ISL_800637 | Lighthouse Lab in Cambridge | Wellcome Sanger Institute for the COVID-19 Genomics UK (COG-UK) Consortium | Rob Howes, The Lighthouse Lab in Cambridge and Alex Alderton, Roberto Amato, Sonia Goncalves, Ewan Harrison, David K. Jackson, Ian Johnston, Dominic Kwiatkowski, Cordelia Langford, John Sillitoe on behalf of the Wellcome Sanger Institute COVID-19 Surveillance Team |
| EPI_ISL_800638, EPI_ISL_800639, EPI_ISL_800640 | Lighthouse Lab in Glasgow | Wellcome Sanger Institute for the COVID-19 Genomics UK (COG-UK) Consortium | Harper VanSteenhouse, Yumi Kasai, David Gray, Carol Clugston, Anna Dominiczak and Alex Alderton, Roberto Amato, Sonia Goncalves, Ewan Harrison, David K. Jackson, Ian Johnston, Dominic Kwiatkowski, Cordelia Langford, John Sillitoe on behalf of the Wellcome Sanger Institute COVID-19 Surveillance Team |
| EPI_ISL_800641 | Lighthouse Lab in Cambridge | Wellcome Sanger Institute for the COVID-19 Genomics UK (COG-UK) Consortium | Rob Howes, The Lighthouse Lab in Cambridge and Alex Alderton, Roberto Amato, Sonia Goncalves, Ewan Harrison, David K. Jackson, Ian Johnston, Dominic Kwiatkowski, Cordelia Langford, John Sillitoe on behalf of the Wellcome Sanger Institute COVID-19 Surveillance Team |
| EPI_ISL_800643, EPI_ISL_800646 | Lighthouse Lab in Glasgow | Wellcome Sanger Institute for the COVID-19 Genomics UK (COG-UK) Consortium | Harper VanSteenhouse, Yumi Kasai, David Gray, Carol Clugston, Anna Dominiczak and Alex Alderton, Roberto Amato, Sonia Goncalves, Ewan Harrison, David K. Jackson, Ian Johnston, Dominic Kwiatkowski, Cordelia Langford, John Sillitoe on behalf of the Wellcome Sanger Institute COVID-19 Surveillance Team |
| EPI_ISL_800647, EPI_ISL_800648, EPI_ISL_800649 | Lighthouse Lab in Cambridge | Wellcome Sanger Institute for the COVID-19 Genomics UK (COG-UK) Consortium | Rob Howes, The Lighthouse Lab in Cambridge and Alex Alderton, Roberto Amato, Sonia Goncalves, Ewan Harrison, David K. Jackson, Ian Johnston, Dominic Kwiatkowski, Cordelia Langford, John Sillitoe on behalf of the Wellcome Sanger Institute COVID-19 Surveillance Team |
| EPI_ISL_800651, EPI_ISL_800653, EPI_ISL_800654, EPI_ISL_800655, EPI_ISL_800656 | Lighthouse Lab in Glasgow | Wellcome Sanger Institute for the COVID-19 Genomics UK (COG-UK) Consortium | Harper VanSteenhouse, Yumi Kasai, David Gray, Carol Clugston, Anna Dominiczak and Alex Alderton, Roberto Amato, Sonia Goncalves, Ewan Harrison, David K. Jackson, Ian Johnston, Dominic Kwiatkowski, Cordelia Langford, John Sillitoe on behalf of the Wellcome Sanger Institute COVID-19 Surveillance Team |
| EPI_ISL_800658 | Lighthouse Lab in Cambridge | Wellcome Sanger Institute for the COVID-19 Genomics UK (COG-UK) Consortium | Rob Howes, The Lighthouse Lab in Cambridge and Alex Alderton, Roberto Amato, Sonia Goncalves, Ewan Harrison, David K. Jackson, Ian Johnston, Dominic Kwiatkowski, Cordelia Langford, John Sillitoe on behalf of the Wellcome Sanger Institute COVID-19 Surveillance Team |
| EPI_ISL_800659, EPI_ISL_800662 | Lighthouse Lab in Glasgow | Wellcome Sanger Institute for the COVID-19 Genomics UK (COG-UK) Consortium | Harper VanSteenhouse, Yumi Kasai, David Gray, Carol Clugston, Anna Dominiczak and Alex Alderton, Roberto Amato, Sonia Goncalves, Ewan Harrison, David K. Jackson, Ian Johnston, Dominic Kwiatkowski, Cordelia Langford, John Sillitoe on behalf of the Wellcome Sanger Institute COVID-19 Surveillance Team |
| EPI_ISL_800663, EPI_ISL_800665 | Lighthouse Lab in Cambridge | Wellcome Sanger Institute for the COVID-19 Genomics UK (COG-UK) Consortium | Rob Howes, The Lighthouse Lab in Cambridge and Alex Alderton, Roberto Amato, Sonia Goncalves, Ewan Harrison, David K. Jackson, Ian Johnston, Dominic Kwiatkowski, Cordelia Langford, John Sillitoe on behalf of the Wellcome Sanger Institute COVID-19 Surveillance Team |
| EPI_ISL_800666, EPI_ISL_800667, EPI_ISL_800668 | Lighthouse Lab in Glasgow | Wellcome Sanger Institute for the COVID-19 Genomics UK (COG-UK) Consortium | Harper VanSteenhouse, Yumi Kasai, David Gray, Carol Clugston, Anna Dominiczak and Alex Alderton, Roberto Amato, Sonia Goncalves, Ewan Harrison, David K. Jackson, Ian Johnston, Dominic Kwiatkowski, Cordelia Langford, John Sillitoe on behalf of the Wellcome Sanger Institute COVID-19 Surveillance Team |
| EPI_ISL_800669 | Lighthouse Lab in Cambridge | Wellcome Sanger Institute for the COVID-19 Genomics UK (COG-UK) Consortium | Rob Howes, The Lighthouse Lab in Cambridge and Alex Alderton, Roberto Amato, Sonia Goncalves, Ewan Harrison, David K. Jackson, Ian Johnston, Dominic Kwiatkowski, Cordelia Langford, John Sillitoe on behalf of the Wellcome Sanger Institute COVID-19 Surveillance Team |
| EPI_ISL_800670, EPI_ISL_800671 | Lighthouse Lab in Glasgow | Wellcome Sanger Institute for the COVID-19 Genomics UK (COG-UK) Consortium | Harper VanSteenhouse, Yumi Kasai, David Gray, Carol Clugston, Anna Dominiczak and Alex Alderton, Roberto Amato, Sonia Goncalves, Ewan Harrison, David K. Jackson, Ian Johnston, Dominic Kwiatkowski, Cordelia Langford, John Sillitoe on behalf of the Wellcome Sanger Institute COVID-19 Surveillance Team |
| EPI_ISL_800672 | Lighthouse Lab in Cambridge | Wellcome Sanger Institute for the COVID-19 Genomics UK (COG-UK) Consortium | Rob Howes, The Lighthouse Lab in Cambridge and Alex Alderton, Roberto Amato, Sonia Goncalves, Ewan Harrison, David K. Jackson, Ian Johnston, Dominic Kwiatkowski, Cordelia Langford, John Sillitoe on behalf of the Wellcome Sanger Institute COVID-19 Surveillance Team |
| EPI_ISL_800675 | Lighthouse Lab in Glasgow | Wellcome Sanger Institute for the COVID-19 Genomics UK (COG-UK) Consortium | Harper VanSteenhouse, Yumi Kasai, David Gray, Carol Clugston, Anna Dominiczak and Alex Alderton, Roberto Amato, Sonia Goncalves, Ewan Harrison, David K. Jackson, Ian Johnston, Dominic Kwiatkowski, Cordelia Langford, John Sillitoe on behalf of the Wellcome Sanger Institute COVID-19 Surveillance Team |
| EPI_ISL_800679, EPI_ISL_800680 | Lighthouse Lab in Cambridge | Wellcome Sanger Institute for the COVID-19 Genomics UK (COG-UK) Consortium | Rob Howes, The Lighthouse Lab in Cambridge and Alex Alderton, Roberto Amato, Sonia Goncalves, Ewan Harrison, David K. Jackson, Ian Johnston, Dominic Kwiatkowski, Cordelia Langford, John Sillitoe on behalf of the Wellcome Sanger Institute COVID-19 Surveillance Team |
| EPI_ISL_800681 | Lighthouse Lab in Milton Keynes | Wellcome Sanger Institute for the COVID-19 Genomics UK (COG-UK) Consortium | The Lighthouse Lab in Milton Keynes and Alex Alderton, Roberto Amato, Sonia Goncalves, Ewan Harrison, David K. Jackson, Ian Johnston, Dominic Kwiatkowski, Cordelia Langford, John Sillitoe on behalf of the Wellcome Sanger Institute COVID-19 Surveillance Team |
| EPI_ISL_800682 | Lighthouse Lab in Cambridge | Wellcome Sanger Institute for the COVID-19 Genomics UK (COG-UK) Consortium | Rob Howes, The Lighthouse Lab in Cambridge and Alex Alderton, Roberto Amato, Sonia Goncalves, Ewan Harrison, David K. Jackson, Ian Johnston, Dominic Kwiatkowski, Cordelia Langford, John Sillitoe on behalf of the Wellcome Sanger Institute COVID-19 Surveillance Team |

| | | | |
|---|---|---|---|
| EPI_ISL_800685 | Lighthouse Lab in Glasgow | Wellcome Sanger Institute for the COVID-19 Genomics UK (COG-UK) Consortium | Harper VanSteenhouse, Yumi Kasai, David Gray, Carol Clugston, Anna Dominiczak and Alex Alderton, Roberto Amato, Sonia Goncalves, Ewan Harrison, David K. Jackson, Ian Johnston, Dominic Kwiatkowski, Cordelia Langford, John Sillitoe on behalf of the Wellcome Sanger Institute COVID-19 Surveillance Team |
| EPI_ISL_800686, EPI_ISL_800687 | Lighthouse Lab in Cambridge | Wellcome Sanger Institute for the COVID-19 Genomics UK (COG-UK) Consortium | Rob Howes, The Lighthouse Lab in Cambridge and Alex Alderton, Roberto Amato, Sonia Goncalves, Ewan Harrison, David K. Jackson, Ian Johnston, Dominic Kwiatkowski, Cordelia Langford, John Sillitoe on behalf of the Wellcome Sanger Institute COVID-19 Surveillance Team |
| EPI_ISL_800688, EPI_ISL_800689 | Lighthouse Lab in Glasgow | Wellcome Sanger Institute for the COVID-19 Genomics UK (COG-UK) Consortium | Harper VanSteenhouse, Yumi Kasai, David Gray, Carol Clugston, Anna Dominiczak and Alex Alderton, Roberto Amato, Sonia Goncalves, Ewan Harrison, David K. Jackson, Ian Johnston, Dominic Kwiatkowski, Cordelia Langford, John Sillitoe on behalf of the Wellcome Sanger Institute COVID-19 Surveillance Team |
| EPI_ISL_800690 | Lighthouse Lab in Cambridge | Wellcome Sanger Institute for the COVID-19 Genomics UK (COG-UK) Consortium | Rob Howes, The Lighthouse Lab in Cambridge and Alex Alderton, Roberto Amato, Sonia Goncalves, Ewan Harrison, David K. Jackson, Ian Johnston, Dominic Kwiatkowski, Cordelia Langford, John Sillitoe on behalf of the Wellcome Sanger Institute COVID-19 Surveillance Team |
| EPI_ISL_800691, EPI_ISL_800692, EPI_ISL_800693, EPI_ISL_800694, EPI_ISL_800695, EPI_ISL_800696, EPI_ISL_800697, EPI_ISL_800699, EPI_ISL_800700 | Lighthouse Lab in Glasgow | Wellcome Sanger Institute for the COVID-19 Genomics UK (COG-UK) Consortium | Harper VanSteenhouse, Yumi Kasai, David Gray, Carol Clugston, Anna Dominiczak and Alex Alderton, Roberto Amato, Sonia Goncalves, Ewan Harrison, David K. Jackson, Ian Johnston, Dominic Kwiatkowski, Cordelia Langford, John Sillitoe on behalf of the Wellcome Sanger Institute COVID-19 Surveillance Team |
| EPI_ISL_800702 | Lighthouse Lab in Cambridge | Wellcome Sanger Institute for the COVID-19 Genomics UK (COG-UK) Consortium | Rob Howes, The Lighthouse Lab in Cambridge and Alex Alderton, Roberto Amato, Sonia Goncalves, Ewan Harrison, David K. Jackson, Ian Johnston, Dominic Kwiatkowski, Cordelia Langford, John Sillitoe on behalf of the Wellcome Sanger Institute COVID-19 Surveillance Team |
| EPI_ISL_800703, EPI_ISL_800704, EPI_ISL_800705, EPI_ISL_800706 | Lighthouse Lab in Glasgow | Wellcome Sanger Institute for the COVID-19 Genomics UK (COG-UK) Consortium | Harper VanSteenhouse, Yumi Kasai, David Gray, Carol Clugston, Anna Dominiczak and Alex Alderton, Roberto Amato, Sonia Goncalves, Ewan Harrison, David K. Jackson, Ian Johnston, Dominic Kwiatkowski, Cordelia Langford, John Sillitoe on behalf of the Wellcome Sanger Institute COVID-19 Surveillance Team |
| EPI_ISL_800707, EPI_ISL_800708, EPI_ISL_800709, EPI_ISL_800712, EPI_ISL_800714 | Lighthouse Lab in Cambridge | Wellcome Sanger Institute for the COVID-19 Genomics UK (COG-UK) Consortium | Rob Howes, The Lighthouse Lab in Cambridge and Alex Alderton, Roberto Amato, Sonia Goncalves, Ewan Harrison, David K. Jackson, Ian Johnston, Dominic Kwiatkowski, Cordelia Langford, John Sillitoe on behalf of the Wellcome Sanger Institute COVID-19 Surveillance Team |
| EPI_ISL_800715 | Lighthouse Lab in Glasgow | Wellcome Sanger Institute for the COVID-19 Genomics UK (COG-UK) Consortium | Harper VanSteenhouse, Yumi Kasai, David Gray, Carol Clugston, Anna Dominiczak and Alex Alderton, Roberto Amato, Sonia Goncalves, Ewan Harrison, David K. Jackson, Ian Johnston, Dominic Kwiatkowski, Cordelia Langford, John Sillitoe on behalf of the Wellcome Sanger Institute COVID-19 Surveillance Team |
| EPI_ISL_800716, EPI_ISL_800717 | Lighthouse Lab in Cambridge | Wellcome Sanger Institute for the COVID-19 Genomics UK (COG-UK) Consortium | Rob Howes, The Lighthouse Lab in Cambridge and Alex Alderton, Roberto Amato, Sonia Goncalves, Ewan Harrison, David K. Jackson, Ian Johnston, Dominic Kwiatkowski, Cordelia Langford, John Sillitoe on behalf of the Wellcome Sanger Institute COVID-19 Surveillance Team |
| EPI_ISL_800719 | Lighthouse Lab in Glasgow | Wellcome Sanger Institute for the COVID-19 Genomics UK (COG-UK) Consortium | Harper VanSteenhouse, Yumi Kasai, David Gray, Carol Clugston, Anna Dominiczak and Alex Alderton, Roberto Amato, Sonia Goncalves, Ewan Harrison, David K. Jackson, Ian Johnston, Dominic Kwiatkowski, Cordelia Langford, John Sillitoe on behalf of the Wellcome Sanger Institute COVID-19 Surveillance Team |
| EPI_ISL_800720, EPI_ISL_800721 | Lighthouse Lab in Cambridge | Wellcome Sanger Institute for the COVID-19 Genomics UK (COG-UK) Consortium | Rob Howes, The Lighthouse Lab in Cambridge and Alex Alderton, Roberto Amato, Sonia Goncalves, Ewan Harrison, David K. Jackson, Ian Johnston, Dominic Kwiatkowski, Cordelia Langford, John Sillitoe on behalf of the Wellcome Sanger Institute COVID-19 Surveillance Team |
| EPI_ISL_800722, EPI_ISL_800723 | Lighthouse Lab in Glasgow | Wellcome Sanger Institute for the COVID-19 Genomics UK (COG-UK) Consortium | Harper VanSteenhouse, Yumi Kasai, David Gray, Carol Clugston, Anna Dominiczak and Alex Alderton, Roberto Amato, Sonia Goncalves, Ewan Harrison, David K. Jackson, Ian Johnston, Dominic Kwiatkowski, Cordelia Langford, John Sillitoe on behalf of the Wellcome Sanger Institute COVID-19 Surveillance Team |
| EPI_ISL_800724 | Lighthouse Lab in Cambridge | Wellcome Sanger Institute for the COVID-19 Genomics UK (COG-UK) Consortium | Rob Howes, The Lighthouse Lab in Cambridge and Alex Alderton, Roberto Amato, Sonia Goncalves, Ewan Harrison, David K. Jackson, Ian Johnston, Dominic Kwiatkowski, Cordelia Langford, John Sillitoe on behalf of the Wellcome Sanger Institute COVID-19 Surveillance Team |
| EPI_ISL_800725, EPI_ISL_800726, EPI_ISL_800727, EPI_ISL_800728 | Lighthouse Lab in Glasgow | Wellcome Sanger Institute for the COVID-19 Genomics UK (COG-UK) Consortium | Harper VanSteenhouse, Yumi Kasai, David Gray, Carol Clugston, Anna Dominiczak and Alex Alderton, Roberto Amato, Sonia Goncalves, Ewan Harrison, David K. Jackson, Ian Johnston, Dominic Kwiatkowski, Cordelia Langford, John Sillitoe on behalf of the Wellcome Sanger Institute COVID-19 Surveillance Team |
| EPI_ISL_800730 | Lighthouse Lab in Cambridge | Wellcome Sanger Institute for the COVID-19 Genomics UK (COG-UK) Consortium | Rob Howes, The Lighthouse Lab in Cambridge and Alex Alderton, Roberto Amato, Sonia Goncalves, Ewan Harrison, David K. Jackson, Ian Johnston, Dominic Kwiatkowski, Cordelia Langford, John Sillitoe on behalf of the Wellcome Sanger Institute COVID-19 Surveillance Team |
| EPI_ISL_800731 | Lighthouse Lab in Glasgow | Wellcome Sanger Institute for the COVID-19 Genomics UK (COG-UK) Consortium | Harper VanSteenhouse, Yumi Kasai, David Gray, Carol Clugston, Anna Dominiczak and Alex Alderton, Roberto Amato, Sonia Goncalves, Ewan Harrison, David K. Jackson, Ian Johnston, Dominic Kwiatkowski, Cordelia Langford, John Sillitoe on behalf of the Wellcome Sanger Institute COVID-19 Surveillance Team |
| EPI_ISL_800733, EPI_ISL_800734 | Lighthouse Lab in Cambridge | Wellcome Sanger Institute for the COVID-19 Genomics UK (COG-UK) Consortium | Rob Howes, The Lighthouse Lab in Cambridge and Alex Alderton, Roberto Amato, Sonia Goncalves, Ewan Harrison, David K. Jackson, Ian Johnston, Dominic Kwiatkowski, Cordelia Langford, John Sillitoe on behalf of the Wellcome Sanger Institute COVID-19 Surveillance Team |
| EPI_ISL_800735, EPI_ISL_800736 | Lighthouse Lab in Glasgow | Wellcome Sanger Institute for the COVID-19 Genomics UK (COG-UK) Consortium | Harper VanSteenhouse, Yumi Kasai, David Gray, Carol Clugston, Anna Dominiczak and Alex Alderton, Roberto Amato, Sonia Goncalves, Ewan Harrison, David K. Jackson, Ian Johnston, Dominic Kwiatkowski, Cordelia Langford, John Sillitoe on behalf of the Wellcome Sanger Institute COVID-19 Surveillance Team |
| EPI_ISL_800737 | Lighthouse Lab in Cambridge | Wellcome Sanger Institute for the COVID-19 Genomics UK (COG-UK) Consortium | Rob Howes, The Lighthouse Lab in Cambridge and Alex Alderton, Roberto Amato, Sonia Goncalves, Ewan Harrison, David K. Jackson, Ian Johnston, Dominic Kwiatkowski, Cordelia Langford, John Sillitoe on behalf of the Wellcome Sanger Institute COVID-19 Surveillance Team |
| EPI_ISL_800739, EPI_ISL_800742 | Lighthouse Lab in Glasgow | Wellcome Sanger Institute for the COVID-19 Genomics UK (COG-UK) Consortium | Harper VanSteenhouse, Yumi Kasai, David Gray, Carol Clugston, Anna Dominiczak and Alex Alderton, Roberto Amato, Sonia Goncalves, Ewan Harrison, David K. Jackson, Ian Johnston, Dominic Kwiatkowski, Cordelia Langford, John Sillitoe on behalf of the Wellcome Sanger Institute COVID-19 Surveillance Team |
| EPI_ISL_800744 | Lighthouse Lab in Cambridge | Wellcome Sanger Institute for the COVID-19 Genomics UK (COG-UK) Consortium | Rob Howes, The Lighthouse Lab in Cambridge and Alex Alderton, Roberto Amato, Sonia Goncalves, Ewan Harrison, David K. Jackson, Ian Johnston, Dominic Kwiatkowski, Cordelia Langford, John Sillitoe on behalf of the Wellcome Sanger Institute COVID-19 Surveillance Team |
| EPI_ISL_800745 | Lighthouse Lab in Glasgow | Wellcome Sanger Institute for the COVID-19 Genomics UK (COG-UK) Consortium | Harper VanSteenhouse, Yumi Kasai, David Gray, Carol Clugston, Anna Dominiczak and Alex Alderton, Roberto Amato, Sonia Goncalves, Ewan Harrison, David K. Jackson, Ian Johnston, Dominic Kwiatkowski, Cordelia Langford, John Sillitoe on behalf of the Wellcome Sanger Institute COVID-19 Surveillance Team |
| EPI_ISL_800746 | Lighthouse Lab in Cambridge | Wellcome Sanger Institute for the COVID-19 Genomics UK (COG-UK) Consortium | Rob Howes, The Lighthouse Lab in Cambridge and Alex Alderton, Roberto Amato, Sonia Goncalves, Ewan Harrison, David K. Jackson, Ian Johnston, Dominic Kwiatkowski, Cordelia Langford, John Sillitoe on behalf of the Wellcome Sanger Institute COVID-19 Surveillance Team |
| EPI_ISL_800748, EPI_ISL_800749, EPI_ISL_800753, EPI_ISL_800754 | Lighthouse Lab in Glasgow | Wellcome Sanger Institute for the COVID-19 Genomics UK (COG-UK) Consortium | Harper VanSteenhouse, Yumi Kasai, David Gray, Carol Clugston, Anna Dominiczak and Alex Alderton, Roberto Amato, Sonia Goncalves, Ewan Harrison, David K. Jackson, Ian Johnston, Dominic Kwiatkowski, Cordelia Langford, John Sillitoe on behalf of the Wellcome Sanger Institute COVID-19 Surveillance Team |
| EPI_ISL_800755 | Lighthouse Lab in Cambridge | Wellcome Sanger Institute for the COVID-19 Genomics UK (COG-UK) Consortium | Rob Howes, The Lighthouse Lab in Cambridge and Alex Alderton, Roberto Amato, Sonia Goncalves, Ewan Harrison, David K. Jackson, Ian Johnston, Dominic Kwiatkowski, Cordelia Langford, John Sillitoe on behalf of the Wellcome Sanger Institute COVID-19 Surveillance Team |
| EPI_ISL_800759, EPI_ISL_800760 | Lighthouse Lab in Glasgow | Wellcome Sanger Institute for the COVID-19 Genomics UK (COG-UK) Consortium | Harper VanSteenhouse, Yumi Kasai, David Gray, Carol Clugston, Anna Dominiczak and Alex Alderton, Roberto Amato, Sonia Goncalves, Ewan Harrison, David K. Jackson, Ian Johnston, Dominic Kwiatkowski, Cordelia Langford, John Sillitoe on behalf of the Wellcome Sanger Institute COVID-19 Surveillance Team |
| EPI_ISL_800763 | Lighthouse Lab in Cambridge | Wellcome Sanger Institute for the COVID-19 Genomics UK (COG-UK) Consortium | Rob Howes, The Lighthouse Lab in Cambridge and Alex Alderton, Roberto Amato, Sonia Goncalves, Ewan Harrison, David K. Jackson, Ian Johnston, Dominic Kwiatkowski, Cordelia Langford, John Sillitoe on behalf of the Wellcome Sanger Institute COVID-19 Surveillance Team |

| | | | |
|---|---|---|---|
| EPI_ISL_800765, EPI_ISL_800766, EPI_ISL_800767 | Lighthouse Lab in Glasgow | Wellcome Sanger Institute for the COVID-19 Genomics UK (COG-UK) Consortium | Harper VanSteenhouse, Yumi Kasai, David Gray, Carol Clugston, Anna Dominiczak and Alex Alderton, Roberto Amato, Sonia Goncalves, Ewan Harrison, David K. Jackson, Ian Johnston, Dominic Kwiatkowski, Cordelia Langford, John Sillitoe on behalf of the Wellcome Sanger Institute COVID-19 Surveillance Team |
| EPI_ISL_800768, EPI_ISL_800770, EPI_ISL_800771 | Lighthouse Lab in Cambridge | Wellcome Sanger Institute for the COVID-19 Genomics UK (COG-UK) Consortium | Rob Howes, The Lighthouse Lab in Cambridge and Alex Alderton, Roberto Amato, Sonia Goncalves, Ewan Harrison, David K. Jackson, Ian Johnston, Dominic Kwiatkowski, Cordelia Langford, John Sillitoe on behalf of the Wellcome Sanger Institute COVID-19 Surveillance Team |
| EPI_ISL_800772, EPI_ISL_800775 | Lighthouse Lab in Glasgow | Wellcome Sanger Institute for the COVID-19 Genomics UK (COG-UK) Consortium | Harper VanSteenhouse, Yumi Kasai, David Gray, Carol Clugston, Anna Dominiczak and Alex Alderton, Roberto Amato, Sonia Goncalves, Ewan Harrison, David K. Jackson, Ian Johnston, Dominic Kwiatkowski, Cordelia Langford, John Sillitoe on behalf of the Wellcome Sanger Institute COVID-19 Surveillance Team |
| EPI_ISL_800776 | Lighthouse Lab in Cambridge | Wellcome Sanger Institute for the COVID-19 Genomics UK (COG-UK) Consortium | Rob Howes, The Lighthouse Lab in Cambridge and Alex Alderton, Roberto Amato, Sonia Goncalves, Ewan Harrison, David K. Jackson, Ian Johnston, Dominic Kwiatkowski, Cordelia Langford, John Sillitoe on behalf of the Wellcome Sanger Institute COVID-19 Surveillance Team |
| EPI_ISL_800779, EPI_ISL_800780, EPI_ISL_800781 | Lighthouse Lab in Glasgow | Wellcome Sanger Institute for the COVID-19 Genomics UK (COG-UK) Consortium | Harper VanSteenhouse, Yumi Kasai, David Gray, Carol Clugston, Anna Dominiczak and Alex Alderton, Roberto Amato, Sonia Goncalves, Ewan Harrison, David K. Jackson, Ian Johnston, Dominic Kwiatkowski, Cordelia Langford, John Sillitoe on behalf of the Wellcome Sanger Institute COVID-19 Surveillance Team |
| EPI_ISL_800784 | Lighthouse Lab in Cambridge | Wellcome Sanger Institute for the COVID-19 Genomics UK (COG-UK) Consortium | Rob Howes, The Lighthouse Lab in Cambridge and Alex Alderton, Roberto Amato, Sonia Goncalves, Ewan Harrison, David K. Jackson, Ian Johnston, Dominic Kwiatkowski, Cordelia Langford, John Sillitoe on behalf of the Wellcome Sanger Institute COVID-19 Surveillance Team |
| EPI_ISL_800785, EPI_ISL_800786, EPI_ISL_800790 | Lighthouse Lab in Glasgow | Wellcome Sanger Institute for the COVID-19 Genomics UK (COG-UK) Consortium | Harper VanSteenhouse, Yumi Kasai, David Gray, Carol Clugston, Anna Dominiczak and Alex Alderton, Roberto Amato, Sonia Goncalves, Ewan Harrison, David K. Jackson, Ian Johnston, Dominic Kwiatkowski, Cordelia Langford, John Sillitoe on behalf of the Wellcome Sanger Institute COVID-19 Surveillance Team |
| EPI_ISL_800791, EPI_ISL_800792, EPI_ISL_800793 | Lighthouse Lab in Cambridge | Wellcome Sanger Institute for the COVID-19 Genomics UK (COG-UK) Consortium | Rob Howes, The Lighthouse Lab in Cambridge and Alex Alderton, Roberto Amato, Sonia Goncalves, Ewan Harrison, David K. Jackson, Ian Johnston, Dominic Kwiatkowski, Cordelia Langford, John Sillitoe on behalf of the Wellcome Sanger Institute COVID-19 Surveillance Team |
| EPI_ISL_800794 | Lighthouse Lab in Glasgow | Wellcome Sanger Institute for the COVID-19 Genomics UK (COG-UK) Consortium | Harper VanSteenhouse, Yumi Kasai, David Gray, Carol Clugston, Anna Dominiczak and Alex Alderton, Roberto Amato, Sonia Goncalves, Ewan Harrison, David K. Jackson, Ian Johnston, Dominic Kwiatkowski, Cordelia Langford, John Sillitoe on behalf of the Wellcome Sanger Institute COVID-19 Surveillance Team |
| EPI_ISL_800795 | Lighthouse Lab in Cambridge | Wellcome Sanger Institute for the COVID-19 Genomics UK (COG-UK) Consortium | Rob Howes, The Lighthouse Lab in Cambridge and Alex Alderton, Roberto Amato, Sonia Goncalves, Ewan Harrison, David K. Jackson, Ian Johnston, Dominic Kwiatkowski, Cordelia Langford, John Sillitoe on behalf of the Wellcome Sanger Institute COVID-19 Surveillance Team |
| EPI_ISL_800796, EPI_ISL_800797 | Lighthouse Lab in Glasgow | Wellcome Sanger Institute for the COVID-19 Genomics UK (COG-UK) Consortium | Harper VanSteenhouse, Yumi Kasai, David Gray, Carol Clugston, Anna Dominiczak and Alex Alderton, Roberto Amato, Sonia Goncalves, Ewan Harrison, David K. Jackson, Ian Johnston, Dominic Kwiatkowski, Cordelia Langford, John Sillitoe on behalf of the Wellcome Sanger Institute COVID-19 Surveillance Team |
| EPI_ISL_800798 | Lighthouse Lab in Cambridge | Wellcome Sanger Institute for the COVID-19 Genomics UK (COG-UK) Consortium | Rob Howes, The Lighthouse Lab in Cambridge and Alex Alderton, Roberto Amato, Sonia Goncalves, Ewan Harrison, David K. Jackson, Ian Johnston, Dominic Kwiatkowski, Cordelia Langford, John Sillitoe on behalf of the Wellcome Sanger Institute COVID-19 Surveillance Team |
| EPI_ISL_800799, EPI_ISL_800800, EPI_ISL_800801 | Lighthouse Lab in Glasgow | Wellcome Sanger Institute for the COVID-19 Genomics UK (COG-UK) Consortium | Harper VanSteenhouse, Yumi Kasai, David Gray, Carol Clugston, Anna Dominiczak and Alex Alderton, Roberto Amato, Sonia Goncalves, Ewan Harrison, David K. Jackson, Ian Johnston, Dominic Kwiatkowski, Cordelia Langford, John Sillitoe on behalf of the Wellcome Sanger Institute COVID-19 Surveillance Team |
| EPI_ISL_800802, EPI_ISL_800803 | Lighthouse Lab in Cambridge | Wellcome Sanger Institute for the COVID-19 Genomics UK (COG-UK) Consortium | Rob Howes, The Lighthouse Lab in Cambridge and Alex Alderton, Roberto Amato, Sonia Goncalves, Ewan Harrison, David K. Jackson, Ian Johnston, Dominic Kwiatkowski, Cordelia Langford, John Sillitoe on behalf of the Wellcome Sanger Institute COVID-19 Surveillance Team |
| EPI_ISL_800804, EPI_ISL_800805 | Lighthouse Lab in Glasgow | Wellcome Sanger Institute for the COVID-19 Genomics UK (COG-UK) Consortium | Harper VanSteenhouse, Yumi Kasai, David Gray, Carol Clugston, Anna Dominiczak and Alex Alderton, Roberto Amato, Sonia Goncalves, Ewan Harrison, David K. Jackson, Ian Johnston, Dominic Kwiatkowski, Cordelia Langford, John Sillitoe on behalf of the Wellcome Sanger Institute COVID-19 Surveillance Team |
| EPI_ISL_800806 | Lighthouse Lab in Cambridge | Wellcome Sanger Institute for the COVID-19 Genomics UK (COG-UK) Consortium | Rob Howes, The Lighthouse Lab in Cambridge and Alex Alderton, Roberto Amato, Sonia Goncalves, Ewan Harrison, David K. Jackson, Ian Johnston, Dominic Kwiatkowski, Cordelia Langford, John Sillitoe on behalf of the Wellcome Sanger Institute COVID-19 Surveillance Team |
| EPI_ISL_800809 | Lighthouse Lab in Glasgow | Wellcome Sanger Institute for the COVID-19 Genomics UK (COG-UK) Consortium | Harper VanSteenhouse, Yumi Kasai, David Gray, Carol Clugston, Anna Dominiczak and Alex Alderton, Roberto Amato, Sonia Goncalves, Ewan Harrison, David K. Jackson, Ian Johnston, Dominic Kwiatkowski, Cordelia Langford, John Sillitoe on behalf of the Wellcome Sanger Institute COVID-19 Surveillance Team |
| EPI_ISL_800810 | Lighthouse Lab in Cambridge | Wellcome Sanger Institute for the COVID-19 Genomics UK (COG-UK) Consortium | Rob Howes, The Lighthouse Lab in Cambridge and Alex Alderton, Roberto Amato, Sonia Goncalves, Ewan Harrison, David K. Jackson, Ian Johnston, Dominic Kwiatkowski, Cordelia Langford, John Sillitoe on behalf of the Wellcome Sanger Institute COVID-19 Surveillance Team |
| EPI_ISL_800812, EPI_ISL_800813, EPI_ISL_800815 | Lighthouse Lab in Glasgow | Wellcome Sanger Institute for the COVID-19 Genomics UK (COG-UK) Consortium | Harper VanSteenhouse, Yumi Kasai, David Gray, Carol Clugston, Anna Dominiczak and Alex Alderton, Roberto Amato, Sonia Goncalves, Ewan Harrison, David K. Jackson, Ian Johnston, Dominic Kwiatkowski, Cordelia Langford, John Sillitoe on behalf of the Wellcome Sanger Institute COVID-19 Surveillance Team |
| EPI_ISL_800816, EPI_ISL_800817 | Lighthouse Lab in Cambridge | Wellcome Sanger Institute for the COVID-19 Genomics UK (COG-UK) Consortium | Rob Howes, The Lighthouse Lab in Cambridge and Alex Alderton, Roberto Amato, Sonia Goncalves, Ewan Harrison, David K. Jackson, Ian Johnston, Dominic Kwiatkowski, Cordelia Langford, John Sillitoe on behalf of the Wellcome Sanger Institute COVID-19 Surveillance Team |
| EPI_ISL_800818 | Lighthouse Lab in Glasgow | Wellcome Sanger Institute for the COVID-19 Genomics UK (COG-UK) Consortium | Harper VanSteenhouse, Yumi Kasai, David Gray, Carol Clugston, Anna Dominiczak and Alex Alderton, Roberto Amato, Sonia Goncalves, Ewan Harrison, David K. Jackson, Ian Johnston, Dominic Kwiatkowski, Cordelia Langford, John Sillitoe on behalf of the Wellcome Sanger Institute COVID-19 Surveillance Team |
| EPI_ISL_800819 | Lighthouse Lab in Cambridge | Wellcome Sanger Institute for the COVID-19 Genomics UK (COG-UK) Consortium | Rob Howes, The Lighthouse Lab in Cambridge and Alex Alderton, Roberto Amato, Sonia Goncalves, Ewan Harrison, David K. Jackson, Ian Johnston, Dominic Kwiatkowski, Cordelia Langford, John Sillitoe on behalf of the Wellcome Sanger Institute COVID-19 Surveillance Team |
| EPI_ISL_800823, EPI_ISL_800825, EPI_ISL_800826, EPI_ISL_800827 | Lighthouse Lab in Glasgow | Wellcome Sanger Institute for the COVID-19 Genomics UK (COG-UK) Consortium | Harper VanSteenhouse, Yumi Kasai, David Gray, Carol Clugston, Anna Dominiczak and Alex Alderton, Roberto Amato, Sonia Goncalves, Ewan Harrison, David K. Jackson, Ian Johnston, Dominic Kwiatkowski, Cordelia Langford, John Sillitoe on behalf of the Wellcome Sanger Institute COVID-19 Surveillance Team |
| EPI_ISL_800828 | Lighthouse Lab in Cambridge | Wellcome Sanger Institute for the COVID-19 Genomics UK (COG-UK) Consortium | Rob Howes, The Lighthouse Lab in Cambridge and Alex Alderton, Roberto Amato, Sonia Goncalves, Ewan Harrison, David K. Jackson, Ian Johnston, Dominic Kwiatkowski, Cordelia Langford, John Sillitoe on behalf of the Wellcome Sanger Institute COVID-19 Surveillance Team |
| EPI_ISL_800829 | Lighthouse Lab in Glasgow | Wellcome Sanger Institute for the COVID-19 Genomics UK (COG-UK) Consortium | Harper VanSteenhouse, Yumi Kasai, David Gray, Carol Clugston, Anna Dominiczak and Alex Alderton, Roberto Amato, Sonia Goncalves, Ewan Harrison, David K. Jackson, Ian Johnston, Dominic Kwiatkowski, Cordelia Langford, John Sillitoe on behalf of the Wellcome Sanger Institute COVID-19 Surveillance Team |
| EPI_ISL_800832 | Lighthouse Lab in Cambridge | Wellcome Sanger Institute for the COVID-19 Genomics UK (COG-UK) Consortium | Rob Howes, The Lighthouse Lab in Cambridge and Alex Alderton, Roberto Amato, Sonia Goncalves, Ewan Harrison, David K. Jackson, Ian Johnston, Dominic Kwiatkowski, Cordelia Langford, John Sillitoe on behalf of the Wellcome Sanger Institute COVID-19 Surveillance Team |
| EPI_ISL_800833, EPI_ISL_800835, EPI_ISL_800838 | Lighthouse Lab in Glasgow | Wellcome Sanger Institute for the COVID-19 Genomics UK (COG-UK) Consortium | Harper VanSteenhouse, Yumi Kasai, David Gray, Carol Clugston, Anna Dominiczak and Alex Alderton, Roberto Amato, Sonia Goncalves, Ewan Harrison, David K. Jackson, Ian Johnston, Dominic Kwiatkowski, Cordelia Langford, John Sillitoe on behalf of the Wellcome Sanger Institute COVID-19 Surveillance Team |
| EPI_ISL_800839 | Lighthouse Lab in Cambridge | Wellcome Sanger Institute for the COVID-19 Genomics UK (COG-UK) Consortium | Rob Howes, The Lighthouse Lab in Cambridge and Alex Alderton, Roberto Amato, Sonia Goncalves, Ewan Harrison, David K. Jackson, Ian Johnston, Dominic Kwiatkowski, Cordelia Langford, John Sillitoe on behalf of the Wellcome Sanger Institute COVID-19 Surveillance Team |
| EPI_ISL_800841 | Lighthouse Lab in Glasgow | Wellcome Sanger Institute for the COVID-19 Genomics UK (COG-UK) Consortium | Harper VanSteenhouse, Yumi Kasai, David Gray, Carol Clugston, Anna Dominiczak and Alex Alderton, Roberto Amato, Sonia Goncalves, Ewan Harrison, David K. Jackson, Ian Johnston, Dominic Kwiatkowski, Cordelia Langford, John Sillitoe on behalf of the Wellcome Sanger Institute COVID-19 Surveillance Team |

| | | | |
|---|---|---|---|
| EPI_ISL_800842 | Lighthouse Lab in Cambridge | Wellcome Sanger Institute for the COVID-19 Genomics UK (COG-UK) Consortium | Rob Howes, The Lighthouse Lab in Cambridge and Alex Alderton, Roberto Amato, Sonia Goncalves, Ewan Harrison, David K. Jackson, Ian Johnston, Dominic Kwiatkowski, Cordelia Langford, John Sillitoe on behalf of the Wellcome Sanger Institute COVID-19 Surveillance Team |
| EPI_ISL_800843, EPI_ISL_800847, EPI_ISL_800848, EPI_ISL_800850, EPI_ISL_800851, EPI_ISL_800852 | Lighthouse Lab in Glasgow | Wellcome Sanger Institute for the COVID-19 Genomics UK (COG-UK) Consortium | Harper VanSteenhouse, Yumi Kasai, David Gray, Carol Clugston, Anna Dominiczak and Alex Alderton, Roberto Amato, Sonia Goncalves, Ewan Harrison, David K. Jackson, Ian Johnston, Dominic Kwiatkowski, Cordelia Langford, John Sillitoe on behalf of the Wellcome Sanger Institute COVID-19 Surveillance Team |
| EPI_ISL_800853, EPI_ISL_800854, EPI_ISL_800855 | Lighthouse Lab in Cambridge | Wellcome Sanger Institute for the COVID-19 Genomics UK (COG-UK) Consortium | Rob Howes, The Lighthouse Lab in Cambridge and Alex Alderton, Roberto Amato, Sonia Goncalves, Ewan Harrison, David K. Jackson, Ian Johnston, Dominic Kwiatkowski, Cordelia Langford, John Sillitoe on behalf of the Wellcome Sanger Institute COVID-19 Surveillance Team |
| EPI_ISL_800856, EPI_ISL_800858 | Lighthouse Lab in Glasgow | Wellcome Sanger Institute for the COVID-19 Genomics UK (COG-UK) Consortium | Harper VanSteenhouse, Yumi Kasai, David Gray, Carol Clugston, Anna Dominiczak and Alex Alderton, Roberto Amato, Sonia Goncalves, Ewan Harrison, David K. Jackson, Ian Johnston, Dominic Kwiatkowski, Cordelia Langford, John Sillitoe on behalf of the Wellcome Sanger Institute COVID-19 Surveillance Team |
| EPI_ISL_800859 | Lighthouse Lab in Cambridge | Wellcome Sanger Institute for the COVID-19 Genomics UK (COG-UK) Consortium | Rob Howes, The Lighthouse Lab in Cambridge and Alex Alderton, Roberto Amato, Sonia Goncalves, Ewan Harrison, David K. Jackson, Ian Johnston, Dominic Kwiatkowski, Cordelia Langford, John Sillitoe on behalf of the Wellcome Sanger Institute COVID-19 Surveillance Team |
| EPI_ISL_800860, EPI_ISL_800861, EPI_ISL_800862 | Lighthouse Lab in Glasgow | Wellcome Sanger Institute for the COVID-19 Genomics UK (COG-UK) Consortium | Harper VanSteenhouse, Yumi Kasai, David Gray, Carol Clugston, Anna Dominiczak and Alex Alderton, Roberto Amato, Sonia Goncalves, Ewan Harrison, David K. Jackson, Ian Johnston, Dominic Kwiatkowski, Cordelia Langford, John Sillitoe on behalf of the Wellcome Sanger Institute COVID-19 Surveillance Team |
| EPI_ISL_800865, EPI_ISL_800870, EPI_ISL_800872, EPI_ISL_800873 | Lighthouse Lab in Cambridge | Wellcome Sanger Institute for the COVID-19 Genomics UK (COG-UK) Consortium | Rob Howes, The Lighthouse Lab in Cambridge and Alex Alderton, Roberto Amato, Sonia Goncalves, Ewan Harrison, David K. Jackson, Ian Johnston, Dominic Kwiatkowski, Cordelia Langford, John Sillitoe on behalf of the Wellcome Sanger Institute COVID-19 Surveillance Team |
| EPI_ISL_800874, EPI_ISL_800875 | Lighthouse Lab in Glasgow | Wellcome Sanger Institute for the COVID-19 Genomics UK (COG-UK) Consortium | Harper VanSteenhouse, Yumi Kasai, David Gray, Carol Clugston, Anna Dominiczak and Alex Alderton, Roberto Amato, Sonia Goncalves, Ewan Harrison, David K. Jackson, Ian Johnston, Dominic Kwiatkowski, Cordelia Langford, John Sillitoe on behalf of the Wellcome Sanger Institute COVID-19 Surveillance Team |
| EPI_ISL_800876 | Lighthouse Lab in Cambridge | Wellcome Sanger Institute for the COVID-19 Genomics UK (COG-UK) Consortium | Rob Howes, The Lighthouse Lab in Cambridge and Alex Alderton, Roberto Amato, Sonia Goncalves, Ewan Harrison, David K. Jackson, Ian Johnston, Dominic Kwiatkowski, Cordelia Langford, John Sillitoe on behalf of the Wellcome Sanger Institute COVID-19 Surveillance Team |
| EPI_ISL_800880, EPI_ISL_800882 | Lighthouse Lab in Glasgow | Wellcome Sanger Institute for the COVID-19 Genomics UK (COG-UK) Consortium | Harper VanSteenhouse, Yumi Kasai, David Gray, Carol Clugston, Anna Dominiczak and Alex Alderton, Roberto Amato, Sonia Goncalves, Ewan Harrison, David K. Jackson, Ian Johnston, Dominic Kwiatkowski, Cordelia Langford, John Sillitoe on behalf of the Wellcome Sanger Institute COVID-19 Surveillance Team |
| EPI_ISL_800883 | Lighthouse Lab in Cambridge | Wellcome Sanger Institute for the COVID-19 Genomics UK (COG-UK) Consortium | Rob Howes, The Lighthouse Lab in Cambridge and Alex Alderton, Roberto Amato, Sonia Goncalves, Ewan Harrison, David K. Jackson, Ian Johnston, Dominic Kwiatkowski, Cordelia Langford, John Sillitoe on behalf of the Wellcome Sanger Institute COVID-19 Surveillance Team |
| EPI_ISL_800884 | Lighthouse Lab in Glasgow | Wellcome Sanger Institute for the COVID-19 Genomics UK (COG-UK) Consortium | Harper VanSteenhouse, Yumi Kasai, David Gray, Carol Clugston, Anna Dominiczak and Alex Alderton, Roberto Amato, Sonia Goncalves, Ewan Harrison, David K. Jackson, Ian Johnston, Dominic Kwiatkowski, Cordelia Langford, John Sillitoe on behalf of the Wellcome Sanger Institute COVID-19 Surveillance Team |
| EPI_ISL_801180, EPI_ISL_801181 | Lighthouse Lab in Milton Keynes | Wellcome Sanger Institute for the COVID-19 Genomics UK (COG-UK) Consortium | The Lighthouse Lab in Milton Keynes and Alex Alderton, Roberto Amato, Sonia Goncalves, Ewan Harrison, David K. Jackson, Ian Johnston, Dominic Kwiatkowski, Cordelia Langford, John Sillitoe on behalf of the Wellcome Sanger Institute COVID-19 Surveillance Team |
| EPI_ISL_801405, EPI_ISL_801409, EPI_ISL_801426, EPI_ISL_801471, EPI_ISL_801473, EPI_ISL_801474, EPI_ISL_801475, EPI_ISL_801476, EPI_ISL_801481 | Dutch COVID-19 response team | Erasmus Medical Center | Bas Oude Munnink, Reina Sikkema, David Nieuwenhuijse, Irina Chestakova, Anne van der Linden, Marjan Boter, Emmanuelle Munger, Corine GeurtsvanKessel, Annemiek van der Eijk, Richard Molenkamp, Marion Koopmans, on behalf of the Dutch national COVID-19 response team. |
| EPI_ISL_802865, EPI_ISL_802866, EPI_ISL_802869, EPI_ISL_802872, EPI_ISL_802876, EPI_ISL_802879, EPI_ISL_802880, EPI_ISL_802886, EPI_ISL_802887, EPI_ISL_802893, EPI_ISL_802894, EPI_ISL_802898, EPI_ISL_802902, EPI_ISL_802906, EPI_ISL_802908, EPI_ISL_802909, EPI_ISL_802911, EPI_ISL_802914, EPI_ISL_802917 | | | |
| see above | Utah Public Health Laboratory | Utah Public Health Laboratory | Erin Young, Kelly Oakeson, Tara Gallagher |
| EPI_ISL_803874, EPI_ISL_803875 | Institute of Medical Microbiology and Hospital Hygiene | Institute of Medical Microbiology and Hospital Hygiene | Prof. Dr. Achim Kaasch |
| EPI_ISL_803911 | BBMP Urban PHC | Department of Neurovirology, National Institute of Mental Health and Neurosciences (NIMHANS) | Chitra Pattabiraman, Pramada Prasad, Risha Rasheed, Anita S Desai, V Ravi |
| EPI_ISL_804609, EPI_ISL_804694, EPI_ISL_804707 | Michigan Department of Health and Human Services, Bureau of Laboratories | Michigan Department of Health and Human Services, Bureau of Laboratories | Blankenship HM, Riner D, Soehnlen MK |
| EPI_ISL_804934 | DC Public Health Lab/ Dept. of Forensic Sciences | DC Public Health Lab/ Dept. of Forensic Sciences | Scott Nguyen, Elizabeth Zelaya, Connie Maza, Monica Mann, Brittany Hamilton, David Payne, Jocelyn Hauser |
| EPI_ISL_806843, EPI_ISL_806844 | Alaska State Virology Laboratory | Alaska State Virology Laboratory | Stephanie DeRonde, Lisa Smith, Ph.D., Devin M. Drown, Ph.D., Jack Chen, Ph.D. |
| EPI_ISL_812402 | Utah Public Health Laboratory | Utah Public Health Laboratory | Erin L. Young, Kelly F. Oakeson, Tara Gallagher |
| EPI_ISL_813030, EPI_ISL_813031, EPI_ISL_813032, EPI_ISL_813033, EPI_ISL_813034, EPI_ISL_813035, EPI_ISL_813036, EPI_ISL_813071 | University of Birmingham | COVID-19 Genomics UK (COG-UK) Consortium | Institute of Microbiology, University of Birmingham: Claire McMurray, Joanne Stockton, Samuel Nicholls, Radoslaw Poplawski, Will Rowe, Josh Quick, Nicholas Loman. University of Birmingham Testing Laboratory: Celina M Whalley, Andrew Bosworth, Charlotte Poxon, Kasun Wanigasooriya, Oliver Pickles, Mike Kidd, Alex Richter, Andrew D Beggs PHE Heartlands Lab: Husam Osman, Andrew Bosworth. Queen Elizabeth Hospital: Anna Casey |
| EPI_ISL_813678, EPI_ISL_813679, EPI_ISL_813681, EPI_ISL_813682, EPI_ISL_813683, EPI_ISL_813684, EPI_ISL_813685, EPI_ISL_813686, EPI_ISL_813688, EPI_ISL_813689, EPI_ISL_813690, EPI_ISL_813691, EPI_ISL_813692, EPI_ISL_813693, EPI_ISL_813694, EPI_ISL_813695, EPI_ISL_813696, EPI_ISL_813697, EPI_ISL_813698, EPI_ISL_813699, EPI_ISL_813700, EPI_ISL_813701, EPI_ISL_813702, EPI_ISL_813703, EPI_ISL_813704, EPI_ISL_813723, EPI_ISL_813724, EPI_ISL_813725, EPI_ISL_813726, EPI_ISL_813727, EPI_ISL_813728, EPI_ISL_813729, EPI_ISL_813730, EPI_ISL_813731 | | | |
| see above | Liverpool Clinical Laboratories | COVID-19 Genomics UK (COG-UK) Consortium | Sam Haldenby, Anita Lucaci, Steve Paterson, Julian Hiscox, Alistair Darby, M Almsaud, A Alrezaihi, Muhannad Alruwaili, Stuart D Armstrong, Jones Benjamin, Eleanor G Bentley, Anu Chawla, Jordan J Clark, Angela Cowell, Richard Eccles, Isabel García-Dorival, Matthew Gemmell, Alessandro Gerada, PKF Gilmore, Richard Gregory, Ximeng Han, Catherine Hartley, Margaret Hughes, Miren Iturriza-Gomara, James Johnson, L Luu, Jenifer Manson, Charlotte Nelson, Elaine O'Toole, Cassie Olateju, Rebekah Penrice-Randal , Lucille Rainbow, N.P Randle, Trevor Ian Robinson, Parul Sharma, Ghada T Shawli, James P Stewart, Neil Swainston, Ecaterina Vamos, Joanne Watts, Mark Whitehead |
| EPI_ISL_814299, EPI_ISL_814307, EPI_ISL_814316, EPI_ISL_814322, EPI_ISL_814327 | Wales Specialist Virology Centre Sequencing lab: Pathogen Genomics Unit | COVID-19 Genomics UK (COG-UK) Consortium | Catherine Moore, Johnathan Evans, Laura Gifford, Malorie Perry, Simon Cottrell, Angela Marchbank, Alec Birchley, Alexander Adams, Amy Gaskin, Bree Gatica-Wilcox, Jason Coombes, Joel Southgate, Lauren Gilbert, Lee Graham, Nicole Pacchiarini, Sara Kumziene-Summerhayes, Sarah Taylor, Sophie Jones, Sara Rey, Matthew Bull, Joanne Watkins, Sally Corden, Tom Connor |
| EPI_ISL_814365 | Virology Department, Royal Infirmary of Edinburgh, NHS Lothian / School of Biological Sciences, University of Edinburgh / Institute of Genetics and Molecular Medicine, University of Edinburgh | COVID-19 Genomics UK (COG-UK) Consortium | McHugh M, Dewar R, Rooke S, Gallagher M, Balcaza C, O'Toole Á, Scher E, Hill V, McCrone JT, Colquhoun R, Yu X, Jackson B, Rambaut A, Williams TC, Templeton K |
| EPI_ISL_814391, EPI_ISL_814409, EPI_ISL_814545, EPI_ISL_814559 | Wales Specialist Virology Centre Sequencing lab: Pathogen Genomics Unit | COVID-19 Genomics UK (COG-UK) Consortium | Catherine Moore, Johnathan Evans, Laura Gifford, Malorie Perry, Simon Cottrell, Angela Marchbank, Alec Birchley, Alexander Adams, Amy Gaskin, Bree Gatica-Wilcox, Jason Coombes, Joel Southgate, Lauren Gilbert, Lee Graham, Nicole Pacchiarini, Sara Kumziene-Summerhayes, Sarah Taylor, Sophie Jones, Sara Rey, Matthew Bull, Joanne Watkins, Sally Corden, Tom Connor |
| EPI_ISL_814590, EPI_ISL_814591, EPI_ISL_814592, EPI_ISL_814593 | West of Scotland Specialist Virology Centre, NHSGGC / MRC-University of Glasgow Centre for Virus Research | COVID-19 Genomics UK (COG-UK) Consortium | Ana da Silva Filipe, Natasha Johnson, Kathy Smollett, Daniel Mair, Stephen Carmichael, Alice Broos, Lily Tong, Jenna Nichols, Kyriaki Nomikou; Sarah McDonald; Richard Orton, Joseph Hughes, Sreenu Vattipally, David L Robertson; Alasdair MacLean, Rory Gunson; Sharif Shaaban, Matthew Holden; Rachel Blacow, Guy Mollett, Kathy Li, James Shepherd, Antonia Ho, Emma Thomson |
| EPI_ISL_814607 | Wales Specialist Virology Centre Sequencing lab: Pathogen | COVID-19 Genomics UK (COG-UK) Consortium | Catherine Moore, Johnathan Evans, Laura Gifford, Malorie Perry, Simon Cottrell, Angela Marchbank, Alec Birchley, Alexander Adams, Amy Gaskin, Bree |

| | | | |
|---|---|---|---|
| | Genomics Unit | | Gatica-Wilcox, Jason Coombes, Joel Southgate, Lauren Gilbert, Lee Graham, Nicole Pacchiarini, Sara Kumziene-Summerhayes, Sarah Taylor, Sophie Jones, Sara Rey, Matthew Bull, Joanne Watkins, Sally Corden, Tom Connor |
| EPI_ISL_814671, EPI_ISL_814672, EPI_ISL_814673, EPI_ISL_814674, EPI_ISL_814676, EPI_ISL_814677 | Virology Department, Royal Infirmary of Edinburgh, NHS Lothian / School of Biological Sciences, University of Edinburgh / Institute of Genetics and Molecular Medicine, University of Edinburgh | COVID-19 Genomics UK (COG-UK) Consortium | McHugh M, Dewar R, Rooke S, Gallagher M, Balcaza C, O'Toole Á, Scher E, Hill V, McCrone JT, Colquhoun R, Yu X, Jackson B, Rambaut A, Williams TC, Templeton K |
| EPI_ISL_814705, EPI_ISL_814706, EPI_ISL_814707, EPI_ISL_814708, EPI_ISL_814709, EPI_ISL_814710, EPI_ISL_814711, EPI_ISL_814712, EPI_ISL_814713, EPI_ISL_814714, EPI_ISL_814715, EPI_ISL_814716, EPI_ISL_815088, EPI_ISL_815089, EPI_ISL_815091, EPI_ISL_815092 | | | |
| see above | Wales Specialist Virology Centre Sequencing lab: Pathogen Genomics Unit | COVID-19 Genomics UK (COG-UK) Consortium | Catherine Moore, Johnathan Evans, Laura Gifford, Malorie Perry, Simon Cottrell, Angela Marchbank, Alec Birchley, Alexander Adams, Amy Gaskin, Bree Gatica-Wilcox, Jason Coombes, Joel Southgate, Lauren Gilbert, Lee Graham, Nicole Pacchiarini, Sara Kumziene-Summerhayes, Sarah Taylor, Sophie Jones, Sara Rey, Matthew Bull, Joanne Watkins, Sally Corden, Tom Connor |
| EPI_ISL_815411, EPI_ISL_815454, EPI_ISL_815455, EPI_ISL_815456, EPI_ISL_815457, EPI_ISL_815458, EPI_ISL_815459, EPI_ISL_815460, EPI_ISL_815461, EPI_ISL_815462, EPI_ISL_815474, EPI_ISL_815480, EPI_ISL_815527, EPI_ISL_815536, EPI_ISL_815540, EPI_ISL_815592, EPI_ISL_815593, EPI_ISL_815594, EPI_ISL_815595, EPI_ISL_815596, EPI_ISL_815607, EPI_ISL_815621, EPI_ISL_816035, EPI_ISL_816036, EPI_ISL_816037, EPI_ISL_816038, EPI_ISL_816039, EPI_ISL_816040, EPI_ISL_816041, EPI_ISL_816042, EPI_ISL_816043, EPI_ISL_816044, EPI_ISL_816045, EPI_ISL_816046, EPI_ISL_816047, EPI_ISL_816048, EPI_ISL_816049, EPI_ISL_816050, EPI_ISL_816051, EPI_ISL_816052, EPI_ISL_816053, EPI_ISL_816054, EPI_ISL_816055, EPI_ISL_816056, EPI_ISL_816057, EPI_ISL_816058, EPI_ISL_816059, EPI_ISL_816060, EPI_ISL_816061, EPI_ISL_816062, EPI_ISL_816063, EPI_ISL_816064, EPI_ISL_816065, EPI_ISL_816066 | | | |
| see above | Department of Virus and Microbiological Special Diagnostics, Statens Serum Institut, Copenhagen, Denmark | Albertsen Lab, Department of Chemistry and Bioscience, Aalborg University, Denmark | Danish Covid-19 Genome Consortium |
| EPI_ISL_816263, EPI_ISL_816270, EPI_ISL_816342, EPI_ISL_816356, EPI_ISL_816397, EPI_ISL_816445, EPI_ISL_816476, EPI_ISL_816485, EPI_ISL_816489, EPI_ISL_816499, EPI_ISL_816516, EPI_ISL_816530, EPI_ISL_816545, EPI_ISL_816570, EPI_ISL_816602 | | | |
| see above | Virology Department, Sheffield Teaching Hospitals NHS Foundation Trust/Department of Infection, Immunity and Cardiovascular Disease, The Medical School, University of Sheffield | COVID-19 Genomics UK (COG-UK) Consortium | Thushan de Silva, Matthew Parker, Nikki Smith, Adri Angyal, Rebecca Brown, Luke Green, Rachel Tucker, Paul Parsons, Danielle Groves, Katie Johnson, Laura Carrilero, Alex Keeley, Dave Partridge, Matthew Wyles, Benjamin Lindsey, Mehmet Yavuz, Mohammad Raza, Cariad Evans |
| EPI_ISL_816861, EPI_ISL_816862, EPI_ISL_816863, EPI_ISL_816864, EPI_ISL_816865, EPI_ISL_816867, EPI_ISL_816868, EPI_ISL_816869, EPI_ISL_816870, EPI_ISL_816871, EPI_ISL_816872, EPI_ISL_816873, EPI_ISL_816874, EPI_ISL_816875, EPI_ISL_816876, EPI_ISL_816878, EPI_ISL_816879, EPI_ISL_816880, EPI_ISL_816881, EPI_ISL_816882, EPI_ISL_816883, EPI_ISL_816884, EPI_ISL_816885, EPI_ISL_816886, EPI_ISL_816887, EPI_ISL_816888, EPI_ISL_816889, EPI_ISL_816890, EPI_ISL_816891, EPI_ISL_816893, EPI_ISL_816895, EPI_ISL_816900, EPI_ISL_816901, EPI_ISL_816902, EPI_ISL_816904, EPI_ISL_816906, EPI_ISL_816909, EPI_ISL_816910, EPI_ISL_816911, EPI_ISL_816913, EPI_ISL_816915, EPI_ISL_816916, EPI_ISL_816918, EPI_ISL_816920, EPI_ISL_816923, EPI_ISL_816924, EPI_ISL_816925, EPI_ISL_816958, EPI_ISL_816960, EPI_ISL_816962, EPI_ISL_816967, EPI_ISL_816968, EPI_ISL_816969, EPI_ISL_816971, EPI_ISL_816973, EPI_ISL_816976, EPI_ISL_816977, EPI_ISL_816978, EPI_ISL_816980, EPI_ISL_816982, EPI_ISL_816983, EPI_ISL_816985, EPI_ISL_816987, EPI_ISL_816990, EPI_ISL_816991, EPI_ISL_816992 | | | |
| see above | Bioinformatics and Biostatistics Lab, Advanced Sequencing Facility | COVID-19 Genomics UK (COG-UK) Consortium | Aengus Stewart,Jerome Nicod,Chelsea Sawyer,Laura Cubitt,Harshil Patel,Margaret Crawford |
| EPI_ISL_819569, EPI_ISL_819570, EPI_ISL_819571, EPI_ISL_819572, EPI_ISL_819573, EPI_ISL_819574 | Northumbria University / South Tees Hospitals NHS Foundation Trust / North Cumbria Integrated Care NHS Foundation Trust / North Tees and Hartlepool NHS Foundation Trust / Newcastle Hospitals NHS Foundation Trust | COVID-19 Genomics UK (COG-UK) Consortium | Darren L Smith,Andrew Nelson,Matthew Bashton,Greg R Young,Joshua Loh,John Allan,Mohammad A Tariq,Giles S Holt,Gary Black,Wen C Yew,Lynn Dover,Paul Baker,Steve Liggett,Sarah Essex,Jane Greenaway,Debra Padgett,Clive Graham,Garren Scott,Edward Barton,Emma Swindells,Brendan Payne,Jennifer Collins,Yusri Taha,Gary Eltringham |
| EPI_ISL_822900, EPI_ISL_823206, EPI_ISL_823207 | Wales Specialist Virology Centre Sequencing lab: Pathogen Genomics Unit | COVID-19 Genomics UK (COG-UK) Consortium | Catherine Moore, Johnathan Evans, Laura Gifford, Malorie Perry, Simon Cottrell, Angela Marchbank, Alec Birchley, Alexander Adams, Amy Gaskin, Bree Gatica-Wilcox, Jason Coombes, Joel Southgate, Lauren Gilbert, Lee Graham, Nicole Pacchiarini, Sara Kumziene-Summerhayes, Sarah Taylor, Sophie Jones, Sara Rey, Matthew Bull, Joanne Watkins, Sally Corden, Tom Connor |
| EPI_ISL_823838, EPI_ISL_823839, EPI_ISL_823840 | DOHMH Central Harlem | New York City Public Health Laboratory | Jade Wang, et al. |
| EPI_ISL_823841 | DOHMH Riverside | New York City Public Health Laboratory | Jade Wang, et al. |
| EPI_ISL_823842 | DOHMH PHL | New York City Public Health Laboratory | Jade Wang, et al. |
| EPI_ISL_823843, EPI_ISL_823844 | DOHMH Central Harlem | New York City Public Health Laboratory | Jade Wang, et al. |
| EPI_ISL_823845 | DOHMH Morrisania | New York City Public Health Laboratory | Jade Wang, et al. |
| EPI_ISL_823846, EPI_ISL_823847, EPI_ISL_823848 | DOHMH Jamaica | New York City Public Health Laboratory | Jade Wang, et al. |
| EPI_ISL_823849 | DOHMH Fort Greene | New York City Public Health Laboratory | Jade Wang, et al. |
| EPI_ISL_823850 | DOHMH Corona | New York City Public Health Laboratory | Jade Wang, et al. |
| EPI_ISL_823851 | DOHMH Fort Greene | New York City Public Health Laboratory | Jade Wang, et al. |
| EPI_ISL_823852, EPI_ISL_823853 | DOHMH Crown Heights | New York City Public Health Laboratory | Jade Wang, et al. |
| EPI_ISL_823854, EPI_ISL_823855, EPI_ISL_823856, EPI_ISL_823857, EPI_ISL_823858, EPI_ISL_823859, EPI_ISL_823860 | DOHMH Morrisania | New York City Public Health Laboratory | Jade Wang, et al. |
| EPI_ISL_823861, EPI_ISL_823862, EPI_ISL_823863 | DOHMH Corona | New York City Public Health Laboratory | Jade Wang, et al. |
| EPI_ISL_823864 | DOHMH Chelsea | New York City Public Health Laboratory | Jade Wang, et al. |
| EPI_ISL_823865 | DOHMH Central Harlem | New York City Public Health Laboratory | Jade Wang, et al. |
| EPI_ISL_823866, EPI_ISL_823867 | DOHMH Fort Greene | New York City Public Health Laboratory | Jade Wang, et al. |
| EPI_ISL_823868 | DOHMH Corona | New York City Public Health Laboratory | Jade Wang, et al. |
| EPI_ISL_823869 | DOHMH Central Harlem | New York City Public Health Laboratory | Jade Wang, et al. |
| EPI_ISL_823870, EPI_ISL_823871 | DOHMH Jamaica | New York City Public Health Laboratory | Jade Wang, et al. |
| EPI_ISL_823872, EPI_ISL_823873, EPI_ISL_823874 | DOHMH Morrisania | New York City Public Health Laboratory | Jade Wang, et al. |
| EPI_ISL_823875 | DOHMH PHL | New York City Public Health Laboratory | Jade Wang, et al. |
| EPI_ISL_823876, EPI_ISL_823877 | DOHMH Jamaica | New York City Public Health Laboratory | Jade Wang, et al. |
| EPI_ISL_823878, EPI_ISL_823879, EPI_ISL_823880, EPI_ISL_823881, EPI_ISL_823882 | DOHMH Corona | New York City Public Health Laboratory | Jade Wang, et al. |
| EPI_ISL_823883 | DOHMH Riverside | New York City Public Health Laboratory | Jade Wang, et al. |
| EPI_ISL_823884, EPI_ISL_823885, EPI_ISL_823886, EPI_ISL_823887 | DOHMH Central Harlem | New York City Public Health Laboratory | Jade Wang, et al. |
| EPI_ISL_823888 | DOHMH Riverside | New York City Public Health Laboratory | Jade Wang, et al. |
| EPI_ISL_823889, EPI_ISL_823890, | DOHMH PHL | New York City Public Health Laboratory | Jade Wang, et al. |

| | | | |
|---|---|---|---|
| EPI_ISL_823891 | | | |
| EPI_ISL_823892, EPI_ISL_823893 | DOHMH Morrisania | New York City Public Health Laboratory | Jade Wang, et al. |
| EPI_ISL_823894, EPI_ISL_823895 | DOHMH Jamaica | New York City Public Health Laboratory | Jade Wang, et al. |
| EPI_ISL_823896, EPI_ISL_823897 | DOHMH PHL | New York City Public Health Laboratory | Jade Wang, et al. |
| EPI_ISL_823898, EPI_ISL_823899 | DOHMH Corona | New York City Public Health Laboratory | Jade Wang, et al. |
| EPI_ISL_823900 | DOHMH Riverside | New York City Public Health Laboratory | Jade Wang, et al. |
| EPI_ISL_823901, EPI_ISL_823902 | DOHMH Central Harlem | New York City Public Health Laboratory | Jade Wang, et al. |
| EPI_ISL_823903 | DOHMH Morrisania | New York City Public Health Laboratory | Jade Wang, et al. |
| EPI_ISL_823904 | DOHMH Fort Greene | New York City Public Health Laboratory | Jade Wang, et al. |
| EPI_ISL_823905, EPI_ISL_823906 | DOHMH Corona | New York City Public Health Laboratory | Jade Wang, et al. |
| EPI_ISL_823907, EPI_ISL_823908 | DOHMH Jamaica | New York City Public Health Laboratory | Jade Wang, et al. |
| EPI_ISL_823909 | DOHMH PHL | New York City Public Health Laboratory | Jade Wang, et al. |
| EPI_ISL_823910, EPI_ISL_823911 | DOHMH Fort Greene | New York City Public Health Laboratory | Jade Wang, et al. |
| EPI_ISL_823912 | DOHMH Central Harlem | New York City Public Health Laboratory | Jade Wang, et al. |
| EPI_ISL_823913, EPI_ISL_823914 | DOHMH PHL | New York City Public Health Laboratory | Jade Wang, et al. |
| EPI_ISL_823915, EPI_ISL_823916, EPI_ISL_823917 | DOHMH Jamaica | New York City Public Health Laboratory | Jade Wang, et al. |
| EPI_ISL_823918, EPI_ISL_823919, EPI_ISL_823920, EPI_ISL_823921, EPI_ISL_823922 | DOHMH Corona | New York City Public Health Laboratory | Jade Wang, et al. |
| EPI_ISL_823923 | DOHMH Fort Greene | New York City Public Health Laboratory | Jade Wang, et al. |
| EPI_ISL_823924 | DOHMH Chelsea | New York City Public Health Laboratory | Jade Wang, et al. |
| EPI_ISL_823925 | DOHMH Crown Heights | New York City Public Health Laboratory | Jade Wang, et al. |
| EPI_ISL_824234, EPI_ISL_824235, EPI_ISL_824236, EPI_ISL_824237, EPI_ISL_824238, EPI_ISL_824239, EPI_ISL_824240 | Dutch COVID-19 response team | National Institute for Public Health and the Environment (RIVM) | Adam Meijer, Harry Vennema, Jeroen Cremer, Sharon van den Brink, Bas van der Veer, AnneMarie van den Brandt, Florian Zwagemaker, Dennis Schmitz, Chantal Reusken, on behalf of the national COVID-19 response team |
| EPI_ISL_824346, EPI_ISL_824362, EPI_ISL_824372, EPI_ISL_824373 | Michigan Department of Health and Human Services, Bureau of Laboratories | Michigan Department of Health and Human Services, Bureau of Laboratories | Blankenship HM, Riner D, Soehnlen MK |
| EPI_ISL_824400 | California Department of Public Health | California Department of Public Health | CDPH IDLB COVIDNet |
| EPI_ISL_824656, EPI_ISL_824666, EPI_ISL_824667, EPI_ISL_824678, EPI_ISL_824679, EPI_ISL_824690, EPI_ISL_824691, EPI_ISL_824702, EPI_ISL_824714, EPI_ISL_824725, EPI_ISL_824726, EPI_ISL_824736, EPI_ISL_824737 | | | |
| see above | Cedars-Sinai Medical Center, Department of Pathology & Laboratory Medicine, Molecular Pathology Laboratory | Cedars-Sinai Medical Center, Molecular Pathology Laboratory of Department of Pathology & Laboratory Medicine and Genomic Core | Wenjuan Zhang, Brian Davis, Stephanie Chen, Jorge Mario Sincuir Martinez, Jasmine T Plummer, Eric Vail |
| EPI_ISL_824765, EPI_ISL_824766, EPI_ISL_824767, EPI_ISL_824768, EPI_ISL_824769, EPI_ISL_824770, EPI_ISL_824771, EPI_ISL_824772, EPI_ISL_824773, EPI_ISL_824774, EPI_ISL_824775, EPI_ISL_824776, EPI_ISL_824777, EPI_ISL_824778, EPI_ISL_824779, EPI_ISL_824780, EPI_ISL_824781, EPI_ISL_824782, EPI_ISL_824783, EPI_ISL_824784, EPI_ISL_824785, EPI_ISL_824786 | | | |
| see above | A. Krumbholz, Labor Dr. Krause und Kollegen MVZ GmbH, Kiel | Charité Universitätsmedizin Berlin, Institut für Virologie | Victor M Corman, Jörn Beheim-Schwarzbach, Julia Schneider, Talitha Veith, Barbara Mühlemann, Terry Jones, Christian Drosten |
| EPI_ISL_825020, EPI_ISL_825022, EPI_ISL_825024, EPI_ISL_825025 | Utah Public Health Laboratory, Utah Public Health Laboratory Infectious Disease submission group | Utah Public Health Laboratory, Utah Public Health Laboratory Infectious Disease submission group | Young,E.L., Oakeson,K.F., Gallagher,T. |
| EPI_ISL_825374 | Hospital Universitari Vall d'Hebron - Vall d'Hebron Institut de Rercerca | Hospital Universitari Vall d'Hebron | Cristina Andrés, Maria Piñana, Josep F Abril, Damir Garcia-Cehic, Ariadna Rando, Juliana Esperalba, Maria Gema Codina, Carla Castillo, Maria Carmen Martin, Tomàs Pumarola, Josep Quer, Andrés Antón |
| EPI_ISL_826539, EPI_ISL_826541, EPI_ISL_826550, EPI_ISL_826551, EPI_ISL_826557, EPI_ISL_826561, EPI_ISL_826566, EPI_ISL_826570, EPI_ISL_826575 | Texas Department of State Health Services | Texas Department of State Health Services | Rashmi Tuladhar, Bonnie Oh, Jenny Zhang, Maliha Rahman, Anita Pokharel, Myong Koag, Chung Wang, Rachel Lee, Grace Kubin, Mayela Pedrueza, James Daniel Bonser |
| EPI_ISL_826806, EPI_ISL_826808, EPI_ISL_826895 | deCODE genetics | deCODE genetics | Daniel F Gudbjartsson; Agnar Helgason; Hakon Jonsson; Olafur T Magnusson; Pall Melsted; Gudmundur L Norddahl; Jona Saemundsdottir; Asgeir Sigurdsson; Patrick Sulem; Arna B Agustsdottir; Hannes Eggertsson; Berglind Eiriksdottir; Run Fridriksdottir; Elisabet E Gardarsdottir; Gudmundur Georgsson; Olafia S Gretarsdottir; Kjartan R Gudmundsson; Thora R Gunnarsdottir; Arnaldur Gylfason; Hilma Holm; Brynjar O Jensson; Aslaug Jonasdottir; Kamilla S Josefsdottir; Thordur Kristjansson; Droplaug N Magnusdottir; Solvi Rognvaldsson; Louise le Roux; Gudrun Sigmundsdottir; Gardar Sveinbjornsson; Kristin E Sveinsdottir; Maney Sveinsdottir; Emil A Thorarensen; Bjarni Thorbjornsson; Gisli Masson; Ingileif Jonsdottir; Alma Moller; Thorolfur Gudnason; Karl G Kristinsson; Unnur Thorsteinsdottir; Kari Stefansson |
| EPI_ISL_826980 | The National University Hospital of Iceland | deCODE genetics | Daniel F Gudbjartsson; Agnar Helgason; Hakon Jonsson; Olafur T Magnusson; Pall Melsted; Gudmundur L Norddahl; Jona Saemundsdottir; Asgeir Sigurdsson; Patrick Sulem; Arna B Agustsdottir; Hannes Eggertsson; Berglind Eiriksdottir; Run Fridriksdottir; Elisabet E Gardarsdottir; Gudmundur Georgsson; Olafia S Gretarsdottir; Kjartan R Gudmundsson; Thora R Gunnarsdottir; Arnaldur Gylfason; Hilma Holm; Brynjar O Jensson; Aslaug Jonasdottir; Kamilla S Josefsdottir; Thordur Kristjansson; Droplaug N Magnusdottir; Solvi Rognvaldsson; Louise le Roux; Gudrun Sigmundsdottir; Gardar Sveinbjornsson; Kristin E Sveinsdottir; Maney Sveinsdottir; Emil A Thorarensen; Bjarni Thorbjornsson; Gisli Masson; Ingileif Jonsdottir; Alma Moller; Thorolfur Gudnason; Karl G Kristinsson; Unnur Thorsteinsdottir; Kari Stefansson |
| EPI_ISL_827089, EPI_ISL_827420, EPI_ISL_827431, EPI_ISL_828088 | deCODE genetics | deCODE genetics | Daniel F Gudbjartsson; Agnar Helgason; Hakon Jonsson; Olafur T Magnusson; Pall Melsted; Gudmundur L Norddahl; Jona Saemundsdottir; Asgeir Sigurdsson; Patrick Sulem; Arna B Agustsdottir; Hannes Eggertsson; Berglind Eiriksdottir; Run Fridriksdottir; Elisabet E Gardarsdottir; Gudmundur Georgsson; Olafia S Gretarsdottir; Kjartan R Gudmundsson; Thora R Gunnarsdottir; Arnaldur Gylfason; Hilma Holm; Brynjar O Jensson; Aslaug Jonasdottir; Kamilla S Josefsdottir; Thordur Kristjansson; Droplaug N Magnusdottir; Solvi Rognvaldsson; Louise le Roux; Gudrun Sigmundsdottir; Gardar Sveinbjornsson; Kristin E Sveinsdottir; Maney Sveinsdottir; Emil A Thorarensen; Bjarni Thorbjornsson; Gisli Masson; Ingileif Jonsdottir; Alma Moller; Thorolfur Gudnason; Karl G Kristinsson; Unnur Thorsteinsdottir; Kari Stefansson |
| EPI_ISL_828363, EPI_ISL_828383 | The National University Hospital of Iceland | deCODE genetics | Daniel F Gudbjartsson; Agnar Helgason; Hakon Jonsson; Olafur T Magnusson; Pall Melsted; Gudmundur L Norddahl; Jona Saemundsdottir; Asgeir Sigurdsson; Patrick Sulem; Arna B Agustsdottir; Hannes Eggertsson; Berglind Eiriksdottir; Run Fridriksdottir; Elisabet E Gardarsdottir; Gudmundur Georgsson; Olafia S Gretarsdottir; Kjartan R Gudmundsson; Thora R Gunnarsdottir; Arnaldur Gylfason; Hilma Holm; Brynjar O Jensson; Aslaug Jonasdottir; Kamilla S Josefsdottir; Thordur Kristjansson; Droplaug N Magnusdottir; Solvi Rognvaldsson; Louise le Roux; Gudrun Sigmundsdottir; Gardar Sveinbjornsson; Kristin E Sveinsdottir; Maney Sveinsdottir; Emil A Thorarensen; Bjarni Thorbjornsson; Gisli Masson; Ingileif Jonsdottir; Alma Moller; Thorolfur Gudnason; Karl G Kristinsson; Unnur Thorsteinsdottir; Kari Stefansson |

| | | | |
|---|---|---|---|
| EPI_ISL_828410, EPI_ISL_829054, EPI_ISL_829058, EPI_ISL_829071, EPI_ISL_829527, EPI_ISL_829528, EPI_ISL_829701, EPI_ISL_829704, EPI_ISL_829794, EPI_ISL_829795, EPI_ISL_829796, EPI_ISL_829797, EPI_ISL_829903, EPI_ISL_830164, EPI_ISL_830165 | | | |
| see above | deCODE genetics | deCODE genetics | Daniel F Gudbjartsson; Agnar Helgason; Hakon Jonsson; Olafur T Magnusson; Pall Melsted; Gudmundur L Norddahl; Jona Saemundsdottir; Asgeir Sigurdsson; Patrick Sulem; Arna B Agustsdottir; Hannes Eggertsson; Berglind Eiriksdottir; Run Fridriksdottir; Elisabet E Gardarsdottir; Gudmundur Georgsson; Olafia S Gretarsdottir; Kjartan R Gudmundsson; Thora R Gunnarsdottir; Arnaldur Gylfason; Hilma Holm; Brynjar O Jensson; Aslaug Jonasdottir; Kamilla S Josefsdottir; Thordur Kristjansson; Droplaug N Magnusdottir; Solvi Rognvaldsson; Louise le Roux; Gudrun Sigmundsdottir; Gardar Sveinbjornsson; Kristin E Sveinsdottir; Maney Sveinsdottir; Emil A Thorarensen; Bjarni Thorbjornsson; Gisli Masson; Ingileif Jonsdottir; Alma Moller; Thorolfur Gudnason; Karl G Kristinsson; Unnur Thorsteinsdottir; Kari Stefansson |
| EPI_ISL_830168, EPI_ISL_830169 | The National University Hospital of Iceland | deCODE genetics | Daniel F Gudbjartsson; Agnar Helgason; Hakon Jonsson; Olafur T Magnusson; Pall Melsted; Gudmundur L Norddahl; Jona Saemundsdottir; Asgeir Sigurdsson; Patrick Sulem; Arna B Agustsdottir; Hannes Eggertsson; Berglind Eiriksdottir; Run Fridriksdottir; Elisabet E Gardarsdottir; Gudmundur Georgsson; Olafia S Gretarsdottir; Kjartan R Gudmundsson; Thora R Gunnarsdottir; Arnaldur Gylfason; Hilma Holm; Brynjar O Jensson; Aslaug Jonasdottir; Kamilla S Josefsdottir; Thordur Kristjansson; Droplaug N Magnusdottir; Solvi Rognvaldsson; Louise le Roux; Gudrun Sigmundsdottir; Gardar Sveinbjornsson; Kristin E Sveinsdottir; Maney Sveinsdottir; Emil A Thorarensen; Bjarni Thorbjornsson; Gisli Masson; Ingileif Jonsdottir; Alma Moller; Thorolfur Gudnason; Karl G Kristinsson; Unnur Thorsteinsdottir; Kari Stefansson |
| EPI_ISL_830499 | deCODE genetics | deCODE genetics | Daniel F Gudbjartsson; Agnar Helgason; Hakon Jonsson; Olafur T Magnusson; Pall Melsted; Gudmundur L Norddahl; Jona Saemundsdottir; Asgeir Sigurdsson; Patrick Sulem; Arna B Agustsdottir; Hannes Eggertsson; Berglind Eiriksdottir; Run Fridriksdottir; Elisabet E Gardarsdottir; Gudmundur Georgsson; Olafia S Gretarsdottir; Kjartan R Gudmundsson; Thora R Gunnarsdottir; Arnaldur Gylfason; Hilma Holm; Brynjar O Jensson; Aslaug Jonasdottir; Kamilla S Josefsdottir; Thordur Kristjansson; Droplaug N Magnusdottir; Solvi Rognvaldsson; Louise le Roux; Gudrun Sigmundsdottir; Gardar Sveinbjornsson; Kristin E Sveinsdottir; Maney Sveinsdottir; Emil A Thorarensen; Bjarni Thorbjornsson; Gisli Masson; Ingileif Jonsdottir; Alma Moller; Thorolfur Gudnason; Karl G Kristinsson; Unnur Thorsteinsdottir; Kari Stefansson |
| EPI_ISL_831247, EPI_ISL_831248, EPI_ISL_831249, EPI_ISL_831250, EPI_ISL_831251, EPI_ISL_831252, EPI_ISL_831253, EPI_ISL_831254, EPI_ISL_831255, EPI_ISL_831257, EPI_ISL_831258 | | | |
| see above | Hospital Universitario La Paz (Madrid) | SeqCOVID-SPAIN consortium/IBV(CSIC) | Fernando Lázaro-Perona, María Rodríguez-Tejedor, Elias Dahdouh, Jesús Mingorance and SeqCOVID-SPAIN consortium |
| EPI_ISL_831389 | Limbach - MVZ Labor Dr. Limbach & Kollegen | Robert Koch Institute, Influenza and respiratory viruses FG17 & Bioinformatics MF1, Berlin, Germany | Dr. Konrad Bode, Stephan Fuchs, Stefan Kroeger, Marianne Wedde, Oliver Drechsel, Aleksandar Radonic, Rene Kmiecinski, Ralf Duerrwald, Thorsten Wolff |
| EPI_ISL_831390 | Labor Krone | Robert Koch Institute, Influenza and respiratory viruses FG17 & Bioinformatics MF1, Berlin, Germany | Dr. Münstermann. Prof. Tiemann , Stephan Fuchs, Stefan Kroeger, Marianne Wedde, Oliver Drechsel, Aleksandar Radonic, Rene Kmiecinski, Ralf Duerrwald, Thorsten Wolff |
| EPI_ISL_831391 | Limbach - MVZ Labor Dr. Limbach & Kollegen | Robert Koch Institute, Influenza and respiratory viruses FG17 & Bioinformatics MF1, Berlin, Germany | Dr. Konrad Bode, Stephan Fuchs, Stefan Kroeger, Marianne Wedde, Oliver Drechsel, Aleksandar Radonic, Rene Kmiecinski, Ralf Duerrwald, Thorsten Wolff |
| EPI_ISL_831392 | Labor Krone | Robert Koch Institute, Influenza and respiratory viruses FG17 & Bioinformatics MF1, Berlin, Germany | Dr. Münstermann. Prof. Tiemann , Stephan Fuchs, Stefan Kroeger, Marianne Wedde, Oliver Drechsel, Aleksandar Radonic, Rene Kmiecinski, Ralf Duerrwald, Thorsten Wolff |
| EPI_ISL_831393 | Limbach - MVZ Labor Dr. Limbach & Kollegen | Robert Koch Institute, Influenza and respiratory viruses FG17 & Bioinformatics MF1, Berlin, Germany | Dr. Konrad Bode, Stephan Fuchs, Stefan Kroeger, Marianne Wedde, Oliver Drechsel, Aleksandar Radonic, Rene Kmiecinski, Ralf Duerrwald, Thorsten Wolff |
| EPI_ISL_831394, EPI_ISL_831395 | Labor Krone | Robert Koch Institute, Influenza and respiratory viruses FG17 & Bioinformatics MF1, Berlin, Germany | Dr. Münstermann. Prof. Tiemann , Stephan Fuchs, Stefan Kroeger, Marianne Wedde, Oliver Drechsel, Aleksandar Radonic, Rene Kmiecinski, Ralf Duerrwald, Thorsten Wolff |
| EPI_ISL_831396 | Limbach - MVZ Labor Dr. Limbach & Kollegen | Robert Koch Institute, Influenza and respiratory viruses FG17 & Bioinformatics MF1, Berlin, Germany | Dr. Konrad Bode, Stephan Fuchs, Stefan Kroeger, Marianne Wedde, Oliver Drechsel, Aleksandar Radonic, Rene Kmiecinski, Ralf Duerrwald, Thorsten Wolff |
| EPI_ISL_831397, EPI_ISL_831398, EPI_ISL_831399 | Labor Krone | Robert Koch Institute, Influenza and respiratory viruses FG17 & Bioinformatics MF1, Berlin, Germany | Dr. Münstermann. Prof. Tiemann , Stephan Fuchs, Stefan Kroeger, Marianne Wedde, Oliver Drechsel, Aleksandar Radonic, Rene Kmiecinski, Ralf Duerrwald, Thorsten Wolff |
| EPI_ISL_831400 | Limbach - MVZ Labor Dr. Limbach & Kollegen | Robert Koch Institute, Influenza and respiratory viruses FG17 & Bioinformatics MF1, Berlin, Germany | Dr. Konrad Bode, Stephan Fuchs, Stefan Kroeger, Marianne Wedde, Oliver Drechsel, Aleksandar Radonic, Rene Kmiecinski, Ralf Duerrwald, Thorsten Wolff |
| EPI_ISL_831401 | Labor Krone | Robert Koch Institute, Influenza and respiratory viruses FG17 & Bioinformatics MF1, Berlin, Germany | Dr. Münstermann. Prof. Tiemann , Stephan Fuchs, Stefan Kroeger, Marianne Wedde, Oliver Drechsel, Aleksandar Radonic, Rene Kmiecinski, Ralf Duerrwald, Thorsten Wolff |
| EPI_ISL_831402 | Labor Dr. Wisplinghoff - Köln | Robert Koch Institute, Influenza and respiratory viruses FG17 & Bioinformatics MF1, Berlin, Germany | Dr. R. Grosser, Stephan Fuchs, Stefan Kroeger, Marianne Wedde, Oliver Drechsel, Aleksandar Radonic, Rene Kmiecinski, Ralf Duerrwald, Thorsten Wolff |
| EPI_ISL_831403 | MVZ Düsseldorf Zentrum GbR | Robert Koch Institute, Influenza and respiratory viruses FG17 & Bioinformatics MF1, Berlin, Germany | Dr. Finzer, Stephan Fuchs, Stefan Kroeger, Marianne Wedde, Oliver Drechsel, Aleksandar Radonic, Rene Kmiecinski, Ralf Duerrwald, Thorsten Wolff |
| EPI_ISL_831404, EPI_ISL_831405 | Labor Dr. Krause & Kollegen MVZ GmbH | Robert Koch Institute, Influenza and respiratory viruses FG17 & Bioinformatics MF1, Berlin, Germany | Dr. Lorentz, Stephan Fuchs, Stefan Kroeger, Marianne Wedde, Oliver Drechsel, Aleksandar Radonic, Rene Kmiecinski, Ralf Duerrwald, Thorsten Wolff |
| EPI_ISL_831406 | MVZ Düsseldorf Zentrum GbR | Robert Koch Institute, Influenza and respiratory viruses FG17 & Bioinformatics MF1, Berlin, Germany | Dr. Finzer, Stephan Fuchs, Stefan Kroeger, Marianne Wedde, Oliver Drechsel, Aleksandar Radonic, Rene Kmiecinski, Ralf Duerrwald, Thorsten Wolff |
| EPI_ISL_831407 | Labor Dr. Krause & Kollegen MVZ GmbH | Robert Koch Institute, Influenza and respiratory viruses FG17 & Bioinformatics MF1, Berlin, Germany | Dr. Lorentz, Stephan Fuchs, Stefan Kroeger, Marianne Wedde, Oliver Drechsel, Aleksandar Radonic, Rene Kmiecinski, Ralf Duerrwald, Thorsten Wolff |
| EPI_ISL_831408 | MVZ Düsseldorf Zentrum GbR | Robert Koch Institute, Influenza and respiratory viruses FG17 & Bioinformatics MF1, Berlin, Germany | Dr. Finzer, Stephan Fuchs, Stefan Kroeger, Marianne Wedde, Oliver Drechsel, Aleksandar Radonic, Rene Kmiecinski, Ralf Duerrwald, Thorsten Wolff |
| EPI_ISL_831409 | Labor Dr. Wisplinghoff - Köln | Robert Koch Institute, Influenza and respiratory viruses FG17 & Bioinformatics MF1, Berlin, Germany | Dr. R. Grosser, Stephan Fuchs, Stefan Kroeger, Marianne Wedde, Oliver Drechsel, Aleksandar Radonic, Rene Kmiecinski, Ralf Duerrwald, Thorsten Wolff |
| EPI_ISL_831410 | MVZ Düsseldorf Zentrum GbR | Robert Koch Institute, Influenza and respiratory viruses FG17 & Bioinformatics MF1, Berlin, Germany | Dr. Finzer, Stephan Fuchs, Stefan Kroeger, Marianne Wedde, Oliver Drechsel, Aleksandar Radonic, Rene Kmiecinski, Ralf Duerrwald, Thorsten Wolff |
| EPI_ISL_831411 | Labor Dr. Wisplinghoff - Köln | Robert Koch Institute, Influenza and respiratory viruses FG17 & Bioinformatics MF1, Berlin, Germany | Dr. R. Grosser, Stephan Fuchs, Stefan Kroeger, Marianne Wedde, Oliver Drechsel, Aleksandar Radonic, Rene Kmiecinski, Ralf Duerrwald, Thorsten Wolff |
| EPI_ISL_831412, EPI_ISL_831413 | Labor Dr. Krause & Kollegen MVZ GmbH | Robert Koch Institute, Influenza and respiratory viruses FG17 & Bioinformatics MF1, Berlin, Germany | Dr. Lorentz, Stephan Fuchs, Stefan Kroeger, Marianne Wedde, Oliver Drechsel, Aleksandar Radonic, Rene Kmiecinski, Ralf Duerrwald, Thorsten Wolff |
| EPI_ISL_831654, EPI_ISL_831655, EPI_ISL_831656 | Institute for Infectious Diseases, University of Bern, Switzerland | Institute for Infectious Diseases, University of Bern, Switzerland | Michel C Koch, Christian Baumann, Miguel A Terrazos Miani, Cora Sägesser, Pascal Bittel, Stephen L Leib, Peter Keller, Franziska Suter-Riniker, Alban Ramette |
| EPI_ISL_831687 | Santa Clara County Public Health Laboratory | Santa Clara County Public Health Laboratory | Santa Clara County Public Health Department |
| EPI_ISL_831698, EPI_ISL_831704, EPI_ISL_831732, EPI_ISL_831762, EPI_ISL_831850, EPI_ISL_831865 | United States Air Force School of Aerospace Medicine | United States Air Force School of Aerospace Medicine | Anthony Fries, Jennifer Meyer, William Gruner, Amanda Javorina, Sarah Purves, Clarise Starr, Elizabeth Macias |
| EPI_ISL_831893, EPI_ISL_831896, EPI_ISL_831897, EPI_ISL_831899 | Santa Clara County Public Health Laboratory | Santa Clara County Public Health Laboratory | Santa Clara County Public Health Department |
| EPI_ISL_831946 | Klinisk mikrobiologi, virus F68 | The Public Health Agency of Sweden | Department of Microbiology, The Public Health Agency of Sweden |
| EPI_ISL_831979 | Klinisk mikrobiologi, Viruslab | The Public Health Agency of Sweden | Department of Microbiology, The Public Health Agency of Sweden |
| EPI_ISL_831991 | Klinisk mikrobiologi | The Public Health Agency of Sweden | Department of Microbiology, The Public Health Agency of Sweden |

| | | | |
|---|---|---|---|
| EPI_ISL_832066 | Santa Clara County Public Health Laboratory | Santa Clara County Public Health Laboratory | Santa Clara County Public Health Department |
| EPI_ISL_832128 | Sentinelles IDF | National Reference Center for Viruses of Respiratory Infections, Institut Pasteur, Paris | Marion Barbet, Sylvie Behillil, Méline Bizard, Angela Brisebarre, Camille Capel, Etienne Simon-Lorière, Vincent Enouf, Maud Vanpeene, Sylvie van der Werf, Patricia Lefebure |
| EPI_ISL_832129 | Sentinelles IDF | National Reference Center for Viruses of Respiratory Infections, Institut Pasteur, Paris | Marion Barbet, Sylvie Behillil, Méline Bizard, Angela Brisebarre, Camille Capel, Etienne Simon-Lorière, Vincent Enouf, Maud Vanpeene, Sylvie van der Werf, Claire Watermez |
| EPI_ISL_832164, EPI_ISL_832165, EPI_ISL_832166 | Hospital | National Reference Center for Viruses of Respiratory Infections, Institut Pasteur, Paris | Marion Barbet, Sylvie Behillil, Méline Bizard, Angela Brisebarre, Camille Capel, Etienne Simon-Lorière, Vincent Enouf, Maud Vanpeene, Sylvie van der Werf, Léa Pilorge |
| EPI_ISL_832369, EPI_ISL_832370, EPI_ISL_832371, EPI_ISL_832372, EPI_ISL_832373, EPI_ISL_832374 | Center of Medical Microbiology, Virology, and Hospital Hygiene, University of Duesseldorf | Center of Medical Microbiology, Virology, and Hospital Hygiene, University of Duesseldorf | Maximilian Damagnez, Alexander Dilthey, Ashley-Jane Duplessis, Torsten Houwaart, Lisanna Hülse, Malte Kohns Vasconcelos, Nadine Lübke, Jessica Nicolai, Klaus Pfeffer, Daniel Strelow, Teresa Tamayo, Jörg Timm, Andreas Walker, Tobias Wienemann |
| EPI_ISL_832402, EPI_ISL_832403 | Santa Clara County Public Health Laboratory | Santa Clara County Public Health Laboratory | Santa Clara County Public Health Department |
| EPI_ISL_833459, EPI_ISL_833464, EPI_ISL_833465, EPI_ISL_833474, EPI_ISL_833480 | CHU Purpan - Laboratoire de Virologie - Institut Fédératif de Biologie | CHU Purpan - Laboratoire de Virologie - Institut Fédératif de Biologie | Latour J., Ranger N., Dubois M., Carcenac R., Harter A., Boyer P., Tremeaux P., Izopet J. |
| EPI_ISL_836143 | Hospital de Campanha COVID-19 de Mairipora | Instituto Adolfo Lutz, Interdiciplinary Procedures Center, Strategic Laboratory | Claudio Tavares Sacchi, Claudia Regina Gonçalves, Erica Valessa Ramos Gomes, Karoline Rodrigues Campos |
| EPI_ISL_836977 | Hospital Municiplal Dr. Jose de Carvalho Florence | Instituto Adolfo Lutz, Interdiciplinary Procedures Center, Strategic Laboratory | Claudio Tavares Sacchi, Claudia Regina Gonçalves, Erica Valessa Ramos Gomes, Karoline Rodrigues Campos |
| EPI_ISL_836978 | Irmandade da Santa Casa de Misericordia de Lorena | Instituto Adolfo Lutz, Interdiciplinary Procedures Center, Strategic Laboratory | Claudio Tavares Sacchi, Claudia Regina Gonçalves, Erica Valessa Ramos Gomes, Karoline Rodrigues Campos |
| EPI_ISL_836979 | Respiratory Virus Unit, National Infection Service, Public Health England | COVID-19 Genomics UK (COG-UK) Consortium | PHE Covid Sequencing Team |
| EPI_ISL_837053 | UBS Darcy Alves e Robalinho | Instituto Adolfo Lutz, Interdiciplinary Procedures Center, Strategic Laboratory | Claudio Tavares Sacchi, Claudia Regina Gonçalves, Erica Valessa Ramos Gomes, Karoline Rodrigues Campos |
| EPI_ISL_837054 | UBS Jose Sabino Ferreira | Instituto Adolfo Lutz, Interdiciplinary Procedures Center, Strategic Laboratory | Claudio Tavares Sacchi, Claudia Regina Gonçalves, Erica Valessa Ramos Gomes, Karoline Rodrigues Campos |
| EPI_ISL_837083, EPI_ISL_837100, EPI_ISL_837112, EPI_ISL_837185, EPI_ISL_837199 | Respiratory Virus Unit, National Infection Service, Public Health England | COVID-19 Genomics UK (COG-UK) Consortium | PHE Covid Sequencing Team |
| EPI_ISL_838072, EPI_ISL_838106, EPI_ISL_838108, EPI_ISL_838109, EPI_ISL_838111, EPI_ISL_838112, EPI_ISL_838113, EPI_ISL_838114, EPI_ISL_838115, EPI_ISL_838209, EPI_ISL_838210, EPI_ISL_838211, EPI_ISL_838212 | | | |
| see above | West of Scotland Specialist Virology Centre, NHSGGC / MRC-University of Glasgow Centre for Virus Research | COVID-19 Genomics UK (COG-UK) Consortium | Ana da Silva Filipe, Natasha Johnson, Kathy Smollett, Daniel Mair, Stephen Carmichael, Alice Broos, Lily Tong, Jenna Nichols, Kyriaki Nomikou; Sarah McDonald; Richard Orton, Joseph Hughes, Sreenu Vattipally, David L Robertson; Alasdair MacLean, Rory Gunson; Sharif Shaaban, Matthew Holden; Rachel Blacow, Guy Mollett, Kathy Li, James Shepherd, Antonia Ho, Emma Thomson |
| EPI_ISL_838216, EPI_ISL_838217, EPI_ISL_838218, EPI_ISL_838219, EPI_ISL_838283 | Virology Department, Royal Infirmary of Edinburgh, NHS Lothian / School of Biological Sciences, University of Edinburgh / Institute of Genetics and Molecular Medicine, University of Edinburgh | COVID-19 Genomics UK (COG-UK) Consortium | McHugh M, Dewar R, Rooke S, Gallagher M, Balcaza C, O'Toole Á, Scher E, Hill V, McCrone JT, Colquhoun R, Yu X, Jackson B, Rambaut A, Williams TC, Templeton K |
| EPI_ISL_838585, EPI_ISL_838664 | Liverpool Clinical Laboratories | COVID-19 Genomics UK (COG-UK) Consortium | Sam Haldenby, Anita Lucaci, Steve Paterson, Julian Hiscox, Alistair Darby, M Almsaud, A Alrezaihi, Muhannad Alruwaili, Stuart D Armstrong, Jones Benjamin, Eleanor G Bentley, Anu Chawla, Jordan J Clark, Angela Cowell, Richard Eccles, Isabel García-Dorival, Matthew Gemmell, Alessandro Gerada, PKF Gilmore, Richard Gregory, Ximeng Han, Catherine Hartley, Margaret Hughes, Miren Iturriza-Gomara, James Johnson, L Luu, Jenifer Manson, Charlotte Nelson, Elaine O'Toole, Cassie Olateju, Rebekah Penrice-Randal , Lucille Rainbow, N.P Randle, Trevor Ian Robinson, Parul Sharma, Ghada T Shawli, James P Stewart, Neil Swainston, Ecaterina Vamos, Joanne Watts, Mark Whitehead |
| EPI_ISL_839331, EPI_ISL_839332, EPI_ISL_839333 | University College London, Great Ormond Street Hospital for Children NHS Foundation Trust, Imperial College Healthcare NHS Trust | COVID-19 Genomics UK (COG-UK) Consortium | Sergi Castellano, Rachel Williams, Mark Kristiansen, Paola Resende Silva, Sunando Roy, Tony Brooks, Helena Tutill, Paola Niola, Patricia Dyal, Charlotte Williams, Leysa Forrest, Yasmin Panchbhaya, Jacqueline Findlay, Samuel Weeks, Julianne Brown, Kathryn Harris, Paul Randell, James Price, Alison Holmes, Judith Breuer |
| EPI_ISL_839520, EPI_ISL_839521, EPI_ISL_839522, EPI_ISL_839523, EPI_ISL_839524, EPI_ISL_839525, EPI_ISL_839526, EPI_ISL_839527, EPI_ISL_839528, EPI_ISL_839529, EPI_ISL_839530, EPI_ISL_839531, EPI_ISL_839532, EPI_ISL_839533, EPI_ISL_839739, EPI_ISL_839740, EPI_ISL_839741, EPI_ISL_839742, EPI_ISL_839743, EPI_ISL_839744, EPI_ISL_839745, EPI_ISL_839746, EPI_ISL_839747, EPI_ISL_839748 | | | |
| see above | Northumbria University / South Tees Hospitals NHS Foundation Trust / North Cumbria Integrated Care NHS Foundation Trust / North Tees and Hartlepool NHS Foundation Trust / Newcastle Hospitals NHS Foundation Trust | COVID-19 Genomics UK (COG-UK) Consortium | Darren L Smith,Andrew Nelson,Matthew Bashton,Greg R Young,Joshua Loh,John Allan,Mohammad A Tariq,Giles S Holt,Gary Black,Wen C Yew,Lynn Dover,Paul Baker,Steve Liggett,Sarah Essex,Jane Greenaway,Debra Padgett,Clive Graham,Garren Scott,Edward Barton,Emma Swindells,Brendan Payne,Jennifer Collins,Yusri Taha,Gary Eltringham |
| EPI_ISL_840037, EPI_ISL_840038, EPI_ISL_840039, EPI_ISL_840040 | Lincolnshire Hospitals and DeepSeq Nottingham | COVID-19 Genomics UK (COG-UK) Consortium | Nichola Duckworth, Tim Sloan, Sarah Walsh, Jonathan Ball, Patrick McClure, Joeseph Chappell, Nadine Holmes, Matthew Carlisle, Christopher Moore, Fei Sang, Johnny Debebe, Victoria Wright, Matthew Loose |
| EPI_ISL_841579 | Originating lab: Wales Specialist Virology Centre Sequencing lab: Pathogen Genomics Unit | Public Health Wales Microbiology Cardiff Wales Specialist Virology Centre | Catherine Moore, Johnathan Evans, Laura Gifford, Malorie Perry, Simon Cottrell, Angela Marchbank, Alec Birchley, Alexander Adams, Amy Gaskin, Bree Gatica-Wilcox, Jason Coombes, Joel Southgate, Lauren Gilbert, Lee Graham, Nicole Pacchiarini, Sara Kumziene-Summerhayes, Sarah Taylor, Sophie Jones, Sara Rey, Matthew Bull, Joanne Watkins, Sally Corden, Tom Connor |
| EPI_ISL_841767 | Centre for Enzyme Innovation, University of Portsmouth / Translational Research Laboratory, Portsmouth Hospitals NHS Trust | COVID-19 Genomics UK (COG-UK) Consortium | Angela Beckett,Yann Bourgeois,Garry Scarlett,Sharon Glaysher,Scott Elliott,Kelly Bicknell,Robert Impey,Allyson Lloyd,Sarah Wyllie,Ethan Butcher,Anoop Chauhan,Samuel Robson |
| EPI_ISL_842355, EPI_ISL_842357, EPI_ISL_842359, EPI_ISL_842364, EPI_ISL_842365, EPI_ISL_842366, EPI_ISL_842368, EPI_ISL_842370, EPI_ISL_842373, EPI_ISL_842374, EPI_ISL_842375, EPI_ISL_842377, EPI_ISL_842379, EPI_ISL_842380, EPI_ISL_842382, EPI_ISL_842384, EPI_ISL_842387, EPI_ISL_842388, EPI_ISL_842389 | | | |
| see above | Bioinformatics and Biostatistics Lab, Advanced Sequencing Facility | COVID-19 Genomics UK (COG-UK) Consortium | Aengus Stewart,Jerome Nicod,Chelsea Sawyer,Laura Cubitt,Harshil Patel,Margaret Crawford |
| EPI_ISL_842653, EPI_ISL_842654, EPI_ISL_842656, EPI_ISL_842657, EPI_ISL_842659, EPI_ISL_842660, EPI_ISL_842661, EPI_ISL_842663, EPI_ISL_842666, EPI_ISL_842667, EPI_ISL_842668, EPI_ISL_842669, EPI_ISL_842697, EPI_ISL_842766, EPI_ISL_842767, EPI_ISL_842783, EPI_ISL_842790 | | | |
| see above | University College London Hospital | COVID-19 Genomics UK (COG-UK) Consortium | Judith Heaney, Matthew Byott, Catherine Houlihan, Dan Frampton, Stuart Kirk, Moira Spyer and Eleni Nastouli |
| EPI_ISL_843006, EPI_ISL_843056, EPI_ISL_843058, EPI_ISL_843059, EPI_ISL_843062, EPI_ISL_843063, EPI_ISL_843088 | Barts Health NHS Trust | COVID-19 Genomics UK (COG-UK) Consortium | CUTINO-MOGUEL, Maria-Teresa; HARRINGTON, David; OWOYEMI, Dola; SHYLINI, Raghavendran; BROAD, Claire; KELE, Beatrix |
| EPI_ISL_843189, EPI_ISL_843190 | Maryland Public Health Laboratory | Maryland Public Health Laboratory | Maryland Department of Health Laboratories Administration |
| EPI_ISL_844205, EPI_ISL_844214, EPI_ISL_844215, EPI_ISL_844216, EPI_ISL_844217, EPI_ISL_844218, EPI_ISL_844219, EPI_ISL_844224, EPI_ISL_844225, EPI_ISL_844227, EPI_ISL_844230, EPI_ISL_844231, EPI_ISL_844237, EPI_ISL_844240, EPI_ISL_844249, EPI_ISL_844250, EPI_ISL_844255, EPI_ISL_844258, EPI_ISL_844261, EPI_ISL_844267, EPI_ISL_844282, EPI_ISL_844283, EPI_ISL_844284, EPI_ISL_844285, EPI_ISL_844286, EPI_ISL_844295, EPI_ISL_844298, EPI_ISL_844308, EPI_ISL_844391, EPI_ISL_844392, EPI_ISL_844393, EPI_ISL_844394, EPI_ISL_844395, EPI_ISL_844396, EPI_ISL_844397, EPI_ISL_844398, | | | |

| | | | |
|---|---|---|---|
| EPI_ISL_844399, EPI_ISL_844400, EPI_ISL_844401, EPI_ISL_844402, EPI_ISL_844403, EPI_ISL_844404, EPI_ISL_844405, EPI_ISL_844406, EPI_ISL_844407, EPI_ISL_844408, EPI_ISL_844409, EPI_ISL_844410, EPI_ISL_844411, EPI_ISL_844412, EPI_ISL_844413, EPI_ISL_844414, EPI_ISL_844415, EPI_ISL_844416, EPI_ISL_844417, EPI_ISL_844418, EPI_ISL_844419, EPI_ISL_844420, EPI_ISL_844421, EPI_ISL_844422, EPI_ISL_844423, EPI_ISL_844424, EPI_ISL_844425, EPI_ISL_844426, EPI_ISL_844427, EPI_ISL_844428, EPI_ISL_844429, EPI_ISL_844430, EPI_ISL_844431, EPI_ISL_844432, EPI_ISL_844433, EPI_ISL_844434, EPI_ISL_844435, EPI_ISL_844436, EPI_ISL_844437, EPI_ISL_844438, EPI_ISL_844439, EPI_ISL_844440, EPI_ISL_844441, EPI_ISL_844442, EPI_ISL_844443, EPI_ISL_844444, EPI_ISL_844445, EPI_ISL_844446, EPI_ISL_844447, EPI_ISL_844448, EPI_ISL_844449, EPI_ISL_844450, EPI_ISL_844451, EPI_ISL_844452, EPI_ISL_844453, EPI_ISL_844454, EPI_ISL_844455, EPI_ISL_844456, EPI_ISL_844457, EPI_ISL_844458, EPI_ISL_844459, EPI_ISL_844460, EPI_ISL_844461 | | | |
| see above | Department of Virus and Microbiological Special Diagnostics, Statens Serum Institut, Copenhagen, Denmark | Albertsen Lab, Department of Chemistry and Bioscience, Aalborg University, Denmark | Danish Covid-19 Genome Consortium |
| EPI_ISL_847525, EPI_ISL_847533, EPI_ISL_847537, EPI_ISL_847542, EPI_ISL_847544, EPI_ISL_847547, EPI_ISL_847562, EPI_ISL_847565, EPI_ISL_847567, EPI_ISL_847581, EPI_ISL_847585, EPI_ISL_847587, EPI_ISL_847649, EPI_ISL_847650, EPI_ISL_847651, EPI_ISL_847652, EPI_ISL_847653, EPI_ISL_847654, EPI_ISL_847655, EPI_ISL_847656, EPI_ISL_847657, EPI_ISL_847658, EPI_ISL_847705, EPI_ISL_847728, EPI_ISL_847733, EPI_ISL_847739, EPI_ISL_847745, EPI_ISL_847752, EPI_ISL_847784, EPI_ISL_847785, EPI_ISL_847786, EPI_ISL_847787, EPI_ISL_847788, EPI_ISL_847789, EPI_ISL_847790, EPI_ISL_847791, EPI_ISL_847792, EPI_ISL_847793 | | | |
| see above | California Department of Public Health | Chiu Laboratory, University of California, San Francisco | Charles Chiu, Xianding (Wayne) Deng, Candace Wang, Brian Bushnell, Scot Federman, Jill Hacker, Debra Wadford |
| EPI_ISL_848006, EPI_ISL_848023, EPI_ISL_848058 | Michigan Department of Health and Human Services, Bureau of Laboratories | Michigan Department of Health and Human Services, Bureau of Laboratories | Blankenship HM, Riner D, Soehnlen MK |
| EPI_ISL_848425 | Illinois Department of Public Health | Gagnon Lab, Southern Illinois University | Keith Gagnon |
| EPI_ISL_848644, EPI_ISL_848645, EPI_ISL_848646, EPI_ISL_848647 | Florida Bureau of Public Health Laboratories | Florida Bureau of Public Health Laboratories | Sarah Schmedes, Jason Blanton |
| EPI_ISL_849385, EPI_ISL_849394, EPI_ISL_849398, EPI_ISL_849403, EPI_ISL_849412, EPI_ISL_849413, EPI_ISL_849415, EPI_ISL_849426, EPI_ISL_849429, EPI_ISL_849436, EPI_ISL_849449, EPI_ISL_849455, EPI_ISL_849460, EPI_ISL_849465, EPI_ISL_849521, EPI_ISL_849524, EPI_ISL_849541, EPI_ISL_849542, EPI_ISL_849543, EPI_ISL_849544, EPI_ISL_849545, EPI_ISL_849546, EPI_ISL_849547, EPI_ISL_849548, EPI_ISL_849549, EPI_ISL_849550, EPI_ISL_849551, EPI_ISL_849559, EPI_ISL_849562, EPI_ISL_849565, EPI_ISL_849566, EPI_ISL_849572, EPI_ISL_849576, EPI_ISL_849578, EPI_ISL_849579, EPI_ISL_849580, EPI_ISL_849581, EPI_ISL_849585, EPI_ISL_849588, EPI_ISL_849616, EPI_ISL_849617, EPI_ISL_849619, EPI_ISL_849628, EPI_ISL_849629, EPI_ISL_849630, EPI_ISL_849631, EPI_ISL_849632, EPI_ISL_849633 | | | |
| see above | Washington State Department of Health | Seattle Flu Study | Deborah A. Nickerson, Chris D. Frazar, Jover Lee, Benjamin Pelle, Matthew Richardson, Amanda Adler, Elisabeth Brandstetter, Peter D. Han, Kairsten Fay, Misja Ilcisin, Kirsten Lacombe, Thomas R. Sibley, Melissa Truong, Caitlin R. Wolf, Romesh Gautom, Geoff Melly, Brian Hiatt, Philip Dykema, Scott Lindquist, Michael Boeckh, Janet A. Englund, Michael Famulare, Barry R. Lutz, Mark J. Rieder, Lea M. Starita, Matthew Thompson, Helen Y. Chu, Jay Shendure, Trevor Bedford |
| EPI_ISL_849731 | unknown | PHV-FSS | Son Nguyen et al. |
| EPI_ISL_849985, EPI_ISL_849986, EPI_ISL_849987, EPI_ISL_849988, EPI_ISL_849989, EPI_ISL_849990, EPI_ISL_849991, EPI_ISL_849992, EPI_ISL_849993, EPI_ISL_849994, EPI_ISL_849995, EPI_ISL_849996, EPI_ISL_849997, EPI_ISL_849998, EPI_ISL_849999, EPI_ISL_850000, EPI_ISL_850001, EPI_ISL_850002, EPI_ISL_850003, EPI_ISL_850004, EPI_ISL_850005, EPI_ISL_850006, EPI_ISL_850007, EPI_ISL_850008, EPI_ISL_850009, EPI_ISL_850010, EPI_ISL_850011 | | | |
| see above | Santa Clara County Public Health Laboratory | Chan-Zuckerberg Biohub | CZB Cliahub Consortium |
| EPI_ISL_850664, EPI_ISL_850665 | The National Institute of Public Health | State Veterinary Institute Prague | Nagy,A;Jirincova,H;Trnka,D;Vecerova,J;Trinklova,M |
| EPI_ISL_852597 | Max von Pettenkofer Institute, Virology, National Reference Center for Retroviruses, LMU München | Laboratory for Functional Genome Analysis, Dept. Genomics, Gene Center of the LMU Munich | Max Muenchhoff, Stefan Krebs, Alexander Graf, Oliver Keppler, Helmut Blum |
| EPI_ISL_853341, EPI_ISL_853342, EPI_ISL_853375, EPI_ISL_853376, EPI_ISL_853381, EPI_ISL_853383, EPI_ISL_853384, EPI_ISL_853385, EPI_ISL_853387, EPI_ISL_853392, EPI_ISL_853393 | | | |
| see above | UPMC Clinical Microbiology Laboratory | Microbial Genome Sequencing Center; Microbial Genomic Epidemiology Laboratory | Mustapha M. Mustapha, Jane W. Marsh, Dan Snyder, Marissa P. Griffith, Stephanie L. Mitchell, Vatsala R. Srinivasa, Kady D. Waggle, Chinelo Ezeonwuku, Vaughn S. Cooper, Lee H. Harrison |
| EPI_ISL_853931, EPI_ISL_853955 | Department of Microbiology, University Innsbruck | Bergthaler laboratory, CeMM Research Center for Molecular Medicine of the Austrian Academy of Sciences | Lukas Endler, Alexandra Popa, Benedikt Agerer, Jakob-Wendelin Genger, Alexander Lercher, Anna Schedl, Thomas Penz, Michael Schuster, Jan Laine, Martin Senekowitsch, Christoph Bock, Andreas Bergthaler |
| EPI_ISL_854438 | SARATOGA HOSPITAL LABORATORY | Wadsworth Center, New York State Department of Health | Kirsten St. George, Daryl M. Lamson, Alexis Russel, Matthew Shudt, Melissa A Leisner, Jonathan Plitnick, Navjot Singh, John Kelly, Erasmus Schneider, Erica Lasek-Nesselquist |
| EPI_ISL_855354 | Hospital | National Reference Center for Viruses of Respiratory Infections, Institut Pasteur, Paris | Marion Barbet, Sylvie Behillil, Méline Bizard, Angela Brisebarre, Camille Capel, Etienne Simon-Lorière, Vincent Enouf, Maud Vanpeene, Sylvie van der Werf |
| EPI_ISL_855410, EPI_ISL_855411, EPI_ISL_855412, EPI_ISL_855413, EPI_ISL_855414, EPI_ISL_855415, EPI_ISL_855416, EPI_ISL_855417, EPI_ISL_855418, EPI_ISL_855419, EPI_ISL_855427 | | | |
| see above | Servicio de Microbiología, Laboratori Clínic Metropolitana Nord. Hospital Universitari Germans Trias i Pujol. Institut d'Investigació en Ciències de la Salut Germans Trias i Pujol (IGTP) | SeqCOVID-SPAIN consortium/IBV(CSIC) | Elisa Martró, Antoni E. Bordoy, Anna Not, Adrián Antuori, Anabel Fernández, Nona Romaní, Verónica Saludes, Cristina Casañ and SeqCOVID-SPAIN consortium |
| EPI_ISL_855515, EPI_ISL_855518, EPI_ISL_855522, EPI_ISL_855535 | KEMRI-Wellcome Trust Research Programme/KEMRI-CGMR-C Kilifi | KEMRI-Wellcome Trust Research Programme/KEMRI-CGMR-C Kilifi | Githinji et al |
| EPI_ISL_856673, EPI_ISL_856674, EPI_ISL_856675, EPI_ISL_856676 | Department of Virus and Microbiological Special Diagnostics, Statens Serum Institut, Copenhagen, Denmark | Aalborg University | Danish Covid-19 Genome Consortium |
| EPI_ISL_856776, EPI_ISL_856780, EPI_ISL_856781, EPI_ISL_856782 | Servicio Virosis Respiratorias-Departamento Virología-INEI | Instituto Nacional Enfermedades Infecciosas C.G.Malbran | Baumeister E., Avaro M., Benedetti E., Russo M., Dattero ME, Pontoriero A., Cisterna D., Molina V., Perandones C., Tuduri E., Lorenzo F., Poklepovich T., Campos J. |
| EPI_ISL_857508, EPI_ISL_857509, EPI_ISL_857510, EPI_ISL_857511, EPI_ISL_857512 | Swiss National Reference Centre for Influenza | Swiss National Reference Centre for Influenza | Ana Rita Goncalves,Samuel Cordey, Laurent Kaiser, Lorenzo Cerutti, Henri Pegeot, Melyssa Elies, Keith Harshman, Ioannis Xenarios, Emmanouil Dermitzakis |
| EPI_ISL_857531, EPI_ISL_857532, EPI_ISL_857533, EPI_ISL_857534, EPI_ISL_857535 | Swiss National Reference Centre for Influenza | Swiss National Reference Centre for Influenza | Tim Roloff, Ana Rita Gonçalves, Madlen Stange, Helena MB Seth-Smith, Alfredo Mari, Karoline Leuzinger, Julia Bielicki, Manuel Battegay, Hans Hirsch, Laurent Kaiser, Adrian Egli |
| EPI_ISL_859469 | Lighthouse Lab in Milton Keynes | Wellcome Sanger Institute for the COVID-19 Genomics UK (COG-UK) Consortium | The Lighthouse Lab in Milton Keynes and Alex Alderton, Roberto Amato, Sonia Goncalves, Ewan Harrison, David K. Jackson, Ian Johnston, Dominic Kwiatkowski, Cordelia Langford, John Sillitoe on behalf of the Wellcome Sanger Institute COVID-19 Surveillance Team |
| EPI_ISL_859541, EPI_ISL_859542, EPI_ISL_859543, EPI_ISL_859544, EPI_ISL_859545, EPI_ISL_859546, EPI_ISL_859547, EPI_ISL_859548, EPI_ISL_859549, EPI_ISL_859550, EPI_ISL_859551, EPI_ISL_859552 | | | |
| see above | Lighthouse Lab in Glasgow | Wellcome Sanger Institute for the COVID-19 Genomics UK (COG-UK) Consortium | Harper VanSteenhouse, Yumi Kasai, David Gray, Carol Clugston, Anna Dominiczak and Alex Alderton, Roberto Amato, Sonia Goncalves, Ewan Harrison, David K. Jackson, Ian Johnston, Dominic Kwiatkowski, Cordelia Langford, John Sillitoe on behalf of the Wellcome Sanger Institute COVID-19 Surveillance Team |
| EPI_ISL_860181 | Keio University School of Medicine | Keio University School of Medicine | Kenjiro Kosaki, Yuka Iwasaki, Hirotsugu Ishizu, Haruhiko Siomi, Kodai Abe |
| EPI_ISL_861470 | UHAS COVID-19 Lab | UHAS COVID-19 Lab | Kwabena O. Duedu, Jones Gyamfi, Reuben Ayivor-Djanie, John O. Gyapong and the UHAS COVID-19 Lab Team |
| EPI_ISL_861584, EPI_ISL_861585 | Instituto Nacional de Saude (INSA) | Instituto Nacional de Saude (INSA) | Borges et al |
| EPI_ISL_861669 | Lab LOC - Itapecerica da Serra | Instituto Adolfo Lutz, Interdiciplinary Procedures Center, Strategic Laboratory | Claudio Tavares Sacchi, Claudia Regina Gonçalves, Erica Valessa Ramos Gomes, Karoline Rodrigues Campos |
| EPI_ISL_861670 | Instituto Adolfo Lutz - Regional de Taubate | Instituto Adolfo Lutz, Interdiciplinary Procedures Center, Strategic Laboratory | Claudio Tavares Sacchi, Claudia Regina Gonçalves, Erica Valessa Ramos Gomes, Karoline Rodrigues Campos |
| EPI_ISL_861773, EPI_ISL_861774, EPI_ISL_861777, EPI_ISL_861789, EPI_ISL_861792, EPI_ISL_861795, EPI_ISL_861797, EPI_ISL_861798, EPI_ISL_861800, EPI_ISL_861801, EPI_ISL_861802, EPI_ISL_861803, EPI_ISL_861808, EPI_ISL_861809, EPI_ISL_861812, EPI_ISL_861815, EPI_ISL_861816, EPI_ISL_861819, EPI_ISL_861820, EPI_ISL_861821, EPI_ISL_861823, EPI_ISL_861830, EPI_ISL_861831, EPI_ISL_861832, EPI_ISL_861833, EPI_ISL_861834, EPI_ISL_861835, EPI_ISL_861836, EPI_ISL_861837, EPI_ISL_861838, EPI_ISL_861839, EPI_ISL_861840, EPI_ISL_861841, EPI_ISL_861842, EPI_ISL_861843, EPI_ISL_861844 | | | |

| | | | |
|---|---|---|---|
| see above | Hospital General Universitario Gregorio Marañón | SeqCOVID-SPAIN consortium/IBV(CSIC) | Darío García de Viedma, Laura Pérez-Lago, Pedro J Sola-Campoy, Sergio Buenestado-Serrano, Marta Herranz, Victor Manuel de la Cueva, Julia Suárez, Pilar Catalán, Patricia Muñoz and SeqCOVID-SPAIN consortium |
| EPI_ISL_862578 | Hospital Clínic | Instituto de Salud Carlos III | Iglesias-Caballero, M.Camarero, S. Molinero Calamita, M. González-Esguevillas, M. Pozo, F. Casas, I. Jiménez, P. Jiménez, M. Zaballos, A. Monzón, S. Varona, S. Juliá, M. Cuesta, I. Marcos, M.A. |
| EPI_ISL_864868, EPI_ISL_864870, EPI_ISL_864888, EPI_ISL_864890, EPI_ISL_864892, EPI_ISL_864894 | Department of Pathology, University of Cambridge | COVID-19 Genomics UK (COG-UK) Consortium | Aminu S. Jahun, Yasmin Chaudhry, Grant Hall, Iliana Georgana, Myra Hosmillo, Martin D. Curran, Malte Pinckert, Surendra Parmar, Ian Goodfellow |
| EPI_ISL_865069, EPI_ISL_865070, EPI_ISL_865071, EPI_ISL_865155, EPI_ISL_865157, EPI_ISL_865161 | Virology Department, Royal Infirmary of Edinburgh, NHS Lothian / School of Biological Sciences, University of Edinburgh / Institute of Genetics and Molecular Medicine, University of Edinburgh | COVID-19 Genomics UK (COG-UK) Consortium | McHugh M, Dewar R, Rooke S, Gallagher M, Balcaza C, O'Toole Á, Scher E, Hill V, McCrone JT, Colquhoun R, Yu X, Jackson B, Rambaut A, Williams TC, Templeton K |
| EPI_ISL_865207, EPI_ISL_865209, EPI_ISL_865211, EPI_ISL_865212, EPI_ISL_865213, EPI_ISL_865214, EPI_ISL_865215, EPI_ISL_865216, EPI_ISL_865261, EPI_ISL_865264, EPI_ISL_865265, EPI_ISL_865268 | | | |
| see above | Liverpool Clinical Laboratories | COVID-19 Genomics UK (COG-UK) Consortium | Sam Haldenby, Anita Lucaci, Steve Paterson, Julian Hiscox, Alistair Darby, M Almsaud, A Alrezaihi, Muhannad Alruwaili, Stuart D Armstrong, Jones Benjamin, Eleanor G Bentley, Anu Chawla, Jordan J Clark, Angela Cowell, Richard Eccles, Isabel García-Dorival, Matthew Gemmell, Alessandro Gerada, PKF Gilmore, Richard Gregory, Ximeng Han, Catherine Hartley, Margaret Hughes, Miren Iturriza-Gomara, James Johnson, L Luu, Jenifer Manson, Charlotte Nelson, Elaine O'Toole, Cassie Olateju, Rebekah Penrice-Randal , Lucille Rainbow, N.P Randle, Trevor Ian Robinson, Parul Sharma, Ghada T Shawli, James P Stewart, Neil Swainston, Ecaterina Vamos, Joanne Watts, Mark Whitehead |
| EPI_ISL_865717 | University College London, Great Ormond Street Hospital for Children NHS Foundation Trust, Imperial College Healthcare NHS Trust | COVID-19 Genomics UK (COG-UK) Consortium | Sergi Castellano, Rachel Williams, Mark Kristiansen, Paola Resende Silva, Sunando Roy, Tony Brooks, Helena Tutill, Paola Niola, Patricia Dyal, Charlotte Williams, Leysa Forrest, Yasmin Panchbhaya, Jacqueline Findlay, Samuel Weeks, Julianne Brown, Kathryn Harris, Paul Randell, James Price, Alison Holmes, Judith Breuer |
| EPI_ISL_866059, EPI_ISL_866060, EPI_ISL_866061, EPI_ISL_866062, EPI_ISL_866063, EPI_ISL_866064 | University College London Hospital | COVID-19 Genomics UK (COG-UK) Consortium | Judith Heaney, Matthew Byott, Catherine Houlihan, Dan Frampton, Stuart Kirk, Moira Spyer and Eleni Nastouli |
| EPI_ISL_866335, EPI_ISL_866341, EPI_ISL_866345 | University College London, Great Ormond Street Hospital for Children NHS Foundation Trust, Imperial College Healthcare NHS Trust | COVID-19 Genomics UK (COG-UK) Consortium | Sergi Castellano, Rachel Williams, Mark Kristiansen, Paola Resende Silva, Sunando Roy, Tony Brooks, Helena Tutill, Paola Niola, Patricia Dyal, Charlotte Williams, Leysa Forrest, Yasmin Panchbhaya, Jacqueline Findlay, Samuel Weeks, Julianne Brown, Kathryn Harris, Paul Randell, James Price, Alison Holmes, Judith Breuer |
| EPI_ISL_866427, EPI_ISL_866430, EPI_ISL_866433, EPI_ISL_866434, EPI_ISL_866436, EPI_ISL_866438, EPI_ISL_866440 | Northumbria University / South Tees Hospitals NHS Foundation Trust / North Cumbria Integrated Care NHS Foundation Trust / North Tees and Hartlepool NHS Foundation Trust / Newcastle Hospitals NHS Foundation Trust | COVID-19 Genomics UK (COG-UK) Consortium | Darren L Smith,Andrew Nelson,Matthew Bashton,Greg R Young,Joshua Loh,John Allan,Mohammad A Tariq,Giles S Holt,Gary Black,Wen C Yew,Lynn Dover,Paul Baker,Steve Liggett,Sarah Essex,Jane Greenaway,Debra Padgett,Clive Graham,Garren Scott,Edward Barton,Emma Swindells,Brendan Payne,Jennifer Collins,Yusri Taha,Gary Eltringham |
| EPI_ISL_866579, EPI_ISL_866809, EPI_ISL_866810, EPI_ISL_866811, EPI_ISL_866816, EPI_ISL_866823 | Quadram Institute Bioscience | COVID-19 Genomics UK (COG-UK) Consortium | Dave J. Baker, Gemma L. Kay, Alp Aydin, Thanh Le-Viet, Steven Rudder, Ana P. Tedim, Anastasia Kolyva, Maria Diaz, Leonardo de Oliveira Martins, Nabil-Fareed Alikhan, Lizzie Meadows, Rachael Stanley, Ngozi Elumogo, Muhammed Yasir, Nicholas M. Thomson, Alexander J Trotter, Rachel Gilroy, Samuel Bloomfield, Claire Stuart, Andrew Bell, Reenesh Prakash, Samir Dervisevic, Alison E. Mather, John Wain, Mark Webber, Andrew J. Page, Justin O'Grady |
| EPI_ISL_867285 | Originating lab: Wales Specialist Virology Centre Sequencing lab: Pathogen Genomics Unit | Public Health Wales Microbiology Cardiff Wales Specialist Virology Centre | Catherine Moore, Johnathan Evans, Laura Gifford, Malorie Perry, Simon Cottrell, Angela Marchbank, Alec Birchley, Alexander Adams, Amy Gaskin, Bree Gatica-Wilcox, Jason Coombes, Joel Southgate, Lauren Gilbert, Lee Graham, Nicole Pacchiarini, Sara Kumziene-Summerhayes, Sarah Taylor, Sophie Jones, Sara Rey, Matthew Bull, Joanne Watkins, Sally Corden, Tom Connor |
| EPI_ISL_867952, EPI_ISL_867953 | Centre for Enzyme Innovation, University of Portsmouth / Translational Research Laboratory, Portsmouth Hospitals NHS Trust | COVID-19 Genomics UK (COG-UK) Consortium | Angela Beckett,Yann Bourgeois,Garry Scarlett,Sharon Glaysher,Scott Elliott,Kelly Bicknell,Robert Impey,Allyson Lloyd,Sarah Wyllie,Ethan Butcher,Anoop Chauhan,Samuel Robson |
| EPI_ISL_868950, EPI_ISL_868951, EPI_ISL_868952, EPI_ISL_868954, EPI_ISL_868955, EPI_ISL_868956, EPI_ISL_868957, EPI_ISL_868958, EPI_ISL_868960, EPI_ISL_868961, EPI_ISL_868962, EPI_ISL_868969, EPI_ISL_868973, EPI_ISL_868979, EPI_ISL_868980 | | | |
| see above | Bioinformatics and Biostatistics Lab, Advanced Sequencing Facility | COVID-19 Genomics UK (COG-UK) Consortium | Aengus Stewart,Jerome Nicod,Chelsea Sawyer,Laura Cubitt,Harshil Patel,Margaret Crawford |
| EPI_ISL_871759, EPI_ISL_871760, EPI_ISL_871761, EPI_ISL_871762, EPI_ISL_871763, EPI_ISL_871764, EPI_ISL_871765, EPI_ISL_871766, EPI_ISL_871767, EPI_ISL_871768, EPI_ISL_871769, EPI_ISL_871770, EPI_ISL_871771, EPI_ISL_871772, EPI_ISL_871773, EPI_ISL_871774, EPI_ISL_871775, EPI_ISL_871776, EPI_ISL_871777, EPI_ISL_871778, EPI_ISL_871779, EPI_ISL_871780 | | | |
| see above | Department of Virus and Microbiological Special Diagnostics, Statens Serum Institut, Copenhagen, Denmark | Aalborg University | Danish Covid-19 Genome Consortium |
| EPI_ISL_872616 | Nigeria Centre for Disease Control (NCDC) | African Centre of Excellence for Genomics of Infectious Diseases (ACEGID), Redeemer's University | Oluniyi P.E. et al |
| EPI_ISL_872732, EPI_ISL_872733, EPI_ISL_872734, EPI_ISL_872735, EPI_ISL_872736, EPI_ISL_872737, EPI_ISL_872738, EPI_ISL_872739, EPI_ISL_872740 | Rhode Island Department of Health | Infectious Disease Program, Broad Institute of Harvard and MIT | Lemieux,J.E., Siddle,K.J., Huard,R., King,E., Azevedo,K., Miller,A., Adams,G., Gladden-Young,A., Lagerborg,K., Rudy,M., DeRuff,K., Carter,A., Normandin,E., Bauer,M., Reilly,S., Tomkins-Tinch,C., Loreth,C., Chaluvadi,S., Birren,B.W., Gallagher,G., Smole,S., Park,D.J., MacInnis,B.L., and Sabeti,P.C. |
| EPI_ISL_872904, EPI_ISL_872906, EPI_ISL_872907, EPI_ISL_872909, EPI_ISL_872912 | WHO National Influenza Centre Russian Federation | WHO National Influenza Centre Russian Federation | Andrey Komissarov, Artem Fadeev, Anna Ivanova, Kseniya Komissarova, Dmitry Bazhenov, Mikhail Bakaev, Daria Danilenko, Ksenia Safina, Elena Nabieva, Georgii Bazykin, Dmitry Lioznov |
| EPI_ISL_873125, EPI_ISL_873161, EPI_ISL_873163 | University of Michigan Clinical Microbiology Laboratory | Lauring Lab, University of Michigan, Department of Microbiology and Immunology | Valesano |
| EPI_ISL_873215 | M Health Fairview | Minnesota Department of Health, Public Health Laboratory | Alexandra Lorentz, Jacob Garfin, Matt Plumb, and Xiong Wang |
| EPI_ISL_876109, EPI_ISL_876126, EPI_ISL_876130, EPI_ISL_876154, EPI_ISL_876156, EPI_ISL_876168, EPI_ISL_876179, EPI_ISL_876201, EPI_ISL_876206, EPI_ISL_876213, EPI_ISL_876305, EPI_ISL_876306, EPI_ISL_876307, EPI_ISL_876308, EPI_ISL_876309, EPI_ISL_876310, EPI_ISL_876311 | | | |
| see above | Massachusetts State Public Health Laboratory | Massachusetts State Public Health Laboratory | Andrew Lang, Timelia Fink, Glen Gallagher, Sandra Smole |
| EPI_ISL_876541, EPI_ISL_876542, EPI_ISL_876543, EPI_ISL_876544, EPI_ISL_876545, EPI_ISL_876546, EPI_ISL_876547, EPI_ISL_876548 | Florida Bureau of Public Health Laboratories | Florida Bureau of Public Health Laboratories | Sarah Schmedes, Jason Blanton |
| EPI_ISL_876590, EPI_ISL_876591, EPI_ISL_876593 | TN Division of Laboratory Services | Pathogen Discovery, Respiratory Viruses Branch, Division of Viral Diseases, Centers for Disease Control and Prevention | Ying Tao, Yan Li, Jing Zhang, Krista Queen, Anna Uehara, Peter Cook, Clinton R. Paden, Haibin Wang, Suxiang Tong |
| EPI_ISL_878152, EPI_ISL_878155, EPI_ISL_878159, EPI_ISL_878162, EPI_ISL_878165, EPI_ISL_878172, EPI_ISL_878175, EPI_ISL_878178, EPI_ISL_878186, EPI_ISL_878189, EPI_ISL_878194, EPI_ISL_878197, EPI_ISL_878199, EPI_ISL_878207, EPI_ISL_878209, EPI_ISL_878217, EPI_ISL_878221, EPI_ISL_878223, EPI_ISL_878234, EPI_ISL_878237, EPI_ISL_878242, EPI_ISL_878245, EPI_ISL_878256, EPI_ISL_878266, EPI_ISL_878268, EPI_ISL_878271, EPI_ISL_878384, EPI_ISL_878389 | | | |
| see above | San Diego County Public Health Laboratory | Andersen lab at Scripps Research | SEARCH Alliance San Diego with Tracy Basler, Jovan Shephard, Brett Austin |
| EPI_ISL_878707 | Rady's Childrens Hospital | Andersen lab at Scripps Research | SEARCH Alliance San Diego with Nanda Radamchar, David Dimmock, Linda Luo, Christina Clarke, Kathryn Bouic, Teresa Mueller, Denise Malicki |
| EPI_ISL_879929, EPI_ISL_879939, | San Diego County Public Health Laboratory | Andersen lab at Scripps Research | SEARCH Alliance San Diego with Tracy Basler, Jovan Shephard, Brett Austin |

| | | | |
|---|---|---|---|
| EPI_ISL_879945, EPI_ISL_880006, EPI_ISL_880024 | | | |
| EPI_ISL_880174, EPI_ISL_880186, EPI_ISL_880187, EPI_ISL_880190, EPI_ISL_880191, EPI_ISL_880194, EPI_ISL_880196 | Rady's Childrens Hospital | Andersen lab at Scripps Research | SEARCH Alliance San Diego with Nanda Radamchar, David Dimmock, Linda Luo, Christina Clarke, Kathryn Bouic, Teresa Mueller, Denise Malicki |
| EPI_ISL_882679, EPI_ISL_882765, EPI_ISL_882766, EPI_ISL_882767 | 1.AO Universitaria 'S. Giovanni di Dio e Ruggi D'Aragona, Scuola Medica Salernitana' Hospital / 2.UOC di Virologia e Microbiologia, Università della Campania 'L. Vanvitelli' / 3.AO Universitaria 'Federico II' Napoli Hospital / 4.AORN 'San Giuseppe Moscati' Avellino Hospital / 5.AO 'San Pio - presidio G. Rummo' Benevento Hospital / 6.AO 'Sant'Anna e San Sebastiano' Caserta Hospital / 7.PO 'Maria Santissima Addolorata' Eboli Hospital / 8.Biogem Istituto di Ricerche Genetiche | 1. Genome Research Center for Health (CRGS) / 2. Laboratory of Molecular Medicine and Genomics(LMMGe) / 3. Center for Research in Pure and Applied Mathematics (CRMPA) | Giorgio Giurato, Francesca Rizzo, Alessandro Weisz, Gianluigi Franci, Giovanni Nassa, Pasquale Pagliano, Roberta Tarallo, Elena Alexandrova, Ylenia D'Agostino, Carlo Ferravante, Jessica Lamberti, Viola Melone, Domenico Memoli, Valeria Mirici Cappa, Domenico Palumbo, Giovanni Pecoraro, Assunta Sellitto, Oriana Strianese, Ilaria Terenzi, Giuseppe Fenza, Aniello Gentile, Antonello Saccomanno, Sonia Amabile, Teresa Rocco, Annamaria Salvati, Emilia Vaccaro, Massimiliano Galdiero, Michele Cennamo, Giuseppe Portella, Maria Grazia Foti, Mariarosaria Ingino, Maria Landi, Maurizio Fumi, Vincenzo Rocco, Rita Greco, Vittoria Letizia, Arnolfo Petruzziello, Maddalena Schioppa, Gregorio Goffredi, Francesca Marciano, Michele Caraglia, Alessia Cossu, Marianna Scrima |
| EPI_ISL_883006, EPI_ISL_883007, EPI_ISL_883008 | Maryland Public Health Laboratory | Maryland Public Health Laboratory | Maryland Department of Health Laboratories Administration |
| EPI_ISL_883867, EPI_ISL_883868, EPI_ISL_883869, EPI_ISL_883870, EPI_ISL_883871, EPI_ISL_883872, EPI_ISL_883873, EPI_ISL_883874, EPI_ISL_883875, EPI_ISL_883876, EPI_ISL_883877, EPI_ISL_883879, EPI_ISL_883880, EPI_ISL_883881, EPI_ISL_883882, EPI_ISL_883883, EPI_ISL_883884, EPI_ISL_883885, EPI_ISL_883886, EPI_ISL_883887, EPI_ISL_883888, EPI_ISL_883889, EPI_ISL_883890, EPI_ISL_883891, EPI_ISL_883892, EPI_ISL_883893, EPI_ISL_883894, EPI_ISL_883895, EPI_ISL_883896, EPI_ISL_883897, EPI_ISL_883898, EPI_ISL_883899, EPI_ISL_883900, EPI_ISL_883901, EPI_ISL_883902, EPI_ISL_883903, EPI_ISL_883904, EPI_ISL_883905, EPI_ISL_883906, EPI_ISL_883907, EPI_ISL_883908, EPI_ISL_883909, EPI_ISL_883910, EPI_ISL_883911, EPI_ISL_883912, EPI_ISL_883913, EPI_ISL_883914, EPI_ISL_883915, EPI_ISL_883916, EPI_ISL_883917, EPI_ISL_883918, EPI_ISL_883919, EPI_ISL_883920, EPI_ISL_883922, EPI_ISL_883923, EPI_ISL_883924, EPI_ISL_883925, EPI_ISL_883926, EPI_ISL_883927, EPI_ISL_883928, EPI_ISL_883929, EPI_ISL_883930, EPI_ISL_883931, EPI_ISL_883932, EPI_ISL_883933, EPI_ISL_883934, EPI_ISL_883935, EPI_ISL_883936, EPI_ISL_883939, EPI_ISL_883940, EPI_ISL_883941, EPI_ISL_883942, EPI_ISL_883943, EPI_ISL_883944, EPI_ISL_883945, EPI_ISL_883946, EPI_ISL_883947, EPI_ISL_883948, EPI_ISL_883949, EPI_ISL_883950, EPI_ISL_883951, EPI_ISL_883952, EPI_ISL_883953, EPI_ISL_883954, EPI_ISL_883955, EPI_ISL_883956 | | | |
| see above | Eurofins Diatherix | Hudsonalpha Genome Sequencing Center | Jane Grimwood, Melissa Williams, Lori H. Handley, Joshua Stough, Leslie Malone, Stefan Brzezinski, Ada Stewart, Teresa Jones, Jenell Webber, John Lovell, Jennifer Cart, and Jeremy Schmutz |
| EPI_ISL_884217, EPI_ISL_884218 | Alaska State Virology Laboratory (Alaska DHHS) | Alaska State Virology Laboratory (Alaska DHHS) | Stephanie DeRonde, Lisa Smith, PhD. Jack Chen, PhD. |
| EPI_ISL_884251, EPI_ISL_884252, EPI_ISL_884253 | Maryland Public Health Laboratory | Maryland Public Health Laboratory | Maryland Department of Health Laboratories Administration |
| EPI_ISL_884890, EPI_ISL_884893, EPI_ISL_884894, EPI_ISL_884895, EPI_ISL_884896, EPI_ISL_884897, EPI_ISL_884902, EPI_ISL_884921, EPI_ISL_884922, EPI_ISL_884957 | Santa Clara County Public Health Laboratory | Chan-Zuckerberg Biohub | CZB Cliahub Consortium |
| EPI_ISL_885131, EPI_ISL_885132 | California Institute of Technology | Chan-Zuckerberg Biohub | CZB Cliahub Consortium |
| EPI_ISL_888844 | Michigan Department of Health and Human Services, Bureau of Laboratories | Michigan Department of Health and Human Services, Bureau of Laboratories | Blankenship HM, Riner D, Soehnlen MK |
| EPI_ISL_888980 | RS Anna | Eijkman Institute for Molecular Biology, Ministry of Research and Technology/National Agency for Research and Innovation | Hidayat Trimarsanto, Iskandar Adnan, Lydia V. Panggalo, Sukma Oktavianthi, Willy Agustine, Edison Johar, Frilasita A Yudhaputri, Safarina G Malik, Khin Saw Myint, Amin Soebandrio |
| EPI_ISL_890189 | Gonoshasthaya-RNA Research Center, Gonoshasthaya-RNA Molecular Diagnostics and Research Center | Gonoshasthaya-RNA Research Center, Gonoshasthaya-RNA Molecular Diagnostics and Research Center | Jamiruddin,M.R., Khondoker,M.U., Sharif,N., Azmuda,N., Ahmed,M.F., Sharmin,S., Akter,S., Mou,T.J., Marzan,M., Liza,S.M., Nahar,S., Jahan,N., Ali,T., Khandker,S.S., Jamiruddin,M., Haq,M.A., Adnan,N., Chaity,M., Oishee,M. |
| EPI_ISL_890949 | Seattle Flu Study | Seattle Flu Study | Deborah A. Nickerson, Chris D. Frazar, Jover Lee, Benjamin Pelle, Erica Ryke, Matthew Richardson, Amanda Adler, Elisabeth Brandstetter, Peter D. Han, Kairsten Fay, Misja Ilcisin, Kirsten Lacombe, Thomas R. Sibley, Melissa Truong, Caitlin R. Wolf, Karen Cowgill, Stephanie Schrag, Jeff Duchin, Michael Boeckh, Janet A. Englund, Michael Famulare, Barry R. Lutz, Mark J. Rieder, Lea M. Starita, Matthew Thompson, Helen Y. Chu, Trevor Bedford, Jay Shendure |
| EPI_ISL_890950 | Seattle Flu Study | Seattle Flu Study | Deborah A. Nickerson, Chris D. Frazar, Jover Lee, Benjamin Pelle, Erica Ryke, Matthew Richardson, Amanda Adler, Elisabeth Brandstetter, Peter D. Han, Kairsten Fay, Misja Ilcisin, Kirsten Lacombe, Thomas R. Sibley, Melissa Truong, Caitlin R. Wolf, Michael Boeckh, Janet A. Englund, Michael Famulare, Barry R. Lutz, Mark J. Rieder, Lea M. Starita, Matthew Thompson, Jay Shendure, Trevor Bedford, Helen Y. Chu |
| EPI_ISL_890958, EPI_ISL_890959 | Seattle Flu Study | Seattle Flu Study | Deborah A. Nickerson, Chris D. Frazar, Jover Lee, Benjamin Pelle, Erica Ryke, Matthew Richardson, Amanda Adler, Elisabeth Brandstetter, Peter D. Han, Kairsten Fay, Misja Ilcisin, Kirsten Lacombe, Thomas R. Sibley, Melissa Truong, Caitlin R. Wolf, Karen Cowgill, Stephanie Schrag, Jeff Duchin, Michael Boeckh, Janet A. Englund, Michael Famulare, Barry R. Lutz, Mark J. Rieder, Lea M. Starita, Matthew Thompson, Helen Y. Chu, Trevor Bedford, Jay Shendure |
| EPI_ISL_892276 | Servicio de Microbiología Clínica (Complejo Hospitalario de Navarra, Pamplona), Instituto de Investigación Sanitaria de Navarra (IdiSNA) | SeqCOVID-SPAIN consortium/IBV(CSIC) | Carmen Ezpeleta Baquedano, Ana Navascués, Ana Miqueleiz and SeqCOVID-SPAIN consortium |
| EPI_ISL_892278, EPI_ISL_892303, EPI_ISL_892304, EPI_ISL_892305, EPI_ISL_892306, EPI_ISL_892307, EPI_ISL_892308, EPI_ISL_892309, EPI_ISL_892310, EPI_ISL_892311, EPI_ISL_892312, EPI_ISL_892313 | | | |
| see above | Servicio de Microbiología. Hospital Universitario Donostia. OSI Donostialdea. Área de Enfermedades Infecciosas, Grupo de Infección Respiratoria y Resistencia Antimicrobiana. Instituto de Investigación Sanitaria Biodonostia | SeqCOVID-SPAIN consortium/IBV(CSIC) | Gustavo Cilla Eguiluz, Milagrosa Montes Ros, Luis Piñeiro Vázquez, Ane Sorrarain, Jose Maria Marimón and SeqCOVID-SPAIN consortium |
| EPI_ISL_892339, EPI_ISL_892340, EPI_ISL_892341, EPI_ISL_892342, EPI_ISL_892343, EPI_ISL_892344 | Servicio de Microbiología Clínica (Complejo Hospitalario de Navarra, Pamplona), Instituto de Investigación Sanitaria de Navarra (IdiSNA) | SeqCOVID-SPAIN consortium/IBV(CSIC) | Carmen Ezpeleta Baquedano, Ana Navascués, Ana Miqueleiz and SeqCOVID-SPAIN consortium |
| EPI_ISL_892364, EPI_ISL_892368, EPI_ISL_893367, EPI_ISL_893740, EPI_ISL_893741, EPI_ISL_893742 | Ohio Department of Health Laboratory | Ohio Department of Health Laboratory | Holmes, Jennifer; Eric Brandt, Keoni Omura, Glen McGillivary, Caitlin McDonnell, Jade Mowery, Stephanie Mccracken, Tyler Payne, Kirtana Ramadugu, Erica Leasure, Brent Lee, Kelsey Florek, Heather Blankenship, Quanta Brown, and Tammy Bannerman |
| EPI_ISL_893785 | Institute of Virology, Medical Center, University of Freiburg, Freiburg, Germany | Institute of Virology, Clinial Virus Genomics, Medical Center, University of Freiburg, Freiburg, Germany | Jonas Fuchs, Lisa Kern, Sandra Reuter, Hajo Grundmann, Marcus Panning |
| EPI_ISL_894170, EPI_ISL_894947, EPI_ISL_895289, EPI_ISL_895727, EPI_ISL_900742 | Ohio Department of Health Laboratory | Ohio Department of Health Laboratory | Holmes, Jennifer; Eric Brandt, Keoni Omura, Glen McGillivary, Caitlin McDonnell, Jade Mowery, Stephanie Mccracken, Tyler Payne, Kirtana Ramadugu, Erica Leasure, Brent Lee, Kelsey Florek, Heather Blankenship, Quanta Brown, and Tammy Bannerman |
| EPI_ISL_903235, EPI_ISL_903253 | M Health Fairview | Minnesota Department of Health, Public Health Laboratory | Alexandra Lorentz, Jacob Garfin, Matt Plumb, and Xiong Wang |
| EPI_ISL_903570 | WVDHHR - Office of Laboratory Services | Genomics and Discovery, Respiratory Viruses Branch, Division of Viral Diseases, Centers for Disease Control and Prevention | Krista Queen, Yan Li, Ying Tao, Jing Zhang, Anna Uehara, Anna Montmayeur, Clinton R. Paden, Peter W. Cook, Rachel Marine, Mili Sheth, Jasmine Padilla, Sarah Nobles, Mark Burroughs, Lori Rowe, Haibin Wang, Ben L. Rambo-Martin, Dhwani Batra, Justin Lee, Suxiang Tong |
| EPI_ISL_903573 | WA State Department of Health | Genomics and Discovery, Respiratory Viruses Branch, Division of Viral Diseases, Centers for Disease Control and | Krista Queen, Yan Li, Ying Tao, Jing Zhang, Anna Uehara, Anna Montmayeur, Clinton R. Paden, Peter W. Cook, Rachel Marine, Mili Sheth, Jasmine Padilla, Sarah Nobles, Mark Burroughs, Lori Rowe, Haibin Wang, Ben L. Rambo-Martin, Dhwani Batra, Justin Lee, Suxiang Tong |

| | | | |
|---|---|---|---|
| | | Prevention | |
| EPI_ISL_903576 | WVDHHR - Office of Laboratory Services | Genomics and Discovery, Respiratory Viruses Branch, Division of Viral Diseases, Centers for Disease Control and Prevention | Krista Queen, Yan Li, Ying Tao, Jing Zhang, Anna Uehara, Anna Montmayeur, Clinton R. Paden, Peter W. Cook, Rachel Marine, Mili Sheth, Jasmine Padilla, Sarah Nobles, Mark Burroughs, Lori Rowe, Haibin Wang, Ben L. Rambo-Martin, Dhwani Batra, Justin Lee, Suxiang Tong |
| EPI_ISL_903590 | PA Department of Health, Bureau of Laboratories | Genomics and Discovery, Respiratory Viruses Branch, Division of Viral Diseases, Centers for Disease Control and Prevention | Krista Queen, Yan Li, Ying Tao, Jing Zhang, Anna Uehara, Anna Montmayeur, Clinton R. Paden, Peter W. Cook, Rachel Marine, Mili Sheth, Jasmine Padilla, Sarah Nobles, Mark Burroughs, Lori Rowe, Haibin Wang, Ben L. Rambo-Martin, Dhwani Batra, Justin Lee, Suxiang Tong |
| EPI_ISL_903601, EPI_ISL_903602, EPI_ISL_903612 | OR State PHL-Virology/Immunology Section | Genomics and Discovery, Respiratory Viruses Branch, Division of Viral Diseases, Centers for Disease Control and Prevention | Krista Queen, Yan Li, Ying Tao, Jing Zhang, Anna Uehara, Anna Montmayeur, Clinton R. Paden, Peter W. Cook, Rachel Marine, Mili Sheth, Jasmine Padilla, Sarah Nobles, Mark Burroughs, Lori Rowe, Haibin Wang, Ben L. Rambo-Martin, Dhwani Batra, Justin Lee, Suxiang Tong |
| EPI_ISL_903643, EPI_ISL_903644 | RI State Health Laboratories | Genomics and Discovery, Respiratory Viruses Branch, Division of Viral Diseases, Centers for Disease Control and Prevention | Krista Queen, Yan Li, Ying Tao, Jing Zhang, Anna Uehara, Anna Montmayeur, Clinton R. Paden, Peter W. Cook, Rachel Marine, Mili Sheth, Jasmine Padilla, Sarah Nobles, Mark Burroughs, Lori Rowe, Haibin Wang, Ben L. Rambo-Martin, Dhwani Batra, Justin Lee, Suxiang Tong |
| EPI_ISL_903653 | WA State Department of Health | Genomics and Discovery, Respiratory Viruses Branch, Division of Viral Diseases, Centers for Disease Control and Prevention | Krista Queen, Yan Li, Ying Tao, Jing Zhang, Anna Uehara, Anna Montmayeur, Clinton R. Paden, Peter W. Cook, Rachel Marine, Mili Sheth, Jasmine Padilla, Sarah Nobles, Mark Burroughs, Lori Rowe, Haibin Wang, Ben L. Rambo-Martin, Dhwani Batra, Justin Lee, Suxiang Tong |
| EPI_ISL_903662 | PA Department of Health, Bureau of Laboratories | Genomics and Discovery, Respiratory Viruses Branch, Division of Viral Diseases, Centers for Disease Control and Prevention | Krista Queen, Yan Li, Ying Tao, Jing Zhang, Anna Uehara, Anna Montmayeur, Clinton R. Paden, Peter W. Cook, Rachel Marine, Mili Sheth, Jasmine Padilla, Sarah Nobles, Mark Burroughs, Lori Rowe, Haibin Wang, Ben L. Rambo-Martin, Dhwani Batra, Justin Lee, Suxiang Tong |
| EPI_ISL_903707, EPI_ISL_903746 | MN PHL Division, Minnesota Department of Health | Genomics and Discovery, Respiratory Viruses Branch, Division of Viral Diseases, Centers for Disease Control and Prevention | Krista Queen, Yan Li, Ying Tao, Jing Zhang, Anna Uehara, Anna Montmayeur, Clinton R. Paden, Peter W. Cook, Rachel Marine, Mili Sheth, Jasmine Padilla, Sarah Nobles, Mark Burroughs, Lori Rowe, Haibin Wang, Ben L. Rambo-Martin, Dhwani Batra, Justin Lee, Suxiang Tong |
| EPI_ISL_903784, EPI_ISL_903786 | WVDHHR - Office of Laboratory Services | Genomics and Discovery, Respiratory Viruses Branch, Division of Viral Diseases, Centers for Disease Control and Prevention | Krista Queen, Yan Li, Ying Tao, Jing Zhang, Anna Uehara, Anna Montmayeur, Clinton R. Paden, Peter W. Cook, Rachel Marine, Mili Sheth, Jasmine Padilla, Sarah Nobles, Mark Burroughs, Lori Rowe, Haibin Wang, Ben L. Rambo-Martin, Dhwani Batra, Justin Lee, Suxiang Tong |
| EPI_ISL_903788 | TX DSHS, Lab Services Section MC 1947 | Genomics and Discovery, Respiratory Viruses Branch, Division of Viral Diseases, Centers for Disease Control and Prevention | Krista Queen, Yan Li, Ying Tao, Jing Zhang, Anna Uehara, Anna Montmayeur, Clinton R. Paden, Peter W. Cook, Rachel Marine, Mili Sheth, Jasmine Padilla, Sarah Nobles, Mark Burroughs, Lori Rowe, Haibin Wang, Ben L. Rambo-Martin, Dhwani Batra, Justin Lee, Suxiang Tong |
| EPI_ISL_903791 | NE Public Health Laboratory | Genomics and Discovery, Respiratory Viruses Branch, Division of Viral Diseases, Centers for Disease Control and Prevention | Krista Queen, Yan Li, Ying Tao, Jing Zhang, Anna Uehara, Anna Montmayeur, Clinton R. Paden, Peter W. Cook, Rachel Marine, Mili Sheth, Jasmine Padilla, Sarah Nobles, Mark Burroughs, Lori Rowe, Haibin Wang, Ben L. Rambo-Martin, Dhwani Batra, Justin Lee, Suxiang Tong |
| EPI_ISL_903795 | WVDHHR - Office of Laboratory Services | Genomics and Discovery, Respiratory Viruses Branch, Division of Viral Diseases, Centers for Disease Control and Prevention | Krista Queen, Yan Li, Ying Tao, Jing Zhang, Anna Uehara, Anna Montmayeur, Clinton R. Paden, Peter W. Cook, Rachel Marine, Mili Sheth, Jasmine Padilla, Sarah Nobles, Mark Burroughs, Lori Rowe, Haibin Wang, Ben L. Rambo-Martin, Dhwani Batra, Justin Lee, Suxiang Tong |
| EPI_ISL_903843 | OR State PHL-Virology/Immunology Section | Genomics and Discovery, Respiratory Viruses Branch, Division of Viral Diseases, Centers for Disease Control and Prevention | Krista Queen, Yan Li, Ying Tao, Jing Zhang, Anna Uehara, Anna Montmayeur, Clinton R. Paden, Peter W. Cook, Rachel Marine, Mili Sheth, Jasmine Padilla, Sarah Nobles, Mark Burroughs, Lori Rowe, Haibin Wang, Ben L. Rambo-Martin, Dhwani Batra, Justin Lee, Suxiang Tong |
| EPI_ISL_903867 | NC State Laboratory of Public Health | Genomics and Discovery, Respiratory Viruses Branch, Division of Viral Diseases, Centers for Disease Control and Prevention | Krista Queen, Yan Li, Ying Tao, Jing Zhang, Anna Uehara, Anna Montmayeur, Clinton R. Paden, Peter W. Cook, Rachel Marine, Mili Sheth, Jasmine Padilla, Sarah Nobles, Mark Burroughs, Lori Rowe, Haibin Wang, Ben L. Rambo-Martin, Dhwani Batra, Justin Lee, Suxiang Tong |
| EPI_ISL_903868 | MN PHL Division, Minnesota Department of Health | Genomics and Discovery, Respiratory Viruses Branch, Division of Viral Diseases, Centers for Disease Control and Prevention | Krista Queen, Yan Li, Ying Tao, Jing Zhang, Anna Uehara, Anna Montmayeur, Clinton R. Paden, Peter W. Cook, Rachel Marine, Mili Sheth, Jasmine Padilla, Sarah Nobles, Mark Burroughs, Lori Rowe, Haibin Wang, Ben L. Rambo-Martin, Dhwani Batra, Justin Lee, Suxiang Tong |
| EPI_ISL_903869 | VA-Division of Consolidated Laboratory Services | Genomics and Discovery, Respiratory Viruses Branch, Division of Viral Diseases, Centers for Disease Control and Prevention | Krista Queen, Yan Li, Ying Tao, Jing Zhang, Anna Uehara, Anna Montmayeur, Clinton R. Paden, Peter W. Cook, Rachel Marine, Mili Sheth, Jasmine Padilla, Sarah Nobles, Mark Burroughs, Lori Rowe, Haibin Wang, Ben L. Rambo-Martin, Dhwani Batra, Justin Lee, Suxiang Tong |
| EPI_ISL_903881 | WVDHHR - Office of Laboratory Services | Genomics and Discovery, Respiratory Viruses Branch, Division of Viral Diseases, Centers for Disease Control and Prevention | Krista Queen, Yan Li, Ying Tao, Jing Zhang, Anna Uehara, Anna Montmayeur, Clinton R. Paden, Peter W. Cook, Rachel Marine, Mili Sheth, Jasmine Padilla, Sarah Nobles, Mark Burroughs, Lori Rowe, Haibin Wang, Ben L. Rambo-Martin, Dhwani Batra, Justin Lee, Suxiang Tong |
| EPI_ISL_903898, EPI_ISL_903916 | MO State Public Health Laboratory | Genomics and Discovery, Respiratory Viruses Branch, Division of Viral Diseases, Centers for Disease Control and Prevention | Krista Queen, Yan Li, Ying Tao, Jing Zhang, Anna Uehara, Anna Montmayeur, Clinton R. Paden, Peter W. Cook, Rachel Marine, Mili Sheth, Jasmine Padilla, Sarah Nobles, Mark Burroughs, Lori Rowe, Haibin Wang, Ben L. Rambo-Martin, Dhwani Batra, Justin Lee, Suxiang Tong |
| EPI_ISL_903917 | PA Department of Health, Bureau of Laboratories | Genomics and Discovery, Respiratory Viruses Branch, Division of Viral Diseases, Centers for Disease Control and Prevention | Krista Queen, Yan Li, Ying Tao, Jing Zhang, Anna Uehara, Anna Montmayeur, Clinton R. Paden, Peter W. Cook, Rachel Marine, Mili Sheth, Jasmine Padilla, Sarah Nobles, Mark Burroughs, Lori Rowe, Haibin Wang, Ben L. Rambo-Martin, Dhwani Batra, Justin Lee, Suxiang Tong |
| EPI_ISL_903932 | AZ SPHL, Arizona Department of Health Services | Genomics and Discovery, Respiratory Viruses Branch, Division of Viral Diseases, Centers for Disease Control and Prevention | Krista Queen, Yan Li, Ying Tao, Jing Zhang, Anna Uehara, Anna Montmayeur, Clinton R. Paden, Peter W. Cook, Rachel Marine, Mili Sheth, Jasmine Padilla, Sarah Nobles, Mark Burroughs, Lori Rowe, Haibin Wang, Ben L. Rambo-Martin, Dhwani Batra, Justin Lee, Suxiang Tong |
| EPI_ISL_903956, EPI_ISL_903974 | WA State Department of Health | Genomics and Discovery, Respiratory Viruses Branch, Division of Viral Diseases, Centers for Disease Control and Prevention | Krista Queen, Yan Li, Ying Tao, Jing Zhang, Anna Uehara, Anna Montmayeur, Clinton R. Paden, Peter W. Cook, Rachel Marine, Mili Sheth, Jasmine Padilla, Sarah Nobles, Mark Burroughs, Lori Rowe, Haibin Wang, Ben L. Rambo-Martin, Dhwani Batra, Justin Lee, Suxiang Tong |
| EPI_ISL_904007, EPI_ISL_904008 | Veterinary Specialized Institute Kraljevo | Veterinary Specialized Institute "Kraljevo", Serbia | Vidanovic,D., Tesovic,B., Knezevic,A., Jovanovic,T., Jankovic,M., Sekler,M., Banovic Djeri,B., Petrovic,T., Volkening,J., Afonso,C. |
| EPI_ISL_904178, EPI_ISL_904242, EPI_ISL_904262, EPI_ISL_904481, EPI_ISL_904482 | Dutch COVID-19 response team | Erasmus Medical Center | Bas Oude Munnink, Reina Sikkema, David Nieuwenhuijse, Irina Chestakova, Anne van der Linden, Marjan Boter, Emmanuelle Munger, Corine GeurtsvanKessel, Annemiek van der Eijk, Richard Molenkamp, Marion Koopmans, on behalf of the Dutch national COVID-19 response team. |
| EPI_ISL_904862, EPI_ISL_904863, EPI_ISL_904865, EPI_ISL_904888, EPI_ISL_904901, EPI_ISL_905085, EPI_ISL_905088, EPI_ISL_905090, EPI_ISL_905535 | Dutch COVID-19 response team | National Institute for Public Health and the Environment (RIVM) | Adam Meijer, Harry Vennema, Dirk Eggink, Jeroen Cremer, Sharon van den Brink, Bas van der Veer, AnneMarie van den Brandt, Florian Zwagemaker, Dennis Schmitz, Chantal Reusken, on behalf of the national COVID-19 response team |
| EPI_ISL_905801, EPI_ISL_905802, EPI_ISL_905803, EPI_ISL_905804, EPI_ISL_905805, EPI_ISL_905806 | OHSU Lab Services Molecular Microbiology Lab | Oregon SARS-CoV-2 Genome Sequencing Center | Brendan L. O'Connell, Sally Grindstaff, Kayla Carter, Ruth V. Nichols, Alec J. Hirsch, Donna Hansel, Guang Fan, Xuan, Qin, Daniel N. Streblow, William B. Messer, Andrew C. Adey, Benjamin N. Bimber, Brian J. O'Roak |
| EPI_ISL_906100 | Child Health Research Foundation | Child Health Research Foundation | Senjuti Saha, Sharmistha Goswami, Afroza Akter Tanni, Syed Muktadir Al Sium, Arif Mohammad Tanmoy, Roly Malaker, Md Hafizur Rahman, Samir K Saha |

| | | | |
|---|---|---|---|
| EPI_ISL_906112 | Child Health Research Foundation | Child Health Research Foundation | Senjuti Saha, Afroza Akter Tanni, Sharmistha Goswami, Syed Muktadir Al Sium, Arif Mohammad Tanmoy, Roly Malaker, Md Hafizur Rahman, Samir K Saha |
| EPI_ISL_906305 | Nigeria Centre for Disease Control (NCDC) | African Centre of Excellence for Genomics of Infectious Diseases (ACEGID), Redeemer's University | Oluniyi P.E. et al |
| EPI_ISL_906549 | Laboratorio de Virologia-Instituto Nacional de Salud | Instituto Nacional de Salud- Dirección de Investigación en Salud Pública, Universidad de los Andes- Applied genomics research group, Vicerrectoria de Investigación y Creación, Universidad de los Andes- Systems and Computing Engineering Department | Katherine Laiton-Donato, Diego A. Álvarez-Díaz, Carlos Franco-Muñoz, Mauricio Pacheco-Montealegre, Héctor Alejandro Ruiz-Moreno, Maria T. Herrera-Sepúlveda, Diego Andrés Prada, Jhonnatan Reales-González, Sheryll Corchuelo, Julian Naizaque, Gerardo Santamaría Jorge Duitama, Laura Natalia Gonzalez, Jorge Ivan Diaz, Silvia Restrepo-Restrepo, Magdalena Wiesner, Martha Lucia Ospina Martinez, Marcela Mercado-Reyes |
| EPI_ISL_909741, EPI_ISL_909743 | Ohio Department of Health Laboratory | Ohio Department of Health Laboratory | Holmes, Jennifer; Eric Brandt, Keoni Omura, Glen McGillivary, Caitlin McDonnell, Jade Mowery, Stephanie Mccracken, Tyler Payne, Kirtana Ramadugu, Erica Leasure, Brent Lee, Kelsey Florek, Heather Blankenship, Quanta Brown, and Tammy Bannerman |
| EPI_ISL_909952 | CUSL/UCLouvain COVID testing federal platform | UCLouvain/IREC/MBLG | Jean Ruelle, Lysa Pinsmaye, Benoit Kabamba Mukadi |
| EPI_ISL_910490, EPI_ISL_910491, EPI_ISL_910492, EPI_ISL_910493, EPI_ISL_910494, EPI_ISL_910495, EPI_ISL_910644, EPI_ISL_910645, EPI_ISL_910646, EPI_ISL_910648, EPI_ISL_910649, EPI_ISL_910650, EPI_ISL_910651, EPI_ISL_910652, EPI_ISL_910653, EPI_ISL_910654, EPI_ISL_910655, EPI_ISL_910656, EPI_ISL_910657, EPI_ISL_910658, EPI_ISL_910659, EPI_ISL_910660, EPI_ISL_910661, EPI_ISL_910662, EPI_ISL_910663, EPI_ISL_910664, EPI_ISL_910665, EPI_ISL_910666, EPI_ISL_910667, EPI_ISL_910668, EPI_ISL_910669, EPI_ISL_910670, EPI_ISL_910671, EPI_ISL_910672, EPI_ISL_910673, EPI_ISL_910674, EPI_ISL_910675, EPI_ISL_910676, EPI_ISL_910677, EPI_ISL_910678, EPI_ISL_910679, EPI_ISL_910680, EPI_ISL_910681, EPI_ISL_910682, EPI_ISL_910683, EPI_ISL_910684, EPI_ISL_910685, EPI_ISL_910686, EPI_ISL_910687, EPI_ISL_910688, EPI_ISL_910689, EPI_ISL_910690, EPI_ISL_910691, EPI_ISL_910692, EPI_ISL_910693, EPI_ISL_910694, EPI_ISL_910695, EPI_ISL_910696, EPI_ISL_910697, EPI_ISL_910698, EPI_ISL_910699, EPI_ISL_910704, EPI_ISL_910706, EPI_ISL_910707, EPI_ISL_910708, EPI_ISL_910709, EPI_ISL_910710, EPI_ISL_910711, EPI_ISL_910712, EPI_ISL_910713, EPI_ISL_910721, EPI_ISL_910722, EPI_ISL_910724, EPI_ISL_910725, EPI_ISL_910726, EPI_ISL_910727, EPI_ISL_910728, EPI_ISL_910729, EPI_ISL_910730, EPI_ISL_910731, EPI_ISL_910732, EPI_ISL_910733, EPI_ISL_910734, EPI_ISL_910735, EPI_ISL_910736, EPI_ISL_910737, EPI_ISL_910738, EPI_ISL_910739, EPI_ISL_910740, EPI_ISL_910741, EPI_ISL_910742, EPI_ISL_910743, EPI_ISL_910744, EPI_ISL_910745, EPI_ISL_910746, EPI_ISL_910747 | | | |
| see above | Laboratoire national de sante, Microbiology, Virology | Laboratoire national de sante, Microbiology, Microbial Genomics Platform | Anke Wienecke-Baldacchino, Catherine Ragimbeau,Jessica Tapp, Fatu Djabi, Lise Pignon, Raoul Salmon, Tamir Abdelrahman |
| EPI_ISL_911514 | Ohio Department of Health Laboratory | Ohio Department of Health Laboratory | Holmes, Jennifer; Eric Brandt, Keoni Omura, Glen McGillivary, Caitlin McDonnell, Jade Mowery, Stephanie Mccracken, Tyler Payne, Kirtana Ramadugu, Erica Leasure, Brent Lee, Kelsey Florek, Heather Blankenship, Quanta Brown, and Tammy Bannerman |
| EPI_ISL_911681, EPI_ISL_911688, EPI_ISL_911691, EPI_ISL_911692 | Alaska State Virology Laboratory | Alaska State Virology Laboratory | Stephanie DeRonde, Lisa Smith, Ph.D., Jack Chen, Ph.D. |
| EPI_ISL_912355, EPI_ISL_912356 | Fondation Congolaise pour la recherche medicale (FCRM), Francine Ntoumi | NGS Competence Center Tuebingen, Institut für Medizinische Mikrobiologie und Hygiene, Universitaetsklinikum Tübingen | Angel Angelov |
| EPI_ISL_913105, EPI_ISL_913106, EPI_ISL_913111 | CHU Purpan - Laboratoire de Virologie - Institut Fédératif de Biologie | CHU Purpan - Laboratoire de Virologie - Institut Fédératif de Biologie | Latour J., Ranger N., Dubois M., Carcenac R., Harter A., Boyer P., Tremeaux P., Izopet J. |
| EPI_ISL_913668, EPI_ISL_913670, EPI_ISL_913672 | Vault Health | Minnesota Department of Health, Public Health Laboratory | Alexandra Lorentz, Jacob Garfin, Matt Plumb, and Xiong Wang |
| EPI_ISL_913940, EPI_ISL_913941, EPI_ISL_913942, EPI_ISL_913943, EPI_ISL_913945, EPI_ISL_913960, EPI_ISL_913961, EPI_ISL_913972, EPI_ISL_913973, EPI_ISL_913974 | Instituto de Diagnostico y Referencia Epidemiologicos INDRE_RNLSP | Instituto de Diagnostico y Referencia Epidemiologicos (INDRE) | Claudia Wong-Arambula, Abril Rodriguez-Maldonado, Fabiola Garces-Ayala, Adnan Araiza-Rodriguez, David Fragoso-Fonseca, Sergio Rangel-Guerrero, Mayra Jimenez-Morales, Nancy Munoz-Hernandez, Natividad Cruz-Ortiz, Tatiana Nunez-Garcia, Gisela Barrera-Badillo, Lucia Hernandez-Rivas, Irma Lopez-Martinez, Ernesto Ramirez-Gonzalez. |
| EPI_ISL_914797 | TAMIZAJE COMUNITARIO - PASO CANOAS | Inciensa, Instituto Costarricense de Investigación y Enseñanza en Nutrición y Salud | Francisco Duarte, Hebleen Porras, Claudio Soto-Garita, Estela Cordero, Adriana Godínez, Melany Calderón & Mariel López |
| EPI_ISL_914883 | Vilnius university hospital Santaros Klinikos, Center of Laboratory Medicine | Vilnius University Hospital Santaros Klinikos | Ingrida Olendraite, Daniel Naumovas, Rimvydas Norvilas, Dovil Ežerskyt, Justinas Šlikas |
| EPI_ISL_915360 | Keio University School of Medicine | Keio University School of Medicine | Kenjiro Kosaki, Yuka Iwasaki, Hirotsugu Ishizu, Haruhiko Siomi, Kodai Abe |
| EPI_ISL_918168 | Department of Infectious Diseases and Immunology, National Hospital Organization Nagoya Medical Center | Clinical Research Center, National Hospital Organization Nagoya Medical Center | Yoshihiro Nakata, Hirotaka Ode, Mai Kubota, Masakazu Matsuda, Kazuhiro Matsuoka, Miho Nakasuji, Mikiko Mori, Mayumi Imahashi, Yoshiyuki Yokomaku, Yasumasa Iwatani |
| EPI_ISL_918186, EPI_ISL_918208 | Innovative Genomics Institute, UC Berkeley | Innovative Genomics Institute, UC Berkeley | Stacia Wyman, Haridha Shivram, Phil Frankino, Liana Lareau, Shana McDevitt, Justin Choi |
| EPI_ISL_918524, EPI_ISL_918525, EPI_ISL_918527, EPI_ISL_918528 | LACEN - Laboratório Central de Saúde Pública do Para | Evandro Chagas Institute | Santos, M.C.; Silva, A.M.; Junior, W.D.C.; Barbagelata, L.S.; Ferreira, J.A.; Sousa, E.M.A.; da Silva, P.S.; Pinheiro, K.C.; L.C.; Sousa Junior, E.C. |
| EPI_ISL_918553 | LACEN - Laboratório Central de Saúde Pública do Amapa | Evandro Chagas Institute | Santos, M.C.; Silva, A.M.; Junior, W.D.C.; Barbagelata, L.S.; Ferreira, J.A.; Sousa, E.M.A.; da Silva, P.S.; Pinheiro, K.C.; L.C.; Sousa Junior, E.C. |
| EPI_ISL_918600, EPI_ISL_918601, EPI_ISL_918604, EPI_ISL_918605, EPI_ISL_918606 | University of Birmingham | COVID-19 Genomics UK (COG-UK) Consortium | Institute of Microbiology, University of Birmingham: Claire McMurray, Joanne Stockton, Samuel Nicholls, Radoslaw Poplawski, Will Rowe, Josh Quick, Nicholas Loman. University of Birmingham Testing Laboratory: Celina M Whalley, Andrew Bosworth, Charlotte Poxon, Kasun Wanigasooriya, Oliver Pickles, Mike Kidd, Alex Richter, Andrew D Beggs PHE Heartlands Lab: Husam Osman, Andrew Bosworth. Queen Elizabeth Hospital: Anna Casey |
| EPI_ISL_919234 | West of Scotland Specialist Virology Centre, NHSGGC / MRC-University of Glasgow Centre for Virus Research | COVID-19 Genomics UK (COG-UK) Consortium | Ana da Silva Filipe, Natasha Johnson, Kathy Smollett, Daniel Mair, Stephen Carmichael, Alice Broos, Lily Tong, Jenna Nichols, Kyriaki Nomikou; Sarah McDonald; Richard Orton, Joseph Hughes, Sreenu Vattipally, David L Robertson; Alasdair MacLean, Rory Gunson; Sharif Shaaban, Matthew Holden; Rachel Blacow, Guy Mollett, Kathy Li, James Shepherd, Antonia Ho, Emma Thomson |
| EPI_ISL_919310, EPI_ISL_919349, EPI_ISL_919350, EPI_ISL_919351, EPI_ISL_919352, EPI_ISL_919353, EPI_ISL_919354, EPI_ISL_919355, EPI_ISL_919356, EPI_ISL_919357, EPI_ISL_919358, EPI_ISL_919382, EPI_ISL_919383, EPI_ISL_919384, EPI_ISL_919385, EPI_ISL_919386, EPI_ISL_919387, EPI_ISL_919402, EPI_ISL_919403 | | | |
| see above | Virology Department, Royal Infirmary of Edinburgh, NHS Lothian / School of Biological Sciences, University of Edinburgh / Institute of Genetics and Molecular Medicine, University of Edinburgh | COVID-19 Genomics UK (COG-UK) Consortium | McHugh M, Dewar R, Rooke S, Gallagher M, Balcaza C, O'Toole Á, Scher E, Hill V, McCrone JT, Colquhoun R, Yu X, Jackson B, Rambaut A, Williams TC, Templeton K |
| EPI_ISL_920377, EPI_ISL_920504 | University College London Hospital | COVID-19 Genomics UK (COG-UK) Consortium | Judith Heaney, Matthew Byott, Catherine Houlihan, Dan Frampton, Stuart Kirk, Moira Spyer and Eleni Nastouli |
| EPI_ISL_922892, EPI_ISL_922893, EPI_ISL_922967, EPI_ISL_922968, EPI_ISL_922969 | Wales Specialist Virology Centre Sequencing lab: Pathogen Genomics Unit | Public Health Wales Microbiology Cardiff Wales Specialist Virology Centre | Catherine Moore, Johnathan Evans, Laura Gifford, Malorie Perry, Simon Cottrell, Angela Marchbank, Alec Birchley, Alexander Adams, Amy Gaskin, Bree Gatica-Wilcox, Jason Coombes, Joel Southgate, Lauren Gilbert, Lee Graham, Nicole Pacchiarini, Sara Kumziene-Summerhayes, Sarah Taylor, Sophie Jones, Sara Rey, Matthew Bull, Joanne Watkins, Sally Corden, Tom Connor |
| EPI_ISL_925942, EPI_ISL_926015, EPI_ISL_926153, EPI_ISL_926187, EPI_ISL_926357, EPI_ISL_926381, EPI_ISL_926396, EPI_ISL_926402, EPI_ISL_926459, EPI_ISL_926517, EPI_ISL_926525, EPI_ISL_926555, EPI_ISL_926564, EPI_ISL_926587, EPI_ISL_926588, EPI_ISL_926594, EPI_ISL_926627, EPI_ISL_926698, EPI_ISL_926717, EPI_ISL_926751, EPI_ISL_926767, EPI_ISL_926873, EPI_ISL_926903, EPI_ISL_926916, EPI_ISL_926964, EPI_ISL_926972, EPI_ISL_926976, EPI_ISL_927101, EPI_ISL_927117, EPI_ISL_927149, EPI_ISL_927215, EPI_ISL_927228, EPI_ISL_927249, EPI_ISL_927262, EPI_ISL_927277, EPI_ISL_927321, EPI_ISL_927342, EPI_ISL_927346, EPI_ISL_927399, EPI_ISL_927456, EPI_ISL_927459, EPI_ISL_927460, EPI_ISL_927569, EPI_ISL_927571, EPI_ISL_927595, EPI_ISL_927651, EPI_ISL_927679, EPI_ISL_927703, EPI_ISL_927818, EPI_ISL_927869, EPI_ISL_927879, EPI_ISL_927882, EPI_ISL_927904, EPI_ISL_928005, EPI_ISL_928068, EPI_ISL_928080, EPI_ISL_928192, EPI_ISL_928211, EPI_ISL_928236, EPI_ISL_928258, EPI_ISL_928320, EPI_ISL_928355, EPI_ISL_928363, EPI_ISL_928372, EPI_ISL_928432, EPI_ISL_928467, EPI_ISL_928509, EPI_ISL_928523, EPI_ISL_928580, EPI_ISL_928585, EPI_ISL_928600, EPI_ISL_928664, EPI_ISL_928676, EPI_ISL_928708, EPI_ISL_928730, EPI_ISL_928794, EPI_ISL_928804, EPI_ISL_928817, EPI_ISL_928869, EPI_ISL_928886, EPI_ISL_928899, EPI_ISL_928909, EPI_ISL_928915, EPI_ISL_928951, EPI_ISL_929018, EPI_ISL_929035, EPI_ISL_929040, EPI_ISL_929047, EPI_ISL_929096, EPI_ISL_929158, EPI_ISL_929269, EPI_ISL_929409, EPI_ISL_929428, EPI_ISL_929433, EPI_ISL_929468, EPI_ISL_929497, EPI_ISL_929598, EPI_ISL_929608, EPI_ISL_929639, EPI_ISL_929647, EPI_ISL_929672, EPI_ISL_929683, EPI_ISL_929701, EPI_ISL_929735, EPI_ISL_929750, EPI_ISL_929757, EPI_ISL_929758, EPI_ISL_929772, EPI_ISL_929803, EPI_ISL_929829, EPI_ISL_929886, EPI_ISL_929911, EPI_ISL_929919, EPI_ISL_929954, EPI_ISL_930068, EPI_ISL_930126, EPI_ISL_930134, EPI_ISL_930151, EPI_ISL_930235, EPI_ISL_930321, EPI_ISL_930375, EPI_ISL_930407, EPI_ISL_930410, EPI_ISL_930444 | | | |
| see above | Department of Virus and Microbiological Special Diagnostics, Statens Serum Institut, Copenhagen, Denmark | Aalborg University | Danish Covid-19 Genome Consortium |
| EPI_ISL_931392, EPI_ISL_931449 | University Hospital Basel, Clinical Virology | University Hospital Basel, Clinical Bacteriology | Tim Roloff, Madlen Stange, Helena MB Seth-Smith, Alfredo Mari, Karoline Leuzinger, Julia Bielicki, Manuel Battegay, Hans Hirsch, Adrian Egli |
| EPI_ISL_933493, EPI_ISL_933494 | Lighthouse Lab in Glasgow | Wellcome Sanger Institute for the COVID-19 Genomics UK | Harper VanSteenhouse, Yumi Kasai, David Gray, Carol Clugston, Anna Dominiczak and Alex Alderton, Roberto Amato, Sonia Goncalves, Ewan Harrison, |

| | | | |
|---|---|---|---|
| | | (COG-UK) Consortium | David K. Jackson, Ian Johnston, Dominic Kwiatkowski, Cordelia Langford, John Sillitoe on behalf of the Wellcome Sanger Institute COVID-19 Surveillance Team |
| EPI_ISL_933533 | Laboratory for HIV and opportunistic infections diagnosis The Republican Research and Practical Center for Epidemiology and Microbiology (RRPCEM) | Laboratory for HIV and opportunistic infections diagnosis The Republican Research and Practical Center for Epidemiology and Microbiology (RRPCEM) | Elena Gasich, Kirill Bulda, Artur Akhremchuk, Leonid Valentovich, Vladimir Gorbunov |
| EPI_ISL_933958, EPI_ISL_933984, EPI_ISL_933985, EPI_ISL_933986, EPI_ISL_933987, EPI_ISL_933988, EPI_ISL_933989, EPI_ISL_933990, EPI_ISL_933992, EPI_ISL_933993, EPI_ISL_933994, EPI_ISL_933995, EPI_ISL_933996, EPI_ISL_933997, EPI_ISL_933998, EPI_ISL_933999, EPI_ISL_934000, EPI_ISL_934001, EPI_ISL_934002, EPI_ISL_934003, EPI_ISL_934004, EPI_ISL_934005, EPI_ISL_934006, EPI_ISL_934007, EPI_ISL_934008, EPI_ISL_934009, EPI_ISL_934010, EPI_ISL_934011, EPI_ISL_934012, EPI_ISL_934013, EPI_ISL_934015, EPI_ISL_934016, EPI_ISL_934017, EPI_ISL_934018, EPI_ISL_934019, EPI_ISL_934020, EPI_ISL_934021, EPI_ISL_934022, EPI_ISL_934023, EPI_ISL_934024, EPI_ISL_934025, EPI_ISL_934026, EPI_ISL_934027, EPI_ISL_934028, EPI_ISL_934029, EPI_ISL_934030, EPI_ISL_934031, EPI_ISL_934032, EPI_ISL_934033, EPI_ISL_934034, EPI_ISL_934035, EPI_ISL_934036, EPI_ISL_934037, EPI_ISL_934038, EPI_ISL_934039, EPI_ISL_934040, EPI_ISL_934041, EPI_ISL_934042, EPI_ISL_934043, EPI_ISL_934044, EPI_ISL_934045, EPI_ISL_934046, EPI_ISL_934047, EPI_ISL_934048, EPI_ISL_934049, EPI_ISL_934050, EPI_ISL_934051, EPI_ISL_934052, EPI_ISL_934053, EPI_ISL_934054, EPI_ISL_934055, EPI_ISL_934056, EPI_ISL_934057, EPI_ISL_934058, EPI_ISL_934059, EPI_ISL_934060 | | | |
| see above | Vilnius university hospital Santaros Klinikos, Center of Laboratory Medicine | Vilnius university hospital Santaros Klinikos, Center of Laboratory Medicine | Ingrida Olendraite, Daniel Naumovas, Rimvydas Norvilas, Dovile Ezerskyte, Justinas Slikas, Gytis Dudas |
| EPI_ISL_935241 | KU Leuven, Rega Institute, Clinical and Epidemiological Virology | KU Leuven, Rega Institute, Clinical and Epidemiological Virology | Tony Wawina-Bokalanga, Bert Vanmechelen, Joan Marti-Carerras, Piet Maes |
| EPI_ISL_936639, EPI_ISL_936640, EPI_ISL_936641 | Northwestern Memorial Hospital | Ozer Lab | Ramon Lorenzo-Redondo, Lacy M. Simons, Chad J. Achenbach, Lawrence J. Jennings, Michael G. Ison, Judd F. Hultquist, Egon A. Ozer |
| EPI_ISL_937338, EPI_ISL_937341, EPI_ISL_937342 | Utah Public Health Laboratory | Utah Public Health Laboratory | Erin L. Young, Kelly F. Oakeson, Tara Gallagher |
| EPI_ISL_940122, EPI_ISL_940123, EPI_ISL_940128, EPI_ISL_940134 | Charlotte Maxeke Johannesburg Academic Hospital, National Health Laboratory Services, Gauteng, South Africa | National Institute for Communicable Diseases of the National Health Laboratory Service | Amoako DG, Mohale T, Ntuli N, Mahlangu B, Allam M, Ismail A, Bhiman JN |
| EPI_ISL_940825, EPI_ISL_940826, EPI_ISL_940832 | Virginia DCLS | Virginia DCLS | Virginia DCLS |
| EPI_ISL_940858 | Vaccines and Infectious Diseases Analytics Research Unit (VIDA) | KRISP, KZN Research Innovation and Sequencing Platform | Baillie Vicky, du Plessis Jeanine, Giandhari Jennifer, Pillay Sureshnee, Naidoo Yeshnee, Tegally Houriiyah, de Oliveira Tulio, Madhi Shabir |
| EPI_ISL_941240, EPI_ISL_941241, EPI_ISL_941242, EPI_ISL_941243, EPI_ISL_941244, EPI_ISL_941249, EPI_ISL_941250 | Virginia DCLS | Virginia DCLS | Virginia DCLS |
| EPI_ISL_942024, EPI_ISL_942025, EPI_ISL_942069, EPI_ISL_942170, EPI_ISL_942171, EPI_ISL_942172, EPI_ISL_942173, EPI_ISL_942174, EPI_ISL_942175, EPI_ISL_942176, EPI_ISL_942177, EPI_ISL_942178, EPI_ISL_942179, EPI_ISL_942180, EPI_ISL_942181, EPI_ISL_942182, EPI_ISL_942183, EPI_ISL_942184, EPI_ISL_942193, EPI_ISL_942194, EPI_ISL_942195, EPI_ISL_942196, EPI_ISL_942197, EPI_ISL_942198, EPI_ISL_942199, EPI_ISL_942200, EPI_ISL_942201, EPI_ISL_942202, EPI_ISL_942203, EPI_ISL_942206, EPI_ISL_942207, EPI_ISL_942208, EPI_ISL_942210, EPI_ISL_942212, EPI_ISL_942214, EPI_ISL_942316 | | | |
| see above | Wisconsin State Laboratory of Hygiene Communicable Disease Division | Wisconsin State Laboratory of Hygiene Communicable Disease Division | Kelsey R. Florek, Abigail C. Shockey |
| EPI_ISL_942629, EPI_ISL_942630, EPI_ISL_942631, EPI_ISL_942632, EPI_ISL_942633, EPI_ISL_942634, EPI_ISL_942635, EPI_ISL_942636, EPI_ISL_942637, EPI_ISL_942638, EPI_ISL_942639, EPI_ISL_942640, EPI_ISL_942641, EPI_ISL_942642, EPI_ISL_942643, EPI_ISL_942644, EPI_ISL_942645, EPI_ISL_942646, EPI_ISL_942647, EPI_ISL_942648, EPI_ISL_942649, EPI_ISL_942650, EPI_ISL_942651, EPI_ISL_942652, EPI_ISL_942653, EPI_ISL_942654, EPI_ISL_942655, EPI_ISL_942656, EPI_ISL_942657, EPI_ISL_942660, EPI_ISL_942661 | | | |
| see above | Gundersen Molecular Diagnostics Laboratory | Kabara Cancer Research Institute | Craig S. Richmond, Paraic A. Kenny |
| EPI_ISL_942963 | General Hospital - Bitola | Research Center for Genetic Engineering and Biotechnology "Georgi D. Efremov" , Macedonian Academy of Sciences and Arts | Aleksandar J.Dimovski, Dijana Plasheska-Karanfilska, Predrag Noveski, Gjorgji Bozinovski, Milena Jakimovska |
| EPI_ISL_942964 | General Hospital - Kumanovo | Research Center for Genetic Engineering and Biotechnology "Georgi D. Efremov" , Macedonian Academy of Sciences and Arts | Aleksandar J.Dimovski, Dijana Plasheska-Karanfilska, Predrag Noveski, Gjorgji Bozinovski, Milen |
| EPI_ISL_942965 | General Hospital - Veles | Research Center for Genetic Engineering and Biotechnology "Georgi D. Efremov" , Macedonian Academy of Sciences and Arts | Aleksandar J.Dimovski, Dijana Plasheska-Karanfilska, Predrag Noveski, Gjorgji Bozinovski, Milena Jakimovska |
| EPI_ISL_943803, EPI_ISL_943804 | Utah Public Health Laboratory | Utah Public Health Laboratory | Erin L. Young, Kelly F. Oakeson, Tara Gallagher |
| EPI_ISL_947293 | Eijkman Institute for Molecular Biology, Ministry of Research and Technology/National Agency for Research and Innovation | Eijkman Institute for Molecular Biology, Ministry of Research and Technology/National Agency for Research and Innovation | Edison Johar, Frilasita A Yudhaputri, Hidayat Trimarsanto, Iskandar Adnan, Lydia V. Panggalo, Sukma Oktavianthi, Willy Agustine, Safarina G Malik, Khin Saw Myint, Amin Soebandrio |
| EPI_ISL_949392, EPI_ISL_949393, EPI_ISL_949394, EPI_ISL_949395, EPI_ISL_949399, EPI_ISL_949400, EPI_ISL_949401 | University of Birmingham | COVID-19 Genomics UK (COG-UK) Consortium | Institute of Microbiology, University of Birmingham: Claire McMurray, Joanne Stockton, Samuel Nicholls, Radoslaw Poplawski, Will Rowe, Josh Quick, Nicholas Loman. University of Birmingham Testing Laboratory: Celina M Whalley, Andrew Bosworth, Charlotte Poxon, Kasun Wanigasooriya, Oliver Pickles, Mike Kidd, Alex Richter, Andrew D Beggs PHE Heartlands Lab: Husam Osman, Andrew Bosworth. Queen Elizabeth Hospital: Anna Casey |
| EPI_ISL_949650 | Queens Medical Centre, Clinical Microbiology Department / DeepSeq Nottingham | COVID-19 Genomics UK (COG-UK) Consortium | Gemma Clark, Wendy Smith, Manjinder Khakh, Vicki M Fleming, Michelle M Lister, Hannah Howson-Wells, Jonathan Ball, Patrick McClure, Joseph Chappell, Theocharis Tsoleridis, Nadine Holmes, Matthew Carlisle, Christopher Moore, Fei Sang, Johnny Debebe, Victoria Wright, Matthew Loose |
| EPI_ISL_949787 | University College London, Great Ormond Street Hospital for Children NHS Foundation Trust, Imperial College Healthcare NHS Trust | COVID-19 Genomics UK (COG-UK) Consortium | Sergi Castellano, Rachel Williams, Mark Kristiansen, Paola Resende Silva, Sunando Roy, Tony Brooks, Helena Tutill, Paola Niola, Patricia Dyal, Charlotte Williams, Leysa Forrest, Yasmin Panchbhaya, Jacqueline Findlay, Samuel Weeks, Julianne Brown, Kathryn Harris, Paul Randell, James Price, Alison Holmes, Judith Breuer |
| EPI_ISL_950577, EPI_ISL_950587 | Quadram Institute Bioscience | COVID-19 Genomics UK (COG-UK) Consortium | Dave J. Baker, Gemma L. Kay, Alp Aydin, Thanh Le-Viet, Steven Rudder, Ana P. Tedim, Anastasia Kolyva, Maria Diaz, Leonardo de Oliveira Martins, Nabil-Fareed Alikhan, Lizzie Meadows, Rachael Stanley, Ngozi Elumogo, Muhammed Yasir, Nicholas M. Thomson, Alexander J Trotter, Rachel Gilroy, Samuel Bloomfield, Claire Stuart, Andrew Bell, Reenesh Prakash, Samir Dervisevic, Alison E. Mather, John Wain, Mark Webber, Andrew J. Page, Justin O'Grady |
| EPI_ISL_954197 | 1.AO Universitaria 'S. Giovanni di Dio e Ruggi D'Aragona, Scuola Medica Salernitana' Hospital / 2.UOC di Virologia e Microbiologia, Università della Campania 'L. Vanvitelli' / 3.AO Universitaria 'Federico II' Napoli Hospital / 4.AORN 'San Giuseppe Moscati' Avellino Hospital / 5.AO 'San Pio - presidio G. Rummo' Benevento Hospital / 6.AO 'Sant'Anna e San Sebastiano' Caserta Hospital / 7.PO 'Maria Santissima Addolorata' Eboli Hospital / 8.Biogem Istituto di Ricerche Genetiche | 1. Genome Research Center for Health (CRGS) / 2. Laboratory of Molecular Medicine and Genomics(LMMGe) / 3. Center for Research in Pure and Applied Mathematics (CRMPA) | Giorgio Giurato, Francesca Rizzo, Alessandro Weisz, Gianluigi Franci, Giovanni Nassa, Pasquale Pagliano, Roberta Tarallo, Elena Alexandrova, Ylenia D'Agostino, Carlo Ferravante, Jessica Lamberti, Viola Melone, Domenico Memoli, Valeria Mirici Cappa, Domenico Palumbo, Giovanni Pecoraro, Assunta Sellitto, Oriana Strianese, Ilaria Terenzi, Giuseppe Fenza, Aniello Gentile, Antonello Saccomanno, Sonia Amabile, Teresa Rocco, Annamaria Salvati, Emilia Vaccaro, Massimiliano Galdiero, Michele Cennamo, Giuseppe Portella, Maria Grazia Foti, Mariarosaria Ingino, Maria Landi, Maurizio Fumi, Vincenzo Rocco, Rita Greco, Vittoria Letizia, Arnolfo Petruzziello, Maddalena Schioppa, Gregorio Goffredi, Francesca Marciano, Michele Caraglia, Alessia Cossu, Marianna Scrima, Edmondo Adorisio, Morena D'Avenia, Michela Iacobellis, Rosanna Piluscio, Giorgio Dirani, Vittorio Sambri, Simona Semprini, Silvia Zanoli, Francesco Curcio, Stefania Marzinotto, Andreina Baj, Fausto Sessa. |
| EPI_ISL_955336 | GA Department of Public Health Laboratory | Pathogen Discovery, Respiratory Viruses Branch, Division of Viral Diseases, Centers for Disease Control and Prevention | Ying Tao, Jing Zhang, Yan Li, Krista Queen, Anna Uehara, Peter Cook, Clinton R. Paden, Haibin Wang, Suxiang Tong |
| EPI_ISL_955353, EPI_ISL_955354, EPI_ISL_955355, EPI_ISL_955356, EPI_ISL_955357, EPI_ISL_955358, EPI_ISL_955359, EPI_ISL_955360, EPI_ISL_955361, EPI_ISL_955371, EPI_ISL_955372, EPI_ISL_955375, EPI_ISL_955376, EPI_ISL_955377 | | | |
| see above | Santa Clara County Public Health Laboratory | Chan-Zuckerberg Biohub | CZB Cliahub Consortium |

| | | | |
|---|---|---|---|
| EPI_ISL_955633, EPI_ISL_955634, EPI_ISL_955635, EPI_ISL_955636, EPI_ISL_955637, EPI_ISL_955638, EPI_ISL_955639, EPI_ISL_955640, EPI_ISL_955641, EPI_ISL_955642, EPI_ISL_955643, EPI_ISL_955644, EPI_ISL_955645, EPI_ISL_955646, EPI_ISL_955647, EPI_ISL_955648, EPI_ISL_955649, EPI_ISL_955650, EPI_ISL_955651, EPI_ISL_955652, EPI_ISL_955653, EPI_ISL_955654, EPI_ISL_955655, EPI_ISL_955656, EPI_ISL_955657, EPI_ISL_955658, EPI_ISL_955659, EPI_ISL_955660, EPI_ISL_955661, EPI_ISL_955662, EPI_ISL_955663, EPI_ISL_955664, EPI_ISL_955666, EPI_ISL_955670 | | | |
| see above | Orange County Public Health Lab | Chan-Zuckerberg Biohub | CZB Cliahub Consortium |
| EPI_ISL_960394 | University of Wisconsin-Madison AIDS Vaccine Research Laboratories | University of Wisconsin-Madison AIDS Vaccine Research Laboratories | Gage Moreno, Katarina Braun, et al. AIDS Vaccine Research Laboratories |
| EPI_ISL_960417 | The National Institute of Public Health | State Veterinary Institute Prague | Nagy,A;Vecerova,J;Cernikova,L;Stara,M;Jirincova,H;Trnka,D |
| EPI_ISL_960446, EPI_ISL_960453, EPI_ISL_960474, EPI_ISL_960503, EPI_ISL_960504, EPI_ISL_960505, EPI_ISL_960506, EPI_ISL_960507, EPI_ISL_960508, EPI_ISL_960509, EPI_ISL_960510, EPI_ISL_960511, EPI_ISL_960512, EPI_ISL_960513, EPI_ISL_960514, EPI_ISL_960515, EPI_ISL_960570, EPI_ISL_960590, EPI_ISL_960593, EPI_ISL_960594, EPI_ISL_960595, EPI_ISL_960596, EPI_ISL_960597, EPI_ISL_960598, EPI_ISL_960599, EPI_ISL_960600, EPI_ISL_960601, EPI_ISL_960602, EPI_ISL_960603, EPI_ISL_960604, EPI_ISL_960605, EPI_ISL_960606 | | | |
| see above | Istituto Zooprofilattico Sperimentale del Mezzogiorno | TIGEM | Patrizia Annunziata, Andrea Ballabio, Valentina Bouche, Davide Cacchiarelli, Pellegrino Cerino, Chiara Colantuono, Maria Concetta Cuomo, Denise Di Concilio, Lucio Di Filippo, Antonio Grimaldi, Antonio Limone, Anna Manfredi, Francesco Panariello, Biancamaria Pierri, Marcello Salvi |
| EPI_ISL_961666 | Hôpital Georges L. Dumont | National Microbiology Laboratory (NML) | Anna Majer, Shari Tyson, Grace Seo, Philip Mabon, Elsie Grudeski, Rhiannon Huzarewich, Russell Mandes, Anneliese Landgraff, Jennifer Tanner, Natalie Knox, Morag Graham, Gary Van Domselaar, Richard Garceau, Guillaume Desnoyers, Nathalie Bastien, Yan Li, Timothy Booth, Darian Hole, Madison Chapel, Kirsten Biggar, CanCOGeN's metadata curation team, Public Health Agency of Canada CanCOGeN team |
| EPI_ISL_961893, EPI_ISL_961928, EPI_ISL_961929, EPI_ISL_961973, EPI_ISL_961984, EPI_ISL_962015, EPI_ISL_962067, EPI_ISL_962068, EPI_ISL_962069, EPI_ISL_962070, EPI_ISL_962071, EPI_ISL_962072, EPI_ISL_962073, EPI_ISL_962074, EPI_ISL_962075, EPI_ISL_962076, EPI_ISL_962077, EPI_ISL_962078, EPI_ISL_962079, EPI_ISL_962082, EPI_ISL_962084, EPI_ISL_962091, EPI_ISL_962158, EPI_ISL_962159 | | | |
| see above | Illinois Department of Public Health | Gagnon Lab, Southern Illinois University | Keith Gagnon |
| EPI_ISL_962506 | UCLA Clinical Micro Lab | Los Angeles County PHL | P. Hemarajata et al. |
| EPI_ISL_962610, EPI_ISL_962768, EPI_ISL_962770, EPI_ISL_962776, EPI_ISL_962777, EPI_ISL_962783, EPI_ISL_962784, EPI_ISL_962787, EPI_ISL_962790, EPI_ISL_962795, EPI_ISL_962799, EPI_ISL_962806 | | | |
| see above | San Diego County Public Health Laboratory | Andersen lab at Scripps Research | SEARCH Alliance San Diego with Tracy Basler, Jovan Shephard, Brett Austin |
| EPI_ISL_962970 | Hospital Universitario de Gran Canaria Dr. Negrín | SeqCOVID-SPAIN consortium/IBV(CSIC) | M. Carmen Pérez González, Francisco J. Chamizo López, Ana Bordes Benítez and SeqCOVID-SPAIN consortium |
| EPI_ISL_965544, EPI_ISL_965574, EPI_ISL_965586, EPI_ISL_965610, EPI_ISL_965616, EPI_ISL_965626, EPI_ISL_965636, EPI_ISL_965647, EPI_ISL_965681, EPI_ISL_965696, EPI_ISL_965725, EPI_ISL_965730, EPI_ISL_965782, EPI_ISL_965807, EPI_ISL_965808 | | | |
| see above | Dutch COVID-19 response team | Medical Microbiology, Maastricht University Medical Centre | Jozef Dingemans*, Brian van der Veer*, Erik Beuken, Carmen Reumkens, Lieke van Alphen, Christian Hoebe, Paul Savelkoul |
| EPI_ISL_967551, EPI_ISL_967552, EPI_ISL_967553, EPI_ISL_967555, EPI_ISL_967556 | State Laboratories Division, Hawaii State Department of Health | State Laboratories Division, Hawaii State Department of Health | Pamela O'Brien, Drew Kuwazaki, Ayana Garnet, Razvan Sultana, Edward Desmond |
| EPI_ISL_968807, EPI_ISL_968813, EPI_ISL_968817, EPI_ISL_968819, EPI_ISL_968820, EPI_ISL_968821, EPI_ISL_968831 | KEMRI-Wellcome Trust Research Programme/KEMRI-CGMR-C Kilifi | KEMRI-Wellcome Trust Research Programme/KEMRI-CGMR-C Kilifi | Githinji et al |
| EPI_ISL_970879, EPI_ISL_971006, EPI_ISL_971143, EPI_ISL_971345, EPI_ISL_971811, EPI_ISL_972238, EPI_ISL_972410, EPI_ISL_972624, EPI_ISL_973276, EPI_ISL_973514 | Department of Virus and Microbiological Special Diagnostics, Statens Serum Institut, Copenhagen, Denmark | Aalborg University | Danish Covid-19 Genome Consortium |
| EPI_ISL_976176, EPI_ISL_976177, EPI_ISL_976178, EPI_ISL_976179, EPI_ISL_976180, EPI_ISL_976181, EPI_ISL_976182, EPI_ISL_976183, EPI_ISL_976184, EPI_ISL_976185, EPI_ISL_976186, EPI_ISL_976187, EPI_ISL_976188, EPI_ISL_976189, EPI_ISL_976190, EPI_ISL_976191, EPI_ISL_976192, EPI_ISL_976193, EPI_ISL_976194, EPI_ISL_976195, EPI_ISL_976196, EPI_ISL_976197, EPI_ISL_976198, EPI_ISL_976199, EPI_ISL_976200, EPI_ISL_976201, EPI_ISL_976202, EPI_ISL_976203, EPI_ISL_976204, EPI_ISL_976205, EPI_ISL_976206, EPI_ISL_976207, EPI_ISL_976208, EPI_ISL_976209, EPI_ISL_976210, EPI_ISL_976211, EPI_ISL_976212, EPI_ISL_976213, EPI_ISL_976214, EPI_ISL_976215, EPI_ISL_976216, EPI_ISL_976217, EPI_ISL_976218, EPI_ISL_976219, EPI_ISL_976220, EPI_ISL_976221, EPI_ISL_976222, EPI_ISL_976223, EPI_ISL_976224, EPI_ISL_976225, EPI_ISL_976226, EPI_ISL_976227, EPI_ISL_976228, EPI_ISL_976229, EPI_ISL_976230, EPI_ISL_976231, EPI_ISL_976232, EPI_ISL_976233, EPI_ISL_976234, EPI_ISL_976235, EPI_ISL_976236, EPI_ISL_976237, EPI_ISL_976238, EPI_ISL_976239, EPI_ISL_976240, EPI_ISL_976241, EPI_ISL_976242, EPI_ISL_976243, EPI_ISL_976244, EPI_ISL_976245, EPI_ISL_976246, EPI_ISL_976247, EPI_ISL_976248, EPI_ISL_976249, EPI_ISL_976250, EPI_ISL_976251, EPI_ISL_976252, EPI_ISL_976253, EPI_ISL_976254, EPI_ISL_976255, EPI_ISL_976256, EPI_ISL_976257, EPI_ISL_976258, EPI_ISL_976259, EPI_ISL_976260, EPI_ISL_976261, EPI_ISL_976262, EPI_ISL_976263, EPI_ISL_976264, EPI_ISL_976265, EPI_ISL_976266, EPI_ISL_976267, EPI_ISL_976268, EPI_ISL_976269, EPI_ISL_976270, EPI_ISL_976271, EPI_ISL_976272, EPI_ISL_976273, EPI_ISL_976274, EPI_ISL_976275, EPI_ISL_976276, EPI_ISL_976277, EPI_ISL_976278, EPI_ISL_976279, EPI_ISL_976280, EPI_ISL_976281, EPI_ISL_976282, EPI_ISL_976283, EPI_ISL_976284, EPI_ISL_976285, EPI_ISL_976286, EPI_ISL_976287, EPI_ISL_976288, EPI_ISL_976289, EPI_ISL_976290, EPI_ISL_976291, EPI_ISL_976292, EPI_ISL_976293, EPI_ISL_976294, EPI_ISL_976295, EPI_ISL_976296, EPI_ISL_976297, EPI_ISL_976298, EPI_ISL_976299, EPI_ISL_976300 | | | |
| see above | BCCDC Public Health Laboratory | BCCDC Public Health Laboratory | Prystajecky Natalie, Linda Hoang, Dan Fornika, John Tyson, Shannon Russell, Kim Macdonald, Kimia Kamelian, Ana Pacagnella, Corrinne Ng, Loretta Janz, Robert Azana Terry Snutch, Mel Krajden |
| EPI_ISL_977024, EPI_ISL_977025, EPI_ISL_977026, EPI_ISL_977027 | Rhode Island Department of Health | Infectious Disease Program, Broad Institute of Harvard and MIT | Lemieux,J.E., Siddle,K.J., Huard,R., King,E., Azevedo,K., Miller,A., Adams,G., Gladden-Young,A., Lagerborg,K., Rudy,M., DeRuff,K., Carter,A., Normandin,E., Bauer,M., Reilly,S., Tomkins-Tinch,C., Loreth,C., Chaluvadi,S., Birren,B.W., Gallagher,G., Smole,S., Park,D.J., MacInnis,B.L., and Sabeti,P.C. |
| EPI_ISL_977200, EPI_ISL_977201, EPI_ISL_977202 | ULSS 7 Pedemontana - Distretto 2 | Istituto Zooprofilattico Sperimentale delle Venezie | Adelaide Milani, Alessia Schivo, Annalisa Salviato, Erika Giorgia Quaranta, Ambra Pastori, Bianca Zecchin, Alice Fusaro, Isabella Monne, Calogero Terregino, Antonia Ricci |
| EPI_ISL_977213, EPI_ISL_977214, EPI_ISL_977215, EPI_ISL_977216, EPI_ISL_977217, EPI_ISL_977218, EPI_ISL_977219, EPI_ISL_977220, EPI_ISL_977221 | ULSS 1 Dolomiti | Istituto Zooprofilattico Sperimentale delle Venezie | Adelaide Milani, Alessia Schivo, Annalisa Salviato, Erika Giorgia Quaranta, Ambra Pastori, Bianca Zecchin, Alice Fusaro, Isabella Monne, Calogero Terregino, Antonia Ricci |
| EPI_ISL_978194, EPI_ISL_978195, EPI_ISL_978196, EPI_ISL_978197, EPI_ISL_978214 | Virginia Division of Consolidated Laboratory Services | Virginia Division of Consolidated Laboratory Services | Virginia DCLS |
| EPI_ISL_978989, EPI_ISL_978990, EPI_ISL_978991, EPI_ISL_978992, EPI_ISL_978993, EPI_ISL_978994, EPI_ISL_978995, EPI_ISL_978996, EPI_ISL_978997, EPI_ISL_978998, EPI_ISL_978999, EPI_ISL_979000, EPI_ISL_979001, EPI_ISL_979002, EPI_ISL_979003, EPI_ISL_979004, EPI_ISL_979005, EPI_ISL_979006, EPI_ISL_979007, EPI_ISL_979008, EPI_ISL_979009, EPI_ISL_979010, EPI_ISL_979011, EPI_ISL_979012, EPI_ISL_979013, EPI_ISL_979014, EPI_ISL_979015, EPI_ISL_979016, EPI_ISL_979019, EPI_ISL_979020, EPI_ISL_979021, EPI_ISL_979022, EPI_ISL_979023, EPI_ISL_979024, EPI_ISL_979025, EPI_ISL_979026, EPI_ISL_979027, EPI_ISL_979028, EPI_ISL_979029, EPI_ISL_979030, EPI_ISL_979031, EPI_ISL_979032, EPI_ISL_979033, EPI_ISL_979034, EPI_ISL_979035, EPI_ISL_979036, EPI_ISL_979037, EPI_ISL_979038, EPI_ISL_979039, EPI_ISL_979040, EPI_ISL_979041, EPI_ISL_979042, EPI_ISL_979043, EPI_ISL_979044, EPI_ISL_979045, EPI_ISL_979046, EPI_ISL_979047, EPI_ISL_979048, EPI_ISL_979049, EPI_ISL_979050, EPI_ISL_979051, EPI_ISL_979052, EPI_ISL_979053, EPI_ISL_979054, EPI_ISL_979055 | | | |
| see above | Santa Clara County Public Health Laboratory | Chan-Zuckerberg Biohub | CZB Cliahub Consortium |
| EPI_ISL_979310 | Cadham Provincial laboratory | National Microbiology Laboratory (NML) | Anna Majer, Shari Tyson, Grace Seo, Philip Mabon, Elsie Grudeski, Rhiannon Huzarewich, Russell Mandes, Anneliese Landgraff, Jennifer Tanner, Natalie Knox, Morag Graham, Gary Van Domselaar, Paul Van Caeseele, Jared Bullard, David Alexander, Kerry Dust, Nathalie Bastien, Yan Li, Timothy Booth, Darian Hole, Madison Chapel, Kirsten Biggar, CanCOGeN's metadata curation team, Public Health Agency of Canada CanCOGeN team |
| EPI_ISL_979545, EPI_ISL_979546, EPI_ISL_979547, EPI_ISL_979548, EPI_ISL_979549, EPI_ISL_979550, EPI_ISL_979551, EPI_ISL_979552, EPI_ISL_979553, EPI_ISL_979554, EPI_ISL_979555, EPI_ISL_979557, EPI_ISL_979558, EPI_ISL_979559, EPI_ISL_979560, EPI_ISL_979561, EPI_ISL_979562, EPI_ISL_979563, EPI_ISL_979564, EPI_ISL_979565, EPI_ISL_979566, EPI_ISL_979567, EPI_ISL_979568, EPI_ISL_979569, EPI_ISL_979570, EPI_ISL_979571, EPI_ISL_979572, EPI_ISL_979573, EPI_ISL_979574, EPI_ISL_979575, EPI_ISL_979576, EPI_ISL_979577, EPI_ISL_979578, EPI_ISL_979579, EPI_ISL_979580, EPI_ISL_979581, EPI_ISL_979582, EPI_ISL_979583, EPI_ISL_979584, EPI_ISL_979585, EPI_ISL_979586, EPI_ISL_979587, EPI_ISL_979588, EPI_ISL_979589, EPI_ISL_979590, EPI_ISL_979591, EPI_ISL_979592, EPI_ISL_979593 | | | |
| see above | Santa Clara County Public Health Laboratory | Chan-Zuckerberg Biohub | CZB Cliahub Consortium |
| EPI_ISL_979756, EPI_ISL_979757, EPI_ISL_979758, EPI_ISL_979759, EPI_ISL_979760, EPI_ISL_979761, EPI_ISL_979762, EPI_ISL_979763, EPI_ISL_979764, EPI_ISL_979765, EPI_ISL_979766, EPI_ISL_979767, EPI_ISL_979768, EPI_ISL_979769, EPI_ISL_979770, EPI_ISL_979771, EPI_ISL_979772, EPI_ISL_979773, EPI_ISL_979774, EPI_ISL_979775, EPI_ISL_979776, EPI_ISL_979777, EPI_ISL_979778, EPI_ISL_979779, EPI_ISL_979780, EPI_ISL_979781, EPI_ISL_979782, EPI_ISL_979783, EPI_ISL_979784, EPI_ISL_979785, EPI_ISL_979786, EPI_ISL_979787, EPI_ISL_979788, EPI_ISL_979789, EPI_ISL_979790, EPI_ISL_979791, EPI_ISL_979792, EPI_ISL_979793, EPI_ISL_979794, EPI_ISL_979795, EPI_ISL_979796 | | | |
| see above | Humboldt County Public Health Laboratory | Chan-Zuckerberg Biohub | CZB Cliahub Consortium |

| | | | |
|---|---|---|---|
| EPI_ISL_981006 | AZ Klina | AZ Klina | Dr. C. Vael |
| EPI_ISL_981882, EPI_ISL_981888, EPI_ISL_981889, EPI_ISL_981890, EPI_ISL_981891, EPI_ISL_981892, EPI_ISL_981893, EPI_ISL_981894, EPI_ISL_981895 | Microbiology Service, Hospital Universitario Clinico San Cecilio, Granada | Microbiology Service, Hospital Universitario Clinico San Cecilio, Granada | Adolfo de Salazar, Natalia Chueca, Laura Viñuela, Ana Fuentes, Federico García |
| EPI_ISL_982585 | Landstuhl Regional Medical Center | US Air Force School of Aerospace Medicine | Anthony Fries, Jennifer Meyer, William Gruner, William Buggele, Amanda Javorina, Sarah Purves, Fritz Castillo, Cole Anderson, Clarise Starr, Elizabeth Macias |
| EPI_ISL_982843 | Kentucky State Public Health Lab | Kentucky State Public Health Lab | Stephanie Lunn, Karim George, Joshua Tobias, William Grooms, Vaneet Arora, Matthew Johnson, Rachel Zinner, Rhonda Lucas |
