## Supplementary material for "Detection and characterization of the SARS-CoV-2 lineage B.1.526 in New York": Supp. Table 3 GISAID Acknowledgment part 2: gisaid_hcov-19_acknowledgement_table_2021_02_13_00-15.pdf

We gratefully acknowledge the following Authors from the Originating laboratories responsible for obtaining the specimens, as well as the Submitting laboratories where the genome data were generated and shared via GISAID, on which this research is based.

All Submitters of data may be contacted directly via www.gisaid.org

Authors are sorted alphabetically.

| Accession ID | Originating Laboratory | Submitting Laboratory | Authors |
|---|---|---|---|
| EPI_ISL_717704, EPI_ISL_717705, EPI_ISL_717706, EPI_ISL_717707 | Area of Virology, Serology and Virology Division (SAViD), New South Wales Health Pathology Randwick | Virology Research Laboratory; Area of Virology, Serology and Virology Division (SAViD), New South Wales Health Pathology Randwick | Foster, C.; Au, J.; Ruiz Silva, M.; Deveson, I.; Bull, R.; Van Hal, S.; Rawlinson, W. |
| EPI_ISL_718318, EPI_ISL_718319, EPI_ISL_718320, EPI_ISL_718321, EPI_ISL_718322, EPI_ISL_718323, EPI_ISL_718324, EPI_ISL_718325, EPI_ISL_718326, EPI_ISL_718327, EPI_ISL_718328, EPI_ISL_718329, EPI_ISL_718330, EPI_ISL_718331, EPI_ISL_718332, EPI_ISL_718333, EPI_ISL_718334, EPI_ISL_718335, EPI_ISL_718336, EPI_ISL_718337, EPI_ISL_718338, EPI_ISL_718339, EPI_ISL_718340, EPI_ISL_718341, EPI_ISL_718342, EPI_ISL_718343, EPI_ISL_718344, EPI_ISL_718345, EPI_ISL_718346, EPI_ISL_718347, EPI_ISL_718348, EPI_ISL_718349, EPI_ISL_718350, EPI_ISL_718351, EPI_ISL_718352, EPI_ISL_718353, EPI_ISL_718354, EPI_ISL_718355, EPI_ISL_718356, EPI_ISL_718357, EPI_ISL_718358, EPI_ISL_718359, EPI_ISL_718360, EPI_ISL_718361, EPI_ISL_718362, EPI_ISL_718363, EPI_ISL_718364, EPI_ISL_718365, EPI_ISL_718366, EPI_ISL_718367, EPI_ISL_718368, EPI_ISL_718369, EPI_ISL_718370, EPI_ISL_718371, EPI_ISL_718372, EPI_ISL_718373, EPI_ISL_718374, EPI_ISL_718375, EPI_ISL_718376, EPI_ISL_718377, EPI_ISL_718378, EPI_ISL_718379, EPI_ISL_718380, EPI_ISL_718381, EPI_ISL_718382, EPI_ISL_718383, EPI_ISL_718384, EPI_ISL_718385, EPI_ISL_718386, EPI_ISL_718387, EPI_ISL_718388, EPI_ISL_718389, EPI_ISL_718390, EPI_ISL_718391, EPI_ISL_718392, EPI_ISL_718393, EPI_ISL_718394, EPI_ISL_718395, EPI_ISL_718396, EPI_ISL_718397, EPI_ISL_718398, EPI_ISL_718399, EPI_ISL_718400, EPI_ISL_718401, EPI_ISL_718402, EPI_ISL_718403, EPI_ISL_718404, EPI_ISL_718405, EPI_ISL_718406, EPI_ISL_718407, EPI_ISL_718408, EPI_ISL_718409, EPI_ISL_718410, EPI_ISL_718411, EPI_ISL_718412, EPI_ISL_718413, EPI_ISL_718414, EPI_ISL_718415, EPI_ISL_718416, EPI_ISL_718417, EPI_ISL_718418, EPI_ISL_718419, EPI_ISL_718420, EPI_ISL_718421, EPI_ISL_718422, EPI_ISL_718423, EPI_ISL_718424, EPI_ISL_718425, EPI_ISL_718426, EPI_ISL_718427, EPI_ISL_718428, EPI_ISL_718429, EPI_ISL_718430, EPI_ISL_718431, EPI_ISL_718432, EPI_ISL_718433, EPI_ISL_718434, EPI_ISL_718435, EPI_ISL_718436, EPI_ISL_718437, EPI_ISL_718438, EPI_ISL_718439, EPI_ISL_718440, EPI_ISL_718441, EPI_ISL_718442, EPI_ISL_718443, EPI_ISL_718444, EPI_ISL_718445, EPI_ISL_718446, EPI_ISL_718447, EPI_ISL_718448, EPI_ISL_718449, EPI_ISL_718450, EPI_ISL_718451, EPI_ISL_718452, EPI_ISL_718453, EPI_ISL_718454, EPI_ISL_718455, EPI_ISL_718456, EPI_ISL_718457, EPI_ISL_718458, EPI_ISL_718459, EPI_ISL_718460, EPI_ISL_718461, EPI_ISL_718462, EPI_ISL_718463, EPI_ISL_718464, EPI_ISL_718465, EPI_ISL_718466, EPI_ISL_718467, EPI_ISL_718468, EPI_ISL_718469, EPI_ISL_718470, EPI_ISL_718471, EPI_ISL_718472, EPI_ISL_718473, EPI_ISL_718474, EPI_ISL_718475, EPI_ISL_718476, EPI_ISL_718477, EPI_ISL_718478, EPI_ISL_718479, EPI_ISL_718480, EPI_ISL_718481, EPI_ISL_718482, EPI_ISL_718483, EPI_ISL_718484, EPI_ISL_718485, EPI_ISL_718486, EPI_ISL_718487, EPI_ISL_718488, EPI_ISL_718489, EPI_ISL_718490, EPI_ISL_718491, EPI_ISL_718492, EPI_ISL_718493, EPI_ISL_718494, EPI_ISL_718495, EPI_ISL_718496, EPI_ISL_718497, EPI_ISL_718498, EPI_ISL_718499, EPI_ISL_718500, EPI_ISL_718501, EPI_ISL_718502, EPI_ISL_718503, EPI_ISL_718504, EPI_ISL_718505, EPI_ISL_718506, EPI_ISL_718507, EPI_ISL_718508, EPI_ISL_718509, EPI_ISL_718510, EPI_ISL_718511, EPI_ISL_718512, EPI_ISL_718513, EPI_ISL_718514, EPI_ISL_718515, EPI_ISL_718516, EPI_ISL_718517, EPI_ISL_718518, EPI_ISL_718519, EPI_ISL_718520, EPI_ISL_718521, EPI_ISL_718522, EPI_ISL_718523, EPI_ISL_718524, EPI_ISL_718525, EPI_ISL_718526, EPI_ISL_718527, EPI_ISL_718528, EPI_ISL_718529, EPI_ISL_718530, EPI_ISL_718531, EPI_ISL_718532, EPI_ISL_718533, EPI_ISL_718534, EPI_ISL_718535, EPI_ISL_718536, EPI_ISL_718537, EPI_ISL_718538, EPI_ISL_718539, EPI_ISL_718540, EPI_ISL_718541, EPI_ISL_718542, EPI_ISL_718543, EPI_ISL_718544, EPI_ISL_718545, EPI_ISL_718546, EPI_ISL_718547, EPI_ISL_718548, EPI_ISL_718549, EPI_ISL_718550, EPI_ISL_718551, EPI_ISL_718552, EPI_ISL_718553, EPI_ISL_718554, EPI_ISL_718555, EPI_ISL_718556, EPI_ISL_718557, EPI_ISL_718558, EPI_ISL_718559, EPI_ISL_718560, EPI_ISL_718561, EPI_ISL_718562, EPI_ISL_718563, EPI_ISL_718564, EPI_ISL_718565, EPI_ISL_718566, EPI_ISL_718567, EPI_ISL_718568, EPI_ISL_718569, EPI_ISL_718570, EPI_ISL_718571, EPI_ISL_718572, EPI_ISL_718573, EPI_ISL_718574, EPI_ISL_718575, EPI_ISL_718576, EPI_ISL_718577, EPI_ISL_718578, EPI_ISL_718579, EPI_ISL_718580, EPI_ISL_718581, EPI_ISL_718582, EPI_ISL_718583, EPI_ISL_718584, EPI_ISL_718585, EPI_ISL_718586, EPI_ISL_718587, EPI_ISL_718588, EPI_ISL_718589, EPI_ISL_718590, EPI_ISL_718591, EPI_ISL_718592, EPI_ISL_718593, EPI_ISL_718594, EPI_ISL_718595, EPI_ISL_718596, EPI_ISL_718597, EPI_ISL_718598, EPI_ISL_718599, EPI_ISL_718600, EPI_ISL_718601, EPI_ISL_718602, EPI_ISL_718603, EPI_ISL_718604, EPI_ISL_718605, EPI_ISL_718606, EPI_ISL_718607, EPI_ISL_718608, EPI_ISL_718609, EPI_ISL_718610, EPI_ISL_718611, EPI_ISL_718612, EPI_ISL_718613, EPI_ISL_718614, EPI_ISL_718615, EPI_ISL_718616, EPI_ISL_718617, EPI_ISL_718618, EPI_ISL_718619, EPI_ISL_718620, EPI_ISL_718621, EPI_ISL_718622, EPI_ISL_718623, EPI_ISL_718624, EPI_ISL_718625, EPI_ISL_718626, EPI_ISL_718627, EPI_ISL_718628 | | | |
| see above | Lighthouse Lab in Cambridge | Wellcome Sanger Institute for the COVID-19 Genomics UK (COG-UK) Consortium | Rob Howes, The Lighthouse Lab in Cambridge and Alex Alderton, Roberto Amato, Sonia Goncalves, Ewan Harrison, David K. Jackson, Ian Johnston, Dominic Kwiatkowski, Cordelia Langford, John Sillitoe on behalf of the Wellcome Sanger Institute COVID-19 Surveillance Team |
| EPI_ISL_718629 | Lighthouse Lab in Alderley Park | Wellcome Sanger Institute for the COVID-19 Genomics UK (COG-UK) Consortium | Jacquelyn Wynn, Mairead Hyland, The Lighthouse Lab in Alderley Park and Alex Alderton, Roberto Amato, Sonia Goncalves, Ewan Harrison, David K. Jackson, Ian Johnston, Dominic Kwiatkowski, Cordelia Langford, John Sillitoe on behalf of the Wellcome Sanger Institute COVID-19 Surveillance Team |
| EPI_ISL_718630 | Lighthouse Lab in Cambridge | Wellcome Sanger Institute for the COVID-19 Genomics UK (COG-UK) Consortium | Rob Howes, The Lighthouse Lab in Cambridge and Alex Alderton, Roberto Amato, Sonia Goncalves, Ewan Harrison, David K. Jackson, Ian Johnston, Dominic Kwiatkowski, Cordelia Langford, John Sillitoe on behalf of the Wellcome Sanger Institute COVID-19 Surveillance Team |
| EPI_ISL_718631 | Lighthouse Lab in Alderley Park | Wellcome Sanger Institute for the COVID-19 Genomics UK (COG-UK) Consortium | Jacquelyn Wynn, Mairead Hyland, The Lighthouse Lab in Alderley Park and Alex Alderton, Roberto Amato, Sonia Goncalves, Ewan Harrison, David K. Jackson, Ian Johnston, Dominic Kwiatkowski, Cordelia Langford, John Sillitoe on behalf of the Wellcome Sanger Institute COVID-19 Surveillance Team |
| EPI_ISL_718632 | Lighthouse Lab in Cambridge | Wellcome Sanger Institute for the COVID-19 Genomics UK (COG-UK) Consortium | Rob Howes, The Lighthouse Lab in Cambridge and Alex Alderton, Roberto Amato, Sonia Goncalves, Ewan Harrison, David K. Jackson, Ian Johnston, Dominic Kwiatkowski, Cordelia Langford, John Sillitoe on behalf of the Wellcome Sanger Institute COVID-19 Surveillance Team |
| EPI_ISL_718633 | Lighthouse Lab in Alderley Park | Wellcome Sanger Institute for the COVID-19 Genomics UK (COG-UK) Consortium | Jacquelyn Wynn, Mairead Hyland, The Lighthouse Lab in Alderley Park and Alex Alderton, Roberto Amato, Sonia Goncalves, Ewan Harrison, David K. Jackson, Ian Johnston, Dominic Kwiatkowski, Cordelia Langford, John Sillitoe on behalf of the Wellcome Sanger Institute COVID-19 Surveillance Team |
| EPI_ISL_718634 | Lighthouse Lab in Cambridge | Wellcome Sanger Institute for the COVID-19 Genomics UK (COG-UK) Consortium | Rob Howes, The Lighthouse Lab in Cambridge and Alex Alderton, Roberto Amato, Sonia Goncalves, Ewan Harrison, David K. Jackson, Ian Johnston, Dominic Kwiatkowski, Cordelia Langford, John Sillitoe on behalf of the Wellcome Sanger Institute COVID-19 Surveillance Team |
| EPI_ISL_718635 | Lighthouse Lab in Alderley Park | Wellcome Sanger Institute for the COVID-19 Genomics UK (COG-UK) Consortium | Jacquelyn Wynn, Mairead Hyland, The Lighthouse Lab in Alderley Park and Alex Alderton, Roberto Amato, Sonia Goncalves, Ewan Harrison, David K. Jackson, Ian Johnston, Dominic Kwiatkowski, Cordelia Langford, John Sillitoe on behalf of the Wellcome Sanger Institute COVID-19 Surveillance Team |
| EPI_ISL_718636 | Lighthouse Lab in Cambridge | Wellcome Sanger Institute for the COVID-19 Genomics UK (COG-UK) Consortium | Rob Howes, The Lighthouse Lab in Cambridge and Alex Alderton, Roberto Amato, Sonia Goncalves, Ewan Harrison, David K. Jackson, Ian Johnston, Dominic Kwiatkowski, Cordelia Langford, John Sillitoe on behalf of the Wellcome Sanger Institute COVID-19 Surveillance Team |
| EPI_ISL_718637 | Lighthouse Lab in Alderley Park | Wellcome Sanger Institute for the COVID-19 Genomics UK (COG-UK) Consortium | Jacquelyn Wynn, Mairead Hyland, The Lighthouse Lab in Alderley Park and Alex Alderton, Roberto Amato, Sonia Goncalves, Ewan Harrison, David K. Jackson, Ian Johnston, Dominic Kwiatkowski, Cordelia Langford, John Sillitoe on behalf of the Wellcome Sanger Institute COVID-19 Surveillance Team |
| EPI_ISL_718638 | Lighthouse Lab in Cambridge | Wellcome Sanger Institute for the COVID-19 Genomics UK (COG-UK) Consortium | Rob Howes, The Lighthouse Lab in Cambridge and Alex Alderton, Roberto Amato, Sonia Goncalves, Ewan Harrison, David K. Jackson, Ian Johnston, Dominic Kwiatkowski, Cordelia Langford, John Sillitoe on behalf of the Wellcome Sanger Institute COVID-19 Surveillance Team |
| EPI_ISL_718639, EPI_ISL_718640, EPI_ISL_718641 | Lighthouse Lab in Alderley Park | Wellcome Sanger Institute for the COVID-19 Genomics UK (COG-UK) Consortium | Jacquelyn Wynn, Mairead Hyland, The Lighthouse Lab in Alderley Park and Alex Alderton, Roberto Amato, Sonia Goncalves, Ewan Harrison, David K. Jackson, Ian Johnston, Dominic Kwiatkowski, Cordelia Langford, John Sillitoe on behalf of the Wellcome Sanger Institute COVID-19 Surveillance Team |
| EPI_ISL_718642 | Lighthouse Lab in Cambridge | Wellcome Sanger Institute for the COVID-19 Genomics UK (COG-UK) Consortium | Rob Howes, The Lighthouse Lab in Cambridge and Alex Alderton, Roberto Amato, Sonia Goncalves, Ewan Harrison, David K. Jackson, Ian Johnston, Dominic Kwiatkowski, Cordelia Langford, John Sillitoe on behalf of the Wellcome Sanger Institute COVID-19 Surveillance Team |
| EPI_ISL_718643, EPI_ISL_718644 | Lighthouse Lab in Alderley Park | Wellcome Sanger Institute for the COVID-19 Genomics UK (COG-UK) Consortium | Jacquelyn Wynn, Mairead Hyland, The Lighthouse Lab in Alderley Park and Alex Alderton, Roberto Amato, Sonia Goncalves, Ewan Harrison, David K. Jackson, Ian Johnston, Dominic Kwiatkowski, Cordelia Langford, John Sillitoe on behalf of the Wellcome Sanger Institute COVID-19 Surveillance Team |
| EPI_ISL_718645, EPI_ISL_718646 | Lighthouse Lab in Cambridge | Wellcome Sanger Institute for the COVID-19 Genomics UK (COG-UK) Consortium | Rob Howes, The Lighthouse Lab in Cambridge and Alex Alderton, Roberto Amato, Sonia Goncalves, Ewan Harrison, David K. Jackson, Ian Johnston, Dominic Kwiatkowski, Cordelia Langford, John Sillitoe on behalf of the Wellcome Sanger Institute COVID-19 Surveillance Team |
| EPI_ISL_718647 | Lighthouse Lab in Alderley Park | Wellcome Sanger Institute for the COVID-19 Genomics UK (COG-UK) Consortium | Jacquelyn Wynn, Mairead Hyland, The Lighthouse Lab in Alderley Park and Alex Alderton, Roberto Amato, Sonia Goncalves, Ewan Harrison, David K. Jackson, Ian Johnston, Dominic Kwiatkowski, Cordelia Langford, John Sillitoe on behalf of the Wellcome Sanger Institute COVID-19 Surveillance Team |
| EPI_ISL_718648, EPI_ISL_718649 | Lighthouse Lab in Cambridge | Wellcome Sanger Institute for the COVID-19 Genomics UK (COG-UK) Consortium | Rob Howes, The Lighthouse Lab in Cambridge and Alex Alderton, Roberto Amato, Sonia Goncalves, Ewan Harrison, David K. Jackson, Ian Johnston, Dominic Kwiatkowski, Cordelia Langford, John Sillitoe on behalf of the Wellcome Sanger Institute COVID-19 Surveillance Team |
| EPI_ISL_718650 | Lighthouse Lab in Alderley Park | Wellcome Sanger Institute for the COVID-19 Genomics UK (COG-UK) Consortium | Jacquelyn Wynn, Mairead Hyland, The Lighthouse Lab in Alderley Park and Alex Alderton, Roberto Amato, Sonia Goncalves, Ewan Harrison, David K. Jackson, Ian Johnston, Dominic Kwiatkowski, Cordelia Langford, John Sillitoe on behalf of the Wellcome Sanger Institute COVID-19 Surveillance Team |
| EPI_ISL_718651, EPI_ISL_718652 | Lighthouse Lab in Cambridge | Wellcome Sanger Institute for the COVID-19 Genomics UK (COG-UK) Consortium | Rob Howes, The Lighthouse Lab in Cambridge and Alex Alderton, Roberto Amato, Sonia Goncalves, Ewan Harrison, David K. Jackson, Ian Johnston, Dominic Kwiatkowski, Cordelia Langford, John Sillitoe on behalf of the Wellcome Sanger Institute COVID-19 Surveillance Team |
| EPI_ISL_718653 | Lighthouse Lab in Alderley Park | Wellcome Sanger Institute for the COVID-19 Genomics UK (COG-UK) Consortium | Jacquelyn Wynn, Mairead Hyland, The Lighthouse Lab in Alderley Park and Alex Alderton, Roberto Amato, Sonia Goncalves, Ewan Harrison, David K. Jackson, Ian Johnston, Dominic Kwiatkowski, Cordelia Langford, John Sillitoe on behalf of the Wellcome Sanger Institute COVID-19 Surveillance Team |
| EPI_ISL_718654, EPI_ISL_718655, EPI_ISL_718656 | Lighthouse Lab in Cambridge | Wellcome Sanger Institute for the COVID-19 Genomics UK (COG-UK) Consortium | Rob Howes, The Lighthouse Lab in Cambridge and Alex Alderton, Roberto Amato, Sonia Goncalves, Ewan Harrison, David K. Jackson, Ian Johnston, Dominic Kwiatkowski, Cordelia Langford, John Sillitoe on behalf of the Wellcome Sanger Institute COVID-19 Surveillance Team |
| EPI_ISL_718657, EPI_ISL_718658 | Lighthouse Lab in Alderley Park | Wellcome Sanger Institute for the COVID-19 Genomics UK | Jacquelyn Wynn, Mairead Hyland, The Lighthouse Lab in Alderley Park and Alex Alderton, Roberto Amato, Sonia Goncalves, Ewan Harrison, David K. |

| | | | |
|---|---|---|---|
| | | (COG-UK) Consortium | Jackson, Ian Johnston, Dominic Kwiatkowski, Cordelia Langford, John Sillitoe on behalf of the Wellcome Sanger Institute COVID-19 Surveillance Team |
| EPI_ISL_718659, EPI_ISL_718660, EPI_ISL_718661 | Lighthouse Lab in Cambridge | Wellcome Sanger Institute for the COVID-19 Genomics UK (COG-UK) Consortium | Rob Howes, The Lighthouse Lab in Cambridge and Alex Alderton, Roberto Amato, Sonia Goncalves, Ewan Harrison, David K. Jackson, Ian Johnston, Dominic Kwiatkowski, Cordelia Langford, John Sillitoe on behalf of the Wellcome Sanger Institute COVID-19 Surveillance Team |
| EPI_ISL_718662, EPI_ISL_718663, EPI_ISL_718664, EPI_ISL_718665, EPI_ISL_718666 | Lighthouse Lab in Alderley Park | Wellcome Sanger Institute for the COVID-19 Genomics UK (COG-UK) Consortium | Jacquelyn Wynn, Mairead Hyland, The Lighthouse Lab in Alderley Park and Alex Alderton, Roberto Amato, Sonia Goncalves, Ewan Harrison, David K. Jackson, Ian Johnston, Dominic Kwiatkowski, Cordelia Langford, John Sillitoe on behalf of the Wellcome Sanger Institute COVID-19 Surveillance Team |
| EPI_ISL_718667, EPI_ISL_718668, EPI_ISL_718669 | Lighthouse Lab in Cambridge | Wellcome Sanger Institute for the COVID-19 Genomics UK (COG-UK) Consortium | Rob Howes, The Lighthouse Lab in Cambridge and Alex Alderton, Roberto Amato, Sonia Goncalves, Ewan Harrison, David K. Jackson, Ian Johnston, Dominic Kwiatkowski, Cordelia Langford, John Sillitoe on behalf of the Wellcome Sanger Institute COVID-19 Surveillance Team |
| EPI_ISL_718670 | Lighthouse Lab in Alderley Park | Wellcome Sanger Institute for the COVID-19 Genomics UK (COG-UK) Consortium | Jacquelyn Wynn, Mairead Hyland, The Lighthouse Lab in Alderley Park and Alex Alderton, Roberto Amato, Sonia Goncalves, Ewan Harrison, David K. Jackson, Ian Johnston, Dominic Kwiatkowski, Cordelia Langford, John Sillitoe on behalf of the Wellcome Sanger Institute COVID-19 Surveillance Team |
| EPI_ISL_718671 | Lighthouse Lab in Cambridge | Wellcome Sanger Institute for the COVID-19 Genomics UK (COG-UK) Consortium | Rob Howes, The Lighthouse Lab in Cambridge and Alex Alderton, Roberto Amato, Sonia Goncalves, Ewan Harrison, David K. Jackson, Ian Johnston, Dominic Kwiatkowski, Cordelia Langford, John Sillitoe on behalf of the Wellcome Sanger Institute COVID-19 Surveillance Team |
| EPI_ISL_718672 | Lighthouse Lab in Alderley Park | Wellcome Sanger Institute for the COVID-19 Genomics UK (COG-UK) Consortium | Jacquelyn Wynn, Mairead Hyland, The Lighthouse Lab in Alderley Park and Alex Alderton, Roberto Amato, Sonia Goncalves, Ewan Harrison, David K. Jackson, Ian Johnston, Dominic Kwiatkowski, Cordelia Langford, John Sillitoe on behalf of the Wellcome Sanger Institute COVID-19 Surveillance Team |
| EPI_ISL_718673 | Lighthouse Lab in Cambridge | Wellcome Sanger Institute for the COVID-19 Genomics UK (COG-UK) Consortium | Rob Howes, The Lighthouse Lab in Cambridge and Alex Alderton, Roberto Amato, Sonia Goncalves, Ewan Harrison, David K. Jackson, Ian Johnston, Dominic Kwiatkowski, Cordelia Langford, John Sillitoe on behalf of the Wellcome Sanger Institute COVID-19 Surveillance Team |
| EPI_ISL_718674, EPI_ISL_718675, EPI_ISL_718676 | Lighthouse Lab in Alderley Park | Wellcome Sanger Institute for the COVID-19 Genomics UK (COG-UK) Consortium | Jacquelyn Wynn, Mairead Hyland, The Lighthouse Lab in Alderley Park and Alex Alderton, Roberto Amato, Sonia Goncalves, Ewan Harrison, David K. Jackson, Ian Johnston, Dominic Kwiatkowski, Cordelia Langford, John Sillitoe on behalf of the Wellcome Sanger Institute COVID-19 Surveillance Team |
| EPI_ISL_718677 | Lighthouse Lab in Cambridge | Wellcome Sanger Institute for the COVID-19 Genomics UK (COG-UK) Consortium | Rob Howes, The Lighthouse Lab in Cambridge and Alex Alderton, Roberto Amato, Sonia Goncalves, Ewan Harrison, David K. Jackson, Ian Johnston, Dominic Kwiatkowski, Cordelia Langford, John Sillitoe on behalf of the Wellcome Sanger Institute COVID-19 Surveillance Team |
| EPI_ISL_718678 | Lighthouse Lab in Alderley Park | Wellcome Sanger Institute for the COVID-19 Genomics UK (COG-UK) Consortium | Jacquelyn Wynn, Mairead Hyland, The Lighthouse Lab in Alderley Park and Alex Alderton, Roberto Amato, Sonia Goncalves, Ewan Harrison, David K. Jackson, Ian Johnston, Dominic Kwiatkowski, Cordelia Langford, John Sillitoe on behalf of the Wellcome Sanger Institute COVID-19 Surveillance Team |
| EPI_ISL_718679, EPI_ISL_718680, EPI_ISL_718681 | Lighthouse Lab in Cambridge | Wellcome Sanger Institute for the COVID-19 Genomics UK (COG-UK) Consortium | Rob Howes, The Lighthouse Lab in Cambridge and Alex Alderton, Roberto Amato, Sonia Goncalves, Ewan Harrison, David K. Jackson, Ian Johnston, Dominic Kwiatkowski, Cordelia Langford, John Sillitoe on behalf of the Wellcome Sanger Institute COVID-19 Surveillance Team |
| EPI_ISL_718682 | Lighthouse Lab in Alderley Park | Wellcome Sanger Institute for the COVID-19 Genomics UK (COG-UK) Consortium | Jacquelyn Wynn, Mairead Hyland, The Lighthouse Lab in Alderley Park and Alex Alderton, Roberto Amato, Sonia Goncalves, Ewan Harrison, David K. Jackson, Ian Johnston, Dominic Kwiatkowski, Cordelia Langford, John Sillitoe on behalf of the Wellcome Sanger Institute COVID-19 Surveillance Team |
| EPI_ISL_718683, EPI_ISL_718684, EPI_ISL_718685 | Lighthouse Lab in Cambridge | Wellcome Sanger Institute for the COVID-19 Genomics UK (COG-UK) Consortium | Rob Howes, The Lighthouse Lab in Cambridge and Alex Alderton, Roberto Amato, Sonia Goncalves, Ewan Harrison, David K. Jackson, Ian Johnston, Dominic Kwiatkowski, Cordelia Langford, John Sillitoe on behalf of the Wellcome Sanger Institute COVID-19 Surveillance Team |
| EPI_ISL_718686, EPI_ISL_718687 | Lighthouse Lab in Alderley Park | Wellcome Sanger Institute for the COVID-19 Genomics UK (COG-UK) Consortium | Jacquelyn Wynn, Mairead Hyland, The Lighthouse Lab in Alderley Park and Alex Alderton, Roberto Amato, Sonia Goncalves, Ewan Harrison, David K. Jackson, Ian Johnston, Dominic Kwiatkowski, Cordelia Langford, John Sillitoe on behalf of the Wellcome Sanger Institute COVID-19 Surveillance Team |
| EPI_ISL_718688, EPI_ISL_718689, EPI_ISL_718690 | Lighthouse Lab in Cambridge | Wellcome Sanger Institute for the COVID-19 Genomics UK (COG-UK) Consortium | Rob Howes, The Lighthouse Lab in Cambridge and Alex Alderton, Roberto Amato, Sonia Goncalves, Ewan Harrison, David K. Jackson, Ian Johnston, Dominic Kwiatkowski, Cordelia Langford, John Sillitoe on behalf of the Wellcome Sanger Institute COVID-19 Surveillance Team |
| EPI_ISL_718691 | Lighthouse Lab in Alderley Park | Wellcome Sanger Institute for the COVID-19 Genomics UK (COG-UK) Consortium | Jacquelyn Wynn, Mairead Hyland, The Lighthouse Lab in Alderley Park and Alex Alderton, Roberto Amato, Sonia Goncalves, Ewan Harrison, David K. Jackson, Ian Johnston, Dominic Kwiatkowski, Cordelia Langford, John Sillitoe on behalf of the Wellcome Sanger Institute COVID-19 Surveillance Team |
| EPI_ISL_718692, EPI_ISL_718693 | Lighthouse Lab in Cambridge | Wellcome Sanger Institute for the COVID-19 Genomics UK (COG-UK) Consortium | Rob Howes, The Lighthouse Lab in Cambridge and Alex Alderton, Roberto Amato, Sonia Goncalves, Ewan Harrison, David K. Jackson, Ian Johnston, Dominic Kwiatkowski, Cordelia Langford, John Sillitoe on behalf of the Wellcome Sanger Institute COVID-19 Surveillance Team |
| EPI_ISL_718694, EPI_ISL_718695 | Lighthouse Lab in Alderley Park | Wellcome Sanger Institute for the COVID-19 Genomics UK (COG-UK) Consortium | Jacquelyn Wynn, Mairead Hyland, The Lighthouse Lab in Alderley Park and Alex Alderton, Roberto Amato, Sonia Goncalves, Ewan Harrison, David K. Jackson, Ian Johnston, Dominic Kwiatkowski, Cordelia Langford, John Sillitoe on behalf of the Wellcome Sanger Institute COVID-19 Surveillance Team |
| EPI_ISL_718696, EPI_ISL_718697, EPI_ISL_718698, EPI_ISL_718699 | Lighthouse Lab in Cambridge | Wellcome Sanger Institute for the COVID-19 Genomics UK (COG-UK) Consortium | Rob Howes, The Lighthouse Lab in Cambridge and Alex Alderton, Roberto Amato, Sonia Goncalves, Ewan Harrison, David K. Jackson, Ian Johnston, Dominic Kwiatkowski, Cordelia Langford, John Sillitoe on behalf of the Wellcome Sanger Institute COVID-19 Surveillance Team |
| EPI_ISL_718700 | Lighthouse Lab in Alderley Park | Wellcome Sanger Institute for the COVID-19 Genomics UK (COG-UK) Consortium | Jacquelyn Wynn, Mairead Hyland, The Lighthouse Lab in Alderley Park and Alex Alderton, Roberto Amato, Sonia Goncalves, Ewan Harrison, David K. Jackson, Ian Johnston, Dominic Kwiatkowski, Cordelia Langford, John Sillitoe on behalf of the Wellcome Sanger Institute COVID-19 Surveillance Team |
| EPI_ISL_718701, EPI_ISL_718702, EPI_ISL_718703, EPI_ISL_718704 | Lighthouse Lab in Cambridge | Wellcome Sanger Institute for the COVID-19 Genomics UK (COG-UK) Consortium | Rob Howes, The Lighthouse Lab in Cambridge and Alex Alderton, Roberto Amato, Sonia Goncalves, Ewan Harrison, David K. Jackson, Ian Johnston, Dominic Kwiatkowski, Cordelia Langford, John Sillitoe on behalf of the Wellcome Sanger Institute COVID-19 Surveillance Team |
| EPI_ISL_718705 | Lighthouse Lab in Alderley Park | Wellcome Sanger Institute for the COVID-19 Genomics UK (COG-UK) Consortium | Jacquelyn Wynn, Mairead Hyland, The Lighthouse Lab in Alderley Park and Alex Alderton, Roberto Amato, Sonia Goncalves, Ewan Harrison, David K. Jackson, Ian Johnston, Dominic Kwiatkowski, Cordelia Langford, John Sillitoe on behalf of the Wellcome Sanger Institute COVID-19 Surveillance Team |
| EPI_ISL_718706 | Lighthouse Lab in Cambridge | Wellcome Sanger Institute for the COVID-19 Genomics UK (COG-UK) Consortium | Rob Howes, The Lighthouse Lab in Cambridge and Alex Alderton, Roberto Amato, Sonia Goncalves, Ewan Harrison, David K. Jackson, Ian Johnston, Dominic Kwiatkowski, Cordelia Langford, John Sillitoe on behalf of the Wellcome Sanger Institute COVID-19 Surveillance Team |
| EPI_ISL_718707 | Lighthouse Lab in Alderley Park | Wellcome Sanger Institute for the COVID-19 Genomics UK (COG-UK) Consortium | Jacquelyn Wynn, Mairead Hyland, The Lighthouse Lab in Alderley Park and Alex Alderton, Roberto Amato, Sonia Goncalves, Ewan Harrison, David K. Jackson, Ian Johnston, Dominic Kwiatkowski, Cordelia Langford, John Sillitoe on behalf of the Wellcome Sanger Institute COVID-19 Surveillance Team |
| EPI_ISL_718708, EPI_ISL_718709 | Lighthouse Lab in Cambridge | Wellcome Sanger Institute for the COVID-19 Genomics UK (COG-UK) Consortium | Rob Howes, The Lighthouse Lab in Cambridge and Alex Alderton, Roberto Amato, Sonia Goncalves, Ewan Harrison, David K. Jackson, Ian Johnston, Dominic Kwiatkowski, Cordelia Langford, John Sillitoe on behalf of the Wellcome Sanger Institute COVID-19 Surveillance Team |
| EPI_ISL_718710, EPI_ISL_718711, EPI_ISL_718712, EPI_ISL_718713, EPI_ISL_718714, EPI_ISL_718715, EPI_ISL_718716 | Lighthouse Lab in Alderley Park | Wellcome Sanger Institute for the COVID-19 Genomics UK (COG-UK) Consortium | Jacquelyn Wynn, Mairead Hyland, The Lighthouse Lab in Alderley Park and Alex Alderton, Roberto Amato, Sonia Goncalves, Ewan Harrison, David K. Jackson, Ian Johnston, Dominic Kwiatkowski, Cordelia Langford, John Sillitoe on behalf of the Wellcome Sanger Institute COVID-19 Surveillance Team |
| EPI_ISL_718717 | Lighthouse Lab in Cambridge | Wellcome Sanger Institute for the COVID-19 Genomics UK (COG-UK) Consortium | Rob Howes, The Lighthouse Lab in Cambridge and Alex Alderton, Roberto Amato, Sonia Goncalves, Ewan Harrison, David K. Jackson, Ian Johnston, Dominic Kwiatkowski, Cordelia Langford, John Sillitoe on behalf of the Wellcome Sanger Institute COVID-19 Surveillance Team |
| EPI_ISL_718718, EPI_ISL_718719 | Lighthouse Lab in Alderley Park | Wellcome Sanger Institute for the COVID-19 Genomics UK (COG-UK) Consortium | Jacquelyn Wynn, Mairead Hyland, The Lighthouse Lab in Alderley Park and Alex Alderton, Roberto Amato, Sonia Goncalves, Ewan Harrison, David K. Jackson, Ian Johnston, Dominic Kwiatkowski, Cordelia Langford, John Sillitoe on behalf of the Wellcome Sanger Institute COVID-19 Surveillance Team |
| EPI_ISL_718720 | Lighthouse Lab in Cambridge | Wellcome Sanger Institute for the COVID-19 Genomics UK (COG-UK) Consortium | Rob Howes, The Lighthouse Lab in Cambridge and Alex Alderton, Roberto Amato, Sonia Goncalves, Ewan Harrison, David K. Jackson, Ian Johnston, Dominic Kwiatkowski, Cordelia Langford, John Sillitoe on behalf of the Wellcome Sanger Institute COVID-19 Surveillance Team |
| EPI_ISL_718721 | Lighthouse Lab in Alderley Park | Wellcome Sanger Institute for the COVID-19 Genomics UK (COG-UK) Consortium | Jacquelyn Wynn, Mairead Hyland, The Lighthouse Lab in Alderley Park and Alex Alderton, Roberto Amato, Sonia Goncalves, Ewan Harrison, David K. Jackson, Ian Johnston, Dominic Kwiatkowski, Cordelia Langford, John Sillitoe on behalf of the Wellcome Sanger Institute COVID-19 Surveillance Team |
| EPI_ISL_718722 | Lighthouse Lab in Cambridge | Wellcome Sanger Institute for the COVID-19 Genomics UK (COG-UK) Consortium | Rob Howes, The Lighthouse Lab in Cambridge and Alex Alderton, Roberto Amato, Sonia Goncalves, Ewan Harrison, David K. Jackson, Ian Johnston, Dominic Kwiatkowski, Cordelia Langford, John Sillitoe on behalf of the Wellcome Sanger Institute COVID-19 Surveillance Team |
| EPI_ISL_718723, EPI_ISL_718724, EPI_ISL_718725 | Lighthouse Lab in Alderley Park | Wellcome Sanger Institute for the COVID-19 Genomics UK (COG-UK) Consortium | Jacquelyn Wynn, Mairead Hyland, The Lighthouse Lab in Alderley Park and Alex Alderton, Roberto Amato, Sonia Goncalves, Ewan Harrison, David K. Jackson, Ian Johnston, Dominic Kwiatkowski, Cordelia Langford, John Sillitoe on behalf of the Wellcome Sanger Institute COVID-19 Surveillance Team |
| EPI_ISL_718726, EPI_ISL_718727 | Lighthouse Lab in Cambridge | Wellcome Sanger Institute for the COVID-19 Genomics UK (COG-UK) Consortium | Rob Howes, The Lighthouse Lab in Cambridge and Alex Alderton, Roberto Amato, Sonia Goncalves, Ewan Harrison, David K. Jackson, Ian Johnston, Dominic Kwiatkowski, Cordelia Langford, John Sillitoe on behalf of the Wellcome Sanger Institute COVID-19 Surveillance Team |
| EPI_ISL_718728, EPI_ISL_718729 | Lighthouse Lab in Alderley Park | Wellcome Sanger Institute for the COVID-19 Genomics UK | Jacquelyn Wynn, Mairead Hyland, The Lighthouse Lab in Alderley Park and Alex Alderton, Roberto Amato, Sonia Goncalves, Ewan Harrison, David K. |

| | | | |
|---|---|---|---|
| | | (COG-UK) Consortium | Jackson, Ian Johnston, Dominic Kwiatkowski, Cordelia Langford, John Sillitoe on behalf of the Wellcome Sanger Institute COVID-19 Surveillance Team |
| EPI_ISL_718730, EPI_ISL_718731, EPI_ISL_718732, EPI_ISL_718733, EPI_ISL_718734 | Lighthouse Lab in Cambridge | Wellcome Sanger Institute for the COVID-19 Genomics UK (COG-UK) Consortium | Rob Howes, The Lighthouse Lab in Cambridge and Alex Alderton, Roberto Amato, Sonia Goncalves, Ewan Harrison, David K. Jackson, Ian Johnston, Dominic Kwiatkowski, Cordelia Langford, John Sillitoe on behalf of the Wellcome Sanger Institute COVID-19 Surveillance Team |
| EPI_ISL_718735 | Lighthouse Lab in Alderley Park | Wellcome Sanger Institute for the COVID-19 Genomics UK (COG-UK) Consortium | Jacquelyn Wynn, Mairead Hyland, The Lighthouse Lab in Alderley Park and Alex Alderton, Roberto Amato, Sonia Goncalves, Ewan Harrison, David K. Jackson, Ian Johnston, Dominic Kwiatkowski, Cordelia Langford, John Sillitoe on behalf of the Wellcome Sanger Institute COVID-19 Surveillance Team |
| EPI_ISL_718736, EPI_ISL_718737 | Lighthouse Lab in Cambridge | Wellcome Sanger Institute for the COVID-19 Genomics UK (COG-UK) Consortium | Rob Howes, The Lighthouse Lab in Cambridge and Alex Alderton, Roberto Amato, Sonia Goncalves, Ewan Harrison, David K. Jackson, Ian Johnston, Dominic Kwiatkowski, Cordelia Langford, John Sillitoe on behalf of the Wellcome Sanger Institute COVID-19 Surveillance Team |
| EPI_ISL_718738, EPI_ISL_718739, EPI_ISL_718740 | Lighthouse Lab in Alderley Park | Wellcome Sanger Institute for the COVID-19 Genomics UK (COG-UK) Consortium | Jacquelyn Wynn, Mairead Hyland, The Lighthouse Lab in Alderley Park and Alex Alderton, Roberto Amato, Sonia Goncalves, Ewan Harrison, David K. Jackson, Ian Johnston, Dominic Kwiatkowski, Cordelia Langford, John Sillitoe on behalf of the Wellcome Sanger Institute COVID-19 Surveillance Team |
| EPI_ISL_718741 | Lighthouse Lab in Cambridge | Wellcome Sanger Institute for the COVID-19 Genomics UK (COG-UK) Consortium | Rob Howes, The Lighthouse Lab in Cambridge and Alex Alderton, Roberto Amato, Sonia Goncalves, Ewan Harrison, David K. Jackson, Ian Johnston, Dominic Kwiatkowski, Cordelia Langford, John Sillitoe on behalf of the Wellcome Sanger Institute COVID-19 Surveillance Team |
| EPI_ISL_718742 | Lighthouse Lab in Alderley Park | Wellcome Sanger Institute for the COVID-19 Genomics UK (COG-UK) Consortium | Jacquelyn Wynn, Mairead Hyland, The Lighthouse Lab in Alderley Park and Alex Alderton, Roberto Amato, Sonia Goncalves, Ewan Harrison, David K. Jackson, Ian Johnston, Dominic Kwiatkowski, Cordelia Langford, John Sillitoe on behalf of the Wellcome Sanger Institute COVID-19 Surveillance Team |
| EPI_ISL_718743 | Lighthouse Lab in Cambridge | Wellcome Sanger Institute for the COVID-19 Genomics UK (COG-UK) Consortium | Rob Howes, The Lighthouse Lab in Cambridge and Alex Alderton, Roberto Amato, Sonia Goncalves, Ewan Harrison, David K. Jackson, Ian Johnston, Dominic Kwiatkowski, Cordelia Langford, John Sillitoe on behalf of the Wellcome Sanger Institute COVID-19 Surveillance Team |
| EPI_ISL_718744, EPI_ISL_718745, EPI_ISL_718746, EPI_ISL_718747 | Lighthouse Lab in Alderley Park | Wellcome Sanger Institute for the COVID-19 Genomics UK (COG-UK) Consortium | Jacquelyn Wynn, Mairead Hyland, The Lighthouse Lab in Alderley Park and Alex Alderton, Roberto Amato, Sonia Goncalves, Ewan Harrison, David K. Jackson, Ian Johnston, Dominic Kwiatkowski, Cordelia Langford, John Sillitoe on behalf of the Wellcome Sanger Institute COVID-19 Surveillance Team |
| EPI_ISL_718748, EPI_ISL_718749 | Lighthouse Lab in Cambridge | Wellcome Sanger Institute for the COVID-19 Genomics UK (COG-UK) Consortium | Rob Howes, The Lighthouse Lab in Cambridge and Alex Alderton, Roberto Amato, Sonia Goncalves, Ewan Harrison, David K. Jackson, Ian Johnston, Dominic Kwiatkowski, Cordelia Langford, John Sillitoe on behalf of the Wellcome Sanger Institute COVID-19 Surveillance Team |
| EPI_ISL_718750 | Lighthouse Lab in Alderley Park | Wellcome Sanger Institute for the COVID-19 Genomics UK (COG-UK) Consortium | Jacquelyn Wynn, Mairead Hyland, The Lighthouse Lab in Alderley Park and Alex Alderton, Roberto Amato, Sonia Goncalves, Ewan Harrison, David K. Jackson, Ian Johnston, Dominic Kwiatkowski, Cordelia Langford, John Sillitoe on behalf of the Wellcome Sanger Institute COVID-19 Surveillance Team |
| EPI_ISL_718751, EPI_ISL_718752, EPI_ISL_718753 | Lighthouse Lab in Cambridge | Wellcome Sanger Institute for the COVID-19 Genomics UK (COG-UK) Consortium | Rob Howes, The Lighthouse Lab in Cambridge and Alex Alderton, Roberto Amato, Sonia Goncalves, Ewan Harrison, David K. Jackson, Ian Johnston, Dominic Kwiatkowski, Cordelia Langford, John Sillitoe on behalf of the Wellcome Sanger Institute COVID-19 Surveillance Team |
| EPI_ISL_718754, EPI_ISL_718755 | Lighthouse Lab in Alderley Park | Wellcome Sanger Institute for the COVID-19 Genomics UK (COG-UK) Consortium | Jacquelyn Wynn, Mairead Hyland, The Lighthouse Lab in Alderley Park and Alex Alderton, Roberto Amato, Sonia Goncalves, Ewan Harrison, David K. Jackson, Ian Johnston, Dominic Kwiatkowski, Cordelia Langford, John Sillitoe on behalf of the Wellcome Sanger Institute COVID-19 Surveillance Team |
| EPI_ISL_718756, EPI_ISL_718757, EPI_ISL_718758 | Lighthouse Lab in Cambridge | Wellcome Sanger Institute for the COVID-19 Genomics UK (COG-UK) Consortium | Rob Howes, The Lighthouse Lab in Cambridge and Alex Alderton, Roberto Amato, Sonia Goncalves, Ewan Harrison, David K. Jackson, Ian Johnston, Dominic Kwiatkowski, Cordelia Langford, John Sillitoe on behalf of the Wellcome Sanger Institute COVID-19 Surveillance Team |
| EPI_ISL_718759 | Lighthouse Lab in Alderley Park | Wellcome Sanger Institute for the COVID-19 Genomics UK (COG-UK) Consortium | Jacquelyn Wynn, Mairead Hyland, The Lighthouse Lab in Alderley Park and Alex Alderton, Roberto Amato, Sonia Goncalves, Ewan Harrison, David K. Jackson, Ian Johnston, Dominic Kwiatkowski, Cordelia Langford, John Sillitoe on behalf of the Wellcome Sanger Institute COVID-19 Surveillance Team |
| EPI_ISL_718760 | Lighthouse Lab in Cambridge | Wellcome Sanger Institute for the COVID-19 Genomics UK (COG-UK) Consortium | Rob Howes, The Lighthouse Lab in Cambridge and Alex Alderton, Roberto Amato, Sonia Goncalves, Ewan Harrison, David K. Jackson, Ian Johnston, Dominic Kwiatkowski, Cordelia Langford, John Sillitoe on behalf of the Wellcome Sanger Institute COVID-19 Surveillance Team |
| EPI_ISL_718761, EPI_ISL_718762 | Lighthouse Lab in Alderley Park | Wellcome Sanger Institute for the COVID-19 Genomics UK (COG-UK) Consortium | Jacquelyn Wynn, Mairead Hyland, The Lighthouse Lab in Alderley Park and Alex Alderton, Roberto Amato, Sonia Goncalves, Ewan Harrison, David K. Jackson, Ian Johnston, Dominic Kwiatkowski, Cordelia Langford, John Sillitoe on behalf of the Wellcome Sanger Institute COVID-19 Surveillance Team |
| EPI_ISL_718763 | Lighthouse Lab in Cambridge | Wellcome Sanger Institute for the COVID-19 Genomics UK (COG-UK) Consortium | Rob Howes, The Lighthouse Lab in Cambridge and Alex Alderton, Roberto Amato, Sonia Goncalves, Ewan Harrison, David K. Jackson, Ian Johnston, Dominic Kwiatkowski, Cordelia Langford, John Sillitoe on behalf of the Wellcome Sanger Institute COVID-19 Surveillance Team |
| EPI_ISL_718764, EPI_ISL_718765, EPI_ISL_718766, EPI_ISL_718767 | Lighthouse Lab in Alderley Park | Wellcome Sanger Institute for the COVID-19 Genomics UK (COG-UK) Consortium | Jacquelyn Wynn, Mairead Hyland, The Lighthouse Lab in Alderley Park and Alex Alderton, Roberto Amato, Sonia Goncalves, Ewan Harrison, David K. Jackson, Ian Johnston, Dominic Kwiatkowski, Cordelia Langford, John Sillitoe on behalf of the Wellcome Sanger Institute COVID-19 Surveillance Team |
| EPI_ISL_718768 | Lighthouse Lab in Cambridge | Wellcome Sanger Institute for the COVID-19 Genomics UK (COG-UK) Consortium | Rob Howes, The Lighthouse Lab in Cambridge and Alex Alderton, Roberto Amato, Sonia Goncalves, Ewan Harrison, David K. Jackson, Ian Johnston, Dominic Kwiatkowski, Cordelia Langford, John Sillitoe on behalf of the Wellcome Sanger Institute COVID-19 Surveillance Team |
| EPI_ISL_718769 | Lighthouse Lab in Alderley Park | Wellcome Sanger Institute for the COVID-19 Genomics UK (COG-UK) Consortium | Jacquelyn Wynn, Mairead Hyland, The Lighthouse Lab in Alderley Park and Alex Alderton, Roberto Amato, Sonia Goncalves, Ewan Harrison, David K. Jackson, Ian Johnston, Dominic Kwiatkowski, Cordelia Langford, John Sillitoe on behalf of the Wellcome Sanger Institute COVID-19 Surveillance Team |
| EPI_ISL_718770, EPI_ISL_718771 | Lighthouse Lab in Cambridge | Wellcome Sanger Institute for the COVID-19 Genomics UK (COG-UK) Consortium | Rob Howes, The Lighthouse Lab in Cambridge and Alex Alderton, Roberto Amato, Sonia Goncalves, Ewan Harrison, David K. Jackson, Ian Johnston, Dominic Kwiatkowski, Cordelia Langford, John Sillitoe on behalf of the Wellcome Sanger Institute COVID-19 Surveillance Team |
| EPI_ISL_718772, EPI_ISL_718773, EPI_ISL_718774 | Lighthouse Lab in Alderley Park | Wellcome Sanger Institute for the COVID-19 Genomics UK (COG-UK) Consortium | Jacquelyn Wynn, Mairead Hyland, The Lighthouse Lab in Alderley Park and Alex Alderton, Roberto Amato, Sonia Goncalves, Ewan Harrison, David K. Jackson, Ian Johnston, Dominic Kwiatkowski, Cordelia Langford, John Sillitoe on behalf of the Wellcome Sanger Institute COVID-19 Surveillance Team |
| EPI_ISL_718775, EPI_ISL_718776 | Lighthouse Lab in Cambridge | Wellcome Sanger Institute for the COVID-19 Genomics UK (COG-UK) Consortium | Rob Howes, The Lighthouse Lab in Cambridge and Alex Alderton, Roberto Amato, Sonia Goncalves, Ewan Harrison, David K. Jackson, Ian Johnston, Dominic Kwiatkowski, Cordelia Langford, John Sillitoe on behalf of the Wellcome Sanger Institute COVID-19 Surveillance Team |
| EPI_ISL_718777 | Lighthouse Lab in Alderley Park | Wellcome Sanger Institute for the COVID-19 Genomics UK (COG-UK) Consortium | Jacquelyn Wynn, Mairead Hyland, The Lighthouse Lab in Alderley Park and Alex Alderton, Roberto Amato, Sonia Goncalves, Ewan Harrison, David K. Jackson, Ian Johnston, Dominic Kwiatkowski, Cordelia Langford, John Sillitoe on behalf of the Wellcome Sanger Institute COVID-19 Surveillance Team |
| EPI_ISL_718778 | Lighthouse Lab in Cambridge | Wellcome Sanger Institute for the COVID-19 Genomics UK (COG-UK) Consortium | Rob Howes, The Lighthouse Lab in Cambridge and Alex Alderton, Roberto Amato, Sonia Goncalves, Ewan Harrison, David K. Jackson, Ian Johnston, Dominic Kwiatkowski, Cordelia Langford, John Sillitoe on behalf of the Wellcome Sanger Institute COVID-19 Surveillance Team |
| EPI_ISL_718779, EPI_ISL_718780, EPI_ISL_718781, EPI_ISL_718782, EPI_ISL_718783 | Lighthouse Lab in Alderley Park | Wellcome Sanger Institute for the COVID-19 Genomics UK (COG-UK) Consortium | Jacquelyn Wynn, Mairead Hyland, The Lighthouse Lab in Alderley Park and Alex Alderton, Roberto Amato, Sonia Goncalves, Ewan Harrison, David K. Jackson, Ian Johnston, Dominic Kwiatkowski, Cordelia Langford, John Sillitoe on behalf of the Wellcome Sanger Institute COVID-19 Surveillance Team |
| EPI_ISL_718784, EPI_ISL_718785, EPI_ISL_718786 | Lighthouse Lab in Cambridge | Wellcome Sanger Institute for the COVID-19 Genomics UK (COG-UK) Consortium | Rob Howes, The Lighthouse Lab in Cambridge and Alex Alderton, Roberto Amato, Sonia Goncalves, Ewan Harrison, David K. Jackson, Ian Johnston, Dominic Kwiatkowski, Cordelia Langford, John Sillitoe on behalf of the Wellcome Sanger Institute COVID-19 Surveillance Team |
| EPI_ISL_718787 | Lighthouse Lab in Alderley Park | Wellcome Sanger Institute for the COVID-19 Genomics UK (COG-UK) Consortium | Jacquelyn Wynn, Mairead Hyland, The Lighthouse Lab in Alderley Park and Alex Alderton, Roberto Amato, Sonia Goncalves, Ewan Harrison, David K. Jackson, Ian Johnston, Dominic Kwiatkowski, Cordelia Langford, John Sillitoe on behalf of the Wellcome Sanger Institute COVID-19 Surveillance Team |
| EPI_ISL_718788 | Lighthouse Lab in Cambridge | Wellcome Sanger Institute for the COVID-19 Genomics UK (COG-UK) Consortium | Rob Howes, The Lighthouse Lab in Cambridge and Alex Alderton, Roberto Amato, Sonia Goncalves, Ewan Harrison, David K. Jackson, Ian Johnston, Dominic Kwiatkowski, Cordelia Langford, John Sillitoe on behalf of the Wellcome Sanger Institute COVID-19 Surveillance Team |
| EPI_ISL_718789 | Lighthouse Lab in Alderley Park | Wellcome Sanger Institute for the COVID-19 Genomics UK (COG-UK) Consortium | Jacquelyn Wynn, Mairead Hyland, The Lighthouse Lab in Alderley Park and Alex Alderton, Roberto Amato, Sonia Goncalves, Ewan Harrison, David K. Jackson, Ian Johnston, Dominic Kwiatkowski, Cordelia Langford, John Sillitoe on behalf of the Wellcome Sanger Institute COVID-19 Surveillance Team |
| EPI_ISL_718790 | Lighthouse Lab in Cambridge | Wellcome Sanger Institute for the COVID-19 Genomics UK (COG-UK) Consortium | Rob Howes, The Lighthouse Lab in Cambridge and Alex Alderton, Roberto Amato, Sonia Goncalves, Ewan Harrison, David K. Jackson, Ian Johnston, Dominic Kwiatkowski, Cordelia Langford, John Sillitoe on behalf of the Wellcome Sanger Institute COVID-19 Surveillance Team |
| EPI_ISL_718791 | Lighthouse Lab in Alderley Park | Wellcome Sanger Institute for the COVID-19 Genomics UK (COG-UK) Consortium | Jacquelyn Wynn, Mairead Hyland, The Lighthouse Lab in Alderley Park and Alex Alderton, Roberto Amato, Sonia Goncalves, Ewan Harrison, David K. Jackson, Ian Johnston, Dominic Kwiatkowski, Cordelia Langford, John Sillitoe on behalf of the Wellcome Sanger Institute COVID-19 Surveillance Team |
| EPI_ISL_718792, EPI_ISL_718793, EPI_ISL_718794 | Lighthouse Lab in Cambridge | Wellcome Sanger Institute for the COVID-19 Genomics UK (COG-UK) Consortium | Rob Howes, The Lighthouse Lab in Cambridge and Alex Alderton, Roberto Amato, Sonia Goncalves, Ewan Harrison, David K. Jackson, Ian Johnston, Dominic Kwiatkowski, Cordelia Langford, John Sillitoe on behalf of the Wellcome Sanger Institute COVID-19 Surveillance Team |
| EPI_ISL_718795, EPI_ISL_718796 | Lighthouse Lab in Alderley Park | Wellcome Sanger Institute for the COVID-19 Genomics UK (COG-UK) Consortium | Jacquelyn Wynn, Mairead Hyland, The Lighthouse Lab in Alderley Park and Alex Alderton, Roberto Amato, Sonia Goncalves, Ewan Harrison, David K. Jackson, Ian Johnston, Dominic Kwiatkowski, Cordelia Langford, John Sillitoe on behalf of the Wellcome Sanger Institute COVID-19 Surveillance Team |

| | | | |
|---|---|---|---|
| EPI_ISL_718797 | Lighthouse Lab in Cambridge | Wellcome Sanger Institute for the COVID-19 Genomics UK (COG-UK) Consortium | Rob Howes, The Lighthouse Lab in Cambridge and Alex Alderton, Roberto Amato, Sonia Goncalves, Ewan Harrison, David K. Jackson, Ian Johnston, Dominic Kwiatkowski, Cordelia Langford, John Sillitoe on behalf of the Wellcome Sanger Institute COVID-19 Surveillance Team |
| EPI_ISL_718798, EPI_ISL_718799 | Lighthouse Lab in Alderley Park | Wellcome Sanger Institute for the COVID-19 Genomics UK (COG-UK) Consortium | Jacquelyn Wynn, Mairead Hyland, The Lighthouse Lab in Alderley Park and Alex Alderton, Roberto Amato, Sonia Goncalves, Ewan Harrison, David K. Jackson, Ian Johnston, Dominic Kwiatkowski, Cordelia Langford, John Sillitoe on behalf of the Wellcome Sanger Institute COVID-19 Surveillance Team |
| EPI_ISL_718800 | Lighthouse Lab in Cambridge | Wellcome Sanger Institute for the COVID-19 Genomics UK (COG-UK) Consortium | Rob Howes, The Lighthouse Lab in Cambridge and Alex Alderton, Roberto Amato, Sonia Goncalves, Ewan Harrison, David K. Jackson, Ian Johnston, Dominic Kwiatkowski, Cordelia Langford, John Sillitoe on behalf of the Wellcome Sanger Institute COVID-19 Surveillance Team |
| EPI_ISL_718801, EPI_ISL_718802, EPI_ISL_718803 | Lighthouse Lab in Alderley Park | Wellcome Sanger Institute for the COVID-19 Genomics UK (COG-UK) Consortium | Jacquelyn Wynn, Mairead Hyland, The Lighthouse Lab in Alderley Park and Alex Alderton, Roberto Amato, Sonia Goncalves, Ewan Harrison, David K. Jackson, Ian Johnston, Dominic Kwiatkowski, Cordelia Langford, John Sillitoe on behalf of the Wellcome Sanger Institute COVID-19 Surveillance Team |
| EPI_ISL_718804, EPI_ISL_718805 | Lighthouse Lab in Cambridge | Wellcome Sanger Institute for the COVID-19 Genomics UK (COG-UK) Consortium | Rob Howes, The Lighthouse Lab in Cambridge and Alex Alderton, Roberto Amato, Sonia Goncalves, Ewan Harrison, David K. Jackson, Ian Johnston, Dominic Kwiatkowski, Cordelia Langford, John Sillitoe on behalf of the Wellcome Sanger Institute COVID-19 Surveillance Team |
| EPI_ISL_718806, EPI_ISL_718807, EPI_ISL_718808 | Lighthouse Lab in Alderley Park | Wellcome Sanger Institute for the COVID-19 Genomics UK (COG-UK) Consortium | Jacquelyn Wynn, Mairead Hyland, The Lighthouse Lab in Alderley Park and Alex Alderton, Roberto Amato, Sonia Goncalves, Ewan Harrison, David K. Jackson, Ian Johnston, Dominic Kwiatkowski, Cordelia Langford, John Sillitoe on behalf of the Wellcome Sanger Institute COVID-19 Surveillance Team |
| EPI_ISL_718809, EPI_ISL_718810 | Lighthouse Lab in Cambridge | Wellcome Sanger Institute for the COVID-19 Genomics UK (COG-UK) Consortium | Rob Howes, The Lighthouse Lab in Cambridge and Alex Alderton, Roberto Amato, Sonia Goncalves, Ewan Harrison, David K. Jackson, Ian Johnston, Dominic Kwiatkowski, Cordelia Langford, John Sillitoe on behalf of the Wellcome Sanger Institute COVID-19 Surveillance Team |
| EPI_ISL_718811, EPI_ISL_718812, EPI_ISL_718813 | Lighthouse Lab in Alderley Park | Wellcome Sanger Institute for the COVID-19 Genomics UK (COG-UK) Consortium | Jacquelyn Wynn, Mairead Hyland, The Lighthouse Lab in Alderley Park and Alex Alderton, Roberto Amato, Sonia Goncalves, Ewan Harrison, David K. Jackson, Ian Johnston, Dominic Kwiatkowski, Cordelia Langford, John Sillitoe on behalf of the Wellcome Sanger Institute COVID-19 Surveillance Team |
| EPI_ISL_718814, EPI_ISL_718815, EPI_ISL_718816, EPI_ISL_718817, EPI_ISL_718818 | Lighthouse Lab in Cambridge | Wellcome Sanger Institute for the COVID-19 Genomics UK (COG-UK) Consortium | Rob Howes, The Lighthouse Lab in Cambridge and Alex Alderton, Roberto Amato, Sonia Goncalves, Ewan Harrison, David K. Jackson, Ian Johnston, Dominic Kwiatkowski, Cordelia Langford, John Sillitoe on behalf of the Wellcome Sanger Institute COVID-19 Surveillance Team |
| EPI_ISL_718819 | Lighthouse Lab in Alderley Park | Wellcome Sanger Institute for the COVID-19 Genomics UK (COG-UK) Consortium | Jacquelyn Wynn, Mairead Hyland, The Lighthouse Lab in Alderley Park and Alex Alderton, Roberto Amato, Sonia Goncalves, Ewan Harrison, David K. Jackson, Ian Johnston, Dominic Kwiatkowski, Cordelia Langford, John Sillitoe on behalf of the Wellcome Sanger Institute COVID-19 Surveillance Team |
| EPI_ISL_718820 | Lighthouse Lab in Cambridge | Wellcome Sanger Institute for the COVID-19 Genomics UK (COG-UK) Consortium | Rob Howes, The Lighthouse Lab in Cambridge and Alex Alderton, Roberto Amato, Sonia Goncalves, Ewan Harrison, David K. Jackson, Ian Johnston, Dominic Kwiatkowski, Cordelia Langford, John Sillitoe on behalf of the Wellcome Sanger Institute COVID-19 Surveillance Team |
| EPI_ISL_718821 | Lighthouse Lab in Alderley Park | Wellcome Sanger Institute for the COVID-19 Genomics UK (COG-UK) Consortium | Jacquelyn Wynn, Mairead Hyland, The Lighthouse Lab in Alderley Park and Alex Alderton, Roberto Amato, Sonia Goncalves, Ewan Harrison, David K. Jackson, Ian Johnston, Dominic Kwiatkowski, Cordelia Langford, John Sillitoe on behalf of the Wellcome Sanger Institute COVID-19 Surveillance Team |
| EPI_ISL_718822 | Lighthouse Lab in Cambridge | Wellcome Sanger Institute for the COVID-19 Genomics UK (COG-UK) Consortium | Rob Howes, The Lighthouse Lab in Cambridge and Alex Alderton, Roberto Amato, Sonia Goncalves, Ewan Harrison, David K. Jackson, Ian Johnston, Dominic Kwiatkowski, Cordelia Langford, John Sillitoe on behalf of the Wellcome Sanger Institute COVID-19 Surveillance Team |
| EPI_ISL_718823 | Lighthouse Lab in Alderley Park | Wellcome Sanger Institute for the COVID-19 Genomics UK (COG-UK) Consortium | Jacquelyn Wynn, Mairead Hyland, The Lighthouse Lab in Alderley Park and Alex Alderton, Roberto Amato, Sonia Goncalves, Ewan Harrison, David K. Jackson, Ian Johnston, Dominic Kwiatkowski, Cordelia Langford, John Sillitoe on behalf of the Wellcome Sanger Institute COVID-19 Surveillance Team |
| EPI_ISL_718824, EPI_ISL_718825, EPI_ISL_718826 | Lighthouse Lab in Cambridge | Wellcome Sanger Institute for the COVID-19 Genomics UK (COG-UK) Consortium | Rob Howes, The Lighthouse Lab in Cambridge and Alex Alderton, Roberto Amato, Sonia Goncalves, Ewan Harrison, David K. Jackson, Ian Johnston, Dominic Kwiatkowski, Cordelia Langford, John Sillitoe on behalf of the Wellcome Sanger Institute COVID-19 Surveillance Team |
| EPI_ISL_718827, EPI_ISL_718828, EPI_ISL_718829, EPI_ISL_718830, EPI_ISL_718831, EPI_ISL_718832, EPI_ISL_718833 | Lighthouse Lab in Alderley Park | Wellcome Sanger Institute for the COVID-19 Genomics UK (COG-UK) Consortium | Jacquelyn Wynn, Mairead Hyland, The Lighthouse Lab in Alderley Park and Alex Alderton, Roberto Amato, Sonia Goncalves, Ewan Harrison, David K. Jackson, Ian Johnston, Dominic Kwiatkowski, Cordelia Langford, John Sillitoe on behalf of the Wellcome Sanger Institute COVID-19 Surveillance Team |
| EPI_ISL_718834, EPI_ISL_718835, EPI_ISL_718836 | Lighthouse Lab in Cambridge | Wellcome Sanger Institute for the COVID-19 Genomics UK (COG-UK) Consortium | Rob Howes, The Lighthouse Lab in Cambridge and Alex Alderton, Roberto Amato, Sonia Goncalves, Ewan Harrison, David K. Jackson, Ian Johnston, Dominic Kwiatkowski, Cordelia Langford, John Sillitoe on behalf of the Wellcome Sanger Institute COVID-19 Surveillance Team |
| EPI_ISL_718837, EPI_ISL_718838, EPI_ISL_718839 | Lighthouse Lab in Alderley Park | Wellcome Sanger Institute for the COVID-19 Genomics UK (COG-UK) Consortium | Jacquelyn Wynn, Mairead Hyland, The Lighthouse Lab in Alderley Park and Alex Alderton, Roberto Amato, Sonia Goncalves, Ewan Harrison, David K. Jackson, Ian Johnston, Dominic Kwiatkowski, Cordelia Langford, John Sillitoe on behalf of the Wellcome Sanger Institute COVID-19 Surveillance Team |
| EPI_ISL_718840 | Lighthouse Lab in Cambridge | Wellcome Sanger Institute for the COVID-19 Genomics UK (COG-UK) Consortium | Rob Howes, The Lighthouse Lab in Cambridge and Alex Alderton, Roberto Amato, Sonia Goncalves, Ewan Harrison, David K. Jackson, Ian Johnston, Dominic Kwiatkowski, Cordelia Langford, John Sillitoe on behalf of the Wellcome Sanger Institute COVID-19 Surveillance Team |
| EPI_ISL_718841 | Lighthouse Lab in Alderley Park | Wellcome Sanger Institute for the COVID-19 Genomics UK (COG-UK) Consortium | Jacquelyn Wynn, Mairead Hyland, The Lighthouse Lab in Alderley Park and Alex Alderton, Roberto Amato, Sonia Goncalves, Ewan Harrison, David K. Jackson, Ian Johnston, Dominic Kwiatkowski, Cordelia Langford, John Sillitoe on behalf of the Wellcome Sanger Institute COVID-19 Surveillance Team |
| EPI_ISL_718842, EPI_ISL_718843 | Lighthouse Lab in Cambridge | Wellcome Sanger Institute for the COVID-19 Genomics UK (COG-UK) Consortium | Rob Howes, The Lighthouse Lab in Cambridge and Alex Alderton, Roberto Amato, Sonia Goncalves, Ewan Harrison, David K. Jackson, Ian Johnston, Dominic Kwiatkowski, Cordelia Langford, John Sillitoe on behalf of the Wellcome Sanger Institute COVID-19 Surveillance Team |
| EPI_ISL_718844 | Lighthouse Lab in Alderley Park | Wellcome Sanger Institute for the COVID-19 Genomics UK (COG-UK) Consortium | Jacquelyn Wynn, Mairead Hyland, The Lighthouse Lab in Alderley Park and Alex Alderton, Roberto Amato, Sonia Goncalves, Ewan Harrison, David K. Jackson, Ian Johnston, Dominic Kwiatkowski, Cordelia Langford, John Sillitoe on behalf of the Wellcome Sanger Institute COVID-19 Surveillance Team |
| EPI_ISL_718845, EPI_ISL_718846 | Lighthouse Lab in Cambridge | Wellcome Sanger Institute for the COVID-19 Genomics UK (COG-UK) Consortium | Rob Howes, The Lighthouse Lab in Cambridge and Alex Alderton, Roberto Amato, Sonia Goncalves, Ewan Harrison, David K. Jackson, Ian Johnston, Dominic Kwiatkowski, Cordelia Langford, John Sillitoe on behalf of the Wellcome Sanger Institute COVID-19 Surveillance Team |
| EPI_ISL_718847 | Lighthouse Lab in Alderley Park | Wellcome Sanger Institute for the COVID-19 Genomics UK (COG-UK) Consortium | Jacquelyn Wynn, Mairead Hyland, The Lighthouse Lab in Alderley Park and Alex Alderton, Roberto Amato, Sonia Goncalves, Ewan Harrison, David K. Jackson, Ian Johnston, Dominic Kwiatkowski, Cordelia Langford, John Sillitoe on behalf of the Wellcome Sanger Institute COVID-19 Surveillance Team |
| EPI_ISL_718848 | Lighthouse Lab in Cambridge | Wellcome Sanger Institute for the COVID-19 Genomics UK (COG-UK) Consortium | Rob Howes, The Lighthouse Lab in Cambridge and Alex Alderton, Roberto Amato, Sonia Goncalves, Ewan Harrison, David K. Jackson, Ian Johnston, Dominic Kwiatkowski, Cordelia Langford, John Sillitoe on behalf of the Wellcome Sanger Institute COVID-19 Surveillance Team |
| EPI_ISL_718849, EPI_ISL_718850 | Lighthouse Lab in Alderley Park | Wellcome Sanger Institute for the COVID-19 Genomics UK (COG-UK) Consortium | Jacquelyn Wynn, Mairead Hyland, The Lighthouse Lab in Alderley Park and Alex Alderton, Roberto Amato, Sonia Goncalves, Ewan Harrison, David K. Jackson, Ian Johnston, Dominic Kwiatkowski, Cordelia Langford, John Sillitoe on behalf of the Wellcome Sanger Institute COVID-19 Surveillance Team |
| EPI_ISL_718851, EPI_ISL_718852 | Lighthouse Lab in Cambridge | Wellcome Sanger Institute for the COVID-19 Genomics UK (COG-UK) Consortium | Rob Howes, The Lighthouse Lab in Cambridge and Alex Alderton, Roberto Amato, Sonia Goncalves, Ewan Harrison, David K. Jackson, Ian Johnston, Dominic Kwiatkowski, Cordelia Langford, John Sillitoe on behalf of the Wellcome Sanger Institute COVID-19 Surveillance Team |
| EPI_ISL_718853, EPI_ISL_718854 | Lighthouse Lab in Alderley Park | Wellcome Sanger Institute for the COVID-19 Genomics UK (COG-UK) Consortium | Jacquelyn Wynn, Mairead Hyland, The Lighthouse Lab in Alderley Park and Alex Alderton, Roberto Amato, Sonia Goncalves, Ewan Harrison, David K. Jackson, Ian Johnston, Dominic Kwiatkowski, Cordelia Langford, John Sillitoe on behalf of the Wellcome Sanger Institute COVID-19 Surveillance Team |
| EPI_ISL_718855, EPI_ISL_718856, EPI_ISL_718857 | Lighthouse Lab in Cambridge | Wellcome Sanger Institute for the COVID-19 Genomics UK (COG-UK) Consortium | Rob Howes, The Lighthouse Lab in Cambridge and Alex Alderton, Roberto Amato, Sonia Goncalves, Ewan Harrison, David K. Jackson, Ian Johnston, Dominic Kwiatkowski, Cordelia Langford, John Sillitoe on behalf of the Wellcome Sanger Institute COVID-19 Surveillance Team |
| EPI_ISL_718858, EPI_ISL_718859 | Lighthouse Lab in Alderley Park | Wellcome Sanger Institute for the COVID-19 Genomics UK (COG-UK) Consortium | Jacquelyn Wynn, Mairead Hyland, The Lighthouse Lab in Alderley Park and Alex Alderton, Roberto Amato, Sonia Goncalves, Ewan Harrison, David K. Jackson, Ian Johnston, Dominic Kwiatkowski, Cordelia Langford, John Sillitoe on behalf of the Wellcome Sanger Institute COVID-19 Surveillance Team |
| EPI_ISL_718860, EPI_ISL_718861, EPI_ISL_718862, EPI_ISL_718863 | Lighthouse Lab in Cambridge | Wellcome Sanger Institute for the COVID-19 Genomics UK (COG-UK) Consortium | Rob Howes, The Lighthouse Lab in Cambridge and Alex Alderton, Roberto Amato, Sonia Goncalves, Ewan Harrison, David K. Jackson, Ian Johnston, Dominic Kwiatkowski, Cordelia Langford, John Sillitoe on behalf of the Wellcome Sanger Institute COVID-19 Surveillance Team |
| EPI_ISL_718864, EPI_ISL_718865, EPI_ISL_718866 | Lighthouse Lab in Alderley Park | Wellcome Sanger Institute for the COVID-19 Genomics UK (COG-UK) Consortium | Jacquelyn Wynn, Mairead Hyland, The Lighthouse Lab in Alderley Park and Alex Alderton, Roberto Amato, Sonia Goncalves, Ewan Harrison, David K. Jackson, Ian Johnston, Dominic Kwiatkowski, Cordelia Langford, John Sillitoe on behalf of the Wellcome Sanger Institute COVID-19 Surveillance Team |
| EPI_ISL_718867, EPI_ISL_718868, EPI_ISL_718869 | Lighthouse Lab in Cambridge | Wellcome Sanger Institute for the COVID-19 Genomics UK (COG-UK) Consortium | Rob Howes, The Lighthouse Lab in Cambridge and Alex Alderton, Roberto Amato, Sonia Goncalves, Ewan Harrison, David K. Jackson, Ian Johnston, Dominic Kwiatkowski, Cordelia Langford, John Sillitoe on behalf of the Wellcome Sanger Institute COVID-19 Surveillance Team |
| EPI_ISL_718870, EPI_ISL_718871, EPI_ISL_718872, EPI_ISL_718873 | Lighthouse Lab in Alderley Park | Wellcome Sanger Institute for the COVID-19 Genomics UK (COG-UK) Consortium | Jacquelyn Wynn, Mairead Hyland, The Lighthouse Lab in Alderley Park and Alex Alderton, Roberto Amato, Sonia Goncalves, Ewan Harrison, David K. Jackson, Ian Johnston, Dominic Kwiatkowski, Cordelia Langford, John Sillitoe on behalf of the Wellcome Sanger Institute COVID-19 Surveillance Team |

| | | | |
|---|---|---|---|
| EPI_ISL_718874, EPI_ISL_718875, EPI_ISL_718876 | Lighthouse Lab in Cambridge | Wellcome Sanger Institute for the COVID-19 Genomics UK (COG-UK) Consortium | Rob Howes, The Lighthouse Lab in Cambridge and Alex Alderton, Roberto Amato, Sonia Goncalves, Ewan Harrison, David K. Jackson, Ian Johnston, Dominic Kwiatkowski, Cordelia Langford, John Sillitoe on behalf of the Wellcome Sanger Institute COVID-19 Surveillance Team |
| EPI_ISL_718877, EPI_ISL_718878 | Lighthouse Lab in Alderley Park | Wellcome Sanger Institute for the COVID-19 Genomics UK (COG-UK) Consortium | Jacquelyn Wynn, Mairead Hyland, The Lighthouse Lab in Alderley Park and Alex Alderton, Roberto Amato, Sonia Goncalves, Ewan Harrison, David K. Jackson, Ian Johnston, Dominic Kwiatkowski, Cordelia Langford, John Sillitoe on behalf of the Wellcome Sanger Institute COVID-19 Surveillance Team |
| EPI_ISL_718879 | Lighthouse Lab in Cambridge | Wellcome Sanger Institute for the COVID-19 Genomics UK (COG-UK) Consortium | Rob Howes, The Lighthouse Lab in Cambridge and Alex Alderton, Roberto Amato, Sonia Goncalves, Ewan Harrison, David K. Jackson, Ian Johnston, Dominic Kwiatkowski, Cordelia Langford, John Sillitoe on behalf of the Wellcome Sanger Institute COVID-19 Surveillance Team |
| EPI_ISL_718880, EPI_ISL_718881 | Lighthouse Lab in Alderley Park | Wellcome Sanger Institute for the COVID-19 Genomics UK (COG-UK) Consortium | Jacquelyn Wynn, Mairead Hyland, The Lighthouse Lab in Alderley Park and Alex Alderton, Roberto Amato, Sonia Goncalves, Ewan Harrison, David K. Jackson, Ian Johnston, Dominic Kwiatkowski, Cordelia Langford, John Sillitoe on behalf of the Wellcome Sanger Institute COVID-19 Surveillance Team |
| EPI_ISL_718882, EPI_ISL_718883, EPI_ISL_718884 | Lighthouse Lab in Cambridge | Wellcome Sanger Institute for the COVID-19 Genomics UK (COG-UK) Consortium | Rob Howes, The Lighthouse Lab in Cambridge and Alex Alderton, Roberto Amato, Sonia Goncalves, Ewan Harrison, David K. Jackson, Ian Johnston, Dominic Kwiatkowski, Cordelia Langford, John Sillitoe on behalf of the Wellcome Sanger Institute COVID-19 Surveillance Team |
| EPI_ISL_718885, EPI_ISL_718886, EPI_ISL_718887, EPI_ISL_718888, EPI_ISL_718889, EPI_ISL_718890, EPI_ISL_718891 | Lighthouse Lab in Alderley Park | Wellcome Sanger Institute for the COVID-19 Genomics UK (COG-UK) Consortium | Jacquelyn Wynn, Mairead Hyland, The Lighthouse Lab in Alderley Park and Alex Alderton, Roberto Amato, Sonia Goncalves, Ewan Harrison, David K. Jackson, Ian Johnston, Dominic Kwiatkowski, Cordelia Langford, John Sillitoe on behalf of the Wellcome Sanger Institute COVID-19 Surveillance Team |
| EPI_ISL_718892, EPI_ISL_718893 | Lighthouse Lab in Cambridge | Wellcome Sanger Institute for the COVID-19 Genomics UK (COG-UK) Consortium | Rob Howes, The Lighthouse Lab in Cambridge and Alex Alderton, Roberto Amato, Sonia Goncalves, Ewan Harrison, David K. Jackson, Ian Johnston, Dominic Kwiatkowski, Cordelia Langford, John Sillitoe on behalf of the Wellcome Sanger Institute COVID-19 Surveillance Team |
| EPI_ISL_718894, EPI_ISL_718895 | Lighthouse Lab in Alderley Park | Wellcome Sanger Institute for the COVID-19 Genomics UK (COG-UK) Consortium | Jacquelyn Wynn, Mairead Hyland, The Lighthouse Lab in Alderley Park and Alex Alderton, Roberto Amato, Sonia Goncalves, Ewan Harrison, David K. Jackson, Ian Johnston, Dominic Kwiatkowski, Cordelia Langford, John Sillitoe on behalf of the Wellcome Sanger Institute COVID-19 Surveillance Team |
| EPI_ISL_718896, EPI_ISL_718897, EPI_ISL_718898, EPI_ISL_718899, EPI_ISL_718900 | Lighthouse Lab in Cambridge | Wellcome Sanger Institute for the COVID-19 Genomics UK (COG-UK) Consortium | Rob Howes, The Lighthouse Lab in Cambridge and Alex Alderton, Roberto Amato, Sonia Goncalves, Ewan Harrison, David K. Jackson, Ian Johnston, Dominic Kwiatkowski, Cordelia Langford, John Sillitoe on behalf of the Wellcome Sanger Institute COVID-19 Surveillance Team |
| EPI_ISL_718901, EPI_ISL_718902, EPI_ISL_718903 | Lighthouse Lab in Alderley Park | Wellcome Sanger Institute for the COVID-19 Genomics UK (COG-UK) Consortium | Jacquelyn Wynn, Mairead Hyland, The Lighthouse Lab in Alderley Park and Alex Alderton, Roberto Amato, Sonia Goncalves, Ewan Harrison, David K. Jackson, Ian Johnston, Dominic Kwiatkowski, Cordelia Langford, John Sillitoe on behalf of the Wellcome Sanger Institute COVID-19 Surveillance Team |
| EPI_ISL_718904, EPI_ISL_718905, EPI_ISL_718906 | Lighthouse Lab in Cambridge | Wellcome Sanger Institute for the COVID-19 Genomics UK (COG-UK) Consortium | Rob Howes, The Lighthouse Lab in Cambridge and Alex Alderton, Roberto Amato, Sonia Goncalves, Ewan Harrison, David K. Jackson, Ian Johnston, Dominic Kwiatkowski, Cordelia Langford, John Sillitoe on behalf of the Wellcome Sanger Institute COVID-19 Surveillance Team |
| EPI_ISL_718907, EPI_ISL_718908 | Lighthouse Lab in Alderley Park | Wellcome Sanger Institute for the COVID-19 Genomics UK (COG-UK) Consortium | Jacquelyn Wynn, Mairead Hyland, The Lighthouse Lab in Alderley Park and Alex Alderton, Roberto Amato, Sonia Goncalves, Ewan Harrison, David K. Jackson, Ian Johnston, Dominic Kwiatkowski, Cordelia Langford, John Sillitoe on behalf of the Wellcome Sanger Institute COVID-19 Surveillance Team |
| EPI_ISL_718909, EPI_ISL_718910 | Lighthouse Lab in Cambridge | Wellcome Sanger Institute for the COVID-19 Genomics UK (COG-UK) Consortium | Rob Howes, The Lighthouse Lab in Cambridge and Alex Alderton, Roberto Amato, Sonia Goncalves, Ewan Harrison, David K. Jackson, Ian Johnston, Dominic Kwiatkowski, Cordelia Langford, John Sillitoe on behalf of the Wellcome Sanger Institute COVID-19 Surveillance Team |
| EPI_ISL_718911 | Lighthouse Lab in Alderley Park | Wellcome Sanger Institute for the COVID-19 Genomics UK (COG-UK) Consortium | Jacquelyn Wynn, Mairead Hyland, The Lighthouse Lab in Alderley Park and Alex Alderton, Roberto Amato, Sonia Goncalves, Ewan Harrison, David K. Jackson, Ian Johnston, Dominic Kwiatkowski, Cordelia Langford, John Sillitoe on behalf of the Wellcome Sanger Institute COVID-19 Surveillance Team |
| EPI_ISL_718912, EPI_ISL_718913 | Lighthouse Lab in Cambridge | Wellcome Sanger Institute for the COVID-19 Genomics UK (COG-UK) Consortium | Rob Howes, The Lighthouse Lab in Cambridge and Alex Alderton, Roberto Amato, Sonia Goncalves, Ewan Harrison, David K. Jackson, Ian Johnston, Dominic Kwiatkowski, Cordelia Langford, John Sillitoe on behalf of the Wellcome Sanger Institute COVID-19 Surveillance Team |
| EPI_ISL_718914, EPI_ISL_718915 | Lighthouse Lab in Alderley Park | Wellcome Sanger Institute for the COVID-19 Genomics UK (COG-UK) Consortium | Jacquelyn Wynn, Mairead Hyland, The Lighthouse Lab in Alderley Park and Alex Alderton, Roberto Amato, Sonia Goncalves, Ewan Harrison, David K. Jackson, Ian Johnston, Dominic Kwiatkowski, Cordelia Langford, John Sillitoe on behalf of the Wellcome Sanger Institute COVID-19 Surveillance Team |
| EPI_ISL_718916 | Lighthouse Lab in Cambridge | Wellcome Sanger Institute for the COVID-19 Genomics UK (COG-UK) Consortium | Rob Howes, The Lighthouse Lab in Cambridge and Alex Alderton, Roberto Amato, Sonia Goncalves, Ewan Harrison, David K. Jackson, Ian Johnston, Dominic Kwiatkowski, Cordelia Langford, John Sillitoe on behalf of the Wellcome Sanger Institute COVID-19 Surveillance Team |
| EPI_ISL_718917 | Lighthouse Lab in Alderley Park | Wellcome Sanger Institute for the COVID-19 Genomics UK (COG-UK) Consortium | Jacquelyn Wynn, Mairead Hyland, The Lighthouse Lab in Alderley Park and Alex Alderton, Roberto Amato, Sonia Goncalves, Ewan Harrison, David K. Jackson, Ian Johnston, Dominic Kwiatkowski, Cordelia Langford, John Sillitoe on behalf of the Wellcome Sanger Institute COVID-19 Surveillance Team |
| EPI_ISL_718918 | Lighthouse Lab in Cambridge | Wellcome Sanger Institute for the COVID-19 Genomics UK (COG-UK) Consortium | Rob Howes, The Lighthouse Lab in Cambridge and Alex Alderton, Roberto Amato, Sonia Goncalves, Ewan Harrison, David K. Jackson, Ian Johnston, Dominic Kwiatkowski, Cordelia Langford, John Sillitoe on behalf of the Wellcome Sanger Institute COVID-19 Surveillance Team |
| EPI_ISL_718919 | Lighthouse Lab in Alderley Park | Wellcome Sanger Institute for the COVID-19 Genomics UK (COG-UK) Consortium | Jacquelyn Wynn, Mairead Hyland, The Lighthouse Lab in Alderley Park and Alex Alderton, Roberto Amato, Sonia Goncalves, Ewan Harrison, David K. Jackson, Ian Johnston, Dominic Kwiatkowski, Cordelia Langford, John Sillitoe on behalf of the Wellcome Sanger Institute COVID-19 Surveillance Team |
| EPI_ISL_718920, EPI_ISL_718921 | Lighthouse Lab in Cambridge | Wellcome Sanger Institute for the COVID-19 Genomics UK (COG-UK) Consortium | Rob Howes, The Lighthouse Lab in Cambridge and Alex Alderton, Roberto Amato, Sonia Goncalves, Ewan Harrison, David K. Jackson, Ian Johnston, Dominic Kwiatkowski, Cordelia Langford, John Sillitoe on behalf of the Wellcome Sanger Institute COVID-19 Surveillance Team |
| EPI_ISL_718922, EPI_ISL_718923 | Lighthouse Lab in Alderley Park | Wellcome Sanger Institute for the COVID-19 Genomics UK (COG-UK) Consortium | Jacquelyn Wynn, Mairead Hyland, The Lighthouse Lab in Alderley Park and Alex Alderton, Roberto Amato, Sonia Goncalves, Ewan Harrison, David K. Jackson, Ian Johnston, Dominic Kwiatkowski, Cordelia Langford, John Sillitoe on behalf of the Wellcome Sanger Institute COVID-19 Surveillance Team |
| EPI_ISL_718924, EPI_ISL_718925 | Lighthouse Lab in Cambridge | Wellcome Sanger Institute for the COVID-19 Genomics UK (COG-UK) Consortium | Rob Howes, The Lighthouse Lab in Cambridge and Alex Alderton, Roberto Amato, Sonia Goncalves, Ewan Harrison, David K. Jackson, Ian Johnston, Dominic Kwiatkowski, Cordelia Langford, John Sillitoe on behalf of the Wellcome Sanger Institute COVID-19 Surveillance Team |
| EPI_ISL_718926 | Lighthouse Lab in Alderley Park | Wellcome Sanger Institute for the COVID-19 Genomics UK (COG-UK) Consortium | Jacquelyn Wynn, Mairead Hyland, The Lighthouse Lab in Alderley Park and Alex Alderton, Roberto Amato, Sonia Goncalves, Ewan Harrison, David K. Jackson, Ian Johnston, Dominic Kwiatkowski, Cordelia Langford, John Sillitoe on behalf of the Wellcome Sanger Institute COVID-19 Surveillance Team |
| EPI_ISL_718927, EPI_ISL_718928 | Lighthouse Lab in Cambridge | Wellcome Sanger Institute for the COVID-19 Genomics UK (COG-UK) Consortium | Rob Howes, The Lighthouse Lab in Cambridge and Alex Alderton, Roberto Amato, Sonia Goncalves, Ewan Harrison, David K. Jackson, Ian Johnston, Dominic Kwiatkowski, Cordelia Langford, John Sillitoe on behalf of the Wellcome Sanger Institute COVID-19 Surveillance Team |
| EPI_ISL_718929, EPI_ISL_718930 | Lighthouse Lab in Alderley Park | Wellcome Sanger Institute for the COVID-19 Genomics UK (COG-UK) Consortium | Jacquelyn Wynn, Mairead Hyland, The Lighthouse Lab in Alderley Park and Alex Alderton, Roberto Amato, Sonia Goncalves, Ewan Harrison, David K. Jackson, Ian Johnston, Dominic Kwiatkowski, Cordelia Langford, John Sillitoe on behalf of the Wellcome Sanger Institute COVID-19 Surveillance Team |
| EPI_ISL_718931 | Lighthouse Lab in Cambridge | Wellcome Sanger Institute for the COVID-19 Genomics UK (COG-UK) Consortium | Rob Howes, The Lighthouse Lab in Cambridge and Alex Alderton, Roberto Amato, Sonia Goncalves, Ewan Harrison, David K. Jackson, Ian Johnston, Dominic Kwiatkowski, Cordelia Langford, John Sillitoe on behalf of the Wellcome Sanger Institute COVID-19 Surveillance Team |
| EPI_ISL_718932 | Lighthouse Lab in Alderley Park | Wellcome Sanger Institute for the COVID-19 Genomics UK (COG-UK) Consortium | Jacquelyn Wynn, Mairead Hyland, The Lighthouse Lab in Alderley Park and Alex Alderton, Roberto Amato, Sonia Goncalves, Ewan Harrison, David K. Jackson, Ian Johnston, Dominic Kwiatkowski, Cordelia Langford, John Sillitoe on behalf of the Wellcome Sanger Institute COVID-19 Surveillance Team |
| EPI_ISL_718933, EPI_ISL_718934 | Lighthouse Lab in Cambridge | Wellcome Sanger Institute for the COVID-19 Genomics UK (COG-UK) Consortium | Rob Howes, The Lighthouse Lab in Cambridge and Alex Alderton, Roberto Amato, Sonia Goncalves, Ewan Harrison, David K. Jackson, Ian Johnston, Dominic Kwiatkowski, Cordelia Langford, John Sillitoe on behalf of the Wellcome Sanger Institute COVID-19 Surveillance Team |
| EPI_ISL_718935, EPI_ISL_718936 | Lighthouse Lab in Alderley Park | Wellcome Sanger Institute for the COVID-19 Genomics UK (COG-UK) Consortium | Jacquelyn Wynn, Mairead Hyland, The Lighthouse Lab in Alderley Park and Alex Alderton, Roberto Amato, Sonia Goncalves, Ewan Harrison, David K. Jackson, Ian Johnston, Dominic Kwiatkowski, Cordelia Langford, John Sillitoe on behalf of the Wellcome Sanger Institute COVID-19 Surveillance Team |
| EPI_ISL_718937, EPI_ISL_718938 | Lighthouse Lab in Cambridge | Wellcome Sanger Institute for the COVID-19 Genomics UK (COG-UK) Consortium | Rob Howes, The Lighthouse Lab in Cambridge and Alex Alderton, Roberto Amato, Sonia Goncalves, Ewan Harrison, David K. Jackson, Ian Johnston, Dominic Kwiatkowski, Cordelia Langford, John Sillitoe on behalf of the Wellcome Sanger Institute COVID-19 Surveillance Team |
| EPI_ISL_718939 | Lighthouse Lab in Alderley Park | Wellcome Sanger Institute for the COVID-19 Genomics UK (COG-UK) Consortium | Jacquelyn Wynn, Mairead Hyland, The Lighthouse Lab in Alderley Park and Alex Alderton, Roberto Amato, Sonia Goncalves, Ewan Harrison, David K. Jackson, Ian Johnston, Dominic Kwiatkowski, Cordelia Langford, John Sillitoe on behalf of the Wellcome Sanger Institute COVID-19 Surveillance Team |
| EPI_ISL_718940, EPI_ISL_718941, EPI_ISL_718942 | Lighthouse Lab in Cambridge | Wellcome Sanger Institute for the COVID-19 Genomics UK (COG-UK) Consortium | Rob Howes, The Lighthouse Lab in Cambridge and Alex Alderton, Roberto Amato, Sonia Goncalves, Ewan Harrison, David K. Jackson, Ian Johnston, Dominic Kwiatkowski, Cordelia Langford, John Sillitoe on behalf of the Wellcome Sanger Institute COVID-19 Surveillance Team |
| EPI_ISL_718943 | Lighthouse Lab in Alderley Park | Wellcome Sanger Institute for the COVID-19 Genomics UK (COG-UK) Consortium | Jacquelyn Wynn, Mairead Hyland, The Lighthouse Lab in Alderley Park and Alex Alderton, Roberto Amato, Sonia Goncalves, Ewan Harrison, David K. Jackson, Ian Johnston, Dominic Kwiatkowski, Cordelia Langford, John Sillitoe on behalf of the Wellcome Sanger Institute COVID-19 Surveillance Team |

| | | | |
|---|---|---|---|
| EPI_ISL_718944 | Lighthouse Lab in Cambridge | Wellcome Sanger Institute for the COVID-19 Genomics UK (COG-UK) Consortium | Rob Howes, The Lighthouse Lab in Cambridge and Alex Alderton, Roberto Amato, Sonia Goncalves, Ewan Harrison, David K. Jackson, Ian Johnston, Dominic Kwiatkowski, Cordelia Langford, John Sillitoe on behalf of the Wellcome Sanger Institute COVID-19 Surveillance Team |
| EPI_ISL_718945 | Lighthouse Lab in Alderley Park | Wellcome Sanger Institute for the COVID-19 Genomics UK (COG-UK) Consortium | Jacquelyn Wynn, Mairead Hyland, The Lighthouse Lab in Alderley Park and Alex Alderton, Roberto Amato, Sonia Goncalves, Ewan Harrison, David K. Jackson, Ian Johnston, Dominic Kwiatkowski, Cordelia Langford, John Sillitoe on behalf of the Wellcome Sanger Institute COVID-19 Surveillance Team |
| EPI_ISL_718946, EPI_ISL_718947, EPI_ISL_718948, EPI_ISL_718949 | Lighthouse Lab in Cambridge | Wellcome Sanger Institute for the COVID-19 Genomics UK (COG-UK) Consortium | Rob Howes, The Lighthouse Lab in Cambridge and Alex Alderton, Roberto Amato, Sonia Goncalves, Ewan Harrison, David K. Jackson, Ian Johnston, Dominic Kwiatkowski, Cordelia Langford, John Sillitoe on behalf of the Wellcome Sanger Institute COVID-19 Surveillance Team |
| EPI_ISL_718950 | Lighthouse Lab in Alderley Park | Wellcome Sanger Institute for the COVID-19 Genomics UK (COG-UK) Consortium | Jacquelyn Wynn, Mairead Hyland, The Lighthouse Lab in Alderley Park and Alex Alderton, Roberto Amato, Sonia Goncalves, Ewan Harrison, David K. Jackson, Ian Johnston, Dominic Kwiatkowski, Cordelia Langford, John Sillitoe on behalf of the Wellcome Sanger Institute COVID-19 Surveillance Team |
| EPI_ISL_718951 | Lighthouse Lab in Cambridge | Wellcome Sanger Institute for the COVID-19 Genomics UK (COG-UK) Consortium | Rob Howes, The Lighthouse Lab in Cambridge and Alex Alderton, Roberto Amato, Sonia Goncalves, Ewan Harrison, David K. Jackson, Ian Johnston, Dominic Kwiatkowski, Cordelia Langford, John Sillitoe on behalf of the Wellcome Sanger Institute COVID-19 Surveillance Team |
| EPI_ISL_718952, EPI_ISL_718953 | Lighthouse Lab in Alderley Park | Wellcome Sanger Institute for the COVID-19 Genomics UK (COG-UK) Consortium | Jacquelyn Wynn, Mairead Hyland, The Lighthouse Lab in Alderley Park and Alex Alderton, Roberto Amato, Sonia Goncalves, Ewan Harrison, David K. Jackson, Ian Johnston, Dominic Kwiatkowski, Cordelia Langford, John Sillitoe on behalf of the Wellcome Sanger Institute COVID-19 Surveillance Team |
| EPI_ISL_718954 | Lighthouse Lab in Cambridge | Wellcome Sanger Institute for the COVID-19 Genomics UK (COG-UK) Consortium | Rob Howes, The Lighthouse Lab in Cambridge and Alex Alderton, Roberto Amato, Sonia Goncalves, Ewan Harrison, David K. Jackson, Ian Johnston, Dominic Kwiatkowski, Cordelia Langford, John Sillitoe on behalf of the Wellcome Sanger Institute COVID-19 Surveillance Team |
| EPI_ISL_718955 | Lighthouse Lab in Alderley Park | Wellcome Sanger Institute for the COVID-19 Genomics UK (COG-UK) Consortium | Jacquelyn Wynn, Mairead Hyland, The Lighthouse Lab in Alderley Park and Alex Alderton, Roberto Amato, Sonia Goncalves, Ewan Harrison, David K. Jackson, Ian Johnston, Dominic Kwiatkowski, Cordelia Langford, John Sillitoe on behalf of the Wellcome Sanger Institute COVID-19 Surveillance Team |
| EPI_ISL_718956 | Lighthouse Lab in Cambridge | Wellcome Sanger Institute for the COVID-19 Genomics UK (COG-UK) Consortium | Rob Howes, The Lighthouse Lab in Cambridge and Alex Alderton, Roberto Amato, Sonia Goncalves, Ewan Harrison, David K. Jackson, Ian Johnston, Dominic Kwiatkowski, Cordelia Langford, John Sillitoe on behalf of the Wellcome Sanger Institute COVID-19 Surveillance Team |
| EPI_ISL_718957 | Lighthouse Lab in Alderley Park | Wellcome Sanger Institute for the COVID-19 Genomics UK (COG-UK) Consortium | Jacquelyn Wynn, Mairead Hyland, The Lighthouse Lab in Alderley Park and Alex Alderton, Roberto Amato, Sonia Goncalves, Ewan Harrison, David K. Jackson, Ian Johnston, Dominic Kwiatkowski, Cordelia Langford, John Sillitoe on behalf of the Wellcome Sanger Institute COVID-19 Surveillance Team |
| EPI_ISL_718958 | Lighthouse Lab in Cambridge | Wellcome Sanger Institute for the COVID-19 Genomics UK (COG-UK) Consortium | Rob Howes, The Lighthouse Lab in Cambridge and Alex Alderton, Roberto Amato, Sonia Goncalves, Ewan Harrison, David K. Jackson, Ian Johnston, Dominic Kwiatkowski, Cordelia Langford, John Sillitoe on behalf of the Wellcome Sanger Institute COVID-19 Surveillance Team |
| EPI_ISL_718959 | Lighthouse Lab in Alderley Park | Wellcome Sanger Institute for the COVID-19 Genomics UK (COG-UK) Consortium | Jacquelyn Wynn, Mairead Hyland, The Lighthouse Lab in Alderley Park and Alex Alderton, Roberto Amato, Sonia Goncalves, Ewan Harrison, David K. Jackson, Ian Johnston, Dominic Kwiatkowski, Cordelia Langford, John Sillitoe on behalf of the Wellcome Sanger Institute COVID-19 Surveillance Team |
| EPI_ISL_718960, EPI_ISL_718961 | Lighthouse Lab in Cambridge | Wellcome Sanger Institute for the COVID-19 Genomics UK (COG-UK) Consortium | Rob Howes, The Lighthouse Lab in Cambridge and Alex Alderton, Roberto Amato, Sonia Goncalves, Ewan Harrison, David K. Jackson, Ian Johnston, Dominic Kwiatkowski, Cordelia Langford, John Sillitoe on behalf of the Wellcome Sanger Institute COVID-19 Surveillance Team |
| EPI_ISL_718962, EPI_ISL_718963, EPI_ISL_718964 | Lighthouse Lab in Alderley Park | Wellcome Sanger Institute for the COVID-19 Genomics UK (COG-UK) Consortium | Jacquelyn Wynn, Mairead Hyland, The Lighthouse Lab in Alderley Park and Alex Alderton, Roberto Amato, Sonia Goncalves, Ewan Harrison, David K. Jackson, Ian Johnston, Dominic Kwiatkowski, Cordelia Langford, John Sillitoe on behalf of the Wellcome Sanger Institute COVID-19 Surveillance Team |
| EPI_ISL_718965 | Lighthouse Lab in Cambridge | Wellcome Sanger Institute for the COVID-19 Genomics UK (COG-UK) Consortium | Rob Howes, The Lighthouse Lab in Cambridge and Alex Alderton, Roberto Amato, Sonia Goncalves, Ewan Harrison, David K. Jackson, Ian Johnston, Dominic Kwiatkowski, Cordelia Langford, John Sillitoe on behalf of the Wellcome Sanger Institute COVID-19 Surveillance Team |
| EPI_ISL_718966, EPI_ISL_718967 | Lighthouse Lab in Alderley Park | Wellcome Sanger Institute for the COVID-19 Genomics UK (COG-UK) Consortium | Jacquelyn Wynn, Mairead Hyland, The Lighthouse Lab in Alderley Park and Alex Alderton, Roberto Amato, Sonia Goncalves, Ewan Harrison, David K. Jackson, Ian Johnston, Dominic Kwiatkowski, Cordelia Langford, John Sillitoe on behalf of the Wellcome Sanger Institute COVID-19 Surveillance Team |
| EPI_ISL_718968, EPI_ISL_718969 | Lighthouse Lab in Cambridge | Wellcome Sanger Institute for the COVID-19 Genomics UK (COG-UK) Consortium | Rob Howes, The Lighthouse Lab in Cambridge and Alex Alderton, Roberto Amato, Sonia Goncalves, Ewan Harrison, David K. Jackson, Ian Johnston, Dominic Kwiatkowski, Cordelia Langford, John Sillitoe on behalf of the Wellcome Sanger Institute COVID-19 Surveillance Team |
| EPI_ISL_718970 | Lighthouse Lab in Alderley Park | Wellcome Sanger Institute for the COVID-19 Genomics UK (COG-UK) Consortium | Jacquelyn Wynn, Mairead Hyland, The Lighthouse Lab in Alderley Park and Alex Alderton, Roberto Amato, Sonia Goncalves, Ewan Harrison, David K. Jackson, Ian Johnston, Dominic Kwiatkowski, Cordelia Langford, John Sillitoe on behalf of the Wellcome Sanger Institute COVID-19 Surveillance Team |
| EPI_ISL_718971, EPI_ISL_718972, EPI_ISL_718973 | Lighthouse Lab in Cambridge | Wellcome Sanger Institute for the COVID-19 Genomics UK (COG-UK) Consortium | Rob Howes, The Lighthouse Lab in Cambridge and Alex Alderton, Roberto Amato, Sonia Goncalves, Ewan Harrison, David K. Jackson, Ian Johnston, Dominic Kwiatkowski, Cordelia Langford, John Sillitoe on behalf of the Wellcome Sanger Institute COVID-19 Surveillance Team |
| EPI_ISL_718974, EPI_ISL_718975, EPI_ISL_718976, EPI_ISL_718977, EPI_ISL_718978, EPI_ISL_718979 | Lighthouse Lab in Alderley Park | Wellcome Sanger Institute for the COVID-19 Genomics UK (COG-UK) Consortium | Jacquelyn Wynn, Mairead Hyland, The Lighthouse Lab in Alderley Park and Alex Alderton, Roberto Amato, Sonia Goncalves, Ewan Harrison, David K. Jackson, Ian Johnston, Dominic Kwiatkowski, Cordelia Langford, John Sillitoe on behalf of the Wellcome Sanger Institute COVID-19 Surveillance Team |
| EPI_ISL_718980, EPI_ISL_718981, EPI_ISL_718982, EPI_ISL_718983, EPI_ISL_718984, EPI_ISL_718985, EPI_ISL_718986, EPI_ISL_718987, EPI_ISL_718988, EPI_ISL_718989, EPI_ISL_718990, EPI_ISL_718991, EPI_ISL_718992, EPI_ISL_718993, EPI_ISL_718994, EPI_ISL_718995, EPI_ISL_718996, EPI_ISL_718997, EPI_ISL_718998, EPI_ISL_718999, EPI_ISL_719000, EPI_ISL_719001, EPI_ISL_719002, EPI_ISL_719003, EPI_ISL_719004, EPI_ISL_719005, EPI_ISL_719006, EPI_ISL_719007, EPI_ISL_719008, EPI_ISL_719009, EPI_ISL_719010, EPI_ISL_719011, EPI_ISL_719012, EPI_ISL_719013, EPI_ISL_719014, EPI_ISL_719015, EPI_ISL_719016, EPI_ISL_719017, EPI_ISL_719018, EPI_ISL_719019, EPI_ISL_719020, EPI_ISL_719021, EPI_ISL_719022, EPI_ISL_719023, EPI_ISL_719024, EPI_ISL_719025, EPI_ISL_719026, EPI_ISL_719027, EPI_ISL_719028, EPI_ISL_719029, EPI_ISL_719030, EPI_ISL_719031, EPI_ISL_719032, EPI_ISL_719033, EPI_ISL_719034, EPI_ISL_719035, EPI_ISL_719036, EPI_ISL_719037, EPI_ISL_719038, EPI_ISL_719039, EPI_ISL_719040, EPI_ISL_719041, EPI_ISL_719042, EPI_ISL_719043, EPI_ISL_719044, EPI_ISL_719045, EPI_ISL_719046, EPI_ISL_719047, EPI_ISL_719048, EPI_ISL_719049, EPI_ISL_719050, EPI_ISL_719051, EPI_ISL_719052, EPI_ISL_719053, EPI_ISL_719054, EPI_ISL_719055, EPI_ISL_719056, EPI_ISL_719057, EPI_ISL_719058, EPI_ISL_719059, EPI_ISL_719060, EPI_ISL_719061, EPI_ISL_719062, EPI_ISL_719063, EPI_ISL_719064, EPI_ISL_719065, EPI_ISL_719066, EPI_ISL_719067, EPI_ISL_719068, EPI_ISL_719069, EPI_ISL_719070, EPI_ISL_719071, EPI_ISL_719072, EPI_ISL_719073, EPI_ISL_719074, EPI_ISL_719075, EPI_ISL_719076, EPI_ISL_719077, EPI_ISL_719078, EPI_ISL_719079, EPI_ISL_719080, EPI_ISL_719081, EPI_ISL_719082, EPI_ISL_719083, EPI_ISL_719084, EPI_ISL_719085, EPI_ISL_719086, EPI_ISL_719087, EPI_ISL_719088, EPI_ISL_719089, EPI_ISL_719090, EPI_ISL_719091, EPI_ISL_719092, EPI_ISL_719093, EPI_ISL_719094, EPI_ISL_719095, EPI_ISL_719096, EPI_ISL_719097, EPI_ISL_719098, EPI_ISL_719099, EPI_ISL_719100, EPI_ISL_719101, EPI_ISL_719102, EPI_ISL_719103, EPI_ISL_719104, EPI_ISL_719105, EPI_ISL_719106, EPI_ISL_719107, EPI_ISL_719108, EPI_ISL_719109, EPI_ISL_719110, EPI_ISL_719111, EPI_ISL_719112, EPI_ISL_719113, EPI_ISL_719114, EPI_ISL_719115, EPI_ISL_719116, EPI_ISL_719117, EPI_ISL_719118, EPI_ISL_719119, EPI_ISL_719120, EPI_ISL_719121, EPI_ISL_719122, EPI_ISL_719123, EPI_ISL_719124, EPI_ISL_719125, EPI_ISL_719126, EPI_ISL_719127, EPI_ISL_719128, EPI_ISL_719129, EPI_ISL_719130, EPI_ISL_719131, EPI_ISL_719132, EPI_ISL_719133, EPI_ISL_719134, EPI_ISL_719135, EPI_ISL_719136, EPI_ISL_719137, EPI_ISL_719138, EPI_ISL_719139, EPI_ISL_719140, EPI_ISL_719141, EPI_ISL_719142, EPI_ISL_719143, EPI_ISL_719144, EPI_ISL_719145, EPI_ISL_719146, EPI_ISL_719147, EPI_ISL_719148, EPI_ISL_719149, EPI_ISL_719150, EPI_ISL_719151, EPI_ISL_719152, EPI_ISL_719153, EPI_ISL_719154, EPI_ISL_719155, EPI_ISL_719156, EPI_ISL_719157, EPI_ISL_719158, EPI_ISL_719159, EPI_ISL_719160, EPI_ISL_719161, EPI_ISL_719162, EPI_ISL_719163, EPI_ISL_719164, EPI_ISL_719165, EPI_ISL_719166, EPI_ISL_719167, EPI_ISL_719168, EPI_ISL_719169, EPI_ISL_719170, EPI_ISL_719171, EPI_ISL_719172, EPI_ISL_719173, EPI_ISL_719174, EPI_ISL_719175, EPI_ISL_719176, EPI_ISL_719177, EPI_ISL_719178, EPI_ISL_719179, EPI_ISL_719180, EPI_ISL_719181, EPI_ISL_719182, EPI_ISL_719183, EPI_ISL_719184, EPI_ISL_719185, EPI_ISL_719186, EPI_ISL_719187, EPI_ISL_719188, EPI_ISL_719189, EPI_ISL_719190, EPI_ISL_719191, EPI_ISL_719192, EPI_ISL_719193, EPI_ISL_719194, EPI_ISL_719195, EPI_ISL_719196, EPI_ISL_719197, EPI_ISL_719198, EPI_ISL_719199, EPI_ISL_719200, EPI_ISL_719201, EPI_ISL_719202, EPI_ISL_719203, EPI_ISL_719204, EPI_ISL_719205, EPI_ISL_719206, EPI_ISL_719207, EPI_ISL_719208, EPI_ISL_719209, EPI_ISL_719210, EPI_ISL_719211, EPI_ISL_719212, EPI_ISL_719213, EPI_ISL_719214, EPI_ISL_719215, EPI_ISL_719216, EPI_ISL_719217, EPI_ISL_719218, EPI_ISL_719219, EPI_ISL_719220, EPI_ISL_719221, EPI_ISL_719222, EPI_ISL_719223, EPI_ISL_719224, EPI_ISL_719225, EPI_ISL_719226, EPI_ISL_719227, EPI_ISL_719228, EPI_ISL_719229, EPI_ISL_719230, EPI_ISL_719231, EPI_ISL_719232, EPI_ISL_719233, EPI_ISL_719234, EPI_ISL_719235, EPI_ISL_719236, EPI_ISL_719237, EPI_ISL_719238, EPI_ISL_719239, EPI_ISL_719240, EPI_ISL_719241, EPI_ISL_719242, EPI_ISL_719243, EPI_ISL_719244, EPI_ISL_719245, EPI_ISL_719246, EPI_ISL_719247, EPI_ISL_719248, EPI_ISL_719249, EPI_ISL_719250, EPI_ISL_719251, EPI_ISL_719252, EPI_ISL_719253, EPI_ISL_719254, EPI_ISL_719255, EPI_ISL_719256, EPI_ISL_719257, EPI_ISL_719258, EPI_ISL_719259, EPI_ISL_719260, EPI_ISL_719261, EPI_ISL_719262, EPI_ISL_719263, EPI_ISL_719264, EPI_ISL_719265, EPI_ISL_719266, EPI_ISL_719267, EPI_ISL_719268, EPI_ISL_719269, EPI_ISL_719270, EPI_ISL_719271, EPI_ISL_719272, EPI_ISL_719273, EPI_ISL_719274, EPI_ISL_719275, EPI_ISL_719276, EPI_ISL_719277, EPI_ISL_719278, EPI_ISL_719279, EPI_ISL_719280, EPI_ISL_719281, EPI_ISL_719282, EPI_ISL_719283, EPI_ISL_719284, EPI_ISL_719285, EPI_ISL_719286, EPI_ISL_719287, EPI_ISL_719288, EPI_ISL_719289, EPI_ISL_719290, EPI_ISL_719291, EPI_ISL_719292, EPI_ISL_719293, EPI_ISL_719294, EPI_ISL_719295, EPI_ISL_719296, EPI_ISL_719297, EPI_ISL_719298, EPI_ISL_719299, EPI_ISL_719300, EPI_ISL_719301, EPI_ISL_719302, EPI_ISL_719303, EPI_ISL_719304, EPI_ISL_719305, EPI_ISL_719306, EPI_ISL_719307, EPI_ISL_719308, EPI_ISL_719309, EPI_ISL_719310, EPI_ISL_719311, EPI_ISL_719312, EPI_ISL_719313, EPI_ISL_719314, EPI_ISL_719315, EPI_ISL_719316, EPI_ISL_719317, EPI_ISL_719318, EPI_ISL_719319, EPI_ISL_719320, EPI_ISL_719321, EPI_ISL_719322, EPI_ISL_719323, EPI_ISL_719324, EPI_ISL_719325, EPI_ISL_719326, EPI_ISL_719327, EPI_ISL_719328, EPI_ISL_719329, EPI_ISL_719330, EPI_ISL_719331, EPI_ISL_719332, EPI_ISL_719333, EPI_ISL_719334, EPI_ISL_719335, EPI_ISL_719336, EPI_ISL_719337, EPI_ISL_719338, EPI_ISL_719339, EPI_ISL_719340, EPI_ISL_719341, EPI_ISL_719342, EPI_ISL_719343, EPI_ISL_719344, EPI_ISL_719345, EPI_ISL_719346, EPI_ISL_719347, EPI_ISL_719348, EPI_ISL_719349, EPI_ISL_719350, EPI_ISL_719351, EPI_ISL_719352, EPI_ISL_719353, EPI_ISL_719354, EPI_ISL_719355, EPI_ISL_719356, EPI_ISL_719357, EPI_ISL_719358, EPI_ISL_719359, EPI_ISL_719360, EPI_ISL_719361, EPI_ISL_719362, EPI_ISL_719363, EPI_ISL_719364, EPI_ISL_719365, EPI_ISL_719366, EPI_ISL_719367, EPI_ISL_719368, EPI_ISL_719369, EPI_ISL_719370, EPI_ISL_719371, EPI_ISL_719372, EPI_ISL_719373, EPI_ISL_719374, EPI_ISL_719375, EPI_ISL_719376, EPI_ISL_719377, EPI_ISL_719378, EPI_ISL_719379, EPI_ISL_719380, EPI_ISL_719381, EPI_ISL_719382, EPI_ISL_719383, EPI_ISL_719384, EPI_ISL_719385, EPI_ISL_719386, EPI_ISL_719387, EPI_ISL_719388, EPI_ISL_719389, EPI_ISL_719390, EPI_ISL_719391, EPI_ISL_719392, EPI_ISL_719393, EPI_ISL_719394, EPI_ISL_719395, EPI_ISL_719396, EPI_ISL_719397, EPI_ISL_719398, EPI_ISL_719399, EPI_ISL_719400, EPI_ISL_719401, EPI_ISL_719402, EPI_ISL_719403, EPI_ISL_719404, EPI_ISL_719405, EPI_ISL_719406, EPI_ISL_719407, EPI_ISL_719408, EPI_ISL_719409, EPI_ISL_719410, EPI_ISL_719411, EPI_ISL_719412, EPI_ISL_719413, EPI_ISL_719414, EPI_ISL_719415, EPI_ISL_719416, EPI_ISL_719417, EPI_ISL_719418, EPI_ISL_719419, EPI_ISL_719420, EPI_ISL_719421, EPI_ISL_719422, EPI_ISL_719423, EPI_ISL_719424, EPI_ISL_719425, EPI_ISL_719426, EPI_ISL_719427, EPI_ISL_719428, EPI_ISL_719429, EPI_ISL_719430, EPI_ISL_719431, EPI_ISL_719432, EPI_ISL_719433, EPI_ISL_719434, EPI_ISL_719435, EPI_ISL_719436, EPI_ISL_719437, EPI_ISL_719438, EPI_ISL_719439, EPI_ISL_719440, EPI_ISL_719441, EPI_ISL_719442, EPI_ISL_719443, EPI_ISL_719444, EPI_ISL_719445, EPI_ISL_719446, EPI_ISL_719447, EPI_ISL_719448, EPI_ISL_719449, EPI_ISL_719450, EPI_ISL_719451, EPI_ISL_719452, EPI_ISL_719453, EPI_ISL_719454, EPI_ISL_719455, EPI_ISL_719456, EPI_ISL_719457, EPI_ISL_719458, EPI_ISL_719459, EPI_ISL_719460, EPI_ISL_719461, EPI_ISL_719462, EPI_ISL_719463, EPI_ISL_719464, EPI_ISL_719465, EPI_ISL_719466, EPI_ISL_719467, EPI_ISL_719468, EPI_ISL_719469, EPI_ISL_719470, EPI_ISL_719471, EPI_ISL_719472, EPI_ISL_719473, EPI_ISL_719474, EPI_ISL_719475, EPI_ISL_719476, EPI_ISL_719477, EPI_ISL_719478, EPI_ISL_719479, EPI_ISL_719480, EPI_ISL_719481, EPI_ISL_719482, EPI_ISL_719483, EPI_ISL_719484, EPI_ISL_719485, EPI_ISL_719486, EPI_ISL_719487, EPI_ISL_719488, EPI_ISL_719489, EPI_ISL_719490, EPI_ISL_719491, EPI_ISL_719492, EPI_ISL_719493, EPI_ISL_719494, EPI_ISL_719495, EPI_ISL_719496, EPI_ISL_719497, EPI_ISL_719498, EPI_ISL_719499, EPI_ISL_719500, EPI_ISL_719501, EPI_ISL_719502, EPI_ISL_719503, EPI_ISL_719504, EPI_ISL_719505, EPI_ISL_719506, EPI_ISL_719507, EPI_ISL_719508, EPI_ISL_719509, EPI_ISL_719510, EPI_ISL_719511, EPI_ISL_719512, EPI_ISL_719513, EPI_ISL_719514, EPI_ISL_719515, EPI_ISL_719516, EPI_ISL_719517, EPI_ISL_719518, EPI_ISL_719519, EPI_ISL_719520, EPI_ISL_719521, EPI_ISL_719522, EPI_ISL_719523, EPI_ISL_719524, EPI_ISL_719525, EPI_ISL_719526, EPI_ISL_719527, EPI_ISL_719528, EPI_ISL_719529, EPI_ISL_719530, EPI_ISL_719531, EPI_ISL_719532, EPI_ISL_719533, EPI_ISL_719534, EPI_ISL_719535, EPI_ISL_719536, EPI_ISL_719537, EPI_ISL_719538, EPI_ISL_719539, EPI_ISL_719540, EPI_ISL_719541, EPI_ISL_719542, EPI_ISL_719543, EPI_ISL_719544, EPI_ISL_719545, EPI_ISL_719546, EPI_ISL_719547, EPI_ISL_719548, EPI_ISL_719549, EPI_ISL_719550, EPI_ISL_719551, EPI_ISL_719552, EPI_ISL_719553, EPI_ISL_719554, EPI_ISL_719555, EPI_ISL_719556, EPI_ISL_719557, EPI_ISL_719558, EPI_ISL_719559, EPI_ISL_719560, EPI_ISL_719561, EPI_ISL_719562, EPI_ISL_719563, EPI_ISL_719564, EPI_ISL_719565, EPI_ISL_719566, EPI_ISL_719567, EPI_ISL_719568, EPI_ISL_719569, EPI_ISL_719570, EPI_ISL_719571, EPI_ISL_719572, EPI_ISL_719573, EPI_ISL_719574, EPI_ISL_719575, EPI_ISL_719576, EPI_ISL_719577, EPI_ISL_719578, EPI_ISL_719579, EPI_ISL_719580, EPI_ISL_719581, EPI_ISL_719582, EPI_ISL_719583, EPI_ISL_719584, EPI_ISL_719585, EPI_ISL_719586, EPI_ISL_719587, EPI_ISL_719588, EPI_ISL_719589, EPI_ISL_719590, EPI_ISL_719591, EPI_ISL_719592, EPI_ISL_719593, EPI_ISL_719594, EPI_ISL_719595, EPI_ISL_719596, EPI_ISL_719597, EPI_ISL_719598, EPI_ISL_719599, EPI_ISL_719600, EPI_ISL_719601, EPI_ISL_719602, EPI_ISL_719603, EPI_ISL_719604 | | | |
| see above | Lighthouse Lab in Cambridge | Wellcome Sanger Institute for the COVID-19 Genomics UK | Rob Howes, The Lighthouse Lab in Cambridge and Alex Alderton, Roberto Amato, Sonia Goncalves, Ewan Harrison, David K. Jackson, Ian Johnston, |

| | | | |
|---|---|---|---|
| | | (COG-UK) Consortium | Dominic Kwiatkowski, Cordelia Langford, John Sillitoe on behalf of the Wellcome Sanger Institute COVID-19 Surveillance Team |
| EPI_ISL_722342, EPI_ISL_722356, EPI_ISL_722381, EPI_ISL_722386, EPI_ISL_722411, EPI_ISL_722433, EPI_ISL_722455, EPI_ISL_722456, EPI_ISL_722457, EPI_ISL_722458, EPI_ISL_722459, EPI_ISL_722460, EPI_ISL_722763, EPI_ISL_722764, EPI_ISL_722765, EPI_ISL_722766, EPI_ISL_722767, EPI_ISL_722768, EPI_ISL_722769, EPI_ISL_722770, EPI_ISL_722771, EPI_ISL_722772, EPI_ISL_722773, EPI_ISL_722774, EPI_ISL_722775, EPI_ISL_722776, EPI_ISL_722777 | | | |
| see above | Dutch COVID-19 response team | Erasmus Medical Center | Bas Oude Munnink, Reina Sikkema, David Nieuwenhuijse, Irina Chestakova, Anne van der Linden, Marjan Boter, Emmanuelle Munger, Corine GeurtsvanKessel, Annemiek van der Eijk, Richard Molenkamp, Marion Koopmans, on behalf of the Dutch national COVID-19 response team. |
| EPI_ISL_722999, EPI_ISL_723000, EPI_ISL_723006, EPI_ISL_723007, EPI_ISL_723015, EPI_ISL_723044 | Respiratory Virus Unit, National Infection Service, Public Health England | COVID-19 Genomics UK (COG-UK) Consortium | PHE Covid Sequencing Team |
| EPI_ISL_723922, EPI_ISL_723923, EPI_ISL_723924, EPI_ISL_723925, EPI_ISL_723926, EPI_ISL_723927, EPI_ISL_723928, EPI_ISL_723929, EPI_ISL_723930, EPI_ISL_723931, EPI_ISL_723932, EPI_ISL_723934, EPI_ISL_723937, EPI_ISL_723938, EPI_ISL_723939, EPI_ISL_723940, EPI_ISL_723941, EPI_ISL_723942, EPI_ISL_723943, EPI_ISL_723944, EPI_ISL_723945, EPI_ISL_723946 | | | |
| see above | Department of Pathology, University of Cambridge | COVID-19 Genomics UK (COG-UK) Consortium | Aminu S. Jahun, Yasmin Chaudhry, Grant Hall, Iliana Georgana, Myra Hosmillo, Martin D. Curran, Malte Pinckert, Surendra Parmar, Ian Goodfellow |
| EPI_ISL_724245, EPI_ISL_724248, EPI_ISL_724249, EPI_ISL_724250, EPI_ISL_724251, EPI_ISL_724252 | Virology Department, Royal Infirmary of Edinburgh, NHS Lothian / School of Biological Sciences, University of Edinburgh / Institute of Genetics and Molecular Medicine, University of Edinburgh | COVID-19 Genomics UK (COG-UK) Consortium | McHugh M, Dewar R, Rooke S, Gallagher M, Balcaza C, O'Toole Á, Scher E, Hill V, McCrone JT, Colquhoun R, Yu X, Jackson B, Rambaut A, Williams TC, Templeton K |
| EPI_ISL_724525, EPI_ISL_724526, EPI_ISL_724527, EPI_ISL_724528, EPI_ISL_724529, EPI_ISL_724530, EPI_ISL_724531, EPI_ISL_724532, EPI_ISL_724533, EPI_ISL_724534, EPI_ISL_724535, EPI_ISL_724536, EPI_ISL_724537, EPI_ISL_724538, EPI_ISL_724539 | | | |
| see above | Liverpool Clinical Laboratories | COVID-19 Genomics UK (COG-UK) Consortium | Sam Haldenby, Anita Lucaci, Steve Paterson, Julian Hiscox, Alistair Darby, M Almsaud, A Alrezaihi, Muhannad Alruwaili, Stuart D Armstrong, Jones Benjamin, Eleanor G Bentley, Anu Chawla, Jordan J Clark, Angela Cowell, Richard Eccles, Isabel García-Dorival, Matthew Gemmell, Alessandro Gerada, PKF Gilmore, Richard Gregory, Ximeng Han, Catherine Hartley, Margaret Hughes, Miren Iturriza-Gomara, James Johnson, L Luu, Jenifer Manson, Charlotte Nelson, Elaine O'Toole, Cassie Olateju, Rebekah Penrice-Randal , Lucille Rainbow, N.P Randle, Trevor Ian Robinson, Parul Sharma, Ghada T Shawli, James P Stewart, Neil Swainston, Ecaterina Vamos, Joanne Watts, Mark Whitehead |
| EPI_ISL_724593, EPI_ISL_724606, EPI_ISL_724607 | University College London, Great Ormond Street Hospital for Children NHS Foundation Trust, Imperial College Healthcare NHS Trust | COVID-19 Genomics UK (COG-UK) Consortium | Sergi Castellano, Rachel Williams, Mark Kristiansen, Paola Resende Silva, Sunando Roy, Tony Brooks, Helena Tutill, Paola Niola, Patricia Dyal, Charlotte Williams, Leysa Forrest, Yasmin Panchbhaya, Jacqueline Findlay, Samuel Weeks, Julianne Brown, Kathryn Harris, Paul Randell, James Price, Alison Holmes, Judith Breuer |
| EPI_ISL_725033, EPI_ISL_725034, EPI_ISL_725035, EPI_ISL_725038, EPI_ISL_725041, EPI_ISL_725055, EPI_ISL_725062, EPI_ISL_725063, EPI_ISL_725064, EPI_ISL_725065, EPI_ISL_725066, EPI_ISL_725067, EPI_ISL_725068, EPI_ISL_725069, EPI_ISL_725070, EPI_ISL_725071, EPI_ISL_725072, EPI_ISL_725073, EPI_ISL_725074, EPI_ISL_725075, EPI_ISL_725076, EPI_ISL_725077, EPI_ISL_725078, EPI_ISL_725079, EPI_ISL_725080, EPI_ISL_725081, EPI_ISL_725082, EPI_ISL_725083, EPI_ISL_725084, EPI_ISL_725085, EPI_ISL_725086, EPI_ISL_725087, EPI_ISL_725088, EPI_ISL_725089, EPI_ISL_725090, EPI_ISL_725091, EPI_ISL_725092, EPI_ISL_725093, EPI_ISL_725094, EPI_ISL_725095, EPI_ISL_725096, EPI_ISL_725097, EPI_ISL_725098, EPI_ISL_725099, EPI_ISL_725100, EPI_ISL_725101, EPI_ISL_725102, EPI_ISL_725103, EPI_ISL_725104, EPI_ISL_725105, EPI_ISL_725151, EPI_ISL_725155, EPI_ISL_725296, EPI_ISL_725297, EPI_ISL_725298, EPI_ISL_725299, EPI_ISL_725300, EPI_ISL_725301, EPI_ISL_725343, EPI_ISL_725344, EPI_ISL_725346, EPI_ISL_725347, EPI_ISL_725348, EPI_ISL_725349, EPI_ISL_725350, EPI_ISL_725351, EPI_ISL_725352, EPI_ISL_725353, EPI_ISL_725354, EPI_ISL_725355, EPI_ISL_725356, EPI_ISL_725357, EPI_ISL_725358, EPI_ISL_725359, EPI_ISL_725360, EPI_ISL_725361, EPI_ISL_725362, EPI_ISL_725363, EPI_ISL_725364, EPI_ISL_725366, EPI_ISL_725367, EPI_ISL_725368, EPI_ISL_725369, EPI_ISL_725417, EPI_ISL_725419, EPI_ISL_725420, EPI_ISL_725422, EPI_ISL_725423, EPI_ISL_725424, EPI_ISL_725428, EPI_ISL_725430, EPI_ISL_725433, EPI_ISL_725438, EPI_ISL_725458, EPI_ISL_725459, EPI_ISL_725463, EPI_ISL_725464, EPI_ISL_725466, EPI_ISL_725467, EPI_ISL_725468, EPI_ISL_725469, EPI_ISL_725471, EPI_ISL_725472, EPI_ISL_725475, EPI_ISL_725476, EPI_ISL_725480, EPI_ISL_725481, EPI_ISL_725487, EPI_ISL_725492, EPI_ISL_725493, EPI_ISL_725494, EPI_ISL_725495 | | | |
| see above | Quadram Institute Bioscience | COVID-19 Genomics UK (COG-UK) Consortium | Dave J. Baker, Gemma L. Kay, Alp Aydin, Thanh Le-Viet, Steven Rudder, Ana P. Tedim, Anastasia Kolyva, Maria Diaz, Leonardo de Oliveira Martins, Nabil-Fareed Alikhan, Lizzie Meadows, Rachael Stanley, Ngozi Elumogo, Muhammed Yasir, Nicholas M. Thomson, Alexander J Trotter, Rachel Gilroy, Samuel Bloomfield, Claire Stuart, Andrew Bell, Reenesh Prakash, Samir Dervisevic, Alison E. Mather, John Wain, Mark Webber, Andrew J. Page, Justin O'Grady |
| EPI_ISL_725643, EPI_ISL_725644, EPI_ISL_725645, EPI_ISL_725646, EPI_ISL_725647, EPI_ISL_725648, EPI_ISL_725649, EPI_ISL_725650, EPI_ISL_725651, EPI_ISL_725652, EPI_ISL_725653, EPI_ISL_725654, EPI_ISL_725655, EPI_ISL_725656, EPI_ISL_725657, EPI_ISL_725658, EPI_ISL_725659, EPI_ISL_725660, EPI_ISL_725661, EPI_ISL_725662, EPI_ISL_725663, EPI_ISL_725664, EPI_ISL_725665, EPI_ISL_725666, EPI_ISL_725667, EPI_ISL_725668, EPI_ISL_725669 | | | |
| see above | Queens Medical Centre, Clinical Microbiology Department / DeepSeq Nottingham | COVID-19 Genomics UK (COG-UK) Consortium | Gemma Clark, Wendy Smith, Manjinder Khakh, Vicki M Fleming, Michelle M Lister, Hannah Howson-Wells, Jonathan Ball, Patrick McClure, Joseph Chappell, Theocharis Tsoleridis, Nadine Holmes, Matthew Carlisle, Christopher Moore, Fei Sang, Johnny Debebe, Victoria Wright, Matthew Loose |
| EPI_ISL_727862, EPI_ISL_727864, EPI_ISL_727866, EPI_ISL_727867, EPI_ISL_727872, EPI_ISL_727873, EPI_ISL_727876, EPI_ISL_727878, EPI_ISL_727881, EPI_ISL_727886, EPI_ISL_727889, EPI_ISL_727893, EPI_ISL_727894, EPI_ISL_727898, EPI_ISL_727899, EPI_ISL_727900, EPI_ISL_727902, EPI_ISL_727904, EPI_ISL_727907, EPI_ISL_727910, EPI_ISL_727912, EPI_ISL_727913, EPI_ISL_727916, EPI_ISL_727921, EPI_ISL_727924, EPI_ISL_727925, EPI_ISL_727926, EPI_ISL_727928, EPI_ISL_727930, EPI_ISL_727933, EPI_ISL_727935, EPI_ISL_727937, EPI_ISL_727944, EPI_ISL_727945, EPI_ISL_727946, EPI_ISL_727952, EPI_ISL_727958, EPI_ISL_727959, EPI_ISL_727962, EPI_ISL_727965, EPI_ISL_727977, EPI_ISL_727984, EPI_ISL_727989 | | | |
| see above | Virology Department, Sheffield Teaching Hospitals NHS Foundation Trust/Department of Infection, Immunity and Cardiovascular Disease, The Medical School, University of Sheffield | COVID-19 Genomics UK (COG-UK) Consortium | Thushan de Silva, Matthew Parker, Nikki Smith, Adri Angyal, Rebecca Brown, Luke Green, Rachel Tucker, Paul Parsons, Danielle Groves, Katie Johnson, Laura Carrilero, Alex Keeley, Dave Partridge, Matthew Wyles, Benjamin Lindsey, Mehmet Yavuz, Mohammad Raza, Cariad Evans |
| EPI_ISL_728192, EPI_ISL_728193, EPI_ISL_728195, EPI_ISL_728196, EPI_ISL_728197, EPI_ISL_728198, EPI_ISL_728199, EPI_ISL_728200 | National Public Health Laboratory, National Centre for Infectious Diseases | National Public Health Laboratory, National Centre for Infectious Diseases | Tze Minn Mak, Sophie Octavia, Zhenyang Zhou, Lin Cui, Raymond Tzer Pin Lin |
| EPI_ISL_728299 | Lighthouse Lab in Cambridge | Wellcome Sanger Institute for the COVID-19 Genomics UK (COG-UK) Consortium | Rob Howes, The Lighthouse Lab in Cambridge and Alex Alderton, Roberto Amato, Sonia Goncalves, Ewan Harrison, David K. Jackson, Ian Johnston, Dominic Kwiatkowski, Cordelia Langford, John Sillitoe on behalf of the Wellcome Sanger Institute COVID-19 Surveillance Team |
| EPI_ISL_728343 | Respiratory Virus Unit, National Infection Service, Public Health England | COVID-19 Genomics UK (COG-UK) Consortium | PHE Covid Sequencing Team |
| EPI_ISL_728568, EPI_ISL_728569, EPI_ISL_728586, EPI_ISL_728592, EPI_ISL_728593, EPI_ISL_728594, EPI_ISL_728614, EPI_ISL_728615, EPI_ISL_728616, EPI_ISL_728617, EPI_ISL_728632, EPI_ISL_728633, EPI_ISL_728634, EPI_ISL_728651, EPI_ISL_728697 | | | |
| see above | Dutch COVID-19 response team | National Institute for Public Health and the Environment (RIVM) | Adam Meijer, Harry Vennema, Jeroen Cremer, Sharon van den Brink, Bas van der Veer, AnneMarie van den Brandt, Florian Zwagemaker, Dennis Schmitz, Chantal Reusken, on behalf of the national COVID-19 response team |
| EPI_ISL_728774 | Viollier AG | Department of Biosystems Science and Engineering, ETH Zürich | Chaoran Chen, Sarah Nadeau, Catharine Aquino, Ivan Topolsky, Pedro Ferreira, Philipp Jablonski, Susana Posada-Céspedes, Andreia Cabral de Gouvea, Maria Domenica Moccia, Simon Grüter, Timothy Sykes, Lennart Opitz, Ralph Schlapbach, Christiane Beckmann, Maurice Redondo, Olivier Kobel, Christoph Noppen, Sophie Seidel, Noemie Santamaria de Souza, Niko Beerenwinkel, Tanja Stadler |
| EPI_ISL_728863, EPI_ISL_728883 | Viollier AG | Department of Biosystems Science and Engineering, ETH Zürich | Christian Beisel, Sarah Nadeau, Chaoran Chen, Ivan Topolsky, Pedro Ferreira, Philipp Jablonski, Susana Posada-Céspedes, Tobias Schär, Ina Nissen, Natascha Santacroce, Elodie Burcklen, Christiane Beckmann, Maurice Redondo, Olivier Kobel, Christoph Noppen, Sophie Seidel, Noemie Santamaria de Souza, Niko Beerenwinkel, Tanja Stadler |
| EPI_ISL_728913, EPI_ISL_728918, EPI_ISL_729007, EPI_ISL_729034, EPI_ISL_729042, EPI_ISL_729047 | Viollier AG | Department of Biosystems Science and Engineering, ETH Zürich | Chaoran Chen, Sarah Nadeau, Catharine Aquino, Ivan Topolsky, Pedro Ferreira, Philipp Jablonski, Susana Posada-Céspedes, Andreia Cabral de Gouvea, Maria Domenica Moccia, Simon Grüter, Timothy Sykes, Lennart Opitz, Ralph Schlapbach, Christiane Beckmann, Maurice Redondo, Olivier Kobel, Christoph Noppen, Sophie Seidel, Noemie Santamaria de Souza, Niko Beerenwinkel, Tanja Stadler |
| EPI_ISL_729049, EPI_ISL_729053, EPI_ISL_729067, EPI_ISL_729088, EPI_ISL_729089, EPI_ISL_729187, EPI_ISL_729188, EPI_ISL_729189 | Viollier AG | Department of Biosystems Science and Engineering, ETH Zürich | Christian Beisel, Sarah Nadeau, Chaoran Chen, Ivan Topolsky, Pedro Ferreira, Philipp Jablonski, Susana Posada-Céspedes, Tobias Schär, Ina Nissen, Natascha Santacroce, Elodie Burcklen, Christiane Beckmann, Maurice Redondo, Olivier Kobel, Christoph Noppen, Sophie Seidel, Noemie Santamaria de Souza, Niko Beerenwinkel, Tanja Stadler |
| EPI_ISL_730655 | Lighthouse Lab in Alderley Park | Wellcome Sanger Institute for the COVID-19 Genomics UK (COG-UK) Consortium | Jacquelyn Wynn, Mairead Hyland, The Lighthouse Lab in Alderley Park and Alex Alderton, Roberto Amato, Sonia Goncalves, Ewan Harrison, David K. Jackson, Ian Johnston, Dominic Kwiatkowski, Cordelia Langford, John Sillitoe on behalf of the Wellcome Sanger Institute COVID-19 Surveillance Team |
| EPI_ISL_730656, EPI_ISL_730657 | Lighthouse Lab in Glasgow | Wellcome Sanger Institute for the COVID-19 Genomics UK (COG-UK) Consortium | Harper VanSteenhouse, Yumi Kasai, David Gray, Carol Clugston, Anna Dominiczak and Alex Alderton, Roberto Amato, Sonia Goncalves, Ewan Harrison, David K. Jackson, Ian Johnston, Dominic Kwiatkowski, Cordelia Langford, John Sillitoe on behalf of the Wellcome Sanger Institute COVID-19 Surveillance |

| | | | |
|---|---|---|---|
| | | | Team |
| EPI_ISL_730658 | Lighthouse Lab in Alderley Park | Wellcome Sanger Institute for the COVID-19 Genomics UK (COG-UK) Consortium | Jacquelyn Wynn, Mairead Hyland, The Lighthouse Lab in Alderley Park and Alex Alderton, Roberto Amato, Sonia Goncalves, Ewan Harrison, David K. Jackson, Ian Johnston, Dominic Kwiatkowski, Cordelia Langford, John Sillitoe on behalf of the Wellcome Sanger Institute COVID-19 Surveillance Team |
| EPI_ISL_730659, EPI_ISL_730664 | Lighthouse Lab in Glasgow | Wellcome Sanger Institute for the COVID-19 Genomics UK (COG-UK) Consortium | Harper VanSteenhouse, Yumi Kasai, David Gray, Carol Clugston, Anna Dominiczak and Alex Alderton, Roberto Amato, Sonia Goncalves, Ewan Harrison, David K. Jackson, Ian Johnston, Dominic Kwiatkowski, Cordelia Langford, John Sillitoe on behalf of the Wellcome Sanger Institute COVID-19 Surveillance Team |
| EPI_ISL_730667, EPI_ISL_730668 | Lighthouse Lab in Alderley Park | Wellcome Sanger Institute for the COVID-19 Genomics UK (COG-UK) Consortium | Jacquelyn Wynn, Mairead Hyland, The Lighthouse Lab in Alderley Park and Alex Alderton, Roberto Amato, Sonia Goncalves, Ewan Harrison, David K. Jackson, Ian Johnston, Dominic Kwiatkowski, Cordelia Langford, John Sillitoe on behalf of the Wellcome Sanger Institute COVID-19 Surveillance Team |
| EPI_ISL_730673, EPI_ISL_730675, EPI_ISL_730676, EPI_ISL_730678 | Lighthouse Lab in Glasgow | Wellcome Sanger Institute for the COVID-19 Genomics UK (COG-UK) Consortium | Harper VanSteenhouse, Yumi Kasai, David Gray, Carol Clugston, Anna Dominiczak and Alex Alderton, Roberto Amato, Sonia Goncalves, Ewan Harrison, David K. Jackson, Ian Johnston, Dominic Kwiatkowski, Cordelia Langford, John Sillitoe on behalf of the Wellcome Sanger Institute COVID-19 Surveillance Team |
| EPI_ISL_730680, EPI_ISL_730682, EPI_ISL_730683 | Lighthouse Lab in Alderley Park | Wellcome Sanger Institute for the COVID-19 Genomics UK (COG-UK) Consortium | Jacquelyn Wynn, Mairead Hyland, The Lighthouse Lab in Alderley Park and Alex Alderton, Roberto Amato, Sonia Goncalves, Ewan Harrison, David K. Jackson, Ian Johnston, Dominic Kwiatkowski, Cordelia Langford, John Sillitoe on behalf of the Wellcome Sanger Institute COVID-19 Surveillance Team |
| EPI_ISL_730685 | Lighthouse Lab in Glasgow | Wellcome Sanger Institute for the COVID-19 Genomics UK (COG-UK) Consortium | Harper VanSteenhouse, Yumi Kasai, David Gray, Carol Clugston, Anna Dominiczak and Alex Alderton, Roberto Amato, Sonia Goncalves, Ewan Harrison, David K. Jackson, Ian Johnston, Dominic Kwiatkowski, Cordelia Langford, John Sillitoe on behalf of the Wellcome Sanger Institute COVID-19 Surveillance Team |
| EPI_ISL_730686, EPI_ISL_730692, EPI_ISL_730695, EPI_ISL_730696 | Lighthouse Lab in Alderley Park | Wellcome Sanger Institute for the COVID-19 Genomics UK (COG-UK) Consortium | Jacquelyn Wynn, Mairead Hyland, The Lighthouse Lab in Alderley Park and Alex Alderton, Roberto Amato, Sonia Goncalves, Ewan Harrison, David K. Jackson, Ian Johnston, Dominic Kwiatkowski, Cordelia Langford, John Sillitoe on behalf of the Wellcome Sanger Institute COVID-19 Surveillance Team |
| EPI_ISL_730697 | Lighthouse Lab in Glasgow | Wellcome Sanger Institute for the COVID-19 Genomics UK (COG-UK) Consortium | Harper VanSteenhouse, Yumi Kasai, David Gray, Carol Clugston, Anna Dominiczak and Alex Alderton, Roberto Amato, Sonia Goncalves, Ewan Harrison, David K. Jackson, Ian Johnston, Dominic Kwiatkowski, Cordelia Langford, John Sillitoe on behalf of the Wellcome Sanger Institute COVID-19 Surveillance Team |
| EPI_ISL_730699, EPI_ISL_730702, EPI_ISL_730704 | Lighthouse Lab in Alderley Park | Wellcome Sanger Institute for the COVID-19 Genomics UK (COG-UK) Consortium | Jacquelyn Wynn, Mairead Hyland, The Lighthouse Lab in Alderley Park and Alex Alderton, Roberto Amato, Sonia Goncalves, Ewan Harrison, David K. Jackson, Ian Johnston, Dominic Kwiatkowski, Cordelia Langford, John Sillitoe on behalf of the Wellcome Sanger Institute COVID-19 Surveillance Team |
| EPI_ISL_730705, EPI_ISL_730709, EPI_ISL_730710 | Lighthouse Lab in Glasgow | Wellcome Sanger Institute for the COVID-19 Genomics UK (COG-UK) Consortium | Harper VanSteenhouse, Yumi Kasai, David Gray, Carol Clugston, Anna Dominiczak and Alex Alderton, Roberto Amato, Sonia Goncalves, Ewan Harrison, David K. Jackson, Ian Johnston, Dominic Kwiatkowski, Cordelia Langford, John Sillitoe on behalf of the Wellcome Sanger Institute COVID-19 Surveillance Team |
| EPI_ISL_730711, EPI_ISL_730712 | Lighthouse Lab in Alderley Park | Wellcome Sanger Institute for the COVID-19 Genomics UK (COG-UK) Consortium | Jacquelyn Wynn, Mairead Hyland, The Lighthouse Lab in Alderley Park and Alex Alderton, Roberto Amato, Sonia Goncalves, Ewan Harrison, David K. Jackson, Ian Johnston, Dominic Kwiatkowski, Cordelia Langford, John Sillitoe on behalf of the Wellcome Sanger Institute COVID-19 Surveillance Team |
| EPI_ISL_730718 | Lighthouse Lab in Glasgow | Wellcome Sanger Institute for the COVID-19 Genomics UK (COG-UK) Consortium | Harper VanSteenhouse, Yumi Kasai, David Gray, Carol Clugston, Anna Dominiczak and Alex Alderton, Roberto Amato, Sonia Goncalves, Ewan Harrison, David K. Jackson, Ian Johnston, Dominic Kwiatkowski, Cordelia Langford, John Sillitoe on behalf of the Wellcome Sanger Institute COVID-19 Surveillance Team |
| EPI_ISL_730722, EPI_ISL_730729, EPI_ISL_730731, EPI_ISL_730732, EPI_ISL_730733, EPI_ISL_730734 | Lighthouse Lab in Alderley Park | Wellcome Sanger Institute for the COVID-19 Genomics UK (COG-UK) Consortium | Jacquelyn Wynn, Mairead Hyland, The Lighthouse Lab in Alderley Park and Alex Alderton, Roberto Amato, Sonia Goncalves, Ewan Harrison, David K. Jackson, Ian Johnston, Dominic Kwiatkowski, Cordelia Langford, John Sillitoe on behalf of the Wellcome Sanger Institute COVID-19 Surveillance Team |
| EPI_ISL_730737 | Lighthouse Lab in Glasgow | Wellcome Sanger Institute for the COVID-19 Genomics UK (COG-UK) Consortium | Harper VanSteenhouse, Yumi Kasai, David Gray, Carol Clugston, Anna Dominiczak and Alex Alderton, Roberto Amato, Sonia Goncalves, Ewan Harrison, David K. Jackson, Ian Johnston, Dominic Kwiatkowski, Cordelia Langford, John Sillitoe on behalf of the Wellcome Sanger Institute COVID-19 Surveillance Team |
| EPI_ISL_730742, EPI_ISL_730743, EPI_ISL_730750, EPI_ISL_730751 | Lighthouse Lab in Alderley Park | Wellcome Sanger Institute for the COVID-19 Genomics UK (COG-UK) Consortium | Jacquelyn Wynn, Mairead Hyland, The Lighthouse Lab in Alderley Park and Alex Alderton, Roberto Amato, Sonia Goncalves, Ewan Harrison, David K. Jackson, Ian Johnston, Dominic Kwiatkowski, Cordelia Langford, John Sillitoe on behalf of the Wellcome Sanger Institute COVID-19 Surveillance Team |
| EPI_ISL_730752 | Lighthouse Lab in Glasgow | Wellcome Sanger Institute for the COVID-19 Genomics UK (COG-UK) Consortium | Harper VanSteenhouse, Yumi Kasai, David Gray, Carol Clugston, Anna Dominiczak and Alex Alderton, Roberto Amato, Sonia Goncalves, Ewan Harrison, David K. Jackson, Ian Johnston, Dominic Kwiatkowski, Cordelia Langford, John Sillitoe on behalf of the Wellcome Sanger Institute COVID-19 Surveillance Team |
| EPI_ISL_730753, EPI_ISL_730754, EPI_ISL_730758 | Lighthouse Lab in Alderley Park | Wellcome Sanger Institute for the COVID-19 Genomics UK (COG-UK) Consortium | Jacquelyn Wynn, Mairead Hyland, The Lighthouse Lab in Alderley Park and Alex Alderton, Roberto Amato, Sonia Goncalves, Ewan Harrison, David K. Jackson, Ian Johnston, Dominic Kwiatkowski, Cordelia Langford, John Sillitoe on behalf of the Wellcome Sanger Institute COVID-19 Surveillance Team |
| EPI_ISL_730759, EPI_ISL_730765 | Lighthouse Lab in Glasgow | Wellcome Sanger Institute for the COVID-19 Genomics UK (COG-UK) Consortium | Harper VanSteenhouse, Yumi Kasai, David Gray, Carol Clugston, Anna Dominiczak and Alex Alderton, Roberto Amato, Sonia Goncalves, Ewan Harrison, David K. Jackson, Ian Johnston, Dominic Kwiatkowski, Cordelia Langford, John Sillitoe on behalf of the Wellcome Sanger Institute COVID-19 Surveillance Team |
| EPI_ISL_730771, EPI_ISL_730773, EPI_ISL_730774 | Lighthouse Lab in Alderley Park | Wellcome Sanger Institute for the COVID-19 Genomics UK (COG-UK) Consortium | Jacquelyn Wynn, Mairead Hyland, The Lighthouse Lab in Alderley Park and Alex Alderton, Roberto Amato, Sonia Goncalves, Ewan Harrison, David K. Jackson, Ian Johnston, Dominic Kwiatkowski, Cordelia Langford, John Sillitoe on behalf of the Wellcome Sanger Institute COVID-19 Surveillance Team |
| EPI_ISL_730777, EPI_ISL_730778, EPI_ISL_730780 | Lighthouse Lab in Glasgow | Wellcome Sanger Institute for the COVID-19 Genomics UK (COG-UK) Consortium | Harper VanSteenhouse, Yumi Kasai, David Gray, Carol Clugston, Anna Dominiczak and Alex Alderton, Roberto Amato, Sonia Goncalves, Ewan Harrison, David K. Jackson, Ian Johnston, Dominic Kwiatkowski, Cordelia Langford, John Sillitoe on behalf of the Wellcome Sanger Institute COVID-19 Surveillance Team |
| EPI_ISL_730784 | Lighthouse Lab in Alderley Park | Wellcome Sanger Institute for the COVID-19 Genomics UK (COG-UK) Consortium | Jacquelyn Wynn, Mairead Hyland, The Lighthouse Lab in Alderley Park and Alex Alderton, Roberto Amato, Sonia Goncalves, Ewan Harrison, David K. Jackson, Ian Johnston, Dominic Kwiatkowski, Cordelia Langford, John Sillitoe on behalf of the Wellcome Sanger Institute COVID-19 Surveillance Team |
| EPI_ISL_730785 | Lighthouse Lab in Glasgow | Wellcome Sanger Institute for the COVID-19 Genomics UK (COG-UK) Consortium | Harper VanSteenhouse, Yumi Kasai, David Gray, Carol Clugston, Anna Dominiczak and Alex Alderton, Roberto Amato, Sonia Goncalves, Ewan Harrison, David K. Jackson, Ian Johnston, Dominic Kwiatkowski, Cordelia Langford, John Sillitoe on behalf of the Wellcome Sanger Institute COVID-19 Surveillance Team |
| EPI_ISL_730786, EPI_ISL_730788 | Lighthouse Lab in Alderley Park | Wellcome Sanger Institute for the COVID-19 Genomics UK (COG-UK) Consortium | Jacquelyn Wynn, Mairead Hyland, The Lighthouse Lab in Alderley Park and Alex Alderton, Roberto Amato, Sonia Goncalves, Ewan Harrison, David K. Jackson, Ian Johnston, Dominic Kwiatkowski, Cordelia Langford, John Sillitoe on behalf of the Wellcome Sanger Institute COVID-19 Surveillance Team |
| EPI_ISL_730789, EPI_ISL_730793 | Lighthouse Lab in Glasgow | Wellcome Sanger Institute for the COVID-19 Genomics UK (COG-UK) Consortium | Harper VanSteenhouse, Yumi Kasai, David Gray, Carol Clugston, Anna Dominiczak and Alex Alderton, Roberto Amato, Sonia Goncalves, Ewan Harrison, David K. Jackson, Ian Johnston, Dominic Kwiatkowski, Cordelia Langford, John Sillitoe on behalf of the Wellcome Sanger Institute COVID-19 Surveillance Team |
| EPI_ISL_730794 | Lighthouse Lab in Alderley Park | Wellcome Sanger Institute for the COVID-19 Genomics UK (COG-UK) Consortium | Jacquelyn Wynn, Mairead Hyland, The Lighthouse Lab in Alderley Park and Alex Alderton, Roberto Amato, Sonia Goncalves, Ewan Harrison, David K. Jackson, Ian Johnston, Dominic Kwiatkowski, Cordelia Langford, John Sillitoe on behalf of the Wellcome Sanger Institute COVID-19 Surveillance Team |
| EPI_ISL_730795, EPI_ISL_730796, EPI_ISL_730801, EPI_ISL_730802, EPI_ISL_730803, EPI_ISL_730804, EPI_ISL_730805, EPI_ISL_730806, EPI_ISL_730809, EPI_ISL_730813 | Lighthouse Lab in Glasgow | Wellcome Sanger Institute for the COVID-19 Genomics UK (COG-UK) Consortium | Harper VanSteenhouse, Yumi Kasai, David Gray, Carol Clugston, Anna Dominiczak and Alex Alderton, Roberto Amato, Sonia Goncalves, Ewan Harrison, David K. Jackson, Ian Johnston, Dominic Kwiatkowski, Cordelia Langford, John Sillitoe on behalf of the Wellcome Sanger Institute COVID-19 Surveillance Team |
| EPI_ISL_730814 | Lighthouse Lab in Alderley Park | Wellcome Sanger Institute for the COVID-19 Genomics UK (COG-UK) Consortium | Jacquelyn Wynn, Mairead Hyland, The Lighthouse Lab in Alderley Park and Alex Alderton, Roberto Amato, Sonia Goncalves, Ewan Harrison, David K. Jackson, Ian Johnston, Dominic Kwiatkowski, Cordelia Langford, John Sillitoe on behalf of the Wellcome Sanger Institute COVID-19 Surveillance Team |
| EPI_ISL_730815 | Lighthouse Lab in Glasgow | Wellcome Sanger Institute for the COVID-19 Genomics UK (COG-UK) Consortium | Harper VanSteenhouse, Yumi Kasai, David Gray, Carol Clugston, Anna Dominiczak and Alex Alderton, Roberto Amato, Sonia Goncalves, Ewan Harrison, David K. Jackson, Ian Johnston, Dominic Kwiatkowski, Cordelia Langford, John Sillitoe on behalf of the Wellcome Sanger Institute COVID-19 Surveillance |

| | | | |
|---|---|---|---|
| | | | Team |
| EPI_ISL_730818, EPI_ISL_730819, EPI_ISL_730823 | Lighthouse Lab in Alderley Park | Wellcome Sanger Institute for the COVID-19 Genomics UK (COG-UK) Consortium | Jacquelyn Wynn, Mairead Hyland, The Lighthouse Lab in Alderley Park and Alex Alderton, Roberto Amato, Sonia Goncalves, Ewan Harrison, David K. Jackson, Ian Johnston, Dominic Kwiatkowski, Cordelia Langford, John Sillitoe on behalf of the Wellcome Sanger Institute COVID-19 Surveillance Team |
| EPI_ISL_730824 | Lighthouse Lab in Glasgow | Wellcome Sanger Institute for the COVID-19 Genomics UK (COG-UK) Consortium | Harper VanSteenhouse, Yumi Kasai, David Gray, Carol Clugston, Anna Dominiczak and Alex Alderton, Roberto Amato, Sonia Goncalves, Ewan Harrison, David K. Jackson, Ian Johnston, Dominic Kwiatkowski, Cordelia Langford, John Sillitoe on behalf of the Wellcome Sanger Institute COVID-19 Surveillance Team |
| EPI_ISL_730825, EPI_ISL_730826 | Lighthouse Lab in Alderley Park | Wellcome Sanger Institute for the COVID-19 Genomics UK (COG-UK) Consortium | Jacquelyn Wynn, Mairead Hyland, The Lighthouse Lab in Alderley Park and Alex Alderton, Roberto Amato, Sonia Goncalves, Ewan Harrison, David K. Jackson, Ian Johnston, Dominic Kwiatkowski, Cordelia Langford, John Sillitoe on behalf of the Wellcome Sanger Institute COVID-19 Surveillance Team |
| EPI_ISL_730828 | Lighthouse Lab in Glasgow | Wellcome Sanger Institute for the COVID-19 Genomics UK (COG-UK) Consortium | Harper VanSteenhouse, Yumi Kasai, David Gray, Carol Clugston, Anna Dominiczak and Alex Alderton, Roberto Amato, Sonia Goncalves, Ewan Harrison, David K. Jackson, Ian Johnston, Dominic Kwiatkowski, Cordelia Langford, John Sillitoe on behalf of the Wellcome Sanger Institute COVID-19 Surveillance Team |
| EPI_ISL_730829, EPI_ISL_730830 | Lighthouse Lab in Alderley Park | Wellcome Sanger Institute for the COVID-19 Genomics UK (COG-UK) Consortium | Jacquelyn Wynn, Mairead Hyland, The Lighthouse Lab in Alderley Park and Alex Alderton, Roberto Amato, Sonia Goncalves, Ewan Harrison, David K. Jackson, Ian Johnston, Dominic Kwiatkowski, Cordelia Langford, John Sillitoe on behalf of the Wellcome Sanger Institute COVID-19 Surveillance Team |
| EPI_ISL_730831, EPI_ISL_730832, EPI_ISL_730833 | Lighthouse Lab in Glasgow | Wellcome Sanger Institute for the COVID-19 Genomics UK (COG-UK) Consortium | Harper VanSteenhouse, Yumi Kasai, David Gray, Carol Clugston, Anna Dominiczak and Alex Alderton, Roberto Amato, Sonia Goncalves, Ewan Harrison, David K. Jackson, Ian Johnston, Dominic Kwiatkowski, Cordelia Langford, John Sillitoe on behalf of the Wellcome Sanger Institute COVID-19 Surveillance Team |
| EPI_ISL_730834, EPI_ISL_730839 | Lighthouse Lab in Alderley Park | Wellcome Sanger Institute for the COVID-19 Genomics UK (COG-UK) Consortium | Jacquelyn Wynn, Mairead Hyland, The Lighthouse Lab in Alderley Park and Alex Alderton, Roberto Amato, Sonia Goncalves, Ewan Harrison, David K. Jackson, Ian Johnston, Dominic Kwiatkowski, Cordelia Langford, John Sillitoe on behalf of the Wellcome Sanger Institute COVID-19 Surveillance Team |
| EPI_ISL_730840 | Lighthouse Lab in Glasgow | Wellcome Sanger Institute for the COVID-19 Genomics UK (COG-UK) Consortium | Harper VanSteenhouse, Yumi Kasai, David Gray, Carol Clugston, Anna Dominiczak and Alex Alderton, Roberto Amato, Sonia Goncalves, Ewan Harrison, David K. Jackson, Ian Johnston, Dominic Kwiatkowski, Cordelia Langford, John Sillitoe on behalf of the Wellcome Sanger Institute COVID-19 Surveillance Team |
| EPI_ISL_730844, EPI_ISL_730845 | Lighthouse Lab in Alderley Park | Wellcome Sanger Institute for the COVID-19 Genomics UK (COG-UK) Consortium | Jacquelyn Wynn, Mairead Hyland, The Lighthouse Lab in Alderley Park and Alex Alderton, Roberto Amato, Sonia Goncalves, Ewan Harrison, David K. Jackson, Ian Johnston, Dominic Kwiatkowski, Cordelia Langford, John Sillitoe on behalf of the Wellcome Sanger Institute COVID-19 Surveillance Team |
| EPI_ISL_730846, EPI_ISL_730847 | Lighthouse Lab in Glasgow | Wellcome Sanger Institute for the COVID-19 Genomics UK (COG-UK) Consortium | Harper VanSteenhouse, Yumi Kasai, David Gray, Carol Clugston, Anna Dominiczak and Alex Alderton, Roberto Amato, Sonia Goncalves, Ewan Harrison, David K. Jackson, Ian Johnston, Dominic Kwiatkowski, Cordelia Langford, John Sillitoe on behalf of the Wellcome Sanger Institute COVID-19 Surveillance Team |
| EPI_ISL_730848 | Lighthouse Lab in Alderley Park | Wellcome Sanger Institute for the COVID-19 Genomics UK (COG-UK) Consortium | Jacquelyn Wynn, Mairead Hyland, The Lighthouse Lab in Alderley Park and Alex Alderton, Roberto Amato, Sonia Goncalves, Ewan Harrison, David K. Jackson, Ian Johnston, Dominic Kwiatkowski, Cordelia Langford, John Sillitoe on behalf of the Wellcome Sanger Institute COVID-19 Surveillance Team |
| EPI_ISL_730849, EPI_ISL_730850 | Lighthouse Lab in Glasgow | Wellcome Sanger Institute for the COVID-19 Genomics UK (COG-UK) Consortium | Harper VanSteenhouse, Yumi Kasai, David Gray, Carol Clugston, Anna Dominiczak and Alex Alderton, Roberto Amato, Sonia Goncalves, Ewan Harrison, David K. Jackson, Ian Johnston, Dominic Kwiatkowski, Cordelia Langford, John Sillitoe on behalf of the Wellcome Sanger Institute COVID-19 Surveillance Team |
| EPI_ISL_730852, EPI_ISL_730854, EPI_ISL_730855 | Lighthouse Lab in Alderley Park | Wellcome Sanger Institute for the COVID-19 Genomics UK (COG-UK) Consortium | Jacquelyn Wynn, Mairead Hyland, The Lighthouse Lab in Alderley Park and Alex Alderton, Roberto Amato, Sonia Goncalves, Ewan Harrison, David K. Jackson, Ian Johnston, Dominic Kwiatkowski, Cordelia Langford, John Sillitoe on behalf of the Wellcome Sanger Institute COVID-19 Surveillance Team |
| EPI_ISL_730858 | Lighthouse Lab in Glasgow | Wellcome Sanger Institute for the COVID-19 Genomics UK (COG-UK) Consortium | Harper VanSteenhouse, Yumi Kasai, David Gray, Carol Clugston, Anna Dominiczak and Alex Alderton, Roberto Amato, Sonia Goncalves, Ewan Harrison, David K. Jackson, Ian Johnston, Dominic Kwiatkowski, Cordelia Langford, John Sillitoe on behalf of the Wellcome Sanger Institute COVID-19 Surveillance Team |
| EPI_ISL_730859 | Lighthouse Lab in Alderley Park | Wellcome Sanger Institute for the COVID-19 Genomics UK (COG-UK) Consortium | Jacquelyn Wynn, Mairead Hyland, The Lighthouse Lab in Alderley Park and Alex Alderton, Roberto Amato, Sonia Goncalves, Ewan Harrison, David K. Jackson, Ian Johnston, Dominic Kwiatkowski, Cordelia Langford, John Sillitoe on behalf of the Wellcome Sanger Institute COVID-19 Surveillance Team |
| EPI_ISL_730860 | Lighthouse Lab in Glasgow | Wellcome Sanger Institute for the COVID-19 Genomics UK (COG-UK) Consortium | Harper VanSteenhouse, Yumi Kasai, David Gray, Carol Clugston, Anna Dominiczak and Alex Alderton, Roberto Amato, Sonia Goncalves, Ewan Harrison, David K. Jackson, Ian Johnston, Dominic Kwiatkowski, Cordelia Langford, John Sillitoe on behalf of the Wellcome Sanger Institute COVID-19 Surveillance Team |
| EPI_ISL_730861 | Lighthouse Lab in Alderley Park | Wellcome Sanger Institute for the COVID-19 Genomics UK (COG-UK) Consortium | Jacquelyn Wynn, Mairead Hyland, The Lighthouse Lab in Alderley Park and Alex Alderton, Roberto Amato, Sonia Goncalves, Ewan Harrison, David K. Jackson, Ian Johnston, Dominic Kwiatkowski, Cordelia Langford, John Sillitoe on behalf of the Wellcome Sanger Institute COVID-19 Surveillance Team |
| EPI_ISL_730862, EPI_ISL_730864 | Lighthouse Lab in Glasgow | Wellcome Sanger Institute for the COVID-19 Genomics UK (COG-UK) Consortium | Harper VanSteenhouse, Yumi Kasai, David Gray, Carol Clugston, Anna Dominiczak and Alex Alderton, Roberto Amato, Sonia Goncalves, Ewan Harrison, David K. Jackson, Ian Johnston, Dominic Kwiatkowski, Cordelia Langford, John Sillitoe on behalf of the Wellcome Sanger Institute COVID-19 Surveillance Team |
| EPI_ISL_730865 | Lighthouse Lab in Alderley Park | Wellcome Sanger Institute for the COVID-19 Genomics UK (COG-UK) Consortium | Jacquelyn Wynn, Mairead Hyland, The Lighthouse Lab in Alderley Park and Alex Alderton, Roberto Amato, Sonia Goncalves, Ewan Harrison, David K. Jackson, Ian Johnston, Dominic Kwiatkowski, Cordelia Langford, John Sillitoe on behalf of the Wellcome Sanger Institute COVID-19 Surveillance Team |
| EPI_ISL_730867, EPI_ISL_730869 | Lighthouse Lab in Glasgow | Wellcome Sanger Institute for the COVID-19 Genomics UK (COG-UK) Consortium | Harper VanSteenhouse, Yumi Kasai, David Gray, Carol Clugston, Anna Dominiczak and Alex Alderton, Roberto Amato, Sonia Goncalves, Ewan Harrison, David K. Jackson, Ian Johnston, Dominic Kwiatkowski, Cordelia Langford, John Sillitoe on behalf of the Wellcome Sanger Institute COVID-19 Surveillance Team |
| EPI_ISL_730870, EPI_ISL_730873 | Lighthouse Lab in Alderley Park | Wellcome Sanger Institute for the COVID-19 Genomics UK (COG-UK) Consortium | Jacquelyn Wynn, Mairead Hyland, The Lighthouse Lab in Alderley Park and Alex Alderton, Roberto Amato, Sonia Goncalves, Ewan Harrison, David K. Jackson, Ian Johnston, Dominic Kwiatkowski, Cordelia Langford, John Sillitoe on behalf of the Wellcome Sanger Institute COVID-19 Surveillance Team |
| EPI_ISL_730875, EPI_ISL_730876, EPI_ISL_730877 | Lighthouse Lab in Glasgow | Wellcome Sanger Institute for the COVID-19 Genomics UK (COG-UK) Consortium | Harper VanSteenhouse, Yumi Kasai, David Gray, Carol Clugston, Anna Dominiczak and Alex Alderton, Roberto Amato, Sonia Goncalves, Ewan Harrison, David K. Jackson, Ian Johnston, Dominic Kwiatkowski, Cordelia Langford, John Sillitoe on behalf of the Wellcome Sanger Institute COVID-19 Surveillance Team |
| EPI_ISL_730878, EPI_ISL_730881, EPI_ISL_730886 | Lighthouse Lab in Alderley Park | Wellcome Sanger Institute for the COVID-19 Genomics UK (COG-UK) Consortium | Jacquelyn Wynn, Mairead Hyland, The Lighthouse Lab in Alderley Park and Alex Alderton, Roberto Amato, Sonia Goncalves, Ewan Harrison, David K. Jackson, Ian Johnston, Dominic Kwiatkowski, Cordelia Langford, John Sillitoe on behalf of the Wellcome Sanger Institute COVID-19 Surveillance Team |
| EPI_ISL_730887, EPI_ISL_730889 | Lighthouse Lab in Glasgow | Wellcome Sanger Institute for the COVID-19 Genomics UK (COG-UK) Consortium | Harper VanSteenhouse, Yumi Kasai, David Gray, Carol Clugston, Anna Dominiczak and Alex Alderton, Roberto Amato, Sonia Goncalves, Ewan Harrison, David K. Jackson, Ian Johnston, Dominic Kwiatkowski, Cordelia Langford, John Sillitoe on behalf of the Wellcome Sanger Institute COVID-19 Surveillance Team |
| EPI_ISL_730890, EPI_ISL_730893 | Lighthouse Lab in Alderley Park | Wellcome Sanger Institute for the COVID-19 Genomics UK (COG-UK) Consortium | Jacquelyn Wynn, Mairead Hyland, The Lighthouse Lab in Alderley Park and Alex Alderton, Roberto Amato, Sonia Goncalves, Ewan Harrison, David K. Jackson, Ian Johnston, Dominic Kwiatkowski, Cordelia Langford, John Sillitoe on behalf of the Wellcome Sanger Institute COVID-19 Surveillance Team |
| EPI_ISL_730894, EPI_ISL_730898 | Lighthouse Lab in Glasgow | Wellcome Sanger Institute for the COVID-19 Genomics UK (COG-UK) Consortium | Harper VanSteenhouse, Yumi Kasai, David Gray, Carol Clugston, Anna Dominiczak and Alex Alderton, Roberto Amato, Sonia Goncalves, Ewan Harrison, David K. Jackson, Ian Johnston, Dominic Kwiatkowski, Cordelia Langford, John Sillitoe on behalf of the Wellcome Sanger Institute COVID-19 Surveillance Team |
| EPI_ISL_730900, EPI_ISL_730902, EPI_ISL_730904 | Lighthouse Lab in Alderley Park | Wellcome Sanger Institute for the COVID-19 Genomics UK (COG-UK) Consortium | Jacquelyn Wynn, Mairead Hyland, The Lighthouse Lab in Alderley Park and Alex Alderton, Roberto Amato, Sonia Goncalves, Ewan Harrison, David K. Jackson, Ian Johnston, Dominic Kwiatkowski, Cordelia Langford, John Sillitoe on behalf of the Wellcome Sanger Institute COVID-19 Surveillance Team |
| EPI_ISL_730908, EPI_ISL_730910, EPI_ISL_730911, EPI_ISL_730913 | Lighthouse Lab in Glasgow | Wellcome Sanger Institute for the COVID-19 Genomics UK (COG-UK) Consortium | Harper VanSteenhouse, Yumi Kasai, David Gray, Carol Clugston, Anna Dominiczak and Alex Alderton, Roberto Amato, Sonia Goncalves, Ewan Harrison, David K. Jackson, Ian Johnston, Dominic Kwiatkowski, Cordelia Langford, John Sillitoe on behalf of the Wellcome Sanger Institute COVID-19 Surveillance Team |

EPI_ISL_730914, EPI_ISL_730917, EPI_ISL_730918, EPI_ISL_730919, EPI_ISL_730920, EPI_ISL_730922, EPI_ISL_730923, EPI_ISL_730925, EPI_ISL_730926, EPI_ISL_730927, EPI_ISL_730928, EPI_ISL_730930, EPI_ISL_730931, EPI_ISL_730932, EPI_ISL_730933, EPI_ISL_730934, EPI_ISL_730936, EPI_ISL_730937, EPI_ISL_730938, EPI_ISL_730940, EPI_ISL_730941, EPI_ISL_730942, EPI_ISL_730943, EPI_ISL_730944, EPI_ISL_730945, EPI_ISL_730946, EPI_ISL_730947, EPI_ISL_730949, EPI_ISL_730950, EPI_ISL_730951, EPI_ISL_730952, EPI_ISL_730953, EPI_ISL_730954, EPI_ISL_730955, EPI_ISL_730956, EPI_ISL_730957,

| | | | |
|---|---|---|---|
| EPI_ISL_730958, EPI_ISL_730960, EPI_ISL_730961, EPI_ISL_730962, EPI_ISL_730963, EPI_ISL_730964, EPI_ISL_730965, EPI_ISL_730968, EPI_ISL_730972, EPI_ISL_730974, EPI_ISL_730975, EPI_ISL_730976, EPI_ISL_730977, EPI_ISL_730978, EPI_ISL_730979, EPI_ISL_730981, EPI_ISL_730982, EPI_ISL_730983, EPI_ISL_730984, EPI_ISL_730985, EPI_ISL_730986, EPI_ISL_730988, EPI_ISL_730990, EPI_ISL_730991, EPI_ISL_730992, EPI_ISL_730993, EPI_ISL_730995, EPI_ISL_730996, EPI_ISL_730997, EPI_ISL_730998, EPI_ISL_730999, EPI_ISL_731001, EPI_ISL_731002, EPI_ISL_731003, EPI_ISL_731004, EPI_ISL_731005, EPI_ISL_731007, EPI_ISL_731008, EPI_ISL_731009, EPI_ISL_731010, EPI_ISL_731011, EPI_ISL_731012, EPI_ISL_731013, EPI_ISL_731014, EPI_ISL_731015, EPI_ISL_731016, EPI_ISL_731018, EPI_ISL_731021, EPI_ISL_731022, EPI_ISL_731023, EPI_ISL_731025, EPI_ISL_731028, EPI_ISL_731029, EPI_ISL_731030, EPI_ISL_731031, EPI_ISL_731032, EPI_ISL_731034, EPI_ISL_731036, EPI_ISL_731037, EPI_ISL_731038, EPI_ISL_731039, EPI_ISL_731041, EPI_ISL_731042, EPI_ISL_731043, EPI_ISL_731044, EPI_ISL_731046, EPI_ISL_731047, EPI_ISL_731048, EPI_ISL_731049, EPI_ISL_731051, EPI_ISL_731053, EPI_ISL_731056, EPI_ISL_731058, EPI_ISL_731060, EPI_ISL_731061, EPI_ISL_731062, EPI_ISL_731063, EPI_ISL_731064, EPI_ISL_731067, EPI_ISL_731068, EPI_ISL_731069, EPI_ISL_731070, EPI_ISL_731071, EPI_ISL_731072, EPI_ISL_731073, EPI_ISL_731075, EPI_ISL_731077, EPI_ISL_731078, EPI_ISL_731081, EPI_ISL_731084, EPI_ISL_731085, EPI_ISL_731089, EPI_ISL_731090, EPI_ISL_731091, EPI_ISL_731092, EPI_ISL_731093, EPI_ISL_731095, EPI_ISL_731097, EPI_ISL_731098, EPI_ISL_731099, EPI_ISL_731100, EPI_ISL_731101, EPI_ISL_731102, EPI_ISL_731103, EPI_ISL_731105, EPI_ISL_731108, EPI_ISL_731109, EPI_ISL_731111, EPI_ISL_731112, EPI_ISL_731113, EPI_ISL_731114, EPI_ISL_731115, EPI_ISL_731117, EPI_ISL_731118, EPI_ISL_731121, EPI_ISL_731122, EPI_ISL_731124, EPI_ISL_731125, EPI_ISL_731127, EPI_ISL_731128, EPI_ISL_731132, EPI_ISL_731134, EPI_ISL_731136, EPI_ISL_731138, EPI_ISL_731139, EPI_ISL_731140, EPI_ISL_731142, EPI_ISL_731144, EPI_ISL_731145, EPI_ISL_731146, EPI_ISL_731148, EPI_ISL_731149, EPI_ISL_731150, EPI_ISL_731151, EPI_ISL_731152, EPI_ISL_731155, EPI_ISL_731156, EPI_ISL_731159, EPI_ISL_731162, EPI_ISL_731163, EPI_ISL_731164, EPI_ISL_731165, EPI_ISL_731166, EPI_ISL_731167, EPI_ISL_731168, EPI_ISL_731170, EPI_ISL_731172, EPI_ISL_731173, EPI_ISL_731174, EPI_ISL_731175, EPI_ISL_731176, EPI_ISL_731177, EPI_ISL_731178, EPI_ISL_731179, EPI_ISL_731180, EPI_ISL_731182, EPI_ISL_731184, EPI_ISL_731185, EPI_ISL_731186, EPI_ISL_731187, EPI_ISL_731189, EPI_ISL_731190, EPI_ISL_731191, EPI_ISL_731192, EPI_ISL_731193, EPI_ISL_731194, EPI_ISL_731195, EPI_ISL_731197, EPI_ISL_731198, EPI_ISL_731201, EPI_ISL_731202, EPI_ISL_731203, EPI_ISL_731205, EPI_ISL_731207, EPI_ISL_731208, EPI_ISL_731209, EPI_ISL_731210, EPI_ISL_731211, EPI_ISL_731212, EPI_ISL_731213, EPI_ISL_731216, EPI_ISL_731217, EPI_ISL_731218, EPI_ISL_731219, EPI_ISL_731220, EPI_ISL_731221, EPI_ISL_731222, EPI_ISL_731223, EPI_ISL_731224, EPI_ISL_731225, EPI_ISL_731226, EPI_ISL_731227, EPI_ISL_731228, EPI_ISL_731229, EPI_ISL_731230, EPI_ISL_731231, EPI_ISL_731232, EPI_ISL_731234, EPI_ISL_731236, EPI_ISL_731237, EPI_ISL_731238, EPI_ISL_731239, EPI_ISL_731241, EPI_ISL_731242, EPI_ISL_731243, EPI_ISL_731244, EPI_ISL_731245, EPI_ISL_731246, EPI_ISL_731249, EPI_ISL_731250 | | | |
| see above | Lighthouse Lab in Alderley Park | Wellcome Sanger Institute for the COVID-19 Genomics UK (COG-UK) Consortium | Jacquelyn Wynn, Mairead Hyland, The Lighthouse Lab in Alderley Park and Alex Alderton, Roberto Amato, Sonia Goncalves, Ewan Harrison, David K. Jackson, Ian Johnston, Dominic Kwiatkowski, Cordelia Langford, John Sillitoe on behalf of the Wellcome Sanger Institute COVID-19 Surveillance Team |
| EPI_ISL_731255, EPI_ISL_731256, EPI_ISL_731257, EPI_ISL_731258, EPI_ISL_731259, EPI_ISL_731267, EPI_ISL_731271, EPI_ISL_731273, EPI_ISL_731280, EPI_ISL_731281, EPI_ISL_731282, EPI_ISL_731286, EPI_ISL_731287, EPI_ISL_731291, EPI_ISL_731295, EPI_ISL_731297, EPI_ISL_731305, EPI_ISL_731308, EPI_ISL_731309, EPI_ISL_731310, EPI_ISL_731314, EPI_ISL_731319, EPI_ISL_731320, EPI_ISL_731321, EPI_ISL_731322, EPI_ISL_731326, EPI_ISL_731327, EPI_ISL_731330, EPI_ISL_731332, EPI_ISL_731337, EPI_ISL_731338, EPI_ISL_731340, EPI_ISL_731341, EPI_ISL_731345, EPI_ISL_731346, EPI_ISL_731354, EPI_ISL_731355, EPI_ISL_731357, EPI_ISL_731361, EPI_ISL_731367, EPI_ISL_731368, EPI_ISL_731372, EPI_ISL_731373, EPI_ISL_731376, EPI_ISL_731379, EPI_ISL_731380, EPI_ISL_731382, EPI_ISL_731383, EPI_ISL_731386, EPI_ISL_731390, EPI_ISL_731392, EPI_ISL_731397, EPI_ISL_731401, EPI_ISL_731402, EPI_ISL_731403, EPI_ISL_731404, EPI_ISL_731406, EPI_ISL_731407, EPI_ISL_731408, EPI_ISL_731411, EPI_ISL_731414, EPI_ISL_731417, EPI_ISL_731420, EPI_ISL_731425, EPI_ISL_731426, EPI_ISL_731429, EPI_ISL_731431, EPI_ISL_731432, EPI_ISL_731435, EPI_ISL_731437, EPI_ISL_731444, EPI_ISL_731447, EPI_ISL_731449, EPI_ISL_731450, EPI_ISL_731452, EPI_ISL_731453, EPI_ISL_731454, EPI_ISL_731458, EPI_ISL_731469, EPI_ISL_731471, EPI_ISL_731472, EPI_ISL_731476, EPI_ISL_731477, EPI_ISL_731482, EPI_ISL_731484, EPI_ISL_731485, EPI_ISL_731486, EPI_ISL_731488, EPI_ISL_731491, EPI_ISL_731493, EPI_ISL_731494, EPI_ISL_731496, EPI_ISL_731497, EPI_ISL_731499, EPI_ISL_731501, EPI_ISL_731505, EPI_ISL_731507, EPI_ISL_731508, EPI_ISL_731513, EPI_ISL_731517, EPI_ISL_731519, EPI_ISL_731523, EPI_ISL_731536, EPI_ISL_731537, EPI_ISL_731544, EPI_ISL_731548, EPI_ISL_731550, EPI_ISL_731551 | | | |
| see above | Lighthouse Lab in Glasgow | Wellcome Sanger Institute for the COVID-19 Genomics UK (COG-UK) Consortium | Harper VanSteenhouse, Yumi Kasai, David Gray, Carol Clugston, Anna Dominiczak and Alex Alderton, Roberto Amato, Sonia Goncalves, Ewan Harrison, David K. Jackson, Ian Johnston, Dominic Kwiatkowski, Cordelia Langford, John Sillitoe on behalf of the Wellcome Sanger Institute COVID-19 Surveillance Team |
| EPI_ISL_731552, EPI_ISL_731553, EPI_ISL_731554, EPI_ISL_731556, EPI_ISL_731557, EPI_ISL_731558, EPI_ISL_731559, EPI_ISL_731560, EPI_ISL_731561, EPI_ISL_731562, EPI_ISL_731563, EPI_ISL_731564, EPI_ISL_731566, EPI_ISL_731567, EPI_ISL_731568, EPI_ISL_731569, EPI_ISL_731572, EPI_ISL_731573, EPI_ISL_731574, EPI_ISL_731575, EPI_ISL_731577, EPI_ISL_731578, EPI_ISL_731579, EPI_ISL_731581, EPI_ISL_731582, EPI_ISL_731583, EPI_ISL_731586, EPI_ISL_731587, EPI_ISL_731589, EPI_ISL_731590, EPI_ISL_731591, EPI_ISL_731593, EPI_ISL_731594, EPI_ISL_731595, EPI_ISL_731596, EPI_ISL_731597, EPI_ISL_731598, EPI_ISL_731600, EPI_ISL_731601, EPI_ISL_731602, EPI_ISL_731603, EPI_ISL_731604, EPI_ISL_731605, EPI_ISL_731606, EPI_ISL_731607, EPI_ISL_731608, EPI_ISL_731609, EPI_ISL_731610, EPI_ISL_731612, EPI_ISL_731613, EPI_ISL_731614, EPI_ISL_731615, EPI_ISL_731616, EPI_ISL_731617, EPI_ISL_731618, EPI_ISL_731619, EPI_ISL_731620, EPI_ISL_731621, EPI_ISL_731622, EPI_ISL_731623, EPI_ISL_731625, EPI_ISL_731627, EPI_ISL_731628, EPI_ISL_731630, EPI_ISL_731631, EPI_ISL_731633, EPI_ISL_731634, EPI_ISL_731636, EPI_ISL_731637, EPI_ISL_731638, EPI_ISL_731642, EPI_ISL_731643, EPI_ISL_731645, EPI_ISL_731646, EPI_ISL_731650, EPI_ISL_731651, EPI_ISL_731652, EPI_ISL_731653, EPI_ISL_731654, EPI_ISL_731657, EPI_ISL_731658, EPI_ISL_731659, EPI_ISL_731660, EPI_ISL_731661, EPI_ISL_731662, EPI_ISL_731663, EPI_ISL_731666, EPI_ISL_731668, EPI_ISL_731669, EPI_ISL_731670, EPI_ISL_731672, EPI_ISL_731673, EPI_ISL_731674, EPI_ISL_731675, EPI_ISL_731676, EPI_ISL_731677, EPI_ISL_731678, EPI_ISL_731679, EPI_ISL_731681, EPI_ISL_731682, EPI_ISL_731683, EPI_ISL_731685, EPI_ISL_731687, EPI_ISL_731689, EPI_ISL_731694, EPI_ISL_731695, EPI_ISL_731696, EPI_ISL_731698, EPI_ISL_731699, EPI_ISL_731700, EPI_ISL_731702, EPI_ISL_731703, EPI_ISL_731704, EPI_ISL_731706, EPI_ISL_731708, EPI_ISL_731710, EPI_ISL_731712, EPI_ISL_731713, EPI_ISL_731715, EPI_ISL_731716, EPI_ISL_731717, EPI_ISL_731718, EPI_ISL_731720, EPI_ISL_731723, EPI_ISL_731725, EPI_ISL_731726, EPI_ISL_731727, EPI_ISL_731728, EPI_ISL_731729, EPI_ISL_731730, EPI_ISL_731732, EPI_ISL_731733, EPI_ISL_731734, EPI_ISL_731735, EPI_ISL_731738, EPI_ISL_731739, EPI_ISL_731740, EPI_ISL_731741, EPI_ISL_731742, EPI_ISL_731743, EPI_ISL_731744, EPI_ISL_731745, EPI_ISL_731746, EPI_ISL_731747, EPI_ISL_731748, EPI_ISL_731749, EPI_ISL_731750, EPI_ISL_731751, EPI_ISL_731752, EPI_ISL_731753, EPI_ISL_731754, EPI_ISL_731755, EPI_ISL_731756, EPI_ISL_731757, EPI_ISL_731758, EPI_ISL_731759, EPI_ISL_731760, EPI_ISL_731761, EPI_ISL_731762, EPI_ISL_731764, EPI_ISL_731765, EPI_ISL_731766, EPI_ISL_731768, EPI_ISL_731769, EPI_ISL_731771, EPI_ISL_731772, EPI_ISL_731773, EPI_ISL_731774, EPI_ISL_731776, EPI_ISL_731777, EPI_ISL_731778, EPI_ISL_731779, EPI_ISL_731780, EPI_ISL_731781, EPI_ISL_731782, EPI_ISL_731783, EPI_ISL_731784, EPI_ISL_731786, EPI_ISL_731789, EPI_ISL_731790, EPI_ISL_731791, EPI_ISL_731793, EPI_ISL_731794, EPI_ISL_731797, EPI_ISL_731798, EPI_ISL_731801, EPI_ISL_731802, EPI_ISL_731803, EPI_ISL_731804, EPI_ISL_731805, EPI_ISL_731807, EPI_ISL_731809, EPI_ISL_731811, EPI_ISL_731812, EPI_ISL_731813, EPI_ISL_731814, EPI_ISL_731815, EPI_ISL_731816, EPI_ISL_731817, EPI_ISL_731818, EPI_ISL_731820, EPI_ISL_731821, EPI_ISL_731822, EPI_ISL_731823, EPI_ISL_731825, EPI_ISL_731826, EPI_ISL_731828, EPI_ISL_731829, EPI_ISL_731830, EPI_ISL_731831, EPI_ISL_731832, EPI_ISL_731833, EPI_ISL_731834, EPI_ISL_731835, EPI_ISL_731837, EPI_ISL_731839, EPI_ISL_731843, EPI_ISL_731844, EPI_ISL_731845, EPI_ISL_731847, EPI_ISL_731848, EPI_ISL_731849, EPI_ISL_731851, EPI_ISL_731852, EPI_ISL_731853, EPI_ISL_731855, EPI_ISL_731856, EPI_ISL_731857, EPI_ISL_731859, EPI_ISL_731860, EPI_ISL_731861, EPI_ISL_731862, EPI_ISL_731864, EPI_ISL_731865, EPI_ISL_731866, EPI_ISL_731867, EPI_ISL_731869, EPI_ISL_731870, EPI_ISL_731871, EPI_ISL_731872, EPI_ISL_731873, EPI_ISL_731876, EPI_ISL_731878, EPI_ISL_731879, EPI_ISL_731880, EPI_ISL_731882, EPI_ISL_731883, EPI_ISL_731886, EPI_ISL_731890 | | | |
| see above | Lighthouse Lab in Alderley Park | Wellcome Sanger Institute for the COVID-19 Genomics UK (COG-UK) Consortium | Jacquelyn Wynn, Mairead Hyland, The Lighthouse Lab in Alderley Park and Alex Alderton, Roberto Amato, Sonia Goncalves, Ewan Harrison, David K. Jackson, Ian Johnston, Dominic Kwiatkowski, Cordelia Langford, John Sillitoe on behalf of the Wellcome Sanger Institute COVID-19 Surveillance Team |
| EPI_ISL_732963 | Canterbury Health Laboratories | Institute of Environmental Science and Research (ESR) | Xiaoyun Ren, Matt Storey, Nikki Freed, Muhammad Faisal, Jing Wang, Hermes Perez, Anja Werno, Antje van der Linden, Arlo Upton, Chris Mansell, David Hammer, Dragana Drinkovic, Gary McAuliffe, Hana Sofia Andersson, James Ussher, Jill Sherwood, Josh Freeman, Julia Howard, Juliet Elvy, Mary DeAlmeida, Matt Blakiston, Matthew Rogers, Max Bloomfield, Michael Addidle, Michelle Balm, Sally Roberts, Sarah Jefferies, Sharmini Muttaiyah, Susan Morpeth, Susan Taylor, Timothy Blackmore, Vani Sathyendran, Veronica Playle, Virginia Hope, Erasmus Smit, Lauren Jelly, Olin Silander, Joep de Ligt |
| EPI_ISL_732964, EPI_ISL_732966 | LabPLUS | Institute of Environmental Science and Research (ESR) | Xiaoyun Ren, Matt Storey, Nikki Freed, Muhammad Faisal, Jing Wang, Hermes Perez, Anja Werno, Antje van der Linden, Arlo Upton, Chris Mansell, David Hammer, Dragana Drinkovic, Gary McAuliffe, Hana Sofia Andersson, James Ussher, Jill Sherwood, Josh Freeman, Julia Howard, Juliet Elvy, Mary DeAlmeida, Matt Blakiston, Matthew Rogers, Max Bloomfield, Michael Addidle, Michelle Balm, Sally Roberts, Sarah Jefferies, Sharmini Muttaiyah, Susan Morpeth, Susan Taylor, Timothy Blackmore, Vani Sathyendran, Veronica Playle, Virginia Hope, Erasmus Smit, Lauren Jelly, Olin Silander, Joep de Ligt |
| EPI_ISL_733568 | Centre for Health Protection | Hong Kong Department of Health | Alan K.L. Tsang, Peter C.W. Yip, Edman T.K. Lam, Rickjason C.W. Chan, Dominic N.C. Tsang |
| EPI_ISL_733572, EPI_ISL_733577 | Temporary Specimen Collection Centre | Hong Kong Department of Health | Alan K.L. Tsang, Peter C.W. Yip, Edman T.K. Lam, Rickjason C.W. Chan, Dominic N.C. Tsang |
| EPI_ISL_733916, EPI_ISL_733917, EPI_ISL_733918, EPI_ISL_733919, EPI_ISL_733920, EPI_ISL_733921, EPI_ISL_733922, EPI_ISL_733923, EPI_ISL_733924, EPI_ISL_733925, EPI_ISL_733926, EPI_ISL_733927, EPI_ISL_733929, EPI_ISL_733930, EPI_ISL_733931, EPI_ISL_733932, EPI_ISL_733933, EPI_ISL_733934, EPI_ISL_733935, EPI_ISL_733936, EPI_ISL_733937, EPI_ISL_733938, EPI_ISL_733939, EPI_ISL_733940, EPI_ISL_733941, EPI_ISL_733942, EPI_ISL_733943, EPI_ISL_733944, EPI_ISL_733945, EPI_ISL_733946, EPI_ISL_733947, EPI_ISL_733948, EPI_ISL_733949, EPI_ISL_733951, EPI_ISL_733952, EPI_ISL_733953, EPI_ISL_733954, EPI_ISL_733955, EPI_ISL_733956, EPI_ISL_733957, EPI_ISL_733958, EPI_ISL_733959, EPI_ISL_733960, EPI_ISL_733961, EPI_ISL_733962, EPI_ISL_733963, EPI_ISL_733964, EPI_ISL_733965, EPI_ISL_733966, EPI_ISL_733967, EPI_ISL_733968, EPI_ISL_733969, EPI_ISL_733970, EPI_ISL_733971, EPI_ISL_733972, EPI_ISL_733973, EPI_ISL_733974, EPI_ISL_733975, EPI_ISL_733976, EPI_ISL_733977, EPI_ISL_733978, EPI_ISL_733979, EPI_ISL_733980, EPI_ISL_733981, EPI_ISL_733982, EPI_ISL_733983, EPI_ISL_733984, EPI_ISL_733985, EPI_ISL_733986, EPI_ISL_733987, EPI_ISL_733988, EPI_ISL_733989, EPI_ISL_733990, EPI_ISL_733991, EPI_ISL_733992, EPI_ISL_733993, EPI_ISL_733994, EPI_ISL_733995, EPI_ISL_733996, EPI_ISL_733997, EPI_ISL_733998, EPI_ISL_733999, EPI_ISL_734000, EPI_ISL_734001, EPI_ISL_734002, EPI_ISL_734003, EPI_ISL_734004, EPI_ISL_734005, EPI_ISL_734006, EPI_ISL_734007, EPI_ISL_734008, EPI_ISL_734009, EPI_ISL_734010, EPI_ISL_734011, EPI_ISL_734012, EPI_ISL_734013, EPI_ISL_734014, EPI_ISL_734015, EPI_ISL_734016, EPI_ISL_734017, EPI_ISL_734018, EPI_ISL_734019, EPI_ISL_734020, EPI_ISL_734021, EPI_ISL_734022, EPI_ISL_734023, EPI_ISL_734024, EPI_ISL_734025, EPI_ISL_734026, EPI_ISL_734027, EPI_ISL_734028, EPI_ISL_734029, EPI_ISL_734030, EPI_ISL_734031, EPI_ISL_734032, EPI_ISL_734033, EPI_ISL_734034, EPI_ISL_734035, EPI_ISL_734036, EPI_ISL_734037, EPI_ISL_734038, EPI_ISL_734039, EPI_ISL_734040, EPI_ISL_734042, EPI_ISL_734043, EPI_ISL_734045, EPI_ISL_734046, EPI_ISL_734047, EPI_ISL_734048, EPI_ISL_734049, EPI_ISL_734050, EPI_ISL_734051, EPI_ISL_734052, EPI_ISL_734053, EPI_ISL_734054, EPI_ISL_734055, EPI_ISL_734056, EPI_ISL_734057, EPI_ISL_734058, EPI_ISL_734059, EPI_ISL_734060, EPI_ISL_734061, EPI_ISL_734062, EPI_ISL_734063, EPI_ISL_734064, EPI_ISL_734065, EPI_ISL_734066, EPI_ISL_734067, EPI_ISL_734068, EPI_ISL_734069, EPI_ISL_734070, EPI_ISL_734071, EPI_ISL_734072, EPI_ISL_734073, EPI_ISL_734074, EPI_ISL_734075, EPI_ISL_734076, EPI_ISL_734077, EPI_ISL_734078, EPI_ISL_734079, EPI_ISL_734080, EPI_ISL_734081, EPI_ISL_734082, EPI_ISL_734083, EPI_ISL_734084, EPI_ISL_734085, EPI_ISL_734086, EPI_ISL_734087, EPI_ISL_734088, EPI_ISL_734089, EPI_ISL_734090, EPI_ISL_734091, EPI_ISL_734092, EPI_ISL_734094, EPI_ISL_734095, EPI_ISL_734096, EPI_ISL_734097, EPI_ISL_734098, EPI_ISL_734099, EPI_ISL_734100, EPI_ISL_734101, EPI_ISL_734102, EPI_ISL_734103, EPI_ISL_734104, EPI_ISL_734105, EPI_ISL_734106, EPI_ISL_734107, EPI_ISL_734108, EPI_ISL_734109, EPI_ISL_734110, EPI_ISL_734111, EPI_ISL_734112, EPI_ISL_734113, EPI_ISL_734114, EPI_ISL_734115, EPI_ISL_734116, EPI_ISL_734118, EPI_ISL_734119, EPI_ISL_734120, EPI_ISL_734121, EPI_ISL_734122, EPI_ISL_734123, EPI_ISL_734124, EPI_ISL_734125, EPI_ISL_734126, EPI_ISL_734127, EPI_ISL_734128, EPI_ISL_734129, EPI_ISL_734130, EPI_ISL_734132, EPI_ISL_734133, EPI_ISL_734134, EPI_ISL_734135, EPI_ISL_734136, EPI_ISL_734137, EPI_ISL_734138, EPI_ISL_734139, EPI_ISL_734140, EPI_ISL_734141, EPI_ISL_734142, EPI_ISL_734143, EPI_ISL_734144, EPI_ISL_734145, EPI_ISL_734146, EPI_ISL_734147, EPI_ISL_734148, EPI_ISL_734149, EPI_ISL_734150, EPI_ISL_734151, EPI_ISL_734152, EPI_ISL_734153, EPI_ISL_734154, EPI_ISL_734155, EPI_ISL_734156, EPI_ISL_734157, EPI_ISL_734158, EPI_ISL_734159, EPI_ISL_734160, EPI_ISL_734161, EPI_ISL_734162, EPI_ISL_734163, EPI_ISL_734164, EPI_ISL_734165 | | | |
| see above | Lighthouse Lab in Glasgow | Wellcome Sanger Institute for the COVID-19 Genomics UK (COG-UK) Consortium | Harper VanSteenhouse, Yumi Kasai, David Gray, Carol Clugston, Anna Dominiczak and Alex Alderton, Roberto Amato, Sonia Goncalves, Ewan Harrison, David K. Jackson, Ian Johnston, Dominic Kwiatkowski, Cordelia Langford, John Sillitoe on behalf of the Wellcome Sanger Institute COVID-19 Surveillance Team |
| EPI_ISL_735512, EPI_ISL_735513, EPI_ISL_735514, EPI_ISL_735515, EPI_ISL_735516, EPI_ISL_735517, EPI_ISL_735518, EPI_ISL_735519, EPI_ISL_735520, EPI_ISL_735521, EPI_ISL_735522, EPI_ISL_735523, EPI_ISL_735524, EPI_ISL_735525, EPI_ISL_735526, EPI_ISL_735527, EPI_ISL_735528, EPI_ISL_735529, EPI_ISL_735530, EPI_ISL_735531, EPI_ISL_735532, EPI_ISL_735533, EPI_ISL_735534, EPI_ISL_735535, EPI_ISL_735536, EPI_ISL_735537, EPI_ISL_735538, EPI_ISL_735539, EPI_ISL_735540, EPI_ISL_735541, EPI_ISL_735542, EPI_ISL_735543, EPI_ISL_735544, EPI_ISL_735545, EPI_ISL_735546, EPI_ISL_735547, EPI_ISL_735548, EPI_ISL_735549, EPI_ISL_735550, EPI_ISL_735551, EPI_ISL_735552, EPI_ISL_735553, EPI_ISL_735554, EPI_ISL_735555, EPI_ISL_735556, EPI_ISL_735557, EPI_ISL_735558, EPI_ISL_735559, EPI_ISL_735560, EPI_ISL_735561, EPI_ISL_735562, EPI_ISL_735563, EPI_ISL_735564, EPI_ISL_735565, EPI_ISL_735566, EPI_ISL_735567, EPI_ISL_735568, EPI_ISL_735569, EPI_ISL_735570, EPI_ISL_735571, EPI_ISL_735572, EPI_ISL_735573, EPI_ISL_735574, EPI_ISL_735575, EPI_ISL_735576, EPI_ISL_735577, EPI_ISL_735578, EPI_ISL_735579, EPI_ISL_735580, EPI_ISL_735581, EPI_ISL_735582, EPI_ISL_735583, EPI_ISL_735584, EPI_ISL_735585, EPI_ISL_735586, EPI_ISL_735587, EPI_ISL_735588, EPI_ISL_735589, EPI_ISL_735590, EPI_ISL_735591, EPI_ISL_735592, EPI_ISL_735593, EPI_ISL_735594, EPI_ISL_735595, EPI_ISL_735596, EPI_ISL_735597, EPI_ISL_735598, EPI_ISL_735599, EPI_ISL_735600, EPI_ISL_735601, EPI_ISL_735602, EPI_ISL_735603, EPI_ISL_735604, EPI_ISL_735605, EPI_ISL_735606, EPI_ISL_735607, EPI_ISL_735608, EPI_ISL_735609, EPI_ISL_735610, EPI_ISL_735611, EPI_ISL_735612, EPI_ISL_735613, EPI_ISL_735614, EPI_ISL_735615, EPI_ISL_735616, EPI_ISL_735617, EPI_ISL_735618, EPI_ISL_735619, EPI_ISL_735620, EPI_ISL_735621, EPI_ISL_735622, EPI_ISL_735623, EPI_ISL_735624, EPI_ISL_735625, EPI_ISL_735626, EPI_ISL_735627, EPI_ISL_735628, EPI_ISL_735629, EPI_ISL_735630, EPI_ISL_735631, EPI_ISL_735632, EPI_ISL_735633, EPI_ISL_735634, EPI_ISL_735635, EPI_ISL_735636, EPI_ISL_735637, EPI_ISL_735638, EPI_ISL_735639, EPI_ISL_735640, EPI_ISL_735641, EPI_ISL_735642, EPI_ISL_735643, EPI_ISL_735644, EPI_ISL_735645, EPI_ISL_735646, EPI_ISL_735647, EPI_ISL_735648, EPI_ISL_735649, EPI_ISL_735650, EPI_ISL_735651, EPI_ISL_735652, EPI_ISL_735653, EPI_ISL_735654, EPI_ISL_735655, EPI_ISL_735656, EPI_ISL_735657, EPI_ISL_735658, EPI_ISL_735659, EPI_ISL_735660, EPI_ISL_735661, EPI_ISL_735662, EPI_ISL_735663, EPI_ISL_735664, EPI_ISL_735665, EPI_ISL_735666, EPI_ISL_735667, EPI_ISL_735668, EPI_ISL_735669, EPI_ISL_735670, EPI_ISL_735671, EPI_ISL_735672, EPI_ISL_735673, EPI_ISL_735674, EPI_ISL_735675, EPI_ISL_735676, EPI_ISL_735677, EPI_ISL_735678, EPI_ISL_735679, EPI_ISL_735680, EPI_ISL_735681, EPI_ISL_735682, EPI_ISL_735683, EPI_ISL_735684, EPI_ISL_735685, EPI_ISL_735686, EPI_ISL_735687, EPI_ISL_735688, EPI_ISL_735689, EPI_ISL_735690, EPI_ISL_735691, EPI_ISL_735692, EPI_ISL_735693, EPI_ISL_735694, EPI_ISL_735695, EPI_ISL_735696, EPI_ISL_735697, EPI_ISL_735698, EPI_ISL_735699, EPI_ISL_735700, EPI_ISL_735701, EPI_ISL_735702, EPI_ISL_735703, EPI_ISL_735704, EPI_ISL_735705, EPI_ISL_735706, EPI_ISL_735707, EPI_ISL_735708, EPI_ISL_735709, EPI_ISL_735710, EPI_ISL_735711, EPI_ISL_735712, EPI_ISL_735713, EPI_ISL_735714, EPI_ISL_735715, EPI_ISL_735716, EPI_ISL_735717, EPI_ISL_735718, EPI_ISL_735719, EPI_ISL_735720, EPI_ISL_735721, EPI_ISL_735722, EPI_ISL_735723, EPI_ISL_735724, EPI_ISL_735725, EPI_ISL_735726, EPI_ISL_735727, EPI_ISL_735728, EPI_ISL_735729, EPI_ISL_735730, EPI_ISL_735731, EPI_ISL_735732, EPI_ISL_735733, EPI_ISL_735734, EPI_ISL_735735, EPI_ISL_735736, EPI_ISL_735737, EPI_ISL_735738, EPI_ISL_735739, EPI_ISL_735740, EPI_ISL_735741, EPI_ISL_735742, EPI_ISL_735743, EPI_ISL_735744, EPI_ISL_735745, EPI_ISL_735746, EPI_ISL_735747, EPI_ISL_735748, EPI_ISL_735749, EPI_ISL_735750, EPI_ISL_735751, EPI_ISL_735752, EPI_ISL_735753, EPI_ISL_735754, EPI_ISL_735755, EPI_ISL_735756, EPI_ISL_735757, EPI_ISL_735758, EPI_ISL_735759, EPI_ISL_735760, EPI_ISL_735761, EPI_ISL_735762, EPI_ISL_735763, EPI_ISL_735764, EPI_ISL_735765, EPI_ISL_735766, EPI_ISL_735767, EPI_ISL_735768, EPI_ISL_735769, EPI_ISL_735770, EPI_ISL_735771, EPI_ISL_735772, EPI_ISL_735773, EPI_ISL_735774, EPI_ISL_735775, EPI_ISL_735776, EPI_ISL_735777, EPI_ISL_735778, EPI_ISL_735779, EPI_ISL_735780, EPI_ISL_735781, | | | |

| | | | |
|---|---|---|---|
| EPI_ISL_735782, EPI_ISL_735783, EPI_ISL_735784, EPI_ISL_735785, EPI_ISL_735786, EPI_ISL_735787, EPI_ISL_735788, EPI_ISL_735789, EPI_ISL_735790, EPI_ISL_735791, EPI_ISL_735792, EPI_ISL_735793, EPI_ISL_735794, EPI_ISL_735795, EPI_ISL_735796, EPI_ISL_735797, EPI_ISL_735798, EPI_ISL_735799, EPI_ISL_735800, EPI_ISL_735801, EPI_ISL_735802, EPI_ISL_735803, EPI_ISL_735804, EPI_ISL_735805, EPI_ISL_735806, EPI_ISL_735807, EPI_ISL_735808, EPI_ISL_735809, EPI_ISL_735810, EPI_ISL_735811, EPI_ISL_735812, EPI_ISL_735813, EPI_ISL_735814, EPI_ISL_735815, EPI_ISL_735816, EPI_ISL_735817, EPI_ISL_735818, EPI_ISL_735819, EPI_ISL_735820, EPI_ISL_735821, EPI_ISL_735822, EPI_ISL_735823, EPI_ISL_735824, EPI_ISL_735825, EPI_ISL_735826, EPI_ISL_735827, EPI_ISL_735828, EPI_ISL_735829, EPI_ISL_735830, EPI_ISL_735831, EPI_ISL_735832, EPI_ISL_735833, EPI_ISL_735834, EPI_ISL_735835, EPI_ISL_735836, EPI_ISL_735837, EPI_ISL_735838, EPI_ISL_735839, EPI_ISL_735840, EPI_ISL_735841, EPI_ISL_735842, EPI_ISL_735843, EPI_ISL_735844, EPI_ISL_735845, EPI_ISL_735846, EPI_ISL_735847, EPI_ISL_735848, EPI_ISL_735849, EPI_ISL_735850, EPI_ISL_735851, EPI_ISL_735852, EPI_ISL_735853, EPI_ISL_735854, EPI_ISL_735855, EPI_ISL_735856, EPI_ISL_735857, EPI_ISL_735858, EPI_ISL_735859 | | | |
| see above | Lighthouse Lab in Milton Keynes | Wellcome Sanger Institute for the COVID-19 Genomics UK (COG-UK) Consortium | The Lighthouse Lab in Milton Keynes and Alex Alderton, Roberto Amato, Sonia Goncalves, Ewan Harrison, David K. Jackson, Ian Johnston, Dominic Kwiatkowski, Cordelia Langford, John Sillitoe on behalf of the Wellcome Sanger Institute COVID-19 Surveillance Team |
| EPI_ISL_735860, EPI_ISL_735862, EPI_ISL_735863, EPI_ISL_735865, EPI_ISL_735866, EPI_ISL_735869, EPI_ISL_735870, EPI_ISL_735871, EPI_ISL_735872, EPI_ISL_735873, EPI_ISL_735874, EPI_ISL_735875, EPI_ISL_735876, EPI_ISL_735879, EPI_ISL_735883, EPI_ISL_735884, EPI_ISL_735885, EPI_ISL_735890, EPI_ISL_735892, EPI_ISL_735893, EPI_ISL_735895, EPI_ISL_735896, EPI_ISL_735898, EPI_ISL_735903, EPI_ISL_735906, EPI_ISL_735910, EPI_ISL_735911, EPI_ISL_735913, EPI_ISL_735914, EPI_ISL_735915, EPI_ISL_735916, EPI_ISL_735918, EPI_ISL_735919, EPI_ISL_735921, EPI_ISL_735922, EPI_ISL_735925, EPI_ISL_735928, EPI_ISL_735929, EPI_ISL_735933, EPI_ISL_735934, EPI_ISL_735935, EPI_ISL_735940, EPI_ISL_735944, EPI_ISL_735946, EPI_ISL_735947, EPI_ISL_735948, EPI_ISL_735950, EPI_ISL_735953, EPI_ISL_735954, EPI_ISL_735955, EPI_ISL_735957, EPI_ISL_735959, EPI_ISL_735960, EPI_ISL_735962, EPI_ISL_735969, EPI_ISL_735970, EPI_ISL_735971, EPI_ISL_735977, EPI_ISL_735979, EPI_ISL_735984, EPI_ISL_735985, EPI_ISL_735988, EPI_ISL_735989, EPI_ISL_735990, EPI_ISL_735991, EPI_ISL_735994, EPI_ISL_735999, EPI_ISL_736002, EPI_ISL_736004, EPI_ISL_736005, EPI_ISL_736006, EPI_ISL_736009, EPI_ISL_736014, EPI_ISL_736020, EPI_ISL_736022, EPI_ISL_736026, EPI_ISL_736031, EPI_ISL_736032, EPI_ISL_736035, EPI_ISL_736036, EPI_ISL_736040, EPI_ISL_736042, EPI_ISL_736043, EPI_ISL_736044, EPI_ISL_736045, EPI_ISL_736049, EPI_ISL_736053, EPI_ISL_736054, EPI_ISL_736056, EPI_ISL_736058, EPI_ISL_736060, EPI_ISL_736061, EPI_ISL_736062, EPI_ISL_736067, EPI_ISL_736068, EPI_ISL_736069, EPI_ISL_736070, EPI_ISL_736071, EPI_ISL_736073, EPI_ISL_736074, EPI_ISL_736075, EPI_ISL_736076, EPI_ISL_736077, EPI_ISL_736081, EPI_ISL_736082, EPI_ISL_736084, EPI_ISL_736085, EPI_ISL_736086, EPI_ISL_736088, EPI_ISL_736089, EPI_ISL_736094, EPI_ISL_736098, EPI_ISL_736103, EPI_ISL_736106, EPI_ISL_736109, EPI_ISL_736111, EPI_ISL_736112, EPI_ISL_736113, EPI_ISL_736114, EPI_ISL_736116, EPI_ISL_736117, EPI_ISL_736119, EPI_ISL_736120, EPI_ISL_736121, EPI_ISL_736122, EPI_ISL_736123, EPI_ISL_736126, EPI_ISL_736128, EPI_ISL_736129, EPI_ISL_736133, EPI_ISL_736134, EPI_ISL_736136, EPI_ISL_736137, EPI_ISL_736138, EPI_ISL_736139, EPI_ISL_736140, EPI_ISL_736144, EPI_ISL_736145, EPI_ISL_736147, EPI_ISL_736148, EPI_ISL_736150, EPI_ISL_736152, EPI_ISL_736154, EPI_ISL_736155, EPI_ISL_736156, EPI_ISL_736157, EPI_ISL_736160, EPI_ISL_736161, EPI_ISL_736163, EPI_ISL_736164, EPI_ISL_736165, EPI_ISL_736166, EPI_ISL_736170, EPI_ISL_736171, EPI_ISL_736173, EPI_ISL_736174, EPI_ISL_736176, EPI_ISL_736177, EPI_ISL_736179, EPI_ISL_736180, EPI_ISL_736181, EPI_ISL_736185, EPI_ISL_736186, EPI_ISL_736188, EPI_ISL_736192, EPI_ISL_736194, EPI_ISL_736195, EPI_ISL_736196, EPI_ISL_736197, EPI_ISL_736198, EPI_ISL_736200 | | | |
| see above | Lighthouse Lab in Alderley Park | Wellcome Sanger Institute for the COVID-19 Genomics UK (COG-UK) Consortium | Jacquelyn Wynn, Mairead Hyland, The Lighthouse Lab in Alderley Park and Alex Alderton, Roberto Amato, Sonia Goncalves, Ewan Harrison, David K. Jackson, Ian Johnston, Dominic Kwiatkowski, Cordelia Langford, John Sillitoe on behalf of the Wellcome Sanger Institute COVID-19 Surveillance Team |
| EPI_ISL_736201, EPI_ISL_736203 | Lighthouse Lab in Milton Keynes | Wellcome Sanger Institute for the COVID-19 Genomics UK (COG-UK) Consortium | The Lighthouse Lab in Milton Keynes and Alex Alderton, Roberto Amato, Sonia Goncalves, Ewan Harrison, David K. Jackson, Ian Johnston, Dominic Kwiatkowski, Cordelia Langford, John Sillitoe on behalf of the Wellcome Sanger Institute COVID-19 Surveillance Team |
| EPI_ISL_736204, EPI_ISL_736205, EPI_ISL_736206 | Lighthouse Lab in Glasgow | Wellcome Sanger Institute for the COVID-19 Genomics UK (COG-UK) Consortium | Harper VanSteenhouse, Yumi Kasai, David Gray, Carol Clugston, Anna Dominiczak and Alex Alderton, Roberto Amato, Sonia Goncalves, Ewan Harrison, David K. Jackson, Ian Johnston, Dominic Kwiatkowski, Cordelia Langford, John Sillitoe on behalf of the Wellcome Sanger Institute COVID-19 Surveillance Team |
| EPI_ISL_736207, EPI_ISL_736208 | Lighthouse Lab in Milton Keynes | Wellcome Sanger Institute for the COVID-19 Genomics UK (COG-UK) Consortium | The Lighthouse Lab in Milton Keynes and Alex Alderton, Roberto Amato, Sonia Goncalves, Ewan Harrison, David K. Jackson, Ian Johnston, Dominic Kwiatkowski, Cordelia Langford, John Sillitoe on behalf of the Wellcome Sanger Institute COVID-19 Surveillance Team |
| EPI_ISL_736209 | Lighthouse Lab in Glasgow | Wellcome Sanger Institute for the COVID-19 Genomics UK (COG-UK) Consortium | Harper VanSteenhouse, Yumi Kasai, David Gray, Carol Clugston, Anna Dominiczak and Alex Alderton, Roberto Amato, Sonia Goncalves, Ewan Harrison, David K. Jackson, Ian Johnston, Dominic Kwiatkowski, Cordelia Langford, John Sillitoe on behalf of the Wellcome Sanger Institute COVID-19 Surveillance Team |
| EPI_ISL_736210, EPI_ISL_736211 | Lighthouse Lab in Milton Keynes | Wellcome Sanger Institute for the COVID-19 Genomics UK (COG-UK) Consortium | The Lighthouse Lab in Milton Keynes and Alex Alderton, Roberto Amato, Sonia Goncalves, Ewan Harrison, David K. Jackson, Ian Johnston, Dominic Kwiatkowski, Cordelia Langford, John Sillitoe on behalf of the Wellcome Sanger Institute COVID-19 Surveillance Team |
| EPI_ISL_736212 | Lighthouse Lab in Glasgow | Wellcome Sanger Institute for the COVID-19 Genomics UK (COG-UK) Consortium | Harper VanSteenhouse, Yumi Kasai, David Gray, Carol Clugston, Anna Dominiczak and Alex Alderton, Roberto Amato, Sonia Goncalves, Ewan Harrison, David K. Jackson, Ian Johnston, Dominic Kwiatkowski, Cordelia Langford, John Sillitoe on behalf of the Wellcome Sanger Institute COVID-19 Surveillance Team |
| EPI_ISL_736213, EPI_ISL_736214, EPI_ISL_736215, EPI_ISL_736216, EPI_ISL_736217, EPI_ISL_736218, EPI_ISL_736219 | Lighthouse Lab in Milton Keynes | Wellcome Sanger Institute for the COVID-19 Genomics UK (COG-UK) Consortium | The Lighthouse Lab in Milton Keynes and Alex Alderton, Roberto Amato, Sonia Goncalves, Ewan Harrison, David K. Jackson, Ian Johnston, Dominic Kwiatkowski, Cordelia Langford, John Sillitoe on behalf of the Wellcome Sanger Institute COVID-19 Surveillance Team |
| EPI_ISL_736220, EPI_ISL_736221, EPI_ISL_736222, EPI_ISL_736223 | Lighthouse Lab in Glasgow | Wellcome Sanger Institute for the COVID-19 Genomics UK (COG-UK) Consortium | Harper VanSteenhouse, Yumi Kasai, David Gray, Carol Clugston, Anna Dominiczak and Alex Alderton, Roberto Amato, Sonia Goncalves, Ewan Harrison, David K. Jackson, Ian Johnston, Dominic Kwiatkowski, Cordelia Langford, John Sillitoe on behalf of the Wellcome Sanger Institute COVID-19 Surveillance Team |
| EPI_ISL_736224, EPI_ISL_736225, EPI_ISL_736226, EPI_ISL_736227, EPI_ISL_736228, EPI_ISL_736229, EPI_ISL_736230 | Lighthouse Lab in Milton Keynes | Wellcome Sanger Institute for the COVID-19 Genomics UK (COG-UK) Consortium | The Lighthouse Lab in Milton Keynes and Alex Alderton, Roberto Amato, Sonia Goncalves, Ewan Harrison, David K. Jackson, Ian Johnston, Dominic Kwiatkowski, Cordelia Langford, John Sillitoe on behalf of the Wellcome Sanger Institute COVID-19 Surveillance Team |
| EPI_ISL_736231 | Lighthouse Lab in Glasgow | Wellcome Sanger Institute for the COVID-19 Genomics UK (COG-UK) Consortium | Harper VanSteenhouse, Yumi Kasai, David Gray, Carol Clugston, Anna Dominiczak and Alex Alderton, Roberto Amato, Sonia Goncalves, Ewan Harrison, David K. Jackson, Ian Johnston, Dominic Kwiatkowski, Cordelia Langford, John Sillitoe on behalf of the Wellcome Sanger Institute COVID-19 Surveillance Team |
| EPI_ISL_736232, EPI_ISL_736233 | Lighthouse Lab in Milton Keynes | Wellcome Sanger Institute for the COVID-19 Genomics UK (COG-UK) Consortium | The Lighthouse Lab in Milton Keynes and Alex Alderton, Roberto Amato, Sonia Goncalves, Ewan Harrison, David K. Jackson, Ian Johnston, Dominic Kwiatkowski, Cordelia Langford, John Sillitoe on behalf of the Wellcome Sanger Institute COVID-19 Surveillance Team |
| EPI_ISL_736234, EPI_ISL_736235, EPI_ISL_736236, EPI_ISL_736237, EPI_ISL_736238, EPI_ISL_736239, EPI_ISL_736241, EPI_ISL_736242, EPI_ISL_736243 | Lighthouse Lab in Glasgow | Wellcome Sanger Institute for the COVID-19 Genomics UK (COG-UK) Consortium | Harper VanSteenhouse, Yumi Kasai, David Gray, Carol Clugston, Anna Dominiczak and Alex Alderton, Roberto Amato, Sonia Goncalves, Ewan Harrison, David K. Jackson, Ian Johnston, Dominic Kwiatkowski, Cordelia Langford, John Sillitoe on behalf of the Wellcome Sanger Institute COVID-19 Surveillance Team |
| EPI_ISL_736244 | Lighthouse Lab in Milton Keynes | Wellcome Sanger Institute for the COVID-19 Genomics UK (COG-UK) Consortium | The Lighthouse Lab in Milton Keynes and Alex Alderton, Roberto Amato, Sonia Goncalves, Ewan Harrison, David K. Jackson, Ian Johnston, Dominic Kwiatkowski, Cordelia Langford, John Sillitoe on behalf of the Wellcome Sanger Institute COVID-19 Surveillance Team |
| EPI_ISL_736245 | Lighthouse Lab in Glasgow | Wellcome Sanger Institute for the COVID-19 Genomics UK (COG-UK) Consortium | Harper VanSteenhouse, Yumi Kasai, David Gray, Carol Clugston, Anna Dominiczak and Alex Alderton, Roberto Amato, Sonia Goncalves, Ewan Harrison, David K. Jackson, Ian Johnston, Dominic Kwiatkowski, Cordelia Langford, John Sillitoe on behalf of the Wellcome Sanger Institute COVID-19 Surveillance Team |
| EPI_ISL_736246, EPI_ISL_736247 | Lighthouse Lab in Milton Keynes | Wellcome Sanger Institute for the COVID-19 Genomics UK (COG-UK) Consortium | The Lighthouse Lab in Milton Keynes and Alex Alderton, Roberto Amato, Sonia Goncalves, Ewan Harrison, David K. Jackson, Ian Johnston, Dominic Kwiatkowski, Cordelia Langford, John Sillitoe on behalf of the Wellcome Sanger Institute COVID-19 Surveillance Team |
| EPI_ISL_736248 | Lighthouse Lab in Glasgow | Wellcome Sanger Institute for the COVID-19 Genomics UK (COG-UK) Consortium | Harper VanSteenhouse, Yumi Kasai, David Gray, Carol Clugston, Anna Dominiczak and Alex Alderton, Roberto Amato, Sonia Goncalves, Ewan Harrison, David K. Jackson, Ian Johnston, Dominic Kwiatkowski, Cordelia Langford, John Sillitoe on behalf of the Wellcome Sanger Institute COVID-19 Surveillance Team |
| EPI_ISL_736249 | Lighthouse Lab in Milton Keynes | Wellcome Sanger Institute for the COVID-19 Genomics UK (COG-UK) Consortium | The Lighthouse Lab in Milton Keynes and Alex Alderton, Roberto Amato, Sonia Goncalves, Ewan Harrison, David K. Jackson, Ian Johnston, Dominic Kwiatkowski, Cordelia Langford, John Sillitoe on behalf of the Wellcome Sanger Institute COVID-19 Surveillance Team |
| EPI_ISL_736250 | Lighthouse Lab in Glasgow | Wellcome Sanger Institute for the COVID-19 Genomics UK (COG-UK) Consortium | Harper VanSteenhouse, Yumi Kasai, David Gray, Carol Clugston, Anna Dominiczak and Alex Alderton, Roberto Amato, Sonia Goncalves, Ewan Harrison, David K. Jackson, Ian Johnston, Dominic Kwiatkowski, Cordelia Langford, John Sillitoe on behalf of the Wellcome Sanger Institute COVID-19 Surveillance Team |
| EPI_ISL_736251, EPI_ISL_736253 | Lighthouse Lab in Milton Keynes | Wellcome Sanger Institute for the COVID-19 Genomics UK (COG-UK) Consortium | The Lighthouse Lab in Milton Keynes and Alex Alderton, Roberto Amato, Sonia Goncalves, Ewan Harrison, David K. Jackson, Ian Johnston, Dominic Kwiatkowski, Cordelia Langford, John Sillitoe on behalf of the Wellcome Sanger Institute COVID-19 Surveillance Team |
| EPI_ISL_736254, EPI_ISL_736255 | Lighthouse Lab in Glasgow | Wellcome Sanger Institute for the COVID-19 Genomics UK (COG-UK) Consortium | Harper VanSteenhouse, Yumi Kasai, David Gray, Carol Clugston, Anna Dominiczak and Alex Alderton, Roberto Amato, Sonia Goncalves, Ewan Harrison, David K. Jackson, Ian Johnston, Dominic Kwiatkowski, Cordelia Langford, John Sillitoe on behalf of the Wellcome Sanger Institute COVID-19 Surveillance Team |

| | | | |
|---|---|---|---|
| EPI_ISL_736256 | Lighthouse Lab in Milton Keynes | Wellcome Sanger Institute for the COVID-19 Genomics UK (COG-UK) Consortium | The Lighthouse Lab in Milton Keynes and Alex Alderton, Roberto Amato, Sonia Goncalves, Ewan Harrison, David K. Jackson, Ian Johnston, Dominic Kwiatkowski, Cordelia Langford, John Sillitoe on behalf of the Wellcome Sanger Institute COVID-19 Surveillance Team |
| EPI_ISL_736257, EPI_ISL_736258 | Lighthouse Lab in Glasgow | Wellcome Sanger Institute for the COVID-19 Genomics UK (COG-UK) Consortium | Harper VanSteenhouse, Yumi Kasai, David Gray, Carol Clugston, Anna Dominiczak and Alex Alderton, Roberto Amato, Sonia Goncalves, Ewan Harrison, David K. Jackson, Ian Johnston, Dominic Kwiatkowski, Cordelia Langford, John Sillitoe on behalf of the Wellcome Sanger Institute COVID-19 Surveillance Team |
| EPI_ISL_736259, EPI_ISL_736260 | Lighthouse Lab in Milton Keynes | Wellcome Sanger Institute for the COVID-19 Genomics UK (COG-UK) Consortium | The Lighthouse Lab in Milton Keynes and Alex Alderton, Roberto Amato, Sonia Goncalves, Ewan Harrison, David K. Jackson, Ian Johnston, Dominic Kwiatkowski, Cordelia Langford, John Sillitoe on behalf of the Wellcome Sanger Institute COVID-19 Surveillance Team |
| EPI_ISL_736261 | Lighthouse Lab in Glasgow | Wellcome Sanger Institute for the COVID-19 Genomics UK (COG-UK) Consortium | Harper VanSteenhouse, Yumi Kasai, David Gray, Carol Clugston, Anna Dominiczak and Alex Alderton, Roberto Amato, Sonia Goncalves, Ewan Harrison, David K. Jackson, Ian Johnston, Dominic Kwiatkowski, Cordelia Langford, John Sillitoe on behalf of the Wellcome Sanger Institute COVID-19 Surveillance Team |
| EPI_ISL_736262, EPI_ISL_736263, EPI_ISL_736265 | Lighthouse Lab in Milton Keynes | Wellcome Sanger Institute for the COVID-19 Genomics UK (COG-UK) Consortium | The Lighthouse Lab in Milton Keynes and Alex Alderton, Roberto Amato, Sonia Goncalves, Ewan Harrison, David K. Jackson, Ian Johnston, Dominic Kwiatkowski, Cordelia Langford, John Sillitoe on behalf of the Wellcome Sanger Institute COVID-19 Surveillance Team |
| EPI_ISL_736266 | Lighthouse Lab in Glasgow | Wellcome Sanger Institute for the COVID-19 Genomics UK (COG-UK) Consortium | Harper VanSteenhouse, Yumi Kasai, David Gray, Carol Clugston, Anna Dominiczak and Alex Alderton, Roberto Amato, Sonia Goncalves, Ewan Harrison, David K. Jackson, Ian Johnston, Dominic Kwiatkowski, Cordelia Langford, John Sillitoe on behalf of the Wellcome Sanger Institute COVID-19 Surveillance Team |
| EPI_ISL_736267, EPI_ISL_736268 | Lighthouse Lab in Milton Keynes | Wellcome Sanger Institute for the COVID-19 Genomics UK (COG-UK) Consortium | The Lighthouse Lab in Milton Keynes and Alex Alderton, Roberto Amato, Sonia Goncalves, Ewan Harrison, David K. Jackson, Ian Johnston, Dominic Kwiatkowski, Cordelia Langford, John Sillitoe on behalf of the Wellcome Sanger Institute COVID-19 Surveillance Team |
| EPI_ISL_736269 | Lighthouse Lab in Glasgow | Wellcome Sanger Institute for the COVID-19 Genomics UK (COG-UK) Consortium | Harper VanSteenhouse, Yumi Kasai, David Gray, Carol Clugston, Anna Dominiczak and Alex Alderton, Roberto Amato, Sonia Goncalves, Ewan Harrison, David K. Jackson, Ian Johnston, Dominic Kwiatkowski, Cordelia Langford, John Sillitoe on behalf of the Wellcome Sanger Institute COVID-19 Surveillance Team |
| EPI_ISL_736270 | Lighthouse Lab in Milton Keynes | Wellcome Sanger Institute for the COVID-19 Genomics UK (COG-UK) Consortium | The Lighthouse Lab in Milton Keynes and Alex Alderton, Roberto Amato, Sonia Goncalves, Ewan Harrison, David K. Jackson, Ian Johnston, Dominic Kwiatkowski, Cordelia Langford, John Sillitoe on behalf of the Wellcome Sanger Institute COVID-19 Surveillance Team |
| EPI_ISL_736271 | Lighthouse Lab in Glasgow | Wellcome Sanger Institute for the COVID-19 Genomics UK (COG-UK) Consortium | Harper VanSteenhouse, Yumi Kasai, David Gray, Carol Clugston, Anna Dominiczak and Alex Alderton, Roberto Amato, Sonia Goncalves, Ewan Harrison, David K. Jackson, Ian Johnston, Dominic Kwiatkowski, Cordelia Langford, John Sillitoe on behalf of the Wellcome Sanger Institute COVID-19 Surveillance Team |
| EPI_ISL_736272, EPI_ISL_736273 | Lighthouse Lab in Milton Keynes | Wellcome Sanger Institute for the COVID-19 Genomics UK (COG-UK) Consortium | The Lighthouse Lab in Milton Keynes and Alex Alderton, Roberto Amato, Sonia Goncalves, Ewan Harrison, David K. Jackson, Ian Johnston, Dominic Kwiatkowski, Cordelia Langford, John Sillitoe on behalf of the Wellcome Sanger Institute COVID-19 Surveillance Team |
| EPI_ISL_736275 | Lighthouse Lab in Glasgow | Wellcome Sanger Institute for the COVID-19 Genomics UK (COG-UK) Consortium | Harper VanSteenhouse, Yumi Kasai, David Gray, Carol Clugston, Anna Dominiczak and Alex Alderton, Roberto Amato, Sonia Goncalves, Ewan Harrison, David K. Jackson, Ian Johnston, Dominic Kwiatkowski, Cordelia Langford, John Sillitoe on behalf of the Wellcome Sanger Institute COVID-19 Surveillance Team |
| EPI_ISL_736276 | Lighthouse Lab in Milton Keynes | Wellcome Sanger Institute for the COVID-19 Genomics UK (COG-UK) Consortium | The Lighthouse Lab in Milton Keynes and Alex Alderton, Roberto Amato, Sonia Goncalves, Ewan Harrison, David K. Jackson, Ian Johnston, Dominic Kwiatkowski, Cordelia Langford, John Sillitoe on behalf of the Wellcome Sanger Institute COVID-19 Surveillance Team |
| EPI_ISL_736277, EPI_ISL_736278 | Lighthouse Lab in Glasgow | Wellcome Sanger Institute for the COVID-19 Genomics UK (COG-UK) Consortium | Harper VanSteenhouse, Yumi Kasai, David Gray, Carol Clugston, Anna Dominiczak and Alex Alderton, Roberto Amato, Sonia Goncalves, Ewan Harrison, David K. Jackson, Ian Johnston, Dominic Kwiatkowski, Cordelia Langford, John Sillitoe on behalf of the Wellcome Sanger Institute COVID-19 Surveillance Team |
| EPI_ISL_736279, EPI_ISL_736280, EPI_ISL_736281, EPI_ISL_736282, EPI_ISL_736283 | Lighthouse Lab in Milton Keynes | Wellcome Sanger Institute for the COVID-19 Genomics UK (COG-UK) Consortium | The Lighthouse Lab in Milton Keynes and Alex Alderton, Roberto Amato, Sonia Goncalves, Ewan Harrison, David K. Jackson, Ian Johnston, Dominic Kwiatkowski, Cordelia Langford, John Sillitoe on behalf of the Wellcome Sanger Institute COVID-19 Surveillance Team |
| EPI_ISL_736284 | Lighthouse Lab in Glasgow | Wellcome Sanger Institute for the COVID-19 Genomics UK (COG-UK) Consortium | Harper VanSteenhouse, Yumi Kasai, David Gray, Carol Clugston, Anna Dominiczak and Alex Alderton, Roberto Amato, Sonia Goncalves, Ewan Harrison, David K. Jackson, Ian Johnston, Dominic Kwiatkowski, Cordelia Langford, John Sillitoe on behalf of the Wellcome Sanger Institute COVID-19 Surveillance Team |
| EPI_ISL_736285, EPI_ISL_736286 | Lighthouse Lab in Milton Keynes | Wellcome Sanger Institute for the COVID-19 Genomics UK (COG-UK) Consortium | The Lighthouse Lab in Milton Keynes and Alex Alderton, Roberto Amato, Sonia Goncalves, Ewan Harrison, David K. Jackson, Ian Johnston, Dominic Kwiatkowski, Cordelia Langford, John Sillitoe on behalf of the Wellcome Sanger Institute COVID-19 Surveillance Team |
| EPI_ISL_736287 | Lighthouse Lab in Glasgow | Wellcome Sanger Institute for the COVID-19 Genomics UK (COG-UK) Consortium | Harper VanSteenhouse, Yumi Kasai, David Gray, Carol Clugston, Anna Dominiczak and Alex Alderton, Roberto Amato, Sonia Goncalves, Ewan Harrison, David K. Jackson, Ian Johnston, Dominic Kwiatkowski, Cordelia Langford, John Sillitoe on behalf of the Wellcome Sanger Institute COVID-19 Surveillance Team |
| EPI_ISL_736288 | Lighthouse Lab in Milton Keynes | Wellcome Sanger Institute for the COVID-19 Genomics UK (COG-UK) Consortium | The Lighthouse Lab in Milton Keynes and Alex Alderton, Roberto Amato, Sonia Goncalves, Ewan Harrison, David K. Jackson, Ian Johnston, Dominic Kwiatkowski, Cordelia Langford, John Sillitoe on behalf of the Wellcome Sanger Institute COVID-19 Surveillance Team |
| EPI_ISL_736290 | Lighthouse Lab in Glasgow | Wellcome Sanger Institute for the COVID-19 Genomics UK (COG-UK) Consortium | Harper VanSteenhouse, Yumi Kasai, David Gray, Carol Clugston, Anna Dominiczak and Alex Alderton, Roberto Amato, Sonia Goncalves, Ewan Harrison, David K. Jackson, Ian Johnston, Dominic Kwiatkowski, Cordelia Langford, John Sillitoe on behalf of the Wellcome Sanger Institute COVID-19 Surveillance Team |
| EPI_ISL_736291, EPI_ISL_736294, EPI_ISL_736295, EPI_ISL_736296, EPI_ISL_736297, EPI_ISL_736298, EPI_ISL_736299, EPI_ISL_736300 | Lighthouse Lab in Milton Keynes | Wellcome Sanger Institute for the COVID-19 Genomics UK (COG-UK) Consortium | The Lighthouse Lab in Milton Keynes and Alex Alderton, Roberto Amato, Sonia Goncalves, Ewan Harrison, David K. Jackson, Ian Johnston, Dominic Kwiatkowski, Cordelia Langford, John Sillitoe on behalf of the Wellcome Sanger Institute COVID-19 Surveillance Team |
| EPI_ISL_736301 | Lighthouse Lab in Glasgow | Wellcome Sanger Institute for the COVID-19 Genomics UK (COG-UK) Consortium | Harper VanSteenhouse, Yumi Kasai, David Gray, Carol Clugston, Anna Dominiczak and Alex Alderton, Roberto Amato, Sonia Goncalves, Ewan Harrison, David K. Jackson, Ian Johnston, Dominic Kwiatkowski, Cordelia Langford, John Sillitoe on behalf of the Wellcome Sanger Institute COVID-19 Surveillance Team |
| EPI_ISL_736302 | Lighthouse Lab in Milton Keynes | Wellcome Sanger Institute for the COVID-19 Genomics UK (COG-UK) Consortium | The Lighthouse Lab in Milton Keynes and Alex Alderton, Roberto Amato, Sonia Goncalves, Ewan Harrison, David K. Jackson, Ian Johnston, Dominic Kwiatkowski, Cordelia Langford, John Sillitoe on behalf of the Wellcome Sanger Institute COVID-19 Surveillance Team |
| EPI_ISL_736304 | Lighthouse Lab in Glasgow | Wellcome Sanger Institute for the COVID-19 Genomics UK (COG-UK) Consortium | Harper VanSteenhouse, Yumi Kasai, David Gray, Carol Clugston, Anna Dominiczak and Alex Alderton, Roberto Amato, Sonia Goncalves, Ewan Harrison, David K. Jackson, Ian Johnston, Dominic Kwiatkowski, Cordelia Langford, John Sillitoe on behalf of the Wellcome Sanger Institute COVID-19 Surveillance Team |
| EPI_ISL_736305 | Lighthouse Lab in Milton Keynes | Wellcome Sanger Institute for the COVID-19 Genomics UK (COG-UK) Consortium | The Lighthouse Lab in Milton Keynes and Alex Alderton, Roberto Amato, Sonia Goncalves, Ewan Harrison, David K. Jackson, Ian Johnston, Dominic Kwiatkowski, Cordelia Langford, John Sillitoe on behalf of the Wellcome Sanger Institute COVID-19 Surveillance Team |
| EPI_ISL_736306 | Lighthouse Lab in Glasgow | Wellcome Sanger Institute for the COVID-19 Genomics UK (COG-UK) Consortium | Harper VanSteenhouse, Yumi Kasai, David Gray, Carol Clugston, Anna Dominiczak and Alex Alderton, Roberto Amato, Sonia Goncalves, Ewan Harrison, David K. Jackson, Ian Johnston, Dominic Kwiatkowski, Cordelia Langford, John Sillitoe on behalf of the Wellcome Sanger Institute COVID-19 Surveillance Team |
| EPI_ISL_736307, EPI_ISL_736308 | Lighthouse Lab in Milton Keynes | Wellcome Sanger Institute for the COVID-19 Genomics UK (COG-UK) Consortium | The Lighthouse Lab in Milton Keynes and Alex Alderton, Roberto Amato, Sonia Goncalves, Ewan Harrison, David K. Jackson, Ian Johnston, Dominic Kwiatkowski, Cordelia Langford, John Sillitoe on behalf of the Wellcome Sanger Institute COVID-19 Surveillance Team |
| EPI_ISL_736309 | Lighthouse Lab in Glasgow | Wellcome Sanger Institute for the COVID-19 Genomics UK (COG-UK) Consortium | Harper VanSteenhouse, Yumi Kasai, David Gray, Carol Clugston, Anna Dominiczak and Alex Alderton, Roberto Amato, Sonia Goncalves, Ewan Harrison, David K. Jackson, Ian Johnston, Dominic Kwiatkowski, Cordelia Langford, John Sillitoe on behalf of the Wellcome Sanger Institute COVID-19 Surveillance Team |

| | | | |
|---|---|---|---|
| EPI_ISL_736310 | Lighthouse Lab in Milton Keynes | Wellcome Sanger Institute for the COVID-19 Genomics UK (COG-UK) Consortium | The Lighthouse Lab in Milton Keynes and Alex Alderton, Roberto Amato, Sonia Goncalves, Ewan Harrison, David K. Jackson, Ian Johnston, Dominic Kwiatkowski, Cordelia Langford, John Sillitoe on behalf of the Wellcome Sanger Institute COVID-19 Surveillance Team |
| EPI_ISL_736311, EPI_ISL_736312, EPI_ISL_736313 | Lighthouse Lab in Glasgow | Wellcome Sanger Institute for the COVID-19 Genomics UK (COG-UK) Consortium | Harper VanSteenhouse, Yumi Kasai, David Gray, Carol Clugston, Anna Dominiczak and Alex Alderton, Roberto Amato, Sonia Goncalves, Ewan Harrison, David K. Jackson, Ian Johnston, Dominic Kwiatkowski, Cordelia Langford, John Sillitoe on behalf of the Wellcome Sanger Institute COVID-19 Surveillance Team |
| EPI_ISL_736314 | Lighthouse Lab in Milton Keynes | Wellcome Sanger Institute for the COVID-19 Genomics UK (COG-UK) Consortium | The Lighthouse Lab in Milton Keynes and Alex Alderton, Roberto Amato, Sonia Goncalves, Ewan Harrison, David K. Jackson, Ian Johnston, Dominic Kwiatkowski, Cordelia Langford, John Sillitoe on behalf of the Wellcome Sanger Institute COVID-19 Surveillance Team |
| EPI_ISL_736315, EPI_ISL_736316 | Lighthouse Lab in Glasgow | Wellcome Sanger Institute for the COVID-19 Genomics UK (COG-UK) Consortium | Harper VanSteenhouse, Yumi Kasai, David Gray, Carol Clugston, Anna Dominiczak and Alex Alderton, Roberto Amato, Sonia Goncalves, Ewan Harrison, David K. Jackson, Ian Johnston, Dominic Kwiatkowski, Cordelia Langford, John Sillitoe on behalf of the Wellcome Sanger Institute COVID-19 Surveillance Team |
| EPI_ISL_736317 | Lighthouse Lab in Milton Keynes | Wellcome Sanger Institute for the COVID-19 Genomics UK (COG-UK) Consortium | The Lighthouse Lab in Milton Keynes and Alex Alderton, Roberto Amato, Sonia Goncalves, Ewan Harrison, David K. Jackson, Ian Johnston, Dominic Kwiatkowski, Cordelia Langford, John Sillitoe on behalf of the Wellcome Sanger Institute COVID-19 Surveillance Team |
| EPI_ISL_736318 | Lighthouse Lab in Glasgow | Wellcome Sanger Institute for the COVID-19 Genomics UK (COG-UK) Consortium | Harper VanSteenhouse, Yumi Kasai, David Gray, Carol Clugston, Anna Dominiczak and Alex Alderton, Roberto Amato, Sonia Goncalves, Ewan Harrison, David K. Jackson, Ian Johnston, Dominic Kwiatkowski, Cordelia Langford, John Sillitoe on behalf of the Wellcome Sanger Institute COVID-19 Surveillance Team |
| EPI_ISL_736319 | Lighthouse Lab in Milton Keynes | Wellcome Sanger Institute for the COVID-19 Genomics UK (COG-UK) Consortium | The Lighthouse Lab in Milton Keynes and Alex Alderton, Roberto Amato, Sonia Goncalves, Ewan Harrison, David K. Jackson, Ian Johnston, Dominic Kwiatkowski, Cordelia Langford, John Sillitoe on behalf of the Wellcome Sanger Institute COVID-19 Surveillance Team |
| EPI_ISL_736320, EPI_ISL_736321 | Lighthouse Lab in Glasgow | Wellcome Sanger Institute for the COVID-19 Genomics UK (COG-UK) Consortium | Harper VanSteenhouse, Yumi Kasai, David Gray, Carol Clugston, Anna Dominiczak and Alex Alderton, Roberto Amato, Sonia Goncalves, Ewan Harrison, David K. Jackson, Ian Johnston, Dominic Kwiatkowski, Cordelia Langford, John Sillitoe on behalf of the Wellcome Sanger Institute COVID-19 Surveillance Team |
| EPI_ISL_736322 | Lighthouse Lab in Milton Keynes | Wellcome Sanger Institute for the COVID-19 Genomics UK (COG-UK) Consortium | The Lighthouse Lab in Milton Keynes and Alex Alderton, Roberto Amato, Sonia Goncalves, Ewan Harrison, David K. Jackson, Ian Johnston, Dominic Kwiatkowski, Cordelia Langford, John Sillitoe on behalf of the Wellcome Sanger Institute COVID-19 Surveillance Team |
| EPI_ISL_736323 | Lighthouse Lab in Glasgow | Wellcome Sanger Institute for the COVID-19 Genomics UK (COG-UK) Consortium | Harper VanSteenhouse, Yumi Kasai, David Gray, Carol Clugston, Anna Dominiczak and Alex Alderton, Roberto Amato, Sonia Goncalves, Ewan Harrison, David K. Jackson, Ian Johnston, Dominic Kwiatkowski, Cordelia Langford, John Sillitoe on behalf of the Wellcome Sanger Institute COVID-19 Surveillance Team |
| EPI_ISL_736324 | Lighthouse Lab in Milton Keynes | Wellcome Sanger Institute for the COVID-19 Genomics UK (COG-UK) Consortium | The Lighthouse Lab in Milton Keynes and Alex Alderton, Roberto Amato, Sonia Goncalves, Ewan Harrison, David K. Jackson, Ian Johnston, Dominic Kwiatkowski, Cordelia Langford, John Sillitoe on behalf of the Wellcome Sanger Institute COVID-19 Surveillance Team |
| EPI_ISL_736325 | Lighthouse Lab in Glasgow | Wellcome Sanger Institute for the COVID-19 Genomics UK (COG-UK) Consortium | Harper VanSteenhouse, Yumi Kasai, David Gray, Carol Clugston, Anna Dominiczak and Alex Alderton, Roberto Amato, Sonia Goncalves, Ewan Harrison, David K. Jackson, Ian Johnston, Dominic Kwiatkowski, Cordelia Langford, John Sillitoe on behalf of the Wellcome Sanger Institute COVID-19 Surveillance Team |
| EPI_ISL_736326 | Lighthouse Lab in Milton Keynes | Wellcome Sanger Institute for the COVID-19 Genomics UK (COG-UK) Consortium | The Lighthouse Lab in Milton Keynes and Alex Alderton, Roberto Amato, Sonia Goncalves, Ewan Harrison, David K. Jackson, Ian Johnston, Dominic Kwiatkowski, Cordelia Langford, John Sillitoe on behalf of the Wellcome Sanger Institute COVID-19 Surveillance Team |
| EPI_ISL_736327 | Lighthouse Lab in Glasgow | Wellcome Sanger Institute for the COVID-19 Genomics UK (COG-UK) Consortium | Harper VanSteenhouse, Yumi Kasai, David Gray, Carol Clugston, Anna Dominiczak and Alex Alderton, Roberto Amato, Sonia Goncalves, Ewan Harrison, David K. Jackson, Ian Johnston, Dominic Kwiatkowski, Cordelia Langford, John Sillitoe on behalf of the Wellcome Sanger Institute COVID-19 Surveillance Team |
| EPI_ISL_736328, EPI_ISL_736329 | Lighthouse Lab in Milton Keynes | Wellcome Sanger Institute for the COVID-19 Genomics UK (COG-UK) Consortium | The Lighthouse Lab in Milton Keynes and Alex Alderton, Roberto Amato, Sonia Goncalves, Ewan Harrison, David K. Jackson, Ian Johnston, Dominic Kwiatkowski, Cordelia Langford, John Sillitoe on behalf of the Wellcome Sanger Institute COVID-19 Surveillance Team |
| EPI_ISL_736330, EPI_ISL_736331, EPI_ISL_736332 | Lighthouse Lab in Glasgow | Wellcome Sanger Institute for the COVID-19 Genomics UK (COG-UK) Consortium | Harper VanSteenhouse, Yumi Kasai, David Gray, Carol Clugston, Anna Dominiczak and Alex Alderton, Roberto Amato, Sonia Goncalves, Ewan Harrison, David K. Jackson, Ian Johnston, Dominic Kwiatkowski, Cordelia Langford, John Sillitoe on behalf of the Wellcome Sanger Institute COVID-19 Surveillance Team |
| EPI_ISL_736333 | Lighthouse Lab in Milton Keynes | Wellcome Sanger Institute for the COVID-19 Genomics UK (COG-UK) Consortium | The Lighthouse Lab in Milton Keynes and Alex Alderton, Roberto Amato, Sonia Goncalves, Ewan Harrison, David K. Jackson, Ian Johnston, Dominic Kwiatkowski, Cordelia Langford, John Sillitoe on behalf of the Wellcome Sanger Institute COVID-19 Surveillance Team |
| EPI_ISL_736334 | Lighthouse Lab in Glasgow | Wellcome Sanger Institute for the COVID-19 Genomics UK (COG-UK) Consortium | Harper VanSteenhouse, Yumi Kasai, David Gray, Carol Clugston, Anna Dominiczak and Alex Alderton, Roberto Amato, Sonia Goncalves, Ewan Harrison, David K. Jackson, Ian Johnston, Dominic Kwiatkowski, Cordelia Langford, John Sillitoe on behalf of the Wellcome Sanger Institute COVID-19 Surveillance Team |
| EPI_ISL_736335, EPI_ISL_736336, EPI_ISL_736337, EPI_ISL_736338, EPI_ISL_736339, EPI_ISL_736340, EPI_ISL_736341, EPI_ISL_736342 | Lighthouse Lab in Milton Keynes | Wellcome Sanger Institute for the COVID-19 Genomics UK (COG-UK) Consortium | The Lighthouse Lab in Milton Keynes and Alex Alderton, Roberto Amato, Sonia Goncalves, Ewan Harrison, David K. Jackson, Ian Johnston, Dominic Kwiatkowski, Cordelia Langford, John Sillitoe on behalf of the Wellcome Sanger Institute COVID-19 Surveillance Team |
| EPI_ISL_736343 | Lighthouse Lab in Glasgow | Wellcome Sanger Institute for the COVID-19 Genomics UK (COG-UK) Consortium | Harper VanSteenhouse, Yumi Kasai, David Gray, Carol Clugston, Anna Dominiczak and Alex Alderton, Roberto Amato, Sonia Goncalves, Ewan Harrison, David K. Jackson, Ian Johnston, Dominic Kwiatkowski, Cordelia Langford, John Sillitoe on behalf of the Wellcome Sanger Institute COVID-19 Surveillance Team |
| EPI_ISL_736344, EPI_ISL_736345 | Lighthouse Lab in Milton Keynes | Wellcome Sanger Institute for the COVID-19 Genomics UK (COG-UK) Consortium | The Lighthouse Lab in Milton Keynes and Alex Alderton, Roberto Amato, Sonia Goncalves, Ewan Harrison, David K. Jackson, Ian Johnston, Dominic Kwiatkowski, Cordelia Langford, John Sillitoe on behalf of the Wellcome Sanger Institute COVID-19 Surveillance Team |
| EPI_ISL_736346, EPI_ISL_736347 | Lighthouse Lab in Glasgow | Wellcome Sanger Institute for the COVID-19 Genomics UK (COG-UK) Consortium | Harper VanSteenhouse, Yumi Kasai, David Gray, Carol Clugston, Anna Dominiczak and Alex Alderton, Roberto Amato, Sonia Goncalves, Ewan Harrison, David K. Jackson, Ian Johnston, Dominic Kwiatkowski, Cordelia Langford, John Sillitoe on behalf of the Wellcome Sanger Institute COVID-19 Surveillance Team |
| EPI_ISL_736348 | Lighthouse Lab in Milton Keynes | Wellcome Sanger Institute for the COVID-19 Genomics UK (COG-UK) Consortium | The Lighthouse Lab in Milton Keynes and Alex Alderton, Roberto Amato, Sonia Goncalves, Ewan Harrison, David K. Jackson, Ian Johnston, Dominic Kwiatkowski, Cordelia Langford, John Sillitoe on behalf of the Wellcome Sanger Institute COVID-19 Surveillance Team |
| EPI_ISL_736350 | Lighthouse Lab in Glasgow | Wellcome Sanger Institute for the COVID-19 Genomics UK (COG-UK) Consortium | Harper VanSteenhouse, Yumi Kasai, David Gray, Carol Clugston, Anna Dominiczak and Alex Alderton, Roberto Amato, Sonia Goncalves, Ewan Harrison, David K. Jackson, Ian Johnston, Dominic Kwiatkowski, Cordelia Langford, John Sillitoe on behalf of the Wellcome Sanger Institute COVID-19 Surveillance Team |
| EPI_ISL_736351 | Lighthouse Lab in Milton Keynes | Wellcome Sanger Institute for the COVID-19 Genomics UK (COG-UK) Consortium | The Lighthouse Lab in Milton Keynes and Alex Alderton, Roberto Amato, Sonia Goncalves, Ewan Harrison, David K. Jackson, Ian Johnston, Dominic Kwiatkowski, Cordelia Langford, John Sillitoe on behalf of the Wellcome Sanger Institute COVID-19 Surveillance Team |
| EPI_ISL_736352 | Lighthouse Lab in Glasgow | Wellcome Sanger Institute for the COVID-19 Genomics UK (COG-UK) Consortium | Harper VanSteenhouse, Yumi Kasai, David Gray, Carol Clugston, Anna Dominiczak and Alex Alderton, Roberto Amato, Sonia Goncalves, Ewan Harrison, David K. Jackson, Ian Johnston, Dominic Kwiatkowski, Cordelia Langford, John Sillitoe on behalf of the Wellcome Sanger Institute COVID-19 Surveillance Team |
| EPI_ISL_736353, EPI_ISL_736354 | Lighthouse Lab in Milton Keynes | Wellcome Sanger Institute for the COVID-19 Genomics UK (COG-UK) Consortium | The Lighthouse Lab in Milton Keynes and Alex Alderton, Roberto Amato, Sonia Goncalves, Ewan Harrison, David K. Jackson, Ian Johnston, Dominic Kwiatkowski, Cordelia Langford, John Sillitoe on behalf of the Wellcome Sanger Institute COVID-19 Surveillance Team |
| EPI_ISL_736355, EPI_ISL_736356 | Lighthouse Lab in Glasgow | Wellcome Sanger Institute for the COVID-19 Genomics UK (COG-UK) Consortium | Harper VanSteenhouse, Yumi Kasai, David Gray, Carol Clugston, Anna Dominiczak and Alex Alderton, Roberto Amato, Sonia Goncalves, Ewan Harrison, David K. Jackson, Ian Johnston, Dominic Kwiatkowski, Cordelia Langford, John Sillitoe on behalf of the Wellcome Sanger Institute COVID-19 Surveillance Team |
| EPI_ISL_736357 | Lighthouse Lab in Milton Keynes | Wellcome Sanger Institute for the COVID-19 Genomics UK | The Lighthouse Lab in Milton Keynes and Alex Alderton, Roberto Amato, Sonia Goncalves, Ewan Harrison, David K. Jackson, Ian Johnston, Dominic |

| | | | |
|---|---|---|---|
| | | (COG-UK) Consortium | Kwiatkowski, Cordelia Langford, John Sillitoe on behalf of the Wellcome Sanger Institute COVID-19 Surveillance Team |
| EPI_ISL_736358 | Lighthouse Lab in Glasgow | Wellcome Sanger Institute for the COVID-19 Genomics UK (COG-UK) Consortium | Harper VanSteenhouse, Yumi Kasai, David Gray, Carol Clugston, Anna Dominiczak and Alex Alderton, Roberto Amato, Sonia Goncalves, Ewan Harrison, David K. Jackson, Ian Johnston, Dominic Kwiatkowski, Cordelia Langford, John Sillitoe on behalf of the Wellcome Sanger Institute COVID-19 Surveillance Team |
| EPI_ISL_736359, EPI_ISL_736360, EPI_ISL_736361 | Lighthouse Lab in Milton Keynes | Wellcome Sanger Institute for the COVID-19 Genomics UK (COG-UK) Consortium | The Lighthouse Lab in Milton Keynes and Alex Alderton, Roberto Amato, Sonia Goncalves, Ewan Harrison, David K. Jackson, Ian Johnston, Dominic Kwiatkowski, Cordelia Langford, John Sillitoe on behalf of the Wellcome Sanger Institute COVID-19 Surveillance Team |
| EPI_ISL_736362 | Lighthouse Lab in Glasgow | Wellcome Sanger Institute for the COVID-19 Genomics UK (COG-UK) Consortium | Harper VanSteenhouse, Yumi Kasai, David Gray, Carol Clugston, Anna Dominiczak and Alex Alderton, Roberto Amato, Sonia Goncalves, Ewan Harrison, David K. Jackson, Ian Johnston, Dominic Kwiatkowski, Cordelia Langford, John Sillitoe on behalf of the Wellcome Sanger Institute COVID-19 Surveillance Team |
| EPI_ISL_736363, EPI_ISL_736364, EPI_ISL_736365, EPI_ISL_736366 | Lighthouse Lab in Milton Keynes | Wellcome Sanger Institute for the COVID-19 Genomics UK (COG-UK) Consortium | The Lighthouse Lab in Milton Keynes and Alex Alderton, Roberto Amato, Sonia Goncalves, Ewan Harrison, David K. Jackson, Ian Johnston, Dominic Kwiatkowski, Cordelia Langford, John Sillitoe on behalf of the Wellcome Sanger Institute COVID-19 Surveillance Team |
| EPI_ISL_736367 | Lighthouse Lab in Glasgow | Wellcome Sanger Institute for the COVID-19 Genomics UK (COG-UK) Consortium | Harper VanSteenhouse, Yumi Kasai, David Gray, Carol Clugston, Anna Dominiczak and Alex Alderton, Roberto Amato, Sonia Goncalves, Ewan Harrison, David K. Jackson, Ian Johnston, Dominic Kwiatkowski, Cordelia Langford, John Sillitoe on behalf of the Wellcome Sanger Institute COVID-19 Surveillance Team |
| EPI_ISL_736368 | Lighthouse Lab in Milton Keynes | Wellcome Sanger Institute for the COVID-19 Genomics UK (COG-UK) Consortium | The Lighthouse Lab in Milton Keynes and Alex Alderton, Roberto Amato, Sonia Goncalves, Ewan Harrison, David K. Jackson, Ian Johnston, Dominic Kwiatkowski, Cordelia Langford, John Sillitoe on behalf of the Wellcome Sanger Institute COVID-19 Surveillance Team |
| EPI_ISL_736370, EPI_ISL_736371 | Lighthouse Lab in Glasgow | Wellcome Sanger Institute for the COVID-19 Genomics UK (COG-UK) Consortium | Harper VanSteenhouse, Yumi Kasai, David Gray, Carol Clugston, Anna Dominiczak and Alex Alderton, Roberto Amato, Sonia Goncalves, Ewan Harrison, David K. Jackson, Ian Johnston, Dominic Kwiatkowski, Cordelia Langford, John Sillitoe on behalf of the Wellcome Sanger Institute COVID-19 Surveillance Team |
| EPI_ISL_736372 | Lighthouse Lab in Milton Keynes | Wellcome Sanger Institute for the COVID-19 Genomics UK (COG-UK) Consortium | The Lighthouse Lab in Milton Keynes and Alex Alderton, Roberto Amato, Sonia Goncalves, Ewan Harrison, David K. Jackson, Ian Johnston, Dominic Kwiatkowski, Cordelia Langford, John Sillitoe on behalf of the Wellcome Sanger Institute COVID-19 Surveillance Team |
| EPI_ISL_736373 | Lighthouse Lab in Glasgow | Wellcome Sanger Institute for the COVID-19 Genomics UK (COG-UK) Consortium | Harper VanSteenhouse, Yumi Kasai, David Gray, Carol Clugston, Anna Dominiczak and Alex Alderton, Roberto Amato, Sonia Goncalves, Ewan Harrison, David K. Jackson, Ian Johnston, Dominic Kwiatkowski, Cordelia Langford, John Sillitoe on behalf of the Wellcome Sanger Institute COVID-19 Surveillance Team |
| EPI_ISL_736374, EPI_ISL_736375 | Lighthouse Lab in Milton Keynes | Wellcome Sanger Institute for the COVID-19 Genomics UK (COG-UK) Consortium | The Lighthouse Lab in Milton Keynes and Alex Alderton, Roberto Amato, Sonia Goncalves, Ewan Harrison, David K. Jackson, Ian Johnston, Dominic Kwiatkowski, Cordelia Langford, John Sillitoe on behalf of the Wellcome Sanger Institute COVID-19 Surveillance Team |
| EPI_ISL_736376 | Lighthouse Lab in Glasgow | Wellcome Sanger Institute for the COVID-19 Genomics UK (COG-UK) Consortium | Harper VanSteenhouse, Yumi Kasai, David Gray, Carol Clugston, Anna Dominiczak and Alex Alderton, Roberto Amato, Sonia Goncalves, Ewan Harrison, David K. Jackson, Ian Johnston, Dominic Kwiatkowski, Cordelia Langford, John Sillitoe on behalf of the Wellcome Sanger Institute COVID-19 Surveillance Team |
| EPI_ISL_736377 | Lighthouse Lab in Milton Keynes | Wellcome Sanger Institute for the COVID-19 Genomics UK (COG-UK) Consortium | The Lighthouse Lab in Milton Keynes and Alex Alderton, Roberto Amato, Sonia Goncalves, Ewan Harrison, David K. Jackson, Ian Johnston, Dominic Kwiatkowski, Cordelia Langford, John Sillitoe on behalf of the Wellcome Sanger Institute COVID-19 Surveillance Team |
| EPI_ISL_736378 | Lighthouse Lab in Glasgow | Wellcome Sanger Institute for the COVID-19 Genomics UK (COG-UK) Consortium | Harper VanSteenhouse, Yumi Kasai, David Gray, Carol Clugston, Anna Dominiczak and Alex Alderton, Roberto Amato, Sonia Goncalves, Ewan Harrison, David K. Jackson, Ian Johnston, Dominic Kwiatkowski, Cordelia Langford, John Sillitoe on behalf of the Wellcome Sanger Institute COVID-19 Surveillance Team |
| EPI_ISL_736379, EPI_ISL_736381 | Lighthouse Lab in Milton Keynes | Wellcome Sanger Institute for the COVID-19 Genomics UK (COG-UK) Consortium | The Lighthouse Lab in Milton Keynes and Alex Alderton, Roberto Amato, Sonia Goncalves, Ewan Harrison, David K. Jackson, Ian Johnston, Dominic Kwiatkowski, Cordelia Langford, John Sillitoe on behalf of the Wellcome Sanger Institute COVID-19 Surveillance Team |
| EPI_ISL_736382, EPI_ISL_736383 | Lighthouse Lab in Glasgow | Wellcome Sanger Institute for the COVID-19 Genomics UK (COG-UK) Consortium | Harper VanSteenhouse, Yumi Kasai, David Gray, Carol Clugston, Anna Dominiczak and Alex Alderton, Roberto Amato, Sonia Goncalves, Ewan Harrison, David K. Jackson, Ian Johnston, Dominic Kwiatkowski, Cordelia Langford, John Sillitoe on behalf of the Wellcome Sanger Institute COVID-19 Surveillance Team |
| EPI_ISL_736384, EPI_ISL_736385 | Lighthouse Lab in Milton Keynes | Wellcome Sanger Institute for the COVID-19 Genomics UK (COG-UK) Consortium | The Lighthouse Lab in Milton Keynes and Alex Alderton, Roberto Amato, Sonia Goncalves, Ewan Harrison, David K. Jackson, Ian Johnston, Dominic Kwiatkowski, Cordelia Langford, John Sillitoe on behalf of the Wellcome Sanger Institute COVID-19 Surveillance Team |
| EPI_ISL_736386 | Lighthouse Lab in Glasgow | Wellcome Sanger Institute for the COVID-19 Genomics UK (COG-UK) Consortium | Harper VanSteenhouse, Yumi Kasai, David Gray, Carol Clugston, Anna Dominiczak and Alex Alderton, Roberto Amato, Sonia Goncalves, Ewan Harrison, David K. Jackson, Ian Johnston, Dominic Kwiatkowski, Cordelia Langford, John Sillitoe on behalf of the Wellcome Sanger Institute COVID-19 Surveillance Team |
| EPI_ISL_736387 | Lighthouse Lab in Milton Keynes | Wellcome Sanger Institute for the COVID-19 Genomics UK (COG-UK) Consortium | The Lighthouse Lab in Milton Keynes and Alex Alderton, Roberto Amato, Sonia Goncalves, Ewan Harrison, David K. Jackson, Ian Johnston, Dominic Kwiatkowski, Cordelia Langford, John Sillitoe on behalf of the Wellcome Sanger Institute COVID-19 Surveillance Team |
| EPI_ISL_736388, EPI_ISL_736389, EPI_ISL_736390, EPI_ISL_736391 | Lighthouse Lab in Glasgow | Wellcome Sanger Institute for the COVID-19 Genomics UK (COG-UK) Consortium | Harper VanSteenhouse, Yumi Kasai, David Gray, Carol Clugston, Anna Dominiczak and Alex Alderton, Roberto Amato, Sonia Goncalves, Ewan Harrison, David K. Jackson, Ian Johnston, Dominic Kwiatkowski, Cordelia Langford, John Sillitoe on behalf of the Wellcome Sanger Institute COVID-19 Surveillance Team |
| EPI_ISL_736393 | Lighthouse Lab in Milton Keynes | Wellcome Sanger Institute for the COVID-19 Genomics UK (COG-UK) Consortium | The Lighthouse Lab in Milton Keynes and Alex Alderton, Roberto Amato, Sonia Goncalves, Ewan Harrison, David K. Jackson, Ian Johnston, Dominic Kwiatkowski, Cordelia Langford, John Sillitoe on behalf of the Wellcome Sanger Institute COVID-19 Surveillance Team |
| EPI_ISL_736394 | Lighthouse Lab in Glasgow | Wellcome Sanger Institute for the COVID-19 Genomics UK (COG-UK) Consortium | Harper VanSteenhouse, Yumi Kasai, David Gray, Carol Clugston, Anna Dominiczak and Alex Alderton, Roberto Amato, Sonia Goncalves, Ewan Harrison, David K. Jackson, Ian Johnston, Dominic Kwiatkowski, Cordelia Langford, John Sillitoe on behalf of the Wellcome Sanger Institute COVID-19 Surveillance Team |
| EPI_ISL_736395, EPI_ISL_736396, EPI_ISL_736397 | Lighthouse Lab in Milton Keynes | Wellcome Sanger Institute for the COVID-19 Genomics UK (COG-UK) Consortium | The Lighthouse Lab in Milton Keynes and Alex Alderton, Roberto Amato, Sonia Goncalves, Ewan Harrison, David K. Jackson, Ian Johnston, Dominic Kwiatkowski, Cordelia Langford, John Sillitoe on behalf of the Wellcome Sanger Institute COVID-19 Surveillance Team |
| EPI_ISL_736398, EPI_ISL_736399, EPI_ISL_736400 | Lighthouse Lab in Glasgow | Wellcome Sanger Institute for the COVID-19 Genomics UK (COG-UK) Consortium | Harper VanSteenhouse, Yumi Kasai, David Gray, Carol Clugston, Anna Dominiczak and Alex Alderton, Roberto Amato, Sonia Goncalves, Ewan Harrison, David K. Jackson, Ian Johnston, Dominic Kwiatkowski, Cordelia Langford, John Sillitoe on behalf of the Wellcome Sanger Institute COVID-19 Surveillance Team |
| EPI_ISL_736401, EPI_ISL_736402, EPI_ISL_736403 | Lighthouse Lab in Milton Keynes | Wellcome Sanger Institute for the COVID-19 Genomics UK (COG-UK) Consortium | The Lighthouse Lab in Milton Keynes and Alex Alderton, Roberto Amato, Sonia Goncalves, Ewan Harrison, David K. Jackson, Ian Johnston, Dominic Kwiatkowski, Cordelia Langford, John Sillitoe on behalf of the Wellcome Sanger Institute COVID-19 Surveillance Team |
| EPI_ISL_736404 | Lighthouse Lab in Glasgow | Wellcome Sanger Institute for the COVID-19 Genomics UK (COG-UK) Consortium | Harper VanSteenhouse, Yumi Kasai, David Gray, Carol Clugston, Anna Dominiczak and Alex Alderton, Roberto Amato, Sonia Goncalves, Ewan Harrison, David K. Jackson, Ian Johnston, Dominic Kwiatkowski, Cordelia Langford, John Sillitoe on behalf of the Wellcome Sanger Institute COVID-19 Surveillance Team |
| EPI_ISL_736405, EPI_ISL_736406, EPI_ISL_736407, EPI_ISL_736409, EPI_ISL_736410 | Lighthouse Lab in Milton Keynes | Wellcome Sanger Institute for the COVID-19 Genomics UK (COG-UK) Consortium | The Lighthouse Lab in Milton Keynes and Alex Alderton, Roberto Amato, Sonia Goncalves, Ewan Harrison, David K. Jackson, Ian Johnston, Dominic Kwiatkowski, Cordelia Langford, John Sillitoe on behalf of the Wellcome Sanger Institute COVID-19 Surveillance Team |
| EPI_ISL_736411, EPI_ISL_736412 | Lighthouse Lab in Glasgow | Wellcome Sanger Institute for the COVID-19 Genomics UK (COG-UK) Consortium | Harper VanSteenhouse, Yumi Kasai, David Gray, Carol Clugston, Anna Dominiczak and Alex Alderton, Roberto Amato, Sonia Goncalves, Ewan Harrison, David K. Jackson, Ian Johnston, Dominic Kwiatkowski, Cordelia Langford, John Sillitoe on behalf of the Wellcome Sanger Institute COVID-19 Surveillance Team |
| EPI_ISL_736413 | Lighthouse Lab in Milton Keynes | Wellcome Sanger Institute for the COVID-19 Genomics UK (COG-UK) Consortium | The Lighthouse Lab in Milton Keynes and Alex Alderton, Roberto Amato, Sonia Goncalves, Ewan Harrison, David K. Jackson, Ian Johnston, Dominic Kwiatkowski, Cordelia Langford, John Sillitoe on behalf of the Wellcome Sanger Institute COVID-19 Surveillance Team |
| EPI_ISL_736414 | Lighthouse Lab in Glasgow | Wellcome Sanger Institute for the COVID-19 Genomics UK | Harper VanSteenhouse, Yumi Kasai, David Gray, Carol Clugston, Anna Dominiczak and Alex Alderton, Roberto Amato, Sonia Goncalves, Ewan Harrison, |

| | | | |
|---|---|---|---|
| | | (COG-UK) Consortium | David K. Jackson, Ian Johnston, Dominic Kwiatkowski, Cordelia Langford, John Sillitoe on behalf of the Wellcome Sanger Institute COVID-19 Surveillance Team |
| EPI_ISL_736415 | Lighthouse Lab in Milton Keynes | Wellcome Sanger Institute for the COVID-19 Genomics UK (COG-UK) Consortium | The Lighthouse Lab in Milton Keynes and Alex Alderton, Roberto Amato, Sonia Goncalves, Ewan Harrison, David K. Jackson, Ian Johnston, Dominic Kwiatkowski, Cordelia Langford, John Sillitoe on behalf of the Wellcome Sanger Institute COVID-19 Surveillance Team |
| EPI_ISL_736416, EPI_ISL_736417 | Lighthouse Lab in Glasgow | Wellcome Sanger Institute for the COVID-19 Genomics UK (COG-UK) Consortium | Harper VanSteenhouse, Yumi Kasai, David Gray, Carol Clugston, Anna Dominiczak and Alex Alderton, Roberto Amato, Sonia Goncalves, Ewan Harrison, David K. Jackson, Ian Johnston, Dominic Kwiatkowski, Cordelia Langford, John Sillitoe on behalf of the Wellcome Sanger Institute COVID-19 Surveillance Team |
| EPI_ISL_736418 | Lighthouse Lab in Milton Keynes | Wellcome Sanger Institute for the COVID-19 Genomics UK (COG-UK) Consortium | The Lighthouse Lab in Milton Keynes and Alex Alderton, Roberto Amato, Sonia Goncalves, Ewan Harrison, David K. Jackson, Ian Johnston, Dominic Kwiatkowski, Cordelia Langford, John Sillitoe on behalf of the Wellcome Sanger Institute COVID-19 Surveillance Team |
| EPI_ISL_736419 | Lighthouse Lab in Glasgow | Wellcome Sanger Institute for the COVID-19 Genomics UK (COG-UK) Consortium | Harper VanSteenhouse, Yumi Kasai, David Gray, Carol Clugston, Anna Dominiczak and Alex Alderton, Roberto Amato, Sonia Goncalves, Ewan Harrison, David K. Jackson, Ian Johnston, Dominic Kwiatkowski, Cordelia Langford, John Sillitoe on behalf of the Wellcome Sanger Institute COVID-19 Surveillance Team |
| EPI_ISL_736420, EPI_ISL_736421, EPI_ISL_736422 | Lighthouse Lab in Milton Keynes | Wellcome Sanger Institute for the COVID-19 Genomics UK (COG-UK) Consortium | The Lighthouse Lab in Milton Keynes and Alex Alderton, Roberto Amato, Sonia Goncalves, Ewan Harrison, David K. Jackson, Ian Johnston, Dominic Kwiatkowski, Cordelia Langford, John Sillitoe on behalf of the Wellcome Sanger Institute COVID-19 Surveillance Team |
| EPI_ISL_736423 | Lighthouse Lab in Glasgow | Wellcome Sanger Institute for the COVID-19 Genomics UK (COG-UK) Consortium | Harper VanSteenhouse, Yumi Kasai, David Gray, Carol Clugston, Anna Dominiczak and Alex Alderton, Roberto Amato, Sonia Goncalves, Ewan Harrison, David K. Jackson, Ian Johnston, Dominic Kwiatkowski, Cordelia Langford, John Sillitoe on behalf of the Wellcome Sanger Institute COVID-19 Surveillance Team |
| EPI_ISL_736424 | Lighthouse Lab in Milton Keynes | Wellcome Sanger Institute for the COVID-19 Genomics UK (COG-UK) Consortium | The Lighthouse Lab in Milton Keynes and Alex Alderton, Roberto Amato, Sonia Goncalves, Ewan Harrison, David K. Jackson, Ian Johnston, Dominic Kwiatkowski, Cordelia Langford, John Sillitoe on behalf of the Wellcome Sanger Institute COVID-19 Surveillance Team |
| EPI_ISL_736425, EPI_ISL_736426 | Lighthouse Lab in Glasgow | Wellcome Sanger Institute for the COVID-19 Genomics UK (COG-UK) Consortium | Harper VanSteenhouse, Yumi Kasai, David Gray, Carol Clugston, Anna Dominiczak and Alex Alderton, Roberto Amato, Sonia Goncalves, Ewan Harrison, David K. Jackson, Ian Johnston, Dominic Kwiatkowski, Cordelia Langford, John Sillitoe on behalf of the Wellcome Sanger Institute COVID-19 Surveillance Team |
| EPI_ISL_736427 | Lighthouse Lab in Milton Keynes | Wellcome Sanger Institute for the COVID-19 Genomics UK (COG-UK) Consortium | The Lighthouse Lab in Milton Keynes and Alex Alderton, Roberto Amato, Sonia Goncalves, Ewan Harrison, David K. Jackson, Ian Johnston, Dominic Kwiatkowski, Cordelia Langford, John Sillitoe on behalf of the Wellcome Sanger Institute COVID-19 Surveillance Team |
| EPI_ISL_736428, EPI_ISL_736429, EPI_ISL_736430, EPI_ISL_736431 | Lighthouse Lab in Glasgow | Wellcome Sanger Institute for the COVID-19 Genomics UK (COG-UK) Consortium | Harper VanSteenhouse, Yumi Kasai, David Gray, Carol Clugston, Anna Dominiczak and Alex Alderton, Roberto Amato, Sonia Goncalves, Ewan Harrison, David K. Jackson, Ian Johnston, Dominic Kwiatkowski, Cordelia Langford, John Sillitoe on behalf of the Wellcome Sanger Institute COVID-19 Surveillance Team |
| EPI_ISL_736432, EPI_ISL_736433, EPI_ISL_736434, EPI_ISL_736435, EPI_ISL_736436 | Lighthouse Lab in Milton Keynes | Wellcome Sanger Institute for the COVID-19 Genomics UK (COG-UK) Consortium | The Lighthouse Lab in Milton Keynes and Alex Alderton, Roberto Amato, Sonia Goncalves, Ewan Harrison, David K. Jackson, Ian Johnston, Dominic Kwiatkowski, Cordelia Langford, John Sillitoe on behalf of the Wellcome Sanger Institute COVID-19 Surveillance Team |
| EPI_ISL_736437, EPI_ISL_736438 | Lighthouse Lab in Glasgow | Wellcome Sanger Institute for the COVID-19 Genomics UK (COG-UK) Consortium | Harper VanSteenhouse, Yumi Kasai, David Gray, Carol Clugston, Anna Dominiczak and Alex Alderton, Roberto Amato, Sonia Goncalves, Ewan Harrison, David K. Jackson, Ian Johnston, Dominic Kwiatkowski, Cordelia Langford, John Sillitoe on behalf of the Wellcome Sanger Institute COVID-19 Surveillance Team |
| EPI_ISL_736439, EPI_ISL_736440, EPI_ISL_736441 | Lighthouse Lab in Milton Keynes | Wellcome Sanger Institute for the COVID-19 Genomics UK (COG-UK) Consortium | The Lighthouse Lab in Milton Keynes and Alex Alderton, Roberto Amato, Sonia Goncalves, Ewan Harrison, David K. Jackson, Ian Johnston, Dominic Kwiatkowski, Cordelia Langford, John Sillitoe on behalf of the Wellcome Sanger Institute COVID-19 Surveillance Team |
| EPI_ISL_736442 | Lighthouse Lab in Glasgow | Wellcome Sanger Institute for the COVID-19 Genomics UK (COG-UK) Consortium | Harper VanSteenhouse, Yumi Kasai, David Gray, Carol Clugston, Anna Dominiczak and Alex Alderton, Roberto Amato, Sonia Goncalves, Ewan Harrison, David K. Jackson, Ian Johnston, Dominic Kwiatkowski, Cordelia Langford, John Sillitoe on behalf of the Wellcome Sanger Institute COVID-19 Surveillance Team |
| EPI_ISL_736443 | Lighthouse Lab in Milton Keynes | Wellcome Sanger Institute for the COVID-19 Genomics UK (COG-UK) Consortium | The Lighthouse Lab in Milton Keynes and Alex Alderton, Roberto Amato, Sonia Goncalves, Ewan Harrison, David K. Jackson, Ian Johnston, Dominic Kwiatkowski, Cordelia Langford, John Sillitoe on behalf of the Wellcome Sanger Institute COVID-19 Surveillance Team |
| EPI_ISL_736444, EPI_ISL_736445 | Lighthouse Lab in Glasgow | Wellcome Sanger Institute for the COVID-19 Genomics UK (COG-UK) Consortium | Harper VanSteenhouse, Yumi Kasai, David Gray, Carol Clugston, Anna Dominiczak and Alex Alderton, Roberto Amato, Sonia Goncalves, Ewan Harrison, David K. Jackson, Ian Johnston, Dominic Kwiatkowski, Cordelia Langford, John Sillitoe on behalf of the Wellcome Sanger Institute COVID-19 Surveillance Team |
| EPI_ISL_736446, EPI_ISL_736447, EPI_ISL_736448, EPI_ISL_736450 | Lighthouse Lab in Milton Keynes | Wellcome Sanger Institute for the COVID-19 Genomics UK (COG-UK) Consortium | The Lighthouse Lab in Milton Keynes and Alex Alderton, Roberto Amato, Sonia Goncalves, Ewan Harrison, David K. Jackson, Ian Johnston, Dominic Kwiatkowski, Cordelia Langford, John Sillitoe on behalf of the Wellcome Sanger Institute COVID-19 Surveillance Team |
| EPI_ISL_736451 | Lighthouse Lab in Glasgow | Wellcome Sanger Institute for the COVID-19 Genomics UK (COG-UK) Consortium | Harper VanSteenhouse, Yumi Kasai, David Gray, Carol Clugston, Anna Dominiczak and Alex Alderton, Roberto Amato, Sonia Goncalves, Ewan Harrison, David K. Jackson, Ian Johnston, Dominic Kwiatkowski, Cordelia Langford, John Sillitoe on behalf of the Wellcome Sanger Institute COVID-19 Surveillance Team |
| EPI_ISL_736452, EPI_ISL_736453, EPI_ISL_736454 | Lighthouse Lab in Milton Keynes | Wellcome Sanger Institute for the COVID-19 Genomics UK (COG-UK) Consortium | The Lighthouse Lab in Milton Keynes and Alex Alderton, Roberto Amato, Sonia Goncalves, Ewan Harrison, David K. Jackson, Ian Johnston, Dominic Kwiatkowski, Cordelia Langford, John Sillitoe on behalf of the Wellcome Sanger Institute COVID-19 Surveillance Team |
| EPI_ISL_736455, EPI_ISL_736456 | Lighthouse Lab in Glasgow | Wellcome Sanger Institute for the COVID-19 Genomics UK (COG-UK) Consortium | Harper VanSteenhouse, Yumi Kasai, David Gray, Carol Clugston, Anna Dominiczak and Alex Alderton, Roberto Amato, Sonia Goncalves, Ewan Harrison, David K. Jackson, Ian Johnston, Dominic Kwiatkowski, Cordelia Langford, John Sillitoe on behalf of the Wellcome Sanger Institute COVID-19 Surveillance Team |
| EPI_ISL_736457 | Lighthouse Lab in Milton Keynes | Wellcome Sanger Institute for the COVID-19 Genomics UK (COG-UK) Consortium | The Lighthouse Lab in Milton Keynes and Alex Alderton, Roberto Amato, Sonia Goncalves, Ewan Harrison, David K. Jackson, Ian Johnston, Dominic Kwiatkowski, Cordelia Langford, John Sillitoe on behalf of the Wellcome Sanger Institute COVID-19 Surveillance Team |
| EPI_ISL_736459, EPI_ISL_736460, EPI_ISL_736461 | Lighthouse Lab in Glasgow | Wellcome Sanger Institute for the COVID-19 Genomics UK (COG-UK) Consortium | Harper VanSteenhouse, Yumi Kasai, David Gray, Carol Clugston, Anna Dominiczak and Alex Alderton, Roberto Amato, Sonia Goncalves, Ewan Harrison, David K. Jackson, Ian Johnston, Dominic Kwiatkowski, Cordelia Langford, John Sillitoe on behalf of the Wellcome Sanger Institute COVID-19 Surveillance Team |
| EPI_ISL_736462, EPI_ISL_736463 | Lighthouse Lab in Milton Keynes | Wellcome Sanger Institute for the COVID-19 Genomics UK (COG-UK) Consortium | The Lighthouse Lab in Milton Keynes and Alex Alderton, Roberto Amato, Sonia Goncalves, Ewan Harrison, David K. Jackson, Ian Johnston, Dominic Kwiatkowski, Cordelia Langford, John Sillitoe on behalf of the Wellcome Sanger Institute COVID-19 Surveillance Team |
| EPI_ISL_736464 | Lighthouse Lab in Glasgow | Wellcome Sanger Institute for the COVID-19 Genomics UK (COG-UK) Consortium | Harper VanSteenhouse, Yumi Kasai, David Gray, Carol Clugston, Anna Dominiczak and Alex Alderton, Roberto Amato, Sonia Goncalves, Ewan Harrison, David K. Jackson, Ian Johnston, Dominic Kwiatkowski, Cordelia Langford, John Sillitoe on behalf of the Wellcome Sanger Institute COVID-19 Surveillance Team |
| EPI_ISL_736465 | Lighthouse Lab in Milton Keynes | Wellcome Sanger Institute for the COVID-19 Genomics UK (COG-UK) Consortium | The Lighthouse Lab in Milton Keynes and Alex Alderton, Roberto Amato, Sonia Goncalves, Ewan Harrison, David K. Jackson, Ian Johnston, Dominic Kwiatkowski, Cordelia Langford, John Sillitoe on behalf of the Wellcome Sanger Institute COVID-19 Surveillance Team |
| EPI_ISL_736466 | Lighthouse Lab in Glasgow | Wellcome Sanger Institute for the COVID-19 Genomics UK (COG-UK) Consortium | Harper VanSteenhouse, Yumi Kasai, David Gray, Carol Clugston, Anna Dominiczak and Alex Alderton, Roberto Amato, Sonia Goncalves, Ewan Harrison, David K. Jackson, Ian Johnston, Dominic Kwiatkowski, Cordelia Langford, John Sillitoe on behalf of the Wellcome Sanger Institute COVID-19 Surveillance Team |
| EPI_ISL_736467 | Lighthouse Lab in Milton Keynes | Wellcome Sanger Institute for the COVID-19 Genomics UK (COG-UK) Consortium | The Lighthouse Lab in Milton Keynes and Alex Alderton, Roberto Amato, Sonia Goncalves, Ewan Harrison, David K. Jackson, Ian Johnston, Dominic Kwiatkowski, Cordelia Langford, John Sillitoe on behalf of the Wellcome Sanger Institute COVID-19 Surveillance Team |
| EPI_ISL_736468, EPI_ISL_736470, EPI_ISL_736471 | Lighthouse Lab in Glasgow | Wellcome Sanger Institute for the COVID-19 Genomics UK (COG-UK) Consortium | Harper VanSteenhouse, Yumi Kasai, David Gray, Carol Clugston, Anna Dominiczak and Alex Alderton, Roberto Amato, Sonia Goncalves, Ewan Harrison, David K. Jackson, Ian Johnston, Dominic Kwiatkowski, Cordelia Langford, John Sillitoe on behalf of the Wellcome Sanger Institute COVID-19 Surveillance Team |

| | | | |
|---|---|---|---|
| EPI_ISL_736472, EPI_ISL_736473 | Lighthouse Lab in Milton Keynes | Wellcome Sanger Institute for the COVID-19 Genomics UK (COG-UK) Consortium | The Lighthouse Lab in Milton Keynes and Alex Alderton, Roberto Amato, Sonia Goncalves, Ewan Harrison, David K. Jackson, Ian Johnston, Dominic Kwiatkowski, Cordelia Langford, John Sillitoe on behalf of the Wellcome Sanger Institute COVID-19 Surveillance Team |
| EPI_ISL_736474 | Lighthouse Lab in Glasgow | Wellcome Sanger Institute for the COVID-19 Genomics UK (COG-UK) Consortium | Harper VanSteenhouse, Yumi Kasai, David Gray, Carol Clugston, Anna Dominiczak and Alex Alderton, Roberto Amato, Sonia Goncalves, Ewan Harrison, David K. Jackson, Ian Johnston, Dominic Kwiatkowski, Cordelia Langford, John Sillitoe on behalf of the Wellcome Sanger Institute COVID-19 Surveillance Team |
| EPI_ISL_736475 | Lighthouse Lab in Milton Keynes | Wellcome Sanger Institute for the COVID-19 Genomics UK (COG-UK) Consortium | The Lighthouse Lab in Milton Keynes and Alex Alderton, Roberto Amato, Sonia Goncalves, Ewan Harrison, David K. Jackson, Ian Johnston, Dominic Kwiatkowski, Cordelia Langford, John Sillitoe on behalf of the Wellcome Sanger Institute COVID-19 Surveillance Team |
| EPI_ISL_736476, EPI_ISL_736478, EPI_ISL_736479 | Lighthouse Lab in Glasgow | Wellcome Sanger Institute for the COVID-19 Genomics UK (COG-UK) Consortium | Harper VanSteenhouse, Yumi Kasai, David Gray, Carol Clugston, Anna Dominiczak and Alex Alderton, Roberto Amato, Sonia Goncalves, Ewan Harrison, David K. Jackson, Ian Johnston, Dominic Kwiatkowski, Cordelia Langford, John Sillitoe on behalf of the Wellcome Sanger Institute COVID-19 Surveillance Team |
| EPI_ISL_736480 | Lighthouse Lab in Milton Keynes | Wellcome Sanger Institute for the COVID-19 Genomics UK (COG-UK) Consortium | The Lighthouse Lab in Milton Keynes and Alex Alderton, Roberto Amato, Sonia Goncalves, Ewan Harrison, David K. Jackson, Ian Johnston, Dominic Kwiatkowski, Cordelia Langford, John Sillitoe on behalf of the Wellcome Sanger Institute COVID-19 Surveillance Team |
| EPI_ISL_736481, EPI_ISL_736482, EPI_ISL_736484, EPI_ISL_736485, EPI_ISL_736486 | Lighthouse Lab in Alderley Park | Wellcome Sanger Institute for the COVID-19 Genomics UK (COG-UK) Consortium | Jacquelyn Wynn, Mairead Hyland, The Lighthouse Lab in Alderley Park and Alex Alderton, Roberto Amato, Sonia Goncalves, Ewan Harrison, David K. Jackson, Ian Johnston, Dominic Kwiatkowski, Cordelia Langford, John Sillitoe on behalf of the Wellcome Sanger Institute COVID-19 Surveillance Team |
| EPI_ISL_736487 | Lighthouse Lab in Milton Keynes | Wellcome Sanger Institute for the COVID-19 Genomics UK (COG-UK) Consortium | The Lighthouse Lab in Milton Keynes and Alex Alderton, Roberto Amato, Sonia Goncalves, Ewan Harrison, David K. Jackson, Ian Johnston, Dominic Kwiatkowski, Cordelia Langford, John Sillitoe on behalf of the Wellcome Sanger Institute COVID-19 Surveillance Team |
| EPI_ISL_736493, EPI_ISL_736494, EPI_ISL_736495, EPI_ISL_736496 | Lighthouse Lab in Alderley Park | Wellcome Sanger Institute for the COVID-19 Genomics UK (COG-UK) Consortium | Jacquelyn Wynn, Mairead Hyland, The Lighthouse Lab in Alderley Park and Alex Alderton, Roberto Amato, Sonia Goncalves, Ewan Harrison, David K. Jackson, Ian Johnston, Dominic Kwiatkowski, Cordelia Langford, John Sillitoe on behalf of the Wellcome Sanger Institute COVID-19 Surveillance Team |
| EPI_ISL_736497 | Lighthouse Lab in Milton Keynes | Wellcome Sanger Institute for the COVID-19 Genomics UK (COG-UK) Consortium | The Lighthouse Lab in Milton Keynes and Alex Alderton, Roberto Amato, Sonia Goncalves, Ewan Harrison, David K. Jackson, Ian Johnston, Dominic Kwiatkowski, Cordelia Langford, John Sillitoe on behalf of the Wellcome Sanger Institute COVID-19 Surveillance Team |
| EPI_ISL_736498, EPI_ISL_736499, EPI_ISL_736500 | Lighthouse Lab in Alderley Park | Wellcome Sanger Institute for the COVID-19 Genomics UK (COG-UK) Consortium | Jacquelyn Wynn, Mairead Hyland, The Lighthouse Lab in Alderley Park and Alex Alderton, Roberto Amato, Sonia Goncalves, Ewan Harrison, David K. Jackson, Ian Johnston, Dominic Kwiatkowski, Cordelia Langford, John Sillitoe on behalf of the Wellcome Sanger Institute COVID-19 Surveillance Team |
| EPI_ISL_736501, EPI_ISL_736502 | Lighthouse Lab in Milton Keynes | Wellcome Sanger Institute for the COVID-19 Genomics UK (COG-UK) Consortium | The Lighthouse Lab in Milton Keynes and Alex Alderton, Roberto Amato, Sonia Goncalves, Ewan Harrison, David K. Jackson, Ian Johnston, Dominic Kwiatkowski, Cordelia Langford, John Sillitoe on behalf of the Wellcome Sanger Institute COVID-19 Surveillance Team |
| EPI_ISL_736503 | Lighthouse Lab in Glasgow | Wellcome Sanger Institute for the COVID-19 Genomics UK (COG-UK) Consortium | Harper VanSteenhouse, Yumi Kasai, David Gray, Carol Clugston, Anna Dominiczak and Alex Alderton, Roberto Amato, Sonia Goncalves, Ewan Harrison, David K. Jackson, Ian Johnston, Dominic Kwiatkowski, Cordelia Langford, John Sillitoe on behalf of the Wellcome Sanger Institute COVID-19 Surveillance Team |
| EPI_ISL_736504 | Lighthouse Lab in Milton Keynes | Wellcome Sanger Institute for the COVID-19 Genomics UK (COG-UK) Consortium | The Lighthouse Lab in Milton Keynes and Alex Alderton, Roberto Amato, Sonia Goncalves, Ewan Harrison, David K. Jackson, Ian Johnston, Dominic Kwiatkowski, Cordelia Langford, John Sillitoe on behalf of the Wellcome Sanger Institute COVID-19 Surveillance Team |
| EPI_ISL_736505, EPI_ISL_736506, EPI_ISL_736507 | Lighthouse Lab in Alderley Park | Wellcome Sanger Institute for the COVID-19 Genomics UK (COG-UK) Consortium | Jacquelyn Wynn, Mairead Hyland, The Lighthouse Lab in Alderley Park and Alex Alderton, Roberto Amato, Sonia Goncalves, Ewan Harrison, David K. Jackson, Ian Johnston, Dominic Kwiatkowski, Cordelia Langford, John Sillitoe on behalf of the Wellcome Sanger Institute COVID-19 Surveillance Team |
| EPI_ISL_736508 | Lighthouse Lab in Milton Keynes | Wellcome Sanger Institute for the COVID-19 Genomics UK (COG-UK) Consortium | The Lighthouse Lab in Milton Keynes and Alex Alderton, Roberto Amato, Sonia Goncalves, Ewan Harrison, David K. Jackson, Ian Johnston, Dominic Kwiatkowski, Cordelia Langford, John Sillitoe on behalf of the Wellcome Sanger Institute COVID-19 Surveillance Team |
| EPI_ISL_736509, EPI_ISL_736510, EPI_ISL_736511, EPI_ISL_736513 | Lighthouse Lab in Alderley Park | Wellcome Sanger Institute for the COVID-19 Genomics UK (COG-UK) Consortium | Jacquelyn Wynn, Mairead Hyland, The Lighthouse Lab in Alderley Park and Alex Alderton, Roberto Amato, Sonia Goncalves, Ewan Harrison, David K. Jackson, Ian Johnston, Dominic Kwiatkowski, Cordelia Langford, John Sillitoe on behalf of the Wellcome Sanger Institute COVID-19 Surveillance Team |
| EPI_ISL_736514, EPI_ISL_736515 | Lighthouse Lab in Milton Keynes | Wellcome Sanger Institute for the COVID-19 Genomics UK (COG-UK) Consortium | The Lighthouse Lab in Milton Keynes and Alex Alderton, Roberto Amato, Sonia Goncalves, Ewan Harrison, David K. Jackson, Ian Johnston, Dominic Kwiatkowski, Cordelia Langford, John Sillitoe on behalf of the Wellcome Sanger Institute COVID-19 Surveillance Team |
| EPI_ISL_736516, EPI_ISL_736517, EPI_ISL_736518, EPI_ISL_736520, EPI_ISL_736521, EPI_ISL_736522 | Lighthouse Lab in Alderley Park | Wellcome Sanger Institute for the COVID-19 Genomics UK (COG-UK) Consortium | Jacquelyn Wynn, Mairead Hyland, The Lighthouse Lab in Alderley Park and Alex Alderton, Roberto Amato, Sonia Goncalves, Ewan Harrison, David K. Jackson, Ian Johnston, Dominic Kwiatkowski, Cordelia Langford, John Sillitoe on behalf of the Wellcome Sanger Institute COVID-19 Surveillance Team |
| EPI_ISL_736523 | Lighthouse Lab in Milton Keynes | Wellcome Sanger Institute for the COVID-19 Genomics UK (COG-UK) Consortium | The Lighthouse Lab in Milton Keynes and Alex Alderton, Roberto Amato, Sonia Goncalves, Ewan Harrison, David K. Jackson, Ian Johnston, Dominic Kwiatkowski, Cordelia Langford, John Sillitoe on behalf of the Wellcome Sanger Institute COVID-19 Surveillance Team |
| EPI_ISL_736524 | Lighthouse Lab in Alderley Park | Wellcome Sanger Institute for the COVID-19 Genomics UK (COG-UK) Consortium | Jacquelyn Wynn, Mairead Hyland, The Lighthouse Lab in Alderley Park and Alex Alderton, Roberto Amato, Sonia Goncalves, Ewan Harrison, David K. Jackson, Ian Johnston, Dominic Kwiatkowski, Cordelia Langford, John Sillitoe on behalf of the Wellcome Sanger Institute COVID-19 Surveillance Team |
| EPI_ISL_736525 | Lighthouse Lab in Milton Keynes | Wellcome Sanger Institute for the COVID-19 Genomics UK (COG-UK) Consortium | The Lighthouse Lab in Milton Keynes and Alex Alderton, Roberto Amato, Sonia Goncalves, Ewan Harrison, David K. Jackson, Ian Johnston, Dominic Kwiatkowski, Cordelia Langford, John Sillitoe on behalf of the Wellcome Sanger Institute COVID-19 Surveillance Team |
| EPI_ISL_736526 | Lighthouse Lab in Alderley Park | Wellcome Sanger Institute for the COVID-19 Genomics UK (COG-UK) Consortium | Jacquelyn Wynn, Mairead Hyland, The Lighthouse Lab in Alderley Park and Alex Alderton, Roberto Amato, Sonia Goncalves, Ewan Harrison, David K. Jackson, Ian Johnston, Dominic Kwiatkowski, Cordelia Langford, John Sillitoe on behalf of the Wellcome Sanger Institute COVID-19 Surveillance Team |
| EPI_ISL_736527, EPI_ISL_736528 | Lighthouse Lab in Milton Keynes | Wellcome Sanger Institute for the COVID-19 Genomics UK (COG-UK) Consortium | The Lighthouse Lab in Milton Keynes and Alex Alderton, Roberto Amato, Sonia Goncalves, Ewan Harrison, David K. Jackson, Ian Johnston, Dominic Kwiatkowski, Cordelia Langford, John Sillitoe on behalf of the Wellcome Sanger Institute COVID-19 Surveillance Team |
| EPI_ISL_736530, EPI_ISL_736531, EPI_ISL_736534, EPI_ISL_736535 | Lighthouse Lab in Alderley Park | Wellcome Sanger Institute for the COVID-19 Genomics UK (COG-UK) Consortium | Jacquelyn Wynn, Mairead Hyland, The Lighthouse Lab in Alderley Park and Alex Alderton, Roberto Amato, Sonia Goncalves, Ewan Harrison, David K. Jackson, Ian Johnston, Dominic Kwiatkowski, Cordelia Langford, John Sillitoe on behalf of the Wellcome Sanger Institute COVID-19 Surveillance Team |
| EPI_ISL_736536 | Lighthouse Lab in Milton Keynes | Wellcome Sanger Institute for the COVID-19 Genomics UK (COG-UK) Consortium | The Lighthouse Lab in Milton Keynes and Alex Alderton, Roberto Amato, Sonia Goncalves, Ewan Harrison, David K. Jackson, Ian Johnston, Dominic Kwiatkowski, Cordelia Langford, John Sillitoe on behalf of the Wellcome Sanger Institute COVID-19 Surveillance Team |
| EPI_ISL_736537 | Lighthouse Lab in Alderley Park | Wellcome Sanger Institute for the COVID-19 Genomics UK (COG-UK) Consortium | Jacquelyn Wynn, Mairead Hyland, The Lighthouse Lab in Alderley Park and Alex Alderton, Roberto Amato, Sonia Goncalves, Ewan Harrison, David K. Jackson, Ian Johnston, Dominic Kwiatkowski, Cordelia Langford, John Sillitoe on behalf of the Wellcome Sanger Institute COVID-19 Surveillance Team |
| EPI_ISL_736538 | Lighthouse Lab in Milton Keynes | Wellcome Sanger Institute for the COVID-19 Genomics UK (COG-UK) Consortium | The Lighthouse Lab in Milton Keynes and Alex Alderton, Roberto Amato, Sonia Goncalves, Ewan Harrison, David K. Jackson, Ian Johnston, Dominic Kwiatkowski, Cordelia Langford, John Sillitoe on behalf of the Wellcome Sanger Institute COVID-19 Surveillance Team |
| EPI_ISL_736539, EPI_ISL_736540, EPI_ISL_736541 | Lighthouse Lab in Alderley Park | Wellcome Sanger Institute for the COVID-19 Genomics UK (COG-UK) Consortium | Jacquelyn Wynn, Mairead Hyland, The Lighthouse Lab in Alderley Park and Alex Alderton, Roberto Amato, Sonia Goncalves, Ewan Harrison, David K. Jackson, Ian Johnston, Dominic Kwiatkowski, Cordelia Langford, John Sillitoe on behalf of the Wellcome Sanger Institute COVID-19 Surveillance Team |
| EPI_ISL_736542 | Lighthouse Lab in Milton Keynes | Wellcome Sanger Institute for the COVID-19 Genomics UK (COG-UK) Consortium | The Lighthouse Lab in Milton Keynes and Alex Alderton, Roberto Amato, Sonia Goncalves, Ewan Harrison, David K. Jackson, Ian Johnston, Dominic Kwiatkowski, Cordelia Langford, John Sillitoe on behalf of the Wellcome Sanger Institute COVID-19 Surveillance Team |
| EPI_ISL_736543, EPI_ISL_736544, EPI_ISL_736545, EPI_ISL_736546, EPI_ISL_736548 | Lighthouse Lab in Alderley Park | Wellcome Sanger Institute for the COVID-19 Genomics UK (COG-UK) Consortium | Jacquelyn Wynn, Mairead Hyland, The Lighthouse Lab in Alderley Park and Alex Alderton, Roberto Amato, Sonia Goncalves, Ewan Harrison, David K. Jackson, Ian Johnston, Dominic Kwiatkowski, Cordelia Langford, John Sillitoe on behalf of the Wellcome Sanger Institute COVID-19 Surveillance Team |
| EPI_ISL_736549, EPI_ISL_736550, EPI_ISL_736551, EPI_ISL_736552 | Lighthouse Lab in Milton Keynes | Wellcome Sanger Institute for the COVID-19 Genomics UK (COG-UK) Consortium | The Lighthouse Lab in Milton Keynes and Alex Alderton, Roberto Amato, Sonia Goncalves, Ewan Harrison, David K. Jackson, Ian Johnston, Dominic Kwiatkowski, Cordelia Langford, John Sillitoe on behalf of the Wellcome Sanger Institute COVID-19 Surveillance Team |
| EPI_ISL_736553, EPI_ISL_736554, EPI_ISL_736555, EPI_ISL_736557 | Lighthouse Lab in Alderley Park | Wellcome Sanger Institute for the COVID-19 Genomics UK (COG-UK) Consortium | Jacquelyn Wynn, Mairead Hyland, The Lighthouse Lab in Alderley Park and Alex Alderton, Roberto Amato, Sonia Goncalves, Ewan Harrison, David K. Jackson, Ian Johnston, Dominic Kwiatkowski, Cordelia Langford, John Sillitoe on behalf of the Wellcome Sanger Institute COVID-19 Surveillance Team |
| EPI_ISL_736558 | Lighthouse Lab in Milton Keynes | Wellcome Sanger Institute for the COVID-19 Genomics UK (COG-UK) Consortium | The Lighthouse Lab in Milton Keynes and Alex Alderton, Roberto Amato, Sonia Goncalves, Ewan Harrison, David K. Jackson, Ian Johnston, Dominic Kwiatkowski, Cordelia Langford, John Sillitoe on behalf of the Wellcome Sanger Institute COVID-19 Surveillance Team |

| | | | |
|---|---|---|---|
| EPI_ISL_736559, EPI_ISL_736560, EPI_ISL_736561, EPI_ISL_736562 | Lighthouse Lab in Alderley Park | Wellcome Sanger Institute for the COVID-19 Genomics UK (COG-UK) Consortium | Jacquelyn Wynn, Mairead Hyland, The Lighthouse Lab in Alderley Park and Alex Alderton, Roberto Amato, Sonia Goncalves, Ewan Harrison, David K. Jackson, Ian Johnston, Dominic Kwiatkowski, Cordelia Langford, John Sillitoe on behalf of the Wellcome Sanger Institute COVID-19 Surveillance Team |
| EPI_ISL_736563 | Lighthouse Lab in Milton Keynes | Wellcome Sanger Institute for the COVID-19 Genomics UK (COG-UK) Consortium | The Lighthouse Lab in Milton Keynes and Alex Alderton, Roberto Amato, Sonia Goncalves, Ewan Harrison, David K. Jackson, Ian Johnston, Dominic Kwiatkowski, Cordelia Langford, John Sillitoe on behalf of the Wellcome Sanger Institute COVID-19 Surveillance Team |
| EPI_ISL_736564 | Lighthouse Lab in Alderley Park | Wellcome Sanger Institute for the COVID-19 Genomics UK (COG-UK) Consortium | Jacquelyn Wynn, Mairead Hyland, The Lighthouse Lab in Alderley Park and Alex Alderton, Roberto Amato, Sonia Goncalves, Ewan Harrison, David K. Jackson, Ian Johnston, Dominic Kwiatkowski, Cordelia Langford, John Sillitoe on behalf of the Wellcome Sanger Institute COVID-19 Surveillance Team |
| EPI_ISL_736565 | Lighthouse Lab in Milton Keynes | Wellcome Sanger Institute for the COVID-19 Genomics UK (COG-UK) Consortium | The Lighthouse Lab in Milton Keynes and Alex Alderton, Roberto Amato, Sonia Goncalves, Ewan Harrison, David K. Jackson, Ian Johnston, Dominic Kwiatkowski, Cordelia Langford, John Sillitoe on behalf of the Wellcome Sanger Institute COVID-19 Surveillance Team |
| EPI_ISL_736566, EPI_ISL_736568, EPI_ISL_736569, EPI_ISL_736570 | Lighthouse Lab in Alderley Park | Wellcome Sanger Institute for the COVID-19 Genomics UK (COG-UK) Consortium | Jacquelyn Wynn, Mairead Hyland, The Lighthouse Lab in Alderley Park and Alex Alderton, Roberto Amato, Sonia Goncalves, Ewan Harrison, David K. Jackson, Ian Johnston, Dominic Kwiatkowski, Cordelia Langford, John Sillitoe on behalf of the Wellcome Sanger Institute COVID-19 Surveillance Team |
| EPI_ISL_736571 | Lighthouse Lab in Milton Keynes | Wellcome Sanger Institute for the COVID-19 Genomics UK (COG-UK) Consortium | The Lighthouse Lab in Milton Keynes and Alex Alderton, Roberto Amato, Sonia Goncalves, Ewan Harrison, David K. Jackson, Ian Johnston, Dominic Kwiatkowski, Cordelia Langford, John Sillitoe on behalf of the Wellcome Sanger Institute COVID-19 Surveillance Team |
| EPI_ISL_736572, EPI_ISL_736574, EPI_ISL_736575, EPI_ISL_736576, EPI_ISL_736577 | Lighthouse Lab in Alderley Park | Wellcome Sanger Institute for the COVID-19 Genomics UK (COG-UK) Consortium | Jacquelyn Wynn, Mairead Hyland, The Lighthouse Lab in Alderley Park and Alex Alderton, Roberto Amato, Sonia Goncalves, Ewan Harrison, David K. Jackson, Ian Johnston, Dominic Kwiatkowski, Cordelia Langford, John Sillitoe on behalf of the Wellcome Sanger Institute COVID-19 Surveillance Team |
| EPI_ISL_736578 | Lighthouse Lab in Milton Keynes | Wellcome Sanger Institute for the COVID-19 Genomics UK (COG-UK) Consortium | The Lighthouse Lab in Milton Keynes and Alex Alderton, Roberto Amato, Sonia Goncalves, Ewan Harrison, David K. Jackson, Ian Johnston, Dominic Kwiatkowski, Cordelia Langford, John Sillitoe on behalf of the Wellcome Sanger Institute COVID-19 Surveillance Team |
| EPI_ISL_736579, EPI_ISL_736581, EPI_ISL_736583, EPI_ISL_736584, EPI_ISL_736585 | Lighthouse Lab in Alderley Park | Wellcome Sanger Institute for the COVID-19 Genomics UK (COG-UK) Consortium | Jacquelyn Wynn, Mairead Hyland, The Lighthouse Lab in Alderley Park and Alex Alderton, Roberto Amato, Sonia Goncalves, Ewan Harrison, David K. Jackson, Ian Johnston, Dominic Kwiatkowski, Cordelia Langford, John Sillitoe on behalf of the Wellcome Sanger Institute COVID-19 Surveillance Team |
| EPI_ISL_736586 | Lighthouse Lab in Milton Keynes | Wellcome Sanger Institute for the COVID-19 Genomics UK (COG-UK) Consortium | The Lighthouse Lab in Milton Keynes and Alex Alderton, Roberto Amato, Sonia Goncalves, Ewan Harrison, David K. Jackson, Ian Johnston, Dominic Kwiatkowski, Cordelia Langford, John Sillitoe on behalf of the Wellcome Sanger Institute COVID-19 Surveillance Team |
| EPI_ISL_736587 | Lighthouse Lab in Alderley Park | Wellcome Sanger Institute for the COVID-19 Genomics UK (COG-UK) Consortium | Jacquelyn Wynn, Mairead Hyland, The Lighthouse Lab in Alderley Park and Alex Alderton, Roberto Amato, Sonia Goncalves, Ewan Harrison, David K. Jackson, Ian Johnston, Dominic Kwiatkowski, Cordelia Langford, John Sillitoe on behalf of the Wellcome Sanger Institute COVID-19 Surveillance Team |
| EPI_ISL_736588 | Lighthouse Lab in Milton Keynes | Wellcome Sanger Institute for the COVID-19 Genomics UK (COG-UK) Consortium | The Lighthouse Lab in Milton Keynes and Alex Alderton, Roberto Amato, Sonia Goncalves, Ewan Harrison, David K. Jackson, Ian Johnston, Dominic Kwiatkowski, Cordelia Langford, John Sillitoe on behalf of the Wellcome Sanger Institute COVID-19 Surveillance Team |
| EPI_ISL_736589 | Lighthouse Lab in Alderley Park | Wellcome Sanger Institute for the COVID-19 Genomics UK (COG-UK) Consortium | Jacquelyn Wynn, Mairead Hyland, The Lighthouse Lab in Alderley Park and Alex Alderton, Roberto Amato, Sonia Goncalves, Ewan Harrison, David K. Jackson, Ian Johnston, Dominic Kwiatkowski, Cordelia Langford, John Sillitoe on behalf of the Wellcome Sanger Institute COVID-19 Surveillance Team |
| EPI_ISL_736590 | Lighthouse Lab in Milton Keynes | Wellcome Sanger Institute for the COVID-19 Genomics UK (COG-UK) Consortium | The Lighthouse Lab in Milton Keynes and Alex Alderton, Roberto Amato, Sonia Goncalves, Ewan Harrison, David K. Jackson, Ian Johnston, Dominic Kwiatkowski, Cordelia Langford, John Sillitoe on behalf of the Wellcome Sanger Institute COVID-19 Surveillance Team |
| EPI_ISL_736591, EPI_ISL_736592 | Lighthouse Lab in Alderley Park | Wellcome Sanger Institute for the COVID-19 Genomics UK (COG-UK) Consortium | Jacquelyn Wynn, Mairead Hyland, The Lighthouse Lab in Alderley Park and Alex Alderton, Roberto Amato, Sonia Goncalves, Ewan Harrison, David K. Jackson, Ian Johnston, Dominic Kwiatkowski, Cordelia Langford, John Sillitoe on behalf of the Wellcome Sanger Institute COVID-19 Surveillance Team |
| EPI_ISL_736593, EPI_ISL_736594 | Lighthouse Lab in Milton Keynes | Wellcome Sanger Institute for the COVID-19 Genomics UK (COG-UK) Consortium | The Lighthouse Lab in Milton Keynes and Alex Alderton, Roberto Amato, Sonia Goncalves, Ewan Harrison, David K. Jackson, Ian Johnston, Dominic Kwiatkowski, Cordelia Langford, John Sillitoe on behalf of the Wellcome Sanger Institute COVID-19 Surveillance Team |
| EPI_ISL_736595, EPI_ISL_736596 | Lighthouse Lab in Alderley Park | Wellcome Sanger Institute for the COVID-19 Genomics UK (COG-UK) Consortium | Jacquelyn Wynn, Mairead Hyland, The Lighthouse Lab in Alderley Park and Alex Alderton, Roberto Amato, Sonia Goncalves, Ewan Harrison, David K. Jackson, Ian Johnston, Dominic Kwiatkowski, Cordelia Langford, John Sillitoe on behalf of the Wellcome Sanger Institute COVID-19 Surveillance Team |
| EPI_ISL_736597, EPI_ISL_736598 | Lighthouse Lab in Milton Keynes | Wellcome Sanger Institute for the COVID-19 Genomics UK (COG-UK) Consortium | The Lighthouse Lab in Milton Keynes and Alex Alderton, Roberto Amato, Sonia Goncalves, Ewan Harrison, David K. Jackson, Ian Johnston, Dominic Kwiatkowski, Cordelia Langford, John Sillitoe on behalf of the Wellcome Sanger Institute COVID-19 Surveillance Team |
| EPI_ISL_736599, EPI_ISL_736600 | Lighthouse Lab in Alderley Park | Wellcome Sanger Institute for the COVID-19 Genomics UK (COG-UK) Consortium | Jacquelyn Wynn, Mairead Hyland, The Lighthouse Lab in Alderley Park and Alex Alderton, Roberto Amato, Sonia Goncalves, Ewan Harrison, David K. Jackson, Ian Johnston, Dominic Kwiatkowski, Cordelia Langford, John Sillitoe on behalf of the Wellcome Sanger Institute COVID-19 Surveillance Team |
| EPI_ISL_736601, EPI_ISL_736602 | Lighthouse Lab in Milton Keynes | Wellcome Sanger Institute for the COVID-19 Genomics UK (COG-UK) Consortium | The Lighthouse Lab in Milton Keynes and Alex Alderton, Roberto Amato, Sonia Goncalves, Ewan Harrison, David K. Jackson, Ian Johnston, Dominic Kwiatkowski, Cordelia Langford, John Sillitoe on behalf of the Wellcome Sanger Institute COVID-19 Surveillance Team |
| EPI_ISL_736603, EPI_ISL_736604, EPI_ISL_736606, EPI_ISL_736607 | Lighthouse Lab in Alderley Park | Wellcome Sanger Institute for the COVID-19 Genomics UK (COG-UK) Consortium | Jacquelyn Wynn, Mairead Hyland, The Lighthouse Lab in Alderley Park and Alex Alderton, Roberto Amato, Sonia Goncalves, Ewan Harrison, David K. Jackson, Ian Johnston, Dominic Kwiatkowski, Cordelia Langford, John Sillitoe on behalf of the Wellcome Sanger Institute COVID-19 Surveillance Team |
| EPI_ISL_736608, EPI_ISL_736609, EPI_ISL_736610 | Lighthouse Lab in Milton Keynes | Wellcome Sanger Institute for the COVID-19 Genomics UK (COG-UK) Consortium | The Lighthouse Lab in Milton Keynes and Alex Alderton, Roberto Amato, Sonia Goncalves, Ewan Harrison, David K. Jackson, Ian Johnston, Dominic Kwiatkowski, Cordelia Langford, John Sillitoe on behalf of the Wellcome Sanger Institute COVID-19 Surveillance Team |
| EPI_ISL_736611, EPI_ISL_736612, EPI_ISL_736613, EPI_ISL_736614, EPI_ISL_736615, EPI_ISL_736616 | Lighthouse Lab in Alderley Park | Wellcome Sanger Institute for the COVID-19 Genomics UK (COG-UK) Consortium | Jacquelyn Wynn, Mairead Hyland, The Lighthouse Lab in Alderley Park and Alex Alderton, Roberto Amato, Sonia Goncalves, Ewan Harrison, David K. Jackson, Ian Johnston, Dominic Kwiatkowski, Cordelia Langford, John Sillitoe on behalf of the Wellcome Sanger Institute COVID-19 Surveillance Team |
| EPI_ISL_736617, EPI_ISL_736618 | Lighthouse Lab in Milton Keynes | Wellcome Sanger Institute for the COVID-19 Genomics UK (COG-UK) Consortium | The Lighthouse Lab in Milton Keynes and Alex Alderton, Roberto Amato, Sonia Goncalves, Ewan Harrison, David K. Jackson, Ian Johnston, Dominic Kwiatkowski, Cordelia Langford, John Sillitoe on behalf of the Wellcome Sanger Institute COVID-19 Surveillance Team |
| EPI_ISL_736619, EPI_ISL_736621 | Lighthouse Lab in Alderley Park | Wellcome Sanger Institute for the COVID-19 Genomics UK (COG-UK) Consortium | Jacquelyn Wynn, Mairead Hyland, The Lighthouse Lab in Alderley Park and Alex Alderton, Roberto Amato, Sonia Goncalves, Ewan Harrison, David K. Jackson, Ian Johnston, Dominic Kwiatkowski, Cordelia Langford, John Sillitoe on behalf of the Wellcome Sanger Institute COVID-19 Surveillance Team |
| EPI_ISL_736622 | Lighthouse Lab in Milton Keynes | Wellcome Sanger Institute for the COVID-19 Genomics UK (COG-UK) Consortium | The Lighthouse Lab in Milton Keynes and Alex Alderton, Roberto Amato, Sonia Goncalves, Ewan Harrison, David K. Jackson, Ian Johnston, Dominic Kwiatkowski, Cordelia Langford, John Sillitoe on behalf of the Wellcome Sanger Institute COVID-19 Surveillance Team |
| EPI_ISL_736623, EPI_ISL_736625, EPI_ISL_736626, EPI_ISL_736629 | Lighthouse Lab in Alderley Park | Wellcome Sanger Institute for the COVID-19 Genomics UK (COG-UK) Consortium | Jacquelyn Wynn, Mairead Hyland, The Lighthouse Lab in Alderley Park and Alex Alderton, Roberto Amato, Sonia Goncalves, Ewan Harrison, David K. Jackson, Ian Johnston, Dominic Kwiatkowski, Cordelia Langford, John Sillitoe on behalf of the Wellcome Sanger Institute COVID-19 Surveillance Team |
| EPI_ISL_736630 | Lighthouse Lab in Milton Keynes | Wellcome Sanger Institute for the COVID-19 Genomics UK (COG-UK) Consortium | The Lighthouse Lab in Milton Keynes and Alex Alderton, Roberto Amato, Sonia Goncalves, Ewan Harrison, David K. Jackson, Ian Johnston, Dominic Kwiatkowski, Cordelia Langford, John Sillitoe on behalf of the Wellcome Sanger Institute COVID-19 Surveillance Team |
| EPI_ISL_736632, EPI_ISL_736634, EPI_ISL_736635, EPI_ISL_736636 | Lighthouse Lab in Alderley Park | Wellcome Sanger Institute for the COVID-19 Genomics UK (COG-UK) Consortium | Jacquelyn Wynn, Mairead Hyland, The Lighthouse Lab in Alderley Park and Alex Alderton, Roberto Amato, Sonia Goncalves, Ewan Harrison, David K. Jackson, Ian Johnston, Dominic Kwiatkowski, Cordelia Langford, John Sillitoe on behalf of the Wellcome Sanger Institute COVID-19 Surveillance Team |
| EPI_ISL_736637 | Lighthouse Lab in Milton Keynes | Wellcome Sanger Institute for the COVID-19 Genomics UK (COG-UK) Consortium | The Lighthouse Lab in Milton Keynes and Alex Alderton, Roberto Amato, Sonia Goncalves, Ewan Harrison, David K. Jackson, Ian Johnston, Dominic Kwiatkowski, Cordelia Langford, John Sillitoe on behalf of the Wellcome Sanger Institute COVID-19 Surveillance Team |
| EPI_ISL_736638, EPI_ISL_736639, EPI_ISL_736640, EPI_ISL_736641, EPI_ISL_736642 | Lighthouse Lab in Alderley Park | Wellcome Sanger Institute for the COVID-19 Genomics UK (COG-UK) Consortium | Jacquelyn Wynn, Mairead Hyland, The Lighthouse Lab in Alderley Park and Alex Alderton, Roberto Amato, Sonia Goncalves, Ewan Harrison, David K. Jackson, Ian Johnston, Dominic Kwiatkowski, Cordelia Langford, John Sillitoe on behalf of the Wellcome Sanger Institute COVID-19 Surveillance Team |
| EPI_ISL_736643 | Lighthouse Lab in Milton Keynes | Wellcome Sanger Institute for the COVID-19 Genomics UK (COG-UK) Consortium | The Lighthouse Lab in Milton Keynes and Alex Alderton, Roberto Amato, Sonia Goncalves, Ewan Harrison, David K. Jackson, Ian Johnston, Dominic Kwiatkowski, Cordelia Langford, John Sillitoe on behalf of the Wellcome Sanger Institute COVID-19 Surveillance Team |
| EPI_ISL_736644, EPI_ISL_736645, EPI_ISL_736647 | Lighthouse Lab in Alderley Park | Wellcome Sanger Institute for the COVID-19 Genomics UK (COG-UK) Consortium | Jacquelyn Wynn, Mairead Hyland, The Lighthouse Lab in Alderley Park and Alex Alderton, Roberto Amato, Sonia Goncalves, Ewan Harrison, David K. Jackson, Ian Johnston, Dominic Kwiatkowski, Cordelia Langford, John Sillitoe on behalf of the Wellcome Sanger Institute COVID-19 Surveillance Team |
| EPI_ISL_736648 | Lighthouse Lab in Milton Keynes | Wellcome Sanger Institute for the COVID-19 Genomics UK | The Lighthouse Lab in Milton Keynes and Alex Alderton, Roberto Amato, Sonia Goncalves, Ewan Harrison, David K. Jackson, Ian Johnston, Dominic |

| | | | |
|---|---|---|---|
| | | (COG-UK) Consortium | Kwiatkowski, Cordelia Langford, John Sillitoe on behalf of the Wellcome Sanger Institute COVID-19 Surveillance Team |
| EPI_ISL_736649, EPI_ISL_736651, EPI_ISL_736652, EPI_ISL_736653, EPI_ISL_736654, EPI_ISL_736655, EPI_ISL_736656, EPI_ISL_736658, EPI_ISL_736660, EPI_ISL_736661, EPI_ISL_736662, EPI_ISL_736663, EPI_ISL_736664, EPI_ISL_736665, EPI_ISL_736666, EPI_ISL_736667, EPI_ISL_736669, EPI_ISL_736670, EPI_ISL_736671, EPI_ISL_736673, EPI_ISL_736674 | | | |
| see above | Lighthouse Lab in Alderley Park | Wellcome Sanger Institute for the COVID-19 Genomics UK (COG-UK) Consortium | Jacquelyn Wynn, Mairead Hyland, The Lighthouse Lab in Alderley Park and Alex Alderton, Roberto Amato, Sonia Goncalves, Ewan Harrison, David K. Jackson, Ian Johnston, Dominic Kwiatkowski, Cordelia Langford, John Sillitoe on behalf of the Wellcome Sanger Institute COVID-19 Surveillance Team |
| EPI_ISL_736675 | Lighthouse Lab in Milton Keynes | Wellcome Sanger Institute for the COVID-19 Genomics UK (COG-UK) Consortium | The Lighthouse Lab in Milton Keynes and Alex Alderton, Roberto Amato, Sonia Goncalves, Ewan Harrison, David K. Jackson, Ian Johnston, Dominic Kwiatkowski, Cordelia Langford, John Sillitoe on behalf of the Wellcome Sanger Institute COVID-19 Surveillance Team |
| EPI_ISL_736676 | Lighthouse Lab in Alderley Park | Wellcome Sanger Institute for the COVID-19 Genomics UK (COG-UK) Consortium | Jacquelyn Wynn, Mairead Hyland, The Lighthouse Lab in Alderley Park and Alex Alderton, Roberto Amato, Sonia Goncalves, Ewan Harrison, David K. Jackson, Ian Johnston, Dominic Kwiatkowski, Cordelia Langford, John Sillitoe on behalf of the Wellcome Sanger Institute COVID-19 Surveillance Team |
| EPI_ISL_736677 | Lighthouse Lab in Milton Keynes | Wellcome Sanger Institute for the COVID-19 Genomics UK (COG-UK) Consortium | The Lighthouse Lab in Milton Keynes and Alex Alderton, Roberto Amato, Sonia Goncalves, Ewan Harrison, David K. Jackson, Ian Johnston, Dominic Kwiatkowski, Cordelia Langford, John Sillitoe on behalf of the Wellcome Sanger Institute COVID-19 Surveillance Team |
| EPI_ISL_736679, EPI_ISL_736680 | Lighthouse Lab in Alderley Park | Wellcome Sanger Institute for the COVID-19 Genomics UK (COG-UK) Consortium | Jacquelyn Wynn, Mairead Hyland, The Lighthouse Lab in Alderley Park and Alex Alderton, Roberto Amato, Sonia Goncalves, Ewan Harrison, David K. Jackson, Ian Johnston, Dominic Kwiatkowski, Cordelia Langford, John Sillitoe on behalf of the Wellcome Sanger Institute COVID-19 Surveillance Team |
| EPI_ISL_736681, EPI_ISL_736682, EPI_ISL_736683 | Lighthouse Lab in Milton Keynes | Wellcome Sanger Institute for the COVID-19 Genomics UK (COG-UK) Consortium | The Lighthouse Lab in Milton Keynes and Alex Alderton, Roberto Amato, Sonia Goncalves, Ewan Harrison, David K. Jackson, Ian Johnston, Dominic Kwiatkowski, Cordelia Langford, John Sillitoe on behalf of the Wellcome Sanger Institute COVID-19 Surveillance Team |
| EPI_ISL_736684 | Lighthouse Lab in Alderley Park | Wellcome Sanger Institute for the COVID-19 Genomics UK (COG-UK) Consortium | Jacquelyn Wynn, Mairead Hyland, The Lighthouse Lab in Alderley Park and Alex Alderton, Roberto Amato, Sonia Goncalves, Ewan Harrison, David K. Jackson, Ian Johnston, Dominic Kwiatkowski, Cordelia Langford, John Sillitoe on behalf of the Wellcome Sanger Institute COVID-19 Surveillance Team |
| EPI_ISL_736685 | Lighthouse Lab in Glasgow | Wellcome Sanger Institute for the COVID-19 Genomics UK (COG-UK) Consortium | Harper VanSteenhouse, Yumi Kasai, David Gray, Carol Clugston, Anna Dominiczak and Alex Alderton, Roberto Amato, Sonia Goncalves, Ewan Harrison, David K. Jackson, Ian Johnston, Dominic Kwiatkowski, Cordelia Langford, John Sillitoe on behalf of the Wellcome Sanger Institute COVID-19 Surveillance Team |
| EPI_ISL_736687, EPI_ISL_736688, EPI_ISL_736690, EPI_ISL_736692 | Lighthouse Lab in Alderley Park | Wellcome Sanger Institute for the COVID-19 Genomics UK (COG-UK) Consortium | Jacquelyn Wynn, Mairead Hyland, The Lighthouse Lab in Alderley Park and Alex Alderton, Roberto Amato, Sonia Goncalves, Ewan Harrison, David K. Jackson, Ian Johnston, Dominic Kwiatkowski, Cordelia Langford, John Sillitoe on behalf of the Wellcome Sanger Institute COVID-19 Surveillance Team |
| EPI_ISL_736693, EPI_ISL_736696, EPI_ISL_736697 | Lighthouse Lab in Milton Keynes | Wellcome Sanger Institute for the COVID-19 Genomics UK (COG-UK) Consortium | The Lighthouse Lab in Milton Keynes and Alex Alderton, Roberto Amato, Sonia Goncalves, Ewan Harrison, David K. Jackson, Ian Johnston, Dominic Kwiatkowski, Cordelia Langford, John Sillitoe on behalf of the Wellcome Sanger Institute COVID-19 Surveillance Team |
| EPI_ISL_736698 | Lighthouse Lab in Alderley Park | Wellcome Sanger Institute for the COVID-19 Genomics UK (COG-UK) Consortium | Jacquelyn Wynn, Mairead Hyland, The Lighthouse Lab in Alderley Park and Alex Alderton, Roberto Amato, Sonia Goncalves, Ewan Harrison, David K. Jackson, Ian Johnston, Dominic Kwiatkowski, Cordelia Langford, John Sillitoe on behalf of the Wellcome Sanger Institute COVID-19 Surveillance Team |
| EPI_ISL_736699 | Lighthouse Lab in Milton Keynes | Wellcome Sanger Institute for the COVID-19 Genomics UK (COG-UK) Consortium | The Lighthouse Lab in Milton Keynes and Alex Alderton, Roberto Amato, Sonia Goncalves, Ewan Harrison, David K. Jackson, Ian Johnston, Dominic Kwiatkowski, Cordelia Langford, John Sillitoe on behalf of the Wellcome Sanger Institute COVID-19 Surveillance Team |
| EPI_ISL_736701, EPI_ISL_736702 | Lighthouse Lab in Alderley Park | Wellcome Sanger Institute for the COVID-19 Genomics UK (COG-UK) Consortium | Jacquelyn Wynn, Mairead Hyland, The Lighthouse Lab in Alderley Park and Alex Alderton, Roberto Amato, Sonia Goncalves, Ewan Harrison, David K. Jackson, Ian Johnston, Dominic Kwiatkowski, Cordelia Langford, John Sillitoe on behalf of the Wellcome Sanger Institute COVID-19 Surveillance Team |
| EPI_ISL_736703 | Lighthouse Lab in Milton Keynes | Wellcome Sanger Institute for the COVID-19 Genomics UK (COG-UK) Consortium | The Lighthouse Lab in Milton Keynes and Alex Alderton, Roberto Amato, Sonia Goncalves, Ewan Harrison, David K. Jackson, Ian Johnston, Dominic Kwiatkowski, Cordelia Langford, John Sillitoe on behalf of the Wellcome Sanger Institute COVID-19 Surveillance Team |
| EPI_ISL_736704, EPI_ISL_736705, EPI_ISL_736706, EPI_ISL_736707, EPI_ISL_736708, EPI_ISL_736709, EPI_ISL_736710, EPI_ISL_736711 | Lighthouse Lab in Alderley Park | Wellcome Sanger Institute for the COVID-19 Genomics UK (COG-UK) Consortium | Jacquelyn Wynn, Mairead Hyland, The Lighthouse Lab in Alderley Park and Alex Alderton, Roberto Amato, Sonia Goncalves, Ewan Harrison, David K. Jackson, Ian Johnston, Dominic Kwiatkowski, Cordelia Langford, John Sillitoe on behalf of the Wellcome Sanger Institute COVID-19 Surveillance Team |
| EPI_ISL_736712, EPI_ISL_736713 | Lighthouse Lab in Milton Keynes | Wellcome Sanger Institute for the COVID-19 Genomics UK (COG-UK) Consortium | The Lighthouse Lab in Milton Keynes and Alex Alderton, Roberto Amato, Sonia Goncalves, Ewan Harrison, David K. Jackson, Ian Johnston, Dominic Kwiatkowski, Cordelia Langford, John Sillitoe on behalf of the Wellcome Sanger Institute COVID-19 Surveillance Team |
| EPI_ISL_736714, EPI_ISL_736715 | Lighthouse Lab in Alderley Park | Wellcome Sanger Institute for the COVID-19 Genomics UK (COG-UK) Consortium | Jacquelyn Wynn, Mairead Hyland, The Lighthouse Lab in Alderley Park and Alex Alderton, Roberto Amato, Sonia Goncalves, Ewan Harrison, David K. Jackson, Ian Johnston, Dominic Kwiatkowski, Cordelia Langford, John Sillitoe on behalf of the Wellcome Sanger Institute COVID-19 Surveillance Team |
| EPI_ISL_736716 | Lighthouse Lab in Milton Keynes | Wellcome Sanger Institute for the COVID-19 Genomics UK (COG-UK) Consortium | The Lighthouse Lab in Milton Keynes and Alex Alderton, Roberto Amato, Sonia Goncalves, Ewan Harrison, David K. Jackson, Ian Johnston, Dominic Kwiatkowski, Cordelia Langford, John Sillitoe on behalf of the Wellcome Sanger Institute COVID-19 Surveillance Team |
| EPI_ISL_736717, EPI_ISL_736718 | Lighthouse Lab in Alderley Park | Wellcome Sanger Institute for the COVID-19 Genomics UK (COG-UK) Consortium | Jacquelyn Wynn, Mairead Hyland, The Lighthouse Lab in Alderley Park and Alex Alderton, Roberto Amato, Sonia Goncalves, Ewan Harrison, David K. Jackson, Ian Johnston, Dominic Kwiatkowski, Cordelia Langford, John Sillitoe on behalf of the Wellcome Sanger Institute COVID-19 Surveillance Team |
| EPI_ISL_736719 | Lighthouse Lab in Milton Keynes | Wellcome Sanger Institute for the COVID-19 Genomics UK (COG-UK) Consortium | The Lighthouse Lab in Milton Keynes and Alex Alderton, Roberto Amato, Sonia Goncalves, Ewan Harrison, David K. Jackson, Ian Johnston, Dominic Kwiatkowski, Cordelia Langford, John Sillitoe on behalf of the Wellcome Sanger Institute COVID-19 Surveillance Team |
| EPI_ISL_736720 | Lighthouse Lab in Alderley Park | Wellcome Sanger Institute for the COVID-19 Genomics UK (COG-UK) Consortium | Jacquelyn Wynn, Mairead Hyland, The Lighthouse Lab in Alderley Park and Alex Alderton, Roberto Amato, Sonia Goncalves, Ewan Harrison, David K. Jackson, Ian Johnston, Dominic Kwiatkowski, Cordelia Langford, John Sillitoe on behalf of the Wellcome Sanger Institute COVID-19 Surveillance Team |
| EPI_ISL_736721 | Lighthouse Lab in Glasgow | Wellcome Sanger Institute for the COVID-19 Genomics UK (COG-UK) Consortium | Harper VanSteenhouse, Yumi Kasai, David Gray, Carol Clugston, Anna Dominiczak and Alex Alderton, Roberto Amato, Sonia Goncalves, Ewan Harrison, David K. Jackson, Ian Johnston, Dominic Kwiatkowski, Cordelia Langford, John Sillitoe on behalf of the Wellcome Sanger Institute COVID-19 Surveillance Team |
| EPI_ISL_736722 | Lighthouse Lab in Alderley Park | Wellcome Sanger Institute for the COVID-19 Genomics UK (COG-UK) Consortium | Jacquelyn Wynn, Mairead Hyland, The Lighthouse Lab in Alderley Park and Alex Alderton, Roberto Amato, Sonia Goncalves, Ewan Harrison, David K. Jackson, Ian Johnston, Dominic Kwiatkowski, Cordelia Langford, John Sillitoe on behalf of the Wellcome Sanger Institute COVID-19 Surveillance Team |
| EPI_ISL_736723, EPI_ISL_736724, EPI_ISL_736725 | Lighthouse Lab in Milton Keynes | Wellcome Sanger Institute for the COVID-19 Genomics UK (COG-UK) Consortium | The Lighthouse Lab in Milton Keynes and Alex Alderton, Roberto Amato, Sonia Goncalves, Ewan Harrison, David K. Jackson, Ian Johnston, Dominic Kwiatkowski, Cordelia Langford, John Sillitoe on behalf of the Wellcome Sanger Institute COVID-19 Surveillance Team |
| EPI_ISL_736726 | Lighthouse Lab in Alderley Park | Wellcome Sanger Institute for the COVID-19 Genomics UK (COG-UK) Consortium | Jacquelyn Wynn, Mairead Hyland, The Lighthouse Lab in Alderley Park and Alex Alderton, Roberto Amato, Sonia Goncalves, Ewan Harrison, David K. Jackson, Ian Johnston, Dominic Kwiatkowski, Cordelia Langford, John Sillitoe on behalf of the Wellcome Sanger Institute COVID-19 Surveillance Team |
| EPI_ISL_736727, EPI_ISL_736728 | Lighthouse Lab in Milton Keynes | Wellcome Sanger Institute for the COVID-19 Genomics UK (COG-UK) Consortium | The Lighthouse Lab in Milton Keynes and Alex Alderton, Roberto Amato, Sonia Goncalves, Ewan Harrison, David K. Jackson, Ian Johnston, Dominic Kwiatkowski, Cordelia Langford, John Sillitoe on behalf of the Wellcome Sanger Institute COVID-19 Surveillance Team |
| EPI_ISL_736729, EPI_ISL_736730 | Lighthouse Lab in Alderley Park | Wellcome Sanger Institute for the COVID-19 Genomics UK (COG-UK) Consortium | Jacquelyn Wynn, Mairead Hyland, The Lighthouse Lab in Alderley Park and Alex Alderton, Roberto Amato, Sonia Goncalves, Ewan Harrison, David K. Jackson, Ian Johnston, Dominic Kwiatkowski, Cordelia Langford, John Sillitoe on behalf of the Wellcome Sanger Institute COVID-19 Surveillance Team |
| EPI_ISL_736731 | Lighthouse Lab in Milton Keynes | Wellcome Sanger Institute for the COVID-19 Genomics UK (COG-UK) Consortium | The Lighthouse Lab in Milton Keynes and Alex Alderton, Roberto Amato, Sonia Goncalves, Ewan Harrison, David K. Jackson, Ian Johnston, Dominic Kwiatkowski, Cordelia Langford, John Sillitoe on behalf of the Wellcome Sanger Institute COVID-19 Surveillance Team |
| EPI_ISL_736732 | Lighthouse Lab in Alderley Park | Wellcome Sanger Institute for the COVID-19 Genomics UK (COG-UK) Consortium | Jacquelyn Wynn, Mairead Hyland, The Lighthouse Lab in Alderley Park and Alex Alderton, Roberto Amato, Sonia Goncalves, Ewan Harrison, David K. Jackson, Ian Johnston, Dominic Kwiatkowski, Cordelia Langford, John Sillitoe on behalf of the Wellcome Sanger Institute COVID-19 Surveillance Team |
| EPI_ISL_736733 | Lighthouse Lab in Milton Keynes | Wellcome Sanger Institute for the COVID-19 Genomics UK (COG-UK) Consortium | The Lighthouse Lab in Milton Keynes and Alex Alderton, Roberto Amato, Sonia Goncalves, Ewan Harrison, David K. Jackson, Ian Johnston, Dominic Kwiatkowski, Cordelia Langford, John Sillitoe on behalf of the Wellcome Sanger Institute COVID-19 Surveillance Team |
| EPI_ISL_736735 | Lighthouse Lab in Alderley Park | Wellcome Sanger Institute for the COVID-19 Genomics UK (COG-UK) Consortium | Jacquelyn Wynn, Mairead Hyland, The Lighthouse Lab in Alderley Park and Alex Alderton, Roberto Amato, Sonia Goncalves, Ewan Harrison, David K. Jackson, Ian Johnston, Dominic Kwiatkowski, Cordelia Langford, John Sillitoe on behalf of the Wellcome Sanger Institute COVID-19 Surveillance Team |
| EPI_ISL_736738 | Lighthouse Lab in Milton Keynes | Wellcome Sanger Institute for the COVID-19 Genomics UK (COG-UK) Consortium | The Lighthouse Lab in Milton Keynes and Alex Alderton, Roberto Amato, Sonia Goncalves, Ewan Harrison, David K. Jackson, Ian Johnston, Dominic Kwiatkowski, Cordelia Langford, John Sillitoe on behalf of the Wellcome Sanger Institute COVID-19 Surveillance Team |

| | | | |
|---|---|---|---|
| EPI_ISL_736739 | Lighthouse Lab in Alderley Park | Wellcome Sanger Institute for the COVID-19 Genomics UK (COG-UK) Consortium | Jacquelyn Wynn, Mairead Hyland, The Lighthouse Lab in Alderley Park and Alex Alderton, Roberto Amato, Sonia Goncalves, Ewan Harrison, David K. Jackson, Ian Johnston, Dominic Kwiatkowski, Cordelia Langford, John Sillitoe on behalf of the Wellcome Sanger Institute COVID-19 Surveillance Team |
| EPI_ISL_736740 | Lighthouse Lab in Milton Keynes | Wellcome Sanger Institute for the COVID-19 Genomics UK (COG-UK) Consortium | The Lighthouse Lab in Milton Keynes and Alex Alderton, Roberto Amato, Sonia Goncalves, Ewan Harrison, David K. Jackson, Ian Johnston, Dominic Kwiatkowski, Cordelia Langford, John Sillitoe on behalf of the Wellcome Sanger Institute COVID-19 Surveillance Team |
| EPI_ISL_736741, EPI_ISL_736742 | Lighthouse Lab in Alderley Park | Wellcome Sanger Institute for the COVID-19 Genomics UK (COG-UK) Consortium | Jacquelyn Wynn, Mairead Hyland, The Lighthouse Lab in Alderley Park and Alex Alderton, Roberto Amato, Sonia Goncalves, Ewan Harrison, David K. Jackson, Ian Johnston, Dominic Kwiatkowski, Cordelia Langford, John Sillitoe on behalf of the Wellcome Sanger Institute COVID-19 Surveillance Team |
| EPI_ISL_736744, EPI_ISL_736746 | Lighthouse Lab in Milton Keynes | Wellcome Sanger Institute for the COVID-19 Genomics UK (COG-UK) Consortium | The Lighthouse Lab in Milton Keynes and Alex Alderton, Roberto Amato, Sonia Goncalves, Ewan Harrison, David K. Jackson, Ian Johnston, Dominic Kwiatkowski, Cordelia Langford, John Sillitoe on behalf of the Wellcome Sanger Institute COVID-19 Surveillance Team |
| EPI_ISL_736747, EPI_ISL_736748, EPI_ISL_736750, EPI_ISL_736751, EPI_ISL_736753, EPI_ISL_736755, EPI_ISL_736756 | Lighthouse Lab in Alderley Park | Wellcome Sanger Institute for the COVID-19 Genomics UK (COG-UK) Consortium | Jacquelyn Wynn, Mairead Hyland, The Lighthouse Lab in Alderley Park and Alex Alderton, Roberto Amato, Sonia Goncalves, Ewan Harrison, David K. Jackson, Ian Johnston, Dominic Kwiatkowski, Cordelia Langford, John Sillitoe on behalf of the Wellcome Sanger Institute COVID-19 Surveillance Team |
| EPI_ISL_736757 | Lighthouse Lab in Milton Keynes | Wellcome Sanger Institute for the COVID-19 Genomics UK (COG-UK) Consortium | The Lighthouse Lab in Milton Keynes and Alex Alderton, Roberto Amato, Sonia Goncalves, Ewan Harrison, David K. Jackson, Ian Johnston, Dominic Kwiatkowski, Cordelia Langford, John Sillitoe on behalf of the Wellcome Sanger Institute COVID-19 Surveillance Team |
| EPI_ISL_736759, EPI_ISL_736760, EPI_ISL_736761, EPI_ISL_736762, EPI_ISL_736763, EPI_ISL_736764, EPI_ISL_736765, EPI_ISL_736766, EPI_ISL_736767, EPI_ISL_736768, EPI_ISL_736769, EPI_ISL_736770, EPI_ISL_736771 | | | |
| see above | Lighthouse Lab in Alderley Park | Wellcome Sanger Institute for the COVID-19 Genomics UK (COG-UK) Consortium | Jacquelyn Wynn, Mairead Hyland, The Lighthouse Lab in Alderley Park and Alex Alderton, Roberto Amato, Sonia Goncalves, Ewan Harrison, David K. Jackson, Ian Johnston, Dominic Kwiatkowski, Cordelia Langford, John Sillitoe on behalf of the Wellcome Sanger Institute COVID-19 Surveillance Team |
| EPI_ISL_736774 | Lighthouse Lab in Glasgow | Wellcome Sanger Institute for the COVID-19 Genomics UK (COG-UK) Consortium | Harper VanSteenhouse, Yumi Kasai, David Gray, Carol Clugston, Anna Dominiczak and Alex Alderton, Roberto Amato, Sonia Goncalves, Ewan Harrison, David K. Jackson, Ian Johnston, Dominic Kwiatkowski, Cordelia Langford, John Sillitoe on behalf of the Wellcome Sanger Institute COVID-19 Surveillance Team |
| EPI_ISL_736972 | NHLS-IALCH | KRISP, KZN Research Innovation and Sequencing Platform | Giandhari J, Pillay S, Lessells R, ChimukangaraB, Mdlalose K, York D, Khan S, Tegally H, Wilkinson E, de Oliveira T |
| EPI_ISL_737400, EPI_ISL_737404, EPI_ISL_737408, EPI_ISL_737410, EPI_ISL_737412, EPI_ISL_737420, EPI_ISL_737426, EPI_ISL_737427, EPI_ISL_737438, EPI_ISL_737444, EPI_ISL_737452, EPI_ISL_737453, EPI_ISL_737465, EPI_ISL_737474, EPI_ISL_737476, EPI_ISL_737487, EPI_ISL_737489, EPI_ISL_737490, EPI_ISL_737493, EPI_ISL_737494, EPI_ISL_737499, EPI_ISL_737516, EPI_ISL_737528, EPI_ISL_737529, EPI_ISL_737540, EPI_ISL_737551, EPI_ISL_737552, EPI_ISL_737555, EPI_ISL_737561, EPI_ISL_737562, EPI_ISL_737572, EPI_ISL_737577, EPI_ISL_737588, EPI_ISL_737592, EPI_ISL_737597, EPI_ISL_737602, EPI_ISL_737621, EPI_ISL_737622, EPI_ISL_737640, EPI_ISL_737641, EPI_ISL_737645, EPI_ISL_737646, EPI_ISL_737656, EPI_ISL_737657, EPI_ISL_737666, EPI_ISL_737667, EPI_ISL_737668, EPI_ISL_737669, EPI_ISL_737710, EPI_ISL_737711, EPI_ISL_737712, EPI_ISL_737713, EPI_ISL_737714, EPI_ISL_737715, EPI_ISL_737716, EPI_ISL_737717, EPI_ISL_737718, EPI_ISL_737719, EPI_ISL_737720, EPI_ISL_737721, EPI_ISL_737722, EPI_ISL_737723, EPI_ISL_737724, EPI_ISL_737739, EPI_ISL_737759, EPI_ISL_737760, EPI_ISL_737765, EPI_ISL_737766, EPI_ISL_737780, EPI_ISL_737781, EPI_ISL_737783, EPI_ISL_737800, EPI_ISL_737801, EPI_ISL_737802, EPI_ISL_737803, EPI_ISL_737804, EPI_ISL_737805, EPI_ISL_737806, EPI_ISL_737807, EPI_ISL_737808, EPI_ISL_737809, EPI_ISL_737810, EPI_ISL_737863, EPI_ISL_737864, EPI_ISL_737865, EPI_ISL_737866, EPI_ISL_737868, EPI_ISL_737902, EPI_ISL_737917 | | | |
| see above | Viollier AG | Department of Biosystems Science and Engineering, ETH Zürich | Chaoran Chen, Sarah Nadeau, Catharine Aquino, Ivan Topolsky, Philipp Jablonski, Lara Fuhrmann, David Dreifuss, Katharina Jahn, Andreia Cabral de Gouvea, Maria Domenica Moccia, Simon Grüter, Timothy Sykes, Lennart Opitz, Griffin White, Laura Neff, Doris Popovic, Andrea Patrignani, Jay Tracy, Ralph Schlapbach, Christiane Beckmann, Maurice Redondo, Olivier Kobel, Christoph Noppen, Sophie Seidel, Noemie Santamaria de Souza, Niko Beerenwinkel, Tanja Stadler |
| EPI_ISL_738314 | Unilabs Laboratory Medicine | Norwegian Institute of Public Health, Department of Virology | Kathrine Stene-Johansen, Kamilla Heddeland Instefjord, Hilde Elshaug, Atiya R Ali,Marie Paulsen Madsen, Rasmus Riis Kopperud, Hilde Vollan, Karoline Bragstad, Olav Hungnes |
| EPI_ISL_738362, EPI_ISL_738364, EPI_ISL_738366, EPI_ISL_738367, EPI_ISL_738368, EPI_ISL_738370, EPI_ISL_738372, EPI_ISL_738373, EPI_ISL_738375, EPI_ISL_738377, EPI_ISL_738378, EPI_ISL_738379, EPI_ISL_738383, EPI_ISL_738384, EPI_ISL_738385, EPI_ISL_738390, EPI_ISL_738391, EPI_ISL_738393, EPI_ISL_738395, EPI_ISL_738396, EPI_ISL_738397, EPI_ISL_738399, EPI_ISL_738401, EPI_ISL_738402, EPI_ISL_738403, EPI_ISL_738404, EPI_ISL_738405, EPI_ISL_738407, EPI_ISL_738408, EPI_ISL_738410, EPI_ISL_738411, EPI_ISL_738416, EPI_ISL_738418, EPI_ISL_738421, EPI_ISL_738422, EPI_ISL_738423, EPI_ISL_738426, EPI_ISL_738428, EPI_ISL_738429, EPI_ISL_738433, EPI_ISL_738434, EPI_ISL_738438, EPI_ISL_738443, EPI_ISL_738444, EPI_ISL_738445, EPI_ISL_738446, EPI_ISL_738449, EPI_ISL_738451, EPI_ISL_738456, EPI_ISL_738457, EPI_ISL_738462, EPI_ISL_738463, EPI_ISL_738465, EPI_ISL_738468, EPI_ISL_738473, EPI_ISL_738478, EPI_ISL_738481, EPI_ISL_738484, EPI_ISL_738487, EPI_ISL_738490, EPI_ISL_738495, EPI_ISL_738496 | | | |
| see above | UZ Leuven, National Reference Laboratory for Coronaviruses, Laboratory Medicine, Leuven, Belgium | KU Leuven, Rega Institute, Clinical and Epidemiological Virology | Tony Wawina-Bokalanga, Joan Marti-Carerras, Bert Vanmechelen, Piet Maes |
| EPI_ISL_739936, EPI_ISL_740160, EPI_ISL_740319 | Laboratoire national de santé, Microbiology, Virology | Laboratoire national de santé, Microbiology, Microbial Genomics Platform | Anke Wienecke-Baldacchino, Catherine Ragimbeau,Jessica Tapp, Fatu Djabi, Lise Pignon, Raoul Salmon, Tamir Abdelrahman |
| EPI_ISL_740869, EPI_ISL_740870 | South Eastern Area Laboratory Services (SEALS) | NSW Health Pathology - Institute of Clinical Pathology and Medical Research; Westmead Hospital; University of Sydney | CIDM-PH et al. |
| EPI_ISL_740995, EPI_ISL_740996, EPI_ISL_740997, EPI_ISL_740998, EPI_ISL_741000, EPI_ISL_741001, EPI_ISL_741002, EPI_ISL_741004, EPI_ISL_741009, EPI_ISL_741011, EPI_ISL_741012 | | | |
| see above | Department of Pathology, University of Cambridge | COVID-19 Genomics UK (COG-UK) Consortium | Aminu S. Jahun, Yasmin Chaudhry, Grant Hall, Iliana Georgana, Myra Hosmillo, Martin D. Curran, Malte Pinckert, Surendra Parmar, Ian Goodfellow |
| EPI_ISL_741186, EPI_ISL_741188, EPI_ISL_741189, EPI_ISL_741190, EPI_ISL_741191, EPI_ISL_741192, EPI_ISL_741193, EPI_ISL_741194, EPI_ISL_741195, EPI_ISL_741196, EPI_ISL_741197, EPI_ISL_741198, EPI_ISL_741199, EPI_ISL_741200, EPI_ISL_741201, EPI_ISL_741202, EPI_ISL_741203, EPI_ISL_741204, EPI_ISL_741205, EPI_ISL_741206, EPI_ISL_741208, EPI_ISL_741209, EPI_ISL_741210, EPI_ISL_741211, EPI_ISL_741212, EPI_ISL_741213, EPI_ISL_741214, EPI_ISL_741215, EPI_ISL_741216, EPI_ISL_741217, EPI_ISL_741218, EPI_ISL_741219, EPI_ISL_741220, EPI_ISL_741221, EPI_ISL_741222, EPI_ISL_741223, EPI_ISL_741224, EPI_ISL_741225, EPI_ISL_741226, EPI_ISL_741227, EPI_ISL_741228, EPI_ISL_741229, EPI_ISL_741230, EPI_ISL_741231, EPI_ISL_741232, EPI_ISL_741233, EPI_ISL_741234, EPI_ISL_741235, EPI_ISL_741236, EPI_ISL_741237, EPI_ISL_741238, EPI_ISL_741239, EPI_ISL_741240, EPI_ISL_741241, EPI_ISL_741242, EPI_ISL_741243, EPI_ISL_741244, EPI_ISL_741245, EPI_ISL_741246, EPI_ISL_741247, EPI_ISL_741248, EPI_ISL_741249, EPI_ISL_741250, EPI_ISL_741251, EPI_ISL_741252, EPI_ISL_741253, EPI_ISL_741254, EPI_ISL_741259, EPI_ISL_741260, EPI_ISL_741261, EPI_ISL_741262, EPI_ISL_741263, EPI_ISL_741270, EPI_ISL_741275, EPI_ISL_741277, EPI_ISL_741287, EPI_ISL_741294, EPI_ISL_741296, EPI_ISL_741297, EPI_ISL_741298, EPI_ISL_741299, EPI_ISL_741301, EPI_ISL_741302, EPI_ISL_741315, EPI_ISL_741316, EPI_ISL_741321 | | | |
| see above | University College London, Great Ormond Street Hospital for Children NHS Foundation Trust, Imperial College Healthcare NHS Trust | COVID-19 Genomics UK (COG-UK) Consortium | Sergi Castellano, Rachel Williams, Mark Kristiansen, Paola Resende Silva, Sunando Roy, Tony Brooks, Helena Tutill, Paola Niola, Patricia Dyal, Charlotte Williams, Leysa Forrest, Yasmin Panchbhaya, Jacqueline Findlay, Samuel Weeks, Julianne Brown, Kathryn Harris, Paul Randell, James Price, Alison Holmes, Judith Breuer |
| EPI_ISL_741485, EPI_ISL_741532 | Quadram Institute Bioscience | COVID-19 Genomics UK (COG-UK) Consortium | Dave J. Baker, Gemma L. Kay, Alp Aydin, Thanh Le-Viet, Steven Rudder, Ana P. Tedim, Anastasia Kolyva, Maria Diaz, Leonardo de Oliveira Martins, Nabil-Fareed Alikhan, Lizzie Meadows, Rachael Stanley, Ngozi Elumogo, Muhammed Yasir, Nicholas M. Thomson, Alexander J Trotter, Rachel Gilroy, Samuel Bloomfield, Claire Stuart, Andrew Bell, Reenesh Prakash, Samir Dervisevic, Alison E. Mather, John Wain, Mark Webber, Andrew J. Page, Justin O'Grady |
| EPI_ISL_741614, EPI_ISL_741615, EPI_ISL_741616, EPI_ISL_741617, EPI_ISL_741618, EPI_ISL_741619 | Queens Medical Centre, Clinical Microbiology Department / DeepSeq Nottingham | COVID-19 Genomics UK (COG-UK) Consortium | Gemma Clark, Wendy Smith, Manjinder Khakh, Vicki M Fleming, Michelle M Lister, Hannah Howson-Wells, Jonathan Ball, Patrick McClure, Joseph Chappell, Theocharis Tsoleridis, Nadine Holmes, Matthew Carlisle, Christopher Moore, Fei Sang, Johnny Debebe, Victoria Wright, Matthew Loose |
| EPI_ISL_741658, EPI_ISL_741731, EPI_ISL_741732, EPI_ISL_741733, EPI_ISL_741748, EPI_ISL_741749 | Centre for Enzyme Innovation, University of Portsmouth / Translational Research Laboratory, Portsmouth Hospitals NHS Trust | COVID-19 Genomics UK (COG-UK) Consortium | Angela Beckett,Yann Bourgeois,Garry Scarlett,Sharon Glaysher,Scott Elliott,Kelly Bicknell,Robert Impey,Allyson Lloyd,Sarah Wyllie,Ethan Butcher,Anoop Chauhan,Samuel Robson |
| EPI_ISL_741945, EPI_ISL_742009, EPI_ISL_742022, EPI_ISL_742023, EPI_ISL_742028, EPI_ISL_742044, EPI_ISL_742047, EPI_ISL_742055, EPI_ISL_742059, EPI_ISL_742065, EPI_ISL_742066, EPI_ISL_742068, EPI_ISL_742071, EPI_ISL_742075, EPI_ISL_742099, EPI_ISL_742103, EPI_ISL_742106 | | | |
| see above | Virology Department, Sheffield Teaching Hospitals NHS Foundation Trust/Department of Infection, Immunity and Cardiovascular Disease, The Medical School, University of Sheffield | COVID-19 Genomics UK (COG-UK) Consortium | Thushan de Silva, Matthew Parker, Nikki Smith, Adri Angyal, Rebecca Brown, Luke Green, Rachel Tucker, Paul Parsons, Danielle Groves, Katie Johnson, Laura Carrilero, Alex Keeley, Dave Partridge, Matthew Wyles, Benjamin Lindsey, Mehmet Yavuz, Mohammad Raza, Cariad Evans |
| EPI_ISL_742177, EPI_ISL_742178, EPI_ISL_742179, EPI_ISL_742181, EPI_ISL_742182, EPI_ISL_742183, EPI_ISL_742184, EPI_ISL_742185, EPI_ISL_742186, EPI_ISL_742187, EPI_ISL_742188, EPI_ISL_742189, EPI_ISL_742190, EPI_ISL_742191, EPI_ISL_742192, EPI_ISL_742193 | | | |
| see above | Virology Department, Royal Infirmary of Edinburgh, NHS Lothian / School of Biological Sciences, University of Edinburgh / Institute of Genetics and Molecular Medicine, University of Edinburgh | COVID-19 Genomics UK (COG-UK) Consortium | McHugh M, Dewar R, Rooke S, Gallagher M, Balcaza C, O'Toole Á, Scher E, Hill V, McCrone JT, Colquhoun R, Yu X, Jackson B, Rambaut A, Williams TC, Templeton K |
| EPI_ISL_742546, EPI_ISL_742555, EPI_ISL_742556, EPI_ISL_742557, EPI_ISL_742558, EPI_ISL_742560, EPI_ISL_742562, EPI_ISL_742566, EPI_ISL_742567, EPI_ISL_742568, EPI_ISL_742569, EPI_ISL_742570, EPI_ISL_742571, EPI_ISL_742572, EPI_ISL_742573, EPI_ISL_742574, EPI_ISL_742575, EPI_ISL_742576, | | | |

EPI_ISL_742577, EPI_ISL_742578, EPI_ISL_742579, EPI_ISL_742580, EPI_ISL_742581, EPI_ISL_742582, EPI_ISL_742583, EPI_ISL_742584, EPI_ISL_742647, EPI_ISL_742648, EPI_ISL_742650, EPI_ISL_742652, EPI_ISL_742680, EPI_ISL_742681, EPI_ISL_742682, EPI_ISL_742692, EPI_ISL_742693, EPI_ISL_742698, EPI_ISL_742699, EPI_ISL_742703, EPI_ISL_742706, EPI_ISL_742708, EPI_ISL_742711, EPI_ISL_742712, EPI_ISL_742715, EPI_ISL_742721, EPI_ISL_742722, EPI_ISL_742723, EPI_ISL_742724, EPI_ISL_742725, EPI_ISL_742726, EPI_ISL_742727, EPI_ISL_742728, EPI_ISL_742729, EPI_ISL_742730, EPI_ISL_742731, EPI_ISL_742732, EPI_ISL_742733, EPI_ISL_742734, EPI_ISL_742735, EPI_ISL_742736, EPI_ISL_742737, EPI_ISL_742738, EPI_ISL_742739, EPI_ISL_742740, EPI_ISL_742741, EPI_ISL_742742, EPI_ISL_742743, EPI_ISL_742744, EPI_ISL_742745, EPI_ISL_742746, EPI_ISL_742747, EPI_ISL_742748, EPI_ISL_742749, EPI_ISL_742750, EPI_ISL_742751, EPI_ISL_742752, EPI_ISL_742753, EPI_ISL_742754, EPI_ISL_742755, EPI_ISL_742756, EPI_ISL_742757, EPI_ISL_742758, EPI_ISL_742759, EPI_ISL_742760, EPI_ISL_742761, EPI_ISL_742762, EPI_ISL_742763, EPI_ISL_742764, EPI_ISL_742765, EPI_ISL_742766, EPI_ISL_742767, EPI_ISL_742768, EPI_ISL_742769, EPI_ISL_742770, EPI_ISL_742771, EPI_ISL_742772, EPI_ISL_742773, EPI_ISL_742774, EPI_ISL_742775, EPI_ISL_742776, EPI_ISL_742777, EPI_ISL_742778, EPI_ISL_742779, EPI_ISL_742780, EPI_ISL_742781, EPI_ISL_742782, EPI_ISL_742783, EPI_ISL_742784, EPI_ISL_742785, EPI_ISL_742786, EPI_ISL_742787, EPI_ISL_742788, EPI_ISL_742789, EPI_ISL_742790, EPI_ISL_742791, EPI_ISL_742792, EPI_ISL_742793, EPI_ISL_742794, EPI_ISL_742795, EPI_ISL_742796, EPI_ISL_742797, EPI_ISL_742798, EPI_ISL_742799, EPI_ISL_742800, EPI_ISL_742801, EPI_ISL_742802, EPI_ISL_742803, EPI_ISL_742804, EPI_ISL_742805, EPI_ISL_742806, EPI_ISL_742807, EPI_ISL_742808, EPI_ISL_742809, EPI_ISL_742810, EPI_ISL_742811, EPI_ISL_742812, EPI_ISL_742813, EPI_ISL_742814, EPI_ISL_742815, EPI_ISL_742816, EPI_ISL_742817, EPI_ISL_742818, EPI_ISL_742819, EPI_ISL_742820, EPI_ISL_742821, EPI_ISL_742822, EPI_ISL_742823, EPI_ISL_742824, EPI_ISL_742825, EPI_ISL_742826, EPI_ISL_742827, EPI_ISL_742828, EPI_ISL_742829, EPI_ISL_742830, EPI_ISL_742831, EPI_ISL_742832, EPI_ISL_742833, EPI_ISL_742834, EPI_ISL_742835, EPI_ISL_742836, EPI_ISL_742837, EPI_ISL_743127, EPI_ISL_743146, EPI_ISL_743147, EPI_ISL_743148, EPI_ISL_743149, EPI_ISL_743150, EPI_ISL_743151, EPI_ISL_743152, EPI_ISL_743153, EPI_ISL_743154, EPI_ISL_743155, EPI_ISL_743156, EPI_ISL_743157, EPI_ISL_743158, EPI_ISL_743321, EPI_ISL_743322, EPI_ISL_743326, EPI_ISL_743341, EPI_ISL_743352, EPI_ISL_743365, EPI_ISL_743366, EPI_ISL_743367, EPI_ISL_743368, EPI_ISL_743369, EPI_ISL_743370, EPI_ISL_743371, EPI_ISL_743373, EPI_ISL_743374, EPI_ISL_743375, EPI_ISL_743376, EPI_ISL_743377, EPI_ISL_743378, EPI_ISL_743379, EPI_ISL_743425, EPI_ISL_743426, EPI_ISL_743427, EPI_ISL_743428, EPI_ISL_743429, EPI_ISL_743430, EPI_ISL_743431, EPI_ISL_743433, EPI_ISL_743434, EPI_ISL_743445, EPI_ISL_743446, EPI_ISL_743447, EPI_ISL_743448, EPI_ISL_743449, EPI_ISL_743509, EPI_ISL_743511, EPI_ISL_743512, EPI_ISL_743513, EPI_ISL_743514, EPI_ISL_743515, EPI_ISL_743516, EPI_ISL_743517, EPI_ISL_743518, EPI_ISL_743519, EPI_ISL_743520, EPI_ISL_743521, EPI_ISL_743522, EPI_ISL_743523, EPI_ISL_743524, EPI_ISL_743525, EPI_ISL_743526, EPI_ISL_743527, EPI_ISL_743528, EPI_ISL_743529, EPI_ISL_743530, EPI_ISL_743531, EPI_ISL_743532, EPI_ISL_743533, EPI_ISL_743534, EPI_ISL_743535, EPI_ISL_743536, EPI_ISL_743537, EPI_ISL_743538, EPI_ISL_743539, EPI_ISL_743540, EPI_ISL_743541, EPI_ISL_743542, EPI_ISL_743543, EPI_ISL_743544, EPI_ISL_743545, EPI_ISL_743546, EPI_ISL_743547, EPI_ISL_743548, EPI_ISL_743549, EPI_ISL_743550, EPI_ISL_743551, EPI_ISL_743552, EPI_ISL_743553, EPI_ISL_743554, EPI_ISL_743555, EPI_ISL_743556, EPI_ISL_743557, EPI_ISL_743558, EPI_ISL_743559, EPI_ISL_743560, EPI_ISL_743561, EPI_ISL_743562, EPI_ISL_743563, EPI_ISL_743564, EPI_ISL_743565, EPI_ISL_743566, EPI_ISL_743567, EPI_ISL_743569, EPI_ISL_743570, EPI_ISL_743571, EPI_ISL_743572, EPI_ISL_743573, EPI_ISL_743574, EPI_ISL_743575, EPI_ISL_743576, EPI_ISL_743577, EPI_ISL_743578, EPI_ISL_743579, EPI_ISL_743581, EPI_ISL_743582, EPI_ISL_743583, EPI_ISL_743584, EPI_ISL_743585, EPI_ISL_743586, EPI_ISL_743587, EPI_ISL_743588, EPI_ISL_743589, EPI_ISL_743590, EPI_ISL_743591, EPI_ISL_743596, EPI_ISL_743597, EPI_ISL_743598, EPI_ISL_743599, EPI_ISL_743600, EPI_ISL_743601, EPI_ISL_743602, EPI_ISL_743641, EPI_ISL_743662, EPI_ISL_743663, EPI_ISL_743664, EPI_ISL_743675, EPI_ISL_743678, EPI_ISL_743680, EPI_ISL_743681, EPI_ISL_743682, EPI_ISL_743683, EPI_ISL_743684, EPI_ISL_743685, EPI_ISL_743686, EPI_ISL_743687, EPI_ISL_743688, EPI_ISL_743689, EPI_ISL_743690, EPI_ISL_743691, EPI_ISL_743692, EPI_ISL_743693, EPI_ISL_743694, EPI_ISL_743695, EPI_ISL_743696, EPI_ISL_743697, EPI_ISL_743698, EPI_ISL_743699, EPI_ISL_743700, EPI_ISL_743701, EPI_ISL_743702, EPI_ISL_743703, EPI_ISL_743704, EPI_ISL_743705, EPI_ISL_743706, EPI_ISL_743707, EPI_ISL_743708, EPI_ISL_743709, EPI_ISL_743710, EPI_ISL_743711, EPI_ISL_743712, EPI_ISL_743713, EPI_ISL_743714, EPI_ISL_743715, EPI_ISL_743716, EPI_ISL_743717, EPI_ISL_743718, EPI_ISL_743719, EPI_ISL_743720, EPI_ISL_743721, EPI_ISL_743722, EPI_ISL_743723, EPI_ISL_743724, EPI_ISL_743725, EPI_ISL_743726, EPI_ISL_743727, EPI_ISL_743728, EPI_ISL_743729, EPI_ISL_743730, EPI_ISL_743731, EPI_ISL_743732, EPI_ISL_743733, EPI_ISL_743734, EPI_ISL_743735, EPI_ISL_743736, EPI_ISL_743737, EPI_ISL_743738, EPI_ISL_743739, EPI_ISL_743740, EPI_ISL_743741, EPI_ISL_743742, EPI_ISL_743743, EPI_ISL_743744, EPI_ISL_743745, EPI_ISL_743746, EPI_ISL_743747, EPI_ISL_743748, EPI_ISL_743749, EPI_ISL_743750, EPI_ISL_743751, EPI_ISL_743752, EPI_ISL_743753, EPI_ISL_743754, EPI_ISL_743755, EPI_ISL_743756, EPI_ISL_743757, EPI_ISL_743758, EPI_ISL_743759, EPI_ISL_743760, EPI_ISL_743761, EPI_ISL_743762, EPI_ISL_743763, EPI_ISL_743764, EPI_ISL_743765, EPI_ISL_743766, EPI_ISL_743767, EPI_ISL_743768, EPI_ISL_743769, EPI_ISL_743770, EPI_ISL_743771, EPI_ISL_743772, EPI_ISL_743773, EPI_ISL_743774, EPI_ISL_743775, EPI_ISL_743776, EPI_ISL_743777, EPI_ISL_743778, EPI_ISL_743779, EPI_ISL_743780, EPI_ISL_743781, EPI_ISL_743783, EPI_ISL_743784, EPI_ISL_743786, EPI_ISL_743787, EPI_ISL_743788, EPI_ISL_743789, EPI_ISL_743790, EPI_ISL_743791, EPI_ISL_743792, EPI_ISL_743793, EPI_ISL_743794, EPI_ISL_743795, EPI_ISL_743796, EPI_ISL_743797, EPI_ISL_743798, EPI_ISL_743799, EPI_ISL_743800, EPI_ISL_743801, EPI_ISL_743802, EPI_ISL_743803, EPI_ISL_743804, EPI_ISL_743807, EPI_ISL_743808, EPI_ISL_743809, EPI_ISL_743810, EPI_ISL_743811, EPI_ISL_743812, EPI_ISL_743813, EPI_ISL_743814, EPI_ISL_743815, EPI_ISL_743816, EPI_ISL_743817, EPI_ISL_743818, EPI_ISL_743819, EPI_ISL_743823, EPI_ISL_743824, EPI_ISL_743829, EPI_ISL_743830, EPI_ISL_743834, EPI_ISL_743837, EPI_ISL_743874, EPI_ISL_743875, EPI_ISL_743876, EPI_ISL_743879, EPI_ISL_743880, EPI_ISL_743881, EPI_ISL_743882, EPI_ISL_743883, EPI_ISL_743884, EPI_ISL_743885, EPI_ISL_743890, EPI_ISL_743891, EPI_ISL_743892, EPI_ISL_743893, EPI_ISL_743894, EPI_ISL_743895, EPI_ISL_743896, EPI_ISL_743897, EPI_ISL_743902, EPI_ISL_743937, EPI_ISL_743942, EPI_ISL_743944, EPI_ISL_743952, EPI_ISL_743958, EPI_ISL_743959, EPI_ISL_743960, EPI_ISL_743961, EPI_ISL_743962, EPI_ISL_743963, EPI_ISL_743964, EPI_ISL_743965, EPI_ISL_743966, EPI_ISL_743967, EPI_ISL_743968, EPI_ISL_743969, EPI_ISL_743970, EPI_ISL_743971, EPI_ISL_743972, EPI_ISL_743973, EPI_ISL_743974, EPI_ISL_743975, EPI_ISL_743976, EPI_ISL_743977, EPI_ISL_743978, EPI_ISL_743979, EPI_ISL_743980, EPI_ISL_743981, EPI_ISL_743982, EPI_ISL_743983, EPI_ISL_743984, EPI_ISL_743985, EPI_ISL_743986, EPI_ISL_743988, EPI_ISL_743989, EPI_ISL_743990, EPI_ISL_743991, EPI_ISL_743992, EPI_ISL_743993, EPI_ISL_743994, EPI_ISL_743995, EPI_ISL_743996, EPI_ISL_743997, EPI_ISL_743998, EPI_ISL_743999

| | | | |
|---|---|---|---|
| see above | Wales Specialist Virology Centre Sequencing lab: Pathogen Genomics Unit | COVID-19 Genomics UK (COG-UK) Consortium | Catherine Moore, Johnathan Evans, Laura Gifford, Malorie Perry, Simon Cottrell, Angela Marchbank, Alec Birchley, Alexander Adams, Amy Gaskin, Bree Gatica-Wilcox, Jason Coombes, Joel Southgate, Lauren Gilbert, Lee Graham, Nicole Pacchiarini, Sara Kumziene-Summerhayes, Sarah Taylor, Sophie Jones, Sara Rey, Matthew Bull, Joanne Watkins, Sally Corden, Tom Connor |
| EPI_ISL_744519, EPI_ISL_744793, EPI_ISL_744946 | Laboratoire national de santé, Microbiology, Virology | Laboratoire national de santé, Microbiology, Microbial Genomics Platform | Anke Wienecke-Baldacchino, Catherine Ragimbeau,Jessica Tapp, Fatu Djabi, Lise Pignon, Raoul Salmon, Tamir Abdelrahman |

EPI_ISL_745056, EPI_ISL_745058, EPI_ISL_745059, EPI_ISL_745060, EPI_ISL_745061, EPI_ISL_745062, EPI_ISL_745063, EPI_ISL_745064, EPI_ISL_745065, EPI_ISL_745066, EPI_ISL_745067, EPI_ISL_745068, EPI_ISL_745069

| | | | |
|---|---|---|---|
| see above | Israel Central Virology laboratory | Israel Central Virology laboratory | Neta Zuckerman, Efrat Dahan Bucris, Oran Erster, Michal Mandelboim, Orna Mor, Ella Mendelson |
| EPI_ISL_745211 | Lab voor klinische biologie | Onderzoeksgroep Virologie | Laurens Lambrechts, Nick Vereecke, Marthe Pauwels, Bruno Verhasselt, Linos Vandekerckhove, Hans Nauwynck, Sebastiaan Theuns |
| EPI_ISL_745213, EPI_ISL_745214, EPI_ISL_745215, EPI_ISL_745216, EPI_ISL_745217, EPI_ISL_745220, EPI_ISL_745221 | Lab voor klinische biologie | Onderzoeksgroep Virologie | Nick Vereecke, Laurens Lambrechts, Marthe Pauwels, Bruno Verhasselt, Linos Vandekerckhove, Hans Nauwynck, Sebastiaan Theuns |

EPI_ISL_745310, EPI_ISL_745331, EPI_ISL_745378, EPI_ISL_745379, EPI_ISL_745380, EPI_ISL_745381, EPI_ISL_745382, EPI_ISL_745383, EPI_ISL_745384, EPI_ISL_745385, EPI_ISL_745386, EPI_ISL_745387, EPI_ISL_745388

| | | | |
|---|---|---|---|
| see above | CNR Virus des Infections Respiratoires - France SUD | CNR Virus des Infections Respiratoires - France SUD | Antonin Bal, Gregory Destras, Claudia Gonzalez, Gwendolyne Burfin, Quentin Semanas, Martine Valette, Bruno Lina, Laurence Josset |
| EPI_ISL_746827 | National Institute for Infectious Diseases, INMI, "L. Spallanzani" IRCCS | National Institute for Infectious Diseases, INMI, "L. Spallanzani" IRCCS | B Bartolini, C.E.M Gruber, M Rueca, F Messina, E Giombini, MR Capobianchi, A Di Caro |

EPI_ISL_747498, EPI_ISL_747499, EPI_ISL_747500, EPI_ISL_747501, EPI_ISL_747502, EPI_ISL_747503, EPI_ISL_747504, EPI_ISL_747505, EPI_ISL_747506, EPI_ISL_747507, EPI_ISL_747508, EPI_ISL_747509, EPI_ISL_747510, EPI_ISL_747511, EPI_ISL_747512, EPI_ISL_747513, EPI_ISL_747514, EPI_ISL_747515, EPI_ISL_747516, EPI_ISL_747517, EPI_ISL_747518, EPI_ISL_747519

| | | | |
|---|---|---|---|
| see above | Respiratory Virus Unit, National Infection Service, Public Health England | COVID-19 Genomics UK (COG-UK) Consortium | PHE Covid Sequencing Team |

EPI_ISL_751802, EPI_ISL_751803, EPI_ISL_751804, EPI_ISL_751805, EPI_ISL_751806, EPI_ISL_751807, EPI_ISL_751808, EPI_ISL_751809, EPI_ISL_751810, EPI_ISL_751811, EPI_ISL_751812, EPI_ISL_751813, EPI_ISL_751814, EPI_ISL_751815, EPI_ISL_751816, EPI_ISL_751817, EPI_ISL_751818, EPI_ISL_751819, EPI_ISL_751820, EPI_ISL_751821, EPI_ISL_751822, EPI_ISL_751823, EPI_ISL_751824, EPI_ISL_751825, EPI_ISL_751826, EPI_ISL_751827, EPI_ISL_751828, EPI_ISL_751829, EPI_ISL_751830, EPI_ISL_751831, EPI_ISL_751832, EPI_ISL_751833, EPI_ISL_751834, EPI_ISL_751835, EPI_ISL_751836, EPI_ISL_751837, EPI_ISL_751838, EPI_ISL_751839, EPI_ISL_751840, EPI_ISL_751841, EPI_ISL_751842, EPI_ISL_751843, EPI_ISL_751844, EPI_ISL_751845, EPI_ISL_751846

| | | | |
|---|---|---|---|
| see above | New Mexico Department of Health Scientific Laboratory | New Mexico Department of Health Scientific Laboratory | D'eldra Malone, Ellie Johnson, Anastacia Griego-Fisher |
| EPI_ISL_752606 | Yale Pathology Lab | Grubaugh Lab - Yale School of Public Health | Tara Alpert, Joseph Fauver, Anderson Brito, Chen Liu, Pei Hui, Jianhui Wang, Kien Pham, Nathan Grubaugh |
| EPI_ISL_753663, EPI_ISL_753669 | Clinical virology Laboratory, Children's Hospital Los Angeles | Center for Personalized Medicine, Children's Hospital Los Angeles | Gai et al |
| EPI_ISL_754142, EPI_ISL_754146, EPI_ISL_754151, EPI_ISL_754167, EPI_ISL_754168, EPI_ISL_754170, EPI_ISL_754171, EPI_ISL_754172 | Laboratoire de Microbiologie | National Reference Center for Viruses of Respiratory Infections, Institut Pasteur, Paris | Marion Barbet, Sylvie Behillil, Méline Bizard, Angela Brisebarre, Camille Capel, Etienne Simon-Lorière, Vincent Enouf, Maud Vanpeene, Sylvie van der Werf, Marie-Sarah Fangous |
| EPI_ISL_754215, EPI_ISL_754216 | Wyoming Public Health Laboratory | Wyoming Public Health Laboratory | Noah Hull, Taylor Fearing, Channing Weber, Ashley Norberg, Bailey Bowcutt, and Wanda Manley |

EPI_ISL_754543, EPI_ISL_754544, EPI_ISL_754550, EPI_ISL_754551, EPI_ISL_754552, EPI_ISL_754553, EPI_ISL_754554, EPI_ISL_754555, EPI_ISL_754556, EPI_ISL_754557, EPI_ISL_754558, EPI_ISL_754559, EPI_ISL_754560, EPI_ISL_754561, EPI_ISL_754562, EPI_ISL_754563, EPI_ISL_754564, EPI_ISL_754565, EPI_ISL_754566, EPI_ISL_754568, EPI_ISL_754569, EPI_ISL_754570, EPI_ISL_754571, EPI_ISL_754573, EPI_ISL_754574

| | | | |
|---|---|---|---|
| see above | Wadsworth Center, New York State Department.of Health | Wadsworth Center, New York State Department.of Health | Kirsten St. George, Daryl M. Lamson, Alexis Russel, Matthew Shudt, Melissa A Leisner, Jonathan Plitnick, Navjot Singh, John Kelly, Sara Griesemer, Erasmus Schneider, Erica Lasek-Nesselquist |
| EPI_ISL_754629, EPI_ISL_754630, EPI_ISL_754631, EPI_ISL_754632, EPI_ISL_754633, EPI_ISL_754634, EPI_ISL_754635 | University of Wisconsin-Madison AIDS Vaccine Research Laboratories | University of Wisconsin-Madison AIDS Vaccine Research Laboratories | Gage Moreno, Katarina Braun, et al. AIDS Vaccine Research Laboratories |
| EPI_ISL_755146, EPI_ISL_755163, EPI_ISL_755164, EPI_ISL_755165, EPI_ISL_755166, EPI_ISL_755167, EPI_ISL_755168, EPI_ISL_755169, EPI_ISL_755170, EPI_ISL_755174 | UCSD EXCITE lab | Andersen lab at Scripps Research | SEARCH Alliance San Diego |
| EPI_ISL_755494, EPI_ISL_755495, EPI_ISL_755497, EPI_ISL_755498, EPI_ISL_755499, EPI_ISL_755561, | Maine Health and Environmental Testing Laboratory | Tewhey Lab, The Jackson Laboratory | Matluk,N., Dewey,H., Iosue,F., Barter,M., Lynch,R., Munger,H. and Tewhey,R. |

| | | | |
|---|---|---|---|
| EPI_ISL_755562, EPI_ISL_755563, EPI_ISL_755564, EPI_ISL_755565 | | | |
| EPI_ISL_755573 | Center of Advanced Studies and Technology, CAST | Center of Advanced Studies and Technology, CAST | Ferrante,R., Mandatori,D., De Fabritiis,S. |
| EPI_ISL_755651, EPI_ISL_755653, EPI_ISL_755654 | Instituto Adolfo Lutz - Central | Instituto Adolfo Lutz, Interdiciplinary Procedures Center, Strategic Laboratory | Claudio Tavares Sacchi, Claudia Regina Gonçalves, Erica Valessa Ramos Gomes, Karoline Rodrigues Campos |
| EPI_ISL_755655 | Instituto Adolfo Lutz - Regional de Campinas | Instituto Adolfo Lutz, Interdiciplinary Procedures Center, Strategic Laboratory | Claudio Tavares Sacchi, Claudia Regina Gonçalves, Erica Valessa Ramos Gomes, Karoline Rodrigues Campos |
| EPI_ISL_756191, EPI_ISL_756192, EPI_ISL_756193, EPI_ISL_756194, EPI_ISL_756195, EPI_ISL_756196, EPI_ISL_756197, EPI_ISL_756198, EPI_ISL_756199, EPI_ISL_756200, EPI_ISL_756201, EPI_ISL_756208, EPI_ISL_756209, EPI_ISL_756210, EPI_ISL_756211, EPI_ISL_756212, EPI_ISL_756213, EPI_ISL_756214, EPI_ISL_756215, EPI_ISL_756217, EPI_ISL_756296, EPI_ISL_756297, EPI_ISL_756299, EPI_ISL_756302 | | | |
| see above | UW Virology Lab | UW Virology Lab | Pavitra Roychoudhury, Hong Xie, Lasata Shrestha, Meei-Li Huang, Keith R Jerome, Alexander Greninger |
| EPI_ISL_756308, EPI_ISL_756309 | The Caribbean Public Health Agency | Carrington Lab, Department of PreClinical Sciences, Faculty of Medical Sciences, The University of the West Indies | Nikita S. D. Sahadeo, Arianne Brown-Jordan, Sarah Hill, Vernie Ramkissoon, Roshan Parasram, Naresh Nandram, Avery Hinds, Jerome Foster, Stanley Giddings, Karla Georges, Marsha Ivey, Rahul Naidu, Risha Singh, SueMin Nathaniel, Rajini Haraksingh, Jaya Jayaraman, Chinna Chinnadurai, Adesh Ramsubhag, Nuno Faria, Oliver Pybus, Christopher Oura, Gabriel Escobar, Christine V. F. Carrington |
| EPI_ISL_756470 | Lighthouse Lab in Alderley Park | Wellcome Sanger Institute for the COVID-19 Genomics UK (COG-UK) Consortium | Jacquelyn Wynn, Mairead Hyland, The Lighthouse Lab in Alderley Park and Alex Alderton, Roberto Amato, Sonia Goncalves, Ewan Harrison, David K. Jackson, Ian Johnston, Dominic Kwiatkowski, Cordelia Langford, John Sillitoe on behalf of the Wellcome Sanger Institute COVID-19 Surveillance Team |
| EPI_ISL_756846, EPI_ISL_756887, EPI_ISL_756903, EPI_ISL_756946, EPI_ISL_757020 | Lighthouse Lab in Milton Keynes | Wellcome Sanger Institute for the COVID-19 Genomics UK (COG-UK) Consortium | The Lighthouse Lab in Milton Keynes and Alex Alderton, Roberto Amato, Sonia Goncalves, Ewan Harrison, David K. Jackson, Ian Johnston, Dominic Kwiatkowski, Cordelia Langford, John Sillitoe on behalf of the Wellcome Sanger Institute COVID-19 Surveillance Team |
| EPI_ISL_757042, EPI_ISL_757043, EPI_ISL_757044, EPI_ISL_757045, EPI_ISL_757046, EPI_ISL_757047, EPI_ISL_757048, EPI_ISL_757049, EPI_ISL_757050, EPI_ISL_757051, EPI_ISL_757052, EPI_ISL_757053, EPI_ISL_757054, EPI_ISL_757055, EPI_ISL_757056, EPI_ISL_757057, EPI_ISL_757058, EPI_ISL_757059, EPI_ISL_757060, EPI_ISL_757061, EPI_ISL_757062, EPI_ISL_757063, EPI_ISL_757064, EPI_ISL_757065, EPI_ISL_757066, EPI_ISL_757067, EPI_ISL_757068, EPI_ISL_757069, EPI_ISL_757070, EPI_ISL_757071, EPI_ISL_757072, EPI_ISL_757073, EPI_ISL_757074, EPI_ISL_757075, EPI_ISL_757076, EPI_ISL_757077, EPI_ISL_757078, EPI_ISL_757079, EPI_ISL_757080, EPI_ISL_757081, EPI_ISL_757082 | | | |
| see above | Lighthouse Lab in Glasgow | Wellcome Sanger Institute for the COVID-19 Genomics UK (COG-UK) Consortium | Harper VanSteenhouse, Yumi Kasai, David Gray, Carol Clugston, Anna Dominiczak and Alex Alderton, Roberto Amato, Sonia Goncalves, Ewan Harrison, David K. Jackson, Ian Johnston, Dominic Kwiatkowski, Cordelia Langford, John Sillitoe on behalf of the Wellcome Sanger Institute COVID-19 Surveillance Team |
| EPI_ISL_757119 | Lighthouse Lab in Cambridge | Wellcome Sanger Institute for the COVID-19 Genomics UK (COG-UK) Consortium | Rob Howes, The Lighthouse Lab in Cambridge and Alex Alderton, Roberto Amato, Sonia Goncalves, Ewan Harrison, David K. Jackson, Ian Johnston, Dominic Kwiatkowski, Cordelia Langford, John Sillitoe on behalf of the Wellcome Sanger Institute COVID-19 Surveillance Team |
| EPI_ISL_757120, EPI_ISL_757121, EPI_ISL_757122, EPI_ISL_757123, EPI_ISL_757124, EPI_ISL_757125, EPI_ISL_757126, EPI_ISL_757127, EPI_ISL_757128, EPI_ISL_757129, EPI_ISL_757130, EPI_ISL_757131, EPI_ISL_757132, EPI_ISL_757133, EPI_ISL_757134, EPI_ISL_757135, EPI_ISL_757136, EPI_ISL_757137, EPI_ISL_757138, EPI_ISL_757139, EPI_ISL_757140, EPI_ISL_757141, EPI_ISL_757142, EPI_ISL_757143, EPI_ISL_757144, EPI_ISL_757145, EPI_ISL_757146, EPI_ISL_757147, EPI_ISL_757148, EPI_ISL_757149, EPI_ISL_757150, EPI_ISL_757151, EPI_ISL_757152, EPI_ISL_757153, EPI_ISL_757154, EPI_ISL_757155, EPI_ISL_757156, EPI_ISL_757157, EPI_ISL_757158, EPI_ISL_757159, EPI_ISL_757160, EPI_ISL_757161, EPI_ISL_757162, EPI_ISL_757163, EPI_ISL_757164, EPI_ISL_757165, EPI_ISL_757166, EPI_ISL_757167, EPI_ISL_757168, EPI_ISL_757169, EPI_ISL_757170, EPI_ISL_757171, EPI_ISL_757172, EPI_ISL_757173, EPI_ISL_757174, EPI_ISL_757175, EPI_ISL_757176, EPI_ISL_757177, EPI_ISL_757178, EPI_ISL_757179, EPI_ISL_757180, EPI_ISL_757181, EPI_ISL_757182, EPI_ISL_757183, EPI_ISL_757184, EPI_ISL_757185, EPI_ISL_757186, EPI_ISL_757225, EPI_ISL_757228, EPI_ISL_757230 | | | |
| see above | Lighthouse Lab in Glasgow | Wellcome Sanger Institute for the COVID-19 Genomics UK (COG-UK) Consortium | Harper VanSteenhouse, Yumi Kasai, David Gray, Carol Clugston, Anna Dominiczak and Alex Alderton, Roberto Amato, Sonia Goncalves, Ewan Harrison, David K. Jackson, Ian Johnston, Dominic Kwiatkowski, Cordelia Langford, John Sillitoe on behalf of the Wellcome Sanger Institute COVID-19 Surveillance Team |
| EPI_ISL_760842 | Lighthouse Lab in Milton Keynes | Wellcome Sanger Institute for the COVID-19 Genomics UK (COG-UK) Consortium | The Lighthouse Lab in Milton Keynes and Alex Alderton, Roberto Amato, Sonia Goncalves, Ewan Harrison, David K. Jackson, Ian Johnston, Dominic Kwiatkowski, Cordelia Langford, John Sillitoe on behalf of the Wellcome Sanger Institute COVID-19 Surveillance Team |
| EPI_ISL_760847, EPI_ISL_760977, EPI_ISL_760980, EPI_ISL_761018 | Lighthouse Lab in Glasgow | Wellcome Sanger Institute for the COVID-19 Genomics UK (COG-UK) Consortium | Harper VanSteenhouse, Yumi Kasai, David Gray, Carol Clugston, Anna Dominiczak and Alex Alderton, Roberto Amato, Sonia Goncalves, Ewan Harrison, David K. Jackson, Ian Johnston, Dominic Kwiatkowski, Cordelia Langford, John Sillitoe on behalf of the Wellcome Sanger Institute COVID-19 Surveillance Team |
| EPI_ISL_761059 | Lighthouse Lab in Milton Keynes | Wellcome Sanger Institute for the COVID-19 Genomics UK (COG-UK) Consortium | The Lighthouse Lab in Milton Keynes and Alex Alderton, Roberto Amato, Sonia Goncalves, Ewan Harrison, David K. Jackson, Ian Johnston, Dominic Kwiatkowski, Cordelia Langford, John Sillitoe on behalf of the Wellcome Sanger Institute COVID-19 Surveillance Team |
| EPI_ISL_761096, EPI_ISL_761727 | Lighthouse Lab in Alderley Park | Wellcome Sanger Institute for the COVID-19 Genomics UK (COG-UK) Consortium | Jacquelyn Wynn, Mairead Hyland, The Lighthouse Lab in Alderley Park and Alex Alderton, Roberto Amato, Sonia Goncalves, Ewan Harrison, David K. Jackson, Ian Johnston, Dominic Kwiatkowski, Cordelia Langford, John Sillitoe on behalf of the Wellcome Sanger Institute COVID-19 Surveillance Team |
| EPI_ISL_762237 | Lighthouse Lab in Milton Keynes | Wellcome Sanger Institute for the COVID-19 Genomics UK (COG-UK) Consortium | The Lighthouse Lab in Milton Keynes and Alex Alderton, Roberto Amato, Sonia Goncalves, Ewan Harrison, David K. Jackson, Ian Johnston, Dominic Kwiatkowski, Cordelia Langford, John Sillitoe on behalf of the Wellcome Sanger Institute COVID-19 Surveillance Team |
| EPI_ISL_763003, EPI_ISL_763022, EPI_ISL_763040, EPI_ISL_763041, EPI_ISL_763042, EPI_ISL_763059 | Unit 17: Influenza & Other Respiratory Viruses, German National Influenza Center | Project group Epidemiology of Highly Pathogenic Microorganisms, Robert Koch-Institute | Ariane Düx, Andreas Sachse, Grit Schubert, Sébastien Calvignac-Spencer, Fabian Leendertz, Thorsten Wolff, Ralf Dürrwald, Djin-Ye Oh, Marianne Wedde |
| EPI_ISL_763079, EPI_ISL_763080 | Microbiologia e Virologia | Istituto Zooprofilattico Sperimentale delle Venezie | Adelaide Milani, Alessia Schivo, Annalisa Salviato, Erika Giorgia Quaranta, Ambra Pastori, Bianca Zecchin, Alice Fusaro, Isabella Monne, Calogero Terregino, Antonia Ricci |
| EPI_ISL_763127, EPI_ISL_763175, EPI_ISL_763177, EPI_ISL_763178, EPI_ISL_763179, EPI_ISL_763180, EPI_ISL_763189, EPI_ISL_763194, EPI_ISL_763195, EPI_ISL_763196, EPI_ISL_763197, EPI_ISL_763198, EPI_ISL_763202, EPI_ISL_763241, EPI_ISL_763285, EPI_ISL_763348, EPI_ISL_763356 | | | |
| see above | Dutch COVID-19 response team | Erasmus Medical Center | Bas Oude Munnink, Reina Sikkema, David Nieuwenhuijse, Irina Chestakova, Anne van der Linden, Marjan Boter, Emmanuelle Munger, Corine GeurtsvanKessel, Annemiek van der Eijk, Richard Molenkamp, Marion Koopmans, on behalf of the Dutch national COVID-19 response team. |
| EPI_ISL_763390, EPI_ISL_763453, EPI_ISL_763457 | University of Exeter | COVID-19 Genomics UK (COG-UK) Consortium | Ben Temperton,Aaron Jeffries,Michelle Michelsen,Joanna Warwick-Dugdale,Audrey Farbos,Robyn Manley,Stephen Michell,Jane Masoli |
| EPI_ISL_763484, EPI_ISL_763487, EPI_ISL_763519, EPI_ISL_763522 | Wales Specialist Virology Centre Sequencing lab: Pathogen Genomics Unit | COVID-19 Genomics UK (COG-UK) Consortium | Catherine Moore, Johnathan Evans, Laura Gifford, Malorie Perry, Simon Cottrell, Angela Marchbank, Alec Birchley, Alexander Adams, Amy Gaskin, Bree Gatica-Wilcox, Jason Coombes, Joel Southgate, Lauren Gilbert, Lee Graham, Nicole Pacchiarini, Sara Kumziene-Summerhayes, Sarah Taylor, Sophie Jones, Sara Rey, Matthew Bull, Joanne Watkins, Sally Corden, Tom Connor |
| EPI_ISL_763542 | Department of Pathology, University of Cambridge | COVID-19 Genomics UK (COG-UK) Consortium | Aminu S. Jahun, Yasmin Chaudhry, Grant Hall, Iliana Georgana, Myra Hosmillo, Martin D. Curran, Malte Pinckert, Surendra Parmar, Ian Goodfellow |
| EPI_ISL_763561, EPI_ISL_763562 | University of Exeter | COVID-19 Genomics UK (COG-UK) Consortium | Ben Temperton,Aaron Jeffries,Michelle Michelsen,Joanna Warwick-Dugdale,Audrey Farbos,Robyn Manley,Stephen Michell,Jane Masoli |
| EPI_ISL_763566, EPI_ISL_763570, EPI_ISL_763571 | Wales Specialist Virology Centre Sequencing lab: Pathogen Genomics Unit | COVID-19 Genomics UK (COG-UK) Consortium | Catherine Moore, Johnathan Evans, Laura Gifford, Malorie Perry, Simon Cottrell, Angela Marchbank, Alec Birchley, Alexander Adams, Amy Gaskin, Bree Gatica-Wilcox, Jason Coombes, Joel Southgate, Lauren Gilbert, Lee Graham, Nicole Pacchiarini, Sara Kumziene-Summerhayes, Sarah Taylor, Sophie Jones, Sara Rey, Matthew Bull, Joanne Watkins, Sally Corden, Tom Connor |
| EPI_ISL_763611 | Centre for Enzyme Innovation, University of Portsmouth / Translational Research Laboratory, Portsmouth Hospitals NHS Trust | COVID-19 Genomics UK (COG-UK) Consortium | Angela Beckett,Yann Bourgeois,Garry Scarlett,Sharon Glaysher,Scott Elliott,Kelly Bicknell,Robert Impey,Allyson Lloyd,Sarah Wyllie,Ethan Butcher,Anoop Chauhan,Samuel Robson |
| EPI_ISL_763634 | Virology Department, Sheffield Teaching Hospitals NHS Foundation Trust/Department of Infection, Immunity and Cardiovascular Disease, The Medical School, University of Sheffield | COVID-19 Genomics UK (COG-UK) Consortium | Thushan de Silva, Matthew Parker, Nikki Smith, Adri Angyal, Rebecca Brown, Luke Green, Rachel Tucker, Paul Parsons, Danielle Groves, Katie Johnson, Laura Carrilero, Alex Keeley, Dave Partridge, Matthew Wyles, Benjamin Lindsey, Mehmet Yavuz, Mohammad Raza, Cariad Evans |
| EPI_ISL_763637 | Department of Pathology, University of Cambridge | COVID-19 Genomics UK (COG-UK) Consortium | Aminu S. Jahun, Yasmin Chaudhry, Grant Hall, Iliana Georgana, Myra Hosmillo, Martin D. Curran, Malte Pinckert, Surendra Parmar, Ian Goodfellow |
| EPI_ISL_763652 | Virology Department, Sheffield Teaching Hospitals NHS Foundation Trust/Department of Infection, Immunity and | COVID-19 Genomics UK (COG-UK) Consortium | Thushan de Silva, Matthew Parker, Nikki Smith, Adri Angyal, Rebecca Brown, Luke Green, Rachel Tucker, Paul Parsons, Danielle Groves, Katie Johnson, Laura Carrilero, Alex Keeley, Dave Partridge, Matthew Wyles, Benjamin Lindsey, Mehmet Yavuz, Mohammad Raza, Cariad Evans |

| | | | |
|---|---|---|---|
| | Cardiovascular Disease, The Medical School, University of Sheffield | | |
| EPI_ISL_763656, EPI_ISL_763664, EPI_ISL_763674, EPI_ISL_763686 | Wales Specialist Virology Centre Sequencing lab: Pathogen Genomics Unit | COVID-19 Genomics UK (COG-UK) Consortium | Catherine Moore, Johnathan Evans, Laura Gifford, Malorie Perry, Simon Cottrell, Angela Marchbank, Alec Birchley, Alexander Adams, Amy Gaskin, Bree Gatica-Wilcox, Jason Coombes, Joel Southgate, Lauren Gilbert, Lee Graham, Nicole Pacchiarini, Sara Kumziene-Summerhayes, Sarah Taylor, Sophie Jones, Sara Rey, Matthew Bull, Joanne Watkins, Sally Corden, Tom Connor |
| EPI_ISL_763696, EPI_ISL_763701, EPI_ISL_763703, EPI_ISL_763704, EPI_ISL_763705, EPI_ISL_763706, EPI_ISL_763707, EPI_ISL_763708 | University of Exeter | COVID-19 Genomics UK (COG-UK) Consortium | Ben Temperton,Aaron Jeffries,Michelle Michelsen,Joanna Warwick-Dugdale,Audrey Farbos,Robyn Manley,Stephen Michell,Jane Masoli |
| EPI_ISL_763732 | Virology Department, Royal Infirmary of Edinburgh, NHS Lothian / School of Biological Sciences, University of Edinburgh / Institute of Genetics and Molecular Medicine, University of Edinburgh | COVID-19 Genomics UK (COG-UK) Consortium | McHugh M, Dewar R, Rooke S, Gallagher M, Balcaza C, O'Toole Á, Scher E, Hill V, McCrone JT, Colquhoun R, Yu X, Jackson B, Rambaut A, Williams TC, Templeton K |
| EPI_ISL_763738, EPI_ISL_763747, EPI_ISL_763768, EPI_ISL_763771, EPI_ISL_763776, EPI_ISL_763786, EPI_ISL_763787 | Wales Specialist Virology Centre Sequencing lab: Pathogen Genomics Unit | COVID-19 Genomics UK (COG-UK) Consortium | Catherine Moore, Johnathan Evans, Laura Gifford, Malorie Perry, Simon Cottrell, Angela Marchbank, Alec Birchley, Alexander Adams, Amy Gaskin, Bree Gatica-Wilcox, Jason Coombes, Joel Southgate, Lauren Gilbert, Lee Graham, Nicole Pacchiarini, Sara Kumziene-Summerhayes, Sarah Taylor, Sophie Jones, Sara Rey, Matthew Bull, Joanne Watkins, Sally Corden, Tom Connor |
| EPI_ISL_763796 | Department of Pathology, University of Cambridge | COVID-19 Genomics UK (COG-UK) Consortium | Aminu S. Jahun, Yasmin Chaudhry, Grant Hall, Iliana Georgana, Myra Hosmillo, Martin D. Curran, Malte Pinckert, Surendra Parmar, Ian Goodfellow |
| EPI_ISL_763820 | Wales Specialist Virology Centre Sequencing lab: Pathogen Genomics Unit | COVID-19 Genomics UK (COG-UK) Consortium | Catherine Moore, Johnathan Evans, Laura Gifford, Malorie Perry, Simon Cottrell, Angela Marchbank, Alec Birchley, Alexander Adams, Amy Gaskin, Bree Gatica-Wilcox, Jason Coombes, Joel Southgate, Lauren Gilbert, Lee Graham, Nicole Pacchiarini, Sara Kumziene-Summerhayes, Sarah Taylor, Sophie Jones, Sara Rey, Matthew Bull, Joanne Watkins, Sally Corden, Tom Connor |
| EPI_ISL_763841, EPI_ISL_763859, EPI_ISL_763861 | Oxford Viromics, NDM, University of Oxford; Oxford University Hospitals; Basingstoke and North Hampshire Hospital | COVID-19 Genomics UK (COG-UK) Consortium | Tanya Golubchik, David Bonsall, George Macintyre, Amy Trebes, Mariateresa de Cesare, Catrin Moore, Alex Mobbs, Anita Justice, Robert Shaw, Monique Andersson, Timothy Peto, Emma Wise, Nathan Moore, Jessica Lynch, Nick Cortes, Matilde Mori, Stephen Kidd, David Buck, John Todd, Christophe Fraser |
| EPI_ISL_763907 | University of Exeter | COVID-19 Genomics UK (COG-UK) Consortium | Ben Temperton,Aaron Jeffries,Michelle Michelsen,Joanna Warwick-Dugdale,Audrey Farbos,Robyn Manley,Stephen Michell,Jane Masoli |
| EPI_ISL_763932, EPI_ISL_763933, EPI_ISL_763942, EPI_ISL_763946, EPI_ISL_763947, EPI_ISL_763948 | Oxford Viromics, NDM, University of Oxford; Oxford University Hospitals; Basingstoke and North Hampshire Hospital | COVID-19 Genomics UK (COG-UK) Consortium | Tanya Golubchik, David Bonsall, George Macintyre, Amy Trebes, Mariateresa de Cesare, Catrin Moore, Alex Mobbs, Anita Justice, Robert Shaw, Monique Andersson, Timothy Peto, Emma Wise, Nathan Moore, Jessica Lynch, Nick Cortes, Matilde Mori, Stephen Kidd, David Buck, John Todd, Christophe Fraser |
| EPI_ISL_763976, EPI_ISL_763977, EPI_ISL_763978, EPI_ISL_763979, EPI_ISL_763980, EPI_ISL_763981, EPI_ISL_763982, EPI_ISL_763989, EPI_ISL_764035 | Wales Specialist Virology Centre Sequencing lab: Pathogen Genomics Unit | COVID-19 Genomics UK (COG-UK) Consortium | Catherine Moore, Johnathan Evans, Laura Gifford, Malorie Perry, Simon Cottrell, Angela Marchbank, Alec Birchley, Alexander Adams, Amy Gaskin, Bree Gatica-Wilcox, Jason Coombes, Joel Southgate, Lauren Gilbert, Lee Graham, Nicole Pacchiarini, Sara Kumziene-Summerhayes, Sarah Taylor, Sophie Jones, Sara Rey, Matthew Bull, Joanne Watkins, Sally Corden, Tom Connor |
| EPI_ISL_764069 | Virology Department, Royal Infirmary of Edinburgh, NHS Lothian / School of Biological Sciences, University of Edinburgh / Institute of Genetics and Molecular Medicine, University of Edinburgh | COVID-19 Genomics UK (COG-UK) Consortium | McHugh M, Dewar R, Rooke S, Gallagher M, Balcaza C, O'Toole Á, Scher E, Hill V, McCrone JT, Colquhoun R, Yu X, Jackson B, Rambaut A, Williams TC, Templeton K |
| EPI_ISL_764077, EPI_ISL_764078, EPI_ISL_764079, EPI_ISL_764086, EPI_ISL_764088, EPI_ISL_764117 | Oxford Viromics, NDM, University of Oxford; Oxford University Hospitals; Basingstoke and North Hampshire Hospital | COVID-19 Genomics UK (COG-UK) Consortium | Tanya Golubchik, David Bonsall, George Macintyre, Amy Trebes, Mariateresa de Cesare, Catrin Moore, Alex Mobbs, Anita Justice, Robert Shaw, Monique Andersson, Timothy Peto, Emma Wise, Nathan Moore, Jessica Lynch, Nick Cortes, Matilde Mori, Stephen Kidd, David Buck, John Todd, Christophe Fraser |
| EPI_ISL_764132, EPI_ISL_764134, EPI_ISL_764135, EPI_ISL_764137 | University of Exeter | COVID-19 Genomics UK (COG-UK) Consortium | Ben Temperton,Aaron Jeffries,Michelle Michelsen,Joanna Warwick-Dugdale,Audrey Farbos,Robyn Manley,Stephen Michell,Jane Masoli |
| EPI_ISL_764147 | Virology Department, Royal Infirmary of Edinburgh, NHS Lothian / School of Biological Sciences, University of Edinburgh / Institute of Genetics and Molecular Medicine, University of Edinburgh | COVID-19 Genomics UK (COG-UK) Consortium | McHugh M, Dewar R, Rooke S, Gallagher M, Balcaza C, O'Toole Á, Scher E, Hill V, McCrone JT, Colquhoun R, Yu X, Jackson B, Rambaut A, Williams TC, Templeton K |
| EPI_ISL_764150, EPI_ISL_764156 | Wales Specialist Virology Centre Sequencing lab: Pathogen Genomics Unit | COVID-19 Genomics UK (COG-UK) Consortium | Catherine Moore, Johnathan Evans, Laura Gifford, Malorie Perry, Simon Cottrell, Angela Marchbank, Alec Birchley, Alexander Adams, Amy Gaskin, Bree Gatica-Wilcox, Jason Coombes, Joel Southgate, Lauren Gilbert, Lee Graham, Nicole Pacchiarini, Sara Kumziene-Summerhayes, Sarah Taylor, Sophie Jones, Sara Rey, Matthew Bull, Joanne Watkins, Sally Corden, Tom Connor |
| EPI_ISL_764165, EPI_ISL_764170, EPI_ISL_764178, EPI_ISL_764182 | University of Exeter | COVID-19 Genomics UK (COG-UK) Consortium | Ben Temperton,Aaron Jeffries,Michelle Michelsen,Joanna Warwick-Dugdale,Audrey Farbos,Robyn Manley,Stephen Michell,Jane Masoli |
| EPI_ISL_764186 | Wales Specialist Virology Centre Sequencing lab: Pathogen Genomics Unit | COVID-19 Genomics UK (COG-UK) Consortium | Catherine Moore, Johnathan Evans, Laura Gifford, Malorie Perry, Simon Cottrell, Angela Marchbank, Alec Birchley, Alexander Adams, Amy Gaskin, Bree Gatica-Wilcox, Jason Coombes, Joel Southgate, Lauren Gilbert, Lee Graham, Nicole Pacchiarini, Sara Kumziene-Summerhayes, Sarah Taylor, Sophie Jones, Sara Rey, Matthew Bull, Joanne Watkins, Sally Corden, Tom Connor |
| EPI_ISL_764198 | Virology Department, Royal Infirmary of Edinburgh, NHS Lothian / School of Biological Sciences, University of Edinburgh / Institute of Genetics and Molecular Medicine, University of Edinburgh | COVID-19 Genomics UK (COG-UK) Consortium | McHugh M, Dewar R, Rooke S, Gallagher M, Balcaza C, O'Toole Á, Scher E, Hill V, McCrone JT, Colquhoun R, Yu X, Jackson B, Rambaut A, Williams TC, Templeton K |
| EPI_ISL_764203 | Wales Specialist Virology Centre Sequencing lab: Pathogen Genomics Unit | COVID-19 Genomics UK (COG-UK) Consortium | Catherine Moore, Johnathan Evans, Laura Gifford, Malorie Perry, Simon Cottrell, Angela Marchbank, Alec Birchley, Alexander Adams, Amy Gaskin, Bree Gatica-Wilcox, Jason Coombes, Joel Southgate, Lauren Gilbert, Lee Graham, Nicole Pacchiarini, Sara Kumziene-Summerhayes, Sarah Taylor, Sophie Jones, Sara Rey, Matthew Bull, Joanne Watkins, Sally Corden, Tom Connor |
| EPI_ISL_764224 | University of Exeter | COVID-19 Genomics UK (COG-UK) Consortium | Ben Temperton,Aaron Jeffries,Michelle Michelsen,Joanna Warwick-Dugdale,Audrey Farbos,Robyn Manley,Stephen Michell,Jane Masoli |
| EPI_ISL_764229, EPI_ISL_764232, EPI_ISL_764234, EPI_ISL_764266 | Wales Specialist Virology Centre Sequencing lab: Pathogen Genomics Unit | COVID-19 Genomics UK (COG-UK) Consortium | Catherine Moore, Johnathan Evans, Laura Gifford, Malorie Perry, Simon Cottrell, Angela Marchbank, Alec Birchley, Alexander Adams, Amy Gaskin, Bree Gatica-Wilcox, Jason Coombes, Joel Southgate, Lauren Gilbert, Lee Graham, Nicole Pacchiarini, Sara Kumziene-Summerhayes, Sarah Taylor, Sophie Jones, Sara Rey, Matthew Bull, Joanne Watkins, Sally Corden, Tom Connor |
| EPI_ISL_764271 | Oxford Viromics, NDM, University of Oxford; Oxford University Hospitals; Basingstoke and North Hampshire Hospital | COVID-19 Genomics UK (COG-UK) Consortium | Tanya Golubchik, David Bonsall, George Macintyre, Amy Trebes, Mariateresa de Cesare, Catrin Moore, Alex Mobbs, Anita Justice, Robert Shaw, Monique Andersson, Timothy Peto, Emma Wise, Nathan Moore, Jessica Lynch, Nick Cortes, Matilde Mori, Stephen Kidd, David Buck, John Todd, Christophe Fraser |
| EPI_ISL_764276, EPI_ISL_764329, EPI_ISL_764330, EPI_ISL_764332, EPI_ISL_764333, EPI_ISL_764334 | Department of Pathology, University of Cambridge | COVID-19 Genomics UK (COG-UK) Consortium | Aminu S. Jahun, Yasmin Chaudhry, Grant Hall, Iliana Georgana, Myra Hosmillo, Martin D. Curran, Malte Pinckert, Surendra Parmar, Ian Goodfellow |
| EPI_ISL_764359 | Virology Department, Royal Infirmary of Edinburgh, NHS Lothian / School of Biological Sciences, University of Edinburgh / Institute of Genetics and Molecular Medicine, University of Edinburgh | COVID-19 Genomics UK (COG-UK) Consortium | McHugh M, Dewar R, Rooke S, Gallagher M, Balcaza C, O'Toole Á, Scher E, Hill V, McCrone JT, Colquhoun R, Yu X, Jackson B, Rambaut A, Williams TC, Templeton K |
| EPI_ISL_764373 | University of Exeter | COVID-19 Genomics UK (COG-UK) Consortium | Ben Temperton,Aaron Jeffries,Michelle Michelsen,Joanna Warwick-Dugdale,Audrey Farbos,Robyn Manley,Stephen Michell,Jane Masoli |

| | | | |
|---|---|---|---|
| EPI_ISL_764534, EPI_ISL_764540, EPI_ISL_764541, EPI_ISL_764542, EPI_ISL_764547 | Oxford Viromics, NDM, University of Oxford; Oxford University Hospitals; Basingstoke and North Hampshire Hospital | COVID-19 Genomics UK (COG-UK) Consortium | Tanya Golubchik, David Bonsall, George Macintyre, Amy Trebes, Mariateresa de Cesare, Catrin Moore, Alex Mobbs, Anita Justice, Robert Shaw, Monique Andersson, Timothy Peto, Emma Wise, Nathan Moore, Jessica Lynch, Nick Cortes, Matilde Mori, Stephen Kidd, David Buck, John Todd, Christophe Fraser |
| EPI_ISL_764573, EPI_ISL_764576, EPI_ISL_764581, EPI_ISL_764585, EPI_ISL_764599, EPI_ISL_764600, EPI_ISL_764601, EPI_ISL_764602, EPI_ISL_764603, EPI_ISL_764604, EPI_ISL_764605, EPI_ISL_764606, EPI_ISL_764607, EPI_ISL_764608, EPI_ISL_764609, EPI_ISL_764610, EPI_ISL_764611, EPI_ISL_764612, EPI_ISL_764613, EPI_ISL_764614, EPI_ISL_764615, EPI_ISL_764616, EPI_ISL_764617, EPI_ISL_764618, EPI_ISL_764619, EPI_ISL_764620, EPI_ISL_764621, EPI_ISL_764622, EPI_ISL_764623, EPI_ISL_764625, EPI_ISL_764626, EPI_ISL_764627, EPI_ISL_764628, EPI_ISL_764629, EPI_ISL_764630, EPI_ISL_764631, EPI_ISL_764632, EPI_ISL_764633, EPI_ISL_764634, EPI_ISL_764635, EPI_ISL_764636, EPI_ISL_764637, EPI_ISL_764638, EPI_ISL_764639, EPI_ISL_764640, EPI_ISL_764641, EPI_ISL_764642, EPI_ISL_764643, EPI_ISL_764644, EPI_ISL_764645, EPI_ISL_764646, EPI_ISL_764647, EPI_ISL_764648, EPI_ISL_764649, EPI_ISL_764650, EPI_ISL_764651, EPI_ISL_764652, EPI_ISL_764739, EPI_ISL_764740, EPI_ISL_764805, EPI_ISL_764806, EPI_ISL_764807, EPI_ISL_764808, EPI_ISL_764809, EPI_ISL_764810, EPI_ISL_764811, EPI_ISL_764812, EPI_ISL_764813, EPI_ISL_764814, EPI_ISL_764815, EPI_ISL_764819, EPI_ISL_764820, EPI_ISL_764822, EPI_ISL_764823, EPI_ISL_764825, EPI_ISL_764952, EPI_ISL_764953, EPI_ISL_764954, EPI_ISL_764955, EPI_ISL_764956, EPI_ISL_764957, EPI_ISL_764958, EPI_ISL_764959, EPI_ISL_764960, EPI_ISL_764961, EPI_ISL_764962, EPI_ISL_764963, EPI_ISL_764964, EPI_ISL_764969 | | | |
| see above | Wales Specialist Virology Centre Sequencing lab: Pathogen Genomics Unit | COVID-19 Genomics UK (COG-UK) Consortium | Catherine Moore, Johnathan Evans, Laura Gifford, Malorie Perry, Simon Cottrell, Angela Marchbank, Alec Birchley, Alexander Adams, Amy Gaskin, Bree Gatica-Wilcox, Jason Coombes, Joel Southgate, Lauren Gilbert, Lee Graham, Nicole Pacchiarini, Sara Kumziene-Summerhayes, Sarah Taylor, Sophie Jones, Sara Rey, Matthew Bull, Joanne Watkins, Sally Corden, Tom Connor |
| EPI_ISL_765517, EPI_ISL_765520 | SARATOGA HOSPITAL LABORATORY | Wadsworth Center, New York State Department.of Health | Kirsten St. George, Daryl M. Lamson, Alexis Russel, Matthew Shudt, Melissa A Leisner, Jonathan Plitnick, Navjot Singh, John Kelly, Sara Griesemer, Erasmus Schneider, Erica Lasek-Nesselquist |
| EPI_ISL_765895 | Unit 17: Influenza & Other Respiratory Viruses, German National Influenza Center | Project group Epidemiology of Highly Pathogenic Microorganisms, Robert Koch-Institute | Ariane Düx, Andreas Sachse, Grit Schubert, Sébastien Calvignac-Spencer, Fabian Leendertz, Thorsten Wolff, Ralf Dürrwald, Djin-Ye Oh, Marianne Wedde |
| EPI_ISL_766070, EPI_ISL_766079, EPI_ISL_766080, EPI_ISL_766106, EPI_ISL_766107, EPI_ISL_766109, EPI_ISL_766117, EPI_ISL_766118, EPI_ISL_766119, EPI_ISL_766120, EPI_ISL_766127, EPI_ISL_766128, EPI_ISL_766131, EPI_ISL_766133, EPI_ISL_766134, EPI_ISL_766144, EPI_ISL_766158, EPI_ISL_766225, EPI_ISL_766226, EPI_ISL_766230, EPI_ISL_766238, EPI_ISL_766248, EPI_ISL_766249, EPI_ISL_766250, EPI_ISL_766260, EPI_ISL_766261, EPI_ISL_766288, EPI_ISL_766290, EPI_ISL_766293, EPI_ISL_766294, EPI_ISL_766295, EPI_ISL_766296, EPI_ISL_766297, EPI_ISL_766298, EPI_ISL_766299, EPI_ISL_766300, EPI_ISL_766302, EPI_ISL_766304, EPI_ISL_766305, EPI_ISL_766306, EPI_ISL_766307, EPI_ISL_766308, EPI_ISL_766309, EPI_ISL_766311, EPI_ISL_766312, EPI_ISL_766313, EPI_ISL_766315, EPI_ISL_766395, EPI_ISL_766396, EPI_ISL_766399, EPI_ISL_766405, EPI_ISL_766409, EPI_ISL_766425, EPI_ISL_766463, EPI_ISL_766508, EPI_ISL_766562, EPI_ISL_766563 | | | |
| see above | Respiratory Virus Unit, National Infection Service, Public Health England | COVID-19 Genomics UK (COG-UK) Consortium | PHE Covid Sequencing Team |
| EPI_ISL_766608 | Klinisk mikrobiologi | The Public Health Agency of Sweden | Department of Microbiology, The Public Health Agency of Sweden |
| EPI_ISL_766632 | Klinisk mikrobiologi, Viruslab | The Public Health Agency of Sweden | Department of Microbiology, The Public Health Agency of Sweden |
| EPI_ISL_766675 | Texas Department of State Health Services | Texas Department of State Health Services | Rashmi Tuladhar, Bonnie Oh, Jenny Zhang, Maliha Rahman, Anita Pokharel, Myong Koag, Chung Wang, Rachel Lee, Grace Kubin, Mayela Pedrueza, James Daniel Bonser |
| EPI_ISL_767121, EPI_ISL_767123, EPI_ISL_767125, EPI_ISL_767128, EPI_ISL_767130, EPI_ISL_767132, EPI_ISL_767133, EPI_ISL_767135, EPI_ISL_767136, EPI_ISL_767137 | Lighthouse Lab in Alderley Park | Wellcome Sanger Institute for the COVID-19 Genomics UK (COG-UK) Consortium | Jacquelyn Wynn, Mairead Hyland, The Lighthouse Lab in Alderley Park and Alex Alderton, Roberto Amato, Sonia Goncalves, Ewan Harrison, David K. Jackson, Ian Johnston, Dominic Kwiatkowski, Cordelia Langford, John Sillitoe on behalf of the Wellcome Sanger Institute COVID-19 Surveillance Team |
| EPI_ISL_767138 | Lighthouse Lab in Cambridge | Wellcome Sanger Institute for the COVID-19 Genomics UK (COG-UK) Consortium | Rob Howes, The Lighthouse Lab in Cambridge and Alex Alderton, Roberto Amato, Sonia Goncalves, Ewan Harrison, David K. Jackson, Ian Johnston, Dominic Kwiatkowski, Cordelia Langford, John Sillitoe on behalf of the Wellcome Sanger Institute COVID-19 Surveillance Team |
| EPI_ISL_767139, EPI_ISL_767140, EPI_ISL_767141, EPI_ISL_767142, EPI_ISL_767143, EPI_ISL_767144, EPI_ISL_767145, EPI_ISL_767146, EPI_ISL_767147, EPI_ISL_767148, EPI_ISL_767149, EPI_ISL_767150, EPI_ISL_767178, EPI_ISL_767216, EPI_ISL_767217, EPI_ISL_767218, EPI_ISL_767219, EPI_ISL_767220, EPI_ISL_767222, EPI_ISL_767223, EPI_ISL_767224, EPI_ISL_767225, EPI_ISL_767226, EPI_ISL_767227, EPI_ISL_767228, EPI_ISL_767229, EPI_ISL_767231, EPI_ISL_767232, EPI_ISL_767233, EPI_ISL_767234, EPI_ISL_767235, EPI_ISL_767236, EPI_ISL_767237, EPI_ISL_767238, EPI_ISL_767277, EPI_ISL_767279, EPI_ISL_767280, EPI_ISL_767281, EPI_ISL_767282, EPI_ISL_767283, EPI_ISL_767285, EPI_ISL_767286, EPI_ISL_767287, EPI_ISL_767288, EPI_ISL_767289, EPI_ISL_767290 | | | |
| see above | Lighthouse Lab in Alderley Park | Wellcome Sanger Institute for the COVID-19 Genomics UK (COG-UK) Consortium | Jacquelyn Wynn, Mairead Hyland, The Lighthouse Lab in Alderley Park and Alex Alderton, Roberto Amato, Sonia Goncalves, Ewan Harrison, David K. Jackson, Ian Johnston, Dominic Kwiatkowski, Cordelia Langford, John Sillitoe on behalf of the Wellcome Sanger Institute COVID-19 Surveillance Team |
| EPI_ISL_767352, EPI_ISL_767355 | Michigan Department of Health and Human Services, Bureau of Laboratories | Michigan Department of Health and Human Services, Bureau of Laboratories | Blankenship HM, Riner D, Soehnlen MK |
| EPI_ISL_767452 | Wadsworth Center, New York State Department.of Health | Wadsworth Center, New York State Department.of Health | Kirsten St. George, Daryl M. Lamson, Alexis Russel, Matthew Shudt, Melissa A Leisner, Jonathan Plitnick, Navjot Singh, John Kelly, Sara Griesemer, Erasmus Schneider, Erica Lasek-Nesselquist |
| EPI_ISL_768419, EPI_ISL_768420, EPI_ISL_768421, EPI_ISL_768422, EPI_ISL_768423, EPI_ISL_768424, EPI_ISL_768425, EPI_ISL_768426, EPI_ISL_768427, EPI_ISL_768428, EPI_ISL_768429, EPI_ISL_768430, EPI_ISL_768431, EPI_ISL_768432, EPI_ISL_768433, EPI_ISL_768434, EPI_ISL_768435, EPI_ISL_768436, EPI_ISL_768437, EPI_ISL_768438, EPI_ISL_768439, EPI_ISL_768440, EPI_ISL_768441, EPI_ISL_768442, EPI_ISL_768443, EPI_ISL_768444, EPI_ISL_768445, EPI_ISL_768446, EPI_ISL_768447, EPI_ISL_768448, EPI_ISL_768449, EPI_ISL_768450, EPI_ISL_768451, EPI_ISL_768452, EPI_ISL_768453, EPI_ISL_768454, EPI_ISL_768455, EPI_ISL_768456, EPI_ISL_768457, EPI_ISL_768458, EPI_ISL_768459, EPI_ISL_768460, EPI_ISL_768461, EPI_ISL_768462, EPI_ISL_768463, EPI_ISL_768464, EPI_ISL_768465, EPI_ISL_768466, EPI_ISL_768467, EPI_ISL_768468, EPI_ISL_768469, EPI_ISL_768470, EPI_ISL_768471, EPI_ISL_768472, EPI_ISL_768473, EPI_ISL_768474 | | | |
| see above | LSUHS Emerging Viral Threat Laboratory | Microbial Genome Sequencing Center | Jeremy P. Kamil, Jennifer L. Carroll, Camille F. Abshire, Maarten Van Diest, Andrew D. Yurochko, Martin J. Sapp, Rona S. Scott, Christopher G. Kevil, Daniel J. Snyder, Vaughn S. Cooper, John A. Vanchiere |
| EPI_ISL_768804, EPI_ISL_768805, EPI_ISL_768806, EPI_ISL_768807, EPI_ISL_768808, EPI_ISL_768809, EPI_ISL_768810, EPI_ISL_768811, EPI_ISL_768812, EPI_ISL_768813, EPI_ISL_768814 | | | |
| see above | Respiratory Virus Unit, National Infection Service, Public Health England | COVID-19 Genomics UK (COG-UK) Consortium | PHE Covid Sequencing Team |
| EPI_ISL_769226, EPI_ISL_769227, EPI_ISL_769228, EPI_ISL_769230, EPI_ISL_769231, EPI_ISL_769232, EPI_ISL_769234, EPI_ISL_769235, EPI_ISL_769237, EPI_ISL_769238, EPI_ISL_769239, EPI_ISL_769240, EPI_ISL_769241, EPI_ISL_769242, EPI_ISL_769247, EPI_ISL_769248, EPI_ISL_769251, EPI_ISL_769253, EPI_ISL_769256, EPI_ISL_769258, EPI_ISL_769262, EPI_ISL_769263, EPI_ISL_769264, EPI_ISL_769265, EPI_ISL_769268, EPI_ISL_769269, EPI_ISL_769270, EPI_ISL_769271, EPI_ISL_769272, EPI_ISL_769274, EPI_ISL_769277, EPI_ISL_769278, EPI_ISL_769279, EPI_ISL_769283, EPI_ISL_769285, EPI_ISL_769287, EPI_ISL_769292, EPI_ISL_769293, EPI_ISL_769295, EPI_ISL_769296, EPI_ISL_769298, EPI_ISL_769299, EPI_ISL_769300, EPI_ISL_769301, EPI_ISL_769302, EPI_ISL_769303, EPI_ISL_769304, EPI_ISL_769305, EPI_ISL_769307, EPI_ISL_769309, EPI_ISL_769311, EPI_ISL_769313, EPI_ISL_769315, EPI_ISL_769317, EPI_ISL_769318, EPI_ISL_769324, EPI_ISL_769326, EPI_ISL_769328, EPI_ISL_769331, EPI_ISL_769332, EPI_ISL_769333, EPI_ISL_769336, EPI_ISL_769337, EPI_ISL_769338, EPI_ISL_769339, EPI_ISL_769340, EPI_ISL_769342, EPI_ISL_769344, EPI_ISL_769346, EPI_ISL_769347, EPI_ISL_769350, EPI_ISL_769351, EPI_ISL_769354, EPI_ISL_769356, EPI_ISL_769357, EPI_ISL_769358, EPI_ISL_769359, EPI_ISL_769362, EPI_ISL_769363, EPI_ISL_769364, EPI_ISL_769366, EPI_ISL_769368, EPI_ISL_769369, EPI_ISL_769371, EPI_ISL_769372, EPI_ISL_769373, EPI_ISL_769375, EPI_ISL_769376, EPI_ISL_769377, EPI_ISL_769378, EPI_ISL_769379, EPI_ISL_769382, EPI_ISL_769386, EPI_ISL_769393, EPI_ISL_769395, EPI_ISL_769397, EPI_ISL_769398, EPI_ISL_769400, EPI_ISL_769403, EPI_ISL_769407, EPI_ISL_769408, EPI_ISL_769409, EPI_ISL_769410, EPI_ISL_769411, EPI_ISL_769414, EPI_ISL_769415, EPI_ISL_769423, EPI_ISL_769424, EPI_ISL_769425, EPI_ISL_769426, EPI_ISL_769429, EPI_ISL_769432, EPI_ISL_769433, EPI_ISL_769435, EPI_ISL_769436, EPI_ISL_769437, EPI_ISL_769438, EPI_ISL_769439, EPI_ISL_769443, EPI_ISL_769444, EPI_ISL_769447, EPI_ISL_769448, EPI_ISL_769451, EPI_ISL_769453, EPI_ISL_769455, EPI_ISL_769456, EPI_ISL_769461, EPI_ISL_769463, EPI_ISL_769464, EPI_ISL_769465, EPI_ISL_769467, EPI_ISL_769472, EPI_ISL_769477, EPI_ISL_769478, EPI_ISL_769479, EPI_ISL_769480, EPI_ISL_769483, EPI_ISL_769484, EPI_ISL_769486, EPI_ISL_769487, EPI_ISL_769488, EPI_ISL_769490, EPI_ISL_769491, EPI_ISL_769492, EPI_ISL_769493, EPI_ISL_769494, EPI_ISL_769495, EPI_ISL_769496, EPI_ISL_769498, EPI_ISL_769505, EPI_ISL_769507, EPI_ISL_769508, EPI_ISL_769509, EPI_ISL_769510, EPI_ISL_769514, EPI_ISL_769515, EPI_ISL_769516, EPI_ISL_769520, EPI_ISL_769521, EPI_ISL_769523, EPI_ISL_769526, EPI_ISL_769527, EPI_ISL_769529, EPI_ISL_769531, EPI_ISL_769532, EPI_ISL_769534, EPI_ISL_769536, EPI_ISL_769537, EPI_ISL_769538, EPI_ISL_769541, EPI_ISL_769544, EPI_ISL_769546, EPI_ISL_769547, EPI_ISL_769548, EPI_ISL_769549, EPI_ISL_769550 | | | |
| see above | Lighthouse Lab in Milton Keynes | Wellcome Sanger Institute for the COVID-19 Genomics UK (COG-UK) Consortium | The Lighthouse Lab in Milton Keynes and Alex Alderton, Roberto Amato, Sonia Goncalves, Ewan Harrison, David K. Jackson, Ian Johnston, Dominic Kwiatkowski, Cordelia Langford, John Sillitoe on behalf of the Wellcome Sanger Institute COVID-19 Surveillance Team |
| EPI_ISL_769551 | Lighthouse Lab in Glasgow | Wellcome Sanger Institute for the COVID-19 Genomics UK (COG-UK) Consortium | Harper VanSteenhouse, Yumi Kasai, David Gray, Carol Clugston, Anna Dominiczak and Alex Alderton, Roberto Amato, Sonia Goncalves, Ewan Harrison, David K. Jackson, Ian Johnston, Dominic Kwiatkowski, Cordelia Langford, John Sillitoe on behalf of the Wellcome Sanger Institute COVID-19 Surveillance Team |
| EPI_ISL_769552, EPI_ISL_769553, EPI_ISL_769554, EPI_ISL_769555 | Lighthouse Lab in Milton Keynes | Wellcome Sanger Institute for the COVID-19 Genomics UK (COG-UK) Consortium | The Lighthouse Lab in Milton Keynes and Alex Alderton, Roberto Amato, Sonia Goncalves, Ewan Harrison, David K. Jackson, Ian Johnston, Dominic Kwiatkowski, Cordelia Langford, John Sillitoe on behalf of the Wellcome Sanger Institute COVID-19 Surveillance Team |
| EPI_ISL_769556 | Lighthouse Lab in Glasgow | Wellcome Sanger Institute for the COVID-19 Genomics UK (COG-UK) Consortium | Harper VanSteenhouse, Yumi Kasai, David Gray, Carol Clugston, Anna Dominiczak and Alex Alderton, Roberto Amato, Sonia Goncalves, Ewan Harrison, David K. Jackson, Ian Johnston, Dominic Kwiatkowski, Cordelia Langford, John Sillitoe on behalf of the Wellcome Sanger Institute COVID-19 Surveillance Team |
| EPI_ISL_769557, EPI_ISL_769558, EPI_ISL_769559, EPI_ISL_769560, EPI_ISL_769561, EPI_ISL_769562, EPI_ISL_769563, EPI_ISL_769564, EPI_ISL_769565, EPI_ISL_769566 | Lighthouse Lab in Milton Keynes | Wellcome Sanger Institute for the COVID-19 Genomics UK (COG-UK) Consortium | The Lighthouse Lab in Milton Keynes and Alex Alderton, Roberto Amato, Sonia Goncalves, Ewan Harrison, David K. Jackson, Ian Johnston, Dominic Kwiatkowski, Cordelia Langford, John Sillitoe on behalf of the Wellcome Sanger Institute COVID-19 Surveillance Team |

| | | | |
|---|---|---|---|
| EPI_ISL_769567, EPI_ISL_769568 | Lighthouse Lab in Glasgow | Wellcome Sanger Institute for the COVID-19 Genomics UK (COG-UK) Consortium | Harper VanSteenhouse, Yumi Kasai, David Gray, Carol Clugston, Anna Dominiczak and Alex Alderton, Roberto Amato, Sonia Goncalves, Ewan Harrison, David K. Jackson, Ian Johnston, Dominic Kwiatkowski, Cordelia Langford, John Sillitoe on behalf of the Wellcome Sanger Institute COVID-19 Surveillance Team |
| EPI_ISL_769569, EPI_ISL_769570, EPI_ISL_769571, EPI_ISL_769572, EPI_ISL_769573, EPI_ISL_769574, EPI_ISL_769575, EPI_ISL_769576, EPI_ISL_769578, EPI_ISL_769579, EPI_ISL_769580, EPI_ISL_769581, EPI_ISL_769582 | | | |
| see above | Lighthouse Lab in Milton Keynes | Wellcome Sanger Institute for the COVID-19 Genomics UK (COG-UK) Consortium | The Lighthouse Lab in Milton Keynes and Alex Alderton, Roberto Amato, Sonia Goncalves, Ewan Harrison, David K. Jackson, Ian Johnston, Dominic Kwiatkowski, Cordelia Langford, John Sillitoe on behalf of the Wellcome Sanger Institute COVID-19 Surveillance Team |
| EPI_ISL_769583 | Lighthouse Lab in Glasgow | Wellcome Sanger Institute for the COVID-19 Genomics UK (COG-UK) Consortium | Harper VanSteenhouse, Yumi Kasai, David Gray, Carol Clugston, Anna Dominiczak and Alex Alderton, Roberto Amato, Sonia Goncalves, Ewan Harrison, David K. Jackson, Ian Johnston, Dominic Kwiatkowski, Cordelia Langford, John Sillitoe on behalf of the Wellcome Sanger Institute COVID-19 Surveillance Team |
| EPI_ISL_769585, EPI_ISL_769586, EPI_ISL_769587, EPI_ISL_769588 | Lighthouse Lab in Milton Keynes | Wellcome Sanger Institute for the COVID-19 Genomics UK (COG-UK) Consortium | The Lighthouse Lab in Milton Keynes and Alex Alderton, Roberto Amato, Sonia Goncalves, Ewan Harrison, David K. Jackson, Ian Johnston, Dominic Kwiatkowski, Cordelia Langford, John Sillitoe on behalf of the Wellcome Sanger Institute COVID-19 Surveillance Team |
| EPI_ISL_769589 | Lighthouse Lab in Glasgow | Wellcome Sanger Institute for the COVID-19 Genomics UK (COG-UK) Consortium | Harper VanSteenhouse, Yumi Kasai, David Gray, Carol Clugston, Anna Dominiczak and Alex Alderton, Roberto Amato, Sonia Goncalves, Ewan Harrison, David K. Jackson, Ian Johnston, Dominic Kwiatkowski, Cordelia Langford, John Sillitoe on behalf of the Wellcome Sanger Institute COVID-19 Surveillance Team |
| EPI_ISL_769591, EPI_ISL_769592, EPI_ISL_769593, EPI_ISL_769595, EPI_ISL_769596, EPI_ISL_769598, EPI_ISL_769599, EPI_ISL_769600, EPI_ISL_769601, EPI_ISL_769602, EPI_ISL_769603, EPI_ISL_769604, EPI_ISL_769605 | | | |
| see above | Lighthouse Lab in Milton Keynes | Wellcome Sanger Institute for the COVID-19 Genomics UK (COG-UK) Consortium | The Lighthouse Lab in Milton Keynes and Alex Alderton, Roberto Amato, Sonia Goncalves, Ewan Harrison, David K. Jackson, Ian Johnston, Dominic Kwiatkowski, Cordelia Langford, John Sillitoe on behalf of the Wellcome Sanger Institute COVID-19 Surveillance Team |
| EPI_ISL_769606, EPI_ISL_769607 | Lighthouse Lab in Glasgow | Wellcome Sanger Institute for the COVID-19 Genomics UK (COG-UK) Consortium | Harper VanSteenhouse, Yumi Kasai, David Gray, Carol Clugston, Anna Dominiczak and Alex Alderton, Roberto Amato, Sonia Goncalves, Ewan Harrison, David K. Jackson, Ian Johnston, Dominic Kwiatkowski, Cordelia Langford, John Sillitoe on behalf of the Wellcome Sanger Institute COVID-19 Surveillance Team |
| EPI_ISL_769608 | Lighthouse Lab in Milton Keynes | Wellcome Sanger Institute for the COVID-19 Genomics UK (COG-UK) Consortium | The Lighthouse Lab in Milton Keynes and Alex Alderton, Roberto Amato, Sonia Goncalves, Ewan Harrison, David K. Jackson, Ian Johnston, Dominic Kwiatkowski, Cordelia Langford, John Sillitoe on behalf of the Wellcome Sanger Institute COVID-19 Surveillance Team |
| EPI_ISL_769609 | Lighthouse Lab in Glasgow | Wellcome Sanger Institute for the COVID-19 Genomics UK (COG-UK) Consortium | Harper VanSteenhouse, Yumi Kasai, David Gray, Carol Clugston, Anna Dominiczak and Alex Alderton, Roberto Amato, Sonia Goncalves, Ewan Harrison, David K. Jackson, Ian Johnston, Dominic Kwiatkowski, Cordelia Langford, John Sillitoe on behalf of the Wellcome Sanger Institute COVID-19 Surveillance Team |
| EPI_ISL_769610 | Lighthouse Lab in Milton Keynes | Wellcome Sanger Institute for the COVID-19 Genomics UK (COG-UK) Consortium | The Lighthouse Lab in Milton Keynes and Alex Alderton, Roberto Amato, Sonia Goncalves, Ewan Harrison, David K. Jackson, Ian Johnston, Dominic Kwiatkowski, Cordelia Langford, John Sillitoe on behalf of the Wellcome Sanger Institute COVID-19 Surveillance Team |
| EPI_ISL_769611, EPI_ISL_769612, EPI_ISL_769613, EPI_ISL_769614 | Lighthouse Lab in Glasgow | Wellcome Sanger Institute for the COVID-19 Genomics UK (COG-UK) Consortium | Harper VanSteenhouse, Yumi Kasai, David Gray, Carol Clugston, Anna Dominiczak and Alex Alderton, Roberto Amato, Sonia Goncalves, Ewan Harrison, David K. Jackson, Ian Johnston, Dominic Kwiatkowski, Cordelia Langford, John Sillitoe on behalf of the Wellcome Sanger Institute COVID-19 Surveillance Team |
| EPI_ISL_769615, EPI_ISL_769616 | Lighthouse Lab in Milton Keynes | Wellcome Sanger Institute for the COVID-19 Genomics UK (COG-UK) Consortium | The Lighthouse Lab in Milton Keynes and Alex Alderton, Roberto Amato, Sonia Goncalves, Ewan Harrison, David K. Jackson, Ian Johnston, Dominic Kwiatkowski, Cordelia Langford, John Sillitoe on behalf of the Wellcome Sanger Institute COVID-19 Surveillance Team |
| EPI_ISL_769617 | Lighthouse Lab in Glasgow | Wellcome Sanger Institute for the COVID-19 Genomics UK (COG-UK) Consortium | Harper VanSteenhouse, Yumi Kasai, David Gray, Carol Clugston, Anna Dominiczak and Alex Alderton, Roberto Amato, Sonia Goncalves, Ewan Harrison, David K. Jackson, Ian Johnston, Dominic Kwiatkowski, Cordelia Langford, John Sillitoe on behalf of the Wellcome Sanger Institute COVID-19 Surveillance Team |
| EPI_ISL_769618 | Lighthouse Lab in Milton Keynes | Wellcome Sanger Institute for the COVID-19 Genomics UK (COG-UK) Consortium | The Lighthouse Lab in Milton Keynes and Alex Alderton, Roberto Amato, Sonia Goncalves, Ewan Harrison, David K. Jackson, Ian Johnston, Dominic Kwiatkowski, Cordelia Langford, John Sillitoe on behalf of the Wellcome Sanger Institute COVID-19 Surveillance Team |
| EPI_ISL_769619 | Lighthouse Lab in Glasgow | Wellcome Sanger Institute for the COVID-19 Genomics UK (COG-UK) Consortium | Harper VanSteenhouse, Yumi Kasai, David Gray, Carol Clugston, Anna Dominiczak and Alex Alderton, Roberto Amato, Sonia Goncalves, Ewan Harrison, David K. Jackson, Ian Johnston, Dominic Kwiatkowski, Cordelia Langford, John Sillitoe on behalf of the Wellcome Sanger Institute COVID-19 Surveillance Team |
| EPI_ISL_769620, EPI_ISL_769621, EPI_ISL_769622, EPI_ISL_769623, EPI_ISL_769624 | Lighthouse Lab in Milton Keynes | Wellcome Sanger Institute for the COVID-19 Genomics UK (COG-UK) Consortium | The Lighthouse Lab in Milton Keynes and Alex Alderton, Roberto Amato, Sonia Goncalves, Ewan Harrison, David K. Jackson, Ian Johnston, Dominic Kwiatkowski, Cordelia Langford, John Sillitoe on behalf of the Wellcome Sanger Institute COVID-19 Surveillance Team |
| EPI_ISL_769625 | Lighthouse Lab in Glasgow | Wellcome Sanger Institute for the COVID-19 Genomics UK (COG-UK) Consortium | Harper VanSteenhouse, Yumi Kasai, David Gray, Carol Clugston, Anna Dominiczak and Alex Alderton, Roberto Amato, Sonia Goncalves, Ewan Harrison, David K. Jackson, Ian Johnston, Dominic Kwiatkowski, Cordelia Langford, John Sillitoe on behalf of the Wellcome Sanger Institute COVID-19 Surveillance Team |
| EPI_ISL_769626, EPI_ISL_769627, EPI_ISL_769628, EPI_ISL_769629, EPI_ISL_769630, EPI_ISL_769631, EPI_ISL_769632, EPI_ISL_769633, EPI_ISL_769634, EPI_ISL_769635, EPI_ISL_769636, EPI_ISL_769637, EPI_ISL_769638, EPI_ISL_769639, EPI_ISL_769640, EPI_ISL_769642, EPI_ISL_769643 | | | |
| see above | Lighthouse Lab in Milton Keynes | Wellcome Sanger Institute for the COVID-19 Genomics UK (COG-UK) Consortium | The Lighthouse Lab in Milton Keynes and Alex Alderton, Roberto Amato, Sonia Goncalves, Ewan Harrison, David K. Jackson, Ian Johnston, Dominic Kwiatkowski, Cordelia Langford, John Sillitoe on behalf of the Wellcome Sanger Institute COVID-19 Surveillance Team |
| EPI_ISL_769644 | Lighthouse Lab in Glasgow | Wellcome Sanger Institute for the COVID-19 Genomics UK (COG-UK) Consortium | Harper VanSteenhouse, Yumi Kasai, David Gray, Carol Clugston, Anna Dominiczak and Alex Alderton, Roberto Amato, Sonia Goncalves, Ewan Harrison, David K. Jackson, Ian Johnston, Dominic Kwiatkowski, Cordelia Langford, John Sillitoe on behalf of the Wellcome Sanger Institute COVID-19 Surveillance Team |
| EPI_ISL_769645, EPI_ISL_769647 | Lighthouse Lab in Milton Keynes | Wellcome Sanger Institute for the COVID-19 Genomics UK (COG-UK) Consortium | The Lighthouse Lab in Milton Keynes and Alex Alderton, Roberto Amato, Sonia Goncalves, Ewan Harrison, David K. Jackson, Ian Johnston, Dominic Kwiatkowski, Cordelia Langford, John Sillitoe on behalf of the Wellcome Sanger Institute COVID-19 Surveillance Team |
| EPI_ISL_769648 | Lighthouse Lab in Glasgow | Wellcome Sanger Institute for the COVID-19 Genomics UK (COG-UK) Consortium | Harper VanSteenhouse, Yumi Kasai, David Gray, Carol Clugston, Anna Dominiczak and Alex Alderton, Roberto Amato, Sonia Goncalves, Ewan Harrison, David K. Jackson, Ian Johnston, Dominic Kwiatkowski, Cordelia Langford, John Sillitoe on behalf of the Wellcome Sanger Institute COVID-19 Surveillance Team |
| EPI_ISL_769649, EPI_ISL_769650, EPI_ISL_769651 | Lighthouse Lab in Milton Keynes | Wellcome Sanger Institute for the COVID-19 Genomics UK (COG-UK) Consortium | The Lighthouse Lab in Milton Keynes and Alex Alderton, Roberto Amato, Sonia Goncalves, Ewan Harrison, David K. Jackson, Ian Johnston, Dominic Kwiatkowski, Cordelia Langford, John Sillitoe on behalf of the Wellcome Sanger Institute COVID-19 Surveillance Team |
| EPI_ISL_769652 | Lighthouse Lab in Glasgow | Wellcome Sanger Institute for the COVID-19 Genomics UK (COG-UK) Consortium | Harper VanSteenhouse, Yumi Kasai, David Gray, Carol Clugston, Anna Dominiczak and Alex Alderton, Roberto Amato, Sonia Goncalves, Ewan Harrison, David K. Jackson, Ian Johnston, Dominic Kwiatkowski, Cordelia Langford, John Sillitoe on behalf of the Wellcome Sanger Institute COVID-19 Surveillance Team |
| EPI_ISL_769653, EPI_ISL_769654, EPI_ISL_769656, EPI_ISL_769657, EPI_ISL_769658, EPI_ISL_769659, EPI_ISL_769660 | Lighthouse Lab in Milton Keynes | Wellcome Sanger Institute for the COVID-19 Genomics UK (COG-UK) Consortium | The Lighthouse Lab in Milton Keynes and Alex Alderton, Roberto Amato, Sonia Goncalves, Ewan Harrison, David K. Jackson, Ian Johnston, Dominic Kwiatkowski, Cordelia Langford, John Sillitoe on behalf of the Wellcome Sanger Institute COVID-19 Surveillance Team |
| EPI_ISL_769662 | Lighthouse Lab in Glasgow | Wellcome Sanger Institute for the COVID-19 Genomics UK (COG-UK) Consortium | Harper VanSteenhouse, Yumi Kasai, David Gray, Carol Clugston, Anna Dominiczak and Alex Alderton, Roberto Amato, Sonia Goncalves, Ewan Harrison, David K. Jackson, Ian Johnston, Dominic Kwiatkowski, Cordelia Langford, John Sillitoe on behalf of the Wellcome Sanger Institute COVID-19 Surveillance Team |
| EPI_ISL_769663, EPI_ISL_769664, EPI_ISL_769665 | Lighthouse Lab in Milton Keynes | Wellcome Sanger Institute for the COVID-19 Genomics UK (COG-UK) Consortium | The Lighthouse Lab in Milton Keynes and Alex Alderton, Roberto Amato, Sonia Goncalves, Ewan Harrison, David K. Jackson, Ian Johnston, Dominic Kwiatkowski, Cordelia Langford, John Sillitoe on behalf of the Wellcome Sanger Institute COVID-19 Surveillance Team |
| EPI_ISL_769666, EPI_ISL_769667 | Lighthouse Lab in Glasgow | Wellcome Sanger Institute for the COVID-19 Genomics UK (COG-UK) Consortium | Harper VanSteenhouse, Yumi Kasai, David Gray, Carol Clugston, Anna Dominiczak and Alex Alderton, Roberto Amato, Sonia Goncalves, Ewan Harrison, David K. Jackson, Ian Johnston, Dominic Kwiatkowski, Cordelia Langford, John Sillitoe on behalf of the Wellcome Sanger Institute COVID-19 Surveillance |

| | | | |
|---|---|---|---|
| | | | Team |
| EPI_ISL_769668, EPI_ISL_769669, EPI_ISL_769670, EPI_ISL_769671, EPI_ISL_769672, EPI_ISL_769673, EPI_ISL_769674, EPI_ISL_769675, EPI_ISL_769676 | Lighthouse Lab in Milton Keynes | Wellcome Sanger Institute for the COVID-19 Genomics UK (COG-UK) Consortium | The Lighthouse Lab in Milton Keynes and Alex Alderton, Roberto Amato, Sonia Goncalves, Ewan Harrison, David K. Jackson, Ian Johnston, Dominic Kwiatkowski, Cordelia Langford, John Sillitoe on behalf of the Wellcome Sanger Institute COVID-19 Surveillance Team |
| EPI_ISL_769677 | Lighthouse Lab in Glasgow | Wellcome Sanger Institute for the COVID-19 Genomics UK (COG-UK) Consortium | Harper VanSteenhouse, Yumi Kasai, David Gray, Carol Clugston, Anna Dominiczak and Alex Alderton, Roberto Amato, Sonia Goncalves, Ewan Harrison, David K. Jackson, Ian Johnston, Dominic Kwiatkowski, Cordelia Langford, John Sillitoe on behalf of the Wellcome Sanger Institute COVID-19 Surveillance Team |
| EPI_ISL_769678, EPI_ISL_769679, EPI_ISL_769680, EPI_ISL_769681, EPI_ISL_769682, EPI_ISL_769683, EPI_ISL_769684, EPI_ISL_769685, EPI_ISL_769686 | Lighthouse Lab in Milton Keynes | Wellcome Sanger Institute for the COVID-19 Genomics UK (COG-UK) Consortium | The Lighthouse Lab in Milton Keynes and Alex Alderton, Roberto Amato, Sonia Goncalves, Ewan Harrison, David K. Jackson, Ian Johnston, Dominic Kwiatkowski, Cordelia Langford, John Sillitoe on behalf of the Wellcome Sanger Institute COVID-19 Surveillance Team |
| EPI_ISL_769687 | Lighthouse Lab in Glasgow | Wellcome Sanger Institute for the COVID-19 Genomics UK (COG-UK) Consortium | Harper VanSteenhouse, Yumi Kasai, David Gray, Carol Clugston, Anna Dominiczak and Alex Alderton, Roberto Amato, Sonia Goncalves, Ewan Harrison, David K. Jackson, Ian Johnston, Dominic Kwiatkowski, Cordelia Langford, John Sillitoe on behalf of the Wellcome Sanger Institute COVID-19 Surveillance Team |
| EPI_ISL_769688 | Lighthouse Lab in Milton Keynes | Wellcome Sanger Institute for the COVID-19 Genomics UK (COG-UK) Consortium | The Lighthouse Lab in Milton Keynes and Alex Alderton, Roberto Amato, Sonia Goncalves, Ewan Harrison, David K. Jackson, Ian Johnston, Dominic Kwiatkowski, Cordelia Langford, John Sillitoe on behalf of the Wellcome Sanger Institute COVID-19 Surveillance Team |
| EPI_ISL_769689 | Lighthouse Lab in Glasgow | Wellcome Sanger Institute for the COVID-19 Genomics UK (COG-UK) Consortium | Harper VanSteenhouse, Yumi Kasai, David Gray, Carol Clugston, Anna Dominiczak and Alex Alderton, Roberto Amato, Sonia Goncalves, Ewan Harrison, David K. Jackson, Ian Johnston, Dominic Kwiatkowski, Cordelia Langford, John Sillitoe on behalf of the Wellcome Sanger Institute COVID-19 Surveillance Team |
| EPI_ISL_769690 | Lighthouse Lab in Milton Keynes | Wellcome Sanger Institute for the COVID-19 Genomics UK (COG-UK) Consortium | The Lighthouse Lab in Milton Keynes and Alex Alderton, Roberto Amato, Sonia Goncalves, Ewan Harrison, David K. Jackson, Ian Johnston, Dominic Kwiatkowski, Cordelia Langford, John Sillitoe on behalf of the Wellcome Sanger Institute COVID-19 Surveillance Team |
| EPI_ISL_769691 | Lighthouse Lab in Glasgow | Wellcome Sanger Institute for the COVID-19 Genomics UK (COG-UK) Consortium | Harper VanSteenhouse, Yumi Kasai, David Gray, Carol Clugston, Anna Dominiczak and Alex Alderton, Roberto Amato, Sonia Goncalves, Ewan Harrison, David K. Jackson, Ian Johnston, Dominic Kwiatkowski, Cordelia Langford, John Sillitoe on behalf of the Wellcome Sanger Institute COVID-19 Surveillance Team |
| EPI_ISL_769692, EPI_ISL_769693, EPI_ISL_769694, EPI_ISL_769695, EPI_ISL_769696, EPI_ISL_769697, EPI_ISL_769698, EPI_ISL_769699, EPI_ISL_769700, EPI_ISL_769701, EPI_ISL_769702 | | | |
| see above | Lighthouse Lab in Milton Keynes | Wellcome Sanger Institute for the COVID-19 Genomics UK (COG-UK) Consortium | The Lighthouse Lab in Milton Keynes and Alex Alderton, Roberto Amato, Sonia Goncalves, Ewan Harrison, David K. Jackson, Ian Johnston, Dominic Kwiatkowski, Cordelia Langford, John Sillitoe on behalf of the Wellcome Sanger Institute COVID-19 Surveillance Team |
| EPI_ISL_769703 | Lighthouse Lab in Glasgow | Wellcome Sanger Institute for the COVID-19 Genomics UK (COG-UK) Consortium | Harper VanSteenhouse, Yumi Kasai, David Gray, Carol Clugston, Anna Dominiczak and Alex Alderton, Roberto Amato, Sonia Goncalves, Ewan Harrison, David K. Jackson, Ian Johnston, Dominic Kwiatkowski, Cordelia Langford, John Sillitoe on behalf of the Wellcome Sanger Institute COVID-19 Surveillance Team |
| EPI_ISL_769704 | Lighthouse Lab in Milton Keynes | Wellcome Sanger Institute for the COVID-19 Genomics UK (COG-UK) Consortium | The Lighthouse Lab in Milton Keynes and Alex Alderton, Roberto Amato, Sonia Goncalves, Ewan Harrison, David K. Jackson, Ian Johnston, Dominic Kwiatkowski, Cordelia Langford, John Sillitoe on behalf of the Wellcome Sanger Institute COVID-19 Surveillance Team |
| EPI_ISL_769705 | Lighthouse Lab in Glasgow | Wellcome Sanger Institute for the COVID-19 Genomics UK (COG-UK) Consortium | Harper VanSteenhouse, Yumi Kasai, David Gray, Carol Clugston, Anna Dominiczak and Alex Alderton, Roberto Amato, Sonia Goncalves, Ewan Harrison, David K. Jackson, Ian Johnston, Dominic Kwiatkowski, Cordelia Langford, John Sillitoe on behalf of the Wellcome Sanger Institute COVID-19 Surveillance Team |
| EPI_ISL_769706, EPI_ISL_769707, EPI_ISL_769708, EPI_ISL_769709, EPI_ISL_769710 | Lighthouse Lab in Milton Keynes | Wellcome Sanger Institute for the COVID-19 Genomics UK (COG-UK) Consortium | The Lighthouse Lab in Milton Keynes and Alex Alderton, Roberto Amato, Sonia Goncalves, Ewan Harrison, David K. Jackson, Ian Johnston, Dominic Kwiatkowski, Cordelia Langford, John Sillitoe on behalf of the Wellcome Sanger Institute COVID-19 Surveillance Team |
| EPI_ISL_769711 | Lighthouse Lab in Glasgow | Wellcome Sanger Institute for the COVID-19 Genomics UK (COG-UK) Consortium | Harper VanSteenhouse, Yumi Kasai, David Gray, Carol Clugston, Anna Dominiczak and Alex Alderton, Roberto Amato, Sonia Goncalves, Ewan Harrison, David K. Jackson, Ian Johnston, Dominic Kwiatkowski, Cordelia Langford, John Sillitoe on behalf of the Wellcome Sanger Institute COVID-19 Surveillance Team |
| EPI_ISL_769712 | Lighthouse Lab in Milton Keynes | Wellcome Sanger Institute for the COVID-19 Genomics UK (COG-UK) Consortium | The Lighthouse Lab in Milton Keynes and Alex Alderton, Roberto Amato, Sonia Goncalves, Ewan Harrison, David K. Jackson, Ian Johnston, Dominic Kwiatkowski, Cordelia Langford, John Sillitoe on behalf of the Wellcome Sanger Institute COVID-19 Surveillance Team |
| EPI_ISL_769713, EPI_ISL_769714, EPI_ISL_769716 | Lighthouse Lab in Glasgow | Wellcome Sanger Institute for the COVID-19 Genomics UK (COG-UK) Consortium | Harper VanSteenhouse, Yumi Kasai, David Gray, Carol Clugston, Anna Dominiczak and Alex Alderton, Roberto Amato, Sonia Goncalves, Ewan Harrison, David K. Jackson, Ian Johnston, Dominic Kwiatkowski, Cordelia Langford, John Sillitoe on behalf of the Wellcome Sanger Institute COVID-19 Surveillance Team |
| EPI_ISL_769717, EPI_ISL_769718, EPI_ISL_769719 | Lighthouse Lab in Milton Keynes | Wellcome Sanger Institute for the COVID-19 Genomics UK (COG-UK) Consortium | The Lighthouse Lab in Milton Keynes and Alex Alderton, Roberto Amato, Sonia Goncalves, Ewan Harrison, David K. Jackson, Ian Johnston, Dominic Kwiatkowski, Cordelia Langford, John Sillitoe on behalf of the Wellcome Sanger Institute COVID-19 Surveillance Team |
| EPI_ISL_769720 | Lighthouse Lab in Glasgow | Wellcome Sanger Institute for the COVID-19 Genomics UK (COG-UK) Consortium | Harper VanSteenhouse, Yumi Kasai, David Gray, Carol Clugston, Anna Dominiczak and Alex Alderton, Roberto Amato, Sonia Goncalves, Ewan Harrison, David K. Jackson, Ian Johnston, Dominic Kwiatkowski, Cordelia Langford, John Sillitoe on behalf of the Wellcome Sanger Institute COVID-19 Surveillance Team |
| EPI_ISL_769721, EPI_ISL_769722, EPI_ISL_769723 | Lighthouse Lab in Milton Keynes | Wellcome Sanger Institute for the COVID-19 Genomics UK (COG-UK) Consortium | The Lighthouse Lab in Milton Keynes and Alex Alderton, Roberto Amato, Sonia Goncalves, Ewan Harrison, David K. Jackson, Ian Johnston, Dominic Kwiatkowski, Cordelia Langford, John Sillitoe on behalf of the Wellcome Sanger Institute COVID-19 Surveillance Team |
| EPI_ISL_769724 | Lighthouse Lab in Glasgow | Wellcome Sanger Institute for the COVID-19 Genomics UK (COG-UK) Consortium | Harper VanSteenhouse, Yumi Kasai, David Gray, Carol Clugston, Anna Dominiczak and Alex Alderton, Roberto Amato, Sonia Goncalves, Ewan Harrison, David K. Jackson, Ian Johnston, Dominic Kwiatkowski, Cordelia Langford, John Sillitoe on behalf of the Wellcome Sanger Institute COVID-19 Surveillance Team |
| EPI_ISL_769725, EPI_ISL_769726, EPI_ISL_769727, EPI_ISL_769728, EPI_ISL_769729, EPI_ISL_769730, EPI_ISL_769731, EPI_ISL_769732, EPI_ISL_769733, EPI_ISL_769734 | Lighthouse Lab in Milton Keynes | Wellcome Sanger Institute for the COVID-19 Genomics UK (COG-UK) Consortium | The Lighthouse Lab in Milton Keynes and Alex Alderton, Roberto Amato, Sonia Goncalves, Ewan Harrison, David K. Jackson, Ian Johnston, Dominic Kwiatkowski, Cordelia Langford, John Sillitoe on behalf of the Wellcome Sanger Institute COVID-19 Surveillance Team |
| EPI_ISL_769735 | Lighthouse Lab in Glasgow | Wellcome Sanger Institute for the COVID-19 Genomics UK (COG-UK) Consortium | Harper VanSteenhouse, Yumi Kasai, David Gray, Carol Clugston, Anna Dominiczak and Alex Alderton, Roberto Amato, Sonia Goncalves, Ewan Harrison, David K. Jackson, Ian Johnston, Dominic Kwiatkowski, Cordelia Langford, John Sillitoe on behalf of the Wellcome Sanger Institute COVID-19 Surveillance Team |
| EPI_ISL_769736, EPI_ISL_769737, EPI_ISL_769738, EPI_ISL_769739, EPI_ISL_769740 | Lighthouse Lab in Milton Keynes | Wellcome Sanger Institute for the COVID-19 Genomics UK (COG-UK) Consortium | The Lighthouse Lab in Milton Keynes and Alex Alderton, Roberto Amato, Sonia Goncalves, Ewan Harrison, David K. Jackson, Ian Johnston, Dominic Kwiatkowski, Cordelia Langford, John Sillitoe on behalf of the Wellcome Sanger Institute COVID-19 Surveillance Team |
| EPI_ISL_769741 | Lighthouse Lab in Glasgow | Wellcome Sanger Institute for the COVID-19 Genomics UK (COG-UK) Consortium | Harper VanSteenhouse, Yumi Kasai, David Gray, Carol Clugston, Anna Dominiczak and Alex Alderton, Roberto Amato, Sonia Goncalves, Ewan Harrison, David K. Jackson, Ian Johnston, Dominic Kwiatkowski, Cordelia Langford, John Sillitoe on behalf of the Wellcome Sanger Institute COVID-19 Surveillance Team |
| EPI_ISL_769742 | Lighthouse Lab in Milton Keynes | Wellcome Sanger Institute for the COVID-19 Genomics UK | The Lighthouse Lab in Milton Keynes and Alex Alderton, Roberto Amato, Sonia Goncalves, Ewan Harrison, David K. Jackson, Ian Johnston, Dominic |

| | | | |
|---|---|---|---|
| | | (COG-UK) Consortium | Kwiatkowski, Cordelia Langford, John Sillitoe on behalf of the Wellcome Sanger Institute COVID-19 Surveillance Team |
| EPI_ISL_769743 | Lighthouse Lab in Glasgow | Wellcome Sanger Institute for the COVID-19 Genomics UK (COG-UK) Consortium | Harper VanSteenhouse, Yumi Kasai, David Gray, Carol Clugston, Anna Dominiczak and Alex Alderton, Roberto Amato, Sonia Goncalves, Ewan Harrison, David K. Jackson, Ian Johnston, Dominic Kwiatkowski, Cordelia Langford, John Sillitoe on behalf of the Wellcome Sanger Institute COVID-19 Surveillance Team |
| EPI_ISL_769745, EPI_ISL_769746, EPI_ISL_769748, EPI_ISL_769749, EPI_ISL_769750 | Lighthouse Lab in Milton Keynes | Wellcome Sanger Institute for the COVID-19 Genomics UK (COG-UK) Consortium | The Lighthouse Lab in Milton Keynes and Alex Alderton, Roberto Amato, Sonia Goncalves, Ewan Harrison, David K. Jackson, Ian Johnston, Dominic Kwiatkowski, Cordelia Langford, John Sillitoe on behalf of the Wellcome Sanger Institute COVID-19 Surveillance Team |
| EPI_ISL_769751, EPI_ISL_769752, EPI_ISL_769753 | Lighthouse Lab in Glasgow | Wellcome Sanger Institute for the COVID-19 Genomics UK (COG-UK) Consortium | Harper VanSteenhouse, Yumi Kasai, David Gray, Carol Clugston, Anna Dominiczak and Alex Alderton, Roberto Amato, Sonia Goncalves, Ewan Harrison, David K. Jackson, Ian Johnston, Dominic Kwiatkowski, Cordelia Langford, John Sillitoe on behalf of the Wellcome Sanger Institute COVID-19 Surveillance Team |
| EPI_ISL_769754, EPI_ISL_769755, EPI_ISL_769756, EPI_ISL_769757 | Lighthouse Lab in Milton Keynes | Wellcome Sanger Institute for the COVID-19 Genomics UK (COG-UK) Consortium | The Lighthouse Lab in Milton Keynes and Alex Alderton, Roberto Amato, Sonia Goncalves, Ewan Harrison, David K. Jackson, Ian Johnston, Dominic Kwiatkowski, Cordelia Langford, John Sillitoe on behalf of the Wellcome Sanger Institute COVID-19 Surveillance Team |
| EPI_ISL_769758, EPI_ISL_769759 | Lighthouse Lab in Glasgow | Wellcome Sanger Institute for the COVID-19 Genomics UK (COG-UK) Consortium | Harper VanSteenhouse, Yumi Kasai, David Gray, Carol Clugston, Anna Dominiczak and Alex Alderton, Roberto Amato, Sonia Goncalves, Ewan Harrison, David K. Jackson, Ian Johnston, Dominic Kwiatkowski, Cordelia Langford, John Sillitoe on behalf of the Wellcome Sanger Institute COVID-19 Surveillance Team |
| EPI_ISL_769760 | Lighthouse Lab in Milton Keynes | Wellcome Sanger Institute for the COVID-19 Genomics UK (COG-UK) Consortium | The Lighthouse Lab in Milton Keynes and Alex Alderton, Roberto Amato, Sonia Goncalves, Ewan Harrison, David K. Jackson, Ian Johnston, Dominic Kwiatkowski, Cordelia Langford, John Sillitoe on behalf of the Wellcome Sanger Institute COVID-19 Surveillance Team |
| EPI_ISL_769761 | Lighthouse Lab in Glasgow | Wellcome Sanger Institute for the COVID-19 Genomics UK (COG-UK) Consortium | Harper VanSteenhouse, Yumi Kasai, David Gray, Carol Clugston, Anna Dominiczak and Alex Alderton, Roberto Amato, Sonia Goncalves, Ewan Harrison, David K. Jackson, Ian Johnston, Dominic Kwiatkowski, Cordelia Langford, John Sillitoe on behalf of the Wellcome Sanger Institute COVID-19 Surveillance Team |
| EPI_ISL_769762, EPI_ISL_769763, EPI_ISL_769765, EPI_ISL_769766, EPI_ISL_769767 | Lighthouse Lab in Milton Keynes | Wellcome Sanger Institute for the COVID-19 Genomics UK (COG-UK) Consortium | The Lighthouse Lab in Milton Keynes and Alex Alderton, Roberto Amato, Sonia Goncalves, Ewan Harrison, David K. Jackson, Ian Johnston, Dominic Kwiatkowski, Cordelia Langford, John Sillitoe on behalf of the Wellcome Sanger Institute COVID-19 Surveillance Team |
| EPI_ISL_769768 | Lighthouse Lab in Glasgow | Wellcome Sanger Institute for the COVID-19 Genomics UK (COG-UK) Consortium | Harper VanSteenhouse, Yumi Kasai, David Gray, Carol Clugston, Anna Dominiczak and Alex Alderton, Roberto Amato, Sonia Goncalves, Ewan Harrison, David K. Jackson, Ian Johnston, Dominic Kwiatkowski, Cordelia Langford, John Sillitoe on behalf of the Wellcome Sanger Institute COVID-19 Surveillance Team |
| EPI_ISL_769769, EPI_ISL_769770, EPI_ISL_769771, EPI_ISL_769772 | Lighthouse Lab in Milton Keynes | Wellcome Sanger Institute for the COVID-19 Genomics UK (COG-UK) Consortium | The Lighthouse Lab in Milton Keynes and Alex Alderton, Roberto Amato, Sonia Goncalves, Ewan Harrison, David K. Jackson, Ian Johnston, Dominic Kwiatkowski, Cordelia Langford, John Sillitoe on behalf of the Wellcome Sanger Institute COVID-19 Surveillance Team |
| EPI_ISL_769773 | Lighthouse Lab in Glasgow | Wellcome Sanger Institute for the COVID-19 Genomics UK (COG-UK) Consortium | Harper VanSteenhouse, Yumi Kasai, David Gray, Carol Clugston, Anna Dominiczak and Alex Alderton, Roberto Amato, Sonia Goncalves, Ewan Harrison, David K. Jackson, Ian Johnston, Dominic Kwiatkowski, Cordelia Langford, John Sillitoe on behalf of the Wellcome Sanger Institute COVID-19 Surveillance Team |
| EPI_ISL_769774, EPI_ISL_769775, EPI_ISL_769777, EPI_ISL_769778, EPI_ISL_769779, EPI_ISL_769780, EPI_ISL_769781, EPI_ISL_769782, EPI_ISL_769783, EPI_ISL_769784, EPI_ISL_769785, EPI_ISL_769786 | | | |
| see above | Lighthouse Lab in Milton Keynes | Wellcome Sanger Institute for the COVID-19 Genomics UK (COG-UK) Consortium | The Lighthouse Lab in Milton Keynes and Alex Alderton, Roberto Amato, Sonia Goncalves, Ewan Harrison, David K. Jackson, Ian Johnston, Dominic Kwiatkowski, Cordelia Langford, John Sillitoe on behalf of the Wellcome Sanger Institute COVID-19 Surveillance Team |
| EPI_ISL_769787, EPI_ISL_769788 | Lighthouse Lab in Glasgow | Wellcome Sanger Institute for the COVID-19 Genomics UK (COG-UK) Consortium | Harper VanSteenhouse, Yumi Kasai, David Gray, Carol Clugston, Anna Dominiczak and Alex Alderton, Roberto Amato, Sonia Goncalves, Ewan Harrison, David K. Jackson, Ian Johnston, Dominic Kwiatkowski, Cordelia Langford, John Sillitoe on behalf of the Wellcome Sanger Institute COVID-19 Surveillance Team |
| EPI_ISL_769789, EPI_ISL_769790, EPI_ISL_769791, EPI_ISL_769792, EPI_ISL_769793 | Lighthouse Lab in Milton Keynes | Wellcome Sanger Institute for the COVID-19 Genomics UK (COG-UK) Consortium | The Lighthouse Lab in Milton Keynes and Alex Alderton, Roberto Amato, Sonia Goncalves, Ewan Harrison, David K. Jackson, Ian Johnston, Dominic Kwiatkowski, Cordelia Langford, John Sillitoe on behalf of the Wellcome Sanger Institute COVID-19 Surveillance Team |
| EPI_ISL_769794 | Lighthouse Lab in Glasgow | Wellcome Sanger Institute for the COVID-19 Genomics UK (COG-UK) Consortium | Harper VanSteenhouse, Yumi Kasai, David Gray, Carol Clugston, Anna Dominiczak and Alex Alderton, Roberto Amato, Sonia Goncalves, Ewan Harrison, David K. Jackson, Ian Johnston, Dominic Kwiatkowski, Cordelia Langford, John Sillitoe on behalf of the Wellcome Sanger Institute COVID-19 Surveillance Team |
| EPI_ISL_769795, EPI_ISL_769796, EPI_ISL_769797, EPI_ISL_769798, EPI_ISL_769799, EPI_ISL_769800, EPI_ISL_769801, EPI_ISL_769803, EPI_ISL_769804, EPI_ISL_769805 | Lighthouse Lab in Milton Keynes | Wellcome Sanger Institute for the COVID-19 Genomics UK (COG-UK) Consortium | The Lighthouse Lab in Milton Keynes and Alex Alderton, Roberto Amato, Sonia Goncalves, Ewan Harrison, David K. Jackson, Ian Johnston, Dominic Kwiatkowski, Cordelia Langford, John Sillitoe on behalf of the Wellcome Sanger Institute COVID-19 Surveillance Team |
| EPI_ISL_769806 | Lighthouse Lab in Glasgow | Wellcome Sanger Institute for the COVID-19 Genomics UK (COG-UK) Consortium | Harper VanSteenhouse, Yumi Kasai, David Gray, Carol Clugston, Anna Dominiczak and Alex Alderton, Roberto Amato, Sonia Goncalves, Ewan Harrison, David K. Jackson, Ian Johnston, Dominic Kwiatkowski, Cordelia Langford, John Sillitoe on behalf of the Wellcome Sanger Institute COVID-19 Surveillance Team |
| EPI_ISL_769807, EPI_ISL_769808, EPI_ISL_769809, EPI_ISL_769810, EPI_ISL_769811, EPI_ISL_769812 | Lighthouse Lab in Milton Keynes | Wellcome Sanger Institute for the COVID-19 Genomics UK (COG-UK) Consortium | The Lighthouse Lab in Milton Keynes and Alex Alderton, Roberto Amato, Sonia Goncalves, Ewan Harrison, David K. Jackson, Ian Johnston, Dominic Kwiatkowski, Cordelia Langford, John Sillitoe on behalf of the Wellcome Sanger Institute COVID-19 Surveillance Team |
| EPI_ISL_769813 | Lighthouse Lab in Glasgow | Wellcome Sanger Institute for the COVID-19 Genomics UK (COG-UK) Consortium | Harper VanSteenhouse, Yumi Kasai, David Gray, Carol Clugston, Anna Dominiczak and Alex Alderton, Roberto Amato, Sonia Goncalves, Ewan Harrison, David K. Jackson, Ian Johnston, Dominic Kwiatkowski, Cordelia Langford, John Sillitoe on behalf of the Wellcome Sanger Institute COVID-19 Surveillance Team |
| EPI_ISL_769814 | Lighthouse Lab in Milton Keynes | Wellcome Sanger Institute for the COVID-19 Genomics UK (COG-UK) Consortium | The Lighthouse Lab in Milton Keynes and Alex Alderton, Roberto Amato, Sonia Goncalves, Ewan Harrison, David K. Jackson, Ian Johnston, Dominic Kwiatkowski, Cordelia Langford, John Sillitoe on behalf of the Wellcome Sanger Institute COVID-19 Surveillance Team |
| EPI_ISL_769815 | Lighthouse Lab in Glasgow | Wellcome Sanger Institute for the COVID-19 Genomics UK (COG-UK) Consortium | Harper VanSteenhouse, Yumi Kasai, David Gray, Carol Clugston, Anna Dominiczak and Alex Alderton, Roberto Amato, Sonia Goncalves, Ewan Harrison, David K. Jackson, Ian Johnston, Dominic Kwiatkowski, Cordelia Langford, John Sillitoe on behalf of the Wellcome Sanger Institute COVID-19 Surveillance Team |
| EPI_ISL_769816, EPI_ISL_769817 | Lighthouse Lab in Milton Keynes | Wellcome Sanger Institute for the COVID-19 Genomics UK (COG-UK) Consortium | The Lighthouse Lab in Milton Keynes and Alex Alderton, Roberto Amato, Sonia Goncalves, Ewan Harrison, David K. Jackson, Ian Johnston, Dominic Kwiatkowski, Cordelia Langford, John Sillitoe on behalf of the Wellcome Sanger Institute COVID-19 Surveillance Team |
| EPI_ISL_769818 | Lighthouse Lab in Glasgow | Wellcome Sanger Institute for the COVID-19 Genomics UK (COG-UK) Consortium | Harper VanSteenhouse, Yumi Kasai, David Gray, Carol Clugston, Anna Dominiczak and Alex Alderton, Roberto Amato, Sonia Goncalves, Ewan Harrison, David K. Jackson, Ian Johnston, Dominic Kwiatkowski, Cordelia Langford, John Sillitoe on behalf of the Wellcome Sanger Institute COVID-19 Surveillance Team |
| EPI_ISL_769819, EPI_ISL_769820, EPI_ISL_769821, EPI_ISL_769822, EPI_ISL_769823, EPI_ISL_769824, EPI_ISL_769825 | Lighthouse Lab in Milton Keynes | Wellcome Sanger Institute for the COVID-19 Genomics UK (COG-UK) Consortium | The Lighthouse Lab in Milton Keynes and Alex Alderton, Roberto Amato, Sonia Goncalves, Ewan Harrison, David K. Jackson, Ian Johnston, Dominic Kwiatkowski, Cordelia Langford, John Sillitoe on behalf of the Wellcome Sanger Institute COVID-19 Surveillance Team |
| EPI_ISL_769826 | Lighthouse Lab in Glasgow | Wellcome Sanger Institute for the COVID-19 Genomics UK (COG-UK) Consortium | Harper VanSteenhouse, Yumi Kasai, David Gray, Carol Clugston, Anna Dominiczak and Alex Alderton, Roberto Amato, Sonia Goncalves, Ewan Harrison, David K. Jackson, Ian Johnston, Dominic Kwiatkowski, Cordelia Langford, John Sillitoe on behalf of the Wellcome Sanger Institute COVID-19 Surveillance Team |

| | | | |
|---|---|---|---|
| EPI_ISL_769827, EPI_ISL_769828, EPI_ISL_769829, EPI_ISL_769830, EPI_ISL_769831, EPI_ISL_769832, EPI_ISL_769833 | Lighthouse Lab in Milton Keynes | Wellcome Sanger Institute for the COVID-19 Genomics UK (COG-UK) Consortium | The Lighthouse Lab in Milton Keynes and Alex Alderton, Roberto Amato, Sonia Goncalves, Ewan Harrison, David K. Jackson, Ian Johnston, Dominic Kwiatkowski, Cordelia Langford, John Sillitoe on behalf of the Wellcome Sanger Institute COVID-19 Surveillance Team |
| EPI_ISL_769834 | Lighthouse Lab in Glasgow | Wellcome Sanger Institute for the COVID-19 Genomics UK (COG-UK) Consortium | Harper VanSteenhouse, Yumi Kasai, David Gray, Carol Clugston, Anna Dominiczak and Alex Alderton, Roberto Amato, Sonia Goncalves, Ewan Harrison, David K. Jackson, Ian Johnston, Dominic Kwiatkowski, Cordelia Langford, John Sillitoe on behalf of the Wellcome Sanger Institute COVID-19 Surveillance Team |
| EPI_ISL_769835 | Lighthouse Lab in Milton Keynes | Wellcome Sanger Institute for the COVID-19 Genomics UK (COG-UK) Consortium | The Lighthouse Lab in Milton Keynes and Alex Alderton, Roberto Amato, Sonia Goncalves, Ewan Harrison, David K. Jackson, Ian Johnston, Dominic Kwiatkowski, Cordelia Langford, John Sillitoe on behalf of the Wellcome Sanger Institute COVID-19 Surveillance Team |
| EPI_ISL_769836, EPI_ISL_769837 | Lighthouse Lab in Glasgow | Wellcome Sanger Institute for the COVID-19 Genomics UK (COG-UK) Consortium | Harper VanSteenhouse, Yumi Kasai, David Gray, Carol Clugston, Anna Dominiczak and Alex Alderton, Roberto Amato, Sonia Goncalves, Ewan Harrison, David K. Jackson, Ian Johnston, Dominic Kwiatkowski, Cordelia Langford, John Sillitoe on behalf of the Wellcome Sanger Institute COVID-19 Surveillance Team |
| EPI_ISL_769838, EPI_ISL_769839, EPI_ISL_769840, EPI_ISL_769841 | Lighthouse Lab in Milton Keynes | Wellcome Sanger Institute for the COVID-19 Genomics UK (COG-UK) Consortium | The Lighthouse Lab in Milton Keynes and Alex Alderton, Roberto Amato, Sonia Goncalves, Ewan Harrison, David K. Jackson, Ian Johnston, Dominic Kwiatkowski, Cordelia Langford, John Sillitoe on behalf of the Wellcome Sanger Institute COVID-19 Surveillance Team |
| EPI_ISL_769842 | Lighthouse Lab in Glasgow | Wellcome Sanger Institute for the COVID-19 Genomics UK (COG-UK) Consortium | Harper VanSteenhouse, Yumi Kasai, David Gray, Carol Clugston, Anna Dominiczak and Alex Alderton, Roberto Amato, Sonia Goncalves, Ewan Harrison, David K. Jackson, Ian Johnston, Dominic Kwiatkowski, Cordelia Langford, John Sillitoe on behalf of the Wellcome Sanger Institute COVID-19 Surveillance Team |
| EPI_ISL_769843, EPI_ISL_769844, EPI_ISL_769846, EPI_ISL_769847, EPI_ISL_769848, EPI_ISL_769849 | Lighthouse Lab in Milton Keynes | Wellcome Sanger Institute for the COVID-19 Genomics UK (COG-UK) Consortium | The Lighthouse Lab in Milton Keynes and Alex Alderton, Roberto Amato, Sonia Goncalves, Ewan Harrison, David K. Jackson, Ian Johnston, Dominic Kwiatkowski, Cordelia Langford, John Sillitoe on behalf of the Wellcome Sanger Institute COVID-19 Surveillance Team |
| EPI_ISL_769850 | Lighthouse Lab in Glasgow | Wellcome Sanger Institute for the COVID-19 Genomics UK (COG-UK) Consortium | Harper VanSteenhouse, Yumi Kasai, David Gray, Carol Clugston, Anna Dominiczak and Alex Alderton, Roberto Amato, Sonia Goncalves, Ewan Harrison, David K. Jackson, Ian Johnston, Dominic Kwiatkowski, Cordelia Langford, John Sillitoe on behalf of the Wellcome Sanger Institute COVID-19 Surveillance Team |
| EPI_ISL_769851, EPI_ISL_769853, EPI_ISL_769854, EPI_ISL_769855, EPI_ISL_769856, EPI_ISL_769858, EPI_ISL_769859, EPI_ISL_769860 | Lighthouse Lab in Milton Keynes | Wellcome Sanger Institute for the COVID-19 Genomics UK (COG-UK) Consortium | The Lighthouse Lab in Milton Keynes and Alex Alderton, Roberto Amato, Sonia Goncalves, Ewan Harrison, David K. Jackson, Ian Johnston, Dominic Kwiatkowski, Cordelia Langford, John Sillitoe on behalf of the Wellcome Sanger Institute COVID-19 Surveillance Team |
| EPI_ISL_769862 | Lighthouse Lab in Glasgow | Wellcome Sanger Institute for the COVID-19 Genomics UK (COG-UK) Consortium | Harper VanSteenhouse, Yumi Kasai, David Gray, Carol Clugston, Anna Dominiczak and Alex Alderton, Roberto Amato, Sonia Goncalves, Ewan Harrison, David K. Jackson, Ian Johnston, Dominic Kwiatkowski, Cordelia Langford, John Sillitoe on behalf of the Wellcome Sanger Institute COVID-19 Surveillance Team |
| EPI_ISL_769869 | Respiratory Virus Unit, National Infection Service, Public Health England | COVID-19 Genomics UK (COG-UK) Consortium | PHE Covid Sequencing Team |
| EPI_ISL_770688, EPI_ISL_770689, EPI_ISL_770690, EPI_ISL_770691, EPI_ISL_770692, EPI_ISL_770693, EPI_ISL_770694, EPI_ISL_770695, EPI_ISL_770696, EPI_ISL_770697, EPI_ISL_770698, EPI_ISL_770699, EPI_ISL_770700, EPI_ISL_770701, EPI_ISL_770702, EPI_ISL_770703, EPI_ISL_770704, EPI_ISL_770705, EPI_ISL_770706, EPI_ISL_770707, EPI_ISL_770708, EPI_ISL_770709, EPI_ISL_770710, EPI_ISL_770711 | | | |
| see above | ZOTZ KLIMAS MVZ Düsseldorf-Centrum GbR ÜBAG für Labormedizin, Genetik, Zytologie, Pathologie | Center of Medical Microbiology, Virology, and Hospital Hygiene, University of Duesseldorf | Maximilian Damagnez, Alexander Dilthey, Ashley-Jane Duplessis, Patrick Finzer, Katrin Hoffmann, Torsten Houwaart, Lisanna Hülse, Malte Kohns Vasconcelos, Marek Korencak, Nadine Lübke, Jessica Nicolai, Klaus Pfeffer, Daniel Strelow, Jörg Timm, Andreas Walker, Tobias Wienemann, Rainer Zotz |
| EPI_ISL_770795, EPI_ISL_770796, EPI_ISL_770798 | Minnesota Department of Health, Public Health Laboratory | Minnesota Department of Health, Public Health Laboratory | Alexandra Lorentz, Jacob Garfin, Matt Plumb, and Xiong Wang |
| EPI_ISL_770815 | Vault Health | Minnesota Department of Health, Public Health Laboratory | Alexandra Lorentz, Jacob Garfin, Matt Plumb, and Xiong Wang |
| EPI_ISL_775270 | Akershus University Hospital, Department for Microbiology and Infectious Disease Control | Norwegian Institute of Public Health, Department of Virology | Kathrine Stene-Johansen, Kamilla Heddeland Instefjord, Hilde Elshaug, Atiya R Ali,Marie Paulsen Madsen, Rasmus Riis Kopperud, Hilde Vollan, Karoline Bragstad, Olav Hungnes |
| EPI_ISL_775410, EPI_ISL_775411, EPI_ISL_775412, EPI_ISL_775413 | Oslo University Hospital, Department of Medical Microbiology | Norwegian Institute of Public Health, Department of Virology | Kathrine Stene-Johansen, Kamilla Heddeland Instefjord, Hilde Elshaug, Atiya R Ali,Marie Paulsen Madsen, Rasmus Riis Kopperud, Hilde Vollan, Karoline Bragstad, Olav Hungnes |
| EPI_ISL_775442 | Medical Microbiology Unit, Department for Laboratory Medicine, Drammen Hospital, Vestre Viken Health Trust, | Norwegian Institute of Public Health, Department of Virology | Kathrine Stene-Johansen, Kamilla Heddeland Instefjord, Hilde Elshaug, Atiya R Ali,Marie Paulsen Madsen, Rasmus Riis Kopperud, Hilde Vollan, Karoline Bragstad, Olav Hungnes |
| EPI_ISL_775518, EPI_ISL_775519, EPI_ISL_775521, EPI_ISL_775522 | Hospital of Southern Norway - Kristiansand, Department of Medical Microbiology | Norwegian Institute of Public Health, Department of Virology | Kathrine Stene-Johansen, Kamilla Heddeland Instefjord, Hilde Elshaug, Atiya R Ali,Marie Paulsen Madsen, Rasmus Riis Kopperud, Hilde Vollan, Karoline Bragstad, Olav Hungnes |
| EPI_ISL_775542, EPI_ISL_775543 | Klinisk mikrobiologi, Viruslab | The Public Health Agency of Sweden | Department of Microbiology, The Public Health Agency of Sweden |
| EPI_ISL_776669, EPI_ISL_776670, EPI_ISL_776708, EPI_ISL_776709, EPI_ISL_776713, EPI_ISL_776715, EPI_ISL_776716, EPI_ISL_776717, EPI_ISL_776718, EPI_ISL_776719, EPI_ISL_776720, EPI_ISL_776721, EPI_ISL_776722, EPI_ISL_776723, EPI_ISL_776724, EPI_ISL_776725, EPI_ISL_776726, EPI_ISL_776727, EPI_ISL_776728, EPI_ISL_776729, EPI_ISL_776742, EPI_ISL_776743, EPI_ISL_776744 | | | |
| see above | UW Virology Lab | UW Virology Lab | Pavitra Roychoudhury, Hong Xie, Lasata Shrestha, Meei-Li Huang, Keith R Jerome, Alexander Greninger |
| EPI_ISL_777033, EPI_ISL_777042, EPI_ISL_777053, EPI_ISL_777063 | Lighthouse Lab in Alderley Park | Wellcome Sanger Institute for the COVID-19 Genomics UK (COG-UK) Consortium | Jacquelyn Wynn, Mairead Hyland, The Lighthouse Lab in Alderley Park and Alex Alderton, Roberto Amato, Sonia Goncalves, Ewan Harrison, David K. Jackson, Ian Johnston, Dominic Kwiatkowski, Cordelia Langford, John Sillitoe on behalf of the Wellcome Sanger Institute COVID-19 Surveillance Team |
| EPI_ISL_777064 | Lighthouse Lab in Milton Keynes | Wellcome Sanger Institute for the COVID-19 Genomics UK (COG-UK) Consortium | The Lighthouse Lab in Milton Keynes and Alex Alderton, Roberto Amato, Sonia Goncalves, Ewan Harrison, David K. Jackson, Ian Johnston, Dominic Kwiatkowski, Cordelia Langford, John Sillitoe on behalf of the Wellcome Sanger Institute COVID-19 Surveillance Team |
| EPI_ISL_777067 | Lighthouse Lab in Alderley Park | Wellcome Sanger Institute for the COVID-19 Genomics UK (COG-UK) Consortium | Jacquelyn Wynn, Mairead Hyland, The Lighthouse Lab in Alderley Park and Alex Alderton, Roberto Amato, Sonia Goncalves, Ewan Harrison, David K. Jackson, Ian Johnston, Dominic Kwiatkowski, Cordelia Langford, John Sillitoe on behalf of the Wellcome Sanger Institute COVID-19 Surveillance Team |
| EPI_ISL_777080 | Lighthouse Lab in Milton Keynes | Wellcome Sanger Institute for the COVID-19 Genomics UK (COG-UK) Consortium | The Lighthouse Lab in Milton Keynes and Alex Alderton, Roberto Amato, Sonia Goncalves, Ewan Harrison, David K. Jackson, Ian Johnston, Dominic Kwiatkowski, Cordelia Langford, John Sillitoe on behalf of the Wellcome Sanger Institute COVID-19 Surveillance Team |
| EPI_ISL_777083 | Lighthouse Lab in Alderley Park | Wellcome Sanger Institute for the COVID-19 Genomics UK (COG-UK) Consortium | Jacquelyn Wynn, Mairead Hyland, The Lighthouse Lab in Alderley Park and Alex Alderton, Roberto Amato, Sonia Goncalves, Ewan Harrison, David K. Jackson, Ian Johnston, Dominic Kwiatkowski, Cordelia Langford, John Sillitoe on behalf of the Wellcome Sanger Institute COVID-19 Surveillance Team |
| EPI_ISL_777084, EPI_ISL_777104 | Lighthouse Lab in Milton Keynes | Wellcome Sanger Institute for the COVID-19 Genomics UK (COG-UK) Consortium | The Lighthouse Lab in Milton Keynes and Alex Alderton, Roberto Amato, Sonia Goncalves, Ewan Harrison, David K. Jackson, Ian Johnston, Dominic Kwiatkowski, Cordelia Langford, John Sillitoe on behalf of the Wellcome Sanger Institute COVID-19 Surveillance Team |
| EPI_ISL_777128 | Lighthouse Lab in Alderley Park | Wellcome Sanger Institute for the COVID-19 Genomics UK (COG-UK) Consortium | Jacquelyn Wynn, Mairead Hyland, The Lighthouse Lab in Alderley Park and Alex Alderton, Roberto Amato, Sonia Goncalves, Ewan Harrison, David K. Jackson, Ian Johnston, Dominic Kwiatkowski, Cordelia Langford, John Sillitoe on behalf of the Wellcome Sanger Institute COVID-19 Surveillance Team |
| EPI_ISL_777133 | Lighthouse Lab in Cambridge | Wellcome Sanger Institute for the COVID-19 Genomics UK (COG-UK) Consortium | Rob Howes, The Lighthouse Lab in Cambridge and Alex Alderton, Roberto Amato, Sonia Goncalves, Ewan Harrison, David K. Jackson, Ian Johnston, Dominic Kwiatkowski, Cordelia Langford, John Sillitoe on behalf of the Wellcome Sanger Institute COVID-19 Surveillance Team |
| EPI_ISL_777141 | Lighthouse Lab in Milton Keynes | Wellcome Sanger Institute for the COVID-19 Genomics UK (COG-UK) Consortium | The Lighthouse Lab in Milton Keynes and Alex Alderton, Roberto Amato, Sonia Goncalves, Ewan Harrison, David K. Jackson, Ian Johnston, Dominic Kwiatkowski, Cordelia Langford, John Sillitoe on behalf of the Wellcome Sanger Institute COVID-19 Surveillance Team |
| EPI_ISL_777153, EPI_ISL_777155 | Lighthouse Lab in Alderley Park | Wellcome Sanger Institute for the COVID-19 Genomics UK (COG-UK) Consortium | Jacquelyn Wynn, Mairead Hyland, The Lighthouse Lab in Alderley Park and Alex Alderton, Roberto Amato, Sonia Goncalves, Ewan Harrison, David K. Jackson, Ian Johnston, Dominic Kwiatkowski, Cordelia Langford, John Sillitoe on behalf of the Wellcome Sanger Institute COVID-19 Surveillance Team |
| EPI_ISL_777158 | Lighthouse Lab in Milton Keynes | Wellcome Sanger Institute for the COVID-19 Genomics UK | The Lighthouse Lab in Milton Keynes and Alex Alderton, Roberto Amato, Sonia Goncalves, Ewan Harrison, David K. Jackson, Ian Johnston, Dominic |

| | | | |
|---|---|---|---|
| | | (COG-UK) Consortium | Kwiatkowski, Cordelia Langford, John Sillitoe on behalf of the Wellcome Sanger Institute COVID-19 Surveillance Team |
| EPI_ISL_777173, EPI_ISL_777181 | Lighthouse Lab in Cambridge | Wellcome Sanger Institute for the COVID-19 Genomics UK (COG-UK) Consortium | Rob Howes, The Lighthouse Lab in Cambridge and Alex Alderton, Roberto Amato, Sonia Goncalves, Ewan Harrison, David K. Jackson, Ian Johnston, Dominic Kwiatkowski, Cordelia Langford, John Sillitoe on behalf of the Wellcome Sanger Institute COVID-19 Surveillance Team |
| EPI_ISL_777183 | Lighthouse Lab in Alderley Park | Wellcome Sanger Institute for the COVID-19 Genomics UK (COG-UK) Consortium | Jacquelyn Wynn, Mairead Hyland, The Lighthouse Lab in Alderley Park and Alex Alderton, Roberto Amato, Sonia Goncalves, Ewan Harrison, David K. Jackson, Ian Johnston, Dominic Kwiatkowski, Cordelia Langford, John Sillitoe on behalf of the Wellcome Sanger Institute COVID-19 Surveillance Team |
| EPI_ISL_777184, EPI_ISL_777200, EPI_ISL_777207 | Lighthouse Lab in Milton Keynes | Wellcome Sanger Institute for the COVID-19 Genomics UK (COG-UK) Consortium | The Lighthouse Lab in Milton Keynes and Alex Alderton, Roberto Amato, Sonia Goncalves, Ewan Harrison, David K. Jackson, Ian Johnston, Dominic Kwiatkowski, Cordelia Langford, John Sillitoe on behalf of the Wellcome Sanger Institute COVID-19 Surveillance Team |
| EPI_ISL_777211 | Lighthouse Lab in Alderley Park | Wellcome Sanger Institute for the COVID-19 Genomics UK (COG-UK) Consortium | Jacquelyn Wynn, Mairead Hyland, The Lighthouse Lab in Alderley Park and Alex Alderton, Roberto Amato, Sonia Goncalves, Ewan Harrison, David K. Jackson, Ian Johnston, Dominic Kwiatkowski, Cordelia Langford, John Sillitoe on behalf of the Wellcome Sanger Institute COVID-19 Surveillance Team |
| EPI_ISL_777218 | Lighthouse Lab in Cambridge | Wellcome Sanger Institute for the COVID-19 Genomics UK (COG-UK) Consortium | Rob Howes, The Lighthouse Lab in Cambridge and Alex Alderton, Roberto Amato, Sonia Goncalves, Ewan Harrison, David K. Jackson, Ian Johnston, Dominic Kwiatkowski, Cordelia Langford, John Sillitoe on behalf of the Wellcome Sanger Institute COVID-19 Surveillance Team |
| EPI_ISL_777224, EPI_ISL_777229 | Lighthouse Lab in Milton Keynes | Wellcome Sanger Institute for the COVID-19 Genomics UK (COG-UK) Consortium | The Lighthouse Lab in Milton Keynes and Alex Alderton, Roberto Amato, Sonia Goncalves, Ewan Harrison, David K. Jackson, Ian Johnston, Dominic Kwiatkowski, Cordelia Langford, John Sillitoe on behalf of the Wellcome Sanger Institute COVID-19 Surveillance Team |
| EPI_ISL_777244 | Lighthouse Lab in Alderley Park | Wellcome Sanger Institute for the COVID-19 Genomics UK (COG-UK) Consortium | Jacquelyn Wynn, Mairead Hyland, The Lighthouse Lab in Alderley Park and Alex Alderton, Roberto Amato, Sonia Goncalves, Ewan Harrison, David K. Jackson, Ian Johnston, Dominic Kwiatkowski, Cordelia Langford, John Sillitoe on behalf of the Wellcome Sanger Institute COVID-19 Surveillance Team |
| EPI_ISL_777245, EPI_ISL_777269 | Lighthouse Lab in Milton Keynes | Wellcome Sanger Institute for the COVID-19 Genomics UK (COG-UK) Consortium | The Lighthouse Lab in Milton Keynes and Alex Alderton, Roberto Amato, Sonia Goncalves, Ewan Harrison, David K. Jackson, Ian Johnston, Dominic Kwiatkowski, Cordelia Langford, John Sillitoe on behalf of the Wellcome Sanger Institute COVID-19 Surveillance Team |
| EPI_ISL_777270 | Lighthouse Lab in Alderley Park | Wellcome Sanger Institute for the COVID-19 Genomics UK (COG-UK) Consortium | Jacquelyn Wynn, Mairead Hyland, The Lighthouse Lab in Alderley Park and Alex Alderton, Roberto Amato, Sonia Goncalves, Ewan Harrison, David K. Jackson, Ian Johnston, Dominic Kwiatkowski, Cordelia Langford, John Sillitoe on behalf of the Wellcome Sanger Institute COVID-19 Surveillance Team |
| EPI_ISL_777275 | Lighthouse Lab in Cambridge | Wellcome Sanger Institute for the COVID-19 Genomics UK (COG-UK) Consortium | Rob Howes, The Lighthouse Lab in Cambridge and Alex Alderton, Roberto Amato, Sonia Goncalves, Ewan Harrison, David K. Jackson, Ian Johnston, Dominic Kwiatkowski, Cordelia Langford, John Sillitoe on behalf of the Wellcome Sanger Institute COVID-19 Surveillance Team |
| EPI_ISL_777286 | Lighthouse Lab in Alderley Park | Wellcome Sanger Institute for the COVID-19 Genomics UK (COG-UK) Consortium | Jacquelyn Wynn, Mairead Hyland, The Lighthouse Lab in Alderley Park and Alex Alderton, Roberto Amato, Sonia Goncalves, Ewan Harrison, David K. Jackson, Ian Johnston, Dominic Kwiatkowski, Cordelia Langford, John Sillitoe on behalf of the Wellcome Sanger Institute COVID-19 Surveillance Team |
| EPI_ISL_777289 | Lighthouse Lab in Cambridge | Wellcome Sanger Institute for the COVID-19 Genomics UK (COG-UK) Consortium | Rob Howes, The Lighthouse Lab in Cambridge and Alex Alderton, Roberto Amato, Sonia Goncalves, Ewan Harrison, David K. Jackson, Ian Johnston, Dominic Kwiatkowski, Cordelia Langford, John Sillitoe on behalf of the Wellcome Sanger Institute COVID-19 Surveillance Team |
| EPI_ISL_777292 | Lighthouse Lab in Alderley Park | Wellcome Sanger Institute for the COVID-19 Genomics UK (COG-UK) Consortium | Jacquelyn Wynn, Mairead Hyland, The Lighthouse Lab in Alderley Park and Alex Alderton, Roberto Amato, Sonia Goncalves, Ewan Harrison, David K. Jackson, Ian Johnston, Dominic Kwiatkowski, Cordelia Langford, John Sillitoe on behalf of the Wellcome Sanger Institute COVID-19 Surveillance Team |
| EPI_ISL_777293 | Lighthouse Lab in Cambridge | Wellcome Sanger Institute for the COVID-19 Genomics UK (COG-UK) Consortium | Rob Howes, The Lighthouse Lab in Cambridge and Alex Alderton, Roberto Amato, Sonia Goncalves, Ewan Harrison, David K. Jackson, Ian Johnston, Dominic Kwiatkowski, Cordelia Langford, John Sillitoe on behalf of the Wellcome Sanger Institute COVID-19 Surveillance Team |
| EPI_ISL_777303, EPI_ISL_777304, EPI_ISL_777310 | Lighthouse Lab in Milton Keynes | Wellcome Sanger Institute for the COVID-19 Genomics UK (COG-UK) Consortium | The Lighthouse Lab in Milton Keynes and Alex Alderton, Roberto Amato, Sonia Goncalves, Ewan Harrison, David K. Jackson, Ian Johnston, Dominic Kwiatkowski, Cordelia Langford, John Sillitoe on behalf of the Wellcome Sanger Institute COVID-19 Surveillance Team |
| EPI_ISL_777314 | Lighthouse Lab in Alderley Park | Wellcome Sanger Institute for the COVID-19 Genomics UK (COG-UK) Consortium | Jacquelyn Wynn, Mairead Hyland, The Lighthouse Lab in Alderley Park and Alex Alderton, Roberto Amato, Sonia Goncalves, Ewan Harrison, David K. Jackson, Ian Johnston, Dominic Kwiatkowski, Cordelia Langford, John Sillitoe on behalf of the Wellcome Sanger Institute COVID-19 Surveillance Team |
| EPI_ISL_777320 | Lighthouse Lab in Cambridge | Wellcome Sanger Institute for the COVID-19 Genomics UK (COG-UK) Consortium | Rob Howes, The Lighthouse Lab in Cambridge and Alex Alderton, Roberto Amato, Sonia Goncalves, Ewan Harrison, David K. Jackson, Ian Johnston, Dominic Kwiatkowski, Cordelia Langford, John Sillitoe on behalf of the Wellcome Sanger Institute COVID-19 Surveillance Team |
| EPI_ISL_778298, EPI_ISL_778300, EPI_ISL_778304, EPI_ISL_778306, EPI_ISL_778307, EPI_ISL_778308, EPI_ISL_778310, EPI_ISL_778312, EPI_ISL_778313, EPI_ISL_778314, EPI_ISL_778316, EPI_ISL_778317, EPI_ISL_778318, EPI_ISL_778319, EPI_ISL_778320, EPI_ISL_778321, EPI_ISL_778323, EPI_ISL_778326, EPI_ISL_778329, EPI_ISL_778338, EPI_ISL_778339, EPI_ISL_778340, EPI_ISL_778341, EPI_ISL_778343, EPI_ISL_778344 | | | |
| see above | Lighthouse Lab in Milton Keynes | Wellcome Sanger Institute for the COVID-19 Genomics UK (COG-UK) Consortium | The Lighthouse Lab in Milton Keynes and Alex Alderton, Roberto Amato, Sonia Goncalves, Ewan Harrison, David K. Jackson, Ian Johnston, Dominic Kwiatkowski, Cordelia Langford, John Sillitoe on behalf of the Wellcome Sanger Institute COVID-19 Surveillance Team |
| EPI_ISL_778347, EPI_ISL_778348 | Lighthouse Lab in Alderley Park | Wellcome Sanger Institute for the COVID-19 Genomics UK (COG-UK) Consortium | Jacquelyn Wynn, Mairead Hyland, The Lighthouse Lab in Alderley Park and Alex Alderton, Roberto Amato, Sonia Goncalves, Ewan Harrison, David K. Jackson, Ian Johnston, Dominic Kwiatkowski, Cordelia Langford, John Sillitoe on behalf of the Wellcome Sanger Institute COVID-19 Surveillance Team |
| EPI_ISL_778349, EPI_ISL_778350, EPI_ISL_778351, EPI_ISL_778354, EPI_ISL_778363, EPI_ISL_778366, EPI_ISL_778367, EPI_ISL_778369, EPI_ISL_778370, EPI_ISL_778371, EPI_ISL_778379, EPI_ISL_778381, EPI_ISL_778383, EPI_ISL_778384, EPI_ISL_778386, EPI_ISL_778387, EPI_ISL_778388, EPI_ISL_778389, EPI_ISL_778391, EPI_ISL_778393, EPI_ISL_778394, EPI_ISL_778395, EPI_ISL_778398, EPI_ISL_778402, EPI_ISL_778405, EPI_ISL_778406, EPI_ISL_778409, EPI_ISL_778413, EPI_ISL_778414, EPI_ISL_778418, EPI_ISL_778425, EPI_ISL_778427, EPI_ISL_778429, EPI_ISL_778431, EPI_ISL_778434, EPI_ISL_778435, EPI_ISL_778436, EPI_ISL_778437, EPI_ISL_778438, EPI_ISL_778439, EPI_ISL_778440, EPI_ISL_778441, EPI_ISL_778442, EPI_ISL_778443, EPI_ISL_778446 | | | |
| see above | Lighthouse Lab in Milton Keynes | Wellcome Sanger Institute for the COVID-19 Genomics UK (COG-UK) Consortium | The Lighthouse Lab in Milton Keynes and Alex Alderton, Roberto Amato, Sonia Goncalves, Ewan Harrison, David K. Jackson, Ian Johnston, Dominic Kwiatkowski, Cordelia Langford, John Sillitoe on behalf of the Wellcome Sanger Institute COVID-19 Surveillance Team |
| EPI_ISL_778448 | Lighthouse Lab in Alderley Park | Wellcome Sanger Institute for the COVID-19 Genomics UK (COG-UK) Consortium | Jacquelyn Wynn, Mairead Hyland, The Lighthouse Lab in Alderley Park and Alex Alderton, Roberto Amato, Sonia Goncalves, Ewan Harrison, David K. Jackson, Ian Johnston, Dominic Kwiatkowski, Cordelia Langford, John Sillitoe on behalf of the Wellcome Sanger Institute COVID-19 Surveillance Team |
| EPI_ISL_778449, EPI_ISL_778452, EPI_ISL_778453, EPI_ISL_778456, EPI_ISL_778458, EPI_ISL_778460, EPI_ISL_778464, EPI_ISL_778466, EPI_ISL_778467, EPI_ISL_778468, EPI_ISL_778469, EPI_ISL_778470, EPI_ISL_778474, EPI_ISL_778475, EPI_ISL_778476, EPI_ISL_778477, EPI_ISL_778479, EPI_ISL_778480, EPI_ISL_778481, EPI_ISL_778483, EPI_ISL_778485, EPI_ISL_778486, EPI_ISL_778489, EPI_ISL_778492, EPI_ISL_778493, EPI_ISL_778495, EPI_ISL_778498, EPI_ISL_778501, EPI_ISL_778502, EPI_ISL_778503, EPI_ISL_778505, EPI_ISL_778507, EPI_ISL_778508, EPI_ISL_778509, EPI_ISL_778511, EPI_ISL_778513, EPI_ISL_778514, EPI_ISL_778515, EPI_ISL_778518 | | | |
| see above | Lighthouse Lab in Milton Keynes | Wellcome Sanger Institute for the COVID-19 Genomics UK (COG-UK) Consortium | The Lighthouse Lab in Milton Keynes and Alex Alderton, Roberto Amato, Sonia Goncalves, Ewan Harrison, David K. Jackson, Ian Johnston, Dominic Kwiatkowski, Cordelia Langford, John Sillitoe on behalf of the Wellcome Sanger Institute COVID-19 Surveillance Team |
| EPI_ISL_778519 | Lighthouse Lab in Alderley Park | Wellcome Sanger Institute for the COVID-19 Genomics UK (COG-UK) Consortium | Jacquelyn Wynn, Mairead Hyland, The Lighthouse Lab in Alderley Park and Alex Alderton, Roberto Amato, Sonia Goncalves, Ewan Harrison, David K. Jackson, Ian Johnston, Dominic Kwiatkowski, Cordelia Langford, John Sillitoe on behalf of the Wellcome Sanger Institute COVID-19 Surveillance Team |
| EPI_ISL_778520, EPI_ISL_778521, EPI_ISL_778523, EPI_ISL_778526, EPI_ISL_778532, EPI_ISL_778533, EPI_ISL_778535, EPI_ISL_778536, EPI_ISL_778537, EPI_ISL_778538, EPI_ISL_778542, EPI_ISL_778544, EPI_ISL_778545, EPI_ISL_778546, EPI_ISL_778549, EPI_ISL_778550, EPI_ISL_778552, EPI_ISL_778554, EPI_ISL_778555, EPI_ISL_778557, EPI_ISL_778559, EPI_ISL_778561, EPI_ISL_778563 | | | |
| see above | Lighthouse Lab in Milton Keynes | Wellcome Sanger Institute for the COVID-19 Genomics UK (COG-UK) Consortium | The Lighthouse Lab in Milton Keynes and Alex Alderton, Roberto Amato, Sonia Goncalves, Ewan Harrison, David K. Jackson, Ian Johnston, Dominic Kwiatkowski, Cordelia Langford, John Sillitoe on behalf of the Wellcome Sanger Institute COVID-19 Surveillance Team |
| EPI_ISL_778564 | Lighthouse Lab in Alderley Park | Wellcome Sanger Institute for the COVID-19 Genomics UK (COG-UK) Consortium | Jacquelyn Wynn, Mairead Hyland, The Lighthouse Lab in Alderley Park and Alex Alderton, Roberto Amato, Sonia Goncalves, Ewan Harrison, David K. Jackson, Ian Johnston, Dominic Kwiatkowski, Cordelia Langford, John Sillitoe on behalf of the Wellcome Sanger Institute COVID-19 Surveillance Team |
| EPI_ISL_778565, EPI_ISL_778566, EPI_ISL_778567, EPI_ISL_778572, EPI_ISL_778573, EPI_ISL_778574, EPI_ISL_778575, EPI_ISL_778576, EPI_ISL_778580, EPI_ISL_778581, EPI_ISL_778582, EPI_ISL_778583, EPI_ISL_778585, EPI_ISL_778587, EPI_ISL_778588, EPI_ISL_778589, EPI_ISL_778590, EPI_ISL_778591, EPI_ISL_778592, EPI_ISL_778595, EPI_ISL_778596, EPI_ISL_778597, EPI_ISL_778600, EPI_ISL_778602, EPI_ISL_778603, EPI_ISL_778604, EPI_ISL_778606, EPI_ISL_778608, EPI_ISL_778609, EPI_ISL_778610, EPI_ISL_778611, EPI_ISL_778613 | | | |
| see above | Lighthouse Lab in Milton Keynes | Wellcome Sanger Institute for the COVID-19 Genomics UK (COG-UK) Consortium | The Lighthouse Lab in Milton Keynes and Alex Alderton, Roberto Amato, Sonia Goncalves, Ewan Harrison, David K. Jackson, Ian Johnston, Dominic Kwiatkowski, Cordelia Langford, John Sillitoe on behalf of the Wellcome Sanger Institute COVID-19 Surveillance Team |
| EPI_ISL_778615 | Lighthouse Lab in Alderley Park | Wellcome Sanger Institute for the COVID-19 Genomics UK (COG-UK) Consortium | Jacquelyn Wynn, Mairead Hyland, The Lighthouse Lab in Alderley Park and Alex Alderton, Roberto Amato, Sonia Goncalves, Ewan Harrison, David K. Jackson, Ian Johnston, Dominic Kwiatkowski, Cordelia Langford, John Sillitoe on behalf of the Wellcome Sanger Institute COVID-19 Surveillance Team |
| EPI_ISL_778622, EPI_ISL_778624, EPI_ISL_778625, EPI_ISL_778626, EPI_ISL_778632, EPI_ISL_778633, EPI_ISL_778634, EPI_ISL_778636, EPI_ISL_778637, EPI_ISL_778639, EPI_ISL_778640, EPI_ISL_778641, EPI_ISL_778642, EPI_ISL_778643 | | | |
| see above | Lighthouse Lab in Milton Keynes | Wellcome Sanger Institute for the COVID-19 Genomics UK (COG-UK) Consortium | The Lighthouse Lab in Milton Keynes and Alex Alderton, Roberto Amato, Sonia Goncalves, Ewan Harrison, David K. Jackson, Ian Johnston, Dominic Kwiatkowski, Cordelia Langford, John Sillitoe on behalf of the Wellcome Sanger Institute COVID-19 Surveillance Team |

| | | | |
|---|---|---|---|
| EPI_ISL_778828, EPI_ISL_778829, EPI_ISL_778831 | AIID | Irish Coronavirus Sequencing Consortium-Teagasc Grange | Matthew McCabe, Aljandro Abner Garcia Leon, Fiona Crispie, Calum Walsh, Michael Carr, John Kenny, Paul Cotter, Patrick Mallon, Gabriel Gonzalez |
| EPI_ISL_778945, EPI_ISL_778946, EPI_ISL_778947, EPI_ISL_778948, EPI_ISL_778949, EPI_ISL_778950, EPI_ISL_778951, EPI_ISL_778952, EPI_ISL_778953, EPI_ISL_778954, EPI_ISL_778955, EPI_ISL_778956, EPI_ISL_778957, EPI_ISL_778958, EPI_ISL_778959, EPI_ISL_778960, EPI_ISL_778961, EPI_ISL_778962, EPI_ISL_778963, EPI_ISL_778964, EPI_ISL_778965, EPI_ISL_778966, EPI_ISL_778967, EPI_ISL_778968, EPI_ISL_778969, EPI_ISL_778970, EPI_ISL_778971, EPI_ISL_778972, EPI_ISL_778973, EPI_ISL_778974, EPI_ISL_778975, EPI_ISL_778976, EPI_ISL_778977, EPI_ISL_778978, EPI_ISL_778979, EPI_ISL_778980, EPI_ISL_778981, EPI_ISL_778982, EPI_ISL_778983, EPI_ISL_778984, EPI_ISL_778985, EPI_ISL_778986, EPI_ISL_778987, EPI_ISL_778988, EPI_ISL_778989, EPI_ISL_778990, EPI_ISL_778991, EPI_ISL_778992, EPI_ISL_778993, EPI_ISL_778994, EPI_ISL_778995, EPI_ISL_778996, EPI_ISL_778997, EPI_ISL_778998, EPI_ISL_778999, EPI_ISL_779000, EPI_ISL_779001, EPI_ISL_779002, EPI_ISL_779003, EPI_ISL_779004, EPI_ISL_779005, EPI_ISL_779006, EPI_ISL_779007, EPI_ISL_779008, EPI_ISL_779009, EPI_ISL_779010, EPI_ISL_779011, EPI_ISL_779012, EPI_ISL_779013, EPI_ISL_779014, EPI_ISL_779015, EPI_ISL_779016, EPI_ISL_779017, EPI_ISL_779018, EPI_ISL_779019, EPI_ISL_779020, EPI_ISL_779021, EPI_ISL_779022, EPI_ISL_779023, EPI_ISL_779024, EPI_ISL_779025, EPI_ISL_779026, EPI_ISL_779027, EPI_ISL_779028, EPI_ISL_779029, EPI_ISL_779030, EPI_ISL_779031, EPI_ISL_779032, EPI_ISL_779033, EPI_ISL_779034, EPI_ISL_779035, EPI_ISL_779036, EPI_ISL_779037, EPI_ISL_779038, EPI_ISL_779039, EPI_ISL_779040, EPI_ISL_779041, EPI_ISL_779042, EPI_ISL_779043, EPI_ISL_779044, EPI_ISL_779045, EPI_ISL_779046, EPI_ISL_779047, EPI_ISL_779048, EPI_ISL_779049, EPI_ISL_779050, EPI_ISL_779051, EPI_ISL_779052, EPI_ISL_779053, EPI_ISL_779054, EPI_ISL_779055, EPI_ISL_779056, EPI_ISL_779057, EPI_ISL_779058, EPI_ISL_779059, EPI_ISL_779060, EPI_ISL_779061, EPI_ISL_779062, EPI_ISL_779063, EPI_ISL_779064, EPI_ISL_779065, EPI_ISL_779066, EPI_ISL_779067, EPI_ISL_779068, EPI_ISL_779069, EPI_ISL_779070, EPI_ISL_779071, EPI_ISL_779072, EPI_ISL_779073, EPI_ISL_779074, EPI_ISL_779075, EPI_ISL_779076, EPI_ISL_779077, EPI_ISL_779078, EPI_ISL_779079, EPI_ISL_779080, EPI_ISL_779081 | | | |
| see above | LSUHS Emerging Viral Threat Laboratory | Microbial Genome Sequencing Center | Jeremy P. Kamil, Jennifer L. Carroll, Camille F. Abshire, Maarten Van Diest, Andrew D. Yurochko, Martin J. Sapp, Rona S. Scott, Christopher G. Kevil, Daniel J. Snyder, Vaughn S. Cooper, John A. Vanchiere |
| EPI_ISL_779194 | Laboratorio Estatal de Salud Pública de Nuevo León | Laboratorio de Infectología Molecular, Departamento de Bioquímica y Medicina Molecular,Facultad de Medicina - Universidad Autónoma de Nuevo León | Kame A. Galán-Huerta, María F. Herrera-Saldivar, Natalia Martínez-Acuña, Sonia A. Lozano-Sepúlveda, Daniel Arellanos-Soto, Ana M. Rivas-Estilla, Samuel Buentello-Wong, Else del Carmen García-García, Gloria A. Jasso-de-la-Peña, Roberto Montes-de-Oca, Consuelo Treviño-Garza, Manuel E. de-la-O-Cavazos |
| EPI_ISL_779605, EPI_ISL_779606 | Victorian Infectious Diseases Reference Laboratory (VIDRL) | VIDRL and MDU-PHL | Caly L., Seemann T., Sait, M.L., Druce J., Sherry, N.L. |
| EPI_ISL_779608 | Microbiological Diagnostic Unit - Public Health Laboratory (MDU-PHL) | MDU-PHL | Seemann T., Sait, M.L., Sherry, N.L. |
| EPI_ISL_779609 | Victorian Infectious Diseases Reference Laboratory (VIDRL) | VIDRL and MDU-PHL | Caly L., Seemann T., Sait, M.L., Druce J., Sherry, N.L. |
| EPI_ISL_779851, EPI_ISL_779853, EPI_ISL_779854, EPI_ISL_779856, EPI_ISL_779857, EPI_ISL_779861, EPI_ISL_779864, EPI_ISL_779868, EPI_ISL_779872, EPI_ISL_779878, EPI_ISL_779882, EPI_ISL_779890, EPI_ISL_779895, EPI_ISL_779901, EPI_ISL_779905, EPI_ISL_779911, EPI_ISL_779919 | | | |
| see above | Servicio de Microbiología, Hospital Universitario Son Espases | SeqCOVID-SPAIN consortium/IBV(CSIC) | Carla López-Causapé, Jordi Reina, Antonio Oliver and SeqCOVID-SPAIN consortium |
| EPI_ISL_782482, EPI_ISL_782483, EPI_ISL_782484, EPI_ISL_782485, EPI_ISL_782487, EPI_ISL_782488, EPI_ISL_782489, EPI_ISL_782490, EPI_ISL_782491, EPI_ISL_782492, EPI_ISL_782493, EPI_ISL_782495, EPI_ISL_782496, EPI_ISL_782497, EPI_ISL_782498, EPI_ISL_782499, EPI_ISL_782501, EPI_ISL_782502, EPI_ISL_782503, EPI_ISL_782505, EPI_ISL_782506, EPI_ISL_782507, EPI_ISL_782508, EPI_ISL_782509, EPI_ISL_782510, EPI_ISL_782511, EPI_ISL_782513, EPI_ISL_782514, EPI_ISL_782515, EPI_ISL_782516, EPI_ISL_782517, EPI_ISL_782518, EPI_ISL_782519, EPI_ISL_782520, EPI_ISL_782521, EPI_ISL_782522, EPI_ISL_782523, EPI_ISL_782524, EPI_ISL_782525, EPI_ISL_782526, EPI_ISL_782527, EPI_ISL_782528, EPI_ISL_782529, EPI_ISL_782530, EPI_ISL_782531, EPI_ISL_782532, EPI_ISL_782533, EPI_ISL_782534, EPI_ISL_782535, EPI_ISL_782537, EPI_ISL_782539, EPI_ISL_782540, EPI_ISL_782541, EPI_ISL_782542, EPI_ISL_782543, EPI_ISL_782544, EPI_ISL_782545, EPI_ISL_782546, EPI_ISL_782547, EPI_ISL_782548, EPI_ISL_782549, EPI_ISL_782550, EPI_ISL_782551, EPI_ISL_782552, EPI_ISL_782553, EPI_ISL_782554, EPI_ISL_782555, EPI_ISL_782556, EPI_ISL_782557, EPI_ISL_782558, EPI_ISL_782559, EPI_ISL_782560, EPI_ISL_782561, EPI_ISL_782562, EPI_ISL_782563, EPI_ISL_782564, EPI_ISL_782565, EPI_ISL_782566, EPI_ISL_782567, EPI_ISL_782569, EPI_ISL_782570, EPI_ISL_782571, EPI_ISL_782572, EPI_ISL_782573, EPI_ISL_782574, EPI_ISL_782575, EPI_ISL_782576, EPI_ISL_782577, EPI_ISL_782578, EPI_ISL_782579, EPI_ISL_782581, EPI_ISL_782582, EPI_ISL_782583, EPI_ISL_782585, EPI_ISL_782586, EPI_ISL_782587, EPI_ISL_782588, EPI_ISL_782589, EPI_ISL_782590, EPI_ISL_782591, EPI_ISL_782592, EPI_ISL_782593, EPI_ISL_782594, EPI_ISL_782595, EPI_ISL_782597, EPI_ISL_782598, EPI_ISL_782599, EPI_ISL_782600, EPI_ISL_782601, EPI_ISL_782602, EPI_ISL_782603, EPI_ISL_782604, EPI_ISL_782606, EPI_ISL_782607, EPI_ISL_782608, EPI_ISL_782609, EPI_ISL_782610, EPI_ISL_782611, EPI_ISL_782612, EPI_ISL_782613, EPI_ISL_782614, EPI_ISL_782616, EPI_ISL_782617, EPI_ISL_782619, EPI_ISL_782620, EPI_ISL_782621, EPI_ISL_782622, EPI_ISL_782623, EPI_ISL_782625, EPI_ISL_782626, EPI_ISL_782627, EPI_ISL_782628, EPI_ISL_782629, EPI_ISL_782630, EPI_ISL_782632, EPI_ISL_782634, EPI_ISL_782635, EPI_ISL_782636, EPI_ISL_782637, EPI_ISL_782638, EPI_ISL_782639, EPI_ISL_782640, EPI_ISL_782641, EPI_ISL_782642, EPI_ISL_782643, EPI_ISL_782644, EPI_ISL_782645, EPI_ISL_782646, EPI_ISL_782647, EPI_ISL_782648, EPI_ISL_782649, EPI_ISL_782650, EPI_ISL_782651, EPI_ISL_782652, EPI_ISL_782653, EPI_ISL_782654, EPI_ISL_782655, EPI_ISL_782656, EPI_ISL_782657, EPI_ISL_782658, EPI_ISL_782659, EPI_ISL_782660, EPI_ISL_782661, EPI_ISL_782662, EPI_ISL_782663, EPI_ISL_782664, EPI_ISL_782665, EPI_ISL_782666, EPI_ISL_782667, EPI_ISL_782668, EPI_ISL_782669, EPI_ISL_782670, EPI_ISL_782671, EPI_ISL_782672, EPI_ISL_782673, EPI_ISL_782675, EPI_ISL_782676, EPI_ISL_782677, EPI_ISL_782678, EPI_ISL_782680, EPI_ISL_782681, EPI_ISL_782682, EPI_ISL_782683, EPI_ISL_782684, EPI_ISL_782685, EPI_ISL_782686, EPI_ISL_782687, EPI_ISL_782688, EPI_ISL_782690, EPI_ISL_782691, EPI_ISL_782692, EPI_ISL_782693, EPI_ISL_782694, EPI_ISL_782695, EPI_ISL_782696, EPI_ISL_782697, EPI_ISL_782698, EPI_ISL_782699, EPI_ISL_782700, EPI_ISL_782701, EPI_ISL_782702, EPI_ISL_782703, EPI_ISL_782704, EPI_ISL_782705, EPI_ISL_782706, EPI_ISL_782707, EPI_ISL_782708, EPI_ISL_782709, EPI_ISL_782711, EPI_ISL_782712, EPI_ISL_782713, EPI_ISL_782714, EPI_ISL_782715, EPI_ISL_782716, EPI_ISL_782717, EPI_ISL_782718, EPI_ISL_782719, EPI_ISL_782720, EPI_ISL_782721, EPI_ISL_782722, EPI_ISL_782723, EPI_ISL_782724, EPI_ISL_782725, EPI_ISL_782726, EPI_ISL_782727, EPI_ISL_782728, EPI_ISL_782729, EPI_ISL_782730, EPI_ISL_782731, EPI_ISL_782732, EPI_ISL_782733 | | | |
| see above | Lighthouse Lab in Glasgow | Wellcome Sanger Institute for the COVID-19 Genomics UK (COG-UK) Consortium | Harper VanSteenhouse, Yumi Kasai, David Gray, Carol Clugston, Anna Dominiczak and Alex Alderton, Roberto Amato, Sonia Goncalves, Ewan Harrison, David K. Jackson, Ian Johnston, Dominic Kwiatkowski, Cordelia Langford, John Sillitoe on behalf of the Wellcome Sanger Institute COVID-19 Surveillance Team |
| EPI_ISL_789045 | Klinisk mikrobiologi | The Public Health Agency of Sweden | Department of Microbiology, The Public Health Agency of Sweden |
| EPI_ISL_790593, EPI_ISL_790596, EPI_ISL_790597, EPI_ISL_790627, EPI_ISL_790648, EPI_ISL_790649, EPI_ISL_790654, EPI_ISL_790655, EPI_ISL_790657, EPI_ISL_790658, EPI_ISL_790659, EPI_ISL_790660, EPI_ISL_790661, EPI_ISL_790662, EPI_ISL_790663, EPI_ISL_790666, EPI_ISL_790668, EPI_ISL_790669, EPI_ISL_790670, EPI_ISL_790671, EPI_ISL_790679, EPI_ISL_790690, EPI_ISL_790691, EPI_ISL_790692, EPI_ISL_790741, EPI_ISL_790792, EPI_ISL_790793, EPI_ISL_790800, EPI_ISL_790810, EPI_ISL_790811, EPI_ISL_790812, EPI_ISL_790818, EPI_ISL_790819, EPI_ISL_790820, EPI_ISL_790821, EPI_ISL_790822, EPI_ISL_790866, EPI_ISL_790867, EPI_ISL_790868, EPI_ISL_790892, EPI_ISL_790900, EPI_ISL_790918, EPI_ISL_790932, EPI_ISL_790952, EPI_ISL_790956, EPI_ISL_790957, EPI_ISL_790958, EPI_ISL_790989, EPI_ISL_790990, EPI_ISL_790991, EPI_ISL_790992, EPI_ISL_790993, EPI_ISL_790994, EPI_ISL_791036, EPI_ISL_791037, EPI_ISL_791042, EPI_ISL_791043, EPI_ISL_791068, EPI_ISL_791069, EPI_ISL_791070, EPI_ISL_791079, EPI_ISL_791082 | | | |
| see above | Dutch COVID-19 response team | National Institute for Public Health and the Environment (RIVM) | Adam Meijer, Harry Vennema, Jeroen Cremer, Sharon van den Brink, Bas van der Veer, AnneMarie van den Brandt, Florian Zwagemaker, Dennis Schmitz, Chantal Reusken, on behalf of the national COVID-19 response team |
| EPI_ISL_791208, EPI_ISL_791220 | Respiratory Virus Unit, National Infection Service, Public Health England | COVID-19 Genomics UK (COG-UK) Consortium | PHE Covid Sequencing Team |
| EPI_ISL_791293, EPI_ISL_791294, EPI_ISL_791307, EPI_ISL_791308, EPI_ISL_791324, EPI_ISL_791325, EPI_ISL_791326 | National Virus Reference Laboratory | Irish Coronavirus Sequencing Consortium - Teagasc Moorepark | Alejandro Abner Garcia Leon, Paul Cotter, Fiona Crispie, John Kenny, Paddy Mallon, Calum Walsh |
| EPI_ISL_791996, EPI_ISL_791997, EPI_ISL_791998 | CHU - Hôpital Cavale Blanche | National Reference Center for Viruses of Respiratory Infections, Institut Pasteur, Paris | Marion Barbet, Sylvie Behillil, Méline Bizard, Angela Brisebarre, Camille Capel, Etienne Simon-Lorière, Vincent Enouf, Maud Vanpeene, Sylvie van der Werf, Léa Pilorgé |
| EPI_ISL_792016 | Plateforme COVID IDF | National Reference Center for Viruses of Respiratory Infections, Institut Pasteur, Paris | Marion Barbet, Sylvie Behillil, Méline Bizard, Angela Brisebarre, Camille Capel, Etienne Simon-Lorière, Vincent Enouf, Maud Vanpeene, Sylvie van der Werf,Jacques Fourgeaud |
| EPI_ISL_794725, EPI_ISL_794728 | PathWest Laboratory Medicine WA | PathWest Laboratory Medicine WA Microbial Surveillance Unit | PathWest Laboratory Medicine WA Microbial Surveillance Unit |
| EPI_ISL_794760, EPI_ISL_794761, EPI_ISL_794762, EPI_ISL_794763, EPI_ISL_794764, EPI_ISL_794765, EPI_ISL_794766, EPI_ISL_794767, EPI_ISL_794768, EPI_ISL_794769, EPI_ISL_794770, EPI_ISL_794771, EPI_ISL_794772, EPI_ISL_794773, EPI_ISL_794791, EPI_ISL_794792 | | | |
| see above | Istituto Zooprofilattico Sperimentale della Puglia e della Basilicata | Istituto Zooprofilattico Sperimentale della Puglia e della Basilicata | Parisi A., Bianco A., Capozzi L., Del Sambro L., Manzulli V, Rondinone V., Pace L., Cipolletta D., Galante D. |
| EPI_ISL_796658 | Dept. of Medical Microbiology, Stavanger University Hospital, Helse Stavanger HF | Norwegian Institute of Public Health, Department of Virology | Kathrine Stene-Johansen, Kamilla Heddeland Instefjord, Hilde Elshaug, Atiya R Ali,Marie Paulsen Madsen, Rasmus Riis Kopperud, Hilde Vollan, Karoline Bragstad, Olav Hungnes |
| EPI_ISL_796684, EPI_ISL_796685 | Norwegian Institute of Public Health, Department of Virology | Norwegian Institute of Public Health, Department of Virology | Kathrine Stene-Johansen, Kamilla Heddeland Instefjord, Hilde Elshaug, Atiya R Ali,Marie Paulsen Madsen, Rasmus Riis Kopperud, Hilde Vollan, Karoline Bragstad, Olav Hungnes |
| EPI_ISL_796689, EPI_ISL_796690 | Nordland Hospital - Bodo, Laboratory Department, Molecular Biology Unit | Norwegian Institute of Public Health, Department of Virology | Kathrine Stene-Johansen, Kamilla Heddeland Instefjord, Hilde Elshaug, Atiya R Ali,Marie Paulsen Madsen, Rasmus Riis Kopperud, Hilde Vollan, Karoline Bragstad, Olav Hungnes |
| EPI_ISL_796706 | Department of Medical Microbiology, St. Olavs hospital | Norwegian Institute of Public Health, Department of Virology | Kathrine Stene-Johansen, Kamilla Heddeland Instefjord, Hilde Elshaug, Atiya R Ali,Marie Paulsen Madsen, Rasmus Riis Kopperud, Hilde Vollan, Karoline Bragstad, Olav Hungnes |
| EPI_ISL_796707, EPI_ISL_796708, EPI_ISL_796709 | University Hospital of Northern Norway, Department for Microbiology and Infectious Disease Control | Norwegian Institute of Public Health, Department of Virology | Kathrine Stene-Johansen, Kamilla Heddeland Instefjord, Hilde Elshaug, Atiya R Ali,Marie Paulsen Madsen, Rasmus Riis Kopperud, Hilde Vollan, Karoline Bragstad, Olav Hungnes |
| EPI_ISL_796710 | Norwegian Institute of Public Health, Department of Virology | Norwegian Institute of Public Health, Department of Virology | Kathrine Stene-Johansen, Kamilla Heddeland Instefjord, Hilde Elshaug, Atiya R Ali,Marie Paulsen Madsen, Rasmus Riis Kopperud, Hilde Vollan, Karoline Bragstad, Olav Hungnes |
| EPI_ISL_796725, EPI_ISL_796727, EPI_ISL_796728 | Department of Medical Microbiology, St. Olavs hospital | Norwegian Institute of Public Health, Department of Virology | Kathrine Stene-Johansen, Kamilla Heddeland Instefjord, Hilde Elshaug, Atiya R Ali,Marie Paulsen Madsen, Rasmus Riis Kopperud, Hilde Vollan, Karoline Bragstad, Olav Hungnes |
| EPI_ISL_796766, EPI_ISL_796770 | Instituto Nacional de Saude (INSA) | Instituto Nacional de Saude (INSA) | Borges et al |

| | | | |
|---|---|---|---|
| EPI_ISL_801179 | Lighthouse Lab in Glasgow | Wellcome Sanger Institute for the COVID-19 Genomics UK (COG-UK) Consortium | Harper VanSteenhouse, Yumi Kasai, David Gray, Carol Clugston, Anna Dominiczak and Alex Alderton, Roberto Amato, Sonia Goncalves, Ewan Harrison, David K. Jackson, Ian Johnston, Dominic Kwiatkowski, Cordelia Langford, John Sillitoe on behalf of the Wellcome Sanger Institute COVID-19 Surveillance Team |
| EPI_ISL_801408, EPI_ISL_801424, EPI_ISL_801425 | Dutch COVID-19 response team | Erasmus Medical Center | Bas Oude Munnink, Reina Sikkema, David Nieuwenhuijse, Irina Chestakova, Anne van der Linden, Marjan Boter, Emmanuelle Munger, Corine GeurtsvanKessel, Annemiek van der Eijk, Richard Molenkamp, Marion Koopmans, on behalf of the Dutch national COVID-19 response team. |
| EPI_ISL_802864, EPI_ISL_802867, EPI_ISL_802868, EPI_ISL_802870, EPI_ISL_802871, EPI_ISL_802873, EPI_ISL_802874, EPI_ISL_802875, EPI_ISL_802877, EPI_ISL_802878, EPI_ISL_802881, EPI_ISL_802882, EPI_ISL_802883, EPI_ISL_802884, EPI_ISL_802885, EPI_ISL_802888, EPI_ISL_802889, EPI_ISL_802890, EPI_ISL_802891, EPI_ISL_802892, EPI_ISL_802895, EPI_ISL_802896, EPI_ISL_802897, EPI_ISL_802899, EPI_ISL_802900, EPI_ISL_802901, EPI_ISL_802903, EPI_ISL_802904, EPI_ISL_802905, EPI_ISL_802907, EPI_ISL_802910, EPI_ISL_802912, EPI_ISL_802913, EPI_ISL_802915, EPI_ISL_802916, EPI_ISL_802918, EPI_ISL_802919, EPI_ISL_802920, EPI_ISL_802922, EPI_ISL_802926, EPI_ISL_802928, EPI_ISL_802930, EPI_ISL_802933, EPI_ISL_802935, EPI_ISL_802937, EPI_ISL_802939, EPI_ISL_802940, EPI_ISL_802942, EPI_ISL_802943, EPI_ISL_802947, EPI_ISL_802950, EPI_ISL_802951, EPI_ISL_802952, EPI_ISL_802955, EPI_ISL_802957, EPI_ISL_802958, EPI_ISL_802962, EPI_ISL_802963, EPI_ISL_802966, EPI_ISL_802969, EPI_ISL_802971, EPI_ISL_802981, EPI_ISL_802982, EPI_ISL_802983, EPI_ISL_802984, EPI_ISL_802985, EPI_ISL_802986, EPI_ISL_802988, EPI_ISL_802989, EPI_ISL_802990, EPI_ISL_802992, EPI_ISL_802993 | | | |
| see above | Utah Public Health Laboratory | Utah Public Health Laboratory | Erin Young, Kelly Oakeson, Tara Gallagher |
| EPI_ISL_803890 | Robert Koch Institute | Robert Koch Institute | Annika Brinkmann, Janine Michel, Livia Schrick, Steven Uddin, Dominique Seifert, Merle Corty, Lars Schaade, Andreas Nitsche |
| EPI_ISL_804373, EPI_ISL_804375, EPI_ISL_804377 | CHU Purpan - Laboratoire de Virologie - Institut Fédératif de Biologie | CHU Purpan - Laboratoire de Virologie - Institut Fédératif de Biologie | Latour J., Ranger N., Dubois M., Carcenac R., Harter A., Boyer P., Tremeaux P., Izopet J. |
| EPI_ISL_804625, EPI_ISL_804654, EPI_ISL_804677, EPI_ISL_804706 | Michigan Department of Health and Human Services, Bureau of Laboratories | Michigan Department of Health and Human Services, Bureau of Laboratories | Blankenship HM, Riner D, Soehnlen MK |
| EPI_ISL_804892 | DC Public Health Lab/ Dept. of Forensic Sciences | DC Public Health Lab/ Dept. of Forensic Sciences | Scott Nguyen, Elizabeth Zelaya, Connie Maza, Monica Mann, Brittany Hamilton, David Payne, Jocelyn Hauser |
| EPI_ISL_806845, EPI_ISL_806848, EPI_ISL_806849, EPI_ISL_806850 | Alaska State Virology Laboratory | Alaska State Virology Laboratory | Stephanie DeRonde, Lisa Smith, Ph.D., Devin M. Drown, Ph.D., Jack Chen, Ph.D. |
| EPI_ISL_812362, EPI_ISL_812363, EPI_ISL_812364, EPI_ISL_812365, EPI_ISL_812366, EPI_ISL_812393, EPI_ISL_812394, EPI_ISL_812395, EPI_ISL_812403, EPI_ISL_812405, EPI_ISL_812406 | | | |
| see above | Utah Public Health Laboratory | Utah Public Health Laboratory | Erin L. Young, Kelly F. Oakeson, Tara Gallagher |
| EPI_ISL_812421, EPI_ISL_812422 | University of Wisconsin-Madison AIDS Vaccine Research Laboratories | University of Wisconsin-Madison AIDS Vaccine Research Laboratories | Gage Moreno, Katarina Braun, et al. AIDS Vaccine Research Laboratories |
| EPI_ISL_812443, EPI_ISL_812444, EPI_ISL_812445, EPI_ISL_812446, EPI_ISL_812447, EPI_ISL_812448, EPI_ISL_812449, EPI_ISL_812450, EPI_ISL_812451, EPI_ISL_812452, EPI_ISL_812453, EPI_ISL_812454, EPI_ISL_812455, EPI_ISL_812456, EPI_ISL_812457, EPI_ISL_812458, EPI_ISL_812459, EPI_ISL_812460, EPI_ISL_812461, EPI_ISL_812462, EPI_ISL_812463, EPI_ISL_812464, EPI_ISL_812465, EPI_ISL_812466, EPI_ISL_812467, EPI_ISL_812468, EPI_ISL_812469, EPI_ISL_812470, EPI_ISL_812471, EPI_ISL_812472, EPI_ISL_812473, EPI_ISL_812474, EPI_ISL_812475, EPI_ISL_812476, EPI_ISL_812477, EPI_ISL_812478, EPI_ISL_812479, EPI_ISL_812480, EPI_ISL_812483, EPI_ISL_812484, EPI_ISL_812485, EPI_ISL_812486, EPI_ISL_812487, EPI_ISL_812489, EPI_ISL_812490, EPI_ISL_812491, EPI_ISL_812492, EPI_ISL_812493, EPI_ISL_812494, EPI_ISL_812495, EPI_ISL_812496, EPI_ISL_812497, EPI_ISL_812498, EPI_ISL_812499, EPI_ISL_812500, EPI_ISL_812501, EPI_ISL_812502, EPI_ISL_812503, EPI_ISL_812504, EPI_ISL_812505, EPI_ISL_812506, EPI_ISL_812507, EPI_ISL_812508, EPI_ISL_812509, EPI_ISL_812510, EPI_ISL_812511, EPI_ISL_812512, EPI_ISL_812513, EPI_ISL_812514, EPI_ISL_812515, EPI_ISL_812516, EPI_ISL_812517, EPI_ISL_812712, EPI_ISL_812713, EPI_ISL_812714, EPI_ISL_812715, EPI_ISL_812716, EPI_ISL_812717, EPI_ISL_812718, EPI_ISL_812719, EPI_ISL_812720, EPI_ISL_812721 | | | |
| see above | Laboratorio de Referencia Nacional de Virus Respiratorios, Instituto Nacional de Salud Peru | Laboratorio de Genómica Microbiana, Universidad Peruana Cayetano Heredia | Pablo Tsukayama, Alejandra Dávila-Barclay, Guillermo Salvatierra, Luis González, Pedro E. Romero, Brenda Ayzanoa, Janet Huancachoque, Pool Marcos, Camila Castillo-Vilcahuamán, Oscar Escalante, Priscila Lope, Nancy Rojas |
| EPI_ISL_812932 | Institute for Urban Disease Control and Prevention | COVID-19 Network Investigations (CONI) Alliance | Kamolthip Atsawawaranunt, Elizabeth Batty, Wasun Chantratita, Thanat Chookajorn, Stefan Fernandez, Angkana Huang, Anthony R. Jones, Khajohn Joonsalak, Chonticha Klungtong, Theerarat Kochakarn, Prayuth Kaewmalang, Amornmas Kongklieng, Namfon Kotanan, Krittikorn Kumpornsin, Duangkamon Loesbanluechai, Wudtichai Manasatienkij, Anek Mungaomklang, Bhakbhoom Panthan, Pukkaporn Parnwijitkul, Ekawat Pasomsub, Kingkan Rakmanee, Insee Sensorn, Janjira Thaipadungpanit, Arporn Wangwiwatsin, Treewat Watthanachockchai |
| EPI_ISL_813037, EPI_ISL_813038, EPI_ISL_813039, EPI_ISL_813040, EPI_ISL_813041, EPI_ISL_813042, EPI_ISL_813043, EPI_ISL_813044, EPI_ISL_813064, EPI_ISL_813065, EPI_ISL_813066, EPI_ISL_813067, EPI_ISL_813068, EPI_ISL_813069, EPI_ISL_813070, EPI_ISL_813072, EPI_ISL_813073, EPI_ISL_813074, EPI_ISL_813075, EPI_ISL_813076, EPI_ISL_813077, EPI_ISL_813078, EPI_ISL_813079, EPI_ISL_813080, EPI_ISL_813081, EPI_ISL_813083, EPI_ISL_813085, EPI_ISL_813086 | | | |
| see above | University of Birmingham | COVID-19 Genomics UK (COG-UK) Consortium | Institute of Microbiology, University of Birmingham: Claire McMurray, Joanne Stockton, Sam Nicholls, Radoslaw Poplawski, Will Rowe, Josh Quick, Nicholas Loman. University of Birmingham Testing Laboratory: Celina M Whalley, Andrew Bosworth, Charlotte Poxon, Kasun Wanigasooriya, Oliver Pickles, Mike Kidd, Alex Richter, Andrew D Beggs PHE Heartlands Lab: Husam Osman, Andrew Bosworth. Queen Elizabeth Hospital: Anna Casey |
| EPI_ISL_813189 | Department of Pathology, University of Cambridge | COVID-19 Genomics UK (COG-UK) Consortium | Aminu S. Jahun, Yasmin Chaudhry, Grant Hall, Iliana Georgana, Myra Hosmillo, Martin D. Curran, Malte Pinckert, Surendra Parmar, Ian Goodfellow |
| EPI_ISL_813655 | Kettering General Hospital | COVID-19 Genomics UK (COG-UK) Consortium | Patrick McClure, Joseph Chappell, Theocharis Tsoleridis, Jonathan Ball, Nadine Holmes, Matthew Carlisle, Christopher Moore, Fei Sang, Johnny Debebe, Victoria Wright, Matthew Loose |
| EPI_ISL_813732, EPI_ISL_813733, EPI_ISL_813734, EPI_ISL_813735, EPI_ISL_813736, EPI_ISL_813737, EPI_ISL_813738, EPI_ISL_813739, EPI_ISL_813740, EPI_ISL_813741, EPI_ISL_813742, EPI_ISL_813743, EPI_ISL_813744, EPI_ISL_813745, EPI_ISL_813746, EPI_ISL_813747, EPI_ISL_813748, EPI_ISL_813749, EPI_ISL_813750, EPI_ISL_813764, EPI_ISL_813765, EPI_ISL_813766, EPI_ISL_813767, EPI_ISL_813768, EPI_ISL_813769, EPI_ISL_813770 | | | |
| see above | Liverpool Clinical Laboratories | COVID-19 Genomics UK (COG-UK) Consortium | Sam Haldenby, Anita Lucaci, Steve Paterson, Julian Hiscox, Alistair Darby, M Almsaud, A Alrezaihi, Muhannad Alruwaili, Stuart D Armstrong, Jones Benjamin, Eleanor G Bentley, Anu Chawla, Jordan J Clark, Angela Cowell, Richard Eccles, Isabel García-Dorival, Matthew Gemmell, Alessandro Gerada, PKF Gilmore, Richard Gregory, Ximeng Han, Catherine Hartley, Margaret Hughes, Miren Iturriza-Gomara, James Johnson, L Luu, Jenifer Manson, Charlotte Nelson, Elaine O'Toole, Cassie Olateju, Rebekah Penrice-Randal , Lucille Rainbow, N.P Randle, Trevor Ian Robinson, Parul Sharma, Ghada T Shawli, James P Stewart, Neil Swainston, Ecaterina Vamos, Joanne Watts, Mark Whitehead |
| EPI_ISL_813978 | University Hospital of Northern Norway, Department for Microbiology and Infectious Disease Control | Norwegian Institute of Public Health, Department of Virology | Kathrine Stene-Johansen, Kamilla Heddeland Instefjord, Hilde Elshaug, Atiya R Ali,Marie Paulsen Madsen, Rasmus Riis Kopperud, Hilde Vollan, Karoline Bragstad, Olav Hungnes |
| EPI_ISL_814285, EPI_ISL_814286 | Liverpool Clinical Laboratories | COVID-19 Genomics UK (COG-UK) Consortium | Sam Haldenby, Anita Lucaci, Steve Paterson, Julian Hiscox, Alistair Darby, M Almsaud, A Alrezaihi, Muhannad Alruwaili, Stuart D Armstrong, Jones Benjamin, Eleanor G Bentley, Anu Chawla, Jordan J Clark, Angela Cowell, Richard Eccles, Isabel García-Dorival, Matthew Gemmell, Alessandro Gerada, PKF Gilmore, Richard Gregory, Ximeng Han, Catherine Hartley, Margaret Hughes, Miren Iturriza-Gomara, James Johnson, L Luu, Jenifer Manson, Charlotte Nelson, Elaine O'Toole, Cassie Olateju, Rebekah Penrice-Randal , Lucille Rainbow, N.P Randle, Trevor Ian Robinson, Parul Sharma, Ghada T Shawli, James P Stewart, Neil Swainston, Ecaterina Vamos, Joanne Watts, Mark Whitehead |
| EPI_ISL_814320, EPI_ISL_814326, EPI_ISL_814404, EPI_ISL_814405, EPI_ISL_814408, EPI_ISL_814468, EPI_ISL_814492, EPI_ISL_814531, EPI_ISL_814536 | Wales Specialist Virology Centre Sequencing lab: Pathogen Genomics Unit | COVID-19 Genomics UK (COG-UK) Consortium | Catherine Moore, Johnathan Evans, Laura Gifford, Malorie Perry, Simon Cottrell, Angela Marchbank, Alec Birchley, Alexander Adams, Amy Gaskin, Bree Gatica-Wilcox, Jason Coombes, Joel Southgate, Lauren Gilbert, Lee Graham, Nicole Pacchiarini, Sara Kumziene-Summerhayes, Sarah Taylor, Sophie Jones, Sara Rey, Matthew Bull, Joanne Watkins, Sally Corden, Tom Connor |
| EPI_ISL_814552 | Liverpool Clinical Laboratories | COVID-19 Genomics UK (COG-UK) Consortium | Sam Haldenby, Anita Lucaci, Steve Paterson, Julian Hiscox, Alistair Darby, M Almsaud, A Alrezaihi, Muhannad Alruwaili, Stuart D Armstrong, Jones Benjamin, Eleanor G Bentley, Anu Chawla, Jordan J Clark, Angela Cowell, Richard Eccles, Isabel García-Dorival, Matthew Gemmell, Alessandro Gerada, PKF Gilmore, Richard Gregory, Ximeng Han, Catherine Hartley, Margaret Hughes, Miren Iturriza-Gomara, James Johnson, L Luu, Jenifer Manson, Charlotte Nelson, Elaine O'Toole, Cassie Olateju, Rebekah Penrice-Randal , Lucille Rainbow, N.P Randle, Trevor Ian Robinson, Parul Sharma, Ghada T Shawli, James P Stewart, Neil Swainston, Ecaterina Vamos, Joanne Watts, Mark Whitehead |
| EPI_ISL_814702, EPI_ISL_814703, EPI_ISL_814704, EPI_ISL_814717, EPI_ISL_814718, EPI_ISL_814719, EPI_ISL_814720, EPI_ISL_814721, EPI_ISL_814722, EPI_ISL_814723, EPI_ISL_814725, EPI_ISL_814726, EPI_ISL_814753, EPI_ISL_814860, EPI_ISL_814861, EPI_ISL_814862, EPI_ISL_814863, EPI_ISL_814864, EPI_ISL_814865, EPI_ISL_814866, EPI_ISL_814867, EPI_ISL_814868, EPI_ISL_814869, EPI_ISL_814870, EPI_ISL_815036, EPI_ISL_815093, EPI_ISL_815094, EPI_ISL_815095, EPI_ISL_815096, EPI_ISL_815097, EPI_ISL_815098, EPI_ISL_815099, EPI_ISL_815100, EPI_ISL_815101, EPI_ISL_815102, EPI_ISL_815103, EPI_ISL_815104, EPI_ISL_815105 | | | |
| see above | Wales Specialist Virology Centre Sequencing lab: Pathogen Genomics Unit | COVID-19 Genomics UK (COG-UK) Consortium | Catherine Moore, Johnathan Evans, Laura Gifford, Malorie Perry, Simon Cottrell, Angela Marchbank, Alec Birchley, Alexander Adams, Amy Gaskin, Bree Gatica-Wilcox, Jason Coombes, Joel Southgate, Lauren Gilbert, Lee Graham, Nicole Pacchiarini, Sara Kumziene-Summerhayes, Sarah Taylor, Sophie Jones, Sara Rey, Matthew Bull, Joanne Watkins, Sally Corden, Tom Connor |
| EPI_ISL_816249, EPI_ISL_816250, EPI_ISL_816254, EPI_ISL_816257, EPI_ISL_816266, EPI_ISL_816278, EPI_ISL_816298, EPI_ISL_816306, EPI_ISL_816346, EPI_ISL_816349, EPI_ISL_816354, EPI_ISL_816360, EPI_ISL_816419, EPI_ISL_816420, EPI_ISL_816422, EPI_ISL_816440, EPI_ISL_816454, EPI_ISL_816459, EPI_ISL_816464, EPI_ISL_816465, EPI_ISL_816466, EPI_ISL_816483, EPI_ISL_816490, EPI_ISL_816515, EPI_ISL_816553, EPI_ISL_816569, EPI_ISL_816579, EPI_ISL_816587, EPI_ISL_816593, EPI_ISL_816594, EPI_ISL_816598, EPI_ISL_816604, EPI_ISL_816610, EPI_ISL_816612, EPI_ISL_816615, EPI_ISL_816628 | | | |

| | | | |
|---|---|---|---|
| see above | Virology Department, Sheffield Teaching Hospitals NHS Foundation Trust/Department of Infection, Immunity and Cardiovascular Disease, The Medical School, University of Sheffield | COVID-19 Genomics UK (COG-UK) Consortium | Thushan de Silva, Matthew Parker, Nikki Smith, Adri Angyal, Rebecca Brown, Luke Green, Rachel Tucker, Paul Parsons, Danielle Groves, Katie Johnson, Laura Carrilero, Alex Keeley, Dave Partridge, Matthew Wyles, Benjamin Lindsey, Mehmet Yavuz, Mohammad Raza, Cariad Evans |
| EPI_ISL_816894, EPI_ISL_816896, EPI_ISL_816899, EPI_ISL_816905, EPI_ISL_816907, EPI_ISL_816908, EPI_ISL_816912, EPI_ISL_816914, EPI_ISL_816917, EPI_ISL_816919, EPI_ISL_816921, EPI_ISL_816922, EPI_ISL_816926, EPI_ISL_816927, EPI_ISL_816928, EPI_ISL_816929, EPI_ISL_816930, EPI_ISL_816931, EPI_ISL_816932, EPI_ISL_816933, EPI_ISL_816934, EPI_ISL_816935, EPI_ISL_816936, EPI_ISL_816937, EPI_ISL_816938, EPI_ISL_816939, EPI_ISL_816940, EPI_ISL_816941, EPI_ISL_816942, EPI_ISL_816943, EPI_ISL_816944, EPI_ISL_816945, EPI_ISL_816946, EPI_ISL_816947, EPI_ISL_816948, EPI_ISL_816949, EPI_ISL_816950, EPI_ISL_816951, EPI_ISL_816952, EPI_ISL_816953, EPI_ISL_816954, EPI_ISL_816955, EPI_ISL_816956, EPI_ISL_816957, EPI_ISL_816961, EPI_ISL_816963, EPI_ISL_816966, EPI_ISL_816972, EPI_ISL_816974, EPI_ISL_816975, EPI_ISL_816979, EPI_ISL_816981, EPI_ISL_816984, EPI_ISL_816986, EPI_ISL_816988, EPI_ISL_816989, EPI_ISL_816993, EPI_ISL_816994, EPI_ISL_816995, EPI_ISL_816996, EPI_ISL_816997, EPI_ISL_816998, EPI_ISL_816999, EPI_ISL_817000, EPI_ISL_817001, EPI_ISL_817002, EPI_ISL_817003, EPI_ISL_817004, EPI_ISL_817005, EPI_ISL_817006, EPI_ISL_817007, EPI_ISL_817008, EPI_ISL_817009, EPI_ISL_817010, EPI_ISL_817011, EPI_ISL_817012, EPI_ISL_817013, EPI_ISL_817014, EPI_ISL_817015, EPI_ISL_817016, EPI_ISL_817017, EPI_ISL_817018, EPI_ISL_817019, EPI_ISL_817020, EPI_ISL_817021, EPI_ISL_817022, EPI_ISL_817023, EPI_ISL_817024, EPI_ISL_817049, EPI_ISL_817060 | | | |
| see above | Bioinformatics and Biostatistics Lab, Advanced Sequencing Facility | COVID-19 Genomics UK (COG-UK) Consortium | Aengus Stewart,Jerome Nicod,Chelsea Sawyer,Laura Cubitt,Harshil Patel,Margaret Crawford |
| EPI_ISL_819575, EPI_ISL_819576 | Northumbria University / South Tees Hospitals NHS Foundation Trust / North Cumbria Integrated Care NHS Foundation Trust / North Tees and Hartlepool NHS Foundation Trust / Newcastle Hospitals NHS Foundation Trust | COVID-19 Genomics UK (COG-UK) Consortium | Darren L Smith,Andrew Nelson,Matthew Bashton,Greg R Young,Joshua Loh,John Allan,Mohammad A Tariq,Giles S Holt,Gary Black,Wen C Yew,Lynn Dover,Paul Baker,Steve Liggett,Sarah Essex,Jane Greenaway,Debra Padgett,Clive Graham,Garren Scott,Edward Barton,Emma Swindells,Brendan Payne,Jennifer Collins,Yusri Taha,Gary Eltringham |
| EPI_ISL_819577 | Queens Medical Centre, Clinical Microbiology Department / DeepSeq Nottingham | COVID-19 Genomics UK (COG-UK) Consortium | Gemma Clark, Wendy Smith, Manjinder Khakh, Vicki M Fleming, Michelle M Lister, Hannah Howson-Wells, Jonathan Ball, Patrick McClure, Joseph Chappell, Theocharis Tsoleridis, Nadine Holmes, Matthew Carlisle, Christopher Moore, Fei Sang, Johnny Debebe, Victoria Wright, Matthew Loose |
| EPI_ISL_820174, EPI_ISL_820346, EPI_ISL_820351, EPI_ISL_820413 | Lighthouse Lab in Alderley Park | Wellcome Sanger Institute for the COVID-19 Genomics UK (COG-UK) Consortium | Jacquelyn Wynn, Mairead Hyland, The Lighthouse Lab in Alderley Park and Alex Alderton, Roberto Amato, Sonia Goncalves, Ewan Harrison, David K. Jackson, Ian Johnston, Dominic Kwiatkowski, Cordelia Langford, John Sillitoe on behalf of the Wellcome Sanger Institute COVID-19 Surveillance Team |
| EPI_ISL_822842, EPI_ISL_822848, EPI_ISL_822849, EPI_ISL_822850, EPI_ISL_822863, EPI_ISL_822869, EPI_ISL_822872, EPI_ISL_822916, EPI_ISL_822927, EPI_ISL_822936, EPI_ISL_822937, EPI_ISL_822977, EPI_ISL_823004, EPI_ISL_823014, EPI_ISL_823057, EPI_ISL_823058, EPI_ISL_823129, EPI_ISL_823162, EPI_ISL_823175, EPI_ISL_823205, EPI_ISL_823208, EPI_ISL_823209, EPI_ISL_823210, EPI_ISL_823211, EPI_ISL_823212, EPI_ISL_823213, EPI_ISL_823214, EPI_ISL_823215, EPI_ISL_823216, EPI_ISL_823217, EPI_ISL_823569, EPI_ISL_823570, EPI_ISL_823571, EPI_ISL_823572, EPI_ISL_823573, EPI_ISL_823574, EPI_ISL_823575, EPI_ISL_823576, EPI_ISL_823577, EPI_ISL_823578, EPI_ISL_823579, EPI_ISL_823580, EPI_ISL_823581, EPI_ISL_823583, EPI_ISL_823584, EPI_ISL_823585, EPI_ISL_823587, EPI_ISL_823588, EPI_ISL_823589, EPI_ISL_823590, EPI_ISL_823591, EPI_ISL_823592, EPI_ISL_823593, EPI_ISL_823594, EPI_ISL_823595, EPI_ISL_823596, EPI_ISL_823597, EPI_ISL_823598, EPI_ISL_823599, EPI_ISL_823600, EPI_ISL_823601, EPI_ISL_823602, EPI_ISL_823603, EPI_ISL_823604, EPI_ISL_823605, EPI_ISL_823606, EPI_ISL_823607, EPI_ISL_823625 | | | |
| see above | Wales Specialist Virology Centre Sequencing lab: Pathogen Genomics Unit | COVID-19 Genomics UK (COG-UK) Consortium | Catherine Moore, Johnathan Evans, Laura Gifford, Malorie Perry, Simon Cottrell, Angela Marchbank, Alec Birchley, Alexander Adams, Amy Gaskin, Bree Gatica-Wilcox, Jason Coombes, Joel Southgate, Lauren Gilbert, Lee Graham, Nicole Pacchiarini, Sara Kumziene-Summerhayes, Sarah Taylor, Sophie Jones, Sara Rey, Matthew Bull, Joanne Watkins, Sally Corden, Tom Connor |
| EPI_ISL_823813 | DOHMH Jamaica | New York City Public Health Laboratory | Jade Wang, et al. |
| EPI_ISL_823814 | DOHMH Riverside | New York City Public Health Laboratory | Jade Wang, et al. |
| EPI_ISL_823815, EPI_ISL_823816 | DOHMH Chelsea | New York City Public Health Laboratory | Jade Wang, et al. |
| EPI_ISL_823817, EPI_ISL_823818, EPI_ISL_823819 | DOHMH Corona | New York City Public Health Laboratory | Jade Wang, et al. |
| EPI_ISL_823820 | DOHMH PHL | New York City Public Health Laboratory | Jade Wang, et al. |
| EPI_ISL_823821, EPI_ISL_823822 | DOHMH Jamaica | New York City Public Health Laboratory | Jade Wang, et al. |
| EPI_ISL_823823, EPI_ISL_823824, EPI_ISL_823825 | DOHMH Riverside | New York City Public Health Laboratory | Jade Wang, et al. |
| EPI_ISL_823826, EPI_ISL_823827, EPI_ISL_823828 | DOHMH Corona | New York City Public Health Laboratory | Jade Wang, et al. |
| EPI_ISL_823829, EPI_ISL_823830, EPI_ISL_823831 | DOHMH Morrisania | New York City Public Health Laboratory | Jade Wang, et al. |
| EPI_ISL_823832, EPI_ISL_823833, EPI_ISL_823834 | DOHMH Central Harlem | New York City Public Health Laboratory | Jade Wang, et al. |
| EPI_ISL_823835, EPI_ISL_823836, EPI_ISL_823837 | DOHMH Corona | New York City Public Health Laboratory | Jade Wang, et al. |
| EPI_ISL_823993, EPI_ISL_824034, EPI_ISL_824055 | Dutch COVID-19 response team | National Institute for Public Health and the Environment (RIVM) | Adam Meijer, Harry Vennema, Jeroen Cremer, Sharon van den Brink, Bas van der Veer, AnneMarie van den Brandt, Florian Zwagemaker, Dennis Schmitz, Chantal Reusken, on behalf of the national COVID-19 response team |
| EPI_ISL_824297 | DOHMH Jamaica | New York City Public Health Laboratory | Jade Wang, et al. |
| EPI_ISL_824299 | DOHMH Morrisania | New York City Public Health Laboratory | Jade Wang, et al. |
| EPI_ISL_824300 | DOHMH Chelsea | New York City Public Health Laboratory | Jade Wang, et al. |
| EPI_ISL_824301 | DOHMH Crown Heights | New York City Public Health Laboratory | Jade Wang, et al. |
| EPI_ISL_824302 | DOHMH Jamaica | New York City Public Health Laboratory | Jade Wang, et al. |
| EPI_ISL_824303 | DOHMH Morrisania | New York City Public Health Laboratory | Jade Wang, et al. |
| EPI_ISL_824304, EPI_ISL_824305 | DOHMH Jamaica | New York City Public Health Laboratory | Jade Wang, et al. |
| EPI_ISL_824306, EPI_ISL_824307 | DOHMH PHL | New York City Public Health Laboratory | Jade Wang, et al. |
| EPI_ISL_824308, EPI_ISL_824309 | DOHMH Jamaica | New York City Public Health Laboratory | Jade Wang, et al. |
| EPI_ISL_824310 | DOHMH Crown Heights | New York City Public Health Laboratory | Jade Wang, et al. |
| EPI_ISL_824347, EPI_ISL_824350, EPI_ISL_824353, EPI_ISL_824354, EPI_ISL_824365, EPI_ISL_824367, EPI_ISL_824368, EPI_ISL_824369, EPI_ISL_824380, EPI_ISL_824383 | Michigan Department of Health and Human Services, Bureau of Laboratories | Michigan Department of Health and Human Services, Bureau of Laboratories | Blankenship HM, Riner D, Soehnlen MK |
| EPI_ISL_824401, EPI_ISL_824403 | California Department of Public Health | California Department of Public Health | CDPH IDLB COVIDNet |
| EPI_ISL_824833 | Department of Clinical Microbiology | GIGA Medical Genomics | Keith Durkin, Maria Artesi, Sébastien Bontems, Raphaël Boreux, Bouchra Boujemla, Cécile Meex, Pierrette Melin, Marie-Pierre Hayette, Vincent Bours |
| EPI_ISL_824982 | Maryland Public Health Laboratory | Maryland Public Health Laboratory | Maryland Department of Health Laboratories Administration |
| EPI_ISL_825017, EPI_ISL_825018, EPI_ISL_825019, EPI_ISL_825021, EPI_ISL_825023 | Utah Public Health Laboratory, Utah Public Health Laboratory Infectious Disease submission group | Utah Public Health Laboratory, Utah Public Health Laboratory Infectious Disease submission group | Young,E.L., Oakeson,K.F., Gallagher,T. |
| EPI_ISL_825131, EPI_ISL_825136 | NHLS-IALCH | KRISP, KZN Research Innovation and Sequencing Platform | Giandhari J, Pillay S, Lessells R, Mdlalose K, York D, Khan S, Tegally H, Wilkinson E, de Oliveira T |
| EPI_ISL_825375, EPI_ISL_825376, EPI_ISL_825377 | Hospital Universitari Vall d'Hebron - Vall d'Hebron Institut de Recerca | Hospital Universitari Vall d'Hebron | Cristina Andrés, Maria Piñana, Josep F Abril, Damir Garcia-Cehic, Ariadna Rando, Juliana Esperalba, Maria Gema Codina, Carla Castillo, Maria Carmen Martin, Tomàs Pumarola, Josep Quer, Andrés Antón |

| | | | |
|---|---|---|---|
| EPI_ISL_825437 | Nigeria Centre For Disease Control | National reference Laboratory, NCDC, Gaduwa, Abuja | Dr Ndodo Nnaemeka, Olusola Anuoluwapo Akanbi, Chimaobi Chukwu, Dr Omoare Adesuyi, Esebanmen Grace, Kingsley Njoku, Anthony Ahumibe, Naidoo Dhamari, Nwando Mba, Dr Chikwe Ihekweazu |
| EPI_ISL_826275 | Sterling Hospital, Memnagar, Ahmedabad | Gujarat Biotechnology Research Centre | Atul K Patel, Nitin Savaliya, Dinesh Kumar, Zuber Saiyed, Labdhi Pandya, Afzal Ansari, Nikha Trivedi, Apurvasinh Puvar, Ramesh Pandit, Janvi Raval, Zarna Patel, Naman Shashtri, Chaitanya Joshi, Madhvi Joshi |
| EPI_ISL_826276 | Sterling Hospital, Memnagar, Ahmedabad | Gujarat Biotechnology Research Centre | Dinesh Kumar, Zuber Saiyed, Labdhi Pandya, Afzal Ansari, Nikha Trivedi, Apurvasinh Puvar, Ramesh Pandit, Janvi Raval, Zarna Patel, Nitin Savaliya, Atul K Patel, Naman Shashtri, Chaitanya Joshi, Madhvi Joshi |
| EPI_ISL_826465 | Lighthouse Lab in Alderley Park | Wellcome Sanger Institute for the COVID-19 Genomics UK (COG-UK) Consortium | Jacquelyn Wynn, Mairead Hyland, The Lighthouse Lab in Alderley Park and Alex Alderton, Roberto Amato, Sonia Goncalves, Ewan Harrison, David K. Jackson, Ian Johnston, Dominic Kwiatkowski, Cordelia Langford, John Sillitoe on behalf of the Wellcome Sanger Institute COVID-19 Surveillance Team |
| EPI_ISL_827183, EPI_ISL_827184 | deCODE genetics | deCODE genetics | Daniel F Gudbjartsson; Agnar Helgason; Hakon Jonsson; Olafur T Magnusson; Pall Melsted; Gudmundur L Norddahl; Jona Saemundsdottir; Asgeir Sigurdsson; Patrick Sulem; Arna B Agustsdottir; Hannes Eggertsson; Berglind Eiriksdottir; Run Fridriksdottir; Elisabet E Gardarsdottir; Gudmundur Georgsson; Olafia S Gretarsdottir; Kjartan R Gudmundsson; Thora R Gunnarsdottir; Arnaldur Gylfason; Hilma Holm; Brynjar O Jensson; Aslaug Jonasdottir; Kamilla S Josefsdottir; Thordur Kristjansson; Droplaug N Magnusdottir; Solvi Rognvaldsson; Louise le Roux; Gudrun Sigmundsdottir; Gardar Sveinbjornsson; Kristin E Sveinsdottir; Maney Sveinsdottir; Emil A Thorarensen; Bjarni Thorbjornsson; Gisli Masson; Ingileif Jonsdottir; Alma Moller; Thorolfur Gudnason; Karl G Kristinsson; Unnur Thorsteinsdottir; Kari Stefansson |
| EPI_ISL_827782 | The National University Hospital of Iceland | deCODE genetics | Daniel F Gudbjartsson; Agnar Helgason; Hakon Jonsson; Olafur T Magnusson; Pall Melsted; Gudmundur L Norddahl; Jona Saemundsdottir; Asgeir Sigurdsson; Patrick Sulem; Arna B Agustsdottir; Hannes Eggertsson; Berglind Eiriksdottir; Run Fridriksdottir; Elisabet E Gardarsdottir; Gudmundur Georgsson; Olafia S Gretarsdottir; Kjartan R Gudmundsson; Thora R Gunnarsdottir; Arnaldur Gylfason; Hilma Holm; Brynjar O Jensson; Aslaug Jonasdottir; Kamilla S Josefsdottir; Thordur Kristjansson; Droplaug N Magnusdottir; Solvi Rognvaldsson; Louise le Roux; Gudrun Sigmundsdottir; Gardar Sveinbjornsson; Kristin E Sveinsdottir; Maney Sveinsdottir; Emil A Thorarensen; Bjarni Thorbjornsson; Gisli Masson; Ingileif Jonsdottir; Alma Moller; Thorolfur Gudnason; Karl G Kristinsson; Unnur Thorsteinsdottir; Kari Stefansson |
| EPI_ISL_828213, EPI_ISL_828248, EPI_ISL_828252, EPI_ISL_829405, EPI_ISL_829406, EPI_ISL_829407, EPI_ISL_829408, EPI_ISL_829409, EPI_ISL_829411, EPI_ISL_829412, EPI_ISL_829661, EPI_ISL_829663, EPI_ISL_829667, EPI_ISL_829668, EPI_ISL_829669, EPI_ISL_829670, EPI_ISL_829892, EPI_ISL_829910, EPI_ISL_829917, EPI_ISL_829929, EPI_ISL_830066 | | | |
| see above | deCODE genetics | deCODE genetics | Daniel F Gudbjartsson; Agnar Helgason; Hakon Jonsson; Olafur T Magnusson; Pall Melsted; Gudmundur L Norddahl; Jona Saemundsdottir; Asgeir Sigurdsson; Patrick Sulem; Arna B Agustsdottir; Hannes Eggertsson; Berglind Eiriksdottir; Run Fridriksdottir; Elisabet E Gardarsdottir; Gudmundur Georgsson; Olafia S Gretarsdottir; Kjartan R Gudmundsson; Thora R Gunnarsdottir; Arnaldur Gylfason; Hilma Holm; Brynjar O Jensson; Aslaug Jonasdottir; Kamilla S Josefsdottir; Thordur Kristjansson; Droplaug N Magnusdottir; Solvi Rognvaldsson; Louise le Roux; Gudrun Sigmundsdottir; Gardar Sveinbjornsson; Kristin E Sveinsdottir; Maney Sveinsdottir; Emil A Thorarensen; Bjarni Thorbjornsson; Gisli Masson; Ingileif Jonsdottir; Alma Moller; Thorolfur Gudnason; Karl G Kristinsson; Unnur Thorsteinsdottir; Kari Stefansson |
| EPI_ISL_830069 | The National University Hospital of Iceland | deCODE genetics | Daniel F Gudbjartsson; Agnar Helgason; Hakon Jonsson; Olafur T Magnusson; Pall Melsted; Gudmundur L Norddahl; Jona Saemundsdottir; Asgeir Sigurdsson; Patrick Sulem; Arna B Agustsdottir; Hannes Eggertsson; Berglind Eiriksdottir; Run Fridriksdottir; Elisabet E Gardarsdottir; Gudmundur Georgsson; Olafia S Gretarsdottir; Kjartan R Gudmundsson; Thora R Gunnarsdottir; Arnaldur Gylfason; Hilma Holm; Brynjar O Jensson; Aslaug Jonasdottir; Kamilla S Josefsdottir; Thordur Kristjansson; Droplaug N Magnusdottir; Solvi Rognvaldsson; Louise le Roux; Gudrun Sigmundsdottir; Gardar Sveinbjornsson; Kristin E Sveinsdottir; Maney Sveinsdottir; Emil A Thorarensen; Bjarni Thorbjornsson; Gisli Masson; Ingileif Jonsdottir; Alma Moller; Thorolfur Gudnason; Karl G Kristinsson; Unnur Thorsteinsdottir; Kari Stefansson |
| EPI_ISL_830071 | deCODE genetics | deCODE genetics | Daniel F Gudbjartsson; Agnar Helgason; Hakon Jonsson; Olafur T Magnusson; Pall Melsted; Gudmundur L Norddahl; Jona Saemundsdottir; Asgeir Sigurdsson; Patrick Sulem; Arna B Agustsdottir; Hannes Eggertsson; Berglind Eiriksdottir; Run Fridriksdottir; Elisabet E Gardarsdottir; Gudmundur Georgsson; Olafia S Gretarsdottir; Kjartan R Gudmundsson; Thora R Gunnarsdottir; Arnaldur Gylfason; Hilma Holm; Brynjar O Jensson; Aslaug Jonasdottir; Kamilla S Josefsdottir; Thordur Kristjansson; Droplaug N Magnusdottir; Solvi Rognvaldsson; Louise le Roux; Gudrun Sigmundsdottir; Gardar Sveinbjornsson; Kristin E Sveinsdottir; Maney Sveinsdottir; Emil A Thorarensen; Bjarni Thorbjornsson; Gisli Masson; Ingileif Jonsdottir; Alma Moller; Thorolfur Gudnason; Karl G Kristinsson; Unnur Thorsteinsdottir; Kari Stefansson |
| EPI_ISL_830072 | The National University Hospital of Iceland | deCODE genetics | Daniel F Gudbjartsson; Agnar Helgason; Hakon Jonsson; Olafur T Magnusson; Pall Melsted; Gudmundur L Norddahl; Jona Saemundsdottir; Asgeir Sigurdsson; Patrick Sulem; Arna B Agustsdottir; Hannes Eggertsson; Berglind Eiriksdottir; Run Fridriksdottir; Elisabet E Gardarsdottir; Gudmundur Georgsson; Olafia S Gretarsdottir; Kjartan R Gudmundsson; Thora R Gunnarsdottir; Arnaldur Gylfason; Hilma Holm; Brynjar O Jensson; Aslaug Jonasdottir; Kamilla S Josefsdottir; Thordur Kristjansson; Droplaug N Magnusdottir; Solvi Rognvaldsson; Louise le Roux; Gudrun Sigmundsdottir; Gardar Sveinbjornsson; Kristin E Sveinsdottir; Maney Sveinsdottir; Emil A Thorarensen; Bjarni Thorbjornsson; Gisli Masson; Ingileif Jonsdottir; Alma Moller; Thorolfur Gudnason; Karl G Kristinsson; Unnur Thorsteinsdottir; Kari Stefansson |
| EPI_ISL_830411, EPI_ISL_830412, EPI_ISL_830416, EPI_ISL_830418, EPI_ISL_830419, EPI_ISL_830420, EPI_ISL_830421, EPI_ISL_830549 | deCODE genetics | deCODE genetics | Daniel F Gudbjartsson; Agnar Helgason; Hakon Jonsson; Olafur T Magnusson; Pall Melsted; Gudmundur L Norddahl; Jona Saemundsdottir; Asgeir Sigurdsson; Patrick Sulem; Arna B Agustsdottir; Hannes Eggertsson; Berglind Eiriksdottir; Run Fridriksdottir; Elisabet E Gardarsdottir; Gudmundur Georgsson; Olafia S Gretarsdottir; Kjartan R Gudmundsson; Thora R Gunnarsdottir; Arnaldur Gylfason; Hilma Holm; Brynjar O Jensson; Aslaug Jonasdottir; Kamilla S Josefsdottir; Thordur Kristjansson; Droplaug N Magnusdottir; Solvi Rognvaldsson; Louise le Roux; Gudrun Sigmundsdottir; Gardar Sveinbjornsson; Kristin E Sveinsdottir; Maney Sveinsdottir; Emil A Thorarensen; Bjarni Thorbjornsson; Gisli Masson; Ingileif Jonsdottir; Alma Moller; Thorolfur Gudnason; Karl G Kristinsson; Unnur Thorsteinsdottir; Kari Stefansson |
| EPI_ISL_831256, EPI_ISL_831280, EPI_ISL_831282, EPI_ISL_831286 | Hospital Universitario La Paz (Madrid) | SeqCOVID-SPAIN consortium/IBV(CSIC) | Fernando Lázaro-Perona, María Rodríguez-Tejedor, Elias Dahdouh, Jesús Mingorance and SeqCOVID-SPAIN consortium |
| EPI_ISL_831648 | Santa Clara County Public Health Laboratory | Santa Clara County Public Health Laboratory | Santa Clara County Public Health Department |
| EPI_ISL_831693, EPI_ISL_831700, EPI_ISL_831754, EPI_ISL_831851, EPI_ISL_831852, EPI_ISL_831853, EPI_ISL_831854, EPI_ISL_831866, EPI_ISL_831867, EPI_ISL_831868, EPI_ISL_831869 | | | |
| see above | United States Air Force School of Aerospace Medicine | United States Air Force School of Aerospace Medicine | Anthony Fries, Jennifer Meyer, William Gruner, Amanda Javorina, Sarah Purves, Clarise Starr, Elizabeth Macias |
| EPI_ISL_831894, EPI_ISL_831895 | Santa Clara County Public Health Laboratory | Santa Clara County Public Health Laboratory | Santa Clara County Public Health Department |
| EPI_ISL_831945 | Klinisk mikrobiologi, virus F68 | The Public Health Agency of Sweden | Department of Microbiology, The Public Health Agency of Sweden |
| EPI_ISL_831994 | TATAA Biocenter | The Public Health Agency of Sweden | Department of Microbiology, The Public Health Agency of Sweden |
| EPI_ISL_832043 | Wyoming Public Health Laboratory | Wyoming Public Health Laboratory | Noah Hull, Taylor Fearing, Lynette Gumbleton, Channing Weber, Ashley Norberg, Bailey Bowcutt, and Wanda Manley |
| EPI_ISL_832099 | Santa Clara County Public Health Laboratory | Santa Clara County Public Health Laboratory | Santa Clara County Public Health Department |
| EPI_ISL_832170 | Hospital | National Reference Center for Viruses of Respiratory Infections, Institut Pasteur, Paris | Marion Barbet, Sylvie Behillil, Méline Bizard, Angela Brisebarre, Camille Capel, Etienne Simon-Lorière, Vincent Enouf, Maud Vanpeene, Sylvie van der Werf, Clémence Guillaume |
| EPI_ISL_832383, EPI_ISL_832391, EPI_ISL_832398, EPI_ISL_832399, EPI_ISL_832400, EPI_ISL_832401 | Santa Clara County Public Health Laboratory | Santa Clara County Public Health Laboratory | Santa Clara County Public Health Department |
| EPI_ISL_833132 | Laboratorio de Ecologia de Doencas Transmissiveis na Amazonia, Instituto Leonidas e Maria Deane - Fiocruz Amazonia | Laboratorio de Ecologia de Doencas Transmissiveis na Amazonia, Instituto Leonidas e Maria Deane - Fiocruz Amazonia | Valdinete Nascimento, Victor Souza, André Corado, Fernanda Nascimento, George Silva, Ágatha Costa, Debora Duarte, Karina Pessoa, Matilde Mejía, Luciana Gonçalves, Maria Júlia Brandão, Michele Jesus, Felipe Naveca |
| EPI_ISL_833159 | Instituto Adolfo Lutz - Central | Instituto Adolfo Lutz, Interdiciplinary Procedures Center, Strategic Laboratory | Claudio Tavares Sacchi, Claudia Regina Gonçalves, Erica Valessa Ramos Gomes, Karoline Rodrigues Campos |
| EPI_ISL_833160 | Instituto Adolfo Lutz - Regional de Santo Andre | Instituto Adolfo Lutz, Interdiciplinary Procedures Center, Strategic Laboratory | Claudio Tavares Sacchi, Claudia Regina Gonçalves, Erica Valessa Ramos Gomes, Karoline Rodrigues Campos |

| | | | |
|---|---|---|---|
| EPI_ISL_837076, EPI_ISL_837186 | Respiratory Virus Unit, National Infection Service, Public Health England | COVID-19 Genomics UK (COG-UK) Consortium | PHE Covid Sequencing Team |
| EPI_ISL_837395, EPI_ISL_837426 | National Virus Reference Laboratory | National Virus Reference Laboratory | Michael Carr, Gabriel Gonzalez, Jonathan Dean, Cillian F De Gascun |
| EPI_ISL_838220, EPI_ISL_838221, EPI_ISL_838284, EPI_ISL_838285, EPI_ISL_838286, EPI_ISL_838287 | Virology Department, Royal Infirmary of Edinburgh, NHS Lothian / School of Biological Sciences, University of Edinburgh / Institute of Genetics and Molecular Medicine, University of Edinburgh | COVID-19 Genomics UK (COG-UK) Consortium | McHugh M, Dewar R, Rooke S, Gallagher M, Balcaza C, O'Toole Á, Scher E, Hill V, McCrone JT, Colquhoun R, Yu X, Jackson B, Rambaut A, Williams TC, Templeton K |
| EPI_ISL_838352, EPI_ISL_838464, EPI_ISL_838465, EPI_ISL_838471, EPI_ISL_838472, EPI_ISL_838481, EPI_ISL_838486, EPI_ISL_838488, EPI_ISL_838490, EPI_ISL_838492, EPI_ISL_838496, EPI_ISL_838665, EPI_ISL_838672 | | | |
| see above | Liverpool Clinical Laboratories | COVID-19 Genomics UK (COG-UK) Consortium | Sam Haldenby, Anita Lucaci, Steve Paterson, Julian Hiscox, Alistair Darby, M Almsaud, A Alrezaihi, Muhannad Alruwaili, Stuart D Armstrong, Jones Benjamin, Eleanor G Bentley, Anu Chawla, Jordan J Clark, Angela Cowell, Richard Eccles, Isabel García-Dorival, Matthew Gemmell, Alessandro Gerada, PKF Gilmore, Richard Gregory, Ximeng Han, Catherine Hartley, Margaret Hughes, Miren Iturriza-Gomara, James Johnson, L Luu, Jenifer Manson, Charlotte Nelson, Elaine O'Toole, Cassie Olateju, Rebekah Penrice-Randal , Lucille Rainbow, N.P Randle, Trevor Ian Robinson, Parul Sharma, Ghada T Shawli, James P Stewart, Neil Swainston, Ecaterina Vamos, Joanne Watts, Mark Whitehead |
| EPI_ISL_838673, EPI_ISL_838674, EPI_ISL_838675, EPI_ISL_838676, EPI_ISL_838677, EPI_ISL_838678, EPI_ISL_838679, EPI_ISL_838680, EPI_ISL_838681, EPI_ISL_838682, EPI_ISL_838683, EPI_ISL_838684, EPI_ISL_838685, EPI_ISL_838686, EPI_ISL_838719 | | | |
| see above | University College London, Great Ormond Street Hospital for Children NHS Foundation Trust, Imperial College Healthcare NHS Trust | COVID-19 Genomics UK (COG-UK) Consortium | Sergi Castellano, Rachel Williams, Mark Kristiansen, Paola Resende Silva, Sunando Roy, Tony Brooks, Helena Tutill, Paola Niola, Patricia Dyal, Charlotte Williams, Leysa Forrest, Yasmin Panchbhaya, Jacqueline Findlay, Samuel Weeks, Julianne Brown, Kathryn Harris, Paul Randell, James Price, Alison Holmes, Judith Breuer |
| EPI_ISL_839534, EPI_ISL_839535, EPI_ISL_839536, EPI_ISL_839537, EPI_ISL_839538, EPI_ISL_839539, EPI_ISL_839540, EPI_ISL_839541, EPI_ISL_839542, EPI_ISL_839749, EPI_ISL_839750, EPI_ISL_839751, EPI_ISL_839752, EPI_ISL_839753, EPI_ISL_839754, EPI_ISL_839755, EPI_ISL_839756, EPI_ISL_839757, EPI_ISL_839758, EPI_ISL_839759, EPI_ISL_839760, EPI_ISL_839761, EPI_ISL_839762, EPI_ISL_839763, EPI_ISL_839764, EPI_ISL_839765, EPI_ISL_839766, EPI_ISL_839767, EPI_ISL_839768, EPI_ISL_839769, EPI_ISL_839770, EPI_ISL_839771, EPI_ISL_839772, EPI_ISL_839773, EPI_ISL_839774, EPI_ISL_839775, EPI_ISL_839776, EPI_ISL_839777, EPI_ISL_839778, EPI_ISL_839779, EPI_ISL_839780, EPI_ISL_839781, EPI_ISL_839782, EPI_ISL_839783, EPI_ISL_839784, EPI_ISL_839785, EPI_ISL_839786, EPI_ISL_839787, EPI_ISL_839788, EPI_ISL_839789, EPI_ISL_839790, EPI_ISL_839791, EPI_ISL_839792, EPI_ISL_839793, EPI_ISL_839794, EPI_ISL_839795, EPI_ISL_839796, EPI_ISL_839797, EPI_ISL_839798, EPI_ISL_839799, EPI_ISL_839800, EPI_ISL_839801, EPI_ISL_839802, EPI_ISL_839803, EPI_ISL_839804, EPI_ISL_839805, EPI_ISL_839806, EPI_ISL_839807, EPI_ISL_839808, EPI_ISL_839809, EPI_ISL_839810, EPI_ISL_839811, EPI_ISL_839812, EPI_ISL_839813, EPI_ISL_839814, EPI_ISL_839815 | | | |
| see above | Northumbria University / South Tees Hospitals NHS Foundation Trust / North Cumbria Integrated Care NHS Foundation Trust / North Tees and Hartlepool NHS Foundation Trust / Newcastle Hospitals NHS Foundation Trust | COVID-19 Genomics UK (COG-UK) Consortium | Darren L Smith,Andrew Nelson,Matthew Bashton,Greg R Young,Joshua Loh,John Allan,Mohammad A Tariq,Giles S Holt,Gary Black,Wen C Yew,Lynn Dover,Paul Baker,Steve Liggett,Sarah Essex,Jane Greenaway,Debra Padgett,Clive Graham,Garren Scott,Edward Barton,Emma Swindells,Brendan Payne,Jennifer Collins,Yusri Taha,Gary Eltringham |
| EPI_ISL_840041, EPI_ISL_840042, EPI_ISL_840043, EPI_ISL_840044, EPI_ISL_840045, EPI_ISL_840046, EPI_ISL_840047 | Lincolnshire Hospitals and DeepSeq Nottingham | COVID-19 Genomics UK (COG-UK) Consortium | Nichola Duckworth, Tim Sloan, Sarah Walsh, Jonathan Ball, Patrick McClure, Joeseph Chappell, Nadine Holmes, Matthew Carlisle, Christopher Moore, Fei Sang, Johnny Debebe, Victoria Wright, Matthew Loose |
| EPI_ISL_840250, EPI_ISL_840252, EPI_ISL_840253, EPI_ISL_840254 | Oxford Viromics, NDM, University of Oxford; Oxford University Hospitals; Basingstoke and North Hampshire Hospital | COVID-19 Genomics UK (COG-UK) Consortium | Tanya Golubchik, David Bonsall, George Macintyre, Amy Trebes, Mariateresa de Cesare, Catrin Moore, Alex Mobbs, Anita Justice, Robert Shaw, Monique Andersson, Timothy Peto, Emma Wise, Nathan Moore, Jessica Lynch, Nick Cortes, Matilde Mori, Stephen Kidd, David Buck, John Todd, Christophe Fraser |
| EPI_ISL_842358, EPI_ISL_842360, EPI_ISL_842363, EPI_ISL_842369, EPI_ISL_842371, EPI_ISL_842372, EPI_ISL_842376, EPI_ISL_842378, EPI_ISL_842381, EPI_ISL_842383, EPI_ISL_842385, EPI_ISL_842386, EPI_ISL_842390, EPI_ISL_842391, EPI_ISL_842392, EPI_ISL_842393, EPI_ISL_842394, EPI_ISL_842395, EPI_ISL_842396, EPI_ISL_842397, EPI_ISL_842398, EPI_ISL_842399, EPI_ISL_842400, EPI_ISL_842401, EPI_ISL_842402, EPI_ISL_842403, EPI_ISL_842404, EPI_ISL_842405, EPI_ISL_842406, EPI_ISL_842407, EPI_ISL_842408, EPI_ISL_842409, EPI_ISL_842410, EPI_ISL_842411, EPI_ISL_842412, EPI_ISL_842413, EPI_ISL_842414, EPI_ISL_842415, EPI_ISL_842416, EPI_ISL_842417, EPI_ISL_842418, EPI_ISL_842419, EPI_ISL_842420, EPI_ISL_842421 | | | |
| see above | Bioinformatics and Biostatistics Lab, Advanced Sequencing Facility | COVID-19 Genomics UK (COG-UK) Consortium | Aengus Stewart,Jerome Nicod,Chelsea Sawyer,Laura Cubitt,Harshil Patel,Margaret Crawford |
| EPI_ISL_842680, EPI_ISL_842681, EPI_ISL_842717, EPI_ISL_842718, EPI_ISL_842720, EPI_ISL_842733, EPI_ISL_842735, EPI_ISL_842736, EPI_ISL_842737, EPI_ISL_842738, EPI_ISL_842739, EPI_ISL_842740, EPI_ISL_842741, EPI_ISL_842742, EPI_ISL_842743, EPI_ISL_842744, EPI_ISL_842746, EPI_ISL_842747, EPI_ISL_842749, EPI_ISL_842750, EPI_ISL_842751, EPI_ISL_842752, EPI_ISL_842753, EPI_ISL_842754, EPI_ISL_842755, EPI_ISL_842756, EPI_ISL_842757, EPI_ISL_842758, EPI_ISL_842759, EPI_ISL_842760, EPI_ISL_842761, EPI_ISL_842762, EPI_ISL_842763, EPI_ISL_842764, EPI_ISL_842765, EPI_ISL_842769, EPI_ISL_842770, EPI_ISL_842771, EPI_ISL_842772, EPI_ISL_842773, EPI_ISL_842774, EPI_ISL_842776, EPI_ISL_842778, EPI_ISL_842779, EPI_ISL_842780, EPI_ISL_842781, EPI_ISL_842782, EPI_ISL_842784, EPI_ISL_842786, EPI_ISL_842787, EPI_ISL_842788, EPI_ISL_842789, EPI_ISL_842791 | | | |
| see above | University College London Hospital | COVID-19 Genomics UK (COG-UK) Consortium | Judith Heaney, Matthew Byott, Catherine Houlihan, Dan Frampton, Stuart Kirk, Moira Spyer and Eleni Nastouli |
| EPI_ISL_843071, EPI_ISL_843072, EPI_ISL_843073, EPI_ISL_843074, EPI_ISL_843075, EPI_ISL_843076 | Barts Health NHS Trust | COVID-19 Genomics UK (COG-UK) Consortium | CUTINO-MOGUEL, Maria-Teresa; HARRINGTON, David; OWOYEMI, Dola; SHYLINI, Raghavendran; BROAD, Claire; KELE, Beatrix |
| EPI_ISL_845757, EPI_ISL_845758, EPI_ISL_845759 | Emory Molecular Diagnostics Laboratory, Emory Healthcare | Piantadosi Lab, Emory Department of Pathology | Ahmed Babiker, Anne Piantadosi |
| EPI_ISL_846591 | Respiratory Virus Unit, National Infection Service, Public Health England | COVID-19 Genomics UK (COG-UK) Consortium | PHE Covid Sequencing Team |
| EPI_ISL_847518, EPI_ISL_847526, EPI_ISL_847645, EPI_ISL_847646, EPI_ISL_847647, EPI_ISL_847648, EPI_ISL_847783 | California Department of Public Health | Chiu Laboratory, University of California, San Francisco | Charles Chiu, Xianding (Wayne) Deng, Candace Wang, Brian Bushnell, Scot Federman, Jill Hacker, Debra Wadford |
| EPI_ISL_847834 | Connecticut Department of Health | Grubaugh Lab - Yale School of Public Health | Tara Alpert, Joseph Fauver, Anderson Brito, Mallery Breban, Anne Wyllie, Chantal Vogels, Mary Petrone, Chaney Kalinich, Isabel Ott, Nathan Grubaugh |
| EPI_ISL_848426 | Illinois Department of Public Health | Gagnon Lab, Southern Illinois University | Keith Gagnon |
| EPI_ISL_849730 | unknown | PHV-FSS | Son Nguyen et al. |
| EPI_ISL_850668, EPI_ISL_850670 | The National Institute of Public Health | State Veterinary Institute Prague | Nagy,A;Jirincova,H;Trnka,D;Vecerova,J;Trinklova,M |
| EPI_ISL_852566, EPI_ISL_852577, EPI_ISL_852578 | Max von Pettenkofer Institute, Virology, National Reference Center for Retroviruses, LMU München | Laboratory for Functional Genome Analysis, Dept. Genomics, Gene Center of the LMU Munich | Max Muenchhoff, Stefan Krebs, Alexander Graf, Oliver Keppler, Helmut Blum |
| EPI_ISL_853343, EPI_ISL_853344, EPI_ISL_853377, EPI_ISL_853378, EPI_ISL_853379, EPI_ISL_853380, EPI_ISL_853382, EPI_ISL_853386, EPI_ISL_853388 | UPMC Clinical Microbiology Laboratory | Microbial Genome Sequencing Center; Microbial Genomic Epidemiology Laboratory | Mustapha M. Mustapha, Jane W. Marsh, Dan Snyder, Marissa P. Griffith, Stephanie L. Mitchell, Vatsala R. Srinivasa, Kady D. Waggle, Chinelo Ezeonwuku, Vaughn S. Cooper, Lee H. Harrison |
| EPI_ISL_853740, EPI_ISL_853935 | Department of Microbiology, University Innsbruck | Bergthaler laboratory, CeMM Research Center for Molecular Medicine of the Austrian Academy of Sciences | Lukas Endler, Alexandra Popa, Benedikt Agerer, Jakob-Wendelin Genger, Alexander Lercher, Anna Schedl, Thomas Penz, Michael Schuster, Jan Laine, Martin Senekowitsch, Christoph Bock, Andreas Bergthaler |
| EPI_ISL_854436, EPI_ISL_854437 | SARATOGA HOSPITAL LABORATORY | Wadsworth Center, New York State Department of Health | Kirsten St. George, Daryl M. Lamson, Alexis Russel, Matthew Shudt, Melissa A Leisner, Jonathan Plitnick, Navjot Singh, John Kelly, Erasmus Schneider, Erica Lasek-Nesselquist |
| EPI_ISL_855392 | California Department of Public Health | Chiu Laboratory, University of California, San Francisco | Charles Chiu, Xianding (Wayne) Deng, Candace Wang, Brian Bushnell, Scot Federman, Jill Hacker, Debra Wadford |
| EPI_ISL_855420, EPI_ISL_855421, EPI_ISL_855422, EPI_ISL_855423, EPI_ISL_855424, EPI_ISL_855425, EPI_ISL_855428, EPI_ISL_855429 | Servicio de Microbiología, Laboratori Clínic Metropolitana Nord. Hospital Universitari Germans Trias i Pujol. Institut d'Investigació en Ciències de la Salut Germans Trias i Pujol (IGTP) | SeqCOVID-SPAIN consortium/IBV(CSIC) | Elisa Martró, Antoni E. Bordoy, Anna Not, Adrián Antuori, Anabel Fernández, Nona Romaní, Verónica Saludes, Cristina Casañ and SeqCOVID-SPAIN consortium |
| EPI_ISL_855542, EPI_ISL_855547 | KEMRI-Wellcome Trust Research | KEMRI-Wellcome Trust Research | Githinji et al |

| | | | |
|---|---|---|---|
| | Programme/KEMRI-CGMR-C Kilifi | Programme/KEMRI-CGMR-C Kilifi | |
| EPI_ISL_856783, EPI_ISL_856784, EPI_ISL_856786 | Servicio Virosis Respiratorias-Departamento Virología-INEI | Instituto Nacional Enfermedades Infecciosas C.G.Malbran | Baumeister E., Avaro M., Benedetti E., Russo M., Dattero ME, Pontoriero A., Cisterna D., Molina V., Perandones C., Tuduri E., Lorenzo F., Poklepovich T., Campos J. |
| EPI_ISL_857536 | Swiss National Reference Centre for Influenza | Swiss National Reference Centre for Influenza | Tim Roloff, Ana Rita Gonçalves, Madlen Stange, Helena MB Seth-Smith, Alfredo Mari, Karoline Leuzinger, Julia Bielicki, Manuel Battegay, Hans Hirsch, Laurent Kaiser, Adrian Egli |
| EPI_ISL_858409, EPI_ISL_858416, EPI_ISL_858994, EPI_ISL_859001, EPI_ISL_859005, EPI_ISL_859008, EPI_ISL_859018, EPI_ISL_859020, EPI_ISL_859024 | Lighthouse Lab in Alderley Park | Wellcome Sanger Institute for the COVID-19 Genomics UK (COG-UK) Consortium | Jacquelyn Wynn, Mairead Hyland, The Lighthouse Lab in Alderley Park and Alex Alderton, Roberto Amato, Sonia Goncalves, Ewan Harrison, David K. Jackson, Ian Johnston, Dominic Kwiatkowski, Cordelia Langford, John Sillitoe on behalf of the Wellcome Sanger Institute COVID-19 Surveillance Team |
| EPI_ISL_859462, EPI_ISL_859463, EPI_ISL_859465, EPI_ISL_859466, EPI_ISL_859467, EPI_ISL_859468, EPI_ISL_859471, EPI_ISL_859472, EPI_ISL_859477, EPI_ISL_859478, EPI_ISL_859479, EPI_ISL_859480, EPI_ISL_859482, EPI_ISL_859483, EPI_ISL_859484, EPI_ISL_859485, EPI_ISL_859486, EPI_ISL_859487, EPI_ISL_859488, EPI_ISL_859489, EPI_ISL_859490, EPI_ISL_859491, EPI_ISL_859492, EPI_ISL_859493, EPI_ISL_859494, EPI_ISL_859495, EPI_ISL_859496, EPI_ISL_859497, EPI_ISL_859498, EPI_ISL_859499, EPI_ISL_859500, EPI_ISL_859501, EPI_ISL_859502, EPI_ISL_859503, EPI_ISL_859504, EPI_ISL_859505, EPI_ISL_859506, EPI_ISL_859507, EPI_ISL_859508, EPI_ISL_859509, EPI_ISL_859510, EPI_ISL_859511, EPI_ISL_859512, EPI_ISL_859513, EPI_ISL_859514, EPI_ISL_859515, EPI_ISL_859516, EPI_ISL_859517, EPI_ISL_859518, EPI_ISL_859519, EPI_ISL_859520, EPI_ISL_859521, EPI_ISL_859522, EPI_ISL_859523, EPI_ISL_859524, EPI_ISL_859525, EPI_ISL_859526, EPI_ISL_859527, EPI_ISL_859528, EPI_ISL_859529, EPI_ISL_859530, EPI_ISL_859531, EPI_ISL_859532 | | | |
| see above | Lighthouse Lab in Glasgow | Wellcome Sanger Institute for the COVID-19 Genomics UK (COG-UK) Consortium | Harper VanSteenhouse, Yumi Kasai, David Gray, Carol Clugston, Anna Dominiczak and Alex Alderton, Roberto Amato, Sonia Goncalves, Ewan Harrison, David K. Jackson, Ian Johnston, Dominic Kwiatkowski, Cordelia Langford, John Sillitoe on behalf of the Wellcome Sanger Institute COVID-19 Surveillance Team |
| EPI_ISL_860174, EPI_ISL_860175, EPI_ISL_860176, EPI_ISL_860177, EPI_ISL_860178, EPI_ISL_860179 | Keio University School of Medicine | Keio University School of Medicine | Kenjiro Kosaki, Yuka Iwasaki, Hirotsugu Ishizu, Haruhiko Siomi, Kodai Abe |
| EPI_ISL_860216, EPI_ISL_860217, EPI_ISL_860237 | Medical Microbiology Unit, Department for Laboratory Medicine, Drammen Hospital, Vestre Viken Health Trust, | Norwegian Institute of Public Health, Department of Virology | Kathrine Stene-Johansen, Kamilla Heddeland Instefjord, Hilde Elshaug, Atiya R Ali,Marie Paulsen Madsen, Rasmus Riis Kopperud, Hilde Vollan, Karoline Bragstad, Olav Hungnes |
| EPI_ISL_860270 | Unilabs Laboratory Medicine | Norwegian Institute of Public Health, Department of Virology | Kathrine Stene-Johansen, Kamilla Heddeland Instefjord, Hilde Elshaug, Atiya R Ali,Marie Paulsen Madsen, Rasmus Riis Kopperud, Hilde Vollan, Karoline Bragstad, Olav Hungnes |
| EPI_ISL_860690 | Respiratory Virus Unit, National Infection Service, Public Health England | COVID-19 Genomics UK (COG-UK) Consortium | PHE Covid Sequencing Team |
| EPI_ISL_860933, EPI_ISL_860934, EPI_ISL_860936 | Johns Hopkins Hospital Department of Pathology | Johns Hopkins Hospital Department of Pathology | C. Paul Morris, Chun Huai Luo, Adannaya Amadi, Nicholas Gallagher, Heba H. Mostafa |
| EPI_ISL_861463, EPI_ISL_861468 | UHAS COVID-19 Lab | UHAS COVID-19 Lab | Kwabena O. Duedu, Jones Gyamfi, Reuben Ayivor-Djanie, John O. Gyapong and the UHAS COVID-19 Lab Team |
| EPI_ISL_861586 | Instituto Nacional de Saude (INSA) | Instituto Nacional de Saude (INSA) | Borges et al |
| EPI_ISL_861781, EPI_ISL_861782, EPI_ISL_861784, EPI_ISL_861785, EPI_ISL_861790, EPI_ISL_861791, EPI_ISL_861796, EPI_ISL_861799, EPI_ISL_861806, EPI_ISL_861810, EPI_ISL_861827, EPI_ISL_861828, EPI_ISL_861829 | | | |
| see above | Hospital General Universitario Gregorio Marañón | SeqCOVID-SPAIN consortium/IBV(CSIC) | Darío García de Viedma, Laura Pérez-Lago, Pedro J Sola-Campoy, Sergio Buenestado-Serrano, Marta Herranz, Victor Manuel de la Cueva, Julia Suárez, Pilar Catalán, Patricia Muñoz and SeqCOVID-SPAIN consortium |
| EPI_ISL_861873 | LATE - Laboratório de Técnicas Especiais - Hospital Israelita Albert Einstein | LATE - Laboratório de Técnicas Especiais - Hospital Israelita Albert Einstein | Deyvid Amgarten, Fernanda de Mello Malta, Raquel Riyuzo, Ana Paula Moreira Salles, Pedro Henrique Sebe Rodrigues, João Renato Rebello Pinho |
| EPI_ISL_862251 | Respiratory Virus Unit, National Infection Service, Public Health England | COVID-19 Genomics UK (COG-UK) Consortium | PHE Covid Sequencing Team |
| EPI_ISL_864578 | Institute of Medical Microbiology and Hospital Hygiene | Institute of Medical Microbiology and Hospital Hygiene | Prof. Dr. Achim Kaasch, Aljoscha Tersteegen |
| EPI_ISL_864786, EPI_ISL_864788, EPI_ISL_864793, EPI_ISL_864802, EPI_ISL_864804, EPI_ISL_864806, EPI_ISL_864808, EPI_ISL_864874, EPI_ISL_864882 | Department of Pathology, University of Cambridge | COVID-19 Genomics UK (COG-UK) Consortium | Aminu S. Jahun, Yasmin Chaudhry, Grant Hall, Iliana Georgana, Myra Hosmillo, Martin D. Curran, Malte Pinckert, Surendra Parmar, Ian Goodfellow |
| EPI_ISL_865208, EPI_ISL_865210, EPI_ISL_865255, EPI_ISL_865256, EPI_ISL_865257, EPI_ISL_865258, EPI_ISL_865259, EPI_ISL_865260, EPI_ISL_865262, EPI_ISL_865263, EPI_ISL_865266, EPI_ISL_865267, EPI_ISL_865269, EPI_ISL_865432 | | | |
| see above | Liverpool Clinical Laboratories | COVID-19 Genomics UK (COG-UK) Consortium | Sam Haldenby, Anita Lucaci, Steve Paterson, Julian Hiscox, Alistair Darby, M Almsaud, A Alrezaihi, Muhannad Alruwaili, Stuart D Armstrong, Jones Benjamin, Eleanor G Bentley, Anu Chawla, Jordan J Clark, Angela Cowell, Richard Eccles, Isabel García-Dorival, Matthew Gemmell, Alessandro Gerada, PKF Gilmore, Richard Gregory, Ximeng Han, Catherine Hartley, Margaret Hughes, Miren Iturriza-Gomara, James Johnson, L Luu, Jenifer Manson, Charlotte Nelson, Elaine O'Toole, Cassie Olateju, Rebekah Penrice-Randal , Lucille Rainbow, N.P Randle, Trevor Ian Robinson, Parul Sharma, Ghada T Shawli, James P Stewart, Neil Swainston, Ecaterina Vamos, Joanne Watts, Mark Whitehead |
| EPI_ISL_865718, EPI_ISL_865719 | University College London, Great Ormond Street Hospital for Children NHS Foundation Trust, Imperial College Healthcare NHS Trust | COVID-19 Genomics UK (COG-UK) Consortium | Sergi Castellano, Rachel Williams, Mark Kristiansen, Paola Resende Silva, Sunando Roy, Tony Brooks, Helena Tutill, Paola Niola, Patricia Dyal, Charlotte Williams, Leysa Forrest, Yasmin Panchbhaya, Jacqueline Findlay, Samuel Weeks, Julianne Brown, Kathryn Harris, Paul Randell, James Price, Alison Holmes, Judith Breuer |
| EPI_ISL_866065, EPI_ISL_866066, EPI_ISL_866067, EPI_ISL_866107, EPI_ISL_866138 | University College London Hospital | COVID-19 Genomics UK (COG-UK) Consortium | Judith Heaney, Matthew Byott, Catherine Houlihan, Dan Frampton, Stuart Kirk, Moira Spyer and Eleni Nastouli |
| EPI_ISL_866400, EPI_ISL_866402, EPI_ISL_866403, EPI_ISL_866404, EPI_ISL_866405, EPI_ISL_866406, EPI_ISL_866407, EPI_ISL_866408, EPI_ISL_866409, EPI_ISL_866410, EPI_ISL_866411, EPI_ISL_866412, EPI_ISL_866413, EPI_ISL_866423, EPI_ISL_866424, EPI_ISL_866469, EPI_ISL_866470, EPI_ISL_866471, EPI_ISL_866472, EPI_ISL_866473, EPI_ISL_866474, EPI_ISL_866475, EPI_ISL_866478, EPI_ISL_866479 | | | |
| see above | Northumbria University / South Tees Hospitals NHS Foundation Trust / North Cumbria Integrated Care NHS Foundation Trust / North Tees and Hartlepool NHS Foundation Trust / Newcastle Hospitals NHS Foundation Trust | COVID-19 Genomics UK (COG-UK) Consortium | Darren L Smith,Andrew Nelson,Matthew Bashton,Greg R Young,Joshua Loh,John Allan,Mohammad A Tariq,Giles S Holt,Gary Black,Wen C Yew,Lynn Dover,Paul Baker,Steve Liggett,Sarah Essex,Jane Greenaway,Debra Padgett,Clive Graham,Garren Scott,Edward Barton,Emma Swindells,Brendan Payne,Jennifer Collins,Yusri Taha,Gary Eltringham |
| EPI_ISL_866581, EPI_ISL_866587, EPI_ISL_866806, EPI_ISL_866815, EPI_ISL_866817 | Quadram Institute Bioscience | COVID-19 Genomics UK (COG-UK) Consortium | Dave J. Baker, Gemma L. Kay, Alp Aydin, Thanh Le-Viet, Steven Rudder, Ana P. Tedim, Anastasia Kolyva, Maria Diaz, Leonardo de Oliveira Martins, Nabil-Fareed Alikhan, Lizzie Meadows, Rachael Stanley, Ngozi Elumogo, Muhammed Yasir, Nicholas M. Thomson, Alexander J Trotter, Rachel Gilroy, Samuel Bloomfield, Claire Stuart, Andrew Bell, Reenesh Prakash, Samir Dervisevic, Alison E. Mather, John Wain, Mark Webber, Andrew J. Page, Justin O'Grady |
| EPI_ISL_866896, EPI_ISL_866897, EPI_ISL_866898, EPI_ISL_866899, EPI_ISL_866900 | Queens Medical Centre, Clinical Microbiology Department / DeepSeq Nottingham | COVID-19 Genomics UK (COG-UK) Consortium | Gemma Clark, Wendy Smith, Manjinder Khakh, Vicki M Fleming, Michelle M Lister, Hannah Howson-Wells, Jonathan Ball, Patrick McClure, Joseph Chappell, Theocharis Tsoleridis, Nadine Holmes, Matthew Carlisle, Christopher Moore, Fei Sang, Johnny Debebe, Victoria Wright, Matthew Loose |
| EPI_ISL_867286, EPI_ISL_867287, EPI_ISL_867288, EPI_ISL_867289, EPI_ISL_867290, EPI_ISL_867291, EPI_ISL_867292, EPI_ISL_867293, EPI_ISL_867294, EPI_ISL_867295, EPI_ISL_867296, EPI_ISL_867297, EPI_ISL_867499 | | | |
| see above | Originating lab: Wales Specialist Virology Centre Sequencing lab: Pathogen Genomics Unit | Public Health Wales Microbiology Cardiff Wales Specialist Virology Centre | Catherine Moore, Johnathan Evans, Laura Gifford, Malorie Perry, Simon Cottrell, Angela Marchbank, Alec Birchley, Alexander Adams, Amy Gaskin, Bree Gatica-Wilcox, Jason Coombes, Joel Southgate, Lauren Gilbert, Lee Graham, Nicole Pacchiarini, Sara Kumziene-Summerhayes, Sarah Taylor, Sophie Jones, Sara Rey, Matthew Bull, Joanne Watkins, Sally Corden, Tom Connor |
| EPI_ISL_871921, EPI_ISL_871929 | Servicio de Microbiología, Laboratori Clínic Metropolitana | SeqCOVID-SPAIN consortium/IBV(CSIC) | Elisa Martró, Antoni E. Bordoy, Anna Not, Adrián Antuori, Anabel Fernández, Nona Romaní, Verónica Saludes, Cristina Casañ and SeqCOVID-SPAIN |

| | | | |
|---|---|---|---|
| | Nord. Hospital Universitari Germans Trias i Pujol. Institut d'Investigació en Ciències de la Salut Germans Trias i Pujol (IGTP) | | consortium |
| EPI_ISL_872596, EPI_ISL_872597 | Department of Laboratory Medicine, National Taiwan University Hospital | Microbial Genomics Core Lab, National Taiwan University Centers of Genomic and Precision Medicine | Shiou-Hwei Yeh, You-Yu Lin, Ya-Yun Lai, Chiao-Ling Li, Shan-Chwen Chang, Pei-Jer Chen, Sui-Yuan Chang |
| EPI_ISL_872617, EPI_ISL_872618, EPI_ISL_872619 | Nigeria Centre for Disease Control (NCDC) | African Centre of Excellence for Genomics of Infectious Diseases (ACEGID), Redeemer's University | Oluniyi P.E. et al |
| EPI_ISL_872920, EPI_ISL_872921, EPI_ISL_872922, EPI_ISL_872923, EPI_ISL_872924, EPI_ISL_872925, EPI_ISL_872926, EPI_ISL_872970 | WHO National Influenza Centre Russian Federation | WHO National Influenza Centre Russian Federation | Andrey Komissarov, Artem Fadeev, Anna Ivanova, Kseniya Komissarova, Dmitry Bazhenov, Mikhail Bakaev, Daria Danilenko, Ksenia Safina, Elena Nabieva, Georgii Bazykin, Dmitry Lioznov |
| EPI_ISL_873214, EPI_ISL_873216, EPI_ISL_873217, EPI_ISL_873218 | M Health Fairview | Minnesota Department of Health, Public Health Laboratory | Alexandra Lorentz, Jacob Garfin, Matt Plumb, and Xiong Wang |
| EPI_ISL_875555, EPI_ISL_875556 | ULSS 2 Treviso | Istituto Zooprofilattico Sperimentale delle Venezie | Adelaide Milani, Alessia Schivo, Annalisa Salviato, Erika Giorgia Quaranta, Ambra Pastori, Bianca Zecchin, Alice Fusaro, Isabella Monne, Calogero Terregino, Antonia Ricci |
| EPI_ISL_876048, EPI_ISL_876089, EPI_ISL_876105, EPI_ISL_876112, EPI_ISL_876157, EPI_ISL_876194, EPI_ISL_876300, EPI_ISL_876301, EPI_ISL_876302, EPI_ISL_876303, EPI_ISL_876304 | | | |
| see above | Massachusetts State Public Health Laboratory | Massachusetts State Public Health Laboratory | Andrew Lang, Timelia Fink, Glen Gallagher, Sandra Smole |
| EPI_ISL_876795, EPI_ISL_876796, EPI_ISL_876797, EPI_ISL_876798, EPI_ISL_876799, EPI_ISL_876800, EPI_ISL_876801, EPI_ISL_876802, EPI_ISL_876803, EPI_ISL_876804, EPI_ISL_876805, EPI_ISL_876806, EPI_ISL_876807, EPI_ISL_876808, EPI_ISL_876809 | | | |
| see above | Istituto Zooprofilattico Sperimentale della Puglia e della Basilicata | Istituto Zooprofilattico Sperimentale della Puglia e della Basilicata | Parisi A., Bianco A., Capozzi L., Del Sambro L., Manzulli V, Rondinone V., Pace L., Cipolletta D., Galante D. |
| EPI_ISL_877419 | Siti Khodijah Hospital | Institute of Tropical Disease, Universitas Airlangga | Krisnoadi Rahardjo, Aldise M Nastri, Jezzy R Dewantari, Rima R Prasetya, Muhammad Hamdan, Gatot Soegiarto, Laksmi Wulandari, Resti Yudhawati, Yasuko Mori, Soetjipto, Kazufumi Shimizu, Maria I Lusida |
| EPI_ISL_877420 | Darus Syifa' Islamic Hospital | Institute of Tropical Disease, Universitas Airlangga | Aldise M Nastri, Jezzy R Dewantari, Rima R Prasetya, Krisnoadi Rahardjo, HM Faiz, Gatot Soegiarto, Laksmi Wulandari, Resti Yudhawati, Soetjipto, Yasuko Mori, Maria I Lusida, Kazufumi Shimizu |
| EPI_ISL_877421 | Institute of Tropical Disease | Institute of Tropical Disease, Universitas Airlangga | Jezzy R Dewantari, Rima R Prasetya, Krisnoadi Rahardjo, Aldise M Nastri, Gatot Soegiarto, Laksmi Wulandari, Resti Yudhawati, Soetjipto, Yasuko Mori, Maria I Lusida, Kazufumi Shimizu |
| EPI_ISL_878584, EPI_ISL_878587 | San Diego County Public Health Laboratory | Andersen lab at Scripps Research | SEARCH Alliance San Diego with Tracy Basler, Jovan Shephard, Brett Austin |
| EPI_ISL_878697, EPI_ISL_878757, EPI_ISL_878763, EPI_ISL_878765, EPI_ISL_878776, EPI_ISL_878781, EPI_ISL_880172, EPI_ISL_880181, EPI_ISL_880189, EPI_ISL_880197 | Rady's Childrens Hospital | Andersen lab at Scripps Research | SEARCH Alliance San Diego with Nanda Radamchar, David Dimmock, Linda Luo, Christina Clarke, Kathryn Bouic, Teresa Mueller, Denise Malicki |
| EPI_ISL_882649 | Swiss National Reference Centre for Influenza Virology laboratory, CNRI | Swiss National Reference Centre for Influenza Virology laboratory, CNRI | Tim Roloff, Ana Rita Gonçalves, Madlen Stange, Helena MB Seth-Smith, Alfredo Mari, Karoline Leuzinger, Julia Bielicki, Manuel Battegay, Hans Hirsch, Laurent Kaiser, Adrian Egli |
| EPI_ISL_883009, EPI_ISL_883010 | Maryland Public Health Laboratory | Maryland Public Health Laboratory | Maryland Department of Health Laboratories Administration |
| EPI_ISL_884214 | Alaska State Virology Laboratory (Alaska DHHS) | Alaska State Virology Laboratory (Alaska DHHS) | Stephanie DeRonde, Lisa Smith, PhD. Jack Chen, PhD. |
| EPI_ISL_884871, EPI_ISL_884872, EPI_ISL_884873, EPI_ISL_884900, EPI_ISL_884901, EPI_ISL_884903, EPI_ISL_884904, EPI_ISL_884905, EPI_ISL_884906, EPI_ISL_884907, EPI_ISL_884908, EPI_ISL_884909, EPI_ISL_884910, EPI_ISL_884911, EPI_ISL_884914, EPI_ISL_884915, EPI_ISL_884916, EPI_ISL_884917, EPI_ISL_884937, EPI_ISL_884947, EPI_ISL_884958, EPI_ISL_884959, EPI_ISL_884961, EPI_ISL_884963 | | | |
| see above | Santa Clara County Public Health Laboratory | Chan-Zuckerberg Biohub | CZB Cliahub Consortium |
| EPI_ISL_885133, EPI_ISL_885134 | California Institute of Technology | Chan-Zuckerberg Biohub | CZB Cliahub Consortium |
| EPI_ISL_888675 | Center of Advanced Studies and Technology, Molecular Genetics Laboratory | Center of Advanced Studies and Technology, Molecular Genetics Laboratory | De Fabritiis Simone, Mandatori Domitilla, Ferrante Rossella |
| EPI_ISL_888683, EPI_ISL_888684, EPI_ISL_888691, EPI_ISL_888752, EPI_ISL_888753, EPI_ISL_888759, EPI_ISL_888773, EPI_ISL_888778, EPI_ISL_888779, EPI_ISL_888788 | KU Leuven, Rega Institute, Clinical and Epidemiological Virology | KU Leuven, Rega Institute, Clinical and Epidemiological Virology | Tony Wawina-Bokalanga, Bert Vanmechelen, Joan Marti-Carerras, Piet Maes |
| EPI_ISL_888993 | RSU Medirossa | Eijkman Institute for Molecular Biology, Ministry of Research and Technology/National Agency for Research and Innovation | Iskandar Adnan, Lydia V. Panggalo, Sukma Oktavianthi, Willy Agustine, Edison Johar, Hidayat Trimarsanto, Frilasita A Yudhaputri, Safarina G Malik, Khin Saw Myint, Amin Soebandrio |
| EPI_ISL_890253, EPI_ISL_890258, EPI_ISL_890285, EPI_ISL_890289 | KU Leuven, Rega Institute, Clinical and Epidemiological Virology | KU Leuven, Rega Institute, Clinical and Epidemiological Virology | Tony Wawina-Bokalanga, Bert Vanmechelen, Joan Marti-Carerras, Piet Maes |
| EPI_ISL_892215, EPI_ISL_892216 | Lighthouse Lab in Alderley Park | Wellcome Sanger Institute for the COVID-19 Genomics UK (COG-UK) Consortium | Jacquelyn Wynn, Mairead Hyland, The Lighthouse Lab in Alderley Park and Alex Alderton, Roberto Amato, Sonia Goncalves, Ewan Harrison, David K. Jackson, Ian Johnston, Dominic Kwiatkowski, Cordelia Langford, John Sillitoe on behalf of the Wellcome Sanger Institute COVID-19 Surveillance Team |
| EPI_ISL_892270, EPI_ISL_892271, EPI_ISL_892275, EPI_ISL_892277 | Servicio de Microbiología Clínica (Complejo Hospitalario de Navarra, Pamplona), Instituto de Investigación Sanitaria de Navarra (IdiSNA) | SeqCOVID-SPAIN consortium/IBV(CSIC) | Carmen Ezpeleta Baquedano, Ana Navascués, Ana Miqueleiz and SeqCOVID-SPAIN consortium |
| EPI_ISL_892279, EPI_ISL_892314, EPI_ISL_892315, EPI_ISL_892316, EPI_ISL_892317, EPI_ISL_892318, EPI_ISL_892319, EPI_ISL_892320, EPI_ISL_892321, EPI_ISL_892322, EPI_ISL_892323, EPI_ISL_892324 | | | |
| see above | Servicio de Microbiología. Hospital Universitario Donostia. OSI Donostialdea. Área de Enfermedades Infecciosas, Grupo de Infección Respiratoria y Resistencia Antimicrobiana. Instituto de Investigación Sanitaria Biodonostia | SeqCOVID-SPAIN consortium/IBV(CSIC) | Gustavo Cilla Eguiluz, Milagrosa Montes Ros, Luis Piñeiro Vázquez, Ane Sorrarain, Jose Maria Marimón and SeqCOVID-SPAIN consortium |
| EPI_ISL_892345, EPI_ISL_892346, EPI_ISL_892347, EPI_ISL_892348, EPI_ISL_892349, EPI_ISL_892350, EPI_ISL_892351, EPI_ISL_892352, EPI_ISL_892353, EPI_ISL_892354, EPI_ISL_892355, EPI_ISL_892356 | | | |
| see above | Servicio de Microbiología Clínica (Complejo Hospitalario de Navarra, Pamplona), Instituto de Investigación Sanitaria de Navarra (IdiSNA) | SeqCOVID-SPAIN consortium/IBV(CSIC) | Carmen Ezpeleta Baquedano, Ana Navascués, Ana Miqueleiz and SeqCOVID-SPAIN consortium |
| EPI_ISL_893736, EPI_ISL_893737, EPI_ISL_893738 | Massachusetts Department of Public Health | Pathogen Discovery, Respiratory Viruses Branch, Division of Viral Diseases, Centers for Disease Control and Prevention | Yan Li, Anna Montmayeur, Ying Tao, Jing Zhang, Krista Queen, Anna Uehara, Brian Lynch, Rachel Marine, Peter Cook, Clinton R. Paden, Haibin Wang, Suxiang Tong |
| EPI_ISL_893739 | Florida Department of Public Health | Pathogen Discovery, Respiratory Viruses Branch, Division of Viral Diseases, Centers for Disease Control and Prevention | Yan Li, Anna Montmayeur, Ying Tao, Jing Zhang, Krista Queen, Anna Uehara, Brian Lynch, Rachel Marine, Peter Cook, Clinton R. Paden, Haibin Wang, Suxiang Tong |
| EPI_ISL_896105, EPI_ISL_896117 | Viollier AG | University Hospital Basel, Clinical Bacteriology | Tim Roloff, Madlen Stange, Helena MB Seth-Smith, Alfredo Mari, Karoline Leuzinger, Julia Bielicki, Christiane Beckmann, Manuel Battegay, Hans Hirsch, Adrian Egli |

| | | | |
|---|---|---|---|
| EPI_ISL_900569 | Hôpital Paris Saint-Joseph | CNR Virus des Infections Respiratoires - France SUD | Antonin Bal, Gregory Destras, Gwendolyne Burfin, Hadrien Règue, Quentin Semanas, Martine Valette, Bruno Lina, Sylvie Larrat, Laurence Josset |
| EPI_ISL_900572 | Hôpital Paris Saint-Joseph | CNR Virus des Infections Respiratoires - France SUD | Antonin Bal, Gregory Destras, Gwendolyne Burfin, Hadrien Règue, Quentin Semanas, Martine Valette, Bruno Lina, Laurence Josset |
| EPI_ISL_900580, EPI_ISL_900606, EPI_ISL_900607, EPI_ISL_900608, EPI_ISL_900609, EPI_ISL_900663 | IZSM | TIGEM | Patrizia Annunziata, Andrea Ballabio, Valentina Bouche, Davide Cacchiarelli (CorrespAuthor), Pellegrino Cerino, Chiara Colantuono, Maria Concetta Cuomo, Denise Di Concilio, Lucio Di Filippo, Antonio Grimaldi, Antonio Limone, Anna Manfredi, Francesco Panariello, Biancamaria Pierri, Marcello Salvi |
| EPI_ISL_903071 | Washington State Department of Health | Seattle Flu Study | Deborah A. Nickerson, Chris D. Frazar, Jover Lee, Benjamin Pelle, Erica Ryke, Matthew Richardson, Amanda Adler, Elisabeth Brandstetter, Peter D. Han, Kairsten Fay, Misja Ilcisin, Kirsten Lacombe, Thomas R. Sibley, Melissa Truong, Caitlin R. Wolf, Romesh Gautom, Geoff Melly, Brian Hiatt, Philip Dykema, Scott Lindquist, Michael Boeckh, Janet A. Englund, Michael Famulare, Barry R. Lutz, Mark J. Rieder, Lea M. Starita, Matthew Thompson, Helen Y. Chu, Jay Shendure, Trevor Bedford |
| EPI_ISL_903230, EPI_ISL_903242, EPI_ISL_903243, EPI_ISL_903247, EPI_ISL_903249, EPI_ISL_903250 | M Health Fairview | Minnesota Department of Health, Public Health Laboratory | Alexandra Lorentz, Jacob Garfin, Matt Plumb, and Xiong Wang |
| EPI_ISL_903618 | MN PHL Division, Minnesota Department of Health | Genomics and Discovery, Respiratory Viruses Branch, Division of Viral Diseases, Centers for Disease Control and Prevention | Krista Queen, Yan Li, Ying Tao, Jing Zhang, Anna Uehara, Anna Montmayeur, Clinton R. Paden, Peter W. Cook, Rachel Marine, Mili Sheth, Jasmine Padilla, Sarah Nobles, Mark Burroughs, Lori Rowe, Haibin Wang, Ben L. Rambo-Martin, Dhwani Batra, Justin Lee, Suxiang Tong |
| EPI_ISL_903621 | IA State Hygienic Laboratory | Genomics and Discovery, Respiratory Viruses Branch, Division of Viral Diseases, Centers for Disease Control and Prevention | Krista Queen, Yan Li, Ying Tao, Jing Zhang, Anna Uehara, Anna Montmayeur, Clinton R. Paden, Peter W. Cook, Rachel Marine, Mili Sheth, Jasmine Padilla, Sarah Nobles, Mark Burroughs, Lori Rowe, Haibin Wang, Ben L. Rambo-Martin, Dhwani Batra, Justin Lee, Suxiang Tong |
| EPI_ISL_903656 | RI State Health Laboratories | Genomics and Discovery, Respiratory Viruses Branch, Division of Viral Diseases, Centers for Disease Control and Prevention | Krista Queen, Yan Li, Ying Tao, Jing Zhang, Anna Uehara, Anna Montmayeur, Clinton R. Paden, Peter W. Cook, Rachel Marine, Mili Sheth, Jasmine Padilla, Sarah Nobles, Mark Burroughs, Lori Rowe, Haibin Wang, Ben L. Rambo-Martin, Dhwani Batra, Justin Lee, Suxiang Tong |
| EPI_ISL_903693, EPI_ISL_903739 | LA Office of Public Health Laboratories | Genomics and Discovery, Respiratory Viruses Branch, Division of Viral Diseases, Centers for Disease Control and Prevention | Krista Queen, Yan Li, Ying Tao, Jing Zhang, Anna Uehara, Anna Montmayeur, Clinton R. Paden, Peter W. Cook, Rachel Marine, Mili Sheth, Jasmine Padilla, Sarah Nobles, Mark Burroughs, Lori Rowe, Haibin Wang, Ben L. Rambo-Martin, Dhwani Batra, Justin Lee, Suxiang Tong |
| EPI_ISL_903747 | NE Public Health Laboratory | Genomics and Discovery, Respiratory Viruses Branch, Division of Viral Diseases, Centers for Disease Control and Prevention | Krista Queen, Yan Li, Ying Tao, Jing Zhang, Anna Uehara, Anna Montmayeur, Clinton R. Paden, Peter W. Cook, Rachel Marine, Mili Sheth, Jasmine Padilla, Sarah Nobles, Mark Burroughs, Lori Rowe, Haibin Wang, Ben L. Rambo-Martin, Dhwani Batra, Justin Lee, Suxiang Tong |
| EPI_ISL_903760 | MN PHL Division, Minnesota Department of Health | Genomics and Discovery, Respiratory Viruses Branch, Division of Viral Diseases, Centers for Disease Control and Prevention | Krista Queen, Yan Li, Ying Tao, Jing Zhang, Anna Uehara, Anna Montmayeur, Clinton R. Paden, Peter W. Cook, Rachel Marine, Mili Sheth, Jasmine Padilla, Sarah Nobles, Mark Burroughs, Lori Rowe, Haibin Wang, Ben L. Rambo-Martin, Dhwani Batra, Justin Lee, Suxiang Tong |
| EPI_ISL_903776 | LA Office of Public Health Laboratories | Genomics and Discovery, Respiratory Viruses Branch, Division of Viral Diseases, Centers for Disease Control and Prevention | Krista Queen, Yan Li, Ying Tao, Jing Zhang, Anna Uehara, Anna Montmayeur, Clinton R. Paden, Peter W. Cook, Rachel Marine, Mili Sheth, Jasmine Padilla, Sarah Nobles, Mark Burroughs, Lori Rowe, Haibin Wang, Ben L. Rambo-Martin, Dhwani Batra, Justin Lee, Suxiang Tong |
| EPI_ISL_903888 | MS Public Health Laboratory | Genomics and Discovery, Respiratory Viruses Branch, Division of Viral Diseases, Centers for Disease Control and Prevention | Krista Queen, Yan Li, Ying Tao, Jing Zhang, Anna Uehara, Anna Montmayeur, Clinton R. Paden, Peter W. Cook, Rachel Marine, Mili Sheth, Jasmine Padilla, Sarah Nobles, Mark Burroughs, Lori Rowe, Haibin Wang, Ben L. Rambo-Martin, Dhwani Batra, Justin Lee, Suxiang Tong |
| EPI_ISL_903889 | TX DSHS, Lab Services Section MC 1947 | Genomics and Discovery, Respiratory Viruses Branch, Division of Viral Diseases, Centers for Disease Control and Prevention | Krista Queen, Yan Li, Ying Tao, Jing Zhang, Anna Uehara, Anna Montmayeur, Clinton R. Paden, Peter W. Cook, Rachel Marine, Mili Sheth, Jasmine Padilla, Sarah Nobles, Mark Burroughs, Lori Rowe, Haibin Wang, Ben L. Rambo-Martin, Dhwani Batra, Justin Lee, Suxiang Tong |
| EPI_ISL_903914, EPI_ISL_903925 | RI State Health Laboratories | Genomics and Discovery, Respiratory Viruses Branch, Division of Viral Diseases, Centers for Disease Control and Prevention | Krista Queen, Yan Li, Ying Tao, Jing Zhang, Anna Uehara, Anna Montmayeur, Clinton R. Paden, Peter W. Cook, Rachel Marine, Mili Sheth, Jasmine Padilla, Sarah Nobles, Mark Burroughs, Lori Rowe, Haibin Wang, Ben L. Rambo-Martin, Dhwani Batra, Justin Lee, Suxiang Tong |
| EPI_ISL_903937 | MS Public Health Laboratory | Genomics and Discovery, Respiratory Viruses Branch, Division of Viral Diseases, Centers for Disease Control and Prevention | Krista Queen, Yan Li, Ying Tao, Jing Zhang, Anna Uehara, Anna Montmayeur, Clinton R. Paden, Peter W. Cook, Rachel Marine, Mili Sheth, Jasmine Padilla, Sarah Nobles, Mark Burroughs, Lori Rowe, Haibin Wang, Ben L. Rambo-Martin, Dhwani Batra, Justin Lee, Suxiang Tong |
| EPI_ISL_903948, EPI_ISL_903961, EPI_ISL_903966 | WA State Department of Health | Genomics and Discovery, Respiratory Viruses Branch, Division of Viral Diseases, Centers for Disease Control and Prevention | Krista Queen, Yan Li, Ying Tao, Jing Zhang, Anna Uehara, Anna Montmayeur, Clinton R. Paden, Peter W. Cook, Rachel Marine, Mili Sheth, Jasmine Padilla, Sarah Nobles, Mark Burroughs, Lori Rowe, Haibin Wang, Ben L. Rambo-Martin, Dhwani Batra, Justin Lee, Suxiang Tong |
| EPI_ISL_904011 | Veterinary Specialized Institute Kraljevo | Veterinary Specialized Institute "Kraljevo", Serbia | Vidanovic,D., Tesovic,B., Knezevic,A., Jovanovic,T., Jankovic,M., Sekler,M., Banovic Djeri,B., Petrovic,T., Volkening,J., Afonso,C. |
| EPI_ISL_904236, EPI_ISL_904240, EPI_ISL_904263, EPI_ISL_904483, EPI_ISL_904484, EPI_ISL_904485, EPI_ISL_904486, EPI_ISL_904487, EPI_ISL_904488, EPI_ISL_904489, EPI_ISL_904490 | | | |
| see above | Dutch COVID-19 response team | Erasmus Medical Center | Bas Oude Munnink, Reina Sikkema, David Nieuwenhuijse, Irina Chestakova, Anne van der Linden, Marjan Boter, Emmanuelle Munger, Corine GeurtsvanKessel, Annemiek van der Eijk, Richard Molenkamp, Marion Koopmans, on behalf of the Dutch national COVID-19 response team. |
| EPI_ISL_904768, EPI_ISL_904795, EPI_ISL_905084, EPI_ISL_905086, EPI_ISL_905089, EPI_ISL_905092, EPI_ISL_905093, EPI_ISL_905094, EPI_ISL_905095, EPI_ISL_905131, EPI_ISL_905134, EPI_ISL_905490, EPI_ISL_905607, EPI_ISL_905651, EPI_ISL_905659, EPI_ISL_905689, EPI_ISL_905690, EPI_ISL_905691, EPI_ISL_905692, EPI_ISL_905693 | | | |
| see above | Dutch COVID-19 response team | National Institute for Public Health and the Environment (RIVM) | Adam Meijer, Harry Vennema, Dirk Eggink, Jeroen Cremer, Sharon van den Brink, Bas van der Veer, AnneMarie van den Brandt, Florian Zwagemaker, Dennis Schmitz, Chantal Reusken, on behalf of the national COVID-19 response team |
| EPI_ISL_905792, EPI_ISL_905793, EPI_ISL_905794, EPI_ISL_905795, EPI_ISL_905796, EPI_ISL_905797, EPI_ISL_905798, EPI_ISL_905807, EPI_ISL_905808, EPI_ISL_905809, EPI_ISL_905810 | | | |
| see above | OHSU Lab Services Molecular Microbiology Lab | Oregon SARS-CoV-2 Genome Sequencing Center | Brendan L. O'Connell, Sally Grindstaff, Kayla Carter, Ruth V. Nichols, Alec J. Hirsch, Donna Hansel, Guang Fan, Xuan, Qin, Daniel N. Streblow, William B. Messer, Andrew C. Adey, Benjamin N. Bimber, Brian J. O'Roak |
| EPI_ISL_906092, EPI_ISL_906097 | Child Health Research Foundation | Child Health Research Foundation | Senjuti Saha, Sharmistha Goswami, Afroza Akter Tanni, Syed Muktadir Al Sium, Arif Mohammad Tanmoy, Roly Malaker, Md Hafizur Rahman, Samir K Saha |
| EPI_ISL_906099 | Child Health Research Foundation | Child Health Research Foundation | Senjuti Saha, Afroza Akter Tanni, Sharmistha Goswami, Syed Muktadir Al Sium, Arif Mohammad Tanmoy, Roly Malaker, Md Hafizur Rahman, Samir K Saha |
| EPI_ISL_906113 | Child Health Research Foundation | Child Health Research Foundation | Senjuti Saha, Syed Muktadir Al Sium, Sharmistha Goswami, Afroza Akter Tanni, Arif Mohammad Tanmoy, Roly Malaker, Md Hafizur Rahman, Samir K Saha |
| EPI_ISL_906114 | Child Health Research Foundation | Child Health Research Foundation | Senjuti Saha, Arif Mohammad Tanmoy, Sharmistha Goswami, Afroza Akter Tanni, Syed Muktadir Al Sium, Roly Malaker, Md Hafizur Rahman, Samir K Saha |
| EPI_ISL_906115 | Child Health Research Foundation | Child Health Research Foundation | Senjuti Saha, Syed Muktadir Al Sium, Sharmistha Goswami, Afroza Akter Tanni, Arif Mohammad Tanmoy, Roly Malaker, Md Hafizur Rahman, Samir K Saha |
| EPI_ISL_906264 | University of Wisconsin-Madison AIDS Vaccine Research Laboratories | University of Wisconsin-Madison AIDS Vaccine Research Laboratories | Gage Moreno, Katarina Braun, et al. AIDS Vaccine Research Laboratories |
| EPI_ISL_906275, EPI_ISL_906276 | Nigeria Centre for Disease Control (NCDC) | African Centre of Excellence for Genomics of Infectious Diseases (ACEGID), Redeemer's University | Oluniyi P.E. et al |

| | | | |
|---|---|---|---|
| EPI_ISL_906562, EPI_ISL_906563, EPI_ISL_906564 | Maine Health and Environmental Testing Laboratory (Maine HETL) | Tewhey Lab, The Jackson Laboratory | Matluk,N., Dewey,H., Iosue,F., Barter,M., Lynch,R., Munger,H. and Tewhey,R. |
| EPI_ISL_909969, EPI_ISL_909978, EPI_ISL_909979, EPI_ISL_909980, EPI_ISL_909981, EPI_ISL_909982 | A. Krumbholz, Labor Dr. Krause und Kollegen MVZ GmbH, Kiel | Charité Universitätsmedizin Berlin, Institut für Virologie | Victor M Corman, Tobias Bleicker, Julia Tesch, Barbara Mühlemann, Jörn Beheim-Schwarzbach, Talitha Veith, Julia Schneider, Cornelia Schlee, Tomasz Zemojtel, Terry Jones, Christian Drosten |
| EPI_ISL_910331 | Department of Infectious Diseases, Istituto Superiore di Sanità, Rome, Italy; AOUP Paolo Giaccone, Palermo, Italy | Istituto Superiore di Sanità (ISS) | Paola Stefanelli, Angela Di Martino, Alessandra Lo Presti, Stefano Fiore, Fabio Tramuto, Francesco Vitale, Carmelo Maida, Daniela Di Naro, Giulia Randazzo, Manuela Marra, Maria Carollo, Marco Crescenzi |
| EPI_ISL_910757, EPI_ISL_910758, EPI_ISL_910759, EPI_ISL_910760, EPI_ISL_910761, EPI_ISL_910762, EPI_ISL_910763, EPI_ISL_910764, EPI_ISL_910765, EPI_ISL_910766, EPI_ISL_910769, EPI_ISL_910770, EPI_ISL_910771, EPI_ISL_910772, EPI_ISL_910773, EPI_ISL_910777, EPI_ISL_910778, EPI_ISL_910779, EPI_ISL_910780, EPI_ISL_910781, EPI_ISL_910782, EPI_ISL_910783, EPI_ISL_910784, EPI_ISL_910785, EPI_ISL_910786, EPI_ISL_910787, EPI_ISL_910788, EPI_ISL_910789, EPI_ISL_910790, EPI_ISL_910791, EPI_ISL_910792, EPI_ISL_910793, EPI_ISL_910794, EPI_ISL_910795, EPI_ISL_910796, EPI_ISL_910797, EPI_ISL_910798, EPI_ISL_910801, EPI_ISL_910802, EPI_ISL_910803, EPI_ISL_910804, EPI_ISL_910805, EPI_ISL_910806, EPI_ISL_910807, EPI_ISL_910808, EPI_ISL_910809, EPI_ISL_910810, EPI_ISL_910811, EPI_ISL_910812, EPI_ISL_910813, EPI_ISL_910814, EPI_ISL_910815, EPI_ISL_910816, EPI_ISL_910817, EPI_ISL_910818, EPI_ISL_910819, EPI_ISL_910820, EPI_ISL_910821, EPI_ISL_910822, EPI_ISL_910823, EPI_ISL_910824, EPI_ISL_910825, EPI_ISL_910826, EPI_ISL_910827, EPI_ISL_910828, EPI_ISL_910829, EPI_ISL_910830, EPI_ISL_910831, EPI_ISL_910832, EPI_ISL_910833, EPI_ISL_910834, EPI_ISL_910835, EPI_ISL_910836, EPI_ISL_910837, EPI_ISL_910838, EPI_ISL_910839, EPI_ISL_910840, EPI_ISL_910841, EPI_ISL_910842, EPI_ISL_910843, EPI_ISL_910844, EPI_ISL_910845, EPI_ISL_910846, EPI_ISL_910850, EPI_ISL_910851, EPI_ISL_910852, EPI_ISL_910853, EPI_ISL_910854, EPI_ISL_910855, EPI_ISL_910856, EPI_ISL_910857, EPI_ISL_910858, EPI_ISL_910859, EPI_ISL_910860, EPI_ISL_910861, EPI_ISL_910862, EPI_ISL_910863, EPI_ISL_910864, EPI_ISL_910865, EPI_ISL_910866, EPI_ISL_910867, EPI_ISL_910868, EPI_ISL_910869, EPI_ISL_910870, EPI_ISL_910871, EPI_ISL_910872, EPI_ISL_910873, EPI_ISL_910874, EPI_ISL_910875, EPI_ISL_910876, EPI_ISL_910877, EPI_ISL_910878, EPI_ISL_910879, EPI_ISL_910880, EPI_ISL_910957, EPI_ISL_910958, EPI_ISL_910959, EPI_ISL_910960, EPI_ISL_910961, EPI_ISL_910962, EPI_ISL_910963, EPI_ISL_910964, EPI_ISL_910965, EPI_ISL_910966, EPI_ISL_910967, EPI_ISL_910968, EPI_ISL_910969, EPI_ISL_910970, EPI_ISL_910971, EPI_ISL_910972, EPI_ISL_910973, EPI_ISL_910974, EPI_ISL_910975, EPI_ISL_910976, EPI_ISL_910977, EPI_ISL_910978, EPI_ISL_910979, EPI_ISL_910980, EPI_ISL_910981, EPI_ISL_910982, EPI_ISL_910983, EPI_ISL_910984, EPI_ISL_910985, EPI_ISL_910986, EPI_ISL_910987, EPI_ISL_910988, EPI_ISL_910989, EPI_ISL_910990, EPI_ISL_910991, EPI_ISL_910992, EPI_ISL_910993, EPI_ISL_910994, EPI_ISL_911003 | | | |
| see above | Laboratoire national de sante, Microbiology, Virology | Laboratoire national de sante, Microbiology, Microbial Genomics Platform | Anke Wienecke-Baldacchino, Catherine Ragimbeau,Jessica Tapp, Fatu Djabi, Lise Pignon, Raoul Salmon, Tamir Abdelrahman |
| EPI_ISL_911471, EPI_ISL_911472, EPI_ISL_911473, EPI_ISL_911474, EPI_ISL_911475, EPI_ISL_911476, EPI_ISL_911477, EPI_ISL_911478, EPI_ISL_911479, EPI_ISL_911480, EPI_ISL_911481, EPI_ISL_911482, EPI_ISL_911483, EPI_ISL_911484, EPI_ISL_911485, EPI_ISL_911486, EPI_ISL_911487, EPI_ISL_911488, EPI_ISL_911489, EPI_ISL_911490, EPI_ISL_911491, EPI_ISL_911492, EPI_ISL_911493 | | | |
| see above | Houston Health Dept. | Houston Health Dept. | Ryker Penn, Pamela Brown, Adolpho Lara |
| EPI_ISL_911515, EPI_ISL_911516, EPI_ISL_911517 | Ohio Department of Health Laboratory | Ohio Department of Health Laboratory | Holmes, Jennifer; Eric Brandt, Keoni Omura, Glen McGillivary, Caitlin McDonnell, Jade Mowery, Stephanie Mccracken, Tyler Payne, Kirtana Ramadugu, Erica Leasure, Brent Lee, Kelsey Florek, Heather Blankenship, Quanta Brown, and Tammy Bannerman |
| EPI_ISL_911694, EPI_ISL_911698 | Alaska State Virology Laboratory | Alaska State Virology Laboratory | Stephanie DeRonde, Lisa Smith, Ph.D., Jack Chen, Ph.D. |
| EPI_ISL_912357, EPI_ISL_912358, EPI_ISL_912359, EPI_ISL_912360, EPI_ISL_912391 | Fondation Congolaise pour la recherche medicale (FCRM), Francine Ntoumi | NGS Competence Center Tuebingen, Institut für Medizinische Mikrobiologie und Hygiene, Universitaetsklinikum Tübingen | Angel Angelov |
| EPI_ISL_912397 | Institute for Medical Research, Infectious Disease Research Centre, National Institutes of Health, Ministry of Health Malaysia | Institute for Medical Research, Infectious Disease Research Centre, National Institutes of Health, Ministry of Health Malaysia | Suppiah J, Kamel K, Azizan MA, Thayan R |
| EPI_ISL_912498, EPI_ISL_912512 | NHLS Universitas Academic | UFS Virology | PA Bester, MM Nyaga, P Nthiga, MT Mogotsi, D Goedhals, T de Oliveira |
| EPI_ISL_913348 | Klinisk mikrobiologi | The Public Health Agency of Sweden | Anna-Malin Linde, Maria Lind Karlberg, Carlo Berg, Oskar Karlsson Lindsjo, Sofia Stamouli, Reza Advani, Mattias Haukland, Petra Holmstrom, Noura Walai, Petra Edquist, Mia Brytting, Anna Risberg, Karin Tegmark-Wisell |
| EPI_ISL_913671, EPI_ISL_913673, EPI_ISL_913674, EPI_ISL_913675 | Vault Health | Minnesota Department of Health, Public Health Laboratory | Alexandra Lorentz, Jacob Garfin, Matt Plumb, and Xiong Wang |
| EPI_ISL_913944, EPI_ISL_913946, EPI_ISL_913947, EPI_ISL_913948, EPI_ISL_913949, EPI_ISL_913951, EPI_ISL_913963, EPI_ISL_913964, EPI_ISL_913976, EPI_ISL_913977, EPI_ISL_913978, EPI_ISL_913981 | | | |
| see above | Instituto de Diagnostico y Referencia Epidemiologicos INDRE_RNLSP | Instituto de Diagnostico y Referencia Epidemiologicos (INDRE) | Claudia Wong-Arambula, Abril Rodriguez-Maldonado, Fabiola Garces-Ayala, Adnan Araiza-Rodriguez, David Fragoso-Fonseca, Sergio Rangel-Guerrero, Mayra Jimenez-Morales, Nancy Munoz-Hernandez, Natividad Cruz-Ortiz, Tatiana Nunez-Garcia, Gisela Barrera-Badillo, Lucia Hernandez-Rivas, Irma Lopez-Martinez, Ernesto Ramirez-Gonzalez. |
| EPI_ISL_914715 | Maryland Public Health Laboratory (MD PHL) | Maryland Public Health Laboratory (MD PHL) | Maryland Department of Health Laboratories Administration |
| EPI_ISL_914799, EPI_ISL_914800 | AREA DE SALUD SAN FRANCISCO-SAN ANTONIO (COOPESANA) | Inciensa, Instituto Costarricense de Investigación y Enseñanza en Nutrición y Salud | Francisco Duarte, Hebleen Porras, Claudio Soto-Garita, Estela Cordero, Adriana Godínez, Melany Calderón & Mariel López |
| EPI_ISL_914885 | Vilnius university hospital Santaros Klinikos, Center of Laboratory Medicine | Vilnius University Hospital Santaros Klinikos | Ingrida Olendraite, Daniel Naumovas, Rimvydas Norvilas, Dovil Ežerskyt, Justinas Šlikas |
| EPI_ISL_915359, EPI_ISL_915379, EPI_ISL_915381 | Keio University School of Medicine | Keio University School of Medicine | Kenjiro Kosaki, Yuka Iwasaki, Hirotsugu Ishizu, Haruhiko Siomi, Kodai Abe |
| EPI_ISL_918169 | Department of Infectious Diseases and Immunology, National Hospital Organization Nagoya Medical Center | Clinical Research Center, National Hospital Organization Nagoya Medical Center | Yoshihiro Nakata, Hirotaka Ode, Mai Kubota, Masakazu Matsuda, Kazuhiro Matsuoka, Miho Nakasuji, Mikiko Mori, Mayumi Imahashi, Yoshiyuki Yokomaku, Yasumasa Iwatani |
| EPI_ISL_918579, EPI_ISL_918580, EPI_ISL_918581, EPI_ISL_918582, EPI_ISL_918599, EPI_ISL_918603 | University of Birmingham | COVID-19 Genomics UK (COG-UK) Consortium | Institute of Microbiology, University of Birmingham: Claire McMurray, Joanne Stockton, Samuel Nicholls, Radoslaw Poplawski, Will Rowe, Josh Quick, Nicholas Loman. University of Birmingham Testing Laboratory: Celina M Whalley, Andrew Bosworth, Charlotte Poxon, Kasun Wanigasooriya, Oliver Pickles, Mike Kidd, Alex Richter, Andrew D Beggs PHE Heartlands Lab: Husam Osman, Andrew Bosworth. Queen Elizabeth Hospital: Anna Casey |
| EPI_ISL_919233, EPI_ISL_919235, EPI_ISL_919237, EPI_ISL_919252, EPI_ISL_919261, EPI_ISL_919266, EPI_ISL_919267 | West of Scotland Specialist Virology Centre, NHSGGC / MRC-University of Glasgow Centre for Virus Research | COVID-19 Genomics UK (COG-UK) Consortium | Ana da Silva Filipe, Natasha Johnson, Kathy Smollett, Daniel Mair, Stephen Carmichael, Alice Broos, Lily Tong, Jenna Nichols, Kyriaki Nomikou; Sarah McDonald; Richard Orton, Joseph Hughes, Sreenu Vattipally, David L Robertson; Alasdair MacLean, Rory Gunson; Sharif Shaaban, Matthew Holden; Rachel Blacow, Guy Mollett, Kathy Li, James Shepherd, Antonia Ho, Emma Thomson |
| EPI_ISL_919311, EPI_ISL_919312, EPI_ISL_919367, EPI_ISL_919380, EPI_ISL_919381, EPI_ISL_919388, EPI_ISL_919389, EPI_ISL_919390, EPI_ISL_919391, EPI_ISL_919392, EPI_ISL_919393, EPI_ISL_919396, EPI_ISL_919398 | | | |
| see above | Virology Department, Royal Infirmary of Edinburgh, NHS Lothian / School of Biological Sciences, University of Edinburgh / Institute of Genetics and Molecular Medicine, University of Edinburgh | COVID-19 Genomics UK (COG-UK) Consortium | McHugh M, Dewar R, Rooke S, Gallagher M, Balcaza C, O'Toole Á, Scher E, Hill V, McCrone JT, Colquhoun R, Yu X, Jackson B, Rambaut A, Williams TC, Templeton K |
| EPI_ISL_920267, EPI_ISL_920277, EPI_ISL_920322, EPI_ISL_920365, EPI_ISL_920372, EPI_ISL_920459, EPI_ISL_920460, EPI_ISL_920516, EPI_ISL_920576, EPI_ISL_920595 | University College London Hospital | COVID-19 Genomics UK (COG-UK) Consortium | Judith Heaney, Matthew Byott, Catherine Houlihan, Dan Frampton, Stuart Kirk, Moira Spyer and Eleni Nastouli |
| EPI_ISL_922894 | Wales Specialist Virology Centre Sequencing lab: Pathogen Genomics Unit | Public Health Wales Microbiology Cardiff Wales Specialist Virology Centre | Catherine Moore, Johnathan Evans, Laura Gifford, Malorie Perry, Simon Cottrell, Angela Marchbank, Alec Birchley, Alexander Adams, Amy Gaskin, Bree Gatica-Wilcox, Jason Coombes, Joel Southgate, Lauren Gilbert, Lee Graham, Nicole Pacchiarini, Sara Kumziene-Summerhayes, Sarah Taylor, Sophie Jones, Sara Rey, Matthew Bull, Joanne Watkins, Sally Corden, Tom Connor |
| EPI_ISL_925853, EPI_ISL_925857, EPI_ISL_925860, EPI_ISL_925861, EPI_ISL_925862, EPI_ISL_925863, EPI_ISL_925864, EPI_ISL_930567 | Nucleic Acid Testing, National Reference Laboratory | GIGA Medical Genomics | Yvan Butera, Keith Durkin, Maria Artesi, Bouchra Boujemla, Robert Rutayisire, Patrick Tuyisenge, Esperence Umumararungu, Sébastien Bontems, Marie-Pierre Hayette, Nathalie Renotte, Swaibu Gatare, Jacob Souopgui, Sabin Nsanzimana, Vincent Bours, Léon Mutesa |
| EPI_ISL_931393, EPI_ISL_931450 | University Hospital Basel, Clinical Virology | University Hospital Basel, Clinical Bacteriology | Tim Roloff, Madlen Stange, Helena MB Seth-Smith, Alfredo Mari, Karoline Leuzinger, Julia Bielicki, Manuel Battegay, Hans Hirsch, Adrian Egli |

| | | | |
|---|---|---|---|
| EPI_ISL_933482, EPI_ISL_933483, EPI_ISL_933484, EPI_ISL_933485, EPI_ISL_933486, EPI_ISL_933487, EPI_ISL_933488, EPI_ISL_933489, EPI_ISL_933490, EPI_ISL_933491 | Lighthouse Lab in Milton Keynes | Wellcome Sanger Institute for the COVID-19 Genomics UK (COG-UK) Consortium | The Lighthouse Lab in Milton Keynes and Alex Alderton, Roberto Amato, Sonia Goncalves, Ewan Harrison, David K. Jackson, Ian Johnston, Dominic Kwiatkowski, Cordelia Langford, John Sillitoe on behalf of the Wellcome Sanger Institute COVID-19 Surveillance Team |
| EPI_ISL_933492 | Lighthouse Lab in Glasgow | Wellcome Sanger Institute for the COVID-19 Genomics UK (COG-UK) Consortium | Harper VanSteenhouse, Yumi Kasai, David Gray, Carol Clugston, Anna Dominiczak and Alex Alderton, Roberto Amato, Sonia Goncalves, Ewan Harrison, David K. Jackson, Ian Johnston, Dominic Kwiatkowski, Cordelia Langford, John Sillitoe on behalf of the Wellcome Sanger Institute COVID-19 Surveillance Team |
| EPI_ISL_933669 | Instituto de Diagnostico y Referencia Epidemiologicos INDRE_RNLSP | Instituto de Diagnostico y Referencia Epidemiologicos (INDRE) | Claudia Wong-Arambula, Abril Rodriguez-Maldonado, Fabiola Garces-Ayala, Adnan Araiza-Rodriguez, David Fragoso-Fonseca, Sergio Rangel-Guerrero, Mayra Jimenez-Morales, Nancy Munoz-Hernandez, Natividad Cruz-Ortiz, Tatiana Nunez-Garcia, Gisela Barrera-Badillo, Lucia Hernandez-Rivas, Irma Lopez-Martinez, Ernesto Ramirez-Gonzalez. |
| EPI_ISL_934061, EPI_ISL_934062, EPI_ISL_934063, EPI_ISL_934064, EPI_ISL_934065, EPI_ISL_934066, EPI_ISL_934067, EPI_ISL_934068, EPI_ISL_934069, EPI_ISL_934070, EPI_ISL_934071, EPI_ISL_934072, EPI_ISL_934073, EPI_ISL_934074, EPI_ISL_934075, EPI_ISL_934076, EPI_ISL_934077, EPI_ISL_934078, EPI_ISL_934079, EPI_ISL_934080, EPI_ISL_934081, EPI_ISL_934082, EPI_ISL_934083, EPI_ISL_934084, EPI_ISL_934085, EPI_ISL_934086, EPI_ISL_934087, EPI_ISL_934088, EPI_ISL_934089, EPI_ISL_934090, EPI_ISL_934091, EPI_ISL_934093, EPI_ISL_934094, EPI_ISL_934095, EPI_ISL_934098, EPI_ISL_934110, EPI_ISL_934111, EPI_ISL_934118, EPI_ISL_934125, EPI_ISL_934126, EPI_ISL_934137, EPI_ISL_934167 | | | |
| see above | Vilnius university hospital Santaros Klinikos, Center of Laboratory Medicine | Vilnius university hospital Santaros Klinikos, Center of Laboratory Medicine | Ingrida Olendraite, Daniel Naumovas, Rimvydas Norvilas, Dovile Ezerskyte, Justinas Slikas, Gytis Dudas |
| EPI_ISL_934325, EPI_ISL_934326, EPI_ISL_934339 | Klinisk mikrobiologi | The Public Health Agency of Sweden | Anna-Malin Linde, Maria Lind Karlberg, Carlo Berg, Oskar Karlsson Lindsjo, Sofia Stamouli, Reza Advani, Mattias Haukland, Petra Holmstrom, Noura Walai, Petra Edquist, Mia Brytting, Anna Risberg, Karin Tegmark-Wisell |
| EPI_ISL_935240 | KU Leuven, Rega Institute, Clinical and Epidemiological Virology | KU Leuven, Rega Institute, Clinical and Epidemiological Virology | Tony Wawina-Bokalanga, Bert Vanmechelen, Joan Marti-Carerras, Piet Maes |
| EPI_ISL_936182, EPI_ISL_936183, EPI_ISL_936184, EPI_ISL_936185, EPI_ISL_936186, EPI_ISL_936187, EPI_ISL_936188, EPI_ISL_936189, EPI_ISL_936190, EPI_ISL_936191, EPI_ISL_936192, EPI_ISL_936193, EPI_ISL_936194, EPI_ISL_936195, EPI_ISL_936196, EPI_ISL_936197, EPI_ISL_936198, EPI_ISL_936199, EPI_ISL_936200, EPI_ISL_936201, EPI_ISL_936202, EPI_ISL_936203, EPI_ISL_936204, EPI_ISL_936205, EPI_ISL_936206, EPI_ISL_936207, EPI_ISL_936208, EPI_ISL_936209, EPI_ISL_936210, EPI_ISL_936211 | | | |
| see above | Houston Health Dept. | Houston Health Dept. | Ryker Penn, Pamela Brown, Adolpho Lara |
| EPI_ISL_936642, EPI_ISL_936643 | Northwestern Memorial Hospital | Ozer Lab | Ramon Lorenzo-Redondo, Lacy M. Simons, Chad J. Achenbach, Lawrence J. Jennings, Michael G. Ison, Judd F. Hultquist, Egon A. Ozer |
| EPI_ISL_937291, EPI_ISL_937292, EPI_ISL_937293, EPI_ISL_937294, EPI_ISL_937295, EPI_ISL_937296, EPI_ISL_937297, EPI_ISL_937298, EPI_ISL_937299, EPI_ISL_937300, EPI_ISL_937301, EPI_ISL_937302, EPI_ISL_937304, EPI_ISL_937305, EPI_ISL_937306, EPI_ISL_937307, EPI_ISL_937308, EPI_ISL_937309, EPI_ISL_937310, EPI_ISL_937319, EPI_ISL_937320, EPI_ISL_937321, EPI_ISL_937322, EPI_ISL_937323, EPI_ISL_937324, EPI_ISL_937325, EPI_ISL_937327, EPI_ISL_937328, EPI_ISL_937329, EPI_ISL_937339, EPI_ISL_937340, EPI_ISL_937343, EPI_ISL_937344 | | | |
| see above | Utah Public Health Laboratory | Utah Public Health Laboratory | Erin L. Young, Kelly F. Oakeson, Tara Gallagher |
| EPI_ISL_940075, EPI_ISL_940076, EPI_ISL_940077, EPI_ISL_940087, EPI_ISL_940093, EPI_ISL_940097, EPI_ISL_940108, EPI_ISL_940109, EPI_ISL_940112, EPI_ISL_940113, EPI_ISL_940116, EPI_ISL_940118, EPI_ISL_940120, EPI_ISL_940127, EPI_ISL_940130, EPI_ISL_940135, EPI_ISL_940136 | | | |
| see above | Charlotte Maxeke Johannesburg Academic Hospital, National Health Laboratory Services, Gauteng, South Africa | National Institute for Communicable Diseases of the National Health Laboratory Service | Amoako DG, Mohale T, Ntuli N, Mahlangu B, Allam M, Ismail A, Bhiman JN |
| EPI_ISL_940879 | Vaccines and Infectious Diseases Analytics Research Unit (VIDA) | KRISP, KZN Research Innovation and Sequencing Platform | Baillie Vicky, du Plessis Jeanine, Giandhari Jennifer, Pillay Sureshnee, Naidoo Yeshnee, Tegally Houriiyah, de Oliveira Tulio, Madhi Shabir |
| EPI_ISL_941209, EPI_ISL_941210 | Servicio de Microbiología. Hospital Clínico Universitario de Valencia | SeqCOVID-SPAIN consortium/IBV(CSIC) | David Navarro Ortega, Eliseo Albert Vicent, Ignacio Torres and SeqCOVID-SPAIN consortium |
| EPI_ISL_941251, EPI_ISL_941252, EPI_ISL_941253, EPI_ISL_941254 | Virginia DCLS | Virginia DCLS | Virginia DCLS |
| EPI_ISL_942185, EPI_ISL_942204, EPI_ISL_942205, EPI_ISL_942209, EPI_ISL_942211, EPI_ISL_942213, EPI_ISL_942215, EPI_ISL_942216, EPI_ISL_942217, EPI_ISL_942218, EPI_ISL_942219, EPI_ISL_942220, EPI_ISL_942221, EPI_ISL_942222, EPI_ISL_942223, EPI_ISL_942224, EPI_ISL_942225, EPI_ISL_942226, EPI_ISL_942227, EPI_ISL_942228, EPI_ISL_942229, EPI_ISL_942231, EPI_ISL_942236, EPI_ISL_942237, EPI_ISL_942238, EPI_ISL_942239, EPI_ISL_942240, EPI_ISL_942241, EPI_ISL_942242, EPI_ISL_942243, EPI_ISL_942244, EPI_ISL_942245, EPI_ISL_942246, EPI_ISL_942247 | | | |
| see above | Wisconsin State Laboratory of Hygiene Communicable Disease Division | Wisconsin State Laboratory of Hygiene Communicable Disease Division | Kelsey R. Florek, Abigail C. Shockey |
| EPI_ISL_942658, EPI_ISL_942659, EPI_ISL_942662, EPI_ISL_942663, EPI_ISL_942664, EPI_ISL_942665, EPI_ISL_942666, EPI_ISL_942667, EPI_ISL_942668, EPI_ISL_942669, EPI_ISL_942670, EPI_ISL_942671, EPI_ISL_942672, EPI_ISL_942673, EPI_ISL_942674, EPI_ISL_942675, EPI_ISL_942676, EPI_ISL_942677, EPI_ISL_942678, EPI_ISL_942679, EPI_ISL_942680, EPI_ISL_942681, EPI_ISL_942682, EPI_ISL_942683, EPI_ISL_942684, EPI_ISL_942685, EPI_ISL_942686, EPI_ISL_942687, EPI_ISL_942688, EPI_ISL_942689, EPI_ISL_942942 | | | |
| see above | Gundersen Molecular Diagnostics Laboratory | Kabara Cancer Research Institute | Craig S. Richmond, Paraic A. Kenny |
| EPI_ISL_942966 | General Hospital - Kumanovo | Research Center for Genetic Engineering and Biotechnology "Georgi D. Efremov" , Macedonian Academy of Sciences and Arts | Aleksandar J.Dimovski, Dijana Plasheska-Karanfilska, Predrag Noveski, Gjorgji Bozinovski, Milena Jakimovska |
| EPI_ISL_942967 | General Hospital - Kumanovo | Research Center for Genetic Engineering and Biotechnology "Georgi D. Efremov" , Macedonian Academy of Sciences and Arts | Aleksandar J. Dimovski, Dijana Plasheska-Karanfilska, Predrag Noveski, Gjorgji Bozinovski, Milena Jakimovska |
| EPI_ISL_943667, EPI_ISL_943756, EPI_ISL_943757, EPI_ISL_943758, EPI_ISL_943759, EPI_ISL_943760, EPI_ISL_943761, EPI_ISL_943762, EPI_ISL_943774, EPI_ISL_943775, EPI_ISL_943776, EPI_ISL_943777, EPI_ISL_943778, EPI_ISL_943779, EPI_ISL_943780, EPI_ISL_943781, EPI_ISL_943782, EPI_ISL_943783, EPI_ISL_943784, EPI_ISL_943785, EPI_ISL_943786, EPI_ISL_943787, EPI_ISL_943788 | | | |
| see above | Utah Public Health Laboratory | Utah Public Health Laboratory | Erin L. Young, Kelly F. Oakeson, Tara Gallagher |
| EPI_ISL_945255 | Lighthouse Lab in Glasgow | Wellcome Sanger Institute for the COVID-19 Genomics UK (COG-UK) Consortium | Harper VanSteenhouse, Yumi Kasai, David Gray, Carol Clugston, Anna Dominiczak and Alex Alderton, Roberto Amato, Sonia Goncalves, Ewan Harrison, David K. Jackson, Ian Johnston, Dominic Kwiatkowski, Cordelia Langford, John Sillitoe on behalf of the Wellcome Sanger Institute COVID-19 Surveillance Team |
| EPI_ISL_947298 | RSU Medirossa | Eijkman Institute for Molecular Biology, Ministry of Research and Technology/National Agency for Research and Innovation | Iskandar Adnan, Lydia V. Panggalo, Sukma Oktavianthi, Willy Agustine, Edison Johar, Hidayat Trimarsanto, Frilasita A Yudhaputri, Safarina G Malik, Khin Saw Myint, Amin Soebandrio |
| EPI_ISL_947301 | RS Mitra Keluarga Cibubur | Eijkman Institute for Molecular Biology, Ministry of Research and Technology/National Agency for Research and Innovation | Iskandar Adnan, Lydia V. Panggalo, Sukma Oktavianthi, Willy Agustine, Edison Johar, Hidayat Trimarsanto, Frilasita A Yudhaputri, Safarina G Malik, Khin Saw Myint, Amin Soebandrio |
| EPI_ISL_947304 | RSU Harapan Bunda | Eijkman Institute for Molecular Biology, Ministry of Research and Technology/National Agency for Research and Innovation | Iskandar Adnan, Lydia V. Panggalo, Sukma Oktavianthi, Willy Agustine, Edison Johar, Hidayat Trimarsanto, Frilasita A Yudhaputri, Safarina G Malik, Khin Saw Myint, Amin Soebandrio |
| EPI_ISL_949396, EPI_ISL_949397, EPI_ISL_949398 | University of Birmingham | COVID-19 Genomics UK (COG-UK) Consortium | Institute of Microbiology, University of Birmingham: Claire McMurray, Joanne Stockton, Samuel Nicholls, Radoslaw Poplawski, Will Rowe, Josh Quick, Nicholas Loman. University of Birmingham Testing Laboratory: Celina M Whalley, Andrew Bosworth, Charlotte Poxon, Kasun Wanigasooriya, Oliver Pickles, Mike Kidd, Alex Richter, Andrew D Beggs PHE Heartlands Lab: Husam Osman, Andrew Bosworth. Queen Elizabeth Hospital: Anna Casey |
| EPI_ISL_949618, EPI_ISL_949622, EPI_ISL_949628 | NHLS Universitas Academic | UFS Virology | PA Bester, MM Nyaga, P Nthiga, MT Mogotsi, Emmanuel Ogunbayo, D Goedhals, T de Oliveira |
| EPI_ISL_949786 | University College London, Great Ormond Street Hospital for Children NHS Foundation Trust, Imperial College Healthcare NHS Trust | COVID-19 Genomics UK (COG-UK) Consortium | Sergi Castellano, Rachel Williams, Mark Kristiansen, Paola Resende Silva, Sunando Roy, Tony Brooks, Helena Tutill, Paola Niola, Patricia Dyal, Charlotte Williams, Leysa Forrest, Yasmin Panchbhaya, Jacqueline Findlay, Samuel Weeks, Julianne Brown, Kathryn Harris, Paul Randell, James Price, Alison Holmes, Judith Breuer |
| EPI_ISL_950699 | Lincolnshire Hospitals and DeepSeq Nottingham | COVID-19 Genomics UK (COG-UK) Consortium | Nichola Duckworth, Tim Sloan, Sarah Walsh, Jonathan Ball, Patrick McClure, Joeseph Chappell, Nadine Holmes, Matthew Carlisle, Christopher Moore, Fei Sang, Johnny Debebe, Victoria Wright, Matthew Loose |

| | | | |
|---|---|---|---|
| EPI_ISL_955362, EPI_ISL_955363, EPI_ISL_955364, EPI_ISL_955365, EPI_ISL_955366, EPI_ISL_955367, EPI_ISL_955368, EPI_ISL_955369, EPI_ISL_955370, EPI_ISL_955373, EPI_ISL_955374, EPI_ISL_955378, EPI_ISL_955379, EPI_ISL_955380, EPI_ISL_955381, EPI_ISL_955382, EPI_ISL_955383, EPI_ISL_955384, EPI_ISL_955385 | | | |
| see above | Santa Clara County Public Health Laboratory | Chan-Zuckerberg Biohub | CZB Cliahub Consortium |
| EPI_ISL_955665, EPI_ISL_955667, EPI_ISL_955668, EPI_ISL_955669, EPI_ISL_955671, EPI_ISL_955672, EPI_ISL_955673 | Orange County Public Health Lab | Chan-Zuckerberg Biohub | CZB Cliahub Consortium |
| EPI_ISL_960414 | The National Institute of Public Health | State Veterinary Institute Prague | Nagy,A;Vecerova,J;Cernikova,L;Stara,M;Jirincova,H;Trnka,D |
| EPI_ISL_960452, EPI_ISL_960456, EPI_ISL_960468, EPI_ISL_960475, EPI_ISL_960476, EPI_ISL_960482, EPI_ISL_960496, EPI_ISL_960497, EPI_ISL_960516, EPI_ISL_960517, EPI_ISL_960518, EPI_ISL_960519, EPI_ISL_960520, EPI_ISL_960521, EPI_ISL_960522, EPI_ISL_960523, EPI_ISL_960524, EPI_ISL_960525, EPI_ISL_960574, EPI_ISL_960575, EPI_ISL_960591, EPI_ISL_960607, EPI_ISL_960608, EPI_ISL_960609, EPI_ISL_960610, EPI_ISL_960611, EPI_ISL_960612, EPI_ISL_960613 | | | |
| see above | Istituto Zooprofilattico Sperimentale del Mezzogiorno | TIGEM | Patrizia Annunziata, Andrea Ballabio, Valentina Bouche, Davide Cacchiarelli, Pellegrino Cerino, Chiara Colantuono, Maria Concetta Cuomo, Denise Di Concilio, Lucio Di Filippo, Antonio Grimaldi, Antonio Limone, Anna Manfredi, Francesco Panariello, Biancamaria Pierri, Marcello Salvi |
| EPI_ISL_961380 | Toronto Invasive Bacterial Diseases Network | McMaster University | Allison McGeer, Patryk Aftanas, Hooman Derakhshani, Angel Li, Kuganya Nirmalarajah, Emily Panousis, Ahmed Draia, Jalees Nasir, Michael Surette, Samira Mubareka, Andrew G. McArthur |
| EPI_ISL_961901, EPI_ISL_961924, EPI_ISL_961939, EPI_ISL_961983, EPI_ISL_962080, EPI_ISL_962081, EPI_ISL_962083, EPI_ISL_962085, EPI_ISL_962086, EPI_ISL_962087, EPI_ISL_962088, EPI_ISL_962089, EPI_ISL_962090, EPI_ISL_962092, EPI_ISL_962093, EPI_ISL_962094, EPI_ISL_962095, EPI_ISL_962096, EPI_ISL_962097, EPI_ISL_962098, EPI_ISL_962099, EPI_ISL_962100 | | | |
| see above | Illinois Department of Public Health | Gagnon Lab, Southern Illinois University | Keith Gagnon |
| EPI_ISL_962766, EPI_ISL_962767, EPI_ISL_962769, EPI_ISL_962772, EPI_ISL_962773, EPI_ISL_962774, EPI_ISL_962778, EPI_ISL_962779, EPI_ISL_962780, EPI_ISL_962785, EPI_ISL_962786, EPI_ISL_962788, EPI_ISL_962791, EPI_ISL_962792, EPI_ISL_962793, EPI_ISL_962796, EPI_ISL_962797, EPI_ISL_962801, EPI_ISL_962802, EPI_ISL_962803, EPI_ISL_962808, EPI_ISL_962809 | | | |
| see above | San Diego County Public Health Laboratory | Andersen lab at Scripps Research | SEARCH Alliance San Diego with Tracy Basler, Jovan Shephard, Brett Austin |
| EPI_ISL_962973 | Hospital Universitario de Gran Canaria Dr. Negrín | SeqCOVID-SPAIN consortium/IBV(CSIC) | M. Carmen Pérez González, Francisco J. Chamizo López, Ana Bordes Benítez and SeqCOVID-SPAIN consortium |
| EPI_ISL_965187, EPI_ISL_965190, EPI_ISL_965221 | Virginia Division of Consolidated Laboratory Services | Virginia Division of Consolidated Laboratory Services | Virginia DCLS |
| EPI_ISL_965538, EPI_ISL_965575, EPI_ISL_965576, EPI_ISL_965599, EPI_ISL_965631, EPI_ISL_965634, EPI_ISL_965635, EPI_ISL_965651, EPI_ISL_965660, EPI_ISL_965664, EPI_ISL_965673, EPI_ISL_965697, EPI_ISL_965698, EPI_ISL_965707, EPI_ISL_965713, EPI_ISL_965740, EPI_ISL_965750, EPI_ISL_965754, EPI_ISL_965789, EPI_ISL_965796, EPI_ISL_965797, EPI_ISL_965803 | | | |
| see above | Dutch COVID-19 response team | Medical Microbiology, Maastricht University Medical Centre | Jozef Dingemans*, Brian van der Veer*, Erik Beuken, Carmen Reumkens, Lieke van Alphen, Christian Hoebe, Paul Savelkoul |
| EPI_ISL_966756 | Maine HETL | Tewhey Lab, The Jackson Laboratory | Matluk,N., Dewey,H., Iosue,F., Barter,M., Lynch,R., Munger,H. and Tewhey,R. |
| EPI_ISL_967525 | State Laboratories Division, Hawaii State Department of Health | State Laboratories Division, Hawaii State Department of Health | Pamela O'Brien, Drew Kuwazaki, Ayana Garnet, Razvan Sultana, Edward Desmond |
| EPI_ISL_968822, EPI_ISL_968825, EPI_ISL_968826, EPI_ISL_968827, EPI_ISL_968828, EPI_ISL_968829, EPI_ISL_968844, EPI_ISL_968846, EPI_ISL_968847, EPI_ISL_968849 | KEMRI-Wellcome Trust Research Programme/KEMRI-CGMR-C Kilifi | KEMRI-Wellcome Trust Research Programme/KEMRI-CGMR-C Kilifi | Githinji et al |
| EPI_ISL_976301, EPI_ISL_976302, EPI_ISL_976303, EPI_ISL_976304, EPI_ISL_976306, EPI_ISL_976307, EPI_ISL_976308, EPI_ISL_976309, EPI_ISL_976310, EPI_ISL_976311, EPI_ISL_976312, EPI_ISL_976313, EPI_ISL_976314, EPI_ISL_976315, EPI_ISL_976316, EPI_ISL_976317, EPI_ISL_976318, EPI_ISL_976319, EPI_ISL_976320, EPI_ISL_976321, EPI_ISL_976322, EPI_ISL_976323, EPI_ISL_976324, EPI_ISL_976325, EPI_ISL_976326, EPI_ISL_976327, EPI_ISL_976328, EPI_ISL_976329, EPI_ISL_976330, EPI_ISL_976331, EPI_ISL_976332, EPI_ISL_976333, EPI_ISL_976334, EPI_ISL_976335, EPI_ISL_976336, EPI_ISL_976337, EPI_ISL_976338, EPI_ISL_976339, EPI_ISL_976340, EPI_ISL_976341, EPI_ISL_976342, EPI_ISL_976343, EPI_ISL_976344, EPI_ISL_976345, EPI_ISL_976346, EPI_ISL_976347, EPI_ISL_976348, EPI_ISL_976349, EPI_ISL_976350, EPI_ISL_976351, EPI_ISL_976352, EPI_ISL_976353, EPI_ISL_976354, EPI_ISL_976355, EPI_ISL_976356, EPI_ISL_976357, EPI_ISL_976358, EPI_ISL_976359, EPI_ISL_976360, EPI_ISL_976361, EPI_ISL_976362, EPI_ISL_976363, EPI_ISL_976364, EPI_ISL_976365, EPI_ISL_976366, EPI_ISL_976367, EPI_ISL_976368, EPI_ISL_976369, EPI_ISL_976370, EPI_ISL_976371, EPI_ISL_976372, EPI_ISL_976373, EPI_ISL_976374, EPI_ISL_976375, EPI_ISL_976376, EPI_ISL_976377, EPI_ISL_976378, EPI_ISL_976379, EPI_ISL_976380, EPI_ISL_976381, EPI_ISL_976382, EPI_ISL_976383, EPI_ISL_976384, EPI_ISL_976385, EPI_ISL_976386, EPI_ISL_976387, EPI_ISL_976388, EPI_ISL_976389, EPI_ISL_976390, EPI_ISL_976391, EPI_ISL_976392, EPI_ISL_976393, EPI_ISL_976394, EPI_ISL_976395, EPI_ISL_976396, EPI_ISL_976397, EPI_ISL_976398, EPI_ISL_976399, EPI_ISL_976400, EPI_ISL_976401, EPI_ISL_976402, EPI_ISL_976403, EPI_ISL_976404, EPI_ISL_976405, EPI_ISL_976406, EPI_ISL_976407, EPI_ISL_976408, EPI_ISL_976409, EPI_ISL_976410, EPI_ISL_976411, EPI_ISL_976412, EPI_ISL_976413, EPI_ISL_976414, EPI_ISL_976415, EPI_ISL_976416, EPI_ISL_976417, EPI_ISL_976418, EPI_ISL_976419, EPI_ISL_976420, EPI_ISL_976421, EPI_ISL_976422, EPI_ISL_976423 | | | |
| see above | BCCDC Public Health Laboratory | BCCDC Public Health Laboratory | Prystajecky Natalie, Linda Hoang, Dan Fornika, John Tyson, Shannon Russell, Kim Macdonald, Kimia Kamelian, Ana Pacagnella, Corrinne Ng, Loretta Janz, Robert Azana Terry Snutch, Mel Krajden |
| EPI_ISL_977028 | Rhode Island Department of Health | Infectious Disease Program, Broad Institute of Harvard and MIT | Lemieux,J.E., Siddle,K.J., Huard,R., King,E., Azevedo,K., Miller,A., Adams,G., Gladden-Young,A., Lagerborg,K., Rudy,M., DeRuff,K., Carter,A., Normandin,E., Bauer,M., Reilly,S., Tomkins-Tinch,C., Loreth,C., Chaluvadi,S., Birren,B.W., Gallagher,G., Smole,S., Park,D.J., MacInnis,B.L., and Sabeti,P.C. |
| EPI_ISL_977162, EPI_ISL_977167, EPI_ISL_977168, EPI_ISL_977169 | Microbiologia e Virologia | Istituto Zooprofilattico Sperimentale delle Venezie | Adelaide Milani, Alessia Schivo, Annalisa Salviato, Erika Giorgia Quaranta, Ambra Pastori, Bianca Zecchin, Alice Fusaro, Isabella Monne, Calogero Terregino, Antonia Ricci |
| EPI_ISL_977222, EPI_ISL_977223 | ULSS 1 Dolomiti | Istituto Zooprofilattico Sperimentale delle Venezie | Adelaide Milani, Alessia Schivo, Annalisa Salviato, Erika Giorgia Quaranta, Ambra Pastori, Bianca Zecchin, Alice Fusaro, Isabella Monne, Calogero Terregino, Antonia Ricci |
| EPI_ISL_977247, EPI_ISL_977248 | ULSS 03 Venezia | Istituto Zooprofilattico Sperimentale delle Venezie | Adelaide Milani, Alessia Schivo, Annalisa Salviato, Erika Giorgia Quaranta, Ambra Pastori, Bianca Zecchin, Alice Fusaro, Isabella Monne, Calogero Terregino, Antonia Ricci |
| EPI_ISL_978193 | Virginia Division of Consolidated Laboratory Services | Virginia Division of Consolidated Laboratory Services | Virginia DCLS |
| EPI_ISL_979017, EPI_ISL_979018 | Santa Clara County Public Health Laboratory | Chan-Zuckerberg Biohub | CZB Cliahub Consortium |
| EPI_ISL_979162, EPI_ISL_979163, EPI_ISL_979164, EPI_ISL_979165, EPI_ISL_979166, EPI_ISL_979167, EPI_ISL_979168, EPI_ISL_979169, EPI_ISL_979170, EPI_ISL_979171, EPI_ISL_979172 | | | |
| see above | Humboldt County Public Health Laboratory | Chan-Zuckerberg Biohub | CZB Cliahub Consortium |
| EPI_ISL_979305, EPI_ISL_979311 | Cadham Provincial laboratory | National Microbiology Laboratory (NML) | Anna Majer, Shari Tyson, Grace Seo, Philip Mabon, Elsie Grudeski, Rhiannon Huzarewich, Russell Mandes, Anneliese Landgraff, Jennifer Tanner, Natalie Knox, Morag Graham, Gary Van Domselaar, Paul Van Caeseele, Jared Bullard, David Alexander, Kerry Dust, Nathalie Bastien, Yan Li, Timothy Booth, Darian Hole, Madison Chapel, Kirsten Biggar, CanCOGeN's metadata curation team, Public Health Agency of Canada CanCOGeN team |
| EPI_ISL_979329 | Laboratorio Estatal de Salud Pública de Nuevo León | Laboratorio de Infectología Molecular, Departamento de Bioquímica y Medicina Molecular,Facultad de Medicina - Universidad Autónoma de Nuevo León | Kame A. Galán-Huerta, María F. Herrera-Saldivar, Natalia Martínez-Acuña, Sonia A. Lozano-Sepúlveda, Daniel Arellanos-Soto, Ana M. Rivas-Estilla, Samuel Buentello-Wong, Else del Carmen García-García, Gloria A. Jasso-de-la-Peña, Roberto Montes-de-Oca, Consuelo Treviño-Garza, Manuel E. de-la-O-Cavazos |
| EPI_ISL_979797 | Humboldt County Public Health Laboratory | Chan-Zuckerberg Biohub | CZB Cliahub Consortium |
| EPI_ISL_981883, EPI_ISL_981884, EPI_ISL_981885, EPI_ISL_981886, EPI_ISL_981887, EPI_ISL_981896, EPI_ISL_981897, EPI_ISL_981898, EPI_ISL_981899, EPI_ISL_981900, EPI_ISL_981901, EPI_ISL_981902 | | | |
| see above | Microbiology Service, Hospital Universitario Clinico San Cecilio, Granada | Microbiology Service, Hospital Universitario Clinico San Cecilio, Granada | Adolfo de Salazar, Natalia Chueca, Laura Viñuela, Ana Fuentes, Federico García |
| EPI_ISL_982844 | Kentucky State Public Health Lab | Kentucky State Public Health Lab | Stephanie Lunn, Karim George, Joshua Tobias, William Grooms, Vaneet Arora, Matthew Johnson, Rachel Zinner, Rhonda Lucas |
| EPI_ISL_982845, EPI_ISL_982846 | UK Healthcare Clinical Microbiology | Kentucky State Public Health Lab | Stephanie Lunn, Karim George, Joshua Tobias, William Grooms, Vaneet Arora, Matthew Johnson, Rachel Zinner, Rhonda Lucas |
