## Supplementary material for "Detection and characterization of the SARS-CoV-2 lineage B.1.526 in New York": Supp. Table 3 GISAID Acknowledgment part 2: gisaid_hcov-19_acknowledgement_table_2021_02_13_00-16.pdf

We gratefully acknowledge the following Authors from the Originating laboratories responsible for obtaining the specimens, as well as the Submitting laboratories where the genome data were generated and shared via GISAID, on which this research is based.

All Submitters of data may be contacted directly via www.gisaid.org

Authors are sorted alphabetically.

| Accession ID | Originating Laboratory | Submitting Laboratory | Authors |
|---|---|---|---|
| EPI_ISL_708798 | Sydney South West Pathology Service (SSWPS) - Royal Prince Alfred Hospital - NSW Health Pathology | NSW Health Pathology - Institute of Clinical Pathology and Medical Research; Westmead Hospital; University of Sydney | CIDM-PH et al. |
| EPI_ISL_708799 | Pathology North - Royal North Shore Hospital - NSW Health Pathology | NSW Health Pathology - Institute of Clinical Pathology and Medical Research; Westmead Hospital; University of Sydney | CIDM-PH et al. |
| EPI_ISL_717701, EPI_ISL_717702 | Area of Virology, Serology and Virology Division (SAViD), New South Wales Health Pathology Randwick | Virology Research Laboratory; Area of Virology, Serology and Virology Division (SAViD), New South Wales Health Pathology Randwick | Foster, C.; Au, J.; Ruiz Silva, M.; Deveson, I.; Bull, R.; Van Hal, S.; Rawlinson, W. |
| EPI_ISL_717978 | Army Medical Center, Scientific Department, Virology Laboratory | Army Medical Center, Scientific Department, Virology Laboratory | Silvia Fillo, Giovanni Faggioni, Riccardo De Santis, Antonella Fortunato, Anna Anselmo, Vanessa Vera Fain, Francesco Giordani, Nino D'Amore, Anella Monte, Marzia Cavalli, Alessandra Amoroso, Stella Lia, Roberta Sorrentino, Rossella Tirelli, Federica Galeano, Annalisa Pelo, Margherita De Santis, Giulia Campoli, Andrea Ciammaruconi, Florigio Lista. |
| EPI_ISL_722357, EPI_ISL_722778, EPI_ISL_722779, EPI_ISL_722780, EPI_ISL_722781, EPI_ISL_722782, EPI_ISL_722783 | Dutch COVID-19 response team | Erasmus Medical Center | Bas Oude Munnink, Reina Sikkema, David Nieuwenhuijse, Irina Chestakova, Anne van der Linden, Marjan Boter, Emmanuelle Munger, Corine GeurtsvanKessel, Annemiek van der Eijk, Richard Molenkamp, Marion Koopmans, on behalf of the Dutch national COVID-19 response team. |
| EPI_ISL_724608, EPI_ISL_724609 | University College London, Great Ormond Street Hospital for Children NHS Foundation Trust, Imperial College Healthcare NHS Trust | COVID-19 Genomics UK (COG-UK) Consortium | Sergi Castellano, Rachel Williams, Mark Kristiansen, Paola Resende Silva, Sunando Roy, Tony Brooks, Helena Tutill, Paola Niola, Patricia Dyal, Charlotte Williams, Leysa Forrest, Yasmin Panchbhaya, Jacqueline Findlay, Samuel Weeks, Julianne Brown, Kathryn Harris, Paul Randell, James Price, Alison Holmes, Judith Breuer |
| EPI_ISL_727865, EPI_ISL_727870, EPI_ISL_727884, EPI_ISL_727891, EPI_ISL_727919, EPI_ISL_727938, EPI_ISL_727953, EPI_ISL_727972, EPI_ISL_727974 | Virology Department, Sheffield Teaching Hospitals NHS Foundation Trust/Department of Infection, Immunity and Cardiovascular Disease, The Medical School, University of Sheffield | COVID-19 Genomics UK (COG-UK) Consortium | Thushan de Silva, Matthew Parker, Nikki Smith, Adri Angyal, Rebecca Brown, Luke Green, Rachel Tucker, Paul Parsons, Danielle Groves, Katie Johnson, Laura Carrilero, Alex Keeley, Dave Partridge, Matthew Wyles, Benjamin Lindsey, Mehmet Yavuz, Mohammad Raza, Cariad Evans |
| EPI_ISL_728642, EPI_ISL_728644, EPI_ISL_728660, EPI_ISL_728661, EPI_ISL_728747 | Dutch COVID-19 response team | National Institute for Public Health and the Environment (RIVM) | Adam Meijer, Harry Vennema, Jeroen Cremer, Sharon van den Brink, Bas van der Veer, AnneMarie van den Brandt, Florian Zwagemaker, Dennis Schmitz, Chantal Reusken, on behalf of the national COVID-19 response team |
| EPI_ISL_732965, EPI_ISL_732967 | LabPLUS | Institute of Environmental Science and Research (ESR) | Xiaoyun Ren, Matt Storey, Nikki Freed, Muhammad Faisal, Jing Wang, Hermes Perez, Anja Werno, Antje van der Linden, Arlo Upton, Chris Mansell, David Hammer, Dragana Drinkovic, Gary McAuliffe, Hana Sofia Andersson, James Ussher, Jill Sherwood, Josh Freeman, Julia Howard, Juliet Elvy, Mary DeAlmeida, Matt Blakiston, Matthew Rogers, Max Bloomfield, Michael Addidle, Michelle Balm, Sally Roberts, Sarah Jefferies, Sharmini Muttaiyah, Susan Morpeth, Susan Taylor, Timothy Blackmore, Vani Sathyendran, Veronica Playle, Virginia Hope, Erasmus Smit, Lauren Jelly, Olin Silander, Joep de Ligt |
| EPI_ISL_733497 | University Hospital Zürich | Institute of Medical Virology, University of Zurich | Stefan Schmutz, Verena Kufner, Maryam Zaheri, Gabriela Ziltener, Thomas Scheier, Jürg Böni, Michael Huber, Alexandra Trkola |
| EPI_ISL_735861, EPI_ISL_735864 | Lighthouse Lab in Glasgow | Wellcome Sanger Institute for the COVID-19 Genomics UK (COG-UK) Consortium | Harper VanSteenhouse, Yumi Kasai, David Gray, Carol Clugston, Anna Dominiczak and Alex Alderton, Roberto Amato, Sonia Goncalves, Ewan Harrison, David K. Jackson, Ian Johnston, Dominic Kwiatkowski, Cordelia Langford, John Sillitoe on behalf of the Wellcome Sanger Institute COVID-19 Surveillance Team |
| EPI_ISL_735867, EPI_ISL_735868, EPI_ISL_735877 | Lighthouse Lab in Alderley Park | Wellcome Sanger Institute for the COVID-19 Genomics UK (COG-UK) Consortium | Jacquelyn Wynn, Mairead Hyland, The Lighthouse Lab in Alderley Park and Alex Alderton, Roberto Amato, Sonia Goncalves, Ewan Harrison, David K. Jackson, Ian Johnston, Dominic Kwiatkowski, Cordelia Langford, John Sillitoe on behalf of the Wellcome Sanger Institute COVID-19 Surveillance Team |
| EPI_ISL_735878, EPI_ISL_735880 | Lighthouse Lab in Glasgow | Wellcome Sanger Institute for the COVID-19 Genomics UK (COG-UK) Consortium | Harper VanSteenhouse, Yumi Kasai, David Gray, Carol Clugston, Anna Dominiczak and Alex Alderton, Roberto Amato, Sonia Goncalves, Ewan Harrison, David K. Jackson, Ian Johnston, Dominic Kwiatkowski, Cordelia Langford, John Sillitoe on behalf of the Wellcome Sanger Institute COVID-19 Surveillance Team |
| EPI_ISL_735881, EPI_ISL_735882 | Lighthouse Lab in Alderley Park | Wellcome Sanger Institute for the COVID-19 Genomics UK (COG-UK) Consortium | Jacquelyn Wynn, Mairead Hyland, The Lighthouse Lab in Alderley Park and Alex Alderton, Roberto Amato, Sonia Goncalves, Ewan Harrison, David K. Jackson, Ian Johnston, Dominic Kwiatkowski, Cordelia Langford, John Sillitoe on behalf of the Wellcome Sanger Institute COVID-19 Surveillance Team |
| EPI_ISL_735886 | Lighthouse Lab in Glasgow | Wellcome Sanger Institute for the COVID-19 Genomics UK (COG-UK) Consortium | Harper VanSteenhouse, Yumi Kasai, David Gray, Carol Clugston, Anna Dominiczak and Alex Alderton, Roberto Amato, Sonia Goncalves, Ewan Harrison, David K. Jackson, Ian Johnston, Dominic Kwiatkowski, Cordelia Langford, John Sillitoe on behalf of the Wellcome Sanger Institute COVID-19 Surveillance Team |
| EPI_ISL_735887, EPI_ISL_735888, EPI_ISL_735889, EPI_ISL_735891 | Lighthouse Lab in Alderley Park | Wellcome Sanger Institute for the COVID-19 Genomics UK (COG-UK) Consortium | Jacquelyn Wynn, Mairead Hyland, The Lighthouse Lab in Alderley Park and Alex Alderton, Roberto Amato, Sonia Goncalves, Ewan Harrison, David K. Jackson, Ian Johnston, Dominic Kwiatkowski, Cordelia Langford, John Sillitoe on behalf of the Wellcome Sanger Institute COVID-19 Surveillance Team |
| EPI_ISL_735894 | Lighthouse Lab in Glasgow | Wellcome Sanger Institute for the COVID-19 Genomics UK (COG-UK) Consortium | Harper VanSteenhouse, Yumi Kasai, David Gray, Carol Clugston, Anna Dominiczak and Alex Alderton, Roberto Amato, Sonia Goncalves, Ewan Harrison, David K. Jackson, Ian Johnston, Dominic Kwiatkowski, Cordelia Langford, John Sillitoe on behalf of the Wellcome Sanger Institute COVID-19 Surveillance Team |
| EPI_ISL_735897, EPI_ISL_735899, EPI_ISL_735900, EPI_ISL_735901, EPI_ISL_735902, EPI_ISL_735904, EPI_ISL_735905, EPI_ISL_735907, EPI_ISL_735908 | Lighthouse Lab in Alderley Park | Wellcome Sanger Institute for the COVID-19 Genomics UK (COG-UK) Consortium | Jacquelyn Wynn, Mairead Hyland, The Lighthouse Lab in Alderley Park and Alex Alderton, Roberto Amato, Sonia Goncalves, Ewan Harrison, David K. Jackson, Ian Johnston, Dominic Kwiatkowski, Cordelia Langford, John Sillitoe on behalf of the Wellcome Sanger Institute COVID-19 Surveillance Team |
| EPI_ISL_735909 | Lighthouse Lab in Glasgow | Wellcome Sanger Institute for the COVID-19 Genomics UK (COG-UK) Consortium | Harper VanSteenhouse, Yumi Kasai, David Gray, Carol Clugston, Anna Dominiczak and Alex Alderton, Roberto Amato, Sonia Goncalves, Ewan Harrison, David K. Jackson, Ian Johnston, Dominic Kwiatkowski, Cordelia Langford, John Sillitoe on behalf of the Wellcome Sanger Institute COVID-19 Surveillance Team |
| EPI_ISL_735912, EPI_ISL_735917 | Lighthouse Lab in Alderley Park | Wellcome Sanger Institute for the COVID-19 Genomics UK (COG-UK) Consortium | Jacquelyn Wynn, Mairead Hyland, The Lighthouse Lab in Alderley Park and Alex Alderton, Roberto Amato, Sonia Goncalves, Ewan Harrison, David K. Jackson, Ian Johnston, Dominic Kwiatkowski, Cordelia Langford, John Sillitoe on behalf of the Wellcome Sanger Institute COVID-19 Surveillance Team |
| EPI_ISL_735920 | Lighthouse Lab in Glasgow | Wellcome Sanger Institute for the COVID-19 Genomics UK (COG-UK) Consortium | Harper VanSteenhouse, Yumi Kasai, David Gray, Carol Clugston, Anna Dominiczak and Alex Alderton, Roberto Amato, Sonia Goncalves, Ewan Harrison, David K. Jackson, Ian Johnston, Dominic Kwiatkowski, Cordelia Langford, John Sillitoe on behalf of the Wellcome Sanger Institute COVID-19 Surveillance Team |
| EPI_ISL_735923, EPI_ISL_735924, EPI_ISL_735926 | Lighthouse Lab in Alderley Park | Wellcome Sanger Institute for the COVID-19 Genomics UK (COG-UK) Consortium | Jacquelyn Wynn, Mairead Hyland, The Lighthouse Lab in Alderley Park and Alex Alderton, Roberto Amato, Sonia Goncalves, Ewan Harrison, David K. Jackson, Ian Johnston, Dominic Kwiatkowski, Cordelia Langford, John Sillitoe on behalf of the Wellcome Sanger Institute COVID-19 Surveillance Team |
| EPI_ISL_735927 | Lighthouse Lab in Glasgow | Wellcome Sanger Institute for the COVID-19 Genomics UK | Harper VanSteenhouse, Yumi Kasai, David Gray, Carol Clugston, Anna Dominiczak and Alex Alderton, Roberto Amato, Sonia Goncalves, Ewan Harrison, |

| | | | |
|---|---|---|---|
| | | (COG-UK) Consortium | David K. Jackson, Ian Johnston, Dominic Kwiatkowski, Cordelia Langford, John Sillitoe on behalf of the Wellcome Sanger Institute COVID-19 Surveillance Team |
| EPI_ISL_735930, EPI_ISL_735931, EPI_ISL_735932, EPI_ISL_735936 | Lighthouse Lab in Alderley Park | Wellcome Sanger Institute for the COVID-19 Genomics UK (COG-UK) Consortium | Jacquelyn Wynn, Mairead Hyland, The Lighthouse Lab in Alderley Park and Alex Alderton, Roberto Amato, Sonia Goncalves, Ewan Harrison, David K. Jackson, Ian Johnston, Dominic Kwiatkowski, Cordelia Langford, John Sillitoe on behalf of the Wellcome Sanger Institute COVID-19 Surveillance Team |
| EPI_ISL_735937 | Lighthouse Lab in Glasgow | Wellcome Sanger Institute for the COVID-19 Genomics UK (COG-UK) Consortium | Harper VanSteenhouse, Yumi Kasai, David Gray, Carol Clugston, Anna Dominiczak and Alex Alderton, Roberto Amato, Sonia Goncalves, Ewan Harrison, David K. Jackson, Ian Johnston, Dominic Kwiatkowski, Cordelia Langford, John Sillitoe on behalf of the Wellcome Sanger Institute COVID-19 Surveillance Team |
| EPI_ISL_735938 | Lighthouse Lab in Alderley Park | Wellcome Sanger Institute for the COVID-19 Genomics UK (COG-UK) Consortium | Jacquelyn Wynn, Mairead Hyland, The Lighthouse Lab in Alderley Park and Alex Alderton, Roberto Amato, Sonia Goncalves, Ewan Harrison, David K. Jackson, Ian Johnston, Dominic Kwiatkowski, Cordelia Langford, John Sillitoe on behalf of the Wellcome Sanger Institute COVID-19 Surveillance Team |
| EPI_ISL_735939 | Lighthouse Lab in Glasgow | Wellcome Sanger Institute for the COVID-19 Genomics UK (COG-UK) Consortium | Harper VanSteenhouse, Yumi Kasai, David Gray, Carol Clugston, Anna Dominiczak and Alex Alderton, Roberto Amato, Sonia Goncalves, Ewan Harrison, David K. Jackson, Ian Johnston, Dominic Kwiatkowski, Cordelia Langford, John Sillitoe on behalf of the Wellcome Sanger Institute COVID-19 Surveillance Team |
| EPI_ISL_735941, EPI_ISL_735942, EPI_ISL_735943 | Lighthouse Lab in Alderley Park | Wellcome Sanger Institute for the COVID-19 Genomics UK (COG-UK) Consortium | Jacquelyn Wynn, Mairead Hyland, The Lighthouse Lab in Alderley Park and Alex Alderton, Roberto Amato, Sonia Goncalves, Ewan Harrison, David K. Jackson, Ian Johnston, Dominic Kwiatkowski, Cordelia Langford, John Sillitoe on behalf of the Wellcome Sanger Institute COVID-19 Surveillance Team |
| EPI_ISL_735945, EPI_ISL_735949 | Lighthouse Lab in Glasgow | Wellcome Sanger Institute for the COVID-19 Genomics UK (COG-UK) Consortium | Harper VanSteenhouse, Yumi Kasai, David Gray, Carol Clugston, Anna Dominiczak and Alex Alderton, Roberto Amato, Sonia Goncalves, Ewan Harrison, David K. Jackson, Ian Johnston, Dominic Kwiatkowski, Cordelia Langford, John Sillitoe on behalf of the Wellcome Sanger Institute COVID-19 Surveillance Team |
| EPI_ISL_735951 | Lighthouse Lab in Alderley Park | Wellcome Sanger Institute for the COVID-19 Genomics UK (COG-UK) Consortium | Jacquelyn Wynn, Mairead Hyland, The Lighthouse Lab in Alderley Park and Alex Alderton, Roberto Amato, Sonia Goncalves, Ewan Harrison, David K. Jackson, Ian Johnston, Dominic Kwiatkowski, Cordelia Langford, John Sillitoe on behalf of the Wellcome Sanger Institute COVID-19 Surveillance Team |
| EPI_ISL_735952, EPI_ISL_735956 | Lighthouse Lab in Glasgow | Wellcome Sanger Institute for the COVID-19 Genomics UK (COG-UK) Consortium | Harper VanSteenhouse, Yumi Kasai, David Gray, Carol Clugston, Anna Dominiczak and Alex Alderton, Roberto Amato, Sonia Goncalves, Ewan Harrison, David K. Jackson, Ian Johnston, Dominic Kwiatkowski, Cordelia Langford, John Sillitoe on behalf of the Wellcome Sanger Institute COVID-19 Surveillance Team |
| EPI_ISL_735958 | Lighthouse Lab in Alderley Park | Wellcome Sanger Institute for the COVID-19 Genomics UK (COG-UK) Consortium | Jacquelyn Wynn, Mairead Hyland, The Lighthouse Lab in Alderley Park and Alex Alderton, Roberto Amato, Sonia Goncalves, Ewan Harrison, David K. Jackson, Ian Johnston, Dominic Kwiatkowski, Cordelia Langford, John Sillitoe on behalf of the Wellcome Sanger Institute COVID-19 Surveillance Team |
| EPI_ISL_735961 | Lighthouse Lab in Glasgow | Wellcome Sanger Institute for the COVID-19 Genomics UK (COG-UK) Consortium | Harper VanSteenhouse, Yumi Kasai, David Gray, Carol Clugston, Anna Dominiczak and Alex Alderton, Roberto Amato, Sonia Goncalves, Ewan Harrison, David K. Jackson, Ian Johnston, Dominic Kwiatkowski, Cordelia Langford, John Sillitoe on behalf of the Wellcome Sanger Institute COVID-19 Surveillance Team |
| EPI_ISL_735963, EPI_ISL_735964 | Lighthouse Lab in Alderley Park | Wellcome Sanger Institute for the COVID-19 Genomics UK (COG-UK) Consortium | Jacquelyn Wynn, Mairead Hyland, The Lighthouse Lab in Alderley Park and Alex Alderton, Roberto Amato, Sonia Goncalves, Ewan Harrison, David K. Jackson, Ian Johnston, Dominic Kwiatkowski, Cordelia Langford, John Sillitoe on behalf of the Wellcome Sanger Institute COVID-19 Surveillance Team |
| EPI_ISL_735965 | Lighthouse Lab in Glasgow | Wellcome Sanger Institute for the COVID-19 Genomics UK (COG-UK) Consortium | Harper VanSteenhouse, Yumi Kasai, David Gray, Carol Clugston, Anna Dominiczak and Alex Alderton, Roberto Amato, Sonia Goncalves, Ewan Harrison, David K. Jackson, Ian Johnston, Dominic Kwiatkowski, Cordelia Langford, John Sillitoe on behalf of the Wellcome Sanger Institute COVID-19 Surveillance Team |
| EPI_ISL_735966, EPI_ISL_735967 | Lighthouse Lab in Alderley Park | Wellcome Sanger Institute for the COVID-19 Genomics UK (COG-UK) Consortium | Jacquelyn Wynn, Mairead Hyland, The Lighthouse Lab in Alderley Park and Alex Alderton, Roberto Amato, Sonia Goncalves, Ewan Harrison, David K. Jackson, Ian Johnston, Dominic Kwiatkowski, Cordelia Langford, John Sillitoe on behalf of the Wellcome Sanger Institute COVID-19 Surveillance Team |
| EPI_ISL_735968, EPI_ISL_735972, EPI_ISL_735973 | Lighthouse Lab in Glasgow | Wellcome Sanger Institute for the COVID-19 Genomics UK (COG-UK) Consortium | Harper VanSteenhouse, Yumi Kasai, David Gray, Carol Clugston, Anna Dominiczak and Alex Alderton, Roberto Amato, Sonia Goncalves, Ewan Harrison, David K. Jackson, Ian Johnston, Dominic Kwiatkowski, Cordelia Langford, John Sillitoe on behalf of the Wellcome Sanger Institute COVID-19 Surveillance Team |
| EPI_ISL_735974, EPI_ISL_735975, EPI_ISL_735976 | Lighthouse Lab in Alderley Park | Wellcome Sanger Institute for the COVID-19 Genomics UK (COG-UK) Consortium | Jacquelyn Wynn, Mairead Hyland, The Lighthouse Lab in Alderley Park and Alex Alderton, Roberto Amato, Sonia Goncalves, Ewan Harrison, David K. Jackson, Ian Johnston, Dominic Kwiatkowski, Cordelia Langford, John Sillitoe on behalf of the Wellcome Sanger Institute COVID-19 Surveillance Team |
| EPI_ISL_735978 | Lighthouse Lab in Glasgow | Wellcome Sanger Institute for the COVID-19 Genomics UK (COG-UK) Consortium | Harper VanSteenhouse, Yumi Kasai, David Gray, Carol Clugston, Anna Dominiczak and Alex Alderton, Roberto Amato, Sonia Goncalves, Ewan Harrison, David K. Jackson, Ian Johnston, Dominic Kwiatkowski, Cordelia Langford, John Sillitoe on behalf of the Wellcome Sanger Institute COVID-19 Surveillance Team |
| EPI_ISL_735980, EPI_ISL_735981 | Lighthouse Lab in Alderley Park | Wellcome Sanger Institute for the COVID-19 Genomics UK (COG-UK) Consortium | Jacquelyn Wynn, Mairead Hyland, The Lighthouse Lab in Alderley Park and Alex Alderton, Roberto Amato, Sonia Goncalves, Ewan Harrison, David K. Jackson, Ian Johnston, Dominic Kwiatkowski, Cordelia Langford, John Sillitoe on behalf of the Wellcome Sanger Institute COVID-19 Surveillance Team |
| EPI_ISL_735982 | Lighthouse Lab in Glasgow | Wellcome Sanger Institute for the COVID-19 Genomics UK (COG-UK) Consortium | Harper VanSteenhouse, Yumi Kasai, David Gray, Carol Clugston, Anna Dominiczak and Alex Alderton, Roberto Amato, Sonia Goncalves, Ewan Harrison, David K. Jackson, Ian Johnston, Dominic Kwiatkowski, Cordelia Langford, John Sillitoe on behalf of the Wellcome Sanger Institute COVID-19 Surveillance Team |
| EPI_ISL_735983, EPI_ISL_735986 | Lighthouse Lab in Alderley Park | Wellcome Sanger Institute for the COVID-19 Genomics UK (COG-UK) Consortium | Jacquelyn Wynn, Mairead Hyland, The Lighthouse Lab in Alderley Park and Alex Alderton, Roberto Amato, Sonia Goncalves, Ewan Harrison, David K. Jackson, Ian Johnston, Dominic Kwiatkowski, Cordelia Langford, John Sillitoe on behalf of the Wellcome Sanger Institute COVID-19 Surveillance Team |
| EPI_ISL_735987 | Lighthouse Lab in Glasgow | Wellcome Sanger Institute for the COVID-19 Genomics UK (COG-UK) Consortium | Harper VanSteenhouse, Yumi Kasai, David Gray, Carol Clugston, Anna Dominiczak and Alex Alderton, Roberto Amato, Sonia Goncalves, Ewan Harrison, David K. Jackson, Ian Johnston, Dominic Kwiatkowski, Cordelia Langford, John Sillitoe on behalf of the Wellcome Sanger Institute COVID-19 Surveillance Team |
| EPI_ISL_735992 | Lighthouse Lab in Alderley Park | Wellcome Sanger Institute for the COVID-19 Genomics UK (COG-UK) Consortium | Jacquelyn Wynn, Mairead Hyland, The Lighthouse Lab in Alderley Park and Alex Alderton, Roberto Amato, Sonia Goncalves, Ewan Harrison, David K. Jackson, Ian Johnston, Dominic Kwiatkowski, Cordelia Langford, John Sillitoe on behalf of the Wellcome Sanger Institute COVID-19 Surveillance Team |
| EPI_ISL_735993, EPI_ISL_735995, EPI_ISL_735996 | Lighthouse Lab in Glasgow | Wellcome Sanger Institute for the COVID-19 Genomics UK (COG-UK) Consortium | Harper VanSteenhouse, Yumi Kasai, David Gray, Carol Clugston, Anna Dominiczak and Alex Alderton, Roberto Amato, Sonia Goncalves, Ewan Harrison, David K. Jackson, Ian Johnston, Dominic Kwiatkowski, Cordelia Langford, John Sillitoe on behalf of the Wellcome Sanger Institute COVID-19 Surveillance Team |
| EPI_ISL_735997, EPI_ISL_735998, EPI_ISL_736000 | Lighthouse Lab in Alderley Park | Wellcome Sanger Institute for the COVID-19 Genomics UK (COG-UK) Consortium | Jacquelyn Wynn, Mairead Hyland, The Lighthouse Lab in Alderley Park and Alex Alderton, Roberto Amato, Sonia Goncalves, Ewan Harrison, David K. Jackson, Ian Johnston, Dominic Kwiatkowski, Cordelia Langford, John Sillitoe on behalf of the Wellcome Sanger Institute COVID-19 Surveillance Team |
| EPI_ISL_736001, EPI_ISL_736003, EPI_ISL_736007 | Lighthouse Lab in Glasgow | Wellcome Sanger Institute for the COVID-19 Genomics UK (COG-UK) Consortium | Harper VanSteenhouse, Yumi Kasai, David Gray, Carol Clugston, Anna Dominiczak and Alex Alderton, Roberto Amato, Sonia Goncalves, Ewan Harrison, David K. Jackson, Ian Johnston, Dominic Kwiatkowski, Cordelia Langford, John Sillitoe on behalf of the Wellcome Sanger Institute COVID-19 Surveillance Team |
| EPI_ISL_736008, EPI_ISL_736010, EPI_ISL_736011, EPI_ISL_736012, EPI_ISL_736013 | Lighthouse Lab in Alderley Park | Wellcome Sanger Institute for the COVID-19 Genomics UK (COG-UK) Consortium | Jacquelyn Wynn, Mairead Hyland, The Lighthouse Lab in Alderley Park and Alex Alderton, Roberto Amato, Sonia Goncalves, Ewan Harrison, David K. Jackson, Ian Johnston, Dominic Kwiatkowski, Cordelia Langford, John Sillitoe on behalf of the Wellcome Sanger Institute COVID-19 Surveillance Team |
| EPI_ISL_736015 | Lighthouse Lab in Glasgow | Wellcome Sanger Institute for the COVID-19 Genomics UK (COG-UK) Consortium | Harper VanSteenhouse, Yumi Kasai, David Gray, Carol Clugston, Anna Dominiczak and Alex Alderton, Roberto Amato, Sonia Goncalves, Ewan Harrison, David K. Jackson, Ian Johnston, Dominic Kwiatkowski, Cordelia Langford, John Sillitoe on behalf of the Wellcome Sanger Institute COVID-19 Surveillance Team |
| EPI_ISL_736016 | Lighthouse Lab in Alderley Park | Wellcome Sanger Institute for the COVID-19 Genomics UK (COG-UK) Consortium | Jacquelyn Wynn, Mairead Hyland, The Lighthouse Lab in Alderley Park and Alex Alderton, Roberto Amato, Sonia Goncalves, Ewan Harrison, David K. Jackson, Ian Johnston, Dominic Kwiatkowski, Cordelia Langford, John Sillitoe on behalf of the Wellcome Sanger Institute COVID-19 Surveillance Team |
| EPI_ISL_736017, EPI_ISL_736018, EPI_ISL_736019 | Lighthouse Lab in Glasgow | Wellcome Sanger Institute for the COVID-19 Genomics UK (COG-UK) Consortium | Harper VanSteenhouse, Yumi Kasai, David Gray, Carol Clugston, Anna Dominiczak and Alex Alderton, Roberto Amato, Sonia Goncalves, Ewan Harrison, David K. Jackson, Ian Johnston, Dominic Kwiatkowski, Cordelia Langford, John Sillitoe on behalf of the Wellcome Sanger Institute COVID-19 Surveillance Team |

| | | | |
|---|---|---|---|
| EPI_ISL_736021, EPI_ISL_736023 | Lighthouse Lab in Alderley Park | Wellcome Sanger Institute for the COVID-19 Genomics UK (COG-UK) Consortium | Jacquelyn Wynn, Mairead Hyland, The Lighthouse Lab in Alderley Park and Alex Alderton, Roberto Amato, Sonia Goncalves, Ewan Harrison, David K. Jackson, Ian Johnston, Dominic Kwiatkowski, Cordelia Langford, John Sillitoe on behalf of the Wellcome Sanger Institute COVID-19 Surveillance Team |
| EPI_ISL_736024, EPI_ISL_736025, EPI_ISL_736027 | Lighthouse Lab in Glasgow | Wellcome Sanger Institute for the COVID-19 Genomics UK (COG-UK) Consortium | Harper VanSteenhouse, Yumi Kasai, David Gray, Carol Clugston, Anna Dominiczak and Alex Alderton, Roberto Amato, Sonia Goncalves, Ewan Harrison, David K. Jackson, Ian Johnston, Dominic Kwiatkowski, Cordelia Langford, John Sillitoe on behalf of the Wellcome Sanger Institute COVID-19 Surveillance Team |
| EPI_ISL_736028, EPI_ISL_736029, EPI_ISL_736030 | Lighthouse Lab in Alderley Park | Wellcome Sanger Institute for the COVID-19 Genomics UK (COG-UK) Consortium | Jacquelyn Wynn, Mairead Hyland, The Lighthouse Lab in Alderley Park and Alex Alderton, Roberto Amato, Sonia Goncalves, Ewan Harrison, David K. Jackson, Ian Johnston, Dominic Kwiatkowski, Cordelia Langford, John Sillitoe on behalf of the Wellcome Sanger Institute COVID-19 Surveillance Team |
| EPI_ISL_736033 | Lighthouse Lab in Glasgow | Wellcome Sanger Institute for the COVID-19 Genomics UK (COG-UK) Consortium | Harper VanSteenhouse, Yumi Kasai, David Gray, Carol Clugston, Anna Dominiczak and Alex Alderton, Roberto Amato, Sonia Goncalves, Ewan Harrison, David K. Jackson, Ian Johnston, Dominic Kwiatkowski, Cordelia Langford, John Sillitoe on behalf of the Wellcome Sanger Institute COVID-19 Surveillance Team |
| EPI_ISL_736034, EPI_ISL_736037 | Lighthouse Lab in Alderley Park | Wellcome Sanger Institute for the COVID-19 Genomics UK (COG-UK) Consortium | Jacquelyn Wynn, Mairead Hyland, The Lighthouse Lab in Alderley Park and Alex Alderton, Roberto Amato, Sonia Goncalves, Ewan Harrison, David K. Jackson, Ian Johnston, Dominic Kwiatkowski, Cordelia Langford, John Sillitoe on behalf of the Wellcome Sanger Institute COVID-19 Surveillance Team |
| EPI_ISL_736038 | Lighthouse Lab in Glasgow | Wellcome Sanger Institute for the COVID-19 Genomics UK (COG-UK) Consortium | Harper VanSteenhouse, Yumi Kasai, David Gray, Carol Clugston, Anna Dominiczak and Alex Alderton, Roberto Amato, Sonia Goncalves, Ewan Harrison, David K. Jackson, Ian Johnston, Dominic Kwiatkowski, Cordelia Langford, John Sillitoe on behalf of the Wellcome Sanger Institute COVID-19 Surveillance Team |
| EPI_ISL_736039 | Lighthouse Lab in Alderley Park | Wellcome Sanger Institute for the COVID-19 Genomics UK (COG-UK) Consortium | Jacquelyn Wynn, Mairead Hyland, The Lighthouse Lab in Alderley Park and Alex Alderton, Roberto Amato, Sonia Goncalves, Ewan Harrison, David K. Jackson, Ian Johnston, Dominic Kwiatkowski, Cordelia Langford, John Sillitoe on behalf of the Wellcome Sanger Institute COVID-19 Surveillance Team |
| EPI_ISL_736041 | Lighthouse Lab in Glasgow | Wellcome Sanger Institute for the COVID-19 Genomics UK (COG-UK) Consortium | Harper VanSteenhouse, Yumi Kasai, David Gray, Carol Clugston, Anna Dominiczak and Alex Alderton, Roberto Amato, Sonia Goncalves, Ewan Harrison, David K. Jackson, Ian Johnston, Dominic Kwiatkowski, Cordelia Langford, John Sillitoe on behalf of the Wellcome Sanger Institute COVID-19 Surveillance Team |
| EPI_ISL_736046, EPI_ISL_736047, EPI_ISL_736048, EPI_ISL_736050, EPI_ISL_736051 | Lighthouse Lab in Alderley Park | Wellcome Sanger Institute for the COVID-19 Genomics UK (COG-UK) Consortium | Jacquelyn Wynn, Mairead Hyland, The Lighthouse Lab in Alderley Park and Alex Alderton, Roberto Amato, Sonia Goncalves, Ewan Harrison, David K. Jackson, Ian Johnston, Dominic Kwiatkowski, Cordelia Langford, John Sillitoe on behalf of the Wellcome Sanger Institute COVID-19 Surveillance Team |
| EPI_ISL_736052 | Lighthouse Lab in Glasgow | Wellcome Sanger Institute for the COVID-19 Genomics UK (COG-UK) Consortium | Harper VanSteenhouse, Yumi Kasai, David Gray, Carol Clugston, Anna Dominiczak and Alex Alderton, Roberto Amato, Sonia Goncalves, Ewan Harrison, David K. Jackson, Ian Johnston, Dominic Kwiatkowski, Cordelia Langford, John Sillitoe on behalf of the Wellcome Sanger Institute COVID-19 Surveillance Team |
| EPI_ISL_736055 | Lighthouse Lab in Alderley Park | Wellcome Sanger Institute for the COVID-19 Genomics UK (COG-UK) Consortium | Jacquelyn Wynn, Mairead Hyland, The Lighthouse Lab in Alderley Park and Alex Alderton, Roberto Amato, Sonia Goncalves, Ewan Harrison, David K. Jackson, Ian Johnston, Dominic Kwiatkowski, Cordelia Langford, John Sillitoe on behalf of the Wellcome Sanger Institute COVID-19 Surveillance Team |
| EPI_ISL_736057, EPI_ISL_736059 | Lighthouse Lab in Glasgow | Wellcome Sanger Institute for the COVID-19 Genomics UK (COG-UK) Consortium | Harper VanSteenhouse, Yumi Kasai, David Gray, Carol Clugston, Anna Dominiczak and Alex Alderton, Roberto Amato, Sonia Goncalves, Ewan Harrison, David K. Jackson, Ian Johnston, Dominic Kwiatkowski, Cordelia Langford, John Sillitoe on behalf of the Wellcome Sanger Institute COVID-19 Surveillance Team |
| EPI_ISL_736063 | Lighthouse Lab in Alderley Park | Wellcome Sanger Institute for the COVID-19 Genomics UK (COG-UK) Consortium | Jacquelyn Wynn, Mairead Hyland, The Lighthouse Lab in Alderley Park and Alex Alderton, Roberto Amato, Sonia Goncalves, Ewan Harrison, David K. Jackson, Ian Johnston, Dominic Kwiatkowski, Cordelia Langford, John Sillitoe on behalf of the Wellcome Sanger Institute COVID-19 Surveillance Team |
| EPI_ISL_736064 | Lighthouse Lab in Glasgow | Wellcome Sanger Institute for the COVID-19 Genomics UK (COG-UK) Consortium | Harper VanSteenhouse, Yumi Kasai, David Gray, Carol Clugston, Anna Dominiczak and Alex Alderton, Roberto Amato, Sonia Goncalves, Ewan Harrison, David K. Jackson, Ian Johnston, Dominic Kwiatkowski, Cordelia Langford, John Sillitoe on behalf of the Wellcome Sanger Institute COVID-19 Surveillance Team |
| EPI_ISL_736065 | Lighthouse Lab in Alderley Park | Wellcome Sanger Institute for the COVID-19 Genomics UK (COG-UK) Consortium | Jacquelyn Wynn, Mairead Hyland, The Lighthouse Lab in Alderley Park and Alex Alderton, Roberto Amato, Sonia Goncalves, Ewan Harrison, David K. Jackson, Ian Johnston, Dominic Kwiatkowski, Cordelia Langford, John Sillitoe on behalf of the Wellcome Sanger Institute COVID-19 Surveillance Team |
| EPI_ISL_736066, EPI_ISL_736072 | Lighthouse Lab in Glasgow | Wellcome Sanger Institute for the COVID-19 Genomics UK (COG-UK) Consortium | Harper VanSteenhouse, Yumi Kasai, David Gray, Carol Clugston, Anna Dominiczak and Alex Alderton, Roberto Amato, Sonia Goncalves, Ewan Harrison, David K. Jackson, Ian Johnston, Dominic Kwiatkowski, Cordelia Langford, John Sillitoe on behalf of the Wellcome Sanger Institute COVID-19 Surveillance Team |
| EPI_ISL_736078, EPI_ISL_736079, EPI_ISL_736080, EPI_ISL_736083, EPI_ISL_736087 | Lighthouse Lab in Alderley Park | Wellcome Sanger Institute for the COVID-19 Genomics UK (COG-UK) Consortium | Jacquelyn Wynn, Mairead Hyland, The Lighthouse Lab in Alderley Park and Alex Alderton, Roberto Amato, Sonia Goncalves, Ewan Harrison, David K. Jackson, Ian Johnston, Dominic Kwiatkowski, Cordelia Langford, John Sillitoe on behalf of the Wellcome Sanger Institute COVID-19 Surveillance Team |
| EPI_ISL_736090, EPI_ISL_736091, EPI_ISL_736092 | Lighthouse Lab in Glasgow | Wellcome Sanger Institute for the COVID-19 Genomics UK (COG-UK) Consortium | Harper VanSteenhouse, Yumi Kasai, David Gray, Carol Clugston, Anna Dominiczak and Alex Alderton, Roberto Amato, Sonia Goncalves, Ewan Harrison, David K. Jackson, Ian Johnston, Dominic Kwiatkowski, Cordelia Langford, John Sillitoe on behalf of the Wellcome Sanger Institute COVID-19 Surveillance Team |
| EPI_ISL_736093, EPI_ISL_736095 | Lighthouse Lab in Alderley Park | Wellcome Sanger Institute for the COVID-19 Genomics UK (COG-UK) Consortium | Jacquelyn Wynn, Mairead Hyland, The Lighthouse Lab in Alderley Park and Alex Alderton, Roberto Amato, Sonia Goncalves, Ewan Harrison, David K. Jackson, Ian Johnston, Dominic Kwiatkowski, Cordelia Langford, John Sillitoe on behalf of the Wellcome Sanger Institute COVID-19 Surveillance Team |
| EPI_ISL_736096 | Lighthouse Lab in Glasgow | Wellcome Sanger Institute for the COVID-19 Genomics UK (COG-UK) Consortium | Harper VanSteenhouse, Yumi Kasai, David Gray, Carol Clugston, Anna Dominiczak and Alex Alderton, Roberto Amato, Sonia Goncalves, Ewan Harrison, David K. Jackson, Ian Johnston, Dominic Kwiatkowski, Cordelia Langford, John Sillitoe on behalf of the Wellcome Sanger Institute COVID-19 Surveillance Team |
| EPI_ISL_736097 | Lighthouse Lab in Alderley Park | Wellcome Sanger Institute for the COVID-19 Genomics UK (COG-UK) Consortium | Jacquelyn Wynn, Mairead Hyland, The Lighthouse Lab in Alderley Park and Alex Alderton, Roberto Amato, Sonia Goncalves, Ewan Harrison, David K. Jackson, Ian Johnston, Dominic Kwiatkowski, Cordelia Langford, John Sillitoe on behalf of the Wellcome Sanger Institute COVID-19 Surveillance Team |
| EPI_ISL_736099, EPI_ISL_736100, EPI_ISL_736101, EPI_ISL_736102, EPI_ISL_736104 | Lighthouse Lab in Glasgow | Wellcome Sanger Institute for the COVID-19 Genomics UK (COG-UK) Consortium | Harper VanSteenhouse, Yumi Kasai, David Gray, Carol Clugston, Anna Dominiczak and Alex Alderton, Roberto Amato, Sonia Goncalves, Ewan Harrison, David K. Jackson, Ian Johnston, Dominic Kwiatkowski, Cordelia Langford, John Sillitoe on behalf of the Wellcome Sanger Institute COVID-19 Surveillance Team |
| EPI_ISL_736105 | Lighthouse Lab in Alderley Park | Wellcome Sanger Institute for the COVID-19 Genomics UK (COG-UK) Consortium | Jacquelyn Wynn, Mairead Hyland, The Lighthouse Lab in Alderley Park and Alex Alderton, Roberto Amato, Sonia Goncalves, Ewan Harrison, David K. Jackson, Ian Johnston, Dominic Kwiatkowski, Cordelia Langford, John Sillitoe on behalf of the Wellcome Sanger Institute COVID-19 Surveillance Team |
| EPI_ISL_736107 | Lighthouse Lab in Glasgow | Wellcome Sanger Institute for the COVID-19 Genomics UK (COG-UK) Consortium | Harper VanSteenhouse, Yumi Kasai, David Gray, Carol Clugston, Anna Dominiczak and Alex Alderton, Roberto Amato, Sonia Goncalves, Ewan Harrison, David K. Jackson, Ian Johnston, Dominic Kwiatkowski, Cordelia Langford, John Sillitoe on behalf of the Wellcome Sanger Institute COVID-19 Surveillance Team |
| EPI_ISL_736108, EPI_ISL_736110 | Lighthouse Lab in Alderley Park | Wellcome Sanger Institute for the COVID-19 Genomics UK (COG-UK) Consortium | Jacquelyn Wynn, Mairead Hyland, The Lighthouse Lab in Alderley Park and Alex Alderton, Roberto Amato, Sonia Goncalves, Ewan Harrison, David K. Jackson, Ian Johnston, Dominic Kwiatkowski, Cordelia Langford, John Sillitoe on behalf of the Wellcome Sanger Institute COVID-19 Surveillance Team |
| EPI_ISL_736115 | Lighthouse Lab in Glasgow | Wellcome Sanger Institute for the COVID-19 Genomics UK (COG-UK) Consortium | Harper VanSteenhouse, Yumi Kasai, David Gray, Carol Clugston, Anna Dominiczak and Alex Alderton, Roberto Amato, Sonia Goncalves, Ewan Harrison, David K. Jackson, Ian Johnston, Dominic Kwiatkowski, Cordelia Langford, John Sillitoe on behalf of the Wellcome Sanger Institute COVID-19 Surveillance Team |
| EPI_ISL_736118 | Lighthouse Lab in Alderley Park | Wellcome Sanger Institute for the COVID-19 Genomics UK (COG-UK) Consortium | Jacquelyn Wynn, Mairead Hyland, The Lighthouse Lab in Alderley Park and Alex Alderton, Roberto Amato, Sonia Goncalves, Ewan Harrison, David K. Jackson, Ian Johnston, Dominic Kwiatkowski, Cordelia Langford, John Sillitoe on behalf of the Wellcome Sanger Institute COVID-19 Surveillance Team |
| EPI_ISL_736124 | Lighthouse Lab in Glasgow | Wellcome Sanger Institute for the COVID-19 Genomics UK (COG-UK) Consortium | Harper VanSteenhouse, Yumi Kasai, David Gray, Carol Clugston, Anna Dominiczak and Alex Alderton, Roberto Amato, Sonia Goncalves, Ewan Harrison, David K. Jackson, Ian Johnston, Dominic Kwiatkowski, Cordelia Langford, John Sillitoe on behalf of the Wellcome Sanger Institute COVID-19 Surveillance Team |
| EPI_ISL_736125, EPI_ISL_736127 | Lighthouse Lab in Alderley Park | Wellcome Sanger Institute for the COVID-19 Genomics UK | Jacquelyn Wynn, Mairead Hyland, The Lighthouse Lab in Alderley Park and Alex Alderton, Roberto Amato, Sonia Goncalves, Ewan Harrison, David K. |

| | | | |
|---|---|---|---|
| | | (COG-UK) Consortium | Jackson, Ian Johnston, Dominic Kwiatkowski, Cordelia Langford, John Sillitoe on behalf of the Wellcome Sanger Institute COVID-19 Surveillance Team |
| EPI_ISL_736130, EPI_ISL_736131, EPI_ISL_736132, EPI_ISL_736135 | Lighthouse Lab in Glasgow | Wellcome Sanger Institute for the COVID-19 Genomics UK (COG-UK) Consortium | Harper VanSteenhouse, Yumi Kasai, David Gray, Carol Clugston, Anna Dominiczak and Alex Alderton, Roberto Amato, Sonia Goncalves, Ewan Harrison, David K. Jackson, Ian Johnston, Dominic Kwiatkowski, Cordelia Langford, John Sillitoe on behalf of the Wellcome Sanger Institute COVID-19 Surveillance Team |
| EPI_ISL_736141, EPI_ISL_736142 | Lighthouse Lab in Alderley Park | Wellcome Sanger Institute for the COVID-19 Genomics UK (COG-UK) Consortium | Jacquelyn Wynn, Mairead Hyland, The Lighthouse Lab in Alderley Park and Alex Alderton, Roberto Amato, Sonia Goncalves, Ewan Harrison, David K. Jackson, Ian Johnston, Dominic Kwiatkowski, Cordelia Langford, John Sillitoe on behalf of the Wellcome Sanger Institute COVID-19 Surveillance Team |
| EPI_ISL_736143, EPI_ISL_736146, EPI_ISL_736149, EPI_ISL_736151 | Lighthouse Lab in Glasgow | Wellcome Sanger Institute for the COVID-19 Genomics UK (COG-UK) Consortium | Harper VanSteenhouse, Yumi Kasai, David Gray, Carol Clugston, Anna Dominiczak and Alex Alderton, Roberto Amato, Sonia Goncalves, Ewan Harrison, David K. Jackson, Ian Johnston, Dominic Kwiatkowski, Cordelia Langford, John Sillitoe on behalf of the Wellcome Sanger Institute COVID-19 Surveillance Team |
| EPI_ISL_736153, EPI_ISL_736158 | Lighthouse Lab in Alderley Park | Wellcome Sanger Institute for the COVID-19 Genomics UK (COG-UK) Consortium | Jacquelyn Wynn, Mairead Hyland, The Lighthouse Lab in Alderley Park and Alex Alderton, Roberto Amato, Sonia Goncalves, Ewan Harrison, David K. Jackson, Ian Johnston, Dominic Kwiatkowski, Cordelia Langford, John Sillitoe on behalf of the Wellcome Sanger Institute COVID-19 Surveillance Team |
| EPI_ISL_736159, EPI_ISL_736162, EPI_ISL_736167, EPI_ISL_736168, EPI_ISL_736169 | Lighthouse Lab in Glasgow | Wellcome Sanger Institute for the COVID-19 Genomics UK (COG-UK) Consortium | Harper VanSteenhouse, Yumi Kasai, David Gray, Carol Clugston, Anna Dominiczak and Alex Alderton, Roberto Amato, Sonia Goncalves, Ewan Harrison, David K. Jackson, Ian Johnston, Dominic Kwiatkowski, Cordelia Langford, John Sillitoe on behalf of the Wellcome Sanger Institute COVID-19 Surveillance Team |
| EPI_ISL_736172 | Lighthouse Lab in Alderley Park | Wellcome Sanger Institute for the COVID-19 Genomics UK (COG-UK) Consortium | Jacquelyn Wynn, Mairead Hyland, The Lighthouse Lab in Alderley Park and Alex Alderton, Roberto Amato, Sonia Goncalves, Ewan Harrison, David K. Jackson, Ian Johnston, Dominic Kwiatkowski, Cordelia Langford, John Sillitoe on behalf of the Wellcome Sanger Institute COVID-19 Surveillance Team |
| EPI_ISL_736175 | Lighthouse Lab in Glasgow | Wellcome Sanger Institute for the COVID-19 Genomics UK (COG-UK) Consortium | Harper VanSteenhouse, Yumi Kasai, David Gray, Carol Clugston, Anna Dominiczak and Alex Alderton, Roberto Amato, Sonia Goncalves, Ewan Harrison, David K. Jackson, Ian Johnston, Dominic Kwiatkowski, Cordelia Langford, John Sillitoe on behalf of the Wellcome Sanger Institute COVID-19 Surveillance Team |
| EPI_ISL_736178 | Lighthouse Lab in Alderley Park | Wellcome Sanger Institute for the COVID-19 Genomics UK (COG-UK) Consortium | Jacquelyn Wynn, Mairead Hyland, The Lighthouse Lab in Alderley Park and Alex Alderton, Roberto Amato, Sonia Goncalves, Ewan Harrison, David K. Jackson, Ian Johnston, Dominic Kwiatkowski, Cordelia Langford, John Sillitoe on behalf of the Wellcome Sanger Institute COVID-19 Surveillance Team |
| EPI_ISL_736182, EPI_ISL_736183, EPI_ISL_736184 | Lighthouse Lab in Glasgow | Wellcome Sanger Institute for the COVID-19 Genomics UK (COG-UK) Consortium | Harper VanSteenhouse, Yumi Kasai, David Gray, Carol Clugston, Anna Dominiczak and Alex Alderton, Roberto Amato, Sonia Goncalves, Ewan Harrison, David K. Jackson, Ian Johnston, Dominic Kwiatkowski, Cordelia Langford, John Sillitoe on behalf of the Wellcome Sanger Institute COVID-19 Surveillance Team |
| EPI_ISL_736187, EPI_ISL_736189 | Lighthouse Lab in Alderley Park | Wellcome Sanger Institute for the COVID-19 Genomics UK (COG-UK) Consortium | Jacquelyn Wynn, Mairead Hyland, The Lighthouse Lab in Alderley Park and Alex Alderton, Roberto Amato, Sonia Goncalves, Ewan Harrison, David K. Jackson, Ian Johnston, Dominic Kwiatkowski, Cordelia Langford, John Sillitoe on behalf of the Wellcome Sanger Institute COVID-19 Surveillance Team |
| EPI_ISL_736190, EPI_ISL_736191 | Lighthouse Lab in Glasgow | Wellcome Sanger Institute for the COVID-19 Genomics UK (COG-UK) Consortium | Harper VanSteenhouse, Yumi Kasai, David Gray, Carol Clugston, Anna Dominiczak and Alex Alderton, Roberto Amato, Sonia Goncalves, Ewan Harrison, David K. Jackson, Ian Johnston, Dominic Kwiatkowski, Cordelia Langford, John Sillitoe on behalf of the Wellcome Sanger Institute COVID-19 Surveillance Team |
| EPI_ISL_736193, EPI_ISL_736199 | Lighthouse Lab in Alderley Park | Wellcome Sanger Institute for the COVID-19 Genomics UK (COG-UK) Consortium | Jacquelyn Wynn, Mairead Hyland, The Lighthouse Lab in Alderley Park and Alex Alderton, Roberto Amato, Sonia Goncalves, Ewan Harrison, David K. Jackson, Ian Johnston, Dominic Kwiatkowski, Cordelia Langford, John Sillitoe on behalf of the Wellcome Sanger Institute COVID-19 Surveillance Team |
| EPI_ISL_736483, EPI_ISL_736488, EPI_ISL_736489, EPI_ISL_736490, EPI_ISL_736491, EPI_ISL_736492, EPI_ISL_736512, EPI_ISL_736519, EPI_ISL_736529, EPI_ISL_736532, EPI_ISL_736533, EPI_ISL_736556, EPI_ISL_736567, EPI_ISL_736573, EPI_ISL_736580, EPI_ISL_736582, EPI_ISL_736605, EPI_ISL_736620, EPI_ISL_736624, EPI_ISL_736627, EPI_ISL_736628, EPI_ISL_736631, EPI_ISL_736633, EPI_ISL_736646, EPI_ISL_736650, EPI_ISL_736657, EPI_ISL_736659, EPI_ISL_736668, EPI_ISL_736672, EPI_ISL_736678, EPI_ISL_736689, EPI_ISL_736691, EPI_ISL_736695, EPI_ISL_736700, EPI_ISL_736734, EPI_ISL_736736, EPI_ISL_736737, EPI_ISL_736743, EPI_ISL_736745, EPI_ISL_736752, EPI_ISL_736754 | | | |
| see above | Lighthouse Lab in Glasgow | Wellcome Sanger Institute for the COVID-19 Genomics UK (COG-UK) Consortium | Harper VanSteenhouse, Yumi Kasai, David Gray, Carol Clugston, Anna Dominiczak and Alex Alderton, Roberto Amato, Sonia Goncalves, Ewan Harrison, David K. Jackson, Ian Johnston, Dominic Kwiatkowski, Cordelia Langford, John Sillitoe on behalf of the Wellcome Sanger Institute COVID-19 Surveillance Team |
| EPI_ISL_737245, EPI_ISL_737246, EPI_ISL_737247, EPI_ISL_737248, EPI_ISL_737249, EPI_ISL_737250, EPI_ISL_737251, EPI_ISL_737252, EPI_ISL_737253 | Los Angeles County PHL | Los Angeles County PHL | P. Hemarajata et al. |
| EPI_ISL_737310, EPI_ISL_737311, EPI_ISL_737312, EPI_ISL_737313, EPI_ISL_737314, EPI_ISL_737315, EPI_ISL_737316, EPI_ISL_737317, EPI_ISL_737318, EPI_ISL_737319, EPI_ISL_737320, EPI_ISL_737321, EPI_ISL_737322, EPI_ISL_737323, EPI_ISL_737324, EPI_ISL_737325, EPI_ISL_737326, EPI_ISL_737327, EPI_ISL_737328, EPI_ISL_737329, EPI_ISL_737330, EPI_ISL_737331, EPI_ISL_737332, EPI_ISL_737333, EPI_ISL_737334, EPI_ISL_737335, EPI_ISL_737336, EPI_ISL_737337, EPI_ISL_737338, EPI_ISL_737339, EPI_ISL_737340, EPI_ISL_737341, EPI_ISL_737342, EPI_ISL_737343, EPI_ISL_737344 | | | |
| see above | Department of Clinical Microbiology | GIGA Medical Genomics | Keith Durkin, Maria Artesi, Sébastien Bontems, Raphaël Boreux, Bouchra Boujemla, Cécile Meex, Pierrette Melin, Marie-Pierre Hayette, Vincent Bours |
| EPI_ISL_737395, EPI_ISL_737396, EPI_ISL_737399, EPI_ISL_737405, EPI_ISL_737406, EPI_ISL_737407, EPI_ISL_737413, EPI_ISL_737416, EPI_ISL_737418, EPI_ISL_737422, EPI_ISL_737424, EPI_ISL_737425, EPI_ISL_737428, EPI_ISL_737430, EPI_ISL_737431, EPI_ISL_737432, EPI_ISL_737435, EPI_ISL_737436, EPI_ISL_737437, EPI_ISL_737441, EPI_ISL_737442, EPI_ISL_737443, EPI_ISL_737445, EPI_ISL_737446, EPI_ISL_737448, EPI_ISL_737450, EPI_ISL_737451, EPI_ISL_737454, EPI_ISL_737455, EPI_ISL_737457, EPI_ISL_737460, EPI_ISL_737461, EPI_ISL_737462, EPI_ISL_737463, EPI_ISL_737466, EPI_ISL_737467, EPI_ISL_737468, EPI_ISL_737470, EPI_ISL_737472, EPI_ISL_737473, EPI_ISL_737475, EPI_ISL_737477, EPI_ISL_737478, EPI_ISL_737479, EPI_ISL_737485, EPI_ISL_737486, EPI_ISL_737488, EPI_ISL_737491, EPI_ISL_737495, EPI_ISL_737496, EPI_ISL_737501, EPI_ISL_737502, EPI_ISL_737503, EPI_ISL_737504, EPI_ISL_737506, EPI_ISL_737507, EPI_ISL_737508, EPI_ISL_737509, EPI_ISL_737510, EPI_ISL_737511, EPI_ISL_737514, EPI_ISL_737517, EPI_ISL_737519, EPI_ISL_737520, EPI_ISL_737521, EPI_ISL_737522, EPI_ISL_737523, EPI_ISL_737524, EPI_ISL_737526, EPI_ISL_737531, EPI_ISL_737532, EPI_ISL_737534, EPI_ISL_737536, EPI_ISL_737537, EPI_ISL_737539, EPI_ISL_737541, EPI_ISL_737542, EPI_ISL_737543, EPI_ISL_737544, EPI_ISL_737545, EPI_ISL_737546, EPI_ISL_737549, EPI_ISL_737554, EPI_ISL_737556, EPI_ISL_737557, EPI_ISL_737558, EPI_ISL_737559, EPI_ISL_737564, EPI_ISL_737565, EPI_ISL_737566, EPI_ISL_737567, EPI_ISL_737570, EPI_ISL_737571, EPI_ISL_737574, EPI_ISL_737579, EPI_ISL_737580, EPI_ISL_737584, EPI_ISL_737586, EPI_ISL_737587, EPI_ISL_737590, EPI_ISL_737594, EPI_ISL_737596, EPI_ISL_737601, EPI_ISL_737604, EPI_ISL_737606, EPI_ISL_737611, EPI_ISL_737616, EPI_ISL_737617, EPI_ISL_737624, EPI_ISL_737627, EPI_ISL_737642, EPI_ISL_737652, EPI_ISL_737658, EPI_ISL_737659, EPI_ISL_737670, EPI_ISL_737671, EPI_ISL_737672, EPI_ISL_737673, EPI_ISL_737674, EPI_ISL_737675, EPI_ISL_737676, EPI_ISL_737677, EPI_ISL_737678, EPI_ISL_737708, EPI_ISL_737725, EPI_ISL_737726, EPI_ISL_737727, EPI_ISL_737731, EPI_ISL_737740, EPI_ISL_737741, EPI_ISL_737742, EPI_ISL_737743, EPI_ISL_737744, EPI_ISL_737745, EPI_ISL_737767, EPI_ISL_737768, EPI_ISL_737769, EPI_ISL_737791, EPI_ISL_737795, EPI_ISL_737796, EPI_ISL_737797, EPI_ISL_737843, EPI_ISL_737845, EPI_ISL_737846, EPI_ISL_737867, EPI_ISL_737869, EPI_ISL_737870, EPI_ISL_737908, EPI_ISL_737909, EPI_ISL_737913, EPI_ISL_737918 | | | |
| see above | Viollier AG | Department of Biosystems Science and Engineering, ETH Zürich | Chaoran Chen, Sarah Nadeau, Catharine Aquino, Ivan Topolsky, Philipp Jablonski, Lara Fuhrmann, David Dreifuss, Katharina Jahn, Andreia Cabral de Gouvea, Maria Domenica Moccia, Simon Grüter, Timothy Sykes, Lennart Opitz, Griffin White, Laura Neff, Doris Popovic, Andrea Patrignani, Jay Tracy, Ralph Schlapbach, Christiane Beckmann, Maurice Redondo, Olivier Kobel, Christoph Noppen, Sophie Seidel, Noemie Santamaria de Souza, Niko Beerenwinkel, Tanja Stadler |
| EPI_ISL_738106 | Instituto Nacional de Saude (INSA) | Instituto Nacional de Saude (INSA) | Borges et al |
| EPI_ISL_738350, EPI_ISL_738351, EPI_ISL_738352, EPI_ISL_738355, EPI_ISL_738357 | Utah Public Health Laboratory | Utah Public Health Laboratory | Erin Young, Kelly Oakeson, Tara Gallagher |
| EPI_ISL_738363, EPI_ISL_738365, EPI_ISL_738369, EPI_ISL_738374, EPI_ISL_738376, EPI_ISL_738380, EPI_ISL_738381, EPI_ISL_738382, EPI_ISL_738386, EPI_ISL_738387, EPI_ISL_738388, EPI_ISL_738392, EPI_ISL_738394, EPI_ISL_738398, EPI_ISL_738400, EPI_ISL_738406, EPI_ISL_738409, EPI_ISL_738412, EPI_ISL_738415, EPI_ISL_738417, EPI_ISL_738420, EPI_ISL_738424, EPI_ISL_738427, EPI_ISL_738430, EPI_ISL_738432, EPI_ISL_738435, EPI_ISL_738439, EPI_ISL_738441, EPI_ISL_738450, EPI_ISL_738452, EPI_ISL_738454, EPI_ISL_738455, EPI_ISL_738458, EPI_ISL_738459, EPI_ISL_738460, EPI_ISL_738461, EPI_ISL_738464, EPI_ISL_738466, EPI_ISL_738467, EPI_ISL_738471, EPI_ISL_738472, EPI_ISL_738474, EPI_ISL_738476, EPI_ISL_738477, EPI_ISL_738479, EPI_ISL_738482, EPI_ISL_738483, EPI_ISL_738485, EPI_ISL_738488, EPI_ISL_738489, EPI_ISL_738491, EPI_ISL_738493, EPI_ISL_738494, EPI_ISL_738497, EPI_ISL_738499 | | | |
| see above | UZ Leuven, National Reference Laboratory for Coronaviruses, Laboratory Medicine, Leuven, Belgium | KU Leuven, Rega Institute, Clinical and Epidemiological Virology | Tony Wawina-Bokalanga, Joan Marti-Carerras, Bert Vanmechelen, Piet Maes |
| EPI_ISL_740025, EPI_ISL_740203, EPI_ISL_740345, EPI_ISL_740489, EPI_ISL_740532 | Laboratoire national de santé, Microbiology, Virology | Laboratoire national de santé, Microbiology, Microbial Genomics Platform | Anke Wienecke-Baldacchino, Catherine Ragimbeau,Jessica Tapp, Fatu Djabi, Lise Pignon, Raoul Salmon, Tamir Abdelrahman |
| EPI_ISL_740867, EPI_ISL_740868 | South Eastern Area Laboratory Services (SEALS) | NSW Health Pathology - Institute of Clinical Pathology and Medical Research; Westmead Hospital; University of Sydney | CIDM-PH et al. |
| EPI_ISL_740894, EPI_ISL_740895 | Sydney South West Pathology Service (SSWPS) - Royal | NSW Health Pathology - Institute of Clinical Pathology and | CIDM-PH et al. |

| | | | |
|---|---|---|---|
| | Prince Alfred Hospital - NSW Health Pathology | Medical Research; Westmead Hospital; University of Sydney | |
| EPI_ISL_740999, EPI_ISL_741005, EPI_ISL_741006, EPI_ISL_741007, EPI_ISL_741008, EPI_ISL_741010, EPI_ISL_741013, EPI_ISL_741014, EPI_ISL_741015, EPI_ISL_741016, EPI_ISL_741017 | | | |
| see above | Department of Pathology, University of Cambridge | COVID-19 Genomics UK (COG-UK) Consortium | Aminu S. Jahun, Yasmin Chaudhry, Grant Hall, Iliana Georgana, Myra Hosmillo, Martin D. Curran, Malte Pinckert, Surendra Parmar, Ian Goodfellow |
| EPI_ISL_741309, EPI_ISL_741310, EPI_ISL_741311, EPI_ISL_741317, EPI_ISL_741318, EPI_ISL_741319, EPI_ISL_741320, EPI_ISL_741322, EPI_ISL_741323, EPI_ISL_741324, EPI_ISL_741325, EPI_ISL_741326, EPI_ISL_741327, EPI_ISL_741328, EPI_ISL_741329, EPI_ISL_741330, EPI_ISL_741331, EPI_ISL_741332, EPI_ISL_741333, EPI_ISL_741334, EPI_ISL_741335, EPI_ISL_741336, EPI_ISL_741340, EPI_ISL_741346 | | | |
| see above | University College London, Great Ormond Street Hospital for Children NHS Foundation Trust, Imperial College Healthcare NHS Trust | COVID-19 Genomics UK (COG-UK) Consortium | Sergi Castellano, Rachel Williams, Mark Kristiansen, Paola Resende Silva, Sunando Roy, Tony Brooks, Helena Tutill, Paola Niola, Patricia Dyal, Charlotte Williams, Leysa Forrest, Yasmin Panchbhaya, Jacqueline Findlay, Samuel Weeks, Julianne Brown, Kathryn Harris, Paul Randell, James Price, Alison Holmes, Judith Breuer |
| EPI_ISL_741453, EPI_ISL_741454, EPI_ISL_741455, EPI_ISL_741456, EPI_ISL_741457, EPI_ISL_741458, EPI_ISL_741459, EPI_ISL_741460, EPI_ISL_741461, EPI_ISL_741462, EPI_ISL_741463, EPI_ISL_741464, EPI_ISL_741465, EPI_ISL_741466, EPI_ISL_741467, EPI_ISL_741468, EPI_ISL_741469, EPI_ISL_741470, EPI_ISL_741471, EPI_ISL_741472, EPI_ISL_741473, EPI_ISL_741474, EPI_ISL_741475, EPI_ISL_741476, EPI_ISL_741477, EPI_ISL_741478, EPI_ISL_741479, EPI_ISL_741480, EPI_ISL_741481, EPI_ISL_741482, EPI_ISL_741483, EPI_ISL_741484, EPI_ISL_741492, EPI_ISL_741583, EPI_ISL_741584, EPI_ISL_741585, EPI_ISL_741586, EPI_ISL_741587, EPI_ISL_741588, EPI_ISL_741589, EPI_ISL_741590, EPI_ISL_741591, EPI_ISL_741592, EPI_ISL_741593, EPI_ISL_741594, EPI_ISL_741595, EPI_ISL_741596, EPI_ISL_741597, EPI_ISL_741598, EPI_ISL_741599, EPI_ISL_741600, EPI_ISL_741601, EPI_ISL_741602, EPI_ISL_741603, EPI_ISL_741604, EPI_ISL_741605, EPI_ISL_741606, EPI_ISL_741607, EPI_ISL_741608 | | | |
| see above | Quadram Institute Bioscience | COVID-19 Genomics UK (COG-UK) Consortium | Dave J. Baker, Gemma L. Kay, Alp Aydin, Thanh Le-Viet, Steven Rudder, Ana P. Tedim, Anastasia Kolyva, Maria Diaz, Leonardo de Oliveira Martins, Nabil-Fareed Alikhan, Lizzie Meadows, Rachael Stanley, Ngozi Elumogo, Muhammed Yasir, Nicholas M. Thomson, Alexander J Trotter, Rachel Gilroy, Samuel Bloomfield, Claire Stuart, Andrew Bell, Reenesh Prakash, Samir Dervisevic, Alison E. Mather, John Wain, Mark Webber, Andrew J. Page, Justin O'Grady |
| EPI_ISL_741620, EPI_ISL_741621, EPI_ISL_741622, EPI_ISL_741623, EPI_ISL_741624, EPI_ISL_741625, EPI_ISL_741626, EPI_ISL_741627, EPI_ISL_741628, EPI_ISL_741629, EPI_ISL_741630 | | | |
| see above | Queens Medical Centre, Clinical Microbiology Department / DeepSeq Nottingham | COVID-19 Genomics UK (COG-UK) Consortium | Gemma Clark, Wendy Smith, Manjinder Khakh, Vicki M Fleming, Michelle M Lister, Hannah Howson-Wells, Jonathan Ball, Patrick McClure, Joseph Chappell, Theocharis Tsoleridis, Nadine Holmes, Matthew Carlisle, Christopher Moore, Fei Sang, Johnny Debebe, Victoria Wright, Matthew Loose |
| EPI_ISL_741675, EPI_ISL_741693, EPI_ISL_741697, EPI_ISL_741730, EPI_ISL_741747 | Centre for Enzyme Innovation, University of Portsmouth / Translational Research Laboratory, Portsmouth Hospitals NHS Trust | COVID-19 Genomics UK (COG-UK) Consortium | Angela Beckett,Yann Bourgeois,Garry Scarlett,Sharon Glaysher,Scott Elliott,Kelly Bicknell,Robert Impey,Allyson Lloyd,Sarah Wyllie,Ethan Butcher,Anoop Chauhan,Samuel Robson |
| EPI_ISL_741966, EPI_ISL_741974, EPI_ISL_742007, EPI_ISL_742017, EPI_ISL_742050, EPI_ISL_742052, EPI_ISL_742064, EPI_ISL_742082, EPI_ISL_742100, EPI_ISL_742105, EPI_ISL_742114 | | | |
| see above | Virology Department, Sheffield Teaching Hospitals NHS Foundation Trust/Department of Infection, Immunity and Cardiovascular Disease, The Medical School, University of Sheffield | COVID-19 Genomics UK (COG-UK) Consortium | Thushan de Silva, Matthew Parker, Nikki Smith, Adri Angyal, Rebecca Brown, Luke Green, Rachel Tucker, Paul Parsons, Danielle Groves, Katie Johnson, Laura Carrilero, Alex Keeley, Dave Partridge, Matthew Wyles, Benjamin Lindsey, Mehmet Yavuz, Mohammad Raza, Cariad Evans |
| EPI_ISL_742194, EPI_ISL_742195, EPI_ISL_742196, EPI_ISL_742197, EPI_ISL_742198, EPI_ISL_742199, EPI_ISL_742200, EPI_ISL_742201 | Virology Department, Royal Infirmary of Edinburgh, NHS Lothian / School of Biological Sciences, University of Edinburgh / Institute of Genetics and Molecular Medicine, University of Edinburgh | COVID-19 Genomics UK (COG-UK) Consortium | McHugh M, Dewar R, Rooke S, Gallagher M, Balcaza C, O'Toole Á, Scher E, Hill V, McCrone JT, Colquhoun R, Yu X, Jackson B, Rambaut A, Williams TC, Templeton K |
| EPI_ISL_743372, EPI_ISL_743603, EPI_ISL_743604, EPI_ISL_743605, EPI_ISL_743782, EPI_ISL_743785, EPI_ISL_743805, EPI_ISL_743806, EPI_ISL_743820, EPI_ISL_743821, EPI_ISL_743822, EPI_ISL_743825, EPI_ISL_743826, EPI_ISL_743827, EPI_ISL_743828, EPI_ISL_743831, EPI_ISL_743832, EPI_ISL_743833, EPI_ISL_743835, EPI_ISL_743836, EPI_ISL_743838, EPI_ISL_743839, EPI_ISL_743840, EPI_ISL_743841, EPI_ISL_743842, EPI_ISL_743843, EPI_ISL_743844, EPI_ISL_743845, EPI_ISL_743846, EPI_ISL_743847, EPI_ISL_743848, EPI_ISL_743849, EPI_ISL_743850, EPI_ISL_743851, EPI_ISL_743852, EPI_ISL_743853, EPI_ISL_743854, EPI_ISL_743855, EPI_ISL_743856, EPI_ISL_743857, EPI_ISL_743858, EPI_ISL_743859, EPI_ISL_743860, EPI_ISL_743861, EPI_ISL_743862, EPI_ISL_743863, EPI_ISL_743864, EPI_ISL_743865, EPI_ISL_743866, EPI_ISL_743867, EPI_ISL_743868, EPI_ISL_743869, EPI_ISL_743870, EPI_ISL_743871, EPI_ISL_743872, EPI_ISL_743873, EPI_ISL_743877, EPI_ISL_743886, EPI_ISL_743887, EPI_ISL_743888, EPI_ISL_743889, EPI_ISL_743898, EPI_ISL_743899, EPI_ISL_743900, EPI_ISL_743903, EPI_ISL_743904, EPI_ISL_743905, EPI_ISL_743906, EPI_ISL_743907, EPI_ISL_743908, EPI_ISL_743909, EPI_ISL_743910, EPI_ISL_743911, EPI_ISL_743912, EPI_ISL_743913, EPI_ISL_743914, EPI_ISL_743915, EPI_ISL_743916, EPI_ISL_743917, EPI_ISL_743918, EPI_ISL_743919, EPI_ISL_743920, EPI_ISL_743921, EPI_ISL_743922, EPI_ISL_743923, EPI_ISL_743924, EPI_ISL_743925, EPI_ISL_743926, EPI_ISL_743927, EPI_ISL_743938, EPI_ISL_743939, EPI_ISL_743940, EPI_ISL_743941, EPI_ISL_743987, EPI_ISL_744000, EPI_ISL_744001, EPI_ISL_744002, EPI_ISL_744003, EPI_ISL_744004, EPI_ISL_744005, EPI_ISL_744006, EPI_ISL_744007, EPI_ISL_744008, EPI_ISL_744009, EPI_ISL_744010, EPI_ISL_744011, EPI_ISL_744012, EPI_ISL_744013, EPI_ISL_744014, EPI_ISL_744015, EPI_ISL_744016, EPI_ISL_744017, EPI_ISL_744018, EPI_ISL_744019, EPI_ISL_744020, EPI_ISL_744021, EPI_ISL_744022, EPI_ISL_744023, EPI_ISL_744024, EPI_ISL_744025, EPI_ISL_744026, EPI_ISL_744027, EPI_ISL_744028, EPI_ISL_744029, EPI_ISL_744030, EPI_ISL_744031, EPI_ISL_744032, EPI_ISL_744033, EPI_ISL_744034, EPI_ISL_744035, EPI_ISL_744036, EPI_ISL_744037, EPI_ISL_744038, EPI_ISL_744039, EPI_ISL_744040, EPI_ISL_744041, EPI_ISL_744042, EPI_ISL_744043, EPI_ISL_744044, EPI_ISL_744045, EPI_ISL_744046, EPI_ISL_744047, EPI_ISL_744048, EPI_ISL_744049, EPI_ISL_744050, EPI_ISL_744051, EPI_ISL_744052, EPI_ISL_744053, EPI_ISL_744054, EPI_ISL_744055, EPI_ISL_744056, EPI_ISL_744057, EPI_ISL_744058, EPI_ISL_744059, EPI_ISL_744060, EPI_ISL_744061, EPI_ISL_744062, EPI_ISL_744063, EPI_ISL_744064, EPI_ISL_744065, EPI_ISL_744066, EPI_ISL_744068, EPI_ISL_744069, EPI_ISL_744070, EPI_ISL_744071, EPI_ISL_744072, EPI_ISL_744073, EPI_ISL_744074, EPI_ISL_744075, EPI_ISL_744076, EPI_ISL_744077, EPI_ISL_744078, EPI_ISL_744079, EPI_ISL_744080, EPI_ISL_744081, EPI_ISL_744082, EPI_ISL_744083, EPI_ISL_744084, EPI_ISL_744085, EPI_ISL_744086, EPI_ISL_744087, EPI_ISL_744088, EPI_ISL_744089, EPI_ISL_744090, EPI_ISL_744091, EPI_ISL_744092, EPI_ISL_744093, EPI_ISL_744094, EPI_ISL_744095, EPI_ISL_744096, EPI_ISL_744097, EPI_ISL_744098, EPI_ISL_744099, EPI_ISL_744100, EPI_ISL_744101, EPI_ISL_744102, EPI_ISL_744103, EPI_ISL_744104, EPI_ISL_744105, EPI_ISL_744106, EPI_ISL_744107, EPI_ISL_744108, EPI_ISL_744109, EPI_ISL_744110, EPI_ISL_744111, EPI_ISL_744112, EPI_ISL_744113, EPI_ISL_744114, EPI_ISL_744115, EPI_ISL_744116, EPI_ISL_744117, EPI_ISL_744118, EPI_ISL_744119, EPI_ISL_744120, EPI_ISL_744121, EPI_ISL_744122, EPI_ISL_744123, EPI_ISL_744124, EPI_ISL_744125, EPI_ISL_744126, EPI_ISL_744127, EPI_ISL_744128, EPI_ISL_744129, EPI_ISL_744130, EPI_ISL_744131, EPI_ISL_744132, EPI_ISL_744133 | | | |
| see above | Wales Specialist Virology Centre Sequencing lab: Pathogen Genomics Unit | COVID-19 Genomics UK (COG-UK) Consortium | Catherine Moore, Johnathan Evans, Laura Gifford, Malorie Perry, Simon Cottrell, Angela Marchbank, Alec Birchley, Alexander Adams, Amy Gaskin, Bree Gatica-Wilcox, Jason Coombes, Joel Southgate, Lauren Gilbert, Lee Graham, Nicole Pacchiarini, Sara Kumziene-Summerhayes, Sarah Taylor, Sophie Jones, Sara Rey, Matthew Bull, Joanne Watkins, Sally Corden, Tom Connor |
| EPI_ISL_744158, EPI_ISL_744206, EPI_ISL_744438, EPI_ISL_744456, EPI_ISL_744769, EPI_ISL_744805, EPI_ISL_744894 | Laboratoire national de santé, Microbiology, Virology | Laboratoire national de santé, Microbiology, Microbial Genomics Platform | Anke Wienecke-Baldacchino, Catherine Ragimbeau,Jessica Tapp, Fatu Djabi, Lise Pignon, Raoul Salmon, Tamir Abdelrahman |
| EPI_ISL_745070, EPI_ISL_745071, EPI_ISL_745072, EPI_ISL_745073, EPI_ISL_745074, EPI_ISL_745075, EPI_ISL_745076, EPI_ISL_745077 | Israel Central Virology laboratory | Israel Central Virology laboratory | Neta Zuckerman, Efrat Dahan Bucris, Oran Erster, Michal Mandelboim, Orna Mor, Ella Mendelson |
| EPI_ISL_745212, EPI_ISL_745218, EPI_ISL_745219 | Lab voor klinische biologie | Onderzoeksgroep Virologie | Nick Vereecke, Laurens Lambrechts, Marthe Pauwels, Bruno Verhasselt, Linos Vandekerckhove, Hans Nauwynck, Sebastiaan Theuns |
| EPI_ISL_745222 | UZ Leuven, National Reference Laboratory for Coronaviruses, Laboratory Medicine, Leuven, Belgium | KU Leuven, Rega Institute, Clinical and Epidemiological Virology | Tony Wawina-Bokalanga, Joan Marti-Carerras, Bert Vanmechelen, Piet Maes |
| EPI_ISL_745260 | CHUM-Site Glen-LAB Microbiologie | Laboratoire de santé publique du Québec | Sandrine Moreira, Ioannis Ragoussis, Guillaume Bourque, Jesse Shapiro, Mark Lathrop and Michel Roger on behalf of the CoVSeQ research group (http://covseq.ca/researchgroup) |
| EPI_ISL_745318, EPI_ISL_745324, EPI_ISL_745332, EPI_ISL_745389 | CNR Virus des Infections Respiratoires - France SUD | CNR Virus des Infections Respiratoires - France SUD | Antonin Bal, Gregory Destras, Claudia Gonzalez, Gwendolyne Burfin, Quentin Semanas, Martine Valette, Bruno Lina, Laurence Josset |
| EPI_ISL_746826 | National Institute for Infectious Diseases, INMI, "L. Spallanzani" IRCCS | National Institute for Infectious Diseases, INMI, "L. Spallanzani" IRCCS | C.E.M Gruber, B Bartolini, M Rueca, F Messina, E Giombini, A Di Caro, MR Capobianchi |
| EPI_ISL_747194 | Clinical Microbiology Laboratory, Faculty of Medicine, Universitas Indonesia | Clinical Microbiology Laboratory, Faculty of Medicine, Universitas Indonesia | Fera Ibrahim, Augustine Natasha, Andi Yasmon, Pratiwi Sudarmono, Yulia Rosa Saharman, Anis Karuniawati, Fithriyah |
| EPI_ISL_749156, EPI_ISL_749157, EPI_ISL_749159, EPI_ISL_749160, EPI_ISL_749161, EPI_ISL_749162, EPI_ISL_749163, EPI_ISL_749164, EPI_ISL_749165, EPI_ISL_749166, EPI_ISL_749168, EPI_ISL_749172, EPI_ISL_749173, EPI_ISL_749174, EPI_ISL_749175, EPI_ISL_749178, EPI_ISL_749179, EPI_ISL_749180, EPI_ISL_749182, EPI_ISL_749183, EPI_ISL_749184, EPI_ISL_749185, EPI_ISL_749186, EPI_ISL_749187, EPI_ISL_749188, EPI_ISL_749189, EPI_ISL_749190, EPI_ISL_749191, EPI_ISL_749192, EPI_ISL_749193, EPI_ISL_749194, EPI_ISL_749195, EPI_ISL_749196, EPI_ISL_749197, EPI_ISL_749198, EPI_ISL_749199, EPI_ISL_749200, EPI_ISL_749201, EPI_ISL_749202, EPI_ISL_749203, EPI_ISL_749204, EPI_ISL_749205, EPI_ISL_749206, EPI_ISL_749207, EPI_ISL_749208, EPI_ISL_749209, EPI_ISL_749210, EPI_ISL_749211, EPI_ISL_749212, EPI_ISL_749213, EPI_ISL_749214, EPI_ISL_749215, EPI_ISL_749216, EPI_ISL_749217, EPI_ISL_749218, EPI_ISL_749219, EPI_ISL_749220, EPI_ISL_749221, EPI_ISL_749222, EPI_ISL_749263, EPI_ISL_749264, EPI_ISL_749265, EPI_ISL_749266, EPI_ISL_749269, EPI_ISL_749270, EPI_ISL_749272, EPI_ISL_749275, EPI_ISL_749277, EPI_ISL_749278, EPI_ISL_749280, EPI_ISL_749281, EPI_ISL_749282, EPI_ISL_749283, EPI_ISL_749284, EPI_ISL_749285, EPI_ISL_749286, EPI_ISL_749288, EPI_ISL_749289, EPI_ISL_749296, EPI_ISL_749297, EPI_ISL_749298, EPI_ISL_749299, EPI_ISL_749300, EPI_ISL_749301, EPI_ISL_749302, EPI_ISL_749303, EPI_ISL_749304, EPI_ISL_749305, EPI_ISL_749306, EPI_ISL_749307, EPI_ISL_749308, EPI_ISL_749309, EPI_ISL_749310, EPI_ISL_749311, EPI_ISL_749312, EPI_ISL_749313, EPI_ISL_749314, EPI_ISL_749315, EPI_ISL_749316, EPI_ISL_749317, EPI_ISL_749318, EPI_ISL_749319, EPI_ISL_749320, EPI_ISL_749355, EPI_ISL_749356, EPI_ISL_749358, EPI_ISL_749359, EPI_ISL_749360, EPI_ISL_749362, EPI_ISL_749363, EPI_ISL_749364, EPI_ISL_749365, EPI_ISL_749366, EPI_ISL_749367, EPI_ISL_749368, EPI_ISL_749370, EPI_ISL_749371, EPI_ISL_749372, EPI_ISL_749373, EPI_ISL_749376, EPI_ISL_749377, EPI_ISL_749378, EPI_ISL_749379, EPI_ISL_749380, EPI_ISL_749381, EPI_ISL_749382, EPI_ISL_749383, EPI_ISL_749384, EPI_ISL_749385, EPI_ISL_749386, EPI_ISL_749387, EPI_ISL_749388, EPI_ISL_749389, EPI_ISL_749407, EPI_ISL_749408, EPI_ISL_749409, EPI_ISL_749410, EPI_ISL_749411, EPI_ISL_749412, EPI_ISL_749413, EPI_ISL_749414, EPI_ISL_749415, EPI_ISL_749416, EPI_ISL_749417, EPI_ISL_749418, EPI_ISL_749419, EPI_ISL_749420, EPI_ISL_749421, EPI_ISL_749422, EPI_ISL_749423, EPI_ISL_749424, EPI_ISL_749425, EPI_ISL_749426, EPI_ISL_749427, EPI_ISL_749428, EPI_ISL_749429, EPI_ISL_749430, EPI_ISL_749431, EPI_ISL_749432, EPI_ISL_749433, EPI_ISL_749434, EPI_ISL_749435, | | | |

EPI_ISL_749436, EPI_ISL_749437, EPI_ISL_749438, EPI_ISL_749439, EPI_ISL_749440, EPI_ISL_749441, EPI_ISL_749442, EPI_ISL_749443, EPI_ISL_749444, EPI_ISL_749445, EPI_ISL_749446, EPI_ISL_749447, EPI_ISL_749448, EPI_ISL_749449, EPI_ISL_749450, EPI_ISL_749451, EPI_ISL_749484, EPI_ISL_749485,
EPI_ISL_749486, EPI_ISL_749487, EPI_ISL_749488, EPI_ISL_749489, EPI_ISL_749490, EPI_ISL_749493, EPI_ISL_749498, EPI_ISL_749500, EPI_ISL_749501, EPI_ISL_749502, EPI_ISL_749503, EPI_ISL_749504, EPI_ISL_749505, EPI_ISL_749507, EPI_ISL_749511, EPI_ISL_749512, EPI_ISL_749515, EPI_ISL_749516,
EPI_ISL_749517, EPI_ISL_749518, EPI_ISL_749519, EPI_ISL_749523, EPI_ISL_749524, EPI_ISL_749528, EPI_ISL_749529, EPI_ISL_749530, EPI_ISL_749531, EPI_ISL_749532, EPI_ISL_749533, EPI_ISL_749534, EPI_ISL_749535, EPI_ISL_749536, EPI_ISL_749537, EPI_ISL_749538, EPI_ISL_749539, EPI_ISL_749542,
EPI_ISL_749543, EPI_ISL_749544, EPI_ISL_749545, EPI_ISL_749546, EPI_ISL_749547, EPI_ISL_749548, EPI_ISL_749549, EPI_ISL_749550, EPI_ISL_749551, EPI_ISL_749552, EPI_ISL_749553, EPI_ISL_749554, EPI_ISL_749555, EPI_ISL_749556, EPI_ISL_749557, EPI_ISL_749558, EPI_ISL_749559, EPI_ISL_749560,
EPI_ISL_749561, EPI_ISL_749562, EPI_ISL_749563, EPI_ISL_749564, EPI_ISL_749565, EPI_ISL_749566, EPI_ISL_749567, EPI_ISL_749568, EPI_ISL_749569, EPI_ISL_749570, EPI_ISL_749571, EPI_ISL_749572, EPI_ISL_749573, EPI_ISL_749574, EPI_ISL_749575, EPI_ISL_749576, EPI_ISL_749577, EPI_ISL_749578,
EPI_ISL_749579, EPI_ISL_749580, EPI_ISL_749581, EPI_ISL_749582, EPI_ISL_749583, EPI_ISL_749584, EPI_ISL_749585, EPI_ISL_749586, EPI_ISL_749587, EPI_ISL_749588, EPI_ISL_749589, EPI_ISL_749590, EPI_ISL_749591, EPI_ISL_749592, EPI_ISL_749593, EPI_ISL_749594, EPI_ISL_749595, EPI_ISL_749596,
EPI_ISL_749597, EPI_ISL_749598, EPI_ISL_749599, EPI_ISL_749600, EPI_ISL_749601, EPI_ISL_749602, EPI_ISL_749603, EPI_ISL_749604, EPI_ISL_749605, EPI_ISL_749606, EPI_ISL_749607, EPI_ISL_749608, EPI_ISL_749609, EPI_ISL_749610, EPI_ISL_749611, EPI_ISL_749612, EPI_ISL_749613, EPI_ISL_749614,
EPI_ISL_749615, EPI_ISL_749616, EPI_ISL_749617, EPI_ISL_749618, EPI_ISL_749619, EPI_ISL_749620, EPI_ISL_749621, EPI_ISL_749622, EPI_ISL_749623, EPI_ISL_749624, EPI_ISL_749625, EPI_ISL_749626, EPI_ISL_749627, EPI_ISL_749628, EPI_ISL_749629, EPI_ISL_749630, EPI_ISL_749631, EPI_ISL_749632,
EPI_ISL_749633, EPI_ISL_749634, EPI_ISL_749635, EPI_ISL_749636, EPI_ISL_749637, EPI_ISL_749638, EPI_ISL_749639, EPI_ISL_749640, EPI_ISL_749641, EPI_ISL_749642, EPI_ISL_749643, EPI_ISL_749644, EPI_ISL_749645, EPI_ISL_749646, EPI_ISL_749647, EPI_ISL_749648, EPI_ISL_749649, EPI_ISL_749650,
EPI_ISL_749651, EPI_ISL_749652, EPI_ISL_749653, EPI_ISL_749654, EPI_ISL_749655, EPI_ISL_749656, EPI_ISL_749657, EPI_ISL_749658, EPI_ISL_749659, EPI_ISL_749660, EPI_ISL_749661, EPI_ISL_749662, EPI_ISL_749663, EPI_ISL_749664, EPI_ISL_749665, EPI_ISL_749666, EPI_ISL_749667, EPI_ISL_749668,
EPI_ISL_749669, EPI_ISL_749670, EPI_ISL_749671, EPI_ISL_749672, EPI_ISL_749673, EPI_ISL_749674, EPI_ISL_749675, EPI_ISL_749676, EPI_ISL_749677, EPI_ISL_749678, EPI_ISL_749679, EPI_ISL_749680, EPI_ISL_749681, EPI_ISL_749682, EPI_ISL_749683, EPI_ISL_749684, EPI_ISL_749685, EPI_ISL_749686,
EPI_ISL_749687, EPI_ISL_749688, EPI_ISL_749689, EPI_ISL_749690, EPI_ISL_749691, EPI_ISL_749692, EPI_ISL_749693, EPI_ISL_749694, EPI_ISL_749695, EPI_ISL_749696, EPI_ISL_749697, EPI_ISL_749698, EPI_ISL_749699, EPI_ISL_749700, EPI_ISL_749701, EPI_ISL_749702, EPI_ISL_749703, EPI_ISL_749704,
EPI_ISL_749705, EPI_ISL_749707, EPI_ISL_749708, EPI_ISL_749709, EPI_ISL_749710, EPI_ISL_749711, EPI_ISL_749712, EPI_ISL_749713, EPI_ISL_749714, EPI_ISL_749715, EPI_ISL_749716, EPI_ISL_749717, EPI_ISL_749718, EPI_ISL_749719, EPI_ISL_749720, EPI_ISL_749721, EPI_ISL_749722, EPI_ISL_749723,
EPI_ISL_749724, EPI_ISL_749725, EPI_ISL_749726, EPI_ISL_749727, EPI_ISL_749728, EPI_ISL_749729, EPI_ISL_749730, EPI_ISL_749731, EPI_ISL_749732, EPI_ISL_749733, EPI_ISL_749734, EPI_ISL_749735, EPI_ISL_749736, EPI_ISL_749737, EPI_ISL_749738, EPI_ISL_749739, EPI_ISL_749740, EPI_ISL_749741,
EPI_ISL_749742, EPI_ISL_749743, EPI_ISL_749744, EPI_ISL_749745, EPI_ISL_749746, EPI_ISL_749747, EPI_ISL_749748, EPI_ISL_749749, EPI_ISL_749750, EPI_ISL_749751, EPI_ISL_749752, EPI_ISL_749753, EPI_ISL_749754, EPI_ISL_749755, EPI_ISL_749756, EPI_ISL_749757, EPI_ISL_749758, EPI_ISL_749759,
EPI_ISL_749760, EPI_ISL_749761, EPI_ISL_749762, EPI_ISL_749763, EPI_ISL_749764, EPI_ISL_749765, EPI_ISL_749766, EPI_ISL_749767, EPI_ISL_749768, EPI_ISL_749769, EPI_ISL_749770, EPI_ISL_749771, EPI_ISL_749772, EPI_ISL_749773, EPI_ISL_749774, EPI_ISL_749775, EPI_ISL_749776, EPI_ISL_749777,
EPI_ISL_749778, EPI_ISL_749779, EPI_ISL_749780, EPI_ISL_749781, EPI_ISL_749782, EPI_ISL_749783, EPI_ISL_749784, EPI_ISL_749785, EPI_ISL_749786, EPI_ISL_749787, EPI_ISL_749788, EPI_ISL_749789, EPI_ISL_749790, EPI_ISL_749791, EPI_ISL_749792, EPI_ISL_749793, EPI_ISL_749794, EPI_ISL_749795,
EPI_ISL_749796, EPI_ISL_749797, EPI_ISL_749798, EPI_ISL_749799, EPI_ISL_749800, EPI_ISL_749801, EPI_ISL_749802, EPI_ISL_749803, EPI_ISL_749804, EPI_ISL_749805, EPI_ISL_749806, EPI_ISL_749807, EPI_ISL_749808, EPI_ISL_749809, EPI_ISL_749810, EPI_ISL_749811, EPI_ISL_749812, EPI_ISL_749813,
EPI_ISL_749814, EPI_ISL_749815, EPI_ISL_749816, EPI_ISL_749817, EPI_ISL_749818, EPI_ISL_749819, EPI_ISL_749820, EPI_ISL_749821, EPI_ISL_749822, EPI_ISL_749823, EPI_ISL_749824, EPI_ISL_749825, EPI_ISL_749826, EPI_ISL_749827, EPI_ISL_749828, EPI_ISL_749829, EPI_ISL_749830, EPI_ISL_749831,
EPI_ISL_749832, EPI_ISL_749833, EPI_ISL_749834, EPI_ISL_749835, EPI_ISL_749836, EPI_ISL_749837, EPI_ISL_749838, EPI_ISL_749839, EPI_ISL_749840, EPI_ISL_749841, EPI_ISL_749842, EPI_ISL_749843, EPI_ISL_749844, EPI_ISL_749845, EPI_ISL_749846, EPI_ISL_749847, EPI_ISL_749848, EPI_ISL_749849,
EPI_ISL_749850, EPI_ISL_749851, EPI_ISL_749852, EPI_ISL_749853, EPI_ISL_749854, EPI_ISL_749855, EPI_ISL_749856, EPI_ISL_749857, EPI_ISL_749858, EPI_ISL_750180, EPI_ISL_750181, EPI_ISL_750182, EPI_ISL_750183, EPI_ISL_750184, EPI_ISL_750185, EPI_ISL_750186, EPI_ISL_750187, EPI_ISL_750188,
EPI_ISL_750189, EPI_ISL_750190, EPI_ISL_750191, EPI_ISL_750192, EPI_ISL_750193, EPI_ISL_750194, EPI_ISL_750195, EPI_ISL_750196, EPI_ISL_750197, EPI_ISL_750198, EPI_ISL_750199, EPI_ISL_750200, EPI_ISL_750201, EPI_ISL_750202, EPI_ISL_750203, EPI_ISL_750204, EPI_ISL_750205, EPI_ISL_750206,
EPI_ISL_750207, EPI_ISL_750208, EPI_ISL_750209, EPI_ISL_750210, EPI_ISL_750211, EPI_ISL_750212, EPI_ISL_750213, EPI_ISL_750214, EPI_ISL_750215, EPI_ISL_750216, EPI_ISL_750217, EPI_ISL_750218, EPI_ISL_750219, EPI_ISL_750220, EPI_ISL_750221, EPI_ISL_750222, EPI_ISL_750223, EPI_ISL_750224,
EPI_ISL_750225, EPI_ISL_750226, EPI_ISL_750227, EPI_ISL_750228, EPI_ISL_750229, EPI_ISL_750230, EPI_ISL_750231, EPI_ISL_750232, EPI_ISL_750233, EPI_ISL_750234, EPI_ISL_750235, EPI_ISL_750236, EPI_ISL_750237, EPI_ISL_750238, EPI_ISL_750239, EPI_ISL_750240, EPI_ISL_750241, EPI_ISL_750242,
EPI_ISL_750243, EPI_ISL_750244, EPI_ISL_750245, EPI_ISL_750246, EPI_ISL_750247, EPI_ISL_750248, EPI_ISL_750249, EPI_ISL_750250, EPI_ISL_750251, EPI_ISL_750252, EPI_ISL_750253, EPI_ISL_750254, EPI_ISL_750255, EPI_ISL_750257, EPI_ISL_750258, EPI_ISL_750259, EPI_ISL_750260, EPI_ISL_750261,
EPI_ISL_750262, EPI_ISL_750263, EPI_ISL_750264, EPI_ISL_750265, EPI_ISL_750266, EPI_ISL_750267, EPI_ISL_750268, EPI_ISL_750269, EPI_ISL_750270, EPI_ISL_750271, EPI_ISL_750272, EPI_ISL_750273, EPI_ISL_750274, EPI_ISL_750275, EPI_ISL_750276, EPI_ISL_750277, EPI_ISL_750278, EPI_ISL_750279,
EPI_ISL_750280, EPI_ISL_750281, EPI_ISL_750282, EPI_ISL_750283, EPI_ISL_750284, EPI_ISL_750285, EPI_ISL_750286, EPI_ISL_750287, EPI_ISL_750288, EPI_ISL_750289, EPI_ISL_750290, EPI_ISL_750291, EPI_ISL_750292, EPI_ISL_750293, EPI_ISL_750294, EPI_ISL_750295, EPI_ISL_750296, EPI_ISL_750297,
EPI_ISL_750298, EPI_ISL_750299, EPI_ISL_750300, EPI_ISL_750301, EPI_ISL_750302, EPI_ISL_750303, EPI_ISL_750304, EPI_ISL_750305, EPI_ISL_750306, EPI_ISL_750307, EPI_ISL_750308, EPI_ISL_750309, EPI_ISL_750310, EPI_ISL_750311, EPI_ISL_750312, EPI_ISL_750313, EPI_ISL_750314, EPI_ISL_750315,
EPI_ISL_750316, EPI_ISL_750317, EPI_ISL_750318, EPI_ISL_750319, EPI_ISL_750320, EPI_ISL_750321, EPI_ISL_750322, EPI_ISL_750323, EPI_ISL_750324, EPI_ISL_750325, EPI_ISL_750326, EPI_ISL_750327, EPI_ISL_750328, EPI_ISL_750329, EPI_ISL_750330, EPI_ISL_750331, EPI_ISL_750332, EPI_ISL_750333,
EPI_ISL_750334, EPI_ISL_750335, EPI_ISL_750336, EPI_ISL_750337, EPI_ISL_750338, EPI_ISL_750339, EPI_ISL_750340, EPI_ISL_750341, EPI_ISL_750342, EPI_ISL_750343, EPI_ISL_750344, EPI_ISL_750345, EPI_ISL_750346, EPI_ISL_750347, EPI_ISL_750348, EPI_ISL_750349, EPI_ISL_750350, EPI_ISL_750351,
EPI_ISL_750352, EPI_ISL_750353, EPI_ISL_750354, EPI_ISL_750355, EPI_ISL_750356, EPI_ISL_750357, EPI_ISL_750358, EPI_ISL_750359, EPI_ISL_750360, EPI_ISL_750361, EPI_ISL_750362, EPI_ISL_750363, EPI_ISL_750364, EPI_ISL_750365, EPI_ISL_750366, EPI_ISL_750367, EPI_ISL_750368, EPI_ISL_750369,
EPI_ISL_750370, EPI_ISL_750371, EPI_ISL_750372, EPI_ISL_750373, EPI_ISL_750374, EPI_ISL_750375, EPI_ISL_750376, EPI_ISL_750377, EPI_ISL_750378, EPI_ISL_750379, EPI_ISL_750380, EPI_ISL_750381, EPI_ISL_750382, EPI_ISL_750383, EPI_ISL_750384, EPI_ISL_750385, EPI_ISL_750386, EPI_ISL_750387,
EPI_ISL_750388, EPI_ISL_750389, EPI_ISL_750390, EPI_ISL_750391, EPI_ISL_750392, EPI_ISL_750393, EPI_ISL_750394, EPI_ISL_750395, EPI_ISL_750396, EPI_ISL_750397, EPI_ISL_750398, EPI_ISL_750399, EPI_ISL_750400, EPI_ISL_750401, EPI_ISL_750402, EPI_ISL_750403, EPI_ISL_750404, EPI_ISL_750405,
EPI_ISL_750406, EPI_ISL_750407, EPI_ISL_750408, EPI_ISL_750409, EPI_ISL_750410, EPI_ISL_750411, EPI_ISL_750412, EPI_ISL_750413, EPI_ISL_750414, EPI_ISL_750415, EPI_ISL_750416, EPI_ISL_750417, EPI_ISL_750418, EPI_ISL_750419, EPI_ISL_750420, EPI_ISL_750421, EPI_ISL_750422, EPI_ISL_750423,
EPI_ISL_750424, EPI_ISL_750425, EPI_ISL_750426, EPI_ISL_750427, EPI_ISL_750428, EPI_ISL_750429, EPI_ISL_750431, EPI_ISL_750432, EPI_ISL_750433, EPI_ISL_750434, EPI_ISL_750435, EPI_ISL_750436, EPI_ISL_750437, EPI_ISL_750438, EPI_ISL_750439, EPI_ISL_750440, EPI_ISL_750441, EPI_ISL_750442,
EPI_ISL_750443, EPI_ISL_750444, EPI_ISL_750445, EPI_ISL_750446, EPI_ISL_750447, EPI_ISL_750448, EPI_ISL_750449, EPI_ISL_750450, EPI_ISL_750451, EPI_ISL_750452, EPI_ISL_750453, EPI_ISL_750454, EPI_ISL_750455, EPI_ISL_750456, EPI_ISL_750457, EPI_ISL_750458, EPI_ISL_750459, EPI_ISL_750460,
EPI_ISL_750461, EPI_ISL_750462, EPI_ISL_750463, EPI_ISL_750464, EPI_ISL_750465, EPI_ISL_750466, EPI_ISL_750467, EPI_ISL_750468, EPI_ISL_750469, EPI_ISL_750470, EPI_ISL_750471, EPI_ISL_750472, EPI_ISL_750473, EPI_ISL_750474, EPI_ISL_750475, EPI_ISL_750476, EPI_ISL_750477, EPI_ISL_750478,
EPI_ISL_750479, EPI_ISL_750480, EPI_ISL_750481, EPI_ISL_750482, EPI_ISL_750483, EPI_ISL_750484, EPI_ISL_750485, EPI_ISL_750486, EPI_ISL_750487, EPI_ISL_750488, EPI_ISL_750489, EPI_ISL_750490, EPI_ISL_750491, EPI_ISL_750492, EPI_ISL_750493, EPI_ISL_750494, EPI_ISL_750495, EPI_ISL_750496,
EPI_ISL_750497, EPI_ISL_750498, EPI_ISL_750499, EPI_ISL_750500, EPI_ISL_750501, EPI_ISL_750502, EPI_ISL_750503, EPI_ISL_750504, EPI_ISL_750505, EPI_ISL_750506, EPI_ISL_750507, EPI_ISL_750508, EPI_ISL_750509, EPI_ISL_750510, EPI_ISL_750511, EPI_ISL_750512, EPI_ISL_750513, EPI_ISL_750514,
EPI_ISL_750515, EPI_ISL_750516, EPI_ISL_750517, EPI_ISL_750518, EPI_ISL_750519, EPI_ISL_750520, EPI_ISL_750521, EPI_ISL_750522, EPI_ISL_750523, EPI_ISL_750524, EPI_ISL_750525, EPI_ISL_750526, EPI_ISL_750527, EPI_ISL_750528, EPI_ISL_750529, EPI_ISL_750530, EPI_ISL_750531, EPI_ISL_750532,
EPI_ISL_750533, EPI_ISL_750534, EPI_ISL_750535, EPI_ISL_750536, EPI_ISL_750537, EPI_ISL_750538, EPI_ISL_750539, EPI_ISL_750540, EPI_ISL_750541, EPI_ISL_750542, EPI_ISL_750543, EPI_ISL_750544, EPI_ISL_750545, EPI_ISL_750546, EPI_ISL_750547, EPI_ISL_750548, EPI_ISL_750549, EPI_ISL_750550,
EPI_ISL_750551, EPI_ISL_750552, EPI_ISL_750553, EPI_ISL_750554, EPI_ISL_750555, EPI_ISL_750556, EPI_ISL_750557, EPI_ISL_750558, EPI_ISL_750559, EPI_ISL_750560, EPI_ISL_750561, EPI_ISL_750562, EPI_ISL_750563, EPI_ISL_750564, EPI_ISL_750565, EPI_ISL_750566, EPI_ISL_750567, EPI_ISL_750568,
EPI_ISL_750569, EPI_ISL_750570, EPI_ISL_750571, EPI_ISL_750572, EPI_ISL_750573, EPI_ISL_750574, EPI_ISL_750575, EPI_ISL_750576, EPI_ISL_750577, EPI_ISL_750578, EPI_ISL_750579, EPI_ISL_750580, EPI_ISL_750581, EPI_ISL_750582, EPI_ISL_750583, EPI_ISL_750584, EPI_ISL_750585, EPI_ISL_750586,
EPI_ISL_750587, EPI_ISL_750588, EPI_ISL_750589, EPI_ISL_750590, EPI_ISL_750591, EPI_ISL_750592, EPI_ISL_750593, EPI_ISL_750594, EPI_ISL_750595, EPI_ISL_750596, EPI_ISL_750597, EPI_ISL_750598, EPI_ISL_750599, EPI_ISL_750600, EPI_ISL_750601, EPI_ISL_750602, EPI_ISL_750603, EPI_ISL_750604,
EPI_ISL_750605, EPI_ISL_750606, EPI_ISL_750607, EPI_ISL_750608, EPI_ISL_750609, EPI_ISL_750610, EPI_ISL_750611, EPI_ISL_750612, EPI_ISL_750613, EPI_ISL_750614, EPI_ISL_750615, EPI_ISL_750616, EPI_ISL_750617, EPI_ISL_750618, EPI_ISL_750619, EPI_ISL_750620, EPI_ISL_750621, EPI_ISL_750622,
EPI_ISL_750623, EPI_ISL_750624, EPI_ISL_750625, EPI_ISL_750626, EPI_ISL_750627, EPI_ISL_750628, EPI_ISL_750629, EPI_ISL_750630, EPI_ISL_750631, EPI_ISL_750632, EPI_ISL_750633, EPI_ISL_750634, EPI_ISL_750635, EPI_ISL_750636, EPI_ISL_750637, EPI_ISL_750638, EPI_ISL_750639, EPI_ISL_750640,
EPI_ISL_750641, EPI_ISL_750642, EPI_ISL_750643, EPI_ISL_750644, EPI_ISL_750645, EPI_ISL_750646, EPI_ISL_750647, EPI_ISL_750648, EPI_ISL_750649, EPI_ISL_750650, EPI_ISL_750651, EPI_ISL_750652, EPI_ISL_750653, EPI_ISL_750654, EPI_ISL_750655, EPI_ISL_750656, EPI_ISL_750657, EPI_ISL_750658,
EPI_ISL_750659, EPI_ISL_750660, EPI_ISL_750661, EPI_ISL_750662, EPI_ISL_750663, EPI_ISL_750664, EPI_ISL_750665, EPI_ISL_750666, EPI_ISL_750667, EPI_ISL_750668, EPI_ISL_750669, EPI_ISL_750670, EPI_ISL_750671, EPI_ISL_750672, EPI_ISL_750673, EPI_ISL_750674, EPI_ISL_750675, EPI_ISL_750676,
EPI_ISL_750677, EPI_ISL_750678, EPI_ISL_750679, EPI_ISL_750680, EPI_ISL_750681, EPI_ISL_750682, EPI_ISL_750683, EPI_ISL_750684, EPI_ISL_750685, EPI_ISL_750686, EPI_ISL_750687, EPI_ISL_750688, EPI_ISL_750689, EPI_ISL_750690, EPI_ISL_750691, EPI_ISL_750692, EPI_ISL_750693, EPI_ISL_750694,
EPI_ISL_750695, EPI_ISL_750696, EPI_ISL_750697, EPI_ISL_750698, EPI_ISL_750699, EPI_ISL_750700, EPI_ISL_750701, EPI_ISL_750702, EPI_ISL_750703, EPI_ISL_750704, EPI_ISL_750705, EPI_ISL_750706, EPI_ISL_750707, EPI_ISL_750708, EPI_ISL_750709, EPI_ISL_750710, EPI_ISL_750711, EPI_ISL_750712,
EPI_ISL_750713, EPI_ISL_750714, EPI_ISL_750715, EPI_ISL_750716, EPI_ISL_750717, EPI_ISL_750718, EPI_ISL_750719, EPI_ISL_750720, EPI_ISL_750721, EPI_ISL_750722, EPI_ISL_750723, EPI_ISL_750724, EPI_ISL_750725, EPI_ISL_750726, EPI_ISL_750727, EPI_ISL_750728, EPI_ISL_750729, EPI_ISL_750730,
EPI_ISL_750731, EPI_ISL_750732, EPI_ISL_750733, EPI_ISL_750734, EPI_ISL_750735, EPI_ISL_750736, EPI_ISL_750737, EPI_ISL_750738, EPI_ISL_750739, EPI_ISL_750740, EPI_ISL_750741, EPI_ISL_750742, EPI_ISL_750743, EPI_ISL_750744, EPI_ISL_750745, EPI_ISL_750746, EPI_ISL_750747, EPI_ISL_750748,
EPI_ISL_750749, EPI_ISL_750750, EPI_ISL_750751, EPI_ISL_750752, EPI_ISL_750753, EPI_ISL_750754, EPI_ISL_750755, EPI_ISL_750756, EPI_ISL_750757, EPI_ISL_750758, EPI_ISL_750759, EPI_ISL_750760, EPI_ISL_750761, EPI_ISL_750762, EPI_ISL_750763, EPI_ISL_750764, EPI_ISL_750765, EPI_ISL_750766,
EPI_ISL_750767, EPI_ISL_750768, EPI_ISL_750769, EPI_ISL_750770, EPI_ISL_750771, EPI_ISL_750772, EPI_ISL_750773, EPI_ISL_750774, EPI_ISL_750775, EPI_ISL_750776, EPI_ISL_750777, EPI_ISL_750778, EPI_ISL_750779, EPI_ISL_750780, EPI_ISL_750781, EPI_ISL_750782, EPI_ISL_750783, EPI_ISL_750784,
EPI_ISL_750785, EPI_ISL_750786, EPI_ISL_750787, EPI_ISL_750788, EPI_ISL_750789, EPI_ISL_750790, EPI_ISL_750791, EPI_ISL_750792, EPI_ISL_750793, EPI_ISL_750794, EPI_ISL_750795, EPI_ISL_750796, EPI_ISL_750797, EPI_ISL_750798, EPI_ISL_750799, EPI_ISL_750800, EPI_ISL_750801, EPI_ISL_750802,
EPI_ISL_750803, EPI_ISL_750804, EPI_ISL_750805, EPI_ISL_750806, EPI_ISL_750807, EPI_ISL_750808, EPI_ISL_750809, EPI_ISL_750810, EPI_ISL_750811, EPI_ISL_750812, EPI_ISL_750813, EPI_ISL_750814, EPI_ISL_750815, EPI_ISL_750816, EPI_ISL_750817, EPI_ISL_750818, EPI_ISL_750819, EPI_ISL_750821,
EPI_ISL_750822, EPI_ISL_750823, EPI_ISL_750824, EPI_ISL_750825, EPI_ISL_750826, EPI_ISL_750827, EPI_ISL_750828, EPI_ISL_750829, EPI_ISL_750830, EPI_ISL_750831, EPI_ISL_750832, EPI_ISL_750833, EPI_ISL_750834, EPI_ISL_750835, EPI_ISL_750836, EPI_ISL_750837, EPI_ISL_750838, EPI_ISL_750839,
EPI_ISL_750840, EPI_ISL_750841, EPI_ISL_750842, EPI_ISL_750843, EPI_ISL_750844, EPI_ISL_750845, EPI_ISL_750846, EPI_ISL_750847, EPI_ISL_750848, EPI_ISL_750849, EPI_ISL_750850, EPI_ISL_750851, EPI_ISL_750852, EPI_ISL_750853, EPI_ISL_750854, EPI_ISL_750855, EPI_ISL_750856, EPI_ISL_750857,
EPI_ISL_750858, EPI_ISL_750859, EPI_ISL_750860, EPI_ISL_750861, EPI_ISL_750862, EPI_ISL_750863, EPI_ISL_750864, EPI_ISL_750865, EPI_ISL_750866, EPI_ISL_750867, EPI_ISL_750868, EPI_ISL_750869, EPI_ISL_750870, EPI_ISL_750871, EPI_ISL_750872, EPI_ISL_750873, EPI_ISL_750874, EPI_ISL_750875,
EPI_ISL_750876, EPI_ISL_750877, EPI_ISL_750878, EPI_ISL_750879, EPI_ISL_750880, EPI_ISL_750881, EPI_ISL_750882, EPI_ISL_750883, EPI_ISL_750884, EPI_ISL_750885, EPI_ISL_750886, EPI_ISL_750887, EPI_ISL_750888, EPI_ISL_750889, EPI_ISL_750890, EPI_ISL_750891, EPI_ISL_750892, EPI_ISL_750893,
EPI_ISL_750894, EPI_ISL_750895, EPI_ISL_750896, EPI_ISL_750897, EPI_ISL_750898, EPI_ISL_750899, EPI_ISL_750900, EPI_ISL_750901, EPI_ISL_750902, EPI_ISL_750903, EPI_ISL_750904, EPI_ISL_750905, EPI_ISL_750906, EPI_ISL_750907, EPI_ISL_750908, EPI_ISL_750909, EPI_ISL_750910, EPI_ISL_750911,
EPI_ISL_750912, EPI_ISL_750913, EPI_ISL_750914, EPI_ISL_750915, EPI_ISL_750916, EPI_ISL_750917, EPI_ISL_750918, EPI_ISL_750919, EPI_ISL_750920, EPI_ISL_750921, EPI_ISL_750922, EPI_ISL_750923, EPI_ISL_750924, EPI_ISL_750925, EPI_ISL_750926, EPI_ISL_750927, EPI_ISL_750928, EPI_ISL_750929,
EPI_ISL_750930, EPI_ISL_750931, EPI_ISL_750932, EPI_ISL_750933, EPI_ISL_750934, EPI_ISL_750935, EPI_ISL_750936, EPI_ISL_750937, EPI_ISL_750938, EPI_ISL_750939, EPI_ISL_750940, EPI_ISL_750941, EPI_ISL_750942, EPI_ISL_750943, EPI_ISL_750944, EPI_ISL_750945, EPI_ISL_750946, EPI_ISL_750947,
EPI_ISL_750948, EPI_ISL_750949, EPI_ISL_750950, EPI_ISL_750951, EPI_ISL_750952, EPI_ISL_750953, EPI_ISL_750954, EPI_ISL_750955, EPI_ISL_750956, EPI_ISL_750957, EPI_ISL_750958, EPI_ISL_750959, EPI_ISL_750960, EPI_ISL_750961, EPI_ISL_750962, EPI_ISL_750963, EPI_ISL_750964, EPI_ISL_750965,
EPI_ISL_750966, EPI_ISL_750967, EPI_ISL_750968, EPI_ISL_750969, EPI_ISL_750970, EPI_ISL_750971, EPI_ISL_750972, EPI_ISL_750973, EPI_ISL_750974, EPI_ISL_750975, EPI_ISL_750976, EPI_ISL_750977, EPI_ISL_750978, EPI_ISL_750979, EPI_ISL_750980, EPI_ISL_750981, EPI_ISL_750982, EPI_ISL_750983,
EPI_ISL_750984, EPI_ISL_750985, EPI_ISL_750986, EPI_ISL_750987, EPI_ISL_750988, EPI_ISL_750989, EPI_ISL_750990, EPI_ISL_750991, EPI_ISL_750992, EPI_ISL_750993, EPI_ISL_750994, EPI_ISL_750995, EPI_ISL_750996, EPI_ISL_750997, EPI_ISL_750998, EPI_ISL_750999, EPI_ISL_751000, EPI_ISL_751001,
EPI_ISL_751002, EPI_ISL_751003, EPI_ISL_751004, EPI_ISL_751005, EPI_ISL_751006, EPI_ISL_751007, EPI_ISL_751008, EPI_ISL_751009, EPI_ISL_751010, EPI_ISL_751012, EPI_ISL_751013, EPI_ISL_751014, EPI_ISL_751015, EPI_ISL_751016, EPI_ISL_751017, EPI_ISL_751018, EPI_ISL_751019, EPI_ISL_751020,
EPI_ISL_751021, EPI_ISL_751022, EPI_ISL_751023, EPI_ISL_751024, EPI_ISL_751025, EPI_ISL_751026, EPI_ISL_751027, EPI_ISL_751028, EPI_ISL_751029, EPI_ISL_751030, EPI_ISL_751031, EPI_ISL_751032, EPI_ISL_751033, EPI_ISL_751034, EPI_ISL_751035, EPI_ISL_751036, EPI_ISL_751037, EPI_ISL_751038,
EPI_ISL_751039, EPI_ISL_751040, EPI_ISL_751041, EPI_ISL_751042, EPI_ISL_751043, EPI_ISL_751044, EPI_ISL_751045, EPI_ISL_751046, EPI_ISL_751047, EPI_ISL_751048, EPI_ISL_751049, EPI_ISL_751050, EPI_ISL_751051, EPI_ISL_751052, EPI_ISL_751053, EPI_ISL_751054, EPI_ISL_751055, EPI_ISL_751056,
EPI_ISL_751057, EPI_ISL_751058, EPI_ISL_751059, EPI_ISL_751060, EPI_ISL_751061, EPI_ISL_751062, EPI_ISL_751063, EPI_ISL_751064, EPI_ISL_751065, EPI_ISL_751066, EPI_ISL_751067, EPI_ISL_751068, EPI_ISL_751069, EPI_ISL_751070, EPI_ISL_751071, EPI_ISL_751072, EPI_ISL_751073, EPI_ISL_751074,
EPI_ISL_751075, EPI_ISL_751076, EPI_ISL_751077, EPI_ISL_751078, EPI_ISL_751079, EPI_ISL_751080, EPI_ISL_751081, EPI_ISL_751082, EPI_ISL_751083, EPI_ISL_751084, EPI_ISL_751085, EPI_ISL_751086, EPI_ISL_751087, EPI_ISL_751088, EPI_ISL_751089, EPI_ISL_751090, EPI_ISL_751091, EPI_ISL_751092,
EPI_ISL_751093, EPI_ISL_751094, EPI_ISL_751095, EPI_ISL_751096, EPI_ISL_751097, EPI_ISL_751098, EPI_ISL_751099, EPI_ISL_751100, EPI_ISL_751101, EPI_ISL_751102, EPI_ISL_751103, EPI_ISL_751104, EPI_ISL_751105, EPI_ISL_751106, EPI_ISL_751107, EPI_ISL_751108, EPI_ISL_751109, EPI_ISL_751110,
EPI_ISL_751111, EPI_ISL_751112, EPI_ISL_751113, EPI_ISL_751114, EPI_ISL_751115, EPI_ISL_751116, EPI_ISL_751117, EPI_ISL_751118, EPI_ISL_751119, EPI_ISL_751120, EPI_ISL_751121, EPI_ISL_751122, EPI_ISL_751123, EPI_ISL_751124, EPI_ISL_751125, EPI_ISL_751126, EPI_ISL_751127, EPI_ISL_751128,
EPI_ISL_751129, EPI_ISL_751130, EPI_ISL_751131, EPI_ISL_751132, EPI_ISL_751133, EPI_ISL_751134, EPI_ISL_751135, EPI_ISL_751136, EPI_ISL_751137, EPI_ISL_751138, EPI_ISL_751139, EPI_ISL_751140, EPI_ISL_751141, EPI_ISL_751142, EPI_ISL_751143, EPI_ISL_751144, EPI_ISL_751145, EPI_ISL_751146,
EPI_ISL_751147, EPI_ISL_751148, EPI_ISL_751149, EPI_ISL_751150, EPI_ISL_751151, EPI_ISL_751152, EPI_ISL_751153, EPI_ISL_751154, EPI_ISL_751155, EPI_ISL_751156, EPI_ISL_751157, EPI_ISL_751158, EPI_ISL_751159, EPI_ISL_751160, EPI_ISL_751161, EPI_ISL_751162, EPI_ISL_751163, EPI_ISL_751164,
EPI_ISL_751165, EPI_ISL_751166, EPI_ISL_751167, EPI_ISL_751168, EPI_ISL_751169, EPI_ISL_751170, EPI_ISL_751171, EPI_ISL_751172, EPI_ISL_751173, EPI_ISL_751174, EPI_ISL_751175, EPI_ISL_751176, EPI_ISL_751177, EPI_ISL_751178, EPI_ISL_751179, EPI_ISL_751180, EPI_ISL_751181, EPI_ISL_751182,
EPI_ISL_751183

| | | | |
|---|---|---|---|
| see above | Department of Virus and Microbiological Special Diagnostics, Statens Serum Institut, Copenhagen, Denmark | Albertsen Lab, Department of Chemistry and Bioscience, Aalborg University, Denmark | Danish Covid-19 Genome Consortium |
| EPI_ISL_753671, EPI_ISL_753672, EPI_ISL_753673, EPI_ISL_753674, EPI_ISL_753689, EPI_ISL_753691 | Clinical virology Laboratory, Children's Hospital Los Angeles | Center for Personalized Medicine, Children's Hospital Los Angeles | Gai et al |
| EPI_ISL_754079, EPI_ISL_754081, EPI_ISL_754085, EPI_ISL_754087, EPI_ISL_754093, EPI_ISL_754096, EPI_ISL_754098, EPI_ISL_754105 | National Public Health Laboratory, National Centre for Infectious Diseases | National Public Health Laboratory, National Centre for Infectious Diseases | Tze Minn Mak, Sophie Octavia, Zhenyang Zhou, Lin Cui, Raymond Tzer Pin Lin |

| | | | |
|---|---|---|---|
| EPI_ISL_754312, EPI_ISL_754367 | Respiratory Virus Unit, National Infection Service, Public Health England | COVID-19 Genomics UK (COG-UK) Consortium | PHE Covid Sequencing Team |
| EPI_ISL_754620, EPI_ISL_754621, EPI_ISL_754622, EPI_ISL_754636, EPI_ISL_754637, EPI_ISL_754638, EPI_ISL_754639, EPI_ISL_754640, EPI_ISL_754641, EPI_ISL_754642, EPI_ISL_754643, EPI_ISL_754644, EPI_ISL_754645, EPI_ISL_754646, EPI_ISL_754647 | | | |
| see above | University of Wisconsin-Madison AIDS Vaccine Research Laboratories | University of Wisconsin-Madison AIDS Vaccine Research Laboratories | Gage Moreno, Katarina Braun, et al. AIDS Vaccine Research Laboratories |
| EPI_ISL_755171, EPI_ISL_755172, EPI_ISL_755173, EPI_ISL_755175, EPI_ISL_755176, EPI_ISL_755177, EPI_ISL_755178, EPI_ISL_755179, EPI_ISL_755180, EPI_ISL_755183 | UCSD EXCITE lab | Andersen lab at Scripps Research | SEARCH Alliance San Diego |
| EPI_ISL_755500, EPI_ISL_755501, EPI_ISL_755502, EPI_ISL_755503, EPI_ISL_755504, EPI_ISL_755505, EPI_ISL_755506, EPI_ISL_755507, EPI_ISL_755508, EPI_ISL_755509, EPI_ISL_755510, EPI_ISL_755554, EPI_ISL_755555, EPI_ISL_755556, EPI_ISL_755557, EPI_ISL_755558, EPI_ISL_755559, EPI_ISL_755566, EPI_ISL_755567 | | | |
| see above | Maine Health and Environmental Testing Laboratory | Tewhey Lab, The Jackson Laboratory | Matluk,N., Dewey,H., Iosue,F., Barter,M., Lynch,R., Munger,H. and Tewhey,R. |
| EPI_ISL_756216, EPI_ISL_756218, EPI_ISL_756219, EPI_ISL_756220, EPI_ISL_756221, EPI_ISL_756222, EPI_ISL_756223, EPI_ISL_756224, EPI_ISL_756225, EPI_ISL_756226, EPI_ISL_756227, EPI_ISL_756231, EPI_ISL_756232, EPI_ISL_756233, EPI_ISL_756300, EPI_ISL_756301, EPI_ISL_756303, EPI_ISL_756304, EPI_ISL_756305 | | | |
| see above | UW Virology Lab | UW Virology Lab | Pavitra Roychoudhury, Hong Xie, Lasata Shrestha, Meei-Li Huang, Keith R Jerome, Alexander Greninger |
| EPI_ISL_756689, EPI_ISL_756691, EPI_ISL_756692, EPI_ISL_756693, EPI_ISL_756694, EPI_ISL_756697, EPI_ISL_756698, EPI_ISL_756699, EPI_ISL_756700, EPI_ISL_756701, EPI_ISL_756703, EPI_ISL_756704, EPI_ISL_756707, EPI_ISL_756708, EPI_ISL_756710, EPI_ISL_756712, EPI_ISL_756713, EPI_ISL_756714, EPI_ISL_756715, EPI_ISL_756717, EPI_ISL_756718, EPI_ISL_756719, EPI_ISL_756720, EPI_ISL_756721, EPI_ISL_756722, EPI_ISL_756724, EPI_ISL_756725, EPI_ISL_756726, EPI_ISL_756727, EPI_ISL_756728, EPI_ISL_756730, EPI_ISL_756731, EPI_ISL_756732, EPI_ISL_756733, EPI_ISL_756734, EPI_ISL_756735, EPI_ISL_756736, EPI_ISL_756738, EPI_ISL_756739, EPI_ISL_756740, EPI_ISL_756741, EPI_ISL_756742, EPI_ISL_756743, EPI_ISL_756746, EPI_ISL_756747, EPI_ISL_756749, EPI_ISL_756751, EPI_ISL_756752, EPI_ISL_756753, EPI_ISL_756754, EPI_ISL_756755, EPI_ISL_756756, EPI_ISL_756757, EPI_ISL_756758, EPI_ISL_756759, EPI_ISL_756760, EPI_ISL_756762, EPI_ISL_756763, EPI_ISL_756764, EPI_ISL_756766, EPI_ISL_756767, EPI_ISL_756768, EPI_ISL_756769, EPI_ISL_756770, EPI_ISL_756771, EPI_ISL_756772, EPI_ISL_756774, EPI_ISL_756776, EPI_ISL_756777, EPI_ISL_756778, EPI_ISL_756779, EPI_ISL_756781, EPI_ISL_756782, EPI_ISL_756783, EPI_ISL_756785, EPI_ISL_756786, EPI_ISL_756787, EPI_ISL_756788, EPI_ISL_756790, EPI_ISL_756791, EPI_ISL_756792, EPI_ISL_756793, EPI_ISL_756794, EPI_ISL_756796, EPI_ISL_756797, EPI_ISL_756798, EPI_ISL_756799, EPI_ISL_756801, EPI_ISL_756802, EPI_ISL_756804, EPI_ISL_756806, EPI_ISL_756807, EPI_ISL_756808, EPI_ISL_756809, EPI_ISL_756810, EPI_ISL_756811, EPI_ISL_756812, EPI_ISL_756813, EPI_ISL_756814, EPI_ISL_756815, EPI_ISL_756816, EPI_ISL_756817, EPI_ISL_756818, EPI_ISL_756819, EPI_ISL_756820, EPI_ISL_756821, EPI_ISL_756822, EPI_ISL_756823, EPI_ISL_756825, EPI_ISL_756826, EPI_ISL_756827, EPI_ISL_756828, EPI_ISL_756829, EPI_ISL_756830, EPI_ISL_756831, EPI_ISL_756832, EPI_ISL_756833, EPI_ISL_756834, EPI_ISL_756836, EPI_ISL_756837, EPI_ISL_756838, EPI_ISL_756839, EPI_ISL_756841, EPI_ISL_756842, EPI_ISL_756843, EPI_ISL_756844, EPI_ISL_756845, EPI_ISL_756847, EPI_ISL_756848, EPI_ISL_756851, EPI_ISL_756853, EPI_ISL_756854, EPI_ISL_756855, EPI_ISL_756856, EPI_ISL_756857, EPI_ISL_756858, EPI_ISL_756859, EPI_ISL_756862, EPI_ISL_756863, EPI_ISL_756864, EPI_ISL_756866, EPI_ISL_756868, EPI_ISL_756869, EPI_ISL_756870, EPI_ISL_756871, EPI_ISL_756872, EPI_ISL_756873, EPI_ISL_756874, EPI_ISL_756875, EPI_ISL_756880, EPI_ISL_756882, EPI_ISL_756883, EPI_ISL_756884, EPI_ISL_756890, EPI_ISL_756891, EPI_ISL_756892, EPI_ISL_756896, EPI_ISL_756898, EPI_ISL_756899, EPI_ISL_756901, EPI_ISL_756902, EPI_ISL_756904, EPI_ISL_756905, EPI_ISL_756906, EPI_ISL_756907, EPI_ISL_756908, EPI_ISL_756911, EPI_ISL_756913, EPI_ISL_756914, EPI_ISL_756915, EPI_ISL_756916, EPI_ISL_756917, EPI_ISL_756919, EPI_ISL_756921, EPI_ISL_756922, EPI_ISL_756924, EPI_ISL_756926, EPI_ISL_756927, EPI_ISL_756929, EPI_ISL_756930, EPI_ISL_756932, EPI_ISL_756933, EPI_ISL_756934, EPI_ISL_756936, EPI_ISL_756937, EPI_ISL_756938, EPI_ISL_756939, EPI_ISL_756941, EPI_ISL_756942, EPI_ISL_756943, EPI_ISL_756944, EPI_ISL_756945, EPI_ISL_756948, EPI_ISL_756949, EPI_ISL_756950, EPI_ISL_756954, EPI_ISL_756956, EPI_ISL_756957, EPI_ISL_756958, EPI_ISL_756960, EPI_ISL_756961, EPI_ISL_756962, EPI_ISL_756963, EPI_ISL_756964, EPI_ISL_756965, EPI_ISL_756966, EPI_ISL_756967, EPI_ISL_756968, EPI_ISL_756972, EPI_ISL_756974, EPI_ISL_756975, EPI_ISL_756976, EPI_ISL_756978, EPI_ISL_756979, EPI_ISL_756980, EPI_ISL_756981, EPI_ISL_756983, EPI_ISL_756984, EPI_ISL_756985, EPI_ISL_756986, EPI_ISL_756987, EPI_ISL_756988, EPI_ISL_756989, EPI_ISL_756990, EPI_ISL_756991, EPI_ISL_756992, EPI_ISL_756993, EPI_ISL_756994, EPI_ISL_756995, EPI_ISL_756996, EPI_ISL_756997, EPI_ISL_756999, EPI_ISL_757000, EPI_ISL_757001, EPI_ISL_757002, EPI_ISL_757003, EPI_ISL_757004, EPI_ISL_757006, EPI_ISL_757010, EPI_ISL_757013, EPI_ISL_757015, EPI_ISL_757016, EPI_ISL_757019, EPI_ISL_757022, EPI_ISL_757023, EPI_ISL_757024, EPI_ISL_757025, EPI_ISL_757026, EPI_ISL_757029, EPI_ISL_757030, EPI_ISL_757031, EPI_ISL_757033 | | | |
| see above | Lighthouse Lab in Milton Keynes | Wellcome Sanger Institute for the COVID-19 Genomics UK (COG-UK) Consortium | The Lighthouse Lab in Milton Keynes and Alex Alderton, Roberto Amato, Sonia Goncalves, Ewan Harrison, David K. Jackson, Ian Johnston, Dominic Kwiatkowski, Cordelia Langford, John Sillitoe on behalf of the Wellcome Sanger Institute COVID-19 Surveillance Team |
| EPI_ISL_757034, EPI_ISL_757035, EPI_ISL_757036, EPI_ISL_757037, EPI_ISL_757038, EPI_ISL_757039, EPI_ISL_757040, EPI_ISL_757041, EPI_ISL_757083, EPI_ISL_757084, EPI_ISL_757085, EPI_ISL_757086, EPI_ISL_757087, EPI_ISL_757088, EPI_ISL_757089, EPI_ISL_757090, EPI_ISL_757091, EPI_ISL_757092, EPI_ISL_757093, EPI_ISL_757094, EPI_ISL_757095, EPI_ISL_757096, EPI_ISL_757097, EPI_ISL_757098, EPI_ISL_757099, EPI_ISL_757100, EPI_ISL_757101, EPI_ISL_757102, EPI_ISL_757103, EPI_ISL_757104, EPI_ISL_757105, EPI_ISL_757106, EPI_ISL_757107, EPI_ISL_757108, EPI_ISL_757109, EPI_ISL_757110, EPI_ISL_757111, EPI_ISL_757112, EPI_ISL_757113, EPI_ISL_757114, EPI_ISL_757115, EPI_ISL_757116, EPI_ISL_757117, EPI_ISL_757118 | | | |
| see above | Lighthouse Lab in Glasgow | Wellcome Sanger Institute for the COVID-19 Genomics UK (COG-UK) Consortium | Harper VanSteenhouse, Yumi Kasai, David Gray, Carol Clugston, Anna Dominiczak and Alex Alderton, Roberto Amato, Sonia Goncalves, Ewan Harrison, David K. Jackson, Ian Johnston, Dominic Kwiatkowski, Cordelia Langford, John Sillitoe on behalf of the Wellcome Sanger Institute COVID-19 Surveillance Team |
| EPI_ISL_757395, EPI_ISL_757396, EPI_ISL_757397, EPI_ISL_757398, EPI_ISL_757399, EPI_ISL_757400, EPI_ISL_757401, EPI_ISL_757402, EPI_ISL_757403, EPI_ISL_757404, EPI_ISL_757405, EPI_ISL_757406, EPI_ISL_757407, EPI_ISL_757408, EPI_ISL_757409, EPI_ISL_757410, EPI_ISL_757411, EPI_ISL_757412, EPI_ISL_757413, EPI_ISL_757414, EPI_ISL_757415, EPI_ISL_757416, EPI_ISL_757417, EPI_ISL_757418, EPI_ISL_757419, EPI_ISL_757420, EPI_ISL_757421, EPI_ISL_757422, EPI_ISL_757423, EPI_ISL_757424, EPI_ISL_757425, EPI_ISL_757426, EPI_ISL_757427, EPI_ISL_757428, EPI_ISL_757429, EPI_ISL_757430, EPI_ISL_757431, EPI_ISL_757432, EPI_ISL_757433, EPI_ISL_757434, EPI_ISL_757435, EPI_ISL_757436, EPI_ISL_757437, EPI_ISL_757438, EPI_ISL_757439, EPI_ISL_757440, EPI_ISL_757441, EPI_ISL_757442, EPI_ISL_757444, EPI_ISL_757445, EPI_ISL_757446, EPI_ISL_757447, EPI_ISL_757449, EPI_ISL_757450, EPI_ISL_757451, EPI_ISL_757452, EPI_ISL_757453, EPI_ISL_757454, EPI_ISL_757455, EPI_ISL_757456, EPI_ISL_757457, EPI_ISL_757458, EPI_ISL_757459, EPI_ISL_757460, EPI_ISL_757461, EPI_ISL_757462, EPI_ISL_757463, EPI_ISL_757464, EPI_ISL_757465, EPI_ISL_757466, EPI_ISL_757467, EPI_ISL_757468, EPI_ISL_757469, EPI_ISL_757470, EPI_ISL_757471, EPI_ISL_757472, EPI_ISL_757473, EPI_ISL_757474, EPI_ISL_757475, EPI_ISL_757476, EPI_ISL_757477, EPI_ISL_757478, EPI_ISL_757479, EPI_ISL_757480, EPI_ISL_757481, EPI_ISL_757482, EPI_ISL_757483, EPI_ISL_757484, EPI_ISL_757485, EPI_ISL_757486, EPI_ISL_757487, EPI_ISL_757488, EPI_ISL_757489, EPI_ISL_757490, EPI_ISL_757492, EPI_ISL_757493, EPI_ISL_757494, EPI_ISL_757495, EPI_ISL_757496, EPI_ISL_757497, EPI_ISL_757499, EPI_ISL_757500, EPI_ISL_757501, EPI_ISL_757502, EPI_ISL_757503, EPI_ISL_757504, EPI_ISL_757505, EPI_ISL_757506, EPI_ISL_757507, EPI_ISL_757508, EPI_ISL_757509, EPI_ISL_757510, EPI_ISL_757511, EPI_ISL_757512, EPI_ISL_757513, EPI_ISL_757514, EPI_ISL_757515, EPI_ISL_757516, EPI_ISL_757517, EPI_ISL_757518, EPI_ISL_757519, EPI_ISL_757520, EPI_ISL_757521, EPI_ISL_757522, EPI_ISL_757523, EPI_ISL_757524, EPI_ISL_757525, EPI_ISL_757526, EPI_ISL_757527, EPI_ISL_757528, EPI_ISL_757529, EPI_ISL_757530, EPI_ISL_757531, EPI_ISL_757532, EPI_ISL_757533, EPI_ISL_757534, EPI_ISL_757535, EPI_ISL_757536, EPI_ISL_757537, EPI_ISL_757538, EPI_ISL_757539, EPI_ISL_757540, EPI_ISL_757541, EPI_ISL_757542, EPI_ISL_757543, EPI_ISL_757544, EPI_ISL_757545, EPI_ISL_757546, EPI_ISL_757547, EPI_ISL_757548, EPI_ISL_757549, EPI_ISL_757550, EPI_ISL_757551, EPI_ISL_757552, EPI_ISL_757553, EPI_ISL_757554, EPI_ISL_757555, EPI_ISL_757556, EPI_ISL_757557, EPI_ISL_757558, EPI_ISL_757559, EPI_ISL_757560, EPI_ISL_757561, EPI_ISL_757562, EPI_ISL_757563, EPI_ISL_757564, EPI_ISL_757565, EPI_ISL_757566, EPI_ISL_757567, EPI_ISL_757568, EPI_ISL_757569, EPI_ISL_757570, EPI_ISL_757571, EPI_ISL_757572, EPI_ISL_757573, EPI_ISL_757574, EPI_ISL_757575, EPI_ISL_757576, EPI_ISL_757577, EPI_ISL_757578, EPI_ISL_757579, EPI_ISL_757580, EPI_ISL_757581, EPI_ISL_757582, EPI_ISL_757583, EPI_ISL_757584, EPI_ISL_757585, EPI_ISL_757586, EPI_ISL_757587, EPI_ISL_757589, EPI_ISL_757590, EPI_ISL_757591, EPI_ISL_757592, EPI_ISL_757593, EPI_ISL_757594, EPI_ISL_757595, EPI_ISL_757596, EPI_ISL_757597, EPI_ISL_757598, EPI_ISL_757599, EPI_ISL_757600, EPI_ISL_757601, EPI_ISL_757602, EPI_ISL_757603, EPI_ISL_757604, EPI_ISL_757605, EPI_ISL_757606, EPI_ISL_757607, EPI_ISL_757608, EPI_ISL_757609, EPI_ISL_757610, EPI_ISL_757611, EPI_ISL_757612, EPI_ISL_757613, EPI_ISL_757614, EPI_ISL_757615, EPI_ISL_757616, EPI_ISL_757617, EPI_ISL_757618, EPI_ISL_757619, EPI_ISL_757620, EPI_ISL_757621, EPI_ISL_757622, EPI_ISL_757623, EPI_ISL_757624, EPI_ISL_757625, EPI_ISL_757626, EPI_ISL_757627, EPI_ISL_757628, EPI_ISL_757629, EPI_ISL_757630, EPI_ISL_757631, EPI_ISL_757632, EPI_ISL_757633, EPI_ISL_757634, EPI_ISL_757635, EPI_ISL_757636, EPI_ISL_757637, EPI_ISL_757638, EPI_ISL_757639, EPI_ISL_757640, EPI_ISL_757641, EPI_ISL_757642, EPI_ISL_757643, EPI_ISL_757644, EPI_ISL_757645, EPI_ISL_757646, EPI_ISL_757647, EPI_ISL_757648, EPI_ISL_757649, EPI_ISL_757650, EPI_ISL_757651, EPI_ISL_757652, EPI_ISL_757653, EPI_ISL_757654, EPI_ISL_757655, EPI_ISL_757656, EPI_ISL_757657, EPI_ISL_757658, EPI_ISL_757659, EPI_ISL_757660, EPI_ISL_757661, EPI_ISL_757662, EPI_ISL_757663, EPI_ISL_757664, EPI_ISL_757665, EPI_ISL_757666, EPI_ISL_757667, EPI_ISL_757668, EPI_ISL_757669, EPI_ISL_757670, EPI_ISL_757671, EPI_ISL_757672, EPI_ISL_757673, EPI_ISL_757674, EPI_ISL_757675, EPI_ISL_757676, EPI_ISL_757677, EPI_ISL_757678, EPI_ISL_757679, EPI_ISL_757680, EPI_ISL_757681, EPI_ISL_757682, EPI_ISL_757683, EPI_ISL_757684, EPI_ISL_757685, EPI_ISL_757686, EPI_ISL_757687, EPI_ISL_757703, EPI_ISL_757704, EPI_ISL_757705, EPI_ISL_757710, EPI_ISL_757712, EPI_ISL_757802, EPI_ISL_757803, EPI_ISL_757804, EPI_ISL_757805, EPI_ISL_757806, EPI_ISL_757807, EPI_ISL_757808, EPI_ISL_757809, EPI_ISL_757810, EPI_ISL_757811, EPI_ISL_757812, EPI_ISL_757813, EPI_ISL_757814, EPI_ISL_757815, EPI_ISL_757816, EPI_ISL_757817, EPI_ISL_757818, EPI_ISL_757819, EPI_ISL_757820, EPI_ISL_757821, EPI_ISL_757822, EPI_ISL_757823, EPI_ISL_757824, EPI_ISL_757825, EPI_ISL_757826, EPI_ISL_757827, EPI_ISL_757828, EPI_ISL_757829, EPI_ISL_757830, EPI_ISL_757831, EPI_ISL_757832, EPI_ISL_757833, EPI_ISL_757834, EPI_ISL_757835, EPI_ISL_757836, EPI_ISL_757837, EPI_ISL_757838, EPI_ISL_757839, EPI_ISL_757840, EPI_ISL_757841, EPI_ISL_757842, EPI_ISL_757843, EPI_ISL_757844, EPI_ISL_757845, EPI_ISL_757846, EPI_ISL_757847, EPI_ISL_757848, EPI_ISL_757849, EPI_ISL_757850, EPI_ISL_757851, EPI_ISL_757852, EPI_ISL_757853, EPI_ISL_757854, EPI_ISL_757859, EPI_ISL_757861, EPI_ISL_757867, EPI_ISL_757872, EPI_ISL_757874, EPI_ISL_757875, EPI_ISL_757876, EPI_ISL_757887, EPI_ISL_757888, EPI_ISL_757889, EPI_ISL_757890, EPI_ISL_757893, EPI_ISL_757894, EPI_ISL_757895, EPI_ISL_757927, EPI_ISL_757928, EPI_ISL_757929, EPI_ISL_757930, EPI_ISL_757931, EPI_ISL_757932, EPI_ISL_757933, EPI_ISL_757934, EPI_ISL_757936, EPI_ISL_757937, EPI_ISL_757938, EPI_ISL_757939, EPI_ISL_757940, EPI_ISL_757941, EPI_ISL_757942, EPI_ISL_757943, EPI_ISL_757944, EPI_ISL_757947, EPI_ISL_757948, EPI_ISL_757949, EPI_ISL_757950, EPI_ISL_757951, EPI_ISL_757953, EPI_ISL_757954, EPI_ISL_757955, EPI_ISL_757956, EPI_ISL_757957, EPI_ISL_757958, EPI_ISL_757967, EPI_ISL_757968, EPI_ISL_757969, EPI_ISL_757970, EPI_ISL_757971, EPI_ISL_757972, EPI_ISL_757973, EPI_ISL_757974, EPI_ISL_758026, EPI_ISL_758027, EPI_ISL_758028, EPI_ISL_758029, EPI_ISL_758030, EPI_ISL_758031, EPI_ISL_758032, EPI_ISL_758033, EPI_ISL_758034, EPI_ISL_758035, EPI_ISL_758036, EPI_ISL_758037, EPI_ISL_758038, EPI_ISL_758039, EPI_ISL_758040, EPI_ISL_758041, EPI_ISL_758042, EPI_ISL_758043, EPI_ISL_758044, EPI_ISL_758045, EPI_ISL_758046, EPI_ISL_758047, EPI_ISL_758048, EPI_ISL_758049, EPI_ISL_758050, EPI_ISL_758051, EPI_ISL_758052, EPI_ISL_758053, EPI_ISL_758054, EPI_ISL_758055, EPI_ISL_758056, EPI_ISL_758057, EPI_ISL_758058, EPI_ISL_758059, EPI_ISL_758060, EPI_ISL_758061, EPI_ISL_758062, EPI_ISL_758063, EPI_ISL_758064, EPI_ISL_758065, EPI_ISL_758066, EPI_ISL_758071, EPI_ISL_758072, EPI_ISL_758082, EPI_ISL_758084, EPI_ISL_758093, EPI_ISL_758101, EPI_ISL_758106, EPI_ISL_758107, EPI_ISL_758108, EPI_ISL_758109, EPI_ISL_758110, EPI_ISL_758111, EPI_ISL_758115, EPI_ISL_758116, EPI_ISL_758122, EPI_ISL_758123, EPI_ISL_758124, EPI_ISL_758125, EPI_ISL_758126, EPI_ISL_758127, EPI_ISL_758484, EPI_ISL_758485, EPI_ISL_758486, EPI_ISL_758487, EPI_ISL_758488, EPI_ISL_758489, EPI_ISL_758490, EPI_ISL_758491, EPI_ISL_758492, EPI_ISL_758493, EPI_ISL_758494, EPI_ISL_758495, EPI_ISL_758496, EPI_ISL_758497, EPI_ISL_758498, EPI_ISL_758499, EPI_ISL_758500, EPI_ISL_758501, EPI_ISL_758502, EPI_ISL_758503, EPI_ISL_758504, EPI_ISL_758505, EPI_ISL_758506, EPI_ISL_758507, EPI_ISL_758508, EPI_ISL_758509, EPI_ISL_758510, EPI_ISL_758511, EPI_ISL_758512, EPI_ISL_758513, EPI_ISL_758514, EPI_ISL_758515, EPI_ISL_758516, EPI_ISL_758517, EPI_ISL_758518, EPI_ISL_758519, EPI_ISL_758520, EPI_ISL_758521, EPI_ISL_758522, EPI_ISL_758523, EPI_ISL_758524, EPI_ISL_758525, EPI_ISL_758526, EPI_ISL_758527, EPI_ISL_758528, EPI_ISL_758529, EPI_ISL_758530, EPI_ISL_758531, EPI_ISL_758532, EPI_ISL_758533, EPI_ISL_758534, EPI_ISL_758535, EPI_ISL_758536, EPI_ISL_758537, EPI_ISL_758538, EPI_ISL_758539, EPI_ISL_758540, EPI_ISL_758541, EPI_ISL_758542, EPI_ISL_758543, EPI_ISL_758544, EPI_ISL_758545, EPI_ISL_758546, EPI_ISL_758547, EPI_ISL_758548, EPI_ISL_758549, EPI_ISL_758550, EPI_ISL_758551, EPI_ISL_758552, EPI_ISL_758553, EPI_ISL_758554, EPI_ISL_758555, EPI_ISL_758556, EPI_ISL_758557, EPI_ISL_758558, EPI_ISL_758559, EPI_ISL_758560, EPI_ISL_758561, EPI_ISL_758562, EPI_ISL_758563, EPI_ISL_758564, EPI_ISL_758565, EPI_ISL_758566, EPI_ISL_758567, EPI_ISL_758568, EPI_ISL_758569, EPI_ISL_758570, EPI_ISL_758571, EPI_ISL_758572, EPI_ISL_758573, EPI_ISL_758574, EPI_ISL_758575, EPI_ISL_758576, EPI_ISL_758577, EPI_ISL_758578, EPI_ISL_758579, EPI_ISL_758580, EPI_ISL_758581, EPI_ISL_758582, EPI_ISL_758583, EPI_ISL_758584, EPI_ISL_758585, EPI_ISL_758586, EPI_ISL_758587, EPI_ISL_758588, EPI_ISL_758589, EPI_ISL_758590, EPI_ISL_758591, EPI_ISL_758592, EPI_ISL_758593, EPI_ISL_758594, EPI_ISL_758595, EPI_ISL_758596, EPI_ISL_758597, EPI_ISL_758598, EPI_ISL_758599, EPI_ISL_758600, EPI_ISL_758601, EPI_ISL_758602, EPI_ISL_758603, EPI_ISL_758604, EPI_ISL_758605, EPI_ISL_758606, EPI_ISL_758607, EPI_ISL_758608, EPI_ISL_758609, EPI_ISL_758610, EPI_ISL_758611, EPI_ISL_758612, EPI_ISL_758613, EPI_ISL_758614, EPI_ISL_758615, EPI_ISL_758616, EPI_ISL_758617, EPI_ISL_758618, EPI_ISL_758619, EPI_ISL_758620, EPI_ISL_758621, EPI_ISL_758622, EPI_ISL_758623, EPI_ISL_758624, EPI_ISL_758625, EPI_ISL_758626, EPI_ISL_758627, EPI_ISL_758628, EPI_ISL_758629, EPI_ISL_758630, EPI_ISL_758631, EPI_ISL_758632, EPI_ISL_758633, EPI_ISL_758634, EPI_ISL_758635, EPI_ISL_758636, EPI_ISL_758637, EPI_ISL_758638, EPI_ISL_758639, EPI_ISL_758640, EPI_ISL_758641, EPI_ISL_758642, EPI_ISL_758643, EPI_ISL_758644, EPI_ISL_758645, EPI_ISL_758646, EPI_ISL_758647, EPI_ISL_758648, EPI_ISL_758649, EPI_ISL_758650, EPI_ISL_758651, EPI_ISL_758652, EPI_ISL_758653, EPI_ISL_758654, EPI_ISL_758655, EPI_ISL_758656, EPI_ISL_758657, EPI_ISL_758658, EPI_ISL_758659, EPI_ISL_758660, EPI_ISL_758661, EPI_ISL_758663, EPI_ISL_758664, EPI_ISL_758665, EPI_ISL_758666, EPI_ISL_758667, EPI_ISL_758668, EPI_ISL_758669, EPI_ISL_758670, EPI_ISL_758671, EPI_ISL_758672, EPI_ISL_758673, EPI_ISL_758674, EPI_ISL_758675, EPI_ISL_758676, EPI_ISL_758677, EPI_ISL_758678, EPI_ISL_758679, EPI_ISL_758680, EPI_ISL_758681, EPI_ISL_758682, EPI_ISL_758683, EPI_ISL_758684, EPI_ISL_758685, EPI_ISL_758686, EPI_ISL_758687, EPI_ISL_758688, EPI_ISL_758689, EPI_ISL_758690, EPI_ISL_758691, EPI_ISL_758692, EPI_ISL_758693, EPI_ISL_758694, EPI_ISL_758695, EPI_ISL_758696, EPI_ISL_758697, EPI_ISL_758698, EPI_ISL_758699, EPI_ISL_758700 | | | |
| see above | Department of Virus and Microbiological Special Diagnostics, Statens Serum Institut, Copenhagen, Denmark | Albertsen Lab, Department of Chemistry and Bioscience, Aalborg University, Denmark | Danish Covid-19 Genome Consortium |
| EPI_ISL_760542, EPI_ISL_760560 | Lighthouse Lab in Glasgow | Wellcome Sanger Institute for the COVID-19 Genomics UK (COG-UK) Consortium | Harper VanSteenhouse, Yumi Kasai, David Gray, Carol Clugston, Anna Dominiczak and Alex Alderton, Roberto Amato, Sonia Goncalves, Ewan Harrison, David K. Jackson, Ian Johnston, Dominic Kwiatkowski, Cordelia Langford, John Sillitoe on behalf of the Wellcome Sanger Institute COVID-19 Surveillance |

| | | | |
|---|---|---|---|
| | | | Team |
| EPI_ISL_760567, EPI_ISL_760600, EPI_ISL_760649 | Lighthouse Lab in Alderley Park | Wellcome Sanger Institute for the COVID-19 Genomics UK (COG-UK) Consortium | Jacquelyn Wynn, Mairead Hyland, The Lighthouse Lab in Alderley Park and Alex Alderton, Roberto Amato, Sonia Goncalves, Ewan Harrison, David K. Jackson, Ian Johnston, Dominic Kwiatkowski, Cordelia Langford, John Sillitoe on behalf of the Wellcome Sanger Institute COVID-19 Surveillance Team |
| EPI_ISL_760658, EPI_ISL_760669, EPI_ISL_760673, EPI_ISL_760739, EPI_ISL_760789 | Lighthouse Lab in Glasgow | Wellcome Sanger Institute for the COVID-19 Genomics UK (COG-UK) Consortium | Harper VanSteenhouse, Yumi Kasai, David Gray, Carol Clugston, Anna Dominiczak and Alex Alderton, Roberto Amato, Sonia Goncalves, Ewan Harrison, David K. Jackson, Ian Johnston, Dominic Kwiatkowski, Cordelia Langford, John Sillitoe on behalf of the Wellcome Sanger Institute COVID-19 Surveillance Team |
| EPI_ISL_760883 | Lighthouse Lab in Cambridge | Wellcome Sanger Institute for the COVID-19 Genomics UK (COG-UK) Consortium | Rob Howes, The Lighthouse Lab in Cambridge and Alex Alderton, Roberto Amato, Sonia Goncalves, Ewan Harrison, David K. Jackson, Ian Johnston, Dominic Kwiatkowski, Cordelia Langford, John Sillitoe on behalf of the Wellcome Sanger Institute COVID-19 Surveillance Team |
| EPI_ISL_760924 | Lighthouse Lab in Alderley Park | Wellcome Sanger Institute for the COVID-19 Genomics UK (COG-UK) Consortium | Jacquelyn Wynn, Mairead Hyland, The Lighthouse Lab in Alderley Park and Alex Alderton, Roberto Amato, Sonia Goncalves, Ewan Harrison, David K. Jackson, Ian Johnston, Dominic Kwiatkowski, Cordelia Langford, John Sillitoe on behalf of the Wellcome Sanger Institute COVID-19 Surveillance Team |
| EPI_ISL_760984 | Lighthouse Lab in Glasgow | Wellcome Sanger Institute for the COVID-19 Genomics UK (COG-UK) Consortium | Harper VanSteenhouse, Yumi Kasai, David Gray, Carol Clugston, Anna Dominiczak and Alex Alderton, Roberto Amato, Sonia Goncalves, Ewan Harrison, David K. Jackson, Ian Johnston, Dominic Kwiatkowski, Cordelia Langford, John Sillitoe on behalf of the Wellcome Sanger Institute COVID-19 Surveillance Team |
| EPI_ISL_760989, EPI_ISL_761036, EPI_ISL_761140, EPI_ISL_761258 | Lighthouse Lab in Alderley Park | Wellcome Sanger Institute for the COVID-19 Genomics UK (COG-UK) Consortium | Jacquelyn Wynn, Mairead Hyland, The Lighthouse Lab in Alderley Park and Alex Alderton, Roberto Amato, Sonia Goncalves, Ewan Harrison, David K. Jackson, Ian Johnston, Dominic Kwiatkowski, Cordelia Langford, John Sillitoe on behalf of the Wellcome Sanger Institute COVID-19 Surveillance Team |
| EPI_ISL_761616 | Lighthouse Lab in Milton Keynes | Wellcome Sanger Institute for the COVID-19 Genomics UK (COG-UK) Consortium | The Lighthouse Lab in Milton Keynes and Alex Alderton, Roberto Amato, Sonia Goncalves, Ewan Harrison, David K. Jackson, Ian Johnston, Dominic Kwiatkowski, Cordelia Langford, John Sillitoe on behalf of the Wellcome Sanger Institute COVID-19 Surveillance Team |
| EPI_ISL_761693, EPI_ISL_761755 | Lighthouse Lab in Alderley Park | Wellcome Sanger Institute for the COVID-19 Genomics UK (COG-UK) Consortium | Jacquelyn Wynn, Mairead Hyland, The Lighthouse Lab in Alderley Park and Alex Alderton, Roberto Amato, Sonia Goncalves, Ewan Harrison, David K. Jackson, Ian Johnston, Dominic Kwiatkowski, Cordelia Langford, John Sillitoe on behalf of the Wellcome Sanger Institute COVID-19 Surveillance Team |
| EPI_ISL_761985, EPI_ISL_761998, EPI_ISL_762013, EPI_ISL_762062, EPI_ISL_762083, EPI_ISL_762110, EPI_ISL_762115, EPI_ISL_762156, EPI_ISL_762169, EPI_ISL_762171 | Lighthouse Lab in Glasgow | Wellcome Sanger Institute for the COVID-19 Genomics UK (COG-UK) Consortium | Harper VanSteenhouse, Yumi Kasai, David Gray, Carol Clugston, Anna Dominiczak and Alex Alderton, Roberto Amato, Sonia Goncalves, Ewan Harrison, David K. Jackson, Ian Johnston, Dominic Kwiatkowski, Cordelia Langford, John Sillitoe on behalf of the Wellcome Sanger Institute COVID-19 Surveillance Team |
| EPI_ISL_762198, EPI_ISL_762204, EPI_ISL_762205, EPI_ISL_762208, EPI_ISL_762227, EPI_ISL_762231, EPI_ISL_762238, EPI_ISL_762244, EPI_ISL_762254, EPI_ISL_762258, EPI_ISL_762271, EPI_ISL_762273, EPI_ISL_762278, EPI_ISL_762281, EPI_ISL_762285, EPI_ISL_762287, EPI_ISL_762293, EPI_ISL_762305, EPI_ISL_762309, EPI_ISL_762324, EPI_ISL_762328, EPI_ISL_762333, EPI_ISL_762340, EPI_ISL_762351, EPI_ISL_762377, EPI_ISL_762378, EPI_ISL_762381, EPI_ISL_762392, EPI_ISL_762395, EPI_ISL_762399, EPI_ISL_762400, EPI_ISL_762408, EPI_ISL_762419, EPI_ISL_762423, EPI_ISL_762434, EPI_ISL_762436, EPI_ISL_762438, EPI_ISL_762440, EPI_ISL_762456, EPI_ISL_762462, EPI_ISL_762463, EPI_ISL_762480, EPI_ISL_762493, EPI_ISL_762494, EPI_ISL_762498, EPI_ISL_762502, EPI_ISL_762505, EPI_ISL_762509, EPI_ISL_762516, EPI_ISL_762519, EPI_ISL_762520, EPI_ISL_762523, EPI_ISL_762864, EPI_ISL_762867, EPI_ISL_762875, EPI_ISL_762880, EPI_ISL_762884, EPI_ISL_762891, EPI_ISL_762895, EPI_ISL_762896, EPI_ISL_762903, EPI_ISL_762908, EPI_ISL_762912, EPI_ISL_762914, EPI_ISL_762917, EPI_ISL_762920, EPI_ISL_762937, EPI_ISL_762943, EPI_ISL_762945, EPI_ISL_762963, EPI_ISL_762965 | | | |
| see above | Lighthouse Lab in Milton Keynes | Wellcome Sanger Institute for the COVID-19 Genomics UK (COG-UK) Consortium | The Lighthouse Lab in Milton Keynes and Alex Alderton, Roberto Amato, Sonia Goncalves, Ewan Harrison, David K. Jackson, Ian Johnston, Dominic Kwiatkowski, Cordelia Langford, John Sillitoe on behalf of the Wellcome Sanger Institute COVID-19 Surveillance Team |
| EPI_ISL_762966 | Lighthouse Lab in Glasgow | Wellcome Sanger Institute for the COVID-19 Genomics UK (COG-UK) Consortium | Harper VanSteenhouse, Yumi Kasai, David Gray, Carol Clugston, Anna Dominiczak and Alex Alderton, Roberto Amato, Sonia Goncalves, Ewan Harrison, David K. Jackson, Ian Johnston, Dominic Kwiatkowski, Cordelia Langford, John Sillitoe on behalf of the Wellcome Sanger Institute COVID-19 Surveillance Team |
| EPI_ISL_762967, EPI_ISL_762968, EPI_ISL_762976, EPI_ISL_762978 | Lighthouse Lab in Milton Keynes | Wellcome Sanger Institute for the COVID-19 Genomics UK (COG-UK) Consortium | The Lighthouse Lab in Milton Keynes and Alex Alderton, Roberto Amato, Sonia Goncalves, Ewan Harrison, David K. Jackson, Ian Johnston, Dominic Kwiatkowski, Cordelia Langford, John Sillitoe on behalf of the Wellcome Sanger Institute COVID-19 Surveillance Team |
| EPI_ISL_762979 | Lighthouse Lab in Glasgow | Wellcome Sanger Institute for the COVID-19 Genomics UK (COG-UK) Consortium | Harper VanSteenhouse, Yumi Kasai, David Gray, Carol Clugston, Anna Dominiczak and Alex Alderton, Roberto Amato, Sonia Goncalves, Ewan Harrison, David K. Jackson, Ian Johnston, Dominic Kwiatkowski, Cordelia Langford, John Sillitoe on behalf of the Wellcome Sanger Institute COVID-19 Surveillance Team |
| EPI_ISL_762981 | Lighthouse Lab in Milton Keynes | Wellcome Sanger Institute for the COVID-19 Genomics UK (COG-UK) Consortium | The Lighthouse Lab in Milton Keynes and Alex Alderton, Roberto Amato, Sonia Goncalves, Ewan Harrison, David K. Jackson, Ian Johnston, Dominic Kwiatkowski, Cordelia Langford, John Sillitoe on behalf of the Wellcome Sanger Institute COVID-19 Surveillance Team |
| EPI_ISL_762986 | Lighthouse Lab in Glasgow | Wellcome Sanger Institute for the COVID-19 Genomics UK (COG-UK) Consortium | Harper VanSteenhouse, Yumi Kasai, David Gray, Carol Clugston, Anna Dominiczak and Alex Alderton, Roberto Amato, Sonia Goncalves, Ewan Harrison, David K. Jackson, Ian Johnston, Dominic Kwiatkowski, Cordelia Langford, John Sillitoe on behalf of the Wellcome Sanger Institute COVID-19 Surveillance Team |
| EPI_ISL_763002, EPI_ISL_763012, EPI_ISL_763013, EPI_ISL_763036, EPI_ISL_763037, EPI_ISL_763039, EPI_ISL_763055, EPI_ISL_763056, EPI_ISL_763057, EPI_ISL_763058 | Unit 17: Influenza & Other Respiratory Viruses, German National Influenza Center | Project group Epidemiology of Highly Pathogenic Microorganisms, Robert Koch-Institute | Ariane Düx, Andreas Sachse, Grit Schubert, Sébastien Calvignac-Spencer, Fabian Leendertz, Thorsten Wolff, Ralf Dürrwald, Djin-Ye Oh, Marianne Wedde |
| EPI_ISL_763078, EPI_ISL_763081, EPI_ISL_763082, EPI_ISL_763083 | Microbiologia e Virologia | Istituto Zooprofilattico Sperimentale delle Venezie | Adelaide Milani, Alessia Schivo, Annalisa Salviato, Erika Giorgia Quaranta, Ambra Pastori, Bianca Zecchin, Alice Fusaro, Isabella Monne, Calogero Terregino, Antonia Ricci |
| EPI_ISL_763181, EPI_ISL_763199, EPI_ISL_763201, EPI_ISL_763206, EPI_ISL_763242, EPI_ISL_763245, EPI_ISL_763246, EPI_ISL_763287, EPI_ISL_763289, EPI_ISL_763290 | Dutch COVID-19 response team | Erasmus Medical Center | Bas Oude Munnink, Reina Sikkema, David Nieuwenhuijse, Irina Chestakova, Anne van der Linden, Marjan Boter, Emmanuelle Munger, Corine GeurtsvanKessel, Annemiek van der Eijk, Richard Molenkamp, Marion Koopmans, on behalf of the Dutch national COVID-19 response team. |
| EPI_ISL_763374 | Wales Specialist Virology Centre Sequencing lab: Pathogen Genomics Unit | COVID-19 Genomics UK (COG-UK) Consortium | Catherine Moore, Johnathan Evans, Laura Gifford, Malorie Perry, Simon Cottrell, Angela Marchbank, Alec Birchley, Alexander Adams, Amy Gaskin, Bree Gatica-Wilcox, Jason Coombes, Joel Southgate, Lauren Gilbert, Lee Graham, Nicole Pacchiarini, Sara Kumziene-Summerhayes, Sarah Taylor, Sophie Jones, Sara Rey, Matthew Bull, Joanne Watkins, Sally Corden, Tom Connor |
| EPI_ISL_763381 | University College London, Great Ormond Street Hospital for Children NHS Foundation Trust, Imperial College Healthcare NHS Trust | COVID-19 Genomics UK (COG-UK) Consortium | Sergi Castellano, Rachel Williams, Mark Kristiansen, Paola Resende Silva, Sunando Roy, Tony Brooks, Helena Tutill, Paola Niola, Patricia Dyal, Charlotte Williams, Leysa Forrest, Yasmin Panchbhaya, Jacqueline Findlay, Samuel Weeks, Julianne Brown, Kathryn Harris, Paul Randell, James Price, Alison Holmes, Judith Breuer |
| EPI_ISL_763382, EPI_ISL_763383, EPI_ISL_763384, EPI_ISL_763385, EPI_ISL_763391 | University of Exeter | COVID-19 Genomics UK (COG-UK) Consortium | Ben Temperton,Aaron Jeffries,Michelle Michelsen,Joanna Warwick-Dugdale,Audrey Farbos,Robyn Manley,Stephen Michell,Jane Masoli |
| EPI_ISL_763399 | University College London, Great Ormond Street Hospital for Children NHS Foundation Trust, Imperial College Healthcare NHS Trust | COVID-19 Genomics UK (COG-UK) Consortium | Sergi Castellano, Rachel Williams, Mark Kristiansen, Paola Resende Silva, Sunando Roy, Tony Brooks, Helena Tutill, Paola Niola, Patricia Dyal, Charlotte Williams, Leysa Forrest, Yasmin Panchbhaya, Jacqueline Findlay, Samuel Weeks, Julianne Brown, Kathryn Harris, Paul Randell, James Price, Alison Holmes, Judith Breuer |
| EPI_ISL_763413 | Wales Specialist Virology Centre Sequencing lab: Pathogen Genomics Unit | COVID-19 Genomics UK (COG-UK) Consortium | Catherine Moore, Johnathan Evans, Laura Gifford, Malorie Perry, Simon Cottrell, Angela Marchbank, Alec Birchley, Alexander Adams, Amy Gaskin, Bree Gatica-Wilcox, Jason Coombes, Joel Southgate, Lauren Gilbert, Lee Graham, Nicole Pacchiarini, Sara Kumziene-Summerhayes, Sarah Taylor, Sophie Jones, Sara Rey, Matthew Bull, Joanne Watkins, Sally Corden, Tom Connor |
| EPI_ISL_763414 | University of Exeter | COVID-19 Genomics UK (COG-UK) Consortium | Ben Temperton,Aaron Jeffries,Michelle Michelsen,Joanna Warwick-Dugdale,Audrey Farbos,Robyn Manley,Stephen Michell,Jane Masoli |
| EPI_ISL_763418 | Wales Specialist Virology Centre Sequencing lab: Pathogen Genomics Unit | COVID-19 Genomics UK (COG-UK) Consortium | Catherine Moore, Johnathan Evans, Laura Gifford, Malorie Perry, Simon Cottrell, Angela Marchbank, Alec Birchley, Alexander Adams, Amy Gaskin, Bree Gatica-Wilcox, Jason Coombes, Joel Southgate, Lauren Gilbert, Lee Graham, Nicole Pacchiarini, Sara Kumziene-Summerhayes, Sarah Taylor, Sophie Jones, Sara Rey, Matthew Bull, Joanne Watkins, Sally Corden, Tom Connor |

| | | | |
|---|---|---|---|
| EPI_ISL_763425 | Centre for Enzyme Innovation, University of Portsmouth / Translational Research Laboratory, Portsmouth Hospitals NHS Trust | COVID-19 Genomics UK (COG-UK) Consortium | Angela Beckett,Yann Bourgeois,Garry Scarlett,Sharon Glaysher,Scott Elliott,Kelly Bicknell,Robert Impey,Allyson Lloyd,Sarah Wyllie,Ethan Butcher,Anoop Chauhan,Samuel Robson |
| EPI_ISL_763432, EPI_ISL_763433 | University of Exeter | COVID-19 Genomics UK (COG-UK) Consortium | Ben Temperton,Aaron Jeffries,Michelle Michelsen,Joanna Warwick-Dugdale,Audrey Farbos,Robyn Manley,Stephen Michell,Jane Masoli |
| EPI_ISL_763441 | Wales Specialist Virology Centre Sequencing lab: Pathogen Genomics Unit | COVID-19 Genomics UK (COG-UK) Consortium | Catherine Moore, Johnathan Evans, Laura Gifford, Malorie Perry, Simon Cottrell, Angela Marchbank, Alec Birchley, Alexander Adams, Amy Gaskin, Bree Gatica-Wilcox, Jason Coombes, Joel Southgate, Lauren Gilbert, Lee Graham, Nicole Pacchiarini, Sara Kumziene-Summerhayes, Sarah Taylor, Sophie Jones, Sara Rey, Matthew Bull, Joanne Watkins, Sally Corden, Tom Connor |
| EPI_ISL_763452 | University of Exeter | COVID-19 Genomics UK (COG-UK) Consortium | Ben Temperton,Aaron Jeffries,Michelle Michelsen,Joanna Warwick-Dugdale,Audrey Farbos,Robyn Manley,Stephen Michell,Jane Masoli |
| EPI_ISL_763455 | University College London, Great Ormond Street Hospital for Children NHS Foundation Trust, Imperial College Healthcare NHS Trust | COVID-19 Genomics UK (COG-UK) Consortium | Sergi Castellano, Rachel Williams, Mark Kristiansen, Paola Resende Silva, Sunando Roy, Tony Brooks, Helena Tutill, Paola Niola, Patricia Dyal, Charlotte Williams, Leysa Forrest, Yasmin Panchbhaya, Jacqueline Findlay, Samuel Weeks, Julianne Brown, Kathryn Harris, Paul Randell, James Price, Alison Holmes, Judith Breuer |
| EPI_ISL_763458 | Wales Specialist Virology Centre Sequencing lab: Pathogen Genomics Unit | COVID-19 Genomics UK (COG-UK) Consortium | Catherine Moore, Johnathan Evans, Laura Gifford, Malorie Perry, Simon Cottrell, Angela Marchbank, Alec Birchley, Alexander Adams, Amy Gaskin, Bree Gatica-Wilcox, Jason Coombes, Joel Southgate, Lauren Gilbert, Lee Graham, Nicole Pacchiarini, Sara Kumziene-Summerhayes, Sarah Taylor, Sophie Jones, Sara Rey, Matthew Bull, Joanne Watkins, Sally Corden, Tom Connor |
| EPI_ISL_763459, EPI_ISL_763460 | University of Exeter | COVID-19 Genomics UK (COG-UK) Consortium | Ben Temperton,Aaron Jeffries,Michelle Michelsen,Joanna Warwick-Dugdale,Audrey Farbos,Robyn Manley,Stephen Michell,Jane Masoli |
| EPI_ISL_763472, EPI_ISL_763493 | Wales Specialist Virology Centre Sequencing lab: Pathogen Genomics Unit | COVID-19 Genomics UK (COG-UK) Consortium | Catherine Moore, Johnathan Evans, Laura Gifford, Malorie Perry, Simon Cottrell, Angela Marchbank, Alec Birchley, Alexander Adams, Amy Gaskin, Bree Gatica-Wilcox, Jason Coombes, Joel Southgate, Lauren Gilbert, Lee Graham, Nicole Pacchiarini, Sara Kumziene-Summerhayes, Sarah Taylor, Sophie Jones, Sara Rey, Matthew Bull, Joanne Watkins, Sally Corden, Tom Connor |
| EPI_ISL_763496 | University of Exeter | COVID-19 Genomics UK (COG-UK) Consortium | Ben Temperton,Aaron Jeffries,Michelle Michelsen,Joanna Warwick-Dugdale,Audrey Farbos,Robyn Manley,Stephen Michell,Jane Masoli |
| EPI_ISL_763497 | Wales Specialist Virology Centre Sequencing lab: Pathogen Genomics Unit | COVID-19 Genomics UK (COG-UK) Consortium | Catherine Moore, Johnathan Evans, Laura Gifford, Malorie Perry, Simon Cottrell, Angela Marchbank, Alec Birchley, Alexander Adams, Amy Gaskin, Bree Gatica-Wilcox, Jason Coombes, Joel Southgate, Lauren Gilbert, Lee Graham, Nicole Pacchiarini, Sara Kumziene-Summerhayes, Sarah Taylor, Sophie Jones, Sara Rey, Matthew Bull, Joanne Watkins, Sally Corden, Tom Connor |
| EPI_ISL_763520 | University College London, Great Ormond Street Hospital for Children NHS Foundation Trust, Imperial College Healthcare NHS Trust | COVID-19 Genomics UK (COG-UK) Consortium | Sergi Castellano, Rachel Williams, Mark Kristiansen, Paola Resende Silva, Sunando Roy, Tony Brooks, Helena Tutill, Paola Niola, Patricia Dyal, Charlotte Williams, Leysa Forrest, Yasmin Panchbhaya, Jacqueline Findlay, Samuel Weeks, Julianne Brown, Kathryn Harris, Paul Randell, James Price, Alison Holmes, Judith Breuer |
| EPI_ISL_763526 | Department of Pathology, University of Cambridge | COVID-19 Genomics UK (COG-UK) Consortium | Aminu S. Jahun, Yasmin Chaudhry, Grant Hall, Iliana Georgana, Myra Hosmillo, Martin D. Curran, Malte Pinckert, Surendra Parmar, Ian Goodfellow |
| EPI_ISL_763527, EPI_ISL_763528, EPI_ISL_763529, EPI_ISL_763530, EPI_ISL_763531, EPI_ISL_763532 | University of Exeter | COVID-19 Genomics UK (COG-UK) Consortium | Ben Temperton,Aaron Jeffries,Michelle Michelsen,Joanna Warwick-Dugdale,Audrey Farbos,Robyn Manley,Stephen Michell,Jane Masoli |
| EPI_ISL_763539 | Wales Specialist Virology Centre Sequencing lab: Pathogen Genomics Unit | COVID-19 Genomics UK (COG-UK) Consortium | Catherine Moore, Johnathan Evans, Laura Gifford, Malorie Perry, Simon Cottrell, Angela Marchbank, Alec Birchley, Alexander Adams, Amy Gaskin, Bree Gatica-Wilcox, Jason Coombes, Joel Southgate, Lauren Gilbert, Lee Graham, Nicole Pacchiarini, Sara Kumziene-Summerhayes, Sarah Taylor, Sophie Jones, Sara Rey, Matthew Bull, Joanne Watkins, Sally Corden, Tom Connor |
| EPI_ISL_763543 | University of Exeter | COVID-19 Genomics UK (COG-UK) Consortium | Ben Temperton,Aaron Jeffries,Michelle Michelsen,Joanna Warwick-Dugdale,Audrey Farbos,Robyn Manley,Stephen Michell,Jane Masoli |
| EPI_ISL_763544 | University College London, Great Ormond Street Hospital for Children NHS Foundation Trust, Imperial College Healthcare NHS Trust | COVID-19 Genomics UK (COG-UK) Consortium | Sergi Castellano, Rachel Williams, Mark Kristiansen, Paola Resende Silva, Sunando Roy, Tony Brooks, Helena Tutill, Paola Niola, Patricia Dyal, Charlotte Williams, Leysa Forrest, Yasmin Panchbhaya, Jacqueline Findlay, Samuel Weeks, Julianne Brown, Kathryn Harris, Paul Randell, James Price, Alison Holmes, Judith Breuer |
| EPI_ISL_763546 | Wales Specialist Virology Centre Sequencing lab: Pathogen Genomics Unit | COVID-19 Genomics UK (COG-UK) Consortium | Catherine Moore, Johnathan Evans, Laura Gifford, Malorie Perry, Simon Cottrell, Angela Marchbank, Alec Birchley, Alexander Adams, Amy Gaskin, Bree Gatica-Wilcox, Jason Coombes, Joel Southgate, Lauren Gilbert, Lee Graham, Nicole Pacchiarini, Sara Kumziene-Summerhayes, Sarah Taylor, Sophie Jones, Sara Rey, Matthew Bull, Joanne Watkins, Sally Corden, Tom Connor |
| EPI_ISL_763553, EPI_ISL_763554, EPI_ISL_763555, EPI_ISL_763556, EPI_ISL_763563, EPI_ISL_763564 | University of Exeter | COVID-19 Genomics UK (COG-UK) Consortium | Ben Temperton,Aaron Jeffries,Michelle Michelsen,Joanna Warwick-Dugdale,Audrey Farbos,Robyn Manley,Stephen Michell,Jane Masoli |
| EPI_ISL_763567 | University College London, Great Ormond Street Hospital for Children NHS Foundation Trust, Imperial College Healthcare NHS Trust | COVID-19 Genomics UK (COG-UK) Consortium | Sergi Castellano, Rachel Williams, Mark Kristiansen, Paola Resende Silva, Sunando Roy, Tony Brooks, Helena Tutill, Paola Niola, Patricia Dyal, Charlotte Williams, Leysa Forrest, Yasmin Panchbhaya, Jacqueline Findlay, Samuel Weeks, Julianne Brown, Kathryn Harris, Paul Randell, James Price, Alison Holmes, Judith Breuer |
| EPI_ISL_763568, EPI_ISL_763574, EPI_ISL_763575, EPI_ISL_763576 | Wales Specialist Virology Centre Sequencing lab: Pathogen Genomics Unit | COVID-19 Genomics UK (COG-UK) Consortium | Catherine Moore, Johnathan Evans, Laura Gifford, Malorie Perry, Simon Cottrell, Angela Marchbank, Alec Birchley, Alexander Adams, Amy Gaskin, Bree Gatica-Wilcox, Jason Coombes, Joel Southgate, Lauren Gilbert, Lee Graham, Nicole Pacchiarini, Sara Kumziene-Summerhayes, Sarah Taylor, Sophie Jones, Sara Rey, Matthew Bull, Joanne Watkins, Sally Corden, Tom Connor |
| EPI_ISL_763578 | University College London, Great Ormond Street Hospital for Children NHS Foundation Trust, Imperial College Healthcare NHS Trust | COVID-19 Genomics UK (COG-UK) Consortium | Sergi Castellano, Rachel Williams, Mark Kristiansen, Paola Resende Silva, Sunando Roy, Tony Brooks, Helena Tutill, Paola Niola, Patricia Dyal, Charlotte Williams, Leysa Forrest, Yasmin Panchbhaya, Jacqueline Findlay, Samuel Weeks, Julianne Brown, Kathryn Harris, Paul Randell, James Price, Alison Holmes, Judith Breuer |
| EPI_ISL_763584, EPI_ISL_763591, EPI_ISL_763594 | Wales Specialist Virology Centre Sequencing lab: Pathogen Genomics Unit | COVID-19 Genomics UK (COG-UK) Consortium | Catherine Moore, Johnathan Evans, Laura Gifford, Malorie Perry, Simon Cottrell, Angela Marchbank, Alec Birchley, Alexander Adams, Amy Gaskin, Bree Gatica-Wilcox, Jason Coombes, Joel Southgate, Lauren Gilbert, Lee Graham, Nicole Pacchiarini, Sara Kumziene-Summerhayes, Sarah Taylor, Sophie Jones, Sara Rey, Matthew Bull, Joanne Watkins, Sally Corden, Tom Connor |
| EPI_ISL_763607, EPI_ISL_763610 | Centre for Enzyme Innovation, University of Portsmouth / Translational Research Laboratory, Portsmouth Hospitals NHS Trust | COVID-19 Genomics UK (COG-UK) Consortium | Angela Beckett,Yann Bourgeois,Garry Scarlett,Sharon Glaysher,Scott Elliott,Kelly Bicknell,Robert Impey,Allyson Lloyd,Sarah Wyllie,Ethan Butcher,Anoop Chauhan,Samuel Robson |
| EPI_ISL_763629, EPI_ISL_763630, EPI_ISL_763631 | University of Exeter | COVID-19 Genomics UK (COG-UK) Consortium | Ben Temperton,Aaron Jeffries,Michelle Michelsen,Joanna Warwick-Dugdale,Audrey Farbos,Robyn Manley,Stephen Michell,Jane Masoli |
| EPI_ISL_763638 | Department of Pathology, University of Cambridge | COVID-19 Genomics UK (COG-UK) Consortium | Aminu S. Jahun, Yasmin Chaudhry, Grant Hall, Iliana Georgana, Myra Hosmillo, Martin D. Curran, Malte Pinckert, Surendra Parmar, Ian Goodfellow |
| EPI_ISL_763668 | Wales Specialist Virology Centre Sequencing lab: Pathogen Genomics Unit | COVID-19 Genomics UK (COG-UK) Consortium | Catherine Moore, Johnathan Evans, Laura Gifford, Malorie Perry, Simon Cottrell, Angela Marchbank, Alec Birchley, Alexander Adams, Amy Gaskin, Bree Gatica-Wilcox, Jason Coombes, Joel Southgate, Lauren Gilbert, Lee Graham, Nicole Pacchiarini, Sara Kumziene-Summerhayes, Sarah Taylor, Sophie Jones, Sara Rey, Matthew Bull, Joanne Watkins, Sally Corden, Tom Connor |
| EPI_ISL_763669 | University of Exeter | COVID-19 Genomics UK (COG-UK) Consortium | Ben Temperton,Aaron Jeffries,Michelle Michelsen,Joanna Warwick-Dugdale,Audrey Farbos,Robyn Manley,Stephen Michell,Jane Masoli |
| EPI_ISL_763679 | Wales Specialist Virology Centre Sequencing lab: Pathogen Genomics Unit | COVID-19 Genomics UK (COG-UK) Consortium | Catherine Moore, Johnathan Evans, Laura Gifford, Malorie Perry, Simon Cottrell, Angela Marchbank, Alec Birchley, Alexander Adams, Amy Gaskin, Bree Gatica-Wilcox, Jason Coombes, Joel Southgate, Lauren Gilbert, Lee Graham, Nicole Pacchiarini, Sara Kumziene-Summerhayes, Sarah Taylor, Sophie Jones, Sara Rey, Matthew Bull, Joanne Watkins, Sally Corden, Tom Connor |
| EPI_ISL_763684 | Quadram Institute Bioscience | COVID-19 Genomics UK (COG-UK) Consortium | Dave J. Baker, Gemma L. Kay, Alp Aydin, Thanh Le-Viet, Steven Rudder, Ana P. Tedim, Anastasia Kolyva, Maria Diaz, Leonardo de Oliveira Martins, Nabil-Fareed Alikhan, Lizzie Meadows, Rachael Stanley, Ngozi Elumogo, Muhammed Yasir, Nicholas M. Thomson, Alexander J Trotter, Rachel Gilroy, Samuel Bloomfield, Claire Stuart, Andrew Bell, Reenesh Prakash, Samir Dervisevic, Alison E. Mather, John Wain, Mark Webber, Andrew J. Page, Justin O'Grady |
| EPI_ISL_763697, EPI_ISL_763698, EPI_ISL_763699 | University of Exeter | COVID-19 Genomics UK (COG-UK) Consortium | Ben Temperton,Aaron Jeffries,Michelle Michelsen,Joanna Warwick-Dugdale,Audrey Farbos,Robyn Manley,Stephen Michell,Jane Masoli |

| | | | |
|---|---|---|---|
| EPI_ISL_763700 | University College London, Great Ormond Street Hospital for Children NHS Foundation Trust, Imperial College Healthcare NHS Trust | COVID-19 Genomics UK (COG-UK) Consortium | Sergi Castellano, Rachel Williams, Mark Kristiansen, Paola Resende Silva, Sunando Roy, Tony Brooks, Helena Tutill, Paola Niola, Patricia Dyal, Charlotte Williams, Leysa Forrest, Yasmin Panchbhaya, Jacqueline Findlay, Samuel Weeks, Julianne Brown, Kathryn Harris, Paul Randell, James Price, Alison Holmes, Judith Breuer |
| EPI_ISL_763709, EPI_ISL_763710, EPI_ISL_763711, EPI_ISL_763712 | University of Exeter | COVID-19 Genomics UK (COG-UK) Consortium | Ben Temperton,Aaron Jeffries,Michelle Michelsen,Joanna Warwick-Dugdale,Audrey Farbos,Robyn Manley,Stephen Michell,Jane Masoli |
| EPI_ISL_763724, EPI_ISL_763733, EPI_ISL_763739 | Wales Specialist Virology Centre Sequencing lab: Pathogen Genomics Unit | COVID-19 Genomics UK (COG-UK) Consortium | Catherine Moore, Johnathan Evans, Laura Gifford, Malorie Perry, Simon Cottrell, Angela Marchbank, Alec Birchley, Alexander Adams, Amy Gaskin, Bree Gatica-Wilcox, Jason Coombes, Joel Southgate, Lauren Gilbert, Lee Graham, Nicole Pacchiarini, Sara Kumziene-Summerhayes, Sarah Taylor, Sophie Jones, Sara Rey, Matthew Bull, Joanne Watkins, Sally Corden, Tom Connor |
| EPI_ISL_763740 | Virology Department, Sheffield Teaching Hospitals NHS Foundation Trust/Department of Infection, Immunity and Cardiovascular Disease, The Medical School, University of Sheffield | COVID-19 Genomics UK (COG-UK) Consortium | Thushan de Silva, Matthew Parker, Nikki Smith, Adri Angyal, Rebecca Brown, Luke Green, Rachel Tucker, Paul Parsons, Danielle Groves, Katie Johnson, Laura Carrilero, Alex Keeley, Dave Partridge, Matthew Wyles, Benjamin Lindsey, Mehmet Yavuz, Mohammad Raza, Cariad Evans |
| EPI_ISL_763751, EPI_ISL_763754 | Wales Specialist Virology Centre Sequencing lab: Pathogen Genomics Unit | COVID-19 Genomics UK (COG-UK) Consortium | Catherine Moore, Johnathan Evans, Laura Gifford, Malorie Perry, Simon Cottrell, Angela Marchbank, Alec Birchley, Alexander Adams, Amy Gaskin, Bree Gatica-Wilcox, Jason Coombes, Joel Southgate, Lauren Gilbert, Lee Graham, Nicole Pacchiarini, Sara Kumziene-Summerhayes, Sarah Taylor, Sophie Jones, Sara Rey, Matthew Bull, Joanne Watkins, Sally Corden, Tom Connor |
| EPI_ISL_763757 | University College London, Great Ormond Street Hospital for Children NHS Foundation Trust, Imperial College Healthcare NHS Trust | COVID-19 Genomics UK (COG-UK) Consortium | Sergi Castellano, Rachel Williams, Mark Kristiansen, Paola Resende Silva, Sunando Roy, Tony Brooks, Helena Tutill, Paola Niola, Patricia Dyal, Charlotte Williams, Leysa Forrest, Yasmin Panchbhaya, Jacqueline Findlay, Samuel Weeks, Julianne Brown, Kathryn Harris, Paul Randell, James Price, Alison Holmes, Judith Breuer |
| EPI_ISL_763762 | Wales Specialist Virology Centre Sequencing lab: Pathogen Genomics Unit | COVID-19 Genomics UK (COG-UK) Consortium | Catherine Moore, Johnathan Evans, Laura Gifford, Malorie Perry, Simon Cottrell, Angela Marchbank, Alec Birchley, Alexander Adams, Amy Gaskin, Bree Gatica-Wilcox, Jason Coombes, Joel Southgate, Lauren Gilbert, Lee Graham, Nicole Pacchiarini, Sara Kumziene-Summerhayes, Sarah Taylor, Sophie Jones, Sara Rey, Matthew Bull, Joanne Watkins, Sally Corden, Tom Connor |
| EPI_ISL_763763 | University of Exeter | COVID-19 Genomics UK (COG-UK) Consortium | Ben Temperton,Aaron Jeffries,Michelle Michelsen,Joanna Warwick-Dugdale,Audrey Farbos,Robyn Manley,Stephen Michell,Jane Masoli |
| EPI_ISL_763764, EPI_ISL_763770, EPI_ISL_763778, EPI_ISL_763788 | Wales Specialist Virology Centre Sequencing lab: Pathogen Genomics Unit | COVID-19 Genomics UK (COG-UK) Consortium | Catherine Moore, Johnathan Evans, Laura Gifford, Malorie Perry, Simon Cottrell, Angela Marchbank, Alec Birchley, Alexander Adams, Amy Gaskin, Bree Gatica-Wilcox, Jason Coombes, Joel Southgate, Lauren Gilbert, Lee Graham, Nicole Pacchiarini, Sara Kumziene-Summerhayes, Sarah Taylor, Sophie Jones, Sara Rey, Matthew Bull, Joanne Watkins, Sally Corden, Tom Connor |
| EPI_ISL_763838, EPI_ISL_763844 | Oxford Viromics, NDM, University of Oxford; Oxford University Hospitals; Basingstoke and North Hampshire Hospital | COVID-19 Genomics UK (COG-UK) Consortium | Tanya Golubchik, David Bonsall, George Macintyre, Amy Trebes, Mariateresa de Cesare, Catrin Moore, Alex Mobbs, Anita Justice, Robert Shaw, Monique Andersson, Timothy Peto, Emma Wise, Nathan Moore, Jessica Lynch, Nick Cortes, Matilde Mori, Stephen Kidd, David Buck, John Todd, Christophe Fraser |
| EPI_ISL_763849 | Department of Pathology, University of Cambridge | COVID-19 Genomics UK (COG-UK) Consortium | Aminu S. Jahun, Yasmin Chaudhry, Grant Hall, Iliana Georgana, Myra Hosmillo, Martin D. Curran, Malte Pinckert, Surendra Parmar, Ian Goodfellow |
| EPI_ISL_763860, EPI_ISL_763862, EPI_ISL_763863 | Oxford Viromics, NDM, University of Oxford; Oxford University Hospitals; Basingstoke and North Hampshire Hospital | COVID-19 Genomics UK (COG-UK) Consortium | Tanya Golubchik, David Bonsall, George Macintyre, Amy Trebes, Mariateresa de Cesare, Catrin Moore, Alex Mobbs, Anita Justice, Robert Shaw, Monique Andersson, Timothy Peto, Emma Wise, Nathan Moore, Jessica Lynch, Nick Cortes, Matilde Mori, Stephen Kidd, David Buck, John Todd, Christophe Fraser |
| EPI_ISL_763874, EPI_ISL_763875, EPI_ISL_763876, EPI_ISL_763877, EPI_ISL_763878, EPI_ISL_763879, EPI_ISL_763880, EPI_ISL_763881 | Department of Pathology, University of Cambridge | COVID-19 Genomics UK (COG-UK) Consortium | Aminu S. Jahun, Yasmin Chaudhry, Grant Hall, Iliana Georgana, Myra Hosmillo, Martin D. Curran, Malte Pinckert, Surendra Parmar, Ian Goodfellow |
| EPI_ISL_763892, EPI_ISL_763902, EPI_ISL_763904 | Virology Department, Royal Infirmary of Edinburgh, NHS Lothian / School of Biological Sciences, University of Edinburgh / Institute of Genetics and Molecular Medicine, University of Edinburgh | COVID-19 Genomics UK (COG-UK) Consortium | McHugh M, Dewar R, Rooke S, Gallagher M, Balcaza C, O'Toole Á, Scher E, Hill V, McCrone JT, Colquhoun R, Yu X, Jackson B, Rambaut A, Williams TC, Templeton K |
| EPI_ISL_763928, EPI_ISL_763929, EPI_ISL_763930, EPI_ISL_763931, EPI_ISL_763936, EPI_ISL_763937, EPI_ISL_763938, EPI_ISL_763939, EPI_ISL_763943, EPI_ISL_763944, EPI_ISL_763949, EPI_ISL_763950, EPI_ISL_763951, EPI_ISL_763952, EPI_ISL_763953, EPI_ISL_763954, EPI_ISL_763955, EPI_ISL_763956, EPI_ISL_763957, EPI_ISL_763958, EPI_ISL_763959, EPI_ISL_763960, EPI_ISL_763961, EPI_ISL_763962, EPI_ISL_763963, EPI_ISL_763964, EPI_ISL_763965, EPI_ISL_763966, EPI_ISL_763967, EPI_ISL_763968, EPI_ISL_763969, EPI_ISL_763970, EPI_ISL_763971, EPI_ISL_763972, EPI_ISL_763973, EPI_ISL_763974 | | | |
| see above | Oxford Viromics, NDM, University of Oxford; Oxford University Hospitals; Basingstoke and North Hampshire Hospital | COVID-19 Genomics UK (COG-UK) Consortium | Tanya Golubchik, David Bonsall, George Macintyre, Amy Trebes, Mariateresa de Cesare, Catrin Moore, Alex Mobbs, Anita Justice, Robert Shaw, Monique Andersson, Timothy Peto, Emma Wise, Nathan Moore, Jessica Lynch, Nick Cortes, Matilde Mori, Stephen Kidd, David Buck, John Todd, Christophe Fraser |
| EPI_ISL_763986, EPI_ISL_763990 | Wales Specialist Virology Centre Sequencing lab: Pathogen Genomics Unit | COVID-19 Genomics UK (COG-UK) Consortium | Catherine Moore, Johnathan Evans, Laura Gifford, Malorie Perry, Simon Cottrell, Angela Marchbank, Alec Birchley, Alexander Adams, Amy Gaskin, Bree Gatica-Wilcox, Jason Coombes, Joel Southgate, Lauren Gilbert, Lee Graham, Nicole Pacchiarini, Sara Kumziene-Summerhayes, Sarah Taylor, Sophie Jones, Sara Rey, Matthew Bull, Joanne Watkins, Sally Corden, Tom Connor |
| EPI_ISL_764046, EPI_ISL_764049 | University College London, Great Ormond Street Hospital for Children NHS Foundation Trust, Imperial College Healthcare NHS Trust | COVID-19 Genomics UK (COG-UK) Consortium | Sergi Castellano, Rachel Williams, Mark Kristiansen, Paola Resende Silva, Sunando Roy, Tony Brooks, Helena Tutill, Paola Niola, Patricia Dyal, Charlotte Williams, Leysa Forrest, Yasmin Panchbhaya, Jacqueline Findlay, Samuel Weeks, Julianne Brown, Kathryn Harris, Paul Randell, James Price, Alison Holmes, Judith Breuer |
| EPI_ISL_764061 | Virology Department, Royal Infirmary of Edinburgh, NHS Lothian / School of Biological Sciences, University of Edinburgh / Institute of Genetics and Molecular Medicine, University of Edinburgh | COVID-19 Genomics UK (COG-UK) Consortium | McHugh M, Dewar R, Rooke S, Gallagher M, Balcaza C, O'Toole Á, Scher E, Hill V, McCrone JT, Colquhoun R, Yu X, Jackson B, Rambaut A, Williams TC, Templeton K |
| EPI_ISL_764062 | Wales Specialist Virology Centre Sequencing lab: Pathogen Genomics Unit | COVID-19 Genomics UK (COG-UK) Consortium | Catherine Moore, Johnathan Evans, Laura Gifford, Malorie Perry, Simon Cottrell, Angela Marchbank, Alec Birchley, Alexander Adams, Amy Gaskin, Bree Gatica-Wilcox, Jason Coombes, Joel Southgate, Lauren Gilbert, Lee Graham, Nicole Pacchiarini, Sara Kumziene-Summerhayes, Sarah Taylor, Sophie Jones, Sara Rey, Matthew Bull, Joanne Watkins, Sally Corden, Tom Connor |
| EPI_ISL_764067, EPI_ISL_764068 | Oxford Viromics, NDM, University of Oxford; Oxford University Hospitals; Basingstoke and North Hampshire Hospital | COVID-19 Genomics UK (COG-UK) Consortium | Tanya Golubchik, David Bonsall, George Macintyre, Amy Trebes, Mariateresa de Cesare, Catrin Moore, Alex Mobbs, Anita Justice, Robert Shaw, Monique Andersson, Timothy Peto, Emma Wise, Nathan Moore, Jessica Lynch, Nick Cortes, Matilde Mori, Stephen Kidd, David Buck, John Todd, Christophe Fraser |
| EPI_ISL_764070, EPI_ISL_764071, EPI_ISL_764072, EPI_ISL_764073, EPI_ISL_764074, EPI_ISL_764075 | Virology Department, Royal Infirmary of Edinburgh, NHS Lothian / School of Biological Sciences, University of Edinburgh / Institute of Genetics and Molecular Medicine, University of Edinburgh | COVID-19 Genomics UK (COG-UK) Consortium | McHugh M, Dewar R, Rooke S, Gallagher M, Balcaza C, O'Toole Á, Scher E, Hill V, McCrone JT, Colquhoun R, Yu X, Jackson B, Rambaut A, Williams TC, Templeton K |
| EPI_ISL_764084 | University of Exeter | COVID-19 Genomics UK (COG-UK) Consortium | Ben Temperton,Aaron Jeffries,Michelle Michelsen,Joanna Warwick-Dugdale,Audrey Farbos,Robyn Manley,Stephen Michell,Jane Masoli |
| EPI_ISL_764085, EPI_ISL_764089, EPI_ISL_764090, EPI_ISL_764091, EPI_ISL_764092, EPI_ISL_764093 | Oxford Viromics, NDM, University of Oxford; Oxford University Hospitals; Basingstoke and North Hampshire Hospital | COVID-19 Genomics UK (COG-UK) Consortium | Tanya Golubchik, David Bonsall, George Macintyre, Amy Trebes, Mariateresa de Cesare, Catrin Moore, Alex Mobbs, Anita Justice, Robert Shaw, Monique Andersson, Timothy Peto, Emma Wise, Nathan Moore, Jessica Lynch, Nick Cortes, Matilde Mori, Stephen Kidd, David Buck, John Todd, Christophe Fraser |
| EPI_ISL_764104, EPI_ISL_764108, EPI_ISL_764109 | Virology Department, Sheffield Teaching Hospitals NHS Foundation Trust/Department of Infection, Immunity and Cardiovascular Disease, The Medical School, University of Sheffield | COVID-19 Genomics UK (COG-UK) Consortium | Thushan de Silva, Matthew Parker, Nikki Smith, Adri Angyal, Rebecca Brown, Luke Green, Rachel Tucker, Paul Parsons, Danielle Groves, Katie Johnson, Laura Carrilero, Alex Keeley, Dave Partridge, Matthew Wyles, Benjamin Lindsey, Mehmet Yavuz, Mohammad Raza, Cariad Evans |
| EPI_ISL_764158 | University of Exeter | COVID-19 Genomics UK (COG-UK) Consortium | Ben Temperton,Aaron Jeffries,Michelle Michelsen,Joanna Warwick-Dugdale,Audrey Farbos,Robyn Manley,Stephen Michell,Jane Masoli |
| EPI_ISL_764161 | Wales Specialist Virology Centre Sequencing lab: Pathogen Genomics Unit | COVID-19 Genomics UK (COG-UK) Consortium | Catherine Moore, Johnathan Evans, Laura Gifford, Malorie Perry, Simon Cottrell, Angela Marchbank, Alec Birchley, Alexander Adams, Amy Gaskin, Bree Gatica-Wilcox, Jason Coombes, Joel Southgate, Lauren Gilbert, Lee Graham, Nicole Pacchiarini, Sara Kumziene-Summerhayes, Sarah Taylor, Sophie Jones, Sara Rey, Matthew Bull, Joanne Watkins, Sally Corden, Tom Connor |
| EPI_ISL_764162 | Department of Pathology, University of Cambridge | COVID-19 Genomics UK (COG-UK) Consortium | Aminu S. Jahun, Yasmin Chaudhry, Grant Hall, Iliana Georgana, Myra Hosmillo, Martin D. Curran, Malte Pinckert, Surendra Parmar, Ian Goodfellow |

| | | | |
|---|---|---|---|
| EPI_ISL_764163, EPI_ISL_764166 | University of Exeter | COVID-19 Genomics UK (COG-UK) Consortium | Ben Temperton,Aaron Jeffries,Michelle Michelsen,Joanna Warwick-Dugdale,Audrey Farbos,Robyn Manley,Stephen Michell,Jane Masoli |
| EPI_ISL_764175 | University College London, Great Ormond Street Hospital for Children NHS Foundation Trust, Imperial College Healthcare NHS Trust | COVID-19 Genomics UK (COG-UK) Consortium | Sergi Castellano, Rachel Williams, Mark Kristiansen, Paola Resende Silva, Sunando Roy, Tony Brooks, Helena Tutill, Paola Niola, Patricia Dyal, Charlotte Williams, Leysa Forrest, Yasmin Panchbhaya, Jacqueline Findlay, Samuel Weeks, Julianne Brown, Kathryn Harris, Paul Randell, James Price, Alison Holmes, Judith Breuer |
| EPI_ISL_764179 | University of Exeter | COVID-19 Genomics UK (COG-UK) Consortium | Ben Temperton,Aaron Jeffries,Michelle Michelsen,Joanna Warwick-Dugdale,Audrey Farbos,Robyn Manley,Stephen Michell,Jane Masoli |
| EPI_ISL_764180 | Wales Specialist Virology Centre Sequencing lab: Pathogen Genomics Unit | COVID-19 Genomics UK (COG-UK) Consortium | Catherine Moore, Johnathan Evans, Laura Gifford, Malorie Perry, Simon Cottrell, Angela Marchbank, Alec Birchley, Alexander Adams, Amy Gaskin, Bree Gatica-Wilcox, Jason Coombes, Joel Southgate, Lauren Gilbert, Lee Graham, Nicole Pacchiarini, Sara Kumziene-Summerhayes, Sarah Taylor, Sophie Jones, Sara Rey, Matthew Bull, Joanne Watkins, Sally Corden, Tom Connor |
| EPI_ISL_764181, EPI_ISL_764183, EPI_ISL_764184, EPI_ISL_764197 | University of Exeter | COVID-19 Genomics UK (COG-UK) Consortium | Ben Temperton,Aaron Jeffries,Michelle Michelsen,Joanna Warwick-Dugdale,Audrey Farbos,Robyn Manley,Stephen Michell,Jane Masoli |
| EPI_ISL_764202 | Department of Pathology, University of Cambridge | COVID-19 Genomics UK (COG-UK) Consortium | Aminu S. Jahun, Yasmin Chaudhry, Grant Hall, Iliana Georgana, Myra Hosmillo, Martin D. Curran, Malte Pinckert, Surendra Parmar, Ian Goodfellow |
| EPI_ISL_764207 | Quadram Institute Bioscience | COVID-19 Genomics UK (COG-UK) Consortium | Dave J. Baker, Gemma L. Kay, Alp Aydin, Thanh Le-Viet, Steven Rudder, Ana P. Tedim, Anastasia Kolyva, Maria Diaz, Leonardo de Oliveira Martins, Nabil-Fareed Alikhan, Lizzie Meadows, Rachael Stanley, Ngozi Elumogo, Muhammed Yasir, Nicholas M. Thomson, Alexander J Trotter, Rachel Gilroy, Samuel Bloomfield, Claire Stuart, Andrew Bell, Reenesh Prakash, Samir Dervisevic, Alison E. Mather, John Wain, Mark Webber, Andrew J. Page, Justin O'Grady |
| EPI_ISL_764211, EPI_ISL_764217, EPI_ISL_764218, EPI_ISL_764219, EPI_ISL_764220, EPI_ISL_764223, EPI_ISL_764240 | University of Exeter | COVID-19 Genomics UK (COG-UK) Consortium | Ben Temperton,Aaron Jeffries,Michelle Michelsen,Joanna Warwick-Dugdale,Audrey Farbos,Robyn Manley,Stephen Michell,Jane Masoli |
| EPI_ISL_764242 | Wales Specialist Virology Centre Sequencing lab: Pathogen Genomics Unit | COVID-19 Genomics UK (COG-UK) Consortium | Catherine Moore, Johnathan Evans, Laura Gifford, Malorie Perry, Simon Cottrell, Angela Marchbank, Alec Birchley, Alexander Adams, Amy Gaskin, Bree Gatica-Wilcox, Jason Coombes, Joel Southgate, Lauren Gilbert, Lee Graham, Nicole Pacchiarini, Sara Kumziene-Summerhayes, Sarah Taylor, Sophie Jones, Sara Rey, Matthew Bull, Joanne Watkins, Sally Corden, Tom Connor |
| EPI_ISL_764243 | Oxford Viromics, NDM, University of Oxford; Oxford University Hospitals; Basingstoke and North Hampshire Hospital | COVID-19 Genomics UK (COG-UK) Consortium | Tanya Golubchik, David Bonsall, George Macintyre, Amy Trebes, Mariateresa de Cesare, Catrin Moore, Alex Mobbs, Anita Justice, Robert Shaw, Monique Andersson, Timothy Peto, Emma Wise, Nathan Moore, Jessica Lynch, Nick Cortes, Matilde Mori, Stephen Kidd, David Buck, John Todd, Christophe Fraser |
| EPI_ISL_764245, EPI_ISL_764247 | Wales Specialist Virology Centre Sequencing lab: Pathogen Genomics Unit | COVID-19 Genomics UK (COG-UK) Consortium | Catherine Moore, Johnathan Evans, Laura Gifford, Malorie Perry, Simon Cottrell, Angela Marchbank, Alec Birchley, Alexander Adams, Amy Gaskin, Bree Gatica-Wilcox, Jason Coombes, Joel Southgate, Lauren Gilbert, Lee Graham, Nicole Pacchiarini, Sara Kumziene-Summerhayes, Sarah Taylor, Sophie Jones, Sara Rey, Matthew Bull, Joanne Watkins, Sally Corden, Tom Connor |
| EPI_ISL_764250 | Oxford Viromics, NDM, University of Oxford; Oxford University Hospitals; Basingstoke and North Hampshire Hospital | COVID-19 Genomics UK (COG-UK) Consortium | Tanya Golubchik, David Bonsall, George Macintyre, Amy Trebes, Mariateresa de Cesare, Catrin Moore, Alex Mobbs, Anita Justice, Robert Shaw, Monique Andersson, Timothy Peto, Emma Wise, Nathan Moore, Jessica Lynch, Nick Cortes, Matilde Mori, Stephen Kidd, David Buck, John Todd, Christophe Fraser |
| EPI_ISL_764254, EPI_ISL_764258 | Wales Specialist Virology Centre Sequencing lab: Pathogen Genomics Unit | COVID-19 Genomics UK (COG-UK) Consortium | Catherine Moore, Johnathan Evans, Laura Gifford, Malorie Perry, Simon Cottrell, Angela Marchbank, Alec Birchley, Alexander Adams, Amy Gaskin, Bree Gatica-Wilcox, Jason Coombes, Joel Southgate, Lauren Gilbert, Lee Graham, Nicole Pacchiarini, Sara Kumziene-Summerhayes, Sarah Taylor, Sophie Jones, Sara Rey, Matthew Bull, Joanne Watkins, Sally Corden, Tom Connor |
| EPI_ISL_764264, EPI_ISL_764265 | Oxford Viromics, NDM, University of Oxford; Oxford University Hospitals; Basingstoke and North Hampshire Hospital | COVID-19 Genomics UK (COG-UK) Consortium | Tanya Golubchik, David Bonsall, George Macintyre, Amy Trebes, Mariateresa de Cesare, Catrin Moore, Alex Mobbs, Anita Justice, Robert Shaw, Monique Andersson, Timothy Peto, Emma Wise, Nathan Moore, Jessica Lynch, Nick Cortes, Matilde Mori, Stephen Kidd, David Buck, John Todd, Christophe Fraser |
| EPI_ISL_764267, EPI_ISL_764275, EPI_ISL_764282 | Wales Specialist Virology Centre Sequencing lab: Pathogen Genomics Unit | COVID-19 Genomics UK (COG-UK) Consortium | Catherine Moore, Johnathan Evans, Laura Gifford, Malorie Perry, Simon Cottrell, Angela Marchbank, Alec Birchley, Alexander Adams, Amy Gaskin, Bree Gatica-Wilcox, Jason Coombes, Joel Southgate, Lauren Gilbert, Lee Graham, Nicole Pacchiarini, Sara Kumziene-Summerhayes, Sarah Taylor, Sophie Jones, Sara Rey, Matthew Bull, Joanne Watkins, Sally Corden, Tom Connor |
| EPI_ISL_764331, EPI_ISL_764335, EPI_ISL_764336, EPI_ISL_764337, EPI_ISL_764338, EPI_ISL_764339, EPI_ISL_764340, EPI_ISL_764341, EPI_ISL_764342 | Department of Pathology, University of Cambridge | COVID-19 Genomics UK (COG-UK) Consortium | Aminu S. Jahun, Yasmin Chaudhry, Grant Hall, Iliana Georgana, Myra Hosmillo, Martin D. Curran, Malte Pinckert, Surendra Parmar, Ian Goodfellow |
| EPI_ISL_764374, EPI_ISL_764375, EPI_ISL_764376, EPI_ISL_764377 | University of Exeter | COVID-19 Genomics UK (COG-UK) Consortium | Ben Temperton,Aaron Jeffries,Michelle Michelsen,Joanna Warwick-Dugdale,Audrey Farbos,Robyn Manley,Stephen Michell,Jane Masoli |
| EPI_ISL_764385 | University College London, Great Ormond Street Hospital for Children NHS Foundation Trust, Imperial College Healthcare NHS Trust | COVID-19 Genomics UK (COG-UK) Consortium | Sergi Castellano, Rachel Williams, Mark Kristiansen, Paola Resende Silva, Sunando Roy, Tony Brooks, Helena Tutill, Paola Niola, Patricia Dyal, Charlotte Williams, Leysa Forrest, Yasmin Panchbhaya, Jacqueline Findlay, Samuel Weeks, Julianne Brown, Kathryn Harris, Paul Randell, James Price, Alison Holmes, Judith Breuer |
| EPI_ISL_764449, EPI_ISL_764451, EPI_ISL_764453, EPI_ISL_764457, EPI_ISL_764477 | Quadram Institute Bioscience | COVID-19 Genomics UK (COG-UK) Consortium | Dave J. Baker, Gemma L. Kay, Alp Aydin, Thanh Le-Viet, Steven Rudder, Ana P. Tedim, Anastasia Kolyva, Maria Diaz, Leonardo de Oliveira Martins, Nabil-Fareed Alikhan, Lizzie Meadows, Rachael Stanley, Ngozi Elumogo, Muhammed Yasir, Nicholas M. Thomson, Alexander J Trotter, Rachel Gilroy, Samuel Bloomfield, Claire Stuart, Andrew Bell, Reenesh Prakash, Samir Dervisevic, Alison E. Mather, John Wain, Mark Webber, Andrew J. Page, Justin O'Grady |
| EPI_ISL_764525, EPI_ISL_764526, EPI_ISL_764527, EPI_ISL_764528, EPI_ISL_764529, EPI_ISL_764530, EPI_ISL_764531, EPI_ISL_764535, EPI_ISL_764536, EPI_ISL_764537, EPI_ISL_764543, EPI_ISL_764544, EPI_ISL_764545, EPI_ISL_764546, EPI_ISL_764548, EPI_ISL_764549, EPI_ISL_764550, EPI_ISL_764551, EPI_ISL_764552, EPI_ISL_764553, EPI_ISL_764554, EPI_ISL_764555, EPI_ISL_764556, EPI_ISL_764557, EPI_ISL_764559, EPI_ISL_764561, EPI_ISL_764564, EPI_ISL_764565, EPI_ISL_764567, EPI_ISL_764568, EPI_ISL_764569, EPI_ISL_764570, EPI_ISL_764572 | | | |
| see above | Oxford Viromics, NDM, University of Oxford; Oxford University Hospitals; Basingstoke and North Hampshire Hospital | COVID-19 Genomics UK (COG-UK) Consortium | Tanya Golubchik, David Bonsall, George Macintyre, Amy Trebes, Mariateresa de Cesare, Catrin Moore, Alex Mobbs, Anita Justice, Robert Shaw, Monique Andersson, Timothy Peto, Emma Wise, Nathan Moore, Jessica Lynch, Nick Cortes, Matilde Mori, Stephen Kidd, David Buck, John Todd, Christophe Fraser |
| EPI_ISL_764574, EPI_ISL_764575, EPI_ISL_764577, EPI_ISL_764583, EPI_ISL_764584, EPI_ISL_764586, EPI_ISL_764587, EPI_ISL_764598, EPI_ISL_764624, EPI_ISL_764653, EPI_ISL_764654, EPI_ISL_764655, EPI_ISL_764656, EPI_ISL_764657, EPI_ISL_764658, EPI_ISL_764660, EPI_ISL_764661, EPI_ISL_764678, EPI_ISL_764679, EPI_ISL_764741, EPI_ISL_764742, EPI_ISL_764743, EPI_ISL_764748, EPI_ISL_764749, EPI_ISL_764750, EPI_ISL_764751, EPI_ISL_764752, EPI_ISL_764756, EPI_ISL_764757, EPI_ISL_764758, EPI_ISL_764759, EPI_ISL_764760, EPI_ISL_764761, EPI_ISL_764762, EPI_ISL_764763, EPI_ISL_764764, EPI_ISL_764768, EPI_ISL_764816, EPI_ISL_764817, EPI_ISL_764818, EPI_ISL_764821, EPI_ISL_764824, EPI_ISL_764826, EPI_ISL_764827, EPI_ISL_764828, EPI_ISL_764829, EPI_ISL_764830, EPI_ISL_764831, EPI_ISL_764832, EPI_ISL_764833, EPI_ISL_764834, EPI_ISL_764835, EPI_ISL_764836, EPI_ISL_764837, EPI_ISL_764838, EPI_ISL_764839, EPI_ISL_764840, EPI_ISL_764841, EPI_ISL_764842, EPI_ISL_764843, EPI_ISL_764844, EPI_ISL_764845, EPI_ISL_764846, EPI_ISL_764847, EPI_ISL_764848, EPI_ISL_764849, EPI_ISL_764850, EPI_ISL_764851, EPI_ISL_764852, EPI_ISL_764853, EPI_ISL_764854, EPI_ISL_764855, EPI_ISL_764856, EPI_ISL_764857, EPI_ISL_764858, EPI_ISL_764859, EPI_ISL_764860, EPI_ISL_764861, EPI_ISL_764862, EPI_ISL_764863, EPI_ISL_764864, EPI_ISL_764865, EPI_ISL_764866, EPI_ISL_764867, EPI_ISL_764868, EPI_ISL_764869, EPI_ISL_764870, EPI_ISL_764871, EPI_ISL_764874, EPI_ISL_764875, EPI_ISL_764879, EPI_ISL_764881, EPI_ISL_764883, EPI_ISL_764884, EPI_ISL_764886, EPI_ISL_764887, EPI_ISL_764897, EPI_ISL_764898, EPI_ISL_764899, EPI_ISL_764900, EPI_ISL_764901, EPI_ISL_764902, EPI_ISL_764903, EPI_ISL_764904, EPI_ISL_764905, EPI_ISL_764906, EPI_ISL_764907, EPI_ISL_764939, EPI_ISL_764940, EPI_ISL_764941, EPI_ISL_764942, EPI_ISL_764943, EPI_ISL_764944, EPI_ISL_764945, EPI_ISL_764946, EPI_ISL_764965, EPI_ISL_764966, EPI_ISL_764967, EPI_ISL_764968, EPI_ISL_764970, EPI_ISL_764971, EPI_ISL_764972, EPI_ISL_764973, EPI_ISL_764974, EPI_ISL_764976, EPI_ISL_764977, EPI_ISL_764978, EPI_ISL_764979, EPI_ISL_764980, EPI_ISL_764981, EPI_ISL_764982, EPI_ISL_764983, EPI_ISL_764984, EPI_ISL_764985, EPI_ISL_764986, EPI_ISL_764987, EPI_ISL_764988, EPI_ISL_764991, EPI_ISL_764992, EPI_ISL_764993, EPI_ISL_764994, EPI_ISL_764995, EPI_ISL_764997, EPI_ISL_765002 | | | |
| see above | Wales Specialist Virology Centre Sequencing lab: Pathogen Genomics Unit | COVID-19 Genomics UK (COG-UK) Consortium | Catherine Moore, Johnathan Evans, Laura Gifford, Malorie Perry, Simon Cottrell, Angela Marchbank, Alec Birchley, Alexander Adams, Amy Gaskin, Bree Gatica-Wilcox, Jason Coombes, Joel Southgate, Lauren Gilbert, Lee Graham, Nicole Pacchiarini, Sara Kumziene-Summerhayes, Sarah Taylor, Sophie Jones, Sara Rey, Matthew Bull, Joanne Watkins, Sally Corden, Tom Connor |
| EPI_ISL_765149, EPI_ISL_765153, EPI_ISL_765165, EPI_ISL_765179 | Virology Department, Sheffield Teaching Hospitals NHS Foundation Trust/Department of Infection, Immunity and Cardiovascular Disease, The Medical School, University of Sheffield | COVID-19 Genomics UK (COG-UK) Consortium | Thushan de Silva, Matthew Parker, Nikki Smith, Adri Angyal, Rebecca Brown, Luke Green, Rachel Tucker, Paul Parsons, Danielle Groves, Katie Johnson, Laura Carrilero, Alex Keeley, Dave Partridge, Matthew Wyles, Benjamin Lindsey, Mehmet Yavuz, Mohammad Raza, Cariad Evans |
| EPI_ISL_765481 | SARATOGA HOSPITAL LABORATORY | Wadsworth Center, New York State Department.of Health | Kirsten St. George, Daryl M. Lamson, Alexis Russel, Matthew Shudt, Melissa A Leisner, Jonathan Plitnick, Navjot Singh, John Kelly, Sara Griesemer, Erasmus Schneider, Erica Lasek-Nesselquist |
| EPI_ISL_765489, EPI_ISL_765490 | MONTEFIORE MEDICAL CENTER LABORATORIES | Wadsworth Center, New York State Department.of Health | Kirsten St. George, Daryl M. Lamson, Alexis Russel, Matthew Shudt, Melissa A Leisner, Jonathan Plitnick, Navjot Singh, John Kelly, Sara Griesemer, Erasmus Schneider, Erica Lasek-Nesselquist |

| | | | |
|---|---|---|---|
| EPI_ISL_765518, EPI_ISL_765519, EPI_ISL_765523, EPI_ISL_765528, EPI_ISL_765529 | SARATOGA HOSPITAL LABORATORY | Wadsworth Center, New York State Department.of Health | Kirsten St. George, Daryl M. Lamson, Alexis Russel, Matthew Shudt, Melissa A Leisner, Jonathan Plitnick, Navjot Singh, John Kelly, Sara Griesemer, Erasmus Schneider, Erica Lasek-Nesselquist |
| EPI_ISL_765530, EPI_ISL_765541, EPI_ISL_765546, EPI_ISL_765548, EPI_ISL_765552, EPI_ISL_765558, EPI_ISL_765563, EPI_ISL_765564 | MONTEFIORE MEDICAL CENTER LABORATORIES | Wadsworth Center, New York State Department.of Health | Kirsten St. George, Daryl M. Lamson, Alexis Russel, Matthew Shudt, Melissa A Leisner, Jonathan Plitnick, Navjot Singh, John Kelly, Sara Griesemer, Erasmus Schneider, Erica Lasek-Nesselquist |
| EPI_ISL_765986 | Massachusetts General Hospital | Infectious Disease Program, Broad Institute of Harvard and MIT | Lemieux,J.E., Siddle,K.J., Shaw,B., Adams,G., Pierce,V., Turbett,S., Anahtar,M., Branda,J., Slater,D., Harris,J., Lin,A.E., Gladden-Young,A., Lagerborg,K., Rudy,M., DeRuff,K., Carter,A., Normandin,E., Bauer,M., Reilly,S., Tomkins-Tinch,C., Loreth,C., Chaluvadi,S., Neumann,A., Cusick,C., Chapman,S.B., Gnirke,A., Flowers,K., Cerrato,F., Birren,B.W., Gallagher,G., Smole,S., Park,D.J., MacInnis,B.L., Ryan,E., LaRocque,R., Rosenberg,E. and Sabeti,P.C. |
| EPI_ISL_766061, EPI_ISL_766072, EPI_ISL_766073, EPI_ISL_766078, EPI_ISL_766081, EPI_ISL_766084, EPI_ISL_766097, EPI_ISL_766102, EPI_ISL_766105, EPI_ISL_766110, EPI_ISL_766121, EPI_ISL_766132, EPI_ISL_766135, EPI_ISL_766136, EPI_ISL_766137, EPI_ISL_766138, EPI_ISL_766139, EPI_ISL_766140, EPI_ISL_766141, EPI_ISL_766142, EPI_ISL_766149, EPI_ISL_766150, EPI_ISL_766151, EPI_ISL_766152, EPI_ISL_766153, EPI_ISL_766202, EPI_ISL_766203, EPI_ISL_766204, EPI_ISL_766223, EPI_ISL_766224, EPI_ISL_766227, EPI_ISL_766237, EPI_ISL_766258, EPI_ISL_766301, EPI_ISL_766303, EPI_ISL_766310, EPI_ISL_766314, EPI_ISL_766316, EPI_ISL_766317, EPI_ISL_766397, EPI_ISL_766400, EPI_ISL_766403, EPI_ISL_766407, EPI_ISL_766410, EPI_ISL_766411, EPI_ISL_766412, EPI_ISL_766415, EPI_ISL_766421, EPI_ISL_766427, EPI_ISL_766437, EPI_ISL_766438, EPI_ISL_766439, EPI_ISL_766555, EPI_ISL_766564 | | | |
| see above | Respiratory Virus Unit, National Infection Service, Public Health England | COVID-19 Genomics UK (COG-UK) Consortium | PHE Covid Sequencing Team |
| EPI_ISL_766575 | ULSS 8 Berica | Istituto Zooprofilattico Sperimentale delle Venezie | Adelaide Milani, Alessia Schivo, Annalisa Salviato, Erika Giorgia Quaranta, Ambra Pastori, Bianca Zecchin, Alice Fusaro, Isabella Monne, Calogero Terregino, Antonia Ricci |
| EPI_ISL_766609, EPI_ISL_766610, EPI_ISL_766637, EPI_ISL_766638, EPI_ISL_766639 | Klinisk mikrobiologi | The Public Health Agency of Sweden | Department of Microbiology, The Public Health Agency of Sweden |
| EPI_ISL_766640, EPI_ISL_766641 | Laboratoriemedicin, Klinisk mikrobiologi | The Public Health Agency of Sweden | Department of Microbiology, The Public Health Agency of Sweden |
| EPI_ISL_766644 | Vardcentralen Brinken | The Public Health Agency of Sweden | Department of Microbiology, The Public Health Agency of Sweden |
| EPI_ISL_766707, EPI_ISL_766711, EPI_ISL_766712, EPI_ISL_766725, EPI_ISL_766726, EPI_ISL_766727, EPI_ISL_766728, EPI_ISL_766729 | Klinisk mikrobiologi | The Public Health Agency of Sweden | Department of Microbiology, The Public Health Agency of Sweden |
| EPI_ISL_767109 | Lighthouse Lab in Milton Keynes | Wellcome Sanger Institute for the COVID-19 Genomics UK (COG-UK) Consortium | The Lighthouse Lab in Milton Keynes and Alex Alderton, Roberto Amato, Sonia Goncalves, Ewan Harrison, David K. Jackson, Ian Johnston, Dominic Kwiatkowski, Cordelia Langford, John Sillitoe on behalf of the Wellcome Sanger Institute COVID-19 Surveillance Team |
| EPI_ISL_767120, EPI_ISL_767122, EPI_ISL_767124, EPI_ISL_767126, EPI_ISL_767127, EPI_ISL_767129, EPI_ISL_767131 | Lighthouse Lab in Alderley Park | Wellcome Sanger Institute for the COVID-19 Genomics UK (COG-UK) Consortium | Jacquelyn Wynn, Mairead Hyland, The Lighthouse Lab in Alderley Park and Alex Alderton, Roberto Amato, Sonia Goncalves, Ewan Harrison, David K. Jackson, Ian Johnston, Dominic Kwiatkowski, Cordelia Langford, John Sillitoe on behalf of the Wellcome Sanger Institute COVID-19 Surveillance Team |
| EPI_ISL_767420, EPI_ISL_767450, EPI_ISL_767451, EPI_ISL_767453, EPI_ISL_767454, EPI_ISL_767455, EPI_ISL_767456, EPI_ISL_767457, EPI_ISL_767458, EPI_ISL_767459, EPI_ISL_767460, EPI_ISL_767461, EPI_ISL_767462, EPI_ISL_767463, EPI_ISL_767464, EPI_ISL_767465, EPI_ISL_767466, EPI_ISL_767467, EPI_ISL_767468, EPI_ISL_767469, EPI_ISL_767470, EPI_ISL_767471, EPI_ISL_767472, EPI_ISL_767473, EPI_ISL_767474, EPI_ISL_767475, EPI_ISL_767476, EPI_ISL_767477, EPI_ISL_767478 | | | |
| see above | Wadsworth Center, New York State Department.of Health | Wadsworth Center, New York State Department.of Health | Kirsten St. George, Daryl M. Lamson, Alexis Russel, Matthew Shudt, Melissa A Leisner, Jonathan Plitnick, Navjot Singh, John Kelly, Sara Griesemer, Erasmus Schneider, Erica Lasek-Nesselquist |
| EPI_ISL_767656 | MEMORIAL SLOAN KETTERING CANCER CENTER | Wadsworth Center, New York State Department.of Health | Kirsten St. George, Daryl M. Lamson, Alexis Russel, Matthew Shudt, Melissa A Leisner, Jonathan Plitnick, Navjot Singh, John Kelly, Sara Griesemer, Erasmus Schneider, Erica Lasek-Nesselquist |
| EPI_ISL_767886 | South Eastern Area Laboratory Services (SEALS) | NSW Health Pathology - Institute of Clinical Pathology and Medical Research; Westmead Hospital; University of Sydney | CIDM-PH et al. |
| EPI_ISL_768475, EPI_ISL_768476, EPI_ISL_768477, EPI_ISL_768478, EPI_ISL_768479, EPI_ISL_768480, EPI_ISL_768481, EPI_ISL_768482, EPI_ISL_768483, EPI_ISL_768484, EPI_ISL_768485, EPI_ISL_768486, EPI_ISL_768487, EPI_ISL_768488, EPI_ISL_768489, EPI_ISL_768490, EPI_ISL_768491, EPI_ISL_768492, EPI_ISL_768493, EPI_ISL_768494, EPI_ISL_768495 | | | |
| see above | LSUHS Emerging Viral Threat Laboratory | Microbial Genome Sequencing Center | Jeremy P. Kamil, Jennifer L. Carroll, Camille F. Abshire, Maarten Van Diest, Andrew D. Yurochko, Martin J. Sapp, Rona S. Scott, Christopher G. Kevil, Daniel J. Snyder, Vaughn S. Cooper, John A. Vanchiere |
| EPI_ISL_768498 | LSUHS Emerging Viral Threat Laboratory | Microbial Genome Sequencing Center | Jennifer L. Carroll, Jeremy P. Kamil, Camille F. Abshire, Maarten Van Diest, Andrew D. Yurochko, Martin J. Sapp, Rona S. Scott, Christopher G. Kevil, Daniel J. Snyder, Vaughn S. Cooper, John A. Vanchiere |
| EPI_ISL_768902, EPI_ISL_768903, EPI_ISL_768904, EPI_ISL_768905, EPI_ISL_768906, EPI_ISL_768907, EPI_ISL_768908, EPI_ISL_768909, EPI_ISL_768910, EPI_ISL_768912, EPI_ISL_768913, EPI_ISL_768914, EPI_ISL_768915, EPI_ISL_768917, EPI_ISL_768918, EPI_ISL_768920, EPI_ISL_768921, EPI_ISL_768922, EPI_ISL_768923, EPI_ISL_768924, EPI_ISL_768926, EPI_ISL_768927, EPI_ISL_768929, EPI_ISL_768930, EPI_ISL_768933, EPI_ISL_768934, EPI_ISL_768935, EPI_ISL_768938, EPI_ISL_768940, EPI_ISL_768941, EPI_ISL_768942, EPI_ISL_768943, EPI_ISL_768944, EPI_ISL_768947, EPI_ISL_768950, EPI_ISL_768951, EPI_ISL_768954, EPI_ISL_768955, EPI_ISL_768956, EPI_ISL_768958, EPI_ISL_768960, EPI_ISL_768961, EPI_ISL_768962, EPI_ISL_768964, EPI_ISL_768967, EPI_ISL_768968, EPI_ISL_768969, EPI_ISL_768970, EPI_ISL_768971, EPI_ISL_768973, EPI_ISL_768974, EPI_ISL_768975, EPI_ISL_768978, EPI_ISL_768979, EPI_ISL_768980, EPI_ISL_768981, EPI_ISL_768982, EPI_ISL_768983, EPI_ISL_768984, EPI_ISL_768985, EPI_ISL_768986, EPI_ISL_768987, EPI_ISL_768988, EPI_ISL_768989, EPI_ISL_768990, EPI_ISL_768991, EPI_ISL_768994, EPI_ISL_768995, EPI_ISL_768996, EPI_ISL_768997, EPI_ISL_768998, EPI_ISL_768999, EPI_ISL_769000, EPI_ISL_769001, EPI_ISL_769003, EPI_ISL_769004, EPI_ISL_769006, EPI_ISL_769007, EPI_ISL_769008, EPI_ISL_769009, EPI_ISL_769010, EPI_ISL_769011, EPI_ISL_769012, EPI_ISL_769013, EPI_ISL_769014, EPI_ISL_769015, EPI_ISL_769018, EPI_ISL_769019, EPI_ISL_769020, EPI_ISL_769021, EPI_ISL_769022, EPI_ISL_769023, EPI_ISL_769024, EPI_ISL_769025, EPI_ISL_769026, EPI_ISL_769028, EPI_ISL_769030, EPI_ISL_769031, EPI_ISL_769032, EPI_ISL_769034, EPI_ISL_769035, EPI_ISL_769037, EPI_ISL_769039, EPI_ISL_769040, EPI_ISL_769041, EPI_ISL_769042, EPI_ISL_769044, EPI_ISL_769046, EPI_ISL_769047, EPI_ISL_769048, EPI_ISL_769050, EPI_ISL_769051, EPI_ISL_769052, EPI_ISL_769053, EPI_ISL_769054, EPI_ISL_769055, EPI_ISL_769056, EPI_ISL_769058, EPI_ISL_769059, EPI_ISL_769061, EPI_ISL_769063, EPI_ISL_769064, EPI_ISL_769065, EPI_ISL_769066, EPI_ISL_769067, EPI_ISL_769068, EPI_ISL_769069, EPI_ISL_769070, EPI_ISL_769071, EPI_ISL_769072, EPI_ISL_769073, EPI_ISL_769074, EPI_ISL_769075, EPI_ISL_769077, EPI_ISL_769078, EPI_ISL_769079, EPI_ISL_769081, EPI_ISL_769082, EPI_ISL_769083, EPI_ISL_769085, EPI_ISL_769086, EPI_ISL_769088, EPI_ISL_769090, EPI_ISL_769091, EPI_ISL_769092, EPI_ISL_769095, EPI_ISL_769096, EPI_ISL_769097, EPI_ISL_769099, EPI_ISL_769100, EPI_ISL_769101, EPI_ISL_769102, EPI_ISL_769103, EPI_ISL_769104, EPI_ISL_769106, EPI_ISL_769107, EPI_ISL_769108, EPI_ISL_769111, EPI_ISL_769112, EPI_ISL_769113, EPI_ISL_769114, EPI_ISL_769115, EPI_ISL_769116, EPI_ISL_769118, EPI_ISL_769120, EPI_ISL_769121, EPI_ISL_769123, EPI_ISL_769124, EPI_ISL_769125, EPI_ISL_769126, EPI_ISL_769127, EPI_ISL_769128, EPI_ISL_769129, EPI_ISL_769130, EPI_ISL_769131, EPI_ISL_769132, EPI_ISL_769133, EPI_ISL_769134, EPI_ISL_769135, EPI_ISL_769136, EPI_ISL_769137, EPI_ISL_769139, EPI_ISL_769140, EPI_ISL_769142, EPI_ISL_769143, EPI_ISL_769144, EPI_ISL_769145, EPI_ISL_769146, EPI_ISL_769147, EPI_ISL_769148, EPI_ISL_769149, EPI_ISL_769150, EPI_ISL_769151, EPI_ISL_769153, EPI_ISL_769154, EPI_ISL_769155, EPI_ISL_769156, EPI_ISL_769158, EPI_ISL_769159, EPI_ISL_769160, EPI_ISL_769161, EPI_ISL_769162, EPI_ISL_769163, EPI_ISL_769164, EPI_ISL_769165, EPI_ISL_769166, EPI_ISL_769167, EPI_ISL_769170, EPI_ISL_769172, EPI_ISL_769174, EPI_ISL_769175, EPI_ISL_769177, EPI_ISL_769178, EPI_ISL_769179, EPI_ISL_769180, EPI_ISL_769181, EPI_ISL_769182, EPI_ISL_769183, EPI_ISL_769184, EPI_ISL_769185, EPI_ISL_769186, EPI_ISL_769187, EPI_ISL_769188, EPI_ISL_769189, EPI_ISL_769190, EPI_ISL_769191, EPI_ISL_769192, EPI_ISL_769197, EPI_ISL_769199, EPI_ISL_769200, EPI_ISL_769201, EPI_ISL_769202, EPI_ISL_769203, EPI_ISL_769204, EPI_ISL_769205, EPI_ISL_769208, EPI_ISL_769209, EPI_ISL_769210, EPI_ISL_769212, EPI_ISL_769214, EPI_ISL_769216, EPI_ISL_769217, EPI_ISL_769220, EPI_ISL_769221, EPI_ISL_769222 | | | |
| see above | Lighthouse Lab in Cambridge | Wellcome Sanger Institute for the COVID-19 Genomics UK (COG-UK) Consortium | Rob Howes, The Lighthouse Lab in Cambridge and Alex Alderton, Roberto Amato, Sonia Goncalves, Ewan Harrison, David K. Jackson, Ian Johnston, Dominic Kwiatkowski, Cordelia Langford, John Sillitoe on behalf of the Wellcome Sanger Institute COVID-19 Surveillance Team |
| EPI_ISL_769223, EPI_ISL_769224, EPI_ISL_769225, EPI_ISL_769229, EPI_ISL_769233, EPI_ISL_769236, EPI_ISL_769243, EPI_ISL_769244, EPI_ISL_769245, EPI_ISL_769246, EPI_ISL_769249, EPI_ISL_769250, EPI_ISL_769252, EPI_ISL_769254, EPI_ISL_769255, EPI_ISL_769257, EPI_ISL_769259, EPI_ISL_769260, EPI_ISL_769261, EPI_ISL_769266, EPI_ISL_769267, EPI_ISL_769273, EPI_ISL_769275, EPI_ISL_769276, EPI_ISL_769280, EPI_ISL_769281, EPI_ISL_769282, EPI_ISL_769284, EPI_ISL_769286, EPI_ISL_769288, EPI_ISL_769289, EPI_ISL_769290, EPI_ISL_769291, EPI_ISL_769294, EPI_ISL_769297, EPI_ISL_769306, EPI_ISL_769308, EPI_ISL_769310, EPI_ISL_769312, EPI_ISL_769314, EPI_ISL_769316, EPI_ISL_769319, EPI_ISL_769320, EPI_ISL_769321, EPI_ISL_769322, EPI_ISL_769323, EPI_ISL_769325, EPI_ISL_769327, EPI_ISL_769329, EPI_ISL_769330, EPI_ISL_769334, EPI_ISL_769335, EPI_ISL_769341, EPI_ISL_769343, EPI_ISL_769345, EPI_ISL_769348, EPI_ISL_769349, EPI_ISL_769352, EPI_ISL_769353, EPI_ISL_769355, EPI_ISL_769360, EPI_ISL_769361, EPI_ISL_769365, EPI_ISL_769367, EPI_ISL_769370, EPI_ISL_769374, EPI_ISL_769380, EPI_ISL_769381, EPI_ISL_769383, EPI_ISL_769384, EPI_ISL_769385, EPI_ISL_769387, EPI_ISL_769388, EPI_ISL_769389, EPI_ISL_769390, EPI_ISL_769391, EPI_ISL_769392, EPI_ISL_769394, EPI_ISL_769396, EPI_ISL_769399, EPI_ISL_769401, EPI_ISL_769402, EPI_ISL_769404, EPI_ISL_769405, EPI_ISL_769406, EPI_ISL_769412, EPI_ISL_769413, EPI_ISL_769416, EPI_ISL_769417, EPI_ISL_769418, EPI_ISL_769419, EPI_ISL_769420, EPI_ISL_769421, EPI_ISL_769422, EPI_ISL_769427, EPI_ISL_769428, EPI_ISL_769430, EPI_ISL_769431, EPI_ISL_769434, EPI_ISL_769440, EPI_ISL_769441, EPI_ISL_769442, EPI_ISL_769445, EPI_ISL_769446, EPI_ISL_769449, EPI_ISL_769450, EPI_ISL_769452, EPI_ISL_769454, EPI_ISL_769457, EPI_ISL_769458, EPI_ISL_769459, EPI_ISL_769460, EPI_ISL_769462, EPI_ISL_769466, EPI_ISL_769468, EPI_ISL_769469, EPI_ISL_769470, EPI_ISL_769471, EPI_ISL_769473, EPI_ISL_769474, EPI_ISL_769475, EPI_ISL_769476, EPI_ISL_769481, EPI_ISL_769482, EPI_ISL_769485, EPI_ISL_769489, EPI_ISL_769497, EPI_ISL_769499, EPI_ISL_769500, EPI_ISL_769501, EPI_ISL_769502, EPI_ISL_769503, EPI_ISL_769504, EPI_ISL_769506, EPI_ISL_769511, EPI_ISL_769512, EPI_ISL_769513, EPI_ISL_769517, EPI_ISL_769518, EPI_ISL_769519, EPI_ISL_769522, EPI_ISL_769524, EPI_ISL_769525, EPI_ISL_769528, EPI_ISL_769530, EPI_ISL_769533, EPI_ISL_769535, EPI_ISL_769539, EPI_ISL_769540, EPI_ISL_769542, EPI_ISL_769543, EPI_ISL_769577, EPI_ISL_769594, EPI_ISL_769641, EPI_ISL_769655, EPI_ISL_769661, EPI_ISL_769802, EPI_ISL_769861 | | | |
| see above | Lighthouse Lab in Glasgow | Wellcome Sanger Institute for the COVID-19 Genomics UK (COG-UK) Consortium | Harper VanSteenhouse, Yumi Kasai, David Gray, Carol Clugston, Anna Dominiczak and Alex Alderton, Roberto Amato, Sonia Goncalves, Ewan Harrison, David K. Jackson, Ian Johnston, Dominic Kwiatkowski, Cordelia Langford, John Sillitoe on behalf of the Wellcome Sanger Institute COVID-19 Surveillance Team |
| EPI_ISL_769863 | Laboratoire Biolife | Laboratoire de Biotechnologie | Mouna Ouadghiri, Tarik Aanniz, Mohammed Walid Chemao Elfihri, Mohamed Chenaoui, Hanae Dakka, Afaf Alaoui, Otmane Touzani, Amina Benouda, Bouchra Belfquih, Lahcen belyamani, Saaid Amzazi and Azeddine Ibrahimi |

| | | | |
|---|---|---|---|
| EPI_ISL_769877 | Respiratory Virus Unit, National Infection Service, Public Health England | COVID-19 Genomics UK (COG-UK) Consortium | PHE Covid Sequencing Team |
| EPI_ISL_770799 | Minnesota Department of Health, Public Health Laboratory | Minnesota Department of Health, Public Health Laboratory | Alexandra Lorentz, Jacob Garfin, Matt Plumb, and Xiong Wang |
| EPI_ISL_775218 | Gonoshasthya-RNA Molecular Research Center | Gonoshasthya-RNA Molecular Research Center | Mohd. Raeed Jamiruddin, Nihad Adnan, Md. Ahsanul Haq, Mohib Ullah Khondoker, Nafisa Azmuda, Firoz Ahmed, Shahana Sharmin, Salma Akter, Taslin Jahan Mou, Mahfuza Marzan, Sayeda Moriam Liza, Nowshin Jahan, Tamanna Ali, Shahad Saif Khandker, Maha Jamiruddin, Mousumi Chaity, Mumtarin Jannat Oishee |
| EPI_ISL_775219, EPI_ISL_775220, EPI_ISL_775221, EPI_ISL_775225 | Laboratoire Biolife | Laboratoire de Biotechnologie | Mouna Ouadghiri, Tarik Aanniz, Mohammed Walid Chemao Elfihri, Mohamed Chenaoui, Hanae Dakka, Afaf Alaoui, Otmane Touzani, Amina Benouda, Bouchra Belfquih, Lahcen belyamani, Saaid Amzazi and Azeddine Ibrahimi |
| EPI_ISL_775268 | National Laboratory for Health, Environment and Food | National Laboratory for Health, Environment and Food | Aleksander Mahnic, Maja Rupnik |
| EPI_ISL_775272 | Dept. of Medical Microbiology, Stavanger University Hospital, Helse Stavanger HF | Norwegian Institute of Public Health, Department of Virology | Kathrine Stene-Johansen, Kamilla Heddeland Instefjord, Hilde Elshaug, Atiya R Ali,Marie Paulsen Madsen, Rasmus Riis Kopperud, Hilde Vollan, Karoline Bragstad, Olav Hungnes |
| EPI_ISL_775529 | Akershus University Hospital, Department for Microbiology and Infectious Disease Control | Norwegian Institute of Public Health, Department of Virology | Kathrine Stene-Johansen, Kamilla Heddeland Instefjord, Hilde Elshaug, Atiya R Ali,Marie Paulsen Madsen, Rasmus Riis Kopperud, Hilde Vollan, Karoline Bragstad, Olav Hungnes |
| EPI_ISL_775583 | Synlab Medilab, Mikrobiologi | The Public Health Agency of Sweden | Department of Microbiology, The Public Health Agency of Sweden |
| EPI_ISL_775584 | Klinisk mikrobiologi | The Public Health Agency of Sweden | Department of Microbiology, The Public Health Agency of Sweden |
| EPI_ISL_775587 | Synlab Medilab, Mikrobiologi | The Public Health Agency of Sweden | Department of Microbiology, The Public Health Agency of Sweden |
| EPI_ISL_775588, EPI_ISL_775590 | Klinisk mikrobiologi | The Public Health Agency of Sweden | Department of Microbiology, The Public Health Agency of Sweden |
| EPI_ISL_776662, EPI_ISL_776667, EPI_ISL_776706, EPI_ISL_776707, EPI_ISL_776710, EPI_ISL_776712, EPI_ISL_776714, EPI_ISL_776741, EPI_ISL_776745 | UW Virology Lab | UW Virology Lab | Pavitra Roychoudhury, Hong Xie, Lasata Shrestha, Meei-Li Huang, Keith R Jerome, Alexander Greninger |
| EPI_ISL_777011, EPI_ISL_777012, EPI_ISL_777014, EPI_ISL_777015, EPI_ISL_777017, EPI_ISL_777018, EPI_ISL_777020 | Lighthouse Lab in Alderley Park | Wellcome Sanger Institute for the COVID-19 Genomics UK (COG-UK) Consortium | Jacquelyn Wynn, Mairead Hyland, The Lighthouse Lab in Alderley Park and Alex Alderton, Roberto Amato, Sonia Goncalves, Ewan Harrison, David K. Jackson, Ian Johnston, Dominic Kwiatkowski, Cordelia Langford, John Sillitoe on behalf of the Wellcome Sanger Institute COVID-19 Surveillance Team |
| EPI_ISL_777022, EPI_ISL_777023 | Lighthouse Lab in Glasgow | Wellcome Sanger Institute for the COVID-19 Genomics UK (COG-UK) Consortium | Harper VanSteenhouse, Yumi Kasai, David Gray, Carol Clugston, Anna Dominiczak and Alex Alderton, Roberto Amato, Sonia Goncalves, Ewan Harrison, David K. Jackson, Ian Johnston, Dominic Kwiatkowski, Cordelia Langford, John Sillitoe on behalf of the Wellcome Sanger Institute COVID-19 Surveillance Team |
| EPI_ISL_777024, EPI_ISL_777025, EPI_ISL_777026, EPI_ISL_777027, EPI_ISL_777028, EPI_ISL_777030, EPI_ISL_777032 | Lighthouse Lab in Alderley Park | Wellcome Sanger Institute for the COVID-19 Genomics UK (COG-UK) Consortium | Jacquelyn Wynn, Mairead Hyland, The Lighthouse Lab in Alderley Park and Alex Alderton, Roberto Amato, Sonia Goncalves, Ewan Harrison, David K. Jackson, Ian Johnston, Dominic Kwiatkowski, Cordelia Langford, John Sillitoe on behalf of the Wellcome Sanger Institute COVID-19 Surveillance Team |
| EPI_ISL_777034 | Lighthouse Lab in Glasgow | Wellcome Sanger Institute for the COVID-19 Genomics UK (COG-UK) Consortium | Harper VanSteenhouse, Yumi Kasai, David Gray, Carol Clugston, Anna Dominiczak and Alex Alderton, Roberto Amato, Sonia Goncalves, Ewan Harrison, David K. Jackson, Ian Johnston, Dominic Kwiatkowski, Cordelia Langford, John Sillitoe on behalf of the Wellcome Sanger Institute COVID-19 Surveillance Team |
| EPI_ISL_777035, EPI_ISL_777036, EPI_ISL_777041, EPI_ISL_777043, EPI_ISL_777044, EPI_ISL_777045, EPI_ISL_777046, EPI_ISL_777048, EPI_ISL_777049, EPI_ISL_777050, EPI_ISL_777051, EPI_ISL_777052, EPI_ISL_777054 | | | |
| see above | Lighthouse Lab in Alderley Park | Wellcome Sanger Institute for the COVID-19 Genomics UK (COG-UK) Consortium | Jacquelyn Wynn, Mairead Hyland, The Lighthouse Lab in Alderley Park and Alex Alderton, Roberto Amato, Sonia Goncalves, Ewan Harrison, David K. Jackson, Ian Johnston, Dominic Kwiatkowski, Cordelia Langford, John Sillitoe on behalf of the Wellcome Sanger Institute COVID-19 Surveillance Team |
| EPI_ISL_777055 | Lighthouse Lab in Glasgow | Wellcome Sanger Institute for the COVID-19 Genomics UK (COG-UK) Consortium | Harper VanSteenhouse, Yumi Kasai, David Gray, Carol Clugston, Anna Dominiczak and Alex Alderton, Roberto Amato, Sonia Goncalves, Ewan Harrison, David K. Jackson, Ian Johnston, Dominic Kwiatkowski, Cordelia Langford, John Sillitoe on behalf of the Wellcome Sanger Institute COVID-19 Surveillance Team |
| EPI_ISL_777057 | Lighthouse Lab in Alderley Park | Wellcome Sanger Institute for the COVID-19 Genomics UK (COG-UK) Consortium | Jacquelyn Wynn, Mairead Hyland, The Lighthouse Lab in Alderley Park and Alex Alderton, Roberto Amato, Sonia Goncalves, Ewan Harrison, David K. Jackson, Ian Johnston, Dominic Kwiatkowski, Cordelia Langford, John Sillitoe on behalf of the Wellcome Sanger Institute COVID-19 Surveillance Team |
| EPI_ISL_777058 | Lighthouse Lab in Glasgow | Wellcome Sanger Institute for the COVID-19 Genomics UK (COG-UK) Consortium | Harper VanSteenhouse, Yumi Kasai, David Gray, Carol Clugston, Anna Dominiczak and Alex Alderton, Roberto Amato, Sonia Goncalves, Ewan Harrison, David K. Jackson, Ian Johnston, Dominic Kwiatkowski, Cordelia Langford, John Sillitoe on behalf of the Wellcome Sanger Institute COVID-19 Surveillance Team |
| EPI_ISL_777060 | Lighthouse Lab in Alderley Park | Wellcome Sanger Institute for the COVID-19 Genomics UK (COG-UK) Consortium | Jacquelyn Wynn, Mairead Hyland, The Lighthouse Lab in Alderley Park and Alex Alderton, Roberto Amato, Sonia Goncalves, Ewan Harrison, David K. Jackson, Ian Johnston, Dominic Kwiatkowski, Cordelia Langford, John Sillitoe on behalf of the Wellcome Sanger Institute COVID-19 Surveillance Team |
| EPI_ISL_777065 | Lighthouse Lab in Glasgow | Wellcome Sanger Institute for the COVID-19 Genomics UK (COG-UK) Consortium | Harper VanSteenhouse, Yumi Kasai, David Gray, Carol Clugston, Anna Dominiczak and Alex Alderton, Roberto Amato, Sonia Goncalves, Ewan Harrison, David K. Jackson, Ian Johnston, Dominic Kwiatkowski, Cordelia Langford, John Sillitoe on behalf of the Wellcome Sanger Institute COVID-19 Surveillance Team |
| EPI_ISL_777069, EPI_ISL_777070, EPI_ISL_777071, EPI_ISL_777072, EPI_ISL_777074, EPI_ISL_777075, EPI_ISL_777076, EPI_ISL_777077, EPI_ISL_777078, EPI_ISL_777079, EPI_ISL_777086, EPI_ISL_777087, EPI_ISL_777088, EPI_ISL_777090, EPI_ISL_777091, EPI_ISL_777092, EPI_ISL_777093, EPI_ISL_777094 | | | |
| see above | Lighthouse Lab in Alderley Park | Wellcome Sanger Institute for the COVID-19 Genomics UK (COG-UK) Consortium | Jacquelyn Wynn, Mairead Hyland, The Lighthouse Lab in Alderley Park and Alex Alderton, Roberto Amato, Sonia Goncalves, Ewan Harrison, David K. Jackson, Ian Johnston, Dominic Kwiatkowski, Cordelia Langford, John Sillitoe on behalf of the Wellcome Sanger Institute COVID-19 Surveillance Team |
| EPI_ISL_777095 | Lighthouse Lab in Glasgow | Wellcome Sanger Institute for the COVID-19 Genomics UK (COG-UK) Consortium | Harper VanSteenhouse, Yumi Kasai, David Gray, Carol Clugston, Anna Dominiczak and Alex Alderton, Roberto Amato, Sonia Goncalves, Ewan Harrison, David K. Jackson, Ian Johnston, Dominic Kwiatkowski, Cordelia Langford, John Sillitoe on behalf of the Wellcome Sanger Institute COVID-19 Surveillance Team |
| EPI_ISL_777096, EPI_ISL_777097 | Lighthouse Lab in Alderley Park | Wellcome Sanger Institute for the COVID-19 Genomics UK (COG-UK) Consortium | Jacquelyn Wynn, Mairead Hyland, The Lighthouse Lab in Alderley Park and Alex Alderton, Roberto Amato, Sonia Goncalves, Ewan Harrison, David K. Jackson, Ian Johnston, Dominic Kwiatkowski, Cordelia Langford, John Sillitoe on behalf of the Wellcome Sanger Institute COVID-19 Surveillance Team |
| EPI_ISL_777098 | Lighthouse Lab in Glasgow | Wellcome Sanger Institute for the COVID-19 Genomics UK (COG-UK) Consortium | Harper VanSteenhouse, Yumi Kasai, David Gray, Carol Clugston, Anna Dominiczak and Alex Alderton, Roberto Amato, Sonia Goncalves, Ewan Harrison, David K. Jackson, Ian Johnston, Dominic Kwiatkowski, Cordelia Langford, John Sillitoe on behalf of the Wellcome Sanger Institute COVID-19 Surveillance Team |
| EPI_ISL_777103, EPI_ISL_777106, EPI_ISL_777107, EPI_ISL_777109, EPI_ISL_777110, EPI_ISL_777111, EPI_ISL_777113, EPI_ISL_777114 | Lighthouse Lab in Alderley Park | Wellcome Sanger Institute for the COVID-19 Genomics UK (COG-UK) Consortium | Jacquelyn Wynn, Mairead Hyland, The Lighthouse Lab in Alderley Park and Alex Alderton, Roberto Amato, Sonia Goncalves, Ewan Harrison, David K. Jackson, Ian Johnston, Dominic Kwiatkowski, Cordelia Langford, John Sillitoe on behalf of the Wellcome Sanger Institute COVID-19 Surveillance Team |
| EPI_ISL_777115 | Lighthouse Lab in Glasgow | Wellcome Sanger Institute for the COVID-19 Genomics UK (COG-UK) Consortium | Harper VanSteenhouse, Yumi Kasai, David Gray, Carol Clugston, Anna Dominiczak and Alex Alderton, Roberto Amato, Sonia Goncalves, Ewan Harrison, David K. Jackson, Ian Johnston, Dominic Kwiatkowski, Cordelia Langford, John Sillitoe on behalf of the Wellcome Sanger Institute COVID-19 Surveillance Team |
| EPI_ISL_777116, EPI_ISL_777119, EPI_ISL_777123, EPI_ISL_777125, EPI_ISL_777126, EPI_ISL_777129, EPI_ISL_777132, EPI_ISL_777134, EPI_ISL_777135, EPI_ISL_777136, EPI_ISL_777137, EPI_ISL_777138, EPI_ISL_777139, EPI_ISL_777143, EPI_ISL_777144, EPI_ISL_777147 | | | |
| see above | Lighthouse Lab in Alderley Park | Wellcome Sanger Institute for the COVID-19 Genomics UK | Jacquelyn Wynn, Mairead Hyland, The Lighthouse Lab in Alderley Park and Alex Alderton, Roberto Amato, Sonia Goncalves, Ewan Harrison, David K. |

| | | | |
|---|---|---|---|
| | | (COG-UK) Consortium | Jackson, Ian Johnston, Dominic Kwiatkowski, Cordelia Langford, John Sillitoe on behalf of the Wellcome Sanger Institute COVID-19 Surveillance Team |
| EPI_ISL_777148, EPI_ISL_777149 | Lighthouse Lab in Glasgow | Wellcome Sanger Institute for the COVID-19 Genomics UK (COG-UK) Consortium | Harper VanSteenhouse, Yumi Kasai, David Gray, Carol Clugston, Anna Dominiczak and Alex Alderton, Roberto Amato, Sonia Goncalves, Ewan Harrison, David K. Jackson, Ian Johnston, Dominic Kwiatkowski, Cordelia Langford, John Sillitoe on behalf of the Wellcome Sanger Institute COVID-19 Surveillance Team |
| EPI_ISL_777150, EPI_ISL_777156, EPI_ISL_777157, EPI_ISL_777159, EPI_ISL_777163, EPI_ISL_777165, EPI_ISL_777166, EPI_ISL_777167, EPI_ISL_777168, EPI_ISL_777170, EPI_ISL_777171, EPI_ISL_777172, EPI_ISL_777174, EPI_ISL_777175, EPI_ISL_777177, EPI_ISL_777178, EPI_ISL_777179, EPI_ISL_777182, EPI_ISL_777185, EPI_ISL_777186 | | | |
| see above | Lighthouse Lab in Alderley Park | Wellcome Sanger Institute for the COVID-19 Genomics UK (COG-UK) Consortium | Jacquelyn Wynn, Mairead Hyland, The Lighthouse Lab in Alderley Park and Alex Alderton, Roberto Amato, Sonia Goncalves, Ewan Harrison, David K. Jackson, Ian Johnston, Dominic Kwiatkowski, Cordelia Langford, John Sillitoe on behalf of the Wellcome Sanger Institute COVID-19 Surveillance Team |
| EPI_ISL_777187 | Lighthouse Lab in Milton Keynes | Wellcome Sanger Institute for the COVID-19 Genomics UK (COG-UK) Consortium | The Lighthouse Lab in Milton Keynes and Alex Alderton, Roberto Amato, Sonia Goncalves, Ewan Harrison, David K. Jackson, Ian Johnston, Dominic Kwiatkowski, Cordelia Langford, John Sillitoe on behalf of the Wellcome Sanger Institute COVID-19 Surveillance Team |
| EPI_ISL_777188, EPI_ISL_777190, EPI_ISL_777193, EPI_ISL_777194, EPI_ISL_777195, EPI_ISL_777197, EPI_ISL_777198, EPI_ISL_777199, EPI_ISL_777202, EPI_ISL_777203, EPI_ISL_777204, EPI_ISL_777206, EPI_ISL_777210, EPI_ISL_777212, EPI_ISL_777213, EPI_ISL_777214, EPI_ISL_777216, EPI_ISL_777217, EPI_ISL_777220, EPI_ISL_777221, EPI_ISL_777226, EPI_ISL_777228, EPI_ISL_777231, EPI_ISL_777234, EPI_ISL_777237, EPI_ISL_777239, EPI_ISL_777240, EPI_ISL_777243, EPI_ISL_777247, EPI_ISL_777248, EPI_ISL_777249 | | | |
| see above | Lighthouse Lab in Alderley Park | Wellcome Sanger Institute for the COVID-19 Genomics UK (COG-UK) Consortium | Jacquelyn Wynn, Mairead Hyland, The Lighthouse Lab in Alderley Park and Alex Alderton, Roberto Amato, Sonia Goncalves, Ewan Harrison, David K. Jackson, Ian Johnston, Dominic Kwiatkowski, Cordelia Langford, John Sillitoe on behalf of the Wellcome Sanger Institute COVID-19 Surveillance Team |
| EPI_ISL_777250 | Lighthouse Lab in Glasgow | Wellcome Sanger Institute for the COVID-19 Genomics UK (COG-UK) Consortium | Harper VanSteenhouse, Yumi Kasai, David Gray, Carol Clugston, Anna Dominiczak and Alex Alderton, Roberto Amato, Sonia Goncalves, Ewan Harrison, David K. Jackson, Ian Johnston, Dominic Kwiatkowski, Cordelia Langford, John Sillitoe on behalf of the Wellcome Sanger Institute COVID-19 Surveillance Team |
| EPI_ISL_777251, EPI_ISL_777254 | Lighthouse Lab in Alderley Park | Wellcome Sanger Institute for the COVID-19 Genomics UK (COG-UK) Consortium | Jacquelyn Wynn, Mairead Hyland, The Lighthouse Lab in Alderley Park and Alex Alderton, Roberto Amato, Sonia Goncalves, Ewan Harrison, David K. Jackson, Ian Johnston, Dominic Kwiatkowski, Cordelia Langford, John Sillitoe on behalf of the Wellcome Sanger Institute COVID-19 Surveillance Team |
| EPI_ISL_777255 | Lighthouse Lab in Glasgow | Wellcome Sanger Institute for the COVID-19 Genomics UK (COG-UK) Consortium | Harper VanSteenhouse, Yumi Kasai, David Gray, Carol Clugston, Anna Dominiczak and Alex Alderton, Roberto Amato, Sonia Goncalves, Ewan Harrison, David K. Jackson, Ian Johnston, Dominic Kwiatkowski, Cordelia Langford, John Sillitoe on behalf of the Wellcome Sanger Institute COVID-19 Surveillance Team |
| EPI_ISL_777256, EPI_ISL_777257, EPI_ISL_777258, EPI_ISL_777259, EPI_ISL_777260 | Lighthouse Lab in Alderley Park | Wellcome Sanger Institute for the COVID-19 Genomics UK (COG-UK) Consortium | Jacquelyn Wynn, Mairead Hyland, The Lighthouse Lab in Alderley Park and Alex Alderton, Roberto Amato, Sonia Goncalves, Ewan Harrison, David K. Jackson, Ian Johnston, Dominic Kwiatkowski, Cordelia Langford, John Sillitoe on behalf of the Wellcome Sanger Institute COVID-19 Surveillance Team |
| EPI_ISL_777261 | Lighthouse Lab in Glasgow | Wellcome Sanger Institute for the COVID-19 Genomics UK (COG-UK) Consortium | Harper VanSteenhouse, Yumi Kasai, David Gray, Carol Clugston, Anna Dominiczak and Alex Alderton, Roberto Amato, Sonia Goncalves, Ewan Harrison, David K. Jackson, Ian Johnston, Dominic Kwiatkowski, Cordelia Langford, John Sillitoe on behalf of the Wellcome Sanger Institute COVID-19 Surveillance Team |
| EPI_ISL_777262, EPI_ISL_777263, EPI_ISL_777264, EPI_ISL_777265, EPI_ISL_777266, EPI_ISL_777267, EPI_ISL_777271, EPI_ISL_777272, EPI_ISL_777273, EPI_ISL_777274, EPI_ISL_777276, EPI_ISL_777277, EPI_ISL_777278, EPI_ISL_777279, EPI_ISL_777280, EPI_ISL_777281, EPI_ISL_777282 | | | |
| see above | Lighthouse Lab in Alderley Park | Wellcome Sanger Institute for the COVID-19 Genomics UK (COG-UK) Consortium | Jacquelyn Wynn, Mairead Hyland, The Lighthouse Lab in Alderley Park and Alex Alderton, Roberto Amato, Sonia Goncalves, Ewan Harrison, David K. Jackson, Ian Johnston, Dominic Kwiatkowski, Cordelia Langford, John Sillitoe on behalf of the Wellcome Sanger Institute COVID-19 Surveillance Team |
| EPI_ISL_777284 | Lighthouse Lab in Glasgow | Wellcome Sanger Institute for the COVID-19 Genomics UK (COG-UK) Consortium | Harper VanSteenhouse, Yumi Kasai, David Gray, Carol Clugston, Anna Dominiczak and Alex Alderton, Roberto Amato, Sonia Goncalves, Ewan Harrison, David K. Jackson, Ian Johnston, Dominic Kwiatkowski, Cordelia Langford, John Sillitoe on behalf of the Wellcome Sanger Institute COVID-19 Surveillance Team |
| EPI_ISL_777285, EPI_ISL_777287, EPI_ISL_777288, EPI_ISL_777291, EPI_ISL_777294, EPI_ISL_777296, EPI_ISL_777297, EPI_ISL_777299, EPI_ISL_777307, EPI_ISL_777308, EPI_ISL_777309, EPI_ISL_777311, EPI_ISL_777312, EPI_ISL_777315, EPI_ISL_777317, EPI_ISL_777318, EPI_ISL_777319, EPI_ISL_777326, EPI_ISL_777327, EPI_ISL_777333, EPI_ISL_777336, EPI_ISL_777338, EPI_ISL_777349, EPI_ISL_777375, EPI_ISL_777380, EPI_ISL_777386, EPI_ISL_777389, EPI_ISL_777394, EPI_ISL_777411, EPI_ISL_777412, EPI_ISL_777416, EPI_ISL_777421, EPI_ISL_777424, EPI_ISL_777425, EPI_ISL_777427, EPI_ISL_777430, EPI_ISL_777435, EPI_ISL_777437, EPI_ISL_777440, EPI_ISL_777448, EPI_ISL_777451, EPI_ISL_777455, EPI_ISL_777458, EPI_ISL_777465, EPI_ISL_777467, EPI_ISL_777470, EPI_ISL_777477, EPI_ISL_777479, EPI_ISL_777481, EPI_ISL_777489, EPI_ISL_777491, EPI_ISL_777495, EPI_ISL_777502, EPI_ISL_777509, EPI_ISL_777515, EPI_ISL_777520, EPI_ISL_777522, EPI_ISL_777524, EPI_ISL_777525, EPI_ISL_777529, EPI_ISL_777530, EPI_ISL_777532, EPI_ISL_777534, EPI_ISL_777535, EPI_ISL_777537, EPI_ISL_777538, EPI_ISL_777539, EPI_ISL_777543, EPI_ISL_777548, EPI_ISL_777550, EPI_ISL_777552, EPI_ISL_777558, EPI_ISL_777561, EPI_ISL_777563, EPI_ISL_777567, EPI_ISL_777568, EPI_ISL_777572, EPI_ISL_777573, EPI_ISL_777576, EPI_ISL_777580, EPI_ISL_777582, EPI_ISL_777584, EPI_ISL_777587, EPI_ISL_777589, EPI_ISL_777591, EPI_ISL_777592, EPI_ISL_777593, EPI_ISL_777595, EPI_ISL_777597, EPI_ISL_777605, EPI_ISL_777621, EPI_ISL_777627, EPI_ISL_777629, EPI_ISL_777633, EPI_ISL_777634, EPI_ISL_777638, EPI_ISL_777639, EPI_ISL_777644, EPI_ISL_777649, EPI_ISL_777650, EPI_ISL_777651, EPI_ISL_777655, EPI_ISL_777659, EPI_ISL_777669, EPI_ISL_777672 | | | |
| see above | Lighthouse Lab in Alderley Park | Wellcome Sanger Institute for the COVID-19 Genomics UK (COG-UK) Consortium | Jacquelyn Wynn, Mairead Hyland, The Lighthouse Lab in Alderley Park and Alex Alderton, Roberto Amato, Sonia Goncalves, Ewan Harrison, David K. Jackson, Ian Johnston, Dominic Kwiatkowski, Cordelia Langford, John Sillitoe on behalf of the Wellcome Sanger Institute COVID-19 Surveillance Team |
| EPI_ISL_778297, EPI_ISL_778299, EPI_ISL_778301, EPI_ISL_778302, EPI_ISL_778303 | Lighthouse Lab in Glasgow | Wellcome Sanger Institute for the COVID-19 Genomics UK (COG-UK) Consortium | Harper VanSteenhouse, Yumi Kasai, David Gray, Carol Clugston, Anna Dominiczak and Alex Alderton, Roberto Amato, Sonia Goncalves, Ewan Harrison, David K. Jackson, Ian Johnston, Dominic Kwiatkowski, Cordelia Langford, John Sillitoe on behalf of the Wellcome Sanger Institute COVID-19 Surveillance Team |
| EPI_ISL_778305, EPI_ISL_778309, EPI_ISL_778311, EPI_ISL_778315 | Lighthouse Lab in Alderley Park | Wellcome Sanger Institute for the COVID-19 Genomics UK (COG-UK) Consortium | Jacquelyn Wynn, Mairead Hyland, The Lighthouse Lab in Alderley Park and Alex Alderton, Roberto Amato, Sonia Goncalves, Ewan Harrison, David K. Jackson, Ian Johnston, Dominic Kwiatkowski, Cordelia Langford, John Sillitoe on behalf of the Wellcome Sanger Institute COVID-19 Surveillance Team |
| EPI_ISL_778322, EPI_ISL_778324, EPI_ISL_778325 | Lighthouse Lab in Glasgow | Wellcome Sanger Institute for the COVID-19 Genomics UK (COG-UK) Consortium | Harper VanSteenhouse, Yumi Kasai, David Gray, Carol Clugston, Anna Dominiczak and Alex Alderton, Roberto Amato, Sonia Goncalves, Ewan Harrison, David K. Jackson, Ian Johnston, Dominic Kwiatkowski, Cordelia Langford, John Sillitoe on behalf of the Wellcome Sanger Institute COVID-19 Surveillance Team |
| EPI_ISL_778327, EPI_ISL_778328 | Lighthouse Lab in Alderley Park | Wellcome Sanger Institute for the COVID-19 Genomics UK (COG-UK) Consortium | Jacquelyn Wynn, Mairead Hyland, The Lighthouse Lab in Alderley Park and Alex Alderton, Roberto Amato, Sonia Goncalves, Ewan Harrison, David K. Jackson, Ian Johnston, Dominic Kwiatkowski, Cordelia Langford, John Sillitoe on behalf of the Wellcome Sanger Institute COVID-19 Surveillance Team |
| EPI_ISL_778330 | Lighthouse Lab in Glasgow | Wellcome Sanger Institute for the COVID-19 Genomics UK (COG-UK) Consortium | Harper VanSteenhouse, Yumi Kasai, David Gray, Carol Clugston, Anna Dominiczak and Alex Alderton, Roberto Amato, Sonia Goncalves, Ewan Harrison, David K. Jackson, Ian Johnston, Dominic Kwiatkowski, Cordelia Langford, John Sillitoe on behalf of the Wellcome Sanger Institute COVID-19 Surveillance Team |
| EPI_ISL_778331 | Lighthouse Lab in Alderley Park | Wellcome Sanger Institute for the COVID-19 Genomics UK (COG-UK) Consortium | Jacquelyn Wynn, Mairead Hyland, The Lighthouse Lab in Alderley Park and Alex Alderton, Roberto Amato, Sonia Goncalves, Ewan Harrison, David K. Jackson, Ian Johnston, Dominic Kwiatkowski, Cordelia Langford, John Sillitoe on behalf of the Wellcome Sanger Institute COVID-19 Surveillance Team |
| EPI_ISL_778332 | Lighthouse Lab in Glasgow | Wellcome Sanger Institute for the COVID-19 Genomics UK (COG-UK) Consortium | Harper VanSteenhouse, Yumi Kasai, David Gray, Carol Clugston, Anna Dominiczak and Alex Alderton, Roberto Amato, Sonia Goncalves, Ewan Harrison, David K. Jackson, Ian Johnston, Dominic Kwiatkowski, Cordelia Langford, John Sillitoe on behalf of the Wellcome Sanger Institute COVID-19 Surveillance Team |
| EPI_ISL_778333 | Lighthouse Lab in Alderley Park | Wellcome Sanger Institute for the COVID-19 Genomics UK (COG-UK) Consortium | Jacquelyn Wynn, Mairead Hyland, The Lighthouse Lab in Alderley Park and Alex Alderton, Roberto Amato, Sonia Goncalves, Ewan Harrison, David K. Jackson, Ian Johnston, Dominic Kwiatkowski, Cordelia Langford, John Sillitoe on behalf of the Wellcome Sanger Institute COVID-19 Surveillance Team |
| EPI_ISL_778334 | Lighthouse Lab in Glasgow | Wellcome Sanger Institute for the COVID-19 Genomics UK (COG-UK) Consortium | Harper VanSteenhouse, Yumi Kasai, David Gray, Carol Clugston, Anna Dominiczak and Alex Alderton, Roberto Amato, Sonia Goncalves, Ewan Harrison, David K. Jackson, Ian Johnston, Dominic Kwiatkowski, Cordelia Langford, John Sillitoe on behalf of the Wellcome Sanger Institute COVID-19 Surveillance Team |
| EPI_ISL_778335 | Lighthouse Lab in Alderley Park | Wellcome Sanger Institute for the COVID-19 Genomics UK (COG-UK) Consortium | Jacquelyn Wynn, Mairead Hyland, The Lighthouse Lab in Alderley Park and Alex Alderton, Roberto Amato, Sonia Goncalves, Ewan Harrison, David K. Jackson, Ian Johnston, Dominic Kwiatkowski, Cordelia Langford, John Sillitoe on behalf of the Wellcome Sanger Institute COVID-19 Surveillance Team |
| EPI_ISL_778336, EPI_ISL_778337 | Lighthouse Lab in Glasgow | Wellcome Sanger Institute for the COVID-19 Genomics UK (COG-UK) Consortium | Harper VanSteenhouse, Yumi Kasai, David Gray, Carol Clugston, Anna Dominiczak and Alex Alderton, Roberto Amato, Sonia Goncalves, Ewan Harrison, David K. Jackson, Ian Johnston, Dominic Kwiatkowski, Cordelia Langford, John Sillitoe on behalf of the Wellcome Sanger Institute COVID-19 Surveillance Team |
| EPI_ISL_778342 | Lighthouse Lab in Alderley Park | Wellcome Sanger Institute for the COVID-19 Genomics UK (COG-UK) Consortium | Jacquelyn Wynn, Mairead Hyland, The Lighthouse Lab in Alderley Park and Alex Alderton, Roberto Amato, Sonia Goncalves, Ewan Harrison, David K. Jackson, Ian Johnston, Dominic Kwiatkowski, Cordelia Langford, John Sillitoe on behalf of the Wellcome Sanger Institute COVID-19 Surveillance Team |
| EPI_ISL_778345 | Lighthouse Lab in Glasgow | Wellcome Sanger Institute for the COVID-19 Genomics UK | Harper VanSteenhouse, Yumi Kasai, David Gray, Carol Clugston, Anna Dominiczak and Alex Alderton, Roberto Amato, Sonia Goncalves, Ewan Harrison, |

| | | | |
|---|---|---|---|
| | | (COG-UK) Consortium | David K. Jackson, Ian Johnston, Dominic Kwiatkowski, Cordelia Langford, John Sillitoe on behalf of the Wellcome Sanger Institute COVID-19 Surveillance Team |
| EPI_ISL_778346 | Lighthouse Lab in Alderley Park | Wellcome Sanger Institute for the COVID-19 Genomics UK (COG-UK) Consortium | Jacquelyn Wynn, Mairead Hyland, The Lighthouse Lab in Alderley Park and Alex Alderton, Roberto Amato, Sonia Goncalves, Ewan Harrison, David K. Jackson, Ian Johnston, Dominic Kwiatkowski, Cordelia Langford, John Sillitoe on behalf of the Wellcome Sanger Institute COVID-19 Surveillance Team |
| EPI_ISL_778352 | Lighthouse Lab in Glasgow | Wellcome Sanger Institute for the COVID-19 Genomics UK (COG-UK) Consortium | Harper VanSteenhouse, Yumi Kasai, David Gray, Carol Clugston, Anna Dominiczak and Alex Alderton, Roberto Amato, Sonia Goncalves, Ewan Harrison, David K. Jackson, Ian Johnston, Dominic Kwiatkowski, Cordelia Langford, John Sillitoe on behalf of the Wellcome Sanger Institute COVID-19 Surveillance Team |
| EPI_ISL_778353, EPI_ISL_778355 | Lighthouse Lab in Alderley Park | Wellcome Sanger Institute for the COVID-19 Genomics UK (COG-UK) Consortium | Jacquelyn Wynn, Mairead Hyland, The Lighthouse Lab in Alderley Park and Alex Alderton, Roberto Amato, Sonia Goncalves, Ewan Harrison, David K. Jackson, Ian Johnston, Dominic Kwiatkowski, Cordelia Langford, John Sillitoe on behalf of the Wellcome Sanger Institute COVID-19 Surveillance Team |
| EPI_ISL_778356 | Lighthouse Lab in Glasgow | Wellcome Sanger Institute for the COVID-19 Genomics UK (COG-UK) Consortium | Harper VanSteenhouse, Yumi Kasai, David Gray, Carol Clugston, Anna Dominiczak and Alex Alderton, Roberto Amato, Sonia Goncalves, Ewan Harrison, David K. Jackson, Ian Johnston, Dominic Kwiatkowski, Cordelia Langford, John Sillitoe on behalf of the Wellcome Sanger Institute COVID-19 Surveillance Team |
| EPI_ISL_778357, EPI_ISL_778358 | Lighthouse Lab in Alderley Park | Wellcome Sanger Institute for the COVID-19 Genomics UK (COG-UK) Consortium | Jacquelyn Wynn, Mairead Hyland, The Lighthouse Lab in Alderley Park and Alex Alderton, Roberto Amato, Sonia Goncalves, Ewan Harrison, David K. Jackson, Ian Johnston, Dominic Kwiatkowski, Cordelia Langford, John Sillitoe on behalf of the Wellcome Sanger Institute COVID-19 Surveillance Team |
| EPI_ISL_778359 | Lighthouse Lab in Glasgow | Wellcome Sanger Institute for the COVID-19 Genomics UK (COG-UK) Consortium | Harper VanSteenhouse, Yumi Kasai, David Gray, Carol Clugston, Anna Dominiczak and Alex Alderton, Roberto Amato, Sonia Goncalves, Ewan Harrison, David K. Jackson, Ian Johnston, Dominic Kwiatkowski, Cordelia Langford, John Sillitoe on behalf of the Wellcome Sanger Institute COVID-19 Surveillance Team |
| EPI_ISL_778360 | Lighthouse Lab in Alderley Park | Wellcome Sanger Institute for the COVID-19 Genomics UK (COG-UK) Consortium | Jacquelyn Wynn, Mairead Hyland, The Lighthouse Lab in Alderley Park and Alex Alderton, Roberto Amato, Sonia Goncalves, Ewan Harrison, David K. Jackson, Ian Johnston, Dominic Kwiatkowski, Cordelia Langford, John Sillitoe on behalf of the Wellcome Sanger Institute COVID-19 Surveillance Team |
| EPI_ISL_778361, EPI_ISL_778362, EPI_ISL_778364 | Lighthouse Lab in Glasgow | Wellcome Sanger Institute for the COVID-19 Genomics UK (COG-UK) Consortium | Harper VanSteenhouse, Yumi Kasai, David Gray, Carol Clugston, Anna Dominiczak and Alex Alderton, Roberto Amato, Sonia Goncalves, Ewan Harrison, David K. Jackson, Ian Johnston, Dominic Kwiatkowski, Cordelia Langford, John Sillitoe on behalf of the Wellcome Sanger Institute COVID-19 Surveillance Team |
| EPI_ISL_778365, EPI_ISL_778368 | Lighthouse Lab in Alderley Park | Wellcome Sanger Institute for the COVID-19 Genomics UK (COG-UK) Consortium | Jacquelyn Wynn, Mairead Hyland, The Lighthouse Lab in Alderley Park and Alex Alderton, Roberto Amato, Sonia Goncalves, Ewan Harrison, David K. Jackson, Ian Johnston, Dominic Kwiatkowski, Cordelia Langford, John Sillitoe on behalf of the Wellcome Sanger Institute COVID-19 Surveillance Team |
| EPI_ISL_778372, EPI_ISL_778373 | Lighthouse Lab in Glasgow | Wellcome Sanger Institute for the COVID-19 Genomics UK (COG-UK) Consortium | Harper VanSteenhouse, Yumi Kasai, David Gray, Carol Clugston, Anna Dominiczak and Alex Alderton, Roberto Amato, Sonia Goncalves, Ewan Harrison, David K. Jackson, Ian Johnston, Dominic Kwiatkowski, Cordelia Langford, John Sillitoe on behalf of the Wellcome Sanger Institute COVID-19 Surveillance Team |
| EPI_ISL_778374 | Lighthouse Lab in Alderley Park | Wellcome Sanger Institute for the COVID-19 Genomics UK (COG-UK) Consortium | Jacquelyn Wynn, Mairead Hyland, The Lighthouse Lab in Alderley Park and Alex Alderton, Roberto Amato, Sonia Goncalves, Ewan Harrison, David K. Jackson, Ian Johnston, Dominic Kwiatkowski, Cordelia Langford, John Sillitoe on behalf of the Wellcome Sanger Institute COVID-19 Surveillance Team |
| EPI_ISL_778375, EPI_ISL_778376, EPI_ISL_778377, EPI_ISL_778378 | Lighthouse Lab in Glasgow | Wellcome Sanger Institute for the COVID-19 Genomics UK (COG-UK) Consortium | Harper VanSteenhouse, Yumi Kasai, David Gray, Carol Clugston, Anna Dominiczak and Alex Alderton, Roberto Amato, Sonia Goncalves, Ewan Harrison, David K. Jackson, Ian Johnston, Dominic Kwiatkowski, Cordelia Langford, John Sillitoe on behalf of the Wellcome Sanger Institute COVID-19 Surveillance Team |
| EPI_ISL_778380, EPI_ISL_778382, EPI_ISL_778385, EPI_ISL_778390, EPI_ISL_778392, EPI_ISL_778396 | Lighthouse Lab in Alderley Park | Wellcome Sanger Institute for the COVID-19 Genomics UK (COG-UK) Consortium | Jacquelyn Wynn, Mairead Hyland, The Lighthouse Lab in Alderley Park and Alex Alderton, Roberto Amato, Sonia Goncalves, Ewan Harrison, David K. Jackson, Ian Johnston, Dominic Kwiatkowski, Cordelia Langford, John Sillitoe on behalf of the Wellcome Sanger Institute COVID-19 Surveillance Team |
| EPI_ISL_778397 | Lighthouse Lab in Glasgow | Wellcome Sanger Institute for the COVID-19 Genomics UK (COG-UK) Consortium | Harper VanSteenhouse, Yumi Kasai, David Gray, Carol Clugston, Anna Dominiczak and Alex Alderton, Roberto Amato, Sonia Goncalves, Ewan Harrison, David K. Jackson, Ian Johnston, Dominic Kwiatkowski, Cordelia Langford, John Sillitoe on behalf of the Wellcome Sanger Institute COVID-19 Surveillance Team |
| EPI_ISL_778399 | Lighthouse Lab in Alderley Park | Wellcome Sanger Institute for the COVID-19 Genomics UK (COG-UK) Consortium | Jacquelyn Wynn, Mairead Hyland, The Lighthouse Lab in Alderley Park and Alex Alderton, Roberto Amato, Sonia Goncalves, Ewan Harrison, David K. Jackson, Ian Johnston, Dominic Kwiatkowski, Cordelia Langford, John Sillitoe on behalf of the Wellcome Sanger Institute COVID-19 Surveillance Team |
| EPI_ISL_778400 | Lighthouse Lab in Glasgow | Wellcome Sanger Institute for the COVID-19 Genomics UK (COG-UK) Consortium | Harper VanSteenhouse, Yumi Kasai, David Gray, Carol Clugston, Anna Dominiczak and Alex Alderton, Roberto Amato, Sonia Goncalves, Ewan Harrison, David K. Jackson, Ian Johnston, Dominic Kwiatkowski, Cordelia Langford, John Sillitoe on behalf of the Wellcome Sanger Institute COVID-19 Surveillance Team |
| EPI_ISL_778401 | Lighthouse Lab in Alderley Park | Wellcome Sanger Institute for the COVID-19 Genomics UK (COG-UK) Consortium | Jacquelyn Wynn, Mairead Hyland, The Lighthouse Lab in Alderley Park and Alex Alderton, Roberto Amato, Sonia Goncalves, Ewan Harrison, David K. Jackson, Ian Johnston, Dominic Kwiatkowski, Cordelia Langford, John Sillitoe on behalf of the Wellcome Sanger Institute COVID-19 Surveillance Team |
| EPI_ISL_778403 | Lighthouse Lab in Glasgow | Wellcome Sanger Institute for the COVID-19 Genomics UK (COG-UK) Consortium | Harper VanSteenhouse, Yumi Kasai, David Gray, Carol Clugston, Anna Dominiczak and Alex Alderton, Roberto Amato, Sonia Goncalves, Ewan Harrison, David K. Jackson, Ian Johnston, Dominic Kwiatkowski, Cordelia Langford, John Sillitoe on behalf of the Wellcome Sanger Institute COVID-19 Surveillance Team |
| EPI_ISL_778404 | Lighthouse Lab in Alderley Park | Wellcome Sanger Institute for the COVID-19 Genomics UK (COG-UK) Consortium | Jacquelyn Wynn, Mairead Hyland, The Lighthouse Lab in Alderley Park and Alex Alderton, Roberto Amato, Sonia Goncalves, Ewan Harrison, David K. Jackson, Ian Johnston, Dominic Kwiatkowski, Cordelia Langford, John Sillitoe on behalf of the Wellcome Sanger Institute COVID-19 Surveillance Team |
| EPI_ISL_778407 | Lighthouse Lab in Glasgow | Wellcome Sanger Institute for the COVID-19 Genomics UK (COG-UK) Consortium | Harper VanSteenhouse, Yumi Kasai, David Gray, Carol Clugston, Anna Dominiczak and Alex Alderton, Roberto Amato, Sonia Goncalves, Ewan Harrison, David K. Jackson, Ian Johnston, Dominic Kwiatkowski, Cordelia Langford, John Sillitoe on behalf of the Wellcome Sanger Institute COVID-19 Surveillance Team |
| EPI_ISL_778408, EPI_ISL_778410 | Lighthouse Lab in Alderley Park | Wellcome Sanger Institute for the COVID-19 Genomics UK (COG-UK) Consortium | Jacquelyn Wynn, Mairead Hyland, The Lighthouse Lab in Alderley Park and Alex Alderton, Roberto Amato, Sonia Goncalves, Ewan Harrison, David K. Jackson, Ian Johnston, Dominic Kwiatkowski, Cordelia Langford, John Sillitoe on behalf of the Wellcome Sanger Institute COVID-19 Surveillance Team |
| EPI_ISL_778411 | Lighthouse Lab in Glasgow | Wellcome Sanger Institute for the COVID-19 Genomics UK (COG-UK) Consortium | Harper VanSteenhouse, Yumi Kasai, David Gray, Carol Clugston, Anna Dominiczak and Alex Alderton, Roberto Amato, Sonia Goncalves, Ewan Harrison, David K. Jackson, Ian Johnston, Dominic Kwiatkowski, Cordelia Langford, John Sillitoe on behalf of the Wellcome Sanger Institute COVID-19 Surveillance Team |
| EPI_ISL_778412, EPI_ISL_778415, EPI_ISL_778416, EPI_ISL_778417 | Lighthouse Lab in Alderley Park | Wellcome Sanger Institute for the COVID-19 Genomics UK (COG-UK) Consortium | Jacquelyn Wynn, Mairead Hyland, The Lighthouse Lab in Alderley Park and Alex Alderton, Roberto Amato, Sonia Goncalves, Ewan Harrison, David K. Jackson, Ian Johnston, Dominic Kwiatkowski, Cordelia Langford, John Sillitoe on behalf of the Wellcome Sanger Institute COVID-19 Surveillance Team |
| EPI_ISL_778419, EPI_ISL_778420, EPI_ISL_778421, EPI_ISL_778422, EPI_ISL_778423, EPI_ISL_778424 | Lighthouse Lab in Glasgow | Wellcome Sanger Institute for the COVID-19 Genomics UK (COG-UK) Consortium | Harper VanSteenhouse, Yumi Kasai, David Gray, Carol Clugston, Anna Dominiczak and Alex Alderton, Roberto Amato, Sonia Goncalves, Ewan Harrison, David K. Jackson, Ian Johnston, Dominic Kwiatkowski, Cordelia Langford, John Sillitoe on behalf of the Wellcome Sanger Institute COVID-19 Surveillance Team |
| EPI_ISL_778426 | Lighthouse Lab in Alderley Park | Wellcome Sanger Institute for the COVID-19 Genomics UK (COG-UK) Consortium | Jacquelyn Wynn, Mairead Hyland, The Lighthouse Lab in Alderley Park and Alex Alderton, Roberto Amato, Sonia Goncalves, Ewan Harrison, David K. Jackson, Ian Johnston, Dominic Kwiatkowski, Cordelia Langford, John Sillitoe on behalf of the Wellcome Sanger Institute COVID-19 Surveillance Team |
| EPI_ISL_778428 | Lighthouse Lab in Glasgow | Wellcome Sanger Institute for the COVID-19 Genomics UK (COG-UK) Consortium | Harper VanSteenhouse, Yumi Kasai, David Gray, Carol Clugston, Anna Dominiczak and Alex Alderton, Roberto Amato, Sonia Goncalves, Ewan Harrison, David K. Jackson, Ian Johnston, Dominic Kwiatkowski, Cordelia Langford, John Sillitoe on behalf of the Wellcome Sanger Institute COVID-19 Surveillance Team |
| EPI_ISL_778430, EPI_ISL_778432 | Lighthouse Lab in Alderley Park | Wellcome Sanger Institute for the COVID-19 Genomics UK (COG-UK) Consortium | Jacquelyn Wynn, Mairead Hyland, The Lighthouse Lab in Alderley Park and Alex Alderton, Roberto Amato, Sonia Goncalves, Ewan Harrison, David K. Jackson, Ian Johnston, Dominic Kwiatkowski, Cordelia Langford, John Sillitoe on behalf of the Wellcome Sanger Institute COVID-19 Surveillance Team |
| EPI_ISL_778433 | Lighthouse Lab in Glasgow | Wellcome Sanger Institute for the COVID-19 Genomics UK (COG-UK) Consortium | Harper VanSteenhouse, Yumi Kasai, David Gray, Carol Clugston, Anna Dominiczak and Alex Alderton, Roberto Amato, Sonia Goncalves, Ewan Harrison, David K. Jackson, Ian Johnston, Dominic Kwiatkowski, Cordelia Langford, John Sillitoe on behalf of the Wellcome Sanger Institute COVID-19 Surveillance Team |

| | | | |
|---|---|---|---|
| EPI_ISL_778444 | Lighthouse Lab in Alderley Park | Wellcome Sanger Institute for the COVID-19 Genomics UK (COG-UK) Consortium | Jacquelyn Wynn, Mairead Hyland, The Lighthouse Lab in Alderley Park and Alex Alderton, Roberto Amato, Sonia Goncalves, Ewan Harrison, David K. Jackson, Ian Johnston, Dominic Kwiatkowski, Cordelia Langford, John Sillitoe on behalf of the Wellcome Sanger Institute COVID-19 Surveillance Team |
| EPI_ISL_778447, EPI_ISL_778450, EPI_ISL_778451 | Lighthouse Lab in Glasgow | Wellcome Sanger Institute for the COVID-19 Genomics UK (COG-UK) Consortium | Harper VanSteenhouse, Yumi Kasai, David Gray, Carol Clugston, Anna Dominiczak and Alex Alderton, Roberto Amato, Sonia Goncalves, Ewan Harrison, David K. Jackson, Ian Johnston, Dominic Kwiatkowski, Cordelia Langford, John Sillitoe on behalf of the Wellcome Sanger Institute COVID-19 Surveillance Team |
| EPI_ISL_778454, EPI_ISL_778455 | Lighthouse Lab in Alderley Park | Wellcome Sanger Institute for the COVID-19 Genomics UK (COG-UK) Consortium | Jacquelyn Wynn, Mairead Hyland, The Lighthouse Lab in Alderley Park and Alex Alderton, Roberto Amato, Sonia Goncalves, Ewan Harrison, David K. Jackson, Ian Johnston, Dominic Kwiatkowski, Cordelia Langford, John Sillitoe on behalf of the Wellcome Sanger Institute COVID-19 Surveillance Team |
| EPI_ISL_778457, EPI_ISL_778459, EPI_ISL_778461, EPI_ISL_778462, EPI_ISL_778463, EPI_ISL_778465 | Lighthouse Lab in Glasgow | Wellcome Sanger Institute for the COVID-19 Genomics UK (COG-UK) Consortium | Harper VanSteenhouse, Yumi Kasai, David Gray, Carol Clugston, Anna Dominiczak and Alex Alderton, Roberto Amato, Sonia Goncalves, Ewan Harrison, David K. Jackson, Ian Johnston, Dominic Kwiatkowski, Cordelia Langford, John Sillitoe on behalf of the Wellcome Sanger Institute COVID-19 Surveillance Team |
| EPI_ISL_778471, EPI_ISL_778472, EPI_ISL_778473, EPI_ISL_778478 | Lighthouse Lab in Alderley Park | Wellcome Sanger Institute for the COVID-19 Genomics UK (COG-UK) Consortium | Jacquelyn Wynn, Mairead Hyland, The Lighthouse Lab in Alderley Park and Alex Alderton, Roberto Amato, Sonia Goncalves, Ewan Harrison, David K. Jackson, Ian Johnston, Dominic Kwiatkowski, Cordelia Langford, John Sillitoe on behalf of the Wellcome Sanger Institute COVID-19 Surveillance Team |
| EPI_ISL_778482 | Lighthouse Lab in Glasgow | Wellcome Sanger Institute for the COVID-19 Genomics UK (COG-UK) Consortium | Harper VanSteenhouse, Yumi Kasai, David Gray, Carol Clugston, Anna Dominiczak and Alex Alderton, Roberto Amato, Sonia Goncalves, Ewan Harrison, David K. Jackson, Ian Johnston, Dominic Kwiatkowski, Cordelia Langford, John Sillitoe on behalf of the Wellcome Sanger Institute COVID-19 Surveillance Team |
| EPI_ISL_778484, EPI_ISL_778487, EPI_ISL_778488, EPI_ISL_778490, EPI_ISL_778491, EPI_ISL_778494 | Lighthouse Lab in Alderley Park | Wellcome Sanger Institute for the COVID-19 Genomics UK (COG-UK) Consortium | Jacquelyn Wynn, Mairead Hyland, The Lighthouse Lab in Alderley Park and Alex Alderton, Roberto Amato, Sonia Goncalves, Ewan Harrison, David K. Jackson, Ian Johnston, Dominic Kwiatkowski, Cordelia Langford, John Sillitoe on behalf of the Wellcome Sanger Institute COVID-19 Surveillance Team |
| EPI_ISL_778496, EPI_ISL_778497, EPI_ISL_778499 | Lighthouse Lab in Glasgow | Wellcome Sanger Institute for the COVID-19 Genomics UK (COG-UK) Consortium | Harper VanSteenhouse, Yumi Kasai, David Gray, Carol Clugston, Anna Dominiczak and Alex Alderton, Roberto Amato, Sonia Goncalves, Ewan Harrison, David K. Jackson, Ian Johnston, Dominic Kwiatkowski, Cordelia Langford, John Sillitoe on behalf of the Wellcome Sanger Institute COVID-19 Surveillance Team |
| EPI_ISL_778500 | Lighthouse Lab in Alderley Park | Wellcome Sanger Institute for the COVID-19 Genomics UK (COG-UK) Consortium | Jacquelyn Wynn, Mairead Hyland, The Lighthouse Lab in Alderley Park and Alex Alderton, Roberto Amato, Sonia Goncalves, Ewan Harrison, David K. Jackson, Ian Johnston, Dominic Kwiatkowski, Cordelia Langford, John Sillitoe on behalf of the Wellcome Sanger Institute COVID-19 Surveillance Team |
| EPI_ISL_778504, EPI_ISL_778506 | Lighthouse Lab in Glasgow | Wellcome Sanger Institute for the COVID-19 Genomics UK (COG-UK) Consortium | Harper VanSteenhouse, Yumi Kasai, David Gray, Carol Clugston, Anna Dominiczak and Alex Alderton, Roberto Amato, Sonia Goncalves, Ewan Harrison, David K. Jackson, Ian Johnston, Dominic Kwiatkowski, Cordelia Langford, John Sillitoe on behalf of the Wellcome Sanger Institute COVID-19 Surveillance Team |
| EPI_ISL_778510, EPI_ISL_778512 | Lighthouse Lab in Alderley Park | Wellcome Sanger Institute for the COVID-19 Genomics UK (COG-UK) Consortium | Jacquelyn Wynn, Mairead Hyland, The Lighthouse Lab in Alderley Park and Alex Alderton, Roberto Amato, Sonia Goncalves, Ewan Harrison, David K. Jackson, Ian Johnston, Dominic Kwiatkowski, Cordelia Langford, John Sillitoe on behalf of the Wellcome Sanger Institute COVID-19 Surveillance Team |
| EPI_ISL_778516 | Lighthouse Lab in Glasgow | Wellcome Sanger Institute for the COVID-19 Genomics UK (COG-UK) Consortium | Harper VanSteenhouse, Yumi Kasai, David Gray, Carol Clugston, Anna Dominiczak and Alex Alderton, Roberto Amato, Sonia Goncalves, Ewan Harrison, David K. Jackson, Ian Johnston, Dominic Kwiatkowski, Cordelia Langford, John Sillitoe on behalf of the Wellcome Sanger Institute COVID-19 Surveillance Team |
| EPI_ISL_778517 | Lighthouse Lab in Alderley Park | Wellcome Sanger Institute for the COVID-19 Genomics UK (COG-UK) Consortium | Jacquelyn Wynn, Mairead Hyland, The Lighthouse Lab in Alderley Park and Alex Alderton, Roberto Amato, Sonia Goncalves, Ewan Harrison, David K. Jackson, Ian Johnston, Dominic Kwiatkowski, Cordelia Langford, John Sillitoe on behalf of the Wellcome Sanger Institute COVID-19 Surveillance Team |
| EPI_ISL_778522 | Lighthouse Lab in Glasgow | Wellcome Sanger Institute for the COVID-19 Genomics UK (COG-UK) Consortium | Harper VanSteenhouse, Yumi Kasai, David Gray, Carol Clugston, Anna Dominiczak and Alex Alderton, Roberto Amato, Sonia Goncalves, Ewan Harrison, David K. Jackson, Ian Johnston, Dominic Kwiatkowski, Cordelia Langford, John Sillitoe on behalf of the Wellcome Sanger Institute COVID-19 Surveillance Team |
| EPI_ISL_778524 | Lighthouse Lab in Alderley Park | Wellcome Sanger Institute for the COVID-19 Genomics UK (COG-UK) Consortium | Jacquelyn Wynn, Mairead Hyland, The Lighthouse Lab in Alderley Park and Alex Alderton, Roberto Amato, Sonia Goncalves, Ewan Harrison, David K. Jackson, Ian Johnston, Dominic Kwiatkowski, Cordelia Langford, John Sillitoe on behalf of the Wellcome Sanger Institute COVID-19 Surveillance Team |
| EPI_ISL_778525, EPI_ISL_778527 | Lighthouse Lab in Glasgow | Wellcome Sanger Institute for the COVID-19 Genomics UK (COG-UK) Consortium | Harper VanSteenhouse, Yumi Kasai, David Gray, Carol Clugston, Anna Dominiczak and Alex Alderton, Roberto Amato, Sonia Goncalves, Ewan Harrison, David K. Jackson, Ian Johnston, Dominic Kwiatkowski, Cordelia Langford, John Sillitoe on behalf of the Wellcome Sanger Institute COVID-19 Surveillance Team |
| EPI_ISL_778528, EPI_ISL_778529 | Lighthouse Lab in Alderley Park | Wellcome Sanger Institute for the COVID-19 Genomics UK (COG-UK) Consortium | Jacquelyn Wynn, Mairead Hyland, The Lighthouse Lab in Alderley Park and Alex Alderton, Roberto Amato, Sonia Goncalves, Ewan Harrison, David K. Jackson, Ian Johnston, Dominic Kwiatkowski, Cordelia Langford, John Sillitoe on behalf of the Wellcome Sanger Institute COVID-19 Surveillance Team |
| EPI_ISL_778530 | Lighthouse Lab in Glasgow | Wellcome Sanger Institute for the COVID-19 Genomics UK (COG-UK) Consortium | Harper VanSteenhouse, Yumi Kasai, David Gray, Carol Clugston, Anna Dominiczak and Alex Alderton, Roberto Amato, Sonia Goncalves, Ewan Harrison, David K. Jackson, Ian Johnston, Dominic Kwiatkowski, Cordelia Langford, John Sillitoe on behalf of the Wellcome Sanger Institute COVID-19 Surveillance Team |
| EPI_ISL_778531 | Lighthouse Lab in Alderley Park | Wellcome Sanger Institute for the COVID-19 Genomics UK (COG-UK) Consortium | Jacquelyn Wynn, Mairead Hyland, The Lighthouse Lab in Alderley Park and Alex Alderton, Roberto Amato, Sonia Goncalves, Ewan Harrison, David K. Jackson, Ian Johnston, Dominic Kwiatkowski, Cordelia Langford, John Sillitoe on behalf of the Wellcome Sanger Institute COVID-19 Surveillance Team |
| EPI_ISL_778534 | Lighthouse Lab in Glasgow | Wellcome Sanger Institute for the COVID-19 Genomics UK (COG-UK) Consortium | Harper VanSteenhouse, Yumi Kasai, David Gray, Carol Clugston, Anna Dominiczak and Alex Alderton, Roberto Amato, Sonia Goncalves, Ewan Harrison, David K. Jackson, Ian Johnston, Dominic Kwiatkowski, Cordelia Langford, John Sillitoe on behalf of the Wellcome Sanger Institute COVID-19 Surveillance Team |
| EPI_ISL_778539 | Lighthouse Lab in Alderley Park | Wellcome Sanger Institute for the COVID-19 Genomics UK (COG-UK) Consortium | Jacquelyn Wynn, Mairead Hyland, The Lighthouse Lab in Alderley Park and Alex Alderton, Roberto Amato, Sonia Goncalves, Ewan Harrison, David K. Jackson, Ian Johnston, Dominic Kwiatkowski, Cordelia Langford, John Sillitoe on behalf of the Wellcome Sanger Institute COVID-19 Surveillance Team |
| EPI_ISL_778540, EPI_ISL_778541 | Lighthouse Lab in Glasgow | Wellcome Sanger Institute for the COVID-19 Genomics UK (COG-UK) Consortium | Harper VanSteenhouse, Yumi Kasai, David Gray, Carol Clugston, Anna Dominiczak and Alex Alderton, Roberto Amato, Sonia Goncalves, Ewan Harrison, David K. Jackson, Ian Johnston, Dominic Kwiatkowski, Cordelia Langford, John Sillitoe on behalf of the Wellcome Sanger Institute COVID-19 Surveillance Team |
| EPI_ISL_778543, EPI_ISL_778547 | Lighthouse Lab in Alderley Park | Wellcome Sanger Institute for the COVID-19 Genomics UK (COG-UK) Consortium | Jacquelyn Wynn, Mairead Hyland, The Lighthouse Lab in Alderley Park and Alex Alderton, Roberto Amato, Sonia Goncalves, Ewan Harrison, David K. Jackson, Ian Johnston, Dominic Kwiatkowski, Cordelia Langford, John Sillitoe on behalf of the Wellcome Sanger Institute COVID-19 Surveillance Team |
| EPI_ISL_778548, EPI_ISL_778551, EPI_ISL_778553 | Lighthouse Lab in Glasgow | Wellcome Sanger Institute for the COVID-19 Genomics UK (COG-UK) Consortium | Harper VanSteenhouse, Yumi Kasai, David Gray, Carol Clugston, Anna Dominiczak and Alex Alderton, Roberto Amato, Sonia Goncalves, Ewan Harrison, David K. Jackson, Ian Johnston, Dominic Kwiatkowski, Cordelia Langford, John Sillitoe on behalf of the Wellcome Sanger Institute COVID-19 Surveillance Team |
| EPI_ISL_778556, EPI_ISL_778558 | Lighthouse Lab in Alderley Park | Wellcome Sanger Institute for the COVID-19 Genomics UK (COG-UK) Consortium | Jacquelyn Wynn, Mairead Hyland, The Lighthouse Lab in Alderley Park and Alex Alderton, Roberto Amato, Sonia Goncalves, Ewan Harrison, David K. Jackson, Ian Johnston, Dominic Kwiatkowski, Cordelia Langford, John Sillitoe on behalf of the Wellcome Sanger Institute COVID-19 Surveillance Team |
| EPI_ISL_778560 | Lighthouse Lab in Glasgow | Wellcome Sanger Institute for the COVID-19 Genomics UK (COG-UK) Consortium | Harper VanSteenhouse, Yumi Kasai, David Gray, Carol Clugston, Anna Dominiczak and Alex Alderton, Roberto Amato, Sonia Goncalves, Ewan Harrison, David K. Jackson, Ian Johnston, Dominic Kwiatkowski, Cordelia Langford, John Sillitoe on behalf of the Wellcome Sanger Institute COVID-19 Surveillance Team |
| EPI_ISL_778562 | Lighthouse Lab in Alderley Park | Wellcome Sanger Institute for the COVID-19 Genomics UK (COG-UK) Consortium | Jacquelyn Wynn, Mairead Hyland, The Lighthouse Lab in Alderley Park and Alex Alderton, Roberto Amato, Sonia Goncalves, Ewan Harrison, David K. Jackson, Ian Johnston, Dominic Kwiatkowski, Cordelia Langford, John Sillitoe on behalf of the Wellcome Sanger Institute COVID-19 Surveillance Team |
| EPI_ISL_778568 | Lighthouse Lab in Glasgow | Wellcome Sanger Institute for the COVID-19 Genomics UK (COG-UK) Consortium | Harper VanSteenhouse, Yumi Kasai, David Gray, Carol Clugston, Anna Dominiczak and Alex Alderton, Roberto Amato, Sonia Goncalves, Ewan Harrison, David K. Jackson, Ian Johnston, Dominic Kwiatkowski, Cordelia Langford, John Sillitoe on behalf of the Wellcome Sanger Institute COVID-19 Surveillance Team |
| EPI_ISL_778569 | Lighthouse Lab in Alderley Park | Wellcome Sanger Institute for the COVID-19 Genomics UK (COG-UK) Consortium | Jacquelyn Wynn, Mairead Hyland, The Lighthouse Lab in Alderley Park and Alex Alderton, Roberto Amato, Sonia Goncalves, Ewan Harrison, David K. Jackson, Ian Johnston, Dominic Kwiatkowski, Cordelia Langford, John Sillitoe on behalf of the Wellcome Sanger Institute COVID-19 Surveillance Team |

| | | | |
|---|---|---|---|
| EPI_ISL_778570 | Lighthouse Lab in Glasgow | Wellcome Sanger Institute for the COVID-19 Genomics UK (COG-UK) Consortium | Harper VanSteenhouse, Yumi Kasai, David Gray, Carol Clugston, Anna Dominiczak and Alex Alderton, Roberto Amato, Sonia Goncalves, Ewan Harrison, David K. Jackson, Ian Johnston, Dominic Kwiatkowski, Cordelia Langford, John Sillitoe on behalf of the Wellcome Sanger Institute COVID-19 Surveillance Team |
| EPI_ISL_778571 | Lighthouse Lab in Alderley Park | Wellcome Sanger Institute for the COVID-19 Genomics UK (COG-UK) Consortium | Jacquelyn Wynn, Mairead Hyland, The Lighthouse Lab in Alderley Park and Alex Alderton, Roberto Amato, Sonia Goncalves, Ewan Harrison, David K. Jackson, Ian Johnston, Dominic Kwiatkowski, Cordelia Langford, John Sillitoe on behalf of the Wellcome Sanger Institute COVID-19 Surveillance Team |
| EPI_ISL_778577 | Lighthouse Lab in Glasgow | Wellcome Sanger Institute for the COVID-19 Genomics UK (COG-UK) Consortium | Harper VanSteenhouse, Yumi Kasai, David Gray, Carol Clugston, Anna Dominiczak and Alex Alderton, Roberto Amato, Sonia Goncalves, Ewan Harrison, David K. Jackson, Ian Johnston, Dominic Kwiatkowski, Cordelia Langford, John Sillitoe on behalf of the Wellcome Sanger Institute COVID-19 Surveillance Team |
| EPI_ISL_778578 | Lighthouse Lab in Alderley Park | Wellcome Sanger Institute for the COVID-19 Genomics UK (COG-UK) Consortium | Jacquelyn Wynn, Mairead Hyland, The Lighthouse Lab in Alderley Park and Alex Alderton, Roberto Amato, Sonia Goncalves, Ewan Harrison, David K. Jackson, Ian Johnston, Dominic Kwiatkowski, Cordelia Langford, John Sillitoe on behalf of the Wellcome Sanger Institute COVID-19 Surveillance Team |
| EPI_ISL_778579 | Lighthouse Lab in Glasgow | Wellcome Sanger Institute for the COVID-19 Genomics UK (COG-UK) Consortium | Harper VanSteenhouse, Yumi Kasai, David Gray, Carol Clugston, Anna Dominiczak and Alex Alderton, Roberto Amato, Sonia Goncalves, Ewan Harrison, David K. Jackson, Ian Johnston, Dominic Kwiatkowski, Cordelia Langford, John Sillitoe on behalf of the Wellcome Sanger Institute COVID-19 Surveillance Team |
| EPI_ISL_778584 | Lighthouse Lab in Alderley Park | Wellcome Sanger Institute for the COVID-19 Genomics UK (COG-UK) Consortium | Jacquelyn Wynn, Mairead Hyland, The Lighthouse Lab in Alderley Park and Alex Alderton, Roberto Amato, Sonia Goncalves, Ewan Harrison, David K. Jackson, Ian Johnston, Dominic Kwiatkowski, Cordelia Langford, John Sillitoe on behalf of the Wellcome Sanger Institute COVID-19 Surveillance Team |
| EPI_ISL_778586 | Lighthouse Lab in Glasgow | Wellcome Sanger Institute for the COVID-19 Genomics UK (COG-UK) Consortium | Harper VanSteenhouse, Yumi Kasai, David Gray, Carol Clugston, Anna Dominiczak and Alex Alderton, Roberto Amato, Sonia Goncalves, Ewan Harrison, David K. Jackson, Ian Johnston, Dominic Kwiatkowski, Cordelia Langford, John Sillitoe on behalf of the Wellcome Sanger Institute COVID-19 Surveillance Team |
| EPI_ISL_778593 | Lighthouse Lab in Alderley Park | Wellcome Sanger Institute for the COVID-19 Genomics UK (COG-UK) Consortium | Jacquelyn Wynn, Mairead Hyland, The Lighthouse Lab in Alderley Park and Alex Alderton, Roberto Amato, Sonia Goncalves, Ewan Harrison, David K. Jackson, Ian Johnston, Dominic Kwiatkowski, Cordelia Langford, John Sillitoe on behalf of the Wellcome Sanger Institute COVID-19 Surveillance Team |
| EPI_ISL_778594 | Lighthouse Lab in Glasgow | Wellcome Sanger Institute for the COVID-19 Genomics UK (COG-UK) Consortium | Harper VanSteenhouse, Yumi Kasai, David Gray, Carol Clugston, Anna Dominiczak and Alex Alderton, Roberto Amato, Sonia Goncalves, Ewan Harrison, David K. Jackson, Ian Johnston, Dominic Kwiatkowski, Cordelia Langford, John Sillitoe on behalf of the Wellcome Sanger Institute COVID-19 Surveillance Team |
| EPI_ISL_778598, EPI_ISL_778599, EPI_ISL_778601, EPI_ISL_778605, EPI_ISL_778607, EPI_ISL_778612, EPI_ISL_778614, EPI_ISL_778616 | Lighthouse Lab in Alderley Park | Wellcome Sanger Institute for the COVID-19 Genomics UK (COG-UK) Consortium | Jacquelyn Wynn, Mairead Hyland, The Lighthouse Lab in Alderley Park and Alex Alderton, Roberto Amato, Sonia Goncalves, Ewan Harrison, David K. Jackson, Ian Johnston, Dominic Kwiatkowski, Cordelia Langford, John Sillitoe on behalf of the Wellcome Sanger Institute COVID-19 Surveillance Team |
| EPI_ISL_778617 | Lighthouse Lab in Glasgow | Wellcome Sanger Institute for the COVID-19 Genomics UK (COG-UK) Consortium | Harper VanSteenhouse, Yumi Kasai, David Gray, Carol Clugston, Anna Dominiczak and Alex Alderton, Roberto Amato, Sonia Goncalves, Ewan Harrison, David K. Jackson, Ian Johnston, Dominic Kwiatkowski, Cordelia Langford, John Sillitoe on behalf of the Wellcome Sanger Institute COVID-19 Surveillance Team |
| EPI_ISL_778618 | Lighthouse Lab in Alderley Park | Wellcome Sanger Institute for the COVID-19 Genomics UK (COG-UK) Consortium | Jacquelyn Wynn, Mairead Hyland, The Lighthouse Lab in Alderley Park and Alex Alderton, Roberto Amato, Sonia Goncalves, Ewan Harrison, David K. Jackson, Ian Johnston, Dominic Kwiatkowski, Cordelia Langford, John Sillitoe on behalf of the Wellcome Sanger Institute COVID-19 Surveillance Team |
| EPI_ISL_778619, EPI_ISL_778620 | Lighthouse Lab in Glasgow | Wellcome Sanger Institute for the COVID-19 Genomics UK (COG-UK) Consortium | Harper VanSteenhouse, Yumi Kasai, David Gray, Carol Clugston, Anna Dominiczak and Alex Alderton, Roberto Amato, Sonia Goncalves, Ewan Harrison, David K. Jackson, Ian Johnston, Dominic Kwiatkowski, Cordelia Langford, John Sillitoe on behalf of the Wellcome Sanger Institute COVID-19 Surveillance Team |
| EPI_ISL_778621, EPI_ISL_778623, EPI_ISL_778627, EPI_ISL_778628 | Lighthouse Lab in Alderley Park | Wellcome Sanger Institute for the COVID-19 Genomics UK (COG-UK) Consortium | Jacquelyn Wynn, Mairead Hyland, The Lighthouse Lab in Alderley Park and Alex Alderton, Roberto Amato, Sonia Goncalves, Ewan Harrison, David K. Jackson, Ian Johnston, Dominic Kwiatkowski, Cordelia Langford, John Sillitoe on behalf of the Wellcome Sanger Institute COVID-19 Surveillance Team |
| EPI_ISL_778629 | Lighthouse Lab in Glasgow | Wellcome Sanger Institute for the COVID-19 Genomics UK (COG-UK) Consortium | Harper VanSteenhouse, Yumi Kasai, David Gray, Carol Clugston, Anna Dominiczak and Alex Alderton, Roberto Amato, Sonia Goncalves, Ewan Harrison, David K. Jackson, Ian Johnston, Dominic Kwiatkowski, Cordelia Langford, John Sillitoe on behalf of the Wellcome Sanger Institute COVID-19 Surveillance Team |
| EPI_ISL_778630 | Lighthouse Lab in Alderley Park | Wellcome Sanger Institute for the COVID-19 Genomics UK (COG-UK) Consortium | Jacquelyn Wynn, Mairead Hyland, The Lighthouse Lab in Alderley Park and Alex Alderton, Roberto Amato, Sonia Goncalves, Ewan Harrison, David K. Jackson, Ian Johnston, Dominic Kwiatkowski, Cordelia Langford, John Sillitoe on behalf of the Wellcome Sanger Institute COVID-19 Surveillance Team |
| EPI_ISL_778631 | Lighthouse Lab in Glasgow | Wellcome Sanger Institute for the COVID-19 Genomics UK (COG-UK) Consortium | Harper VanSteenhouse, Yumi Kasai, David Gray, Carol Clugston, Anna Dominiczak and Alex Alderton, Roberto Amato, Sonia Goncalves, Ewan Harrison, David K. Jackson, Ian Johnston, Dominic Kwiatkowski, Cordelia Langford, John Sillitoe on behalf of the Wellcome Sanger Institute COVID-19 Surveillance Team |
| EPI_ISL_778635, EPI_ISL_778638 | Lighthouse Lab in Alderley Park | Wellcome Sanger Institute for the COVID-19 Genomics UK (COG-UK) Consortium | Jacquelyn Wynn, Mairead Hyland, The Lighthouse Lab in Alderley Park and Alex Alderton, Roberto Amato, Sonia Goncalves, Ewan Harrison, David K. Jackson, Ian Johnston, Dominic Kwiatkowski, Cordelia Langford, John Sillitoe on behalf of the Wellcome Sanger Institute COVID-19 Surveillance Team |
| EPI_ISL_778822 | AIID | Irish Coronavirus Sequencing Consortium-Teagasc Grange | Matthew McCabe, Aljandro Abner Garcia Leon, Fiona Crispie, Calum Walsh, Michael Carr, John Kenny, Paul Cotter, Patrick Mallon, Gabriel Gonzalez |
| EPI_ISL_779082, EPI_ISL_779083, EPI_ISL_779084, EPI_ISL_779085, EPI_ISL_779086, EPI_ISL_779087, EPI_ISL_779088, EPI_ISL_779089, EPI_ISL_779090, EPI_ISL_779091, EPI_ISL_779092, EPI_ISL_779093, EPI_ISL_779094, EPI_ISL_779095, EPI_ISL_779096, EPI_ISL_779097, EPI_ISL_779098, EPI_ISL_779099, EPI_ISL_779100, EPI_ISL_779101, EPI_ISL_779102, EPI_ISL_779103, EPI_ISL_779104 | | | |
| see above | LSUHS Emerging Viral Threat Laboratory | Microbial Genome Sequencing Center | Jeremy P. Kamil, Jennifer L. Carroll, Camille F. Abshire, Maarten Van Diest, Andrew D. Yurochko, Martin J. Sapp, Rona S. Scott, Christopher G. Kevil, Daniel J. Snyder, Vaughn S. Cooper, John A. Vanchiere |
| EPI_ISL_779105, EPI_ISL_779106, EPI_ISL_779107, EPI_ISL_779108 | LSUHS Emerging Viral Threat Laboratory | Microbial Genome Sequencing Center | Jennifer L. Carroll, Jeremy P. Kamil, Camille F. Abshire, Maarten Van Diest, Andrew D. Yurochko, Martin J. Sapp, Rona S. Scott, Christopher G. Kevil, Daniel J. Snyder, Vaughn S. Cooper, John A. Vanchiere |
| EPI_ISL_779847 | Centre Hospotalier Cannes | CNR Virus des Infections Respiratoires - France SUD | Antonin Bal, Gregory Destras, Gwendolyne Burfin, Hadrien Règue, Quentin Semanas, Martine Valette, Bruno Lina, Laurence Josset |
| EPI_ISL_779855, EPI_ISL_779858, EPI_ISL_779859, EPI_ISL_779860, EPI_ISL_779869, EPI_ISL_779880, EPI_ISL_779884, EPI_ISL_779896, EPI_ISL_779897, EPI_ISL_779907, EPI_ISL_779912, EPI_ISL_779913 | | | |
| see above | Servicio de Microbiología, Hospital Universitario Son Espases | SeqCOVID-SPAIN consortium/IBV(CSIC) | Carla López-Causapé, Jordi Reina, Antonio Oliver and SeqCOVID-SPAIN consortium |
| EPI_ISL_779922, EPI_ISL_779924, EPI_ISL_779927, EPI_ISL_779932, EPI_ISL_779939, EPI_ISL_779940, EPI_ISL_779941, EPI_ISL_779942, EPI_ISL_779943, EPI_ISL_779944, EPI_ISL_779945, EPI_ISL_779946, EPI_ISL_779947, EPI_ISL_779948, EPI_ISL_779949, EPI_ISL_779950, EPI_ISL_779951 | | | |
| see above | Center of Medical Microbiology, Virology, and Hospital Hygiene, University of Duesseldorf | Center of Medical Microbiology, Virology, and Hospital Hygiene, University of Duesseldorf | Maximilian Damagnez, Alexander Dilthey, Ashley-Jane Duplessis, Torsten Houwaart, Lisanna Hülse, Malte Kohns Vasconcelos, Nadine Lübke, Jessica Nicolai, Klaus Pfeffer, Daniel Strelow, Teresa Tamayo, Jörg Timm, Andreas Walker, Tobias Wienemann |
| EPI_ISL_781551, EPI_ISL_781571, EPI_ISL_781625, EPI_ISL_781671, EPI_ISL_781690, EPI_ISL_781710, EPI_ISL_781824, EPI_ISL_781863 | Lighthouse Lab in Cambridge | Wellcome Sanger Institute for the COVID-19 Genomics UK (COG-UK) Consortium | Rob Howes, The Lighthouse Lab in Cambridge and Alex Alderton, Roberto Amato, Sonia Goncalves, Ewan Harrison, David K. Jackson, Ian Johnston, Dominic Kwiatkowski, Cordelia Langford, John Sillitoe on behalf of the Wellcome Sanger Institute COVID-19 Surveillance Team |
| EPI_ISL_781957, EPI_ISL_782123 | Lighthouse Lab in Alderley Park | Wellcome Sanger Institute for the COVID-19 Genomics UK (COG-UK) Consortium | Jacquelyn Wynn, Mairead Hyland, The Lighthouse Lab in Alderley Park and Alex Alderton, Roberto Amato, Sonia Goncalves, Ewan Harrison, David K. Jackson, Ian Johnston, Dominic Kwiatkowski, Cordelia Langford, John Sillitoe on behalf of the Wellcome Sanger Institute COVID-19 Surveillance Team |
| EPI_ISL_789038, EPI_ISL_789040, EPI_ISL_789041, EPI_ISL_789042, EPI_ISL_789043 | Klinisk mikrobiologi | The Public Health Agency of Sweden | Department of Microbiology, The Public Health Agency of Sweden |
| EPI_ISL_789046 | unknown | The Public Health Agency of Sweden | Department of Microbiology, The Public Health Agency of Sweden |
| EPI_ISL_789047 | Klinisk mikrobiologi | The Public Health Agency of Sweden | Department of Microbiology, The Public Health Agency of Sweden |
| EPI_ISL_790585, EPI_ISL_790592, EPI_ISL_790598, EPI_ISL_790611, EPI_ISL_790624, EPI_ISL_790625, EPI_ISL_790629, EPI_ISL_790633, EPI_ISL_790635, EPI_ISL_790636, EPI_ISL_790637, EPI_ISL_790638, EPI_ISL_790639, EPI_ISL_790640, EPI_ISL_790641, EPI_ISL_790642, EPI_ISL_790643, EPI_ISL_790644, | | | |

| | | | |
|---|---|---|---|
| EPI_ISL_790645, EPI_ISL_790646, EPI_ISL_790650, EPI_ISL_790667, EPI_ISL_790672, EPI_ISL_790673, EPI_ISL_790703, EPI_ISL_790709, EPI_ISL_790714, EPI_ISL_790715, EPI_ISL_790716, EPI_ISL_790717, EPI_ISL_790718, EPI_ISL_790719, EPI_ISL_790720, EPI_ISL_790721, EPI_ISL_790729, EPI_ISL_790734, EPI_ISL_790735, EPI_ISL_790736, EPI_ISL_790737, EPI_ISL_790742, EPI_ISL_790765, EPI_ISL_790770, EPI_ISL_790772, EPI_ISL_790773, EPI_ISL_790777, EPI_ISL_790784, EPI_ISL_790788, EPI_ISL_790794, EPI_ISL_790795, EPI_ISL_790796, EPI_ISL_790797, EPI_ISL_790807, EPI_ISL_790813, EPI_ISL_790816, EPI_ISL_790832, EPI_ISL_790837, EPI_ISL_790850, EPI_ISL_790876, EPI_ISL_790877, EPI_ISL_790878, EPI_ISL_790889, EPI_ISL_790890, EPI_ISL_790891, EPI_ISL_790901, EPI_ISL_790920, EPI_ISL_790935, EPI_ISL_790936, EPI_ISL_790937, EPI_ISL_790955, EPI_ISL_791081, EPI_ISL_791083 | | | |
| see above | Dutch COVID-19 response team | National Institute for Public Health and the Environment (RIVM) | Adam Meijer, Harry Vennema, Jeroen Cremer, Sharon van den Brink, Bas van der Veer, AnneMarie van den Brandt, Florian Zwagemaker, Dennis Schmitz, Chantal Reusken, on behalf of the national COVID-19 response team |
| EPI_ISL_791085 | Instituto Nacional de Salud - Unidad de Secuenciación y Análisis Genómico | Instituto Nacional de Salud - Dirección de Investigación en Salud Pública | Katherine Laiton-Donato, Diego A. Álvarez-Díaz, Carlos Franco-Muñoz, Mauricio Pacheco-Montealegre, Jeadran Malagon-Rojas, Jesith Toloza, Julia Almentero, Ronald Lopez, Jonathan Reales, Diego Andrés Prada, Magdalena Wiesner, Martha Lucia Ospina Martinez, Marcela Mercado-Reyes |
| EPI_ISL_791184, EPI_ISL_791185, EPI_ISL_791186 | Respiratory Virus Unit, National Infection Service, Public Health England | COVID-19 Genomics UK (COG-UK) Consortium | PHE Covid Sequencing Team |
| EPI_ISL_791287, EPI_ISL_791301, EPI_ISL_791306, EPI_ISL_791321 | National Virus Reference Laboratory | Irish Coronavirus Sequencing Consortium - Teagasc Moorepark | Alejandro Abner Garcia Leon, Paul Cotter, Fiona Crispie, John Kenny, Paddy Mallon, Calum Walsh |
| EPI_ISL_791417, EPI_ISL_791484 | Johns Hopkins Hospital Department of Pathology | Johns Hopkins Hospital Department of Pathology | C. Paul Morris, Chun Huai Luo, Adannaya Amadi, Nicholas Gallagher, Heba H. Mostafa |
| EPI_ISL_792522 | Laboratorio de Virología del Hospital de Niños Dr. Ricardo Gutierrez | Grupo de Genómica y Bioinformática del Instituto de Investigación de la Cadena Láctea CONICET-INTA on behalf of 'Proyecto Argentino Interinstitucional de genomica de SARS-CoV-2' (PAIS Consortium) | Amadio, AF, Eberhardt, MF; Irazoqui, M; Torres, C; Aulicino, P; König, G; Acevedo, ME; Alvarez Lopez, C; Alexay, S; Jacques, O; Mistchenko, AS, Goya, S; Nabaes Jodar, MS; Viegas, M. |
| EPI_ISL_792557 | Connecticut State Department of Public Health | Grubaugh Lab - Yale School of Public Health | Joseph Fauver, Tara Alpert, Chantal Vogels, Anderson Brito, Nagarjuna Cheemarla, Ellen Foxman, Anthony Muyombwe, Jafar Razeq, Nathan Grubaugh |
| EPI_ISL_792708, EPI_ISL_792709, EPI_ISL_792711, EPI_ISL_792712, EPI_ISL_792714, EPI_ISL_792715, EPI_ISL_792716, EPI_ISL_792717, EPI_ISL_792718, EPI_ISL_792719, EPI_ISL_792720, EPI_ISL_792721, EPI_ISL_792722, EPI_ISL_792723, EPI_ISL_792724, EPI_ISL_792725, EPI_ISL_792726, EPI_ISL_792727, EPI_ISL_792728, EPI_ISL_792729, EPI_ISL_792730, EPI_ISL_792731, EPI_ISL_792732, EPI_ISL_792733, EPI_ISL_792734, EPI_ISL_792735, EPI_ISL_792736, EPI_ISL_792737, EPI_ISL_792738, EPI_ISL_792739, EPI_ISL_792749, EPI_ISL_792750, EPI_ISL_792751, EPI_ISL_792752, EPI_ISL_792753, EPI_ISL_792755, EPI_ISL_792756, EPI_ISL_792757, EPI_ISL_792758, EPI_ISL_792760, EPI_ISL_792761, EPI_ISL_792762, EPI_ISL_792763, EPI_ISL_792764, EPI_ISL_792765, EPI_ISL_792766, EPI_ISL_792767, EPI_ISL_792768, EPI_ISL_792769, EPI_ISL_792770, EPI_ISL_792771, EPI_ISL_792772, EPI_ISL_792773, EPI_ISL_792774, EPI_ISL_792775, EPI_ISL_792776, EPI_ISL_792777, EPI_ISL_792778, EPI_ISL_792779, EPI_ISL_792781, EPI_ISL_792782, EPI_ISL_792785, EPI_ISL_792786, EPI_ISL_792787, EPI_ISL_792788, EPI_ISL_792789, EPI_ISL_792790, EPI_ISL_792791, EPI_ISL_792792, EPI_ISL_792794, EPI_ISL_792795, EPI_ISL_792796, EPI_ISL_792797, EPI_ISL_792798, EPI_ISL_792799, EPI_ISL_792800, EPI_ISL_792801, EPI_ISL_792802, EPI_ISL_792803, EPI_ISL_792804, EPI_ISL_792805, EPI_ISL_792806, EPI_ISL_792807, EPI_ISL_792808, EPI_ISL_792809, EPI_ISL_792810, EPI_ISL_792811, EPI_ISL_792812, EPI_ISL_792813, EPI_ISL_792814, EPI_ISL_792815, EPI_ISL_792816, EPI_ISL_792817, EPI_ISL_792818, EPI_ISL_792819, EPI_ISL_792820, EPI_ISL_792821, EPI_ISL_792822, EPI_ISL_792823, EPI_ISL_792824, EPI_ISL_792829, EPI_ISL_792830, EPI_ISL_792834, EPI_ISL_792835, EPI_ISL_792840, EPI_ISL_792841, EPI_ISL_792842, EPI_ISL_792843, EPI_ISL_792844, EPI_ISL_792845, EPI_ISL_792846, EPI_ISL_792847, EPI_ISL_792848, EPI_ISL_792849, EPI_ISL_792850, EPI_ISL_792851, EPI_ISL_792852, EPI_ISL_792853, EPI_ISL_792854, EPI_ISL_792855, EPI_ISL_792856, EPI_ISL_792857, EPI_ISL_792858, EPI_ISL_792859, EPI_ISL_792860, EPI_ISL_792861, EPI_ISL_792862, EPI_ISL_792863, EPI_ISL_792864, EPI_ISL_792865, EPI_ISL_792866, EPI_ISL_792867, EPI_ISL_792868, EPI_ISL_792869, EPI_ISL_792870, EPI_ISL_792871, EPI_ISL_792872, EPI_ISL_792873, EPI_ISL_792874, EPI_ISL_792875, EPI_ISL_792876, EPI_ISL_792877, EPI_ISL_792878, EPI_ISL_792879, EPI_ISL_792880, EPI_ISL_792881, EPI_ISL_792882, EPI_ISL_792883, EPI_ISL_792884, EPI_ISL_792885, EPI_ISL_792886, EPI_ISL_792887, EPI_ISL_792888, EPI_ISL_792889, EPI_ISL_792890, EPI_ISL_792891, EPI_ISL_792892, EPI_ISL_792893, EPI_ISL_792894, EPI_ISL_792895, EPI_ISL_792896, EPI_ISL_792897, EPI_ISL_792898, EPI_ISL_792899, EPI_ISL_792900, EPI_ISL_792901, EPI_ISL_792902, EPI_ISL_792903, EPI_ISL_792904, EPI_ISL_792905, EPI_ISL_792906, EPI_ISL_792907, EPI_ISL_792908, EPI_ISL_792909, EPI_ISL_792910, EPI_ISL_792911, EPI_ISL_792912, EPI_ISL_792913, EPI_ISL_792914, EPI_ISL_792915, EPI_ISL_792916, EPI_ISL_792917, EPI_ISL_792918, EPI_ISL_792919, EPI_ISL_792920, EPI_ISL_792921, EPI_ISL_792922, EPI_ISL_792923, EPI_ISL_792924, EPI_ISL_792925, EPI_ISL_792926, EPI_ISL_792927, EPI_ISL_792928, EPI_ISL_792929, EPI_ISL_792930, EPI_ISL_792931, EPI_ISL_792932, EPI_ISL_792933, EPI_ISL_792934, EPI_ISL_792935, EPI_ISL_792936, EPI_ISL_792937, EPI_ISL_792938, EPI_ISL_792939, EPI_ISL_792940, EPI_ISL_792941, EPI_ISL_792942, EPI_ISL_792943, EPI_ISL_792944, EPI_ISL_792945, EPI_ISL_792946, EPI_ISL_792947, EPI_ISL_792948, EPI_ISL_792949, EPI_ISL_792950, EPI_ISL_792951, EPI_ISL_792952, EPI_ISL_792953, EPI_ISL_792954, EPI_ISL_792955, EPI_ISL_792956, EPI_ISL_792957, EPI_ISL_792958, EPI_ISL_792959, EPI_ISL_792960, EPI_ISL_792961, EPI_ISL_792962, EPI_ISL_792963, EPI_ISL_792964, EPI_ISL_792965, EPI_ISL_792966, EPI_ISL_792967, EPI_ISL_792968, EPI_ISL_792969, EPI_ISL_792970, EPI_ISL_792971, EPI_ISL_792972, EPI_ISL_792973, EPI_ISL_792974, EPI_ISL_792975, EPI_ISL_792976, EPI_ISL_792977, EPI_ISL_792978, EPI_ISL_792979, EPI_ISL_793111, EPI_ISL_793112, EPI_ISL_793113, EPI_ISL_793114, EPI_ISL_793146, EPI_ISL_793155, EPI_ISL_793156, EPI_ISL_793157, EPI_ISL_793162, EPI_ISL_793178, EPI_ISL_793201, EPI_ISL_793248, EPI_ISL_793259, EPI_ISL_793268, EPI_ISL_793278, EPI_ISL_793292, EPI_ISL_793315, EPI_ISL_793368, EPI_ISL_793369, EPI_ISL_793380, EPI_ISL_793990, EPI_ISL_793991, EPI_ISL_793992, EPI_ISL_793993, EPI_ISL_793994, EPI_ISL_793995, EPI_ISL_793996, EPI_ISL_793997, EPI_ISL_793998, EPI_ISL_793999, EPI_ISL_794000, EPI_ISL_794001, EPI_ISL_794002, EPI_ISL_794003, EPI_ISL_794004, EPI_ISL_794005, EPI_ISL_794006, EPI_ISL_794007, EPI_ISL_794008, EPI_ISL_794009, EPI_ISL_794010, EPI_ISL_794011, EPI_ISL_794012, EPI_ISL_794013 | | | |
| see above | Department of Virus and Microbiological Special Diagnostics, Statens Serum Institut, Copenhagen, Denmark | Albertsen Lab, Department of Chemistry and Bioscience, Aalborg University, Denmark | Danish Covid-19 Genome Consortium |
| EPI_ISL_794297 | Columbia University Irving Medical Center | Wadsworth Center, New York State Department.of Health | Kirsten St. George, Daryl M. Lamson, Alexis Russel, Matthew Shudt, Melissa A Leisner, Jonathan Plitnick, Navjot Singh, John Kelly, Sara Griesemer, Erasmus Schneider, Erica Lasek-Nesselquist |
| EPI_ISL_794326, EPI_ISL_794327, EPI_ISL_794328, EPI_ISL_794339, EPI_ISL_794348, EPI_ISL_794354, EPI_ISL_794355, EPI_ISL_794356, EPI_ISL_794357, EPI_ISL_794358, EPI_ISL_794359, EPI_ISL_794360, EPI_ISL_794368, EPI_ISL_794379, EPI_ISL_794380, EPI_ISL_794381, EPI_ISL_794382, EPI_ISL_794390, EPI_ISL_794391, EPI_ISL_794392, EPI_ISL_794393, EPI_ISL_794394, EPI_ISL_794395, EPI_ISL_794396, EPI_ISL_794397, EPI_ISL_794422, EPI_ISL_794428, EPI_ISL_794429, EPI_ISL_794432, EPI_ISL_794433, EPI_ISL_794436, EPI_ISL_794438, EPI_ISL_794439, EPI_ISL_794449, EPI_ISL_794450, EPI_ISL_794451, EPI_ISL_794452, EPI_ISL_794453, EPI_ISL_794454, EPI_ISL_794455, EPI_ISL_794456, EPI_ISL_794457, EPI_ISL_794458, EPI_ISL_794459, EPI_ISL_794460, EPI_ISL_794461, EPI_ISL_794462, EPI_ISL_794463, EPI_ISL_794464, EPI_ISL_794465, EPI_ISL_794466, EPI_ISL_794467, EPI_ISL_794468, EPI_ISL_794469, EPI_ISL_794470, EPI_ISL_794471, EPI_ISL_794472, EPI_ISL_794473, EPI_ISL_794474, EPI_ISL_794475, EPI_ISL_794476, EPI_ISL_794477, EPI_ISL_794478, EPI_ISL_794479, EPI_ISL_794480, EPI_ISL_794481, EPI_ISL_794482, EPI_ISL_794483, EPI_ISL_794484, EPI_ISL_794485, EPI_ISL_794486, EPI_ISL_794487, EPI_ISL_794488, EPI_ISL_794489, EPI_ISL_794490, EPI_ISL_794491, EPI_ISL_794492, EPI_ISL_794493, EPI_ISL_794494, EPI_ISL_794495 | | | |
| see above | Department of Virus and Microbiological Special Diagnostics, Statens Serum Institut, Copenhagen, Denmark | Albertsen Lab, Department of Chemistry and Bioscience, Aalborg University, Denmark | Danish Covid-19 Genome Consortium |
| EPI_ISL_794726, EPI_ISL_794734 | PathWest Laboratory Medicine WA | PathWest Laboratory Medicine WA Microbial Surveillance Unit | PathWest Laboratory Medicine WA Microbial Surveillance Unit |
| EPI_ISL_794788, EPI_ISL_794789, EPI_ISL_794790, EPI_ISL_794793, EPI_ISL_794803, EPI_ISL_794804, EPI_ISL_794805, EPI_ISL_794806, EPI_ISL_794807, EPI_ISL_794808 | Istituto Zooprofilattico Sperimentale della Puglia e della Basilicata | Istituto Zooprofilattico Sperimentale della Puglia e della Basilicata | Parisi A., Bianco A., Capozzi L., Del Sambro L., Manzulli V, Rondinone V., Pace L., Cipolletta D., Galante D. |
| EPI_ISL_794831, EPI_ISL_794832, EPI_ISL_794837, EPI_ISL_794838, EPI_ISL_794839, EPI_ISL_794840, EPI_ISL_794841, EPI_ISL_794842, EPI_ISL_794843, EPI_ISL_794844, EPI_ISL_794845, EPI_ISL_794846, EPI_ISL_794847, EPI_ISL_794848, EPI_ISL_794849, EPI_ISL_794861, EPI_ISL_794863, EPI_ISL_794865, EPI_ISL_794866, EPI_ISL_794867, EPI_ISL_794869, EPI_ISL_794873, EPI_ISL_794875, EPI_ISL_794877, EPI_ISL_794879, EPI_ISL_794881, EPI_ISL_794882, EPI_ISL_794884, EPI_ISL_794889, EPI_ISL_794890, EPI_ISL_794891, EPI_ISL_794892, EPI_ISL_794893, EPI_ISL_794894, EPI_ISL_794904, EPI_ISL_794930, EPI_ISL_794931, EPI_ISL_794932, EPI_ISL_794933, EPI_ISL_794934, EPI_ISL_794935, EPI_ISL_794936, EPI_ISL_794937, EPI_ISL_794938, EPI_ISL_794939, EPI_ISL_794940, EPI_ISL_794941, EPI_ISL_794942, EPI_ISL_794943, EPI_ISL_794944, EPI_ISL_794945, EPI_ISL_794946, EPI_ISL_794947, EPI_ISL_794948, EPI_ISL_794949, EPI_ISL_794950, EPI_ISL_794951, EPI_ISL_794952, EPI_ISL_794953, EPI_ISL_794954, EPI_ISL_794955, EPI_ISL_794956, EPI_ISL_794957, EPI_ISL_794958, EPI_ISL_794959, EPI_ISL_794960, EPI_ISL_794961, EPI_ISL_794962, EPI_ISL_794963, EPI_ISL_794964, EPI_ISL_794965, EPI_ISL_794966, EPI_ISL_794967, EPI_ISL_794968, EPI_ISL_794969, EPI_ISL_794970, EPI_ISL_794971, EPI_ISL_794972, EPI_ISL_794973, EPI_ISL_794974, EPI_ISL_794975, EPI_ISL_794976, EPI_ISL_794977, EPI_ISL_794978, EPI_ISL_794979, EPI_ISL_794980, EPI_ISL_794981, EPI_ISL_794982 | | | |
| see above | Department of Virus and Microbiological Special Diagnostics, Statens Serum Institut, Copenhagen, Denmark | Albertsen Lab, Department of Chemistry and Bioscience, Aalborg University, Denmark | Danish Covid-19 Genome Consortium |
| EPI_ISL_796651 | Medical Microbiology Unit, Department for Laboratory Medicine, Drammen Hospital, Vestre Viken Health Trust, | Norwegian Institute of Public Health, Department of Virology | Kathrine Stene-Johansen, Kamilla Heddeland Instefjord, Hilde Elshaug, Atiya R Ali,Marie Paulsen Madsen, Rasmus Riis Kopperud, Hilde Vollan, Karoline Bragstad, Olav Hungnes |
| EPI_ISL_796683 | Department of Medical Microbiology - section Molde, Molde Hospital | Norwegian Institute of Public Health, Department of Virology | Kathrine Stene-Johansen, Kamilla Heddeland Instefjord, Hilde Elshaug, Atiya R Ali,Marie Paulsen Madsen, Rasmus Riis Kopperud, Hilde Vollan, Karoline Bragstad, Olav Hungnes |
| EPI_ISL_796686 | Unilabs Laboratory Medicine | Norwegian Institute of Public Health, Department of Virology | Kathrine Stene-Johansen, Kamilla Heddeland Instefjord, Hilde Elshaug, Atiya R Ali,Marie Paulsen Madsen, Rasmus Riis Kopperud, Hilde Vollan, Karoline Bragstad, Olav Hungnes |
| EPI_ISL_796688 | Hospital of Southern Norway - Kristiansand, Department of Medical Microbiology | Norwegian Institute of Public Health, Department of Virology | Kathrine Stene-Johansen, Kamilla Heddeland Instefjord, Hilde Elshaug, Atiya R Ali,Marie Paulsen Madsen, Rasmus Riis Kopperud, Hilde Vollan, Karoline Bragstad, Olav Hungnes |
| EPI_ISL_796700, EPI_ISL_796701 | Unilabs Laboratory Medicine | Norwegian Institute of Public Health, Department of Virology | Kathrine Stene-Johansen, Kamilla Heddeland Instefjord, Hilde Elshaug, Atiya R Ali,Marie Paulsen Madsen, Rasmus Riis Kopperud, Hilde Vollan, Karoline Bragstad, Olav Hungnes |
| EPI_ISL_796715 | Hospital of Southern Norway - Kristiansand, Department of Medical Microbiology | Norwegian Institute of Public Health, Department of Virology | Kathrine Stene-Johansen, Kamilla Heddeland Instefjord, Hilde Elshaug, Atiya R Ali,Marie Paulsen Madsen, Rasmus Riis Kopperud, Hilde Vollan, Karoline Bragstad, Olav Hungnes |
| EPI_ISL_796717, EPI_ISL_796718 | Department of Medical Microbiology, St. Olavs hospital | Norwegian Institute of Public Health, Department of Virology | Kathrine Stene-Johansen, Kamilla Heddeland Instefjord, Hilde Elshaug, Atiya R Ali,Marie Paulsen Madsen, Rasmus Riis Kopperud, Hilde Vollan, Karoline Bragstad, Olav Hungnes |
| EPI_ISL_796722, EPI_ISL_796723 | Furst Medical Laboratory | Norwegian Institute of Public Health, Department of Virology | Kathrine Stene-Johansen, Kamilla Heddeland Instefjord, Hilde Elshaug, Atiya R Ali,Marie Paulsen Madsen, Rasmus Riis Kopperud, Hilde Vollan, Karoline Bragstad, Olav Hungnes |
| EPI_ISL_796729 | Hospital of Southern Norway - Kristiansand, Department of Medical Microbiology | Norwegian Institute of Public Health, Department of Virology | Kathrine Stene-Johansen, Kamilla Heddeland Instefjord, Hilde Elshaug, Atiya R Ali,Marie Paulsen Madsen, Rasmus Riis Kopperud, Hilde Vollan, Karoline Bragstad, Olav Hungnes |

| | | | |
|---|---|---|---|
| EPI_ISL_796780 | Instituto Nacional de Saude (INSA) | Instituto Nacional de Saude (INSA) | Borges et al |
| EPI_ISL_799710, EPI_ISL_799900 | Lighthouse Lab in Milton Keynes | Wellcome Sanger Institute for the COVID-19 Genomics UK (COG-UK) Consortium | The Lighthouse Lab in Milton Keynes and Alex Alderton, Roberto Amato, Sonia Goncalves, Ewan Harrison, David K. Jackson, Ian Johnston, Dominic Kwiatkowski, Cordelia Langford, John Sillitoe on behalf of the Wellcome Sanger Institute COVID-19 Surveillance Team |
| EPI_ISL_799936, EPI_ISL_799971, EPI_ISL_799972, EPI_ISL_799993, EPI_ISL_799994, EPI_ISL_800054, EPI_ISL_800100, EPI_ISL_800139, EPI_ISL_800192, EPI_ISL_800225, EPI_ISL_800244 | | | |
| see above | Lighthouse Lab in Cambridge | Wellcome Sanger Institute for the COVID-19 Genomics UK (COG-UK) Consortium | Rob Howes, The Lighthouse Lab in Cambridge and Alex Alderton, Roberto Amato, Sonia Goncalves, Ewan Harrison, David K. Jackson, Ian Johnston, Dominic Kwiatkowski, Cordelia Langford, John Sillitoe on behalf of the Wellcome Sanger Institute COVID-19 Surveillance Team |
| EPI_ISL_801443, EPI_ISL_801448 | Dutch COVID-19 response team | Erasmus Medical Center | Bas Oude Munnink, Reina Sikkema, David Nieuwenhuijse, Irina Chestakova, Anne van der Linden, Marjan Boter, Emmanuelle Munger, Corine GeurtsvanKessel, Annemiek van der Eijk, Richard Molenkamp, Marion Koopmans, on behalf of the Dutch national COVID-19 response team. |
| EPI_ISL_801524, EPI_ISL_801525, EPI_ISL_801526, EPI_ISL_801527, EPI_ISL_801528 | Sonic Reference Laboratory | Pathogen Discovery, Respiratory Viruses Branch, Division of Viral Diseases, Centers for Disease Control and Prevention | Ying Tao, Yan Li, Jing Zhang, Krista Queen, Anna Uehara, Peter Cook, Clinton R. Paden, Haibin Wang, Suxiang Tong |
| EPI_ISL_802512 | Dutch COVID-19 response team | Erasmus Medical Center | Bas Oude Munnink, Reina Sikkema, David Nieuwenhuijse, Irina Chestakova, Anne van der Linden, Marjan Boter, Emmanuelle Munger, Corine GeurtsvanKessel, Annemiek van der Eijk, Richard Molenkamp, Marion Koopmans, on behalf of the Dutch national COVID-19 response team. |
| EPI_ISL_802921, EPI_ISL_802923, EPI_ISL_802924, EPI_ISL_802925, EPI_ISL_802927, EPI_ISL_802929, EPI_ISL_802931, EPI_ISL_802932, EPI_ISL_802934, EPI_ISL_802936, EPI_ISL_802938, EPI_ISL_802941, EPI_ISL_802944, EPI_ISL_802945, EPI_ISL_802946, EPI_ISL_802948, EPI_ISL_802949, EPI_ISL_802953, EPI_ISL_802954, EPI_ISL_802956, EPI_ISL_802959, EPI_ISL_802960, EPI_ISL_802961, EPI_ISL_802964, EPI_ISL_802965, EPI_ISL_802967, EPI_ISL_802968, EPI_ISL_802970, EPI_ISL_802972, EPI_ISL_802973, EPI_ISL_802974, EPI_ISL_802975, EPI_ISL_802976, EPI_ISL_802978, EPI_ISL_802979, EPI_ISL_802980, EPI_ISL_802987, EPI_ISL_802991 | | | |
| see above | Utah Public Health Laboratory | Utah Public Health Laboratory | Erin Young, Kelly Oakeson, Tara Gallagher |
| EPI_ISL_803899 | Genomic Medicine Laboratory, IRCCS Santa Lucia Foundation | National Institute for Infectious Diseases, INMI, "L. Spallanzani" IRCCS | E Giombini, M. Rueca, B Bartolini, O Butera, C.E.M Gruber, F Messina, E Giardina, MR Capobianchi, A Di Caro |
| EPI_ISL_804214, EPI_ISL_804215 | Respiratory Virus Unit, National Infection Service, Public Health England | COVID-19 Genomics UK (COG-UK) Consortium | PHE Covid Sequencing Team |
| EPI_ISL_804604, EPI_ISL_804636, EPI_ISL_804642 | Michigan Department of Health and Human Services, Bureau of Laboratories | Michigan Department of Health and Human Services, Bureau of Laboratories | Blankenship HM, Riner D, Soehnlen MK |
| EPI_ISL_804825, EPI_ISL_804827 | DB Diagnosticos do Brasil | Laboratório de Parasitologia Médica - Instituto de Medicina Tropical - Universidade de São Paulo | Nuno Faria, Ingra Morales Claro, Darlan Candido, Lucas A. Moyses Franco, Pamela dos Santos Andrade, Thais de Moura Coletti, Camila A. Maia da Silva, Flavia Cristina Sales, Erika Regina Manuli, Renato A. Santana, Nelson Gaburo, Cecília da Cunha Camilo, Nelson Abrahim Fraiji, Myuki Alfaia Esashika Crispim, Maria do Perpétuo Socorro Sampaio Carvalho, Andrew Rambaut, Nick Loman, Oliver G. Pybus, Ester C. Sabino; DB; HEMOAM; CDL; CADDE Genomic Network. |
| EPI_ISL_804847, EPI_ISL_804870, EPI_ISL_804883, EPI_ISL_804889, EPI_ISL_804935, EPI_ISL_804936, EPI_ISL_804937, EPI_ISL_804938, EPI_ISL_804939, EPI_ISL_804940 | DC Public Health Lab/ Dept. of Forensic Sciences | DC Public Health Lab/ Dept. of Forensic Sciences | Scott Nguyen, Elizabeth Zelaya, Connie Maza, Monica Mann, Brittany Hamilton, David Payne, Jocelyn Hauser |
| EPI_ISL_806851 | Alaska State Virology Laboratory | Alaska State Virology Laboratory | Stephanie DeRonde, Lisa Smith, Ph.D., Devin M. Drown, Ph.D., Jack Chen, Ph.D. |
| EPI_ISL_812360, EPI_ISL_812361, EPI_ISL_812367, EPI_ISL_812368, EPI_ISL_812369, EPI_ISL_812370, EPI_ISL_812371, EPI_ISL_812372, EPI_ISL_812373, EPI_ISL_812374, EPI_ISL_812375, EPI_ISL_812376, EPI_ISL_812377, EPI_ISL_812378, EPI_ISL_812379, EPI_ISL_812380, EPI_ISL_812381, EPI_ISL_812384, EPI_ISL_812385, EPI_ISL_812386, EPI_ISL_812387, EPI_ISL_812388, EPI_ISL_812389, EPI_ISL_812390, EPI_ISL_812391, EPI_ISL_812392, EPI_ISL_812397, EPI_ISL_812398, EPI_ISL_812399, EPI_ISL_812400, EPI_ISL_812401, EPI_ISL_812407, EPI_ISL_812408, EPI_ISL_812409, EPI_ISL_812410, EPI_ISL_812411, EPI_ISL_812412, EPI_ISL_812413, EPI_ISL_812414, EPI_ISL_812415, EPI_ISL_812416, EPI_ISL_812417, EPI_ISL_812418, EPI_ISL_812419, EPI_ISL_812420 | | | |
| see above | Utah Public Health Laboratory | Utah Public Health Laboratory | Erin L. Young, Kelly F. Oakeson, Tara Gallagher |
| EPI_ISL_812722, EPI_ISL_812723, EPI_ISL_812724 | DOHMH PHL | New York City Public Health Laboratory | Jade Wang, et al. |
| EPI_ISL_812725 | DOHMH Crown Heights | New York City Public Health Laboratory | Jade Wang, et al. |
| EPI_ISL_812726 | DOHMH Corona | New York City Public Health Laboratory | Jade Wang, et al. |
| EPI_ISL_812727 | DOHMH Jamaica | New York City Public Health Laboratory | Jade Wang, et al. |
| EPI_ISL_812728 | DOHMH Riverside | New York City Public Health Laboratory | Jade Wang, et al. |
| EPI_ISL_812729 | DOHMH Central Harlem | New York City Public Health Laboratory | Jade Wang, et al. |
| EPI_ISL_812730 | DOHMH Crown Heights | New York City Public Health Laboratory | Jade Wang, et al. |
| EPI_ISL_812731 | DOHMH Jamaica | New York City Public Health Laboratory | Jade Wang, et al. |
| EPI_ISL_812732, EPI_ISL_812733 | DOHMH Corona | New York City Public Health Laboratory | Jade Wang, et al. |
| EPI_ISL_812734 | DOHMH Crown Heights | New York City Public Health Laboratory | Jade Wang, et al. |
| EPI_ISL_812735, EPI_ISL_812736, EPI_ISL_812737 | DOHMH Morrisania | New York City Public Health Laboratory | Jade Wang, et al. |
| EPI_ISL_812738 | DOHMH Crown Heights | New York City Public Health Laboratory | Jade Wang, et al. |
| EPI_ISL_812739 | DOHMH Central Harlem | New York City Public Health Laboratory | Jade Wang, et al. |
| EPI_ISL_812740 | DOHMH Corona | New York City Public Health Laboratory | Jade Wang, et al. |
| EPI_ISL_812741 | DOHMH Central Harlem | New York City Public Health Laboratory | Jade Wang, et al. |
| EPI_ISL_812742, EPI_ISL_812743, EPI_ISL_812744 | DOHMH Corona | New York City Public Health Laboratory | Jade Wang, et al. |
| EPI_ISL_813060, EPI_ISL_813061, EPI_ISL_813063, EPI_ISL_813082, EPI_ISL_813084, EPI_ISL_813088, EPI_ISL_813089, EPI_ISL_813090, EPI_ISL_813153 | University of Birmingham | COVID-19 Genomics UK (COG-UK) Consortium | Institute of Microbiology, University of Birmingham: Claire McMurray, Joanne Stockton, Samuel Nicholls, Radoslaw Poplawski, Will Rowe, Josh Quick, Nicholas Loman. University of Birmingham Testing Laboratory: Celina M Whalley, Andrew Bosworth, Charlotte Poxon, Kasun Wanigasooriya, Oliver Pickles, Mike Kidd, Alex Richter, Andrew D Beggs PHE Heartlands Lab: Husam Osman, Andrew Bosworth. Queen Elizabeth Hospital: Anna Casey |
| EPI_ISL_813225, EPI_ISL_813226, EPI_ISL_813239, EPI_ISL_813243, EPI_ISL_813246, EPI_ISL_813252, EPI_ISL_813254, EPI_ISL_813255, EPI_ISL_813262, EPI_ISL_813264, EPI_ISL_813273, EPI_ISL_813274, EPI_ISL_813275, EPI_ISL_813276, EPI_ISL_813277, EPI_ISL_813279, EPI_ISL_813286, EPI_ISL_813288, EPI_ISL_813290, EPI_ISL_813292 | | | |
| see above | Department of Pathology, University of Cambridge | COVID-19 Genomics UK (COG-UK) Consortium | Aminu S. Jahun, Yasmin Chaudhry, Grant Hall, Iliana Georgana, Myra Hosmillo, Martin D. Curran, Malte Pinckert, Surendra Parmar, Ian Goodfellow |
| EPI_ISL_813771, EPI_ISL_813772, EPI_ISL_813809 | Liverpool Clinical Laboratories | COVID-19 Genomics UK (COG-UK) Consortium | Sam Haldenby, Anita Lucaci, Steve Paterson, Julian Hiscox, Alistair Darby, M Almsaud, A Alrezaihi, Muhannad Alruwaili, Stuart D Armstrong, Jones Benjamin, Eleanor G Bentley, Anu Chawla, Jordan J Clark, Angela Cowell, Richard Eccles, Isabel García-Dorival, Matthew Gemmell, Alessandro Gerada, PKF Gilmore, Richard Gregory, Ximeng Han, Catherine Hartley, Margaret Hughes, Miren Iturriza-Gomara, James Johnson, L Luu, Jenifer Manson, Charlotte Nelson, Elaine O'Toole, Cassie Olateju, Rebekah Penrice-Randal , Lucille Rainbow, N.P Randle, Trevor Ian Robinson, Parul Sharma, Ghada T Shawli, James P Stewart, Neil Swainston, Ecaterina Vamos, Joanne Watts, Mark Whitehead |

| | | | |
|---|---|---|---|
| EPI_ISL_814274, EPI_ISL_814277, EPI_ISL_814278, EPI_ISL_814279, EPI_ISL_814282 | Wales Specialist Virology Centre Sequencing lab: Pathogen Genomics Unit | COVID-19 Genomics UK (COG-UK) Consortium | Catherine Moore, Johnathan Evans, Laura Gifford, Malorie Perry, Simon Cottrell, Angela Marchbank, Alec Birchley, Alexander Adams, Amy Gaskin, Bree Gatica-Wilcox, Jason Coombes, Joel Southgate, Lauren Gilbert, Lee Graham, Nicole Pacchiarini, Sara Kumziene-Summerhayes, Sarah Taylor, Sophie Jones, Sara Rey, Matthew Bull, Joanne Watkins, Sally Corden, Tom Connor |
| EPI_ISL_814293 | Liverpool Clinical Laboratories | COVID-19 Genomics UK (COG-UK) Consortium | Sam Haldenby, Anita Lucaci, Steve Paterson, Julian Hiscox, Alistair Darby, M Almsaud, A Alrezaihi, Muhannad Alruwaili, Stuart D Armstrong, Jones Benjamin, Eleanor G Bentley, Anu Chawla, Jordan J Clark, Angela Cowell, Richard Eccles, Isabel García-Dorival, Matthew Gemmell, Alessandro Gerada, PKF Gilmore, Richard Gregory, Ximeng Han, Catherine Hartley, Margaret Hughes, Miren Iturriza-Gomara, James Johnson, L Luu, Jenifer Manson, Charlotte Nelson, Elaine O'Toole, Cassie Olateju, Rebekah Penrice-Randal , Lucille Rainbow, N.P Randle, Trevor Ian Robinson, Parul Sharma, Ghada T Shawli, James P Stewart, Neil Swainston, Ecaterina Vamos, Joanne Watts, Mark Whitehead |
| EPI_ISL_814294, EPI_ISL_814295, EPI_ISL_814296, EPI_ISL_814300, EPI_ISL_814304, EPI_ISL_814306, EPI_ISL_814309 | Wales Specialist Virology Centre Sequencing lab: Pathogen Genomics Unit | COVID-19 Genomics UK (COG-UK) Consortium | Catherine Moore, Johnathan Evans, Laura Gifford, Malorie Perry, Simon Cottrell, Angela Marchbank, Alec Birchley, Alexander Adams, Amy Gaskin, Bree Gatica-Wilcox, Jason Coombes, Joel Southgate, Lauren Gilbert, Lee Graham, Nicole Pacchiarini, Sara Kumziene-Summerhayes, Sarah Taylor, Sophie Jones, Sara Rey, Matthew Bull, Joanne Watkins, Sally Corden, Tom Connor |
| EPI_ISL_814312 | West of Scotland Specialist Virology Centre, NHSGGC / MRC-University of Glasgow Centre for Virus Research | COVID-19 Genomics UK (COG-UK) Consortium | Ana da Silva Filipe, Natasha Johnson, Kathy Smollett, Daniel Mair, Stephen Carmichael, Alice Broos, Lily Tong, Jenna Nichols, Kyriaki Nomikou; Sarah McDonald; Richard Orton, Joseph Hughes, Sreenu Vattipally, David L Robertson; Alasdair MacLean, Rory Gunson; Sharif Shaaban, Matthew Holden; Rachel Blacow, Guy Mollett, Kathy Li, James Shepherd, Antonia Ho, Emma Thomson |
| EPI_ISL_814319, EPI_ISL_814328, EPI_ISL_814337, EPI_ISL_814356, EPI_ISL_814363, EPI_ISL_814367, EPI_ISL_814369, EPI_ISL_814371, EPI_ISL_814372, EPI_ISL_814373, EPI_ISL_814377, EPI_ISL_814383, EPI_ISL_814386, EPI_ISL_814387, EPI_ISL_814389, EPI_ISL_814394, EPI_ISL_814395, EPI_ISL_814400, EPI_ISL_814411, EPI_ISL_814413, EPI_ISL_814415 | | | |
| see above | Wales Specialist Virology Centre Sequencing lab: Pathogen Genomics Unit | COVID-19 Genomics UK (COG-UK) Consortium | Catherine Moore, Johnathan Evans, Laura Gifford, Malorie Perry, Simon Cottrell, Angela Marchbank, Alec Birchley, Alexander Adams, Amy Gaskin, Bree Gatica-Wilcox, Jason Coombes, Joel Southgate, Lauren Gilbert, Lee Graham, Nicole Pacchiarini, Sara Kumziene-Summerhayes, Sarah Taylor, Sophie Jones, Sara Rey, Matthew Bull, Joanne Watkins, Sally Corden, Tom Connor |
| EPI_ISL_814449, EPI_ISL_814457 | West of Scotland Specialist Virology Centre, NHSGGC / MRC-University of Glasgow Centre for Virus Research | COVID-19 Genomics UK (COG-UK) Consortium | Ana da Silva Filipe, Natasha Johnson, Kathy Smollett, Daniel Mair, Stephen Carmichael, Alice Broos, Lily Tong, Jenna Nichols, Kyriaki Nomikou; Sarah McDonald; Richard Orton, Joseph Hughes, Sreenu Vattipally, David L Robertson; Alasdair MacLean, Rory Gunson; Sharif Shaaban, Matthew Holden; Rachel Blacow, Guy Mollett, Kathy Li, James Shepherd, Antonia Ho, Emma Thomson |
| EPI_ISL_814473, EPI_ISL_814474, EPI_ISL_814475, EPI_ISL_814476, EPI_ISL_814477, EPI_ISL_814478, EPI_ISL_814479, EPI_ISL_814511, EPI_ISL_814523, EPI_ISL_814529 | Wales Specialist Virology Centre Sequencing lab: Pathogen Genomics Unit | COVID-19 Genomics UK (COG-UK) Consortium | Catherine Moore, Johnathan Evans, Laura Gifford, Malorie Perry, Simon Cottrell, Angela Marchbank, Alec Birchley, Alexander Adams, Amy Gaskin, Bree Gatica-Wilcox, Jason Coombes, Joel Southgate, Lauren Gilbert, Lee Graham, Nicole Pacchiarini, Sara Kumziene-Summerhayes, Sarah Taylor, Sophie Jones, Sara Rey, Matthew Bull, Joanne Watkins, Sally Corden, Tom Connor |
| EPI_ISL_814530 | Bioinformatics and Biostatistics Lab, Advanced Sequencing Facility | COVID-19 Genomics UK (COG-UK) Consortium | Aengus Stewart,Jerome Nicod,Chelsea Sawyer,Laura Cubitt,Harshil Patel,Margaret Crawford |
| EPI_ISL_814537 | Wales Specialist Virology Centre Sequencing lab: Pathogen Genomics Unit | COVID-19 Genomics UK (COG-UK) Consortium | Catherine Moore, Johnathan Evans, Laura Gifford, Malorie Perry, Simon Cottrell, Angela Marchbank, Alec Birchley, Alexander Adams, Amy Gaskin, Bree Gatica-Wilcox, Jason Coombes, Joel Southgate, Lauren Gilbert, Lee Graham, Nicole Pacchiarini, Sara Kumziene-Summerhayes, Sarah Taylor, Sophie Jones, Sara Rey, Matthew Bull, Joanne Watkins, Sally Corden, Tom Connor |
| EPI_ISL_814547 | West of Scotland Specialist Virology Centre, NHSGGC / MRC-University of Glasgow Centre for Virus Research | COVID-19 Genomics UK (COG-UK) Consortium | Ana da Silva Filipe, Natasha Johnson, Kathy Smollett, Daniel Mair, Stephen Carmichael, Alice Broos, Lily Tong, Jenna Nichols, Kyriaki Nomikou; Sarah McDonald; Richard Orton, Joseph Hughes, Sreenu Vattipally, David L Robertson; Alasdair MacLean, Rory Gunson; Sharif Shaaban, Matthew Holden; Rachel Blacow, Guy Mollett, Kathy Li, James Shepherd, Antonia Ho, Emma Thomson |
| EPI_ISL_814551, EPI_ISL_814556, EPI_ISL_814561, EPI_ISL_814573, EPI_ISL_814578, EPI_ISL_814584, EPI_ISL_814595, EPI_ISL_814606 | Wales Specialist Virology Centre Sequencing lab: Pathogen Genomics Unit | COVID-19 Genomics UK (COG-UK) Consortium | Catherine Moore, Johnathan Evans, Laura Gifford, Malorie Perry, Simon Cottrell, Angela Marchbank, Alec Birchley, Alexander Adams, Amy Gaskin, Bree Gatica-Wilcox, Jason Coombes, Joel Southgate, Lauren Gilbert, Lee Graham, Nicole Pacchiarini, Sara Kumziene-Summerhayes, Sarah Taylor, Sophie Jones, Sara Rey, Matthew Bull, Joanne Watkins, Sally Corden, Tom Connor |
| EPI_ISL_814629, EPI_ISL_814630, EPI_ISL_814648, EPI_ISL_814652, EPI_ISL_814659, EPI_ISL_814660, EPI_ISL_814662 | West of Scotland Specialist Virology Centre, NHSGGC / MRC-University of Glasgow Centre for Virus Research | COVID-19 Genomics UK (COG-UK) Consortium | Ana da Silva Filipe, Natasha Johnson, Kathy Smollett, Daniel Mair, Stephen Carmichael, Alice Broos, Lily Tong, Jenna Nichols, Kyriaki Nomikou; Sarah McDonald; Richard Orton, Joseph Hughes, Sreenu Vattipally, David L Robertson; Alasdair MacLean, Rory Gunson; Sharif Shaaban, Matthew Holden; Rachel Blacow, Guy Mollett, Kathy Li, James Shepherd, Antonia Ho, Emma Thomson |
| EPI_ISL_814689 | Oxford Viromics, NDM, University of Oxford; Oxford University Hospitals; Basingstoke and North Hampshire Hospital | COVID-19 Genomics UK (COG-UK) Consortium | Tanya Golubchik, David Bonsall, George Macintyre, Amy Trebes, Mariateresa de Cesare, Catrin Moore, Alex Mobbs, Anita Justice, Robert Shaw, Monique Andersson, Timothy Peto, Emma Wise, Nathan Moore, Jessica Lynch, Nick Cortes, Matilde Mori, Stephen Kidd, David Buck, John Todd, Christophe Fraser |
| EPI_ISL_814724, EPI_ISL_814727, EPI_ISL_814728, EPI_ISL_814729, EPI_ISL_814730, EPI_ISL_814731, EPI_ISL_814732, EPI_ISL_814733, EPI_ISL_814734, EPI_ISL_814735, EPI_ISL_814736, EPI_ISL_814737, EPI_ISL_814738, EPI_ISL_814739, EPI_ISL_814740, EPI_ISL_814741, EPI_ISL_814742, EPI_ISL_814743, EPI_ISL_814744, EPI_ISL_814745, EPI_ISL_814746, EPI_ISL_814747, EPI_ISL_814748, EPI_ISL_814749, EPI_ISL_814750, EPI_ISL_814751, EPI_ISL_814752, EPI_ISL_814798, EPI_ISL_814799, EPI_ISL_814871, EPI_ISL_814872, EPI_ISL_814873, EPI_ISL_814874, EPI_ISL_814875, EPI_ISL_814876, EPI_ISL_814877, EPI_ISL_814878, EPI_ISL_814879, EPI_ISL_814880, EPI_ISL_814881, EPI_ISL_814882, EPI_ISL_814883, EPI_ISL_814884, EPI_ISL_814885, EPI_ISL_814886, EPI_ISL_814887, EPI_ISL_814888, EPI_ISL_814889, EPI_ISL_814890, EPI_ISL_814891, EPI_ISL_814892, EPI_ISL_814893, EPI_ISL_814894, EPI_ISL_814895, EPI_ISL_814896, EPI_ISL_814897, EPI_ISL_814898, EPI_ISL_814899, EPI_ISL_814900, EPI_ISL_814901, EPI_ISL_814902, EPI_ISL_814903, EPI_ISL_814904, EPI_ISL_814905, EPI_ISL_814906, EPI_ISL_814907, EPI_ISL_814908, EPI_ISL_814909, EPI_ISL_814910, EPI_ISL_814911, EPI_ISL_814912, EPI_ISL_814913, EPI_ISL_814914, EPI_ISL_814915, EPI_ISL_814916, EPI_ISL_814917, EPI_ISL_814918, EPI_ISL_814919, EPI_ISL_814920, EPI_ISL_814921, EPI_ISL_814922, EPI_ISL_814923, EPI_ISL_814924, EPI_ISL_814925, EPI_ISL_814926, EPI_ISL_814927, EPI_ISL_814928, EPI_ISL_814929, EPI_ISL_814930, EPI_ISL_814931, EPI_ISL_814932, EPI_ISL_814933, EPI_ISL_814934, EPI_ISL_814935, EPI_ISL_814936, EPI_ISL_814937, EPI_ISL_814938, EPI_ISL_814939, EPI_ISL_814940, EPI_ISL_814942, EPI_ISL_814943, EPI_ISL_814944, EPI_ISL_814945, EPI_ISL_814946, EPI_ISL_814947, EPI_ISL_814948, EPI_ISL_814949, EPI_ISL_814950, EPI_ISL_814951, EPI_ISL_814952, EPI_ISL_814953, EPI_ISL_815035, EPI_ISL_815037, EPI_ISL_815048, EPI_ISL_815049, EPI_ISL_815050, EPI_ISL_815052, EPI_ISL_815053, EPI_ISL_815054, EPI_ISL_815055, EPI_ISL_815056, EPI_ISL_815057, EPI_ISL_815058, EPI_ISL_815059, EPI_ISL_815060, EPI_ISL_815061, EPI_ISL_815072, EPI_ISL_815073, EPI_ISL_815074, EPI_ISL_815075, EPI_ISL_815106, EPI_ISL_815107, EPI_ISL_815108, EPI_ISL_815109, EPI_ISL_815110, EPI_ISL_815111, EPI_ISL_815112, EPI_ISL_815113, EPI_ISL_815114, EPI_ISL_815115, EPI_ISL_815116, EPI_ISL_815117, EPI_ISL_815118, EPI_ISL_815119, EPI_ISL_815120, EPI_ISL_815121, EPI_ISL_815122, EPI_ISL_815123, EPI_ISL_815124, EPI_ISL_815125, EPI_ISL_815126, EPI_ISL_815127, EPI_ISL_815128, EPI_ISL_815129, EPI_ISL_815130, EPI_ISL_815131, EPI_ISL_815132, EPI_ISL_815133, EPI_ISL_815134, EPI_ISL_815135, EPI_ISL_815136, EPI_ISL_815247, EPI_ISL_815248, EPI_ISL_815249, EPI_ISL_815250, EPI_ISL_815251, EPI_ISL_815252, EPI_ISL_815253, EPI_ISL_815254 | | | |
| see above | Wales Specialist Virology Centre Sequencing lab: Pathogen Genomics Unit | COVID-19 Genomics UK (COG-UK) Consortium | Catherine Moore, Johnathan Evans, Laura Gifford, Malorie Perry, Simon Cottrell, Angela Marchbank, Alec Birchley, Alexander Adams, Amy Gaskin, Bree Gatica-Wilcox, Jason Coombes, Joel Southgate, Lauren Gilbert, Lee Graham, Nicole Pacchiarini, Sara Kumziene-Summerhayes, Sarah Taylor, Sophie Jones, Sara Rey, Matthew Bull, Joanne Watkins, Sally Corden, Tom Connor |
| EPI_ISL_815401, EPI_ISL_815407, EPI_ISL_815410, EPI_ISL_815441, EPI_ISL_815442, EPI_ISL_815443, EPI_ISL_815444, EPI_ISL_815445, EPI_ISL_815446, EPI_ISL_815447, EPI_ISL_815448, EPI_ISL_815449, EPI_ISL_815450, EPI_ISL_815451, EPI_ISL_815452, EPI_ISL_815453, EPI_ISL_815468, EPI_ISL_815469, EPI_ISL_815471, EPI_ISL_815475, EPI_ISL_815501, EPI_ISL_815523, EPI_ISL_815524, EPI_ISL_815526, EPI_ISL_815534, EPI_ISL_815535, EPI_ISL_815539, EPI_ISL_815544, EPI_ISL_815581, EPI_ISL_815582, EPI_ISL_815583, EPI_ISL_815584, EPI_ISL_815585, EPI_ISL_815586, EPI_ISL_815587, EPI_ISL_815588, EPI_ISL_815589, EPI_ISL_815590, EPI_ISL_815591, EPI_ISL_815606, EPI_ISL_815650, EPI_ISL_815656, EPI_ISL_815659, EPI_ISL_815962, EPI_ISL_815963, EPI_ISL_815964, EPI_ISL_815965, EPI_ISL_815966, EPI_ISL_815967, EPI_ISL_815968, EPI_ISL_815969, EPI_ISL_815970, EPI_ISL_815971, EPI_ISL_815972, EPI_ISL_815973, EPI_ISL_815974, EPI_ISL_815975, EPI_ISL_815976, EPI_ISL_815977, EPI_ISL_815978, EPI_ISL_815979, EPI_ISL_815980, EPI_ISL_815981, EPI_ISL_815982, EPI_ISL_815983, EPI_ISL_815984, EPI_ISL_815985, EPI_ISL_815986, EPI_ISL_815987, EPI_ISL_815988, EPI_ISL_815989, EPI_ISL_815990, EPI_ISL_815991, EPI_ISL_815992, EPI_ISL_815993, EPI_ISL_815994, EPI_ISL_815995, EPI_ISL_815996, EPI_ISL_815997, EPI_ISL_815998, EPI_ISL_815999, EPI_ISL_816000, EPI_ISL_816001, EPI_ISL_816002, EPI_ISL_816003, EPI_ISL_816004, EPI_ISL_816005, EPI_ISL_816006, EPI_ISL_816007, EPI_ISL_816008, EPI_ISL_816009, EPI_ISL_816010, EPI_ISL_816011, EPI_ISL_816012, EPI_ISL_816013, EPI_ISL_816014, EPI_ISL_816015, EPI_ISL_816016, EPI_ISL_816017, EPI_ISL_816018, EPI_ISL_816019, EPI_ISL_816020, EPI_ISL_816021, EPI_ISL_816022, EPI_ISL_816023, EPI_ISL_816024, EPI_ISL_816025, EPI_ISL_816026, EPI_ISL_816027, EPI_ISL_816028, EPI_ISL_816029, EPI_ISL_816030, EPI_ISL_816031, EPI_ISL_816032, EPI_ISL_816033, EPI_ISL_816034 | | | |
| see above | Department of Virus and Microbiological Special Diagnostics, Statens Serum Institut, Copenhagen, Denmark | Albertsen Lab, Department of Chemistry and Bioscience, Aalborg University, Denmark | Danish Covid-19 Genome Consortium |
| EPI_ISL_816234, EPI_ISL_816240, EPI_ISL_816245, EPI_ISL_816259, EPI_ISL_816265, EPI_ISL_816269, EPI_ISL_816274, EPI_ISL_816282, EPI_ISL_816300, EPI_ISL_816301, EPI_ISL_816304, EPI_ISL_816325, EPI_ISL_816329, EPI_ISL_816344, EPI_ISL_816347, EPI_ISL_816375, EPI_ISL_816385, EPI_ISL_816388, EPI_ISL_816400, EPI_ISL_816404, EPI_ISL_816410, EPI_ISL_816418, EPI_ISL_816460, EPI_ISL_816482, EPI_ISL_816486, EPI_ISL_816487, EPI_ISL_816504, EPI_ISL_816514, EPI_ISL_816528, EPI_ISL_816529, EPI_ISL_816535, EPI_ISL_816538, EPI_ISL_816541, EPI_ISL_816544, EPI_ISL_816547, EPI_ISL_816554, EPI_ISL_816556, EPI_ISL_816568, EPI_ISL_816595, EPI_ISL_816617, EPI_ISL_816626, EPI_ISL_816630, EPI_ISL_816640, EPI_ISL_816644, EPI_ISL_816650, EPI_ISL_816654, EPI_ISL_816656 | | | |
| see above | Virology Department, Sheffield Teaching Hospitals NHS Foundation Trust/Department of Infection, Immunity and Cardiovascular Disease, The Medical School, University of Sheffield | COVID-19 Genomics UK (COG-UK) Consortium | Thushan de Silva, Matthew Parker, Nikki Smith, Adri Angyal, Rebecca Brown, Luke Green, Rachel Tucker, Paul Parsons, Danielle Groves, Katie Johnson, Laura Carrilero, Alex Keeley, Dave Partridge, Matthew Wyles, Benjamin Lindsey, Mehmet Yavuz, Mohammad Raza, Cariad Evans |
| EPI_ISL_817025, EPI_ISL_817026, EPI_ISL_817027, EPI_ISL_817028, EPI_ISL_817029, EPI_ISL_817030, EPI_ISL_817031, EPI_ISL_817032, EPI_ISL_817033, EPI_ISL_817034, EPI_ISL_817035, EPI_ISL_817036, EPI_ISL_817037, EPI_ISL_817038, EPI_ISL_817040, EPI_ISL_817041, EPI_ISL_817042, EPI_ISL_817043, EPI_ISL_817045, EPI_ISL_817046, EPI_ISL_817047, EPI_ISL_817048, EPI_ISL_817052, EPI_ISL_817055, EPI_ISL_817056, EPI_ISL_817058, EPI_ISL_817059, EPI_ISL_817061, EPI_ISL_817064, EPI_ISL_817065, EPI_ISL_817122 | | | |

| | | | |
|---|---|---|---|
| see above | Bioinformatics and Biostatistics Lab, Advanced Sequencing Facility | COVID-19 Genomics UK (COG-UK) Consortium | Aengus Stewart,Jerome Nicod,Chelsea Sawyer,Laura Cubitt,Harshil Patel,Margaret Crawford |
| EPI_ISL_819578 | Oxford Viromics, NDM, University of Oxford; Oxford University Hospitals; Basingstoke and North Hampshire Hospital | COVID-19 Genomics UK (COG-UK) Consortium | Tanya Golubchik, David Bonsall, George Macintyre, Amy Trebes, Mariateresa de Cesare, Catrin Moore, Alex Mobbs, Anita Justice, Robert Shaw, Monique Andersson, Timothy Peto, Emma Wise, Nathan Moore, Jessica Lynch, Nick Cortes, Matilde Mori, Stephen Kidd, David Buck, John Todd, Christophe Fraser |
| EPI_ISL_819579, EPI_ISL_819580, EPI_ISL_819581, EPI_ISL_819582, EPI_ISL_819583, EPI_ISL_819584 | Queens Medical Centre, Clinical Microbiology Department / DeepSeq Nottingham | COVID-19 Genomics UK (COG-UK) Consortium | Gemma Clark, Wendy Smith, Manjinder Khakh, Vicki M Fleming, Michelle M Lister, Hannah Howson-Wells, Jonathan Ball, Patrick McClure, Joseph Chappell, Theocharis Tsoleridis, Nadine Holmes, Matthew Carlisle, Christopher Moore, Fei Sang, Johnny Debebe, Victoria Wright, Matthew Loose |
| EPI_ISL_819585, EPI_ISL_819586, EPI_ISL_819587, EPI_ISL_819588, EPI_ISL_819589, EPI_ISL_819590, EPI_ISL_819591, EPI_ISL_819592, EPI_ISL_819593 | Oxford Viromics, NDM, University of Oxford; Oxford University Hospitals; Basingstoke and North Hampshire Hospital | COVID-19 Genomics UK (COG-UK) Consortium | Tanya Golubchik, David Bonsall, George Macintyre, Amy Trebes, Mariateresa de Cesare, Catrin Moore, Alex Mobbs, Anita Justice, Robert Shaw, Monique Andersson, Timothy Peto, Emma Wise, Nathan Moore, Jessica Lynch, Nick Cortes, Matilde Mori, Stephen Kidd, David Buck, John Todd, Christophe Fraser |
| EPI_ISL_819594, EPI_ISL_819595, EPI_ISL_819596, EPI_ISL_819597, EPI_ISL_819598, EPI_ISL_819599, EPI_ISL_819600, EPI_ISL_819601, EPI_ISL_819602, EPI_ISL_819603, EPI_ISL_819604 | | | |
| see above | Queens Medical Centre, Clinical Microbiology Department / DeepSeq Nottingham | COVID-19 Genomics UK (COG-UK) Consortium | Gemma Clark, Wendy Smith, Manjinder Khakh, Vicki M Fleming, Michelle M Lister, Hannah Howson-Wells, Jonathan Ball, Patrick McClure, Joseph Chappell, Theocharis Tsoleridis, Nadine Holmes, Matthew Carlisle, Christopher Moore, Fei Sang, Johnny Debebe, Victoria Wright, Matthew Loose |
| EPI_ISL_819605, EPI_ISL_819606, EPI_ISL_819607, EPI_ISL_819608 | Oxford Viromics, NDM, University of Oxford; Oxford University Hospitals; Basingstoke and North Hampshire Hospital | COVID-19 Genomics UK (COG-UK) Consortium | Tanya Golubchik, David Bonsall, George Macintyre, Amy Trebes, Mariateresa de Cesare, Catrin Moore, Alex Mobbs, Anita Justice, Robert Shaw, Monique Andersson, Timothy Peto, Emma Wise, Nathan Moore, Jessica Lynch, Nick Cortes, Matilde Mori, Stephen Kidd, David Buck, John Todd, Christophe Fraser |
| EPI_ISL_820129, EPI_ISL_820130, EPI_ISL_820132, EPI_ISL_820133, EPI_ISL_820135, EPI_ISL_820137, EPI_ISL_820139, EPI_ISL_820140, EPI_ISL_820142, EPI_ISL_820143, EPI_ISL_820145 | | | |
| see above | Lighthouse Lab in Alderley Park | Wellcome Sanger Institute for the COVID-19 Genomics UK (COG-UK) Consortium | Jacquelyn Wynn, Mairead Hyland, The Lighthouse Lab in Alderley Park and Alex Alderton, Roberto Amato, Sonia Goncalves, Ewan Harrison, David K. Jackson, Ian Johnston, Dominic Kwiatkowski, Cordelia Langford, John Sillitoe on behalf of the Wellcome Sanger Institute COVID-19 Surveillance Team |
| EPI_ISL_820147 | Lighthouse Lab in Glasgow | Wellcome Sanger Institute for the COVID-19 Genomics UK (COG-UK) Consortium | Harper VanSteenhouse, Yumi Kasai, David Gray, Carol Clugston, Anna Dominiczak and Alex Alderton, Roberto Amato, Sonia Goncalves, Ewan Harrison, David K. Jackson, Ian Johnston, Dominic Kwiatkowski, Cordelia Langford, John Sillitoe on behalf of the Wellcome Sanger Institute COVID-19 Surveillance Team |
| EPI_ISL_820148 | Lighthouse Lab in Alderley Park | Wellcome Sanger Institute for the COVID-19 Genomics UK (COG-UK) Consortium | Jacquelyn Wynn, Mairead Hyland, The Lighthouse Lab in Alderley Park and Alex Alderton, Roberto Amato, Sonia Goncalves, Ewan Harrison, David K. Jackson, Ian Johnston, Dominic Kwiatkowski, Cordelia Langford, John Sillitoe on behalf of the Wellcome Sanger Institute COVID-19 Surveillance Team |
| EPI_ISL_820150, EPI_ISL_820152 | Lighthouse Lab in Glasgow | Wellcome Sanger Institute for the COVID-19 Genomics UK (COG-UK) Consortium | Harper VanSteenhouse, Yumi Kasai, David Gray, Carol Clugston, Anna Dominiczak and Alex Alderton, Roberto Amato, Sonia Goncalves, Ewan Harrison, David K. Jackson, Ian Johnston, Dominic Kwiatkowski, Cordelia Langford, John Sillitoe on behalf of the Wellcome Sanger Institute COVID-19 Surveillance Team |
| EPI_ISL_820153, EPI_ISL_820155, EPI_ISL_820157, EPI_ISL_820158, EPI_ISL_820160, EPI_ISL_820162, EPI_ISL_820163, EPI_ISL_820165, EPI_ISL_820166, EPI_ISL_820168, EPI_ISL_820170 | | | |
| see above | Lighthouse Lab in Alderley Park | Wellcome Sanger Institute for the COVID-19 Genomics UK (COG-UK) Consortium | Jacquelyn Wynn, Mairead Hyland, The Lighthouse Lab in Alderley Park and Alex Alderton, Roberto Amato, Sonia Goncalves, Ewan Harrison, David K. Jackson, Ian Johnston, Dominic Kwiatkowski, Cordelia Langford, John Sillitoe on behalf of the Wellcome Sanger Institute COVID-19 Surveillance Team |
| EPI_ISL_820171 | Lighthouse Lab in Glasgow | Wellcome Sanger Institute for the COVID-19 Genomics UK (COG-UK) Consortium | Harper VanSteenhouse, Yumi Kasai, David Gray, Carol Clugston, Anna Dominiczak and Alex Alderton, Roberto Amato, Sonia Goncalves, Ewan Harrison, David K. Jackson, Ian Johnston, Dominic Kwiatkowski, Cordelia Langford, John Sillitoe on behalf of the Wellcome Sanger Institute COVID-19 Surveillance Team |
| EPI_ISL_820173 | Lighthouse Lab in Alderley Park | Wellcome Sanger Institute for the COVID-19 Genomics UK (COG-UK) Consortium | Jacquelyn Wynn, Mairead Hyland, The Lighthouse Lab in Alderley Park and Alex Alderton, Roberto Amato, Sonia Goncalves, Ewan Harrison, David K. Jackson, Ian Johnston, Dominic Kwiatkowski, Cordelia Langford, John Sillitoe on behalf of the Wellcome Sanger Institute COVID-19 Surveillance Team |
| EPI_ISL_820176, EPI_ISL_820177 | Lighthouse Lab in Glasgow | Wellcome Sanger Institute for the COVID-19 Genomics UK (COG-UK) Consortium | Harper VanSteenhouse, Yumi Kasai, David Gray, Carol Clugston, Anna Dominiczak and Alex Alderton, Roberto Amato, Sonia Goncalves, Ewan Harrison, David K. Jackson, Ian Johnston, Dominic Kwiatkowski, Cordelia Langford, John Sillitoe on behalf of the Wellcome Sanger Institute COVID-19 Surveillance Team |
| EPI_ISL_820179, EPI_ISL_820181 | Lighthouse Lab in Alderley Park | Wellcome Sanger Institute for the COVID-19 Genomics UK (COG-UK) Consortium | Jacquelyn Wynn, Mairead Hyland, The Lighthouse Lab in Alderley Park and Alex Alderton, Roberto Amato, Sonia Goncalves, Ewan Harrison, David K. Jackson, Ian Johnston, Dominic Kwiatkowski, Cordelia Langford, John Sillitoe on behalf of the Wellcome Sanger Institute COVID-19 Surveillance Team |
| EPI_ISL_820182, EPI_ISL_820184 | Lighthouse Lab in Glasgow | Wellcome Sanger Institute for the COVID-19 Genomics UK (COG-UK) Consortium | Harper VanSteenhouse, Yumi Kasai, David Gray, Carol Clugston, Anna Dominiczak and Alex Alderton, Roberto Amato, Sonia Goncalves, Ewan Harrison, David K. Jackson, Ian Johnston, Dominic Kwiatkowski, Cordelia Langford, John Sillitoe on behalf of the Wellcome Sanger Institute COVID-19 Surveillance Team |
| EPI_ISL_820185, EPI_ISL_820187, EPI_ISL_820189 | Lighthouse Lab in Alderley Park | Wellcome Sanger Institute for the COVID-19 Genomics UK (COG-UK) Consortium | Jacquelyn Wynn, Mairead Hyland, The Lighthouse Lab in Alderley Park and Alex Alderton, Roberto Amato, Sonia Goncalves, Ewan Harrison, David K. Jackson, Ian Johnston, Dominic Kwiatkowski, Cordelia Langford, John Sillitoe on behalf of the Wellcome Sanger Institute COVID-19 Surveillance Team |
| EPI_ISL_820190 | Lighthouse Lab in Glasgow | Wellcome Sanger Institute for the COVID-19 Genomics UK (COG-UK) Consortium | Harper VanSteenhouse, Yumi Kasai, David Gray, Carol Clugston, Anna Dominiczak and Alex Alderton, Roberto Amato, Sonia Goncalves, Ewan Harrison, David K. Jackson, Ian Johnston, Dominic Kwiatkowski, Cordelia Langford, John Sillitoe on behalf of the Wellcome Sanger Institute COVID-19 Surveillance Team |
| EPI_ISL_820192, EPI_ISL_820194, EPI_ISL_820195 | Lighthouse Lab in Alderley Park | Wellcome Sanger Institute for the COVID-19 Genomics UK (COG-UK) Consortium | Jacquelyn Wynn, Mairead Hyland, The Lighthouse Lab in Alderley Park and Alex Alderton, Roberto Amato, Sonia Goncalves, Ewan Harrison, David K. Jackson, Ian Johnston, Dominic Kwiatkowski, Cordelia Langford, John Sillitoe on behalf of the Wellcome Sanger Institute COVID-19 Surveillance Team |
| EPI_ISL_820197 | Lighthouse Lab in Glasgow | Wellcome Sanger Institute for the COVID-19 Genomics UK (COG-UK) Consortium | Harper VanSteenhouse, Yumi Kasai, David Gray, Carol Clugston, Anna Dominiczak and Alex Alderton, Roberto Amato, Sonia Goncalves, Ewan Harrison, David K. Jackson, Ian Johnston, Dominic Kwiatkowski, Cordelia Langford, John Sillitoe on behalf of the Wellcome Sanger Institute COVID-19 Surveillance Team |
| EPI_ISL_820198 | Lighthouse Lab in Alderley Park | Wellcome Sanger Institute for the COVID-19 Genomics UK (COG-UK) Consortium | Jacquelyn Wynn, Mairead Hyland, The Lighthouse Lab in Alderley Park and Alex Alderton, Roberto Amato, Sonia Goncalves, Ewan Harrison, David K. Jackson, Ian Johnston, Dominic Kwiatkowski, Cordelia Langford, John Sillitoe on behalf of the Wellcome Sanger Institute COVID-19 Surveillance Team |
| EPI_ISL_820200 | Lighthouse Lab in Glasgow | Wellcome Sanger Institute for the COVID-19 Genomics UK (COG-UK) Consortium | Harper VanSteenhouse, Yumi Kasai, David Gray, Carol Clugston, Anna Dominiczak and Alex Alderton, Roberto Amato, Sonia Goncalves, Ewan Harrison, David K. Jackson, Ian Johnston, Dominic Kwiatkowski, Cordelia Langford, John Sillitoe on behalf of the Wellcome Sanger Institute COVID-19 Surveillance Team |
| EPI_ISL_820201, EPI_ISL_820203, EPI_ISL_820205, EPI_ISL_820207, EPI_ISL_820208, EPI_ISL_820210 | Lighthouse Lab in Alderley Park | Wellcome Sanger Institute for the COVID-19 Genomics UK (COG-UK) Consortium | Jacquelyn Wynn, Mairead Hyland, The Lighthouse Lab in Alderley Park and Alex Alderton, Roberto Amato, Sonia Goncalves, Ewan Harrison, David K. Jackson, Ian Johnston, Dominic Kwiatkowski, Cordelia Langford, John Sillitoe on behalf of the Wellcome Sanger Institute COVID-19 Surveillance Team |
| EPI_ISL_820212 | Lighthouse Lab in Glasgow | Wellcome Sanger Institute for the COVID-19 Genomics UK (COG-UK) Consortium | Harper VanSteenhouse, Yumi Kasai, David Gray, Carol Clugston, Anna Dominiczak and Alex Alderton, Roberto Amato, Sonia Goncalves, Ewan Harrison, David K. Jackson, Ian Johnston, Dominic Kwiatkowski, Cordelia Langford, John Sillitoe on behalf of the Wellcome Sanger Institute COVID-19 Surveillance Team |
| EPI_ISL_820214 | Lighthouse Lab in Alderley Park | Wellcome Sanger Institute for the COVID-19 Genomics UK (COG-UK) Consortium | Jacquelyn Wynn, Mairead Hyland, The Lighthouse Lab in Alderley Park and Alex Alderton, Roberto Amato, Sonia Goncalves, Ewan Harrison, David K. Jackson, Ian Johnston, Dominic Kwiatkowski, Cordelia Langford, John Sillitoe on behalf of the Wellcome Sanger Institute COVID-19 Surveillance Team |
| EPI_ISL_820215 | Lighthouse Lab in Glasgow | Wellcome Sanger Institute for the COVID-19 Genomics UK (COG-UK) Consortium | Harper VanSteenhouse, Yumi Kasai, David Gray, Carol Clugston, Anna Dominiczak and Alex Alderton, Roberto Amato, Sonia Goncalves, Ewan Harrison, David K. Jackson, Ian Johnston, Dominic Kwiatkowski, Cordelia Langford, John Sillitoe on behalf of the Wellcome Sanger Institute COVID-19 Surveillance Team |
| EPI_ISL_820217, EPI_ISL_820219 | Lighthouse Lab in Alderley Park | Wellcome Sanger Institute for the COVID-19 Genomics UK (COG-UK) Consortium | Jacquelyn Wynn, Mairead Hyland, The Lighthouse Lab in Alderley Park and Alex Alderton, Roberto Amato, Sonia Goncalves, Ewan Harrison, David K. Jackson, Ian Johnston, Dominic Kwiatkowski, Cordelia Langford, John Sillitoe on behalf of the Wellcome Sanger Institute COVID-19 Surveillance Team |
| EPI_ISL_820220 | Lighthouse Lab in Glasgow | Wellcome Sanger Institute for the COVID-19 Genomics UK | Harper VanSteenhouse, Yumi Kasai, David Gray, Carol Clugston, Anna Dominiczak and Alex Alderton, Roberto Amato, Sonia Goncalves, Ewan Harrison, |

| | | | |
|---|---|---|---|
| | | (COG-UK) Consortium | David K. Jackson, Ian Johnston, Dominic Kwiatkowski, Cordelia Langford, John Sillitoe on behalf of the Wellcome Sanger Institute COVID-19 Surveillance Team |
| EPI_ISL_820222, EPI_ISL_820224 | Lighthouse Lab in Alderley Park | Wellcome Sanger Institute for the COVID-19 Genomics UK (COG-UK) Consortium | Jacquelyn Wynn, Mairead Hyland, The Lighthouse Lab in Alderley Park and Alex Alderton, Roberto Amato, Sonia Goncalves, Ewan Harrison, David K. Jackson, Ian Johnston, Dominic Kwiatkowski, Cordelia Langford, John Sillitoe on behalf of the Wellcome Sanger Institute COVID-19 Surveillance Team |
| EPI_ISL_820225, EPI_ISL_820227 | Lighthouse Lab in Glasgow | Wellcome Sanger Institute for the COVID-19 Genomics UK (COG-UK) Consortium | Harper VanSteenhouse, Yumi Kasai, David Gray, Carol Clugston, Anna Dominiczak and Alex Alderton, Roberto Amato, Sonia Goncalves, Ewan Harrison, David K. Jackson, Ian Johnston, Dominic Kwiatkowski, Cordelia Langford, John Sillitoe on behalf of the Wellcome Sanger Institute COVID-19 Surveillance Team |
| EPI_ISL_820228, EPI_ISL_820230, EPI_ISL_820232 | Lighthouse Lab in Alderley Park | Wellcome Sanger Institute for the COVID-19 Genomics UK (COG-UK) Consortium | Jacquelyn Wynn, Mairead Hyland, The Lighthouse Lab in Alderley Park and Alex Alderton, Roberto Amato, Sonia Goncalves, Ewan Harrison, David K. Jackson, Ian Johnston, Dominic Kwiatkowski, Cordelia Langford, John Sillitoe on behalf of the Wellcome Sanger Institute COVID-19 Surveillance Team |
| EPI_ISL_820233 | Lighthouse Lab in Glasgow | Wellcome Sanger Institute for the COVID-19 Genomics UK (COG-UK) Consortium | Harper VanSteenhouse, Yumi Kasai, David Gray, Carol Clugston, Anna Dominiczak and Alex Alderton, Roberto Amato, Sonia Goncalves, Ewan Harrison, David K. Jackson, Ian Johnston, Dominic Kwiatkowski, Cordelia Langford, John Sillitoe on behalf of the Wellcome Sanger Institute COVID-19 Surveillance Team |
| EPI_ISL_820235, EPI_ISL_820237, EPI_ISL_820239, EPI_ISL_820240, EPI_ISL_820242 | Lighthouse Lab in Alderley Park | Wellcome Sanger Institute for the COVID-19 Genomics UK (COG-UK) Consortium | Jacquelyn Wynn, Mairead Hyland, The Lighthouse Lab in Alderley Park and Alex Alderton, Roberto Amato, Sonia Goncalves, Ewan Harrison, David K. Jackson, Ian Johnston, Dominic Kwiatkowski, Cordelia Langford, John Sillitoe on behalf of the Wellcome Sanger Institute COVID-19 Surveillance Team |
| EPI_ISL_820243 | Lighthouse Lab in Glasgow | Wellcome Sanger Institute for the COVID-19 Genomics UK (COG-UK) Consortium | Harper VanSteenhouse, Yumi Kasai, David Gray, Carol Clugston, Anna Dominiczak and Alex Alderton, Roberto Amato, Sonia Goncalves, Ewan Harrison, David K. Jackson, Ian Johnston, Dominic Kwiatkowski, Cordelia Langford, John Sillitoe on behalf of the Wellcome Sanger Institute COVID-19 Surveillance Team |
| EPI_ISL_820245, EPI_ISL_820247, EPI_ISL_820248, EPI_ISL_820250, EPI_ISL_820252, EPI_ISL_820254, EPI_ISL_820255, EPI_ISL_820257, EPI_ISL_820258 | Lighthouse Lab in Alderley Park | Wellcome Sanger Institute for the COVID-19 Genomics UK (COG-UK) Consortium | Jacquelyn Wynn, Mairead Hyland, The Lighthouse Lab in Alderley Park and Alex Alderton, Roberto Amato, Sonia Goncalves, Ewan Harrison, David K. Jackson, Ian Johnston, Dominic Kwiatkowski, Cordelia Langford, John Sillitoe on behalf of the Wellcome Sanger Institute COVID-19 Surveillance Team |
| EPI_ISL_820260, EPI_ISL_820262 | Lighthouse Lab in Glasgow | Wellcome Sanger Institute for the COVID-19 Genomics UK (COG-UK) Consortium | Harper VanSteenhouse, Yumi Kasai, David Gray, Carol Clugston, Anna Dominiczak and Alex Alderton, Roberto Amato, Sonia Goncalves, Ewan Harrison, David K. Jackson, Ian Johnston, Dominic Kwiatkowski, Cordelia Langford, John Sillitoe on behalf of the Wellcome Sanger Institute COVID-19 Surveillance Team |
| EPI_ISL_820263, EPI_ISL_820265, EPI_ISL_820267, EPI_ISL_820269 | Lighthouse Lab in Alderley Park | Wellcome Sanger Institute for the COVID-19 Genomics UK (COG-UK) Consortium | Jacquelyn Wynn, Mairead Hyland, The Lighthouse Lab in Alderley Park and Alex Alderton, Roberto Amato, Sonia Goncalves, Ewan Harrison, David K. Jackson, Ian Johnston, Dominic Kwiatkowski, Cordelia Langford, John Sillitoe on behalf of the Wellcome Sanger Institute COVID-19 Surveillance Team |
| EPI_ISL_820270, EPI_ISL_820272 | Lighthouse Lab in Glasgow | Wellcome Sanger Institute for the COVID-19 Genomics UK (COG-UK) Consortium | Harper VanSteenhouse, Yumi Kasai, David Gray, Carol Clugston, Anna Dominiczak and Alex Alderton, Roberto Amato, Sonia Goncalves, Ewan Harrison, David K. Jackson, Ian Johnston, Dominic Kwiatkowski, Cordelia Langford, John Sillitoe on behalf of the Wellcome Sanger Institute COVID-19 Surveillance Team |
| EPI_ISL_820274, EPI_ISL_820275, EPI_ISL_820277, EPI_ISL_820278, EPI_ISL_820280, EPI_ISL_820281 | Lighthouse Lab in Alderley Park | Wellcome Sanger Institute for the COVID-19 Genomics UK (COG-UK) Consortium | Jacquelyn Wynn, Mairead Hyland, The Lighthouse Lab in Alderley Park and Alex Alderton, Roberto Amato, Sonia Goncalves, Ewan Harrison, David K. Jackson, Ian Johnston, Dominic Kwiatkowski, Cordelia Langford, John Sillitoe on behalf of the Wellcome Sanger Institute COVID-19 Surveillance Team |
| EPI_ISL_820283 | Lighthouse Lab in Glasgow | Wellcome Sanger Institute for the COVID-19 Genomics UK (COG-UK) Consortium | Harper VanSteenhouse, Yumi Kasai, David Gray, Carol Clugston, Anna Dominiczak and Alex Alderton, Roberto Amato, Sonia Goncalves, Ewan Harrison, David K. Jackson, Ian Johnston, Dominic Kwiatkowski, Cordelia Langford, John Sillitoe on behalf of the Wellcome Sanger Institute COVID-19 Surveillance Team |
| EPI_ISL_820285, EPI_ISL_820286, EPI_ISL_820288, EPI_ISL_820290, EPI_ISL_820291, EPI_ISL_820293 | Lighthouse Lab in Alderley Park | Wellcome Sanger Institute for the COVID-19 Genomics UK (COG-UK) Consortium | Jacquelyn Wynn, Mairead Hyland, The Lighthouse Lab in Alderley Park and Alex Alderton, Roberto Amato, Sonia Goncalves, Ewan Harrison, David K. Jackson, Ian Johnston, Dominic Kwiatkowski, Cordelia Langford, John Sillitoe on behalf of the Wellcome Sanger Institute COVID-19 Surveillance Team |
| EPI_ISL_820295 | Lighthouse Lab in Glasgow | Wellcome Sanger Institute for the COVID-19 Genomics UK (COG-UK) Consortium | Harper VanSteenhouse, Yumi Kasai, David Gray, Carol Clugston, Anna Dominiczak and Alex Alderton, Roberto Amato, Sonia Goncalves, Ewan Harrison, David K. Jackson, Ian Johnston, Dominic Kwiatkowski, Cordelia Langford, John Sillitoe on behalf of the Wellcome Sanger Institute COVID-19 Surveillance Team |
| EPI_ISL_820296, EPI_ISL_820298, EPI_ISL_820299, EPI_ISL_820301 | Lighthouse Lab in Alderley Park | Wellcome Sanger Institute for the COVID-19 Genomics UK (COG-UK) Consortium | Jacquelyn Wynn, Mairead Hyland, The Lighthouse Lab in Alderley Park and Alex Alderton, Roberto Amato, Sonia Goncalves, Ewan Harrison, David K. Jackson, Ian Johnston, Dominic Kwiatkowski, Cordelia Langford, John Sillitoe on behalf of the Wellcome Sanger Institute COVID-19 Surveillance Team |
| EPI_ISL_820303 | Lighthouse Lab in Glasgow | Wellcome Sanger Institute for the COVID-19 Genomics UK (COG-UK) Consortium | Harper VanSteenhouse, Yumi Kasai, David Gray, Carol Clugston, Anna Dominiczak and Alex Alderton, Roberto Amato, Sonia Goncalves, Ewan Harrison, David K. Jackson, Ian Johnston, Dominic Kwiatkowski, Cordelia Langford, John Sillitoe on behalf of the Wellcome Sanger Institute COVID-19 Surveillance Team |
| EPI_ISL_820305, EPI_ISL_820307, EPI_ISL_820308, EPI_ISL_820310 | Lighthouse Lab in Alderley Park | Wellcome Sanger Institute for the COVID-19 Genomics UK (COG-UK) Consortium | Jacquelyn Wynn, Mairead Hyland, The Lighthouse Lab in Alderley Park and Alex Alderton, Roberto Amato, Sonia Goncalves, Ewan Harrison, David K. Jackson, Ian Johnston, Dominic Kwiatkowski, Cordelia Langford, John Sillitoe on behalf of the Wellcome Sanger Institute COVID-19 Surveillance Team |
| EPI_ISL_820312 | Lighthouse Lab in Glasgow | Wellcome Sanger Institute for the COVID-19 Genomics UK (COG-UK) Consortium | Harper VanSteenhouse, Yumi Kasai, David Gray, Carol Clugston, Anna Dominiczak and Alex Alderton, Roberto Amato, Sonia Goncalves, Ewan Harrison, David K. Jackson, Ian Johnston, Dominic Kwiatkowski, Cordelia Langford, John Sillitoe on behalf of the Wellcome Sanger Institute COVID-19 Surveillance Team |
| EPI_ISL_820313, EPI_ISL_820315, EPI_ISL_820317 | Lighthouse Lab in Alderley Park | Wellcome Sanger Institute for the COVID-19 Genomics UK (COG-UK) Consortium | Jacquelyn Wynn, Mairead Hyland, The Lighthouse Lab in Alderley Park and Alex Alderton, Roberto Amato, Sonia Goncalves, Ewan Harrison, David K. Jackson, Ian Johnston, Dominic Kwiatkowski, Cordelia Langford, John Sillitoe on behalf of the Wellcome Sanger Institute COVID-19 Surveillance Team |
| EPI_ISL_820319, EPI_ISL_820320 | Lighthouse Lab in Glasgow | Wellcome Sanger Institute for the COVID-19 Genomics UK (COG-UK) Consortium | Harper VanSteenhouse, Yumi Kasai, David Gray, Carol Clugston, Anna Dominiczak and Alex Alderton, Roberto Amato, Sonia Goncalves, Ewan Harrison, David K. Jackson, Ian Johnston, Dominic Kwiatkowski, Cordelia Langford, John Sillitoe on behalf of the Wellcome Sanger Institute COVID-19 Surveillance Team |
| EPI_ISL_820322, EPI_ISL_820324, EPI_ISL_820325, EPI_ISL_820327, EPI_ISL_820329, EPI_ISL_820330, EPI_ISL_820332, EPI_ISL_820334, EPI_ISL_820336, EPI_ISL_820338, EPI_ISL_820339, EPI_ISL_820341, EPI_ISL_820342, EPI_ISL_820344, EPI_ISL_820348 | | | |
| see above | Lighthouse Lab in Alderley Park | Wellcome Sanger Institute for the COVID-19 Genomics UK (COG-UK) Consortium | Jacquelyn Wynn, Mairead Hyland, The Lighthouse Lab in Alderley Park and Alex Alderton, Roberto Amato, Sonia Goncalves, Ewan Harrison, David K. Jackson, Ian Johnston, Dominic Kwiatkowski, Cordelia Langford, John Sillitoe on behalf of the Wellcome Sanger Institute COVID-19 Surveillance Team |
| EPI_ISL_820349 | Lighthouse Lab in Glasgow | Wellcome Sanger Institute for the COVID-19 Genomics UK (COG-UK) Consortium | Harper VanSteenhouse, Yumi Kasai, David Gray, Carol Clugston, Anna Dominiczak and Alex Alderton, Roberto Amato, Sonia Goncalves, Ewan Harrison, David K. Jackson, Ian Johnston, Dominic Kwiatkowski, Cordelia Langford, John Sillitoe on behalf of the Wellcome Sanger Institute COVID-19 Surveillance Team |
| EPI_ISL_820352, EPI_ISL_820354 | Lighthouse Lab in Alderley Park | Wellcome Sanger Institute for the COVID-19 Genomics UK (COG-UK) Consortium | Jacquelyn Wynn, Mairead Hyland, The Lighthouse Lab in Alderley Park and Alex Alderton, Roberto Amato, Sonia Goncalves, Ewan Harrison, David K. Jackson, Ian Johnston, Dominic Kwiatkowski, Cordelia Langford, John Sillitoe on behalf of the Wellcome Sanger Institute COVID-19 Surveillance Team |
| EPI_ISL_820356 | Lighthouse Lab in Glasgow | Wellcome Sanger Institute for the COVID-19 Genomics UK (COG-UK) Consortium | Harper VanSteenhouse, Yumi Kasai, David Gray, Carol Clugston, Anna Dominiczak and Alex Alderton, Roberto Amato, Sonia Goncalves, Ewan Harrison, David K. Jackson, Ian Johnston, Dominic Kwiatkowski, Cordelia Langford, John Sillitoe on behalf of the Wellcome Sanger Institute COVID-19 Surveillance Team |
| EPI_ISL_820357, EPI_ISL_820359, EPI_ISL_820361, EPI_ISL_820362, EPI_ISL_820364, EPI_ISL_820366, EPI_ISL_820367, EPI_ISL_820369, EPI_ISL_820370, EPI_ISL_820372, EPI_ISL_820374, EPI_ISL_820375, EPI_ISL_820377, EPI_ISL_820379, EPI_ISL_820380, EPI_ISL_820382 | | | |
| see above | Lighthouse Lab in Alderley Park | Wellcome Sanger Institute for the COVID-19 Genomics UK (COG-UK) Consortium | Jacquelyn Wynn, Mairead Hyland, The Lighthouse Lab in Alderley Park and Alex Alderton, Roberto Amato, Sonia Goncalves, Ewan Harrison, David K. Jackson, Ian Johnston, Dominic Kwiatkowski, Cordelia Langford, John Sillitoe on behalf of the Wellcome Sanger Institute COVID-19 Surveillance Team |
| EPI_ISL_820384, EPI_ISL_820386, | Lighthouse Lab in Glasgow | Wellcome Sanger Institute for the COVID-19 Genomics UK | Harper VanSteenhouse, Yumi Kasai, David Gray, Carol Clugston, Anna Dominiczak and Alex Alderton, Roberto Amato, Sonia Goncalves, Ewan Harrison, |

| | | | |
|---|---|---|---|
| EPI_ISL_820387 | | (COG-UK) Consortium | David K. Jackson, Ian Johnston, Dominic Kwiatkowski, Cordelia Langford, John Sillitoe on behalf of the Wellcome Sanger Institute COVID-19 Surveillance Team |
| EPI_ISL_820389, EPI_ISL_820390, EPI_ISL_820392, EPI_ISL_820393, EPI_ISL_820395 | Lighthouse Lab in Alderley Park | Wellcome Sanger Institute for the COVID-19 Genomics UK (COG-UK) Consortium | Jacquelyn Wynn, Mairead Hyland, The Lighthouse Lab in Alderley Park and Alex Alderton, Roberto Amato, Sonia Goncalves, Ewan Harrison, David K. Jackson, Ian Johnston, Dominic Kwiatkowski, Cordelia Langford, John Sillitoe on behalf of the Wellcome Sanger Institute COVID-19 Surveillance Team |
| EPI_ISL_820397 | Lighthouse Lab in Glasgow | Wellcome Sanger Institute for the COVID-19 Genomics UK (COG-UK) Consortium | Harper VanSteenhouse, Yumi Kasai, David Gray, Carol Clugston, Anna Dominiczak and Alex Alderton, Roberto Amato, Sonia Goncalves, Ewan Harrison, David K. Jackson, Ian Johnston, Dominic Kwiatkowski, Cordelia Langford, John Sillitoe on behalf of the Wellcome Sanger Institute COVID-19 Surveillance Team |
| EPI_ISL_820398, EPI_ISL_820400, EPI_ISL_820401, EPI_ISL_820403, EPI_ISL_820405, EPI_ISL_820406, EPI_ISL_820408 | Lighthouse Lab in Alderley Park | Wellcome Sanger Institute for the COVID-19 Genomics UK (COG-UK) Consortium | Jacquelyn Wynn, Mairead Hyland, The Lighthouse Lab in Alderley Park and Alex Alderton, Roberto Amato, Sonia Goncalves, Ewan Harrison, David K. Jackson, Ian Johnston, Dominic Kwiatkowski, Cordelia Langford, John Sillitoe on behalf of the Wellcome Sanger Institute COVID-19 Surveillance Team |
| EPI_ISL_820409 | Lighthouse Lab in Glasgow | Wellcome Sanger Institute for the COVID-19 Genomics UK (COG-UK) Consortium | Harper VanSteenhouse, Yumi Kasai, David Gray, Carol Clugston, Anna Dominiczak and Alex Alderton, Roberto Amato, Sonia Goncalves, Ewan Harrison, David K. Jackson, Ian Johnston, Dominic Kwiatkowski, Cordelia Langford, John Sillitoe on behalf of the Wellcome Sanger Institute COVID-19 Surveillance Team |
| EPI_ISL_820411, EPI_ISL_820414, EPI_ISL_820416 | Lighthouse Lab in Alderley Park | Wellcome Sanger Institute for the COVID-19 Genomics UK (COG-UK) Consortium | Jacquelyn Wynn, Mairead Hyland, The Lighthouse Lab in Alderley Park and Alex Alderton, Roberto Amato, Sonia Goncalves, Ewan Harrison, David K. Jackson, Ian Johnston, Dominic Kwiatkowski, Cordelia Langford, John Sillitoe on behalf of the Wellcome Sanger Institute COVID-19 Surveillance Team |
| EPI_ISL_820417 | Lighthouse Lab in Glasgow | Wellcome Sanger Institute for the COVID-19 Genomics UK (COG-UK) Consortium | Harper VanSteenhouse, Yumi Kasai, David Gray, Carol Clugston, Anna Dominiczak and Alex Alderton, Roberto Amato, Sonia Goncalves, Ewan Harrison, David K. Jackson, Ian Johnston, Dominic Kwiatkowski, Cordelia Langford, John Sillitoe on behalf of the Wellcome Sanger Institute COVID-19 Surveillance Team |
| EPI_ISL_820419 | Lighthouse Lab in Alderley Park | Wellcome Sanger Institute for the COVID-19 Genomics UK (COG-UK) Consortium | Jacquelyn Wynn, Mairead Hyland, The Lighthouse Lab in Alderley Park and Alex Alderton, Roberto Amato, Sonia Goncalves, Ewan Harrison, David K. Jackson, Ian Johnston, Dominic Kwiatkowski, Cordelia Langford, John Sillitoe on behalf of the Wellcome Sanger Institute COVID-19 Surveillance Team |
| EPI_ISL_820421 | Lighthouse Lab in Glasgow | Wellcome Sanger Institute for the COVID-19 Genomics UK (COG-UK) Consortium | Harper VanSteenhouse, Yumi Kasai, David Gray, Carol Clugston, Anna Dominiczak and Alex Alderton, Roberto Amato, Sonia Goncalves, Ewan Harrison, David K. Jackson, Ian Johnston, Dominic Kwiatkowski, Cordelia Langford, John Sillitoe on behalf of the Wellcome Sanger Institute COVID-19 Surveillance Team |
| EPI_ISL_820422, EPI_ISL_820424, EPI_ISL_820425, EPI_ISL_820427, EPI_ISL_820429, EPI_ISL_820430, EPI_ISL_820432, EPI_ISL_820434, EPI_ISL_820435 | Lighthouse Lab in Alderley Park | Wellcome Sanger Institute for the COVID-19 Genomics UK (COG-UK) Consortium | Jacquelyn Wynn, Mairead Hyland, The Lighthouse Lab in Alderley Park and Alex Alderton, Roberto Amato, Sonia Goncalves, Ewan Harrison, David K. Jackson, Ian Johnston, Dominic Kwiatkowski, Cordelia Langford, John Sillitoe on behalf of the Wellcome Sanger Institute COVID-19 Surveillance Team |
| EPI_ISL_820437 | Lighthouse Lab in Glasgow | Wellcome Sanger Institute for the COVID-19 Genomics UK (COG-UK) Consortium | Harper VanSteenhouse, Yumi Kasai, David Gray, Carol Clugston, Anna Dominiczak and Alex Alderton, Roberto Amato, Sonia Goncalves, Ewan Harrison, David K. Jackson, Ian Johnston, Dominic Kwiatkowski, Cordelia Langford, John Sillitoe on behalf of the Wellcome Sanger Institute COVID-19 Surveillance Team |
| EPI_ISL_820438, EPI_ISL_820440, EPI_ISL_820442, EPI_ISL_820443, EPI_ISL_820445, EPI_ISL_820447 | Lighthouse Lab in Alderley Park | Wellcome Sanger Institute for the COVID-19 Genomics UK (COG-UK) Consortium | Jacquelyn Wynn, Mairead Hyland, The Lighthouse Lab in Alderley Park and Alex Alderton, Roberto Amato, Sonia Goncalves, Ewan Harrison, David K. Jackson, Ian Johnston, Dominic Kwiatkowski, Cordelia Langford, John Sillitoe on behalf of the Wellcome Sanger Institute COVID-19 Surveillance Team |
| EPI_ISL_820448 | Lighthouse Lab in Glasgow | Wellcome Sanger Institute for the COVID-19 Genomics UK (COG-UK) Consortium | Harper VanSteenhouse, Yumi Kasai, David Gray, Carol Clugston, Anna Dominiczak and Alex Alderton, Roberto Amato, Sonia Goncalves, Ewan Harrison, David K. Jackson, Ian Johnston, Dominic Kwiatkowski, Cordelia Langford, John Sillitoe on behalf of the Wellcome Sanger Institute COVID-19 Surveillance Team |
| EPI_ISL_820450, EPI_ISL_820452, EPI_ISL_820453, EPI_ISL_820455, EPI_ISL_820457, EPI_ISL_820458, EPI_ISL_820460 | Lighthouse Lab in Alderley Park | Wellcome Sanger Institute for the COVID-19 Genomics UK (COG-UK) Consortium | Jacquelyn Wynn, Mairead Hyland, The Lighthouse Lab in Alderley Park and Alex Alderton, Roberto Amato, Sonia Goncalves, Ewan Harrison, David K. Jackson, Ian Johnston, Dominic Kwiatkowski, Cordelia Langford, John Sillitoe on behalf of the Wellcome Sanger Institute COVID-19 Surveillance Team |
| EPI_ISL_820461 | Lighthouse Lab in Glasgow | Wellcome Sanger Institute for the COVID-19 Genomics UK (COG-UK) Consortium | Harper VanSteenhouse, Yumi Kasai, David Gray, Carol Clugston, Anna Dominiczak and Alex Alderton, Roberto Amato, Sonia Goncalves, Ewan Harrison, David K. Jackson, Ian Johnston, Dominic Kwiatkowski, Cordelia Langford, John Sillitoe on behalf of the Wellcome Sanger Institute COVID-19 Surveillance Team |
| EPI_ISL_820463 | Lighthouse Lab in Alderley Park | Wellcome Sanger Institute for the COVID-19 Genomics UK (COG-UK) Consortium | Jacquelyn Wynn, Mairead Hyland, The Lighthouse Lab in Alderley Park and Alex Alderton, Roberto Amato, Sonia Goncalves, Ewan Harrison, David K. Jackson, Ian Johnston, Dominic Kwiatkowski, Cordelia Langford, John Sillitoe on behalf of the Wellcome Sanger Institute COVID-19 Surveillance Team |
| EPI_ISL_820465 | Lighthouse Lab in Glasgow | Wellcome Sanger Institute for the COVID-19 Genomics UK (COG-UK) Consortium | Harper VanSteenhouse, Yumi Kasai, David Gray, Carol Clugston, Anna Dominiczak and Alex Alderton, Roberto Amato, Sonia Goncalves, Ewan Harrison, David K. Jackson, Ian Johnston, Dominic Kwiatkowski, Cordelia Langford, John Sillitoe on behalf of the Wellcome Sanger Institute COVID-19 Surveillance Team |
| EPI_ISL_820466 | Lighthouse Lab in Alderley Park | Wellcome Sanger Institute for the COVID-19 Genomics UK (COG-UK) Consortium | Jacquelyn Wynn, Mairead Hyland, The Lighthouse Lab in Alderley Park and Alex Alderton, Roberto Amato, Sonia Goncalves, Ewan Harrison, David K. Jackson, Ian Johnston, Dominic Kwiatkowski, Cordelia Langford, John Sillitoe on behalf of the Wellcome Sanger Institute COVID-19 Surveillance Team |
| EPI_ISL_820468 | Lighthouse Lab in Glasgow | Wellcome Sanger Institute for the COVID-19 Genomics UK (COG-UK) Consortium | Harper VanSteenhouse, Yumi Kasai, David Gray, Carol Clugston, Anna Dominiczak and Alex Alderton, Roberto Amato, Sonia Goncalves, Ewan Harrison, David K. Jackson, Ian Johnston, Dominic Kwiatkowski, Cordelia Langford, John Sillitoe on behalf of the Wellcome Sanger Institute COVID-19 Surveillance Team |
| EPI_ISL_820469, EPI_ISL_820471, EPI_ISL_820473, EPI_ISL_820474, EPI_ISL_820476 | Lighthouse Lab in Alderley Park | Wellcome Sanger Institute for the COVID-19 Genomics UK (COG-UK) Consortium | Jacquelyn Wynn, Mairead Hyland, The Lighthouse Lab in Alderley Park and Alex Alderton, Roberto Amato, Sonia Goncalves, Ewan Harrison, David K. Jackson, Ian Johnston, Dominic Kwiatkowski, Cordelia Langford, John Sillitoe on behalf of the Wellcome Sanger Institute COVID-19 Surveillance Team |
| EPI_ISL_820477, EPI_ISL_820479, EPI_ISL_820481 | Lighthouse Lab in Glasgow | Wellcome Sanger Institute for the COVID-19 Genomics UK (COG-UK) Consortium | Harper VanSteenhouse, Yumi Kasai, David Gray, Carol Clugston, Anna Dominiczak and Alex Alderton, Roberto Amato, Sonia Goncalves, Ewan Harrison, David K. Jackson, Ian Johnston, Dominic Kwiatkowski, Cordelia Langford, John Sillitoe on behalf of the Wellcome Sanger Institute COVID-19 Surveillance Team |
| EPI_ISL_820482 | Lighthouse Lab in Alderley Park | Wellcome Sanger Institute for the COVID-19 Genomics UK (COG-UK) Consortium | Jacquelyn Wynn, Mairead Hyland, The Lighthouse Lab in Alderley Park and Alex Alderton, Roberto Amato, Sonia Goncalves, Ewan Harrison, David K. Jackson, Ian Johnston, Dominic Kwiatkowski, Cordelia Langford, John Sillitoe on behalf of the Wellcome Sanger Institute COVID-19 Surveillance Team |
| EPI_ISL_820484 | Lighthouse Lab in Glasgow | Wellcome Sanger Institute for the COVID-19 Genomics UK (COG-UK) Consortium | Harper VanSteenhouse, Yumi Kasai, David Gray, Carol Clugston, Anna Dominiczak and Alex Alderton, Roberto Amato, Sonia Goncalves, Ewan Harrison, David K. Jackson, Ian Johnston, Dominic Kwiatkowski, Cordelia Langford, John Sillitoe on behalf of the Wellcome Sanger Institute COVID-19 Surveillance Team |
| EPI_ISL_820486, EPI_ISL_820487 | Lighthouse Lab in Alderley Park | Wellcome Sanger Institute for the COVID-19 Genomics UK (COG-UK) Consortium | Jacquelyn Wynn, Mairead Hyland, The Lighthouse Lab in Alderley Park and Alex Alderton, Roberto Amato, Sonia Goncalves, Ewan Harrison, David K. Jackson, Ian Johnston, Dominic Kwiatkowski, Cordelia Langford, John Sillitoe on behalf of the Wellcome Sanger Institute COVID-19 Surveillance Team |
| EPI_ISL_820489 | Lighthouse Lab in Glasgow | Wellcome Sanger Institute for the COVID-19 Genomics UK (COG-UK) Consortium | Harper VanSteenhouse, Yumi Kasai, David Gray, Carol Clugston, Anna Dominiczak and Alex Alderton, Roberto Amato, Sonia Goncalves, Ewan Harrison, David K. Jackson, Ian Johnston, Dominic Kwiatkowski, Cordelia Langford, John Sillitoe on behalf of the Wellcome Sanger Institute COVID-19 Surveillance Team |
| EPI_ISL_820491, EPI_ISL_820492, EPI_ISL_820494, EPI_ISL_820495, EPI_ISL_820497, EPI_ISL_820499, | Lighthouse Lab in Alderley Park | Wellcome Sanger Institute for the COVID-19 Genomics UK (COG-UK) Consortium | Jacquelyn Wynn, Mairead Hyland, The Lighthouse Lab in Alderley Park and Alex Alderton, Roberto Amato, Sonia Goncalves, Ewan Harrison, David K. Jackson, Ian Johnston, Dominic Kwiatkowski, Cordelia Langford, John Sillitoe on behalf of the Wellcome Sanger Institute COVID-19 Surveillance Team |

| | | | |
|---|---|---|---|
| EPI_ISL_820500, EPI_ISL_820502, EPI_ISL_820503 | | | |
| EPI_ISL_820505, EPI_ISL_820507 | Lighthouse Lab in Glasgow | Wellcome Sanger Institute for the COVID-19 Genomics UK (COG-UK) Consortium | Harper VanSteenhouse, Yumi Kasai, David Gray, Carol Clugston, Anna Dominiczak and Alex Alderton, Roberto Amato, Sonia Goncalves, Ewan Harrison, David K. Jackson, Ian Johnston, Dominic Kwiatkowski, Cordelia Langford, John Sillitoe on behalf of the Wellcome Sanger Institute COVID-19 Surveillance Team |
| EPI_ISL_820508 | Lighthouse Lab in Alderley Park | Wellcome Sanger Institute for the COVID-19 Genomics UK (COG-UK) Consortium | Jacquelyn Wynn, Mairead Hyland, The Lighthouse Lab in Alderley Park and Alex Alderton, Roberto Amato, Sonia Goncalves, Ewan Harrison, David K. Jackson, Ian Johnston, Dominic Kwiatkowski, Cordelia Langford, John Sillitoe on behalf of the Wellcome Sanger Institute COVID-19 Surveillance Team |
| EPI_ISL_820510, EPI_ISL_820512, EPI_ISL_820513 | Lighthouse Lab in Glasgow | Wellcome Sanger Institute for the COVID-19 Genomics UK (COG-UK) Consortium | Harper VanSteenhouse, Yumi Kasai, David Gray, Carol Clugston, Anna Dominiczak and Alex Alderton, Roberto Amato, Sonia Goncalves, Ewan Harrison, David K. Jackson, Ian Johnston, Dominic Kwiatkowski, Cordelia Langford, John Sillitoe on behalf of the Wellcome Sanger Institute COVID-19 Surveillance Team |
| EPI_ISL_820515, EPI_ISL_820517 | Lighthouse Lab in Alderley Park | Wellcome Sanger Institute for the COVID-19 Genomics UK (COG-UK) Consortium | Jacquelyn Wynn, Mairead Hyland, The Lighthouse Lab in Alderley Park and Alex Alderton, Roberto Amato, Sonia Goncalves, Ewan Harrison, David K. Jackson, Ian Johnston, Dominic Kwiatkowski, Cordelia Langford, John Sillitoe on behalf of the Wellcome Sanger Institute COVID-19 Surveillance Team |
| EPI_ISL_820518 | Lighthouse Lab in Glasgow | Wellcome Sanger Institute for the COVID-19 Genomics UK (COG-UK) Consortium | Harper VanSteenhouse, Yumi Kasai, David Gray, Carol Clugston, Anna Dominiczak and Alex Alderton, Roberto Amato, Sonia Goncalves, Ewan Harrison, David K. Jackson, Ian Johnston, Dominic Kwiatkowski, Cordelia Langford, John Sillitoe on behalf of the Wellcome Sanger Institute COVID-19 Surveillance Team |
| EPI_ISL_820520, EPI_ISL_820522, EPI_ISL_820523, EPI_ISL_820525, EPI_ISL_820527, EPI_ISL_820529, EPI_ISL_820530, EPI_ISL_820532, EPI_ISL_820534, EPI_ISL_820535, EPI_ISL_820537, EPI_ISL_820538, EPI_ISL_820540, EPI_ISL_820542, EPI_ISL_820544, EPI_ISL_820545 | | | |
| see above | Lighthouse Lab in Alderley Park | Wellcome Sanger Institute for the COVID-19 Genomics UK (COG-UK) Consortium | Jacquelyn Wynn, Mairead Hyland, The Lighthouse Lab in Alderley Park and Alex Alderton, Roberto Amato, Sonia Goncalves, Ewan Harrison, David K. Jackson, Ian Johnston, Dominic Kwiatkowski, Cordelia Langford, John Sillitoe on behalf of the Wellcome Sanger Institute COVID-19 Surveillance Team |
| EPI_ISL_820547 | Lighthouse Lab in Glasgow | Wellcome Sanger Institute for the COVID-19 Genomics UK (COG-UK) Consortium | Harper VanSteenhouse, Yumi Kasai, David Gray, Carol Clugston, Anna Dominiczak and Alex Alderton, Roberto Amato, Sonia Goncalves, Ewan Harrison, David K. Jackson, Ian Johnston, Dominic Kwiatkowski, Cordelia Langford, John Sillitoe on behalf of the Wellcome Sanger Institute COVID-19 Surveillance Team |
| EPI_ISL_820548, EPI_ISL_820550, EPI_ISL_820551, EPI_ISL_820553, EPI_ISL_820555, EPI_ISL_820556 | Lighthouse Lab in Alderley Park | Wellcome Sanger Institute for the COVID-19 Genomics UK (COG-UK) Consortium | Jacquelyn Wynn, Mairead Hyland, The Lighthouse Lab in Alderley Park and Alex Alderton, Roberto Amato, Sonia Goncalves, Ewan Harrison, David K. Jackson, Ian Johnston, Dominic Kwiatkowski, Cordelia Langford, John Sillitoe on behalf of the Wellcome Sanger Institute COVID-19 Surveillance Team |
| EPI_ISL_820558 | Lighthouse Lab in Glasgow | Wellcome Sanger Institute for the COVID-19 Genomics UK (COG-UK) Consortium | Harper VanSteenhouse, Yumi Kasai, David Gray, Carol Clugston, Anna Dominiczak and Alex Alderton, Roberto Amato, Sonia Goncalves, Ewan Harrison, David K. Jackson, Ian Johnston, Dominic Kwiatkowski, Cordelia Langford, John Sillitoe on behalf of the Wellcome Sanger Institute COVID-19 Surveillance Team |
| EPI_ISL_820559, EPI_ISL_820561, EPI_ISL_820562, EPI_ISL_820564, EPI_ISL_820566, EPI_ISL_820567, EPI_ISL_820569, EPI_ISL_820571, EPI_ISL_820572, EPI_ISL_820574, EPI_ISL_820576 | | | |
| see above | Lighthouse Lab in Alderley Park | Wellcome Sanger Institute for the COVID-19 Genomics UK (COG-UK) Consortium | Jacquelyn Wynn, Mairead Hyland, The Lighthouse Lab in Alderley Park and Alex Alderton, Roberto Amato, Sonia Goncalves, Ewan Harrison, David K. Jackson, Ian Johnston, Dominic Kwiatkowski, Cordelia Langford, John Sillitoe on behalf of the Wellcome Sanger Institute COVID-19 Surveillance Team |
| EPI_ISL_820577 | Lighthouse Lab in Glasgow | Wellcome Sanger Institute for the COVID-19 Genomics UK (COG-UK) Consortium | Harper VanSteenhouse, Yumi Kasai, David Gray, Carol Clugston, Anna Dominiczak and Alex Alderton, Roberto Amato, Sonia Goncalves, Ewan Harrison, David K. Jackson, Ian Johnston, Dominic Kwiatkowski, Cordelia Langford, John Sillitoe on behalf of the Wellcome Sanger Institute COVID-19 Surveillance Team |
| EPI_ISL_820579, EPI_ISL_820581, EPI_ISL_820582, EPI_ISL_820584, EPI_ISL_820585, EPI_ISL_820587, EPI_ISL_820589 | Lighthouse Lab in Alderley Park | Wellcome Sanger Institute for the COVID-19 Genomics UK (COG-UK) Consortium | Jacquelyn Wynn, Mairead Hyland, The Lighthouse Lab in Alderley Park and Alex Alderton, Roberto Amato, Sonia Goncalves, Ewan Harrison, David K. Jackson, Ian Johnston, Dominic Kwiatkowski, Cordelia Langford, John Sillitoe on behalf of the Wellcome Sanger Institute COVID-19 Surveillance Team |
| EPI_ISL_820590 | Lighthouse Lab in Glasgow | Wellcome Sanger Institute for the COVID-19 Genomics UK (COG-UK) Consortium | Harper VanSteenhouse, Yumi Kasai, David Gray, Carol Clugston, Anna Dominiczak and Alex Alderton, Roberto Amato, Sonia Goncalves, Ewan Harrison, David K. Jackson, Ian Johnston, Dominic Kwiatkowski, Cordelia Langford, John Sillitoe on behalf of the Wellcome Sanger Institute COVID-19 Surveillance Team |
| EPI_ISL_820592, EPI_ISL_820594, EPI_ISL_820595 | Lighthouse Lab in Alderley Park | Wellcome Sanger Institute for the COVID-19 Genomics UK (COG-UK) Consortium | Jacquelyn Wynn, Mairead Hyland, The Lighthouse Lab in Alderley Park and Alex Alderton, Roberto Amato, Sonia Goncalves, Ewan Harrison, David K. Jackson, Ian Johnston, Dominic Kwiatkowski, Cordelia Langford, John Sillitoe on behalf of the Wellcome Sanger Institute COVID-19 Surveillance Team |
| EPI_ISL_820597 | Lighthouse Lab in Glasgow | Wellcome Sanger Institute for the COVID-19 Genomics UK (COG-UK) Consortium | Harper VanSteenhouse, Yumi Kasai, David Gray, Carol Clugston, Anna Dominiczak and Alex Alderton, Roberto Amato, Sonia Goncalves, Ewan Harrison, David K. Jackson, Ian Johnston, Dominic Kwiatkowski, Cordelia Langford, John Sillitoe on behalf of the Wellcome Sanger Institute COVID-19 Surveillance Team |
| EPI_ISL_820599, EPI_ISL_820600, EPI_ISL_820602, EPI_ISL_820604, EPI_ISL_820606, EPI_ISL_820608 | Lighthouse Lab in Alderley Park | Wellcome Sanger Institute for the COVID-19 Genomics UK (COG-UK) Consortium | Jacquelyn Wynn, Mairead Hyland, The Lighthouse Lab in Alderley Park and Alex Alderton, Roberto Amato, Sonia Goncalves, Ewan Harrison, David K. Jackson, Ian Johnston, Dominic Kwiatkowski, Cordelia Langford, John Sillitoe on behalf of the Wellcome Sanger Institute COVID-19 Surveillance Team |
| EPI_ISL_820609 | Lighthouse Lab in Glasgow | Wellcome Sanger Institute for the COVID-19 Genomics UK (COG-UK) Consortium | Harper VanSteenhouse, Yumi Kasai, David Gray, Carol Clugston, Anna Dominiczak and Alex Alderton, Roberto Amato, Sonia Goncalves, Ewan Harrison, David K. Jackson, Ian Johnston, Dominic Kwiatkowski, Cordelia Langford, John Sillitoe on behalf of the Wellcome Sanger Institute COVID-19 Surveillance Team |
| EPI_ISL_820611, EPI_ISL_820612 | Lighthouse Lab in Alderley Park | Wellcome Sanger Institute for the COVID-19 Genomics UK (COG-UK) Consortium | Jacquelyn Wynn, Mairead Hyland, The Lighthouse Lab in Alderley Park and Alex Alderton, Roberto Amato, Sonia Goncalves, Ewan Harrison, David K. Jackson, Ian Johnston, Dominic Kwiatkowski, Cordelia Langford, John Sillitoe on behalf of the Wellcome Sanger Institute COVID-19 Surveillance Team |
| EPI_ISL_820613 | Lighthouse Lab in Glasgow | Wellcome Sanger Institute for the COVID-19 Genomics UK (COG-UK) Consortium | Harper VanSteenhouse, Yumi Kasai, David Gray, Carol Clugston, Anna Dominiczak and Alex Alderton, Roberto Amato, Sonia Goncalves, Ewan Harrison, David K. Jackson, Ian Johnston, Dominic Kwiatkowski, Cordelia Langford, John Sillitoe on behalf of the Wellcome Sanger Institute COVID-19 Surveillance Team |
| EPI_ISL_820614, EPI_ISL_820615, EPI_ISL_820616 | Lighthouse Lab in Alderley Park | Wellcome Sanger Institute for the COVID-19 Genomics UK (COG-UK) Consortium | Jacquelyn Wynn, Mairead Hyland, The Lighthouse Lab in Alderley Park and Alex Alderton, Roberto Amato, Sonia Goncalves, Ewan Harrison, David K. Jackson, Ian Johnston, Dominic Kwiatkowski, Cordelia Langford, John Sillitoe on behalf of the Wellcome Sanger Institute COVID-19 Surveillance Team |
| EPI_ISL_820617, EPI_ISL_820618 | Lighthouse Lab in Glasgow | Wellcome Sanger Institute for the COVID-19 Genomics UK (COG-UK) Consortium | Harper VanSteenhouse, Yumi Kasai, David Gray, Carol Clugston, Anna Dominiczak and Alex Alderton, Roberto Amato, Sonia Goncalves, Ewan Harrison, David K. Jackson, Ian Johnston, Dominic Kwiatkowski, Cordelia Langford, John Sillitoe on behalf of the Wellcome Sanger Institute COVID-19 Surveillance Team |
| EPI_ISL_820619 | Lighthouse Lab in Alderley Park | Wellcome Sanger Institute for the COVID-19 Genomics UK (COG-UK) Consortium | Jacquelyn Wynn, Mairead Hyland, The Lighthouse Lab in Alderley Park and Alex Alderton, Roberto Amato, Sonia Goncalves, Ewan Harrison, David K. Jackson, Ian Johnston, Dominic Kwiatkowski, Cordelia Langford, John Sillitoe on behalf of the Wellcome Sanger Institute COVID-19 Surveillance Team |
| EPI_ISL_820620 | Lighthouse Lab in Glasgow | Wellcome Sanger Institute for the COVID-19 Genomics UK (COG-UK) Consortium | Harper VanSteenhouse, Yumi Kasai, David Gray, Carol Clugston, Anna Dominiczak and Alex Alderton, Roberto Amato, Sonia Goncalves, Ewan Harrison, David K. Jackson, Ian Johnston, Dominic Kwiatkowski, Cordelia Langford, John Sillitoe on behalf of the Wellcome Sanger Institute COVID-19 Surveillance Team |
| EPI_ISL_820621, EPI_ISL_820622, EPI_ISL_820623, EPI_ISL_820624, EPI_ISL_820625, EPI_ISL_820626, EPI_ISL_820627 | Lighthouse Lab in Alderley Park | Wellcome Sanger Institute for the COVID-19 Genomics UK (COG-UK) Consortium | Jacquelyn Wynn, Mairead Hyland, The Lighthouse Lab in Alderley Park and Alex Alderton, Roberto Amato, Sonia Goncalves, Ewan Harrison, David K. Jackson, Ian Johnston, Dominic Kwiatkowski, Cordelia Langford, John Sillitoe on behalf of the Wellcome Sanger Institute COVID-19 Surveillance Team |
| EPI_ISL_820628 | Lighthouse Lab in Glasgow | Wellcome Sanger Institute for the COVID-19 Genomics UK (COG-UK) Consortium | Harper VanSteenhouse, Yumi Kasai, David Gray, Carol Clugston, Anna Dominiczak and Alex Alderton, Roberto Amato, Sonia Goncalves, Ewan Harrison, David K. Jackson, Ian Johnston, Dominic Kwiatkowski, Cordelia Langford, John Sillitoe on behalf of the Wellcome Sanger Institute COVID-19 Surveillance Team |

| | | | |
|---|---|---|---|
| EPI_ISL_820629, EPI_ISL_820630, EPI_ISL_820631, EPI_ISL_820632 | Lighthouse Lab in Alderley Park | Wellcome Sanger Institute for the COVID-19 Genomics UK (COG-UK) Consortium | Jacquelyn Wynn, Mairead Hyland, The Lighthouse Lab in Alderley Park and Alex Alderton, Roberto Amato, Sonia Goncalves, Ewan Harrison, David K. Jackson, Ian Johnston, Dominic Kwiatkowski, Cordelia Langford, John Sillitoe on behalf of the Wellcome Sanger Institute COVID-19 Surveillance Team |
| EPI_ISL_820633, EPI_ISL_820634 | Lighthouse Lab in Glasgow | Wellcome Sanger Institute for the COVID-19 Genomics UK (COG-UK) Consortium | Harper VanSteenhouse, Yumi Kasai, David Gray, Carol Clugston, Anna Dominiczak and Alex Alderton, Roberto Amato, Sonia Goncalves, Ewan Harrison, David K. Jackson, Ian Johnston, Dominic Kwiatkowski, Cordelia Langford, John Sillitoe on behalf of the Wellcome Sanger Institute COVID-19 Surveillance Team |
| EPI_ISL_820635, EPI_ISL_820636, EPI_ISL_820637, EPI_ISL_820638 | Lighthouse Lab in Alderley Park | Wellcome Sanger Institute for the COVID-19 Genomics UK (COG-UK) Consortium | Jacquelyn Wynn, Mairead Hyland, The Lighthouse Lab in Alderley Park and Alex Alderton, Roberto Amato, Sonia Goncalves, Ewan Harrison, David K. Jackson, Ian Johnston, Dominic Kwiatkowski, Cordelia Langford, John Sillitoe on behalf of the Wellcome Sanger Institute COVID-19 Surveillance Team |
| EPI_ISL_820639 | Lighthouse Lab in Glasgow | Wellcome Sanger Institute for the COVID-19 Genomics UK (COG-UK) Consortium | Harper VanSteenhouse, Yumi Kasai, David Gray, Carol Clugston, Anna Dominiczak and Alex Alderton, Roberto Amato, Sonia Goncalves, Ewan Harrison, David K. Jackson, Ian Johnston, Dominic Kwiatkowski, Cordelia Langford, John Sillitoe on behalf of the Wellcome Sanger Institute COVID-19 Surveillance Team |
| EPI_ISL_820640 | Lighthouse Lab in Alderley Park | Wellcome Sanger Institute for the COVID-19 Genomics UK (COG-UK) Consortium | Jacquelyn Wynn, Mairead Hyland, The Lighthouse Lab in Alderley Park and Alex Alderton, Roberto Amato, Sonia Goncalves, Ewan Harrison, David K. Jackson, Ian Johnston, Dominic Kwiatkowski, Cordelia Langford, John Sillitoe on behalf of the Wellcome Sanger Institute COVID-19 Surveillance Team |
| EPI_ISL_820641 | Lighthouse Lab in Glasgow | Wellcome Sanger Institute for the COVID-19 Genomics UK (COG-UK) Consortium | Harper VanSteenhouse, Yumi Kasai, David Gray, Carol Clugston, Anna Dominiczak and Alex Alderton, Roberto Amato, Sonia Goncalves, Ewan Harrison, David K. Jackson, Ian Johnston, Dominic Kwiatkowski, Cordelia Langford, John Sillitoe on behalf of the Wellcome Sanger Institute COVID-19 Surveillance Team |
| EPI_ISL_820642, EPI_ISL_820643, EPI_ISL_820644 | Lighthouse Lab in Alderley Park | Wellcome Sanger Institute for the COVID-19 Genomics UK (COG-UK) Consortium | Jacquelyn Wynn, Mairead Hyland, The Lighthouse Lab in Alderley Park and Alex Alderton, Roberto Amato, Sonia Goncalves, Ewan Harrison, David K. Jackson, Ian Johnston, Dominic Kwiatkowski, Cordelia Langford, John Sillitoe on behalf of the Wellcome Sanger Institute COVID-19 Surveillance Team |
| EPI_ISL_820645 | Lighthouse Lab in Glasgow | Wellcome Sanger Institute for the COVID-19 Genomics UK (COG-UK) Consortium | Harper VanSteenhouse, Yumi Kasai, David Gray, Carol Clugston, Anna Dominiczak and Alex Alderton, Roberto Amato, Sonia Goncalves, Ewan Harrison, David K. Jackson, Ian Johnston, Dominic Kwiatkowski, Cordelia Langford, John Sillitoe on behalf of the Wellcome Sanger Institute COVID-19 Surveillance Team |
| EPI_ISL_820646 | Lighthouse Lab in Alderley Park | Wellcome Sanger Institute for the COVID-19 Genomics UK (COG-UK) Consortium | Jacquelyn Wynn, Mairead Hyland, The Lighthouse Lab in Alderley Park and Alex Alderton, Roberto Amato, Sonia Goncalves, Ewan Harrison, David K. Jackson, Ian Johnston, Dominic Kwiatkowski, Cordelia Langford, John Sillitoe on behalf of the Wellcome Sanger Institute COVID-19 Surveillance Team |
| EPI_ISL_820647 | Lighthouse Lab in Glasgow | Wellcome Sanger Institute for the COVID-19 Genomics UK (COG-UK) Consortium | Harper VanSteenhouse, Yumi Kasai, David Gray, Carol Clugston, Anna Dominiczak and Alex Alderton, Roberto Amato, Sonia Goncalves, Ewan Harrison, David K. Jackson, Ian Johnston, Dominic Kwiatkowski, Cordelia Langford, John Sillitoe on behalf of the Wellcome Sanger Institute COVID-19 Surveillance Team |
| EPI_ISL_822346, EPI_ISL_822853, EPI_ISL_822864, EPI_ISL_822868, EPI_ISL_822891, EPI_ISL_822896, EPI_ISL_822908, EPI_ISL_822942, EPI_ISL_822988, EPI_ISL_823007, EPI_ISL_823008, EPI_ISL_823012, EPI_ISL_823015, EPI_ISL_823021, EPI_ISL_823022, EPI_ISL_823023, EPI_ISL_823024, EPI_ISL_823056, EPI_ISL_823059, EPI_ISL_823060, EPI_ISL_823061, EPI_ISL_823062, EPI_ISL_823063, EPI_ISL_823116, EPI_ISL_823131, EPI_ISL_823137, EPI_ISL_823143, EPI_ISL_823156, EPI_ISL_823170, EPI_ISL_823171, EPI_ISL_823176, EPI_ISL_823177, EPI_ISL_823218, EPI_ISL_823219, EPI_ISL_823220, EPI_ISL_823221, EPI_ISL_823222, EPI_ISL_823223, EPI_ISL_823224, EPI_ISL_823225, EPI_ISL_823226, EPI_ISL_823227, EPI_ISL_823228, EPI_ISL_823229, EPI_ISL_823230, EPI_ISL_823231, EPI_ISL_823232, EPI_ISL_823233, EPI_ISL_823236, EPI_ISL_823237, EPI_ISL_823238, EPI_ISL_823239, EPI_ISL_823240, EPI_ISL_823241, EPI_ISL_823242, EPI_ISL_823243, EPI_ISL_823244, EPI_ISL_823245, EPI_ISL_823397, EPI_ISL_823398, EPI_ISL_823399, EPI_ISL_823400, EPI_ISL_823401, EPI_ISL_823402, EPI_ISL_823403, EPI_ISL_823582, EPI_ISL_823586, EPI_ISL_823608, EPI_ISL_823609, EPI_ISL_823610, EPI_ISL_823611, EPI_ISL_823612, EPI_ISL_823613, EPI_ISL_823614, EPI_ISL_823615, EPI_ISL_823616, EPI_ISL_823617, EPI_ISL_823618, EPI_ISL_823619, EPI_ISL_823620, EPI_ISL_823621, EPI_ISL_823622, EPI_ISL_823623, EPI_ISL_823624, EPI_ISL_823626, EPI_ISL_823627, EPI_ISL_823628, EPI_ISL_823629, EPI_ISL_823630, EPI_ISL_823631, EPI_ISL_823632, EPI_ISL_823633 | | | |
| see above | Wales Specialist Virology Centre Sequencing lab: Pathogen Genomics Unit | COVID-19 Genomics UK (COG-UK) Consortium | Catherine Moore, Johnathan Evans, Laura Gifford, Malorie Perry, Simon Cottrell, Angela Marchbank, Alec Birchley, Alexander Adams, Amy Gaskin, Bree Gatica-Wilcox, Jason Coombes, Joel Southgate, Lauren Gilbert, Lee Graham, Nicole Pacchiarini, Sara Kumziene-Summerhayes, Sarah Taylor, Sophie Jones, Sara Rey, Matthew Bull, Joanne Watkins, Sally Corden, Tom Connor |
| EPI_ISL_824284, EPI_ISL_824285 | Institute of Microbiology, Universidad San Francisco de Quito | Institute of Microbiology, Universidad San Francisco de Quito | Belén Prado-Vivar, Sully Márquez, Juan José Guadalupe, Monica Becerra-Wong, Bernardo Gutiérrez, Tania Guayasamin, Patricio Reyes, Verónica Barragán, Patricio Rojas-Silva, Gabriel Trueba, Michelle Grunauer, Paúl Cárdenas |
| EPI_ISL_824295 | DOHMH Morrisania | New York City Public Health Laboratory | Jade Wang, et al. |
| EPI_ISL_824311, EPI_ISL_824312 | DOHMH Fort Greene | New York City Public Health Laboratory | Jade Wang, et al. |
| EPI_ISL_824313 | DOHMH Jamaica | New York City Public Health Laboratory | Jade Wang, et al. |
| EPI_ISL_824314, EPI_ISL_824315 | DOHMH Corona | New York City Public Health Laboratory | Jade Wang, et al. |
| EPI_ISL_824316 | DOHMH Crown Heights | New York City Public Health Laboratory | Jade Wang, et al. |
| EPI_ISL_824317, EPI_ISL_824318 | DOHMH Central Harlem | New York City Public Health Laboratory | Jade Wang, et al. |
| EPI_ISL_824319 | DOHMH Chelsea | New York City Public Health Laboratory | Jade Wang, et al. |
| EPI_ISL_824320 | DOHMH Corona | New York City Public Health Laboratory | Jade Wang, et al. |
| EPI_ISL_824321, EPI_ISL_824322 | DOHMH Fort Greene | New York City Public Health Laboratory | Jade Wang, et al. |
| EPI_ISL_824323, EPI_ISL_824324, EPI_ISL_824325, EPI_ISL_824326, EPI_ISL_824327 | DOHMH Morrisania | New York City Public Health Laboratory | Jade Wang, et al. |
| EPI_ISL_824328 | DOHMH Corona | New York City Public Health Laboratory | Jade Wang, et al. |
| EPI_ISL_824329, EPI_ISL_824330 | DOHMH Jamaica | New York City Public Health Laboratory | Jade Wang, et al. |
| EPI_ISL_824331 | DOHMH Corona | New York City Public Health Laboratory | Jade Wang, et al. |
| EPI_ISL_824343, EPI_ISL_824345, EPI_ISL_824358, EPI_ISL_824359, EPI_ISL_824360, EPI_ISL_824361, EPI_ISL_824363, EPI_ISL_824377, EPI_ISL_824378 | Michigan Department of Health and Human Services, Bureau of Laboratories | Michigan Department of Health and Human Services, Bureau of Laboratories | Blankenship HM, Riner D, Soehnlen MK |
| EPI_ISL_824402 | California Department of Public Health | California Department of Public Health | CDPH IDLB COVIDNet |
| EPI_ISL_824983, EPI_ISL_824984 | Maryland Public Health Laboratory | Maryland Public Health Laboratory | Maryland Department of Health Laboratories Administration |
| EPI_ISL_825026, EPI_ISL_825027 | Utah Public Health Laboratory, Utah Public Health Laboratory Infectious Disease submission group | Utah Public Health Laboratory, Utah Public Health Laboratory Infectious Disease submission group | Young,E.L., Oakeson,K.F., Gallagher,T. |
| EPI_ISL_825378 | Hospital Universitari Vall d'Hebron - Vall d'Hebron Institut de Rercerca | Hospital Universitari Vall d'Hebron | Cristina Andrés, Maria Piñana, Josep F Abril, Damir Garcia-Cehic, Ariadna Rando, Juliana Esperalba, Maria Gema Codina, Carla Castillo, Maria Carmen Martin, Tomàs Pumarola, Josep Quer, Andrés Antón |
| EPI_ISL_825385 | Nigeria Centre For Disease Control | National reference Laboratory, NCDC, Gaduwa, Abuja | Dr Ndodo Nnaemeka, Olusola Anuoluwapo Akanbi, Chimaobi Chukwu, Dr Omoare Adesuyi, Kingsley Madubuike, Anthony Ahumibe, Naidoo Dhamari, Nwando Mba, Dr Chikwe Ihekweazu |
| EPI_ISL_825446, EPI_ISL_825447, EPI_ISL_825448, EPI_ISL_825449, EPI_ISL_825450, EPI_ISL_825451, EPI_ISL_825452, EPI_ISL_825453, EPI_ISL_825454, EPI_ISL_825455, EPI_ISL_825456, EPI_ISL_825457, EPI_ISL_825458, EPI_ISL_825459, EPI_ISL_825460, EPI_ISL_825465, EPI_ISL_825466, EPI_ISL_825467, EPI_ISL_825468, EPI_ISL_825469, EPI_ISL_825470, EPI_ISL_825471, EPI_ISL_825472, EPI_ISL_825473, EPI_ISL_825474, EPI_ISL_825475, EPI_ISL_825476, EPI_ISL_825477, EPI_ISL_825478, EPI_ISL_825479, EPI_ISL_825480, EPI_ISL_825481, EPI_ISL_825482, EPI_ISL_825483, EPI_ISL_825484, EPI_ISL_825485, EPI_ISL_825486, EPI_ISL_825487, EPI_ISL_825488, EPI_ISL_825489 | | | |
| see above | NHLS-IALCH | KRISP, KZN Research Innovation and Sequencing Platform | Giandhari J, Pillay S, Lessells R, Mdlalose K, York D, Khan S, Tegally H, Wilkinson E, de Oliveira T |
| EPI_ISL_825617 | Nigeria Centre For Disease Control | National reference Laboratory, NCDC, Gaduwa, Abuja | Dr Ndodo Nnaemeka, Olusola Anuoluwapo Akanbi, Chimaobi Chukwu, Dr Omoare Adesuyi, Esebanmen Grace, Anthony Ahumibe, Naidoo Dhamari, Nwando Mba, Dr Chikwe Ihekweazu |

| | | | |
|---|---|---|---|
| EPI_ISL_826464 | Lighthouse Lab in Alderley Park | Wellcome Sanger Institute for the COVID-19 Genomics UK (COG-UK) Consortium | Jacquelyn Wynn, Mairead Hyland, The Lighthouse Lab in Alderley Park and Alex Alderton, Roberto Amato, Sonia Goncalves, Ewan Harrison, David K. Jackson, Ian Johnston, Dominic Kwiatkowski, Cordelia Langford, John Sillitoe on behalf of the Wellcome Sanger Institute COVID-19 Surveillance Team |
| EPI_ISL_827039, EPI_ISL_827778, EPI_ISL_827779, EPI_ISL_828069, EPI_ISL_828071, EPI_ISL_828073, EPI_ISL_829954, EPI_ISL_830219, EPI_ISL_830222, EPI_ISL_830226, EPI_ISL_830228 | | | |
| see above | deCODE genetics | deCODE genetics | Daniel F Gudbjartsson; Agnar Helgason; Hakon Jonsson; Olafur T Magnusson; Pall Melsted; Gudmundur L Norddahl; Jona Saemundsdottir; Asgeir Sigurdsson; Patrick Sulem; Arna B Agustsdottir; Hannes Eggertsson; Berglind Eiriksdottir; Run Fridriksdottir; Elisabet E Gardarsdottir; Gudmundur Georgsson; Olafia S Gretarsdottir; Kjartan R Gudmundsson; Thora R Gunnarsdottir; Arnaldur Gylfason; Hilma Holm; Brynjar O Jensson; Aslaug Jonasdottir; Kamilla S Josefsdottir; Thordur Kristjansson; Droplaug N Magnusdottir; Solvi Rognvaldsson; Louise le Roux; Gudrun Sigmundsdottir; Gardar Sveinbjornsson; Kristin E Sveinsdottir; Maney Sveinsdottir; Emil A Thorarensen; Bjarni Thorbjornsson; Gisli Masson; Ingileif Jonsdottir; Alma Moller; Thorolfur Gudnason; Karl G Kristinsson; Unnur Thorsteinsdottir; Kari Stefansson |
| EPI_ISL_830519 | The National University Hospital of Iceland | deCODE genetics | Daniel F Gudbjartsson; Agnar Helgason; Hakon Jonsson; Olafur T Magnusson; Pall Melsted; Gudmundur L Norddahl; Jona Saemundsdottir; Asgeir Sigurdsson; Patrick Sulem; Arna B Agustsdottir; Hannes Eggertsson; Berglind Eiriksdottir; Run Fridriksdottir; Elisabet E Gardarsdottir; Gudmundur Georgsson; Olafia S Gretarsdottir; Kjartan R Gudmundsson; Thora R Gunnarsdottir; Arnaldur Gylfason; Hilma Holm; Brynjar O Jensson; Aslaug Jonasdottir; Kamilla S Josefsdottir; Thordur Kristjansson; Droplaug N Magnusdottir; Solvi Rognvaldsson; Louise le Roux; Gudrun Sigmundsdottir; Gardar Sveinbjornsson; Kristin E Sveinsdottir; Maney Sveinsdottir; Emil A Thorarensen; Bjarni Thorbjornsson; Gisli Masson; Ingileif Jonsdottir; Alma Moller; Thorolfur Gudnason; Karl G Kristinsson; Unnur Thorsteinsdottir; Kari Stefansson |
| EPI_ISL_831027, EPI_ISL_831030, EPI_ISL_831275, EPI_ISL_831276, EPI_ISL_831277, EPI_ISL_831278, EPI_ISL_831279, EPI_ISL_831285, EPI_ISL_831287, EPI_ISL_831288 | Hospital Universitario La Paz (Madrid) | SeqCOVID-SPAIN consortium/IBV(CSIC) | Fernando Lázaro-Perona, María Rodríguez-Tejedor, Elias Dahdouh, Jesús Mingorance and SeqCOVID-SPAIN consortium |
| EPI_ISL_831379 | Labor Krone | Robert Koch Institute, Influenza and respiratory viruses FG17 & Bioinformatics MF1, Berlin, Germany | Dr. Münstermann. Prof. Tiemann , Stephan Fuchs, Stefan Kroeger, Marianne Wedde, Oliver Drechsel, Aleksandar Radonic, Rene Kmiecinski, Ralf Duerrwald, Thorsten Wolff |
| EPI_ISL_831380, EPI_ISL_831381 | Limbach - MVZ Labor Dr. Limbach & Kollegen | Robert Koch Institute, Influenza and respiratory viruses FG17 & Bioinformatics MF1, Berlin, Germany | Dr. Konrad Bode, Stephan Fuchs, Stefan Kroeger, Marianne Wedde, Oliver Drechsel, Aleksandar Radonic, Rene Kmiecinski, Ralf Duerrwald, Thorsten Wolff |
| EPI_ISL_831382, EPI_ISL_831383, EPI_ISL_831384, EPI_ISL_831385 | Labor Krone | Robert Koch Institute, Influenza and respiratory viruses FG17 & Bioinformatics MF1, Berlin, Germany | Dr. Münstermann. Prof. Tiemann , Stephan Fuchs, Stefan Kroeger, Marianne Wedde, Oliver Drechsel, Aleksandar Radonic, Rene Kmiecinski, Ralf Duerrwald, Thorsten Wolff |
| EPI_ISL_831386, EPI_ISL_831387, EPI_ISL_831388 | Limbach - MVZ Labor Dr. Limbach & Kollegen | Robert Koch Institute, Influenza and respiratory viruses FG17 & Bioinformatics MF1, Berlin, Germany | Dr. Konrad Bode, Stephan Fuchs, Stefan Kroeger, Marianne Wedde, Oliver Drechsel, Aleksandar Radonic, Rene Kmiecinski, Ralf Duerrwald, Thorsten Wolff |
| EPI_ISL_831651 | Institute for Infectious Diseases, University of Bern, Switzerland | Institute for Infectious Diseases, University of Bern, Switzerland | Michel C Koch, Christian Baumann, Miguel A Terrazos Miani, Cora Sägesser, Pascal Bittel, Stephen L Leib, Peter Keller, Franziska Suter-Riniker, Alban Ramette |
| EPI_ISL_831695, EPI_ISL_831703, EPI_ISL_831705, EPI_ISL_831712, EPI_ISL_831716, EPI_ISL_831718, EPI_ISL_831737, EPI_ISL_831739, EPI_ISL_831742, EPI_ISL_831749, EPI_ISL_831750, EPI_ISL_831757, EPI_ISL_831758, EPI_ISL_831870, EPI_ISL_831871, EPI_ISL_831872, EPI_ISL_831873, EPI_ISL_831877, EPI_ISL_831878, EPI_ISL_831879, EPI_ISL_831880, EPI_ISL_831881 | | | |
| see above | United States Air Force School of Aerospace Medicine | United States Air Force School of Aerospace Medicine | Anthony Fries, Jennifer Meyer, William Gruner, Amanda Javorina, Sarah Purves, Clarise Starr, Elizabeth Macias |
| EPI_ISL_831986, EPI_ISL_831987, EPI_ISL_831988 | Synlab Medilab, Mikrobiologi | The Public Health Agency of Sweden | Department of Microbiology, The Public Health Agency of Sweden |
| EPI_ISL_831989 | Laboratorium for klinisk mikrobiologi | The Public Health Agency of Sweden | Department of Microbiology, The Public Health Agency of Sweden |
| EPI_ISL_831990, EPI_ISL_831992, EPI_ISL_831996, EPI_ISL_831997 | Klinisk mikrobiologi | The Public Health Agency of Sweden | Department of Microbiology, The Public Health Agency of Sweden |
| EPI_ISL_832068 | Santa Clara County Public Health Laboratory | Santa Clara County Public Health Laboratory | Santa Clara County Public Health Department |
| EPI_ISL_832106 | MD Laboratories | Los Angeles County PHL | P. Hemarajata et al. |
| EPI_ISL_832392, EPI_ISL_832394, EPI_ISL_832395, EPI_ISL_832396 | Santa Clara County Public Health Laboratory | Santa Clara County Public Health Laboratory | Santa Clara County Public Health Department |
| EPI_ISL_833134 | Laboratorio de Ecologia de Doencas Transmissiveis na Amazonia, Instituto Leonidas e Maria Deane - Fiocruz Amazonia | Laboratorio de Ecologia de Doencas Transmissiveis na Amazonia, Instituto Leonidas e Maria Deane - Fiocruz Amazonia | Valdinete Nascimento, Victor Souza, André Corado, Fernanda Nascimento, George Silva, Ágatha Costa, Debora Duarte, Karina Pessoa, Matilde Mejía, Luciana Gonçalves, Maria Júlia Brandão, Michele Jesus, Felipe Naveca |
| EPI_ISL_833456 | UZA/UAntwerpen (BIS lab), University Hospital Antwerp, Edegem, Belgium | Laboratory of Medical Microbiology, University of Antwerp, Campus Drie Eiken, S6.26, Universiteitsplein 1, 2610, Wilrijk, Antwerp, Belgium | Basil Britto Xavier, Jasmine Coppens, Christine Lammens, Veerle Matheeussen, Herman Goossens |
| EPI_ISL_836980, EPI_ISL_837113, EPI_ISL_837114, EPI_ISL_837115, EPI_ISL_837200, EPI_ISL_837201, EPI_ISL_837202 | Respiratory Virus Unit, National Infection Service, Public Health England | COVID-19 Genomics UK (COG-UK) Consortium | PHE Covid Sequencing Team |
| EPI_ISL_837342, EPI_ISL_837350 | National Virus Reference Laboratory | National Virus Reference Laboratory | Michael Carr, Gabriel Gonzalez, Jonathan Dean, Cillian F De Gascun |
| EPI_ISL_837444 | Istituto Zooprofilattico Sperimentale del Mezzogiorno | TIGEM | Patrizia Annunziata, Andrea Ballabio, Valentina Bouche, Davide Cacchiarelli (CorrespAuthor), Pellegrino Cerino, Chiara Colantuono, Lucio Di Filippo, Antonio Grimaldi, Antonio Limone, Gabriella Loconte, Anna Manfredi, Francesco Panariello, Biancamaria Pierri, Marcello Salvi, Lucia Vassallo |
| EPI_ISL_838117, EPI_ISL_838119, EPI_ISL_838120, EPI_ISL_838123, EPI_ISL_838127, EPI_ISL_838129, EPI_ISL_838130, EPI_ISL_838131 | West of Scotland Specialist Virology Centre, NHSGGC / MRC-University of Glasgow Centre for Virus Research | COVID-19 Genomics UK (COG-UK) Consortium | Ana da Silva Filipe, Natasha Johnson, Kathy Smollett, Daniel Mair, Stephen Carmichael, Alice Broos, Lily Tong, Jenna Nichols, Kyriaki Nomikou; Sarah McDonald; Richard Orton, Joseph Hughes, Sreenu Vattipally, David L Robertson; Alasdair MacLean, Rory Gunson; Sharif Shaaban, Matthew Holden; Rachel Blacow, Guy Mollett, Kathy Li, James Shepherd, Antonia Ho, Emma Thomson |
| EPI_ISL_838222, EPI_ISL_838223, EPI_ISL_838224, EPI_ISL_838225, EPI_ISL_838226, EPI_ISL_838234, EPI_ISL_838288, EPI_ISL_838289 | Virology Department, Royal Infirmary of Edinburgh, NHS Lothian / School of Biological Sciences, University of Edinburgh / Institute of Genetics and Molecular Medicine, University of Edinburgh | COVID-19 Genomics UK (COG-UK) Consortium | McHugh M, Dewar R, Rooke S, Gallagher M, Balcaza C, O'Toole Á, Scher E, Hill V, McCrone JT, Colquhoun R, Yu X, Jackson B, Rambaut A, Williams TC, Templeton K |
| EPI_ISL_838458, EPI_ISL_838459, EPI_ISL_838460, EPI_ISL_838461, EPI_ISL_838462, EPI_ISL_838463, EPI_ISL_838466, EPI_ISL_838467, EPI_ISL_838468, EPI_ISL_838469, EPI_ISL_838470, EPI_ISL_838473, EPI_ISL_838475, EPI_ISL_838476, EPI_ISL_838477, EPI_ISL_838478, EPI_ISL_838479, EPI_ISL_838480, EPI_ISL_838482, EPI_ISL_838483, EPI_ISL_838484, EPI_ISL_838485, EPI_ISL_838487, EPI_ISL_838489, EPI_ISL_838491, EPI_ISL_838493, EPI_ISL_838494, EPI_ISL_838495, EPI_ISL_838497, EPI_ISL_838498, EPI_ISL_838499, EPI_ISL_838500, EPI_ISL_838501, EPI_ISL_838502, EPI_ISL_838503, EPI_ISL_838504, EPI_ISL_838505, EPI_ISL_838506, EPI_ISL_838507, EPI_ISL_838508, EPI_ISL_838509, EPI_ISL_838510, EPI_ISL_838511, EPI_ISL_838512, EPI_ISL_838513, EPI_ISL_838514, EPI_ISL_838516, EPI_ISL_838517, EPI_ISL_838518, EPI_ISL_838519, EPI_ISL_838520, EPI_ISL_838521, EPI_ISL_838522, EPI_ISL_838523, EPI_ISL_838524, EPI_ISL_838525, EPI_ISL_838526, EPI_ISL_838527, EPI_ISL_838528, EPI_ISL_838530, EPI_ISL_838538, EPI_ISL_838539, EPI_ISL_838540, EPI_ISL_838543, EPI_ISL_838544, EPI_ISL_838546, EPI_ISL_838548, EPI_ISL_838550, EPI_ISL_838557, EPI_ISL_838560, EPI_ISL_838561, EPI_ISL_838562, EPI_ISL_838563, EPI_ISL_838565, EPI_ISL_838566, EPI_ISL_838567 | | | |
| see above | Liverpool Clinical Laboratories | COVID-19 Genomics UK (COG-UK) Consortium | Sam Haldenby, Anita Lucaci, Steve Paterson, Julian Hiscox, Alistair Darby, M Almsaud, A Alrezaihi, Muhannad Alruwaili, Stuart D Armstrong, Jones Benjamin, Eleanor G Bentley, Anu Chawla, Jordan J Clark, Angela Cowell, Richard Eccles, Isabel García-Dorival, Matthew Gemmell, Alessandro Gerada, PKF Gilmore, Richard Gregory, Ximeng Han, Catherine Hartley, Margaret Hughes, Miren Iturriza-Gomara, James Johnson, L Luu, Jenifer Manson, Charlotte Nelson, Elaine O'Toole, Cassie Olateju, Rebekah Penrice-Randal , Lucille Rainbow, N.P Randle, Trevor Ian Robinson, Parul Sharma, Ghada T Shawli, James P Stewart, Neil Swainston, Ecaterina Vamos, Joanne Watts, Mark Whitehead |
| EPI_ISL_838687, EPI_ISL_838688, EPI_ISL_838689, EPI_ISL_838690, EPI_ISL_838691, EPI_ISL_838692, EPI_ISL_838693, EPI_ISL_838694, EPI_ISL_838695, EPI_ISL_838696, EPI_ISL_838697, EPI_ISL_838698, EPI_ISL_838699, EPI_ISL_838700, EPI_ISL_838701, EPI_ISL_838702, EPI_ISL_838703, EPI_ISL_838704, | | | |

| | | | |
|---|---|---|---|
| EPI_ISL_838706, EPI_ISL_838707, EPI_ISL_838716, EPI_ISL_838720, EPI_ISL_838722, EPI_ISL_838728, EPI_ISL_838729 | | | |
| see above | University College London, Great Ormond Street Hospital for Children NHS Foundation Trust, Imperial College Healthcare NHS Trust | COVID-19 Genomics UK (COG-UK) Consortium | Sergi Castellano, Rachel Williams, Mark Kristiansen, Paola Resende Silva, Sunando Roy, Tony Brooks, Helena Tutill, Paola Niola, Patricia Dyal, Charlotte Williams, Leysa Forrest, Yasmin Panchbhaya, Jacqueline Findlay, Samuel Weeks, Julianne Brown, Kathryn Harris, Paul Randell, James Price, Alison Holmes, Judith Breuer |
| EPI_ISL_839485, EPI_ISL_839488, EPI_ISL_839490, EPI_ISL_839491, EPI_ISL_839493, EPI_ISL_839494, EPI_ISL_839497, EPI_ISL_839500, EPI_ISL_839507, EPI_ISL_839544, EPI_ISL_839545, EPI_ISL_839546, EPI_ISL_839547, EPI_ISL_839548, EPI_ISL_839549, EPI_ISL_839550, EPI_ISL_839551, EPI_ISL_839552, EPI_ISL_839553, EPI_ISL_839554, EPI_ISL_839555, EPI_ISL_839556, EPI_ISL_839558, EPI_ISL_839559, EPI_ISL_839560, EPI_ISL_839561, EPI_ISL_839562, EPI_ISL_839563, EPI_ISL_839564, EPI_ISL_839565, EPI_ISL_839566, EPI_ISL_839567, EPI_ISL_839568, EPI_ISL_839569, EPI_ISL_839570, EPI_ISL_839571, EPI_ISL_839572, EPI_ISL_839573, EPI_ISL_839574, EPI_ISL_839575, EPI_ISL_839576, EPI_ISL_839577, EPI_ISL_839578, EPI_ISL_839579, EPI_ISL_839580, EPI_ISL_839581, EPI_ISL_839590, EPI_ISL_839632, EPI_ISL_839634, EPI_ISL_839635, EPI_ISL_839637, EPI_ISL_839642, EPI_ISL_839661, EPI_ISL_839662, EPI_ISL_839665 | | | |
| see above | Northumbria University / South Tees Hospitals NHS Foundation Trust / North Cumbria Integrated Care NHS Foundation Trust / North Tees and Hartlepool NHS Foundation Trust / Newcastle Hospitals NHS Foundation Trust | COVID-19 Genomics UK (COG-UK) Consortium | Darren L Smith,Andrew Nelson,Matthew Bashton,Greg R Young,Joshua Loh,John Allan,Mohammad A Tariq,Giles S Holt,Gary Black,Wen C Yew,Lynn Dover,Paul Baker,Steve Liggett,Sarah Essex,Jane Greenaway,Debra Padgett,Clive Graham,Garren Scott,Edward Barton,Emma Swindells,Brendan Payne,Jennifer Collins,Yusri Taha,Gary Eltringham |
| EPI_ISL_840048, EPI_ISL_840049, EPI_ISL_840050, EPI_ISL_840146, EPI_ISL_840147, EPI_ISL_840148, EPI_ISL_840149, EPI_ISL_840150, EPI_ISL_840151, EPI_ISL_840152, EPI_ISL_840153, EPI_ISL_840154, EPI_ISL_840155, EPI_ISL_840156, EPI_ISL_840157, EPI_ISL_840158, EPI_ISL_840159, EPI_ISL_840160, EPI_ISL_840161, EPI_ISL_840162, EPI_ISL_840163, EPI_ISL_840164, EPI_ISL_840165, EPI_ISL_840166, EPI_ISL_840167, EPI_ISL_840168, EPI_ISL_840169, EPI_ISL_840170, EPI_ISL_840171, EPI_ISL_840172, EPI_ISL_840173, EPI_ISL_840174, EPI_ISL_840175, EPI_ISL_840176, EPI_ISL_840177, EPI_ISL_840178, EPI_ISL_840179 | | | |
| see above | Lincolnshire Hospitals and DeepSeq Nottingham | COVID-19 Genomics UK (COG-UK) Consortium | Nichola Duckworth, Tim Sloan, Sarah Walsh, Jonathan Ball, Patrick McClure, Joeseph Chappell, Nadine Holmes, Matthew Carlisle, Christopher Moore, Fei Sang, Johnny Debebe, Victoria Wright, Matthew Loose |
| EPI_ISL_840259, EPI_ISL_840265, EPI_ISL_840266, EPI_ISL_840281, EPI_ISL_840282, EPI_ISL_840283, EPI_ISL_840284, EPI_ISL_840286, EPI_ISL_840351, EPI_ISL_840358 | Oxford Viromics, NDM, University of Oxford; Oxford University Hospitals; Basingstoke and North Hampshire Hospital | COVID-19 Genomics UK (COG-UK) Consortium | Tanya Golubchik, David Bonsall, George Macintyre, Amy Trebes, Mariateresa de Cesare, Catrin Moore, Alex Mobbs, Anita Justice, Robert Shaw, Monique Andersson, Timothy Peto, Emma Wise, Nathan Moore, Jessica Lynch, Nick Cortes, Matilde Mori, Stephen Kidd, David Buck, John Todd, Christophe Fraser |
| EPI_ISL_840835, EPI_ISL_840885, EPI_ISL_840886 | Wales Specialist Virology Centre Sequencing lab: Pathogen Genomics Unit | Public Health Wales Microbiology Cardiff Wales Specialist Virology Centre | Catherine Moore, Johnathan Evans, Laura Gifford, Malorie Perry, Simon Cottrell, Angela Marchbank, Alec Birchley, Alexander Adams, Amy Gaskin, Bree Gatica-Wilcox, Jason Coombes, Joel Southgate, Lauren Gilbert, Lee Graham, Nicole Pacchiarini, Sara Kumziene-Summerhayes, Sarah Taylor, Sophie Jones, Sara Rey, Matthew Bull, Joanne Watkins, Sally Corden, Tom Connor |
| EPI_ISL_841769, EPI_ISL_841774, EPI_ISL_841780, EPI_ISL_841782, EPI_ISL_841786, EPI_ISL_841788, EPI_ISL_841794, EPI_ISL_841796, EPI_ISL_841812, EPI_ISL_841814, EPI_ISL_841815, EPI_ISL_841816, EPI_ISL_841817, EPI_ISL_841818, EPI_ISL_841819, EPI_ISL_841821, EPI_ISL_841822, EPI_ISL_841823, EPI_ISL_841824, EPI_ISL_841825, EPI_ISL_841826, EPI_ISL_841827, EPI_ISL_841828, EPI_ISL_841829, EPI_ISL_841848, EPI_ISL_841850, EPI_ISL_841852, EPI_ISL_841853, EPI_ISL_841854, EPI_ISL_841861 | | | |
| see above | Centre for Enzyme Innovation, University of Portsmouth / Translational Research Laboratory, Portsmouth Hospitals NHS Trust | COVID-19 Genomics UK (COG-UK) Consortium | Angela Beckett,Yann Bourgeois,Garry Scarlett,Sharon Glaysher,Scott Elliott,Kelly Bicknell,Robert Impey,Allyson Lloyd,Sarah Wyllie,Ethan Butcher,Anoop Chauhan,Samuel Robson |
| EPI_ISL_842670, EPI_ISL_842671, EPI_ISL_842672, EPI_ISL_842674, EPI_ISL_842675, EPI_ISL_842676, EPI_ISL_842677, EPI_ISL_842678, EPI_ISL_842679, EPI_ISL_842682, EPI_ISL_842683, EPI_ISL_842684, EPI_ISL_842685, EPI_ISL_842688, EPI_ISL_842691, EPI_ISL_842693, EPI_ISL_842694, EPI_ISL_842695, EPI_ISL_842705, EPI_ISL_842706, EPI_ISL_842707, EPI_ISL_842708, EPI_ISL_842710, EPI_ISL_842711, EPI_ISL_842712, EPI_ISL_842713, EPI_ISL_842714, EPI_ISL_842715, EPI_ISL_842716, EPI_ISL_842726, EPI_ISL_842727, EPI_ISL_842728, EPI_ISL_842729, EPI_ISL_842730, EPI_ISL_842731, EPI_ISL_842732, EPI_ISL_842734 | | | |
| see above | University College London Hospital | COVID-19 Genomics UK (COG-UK) Consortium | Judith Heaney, Matthew Byott, Catherine Houlihan, Dan Frampton, Stuart Kirk, Moira Spyer and Eleni Nastouli |
| EPI_ISL_843078, EPI_ISL_843089, EPI_ISL_843090 | Barts Health NHS Trust | COVID-19 Genomics UK (COG-UK) Consortium | CUTINO-MOGUEL, Maria-Teresa; HARRINGTON, David; OWOYEMI, Dola; SHYLINI, Raghavendran; BROAD, Claire; KELE, Beatrix |
| EPI_ISL_844202, EPI_ISL_844203, EPI_ISL_844206, EPI_ISL_844207, EPI_ISL_844208, EPI_ISL_844209, EPI_ISL_844210, EPI_ISL_844211, EPI_ISL_844212, EPI_ISL_844213, EPI_ISL_844226, EPI_ISL_844235, EPI_ISL_844242, EPI_ISL_844243, EPI_ISL_844244, EPI_ISL_844245, EPI_ISL_844246, EPI_ISL_844247, EPI_ISL_844248, EPI_ISL_844253, EPI_ISL_844256, EPI_ISL_844257, EPI_ISL_844259, EPI_ISL_844260, EPI_ISL_844265, EPI_ISL_844268, EPI_ISL_844270, EPI_ISL_844271, EPI_ISL_844272, EPI_ISL_844273, EPI_ISL_844274, EPI_ISL_844275, EPI_ISL_844276, EPI_ISL_844277, EPI_ISL_844278, EPI_ISL_844279, EPI_ISL_844280, EPI_ISL_844281, EPI_ISL_844301, EPI_ISL_844306, EPI_ISL_844307, EPI_ISL_844313, EPI_ISL_844314, EPI_ISL_844315, EPI_ISL_844316, EPI_ISL_844317, EPI_ISL_844318, EPI_ISL_844319, EPI_ISL_844320, EPI_ISL_844321, EPI_ISL_844322, EPI_ISL_844323, EPI_ISL_844324, EPI_ISL_844325, EPI_ISL_844326, EPI_ISL_844327, EPI_ISL_844328, EPI_ISL_844329, EPI_ISL_844330, EPI_ISL_844331, EPI_ISL_844332, EPI_ISL_844333, EPI_ISL_844334, EPI_ISL_844335, EPI_ISL_844336, EPI_ISL_844337, EPI_ISL_844338, EPI_ISL_844339, EPI_ISL_844340, EPI_ISL_844341, EPI_ISL_844342, EPI_ISL_844343, EPI_ISL_844344, EPI_ISL_844345, EPI_ISL_844346, EPI_ISL_844347, EPI_ISL_844348, EPI_ISL_844349, EPI_ISL_844350, EPI_ISL_844351, EPI_ISL_844352, EPI_ISL_844353, EPI_ISL_844354, EPI_ISL_844355, EPI_ISL_844356, EPI_ISL_844357, EPI_ISL_844358, EPI_ISL_844359, EPI_ISL_844360, EPI_ISL_844361, EPI_ISL_844362, EPI_ISL_844363, EPI_ISL_844364, EPI_ISL_844365, EPI_ISL_844366, EPI_ISL_844367, EPI_ISL_844368, EPI_ISL_844369, EPI_ISL_844370, EPI_ISL_844371, EPI_ISL_844372, EPI_ISL_844373, EPI_ISL_844374, EPI_ISL_844375, EPI_ISL_844376, EPI_ISL_844377, EPI_ISL_844378, EPI_ISL_844379, EPI_ISL_844380, EPI_ISL_844381, EPI_ISL_844382, EPI_ISL_844383, EPI_ISL_844384, EPI_ISL_844385, EPI_ISL_844386, EPI_ISL_844387, EPI_ISL_844388, EPI_ISL_844389, EPI_ISL_844390 | | | |
| see above | Department of Virus and Microbiological Special Diagnostics, Statens Serum Institut, Copenhagen, Denmark | Albertsen Lab, Department of Chemistry and Bioscience, Aalborg University, Denmark | Danish Covid-19 Genome Consortium |
| EPI_ISL_846582 | National Laboratory for Health, Environment and Food | National Laboratory for Health, Environment and Food | Aleksander Mahnic, Sandra Janezic, Maja Rupnik |
| EPI_ISL_847260, EPI_ISL_847514 | Department of Virus and Microbiological Special Diagnostics, Statens Serum Institut, Copenhagen, Denmark | Albertsen Lab, Department of Chemistry and Bioscience, Aalborg University, Denmark | Danish Covid-19 Genome Consortium |
| EPI_ISL_847517 | Chiu Laboratory, University of California, San Francisco | Chiu Laboratory, University of California, San Francisco | Charles Chiu, Xianding (Wayne) Deng, Candace Wang, Brian Bushnell, Scot Federman, Jill Hacker, Debra Wadford |
| EPI_ISL_847528 | California Department of Public Health | Chiu Laboratory, University of California, San Francisco | Charles Chiu, Xianding (Wayne) Deng, Candace Wang, Brian Bushnell, Scot Federman, Jill Hacker, Debra Wadford |
| EPI_ISL_847540, EPI_ISL_847553 | Chiu Laboratory, University of California, San Francisco | Chiu Laboratory, University of California, San Francisco | Charles Chiu, Xianding (Wayne) Deng, Candace Wang, Brian Bushnell, Scot Federman, Jill Hacker, Debra Wadford |
| EPI_ISL_847555, EPI_ISL_847559, EPI_ISL_847569 | California Department of Public Health | Chiu Laboratory, University of California, San Francisco | Charles Chiu, Xianding (Wayne) Deng, Candace Wang, Brian Bushnell, Scot Federman, Jill Hacker, Debra Wadford |
| EPI_ISL_847579 | Chiu Laboratory, University of California, San Francisco | Chiu Laboratory, University of California, San Francisco | Charles Chiu, Xianding (Wayne) Deng, Candace Wang, Brian Bushnell, Scot Federman, Jill Hacker, Debra Wadford |
| EPI_ISL_847638, EPI_ISL_847639 | California Department of Public Health | Chiu Laboratory, University of California, San Francisco | Charles Chiu, Xianding (Wayne) Deng, Candace Wang, Brian Bushnell, Scot Federman, Jill Hacker, Debra Wadford |
| EPI_ISL_847640, EPI_ISL_847641, EPI_ISL_847642, EPI_ISL_847643, EPI_ISL_847644 | Chiu Laboratory, University of California, San Francisco | Chiu Laboratory, University of California, San Francisco | Charles Chiu, Xianding (Wayne) Deng, Candace Wang, Brian Bushnell, Scot Federman, Jill Hacker, Debra Wadford |
| EPI_ISL_847713 | California Department of Public Health | Chiu Laboratory, University of California, San Francisco | Charles Chiu, Xianding (Wayne) Deng, Candace Wang, Brian Bushnell, Scot Federman, Jill Hacker, Debra Wadford |
| EPI_ISL_847716, EPI_ISL_847720 | Chiu Laboratory, University of California, San Francisco | Chiu Laboratory, University of California, San Francisco | Charles Chiu, Xianding (Wayne) Deng, Candace Wang, Brian Bushnell, Scot Federman, Jill Hacker, Debra Wadford |
| EPI_ISL_847724, EPI_ISL_847740, EPI_ISL_847781 | California Department of Public Health | Chiu Laboratory, University of California, San Francisco | Charles Chiu, Xianding (Wayne) Deng, Candace Wang, Brian Bushnell, Scot Federman, Jill Hacker, Debra Wadford |
| EPI_ISL_847782 | Chiu Laboratory, University of California, San Francisco | Chiu Laboratory, University of California, San Francisco | Charles Chiu, Xianding (Wayne) Deng, Candace Wang, Brian Bushnell, Scot Federman, Jill Hacker, Debra Wadford |
| EPI_ISL_847979, EPI_ISL_847982, EPI_ISL_847990, EPI_ISL_848009, EPI_ISL_848016, EPI_ISL_848021, EPI_ISL_848049, EPI_ISL_848051, EPI_ISL_848053, EPI_ISL_848057 | Michigan Department of Health and Human Services, Bureau of Laboratories | Michigan Department of Health and Human Services, Bureau of Laboratories | Blankenship HM, Riner D, Soehnlen MK |
| EPI_ISL_848260, EPI_ISL_848268, EPI_ISL_848427, EPI_ISL_848428, EPI_ISL_848429, EPI_ISL_848430, EPI_ISL_848441, EPI_ISL_848458, EPI_ISL_848535, EPI_ISL_848536, EPI_ISL_848537 | | | |
| see above | Illinois Department of Public Health | Gagnon Lab, Southern Illinois University | Keith Gagnon |
| EPI_ISL_849208, EPI_ISL_849209, EPI_ISL_849212, EPI_ISL_849213, EPI_ISL_849214, EPI_ISL_849216, EPI_ISL_849218, EPI_ISL_849225, EPI_ISL_849226, EPI_ISL_849227, EPI_ISL_849231, EPI_ISL_849233, EPI_ISL_849234, EPI_ISL_849235, EPI_ISL_849236, EPI_ISL_849239, EPI_ISL_849247, EPI_ISL_849248, EPI_ISL_849249, EPI_ISL_849250, EPI_ISL_849251, EPI_ISL_849252, EPI_ISL_849253, EPI_ISL_849254, EPI_ISL_849255, EPI_ISL_849258, EPI_ISL_849260, EPI_ISL_849263, EPI_ISL_849266, EPI_ISL_849270, EPI_ISL_849273, EPI_ISL_849274, EPI_ISL_849277, EPI_ISL_849278 | | | |

| | | | |
|---|---|---|---|
| see above | Utah Public Health Laboratory | Utah Public Health Laboratory | Erin L. Young, Kelly F. Oakeson, Tara Gallagher |
| EPI_ISL_850658 | Division of Emerging Infectious Diseases, Bureau of Infectious Diseases Diagnosis Control, Korea Disease Control and Prevention Agency | Division of Emerging Infectious Diseases, Bureau of Infectious Diseases Diagnosis Control, Korea Disease Control and Prevention Agency | Ae Kyung Park, Il-Hwan Kim, Heui Man Kim, Jeong-Min Kim, Namjoo Lee, Chaeyoung Lee, Sang Hee Woo, Eun-Jin Kim |
| EPI_ISL_850667, EPI_ISL_850674 | The National Institute of Public Health | State Veterinary Institute Prague | Nagy,A;Jirincova,H;Trnka,D;Vecerova,J;Trinklova,M |
| EPI_ISL_852571, EPI_ISL_852572 | Max von Pettenkofer Institute, Virology, National Reference Center for Retroviruses, LMU München | Laboratory for Functional Genome Analysis, Dept. Genomics, Gene Center of the LMU Munich | Max Muenchhoff, Stefan Krebs, Alexander Graf, Oliver Keppler, Helmut Blum |
| EPI_ISL_852616 | National Laboratory for Health, Environment and Food | National Laboratory for Health, Environment and Food | Aleksander Mahnic, Sandra Janezic, Maja Rupnik |
| EPI_ISL_852817 | CHU Purpan - Laboratoire de Virologie - Institut Fédératif de Biologie | CHU Purpan - Laboratoire de Virologie - Institut Fédératif de Biologie | Latour J., Ranger N., Dubois M., Carcenac R., Harter A., Boyer P., Tremeaux P., Izopet J. |
| EPI_ISL_852958, EPI_ISL_852989 | Hospital General Universitario Gregorio Marañón | SeqCOVID-SPAIN consortium/IBV(CSIC) | Darío García de Viedma, Laura Pérez-Lago, Pedro J Sola-Campoy, Sergio Buenestado-Serrano, Marta Herranz, Victor Manuel de la Cueva, Julia Suárez, Pilar Catalán, Patricia Muñoz and SeqCOVID-SPAIN consortium |
| EPI_ISL_853345 | UPMC Clinical Microbiology Laboratory | Microbial Genome Sequencing Center; Microbial Genomic Epidemiology Laboratory | Mustapha M. Mustapha, Jane W. Marsh, Dan Snyder, Marissa P. Griffith, Stephanie L. Mitchell, Vatsala R. Srinivasa, Kady D. Waggle, Chinelo Ezeonwuku, Vaughn S. Cooper, Lee H. Harrison |
| EPI_ISL_853725, EPI_ISL_853739, EPI_ISL_853744, EPI_ISL_853747, EPI_ISL_853765 | Department of Microbiology, University Innsbruck | Bergthaler laboratory, CeMM Research Center for Molecular Medicine of the Austrian Academy of Sciences | Lukas Endler, Alexandra Popa, Benedikt Agerer, Jakob-Wendelin Genger, Alexander Lercher, Anna Schedl, Thomas Penz, Michael Schuster, Jan Laine, Martin Senekowitsch, Christoph Bock, Andreas Bergthaler |
| EPI_ISL_855409 | Servicio de Microbiología, Laboratori Clínic Metropolitana Nord. Hospital Universitari Germans Trias i Pujol. Institut d'Investigació en Ciències de la Salut Germans Trias i Pujol (IGTP) | SeqCOVID-SPAIN consortium/IBV(CSIC) | Elisa Martró, Antoni E. Bordoy, Anna Not, Adrián Antuori, Anabel Fernández, Nona Romaní, Verónica Saludes, Cristina Casañ and SeqCOVID-SPAIN consortium |
| EPI_ISL_855517, EPI_ISL_855519, EPI_ISL_855531 | KEMRI-Wellcome Trust Research Programme/KEMRI-CGMR-C Kilifi | KEMRI-Wellcome Trust Research Programme/KEMRI-CGMR-C Kilifi | Githinji et al |
| EPI_ISL_856663, EPI_ISL_856664, EPI_ISL_856665, EPI_ISL_856666, EPI_ISL_856667, EPI_ISL_856668, EPI_ISL_856669, EPI_ISL_856670, EPI_ISL_856671, EPI_ISL_856672 | Department of Virus and Microbiological Special Diagnostics, Statens Serum Institut, Copenhagen, Denmark | Aalborg University | Danish Covid-19 Genome Consortium |
| EPI_ISL_856696, EPI_ISL_856697, EPI_ISL_856699, EPI_ISL_856701 | Ohio Department of Health Laboratory | Ohio Department of Health Laboratory | Holmes, Jennifer; Eric Brandt, Keoni Omura, Glen McGillivary, Caitlin McDonnell, Kirtana Ramadugu, Erica Leasure, Kelsey Florek, Heather Blankenship, Quanta Brown, and Tammy Bannerman |
| EPI_ISL_856785, EPI_ISL_856787 | Servicio Virosis Respiratorias-Departamento Virología-INEI | Instituto Nacional Enfermedades Infecciosas C.G.Malbran | Baumeister E., Avaro M., Benedetti E., Russo M., Dattero ME, Pontoriero A., Cisterna D., Molina V., Perandones C., Tuduri E., Lorenzo F., Poklepovich T., Campos J. |
| EPI_ISL_856810, EPI_ISL_856811, EPI_ISL_856814, EPI_ISL_856821, EPI_ISL_856823, EPI_ISL_856827, EPI_ISL_856831, EPI_ISL_856832, EPI_ISL_856835, EPI_ISL_856836, EPI_ISL_856840, EPI_ISL_856843, EPI_ISL_856846, EPI_ISL_856847, EPI_ISL_856851, EPI_ISL_856854, EPI_ISL_856857 | | | |
| see above | Utah Public Health Laboratory | Utah Public Health Laboratory | Erin L. Young, Kelly F. Oakeson, Tara Gallagher |
| EPI_ISL_857514 | Swiss National Reference Centre for Influenza | Swiss National Reference Centre for Influenza | Ana Rita Goncalves,Samuel Cordey, Laurent Kaiser, Lorenzo Cerutti, Henri Pegeot, Melyssa Elies, Keith Harshman, Ioannis Xenarios, Emmanouil Dermitzakis |
| EPI_ISL_857537, EPI_ISL_857538, EPI_ISL_857539 | Swiss National Reference Centre for Influenza | Swiss National Reference Centre for Influenza | Tim Roloff, Ana Rita Gonçalves, Madlen Stange, Helena MB Seth-Smith, Alfredo Mari, Karoline Leuzinger, Julia Bielicki, Manuel Battegay, Hans Hirsch, Laurent Kaiser, Adrian Egli |
| EPI_ISL_858408 | Lighthouse Lab in Alderley Park | Wellcome Sanger Institute for the COVID-19 Genomics UK (COG-UK) Consortium | Jacquelyn Wynn, Mairead Hyland, The Lighthouse Lab in Alderley Park and Alex Alderton, Roberto Amato, Sonia Goncalves, Ewan Harrison, David K. Jackson, Ian Johnston, Dominic Kwiatkowski, Cordelia Langford, John Sillitoe on behalf of the Wellcome Sanger Institute COVID-19 Surveillance Team |
| EPI_ISL_858410 | Lighthouse Lab in Glasgow | Wellcome Sanger Institute for the COVID-19 Genomics UK (COG-UK) Consortium | Harper VanSteenhouse, Yumi Kasai, David Gray, Carol Clugston, Anna Dominiczak and Alex Alderton, Roberto Amato, Sonia Goncalves, Ewan Harrison, David K. Jackson, Ian Johnston, Dominic Kwiatkowski, Cordelia Langford, John Sillitoe on behalf of the Wellcome Sanger Institute COVID-19 Surveillance Team |
| EPI_ISL_858411 | Lighthouse Lab in Alderley Park | Wellcome Sanger Institute for the COVID-19 Genomics UK (COG-UK) Consortium | Jacquelyn Wynn, Mairead Hyland, The Lighthouse Lab in Alderley Park and Alex Alderton, Roberto Amato, Sonia Goncalves, Ewan Harrison, David K. Jackson, Ian Johnston, Dominic Kwiatkowski, Cordelia Langford, John Sillitoe on behalf of the Wellcome Sanger Institute COVID-19 Surveillance Team |
| EPI_ISL_858412 | Lighthouse Lab in Glasgow | Wellcome Sanger Institute for the COVID-19 Genomics UK (COG-UK) Consortium | Harper VanSteenhouse, Yumi Kasai, David Gray, Carol Clugston, Anna Dominiczak and Alex Alderton, Roberto Amato, Sonia Goncalves, Ewan Harrison, David K. Jackson, Ian Johnston, Dominic Kwiatkowski, Cordelia Langford, John Sillitoe on behalf of the Wellcome Sanger Institute COVID-19 Surveillance Team |
| EPI_ISL_858413, EPI_ISL_858414 | Lighthouse Lab in Alderley Park | Wellcome Sanger Institute for the COVID-19 Genomics UK (COG-UK) Consortium | Jacquelyn Wynn, Mairead Hyland, The Lighthouse Lab in Alderley Park and Alex Alderton, Roberto Amato, Sonia Goncalves, Ewan Harrison, David K. Jackson, Ian Johnston, Dominic Kwiatkowski, Cordelia Langford, John Sillitoe on behalf of the Wellcome Sanger Institute COVID-19 Surveillance Team |
| EPI_ISL_858415, EPI_ISL_858417 | Lighthouse Lab in Glasgow | Wellcome Sanger Institute for the COVID-19 Genomics UK (COG-UK) Consortium | Harper VanSteenhouse, Yumi Kasai, David Gray, Carol Clugston, Anna Dominiczak and Alex Alderton, Roberto Amato, Sonia Goncalves, Ewan Harrison, David K. Jackson, Ian Johnston, Dominic Kwiatkowski, Cordelia Langford, John Sillitoe on behalf of the Wellcome Sanger Institute COVID-19 Surveillance Team |
| EPI_ISL_858418 | Lighthouse Lab in Alderley Park | Wellcome Sanger Institute for the COVID-19 Genomics UK (COG-UK) Consortium | Jacquelyn Wynn, Mairead Hyland, The Lighthouse Lab in Alderley Park and Alex Alderton, Roberto Amato, Sonia Goncalves, Ewan Harrison, David K. Jackson, Ian Johnston, Dominic Kwiatkowski, Cordelia Langford, John Sillitoe on behalf of the Wellcome Sanger Institute COVID-19 Surveillance Team |
| EPI_ISL_858419, EPI_ISL_858420, EPI_ISL_858421 | Lighthouse Lab in Glasgow | Wellcome Sanger Institute for the COVID-19 Genomics UK (COG-UK) Consortium | Harper VanSteenhouse, Yumi Kasai, David Gray, Carol Clugston, Anna Dominiczak and Alex Alderton, Roberto Amato, Sonia Goncalves, Ewan Harrison, David K. Jackson, Ian Johnston, Dominic Kwiatkowski, Cordelia Langford, John Sillitoe on behalf of the Wellcome Sanger Institute COVID-19 Surveillance Team |
| EPI_ISL_858422 | Lighthouse Lab in Alderley Park | Wellcome Sanger Institute for the COVID-19 Genomics UK (COG-UK) Consortium | Jacquelyn Wynn, Mairead Hyland, The Lighthouse Lab in Alderley Park and Alex Alderton, Roberto Amato, Sonia Goncalves, Ewan Harrison, David K. Jackson, Ian Johnston, Dominic Kwiatkowski, Cordelia Langford, John Sillitoe on behalf of the Wellcome Sanger Institute COVID-19 Surveillance Team |
| EPI_ISL_858423, EPI_ISL_858424 | Lighthouse Lab in Glasgow | Wellcome Sanger Institute for the COVID-19 Genomics UK (COG-UK) Consortium | Harper VanSteenhouse, Yumi Kasai, David Gray, Carol Clugston, Anna Dominiczak and Alex Alderton, Roberto Amato, Sonia Goncalves, Ewan Harrison, David K. Jackson, Ian Johnston, Dominic Kwiatkowski, Cordelia Langford, John Sillitoe on behalf of the Wellcome Sanger Institute COVID-19 Surveillance Team |
| EPI_ISL_858929, EPI_ISL_858973, EPI_ISL_858992, EPI_ISL_858993, EPI_ISL_858995, EPI_ISL_858996, EPI_ISL_858998, EPI_ISL_858999, EPI_ISL_859000, EPI_ISL_859002, EPI_ISL_859004, EPI_ISL_859006, EPI_ISL_859007, EPI_ISL_859009, EPI_ISL_859010, EPI_ISL_859011, EPI_ISL_859013, EPI_ISL_859014, EPI_ISL_859015, EPI_ISL_859016, EPI_ISL_859017, EPI_ISL_859019, EPI_ISL_859021, EPI_ISL_859022, EPI_ISL_859023, EPI_ISL_859025, EPI_ISL_859026, EPI_ISL_859027 | | | |
| see above | Lighthouse Lab in Alderley Park | Wellcome Sanger Institute for the COVID-19 Genomics UK (COG-UK) Consortium | Jacquelyn Wynn, Mairead Hyland, The Lighthouse Lab in Alderley Park and Alex Alderton, Roberto Amato, Sonia Goncalves, Ewan Harrison, David K. Jackson, Ian Johnston, Dominic Kwiatkowski, Cordelia Langford, John Sillitoe on behalf of the Wellcome Sanger Institute COVID-19 Surveillance Team |
| EPI_ISL_859326 | Lighthouse Lab in Glasgow | Wellcome Sanger Institute for the COVID-19 Genomics UK (COG-UK) Consortium | Harper VanSteenhouse, Yumi Kasai, David Gray, Carol Clugston, Anna Dominiczak and Alex Alderton, Roberto Amato, Sonia Goncalves, Ewan Harrison, David K. Jackson, Ian Johnston, Dominic Kwiatkowski, Cordelia Langford, John Sillitoe on behalf of the Wellcome Sanger Institute COVID-19 Surveillance Team |
| EPI_ISL_859401, EPI_ISL_859427, EPI_ISL_859432, EPI_ISL_859433 | Lighthouse Lab in Alderley Park | Wellcome Sanger Institute for the COVID-19 Genomics UK (COG-UK) Consortium | Jacquelyn Wynn, Mairead Hyland, The Lighthouse Lab in Alderley Park and Alex Alderton, Roberto Amato, Sonia Goncalves, Ewan Harrison, David K. Jackson, Ian Johnston, Dominic Kwiatkowski, Cordelia Langford, John Sillitoe on behalf of the Wellcome Sanger Institute COVID-19 Surveillance Team |

| | | | |
|---|---|---|---|
| EPI_ISL_859456, EPI_ISL_859457, EPI_ISL_859458, EPI_ISL_859459, EPI_ISL_859460, EPI_ISL_859461, EPI_ISL_859464, EPI_ISL_859470, EPI_ISL_859473, EPI_ISL_859474, EPI_ISL_859475, EPI_ISL_859476, EPI_ISL_859481 | | | |
| see above | Lighthouse Lab in Glasgow | Wellcome Sanger Institute for the COVID-19 Genomics UK (COG-UK) Consortium | Harper VanSteenhouse, Yumi Kasai, David Gray, Carol Clugston, Anna Dominiczak and Alex Alderton, Roberto Amato, Sonia Goncalves, Ewan Harrison, David K. Jackson, Ian Johnston, Dominic Kwiatkowski, Cordelia Langford, John Sillitoe on behalf of the Wellcome Sanger Institute COVID-19 Surveillance Team |
| EPI_ISL_860114, EPI_ISL_860115 | Medical Microbiology Unit, Department for Laboratory Medicine, Drammen Hospital, Vestre Viken Health Trust, | Norwegian Institute of Public Health, Department of Virology | Kathrine Stene-Johansen, Kamilla Heddeland Instefjord, Hilde Elshaug, Atiya R Ali,Marie Paulsen Madsen, Rasmus Riis Kopperud, Hilde Vollan, Karoline Bragstad, Olav Hungnes |
| EPI_ISL_860218 | Ostfold Hospital Trust - Kalnes, Centre for Laboratory Medicine, Section for gene technology and infection serology | Norwegian Institute of Public Health, Department of Virology | Kathrine Stene-Johansen, Kamilla Heddeland Instefjord, Hilde Elshaug, Atiya R Ali,Marie Paulsen Madsen, Rasmus Riis Kopperud, Hilde Vollan, Karoline Bragstad, Olav Hungnes |
| EPI_ISL_860292 | Innlandet Hospital Trust, Division Lillehammer, Department for Medical Microbiology | Norwegian Institute of Public Health, Department of Virology | Kathrine Stene-Johansen, Kamilla Heddeland Instefjord, Hilde Elshaug, Atiya R Ali,Marie Paulsen Madsen, Rasmus Riis Kopperud, Hilde Vollan, Karoline Bragstad, Olav Hungnes |
| EPI_ISL_860932 | Johns Hopkins Hospital Department of Pathology | Johns Hopkins Hospital Department of Pathology | C. Paul Morris, Chun Huai Luo, Adannaya Amadi, Nicholas Gallagher, Heba H. Mostafa |
| EPI_ISL_861465, EPI_ISL_861469, EPI_ISL_861471 | UHAS COVID-19 Lab | UHAS COVID-19 Lab | Kwabena O. Duedu, Jones Gyamfi, Reuben Ayivor-Djanie, John O. Gyapong and the UHAS COVID-19 Lab Team |
| EPI_ISL_861531, EPI_ISL_861590 | Instituto Nacional de Saude (INSA) | Instituto Nacional de Saude (INSA) | Borges et al |
| EPI_ISL_861775, EPI_ISL_861776, EPI_ISL_861783, EPI_ISL_861786, EPI_ISL_861805, EPI_ISL_861825, EPI_ISL_861826 | Hospital General Universitario Gregorio Marañón | SeqCOVID-SPAIN consortium/IBV(CSIC) | Darío García de Viedma, Laura Pérez-Lago, Pedro J Sola-Campoy, Sergio Buenestado-Serrano, Marta Herranz, Victor Manuel de la Cueva, Julia Suárez, Pilar Catalán, Patricia Muñoz and SeqCOVID-SPAIN consortium |
| EPI_ISL_862190 | Respiratory Virus Unit, National Infection Service, Public Health England | COVID-19 Genomics UK (COG-UK) Consortium | PHE Covid Sequencing Team |
| EPI_ISL_862581, EPI_ISL_862582, EPI_ISL_862583, EPI_ISL_862584 | Hospital Clínic | Instituto de Salud Carlos III | Iglesias-Caballero, M.Camarero, S. Molinero Calamita, M. González-Esguevillas, M. Pozo, F. Casas, I. Jiménez, P. Jiménez, M. Zaballos, A. Monzón, S. Varona, S. Juliá, M. Cuesta, I. Marcos, M.A. |
| EPI_ISL_862780, EPI_ISL_862782 | Utah Public Health Laboratory, Utah Public Health Laboratory Infectious Disease submission group | Utah Public Health Laboratory, Utah Public Health Laboratory Infectious Disease submission group | Young,E.L., Oakeson,K.F., Gallagher,T. |
| EPI_ISL_864574, EPI_ISL_864576 | Institute of Medical Microbiology and Hospital Hygiene | Institute of Medical Microbiology and Hospital Hygiene | Prof. Dr. Achim Kaasch, Aljoscha Tersteegen |
| EPI_ISL_864580 | National Institute of Public Health - National Institute of Hygiene | National Institute of Public Health - National Institute of Hygiene, DNA Sequencing and Synthesis Facility (oligo.pl) - Institute of Biochemistry and Biophysics PAS | Wokowicz Tomasz, Zacharczuk Katarzyna, Gawor Jan |
| EPI_ISL_864796, EPI_ISL_864847, EPI_ISL_864849, EPI_ISL_864851, EPI_ISL_864852 | Department of Pathology, University of Cambridge | COVID-19 Genomics UK (COG-UK) Consortium | Aminu S. Jahun, Yasmin Chaudhry, Grant Hall, Iliana Georgana, Myra Hosmillo, Martin D. Curran, Malte Pinckert, Surendra Parmar, Ian Goodfellow |
| EPI_ISL_865411, EPI_ISL_865412, EPI_ISL_865413, EPI_ISL_865414, EPI_ISL_865415, EPI_ISL_865416, EPI_ISL_865417, EPI_ISL_865418, EPI_ISL_865419, EPI_ISL_865420, EPI_ISL_865422, EPI_ISL_865427, EPI_ISL_865428, EPI_ISL_865433, EPI_ISL_865436, EPI_ISL_865437 | | | |
| see above | Liverpool Clinical Laboratories | COVID-19 Genomics UK (COG-UK) Consortium | Sam Haldenby, Anita Lucaci, Steve Paterson, Julian Hiscox, Alistair Darby, M Almsaud, A Alrezaihi, Muhannad Alruwaili, Stuart D Armstrong, Jones Benjamin, Eleanor G Bentley, Anu Chawla, Jordan J Clark, Angela Cowell, Richard Eccles, Isabel García-Dorival, Matthew Gemmell, Alessandro Gerada, PKF Gilmore, Richard Gregory, Ximeng Han, Catherine Hartley, Margaret Hughes, Miren Iturriza-Gomara, James Johnson, L Luu, Jenifer Manson, Charlotte Nelson, Elaine O'Toole, Cassie Olateju, Rebekah Penrice-Randal , Lucille Rainbow, N.P Randle, Trevor Ian Robinson, Parul Sharma, Ghada T Shawli, James P Stewart, Neil Swainston, Ecaterina Vamos, Joanne Watts, Mark Whitehead |
| EPI_ISL_865720 | University College London, Great Ormond Street Hospital for Children NHS Foundation Trust, Imperial College Healthcare NHS Trust | COVID-19 Genomics UK (COG-UK) Consortium | Sergi Castellano, Rachel Williams, Mark Kristiansen, Paola Resende Silva, Sunando Roy, Tony Brooks, Helena Tutill, Paola Niola, Patricia Dyal, Charlotte Williams, Leysa Forrest, Yasmin Panchbhaya, Jacqueline Findlay, Samuel Weeks, Julianne Brown, Kathryn Harris, Paul Randell, James Price, Alison Holmes, Judith Breuer |
| EPI_ISL_866049, EPI_ISL_866106, EPI_ISL_866121 | University College London Hospital | COVID-19 Genomics UK (COG-UK) Consortium | Judith Heaney, Matthew Byott, Catherine Houlihan, Dan Frampton, Stuart Kirk, Moira Spyer and Eleni Nastouli |
| EPI_ISL_866414, EPI_ISL_866415, EPI_ISL_866416, EPI_ISL_866417, EPI_ISL_866418, EPI_ISL_866419, EPI_ISL_866420, EPI_ISL_866421, EPI_ISL_866422, EPI_ISL_866463, EPI_ISL_866464, EPI_ISL_866465, EPI_ISL_866466, EPI_ISL_866467, EPI_ISL_866468, EPI_ISL_866476, EPI_ISL_866477, EPI_ISL_866480, EPI_ISL_866481, EPI_ISL_866482, EPI_ISL_866483, EPI_ISL_866485 | | | |
| see above | Northumbria University / South Tees Hospitals NHS Foundation Trust / North Cumbria Integrated Care NHS Foundation Trust / North Tees and Hartlepool NHS Foundation Trust / Newcastle Hospitals NHS Foundation Trust | COVID-19 Genomics UK (COG-UK) Consortium | Darren L Smith,Andrew Nelson,Matthew Bashton,Greg R Young,Joshua Loh,John Allan,Mohammad A Tariq,Giles S Holt,Gary Black,Wen C Yew,Lynn Dover,Paul Baker,Steve Liggett,Sarah Essex,Jane Greenaway,Debra Padgett,Clive Graham,Garren Scott,Edward Barton,Emma Swindells,Brendan Payne,Jennifer Collins,Yusri Taha,Gary Eltringham |
| EPI_ISL_866584, EPI_ISL_866828, EPI_ISL_866830 | Quadram Institute Bioscience | COVID-19 Genomics UK (COG-UK) Consortium | Dave J. Baker, Gemma L. Kay, Alp Aydin, Thanh Le-Viet, Steven Rudder, Ana P. Tedim, Anastasia Kolyva, Maria Diaz, Leonardo de Oliveira Martins, Nabil-Fareed Alikhan, Lizzie Meadows, Rachael Stanley, Ngozi Elumogo, Muhammed Yasir, Nicholas M. Thomson, Alexander J Trotter, Rachel Gilroy, Samuel Bloomfield, Claire Stuart, Andrew Bell, Reenesh Prakash, Samir Dervisevic, Alison E. Mather, John Wain, Mark Webber, Andrew J. Page, Justin O'Grady |
| EPI_ISL_866990, EPI_ISL_866991 | Queens Medical Centre, Clinical Microbiology Department / DeepSeq Nottingham | COVID-19 Genomics UK (COG-UK) Consortium | Gemma Clark, Wendy Smith, Manjinder Khakh, Vicki M Fleming, Michelle M Lister, Hannah Howson-Wells, Jonathan Ball, Patrick McClure, Joseph Chappell, Theocharis Tsoleridis, Nadine Holmes, Matthew Carlisle, Christopher Moore, Fei Sang, Johnny Debebe, Victoria Wright, Matthew Loose |
| EPI_ISL_871716, EPI_ISL_871717, EPI_ISL_871718, EPI_ISL_871719, EPI_ISL_871720, EPI_ISL_871721, EPI_ISL_871722, EPI_ISL_871723, EPI_ISL_871724, EPI_ISL_871725, EPI_ISL_871726, EPI_ISL_871727, EPI_ISL_871728, EPI_ISL_871729, EPI_ISL_871730, EPI_ISL_871731, EPI_ISL_871732, EPI_ISL_871733, EPI_ISL_871734, EPI_ISL_871735, EPI_ISL_871736, EPI_ISL_871737, EPI_ISL_871738, EPI_ISL_871739, EPI_ISL_871740, EPI_ISL_871741, EPI_ISL_871742, EPI_ISL_871743, EPI_ISL_871744, EPI_ISL_871745, EPI_ISL_871746, EPI_ISL_871747, EPI_ISL_871748, EPI_ISL_871749, EPI_ISL_871750, EPI_ISL_871751, EPI_ISL_871752, EPI_ISL_871753, EPI_ISL_871754, EPI_ISL_871755, EPI_ISL_871756, EPI_ISL_871757, EPI_ISL_871758 | | | |
| see above | Department of Virus and Microbiological Special Diagnostics, Statens Serum Institut, Copenhagen, Denmark | Aalborg University | Danish Covid-19 Genome Consortium |
| EPI_ISL_871924, EPI_ISL_871932, EPI_ISL_871936 | Servicio de Microbiología, Laboratori Clínic Metropolitana Nord. Hospital Universitari Germans Trias i Pujol. Institut d'Investigació en Ciències de la Salut Germans Trias i Pujol (IGTP) | SeqCOVID-SPAIN consortium/IBV(CSIC) | Elisa Martró, Antoni E. Bordoy, Anna Not, Adrián Antuori, Anabel Fernández, Nona Romaní, Verónica Saludes, Cristina Casañ and SeqCOVID-SPAIN consortium |
| EPI_ISL_872363, EPI_ISL_872364, EPI_ISL_872365, EPI_ISL_872366, EPI_ISL_872367, EPI_ISL_872368 | Texas Department of State Health Services (TXDSHS) | Texas Department of State Health Services (TXDSHS) | Bonnie Oh, Anita Pokharel, James Daniel Bonser, Myong Koag, Chung Wang, Rachel Lee, Grace Kubin, Rashmi Tuladhar, Mayela Pedrueza, Maliha Rahman, Jenny Zhang |
| EPI_ISL_872927, EPI_ISL_872928, EPI_ISL_872929, EPI_ISL_872930, EPI_ISL_872931, EPI_ISL_872932, EPI_ISL_872933, EPI_ISL_872934, EPI_ISL_872935, EPI_ISL_872936, EPI_ISL_872938, EPI_ISL_872939, EPI_ISL_872940, EPI_ISL_872941, EPI_ISL_872942, EPI_ISL_872944, EPI_ISL_872945, EPI_ISL_872947, EPI_ISL_872948, EPI_ISL_872951, EPI_ISL_872959, EPI_ISL_872960, EPI_ISL_872961, EPI_ISL_872962, EPI_ISL_872963, EPI_ISL_872964, EPI_ISL_872965, EPI_ISL_872966, EPI_ISL_872967, EPI_ISL_872968, EPI_ISL_872969, EPI_ISL_872971 | | | |
| see above | WHO National Influenza Centre Russian Federation | WHO National Influenza Centre Russian Federation | Andrey Komissarov, Artem Fadeev, Anna Ivanova, Kseniya Komissarova, Dmitry Bazhenov, Mikhail Bakaev, Daria Danilenko, Ksenia Safina, Elena Nabieva, Georgii Bazykin, Dmitry Lioznov |
| EPI_ISL_873219, EPI_ISL_873220 | M Health Fairview | Minnesota Department of Health, Public Health Laboratory | Alexandra Lorentz, Jacob Garfin, Matt Plumb, and Xiong Wang |
| EPI_ISL_876075, EPI_ISL_876100, EPI_ISL_876108, EPI_ISL_876152, EPI_ISL_876166, EPI_ISL_876183, EPI_ISL_876211, EPI_ISL_876212, EPI_ISL_876217, EPI_ISL_876297, EPI_ISL_876298, EPI_ISL_876299 | | | |

| | | | |
|---|---|---|---|
| see above | Massachusetts State Public Health Laboratory | Massachusetts State Public Health Laboratory | Andrew Lang, Timelia Fink, Glen Gallagher, Sandra Smole |
| EPI_ISL_876520, EPI_ISL_876531, EPI_ISL_876534, EPI_ISL_876535, EPI_ISL_876536, EPI_ISL_876537, EPI_ISL_876538, EPI_ISL_876539, EPI_ISL_876540 | Florida Bureau of Public Health Laboratories | Florida Bureau of Public Health Laboratories | Sarah Schmedes, Jason Blanton |
| EPI_ISL_876784, EPI_ISL_876785, EPI_ISL_876786, EPI_ISL_876787, EPI_ISL_876788, EPI_ISL_876789, EPI_ISL_876790, EPI_ISL_876791, EPI_ISL_876792, EPI_ISL_876793, EPI_ISL_876794 | | | |
| see above | Istituto Zooprofilattico Sperimentale della Puglia e della Basilicata | Istituto Zooprofilattico Sperimentale della Puglia e della Basilicata | Parisi A., Bianco A., Capozzi L., Del Sambro L., Manzulli V, Rondinone V., Pace L., Cipolletta D., Galante D. |
| EPI_ISL_877422 | Citra Medika Hospital | Institute of Tropical Disease, Universitas Airlangga | Rima R Prasetya, Krisnoadi Rahardjo, Aldise M Nastri, Jezzy R Dewantari, Wahidah Rachmaniyah, Gatot Soegiarto, Laksmi Wulandari, Resti Yudhawati, Soetjipto, Yasuko Mori, Maria I Lusida, Kazufumi Shimizu |
| EPI_ISL_877437, EPI_ISL_877438, EPI_ISL_877439, EPI_ISL_877440, EPI_ISL_877441, EPI_ISL_877442, EPI_ISL_877443, EPI_ISL_877444, EPI_ISL_877445, EPI_ISL_877446, EPI_ISL_877447, EPI_ISL_877448, EPI_ISL_877449, EPI_ISL_877450 | | | |
| see above | Institute of Microbiology and Immunology, Faculty of Medicine, University of Ljubljana | Institute of Microbiology and Immunology, Faculty of Medicine, University of Ljubljana | Samo Zakotnik, Tomaž Mark Zorec, Matic Brvar, Miša Korva, Mario Poljak, Tatjana Avši - Županc |
| EPI_ISL_878721, EPI_ISL_880180, EPI_ISL_880199, EPI_ISL_880200 | Rady's Childrens Hospital | Andersen lab at Scripps Research | SEARCH Alliance San Diego with Nanda Radamchar, David Dimmock, Linda Luo, Christina Clarke, Kathryn Bouic, Teresa Mueller, Denise Malicki |
| EPI_ISL_882958, EPI_ISL_882959 | National Institute of Public Health - National Institute of Hygiene | National Institute of Public Health - National Institute of Hygiene | Wokowicz Tomasz, Zacharczuk Katarzyna, Gawor Jan |
| EPI_ISL_884874, EPI_ISL_884875, EPI_ISL_884876, EPI_ISL_884877, EPI_ISL_884878, EPI_ISL_884879, EPI_ISL_884880, EPI_ISL_884881, EPI_ISL_884882, EPI_ISL_884883, EPI_ISL_884918, EPI_ISL_884919, EPI_ISL_884938, EPI_ISL_884960, EPI_ISL_884962, EPI_ISL_884964, EPI_ISL_884965, EPI_ISL_884966, EPI_ISL_884967, EPI_ISL_884968, EPI_ISL_884969, EPI_ISL_884970, EPI_ISL_884971, EPI_ISL_884973, EPI_ISL_884974, EPI_ISL_884975, EPI_ISL_884976, EPI_ISL_884977, EPI_ISL_884979, EPI_ISL_884980 | | | |
| see above | Santa Clara County Public Health Laboratory | Chan-Zuckerberg Biohub | CZB Cliahub Consortium |
| EPI_ISL_887445, EPI_ISL_887469, EPI_ISL_887470, EPI_ISL_887471, EPI_ISL_887474, EPI_ISL_887475 | Instituto Nacional de Saude (INS), Mozambique | KRISP, KZN Research Innovation and Sequencing Platform | Nalia Ismael, Nadia Sitoe, Paulo Arnaldo, Nedio Mabunda, Giandhari J, Pillay S, Tegally H, Wilkinson E, de Oliveira T |
| EPI_ISL_888677 | Laboratory for Respiratory Viruses, Cantacuzino National Military-Medical Institute for Research and Development | Cantacuzino Institute Virology | Mihaela Lazar, Luiza Ustea, Nicoleta Paraschiv, Tim Durfee |
| EPI_ISL_888686, EPI_ISL_888687, EPI_ISL_888688, EPI_ISL_888689, EPI_ISL_888690, EPI_ISL_888693, EPI_ISL_888694, EPI_ISL_888695, EPI_ISL_888696, EPI_ISL_888697, EPI_ISL_888698, EPI_ISL_888699, EPI_ISL_888748, EPI_ISL_888751, EPI_ISL_888772, EPI_ISL_888774, EPI_ISL_888775, EPI_ISL_888777, EPI_ISL_888780, EPI_ISL_888781, EPI_ISL_888782, EPI_ISL_888783, EPI_ISL_888784, EPI_ISL_888787 | | | |
| see above | KU Leuven, Rega Institute, Clinical and Epidemiological Virology | KU Leuven, Rega Institute, Clinical and Epidemiological Virology | Tony Wawina-Bokalanga, Bert Vanmechelen, Joan Marti-Carerras, Piet Maes |
| EPI_ISL_888857 | Michigan Department of Health and Human Services, Bureau of Laboratories | Michigan Department of Health and Human Services, Bureau of Laboratories | Blankenship HM, Riner D, Soehnlen MK |
| EPI_ISL_889013 | RS Rawa Lumbu | Eijkman Institute for Molecular Biology, Ministry of Research and Technology/National Agency for Research and Innovation | Iskandar Adnan, Lydia V. Panggalo, Sukma Oktavianthi, Willy Agustine, Edison Johar, Hidayat Trimarsanto, Frilasita A Yudhaputri, Safarina G Malik, Khin Saw Myint, Amin Soebandrio |
| EPI_ISL_890194 | Gonoshasthaya-RNA Research Center, Gonoshasthaya-RNA Molecular Diagnostics and Research Center | Gonoshasthaya-RNA Research Center, Gonoshasthaya-RNA Molecular Diagnostics and Research Center | Jamiruddin,M.R., Khondoker,M.U., Sharif,N., Azmuda,N., Ahmed,M.F., Sharmin,S., Akter,S., Mou,T.J., Marzan,M., Liza,S.M., Nahar,S., Jahan,N., Ali,T., Khandker,S.S., Jamiruddin,M., Haq,M.A., Adnan,N., Chaity,M., Oishee,M. |
| EPI_ISL_892274, EPI_ISL_892357, EPI_ISL_892358, EPI_ISL_892359, EPI_ISL_892360, EPI_ISL_892361, EPI_ISL_892362 | Servicio de Microbiología Clínica (Complejo Hospitalario de Navarra, Pamplona), Instituto de Investigación Sanitaria de Navarra (IdiSNA) | SeqCOVID-SPAIN consortium/IBV(CSIC) | Carmen Ezpeleta Baquedano, Ana Navascués, Ana Miqueleiz and SeqCOVID-SPAIN consortium |
| EPI_ISL_896094 | Viollier AG | University Hospital Basel, Clinical Bacteriology | Tim Roloff, Madlen Stange, Helena MB Seth-Smith, Alfredo Mari, Karoline Leuzinger, Julia Bielicki, Christiane Beckmann, Manuel Battegay, Hans Hirsch, Adrian Egli |
| EPI_ISL_896128, EPI_ISL_896160, EPI_ISL_896187, EPI_ISL_896188, EPI_ISL_896193 | MEPHI, Aix Marseille University | MEPHI, Aix Marseille University | Anthony LEVASSEUR |
| EPI_ISL_896299, EPI_ISL_896310, EPI_ISL_896311, EPI_ISL_896312, EPI_ISL_896313, EPI_ISL_896314, EPI_ISL_896315, EPI_ISL_896316, EPI_ISL_896317, EPI_ISL_896318, EPI_ISL_896319, EPI_ISL_896320, EPI_ISL_896321, EPI_ISL_896322, EPI_ISL_896323, EPI_ISL_896324, EPI_ISL_896325 | | | |
| see above | MEMORIAL SLOAN KETTERING CANCER CENTER | Wadsworth Center, New York State Department of Health | Kirsten St. George, Daryl M. Lamson, Alexis Russel, Matthew Shudt, Melissa A Leisner, Jonathan Plitnick, Navjot Singh, John Kelly, Erasmus Schneider, Erica Lasek-Nesselquist |
| EPI_ISL_900086, EPI_ISL_900092, EPI_ISL_900162, EPI_ISL_900236, EPI_ISL_900342, EPI_ISL_900394 | MEPHI, Aix Marseille University | MEPHI, Aix Marseille University | Anthony LEVASSEUR |
| EPI_ISL_900503, EPI_ISL_900571 | Hôpital Paris Saint-Joseph | CNR Virus des Infections Respiratoires - France SUD | Antonin Bal, Gregory Destras, Gwendolyne Burfin, Hadrien Règue, Quentin Semanas, Martine Valette, Bruno Lina, Pantxika Bellecave, Camille Ciccone, Isabelle Garrigue, Marie-Edith Lafon, Pascale Trimoulet, Laurence Josset |
| EPI_ISL_900574, EPI_ISL_900576, EPI_ISL_900579, EPI_ISL_900585, EPI_ISL_900593, EPI_ISL_900610, EPI_ISL_900611, EPI_ISL_900612, EPI_ISL_900613, EPI_ISL_900614, EPI_ISL_900615, EPI_ISL_900616, EPI_ISL_900664, EPI_ISL_900665, EPI_ISL_900666, EPI_ISL_900667, EPI_ISL_900668, EPI_ISL_900669, EPI_ISL_900670 | | | |
| see above | IZSM | TIGEM | Patrizia Annunziata, Andrea Ballabio, Valentina Bouche, Davide Cacchiarelli (CorrespAuthor), Pellegrino Cerino, Chiara Colantuono, Maria Concetta Cuomo, Denise Di Concilio, Lucio Di Filippo, Antonio Grimaldi, Antonio Limone, Anna Manfredi, Francesco Panariello, Biancamaria Pierri, Marcello Salvi |
| EPI_ISL_903251, EPI_ISL_903252 | M Health Fairview | Minnesota Department of Health, Public Health Laboratory | Alexandra Lorentz, Jacob Garfin, Matt Plumb, and Xiong Wang |
| EPI_ISL_903566 | MO State Public Health Laboratory | Genomics and Discovery, Respiratory Viruses Branch, Division of Viral Diseases, Centers for Disease Control and Prevention | Krista Queen, Yan Li, Ying Tao, Jing Zhang, Anna Uehara, Anna Montmayeur, Clinton R. Paden, Peter W. Cook, Rachel Marine, Mili Sheth, Jasmine Padilla, Sarah Nobles, Mark Burroughs, Lori Rowe, Haibin Wang, Ben L. Rambo-Martin, Dhwani Batra, Justin Lee, Suxiang Tong |
| EPI_ISL_903641, EPI_ISL_903647 | RI State Health Laboratories | Genomics and Discovery, Respiratory Viruses Branch, Division of Viral Diseases, Centers for Disease Control and Prevention | Krista Queen, Yan Li, Ying Tao, Jing Zhang, Anna Uehara, Anna Montmayeur, Clinton R. Paden, Peter W. Cook, Rachel Marine, Mili Sheth, Jasmine Padilla, Sarah Nobles, Mark Burroughs, Lori Rowe, Haibin Wang, Ben L. Rambo-Martin, Dhwani Batra, Justin Lee, Suxiang Tong |
| EPI_ISL_903652 | NC State Laboratory of Public Health | Genomics and Discovery, Respiratory Viruses Branch, Division of Viral Diseases, Centers for Disease Control and Prevention | Krista Queen, Yan Li, Ying Tao, Jing Zhang, Anna Uehara, Anna Montmayeur, Clinton R. Paden, Peter W. Cook, Rachel Marine, Mili Sheth, Jasmine Padilla, Sarah Nobles, Mark Burroughs, Lori Rowe, Haibin Wang, Ben L. Rambo-Martin, Dhwani Batra, Justin Lee, Suxiang Tong |
| EPI_ISL_903701 | MN PHL Division, Minnesota Department of Health | Genomics and Discovery, Respiratory Viruses Branch, Division of Viral Diseases, Centers for Disease Control and Prevention | Krista Queen, Yan Li, Ying Tao, Jing Zhang, Anna Uehara, Anna Montmayeur, Clinton R. Paden, Peter W. Cook, Rachel Marine, Mili Sheth, Jasmine Padilla, Sarah Nobles, Mark Burroughs, Lori Rowe, Haibin Wang, Ben L. Rambo-Martin, Dhwani Batra, Justin Lee, Suxiang Tong |
| EPI_ISL_903725 | SD Public Health Laboratory | Genomics and Discovery, Respiratory Viruses Branch, Division of Viral Diseases, Centers for Disease Control and | Krista Queen, Yan Li, Ying Tao, Jing Zhang, Anna Uehara, Anna Montmayeur, Clinton R. Paden, Peter W. Cook, Rachel Marine, Mili Sheth, Jasmine Padilla, Sarah Nobles, Mark Burroughs, Lori Rowe, Haibin Wang, Ben L. Rambo-Martin, Dhwani Batra, Justin Lee, Suxiang Tong |

| | | | |
|---|---|---|---|
| | | Prevention | |
| EPI_ISL_903731, EPI_ISL_903732, EPI_ISL_903735 | NV State Public Health Laboratory | Genomics and Discovery, Respiratory Viruses Branch, Division of Viral Diseases, Centers for Disease Control and Prevention | Krista Queen, Yan Li, Ying Tao, Jing Zhang, Anna Uehara, Anna Montmayeur, Clinton R. Paden, Peter W. Cook, Rachel Marine, Mili Sheth, Jasmine Padilla, Sarah Nobles, Mark Burroughs, Lori Rowe, Haibin Wang, Ben L. Rambo-Martin, Dhwani Batra, Justin Lee, Suxiang Tong |
| EPI_ISL_903738 | MN PHL Division, Minnesota Department of Health | Genomics and Discovery, Respiratory Viruses Branch, Division of Viral Diseases, Centers for Disease Control and Prevention | Krista Queen, Yan Li, Ying Tao, Jing Zhang, Anna Uehara, Anna Montmayeur, Clinton R. Paden, Peter W. Cook, Rachel Marine, Mili Sheth, Jasmine Padilla, Sarah Nobles, Mark Burroughs, Lori Rowe, Haibin Wang, Ben L. Rambo-Martin, Dhwani Batra, Justin Lee, Suxiang Tong |
| EPI_ISL_903744 | SD Public Health Laboratory | Genomics and Discovery, Respiratory Viruses Branch, Division of Viral Diseases, Centers for Disease Control and Prevention | Krista Queen, Yan Li, Ying Tao, Jing Zhang, Anna Uehara, Anna Montmayeur, Clinton R. Paden, Peter W. Cook, Rachel Marine, Mili Sheth, Jasmine Padilla, Sarah Nobles, Mark Burroughs, Lori Rowe, Haibin Wang, Ben L. Rambo-Martin, Dhwani Batra, Justin Lee, Suxiang Tong |
| EPI_ISL_903772 | IA State Hygienic Laboratory | Genomics and Discovery, Respiratory Viruses Branch, Division of Viral Diseases, Centers for Disease Control and Prevention | Krista Queen, Yan Li, Ying Tao, Jing Zhang, Anna Uehara, Anna Montmayeur, Clinton R. Paden, Peter W. Cook, Rachel Marine, Mili Sheth, Jasmine Padilla, Sarah Nobles, Mark Burroughs, Lori Rowe, Haibin Wang, Ben L. Rambo-Martin, Dhwani Batra, Justin Lee, Suxiang Tong |
| EPI_ISL_903794 | AZ SPHL, Arizona Department of Health Services | Genomics and Discovery, Respiratory Viruses Branch, Division of Viral Diseases, Centers for Disease Control and Prevention | Krista Queen, Yan Li, Ying Tao, Jing Zhang, Anna Uehara, Anna Montmayeur, Clinton R. Paden, Peter W. Cook, Rachel Marine, Mili Sheth, Jasmine Padilla, Sarah Nobles, Mark Burroughs, Lori Rowe, Haibin Wang, Ben L. Rambo-Martin, Dhwani Batra, Justin Lee, Suxiang Tong |
| EPI_ISL_903815 | IA State Hygienic Laboratory | Genomics and Discovery, Respiratory Viruses Branch, Division of Viral Diseases, Centers for Disease Control and Prevention | Krista Queen, Yan Li, Ying Tao, Jing Zhang, Anna Uehara, Anna Montmayeur, Clinton R. Paden, Peter W. Cook, Rachel Marine, Mili Sheth, Jasmine Padilla, Sarah Nobles, Mark Burroughs, Lori Rowe, Haibin Wang, Ben L. Rambo-Martin, Dhwani Batra, Justin Lee, Suxiang Tong |
| EPI_ISL_903831 | MO State Public Health Laboratory | Genomics and Discovery, Respiratory Viruses Branch, Division of Viral Diseases, Centers for Disease Control and Prevention | Krista Queen, Yan Li, Ying Tao, Jing Zhang, Anna Uehara, Anna Montmayeur, Clinton R. Paden, Peter W. Cook, Rachel Marine, Mili Sheth, Jasmine Padilla, Sarah Nobles, Mark Burroughs, Lori Rowe, Haibin Wang, Ben L. Rambo-Martin, Dhwani Batra, Justin Lee, Suxiang Tong |
| EPI_ISL_903836 | SD Public Health Laboratory | Genomics and Discovery, Respiratory Viruses Branch, Division of Viral Diseases, Centers for Disease Control and Prevention | Krista Queen, Yan Li, Ying Tao, Jing Zhang, Anna Uehara, Anna Montmayeur, Clinton R. Paden, Peter W. Cook, Rachel Marine, Mili Sheth, Jasmine Padilla, Sarah Nobles, Mark Burroughs, Lori Rowe, Haibin Wang, Ben L. Rambo-Martin, Dhwani Batra, Justin Lee, Suxiang Tong |
| EPI_ISL_903842 | AZ SPHL, Arizona Department of Health Services | Genomics and Discovery, Respiratory Viruses Branch, Division of Viral Diseases, Centers for Disease Control and Prevention | Krista Queen, Yan Li, Ying Tao, Jing Zhang, Anna Uehara, Anna Montmayeur, Clinton R. Paden, Peter W. Cook, Rachel Marine, Mili Sheth, Jasmine Padilla, Sarah Nobles, Mark Burroughs, Lori Rowe, Haibin Wang, Ben L. Rambo-Martin, Dhwani Batra, Justin Lee, Suxiang Tong |
| EPI_ISL_903861 | CO Dept. of Public Health and Environment, Lab Services Division | Genomics and Discovery, Respiratory Viruses Branch, Division of Viral Diseases, Centers for Disease Control and Prevention | Krista Queen, Yan Li, Ying Tao, Jing Zhang, Anna Uehara, Anna Montmayeur, Clinton R. Paden, Peter W. Cook, Rachel Marine, Mili Sheth, Jasmine Padilla, Sarah Nobles, Mark Burroughs, Lori Rowe, Haibin Wang, Ben L. Rambo-Martin, Dhwani Batra, Justin Lee, Suxiang Tong |
| EPI_ISL_903873 | AZ SPHL, Arizona Department of Health Services | Genomics and Discovery, Respiratory Viruses Branch, Division of Viral Diseases, Centers for Disease Control and Prevention | Krista Queen, Yan Li, Ying Tao, Jing Zhang, Anna Uehara, Anna Montmayeur, Clinton R. Paden, Peter W. Cook, Rachel Marine, Mili Sheth, Jasmine Padilla, Sarah Nobles, Mark Burroughs, Lori Rowe, Haibin Wang, Ben L. Rambo-Martin, Dhwani Batra, Justin Lee, Suxiang Tong |
| EPI_ISL_903884 | TX DSHS, Lab Services Section MC 1947 | Genomics and Discovery, Respiratory Viruses Branch, Division of Viral Diseases, Centers for Disease Control and Prevention | Krista Queen, Yan Li, Ying Tao, Jing Zhang, Anna Uehara, Anna Montmayeur, Clinton R. Paden, Peter W. Cook, Rachel Marine, Mili Sheth, Jasmine Padilla, Sarah Nobles, Mark Burroughs, Lori Rowe, Haibin Wang, Ben L. Rambo-Martin, Dhwani Batra, Justin Lee, Suxiang Tong |
| EPI_ISL_903894 | SD Public Health Laboratory | Genomics and Discovery, Respiratory Viruses Branch, Division of Viral Diseases, Centers for Disease Control and Prevention | Krista Queen, Yan Li, Ying Tao, Jing Zhang, Anna Uehara, Anna Montmayeur, Clinton R. Paden, Peter W. Cook, Rachel Marine, Mili Sheth, Jasmine Padilla, Sarah Nobles, Mark Burroughs, Lori Rowe, Haibin Wang, Ben L. Rambo-Martin, Dhwani Batra, Justin Lee, Suxiang Tong |
| EPI_ISL_903897 | TX DSHS, Lab Services Section MC 1947 | Genomics and Discovery, Respiratory Viruses Branch, Division of Viral Diseases, Centers for Disease Control and Prevention | Krista Queen, Yan Li, Ying Tao, Jing Zhang, Anna Uehara, Anna Montmayeur, Clinton R. Paden, Peter W. Cook, Rachel Marine, Mili Sheth, Jasmine Padilla, Sarah Nobles, Mark Burroughs, Lori Rowe, Haibin Wang, Ben L. Rambo-Martin, Dhwani Batra, Justin Lee, Suxiang Tong |
| EPI_ISL_903934 | SD Public Health Laboratory | Genomics and Discovery, Respiratory Viruses Branch, Division of Viral Diseases, Centers for Disease Control and Prevention | Krista Queen, Yan Li, Ying Tao, Jing Zhang, Anna Uehara, Anna Montmayeur, Clinton R. Paden, Peter W. Cook, Rachel Marine, Mili Sheth, Jasmine Padilla, Sarah Nobles, Mark Burroughs, Lori Rowe, Haibin Wang, Ben L. Rambo-Martin, Dhwani Batra, Justin Lee, Suxiang Tong |
| EPI_ISL_903950 | NYSDOH Wadsworth Center, Virology Lab | Genomics and Discovery, Respiratory Viruses Branch, Division of Viral Diseases, Centers for Disease Control and Prevention | Krista Queen, Yan Li, Ying Tao, Jing Zhang, Anna Uehara, Anna Montmayeur, Clinton R. Paden, Peter W. Cook, Rachel Marine, Mili Sheth, Jasmine Padilla, Sarah Nobles, Mark Burroughs, Lori Rowe, Haibin Wang, Ben L. Rambo-Martin, Dhwani Batra, Justin Lee, Suxiang Tong |
| EPI_ISL_903962 | WA State Department of Health | Genomics and Discovery, Respiratory Viruses Branch, Division of Viral Diseases, Centers for Disease Control and Prevention | Krista Queen, Yan Li, Ying Tao, Jing Zhang, Anna Uehara, Anna Montmayeur, Clinton R. Paden, Peter W. Cook, Rachel Marine, Mili Sheth, Jasmine Padilla, Sarah Nobles, Mark Burroughs, Lori Rowe, Haibin Wang, Ben L. Rambo-Martin, Dhwani Batra, Justin Lee, Suxiang Tong |
| EPI_ISL_904174, EPI_ISL_904234, EPI_ISL_904244, EPI_ISL_904491, EPI_ISL_904492, EPI_ISL_904493, EPI_ISL_904494, EPI_ISL_904612, EPI_ISL_904613 | Dutch COVID-19 response team | Erasmus Medical Center | Bas Oude Munnink, Reina Sikkema, David Nieuwenhuijse, Irina Chestakova, Anne van der Linden, Marjan Boter, Emmanuelle Munger, Corine GeurtsvanKessel, Annemiek van der Eijk, Richard Molenkamp, Marion Koopmans, on behalf of the Dutch national COVID-19 response team. |
| EPI_ISL_905489, EPI_ISL_905532 | Dutch COVID-19 response team | National Institute for Public Health and the Environment (RIVM) | Adam Meijer, Harry Vennema, Dirk Eggink, Jeroen Cremer, Sharon van den Brink, Bas van der Veer, AnneMarie van den Brandt, Florian Zwagemaker, Dennis Schmitz, Chantal Reusken, on behalf of the national COVID-19 response team |
| EPI_ISL_905799, EPI_ISL_905811, EPI_ISL_905812, EPI_ISL_905813, EPI_ISL_905814, EPI_ISL_905815, EPI_ISL_905816, EPI_ISL_905817, EPI_ISL_905818, EPI_ISL_905819, EPI_ISL_905820 | | | |
| see above | OHSU Lab Services Molecular Microbiology Lab | Oregon SARS-CoV-2 Genome Sequencing Center | Brendan L. O'Connell, Sally Grindstaff, Kayla Carter, Ruth V. Nichols, Alec J. Hirsch, Donna Hansel, Guang Fan, Xuan, Qin, Daniel N. Streblow, William B. Messer, Andrew C. Adey, Benjamin N. Bimber, Brian J. O'Roak |
| EPI_ISL_906057 | Tilia Laboratories s.r.o. | Tilia Laboratories s.r.o. | Sona Pekova, MD, PhD. |
| EPI_ISL_906120, EPI_ISL_906121, EPI_ISL_906122, EPI_ISL_906123, EPI_ISL_906124, EPI_ISL_906125, EPI_ISL_906126, EPI_ISL_906127 | Institute of Microbiology and Immunology, Faculty of Medicine, University of Ljubljana | Institute of Microbiology and Immunology, Faculty of Medicine, University of Ljubljana | Samo Zakotnik, Tomaž Mark Zorec, Matic Brvar, Miša Korva, Mario Poljak, Tatjana Avši - Županc |
| EPI_ISL_906294 | Nigeria Centre for Disease Control (NCDC) | African Centre of Excellence for Genomics of Infectious Diseases (ACEGID), Redeemer's University | Oluniyi P.E. et al |
| EPI_ISL_909951 | CUSL/UCLouvain COVID testing federal platform | UCLouvain/IREC/MBLG | Jean Ruelle, Lysa Pinsmaye, Benoit Kabamba Mukadi |
| EPI_ISL_910995, EPI_ISL_910996, EPI_ISL_910997, EPI_ISL_910998, EPI_ISL_910999, EPI_ISL_911000, EPI_ISL_911004, EPI_ISL_911005, EPI_ISL_911006, EPI_ISL_911007, EPI_ISL_911008, EPI_ISL_911009, EPI_ISL_911010, EPI_ISL_911011, EPI_ISL_911012, EPI_ISL_911013, EPI_ISL_911014, EPI_ISL_911015, EPI_ISL_911016, EPI_ISL_911017, EPI_ISL_911018, EPI_ISL_911019, EPI_ISL_911020, EPI_ISL_911021, EPI_ISL_911022, EPI_ISL_911023, EPI_ISL_911024, EPI_ISL_911025, EPI_ISL_911026, EPI_ISL_911027, EPI_ISL_911028, EPI_ISL_911029, EPI_ISL_911030, EPI_ISL_911031, EPI_ISL_911032, EPI_ISL_911033, EPI_ISL_911034, EPI_ISL_911035, EPI_ISL_911036, EPI_ISL_911037, EPI_ISL_911038, EPI_ISL_911039, EPI_ISL_911040, EPI_ISL_911041, EPI_ISL_911042, EPI_ISL_911043, EPI_ISL_911044, EPI_ISL_911045, EPI_ISL_911046, EPI_ISL_911047, EPI_ISL_911048, EPI_ISL_911049, EPI_ISL_911050, EPI_ISL_911051, EPI_ISL_911052, EPI_ISL_911053, EPI_ISL_911054, EPI_ISL_911055, EPI_ISL_911056, EPI_ISL_911057, EPI_ISL_911058, EPI_ISL_911059, EPI_ISL_911060, EPI_ISL_911061, EPI_ISL_911062, EPI_ISL_911063, EPI_ISL_911064, EPI_ISL_911065, EPI_ISL_911066 | | | |

| | | | |
|---|---|---|---|
| see above | Laboratoire national de sante, Microbiology, Virology | Laboratoire national de sante, Microbiology, Microbial Genomics Platform | Anke Wienecke-Baldacchino, Catherine Ragimbeau,Jessica Tapp, Fatu Djabi, Lise Pignon, Raoul Salmon, Tamir Abdelrahman |
| EPI_ISL_911684, EPI_ISL_911693, EPI_ISL_911695, EPI_ISL_911699 | Alaska State Virology Laboratory | Alaska State Virology Laboratory | Stephanie DeRonde, Lisa Smith, Ph.D., Jack Chen, Ph.D. |
| EPI_ISL_912369, EPI_ISL_912392, EPI_ISL_912393 | Fondation Congolaise pour la recherche medicale (FCRM), Francine Ntoumi | NGS Competence Center Tuebingen, Institut für Medizinische Mikrobiologie und Hygiene, Universitaetsklinikum Tübingen | Angel Angelov |
| EPI_ISL_912399, EPI_ISL_912400 | Institute for Medical Research, Infectious Disease Research Centre, National Institutes of Health, Ministry of Health Malaysia | Institute for Medical Research, Infectious Disease Research Centre, National Institutes of Health, Ministry of Health Malaysia | Suppiah J, Kamel K, Azizan MA, Thayan R |
| EPI_ISL_912459, EPI_ISL_912468, EPI_ISL_912501, EPI_ISL_912520 | NHLS Universitas Academic | UFS Virology | PA Bester, MM Nyaga, P Nthiga, MT Mogotsi, D Goedhals, T de Oliveira |
| EPI_ISL_913113 | CHU Purpan - Laboratoire de Virologie - Institut Fédératif de Biologie | CHU Purpan - Laboratoire de Virologie - Institut Fédératif de Biologie | Latour J., Ranger N., Dubois M., Carcenac R., Harter A., Boyer P., Tremeaux P., Izopet J. |
| EPI_ISL_913310, EPI_ISL_913349, EPI_ISL_913350, EPI_ISL_913481, EPI_ISL_913482, EPI_ISL_913483 | Klinisk mikrobiologi | The Public Health Agency of Sweden | Anna-Malin Linde, Maria Lind Karlberg, Carlo Berg, Oskar Karlsson Lindsjo, Sofia Stamouli, Reza Advani, Mattias Haukland, Petra Holmstrom, Noura Walai, Petra Edquist, Mia Brytting, Anna Risberg, Karin Tegmark-Wisell |
| EPI_ISL_913950, EPI_ISL_913952, EPI_ISL_913962, EPI_ISL_913965, EPI_ISL_913966, EPI_ISL_913979, EPI_ISL_913980, EPI_ISL_913982, EPI_ISL_913984 | Instituto de Diagnostico y Referencia Epidemiologicos INDRE_RNLSP | Instituto de Diagnostico y Referencia Epidemiologicos (INDRE) | Claudia Wong-Arambula, Abril Rodriguez-Maldonado, Fabiola Garces-Ayala, Adnan Araiza-Rodriguez, David Fragoso-Fonseca, Sergio Rangel-Guerrero, Mayra Jimenez-Morales, Nancy Munoz-Hernandez, Natividad Cruz-Ortiz, Tatiana Nunez-Garcia, Gisela Barrera-Badillo, Lucia Hernandez-Rivas, Irma Lopez-Martinez, Ernesto Ramirez-Gonzalez. |
| EPI_ISL_914801 | HOSPITAL DR. TOMAS CASAS CASAJUS | Inciensa, Instituto Costarricense de Investigación y Enseñanza en Nutrición y Salud | Francisco Duarte, Hebleen Porras, Claudio Soto-Garita, Estela Cordero, Adriana Godínez, Melany Calderón & Mariel López |
| EPI_ISL_914804 | AREA DE SALUD TURRIALBA-JIMENEZ | Inciensa, Instituto Costarricense de Investigación y Enseñanza en Nutrición y Salud | Francisco Duarte, Hebleen Porras, Claudio Soto-Garita, Estela Cordero, Adriana Godínez, Melany Calderón & Mónica Charpentier-Artavia |
| EPI_ISL_914880 | Instituto de Diagnostico y Referencia Epidemiologicos INDRE_RNLSP | Instituto de Diagnostico y Referencia Epidemiologicos (INDRE) | Claudia Wong-Arambula, Abril Rodriguez-Maldonado, Fabiola Garces-Ayala, Adnan Araiza-Rodriguez, David Fragoso-Fonseca, Sergio Rangel-Guerrero, Mayra Jimenez-Morales, Nancy Munoz-Hernandez, Natividad Cruz-Ortiz, Tatiana Nunez-Garcia, Gisela Barrera-Badillo, Lucia Hernandez-Rivas, Irma Lopez-Martinez, Ernesto Ramirez-Gonzalez. |
| EPI_ISL_915358, EPI_ISL_915380, EPI_ISL_915382, EPI_ISL_915383, EPI_ISL_915417, EPI_ISL_915418, EPI_ISL_915419, EPI_ISL_915420 | Keio University School of Medicine | Keio University School of Medicine | Kenjiro Kosaki, Yuka Iwasaki, Hirotsugu Ishizu, Haruhiko Siomi, Kodai Abe |
| EPI_ISL_918485, EPI_ISL_918491, EPI_ISL_918494, EPI_ISL_918498 | Ohio Department of Health Laboratory | Ohio Department of Health Laboratory | Holmes, Jennifer; Eric Brandt, Keoni Omura, Glen McGillivary, Caitlin McDonnell, Jade Mowery, Stephanie Mccracken, Tyler Payne, Kirtana Ramadugu, Erica Leasure, Brent Lee, Kelsey Florek, Heather Blankenship, Quanta Brown, and Tammy Bannerman |
| EPI_ISL_918518, EPI_ISL_918519, EPI_ISL_918520 | Evandro Chagas Institute | Evandro Chagas Institute | Santos, M.C.; Silva, A.M.; Junior, W.D.C.; Barbagelata, L.S.; Ferreira, J.A.; Sousa, E.M.A.; da Silva, P.S.; Pinheiro, K.C.; L.C.; Sousa Junior, E.C. |
| EPI_ISL_918540 | LACEN - Laboratório Central de Saúde Pública do Ceara | Evandro Chagas Institute | Santos, M.C.; Silva, A.M.; Junior, W.D.C.; Barbagelata, L.S.; Ferreira, J.A.; Sousa, E.M.A.; da Silva, P.S.; Pinheiro, K.C.; L.C.; Sousa Junior, E.C. |
| EPI_ISL_918562 | Ohio Department of Health Laboratory | Ohio Department of Health Laboratory | Holmes, Jennifer; Eric Brandt, Keoni Omura, Glen McGillivary, Caitlin McDonnell, Jade Mowery, Stephanie Mccracken, Tyler Payne, Kirtana Ramadugu, Erica Leasure, Brent Lee, Kelsey Florek, Heather Blankenship, Quanta Brown, and Tammy Bannerman |
| EPI_ISL_918841, EPI_ISL_918957 | University of Birmingham | COVID-19 Genomics UK (COG-UK) Consortium | Institute of Microbiology, University of Birmingham: Claire McMurray, Joanne Stockton, Samuel Nicholls, Radoslaw Poplawski, Will Rowe, Josh Quick, Nicholas Loman. University of Birmingham Testing Laboratory: Celina M Whalley, Andrew Bosworth, Charlotte Poxon, Kasun Wanigasooriya, Oliver Pickles, Mike Kidd, Alex Richter, Andrew D Beggs PHE Heartlands Lab: Husam Osman, Andrew Bosworth. Queen Elizabeth Hospital: Anna Casey |
| EPI_ISL_919232, EPI_ISL_919236, EPI_ISL_919238, EPI_ISL_919240, EPI_ISL_919244, EPI_ISL_919249, EPI_ISL_919259, EPI_ISL_919263, EPI_ISL_919268 | West of Scotland Specialist Virology Centre, NHSGGC / MRC-University of Glasgow Centre for Virus Research | COVID-19 Genomics UK (COG-UK) Consortium | Ana da Silva Filipe, Natasha Johnson, Kathy Smollett, Daniel Mair, Stephen Carmichael, Alice Broos, Lily Tong, Jenna Nichols, Kyriaki Nomikou; Sarah McDonald; Richard Orton, Joseph Hughes, Sreenu Vattipally, David L Robertson; Alasdair MacLean, Rory Gunson; Sharif Shaaban, Matthew Holden; Rachel Blacow, Guy Mollett, Kathy Li, James Shepherd, Antonia Ho, Emma Thomson |
| EPI_ISL_919313, EPI_ISL_919365, EPI_ISL_919366, EPI_ISL_919368, EPI_ISL_919369, EPI_ISL_919370, EPI_ISL_919394, EPI_ISL_919395, EPI_ISL_919397 | Virology Department, Royal Infirmary of Edinburgh, NHS Lothian / School of Biological Sciences, University of Edinburgh / Institute of Genetics and Molecular Medicine, University of Edinburgh | COVID-19 Genomics UK (COG-UK) Consortium | McHugh M, Dewar R, Rooke S, Gallagher M, Balcaza C, O'Toole Á, Scher E, Hill V, McCrone JT, Colquhoun R, Yu X, Jackson B, Rambaut A, Williams TC, Templeton K |
| EPI_ISL_920199, EPI_ISL_920216, EPI_ISL_920220, EPI_ISL_920256, EPI_ISL_920257, EPI_ISL_920290, EPI_ISL_920331, EPI_ISL_920356, EPI_ISL_920373, EPI_ISL_920378, EPI_ISL_920494, EPI_ISL_920500, EPI_ISL_920503, EPI_ISL_920510, EPI_ISL_920511 | | | |
| see above | University College London Hospital | COVID-19 Genomics UK (COG-UK) Consortium | Judith Heaney, Matthew Byott, Catherine Houlihan, Dan Frampton, Stuart Kirk, Moira Spyer and Eleni Nastouli |
| EPI_ISL_925141, EPI_ISL_925167, EPI_ISL_925168, EPI_ISL_925169, EPI_ISL_925170, EPI_ISL_925171, EPI_ISL_925172 | Virginia DCLS | Virginia DCLS | Virginia DCLS |
| EPI_ISL_925387, EPI_ISL_925388, EPI_ISL_925389 | Department of Clinical Microbiology | GIGA Medical Genomics | Keith Durkin, Maria Artesi, Sébastien Bontems, Raphaël Boreux, Bouchra Boujemla, Cécile Meex, Pierrette Melin, Marie-Pierre Hayette, Vincent Bours |
| EPI_ISL_925848, EPI_ISL_925851, EPI_ISL_925852, EPI_ISL_925854, EPI_ISL_925858, EPI_ISL_925859, EPI_ISL_925894, EPI_ISL_925895, EPI_ISL_925896 | Nucleic Acid Testing, National Reference Laboratory | GIGA Medical Genomics | Yvan Butera, Keith Durkin, Maria Artesi, Bouchra Boujemla, Robert Rutayisire, Patrick Tuyisenge, Esperence Umumararungu, Sébastien Bontems, Marie-Pierre Hayette, Nathalie Renotte, Swaibu Gatare, Jacob Souopgui, Sabin Nsanzimana, Vincent Bours, Léon Mutesa |
| EPI_ISL_925952, EPI_ISL_925969, EPI_ISL_925971, EPI_ISL_925983, EPI_ISL_925988, EPI_ISL_926002, EPI_ISL_926005, EPI_ISL_926007, EPI_ISL_926050, EPI_ISL_926053, EPI_ISL_926055, EPI_ISL_926070, EPI_ISL_926093, EPI_ISL_926095, EPI_ISL_926124, EPI_ISL_926128, EPI_ISL_926178, EPI_ISL_926201, EPI_ISL_926209, EPI_ISL_926233, EPI_ISL_926235, EPI_ISL_926255, EPI_ISL_926275, EPI_ISL_926283, EPI_ISL_926315, EPI_ISL_926316, EPI_ISL_926324, EPI_ISL_926345, EPI_ISL_926404, EPI_ISL_926441, EPI_ISL_926469, EPI_ISL_926484, EPI_ISL_926485, EPI_ISL_926528, EPI_ISL_926537, EPI_ISL_926541, EPI_ISL_926547, EPI_ISL_926566, EPI_ISL_926573, EPI_ISL_926576, EPI_ISL_926607, EPI_ISL_926615, EPI_ISL_926617, EPI_ISL_926648, EPI_ISL_926671, EPI_ISL_926679, EPI_ISL_926681, EPI_ISL_926727, EPI_ISL_926773, EPI_ISL_926775, EPI_ISL_926826, EPI_ISL_926838, EPI_ISL_926851, EPI_ISL_926852, EPI_ISL_926859, EPI_ISL_926862, EPI_ISL_926866, EPI_ISL_926870, EPI_ISL_926881, EPI_ISL_926932, EPI_ISL_926941, EPI_ISL_926949, EPI_ISL_926965, EPI_ISL_926970, EPI_ISL_926980, EPI_ISL_926981, EPI_ISL_926990, EPI_ISL_927003, EPI_ISL_927039, EPI_ISL_927047, EPI_ISL_927060, EPI_ISL_927072, EPI_ISL_927089, EPI_ISL_927092, EPI_ISL_927095, EPI_ISL_927119, EPI_ISL_927123, EPI_ISL_927145, EPI_ISL_927151, EPI_ISL_927161, EPI_ISL_927162, EPI_ISL_927164, EPI_ISL_927186, EPI_ISL_927247, EPI_ISL_927276, EPI_ISL_927283, EPI_ISL_927299, EPI_ISL_927311, EPI_ISL_927348, EPI_ISL_927351, EPI_ISL_927353, EPI_ISL_927364, EPI_ISL_927371, EPI_ISL_927375, EPI_ISL_927411, EPI_ISL_927422, EPI_ISL_927428, EPI_ISL_927439, EPI_ISL_927444, EPI_ISL_927474, EPI_ISL_927476, EPI_ISL_927483, EPI_ISL_927501, EPI_ISL_927504, EPI_ISL_927519, EPI_ISL_927558, EPI_ISL_927593, EPI_ISL_927624, EPI_ISL_927627, EPI_ISL_927637, EPI_ISL_927660, EPI_ISL_927665, EPI_ISL_927673, EPI_ISL_927685, EPI_ISL_927689, EPI_ISL_927695, EPI_ISL_927705, EPI_ISL_927707, EPI_ISL_927708, EPI_ISL_927730, EPI_ISL_927744, EPI_ISL_927748, EPI_ISL_927756, EPI_ISL_927771, EPI_ISL_927781, EPI_ISL_927787, EPI_ISL_927789, EPI_ISL_927840, EPI_ISL_927855, EPI_ISL_927859, EPI_ISL_927860, EPI_ISL_927863, EPI_ISL_927871, EPI_ISL_927881, EPI_ISL_927888, EPI_ISL_927891, EPI_ISL_927946, EPI_ISL_927955, EPI_ISL_927969, EPI_ISL_927996, EPI_ISL_928012, EPI_ISL_928032, EPI_ISL_928051, EPI_ISL_928061, EPI_ISL_928066, EPI_ISL_928067, EPI_ISL_928074, EPI_ISL_928076, EPI_ISL_928155, EPI_ISL_928171, EPI_ISL_928221, EPI_ISL_928224, EPI_ISL_928229, EPI_ISL_928231, EPI_ISL_928250, EPI_ISL_928262, EPI_ISL_928307, EPI_ISL_928327, EPI_ISL_928333, EPI_ISL_928344, EPI_ISL_928356, EPI_ISL_928386, EPI_ISL_928389, EPI_ISL_928397, EPI_ISL_928403, EPI_ISL_928420, EPI_ISL_928427, EPI_ISL_928464, EPI_ISL_928490, EPI_ISL_928493, EPI_ISL_928503, EPI_ISL_928522, EPI_ISL_928527, EPI_ISL_928539, EPI_ISL_928563, EPI_ISL_928581, EPI_ISL_928617, EPI_ISL_928631, EPI_ISL_928638, EPI_ISL_928665, | | | |

| | | | |
|---|---|---|---|
| EPI_ISL_928683, EPI_ISL_928751, EPI_ISL_928777, EPI_ISL_928778, EPI_ISL_928791, EPI_ISL_928821, EPI_ISL_928850, EPI_ISL_928864, EPI_ISL_928876, EPI_ISL_928916, EPI_ISL_928919, EPI_ISL_928924, EPI_ISL_928934, EPI_ISL_928942, EPI_ISL_928967, EPI_ISL_928976, EPI_ISL_929005, EPI_ISL_929057, EPI_ISL_929063, EPI_ISL_929082, EPI_ISL_929136, EPI_ISL_929157, EPI_ISL_929171, EPI_ISL_929185, EPI_ISL_929199, EPI_ISL_929215, EPI_ISL_929219, EPI_ISL_929228, EPI_ISL_929241, EPI_ISL_929253, EPI_ISL_929265, EPI_ISL_929280, EPI_ISL_929281, EPI_ISL_929292, EPI_ISL_929307, EPI_ISL_929339, EPI_ISL_929353, EPI_ISL_929366, EPI_ISL_929383, EPI_ISL_929388, EPI_ISL_929422, EPI_ISL_929438, EPI_ISL_929446, EPI_ISL_929464, EPI_ISL_929466, EPI_ISL_929488, EPI_ISL_929494, EPI_ISL_929501, EPI_ISL_929519, EPI_ISL_929523, EPI_ISL_929534, EPI_ISL_929535, EPI_ISL_929559, EPI_ISL_929583, EPI_ISL_929619, EPI_ISL_929628, EPI_ISL_929655, EPI_ISL_929659, EPI_ISL_929676, EPI_ISL_929694, EPI_ISL_929702, EPI_ISL_929730, EPI_ISL_929740, EPI_ISL_929746, EPI_ISL_929773, EPI_ISL_929782, EPI_ISL_929796, EPI_ISL_929811, EPI_ISL_929832, EPI_ISL_929833, EPI_ISL_929845, EPI_ISL_929854, EPI_ISL_929864, EPI_ISL_929870, EPI_ISL_929880, EPI_ISL_929882, EPI_ISL_929913, EPI_ISL_929926, EPI_ISL_929939, EPI_ISL_930003, EPI_ISL_930007, EPI_ISL_930017, EPI_ISL_930029, EPI_ISL_930030, EPI_ISL_930031, EPI_ISL_930045, EPI_ISL_930058, EPI_ISL_930063, EPI_ISL_930081, EPI_ISL_930082, EPI_ISL_930093, EPI_ISL_930104, EPI_ISL_930129, EPI_ISL_930154, EPI_ISL_930161, EPI_ISL_930170, EPI_ISL_930174, EPI_ISL_930200, EPI_ISL_930206, EPI_ISL_930223, EPI_ISL_930227, EPI_ISL_930252, EPI_ISL_930253, EPI_ISL_930273, EPI_ISL_930275, EPI_ISL_930284, EPI_ISL_930285, EPI_ISL_930307, EPI_ISL_930338, EPI_ISL_930392, EPI_ISL_930394, EPI_ISL_930399, EPI_ISL_930414, EPI_ISL_930430, EPI_ISL_930432, EPI_ISL_930457, EPI_ISL_930464, EPI_ISL_930476, EPI_ISL_930486, EPI_ISL_930503, EPI_ISL_930516, EPI_ISL_930559 | | | |
| see above | Department of Virus and Microbiological Special Diagnostics, Statens Serum Institut, Copenhagen, Denmark | Aalborg University | Danish Covid-19 Genome Consortium |
| EPI_ISL_931448 | University Hospital Basel, Clinical Virology | University Hospital Basel, Clinical Bacteriology | Tim Roloff, Madlen Stange, Helena MB Seth-Smith, Alfredo Mari, Karoline Leuzinger, Julia Bielicki, Manuel Battegay, Hans Hirsch, Adrian Egli |
| EPI_ISL_933693, EPI_ISL_933703 | Instituto de Diagnostico y Referencia Epidemiologicos INDRE_RNLSP | Instituto de Diagnostico y Referencia Epidemiologicos (INDRE) | Claudia Wong-Arambula, Abril Rodriguez-Maldonado, Fabiola Garces-Ayala, Adnan Araiza-Rodriguez, David Fragoso-Fonseca, Sergio Rangel-Guerrero, Mayra Jimenez-Morales, Nancy Munoz-Hernandez, Natividad Cruz-Ortiz, Tatiana Nunez-Garcia, Gisela Barrera-Badillo, Lucia Hernandez-Rivas, Irma Lopez-Martinez, Ernesto Ramirez-Gonzalez. |
| EPI_ISL_934096, EPI_ISL_934097, EPI_ISL_934099, EPI_ISL_934100, EPI_ISL_934101, EPI_ISL_934102, EPI_ISL_934103, EPI_ISL_934104, EPI_ISL_934105, EPI_ISL_934106, EPI_ISL_934107, EPI_ISL_934108, EPI_ISL_934109, EPI_ISL_934112, EPI_ISL_934113, EPI_ISL_934114, EPI_ISL_934115, EPI_ISL_934116, EPI_ISL_934117, EPI_ISL_934119, EPI_ISL_934120, EPI_ISL_934121, EPI_ISL_934122, EPI_ISL_934123, EPI_ISL_934124, EPI_ISL_934127, EPI_ISL_934128, EPI_ISL_934129, EPI_ISL_934130, EPI_ISL_934131, EPI_ISL_934132, EPI_ISL_934133, EPI_ISL_934134, EPI_ISL_934135, EPI_ISL_934136, EPI_ISL_934138, EPI_ISL_934139, EPI_ISL_934140, EPI_ISL_934141, EPI_ISL_934142, EPI_ISL_934143, EPI_ISL_934144, EPI_ISL_934145, EPI_ISL_934146, EPI_ISL_934147, EPI_ISL_934148, EPI_ISL_934149, EPI_ISL_934150, EPI_ISL_934151, EPI_ISL_934152, EPI_ISL_934153, EPI_ISL_934154, EPI_ISL_934155, EPI_ISL_934156, EPI_ISL_934157, EPI_ISL_934158, EPI_ISL_934159, EPI_ISL_934160, EPI_ISL_934161, EPI_ISL_934162, EPI_ISL_934163, EPI_ISL_934164, EPI_ISL_934165, EPI_ISL_934166, EPI_ISL_934168, EPI_ISL_934169, EPI_ISL_934170 | | | |
| see above | Vilnius university hospital Santaros Klinikos, Center of Laboratory Medicine | Vilnius university hospital Santaros Klinikos, Center of Laboratory Medicine | Ingrida Olendraite, Daniel Naumovas, Rimvydas Norvilas, Dovile Ezerskyte, Justinas Slikas, Gytis Dudas |
| EPI_ISL_934316, EPI_ISL_934317, EPI_ISL_934318, EPI_ISL_934319, EPI_ISL_934320, EPI_ISL_934321 | Laboratoriemedicin, Klinisk mikrobiologi | The Public Health Agency of Sweden | Anna-Malin Linde, Maria Lind Karlberg, Carlo Berg, Oskar Karlsson Lindsjo, Sofia Stamouli, Reza Advani, Mattias Haukland, Petra Holmstrom, Noura Walai, Petra Edquist, Mia Brytting, Anna Risberg, Karin Tegmark-Wisell |
| EPI_ISL_934327, EPI_ISL_934328, EPI_ISL_934329, EPI_ISL_934354 | Klinisk mikrobiologi | The Public Health Agency of Sweden | Anna-Malin Linde, Maria Lind Karlberg, Carlo Berg, Oskar Karlsson Lindsjo, Sofia Stamouli, Reza Advani, Mattias Haukland, Petra Holmstrom, Noura Walai, Petra Edquist, Mia Brytting, Anna Risberg, Karin Tegmark-Wisell |
| EPI_ISL_934414, EPI_ISL_934417 | Ohio Department of Health Laboratory | Ohio Department of Health Laboratory | Holmes, Jennifer; Eric Brandt, Keoni Omura, Glen McGillivary, Caitlin McDonnell, Jade Mowery, Stephanie Mccracken, Tyler Payne, Kirtana Ramadugu, Erica Leasure, Brent Lee, Kelsey Florek, Heather Blankenship, Quanta Brown, and Tammy Bannerman |
| EPI_ISL_934570, EPI_ISL_934571 | Department of Laboratory Medicine, Division of Clinical Virology, University of Medicine, Vienna | Bergthaler laboratory, CeMM Research Center for Molecular Medicine of the Austrian Academy of Sciences | Lukas Endler, Anna Schedl, Thomas Penz, Benedikt Agerer, Maelle Le Moing, Michael Schuster, Bekir Erguner, Jan Laine, Martin Senekowitsch, Christoph Bock, Andreas Bergthaler |
| EPI_ISL_934973, EPI_ISL_934975, EPI_ISL_935020, EPI_ISL_935023, EPI_ISL_935024 | Ohio Department of Health Laboratory | Ohio Department of Health Laboratory | Holmes, Jennifer; Eric Brandt, Keoni Omura, Glen McGillivary, Caitlin McDonnell, Jade Mowery, Stephanie Mccracken, Tyler Payne, Kirtana Ramadugu, Erica Leasure, Brent Lee, Kelsey Florek, Heather Blankenship, Quanta Brown, and Tammy Bannerman |
| EPI_ISL_936491 | Institute for Medical Research, Infectious Disease Research Centre, National Institutes of Health, Ministry of Health Malaysia | Institute for Medical Research, Infectious Disease Research Centre, National Institutes of Health, Ministry of Health Malaysia | Suppiah J, Kamel K, Azizan MA, Thayan R |
| EPI_ISL_936644, EPI_ISL_936645, EPI_ISL_936653, EPI_ISL_936654, EPI_ISL_936655, EPI_ISL_936656, EPI_ISL_936657, EPI_ISL_936658, EPI_ISL_936659 | Northwestern Memorial Hospital | Ozer Lab | Ramon Lorenzo-Redondo, Lacy M. Simons, Chad J. Achenbach, Lawrence J. Jennings, Michael G. Ison, Judd F. Hultquist, Egon A. Ozer |
| EPI_ISL_940078, EPI_ISL_940079, EPI_ISL_940080, EPI_ISL_940083, EPI_ISL_940084, EPI_ISL_940094, EPI_ISL_940095, EPI_ISL_940098, EPI_ISL_940099, EPI_ISL_940101, EPI_ISL_940102, EPI_ISL_940104, EPI_ISL_940105, EPI_ISL_940106, EPI_ISL_940107, EPI_ISL_940114, EPI_ISL_940115, EPI_ISL_940117, EPI_ISL_940121, EPI_ISL_940125, EPI_ISL_940129, EPI_ISL_940132, EPI_ISL_940133 | | | |
| see above | Charlotte Maxeke Johannesburg Academic Hospital, National Health Laboratory Services, Gauteng, South Africa | National Institute for Communicable Diseases of the National Health Laboratory Service | Amoako DG, Mohale T, Ntuli N, Mahlangu B, Allam M, Ismail A, Bhiman JN |
| EPI_ISL_940823, EPI_ISL_940824 | Virginia DCLS | Virginia DCLS | Virginia DCLS |
| EPI_ISL_940859 | Vaccines and Infectious Diseases Analytics Research Unit (VIDA) | KRISP, KZN Research Innovation and Sequencing Platform | Baillie Vicky, du Plessis Jeanine, Giandhari Jennifer, Pillay Sureshnee, Naidoo Yeshnee, Tegally Houriiyah, de Oliveira Tulio, Madhi Shabir |
| EPI_ISL_941255, EPI_ISL_941256, EPI_ISL_941257, EPI_ISL_941258, EPI_ISL_941259, EPI_ISL_941260, EPI_ISL_941261, EPI_ISL_941262, EPI_ISL_941263, EPI_ISL_941264, EPI_ISL_941265, EPI_ISL_941266, EPI_ISL_941267, EPI_ISL_941268 | | | |
| see above | Virginia DCLS | Virginia DCLS | Virginia DCLS |
| EPI_ISL_942070, EPI_ISL_942071, EPI_ISL_942072, EPI_ISL_942230, EPI_ISL_942232, EPI_ISL_942233, EPI_ISL_942234, EPI_ISL_942235, EPI_ISL_942248, EPI_ISL_942249, EPI_ISL_942250, EPI_ISL_942251, EPI_ISL_942252, EPI_ISL_942253, EPI_ISL_942254, EPI_ISL_942255, EPI_ISL_942256, EPI_ISL_942257, EPI_ISL_942258, EPI_ISL_942259, EPI_ISL_942260, EPI_ISL_942261, EPI_ISL_942264, EPI_ISL_942265, EPI_ISL_942287, EPI_ISL_942288, EPI_ISL_942289, EPI_ISL_942290, EPI_ISL_942291, EPI_ISL_942292, EPI_ISL_942293, EPI_ISL_942294, EPI_ISL_942295, EPI_ISL_942296, EPI_ISL_942297, EPI_ISL_942298, EPI_ISL_942299, EPI_ISL_942300, EPI_ISL_942301, EPI_ISL_942302, EPI_ISL_942303, EPI_ISL_942304, EPI_ISL_942305, EPI_ISL_942306, EPI_ISL_942307, EPI_ISL_942313 | | | |
| see above | Wisconsin State Laboratory of Hygiene Communicable Disease Division | Wisconsin State Laboratory of Hygiene Communicable Disease Division | Kelsey R. Florek, Abigail C. Shockey |
| EPI_ISL_942690, EPI_ISL_942691, EPI_ISL_942692, EPI_ISL_942693, EPI_ISL_942694, EPI_ISL_942695, EPI_ISL_942696, EPI_ISL_942697, EPI_ISL_942698, EPI_ISL_942699, EPI_ISL_942700, EPI_ISL_942701, EPI_ISL_942702, EPI_ISL_942703, EPI_ISL_942704, EPI_ISL_942705, EPI_ISL_942706, EPI_ISL_942943 | | | |
| see above | Gundersen Molecular Diagnostics Laboratory | Kabara Cancer Research Institute | Craig S. Richmond, Paraic A. Kenny |
| EPI_ISL_942968 | General Hospital - Kumanovo | Research Center for Genetic Engineering and Biotechnology "Georgi D. Efremov" , Macedonian Academy of Sciences and Arts | Aleksandar J. Dimovski, Dijana Plasheska-Karanfilska, Predrag Noveski, Gjorgji Bozinovski, Milena Jakimovska |
| EPI_ISL_942969 | General Hospital - Veles | Research Center for Genetic Engineering and Biotechnology "Georgi D. Efremov" , Macedonian Academy of Sciences and Arts | Aleksandar J. Dimovski, Dijana Plasheska-Karanfilska, Predrag Noveski, Gjorgji Bozinovski, Milena Jakimovska |
| EPI_ISL_942970 | General Hospital - Bitola | Research Center for Genetic Engineering and Biotechnology "Georgi D. Efremov" , Macedonian Academy of Sciences and Arts | Aleksandar J. Dimovski, Dijana Plasheska-Karanfilska, Predrag Noveski, Gjorgji Bozinovski, Milena Jakimovska |
| EPI_ISL_943993 | General Hospital - Strumica | Research Center for Genetic Engineering and Biotechnology "Georgi D. Efremov" , Macedon | Aleksandar J. Dimovski, Dijana Plasheska-Karanfilska, Predrag Noveski, Gjorgji Bozinovski |
| EPI_ISL_943994 | General Hospital - Strumica | Research Center for Genetic Engineering and Biotechnology "Georgi D. Efremov" , Macedonian Academy of Sciences and Arts | Aleksandar J. Dimovski, Dijana Plasheska-Karanfilska, Predrag Noveski, Gjorgji Bozinovski, Mi |
| EPI_ISL_944094, EPI_ISL_944095, EPI_ISL_944096 | Institute for Medical Research, Infectious Disease Research Centre, National Institutes of Health, Ministry of Health Malaysia | Institute for Medical Research, Infectious Disease Research Centre, National Institutes of Health, Ministry of Health Malaysia | Suppiah J, Kamel K, Azizan MA, Thayan R |
| EPI_ISL_945073 | Lighthouse Lab in Glasgow | Wellcome Sanger Institute for the COVID-19 Genomics UK | Harper VanSteenhouse, Yumi Kasai, David Gray, Carol Clugston, Anna Dominiczak and Alex Alderton, Roberto Amato, Sonia Goncalves, Ewan Harrison, |

| | | | |
|---|---|---|---|
| | | (COG-UK) Consortium | David K. Jackson, Ian Johnston, Dominic Kwiatkowski, Cordelia Langford, John Sillitoe on behalf of the Wellcome Sanger Institute COVID-19 Surveillance Team |
| EPI_ISL_945140, EPI_ISL_945184 | Lighthouse Lab in Milton Keynes | Wellcome Sanger Institute for the COVID-19 Genomics UK (COG-UK) Consortium | The Lighthouse Lab in Milton Keynes and Alex Alderton, Roberto Amato, Sonia Goncalves, Ewan Harrison, David K. Jackson, Ian Johnston, Dominic Kwiatkowski, Cordelia Langford, John Sillitoe on behalf of the Wellcome Sanger Institute COVID-19 Surveillance Team |
| EPI_ISL_945240, EPI_ISL_945252, EPI_ISL_945289 | Lighthouse Lab in Alderley Park | Wellcome Sanger Institute for the COVID-19 Genomics UK (COG-UK) Consortium | Jacquelyn Wynn, Mairead Hyland, The Lighthouse Lab in Alderley Park and Alex Alderton, Roberto Amato, Sonia Goncalves, Ewan Harrison, David K. Jackson, Ian Johnston, Dominic Kwiatkowski, Cordelia Langford, John Sillitoe on behalf of the Wellcome Sanger Institute COVID-19 Surveillance Team |
| EPI_ISL_945307, EPI_ISL_945338 | Lighthouse Lab in Milton Keynes | Wellcome Sanger Institute for the COVID-19 Genomics UK (COG-UK) Consortium | The Lighthouse Lab in Milton Keynes and Alex Alderton, Roberto Amato, Sonia Goncalves, Ewan Harrison, David K. Jackson, Ian Johnston, Dominic Kwiatkowski, Cordelia Langford, John Sillitoe on behalf of the Wellcome Sanger Institute COVID-19 Surveillance Team |
| EPI_ISL_945357 | Lighthouse Lab in Glasgow | Wellcome Sanger Institute for the COVID-19 Genomics UK (COG-UK) Consortium | Harper VanSteenhouse, Yumi Kasai, David Gray, Carol Clugston, Anna Dominiczak and Alex Alderton, Roberto Amato, Sonia Goncalves, Ewan Harrison, David K. Jackson, Ian Johnston, Dominic Kwiatkowski, Cordelia Langford, John Sillitoe on behalf of the Wellcome Sanger Institute COVID-19 Surveillance Team |
| EPI_ISL_947307, EPI_ISL_947309 | RS Sentra Medika Cikarang | Eijkman Institute for Molecular Biology, Ministry of Research and Technology/National Agency for Research and Innovation | Lydia V. Panggalo, Sukma Oktavianthi, Willy Agustine, Edison Johar, Hidayat Trimarsanto, Iskandar Adnan, Frilasita A Yudhaputri, Safarina G Malik, Khin Saw Myint, Amin Soebandrio |
| EPI_ISL_949403, EPI_ISL_949404 | University of Birmingham | COVID-19 Genomics UK (COG-UK) Consortium | Institute of Microbiology, University of Birmingham: Claire McMurray, Joanne Stockton, Sam Nicholls, Radoslaw Poplawski, Will Rowe, Josh Quick, Nicholas Loman. University of Birmingham Testing Laboratory: Celina M Whalley, Andrew Bosworth, Charlotte Poxon, Kasun Wanigasooriya, Oliver Pickles, Mike Kidd, Alex Richter, Andrew D Beggs PHE Heartlands Lab: Husam Osman, Andrew Bosworth. Queen Elizabeth Hospital: Anna Casey |
| EPI_ISL_949651 | Queens Medical Centre, Clinical Microbiology Department / DeepSeq Nottingham | COVID-19 Genomics UK (COG-UK) Consortium | Gemma Clark, Wendy Smith, Manjinder Khakh, Vicki M Fleming, Michelle M Lister, Hannah Howson-Wells, Jonathan Ball, Patrick McClure, Joseph Chappell, Theocharis Tsoleridis, Nadine Holmes, Matthew Carlisle, Christopher Moore, Fei Sang, Johnny Debebe, Victoria Wright, Matthew Loose |
| EPI_ISL_950166 | University College London, Great Ormond Street Hospital for Children NHS Foundation Trust, Imperial College Healthcare NHS Trust | COVID-19 Genomics UK (COG-UK) Consortium | Sergi Castellano, Rachel Williams, Mark Kristiansen, Paola Resende Silva, Sunando Roy, Tony Brooks, Helena Tutill, Paola Niola, Patricia Dyal, Charlotte Williams, Leysa Forrest, Yasmin Panchbhaya, Jacqueline Findlay, Samuel Weeks, Julianne Brown, Kathryn Harris, Paul Randell, James Price, Alison Holmes, Judith Breuer |
| EPI_ISL_950698, EPI_ISL_950700, EPI_ISL_950701, EPI_ISL_950702, EPI_ISL_950703, EPI_ISL_950704 | Lincolnshire Hospitals and DeepSeq Nottingham | COVID-19 Genomics UK (COG-UK) Consortium | Nichola Duckworth, Tim Sloan, Sarah Walsh, Jonathan Ball, Patrick McClure, Joeseph Chappell, Nadine Holmes, Matthew Carlisle, Christopher Moore, Fei Sang, Johnny Debebe, Victoria Wright, Matthew Loose |
| EPI_ISL_950903, EPI_ISL_950904, EPI_ISL_950911, EPI_ISL_950914, EPI_ISL_950915, EPI_ISL_950916 | Oxford Viromics, NDM, University of Oxford; Oxford University Hospitals; Basingstoke and North Hampshire Hospital | COVID-19 Genomics UK (COG-UK) Consortium | Tanya Golubchik, David Bonsall, George Macintyre, Amy Trebes, Mariateresa de Cesare, Catrin Moore, Alex Mobbs, Anita Justice, Robert Shaw, Monique Andersson, Timothy Peto, Emma Wise, Nathan Moore, Jessica Lynch, Nick Cortes, Matilde Mori, Stephen Kidd, David Buck, John Todd, Christophe Fraser |
| EPI_ISL_959857, EPI_ISL_959859 | National Virus Reference Laboratory | National Virus Reference Laboratory | Michael Carr, Gabriel Gonzalez, Jonathan Dean, Cillian F De Gascun |
| EPI_ISL_960189 | Virginia Division of Consolidated Laboratory Services | Virginia Division of Consolidated Laboratory Services | Virginia DCLS |
| EPI_ISL_960442, EPI_ISL_960472, EPI_ISL_960478, EPI_ISL_960526, EPI_ISL_960527, EPI_ISL_960528, EPI_ISL_960529, EPI_ISL_960530, EPI_ISL_960614, EPI_ISL_960615, EPI_ISL_960616, EPI_ISL_960617, EPI_ISL_960618 | | | |
| see above | Istituto Zooprofilattico Sperimentale del Mezzogiorno | TIGEM | Patrizia Annunziata, Andrea Ballabio, Valentina Bouche, Davide Cacchiarelli, Pellegrino Cerino, Chiara Colantuono, Maria Concetta Cuomo, Denise Di Concilio, Lucio Di Filippo, Antonio Grimaldi, Antonio Limone, Anna Manfredi, Francesco Panariello, Biancamaria Pierri, Marcello Salvi |
| EPI_ISL_962123 | Illinois Department of Public Health | Gagnon Lab, Southern Illinois University | Keith Gagnon |
| EPI_ISL_962800, EPI_ISL_962807 | San Diego County Public Health Laboratory | Andersen lab at Scripps Research | SEARCH Alliance San Diego with Tracy Basler, Jovan Shephard, Brett Austin |
| EPI_ISL_965220 | Virginia Division of Consolidated Laboratory Services | Virginia Division of Consolidated Laboratory Services | Virginia DCLS |
| EPI_ISL_965388 | University of Liège COVID-19 testing center | GIGA Medical Genomics | Keith Durkin, Maria Artesi, Bouchra Boujemla, Emmanuel André, Marc Van Ranst, Fabrice Bureau, Laurent Gillet, Wouter Coppieters, Vincent Bours |
| EPI_ISL_965551, EPI_ISL_965569, EPI_ISL_965683, EPI_ISL_965751, EPI_ISL_965804 | Dutch COVID-19 response team | Medical Microbiology, Maastricht University Medical Centre | Jozef Dingemans*, Brian van der Veer*, Erik Beuken, Carmen Reumkens, Lieke van Alphen, Christian Hoebe, Paul Savelkoul |
| EPI_ISL_966757, EPI_ISL_966758, EPI_ISL_966759, EPI_ISL_966760 | Maine HETL | Tewhey Lab, The Jackson Laboratory | Matluk,N., Dewey,H., Iosue,F., Barter,M., Lynch,R., Munger,H. and Tewhey,R. |
| EPI_ISL_967523, EPI_ISL_967526, EPI_ISL_967538 | State Laboratories Division, Hawaii State Department of Health | State Laboratories Division, Hawaii State Department of Health | Pamela O'Brien, Drew Kuwazaki, Ayana Garnet, Razvan Sultana, Edward Desmond |
| EPI_ISL_968832, EPI_ISL_968850 | KEMRI-Wellcome Trust Research Programme/KEMRI-CGMR-C Kilifi | KEMRI-Wellcome Trust Research Programme/KEMRI-CGMR-C Kilifi | Githinji et al |
| EPI_ISL_970834, EPI_ISL_970982, EPI_ISL_971085, EPI_ISL_971151, EPI_ISL_971395, EPI_ISL_971550, EPI_ISL_971608, EPI_ISL_971643, EPI_ISL_971675, EPI_ISL_971749, EPI_ISL_971767, EPI_ISL_971798, EPI_ISL_971846, EPI_ISL_971875, EPI_ISL_971936, EPI_ISL_972025, EPI_ISL_972050, EPI_ISL_972070, EPI_ISL_972082, EPI_ISL_972089, EPI_ISL_972111, EPI_ISL_972135, EPI_ISL_972176, EPI_ISL_972282, EPI_ISL_972398, EPI_ISL_972541, EPI_ISL_972566, EPI_ISL_972585, EPI_ISL_972634, EPI_ISL_972816, EPI_ISL_972877, EPI_ISL_972908, EPI_ISL_972916, EPI_ISL_972917, EPI_ISL_972963, EPI_ISL_972970, EPI_ISL_972973, EPI_ISL_973002, EPI_ISL_973022, EPI_ISL_973088, EPI_ISL_973101, EPI_ISL_973113, EPI_ISL_973178, EPI_ISL_973194, EPI_ISL_973233, EPI_ISL_973357, EPI_ISL_973478, EPI_ISL_973644, EPI_ISL_973701, EPI_ISL_973774 | | | |
| see above | Department of Virus and Microbiological Special Diagnostics, Statens Serum Institut, Copenhagen, Denmark | Aalborg University | Danish Covid-19 Genome Consortium |
| EPI_ISL_976424, EPI_ISL_976425, EPI_ISL_976426, EPI_ISL_976427, EPI_ISL_976428, EPI_ISL_976429, EPI_ISL_976430, EPI_ISL_976431, EPI_ISL_976432, EPI_ISL_976433, EPI_ISL_976434, EPI_ISL_976435, EPI_ISL_976436, EPI_ISL_976437, EPI_ISL_976438, EPI_ISL_976439, EPI_ISL_976440, EPI_ISL_976441, EPI_ISL_976442, EPI_ISL_976443, EPI_ISL_976444, EPI_ISL_976445, EPI_ISL_976446, EPI_ISL_976447, EPI_ISL_976448, EPI_ISL_976449, EPI_ISL_976450, EPI_ISL_976451, EPI_ISL_976452, EPI_ISL_976453, EPI_ISL_976454, EPI_ISL_976455, EPI_ISL_976456, EPI_ISL_976457, EPI_ISL_976458, EPI_ISL_976459, EPI_ISL_976460, EPI_ISL_976461, EPI_ISL_976462, EPI_ISL_976463, EPI_ISL_976464, EPI_ISL_976465, EPI_ISL_976466, EPI_ISL_976467, EPI_ISL_976468, EPI_ISL_976469, EPI_ISL_976470, EPI_ISL_976471, EPI_ISL_976472, EPI_ISL_976473, EPI_ISL_976474, EPI_ISL_976475, EPI_ISL_976476, EPI_ISL_976477, EPI_ISL_976478, EPI_ISL_976479, EPI_ISL_976480, EPI_ISL_976481, EPI_ISL_976482, EPI_ISL_976483, EPI_ISL_976484, EPI_ISL_976485, EPI_ISL_976486, EPI_ISL_976487, EPI_ISL_976488, EPI_ISL_976489, EPI_ISL_976490, EPI_ISL_976491, EPI_ISL_976492, EPI_ISL_976493, EPI_ISL_976494, EPI_ISL_976495, EPI_ISL_976496, EPI_ISL_976497, EPI_ISL_976498, EPI_ISL_976499, EPI_ISL_976500, EPI_ISL_976501, EPI_ISL_976502, EPI_ISL_976503, EPI_ISL_976504, EPI_ISL_976505, EPI_ISL_976506, EPI_ISL_976507, EPI_ISL_976508, EPI_ISL_976509, EPI_ISL_976510, EPI_ISL_976511, EPI_ISL_976512, EPI_ISL_976513, EPI_ISL_976514, EPI_ISL_976515, EPI_ISL_976516, EPI_ISL_976517, EPI_ISL_976518, EPI_ISL_976519, EPI_ISL_976520 | | | |
| see above | BCCDC Public Health Laboratory | BCCDC Public Health Laboratory | Prystajecky Natalie, Linda Hoang, Dan Fornika, John Tyson, Shannon Russell, Kim Macdonald, Kimia Kamelian, Ana Pacagnella, Corrinne Ng, Loretta Janz, Robert Azana Terry Snutch, Mel Krajden |
| EPI_ISL_977029, EPI_ISL_977030, EPI_ISL_977031, EPI_ISL_977032 | Rhode Island Department of Health | Infectious Disease Program, Broad Institute of Harvard and MIT | Lemieux,J.E., Siddle,K.J., Huard,R., King,E., Azevedo,K., Miller,A., Adams,G., Gladden-Young,A., Lagerborg,K., Rudy,M., DeRuff,K., Carter,A., Normandin,E., Bauer,M., Reilly,S., Tomkins-Tinch,C., Loreth,C., Chaluvadi,S., Birren,B.W., Gallagher,G., Smole,S., Park,D.J., MacInnis,B.L., and Sabeti,P.C. |
| EPI_ISL_977164, EPI_ISL_977170 | Microbiologia e Virologia | Istituto Zooprofilattico Sperimentale delle Venezie | Adelaide Milani, Alessia Schivo, Annalisa Salviato, Erika Giorgia Quaranta, Ambra Pastori, Bianca Zecchin, Alice Fusaro, Isabella Monne, Calogero Terregino, Antonia Ricci |
| EPI_ISL_977336 | University of Zambia, School of Veterinary Medicine | UNZAVET and PATH | Mulenga Mwenda-Chimfwembe, Ngonda Saasa, Daniel Bridges |
| EPI_ISL_978010 | Chiu Laboratory, University of California, San Francisco | Chiu Laboratory, University of California, San Francisco | Charles Chiu, Xianding (Wayne) Deng, Candace Wang, Venice Servellita, Jill Hacker, Debra Wadford |
| EPI_ISL_978198, EPI_ISL_978199, EPI_ISL_978200, EPI_ISL_978213 | Virginia Division of Consolidated Laboratory Services | Virginia Division of Consolidated Laboratory Services | Virginia DCLS |
| EPI_ISL_979173, EPI_ISL_979174, EPI_ISL_979175, EPI_ISL_979176, EPI_ISL_979177, EPI_ISL_979178, EPI_ISL_979179, EPI_ISL_979180, EPI_ISL_979181, EPI_ISL_979182, EPI_ISL_979185, EPI_ISL_979186, EPI_ISL_979192, EPI_ISL_979193, EPI_ISL_979194 | | | |
| see above | Humboldt County Public Health Laboratory | Chan-Zuckerberg Biohub | CZB Cliahub Consortium |
| EPI_ISL_979248 | Institute of Microbiology and Immunology, Faculty of Medicine, University of Ljubljana | Institute of Microbiology and Immunology, Faculty of Medicine, University of Ljubljana | Samo Zakotnik, Tomaž Mark Zorec, Matic Brvar, Doroteja Vlaj, Patricija Pozvek,Špela Pleh, Miša Korva, Mario Poljak, Tatjana Avši - Županc |

| | | | |
|---|---|---|---|
| EPI_ISL_979295, EPI_ISL_979296, EPI_ISL_979297, EPI_ISL_979298, EPI_ISL_979303 | Cadham Provincial laboratory | National Microbiology Laboratory (NML) | Anna Majer, Shari Tyson, Grace Seo, Philip Mabon, Elsie Grudeski, Rhiannon Huzarewich, Russell Mandes, Anneliese Landgraff, Jennifer Tanner, Natalie Knox, Morag Graham, Gary Van Domselaar, Paul Van Caeseele, Jared Bullard, David Alexander, Kerry Dust, Nathalie Bastien, Yan Li, Timothy Booth, Darian Hole, Madison Chapel, Kirsten Biggar, CanCOGeN's metadata curation team, Public Health Agency of Canada CanCOGeN team |
| EPI_ISL_981903, EPI_ISL_981904, EPI_ISL_981905, EPI_ISL_981906, EPI_ISL_981907, EPI_ISL_981908, EPI_ISL_981909, EPI_ISL_981910, EPI_ISL_981911, EPI_ISL_981912, EPI_ISL_981913, EPI_ISL_981914, EPI_ISL_981915, EPI_ISL_981916, EPI_ISL_981917, EPI_ISL_981918, EPI_ISL_981919 | | | |
| see above | Microbiology Service, Hospital Universitario Clinico San Cecilio, Granada | Microbiology Service, Hospital Universitario Clinico San Cecilio, Granada | Adolfo de Salazar, Natalia Chueca, Laura Viñuela, Ana Fuentes, Federico García |
| EPI_ISL_982847, EPI_ISL_982848, EPI_ISL_982849, EPI_ISL_982850 | UK Healthcare Clinical Microbiology | Kentucky State Public Health Lab | Stephanie Lunn, Karim George, Joshua Tobias, William Grooms, Vaneet Arora, Matthew Johnson, Rachel Zinner, Rhonda Lucas |
