## Supplementary material for "Detection and characterization of the SARS-CoV-2 lineage B.1.526 in New York": Supp. Table 3 GISAID Acknowledgment part 2: gisaid_hcov-19_acknowledgement_table_2021_02_13_00.pdf

We gratefully acknowledge the following Authors from the Originating laboratories responsible for obtaining the specimens, as well as the Submitting laboratories where the genome data were generated and shared via GISAID, on which this research is based.

All Submitters of data may be contacted directly via www.gisaid.org

Authors are sorted alphabetically.

| Accession ID | Originating Laboratory | Submitting Laboratory | Authors |
|---|---|---|---|
| EPI_ISL_584788, EPI_ISL_584789, EPI_ISL_584790, EPI_ISL_584791, EPI_ISL_584792 | Queens Medical Centre, Clinical Microbiology Department / DeepSeq Nottingham | COVID-19 Genomics UK (COG-UK) Consortium | Gemma Clark, Wendy Smith, Manjinder Khakh, Vicki M Fleming, Michelle M Lister, Hannah Howson-Wells, Jonathan Ball, Patrick McClure, Joseph Chappell, Theocharis Tsoleridis, Nadine Holmes, Matthew Carlisle, Christopher Moore, Fei Sang, Johnny Debebe, Victoria Wright, Matthew Loose |
| EPI_ISL_591486 | Histopath | NSW Health Pathology - Institute of Clinical Pathology and Medical Research; Westmead Hospital; University of Sydney | CIDM-PH et al. |
| EPI_ISL_591516 | Sydney South West Pathology Service (SSWPS) - Liverpool Hospital - NSW Health Pathology | NSW Health Pathology - Institute of Clinical Pathology and Medical Research; Westmead Hospital; University of Sydney | CIDM-PH et al. |
| EPI_ISL_593998 | Respiratory Virus Unit, Microbiology Services Colindale, Public Health England | Respiratory Virus Unit, Microbiology Services Colindale, Public Health England | PHE Covid Sequencing Team |
| EPI_ISL_594846, EPI_ISL_594847, EPI_ISL_594848 | Virology Department, Royal Infirmary of Edinburgh, NHS Lothian / School of Biological Sciences, University of Edinburgh / Institute of Genetics and Molecular Medicine, University of Edinburgh | COVID-19 Genomics UK (COG-UK) Consortium | McHugh M, Dewar R, Rooke S, Gallagher M, Balcaza C, O'Toole Á, Scher E, Hill V, McCrone JT, Colquhoun R, Yu X, Jackson B, Rambaut A, Williams TC, Templeton K |
| EPI_ISL_595180, EPI_ISL_595181, EPI_ISL_595182, EPI_ISL_595183, EPI_ISL_595184, EPI_ISL_595185, EPI_ISL_595186, EPI_ISL_595188, EPI_ISL_595189, EPI_ISL_595191, EPI_ISL_595192, EPI_ISL_595193, EPI_ISL_595194, EPI_ISL_595195, EPI_ISL_595196, EPI_ISL_595197, EPI_ISL_595198, EPI_ISL_595199, EPI_ISL_595201, EPI_ISL_595202, EPI_ISL_595203, EPI_ISL_595204, EPI_ISL_595205, EPI_ISL_595206, EPI_ISL_595207, EPI_ISL_595208, EPI_ISL_595210, EPI_ISL_595211, EPI_ISL_595212, EPI_ISL_595213, EPI_ISL_595214, EPI_ISL_595215, EPI_ISL_595216, EPI_ISL_595217, EPI_ISL_595218, EPI_ISL_595219, EPI_ISL_595220, EPI_ISL_595222, EPI_ISL_595224, EPI_ISL_595227, EPI_ISL_595228, EPI_ISL_595230, EPI_ISL_595231 | | | |
| see above | Quadram Institute Bioscience | COVID-19 Genomics UK (COG-UK) Consortium | Dave J. Baker, Gemma L. Kay, Alp Aydin, Thanh Le-Viet, Steven Rudder, Ana P. Tedim, Anastasia Kolyva, Maria Diaz, Leonardo de Oliveira Martins, Nabil-Fareed Alikhan, Lizzie Meadows, Rachael Stanley, Ngozi Elumogo, Muhammed Yasir, Nicholas M. Thomson, Alexander J Trotter, Rachel Gilroy, Samuel Bloomfield, Claire Stuart, Andrew Bell, Reenesh Prakash, Samir Dervisevic, Alison E. Mather, John Wain, Mark Webber, Andrew J. Page, Justin O'Grady |
| EPI_ISL_595237, EPI_ISL_595238, EPI_ISL_595239, EPI_ISL_595240, EPI_ISL_595241, EPI_ISL_595242, EPI_ISL_595243, EPI_ISL_595244, EPI_ISL_595245, EPI_ISL_595246, EPI_ISL_595247, EPI_ISL_595248, EPI_ISL_595249, EPI_ISL_595250, EPI_ISL_595251, EPI_ISL_595252, EPI_ISL_595253, EPI_ISL_595254, EPI_ISL_595255, EPI_ISL_595256, EPI_ISL_595257, EPI_ISL_595258, EPI_ISL_595259, EPI_ISL_595260, EPI_ISL_595261, EPI_ISL_595262, EPI_ISL_595263, EPI_ISL_595264, EPI_ISL_595265, EPI_ISL_595266, EPI_ISL_595267, EPI_ISL_595268, EPI_ISL_595269, EPI_ISL_595270, EPI_ISL_595271, EPI_ISL_595272, EPI_ISL_595273, EPI_ISL_595274, EPI_ISL_595275, EPI_ISL_595276, EPI_ISL_595277, EPI_ISL_595278, EPI_ISL_595279, EPI_ISL_595280, EPI_ISL_595281, EPI_ISL_595282, EPI_ISL_595283, EPI_ISL_595284, EPI_ISL_595285, EPI_ISL_595286, EPI_ISL_595287, EPI_ISL_595288, EPI_ISL_595289, EPI_ISL_595290, EPI_ISL_595291, EPI_ISL_595292, EPI_ISL_595293, EPI_ISL_595294, EPI_ISL_595295, EPI_ISL_595296, EPI_ISL_595297, EPI_ISL_595298, EPI_ISL_595299, EPI_ISL_595300, EPI_ISL_595301, EPI_ISL_595302, EPI_ISL_595303, EPI_ISL_595304, EPI_ISL_595305, EPI_ISL_595306, EPI_ISL_595317, EPI_ISL_595318, EPI_ISL_595319, EPI_ISL_595320, EPI_ISL_595321, EPI_ISL_595322 | | | |
| see above | Queens Medical Centre, Clinical Microbiology Department / DeepSeq Nottingham | COVID-19 Genomics UK (COG-UK) Consortium | Gemma Clark, Wendy Smith, Manjinder Khakh, Vicki M Fleming, Michelle M Lister, Hannah Howson-Wells, Jonathan Ball, Patrick McClure, Joseph Chappell, Theocharis Tsoleridis, Nadine Holmes, Matthew Carlisle, Christopher Moore, Fei Sang, Johnny Debebe, Victoria Wright, Matthew Loose |
| EPI_ISL_595331, EPI_ISL_595333, EPI_ISL_595342, EPI_ISL_595344, EPI_ISL_595349, EPI_ISL_595350, EPI_ISL_595359, EPI_ISL_595392, EPI_ISL_595401, EPI_ISL_595409, EPI_ISL_595425, EPI_ISL_595428, EPI_ISL_595445, EPI_ISL_595453, EPI_ISL_595456, EPI_ISL_595484, EPI_ISL_595485, EPI_ISL_595506, EPI_ISL_595510, EPI_ISL_595518, EPI_ISL_595526, EPI_ISL_595529, EPI_ISL_595533, EPI_ISL_595539, EPI_ISL_595560, EPI_ISL_595570, EPI_ISL_595577, EPI_ISL_595582, EPI_ISL_595587, EPI_ISL_595613 | | | |
| see above | Wales Specialist Virology Centre Sequencing lab: Pathogen Genomics Unit | COVID-19 Genomics UK (COG-UK) Consortium | Catherine Moore, Johnathan Evans, Laura Gifford, Malorie Perry, Simon Cottrell, Angela Marchbank, Alec Birchley, Alexander Adams, Amy Gaskin, Bree Gatica-Wilcox, Jason Coombes, Joel Southgate, Lauren Gilbert, Lee Graham, Nicole Pacchiarini, Sara Kumziene-Summerhayes, Sarah Taylor, Sophie Jones, Sara Rey, Matthew Bull, Joanne Watkins, Sally Corden, Tom Connor |
| EPI_ISL_596128, EPI_ISL_596183 | Quadram Institute Bioscience | COVID-19 Genomics UK (COG-UK) Consortium | Dave J. Baker, Gemma L. Kay, Alp Aydin, Thanh Le-Viet, Steven Rudder, Ana P. Tedim, Anastasia Kolyva, Maria Diaz, Leonardo de Oliveira Martins, Nabil-Fareed Alikhan, Lizzie Meadows, Rachael Stanley, Ngozi Elumogo, Muhammed Yasir, Nicholas M. Thomson, Alexander J Trotter, Rachel Gilroy, Samuel Bloomfield, Claire Stuart, Andrew Bell, Reenesh Prakash, Samir Dervisevic, Alison E. Mather, John Wain, Mark Webber, Andrew J. Page, Justin O'Grady |
| EPI_ISL_596217 | Virology Department, Sheffield Teaching Hospitals NHS Foundation Trust/Department of Infection, Immunity and Cardiovascular Disease, The Medical School, University of Sheffield | COVID-19 Genomics UK (COG-UK) Consortium | Thushan de Silva, Matthew Parker, Nikki Smith, Adri Angyal, Rebecca Brown, Luke Green, Rachel Tucker, Paul Parsons, Danielle Groves, Katie Johnson, Laura Carrilero, Alex Keeley, Dave Partridge, Matthew Wyles, Benjamin Lindsey, Mehmet Yavuz, Mohammad Raza, Cariad Evans |
| EPI_ISL_596486, EPI_ISL_596487, EPI_ISL_596488, EPI_ISL_596489, EPI_ISL_596491, EPI_ISL_596493, EPI_ISL_596494, EPI_ISL_596495 | National Public Health Laboratory, National Centre for Infectious Diseases | National Public Health Laboratory, National Centre for Infectious Diseases | Tze Minn Mak, Sophie Octavia, Zhenyang Zhou, Lin Cui, Raymond Tzer Pin Lin |
| EPI_ISL_596624, EPI_ISL_596625, EPI_ISL_596626 | University of Michigan Clinical Microbiology Laboratory | Lauring Lab, University of Michigan, Department of Microbiology and Immunology | Valesano |
| EPI_ISL_597264, EPI_ISL_597354 | Lighthouse Lab in Cambridge | Wellcome Sanger Institute for the COVID-19 Genomics UK (COG-UK) consortium | Rob Howes, The Lighthouse Lab in Cambridge and Alex Alderton, Roberto Amato, Sonia Goncalves, Ewan Harrison, David K. Jackson, Ian Johnston, Dominic Kwiatkowski, Cordelia Langford, John Sillitoe on behalf of the Wellcome Sanger Institute COVID-19 Surveillance Team (http://www.sanger.ac.uk/covid-team) |
| EPI_ISL_602572, EPI_ISL_602573 | Centre for Dengue Research, Department of Immunology and Molecular Medicine | Centre for Dengue Research, Department of Immunology and Molecular Medicine | Chandima Jeewandara, Deshni Jayathilaka, Dinuka Ariyaratne, Diyanath Ranasinghe, Laksiri Gomes, Dinuka Guruge, Ruwan Wiajayamuni, Gathsaurie Neelika Malavige |
| EPI_ISL_603006, EPI_ISL_603007, EPI_ISL_603008, EPI_ISL_603009, EPI_ISL_603015, EPI_ISL_603018, EPI_ISL_603019, EPI_ISL_603020 | Minnesota Department of Health, Public Health Laboratory | Minnesota Department of Health, Public Health Laboratory | Matt Plumb, Jacob Garfin, Alexandra Lorentz, and Xiong Wang |
| EPI_ISL_603205, EPI_ISL_603207, EPI_ISL_603208, EPI_ISL_603209, EPI_ISL_603210, EPI_ISL_603211, EPI_ISL_603213, EPI_ISL_603214 | Respiratory Virus Unit, Microbiology Services Colindale, Public Health England | Respiratory Virus Unit, Microbiology Services Colindale, Public Health England | PHE Covid Sequencing Team |
| EPI_ISL_603221 | National Institute of Laboratory Medicine and Referral Center | Genomic Research Lab, BCSIR | Abu Sayeed Mohammad Mahmud, Mohammad Samir Uzzaman, Eshrar Osman, Md. Ahashan Habib, Shahina Akter, Tanjina Akhter Banu, Md. Murshed Hasan Sarkar, Barna Goswami, Iffat Jahan, Md. Saddam Hossain, Tasnim Nafisa, Md. Maruf Ahmed Molla, Mahmuda Yeasmin, Asish Kumar Ghosh, A. K. M. Shamsuzzaman, Monira Parveen, Md. Masum Hossain Arif, Md. Salim Khan |
| EPI_ISL_603238 | National Institute of Laboratory Medicine and Referral Center | Genomic Research Lab, BCSIR | Shahina Akter, Abu Sayeed Mohammad Mahmud, Mohammad Samir Uzzaman, Eshrar Osman, Md. Ahashan Habib, Tanjina Akhter Banu, Md. Murshed Hasan Sarkar, Barna Goswami, Iffat Jahan, Md. Saddam Hossain, Tasnim Nafisa, Md. Maruf Ahmed Molla, Mahmuda Yeasmin, Asish Kumar Ghosh, A. K. M. Shamsuzzaman, Monira Parveen, Md. Masum Hossain Arif, Md. Salim Khan |

| | | | |
|---|---|---|---|
| EPI_ISL_603241 | National Institute of Laboratory Medicine and Referral Center | Genomic Research Lab, BCSIR | Tanjina Akhter Banu, Abu Sayeed Mohammad Mahmud, Mohammad Samir Uzzaman, Eshrar Osman, Md. Ahashan Habib, Shahina Akter, Md. Murshed Hasan Sarkar, Barna Goswami, Iffat Jahan, Md. Saddam Hossain, Tasnim Nafisa, Md. Maruf Ahmed Molla, Mahmuda Yeasmin, Asish Kumar Ghosh, A. K. M. Shamsuzzaman, Monira Parveen, Md. Masum Hossain Arif, Md. Salim Khan |
| EPI_ISL_603600, EPI_ISL_603601, EPI_ISL_603602, EPI_ISL_603603, EPI_ISL_603604, EPI_ISL_603605, EPI_ISL_603606, EPI_ISL_603607, EPI_ISL_603608, EPI_ISL_603609, EPI_ISL_603610, EPI_ISL_603611, EPI_ISL_603612, EPI_ISL_603613, EPI_ISL_603614, EPI_ISL_603615, EPI_ISL_603616, EPI_ISL_603617, EPI_ISL_603618, EPI_ISL_603619, EPI_ISL_603620, EPI_ISL_603621, EPI_ISL_603622, EPI_ISL_603623, EPI_ISL_603624, EPI_ISL_603625, EPI_ISL_603626, EPI_ISL_603627, EPI_ISL_603628, EPI_ISL_603629, EPI_ISL_603630, EPI_ISL_603631, EPI_ISL_603632, EPI_ISL_603633, EPI_ISL_603634, EPI_ISL_603635, EPI_ISL_603636, EPI_ISL_603637, EPI_ISL_603638, EPI_ISL_603639, EPI_ISL_603640, EPI_ISL_603641, EPI_ISL_603642, EPI_ISL_603643, EPI_ISL_603644, EPI_ISL_603645, EPI_ISL_603646, EPI_ISL_603647, EPI_ISL_603648, EPI_ISL_603649, EPI_ISL_603650, EPI_ISL_603651, EPI_ISL_603652, EPI_ISL_603653, EPI_ISL_603654, EPI_ISL_603655, EPI_ISL_603656, EPI_ISL_603657, EPI_ISL_603658, EPI_ISL_603659, EPI_ISL_603660, EPI_ISL_603661, EPI_ISL_603662, EPI_ISL_603663, EPI_ISL_603664, EPI_ISL_603665, EPI_ISL_603666, EPI_ISL_603667, EPI_ISL_603668, EPI_ISL_603669, EPI_ISL_603670, EPI_ISL_603671, EPI_ISL_603672, EPI_ISL_603673, EPI_ISL_603674, EPI_ISL_603675, EPI_ISL_603676, EPI_ISL_603677, EPI_ISL_603678, EPI_ISL_603679, EPI_ISL_603680, EPI_ISL_603681, EPI_ISL_603682, EPI_ISL_603683, EPI_ISL_603684, EPI_ISL_603685, EPI_ISL_603686, EPI_ISL_603687, EPI_ISL_603688, EPI_ISL_603689, EPI_ISL_603690, EPI_ISL_603691, EPI_ISL_603692, EPI_ISL_603693, EPI_ISL_603694, EPI_ISL_603695, EPI_ISL_603696, EPI_ISL_603697, EPI_ISL_603698, EPI_ISL_603699, EPI_ISL_603700, EPI_ISL_603701, EPI_ISL_603702, EPI_ISL_603703, EPI_ISL_603704, EPI_ISL_603705, EPI_ISL_603706, EPI_ISL_603707, EPI_ISL_603708, EPI_ISL_603709, EPI_ISL_603710, EPI_ISL_603711, EPI_ISL_603712, EPI_ISL_603713, EPI_ISL_603714, EPI_ISL_603715, EPI_ISL_603716, EPI_ISL_603717, EPI_ISL_603718, EPI_ISL_603719, EPI_ISL_603720, EPI_ISL_603721, EPI_ISL_603722, EPI_ISL_603723, EPI_ISL_603724, EPI_ISL_603725, EPI_ISL_603726, EPI_ISL_603727, EPI_ISL_603728, EPI_ISL_603729, EPI_ISL_603730, EPI_ISL_603731, EPI_ISL_603732, EPI_ISL_603733, EPI_ISL_603734, EPI_ISL_603735, EPI_ISL_603736, EPI_ISL_603737, EPI_ISL_603738, EPI_ISL_603739, EPI_ISL_603740, EPI_ISL_603741, EPI_ISL_603742, EPI_ISL_603743, EPI_ISL_603744, EPI_ISL_603745, EPI_ISL_603746, EPI_ISL_603747, EPI_ISL_603748, EPI_ISL_603749, EPI_ISL_603750, EPI_ISL_603751, EPI_ISL_603752, EPI_ISL_603753, EPI_ISL_603754, EPI_ISL_603755, EPI_ISL_603756, EPI_ISL_603757, EPI_ISL_603758 | | | |
| see above | Viollier AG | Department of Biosystems Science and Engineering, ETH Zürich | Christian Beisel, Sarah Nadeau, Ivan Topolsky, Pedro Ferreira, Philipp Jablonski, Susana Posada-Céspedes, Tobias Schär, Ina Nissen, Natascha Santacroce, Elodie Burcklen, Christiane Beckmann, Maurice Redondo, Olivier Kobel, Christoph Noppen, Sophie Seidel, Noemie Santamaria de Souza, Niko Beerenwinkel, Tanja Stadler |
| EPI_ISL_605791 | Brotman Baty Institute for Precision Medicine | Brotman Baty Institute for Precision Medicine | Trevor Bedford, Jennifer K. Logue, Peter D. Han, Caitlin R. Wolf, Chris D. Frazar, Benjamin Pelle, Erica Ryke, Jover Lee, Mark J. Rieder, Deborah A. Nickerson, Christina M. Lockwood, Lea M. Starita, Helen Y. Chu, Jay Shendure |
| EPI_ISL_605826, EPI_ISL_605828, EPI_ISL_605829, EPI_ISL_605830, EPI_ISL_605831, EPI_ISL_605832, EPI_ISL_605834, EPI_ISL_605835, EPI_ISL_605839, EPI_ISL_605840, EPI_ISL_605841, EPI_ISL_605845, EPI_ISL_605846, EPI_ISL_605847, EPI_ISL_605848, EPI_ISL_605855, EPI_ISL_605860, EPI_ISL_605863, EPI_ISL_605866, EPI_ISL_605871, EPI_ISL_605873, EPI_ISL_605874, EPI_ISL_605875, EPI_ISL_605879 | | | |
| see above | PathWest Laboratory Medicine WA | PathWest Laboratory Medicine WA Microbial Surveillance Unit | PathWest Laboratory Medicine WA Microbial Surveillance Unit |
| EPI_ISL_605932, EPI_ISL_605933, EPI_ISL_605934, EPI_ISL_605935, EPI_ISL_605936, EPI_ISL_605937, EPI_ISL_605938, EPI_ISL_605939, EPI_ISL_605940, EPI_ISL_605941, EPI_ISL_605942, EPI_ISL_605943, EPI_ISL_605944, EPI_ISL_605945, EPI_ISL_605946, EPI_ISL_605947, EPI_ISL_605948, EPI_ISL_605949, EPI_ISL_605950, EPI_ISL_605951, EPI_ISL_605952, EPI_ISL_605953, EPI_ISL_605954, EPI_ISL_605955, EPI_ISL_605956, EPI_ISL_605957, EPI_ISL_605958, EPI_ISL_605959, EPI_ISL_605960, EPI_ISL_605961, EPI_ISL_605962, EPI_ISL_605963, EPI_ISL_605964, EPI_ISL_605965, EPI_ISL_605966, EPI_ISL_605967, EPI_ISL_605968, EPI_ISL_605969, EPI_ISL_605970, EPI_ISL_605971, EPI_ISL_605972, EPI_ISL_605973, EPI_ISL_605974, EPI_ISL_605975, EPI_ISL_605976, EPI_ISL_605977, EPI_ISL_605978, EPI_ISL_605979, EPI_ISL_605980, EPI_ISL_605981, EPI_ISL_605982, EPI_ISL_605983, EPI_ISL_605984, EPI_ISL_605985, EPI_ISL_605986, EPI_ISL_605987, EPI_ISL_605988, EPI_ISL_605989, EPI_ISL_605990, EPI_ISL_605991, EPI_ISL_605992, EPI_ISL_605993, EPI_ISL_605994, EPI_ISL_605995, EPI_ISL_605996, EPI_ISL_605997, EPI_ISL_605998, EPI_ISL_605999, EPI_ISL_606000, EPI_ISL_606001, EPI_ISL_606002, EPI_ISL_606003, EPI_ISL_606004, EPI_ISL_606005, EPI_ISL_606006, EPI_ISL_606007, EPI_ISL_606008, EPI_ISL_606009, EPI_ISL_606010, EPI_ISL_606011, EPI_ISL_606012, EPI_ISL_606013, EPI_ISL_606014, EPI_ISL_606015, EPI_ISL_606016, EPI_ISL_606017, EPI_ISL_606018, EPI_ISL_606019, EPI_ISL_606020, EPI_ISL_606021, EPI_ISL_606022, EPI_ISL_606023, EPI_ISL_606024, EPI_ISL_606025, EPI_ISL_606026, EPI_ISL_606027, EPI_ISL_606028, EPI_ISL_606029, EPI_ISL_606030, EPI_ISL_606031, EPI_ISL_606032, EPI_ISL_606033, EPI_ISL_606034, EPI_ISL_606035, EPI_ISL_606036, EPI_ISL_606037, EPI_ISL_606038, EPI_ISL_606039, EPI_ISL_606040, EPI_ISL_606041, EPI_ISL_606042, EPI_ISL_606043, EPI_ISL_606044, EPI_ISL_606045, EPI_ISL_606046, EPI_ISL_606047, EPI_ISL_606048, EPI_ISL_606049, EPI_ISL_606050, EPI_ISL_606051, EPI_ISL_606052, EPI_ISL_606053, EPI_ISL_606054, EPI_ISL_606055, EPI_ISL_606056, EPI_ISL_606057, EPI_ISL_606058, EPI_ISL_606059, EPI_ISL_606060, EPI_ISL_606061, EPI_ISL_606062, EPI_ISL_606063, EPI_ISL_606064, EPI_ISL_606065, EPI_ISL_606066, EPI_ISL_606067, EPI_ISL_606068, EPI_ISL_606069, EPI_ISL_606070, EPI_ISL_606071, EPI_ISL_606072, EPI_ISL_606073, EPI_ISL_606074, EPI_ISL_606075, EPI_ISL_606076, EPI_ISL_606077, EPI_ISL_606078, EPI_ISL_606079, EPI_ISL_606080, EPI_ISL_606081, EPI_ISL_606082, EPI_ISL_606083, EPI_ISL_606084, EPI_ISL_606085, EPI_ISL_606086, EPI_ISL_606087, EPI_ISL_606088, EPI_ISL_606089, EPI_ISL_606090, EPI_ISL_606091, EPI_ISL_606092, EPI_ISL_606093, EPI_ISL_606094, EPI_ISL_606095, EPI_ISL_606096, EPI_ISL_606097, EPI_ISL_606098, EPI_ISL_606099, EPI_ISL_606100, EPI_ISL_606101, EPI_ISL_606102, EPI_ISL_606103, EPI_ISL_606104, EPI_ISL_606105, EPI_ISL_606106, EPI_ISL_606107, EPI_ISL_606108, EPI_ISL_606109, EPI_ISL_606110, EPI_ISL_606111, EPI_ISL_606112, EPI_ISL_606113, EPI_ISL_606114, EPI_ISL_606115, EPI_ISL_606116, EPI_ISL_606117, EPI_ISL_606118, EPI_ISL_606119, EPI_ISL_606120, EPI_ISL_606121, EPI_ISL_606122, EPI_ISL_606123, EPI_ISL_606124, EPI_ISL_606125, EPI_ISL_606126, EPI_ISL_606127, EPI_ISL_606128, EPI_ISL_606129, EPI_ISL_606130, EPI_ISL_606131, EPI_ISL_606132, EPI_ISL_606133, EPI_ISL_606134, EPI_ISL_606135, EPI_ISL_606136, EPI_ISL_606137, EPI_ISL_606138, EPI_ISL_606139, EPI_ISL_606140, EPI_ISL_606141, EPI_ISL_606142, EPI_ISL_606143, EPI_ISL_606144, EPI_ISL_606145, EPI_ISL_606146, EPI_ISL_606147, EPI_ISL_606148, EPI_ISL_606149, EPI_ISL_606150, EPI_ISL_606151, EPI_ISL_606152, EPI_ISL_606153, EPI_ISL_606154, EPI_ISL_606155, EPI_ISL_606156, EPI_ISL_606157, EPI_ISL_606158, EPI_ISL_606159, EPI_ISL_606160, EPI_ISL_606161, EPI_ISL_606162, EPI_ISL_606163, EPI_ISL_606164, EPI_ISL_606165, EPI_ISL_606166, EPI_ISL_606167, EPI_ISL_606168, EPI_ISL_606169, EPI_ISL_606170, EPI_ISL_606171, EPI_ISL_606172, EPI_ISL_606173, EPI_ISL_606174, EPI_ISL_606175, EPI_ISL_606176, EPI_ISL_606177, EPI_ISL_606178, EPI_ISL_606179, EPI_ISL_606180, EPI_ISL_606181, EPI_ISL_606182, EPI_ISL_606183, EPI_ISL_606184, EPI_ISL_606185, EPI_ISL_606186, EPI_ISL_606187, EPI_ISL_606188, EPI_ISL_606189, EPI_ISL_606190, EPI_ISL_606191, EPI_ISL_606192, EPI_ISL_606193, EPI_ISL_606194, EPI_ISL_606195, EPI_ISL_606196, EPI_ISL_606197, EPI_ISL_606198, EPI_ISL_606199, EPI_ISL_606200, EPI_ISL_606201, EPI_ISL_606202, EPI_ISL_606203, EPI_ISL_606204, EPI_ISL_606205, EPI_ISL_606206, EPI_ISL_606207, EPI_ISL_606208, EPI_ISL_606209, EPI_ISL_606210, EPI_ISL_606211, EPI_ISL_606212, EPI_ISL_606213, EPI_ISL_606214, EPI_ISL_606215, EPI_ISL_606216, EPI_ISL_606217, EPI_ISL_606218, EPI_ISL_606219, EPI_ISL_606220, EPI_ISL_606221, EPI_ISL_606222, EPI_ISL_606223, EPI_ISL_606224, EPI_ISL_606225, EPI_ISL_606226, EPI_ISL_606227, EPI_ISL_606228, EPI_ISL_606229, EPI_ISL_606230, EPI_ISL_606231, EPI_ISL_606232, EPI_ISL_606233, EPI_ISL_606234, EPI_ISL_606235, EPI_ISL_606236, EPI_ISL_606237, EPI_ISL_606238, EPI_ISL_606239, EPI_ISL_606240, EPI_ISL_606241, EPI_ISL_606242, EPI_ISL_606243, EPI_ISL_606244, EPI_ISL_606245, EPI_ISL_606246, EPI_ISL_606247, EPI_ISL_606248, EPI_ISL_606249, EPI_ISL_606250, EPI_ISL_606251, EPI_ISL_606252, EPI_ISL_606253, EPI_ISL_606254, EPI_ISL_606255, EPI_ISL_606256, EPI_ISL_606257, EPI_ISL_606258, EPI_ISL_606259, EPI_ISL_606260, EPI_ISL_606261, EPI_ISL_606262, EPI_ISL_606263, EPI_ISL_606264, EPI_ISL_606265, EPI_ISL_606266, EPI_ISL_606267, EPI_ISL_606268, EPI_ISL_606269, EPI_ISL_606270, EPI_ISL_606271, EPI_ISL_606272, EPI_ISL_606273, EPI_ISL_606274, EPI_ISL_606275, EPI_ISL_606276, EPI_ISL_606277, EPI_ISL_606278, EPI_ISL_606279, EPI_ISL_606280, EPI_ISL_606281, EPI_ISL_606282, EPI_ISL_606283, EPI_ISL_606284, EPI_ISL_606285, EPI_ISL_606286, EPI_ISL_606287, EPI_ISL_606288, EPI_ISL_606289, EPI_ISL_606290, EPI_ISL_606291, EPI_ISL_606292, EPI_ISL_606293, EPI_ISL_606294, EPI_ISL_606295 | | | |
| see above | Lighthouse Lab in Alderley Park | Wellcome Sanger Institute for the COVID-19 Genomics UK (COG-UK) consortium | Jacquelyn Wynn, Mairead Hyland, The Lighthouse Lab in Alderley Park and Alex Alderton, Roberto Amato, Sonia Goncalves, Ewan Harrison, David K. Jackson, Ian Johnston, Dominic Kwiatkowski, Cordelia Langford, John Sillitoe on behalf of the Wellcome Sanger Institute COVID-19 Surveillance Team |
| EPI_ISL_606297, EPI_ISL_606298, EPI_ISL_606299, EPI_ISL_606300, EPI_ISL_606301, EPI_ISL_606302, EPI_ISL_606303, EPI_ISL_606304, EPI_ISL_606305, EPI_ISL_606307, EPI_ISL_606308, EPI_ISL_606309, EPI_ISL_606310 | | | |
| see above | Lighthouse Lab in Milton Keynes | Wellcome Sanger Institute for the COVID-19 Genomics UK (COG-UK) consortium | The Lighthouse Lab in Milton Keynes and Alex Alderton, Roberto Amato, Sonia Goncalves, Ewan Harrison, David K. Jackson, Ian Johnston, Dominic Kwiatkowski, Cordelia Langford, John Sillitoe on behalf of the Wellcome Sanger Institute COVID-19 Surveillance Team |
| EPI_ISL_606311 | Lighthouse Lab in Milton Keynes | Wellcome Sanger Institute for the COVID-19 Genomics UK (COG-UK) Consortium | The Lighthouse Lab in Milton Keynes and Alex Alderton, Roberto Amato, Sonia Goncalves, Ewan Harrison, David K. Jackson, Ian Johnston, Dominic Kwiatkowski, Cordelia Langford, John Sillitoe on behalf of the Wellcome Sanger Institute COVID-19 Surveillance Team |
| EPI_ISL_606312, EPI_ISL_606313 | Lighthouse Lab in Milton Keynes | Wellcome Sanger Institute for the COVID-19 Genomics UK (COG-UK) consortium | The Lighthouse Lab in Milton Keynes and Alex Alderton, Roberto Amato, Sonia Goncalves, Ewan Harrison, David K. Jackson, Ian Johnston, Dominic Kwiatkowski, Cordelia Langford, John Sillitoe on behalf of the Wellcome Sanger Institute COVID-19 Surveillance Team |
| EPI_ISL_606314 | Lighthouse Lab in Milton Keynes | Wellcome Sanger Institute for the COVID-19 Genomics UK (COG-UK) Consortium | The Lighthouse Lab in Milton Keynes and Alex Alderton, Roberto Amato, Sonia Goncalves, Ewan Harrison, David K. Jackson, Ian Johnston, Dominic Kwiatkowski, Cordelia Langford, John Sillitoe on behalf of the Wellcome Sanger Institute COVID-19 Surveillance Team |
| EPI_ISL_606315, EPI_ISL_606316, EPI_ISL_606317, EPI_ISL_606318, EPI_ISL_606319, EPI_ISL_606320, EPI_ISL_606322, EPI_ISL_606323, EPI_ISL_606324, EPI_ISL_606325, EPI_ISL_606326, EPI_ISL_606327, EPI_ISL_606329, EPI_ISL_606330, EPI_ISL_606331, EPI_ISL_606332, EPI_ISL_606333, EPI_ISL_606334, EPI_ISL_606335, EPI_ISL_606336, EPI_ISL_606337, EPI_ISL_606338, EPI_ISL_606339, EPI_ISL_606340, EPI_ISL_606341, EPI_ISL_606342, EPI_ISL_606343, EPI_ISL_606345, EPI_ISL_606346, EPI_ISL_606347, EPI_ISL_606348, EPI_ISL_606349, EPI_ISL_606350, EPI_ISL_606351, EPI_ISL_606352 | | | |
| see above | Lighthouse Lab in Milton Keynes | Wellcome Sanger Institute for the COVID-19 Genomics UK (COG-UK) consortium | The Lighthouse Lab in Milton Keynes and Alex Alderton, Roberto Amato, Sonia Goncalves, Ewan Harrison, David K. Jackson, Ian Johnston, Dominic Kwiatkowski, Cordelia Langford, John Sillitoe on behalf of the Wellcome Sanger Institute COVID-19 Surveillance Team |
| EPI_ISL_606353 | Lighthouse Lab in Milton Keynes | Wellcome Sanger Institute for the COVID-19 Genomics UK (COG-UK) Consortium | The Lighthouse Lab in Milton Keynes and Alex Alderton, Roberto Amato, Sonia Goncalves, Ewan Harrison, David K. Jackson, Ian Johnston, Dominic Kwiatkowski, Cordelia Langford, John Sillitoe on behalf of the Wellcome Sanger Institute COVID-19 Surveillance Team |
| EPI_ISL_606354, EPI_ISL_606355, EPI_ISL_606356, EPI_ISL_606357, EPI_ISL_606358, EPI_ISL_606359, EPI_ISL_606360, EPI_ISL_606361, EPI_ISL_606362, EPI_ISL_606363, EPI_ISL_606364, EPI_ISL_606365, EPI_ISL_606367, EPI_ISL_606368, EPI_ISL_606369, EPI_ISL_606370, EPI_ISL_606371, EPI_ISL_606372, EPI_ISL_606373, EPI_ISL_606374, EPI_ISL_606375, EPI_ISL_606376, EPI_ISL_606377, EPI_ISL_606378, EPI_ISL_606379, EPI_ISL_606380, EPI_ISL_606381, EPI_ISL_606382, EPI_ISL_606383, EPI_ISL_606384, EPI_ISL_606385, EPI_ISL_606387, EPI_ISL_606389, EPI_ISL_606390, EPI_ISL_606391, EPI_ISL_606392, EPI_ISL_606393, EPI_ISL_606394, EPI_ISL_606395, EPI_ISL_606396, EPI_ISL_606397, EPI_ISL_606398, EPI_ISL_606399, EPI_ISL_606400, EPI_ISL_606401 | | | |
| see above | Lighthouse Lab in Milton Keynes | Wellcome Sanger Institute for the COVID-19 Genomics UK (COG-UK) consortium | The Lighthouse Lab in Milton Keynes and Alex Alderton, Roberto Amato, Sonia Goncalves, Ewan Harrison, David K. Jackson, Ian Johnston, Dominic Kwiatkowski, Cordelia Langford, John Sillitoe on behalf of the Wellcome Sanger Institute COVID-19 Surveillance Team |
| EPI_ISL_606402 | Lighthouse Lab in Milton Keynes | Wellcome Sanger Institute for the COVID-19 Genomics UK (COG-UK) Consortium | The Lighthouse Lab in Milton Keynes and Alex Alderton, Roberto Amato, Sonia Goncalves, Ewan Harrison, David K. Jackson, Ian Johnston, Dominic Kwiatkowski, Cordelia Langford, John Sillitoe on behalf of the Wellcome Sanger Institute COVID-19 Surveillance Team |
| EPI_ISL_606403, EPI_ISL_606404, EPI_ISL_606405, EPI_ISL_606406, EPI_ISL_606407, EPI_ISL_606408, EPI_ISL_606409, EPI_ISL_606410, EPI_ISL_606411, EPI_ISL_606412, EPI_ISL_606413, EPI_ISL_606414 | | | |
| see above | Lighthouse Lab in Milton Keynes | Wellcome Sanger Institute for the COVID-19 Genomics UK (COG-UK) consortium | The Lighthouse Lab in Milton Keynes and Alex Alderton, Roberto Amato, Sonia Goncalves, Ewan Harrison, David K. Jackson, Ian Johnston, Dominic Kwiatkowski, Cordelia Langford, John Sillitoe on behalf of the Wellcome Sanger Institute COVID-19 Surveillance Team |
| EPI_ISL_606415 | Lighthouse Lab in Milton Keynes | Wellcome Sanger Institute for the COVID-19 Genomics UK (COG-UK) Consortium | The Lighthouse Lab in Milton Keynes and Alex Alderton, Roberto Amato, Sonia Goncalves, Ewan Harrison, David K. Jackson, Ian Johnston, Dominic Kwiatkowski, Cordelia Langford, John Sillitoe on behalf of the Wellcome Sanger Institute COVID-19 Surveillance Team |
| EPI_ISL_606416, EPI_ISL_606417, EPI_ISL_606418, EPI_ISL_606419, EPI_ISL_606420, EPI_ISL_606421, EPI_ISL_606422, EPI_ISL_606423, EPI_ISL_606424, EPI_ISL_606425, EPI_ISL_606426, EPI_ISL_606427, EPI_ISL_606428, EPI_ISL_606429, EPI_ISL_606430, EPI_ISL_606431, EPI_ISL_606432, EPI_ISL_606433, EPI_ISL_606434, EPI_ISL_606435, EPI_ISL_606436, EPI_ISL_606437, EPI_ISL_606438, EPI_ISL_606439, EPI_ISL_606440, EPI_ISL_606441, EPI_ISL_606442, EPI_ISL_606443, EPI_ISL_606444, EPI_ISL_606445, EPI_ISL_606446, EPI_ISL_606447, EPI_ISL_606448, EPI_ISL_606449, EPI_ISL_606450, EPI_ISL_606451, EPI_ISL_606452, EPI_ISL_606453, EPI_ISL_606454, EPI_ISL_606455, EPI_ISL_606456, EPI_ISL_606457, EPI_ISL_606458, EPI_ISL_606459, EPI_ISL_606460, EPI_ISL_606461 | | | |
| see above | Lighthouse Lab in Milton Keynes | Wellcome Sanger Institute for the COVID-19 Genomics UK (COG-UK) consortium | The Lighthouse Lab in Milton Keynes and Alex Alderton, Roberto Amato, Sonia Goncalves, Ewan Harrison, David K. Jackson, Ian Johnston, Dominic Kwiatkowski, Cordelia Langford, John Sillitoe on behalf of the Wellcome Sanger Institute COVID-19 Surveillance Team |

| | | | |
|---|---|---|---|
| EPI_ISL_606462 | Lighthouse Lab in Milton Keynes | Wellcome Sanger Institute for the COVID-19 Genomics UK (COG-UK) Consortium | The Lighthouse Lab in Milton Keynes and Alex Alderton, Roberto Amato, Sonia Goncalves, Ewan Harrison, David K. Jackson, Ian Johnston, Dominic Kwiatkowski, Cordelia Langford, John Sillitoe on behalf of the Wellcome Sanger Institute COVID-19 Surveillance Team |
| EPI_ISL_606463, EPI_ISL_606464, EPI_ISL_606465, EPI_ISL_606466, EPI_ISL_606467, EPI_ISL_606468, EPI_ISL_606469, EPI_ISL_606470, EPI_ISL_606471, EPI_ISL_606472, EPI_ISL_606473, EPI_ISL_606474, EPI_ISL_606475, EPI_ISL_606476, EPI_ISL_606477, EPI_ISL_606478, EPI_ISL_606479, EPI_ISL_606480, EPI_ISL_606481, EPI_ISL_606482, EPI_ISL_606483, EPI_ISL_606484, EPI_ISL_606485, EPI_ISL_606486, EPI_ISL_606487, EPI_ISL_606488, EPI_ISL_606489, EPI_ISL_606490, EPI_ISL_606491, EPI_ISL_606492, EPI_ISL_606493, EPI_ISL_606494, EPI_ISL_606495, EPI_ISL_606496, EPI_ISL_606497, EPI_ISL_606498, EPI_ISL_606499, EPI_ISL_606500, EPI_ISL_606501, EPI_ISL_606502, EPI_ISL_606503, EPI_ISL_606504, EPI_ISL_606505, EPI_ISL_606506, EPI_ISL_606507, EPI_ISL_606508, EPI_ISL_606509, EPI_ISL_606510, EPI_ISL_606511, EPI_ISL_606512, EPI_ISL_606513, EPI_ISL_606514, EPI_ISL_606515 | | | |
| see above | Lighthouse Lab in Milton Keynes | Wellcome Sanger Institute for the COVID-19 Genomics UK (COG-UK) consortium | The Lighthouse Lab in Milton Keynes and Alex Alderton, Roberto Amato, Sonia Goncalves, Ewan Harrison, David K. Jackson, Ian Johnston, Dominic Kwiatkowski, Cordelia Langford, John Sillitoe on behalf of the Wellcome Sanger Institute COVID-19 Surveillance Team |
| EPI_ISL_606516 | Lighthouse Lab in Milton Keynes | Wellcome Sanger Institute for the COVID-19 Genomics UK (COG-UK) Consortium | The Lighthouse Lab in Milton Keynes and Alex Alderton, Roberto Amato, Sonia Goncalves, Ewan Harrison, David K. Jackson, Ian Johnston, Dominic Kwiatkowski, Cordelia Langford, John Sillitoe on behalf of the Wellcome Sanger Institute COVID-19 Surveillance Team |
| EPI_ISL_606517, EPI_ISL_606518, EPI_ISL_606519, EPI_ISL_606520, EPI_ISL_606521, EPI_ISL_606522, EPI_ISL_606523 | Lighthouse Lab in Milton Keynes | Wellcome Sanger Institute for the COVID-19 Genomics UK (COG-UK) consortium | The Lighthouse Lab in Milton Keynes and Alex Alderton, Roberto Amato, Sonia Goncalves, Ewan Harrison, David K. Jackson, Ian Johnston, Dominic Kwiatkowski, Cordelia Langford, John Sillitoe on behalf of the Wellcome Sanger Institute COVID-19 Surveillance Team |
| EPI_ISL_606524 | Lighthouse Lab in Milton Keynes | Wellcome Sanger Institute for the COVID-19 Genomics UK (COG-UK) Consortium | The Lighthouse Lab in Milton Keynes and Alex Alderton, Roberto Amato, Sonia Goncalves, Ewan Harrison, David K. Jackson, Ian Johnston, Dominic Kwiatkowski, Cordelia Langford, John Sillitoe on behalf of the Wellcome Sanger Institute COVID-19 Surveillance Team |
| EPI_ISL_606525, EPI_ISL_606526, EPI_ISL_606527, EPI_ISL_606528, EPI_ISL_606529, EPI_ISL_606531, EPI_ISL_606533, EPI_ISL_606534, EPI_ISL_606535, EPI_ISL_606536, EPI_ISL_606537, EPI_ISL_606538, EPI_ISL_606539, EPI_ISL_606540, EPI_ISL_606541, EPI_ISL_606542, EPI_ISL_606543, EPI_ISL_606544, EPI_ISL_606545, EPI_ISL_606546, EPI_ISL_606547, EPI_ISL_606548, EPI_ISL_606549, EPI_ISL_606550, EPI_ISL_606551, EPI_ISL_606552, EPI_ISL_606553, EPI_ISL_606554, EPI_ISL_606555, EPI_ISL_606556, EPI_ISL_606557, EPI_ISL_606558, EPI_ISL_606559, EPI_ISL_606560, EPI_ISL_606562, EPI_ISL_606563, EPI_ISL_606564, EPI_ISL_606565, EPI_ISL_606566, EPI_ISL_606567, EPI_ISL_606568, EPI_ISL_606569, EPI_ISL_606570, EPI_ISL_606571, EPI_ISL_606572, EPI_ISL_606573, EPI_ISL_606574, EPI_ISL_606575, EPI_ISL_606577, EPI_ISL_606578, EPI_ISL_606579, EPI_ISL_606580, EPI_ISL_606581, EPI_ISL_606582, EPI_ISL_606583, EPI_ISL_606585, EPI_ISL_606586, EPI_ISL_606587, EPI_ISL_606589, EPI_ISL_606590, EPI_ISL_606591, EPI_ISL_606592, EPI_ISL_606593, EPI_ISL_606594, EPI_ISL_606595, EPI_ISL_606596, EPI_ISL_606597, EPI_ISL_606598, EPI_ISL_606599, EPI_ISL_606600, EPI_ISL_606601, EPI_ISL_606602, EPI_ISL_606603, EPI_ISL_606604, EPI_ISL_606605, EPI_ISL_606606, EPI_ISL_606607, EPI_ISL_606609, EPI_ISL_606610, EPI_ISL_606612, EPI_ISL_606614, EPI_ISL_606615, EPI_ISL_606616, EPI_ISL_606617, EPI_ISL_606618, EPI_ISL_606619, EPI_ISL_606620, EPI_ISL_606621, EPI_ISL_606622, EPI_ISL_606623, EPI_ISL_606624, EPI_ISL_606625, EPI_ISL_606626, EPI_ISL_606627, EPI_ISL_606628, EPI_ISL_606629, EPI_ISL_606630, EPI_ISL_606631, EPI_ISL_606632, EPI_ISL_606633, EPI_ISL_606634, EPI_ISL_606635, EPI_ISL_606637, EPI_ISL_606638, EPI_ISL_606639, EPI_ISL_606640, EPI_ISL_606641, EPI_ISL_606642, EPI_ISL_606643, EPI_ISL_606644, EPI_ISL_606645, EPI_ISL_606646, EPI_ISL_606647, EPI_ISL_606648, EPI_ISL_606649, EPI_ISL_606650, EPI_ISL_606651, EPI_ISL_606652, EPI_ISL_606653 | | | |
| see above | Lighthouse Lab in Milton Keynes | Wellcome Sanger Institute for the COVID-19 Genomics UK (COG-UK) consortium | The Lighthouse Lab in Milton Keynes and Alex Alderton, Roberto Amato, Sonia Goncalves, Ewan Harrison, David K. Jackson, Ian Johnston, Dominic Kwiatkowski, Cordelia Langford, John Sillitoe on behalf of the Wellcome Sanger Institute COVID-19 Surveillance Team |
| EPI_ISL_606654 | Lighthouse Lab in Milton Keynes | Wellcome Sanger Institute for the COVID-19 Genomics UK (COG-UK) Consortium | The Lighthouse Lab in Milton Keynes and Alex Alderton, Roberto Amato, Sonia Goncalves, Ewan Harrison, David K. Jackson, Ian Johnston, Dominic Kwiatkowski, Cordelia Langford, John Sillitoe on behalf of the Wellcome Sanger Institute COVID-19 Surveillance Team |
| EPI_ISL_606655, EPI_ISL_606656, EPI_ISL_606657, EPI_ISL_606658, EPI_ISL_606659, EPI_ISL_606661, EPI_ISL_606662, EPI_ISL_606663, EPI_ISL_606664, EPI_ISL_606665, EPI_ISL_606666, EPI_ISL_606667, EPI_ISL_606668, EPI_ISL_606669, EPI_ISL_606670, EPI_ISL_606671, EPI_ISL_606672, EPI_ISL_606673, EPI_ISL_606674, EPI_ISL_606675, EPI_ISL_606676, EPI_ISL_606677, EPI_ISL_606678, EPI_ISL_606679, EPI_ISL_606680, EPI_ISL_606681, EPI_ISL_606682, EPI_ISL_606683, EPI_ISL_606684, EPI_ISL_606685, EPI_ISL_606686, EPI_ISL_606687, EPI_ISL_606688, EPI_ISL_606690, EPI_ISL_606691, EPI_ISL_606692, EPI_ISL_606693, EPI_ISL_606694, EPI_ISL_606695, EPI_ISL_606696, EPI_ISL_606697, EPI_ISL_606698, EPI_ISL_606699, EPI_ISL_606700, EPI_ISL_606701, EPI_ISL_606702, EPI_ISL_606703, EPI_ISL_606704, EPI_ISL_606705, EPI_ISL_606706, EPI_ISL_606707, EPI_ISL_606708, EPI_ISL_606709, EPI_ISL_606710, EPI_ISL_606711, EPI_ISL_606712, EPI_ISL_606713, EPI_ISL_606714, EPI_ISL_606715, EPI_ISL_606716, EPI_ISL_606717, EPI_ISL_606718, EPI_ISL_606719, EPI_ISL_606720, EPI_ISL_606721, EPI_ISL_606722, EPI_ISL_606723, EPI_ISL_606724, EPI_ISL_606726, EPI_ISL_606728, EPI_ISL_606729, EPI_ISL_606730, EPI_ISL_606731, EPI_ISL_606732, EPI_ISL_606733, EPI_ISL_606734, EPI_ISL_606735, EPI_ISL_606736, EPI_ISL_606737, EPI_ISL_606738, EPI_ISL_606739, EPI_ISL_606740, EPI_ISL_606741, EPI_ISL_606742, EPI_ISL_606743, EPI_ISL_606744, EPI_ISL_606745, EPI_ISL_606746, EPI_ISL_606748, EPI_ISL_606749, EPI_ISL_606750, EPI_ISL_606752, EPI_ISL_606753, EPI_ISL_606754, EPI_ISL_606755, EPI_ISL_606756, EPI_ISL_606757, EPI_ISL_606758, EPI_ISL_606759, EPI_ISL_606760, EPI_ISL_606762, EPI_ISL_606763, EPI_ISL_606764, EPI_ISL_606765, EPI_ISL_606766, EPI_ISL_606767, EPI_ISL_606768, EPI_ISL_606770, EPI_ISL_606771, EPI_ISL_606772, EPI_ISL_606773, EPI_ISL_606774, EPI_ISL_606775, EPI_ISL_606776, EPI_ISL_606777, EPI_ISL_606778, EPI_ISL_606779, EPI_ISL_606780, EPI_ISL_606781, EPI_ISL_606782, EPI_ISL_606783, EPI_ISL_606784, EPI_ISL_606785, EPI_ISL_606786, EPI_ISL_606787, EPI_ISL_606788, EPI_ISL_606789, EPI_ISL_606790, EPI_ISL_606791, EPI_ISL_606792, EPI_ISL_606793, EPI_ISL_606794, EPI_ISL_606795, EPI_ISL_606796 | | | |
| see above | Lighthouse Lab in Milton Keynes | Wellcome Sanger Institute for the COVID-19 Genomics UK (COG-UK) consortium | The Lighthouse Lab in Milton Keynes and Alex Alderton, Roberto Amato, Sonia Goncalves, Ewan Harrison, David K. Jackson, Ian Johnston, Dominic Kwiatkowski, Cordelia Langford, John Sillitoe on behalf of the Wellcome Sanger Institute COVID-19 Surveillance Team |
| EPI_ISL_606797 | Lighthouse Lab in Milton Keynes | Wellcome Sanger Institute for the COVID-19 Genomics UK (COG-UK) Consortium | The Lighthouse Lab in Milton Keynes and Alex Alderton, Roberto Amato, Sonia Goncalves, Ewan Harrison, David K. Jackson, Ian Johnston, Dominic Kwiatkowski, Cordelia Langford, John Sillitoe on behalf of the Wellcome Sanger Institute COVID-19 Surveillance Team |
| EPI_ISL_606798, EPI_ISL_606799, EPI_ISL_606800, EPI_ISL_606801, EPI_ISL_606802, EPI_ISL_606803, EPI_ISL_606804, EPI_ISL_606805, EPI_ISL_606806, EPI_ISL_606808, EPI_ISL_606809, EPI_ISL_606810, EPI_ISL_606811, EPI_ISL_606812, EPI_ISL_606813, EPI_ISL_606814, EPI_ISL_606815, EPI_ISL_606816, EPI_ISL_606817, EPI_ISL_606818, EPI_ISL_606819, EPI_ISL_606820, EPI_ISL_606821, EPI_ISL_606823, EPI_ISL_606824, EPI_ISL_606825, EPI_ISL_606826, EPI_ISL_606828, EPI_ISL_606829, EPI_ISL_606830, EPI_ISL_606831, EPI_ISL_606832, EPI_ISL_606833, EPI_ISL_606834, EPI_ISL_606836, EPI_ISL_606837, EPI_ISL_606838, EPI_ISL_606839, EPI_ISL_606840, EPI_ISL_606841, EPI_ISL_606842, EPI_ISL_606843, EPI_ISL_606844, EPI_ISL_606845, EPI_ISL_606846, EPI_ISL_606847, EPI_ISL_606848, EPI_ISL_606849, EPI_ISL_606850, EPI_ISL_606851, EPI_ISL_606852, EPI_ISL_606853, EPI_ISL_606854, EPI_ISL_606855, EPI_ISL_606856, EPI_ISL_606857, EPI_ISL_606858, EPI_ISL_606859, EPI_ISL_606860, EPI_ISL_606861, EPI_ISL_606862, EPI_ISL_606864, EPI_ISL_606866, EPI_ISL_606867, EPI_ISL_606868, EPI_ISL_606869, EPI_ISL_606870, EPI_ISL_606871, EPI_ISL_606872, EPI_ISL_606873, EPI_ISL_606874, EPI_ISL_606875, EPI_ISL_606876, EPI_ISL_606877, EPI_ISL_606878, EPI_ISL_606879, EPI_ISL_606880, EPI_ISL_606881, EPI_ISL_606882, EPI_ISL_606883, EPI_ISL_606884, EPI_ISL_606885, EPI_ISL_606886, EPI_ISL_606887, EPI_ISL_606888, EPI_ISL_606889, EPI_ISL_606890, EPI_ISL_606891, EPI_ISL_606892, EPI_ISL_606893, EPI_ISL_606894, EPI_ISL_606895, EPI_ISL_606896, EPI_ISL_606897, EPI_ISL_606898, EPI_ISL_606899, EPI_ISL_606901, EPI_ISL_606903, EPI_ISL_606904, EPI_ISL_606905, EPI_ISL_606906, EPI_ISL_606907, EPI_ISL_606908, EPI_ISL_606909, EPI_ISL_606910, EPI_ISL_606911, EPI_ISL_606912, EPI_ISL_606913, EPI_ISL_606914, EPI_ISL_606915, EPI_ISL_606916, EPI_ISL_606917, EPI_ISL_606918, EPI_ISL_606919, EPI_ISL_606920, EPI_ISL_606921, EPI_ISL_606922, EPI_ISL_606923 | | | |
| see above | Lighthouse Lab in Milton Keynes | Wellcome Sanger Institute for the COVID-19 Genomics UK (COG-UK) consortium | The Lighthouse Lab in Milton Keynes and Alex Alderton, Roberto Amato, Sonia Goncalves, Ewan Harrison, David K. Jackson, Ian Johnston, Dominic Kwiatkowski, Cordelia Langford, John Sillitoe on behalf of the Wellcome Sanger Institute COVID-19 Surveillance Team |
| EPI_ISL_606924, EPI_ISL_606925, EPI_ISL_606926, EPI_ISL_606927, EPI_ISL_606928, EPI_ISL_606929, EPI_ISL_606930, EPI_ISL_606931, EPI_ISL_606932, EPI_ISL_606933, EPI_ISL_606934, EPI_ISL_606935, EPI_ISL_606936, EPI_ISL_606937, EPI_ISL_606938, EPI_ISL_606939, EPI_ISL_606940, EPI_ISL_606941, EPI_ISL_606942, EPI_ISL_606943, EPI_ISL_606944, EPI_ISL_606945, EPI_ISL_606946, EPI_ISL_606947 | | | |
| see above | Lighthouse Lab in Alderley Park | Wellcome Sanger Institute for the COVID-19 Genomics UK (COG-UK) consortium | Jacquelyn Wynn, Mairead Hyland, The Lighthouse Lab in Alderley Park and Alex Alderton, Roberto Amato, Sonia Goncalves, Ewan Harrison, David K. Jackson, Ian Johnston, Dominic Kwiatkowski, Cordelia Langford, John Sillitoe on behalf of the Wellcome Sanger Institute COVID-19 Surveillance Team |
| EPI_ISL_606948 | Lighthouse Lab in Alderley Park | Wellcome Sanger Institute for the COVID-19 Genomics UK (COG-UK) Consortium | Jacquelyn Wynn, Mairead Hyland, The Lighthouse Lab in Alderley Park and Alex Alderton, Roberto Amato, Sonia Goncalves, Ewan Harrison, David K. Jackson, Ian Johnston, Dominic Kwiatkowski, Cordelia Langford, John Sillitoe on behalf of the Wellcome Sanger Institute COVID-19 Surveillance Team |
| EPI_ISL_606949, EPI_ISL_606950, EPI_ISL_606951, EPI_ISL_606952, EPI_ISL_606953, EPI_ISL_606954, EPI_ISL_606955, EPI_ISL_606956, EPI_ISL_606957, EPI_ISL_606958, EPI_ISL_606959 | | | |
| see above | Lighthouse Lab in Alderley Park | Wellcome Sanger Institute for the COVID-19 Genomics UK (COG-UK) consortium | Jacquelyn Wynn, Mairead Hyland, The Lighthouse Lab in Alderley Park and Alex Alderton, Roberto Amato, Sonia Goncalves, Ewan Harrison, David K. Jackson, Ian Johnston, Dominic Kwiatkowski, Cordelia Langford, John Sillitoe on behalf of the Wellcome Sanger Institute COVID-19 Surveillance Team |
| EPI_ISL_606960 | Lighthouse Lab in Alderley Park | Wellcome Sanger Institute for the COVID-19 Genomics UK (COG-UK) Consortium | Jacquelyn Wynn, Mairead Hyland, The Lighthouse Lab in Alderley Park and Alex Alderton, Roberto Amato, Sonia Goncalves, Ewan Harrison, David K. Jackson, Ian Johnston, Dominic Kwiatkowski, Cordelia Langford, John Sillitoe on behalf of the Wellcome Sanger Institute COVID-19 Surveillance Team |
| EPI_ISL_606961, EPI_ISL_606962, EPI_ISL_606963, EPI_ISL_606964, EPI_ISL_606965, EPI_ISL_606966, EPI_ISL_606967, EPI_ISL_606968, EPI_ISL_606969, EPI_ISL_606970, EPI_ISL_606971, EPI_ISL_606972, EPI_ISL_606973, EPI_ISL_606974, EPI_ISL_606975, EPI_ISL_606976, EPI_ISL_606977, EPI_ISL_606978, EPI_ISL_606979, EPI_ISL_606980, EPI_ISL_606981, EPI_ISL_606982, EPI_ISL_606983, EPI_ISL_606984, EPI_ISL_606985, EPI_ISL_606986, EPI_ISL_606987, EPI_ISL_606988, EPI_ISL_606989, EPI_ISL_606990, EPI_ISL_606991, EPI_ISL_606992, EPI_ISL_606993, EPI_ISL_606994, EPI_ISL_606995, EPI_ISL_606996, EPI_ISL_606997, EPI_ISL_606998, EPI_ISL_606999, EPI_ISL_607000, EPI_ISL_607001, EPI_ISL_607002, EPI_ISL_607003, EPI_ISL_607004, EPI_ISL_607005, EPI_ISL_607006, EPI_ISL_607007, EPI_ISL_607008, EPI_ISL_607009, EPI_ISL_607010, EPI_ISL_607011, EPI_ISL_607012, EPI_ISL_607013, EPI_ISL_607014, EPI_ISL_607015, EPI_ISL_607016, EPI_ISL_607017, EPI_ISL_607018, EPI_ISL_607019, EPI_ISL_607020, EPI_ISL_607021, EPI_ISL_607022, EPI_ISL_607023, EPI_ISL_607024, EPI_ISL_607025, EPI_ISL_607026, EPI_ISL_607027, EPI_ISL_607028, EPI_ISL_607029, EPI_ISL_607030, EPI_ISL_607031, EPI_ISL_607032, EPI_ISL_607033, EPI_ISL_607034, EPI_ISL_607035, EPI_ISL_607036, EPI_ISL_607037, EPI_ISL_607038, EPI_ISL_607039, EPI_ISL_607040, EPI_ISL_607041, EPI_ISL_607042, EPI_ISL_607043, EPI_ISL_607044, EPI_ISL_607045, EPI_ISL_607046, EPI_ISL_607047, EPI_ISL_607048, EPI_ISL_607049, EPI_ISL_607050, EPI_ISL_607051, EPI_ISL_607052, EPI_ISL_607053, EPI_ISL_607054, EPI_ISL_607055, EPI_ISL_607056, EPI_ISL_607057, EPI_ISL_607058, EPI_ISL_607059, EPI_ISL_607060, EPI_ISL_607061, EPI_ISL_607062, EPI_ISL_607063, EPI_ISL_607064, EPI_ISL_607065, EPI_ISL_607066, EPI_ISL_607067, EPI_ISL_607068, EPI_ISL_607069, EPI_ISL_607070, EPI_ISL_607071, EPI_ISL_607072, EPI_ISL_607073, EPI_ISL_607074, EPI_ISL_607075, EPI_ISL_607076, EPI_ISL_607077, EPI_ISL_607078, EPI_ISL_607079, EPI_ISL_607080, EPI_ISL_607081, EPI_ISL_607082, EPI_ISL_607083, EPI_ISL_607084, EPI_ISL_607085, EPI_ISL_607086, EPI_ISL_607087, EPI_ISL_607088, EPI_ISL_607089, EPI_ISL_607090, EPI_ISL_607091, EPI_ISL_607092, EPI_ISL_607093, EPI_ISL_607094, EPI_ISL_607095, EPI_ISL_607096, EPI_ISL_607097, EPI_ISL_607098, EPI_ISL_607099, EPI_ISL_607100, EPI_ISL_607101, EPI_ISL_607102, EPI_ISL_607103, EPI_ISL_607104, EPI_ISL_607105, EPI_ISL_607106, EPI_ISL_607107, EPI_ISL_607108, EPI_ISL_607109, EPI_ISL_607110, EPI_ISL_607111, EPI_ISL_607112, EPI_ISL_607113, EPI_ISL_607114, EPI_ISL_607115, EPI_ISL_607116, EPI_ISL_607117, EPI_ISL_607118, EPI_ISL_607119, EPI_ISL_607120, EPI_ISL_607121, EPI_ISL_607122, EPI_ISL_607123, EPI_ISL_607124, EPI_ISL_607125, EPI_ISL_607126, EPI_ISL_607127, EPI_ISL_607128, EPI_ISL_607129, EPI_ISL_607130, EPI_ISL_607131, EPI_ISL_607132, EPI_ISL_607133, EPI_ISL_607134, EPI_ISL_607135, EPI_ISL_607136, EPI_ISL_607137, EPI_ISL_607138, EPI_ISL_607139, EPI_ISL_607140, EPI_ISL_607141, EPI_ISL_607142, EPI_ISL_607143, EPI_ISL_607144, EPI_ISL_607145, EPI_ISL_607146, EPI_ISL_607147, EPI_ISL_607148, EPI_ISL_607149, EPI_ISL_607150, EPI_ISL_607151, EPI_ISL_607152, EPI_ISL_607153, EPI_ISL_607154, EPI_ISL_607155, EPI_ISL_607156, EPI_ISL_607157, EPI_ISL_607158, EPI_ISL_607159, EPI_ISL_607160, EPI_ISL_607161, EPI_ISL_607162, EPI_ISL_607163, EPI_ISL_607164, EPI_ISL_607165, EPI_ISL_607166, EPI_ISL_607167, EPI_ISL_607168, EPI_ISL_607169, EPI_ISL_607170, EPI_ISL_607171, EPI_ISL_607172, EPI_ISL_607173, EPI_ISL_607174, EPI_ISL_607175, EPI_ISL_607176, EPI_ISL_607177, EPI_ISL_607178, EPI_ISL_607179, EPI_ISL_607180, EPI_ISL_607181, EPI_ISL_607182, EPI_ISL_607183, EPI_ISL_607184, EPI_ISL_607185, EPI_ISL_607186, EPI_ISL_607187, EPI_ISL_607188 | | | |
| see above | Lighthouse Lab in Alderley Park | Wellcome Sanger Institute for the COVID-19 Genomics UK (COG-UK) consortium | Jacquelyn Wynn, Mairead Hyland, The Lighthouse Lab in Alderley Park and Alex Alderton, Roberto Amato, Sonia Goncalves, Ewan Harrison, David K. Jackson, Ian Johnston, Dominic Kwiatkowski, Cordelia Langford, John Sillitoe on behalf of the Wellcome Sanger Institute COVID-19 Surveillance Team |
| EPI_ISL_607189 | Lighthouse Lab in Alderley Park | Wellcome Sanger Institute for the COVID-19 Genomics UK (COG-UK) Consortium | Jacquelyn Wynn, Mairead Hyland, The Lighthouse Lab in Alderley Park and Alex Alderton, Roberto Amato, Sonia Goncalves, Ewan Harrison, David K. Jackson, Ian Johnston, Dominic Kwiatkowski, Cordelia Langford, John Sillitoe on behalf of the Wellcome Sanger Institute COVID-19 Surveillance Team |
| EPI_ISL_607190, EPI_ISL_607191, | Lighthouse Lab in Alderley Park | Wellcome Sanger Institute for the COVID-19 Genomics UK | Jacquelyn Wynn, Mairead Hyland, The Lighthouse Lab in Alderley Park and Alex Alderton, Roberto Amato, Sonia Goncalves, Ewan Harrison, David K. |

| | | | |
|---|---|---|---|
| EPI_ISL_607192, EPI_ISL_607193, EPI_ISL_607194, EPI_ISL_607195, EPI_ISL_607196, EPI_ISL_607197, EPI_ISL_607198, EPI_ISL_607199 | | (COG-UK) consortium | Jackson, Ian Johnston, Dominic Kwiatkowski, Cordelia Langford, John Sillitoe on behalf of the Wellcome Sanger Institute COVID-19 Surveillance Team |
| EPI_ISL_607200 | Lighthouse Lab in Alderley Park | Wellcome Sanger Institute for the COVID-19 Genomics UK (COG-UK) Consortium | Jacquelyn Wynn, Mairead Hyland, The Lighthouse Lab in Alderley Park and Alex Alderton, Roberto Amato, Sonia Goncalves, Ewan Harrison, David K. Jackson, Ian Johnston, Dominic Kwiatkowski, Cordelia Langford, John Sillitoe on behalf of the Wellcome Sanger Institute COVID-19 Surveillance Team |
| EPI_ISL_607201, EPI_ISL_607202, EPI_ISL_607203, EPI_ISL_607204, EPI_ISL_607205, EPI_ISL_607206, EPI_ISL_607207, EPI_ISL_607208, EPI_ISL_607209, EPI_ISL_607210, EPI_ISL_607211, EPI_ISL_607212, EPI_ISL_607213, EPI_ISL_607214, EPI_ISL_607215, EPI_ISL_607216, EPI_ISL_607217, EPI_ISL_607218, EPI_ISL_607219, EPI_ISL_607220, EPI_ISL_607221, EPI_ISL_607222, EPI_ISL_607223, EPI_ISL_607224, EPI_ISL_607225, EPI_ISL_607226, EPI_ISL_607227, EPI_ISL_607228, EPI_ISL_607229, EPI_ISL_607230, EPI_ISL_607231, EPI_ISL_607232, EPI_ISL_607233, EPI_ISL_607234, EPI_ISL_607235, EPI_ISL_607236, EPI_ISL_607237, EPI_ISL_607238, EPI_ISL_607239, EPI_ISL_607240, EPI_ISL_607241, EPI_ISL_607242, EPI_ISL_607243, EPI_ISL_607244, EPI_ISL_607245, EPI_ISL_607246, EPI_ISL_607247, EPI_ISL_607248, EPI_ISL_607249, EPI_ISL_607250, EPI_ISL_607251, EPI_ISL_607252, EPI_ISL_607253, EPI_ISL_607254, EPI_ISL_607255, EPI_ISL_607256, EPI_ISL_607257, EPI_ISL_607258, EPI_ISL_607259, EPI_ISL_607260, EPI_ISL_607261, EPI_ISL_607262, EPI_ISL_607263, EPI_ISL_607264, EPI_ISL_607265, EPI_ISL_607266, EPI_ISL_607267, EPI_ISL_607268, EPI_ISL_607269, EPI_ISL_607270, EPI_ISL_607271, EPI_ISL_607272, EPI_ISL_607273, EPI_ISL_607274, EPI_ISL_607275, EPI_ISL_607276, EPI_ISL_607277, EPI_ISL_607278, EPI_ISL_607279, EPI_ISL_607280, EPI_ISL_607281, EPI_ISL_607282, EPI_ISL_607283, EPI_ISL_607284, EPI_ISL_607285, EPI_ISL_607286, EPI_ISL_607287, EPI_ISL_607288, EPI_ISL_607289, EPI_ISL_607290 | | | |
| see above | Lighthouse Lab in Alderley Park | Wellcome Sanger Institute for the COVID-19 Genomics UK (COG-UK) consortium | Jacquelyn Wynn, Mairead Hyland, The Lighthouse Lab in Alderley Park and Alex Alderton, Roberto Amato, Sonia Goncalves, Ewan Harrison, David K. Jackson, Ian Johnston, Dominic Kwiatkowski, Cordelia Langford, John Sillitoe on behalf of the Wellcome Sanger Institute COVID-19 Surveillance Team |
| EPI_ISL_607291, EPI_ISL_607292, EPI_ISL_607293, EPI_ISL_607294, EPI_ISL_607295, EPI_ISL_607296, EPI_ISL_607297, EPI_ISL_607298, EPI_ISL_607299, EPI_ISL_607300, EPI_ISL_607301, EPI_ISL_607302, EPI_ISL_607303, EPI_ISL_607304, EPI_ISL_607305, EPI_ISL_607306, EPI_ISL_607307, EPI_ISL_607308, EPI_ISL_607309, EPI_ISL_607310, EPI_ISL_607311, EPI_ISL_607312, EPI_ISL_607313, EPI_ISL_607314, EPI_ISL_607315, EPI_ISL_607316, EPI_ISL_607317, EPI_ISL_607318, EPI_ISL_607319, EPI_ISL_607320, EPI_ISL_607321, EPI_ISL_607322, EPI_ISL_607323, EPI_ISL_607324, EPI_ISL_607325, EPI_ISL_607326, EPI_ISL_607327, EPI_ISL_607328, EPI_ISL_607329, EPI_ISL_607330, EPI_ISL_607331, EPI_ISL_607332, EPI_ISL_607333, EPI_ISL_607334, EPI_ISL_607335, EPI_ISL_607336, EPI_ISL_607337, EPI_ISL_607338, EPI_ISL_607339, EPI_ISL_607340, EPI_ISL_607341, EPI_ISL_607342, EPI_ISL_607343, EPI_ISL_607344, EPI_ISL_607345, EPI_ISL_607346, EPI_ISL_607347, EPI_ISL_607348, EPI_ISL_607349, EPI_ISL_607350, EPI_ISL_607351, EPI_ISL_607352, EPI_ISL_607353, EPI_ISL_607354, EPI_ISL_607355, EPI_ISL_607356, EPI_ISL_607357, EPI_ISL_607358, EPI_ISL_607359, EPI_ISL_607360, EPI_ISL_607361, EPI_ISL_607362, EPI_ISL_607363, EPI_ISL_607364, EPI_ISL_607365, EPI_ISL_607366, EPI_ISL_607367, EPI_ISL_607368, EPI_ISL_607369, EPI_ISL_607370, EPI_ISL_607371, EPI_ISL_607372, EPI_ISL_607373, EPI_ISL_607374, EPI_ISL_607375, EPI_ISL_607376, EPI_ISL_607377, EPI_ISL_607378, EPI_ISL_607379, EPI_ISL_607380, EPI_ISL_607381, EPI_ISL_607382, EPI_ISL_607383, EPI_ISL_607384, EPI_ISL_607385, EPI_ISL_607386, EPI_ISL_607387, EPI_ISL_607388, EPI_ISL_607389, EPI_ISL_607390, EPI_ISL_607391, EPI_ISL_607392, EPI_ISL_607393, EPI_ISL_607394, EPI_ISL_607395, EPI_ISL_607396, EPI_ISL_607397, EPI_ISL_607398, EPI_ISL_607399, EPI_ISL_607400, EPI_ISL_607401, EPI_ISL_607402, EPI_ISL_607403, EPI_ISL_607404, EPI_ISL_607405, EPI_ISL_607406, EPI_ISL_607407, EPI_ISL_607408, EPI_ISL_607409, EPI_ISL_607410, EPI_ISL_607411, EPI_ISL_607412, EPI_ISL_607413, EPI_ISL_607414, EPI_ISL_607415, EPI_ISL_607416, EPI_ISL_607417, EPI_ISL_607418, EPI_ISL_607419, EPI_ISL_607420, EPI_ISL_607421, EPI_ISL_607422, EPI_ISL_607423, EPI_ISL_607424, EPI_ISL_607425, EPI_ISL_607426, EPI_ISL_607427, EPI_ISL_607428, EPI_ISL_607429, EPI_ISL_607430, EPI_ISL_607431, EPI_ISL_607432, EPI_ISL_607433, EPI_ISL_607434, EPI_ISL_607435, EPI_ISL_607436, EPI_ISL_607437, EPI_ISL_607438, EPI_ISL_607439, EPI_ISL_607440, EPI_ISL_607441, EPI_ISL_607442, EPI_ISL_607443, EPI_ISL_607444, EPI_ISL_607445, EPI_ISL_607446, EPI_ISL_607447 | | | |
| see above | Lighthouse Lab in Milton Keynes | Wellcome Sanger Institute for the COVID-19 Genomics UK (COG-UK) consortium | The Lighthouse Lab in Milton Keynes and Alex Alderton, Roberto Amato, Sonia Goncalves, Ewan Harrison, David K. Jackson, Ian Johnston, Dominic Kwiatkowski, Cordelia Langford, John Sillitoe on behalf of the Wellcome Sanger Institute COVID-19 Surveillance Team |
| EPI_ISL_607448 | Lighthouse Lab in Milton Keynes | Wellcome Sanger Institute for the COVID-19 Genomics UK (COG-UK) Consortium | The Lighthouse Lab in Milton Keynes and Alex Alderton, Roberto Amato, Sonia Goncalves, Ewan Harrison, David K. Jackson, Ian Johnston, Dominic Kwiatkowski, Cordelia Langford, John Sillitoe on behalf of the Wellcome Sanger Institute COVID-19 Surveillance Team |
| EPI_ISL_607449, EPI_ISL_607450, EPI_ISL_607451, EPI_ISL_607452, EPI_ISL_607453, EPI_ISL_607454, EPI_ISL_607455, EPI_ISL_607456, EPI_ISL_607457, EPI_ISL_607458, EPI_ISL_607459, EPI_ISL_607460 | | | |
| see above | Lighthouse Lab in Milton Keynes | Wellcome Sanger Institute for the COVID-19 Genomics UK (COG-UK) consortium | The Lighthouse Lab in Milton Keynes and Alex Alderton, Roberto Amato, Sonia Goncalves, Ewan Harrison, David K. Jackson, Ian Johnston, Dominic Kwiatkowski, Cordelia Langford, John Sillitoe on behalf of the Wellcome Sanger Institute COVID-19 Surveillance Team |
| EPI_ISL_607461 | Lighthouse Lab in Milton Keynes | Wellcome Sanger Institute for the COVID-19 Genomics UK (COG-UK) Consortium | The Lighthouse Lab in Milton Keynes and Alex Alderton, Roberto Amato, Sonia Goncalves, Ewan Harrison, David K. Jackson, Ian Johnston, Dominic Kwiatkowski, Cordelia Langford, John Sillitoe on behalf of the Wellcome Sanger Institute COVID-19 Surveillance Team |
| EPI_ISL_607462, EPI_ISL_607463, EPI_ISL_607464, EPI_ISL_607465, EPI_ISL_607466, EPI_ISL_607467, EPI_ISL_607468, EPI_ISL_607469, EPI_ISL_607470, EPI_ISL_607471, EPI_ISL_607472, EPI_ISL_607473, EPI_ISL_607474, EPI_ISL_607475, EPI_ISL_607476, EPI_ISL_607477, EPI_ISL_607478, EPI_ISL_607479, EPI_ISL_607480, EPI_ISL_607481, EPI_ISL_607482, EPI_ISL_607483, EPI_ISL_607484, EPI_ISL_607485, EPI_ISL_607486, EPI_ISL_607487, EPI_ISL_607488, EPI_ISL_607489, EPI_ISL_607490, EPI_ISL_607491, EPI_ISL_607492, EPI_ISL_607493, EPI_ISL_607494, EPI_ISL_607495, EPI_ISL_607496, EPI_ISL_607497, EPI_ISL_607498, EPI_ISL_607499, EPI_ISL_607500, EPI_ISL_607501, EPI_ISL_607502, EPI_ISL_607503, EPI_ISL_607504, EPI_ISL_607505, EPI_ISL_607506, EPI_ISL_607507, EPI_ISL_607508, EPI_ISL_607509, EPI_ISL_607510, EPI_ISL_607511, EPI_ISL_607512, EPI_ISL_607513, EPI_ISL_607514, EPI_ISL_607515 | | | |
| see above | Lighthouse Lab in Milton Keynes | Wellcome Sanger Institute for the COVID-19 Genomics UK (COG-UK) consortium | The Lighthouse Lab in Milton Keynes and Alex Alderton, Roberto Amato, Sonia Goncalves, Ewan Harrison, David K. Jackson, Ian Johnston, Dominic Kwiatkowski, Cordelia Langford, John Sillitoe on behalf of the Wellcome Sanger Institute COVID-19 Surveillance Team |
| EPI_ISL_607516 | Lighthouse Lab in Milton Keynes | Wellcome Sanger Institute for the COVID-19 Genomics UK (COG-UK) Consortium | The Lighthouse Lab in Milton Keynes and Alex Alderton, Roberto Amato, Sonia Goncalves, Ewan Harrison, David K. Jackson, Ian Johnston, Dominic Kwiatkowski, Cordelia Langford, John Sillitoe on behalf of the Wellcome Sanger Institute COVID-19 Surveillance Team |
| EPI_ISL_607517, EPI_ISL_607518, EPI_ISL_607519, EPI_ISL_607520, EPI_ISL_607521, EPI_ISL_607522, EPI_ISL_607523, EPI_ISL_607524, EPI_ISL_607525, EPI_ISL_607526, EPI_ISL_607527, EPI_ISL_607528, EPI_ISL_607529, EPI_ISL_607530, EPI_ISL_607531, EPI_ISL_607532, EPI_ISL_607533, EPI_ISL_607534, EPI_ISL_607535, EPI_ISL_607536, EPI_ISL_607537, EPI_ISL_607538, EPI_ISL_607539, EPI_ISL_607540, EPI_ISL_607541, EPI_ISL_607542, EPI_ISL_607543, EPI_ISL_607544, EPI_ISL_607545, EPI_ISL_607546, EPI_ISL_607547, EPI_ISL_607548, EPI_ISL_607549 | | | |
| see above | Lighthouse Lab in Milton Keynes | Wellcome Sanger Institute for the COVID-19 Genomics UK (COG-UK) consortium | The Lighthouse Lab in Milton Keynes and Alex Alderton, Roberto Amato, Sonia Goncalves, Ewan Harrison, David K. Jackson, Ian Johnston, Dominic Kwiatkowski, Cordelia Langford, John Sillitoe on behalf of the Wellcome Sanger Institute COVID-19 Surveillance Team |
| EPI_ISL_607550 | Lighthouse Lab in Milton Keynes | Wellcome Sanger Institute for the COVID-19 Genomics UK (COG-UK) Consortium | The Lighthouse Lab in Milton Keynes and Alex Alderton, Roberto Amato, Sonia Goncalves, Ewan Harrison, David K. Jackson, Ian Johnston, Dominic Kwiatkowski, Cordelia Langford, John Sillitoe on behalf of the Wellcome Sanger Institute COVID-19 Surveillance Team |
| EPI_ISL_607551, EPI_ISL_607552, EPI_ISL_607553, EPI_ISL_607554, EPI_ISL_607555, EPI_ISL_607556, EPI_ISL_607557, EPI_ISL_607558, EPI_ISL_607559, EPI_ISL_607560, EPI_ISL_607561, EPI_ISL_607562, EPI_ISL_607563, EPI_ISL_607564, EPI_ISL_607565, EPI_ISL_607566, EPI_ISL_607567, EPI_ISL_607568, EPI_ISL_607569, EPI_ISL_607570, EPI_ISL_607571 | | | |
| see above | Lighthouse Lab in Milton Keynes | Wellcome Sanger Institute for the COVID-19 Genomics UK (COG-UK) consortium | The Lighthouse Lab in Milton Keynes and Alex Alderton, Roberto Amato, Sonia Goncalves, Ewan Harrison, David K. Jackson, Ian Johnston, Dominic Kwiatkowski, Cordelia Langford, John Sillitoe on behalf of the Wellcome Sanger Institute COVID-19 Surveillance Team |
| EPI_ISL_607572 | Lighthouse Lab in Milton Keynes | Wellcome Sanger Institute for the COVID-19 Genomics UK (COG-UK) Consortium | The Lighthouse Lab in Milton Keynes and Alex Alderton, Roberto Amato, Sonia Goncalves, Ewan Harrison, David K. Jackson, Ian Johnston, Dominic Kwiatkowski, Cordelia Langford, John Sillitoe on behalf of the Wellcome Sanger Institute COVID-19 Surveillance Team |
| EPI_ISL_607573, EPI_ISL_607574, EPI_ISL_607575, EPI_ISL_607576, EPI_ISL_607577, EPI_ISL_607578 | Lighthouse Lab in Milton Keynes | Wellcome Sanger Institute for the COVID-19 Genomics UK (COG-UK) consortium | The Lighthouse Lab in Milton Keynes and Alex Alderton, Roberto Amato, Sonia Goncalves, Ewan Harrison, David K. Jackson, Ian Johnston, Dominic Kwiatkowski, Cordelia Langford, John Sillitoe on behalf of the Wellcome Sanger Institute COVID-19 Surveillance Team |
| EPI_ISL_607579 | Lighthouse Lab in Milton Keynes | Wellcome Sanger Institute for the COVID-19 Genomics UK (COG-UK) Consortium | The Lighthouse Lab in Milton Keynes and Alex Alderton, Roberto Amato, Sonia Goncalves, Ewan Harrison, David K. Jackson, Ian Johnston, Dominic Kwiatkowski, Cordelia Langford, John Sillitoe on behalf of the Wellcome Sanger Institute COVID-19 Surveillance Team |
| EPI_ISL_607580, EPI_ISL_607581, EPI_ISL_607582, EPI_ISL_607583, EPI_ISL_607584, EPI_ISL_607586 | Lighthouse Lab in Milton Keynes | Wellcome Sanger Institute for the COVID-19 Genomics UK (COG-UK) consortium | The Lighthouse Lab in Milton Keynes and Alex Alderton, Roberto Amato, Sonia Goncalves, Ewan Harrison, David K. Jackson, Ian Johnston, Dominic Kwiatkowski, Cordelia Langford, John Sillitoe on behalf of the Wellcome Sanger Institute COVID-19 Surveillance Team |
| EPI_ISL_607587 | Lighthouse Lab in Milton Keynes | Wellcome Sanger Institute for the COVID-19 Genomics UK (COG-UK) Consortium | The Lighthouse Lab in Milton Keynes and Alex Alderton, Roberto Amato, Sonia Goncalves, Ewan Harrison, David K. Jackson, Ian Johnston, Dominic Kwiatkowski, Cordelia Langford, John Sillitoe on behalf of the Wellcome Sanger Institute COVID-19 Surveillance Team |
| EPI_ISL_607588, EPI_ISL_607589, EPI_ISL_607590, EPI_ISL_607591, EPI_ISL_607592, EPI_ISL_607593, EPI_ISL_607594, EPI_ISL_607595, EPI_ISL_607596, EPI_ISL_607597, EPI_ISL_607598, EPI_ISL_607599, EPI_ISL_607600, EPI_ISL_607601, EPI_ISL_607602, EPI_ISL_607603, EPI_ISL_607604, EPI_ISL_607605, EPI_ISL_607606, EPI_ISL_607607, EPI_ISL_607608, EPI_ISL_607609, EPI_ISL_607610, EPI_ISL_607611, EPI_ISL_607612, EPI_ISL_607613, EPI_ISL_607614, EPI_ISL_607615, EPI_ISL_607616, EPI_ISL_607617, EPI_ISL_607618, EPI_ISL_607619, EPI_ISL_607620, EPI_ISL_607621, EPI_ISL_607622 | | | |
| see above | Lighthouse Lab in Milton Keynes | Wellcome Sanger Institute for the COVID-19 Genomics UK (COG-UK) consortium | The Lighthouse Lab in Milton Keynes and Alex Alderton, Roberto Amato, Sonia Goncalves, Ewan Harrison, David K. Jackson, Ian Johnston, Dominic Kwiatkowski, Cordelia Langford, John Sillitoe on behalf of the Wellcome Sanger Institute COVID-19 Surveillance Team |
| EPI_ISL_607623, EPI_ISL_607624, EPI_ISL_607625, EPI_ISL_607626, EPI_ISL_607627, EPI_ISL_607628, EPI_ISL_607629, EPI_ISL_607630, EPI_ISL_607631, EPI_ISL_607632, EPI_ISL_607633, EPI_ISL_607634, EPI_ISL_607635, EPI_ISL_607636, EPI_ISL_607637, EPI_ISL_607638, EPI_ISL_607639, EPI_ISL_607640, EPI_ISL_607641, EPI_ISL_607642, EPI_ISL_607643, EPI_ISL_607644, EPI_ISL_607645, EPI_ISL_607646, EPI_ISL_607647, EPI_ISL_607648, EPI_ISL_607649, EPI_ISL_607650, EPI_ISL_607651, EPI_ISL_607652, EPI_ISL_607653, EPI_ISL_607654, EPI_ISL_607655, EPI_ISL_607656, EPI_ISL_607657, EPI_ISL_607658, EPI_ISL_607659, EPI_ISL_607660, EPI_ISL_607661, EPI_ISL_607662, EPI_ISL_607663, EPI_ISL_607664, EPI_ISL_607665, EPI_ISL_607666, EPI_ISL_607667, EPI_ISL_607668, EPI_ISL_607669, EPI_ISL_607670, EPI_ISL_607671, EPI_ISL_607672, EPI_ISL_607673, EPI_ISL_607674, EPI_ISL_607675, EPI_ISL_607676, EPI_ISL_607677, EPI_ISL_607678, EPI_ISL_607679, EPI_ISL_607680, EPI_ISL_607681, EPI_ISL_607682, EPI_ISL_607683, EPI_ISL_607684, EPI_ISL_607685, EPI_ISL_607686, EPI_ISL_607687, EPI_ISL_607688, EPI_ISL_607689, EPI_ISL_607690, EPI_ISL_607691, EPI_ISL_607692, EPI_ISL_607693, EPI_ISL_607694, EPI_ISL_607695, EPI_ISL_607696, EPI_ISL_607697, EPI_ISL_607698, EPI_ISL_607699, EPI_ISL_607700, EPI_ISL_607701, EPI_ISL_607702, EPI_ISL_607703, EPI_ISL_607704, EPI_ISL_607705, EPI_ISL_607706, EPI_ISL_607707, EPI_ISL_607708, EPI_ISL_607709, EPI_ISL_607710, EPI_ISL_607711, EPI_ISL_607712, EPI_ISL_607713, EPI_ISL_607714, EPI_ISL_607715, EPI_ISL_607716, EPI_ISL_607717, EPI_ISL_607718, EPI_ISL_607719, EPI_ISL_607720, EPI_ISL_607721, EPI_ISL_607722, EPI_ISL_607723, EPI_ISL_607724, EPI_ISL_607725, EPI_ISL_607726, EPI_ISL_607727, EPI_ISL_607728, EPI_ISL_607729, EPI_ISL_607730, EPI_ISL_607731, EPI_ISL_607732, EPI_ISL_607733, EPI_ISL_607734, EPI_ISL_607735, EPI_ISL_607736, EPI_ISL_607737, EPI_ISL_607738, EPI_ISL_607739, EPI_ISL_607740, EPI_ISL_607741, EPI_ISL_607742, EPI_ISL_607743, EPI_ISL_607744, EPI_ISL_607745, EPI_ISL_607746, EPI_ISL_607747, EPI_ISL_607748, EPI_ISL_607749, EPI_ISL_607750, EPI_ISL_607751, EPI_ISL_607752, EPI_ISL_607753, EPI_ISL_607754, EPI_ISL_607755, EPI_ISL_607756, EPI_ISL_607757, EPI_ISL_607758, EPI_ISL_607759, EPI_ISL_607760, EPI_ISL_607761, EPI_ISL_607762, EPI_ISL_607763, EPI_ISL_607764, EPI_ISL_607765, EPI_ISL_607766, EPI_ISL_607767, EPI_ISL_607768, EPI_ISL_607769, EPI_ISL_607770, EPI_ISL_607771, EPI_ISL_607772, EPI_ISL_607773, EPI_ISL_607774, EPI_ISL_607775, EPI_ISL_607776, EPI_ISL_607777, EPI_ISL_607778, EPI_ISL_607779, EPI_ISL_607780, EPI_ISL_607781, EPI_ISL_607782, EPI_ISL_607783, EPI_ISL_607784, EPI_ISL_607785, EPI_ISL_607786, EPI_ISL_607787, EPI_ISL_607788, EPI_ISL_607789, EPI_ISL_607790, EPI_ISL_607791, EPI_ISL_607792, EPI_ISL_607793, EPI_ISL_607794, EPI_ISL_607795, EPI_ISL_607796, EPI_ISL_607797, EPI_ISL_607798, EPI_ISL_607799, EPI_ISL_607800, EPI_ISL_607801, EPI_ISL_607802, EPI_ISL_607803, EPI_ISL_607804, EPI_ISL_607805, EPI_ISL_607806, EPI_ISL_607807, EPI_ISL_607808, EPI_ISL_607809, EPI_ISL_607810, EPI_ISL_607811, EPI_ISL_607812, EPI_ISL_607813, EPI_ISL_607814, EPI_ISL_607815, EPI_ISL_607816, EPI_ISL_607817, EPI_ISL_607818, EPI_ISL_607819, EPI_ISL_607820, | | | |

| | | | |
|---|---|---|---|
| EPI_ISL_607821, EPI_ISL_607822, EPI_ISL_607823, EPI_ISL_607824, EPI_ISL_607825, EPI_ISL_607826, EPI_ISL_607827, EPI_ISL_607828, EPI_ISL_607829, EPI_ISL_607830, EPI_ISL_607831, EPI_ISL_607832, EPI_ISL_607833, EPI_ISL_607834, EPI_ISL_607835, EPI_ISL_607836, EPI_ISL_607837, EPI_ISL_607838, EPI_ISL_607839, EPI_ISL_607840, EPI_ISL_607841, EPI_ISL_607842, EPI_ISL_607843, EPI_ISL_607844, EPI_ISL_607845, EPI_ISL_607846, EPI_ISL_607847, EPI_ISL_607848, EPI_ISL_607849, EPI_ISL_607850, EPI_ISL_607851, EPI_ISL_607852, EPI_ISL_607853, EPI_ISL_607854, EPI_ISL_607855, EPI_ISL_607856, EPI_ISL_607857, EPI_ISL_607858, EPI_ISL_607859, EPI_ISL_607860, EPI_ISL_607861, EPI_ISL_607862, EPI_ISL_607863, EPI_ISL_607864, EPI_ISL_607865, EPI_ISL_607866, EPI_ISL_607867, EPI_ISL_607868, EPI_ISL_607869, EPI_ISL_607870, EPI_ISL_607871, EPI_ISL_607872, EPI_ISL_607873, EPI_ISL_607874, EPI_ISL_607875, EPI_ISL_607876, EPI_ISL_607877, EPI_ISL_607878, EPI_ISL_607879, EPI_ISL_607880, EPI_ISL_607881, EPI_ISL_607882, EPI_ISL_607883, EPI_ISL_607884, EPI_ISL_607885, EPI_ISL_607886, EPI_ISL_607887, EPI_ISL_607888, EPI_ISL_607889, EPI_ISL_607890, EPI_ISL_607891, EPI_ISL_607892, EPI_ISL_607893 | | | |
| see above | Lighthouse Lab in Alderley Park | Wellcome Sanger Institute for the COVID-19 Genomics UK (COG-UK) consortium | Jacquelyn Wynn, Mairead Hyland, The Lighthouse Lab in Alderley Park and Alex Alderton, Roberto Amato, Sonia Goncalves, Ewan Harrison, David K. Jackson, Ian Johnston, Dominic Kwiatkowski, Cordelia Langford, John Sillitoe on behalf of the Wellcome Sanger Institute COVID-19 Surveillance Team |
| EPI_ISL_607894 | Lighthouse Lab in Alderley Park | Wellcome Sanger Institute for the COVID-19 Genomics UK (COG-UK) Consortium | Jacquelyn Wynn, Mairead Hyland, The Lighthouse Lab in Alderley Park and Alex Alderton, Roberto Amato, Sonia Goncalves, Ewan Harrison, David K. Jackson, Ian Johnston, Dominic Kwiatkowski, Cordelia Langford, John Sillitoe on behalf of the Wellcome Sanger Institute COVID-19 Surveillance Team |
| EPI_ISL_607895, EPI_ISL_607896, EPI_ISL_607897, EPI_ISL_607898, EPI_ISL_607899, EPI_ISL_607900, EPI_ISL_607901, EPI_ISL_607902, EPI_ISL_607903, EPI_ISL_607904, EPI_ISL_607905, EPI_ISL_607906, EPI_ISL_607907, EPI_ISL_607908, EPI_ISL_607909, EPI_ISL_607910, EPI_ISL_607911, EPI_ISL_607912, EPI_ISL_607913, EPI_ISL_607914, EPI_ISL_607915, EPI_ISL_607916, EPI_ISL_607917, EPI_ISL_607918, EPI_ISL_607919, EPI_ISL_607920, EPI_ISL_607921, EPI_ISL_607922, EPI_ISL_607923, EPI_ISL_607924, EPI_ISL_607925, EPI_ISL_607926, EPI_ISL_607927, EPI_ISL_607928, EPI_ISL_607929, EPI_ISL_607930, EPI_ISL_607931, EPI_ISL_607932, EPI_ISL_607933, EPI_ISL_607934, EPI_ISL_607935, EPI_ISL_607936, EPI_ISL_607937, EPI_ISL_607938, EPI_ISL_607939, EPI_ISL_607940, EPI_ISL_607941, EPI_ISL_607942, EPI_ISL_607943, EPI_ISL_607944, EPI_ISL_607945, EPI_ISL_607946, EPI_ISL_607947, EPI_ISL_607948, EPI_ISL_607949, EPI_ISL_607950, EPI_ISL_607951, EPI_ISL_607952, EPI_ISL_607953, EPI_ISL_607954, EPI_ISL_607955, EPI_ISL_607956, EPI_ISL_607957, EPI_ISL_607958, EPI_ISL_607959, EPI_ISL_607960, EPI_ISL_607961, EPI_ISL_607962, EPI_ISL_607963, EPI_ISL_607964, EPI_ISL_607965, EPI_ISL_607966, EPI_ISL_607967, EPI_ISL_607968, EPI_ISL_607969, EPI_ISL_607970, EPI_ISL_607971, EPI_ISL_607972, EPI_ISL_607973, EPI_ISL_607974, EPI_ISL_607975, EPI_ISL_607976, EPI_ISL_607977, EPI_ISL_607978, EPI_ISL_607979, EPI_ISL_607980, EPI_ISL_607981, EPI_ISL_607982, EPI_ISL_607983, EPI_ISL_607984, EPI_ISL_607985, EPI_ISL_607986, EPI_ISL_607987, EPI_ISL_607988, EPI_ISL_607989, EPI_ISL_607990, EPI_ISL_607991, EPI_ISL_607992, EPI_ISL_607993, EPI_ISL_607994, EPI_ISL_607995, EPI_ISL_607996, EPI_ISL_607997, EPI_ISL_607998, EPI_ISL_607999, EPI_ISL_608000, EPI_ISL_608001, EPI_ISL_608002, EPI_ISL_608003, EPI_ISL_608004, EPI_ISL_608005, EPI_ISL_608006, EPI_ISL_608007, EPI_ISL_608008, EPI_ISL_608009, EPI_ISL_608010, EPI_ISL_608011, EPI_ISL_608012, EPI_ISL_608013, EPI_ISL_608014, EPI_ISL_608015, EPI_ISL_608016, EPI_ISL_608017, EPI_ISL_608018, EPI_ISL_608019, EPI_ISL_608020, EPI_ISL_608021, EPI_ISL_608022, EPI_ISL_608023, EPI_ISL_608024, EPI_ISL_608025, EPI_ISL_608026, EPI_ISL_608027, EPI_ISL_608028, EPI_ISL_608029, EPI_ISL_608030, EPI_ISL_608031, EPI_ISL_608032, EPI_ISL_608033, EPI_ISL_608034, EPI_ISL_608035, EPI_ISL_608036, EPI_ISL_608037, EPI_ISL_608038, EPI_ISL_608039, EPI_ISL_608040, EPI_ISL_608041, EPI_ISL_608042, EPI_ISL_608043, EPI_ISL_608044, EPI_ISL_608045, EPI_ISL_608046, EPI_ISL_608047, EPI_ISL_608048, EPI_ISL_608049, EPI_ISL_608050, EPI_ISL_608051, EPI_ISL_608052, EPI_ISL_608053, EPI_ISL_608054, EPI_ISL_608055, EPI_ISL_608056, EPI_ISL_608057, EPI_ISL_608058, EPI_ISL_608059, EPI_ISL_608060, EPI_ISL_608061, EPI_ISL_608062, EPI_ISL_608063, EPI_ISL_608064, EPI_ISL_608065, EPI_ISL_608066, EPI_ISL_608067, EPI_ISL_608068, EPI_ISL_608069, EPI_ISL_608070, EPI_ISL_608071, EPI_ISL_608072, EPI_ISL_608073, EPI_ISL_608074, EPI_ISL_608075, EPI_ISL_608076, EPI_ISL_608077, EPI_ISL_608078, EPI_ISL_608079, EPI_ISL_608080, EPI_ISL_608081, EPI_ISL_608082, EPI_ISL_608083, EPI_ISL_608084, EPI_ISL_608085, EPI_ISL_608086, EPI_ISL_608087, EPI_ISL_608088, EPI_ISL_608089, EPI_ISL_608090, EPI_ISL_608091, EPI_ISL_608092, EPI_ISL_608093, EPI_ISL_608094, EPI_ISL_608095, EPI_ISL_608096, EPI_ISL_608097, EPI_ISL_608098, EPI_ISL_608099, EPI_ISL_608100, EPI_ISL_608101, EPI_ISL_608102, EPI_ISL_608103, EPI_ISL_608104, EPI_ISL_608105, EPI_ISL_608106, EPI_ISL_608107, EPI_ISL_608108, EPI_ISL_608109, EPI_ISL_608110, EPI_ISL_608111, EPI_ISL_608112, EPI_ISL_608113, EPI_ISL_608114, EPI_ISL_608115, EPI_ISL_608116, EPI_ISL_608117, EPI_ISL_608118, EPI_ISL_608119, EPI_ISL_608120, EPI_ISL_608121, EPI_ISL_608122, EPI_ISL_608123, EPI_ISL_608124, EPI_ISL_608125, EPI_ISL_608126, EPI_ISL_608127, EPI_ISL_608128, EPI_ISL_608129, EPI_ISL_608130, EPI_ISL_608131, EPI_ISL_608132, EPI_ISL_608133, EPI_ISL_608134, EPI_ISL_608135, EPI_ISL_608136, EPI_ISL_608137, EPI_ISL_608138, EPI_ISL_608139, EPI_ISL_608140, EPI_ISL_608141, EPI_ISL_608142, EPI_ISL_608143, EPI_ISL_608144, EPI_ISL_608145, EPI_ISL_608146 | | | |
| see above | Lighthouse Lab in Alderley Park | Wellcome Sanger Institute for the COVID-19 Genomics UK (COG-UK) consortium | Jacquelyn Wynn, Mairead Hyland, The Lighthouse Lab in Alderley Park and Alex Alderton, Roberto Amato, Sonia Goncalves, Ewan Harrison, David K. Jackson, Ian Johnston, Dominic Kwiatkowski, Cordelia Langford, John Sillitoe on behalf of the Wellcome Sanger Institute COVID-19 Surveillance Team |
| EPI_ISL_608147 | Lighthouse Lab in Alderley Park | Wellcome Sanger Institute for the COVID-19 Genomics UK (COG-UK) Consortium | Jacquelyn Wynn, Mairead Hyland, The Lighthouse Lab in Alderley Park and Alex Alderton, Roberto Amato, Sonia Goncalves, Ewan Harrison, David K. Jackson, Ian Johnston, Dominic Kwiatkowski, Cordelia Langford, John Sillitoe on behalf of the Wellcome Sanger Institute COVID-19 Surveillance Team |
| EPI_ISL_608148, EPI_ISL_608149, EPI_ISL_608150, EPI_ISL_608151, EPI_ISL_608152, EPI_ISL_608153, EPI_ISL_608154, EPI_ISL_608155, EPI_ISL_608156, EPI_ISL_608157, EPI_ISL_608158, EPI_ISL_608159, EPI_ISL_608160, EPI_ISL_608161, EPI_ISL_608162, EPI_ISL_608163, EPI_ISL_608164, EPI_ISL_608165, EPI_ISL_608166, EPI_ISL_608167, EPI_ISL_608168, EPI_ISL_608169, EPI_ISL_608170, EPI_ISL_608171, EPI_ISL_608172, EPI_ISL_608173, EPI_ISL_608174, EPI_ISL_608175, EPI_ISL_608176, EPI_ISL_608177, EPI_ISL_608178, EPI_ISL_608179, EPI_ISL_608180, EPI_ISL_608181, EPI_ISL_608182, EPI_ISL_608183, EPI_ISL_608184, EPI_ISL_608185, EPI_ISL_608186, EPI_ISL_608187, EPI_ISL_608188, EPI_ISL_608189, EPI_ISL_608190, EPI_ISL_608191, EPI_ISL_608192, EPI_ISL_608193, EPI_ISL_608194, EPI_ISL_608195, EPI_ISL_608196, EPI_ISL_608197, EPI_ISL_608198, EPI_ISL_608199, EPI_ISL_608200, EPI_ISL_608201, EPI_ISL_608202, EPI_ISL_608203, EPI_ISL_608204, EPI_ISL_608205, EPI_ISL_608206, EPI_ISL_608207, EPI_ISL_608208, EPI_ISL_608209, EPI_ISL_608210, EPI_ISL_608211, EPI_ISL_608212, EPI_ISL_608213, EPI_ISL_608214, EPI_ISL_608215 | | | |
| see above | Lighthouse Lab in Alderley Park | Wellcome Sanger Institute for the COVID-19 Genomics UK (COG-UK) consortium | Jacquelyn Wynn, Mairead Hyland, The Lighthouse Lab in Alderley Park and Alex Alderton, Roberto Amato, Sonia Goncalves, Ewan Harrison, David K. Jackson, Ian Johnston, Dominic Kwiatkowski, Cordelia Langford, John Sillitoe on behalf of the Wellcome Sanger Institute COVID-19 Surveillance Team |
| EPI_ISL_608216 | Lighthouse Lab in Alderley Park | Wellcome Sanger Institute for the COVID-19 Genomics UK (COG-UK) Consortium | Jacquelyn Wynn, Mairead Hyland, The Lighthouse Lab in Alderley Park and Alex Alderton, Roberto Amato, Sonia Goncalves, Ewan Harrison, David K. Jackson, Ian Johnston, Dominic Kwiatkowski, Cordelia Langford, John Sillitoe on behalf of the Wellcome Sanger Institute COVID-19 Surveillance Team |
| EPI_ISL_608217, EPI_ISL_608218, EPI_ISL_608219, EPI_ISL_608220, EPI_ISL_608221, EPI_ISL_608222, EPI_ISL_608223, EPI_ISL_608224, EPI_ISL_608225, EPI_ISL_608226, EPI_ISL_608227, EPI_ISL_608228, EPI_ISL_608229, EPI_ISL_608230, EPI_ISL_608231, EPI_ISL_608232, EPI_ISL_608233, EPI_ISL_608234, EPI_ISL_608235, EPI_ISL_608236, EPI_ISL_608237, EPI_ISL_608238, EPI_ISL_608239, EPI_ISL_608240, EPI_ISL_608241, EPI_ISL_608242, EPI_ISL_608243, EPI_ISL_608244, EPI_ISL_608245, EPI_ISL_608246, EPI_ISL_608247, EPI_ISL_608248, EPI_ISL_608249, EPI_ISL_608250, EPI_ISL_608251, EPI_ISL_608252, EPI_ISL_608253, EPI_ISL_608254, EPI_ISL_608255, EPI_ISL_608256, EPI_ISL_608257, EPI_ISL_608258, EPI_ISL_608259, EPI_ISL_608260, EPI_ISL_608261, EPI_ISL_608262, EPI_ISL_608263, EPI_ISL_608264, EPI_ISL_608265, EPI_ISL_608266, EPI_ISL_608267, EPI_ISL_608268, EPI_ISL_608269, EPI_ISL_608270, EPI_ISL_608271, EPI_ISL_608272, EPI_ISL_608273, EPI_ISL_608274, EPI_ISL_608275, EPI_ISL_608276, EPI_ISL_608277, EPI_ISL_608278, EPI_ISL_608279, EPI_ISL_608280, EPI_ISL_608281, EPI_ISL_608282, EPI_ISL_608283, EPI_ISL_608284, EPI_ISL_608285, EPI_ISL_608286, EPI_ISL_608287, EPI_ISL_608288, EPI_ISL_608289, EPI_ISL_608290, EPI_ISL_608291, EPI_ISL_608292, EPI_ISL_608293, EPI_ISL_608294, EPI_ISL_608295, EPI_ISL_608296, EPI_ISL_608297, EPI_ISL_608298, EPI_ISL_608299, EPI_ISL_608300, EPI_ISL_608301, EPI_ISL_608302, EPI_ISL_608303, EPI_ISL_608304, EPI_ISL_608305, EPI_ISL_608306, EPI_ISL_608307, EPI_ISL_608308, EPI_ISL_608309 | | | |
| see above | Lighthouse Lab in Alderley Park | Wellcome Sanger Institute for the COVID-19 Genomics UK (COG-UK) consortium | Jacquelyn Wynn, Mairead Hyland, The Lighthouse Lab in Alderley Park and Alex Alderton, Roberto Amato, Sonia Goncalves, Ewan Harrison, David K. Jackson, Ian Johnston, Dominic Kwiatkowski, Cordelia Langford, John Sillitoe on behalf of the Wellcome Sanger Institute COVID-19 Surveillance Team |
| EPI_ISL_608310 | Lighthouse Lab in Alderley Park | Wellcome Sanger Institute for the COVID-19 Genomics UK (COG-UK) Consortium | Jacquelyn Wynn, Mairead Hyland, The Lighthouse Lab in Alderley Park and Alex Alderton, Roberto Amato, Sonia Goncalves, Ewan Harrison, David K. Jackson, Ian Johnston, Dominic Kwiatkowski, Cordelia Langford, John Sillitoe on behalf of the Wellcome Sanger Institute COVID-19 Surveillance Team |
| EPI_ISL_608311, EPI_ISL_608312, EPI_ISL_608313, EPI_ISL_608314, EPI_ISL_608315, EPI_ISL_608316, EPI_ISL_608317, EPI_ISL_608318, EPI_ISL_608319, EPI_ISL_608320, EPI_ISL_608321, EPI_ISL_608322, EPI_ISL_608323, EPI_ISL_608324, EPI_ISL_608325, EPI_ISL_608326, EPI_ISL_608327, EPI_ISL_608328, EPI_ISL_608329, EPI_ISL_608330, EPI_ISL_608331, EPI_ISL_608332, EPI_ISL_608333, EPI_ISL_608334, EPI_ISL_608335, EPI_ISL_608336, EPI_ISL_608337, EPI_ISL_608338, EPI_ISL_608339, EPI_ISL_608340, EPI_ISL_608341, EPI_ISL_608342, EPI_ISL_608343, EPI_ISL_608344, EPI_ISL_608345, EPI_ISL_608346, EPI_ISL_608445, EPI_ISL_608505, EPI_ISL_608512, EPI_ISL_608539, EPI_ISL_608569, EPI_ISL_608639, EPI_ISL_608649, EPI_ISL_608658, EPI_ISL_608880, EPI_ISL_608948 | | | |
| see above | Lighthouse Lab in Alderley Park | Wellcome Sanger Institute for the COVID-19 Genomics UK (COG-UK) consortium | Jacquelyn Wynn, Mairead Hyland, The Lighthouse Lab in Alderley Park and Alex Alderton, Roberto Amato, Sonia Goncalves, Ewan Harrison, David K. Jackson, Ian Johnston, Dominic Kwiatkowski, Cordelia Langford, John Sillitoe on behalf of the Wellcome Sanger Institute COVID-19 Surveillance Team |
| EPI_ISL_609890, EPI_ISL_609891, EPI_ISL_609892, EPI_ISL_609893, EPI_ISL_609894, EPI_ISL_609895, EPI_ISL_609896, EPI_ISL_609898, EPI_ISL_609899, EPI_ISL_609900, EPI_ISL_609901, EPI_ISL_609902, EPI_ISL_609903, EPI_ISL_609905, EPI_ISL_609906, EPI_ISL_609907, EPI_ISL_609908 | | | |
| see above | Respiratory Virus Unit, Microbiology Services Colindale, Public Health England | Respiratory Virus Unit, Microbiology Services Colindale, Public Health England | PHE Covid Sequencing Team |
| EPI_ISL_609994 | INMI Lazzaro Spallanzani IRCCS | INMI Lazzaro Spallanzani IRCCS | C.E.M Gruber, F Messina, M Rueca, B Bartolini, E Giombini, MR Capobianchi, A Di Caro |
| EPI_ISL_609995 | INMI Lazzaro Spallanzani IRCCS | INMI Lazzaro Spallanzani IRCCS | E Giombini, C.E.M Gruber, M Rueca, B Bartolini, F Messina, A Di Caro, MR Capobianchi |
| EPI_ISL_609996 | INMI Lazzaro Spallanzani IRCCS | INMI Lazzaro Spallanzani IRCCS | F Messina, M Rueca, B Bartolini, C.E.M Gruber, E Giombini, MR Capobianchi, A Di Caro |
| EPI_ISL_609997 | INMI Lazzaro Spallanzani IRCCS | INMI Lazzaro Spallanzani IRCCS | M Rueca, B Bartolini, C.E.M Gruber, F Messina, E Giombini, A Di Caro, MR Capobianchi |
| EPI_ISL_610049, EPI_ISL_610051, EPI_ISL_610052, EPI_ISL_610060, EPI_ISL_610074, EPI_ISL_610077, EPI_ISL_610079, EPI_ISL_610081, EPI_ISL_610083, EPI_ISL_610086, EPI_ISL_610089, EPI_ISL_610090, EPI_ISL_610093, EPI_ISL_610094, EPI_ISL_610095, EPI_ISL_610096, EPI_ISL_610097, EPI_ISL_610098, EPI_ISL_610099, EPI_ISL_610101, EPI_ISL_610102, EPI_ISL_610103, EPI_ISL_610104, EPI_ISL_610105, EPI_ISL_610106, EPI_ISL_610107, EPI_ISL_610108, EPI_ISL_610109, EPI_ISL_610110, EPI_ISL_610111, EPI_ISL_610112, EPI_ISL_610113, EPI_ISL_610114, EPI_ISL_610115, EPI_ISL_610116, EPI_ISL_610117, EPI_ISL_610246, EPI_ISL_610248 | | | |
| see above | University of Michigan Clinical Microbiology Laboratory | Lauring Lab, University of Michigan, Department of Microbiology and Immunology | Valesano |
| EPI_ISL_610286, EPI_ISL_610287, EPI_ISL_610288, EPI_ISL_610289, EPI_ISL_610290, EPI_ISL_610291, EPI_ISL_610292, EPI_ISL_610293, EPI_ISL_610294, EPI_ISL_610295, EPI_ISL_610296, EPI_ISL_610297, EPI_ISL_610298, EPI_ISL_610299, EPI_ISL_610300, EPI_ISL_610301, EPI_ISL_610302, EPI_ISL_610303, EPI_ISL_610304, EPI_ISL_610305, EPI_ISL_610306, EPI_ISL_610307, EPI_ISL_610308, EPI_ISL_610309, EPI_ISL_610310, EPI_ISL_610311, EPI_ISL_610312, EPI_ISL_610313, EPI_ISL_610314, EPI_ISL_610315, EPI_ISL_610316, EPI_ISL_610317, EPI_ISL_610318, EPI_ISL_610319, EPI_ISL_610320, EPI_ISL_610321, EPI_ISL_610322, EPI_ISL_610323, EPI_ISL_610324, EPI_ISL_610325, EPI_ISL_610326, EPI_ISL_610327, EPI_ISL_610328, EPI_ISL_610329, EPI_ISL_610330, EPI_ISL_610331 | | | |
| see above | Lighthouse Lab in Alderley Park | Wellcome Sanger Institute for the COVID-19 Genomics UK (COG-UK) consortium | Jacquelyn Wynn, Mairead Hyland, The Lighthouse Lab in Alderley Park and Alex Alderton, Roberto Amato, Sonia Goncalves, Ewan Harrison, David K. Jackson, Ian Johnston, Dominic Kwiatkowski, Cordelia Langford, John Sillitoe on behalf of the Wellcome Sanger Institute COVID-19 Surveillance Team |
| EPI_ISL_610332 | Lighthouse Lab in Alderley Park | Wellcome Sanger Institute for the COVID-19 Genomics UK (COG-UK) Consortium | Jacquelyn Wynn, Mairead Hyland, The Lighthouse Lab in Alderley Park and Alex Alderton, Roberto Amato, Sonia Goncalves, Ewan Harrison, David K. Jackson, Ian Johnston, Dominic Kwiatkowski, Cordelia Langford, John Sillitoe on behalf of the Wellcome Sanger Institute COVID-19 Surveillance Team |
| EPI_ISL_610333, EPI_ISL_610334, EPI_ISL_610335, EPI_ISL_610336, EPI_ISL_610337, EPI_ISL_610338, EPI_ISL_610339, EPI_ISL_610340, EPI_ISL_610341, EPI_ISL_610342, EPI_ISL_610343, EPI_ISL_610344, EPI_ISL_610345, EPI_ISL_610346, EPI_ISL_610347, EPI_ISL_610348, EPI_ISL_610349, EPI_ISL_610350, EPI_ISL_610351, EPI_ISL_610352, EPI_ISL_610353, EPI_ISL_610354, EPI_ISL_610355, EPI_ISL_610356, EPI_ISL_610357, EPI_ISL_610358, EPI_ISL_610359, EPI_ISL_610360, EPI_ISL_610361, EPI_ISL_610362, EPI_ISL_610363, EPI_ISL_610364, EPI_ISL_610365, EPI_ISL_610366, EPI_ISL_610367, EPI_ISL_610368, EPI_ISL_610369, EPI_ISL_610370, EPI_ISL_610371, EPI_ISL_610372, EPI_ISL_610373, EPI_ISL_610374, EPI_ISL_610375, EPI_ISL_610376, EPI_ISL_610377, EPI_ISL_610378, EPI_ISL_610379, EPI_ISL_610380, EPI_ISL_610381, EPI_ISL_610382, EPI_ISL_610383, EPI_ISL_610384, EPI_ISL_610385, EPI_ISL_610386, EPI_ISL_610387, EPI_ISL_610388, EPI_ISL_610389, EPI_ISL_610390, EPI_ISL_610391, EPI_ISL_610392, EPI_ISL_610393, EPI_ISL_610394, EPI_ISL_610395, EPI_ISL_610396, EPI_ISL_610397, EPI_ISL_610398, EPI_ISL_610399, EPI_ISL_610400, EPI_ISL_610401, EPI_ISL_610402, EPI_ISL_610403, EPI_ISL_610404, EPI_ISL_610405, EPI_ISL_610406, EPI_ISL_610407, EPI_ISL_610408, EPI_ISL_610409, EPI_ISL_610410, EPI_ISL_610411, EPI_ISL_610412, EPI_ISL_610413, EPI_ISL_610414, EPI_ISL_610415, EPI_ISL_610416, EPI_ISL_610417, EPI_ISL_610418, EPI_ISL_610419, EPI_ISL_610420, EPI_ISL_610421, EPI_ISL_610422, EPI_ISL_610423, EPI_ISL_610424, EPI_ISL_610425, EPI_ISL_610426, EPI_ISL_610427, EPI_ISL_610428, EPI_ISL_610429, EPI_ISL_610430, EPI_ISL_610431, EPI_ISL_610432, EPI_ISL_610433, EPI_ISL_610434, EPI_ISL_610435, EPI_ISL_610436, EPI_ISL_610437, EPI_ISL_610438, EPI_ISL_610439, EPI_ISL_610440, | | | |

| | | | |
|---|---|---|---|
| EPI_ISL_610441, EPI_ISL_610442, EPI_ISL_610443, EPI_ISL_610444, EPI_ISL_610445, EPI_ISL_610446, EPI_ISL_610447, EPI_ISL_610448, EPI_ISL_610449, EPI_ISL_610450, EPI_ISL_610451, EPI_ISL_610452, EPI_ISL_610453, EPI_ISL_610454, EPI_ISL_610455, EPI_ISL_610456, EPI_ISL_610457, EPI_ISL_610458, EPI_ISL_610459, EPI_ISL_610460, EPI_ISL_610461, EPI_ISL_610462, EPI_ISL_610463, EPI_ISL_610464, EPI_ISL_610465, EPI_ISL_610466, EPI_ISL_610467, EPI_ISL_610468, EPI_ISL_610469, EPI_ISL_610470, EPI_ISL_610471, EPI_ISL_610472, EPI_ISL_610473, EPI_ISL_610474, EPI_ISL_610475, EPI_ISL_610476, EPI_ISL_610477, EPI_ISL_610478, EPI_ISL_610479, EPI_ISL_610480, EPI_ISL_610481, EPI_ISL_610482, EPI_ISL_610483, EPI_ISL_610484, EPI_ISL_610485, EPI_ISL_610486, EPI_ISL_610487, EPI_ISL_610488, EPI_ISL_610489, EPI_ISL_610490, EPI_ISL_610491, EPI_ISL_610492, EPI_ISL_610493, EPI_ISL_610494, EPI_ISL_610495, EPI_ISL_610496, EPI_ISL_610497, EPI_ISL_610498, EPI_ISL_610499, EPI_ISL_610500, EPI_ISL_610501 | | | |
| see above | Lighthouse Lab in Alderley Park | Wellcome Sanger Institute for the COVID-19 Genomics UK (COG-UK) consortium | Jacquelyn Wynn, Mairead Hyland, The Lighthouse Lab in Alderley Park and Alex Alderton, Roberto Amato, Sonia Goncalves, Ewan Harrison, David K. Jackson, Ian Johnston, Dominic Kwiatkowski, Cordelia Langford, John Sillitoe on behalf of the Wellcome Sanger Institute COVID-19 Surveillance Team |
| EPI_ISL_610502, EPI_ISL_610503 | Lighthouse Lab in Alderley Park | Wellcome Sanger Institute for the COVID-19 Genomics UK (COG-UK) Consortium | Jacquelyn Wynn, Mairead Hyland, The Lighthouse Lab in Alderley Park and Alex Alderton, Roberto Amato, Sonia Goncalves, Ewan Harrison, David K. Jackson, Ian Johnston, Dominic Kwiatkowski, Cordelia Langford, John Sillitoe on behalf of the Wellcome Sanger Institute COVID-19 Surveillance Team |
| EPI_ISL_610504, EPI_ISL_610505, EPI_ISL_610506, EPI_ISL_610507, EPI_ISL_610508, EPI_ISL_610509, EPI_ISL_610510, EPI_ISL_610511, EPI_ISL_610512, EPI_ISL_610513, EPI_ISL_610514, EPI_ISL_610515, EPI_ISL_610516, EPI_ISL_610517, EPI_ISL_610518, EPI_ISL_610519, EPI_ISL_610520, EPI_ISL_610521, EPI_ISL_610522, EPI_ISL_610523, EPI_ISL_610524, EPI_ISL_610525, EPI_ISL_610526, EPI_ISL_610527, EPI_ISL_610528, EPI_ISL_610529, EPI_ISL_610530, EPI_ISL_610531, EPI_ISL_610532, EPI_ISL_610533, EPI_ISL_610534, EPI_ISL_610535, EPI_ISL_610536, EPI_ISL_610537, EPI_ISL_610538, EPI_ISL_610539, EPI_ISL_610540, EPI_ISL_610541, EPI_ISL_610542, EPI_ISL_610543, EPI_ISL_610544, EPI_ISL_610545, EPI_ISL_610546, EPI_ISL_610547, EPI_ISL_610548, EPI_ISL_610549, EPI_ISL_610550, EPI_ISL_610551, EPI_ISL_610552, EPI_ISL_610553, EPI_ISL_610554, EPI_ISL_610555, EPI_ISL_610556, EPI_ISL_610557, EPI_ISL_610558, EPI_ISL_610559, EPI_ISL_610560, EPI_ISL_610561, EPI_ISL_610562, EPI_ISL_610563, EPI_ISL_610564, EPI_ISL_610565, EPI_ISL_610566, EPI_ISL_610567, EPI_ISL_610568, EPI_ISL_610569, EPI_ISL_610570, EPI_ISL_610571, EPI_ISL_610572, EPI_ISL_610573, EPI_ISL_610574, EPI_ISL_610575, EPI_ISL_610576, EPI_ISL_610577, EPI_ISL_610578, EPI_ISL_610579, EPI_ISL_610580, EPI_ISL_610581, EPI_ISL_610582, EPI_ISL_610583, EPI_ISL_610584, EPI_ISL_610585, EPI_ISL_610586, EPI_ISL_610587, EPI_ISL_610588, EPI_ISL_610589, EPI_ISL_610590, EPI_ISL_610591, EPI_ISL_610592, EPI_ISL_610593, EPI_ISL_610594, EPI_ISL_610595, EPI_ISL_610596, EPI_ISL_610597, EPI_ISL_610598, EPI_ISL_610599, EPI_ISL_610600, EPI_ISL_610601, EPI_ISL_610602, EPI_ISL_610603, EPI_ISL_610604, EPI_ISL_610605, EPI_ISL_610606, EPI_ISL_610607, EPI_ISL_610608, EPI_ISL_610609, EPI_ISL_610610, EPI_ISL_610611, EPI_ISL_610612, EPI_ISL_610613, EPI_ISL_610614, EPI_ISL_610615, EPI_ISL_610616, EPI_ISL_610617, EPI_ISL_610618, EPI_ISL_610619, EPI_ISL_610620, EPI_ISL_610621, EPI_ISL_610622, EPI_ISL_610623, EPI_ISL_610624, EPI_ISL_610625, EPI_ISL_610626, EPI_ISL_610627, EPI_ISL_610628 | | | |
| see above | Lighthouse Lab in Alderley Park | Wellcome Sanger Institute for the COVID-19 Genomics UK (COG-UK) consortium | Jacquelyn Wynn, Mairead Hyland, The Lighthouse Lab in Alderley Park and Alex Alderton, Roberto Amato, Sonia Goncalves, Ewan Harrison, David K. Jackson, Ian Johnston, Dominic Kwiatkowski, Cordelia Langford, John Sillitoe on behalf of the Wellcome Sanger Institute COVID-19 Surveillance Team |
| EPI_ISL_610629, EPI_ISL_610630, EPI_ISL_610631, EPI_ISL_610634, EPI_ISL_610636, EPI_ISL_610640, EPI_ISL_610641, EPI_ISL_610642, EPI_ISL_610644, EPI_ISL_610647, EPI_ISL_610649, EPI_ISL_610651, EPI_ISL_610652, EPI_ISL_610653, EPI_ISL_610655, EPI_ISL_610658, EPI_ISL_610660, EPI_ISL_610662, EPI_ISL_610663, EPI_ISL_610664, EPI_ISL_610665, EPI_ISL_610666, EPI_ISL_610667, EPI_ISL_610668, EPI_ISL_610670, EPI_ISL_610672, EPI_ISL_610674, EPI_ISL_610675, EPI_ISL_610676, EPI_ISL_610677, EPI_ISL_610679, EPI_ISL_610681, EPI_ISL_610683, EPI_ISL_610684, EPI_ISL_610685, EPI_ISL_610686, EPI_ISL_610687, EPI_ISL_610689, EPI_ISL_610691, EPI_ISL_610692, EPI_ISL_610693, EPI_ISL_610694, EPI_ISL_610695, EPI_ISL_610697, EPI_ISL_610698, EPI_ISL_610699 | | | |
| see above | Lighthouse Lab in Cambridge | Wellcome Sanger Institute for the COVID-19 Genomics UK (COG-UK) consortium | Rob Howes, The Lighthouse Lab in Cambridge and Alex Alderton, Roberto Amato, Sonia Goncalves, Ewan Harrison, David K. Jackson, Ian Johnston, Dominic Kwiatkowski, Cordelia Langford, John Sillitoe on behalf of the Wellcome Sanger Institute COVID-19 Surveillance Team |
| EPI_ISL_610702 | Lighthouse Lab in Cambridge | Wellcome Sanger Institute for the COVID-19 Genomics UK (COG-UK) Consortium | Rob Howes, The Lighthouse Lab in Cambridge and Alex Alderton, Roberto Amato, Sonia Goncalves, Ewan Harrison, David K. Jackson, Ian Johnston, Dominic Kwiatkowski, Cordelia Langford, John Sillitoe on behalf of the Wellcome Sanger Institute COVID-19 Surveillance Team |
| EPI_ISL_610705, EPI_ISL_610708, EPI_ISL_610709, EPI_ISL_610710, EPI_ISL_610712, EPI_ISL_610713, EPI_ISL_610714, EPI_ISL_610715, EPI_ISL_610716, EPI_ISL_610717, EPI_ISL_610719, EPI_ISL_610720, EPI_ISL_610721, EPI_ISL_610722, EPI_ISL_610723, EPI_ISL_610725, EPI_ISL_610727, EPI_ISL_610728, EPI_ISL_610729, EPI_ISL_610730, EPI_ISL_610734, EPI_ISL_610736, EPI_ISL_610738, EPI_ISL_610740, EPI_ISL_610741, EPI_ISL_610743, EPI_ISL_610744, EPI_ISL_610745, EPI_ISL_610746, EPI_ISL_610747, EPI_ISL_610748, EPI_ISL_610749, EPI_ISL_610756, EPI_ISL_610757, EPI_ISL_610758, EPI_ISL_610759, EPI_ISL_610761, EPI_ISL_610762, EPI_ISL_610763, EPI_ISL_610768, EPI_ISL_610769, EPI_ISL_610770, EPI_ISL_610771, EPI_ISL_610772, EPI_ISL_610775, EPI_ISL_610776 | | | |
| see above | Lighthouse Lab in Cambridge | Wellcome Sanger Institute for the COVID-19 Genomics UK (COG-UK) consortium | Rob Howes, The Lighthouse Lab in Cambridge and Alex Alderton, Roberto Amato, Sonia Goncalves, Ewan Harrison, David K. Jackson, Ian Johnston, Dominic Kwiatkowski, Cordelia Langford, John Sillitoe on behalf of the Wellcome Sanger Institute COVID-19 Surveillance Team |
| EPI_ISL_610779 | Lighthouse Lab in Cambridge | Wellcome Sanger Institute for the COVID-19 Genomics UK (COG-UK) Consortium | Rob Howes, The Lighthouse Lab in Cambridge and Alex Alderton, Roberto Amato, Sonia Goncalves, Ewan Harrison, David K. Jackson, Ian Johnston, Dominic Kwiatkowski, Cordelia Langford, John Sillitoe on behalf of the Wellcome Sanger Institute COVID-19 Surveillance Team |
| EPI_ISL_610780, EPI_ISL_610781, EPI_ISL_610783, EPI_ISL_610784 | Lighthouse Lab in Cambridge | Wellcome Sanger Institute for the COVID-19 Genomics UK (COG-UK) consortium | Rob Howes, The Lighthouse Lab in Cambridge and Alex Alderton, Roberto Amato, Sonia Goncalves, Ewan Harrison, David K. Jackson, Ian Johnston, Dominic Kwiatkowski, Cordelia Langford, John Sillitoe on behalf of the Wellcome Sanger Institute COVID-19 Surveillance Team |
| EPI_ISL_610787, EPI_ISL_610789, EPI_ISL_610790, EPI_ISL_610791, EPI_ISL_610793, EPI_ISL_610794, EPI_ISL_610795, EPI_ISL_610797, EPI_ISL_610798, EPI_ISL_610800, EPI_ISL_610802, EPI_ISL_610803, EPI_ISL_610804, EPI_ISL_610807, EPI_ISL_610808, EPI_ISL_610809, EPI_ISL_610812, EPI_ISL_610813, EPI_ISL_610817, EPI_ISL_610818, EPI_ISL_610821, EPI_ISL_610824, EPI_ISL_610825, EPI_ISL_610828, EPI_ISL_610831, EPI_ISL_610832, EPI_ISL_610835, EPI_ISL_610838, EPI_ISL_610840, EPI_ISL_610841 | | | |
| see above | Lighthouse Lab in Cambridge | Wellcome Sanger Institute for the COVID-19 Genomics UK (COG-UK) consortium | Rob Howes, The Lighthouse Lab in Cambridge and Alex Alderton, Roberto Amato, Sonia Goncalves, Ewan Harrison, David K. Jackson, Ian Johnston, Dominic Kwiatkowski, Cordelia Langford, John Sillitoe on behalf of the Wellcome Sanger Institute COVID-19 Surveillance Team (http://www.sanger.ac.uk/covid-team) |
| EPI_ISL_610843 | Lighthouse Lab in Cambridge | Wellcome Sanger Institute for the COVID-19 Genomics UK (COG-UK) Consortium | Rob Howes, The Lighthouse Lab in Cambridge and Alex Alderton, Roberto Amato, Sonia Goncalves, Ewan Harrison, David K. Jackson, Ian Johnston, Dominic Kwiatkowski, Cordelia Langford, John Sillitoe on behalf of the Wellcome Sanger Institute COVID-19 Surveillance Team (http://www.sanger.ac.uk/covid-team) |
| EPI_ISL_610844, EPI_ISL_610845, EPI_ISL_610846, EPI_ISL_610848, EPI_ISL_610849, EPI_ISL_610850, EPI_ISL_610851, EPI_ISL_610854, EPI_ISL_610855, EPI_ISL_610858, EPI_ISL_610859, EPI_ISL_610860, EPI_ISL_610862, EPI_ISL_610863, EPI_ISL_610864, EPI_ISL_610865, EPI_ISL_610866, EPI_ISL_610868, EPI_ISL_610869, EPI_ISL_610870, EPI_ISL_610872, EPI_ISL_610873, EPI_ISL_610874, EPI_ISL_610876, EPI_ISL_610878, EPI_ISL_610880, EPI_ISL_610882, EPI_ISL_610883, EPI_ISL_610886, EPI_ISL_610891, EPI_ISL_610893, EPI_ISL_610894, EPI_ISL_610897, EPI_ISL_610898, EPI_ISL_610901, EPI_ISL_610902, EPI_ISL_610903, EPI_ISL_610904, EPI_ISL_610908, EPI_ISL_610909, EPI_ISL_610910, EPI_ISL_610911, EPI_ISL_610912, EPI_ISL_610918, EPI_ISL_610919, EPI_ISL_610921 | | | |
| see above | Lighthouse Lab in Cambridge | Wellcome Sanger Institute for the COVID-19 Genomics UK (COG-UK) consortium | Rob Howes, The Lighthouse Lab in Cambridge and Alex Alderton, Roberto Amato, Sonia Goncalves, Ewan Harrison, David K. Jackson, Ian Johnston, Dominic Kwiatkowski, Cordelia Langford, John Sillitoe on behalf of the Wellcome Sanger Institute COVID-19 Surveillance Team (http://www.sanger.ac.uk/covid-team) |
| EPI_ISL_611422, EPI_ISL_611426 | Lighthouse Lab in Glasgow | Wellcome Sanger Institute for the COVID-19 Genomics UK (COG-UK) consortium | Harper VanSteenhouse, Yumi Kasai, David Gray, Carol Clugston, Anna Dominiczak and Alex Alderton, Roberto Amato, Sonia Goncalves, Ewan Harrison, David K. Jackson, Ian Johnston, Dominic Kwiatkowski, Cordelia Langford, John Sillitoe on behalf of the Wellcome Sanger Institute COVID-19 Surveillance Team (http://www.sanger.ac.uk/covid-team) |
| EPI_ISL_611518 | Wales Specialist Virology Centre Sequencing lab: Pathogen Genomics Unit | COVID-19 Genomics UK (COG-UK) Consortium | Catherine Moore, Johnathan Evans, Laura Gifford, Malorie Perry, Simon Cottrell, Angela Marchbank, Alec Birchley, Alexander Adams, Amy Gaskin, Bree Gatica-Wilcox, Jason Coombes, Joel Southgate, Lauren Gilbert, Lee Graham, Nicole Pacchiarini, Sara Kumziene-Summerhayes, Sarah Taylor, Sophie Jones, Sara Rey, Matthew Bull, Joanne Watkins, Sally Corden, Tom Connor |
| EPI_ISL_611523 | University College London, Great Ormond Street Hospital for Children NHS Foundation Trust, Imperial College Healthcare NHS Trust | COVID-19 Genomics UK (COG-UK) Consortium | Sergi Castellano, Rachel Williams, Mark Kristiansen, Paola Resende Silva, Sunando Roy, Tony Brooks, Helena Tutill, Paola Niola, Patricia Dyal, Charlotte Williams, Leysa Forrest, Yasmin Panchbhaya, Jacqueline Findlay, Samuel Weeks, Julianne Brown, Kathryn Harris, Paul Randell, James Price, Alison Holmes, Judith Breuer |
| EPI_ISL_611525 | Virology Department, Sheffield Teaching Hospitals NHS Foundation Trust/Department of Infection, Immunity and Cardiovascular Disease, The Medical School, University of Sheffield | COVID-19 Genomics UK (COG-UK) Consortium | Thushan de Silva, Matthew Parker, Nikki Smith, Adri Angyal, Rebecca Brown, Luke Green, Rachel Tucker, Paul Parsons, Danielle Groves, Katie Johnson, Laura Carrilero, Alex Keeley, Dave Partridge, Matthew Wyles, Benjamin Lindsey, Mehmet Yavuz, Mohammad Raza, Cariad Evans |
| EPI_ISL_611526 | Centre for Enzyme Innovation, University of Portsmouth / Translational Research Laboratory, Portsmouth Hospitals NHS Trust | COVID-19 Genomics UK (COG-UK) Consortium | Angela Beckett,Yann Bourgeois,Garry Scarlett,Sharon Glaysher,Scott Elliott,Kelly Bicknell,Robert Impey,Allyson Lloyd,Sarah Wyllie,Ethan Butcher,Anoop Chauhan,Samuel Robson |
| EPI_ISL_611528 | Virology Department, Sheffield Teaching Hospitals NHS Foundation Trust/Department of Infection, Immunity and Cardiovascular Disease, The Medical School, University of Sheffield | COVID-19 Genomics UK (COG-UK) Consortium | Thushan de Silva, Matthew Parker, Nikki Smith, Adri Angyal, Rebecca Brown, Luke Green, Rachel Tucker, Paul Parsons, Danielle Groves, Katie Johnson, Laura Carrilero, Alex Keeley, Dave Partridge, Matthew Wyles, Benjamin Lindsey, Mehmet Yavuz, Mohammad Raza, Cariad Evans |
| EPI_ISL_611532, EPI_ISL_611535, EPI_ISL_611537, EPI_ISL_611538 | Centre for Enzyme Innovation, University of Portsmouth / Translational Research Laboratory, Portsmouth Hospitals NHS Trust | COVID-19 Genomics UK (COG-UK) Consortium | Angela Beckett,Yann Bourgeois,Garry Scarlett,Sharon Glaysher,Scott Elliott,Kelly Bicknell,Robert Impey,Allyson Lloyd,Sarah Wyllie,Ethan Butcher,Anoop Chauhan,Samuel Robson |
| EPI_ISL_611544, EPI_ISL_611549 | Virology Department, Royal Infirmary of Edinburgh, NHS Lothian / School of Biological Sciences, University of Edinburgh / Institute of Genetics and Molecular Medicine, University of Edinburgh | COVID-19 Genomics UK (COG-UK) Consortium | McHugh M, Dewar R, Rooke S, Gallagher M, Balcaza C, O'Toole Á, Scher E, Hill V, McCrone JT, Colquhoun R, Yu X, Jackson B, Rambaut A, Williams TC, Templeton K |
| EPI_ISL_611553 | Virology Department, Sheffield Teaching Hospitals NHS | COVID-19 Genomics UK (COG-UK) Consortium | Thushan de Silva, Matthew Parker, Nikki Smith, Adri Angyal, Rebecca Brown, Luke Green, Rachel Tucker, Paul Parsons, Danielle Groves, Katie Johnson, |

| | | | |
|---|---|---|---|
| | Foundation Trust/Department of Infection, Immunity and Cardiovascular Disease, The Medical School, University of Sheffield | | Laura Carrilero, Alex Keeley, Dave Partridge, Matthew Wyles, Benjamin Lindsey, Mehmet Yavuz, Mohammad Raza, Cariad Evans |
| EPI_ISL_611559 | University College London, Great Ormond Street Hospital for Children NHS Foundation Trust, Imperial College Healthcare NHS Trust | COVID-19 Genomics UK (COG-UK) Consortium | Sergi Castellano, Rachel Williams, Mark Kristiansen, Paola Resende Silva, Sunando Roy, Tony Brooks, Helena Tutill, Paola Niola, Patricia Dyal, Charlotte Williams, Leysa Forrest, Yasmin Panchbhaya, Jacqueline Findlay, Samuel Weeks, Julianne Brown, Kathryn Harris, Paul Randell, James Price, Alison Holmes, Judith Breuer |
| EPI_ISL_611560, EPI_ISL_611562, EPI_ISL_611564 | Wales Specialist Virology Centre Sequencing lab: Pathogen Genomics Unit | COVID-19 Genomics UK (COG-UK) Consortium | Catherine Moore, Johnathan Evans, Laura Gifford, Malorie Perry, Simon Cottrell, Angela Marchbank, Alec Birchley, Alexander Adams, Amy Gaskin, Bree Gatica-Wilcox, Jason Coombes, Joel Southgate, Lauren Gilbert, Lee Graham, Nicole Pacchiarini, Sara Kumziene-Summerhayes, Sarah Taylor, Sophie Jones, Sara Rey, Matthew Bull, Joanne Watkins, Sally Corden, Tom Connor |
| EPI_ISL_611570 | Liverpool Clinical Laboratories | COVID-19 Genomics UK (COG-UK) Consortium | Sam Haldenby, Anita Lucaci, Steve Paterson, Julian Hiscox, Alistair Darby, M Almsaud, A Alrezaihi, Muhannad Alruwaili, Stuart D Armstrong, Jones Benjamin, Eleanor G Bentley, Anu Chawla, Jordan J Clark, Angela Cowell, Richard Eccles, Isabel García-Dorival, Matthew Gemmell, Alessandro Gerada, PKF Gilmore, Richard Gregory, Ximeng Han, Catherine Hartley, Margaret Hughes, Miren Iturriza-Gomara, James Johnson, L Luu, Jenifer Manson, Charlotte Nelson, Elaine O'Toole, Cassie Olateju, Rebekah Penrice-Randal , Lucille Rainbow, N.P Randle, Trevor Ian Robinson, Parul Sharma, Ghada T Shawli, James P Stewart, Neil Swainston, Ecaterina Vamos, Joanne Watts, Mark Whitehead |
| EPI_ISL_611573, EPI_ISL_611578 | Wales Specialist Virology Centre Sequencing lab: Pathogen Genomics Unit | COVID-19 Genomics UK (COG-UK) Consortium | Catherine Moore, Johnathan Evans, Laura Gifford, Malorie Perry, Simon Cottrell, Angela Marchbank, Alec Birchley, Alexander Adams, Amy Gaskin, Bree Gatica-Wilcox, Jason Coombes, Joel Southgate, Lauren Gilbert, Lee Graham, Nicole Pacchiarini, Sara Kumziene-Summerhayes, Sarah Taylor, Sophie Jones, Sara Rey, Matthew Bull, Joanne Watkins, Sally Corden, Tom Connor |
| EPI_ISL_611583, EPI_ISL_611584 | Virology Department, Sheffield Teaching Hospitals NHS Foundation Trust/Department of Infection, Immunity and Cardiovascular Disease, The Medical School, University of Sheffield | COVID-19 Genomics UK (COG-UK) Consortium | Thushan de Silva, Matthew Parker, Nikki Smith, Adri Angyal, Rebecca Brown, Luke Green, Rachel Tucker, Paul Parsons, Danielle Groves, Katie Johnson, Laura Carrilero, Alex Keeley, Dave Partridge, Matthew Wyles, Benjamin Lindsey, Mehmet Yavuz, Mohammad Raza, Cariad Evans |
| EPI_ISL_611585 | Centre for Enzyme Innovation, University of Portsmouth / Translational Research Laboratory, Portsmouth Hospitals NHS Trust | COVID-19 Genomics UK (COG-UK) Consortium | Angela Beckett,Yann Bourgeois,Garry Scarlett,Sharon Glaysher,Scott Elliott,Kelly Bicknell,Robert Impey,Allyson Lloyd,Sarah Wyllie,Ethan Butcher,Anoop Chauhan,Samuel Robson |
| EPI_ISL_611588 | Wales Specialist Virology Centre Sequencing lab: Pathogen Genomics Unit | COVID-19 Genomics UK (COG-UK) Consortium | Catherine Moore, Johnathan Evans, Laura Gifford, Malorie Perry, Simon Cottrell, Angela Marchbank, Alec Birchley, Alexander Adams, Amy Gaskin, Bree Gatica-Wilcox, Jason Coombes, Joel Southgate, Lauren Gilbert, Lee Graham, Nicole Pacchiarini, Sara Kumziene-Summerhayes, Sarah Taylor, Sophie Jones, Sara Rey, Matthew Bull, Joanne Watkins, Sally Corden, Tom Connor |
| EPI_ISL_611590, EPI_ISL_611592 | Centre for Enzyme Innovation, University of Portsmouth / Translational Research Laboratory, Portsmouth Hospitals NHS Trust | COVID-19 Genomics UK (COG-UK) Consortium | Angela Beckett,Yann Bourgeois,Garry Scarlett,Sharon Glaysher,Scott Elliott,Kelly Bicknell,Robert Impey,Allyson Lloyd,Sarah Wyllie,Ethan Butcher,Anoop Chauhan,Samuel Robson |
| EPI_ISL_611593 | Department of Pathology, University of Cambridge | COVID-19 Genomics UK (COG-UK) Consortium | Aminu S. Jahun, Yasmin Chaudhry, Grant Hall, Iliana Georgana, Myra Hosmillo, Martin D. Curran, Malte Pinckert, Surendra Parmar, Ian Goodfellow |
| EPI_ISL_611595, EPI_ISL_611596, EPI_ISL_611597 | University College London, Great Ormond Street Hospital for Children NHS Foundation Trust, Imperial College Healthcare NHS Trust | COVID-19 Genomics UK (COG-UK) Consortium | Sergi Castellano, Rachel Williams, Mark Kristiansen, Paola Resende Silva, Sunando Roy, Tony Brooks, Helena Tutill, Paola Niola, Patricia Dyal, Charlotte Williams, Leysa Forrest, Yasmin Panchbhaya, Jacqueline Findlay, Samuel Weeks, Julianne Brown, Kathryn Harris, Paul Randell, James Price, Alison Holmes, Judith Breuer |
| EPI_ISL_611602 | Virology Department, Sheffield Teaching Hospitals NHS Foundation Trust/Department of Infection, Immunity and Cardiovascular Disease, The Medical School, University of Sheffield | COVID-19 Genomics UK (COG-UK) Consortium | Thushan de Silva, Matthew Parker, Nikki Smith, Adri Angyal, Rebecca Brown, Luke Green, Rachel Tucker, Paul Parsons, Danielle Groves, Katie Johnson, Laura Carrilero, Alex Keeley, Dave Partridge, Matthew Wyles, Benjamin Lindsey, Mehmet Yavuz, Mohammad Raza, Cariad Evans |
| EPI_ISL_611604 | Queens Medical Centre, Clinical Microbiology Department / DeepSeq Nottingham | COVID-19 Genomics UK (COG-UK) Consortium | Gemma Clark, Wendy Smith, Manjinder Khakh, Vicki M Fleming, Michelle M Lister, Hannah Howson-Wells, Jonathan Ball, Patrick McClure, Joseph Chappell, Theocharis Tsoleridis, Nadine Holmes, Matthew Carlisle, Christopher Moore, Fei Sang, Johnny Debebe, Victoria Wright, Matthew Loose |
| EPI_ISL_611609 | Wales Specialist Virology Centre Sequencing lab: Pathogen Genomics Unit | COVID-19 Genomics UK (COG-UK) Consortium | Catherine Moore, Johnathan Evans, Laura Gifford, Malorie Perry, Simon Cottrell, Angela Marchbank, Alec Birchley, Alexander Adams, Amy Gaskin, Bree Gatica-Wilcox, Jason Coombes, Joel Southgate, Lauren Gilbert, Lee Graham, Nicole Pacchiarini, Sara Kumziene-Summerhayes, Sarah Taylor, Sophie Jones, Sara Rey, Matthew Bull, Joanne Watkins, Sally Corden, Tom Connor |
| EPI_ISL_611623 | University College London, Great Ormond Street Hospital for Children NHS Foundation Trust, Imperial College Healthcare NHS Trust | COVID-19 Genomics UK (COG-UK) Consortium | Sergi Castellano, Rachel Williams, Mark Kristiansen, Paola Resende Silva, Sunando Roy, Tony Brooks, Helena Tutill, Paola Niola, Patricia Dyal, Charlotte Williams, Leysa Forrest, Yasmin Panchbhaya, Jacqueline Findlay, Samuel Weeks, Julianne Brown, Kathryn Harris, Paul Randell, James Price, Alison Holmes, Judith Breuer |
| EPI_ISL_611626, EPI_ISL_611637 | Wales Specialist Virology Centre Sequencing lab: Pathogen Genomics Unit | COVID-19 Genomics UK (COG-UK) Consortium | Catherine Moore, Johnathan Evans, Laura Gifford, Malorie Perry, Simon Cottrell, Angela Marchbank, Alec Birchley, Alexander Adams, Amy Gaskin, Bree Gatica-Wilcox, Jason Coombes, Joel Southgate, Lauren Gilbert, Lee Graham, Nicole Pacchiarini, Sara Kumziene-Summerhayes, Sarah Taylor, Sophie Jones, Sara Rey, Matthew Bull, Joanne Watkins, Sally Corden, Tom Connor |
| EPI_ISL_611638 | Centre for Enzyme Innovation, University of Portsmouth / Translational Research Laboratory, Portsmouth Hospitals NHS Trust | COVID-19 Genomics UK (COG-UK) Consortium | Angela Beckett,Yann Bourgeois,Garry Scarlett,Sharon Glaysher,Scott Elliott,Kelly Bicknell,Robert Impey,Allyson Lloyd,Sarah Wyllie,Ethan Butcher,Anoop Chauhan,Samuel Robson |
| EPI_ISL_611650, EPI_ISL_611651, EPI_ISL_611656, EPI_ISL_611669 | Virology Department, Sheffield Teaching Hospitals NHS Foundation Trust/Department of Infection, Immunity and Cardiovascular Disease, The Medical School, University of Sheffield | COVID-19 Genomics UK (COG-UK) Consortium | Thushan de Silva, Matthew Parker, Nikki Smith, Adri Angyal, Rebecca Brown, Luke Green, Rachel Tucker, Paul Parsons, Danielle Groves, Katie Johnson, Laura Carrilero, Alex Keeley, Dave Partridge, Matthew Wyles, Benjamin Lindsey, Mehmet Yavuz, Mohammad Raza, Cariad Evans |
| EPI_ISL_611673, EPI_ISL_611674 | Wales Specialist Virology Centre Sequencing lab: Pathogen Genomics Unit | COVID-19 Genomics UK (COG-UK) Consortium | Catherine Moore, Johnathan Evans, Laura Gifford, Malorie Perry, Simon Cottrell, Angela Marchbank, Alec Birchley, Alexander Adams, Amy Gaskin, Bree Gatica-Wilcox, Jason Coombes, Joel Southgate, Lauren Gilbert, Lee Graham, Nicole Pacchiarini, Sara Kumziene-Summerhayes, Sarah Taylor, Sophie Jones, Sara Rey, Matthew Bull, Joanne Watkins, Sally Corden, Tom Connor |
| EPI_ISL_611680 | Queens Medical Centre, Clinical Microbiology Department / DeepSeq Nottingham | COVID-19 Genomics UK (COG-UK) Consortium | Gemma Clark, Wendy Smith, Manjinder Khakh, Vicki M Fleming, Michelle M Lister, Hannah Howson-Wells, Jonathan Ball, Patrick McClure, Joseph Chappell, Theocharis Tsoleridis, Nadine Holmes, Matthew Carlisle, Christopher Moore, Fei Sang, Johnny Debebe, Victoria Wright, Matthew Loose |
| EPI_ISL_611681, EPI_ISL_611682 | Virology Department, Sheffield Teaching Hospitals NHS Foundation Trust/Department of Infection, Immunity and Cardiovascular Disease, The Medical School, University of Sheffield | COVID-19 Genomics UK (COG-UK) Consortium | Thushan de Silva, Matthew Parker, Nikki Smith, Adri Angyal, Rebecca Brown, Luke Green, Rachel Tucker, Paul Parsons, Danielle Groves, Katie Johnson, Laura Carrilero, Alex Keeley, Dave Partridge, Matthew Wyles, Benjamin Lindsey, Mehmet Yavuz, Mohammad Raza, Cariad Evans |
| EPI_ISL_611686, EPI_ISL_611688 | Wales Specialist Virology Centre Sequencing lab: Pathogen Genomics Unit | COVID-19 Genomics UK (COG-UK) Consortium | Catherine Moore, Johnathan Evans, Laura Gifford, Malorie Perry, Simon Cottrell, Angela Marchbank, Alec Birchley, Alexander Adams, Amy Gaskin, Bree Gatica-Wilcox, Jason Coombes, Joel Southgate, Lauren Gilbert, Lee Graham, Nicole Pacchiarini, Sara Kumziene-Summerhayes, Sarah Taylor, Sophie Jones, Sara Rey, Matthew Bull, Joanne Watkins, Sally Corden, Tom Connor |
| EPI_ISL_611690 | Virology Department, Sheffield Teaching Hospitals NHS Foundation Trust/Department of Infection, Immunity and Cardiovascular Disease, The Medical School, University of Sheffield | COVID-19 Genomics UK (COG-UK) Consortium | Thushan de Silva, Matthew Parker, Nikki Smith, Adri Angyal, Rebecca Brown, Luke Green, Rachel Tucker, Paul Parsons, Danielle Groves, Katie Johnson, Laura Carrilero, Alex Keeley, Dave Partridge, Matthew Wyles, Benjamin Lindsey, Mehmet Yavuz, Mohammad Raza, Cariad Evans |
| EPI_ISL_611691 | Wales Specialist Virology Centre Sequencing lab: Pathogen Genomics Unit | COVID-19 Genomics UK (COG-UK) Consortium | Catherine Moore, Johnathan Evans, Laura Gifford, Malorie Perry, Simon Cottrell, Angela Marchbank, Alec Birchley, Alexander Adams, Amy Gaskin, Bree Gatica-Wilcox, Jason Coombes, Joel Southgate, Lauren Gilbert, Lee Graham, Nicole Pacchiarini, Sara Kumziene-Summerhayes, Sarah Taylor, Sophie Jones, Sara Rey, Matthew Bull, Joanne Watkins, Sally Corden, Tom Connor |

| | | | |
|---|---|---|---|
| EPI_ISL_611698, EPI_ISL_611699 | Virology Department, Sheffield Teaching Hospitals NHS Foundation Trust/Department of Infection, Immunity and Cardiovascular Disease, The Medical School, University of Sheffield | COVID-19 Genomics UK (COG-UK) Consortium | Thushan de Silva, Matthew Parker, Nikki Smith, Adri Angyal, Rebecca Brown, Luke Green, Rachel Tucker, Paul Parsons, Danielle Groves, Katie Johnson, Laura Carrilero, Alex Keeley, Dave Partridge, Matthew Wyles, Benjamin Lindsey, Mehmet Yavuz, Mohammad Raza, Cariad Evans |
| EPI_ISL_611704, EPI_ISL_611711, EPI_ISL_611738, EPI_ISL_611740, EPI_ISL_611742 | Wales Specialist Virology Centre Sequencing lab: Pathogen Genomics Unit | COVID-19 Genomics UK (COG-UK) Consortium | Catherine Moore, Johnathan Evans, Laura Gifford, Malorie Perry, Simon Cottrell, Angela Marchbank, Alec Birchley, Alexander Adams, Amy Gaskin, Bree Gatica-Wilcox, Jason Coombes, Joel Southgate, Lauren Gilbert, Lee Graham, Nicole Pacchiarini, Sara Kumziene-Summerhayes, Sarah Taylor, Sophie Jones, Sara Rey, Matthew Bull, Joanne Watkins, Sally Corden, Tom Connor |
| EPI_ISL_611745 | Virology Department, Sheffield Teaching Hospitals NHS Foundation Trust/Department of Infection, Immunity and Cardiovascular Disease, The Medical School, University of Sheffield | COVID-19 Genomics UK (COG-UK) Consortium | Thushan de Silva, Matthew Parker, Nikki Smith, Adri Angyal, Rebecca Brown, Luke Green, Rachel Tucker, Paul Parsons, Danielle Groves, Katie Johnson, Laura Carrilero, Alex Keeley, Dave Partridge, Matthew Wyles, Benjamin Lindsey, Mehmet Yavuz, Mohammad Raza, Cariad Evans |
| EPI_ISL_611748, EPI_ISL_611749 | Wales Specialist Virology Centre Sequencing lab: Pathogen Genomics Unit | COVID-19 Genomics UK (COG-UK) Consortium | Catherine Moore, Johnathan Evans, Laura Gifford, Malorie Perry, Simon Cottrell, Angela Marchbank, Alec Birchley, Alexander Adams, Amy Gaskin, Bree Gatica-Wilcox, Jason Coombes, Joel Southgate, Lauren Gilbert, Lee Graham, Nicole Pacchiarini, Sara Kumziene-Summerhayes, Sarah Taylor, Sophie Jones, Sara Rey, Matthew Bull, Joanne Watkins, Sally Corden, Tom Connor |
| EPI_ISL_611753 | Queens Medical Centre, Clinical Microbiology Department / DeepSeq Nottingham | COVID-19 Genomics UK (COG-UK) Consortium | Gemma Clark, Wendy Smith, Manjinder Khakh, Vicki M Fleming, Michelle M Lister, Hannah Howson-Wells, Jonathan Ball, Patrick McClure, Joseph Chappell, Theocharis Tsoleridis, Nadine Holmes, Matthew Carlisle, Christopher Moore, Fei Sang, Johnny Debebe, Victoria Wright, Matthew Loose |
| EPI_ISL_611754, EPI_ISL_611755, EPI_ISL_611761, EPI_ISL_611772, EPI_ISL_611773 | Centre for Enzyme Innovation, University of Portsmouth / Translational Research Laboratory, Portsmouth Hospitals NHS Trust | COVID-19 Genomics UK (COG-UK) Consortium | Angela Beckett,Yann Bourgeois,Garry Scarlett,Sharon Glaysher,Scott Elliott,Kelly Bicknell,Robert Impey,Allyson Lloyd,Sarah Wyllie,Ethan Butcher,Anoop Chauhan,Samuel Robson |
| EPI_ISL_611774, EPI_ISL_611775, EPI_ISL_611777, EPI_ISL_611778, EPI_ISL_611779, EPI_ISL_611780, EPI_ISL_611781, EPI_ISL_611782, EPI_ISL_611783, EPI_ISL_611788 | Virology Department, Sheffield Teaching Hospitals NHS Foundation Trust/Department of Infection, Immunity and Cardiovascular Disease, The Medical School, University of Sheffield | COVID-19 Genomics UK (COG-UK) Consortium | Thushan de Silva, Matthew Parker, Nikki Smith, Adri Angyal, Rebecca Brown, Luke Green, Rachel Tucker, Paul Parsons, Danielle Groves, Katie Johnson, Laura Carrilero, Alex Keeley, Dave Partridge, Matthew Wyles, Benjamin Lindsey, Mehmet Yavuz, Mohammad Raza, Cariad Evans |
| EPI_ISL_611792 | Wales Specialist Virology Centre Sequencing lab: Pathogen Genomics Unit | COVID-19 Genomics UK (COG-UK) Consortium | Catherine Moore, Johnathan Evans, Laura Gifford, Malorie Perry, Simon Cottrell, Angela Marchbank, Alec Birchley, Alexander Adams, Amy Gaskin, Bree Gatica-Wilcox, Jason Coombes, Joel Southgate, Lauren Gilbert, Lee Graham, Nicole Pacchiarini, Sara Kumziene-Summerhayes, Sarah Taylor, Sophie Jones, Sara Rey, Matthew Bull, Joanne Watkins, Sally Corden, Tom Connor |
| EPI_ISL_611793, EPI_ISL_611797, EPI_ISL_611798, EPI_ISL_611801, EPI_ISL_611802, EPI_ISL_611804 | Virology Department, Sheffield Teaching Hospitals NHS Foundation Trust/Department of Infection, Immunity and Cardiovascular Disease, The Medical School, University of Sheffield | COVID-19 Genomics UK (COG-UK) Consortium | Thushan de Silva, Matthew Parker, Nikki Smith, Adri Angyal, Rebecca Brown, Luke Green, Rachel Tucker, Paul Parsons, Danielle Groves, Katie Johnson, Laura Carrilero, Alex Keeley, Dave Partridge, Matthew Wyles, Benjamin Lindsey, Mehmet Yavuz, Mohammad Raza, Cariad Evans |
| EPI_ISL_611809 | Wales Specialist Virology Centre Sequencing lab: Pathogen Genomics Unit | COVID-19 Genomics UK (COG-UK) Consortium | Catherine Moore, Johnathan Evans, Laura Gifford, Malorie Perry, Simon Cottrell, Angela Marchbank, Alec Birchley, Alexander Adams, Amy Gaskin, Bree Gatica-Wilcox, Jason Coombes, Joel Southgate, Lauren Gilbert, Lee Graham, Nicole Pacchiarini, Sara Kumziene-Summerhayes, Sarah Taylor, Sophie Jones, Sara Rey, Matthew Bull, Joanne Watkins, Sally Corden, Tom Connor |
| EPI_ISL_611810, EPI_ISL_611813 | Queens Medical Centre, Clinical Microbiology Department / DeepSeq Nottingham | COVID-19 Genomics UK (COG-UK) Consortium | Gemma Clark, Wendy Smith, Manjinder Khakh, Vicki M Fleming, Michelle M Lister, Hannah Howson-Wells, Jonathan Ball, Patrick McClure, Joseph Chappell, Theocharis Tsoleridis, Nadine Holmes, Matthew Carlisle, Christopher Moore, Fei Sang, Johnny Debebe, Victoria Wright, Matthew Loose |
| EPI_ISL_611817 | Virology Department, Sheffield Teaching Hospitals NHS Foundation Trust/Department of Infection, Immunity and Cardiovascular Disease, The Medical School, University of Sheffield | COVID-19 Genomics UK (COG-UK) Consortium | Thushan de Silva, Matthew Parker, Nikki Smith, Adri Angyal, Rebecca Brown, Luke Green, Rachel Tucker, Paul Parsons, Danielle Groves, Katie Johnson, Laura Carrilero, Alex Keeley, Dave Partridge, Matthew Wyles, Benjamin Lindsey, Mehmet Yavuz, Mohammad Raza, Cariad Evans |
| EPI_ISL_611822 | University College London, Great Ormond Street Hospital for Children NHS Foundation Trust, Imperial College Healthcare NHS Trust | COVID-19 Genomics UK (COG-UK) Consortium | Sergi Castellano, Rachel Williams, Mark Kristiansen, Paola Resende Silva, Sunando Roy, Tony Brooks, Helena Tutill, Paola Niola, Patricia Dyal, Charlotte Williams, Leysa Forrest, Yasmin Panchbhaya, Jacqueline Findlay, Samuel Weeks, Julianne Brown, Kathryn Harris, Paul Randell, James Price, Alison Holmes, Judith Breuer |
| EPI_ISL_611824, EPI_ISL_611828 | Wales Specialist Virology Centre Sequencing lab: Pathogen Genomics Unit | COVID-19 Genomics UK (COG-UK) Consortium | Catherine Moore, Johnathan Evans, Laura Gifford, Malorie Perry, Simon Cottrell, Angela Marchbank, Alec Birchley, Alexander Adams, Amy Gaskin, Bree Gatica-Wilcox, Jason Coombes, Joel Southgate, Lauren Gilbert, Lee Graham, Nicole Pacchiarini, Sara Kumziene-Summerhayes, Sarah Taylor, Sophie Jones, Sara Rey, Matthew Bull, Joanne Watkins, Sally Corden, Tom Connor |
| EPI_ISL_611838 | University of Exeter | COVID-19 Genomics UK (COG-UK) Consortium | Ben Temperton,Aaron Jeffries,Michelle Michelsen,Joanna Warwick-Dugdale,Audrey Farbos,Robyn Manley,Stephen Michell,Jane Masoli |
| EPI_ISL_611842, EPI_ISL_611850 | Virology Department, Royal Infirmary of Edinburgh, NHS Lothian / School of Biological Sciences, University of Edinburgh / Institute of Genetics and Molecular Medicine, University of Edinburgh | COVID-19 Genomics UK (COG-UK) Consortium | McHugh M, Dewar R, Rooke S, Gallagher M, Balcaza C, O'Toole Á, Scher E, Hill V, McCrone JT, Colquhoun R, Yu X, Jackson B, Rambaut A, Williams TC, Templeton K |
| EPI_ISL_611854 | Wales Specialist Virology Centre Sequencing lab: Pathogen Genomics Unit | COVID-19 Genomics UK (COG-UK) Consortium | Catherine Moore, Johnathan Evans, Laura Gifford, Malorie Perry, Simon Cottrell, Angela Marchbank, Alec Birchley, Alexander Adams, Amy Gaskin, Bree Gatica-Wilcox, Jason Coombes, Joel Southgate, Lauren Gilbert, Lee Graham, Nicole Pacchiarini, Sara Kumziene-Summerhayes, Sarah Taylor, Sophie Jones, Sara Rey, Matthew Bull, Joanne Watkins, Sally Corden, Tom Connor |
| EPI_ISL_611861 | Virology Department, Sheffield Teaching Hospitals NHS Foundation Trust/Department of Infection, Immunity and Cardiovascular Disease, The Medical School, University of Sheffield | COVID-19 Genomics UK (COG-UK) Consortium | Thushan de Silva, Matthew Parker, Nikki Smith, Adri Angyal, Rebecca Brown, Luke Green, Rachel Tucker, Paul Parsons, Danielle Groves, Katie Johnson, Laura Carrilero, Alex Keeley, Dave Partridge, Matthew Wyles, Benjamin Lindsey, Mehmet Yavuz, Mohammad Raza, Cariad Evans |
| EPI_ISL_611863, EPI_ISL_611887, EPI_ISL_611888 | Wales Specialist Virology Centre Sequencing lab: Pathogen Genomics Unit | COVID-19 Genomics UK (COG-UK) Consortium | Catherine Moore, Johnathan Evans, Laura Gifford, Malorie Perry, Simon Cottrell, Angela Marchbank, Alec Birchley, Alexander Adams, Amy Gaskin, Bree Gatica-Wilcox, Jason Coombes, Joel Southgate, Lauren Gilbert, Lee Graham, Nicole Pacchiarini, Sara Kumziene-Summerhayes, Sarah Taylor, Sophie Jones, Sara Rey, Matthew Bull, Joanne Watkins, Sally Corden, Tom Connor |
| EPI_ISL_611894 | Virology Department, Royal Infirmary of Edinburgh, NHS Lothian / School of Biological Sciences, University of Edinburgh / Institute of Genetics and Molecular Medicine, University of Edinburgh | COVID-19 Genomics UK (COG-UK) Consortium | McHugh M, Dewar R, Rooke S, Gallagher M, Balcaza C, O'Toole Á, Scher E, Hill V, McCrone JT, Colquhoun R, Yu X, Jackson B, Rambaut A, Williams TC, Templeton K |
| EPI_ISL_611901, EPI_ISL_611902, EPI_ISL_611909, EPI_ISL_611913 | Wales Specialist Virology Centre Sequencing lab: Pathogen Genomics Unit | COVID-19 Genomics UK (COG-UK) Consortium | Catherine Moore, Johnathan Evans, Laura Gifford, Malorie Perry, Simon Cottrell, Angela Marchbank, Alec Birchley, Alexander Adams, Amy Gaskin, Bree Gatica-Wilcox, Jason Coombes, Joel Southgate, Lauren Gilbert, Lee Graham, Nicole Pacchiarini, Sara Kumziene-Summerhayes, Sarah Taylor, Sophie Jones, Sara Rey, Matthew Bull, Joanne Watkins, Sally Corden, Tom Connor |
| EPI_ISL_611920 | Virology Department, Sheffield Teaching Hospitals NHS Foundation Trust/Department of Infection, Immunity and Cardiovascular Disease, The Medical School, University of Sheffield | COVID-19 Genomics UK (COG-UK) Consortium | Thushan de Silva, Matthew Parker, Nikki Smith, Adri Angyal, Rebecca Brown, Luke Green, Rachel Tucker, Paul Parsons, Danielle Groves, Katie Johnson, Laura Carrilero, Alex Keeley, Dave Partridge, Matthew Wyles, Benjamin Lindsey, Mehmet Yavuz, Mohammad Raza, Cariad Evans |
| EPI_ISL_611922 | Wales Specialist Virology Centre Sequencing lab: Pathogen Genomics Unit | COVID-19 Genomics UK (COG-UK) Consortium | Catherine Moore, Johnathan Evans, Laura Gifford, Malorie Perry, Simon Cottrell, Angela Marchbank, Alec Birchley, Alexander Adams, Amy Gaskin, Bree Gatica-Wilcox, Jason Coombes, Joel Southgate, Lauren Gilbert, Lee Graham, Nicole Pacchiarini, Sara Kumziene-Summerhayes, Sarah Taylor, Sophie Jones, Sara Rey, Matthew Bull, Joanne Watkins, Sally Corden, Tom Connor |
| EPI_ISL_611924 | Virology Department, Royal Infirmary of Edinburgh, NHS | COVID-19 Genomics UK (COG-UK) Consortium | McHugh M, Dewar R, Rooke S, Gallagher M, Balcaza C, O'Toole Á, Scher E, Hill V, McCrone JT, Colquhoun R, Yu X, Jackson B, Rambaut A, Williams TC, |

| | | | |
|---|---|---|---|
| | Lothian / School of Biological Sciences, University of Edinburgh / Institute of Genetics and Molecular Medicine, University of Edinburgh | | Templeton K |
| EPI_ISL_611934 | Centre for Enzyme Innovation, University of Portsmouth / Translational Research Laboratory, Portsmouth Hospitals NHS Trust | COVID-19 Genomics UK (COG-UK) Consortium | Angela Beckett,Yann Bourgeois,Garry Scarlett,Sharon Glaysher,Scott Elliott,Kelly Bicknell,Robert Impey,Allyson Lloyd,Sarah Wyllie,Ethan Butcher,Anoop Chauhan,Samuel Robson |
| EPI_ISL_611938 | Department of Pathology, University of Cambridge | COVID-19 Genomics UK (COG-UK) Consortium | Aminu S. Jahun, Yasmin Chaudhry, Grant Hall, Iliana Georgana, Myra Hosmillo, Martin D. Curran, Malte Pinckert, Surendra Parmar, Ian Goodfellow |
| EPI_ISL_611960 | Centre for Enzyme Innovation, University of Portsmouth / Translational Research Laboratory, Portsmouth Hospitals NHS Trust | COVID-19 Genomics UK (COG-UK) Consortium | Angela Beckett,Yann Bourgeois,Garry Scarlett,Sharon Glaysher,Scott Elliott,Kelly Bicknell,Robert Impey,Allyson Lloyd,Sarah Wyllie,Ethan Butcher,Anoop Chauhan,Samuel Robson |
| EPI_ISL_611964, EPI_ISL_611965, EPI_ISL_611966 | Department of Pathology, University of Cambridge | COVID-19 Genomics UK (COG-UK) Consortium | Aminu S. Jahun, Yasmin Chaudhry, Grant Hall, Iliana Georgana, Myra Hosmillo, Martin D. Curran, Malte Pinckert, Surendra Parmar, Ian Goodfellow |
| EPI_ISL_611972, EPI_ISL_611975 | Wales Specialist Virology Centre Sequencing lab: Pathogen Genomics Unit | COVID-19 Genomics UK (COG-UK) Consortium | Catherine Moore, Johnathan Evans, Laura Gifford, Malorie Perry, Simon Cottrell, Angela Marchbank, Alec Birchley, Alexander Adams, Amy Gaskin, Bree Gatica-Wilcox, Jason Coombes, Joel Southgate, Lauren Gilbert, Lee Graham, Nicole Pacchiarini, Sara Kumziene-Summerhayes, Sarah Taylor, Sophie Jones, Sara Rey, Matthew Bull, Joanne Watkins, Sally Corden, Tom Connor |
| EPI_ISL_611979 | Virology Department, Sheffield Teaching Hospitals NHS Foundation Trust/Department of Infection, Immunity and Cardiovascular Disease, The Medical School, University of Sheffield | COVID-19 Genomics UK (COG-UK) Consortium | Thushan de Silva, Matthew Parker, Nikki Smith, Adri Angyal, Rebecca Brown, Luke Green, Rachel Tucker, Paul Parsons, Danielle Groves, Katie Johnson, Laura Carrilero, Alex Keeley, Dave Partridge, Matthew Wyles, Benjamin Lindsey, Mehmet Yavuz, Mohammad Raza, Cariad Evans |
| EPI_ISL_611983 | Centre for Enzyme Innovation, University of Portsmouth / Translational Research Laboratory, Portsmouth Hospitals NHS Trust | COVID-19 Genomics UK (COG-UK) Consortium | Angela Beckett,Yann Bourgeois,Garry Scarlett,Sharon Glaysher,Scott Elliott,Kelly Bicknell,Robert Impey,Allyson Lloyd,Sarah Wyllie,Ethan Butcher,Anoop Chauhan,Samuel Robson |
| EPI_ISL_611986 | Wales Specialist Virology Centre Sequencing lab: Pathogen Genomics Unit | COVID-19 Genomics UK (COG-UK) Consortium | Catherine Moore, Johnathan Evans, Laura Gifford, Malorie Perry, Simon Cottrell, Angela Marchbank, Alec Birchley, Alexander Adams, Amy Gaskin, Bree Gatica-Wilcox, Jason Coombes, Joel Southgate, Lauren Gilbert, Lee Graham, Nicole Pacchiarini, Sara Kumziene-Summerhayes, Sarah Taylor, Sophie Jones, Sara Rey, Matthew Bull, Joanne Watkins, Sally Corden, Tom Connor |
| EPI_ISL_611987 | University of Exeter | COVID-19 Genomics UK (COG-UK) Consortium | Ben Temperton,Aaron Jeffries,Michelle Michelsen,Joanna Warwick-Dugdale,Audrey Farbos,Robyn Manley,Stephen Michell,Jane Masoli |
| EPI_ISL_611992, EPI_ISL_611993 | Wales Specialist Virology Centre Sequencing lab: Pathogen Genomics Unit | COVID-19 Genomics UK (COG-UK) Consortium | Catherine Moore, Johnathan Evans, Laura Gifford, Malorie Perry, Simon Cottrell, Angela Marchbank, Alec Birchley, Alexander Adams, Amy Gaskin, Bree Gatica-Wilcox, Jason Coombes, Joel Southgate, Lauren Gilbert, Lee Graham, Nicole Pacchiarini, Sara Kumziene-Summerhayes, Sarah Taylor, Sophie Jones, Sara Rey, Matthew Bull, Joanne Watkins, Sally Corden, Tom Connor |
| EPI_ISL_611996 | University College London, Great Ormond Street Hospital for Children NHS Foundation Trust, Imperial College Healthcare NHS Trust | COVID-19 Genomics UK (COG-UK) Consortium | Sergi Castellano, Rachel Williams, Mark Kristiansen, Paola Resende Silva, Sunando Roy, Tony Brooks, Helena Tutill, Paola Niola, Patricia Dyal, Charlotte Williams, Leysa Forrest, Yasmin Panchbhaya, Jacqueline Findlay, Samuel Weeks, Julianne Brown, Kathryn Harris, Paul Randell, James Price, Alison Holmes, Judith Breuer |
| EPI_ISL_611997 | Virology Department, Sheffield Teaching Hospitals NHS Foundation Trust/Department of Infection, Immunity and Cardiovascular Disease, The Medical School, University of Sheffield | COVID-19 Genomics UK (COG-UK) Consortium | Thushan de Silva, Matthew Parker, Nikki Smith, Adri Angyal, Rebecca Brown, Luke Green, Rachel Tucker, Paul Parsons, Danielle Groves, Katie Johnson, Laura Carrilero, Alex Keeley, Dave Partridge, Matthew Wyles, Benjamin Lindsey, Mehmet Yavuz, Mohammad Raza, Cariad Evans |
| EPI_ISL_611998 | Centre for Enzyme Innovation, University of Portsmouth / Translational Research Laboratory, Portsmouth Hospitals NHS Trust | COVID-19 Genomics UK (COG-UK) Consortium | Angela Beckett,Yann Bourgeois,Garry Scarlett,Sharon Glaysher,Scott Elliott,Kelly Bicknell,Robert Impey,Allyson Lloyd,Sarah Wyllie,Ethan Butcher,Anoop Chauhan,Samuel Robson |
| EPI_ISL_612002 | Wales Specialist Virology Centre Sequencing lab: Pathogen Genomics Unit | COVID-19 Genomics UK (COG-UK) Consortium | Catherine Moore, Johnathan Evans, Laura Gifford, Malorie Perry, Simon Cottrell, Angela Marchbank, Alec Birchley, Alexander Adams, Amy Gaskin, Bree Gatica-Wilcox, Jason Coombes, Joel Southgate, Lauren Gilbert, Lee Graham, Nicole Pacchiarini, Sara Kumziene-Summerhayes, Sarah Taylor, Sophie Jones, Sara Rey, Matthew Bull, Joanne Watkins, Sally Corden, Tom Connor |
| EPI_ISL_612005 | Virology Department, Sheffield Teaching Hospitals NHS Foundation Trust/Department of Infection, Immunity and Cardiovascular Disease, The Medical School, University of Sheffield | COVID-19 Genomics UK (COG-UK) Consortium | Thushan de Silva, Matthew Parker, Nikki Smith, Adri Angyal, Rebecca Brown, Luke Green, Rachel Tucker, Paul Parsons, Danielle Groves, Katie Johnson, Laura Carrilero, Alex Keeley, Dave Partridge, Matthew Wyles, Benjamin Lindsey, Mehmet Yavuz, Mohammad Raza, Cariad Evans |
| EPI_ISL_612006 | Wales Specialist Virology Centre Sequencing lab: Pathogen Genomics Unit | COVID-19 Genomics UK (COG-UK) Consortium | Catherine Moore, Johnathan Evans, Laura Gifford, Malorie Perry, Simon Cottrell, Angela Marchbank, Alec Birchley, Alexander Adams, Amy Gaskin, Bree Gatica-Wilcox, Jason Coombes, Joel Southgate, Lauren Gilbert, Lee Graham, Nicole Pacchiarini, Sara Kumziene-Summerhayes, Sarah Taylor, Sophie Jones, Sara Rey, Matthew Bull, Joanne Watkins, Sally Corden, Tom Connor |
| EPI_ISL_612014 | University College London, Great Ormond Street Hospital for Children NHS Foundation Trust, Imperial College Healthcare NHS Trust | COVID-19 Genomics UK (COG-UK) Consortium | Sergi Castellano, Rachel Williams, Mark Kristiansen, Paola Resende Silva, Sunando Roy, Tony Brooks, Helena Tutill, Paola Niola, Patricia Dyal, Charlotte Williams, Leysa Forrest, Yasmin Panchbhaya, Jacqueline Findlay, Samuel Weeks, Julianne Brown, Kathryn Harris, Paul Randell, James Price, Alison Holmes, Judith Breuer |
| EPI_ISL_612022, EPI_ISL_612023 | Virology Department, Sheffield Teaching Hospitals NHS Foundation Trust/Department of Infection, Immunity and Cardiovascular Disease, The Medical School, University of Sheffield | COVID-19 Genomics UK (COG-UK) Consortium | Thushan de Silva, Matthew Parker, Nikki Smith, Adri Angyal, Rebecca Brown, Luke Green, Rachel Tucker, Paul Parsons, Danielle Groves, Katie Johnson, Laura Carrilero, Alex Keeley, Dave Partridge, Matthew Wyles, Benjamin Lindsey, Mehmet Yavuz, Mohammad Raza, Cariad Evans |
| EPI_ISL_612025, EPI_ISL_612031, EPI_ISL_612038, EPI_ISL_612045, EPI_ISL_612049 | Wales Specialist Virology Centre Sequencing lab: Pathogen Genomics Unit | COVID-19 Genomics UK (COG-UK) Consortium | Catherine Moore, Johnathan Evans, Laura Gifford, Malorie Perry, Simon Cottrell, Angela Marchbank, Alec Birchley, Alexander Adams, Amy Gaskin, Bree Gatica-Wilcox, Jason Coombes, Joel Southgate, Lauren Gilbert, Lee Graham, Nicole Pacchiarini, Sara Kumziene-Summerhayes, Sarah Taylor, Sophie Jones, Sara Rey, Matthew Bull, Joanne Watkins, Sally Corden, Tom Connor |
| EPI_ISL_612050 | Virology Department, Sheffield Teaching Hospitals NHS Foundation Trust/Department of Infection, Immunity and Cardiovascular Disease, The Medical School, University of Sheffield | COVID-19 Genomics UK (COG-UK) Consortium | Thushan de Silva, Matthew Parker, Nikki Smith, Adri Angyal, Rebecca Brown, Luke Green, Rachel Tucker, Paul Parsons, Danielle Groves, Katie Johnson, Laura Carrilero, Alex Keeley, Dave Partridge, Matthew Wyles, Benjamin Lindsey, Mehmet Yavuz, Mohammad Raza, Cariad Evans |
| EPI_ISL_612052 | Wales Specialist Virology Centre Sequencing lab: Pathogen Genomics Unit | COVID-19 Genomics UK (COG-UK) Consortium | Catherine Moore, Johnathan Evans, Laura Gifford, Malorie Perry, Simon Cottrell, Angela Marchbank, Alec Birchley, Alexander Adams, Amy Gaskin, Bree Gatica-Wilcox, Jason Coombes, Joel Southgate, Lauren Gilbert, Lee Graham, Nicole Pacchiarini, Sara Kumziene-Summerhayes, Sarah Taylor, Sophie Jones, Sara Rey, Matthew Bull, Joanne Watkins, Sally Corden, Tom Connor |
| EPI_ISL_612055, EPI_ISL_612058, EPI_ISL_612062, EPI_ISL_612067 | Virology Department, Sheffield Teaching Hospitals NHS Foundation Trust/Department of Infection, Immunity and Cardiovascular Disease, The Medical School, University of Sheffield | COVID-19 Genomics UK (COG-UK) Consortium | Thushan de Silva, Matthew Parker, Nikki Smith, Adri Angyal, Rebecca Brown, Luke Green, Rachel Tucker, Paul Parsons, Danielle Groves, Katie Johnson, Laura Carrilero, Alex Keeley, Dave Partridge, Matthew Wyles, Benjamin Lindsey, Mehmet Yavuz, Mohammad Raza, Cariad Evans |
| EPI_ISL_612069 | Centre for Enzyme Innovation, University of Portsmouth / Translational Research Laboratory, Portsmouth Hospitals NHS Trust | COVID-19 Genomics UK (COG-UK) Consortium | Angela Beckett,Yann Bourgeois,Garry Scarlett,Sharon Glaysher,Scott Elliott,Kelly Bicknell,Robert Impey,Allyson Lloyd,Sarah Wyllie,Ethan Butcher,Anoop Chauhan,Samuel Robson |
| EPI_ISL_612074 | Virology Department, Royal Infirmary of Edinburgh, NHS Lothian / School of Biological Sciences, University of | COVID-19 Genomics UK (COG-UK) Consortium | McHugh M, Dewar R, Rooke S, Gallagher M, Balcaza C, O'Toole Á, Scher E, Hill V, McCrone JT, Colquhoun R, Yu X, Jackson B, Rambaut A, Williams TC, Templeton K |

| | | | |
|---|---|---|---|
| | Edinburgh / Institute of Genetics and Molecular Medicine, University of Edinburgh | | |
| EPI_ISL_612076 | Wales Specialist Virology Centre Sequencing lab: Pathogen Genomics Unit | COVID-19 Genomics UK (COG-UK) Consortium | Catherine Moore, Johnathan Evans, Laura Gifford, Malorie Perry, Simon Cottrell, Angela Marchbank, Alec Birchley, Alexander Adams, Amy Gaskin, Bree Gatica-Wilcox, Jason Coombes, Joel Southgate, Lauren Gilbert, Lee Graham, Nicole Pacchiarini, Sara Kumziene-Summerhayes, Sarah Taylor, Sophie Jones, Sara Rey, Matthew Bull, Joanne Watkins, Sally Corden, Tom Connor |
| EPI_ISL_612077 | Queens Medical Centre, Clinical Microbiology Department / DeepSeq Nottingham | COVID-19 Genomics UK (COG-UK) Consortium | Gemma Clark, Wendy Smith, Manjinder Khakh, Vicki M Fleming, Michelle M Lister, Hannah Howson-Wells, Jonathan Ball, Patrick McClure, Joseph Chappell, Theocharis Tsoleridis, Nadine Holmes, Matthew Carlisle, Christopher Moore, Fei Sang, Johnny Debebe, Victoria Wright, Matthew Loose |
| EPI_ISL_612078, EPI_ISL_612079 | University College London, Great Ormond Street Hospital for Children NHS Foundation Trust, Imperial College Healthcare NHS Trust | COVID-19 Genomics UK (COG-UK) Consortium | Sergi Castellano, Rachel Williams, Mark Kristiansen, Paola Resende Silva, Sunando Roy, Tony Brooks, Helena Tutill, Paola Niola, Patricia Dyal, Charlotte Williams, Leysa Forrest, Yasmin Panchbhaya, Jacqueline Findlay, Samuel Weeks, Julianne Brown, Kathryn Harris, Paul Randell, James Price, Alison Holmes, Judith Breuer |
| EPI_ISL_612087 | Wales Specialist Virology Centre Sequencing lab: Pathogen Genomics Unit | COVID-19 Genomics UK (COG-UK) Consortium | Catherine Moore, Johnathan Evans, Laura Gifford, Malorie Perry, Simon Cottrell, Angela Marchbank, Alec Birchley, Alexander Adams, Amy Gaskin, Bree Gatica-Wilcox, Jason Coombes, Joel Southgate, Lauren Gilbert, Lee Graham, Nicole Pacchiarini, Sara Kumziene-Summerhayes, Sarah Taylor, Sophie Jones, Sara Rey, Matthew Bull, Joanne Watkins, Sally Corden, Tom Connor |
| EPI_ISL_612089, EPI_ISL_612092, EPI_ISL_612094 | Centre for Enzyme Innovation, University of Portsmouth / Translational Research Laboratory, Portsmouth Hospitals NHS Trust | COVID-19 Genomics UK (COG-UK) Consortium | Angela Beckett,Yann Bourgeois,Garry Scarlett,Sharon Glaysher,Scott Elliott,Kelly Bicknell,Robert Impey,Allyson Lloyd,Sarah Wyllie,Ethan Butcher,Anoop Chauhan,Samuel Robson |
| EPI_ISL_612098 | Wales Specialist Virology Centre Sequencing lab: Pathogen Genomics Unit | COVID-19 Genomics UK (COG-UK) Consortium | Catherine Moore, Johnathan Evans, Laura Gifford, Malorie Perry, Simon Cottrell, Angela Marchbank, Alec Birchley, Alexander Adams, Amy Gaskin, Bree Gatica-Wilcox, Jason Coombes, Joel Southgate, Lauren Gilbert, Lee Graham, Nicole Pacchiarini, Sara Kumziene-Summerhayes, Sarah Taylor, Sophie Jones, Sara Rey, Matthew Bull, Joanne Watkins, Sally Corden, Tom Connor |
| EPI_ISL_612099 | Department of Pathology, University of Cambridge | COVID-19 Genomics UK (COG-UK) Consortium | Aminu S. Jahun, Yasmin Chaudhry, Grant Hall, Iliana Georgana, Myra Hosmillo, Martin D. Curran, Malte Pinckert, Surendra Parmar, Ian Goodfellow |
| EPI_ISL_612102 | Virology Department, Sheffield Teaching Hospitals NHS Foundation Trust/Department of Infection, Immunity and Cardiovascular Disease, The Medical School, University of Sheffield | COVID-19 Genomics UK (COG-UK) Consortium | Thushan de Silva, Matthew Parker, Nikki Smith, Adri Angyal, Rebecca Brown, Luke Green, Rachel Tucker, Paul Parsons, Danielle Groves, Katie Johnson, Laura Carrilero, Alex Keeley, Dave Partridge, Matthew Wyles, Benjamin Lindsey, Mehmet Yavuz, Mohammad Raza, Cariad Evans |
| EPI_ISL_612109, EPI_ISL_612119 | Wales Specialist Virology Centre Sequencing lab: Pathogen Genomics Unit | COVID-19 Genomics UK (COG-UK) Consortium | Catherine Moore, Johnathan Evans, Laura Gifford, Malorie Perry, Simon Cottrell, Angela Marchbank, Alec Birchley, Alexander Adams, Amy Gaskin, Bree Gatica-Wilcox, Jason Coombes, Joel Southgate, Lauren Gilbert, Lee Graham, Nicole Pacchiarini, Sara Kumziene-Summerhayes, Sarah Taylor, Sophie Jones, Sara Rey, Matthew Bull, Joanne Watkins, Sally Corden, Tom Connor |
| EPI_ISL_612123 | Department of Pathology, University of Cambridge | COVID-19 Genomics UK (COG-UK) Consortium | Aminu S. Jahun, Yasmin Chaudhry, Grant Hall, Iliana Georgana, Myra Hosmillo, Martin D. Curran, Malte Pinckert, Surendra Parmar, Ian Goodfellow |
| EPI_ISL_612127, EPI_ISL_612128, EPI_ISL_612129 | Virology Department, Royal Infirmary of Edinburgh, NHS Lothian / School of Biological Sciences, University of Edinburgh / Institute of Genetics and Molecular Medicine, University of Edinburgh | COVID-19 Genomics UK (COG-UK) Consortium | McHugh M, Dewar R, Rooke S, Gallagher M, Balcaza C, O'Toole Á, Scher E, Hill V, McCrone JT, Colquhoun R, Yu X, Jackson B, Rambaut A, Williams TC, Templeton K |
| EPI_ISL_612136, EPI_ISL_612140, EPI_ISL_612143, EPI_ISL_612148, EPI_ISL_612150, EPI_ISL_612151 | Wales Specialist Virology Centre Sequencing lab: Pathogen Genomics Unit | COVID-19 Genomics UK (COG-UK) Consortium | Catherine Moore, Johnathan Evans, Laura Gifford, Malorie Perry, Simon Cottrell, Angela Marchbank, Alec Birchley, Alexander Adams, Amy Gaskin, Bree Gatica-Wilcox, Jason Coombes, Joel Southgate, Lauren Gilbert, Lee Graham, Nicole Pacchiarini, Sara Kumziene-Summerhayes, Sarah Taylor, Sophie Jones, Sara Rey, Matthew Bull, Joanne Watkins, Sally Corden, Tom Connor |
| EPI_ISL_612158, EPI_ISL_612159 | Queens Medical Centre, Clinical Microbiology Department / DeepSeq Nottingham | COVID-19 Genomics UK (COG-UK) Consortium | Gemma Clark, Wendy Smith, Manjinder Khakh, Vicki M Fleming, Michelle M Lister, Hannah Howson-Wells, Jonathan Ball, Patrick McClure, Joseph Chappell, Theocharis Tsoleridis, Nadine Holmes, Matthew Carlisle, Christopher Moore, Fei Sang, Johnny Debebe, Victoria Wright, Matthew Loose |
| EPI_ISL_612160, EPI_ISL_612161, EPI_ISL_612172, EPI_ISL_612173, EPI_ISL_612174, EPI_ISL_612175, EPI_ISL_612176, EPI_ISL_612178, EPI_ISL_612179, EPI_ISL_612180 | Wales Specialist Virology Centre Sequencing lab: Pathogen Genomics Unit | COVID-19 Genomics UK (COG-UK) Consortium | Catherine Moore, Johnathan Evans, Laura Gifford, Malorie Perry, Simon Cottrell, Angela Marchbank, Alec Birchley, Alexander Adams, Amy Gaskin, Bree Gatica-Wilcox, Jason Coombes, Joel Southgate, Lauren Gilbert, Lee Graham, Nicole Pacchiarini, Sara Kumziene-Summerhayes, Sarah Taylor, Sophie Jones, Sara Rey, Matthew Bull, Joanne Watkins, Sally Corden, Tom Connor |
| EPI_ISL_612181, EPI_ISL_612182, EPI_ISL_612183 | Queens Medical Centre, Clinical Microbiology Department / DeepSeq Nottingham | COVID-19 Genomics UK (COG-UK) Consortium | Gemma Clark, Wendy Smith, Manjinder Khakh, Vicki M Fleming, Michelle M Lister, Hannah Howson-Wells, Jonathan Ball, Patrick McClure, Joseph Chappell, Theocharis Tsoleridis, Nadine Holmes, Matthew Carlisle, Christopher Moore, Fei Sang, Johnny Debebe, Victoria Wright, Matthew Loose |
| EPI_ISL_612186 | Department of Pathology, University of Cambridge | COVID-19 Genomics UK (COG-UK) Consortium | Aminu S. Jahun, Yasmin Chaudhry, Grant Hall, Iliana Georgana, Myra Hosmillo, Martin D. Curran, Malte Pinckert, Surendra Parmar, Ian Goodfellow |
| EPI_ISL_612187, EPI_ISL_612188, EPI_ISL_612189 | Virology Department, Royal Infirmary of Edinburgh, NHS Lothian / School of Biological Sciences, University of Edinburgh / Institute of Genetics and Molecular Medicine, University of Edinburgh | COVID-19 Genomics UK (COG-UK) Consortium | McHugh M, Dewar R, Rooke S, Gallagher M, Balcaza C, O'Toole Á, Scher E, Hill V, McCrone JT, Colquhoun R, Yu X, Jackson B, Rambaut A, Williams TC, Templeton K |
| EPI_ISL_612194, EPI_ISL_612195 | Wales Specialist Virology Centre Sequencing lab: Pathogen Genomics Unit | COVID-19 Genomics UK (COG-UK) Consortium | Catherine Moore, Johnathan Evans, Laura Gifford, Malorie Perry, Simon Cottrell, Angela Marchbank, Alec Birchley, Alexander Adams, Amy Gaskin, Bree Gatica-Wilcox, Jason Coombes, Joel Southgate, Lauren Gilbert, Lee Graham, Nicole Pacchiarini, Sara Kumziene-Summerhayes, Sarah Taylor, Sophie Jones, Sara Rey, Matthew Bull, Joanne Watkins, Sally Corden, Tom Connor |
| EPI_ISL_612197 | Queens Medical Centre, Clinical Microbiology Department / DeepSeq Nottingham | COVID-19 Genomics UK (COG-UK) Consortium | Gemma Clark, Wendy Smith, Manjinder Khakh, Vicki M Fleming, Michelle M Lister, Hannah Howson-Wells, Jonathan Ball, Patrick McClure, Joseph Chappell, Theocharis Tsoleridis, Nadine Holmes, Matthew Carlisle, Christopher Moore, Fei Sang, Johnny Debebe, Victoria Wright, Matthew Loose |
| EPI_ISL_612277, EPI_ISL_612278, EPI_ISL_612279, EPI_ISL_612281, EPI_ISL_612282, EPI_ISL_612283 | Department of Pathology, University of Cambridge | COVID-19 Genomics UK (COG-UK) Consortium | Aminu S. Jahun, Yasmin Chaudhry, Grant Hall, Iliana Georgana, Myra Hosmillo, Martin D. Curran, Malte Pinckert, Surendra Parmar, Ian Goodfellow |
| EPI_ISL_612341, EPI_ISL_612342, EPI_ISL_612343, EPI_ISL_612345, EPI_ISL_612346, EPI_ISL_612348, EPI_ISL_612349, EPI_ISL_612350, EPI_ISL_612351, EPI_ISL_612352, EPI_ISL_612353, EPI_ISL_612354, EPI_ISL_612355, EPI_ISL_612356, EPI_ISL_612357, EPI_ISL_612358, EPI_ISL_612359, EPI_ISL_612360, EPI_ISL_612361, EPI_ISL_612362, EPI_ISL_612363, EPI_ISL_612364, EPI_ISL_612365, EPI_ISL_612366, EPI_ISL_612367, EPI_ISL_612368, EPI_ISL_612369, EPI_ISL_612370, EPI_ISL_612371, EPI_ISL_612372, EPI_ISL_612374 | | | |
| see above | Virology Department, Royal Infirmary of Edinburgh, NHS Lothian / School of Biological Sciences, University of Edinburgh / Institute of Genetics and Molecular Medicine, University of Edinburgh | COVID-19 Genomics UK (COG-UK) Consortium | McHugh M, Dewar R, Rooke S, Gallagher M, Balcaza C, O'Toole Á, Scher E, Hill V, McCrone JT, Colquhoun R, Yu X, Jackson B, Rambaut A, Williams TC, Templeton K |
| EPI_ISL_612400 | University of Exeter | COVID-19 Genomics UK (COG-UK) Consortium | Ben Temperton,Aaron Jeffries,Michelle Michelsen,Joanna Warwick-Dugdale,Audrey Farbos,Robyn Manley,Stephen Michell,Jane Masoli |
| EPI_ISL_612450, EPI_ISL_612452, EPI_ISL_612453 | University College London, Great Ormond Street Hospital for Children NHS Foundation Trust, Imperial College Healthcare NHS Trust | COVID-19 Genomics UK (COG-UK) Consortium | Sergi Castellano, Rachel Williams, Mark Kristiansen, Paola Resende Silva, Sunando Roy, Tony Brooks, Helena Tutill, Paola Niola, Patricia Dyal, Charlotte Williams, Leysa Forrest, Yasmin Panchbhaya, Jacqueline Findlay, Samuel Weeks, Julianne Brown, Kathryn Harris, Paul Randell, James Price, Alison Holmes, Judith Breuer |
| EPI_ISL_612571, EPI_ISL_612572, EPI_ISL_612573 | Queens Medical Centre, Clinical Microbiology Department / DeepSeq Nottingham | COVID-19 Genomics UK (COG-UK) Consortium | Gemma Clark, Wendy Smith, Manjinder Khakh, Vicki M Fleming, Michelle M Lister, Hannah Howson-Wells, Jonathan Ball, Patrick McClure, Joseph Chappell, Theocharis Tsoleridis, Nadine Holmes, Matthew Carlisle, Christopher Moore, Fei Sang, Johnny Debebe, Victoria Wright, Matthew Loose |
| EPI_ISL_612611, EPI_ISL_612614, EPI_ISL_612618, EPI_ISL_612619, EPI_ISL_612623, EPI_ISL_612626, EPI_ISL_612630, EPI_ISL_612633, EPI_ISL_612634, EPI_ISL_612638, EPI_ISL_612640, EPI_ISL_612645, EPI_ISL_612648, EPI_ISL_612649, EPI_ISL_612652, EPI_ISL_612653, EPI_ISL_612654, EPI_ISL_612655, EPI_ISL_612659, EPI_ISL_612660, EPI_ISL_612667, EPI_ISL_612673, EPI_ISL_612674, EPI_ISL_612675, EPI_ISL_612676, EPI_ISL_612677, EPI_ISL_612683, EPI_ISL_612685, EPI_ISL_612686, EPI_ISL_612689, EPI_ISL_612692, EPI_ISL_612694, EPI_ISL_612698, EPI_ISL_612699, EPI_ISL_612705, EPI_ISL_612723, EPI_ISL_612724, EPI_ISL_612729, EPI_ISL_612730, EPI_ISL_612734, EPI_ISL_612735, EPI_ISL_612736, EPI_ISL_612741, EPI_ISL_612742, EPI_ISL_612743, EPI_ISL_612744, EPI_ISL_612758, EPI_ISL_612761, EPI_ISL_612772, EPI_ISL_612775, EPI_ISL_612776, EPI_ISL_612778, EPI_ISL_612780, EPI_ISL_612783, EPI_ISL_612792, EPI_ISL_612800, EPI_ISL_612801, EPI_ISL_612805, EPI_ISL_612806, EPI_ISL_612807, EPI_ISL_612810, EPI_ISL_612815, EPI_ISL_612820, EPI_ISL_612822, EPI_ISL_612823, EPI_ISL_612825, EPI_ISL_612827, EPI_ISL_612832, EPI_ISL_612837, EPI_ISL_612841, EPI_ISL_612842, EPI_ISL_612846, EPI_ISL_612847, EPI_ISL_612853, EPI_ISL_612856, EPI_ISL_612866, EPI_ISL_612871, EPI_ISL_612872, EPI_ISL_612873, EPI_ISL_612874, EPI_ISL_612876, EPI_ISL_612879, EPI_ISL_612884, EPI_ISL_612886, EPI_ISL_612888, EPI_ISL_612889, EPI_ISL_612890, EPI_ISL_612894, EPI_ISL_612896, EPI_ISL_612898, | | | |

| | | | |
|---|---|---|---|
| EPI_ISL_612900, EPI_ISL_612906, EPI_ISL_612909, EPI_ISL_612912, EPI_ISL_612914, EPI_ISL_612916, EPI_ISL_612919, EPI_ISL_612921, EPI_ISL_612926, EPI_ISL_612929, EPI_ISL_612930, EPI_ISL_612931, EPI_ISL_612945, EPI_ISL_612947, EPI_ISL_612948, EPI_ISL_612949, EPI_ISL_612955, EPI_ISL_612965, EPI_ISL_612966, EPI_ISL_612970, EPI_ISL_612973, EPI_ISL_612980, EPI_ISL_612981, EPI_ISL_612996, EPI_ISL_613000, EPI_ISL_613001, EPI_ISL_613002, EPI_ISL_613004, EPI_ISL_613005, EPI_ISL_613006, EPI_ISL_613007, EPI_ISL_613008, EPI_ISL_613009, EPI_ISL_613011, EPI_ISL_613012, EPI_ISL_613013, EPI_ISL_613015, EPI_ISL_613016, EPI_ISL_613017, EPI_ISL_613022, EPI_ISL_613023, EPI_ISL_613024, EPI_ISL_613027, EPI_ISL_613028, EPI_ISL_613029, EPI_ISL_613030, EPI_ISL_613031, EPI_ISL_613033, EPI_ISL_613034, EPI_ISL_613036, EPI_ISL_613037, EPI_ISL_613038, EPI_ISL_613039, EPI_ISL_613040, EPI_ISL_613041, EPI_ISL_613047, EPI_ISL_613050, EPI_ISL_613052, EPI_ISL_613053, EPI_ISL_613054, EPI_ISL_613055, EPI_ISL_613058, EPI_ISL_613060, EPI_ISL_613062, EPI_ISL_613063, EPI_ISL_613064, EPI_ISL_613065, EPI_ISL_613069, EPI_ISL_613070, EPI_ISL_613071, EPI_ISL_613072, EPI_ISL_613079, EPI_ISL_613081, EPI_ISL_613082, EPI_ISL_613083, EPI_ISL_613084, EPI_ISL_613085, EPI_ISL_613086, EPI_ISL_613087, EPI_ISL_613088, EPI_ISL_613092, EPI_ISL_613093, EPI_ISL_613095, EPI_ISL_613097, EPI_ISL_613099, EPI_ISL_613104, EPI_ISL_613106, EPI_ISL_613108, EPI_ISL_613109, EPI_ISL_613110, EPI_ISL_613111, EPI_ISL_613113, EPI_ISL_613115, EPI_ISL_613117, EPI_ISL_613118, EPI_ISL_613119, EPI_ISL_613120, EPI_ISL_613121, EPI_ISL_613122, EPI_ISL_613123, EPI_ISL_613124, EPI_ISL_613126, EPI_ISL_613127, EPI_ISL_613128, EPI_ISL_613129, EPI_ISL_613130, EPI_ISL_613131, EPI_ISL_613132, EPI_ISL_613133, EPI_ISL_613140, EPI_ISL_613142, EPI_ISL_613143, EPI_ISL_613144, EPI_ISL_613145, EPI_ISL_613146, EPI_ISL_613150, EPI_ISL_613152, EPI_ISL_613154, EPI_ISL_613156, EPI_ISL_613157, EPI_ISL_613158, EPI_ISL_613159, EPI_ISL_613160, EPI_ISL_613161, EPI_ISL_613165, EPI_ISL_613167, EPI_ISL_613169, EPI_ISL_613171, EPI_ISL_613172, EPI_ISL_613174, EPI_ISL_613175, EPI_ISL_613176, EPI_ISL_613179, EPI_ISL_613181, EPI_ISL_613182, EPI_ISL_613183, EPI_ISL_613184, EPI_ISL_613187, EPI_ISL_613188, EPI_ISL_613216, EPI_ISL_613265, EPI_ISL_613271, EPI_ISL_613275, EPI_ISL_613278, EPI_ISL_613280 | | | |
| see above | Wales Specialist Virology Centre Sequencing lab: Pathogen Genomics Unit | COVID-19 Genomics UK (COG-UK) Consortium | Catherine Moore, Johnathan Evans, Laura Gifford, Malorie Perry, Simon Cottrell, Angela Marchbank, Alec Birchley, Alexander Adams, Amy Gaskin, Bree Gatica-Wilcox, Jason Coombes, Joel Southgate, Lauren Gilbert, Lee Graham, Nicole Pacchiarini, Sara Kumziene-Summerhayes, Sarah Taylor, Sophie Jones, Sara Rey, Matthew Bull, Joanne Watkins, Sally Corden, Tom Connor |
| EPI_ISL_613296, EPI_ISL_613297, EPI_ISL_613298 | Centre for Enzyme Innovation, University of Portsmouth / Translational Research Laboratory, Portsmouth Hospitals NHS Trust | COVID-19 Genomics UK (COG-UK) Consortium | Angela Beckett,Yann Bourgeois,Garry Scarlett,Sharon Glaysher,Scott Elliott,Kelly Bicknell,Robert Impey,Allyson Lloyd,Sarah Wyllie,Ethan Butcher,Anoop Chauhan,Samuel Robson |
| EPI_ISL_613307, EPI_ISL_613308, EPI_ISL_613309, EPI_ISL_613310, EPI_ISL_613311, EPI_ISL_613312, EPI_ISL_613313, EPI_ISL_613314, EPI_ISL_613316, EPI_ISL_613317, EPI_ISL_613319, EPI_ISL_613320, EPI_ISL_613321, EPI_ISL_613322, EPI_ISL_613323, EPI_ISL_613324, EPI_ISL_613325, EPI_ISL_613326, EPI_ISL_613327, EPI_ISL_613328, EPI_ISL_613330, EPI_ISL_613331, EPI_ISL_613332, EPI_ISL_613333, EPI_ISL_613334, EPI_ISL_613335, EPI_ISL_613336, EPI_ISL_613337, EPI_ISL_613338, EPI_ISL_613339, EPI_ISL_613340, EPI_ISL_613341, EPI_ISL_613342, EPI_ISL_613343, EPI_ISL_613344, EPI_ISL_613345, EPI_ISL_613346, EPI_ISL_613348, EPI_ISL_613349, EPI_ISL_613350, EPI_ISL_613351, EPI_ISL_613352, EPI_ISL_613353, EPI_ISL_613354, EPI_ISL_613355 | | | |
| see above | Queens Medical Centre, Clinical Microbiology Department / DeepSeq Nottingham | COVID-19 Genomics UK (COG-UK) Consortium | Gemma Clark, Wendy Smith, Manjinder Khakh, Vicki M Fleming, Michelle M Lister, Hannah Howson-Wells, Jonathan Ball, Patrick McClure, Joseph Chappell, Theocharis Tsoleridis, Nadine Holmes, Matthew Carlisle, Christopher Moore, Fei Sang, Johnny Debebe, Victoria Wright, Matthew Loose |
| EPI_ISL_613356, EPI_ISL_613357, EPI_ISL_613359, EPI_ISL_613361, EPI_ISL_613363, EPI_ISL_613365, EPI_ISL_613367, EPI_ISL_613369, EPI_ISL_613370, EPI_ISL_613373, EPI_ISL_613374, EPI_ISL_613376, EPI_ISL_613378, EPI_ISL_613382, EPI_ISL_613385, EPI_ISL_613388, EPI_ISL_613390, EPI_ISL_613394, EPI_ISL_613395, EPI_ISL_613397, EPI_ISL_613398, EPI_ISL_613401, EPI_ISL_613402, EPI_ISL_613404, EPI_ISL_613406 | | | |
| see above | Virology Department, Sheffield Teaching Hospitals NHS Foundation Trust/Department of Infection, Immunity and Cardiovascular Disease, The Medical School, University of Sheffield | COVID-19 Genomics UK (COG-UK) Consortium | Thushan de Silva, Matthew Parker, Nikki Smith, Adri Angyal, Rebecca Brown, Luke Green, Rachel Tucker, Paul Parsons, Danielle Groves, Katie Johnson, Laura Carrilero, Alex Keeley, Dave Partridge, Matthew Wyles, Benjamin Lindsey, Mehmet Yavuz, Mohammad Raza, Cariad Evans |
| EPI_ISL_613503, EPI_ISL_613504, EPI_ISL_613505, EPI_ISL_613506, EPI_ISL_613507, EPI_ISL_613508, EPI_ISL_613509, EPI_ISL_613510, EPI_ISL_613511, EPI_ISL_613512, EPI_ISL_613513, EPI_ISL_613514, EPI_ISL_613515, EPI_ISL_613516, EPI_ISL_613517, EPI_ISL_613518, EPI_ISL_613519, EPI_ISL_613520, EPI_ISL_613521, EPI_ISL_613522, EPI_ISL_613524, EPI_ISL_613525, EPI_ISL_613526, EPI_ISL_613527 | | | |
| see above | Public Health Laboratory - Infectious Disease Lab, Minnesota Department of Health Infectious Disease Laboratory Submission Group | Minnesota Department of Health, Public Health Laboratory | Plumb,M., Garfin,J., Lorentz,A., Wang,X. |
| EPI_ISL_613565, EPI_ISL_613568, EPI_ISL_613569, EPI_ISL_613571, EPI_ISL_613572, EPI_ISL_613573, EPI_ISL_613574, EPI_ISL_613575, EPI_ISL_613576, EPI_ISL_613577, EPI_ISL_613578, EPI_ISL_613579, EPI_ISL_613580, EPI_ISL_613581, EPI_ISL_613582, EPI_ISL_613583, EPI_ISL_613584, EPI_ISL_613585, EPI_ISL_613586, EPI_ISL_613590, EPI_ISL_613591, EPI_ISL_613592, EPI_ISL_613593, EPI_ISL_613594, EPI_ISL_613595, EPI_ISL_613596, EPI_ISL_613597, EPI_ISL_613598, EPI_ISL_613599, EPI_ISL_613600, EPI_ISL_613601, EPI_ISL_613602, EPI_ISL_613603, EPI_ISL_613604, EPI_ISL_613605, EPI_ISL_613606, EPI_ISL_613607, EPI_ISL_613608, EPI_ISL_613609, EPI_ISL_613610, EPI_ISL_613611, EPI_ISL_613612, EPI_ISL_613613, EPI_ISL_613614, EPI_ISL_613615, EPI_ISL_613616, EPI_ISL_613617, EPI_ISL_613618, EPI_ISL_613619, EPI_ISL_613620, EPI_ISL_613621, EPI_ISL_613622, EPI_ISL_613623, EPI_ISL_613624, EPI_ISL_613625, EPI_ISL_613626, EPI_ISL_613627, EPI_ISL_613628, EPI_ISL_613629, EPI_ISL_613630, EPI_ISL_613631, EPI_ISL_613632, EPI_ISL_613957 | | | |
| see above | Respiratory Virus Unit, Microbiology Services Colindale, Public Health England | Respiratory Virus Unit, Microbiology Services Colindale, Public Health England | PHE Covid Sequencing Team |
| EPI_ISL_614158, EPI_ISL_614162, EPI_ISL_614164, EPI_ISL_614169, EPI_ISL_614170, EPI_ISL_614172, EPI_ISL_614173, EPI_ISL_614177, EPI_ISL_614183, EPI_ISL_614184, EPI_ISL_614186, EPI_ISL_614197, EPI_ISL_614200, EPI_ISL_614202, EPI_ISL_614203, EPI_ISL_614204, EPI_ISL_614208, EPI_ISL_614211, EPI_ISL_614212, EPI_ISL_614214, EPI_ISL_614217, EPI_ISL_614220, EPI_ISL_614224, EPI_ISL_614231, EPI_ISL_614233, EPI_ISL_614235, EPI_ISL_614236, EPI_ISL_614238, EPI_ISL_614241, EPI_ISL_614242, EPI_ISL_614243 | | | |
| see above | Michigan Department of Health and Human Services, Bureau of Laboratories | Michigan Department of Health and Human Services, Bureau of Laboratories | Blankenship HM, Riner D, Soehnlen MK |
| EPI_ISL_614286, EPI_ISL_614290 | General practitioner | National Reference Center for Viruses of Respiratory Infections, Institut Pasteur, Paris | Marion Barbet, Sylvie Behillil, Méline Bizard, Angela Brisebarre, Camille Capel, Etienne Simon-Lorière, Vincent Enouf, Maud Vanpeene, Sylvie van der Werf |
| EPI_ISL_614895, EPI_ISL_614896, EPI_ISL_614897, EPI_ISL_614898, EPI_ISL_614899, EPI_ISL_614900, EPI_ISL_614901, EPI_ISL_614902, EPI_ISL_614903, EPI_ISL_614904, EPI_ISL_614905, EPI_ISL_614906, EPI_ISL_614907, EPI_ISL_614908, EPI_ISL_614909, EPI_ISL_614910, EPI_ISL_614911, EPI_ISL_614934, EPI_ISL_614935, EPI_ISL_614936, EPI_ISL_614947, EPI_ISL_614948, EPI_ISL_614949, EPI_ISL_614951, EPI_ISL_614952, EPI_ISL_614953, EPI_ISL_614954, EPI_ISL_614955, EPI_ISL_614956, EPI_ISL_614957, EPI_ISL_614958, EPI_ISL_614959, EPI_ISL_614960, EPI_ISL_614961, EPI_ISL_614962, EPI_ISL_614964 | | | |
| see above | Viollier AG | Department of Biosystems Science and Engineering, ETH Zürich | Christian Beisel, Sarah Nadeau, Ivan Topolsky, Pedro Ferreira, Philipp Jablonski, Susana Posada-Céspedes, Tobias Schär, Ina Nissen, Natascha Santacroce, Elodie Burcklen, Christiane Beckmann, Maurice Redondo, Olivier Kobel, Christoph Noppen, Sophie Seidel, Noemie Santamaria de Souza, Chaoran Chen, Niko Beerenwinkel, Tanja Stadler |
| EPI_ISL_620409 | Utah Public Health Laboratory, Utah Public Health Laboratory Infectious Disease submission group | Utah Public Health Laboratory, Utah Public Health Laboratory Infectious Disease submission group | Young,E.L., Oakeson,K.F. |
| EPI_ISL_622761 | SA Pathology | SA Pathology | Lex Leong, Julien Soubrier, Chuan Kok Lim, Song Gao, Mark Turra, Karin Kassahn, Ivan Bastian, Geoff Higgins |
| EPI_ISL_622777, EPI_ISL_622779 | LabPLUS | Institute of Environmental Science and Research (ESR) | Xiaoyun Ren, Matt Storey, Nikki Freed, Muhammad Faisal, Jing Wang, Hermes Perez, Anja Werno, Antje van der Linden, Arlo Upton, Chris Mansell, David Hammer, Dragana Drinkovic, Gary McAuliffe, Hana Sofia Andersson, James Ussher, Jill Sherwood, Josh Freeman, Julia Howard, Juliet Elvy, Mary DeAlmeida, Matt Blakiston, Matthew Rogers, Max Bloomfield, Michael Addidle, Michelle Balm, Sally Roberts, Sarah Jefferies, Sharmini Muttaiyah, Susan Morpeth, Susan Taylor, Timothy Blackmore, Vani Sathyendran, Veronica Playle, Virginia Hope, Erasmus Smit, Lauren Jelly, Olin Silander, Joep de Ligt |
| EPI_ISL_622793 | LabTests | Institute of Environmental Science and Research (ESR) | Xiaoyun Ren, Matt Storey, Nikki Freed, Muhammad Faisal, Jing Wang, Hermes Perez, Anja Werno, Antje van der Linden, Arlo Upton, Chris Mansell, David Hammer, Dragana Drinkovic, Gary McAuliffe, Hana Sofia Andersson, James Ussher, Jill Sherwood, Josh Freeman, Julia Howard, Juliet Elvy, Mary DeAlmeida, Matt Blakiston, Matthew Rogers, Max Bloomfield, Michael Addidle, Michelle Balm, Sally Roberts, Sarah Jefferies, Sharmini Muttaiyah, Susan Morpeth, Susan Taylor, Timothy Blackmore, Vani Sathyendran, Veronica Playle, Virginia Hope, Erasmus Smit, Lauren Jelly, Olin Silander, Joep de Ligt |
| EPI_ISL_622808 | Canterbury Health Laboratories | Institute of Environmental Science and Research (ESR) | Xiaoyun Ren, Matt Storey, Nikki Freed, Muhammad Faisal, Jing Wang, Hermes Perez, Anja Werno, Antje van der Linden, Arlo Upton, Chris Mansell, David Hammer, Dragana Drinkovic, Gary McAuliffe, Hana Sofia Andersson, James Ussher, Jill Sherwood, Josh Freeman, Julia Howard, Juliet Elvy, Mary DeAlmeida, Matt Blakiston, Matthew Rogers, Max Bloomfield, Michael Addidle, Michelle Balm, Sally Roberts, Sarah Jefferies, Sharmini Muttaiyah, Susan Morpeth, Susan Taylor, Timothy Blackmore, Vani Sathyendran, Veronica Playle, Virginia Hope, Erasmus Smit, Lauren Jelly, Olin Silander, Joep de Ligt |
| EPI_ISL_622844, EPI_ISL_622845, EPI_ISL_622847, EPI_ISL_622849, EPI_ISL_622850, EPI_ISL_622852, EPI_ISL_622853, EPI_ISL_622854, EPI_ISL_622855, EPI_ISL_622856, EPI_ISL_622857, EPI_ISL_622858, EPI_ISL_622859, EPI_ISL_622860, EPI_ISL_622861, EPI_ISL_622862, EPI_ISL_622863, EPI_ISL_622864, EPI_ISL_622865, EPI_ISL_622866, EPI_ISL_622867, EPI_ISL_622868, EPI_ISL_622869, EPI_ISL_622870, EPI_ISL_622871, EPI_ISL_622873, EPI_ISL_622874, EPI_ISL_622875, EPI_ISL_622876, EPI_ISL_622877, EPI_ISL_622884, EPI_ISL_622885, EPI_ISL_622888, EPI_ISL_622890 | | | |
| see above | Respiratory Virus Unit, Microbiology Services Colindale, Public Health England | Respiratory Virus Unit, Microbiology Services Colindale, Public Health England | PHE Covid Sequencing Team |
| EPI_ISL_623103 | General practitioner | National Reference Center for Viruses of Respiratory Infections, Institut Pasteur, Paris | Marion Barbet, Sylvie Behillil, Méline Bizard, Angela Brisebarre, Camille Capel, Etienne Simon-Lorière, Vincent Enouf, Maud Vanpeene, Sylvie van der Werf |
| EPI_ISL_623377, EPI_ISL_623421, EPI_ISL_623486, EPI_ISL_623496 | Lighthouse Lab in Milton Keynes | Wellcome Sanger Institute for the COVID-19 Genomics UK (COG-UK) consortium | The Lighthouse Lab in Milton Keynes and Alex Alderton, Roberto Amato, Sonia Goncalves, Ewan Harrison, David K. Jackson, Ian Johnston, Dominic Kwiatkowski, Cordelia Langford, John Sillitoe on behalf of the Wellcome Sanger Institute COVID-19 Surveillance Team (http://www.sanger.ac.uk/covid-team) |
| EPI_ISL_623895 | Lighthouse Lab in Glasgow | Wellcome Sanger Institute for the COVID-19 Genomics UK (COG-UK) consortium | Harper VanSteenhouse, Yumi Kasai, David Gray, Carol Clugston, Anna Dominiczak and Alex Alderton, Roberto Amato, Sonia Goncalves, Ewan Harrison, David K. Jackson, Ian Johnston, Dominic Kwiatkowski, Cordelia Langford, John Sillitoe on behalf of the Wellcome Sanger Institute COVID-19 Surveillance Team (http://www.sanger.ac.uk/covid-team) |

| | | | |
|---|---|---|---|
| EPI_ISL_625267, EPI_ISL_625268, EPI_ISL_625269, EPI_ISL_625270, EPI_ISL_625271, EPI_ISL_625272, EPI_ISL_625273, EPI_ISL_625274, EPI_ISL_625275, EPI_ISL_625276, EPI_ISL_625277, EPI_ISL_625278, EPI_ISL_625279, EPI_ISL_625280 | | | |
| see above | Lighthouse Lab in Alderley Park | Wellcome Sanger Institute for the COVID-19 Genomics UK (COG-UK) consortium | Jacquelyn Wynn, Mairead Hyland, The Lighthouse Lab in Alderley Park and Alex Alderton, Roberto Amato, Sonia Goncalves, Ewan Harrison, David K. Jackson, Ian Johnston, Dominic Kwiatkowski, Cordelia Langford, John Sillitoe on behalf of the Wellcome Sanger Institute COVID-19 Surveillance Team (http://www.sanger.ac.uk/covid-team) |
| EPI_ISL_625283, EPI_ISL_625287, EPI_ISL_625288, EPI_ISL_625291, EPI_ISL_625294, EPI_ISL_625295, EPI_ISL_625296, EPI_ISL_625304, EPI_ISL_625305, EPI_ISL_625306, EPI_ISL_625310, EPI_ISL_625311 | | | |
| see above | Lighthouse Lab in Cambridge | Wellcome Sanger Institute for the COVID-19 Genomics UK (COG-UK) consortium | Rob Howes, The Lighthouse Lab in Cambridge and Alex Alderton, Roberto Amato, Sonia Goncalves, Ewan Harrison, David K. Jackson, Ian Johnston, Dominic Kwiatkowski, Cordelia Langford, John Sillitoe on behalf of the Wellcome Sanger Institute COVID-19 Surveillance Team (http://www.sanger.ac.uk/covid-team) |
| EPI_ISL_625312 | Lighthouse Lab in Cambridge | Wellcome Sanger Institute for the COVID-19 Genomics UK (COG-UK) Consortium | Rob Howes, The Lighthouse Lab in Cambridge and Alex Alderton, Roberto Amato, Sonia Goncalves, Ewan Harrison, David K. Jackson, Ian Johnston, Dominic Kwiatkowski, Cordelia Langford, John Sillitoe on behalf of the Wellcome Sanger Institute COVID-19 Surveillance Team (http://www.sanger.ac.uk/covid-team) |
| EPI_ISL_625396, EPI_ISL_625397, EPI_ISL_625398, EPI_ISL_625399, EPI_ISL_625400, EPI_ISL_625401, EPI_ISL_625402, EPI_ISL_625403, EPI_ISL_625404 | Lighthouse Lab in Alderley Park | Wellcome Sanger Institute for the COVID-19 Genomics UK (COG-UK) consortium | Jacquelyn Wynn, Mairead Hyland, The Lighthouse Lab in Alderley Park and Alex Alderton, Roberto Amato, Sonia Goncalves, Ewan Harrison, David K. Jackson, Ian Johnston, Dominic Kwiatkowski, Cordelia Langford, John Sillitoe on behalf of the Wellcome Sanger Institute COVID-19 Surveillance Team (http://www.sanger.ac.uk/covid-team) |
| EPI_ISL_626623, EPI_ISL_626625 | The National Institute of Public Health | State Veterinary Institute Prague | Nagy,A;Jirincova,H;Novakova,L;Trnka,D;Vecerova,J |
| EPI_ISL_626656, EPI_ISL_626657, EPI_ISL_626660, EPI_ISL_626681 | Quadram Institute Bioscience | COVID-19 Genomics UK (COG-UK) Consortium | Dave J. Baker, Gemma L. Kay, Alp Aydin, Thanh Le-Viet, Steven Rudder, Ana P. Tedim, Anastasia Kolyva, Maria Diaz, Leonardo de Oliveira Martins, Nabil-Fareed Alikhan, Lizzie Meadows, Rachael Stanley, Ngozi Elumogo, Muhammed Yasir, Nicholas M. Thomson, Alexander J Trotter, Rachel Gilroy, Samuel Bloomfield, Claire Stuart, Andrew Bell, Reenesh Prakash, Samir Dervisevic, Alison E. Mather, John Wain, Mark Webber, Andrew J. Page, Justin O'Grady |
| EPI_ISL_626683, EPI_ISL_626685, EPI_ISL_626686 | University of Exeter | COVID-19 Genomics UK (COG-UK) Consortium | Ben Temperton,Aaron Jeffries,Michelle Michelsen,Joanna Warwick-Dugdale,Audrey Farbos,Robyn Manley,Stephen Michell,Jane Masoli |
| EPI_ISL_626687 | West of Scotland Specialist Virology Centre, NHSGGC / MRC-University of Glasgow Centre for Virus Research | COVID-19 Genomics UK (COG-UK) Consortium | Ana da Silva Filipe, Natasha Johnson, Kathy Smollett, Daniel Mair, Stephen Carmichael, Lily Tong, Jenna Nichols, Elihu Aranday-Cortes, Kyriaki Nomikou; Sarah McDonald, Marc Niebel, Patawee Asamaphan; Richard Orton, Joseph Hughes, Sreenu Vattipally, David L Robertson; Alasdair MacLean, Rory Gunson; Kathy Li, Igor Starinskij, Natasha Jesudason, Rajiv Shah, James Shepherd, Antonia Ho, Emma Thomson |
| EPI_ISL_626694 | Liverpool Clinical Laboratories | COVID-19 Genomics UK (COG-UK) Consortium | Sam Haldenby, Anita Lucaci, Steve Paterson, Julian Hiscox, Alistair Darby, M Almsaud, A Alrezaihi, Muhannad Alruwaili, Stuart D Armstrong, Jones Benjamin, Eleanor G Bentley, Anu Chawla, Jordan J Clark, Angela Cowell, Richard Eccles, Isabel García-Dorival, Matthew Gemmell, Alessandro Gerada, PKF Gilmore, Richard Gregory, Ximeng Han, Catherine Hartley, Margaret Hughes, Miren Iturriza-Gomara, James Johnson, L Luu, Jenifer Manson, Charlotte Nelson, Elaine O'Toole, Cassie Olateju, Rebekah Penrice-Randal , Lucille Rainbow, N.P Randle, Trevor Ian Robinson, Parul Sharma, Ghada T Shawli, James P Stewart, Neil Swainston, Ecaterina Vamos, Joanne Watts, Mark Whitehead |
| EPI_ISL_626698, EPI_ISL_626700, EPI_ISL_626701, EPI_ISL_626704, EPI_ISL_626705 | Quadram Institute Bioscience | COVID-19 Genomics UK (COG-UK) Consortium | Dave J. Baker, Gemma L. Kay, Alp Aydin, Thanh Le-Viet, Steven Rudder, Ana P. Tedim, Anastasia Kolyva, Maria Diaz, Leonardo de Oliveira Martins, Nabil-Fareed Alikhan, Lizzie Meadows, Rachael Stanley, Ngozi Elumogo, Muhammed Yasir, Nicholas M. Thomson, Alexander J Trotter, Rachel Gilroy, Samuel Bloomfield, Claire Stuart, Andrew Bell, Reenesh Prakash, Samir Dervisevic, Alison E. Mather, John Wain, Mark Webber, Andrew J. Page, Justin O'Grady |
| EPI_ISL_626726, EPI_ISL_626727, EPI_ISL_626728 | University of Exeter | COVID-19 Genomics UK (COG-UK) Consortium | Ben Temperton,Aaron Jeffries,Michelle Michelsen,Joanna Warwick-Dugdale,Audrey Farbos,Robyn Manley,Stephen Michell,Jane Masoli |
| EPI_ISL_626736 | Liverpool Clinical Laboratories | COVID-19 Genomics UK (COG-UK) Consortium | Sam Haldenby, Anita Lucaci, Steve Paterson, Julian Hiscox, Alistair Darby, M Almsaud, A Alrezaihi, Muhannad Alruwaili, Stuart D Armstrong, Jones Benjamin, Eleanor G Bentley, Anu Chawla, Jordan J Clark, Angela Cowell, Richard Eccles, Isabel García-Dorival, Matthew Gemmell, Alessandro Gerada, PKF Gilmore, Richard Gregory, Ximeng Han, Catherine Hartley, Margaret Hughes, Miren Iturriza-Gomara, James Johnson, L Luu, Jenifer Manson, Charlotte Nelson, Elaine O'Toole, Cassie Olateju, Rebekah Penrice-Randal , Lucille Rainbow, N.P Randle, Trevor Ian Robinson, Parul Sharma, Ghada T Shawli, James P Stewart, Neil Swainston, Ecaterina Vamos, Joanne Watts, Mark Whitehead |
| EPI_ISL_626737, EPI_ISL_626741, EPI_ISL_626744, EPI_ISL_626745, EPI_ISL_626751, EPI_ISL_626753 | Quadram Institute Bioscience | COVID-19 Genomics UK (COG-UK) Consortium | Dave J. Baker, Gemma L. Kay, Alp Aydin, Thanh Le-Viet, Steven Rudder, Ana P. Tedim, Anastasia Kolyva, Maria Diaz, Leonardo de Oliveira Martins, Nabil-Fareed Alikhan, Lizzie Meadows, Rachael Stanley, Ngozi Elumogo, Muhammed Yasir, Nicholas M. Thomson, Alexander J Trotter, Rachel Gilroy, Samuel Bloomfield, Claire Stuart, Andrew Bell, Reenesh Prakash, Samir Dervisevic, Alison E. Mather, John Wain, Mark Webber, Andrew J. Page, Justin O'Grady |
| EPI_ISL_626758 | West of Scotland Specialist Virology Centre, NHSGGC / MRC-University of Glasgow Centre for Virus Research | COVID-19 Genomics UK (COG-UK) Consortium | Ana da Silva Filipe, Natasha Johnson, Kathy Smollett, Daniel Mair, Stephen Carmichael, Lily Tong, Jenna Nichols, Elihu Aranday-Cortes, Kyriaki Nomikou; Sarah McDonald, Marc Niebel, Patawee Asamaphan; Richard Orton, Joseph Hughes, Sreenu Vattipally, David L Robertson; Alasdair MacLean, Rory Gunson; Kathy Li, Igor Starinskij, Natasha Jesudason, Rajiv Shah, James Shepherd, Antonia Ho, Emma Thomson |
| EPI_ISL_626760 | Liverpool Clinical Laboratories | COVID-19 Genomics UK (COG-UK) Consortium | Sam Haldenby, Anita Lucaci, Steve Paterson, Julian Hiscox, Alistair Darby, M Almsaud, A Alrezaihi, Muhannad Alruwaili, Stuart D Armstrong, Jones Benjamin, Eleanor G Bentley, Anu Chawla, Jordan J Clark, Angela Cowell, Richard Eccles, Isabel García-Dorival, Matthew Gemmell, Alessandro Gerada, PKF Gilmore, Richard Gregory, Ximeng Han, Catherine Hartley, Margaret Hughes, Miren Iturriza-Gomara, James Johnson, L Luu, Jenifer Manson, Charlotte Nelson, Elaine O'Toole, Cassie Olateju, Rebekah Penrice-Randal , Lucille Rainbow, N.P Randle, Trevor Ian Robinson, Parul Sharma, Ghada T Shawli, James P Stewart, Neil Swainston, Ecaterina Vamos, Joanne Watts, Mark Whitehead |
| EPI_ISL_626765 | Quadram Institute Bioscience | COVID-19 Genomics UK (COG-UK) Consortium | Dave J. Baker, Gemma L. Kay, Alp Aydin, Thanh Le-Viet, Steven Rudder, Ana P. Tedim, Anastasia Kolyva, Maria Diaz, Leonardo de Oliveira Martins, Nabil-Fareed Alikhan, Lizzie Meadows, Rachael Stanley, Ngozi Elumogo, Muhammed Yasir, Nicholas M. Thomson, Alexander J Trotter, Rachel Gilroy, Samuel Bloomfield, Claire Stuart, Andrew Bell, Reenesh Prakash, Samir Dervisevic, Alison E. Mather, John Wain, Mark Webber, Andrew J. Page, Justin O'Grady |
| EPI_ISL_626776 | University of Exeter | COVID-19 Genomics UK (COG-UK) Consortium | Ben Temperton,Aaron Jeffries,Michelle Michelsen,Joanna Warwick-Dugdale,Audrey Farbos,Robyn Manley,Stephen Michell,Jane Masoli |
| EPI_ISL_626779, EPI_ISL_626790, EPI_ISL_626795 | Quadram Institute Bioscience | COVID-19 Genomics UK (COG-UK) Consortium | Dave J. Baker, Gemma L. Kay, Alp Aydin, Thanh Le-Viet, Steven Rudder, Ana P. Tedim, Anastasia Kolyva, Maria Diaz, Leonardo de Oliveira Martins, Nabil-Fareed Alikhan, Lizzie Meadows, Rachael Stanley, Ngozi Elumogo, Muhammed Yasir, Nicholas M. Thomson, Alexander J Trotter, Rachel Gilroy, Samuel Bloomfield, Claire Stuart, Andrew Bell, Reenesh Prakash, Samir Dervisevic, Alison E. Mather, John Wain, Mark Webber, Andrew J. Page, Justin O'Grady |
| EPI_ISL_626797, EPI_ISL_626801, EPI_ISL_626803, EPI_ISL_626805 | University of Exeter | COVID-19 Genomics UK (COG-UK) Consortium | Ben Temperton,Aaron Jeffries,Michelle Michelsen,Joanna Warwick-Dugdale,Audrey Farbos,Robyn Manley,Stephen Michell,Jane Masoli |
| EPI_ISL_626807 | Liverpool Clinical Laboratories | COVID-19 Genomics UK (COG-UK) Consortium | Sam Haldenby, Anita Lucaci, Steve Paterson, Julian Hiscox, Alistair Darby, M Almsaud, A Alrezaihi, Muhannad Alruwaili, Stuart D Armstrong, Jones Benjamin, Eleanor G Bentley, Anu Chawla, Jordan J Clark, Angela Cowell, Richard Eccles, Isabel García-Dorival, Matthew Gemmell, Alessandro Gerada, PKF Gilmore, Richard Gregory, Ximeng Han, Catherine Hartley, Margaret Hughes, Miren Iturriza-Gomara, James Johnson, L Luu, Jenifer Manson, Charlotte Nelson, Elaine O'Toole, Cassie Olateju, Rebekah Penrice-Randal , Lucille Rainbow, N.P Randle, Trevor Ian Robinson, Parul Sharma, Ghada T Shawli, James P Stewart, Neil Swainston, Ecaterina Vamos, Joanne Watts, Mark Whitehead |
| EPI_ISL_626812, EPI_ISL_626818 | West of Scotland Specialist Virology Centre, NHSGGC / MRC-University of Glasgow Centre for Virus Research | COVID-19 Genomics UK (COG-UK) Consortium | Ana da Silva Filipe, Natasha Johnson, Kathy Smollett, Daniel Mair, Stephen Carmichael, Lily Tong, Jenna Nichols, Elihu Aranday-Cortes, Kyriaki Nomikou; Sarah McDonald, Marc Niebel, Patawee Asamaphan; Richard Orton, Joseph Hughes, Sreenu Vattipally, David L Robertson; Alasdair MacLean, Rory Gunson; Kathy Li, Igor Starinskij, Natasha Jesudason, Rajiv Shah, James Shepherd, Antonia Ho, Emma Thomson |
| EPI_ISL_626819 | Quadram Institute Bioscience | COVID-19 Genomics UK (COG-UK) Consortium | Dave J. Baker, Gemma L. Kay, Alp Aydin, Thanh Le-Viet, Steven Rudder, Ana P. Tedim, Anastasia Kolyva, Maria Diaz, Leonardo de Oliveira Martins, Nabil-Fareed Alikhan, Lizzie Meadows, Rachael Stanley, Ngozi Elumogo, Muhammed Yasir, Nicholas M. Thomson, Alexander J Trotter, Rachel Gilroy, Samuel Bloomfield, Claire Stuart, Andrew Bell, Reenesh Prakash, Samir Dervisevic, Alison E. Mather, John Wain, Mark Webber, Andrew J. Page, Justin |

| | | | |
|---|---|---|---|
| | | | O'Grady |
| EPI_ISL_626820 | Wales Specialist Virology Centre Sequencing lab: Pathogen Genomics Unit | COVID-19 Genomics UK (COG-UK) Consortium | Catherine Moore, Johnathan Evans, Laura Gifford, Malorie Perry, Simon Cottrell, Angela Marchbank, Alec Birchley, Alexander Adams, Amy Gaskin, Bree Gatica-Wilcox, Jason Coombes, Joel Southgate, Lauren Gilbert, Lee Graham, Nicole Pacchiarini, Sara Kumziene-Summerhayes, Sarah Taylor, Sophie Jones, Sara Rey, Matthew Bull, Joanne Watkins, Sally Corden, Tom Connor |
| EPI_ISL_626822, EPI_ISL_626837, EPI_ISL_626838, EPI_ISL_626839, EPI_ISL_626840, EPI_ISL_626841, EPI_ISL_626842, EPI_ISL_626844, EPI_ISL_626845, EPI_ISL_626847, EPI_ISL_626848 | | | |
| see above | West of Scotland Specialist Virology Centre, NHSGGC / MRC-University of Glasgow Centre for Virus Research | COVID-19 Genomics UK (COG-UK) Consortium | Ana da Silva Filipe, Natasha Johnson, Kathy Smollett, Daniel Mair, Stephen Carmichael, Lily Tong, Jenna Nichols, Elihu Aranday-Cortes, Kyriaki Nomikou; Sarah McDonald, Marc Niebel, Patawee Asamaphan; Richard Orton, Joseph Hughes, Sreenu Vattipally, David L Robertson; Alasdair MacLean, Rory Gunson; Kathy Li, Igor Starinskij, Natasha Jesudason, Rajiv Shah, James Shepherd, Antonia Ho, Emma Thomson |
| EPI_ISL_626860 | Quadram Institute Bioscience | COVID-19 Genomics UK (COG-UK) Consortium | Dave J. Baker, Gemma L. Kay, Alp Aydin, Thanh Le-Viet, Steven Rudder, Ana P. Tedim, Anastasia Kolyva, Maria Diaz, Leonardo de Oliveira Martins, Nabil-Fareed Alikhan, Lizzie Meadows, Rachael Stanley, Ngozi Elumogo, Muhammed Yasir, Nicholas M. Thomson, Alexander J Trotter, Rachel Gilroy, Samuel Bloomfield, Claire Stuart, Andrew Bell, Reenesh Prakash, Samir Dervisevic, Alison E. Mather, John Wain, Mark Webber, Andrew J. Page, Justin O'Grady |
| EPI_ISL_626862 | Wales Specialist Virology Centre Sequencing lab: Pathogen Genomics Unit | COVID-19 Genomics UK (COG-UK) Consortium | Catherine Moore, Johnathan Evans, Laura Gifford, Malorie Perry, Simon Cottrell, Angela Marchbank, Alec Birchley, Alexander Adams, Amy Gaskin, Bree Gatica-Wilcox, Jason Coombes, Joel Southgate, Lauren Gilbert, Lee Graham, Nicole Pacchiarini, Sara Kumziene-Summerhayes, Sarah Taylor, Sophie Jones, Sara Rey, Matthew Bull, Joanne Watkins, Sally Corden, Tom Connor |
| EPI_ISL_626866 | Liverpool Clinical Laboratories | COVID-19 Genomics UK (COG-UK) Consortium | Sam Haldenby, Anita Lucaci, Steve Paterson, Julian Hiscox, Alistair Darby, M Almsaud, A Alrezaihi, Muhannad Alruwaili, Stuart D Armstrong, Jones Benjamin, Eleanor G Bentley, Anu Chawla, Jordan J Clark, Angela Cowell, Richard Eccles, Isabel García-Dorival, Matthew Gemmell, Alessandro Gerada, PKF Gilmore, Richard Gregory, Ximeng Han, Catherine Hartley, Margaret Hughes, Miren Iturriza-Gomara, James Johnson, L Luu, Jenifer Manson, Charlotte Nelson, Elaine O'Toole, Cassie Olateju, Rebekah Penrice-Randal , Lucille Rainbow, N.P Randle, Trevor Ian Robinson, Parul Sharma, Ghada T Shawli, James P Stewart, Neil Swainston, Ecaterina Vamos, Joanne Watts, Mark Whitehead |
| EPI_ISL_626868 | Quadram Institute Bioscience | COVID-19 Genomics UK (COG-UK) Consortium | Dave J. Baker, Gemma L. Kay, Alp Aydin, Thanh Le-Viet, Steven Rudder, Ana P. Tedim, Anastasia Kolyva, Maria Diaz, Leonardo de Oliveira Martins, Nabil-Fareed Alikhan, Lizzie Meadows, Rachael Stanley, Ngozi Elumogo, Muhammed Yasir, Nicholas M. Thomson, Alexander J Trotter, Rachel Gilroy, Samuel Bloomfield, Claire Stuart, Andrew Bell, Reenesh Prakash, Samir Dervisevic, Alison E. Mather, John Wain, Mark Webber, Andrew J. Page, Justin O'Grady |
| EPI_ISL_626873 | West of Scotland Specialist Virology Centre, NHSGGC / MRC-University of Glasgow Centre for Virus Research | COVID-19 Genomics UK (COG-UK) Consortium | Ana da Silva Filipe, Natasha Johnson, Kathy Smollett, Daniel Mair, Stephen Carmichael, Lily Tong, Jenna Nichols, Elihu Aranday-Cortes, Kyriaki Nomikou; Sarah McDonald, Marc Niebel, Patawee Asamaphan; Richard Orton, Joseph Hughes, Sreenu Vattipally, David L Robertson; Alasdair MacLean, Rory Gunson; Kathy Li, Igor Starinskij, Natasha Jesudason, Rajiv Shah, James Shepherd, Antonia Ho, Emma Thomson |
| EPI_ISL_626894 | Quadram Institute Bioscience | COVID-19 Genomics UK (COG-UK) Consortium | Dave J. Baker, Gemma L. Kay, Alp Aydin, Thanh Le-Viet, Steven Rudder, Ana P. Tedim, Anastasia Kolyva, Maria Diaz, Leonardo de Oliveira Martins, Nabil-Fareed Alikhan, Lizzie Meadows, Rachael Stanley, Ngozi Elumogo, Muhammed Yasir, Nicholas M. Thomson, Alexander J Trotter, Rachel Gilroy, Samuel Bloomfield, Claire Stuart, Andrew Bell, Reenesh Prakash, Samir Dervisevic, Alison E. Mather, John Wain, Mark Webber, Andrew J. Page, Justin O'Grady |
| EPI_ISL_626897, EPI_ISL_626904 | West of Scotland Specialist Virology Centre, NHSGGC / MRC-University of Glasgow Centre for Virus Research | COVID-19 Genomics UK (COG-UK) Consortium | Ana da Silva Filipe, Natasha Johnson, Kathy Smollett, Daniel Mair, Stephen Carmichael, Lily Tong, Jenna Nichols, Elihu Aranday-Cortes, Kyriaki Nomikou; Sarah McDonald, Marc Niebel, Patawee Asamaphan; Richard Orton, Joseph Hughes, Sreenu Vattipally, David L Robertson; Alasdair MacLean, Rory Gunson; Kathy Li, Igor Starinskij, Natasha Jesudason, Rajiv Shah, James Shepherd, Antonia Ho, Emma Thomson |
| EPI_ISL_626914 | Quadram Institute Bioscience | COVID-19 Genomics UK (COG-UK) Consortium | Dave J. Baker, Gemma L. Kay, Alp Aydin, Thanh Le-Viet, Steven Rudder, Ana P. Tedim, Anastasia Kolyva, Maria Diaz, Leonardo de Oliveira Martins, Nabil-Fareed Alikhan, Lizzie Meadows, Rachael Stanley, Ngozi Elumogo, Muhammed Yasir, Nicholas M. Thomson, Alexander J Trotter, Rachel Gilroy, Samuel Bloomfield, Claire Stuart, Andrew Bell, Reenesh Prakash, Samir Dervisevic, Alison E. Mather, John Wain, Mark Webber, Andrew J. Page, Justin O'Grady |
| EPI_ISL_626917, EPI_ISL_626927, EPI_ISL_626935 | West of Scotland Specialist Virology Centre, NHSGGC / MRC-University of Glasgow Centre for Virus Research | COVID-19 Genomics UK (COG-UK) Consortium | Ana da Silva Filipe, Natasha Johnson, Kathy Smollett, Daniel Mair, Stephen Carmichael, Lily Tong, Jenna Nichols, Elihu Aranday-Cortes, Kyriaki Nomikou; Sarah McDonald, Marc Niebel, Patawee Asamaphan; Richard Orton, Joseph Hughes, Sreenu Vattipally, David L Robertson; Alasdair MacLean, Rory Gunson; Kathy Li, Igor Starinskij, Natasha Jesudason, Rajiv Shah, James Shepherd, Antonia Ho, Emma Thomson |
| EPI_ISL_626941, EPI_ISL_626942 | University of Exeter | COVID-19 Genomics UK (COG-UK) Consortium | Ben Temperton,Aaron Jeffries,Michelle Michelsen,Joanna Warwick-Dugdale,Audrey Farbos,Robyn Manley,Stephen Michell,Jane Masoli |
| EPI_ISL_626954, EPI_ISL_626956 | Quadram Institute Bioscience | COVID-19 Genomics UK (COG-UK) Consortium | Dave J. Baker, Gemma L. Kay, Alp Aydin, Thanh Le-Viet, Steven Rudder, Ana P. Tedim, Anastasia Kolyva, Maria Diaz, Leonardo de Oliveira Martins, Nabil-Fareed Alikhan, Lizzie Meadows, Rachael Stanley, Ngozi Elumogo, Muhammed Yasir, Nicholas M. Thomson, Alexander J Trotter, Rachel Gilroy, Samuel Bloomfield, Claire Stuart, Andrew Bell, Reenesh Prakash, Samir Dervisevic, Alison E. Mather, John Wain, Mark Webber, Andrew J. Page, Justin O'Grady |
| EPI_ISL_626958, EPI_ISL_626962, EPI_ISL_626963 | West of Scotland Specialist Virology Centre, NHSGGC / MRC-University of Glasgow Centre for Virus Research | COVID-19 Genomics UK (COG-UK) Consortium | Ana da Silva Filipe, Natasha Johnson, Kathy Smollett, Daniel Mair, Stephen Carmichael, Lily Tong, Jenna Nichols, Elihu Aranday-Cortes, Kyriaki Nomikou; Sarah McDonald, Marc Niebel, Patawee Asamaphan; Richard Orton, Joseph Hughes, Sreenu Vattipally, David L Robertson; Alasdair MacLean, Rory Gunson; Kathy Li, Igor Starinskij, Natasha Jesudason, Rajiv Shah, James Shepherd, Antonia Ho, Emma Thomson |
| EPI_ISL_626964 | Quadram Institute Bioscience | COVID-19 Genomics UK (COG-UK) Consortium | Dave J. Baker, Gemma L. Kay, Alp Aydin, Thanh Le-Viet, Steven Rudder, Ana P. Tedim, Anastasia Kolyva, Maria Diaz, Leonardo de Oliveira Martins, Nabil-Fareed Alikhan, Lizzie Meadows, Rachael Stanley, Ngozi Elumogo, Muhammed Yasir, Nicholas M. Thomson, Alexander J Trotter, Rachel Gilroy, Samuel Bloomfield, Claire Stuart, Andrew Bell, Reenesh Prakash, Samir Dervisevic, Alison E. Mather, John Wain, Mark Webber, Andrew J. Page, Justin O'Grady |
| EPI_ISL_626967, EPI_ISL_626968, EPI_ISL_626971, EPI_ISL_626973, EPI_ISL_626974, EPI_ISL_626975, EPI_ISL_626976, EPI_ISL_626977, EPI_ISL_626979, EPI_ISL_626980, EPI_ISL_626981, EPI_ISL_626984, EPI_ISL_626985, EPI_ISL_626986, EPI_ISL_626987, EPI_ISL_626989, EPI_ISL_626990, EPI_ISL_626992 | | | |
| see above | West of Scotland Specialist Virology Centre, NHSGGC / MRC-University of Glasgow Centre for Virus Research | COVID-19 Genomics UK (COG-UK) Consortium | Ana da Silva Filipe, Natasha Johnson, Kathy Smollett, Daniel Mair, Stephen Carmichael, Lily Tong, Jenna Nichols, Elihu Aranday-Cortes, Kyriaki Nomikou; Sarah McDonald, Marc Niebel, Patawee Asamaphan; Richard Orton, Joseph Hughes, Sreenu Vattipally, David L Robertson; Alasdair MacLean, Rory Gunson; Kathy Li, Igor Starinskij, Natasha Jesudason, Rajiv Shah, James Shepherd, Antonia Ho, Emma Thomson |
| EPI_ISL_627002 | Wales Specialist Virology Centre Sequencing lab: Pathogen Genomics Unit | COVID-19 Genomics UK (COG-UK) Consortium | Catherine Moore, Johnathan Evans, Laura Gifford, Malorie Perry, Simon Cottrell, Angela Marchbank, Alec Birchley, Alexander Adams, Amy Gaskin, Bree Gatica-Wilcox, Jason Coombes, Joel Southgate, Lauren Gilbert, Lee Graham, Nicole Pacchiarini, Sara Kumziene-Summerhayes, Sarah Taylor, Sophie Jones, Sara Rey, Matthew Bull, Joanne Watkins, Sally Corden, Tom Connor |
| EPI_ISL_627009 | Quadram Institute Bioscience | COVID-19 Genomics UK (COG-UK) Consortium | Dave J. Baker, Gemma L. Kay, Alp Aydin, Thanh Le-Viet, Steven Rudder, Ana P. Tedim, Anastasia Kolyva, Maria Diaz, Leonardo de Oliveira Martins, Nabil-Fareed Alikhan, Lizzie Meadows, Rachael Stanley, Ngozi Elumogo, Muhammed Yasir, Nicholas M. Thomson, Alexander J Trotter, Rachel Gilroy, Samuel Bloomfield, Claire Stuart, Andrew Bell, Reenesh Prakash, Samir Dervisevic, Alison E. Mather, John Wain, Mark Webber, Andrew J. Page, Justin O'Grady |
| EPI_ISL_627010, EPI_ISL_627027, EPI_ISL_627032 | Wales Specialist Virology Centre Sequencing lab: Pathogen Genomics Unit | COVID-19 Genomics UK (COG-UK) Consortium | Catherine Moore, Johnathan Evans, Laura Gifford, Malorie Perry, Simon Cottrell, Angela Marchbank, Alec Birchley, Alexander Adams, Amy Gaskin, Bree Gatica-Wilcox, Jason Coombes, Joel Southgate, Lauren Gilbert, Lee Graham, Nicole Pacchiarini, Sara Kumziene-Summerhayes, Sarah Taylor, Sophie Jones, Sara Rey, Matthew Bull, Joanne Watkins, Sally Corden, Tom Connor |
| EPI_ISL_627045 | West of Scotland Specialist Virology Centre, NHSGGC / MRC-University of Glasgow Centre for Virus Research | COVID-19 Genomics UK (COG-UK) Consortium | Ana da Silva Filipe, Natasha Johnson, Kathy Smollett, Daniel Mair, Stephen Carmichael, Lily Tong, Jenna Nichols, Elihu Aranday-Cortes, Kyriaki Nomikou; Sarah McDonald, Marc Niebel, Patawee Asamaphan; Richard Orton, Joseph Hughes, Sreenu Vattipally, David L Robertson; Alasdair MacLean, Rory Gunson; Kathy Li, Igor Starinskij, Natasha Jesudason, Rajiv Shah, James Shepherd, Antonia Ho, Emma Thomson |
| EPI_ISL_627048 | Wales Specialist Virology Centre Sequencing lab: Pathogen Genomics Unit | COVID-19 Genomics UK (COG-UK) Consortium | Catherine Moore, Johnathan Evans, Laura Gifford, Malorie Perry, Simon Cottrell, Angela Marchbank, Alec Birchley, Alexander Adams, Amy Gaskin, Bree Gatica-Wilcox, Jason Coombes, Joel Southgate, Lauren Gilbert, Lee Graham, Nicole Pacchiarini, Sara Kumziene-Summerhayes, Sarah Taylor, Sophie Jones, Sara Rey, Matthew Bull, Joanne Watkins, Sally Corden, Tom Connor |
| EPI_ISL_627049 | West of Scotland Specialist Virology Centre, NHSGGC / MRC-University of Glasgow Centre for Virus Research | COVID-19 Genomics UK (COG-UK) Consortium | Ana da Silva Filipe, Natasha Johnson, Kathy Smollett, Daniel Mair, Stephen Carmichael, Lily Tong, Jenna Nichols, Elihu Aranday-Cortes, Kyriaki Nomikou; Sarah McDonald, Marc Niebel, Patawee Asamaphan; Richard Orton, Joseph Hughes, Sreenu Vattipally, David L Robertson; Alasdair MacLean, Rory Gunson; |

| | | | |
|---|---|---|---|
| | | | Kathy Li, Igor Starinskij, Natasha Jesudason, Rajiv Shah, James Shepherd, Antonia Ho, Emma Thomson |
| EPI_ISL_627078 | Wales Specialist Virology Centre Sequencing lab: Pathogen Genomics Unit | COVID-19 Genomics UK (COG-UK) Consortium | Catherine Moore, Johnathan Evans, Laura Gifford, Malorie Perry, Simon Cottrell, Angela Marchbank, Alec Birchley, Alexander Adams, Amy Gaskin, Bree Gatica-Wilcox, Jason Coombes, Joel Southgate, Lauren Gilbert, Lee Graham, Nicole Pacchiarini, Sara Kumziene-Summerhayes, Sarah Taylor, Sophie Jones, Sara Rey, Matthew Bull, Joanne Watkins, Sally Corden, Tom Connor |
| EPI_ISL_627082 | University College London, Great Ormond Street Hospital for Children NHS Foundation Trust, Imperial College Healthcare NHS Trust | COVID-19 Genomics UK (COG-UK) Consortium | Sergi Castellano, Rachel Williams, Mark Kristiansen, Paola Resende Silva, Sunando Roy, Tony Brooks, Helena Tutill, Paola Niola, Patricia Dyal, Charlotte Williams, Leysa Forrest, Yasmin Panchbhaya, Jacqueline Findlay, Samuel Weeks, Julianne Brown, Kathryn Harris, Paul Randell, James Price, Alison Holmes, Judith Breuer |
| EPI_ISL_627088, EPI_ISL_627089, EPI_ISL_627090, EPI_ISL_627092, EPI_ISL_627093, EPI_ISL_627094 | Wales Specialist Virology Centre Sequencing lab: Pathogen Genomics Unit | COVID-19 Genomics UK (COG-UK) Consortium | Catherine Moore, Johnathan Evans, Laura Gifford, Malorie Perry, Simon Cottrell, Angela Marchbank, Alec Birchley, Alexander Adams, Amy Gaskin, Bree Gatica-Wilcox, Jason Coombes, Joel Southgate, Lauren Gilbert, Lee Graham, Nicole Pacchiarini, Sara Kumziene-Summerhayes, Sarah Taylor, Sophie Jones, Sara Rey, Matthew Bull, Joanne Watkins, Sally Corden, Tom Connor |
| EPI_ISL_627099 | Quadram Institute Bioscience | COVID-19 Genomics UK (COG-UK) Consortium | Dave J. Baker, Gemma L. Kay, Alp Aydin, Thanh Le-Viet, Steven Rudder, Ana P. Tedim, Anastasia Kolyva, Maria Diaz, Leonardo de Oliveira Martins, Nabil-Fareed Alikhan, Lizzie Meadows, Rachael Stanley, Ngozi Elumogo, Muhammed Yasir, Nicholas M. Thomson, Alexander J Trotter, Rachel Gilroy, Samuel Bloomfield, Claire Stuart, Andrew Bell, Reenesh Prakash, Samir Dervisevic, Alison E. Mather, John Wain, Mark Webber, Andrew J. Page, Justin O'Grady |
| EPI_ISL_627106, EPI_ISL_627107 | Liverpool Clinical Laboratories | COVID-19 Genomics UK (COG-UK) Consortium | Sam Haldenby, Anita Lucaci, Steve Paterson, Julian Hiscox, Alistair Darby, M Almsaud, A Alrezaihi, Muhannad Alruwaili, Stuart D Armstrong, Jones Benjamin, Eleanor G Bentley, Anu Chawla, Jordan J Clark, Angela Cowell, Richard Eccles, Isabel García-Dorival, Matthew Gemmell, Alessandro Gerada, PKF Gilmore, Richard Gregory, Ximeng Han, Catherine Hartley, Margaret Hughes, Miren Iturriza-Gomara, James Johnson, L Luu, Jenifer Manson, Charlotte Nelson, Elaine O'Toole, Cassie Olateju, Rebekah Penrice-Randal , Lucille Rainbow, N.P Randle, Trevor Ian Robinson, Parul Sharma, Ghada T Shawli, James P Stewart, Neil Swainston, Ecaterina Vamos, Joanne Watts, Mark Whitehead |
| EPI_ISL_627111 | University of Exeter | COVID-19 Genomics UK (COG-UK) Consortium | Ben Temperton,Aaron Jeffries,Michelle Michelsen,Joanna Warwick-Dugdale,Audrey Farbos,Robyn Manley,Stephen Michell,Jane Masoli |
| EPI_ISL_627113 | Wales Specialist Virology Centre Sequencing lab: Pathogen Genomics Unit | COVID-19 Genomics UK (COG-UK) Consortium | Catherine Moore, Johnathan Evans, Laura Gifford, Malorie Perry, Simon Cottrell, Angela Marchbank, Alec Birchley, Alexander Adams, Amy Gaskin, Bree Gatica-Wilcox, Jason Coombes, Joel Southgate, Lauren Gilbert, Lee Graham, Nicole Pacchiarini, Sara Kumziene-Summerhayes, Sarah Taylor, Sophie Jones, Sara Rey, Matthew Bull, Joanne Watkins, Sally Corden, Tom Connor |
| EPI_ISL_627114, EPI_ISL_627125, EPI_ISL_627132, EPI_ISL_627141, EPI_ISL_627156 | Liverpool Clinical Laboratories | COVID-19 Genomics UK (COG-UK) Consortium | Sam Haldenby, Anita Lucaci, Steve Paterson, Julian Hiscox, Alistair Darby, M Almsaud, A Alrezaihi, Muhannad Alruwaili, Stuart D Armstrong, Jones Benjamin, Eleanor G Bentley, Anu Chawla, Jordan J Clark, Angela Cowell, Richard Eccles, Isabel García-Dorival, Matthew Gemmell, Alessandro Gerada, PKF Gilmore, Richard Gregory, Ximeng Han, Catherine Hartley, Margaret Hughes, Miren Iturriza-Gomara, James Johnson, L Luu, Jenifer Manson, Charlotte Nelson, Elaine O'Toole, Cassie Olateju, Rebekah Penrice-Randal , Lucille Rainbow, N.P Randle, Trevor Ian Robinson, Parul Sharma, Ghada T Shawli, James P Stewart, Neil Swainston, Ecaterina Vamos, Joanne Watts, Mark Whitehead |
| EPI_ISL_627163 | University of Exeter | COVID-19 Genomics UK (COG-UK) Consortium | Ben Temperton,Aaron Jeffries,Michelle Michelsen,Joanna Warwick-Dugdale,Audrey Farbos,Robyn Manley,Stephen Michell,Jane Masoli |
| EPI_ISL_627168 | Wales Specialist Virology Centre Sequencing lab: Pathogen Genomics Unit | COVID-19 Genomics UK (COG-UK) Consortium | Catherine Moore, Johnathan Evans, Laura Gifford, Malorie Perry, Simon Cottrell, Angela Marchbank, Alec Birchley, Alexander Adams, Amy Gaskin, Bree Gatica-Wilcox, Jason Coombes, Joel Southgate, Lauren Gilbert, Lee Graham, Nicole Pacchiarini, Sara Kumziene-Summerhayes, Sarah Taylor, Sophie Jones, Sara Rey, Matthew Bull, Joanne Watkins, Sally Corden, Tom Connor |
| EPI_ISL_627171 | Liverpool Clinical Laboratories | COVID-19 Genomics UK (COG-UK) Consortium | Sam Haldenby, Anita Lucaci, Steve Paterson, Julian Hiscox, Alistair Darby, M Almsaud, A Alrezaihi, Muhannad Alruwaili, Stuart D Armstrong, Jones Benjamin, Eleanor G Bentley, Anu Chawla, Jordan J Clark, Angela Cowell, Richard Eccles, Isabel García-Dorival, Matthew Gemmell, Alessandro Gerada, PKF Gilmore, Richard Gregory, Ximeng Han, Catherine Hartley, Margaret Hughes, Miren Iturriza-Gomara, James Johnson, L Luu, Jenifer Manson, Charlotte Nelson, Elaine O'Toole, Cassie Olateju, Rebekah Penrice-Randal , Lucille Rainbow, N.P Randle, Trevor Ian Robinson, Parul Sharma, Ghada T Shawli, James P Stewart, Neil Swainston, Ecaterina Vamos, Joanne Watts, Mark Whitehead |
| EPI_ISL_627173 | West of Scotland Specialist Virology Centre, NHSGGC / MRC-University of Glasgow Centre for Virus Research | COVID-19 Genomics UK (COG-UK) Consortium | Ana da Silva Filipe, Natasha Johnson, Kathy Smollett, Daniel Mair, Stephen Carmichael, Lily Tong, Jenna Nichols, Elihu Aranday-Cortes, Kyriaki Nomikou; Sarah McDonald, Marc Niebel, Patawee Asamaphan; Richard Orton, Joseph Hughes, Sreenu Vattipally, David L Robertson; Alasdair MacLean, Rory Gunson; Kathy Li, Igor Starinskij, Natasha Jesudason, Rajiv Shah, James Shepherd, Antonia Ho, Emma Thomson |
| EPI_ISL_627175 | University College London, Great Ormond Street Hospital for Children NHS Foundation Trust, Imperial College Healthcare NHS Trust | COVID-19 Genomics UK (COG-UK) Consortium | Sergi Castellano, Rachel Williams, Mark Kristiansen, Paola Resende Silva, Sunando Roy, Tony Brooks, Helena Tutill, Paola Niola, Patricia Dyal, Charlotte Williams, Leysa Forrest, Yasmin Panchbhaya, Jacqueline Findlay, Samuel Weeks, Julianne Brown, Kathryn Harris, Paul Randell, James Price, Alison Holmes, Judith Breuer |
| EPI_ISL_627180 | Quadram Institute Bioscience | COVID-19 Genomics UK (COG-UK) Consortium | Dave J. Baker, Gemma L. Kay, Alp Aydin, Thanh Le-Viet, Steven Rudder, Ana P. Tedim, Anastasia Kolyva, Maria Diaz, Leonardo de Oliveira Martins, Nabil-Fareed Alikhan, Lizzie Meadows, Rachael Stanley, Ngozi Elumogo, Muhammed Yasir, Nicholas M. Thomson, Alexander J Trotter, Rachel Gilroy, Samuel Bloomfield, Claire Stuart, Andrew Bell, Reenesh Prakash, Samir Dervisevic, Alison E. Mather, John Wain, Mark Webber, Andrew J. Page, Justin O'Grady |
| EPI_ISL_627181, EPI_ISL_627185 | Liverpool Clinical Laboratories | COVID-19 Genomics UK (COG-UK) Consortium | Sam Haldenby, Anita Lucaci, Steve Paterson, Julian Hiscox, Alistair Darby, M Almsaud, A Alrezaihi, Muhannad Alruwaili, Stuart D Armstrong, Jones Benjamin, Eleanor G Bentley, Anu Chawla, Jordan J Clark, Angela Cowell, Richard Eccles, Isabel García-Dorival, Matthew Gemmell, Alessandro Gerada, PKF Gilmore, Richard Gregory, Ximeng Han, Catherine Hartley, Margaret Hughes, Miren Iturriza-Gomara, James Johnson, L Luu, Jenifer Manson, Charlotte Nelson, Elaine O'Toole, Cassie Olateju, Rebekah Penrice-Randal , Lucille Rainbow, N.P Randle, Trevor Ian Robinson, Parul Sharma, Ghada T Shawli, James P Stewart, Neil Swainston, Ecaterina Vamos, Joanne Watts, Mark Whitehead |
| EPI_ISL_627186, EPI_ISL_627190 | University of Exeter | COVID-19 Genomics UK (COG-UK) Consortium | Ben Temperton,Aaron Jeffries,Michelle Michelsen,Joanna Warwick-Dugdale,Audrey Farbos,Robyn Manley,Stephen Michell,Jane Masoli |
| EPI_ISL_627195 | West of Scotland Specialist Virology Centre, NHSGGC / MRC-University of Glasgow Centre for Virus Research | COVID-19 Genomics UK (COG-UK) Consortium | Ana da Silva Filipe, Natasha Johnson, Kathy Smollett, Daniel Mair, Stephen Carmichael, Lily Tong, Jenna Nichols, Elihu Aranday-Cortes, Kyriaki Nomikou; Sarah McDonald, Marc Niebel, Patawee Asamaphan; Richard Orton, Joseph Hughes, Sreenu Vattipally, David L Robertson; Alasdair MacLean, Rory Gunson; Kathy Li, Igor Starinskij, Natasha Jesudason, Rajiv Shah, James Shepherd, Antonia Ho, Emma Thomson |
| EPI_ISL_627198 | Quadram Institute Bioscience | COVID-19 Genomics UK (COG-UK) Consortium | Dave J. Baker, Gemma L. Kay, Alp Aydin, Thanh Le-Viet, Steven Rudder, Ana P. Tedim, Anastasia Kolyva, Maria Diaz, Leonardo de Oliveira Martins, Nabil-Fareed Alikhan, Lizzie Meadows, Rachael Stanley, Ngozi Elumogo, Muhammed Yasir, Nicholas M. Thomson, Alexander J Trotter, Rachel Gilroy, Samuel Bloomfield, Claire Stuart, Andrew Bell, Reenesh Prakash, Samir Dervisevic, Alison E. Mather, John Wain, Mark Webber, Andrew J. Page, Justin O'Grady |
| EPI_ISL_627200, EPI_ISL_627201, EPI_ISL_627202 | West of Scotland Specialist Virology Centre, NHSGGC / MRC-University of Glasgow Centre for Virus Research | COVID-19 Genomics UK (COG-UK) Consortium | Ana da Silva Filipe, Natasha Johnson, Kathy Smollett, Daniel Mair, Stephen Carmichael, Lily Tong, Jenna Nichols, Elihu Aranday-Cortes, Kyriaki Nomikou; Sarah McDonald, Marc Niebel, Patawee Asamaphan; Richard Orton, Joseph Hughes, Sreenu Vattipally, David L Robertson; Alasdair MacLean, Rory Gunson; Kathy Li, Igor Starinskij, Natasha Jesudason, Rajiv Shah, James Shepherd, Antonia Ho, Emma Thomson |
| EPI_ISL_627260 | Oxford Viromics, NDM, University of Oxford; Oxford University Hospitals; Basingstoke and North Hampshire Hospital | COVID-19 Genomics UK (COG-UK) Consortium | Tanya Golubchik, David Bonsall, George Macintyre, Amy Trebes, Mariateresa de Cesare, Catrin Moore, Alex Mobbs, Anita Justice, Robert Shaw, Monique Andersson, Timothy Peto, Emma Wise, Nathan Moore, Jessica Lynch, Nick Cortes, Matilde Mori, Stephen Kidd, David Buck, John Todd, Christophe Fraser |
| EPI_ISL_627275, EPI_ISL_627276, EPI_ISL_627293 | Wales Specialist Virology Centre Sequencing lab: Pathogen Genomics Unit | COVID-19 Genomics UK (COG-UK) Consortium | Catherine Moore, Johnathan Evans, Laura Gifford, Malorie Perry, Simon Cottrell, Angela Marchbank, Alec Birchley, Alexander Adams, Amy Gaskin, Bree Gatica-Wilcox, Jason Coombes, Joel Southgate, Lauren Gilbert, Lee Graham, Nicole Pacchiarini, Sara Kumziene-Summerhayes, Sarah Taylor, Sophie Jones, Sara Rey, Matthew Bull, Joanne Watkins, Sally Corden, Tom Connor |
| EPI_ISL_627300 | Quadram Institute Bioscience | COVID-19 Genomics UK (COG-UK) Consortium | Dave J. Baker, Gemma L. Kay, Alp Aydin, Thanh Le-Viet, Steven Rudder, Ana P. Tedim, Anastasia Kolyva, Maria Diaz, Leonardo de Oliveira Martins, Nabil-Fareed Alikhan, Lizzie Meadows, Rachael Stanley, Ngozi Elumogo, Muhammed Yasir, Nicholas M. Thomson, Alexander J Trotter, Rachel Gilroy, Samuel Bloomfield, Claire Stuart, Andrew Bell, Reenesh Prakash, Samir Dervisevic, Alison E. Mather, John Wain, Mark Webber, Andrew J. Page, Justin O'Grady |
| EPI_ISL_627368, EPI_ISL_627369, EPI_ISL_627370, EPI_ISL_627371, EPI_ISL_627372, EPI_ISL_627378, EPI_ISL_627381, EPI_ISL_627382, EPI_ISL_627383, EPI_ISL_627385, EPI_ISL_627386, EPI_ISL_627387, EPI_ISL_627389, EPI_ISL_627390, EPI_ISL_627392, EPI_ISL_627393, EPI_ISL_627394, EPI_ISL_627395, EPI_ISL_627399, EPI_ISL_627400, EPI_ISL_627401, EPI_ISL_627402, EPI_ISL_627403, EPI_ISL_627405, EPI_ISL_627406, EPI_ISL_627407, EPI_ISL_627408, EPI_ISL_627411, EPI_ISL_627412, EPI_ISL_627413, EPI_ISL_627414, EPI_ISL_627415, EPI_ISL_627416, EPI_ISL_627417, EPI_ISL_627420, EPI_ISL_627421, EPI_ISL_627422, EPI_ISL_627423 | | | |

| | | | |
|---|---|---|---|
| see above | West of Scotland Specialist Virology Centre, NHSGGC / MRC-University of Glasgow Centre for Virus Research | COVID-19 Genomics UK (COG-UK) Consortium | Ana da Silva Filipe, Natasha Johnson, Kathy Smollett, Daniel Mair, Stephen Carmichael, Lily Tong, Jenna Nichols, Elihu Aranday-Cortes, Kyriaki Nomikou; Sarah McDonald, Marc Niebel, Patawee Asamaphan; Richard Orton, Joseph Hughes, Sreenu Vattipally, David L Robertson; Alasdair MacLean, Rory Gunson; Kathy Li, Igor Starinskij, Natasha Jesudason, Rajiv Shah, James Shepherd, Antonia Ho, Emma Thomson |
| EPI_ISL_627425, EPI_ISL_627426, EPI_ISL_627427 | University of Exeter | COVID-19 Genomics UK (COG-UK) Consortium | Ben Temperton,Aaron Jeffries,Michelle Michelsen,Joanna Warwick-Dugdale,Audrey Farbos,Robyn Manley,Stephen Michell,Jane Masoli |
| EPI_ISL_627447, EPI_ISL_627448, EPI_ISL_627449 | Liverpool Clinical Laboratories | COVID-19 Genomics UK (COG-UK) Consortium | Sam Haldenby, Anita Lucaci, Steve Paterson, Julian Hiscox, Alistair Darby, M Almsaud, A Alrezaihi, Muhannad Alruwaili, Stuart D Armstrong, Jones Benjamin, Eleanor G Bentley, Anu Chawla, Jordan J Clark, Angela Cowell, Richard Eccles, Isabel García-Dorival, Matthew Gemmell, Alessandro Gerada, PKF Gilmore, Richard Gregory, Ximeng Han, Catherine Hartley, Margaret Hughes, Miren Iturriza-Gomara, James Johnson, L Luu, Jenifer Manson, Charlotte Nelson, Elaine O'Toole, Cassie Olateju, Rebekah Penrice-Randal , Lucille Rainbow, N.P Randle, Trevor Ian Robinson, Parul Sharma, Ghada T Shawli, James P Stewart, Neil Swainston, Ecaterina Vamos, Joanne Watts, Mark Whitehead |
| EPI_ISL_627450, EPI_ISL_627451, EPI_ISL_627452, EPI_ISL_627453, EPI_ISL_627454, EPI_ISL_627455, EPI_ISL_627456, EPI_ISL_627457, EPI_ISL_627458, EPI_ISL_627459, EPI_ISL_627460, EPI_ISL_627461, EPI_ISL_627462, EPI_ISL_627463, EPI_ISL_627464 | | | |
| see above | University College London, Great Ormond Street Hospital for Children NHS Foundation Trust, Imperial College Healthcare NHS Trust | COVID-19 Genomics UK (COG-UK) Consortium | Sergi Castellano, Rachel Williams, Mark Kristiansen, Paola Resende Silva, Sunando Roy, Tony Brooks, Helena Tutill, Paola Niola, Patricia Dyal, Charlotte Williams, Leysa Forrest, Yasmin Panchbhaya, Jacqueline Findlay, Samuel Weeks, Julianne Brown, Kathryn Harris, Paul Randell, James Price, Alison Holmes, Judith Breuer |
| EPI_ISL_627570, EPI_ISL_627571, EPI_ISL_627572, EPI_ISL_627573, EPI_ISL_627574, EPI_ISL_627575, EPI_ISL_627576, EPI_ISL_627577, EPI_ISL_627578, EPI_ISL_627579, EPI_ISL_627580, EPI_ISL_627581, EPI_ISL_627582, EPI_ISL_627583, EPI_ISL_627587, EPI_ISL_627588, EPI_ISL_627589, EPI_ISL_627590, EPI_ISL_627591, EPI_ISL_627592, EPI_ISL_627593, EPI_ISL_627594, EPI_ISL_627595, EPI_ISL_627596, EPI_ISL_627602, EPI_ISL_627616, EPI_ISL_627631, EPI_ISL_627635, EPI_ISL_627640 | | | |
| see above | Quadram Institute Bioscience | COVID-19 Genomics UK (COG-UK) Consortium | Dave J. Baker, Gemma L. Kay, Alp Aydin, Thanh Le-Viet, Steven Rudder, Ana P. Tedim, Anastasia Kolyva, Maria Diaz, Leonardo de Oliveira Martins, Nabil-Fareed Alikhan, Lizzie Meadows, Rachael Stanley, Ngozi Elumogo, Muhammed Yasir, Nicholas M. Thomson, Alexander J Trotter, Rachel Gilroy, Samuel Bloomfield, Claire Stuart, Andrew Bell, Reenesh Prakash, Samir Dervisevic, Alison E. Mather, John Wain, Mark Webber, Andrew J. Page, Justin O'Grady |
| EPI_ISL_627666 | Queens Medical Centre, Clinical Microbiology Department / DeepSeq Nottingham | COVID-19 Genomics UK (COG-UK) Consortium | Gemma Clark, Wendy Smith, Manjinder Khakh, Vicki M Fleming, Michelle M Lister, Hannah Howson-Wells, Jonathan Ball, Patrick McClure, Joseph Chappell, Theocharis Tsoleridis, Nadine Holmes, Matthew Carlisle, Christopher Moore, Fei Sang, Johnny Debebe, Victoria Wright, Matthew Loose |
| EPI_ISL_627738, EPI_ISL_627739, EPI_ISL_627740, EPI_ISL_627746, EPI_ISL_627749, EPI_ISL_627752, EPI_ISL_627754, EPI_ISL_627755, EPI_ISL_627760, EPI_ISL_627767, EPI_ISL_627769, EPI_ISL_627778, EPI_ISL_627786, EPI_ISL_627788, EPI_ISL_627792, EPI_ISL_627794, EPI_ISL_627809, EPI_ISL_627810, EPI_ISL_627817, EPI_ISL_627820, EPI_ISL_627821, EPI_ISL_627822, EPI_ISL_627826, EPI_ISL_627836, EPI_ISL_627839, EPI_ISL_627840, EPI_ISL_627843, EPI_ISL_627845, EPI_ISL_627846, EPI_ISL_627847, EPI_ISL_627858, EPI_ISL_627868, EPI_ISL_627869, EPI_ISL_627878, EPI_ISL_627887, EPI_ISL_627888, EPI_ISL_627897, EPI_ISL_627902, EPI_ISL_627907, EPI_ISL_627918, EPI_ISL_627932, EPI_ISL_627933, EPI_ISL_627934, EPI_ISL_627936, EPI_ISL_627942, EPI_ISL_627976, EPI_ISL_627977, EPI_ISL_627978, EPI_ISL_627983, EPI_ISL_628001, EPI_ISL_628012, EPI_ISL_628020, EPI_ISL_628030, EPI_ISL_628034, EPI_ISL_628037, EPI_ISL_628059, EPI_ISL_628064, EPI_ISL_628119, EPI_ISL_628128 | | | |
| see above | Wales Specialist Virology Centre Sequencing lab: Pathogen Genomics Unit | COVID-19 Genomics UK (COG-UK) Consortium | Catherine Moore, Johnathan Evans, Laura Gifford, Malorie Perry, Simon Cottrell, Angela Marchbank, Alec Birchley, Alexander Adams, Amy Gaskin, Bree Gatica-Wilcox, Jason Coombes, Joel Southgate, Lauren Gilbert, Lee Graham, Nicole Pacchiarini, Sara Kumziene-Summerhayes, Sarah Taylor, Sophie Jones, Sara Rey, Matthew Bull, Joanne Watkins, Sally Corden, Tom Connor |
| EPI_ISL_628628, EPI_ISL_628629, EPI_ISL_628630, EPI_ISL_628631, EPI_ISL_628632, EPI_ISL_628633, EPI_ISL_628646, EPI_ISL_628648, EPI_ISL_628649, EPI_ISL_628650, EPI_ISL_628651, EPI_ISL_628652 | | | |
| see above | Oxford Viromics, NDM, University of Oxford; Oxford University Hospitals; Basingstoke and North Hampshire Hospital | COVID-19 Genomics UK (COG-UK) Consortium | Tanya Golubchik, David Bonsall, George Macintyre, Amy Trebes, Mariateresa de Cesare, Catrin Moore, Alex Mobbs, Anita Justice, Robert Shaw, Monique Andersson, Timothy Peto, Emma Wise, Nathan Moore, Jessica Lynch, Nick Cortes, Matilde Mori, Stephen Kidd, David Buck, John Todd, Christophe Fraser |
| EPI_ISL_628804, EPI_ISL_628805, EPI_ISL_628807, EPI_ISL_628808, EPI_ISL_628809, EPI_ISL_628810, EPI_ISL_628811, EPI_ISL_628812, EPI_ISL_628813, EPI_ISL_628814, EPI_ISL_628815, EPI_ISL_628816, EPI_ISL_628817, EPI_ISL_628818, EPI_ISL_628819, EPI_ISL_628820, EPI_ISL_628821, EPI_ISL_628822, EPI_ISL_628823, EPI_ISL_628824, EPI_ISL_628825, EPI_ISL_628826, EPI_ISL_628827, EPI_ISL_628828, EPI_ISL_628829, EPI_ISL_628830, EPI_ISL_628831, EPI_ISL_628832, EPI_ISL_628833, EPI_ISL_628834, EPI_ISL_628835, EPI_ISL_628836, EPI_ISL_628837, EPI_ISL_628838, EPI_ISL_628839, EPI_ISL_628840, EPI_ISL_628841, EPI_ISL_628842, EPI_ISL_628843, EPI_ISL_628844, EPI_ISL_628845, EPI_ISL_628846, EPI_ISL_628847, EPI_ISL_628848, EPI_ISL_628849, EPI_ISL_628850, EPI_ISL_628851, EPI_ISL_628852, EPI_ISL_628853, EPI_ISL_628854, EPI_ISL_628855, EPI_ISL_628856, EPI_ISL_628857, EPI_ISL_628859, EPI_ISL_628860, EPI_ISL_628861, EPI_ISL_628862, EPI_ISL_628863, EPI_ISL_628864, EPI_ISL_628865, EPI_ISL_628866 | | | |
| see above | University of Michigan Clinical Microbiology Laboratory | Lauring Lab, University of Michigan, Department of Microbiology and Immunology | Valesano |
| EPI_ISL_628942 | Utah Public Health Laboratory | Utah Public Health Laboratory | Erin Young, Kelly Oakeson |
| EPI_ISL_629002 | Sydney South West Pathology Service (SSWPS) - Liverpool Hospital - NSW Health Pathology | NSW Health Pathology - Institute of Clinical Pathology and Medical Research; Westmead Hospital; University of Sydney | CIDM-PH et al. |
| EPI_ISL_629010 | Sydney South West Pathology Service (SSWPS) - Royal Prince Alfred Hospital - NSW Health Pathology | NSW Health Pathology - Institute of Clinical Pathology and Medical Research; Westmead Hospital; University of Sydney | CIDM-PH et al. |
| EPI_ISL_629041, EPI_ISL_629042, EPI_ISL_629043, EPI_ISL_629044, EPI_ISL_629045, EPI_ISL_629046, EPI_ISL_629047, EPI_ISL_629048, EPI_ISL_629049, EPI_ISL_629050, EPI_ISL_629051, EPI_ISL_629052, EPI_ISL_629053, EPI_ISL_629054, EPI_ISL_629055, EPI_ISL_629056, EPI_ISL_629057, EPI_ISL_629058, EPI_ISL_629059, EPI_ISL_629060, EPI_ISL_629061, EPI_ISL_629062, EPI_ISL_629063, EPI_ISL_629066, EPI_ISL_629067, EPI_ISL_629070, EPI_ISL_629071, EPI_ISL_629072, EPI_ISL_629073, EPI_ISL_629074, EPI_ISL_629075, EPI_ISL_629077, EPI_ISL_629078, EPI_ISL_629079, EPI_ISL_629080 | | | |
| see above | Maryland Public Health Laboratory | Maryland Public Health Laboratory | Maryland Department of Health Laboratories Administration |
| EPI_ISL_629563, EPI_ISL_630104, EPI_ISL_630174, EPI_ISL_630272 | Lighthouse Lab in Milton Keynes | Wellcome Sanger Institute for the COVID-19 Genomics UK (COG-UK) consortium | The Lighthouse Lab in Milton Keynes and Alex Alderton, Roberto Amato, Sonia Goncalves, Ewan Harrison, David K. Jackson, Ian Johnston, Dominic Kwiatkowski, Cordelia Langford, John Sillitoe on behalf of the Wellcome Sanger Institute COVID-19 Surveillance Team |
| EPI_ISL_630983, EPI_ISL_630999, EPI_ISL_631012, EPI_ISL_631014, EPI_ISL_631018, EPI_ISL_631027, EPI_ISL_631033, EPI_ISL_631035, EPI_ISL_631041, EPI_ISL_631046, EPI_ISL_631056, EPI_ISL_631081, EPI_ISL_631093, EPI_ISL_631099, EPI_ISL_631107, EPI_ISL_631131, EPI_ISL_631133, EPI_ISL_631150, EPI_ISL_631189, EPI_ISL_631215, EPI_ISL_631217, EPI_ISL_631250, EPI_ISL_631265 | | | |
| see above | Lighthouse Lab in Alderley Park | Wellcome Sanger Institute for the COVID-19 Genomics UK (COG-UK) consortium | Jacquelyn Wynn, Mairead Hyland, The Lighthouse Lab in Alderley Park and Alex Alderton, Roberto Amato, Sonia Goncalves, Ewan Harrison, David K. Jackson, Ian Johnston, Dominic Kwiatkowski, Cordelia Langford, John Sillitoe on behalf of the Wellcome Sanger Institute COVID-19 Surveillance Team |
| EPI_ISL_632250, EPI_ISL_632251, EPI_ISL_632252, EPI_ISL_632253, EPI_ISL_632254, EPI_ISL_632255 | Communicable Disease Laboratory, Public Health Directorate | Communicable Disease Laboratory, Public Health Directorate | AlWasti,H., Altaif,Z., AlHujairi,Z., AlAbbas,Z. |
| EPI_ISL_632256, EPI_ISL_632257, EPI_ISL_632258 | Communicable Disease Laboratory, Public Health Directorate | Communicable Disease Laboratory, Public Health Directorate | AlHujairi,Z., Altaif,Z., AlWasti,H., AlAbbas,Z. |
| EPI_ISL_632259, EPI_ISL_632260 | Communicable Disease Laboratory, Public Health Directorate | Communicable Disease Laboratory, Public Health Directorate | AlTaif,Z., AlWasti,H., AlHujairi,Z., AlAbbas,Z. |
| EPI_ISL_632268, EPI_ISL_632269, EPI_ISL_632270, EPI_ISL_632271, EPI_ISL_632272, EPI_ISL_632273, EPI_ISL_632274, EPI_ISL_632275, EPI_ISL_632276, EPI_ISL_632277, EPI_ISL_632278, EPI_ISL_632279, EPI_ISL_632280, EPI_ISL_632281, EPI_ISL_632282 | | | |
| see above | Communicable Disease Laboratory, Public Health Directorate | Communicable Disease Laboratory, Public Health Directorate | AlWasti,H., AlTaif,Z., AlHujairi,Z., AlAbbas,Z. |
| EPI_ISL_632324 | Dutch COVID-19 response team | Erasmus Medical Center | OH consortium |
| EPI_ISL_632328, EPI_ISL_632330, EPI_ISL_632338, EPI_ISL_632340, EPI_ISL_632343 | Dutch COVID-19 response team | Erasmus Medical Center | Bas Oude Munnink, David Nieuwenhuijse, Reina Sikkema, Claudia Schapendonk, Irina Chestakova, Anne van der Linden, Theo Bestebroer, Stefan van Nieuwkoop, Mark Pronk, Pascal Lexmond, Corien Swaan, Manon Haverkate, Madelief Mollers, Mart Stein, Sandra Kengne Kamga Mobou, Jeroen van Kampen, Jolanda Voermans, Aura Timen, Corine GeurtsvanKessel, Annemiek van der Eijk, Richard Molenkamp, Marion Koopmans, on behalf of the Dutch national COVID-19 response team. |
| EPI_ISL_632348 | Dutch COVID-19 response team | Erasmus Medical Center | OH consortium |
| EPI_ISL_632349, EPI_ISL_632350 | Dutch COVID-19 response team | Erasmus Medical Center | Bas Oude Munnink, David Nieuwenhuijse, Reina Sikkema, Claudia Schapendonk, Irina Chestakova, Anne van der Linden, Theo Bestebroer, Stefan van Nieuwkoop, Mark Pronk, Pascal Lexmond, Corien Swaan, Manon Haverkate, Madelief Mollers, Mart Stein, Sandra Kengne Kamga Mobou, Jeroen van Kampen, Jolanda Voermans, Aura Timen, Corine GeurtsvanKessel, Annemiek van der Eijk, Richard Molenkamp, Marion Koopmans, on behalf of the Dutch national COVID-19 response team. |
| EPI_ISL_632353 | Dutch COVID-19 response team | Erasmus Medical Center | OH consortium |

| | | | |
|---|---|---|---|
| EPI_ISL_632362, EPI_ISL_632374, EPI_ISL_632376, EPI_ISL_632383, EPI_ISL_632393, EPI_ISL_632394, EPI_ISL_632395, EPI_ISL_632404, EPI_ISL_632407, EPI_ISL_632412, EPI_ISL_632416, EPI_ISL_632424 | | | |
| see above | Dutch COVID-19 response team | Erasmus Medical Center | Bas Oude Munnink, David Nieuwenhuijse, Reina Sikkema, Claudia Schapendonk, Irina Chestakova, Anne van der Linden, Theo Bestebroer, Stefan van Nieuwkoop, Mark Pronk, Pascal Lexmond, Corien Swaan, Manon Haverkate, Madelief Mollers, Mart Stein, Sandra Kengne Kamga Mobou, Jeroen van Kampen, Jolanda Voermans, Aura Timen, Corine GeurtsvanKessel, Annemiek van der Eijk, Richard Molenkamp, Marion Koopmans, on behalf of the Dutch national COVID-19 response team. |
| EPI_ISL_632439 | Dutch COVID-19 response team | Erasmus Medical Center | OH consortium |
| EPI_ISL_632441, EPI_ISL_632442, EPI_ISL_632443, EPI_ISL_632444, EPI_ISL_632448 | Dutch COVID-19 response team | Erasmus Medical Center | Bas Oude Munnink, David Nieuwenhuijse, Reina Sikkema, Claudia Schapendonk, Irina Chestakova, Anne van der Linden, Theo Bestebroer, Stefan van Nieuwkoop, Mark Pronk, Pascal Lexmond, Corien Swaan, Manon Haverkate, Madelief Mollers, Mart Stein, Sandra Kengne Kamga Mobou, Jeroen van Kampen, Jolanda Voermans, Aura Timen, Corine GeurtsvanKessel, Annemiek van der Eijk, Richard Molenkamp, Marion Koopmans, on behalf of the Dutch national COVID-19 response team. |
| EPI_ISL_632486, EPI_ISL_632487, EPI_ISL_632488, EPI_ISL_632489, EPI_ISL_632490, EPI_ISL_632491 | Dutch COVID-19 response team | Erasmus Medical Center | OH consortium |
| EPI_ISL_632656, EPI_ISL_632657, EPI_ISL_632675, EPI_ISL_632676, EPI_ISL_632677, EPI_ISL_632678, EPI_ISL_632679, EPI_ISL_632680, EPI_ISL_632681, EPI_ISL_632682, EPI_ISL_632683, EPI_ISL_632684, EPI_ISL_632685, EPI_ISL_632686, EPI_ISL_632687, EPI_ISL_632688, EPI_ISL_632689, EPI_ISL_632690, EPI_ISL_632691, EPI_ISL_632692, EPI_ISL_632693, EPI_ISL_632694, EPI_ISL_632695, EPI_ISL_632701, EPI_ISL_632705, EPI_ISL_632706, EPI_ISL_632711, EPI_ISL_632712, EPI_ISL_632713, EPI_ISL_632720, EPI_ISL_632726, EPI_ISL_632727, EPI_ISL_632728, EPI_ISL_632729, EPI_ISL_632730, EPI_ISL_632731, EPI_ISL_632736, EPI_ISL_632737, EPI_ISL_632739, EPI_ISL_632740, EPI_ISL_632741, EPI_ISL_632747, EPI_ISL_632750, EPI_ISL_632751, EPI_ISL_632752, EPI_ISL_632753, EPI_ISL_632759, EPI_ISL_632760, EPI_ISL_632776, EPI_ISL_632778, EPI_ISL_632779, EPI_ISL_632780, EPI_ISL_632781, EPI_ISL_632782, EPI_ISL_632783, EPI_ISL_632784, EPI_ISL_632787 | | | |
| see above | Dutch COVID-19 response team | Erasmus Medical Center | Bas Oude Munnink, David Nieuwenhuijse, Reina Sikkema, Claudia Schapendonk, Irina Chestakova, Anne van der Linden, Theo Bestebroer, Stefan van Nieuwkoop, Mark Pronk, Pascal Lexmond, Corien Swaan, Manon Haverkate, Madelief Mollers, Mart Stein, Sandra Kengne Kamga Mobou, Jeroen van Kampen, Jolanda Voermans, Aura Timen, Corine GeurtsvanKessel, Annemiek van der Eijk, Richard Molenkamp, Marion Koopmans, on behalf of the Dutch national COVID-19 response team. |
| EPI_ISL_632899, EPI_ISL_632900, EPI_ISL_632901, EPI_ISL_632902, EPI_ISL_632903 | Communicable Disease Laboratory, Public Health Directorate | Communicable Disease Laboratory, Public Health Directorate | AlAbbas,Z., Altaif,Z., AlWasti,H., Alhujairi,Z. |
| EPI_ISL_632905 | Communicable Disease Laboratory, Public Health Directorate | Communicable Disease Laboratory, Public Health Directorate | AlHujairi,Z., Altaif,Z., AlWasti,H., AlAbbas,Z. |
| EPI_ISL_632906, EPI_ISL_632907 | Communicable Disease Laboratory, Public Health Directorate | Communicable Disease Laboratory, Public Health Directorate | AlAbbas,Z., Altaif,Z., AlWasti,H., Alhujairi,Z. |
| EPI_ISL_633061, EPI_ISL_633062 | DOHMH Jamaica | New York City Public Health Laboratory | Jade Wang, et al. |
| EPI_ISL_633063 | DOHMH Crown Heights | New York City Public Health Laboratory | Jade Wang, et al. |
| EPI_ISL_633064 | DOHMH Jamaica | New York City Public Health Laboratory | Jade Wang, et al. |
| EPI_ISL_633067, EPI_ISL_633068 | DOHMH Morrisania | New York City Public Health Laboratory | Jade Wang, et al. |
| EPI_ISL_633069 | DOHMH Chelsea | New York City Public Health Laboratory | Jade Wang, et al. |
| EPI_ISL_633071 | DOHMH PHL | New York City Public Health Laboratory | Jade Wang, et al. |
| EPI_ISL_633072 | DOHMH Riverside | New York City Public Health Laboratory | Jade Wang, et al. |
| EPI_ISL_633073 | DOHMH Fort Greene | New York City Public Health Laboratory | Jade Wang, et al. |
| EPI_ISL_633074 | DOHMH Corona | New York City Public Health Laboratory | Jade Wang, et al. |
| EPI_ISL_633075 | DOHMH Morrisania | New York City Public Health Laboratory | Jade Wang, et al. |
| EPI_ISL_633076, EPI_ISL_633077, EPI_ISL_633078 | DOHMH Chelsea | New York City Public Health Laboratory | Jade Wang, et al. |
| EPI_ISL_633079 | DOHMH PHL | New York City Public Health Laboratory | Jade Wang, et al. |
| EPI_ISL_633080, EPI_ISL_633081, EPI_ISL_633082 | DOHMH Morrisania | New York City Public Health Laboratory | Jade Wang, et al. |
| EPI_ISL_633085, EPI_ISL_633086, EPI_ISL_633087, EPI_ISL_633088, EPI_ISL_633089, EPI_ISL_633090, EPI_ISL_633094 | MD PHL | MD PHL | Maryland Department of Health Laboratories Administration |
| EPI_ISL_634839 | Minnesota Department of Health, Public Health Laboratory | Minnesota Department of Health, Public Health Laboratory | Matt Plumb, Jacob Garfin, Alexandra Lorentz, and Xiong Wang |
| EPI_ISL_634890, EPI_ISL_634891, EPI_ISL_634893, EPI_ISL_634894 | Lab voor klinische biologie | Onderzoeksgroep Virologie | Nick Vereecke, Laurens Lambrechts, Marthe Pauwels, Bruno Verhasselt, Linos Vandekerckhove, Hans Nauwynck, Sebastiaan Theuns |
| EPI_ISL_635001, EPI_ISL_635003, EPI_ISL_635004, EPI_ISL_635009, EPI_ISL_635015, EPI_ISL_635016, EPI_ISL_635017, EPI_ISL_635018, EPI_ISL_635019, EPI_ISL_635020, EPI_ISL_635021, EPI_ISL_635022, EPI_ISL_635023, EPI_ISL_635024, EPI_ISL_635025, EPI_ISL_635032, EPI_ISL_635036, EPI_ISL_635037, EPI_ISL_635038, EPI_ISL_635040, EPI_ISL_635052, EPI_ISL_635053 | | | |
| see above | National Health Laboratory Service - Inkosi Albert Luthuli Central Hospital (NHLS-IALCH) | KRISP, KZN Research Innovation and Sequencing Platform | Giandhari J, Pillay S, Lessells R, Mdlalose K, York D, Khan S, Tegally H, Wilkinson E, de Oliveira T |
| EPI_ISL_635068, EPI_ISL_635069, EPI_ISL_635070 | University Hospital of Northern Norway, Department for Microbiology and Infectious Disease Control | Norwegian Institute of Public Health, Department of Virology | Kathrine Stene-Johansen, Kamilla Heddeland Instefjord, Hilde Elshaug, Marie Paulsen Madsen, Rasmus Riis Kopperud, Hilde Vollan, Karoline Bragstad, Olav Hungnes |
| EPI_ISL_635100 | Department of Medical Microbiology, St. Olavs hospital | Norwegian Institute of Public Health, Department of Virology | Kathrine Stene-Johansen, Kamilla Heddeland Instefjord, Hilde Elshaug, Marie Paulsen Madsen, Rasmus Riis Kopperud, Hilde Vollan, Karoline Bragstad, Olav Hungnes |
| EPI_ISL_635101 | Medical Microbiology Unit, Department for Laboratory Medicine, Drammen Hospital, Vestre Viken Health Trust, | Norwegian Institute of Public Health, Department of Virology | Kathrine Stene-Johansen, Kamilla Heddeland Instefjord, Hilde Elshaug, Marie Paulsen Madsen, Rasmus Riis Kopperud, Hilde Vollan, Karoline Bragstad, Olav Hungnes |
| EPI_ISL_635103 | Vestfold Hospital, Toensberg Department of Microbiology | Norwegian Institute of Public Health, Department of Virology | Kathrine Stene-Johansen, Kamilla Heddeland Instefjord, Hilde Elshaug, Marie Paulsen Madsen, Rasmus Riis Kopperud, Hilde Vollan, Karoline Bragstad, Olav Hungnes |
| EPI_ISL_635161 | Hospital of Southern Norway - Kristiansand, Department of Medical Microbiology | Norwegian Institute of Public Health, Department of Virology | Kathrine Stene-Johansen, Kamilla Heddeland Instefjord, Hilde Elshaug, Marie Paulsen Madsen, Rasmus Riis Kopperud, Hilde Vollan, Karoline Bragstad, Olav Hungnes |
| EPI_ISL_635164 | Ostfold Hospital Trust - Kalnes, Centre for Laboratory Medicine, Section for gene technology and infection serology | Norwegian Institute of Public Health, Department of Virology | Kathrine Stene-Johansen, Kamilla Heddeland Instefjord, Hilde Elshaug, Marie Paulsen Madsen, Rasmus Riis Kopperud, Hilde Vollan, Karoline Bragstad, Olav Hungnes |
| EPI_ISL_635165 | Department of Medical Microbiology, St. Olavs hospital | Norwegian Institute of Public Health, Department of Virology | Kathrine Stene-Johansen, Kamilla Heddeland Instefjord, Hilde Elshaug, Marie Paulsen Madsen, Rasmus Riis Kopperud, Hilde Vollan, Karoline Bragstad, Olav Hungnes |
| EPI_ISL_635168 | Vestfold Hospital, Toensberg Department of Microbiology | Norwegian Institute of Public Health, Department of Virology | Kathrine Stene-Johansen, Kamilla Heddeland Instefjord, Hilde Elshaug, Marie Paulsen Madsen, Rasmus Riis Kopperud, Hilde Vollan, Karoline Bragstad, Olav Hungnes |
| EPI_ISL_635169 | Medical Microbiology Unit, Department for Laboratory Medicine, Drammen Hospital, Vestre Viken Health Trust, | Norwegian Institute of Public Health, Department of Virology | Kathrine Stene-Johansen, Kamilla Heddeland Instefjord, Hilde Elshaug, Marie Paulsen Madsen, Rasmus Riis Kopperud, Hilde Vollan, Karoline Bragstad, Olav Hungnes |

| | | | |
|---|---|---|---|
| EPI_ISL_635174, EPI_ISL_635175 | Ostfold Hospital Trust - Kalnes, Centre for Laboratory Medicine, Section for gene technology and infection serology | Norwegian Institute of Public Health, Department of Virology | Kathrine Stene-Johansen, Kamilla Heddeland Instefjord, Hilde Elshaug, Marie Paulsen Madsen, Rasmus Riis Kopperud, Hilde Vollan, Karoline Bragstad, Olav Hungnes |
| EPI_ISL_635179, EPI_ISL_635180 | University Hospital of Northern Norway, Department for Microbiology and Infectious Disease Control | Norwegian Institute of Public Health, Department of Virology | Kathrine Stene-Johansen, Kamilla Heddeland Instefjord, Hilde Elshaug, Marie Paulsen Madsen, Rasmus Riis Kopperud, Hilde Vollan, Karoline Bragstad, Olav Hungnes |
| EPI_ISL_635192, EPI_ISL_635193 | Medical Microbiology Unit, Department for Laboratory Medicine, Drammen Hospital, Vestre Viken Health Trust, | Norwegian Institute of Public Health, Department of Virology | Kathrine Stene-Johansen, Kamilla Heddeland Instefjord, Hilde Elshaug, Marie Paulsen Madsen, Rasmus Riis Kopperud, Hilde Vollan, Karoline Bragstad, Olav Hungnes |
| EPI_ISL_635195 | Vestfold Hospital, Toensberg Department of Microbiology | Norwegian Institute of Public Health, Department of Virology | Kathrine Stene-Johansen, Kamilla Heddeland Instefjord, Hilde Elshaug, Marie Paulsen Madsen, Rasmus Riis Kopperud, Hilde Vollan, Karoline Bragstad, Olav Hungnes |
| EPI_ISL_636491, EPI_ISL_636550, EPI_ISL_636568, EPI_ISL_636569, EPI_ISL_636570, EPI_ISL_636571, EPI_ISL_636585 | Dutch COVID-19 response team | National Institute for Public Health and the Environment (RIVM) | Adam Meijer, Harry Vennema, Jeroen Cremer, Sharon van den Brink, Bas van der Veer, AnneMarie van den Brandt, Florian Zwagemaker, Dennis Schmitz, Chantal Reusken, on behalf of the national COVID-19 response team |
| EPI_ISL_636705, EPI_ISL_636706, EPI_ISL_636707, EPI_ISL_636708, EPI_ISL_636709, EPI_ISL_636710, EPI_ISL_636711, EPI_ISL_636712, EPI_ISL_636713, EPI_ISL_636714, EPI_ISL_636715, EPI_ISL_636736 | | | |
| see above | Respiratory Virus Unit, Microbiology Services Colindale, Public Health England | Respiratory Virus Unit, Microbiology Services Colindale, Public Health England | PHE Covid Sequencing Team |
| EPI_ISL_637088 | PathLab Bay of Plenty | Institute of Environmental Science and Research (ESR) | Xiaoyun Ren, Matt Storey, Nikki Freed, Muhammad Faisal, Jing Wang, Hermes Perez, Anja Werno, Antje van der Linden, Arlo Upton, Chris Mansell, David Hammer, Dragana Drinkovic, Gary McAuliffe, Hana Sofia Andersson, James Ussher, Jill Sherwood, Josh Freeman, Julia Howard, Juliet Elvy, Mary DeAlmeida, Matt Blakiston, Matthew Rogers, Max Bloomfield, Michael Addidle, Michelle Balm, Sally Roberts, Sarah Jefferies, Sharmini Muttaiyah, Susan Morpeth, Susan Taylor, Timothy Blackmore, Vani Sathyendran, Veronica Playle, Virginia Hope, Erasmus Smit, Lauren Jelly, Olin Silander, Joep de Ligt |
| EPI_ISL_637155, EPI_ISL_637156, EPI_ISL_637157 | Respiratory Virus Unit, Microbiology Services Colindale, Public Health England | COVID-19 Genomics UK (COG-UK) Consortium | PHE Covid Sequencing Team |
| EPI_ISL_637258 | Wales Specialist Virology Centre Sequencing lab: Pathogen Genomics Unit | COVID-19 Genomics UK (COG-UK) Consortium | Catherine Moore, Johnathan Evans, Laura Gifford, Malorie Perry, Simon Cottrell, Angela Marchbank, Alec Birchley, Alexander Adams, Amy Gaskin, Bree Gatica-Wilcox, Jason Coombes, Joel Southgate, Lauren Gilbert, Lee Graham, Nicole Pacchiarini, Sara Kumziene-Summerhayes, Sarah Taylor, Sophie Jones, Sara Rey, Matthew Bull, Joanne Watkins, Sally Corden, Tom Connor |
| EPI_ISL_637267, EPI_ISL_637268 | Department of Pathology, University of Cambridge | COVID-19 Genomics UK (COG-UK) Consortium | Aminu S. Jahun, Yasmin Chaudhry, Grant Hall, Iliana Georgana, Myra Hosmillo, Martin D. Curran, Malte Pinckert, Surendra Parmar, Ian Goodfellow |
| EPI_ISL_637290 | Virology Department, Sheffield Teaching Hospitals NHS Foundation Trust/Department of Infection, Immunity and Cardiovascular Disease, The Medical School, University of Sheffield | COVID-19 Genomics UK (COG-UK) Consortium | Thushan de Silva, Matthew Parker, Nikki Smith, Adri Angyal, Rebecca Brown, Luke Green, Rachel Tucker, Paul Parsons, Danielle Groves, Katie Johnson, Laura Carrilero, Alex Keeley, Dave Partridge, Matthew Wyles, Benjamin Lindsey, Mehmet Yavuz, Mohammad Raza, Cariad Evans |
| EPI_ISL_637322 | Northumbria University / South Tees Hospitals NHS Foundation Trust / North Cumbria Integrated Care NHS Foundation Trust / North Tees and Hartlepool NHS Foundation Trust / Newcastle Hospitals NHS Foundation Trust | COVID-19 Genomics UK (COG-UK) Consortium | Darren L Smith,Andrew Nelson,Matthew Bashton,Greg R Young,Joshua Loh,John Allan,Mohammad A Tariq,Giles S Holt,Gary Black,Wen C Yew,Lynn Dover,Paul Baker,Steve Liggett,Sarah Essex,Jane Greenaway,Debra Padgett,Clive Graham,Garren Scott,Edward Barton,Emma Swindells,Brendan Payne,Jennifer Collins,Yusri Taha,Gary Eltringham |
| EPI_ISL_637327 | Wales Specialist Virology Centre Sequencing lab: Pathogen Genomics Unit | COVID-19 Genomics UK (COG-UK) Consortium | Catherine Moore, Johnathan Evans, Laura Gifford, Malorie Perry, Simon Cottrell, Angela Marchbank, Alec Birchley, Alexander Adams, Amy Gaskin, Bree Gatica-Wilcox, Jason Coombes, Joel Southgate, Lauren Gilbert, Lee Graham, Nicole Pacchiarini, Sara Kumziene-Summerhayes, Sarah Taylor, Sophie Jones, Sara Rey, Matthew Bull, Joanne Watkins, Sally Corden, Tom Connor |
| EPI_ISL_637352 | Northumbria University / South Tees Hospitals NHS Foundation Trust / North Cumbria Integrated Care NHS Foundation Trust / North Tees and Hartlepool NHS Foundation Trust / Newcastle Hospitals NHS Foundation Trust | COVID-19 Genomics UK (COG-UK) Consortium | Darren L Smith,Andrew Nelson,Matthew Bashton,Greg R Young,Joshua Loh,John Allan,Mohammad A Tariq,Giles S Holt,Gary Black,Wen C Yew,Lynn Dover,Paul Baker,Steve Liggett,Sarah Essex,Jane Greenaway,Debra Padgett,Clive Graham,Garren Scott,Edward Barton,Emma Swindells,Brendan Payne,Jennifer Collins,Yusri Taha,Gary Eltringham |
| EPI_ISL_637357, EPI_ISL_637388, EPI_ISL_637389, EPI_ISL_637404, EPI_ISL_637405, EPI_ISL_637406, EPI_ISL_637426, EPI_ISL_637435 | Department of Pathology, University of Cambridge | COVID-19 Genomics UK (COG-UK) Consortium | Aminu S. Jahun, Yasmin Chaudhry, Grant Hall, Iliana Georgana, Myra Hosmillo, Martin D. Curran, Malte Pinckert, Surendra Parmar, Ian Goodfellow |
| EPI_ISL_637445, EPI_ISL_637448, EPI_ISL_637449 | Wales Specialist Virology Centre Sequencing lab: Pathogen Genomics Unit | COVID-19 Genomics UK (COG-UK) Consortium | Catherine Moore, Johnathan Evans, Laura Gifford, Malorie Perry, Simon Cottrell, Angela Marchbank, Alec Birchley, Alexander Adams, Amy Gaskin, Bree Gatica-Wilcox, Jason Coombes, Joel Southgate, Lauren Gilbert, Lee Graham, Nicole Pacchiarini, Sara Kumziene-Summerhayes, Sarah Taylor, Sophie Jones, Sara Rey, Matthew Bull, Joanne Watkins, Sally Corden, Tom Connor |
| EPI_ISL_637460, EPI_ISL_637472, EPI_ISL_637473, EPI_ISL_637474, EPI_ISL_637503, EPI_ISL_637508 | Department of Pathology, University of Cambridge | COVID-19 Genomics UK (COG-UK) Consortium | Aminu S. Jahun, Yasmin Chaudhry, Grant Hall, Iliana Georgana, Myra Hosmillo, Martin D. Curran, Malte Pinckert, Surendra Parmar, Ian Goodfellow |
| EPI_ISL_637520 | Northumbria University / South Tees Hospitals NHS Foundation Trust / North Cumbria Integrated Care NHS Foundation Trust / North Tees and Hartlepool NHS Foundation Trust / Newcastle Hospitals NHS Foundation Trust | COVID-19 Genomics UK (COG-UK) Consortium | Darren L Smith,Andrew Nelson,Matthew Bashton,Greg R Young,Joshua Loh,John Allan,Mohammad A Tariq,Giles S Holt,Gary Black,Wen C Yew,Lynn Dover,Paul Baker,Steve Liggett,Sarah Essex,Jane Greenaway,Debra Padgett,Clive Graham,Garren Scott,Edward Barton,Emma Swindells,Brendan Payne,Jennifer Collins,Yusri Taha,Gary Eltringham |
| EPI_ISL_637527 | Department of Pathology, University of Cambridge | COVID-19 Genomics UK (COG-UK) Consortium | Aminu S. Jahun, Yasmin Chaudhry, Grant Hall, Iliana Georgana, Myra Hosmillo, Martin D. Curran, Malte Pinckert, Surendra Parmar, Ian Goodfellow |
| EPI_ISL_637538 | Virology Department, Sheffield Teaching Hospitals NHS Foundation Trust/Department of Infection, Immunity and Cardiovascular Disease, The Medical School, University of Sheffield | COVID-19 Genomics UK (COG-UK) Consortium | Thushan de Silva, Matthew Parker, Nikki Smith, Adri Angyal, Rebecca Brown, Luke Green, Rachel Tucker, Paul Parsons, Danielle Groves, Katie Johnson, Laura Carrilero, Alex Keeley, Dave Partridge, Matthew Wyles, Benjamin Lindsey, Mehmet Yavuz, Mohammad Raza, Cariad Evans |
| EPI_ISL_637544, EPI_ISL_637546 | Wales Specialist Virology Centre Sequencing lab: Pathogen Genomics Unit | COVID-19 Genomics UK (COG-UK) Consortium | Catherine Moore, Johnathan Evans, Laura Gifford, Malorie Perry, Simon Cottrell, Angela Marchbank, Alec Birchley, Alexander Adams, Amy Gaskin, Bree Gatica-Wilcox, Jason Coombes, Joel Southgate, Lauren Gilbert, Lee Graham, Nicole Pacchiarini, Sara Kumziene-Summerhayes, Sarah Taylor, Sophie Jones, Sara Rey, Matthew Bull, Joanne Watkins, Sally Corden, Tom Connor |
| EPI_ISL_637560 | Department of Pathology, University of Cambridge | COVID-19 Genomics UK (COG-UK) Consortium | Aminu S. Jahun, Yasmin Chaudhry, Grant Hall, Iliana Georgana, Myra Hosmillo, Martin D. Curran, Malte Pinckert, Surendra Parmar, Ian Goodfellow |
| EPI_ISL_637561 | Virology Department, Sheffield Teaching Hospitals NHS Foundation Trust/Department of Infection, Immunity and Cardiovascular Disease, The Medical School, University of Sheffield | COVID-19 Genomics UK (COG-UK) Consortium | Thushan de Silva, Matthew Parker, Nikki Smith, Adri Angyal, Rebecca Brown, Luke Green, Rachel Tucker, Paul Parsons, Danielle Groves, Katie Johnson, Laura Carrilero, Alex Keeley, Dave Partridge, Matthew Wyles, Benjamin Lindsey, Mehmet Yavuz, Mohammad Raza, Cariad Evans |
| EPI_ISL_637601, EPI_ISL_637606 | Wales Specialist Virology Centre Sequencing lab: Pathogen Genomics Unit | COVID-19 Genomics UK (COG-UK) Consortium | Catherine Moore, Johnathan Evans, Laura Gifford, Malorie Perry, Simon Cottrell, Angela Marchbank, Alec Birchley, Alexander Adams, Amy Gaskin, Bree Gatica-Wilcox, Jason Coombes, Joel Southgate, Lauren Gilbert, Lee Graham, Nicole Pacchiarini, Sara Kumziene-Summerhayes, Sarah Taylor, Sophie Jones, Sara Rey, Matthew Bull, Joanne Watkins, Sally Corden, Tom Connor |
| EPI_ISL_637613 | Department of Pathology, University of Cambridge | COVID-19 Genomics UK (COG-UK) Consortium | Aminu S. Jahun, Yasmin Chaudhry, Grant Hall, Iliana Georgana, Myra Hosmillo, Martin D. Curran, Malte Pinckert, Surendra Parmar, Ian Goodfellow |

| | | | |
|---|---|---|---|
| EPI_ISL_637616, EPI_ISL_637618, EPI_ISL_637619 | Wales Specialist Virology Centre Sequencing lab: Pathogen Genomics Unit | COVID-19 Genomics UK (COG-UK) Consortium | Catherine Moore, Johnathan Evans, Laura Gifford, Malorie Perry, Simon Cottrell, Angela Marchbank, Alec Birchley, Alexander Adams, Amy Gaskin, Bree Gatica-Wilcox, Jason Coombes, Joel Southgate, Lauren Gilbert, Lee Graham, Nicole Pacchiarini, Sara Kumziene-Summerhayes, Sarah Taylor, Sophie Jones, Sara Rey, Matthew Bull, Joanne Watkins, Sally Corden, Tom Connor |
| EPI_ISL_637621 | Virology Department, Sheffield Teaching Hospitals NHS Foundation Trust/Department of Infection, Immunity and Cardiovascular Disease, The Medical School, University of Sheffield | COVID-19 Genomics UK (COG-UK) Consortium | Thushan de Silva, Matthew Parker, Nikki Smith, Adri Angyal, Rebecca Brown, Luke Green, Rachel Tucker, Paul Parsons, Danielle Groves, Katie Johnson, Laura Carrilero, Alex Keeley, Dave Partridge, Matthew Wyles, Benjamin Lindsey, Mehmet Yavuz, Mohammad Raza, Cariad Evans |
| EPI_ISL_637638, EPI_ISL_637641, EPI_ISL_637643, EPI_ISL_637685, EPI_ISL_637686, EPI_ISL_637687, EPI_ISL_637688, EPI_ISL_637689, EPI_ISL_637690, EPI_ISL_637691, EPI_ISL_637692, EPI_ISL_637693, EPI_ISL_637704, EPI_ISL_637706, EPI_ISL_637707, EPI_ISL_637721, EPI_ISL_637722, EPI_ISL_637728, EPI_ISL_637729, EPI_ISL_637730, EPI_ISL_637731, EPI_ISL_637732, EPI_ISL_637733, EPI_ISL_637734, EPI_ISL_637735, EPI_ISL_637736, EPI_ISL_637748 | | | |
| see above | Department of Pathology, University of Cambridge | COVID-19 Genomics UK (COG-UK) Consortium | Aminu S. Jahun, Yasmin Chaudhry, Grant Hall, Iliana Georgana, Myra Hosmillo, Martin D. Curran, Malte Pinckert, Surendra Parmar, Ian Goodfellow |
| EPI_ISL_637785, EPI_ISL_637786, EPI_ISL_637787, EPI_ISL_637795, EPI_ISL_637796 | Northumbria University / South Tees Hospitals NHS Foundation Trust / North Cumbria Integrated Care NHS Foundation Trust / North Tees and Hartlepool NHS Foundation Trust / Newcastle Hospitals NHS Foundation Trust | COVID-19 Genomics UK (COG-UK) Consortium | Darren L Smith,Andrew Nelson,Matthew Bashton,Greg R Young,Joshua Loh,John Allan,Mohammad A Tariq,Giles S Holt,Gary Black,Wen C Yew,Lynn Dover,Paul Baker,Steve Liggett,Sarah Essex,Jane Greenaway,Debra Padgett,Clive Graham,Garren Scott,Edward Barton,Emma Swindells,Brendan Payne,Jennifer Collins,Yusri Taha,Gary Eltringham |
| EPI_ISL_637858 | Oxford Viromics, NDM, University of Oxford; Oxford University Hospitals; Basingstoke and North Hampshire Hospital | COVID-19 Genomics UK (COG-UK) Consortium | Tanya Golubchik, David Bonsall, George Macintyre, Amy Trebes, Mariateresa de Cesare, Catrin Moore, Alex Mobbs, Anita Justice, Robert Shaw, Monique Andersson, Timothy Peto, Emma Wise, Nathan Moore, Jessica Lynch, Nick Cortes, Matilde Mori, Stephen Kidd, David Buck, John Todd, Christophe Fraser |
| EPI_ISL_637859 | Department of Pathology, University of Cambridge | COVID-19 Genomics UK (COG-UK) Consortium | Aminu S. Jahun, Yasmin Chaudhry, Grant Hall, Iliana Georgana, Myra Hosmillo, Martin D. Curran, Malte Pinckert, Surendra Parmar, Ian Goodfellow |
| EPI_ISL_637861, EPI_ISL_637871 | Oxford Viromics, NDM, University of Oxford; Oxford University Hospitals; Basingstoke and North Hampshire Hospital | COVID-19 Genomics UK (COG-UK) Consortium | Tanya Golubchik, David Bonsall, George Macintyre, Amy Trebes, Mariateresa de Cesare, Catrin Moore, Alex Mobbs, Anita Justice, Robert Shaw, Monique Andersson, Timothy Peto, Emma Wise, Nathan Moore, Jessica Lynch, Nick Cortes, Matilde Mori, Stephen Kidd, David Buck, John Todd, Christophe Fraser |
| EPI_ISL_637882 | Wales Specialist Virology Centre Sequencing lab: Pathogen Genomics Unit | COVID-19 Genomics UK (COG-UK) Consortium | Catherine Moore, Johnathan Evans, Laura Gifford, Malorie Perry, Simon Cottrell, Angela Marchbank, Alec Birchley, Alexander Adams, Amy Gaskin, Bree Gatica-Wilcox, Jason Coombes, Joel Southgate, Lauren Gilbert, Lee Graham, Nicole Pacchiarini, Sara Kumziene-Summerhayes, Sarah Taylor, Sophie Jones, Sara Rey, Matthew Bull, Joanne Watkins, Sally Corden, Tom Connor |
| EPI_ISL_637887, EPI_ISL_637901 | Virology Department, Sheffield Teaching Hospitals NHS Foundation Trust/Department of Infection, Immunity and Cardiovascular Disease, The Medical School, University of Sheffield | COVID-19 Genomics UK (COG-UK) Consortium | Thushan de Silva, Matthew Parker, Nikki Smith, Adri Angyal, Rebecca Brown, Luke Green, Rachel Tucker, Paul Parsons, Danielle Groves, Katie Johnson, Laura Carrilero, Alex Keeley, Dave Partridge, Matthew Wyles, Benjamin Lindsey, Mehmet Yavuz, Mohammad Raza, Cariad Evans |
| EPI_ISL_637936, EPI_ISL_637939, EPI_ISL_637942 | Department of Pathology, University of Cambridge | COVID-19 Genomics UK (COG-UK) Consortium | Aminu S. Jahun, Yasmin Chaudhry, Grant Hall, Iliana Georgana, Myra Hosmillo, Martin D. Curran, Malte Pinckert, Surendra Parmar, Ian Goodfellow |
| EPI_ISL_637946 | Virology Department, Sheffield Teaching Hospitals NHS Foundation Trust/Department of Infection, Immunity and Cardiovascular Disease, The Medical School, University of Sheffield | COVID-19 Genomics UK (COG-UK) Consortium | Thushan de Silva, Matthew Parker, Nikki Smith, Adri Angyal, Rebecca Brown, Luke Green, Rachel Tucker, Paul Parsons, Danielle Groves, Katie Johnson, Laura Carrilero, Alex Keeley, Dave Partridge, Matthew Wyles, Benjamin Lindsey, Mehmet Yavuz, Mohammad Raza, Cariad Evans |
| EPI_ISL_637974 | Department of Pathology, University of Cambridge | COVID-19 Genomics UK (COG-UK) Consortium | Aminu S. Jahun, Yasmin Chaudhry, Grant Hall, Iliana Georgana, Myra Hosmillo, Martin D. Curran, Malte Pinckert, Surendra Parmar, Ian Goodfellow |
| EPI_ISL_637979 | Northumbria University / South Tees Hospitals NHS Foundation Trust / North Cumbria Integrated Care NHS Foundation Trust / North Tees and Hartlepool NHS Foundation Trust / Newcastle Hospitals NHS Foundation Trust | COVID-19 Genomics UK (COG-UK) Consortium | Darren L Smith,Andrew Nelson,Matthew Bashton,Greg R Young,Joshua Loh,John Allan,Mohammad A Tariq,Giles S Holt,Gary Black,Wen C Yew,Lynn Dover,Paul Baker,Steve Liggett,Sarah Essex,Jane Greenaway,Debra Padgett,Clive Graham,Garren Scott,Edward Barton,Emma Swindells,Brendan Payne,Jennifer Collins,Yusri Taha,Gary Eltringham |
| EPI_ISL_638019 | Department of Pathology, University of Cambridge | COVID-19 Genomics UK (COG-UK) Consortium | Aminu S. Jahun, Yasmin Chaudhry, Grant Hall, Iliana Georgana, Myra Hosmillo, Martin D. Curran, Malte Pinckert, Surendra Parmar, Ian Goodfellow |
| EPI_ISL_638037 | Oxford Viromics, NDM, University of Oxford; Oxford University Hospitals; Basingstoke and North Hampshire Hospital | COVID-19 Genomics UK (COG-UK) Consortium | Tanya Golubchik, David Bonsall, George Macintyre, Amy Trebes, Mariateresa de Cesare, Catrin Moore, Alex Mobbs, Anita Justice, Robert Shaw, Monique Andersson, Timothy Peto, Emma Wise, Nathan Moore, Jessica Lynch, Nick Cortes, Matilde Mori, Stephen Kidd, David Buck, John Todd, Christophe Fraser |
| EPI_ISL_638038, EPI_ISL_638039 | Department of Pathology, University of Cambridge | COVID-19 Genomics UK (COG-UK) Consortium | Aminu S. Jahun, Yasmin Chaudhry, Grant Hall, Iliana Georgana, Myra Hosmillo, Martin D. Curran, Malte Pinckert, Surendra Parmar, Ian Goodfellow |
| EPI_ISL_638051 | Wales Specialist Virology Centre Sequencing lab: Pathogen Genomics Unit | COVID-19 Genomics UK (COG-UK) Consortium | Catherine Moore, Johnathan Evans, Laura Gifford, Malorie Perry, Simon Cottrell, Angela Marchbank, Alec Birchley, Alexander Adams, Amy Gaskin, Bree Gatica-Wilcox, Jason Coombes, Joel Southgate, Lauren Gilbert, Lee Graham, Nicole Pacchiarini, Sara Kumziene-Summerhayes, Sarah Taylor, Sophie Jones, Sara Rey, Matthew Bull, Joanne Watkins, Sally Corden, Tom Connor |
| EPI_ISL_638067, EPI_ISL_638068 | Department of Pathology, University of Cambridge | COVID-19 Genomics UK (COG-UK) Consortium | Aminu S. Jahun, Yasmin Chaudhry, Grant Hall, Iliana Georgana, Myra Hosmillo, Martin D. Curran, Malte Pinckert, Surendra Parmar, Ian Goodfellow |
| EPI_ISL_638073, EPI_ISL_638074, EPI_ISL_638075, EPI_ISL_638076, EPI_ISL_638077, EPI_ISL_638078, EPI_ISL_638088, EPI_ISL_638089 | Oxford Viromics, NDM, University of Oxford; Oxford University Hospitals; Basingstoke and North Hampshire Hospital | COVID-19 Genomics UK (COG-UK) Consortium | Tanya Golubchik, David Bonsall, George Macintyre, Amy Trebes, Mariateresa de Cesare, Catrin Moore, Alex Mobbs, Anita Justice, Robert Shaw, Monique Andersson, Timothy Peto, Emma Wise, Nathan Moore, Jessica Lynch, Nick Cortes, Matilde Mori, Stephen Kidd, David Buck, John Todd, Christophe Fraser |
| EPI_ISL_638096, EPI_ISL_638099 | Wales Specialist Virology Centre Sequencing lab: Pathogen Genomics Unit | COVID-19 Genomics UK (COG-UK) Consortium | Catherine Moore, Johnathan Evans, Laura Gifford, Malorie Perry, Simon Cottrell, Angela Marchbank, Alec Birchley, Alexander Adams, Amy Gaskin, Bree Gatica-Wilcox, Jason Coombes, Joel Southgate, Lauren Gilbert, Lee Graham, Nicole Pacchiarini, Sara Kumziene-Summerhayes, Sarah Taylor, Sophie Jones, Sara Rey, Matthew Bull, Joanne Watkins, Sally Corden, Tom Connor |
| EPI_ISL_638138 | Department of Pathology, University of Cambridge | COVID-19 Genomics UK (COG-UK) Consortium | Aminu S. Jahun, Yasmin Chaudhry, Grant Hall, Iliana Georgana, Myra Hosmillo, Martin D. Curran, Malte Pinckert, Surendra Parmar, Ian Goodfellow |
| EPI_ISL_638144, EPI_ISL_638164, EPI_ISL_638170, EPI_ISL_638171, EPI_ISL_638173, EPI_ISL_638175, EPI_ISL_638176 | Oxford Viromics, NDM, University of Oxford; Oxford University Hospitals; Basingstoke and North Hampshire Hospital | COVID-19 Genomics UK (COG-UK) Consortium | Tanya Golubchik, David Bonsall, George Macintyre, Amy Trebes, Mariateresa de Cesare, Catrin Moore, Alex Mobbs, Anita Justice, Robert Shaw, Monique Andersson, Timothy Peto, Emma Wise, Nathan Moore, Jessica Lynch, Nick Cortes, Matilde Mori, Stephen Kidd, David Buck, John Todd, Christophe Fraser |
| EPI_ISL_638177, EPI_ISL_638178, EPI_ISL_638179, EPI_ISL_638180, EPI_ISL_638181, EPI_ISL_638182, EPI_ISL_638183, EPI_ISL_638184, EPI_ISL_638185, EPI_ISL_638186, EPI_ISL_638190, EPI_ISL_638191, EPI_ISL_638192, EPI_ISL_638202, EPI_ISL_638204, EPI_ISL_638205, EPI_ISL_638293, EPI_ISL_638294, EPI_ISL_638295, EPI_ISL_638296, EPI_ISL_638297, EPI_ISL_638298, EPI_ISL_638299, EPI_ISL_638302, EPI_ISL_638303, EPI_ISL_638304, EPI_ISL_638305, EPI_ISL_638306, EPI_ISL_638307, EPI_ISL_638308, EPI_ISL_638309, EPI_ISL_638310, EPI_ISL_638311, EPI_ISL_638312, EPI_ISL_638317, EPI_ISL_638318, EPI_ISL_638319, EPI_ISL_638372, EPI_ISL_638373, EPI_ISL_638374, EPI_ISL_638375, EPI_ISL_638376, EPI_ISL_638377, EPI_ISL_638378, EPI_ISL_638379, EPI_ISL_638380, EPI_ISL_638381, EPI_ISL_638382, EPI_ISL_638389, EPI_ISL_638391, EPI_ISL_638393, EPI_ISL_638395, EPI_ISL_638396, EPI_ISL_638397, EPI_ISL_638398, EPI_ISL_638399, EPI_ISL_638400 | | | |
| see above | Department of Pathology, University of Cambridge | COVID-19 Genomics UK (COG-UK) Consortium | Aminu S. Jahun, Yasmin Chaudhry, Grant Hall, Iliana Georgana, Myra Hosmillo, Martin D. Curran, Malte Pinckert, Surendra Parmar, Ian Goodfellow |
| EPI_ISL_638459, EPI_ISL_638460, EPI_ISL_638461, EPI_ISL_638462, EPI_ISL_638463, EPI_ISL_638464, EPI_ISL_638465, EPI_ISL_638466, EPI_ISL_638467, EPI_ISL_638468, EPI_ISL_638469, EPI_ISL_638470, EPI_ISL_638471, EPI_ISL_638472, EPI_ISL_638473, EPI_ISL_638474, EPI_ISL_638475, EPI_ISL_638476, EPI_ISL_638477, EPI_ISL_638478, EPI_ISL_638479, EPI_ISL_638480, EPI_ISL_638481, EPI_ISL_638483, EPI_ISL_638484, EPI_ISL_638485, EPI_ISL_638486, EPI_ISL_638487, EPI_ISL_638488, EPI_ISL_638489, EPI_ISL_638490, EPI_ISL_638491, EPI_ISL_638492, EPI_ISL_638493, EPI_ISL_638494, EPI_ISL_638495, EPI_ISL_638496, EPI_ISL_638497, EPI_ISL_638498, EPI_ISL_638499, EPI_ISL_638500, EPI_ISL_638501, EPI_ISL_638502, EPI_ISL_638503, EPI_ISL_638504, EPI_ISL_638505, EPI_ISL_638506, EPI_ISL_638507, EPI_ISL_638508, EPI_ISL_638509, EPI_ISL_638510, EPI_ISL_638511, EPI_ISL_638512, EPI_ISL_638513, EPI_ISL_638514, EPI_ISL_638515, EPI_ISL_638516, EPI_ISL_638517, EPI_ISL_638518, EPI_ISL_638519, EPI_ISL_638520, EPI_ISL_638521, EPI_ISL_638522, EPI_ISL_638523, EPI_ISL_638524, EPI_ISL_638525, EPI_ISL_638526, EPI_ISL_638527, EPI_ISL_638528, EPI_ISL_638529, EPI_ISL_638530 | | | |
| see above | Oxford Viromics, NDM, University of Oxford; Oxford University Hospitals; Basingstoke and North Hampshire Hospital | COVID-19 Genomics UK (COG-UK) Consortium | Tanya Golubchik, David Bonsall, George Macintyre, Amy Trebes, Mariateresa de Cesare, Catrin Moore, Alex Mobbs, Anita Justice, Robert Shaw, Monique Andersson, Timothy Peto, Emma Wise, Nathan Moore, Jessica Lynch, Nick Cortes, Matilde Mori, Stephen Kidd, David Buck, John Todd, Christophe Fraser |

| | | | |
|---|---|---|---|
| EPI_ISL_638555, EPI_ISL_638556, EPI_ISL_638571, EPI_ISL_638572, EPI_ISL_638573, EPI_ISL_638574, EPI_ISL_638575, EPI_ISL_638576, EPI_ISL_638577, EPI_ISL_638580 | Northumbria University / South Tees Hospitals NHS Foundation Trust / North Cumbria Integrated Care NHS Foundation Trust / North Tees and Hartlepool NHS Foundation Trust / Newcastle Hospitals NHS Foundation Trust | COVID-19 Genomics UK (COG-UK) Consortium | Darren L Smith,Andrew Nelson,Matthew Bashton,Greg R Young,Joshua Loh,John Allan,Mohammad A Tariq,Giles S Holt,Gary Black,Wen C Yew,Lynn Dover,Paul Baker,Steve Liggett,Sarah Essex,Jane Greenaway,Debra Padgett,Clive Graham,Garren Scott,Edward Barton,Emma Swindells,Brendan Payne,Jennifer Collins,Yusri Taha,Gary Eltringham |
| EPI_ISL_638739, EPI_ISL_638740, EPI_ISL_638746, EPI_ISL_638747, EPI_ISL_638748, EPI_ISL_638761, EPI_ISL_638765, EPI_ISL_638772, EPI_ISL_638773, EPI_ISL_638774, EPI_ISL_638775, EPI_ISL_638800, EPI_ISL_638801, EPI_ISL_638802, EPI_ISL_638803, EPI_ISL_638804, EPI_ISL_638810, EPI_ISL_638811, EPI_ISL_638812, EPI_ISL_638813, EPI_ISL_638814, EPI_ISL_638815, EPI_ISL_638816, EPI_ISL_638817, EPI_ISL_638818, EPI_ISL_638819, EPI_ISL_638820, EPI_ISL_638821, EPI_ISL_638822, EPI_ISL_638823, EPI_ISL_638824, EPI_ISL_638825, EPI_ISL_638826, EPI_ISL_638827, EPI_ISL_638828, EPI_ISL_638829, EPI_ISL_638830, EPI_ISL_638831, EPI_ISL_638832, EPI_ISL_638833, EPI_ISL_638834, EPI_ISL_638835, EPI_ISL_638836, EPI_ISL_638837, EPI_ISL_638838, EPI_ISL_638839, EPI_ISL_638840, EPI_ISL_638841, EPI_ISL_638842, EPI_ISL_638843, EPI_ISL_638844, EPI_ISL_638845, EPI_ISL_638846, EPI_ISL_638847, EPI_ISL_638848, EPI_ISL_638849, EPI_ISL_638850, EPI_ISL_638851, EPI_ISL_638852, EPI_ISL_638853, EPI_ISL_638854, EPI_ISL_638855, EPI_ISL_638856, EPI_ISL_638857, EPI_ISL_638858, EPI_ISL_638859, EPI_ISL_638860, EPI_ISL_638861, EPI_ISL_638862, EPI_ISL_638863, EPI_ISL_638864, EPI_ISL_638865, EPI_ISL_638866, EPI_ISL_638867, EPI_ISL_638868, EPI_ISL_638869, EPI_ISL_638870, EPI_ISL_638871, EPI_ISL_638872, EPI_ISL_638873, EPI_ISL_638874, EPI_ISL_638875, EPI_ISL_638876 | | | |
| see above | Oxford Viromics, NDM, University of Oxford; Oxford University Hospitals; Basingstoke and North Hampshire Hospital | COVID-19 Genomics UK (COG-UK) Consortium | Tanya Golubchik, David Bonsall, George Macintyre, Amy Trebes, Mariateresa de Cesare, Catrin Moore, Alex Mobbs, Anita Justice, Robert Shaw, Monique Andersson, Timothy Peto, Emma Wise, Nathan Moore, Jessica Lynch, Nick Cortes, Matilde Mori, Stephen Kidd, David Buck, John Todd, Christophe Fraser |
| EPI_ISL_638892 | Virology Department, Sheffield Teaching Hospitals NHS Foundation Trust/Department of Infection, Immunity and Cardiovascular Disease, The Medical School, University of Sheffield | COVID-19 Genomics UK (COG-UK) Consortium | Thushan de Silva, Matthew Parker, Nikki Smith, Adri Angyal, Rebecca Brown, Luke Green, Rachel Tucker, Paul Parsons, Danielle Groves, Katie Johnson, Laura Carrilero, Alex Keeley, Dave Partridge, Matthew Wyles, Benjamin Lindsey, Mehmet Yavuz, Mohammad Raza, Cariad Evans |
| EPI_ISL_638894, EPI_ISL_638944, EPI_ISL_638945, EPI_ISL_638946, EPI_ISL_638947, EPI_ISL_638948, EPI_ISL_638949, EPI_ISL_638950, EPI_ISL_638951, EPI_ISL_638952, EPI_ISL_638953, EPI_ISL_638954, EPI_ISL_638955, EPI_ISL_638956, EPI_ISL_638957, EPI_ISL_638958, EPI_ISL_638959, EPI_ISL_638960, EPI_ISL_638961, EPI_ISL_638962, EPI_ISL_638963, EPI_ISL_638964, EPI_ISL_638965, EPI_ISL_638966, EPI_ISL_638967, EPI_ISL_638968, EPI_ISL_638969, EPI_ISL_638970, EPI_ISL_638971, EPI_ISL_638972, EPI_ISL_638973, EPI_ISL_638974, EPI_ISL_638975, EPI_ISL_638976, EPI_ISL_638977, EPI_ISL_638978, EPI_ISL_638979, EPI_ISL_638980, EPI_ISL_638981, EPI_ISL_638982, EPI_ISL_638983, EPI_ISL_638984, EPI_ISL_638986, EPI_ISL_638987, EPI_ISL_638988, EPI_ISL_638989, EPI_ISL_638990, EPI_ISL_638991, EPI_ISL_638992, EPI_ISL_638993, EPI_ISL_638994, EPI_ISL_638995, EPI_ISL_638996, EPI_ISL_638997, EPI_ISL_638998, EPI_ISL_638999, EPI_ISL_639000, EPI_ISL_639001 | | | |
| see above | Oxford Viromics, NDM, University of Oxford; Oxford University Hospitals; Basingstoke and North Hampshire Hospital | COVID-19 Genomics UK (COG-UK) Consortium | Tanya Golubchik, David Bonsall, George Macintyre, Amy Trebes, Mariateresa de Cesare, Catrin Moore, Alex Mobbs, Anita Justice, Robert Shaw, Monique Andersson, Timothy Peto, Emma Wise, Nathan Moore, Jessica Lynch, Nick Cortes, Matilde Mori, Stephen Kidd, David Buck, John Todd, Christophe Fraser |
| EPI_ISL_639012, EPI_ISL_639013, EPI_ISL_639014, EPI_ISL_639023, EPI_ISL_639030, EPI_ISL_639046, EPI_ISL_639050, EPI_ISL_639052, EPI_ISL_639064, EPI_ISL_639065, EPI_ISL_639071, EPI_ISL_639072, EPI_ISL_639073, EPI_ISL_639081, EPI_ISL_639083, EPI_ISL_639086, EPI_ISL_639087, EPI_ISL_639094, EPI_ISL_639095, EPI_ISL_639097, EPI_ISL_639100, EPI_ISL_639104, EPI_ISL_639107, EPI_ISL_639130, EPI_ISL_639134, EPI_ISL_639135, EPI_ISL_639136, EPI_ISL_639141, EPI_ISL_639143, EPI_ISL_639147, EPI_ISL_639149, EPI_ISL_639150, EPI_ISL_639151, EPI_ISL_639153, EPI_ISL_639163, EPI_ISL_639165, EPI_ISL_639168, EPI_ISL_639169, EPI_ISL_639170, EPI_ISL_639172, EPI_ISL_639177, EPI_ISL_639184, EPI_ISL_639187 | | | |
| see above | Wales Specialist Virology Centre Sequencing lab: Pathogen Genomics Unit | COVID-19 Genomics UK (COG-UK) Consortium | Catherine Moore, Johnathan Evans, Laura Gifford, Malorie Perry, Simon Cottrell, Angela Marchbank, Alec Birchley, Alexander Adams, Amy Gaskin, Bree Gatica-Wilcox, Jason Coombes, Joel Southgate, Lauren Gilbert, Lee Graham, Nicole Pacchiarini, Sara Kumziene-Summerhayes, Sarah Taylor, Sophie Jones, Sara Rey, Matthew Bull, Joanne Watkins, Sally Corden, Tom Connor |
| EPI_ISL_639628, EPI_ISL_639629, EPI_ISL_639630 | Oxford Viromics, NDM, University of Oxford; Oxford University Hospitals; Basingstoke and North Hampshire Hospital | COVID-19 Genomics UK (COG-UK) Consortium | Tanya Golubchik, David Bonsall, George Macintyre, Amy Trebes, Mariateresa de Cesare, Catrin Moore, Alex Mobbs, Anita Justice, Robert Shaw, Monique Andersson, Timothy Peto, Emma Wise, Nathan Moore, Jessica Lynch, Nick Cortes, Matilde Mori, Stephen Kidd, David Buck, John Todd, Christophe Fraser |
| EPI_ISL_639674, EPI_ISL_639686, EPI_ISL_639691 | E. Gulbja Laboratorija | Latvian Biomedical Research and Study Centre | Ivars Silamielis, Kaspars Megnis, Monta Ustinova, ikita Zrelovs, Vita Rovte, Mikus Gavars, Dmitrijs Perminovs, Uga Dumpis, Jnis Kloviš |
| EPI_ISL_639732 | Sydney South West Pathology Service (SSWPS) - Liverpool Hospital - NSW Health Pathology | NSW Health Pathology - Institute of Clinical Pathology and Medical Research; Westmead Hospital; University of Sydney | CIDM-PH et al. |
| EPI_ISL_639832, EPI_ISL_639840, EPI_ISL_639842, EPI_ISL_639885, EPI_ISL_639910 | National Virus Reference Laboratory | National Virus Reference Laboratory | Michael Carr, Gabriel Gonzalez, Jonathan Dean, Daniel Hare, Cillian F De Gascun |
| EPI_ISL_639977, EPI_ISL_639981, EPI_ISL_640006 | CNR Virus des Infections Respiratoires - France SUD | CNR Virus des Infections Respiratoires - France SUD | Antonin Bal, Gregory Destras, Gwendolyne Burfin, Hadrien Règue, Alexandre Gaymard, Maude Bouscambert-Duchamp, Florence Morfin-Sherpa, Martine Valette, Bruno Lina, Laurence Josset |
| EPI_ISL_640161 | University of Michigan Clinical Microbiology Laboratory | Lauring Lab, University of Michigan, Department of Microbiology and Immunology | Valesano |
| EPI_ISL_640872 | Microbiological Diagnostic Unit - Public Health Laboratory (MDU-PHL) | MDU-PHL | Seemann T., Schultz M.B., Sait, M.L., Sherry, N.L. |
| EPI_ISL_641012, EPI_ISL_641017 | Victorian Infectious Diseases Reference Laboratory (VIDRL) | VIDRL and MDU-PHL | Caly L., Seemann T., Sait, M.L., Schultz M.B., Druce J., Sherry, N.L. |
| EPI_ISL_641214 | Microbiological Diagnostic Unit - Public Health Laboratory (MDU-PHL) | MDU-PHL | Seemann T., Schultz M.B., Sait, M.L., Sherry, N.L. |
| EPI_ISL_641417, EPI_ISL_641418, EPI_ISL_641419, EPI_ISL_641420, EPI_ISL_641421, EPI_ISL_641422, EPI_ISL_641423 | Department of Virus and Microbiological Special Diagnostics, Statens Serum Institut, Copenhagen, Denmark | Albertsen lab, Department of Chemistry and Bioscience, Aalborg University, Denmark | Thomas Bruun Rasmussen, Jannik Fonager, Morten Rasmussen |
| EPI_ISL_643044, EPI_ISL_643045, EPI_ISL_643046, EPI_ISL_643047, EPI_ISL_643048, EPI_ISL_643049, EPI_ISL_643050, EPI_ISL_643051, EPI_ISL_643052, EPI_ISL_643053, EPI_ISL_643054, EPI_ISL_643055, EPI_ISL_643056, EPI_ISL_643057, EPI_ISL_643058, EPI_ISL_643059, EPI_ISL_643060, EPI_ISL_643061, EPI_ISL_643062, EPI_ISL_643063, EPI_ISL_643064, EPI_ISL_643065, EPI_ISL_643066, EPI_ISL_643067, EPI_ISL_643068, EPI_ISL_643069, EPI_ISL_643070, EPI_ISL_643071, EPI_ISL_643072, EPI_ISL_643073, EPI_ISL_643074, EPI_ISL_643075, EPI_ISL_643076, EPI_ISL_643077, EPI_ISL_643078, EPI_ISL_643079, EPI_ISL_643080, EPI_ISL_643081, EPI_ISL_643082, EPI_ISL_643083, EPI_ISL_643084, EPI_ISL_643085, EPI_ISL_643086, EPI_ISL_643087, EPI_ISL_643088, EPI_ISL_643089, EPI_ISL_643090, EPI_ISL_643091, EPI_ISL_643092, EPI_ISL_643093, EPI_ISL_643094, EPI_ISL_643095, EPI_ISL_643096, EPI_ISL_643097, EPI_ISL_643098, EPI_ISL_643099, EPI_ISL_643100, EPI_ISL_643101, EPI_ISL_643102, EPI_ISL_643103, EPI_ISL_643104, EPI_ISL_643105, EPI_ISL_643106, EPI_ISL_643107, EPI_ISL_643108, EPI_ISL_643109, EPI_ISL_643110, EPI_ISL_643111, EPI_ISL_643112, EPI_ISL_643113, EPI_ISL_643114, EPI_ISL_643115, EPI_ISL_643116, EPI_ISL_643118, EPI_ISL_643119, EPI_ISL_643120, EPI_ISL_643121, EPI_ISL_643122, EPI_ISL_643123, EPI_ISL_643124, EPI_ISL_643125, EPI_ISL_643126, EPI_ISL_643127, EPI_ISL_643128, EPI_ISL_643129, EPI_ISL_643130, EPI_ISL_643131, EPI_ISL_643132, EPI_ISL_643133, EPI_ISL_643134, EPI_ISL_643135, EPI_ISL_643136, EPI_ISL_643137, EPI_ISL_643138, EPI_ISL_643139, EPI_ISL_643140, EPI_ISL_643141, EPI_ISL_643142, EPI_ISL_643143, EPI_ISL_643144, EPI_ISL_643145, EPI_ISL_643146, EPI_ISL_643147, EPI_ISL_643148, EPI_ISL_643149, EPI_ISL_643150, EPI_ISL_643151, EPI_ISL_643152, EPI_ISL_643153, EPI_ISL_643154, EPI_ISL_643155, EPI_ISL_643156, EPI_ISL_643157, EPI_ISL_643158, EPI_ISL_643159, EPI_ISL_643160, EPI_ISL_643161, EPI_ISL_643162, EPI_ISL_643163, EPI_ISL_643164, EPI_ISL_643165, EPI_ISL_643166, EPI_ISL_643167, EPI_ISL_643168, EPI_ISL_643169, EPI_ISL_643170, EPI_ISL_643171, EPI_ISL_643172, EPI_ISL_643173, EPI_ISL_643174, EPI_ISL_643175, EPI_ISL_643176, EPI_ISL_643177, EPI_ISL_643178, EPI_ISL_643179, EPI_ISL_643180, EPI_ISL_643181, EPI_ISL_643182, EPI_ISL_643183, EPI_ISL_643184, EPI_ISL_643185, EPI_ISL_643186, EPI_ISL_643187, EPI_ISL_643188, EPI_ISL_643189, EPI_ISL_643190, EPI_ISL_643191, EPI_ISL_643192, EPI_ISL_643193, EPI_ISL_643194, EPI_ISL_643195, EPI_ISL_643196, EPI_ISL_643197, EPI_ISL_643198, EPI_ISL_643199, EPI_ISL_643200, EPI_ISL_643201, EPI_ISL_643202, EPI_ISL_643203, EPI_ISL_643204, EPI_ISL_643205, EPI_ISL_643206, EPI_ISL_643207, EPI_ISL_643208, EPI_ISL_643209, EPI_ISL_643210, EPI_ISL_643211, EPI_ISL_643212, EPI_ISL_643213, EPI_ISL_643214, EPI_ISL_643215, EPI_ISL_643216, EPI_ISL_643217, EPI_ISL_643218, EPI_ISL_643219, EPI_ISL_643220, EPI_ISL_643221, EPI_ISL_643222, EPI_ISL_643223, EPI_ISL_643224, EPI_ISL_643225, EPI_ISL_643226, EPI_ISL_643227, EPI_ISL_643228, EPI_ISL_643229, EPI_ISL_643230, EPI_ISL_643231, EPI_ISL_643232, EPI_ISL_643233, EPI_ISL_643234, EPI_ISL_643235, EPI_ISL_643236, EPI_ISL_643237, EPI_ISL_643238, EPI_ISL_643239, EPI_ISL_643240, EPI_ISL_643241, EPI_ISL_643242, EPI_ISL_643243, EPI_ISL_643244, EPI_ISL_643245, EPI_ISL_643246, EPI_ISL_643247, EPI_ISL_643248, EPI_ISL_643249, EPI_ISL_643250, EPI_ISL_643251, EPI_ISL_643252, EPI_ISL_643253, EPI_ISL_643254, EPI_ISL_643255, EPI_ISL_643256, EPI_ISL_643257, EPI_ISL_643258, EPI_ISL_643259, EPI_ISL_643260, EPI_ISL_643261, EPI_ISL_643262, EPI_ISL_643263, EPI_ISL_643264, EPI_ISL_643265, EPI_ISL_643266, EPI_ISL_643267, EPI_ISL_643268, EPI_ISL_643269, EPI_ISL_643270, EPI_ISL_643271, EPI_ISL_643272, EPI_ISL_643273, EPI_ISL_643274, EPI_ISL_643275, EPI_ISL_643276, EPI_ISL_643277, EPI_ISL_643278, EPI_ISL_643279, EPI_ISL_643280, EPI_ISL_643281, EPI_ISL_643282, EPI_ISL_643283, EPI_ISL_643284, EPI_ISL_643285, EPI_ISL_643286, EPI_ISL_643287, EPI_ISL_643288, EPI_ISL_643289, EPI_ISL_643290, EPI_ISL_643291, EPI_ISL_643292, EPI_ISL_643293, EPI_ISL_643294, EPI_ISL_643295, EPI_ISL_643296, EPI_ISL_643297, EPI_ISL_643298, EPI_ISL_643299, EPI_ISL_643300, EPI_ISL_643301, EPI_ISL_643302, EPI_ISL_643303, EPI_ISL_643304, EPI_ISL_643305, EPI_ISL_643306, EPI_ISL_643307, EPI_ISL_643308, EPI_ISL_643309, EPI_ISL_643310, EPI_ISL_643311, EPI_ISL_643312, EPI_ISL_643313, EPI_ISL_643314 | | | |
| see above | Lighthouse Lab in Glasgow | Wellcome Sanger Institute for the COVID-19 Genomics UK (COG-UK) Consortium | Harper VanSteenhouse, Yumi Kasai, David Gray, Carol Clugston, Anna Dominiczak and Alex Alderton, Roberto Amato, Sonia Goncalves, Ewan Harrison, David K. Jackson, Ian Johnston, Dominic Kwiatkowski, Cordelia Langford, John Sillitoe on behalf of the Wellcome Sanger Institute COVID-19 Surveillance Team |
| EPI_ISL_644354, EPI_ISL_644355 | Michigan Department of Health and Human Services, Bureau of Laboratories | Michigan Department of Health and Human Services, Bureau of Laboratories | Blankenship HM, Riner D, Soehnlen MK |
| EPI_ISL_644556, EPI_ISL_644557, EPI_ISL_644558, EPI_ISL_644559, | MEPHI, Aix Marseille University | MEPHI, Aix Marseille University | Anthony LEVASSEUR |

| | | | |
|---|---|---|---|
| EPI_ISL_644560, EPI_ISL_644561, EPI_ISL_644562, EPI_ISL_644563, EPI_ISL_644564 | | | |
| EPI_ISL_644902, EPI_ISL_644903, EPI_ISL_644904, EPI_ISL_644905, EPI_ISL_644906, EPI_ISL_644907, EPI_ISL_644908, EPI_ISL_644909, EPI_ISL_644910 | Virginia DCLS | Virginia DCLS | Virginia DCLS |
| EPI_ISL_644934, EPI_ISL_644935 | Essentia Health-St. Mary's Medical Center | Minnesota Department of Health, Public Health Laboratory | Matt Plumb, Jacob Garfin, Alexandra Lorentz, and Xiong Wang |
| EPI_ISL_647697 | Lighthouse Lab in Cambridge | Wellcome Sanger Institute for the COVID-19 Genomics UK (COG-UK) Consortium | Rob Howes, The Lighthouse Lab in Cambridge and Alex Alderton, Roberto Amato, Sonia Goncalves, Ewan Harrison, David K. Jackson, Ian Johnston, Dominic Kwiatkowski, Cordelia Langford, John Sillitoe on behalf of the Wellcome Sanger Institute COVID-19 Surveillance Team |
| EPI_ISL_647854, EPI_ISL_647855, EPI_ISL_647856, EPI_ISL_647857, EPI_ISL_647858, EPI_ISL_647859, EPI_ISL_647860 | Lighthouse Lab in Glasgow | Wellcome Sanger Institute for the COVID-19 Genomics UK (COG-UK) Consortium | Harper VanSteenhouse, Yumi Kasai, David Gray, Carol Clugston, Anna Dominiczak and Alex Alderton, Roberto Amato, Sonia Goncalves, Ewan Harrison, David K. Jackson, Ian Johnston, Dominic Kwiatkowski, Cordelia Langford, John Sillitoe on behalf of the Wellcome Sanger Institute COVID-19 Surveillance Team |
| EPI_ISL_648152, EPI_ISL_648154, EPI_ISL_648155 | Stockholm | The Public Health Agency of Sweden | Anna-Malin Linde, Maria Lind Karlberg, Mattias Haukland, Reza Advani, Olov Svartstrom, Oskar Karlsson Lindsjo, Sandra Broddesson, Petra Edquist, Mia Brytting, Anna Risberg, Karin Tegmark-Wisell |
| EPI_ISL_648168 | Eskilstuna | The Public Health Agency of Sweden | Anna-Malin Linde, Maria Lind Karlberg, Mattias Haukland, Reza Advani, Olov Svartstrom, Oskar Karlsson Lindsjo, Sandra Broddesson, Petra Edquist, Mia Brytting, Anna Risberg, Karin Tegmark-Wisell |
| EPI_ISL_648271, EPI_ISL_648272, EPI_ISL_648273, EPI_ISL_648274, EPI_ISL_648275, EPI_ISL_648276, EPI_ISL_648277, EPI_ISL_648278, EPI_ISL_648279, EPI_ISL_648280, EPI_ISL_648281, EPI_ISL_648282, EPI_ISL_648283, EPI_ISL_648284, EPI_ISL_648285, EPI_ISL_648286 | | | |
| see above | MD PHL | MD PHL | Maryland Department of Health Laboratories Administration |
| EPI_ISL_648424, EPI_ISL_648438, EPI_ISL_648439, EPI_ISL_648440, EPI_ISL_648441, EPI_ISL_648442, EPI_ISL_648443, EPI_ISL_648444, EPI_ISL_648445, EPI_ISL_648446, EPI_ISL_648447, EPI_ISL_648448, EPI_ISL_648449, EPI_ISL_648450, EPI_ISL_648451, EPI_ISL_648452, EPI_ISL_648453, EPI_ISL_648454, EPI_ISL_648455, EPI_ISL_648456, EPI_ISL_648457, EPI_ISL_648458, EPI_ISL_648459, EPI_ISL_648476, EPI_ISL_648477, EPI_ISL_648478, EPI_ISL_648479 | | | |
| see above | Santa Clara County Public Health Laboratory | Chan-Zuckerberg Biohub | CZB Cliahub Consortium |
| EPI_ISL_648513, EPI_ISL_648514, EPI_ISL_648516, EPI_ISL_648517, EPI_ISL_648518, EPI_ISL_648519, EPI_ISL_648520, EPI_ISL_648521, EPI_ISL_648522, EPI_ISL_648523, EPI_ISL_648524, EPI_ISL_648525, EPI_ISL_648526, EPI_ISL_648527, EPI_ISL_648528, EPI_ISL_648529, EPI_ISL_648530 | | | |
| see above | Orange County Public Health Lab | Chan-Zuckerberg Biohub | CZB Cliahub Consortium |
| EPI_ISL_648927, EPI_ISL_648928, EPI_ISL_648934, EPI_ISL_648939, EPI_ISL_648966, EPI_ISL_648970, EPI_ISL_648975, EPI_ISL_649021, EPI_ISL_649024 | San Diego County Public Health Laboratory | Andersen lab at Scripps Research | SEARCH Alliance San Diego with Tracy Basler, Jovan Shephard, Brett Austin |
| EPI_ISL_649426, EPI_ISL_649457 | Lighthouse Lab in Alderley Park | Wellcome Sanger Institute for the COVID-19 Genomics UK (COG-UK) Consortium | Jacquelyn Wynn, Mairead Hyland, The Lighthouse Lab in Alderley Park and Alex Alderton, Roberto Amato, Sonia Goncalves, Ewan Harrison, David K. Jackson, Ian Johnston, Dominic Kwiatkowski, Cordelia Langford, John Sillitoe on behalf of the Wellcome Sanger Institute COVID-19 Surveillance Team (http://www.sanger.ac.uk/covid-team) |
| EPI_ISL_649556, EPI_ISL_649557, EPI_ISL_649558, EPI_ISL_649559, EPI_ISL_649560, EPI_ISL_649561, EPI_ISL_649562, EPI_ISL_649563, EPI_ISL_649564, EPI_ISL_649565, EPI_ISL_649566, EPI_ISL_649567, EPI_ISL_649568 | | | |
| see above | Lighthouse Lab in Glasgow | Wellcome Sanger Institute for the COVID-19 Genomics UK (COG-UK) Consortium | Harper VanSteenhouse, Yumi Kasai, David Gray, Carol Clugston, Anna Dominiczak and Alex Alderton, Roberto Amato, Sonia Goncalves, Ewan Harrison, David K. Jackson, Ian Johnston, Dominic Kwiatkowski, Cordelia Langford, John Sillitoe on behalf of the Wellcome Sanger Institute COVID-19 Surveillance Team (http://www.sanger.ac.uk/covid-team) |
| EPI_ISL_649808, EPI_ISL_649809, EPI_ISL_649810, EPI_ISL_649811, EPI_ISL_649812, EPI_ISL_649821, EPI_ISL_649822 | Respiratory Virus Unit, Microbiology Services Colindale, Public Health England | COVID-19 Genomics UK (COG-UK) Consortium | PHE Covid Sequencing Team |
| EPI_ISL_650125, EPI_ISL_650147, EPI_ISL_650148 | Northumbria University / South Tees Hospitals NHS Foundation Trust / North Cumbria Integrated Care NHS Foundation Trust / North Tees and Hartlepool NHS Foundation Trust / Newcastle Hospitals NHS Foundation Trust | COVID-19 Genomics UK (COG-UK) Consortium | Darren L Smith,Andrew Nelson,Matthew Bashton,Greg R Young,Joshua Loh,John Allan,Mohammad A Tariq,Giles S Holt,Gary Black,Wen C Yew,Lynn Dover,Paul Baker,Steve Liggett,Sarah Essex,Jane Greenaway,Debra Padgett,Clive Graham,Garren Scott,Edward Barton,Emma Swindells,Brendan Payne,Jennifer Collins,Yusri Taha,Gary Eltringham |
| EPI_ISL_650183 | Liverpool Clinical Laboratories | COVID-19 Genomics UK (COG-UK) Consortium | Sam Haldenby, Anita Lucaci, Steve Paterson, Julian Hiscox, Alistair Darby, M Almsaud, A Alrezaihi, Muhannad Alruwaili, Stuart D Armstrong, Jones Benjamin, Eleanor G Bentley, Anu Chawla, Jordan J Clark, Angela Cowell, Richard Eccles, Isabel García-Dorival, Matthew Gemmell, Alessandro Gerada, PKF Gilmore, Richard Gregory, Ximeng Han, Catherine Hartley, Margaret Hughes, Miren Iturriza-Gomara, James Johnson, L Luu, Jenifer Manson, Charlotte Nelson, Elaine O'Toole, Cassie Olateju, Rebekah Penrice-Randal , Lucille Rainbow, N.P Randle, Trevor Ian Robinson, Parul Sharma, Ghada T Shawli, James P Stewart, Neil Swainston, Ecaterina Vamos, Joanne Watts, Mark Whitehead |
| EPI_ISL_650246, EPI_ISL_650247, EPI_ISL_650279, EPI_ISL_650282 | Northumbria University / South Tees Hospitals NHS Foundation Trust / North Cumbria Integrated Care NHS Foundation Trust / North Tees and Hartlepool NHS Foundation Trust / Newcastle Hospitals NHS Foundation Trust | COVID-19 Genomics UK (COG-UK) Consortium | Darren L Smith,Andrew Nelson,Matthew Bashton,Greg R Young,Joshua Loh,John Allan,Mohammad A Tariq,Giles S Holt,Gary Black,Wen C Yew,Lynn Dover,Paul Baker,Steve Liggett,Sarah Essex,Jane Greenaway,Debra Padgett,Clive Graham,Garren Scott,Edward Barton,Emma Swindells,Brendan Payne,Jennifer Collins,Yusri Taha,Gary Eltringham |
| EPI_ISL_650336 | Liverpool Clinical Laboratories | COVID-19 Genomics UK (COG-UK) Consortium | Sam Haldenby, Anita Lucaci, Steve Paterson, Julian Hiscox, Alistair Darby, M Almsaud, A Alrezaihi, Muhannad Alruwaili, Stuart D Armstrong, Jones Benjamin, Eleanor G Bentley, Anu Chawla, Jordan J Clark, Angela Cowell, Richard Eccles, Isabel García-Dorival, Matthew Gemmell, Alessandro Gerada, PKF Gilmore, Richard Gregory, Ximeng Han, Catherine Hartley, Margaret Hughes, Miren Iturriza-Gomara, James Johnson, L Luu, Jenifer Manson, Charlotte Nelson, Elaine O'Toole, Cassie Olateju, Rebekah Penrice-Randal , Lucille Rainbow, N.P Randle, Trevor Ian Robinson, Parul Sharma, Ghada T Shawli, James P Stewart, Neil Swainston, Ecaterina Vamos, Joanne Watts, Mark Whitehead |
| EPI_ISL_650355 | Northumbria University / South Tees Hospitals NHS Foundation Trust / North Cumbria Integrated Care NHS Foundation Trust / North Tees and Hartlepool NHS Foundation Trust / Newcastle Hospitals NHS Foundation Trust | COVID-19 Genomics UK (COG-UK) Consortium | Darren L Smith,Andrew Nelson,Matthew Bashton,Greg R Young,Joshua Loh,John Allan,Mohammad A Tariq,Giles S Holt,Gary Black,Wen C Yew,Lynn Dover,Paul Baker,Steve Liggett,Sarah Essex,Jane Greenaway,Debra Padgett,Clive Graham,Garren Scott,Edward Barton,Emma Swindells,Brendan Payne,Jennifer Collins,Yusri Taha,Gary Eltringham |
| EPI_ISL_650415, EPI_ISL_650535, EPI_ISL_650543 | Regional Virus Laboratory, Belfast Health and Social Care Trust | COVID-19 Genomics UK (COG-UK) Consortium | Conall McCaughey, James McKenna, Tanya Curran, Susan Feeney, Alison Watt, Ciara Cox, Mairead Connor, Zoltan Molnar, David Simpson, Derek Fairley |
| EPI_ISL_650572, EPI_ISL_650587, EPI_ISL_650588 | Northumbria University / South Tees Hospitals NHS Foundation Trust / North Cumbria Integrated Care NHS Foundation Trust / North Tees and Hartlepool NHS Foundation Trust / Newcastle Hospitals NHS Foundation Trust | COVID-19 Genomics UK (COG-UK) Consortium | Darren L Smith,Andrew Nelson,Matthew Bashton,Greg R Young,Joshua Loh,John Allan,Mohammad A Tariq,Giles S Holt,Gary Black,Wen C Yew,Lynn Dover,Paul Baker,Steve Liggett,Sarah Essex,Jane Greenaway,Debra Padgett,Clive Graham,Garren Scott,Edward Barton,Emma Swindells,Brendan Payne,Jennifer Collins,Yusri Taha,Gary Eltringham |
| EPI_ISL_650598 | Regional Virus Laboratory, Belfast Health and Social Care | COVID-19 Genomics UK (COG-UK) Consortium | Conall McCaughey, James McKenna, Tanya Curran, Susan Feeney, Alison Watt, Ciara Cox, Mairead Connor, Zoltan Molnar, David Simpson, Derek Fairley |

| | | | |
|---|---|---|---|
| | Trust | | |
| EPI_ISL_650624 | Northumbria University / South Tees Hospitals NHS Foundation Trust / North Cumbria Integrated Care NHS Foundation Trust / North Tees and Hartlepool NHS Foundation Trust / Newcastle Hospitals NHS Foundation Trust | COVID-19 Genomics UK (COG-UK) Consortium | Darren L Smith,Andrew Nelson,Matthew Bashton,Greg R Young,Joshua Loh,John Allan,Mohammad A Tariq,Giles S Holt,Gary Black,Wen C Yew,Lynn Dover,Paul Baker,Steve Liggett,Sarah Essex,Jane Greenaway,Debra Padgett,Clive Graham,Garren Scott,Edward Barton,Emma Swindells,Brendan Payne,Jennifer Collins,Yusri Taha,Gary Eltringham |
| EPI_ISL_650672 | Liverpool Clinical Laboratories | COVID-19 Genomics UK (COG-UK) Consortium | Sam Haldenby, Anita Lucaci, Steve Paterson, Julian Hiscox, Alistair Darby, M Almsaud, A Alrezaihi, Muhannad Alruwaili, Stuart D Armstrong, Jones Benjamin, Eleanor G Bentley, Anu Chawla, Jordan J Clark, Angela Cowell, Richard Eccles, Isabel García-Dorival, Matthew Gemmell, Alessandro Gerada, PKF Gilmore, Richard Gregory, Ximeng Han, Catherine Hartley, Margaret Hughes, Miren Iturriza-Gomara, James Johnson, L Luu, Jenifer Manson, Charlotte Nelson, Elaine O'Toole, Cassie Olateju, Rebekah Penrice-Randal , Lucille Rainbow, N.P Randle, Trevor Ian Robinson, Parul Sharma, Ghada T Shawli, James P Stewart, Neil Swainston, Ecaterina Vamos, Joanne Watts, Mark Whitehead |
| EPI_ISL_650680 | Virology Department, Sheffield Teaching Hospitals NHS Foundation Trust/Department of Infection, Immunity and Cardiovascular Disease, The Medical School, University of Sheffield | COVID-19 Genomics UK (COG-UK) Consortium | Thushan de Silva, Matthew Parker, Nikki Smith, Adri Angyal, Rebecca Brown, Luke Green, Rachel Tucker, Paul Parsons, Danielle Groves, Katie Johnson, Laura Carrilero, Alex Keeley, Dave Partridge, Matthew Wyles, Benjamin Lindsey, Mehmet Yavuz, Mohammad Raza, Cariad Evans |
| EPI_ISL_650730 | West of Scotland Specialist Virology Centre, NHSGGC / MRC-University of Glasgow Centre for Virus Research | COVID-19 Genomics UK (COG-UK) Consortium | Ana da Silva Filipe, Natasha Johnson, Kathy Smollett, Daniel Mair, Stephen Carmichael, Alice Broos, Lily Tong, Jenna Nichols, Kyriaki Nomikou; Sarah McDonald; Richard Orton, Joseph Hughes, Sreenu Vattipally, David L Robertson; Alasdair MacLean, Rory Gunson; Sharif Shaaban, Matthew Holden; Rachel Blacow, Guy Mollett, Kathy Li, James Shepherd, Antonia Ho, Emma Thomson |
| EPI_ISL_650810, EPI_ISL_650888, EPI_ISL_650901 | Regional Virus Laboratory, Belfast Health and Social Care Trust | COVID-19 Genomics UK (COG-UK) Consortium | Conall McCaughey, James McKenna, Tanya Curran, Susan Feeney, Alison Watt, Ciara Cox, Mairead Connor, Zoltan Molnar, David Simpson, Derek Fairley |
| EPI_ISL_650950 | Northumbria University / South Tees Hospitals NHS Foundation Trust / North Cumbria Integrated Care NHS Foundation Trust / North Tees and Hartlepool NHS Foundation Trust / Newcastle Hospitals NHS Foundation Trust | COVID-19 Genomics UK (COG-UK) Consortium | Darren L Smith,Andrew Nelson,Matthew Bashton,Greg R Young,Joshua Loh,John Allan,Mohammad A Tariq,Giles S Holt,Gary Black,Wen C Yew,Lynn Dover,Paul Baker,Steve Liggett,Sarah Essex,Jane Greenaway,Debra Padgett,Clive Graham,Garren Scott,Edward Barton,Emma Swindells,Brendan Payne,Jennifer Collins,Yusri Taha,Gary Eltringham |
| EPI_ISL_650957 | Centre for Enzyme Innovation, University of Portsmouth / Translational Research Laboratory, Portsmouth Hospitals NHS Trust | COVID-19 Genomics UK (COG-UK) Consortium | Angela Beckett,Yann Bourgeois,Garry Scarlett,Sharon Glaysher,Scott Elliott,Kelly Bicknell,Robert Impey,Allyson Lloyd,Sarah Wyllie,Ethan Butcher,Anoop Chauhan,Samuel Robson |
| EPI_ISL_650967, EPI_ISL_650968, EPI_ISL_651013 | Northumbria University / South Tees Hospitals NHS Foundation Trust / North Cumbria Integrated Care NHS Foundation Trust / North Tees and Hartlepool NHS Foundation Trust / Newcastle Hospitals NHS Foundation Trust | COVID-19 Genomics UK (COG-UK) Consortium | Darren L Smith,Andrew Nelson,Matthew Bashton,Greg R Young,Joshua Loh,John Allan,Mohammad A Tariq,Giles S Holt,Gary Black,Wen C Yew,Lynn Dover,Paul Baker,Steve Liggett,Sarah Essex,Jane Greenaway,Debra Padgett,Clive Graham,Garren Scott,Edward Barton,Emma Swindells,Brendan Payne,Jennifer Collins,Yusri Taha,Gary Eltringham |
| EPI_ISL_651128 | Regional Virus Laboratory, Belfast Health and Social Care Trust | COVID-19 Genomics UK (COG-UK) Consortium | Conall McCaughey, James McKenna, Tanya Curran, Susan Feeney, Alison Watt, Ciara Cox, Mairead Connor, Zoltan Molnar, David Simpson, Derek Fairley |
| EPI_ISL_651132 | University College London, Great Ormond Street Hospital for Children NHS Foundation Trust, Imperial College Healthcare NHS Trust | COVID-19 Genomics UK (COG-UK) Consortium | Sergi Castellano, Rachel Williams, Mark Kristiansen, Paola Resende Silva, Sunando Roy, Tony Brooks, Helena Tutill, Paola Niola, Patricia Dyal, Charlotte Williams, Leysa Forrest, Yasmin Panchbhaya, Jacqueline Findlay, Samuel Weeks, Julianne Brown, Kathryn Harris, Paul Randell, James Price, Alison Holmes, Judith Breuer |
| EPI_ISL_651215 | Regional Virus Laboratory, Belfast Health and Social Care Trust | COVID-19 Genomics UK (COG-UK) Consortium | Conall McCaughey, James McKenna, Tanya Curran, Susan Feeney, Alison Watt, Ciara Cox, Mairead Connor, Zoltan Molnar, David Simpson, Derek Fairley |
| EPI_ISL_651295 | Liverpool Clinical Laboratories | COVID-19 Genomics UK (COG-UK) Consortium | Sam Haldenby, Anita Lucaci, Steve Paterson, Julian Hiscox, Alistair Darby, M Almsaud, A Alrezaihi, Muhannad Alruwaili, Stuart D Armstrong, Jones Benjamin, Eleanor G Bentley, Anu Chawla, Jordan J Clark, Angela Cowell, Richard Eccles, Isabel García-Dorival, Matthew Gemmell, Alessandro Gerada, PKF Gilmore, Richard Gregory, Ximeng Han, Catherine Hartley, Margaret Hughes, Miren Iturriza-Gomara, James Johnson, L Luu, Jenifer Manson, Charlotte Nelson, Elaine O'Toole, Cassie Olateju, Rebekah Penrice-Randal , Lucille Rainbow, N.P Randle, Trevor Ian Robinson, Parul Sharma, Ghada T Shawli, James P Stewart, Neil Swainston, Ecaterina Vamos, Joanne Watts, Mark Whitehead |
| EPI_ISL_651329 | Regional Virus Laboratory, Belfast Health and Social Care Trust | COVID-19 Genomics UK (COG-UK) Consortium | Conall McCaughey, James McKenna, Tanya Curran, Susan Feeney, Alison Watt, Ciara Cox, Mairead Connor, Zoltan Molnar, David Simpson, Derek Fairley |
| EPI_ISL_651388 | Northumbria University / South Tees Hospitals NHS Foundation Trust / North Cumbria Integrated Care NHS Foundation Trust / North Tees and Hartlepool NHS Foundation Trust / Newcastle Hospitals NHS Foundation Trust | COVID-19 Genomics UK (COG-UK) Consortium | Darren L Smith,Andrew Nelson,Matthew Bashton,Greg R Young,Joshua Loh,John Allan,Mohammad A Tariq,Giles S Holt,Gary Black,Wen C Yew,Lynn Dover,Paul Baker,Steve Liggett,Sarah Essex,Jane Greenaway,Debra Padgett,Clive Graham,Garren Scott,Edward Barton,Emma Swindells,Brendan Payne,Jennifer Collins,Yusri Taha,Gary Eltringham |
| EPI_ISL_651501 | Regional Virus Laboratory, Belfast Health and Social Care Trust | COVID-19 Genomics UK (COG-UK) Consortium | Conall McCaughey, James McKenna, Tanya Curran, Susan Feeney, Alison Watt, Ciara Cox, Mairead Connor, Zoltan Molnar, David Simpson, Derek Fairley |
| EPI_ISL_651814, EPI_ISL_651815, EPI_ISL_651816 | West of Scotland Specialist Virology Centre, NHSGGC / MRC-University of Glasgow Centre for Virus Research | COVID-19 Genomics UK (COG-UK) Consortium | Ana da Silva Filipe, Natasha Johnson, Kathy Smollett, Daniel Mair, Stephen Carmichael, Alice Broos, Lily Tong, Jenna Nichols, Kyriaki Nomikou; Sarah McDonald; Richard Orton, Joseph Hughes, Sreenu Vattipally, David L Robertson; Alasdair MacLean, Rory Gunson; Sharif Shaaban, Matthew Holden; Rachel Blacow, Guy Mollett, Kathy Li, James Shepherd, Antonia Ho, Emma Thomson |
| EPI_ISL_651958, EPI_ISL_651959 | Virology Department, Royal Infirmary of Edinburgh, NHS Lothian / School of Biological Sciences, University of Edinburgh / Institute of Genetics and Molecular Medicine, University of Edinburgh | COVID-19 Genomics UK (COG-UK) Consortium | McHugh M, Dewar R, Rooke S, Gallagher M, Balcaza C, O'Toole Á, Scher E, Hill V, McCrone JT, Colquhoun R, Yu X, Jackson B, Rambaut A, Williams TC, Templeton K |
| EPI_ISL_652084 | Liverpool Clinical Laboratories | COVID-19 Genomics UK (COG-UK) Consortium | Sam Haldenby, Anita Lucaci, Steve Paterson, Julian Hiscox, Alistair Darby, M Almsaud, A Alrezaihi, Muhannad Alruwaili, Stuart D Armstrong, Jones Benjamin, Eleanor G Bentley, Anu Chawla, Jordan J Clark, Angela Cowell, Richard Eccles, Isabel García-Dorival, Matthew Gemmell, Alessandro Gerada, PKF Gilmore, Richard Gregory, Ximeng Han, Catherine Hartley, Margaret Hughes, Miren Iturriza-Gomara, James Johnson, L Luu, Jenifer Manson, Charlotte Nelson, Elaine O'Toole, Cassie Olateju, Rebekah Penrice-Randal , Lucille Rainbow, N.P Randle, Trevor Ian Robinson, Parul Sharma, Ghada T Shawli, James P Stewart, Neil Swainston, Ecaterina Vamos, Joanne Watts, Mark Whitehead |
| EPI_ISL_652112, EPI_ISL_652114, EPI_ISL_652115, EPI_ISL_652116 | Northumbria University / South Tees Hospitals NHS Foundation Trust / North Cumbria Integrated Care NHS Foundation Trust / North Tees and Hartlepool NHS Foundation Trust / Newcastle Hospitals NHS Foundation Trust | COVID-19 Genomics UK (COG-UK) Consortium | Darren L Smith,Andrew Nelson,Matthew Bashton,Greg R Young,Joshua Loh,John Allan,Mohammad A Tariq,Giles S Holt,Gary Black,Wen C Yew,Lynn Dover,Paul Baker,Steve Liggett,Sarah Essex,Jane Greenaway,Debra Padgett,Clive Graham,Garren Scott,Edward Barton,Emma Swindells,Brendan Payne,Jennifer Collins,Yusri Taha,Gary Eltringham |
| EPI_ISL_652121, EPI_ISL_652122 | Oxford Viromics, NDM, University of Oxford; Oxford University Hospitals; Basingstoke and North Hampshire Hospital | COVID-19 Genomics UK (COG-UK) Consortium | Tanya Golubchik, David Bonsall, George Macintyre, Amy Trebes, Mariateresa de Cesare, Catrin Moore, Alex Mobbs, Anita Justice, Robert Shaw, Monique Andersson, Timothy Peto, Emma Wise, Nathan Moore, Jessica Lynch, Nick Cortes, Matilde Mori, Stephen Kidd, David Buck, John Todd, Christophe Fraser |
| EPI_ISL_652305, EPI_ISL_652307, EPI_ISL_652309, EPI_ISL_652310, EPI_ISL_652330, EPI_ISL_652331, EPI_ISL_652332, EPI_ISL_652333, EPI_ISL_652334, EPI_ISL_652335, EPI_ISL_652336 | | | |
| see above | Regional Virus Laboratory, Belfast Health and Social Care | COVID-19 Genomics UK (COG-UK) Consortium | Conall McCaughey, James McKenna, Tanya Curran, Susan Feeney, Alison Watt, Ciara Cox, Mairead Connor, Zoltan Molnar, David Simpson, Derek Fairley |

| | | | |
|---|---|---|---|
| | Trust | | |
| EPI_ISL_652622 | Centre for Enzyme Innovation, University of Portsmouth / Translational Research Laboratory, Portsmouth Hospitals NHS Trust | COVID-19 Genomics UK (COG-UK) Consortium | Angela Beckett,Yann Bourgeois,Garry Scarlett,Sharon Glayshser,Scott Elliott,Kelly Bicknell,Robert Impey,Allyson Lloyd,Sarah Wyllie,Ethan Butcher,Anoop Chauhan,Samuel Robson |
| EPI_ISL_652627 | Queens Medical Centre, Clinical Microbiology Department / DeepSeq Nottingham | COVID-19 Genomics UK (COG-UK) Consortium | Gemma Clark, Wendy Smith, Manjinder Khakh, Vicki M Fleming, Michelle M Lister, Hannah Howson-Wells, Jonathan Ball, Patrick McClure, Joseph Chappell, Theocharis Tsoleridis, Nadine Holmes, Matthew Carlisle, Christopher Moore, Fei Sang, Johnny Debebe, Victoria Wright, Matthew Loose |
| EPI_ISL_652701 | University of Exeter | COVID-19 Genomics UK (COG-UK) Consortium | Ben Temperton,Aaron Jeffries,Michelle Michelsen,Joanna Warwick-Dugdale,Audrey Farbos,Robyn Manley,Stephen Michell,Jane Masoli |
| EPI_ISL_653026 | Oxford Viromics, NDM, University of Oxford; Oxford University Hospitals; Basingstoke and North Hampshire Hospital | COVID-19 Genomics UK (COG-UK) Consortium | Tanya Golubchik, David Bonsall, George Macintyre, Amy Trebes, Mariateresa de Cesare, Catrin Moore, Alex Mobbs, Anita Justice, Robert Shaw, Monique Andersson, Timothy Peto, Emma Wise, Nathan Moore, Jessica Lynch, Nick Cortes, Matilde Mori, Stephen Kidd, David Buck, John Todd, Christophe Fraser |
| EPI_ISL_653077, EPI_ISL_653084, EPI_ISL_653092 | Virology Department, Sheffield Teaching Hospitals NHS Foundation Trust/Department of Infection, Immunity and Cardiovascular Disease, The Medical School, University of Sheffield | COVID-19 Genomics UK (COG-UK) Consortium | Thushan de Silva, Matthew Parker, Nikki Smith, Adri Angyal, Rebecca Brown, Luke Green, Rachel Tucker, Paul Parsons, Danielle Groves, Katie Johnson, Laura Carrilero, Alex Keeley, Dave Partridge, Matthew Wyles, Benjamin Lindsey, Mehmet Yavuz, Mohammad Raza, Cariad Evans |
| EPI_ISL_654250, EPI_ISL_654254, EPI_ISL_654255, EPI_ISL_654258, EPI_ISL_654259, EPI_ISL_654261, EPI_ISL_654263, EPI_ISL_654264, EPI_ISL_654265, EPI_ISL_654266, EPI_ISL_654268, EPI_ISL_654276, EPI_ISL_654277, EPI_ISL_654295, EPI_ISL_654296, EPI_ISL_654297, EPI_ISL_654298, EPI_ISL_654299, EPI_ISL_654300, EPI_ISL_654301, EPI_ISL_654302, EPI_ISL_654303, EPI_ISL_654304, EPI_ISL_654305, EPI_ISL_654306, EPI_ISL_654307, EPI_ISL_654308, EPI_ISL_654309, EPI_ISL_654310, EPI_ISL_654311, EPI_ISL_654351, EPI_ISL_654353, EPI_ISL_654358, EPI_ISL_654361, EPI_ISL_654363 | | | |
| see above | Hospital General Universitario Gregorio Marañón | SeqCOVID-SPAIN consortium/IBV(CSIC) | Darío García de Viedma, Laura Pérez-Lago, Marta Herranz, Jon Sicilia, Julia Suárez, Pilar Catalán, Patricia Muñoz and SeqCOVID-SPAIN consortium |
| EPI_ISL_654403, EPI_ISL_654405, EPI_ISL_654410, EPI_ISL_654411, EPI_ISL_654417, EPI_ISL_654418, EPI_ISL_654419, EPI_ISL_654420, EPI_ISL_654426, EPI_ISL_654427, EPI_ISL_654428, EPI_ISL_654429, EPI_ISL_654430, EPI_ISL_654431, EPI_ISL_654432, EPI_ISL_654457, EPI_ISL_654458, EPI_ISL_654459, EPI_ISL_654460, EPI_ISL_654461, EPI_ISL_654462, EPI_ISL_654463, EPI_ISL_654464, EPI_ISL_654465, EPI_ISL_654466, EPI_ISL_654467 | | | |
| see above | Servicio de Microbiología, Hospital Miguel Servet, Zaragoza | SeqCOVID-SPAIN consortium/IBV(CSIC) | Antonio Rezusta López, Alexander Tristancho Baró, Ana Milagro, Yolanda Gracia Grataloup, Nieves Martínez Cameo and SeqCOVID-SPAIN consortium |
| EPI_ISL_654493, EPI_ISL_654494, EPI_ISL_654495, EPI_ISL_654496, EPI_ISL_654497 | Servicio de Microbiología. Hospital Clínico Universitario de Valencia | SeqCOVID-SPAIN consortium/IBV(CSIC) | David Navarro Ortega, Eliseo Albert Vicent, Ignacio Torres and SeqCOVID-SPAIN consortium |
| EPI_ISL_654540, EPI_ISL_654544, EPI_ISL_654545, EPI_ISL_654574, EPI_ISL_654575, EPI_ISL_654576, EPI_ISL_654577, EPI_ISL_654578, EPI_ISL_654579, EPI_ISL_654580, EPI_ISL_654581, EPI_ISL_654582, EPI_ISL_654583, EPI_ISL_654584, EPI_ISL_654585, EPI_ISL_654587, EPI_ISL_654588, EPI_ISL_654589, EPI_ISL_654590, EPI_ISL_654591 | | | |
| see above | Servicio de Microbiología. Hospital Universitario Donostia. OSI Donostialdea. Área de Enfermedades Infecciosas, Grupo de Infección Respiratoria y Resistencia Antimicrobiana. Instituto de Investigación Sanitaria Biodonostia | SeqCOVID-SPAIN consortium/IBV(CSIC) | Gustavo Cilla Eguiluz, Milagrosa Montes Ros, Luis Piñeiro Vázquez, Ane Sorrarain, Jose Maria Marimón and SeqCOVID-SPAIN consortium |
| EPI_ISL_660153, EPI_ISL_660155, EPI_ISL_660157 | PathCare | National Health Laboratory Service (NHLS), Tygerberg | Susan Engelbrecht, Draper C, Davis M-A, Siegfried N, Williamson C, Hsiao M, Kayla Delaney, Bronwyn Kleinhans, Houriiyah Tegally, Eduan Wilkindon, Gert van Zyl, Wolfgang Preiser, Tulio de Oliveira |
| EPI_ISL_660234, EPI_ISL_660235, EPI_ISL_660236, EPI_ISL_660237, EPI_ISL_660238 | NHLS-IALCH | KRISP, KZN Research Innovation and Sequencing Platform | Giandhari J, Pillay S, Lessells R, Mdlalose K, York D, Khan S, Tegally H, Wilkinson E, de Oliveira T |
| EPI_ISL_660442 | Centre Muraz | Project group Epidemology of Highly Pathogenic Microorganisms, Robert Koch Institut | Soumeya Ouangraoua, Abdoul-Salam Ouedraogo, Arsène Zongo, Yacouba Sawadogo, Essia Belarbi, Grit Schubert, Fabian Leendertz |
| EPI_ISL_660559, EPI_ISL_660560, EPI_ISL_660561, EPI_ISL_660562, EPI_ISL_660564, EPI_ISL_660579, EPI_ISL_660581, EPI_ISL_660582, EPI_ISL_660589, EPI_ISL_660590, EPI_ISL_660598, EPI_ISL_660599 | | | |
| see above | The National Institute of Public Health | State Veterinary Institute Prague | Nagy,A;Jirincova,H;Novakova,L;Trnka,D;Vecerova,J |
| EPI_ISL_660742, EPI_ISL_660743, EPI_ISL_660744, EPI_ISL_660745 | Respiratory Virus Unit, Microbiology Services Colindale, Public Health England | COVID-19 Genomics UK (COG-UK) Consortium | PHE Covid Sequencing Team |
| EPI_ISL_660932, EPI_ISL_660933, EPI_ISL_660934, EPI_ISL_660935, EPI_ISL_660936, EPI_ISL_660937, EPI_ISL_660938, EPI_ISL_660939, EPI_ISL_660940, EPI_ISL_660941, EPI_ISL_660942, EPI_ISL_660943, EPI_ISL_660944, EPI_ISL_660945, EPI_ISL_660946, EPI_ISL_660947, EPI_ISL_660948, EPI_ISL_660949, EPI_ISL_660950, EPI_ISL_660951, EPI_ISL_660952, EPI_ISL_660953, EPI_ISL_660954, EPI_ISL_660955 | | | |
| see above | Gundersen Molecular Diagnostics Laboratory | Kabara Cancer Research Institute | Craig S. Richmond, Paraic A. Kenny |
| EPI_ISL_661267 | Russian Academy of Sciences, Federal Research Center for Virology and Microbiology | Russian Academy of Sciences, Federal Research Center for Virology and Microbiology | Titov,I., Nefedeva,M., Egorova,I., Malogolovkin,A. |
| EPI_ISL_664340, EPI_ISL_664498, EPI_ISL_664507, EPI_ISL_665199, EPI_ISL_665200, EPI_ISL_665201 | University College London Hospital | COVID-19 Genomics UK (COG-UK) Consortium | Judith Heaney, Matthew Byott, Catherine Houlihan, Dan Frampton, Stuart Kirk, Moira Spyer and Eleni Nastouli |
| EPI_ISL_665262, EPI_ISL_665264, EPI_ISL_665303, EPI_ISL_665308, EPI_ISL_665311, EPI_ISL_665313, EPI_ISL_665314, EPI_ISL_665319, EPI_ISL_665323, EPI_ISL_665379, EPI_ISL_665387, EPI_ISL_665402, EPI_ISL_665403, EPI_ISL_665404, EPI_ISL_665412, EPI_ISL_665432, EPI_ISL_665490, EPI_ISL_665491, EPI_ISL_665499, EPI_ISL_665500, EPI_ISL_665501, EPI_ISL_665502, EPI_ISL_665600, EPI_ISL_665618, EPI_ISL_665630, EPI_ISL_665735 | | | |
| see above | Northumbria University / South Tees Hospitals NHS Foundation Trust / North Cumbria Integrated Care NHS Foundation Trust / North Tees and Hartlepool NHS Foundation Trust / Newcastle Hospitals NHS Foundation Trust | COVID-19 Genomics UK (COG-UK) Consortium | Darren L Smith,Andrew Nelson,Matthew Bashton,Greg R Young,Joshua Loh,John Allan,Mohammad A Tariq,Giles S Holt,Gary Black,Wen C Yew,Lynn Dover,Paul Baker,Steve Liggett,Sarah Essex,Jane Greenaway,Debra Padgett,Clive Graham,Garren Scott,Edward Barton,Emma Swindells,Brendan Payne,Jennifer Collins,Yusri Taha,Gary Eltringham |
| EPI_ISL_665822 | University College London Hospital | COVID-19 Genomics UK (COG-UK) Consortium | Judith Heaney, Matthew Byott, Catherine Houlihan, Dan Frampton, Stuart Kirk, Moira Spyer and Eleni Nastouli |
| EPI_ISL_665965 | Northumbria University / South Tees Hospitals NHS Foundation Trust / North Cumbria Integrated Care NHS Foundation Trust / North Tees and Hartlepool NHS Foundation Trust / Newcastle Hospitals NHS Foundation Trust | COVID-19 Genomics UK (COG-UK) Consortium | Darren L Smith,Andrew Nelson,Matthew Bashton,Greg R Young,Joshua Loh,John Allan,Mohammad A Tariq,Giles S Holt,Gary Black,Wen C Yew,Lynn Dover,Paul Baker,Steve Liggett,Sarah Essex,Jane Greenaway,Debra Padgett,Clive Graham,Garren Scott,Edward Barton,Emma Swindells,Brendan Payne,Jennifer Collins,Yusri Taha,Gary Eltringham |
| EPI_ISL_665973, EPI_ISL_665974, EPI_ISL_665975 | Queens Medical Centre, Clinical Microbiology Department / DeepSeq Nottingham | COVID-19 Genomics UK (COG-UK) Consortium | Gemma Clark, Wendy Smith, Manjinder Khakh, Vicki M Fleming, Michelle M Lister, Hannah Howson-Wells, Jonathan Ball, Patrick McClure, Joseph Chappell, Theocharis Tsoleridis, Nadine Holmes, Matthew Carlisle, Christopher Moore, Fei Sang, Johnny Debebe, Victoria Wright, Matthew Loose |
| EPI_ISL_666084, EPI_ISL_666085, EPI_ISL_666086, EPI_ISL_666087 | Northumbria University / South Tees Hospitals NHS Foundation Trust / North Cumbria Integrated Care NHS Foundation Trust / North Tees and Hartlepool NHS Foundation Trust / Newcastle Hospitals NHS Foundation Trust | COVID-19 Genomics UK (COG-UK) Consortium | Darren L Smith,Andrew Nelson,Matthew Bashton,Greg R Young,Joshua Loh,John Allan,Mohammad A Tariq,Giles S Holt,Gary Black,Wen C Yew,Lynn Dover,Paul Baker,Steve Liggett,Sarah Essex,Jane Greenaway,Debra Padgett,Clive Graham,Garren Scott,Edward Barton,Emma Swindells,Brendan Payne,Jennifer Collins,Yusri Taha,Gary Eltringham |
| EPI_ISL_666195, EPI_ISL_666205, EPI_ISL_666288, EPI_ISL_666323 | Wales Specialist Virology Centre Sequencing lab: Pathogen Genomics Unit | COVID-19 Genomics UK (COG-UK) Consortium | Catherine Moore, Johnathan Evans, Laura Gifford, Malorie Perry, Simon Cottrell, Angela Marchbank, Alec Birchley, Alexander Adams, Amy Gaskin, Bree Gatica-Wilcox, Jason Coombes, Joel Southgate, Lauren Gilbert, Lee Graham, Nicole Pacchiarini, Sara Kumziene-Summerhayes, Sarah Taylor, Sophie Jones, Sara Rey, Matthew Bull, Joanne Watkins, Sally Corden, Tom Connor |
| EPI_ISL_666618, EPI_ISL_666619, EPI_ISL_666620, EPI_ISL_666621 | Environmental and Global Health, University of Florida | Environmental and Global Health, University of Florida | Loeb,J.C., Silva,L.O., Elbadry,M.A., Stephenson,C.J., Morris,J.G., Lednicky,J.A. |

| | | | |
|---|---|---|---|
| EPI_ISL_666628, EPI_ISL_666629 | ZOTZ KLIMAS MVZ Düsseldorf-Centrum GbR ÜBAG für Labormedizin, Genetik, Zytologie, Pathologie | Center of Medical Microbiology, Virology, and Hospital Hygiene, University of Duesseldorf | Maximilian Damagnez, Alexander Dilthey, Ashley-Jane Duplessis, Patrick Finzer, Katrin Hoffmann, Torsten Houwaart, Lisanna Hülse, Malte Kohns Vasconcelos, Marek Korencak, Nadine Lübke, Jessica Nicolai, Klaus Pfeffer, Daniel Strelow, Jörg Timm, Andreas Walker, Tobias Wienemann, Rainer Zotz |
| EPI_ISL_666769 | Respiratory Virus Unit, Microbiology Services Colindale, Public Health England | COVID-19 Genomics UK (COG-UK) Consortium | PHE Covid Sequencing Team |
| EPI_ISL_666771, EPI_ISL_666772, EPI_ISL_666773, EPI_ISL_666774, EPI_ISL_666775, EPI_ISL_666776, EPI_ISL_666794, EPI_ISL_666795, EPI_ISL_666796, EPI_ISL_666797, EPI_ISL_666798 | | | |
| see above | Maryland Public Health Laboratory | Maryland Public Health Laboratory | Maryland Department of Health Laboratories Administration |
| EPI_ISL_666923 | Michigan Department of Health and Human Services, Bureau of Laboratories | Michigan Department of Health and Human Services, Bureau of Laboratories | Blankenship HM, Riner D, Soehnlen MK |
| EPI_ISL_666977, EPI_ISL_666978, EPI_ISL_666981 | San Diego County Public Health Laboratory | Andersen lab at Scripps Research | SEARCH Alliance San Diego with Tracy Basler, Jovan Shephard, Brett Austin |
| EPI_ISL_667504, EPI_ISL_667508 | OHSU Lab Services Molecular Microbiology Lab | Oregon SARS-CoV-2 Genome Sequencing Center | Brendan L. O'Connell, Ruth V. Nichols, Sally Grindstaff, Alec J. Hirsch, Donna Hansel, Guang Fan, Daniel N. Streblow, William B. Messer, Andrew C. Adey, Benjamin N. Bimber, Brian J. O'Roak |
| EPI_ISL_667779, EPI_ISL_667797 | South Eastern Area Laboratory Services (SEALS) | NSW Health Pathology - Institute of Clinical Pathology and Medical Research; Westmead Hospital; University of Sydney | CIDM-PH et al. |
| EPI_ISL_668392 | Haukeland University Hospital, Dept. of Microbiology | Norwegian Institute of Public Health, Department of Virology | Kathrine Stene-Johansen, Kamilla Heddeland Instefjord, Hilde Elshaug, Marie Paulsen Madsen, Rasmus Riis Kopperud, Hilde Vollan, Karoline Bragstad, Olav Hungnes |
| EPI_ISL_668398 | Nordland Hospital - Bodo, Laboratory Department, Molecular Biology Unit | Norwegian Institute of Public Health, Department of Virology | Kathrine Stene-Johansen, Kamilla Heddeland Instefjord, Hilde Elshaug, Marie Paulsen Madsen, Rasmus Riis Kopperud, Hilde Vollan, Karoline Bragstad, Olav Hungnes |
| EPI_ISL_668402 | Oslo University Hospital, Department of Medical Microbiology | Norwegian Institute of Public Health, Department of Virology | Kathrine Stene-Johansen, Kamilla Heddeland Instefjord, Hilde Elshaug, Marie Paulsen Madsen, Rasmus Riis Kopperud, Hilde Vollan, Karoline Bragstad, Olav Hungnes |
| EPI_ISL_668418, EPI_ISL_668441 | Department of Medical Microbiology, St. Olavs hospital | Norwegian Institute of Public Health, Department of Virology | Kathrine Stene-Johansen, Kamilla Heddeland Instefjord, Hilde Elshaug, Marie Paulsen Madsen, Rasmus Riis Kopperud, Hilde Vollan, Karoline Bragstad, Olav Hungnes |
| EPI_ISL_668444 | Respiratory Virus Unit, Microbiology Services Colindale, Public Health England | COVID-19 Genomics UK (COG-UK) Consortium | PHE Covid Sequencing Team |
| EPI_ISL_671676 | DOHMH PHL | New York City Public Health Laboratory | Jade Wang, et al. |
| EPI_ISL_671686, EPI_ISL_671687, EPI_ISL_671688 | DOHMH Morrisania | New York City Public Health Laboratory | Jade Wang, et al. |
| EPI_ISL_671865, EPI_ISL_671881, EPI_ISL_671882, EPI_ISL_671890, EPI_ISL_671891, EPI_ISL_671893, EPI_ISL_671894, EPI_ISL_671895, EPI_ISL_671896, EPI_ISL_671897, EPI_ISL_671898, EPI_ISL_671899, EPI_ISL_671900, EPI_ISL_671901, EPI_ISL_671903, EPI_ISL_671904, EPI_ISL_671905, EPI_ISL_671906, EPI_ISL_671907, EPI_ISL_671910, EPI_ISL_671922, EPI_ISL_671923, EPI_ISL_671927, EPI_ISL_671929, EPI_ISL_671936, EPI_ISL_671937, EPI_ISL_671940 | | | |
| see above | National Virus Reference Laboratory | National Virus Reference Laboratory | Michael Carr, Gabriel Gonzalez, Jonathan Dean, Daniel Hare, Cillian F De Gascun |
| EPI_ISL_671966, EPI_ISL_671967 | CHU Purpan - Laboratoire de Virologie - Institut Fédératif de Biologie | CHU Purpan - Laboratoire de Virologie - Institut Fédératif de Biologie | Latour J., Ranger N., Dubois M., Carcenac R., Harter A., Boyer P., Tremeaux P., Izopet J. |
| EPI_ISL_672075, EPI_ISL_672084, EPI_ISL_672103, EPI_ISL_672361, EPI_ISL_672362, EPI_ISL_672363, EPI_ISL_672364, EPI_ISL_672365 | Orange County Public Health Lab | Chan-Zuckerberg Biohub | CZB Cliahub Consortium |
| EPI_ISL_676515 | Klinisk mikrobiologi, Viruslab | The Public Health Agency of Sweden | Department of Microbiology, The Public Health Agency of Sweden |
| EPI_ISL_677143, EPI_ISL_677145, EPI_ISL_677164, EPI_ISL_677165, EPI_ISL_677166, EPI_ISL_677167, EPI_ISL_677168, EPI_ISL_677169, EPI_ISL_677170, EPI_ISL_677171, EPI_ISL_677172, EPI_ISL_677173, EPI_ISL_677174, EPI_ISL_677175, EPI_ISL_677176, EPI_ISL_677177, EPI_ISL_677178, EPI_ISL_677215, EPI_ISL_677216, EPI_ISL_677234, EPI_ISL_677238, EPI_ISL_677240 | | | |
| see above | Virginia Division of Consolidated Laboratory Services | Virginia Division of Consolidated Laboratory Services | Virginia DCLS |
| EPI_ISL_677460, EPI_ISL_677463, EPI_ISL_677472, EPI_ISL_677475, EPI_ISL_677476, EPI_ISL_677480, EPI_ISL_677481, EPI_ISL_677482, EPI_ISL_677483, EPI_ISL_677486, EPI_ISL_677487, EPI_ISL_677488, EPI_ISL_677489, EPI_ISL_677539, EPI_ISL_677540, EPI_ISL_677541, EPI_ISL_677542, EPI_ISL_677543, EPI_ISL_677544, EPI_ISL_677545, EPI_ISL_677546, EPI_ISL_677547, EPI_ISL_677548, EPI_ISL_677549, EPI_ISL_677550, EPI_ISL_677551, EPI_ISL_677552, EPI_ISL_677553, EPI_ISL_677557, EPI_ISL_677558, EPI_ISL_677559, EPI_ISL_677560, EPI_ISL_677561, EPI_ISL_677562, EPI_ISL_677563, EPI_ISL_677564, EPI_ISL_677565, EPI_ISL_677566, EPI_ISL_677567, EPI_ISL_677568, EPI_ISL_677569, EPI_ISL_677570, EPI_ISL_677571, EPI_ISL_677572, EPI_ISL_677573, EPI_ISL_677574, EPI_ISL_677575, EPI_ISL_677576, EPI_ISL_677627 | | | |
| see above | University of Wisconsin-Madison AIDS Vaccine Research Laboratories | University of Wisconsin-Madison AIDS Vaccine Research Laboratories | Gage Moreno, Katarina Braun, et al. AIDS Vaccine Research Laboratories |
| EPI_ISL_678371, EPI_ISL_678372, EPI_ISL_678373, EPI_ISL_678374 | Area of Virology, Serology and Virology Division (SAViD), New South Wales Health Pathology Randwick | Virology Research Laboratory; Area of Virology, Serology and Virology Division (SAViD), New South Wales Health Pathology Randwick | Foster, C.; Au, J.; Ruiz Silva, M.; Deveson, I.; Bull, R.; Van Hal, S.; Rawlinson, W. |
| EPI_ISL_678751 | Respiratory Virus Unit, Microbiology Services Colindale, Public Health England | COVID-19 Genomics UK (COG-UK) Consortium | PHE Covid Sequencing Team |
| EPI_ISL_679132 | University of Birmingham | COVID-19 Genomics UK (COG-UK) Consortium | Institute of Microbiology, University of Birmingham: Claire McMurray, Joanne Stockton, Samuel Nicholls, Radoslaw Poplawski, Will Rowe, Josh Quick, Nicholas Loman. University of Birmingham Testing Laboratory: Celina M Whalley, Andrew Bosworth, Charlotte Poxon, Kasun Wanigasooriya, Oliver Pickles, Mike Kidd, Alex Richter, Andrew D Beggs PHE Heartlands Lab: Husam Osman, Andrew Bosworth. Queen Elizabeth Hospital: Anna Casey |
| EPI_ISL_680206, EPI_ISL_680208, EPI_ISL_680210, EPI_ISL_680212, EPI_ISL_680214, EPI_ISL_680220, EPI_ISL_680221, EPI_ISL_680222, EPI_ISL_680225, EPI_ISL_680329, EPI_ISL_680331, EPI_ISL_680332, EPI_ISL_680333, EPI_ISL_680334, EPI_ISL_680335, EPI_ISL_680336, EPI_ISL_680337, EPI_ISL_680338, EPI_ISL_680339, EPI_ISL_680342, EPI_ISL_680343, EPI_ISL_680344, EPI_ISL_680345, EPI_ISL_680346, EPI_ISL_680347, EPI_ISL_680348, EPI_ISL_680349, EPI_ISL_680350, EPI_ISL_680351, EPI_ISL_680352, EPI_ISL_680353, EPI_ISL_680354, EPI_ISL_680355, EPI_ISL_680356, EPI_ISL_680357, EPI_ISL_680358, EPI_ISL_680359, EPI_ISL_680360, EPI_ISL_680361, EPI_ISL_680362, EPI_ISL_680363, EPI_ISL_680364, EPI_ISL_680365, EPI_ISL_680366, EPI_ISL_680367, EPI_ISL_680368, EPI_ISL_680369, EPI_ISL_680370, EPI_ISL_680371, EPI_ISL_680372, EPI_ISL_680373, EPI_ISL_680374, EPI_ISL_680375, EPI_ISL_680376, EPI_ISL_680377, EPI_ISL_680378, EPI_ISL_680379, EPI_ISL_680380, EPI_ISL_680381, EPI_ISL_680382, EPI_ISL_680383, EPI_ISL_680384, EPI_ISL_680385, EPI_ISL_680386, EPI_ISL_680387, EPI_ISL_680409, EPI_ISL_680410, EPI_ISL_680411, EPI_ISL_680412, EPI_ISL_680413, EPI_ISL_680414, EPI_ISL_680415, EPI_ISL_680416, EPI_ISL_680417, EPI_ISL_680418, EPI_ISL_680419, EPI_ISL_680420, EPI_ISL_680421, EPI_ISL_680422, EPI_ISL_680423, EPI_ISL_680424, EPI_ISL_680425, EPI_ISL_680426, EPI_ISL_680427, EPI_ISL_680428, EPI_ISL_680429, EPI_ISL_680430, EPI_ISL_680431, EPI_ISL_680432, EPI_ISL_680433, EPI_ISL_680434, EPI_ISL_680435 | | | |
| see above | Regional Virus Laboratory, Belfast Health and Social Care Trust | COVID-19 Genomics UK (COG-UK) Consortium | Conall McCaughey, James McKenna, Tanya Curran, Susan Feeney, Alison Watt, Ciara Cox, Mairead Connor, Zoltan Molnar, David Simpson, Derek Fairley |
| EPI_ISL_680477, EPI_ISL_680478, EPI_ISL_680479, EPI_ISL_680480, EPI_ISL_680481, EPI_ISL_680483, EPI_ISL_680484, EPI_ISL_680511, EPI_ISL_680512 | Virology Department, Royal Infirmary of Edinburgh, NHS Lothian / School of Biological Sciences, University of Edinburgh / Institute of Genetics and Molecular Medicine, University of Edinburgh | COVID-19 Genomics UK (COG-UK) Consortium | McHugh M, Dewar R, Rooke S, Gallagher M, Balcaza C, O'Toole Á, Scher E, Hill V, McCrone JT, Colquhoun R, Yu X, Jackson B, Rambaut A, Williams TC, Templeton K |
| EPI_ISL_680580, EPI_ISL_680581, EPI_ISL_680589, EPI_ISL_680611, EPI_ISL_680613, EPI_ISL_680615 | Wales Specialist Virology Centre Sequencing lab: Pathogen Genomics Unit | COVID-19 Genomics UK (COG-UK) Consortium | Catherine Moore, Johnathan Evans, Laura Gifford, Malorie Perry, Simon Cottrell, Angela Marchbank, Alec Birchley, Alexander Adams, Amy Gaskin, Bree Gatica-Wilcox, Jason Coombes, Joel Southgate, Lauren Gilbert, Lee Graham, Nicole Pacchiarini, Sara Kumziene-Summerhayes, Sarah Taylor, Sophie Jones, Sara Rey, Matthew Bull, Joanne Watkins, Sally Corden, Tom Connor |
| EPI_ISL_681298, EPI_ISL_681299, EPI_ISL_681300, EPI_ISL_681304, EPI_ISL_681305 | Communicable Disease Laboratory, Public Health Directorate | Communicable Disease Laboratory, Public Health Directorate | Alwasti,H., Altaif,Z., AlHujairi,Z., AlAbbas,Z. |

| | | | |
|---|---|---|---|
| EPI_ISL_682041, EPI_ISL_682053, EPI_ISL_682054 | UPMC Clinical Microbiology Laboratory | Microbial Genomic Epidemiology Laboratory, University of Pittsburgh | Mustapha M. Mustapha, Jane W. Marsh, Dan Snyder, Marissa P. Griffith, Stephanie L. Mitchell, Vatsala R. Srinivasa, Kady D. Waggle, Chinelo Ezeonwuku, Vaughn S. Cooper, Lee H. Harrison |
| EPI_ISL_682304, EPI_ISL_682305, EPI_ISL_682310, EPI_ISL_682311, EPI_ISL_682312, EPI_ISL_682313 | Communicable Disease Laboratory, Public Health Directorate | Communicable Disease Laboratory, Public Health Directorate | Alwasti,H., Altaif,Z., AlHujairi,Z., AlAbbas,Z. |
| EPI_ISL_682323, EPI_ISL_682324, EPI_ISL_682325, EPI_ISL_682326, EPI_ISL_682328, EPI_ISL_682331, EPI_ISL_682333, EPI_ISL_682336, EPI_ISL_682340, EPI_ISL_682345, EPI_ISL_682346, EPI_ISL_682348, EPI_ISL_682349, EPI_ISL_682350, EPI_ISL_682351, EPI_ISL_682352 | | | |
| see above | NHLS Universitas Academic | UFS Virology | PA Bester, MM Nyaga, P Nthiga, MT Mogotsi, D Goedhals, T de Oliveira |
| EPI_ISL_683267, EPI_ISL_683268, EPI_ISL_683269, EPI_ISL_683270, EPI_ISL_683271, EPI_ISL_683275, EPI_ISL_683276, EPI_ISL_683279, EPI_ISL_683280, EPI_ISL_683281, EPI_ISL_683282, EPI_ISL_683283, EPI_ISL_683298, EPI_ISL_683299, EPI_ISL_683300, EPI_ISL_683301, EPI_ISL_683302, EPI_ISL_683303, EPI_ISL_683304, EPI_ISL_683305, EPI_ISL_683306, EPI_ISL_683307, EPI_ISL_683308, EPI_ISL_683309, EPI_ISL_683310, EPI_ISL_683311, EPI_ISL_683312, EPI_ISL_683313, EPI_ISL_683314, EPI_ISL_683315, EPI_ISL_683316, EPI_ISL_683317, EPI_ISL_683318, EPI_ISL_683319, EPI_ISL_683320, EPI_ISL_683321, EPI_ISL_683322, EPI_ISL_683323, EPI_ISL_683324, EPI_ISL_683325, EPI_ISL_683326, EPI_ISL_683327, EPI_ISL_683328 | | | |
| see above | Department of Virus and Microbiological Special Diagnostics, Statens Serum Institut, Copenhagen, Denmark | Albertsen Lab, Department of Chemistry and Bioscience, Aalborg University, Denmark | Danish Covid-19 Genome Consortium |
| EPI_ISL_683638 | Servicio de Microbiología, Laboratori Clínic Metropolitana Nord. Hospital Universitari Germans Trias i Pujol. Institut d'Investigació en Ciències de la Salut Germans Trias i Pujol (IGTP) | SeqCOVID-SPAIN consortium/IBV(CSIC) | Elisa Martró, Antoni E. Bordoy, Anna Not, Adrián Antuori, Anabel Fernández, Nona Romaní, Verónica Saludes, Cristina Casañ and SeqCOVID-SPAIN consortium |
| EPI_ISL_683727 | Mayo Clinic & Mayo Clinic Laboratories | Minnesota Department of Health, Public Health Laboratory | Alexandra Lorentz, Jacob Garfin, Matt Plumb, and Xiong Wang |
| EPI_ISL_684047 | NHLS Universitas Academic | UFS Virology | PA Bester, MM Nyaga, P Nthiga, MT Mogotsi, D Goedhals, T de Oliveira |
| EPI_ISL_691615, EPI_ISL_691625, EPI_ISL_691657, EPI_ISL_691660, EPI_ISL_691674, EPI_ISL_691675, EPI_ISL_691676, EPI_ISL_691679 | Servicio de Microbiología, Hospital Universitario Son Espases | SeqCOVID-SPAIN consortium/IBV(CSIC) | Carla López-Causapé, Jordi Reina, Antonio Oliver and SeqCOVID-SPAIN consortium |
| EPI_ISL_691686, EPI_ISL_691687, EPI_ISL_691688, EPI_ISL_691689, EPI_ISL_691690, EPI_ISL_691691, EPI_ISL_691692, EPI_ISL_691693, EPI_ISL_691694, EPI_ISL_691695, EPI_ISL_691696, EPI_ISL_691697, EPI_ISL_691698, EPI_ISL_691699 | | | |
| see above | Hospital Universitario de Ceuta | Instituto de Salud Carlos III | Iglesias-Caballero, M. Camarero, S. Molinero Calamita, M. González-Esguevillas, M. Pozo, F. Casas, I. Jiménez, P. Jiménez, M. Zaballos, A. Monzón, S. Varona, S. Juliá, M. Cuesta, I. Hijano, S. |
| EPI_ISL_692848, EPI_ISL_692849, EPI_ISL_692850, EPI_ISL_692851, EPI_ISL_692852, EPI_ISL_692853, EPI_ISL_692854, EPI_ISL_692855, EPI_ISL_692869, EPI_ISL_692870 | Massachusetts State Public Health Laboratory | Massachusetts State Public Health Laboratory | Andrew Lang, Timelia Fink, Glen Gallagher, Sandra Smole |
| EPI_ISL_693662 | The National Institute of Public Health | State Veterinary Institute Prague | Nagy,A;Jirincova,H;Trnka,D;Vecerova,J |
| EPI_ISL_693764 | Hospital | National Reference Center for Viruses of Respiratory Infections, Institut Pasteur, Paris | Marion Barbet, Sylvie Behillil, Méline Bizard, Angela Brisebarre, Camille Capel, Etienne Simon-Lorière, Vincent Enouf, Maud Vanpeene, Sylvie van der Werf, Gisèle Lagathu |
| EPI_ISL_693765 | hospital | National Reference Center for Viruses of Respiratory Infections, Institut Pasteur, Paris | Marion Barbet, Sylvie Behillil, Méline Bizard, Angela Brisebarre, Camille Capel, Etienne Simon-Lorière, Vincent Enouf, Maud Vanpeene, Sylvie van der Werf, Gisèle Lagathu |
| EPI_ISL_699570, EPI_ISL_699571 | Group of Genetic Engeneering and Biotechnology, Federal Budget Institution of Science 'Central Research Institute of Epidemiology' of The Federal Service on Customers' Rights Protection and Human Well-being Surveillance | Group of Genetic Engeneering and Biotechnology, Federal Budget Institution of Science 'Central Research Institute of Epidemiology' of The Federal Service on Customers' Rights Protection and Human Well-being Surveillance | Cherkashina,A.S., Golubeva,A.G., Soloviova,E.D., Zotova,M.I., Berlina,Y.Y., Valdokhina,A.V., Bulanenko,V.P., Speranskaya,A.S., Tivanova,E.V., Shipulina,O.Y., Akimkin,V.G. |
| EPI_ISL_700441, EPI_ISL_700492 | Conville CDC wc CVC | NHLS/UCT | Houriiyah Tegally, Arash Iranzadeh, Deelan Doolabh, Lynn Tyers, Bruna Galvao, Innocent Mudau, Marvin Hsiao, Kruger Marais, Diana Hardie, Stephen Korsman, Carolyn Williamson |
| EPI_ISL_700518 | Plettenberg Bay Clinic wc PLC | NHLS/UCT | Arash Iranzadeh, Deelan Doolabh, Lynn Tyers, Bruna Galvao, Innocent Mudau, Marvin Hsiao, Kruger Marais, Diana Hardie, Stephen Korsman, Carolyn Williamson |
| EPI_ISL_700519 | Clinic-in-Asla | NHLS/UCT | Arash Iranzadeh, Deelan Doolabh, Lynn Tyers, Bruna Galvao, Innocent Mudau, Marvin Hsiao, Kruger Marais, Diana Hardie, Stephen Korsman, Carolyn Williamson |
| EPI_ISL_700543 | Great Brak River Clinic wc GBC | NHLS/UCT | Arash Iranzadeh, Deelan Doolabh, Lynn Tyers, Bruna Galvao, Innocent Mudau, Marvin Hsiao, Kruger Marais, Diana Hardie, Stephen Korsman, Carolyn Williamson |
| EPI_ISL_700551 | Pacaltsdorp Clinic wc PAC | NHLS/UCT | Houriiyah Tegally, Arash Iranzadeh, Deelan Doolabh, Lynn Tyers, Bruna Galvao, Innocent Mudau, Marvin Hsiao, Kruger Marais, Diana Hardie, Stephen Korsman, Carolyn Williamson |
| EPI_ISL_700569 | D'Almeida Clinic wc DAL | NHLS/UCT | Arash Iranzadeh, Deelan Doolabh, Lynn Tyers, Bruna Galvao, Innocent Mudau, Marvin Hsiao, Kruger Marais, Diana Hardie, Stephen Korsman, Carolyn Williamson |
| EPI_ISL_700705, EPI_ISL_700710, EPI_ISL_700738 | Texas Department of State Health Services | Texas Department of State Health Services | Rashmi Tuladhar, Bonnie Oh, Jenny Zhang, Maliha Rahman, Anita Pokharel, Myong Koag, Chung Wang, Rachel Lee, Grace Kubin, Mayela Pedrueza, James Daniel Bonser |
| EPI_ISL_703198 | Virology Department, Royal Infirmary of Edinburgh, NHS Lothian / School of Biological Sciences, University of Edinburgh / Institute of Genetics and Molecular Medicine, University of Edinburgh | COVID-19 Genomics UK (COG-UK) Consortium | McHugh M, Dewar R, Rooke S, Gallagher M, Balcaza C, O'Toole Á, Scher E, Hill V, McCrone JT, Colquhoun R, Yu X, Jackson B, Rambaut A, Williams TC, Templeton K |
| EPI_ISL_704812 | Oxford Viromics, NDM, University of Oxford; Oxford University Hospitals; Basingstoke and North Hampshire Hospital | COVID-19 Genomics UK (COG-UK) Consortium | Tanya Golubchik, David Bonsall, George Macintyre, Amy Trebes, Mariateresa de Cesare, Catrin Moore, Alex Mobbs, Anita Justice, Robert Shaw, Monique Andersson, Timothy Peto, Emma Wise, Nathan Moore, Jessica Lynch, Nick Cortes, Matilde Mori, Stephen Kidd, David Buck, John Todd, Christophe Fraser |
| EPI_ISL_704918, EPI_ISL_705229 | Wales Specialist Virology Centre Sequencing lab: Pathogen Genomics Unit | COVID-19 Genomics UK (COG-UK) Consortium | Catherine Moore, Johnathan Evans, Laura Gifford, Malorie Perry, Simon Cottrell, Angela Marchbank, Alec Birchley, Alexander Adams, Amy Gaskin, Bree Gatica-Wilcox, Jason Coombes, Joel Southgate, Lauren Gilbert, Lee Graham, Nicole Pacchiarini, Sara Kumziene-Summerhayes, Sarah Taylor, Sophie Jones, Sara Rey, Matthew Bull, Joanne Watkins, Sally Corden, Tom Connor |
| EPI_ISL_705391 | Virology Department, Royal Infirmary of Edinburgh, NHS Lothian / School of Biological Sciences, University of Edinburgh / Institute of Genetics and Molecular Medicine, University of Edinburgh | COVID-19 Genomics UK (COG-UK) Consortium | McHugh M, Dewar R, Rooke S, Gallagher M, Balcaza C, O'Toole Á, Scher E, Hill V, McCrone JT, Colquhoun R, Yu X, Jackson B, Rambaut A, Williams TC, Templeton K |
| EPI_ISL_705433, EPI_ISL_705463 | Wales Specialist Virology Centre Sequencing lab: Pathogen Genomics Unit | COVID-19 Genomics UK (COG-UK) Consortium | Catherine Moore, Johnathan Evans, Laura Gifford, Malorie Perry, Simon Cottrell, Angela Marchbank, Alec Birchley, Alexander Adams, Amy Gaskin, Bree Gatica-Wilcox, Jason Coombes, Joel Southgate, Lauren Gilbert, Lee Graham, Nicole Pacchiarini, Sara Kumziene-Summerhayes, Sarah Taylor, Sophie Jones, Sara Rey, Matthew Bull, Joanne Watkins, Sally Corden, Tom Connor |
| EPI_ISL_705473 | Oxford Viromics, NDM, University of Oxford; Oxford University Hospitals; Basingstoke and North Hampshire Hospital | COVID-19 Genomics UK (COG-UK) Consortium | Tanya Golubchik, David Bonsall, George Macintyre, Amy Trebes, Mariateresa de Cesare, Catrin Moore, Alex Mobbs, Anita Justice, Robert Shaw, Monique Andersson, Timothy Peto, Emma Wise, Nathan Moore, Jessica Lynch, Nick Cortes, Matilde Mori, Stephen Kidd, David Buck, John Todd, Christophe Fraser |

| | | | |
|---|---|---|---|
| EPI_ISL_705488 | Wales Specialist Virology Centre Sequencing lab: Pathogen Genomics Unit | COVID-19 Genomics UK (COG-UK) Consortium | Catherine Moore, Johnathan Evans, Laura Gifford, Malorie Perry, Simon Cottrell, Angela Marchbank, Alec Birchley, Alexander Adams, Amy Gaskin, Bree Gatica-Wilcox, Jason Coombes, Joel Southgate, Lauren Gilbert, Lee Graham, Nicole Pacchiarini, Sara Kumziene-Summerhayes, Sarah Taylor, Sophie Jones, Sara Rey, Matthew Bull, Joanne Watkins, Sally Corden, Tom Connor |
| EPI_ISL_705762, EPI_ISL_705779 | Virology Department, Royal Infirmary of Edinburgh, NHS Lothian / School of Biological Sciences, University of Edinburgh / Institute of Genetics and Molecular Medicine, University of Edinburgh | COVID-19 Genomics UK (COG-UK) Consortium | McHugh M, Dewar R, Rooke S, Gallagher M, Balcaza C, O'Toole Á, Scher E, Hill V, McCrone JT, Colquhoun R, Yu X, Jackson B, Rambaut A, Williams TC, Templeton K |
| EPI_ISL_705972, EPI_ISL_705973, EPI_ISL_705974, EPI_ISL_705975 | Oxford Viromics, NDM, University of Oxford; Oxford University Hospitals; Basingstoke and North Hampshire Hospital | COVID-19 Genomics UK (COG-UK) Consortium | Tanya Golubchik, David Bonsall, George Macintyre, Amy Trebes, Mariateresa de Cesare, Catrin Moore, Alex Mobbs, Anita Justice, Robert Shaw, Monique Andersson, Timothy Peto, Emma Wise, Nathan Moore, Jessica Lynch, Nick Cortes, Matilde Mori, Stephen Kidd, David Buck, John Todd, Christophe Fraser |
| EPI_ISL_706470, EPI_ISL_706603, EPI_ISL_706660, EPI_ISL_706661 | Wales Specialist Virology Centre Sequencing lab: Pathogen Genomics Unit | COVID-19 Genomics UK (COG-UK) Consortium | Catherine Moore, Johnathan Evans, Laura Gifford, Malorie Perry, Simon Cottrell, Angela Marchbank, Alec Birchley, Alexander Adams, Amy Gaskin, Bree Gatica-Wilcox, Jason Coombes, Joel Southgate, Lauren Gilbert, Lee Graham, Nicole Pacchiarini, Sara Kumziene-Summerhayes, Sarah Taylor, Sophie Jones, Sara Rey, Matthew Bull, Joanne Watkins, Sally Corden, Tom Connor |
| EPI_ISL_706930, EPI_ISL_706931, EPI_ISL_706932, EPI_ISL_706933, EPI_ISL_706934, EPI_ISL_706935, EPI_ISL_706936, EPI_ISL_706937, EPI_ISL_706974 | Oxford Viromics, NDM, University of Oxford; Oxford University Hospitals; Basingstoke and North Hampshire Hospital | COVID-19 Genomics UK (COG-UK) Consortium | Tanya Golubchik, David Bonsall, George Macintyre, Amy Trebes, Mariateresa de Cesare, Catrin Moore, Alex Mobbs, Anita Justice, Robert Shaw, Monique Andersson, Timothy Peto, Emma Wise, Nathan Moore, Jessica Lynch, Nick Cortes, Matilde Mori, Stephen Kidd, David Buck, John Todd, Christophe Fraser |
| EPI_ISL_707696 | Medical Research Center, Faculty of Medicine, Syarif Hidayatullah State Islamic University Jakarta | Medical Research Center, Faculty of Medicine, Syarif Hidayatullah State Islamic University Jakarta | Laifa Hendarmin, Chris Adhiyanto, Erike Suwarsono, Zeti Harriyati, Rini Puspitaningrum, Ferania Mela, Dennis Nurjadi |
| EPI_ISL_707976, EPI_ISL_707977, EPI_ISL_707978, EPI_ISL_707979, EPI_ISL_707980, EPI_ISL_707981, EPI_ISL_707982, EPI_ISL_707983, EPI_ISL_707984, EPI_ISL_707985, EPI_ISL_707986, EPI_ISL_707987, EPI_ISL_707988, EPI_ISL_707989, EPI_ISL_707990, EPI_ISL_707991, EPI_ISL_707992, EPI_ISL_707993, EPI_ISL_707994, EPI_ISL_707995, EPI_ISL_707996, EPI_ISL_707997, EPI_ISL_707998, EPI_ISL_708013, EPI_ISL_708014 | | | |
| see above | Virology, Universitätsklinikum des Saarlandes | Epigenetics, Saarland University | Kathrin Kattler, Markus Vogelgesang, Sascha Lohse, Sascha Tierling, Sigrun Smola, Jörn Walter |
| EPI_ISL_708028, EPI_ISL_708029 | Norwegian Institute of Public Health, Department of Virology | Norwegian Institute of Public Health, Department of Virology | Kathrine Stene-Johansen, Kamilla Heddeland Instefjord, Hilde Elshaug, Marie Paulsen Madsen, Rasmus Riis Kopperud, Hilde Vollan, Karoline Bragstad, Olav Hungnes |
| EPI_ISL_708030, EPI_ISL_708031 | Innlandet Hospital Trust, Division Lillehammer, Department for Medical Microbiology | Norwegian Institute of Public Health, Department of Virology | Kathrine Stene-Johansen, Kamilla Heddeland Instefjord, Hilde Elshaug, Marie Paulsen Madsen, Rasmus Riis Kopperud, Hilde Vollan, Karoline Bragstad, Olav Hungnes |
| EPI_ISL_708032 | Ostfold Hospital Trust - Kalnes, Centre for Laboratory Medicine, Section for gene technology and infection serology | Norwegian Institute of Public Health, Department of Virology | Kathrine Stene-Johansen, Kamilla Heddeland Instefjord, Hilde Elshaug, Marie Paulsen Madsen, Rasmus Riis Kopperud, Hilde Vollan, Karoline Bragstad, Olav Hungnes |
| EPI_ISL_708383, EPI_ISL_708388, EPI_ISL_708392, EPI_ISL_708393, EPI_ISL_708399, EPI_ISL_708417 | Delaware Public Health Lab | Delaware Public Health Lab | Gregory Hovan |
| EPI_ISL_708528 | Los Angeles County PHL | Los Angeles County PHL | P. Hemarajata et al. |
| EPI_ISL_708652 | Medical Research Center, Faculty of Medicine, Syarif Hidayatullah State Islamic University Jakarta | Medical Research Center, Faculty of Medicine, Syarif Hidayatullah State Islamic University Jakarta | Chris Adhiyanto, Laifa Hendarmin, Erike A Suwarsono, Zeti Harriyati, Endah Wulandari, Flori R Sari, Hari Hendarto |
| EPI_ISL_708724 | Los Angeles County Public Health Laboratory | Los Angeles County Public Health Laboratory | P. Hemarajata et al. |
| EPI_ISL_708783 | PathWest Laboratory Medicine WA | PathWest Laboratory Medicine WA Microbial Surveillance Unit | PathWest Laboratory Medicine WA Microbial Surveillance Unit |
| EPI_ISL_708807 | Regional medical sciences center 6 chonburi | National Institute of Health, Department of Medical Sciences, Ministry of Public Health, Thailand | Pilailuk Okada; Siripaporn Phuygun; Thanutsapa Thanadachakul; Sittiporn Parnmen; Pakorn Piromtong; Warawan Wongboot; Sunthareeya Waicharoen; Malinee Chittaganpitch |
| EPI_ISL_708822 | Emergency Operation Center, (EOC) | National Institute of Health, Department of Medical Sciences, Ministry of Public Health, Thailand | Pilailuk Okada; Siripaporn Phuygun; Thanutsapa Thanadachakul; Sittiporn Parnmen; Pakorn Piromtong; Warawan Wongboot; Sunthareeya Waicharoen; Malinee Chittaganpitch |
| EPI_ISL_709542 | National Institute of Blood Diseases (NIBD), Molecular Biology Lab | Genomics Lab NIBD | Samina Naz Mukry, Sayed Ali Raza, Shariq Ahmed, Aneeta Shahni, Gul Sufaida, Arshi Naz , Tahir Sultan Shamsi |
| EPI_ISL_709911, EPI_ISL_709936, EPI_ISL_709937, EPI_ISL_709938, EPI_ISL_709939, EPI_ISL_709940, EPI_ISL_709941 | Lighthouse Lab in Milton Keynes | Wellcome Sanger Institute for the COVID-19 Genomics UK (COG-UK) Consortium | The Lighthouse Lab in Milton Keynes and Alex Alderton, Roberto Amato, Sonia Goncalves, Ewan Harrison, David K. Jackson, Ian Johnston, Dominic Kwiatkowski, Cordelia Langford, John Sillitoe on behalf of the Wellcome Sanger Institute COVID-19 Surveillance Team |
| EPI_ISL_710354 | Colorado Department of Public Health and Environment | Colorado Department of Puplic Health and Environment | Laura Bankers, Molly C. Hetherington-Rauth, Shannon Ely, Shannon R. Matzinger, Sarah Elizabeth Totten, Emily A. Travanty |
| EPI_ISL_710416, EPI_ISL_710417, EPI_ISL_710421, EPI_ISL_710423, EPI_ISL_710425, EPI_ISL_710429, EPI_ISL_710430, EPI_ISL_710432 | Los Angeles County PHL | Los Angeles County PHL | P. Hemarajata et al. |
| EPI_ISL_710534, EPI_ISL_710537, EPI_ISL_710540, EPI_ISL_710541, EPI_ISL_710575 | Hôpital Fattouma-Bourguiba de Monastir | Laboratoire des Procédés de Criblage Moléculaire et Cellulaire-Centre de Biotechnologie de Sfax | Souissi,A., Abid,N., Ben Ayed,I., Gargouri,S., Abdelmoulah,F.,Elargoubi,A., Smeti,I., Bensaid,M., Stambouli,N., Kharat,N., Ajili,F., Fki-berrajah,L., Mhalla,S., Chtourou,A., Gaaloul,I., Nabli,A., Turki,M., Aouni,M., Hammami,A., Mastouri,M., Karray Hakim,H., Kamoun,S., Rebai,A. and Masmoudi,S. |
| EPI_ISL_718027, EPI_ISL_718029, EPI_ISL_718034, EPI_ISL_718035, EPI_ISL_718036, EPI_ISL_718037, EPI_ISL_718038, EPI_ISL_718039 | ZOTZ KLIMAS MVZ Düsseldorf-Centrum GbR ÜBAG für Labormedizin, Genetik, Zytologie, Pathologie | Center of Medical Microbiology, Virology, and Hospital Hygiene, University of Duesseldorf | Maximilian Damagnez, Alexander Dilthey, Ashley-Jane Duplessis, Patrick Finzer, Katrin Hoffmann, Torsten Houwaart, Lisanna Hülse, Malte Kohns Vasconcelos, Marek Korencak, Nadine Lübke, Jessica Nicolai, Klaus Pfeffer, Daniel Strelow, Jörg Timm, Andreas Walker, Tobias Wienemann, Rainer Zotz |
| EPI_ISL_718168 | Ministry of Health Hospitals | Institute of Health and Community Medicine | David Perera, Ooi Mong How, Chua Hock Hin, Tonnii Sia Loong Loong, Wong Jyn Shan, Wong Kiing Aik, Chan Chia Jui |
| EPI_ISL_718238, EPI_ISL_718242, EPI_ISL_718245 | Hospital | National Reference Center for Viruses of Respiratory Infections, Institut Pasteur, Paris | Marion Barbet, Sylvie Behillil, Méline Bizard, Angela Brisebarre, Camille Capel, Etienne Simon-Lorière, Vincent Enouf, Maud Vanpeene, Sylvie van der Werf, Gisèle Lagathu |
| EPI_ISL_718307, EPI_ISL_718308, EPI_ISL_718309 | Institute for Medical Research, Infectious Disease Research Centre, National Institutes of Health, Ministry of Health Malaysia | Institute for Medical Research, Infectious Disease Research Centre, National Institutes of Health, Ministry of Health Malaysia | Suppiah J, Kamel K, Mohd-Zawawi Z, Thayan R |
| EPI_ISL_721536, EPI_ISL_721537, EPI_ISL_721538, EPI_ISL_721539, EPI_ISL_721540, EPI_ISL_721541, EPI_ISL_721544 | Lighthouse Lab in Glasgow | Wellcome Sanger Institute for the COVID-19 Genomics UK (COG-UK) Consortium | Harper VanSteenhouse, Yumi Kasai, David Gray, Carol Clugston, Anna Dominiczak and Alex Alderton, Roberto Amato, Sonia Goncalves, Ewan Harrison, David K. Jackson, Ian Johnston, Dominic Kwiatkowski, Cordelia Langford, John Sillitoe on behalf of the Wellcome Sanger Institute COVID-19 Surveillance Team |
| EPI_ISL_722285, EPI_ISL_722298, EPI_ISL_722355, EPI_ISL_722366, EPI_ISL_722369, EPI_ISL_722383, EPI_ISL_722420, EPI_ISL_722421, EPI_ISL_722422, EPI_ISL_722423, EPI_ISL_722437, EPI_ISL_722438, EPI_ISL_722439, EPI_ISL_722440, EPI_ISL_722441, EPI_ISL_722500, EPI_ISL_722501, EPI_ISL_722502, EPI_ISL_722503, EPI_ISL_722504, EPI_ISL_722505, EPI_ISL_722506, EPI_ISL_722507, EPI_ISL_722508, EPI_ISL_722509, EPI_ISL_722510, EPI_ISL_722511, EPI_ISL_722512, EPI_ISL_722513 | | | |
| see above | Dutch COVID-19 response team | Erasmus Medical Center | Bas Oude Munnink, Reina Sikkema, David Nieuwenhuijse, Irina Chestakova, Anne van der Linden, Marjan Boter, Emmanuelle Munger, Corine |

| | | | |
|---|---|---|---|
| | | | GeurtsvanKessel, Annemiek van der Eijk, Richard Molenkamp, Marion Koopmans, on behalf of the Dutch national COVID-19 response team. |
| EPI_ISL_722878, EPI_ISL_722879, EPI_ISL_722880, EPI_ISL_722881, EPI_ISL_722904, EPI_ISL_722915, EPI_ISL_722916, EPI_ISL_722917, EPI_ISL_722918, EPI_ISL_722919 | Istituto Zooprofilattico Sperimentale della Puglia e della Basilicata | Istituto Zooprofilattico Sperimentale della Puglia e della Basilicata | Parisi A., Bianco A., Capozzi L., Del Sambro L., Manzulli V, Rondinone V., Pace L., Cipolletta D., Galante D. |
| EPI_ISL_723517 | Virginia Division of Consolidated Laboratory Services (DCLS) | Virginia Division of Consolidated Laboratory Services (DCLS) | Virginia DCLS |
| EPI_ISL_724977 | Northumbria University / South Tees Hospitals NHS Foundation Trust / North Cumbria Integrated Care NHS Foundation Trust / North Tees and Hartlepool NHS Foundation Trust / Newcastle Hospitals NHS Foundation Trust | COVID-19 Genomics UK (COG-UK) Consortium | Darren L Smith,Andrew Nelson,Matthew Bashton,Greg R Young,Joshua Loh,John Allan,Mohammad A Tariq,Giles S Holt,Gary Black,Wen C Yew,Lynn Dover,Paul Baker,Steve Liggett,Sarah Essex,Jane Greenaway,Debra Padgett,Clive Graham,Garren Scott,Edward Barton,Emma Swindells,Brendan Payne,Jennifer Collins,Yusri Taha,Gary Eltringham |
| EPI_ISL_727180 | Wales Specialist Virology Centre Sequencing lab: Pathogen Genomics Unit | COVID-19 Genomics UK (COG-UK) Consortium | Catherine Moore, Johnathan Evans, Laura Gifford, Malorie Perry, Simon Cottrell, Angela Marchbank, Alec Birchley, Alexander Adams, Amy Gaskin, Bree Gatica-Wilcox, Jason Coombes, Joel Southgate, Lauren Gilbert, Lee Graham, Nicole Pacchiarini, Sara Kumziene-Summerhayes, Sarah Taylor, Sophie Jones, Sara Rey, Matthew Bull, Joanne Watkins, Sally Corden, Tom Connor |
| EPI_ISL_727706, EPI_ISL_727709 | Centre for Enzyme Innovation, University of Portsmouth / Translational Research Laboratory, Portsmouth Hospitals NHS Trust | COVID-19 Genomics UK (COG-UK) Consortium | Angela Beckett,Yann Bourgeois,Garry Scarlett,Sharon Glaysher,Scott Elliott,Kelly Bicknell,Robert Impey,Allyson Lloyd,Sarah Wyllie,Ethan Butcher,Anoop Chauhan,Samuel Robson |
| EPI_ISL_728050, EPI_ISL_728076, EPI_ISL_728555 | University of Wisconsin-Madison AIDS Vaccine Research Laboratories | University of Wisconsin-Madison AIDS Vaccine Research Laboratories | Gage Moreno, Katarina Braun, et al. AIDS Vaccine Research Laboratories |
| EPI_ISL_728731 | Dutch COVID-19 response team | National Institute for Public Health and the Environment (RIVM) | Adam Meijer, Harry Vennema, Jeroen Cremer, Sharon van den Brink, Bas van der Veer, AnneMarie van den Brandt, Florian Zwagemaker, Dennis Schmitz, Chantal Reusken, on behalf of the national COVID-19 response team |
| EPI_ISL_729339, EPI_ISL_729361, EPI_ISL_729363, EPI_ISL_729367, EPI_ISL_729383, EPI_ISL_729399, EPI_ISL_729422, EPI_ISL_729423, EPI_ISL_729619 | A. Krumbholz, Labor Dr. Krause und Kollegen MVZ GmbH, Kiel | Charité Universitätsmedizin Berlin, Institut für Virologie | Victor M Corman, Barbara Mühlemann, Jörn Beheim-Schwarzbach, Talitha Veith, Julia Schneider, Terry Jones, Christian Drosten |
| EPI_ISL_729738, EPI_ISL_729757, EPI_ISL_729761, EPI_ISL_729762, EPI_ISL_729763, EPI_ISL_729769 | Yale Pathology Lab | Grubaugh Lab - Yale School of Public Health | Joseph Fauver, Tara Alpert, Anderson Brito, Annie Watkins, Anne Wyllie, Chantal Vogels, Mary Petrone, Chaney Kalinich, Isabel Ott, Arnau Casanovas, Catherine Muenker, Adam Moore, Alice Lu, Maria Tokuyama, Patrick Wong, Peiwen Lu, Saad Omer, Richard Martinello, Allison Nelson, Shelli Farhadian, Akiko Iwasaki, Charlese Dela Cruz, Albert Ko, Nathan Grubaugh |
| EPI_ISL_729784, EPI_ISL_729787, EPI_ISL_729788 | Grubaugh Lab - Yale School of Public Health | Grubaugh Lab - Yale School of Public Health | Joseph Fauver, Tara Alpert, Anderson Brito, Annie Watkins, Anne Wyllie, Chantal Vogels, Mary Petrone, Chaney Kalinich, Isabel Ott, Arnau Casanovas, Catherine Muenker, Adam Moore, Alice Lu, Maria Tokuyama, Patrick Wong, Peiwen Lu, Saad Omer, Richard Martinello, Allison Nelson, Shelli Farhadian, Akiko Iwasaki, Charlese Dela Cruz, Albert Ko, Nathan Grubaugh |
| EPI_ISL_729862, EPI_ISL_729863, EPI_ISL_729864, EPI_ISL_729865, EPI_ISL_729866, EPI_ISL_729867, EPI_ISL_729868, EPI_ISL_729872, EPI_ISL_729873, EPI_ISL_729874, EPI_ISL_729875, EPI_ISL_729877, EPI_ISL_729878, EPI_ISL_729879, EPI_ISL_729880, EPI_ISL_729881, EPI_ISL_729882, EPI_ISL_729883, EPI_ISL_729884, EPI_ISL_729885, EPI_ISL_729886, EPI_ISL_729887, EPI_ISL_729888, EPI_ISL_729889, EPI_ISL_729890, EPI_ISL_729891, EPI_ISL_729892, EPI_ISL_729893, EPI_ISL_729894, EPI_ISL_729895, EPI_ISL_729896, EPI_ISL_729897, EPI_ISL_729898, EPI_ISL_729899, EPI_ISL_729900, EPI_ISL_729901 | | | |
| see above | Laboratorio de Referencia Nacional de Virus Respiratorios, Instituto Nacional de Salud Peru | Laboratorio de Genómica Microbiana, Universidad Peruana Cayetano Heredia | Pablo Tsukayama, Alejandra Dávila-Barclay, Luis González, Guillermo Salvatierra, Pedro E. Romero, Brenda Ayzanoa, Janet Huancachoque, Pool Marcos, Marco Galarza, Priscila Lope, Nancy Rojas |
| EPI_ISL_730089, EPI_ISL_730114 | San Diego County Public Health Laboratory | Andersen lab at Scripps Research | SEARCH Alliance San Diego with Tracy Basler, Jovan Shephard, Brett Austin |
| EPI_ISL_730373, EPI_ISL_730374, EPI_ISL_730375, EPI_ISL_730377, EPI_ISL_730381, EPI_ISL_730383, EPI_ISL_730384, EPI_ISL_730385, EPI_ISL_730386, EPI_ISL_730387, EPI_ISL_730388, EPI_ISL_730392, EPI_ISL_730393, EPI_ISL_730395, EPI_ISL_730396, EPI_ISL_730397, EPI_ISL_730398, EPI_ISL_730400, EPI_ISL_730401, EPI_ISL_730402, EPI_ISL_730405, EPI_ISL_730406, EPI_ISL_730407, EPI_ISL_730408, EPI_ISL_730409, EPI_ISL_730410, EPI_ISL_730413, EPI_ISL_730414, EPI_ISL_730415, EPI_ISL_730416, EPI_ISL_730420, EPI_ISL_730421, EPI_ISL_730423, EPI_ISL_730424, EPI_ISL_730425, EPI_ISL_730426, EPI_ISL_730427, EPI_ISL_730435, EPI_ISL_730438, EPI_ISL_730447, EPI_ISL_730453, EPI_ISL_730454, EPI_ISL_730459, EPI_ISL_730460, EPI_ISL_730465, EPI_ISL_730469, EPI_ISL_730470, EPI_ISL_730472, EPI_ISL_730477, EPI_ISL_730478, EPI_ISL_730481, EPI_ISL_730485, EPI_ISL_730489, EPI_ISL_730490, EPI_ISL_730496, EPI_ISL_730497, EPI_ISL_730502, EPI_ISL_730507, EPI_ISL_730514 | | | |
| see above | Biolab Diagnostic Laboratories | Andersen lab at Scripps Research | Issa Abu-Dayyeh, Ahmad Tibi, Lama Hussein, Lina Mohammad, Zein Naber, Amid Abdelnour with SEARCH Alliance San Diego |
| EPI_ISL_730600 | Queen Mary Hospital | Hong Kong Department of Health | Mak Gannon C.K., Lam Edman T.K., Chan Rickjason C.W., Tsang Dominic N.C. |
| EPI_ISL_730615 | United Christian Hospital | Hong Kong Department of Health | Mak Gannon C.K., Lam Edman T.K., Chan Rickjason C.W., Tsang Dominic N.C. |
| EPI_ISL_732425, EPI_ISL_732439, EPI_ISL_732458, EPI_ISL_732459, EPI_ISL_732460, EPI_ISL_732461, EPI_ISL_732462, EPI_ISL_732463, EPI_ISL_732464, EPI_ISL_732465, EPI_ISL_732466, EPI_ISL_732467, EPI_ISL_732468, EPI_ISL_732469, EPI_ISL_732470, EPI_ISL_732471, EPI_ISL_732473, EPI_ISL_732474, EPI_ISL_732475, EPI_ISL_732476, EPI_ISL_732477, EPI_ISL_732478, EPI_ISL_732479, EPI_ISL_732480, EPI_ISL_732481, EPI_ISL_732483, EPI_ISL_732485, EPI_ISL_732486, EPI_ISL_732487, EPI_ISL_732488, EPI_ISL_732490, EPI_ISL_732491, EPI_ISL_732492, EPI_ISL_732494, EPI_ISL_732496, EPI_ISL_732518, EPI_ISL_732519, EPI_ISL_732520, EPI_ISL_732521, EPI_ISL_732523 | | | |
| see above | National Virus Reference Laboratory | National Virus Reference Laboratory | Michael Carr, Gabriel Gonzalez, Jonathan Dean, Daniel Hare, Cillian F De Gascun |
| EPI_ISL_732696, EPI_ISL_732697, EPI_ISL_732698 | CNR Virus des Infections Respiratoires - France SUD | CNR Virus des Infections Respiratoires - France SUD | Antonin Bal, Gregory Destras, Claudia Gonzalez, Gwendolyne Burfin, Quentin Semanas, Martine Valette, Bruno Lina, Laurence Josset |
| EPI_ISL_732813, EPI_ISL_732814, EPI_ISL_732818 | Centro de Investigación Biomédica de La Rioja - Hospital San Pedro Logroño | SeqCOVID-SPAIN consortium/IBV(CSIC) | María de Toro, José Manuel Azcona Gutiérrez, María Pilar Bea Escudero, Miriam Blasco Alberdi and SeqCOVID-SPAIN consortium |
| EPI_ISL_733154, EPI_ISL_733155 | WHO National Influenza Centre Russian Federation | WHO National Influenza Centre Russian Federation | Andrey Komissarov, Artem Fadeev, Anna Ivanova, Kseniya Komissarova, Dmitry Bazhenov, Daria Danilenko, Ksenia Safina, Elena Nabieva, Georgii Bazykin, Dmitry Lioznov |
| EPI_ISL_733237 | UMMC-Health | WHO National Influenza Centre Russian Federation | Andrey Komissarov, Artem Fadeev, Anna Ivanova, Kseniya Komissarova, Dmitry Bazhenov, Tatiana Platonova, Daria Danilenko, Ksenia Safina, Elena Nabieva, Georgii Bazykin, Dmitry Lioznov |
| EPI_ISL_733248, EPI_ISL_733249, EPI_ISL_733250, EPI_ISL_733251, EPI_ISL_733252, EPI_ISL_733253, EPI_ISL_733272, EPI_ISL_733273, EPI_ISL_733274, EPI_ISL_733275, EPI_ISL_733276, EPI_ISL_733282, EPI_ISL_733283, EPI_ISL_733284, EPI_ISL_733285, EPI_ISL_733286, EPI_ISL_733287, EPI_ISL_733288, EPI_ISL_733289, EPI_ISL_733290, EPI_ISL_733291, EPI_ISL_733292 | | | |
| see above | WHO National Influenza Centre Russian Federation | WHO National Influenza Centre Russian Federation | Andrey Komissarov, Artem Fadeev, Anna Ivanova, Kseniya Komissarova, Dmitry Bazhenov, Daria Danilenko, Ksenia Safina, Elena Nabieva, Georgii Bazykin, Dmitry Lioznov |
| EPI_ISL_733417, EPI_ISL_733418, EPI_ISL_733422, EPI_ISL_733423, EPI_ISL_733426, EPI_ISL_733427, EPI_ISL_733428, EPI_ISL_733429, EPI_ISL_733431, EPI_ISL_733432, EPI_ISL_733433, EPI_ISL_733434, EPI_ISL_733438, EPI_ISL_733441, EPI_ISL_733443, EPI_ISL_733446, EPI_ISL_733449, EPI_ISL_733450, EPI_ISL_733451, EPI_ISL_733452, EPI_ISL_733453, EPI_ISL_733455, EPI_ISL_733456 | | | |
| see above | HELIX LLC | WHO National Influenza Centre Russian Federation | Andrey Komissarov, Artem Fadeev, Anna Ivanova, Kseniya Komissarova, Dmitry Bazhenov, Daria Danilenko, Ksenia Safina, Elena Nabieva, Georgii Bazykin, Dmitry Lioznov |
| EPI_ISL_734289, EPI_ISL_734290, EPI_ISL_734349, EPI_ISL_734351, EPI_ISL_734352, EPI_ISL_734353, EPI_ISL_734354, EPI_ISL_734355, EPI_ISL_734356, EPI_ISL_734357, EPI_ISL_734358, EPI_ISL_734359, EPI_ISL_734360, EPI_ISL_734361, EPI_ISL_734362, EPI_ISL_734363, EPI_ISL_734364, EPI_ISL_734365, EPI_ISL_734366, EPI_ISL_734367, EPI_ISL_734368 | | | |
| see above | Wadsworth Center, New York State Department.of Health | Wadsworth Center, New York State Department.of Health | Kirsten St. George, Daryl M. Lamson, Alexis Russel, Jonathan Plitnick, Navjot Singh, John Kelly, Sara Griesemer, Erasmus Schneider, Erica Lasek-Nesselquist |
| EPI_ISL_734449, EPI_ISL_734450, EPI_ISL_734451, EPI_ISL_734452 | Masonic Medical Research Institute | Wadsworth Center, New York State Department.of Health | Kirsten St. George, Nathan Tucker, Ryan D. Pfeiffer, Daryl M. Lamson, Alexis Russel, Jonathan Plitnick, Navjot Singh, John Kelly, Sara Griesemer, Erasmus Schneider, Erica Lasek-Nesselquist |
| EPI_ISL_734495, EPI_ISL_735095, EPI_ISL_735096, EPI_ISL_735097, EPI_ISL_735098, EPI_ISL_735099, EPI_ISL_735100, EPI_ISL_735101, EPI_ISL_735102, EPI_ISL_735103, EPI_ISL_735104, EPI_ISL_735105, EPI_ISL_735106, EPI_ISL_735107, EPI_ISL_735108, EPI_ISL_735109, EPI_ISL_735110, EPI_ISL_735111, | | | |

| | | | |
|---|---|---|---|
| EPI_ISL_735112, EPI_ISL_735113, EPI_ISL_735114, EPI_ISL_735115, EPI_ISL_735116, EPI_ISL_735117, EPI_ISL_735118, EPI_ISL_735119, EPI_ISL_735120, EPI_ISL_735121, EPI_ISL_735122, EPI_ISL_735123, EPI_ISL_735124, EPI_ISL_735125, EPI_ISL_735126, EPI_ISL_735127, EPI_ISL_735128, EPI_ISL_735129, EPI_ISL_735130, EPI_ISL_735131, EPI_ISL_735132, EPI_ISL_735133, EPI_ISL_735134, EPI_ISL_735135, EPI_ISL_735136, EPI_ISL_735137, EPI_ISL_735138, EPI_ISL_735139, EPI_ISL_735161 | | | |
| see above | UZ Leuven, National Reference Laboratory for Coronaviruses, Laboratory Medicine, Leuven, Belgium | KU Leuven, Rega Institute, Clinical and Epidemiological Virology | Tony Wawina-Bokalanga, Joan Marti-Carerras, Bert Vanmechelen, Piet Maes |
| EPI_ISL_735257 | Group of Genetic Engeneering and Biotechnology, Federal Budget Institution of Science 'Central Research Institute of Epidemiology' of The Federal Service on Customers' Rights Protection and Human Well-being Surveillance | Group of Genetic Engeneering and Biotechnology, Federal Budget Institution of Science 'Central Research Institute of Epidemiology' of The Federal Service on Customers' Rights Protection and Human Well-being Surveillance | Cherkashina,A.S., Golubeva,A.G., Solovyova,E.D., Valdokhina,A.V., Bulanenko,V.P., Zotova,M.I., Berlina,Y.Y., Speranskaya,A.S., Tivanova,E.V., Shipulina,O.Y., Akimkin,V.G. |
| EPI_ISL_735446, EPI_ISL_735447 | Nucleic Acid Testing - Rwanda National Reference Laboratory | GIGA Medical Genomics | Yvan Butera, Keith Durkin, Maria Artesi, Bouchra Boujemla, Robert Rutayisire, Patrick Tuyisenge, Esperence Umumararungu, Sébastien Bontems, Marie-Pierre Hayette, Swaibu Gatare, Jacob Souopgui, Sabin Nsanzimana, Vincent Bours, Léon Mutesa |
| EPI_ISL_735452, EPI_ISL_735453, EPI_ISL_735457, EPI_ISL_735459, EPI_ISL_735460, EPI_ISL_735461, EPI_ISL_735462, EPI_ISL_735463, EPI_ISL_735466, EPI_ISL_735469, EPI_ISL_735471, EPI_ISL_735475, EPI_ISL_735476, EPI_ISL_735477, EPI_ISL_735478, EPI_ISL_735479, EPI_ISL_735480, EPI_ISL_735481, EPI_ISL_735482, EPI_ISL_735483, EPI_ISL_735484, EPI_ISL_735485, EPI_ISL_735486, EPI_ISL_735489 | | | |
| see above | UW Virology Lab | UW Virology Lab | Pavitra Roychoudhury, Hong Xie, Lasata Shrestha, Meei-Li Huang, Keith R Jerome, Alexander Greninger |
| EPI_ISL_735492 | Kumilla Medical College | Central Biological Research Laboratory and Department of Biochemistry and Molecular Biology | Robiul Hasan Bhuiyan, Md. Imranul Hoq, Md. Khondakar Raziur Rahman, Imam Hossen, Sajib Rudra, Md. Arif Hossain, Shanta Paul, Md. Omer Faruq, H. M. Abdullah Al Masud, Mohammad Omar Faruque |
| EPI_ISL_736776, EPI_ISL_737088, EPI_ISL_737089, EPI_ISL_737090, EPI_ISL_737091, EPI_ISL_737092, EPI_ISL_737093, EPI_ISL_737094, EPI_ISL_737095, EPI_ISL_737096, EPI_ISL_737097, EPI_ISL_737098, EPI_ISL_737099 | | | |
| see above | UW Virology Lab | UW Virology Lab | Pavitra Roychoudhury, Hong Xie, Lasata Shrestha, Meei-Li Huang, Keith R Jerome, Alexander Greninger |
| EPI_ISL_737212, EPI_ISL_737213, EPI_ISL_737214, EPI_ISL_737215, EPI_ISL_737216, EPI_ISL_737217, EPI_ISL_737218, EPI_ISL_737219, EPI_ISL_737220, EPI_ISL_737221, EPI_ISL_737222, EPI_ISL_737223, EPI_ISL_737224, EPI_ISL_737225, EPI_ISL_737226, EPI_ISL_737227, EPI_ISL_737228, EPI_ISL_737229, EPI_ISL_737230, EPI_ISL_737231, EPI_ISL_737232, EPI_ISL_737233 | | | |
| see above | Department of Virology and Immunology, University of Helsinki and Helsinki University Hospital, Huslab Finland | Department of Virology, Faculty of Medicine, University of Helsinki, Helsinki, Finland | Teemu Smura, Ravi Kant, Phuoc Truong, Hussein Alburkat, Hannimari Kallio-Kokko, Jenni Virtanen, Maija Suvanto, Sari Hannula, Harri Kangas, Pekka Ellonen, Olli Vapalahti |
| EPI_ISL_737309 | UW Virology Lab | UW Virology Lab | Pavitra Roychoudhury, Hong Xie, Lasata Shrestha, Meei-Li Huang, Keith R Jerome, Alexander Greninger |
| EPI_ISL_737593, EPI_ISL_737607, EPI_ISL_737692, EPI_ISL_737693, EPI_ISL_737694, EPI_ISL_737695, EPI_ISL_737696, EPI_ISL_737697, EPI_ISL_737698, EPI_ISL_737699, EPI_ISL_737700, EPI_ISL_737701, EPI_ISL_737702, EPI_ISL_737703, EPI_ISL_737704, EPI_ISL_737705, EPI_ISL_737706, EPI_ISL_737763, EPI_ISL_737764, EPI_ISL_737892, EPI_ISL_737893, EPI_ISL_737894, EPI_ISL_737895, EPI_ISL_737896, EPI_ISL_737897, EPI_ISL_737898, EPI_ISL_737900 | | | |
| see above | Viollier AG | Department of Biosystems Science and Engineering, ETH Zürich | Chaoran Chen, Sarah Nadeau, Ivan Topolsky, Emmanouil Dermitzakis, Keith Harshman, Ioannis Xenarios, Henri Pegeot, Lorenzo Cerutti, Deborah Penet, Philipp Jablonski, Lara Fuhrmann, David Dreifuss, Katharina Jahn, Christiane Beckmann, Maurice Redondo, Olivier Kobel, Christoph Noppen, Sophie Seidel, Noemie Santamaria de Souza, Niko Beerenwinkel, Tanja Stadler |
| EPI_ISL_738360, EPI_ISL_738413, EPI_ISL_738419, EPI_ISL_738425, EPI_ISL_738436, EPI_ISL_738442, EPI_ISL_738447, EPI_ISL_738448, EPI_ISL_738453, EPI_ISL_738470, EPI_ISL_738475, EPI_ISL_738480, EPI_ISL_738486, EPI_ISL_738492 | | | |
| see above | UZ Leuven, National Reference Laboratory for Coronaviruses, Laboratory Medicine, Leuven, Belgium | KU Leuven, Rega Institute, Clinical and Epidemiological Virology | Tony Wawina-Bokalanga, Joan Marti-Carerras, Bert Vanmechelen, Piet Maes |
| EPI_ISL_738530 | Humboldt County Public Health Laboratory | Chan-Zuckerberg Biohub | CZB Cliahub Consortium |
| EPI_ISL_738707, EPI_ISL_738708 | Madera County Department of Public Health | Chan-Zuckerberg Biohub | CZB Cliahub Consortium |
| EPI_ISL_738961 | Humboldt County Public Health Laboratory | Chan-Zuckerberg Biohub | CZB Cliahub Consortium |
| EPI_ISL_738964 | Madera County Department of Public Health | Chan-Zuckerberg Biohub | CZB Cliahub Consortium |
| EPI_ISL_739004 | UCSF Clinical Microbiology Laboratory | Chan-Zuckerberg Biohub | CZB Cliahub Consortium |
| EPI_ISL_739061, EPI_ISL_739133 | Madera County Department of Public Health | Chan-Zuckerberg Biohub | CZB Cliahub Consortium |
| EPI_ISL_739177 | Humboldt County Public Health Laboratory | Chan-Zuckerberg Biohub | CZB Cliahub Consortium |
| EPI_ISL_739197 | Madera County Department of Public Health | Chan-Zuckerberg Biohub | CZB Cliahub Consortium |
| EPI_ISL_739295 | UCSF Clinical Microbiology Laboratory | Chan-Zuckerberg Biohub | CZB Cliahub Consortium |
| EPI_ISL_739390, EPI_ISL_739429, EPI_ISL_739539 | Madera County Department of Public Health | Chan-Zuckerberg Biohub | CZB Cliahub Consortium |
| EPI_ISL_739558 | Humboldt County Public Health Laboratory | Chan-Zuckerberg Biohub | CZB Cliahub Consortium |
| EPI_ISL_739613, EPI_ISL_739632, EPI_ISL_739633 | UCSF Clinical Microbiology Laboratory | Chan-Zuckerberg Biohub | CZB Cliahub Consortium |
| EPI_ISL_739679 | Instituto Nacional de Salud, Bogotá, Colombia | Instituto Nacional de Salud, Bogotá, Colombia | Katherine Laiton-Donato, Diego A. Álvarez-Díaz, Carlos Franco-Muñoz, Mauricio Pacheco-Montealegre, Jonathan Reales, Diego Andrés Prada, Sheryl Corchuelo, Magdalena Weisner, Martha Lucia Ospina Martinez, Marcela Mercado-Reyes |
| EPI_ISL_739705, EPI_ISL_739716, EPI_ISL_739750, EPI_ISL_739757, EPI_ISL_739875, EPI_ISL_739912, EPI_ISL_740022, EPI_ISL_740081, EPI_ISL_740122, EPI_ISL_740126, EPI_ISL_740176, EPI_ISL_740322, EPI_ISL_740326, EPI_ISL_740447, EPI_ISL_740479, EPI_ISL_740519 | | | |
| see above | Laboratoire national de santé, Microbiology, Virology | Laboratoire national de santé, Microbiology, Microbial Genomics Platform | Anke Wienecke-Baldacchino, Catherine Ragimbeau,Jessica Tapp, Fatu Djabi, Lise Pignon, Raoul Salmon, Tamir Abdelrahman |
| EPI_ISL_741349, EPI_ISL_741350, EPI_ISL_741351, EPI_ISL_741352, EPI_ISL_741353, EPI_ISL_741354, EPI_ISL_741355, EPI_ISL_741770, EPI_ISL_741771 | Oxford Viromics, NDM, University of Oxford; Oxford University Hospitals; Basingstoke and North Hampshire Hospital | COVID-19 Genomics UK (COG-UK) Consortium | Tanya Golubchik, David Bonsall, George Macintyre, Amy Trebes, Mariateresa de Cesare, Catrin Moore, Alex Mobbs, Anita Justice, Robert Shaw, Monique Andersson, Timothy Peto, Emma Wise, Nathan Moore, Jessica Lynch, Nick Cortes, Matilde Mori, Stephen Kidd, David Buck, John Todd, Christophe Fraser |
| EPI_ISL_744154, EPI_ISL_744298, EPI_ISL_744357, EPI_ISL_744466, EPI_ISL_744495, EPI_ISL_744514, EPI_ISL_744599, EPI_ISL_744653, EPI_ISL_744663, EPI_ISL_744833, EPI_ISL_744899, EPI_ISL_744967, EPI_ISL_745006 | | | |
| see above | Laboratoire national de santé, Microbiology, Virology | Laboratoire national de santé, Microbiology, Microbial Genomics Platform | Anke Wienecke-Baldacchino, Catherine Ragimbeau,Jessica Tapp, Fatu Djabi, Lise Pignon, Raoul Salmon, Tamir Abdelrahman |
| EPI_ISL_745507, EPI_ISL_746302, EPI_ISL_746312 | Ginkgo Bioworks Clinical Laboratory | Utah Public Health Laboratory | Erin L. Young, Kelly Oakeson, Tara Gallagher, Michael T. Pyne, E. Susan Slechta, Melanie A. Mallory, Jeffrey B. Stevenson, Salika M. Shakir, David R. Hillyard, Malaika McKenzie-Bennett, James McGann, Jim Griffin, Keith Robison, Alex Plocik, Becky Schilling, Martha Pierson, Rebecca Littlefield, Michelle Spencer, Birgitte Simen |
| EPI_ISL_746503, EPI_ISL_746504, EPI_ISL_746505, EPI_ISL_746517, EPI_ISL_746518, EPI_ISL_746519, EPI_ISL_746543, EPI_ISL_746544, EPI_ISL_746765, EPI_ISL_746766, EPI_ISL_746767, EPI_ISL_746768, EPI_ISL_746769, EPI_ISL_746770, EPI_ISL_746771, EPI_ISL_746772, EPI_ISL_746773 | | | |
| see above | Genetica Molecular and Subdepartamento de Virologia ISP Chile | Instituto de Salud Publica de Chile | Javier Tognarelli, Barbara Parra, Loredana Arata, Jaime Lagos, Gisselle Barra, Patricia Bustos, Rodrigo Fasce, Andres Castillo, Jorge Fernandez |
| EPI_ISL_747042, EPI_ISL_747116 | Respiratory Viruses Branch, Centers for Disease Control and Prevention | Respiratory Viruses Branch, Centers for Disease Control and Prevention | Queen,K., Li,Y., Tao,Y., Uehara,A., Montmayeur,A., Paden,C.R., Cook,P.W., Marine,R., Sheth,M., Wang,H., Lee,J., Tong,S. |
| EPI_ISL_747240 | Al Islam Hospital | West Java Health Laboratory; School of Life Sciences and Technology, Institut Teknologi Bandung | Azzania Fibriani, Ema Rahmawati, Ryan Bayusantika Ristandi, Rifky Waluyajati Rachman, Cut Nur Cinthia Alamanda, Isak Solihin, Rini Robiani, Miftahul Faridl, Karimatu Khoirunnisa |
| EPI_ISL_747241 | Subang Public Health Office | West Java Health Laboratory; School of Life Sciences and | Azzania Fibriani, Ema Rahmawati, Ryan Bayusantika Ristandi, Rifky Waluyajati Rachman, Cut Nur Cinthia Alamanda, Isak Solihin, Rini Robiani, Miftahul |

| | | | |
|---|---|---|---|
| | | Technology, Institut Teknologi Bandung | Faridl, Karimatu Khoirunnisa |
| EPI_ISL_747369, EPI_ISL_747370, EPI_ISL_747371, EPI_ISL_747372, EPI_ISL_747373, EPI_ISL_747377, EPI_ISL_747378 | Division of Emerging Infectious Diseases, Bureau of Infectious Diseases Diagnosis Control, Korea Disease Control and Prevention Agency | Division of Emerging Infectious Diseases, Bureau of Infectious Diseases Diagnosis Control, Korea Disease Control and Prevention Agency | Ae Kyung Park, Il-Hwan Kim, Heui Man Kim, Jeong-Min Kim, Namjoo Lee, Chaeyoung Lee, Sang Hee Woo, Eun-Jin Kim |
| EPI_ISL_751477, EPI_ISL_751487 | CHU Purpan - Laboratoire de Virologie - Institut Fédératif de Biologie | CHU Purpan - Laboratoire de Virologie - Institut Fédératif de Biologie | Latour J., Ranger N., Dubois M., Carcenac R., Harter A., Boyer P., Tremeaux P., Izopet J. |
| EPI_ISL_751601 | MI - Michigan Department of Health and Human Services - Bureau of Laboratories | Genomics and Discovery, Respiratory Viruses Branch, Division of Viral Diseases, Centers for Disease Control and Prevention | Krista Queen, Yan Li, Ying Tao, Jing Zhang, Anna Uehara, Anna Montmayeur, Clinton R. Paden, Peter W. Cook,Rachel Marine, Mili Sheth, Haibin Wang, Justin Lee, Suxiang Tong |
| EPI_ISL_753038, EPI_ISL_753105, EPI_ISL_753106, EPI_ISL_753107 | State Laboratories Division, Hawaii State Department of Health | State Laboratories Division, Hawaii State Department of Health | Pamela O'Brien, Sabrina Diemert, Drew Kuwazaki, Razvan Sultana, Edward Desmond |
| EPI_ISL_753711, EPI_ISL_753715, EPI_ISL_753719, EPI_ISL_753776, EPI_ISL_753791, EPI_ISL_753792, EPI_ISL_753793, EPI_ISL_753888, EPI_ISL_753891, EPI_ISL_753892, EPI_ISL_753893, EPI_ISL_753895, EPI_ISL_753896, EPI_ISL_753897, EPI_ISL_753898, EPI_ISL_753899, EPI_ISL_753900, EPI_ISL_753901, EPI_ISL_753909, EPI_ISL_753910, EPI_ISL_753912, EPI_ISL_754018, EPI_ISL_754019, EPI_ISL_754020, EPI_ISL_754186, EPI_ISL_754190, EPI_ISL_754191 | | | |
| see above | Charité Universitätsmedizin Berlin, Institut für Virologie/Labor Berlin | Charité Universitätsmedizin Berlin, Institut für Virologie | Victor M Corman, Jörn Beheim-Schwarzbach, Barbara Mühlemann, Julia Schneider, Talitha Veith, Terry Jones, Christian Drosten |
| EPI_ISL_754677, EPI_ISL_754678, EPI_ISL_754679, EPI_ISL_754680, EPI_ISL_754681, EPI_ISL_754682, EPI_ISL_754683, EPI_ISL_754684, EPI_ISL_754685, EPI_ISL_754686, EPI_ISL_754687, EPI_ISL_754688, EPI_ISL_754689, EPI_ISL_754690, EPI_ISL_754695, EPI_ISL_754696, EPI_ISL_754697, EPI_ISL_754698, EPI_ISL_754699, EPI_ISL_754700, EPI_ISL_754701, EPI_ISL_754702, EPI_ISL_754703, EPI_ISL_754704, EPI_ISL_754705, EPI_ISL_754706, EPI_ISL_754707, EPI_ISL_754708, EPI_ISL_754709, EPI_ISL_754710, EPI_ISL_754711, EPI_ISL_754712 | | | |
| see above | Wadsworth Center, New York State Department.of Health | Wadsworth Center, New York State Department.of Health | Kirsten St. George, Daryl M. Lamson, Alexis Russel, Matthew Shudt, Melissa A Leisner, Jonathan Plitnick, Navjot Singh, John Kelly, Sara Griesemer, Erasmus Schneider, Erica Lasek-Nesselquist |
| EPI_ISL_755215 | UCSD EXCITE lab | Andersen lab at Scripps Research | SEARCH Alliance San Diego |
| EPI_ISL_755745, EPI_ISL_755811 | Toronto Invasive Bacterial Diseases Network | McMaster University | Allison McGeer, Patryk Aftanas, Hooman Derakhshani, Angel Li, Kuganya Nirmalarajah, Emily Panousis, Ahmed Draia, Jalees Nasir, Michael Surette, Samira Mubareka, Andrew G. McArthur |
| EPI_ISL_756167, EPI_ISL_756168, EPI_ISL_756169, EPI_ISL_756170, EPI_ISL_756171, EPI_ISL_756172, EPI_ISL_756173, EPI_ISL_756174, EPI_ISL_756175, EPI_ISL_756176, EPI_ISL_756177, EPI_ISL_756178, EPI_ISL_756179, EPI_ISL_756180, EPI_ISL_756285, EPI_ISL_756286, EPI_ISL_756287 | | | |
| see above | Department of Virology and Immunology, University of Helsinki and Helsinki University Hospital, Huslab Finland | Department of Virology, Faculty of Medicine, University of Helsinki, Helsinki, Finland | Teemu Smura, Ravi Kant, Phuoc Truong, Hussein Alburkat, Hannimari Kallio-Kokko, Jenni Virtanen, Maija Suvanto, Sari Hannula, Harri Kangas, Pekka Ellonen, Olli Vapalahti |
| EPI_ISL_759701 | RS UNS, Surakarta | UNS/RS-UNS, Faculty of Medicine | Betty Suryawati, Yulia Sari, Hartono, Maryani, Afif Avicenna Gufron, HanaHana Apsari Pawestri, Kartika Dewi Puspa, Hartanti Dian Ika, Arie Ardiansyah Nugraha, Vivi Setiawaty. |
| EPI_ISL_759969 | Department of Medical Microbiology, St. Olavs hospital | Norwegian Institute of Public Health, Department of Virology | Kathrine Stene-Johansen, Kamilla Heddeland Instefjord, Hilde Elshaug, Marie Paulsen Madsen, Rasmus Riis Kopperud, Hilde Vollan, Karoline Bragstad, Olav Hungnes |
| EPI_ISL_759992, EPI_ISL_759993, EPI_ISL_759994 | Division of Emerging Infectious Diseases, Bureau of Infectious Diseases Diagnosis Control, Korea Disease Control and Prevention Agency | Division of Emerging Infectious Diseases, Bureau of Infectious Diseases Diagnosis Control, Korea Disease Control and Prevention Agency | Ae Kyung Park, Il-Hwan Kim, Heui Man Kim, Jeong-Min Kim, Namjoo Lee, Chaeyoung Lee, Sang Hee Woo, Eun-Jin Kim |
| EPI_ISL_760057, EPI_ISL_760058 | Hong Kong Department of Health | School of Public Health, The University of Hong Kong | Daniel Chu, Haogao Gu, Pavithra Krishnan, Daisy Ng, Gigi Liu, Carrie Wan, Malik Peiris, Leo Poon |
| EPI_ISL_760064, EPI_ISL_760065, EPI_ISL_760104, EPI_ISL_760105, EPI_ISL_760112, EPI_ISL_760113, EPI_ISL_760119, EPI_ISL_760120, EPI_ISL_760203, EPI_ISL_760204 | Division of Emerging Infectious Diseases, Bureau of Infectious Diseases Diagnosis Control, Korea Disease Control and Prevention Agency | Division of Emerging Infectious Diseases, Bureau of Infectious Diseases Diagnosis Control, Korea Disease Control and Prevention Agency | Ae Kyung Park, Il-Hwan Kim, Heui Man Kim, Jeong-Min Kim, Namjoo Lee, Chaeyoung Lee, Sang Hee Woo, Eun-Jin Kim |
| EPI_ISL_763109 | Dutch COVID-19 response team | Erasmus Medical Center | Bas Oude Munnink, Reina Sikkema, David Nieuwenhuijse, Irina Chestakova, Anne van der Linden, Marjan Boter, Emmanuelle Munger, Corine GeurtsvanKessel, Annemiek van der Eijk, Richard Molenkamp, Marion Koopmans, on behalf of the Dutch national COVID-19 response team. |
| EPI_ISL_763388, EPI_ISL_763405, EPI_ISL_763408, EPI_ISL_763417, EPI_ISL_763446, EPI_ISL_763573, EPI_ISL_763583, EPI_ISL_763667, EPI_ISL_763729, EPI_ISL_764154, EPI_ISL_764173, EPI_ISL_764206, EPI_ISL_764262, EPI_ISL_764387, EPI_ISL_764388, EPI_ISL_764389, EPI_ISL_764390, EPI_ISL_764391, EPI_ISL_764392, EPI_ISL_764393, EPI_ISL_764394, EPI_ISL_764403, EPI_ISL_764404, EPI_ISL_764405, EPI_ISL_764406, EPI_ISL_764407, EPI_ISL_764408, EPI_ISL_764409, EPI_ISL_764410, EPI_ISL_764411, EPI_ISL_764412, EPI_ISL_764413, EPI_ISL_764414, EPI_ISL_764415, EPI_ISL_764416, EPI_ISL_764417, EPI_ISL_764418, EPI_ISL_764419, EPI_ISL_764420, EPI_ISL_764421, EPI_ISL_764422, EPI_ISL_764423, EPI_ISL_764424, EPI_ISL_764425 | | | |
| see above | Regional Virus Laboratory, Belfast Health and Social Care Trust | COVID-19 Genomics UK (COG-UK) Consortium | Conall McCaughey, James McKenna, Tanya Curran, Susan Feeney, Alison Watt, Ciara Cox, Mairead Connor, Zoltan Molnar, David Simpson, Derek Fairley |
| EPI_ISL_765650, EPI_ISL_765651, EPI_ISL_765654, EPI_ISL_765656, EPI_ISL_765756, EPI_ISL_765757, EPI_ISL_765758, EPI_ISL_765759, EPI_ISL_765760, EPI_ISL_765761 | Massachusetts General Hospital | Infectious Disease Program, Broad Institute of Harvard and MIT | Lemieux,J.E., Siddle,K.J., Shaw,B., Adams,G., Pierce,V., Turbett,S., Anahtar,M., Branda,J., Slater,D., Harris,J., Lin,A.E., Gladden-Young,A., Lagerborg,K., Rudy,M., DeRuff,K., Carter,A., Normandin,E., Bauer,M., Reilly,S., Tomkins-Tinch,C., Loreth,C., Chaluvadi,S., Neumann,A., Cusick,C., Chapman,S.B., Gnirke,A., Flowers,K., Cerrato,F., Birren,B.W., Gallagher,G., Smole,S., Park,D.J., MacInnis,B.L., Ryan,E., LaRocque,R., Rosenberg,E. and Sabeti,P.C. |
| EPI_ISL_765956, EPI_ISL_765957 | Worobey Lab, Department of Ecology and Evolutionary Biology, University of Arizona | Worobey Lab, Department of Ecology and Evolutionary Biology, University of Arizona | Brendan Larsen, Grace Quirk, Thomas Watts, David Baltrus, Michael Worobey |
| EPI_ISL_768756 | AIID | Irish Coronavirus Sequencing Consortium - National Virus Reference Laboratory | Michael Carr, Gabriel Gonzalez, Alejandro Abner Garcia Leon, Patrick Mallon |
| EPI_ISL_769987 | Laboratorio Clinico Labin | Inciensa, Instituto Costarricense de Investigación y Enseñanza en Nutrición y Salud | Francisco Duarte, Hebleen Porras, Claudio Soto-Garita, Estela Cordero, Adriana Godinez, Melany Calderon & Pei Chan Ma |
| EPI_ISL_769996 | Area De Salud San Juan-San Diego-Concepcion 2 | Inciensa, Instituto Costarricense de Investigación y Enseñanza en Nutrición y Salud | Francisco Duarte, Hebleen Porras, Claudio Soto-Garita, Estela Cordero, Adriana Godínez, Melany Calderón & Mariel López |
| EPI_ISL_770015 | Area De Salud San Francisco-San Antonio (Coopesana) | Inciensa, Instituto Costarricense de Investigación y Enseñanza en Nutrición y Salud | Francisco Duarte, Hebleen Porras, Claudio Soto-Garita, Estela Cordero, Adriana Godínez, Melany Calderón & Mariel López |
| EPI_ISL_775216 | Gonoshasthya-RNA Molecular Research Center | Gonoshasthya-RNA Molecular Research Center | Mohd. Raeed Jamiruddin, Nihad Adnan, Md. Ahsanul Haq, Mohib Ullah Khondoker, Nafisa Azmuda, Firoz Ahmed, Shahana Sharmin, Salma Akter, Taslin Jahan Mou, Mahfuza Marzan, Sayeda Moriam Liza, Nowshin Jahan, Tamanna Ali, Shahad Saif Khandker, Maha Jamiruddin, Mousumi Chaity, Mumtarin Jannat Oishee |
| EPI_ISL_775280, EPI_ISL_775314 | Dept. of Medical Microbiology, Stavanger University Hospital, Helse Stavanger HF | Norwegian Institute of Public Health, Department of Virology | Kathrine Stene-Johansen, Kamilla Heddeland Instefjord, Hilde Elshaug, Atiya R Ali,Marie Paulsen Madsen, Rasmus Riis Kopperud, Hilde Vollan, Karoline Bragstad, Olav Hungnes |
| EPI_ISL_775315 | Ostfold Hospital Trust - Kalnes, Centre for Laboratory Medicine, Section for gene technology and infection serology | Norwegian Institute of Public Health, Department of Virology | Kathrine Stene-Johansen, Kamilla Heddeland Instefjord, Hilde Elshaug, Atiya R Ali,Marie Paulsen Madsen, Rasmus Riis Kopperud, Hilde Vollan, Karoline Bragstad, Olav Hungnes |
| EPI_ISL_775450, EPI_ISL_775451, EPI_ISL_775474 | Vestfold Hospital, Toensberg Department of Microbiology | Norwegian Institute of Public Health, Department of Virology | Kathrine Stene-Johansen, Kamilla Heddeland Instefjord, Hilde Elshaug, Atiya R Ali,Marie Paulsen Madsen, Rasmus Riis Kopperud, Hilde Vollan, Karoline Bragstad, Olav Hungnes |
| EPI_ISL_775475 | Department of Medical Microbiology, St. Olavs hospital | Norwegian Institute of Public Health, Department of Virology | Kathrine Stene-Johansen, Kamilla Heddeland Instefjord, Hilde Elshaug, Atiya R Ali,Marie Paulsen Madsen, Rasmus Riis Kopperud, Hilde Vollan, Karoline Bragstad, Olav Hungnes |

| | | | |
|---|---|---|---|
| EPI_ISL_775478 | Unilabs Laboratory Medicine | Norwegian Institute of Public Health, Department of Virology | Kathrine Stene-Johansen, Kamilla Heddeland Instefjord, Hilde Elshaug, Atiya R Ali,Marie Paulsen Madsen, Rasmus Riis Kopperud, Hilde Vollan, Karoline Bragstad, Olav Hungnes |
| EPI_ISL_780073 | Hospital General Universitario Gregorio Marañón | SeqCOVID-SPAIN consortium/IBV(CSIC) | Darío García de Viedma, Laura Pérez-Lago, Marta Herranz, Jon Sicilia, Julia Suárez, Pilar Catalán, Patricia Muñoz and SeqCOVID-SPAIN consortium |
| EPI_ISL_785718, EPI_ISL_785854, EPI_ISL_786265, EPI_ISL_786266, EPI_ISL_786267, EPI_ISL_786268, EPI_ISL_786269, EPI_ISL_786270, EPI_ISL_786271, EPI_ISL_786277, EPI_ISL_786278, EPI_ISL_786279, EPI_ISL_786280, EPI_ISL_786281, EPI_ISL_786282, EPI_ISL_786283, EPI_ISL_786288, EPI_ISL_786289, EPI_ISL_786290, EPI_ISL_786291, EPI_ISL_786292, EPI_ISL_786293, EPI_ISL_786294, EPI_ISL_786295, EPI_ISL_786299, EPI_ISL_786300, EPI_ISL_786301, EPI_ISL_786302, EPI_ISL_786303, EPI_ISL_786305, EPI_ISL_786306, EPI_ISL_786311, EPI_ISL_786312, EPI_ISL_786314, EPI_ISL_786315, EPI_ISL_786316, EPI_ISL_786317, EPI_ISL_786318, EPI_ISL_786323, EPI_ISL_786324, EPI_ISL_786325, EPI_ISL_786326, EPI_ISL_786327, EPI_ISL_786328, EPI_ISL_786329, EPI_ISL_786330, EPI_ISL_786335, EPI_ISL_786336, EPI_ISL_786337, EPI_ISL_786338, EPI_ISL_786339, EPI_ISL_786340, EPI_ISL_786341, EPI_ISL_786342, EPI_ISL_786347, EPI_ISL_786348, EPI_ISL_786350, EPI_ISL_786351, EPI_ISL_786352, EPI_ISL_786353, EPI_ISL_786354, EPI_ISL_786355, EPI_ISL_786356, EPI_ISL_786357, EPI_ISL_786358, EPI_ISL_786359, EPI_ISL_786360, EPI_ISL_786361, EPI_ISL_786362, EPI_ISL_786363, EPI_ISL_786364, EPI_ISL_786365, EPI_ISL_786366, EPI_ISL_786367, EPI_ISL_786368, EPI_ISL_786369, EPI_ISL_786370, EPI_ISL_786371, EPI_ISL_786372, EPI_ISL_786373, EPI_ISL_786374, EPI_ISL_786375, EPI_ISL_786376, EPI_ISL_786377, EPI_ISL_786378, EPI_ISL_786379, EPI_ISL_786380, EPI_ISL_786381, EPI_ISL_786382, EPI_ISL_786383, EPI_ISL_786384, EPI_ISL_786386, EPI_ISL_786387, EPI_ISL_786388, EPI_ISL_786389, EPI_ISL_786390, EPI_ISL_786392, EPI_ISL_786393, EPI_ISL_786394, EPI_ISL_786395, EPI_ISL_786396, EPI_ISL_786397, EPI_ISL_786400, EPI_ISL_786401, EPI_ISL_786402, EPI_ISL_786403, EPI_ISL_786404, EPI_ISL_786405, EPI_ISL_786406, EPI_ISL_786407, EPI_ISL_786408, EPI_ISL_786409, EPI_ISL_786410, EPI_ISL_786411, EPI_ISL_786412, EPI_ISL_786413, EPI_ISL_786414, EPI_ISL_786415, EPI_ISL_786416, EPI_ISL_786417, EPI_ISL_786418, EPI_ISL_786419, EPI_ISL_786420, EPI_ISL_786421, EPI_ISL_786422, EPI_ISL_786423, EPI_ISL_786424, EPI_ISL_786425, EPI_ISL_786426, EPI_ISL_786427, EPI_ISL_786428, EPI_ISL_786429, EPI_ISL_786430, EPI_ISL_786431, EPI_ISL_786432, EPI_ISL_786433, EPI_ISL_786434, EPI_ISL_786435, EPI_ISL_786436, EPI_ISL_786437, EPI_ISL_786438, EPI_ISL_786439, EPI_ISL_786440, EPI_ISL_786441, EPI_ISL_786442, EPI_ISL_786443, EPI_ISL_786444, EPI_ISL_786445, EPI_ISL_786446, EPI_ISL_786447, EPI_ISL_786448, EPI_ISL_786449, EPI_ISL_786450, EPI_ISL_786451, EPI_ISL_786452, EPI_ISL_786453, EPI_ISL_786454, EPI_ISL_786455, EPI_ISL_786456, EPI_ISL_786457, EPI_ISL_786458, EPI_ISL_786459, EPI_ISL_786460, EPI_ISL_786461, EPI_ISL_786462, EPI_ISL_786463, EPI_ISL_786464, EPI_ISL_786465, EPI_ISL_786466, EPI_ISL_786467, EPI_ISL_786468, EPI_ISL_786469, EPI_ISL_786470, EPI_ISL_786471, EPI_ISL_786472, EPI_ISL_786473, EPI_ISL_786474, EPI_ISL_786475, EPI_ISL_786476, EPI_ISL_786477, EPI_ISL_786478, EPI_ISL_786479, EPI_ISL_786480, EPI_ISL_786481, EPI_ISL_786482, EPI_ISL_786483, EPI_ISL_786484, EPI_ISL_786485, EPI_ISL_786486, EPI_ISL_786487, EPI_ISL_786488, EPI_ISL_786489, EPI_ISL_786490, EPI_ISL_786491, EPI_ISL_786492, EPI_ISL_786493, EPI_ISL_786494, EPI_ISL_786495, EPI_ISL_786496, EPI_ISL_786497, EPI_ISL_786498, EPI_ISL_786499, EPI_ISL_786500, EPI_ISL_786501, EPI_ISL_786502, EPI_ISL_786503, EPI_ISL_786504, EPI_ISL_786505, EPI_ISL_786506, EPI_ISL_786507, EPI_ISL_786508, EPI_ISL_786509, EPI_ISL_786510, EPI_ISL_786511, EPI_ISL_786512, EPI_ISL_786513, EPI_ISL_786514, EPI_ISL_786515, EPI_ISL_786516, EPI_ISL_786517, EPI_ISL_786518, EPI_ISL_786519, EPI_ISL_786520, EPI_ISL_786521, EPI_ISL_786522, EPI_ISL_786523, EPI_ISL_786524, EPI_ISL_786525, EPI_ISL_786526, EPI_ISL_786527, EPI_ISL_786528, EPI_ISL_786529, EPI_ISL_786530, EPI_ISL_786531, EPI_ISL_786532, EPI_ISL_786533, EPI_ISL_786534, EPI_ISL_786535, EPI_ISL_786536, EPI_ISL_786537, EPI_ISL_786538, EPI_ISL_786539, EPI_ISL_786540, EPI_ISL_786541, EPI_ISL_786542, EPI_ISL_786543, EPI_ISL_786544, EPI_ISL_786545, EPI_ISL_786546, EPI_ISL_786547, EPI_ISL_786548, EPI_ISL_786549, EPI_ISL_786550, EPI_ISL_786551, EPI_ISL_786552, EPI_ISL_786553, EPI_ISL_786554, EPI_ISL_786555, EPI_ISL_786556, EPI_ISL_786559, EPI_ISL_786560, EPI_ISL_786561, EPI_ISL_786562, EPI_ISL_786563, EPI_ISL_786564, EPI_ISL_786565, EPI_ISL_786566, EPI_ISL_786567, EPI_ISL_786568, EPI_ISL_786571, EPI_ISL_786572, EPI_ISL_786573, EPI_ISL_786574, EPI_ISL_786575, EPI_ISL_786576, EPI_ISL_786577, EPI_ISL_786578, EPI_ISL_786579, EPI_ISL_786580, EPI_ISL_786583, EPI_ISL_786584, EPI_ISL_786585, EPI_ISL_786586, EPI_ISL_786587, EPI_ISL_786588, EPI_ISL_786589, EPI_ISL_786590, EPI_ISL_786591, EPI_ISL_786592, EPI_ISL_786595, EPI_ISL_786596, EPI_ISL_786597, EPI_ISL_786598, EPI_ISL_786599, EPI_ISL_786600, EPI_ISL_786601, EPI_ISL_786602, EPI_ISL_786603, EPI_ISL_786604, EPI_ISL_786607, EPI_ISL_786608, EPI_ISL_786609, EPI_ISL_786610, EPI_ISL_786611, EPI_ISL_786612, EPI_ISL_786613, EPI_ISL_786614, EPI_ISL_786615, EPI_ISL_786616, EPI_ISL_786619, EPI_ISL_786620, EPI_ISL_786621, EPI_ISL_786622, EPI_ISL_786624, EPI_ISL_786625, EPI_ISL_786626, EPI_ISL_786627, EPI_ISL_786628, EPI_ISL_786631, EPI_ISL_786632, EPI_ISL_786633, EPI_ISL_786634, EPI_ISL_786635, EPI_ISL_786636, EPI_ISL_786637, EPI_ISL_786638, EPI_ISL_786639 | | | |
| see above | Houston Methodist Hospital | Houston Methodist Hospital | S. Wesley Long, Randall J. Olsen, Paul A. Christensen, David W. Bernard, James J. Davis, Maulik Shukla, Marcus Nguyen, Matthew Ojeda Saavedra, Prasanti Yerramilli, Layne Pruitt, Sishir Subedi, Heather Hendrickson, and James M. Musser |
| EPI_ISL_791988 | Balai Besar POM Semarang | National Institute of Health Research and Development | Subangkit;Pawestri,HA;Ikawati,HD;Nugraha,AA;Puspa,KD;Aryanti;Pangesti,KNA;Soekarso,T;Puspandari,N;Setiawaty,V |
| EPI_ISL_792562, EPI_ISL_792624 | LACEN-PB | Laboratory of Respiratory Viruses and Measles, Oswaldo Cruz Institute, FIOCRUZ | Paola Resende, Luciana Appolinario, Fernando Motta, Anna Carolina Paixao, Ana Carolina Mendonca, João Felipe Bezerra, Romero Henrique Teixeira de Vasconcelos, Dalane Loudal Florentino Teixeira, Thiago Franco de Oliveira Carneiro, Marilda Siqueira |
| EPI_ISL_794722, EPI_ISL_794723 | PathWest Laboratory Medicine WA | PathWest Laboratory Medicine WA Microbial Surveillance Unit | PathWest Laboratory Medicine WA Microbial Surveillance Unit |
| EPI_ISL_801431, EPI_ISL_801484, EPI_ISL_801485 | Dutch COVID-19 response team | Erasmus Medical Center | Bas Oude Munnink, Reina Sikkema, David Nieuwenhuijse, Irina Chestakova, Anne van der Linden, Marjan Boter, Emmanuelle Munger, Corine GeurtsvanKessel, Annemiek van der Eijk, Richard Molenkamp, Marion Koopmans, on behalf of the Dutch national COVID-19 response team. |
| EPI_ISL_802417, EPI_ISL_802418, EPI_ISL_802419, EPI_ISL_802420, EPI_ISL_802421, EPI_ISL_802422, EPI_ISL_802423, EPI_ISL_802424, EPI_ISL_802425, EPI_ISL_802426, EPI_ISL_802427 | | | |
| see above | MSHS Clinical Microbiology Laboratories | MSHS Pathogen Surveillance Program | Ana S. Gonzalez-Reiche, Hala Alshammary, Mitchell J. Sullivan, Brianne Ciferri, Ajay Obla, Angela Amoako, Mahmoud Awawda, Elena Hirsch, Ashley S. Salimbangon, Levy Sominsky, Katherine Beach, Kayla Russo, Charles Gleason, Shelcie Fabre, Giulio Kleiner, Zenab Khan, Bremy Alburquerque, Adriana van de Guchte, Komal Srivastava, Matthew M. Hernandez, Jayeeta Dutta, Denise Jurczyszak, Emily Ferreri, Rachel Chernet, Nancy Francoeur, Betsaida Salom Melo, Irina Oussenko, Gintaras Deikus, Juan Soto, Shwetha Hara Sridhar, Ying-Chih Wang, Kathryn Twyman, Andrew Kasarskis, Deena R. Altman, Robert Sebra, Adolfo García-Sastre, Marta Luksza, Gopi Patel, Sarah Schaefer, Melissa Gitman, Michael D. Nowak, Alberto Paniz-Mondolfi, Emilia Mia Sordillo, Viviana Simon, Harm van Bakel |
| EPI_ISL_802861 | Vilnius University Hospital Santaros Klinikos, Vilnius University | Institute of Biotechnology, Life Sciences Center, Vilnius University | Emilija Vasiliunaite, Milda Norkiene, Albertas Timinskas, Alma Gedvilaite, Aurelija Zvirbliene, Daniel Naumovas, Laimonas Griskevicius |
| EPI_ISL_803369, EPI_ISL_803370, EPI_ISL_803371, EPI_ISL_803372, EPI_ISL_803373, EPI_ISL_803374, EPI_ISL_803375, EPI_ISL_803376, EPI_ISL_803377, EPI_ISL_803392, EPI_ISL_803393, EPI_ISL_803394, EPI_ISL_803395, EPI_ISL_803396, EPI_ISL_803397, EPI_ISL_803399, EPI_ISL_803400, EPI_ISL_803437, EPI_ISL_803438, EPI_ISL_803439, EPI_ISL_803658, EPI_ISL_803659, EPI_ISL_803660, EPI_ISL_803661, EPI_ISL_803662, EPI_ISL_803663, EPI_ISL_803664, EPI_ISL_803665, EPI_ISL_803666, EPI_ISL_803667, EPI_ISL_803668, EPI_ISL_803669, EPI_ISL_803670, EPI_ISL_803671, EPI_ISL_803672, EPI_ISL_803688, EPI_ISL_803689, EPI_ISL_803690, EPI_ISL_803691, EPI_ISL_803692, EPI_ISL_803693, EPI_ISL_803694, EPI_ISL_803695, EPI_ISL_803696, EPI_ISL_803697, EPI_ISL_803698, EPI_ISL_803768, EPI_ISL_803769, EPI_ISL_803770, EPI_ISL_803771, EPI_ISL_803772, EPI_ISL_803773, EPI_ISL_803774, EPI_ISL_803775, EPI_ISL_803776, EPI_ISL_803777, EPI_ISL_803779, EPI_ISL_803780, EPI_ISL_803781, EPI_ISL_803782, EPI_ISL_803783, EPI_ISL_803784, EPI_ISL_803785, EPI_ISL_803786, EPI_ISL_803787, EPI_ISL_803788, EPI_ISL_803791, EPI_ISL_803792, EPI_ISL_803793, EPI_ISL_803794, EPI_ISL_803795, EPI_ISL_803796, EPI_ISL_803797, EPI_ISL_803798, EPI_ISL_803799, EPI_ISL_803800, EPI_ISL_803801, EPI_ISL_803802, EPI_ISL_803803, EPI_ISL_803804, EPI_ISL_803805, EPI_ISL_803806, EPI_ISL_803807, EPI_ISL_803808, EPI_ISL_803829, EPI_ISL_803830 | | | |
| see above | Wisconsin State Laboratory of Hygiene Communicable Disease Division | Wisconsin State Laboratory of Hygiene Communicable Disease Division | Kelsey R. Florek, Abigail C. Shockey |
| EPI_ISL_804376 | CHU Purpan - Laboratoire de Virologie - Institut Fédératif de Biologie | CHU Purpan - Laboratoire de Virologie - Institut Fédératif de Biologie | Latour J., Ranger N., Dubois M., Carcenac R., Harter A., Boyer P., Tremeaux P., Izopet J. |
| EPI_ISL_804479, EPI_ISL_804482, EPI_ISL_804493, EPI_ISL_804494, EPI_ISL_804495, EPI_ISL_804496, EPI_ISL_804497, EPI_ISL_804501, EPI_ISL_804502, EPI_ISL_804503, EPI_ISL_804504, EPI_ISL_804505, EPI_ISL_804506, EPI_ISL_804507, EPI_ISL_804508, EPI_ISL_804509, EPI_ISL_804510, EPI_ISL_804511, EPI_ISL_804512, EPI_ISL_804513, EPI_ISL_804537, EPI_ISL_804538 | | | |
| see above | MEPHI, Aix Marseille University | MEPHI, Aix Marseille University | Anthony LEVASSEUR |
| EPI_ISL_804592, EPI_ISL_804593, EPI_ISL_804594, EPI_ISL_804595, EPI_ISL_804596, EPI_ISL_804597 | Maryland Public Health Laboratory | Maryland Public Health Laboratory | Maryland Department of Health Laboratories Administration |
| EPI_ISL_819449 | Northumbria University / South Tees Hospitals NHS Foundation Trust / North Cumbria Integrated Care NHS Foundation Trust / North Tees and Hartlepool NHS Foundation Trust / Newcastle Hospitals NHS Foundation Trust | COVID-19 Genomics UK (COG-UK) Consortium | Darren L Smith,Andrew Nelson,Matthew Bashton,Greg R Young,Joshua Loh,John Allan,Mohammad A Tariq,Giles S Holt,Gary Black,Wen C Yew,Lynn Dover,Paul Baker,Steve Liggett,Sarah Essex,Jane Greenaway,Debra Padgett,Clive Graham,Garren Scott,Edward Barton,Emma Swindells,Brendan Payne,Jennifer Collins,Yusri Taha,Gary Eltringham |
| EPI_ISL_824411, EPI_ISL_824511, EPI_ISL_824512 | Hospital Universitari Vall d'Hebron - Vall d'Hebron Institut de Rercerca | Hospital Universitari Vall d'Hebron | Cristina Andrés, Maria Piñana, Josep F Abril, Damir Garcia-Cehic, Ariadna Rando, Juliana Esperalba, Maria Gema Codina, Carla Castillo, Maria Carmen Martin, Tomàs Pumarola, Josep Quer, Andrés Antón |
| EPI_ISL_824514 | New Mexico Department of Health Scientific Laboratory | New Mexico Department of Health Scientific Laboratory | Ellie Johnson, Anastacia Griego-Fisher, D'eldra Malone |
| EPI_ISL_825492, EPI_ISL_825495, EPI_ISL_825496, EPI_ISL_825580, EPI_ISL_825581, EPI_ISL_825582 | Respiratory Virus Unit, National Infection Service, Public Health England | COVID-19 Genomics UK (COG-UK) Consortium | PHE Covid Sequencing Team |
| EPI_ISL_826517 | Lighthouse Lab in Alderley Park | Wellcome Sanger Institute for the COVID-19 Genomics UK (COG-UK) Consortium | Jacquelyn Wynn, Mairead Hyland, The Lighthouse Lab in Alderley Park and Alex Alderton, Roberto Amato, Sonia Goncalves, Ewan Harrison, David K. Jackson, Ian Johnston, Dominic Kwiatkowski, Cordelia Langford, John Sillitoe on behalf of the Wellcome Sanger Institute COVID-19 Surveillance Team |
| EPI_ISL_826518 | Lighthouse Lab in Cambridge | Wellcome Sanger Institute for the COVID-19 Genomics UK (COG-UK) Consortium | Rob Howes, The Lighthouse Lab in Cambridge and Alex Alderton, Roberto Amato, Sonia Goncalves, Ewan Harrison, David K. Jackson, Ian Johnston, Dominic Kwiatkowski, Cordelia Langford, John Sillitoe on behalf of the Wellcome Sanger Institute COVID-19 Surveillance Team |
| EPI_ISL_826519 | Lighthouse Lab in Alderley Park | Wellcome Sanger Institute for the COVID-19 Genomics UK (COG-UK) Consortium | Jacquelyn Wynn, Mairead Hyland, The Lighthouse Lab in Alderley Park and Alex Alderton, Roberto Amato, Sonia Goncalves, Ewan Harrison, David K. Jackson, Ian Johnston, Dominic Kwiatkowski, Cordelia Langford, John Sillitoe on behalf of the Wellcome Sanger Institute COVID-19 Surveillance Team |
| EPI_ISL_826667, EPI_ISL_826668 | Montefiore Medical Center | Albert Einstein College of Medicine, Dept. of Microbiology & Immunology, Chandran lab | J. Maximilian Fels, Saad Khan, Ryan Forster, Karin A. Skalina, Surksha Sirichand, Amy S. Fox, Aviv Bergman, William B. Mitchell, Lucia R. Wolgast, Wendy Szymczak, Robert H. Bortz III, M. Eugenia Dieterle, Catalina Florez, Denise Haslwanter, Rohit K. Jangra, Ethan Laudermilch, Ariel S. Wirchnianski, Jason Barnhill, David L. Goldman, Hnin Khine, D. Yitzchak Goldstein, Johanna P. Daily, Kartik Chandran, Libusha Kelly |

| | | | |
|---|---|---|---|
| EPI_ISL_826679, EPI_ISL_826745, EPI_ISL_826746, EPI_ISL_826749, EPI_ISL_826750, EPI_ISL_826751, EPI_ISL_826752, EPI_ISL_826762, EPI_ISL_826768, EPI_ISL_826774, EPI_ISL_826789, EPI_ISL_826790, EPI_ISL_826791, EPI_ISL_826841, EPI_ISL_826843, EPI_ISL_826844 | | | |
| see above | deCODE genetics | deCODE genetics | Daniel F Gudbjartsson; Agnar Helgason; Hakon Jonsson; Olafur T Magnusson; Pall Melsted; Gudmundur L Norddahl; Jona Saemundsdottir; Asgeir Sigurdsson; Patrick Sulem; Arna B Agustsdottir; Hannes Eggertsson; Berglind Eiriksdottir; Run Fridriksdottir; Elisabet E Gardarsdottir; Gudmundur Georgsson; Olafia S Gretarsdottir; Kjartan R Gudmundsson; Thora R Gunnarsdottir; Arnaldur Gylfason; Hilma Holm; Brynjar O Jensson; Aslaug Jonasdottir; Kamilla S Josefsdottir; Thordur Kristjansson; Droplaug N Magnusdottir; Solvi Rognvaldsson; Louise le Roux; Gudrun Sigmundsdottir; Gardar Sveinbjornsson; Kristin E Sveinsdottir; Maney Sveinsdottir; Emil A Thorarensen; Bjarni Thorbjornsson; Gisli Masson; Ingileif Jonsdottir; Alma Moller; Thorolfur Gudnason; Karl G Kristinsson; Unnur Thorsteinsdottir; Kari Stefansson |
| EPI_ISL_826897 | The National University Hospital of Iceland | deCODE genetics | Daniel F Gudbjartsson; Agnar Helgason; Hakon Jonsson; Olafur T Magnusson; Pall Melsted; Gudmundur L Norddahl; Jona Saemundsdottir; Asgeir Sigurdsson; Patrick Sulem; Arna B Agustsdottir; Hannes Eggertsson; Berglind Eiriksdottir; Run Fridriksdottir; Elisabet E Gardarsdottir; Gudmundur Georgsson; Olafia S Gretarsdottir; Kjartan R Gudmundsson; Thora R Gunnarsdottir; Arnaldur Gylfason; Hilma Holm; Brynjar O Jensson; Aslaug Jonasdottir; Kamilla S Josefsdottir; Thordur Kristjansson; Droplaug N Magnusdottir; Solvi Rognvaldsson; Louise le Roux; Gudrun Sigmundsdottir; Gardar Sveinbjornsson; Kristin E Sveinsdottir; Maney Sveinsdottir; Emil A Thorarensen; Bjarni Thorbjornsson; Gisli Masson; Ingileif Jonsdottir; Alma Moller; Thorolfur Gudnason; Karl G Kristinsson; Unnur Thorsteinsdottir; Kari Stefansson |
| EPI_ISL_826903, EPI_ISL_826904, EPI_ISL_826905, EPI_ISL_826906, EPI_ISL_826907, EPI_ISL_826908, EPI_ISL_826909, EPI_ISL_826910, EPI_ISL_826911, EPI_ISL_826912, EPI_ISL_826920, EPI_ISL_826921, EPI_ISL_826925, EPI_ISL_826926, EPI_ISL_826928, EPI_ISL_826929, EPI_ISL_827047, EPI_ISL_827049, EPI_ISL_827050, EPI_ISL_827052, EPI_ISL_827053, EPI_ISL_827054, EPI_ISL_827055, EPI_ISL_827056, EPI_ISL_827099, EPI_ISL_827100, EPI_ISL_827101, EPI_ISL_827102, EPI_ISL_827105, EPI_ISL_827106, EPI_ISL_827143, EPI_ISL_827147, EPI_ISL_827148, EPI_ISL_827150, EPI_ISL_827153, EPI_ISL_827154, EPI_ISL_827155, EPI_ISL_827156, EPI_ISL_827157, EPI_ISL_827158, EPI_ISL_827159, EPI_ISL_827160, EPI_ISL_827161, EPI_ISL_827162, EPI_ISL_827165, EPI_ISL_827169, EPI_ISL_827174, EPI_ISL_827185, EPI_ISL_827186, EPI_ISL_827189, EPI_ISL_827190, EPI_ISL_827191, EPI_ISL_827192 | | | |
| see above | deCODE genetics | deCODE genetics | Daniel F Gudbjartsson; Agnar Helgason; Hakon Jonsson; Olafur T Magnusson; Pall Melsted; Gudmundur L Norddahl; Jona Saemundsdottir; Asgeir Sigurdsson; Patrick Sulem; Arna B Agustsdottir; Hannes Eggertsson; Berglind Eiriksdottir; Run Fridriksdottir; Elisabet E Gardarsdottir; Gudmundur Georgsson; Olafia S Gretarsdottir; Kjartan R Gudmundsson; Thora R Gunnarsdottir; Arnaldur Gylfason; Hilma Holm; Brynjar O Jensson; Aslaug Jonasdottir; Kamilla S Josefsdottir; Thordur Kristjansson; Droplaug N Magnusdottir; Solvi Rognvaldsson; Louise le Roux; Gudrun Sigmundsdottir; Gardar Sveinbjornsson; Kristin E Sveinsdottir; Maney Sveinsdottir; Emil A Thorarensen; Bjarni Thorbjornsson; Gisli Masson; Ingileif Jonsdottir; Alma Moller; Thorolfur Gudnason; Karl G Kristinsson; Unnur Thorsteinsdottir; Kari Stefansson |
| EPI_ISL_827238, EPI_ISL_827242 | The National University Hospital of Iceland | deCODE genetics | Daniel F Gudbjartsson; Agnar Helgason; Hakon Jonsson; Olafur T Magnusson; Pall Melsted; Gudmundur L Norddahl; Jona Saemundsdottir; Asgeir Sigurdsson; Patrick Sulem; Arna B Agustsdottir; Hannes Eggertsson; Berglind Eiriksdottir; Run Fridriksdottir; Elisabet E Gardarsdottir; Gudmundur Georgsson; Olafia S Gretarsdottir; Kjartan R Gudmundsson; Thora R Gunnarsdottir; Arnaldur Gylfason; Hilma Holm; Brynjar O Jensson; Aslaug Jonasdottir; Kamilla S Josefsdottir; Thordur Kristjansson; Droplaug N Magnusdottir; Solvi Rognvaldsson; Louise le Roux; Gudrun Sigmundsdottir; Gardar Sveinbjornsson; Kristin E Sveinsdottir; Maney Sveinsdottir; Emil A Thorarensen; Bjarni Thorbjornsson; Gisli Masson; Ingileif Jonsdottir; Alma Moller; Thorolfur Gudnason; Karl G Kristinsson; Unnur Thorsteinsdottir; Kari Stefansson |
| EPI_ISL_827243, EPI_ISL_827274, EPI_ISL_827276, EPI_ISL_827277, EPI_ISL_827336, EPI_ISL_827346, EPI_ISL_827347, EPI_ISL_827348, EPI_ISL_827349, EPI_ISL_827350, EPI_ISL_827351, EPI_ISL_827352, EPI_ISL_827353, EPI_ISL_827355, EPI_ISL_827356, EPI_ISL_827414, EPI_ISL_827421, EPI_ISL_827423, EPI_ISL_827424, EPI_ISL_827425, EPI_ISL_827426, EPI_ISL_827428, EPI_ISL_827432, EPI_ISL_827433 | | | |
| see above | deCODE genetics | deCODE genetics | Daniel F Gudbjartsson; Agnar Helgason; Hakon Jonsson; Olafur T Magnusson; Pall Melsted; Gudmundur L Norddahl; Jona Saemundsdottir; Asgeir Sigurdsson; Patrick Sulem; Arna B Agustsdottir; Hannes Eggertsson; Berglind Eiriksdottir; Run Fridriksdottir; Elisabet E Gardarsdottir; Gudmundur Georgsson; Olafia S Gretarsdottir; Kjartan R Gudmundsson; Thora R Gunnarsdottir; Arnaldur Gylfason; Hilma Holm; Brynjar O Jensson; Aslaug Jonasdottir; Kamilla S Josefsdottir; Thordur Kristjansson; Droplaug N Magnusdottir; Solvi Rognvaldsson; Louise le Roux; Gudrun Sigmundsdottir; Gardar Sveinbjornsson; Kristin E Sveinsdottir; Maney Sveinsdottir; Emil A Thorarensen; Bjarni Thorbjornsson; Gisli Masson; Ingileif Jonsdottir; Alma Moller; Thorolfur Gudnason; Karl G Kristinsson; Unnur Thorsteinsdottir; Kari Stefansson |
| EPI_ISL_827435 | The National University Hospital of Iceland | deCODE genetics | Daniel F Gudbjartsson; Agnar Helgason; Hakon Jonsson; Olafur T Magnusson; Pall Melsted; Gudmundur L Norddahl; Jona Saemundsdottir; Asgeir Sigurdsson; Patrick Sulem; Arna B Agustsdottir; Hannes Eggertsson; Berglind Eiriksdottir; Run Fridriksdottir; Elisabet E Gardarsdottir; Gudmundur Georgsson; Olafia S Gretarsdottir; Kjartan R Gudmundsson; Thora R Gunnarsdottir; Arnaldur Gylfason; Hilma Holm; Brynjar O Jensson; Aslaug Jonasdottir; Kamilla S Josefsdottir; Thordur Kristjansson; Droplaug N Magnusdottir; Solvi Rognvaldsson; Louise le Roux; Gudrun Sigmundsdottir; Gardar Sveinbjornsson; Kristin E Sveinsdottir; Maney Sveinsdottir; Emil A Thorarensen; Bjarni Thorbjornsson; Gisli Masson; Ingileif Jonsdottir; Alma Moller; Thorolfur Gudnason; Karl G Kristinsson; Unnur Thorsteinsdottir; Kari Stefansson |
| EPI_ISL_827436, EPI_ISL_827440, EPI_ISL_827447, EPI_ISL_827448, EPI_ISL_827450, EPI_ISL_827451, EPI_ISL_827452, EPI_ISL_827453, EPI_ISL_827454, EPI_ISL_827455, EPI_ISL_827456, EPI_ISL_827457, EPI_ISL_827466, EPI_ISL_827467, EPI_ISL_827469, EPI_ISL_827470, EPI_ISL_827471, EPI_ISL_827472, EPI_ISL_827478, EPI_ISL_827480 | | | |
| see above | deCODE genetics | deCODE genetics | Daniel F Gudbjartsson; Agnar Helgason; Hakon Jonsson; Olafur T Magnusson; Pall Melsted; Gudmundur L Norddahl; Jona Saemundsdottir; Asgeir Sigurdsson; Patrick Sulem; Arna B Agustsdottir; Hannes Eggertsson; Berglind Eiriksdottir; Run Fridriksdottir; Elisabet E Gardarsdottir; Gudmundur Georgsson; Olafia S Gretarsdottir; Kjartan R Gudmundsson; Thora R Gunnarsdottir; Arnaldur Gylfason; Hilma Holm; Brynjar O Jensson; Aslaug Jonasdottir; Kamilla S Josefsdottir; Thordur Kristjansson; Droplaug N Magnusdottir; Solvi Rognvaldsson; Louise le Roux; Gudrun Sigmundsdottir; Gardar Sveinbjornsson; Kristin E Sveinsdottir; Maney Sveinsdottir; Emil A Thorarensen; Bjarni Thorbjornsson; Gisli Masson; Ingileif Jonsdottir; Alma Moller; Thorolfur Gudnason; Karl G Kristinsson; Unnur Thorsteinsdottir; Kari Stefansson |
| EPI_ISL_827539, EPI_ISL_827540 | The National University Hospital of Iceland | deCODE genetics | Daniel F Gudbjartsson; Agnar Helgason; Hakon Jonsson; Olafur T Magnusson; Pall Melsted; Gudmundur L Norddahl; Jona Saemundsdottir; Asgeir Sigurdsson; Patrick Sulem; Arna B Agustsdottir; Hannes Eggertsson; Berglind Eiriksdottir; Run Fridriksdottir; Elisabet E Gardarsdottir; Gudmundur Georgsson; Olafia S Gretarsdottir; Kjartan R Gudmundsson; Thora R Gunnarsdottir; Arnaldur Gylfason; Hilma Holm; Brynjar O Jensson; Aslaug Jonasdottir; Kamilla S Josefsdottir; Thordur Kristjansson; Droplaug N Magnusdottir; Solvi Rognvaldsson; Louise le Roux; Gudrun Sigmundsdottir; Gardar Sveinbjornsson; Kristin E Sveinsdottir; Maney Sveinsdottir; Emil A Thorarensen; Bjarni Thorbjornsson; Gisli Masson; Ingileif Jonsdottir; Alma Moller; Thorolfur Gudnason; Karl G Kristinsson; Unnur Thorsteinsdottir; Kari Stefansson |
| EPI_ISL_827565, EPI_ISL_827566, EPI_ISL_827567, EPI_ISL_827568, EPI_ISL_827569, EPI_ISL_827570, EPI_ISL_827573, EPI_ISL_827574, EPI_ISL_827661, EPI_ISL_827768, EPI_ISL_827769, EPI_ISL_827770, EPI_ISL_827771, EPI_ISL_827772, EPI_ISL_827773, EPI_ISL_827774, EPI_ISL_827775, EPI_ISL_827776, EPI_ISL_827777, EPI_ISL_827788, EPI_ISL_827792, EPI_ISL_827793, EPI_ISL_827794, EPI_ISL_827795, EPI_ISL_827796, EPI_ISL_827797, EPI_ISL_827798, EPI_ISL_827799, EPI_ISL_827806, EPI_ISL_827807, EPI_ISL_827810, EPI_ISL_827811, EPI_ISL_827812, EPI_ISL_827813, EPI_ISL_827884, EPI_ISL_827885, EPI_ISL_827887, EPI_ISL_827888, EPI_ISL_827889, EPI_ISL_827890, EPI_ISL_827892, EPI_ISL_827893 | | | |
| see above | deCODE genetics | deCODE genetics | Daniel F Gudbjartsson; Agnar Helgason; Hakon Jonsson; Olafur T Magnusson; Pall Melsted; Gudmundur L Norddahl; Jona Saemundsdottir; Asgeir Sigurdsson; Patrick Sulem; Arna B Agustsdottir; Hannes Eggertsson; Berglind Eiriksdottir; Run Fridriksdottir; Elisabet E Gardarsdottir; Gudmundur Georgsson; Olafia S Gretarsdottir; Kjartan R Gudmundsson; Thora R Gunnarsdottir; Arnaldur Gylfason; Hilma Holm; Brynjar O Jensson; Aslaug Jonasdottir; Kamilla S Josefsdottir; Thordur Kristjansson; Droplaug N Magnusdottir; Solvi Rognvaldsson; Louise le Roux; Gudrun Sigmundsdottir; Gardar Sveinbjornsson; Kristin E Sveinsdottir; Maney Sveinsdottir; Emil A Thorarensen; Bjarni Thorbjornsson; Gisli Masson; Ingileif Jonsdottir; Alma Moller; Thorolfur Gudnason; Karl G Kristinsson; Unnur Thorsteinsdottir; Kari Stefansson |
| EPI_ISL_827913 | The National University Hospital of Iceland | deCODE genetics | Daniel F Gudbjartsson; Agnar Helgason; Hakon Jonsson; Olafur T Magnusson; Pall Melsted; Gudmundur L Norddahl; Jona Saemundsdottir; Asgeir Sigurdsson; Patrick Sulem; Arna B Agustsdottir; Hannes Eggertsson; Berglind Eiriksdottir; Run Fridriksdottir; Elisabet E Gardarsdottir; Gudmundur Georgsson; Olafia S Gretarsdottir; Kjartan R Gudmundsson; Thora R Gunnarsdottir; Arnaldur Gylfason; Hilma Holm; Brynjar O Jensson; Aslaug Jonasdottir; Kamilla S Josefsdottir; Thordur Kristjansson; Droplaug N Magnusdottir; Solvi Rognvaldsson; Louise le Roux; Gudrun Sigmundsdottir; Gardar Sveinbjornsson; Kristin E Sveinsdottir; Maney Sveinsdottir; Emil A Thorarensen; Bjarni Thorbjornsson; Gisli Masson; Ingileif Jonsdottir; Alma Moller; Thorolfur Gudnason; Karl G Kristinsson; Unnur Thorsteinsdottir; Kari Stefansson |
| EPI_ISL_827974, EPI_ISL_827991, EPI_ISL_828063, EPI_ISL_828074, EPI_ISL_828075, EPI_ISL_828076, EPI_ISL_828077, EPI_ISL_828078, EPI_ISL_828079, EPI_ISL_828080, EPI_ISL_828081, EPI_ISL_828082, EPI_ISL_828083, EPI_ISL_828112, EPI_ISL_828134, EPI_ISL_828149, EPI_ISL_828172, EPI_ISL_828194, EPI_ISL_828195, EPI_ISL_828196, EPI_ISL_828197, EPI_ISL_828198, EPI_ISL_828199, EPI_ISL_828201, EPI_ISL_828202, EPI_ISL_828203, EPI_ISL_828290, EPI_ISL_828309, EPI_ISL_828337, EPI_ISL_828338, EPI_ISL_828339, EPI_ISL_828340, EPI_ISL_828341, EPI_ISL_828342, EPI_ISL_828343, EPI_ISL_828344, EPI_ISL_828374, EPI_ISL_828375, EPI_ISL_828376, EPI_ISL_828377, EPI_ISL_828378, EPI_ISL_828379, EPI_ISL_828384, EPI_ISL_828391 | | | |
| see above | deCODE genetics | deCODE genetics | Daniel F Gudbjartsson; Agnar Helgason; Hakon Jonsson; Olafur T Magnusson; Pall Melsted; Gudmundur L Norddahl; Jona Saemundsdottir; Asgeir Sigurdsson; Patrick Sulem; Arna B Agustsdottir; Hannes Eggertsson; Berglind Eiriksdottir; Run Fridriksdottir; Elisabet E Gardarsdottir; Gudmundur Georgsson; Olafia S Gretarsdottir; Kjartan R Gudmundsson; Thora R Gunnarsdottir; Arnaldur Gylfason; Hilma Holm; Brynjar O Jensson; Aslaug Jonasdottir; |

| | | | |
|---|---|---|---|
| | | | Kamilla S Josefsdottir; Thordur Kristjansson; Droplaug N Magnusdottir; Solvi Rognvaldsson; Louise le Roux; Gudrun Sigmundsdottir; Gardar Sveinbjornsson; Kristin E Sveinsdottir; Maney Sveinsdottir; Emil A Thorarensen; Bjarni Thorbjornsson; Gisli Masson; Ingileif Jonsdottir; Alma Moller; Thorolfur Gudnason; Karl G Kristinsson; Unnur Thorsteinsdottir; Kari Stefansson |
| EPI_ISL_828397, EPI_ISL_828398 | The National University Hospital of Iceland | deCODE genetics | Daniel F Gudbjartsson; Agnar Helgason; Hakon Jonsson; Olafur T Magnusson; Pall Melsted; Gudmundur L Norddahl; Jona Saemundsdottir; Asgeir Sigurdsson; Patrick Sulem; Arna B Agustsdottir; Hannes Eggertsson; Berglind Eiriksdottir; Run Fridriksdottir; Elisabet E Gardarsdottir; Gudmundur Georgsson; Olafia S Gretarsdottir; Kjartan R Gudmundsson; Thora R Gunnarsdottir; Arnaldur Gylfason; Hilma Holm; Brynjar O Jensson; Aslaug Jonasdottir; Kamilla S Josefsdottir; Thordur Kristjansson; Droplaug N Magnusdottir; Solvi Rognvaldsson; Louise le Roux; Gudrun Sigmundsdottir; Gardar Sveinbjornsson; Kristin E Sveinsdottir; Maney Sveinsdottir; Emil A Thorarensen; Bjarni Thorbjornsson; Gisli Masson; Ingileif Jonsdottir; Alma Moller; Thorolfur Gudnason; Karl G Kristinsson; Unnur Thorsteinsdottir; Kari Stefansson |
| EPI_ISL_828407 | deCODE genetics | deCODE genetics | Daniel F Gudbjartsson; Agnar Helgason; Hakon Jonsson; Olafur T Magnusson; Pall Melsted; Gudmundur L Norddahl; Jona Saemundsdottir; Asgeir Sigurdsson; Patrick Sulem; Arna B Agustsdottir; Hannes Eggertsson; Berglind Eiriksdottir; Run Fridriksdottir; Elisabet E Gardarsdottir; Gudmundur Georgsson; Olafia S Gretarsdottir; Kjartan R Gudmundsson; Thora R Gunnarsdottir; Arnaldur Gylfason; Hilma Holm; Brynjar O Jensson; Aslaug Jonasdottir; Kamilla S Josefsdottir; Thordur Kristjansson; Droplaug N Magnusdottir; Solvi Rognvaldsson; Louise le Roux; Gudrun Sigmundsdottir; Gardar Sveinbjornsson; Kristin E Sveinsdottir; Maney Sveinsdottir; Emil A Thorarensen; Bjarni Thorbjornsson; Gisli Masson; Ingileif Jonsdottir; Alma Moller; Thorolfur Gudnason; Karl G Kristinsson; Unnur Thorsteinsdottir; Kari Stefansson |
| EPI_ISL_828413 | The National University Hospital of Iceland | deCODE genetics | Daniel F Gudbjartsson; Agnar Helgason; Hakon Jonsson; Olafur T Magnusson; Pall Melsted; Gudmundur L Norddahl; Jona Saemundsdottir; Asgeir Sigurdsson; Patrick Sulem; Arna B Agustsdottir; Hannes Eggertsson; Berglind Eiriksdottir; Run Fridriksdottir; Elisabet E Gardarsdottir; Gudmundur Georgsson; Olafia S Gretarsdottir; Kjartan R Gudmundsson; Thora R Gunnarsdottir; Arnaldur Gylfason; Hilma Holm; Brynjar O Jensson; Aslaug Jonasdottir; Kamilla S Josefsdottir; Thordur Kristjansson; Droplaug N Magnusdottir; Solvi Rognvaldsson; Louise le Roux; Gudrun Sigmundsdottir; Gardar Sveinbjornsson; Kristin E Sveinsdottir; Maney Sveinsdottir; Emil A Thorarensen; Bjarni Thorbjornsson; Gisli Masson; Ingileif Jonsdottir; Alma Moller; Thorolfur Gudnason; Karl G Kristinsson; Unnur Thorsteinsdottir; Kari Stefansson |
| EPI_ISL_828471, EPI_ISL_828472, EPI_ISL_828473, EPI_ISL_828474, EPI_ISL_828475, EPI_ISL_828476, EPI_ISL_828478, EPI_ISL_828479, EPI_ISL_828480 | deCODE genetics | deCODE genetics | Daniel F Gudbjartsson; Agnar Helgason; Hakon Jonsson; Olafur T Magnusson; Pall Melsted; Gudmundur L Norddahl; Jona Saemundsdottir; Asgeir Sigurdsson; Patrick Sulem; Arna B Agustsdottir; Hannes Eggertsson; Berglind Eiriksdottir; Run Fridriksdottir; Elisabet E Gardarsdottir; Gudmundur Georgsson; Olafia S Gretarsdottir; Kjartan R Gudmundsson; Thora R Gunnarsdottir; Arnaldur Gylfason; Hilma Holm; Brynjar O Jensson; Aslaug Jonasdottir; Kamilla S Josefsdottir; Thordur Kristjansson; Droplaug N Magnusdottir; Solvi Rognvaldsson; Louise le Roux; Gudrun Sigmundsdottir; Gardar Sveinbjornsson; Kristin E Sveinsdottir; Maney Sveinsdottir; Emil A Thorarensen; Bjarni Thorbjornsson; Gisli Masson; Ingileif Jonsdottir; Alma Moller; Thorolfur Gudnason; Karl G Kristinsson; Unnur Thorsteinsdottir; Kari Stefansson |
| EPI_ISL_828484, EPI_ISL_828485, EPI_ISL_828486, EPI_ISL_828487 | The National University Hospital of Iceland | deCODE genetics | Daniel F Gudbjartsson; Agnar Helgason; Hakon Jonsson; Olafur T Magnusson; Pall Melsted; Gudmundur L Norddahl; Jona Saemundsdottir; Asgeir Sigurdsson; Patrick Sulem; Arna B Agustsdottir; Hannes Eggertsson; Berglind Eiriksdottir; Run Fridriksdottir; Elisabet E Gardarsdottir; Gudmundur Georgsson; Olafia S Gretarsdottir; Kjartan R Gudmundsson; Thora R Gunnarsdottir; Arnaldur Gylfason; Hilma Holm; Brynjar O Jensson; Aslaug Jonasdottir; Kamilla S Josefsdottir; Thordur Kristjansson; Droplaug N Magnusdottir; Solvi Rognvaldsson; Louise le Roux; Gudrun Sigmundsdottir; Gardar Sveinbjornsson; Kristin E Sveinsdottir; Maney Sveinsdottir; Emil A Thorarensen; Bjarni Thorbjornsson; Gisli Masson; Ingileif Jonsdottir; Alma Moller; Thorolfur Gudnason; Karl G Kristinsson; Unnur Thorsteinsdottir; Kari Stefansson |
| EPI_ISL_828488, EPI_ISL_828489, EPI_ISL_828490 | deCODE genetics | deCODE genetics | Daniel F Gudbjartsson; Agnar Helgason; Hakon Jonsson; Olafur T Magnusson; Pall Melsted; Gudmundur L Norddahl; Jona Saemundsdottir; Asgeir Sigurdsson; Patrick Sulem; Arna B Agustsdottir; Hannes Eggertsson; Berglind Eiriksdottir; Run Fridriksdottir; Elisabet E Gardarsdottir; Gudmundur Georgsson; Olafia S Gretarsdottir; Kjartan R Gudmundsson; Thora R Gunnarsdottir; Arnaldur Gylfason; Hilma Holm; Brynjar O Jensson; Aslaug Jonasdottir; Kamilla S Josefsdottir; Thordur Kristjansson; Droplaug N Magnusdottir; Solvi Rognvaldsson; Louise le Roux; Gudrun Sigmundsdottir; Gardar Sveinbjornsson; Kristin E Sveinsdottir; Maney Sveinsdottir; Emil A Thorarensen; Bjarni Thorbjornsson; Gisli Masson; Ingileif Jonsdottir; Alma Moller; Thorolfur Gudnason; Karl G Kristinsson; Unnur Thorsteinsdottir; Kari Stefansson |
| EPI_ISL_828494 | The National University Hospital of Iceland | deCODE genetics | Daniel F Gudbjartsson; Agnar Helgason; Hakon Jonsson; Olafur T Magnusson; Pall Melsted; Gudmundur L Norddahl; Jona Saemundsdottir; Asgeir Sigurdsson; Patrick Sulem; Arna B Agustsdottir; Hannes Eggertsson; Berglind Eiriksdottir; Run Fridriksdottir; Elisabet E Gardarsdottir; Gudmundur Georgsson; Olafia S Gretarsdottir; Kjartan R Gudmundsson; Thora R Gunnarsdottir; Arnaldur Gylfason; Hilma Holm; Brynjar O Jensson; Aslaug Jonasdottir; Kamilla S Josefsdottir; Thordur Kristjansson; Droplaug N Magnusdottir; Solvi Rognvaldsson; Louise le Roux; Gudrun Sigmundsdottir; Gardar Sveinbjornsson; Kristin E Sveinsdottir; Maney Sveinsdottir; Emil A Thorarensen; Bjarni Thorbjornsson; Gisli Masson; Ingileif Jonsdottir; Alma Moller; Thorolfur Gudnason; Karl G Kristinsson; Unnur Thorsteinsdottir; Kari Stefansson |
| EPI_ISL_828499, EPI_ISL_828526, EPI_ISL_828577, EPI_ISL_828578, EPI_ISL_828603, EPI_ISL_828615, EPI_ISL_828616, EPI_ISL_828617, EPI_ISL_828639 | deCODE genetics | deCODE genetics | Daniel F Gudbjartsson; Agnar Helgason; Hakon Jonsson; Olafur T Magnusson; Pall Melsted; Gudmundur L Norddahl; Jona Saemundsdottir; Asgeir Sigurdsson; Patrick Sulem; Arna B Agustsdottir; Hannes Eggertsson; Berglind Eiriksdottir; Run Fridriksdottir; Elisabet E Gardarsdottir; Gudmundur Georgsson; Olafia S Gretarsdottir; Kjartan R Gudmundsson; Thora R Gunnarsdottir; Arnaldur Gylfason; Hilma Holm; Brynjar O Jensson; Aslaug Jonasdottir; Kamilla S Josefsdottir; Thordur Kristjansson; Droplaug N Magnusdottir; Solvi Rognvaldsson; Louise le Roux; Gudrun Sigmundsdottir; Gardar Sveinbjornsson; Kristin E Sveinsdottir; Maney Sveinsdottir; Emil A Thorarensen; Bjarni Thorbjornsson; Gisli Masson; Ingileif Jonsdottir; Alma Moller; Thorolfur Gudnason; Karl G Kristinsson; Unnur Thorsteinsdottir; Kari Stefansson |
| EPI_ISL_828643, EPI_ISL_828644, EPI_ISL_828646 | The National University Hospital of Iceland | deCODE genetics | Daniel F Gudbjartsson; Agnar Helgason; Hakon Jonsson; Olafur T Magnusson; Pall Melsted; Gudmundur L Norddahl; Jona Saemundsdottir; Asgeir Sigurdsson; Patrick Sulem; Arna B Agustsdottir; Hannes Eggertsson; Berglind Eiriksdottir; Run Fridriksdottir; Elisabet E Gardarsdottir; Gudmundur Georgsson; Olafia S Gretarsdottir; Kjartan R Gudmundsson; Thora R Gunnarsdottir; Arnaldur Gylfason; Hilma Holm; Brynjar O Jensson; Aslaug Jonasdottir; Kamilla S Josefsdottir; Thordur Kristjansson; Droplaug N Magnusdottir; Solvi Rognvaldsson; Louise le Roux; Gudrun Sigmundsdottir; Gardar Sveinbjornsson; Kristin E Sveinsdottir; Maney Sveinsdottir; Emil A Thorarensen; Bjarni Thorbjornsson; Gisli Masson; Ingileif Jonsdottir; Alma Moller; Thorolfur Gudnason; Karl G Kristinsson; Unnur Thorsteinsdottir; Kari Stefansson |
| EPI_ISL_828681, EPI_ISL_828682, EPI_ISL_828683, EPI_ISL_828684, EPI_ISL_828685, EPI_ISL_828686, EPI_ISL_828687, EPI_ISL_828688, EPI_ISL_828689 | deCODE genetics | deCODE genetics | Daniel F Gudbjartsson; Agnar Helgason; Hakon Jonsson; Olafur T Magnusson; Pall Melsted; Gudmundur L Norddahl; Jona Saemundsdottir; Asgeir Sigurdsson; Patrick Sulem; Arna B Agustsdottir; Hannes Eggertsson; Berglind Eiriksdottir; Run Fridriksdottir; Elisabet E Gardarsdottir; Gudmundur Georgsson; Olafia S Gretarsdottir; Kjartan R Gudmundsson; Thora R Gunnarsdottir; Arnaldur Gylfason; Hilma Holm; Brynjar O Jensson; Aslaug Jonasdottir; Kamilla S Josefsdottir; Thordur Kristjansson; Droplaug N Magnusdottir; Solvi Rognvaldsson; Louise le Roux; Gudrun Sigmundsdottir; Gardar Sveinbjornsson; Kristin E Sveinsdottir; Maney Sveinsdottir; Emil A Thorarensen; Bjarni Thorbjornsson; Gisli Masson; Ingileif Jonsdottir; Alma Moller; Thorolfur Gudnason; Karl G Kristinsson; Unnur Thorsteinsdottir; Kari Stefansson |
| EPI_ISL_828709 | The National University Hospital of Iceland | deCODE genetics | Daniel F Gudbjartsson; Agnar Helgason; Hakon Jonsson; Olafur T Magnusson; Pall Melsted; Gudmundur L Norddahl; Jona Saemundsdottir; Asgeir Sigurdsson; Patrick Sulem; Arna B Agustsdottir; Hannes Eggertsson; Berglind Eiriksdottir; Run Fridriksdottir; Elisabet E Gardarsdottir; Gudmundur Georgsson; Olafia S Gretarsdottir; Kjartan R Gudmundsson; Thora R Gunnarsdottir; Arnaldur Gylfason; Hilma Holm; Brynjar O Jensson; Aslaug Jonasdottir; Kamilla S Josefsdottir; Thordur Kristjansson; Droplaug N Magnusdottir; Solvi Rognvaldsson; Louise le Roux; Gudrun Sigmundsdottir; Gardar Sveinbjornsson; Kristin E Sveinsdottir; Maney Sveinsdottir; Emil A Thorarensen; Bjarni Thorbjornsson; Gisli Masson; Ingileif Jonsdottir; Alma Moller; Thorolfur Gudnason; Karl G Kristinsson; Unnur Thorsteinsdottir; Kari Stefansson |
| EPI_ISL_828861, EPI_ISL_828871, EPI_ISL_828872, EPI_ISL_828879, EPI_ISL_828882, EPI_ISL_828884, EPI_ISL_828885, EPI_ISL_828914, EPI_ISL_828918 | deCODE genetics | deCODE genetics | Daniel F Gudbjartsson; Agnar Helgason; Hakon Jonsson; Olafur T Magnusson; Pall Melsted; Gudmundur L Norddahl; Jona Saemundsdottir; Asgeir Sigurdsson; Patrick Sulem; Arna B Agustsdottir; Hannes Eggertsson; Berglind Eiriksdottir; Run Fridriksdottir; Elisabet E Gardarsdottir; Gudmundur Georgsson; Olafia S Gretarsdottir; Kjartan R Gudmundsson; Thora R Gunnarsdottir; Arnaldur Gylfason; Hilma Holm; Brynjar O Jensson; Aslaug Jonasdottir; Kamilla S Josefsdottir; Thordur Kristjansson; Droplaug N Magnusdottir; Solvi Rognvaldsson; Louise le Roux; Gudrun Sigmundsdottir; Gardar Sveinbjornsson; Kristin E Sveinsdottir; Maney Sveinsdottir; Emil A Thorarensen; Bjarni Thorbjornsson; Gisli Masson; Ingileif Jonsdottir; Alma Moller; Thorolfur Gudnason; Karl G Kristinsson; Unnur Thorsteinsdottir; Kari Stefansson |
| EPI_ISL_828931, EPI_ISL_828932, EPI_ISL_828934 | The National University Hospital of Iceland | deCODE genetics | Daniel F Gudbjartsson; Agnar Helgason; Hakon Jonsson; Olafur T Magnusson; Pall Melsted; Gudmundur L Norddahl; Jona Saemundsdottir; Asgeir Sigurdsson; Patrick Sulem; Arna B Agustsdottir; Hannes Eggertsson; Berglind Eiriksdottir; Run Fridriksdottir; Elisabet E Gardarsdottir; Gudmundur Georgsson; Olafia S Gretarsdottir; Kjartan R Gudmundsson; Thora R Gunnarsdottir; Arnaldur Gylfason; Hilma Holm; Brynjar O Jensson; Aslaug Jonasdottir; Kamilla S Josefsdottir; Thordur Kristjansson; Droplaug N Magnusdottir; Solvi Rognvaldsson; Louise le Roux; Gudrun Sigmundsdottir; Gardar Sveinbjornsson; Kristin E Sveinsdottir; Maney Sveinsdottir; Emil A Thorarensen; Bjarni Thorbjornsson; Gisli Masson; Ingileif Jonsdottir; Alma Moller; Thorolfur Gudnason; Karl G Kristinsson; Unnur Thorsteinsdottir; Kari Stefansson |

| | | | |
|---|---|---|---|
| EPI_ISL_829037, EPI_ISL_829047, EPI_ISL_829048, EPI_ISL_829072, EPI_ISL_829074, EPI_ISL_829075, EPI_ISL_829106, EPI_ISL_829108, EPI_ISL_829109, EPI_ISL_829110, EPI_ISL_829111, EPI_ISL_829112, EPI_ISL_829113, EPI_ISL_829114, EPI_ISL_829115, EPI_ISL_829118, EPI_ISL_829119, EPI_ISL_829122, EPI_ISL_829123, EPI_ISL_829124 | | | |
| see above | deCODE genetics | deCODE genetics | Daniel F Gudbjartsson; Agnar Helgason; Hakon Jonsson; Olafur T Magnusson; Pall Melsted; Gudmundur L Norddahl; Jona Saemundsdottir; Asgeir Sigurdsson; Patrick Sulem; Arna B Agustsdottir; Hannes Eggertsson; Berglind Eiriksdottir; Run Fridriksdottir; Elisabet E Gardarsdottir; Gudmundur Georgsson; Olafia S Gretarsdottir; Kjartan R Gudmundsson; Thora R Gunnarsdottir; Arnaldur Gylfason; Hilma Holm; Brynjar O Jensson; Aslaug Jonasdottir; Kamilla S Josefsdottir; Thordur Kristjansson; Droplaug N Magnusdottir; Solvi Rognvaldsson; Louise le Roux; Gudrun Sigmundsdottir; Gardar Sveinbjornsson; Kristin E Sveinsdottir; Maney Sveinsdottir; Emil A Thorarensen; Bjarni Thorbjornsson; Gisli Masson; Ingileif Jonsdottir; Alma Moller; Thorolfur Gudnason; Karl G Kristinsson; Unnur Thorsteinsdottir; Kari Stefansson |
| EPI_ISL_829133 | The National University Hospital of Iceland | deCODE genetics | Daniel F Gudbjartsson; Agnar Helgason; Hakon Jonsson; Olafur T Magnusson; Pall Melsted; Gudmundur L Norddahl; Jona Saemundsdottir; Asgeir Sigurdsson; Patrick Sulem; Arna B Agustsdottir; Hannes Eggertsson; Berglind Eiriksdottir; Run Fridriksdottir; Elisabet E Gardarsdottir; Gudmundur Georgsson; Olafia S Gretarsdottir; Kjartan R Gudmundsson; Thora R Gunnarsdottir; Arnaldur Gylfason; Hilma Holm; Brynjar O Jensson; Aslaug Jonasdottir; Kamilla S Josefsdottir; Thordur Kristjansson; Droplaug N Magnusdottir; Solvi Rognvaldsson; Louise le Roux; Gudrun Sigmundsdottir; Gardar Sveinbjornsson; Kristin E Sveinsdottir; Maney Sveinsdottir; Emil A Thorarensen; Bjarni Thorbjornsson; Gisli Masson; Ingileif Jonsdottir; Alma Moller; Thorolfur Gudnason; Karl G Kristinsson; Unnur Thorsteinsdottir; Kari Stefansson |
| EPI_ISL_829181 | deCODE genetics | deCODE genetics | Daniel F Gudbjartsson; Agnar Helgason; Hakon Jonsson; Olafur T Magnusson; Pall Melsted; Gudmundur L Norddahl; Jona Saemundsdottir; Asgeir Sigurdsson; Patrick Sulem; Arna B Agustsdottir; Hannes Eggertsson; Berglind Eiriksdottir; Run Fridriksdottir; Elisabet E Gardarsdottir; Gudmundur Georgsson; Olafia S Gretarsdottir; Kjartan R Gudmundsson; Thora R Gunnarsdottir; Arnaldur Gylfason; Hilma Holm; Brynjar O Jensson; Aslaug Jonasdottir; Kamilla S Josefsdottir; Thordur Kristjansson; Droplaug N Magnusdottir; Solvi Rognvaldsson; Louise le Roux; Gudrun Sigmundsdottir; Gardar Sveinbjornsson; Kristin E Sveinsdottir; Maney Sveinsdottir; Emil A Thorarensen; Bjarni Thorbjornsson; Gisli Masson; Ingileif Jonsdottir; Alma Moller; Thorolfur Gudnason; Karl G Kristinsson; Unnur Thorsteinsdottir; Kari Stefansson |
| EPI_ISL_829188, EPI_ISL_829189, EPI_ISL_829190 | The National University Hospital of Iceland | deCODE genetics | Daniel F Gudbjartsson; Agnar Helgason; Hakon Jonsson; Olafur T Magnusson; Pall Melsted; Gudmundur L Norddahl; Jona Saemundsdottir; Asgeir Sigurdsson; Patrick Sulem; Arna B Agustsdottir; Hannes Eggertsson; Berglind Eiriksdottir; Run Fridriksdottir; Elisabet E Gardarsdottir; Gudmundur Georgsson; Olafia S Gretarsdottir; Kjartan R Gudmundsson; Thora R Gunnarsdottir; Arnaldur Gylfason; Hilma Holm; Brynjar O Jensson; Aslaug Jonasdottir; Kamilla S Josefsdottir; Thordur Kristjansson; Droplaug N Magnusdottir; Solvi Rognvaldsson; Louise le Roux; Gudrun Sigmundsdottir; Gardar Sveinbjornsson; Kristin E Sveinsdottir; Maney Sveinsdottir; Emil A Thorarensen; Bjarni Thorbjornsson; Gisli Masson; Ingileif Jonsdottir; Alma Moller; Thorolfur Gudnason; Karl G Kristinsson; Unnur Thorsteinsdottir; Kari Stefansson |
| EPI_ISL_829191 | deCODE genetics | deCODE genetics | Daniel F Gudbjartsson; Agnar Helgason; Hakon Jonsson; Olafur T Magnusson; Pall Melsted; Gudmundur L Norddahl; Jona Saemundsdottir; Asgeir Sigurdsson; Patrick Sulem; Arna B Agustsdottir; Hannes Eggertsson; Berglind Eiriksdottir; Run Fridriksdottir; Elisabet E Gardarsdottir; Gudmundur Georgsson; Olafia S Gretarsdottir; Kjartan R Gudmundsson; Thora R Gunnarsdottir; Arnaldur Gylfason; Hilma Holm; Brynjar O Jensson; Aslaug Jonasdottir; Kamilla S Josefsdottir; Thordur Kristjansson; Droplaug N Magnusdottir; Solvi Rognvaldsson; Louise le Roux; Gudrun Sigmundsdottir; Gardar Sveinbjornsson; Kristin E Sveinsdottir; Maney Sveinsdottir; Emil A Thorarensen; Bjarni Thorbjornsson; Gisli Masson; Ingileif Jonsdottir; Alma Moller; Thorolfur Gudnason; Karl G Kristinsson; Unnur Thorsteinsdottir; Kari Stefansson |
| EPI_ISL_829192, EPI_ISL_829193, EPI_ISL_829194, EPI_ISL_829195, EPI_ISL_829196, EPI_ISL_829197, EPI_ISL_829224, EPI_ISL_829361, EPI_ISL_829366, EPI_ISL_829377, EPI_ISL_829381 | | | |
| see above | The National University Hospital of Iceland | deCODE genetics | Daniel F Gudbjartsson; Agnar Helgason; Hakon Jonsson; Olafur T Magnusson; Pall Melsted; Gudmundur L Norddahl; Jona Saemundsdottir; Asgeir Sigurdsson; Patrick Sulem; Arna B Agustsdottir; Hannes Eggertsson; Berglind Eiriksdottir; Run Fridriksdottir; Elisabet E Gardarsdottir; Gudmundur Georgsson; Olafia S Gretarsdottir; Kjartan R Gudmundsson; Thora R Gunnarsdottir; Arnaldur Gylfason; Hilma Holm; Brynjar O Jensson; Aslaug Jonasdottir; Kamilla S Josefsdottir; Thordur Kristjansson; Droplaug N Magnusdottir; Solvi Rognvaldsson; Louise le Roux; Gudrun Sigmundsdottir; Gardar Sveinbjornsson; Kristin E Sveinsdottir; Maney Sveinsdottir; Emil A Thorarensen; Bjarni Thorbjornsson; Gisli Masson; Ingileif Jonsdottir; Alma Moller; Thorolfur Gudnason; Karl G Kristinsson; Unnur Thorsteinsdottir; Kari Stefansson |
| EPI_ISL_829394, EPI_ISL_829395, EPI_ISL_829396, EPI_ISL_829397, EPI_ISL_829398, EPI_ISL_829399, EPI_ISL_829400, EPI_ISL_829401, EPI_ISL_829402, EPI_ISL_829403 | deCODE genetics | deCODE genetics | Daniel F Gudbjartsson; Agnar Helgason; Hakon Jonsson; Olafur T Magnusson; Pall Melsted; Gudmundur L Norddahl; Jona Saemundsdottir; Asgeir Sigurdsson; Patrick Sulem; Arna B Agustsdottir; Hannes Eggertsson; Berglind Eiriksdottir; Run Fridriksdottir; Elisabet E Gardarsdottir; Gudmundur Georgsson; Olafia S Gretarsdottir; Kjartan R Gudmundsson; Thora R Gunnarsdottir; Arnaldur Gylfason; Hilma Holm; Brynjar O Jensson; Aslaug Jonasdottir; Kamilla S Josefsdottir; Thordur Kristjansson; Droplaug N Magnusdottir; Solvi Rognvaldsson; Louise le Roux; Gudrun Sigmundsdottir; Gardar Sveinbjornsson; Kristin E Sveinsdottir; Maney Sveinsdottir; Emil A Thorarensen; Bjarni Thorbjornsson; Gisli Masson; Ingileif Jonsdottir; Alma Moller; Thorolfur Gudnason; Karl G Kristinsson; Unnur Thorsteinsdottir; Kari Stefansson |
| EPI_ISL_829414, EPI_ISL_829426, EPI_ISL_829449 | The National University Hospital of Iceland | deCODE genetics | Daniel F Gudbjartsson; Agnar Helgason; Hakon Jonsson; Olafur T Magnusson; Pall Melsted; Gudmundur L Norddahl; Jona Saemundsdottir; Asgeir Sigurdsson; Patrick Sulem; Arna B Agustsdottir; Hannes Eggertsson; Berglind Eiriksdottir; Run Fridriksdottir; Elisabet E Gardarsdottir; Gudmundur Georgsson; Olafia S Gretarsdottir; Kjartan R Gudmundsson; Thora R Gunnarsdottir; Arnaldur Gylfason; Hilma Holm; Brynjar O Jensson; Aslaug Jonasdottir; Kamilla S Josefsdottir; Thordur Kristjansson; Droplaug N Magnusdottir; Solvi Rognvaldsson; Louise le Roux; Gudrun Sigmundsdottir; Gardar Sveinbjornsson; Kristin E Sveinsdottir; Maney Sveinsdottir; Emil A Thorarensen; Bjarni Thorbjornsson; Gisli Masson; Ingileif Jonsdottir; Alma Moller; Thorolfur Gudnason; Karl G Kristinsson; Unnur Thorsteinsdottir; Kari Stefansson |
| EPI_ISL_829457, EPI_ISL_829461, EPI_ISL_829479, EPI_ISL_829531, EPI_ISL_829532, EPI_ISL_829542, EPI_ISL_829548, EPI_ISL_829552, EPI_ISL_829555, EPI_ISL_829640, EPI_ISL_829641, EPI_ISL_829643, EPI_ISL_829644, EPI_ISL_829645, EPI_ISL_829646, EPI_ISL_829647, EPI_ISL_829648, EPI_ISL_829649, EPI_ISL_829651, EPI_ISL_829652, EPI_ISL_829653, EPI_ISL_829655, EPI_ISL_829657, EPI_ISL_829658, EPI_ISL_829659, EPI_ISL_829660 | | | |
| see above | deCODE genetics | deCODE genetics | Daniel F Gudbjartsson; Agnar Helgason; Hakon Jonsson; Olafur T Magnusson; Pall Melsted; Gudmundur L Norddahl; Jona Saemundsdottir; Asgeir Sigurdsson; Patrick Sulem; Arna B Agustsdottir; Hannes Eggertsson; Berglind Eiriksdottir; Run Fridriksdottir; Elisabet E Gardarsdottir; Gudmundur Georgsson; Olafia S Gretarsdottir; Kjartan R Gudmundsson; Thora R Gunnarsdottir; Arnaldur Gylfason; Hilma Holm; Brynjar O Jensson; Aslaug Jonasdottir; Kamilla S Josefsdottir; Thordur Kristjansson; Droplaug N Magnusdottir; Solvi Rognvaldsson; Louise le Roux; Gudrun Sigmundsdottir; Gardar Sveinbjornsson; Kristin E Sveinsdottir; Maney Sveinsdottir; Emil A Thorarensen; Bjarni Thorbjornsson; Gisli Masson; Ingileif Jonsdottir; Alma Moller; Thorolfur Gudnason; Karl G Kristinsson; Unnur Thorsteinsdottir; Kari Stefansson |
| EPI_ISL_829734 | The National University Hospital of Iceland | deCODE genetics | Daniel F Gudbjartsson; Agnar Helgason; Hakon Jonsson; Olafur T Magnusson; Pall Melsted; Gudmundur L Norddahl; Jona Saemundsdottir; Asgeir Sigurdsson; Patrick Sulem; Arna B Agustsdottir; Hannes Eggertsson; Berglind Eiriksdottir; Run Fridriksdottir; Elisabet E Gardarsdottir; Gudmundur Georgsson; Olafia S Gretarsdottir; Kjartan R Gudmundsson; Thora R Gunnarsdottir; Arnaldur Gylfason; Hilma Holm; Brynjar O Jensson; Aslaug Jonasdottir; Kamilla S Josefsdottir; Thordur Kristjansson; Droplaug N Magnusdottir; Solvi Rognvaldsson; Louise le Roux; Gudrun Sigmundsdottir; Gardar Sveinbjornsson; Kristin E Sveinsdottir; Maney Sveinsdottir; Emil A Thorarensen; Bjarni Thorbjornsson; Gisli Masson; Ingileif Jonsdottir; Alma Moller; Thorolfur Gudnason; Karl G Kristinsson; Unnur Thorsteinsdottir; Kari Stefansson |
| EPI_ISL_829876, EPI_ISL_829877, EPI_ISL_829878, EPI_ISL_829879, EPI_ISL_829880, EPI_ISL_829883, EPI_ISL_829893, EPI_ISL_829894, EPI_ISL_829895, EPI_ISL_829896, EPI_ISL_829897, EPI_ISL_829898, EPI_ISL_829899, EPI_ISL_829900, EPI_ISL_829901 | | | |
| see above | deCODE genetics | deCODE genetics | Daniel F Gudbjartsson; Agnar Helgason; Hakon Jonsson; Olafur T Magnusson; Pall Melsted; Gudmundur L Norddahl; Jona Saemundsdottir; Asgeir Sigurdsson; Patrick Sulem; Arna B Agustsdottir; Hannes Eggertsson; Berglind Eiriksdottir; Run Fridriksdottir; Elisabet E Gardarsdottir; Gudmundur Georgsson; Olafia S Gretarsdottir; Kjartan R Gudmundsson; Thora R Gunnarsdottir; Arnaldur Gylfason; Hilma Holm; Brynjar O Jensson; Aslaug Jonasdottir; Kamilla S Josefsdottir; Thordur Kristjansson; Droplaug N Magnusdottir; Solvi Rognvaldsson; Louise le Roux; Gudrun Sigmundsdottir; Gardar Sveinbjornsson; Kristin E Sveinsdottir; Maney Sveinsdottir; Emil A Thorarensen; Bjarni Thorbjornsson; Gisli Masson; Ingileif Jonsdottir; Alma Moller; Thorolfur Gudnason; Karl G Kristinsson; Unnur Thorsteinsdottir; Kari Stefansson |
| EPI_ISL_829930, EPI_ISL_829934 | The National University Hospital of Iceland | deCODE genetics | Daniel F Gudbjartsson; Agnar Helgason; Hakon Jonsson; Olafur T Magnusson; Pall Melsted; Gudmundur L Norddahl; Jona Saemundsdottir; Asgeir Sigurdsson; Patrick Sulem; Arna B Agustsdottir; Hannes Eggertsson; Berglind Eiriksdottir; Run Fridriksdottir; Elisabet E Gardarsdottir; Gudmundur Georgsson; Olafia S Gretarsdottir; Kjartan R Gudmundsson; Thora R Gunnarsdottir; Arnaldur Gylfason; Hilma Holm; Brynjar O Jensson; Aslaug Jonasdottir; Kamilla S Josefsdottir; Thordur Kristjansson; Droplaug N Magnusdottir; Solvi Rognvaldsson; Louise le Roux; Gudrun Sigmundsdottir; Gardar Sveinbjornsson; Kristin E Sveinsdottir; Maney Sveinsdottir; Emil A Thorarensen; Bjarni Thorbjornsson; Gisli Masson; Ingileif Jonsdottir; Alma Moller; Thorolfur Gudnason; Karl G Kristinsson; Unnur Thorsteinsdottir; Kari Stefansson |
| EPI_ISL_829935, EPI_ISL_829936 | deCODE genetics | deCODE genetics | Daniel F Gudbjartsson; Agnar Helgason; Hakon Jonsson; Olafur T Magnusson; Pall Melsted; Gudmundur L Norddahl; Jona Saemundsdottir; Asgeir |

| | | | |
|---|---|---|---|
| | | | Sigurdsson; Patrick Sulem; Arna B Agustsdottir; Hannes Eggertsson; Berglind Eiriksdottir; Run Fridriksdottir; Elisabet E Gardarsdottir; Gudmundur Georgsson; Olafia S Gretarsdottir; Kjartan R Gudmundsson; Thora R Gunnarsdottir; Arnaldur Gylfason; Hilma Holm; Brynjar O Jensson; Aslaug Jonasdottir; Kamilla S Josefsdottir; Thordur Kristjansson; Droplaug N Magnusdottir; Solvi Rognvaldsson; Louise le Roux; Gudrun Sigmundsdottir; Gardar Sveinbjornsson; Kristin E Sveinsdottir; Maney Sveinsdottir; Emil A Thorarensen; Bjarni Thorbjornsson; Gisli Masson; Ingileif Jonsdottir; Alma Moller; Thorolfur Gudnason; Karl G Kristinsson; Unnur Thorsteinsdottir; Kari Stefansson |
| EPI_ISL_829938, EPI_ISL_829939 | The National University Hospital of Iceland | deCODE genetics | Daniel F Gudbjartsson; Agnar Helgason; Hakon Jonsson; Olafur T Magnusson; Pall Melsted; Gudmundur L Norddahl; Jona Saemundsdottir; Asgeir Sigurdsson; Patrick Sulem; Arna B Agustsdottir; Hannes Eggertsson; Berglind Eiriksdottir; Run Fridriksdottir; Elisabet E Gardarsdottir; Gudmundur Georgsson; Olafia S Gretarsdottir; Kjartan R Gudmundsson; Thora R Gunnarsdottir; Arnaldur Gylfason; Hilma Holm; Brynjar O Jensson; Aslaug Jonasdottir; Kamilla S Josefsdottir; Thordur Kristjansson; Droplaug N Magnusdottir; Solvi Rognvaldsson; Louise le Roux; Gudrun Sigmundsdottir; Gardar Sveinbjornsson; Kristin E Sveinsdottir; Maney Sveinsdottir; Emil A Thorarensen; Bjarni Thorbjornsson; Gisli Masson; Ingileif Jonsdottir; Alma Moller; Thorolfur Gudnason; Karl G Kristinsson; Unnur Thorsteinsdottir; Kari Stefansson |
| EPI_ISL_829946, EPI_ISL_830052 | deCODE genetics | deCODE genetics | Daniel F Gudbjartsson; Agnar Helgason; Hakon Jonsson; Olafur T Magnusson; Pall Melsted; Gudmundur L Norddahl; Jona Saemundsdottir; Asgeir Sigurdsson; Patrick Sulem; Arna B Agustsdottir; Hannes Eggertsson; Berglind Eiriksdottir; Run Fridriksdottir; Elisabet E Gardarsdottir; Gudmundur Georgsson; Olafia S Gretarsdottir; Kjartan R Gudmundsson; Thora R Gunnarsdottir; Arnaldur Gylfason; Hilma Holm; Brynjar O Jensson; Aslaug Jonasdottir; Kamilla S Josefsdottir; Thordur Kristjansson; Droplaug N Magnusdottir; Solvi Rognvaldsson; Louise le Roux; Gudrun Sigmundsdottir; Gardar Sveinbjornsson; Kristin E Sveinsdottir; Maney Sveinsdottir; Emil A Thorarensen; Bjarni Thorbjornsson; Gisli Masson; Ingileif Jonsdottir; Alma Moller; Thorolfur Gudnason; Karl G Kristinsson; Unnur Thorsteinsdottir; Kari Stefansson |
| EPI_ISL_830085, EPI_ISL_830086, EPI_ISL_830093, EPI_ISL_830094, EPI_ISL_830095, EPI_ISL_830096, EPI_ISL_830120, EPI_ISL_830128 | The National University Hospital of Iceland | deCODE genetics | Daniel F Gudbjartsson; Agnar Helgason; Hakon Jonsson; Olafur T Magnusson; Pall Melsted; Gudmundur L Norddahl; Jona Saemundsdottir; Asgeir Sigurdsson; Patrick Sulem; Arna B Agustsdottir; Hannes Eggertsson; Berglind Eiriksdottir; Run Fridriksdottir; Elisabet E Gardarsdottir; Gudmundur Georgsson; Olafia S Gretarsdottir; Kjartan R Gudmundsson; Thora R Gunnarsdottir; Arnaldur Gylfason; Hilma Holm; Brynjar O Jensson; Aslaug Jonasdottir; Kamilla S Josefsdottir; Thordur Kristjansson; Droplaug N Magnusdottir; Solvi Rognvaldsson; Louise le Roux; Gudrun Sigmundsdottir; Gardar Sveinbjornsson; Kristin E Sveinsdottir; Maney Sveinsdottir; Emil A Thorarensen; Bjarni Thorbjornsson; Gisli Masson; Ingileif Jonsdottir; Alma Moller; Thorolfur Gudnason; Karl G Kristinsson; Unnur Thorsteinsdottir; Kari Stefansson |
| EPI_ISL_830130, EPI_ISL_830139 | deCODE genetics | deCODE genetics | Daniel F Gudbjartsson; Agnar Helgason; Hakon Jonsson; Olafur T Magnusson; Pall Melsted; Gudmundur L Norddahl; Jona Saemundsdottir; Asgeir Sigurdsson; Patrick Sulem; Arna B Agustsdottir; Hannes Eggertsson; Berglind Eiriksdottir; Run Fridriksdottir; Elisabet E Gardarsdottir; Gudmundur Georgsson; Olafia S Gretarsdottir; Kjartan R Gudmundsson; Thora R Gunnarsdottir; Arnaldur Gylfason; Hilma Holm; Brynjar O Jensson; Aslaug Jonasdottir; Kamilla S Josefsdottir; Thordur Kristjansson; Droplaug N Magnusdottir; Solvi Rognvaldsson; Louise le Roux; Gudrun Sigmundsdottir; Gardar Sveinbjornsson; Kristin E Sveinsdottir; Maney Sveinsdottir; Emil A Thorarensen; Bjarni Thorbjornsson; Gisli Masson; Ingileif Jonsdottir; Alma Moller; Thorolfur Gudnason; Karl G Kristinsson; Unnur Thorsteinsdottir; Kari Stefansson |
| EPI_ISL_830140, EPI_ISL_830145 | The National University Hospital of Iceland | deCODE genetics | Daniel F Gudbjartsson; Agnar Helgason; Hakon Jonsson; Olafur T Magnusson; Pall Melsted; Gudmundur L Norddahl; Jona Saemundsdottir; Asgeir Sigurdsson; Patrick Sulem; Arna B Agustsdottir; Hannes Eggertsson; Berglind Eiriksdottir; Run Fridriksdottir; Elisabet E Gardarsdottir; Gudmundur Georgsson; Olafia S Gretarsdottir; Kjartan R Gudmundsson; Thora R Gunnarsdottir; Arnaldur Gylfason; Hilma Holm; Brynjar O Jensson; Aslaug Jonasdottir; Kamilla S Josefsdottir; Thordur Kristjansson; Droplaug N Magnusdottir; Solvi Rognvaldsson; Louise le Roux; Gudrun Sigmundsdottir; Gardar Sveinbjornsson; Kristin E Sveinsdottir; Maney Sveinsdottir; Emil A Thorarensen; Bjarni Thorbjornsson; Gisli Masson; Ingileif Jonsdottir; Alma Moller; Thorolfur Gudnason; Karl G Kristinsson; Unnur Thorsteinsdottir; Kari Stefansson |
| EPI_ISL_830150, EPI_ISL_830199, EPI_ISL_830200, EPI_ISL_830201, EPI_ISL_830231, EPI_ISL_830422, EPI_ISL_830423 | deCODE genetics | deCODE genetics | Daniel F Gudbjartsson; Agnar Helgason; Hakon Jonsson; Olafur T Magnusson; Pall Melsted; Gudmundur L Norddahl; Jona Saemundsdottir; Asgeir Sigurdsson; Patrick Sulem; Arna B Agustsdottir; Hannes Eggertsson; Berglind Eiriksdottir; Run Fridriksdottir; Elisabet E Gardarsdottir; Gudmundur Georgsson; Olafia S Gretarsdottir; Kjartan R Gudmundsson; Thora R Gunnarsdottir; Arnaldur Gylfason; Hilma Holm; Brynjar O Jensson; Aslaug Jonasdottir; Kamilla S Josefsdottir; Thordur Kristjansson; Droplaug N Magnusdottir; Solvi Rognvaldsson; Louise le Roux; Gudrun Sigmundsdottir; Gardar Sveinbjornsson; Kristin E Sveinsdottir; Maney Sveinsdottir; Emil A Thorarensen; Bjarni Thorbjornsson; Gisli Masson; Ingileif Jonsdottir; Alma Moller; Thorolfur Gudnason; Karl G Kristinsson; Unnur Thorsteinsdottir; Kari Stefansson |
| EPI_ISL_830424, EPI_ISL_830425, EPI_ISL_830428, EPI_ISL_830429 | The National University Hospital of Iceland | deCODE genetics | Daniel F Gudbjartsson; Agnar Helgason; Hakon Jonsson; Olafur T Magnusson; Pall Melsted; Gudmundur L Norddahl; Jona Saemundsdottir; Asgeir Sigurdsson; Patrick Sulem; Arna B Agustsdottir; Hannes Eggertsson; Berglind Eiriksdottir; Run Fridriksdottir; Elisabet E Gardarsdottir; Gudmundur Georgsson; Olafia S Gretarsdottir; Kjartan R Gudmundsson; Thora R Gunnarsdottir; Arnaldur Gylfason; Hilma Holm; Brynjar O Jensson; Aslaug Jonasdottir; Kamilla S Josefsdottir; Thordur Kristjansson; Droplaug N Magnusdottir; Solvi Rognvaldsson; Louise le Roux; Gudrun Sigmundsdottir; Gardar Sveinbjornsson; Kristin E Sveinsdottir; Maney Sveinsdottir; Emil A Thorarensen; Bjarni Thorbjornsson; Gisli Masson; Ingileif Jonsdottir; Alma Moller; Thorolfur Gudnason; Karl G Kristinsson; Unnur Thorsteinsdottir; Kari Stefansson |
| EPI_ISL_830493, EPI_ISL_830494, EPI_ISL_830497, EPI_ISL_830498, EPI_ISL_830500, EPI_ISL_830505, EPI_ISL_830521, EPI_ISL_830522, EPI_ISL_830523, EPI_ISL_830524, EPI_ISL_830525, EPI_ISL_830526, EPI_ISL_830527, EPI_ISL_830528, EPI_ISL_830531, EPI_ISL_830538, EPI_ISL_830540, EPI_ISL_830544, EPI_ISL_830562 | | | |
| see above | deCODE genetics | deCODE genetics | Daniel F Gudbjartsson; Agnar Helgason; Hakon Jonsson; Olafur T Magnusson; Pall Melsted; Gudmundur L Norddahl; Jona Saemundsdottir; Asgeir Sigurdsson; Patrick Sulem; Arna B Agustsdottir; Hannes Eggertsson; Berglind Eiriksdottir; Run Fridriksdottir; Elisabet E Gardarsdottir; Gudmundur Georgsson; Olafia S Gretarsdottir; Kjartan R Gudmundsson; Thora R Gunnarsdottir; Arnaldur Gylfason; Hilma Holm; Brynjar O Jensson; Aslaug Jonasdottir; Kamilla S Josefsdottir; Thordur Kristjansson; Droplaug N Magnusdottir; Solvi Rognvaldsson; Louise le Roux; Gudrun Sigmundsdottir; Gardar Sveinbjornsson; Kristin E Sveinsdottir; Maney Sveinsdottir; Emil A Thorarensen; Bjarni Thorbjornsson; Gisli Masson; Ingileif Jonsdottir; Alma Moller; Thorolfur Gudnason; Karl G Kristinsson; Unnur Thorsteinsdottir; Kari Stefansson |
| EPI_ISL_830729, EPI_ISL_830733, EPI_ISL_830734, EPI_ISL_830741, EPI_ISL_830742, EPI_ISL_830750, EPI_ISL_830751, EPI_ISL_830752, EPI_ISL_830758, EPI_ISL_830759, EPI_ISL_830760, EPI_ISL_830762, EPI_ISL_830766, EPI_ISL_830767, EPI_ISL_830774, EPI_ISL_830776, EPI_ISL_830777, EPI_ISL_830778, EPI_ISL_830910, EPI_ISL_830911, EPI_ISL_830912, EPI_ISL_830913, EPI_ISL_830914, EPI_ISL_830915, EPI_ISL_830916, EPI_ISL_830917, EPI_ISL_830918, EPI_ISL_830919, EPI_ISL_830920, EPI_ISL_830921, EPI_ISL_830922, EPI_ISL_830923, EPI_ISL_830924, EPI_ISL_830925, EPI_ISL_830926, EPI_ISL_830927, EPI_ISL_830928, EPI_ISL_830929, EPI_ISL_830930, EPI_ISL_830931, EPI_ISL_830932, EPI_ISL_830933, EPI_ISL_830934, EPI_ISL_830935, EPI_ISL_830936, EPI_ISL_830937, EPI_ISL_830938, EPI_ISL_830939, EPI_ISL_830940, EPI_ISL_830941, EPI_ISL_830942, EPI_ISL_830943, EPI_ISL_830944, EPI_ISL_830945, EPI_ISL_830946, EPI_ISL_830947, EPI_ISL_830948, EPI_ISL_830949, EPI_ISL_830950, EPI_ISL_830951, EPI_ISL_830952, EPI_ISL_830953, EPI_ISL_830954, EPI_ISL_830955, EPI_ISL_830956, EPI_ISL_830957, EPI_ISL_830958, EPI_ISL_830959, EPI_ISL_830960, EPI_ISL_830961, EPI_ISL_830962, EPI_ISL_830963, EPI_ISL_830964, EPI_ISL_830965, EPI_ISL_830966, EPI_ISL_830967, EPI_ISL_830968, EPI_ISL_830969, EPI_ISL_830970, EPI_ISL_830971, EPI_ISL_830972, EPI_ISL_830973, EPI_ISL_830974, EPI_ISL_830975, EPI_ISL_830976, EPI_ISL_830977, EPI_ISL_830980, EPI_ISL_830981, EPI_ISL_830982, EPI_ISL_830983, EPI_ISL_830992, EPI_ISL_830993, EPI_ISL_830994, EPI_ISL_830995, EPI_ISL_830996, EPI_ISL_830997, EPI_ISL_830998, EPI_ISL_830999, EPI_ISL_831000, EPI_ISL_831006, EPI_ISL_831007, EPI_ISL_831008, EPI_ISL_831009, EPI_ISL_831011, EPI_ISL_831014, EPI_ISL_831016 | | | |
| see above | University Hospital Basel, Clinical Virology | University Hospital Basel, Clinical Bacteriology | Tim Roloff, Madlen Stange, Helena MB Seth-Smith, Alfredo Mari, Karoline Leuzinger, Julia Bielicki, Manuel Battegay, Hans Hirsch, Adrian Egli |
| EPI_ISL_831100, EPI_ISL_831101, EPI_ISL_831102, EPI_ISL_831103, EPI_ISL_831104, EPI_ISL_831105, EPI_ISL_831231 | Hospital Universitario La Paz (Madrid) | SeqCOVID-SPAIN consortium/IBV(CSIC) | María Rodríguez-Tejedor, Elias Dahdouh, Fernando Lázaro-Perona, Jesús Mingorance and SeqCOVID-SPAIN consortium |
| EPI_ISL_831770, EPI_ISL_831771, EPI_ISL_831772, EPI_ISL_831773, EPI_ISL_831774, EPI_ISL_831775, EPI_ISL_831776, EPI_ISL_831777 | United States Air Force School of Aerospace Medicine | United States Air Force School of Aerospace Medicine | Anthony Fries, Jennifer Meyer, William Gruner, Amanda Javorina, Sarah Purves, Clarise Starr, Elizabeth Macias |
| EPI_ISL_832010, EPI_ISL_832011, EPI_ISL_832012, EPI_ISL_832013 | Laboratório de Microbiologia Molecular - Universidade FEEVALE | Universidade Federal de Ciências da Saúde de Porto Alegre | Vinícius Bonetti Franceschi, Amanda de Menezes Mayer, Gabriel Dickin Caldana, Carla Andretta Moreira Neves, Patrícia Aline Gröhs Ferrareze, Gabriela Bettella Cybis, Ricardo Ariel Zimerman, Lívia Kmetzsch, Fernando Rosado Spilki, Claudia Elizabeth Thompson |
| EPI_ISL_832468, EPI_ISL_832469, EPI_ISL_832470, EPI_ISL_832471, EPI_ISL_832472, EPI_ISL_832473, EPI_ISL_832474, EPI_ISL_832475, EPI_ISL_832476, EPI_ISL_832477, EPI_ISL_832478, EPI_ISL_832479, EPI_ISL_832480, EPI_ISL_832481, EPI_ISL_832482, EPI_ISL_832483, EPI_ISL_832484, EPI_ISL_832485, EPI_ISL_832486, EPI_ISL_832487, EPI_ISL_832488, EPI_ISL_832489, EPI_ISL_832490, EPI_ISL_832491, EPI_ISL_832492, EPI_ISL_832493, EPI_ISL_832494, EPI_ISL_832495, EPI_ISL_832496, EPI_ISL_832547, EPI_ISL_832548, EPI_ISL_832549 | | | |
| see above | OHSU Lab Services Molecular Microbiology Lab | Oregon SARS-CoV-2 Genome Sequencing Center | Brendan L. O'Connell, Ruth V. Nichols, Sally Grindstaff, Alec J. Hirsch, Donna Hansel, Guang Fan, Daniel N. Streblow, William B. Messer, Andrew C. Adey, Benjamin N. Bimber, Brian J. O'Roak |
| EPI_ISL_837078 | Respiratory Virus Unit, National Infection Service, Public Health England | COVID-19 Genomics UK (COG-UK) Consortium | PHE Covid Sequencing Team |

| | | | |
|---|---|---|---|
| EPI_ISL_839981 | Queens Medical Centre, Clinical Microbiology Department / DeepSeq Nottingham | COVID-19 Genomics UK (COG-UK) Consortium | Gemma Clark, Wendy Smith, Manjinder Khakh, Vicki M Fleming, Michelle M Lister, Hannah Howson-Wells, Jonathan Ball, Patrick McClure, Joseph Chappell, Theocharis Tsoleridis, Nadine Holmes, Matthew Carlisle, Christopher Moore, Fei Sang, Johnny Debebe, Victoria Wright, Matthew Loose |
| EPI_ISL_842175, EPI_ISL_842178, EPI_ISL_842179, EPI_ISL_842180, EPI_ISL_842181, EPI_ISL_842184, EPI_ISL_842186, EPI_ISL_842187, EPI_ISL_842188, EPI_ISL_842190, EPI_ISL_842191, EPI_ISL_842192, EPI_ISL_842194, EPI_ISL_842195, EPI_ISL_842196, EPI_ISL_842197, EPI_ISL_842198, EPI_ISL_842199, EPI_ISL_842200 | | | |
| see above | Oxford Viromics, NDM, University of Oxford; Oxford University Hospitals; Basingstoke and North Hampshire Hospital | COVID-19 Genomics UK (COG-UK) Consortium | Tanya Golubchik, David Bonsall, George Macintyre, Amy Trebes, Mariateresa de Cesare, Catrin Moore, Alex Mobbs, Anita Justice, Robert Shaw, Monique Andersson, Timothy Peto, Emma Wise, Nathan Moore, Jessica Lynch, Nick Cortes, Matilde Mori, Stephen Kidd, David Buck, John Todd, Christophe Fraser |
| EPI_ISL_842862 | Barts Health NHS Trust | COVID-19 Genomics UK (COG-UK) Consortium | CUTINO-MOGUEL, Maria-Teresa; HARRINGTON, David; OWOYEMI, Dola; SHYLINI, Raghavendran; BROAD, Claire; KELE, Beatrix |
| EPI_ISL_845891, EPI_ISL_845892 | Benaroya Research Institute | UW Virology Lab | Pavitra Roychoudhury, Hong Xie, Lasata Shrestha, Michelle Lin, Meei-Li Huang, Keith R Jerome, Alexander Greninger |
| EPI_ISL_848224, EPI_ISL_848235, EPI_ISL_848238, EPI_ISL_848239, EPI_ISL_848248, EPI_ISL_848255, EPI_ISL_848259, EPI_ISL_848263, EPI_ISL_848267, EPI_ISL_848272, EPI_ISL_848356, EPI_ISL_848357, EPI_ISL_848358, EPI_ISL_848359, EPI_ISL_848360, EPI_ISL_848361, EPI_ISL_848362, EPI_ISL_848363, EPI_ISL_848364, EPI_ISL_848365, EPI_ISL_848366, EPI_ISL_848367, EPI_ISL_848368, EPI_ISL_848369, EPI_ISL_848370, EPI_ISL_848371, EPI_ISL_848372, EPI_ISL_848373, EPI_ISL_848374, EPI_ISL_848375, EPI_ISL_848455, EPI_ISL_848510, EPI_ISL_848511, EPI_ISL_848512, EPI_ISL_848513, EPI_ISL_848514, EPI_ISL_848515, EPI_ISL_848516, EPI_ISL_848517, EPI_ISL_848518, EPI_ISL_848519, EPI_ISL_848520, EPI_ISL_848521 | | | |
| see above | Illinois Department of Public Health | Gagnon Lab, Southern Illinois University | Keith Gagnon |
| EPI_ISL_848555, EPI_ISL_848607 | Evandro Chagas Institute | Evandro Chagas Institute | Santos, M.C.; Silva, A.M.; Junior, W.D.C.; Barbagelata, L.S.; Ferreira, J.A.; Sousa, E.M.A.; da Silva, P.S.; Pinheiro, K.C.; L.C.; Sousa Junior, E.C. |
| EPI_ISL_848932, EPI_ISL_848933, EPI_ISL_848934, EPI_ISL_848935, EPI_ISL_848936, EPI_ISL_848937, EPI_ISL_848938, EPI_ISL_848939, EPI_ISL_848940, EPI_ISL_848941, EPI_ISL_848942, EPI_ISL_848943, EPI_ISL_848944, EPI_ISL_848945, EPI_ISL_848946, EPI_ISL_848947, EPI_ISL_848948, EPI_ISL_848949, EPI_ISL_848950 | | | |
| see above | Florida Bureau of Public Health Laboratories | Florida Bureau of Public Health Laboratories | Sarah Schmedes, Jason Blanton |
| EPI_ISL_849350, EPI_ISL_849351, EPI_ISL_849353, EPI_ISL_849354, EPI_ISL_849357, EPI_ISL_849360, EPI_ISL_849361, EPI_ISL_849362, EPI_ISL_849363, EPI_ISL_849367 | Servicio Virosis Respiratorias-Departamento Virología-INEI | Instituto Nacional Enfermedades Infecciosas C.G.Malbran | Baumeister E., Avaro M., Benedetti E., Russo M., Dattero ME, Pontoriero A., Cisterna D., Molina V., Perandones C., Tuduri E., Lorenzo F., Poklepovich T., Campos J. |
| EPI_ISL_849408, EPI_ISL_849490, EPI_ISL_849491, EPI_ISL_849599, EPI_ISL_849600, EPI_ISL_849601, EPI_ISL_849602 | Seattle Flu Study | Seattle Flu Study | Deborah A. Nickerson, Chris D. Frazar, Jover Lee, Benjamin Pelle, Matthew Richardson, Amanda Adler, Elisabeth Brandstetter, Peter D. Han, Kairsten Fay, Misja Ilcisin, Kirsten Lacombe, Thomas R. Sibley, Melissa Truong, Caitlin R. Wolf, Michael Boeckh, Janet A. Englund, Michael Famulare, Barry R. Lutz, Mark J. Rieder, Lea M. Starita, Matthew Thompson, Jay Shendure, Trevor Bedford, Helen Y. Chu |
| EPI_ISL_852653 | Institute of Virology, Medical Center, University of Freiburg, Freiburg, Germany | Institute of Virology, Clinial Virus Genomics, Medical Center, University of Freiburg, Freiburg, Germany | Jonas Fuchs, Lisa Kern, Sandra Reuter, Hajo Grundmann, Marcus Panning |
| EPI_ISL_853312, EPI_ISL_853319, EPI_ISL_853320, EPI_ISL_853321, EPI_ISL_853322, EPI_ISL_853391 | UPMC Clinical Microbiology Laboratory | Microbial Genome Sequencing Center; Microbial Genomic Epidemiology Laboratory | Mustapha M. Mustapha, Jane W. Marsh, Dan Snyder, Marissa P. Griffith, Stephanie L. Mitchell, Vatsala R. Srinivasa, Kady D. Waggle, Chinelo Ezeonwuku, Vaughn S. Cooper, Lee H. Harrison |
| EPI_ISL_853720, EPI_ISL_853732, EPI_ISL_853755, EPI_ISL_853783 | Department of Microbiology, University Innsbruck | Bergthaler laboratory, CeMM Research Center for Molecular Medicine of the Austrian Academy of Sciences | Lukas Endler, Alexandra Popa, Benedikt Agerer, Jakob-Wendelin Genger, Alexander Lercher, Anna Schedl, Thomas Penz, Michael Schuster, Jan Laine, Martin Senekowitsch, Christoph Bock, Andreas Bergthaler |
| EPI_ISL_853866, EPI_ISL_853867, EPI_ISL_853868, EPI_ISL_853869 | Institute for Laboratory Diagnostics and Microbiology, Klinikum Klagenfurt am Worthersee | Bergthaler laboratory, CeMM Research Center for Molecular Medicine of the Austrian Academy of Sciences | Lukas Endler, Alexandra Popa, Benedikt Agerer, Jakob-Wendelin Genger, Alexander Lercher, Anna Schedl, Thomas Penz, Michael Schuster, Jan Laine, Martin Senekowitsch, Christoph Bock, Andreas Bergthaler |
| EPI_ISL_854220, EPI_ISL_854221 | Department of Microbiology, University Innsbruck | Bergthaler laboratory, CeMM Research Center for Molecular Medicine of the Austrian Academy of Sciences | Lukas Endler, Alexandra Popa, Benedikt Agerer, Jakob-Wendelin Genger, Alexander Lercher, Anna Schedl, Thomas Penz, Michael Schuster, Jan Laine, Martin Senekowitsch, Christoph Bock, Andreas Bergthaler |
| EPI_ISL_854233 | Institute for Water Quality and Resource Management, Technical University Vienna | Bergthaler laboratory, CeMM Research Center for Molecular Medicine of the Austrian Academy of Sciences | Lukas Endler, Alexandra Popa, Benedikt Agerer, Jakob-Wendelin Genger, Alexander Lercher, Anna Schedl, Thomas Penz, Michael Schuster, Jan Laine, Martin Senekowitsch, Christoph Bock, Andreas Bergthaler |
| EPI_ISL_855355 | Hospital | National Reference Center for Viruses of Respiratory Infections, Institut Pasteur, Paris | Marion Barbet, Sylvie Behillil, Méline Bizard, Angela Brisebarre, Camille Capel, Etienne Simon-Lorière, Vincent Enouf, Maud Vanpeene, Sylvie van der Werf |
| EPI_ISL_855534 | KEMRI-Wellcome Trust Research Programme/KEMRI-CGMR-C Kilifi | KEMRI-Wellcome Trust Research Programme/KEMRI-CGMR-C Kilifi | Githinji et al |
| EPI_ISL_856679, EPI_ISL_856698 | Lab. Microbiologia e Virologia, Cotugno, A.O. dei Colli | Lab. Microbiologia e Virologia, Cotugno, A.O. dei Colli | Luigi Atripaldi, Claudia Tiberio, Anna Perfetti |
| EPI_ISL_859789, EPI_ISL_859790, EPI_ISL_859819, EPI_ISL_859823, EPI_ISL_859825, EPI_ISL_859827, EPI_ISL_859831, EPI_ISL_859833, EPI_ISL_859891, EPI_ISL_859892, EPI_ISL_859893, EPI_ISL_859894 | | | |
| see above | BTC, Khalifa University | BTC, Khalifa University | Al Safar et al |
| EPI_ISL_860122 | Keio University School of Medicine | Keio University School of Medicine | Kenjiro Kosaki, Yuka Iwasaki, Hirotsugu Ishizu, Haruhiko Siomi, Kodai Abe |
| EPI_ISL_860747, EPI_ISL_860748, EPI_ISL_860749, EPI_ISL_860750, EPI_ISL_860751 | Swiss National Reference Centre for Influenza | Swiss National Reference Centre for Influenza | Ana Rita Gonçalves Cabecinhas, Samuel Cordey, Florian Laubscher,Christoph Grünig, Laurent Kaiser |
| EPI_ISL_866820, EPI_ISL_866821 | Quadram Institute Bioscience | COVID-19 Genomics UK (COG-UK) Consortium | Dave J. Baker, Gemma L. Kay, Alp Aydin, Thanh Le-Viet, Steven Rudder, Ana P. Tedim, Anastasia Kolyva, Maria Diaz, Leonardo de Oliveira Martins, Nabil-Fareed Alikhan, Lizzie Meadows, Rachael Stanley, Ngozi Elumogo, Muhammed Yasir, Nicholas M. Thomson, Alexander J Trotter, Rachel Gilroy, Samuel Bloomfield, Claire Stuart, Andrew Bell, Reenesh Prakash, Samir Dervisevic, Alison E. Mather, John Wain, Mark Webber, Andrew J. Page, Justin O'Grady |
| EPI_ISL_867933, EPI_ISL_867935, EPI_ISL_867955, EPI_ISL_867956, EPI_ISL_867957, EPI_ISL_867969, EPI_ISL_867970, EPI_ISL_867975 | Centre for Enzyme Innovation, University of Portsmouth / Translational Research Laboratory, Portsmouth Hospitals NHS Trust | COVID-19 Genomics UK (COG-UK) Consortium | Angela Beckett,Yann Bourgeois,Garry Scarlett,Sharon Glaysher,Scott Elliott,Kelly Bicknell,Robert Impey,Allyson Lloyd,Sarah Wyllie,Ethan Butcher,Anoop Chauhan,Samuel Robson |
| EPI_ISL_868789, EPI_ISL_868790, EPI_ISL_868791, EPI_ISL_868793, EPI_ISL_868795, EPI_ISL_868798, EPI_ISL_868799, EPI_ISL_868800, EPI_ISL_868801, EPI_ISL_868802, EPI_ISL_868804, EPI_ISL_868805, EPI_ISL_868808, EPI_ISL_868810, EPI_ISL_868813, EPI_ISL_868815, EPI_ISL_868816, EPI_ISL_868817, EPI_ISL_868820, EPI_ISL_868822, EPI_ISL_868829, EPI_ISL_868830, EPI_ISL_868832, EPI_ISL_868834, EPI_ISL_868837, EPI_ISL_868838, EPI_ISL_868841 | | | |
| see above | Bioinformatics and Biostatistics Lab, Advanced Sequencing Facility | COVID-19 Genomics UK (COG-UK) Consortium | Aengus Stewart,Jerome Nicod,Chelsea Sawyer,Laura Cubitt,Harshil Patel,Margaret Crawford |
| EPI_ISL_871995 | Hospital Universitario de Ceuta | Instituto de Salud Carlos III | Iglesias-Caballero, M. Camarero, S. Molinero Calamita, M. González-Esguevillas, M. Pozo, F. Casas, I. Jiménez, P. Jiménez, M. Zaballos, A. Monzón, S. Varona, S. Juliá, M. Cuesta, I. Hijano, S. |
| EPI_ISL_872458, EPI_ISL_872459, EPI_ISL_872460, EPI_ISL_872461, EPI_ISL_872462, EPI_ISL_872463, EPI_ISL_872464, EPI_ISL_872465, EPI_ISL_872466, EPI_ISL_872467, EPI_ISL_872468 | | | |
| see above | Eurofins Diatherix | Hudsonalpha Genome Sequencing Center | Jane Grimwood, Melissa Williams, Lori H. Handley, Joshua Stough, Leslie Malone, Stefan Brzezinski, Ada Stewart, Teresa Jones, Jenell Webber, John Lovell, Jennifer Cart, and Jeremy Schmutz |
| EPI_ISL_872680 | Rhode Island Department of Health | Infectious Disease Program, Broad Institute of Harvard and MIT | Lemieux,J.E., Siddle,K.J., Huard,R., King,E., Azevedo,K., Miller,A., Adams,G., Gladden-Young,A., Lagerborg,K., Rudy,M., DeRuff,K., Carter,A., Normandin,E., Bauer,M., Reilly,S., Tomkins-Tinch,C., Loreth,C., Chaluvadi,S., Birren,B.W., Gallagher,G., Smole,S., Park,D.J., MacInnis,B.L., and Sabeti,P.C. |
| EPI_ISL_872898, EPI_ISL_872899, EPI_ISL_872900, EPI_ISL_872901, | WHO National Influenza Centre Russian Federation | WHO National Influenza Centre Russian Federation | Andrey Komissarov, Artem Fadeev, Anna Ivanova, Kseniya Komissarova, Dmitry Bazhenov, Mikhail Bakaev, Daria Danilenko, Ksenia Safina, Elena Nabieva, Georgii Bazykin, Dmitry Lioznov |

| | | | |
|---|---|---|---|
| EPI_ISL_872902 | | | |
| EPI_ISL_876570, EPI_ISL_876571 | Florida Bureau of Public Health Laboratories | Florida Bureau of Public Health Laboratories | Sarah Schmedes, Jason Blanton |
| EPI_ISL_876823, EPI_ISL_876831, EPI_ISL_876869, EPI_ISL_876870, EPI_ISL_876871, EPI_ISL_876872, EPI_ISL_876873, EPI_ISL_876874, EPI_ISL_876875, EPI_ISL_876876 | Quest Diagnostics | Quest Diagnostics | Rosenthal,S.H., Gerasimova,A., Kagan,R.M., Anderson, B., Hua, M., Liu Y., Bernstein, L.E., Livingston, K.E., Perez, A., Shalhout, D.F., Shlyakhter, I.A., Owen, R., Tanpaiboon, P., Lacbawan, F. |
| EPI_ISL_877425, EPI_ISL_877426, EPI_ISL_877455 | Institute of Microbiology and Immunology, Faculty of Medicine, University of Ljubljana | Institute of Microbiology and Immunology, Faculty of Medicine, University of Ljubljana | Samo Zakotnik, Tomaž Mark Zorec, Matic Brvar, Miša Korva, Mario Poljak, Tatjana Avši - Županc |
| EPI_ISL_878656, EPI_ISL_878671 | Rady's Childrens Hospital | Andersen lab at Scripps Research | SEARCH Alliance San Diego with Nanda Radamchar, David Dimmock, Linda Luo, Christina Clarke, Kathryn Bouic, Teresa Mueller, Denise Malicki |
| EPI_ISL_882675, EPI_ISL_882676, EPI_ISL_882681, EPI_ISL_882685, EPI_ISL_882691, EPI_ISL_882694, EPI_ISL_882695, EPI_ISL_882717, EPI_ISL_882718, EPI_ISL_882719, EPI_ISL_882720, EPI_ISL_882721, EPI_ISL_882722, EPI_ISL_882723, EPI_ISL_882724, EPI_ISL_882740, EPI_ISL_882741, EPI_ISL_882743, EPI_ISL_882746, EPI_ISL_882749, EPI_ISL_882750 | | | |
| see above | 1.AO Universitaria 'S. Giovanni di Dio e Ruggi D'Aragona, Scuola Medica Salernitana' Hospital / 2.UOC di Virologia e Microbiologia, Università della Campania 'L. Vanvitelli' / 3.AO Universitaria 'Federico II' Napoli Hospital / 4.AORN 'San Giuseppe Moscati' Avellino Hospital / 5.AO 'San Pio - presidio G. Rummo' Benevento Hospital / 6.AO 'Sant'Anna e San Sebastiano' Caserta Hospital / 7.PO 'Maria Santissima Addolorata' Eboli Hospital / 8.Biogem Istituto di Ricerche Genetiche | 1. Genome Research Center for Health (CRGS) / 2. Laboratory of Molecular Medicine and Genomics(LMMGe) / 3. Center for Research in Pure and Applied Mathematics (CRMPA) | Giorgio Giurato, Francesca Rizzo, Alessandro Weisz, Gianluigi Franci, Giovanni Nassa, Pasquale Pagliano, Roberta Tarallo, Elena Alexandrova, Ylenia D'Agostino, Carlo Ferravante, Jessica Lamberti, Viola Melone, Domenico Memoli, Valeria Mirici Cappa, Domenico Palumbo, Giovanni Pecoraro, Assunta Sellitto, Oriana Strianese, Ilaria Terenzi, Giuseppe Fenza, Aniello Gentile, Antonello Saccomanno, Sonia Amabile, Teresa Rocco, Annamaria Salvati, Emilia Vaccaro, Massimiliano Galdiero, Michele Cennamo, Giuseppe Portella, Maria Grazia Foti, Mariarosaria Ingino, Maria Landi, Maurizio Fumi, Vincenzo Rocco, Rita Greco, Vittoria Letizia, Arnolfo Petruzziello, Maddalena Schioppa, Gregorio Goffredi, Francesca Marciano, Michele Caraglia, Alessia Cossu, Marianna Scrima |
| EPI_ISL_884320, EPI_ISL_884366, EPI_ISL_884367, EPI_ISL_884376, EPI_ISL_884377, EPI_ISL_884393, EPI_ISL_884441, EPI_ISL_884443 | Infectious Diseases, Quest Diagnostics | Infectious Diseases, Quest Diagnostics | Rosenthal,S.H., Gerasimova,A., Kagan,R.M., Anderson,B., Bernstein,L.E., Livingston,K.E., Hua,M., Liu,Y., Shalhout,D.F., Owen,R., Lacbawan,F. |
| EPI_ISL_888692, EPI_ISL_888700, EPI_ISL_888705, EPI_ISL_888714 | KU Leuven, Rega Institute, Clinical and Epidemiological Virology | KU Leuven, Rega Institute, Clinical and Epidemiological Virology | Tony Wawina-Bokalanga, Bert Vanmechelen, Joan Marti-Carerras, Piet Maes |
| EPI_ISL_890192 | Gonoshasthaya-RNA Research Center, Gonoshasthaya-RNA Molecular Diagnostics and Research Center | Gonoshasthaya-RNA Research Center, Gonoshasthaya-RNA Molecular Diagnostics and Research Center | Jamiruddin,M.R., Khondoker,M.U., Sharif,N., Azmuda,N., Ahmed,M.F., Sharmin,S., Akter,S., Mou,T.J., Marzan,M., Liza,S.M., Nahar,S., Jahan,N., Ali,T., Khandker,S.S., Jamiruddin,M., Haq,M.A., Adnan,N., Chaity,M., Oishee,M. |
| EPI_ISL_892224 | Lighthouse Lab in Milton Keynes | Wellcome Sanger Institute for the COVID-19 Genomics UK (COG-UK) Consortium | The Lighthouse Lab in Milton Keynes and Alex Alderton, Roberto Amato, Sonia Goncalves, Ewan Harrison, David K. Jackson, Ian Johnston, Dominic Kwiatkowski, Cordelia Langford, John Sillitoe on behalf of the Wellcome Sanger Institute COVID-19 Surveillance Team |
| EPI_ISL_896134, EPI_ISL_896138, EPI_ISL_896170, EPI_ISL_900083, EPI_ISL_900106, EPI_ISL_900119, EPI_ISL_900147, EPI_ISL_900171, EPI_ISL_900191, EPI_ISL_900243, EPI_ISL_900291, EPI_ISL_900321, EPI_ISL_900436, EPI_ISL_900438, EPI_ISL_900451, EPI_ISL_900458, EPI_ISL_900476 | | | |
| see above | MEPHI, Aix Marseille University | MEPHI, Aix Marseille University | Anthony LEVASSEUR |
| EPI_ISL_902921 | Tanjungpura University Hospital | Tanjungpura University Hospital | Puji Astuti; Andriani; Mahyarudin; Delima Fajar Liana; Virhan Novianry ; Sofi Siti Shofiyah |
| EPI_ISL_904137, EPI_ISL_904230, EPI_ISL_904281, EPI_ISL_904431, EPI_ISL_904432, EPI_ISL_904433, EPI_ISL_904434, EPI_ISL_904435, EPI_ISL_904436 | Dutch COVID-19 response team | Erasmus Medical Center | Bas Oude Munnink, Reina Sikkema, David Nieuwenhuijse, Irina Chestakova, Anne van der Linden, Marjan Boter, Emmanuelle Munger, Corine GeurtsvanKessel, Annemiek van der Eijk, Richard Molenkamp, Marion Koopmans, on behalf of the Dutch national COVID-19 response team. |
| EPI_ISL_904945 | Vilnius University Hospital Santaros Klinikos, Vilnius University | Institute of Biotechnology, Life Sciences Center, Vilnius University | Emilija Vasiliunaite, Milda Norkiene, Albertas Timinskas, Alma Gedvilaite, Aurelija Zvirbliene, Daniel Naumovas, Laimonas Griskevicius |
| EPI_ISL_905604, EPI_ISL_905652, EPI_ISL_905653, EPI_ISL_905654, EPI_ISL_905655, EPI_ISL_905656, EPI_ISL_905687 | Dutch COVID-19 response team | National Institute for Public Health and the Environment (RIVM) | Adam Meijer, Harry Vennema, Dirk Eggink, Jeroen Cremer, Sharon van den Brink, Bas van der Veer, AnneMarie van den Brandt, Florian Zwagemaker, Dennis Schmitz, Chantal Reusken, on behalf of the national COVID-19 response team |
| EPI_ISL_910326 | General Hospital - Strumica | Research Center for Genetic Engineering and Biotechnology "Georgi D. Efremov" , Macedonian Academ | RCGEB - MASA |
| EPI_ISL_910327 | General Hospital - Kumanovo | Research Center for Genetic Engineering and Biotechnology "Georgi D. Efremov" , Macedonian Academy of Sciences and Arts | RCGEB - MASA |
| EPI_ISL_910328 | General Hospital - Tetovo | Research Center for Genetic Engineering and Biotechnology "Georgi D. Efremov" , Macedonian Academy of Sciences and Arts | RCGEB - MASA |
| EPI_ISL_913071, EPI_ISL_913072, EPI_ISL_913086, EPI_ISL_913090, EPI_ISL_913095 | Center for Virology | Center for Virology | Jeremy V. Camp, Irene Goerzer, Monika Redlberger-Fritz, Stephan W. Aberle |
| EPI_ISL_913104, EPI_ISL_913107 | CHU Purpan - Laboratoire de Virologie - Institut Fédératif de Biologie | CHU Purpan - Laboratoire de Virologie - Institut Fédératif de Biologie | Latour J., Ranger N., Dubois M., Carcenac R., Harter A., Boyer P., Tremeaux P., Izopet J. |
| EPI_ISL_913920, EPI_ISL_913921, EPI_ISL_913922, EPI_ISL_913923, EPI_ISL_913924, EPI_ISL_913926, EPI_ISL_913967 | Instituto de Diagnostico y Referencia Epidemiologicos INDRE_RNLSP | Instituto de Diagnostico y Referencia Epidemiologicos (INDRE) | Claudia Wong-Arambula, Abril Rodriguez-Maldonado, Fabiola Garces-Ayala, Adnan Araiza-Rodriguez, David Fragoso-Fonseca, Sergio Rangel-Guerrero, Mayra Jimenez-Morales, Nancy Munoz-Hernandez, Natividad Cruz-Ortiz, Tatiana Nunez-Garcia, Gisela Barrera-Badillo, Lucia Hernandez-Rivas, Irma Lopez-Martinez, Ernesto Ramirez-Gonzalez. |
| EPI_ISL_918488 | General Hospital - Strumica | Research Center for Genetic Engineering and Biotechnology "Georgi D. Efremov" , Macedonian Academy of Sciences and Arts | RCGEB - MASA |
| EPI_ISL_918514 | Evandro Chagas Institute | Evandro Chagas Institute | Santos, M.C.; Silva, A.M.; Junior, W.D.C.; Barbagelata, L.S.; Ferreira, J.A.; Sousa, E.M.A.; da Silva, P.S.; Pinheiro, K.C.; L.C.; Sousa Junior, E.C. |
| EPI_ISL_923228, EPI_ISL_923259, EPI_ISL_923264, EPI_ISL_923269, EPI_ISL_923270, EPI_ISL_923271, EPI_ISL_923274, EPI_ISL_923282, EPI_ISL_923283, EPI_ISL_923285, EPI_ISL_923286, EPI_ISL_923288, EPI_ISL_923299 | | | |
| see above | Centre for Enzyme Innovation, University of Portsmouth / Translational Research Laboratory, Portsmouth Hospitals NHS Trust | COVID-19 Genomics UK (COG-UK) Consortium | Angela Beckett,Salman Goudarzi,Christopher Fearn,Kate Cook,Katie Loveson,Sharon Glaysher,Scott Elliott,Samuel Robson |
| EPI_ISL_924420, EPI_ISL_924424 | Virology Department, Sheffield Teaching Hospitals NHS Foundation Trust/Department of Infection, Immunity and Cardiovascular Disease, The Medical School, University of Sheffield | COVID-19 Genomics UK (COG-UK) Consortium | Thushan de Silva, Matthew Parker, Nikki Smith, Adri Angyal, Rebecca Brown, Luke Green, Rachel Tucker, Paul Parsons, Danielle Groves, Katie Johnson, Laura Carrilero, Alex Keeley, Dave Partridge, Matthew Wyles, Benjamin Lindsey, Mehmet Yavuz, Mohammad Raza, Cariad Evans |

| | | | |
|---|---|---|---|
| EPI_ISL_925099, EPI_ISL_925100, EPI_ISL_925101, EPI_ISL_925121, EPI_ISL_925122, EPI_ISL_925123 | 1.AO Universitaria 'S. Giovanni di Dio e Ruggi D'Aragona, Scuola Medica Salernitana' Hospital / 2.UOC di Virologia e Microbiologia, Università della Campania 'L. Vanvitelli' / 3.AO Universitaria 'Federico II' Napoli Hospital / 4.AORN 'San Giuseppe Moscati' Avellino Hospital / 5.AO 'San Pio - presidio G. Rummo' Benevento Hospital / 6.AO 'Sant'Anna e San Sebastiano' Caserta Hospital / 7.PO 'Maria Santissima Addolorata' Eboli Hospital / 8.Biogem Istituto di Ricerche Genetiche | 1. Genome Research Center for Health (CRGS) / 2. Laboratory of Molecular Medicine and Genomics(LMMGe) / 3. Center for Research in Pure and Applied Mathematics (CRMPA) | Giorgio Giurato, Francesca Rizzo, Alessandro Weisz, Gianluigi Franci, Giovanni Nassa, Pasquale Pagliano, Roberta Tarallo, Elena Alexandrova, Ylenia D'Agostino, Carlo Ferravante, Jessica Lamberti, Viola Melone, Domenico Memoli, Valeria Mirici Cappa, Domenico Palumbo, Giovanni Pecoraro, Assunta Sellitto, Oriana Strianese, Ilaria Terenzi, Giuseppe Fenza, Aniello Gentile, Antonello Saccomanno, Sonia Amabile, Teresa Rocco, Annamaria Salvati, Emilia Vaccaro, Massimiliano Galdiero, Michele Cennamo, Giuseppe Portella, Maria Grazia Foti, Mariarosaria Ingino, Maria Landi, Maurizio Fumi, Vincenzo Rocco, Rita Greco, Vittoria Letizia, Arnolfo Petruzziello, Maddalena Schioppa, Gregorio Goffredi, Francesca Marciano, Michele Caraglia, Alessia Cossu, Marianna Scrima |
| EPI_ISL_925307, EPI_ISL_925308, EPI_ISL_925309, EPI_ISL_925310, EPI_ISL_925311, EPI_ISL_925312, EPI_ISL_925313, EPI_ISL_925314, EPI_ISL_925315, EPI_ISL_925316, EPI_ISL_925317, EPI_ISL_925318, EPI_ISL_925319, EPI_ISL_925320, EPI_ISL_925321, EPI_ISL_925322, EPI_ISL_925323, EPI_ISL_925324, EPI_ISL_925325, EPI_ISL_925326, EPI_ISL_925327, EPI_ISL_925328, EPI_ISL_925329, EPI_ISL_925330, EPI_ISL_925331, EPI_ISL_925332, EPI_ISL_925333, EPI_ISL_925334, EPI_ISL_925335, EPI_ISL_925336, EPI_ISL_925337, EPI_ISL_925338, EPI_ISL_925339, EPI_ISL_925340, EPI_ISL_925341, EPI_ISL_925342, EPI_ISL_925343, EPI_ISL_925344, EPI_ISL_925345, EPI_ISL_925346, EPI_ISL_925347, EPI_ISL_925348, EPI_ISL_925349, EPI_ISL_925350, EPI_ISL_925351, EPI_ISL_925352, EPI_ISL_925353, EPI_ISL_925354, EPI_ISL_925355, EPI_ISL_925356, EPI_ISL_925357, EPI_ISL_925358, EPI_ISL_925359, EPI_ISL_925360 | | | |
| see above | Michigan Department of Agriculture and Rural Development | Pathogen Discovery, Respiratory Viruses Branch, Division of Viral Diseases, Centers for Disease Control and Prevention | Yan Li, Jing Zhang, Ying Tao, Anna montmayeur, Brian Lynch, Krista Queen, Anna Uehara, Rachel Marine, Peter Cook, Clinton R. Paden, Haibin Wang, Suxiang Tong |
| EPI_ISL_925476, EPI_ISL_925477, EPI_ISL_925478 | Department of Clinical Microbiology | GIGA Medical Genomics | Keith Durkin, Maria Artesi, Sébastien Bontems, Raphaël Boreux, Bouchra Boujemla, Cécile Meex, Pierrette Melin, Marie-Pierre Hayette, Vincent Bours |
| EPI_ISL_930859, EPI_ISL_930860, EPI_ISL_930861, EPI_ISL_930862, EPI_ISL_930863, EPI_ISL_930864, EPI_ISL_930865, EPI_ISL_930866, EPI_ISL_930867, EPI_ISL_930868, EPI_ISL_930869, EPI_ISL_930870, EPI_ISL_930871, EPI_ISL_930872, EPI_ISL_930873, EPI_ISL_930874, EPI_ISL_930875, EPI_ISL_930876, EPI_ISL_930877, EPI_ISL_930878, EPI_ISL_930879, EPI_ISL_930880, EPI_ISL_930881, EPI_ISL_930882, EPI_ISL_930883, EPI_ISL_930884, EPI_ISL_930885, EPI_ISL_930886, EPI_ISL_930887, EPI_ISL_930888, EPI_ISL_930889, EPI_ISL_930890, EPI_ISL_930891, EPI_ISL_930892, EPI_ISL_930893, EPI_ISL_930894, EPI_ISL_930895, EPI_ISL_930896, EPI_ISL_930897, EPI_ISL_930898, EPI_ISL_930899, EPI_ISL_930900, EPI_ISL_930901, EPI_ISL_930902, EPI_ISL_930903, EPI_ISL_930904, EPI_ISL_930907, EPI_ISL_930908, EPI_ISL_930909, EPI_ISL_930910, EPI_ISL_930911, EPI_ISL_930912, EPI_ISL_930913, EPI_ISL_930914, EPI_ISL_930915, EPI_ISL_930916, EPI_ISL_930917, EPI_ISL_930918, EPI_ISL_930919, EPI_ISL_930930, EPI_ISL_930932, EPI_ISL_930933, EPI_ISL_930965 | | | |
| see above | University Hospital Basel, Clinical Virology | University Hospital Basel, Clinical Bacteriology | Tim Roloff, Madlen Stange, Helena MB Seth-Smith, Alfredo Mari, Karoline Leuzinger, Julia Bielicki, Manuel Battegay, Hans Hirsch, Adrian Egli |
| EPI_ISL_934405, EPI_ISL_934987 | ADMED Microbiologie | Genomics and Transcriptomics, Philip Morris International | Reto Lienhard, Marie-Lise Tritten, Emmanuel Guedj, Nicolas Sierro, Rémi Dulize, David Bornand, Mehdi Auberson, Maxime Berthouzoz, Nikolai Ivanov, Manuel Peitsch |
| EPI_ISL_935132, EPI_ISL_935133, EPI_ISL_935134, EPI_ISL_935135, EPI_ISL_935136, EPI_ISL_935137, EPI_ISL_935138, EPI_ISL_935139, EPI_ISL_935140, EPI_ISL_935141, EPI_ISL_935142 | | | |
| see above | 1.AO Universitaria 'S. Giovanni di Dio e Ruggi D'Aragona, Scuola Medica Salernitana' Hospital / 2.UOC di Virologia e Microbiologia, Università della Campania 'L. Vanvitelli' / 3.AO Universitaria 'Federico II' Napoli Hospital / 4.AORN 'San Giuseppe Moscati' Avellino Hospital / 5.AO 'San Pio - presidio G. Rummo' Benevento Hospital / 6.AO 'Sant'Anna e San Sebastiano' Caserta Hospital / 7.PO 'Maria Santissima Addolorata' Eboli Hospital / 8.Biogem Istituto di Ricerche Genetiche | 1. Genome Research Center for Health (CRGS) / 2. Laboratory of Molecular Medicine and Genomics(LMMGe) / 3. Center for Research in Pure and Applied Mathematics (CRMPA) | Giorgio Giurato (Corresponding Author), Francesca Rizzo (Corresponding Author), Alessandro Weisz (Corresponding Author), Gianluigi Franci, Giovanni Nassa, Pasquale Pagliano, Roberta Tarallo, Elena Alexandrova, Ylenia D'Agostino, Carlo Ferravante, Jessica Lamberti, Viola Melone, Domenico Memoli, Valeria Mirici Cappa, Domenico Palumbo, Giovanni Pecoraro, Assunta Sellitto, Oriana Strianese, Ilaria Terenzi, Giuseppe Fenza, Aniello Gentile, Antonello Saccomanno, Sonia Amabile, Teresa Rocco, Annamaria Salvati, Emilia Vaccaro, Massimiliano Galdiero, Michele Cennamo, Giuseppe Portella, Maria Grazia Foti, Mariarosaria Ingino, Maria Landi, Maurizio Fumi, Vincenzo Rocco, Rita Greco, Vittoria Letizia, Arnolfo Petruzziello, Maddalena Schioppa, Gregorio Goffredi, Francesca Marciano, Michele Caraglia, Alessia Cossu, Marianna Scrima |
| EPI_ISL_936588, EPI_ISL_936858, EPI_ISL_936859, EPI_ISL_936860 | Northwestern Memorial Hospital | Ozer Lab | Ramon Lorenzo-Redondo, Lacy M. Simons, Chad J. Achenbach, Lawrence J. Jennings, Michael G. Ison, Judd F. Hultquist, Egon A. Ozer |
| EPI_ISL_937068, EPI_ISL_937083, EPI_ISL_937108, EPI_ISL_937118 | Quest Diagnostics | Quest Diagnostics | Rosenthal,S.H., Gerasimova,A., Kagan,R.M., Anderson, B., Livingston, K.E., Hua, M., Liu Y., Shalhout, D.F., Owen, R., Lacbawan, F. |
| EPI_ISL_940936, EPI_ISL_940937, EPI_ISL_940938, EPI_ISL_940939, EPI_ISL_940940, EPI_ISL_940941, EPI_ISL_940980 | Centers for Disease Control and Prevention, Dengue Branch | Centers for Disease Control and Prevention, Dengue Branch | Gilberto A. Santiago, Glenda Gonzalez, Betzabel Flores, Keyla Charriez, Gabriela Paz-Bailey, Jorge L. Munoz-Jordan |
| EPI_ISL_941185, EPI_ISL_941186, EPI_ISL_941187, EPI_ISL_941188, EPI_ISL_941189, EPI_ISL_941190, EPI_ISL_941191, EPI_ISL_941192, EPI_ISL_941193, EPI_ISL_941194, EPI_ISL_941195, EPI_ISL_941196, EPI_ISL_941197, EPI_ISL_941198, EPI_ISL_941211 | | | |
| see above | Servicio de Microbiología. Hospital Clínico Universitario de Valencia | SeqCOVID-SPAIN consortium/IBV(CSIC) | David Navarro Ortega, Eliseo Albert Vicent, Ignacio Torres and SeqCOVID-SPAIN consortium |
| EPI_ISL_943949 | Utah Public Health Laboratory | Utah Public Health Laboratory | Erin L. Young, Kelly F. Oakeson, Tara Gallagher |
| EPI_ISL_943981, EPI_ISL_943982, EPI_ISL_943983, EPI_ISL_943991 | LACEN do Estado de Tocantins | Instituto Adolfo Lutz, Interdiciplinary Procedures Center, Strategic Laboratory | Claudio Tavares Sacchi, Claudia Regina Gonçalves, Erica Valessa Ramos Gomes, Karoline Rodrigues Campos |
| EPI_ISL_949230 | Departamento de Microbiología, CDB, Hospital Clínic, Barcelona | SeqCOVID-SPAIN consortium/IBV(CSIC) | Andrea Vergara, Mikel Martínez, Elisa Rubio, Jéssica Navero, Aida Peiró and SeqCOVID-SPAIN consortium |
| EPI_ISL_954186 | 1.AO Universitaria 'S. Giovanni di Dio e Ruggi D'Aragona, Scuola Medica Salernitana' Hospital / 2.UOC di Virologia e Microbiologia, Università della Campania 'L. Vanvitelli' / 3.AO Universitaria 'Federico II' Napoli Hospital / 4.AORN 'San Giuseppe Moscati' Avellino Hospital / 5.AO 'San Pio - presidio G. Rummo' Benevento Hospital / 6.AO 'Sant'Anna e San Sebastiano' Caserta Hospital / 7.PO 'Maria Santissima Addolorata' Eboli Hospital / 8.Biogem Istituto di Ricerche Genetiche | 1. Genome Research Center for Health (CRGS) / 2. Laboratory of Molecular Medicine and Genomics(LMMGe) / 3. Center for Research in Pure and Applied Mathematics (CRMPA) | Giorgio Giurato, Francesca Rizzo, Alessandro Weisz, Gianluigi Franci, Giovanni Nassa, Pasquale Pagliano, Roberta Tarallo, Elena Alexandrova, Ylenia D'Agostino, Carlo Ferravante, Jessica Lamberti, Viola Melone, Domenico Memoli, Valeria Mirici Cappa, Domenico Palumbo, Giovanni Pecoraro, Assunta Sellitto, Oriana Strianese, Ilaria Terenzi, Giuseppe Fenza, Aniello Gentile, Antonello Saccomanno, Sonia Amabile, Teresa Rocco, Annamaria Salvati, Emilia Vaccaro, Massimiliano Galdiero, Michele Cennamo, Giuseppe Portella, Maria Grazia Foti, Mariarosaria Ingino, Maria Landi, Maurizio Fumi, Vincenzo Rocco, Rita Greco, Vittoria Letizia, Arnolfo Petruzziello, Maddalena Schioppa, Gregorio Goffredi, Francesca Marciano, Michele Caraglia, Alessia Cossu, Marianna Scrima, Edmondo Adorisio, Morena D'Avenia, Michela Iacobellis, Rosanna Piluscio, Giorgio Dirani, Vittorio Sambri, Simona Semprini, Silvia Zanoli, Francesco Curcio, Stefania Marzinotto, Andreina Baj, Fausto Sessa. |
| EPI_ISL_960136 | Groote Schuur Hospital wc GSH | National Health Laboratory Service/UCT | Arash Iranzadeh, Deelan Doolabh, Lynn Tyers, Bruna Galvao, Innocent Mudau, Marvin Hsiao, Kruger Marais, Diana Hardie, Stephen Korsman, Carolyn Williamson |
| EPI_ISL_960137 | Mitchells Plain Hospital wc MPH | National Health Laboratory Service/UCT | Arash Iranzadeh, Deelan Doolabh, Lynn Tyers, Bruna Galvao, Innocent Mudau, Marvin Hsiao, Kruger Marais, Diana Hardie, Stephen Korsman, Carolyn Williamson |
| EPI_ISL_960138 | Victoria Hospital wc VHW | National Health Laboratory Service/UCT | Arash Iranzadeh, Deelan Doolabh, Lynn Tyers, Bruna Galvao, Innocent Mudau, Marvin Hsiao, Kruger Marais, Diana Hardie, Stephen Korsman, Carolyn Williamson |
| EPI_ISL_960139 | Thembalethu CDC wc THC | National Health Laboratory Service/UCT | Arash Iranzadeh, Deelan Doolabh, Lynn Tyers, Bruna Galvao, Innocent Mudau, Marvin Hsiao, Kruger Marais, Diana Hardie, Stephen Korsman, Carolyn Williamson |
| EPI_ISL_960140 | Groote Schuur Hospital wc GSH | National Health Laboratory Service/UCT | Arash Iranzadeh, Deelan Doolabh, Lynn Tyers, Bruna Galvao, Innocent Mudau, Marvin Hsiao, Kruger Marais, Diana Hardie, Stephen Korsman, Carolyn Williamson |
| EPI_ISL_960353, EPI_ISL_960388, EPI_ISL_960390 | University of Wisconsin-Madison AIDS Vaccine Research Laboratories | University of Wisconsin-Madison AIDS Vaccine Research Laboratories | Gage Moreno, Katarina Braun, et al. AIDS Vaccine Research Laboratories |
| EPI_ISL_961890 | Illinois Department of Public Health | Gagnon Lab, Southern Illinois University | Keith Gagnon |
| EPI_ISL_965179 | Laboratorio Biologia Molecolare SARS-COV2- UOC Laboratorio Analisi- Servizio Medicina di Laboratorio, Ospedale San Francesco - ATS-ASSLNUoro | LABORATORIO SPECIALISTICO UOC EMATOLOGIA-Ospedale San Francesco - ATS-ASSLNUoro | SULIS VINCENZO, PIRAS GIOVANNA, ASPRONI ROSANNA, FANCELLO TATIANA, MONNE MARIA ITRIA, FIAMMA MAURA, TOJA ALESSANDRO, SANNA FILOMENA, FLORE ANNA RITA. PALMAS ANGELO DOMENICO, IANA LO MAGLIO, MAMELI GIUSEPPE |

| | | | |
|---|---|---|---|
| EPI_ISL_965303 | NYU Langone Health | Departments of Pathology and Medicine, New York University School of Medicine | Adriana Heguy, Dacia Dimartino, Emily Guzman, Christian Marier, Peter Meyn, Sitharam Ramaswami, Gael Westby, Paul Zappile, Yutong Zhang, Paolo Cotzia, Guiqing Wang |
| EPI_ISL_965687, EPI_ISL_965717, EPI_ISL_965719 | Dutch COVID-19 response team | Medical Microbiology, Maastricht University Medical Centre | Jozef Dingemans*, Brian van der Veer*, Erik Beuken, Carmen Reumkens, Lieke van Alphen, Christian Hoebe, Paul Savelkoul |
| EPI_ISL_966327, EPI_ISL_966328, EPI_ISL_966329, EPI_ISL_966330, EPI_ISL_966331, EPI_ISL_966332 | Kentucky State Public Health Lab | Kentucky State Public Health Lab | Stephanie Lunn, Karim George, Joshua Tobias, William Grooms, Vaneet Arora, Matthew Johnson, Rachel Zinner, Rhonda Lucas |
| EPI_ISL_968883, EPI_ISL_968884, EPI_ISL_968885, EPI_ISL_968887, EPI_ISL_968889, EPI_ISL_968890, EPI_ISL_968891, EPI_ISL_968892 | KEMRI-Wellcome Trust Research Programme/KEMRI-CGMR-C Kilifi | KEMRI-Wellcome Trust Research Programme/KEMRI-CGMR-C Kilifi | Githinji et al |
| EPI_ISL_974284, EPI_ISL_974285, EPI_ISL_974286, EPI_ISL_974287, EPI_ISL_974288, EPI_ISL_974289, EPI_ISL_974290, EPI_ISL_974291, EPI_ISL_974292, EPI_ISL_974293, EPI_ISL_974294, EPI_ISL_974295, EPI_ISL_974296, EPI_ISL_974297, EPI_ISL_974298, EPI_ISL_974299, EPI_ISL_974300, EPI_ISL_974301, EPI_ISL_974302, EPI_ISL_974303, EPI_ISL_974304, EPI_ISL_974305, EPI_ISL_974306, EPI_ISL_974307, EPI_ISL_974308, EPI_ISL_974309, EPI_ISL_974310, EPI_ISL_974311, EPI_ISL_974312, EPI_ISL_974313, EPI_ISL_974314, EPI_ISL_974315, EPI_ISL_974316, EPI_ISL_974317, EPI_ISL_974318, EPI_ISL_974319, EPI_ISL_974320, EPI_ISL_974321, EPI_ISL_974322, EPI_ISL_974323, EPI_ISL_974324, EPI_ISL_974325, EPI_ISL_974326, EPI_ISL_974327, EPI_ISL_974328, EPI_ISL_974329, EPI_ISL_974330, EPI_ISL_974331, EPI_ISL_974332, EPI_ISL_974333, EPI_ISL_974334, EPI_ISL_974335, EPI_ISL_974336, EPI_ISL_974337, EPI_ISL_974338, EPI_ISL_974339, EPI_ISL_974340, EPI_ISL_974341, EPI_ISL_974342, EPI_ISL_974343, EPI_ISL_974344, EPI_ISL_974345, EPI_ISL_974346, EPI_ISL_974347, EPI_ISL_974348, EPI_ISL_974349, EPI_ISL_974350, EPI_ISL_974351, EPI_ISL_974352, EPI_ISL_974353, EPI_ISL_974354, EPI_ISL_974355, EPI_ISL_974356, EPI_ISL_974357, EPI_ISL_974358, EPI_ISL_974359, EPI_ISL_974360, EPI_ISL_974361, EPI_ISL_974362, EPI_ISL_974363, EPI_ISL_974364, EPI_ISL_974365, EPI_ISL_974366, EPI_ISL_974367, EPI_ISL_974368, EPI_ISL_974369, EPI_ISL_974370, EPI_ISL_974371, EPI_ISL_974372, EPI_ISL_974373, EPI_ISL_974374, EPI_ISL_974375, EPI_ISL_974376, EPI_ISL_974377, EPI_ISL_974378, EPI_ISL_974379, EPI_ISL_974380, EPI_ISL_974381, EPI_ISL_974382, EPI_ISL_974383, EPI_ISL_974384, EPI_ISL_974385, EPI_ISL_974386, EPI_ISL_974387, EPI_ISL_974388, EPI_ISL_974389, EPI_ISL_974390, EPI_ISL_974391, EPI_ISL_974392, EPI_ISL_974393, EPI_ISL_974394, EPI_ISL_974395, EPI_ISL_974396, EPI_ISL_974397, EPI_ISL_974398, EPI_ISL_974399, EPI_ISL_974400, EPI_ISL_974401, EPI_ISL_974402, EPI_ISL_974403, EPI_ISL_974404, EPI_ISL_974405, EPI_ISL_974406, EPI_ISL_974407, EPI_ISL_974408, EPI_ISL_974409, EPI_ISL_974410, EPI_ISL_974411, EPI_ISL_974412, EPI_ISL_974413, EPI_ISL_974414, EPI_ISL_974415, EPI_ISL_974416, EPI_ISL_974417, EPI_ISL_974418, EPI_ISL_974419, EPI_ISL_974420, EPI_ISL_974421, EPI_ISL_974422, EPI_ISL_974423, EPI_ISL_974424, EPI_ISL_974425, EPI_ISL_974426, EPI_ISL_974427, EPI_ISL_974428, EPI_ISL_974429, EPI_ISL_974430, EPI_ISL_974431, EPI_ISL_974432, EPI_ISL_974433, EPI_ISL_974434, EPI_ISL_974435, EPI_ISL_974436, EPI_ISL_974437, EPI_ISL_974438, EPI_ISL_974439, EPI_ISL_974440, EPI_ISL_974441, EPI_ISL_974442, EPI_ISL_974443, EPI_ISL_974444, EPI_ISL_974445, EPI_ISL_974446, EPI_ISL_974447, EPI_ISL_974448, EPI_ISL_974449, EPI_ISL_974450, EPI_ISL_974451, EPI_ISL_974452, EPI_ISL_974453, EPI_ISL_974454, EPI_ISL_974455, EPI_ISL_974456, EPI_ISL_974457, EPI_ISL_974458, EPI_ISL_974459, EPI_ISL_974460, EPI_ISL_974461, EPI_ISL_974462, EPI_ISL_974463, EPI_ISL_974464 | | | |
| see above | BCCDC Public Health Laboratory | BCCDC Public Health Laboratory | Prystajecky Natalie, Linda Hoang, Dan Fornika, John Tyson, Shannon Russell, Kim Macdonald, Kimia Kamelian, Ana Pacagnella, Corrinne Ng, Loretta Janz, Robert Azana Terry Snutch, Mel Krajden |
| EPI_ISL_977175 | Microbiologia e Virologia | Istituto Zooprofilattico Sperimentale delle Venezie | Adelaide Milani, Alessia Schivo, Annalisa Salviato, Erika Giorgia Quaranta, Ambra Pastori, Bianca Zecchin, Alice Fusaro, Isabella Monne, Calogero Terregino, Antonia Ricci |
| EPI_ISL_977910 | Chiu Laboratory, University of California, San Francisco | Chiu Laboratory, University of California, San Francisco | Charles Chiu, Xianding (Wayne) Deng, Candace Wang, Venice Servellita, Jill Hacker, Debra Wadford |
