## Supplementary material for "Detection and characterization of the SARS-CoV-2 lineage B.1.526 in New York": Supp. Table 3 GISAID Acknowledgment part 2: gisaid_hcov-19_acknowledgement_table_2021_02_13_00_.pdf

We gratefully acknowledge the following Authors from the Originating laboratories responsible for obtaining the specimens, as well as the Submitting laboratories where the genome data were generated and shared via GISAID, on which this research is based.

All Submitters of data may be contacted directly via www.gisaid.org

Authors are sorted alphabetically.

| Accession ID | Originating Laboratory | Submitting Laboratory | Authors |
|---|---|---|---|
| EPI_ISL_595221, EPI_ISL_595223, EPI_ISL_595225, EPI_ISL_595226, EPI_ISL_595229 | Quadram Institute Bioscience | COVID-19 Genomics UK (COG-UK) Consortium | Dave J. Baker, Gemma L. Kay, Alp Aydin, Thanh Le-Viet, Steven Rudder, Ana P. Tedim, Anastasia Kolyva, Maria Diaz, Leonardo de Oliveira Martins, Nabil-Fareed Alikhan, Lizzie Meadows, Rachael Stanley, Ngozi Elumogo, Muhammed Yasir, Nicholas M. Thomson, Alexander J Trotter, Rachel Gilroy, Samuel Bloomfield, Claire Stuart, Andrew Bell, Reenesh Prakash, Samir Dervisevic, Alison E. Mather, John Wain, Mark Webber, Andrew J. Page, Justin O'Grady |
| EPI_ISL_595307, EPI_ISL_595308, EPI_ISL_595309, EPI_ISL_595310, EPI_ISL_595311, EPI_ISL_595312, EPI_ISL_595313, EPI_ISL_595314, EPI_ISL_595315, EPI_ISL_595316, EPI_ISL_595323, EPI_ISL_595324, EPI_ISL_595325 | | | |
| see above | Queens Medical Centre, Clinical Microbiology Department / DeepSeq Nottingham | COVID-19 Genomics UK (COG-UK) Consortium | Gemma Clark, Wendy Smith, Manjinder Khakh, Vicki M Fleming, Michelle M Lister, Hannah Howson-Wells, Jonathan Ball, Patrick McClure, Joseph Chappell, Theocharis Tsoleridis, Nadine Holmes, Matthew Carlisle, Christopher Moore, Fei Sang, Johnny Debebe, Victoria Wright, Matthew Loose |
| EPI_ISL_605818, EPI_ISL_605820, EPI_ISL_605821, EPI_ISL_605822, EPI_ISL_605823, EPI_ISL_605824 | National Public Health Laboratory, National Centre for Infectious Diseases | National Public Health Laboratory, National Centre for Infectious Diseases | Tze Minn Mak, Sophie Octavia, Zhenyang Zhou, Lin Cui, Raymond Tzer Pin Lin |
| EPI_ISL_605825, EPI_ISL_605836, EPI_ISL_605842, EPI_ISL_605850, EPI_ISL_605853, EPI_ISL_605854, EPI_ISL_605857, EPI_ISL_605867 | PathWest Laboratory Medicine WA | PathWest Laboratory Medicine WA Microbial Surveillance Unit | PathWest Laboratory Medicine WA Microbial Surveillance Unit |
| EPI_ISL_609904 | Respiratory Virus Unit, Microbiology Services Colindale, Public Health England | Respiratory Virus Unit, Microbiology Services Colindale, Public Health England | PHE Covid Sequencing Team |
| EPI_ISL_609998 | INMI Lazzaro Spallanzani IRCCS | INMI Lazzaro Spallanzani IRCCS | F Messina, B Bartolini, M Rueca, C.E.M Gruber, E Giombini, A Di Caro, MR Capobianchi |
| EPI_ISL_609999 | INMI Lazzaro Spallanzani IRCCS | INMI Lazzaro Spallanzani IRCCS | B Bartolini, M Rueca, C.E.M Gruber, F Messina, E Giombini, MR Capobianchi, A Di Caro |
| EPI_ISL_610922, EPI_ISL_610923, EPI_ISL_610924, EPI_ISL_610925, EPI_ISL_610926, EPI_ISL_610927, EPI_ISL_610928, EPI_ISL_610929, EPI_ISL_610930, EPI_ISL_610931, EPI_ISL_610932, EPI_ISL_610933, EPI_ISL_610934, EPI_ISL_610935, EPI_ISL_610936, EPI_ISL_610937, EPI_ISL_610938, EPI_ISL_610939, EPI_ISL_610940, EPI_ISL_610941, EPI_ISL_610942, EPI_ISL_610943, EPI_ISL_610944, EPI_ISL_610945, EPI_ISL_610946, EPI_ISL_610947, EPI_ISL_610948, EPI_ISL_610949, EPI_ISL_610950, EPI_ISL_610951, EPI_ISL_610952, EPI_ISL_610953, EPI_ISL_610954, EPI_ISL_610955, EPI_ISL_610956, EPI_ISL_610957, EPI_ISL_610958, EPI_ISL_610959, EPI_ISL_610960, EPI_ISL_610961, EPI_ISL_610962, EPI_ISL_610963, EPI_ISL_610964, EPI_ISL_610965, EPI_ISL_610966, EPI_ISL_610967, EPI_ISL_610968, EPI_ISL_610969, EPI_ISL_610970, EPI_ISL_610971, EPI_ISL_610972, EPI_ISL_610973, EPI_ISL_610974, EPI_ISL_610975, EPI_ISL_610976, EPI_ISL_610977, EPI_ISL_610978, EPI_ISL_610979, EPI_ISL_610980, EPI_ISL_610981, EPI_ISL_610982, EPI_ISL_610983, EPI_ISL_610984, EPI_ISL_610985, EPI_ISL_610986, EPI_ISL_610987, EPI_ISL_610988, EPI_ISL_610989, EPI_ISL_610990, EPI_ISL_610991, EPI_ISL_610992, EPI_ISL_610993, EPI_ISL_610994, EPI_ISL_610995, EPI_ISL_610996, EPI_ISL_610997, EPI_ISL_610998, EPI_ISL_610999, EPI_ISL_611000, EPI_ISL_611001, EPI_ISL_611002, EPI_ISL_611003, EPI_ISL_611004, EPI_ISL_611005, EPI_ISL_611006, EPI_ISL_611007, EPI_ISL_611008, EPI_ISL_611009, EPI_ISL_611010, EPI_ISL_611011, EPI_ISL_611012, EPI_ISL_611013, EPI_ISL_611014, EPI_ISL_611015, EPI_ISL_611016, EPI_ISL_611017, EPI_ISL_611018, EPI_ISL_611019, EPI_ISL_611020, EPI_ISL_611021, EPI_ISL_611022, EPI_ISL_611023, EPI_ISL_611024, EPI_ISL_611025, EPI_ISL_611026, EPI_ISL_611027, EPI_ISL_611028, EPI_ISL_611029, EPI_ISL_611030, EPI_ISL_611031, EPI_ISL_611032, EPI_ISL_611033, EPI_ISL_611034, EPI_ISL_611035, EPI_ISL_611036, EPI_ISL_611037, EPI_ISL_611038, EPI_ISL_611039, EPI_ISL_611040, EPI_ISL_611041, EPI_ISL_611042, EPI_ISL_611043, EPI_ISL_611044, EPI_ISL_611045, EPI_ISL_611046, EPI_ISL_611047, EPI_ISL_611048, EPI_ISL_611049, EPI_ISL_611050, EPI_ISL_611051, EPI_ISL_611052, EPI_ISL_611053, EPI_ISL_611054, EPI_ISL_611055, EPI_ISL_611056, EPI_ISL_611057, EPI_ISL_611058, EPI_ISL_611059, EPI_ISL_611060, EPI_ISL_611061, EPI_ISL_611062, EPI_ISL_611063, EPI_ISL_611064, EPI_ISL_611065, EPI_ISL_611066, EPI_ISL_611067, EPI_ISL_611068, EPI_ISL_611069, EPI_ISL_611070, EPI_ISL_611071, EPI_ISL_611072, EPI_ISL_611073, EPI_ISL_611074, EPI_ISL_611075, EPI_ISL_611076, EPI_ISL_611077, EPI_ISL_611078, EPI_ISL_611079, EPI_ISL_611080, EPI_ISL_611081, EPI_ISL_611082, EPI_ISL_611083, EPI_ISL_611084, EPI_ISL_611085, EPI_ISL_611086, EPI_ISL_611087, EPI_ISL_611088, EPI_ISL_611089, EPI_ISL_611090, EPI_ISL_611091, EPI_ISL_611092 | | | |
| see above | Lighthouse Lab in Alderley Park | Wellcome Sanger Institute for the COVID-19 Genomics UK (COG-UK) consortium | Jacquelyn Wynn, Mairead Hyland, The Lighthouse Lab in Alderley Park and Alex Alderton, Roberto Amato, Sonia Goncalves, Ewan Harrison, David K. Jackson, Ian Johnston, Dominic Kwiatkowski, Cordelia Langford, John Sillitoe on behalf of the Wellcome Sanger Institute COVID-19 Surveillance Team (http://www.sanger.ac.uk/covid-team) |
| EPI_ISL_611093 | Lighthouse Lab in Alderley Park | Wellcome Sanger Institute for the COVID-19 Genomics UK (COG-UK) Consortium | Jacquelyn Wynn, Mairead Hyland, The Lighthouse Lab in Alderley Park and Alex Alderton, Roberto Amato, Sonia Goncalves, Ewan Harrison, David K. Jackson, Ian Johnston, Dominic Kwiatkowski, Cordelia Langford, John Sillitoe on behalf of the Wellcome Sanger Institute COVID-19 Surveillance Team (http://www.sanger.ac.uk/covid-team) |
| EPI_ISL_611094, EPI_ISL_611095, EPI_ISL_611096, EPI_ISL_611097, EPI_ISL_611098, EPI_ISL_611099, EPI_ISL_611100, EPI_ISL_611101, EPI_ISL_611102, EPI_ISL_611103, EPI_ISL_611104, EPI_ISL_611105, EPI_ISL_611106, EPI_ISL_611107, EPI_ISL_611108, EPI_ISL_611109, EPI_ISL_611110, EPI_ISL_611111, EPI_ISL_611112, EPI_ISL_611113, EPI_ISL_611114, EPI_ISL_611115, EPI_ISL_611116, EPI_ISL_611117, EPI_ISL_611118, EPI_ISL_611119, EPI_ISL_611120, EPI_ISL_611121, EPI_ISL_611122, EPI_ISL_611123, EPI_ISL_611124, EPI_ISL_611125, EPI_ISL_611126, EPI_ISL_611127, EPI_ISL_611128, EPI_ISL_611129, EPI_ISL_611130, EPI_ISL_611131, EPI_ISL_611132, EPI_ISL_611133, EPI_ISL_611134, EPI_ISL_611135, EPI_ISL_611136, EPI_ISL_611137, EPI_ISL_611138, EPI_ISL_611139, EPI_ISL_611140, EPI_ISL_611141, EPI_ISL_611142, EPI_ISL_611143, EPI_ISL_611144, EPI_ISL_611145, EPI_ISL_611146, EPI_ISL_611147, EPI_ISL_611148, EPI_ISL_611149, EPI_ISL_611150, EPI_ISL_611151, EPI_ISL_611152, EPI_ISL_611153, EPI_ISL_611154, EPI_ISL_611155, EPI_ISL_611156, EPI_ISL_611157, EPI_ISL_611158, EPI_ISL_611159, EPI_ISL_611160, EPI_ISL_611161, EPI_ISL_611162, EPI_ISL_611163, EPI_ISL_611164, EPI_ISL_611165, EPI_ISL_611166, EPI_ISL_611167, EPI_ISL_611168, EPI_ISL_611169, EPI_ISL_611170, EPI_ISL_611171, EPI_ISL_611172, EPI_ISL_611173, EPI_ISL_611174, EPI_ISL_611175, EPI_ISL_611176, EPI_ISL_611177, EPI_ISL_611178, EPI_ISL_611179, EPI_ISL_611180, EPI_ISL_611181, EPI_ISL_611182, EPI_ISL_611183, EPI_ISL_611184, EPI_ISL_611185, EPI_ISL_611186, EPI_ISL_611187, EPI_ISL_611188, EPI_ISL_611189, EPI_ISL_611190, EPI_ISL_611191, EPI_ISL_611192, EPI_ISL_611193, EPI_ISL_611194, EPI_ISL_611195, EPI_ISL_611196, EPI_ISL_611197, EPI_ISL_611198, EPI_ISL_611199, EPI_ISL_611200, EPI_ISL_611201, EPI_ISL_611202, EPI_ISL_611203, EPI_ISL_611204, EPI_ISL_611205, EPI_ISL_611206, EPI_ISL_611207, EPI_ISL_611208, EPI_ISL_611209, EPI_ISL_611210, EPI_ISL_611211, EPI_ISL_611212, EPI_ISL_611213, EPI_ISL_611214, EPI_ISL_611215, EPI_ISL_611216, EPI_ISL_611217, EPI_ISL_611218, EPI_ISL_611219, EPI_ISL_611220, EPI_ISL_611221, EPI_ISL_611222, EPI_ISL_611223, EPI_ISL_611224, EPI_ISL_611225, EPI_ISL_611226, EPI_ISL_611227, EPI_ISL_611228, EPI_ISL_611229, EPI_ISL_611230, EPI_ISL_611231, EPI_ISL_611232, EPI_ISL_611233, EPI_ISL_611234, EPI_ISL_611235, EPI_ISL_611236, EPI_ISL_611237, EPI_ISL_611238, EPI_ISL_611239, EPI_ISL_611240, EPI_ISL_611241, EPI_ISL_611242, EPI_ISL_611243, EPI_ISL_611244, EPI_ISL_611245, EPI_ISL_611246, EPI_ISL_611247, EPI_ISL_611248, EPI_ISL_611249, EPI_ISL_611250, EPI_ISL_611251, EPI_ISL_611252, EPI_ISL_611253, EPI_ISL_611254, EPI_ISL_611255, EPI_ISL_611256, EPI_ISL_611257, EPI_ISL_611258, EPI_ISL_611259, EPI_ISL_611260, EPI_ISL_611261, EPI_ISL_611262, EPI_ISL_611263, EPI_ISL_611264, EPI_ISL_611265, EPI_ISL_611266, EPI_ISL_611267, EPI_ISL_611268, EPI_ISL_611269, EPI_ISL_611270, EPI_ISL_611271, EPI_ISL_611272, EPI_ISL_611273, EPI_ISL_611274, EPI_ISL_611275, EPI_ISL_611276, EPI_ISL_611277, EPI_ISL_611278, EPI_ISL_611279, EPI_ISL_611280, EPI_ISL_611281, EPI_ISL_611282, EPI_ISL_611283, EPI_ISL_611284, EPI_ISL_611285, EPI_ISL_611286 | | | |
| see above | Lighthouse Lab in Alderley Park | Wellcome Sanger Institute for the COVID-19 Genomics UK (COG-UK) consortium | Jacquelyn Wynn, Mairead Hyland, The Lighthouse Lab in Alderley Park and Alex Alderton, Roberto Amato, Sonia Goncalves, Ewan Harrison, David K. Jackson, Ian Johnston, Dominic Kwiatkowski, Cordelia Langford, John Sillitoe on behalf of the Wellcome Sanger Institute COVID-19 Surveillance Team (http://www.sanger.ac.uk/covid-team) |
| EPI_ISL_611287, EPI_ISL_611288, EPI_ISL_611289, EPI_ISL_611290, EPI_ISL_611291, EPI_ISL_611292, EPI_ISL_611293, EPI_ISL_611294, EPI_ISL_611295, EPI_ISL_611296, EPI_ISL_611297, EPI_ISL_611298, EPI_ISL_611299, EPI_ISL_611300, EPI_ISL_611301, EPI_ISL_611302, EPI_ISL_611303 | | | |
| see above | Lighthouse Lab in Glasgow | Wellcome Sanger Institute for the COVID-19 Genomics UK (COG-UK) consortium | Harper VanSteenhouse, Yumi Kasai, David Gray, Carol Clugston, Anna Dominiczak and Alex Alderton, Roberto Amato, Sonia Goncalves, Ewan Harrison, David K. Jackson, Ian Johnston, Dominic Kwiatkowski, Cordelia Langford, John Sillitoe on behalf of the Wellcome Sanger Institute COVID-19 Surveillance Team (http://www.sanger.ac.uk/covid-team) |
| EPI_ISL_611304 | Lighthouse Lab in Glasgow | Wellcome Sanger Institute for the COVID-19 Genomics UK (COG-UK) Consortium | Harper VanSteenhouse, Yumi Kasai, David Gray, Carol Clugston, Anna Dominiczak and Alex Alderton, Roberto Amato, Sonia Goncalves, Ewan Harrison, David K. Jackson, Ian Johnston, Dominic Kwiatkowski, Cordelia Langford, John Sillitoe on behalf of the Wellcome Sanger Institute COVID-19 Surveillance Team (http://www.sanger.ac.uk/covid-team) |
| EPI_ISL_611305, EPI_ISL_611306, EPI_ISL_611307, EPI_ISL_611308, EPI_ISL_611309, EPI_ISL_611310, EPI_ISL_611311, EPI_ISL_611312, EPI_ISL_611313, EPI_ISL_611314, EPI_ISL_611315, EPI_ISL_611316, EPI_ISL_611317, EPI_ISL_611318, EPI_ISL_611319, EPI_ISL_611320, EPI_ISL_611321, EPI_ISL_611322, EPI_ISL_611323, EPI_ISL_611324, EPI_ISL_611325, EPI_ISL_611326, EPI_ISL_611327, EPI_ISL_611328, EPI_ISL_611329, EPI_ISL_611330, EPI_ISL_611331, EPI_ISL_611332, EPI_ISL_611333, EPI_ISL_611334, EPI_ISL_611335, EPI_ISL_611336, EPI_ISL_611337, EPI_ISL_611338, EPI_ISL_611339, EPI_ISL_611340, EPI_ISL_611341, EPI_ISL_611342, EPI_ISL_611343, EPI_ISL_611344, EPI_ISL_611345, EPI_ISL_611346, EPI_ISL_611347, EPI_ISL_611348, EPI_ISL_611349, EPI_ISL_611350, EPI_ISL_611351, EPI_ISL_611352, EPI_ISL_611353, EPI_ISL_611354, EPI_ISL_611355, EPI_ISL_611356, EPI_ISL_611357, EPI_ISL_611358, EPI_ISL_611359, EPI_ISL_611360, EPI_ISL_611361, EPI_ISL_611362, EPI_ISL_611363, EPI_ISL_611364, EPI_ISL_611365, EPI_ISL_611366, EPI_ISL_611367, EPI_ISL_611368, EPI_ISL_611369, EPI_ISL_611370, EPI_ISL_611371, EPI_ISL_611372, EPI_ISL_611373, EPI_ISL_611374, EPI_ISL_611375, EPI_ISL_611376, EPI_ISL_611377 | | | |
| see above | Lighthouse Lab in Glasgow | Wellcome Sanger Institute for the COVID-19 Genomics UK (COG-UK) consortium | Harper VanSteenhouse, Yumi Kasai, David Gray, Carol Clugston, Anna Dominiczak and Alex Alderton, Roberto Amato, Sonia Goncalves, Ewan Harrison, David K. Jackson, Ian Johnston, Dominic Kwiatkowski, Cordelia Langford, John Sillitoe on behalf of the Wellcome Sanger Institute COVID-19 Surveillance Team (http://www.sanger.ac.uk/covid-team) |
| EPI_ISL_611378 | Lighthouse Lab in Glasgow | Wellcome Sanger Institute for the COVID-19 Genomics UK (COG-UK) Consortium | Harper VanSteenhouse, Yumi Kasai, David Gray, Carol Clugston, Anna Dominiczak and Alex Alderton, Roberto Amato, Sonia Goncalves, Ewan Harrison, David K. Jackson, Ian Johnston, Dominic Kwiatkowski, Cordelia Langford, John Sillitoe on behalf of the Wellcome Sanger Institute COVID-19 Surveillance Team (http://www.sanger.ac.uk/covid-team) |

EPI_ISL_611379, EPI_ISL_611380, EPI_ISL_611381, EPI_ISL_611382, EPI_ISL_611383, EPI_ISL_611384, EPI_ISL_611385, EPI_ISL_611386, EPI_ISL_611387, EPI_ISL_611388, EPI_ISL_611389, EPI_ISL_611390, EPI_ISL_611391, EPI_ISL_611392, EPI_ISL_611393, EPI_ISL_611394, EPI_ISL_611395, EPI_ISL_611396, EPI_ISL_611397, EPI_ISL_611398, EPI_ISL_611399, EPI_ISL_611400, EPI_ISL_611401, EPI_ISL_611402, EPI_ISL_611403, EPI_ISL_611404, EPI_ISL_611405, EPI_ISL_611406, EPI_ISL_611407, EPI_ISL_611408, EPI_ISL_611409, EPI_ISL_611410, EPI_ISL_611411, EPI_ISL_611412, EPI_ISL_611413, EPI_ISL_611414, EPI_ISL_611415, EPI_ISL_611416, EPI_ISL_611417, EPI_ISL_611418, EPI_ISL_611419, EPI_ISL_611420, EPI_ISL_611421, EPI_ISL_611423, EPI_ISL_611424, EPI_ISL_611425, EPI_ISL_611427, EPI_ISL_611428, EPI_ISL_611429, EPI_ISL_611430, EPI_ISL_611431, EPI_ISL_611432, EPI_ISL_611433, EPI_ISL_611434, EPI_ISL_611435, EPI_ISL_611436, EPI_ISL_611437, EPI_ISL_611438, EPI_ISL_611439, EPI_ISL_611440, EPI_ISL_611441, EPI_ISL_611442, EPI_ISL_611443, EPI_ISL_611444, EPI_ISL_611445, EPI_ISL_611446, EPI_ISL_611447, EPI_ISL_611448, EPI_ISL_611449, EPI_ISL_611450, EPI_ISL_611451, EPI_ISL_611452, EPI_ISL_611453, EPI_ISL_611454, EPI_ISL_611455, EPI_ISL_611456, EPI_ISL_611457, EPI_ISL_611458, EPI_ISL_611459, EPI_ISL_611460, EPI_ISL_611461, EPI_ISL_611462, EPI_ISL_611463, EPI_ISL_611464, EPI_ISL_611465, EPI_ISL_611466, EPI_ISL_611467, EPI_ISL_611468, EPI_ISL_611469, EPI_ISL_611470, EPI_ISL_611471, EPI_ISL_611472, EPI_ISL_611473, EPI_ISL_611474, EPI_ISL_611475, EPI_ISL_611476, EPI_ISL_611477, EPI_ISL_611478, EPI_ISL_611479, EPI_ISL_611480, EPI_ISL_611481, EPI_ISL_611482, EPI_ISL_611483, EPI_ISL_611484

| | | | |
|---|---|---|---|
| see above | Lighthouse Lab in Glasgow | Wellcome Sanger Institute for the COVID-19 Genomics UK (COG-UK) consortium | Harper VanSteenhouse, Yumi Kasai, David Gray, Carol Clugston, Anna Dominiczak and Alex Alderton, Roberto Amato, Sonia Goncalves, Ewan Harrison, David K. Jackson, Ian Johnston, Dominic Kwiatkowski, Cordelia Langford, John Sillitoe on behalf of the Wellcome Sanger Institute COVID-19 Surveillance Team (http://www.sanger.ac.uk/covid-team) |
| EPI_ISL_611519 | Wales Specialist Virology Centre Sequencing lab: Pathogen Genomics Unit | COVID-19 Genomics UK (COG-UK) Consortium | Catherine Moore, Johnathan Evans, Laura Gifford, Malorie Perry, Simon Cottrell, Angela Marchbank, Alec Birchley, Alexander Adams, Amy Gaskin, Bree Gatica-Wilcox, Jason Coombes, Joel Southgate, Lauren Gilbert, Lee Graham, Nicole Pacchiarini, Sara Kumziene-Summerhayes, Sarah Taylor, Sophie Jones, Sara Rey, Matthew Bull, Joanne Watkins, Sally Corden, Tom Connor |
| EPI_ISL_611520 | Virology Department, Sheffield Teaching Hospitals NHS Foundation Trust/Department of Infection, Immunity and Cardiovascular Disease, The Medical School, University of Sheffield | COVID-19 Genomics UK (COG-UK) Consortium | Thushan de Silva, Matthew Parker, Nikki Smith, Adri Angyal, Rebecca Brown, Luke Green, Rachel Tucker, Paul Parsons, Danielle Groves, Katie Johnson, Laura Carrilero, Alex Keeley, Dave Partridge, Matthew Wyles, Benjamin Lindsey, Mehmet Yavuz, Mohammad Raza, Cariad Evans |
| EPI_ISL_611527 | University College London, Great Ormond Street Hospital for Children NHS Foundation Trust, Imperial College Healthcare NHS Trust | COVID-19 Genomics UK (COG-UK) Consortium | Sergi Castellano, Rachel Williams, Mark Kristiansen, Paola Resende Silva, Sunando Roy, Tony Brooks, Helena Tutill, Paola Niola, Patricia Dyal, Charlotte Williams, Leysa Forrest, Yasmin Panchbhaya, Jacqueline Findlay, Samuel Weeks, Julianne Brown, Kathryn Harris, Paul Randell, James Price, Alison Holmes, Judith Breuer |
| EPI_ISL_611529, EPI_ISL_611530, EPI_ISL_611534 | Wales Specialist Virology Centre Sequencing lab: Pathogen Genomics Unit | COVID-19 Genomics UK (COG-UK) Consortium | Catherine Moore, Johnathan Evans, Laura Gifford, Malorie Perry, Simon Cottrell, Angela Marchbank, Alec Birchley, Alexander Adams, Amy Gaskin, Bree Gatica-Wilcox, Jason Coombes, Joel Southgate, Lauren Gilbert, Lee Graham, Nicole Pacchiarini, Sara Kumziene-Summerhayes, Sarah Taylor, Sophie Jones, Sara Rey, Matthew Bull, Joanne Watkins, Sally Corden, Tom Connor |
| EPI_ISL_611546 | Queens Medical Centre, Clinical Microbiology Department / DeepSeq Nottingham | COVID-19 Genomics UK (COG-UK) Consortium | Gemma Clark, Wendy Smith, Manjinder Khakh, Vicki M Fleming, Michelle M Lister, Hannah Howson-Wells, Jonathan Ball, Patrick McClure, Joseph Chappell, Theocharis Tsoleridis, Nadine Holmes, Matthew Carlisle, Christopher Moore, Fei Sang, Johnny Debebe, Victoria Wright, Matthew Loose |
| EPI_ISL_611558 | University College London, Great Ormond Street Hospital for Children NHS Foundation Trust, Imperial College Healthcare NHS Trust | COVID-19 Genomics UK (COG-UK) Consortium | Sergi Castellano, Rachel Williams, Mark Kristiansen, Paola Resende Silva, Sunando Roy, Tony Brooks, Helena Tutill, Paola Niola, Patricia Dyal, Charlotte Williams, Leysa Forrest, Yasmin Panchbhaya, Jacqueline Findlay, Samuel Weeks, Julianne Brown, Kathryn Harris, Paul Randell, James Price, Alison Holmes, Judith Breuer |
| EPI_ISL_611561, EPI_ISL_611563, EPI_ISL_611566, EPI_ISL_611581 | Wales Specialist Virology Centre Sequencing lab: Pathogen Genomics Unit | COVID-19 Genomics UK (COG-UK) Consortium | Catherine Moore, Johnathan Evans, Laura Gifford, Malorie Perry, Simon Cottrell, Angela Marchbank, Alec Birchley, Alexander Adams, Amy Gaskin, Bree Gatica-Wilcox, Jason Coombes, Joel Southgate, Lauren Gilbert, Lee Graham, Nicole Pacchiarini, Sara Kumziene-Summerhayes, Sarah Taylor, Sophie Jones, Sara Rey, Matthew Bull, Joanne Watkins, Sally Corden, Tom Connor |
| EPI_ISL_611587, EPI_ISL_611598 | Virology Department, Sheffield Teaching Hospitals NHS Foundation Trust/Department of Infection, Immunity and Cardiovascular Disease, The Medical School, University of Sheffield | COVID-19 Genomics UK (COG-UK) Consortium | Thushan de Silva, Matthew Parker, Nikki Smith, Adri Angyal, Rebecca Brown, Luke Green, Rachel Tucker, Paul Parsons, Danielle Groves, Katie Johnson, Laura Carrilero, Alex Keeley, Dave Partridge, Matthew Wyles, Benjamin Lindsey, Mehmet Yavuz, Mohammad Raza, Cariad Evans |
| EPI_ISL_611601 | Wales Specialist Virology Centre Sequencing lab: Pathogen Genomics Unit | COVID-19 Genomics UK (COG-UK) Consortium | Catherine Moore, Johnathan Evans, Laura Gifford, Malorie Perry, Simon Cottrell, Angela Marchbank, Alec Birchley, Alexander Adams, Amy Gaskin, Bree Gatica-Wilcox, Jason Coombes, Joel Southgate, Lauren Gilbert, Lee Graham, Nicole Pacchiarini, Sara Kumziene-Summerhayes, Sarah Taylor, Sophie Jones, Sara Rey, Matthew Bull, Joanne Watkins, Sally Corden, Tom Connor |
| EPI_ISL_611606 | Virology Department, Sheffield Teaching Hospitals NHS Foundation Trust/Department of Infection, Immunity and Cardiovascular Disease, The Medical School, University of Sheffield | COVID-19 Genomics UK (COG-UK) Consortium | Thushan de Silva, Matthew Parker, Nikki Smith, Adri Angyal, Rebecca Brown, Luke Green, Rachel Tucker, Paul Parsons, Danielle Groves, Katie Johnson, Laura Carrilero, Alex Keeley, Dave Partridge, Matthew Wyles, Benjamin Lindsey, Mehmet Yavuz, Mohammad Raza, Cariad Evans |
| EPI_ISL_611624, EPI_ISL_611625, EPI_ISL_611640, EPI_ISL_611652 | Wales Specialist Virology Centre Sequencing lab: Pathogen Genomics Unit | COVID-19 Genomics UK (COG-UK) Consortium | Catherine Moore, Johnathan Evans, Laura Gifford, Malorie Perry, Simon Cottrell, Angela Marchbank, Alec Birchley, Alexander Adams, Amy Gaskin, Bree Gatica-Wilcox, Jason Coombes, Joel Southgate, Lauren Gilbert, Lee Graham, Nicole Pacchiarini, Sara Kumziene-Summerhayes, Sarah Taylor, Sophie Jones, Sara Rey, Matthew Bull, Joanne Watkins, Sally Corden, Tom Connor |
| EPI_ISL_611665 | University College London, Great Ormond Street Hospital for Children NHS Foundation Trust, Imperial College Healthcare NHS Trust | COVID-19 Genomics UK (COG-UK) Consortium | Sergi Castellano, Rachel Williams, Mark Kristiansen, Paola Resende Silva, Sunando Roy, Tony Brooks, Helena Tutill, Paola Niola, Patricia Dyal, Charlotte Williams, Leysa Forrest, Yasmin Panchbhaya, Jacqueline Findlay, Samuel Weeks, Julianne Brown, Kathryn Harris, Paul Randell, James Price, Alison Holmes, Judith Breuer |
| EPI_ISL_611666 | Wales Specialist Virology Centre Sequencing lab: Pathogen Genomics Unit | COVID-19 Genomics UK (COG-UK) Consortium | Catherine Moore, Johnathan Evans, Laura Gifford, Malorie Perry, Simon Cottrell, Angela Marchbank, Alec Birchley, Alexander Adams, Amy Gaskin, Bree Gatica-Wilcox, Jason Coombes, Joel Southgate, Lauren Gilbert, Lee Graham, Nicole Pacchiarini, Sara Kumziene-Summerhayes, Sarah Taylor, Sophie Jones, Sara Rey, Matthew Bull, Joanne Watkins, Sally Corden, Tom Connor |
| EPI_ISL_611676, EPI_ISL_611677 | Queens Medical Centre, Clinical Microbiology Department / DeepSeq Nottingham | COVID-19 Genomics UK (COG-UK) Consortium | Gemma Clark, Wendy Smith, Manjinder Khakh, Vicki M Fleming, Michelle M Lister, Hannah Howson-Wells, Jonathan Ball, Patrick McClure, Joseph Chappell, Theocharis Tsoleridis, Nadine Holmes, Matthew Carlisle, Christopher Moore, Fei Sang, Johnny Debebe, Victoria Wright, Matthew Loose |
| EPI_ISL_611678 | Virology Department, Sheffield Teaching Hospitals NHS Foundation Trust/Department of Infection, Immunity and Cardiovascular Disease, The Medical School, University of Sheffield | COVID-19 Genomics UK (COG-UK) Consortium | Thushan de Silva, Matthew Parker, Nikki Smith, Adri Angyal, Rebecca Brown, Luke Green, Rachel Tucker, Paul Parsons, Danielle Groves, Katie Johnson, Laura Carrilero, Alex Keeley, Dave Partridge, Matthew Wyles, Benjamin Lindsey, Mehmet Yavuz, Mohammad Raza, Cariad Evans |
| EPI_ISL_611687, EPI_ISL_611697, EPI_ISL_611706, EPI_ISL_611722, EPI_ISL_611726, EPI_ISL_611732 | Wales Specialist Virology Centre Sequencing lab: Pathogen Genomics Unit | COVID-19 Genomics UK (COG-UK) Consortium | Catherine Moore, Johnathan Evans, Laura Gifford, Malorie Perry, Simon Cottrell, Angela Marchbank, Alec Birchley, Alexander Adams, Amy Gaskin, Bree Gatica-Wilcox, Jason Coombes, Joel Southgate, Lauren Gilbert, Lee Graham, Nicole Pacchiarini, Sara Kumziene-Summerhayes, Sarah Taylor, Sophie Jones, Sara Rey, Matthew Bull, Joanne Watkins, Sally Corden, Tom Connor |
| EPI_ISL_611744, EPI_ISL_611794, EPI_ISL_611799, EPI_ISL_611803 | Virology Department, Sheffield Teaching Hospitals NHS Foundation Trust/Department of Infection, Immunity and Cardiovascular Disease, The Medical School, University of Sheffield | COVID-19 Genomics UK (COG-UK) Consortium | Thushan de Silva, Matthew Parker, Nikki Smith, Adri Angyal, Rebecca Brown, Luke Green, Rachel Tucker, Paul Parsons, Danielle Groves, Katie Johnson, Laura Carrilero, Alex Keeley, Dave Partridge, Matthew Wyles, Benjamin Lindsey, Mehmet Yavuz, Mohammad Raza, Cariad Evans |
| EPI_ISL_611823 | University College London, Great Ormond Street Hospital for Children NHS Foundation Trust, Imperial College Healthcare NHS Trust | COVID-19 Genomics UK (COG-UK) Consortium | Sergi Castellano, Rachel Williams, Mark Kristiansen, Paola Resende Silva, Sunando Roy, Tony Brooks, Helena Tutill, Paola Niola, Patricia Dyal, Charlotte Williams, Leysa Forrest, Yasmin Panchbhaya, Jacqueline Findlay, Samuel Weeks, Julianne Brown, Kathryn Harris, Paul Randell, James Price, Alison Holmes, Judith Breuer |
| EPI_ISL_611830 | Virology Department, Sheffield Teaching Hospitals NHS Foundation Trust/Department of Infection, Immunity and Cardiovascular Disease, The Medical School, University of Sheffield | COVID-19 Genomics UK (COG-UK) Consortium | Thushan de Silva, Matthew Parker, Nikki Smith, Adri Angyal, Rebecca Brown, Luke Green, Rachel Tucker, Paul Parsons, Danielle Groves, Katie Johnson, Laura Carrilero, Alex Keeley, Dave Partridge, Matthew Wyles, Benjamin Lindsey, Mehmet Yavuz, Mohammad Raza, Cariad Evans |
| EPI_ISL_611865 | Wales Specialist Virology Centre Sequencing lab: Pathogen Genomics Unit | COVID-19 Genomics UK (COG-UK) Consortium | Catherine Moore, Johnathan Evans, Laura Gifford, Malorie Perry, Simon Cottrell, Angela Marchbank, Alec Birchley, Alexander Adams, Amy Gaskin, Bree Gatica-Wilcox, Jason Coombes, Joel Southgate, Lauren Gilbert, Lee Graham, Nicole Pacchiarini, Sara Kumziene-Summerhayes, Sarah Taylor, Sophie Jones, Sara Rey, Matthew Bull, Joanne Watkins, Sally Corden, Tom Connor |
| EPI_ISL_611869 | Virology Department, Sheffield Teaching Hospitals NHS Foundation Trust/Department of Infection, Immunity and Cardiovascular Disease, The Medical School, University of | COVID-19 Genomics UK (COG-UK) Consortium | Thushan de Silva, Matthew Parker, Nikki Smith, Adri Angyal, Rebecca Brown, Luke Green, Rachel Tucker, Paul Parsons, Danielle Groves, Katie Johnson, Laura Carrilero, Alex Keeley, Dave Partridge, Matthew Wyles, Benjamin Lindsey, Mehmet Yavuz, Mohammad Raza, Cariad Evans |

| | | | |
|---|---|---|---|
| | Sheffield | | |
| EPI_ISL_611903, EPI_ISL_611904, EPI_ISL_611907, EPI_ISL_611923, EPI_ISL_611932 | Wales Specialist Virology Centre Sequencing lab: Pathogen Genomics Unit | COVID-19 Genomics UK (COG-UK) Consortium | Catherine Moore, Johnathan Evans, Laura Gifford, Malorie Perry, Simon Cottrell, Angela Marchbank, Alec Birchley, Alexander Adams, Amy Gaskin, Bree Gatica-Wilcox, Jason Coombes, Joel Southgate, Lauren Gilbert, Lee Graham, Nicole Pacchiarini, Sara Kumziene-Summerhayes, Sarah Taylor, Sophie Jones, Sara Rey, Matthew Bull, Joanne Watkins, Sally Corden, Tom Connor |
| EPI_ISL_611936 | Virology Department, Sheffield Teaching Hospitals NHS Foundation Trust/Department of Infection, Immunity and Cardiovascular Disease, The Medical School, University of Sheffield | COVID-19 Genomics UK (COG-UK) Consortium | Thushan de Silva, Matthew Parker, Nikki Smith, Adri Angyal, Rebecca Brown, Luke Green, Rachel Tucker, Paul Parsons, Danielle Groves, Katie Johnson, Laura Carrilero, Alex Keeley, Dave Partridge, Matthew Wyles, Benjamin Lindsey, Mehmet Yavuz, Mohammad Raza, Cariad Evans |
| EPI_ISL_611954 | Queens Medical Centre, Clinical Microbiology Department / DeepSeq Nottingham | COVID-19 Genomics UK (COG-UK) Consortium | Gemma Clark, Wendy Smith, Manjinder Khakh, Vicki M Fleming, Michelle M Lister, Hannah Howson-Wells, Jonathan Ball, Patrick McClure, Joseph Chappell, Theocharis Tsoleridis, Nadine Holmes, Matthew Carlisle, Christopher Moore, Fei Sang, Johnny Debebe, Victoria Wright, Matthew Loose |
| EPI_ISL_611973, EPI_ISL_611982, EPI_ISL_612013, EPI_ISL_612081, EPI_ISL_612083, EPI_ISL_612093 | Wales Specialist Virology Centre Sequencing lab: Pathogen Genomics Unit | COVID-19 Genomics UK (COG-UK) Consortium | Catherine Moore, Johnathan Evans, Laura Gifford, Malorie Perry, Simon Cottrell, Angela Marchbank, Alec Birchley, Alexander Adams, Amy Gaskin, Bree Gatica-Wilcox, Jason Coombes, Joel Southgate, Lauren Gilbert, Lee Graham, Nicole Pacchiarini, Sara Kumziene-Summerhayes, Sarah Taylor, Sophie Jones, Sara Rey, Matthew Bull, Joanne Watkins, Sally Corden, Tom Connor |
| EPI_ISL_612105 | University College London, Great Ormond Street Hospital for Children NHS Foundation Trust, Imperial College Healthcare NHS Trust | COVID-19 Genomics UK (COG-UK) Consortium | Sergi Castellano, Rachel Williams, Mark Kristiansen, Paola Resende Silva, Sunando Roy, Tony Brooks, Helena Tutill, Paola Niola, Patricia Dyal, Charlotte Williams, Leysa Forrest, Yasmin Panchbhaya, Jacqueline Findlay, Samuel Weeks, Julianne Brown, Kathryn Harris, Paul Randell, James Price, Alison Holmes, Judith Breuer |
| EPI_ISL_612116, EPI_ISL_612120, EPI_ISL_612146, EPI_ISL_612147, EPI_ISL_612152, EPI_ISL_612153, EPI_ISL_612154, EPI_ISL_612155, EPI_ISL_612156, EPI_ISL_612157, EPI_ISL_612177, EPI_ISL_612193, EPI_ISL_612199, EPI_ISL_612202 | | | |
| see above | Wales Specialist Virology Centre Sequencing lab: Pathogen Genomics Unit | COVID-19 Genomics UK (COG-UK) Consortium | Catherine Moore, Johnathan Evans, Laura Gifford, Malorie Perry, Simon Cottrell, Angela Marchbank, Alec Birchley, Alexander Adams, Amy Gaskin, Bree Gatica-Wilcox, Jason Coombes, Joel Southgate, Lauren Gilbert, Lee Graham, Nicole Pacchiarini, Sara Kumziene-Summerhayes, Sarah Taylor, Sophie Jones, Sara Rey, Matthew Bull, Joanne Watkins, Sally Corden, Tom Connor |
| EPI_ISL_612373 | Virology Department, Royal Infirmary of Edinburgh, NHS Lothian / School of Biological Sciences, University of Edinburgh / Institute of Genetics and Molecular Medicine, University of Edinburgh | COVID-19 Genomics UK (COG-UK) Consortium | McHugh M, Dewar R, Rooke S, Gallagher M, Balcaza C, O'Toole Á, Scher E, Hill V, McCrone JT, Colquhoun R, Yu X, Jackson B, Rambaut A, Williams TC, Templeton K |
| EPI_ISL_612451, EPI_ISL_612454 | University College London, Great Ormond Street Hospital for Children NHS Foundation Trust, Imperial College Healthcare NHS Trust | COVID-19 Genomics UK (COG-UK) Consortium | Sergi Castellano, Rachel Williams, Mark Kristiansen, Paola Resende Silva, Sunando Roy, Tony Brooks, Helena Tutill, Paola Niola, Patricia Dyal, Charlotte Williams, Leysa Forrest, Yasmin Panchbhaya, Jacqueline Findlay, Samuel Weeks, Julianne Brown, Kathryn Harris, Paul Randell, James Price, Alison Holmes, Judith Breuer |
| EPI_ISL_612574, EPI_ISL_612575, EPI_ISL_612576 | Queens Medical Centre, Clinical Microbiology Department / DeepSeq Nottingham | COVID-19 Genomics UK (COG-UK) Consortium | Gemma Clark, Wendy Smith, Manjinder Khakh, Vicki M Fleming, Michelle M Lister, Hannah Howson-Wells, Jonathan Ball, Patrick McClure, Joseph Chappell, Theocharis Tsoleridis, Nadine Holmes, Matthew Carlisle, Christopher Moore, Fei Sang, Johnny Debebe, Victoria Wright, Matthew Loose |
| EPI_ISL_612849, EPI_ISL_612861, EPI_ISL_612864, EPI_ISL_612865, EPI_ISL_612893, EPI_ISL_612904, EPI_ISL_612917, EPI_ISL_612918, EPI_ISL_612946, EPI_ISL_612951, EPI_ISL_612956, EPI_ISL_612971, EPI_ISL_612987, EPI_ISL_612999, EPI_ISL_613018, EPI_ISL_613019, EPI_ISL_613021, EPI_ISL_613025, EPI_ISL_613026, EPI_ISL_613032, EPI_ISL_613035, EPI_ISL_613042, EPI_ISL_613043, EPI_ISL_613044, EPI_ISL_613046, EPI_ISL_613048, EPI_ISL_613049, EPI_ISL_613057, EPI_ISL_613059, EPI_ISL_613066, EPI_ISL_613067, EPI_ISL_613068, EPI_ISL_613076, EPI_ISL_613077, EPI_ISL_613089, EPI_ISL_613090, EPI_ISL_613091, EPI_ISL_613094, EPI_ISL_613101, EPI_ISL_613107, EPI_ISL_613116, EPI_ISL_613125, EPI_ISL_613134, EPI_ISL_613135, EPI_ISL_613137, EPI_ISL_613138, EPI_ISL_613139, EPI_ISL_613141, EPI_ISL_613147, EPI_ISL_613148, EPI_ISL_613153, EPI_ISL_613155, EPI_ISL_613162, EPI_ISL_613163, EPI_ISL_613164, EPI_ISL_613166, EPI_ISL_613173, EPI_ISL_613178, EPI_ISL_613185, EPI_ISL_613186, EPI_ISL_613189, EPI_ISL_613190, EPI_ISL_613191, EPI_ISL_613192, EPI_ISL_613193, EPI_ISL_613194, EPI_ISL_613195, EPI_ISL_613196, EPI_ISL_613197, EPI_ISL_613198, EPI_ISL_613199, EPI_ISL_613200, EPI_ISL_613201, EPI_ISL_613202, EPI_ISL_613203, EPI_ISL_613204, EPI_ISL_613205, EPI_ISL_613206, EPI_ISL_613207, EPI_ISL_613208, EPI_ISL_613209, EPI_ISL_613210, EPI_ISL_613211, EPI_ISL_613212, EPI_ISL_613213, EPI_ISL_613214, EPI_ISL_613215, EPI_ISL_613217, EPI_ISL_613218, EPI_ISL_613219, EPI_ISL_613220, EPI_ISL_613221, EPI_ISL_613222, EPI_ISL_613223, EPI_ISL_613224, EPI_ISL_613225, EPI_ISL_613226, EPI_ISL_613227, EPI_ISL_613228, EPI_ISL_613229, EPI_ISL_613230, EPI_ISL_613231, EPI_ISL_613232, EPI_ISL_613233, EPI_ISL_613234, EPI_ISL_613235, EPI_ISL_613236, EPI_ISL_613237, EPI_ISL_613238, EPI_ISL_613239, EPI_ISL_613240, EPI_ISL_613241, EPI_ISL_613242, EPI_ISL_613243, EPI_ISL_613244, EPI_ISL_613245, EPI_ISL_613246, EPI_ISL_613247, EPI_ISL_613248, EPI_ISL_613249, EPI_ISL_613250, EPI_ISL_613251, EPI_ISL_613252, EPI_ISL_613253, EPI_ISL_613254, EPI_ISL_613255, EPI_ISL_613256, EPI_ISL_613257, EPI_ISL_613258, EPI_ISL_613259, EPI_ISL_613260, EPI_ISL_613261, EPI_ISL_613262, EPI_ISL_613263, EPI_ISL_613264, EPI_ISL_613266, EPI_ISL_613267, EPI_ISL_613268, EPI_ISL_613269, EPI_ISL_613270, EPI_ISL_613272, EPI_ISL_613273, EPI_ISL_613274, EPI_ISL_613276, EPI_ISL_613277, EPI_ISL_613279, EPI_ISL_613281 | | | |
| see above | Wales Specialist Virology Centre Sequencing lab: Pathogen Genomics Unit | COVID-19 Genomics UK (COG-UK) Consortium | Catherine Moore, Johnathan Evans, Laura Gifford, Malorie Perry, Simon Cottrell, Angela Marchbank, Alec Birchley, Alexander Adams, Amy Gaskin, Bree Gatica-Wilcox, Jason Coombes, Joel Southgate, Lauren Gilbert, Lee Graham, Nicole Pacchiarini, Sara Kumziene-Summerhayes, Sarah Taylor, Sophie Jones, Sara Rey, Matthew Bull, Joanne Watkins, Sally Corden, Tom Connor |
| EPI_ISL_613366, EPI_ISL_613368, EPI_ISL_613383, EPI_ISL_613391, EPI_ISL_613400 | Virology Department, Sheffield Teaching Hospitals NHS Foundation Trust/Department of Infection, Immunity and Cardiovascular Disease, The Medical School, University of Sheffield | COVID-19 Genomics UK (COG-UK) Consortium | Thushan de Silva, Matthew Parker, Nikki Smith, Adri Angyal, Rebecca Brown, Luke Green, Rachel Tucker, Paul Parsons, Danielle Groves, Katie Johnson, Laura Carrilero, Alex Keeley, Dave Partridge, Matthew Wyles, Benjamin Lindsey, Mehmet Yavuz, Mohammad Raza, Cariad Evans |
| EPI_ISL_613523 | Public Health Laboratory - Infectious Disease Lab, Minnesota Department of Health Infectious Disease Laboratory Submission Group | Minnesota Department of Health, Public Health Laboratory | Plumb,M., Garfin,J., Lorentz,A., Wang,X. |
| EPI_ISL_613634 | Respiratory Virus Unit, Microbiology Services Colindale, Public Health England | Respiratory Virus Unit, Microbiology Services Colindale, Public Health England | PHE Covid Sequencing Team |
| EPI_ISL_614182, EPI_ISL_614185, EPI_ISL_614188, EPI_ISL_614189, EPI_ISL_614192, EPI_ISL_614194, EPI_ISL_614195, EPI_ISL_614201, EPI_ISL_614205, EPI_ISL_614207, EPI_ISL_614209, EPI_ISL_614215, EPI_ISL_614221, EPI_ISL_614222, EPI_ISL_614226, EPI_ISL_614230, EPI_ISL_614237, EPI_ISL_614239, EPI_ISL_614240, EPI_ISL_614244, EPI_ISL_614245, EPI_ISL_614246 | | | |
| see above | Michigan Department of Health and Human Services, Bureau of Laboratories | Michigan Department of Health and Human Services, Bureau of Laboratories | Blankenship HM, Riner D, Soehnlen MK |
| EPI_ISL_614287, EPI_ISL_614288, EPI_ISL_614289, EPI_ISL_614291 | General practitioner | National Reference Center for Viruses of Respiratory Infections, Institut Pasteur, Paris | Marion Barbet, Sylvie Behillil, Méline Bizard, Angela Brisebarre, Camille Capel, Etienne Simon-Lorière, Vincent Enouf, Maud Vanpeene, Sylvie van der Werf |
| EPI_ISL_614893, EPI_ISL_614894, EPI_ISL_614912, EPI_ISL_614913, EPI_ISL_614914, EPI_ISL_614915, EPI_ISL_614916, EPI_ISL_614917, EPI_ISL_614918, EPI_ISL_614919, EPI_ISL_614920, EPI_ISL_614921, EPI_ISL_614922, EPI_ISL_614923, EPI_ISL_614924, EPI_ISL_614925, EPI_ISL_614926, EPI_ISL_614927, EPI_ISL_614937, EPI_ISL_614938, EPI_ISL_614939, EPI_ISL_614940, EPI_ISL_614941, EPI_ISL_614942, EPI_ISL_614943, EPI_ISL_614944, EPI_ISL_614945, EPI_ISL_614946, EPI_ISL_614965, EPI_ISL_614966, EPI_ISL_614967, EPI_ISL_614968, EPI_ISL_614969, EPI_ISL_614970, EPI_ISL_614971, EPI_ISL_614972, EPI_ISL_614973, EPI_ISL_614974, EPI_ISL_614975, EPI_ISL_614976, EPI_ISL_614977 | | | |
| see above | Viollier AG | Department of Biosystems Science and Engineering, ETH Zürich | Christian Beisel, Sarah Nadeau, Ivan Topolsky, Pedro Ferreira, Philipp Jablonski, Susana Posada-Céspedes, Tobias Schär, Ina Nissen, Natascha Santacroce, Elodie Burcklen, Christiane Beckmann, Maurice Redondo, Olivier Kobel, Christoph Noppen, Sophie Seidel, Noemie Santamaria de Souza, Chaoran Chen, Niko Beerenwinkel, Tanja Stadler |
| EPI_ISL_617026, EPI_ISL_617251, EPI_ISL_617257, EPI_ISL_618499, EPI_ISL_618500, EPI_ISL_618501, EPI_ISL_618502, EPI_ISL_618503, EPI_ISL_618504, EPI_ISL_618505, EPI_ISL_618506, EPI_ISL_618507, EPI_ISL_618508, EPI_ISL_618509, EPI_ISL_618510, EPI_ISL_618512, EPI_ISL_618513, EPI_ISL_618514, EPI_ISL_618515, EPI_ISL_618516, EPI_ISL_618518, EPI_ISL_618519, EPI_ISL_618520, EPI_ISL_618521, EPI_ISL_618522, EPI_ISL_618523, EPI_ISL_618524, EPI_ISL_618526, EPI_ISL_618527, EPI_ISL_618528, EPI_ISL_618529, EPI_ISL_618530, EPI_ISL_618531, EPI_ISL_618532, EPI_ISL_618533, EPI_ISL_618534, EPI_ISL_618535, EPI_ISL_618536, EPI_ISL_618537, EPI_ISL_618538, EPI_ISL_618539, EPI_ISL_618540, EPI_ISL_618541, EPI_ISL_618542, EPI_ISL_618543, EPI_ISL_618544, EPI_ISL_618545, EPI_ISL_618546, EPI_ISL_618547, EPI_ISL_618548, EPI_ISL_618549, EPI_ISL_618550, EPI_ISL_618551, EPI_ISL_618552, EPI_ISL_618553, EPI_ISL_618554, EPI_ISL_618555, EPI_ISL_618556, EPI_ISL_618557, EPI_ISL_618558, EPI_ISL_618559, EPI_ISL_618560, EPI_ISL_618561, EPI_ISL_618562, EPI_ISL_618563, EPI_ISL_618564, EPI_ISL_618565, EPI_ISL_618566, EPI_ISL_618567, EPI_ISL_618568, EPI_ISL_618569, EPI_ISL_618570, EPI_ISL_618571, EPI_ISL_618572, EPI_ISL_618573, EPI_ISL_618574, EPI_ISL_618575, EPI_ISL_618576, EPI_ISL_618577, EPI_ISL_618578, EPI_ISL_618579, EPI_ISL_618580, EPI_ISL_618581, EPI_ISL_618582, EPI_ISL_618583, EPI_ISL_618584, EPI_ISL_618585, EPI_ISL_618586, EPI_ISL_618587, EPI_ISL_618588, EPI_ISL_618589, EPI_ISL_618590, EPI_ISL_618591, EPI_ISL_618592, EPI_ISL_618593, EPI_ISL_618594, EPI_ISL_618596, EPI_ISL_618597, EPI_ISL_618598, EPI_ISL_618599, EPI_ISL_618600, EPI_ISL_618601, EPI_ISL_618602, EPI_ISL_618603, EPI_ISL_618604, EPI_ISL_618605, EPI_ISL_618606, EPI_ISL_618607, EPI_ISL_618608, EPI_ISL_618610, EPI_ISL_618611, EPI_ISL_618612, EPI_ISL_618613, EPI_ISL_618614, EPI_ISL_618615, EPI_ISL_618616, EPI_ISL_618617, EPI_ISL_618618, EPI_ISL_618619, EPI_ISL_618620, EPI_ISL_618621, EPI_ISL_618622, EPI_ISL_618623, EPI_ISL_618624, EPI_ISL_618625, EPI_ISL_618663, EPI_ISL_618664, EPI_ISL_618665, EPI_ISL_618666, EPI_ISL_618667, EPI_ISL_618668, EPI_ISL_618669, EPI_ISL_618670, EPI_ISL_618671, EPI_ISL_618672, EPI_ISL_618673, EPI_ISL_618674, EPI_ISL_618675, EPI_ISL_618676, EPI_ISL_618677, EPI_ISL_618678, EPI_ISL_618679, EPI_ISL_618680, EPI_ISL_618681, EPI_ISL_618682, EPI_ISL_618683, EPI_ISL_618684, EPI_ISL_618685, EPI_ISL_618686, EPI_ISL_618687, EPI_ISL_618688, EPI_ISL_618689, EPI_ISL_618690, EPI_ISL_618691, EPI_ISL_618692, EPI_ISL_618693, EPI_ISL_618694, EPI_ISL_618695, EPI_ISL_618696, EPI_ISL_618697, EPI_ISL_618698, EPI_ISL_618699, EPI_ISL_618700, EPI_ISL_618701, EPI_ISL_618702, EPI_ISL_618703, EPI_ISL_618704, EPI_ISL_618705, EPI_ISL_618706, EPI_ISL_618707, EPI_ISL_618708, EPI_ISL_618709, EPI_ISL_618710, EPI_ISL_618711, EPI_ISL_618712, EPI_ISL_618713, EPI_ISL_618714, EPI_ISL_618715, EPI_ISL_618716, EPI_ISL_618717, EPI_ISL_618718, EPI_ISL_618719, EPI_ISL_618721, EPI_ISL_618722, EPI_ISL_618723, EPI_ISL_618724, EPI_ISL_618725, EPI_ISL_618726, EPI_ISL_618727, EPI_ISL_618728, EPI_ISL_618729, EPI_ISL_618731, EPI_ISL_618732, EPI_ISL_618733, EPI_ISL_618734, EPI_ISL_618735, EPI_ISL_618736, EPI_ISL_618737, EPI_ISL_618738, EPI_ISL_618739, EPI_ISL_618741, EPI_ISL_618742, EPI_ISL_618743, EPI_ISL_618744, EPI_ISL_618745, EPI_ISL_618746, EPI_ISL_618747, EPI_ISL_618748, EPI_ISL_618749, EPI_ISL_618750, EPI_ISL_618751, EPI_ISL_618752, EPI_ISL_618753, EPI_ISL_618754, EPI_ISL_618755, EPI_ISL_618756, EPI_ISL_618757, EPI_ISL_618758, EPI_ISL_618759, EPI_ISL_618760, EPI_ISL_618761, EPI_ISL_618762, EPI_ISL_618763, EPI_ISL_618764, EPI_ISL_618765, EPI_ISL_618766, EPI_ISL_618767, EPI_ISL_618769, EPI_ISL_618770, EPI_ISL_618771, EPI_ISL_618772, EPI_ISL_618773, EPI_ISL_618774, EPI_ISL_618775, EPI_ISL_618776, EPI_ISL_618777, EPI_ISL_618778, EPI_ISL_618779, EPI_ISL_618780, EPI_ISL_618781, EPI_ISL_618782, EPI_ISL_618783, EPI_ISL_618784, EPI_ISL_618785, EPI_ISL_618786, EPI_ISL_618787, EPI_ISL_618788, EPI_ISL_618789, EPI_ISL_618790, EPI_ISL_618791, EPI_ISL_618792, EPI_ISL_618793, EPI_ISL_618794, EPI_ISL_618795, EPI_ISL_618796, EPI_ISL_618797, EPI_ISL_618798, EPI_ISL_618799, EPI_ISL_618813, EPI_ISL_618822, EPI_ISL_618830, EPI_ISL_618833, EPI_ISL_618837, EPI_ISL_618844, EPI_ISL_618846, EPI_ISL_618847, EPI_ISL_618852, EPI_ISL_618855, EPI_ISL_618857, EPI_ISL_618872 | | | |

| | | | |
|---|---|---|---|
| see above | Department of Virus and Microbiological Special Diagnostics, Statens Serum Institut, Denmark | Albertsen lab, Department of Chemistry and Bioscience, Aalborg University, Denmark | Danish Covid-19 Genome Consortia |
| EPI_ISL_622769, EPI_ISL_622770, EPI_ISL_622771 | Canterbury Health Laboratories | Institute of Environmental Science and Research (ESR) | Xiaoyun Ren, Matt Storey, Nikki Freed, Muhammad Faisal, Jing Wang, Hermes Perez, Anja Werno, Antje van der Linden, Arlo Upton, Chris Mansell, David Hammer, Dragana Drinkovic, Gary McAuliffe, Hana Sofia Andersson, James Ussher, Jill Sherwood, Josh Freeman, Julia Howard, Juliet Elvy, Mary DeAlmeida, Matt Blakiston, Matthew Rogers, Max Bloomfield, Michael Addidle, Michelle Balm, Sally Roberts, Sarah Jefferies, Sharmini Muttaiyah, Susan Morpeth, Susan Taylor, Timothy Blackmore, Vani Sathyendran, Veronica Playle, Virginia Hope, Erasmus Smit, Lauren Jelly, Olin Silander, Joep de Ligt |
| EPI_ISL_622772 | Middlemore Hospital | Institute of Environmental Science and Research (ESR) | Xiaoyun Ren, Matt Storey, Nikki Freed, Muhammad Faisal, Jing Wang, Hermes Perez, Anja Werno, Antje van der Linden, Arlo Upton, Chris Mansell, David Hammer, Dragana Drinkovic, Gary McAuliffe, Hana Sofia Andersson, James Ussher, Jill Sherwood, Josh Freeman, Julia Howard, Juliet Elvy, Mary DeAlmeida, Matt Blakiston, Matthew Rogers, Max Bloomfield, Michael Addidle, Michelle Balm, Sally Roberts, Sarah Jefferies, Sharmini Muttaiyah, Susan Morpeth, Susan Taylor, Timothy Blackmore, Vani Sathyendran, Veronica Playle, Virginia Hope, Erasmus Smit, Lauren Jelly, Olin Silander, Joep de Ligt |
| EPI_ISL_622774 | LabPLUS | Institute of Environmental Science and Research (ESR) | Xiaoyun Ren, Matt Storey, Nikki Freed, Muhammad Faisal, Jing Wang, Hermes Perez, Anja Werno, Antje van der Linden, Arlo Upton, Chris Mansell, David Hammer, Dragana Drinkovic, Gary McAuliffe, Hana Sofia Andersson, James Ussher, Jill Sherwood, Josh Freeman, Julia Howard, Juliet Elvy, Mary DeAlmeida, Matt Blakiston, Matthew Rogers, Max Bloomfield, Michael Addidle, Michelle Balm, Sally Roberts, Sarah Jefferies, Sharmini Muttaiyah, Susan Morpeth, Susan Taylor, Timothy Blackmore, Vani Sathyendran, Veronica Playle, Virginia Hope, Erasmus Smit, Lauren Jelly, Olin Silander, Joep de Ligt |
| EPI_ISL_622775 | Canterbury Health Laboratories | Institute of Environmental Science and Research (ESR) | Xiaoyun Ren, Matt Storey, Nikki Freed, Muhammad Faisal, Jing Wang, Hermes Perez, Anja Werno, Antje van der Linden, Arlo Upton, Chris Mansell, David Hammer, Dragana Drinkovic, Gary McAuliffe, Hana Sofia Andersson, James Ussher, Jill Sherwood, Josh Freeman, Julia Howard, Juliet Elvy, Mary DeAlmeida, Matt Blakiston, Matthew Rogers, Max Bloomfield, Michael Addidle, Michelle Balm, Sally Roberts, Sarah Jefferies, Sharmini Muttaiyah, Susan Morpeth, Susan Taylor, Timothy Blackmore, Vani Sathyendran, Veronica Playle, Virginia Hope, Erasmus Smit, Lauren Jelly, Olin Silander, Joep de Ligt |
| EPI_ISL_622776 | Middlemore Hospital | Institute of Environmental Science and Research (ESR) | Xiaoyun Ren, Matt Storey, Nikki Freed, Muhammad Faisal, Jing Wang, Hermes Perez, Anja Werno, Antje van der Linden, Arlo Upton, Chris Mansell, David Hammer, Dragana Drinkovic, Gary McAuliffe, Hana Sofia Andersson, James Ussher, Jill Sherwood, Josh Freeman, Julia Howard, Juliet Elvy, Mary DeAlmeida, Matt Blakiston, Matthew Rogers, Max Bloomfield, Michael Addidle, Michelle Balm, Sally Roberts, Sarah Jefferies, Sharmini Muttaiyah, Susan Morpeth, Susan Taylor, Timothy Blackmore, Vani Sathyendran, Veronica Playle, Virginia Hope, Erasmus Smit, Lauren Jelly, Olin Silander, Joep de Ligt |
| EPI_ISL_622778 | LabTests | Institute of Environmental Science and Research (ESR) | Xiaoyun Ren, Matt Storey, Nikki Freed, Muhammad Faisal, Jing Wang, Hermes Perez, Anja Werno, Antje van der Linden, Arlo Upton, Chris Mansell, David Hammer, Dragana Drinkovic, Gary McAuliffe, Hana Sofia Andersson, James Ussher, Jill Sherwood, Josh Freeman, Julia Howard, Juliet Elvy, Mary DeAlmeida, Matt Blakiston, Matthew Rogers, Max Bloomfield, Michael Addidle, Michelle Balm, Sally Roberts, Sarah Jefferies, Sharmini Muttaiyah, Susan Morpeth, Susan Taylor, Timothy Blackmore, Vani Sathyendran, Veronica Playle, Virginia Hope, Erasmus Smit, Lauren Jelly, Olin Silander, Joep de Ligt |
| EPI_ISL_622781, EPI_ISL_622782, EPI_ISL_622789, EPI_ISL_622791, EPI_ISL_622809 | Canterbury Health Laboratories | Institute of Environmental Science and Research (ESR) | Xiaoyun Ren, Matt Storey, Nikki Freed, Muhammad Faisal, Jing Wang, Hermes Perez, Anja Werno, Antje van der Linden, Arlo Upton, Chris Mansell, David Hammer, Dragana Drinkovic, Gary McAuliffe, Hana Sofia Andersson, James Ussher, Jill Sherwood, Josh Freeman, Julia Howard, Juliet Elvy, Mary DeAlmeida, Matt Blakiston, Matthew Rogers, Max Bloomfield, Michael Addidle, Michelle Balm, Sally Roberts, Sarah Jefferies, Sharmini Muttaiyah, Susan Morpeth, Susan Taylor, Timothy Blackmore, Vani Sathyendran, Veronica Playle, Virginia Hope, Erasmus Smit, Lauren Jelly, Olin Silander, Joep de Ligt |
| EPI_ISL_622818 | Middlemore Hospital | Institute of Environmental Science and Research (ESR) | Xiaoyun Ren, Matt Storey, Nikki Freed, Muhammad Faisal, Jing Wang, Hermes Perez, Anja Werno, Antje van der Linden, Arlo Upton, Chris Mansell, David Hammer, Dragana Drinkovic, Gary McAuliffe, Hana Sofia Andersson, James Ussher, Jill Sherwood, Josh Freeman, Julia Howard, Juliet Elvy, Mary DeAlmeida, Matt Blakiston, Matthew Rogers, Max Bloomfield, Michael Addidle, Michelle Balm, Sally Roberts, Sarah Jefferies, Sharmini Muttaiyah, Susan Morpeth, Susan Taylor, Timothy Blackmore, Vani Sathyendran, Veronica Playle, Virginia Hope, Erasmus Smit, Lauren Jelly, Olin Silander, Joep de Ligt |
| EPI_ISL_622879, EPI_ISL_622880, EPI_ISL_622881, EPI_ISL_622882, EPI_ISL_622883, EPI_ISL_622886, EPI_ISL_622887, EPI_ISL_622889 | Respiratory Virus Unit, Microbiology Services Colindale, Public Health England | Respiratory Virus Unit, Microbiology Services Colindale, Public Health England | PHE Covid Sequencing Team |
| EPI_ISL_623097 | General practitioner | National Reference Center for Viruses of Respiratory Infections, Institut Pasteur, Paris | Marion Barbet, Sylvie Behillil, Méline Bizard, Angela Brisebarre, Camille Capel, Etienne Simon-Lorière, Vincent Enouf, Maud Vanpeene, Sylvie van der Werf |
| EPI_ISL_623211, EPI_ISL_623212 | Lighthouse Lab in Milton Keynes | Wellcome Sanger Institute for the COVID-19 Genomics UK (COG-UK) consortium | The Lighthouse Lab in Milton Keynes and Alex Alderton, Roberto Amato, Sonia Goncalves, Ewan Harrison, David K. Jackson, Ian Johnston, Dominic Kwiatkowski, Cordelia Langford, John Sillitoe on behalf of the Wellcome Sanger Institute COVID-19 Surveillance Team (http://www.sanger.ac.uk/covid-team) |
| EPI_ISL_623214 | Lighthouse Lab in Milton Keynes | Wellcome Sanger Institute for the COVID-19 Genomics UK (COG-UK) Consortium | The Lighthouse Lab in Milton Keynes and Alex Alderton, Roberto Amato, Sonia Goncalves, Ewan Harrison, David K. Jackson, Ian Johnston, Dominic Kwiatkowski, Cordelia Langford, John Sillitoe on behalf of the Wellcome Sanger Institute COVID-19 Surveillance Team (http://www.sanger.ac.uk/covid-team) |
| EPI_ISL_623215, EPI_ISL_623216, EPI_ISL_623217, EPI_ISL_623218, EPI_ISL_623219, EPI_ISL_623221 | Lighthouse Lab in Milton Keynes | Wellcome Sanger Institute for the COVID-19 Genomics UK (COG-UK) consortium | The Lighthouse Lab in Milton Keynes and Alex Alderton, Roberto Amato, Sonia Goncalves, Ewan Harrison, David K. Jackson, Ian Johnston, Dominic Kwiatkowski, Cordelia Langford, John Sillitoe on behalf of the Wellcome Sanger Institute COVID-19 Surveillance Team (http://www.sanger.ac.uk/covid-team) |
| EPI_ISL_623222 | Lighthouse Lab in Cambridge | Wellcome Sanger Institute for the COVID-19 Genomics UK (COG-UK) consortium | Rob Howes, The Lighthouse Lab in Cambridge and Alex Alderton, Roberto Amato, Sonia Goncalves, Ewan Harrison, David K. Jackson, Ian Johnston, Dominic Kwiatkowski, Cordelia Langford, John Sillitoe on behalf of the Wellcome Sanger Institute COVID-19 Surveillance Team (http://www.sanger.ac.uk/covid-team) |
| EPI_ISL_623223, EPI_ISL_623224, EPI_ISL_623225, EPI_ISL_623226, EPI_ISL_623227, EPI_ISL_623228, EPI_ISL_623229, EPI_ISL_623231, EPI_ISL_623232, EPI_ISL_623233 | Lighthouse Lab in Milton Keynes | Wellcome Sanger Institute for the COVID-19 Genomics UK (COG-UK) consortium | The Lighthouse Lab in Milton Keynes and Alex Alderton, Roberto Amato, Sonia Goncalves, Ewan Harrison, David K. Jackson, Ian Johnston, Dominic Kwiatkowski, Cordelia Langford, John Sillitoe on behalf of the Wellcome Sanger Institute COVID-19 Surveillance Team (http://www.sanger.ac.uk/covid-team) |
| EPI_ISL_623234 | Lighthouse Lab in Cambridge | Wellcome Sanger Institute for the COVID-19 Genomics UK (COG-UK) consortium | Rob Howes, The Lighthouse Lab in Cambridge and Alex Alderton, Roberto Amato, Sonia Goncalves, Ewan Harrison, David K. Jackson, Ian Johnston, Dominic Kwiatkowski, Cordelia Langford, John Sillitoe on behalf of the Wellcome Sanger Institute COVID-19 Surveillance Team (http://www.sanger.ac.uk/covid-team) |
| EPI_ISL_623236, EPI_ISL_623237, EPI_ISL_623238, EPI_ISL_623239, EPI_ISL_623240, EPI_ISL_623241, EPI_ISL_623243 | Lighthouse Lab in Milton Keynes | Wellcome Sanger Institute for the COVID-19 Genomics UK (COG-UK) consortium | The Lighthouse Lab in Milton Keynes and Alex Alderton, Roberto Amato, Sonia Goncalves, Ewan Harrison, David K. Jackson, Ian Johnston, Dominic Kwiatkowski, Cordelia Langford, John Sillitoe on behalf of the Wellcome Sanger Institute COVID-19 Surveillance Team (http://www.sanger.ac.uk/covid-team) |
| EPI_ISL_623247 | Lighthouse Lab in Cambridge | Wellcome Sanger Institute for the COVID-19 Genomics UK (COG-UK) consortium | Rob Howes, The Lighthouse Lab in Cambridge and Alex Alderton, Roberto Amato, Sonia Goncalves, Ewan Harrison, David K. Jackson, Ian Johnston, Dominic Kwiatkowski, Cordelia Langford, John Sillitoe on behalf of the Wellcome Sanger Institute COVID-19 Surveillance Team (http://www.sanger.ac.uk/covid-team) |
| EPI_ISL_623249, EPI_ISL_623250, EPI_ISL_623251, EPI_ISL_623253, EPI_ISL_623254, EPI_ISL_623255 | Lighthouse Lab in Milton Keynes | Wellcome Sanger Institute for the COVID-19 Genomics UK (COG-UK) consortium | The Lighthouse Lab in Milton Keynes and Alex Alderton, Roberto Amato, Sonia Goncalves, Ewan Harrison, David K. Jackson, Ian Johnston, Dominic Kwiatkowski, Cordelia Langford, John Sillitoe on behalf of the Wellcome Sanger Institute COVID-19 Surveillance Team (http://www.sanger.ac.uk/covid-team) |
| EPI_ISL_623256 | Lighthouse Lab in Cambridge | Wellcome Sanger Institute for the COVID-19 Genomics UK (COG-UK) consortium | Rob Howes, The Lighthouse Lab in Cambridge and Alex Alderton, Roberto Amato, Sonia Goncalves, Ewan Harrison, David K. Jackson, Ian Johnston, Dominic Kwiatkowski, Cordelia Langford, John Sillitoe on behalf of the Wellcome Sanger Institute COVID-19 Surveillance Team (http://www.sanger.ac.uk/covid-team) |
| EPI_ISL_623257, EPI_ISL_623258 | Lighthouse Lab in Milton Keynes | Wellcome Sanger Institute for the COVID-19 Genomics UK (COG-UK) consortium | The Lighthouse Lab in Milton Keynes and Alex Alderton, Roberto Amato, Sonia Goncalves, Ewan Harrison, David K. Jackson, Ian Johnston, Dominic Kwiatkowski, Cordelia Langford, John Sillitoe on behalf of the Wellcome Sanger Institute COVID-19 Surveillance Team (http://www.sanger.ac.uk/covid-team) |
| EPI_ISL_623259 | Lighthouse Lab in Milton Keynes | Wellcome Sanger Institute for the COVID-19 Genomics UK (COG-UK) Consortium | The Lighthouse Lab in Milton Keynes and Alex Alderton, Roberto Amato, Sonia Goncalves, Ewan Harrison, David K. Jackson, Ian Johnston, Dominic Kwiatkowski, Cordelia Langford, John Sillitoe on behalf of the Wellcome Sanger Institute COVID-19 Surveillance Team (http://www.sanger.ac.uk/covid-team) |

EPI_ISL_623260, EPI_ISL_623261, EPI_ISL_623262, EPI_ISL_623264, EPI_ISL_623266, EPI_ISL_623267, EPI_ISL_623269, EPI_ISL_623270, EPI_ISL_623271, EPI_ISL_623272, EPI_ISL_623274, EPI_ISL_623275

| | | | |
|---|---|---|---|
| see above | Lighthouse Lab in Milton Keynes | Wellcome Sanger Institute for the COVID-19 Genomics UK (COG-UK) consortium | The Lighthouse Lab in Milton Keynes and Alex Alderton, Roberto Amato, Sonia Goncalves, Ewan Harrison, David K. Jackson, Ian Johnston, Dominic Kwiatkowski, Cordelia Langford, John Sillitoe on behalf of the Wellcome Sanger Institute COVID-19 Surveillance Team (http://www.sanger.ac.uk/covid-team) |
| EPI_ISL_623276 | Lighthouse Lab in Cambridge | Wellcome Sanger Institute for the COVID-19 Genomics UK (COG-UK) consortium | Rob Howes, The Lighthouse Lab in Cambridge and Alex Alderton, Roberto Amato, Sonia Goncalves, Ewan Harrison, David K. Jackson, Ian Johnston, Dominic Kwiatkowski, Cordelia Langford, John Sillitoe on behalf of the Wellcome Sanger Institute COVID-19 Surveillance Team (http://www.sanger.ac.uk/covid-team) |
| EPI_ISL_623277, EPI_ISL_623278, EPI_ISL_623279, EPI_ISL_623280, EPI_ISL_623282, EPI_ISL_623283, EPI_ISL_623284, EPI_ISL_623285 | Lighthouse Lab in Milton Keynes | Wellcome Sanger Institute for the COVID-19 Genomics UK (COG-UK) consortium | The Lighthouse Lab in Milton Keynes and Alex Alderton, Roberto Amato, Sonia Goncalves, Ewan Harrison, David K. Jackson, Ian Johnston, Dominic Kwiatkowski, Cordelia Langford, John Sillitoe on behalf of the Wellcome Sanger Institute COVID-19 Surveillance Team (http://www.sanger.ac.uk/covid-team) |
| EPI_ISL_623286 | Lighthouse Lab in Cambridge | Wellcome Sanger Institute for the COVID-19 Genomics UK (COG-UK) consortium | Rob Howes, The Lighthouse Lab in Cambridge and Alex Alderton, Roberto Amato, Sonia Goncalves, Ewan Harrison, David K. Jackson, Ian Johnston, Dominic Kwiatkowski, Cordelia Langford, John Sillitoe on behalf of the Wellcome Sanger Institute COVID-19 Surveillance Team (http://www.sanger.ac.uk/covid-team) |
| EPI_ISL_623287 | Lighthouse Lab in Milton Keynes | Wellcome Sanger Institute for the COVID-19 Genomics UK (COG-UK) consortium | The Lighthouse Lab in Milton Keynes and Alex Alderton, Roberto Amato, Sonia Goncalves, Ewan Harrison, David K. Jackson, Ian Johnston, Dominic Kwiatkowski, Cordelia Langford, John Sillitoe on behalf of the Wellcome Sanger Institute COVID-19 Surveillance Team (http://www.sanger.ac.uk/covid-team) |
| EPI_ISL_623288 | Lighthouse Lab in Cambridge | Wellcome Sanger Institute for the COVID-19 Genomics UK (COG-UK) consortium | Rob Howes, The Lighthouse Lab in Cambridge and Alex Alderton, Roberto Amato, Sonia Goncalves, Ewan Harrison, David K. Jackson, Ian Johnston, Dominic Kwiatkowski, Cordelia Langford, John Sillitoe on behalf of the Wellcome Sanger Institute COVID-19 Surveillance Team (http://www.sanger.ac.uk/covid-team) |
| EPI_ISL_623289, EPI_ISL_623290, EPI_ISL_623291, EPI_ISL_623292, EPI_ISL_623293, EPI_ISL_623294, EPI_ISL_623295, EPI_ISL_623296 | Lighthouse Lab in Milton Keynes | Wellcome Sanger Institute for the COVID-19 Genomics UK (COG-UK) consortium | The Lighthouse Lab in Milton Keynes and Alex Alderton, Roberto Amato, Sonia Goncalves, Ewan Harrison, David K. Jackson, Ian Johnston, Dominic Kwiatkowski, Cordelia Langford, John Sillitoe on behalf of the Wellcome Sanger Institute COVID-19 Surveillance Team (http://www.sanger.ac.uk/covid-team) |
| EPI_ISL_623298 | Lighthouse Lab in Milton Keynes | Wellcome Sanger Institute for the COVID-19 Genomics UK (COG-UK) Consortium | The Lighthouse Lab in Milton Keynes and Alex Alderton, Roberto Amato, Sonia Goncalves, Ewan Harrison, David K. Jackson, Ian Johnston, Dominic Kwiatkowski, Cordelia Langford, John Sillitoe on behalf of the Wellcome Sanger Institute COVID-19 Surveillance Team (http://www.sanger.ac.uk/covid-team) |
| EPI_ISL_623299 | Lighthouse Lab in Milton Keynes | Wellcome Sanger Institute for the COVID-19 Genomics UK (COG-UK) consortium | The Lighthouse Lab in Milton Keynes and Alex Alderton, Roberto Amato, Sonia Goncalves, Ewan Harrison, David K. Jackson, Ian Johnston, Dominic Kwiatkowski, Cordelia Langford, John Sillitoe on behalf of the Wellcome Sanger Institute COVID-19 Surveillance Team (http://www.sanger.ac.uk/covid-team) |
| EPI_ISL_623301 | Lighthouse Lab in Cambridge | Wellcome Sanger Institute for the COVID-19 Genomics UK (COG-UK) consortium | Rob Howes, The Lighthouse Lab in Cambridge and Alex Alderton, Roberto Amato, Sonia Goncalves, Ewan Harrison, David K. Jackson, Ian Johnston, Dominic Kwiatkowski, Cordelia Langford, John Sillitoe on behalf of the Wellcome Sanger Institute COVID-19 Surveillance Team (http://www.sanger.ac.uk/covid-team) |
| EPI_ISL_623302, EPI_ISL_623303, EPI_ISL_623304, EPI_ISL_623305, EPI_ISL_623306, EPI_ISL_623307, EPI_ISL_623309, EPI_ISL_623310 | Lighthouse Lab in Milton Keynes | Wellcome Sanger Institute for the COVID-19 Genomics UK (COG-UK) consortium | The Lighthouse Lab in Milton Keynes and Alex Alderton, Roberto Amato, Sonia Goncalves, Ewan Harrison, David K. Jackson, Ian Johnston, Dominic Kwiatkowski, Cordelia Langford, John Sillitoe on behalf of the Wellcome Sanger Institute COVID-19 Surveillance Team (http://www.sanger.ac.uk/covid-team) |
| EPI_ISL_623313 | Lighthouse Lab in Milton Keynes | Wellcome Sanger Institute for the COVID-19 Genomics UK (COG-UK) Consortium | The Lighthouse Lab in Milton Keynes and Alex Alderton, Roberto Amato, Sonia Goncalves, Ewan Harrison, David K. Jackson, Ian Johnston, Dominic Kwiatkowski, Cordelia Langford, John Sillitoe on behalf of the Wellcome Sanger Institute COVID-19 Surveillance Team (http://www.sanger.ac.uk/covid-team) |
| EPI_ISL_623314, EPI_ISL_623315, EPI_ISL_623316, EPI_ISL_623317, EPI_ISL_623318, EPI_ISL_623319, EPI_ISL_623320 | Lighthouse Lab in Milton Keynes | Wellcome Sanger Institute for the COVID-19 Genomics UK (COG-UK) consortium | The Lighthouse Lab in Milton Keynes and Alex Alderton, Roberto Amato, Sonia Goncalves, Ewan Harrison, David K. Jackson, Ian Johnston, Dominic Kwiatkowski, Cordelia Langford, John Sillitoe on behalf of the Wellcome Sanger Institute COVID-19 Surveillance Team (http://www.sanger.ac.uk/covid-team) |
| EPI_ISL_623321 | Lighthouse Lab in Cambridge | Wellcome Sanger Institute for the COVID-19 Genomics UK (COG-UK) consortium | Rob Howes, The Lighthouse Lab in Cambridge and Alex Alderton, Roberto Amato, Sonia Goncalves, Ewan Harrison, David K. Jackson, Ian Johnston, Dominic Kwiatkowski, Cordelia Langford, John Sillitoe on behalf of the Wellcome Sanger Institute COVID-19 Surveillance Team (http://www.sanger.ac.uk/covid-team) |
| EPI_ISL_623322 | Lighthouse Lab in Milton Keynes | Wellcome Sanger Institute for the COVID-19 Genomics UK (COG-UK) consortium | The Lighthouse Lab in Milton Keynes and Alex Alderton, Roberto Amato, Sonia Goncalves, Ewan Harrison, David K. Jackson, Ian Johnston, Dominic Kwiatkowski, Cordelia Langford, John Sillitoe on behalf of the Wellcome Sanger Institute COVID-19 Surveillance Team (http://www.sanger.ac.uk/covid-team) |
| EPI_ISL_623323 | Lighthouse Lab in Cambridge | Wellcome Sanger Institute for the COVID-19 Genomics UK (COG-UK) consortium | Rob Howes, The Lighthouse Lab in Cambridge and Alex Alderton, Roberto Amato, Sonia Goncalves, Ewan Harrison, David K. Jackson, Ian Johnston, Dominic Kwiatkowski, Cordelia Langford, John Sillitoe on behalf of the Wellcome Sanger Institute COVID-19 Surveillance Team (http://www.sanger.ac.uk/covid-team) |
| EPI_ISL_623324, EPI_ISL_623325, EPI_ISL_623326, EPI_ISL_623327, EPI_ISL_623328, EPI_ISL_623329, EPI_ISL_623331, EPI_ISL_623332 | Lighthouse Lab in Milton Keynes | Wellcome Sanger Institute for the COVID-19 Genomics UK (COG-UK) consortium | The Lighthouse Lab in Milton Keynes and Alex Alderton, Roberto Amato, Sonia Goncalves, Ewan Harrison, David K. Jackson, Ian Johnston, Dominic Kwiatkowski, Cordelia Langford, John Sillitoe on behalf of the Wellcome Sanger Institute COVID-19 Surveillance Team (http://www.sanger.ac.uk/covid-team) |
| EPI_ISL_623334 | Lighthouse Lab in Cambridge | Wellcome Sanger Institute for the COVID-19 Genomics UK (COG-UK) consortium | Rob Howes, The Lighthouse Lab in Cambridge and Alex Alderton, Roberto Amato, Sonia Goncalves, Ewan Harrison, David K. Jackson, Ian Johnston, Dominic Kwiatkowski, Cordelia Langford, John Sillitoe on behalf of the Wellcome Sanger Institute COVID-19 Surveillance Team (http://www.sanger.ac.uk/covid-team) |
| EPI_ISL_623335, EPI_ISL_623336, EPI_ISL_623338, EPI_ISL_623339, EPI_ISL_623340, EPI_ISL_623341, EPI_ISL_623342 | Lighthouse Lab in Milton Keynes | Wellcome Sanger Institute for the COVID-19 Genomics UK (COG-UK) consortium | The Lighthouse Lab in Milton Keynes and Alex Alderton, Roberto Amato, Sonia Goncalves, Ewan Harrison, David K. Jackson, Ian Johnston, Dominic Kwiatkowski, Cordelia Langford, John Sillitoe on behalf of the Wellcome Sanger Institute COVID-19 Surveillance Team (http://www.sanger.ac.uk/covid-team) |
| EPI_ISL_623344, EPI_ISL_623346 | Lighthouse Lab in Cambridge | Wellcome Sanger Institute for the COVID-19 Genomics UK (COG-UK) consortium | Rob Howes, The Lighthouse Lab in Cambridge and Alex Alderton, Roberto Amato, Sonia Goncalves, Ewan Harrison, David K. Jackson, Ian Johnston, Dominic Kwiatkowski, Cordelia Langford, John Sillitoe on behalf of the Wellcome Sanger Institute COVID-19 Surveillance Team (http://www.sanger.ac.uk/covid-team) |
| EPI_ISL_623348, EPI_ISL_623349, EPI_ISL_623350, EPI_ISL_623351, EPI_ISL_623352, EPI_ISL_623353, EPI_ISL_623355, EPI_ISL_623356, EPI_ISL_623357, EPI_ISL_623358 | Lighthouse Lab in Milton Keynes | Wellcome Sanger Institute for the COVID-19 Genomics UK (COG-UK) consortium | The Lighthouse Lab in Milton Keynes and Alex Alderton, Roberto Amato, Sonia Goncalves, Ewan Harrison, David K. Jackson, Ian Johnston, Dominic Kwiatkowski, Cordelia Langford, John Sillitoe on behalf of the Wellcome Sanger Institute COVID-19 Surveillance Team (http://www.sanger.ac.uk/covid-team) |
| EPI_ISL_623360 | Lighthouse Lab in Cambridge | Wellcome Sanger Institute for the COVID-19 Genomics UK (COG-UK) consortium | Rob Howes, The Lighthouse Lab in Cambridge and Alex Alderton, Roberto Amato, Sonia Goncalves, Ewan Harrison, David K. Jackson, Ian Johnston, Dominic Kwiatkowski, Cordelia Langford, John Sillitoe on behalf of the Wellcome Sanger Institute COVID-19 Surveillance Team (http://www.sanger.ac.uk/covid-team) |
| EPI_ISL_623361, EPI_ISL_623362 | Lighthouse Lab in Milton Keynes | Wellcome Sanger Institute for the COVID-19 Genomics UK (COG-UK) consortium | The Lighthouse Lab in Milton Keynes and Alex Alderton, Roberto Amato, Sonia Goncalves, Ewan Harrison, David K. Jackson, Ian Johnston, Dominic Kwiatkowski, Cordelia Langford, John Sillitoe on behalf of the Wellcome Sanger Institute COVID-19 Surveillance Team (http://www.sanger.ac.uk/covid-team) |
| EPI_ISL_623364 | Lighthouse Lab in Cambridge | Wellcome Sanger Institute for the COVID-19 Genomics UK (COG-UK) consortium | Rob Howes, The Lighthouse Lab in Cambridge and Alex Alderton, Roberto Amato, Sonia Goncalves, Ewan Harrison, David K. Jackson, Ian Johnston, Dominic Kwiatkowski, Cordelia Langford, John Sillitoe on behalf of the Wellcome Sanger Institute COVID-19 Surveillance Team (http://www.sanger.ac.uk/covid-team) |
| EPI_ISL_623365 | Lighthouse Lab in Milton Keynes | Wellcome Sanger Institute for the COVID-19 Genomics UK (COG-UK) consortium | The Lighthouse Lab in Milton Keynes and Alex Alderton, Roberto Amato, Sonia Goncalves, Ewan Harrison, David K. Jackson, Ian Johnston, Dominic Kwiatkowski, Cordelia Langford, John Sillitoe on behalf of the Wellcome Sanger Institute COVID-19 Surveillance Team (http://www.sanger.ac.uk/covid-team) |

| | | | |
|---|---|---|---|
| EPI_ISL_623366 | Lighthouse Lab in Cambridge | Wellcome Sanger Institute for the COVID-19 Genomics UK (COG-UK) consortium | Rob Howes, The Lighthouse Lab in Cambridge and Alex Alderton, Roberto Amato, Sonia Goncalves, Ewan Harrison, David K. Jackson, Ian Johnston, Dominic Kwiatkowski, Cordelia Langford, John Sillitoe on behalf of the Wellcome Sanger Institute COVID-19 Surveillance Team (http://www.sanger.ac.uk/covid-team) |
| EPI_ISL_623367, EPI_ISL_623368, EPI_ISL_623369, EPI_ISL_623371, EPI_ISL_623373, EPI_ISL_623374, EPI_ISL_623375, EPI_ISL_623376, EPI_ISL_623378, EPI_ISL_623379, EPI_ISL_623380, EPI_ISL_623381, EPI_ISL_623382, EPI_ISL_623383 | | | |
| see above | Lighthouse Lab in Milton Keynes | Wellcome Sanger Institute for the COVID-19 Genomics UK (COG-UK) consortium | The Lighthouse Lab in Milton Keynes and Alex Alderton, Roberto Amato, Sonia Goncalves, Ewan Harrison, David K. Jackson, Ian Johnston, Dominic Kwiatkowski, Cordelia Langford, John Sillitoe on behalf of the Wellcome Sanger Institute COVID-19 Surveillance Team (http://www.sanger.ac.uk/covid-team) |
| EPI_ISL_623385, EPI_ISL_623386 | Lighthouse Lab in Cambridge | Wellcome Sanger Institute for the COVID-19 Genomics UK (COG-UK) consortium | Rob Howes, The Lighthouse Lab in Cambridge and Alex Alderton, Roberto Amato, Sonia Goncalves, Ewan Harrison, David K. Jackson, Ian Johnston, Dominic Kwiatkowski, Cordelia Langford, John Sillitoe on behalf of the Wellcome Sanger Institute COVID-19 Surveillance Team (http://www.sanger.ac.uk/covid-team) |
| EPI_ISL_623387, EPI_ISL_623388, EPI_ISL_623389, EPI_ISL_623391 | Lighthouse Lab in Milton Keynes | Wellcome Sanger Institute for the COVID-19 Genomics UK (COG-UK) consortium | The Lighthouse Lab in Milton Keynes and Alex Alderton, Roberto Amato, Sonia Goncalves, Ewan Harrison, David K. Jackson, Ian Johnston, Dominic Kwiatkowski, Cordelia Langford, John Sillitoe on behalf of the Wellcome Sanger Institute COVID-19 Surveillance Team (http://www.sanger.ac.uk/covid-team) |
| EPI_ISL_623392, EPI_ISL_623393 | Lighthouse Lab in Cambridge | Wellcome Sanger Institute for the COVID-19 Genomics UK (COG-UK) consortium | Rob Howes, The Lighthouse Lab in Cambridge and Alex Alderton, Roberto Amato, Sonia Goncalves, Ewan Harrison, David K. Jackson, Ian Johnston, Dominic Kwiatkowski, Cordelia Langford, John Sillitoe on behalf of the Wellcome Sanger Institute COVID-19 Surveillance Team (http://www.sanger.ac.uk/covid-team) |
| EPI_ISL_623394, EPI_ISL_623396 | Lighthouse Lab in Milton Keynes | Wellcome Sanger Institute for the COVID-19 Genomics UK (COG-UK) consortium | The Lighthouse Lab in Milton Keynes and Alex Alderton, Roberto Amato, Sonia Goncalves, Ewan Harrison, David K. Jackson, Ian Johnston, Dominic Kwiatkowski, Cordelia Langford, John Sillitoe on behalf of the Wellcome Sanger Institute COVID-19 Surveillance Team (http://www.sanger.ac.uk/covid-team) |
| EPI_ISL_623398 | Lighthouse Lab in Cambridge | Wellcome Sanger Institute for the COVID-19 Genomics UK (COG-UK) consortium | Rob Howes, The Lighthouse Lab in Cambridge and Alex Alderton, Roberto Amato, Sonia Goncalves, Ewan Harrison, David K. Jackson, Ian Johnston, Dominic Kwiatkowski, Cordelia Langford, John Sillitoe on behalf of the Wellcome Sanger Institute COVID-19 Surveillance Team (http://www.sanger.ac.uk/covid-team) |
| EPI_ISL_623399, EPI_ISL_623401, EPI_ISL_623402, EPI_ISL_623403 | Lighthouse Lab in Milton Keynes | Wellcome Sanger Institute for the COVID-19 Genomics UK (COG-UK) consortium | The Lighthouse Lab in Milton Keynes and Alex Alderton, Roberto Amato, Sonia Goncalves, Ewan Harrison, David K. Jackson, Ian Johnston, Dominic Kwiatkowski, Cordelia Langford, John Sillitoe on behalf of the Wellcome Sanger Institute COVID-19 Surveillance Team (http://www.sanger.ac.uk/covid-team) |
| EPI_ISL_623404 | Lighthouse Lab in Cambridge | Wellcome Sanger Institute for the COVID-19 Genomics UK (COG-UK) consortium | Rob Howes, The Lighthouse Lab in Cambridge and Alex Alderton, Roberto Amato, Sonia Goncalves, Ewan Harrison, David K. Jackson, Ian Johnston, Dominic Kwiatkowski, Cordelia Langford, John Sillitoe on behalf of the Wellcome Sanger Institute COVID-19 Surveillance Team (http://www.sanger.ac.uk/covid-team) |
| EPI_ISL_623405, EPI_ISL_623406, EPI_ISL_623407 | Lighthouse Lab in Milton Keynes | Wellcome Sanger Institute for the COVID-19 Genomics UK (COG-UK) consortium | The Lighthouse Lab in Milton Keynes and Alex Alderton, Roberto Amato, Sonia Goncalves, Ewan Harrison, David K. Jackson, Ian Johnston, Dominic Kwiatkowski, Cordelia Langford, John Sillitoe on behalf of the Wellcome Sanger Institute COVID-19 Surveillance Team (http://www.sanger.ac.uk/covid-team) |
| EPI_ISL_623409 | Lighthouse Lab in Cambridge | Wellcome Sanger Institute for the COVID-19 Genomics UK (COG-UK) consortium | Rob Howes, The Lighthouse Lab in Cambridge and Alex Alderton, Roberto Amato, Sonia Goncalves, Ewan Harrison, David K. Jackson, Ian Johnston, Dominic Kwiatkowski, Cordelia Langford, John Sillitoe on behalf of the Wellcome Sanger Institute COVID-19 Surveillance Team (http://www.sanger.ac.uk/covid-team) |
| EPI_ISL_623410 | Lighthouse Lab in Milton Keynes | Wellcome Sanger Institute for the COVID-19 Genomics UK (COG-UK) consortium | The Lighthouse Lab in Milton Keynes and Alex Alderton, Roberto Amato, Sonia Goncalves, Ewan Harrison, David K. Jackson, Ian Johnston, Dominic Kwiatkowski, Cordelia Langford, John Sillitoe on behalf of the Wellcome Sanger Institute COVID-19 Surveillance Team (http://www.sanger.ac.uk/covid-team) |
| EPI_ISL_623411 | Lighthouse Lab in Cambridge | Wellcome Sanger Institute for the COVID-19 Genomics UK (COG-UK) consortium | Rob Howes, The Lighthouse Lab in Cambridge and Alex Alderton, Roberto Amato, Sonia Goncalves, Ewan Harrison, David K. Jackson, Ian Johnston, Dominic Kwiatkowski, Cordelia Langford, John Sillitoe on behalf of the Wellcome Sanger Institute COVID-19 Surveillance Team (http://www.sanger.ac.uk/covid-team) |
| EPI_ISL_623412, EPI_ISL_623413, EPI_ISL_623414, EPI_ISL_623416, EPI_ISL_623417, EPI_ISL_623418, EPI_ISL_623419, EPI_ISL_623420, EPI_ISL_623422, EPI_ISL_623423 | Lighthouse Lab in Milton Keynes | Wellcome Sanger Institute for the COVID-19 Genomics UK (COG-UK) consortium | The Lighthouse Lab in Milton Keynes and Alex Alderton, Roberto Amato, Sonia Goncalves, Ewan Harrison, David K. Jackson, Ian Johnston, Dominic Kwiatkowski, Cordelia Langford, John Sillitoe on behalf of the Wellcome Sanger Institute COVID-19 Surveillance Team (http://www.sanger.ac.uk/covid-team) |
| EPI_ISL_623425 | Lighthouse Lab in Cambridge | Wellcome Sanger Institute for the COVID-19 Genomics UK (COG-UK) consortium | Rob Howes, The Lighthouse Lab in Cambridge and Alex Alderton, Roberto Amato, Sonia Goncalves, Ewan Harrison, David K. Jackson, Ian Johnston, Dominic Kwiatkowski, Cordelia Langford, John Sillitoe on behalf of the Wellcome Sanger Institute COVID-19 Surveillance Team (http://www.sanger.ac.uk/covid-team) |
| EPI_ISL_623428 | Lighthouse Lab in Milton Keynes | Wellcome Sanger Institute for the COVID-19 Genomics UK (COG-UK) Consortium | The Lighthouse Lab in Milton Keynes and Alex Alderton, Roberto Amato, Sonia Goncalves, Ewan Harrison, David K. Jackson, Ian Johnston, Dominic Kwiatkowski, Cordelia Langford, John Sillitoe on behalf of the Wellcome Sanger Institute COVID-19 Surveillance Team (http://www.sanger.ac.uk/covid-team) |
| EPI_ISL_623429, EPI_ISL_623430, EPI_ISL_623431, EPI_ISL_623432, EPI_ISL_623434, EPI_ISL_623435, EPI_ISL_623436, EPI_ISL_623438, EPI_ISL_623439, EPI_ISL_623441, EPI_ISL_623446, EPI_ISL_623447 | | | |
| see above | Lighthouse Lab in Milton Keynes | Wellcome Sanger Institute for the COVID-19 Genomics UK (COG-UK) consortium | The Lighthouse Lab in Milton Keynes and Alex Alderton, Roberto Amato, Sonia Goncalves, Ewan Harrison, David K. Jackson, Ian Johnston, Dominic Kwiatkowski, Cordelia Langford, John Sillitoe on behalf of the Wellcome Sanger Institute COVID-19 Surveillance Team (http://www.sanger.ac.uk/covid-team) |
| EPI_ISL_623448, EPI_ISL_623449 | Lighthouse Lab in Cambridge | Wellcome Sanger Institute for the COVID-19 Genomics UK (COG-UK) consortium | Rob Howes, The Lighthouse Lab in Cambridge and Alex Alderton, Roberto Amato, Sonia Goncalves, Ewan Harrison, David K. Jackson, Ian Johnston, Dominic Kwiatkowski, Cordelia Langford, John Sillitoe on behalf of the Wellcome Sanger Institute COVID-19 Surveillance Team (http://www.sanger.ac.uk/covid-team) |
| EPI_ISL_623450 | Lighthouse Lab in Milton Keynes | Wellcome Sanger Institute for the COVID-19 Genomics UK (COG-UK) consortium | The Lighthouse Lab in Milton Keynes and Alex Alderton, Roberto Amato, Sonia Goncalves, Ewan Harrison, David K. Jackson, Ian Johnston, Dominic Kwiatkowski, Cordelia Langford, John Sillitoe on behalf of the Wellcome Sanger Institute COVID-19 Surveillance Team (http://www.sanger.ac.uk/covid-team) |
| EPI_ISL_623451 | Lighthouse Lab in Cambridge | Wellcome Sanger Institute for the COVID-19 Genomics UK (COG-UK) consortium | Rob Howes, The Lighthouse Lab in Cambridge and Alex Alderton, Roberto Amato, Sonia Goncalves, Ewan Harrison, David K. Jackson, Ian Johnston, Dominic Kwiatkowski, Cordelia Langford, John Sillitoe on behalf of the Wellcome Sanger Institute COVID-19 Surveillance Team (http://www.sanger.ac.uk/covid-team) |
| EPI_ISL_623452, EPI_ISL_623453, EPI_ISL_623454, EPI_ISL_623455 | Lighthouse Lab in Milton Keynes | Wellcome Sanger Institute for the COVID-19 Genomics UK (COG-UK) consortium | The Lighthouse Lab in Milton Keynes and Alex Alderton, Roberto Amato, Sonia Goncalves, Ewan Harrison, David K. Jackson, Ian Johnston, Dominic Kwiatkowski, Cordelia Langford, John Sillitoe on behalf of the Wellcome Sanger Institute COVID-19 Surveillance Team (http://www.sanger.ac.uk/covid-team) |
| EPI_ISL_623456 | Lighthouse Lab in Milton Keynes | Wellcome Sanger Institute for the COVID-19 Genomics UK (COG-UK) Consortium | The Lighthouse Lab in Milton Keynes and Alex Alderton, Roberto Amato, Sonia Goncalves, Ewan Harrison, David K. Jackson, Ian Johnston, Dominic Kwiatkowski, Cordelia Langford, John Sillitoe on behalf of the Wellcome Sanger Institute COVID-19 Surveillance Team (http://www.sanger.ac.uk/covid-team) |
| EPI_ISL_623458, EPI_ISL_623459, EPI_ISL_623460, EPI_ISL_623461, EPI_ISL_623462, EPI_ISL_623463, EPI_ISL_623464, EPI_ISL_623465, EPI_ISL_623466, EPI_ISL_623467, EPI_ISL_623469, EPI_ISL_623470, EPI_ISL_623471, EPI_ISL_623472, EPI_ISL_623473 | | | |
| see above | Lighthouse Lab in Milton Keynes | Wellcome Sanger Institute for the COVID-19 Genomics UK (COG-UK) consortium | The Lighthouse Lab in Milton Keynes and Alex Alderton, Roberto Amato, Sonia Goncalves, Ewan Harrison, David K. Jackson, Ian Johnston, Dominic Kwiatkowski, Cordelia Langford, John Sillitoe on behalf of the Wellcome Sanger Institute COVID-19 Surveillance Team (http://www.sanger.ac.uk/covid-team) |
| EPI_ISL_623474 | Lighthouse Lab in Cambridge | Wellcome Sanger Institute for the COVID-19 Genomics UK (COG-UK) consortium | Rob Howes, The Lighthouse Lab in Cambridge and Alex Alderton, Roberto Amato, Sonia Goncalves, Ewan Harrison, David K. Jackson, Ian Johnston, Dominic Kwiatkowski, Cordelia Langford, John Sillitoe on behalf of the Wellcome Sanger Institute COVID-19 Surveillance Team (http://www.sanger.ac.uk/covid-team) |
| EPI_ISL_623476, EPI_ISL_623478, EPI_ISL_623479, EPI_ISL_623480, EPI_ISL_623481 | Lighthouse Lab in Milton Keynes | Wellcome Sanger Institute for the COVID-19 Genomics UK (COG-UK) consortium | The Lighthouse Lab in Milton Keynes and Alex Alderton, Roberto Amato, Sonia Goncalves, Ewan Harrison, David K. Jackson, Ian Johnston, Dominic Kwiatkowski, Cordelia Langford, John Sillitoe on behalf of the Wellcome Sanger Institute COVID-19 Surveillance Team (http://www.sanger.ac.uk/covid-team) |
| EPI_ISL_623482 | Lighthouse Lab in Cambridge | Wellcome Sanger Institute for the COVID-19 Genomics UK (COG-UK) consortium | Rob Howes, The Lighthouse Lab in Cambridge and Alex Alderton, Roberto Amato, Sonia Goncalves, Ewan Harrison, David K. Jackson, Ian Johnston, Dominic Kwiatkowski, Cordelia Langford, John Sillitoe on behalf of the Wellcome Sanger Institute COVID-19 Surveillance Team (http://www.sanger.ac.uk/covid-team) |
| EPI_ISL_623483 | Lighthouse Lab in Milton Keynes | Wellcome Sanger Institute for the COVID-19 Genomics UK (COG-UK) consortium | The Lighthouse Lab in Milton Keynes and Alex Alderton, Roberto Amato, Sonia Goncalves, Ewan Harrison, David K. Jackson, Ian Johnston, Dominic Kwiatkowski, Cordelia Langford, John Sillitoe on behalf of the Wellcome Sanger Institute COVID-19 Surveillance Team (http://www.sanger.ac.uk/covid-team) |

| | | | |
|---|---|---|---|
| EPI_ISL_623484 | Lighthouse Lab in Milton Keynes | Wellcome Sanger Institute for the COVID-19 Genomics UK (COG-UK) Consortium | The Lighthouse Lab in Milton Keynes and Alex Alderton, Roberto Amato, Sonia Goncalves, Ewan Harrison, David K. Jackson, Ian Johnston, Dominic Kwiatkowski, Cordelia Langford, John Sillitoe on behalf of the Wellcome Sanger Institute COVID-19 Surveillance Team (http://www.sanger.ac.uk/covid-team) |
| EPI_ISL_623485 | Lighthouse Lab in Cambridge | Wellcome Sanger Institute for the COVID-19 Genomics UK (COG-UK) consortium | Rob Howes, The Lighthouse Lab in Cambridge and Alex Alderton, Roberto Amato, Sonia Goncalves, Ewan Harrison, David K. Jackson, Ian Johnston, Dominic Kwiatkowski, Cordelia Langford, John Sillitoe on behalf of the Wellcome Sanger Institute COVID-19 Surveillance Team (http://www.sanger.ac.uk/covid-team) |
| EPI_ISL_623487 | Lighthouse Lab in Milton Keynes | Wellcome Sanger Institute for the COVID-19 Genomics UK (COG-UK) consortium | The Lighthouse Lab in Milton Keynes and Alex Alderton, Roberto Amato, Sonia Goncalves, Ewan Harrison, David K. Jackson, Ian Johnston, Dominic Kwiatkowski, Cordelia Langford, John Sillitoe on behalf of the Wellcome Sanger Institute COVID-19 Surveillance Team (http://www.sanger.ac.uk/covid-team) |
| EPI_ISL_623488, EPI_ISL_623489 | Lighthouse Lab in Cambridge | Wellcome Sanger Institute for the COVID-19 Genomics UK (COG-UK) consortium | Rob Howes, The Lighthouse Lab in Cambridge and Alex Alderton, Roberto Amato, Sonia Goncalves, Ewan Harrison, David K. Jackson, Ian Johnston, Dominic Kwiatkowski, Cordelia Langford, John Sillitoe on behalf of the Wellcome Sanger Institute COVID-19 Surveillance Team (http://www.sanger.ac.uk/covid-team) |
| EPI_ISL_623491, EPI_ISL_623492 | Lighthouse Lab in Milton Keynes | Wellcome Sanger Institute for the COVID-19 Genomics UK (COG-UK) consortium | The Lighthouse Lab in Milton Keynes and Alex Alderton, Roberto Amato, Sonia Goncalves, Ewan Harrison, David K. Jackson, Ian Johnston, Dominic Kwiatkowski, Cordelia Langford, John Sillitoe on behalf of the Wellcome Sanger Institute COVID-19 Surveillance Team (http://www.sanger.ac.uk/covid-team) |
| EPI_ISL_623494 | Lighthouse Lab in Cambridge | Wellcome Sanger Institute for the COVID-19 Genomics UK (COG-UK) consortium | Rob Howes, The Lighthouse Lab in Cambridge and Alex Alderton, Roberto Amato, Sonia Goncalves, Ewan Harrison, David K. Jackson, Ian Johnston, Dominic Kwiatkowski, Cordelia Langford, John Sillitoe on behalf of the Wellcome Sanger Institute COVID-19 Surveillance Team (http://www.sanger.ac.uk/covid-team) |
| EPI_ISL_623497, EPI_ISL_623498, EPI_ISL_623499, EPI_ISL_623500, EPI_ISL_623501, EPI_ISL_623502, EPI_ISL_623503, EPI_ISL_623504, EPI_ISL_623505, EPI_ISL_623506, EPI_ISL_623507, EPI_ISL_623508, EPI_ISL_623509, EPI_ISL_623510, EPI_ISL_623511, EPI_ISL_623513, EPI_ISL_623514 | | | |
| see above | Lighthouse Lab in Milton Keynes | Wellcome Sanger Institute for the COVID-19 Genomics UK (COG-UK) consortium | The Lighthouse Lab in Milton Keynes and Alex Alderton, Roberto Amato, Sonia Goncalves, Ewan Harrison, David K. Jackson, Ian Johnston, Dominic Kwiatkowski, Cordelia Langford, John Sillitoe on behalf of the Wellcome Sanger Institute COVID-19 Surveillance Team (http://www.sanger.ac.uk/covid-team) |
| EPI_ISL_623515 | Lighthouse Lab in Milton Keynes | Wellcome Sanger Institute for the COVID-19 Genomics UK (COG-UK) Consortium | The Lighthouse Lab in Milton Keynes and Alex Alderton, Roberto Amato, Sonia Goncalves, Ewan Harrison, David K. Jackson, Ian Johnston, Dominic Kwiatkowski, Cordelia Langford, John Sillitoe on behalf of the Wellcome Sanger Institute COVID-19 Surveillance Team (http://www.sanger.ac.uk/covid-team) |
| EPI_ISL_623516, EPI_ISL_623517, EPI_ISL_623518, EPI_ISL_623519 | Lighthouse Lab in Milton Keynes | Wellcome Sanger Institute for the COVID-19 Genomics UK (COG-UK) consortium | The Lighthouse Lab in Milton Keynes and Alex Alderton, Roberto Amato, Sonia Goncalves, Ewan Harrison, David K. Jackson, Ian Johnston, Dominic Kwiatkowski, Cordelia Langford, John Sillitoe on behalf of the Wellcome Sanger Institute COVID-19 Surveillance Team (http://www.sanger.ac.uk/covid-team) |
| EPI_ISL_623520 | Lighthouse Lab in Cambridge | Wellcome Sanger Institute for the COVID-19 Genomics UK (COG-UK) consortium | Rob Howes, The Lighthouse Lab in Cambridge and Alex Alderton, Roberto Amato, Sonia Goncalves, Ewan Harrison, David K. Jackson, Ian Johnston, Dominic Kwiatkowski, Cordelia Langford, John Sillitoe on behalf of the Wellcome Sanger Institute COVID-19 Surveillance Team (http://www.sanger.ac.uk/covid-team) |
| EPI_ISL_623521 | Lighthouse Lab in Milton Keynes | Wellcome Sanger Institute for the COVID-19 Genomics UK (COG-UK) consortium | The Lighthouse Lab in Milton Keynes and Alex Alderton, Roberto Amato, Sonia Goncalves, Ewan Harrison, David K. Jackson, Ian Johnston, Dominic Kwiatkowski, Cordelia Langford, John Sillitoe on behalf of the Wellcome Sanger Institute COVID-19 Surveillance Team (http://www.sanger.ac.uk/covid-team) |
| EPI_ISL_623522 | Lighthouse Lab in Milton Keynes | Wellcome Sanger Institute for the COVID-19 Genomics UK (COG-UK) Consortium | The Lighthouse Lab in Milton Keynes and Alex Alderton, Roberto Amato, Sonia Goncalves, Ewan Harrison, David K. Jackson, Ian Johnston, Dominic Kwiatkowski, Cordelia Langford, John Sillitoe on behalf of the Wellcome Sanger Institute COVID-19 Surveillance Team (http://www.sanger.ac.uk/covid-team) |
| EPI_ISL_623523, EPI_ISL_623524, EPI_ISL_623526 | Lighthouse Lab in Milton Keynes | Wellcome Sanger Institute for the COVID-19 Genomics UK (COG-UK) consortium | The Lighthouse Lab in Milton Keynes and Alex Alderton, Roberto Amato, Sonia Goncalves, Ewan Harrison, David K. Jackson, Ian Johnston, Dominic Kwiatkowski, Cordelia Langford, John Sillitoe on behalf of the Wellcome Sanger Institute COVID-19 Surveillance Team (http://www.sanger.ac.uk/covid-team) |
| EPI_ISL_623527 | Lighthouse Lab in Cambridge | Wellcome Sanger Institute for the COVID-19 Genomics UK (COG-UK) consortium | Rob Howes, The Lighthouse Lab in Cambridge and Alex Alderton, Roberto Amato, Sonia Goncalves, Ewan Harrison, David K. Jackson, Ian Johnston, Dominic Kwiatkowski, Cordelia Langford, John Sillitoe on behalf of the Wellcome Sanger Institute COVID-19 Surveillance Team (http://www.sanger.ac.uk/covid-team) |
| EPI_ISL_623530, EPI_ISL_623533, EPI_ISL_623534, EPI_ISL_623535, EPI_ISL_623536 | Lighthouse Lab in Milton Keynes | Wellcome Sanger Institute for the COVID-19 Genomics UK (COG-UK) consortium | The Lighthouse Lab in Milton Keynes and Alex Alderton, Roberto Amato, Sonia Goncalves, Ewan Harrison, David K. Jackson, Ian Johnston, Dominic Kwiatkowski, Cordelia Langford, John Sillitoe on behalf of the Wellcome Sanger Institute COVID-19 Surveillance Team (http://www.sanger.ac.uk/covid-team) |
| EPI_ISL_623537 | Lighthouse Lab in Cambridge | Wellcome Sanger Institute for the COVID-19 Genomics UK (COG-UK) consortium | Rob Howes, The Lighthouse Lab in Cambridge and Alex Alderton, Roberto Amato, Sonia Goncalves, Ewan Harrison, David K. Jackson, Ian Johnston, Dominic Kwiatkowski, Cordelia Langford, John Sillitoe on behalf of the Wellcome Sanger Institute COVID-19 Surveillance Team (http://www.sanger.ac.uk/covid-team) |
| EPI_ISL_623538 | Lighthouse Lab in Milton Keynes | Wellcome Sanger Institute for the COVID-19 Genomics UK (COG-UK) consortium | The Lighthouse Lab in Milton Keynes and Alex Alderton, Roberto Amato, Sonia Goncalves, Ewan Harrison, David K. Jackson, Ian Johnston, Dominic Kwiatkowski, Cordelia Langford, John Sillitoe on behalf of the Wellcome Sanger Institute COVID-19 Surveillance Team (http://www.sanger.ac.uk/covid-team) |
| EPI_ISL_623539 | Lighthouse Lab in Cambridge | Wellcome Sanger Institute for the COVID-19 Genomics UK (COG-UK) consortium | Rob Howes, The Lighthouse Lab in Cambridge and Alex Alderton, Roberto Amato, Sonia Goncalves, Ewan Harrison, David K. Jackson, Ian Johnston, Dominic Kwiatkowski, Cordelia Langford, John Sillitoe on behalf of the Wellcome Sanger Institute COVID-19 Surveillance Team (http://www.sanger.ac.uk/covid-team) |
| EPI_ISL_623540, EPI_ISL_623541, EPI_ISL_623543 | Lighthouse Lab in Milton Keynes | Wellcome Sanger Institute for the COVID-19 Genomics UK (COG-UK) consortium | The Lighthouse Lab in Milton Keynes and Alex Alderton, Roberto Amato, Sonia Goncalves, Ewan Harrison, David K. Jackson, Ian Johnston, Dominic Kwiatkowski, Cordelia Langford, John Sillitoe on behalf of the Wellcome Sanger Institute COVID-19 Surveillance Team (http://www.sanger.ac.uk/covid-team) |
| EPI_ISL_623545 | Lighthouse Lab in Cambridge | Wellcome Sanger Institute for the COVID-19 Genomics UK (COG-UK) consortium | Rob Howes, The Lighthouse Lab in Cambridge and Alex Alderton, Roberto Amato, Sonia Goncalves, Ewan Harrison, David K. Jackson, Ian Johnston, Dominic Kwiatkowski, Cordelia Langford, John Sillitoe on behalf of the Wellcome Sanger Institute COVID-19 Surveillance Team (http://www.sanger.ac.uk/covid-team) |
| EPI_ISL_623546, EPI_ISL_623549, EPI_ISL_623552, EPI_ISL_623554 | Lighthouse Lab in Milton Keynes | Wellcome Sanger Institute for the COVID-19 Genomics UK (COG-UK) consortium | The Lighthouse Lab in Milton Keynes and Alex Alderton, Roberto Amato, Sonia Goncalves, Ewan Harrison, David K. Jackson, Ian Johnston, Dominic Kwiatkowski, Cordelia Langford, John Sillitoe on behalf of the Wellcome Sanger Institute COVID-19 Surveillance Team (http://www.sanger.ac.uk/covid-team) |
| EPI_ISL_623558 | Lighthouse Lab in Cambridge | Wellcome Sanger Institute for the COVID-19 Genomics UK (COG-UK) consortium | Rob Howes, The Lighthouse Lab in Cambridge and Alex Alderton, Roberto Amato, Sonia Goncalves, Ewan Harrison, David K. Jackson, Ian Johnston, Dominic Kwiatkowski, Cordelia Langford, John Sillitoe on behalf of the Wellcome Sanger Institute COVID-19 Surveillance Team (http://www.sanger.ac.uk/covid-team) |
| EPI_ISL_623561, EPI_ISL_623563, EPI_ISL_623564, EPI_ISL_623565, EPI_ISL_623566, EPI_ISL_623571, EPI_ISL_623573, EPI_ISL_623577, EPI_ISL_623579, EPI_ISL_623580, EPI_ISL_623582, EPI_ISL_623583, EPI_ISL_623586, EPI_ISL_623588, EPI_ISL_623589, EPI_ISL_623590, EPI_ISL_623594, EPI_ISL_623596, EPI_ISL_623598, EPI_ISL_623599 | | | |
| see above | Lighthouse Lab in Milton Keynes | Wellcome Sanger Institute for the COVID-19 Genomics UK (COG-UK) consortium | The Lighthouse Lab in Milton Keynes and Alex Alderton, Roberto Amato, Sonia Goncalves, Ewan Harrison, David K. Jackson, Ian Johnston, Dominic Kwiatkowski, Cordelia Langford, John Sillitoe on behalf of the Wellcome Sanger Institute COVID-19 Surveillance Team (http://www.sanger.ac.uk/covid-team) |
| EPI_ISL_623602 | Lighthouse Lab in Milton Keynes | Wellcome Sanger Institute for the COVID-19 Genomics UK (COG-UK) Consortium | The Lighthouse Lab in Milton Keynes and Alex Alderton, Roberto Amato, Sonia Goncalves, Ewan Harrison, David K. Jackson, Ian Johnston, Dominic Kwiatkowski, Cordelia Langford, John Sillitoe on behalf of the Wellcome Sanger Institute COVID-19 Surveillance Team (http://www.sanger.ac.uk/covid-team) |
| EPI_ISL_623603, EPI_ISL_623604, EPI_ISL_623606, EPI_ISL_623607 | Lighthouse Lab in Milton Keynes | Wellcome Sanger Institute for the COVID-19 Genomics UK (COG-UK) consortium | The Lighthouse Lab in Milton Keynes and Alex Alderton, Roberto Amato, Sonia Goncalves, Ewan Harrison, David K. Jackson, Ian Johnston, Dominic Kwiatkowski, Cordelia Langford, John Sillitoe on behalf of the Wellcome Sanger Institute COVID-19 Surveillance Team (http://www.sanger.ac.uk/covid-team) |
| EPI_ISL_623609 | Lighthouse Lab in Cambridge | Wellcome Sanger Institute for the COVID-19 Genomics UK (COG-UK) consortium | Rob Howes, The Lighthouse Lab in Cambridge and Alex Alderton, Roberto Amato, Sonia Goncalves, Ewan Harrison, David K. Jackson, Ian Johnston, Dominic Kwiatkowski, Cordelia Langford, John Sillitoe on behalf of the Wellcome Sanger Institute COVID-19 Surveillance Team (http://www.sanger.ac.uk/covid-team) |
| EPI_ISL_623610, EPI_ISL_623614, EPI_ISL_623615, EPI_ISL_623618, EPI_ISL_623621, EPI_ISL_623622, EPI_ISL_623623, EPI_ISL_623624, EPI_ISL_623625, EPI_ISL_623627, EPI_ISL_623628, EPI_ISL_623629, EPI_ISL_623631, EPI_ISL_623632, EPI_ISL_623634, EPI_ISL_623636, EPI_ISL_623637, EPI_ISL_623638 | | | |
| see above | Lighthouse Lab in Milton Keynes | Wellcome Sanger Institute for the COVID-19 Genomics UK (COG-UK) consortium | The Lighthouse Lab in Milton Keynes and Alex Alderton, Roberto Amato, Sonia Goncalves, Ewan Harrison, David K. Jackson, Ian Johnston, Dominic Kwiatkowski, Cordelia Langford, John Sillitoe on behalf of the Wellcome Sanger Institute COVID-19 Surveillance Team (http://www.sanger.ac.uk/covid-team) |
| EPI_ISL_623639 | Lighthouse Lab in Cambridge | Wellcome Sanger Institute for the COVID-19 Genomics UK (COG-UK) consortium | Rob Howes, The Lighthouse Lab in Cambridge and Alex Alderton, Roberto Amato, Sonia Goncalves, Ewan Harrison, David K. Jackson, Ian Johnston, Dominic Kwiatkowski, Cordelia Langford, John Sillitoe on behalf of the Wellcome Sanger Institute COVID-19 Surveillance Team (http://www.sanger.ac.uk/covid-team) |

| | | | |
|---|---|---|---|
| EPI_ISL_623642, EPI_ISL_623646, EPI_ISL_623648, EPI_ISL_623649, EPI_ISL_623656, EPI_ISL_623659, EPI_ISL_623660, EPI_ISL_623668, EPI_ISL_623669, EPI_ISL_623670, EPI_ISL_623671, EPI_ISL_623676, EPI_ISL_623678, EPI_ISL_623681, EPI_ISL_623682, EPI_ISL_623684, EPI_ISL_623685, EPI_ISL_623687, EPI_ISL_623689, EPI_ISL_623692 | | | |
| see above | Lighthouse Lab in Milton Keynes | Wellcome Sanger Institute for the COVID-19 Genomics UK (COG-UK) consortium | The Lighthouse Lab in Milton Keynes and Alex Alderton, Roberto Amato, Sonia Goncalves, Ewan Harrison, David K. Jackson, Ian Johnston, Dominic Kwiatkowski, Cordelia Langford, John Sillitoe on behalf of the Wellcome Sanger Institute COVID-19 Surveillance Team (http://www.sanger.ac.uk/covid-team) |
| EPI_ISL_623694 | Lighthouse Lab in Cambridge | Wellcome Sanger Institute for the COVID-19 Genomics UK (COG-UK) consortium | Rob Howes, The Lighthouse Lab in Cambridge and Alex Alderton, Roberto Amato, Sonia Goncalves, Ewan Harrison, David K. Jackson, Ian Johnston, Dominic Kwiatkowski, Cordelia Langford, John Sillitoe on behalf of the Wellcome Sanger Institute COVID-19 Surveillance Team (http://www.sanger.ac.uk/covid-team) |
| EPI_ISL_623695, EPI_ISL_623696, EPI_ISL_623698, EPI_ISL_623701, EPI_ISL_623707, EPI_ISL_623709, EPI_ISL_623711 | Lighthouse Lab in Milton Keynes | Wellcome Sanger Institute for the COVID-19 Genomics UK (COG-UK) consortium | The Lighthouse Lab in Milton Keynes and Alex Alderton, Roberto Amato, Sonia Goncalves, Ewan Harrison, David K. Jackson, Ian Johnston, Dominic Kwiatkowski, Cordelia Langford, John Sillitoe on behalf of the Wellcome Sanger Institute COVID-19 Surveillance Team (http://www.sanger.ac.uk/covid-team) |
| EPI_ISL_623712 | Lighthouse Lab in Cambridge | Wellcome Sanger Institute for the COVID-19 Genomics UK (COG-UK) consortium | Rob Howes, The Lighthouse Lab in Cambridge and Alex Alderton, Roberto Amato, Sonia Goncalves, Ewan Harrison, David K. Jackson, Ian Johnston, Dominic Kwiatkowski, Cordelia Langford, John Sillitoe on behalf of the Wellcome Sanger Institute COVID-19 Surveillance Team (http://www.sanger.ac.uk/covid-team) |
| EPI_ISL_623714, EPI_ISL_623715, EPI_ISL_623718, EPI_ISL_623719, EPI_ISL_623720, EPI_ISL_623722, EPI_ISL_623727, EPI_ISL_623728, EPI_ISL_623729, EPI_ISL_623730, EPI_ISL_623732, EPI_ISL_623734, EPI_ISL_623737, EPI_ISL_623738, EPI_ISL_623739, EPI_ISL_623744, EPI_ISL_623745, EPI_ISL_623748, EPI_ISL_623753, EPI_ISL_623755, EPI_ISL_623756, EPI_ISL_623757, EPI_ISL_623760, EPI_ISL_623761, EPI_ISL_623766, EPI_ISL_623767, EPI_ISL_623769, EPI_ISL_623771, EPI_ISL_623778, EPI_ISL_623779, EPI_ISL_623780, EPI_ISL_623782, EPI_ISL_623783, EPI_ISL_623784, EPI_ISL_623786, EPI_ISL_623787 | | | |
| see above | Lighthouse Lab in Milton Keynes | Wellcome Sanger Institute for the COVID-19 Genomics UK (COG-UK) consortium | The Lighthouse Lab in Milton Keynes and Alex Alderton, Roberto Amato, Sonia Goncalves, Ewan Harrison, David K. Jackson, Ian Johnston, Dominic Kwiatkowski, Cordelia Langford, John Sillitoe on behalf of the Wellcome Sanger Institute COVID-19 Surveillance Team (http://www.sanger.ac.uk/covid-team) |
| EPI_ISL_623790 | Lighthouse Lab in Cambridge | Wellcome Sanger Institute for the COVID-19 Genomics UK (COG-UK) consortium | Rob Howes, The Lighthouse Lab in Cambridge and Alex Alderton, Roberto Amato, Sonia Goncalves, Ewan Harrison, David K. Jackson, Ian Johnston, Dominic Kwiatkowski, Cordelia Langford, John Sillitoe on behalf of the Wellcome Sanger Institute COVID-19 Surveillance Team (http://www.sanger.ac.uk/covid-team) |
| EPI_ISL_623792, EPI_ISL_623795, EPI_ISL_623801, EPI_ISL_623803, EPI_ISL_623804, EPI_ISL_623805, EPI_ISL_623807, EPI_ISL_623814, EPI_ISL_623817, EPI_ISL_623818, EPI_ISL_623819, EPI_ISL_623821, EPI_ISL_623826, EPI_ISL_623827, EPI_ISL_623830, EPI_ISL_623836, EPI_ISL_623837, EPI_ISL_623842, EPI_ISL_623853 | | | |
| see above | Lighthouse Lab in Milton Keynes | Wellcome Sanger Institute for the COVID-19 Genomics UK (COG-UK) consortium | The Lighthouse Lab in Milton Keynes and Alex Alderton, Roberto Amato, Sonia Goncalves, Ewan Harrison, David K. Jackson, Ian Johnston, Dominic Kwiatkowski, Cordelia Langford, John Sillitoe on behalf of the Wellcome Sanger Institute COVID-19 Surveillance Team (http://www.sanger.ac.uk/covid-team) |
| EPI_ISL_623856 | Lighthouse Lab in Cambridge | Wellcome Sanger Institute for the COVID-19 Genomics UK (COG-UK) consortium | Rob Howes, The Lighthouse Lab in Cambridge and Alex Alderton, Roberto Amato, Sonia Goncalves, Ewan Harrison, David K. Jackson, Ian Johnston, Dominic Kwiatkowski, Cordelia Langford, John Sillitoe on behalf of the Wellcome Sanger Institute COVID-19 Surveillance Team (http://www.sanger.ac.uk/covid-team) |
| EPI_ISL_623862 | Lighthouse Lab in Alderley Park | Wellcome Sanger Institute for the COVID-19 Genomics UK (COG-UK) consortium | Jacquelyn Wynn, Mairead Hyland, The Lighthouse Lab in Alderley Park and Alex Alderton, Roberto Amato, Sonia Goncalves, Ewan Harrison, David K. Jackson, Ian Johnston, Dominic Kwiatkowski, Cordelia Langford, John Sillitoe on behalf of the Wellcome Sanger Institute COVID-19 Surveillance Team (http://www.sanger.ac.uk/covid-team) |
| EPI_ISL_623863, EPI_ISL_623873 | Lighthouse Lab in Glasgow | Wellcome Sanger Institute for the COVID-19 Genomics UK (COG-UK) consortium | Harper VanSteenhouse, Yumi Kasai, David Gray, Carol Clugston, Anna Dominiczak and Alex Alderton, Roberto Amato, Sonia Goncalves, Ewan Harrison, David K. Jackson, Ian Johnston, Dominic Kwiatkowski, Cordelia Langford, John Sillitoe on behalf of the Wellcome Sanger Institute COVID-19 Surveillance Team (http://www.sanger.ac.uk/covid-team) |
| EPI_ISL_623879 | Lighthouse Lab in Alderley Park | Wellcome Sanger Institute for the COVID-19 Genomics UK (COG-UK) consortium | Jacquelyn Wynn, Mairead Hyland, The Lighthouse Lab in Alderley Park and Alex Alderton, Roberto Amato, Sonia Goncalves, Ewan Harrison, David K. Jackson, Ian Johnston, Dominic Kwiatkowski, Cordelia Langford, John Sillitoe on behalf of the Wellcome Sanger Institute COVID-19 Surveillance Team (http://www.sanger.ac.uk/covid-team) |
| EPI_ISL_623908 | Lighthouse Lab in Glasgow | Wellcome Sanger Institute for the COVID-19 Genomics UK (COG-UK) consortium | Harper VanSteenhouse, Yumi Kasai, David Gray, Carol Clugston, Anna Dominiczak and Alex Alderton, Roberto Amato, Sonia Goncalves, Ewan Harrison, David K. Jackson, Ian Johnston, Dominic Kwiatkowski, Cordelia Langford, John Sillitoe on behalf of the Wellcome Sanger Institute COVID-19 Surveillance Team (http://www.sanger.ac.uk/covid-team) |
| EPI_ISL_623910 | Lighthouse Lab in Alderley Park | Wellcome Sanger Institute for the COVID-19 Genomics UK (COG-UK) consortium | Jacquelyn Wynn, Mairead Hyland, The Lighthouse Lab in Alderley Park and Alex Alderton, Roberto Amato, Sonia Goncalves, Ewan Harrison, David K. Jackson, Ian Johnston, Dominic Kwiatkowski, Cordelia Langford, John Sillitoe on behalf of the Wellcome Sanger Institute COVID-19 Surveillance Team (http://www.sanger.ac.uk/covid-team) |
| EPI_ISL_623911 | Lighthouse Lab in Glasgow | Wellcome Sanger Institute for the COVID-19 Genomics UK (COG-UK) consortium | Harper VanSteenhouse, Yumi Kasai, David Gray, Carol Clugston, Anna Dominiczak and Alex Alderton, Roberto Amato, Sonia Goncalves, Ewan Harrison, David K. Jackson, Ian Johnston, Dominic Kwiatkowski, Cordelia Langford, John Sillitoe on behalf of the Wellcome Sanger Institute COVID-19 Surveillance Team (http://www.sanger.ac.uk/covid-team) |
| EPI_ISL_623918 | Lighthouse Lab in Alderley Park | Wellcome Sanger Institute for the COVID-19 Genomics UK (COG-UK) consortium | Jacquelyn Wynn, Mairead Hyland, The Lighthouse Lab in Alderley Park and Alex Alderton, Roberto Amato, Sonia Goncalves, Ewan Harrison, David K. Jackson, Ian Johnston, Dominic Kwiatkowski, Cordelia Langford, John Sillitoe on behalf of the Wellcome Sanger Institute COVID-19 Surveillance Team (http://www.sanger.ac.uk/covid-team) |
| EPI_ISL_623924 | Lighthouse Lab in Glasgow | Wellcome Sanger Institute for the COVID-19 Genomics UK (COG-UK) consortium | Harper VanSteenhouse, Yumi Kasai, David Gray, Carol Clugston, Anna Dominiczak and Alex Alderton, Roberto Amato, Sonia Goncalves, Ewan Harrison, David K. Jackson, Ian Johnston, Dominic Kwiatkowski, Cordelia Langford, John Sillitoe on behalf of the Wellcome Sanger Institute COVID-19 Surveillance Team (http://www.sanger.ac.uk/covid-team) |
| EPI_ISL_623925, EPI_ISL_623935, EPI_ISL_623939 | Lighthouse Lab in Alderley Park | Wellcome Sanger Institute for the COVID-19 Genomics UK (COG-UK) consortium | Jacquelyn Wynn, Mairead Hyland, The Lighthouse Lab in Alderley Park and Alex Alderton, Roberto Amato, Sonia Goncalves, Ewan Harrison, David K. Jackson, Ian Johnston, Dominic Kwiatkowski, Cordelia Langford, John Sillitoe on behalf of the Wellcome Sanger Institute COVID-19 Surveillance Team (http://www.sanger.ac.uk/covid-team) |
| EPI_ISL_623942, EPI_ISL_623954 | Lighthouse Lab in Glasgow | Wellcome Sanger Institute for the COVID-19 Genomics UK (COG-UK) consortium | Harper VanSteenhouse, Yumi Kasai, David Gray, Carol Clugston, Anna Dominiczak and Alex Alderton, Roberto Amato, Sonia Goncalves, Ewan Harrison, David K. Jackson, Ian Johnston, Dominic Kwiatkowski, Cordelia Langford, John Sillitoe on behalf of the Wellcome Sanger Institute COVID-19 Surveillance Team (http://www.sanger.ac.uk/covid-team) |
| EPI_ISL_623955 | Lighthouse Lab in Alderley Park | Wellcome Sanger Institute for the COVID-19 Genomics UK (COG-UK) consortium | Jacquelyn Wynn, Mairead Hyland, The Lighthouse Lab in Alderley Park and Alex Alderton, Roberto Amato, Sonia Goncalves, Ewan Harrison, David K. Jackson, Ian Johnston, Dominic Kwiatkowski, Cordelia Langford, John Sillitoe on behalf of the Wellcome Sanger Institute COVID-19 Surveillance Team (http://www.sanger.ac.uk/covid-team) |
| EPI_ISL_623956 | Lighthouse Lab in Glasgow | Wellcome Sanger Institute for the COVID-19 Genomics UK (COG-UK) consortium | Harper VanSteenhouse, Yumi Kasai, David Gray, Carol Clugston, Anna Dominiczak and Alex Alderton, Roberto Amato, Sonia Goncalves, Ewan Harrison, David K. Jackson, Ian Johnston, Dominic Kwiatkowski, Cordelia Langford, John Sillitoe on behalf of the Wellcome Sanger Institute COVID-19 Surveillance Team (http://www.sanger.ac.uk/covid-team) |
| EPI_ISL_623965, EPI_ISL_623971, EPI_ISL_623981 | Lighthouse Lab in Alderley Park | Wellcome Sanger Institute for the COVID-19 Genomics UK (COG-UK) consortium | Jacquelyn Wynn, Mairead Hyland, The Lighthouse Lab in Alderley Park and Alex Alderton, Roberto Amato, Sonia Goncalves, Ewan Harrison, David K. Jackson, Ian Johnston, Dominic Kwiatkowski, Cordelia Langford, John Sillitoe on behalf of the Wellcome Sanger Institute COVID-19 Surveillance Team (http://www.sanger.ac.uk/covid-team) |
| EPI_ISL_623991 | Lighthouse Lab in Glasgow | Wellcome Sanger Institute for the COVID-19 Genomics UK (COG-UK) consortium | Harper VanSteenhouse, Yumi Kasai, David Gray, Carol Clugston, Anna Dominiczak and Alex Alderton, Roberto Amato, Sonia Goncalves, Ewan Harrison, David K. Jackson, Ian Johnston, Dominic Kwiatkowski, Cordelia Langford, John Sillitoe on behalf of the Wellcome Sanger Institute COVID-19 Surveillance Team (http://www.sanger.ac.uk/covid-team) |
| EPI_ISL_624013 | Lighthouse Lab in Alderley Park | Wellcome Sanger Institute for the COVID-19 Genomics UK (COG-UK) consortium | Jacquelyn Wynn, Mairead Hyland, The Lighthouse Lab in Alderley Park and Alex Alderton, Roberto Amato, Sonia Goncalves, Ewan Harrison, David K. Jackson, Ian Johnston, Dominic Kwiatkowski, Cordelia Langford, John Sillitoe on behalf of the Wellcome Sanger Institute COVID-19 Surveillance Team (http://www.sanger.ac.uk/covid-team) |
| EPI_ISL_624019, EPI_ISL_624027, | Lighthouse Lab in Glasgow | Wellcome Sanger Institute for the COVID-19 Genomics UK | Harper VanSteenhouse, Yumi Kasai, David Gray, Carol Clugston, Anna Dominiczak and Alex Alderton, Roberto Amato, Sonia Goncalves, Ewan Harrison, |

| | | | |
|---|---|---|---|
| EPI_ISL_624028 | | (COG-UK) consortium | David K. Jackson, Ian Johnston, Dominic Kwiatkowski, Cordelia Langford, John Sillitoe on behalf of the Wellcome Sanger Institute COVID-19 Surveillance Team (http://www.sanger.ac.uk/covid-team) |
| EPI_ISL_624037, EPI_ISL_624038, EPI_ISL_624047, EPI_ISL_624048, EPI_ISL_624059, EPI_ISL_624062, EPI_ISL_624066 | Lighthouse Lab in Alderley Park | Wellcome Sanger Institute for the COVID-19 Genomics UK (COG-UK) consortium | Jacquelyn Wynn, Mairead Hyland, The Lighthouse Lab in Alderley Park and Alex Alderton, Roberto Amato, Sonia Goncalves, Ewan Harrison, David K. Jackson, Ian Johnston, Dominic Kwiatkowski, Cordelia Langford, John Sillitoe on behalf of the Wellcome Sanger Institute COVID-19 Surveillance Team (http://www.sanger.ac.uk/covid-team) |
| EPI_ISL_624073 | Lighthouse Lab in Glasgow | Wellcome Sanger Institute for the COVID-19 Genomics UK (COG-UK) consortium | Harper VanSteenhouse, Yumi Kasai, David Gray, Carol Clugston, Anna Dominiczak and Alex Alderton, Roberto Amato, Sonia Goncalves, Ewan Harrison, David K. Jackson, Ian Johnston, Dominic Kwiatkowski, Cordelia Langford, John Sillitoe on behalf of the Wellcome Sanger Institute COVID-19 Surveillance Team (http://www.sanger.ac.uk/covid-team) |
| EPI_ISL_624074 | Lighthouse Lab in Alderley Park | Wellcome Sanger Institute for the COVID-19 Genomics UK (COG-UK) consortium | Jacquelyn Wynn, Mairead Hyland, The Lighthouse Lab in Alderley Park and Alex Alderton, Roberto Amato, Sonia Goncalves, Ewan Harrison, David K. Jackson, Ian Johnston, Dominic Kwiatkowski, Cordelia Langford, John Sillitoe on behalf of the Wellcome Sanger Institute COVID-19 Surveillance Team (http://www.sanger.ac.uk/covid-team) |
| EPI_ISL_624077 | Lighthouse Lab in Glasgow | Wellcome Sanger Institute for the COVID-19 Genomics UK (COG-UK) consortium | Harper VanSteenhouse, Yumi Kasai, David Gray, Carol Clugston, Anna Dominiczak and Alex Alderton, Roberto Amato, Sonia Goncalves, Ewan Harrison, David K. Jackson, Ian Johnston, Dominic Kwiatkowski, Cordelia Langford, John Sillitoe on behalf of the Wellcome Sanger Institute COVID-19 Surveillance Team (http://www.sanger.ac.uk/covid-team) |
| EPI_ISL_624081, EPI_ISL_624083, EPI_ISL_624106, EPI_ISL_624117, EPI_ISL_624125, EPI_ISL_624126 | Lighthouse Lab in Alderley Park | Wellcome Sanger Institute for the COVID-19 Genomics UK (COG-UK) consortium | Jacquelyn Wynn, Mairead Hyland, The Lighthouse Lab in Alderley Park and Alex Alderton, Roberto Amato, Sonia Goncalves, Ewan Harrison, David K. Jackson, Ian Johnston, Dominic Kwiatkowski, Cordelia Langford, John Sillitoe on behalf of the Wellcome Sanger Institute COVID-19 Surveillance Team (http://www.sanger.ac.uk/covid-team) |
| EPI_ISL_624128 | Lighthouse Lab in Glasgow | Wellcome Sanger Institute for the COVID-19 Genomics UK (COG-UK) consortium | Harper VanSteenhouse, Yumi Kasai, David Gray, Carol Clugston, Anna Dominiczak and Alex Alderton, Roberto Amato, Sonia Goncalves, Ewan Harrison, David K. Jackson, Ian Johnston, Dominic Kwiatkowski, Cordelia Langford, John Sillitoe on behalf of the Wellcome Sanger Institute COVID-19 Surveillance Team (http://www.sanger.ac.uk/covid-team) |
| EPI_ISL_624134, EPI_ISL_624136 | Lighthouse Lab in Alderley Park | Wellcome Sanger Institute for the COVID-19 Genomics UK (COG-UK) consortium | Jacquelyn Wynn, Mairead Hyland, The Lighthouse Lab in Alderley Park and Alex Alderton, Roberto Amato, Sonia Goncalves, Ewan Harrison, David K. Jackson, Ian Johnston, Dominic Kwiatkowski, Cordelia Langford, John Sillitoe on behalf of the Wellcome Sanger Institute COVID-19 Surveillance Team (http://www.sanger.ac.uk/covid-team) |
| EPI_ISL_624147 | Lighthouse Lab in Glasgow | Wellcome Sanger Institute for the COVID-19 Genomics UK (COG-UK) consortium | Harper VanSteenhouse, Yumi Kasai, David Gray, Carol Clugston, Anna Dominiczak and Alex Alderton, Roberto Amato, Sonia Goncalves, Ewan Harrison, David K. Jackson, Ian Johnston, Dominic Kwiatkowski, Cordelia Langford, John Sillitoe on behalf of the Wellcome Sanger Institute COVID-19 Surveillance Team (http://www.sanger.ac.uk/covid-team) |
| EPI_ISL_624153 | Lighthouse Lab in Alderley Park | Wellcome Sanger Institute for the COVID-19 Genomics UK (COG-UK) consortium | Jacquelyn Wynn, Mairead Hyland, The Lighthouse Lab in Alderley Park and Alex Alderton, Roberto Amato, Sonia Goncalves, Ewan Harrison, David K. Jackson, Ian Johnston, Dominic Kwiatkowski, Cordelia Langford, John Sillitoe on behalf of the Wellcome Sanger Institute COVID-19 Surveillance Team (http://www.sanger.ac.uk/covid-team) |
| EPI_ISL_624158 | Lighthouse Lab in Glasgow | Wellcome Sanger Institute for the COVID-19 Genomics UK (COG-UK) consortium | Harper VanSteenhouse, Yumi Kasai, David Gray, Carol Clugston, Anna Dominiczak and Alex Alderton, Roberto Amato, Sonia Goncalves, Ewan Harrison, David K. Jackson, Ian Johnston, Dominic Kwiatkowski, Cordelia Langford, John Sillitoe on behalf of the Wellcome Sanger Institute COVID-19 Surveillance Team (http://www.sanger.ac.uk/covid-team) |
| EPI_ISL_624161 | Lighthouse Lab in Alderley Park | Wellcome Sanger Institute for the COVID-19 Genomics UK (COG-UK) consortium | Jacquelyn Wynn, Mairead Hyland, The Lighthouse Lab in Alderley Park and Alex Alderton, Roberto Amato, Sonia Goncalves, Ewan Harrison, David K. Jackson, Ian Johnston, Dominic Kwiatkowski, Cordelia Langford, John Sillitoe on behalf of the Wellcome Sanger Institute COVID-19 Surveillance Team (http://www.sanger.ac.uk/covid-team) |
| EPI_ISL_624163 | Lighthouse Lab in Glasgow | Wellcome Sanger Institute for the COVID-19 Genomics UK (COG-UK) consortium | Harper VanSteenhouse, Yumi Kasai, David Gray, Carol Clugston, Anna Dominiczak and Alex Alderton, Roberto Amato, Sonia Goncalves, Ewan Harrison, David K. Jackson, Ian Johnston, Dominic Kwiatkowski, Cordelia Langford, John Sillitoe on behalf of the Wellcome Sanger Institute COVID-19 Surveillance Team (http://www.sanger.ac.uk/covid-team) |
| EPI_ISL_624166, EPI_ISL_624170, EPI_ISL_624172, EPI_ISL_624177, EPI_ISL_624179, EPI_ISL_624182 | Lighthouse Lab in Alderley Park | Wellcome Sanger Institute for the COVID-19 Genomics UK (COG-UK) consortium | Jacquelyn Wynn, Mairead Hyland, The Lighthouse Lab in Alderley Park and Alex Alderton, Roberto Amato, Sonia Goncalves, Ewan Harrison, David K. Jackson, Ian Johnston, Dominic Kwiatkowski, Cordelia Langford, John Sillitoe on behalf of the Wellcome Sanger Institute COVID-19 Surveillance Team (http://www.sanger.ac.uk/covid-team) |
| EPI_ISL_624188, EPI_ISL_624189 | Lighthouse Lab in Glasgow | Wellcome Sanger Institute for the COVID-19 Genomics UK (COG-UK) consortium | Harper VanSteenhouse, Yumi Kasai, David Gray, Carol Clugston, Anna Dominiczak and Alex Alderton, Roberto Amato, Sonia Goncalves, Ewan Harrison, David K. Jackson, Ian Johnston, Dominic Kwiatkowski, Cordelia Langford, John Sillitoe on behalf of the Wellcome Sanger Institute COVID-19 Surveillance Team (http://www.sanger.ac.uk/covid-team) |
| EPI_ISL_624197 | Lighthouse Lab in Glasgow | Wellcome Sanger Institute for the COVID-19 Genomics UK (COG-UK) Consortium | Harper VanSteenhouse, Yumi Kasai, David Gray, Carol Clugston, Anna Dominiczak and Alex Alderton, Roberto Amato, Sonia Goncalves, Ewan Harrison, David K. Jackson, Ian Johnston, Dominic Kwiatkowski, Cordelia Langford, John Sillitoe on behalf of the Wellcome Sanger Institute COVID-19 Surveillance Team (http://www.sanger.ac.uk/covid-team) |
| EPI_ISL_624206, EPI_ISL_624208, EPI_ISL_624221, EPI_ISL_624230, EPI_ISL_624231, EPI_ISL_624234, EPI_ISL_624241, EPI_ISL_624243 | Lighthouse Lab in Alderley Park | Wellcome Sanger Institute for the COVID-19 Genomics UK (COG-UK) consortium | Jacquelyn Wynn, Mairead Hyland, The Lighthouse Lab in Alderley Park and Alex Alderton, Roberto Amato, Sonia Goncalves, Ewan Harrison, David K. Jackson, Ian Johnston, Dominic Kwiatkowski, Cordelia Langford, John Sillitoe on behalf of the Wellcome Sanger Institute COVID-19 Surveillance Team (http://www.sanger.ac.uk/covid-team) |
| EPI_ISL_624244 | Lighthouse Lab in Alderley Park | Wellcome Sanger Institute for the COVID-19 Genomics UK (COG-UK) Consortium | Jacquelyn Wynn, Mairead Hyland, The Lighthouse Lab in Alderley Park and Alex Alderton, Roberto Amato, Sonia Goncalves, Ewan Harrison, David K. Jackson, Ian Johnston, Dominic Kwiatkowski, Cordelia Langford, John Sillitoe on behalf of the Wellcome Sanger Institute COVID-19 Surveillance Team (http://www.sanger.ac.uk/covid-team) |
| EPI_ISL_624256, EPI_ISL_624261, EPI_ISL_624268, EPI_ISL_624278, EPI_ISL_624280, EPI_ISL_624284, EPI_ISL_624290, EPI_ISL_624295, EPI_ISL_624316, EPI_ISL_624319, EPI_ISL_624323, EPI_ISL_624328, EPI_ISL_624329, EPI_ISL_624334, EPI_ISL_624337, EPI_ISL_624344, EPI_ISL_624348, EPI_ISL_624352, EPI_ISL_624358, EPI_ISL_624359, EPI_ISL_624362, EPI_ISL_624364, EPI_ISL_624370, EPI_ISL_624372, EPI_ISL_624381, EPI_ISL_624385, EPI_ISL_624386, EPI_ISL_624397, EPI_ISL_624399, EPI_ISL_624403, EPI_ISL_624405, EPI_ISL_624416, EPI_ISL_624426, EPI_ISL_624427, EPI_ISL_624431, EPI_ISL_624439, EPI_ISL_624443, EPI_ISL_624447, EPI_ISL_624451, EPI_ISL_624465, EPI_ISL_624472, EPI_ISL_624473, EPI_ISL_624476, EPI_ISL_624479, EPI_ISL_624484, EPI_ISL_624488, EPI_ISL_624489, EPI_ISL_624492, EPI_ISL_624493, EPI_ISL_624501, EPI_ISL_624504, EPI_ISL_624509, EPI_ISL_624515, EPI_ISL_624517, EPI_ISL_624521, EPI_ISL_624522, EPI_ISL_624523, EPI_ISL_624536, EPI_ISL_624540, EPI_ISL_624548, EPI_ISL_624554, EPI_ISL_624559, EPI_ISL_624560, EPI_ISL_624561, EPI_ISL_624562, EPI_ISL_624563, EPI_ISL_624564, EPI_ISL_624565, EPI_ISL_624566, EPI_ISL_624567, EPI_ISL_624568, EPI_ISL_624569, EPI_ISL_624570, EPI_ISL_624571, EPI_ISL_624572, EPI_ISL_624573, EPI_ISL_624574, EPI_ISL_624575, EPI_ISL_624576, EPI_ISL_624577, EPI_ISL_624578, EPI_ISL_624579, EPI_ISL_624580, EPI_ISL_624581, EPI_ISL_624582, EPI_ISL_624583, EPI_ISL_624584, EPI_ISL_624585, EPI_ISL_624586, EPI_ISL_624587, EPI_ISL_624588, EPI_ISL_624589, EPI_ISL_624590, EPI_ISL_624591, EPI_ISL_624592, EPI_ISL_624593, EPI_ISL_624594, EPI_ISL_624595, EPI_ISL_624596, EPI_ISL_624597, EPI_ISL_624598, EPI_ISL_624599, EPI_ISL_624600, EPI_ISL_624601, EPI_ISL_624602, EPI_ISL_624603, EPI_ISL_624604, EPI_ISL_624605, EPI_ISL_624606, EPI_ISL_624607, EPI_ISL_624608, EPI_ISL_624609, EPI_ISL_624610, EPI_ISL_624611, EPI_ISL_624612, EPI_ISL_624613, EPI_ISL_624614, EPI_ISL_624615, EPI_ISL_624616, EPI_ISL_624617, EPI_ISL_624618, EPI_ISL_624619, EPI_ISL_624620, EPI_ISL_624621, EPI_ISL_624622, EPI_ISL_624623, EPI_ISL_624624, EPI_ISL_624625, EPI_ISL_624626, EPI_ISL_624627, EPI_ISL_624628, EPI_ISL_624629, EPI_ISL_624630, EPI_ISL_624631, EPI_ISL_624632, EPI_ISL_624633, EPI_ISL_624634, EPI_ISL_624635, EPI_ISL_624636, EPI_ISL_624637, EPI_ISL_624638, EPI_ISL_624639, EPI_ISL_624640, EPI_ISL_624641, EPI_ISL_624642, EPI_ISL_624643, EPI_ISL_624644, EPI_ISL_624645, EPI_ISL_624646, EPI_ISL_624647, EPI_ISL_624648, EPI_ISL_624649, EPI_ISL_624650, EPI_ISL_624651, EPI_ISL_624652, EPI_ISL_624653, EPI_ISL_624654, EPI_ISL_624655, EPI_ISL_624656, EPI_ISL_624657, EPI_ISL_624658, EPI_ISL_624659, EPI_ISL_624660, EPI_ISL_624661, EPI_ISL_624662, EPI_ISL_624663, EPI_ISL_624664, EPI_ISL_624665, EPI_ISL_624666, EPI_ISL_624667, EPI_ISL_624668, EPI_ISL_624669, EPI_ISL_624670, EPI_ISL_624671, EPI_ISL_624672, EPI_ISL_624673, EPI_ISL_624674, EPI_ISL_624675, EPI_ISL_624676, EPI_ISL_624677, EPI_ISL_624678, EPI_ISL_624679, EPI_ISL_624680, EPI_ISL_624681, EPI_ISL_624682, EPI_ISL_624683, EPI_ISL_624684, EPI_ISL_624685, EPI_ISL_624686, EPI_ISL_624687, EPI_ISL_624688, EPI_ISL_624689, EPI_ISL_624690, EPI_ISL_624691, EPI_ISL_624692, EPI_ISL_624693, EPI_ISL_624694, EPI_ISL_624695, EPI_ISL_624696, EPI_ISL_624697, EPI_ISL_624698, EPI_ISL_624699, EPI_ISL_624700, EPI_ISL_624701, EPI_ISL_624702, EPI_ISL_624703, EPI_ISL_624704, EPI_ISL_624705, EPI_ISL_624706, EPI_ISL_624707, EPI_ISL_624708, EPI_ISL_624709, EPI_ISL_624710, EPI_ISL_624711, EPI_ISL_624712, EPI_ISL_624713, EPI_ISL_624714, EPI_ISL_624715, EPI_ISL_624716, EPI_ISL_624717, EPI_ISL_624718, EPI_ISL_624719, EPI_ISL_624720, EPI_ISL_624721, EPI_ISL_624722, EPI_ISL_624723, EPI_ISL_624724, EPI_ISL_624725, EPI_ISL_624726, EPI_ISL_624727, EPI_ISL_624728, EPI_ISL_624729, EPI_ISL_624730, EPI_ISL_624731, EPI_ISL_624732, EPI_ISL_624733, EPI_ISL_624734, EPI_ISL_624735, EPI_ISL_624736, EPI_ISL_624737, EPI_ISL_624738, EPI_ISL_624739, EPI_ISL_624740, EPI_ISL_624741, EPI_ISL_624742, EPI_ISL_624743, EPI_ISL_624744, EPI_ISL_624745, EPI_ISL_624746, EPI_ISL_624747, EPI_ISL_624748, EPI_ISL_624749, EPI_ISL_624750, EPI_ISL_624751, EPI_ISL_624752, EPI_ISL_624753, EPI_ISL_624754, EPI_ISL_624755, EPI_ISL_624756, EPI_ISL_624757, EPI_ISL_624758, EPI_ISL_624759, EPI_ISL_624760, EPI_ISL_624761, EPI_ISL_624762, EPI_ISL_624763, EPI_ISL_624764, EPI_ISL_624765, EPI_ISL_624766, EPI_ISL_624767, EPI_ISL_624768, EPI_ISL_624769, EPI_ISL_624770, EPI_ISL_624771, EPI_ISL_624772, EPI_ISL_624773, EPI_ISL_624774, EPI_ISL_624775, EPI_ISL_624776, EPI_ISL_624777, EPI_ISL_624778, EPI_ISL_624779, EPI_ISL_624780, EPI_ISL_624781, EPI_ISL_624782, EPI_ISL_624783, EPI_ISL_624784, EPI_ISL_624785, EPI_ISL_624786, EPI_ISL_624787, EPI_ISL_624788, EPI_ISL_624789, EPI_ISL_624790, EPI_ISL_624791, EPI_ISL_624792, EPI_ISL_624793, EPI_ISL_624794, EPI_ISL_624795, EPI_ISL_624796, EPI_ISL_624797, EPI_ISL_624798, EPI_ISL_624799, EPI_ISL_624800, EPI_ISL_624801, EPI_ISL_624802, EPI_ISL_624803, EPI_ISL_624804, EPI_ISL_624805, EPI_ISL_624806, EPI_ISL_624807, EPI_ISL_624808, EPI_ISL_624809, EPI_ISL_624810, EPI_ISL_624811, EPI_ISL_624812, EPI_ISL_624813, EPI_ISL_624814, EPI_ISL_624815, EPI_ISL_624816, EPI_ISL_624817, EPI_ISL_624818, EPI_ISL_624819, EPI_ISL_624820, EPI_ISL_624821, EPI_ISL_624822, EPI_ISL_624823, EPI_ISL_624824, EPI_ISL_624825, EPI_ISL_624826, EPI_ISL_624827, EPI_ISL_624828, EPI_ISL_624829, EPI_ISL_624830, EPI_ISL_624831, EPI_ISL_624832, EPI_ISL_624833, EPI_ISL_624834, EPI_ISL_624835, EPI_ISL_624836, EPI_ISL_624837, EPI_ISL_624838, EPI_ISL_624839, EPI_ISL_624840, EPI_ISL_624841, EPI_ISL_624842, EPI_ISL_624843, EPI_ISL_624844, EPI_ISL_624845, EPI_ISL_624846, EPI_ISL_624847, EPI_ISL_624848, EPI_ISL_624849, EPI_ISL_624850, EPI_ISL_624851, EPI_ISL_624852, EPI_ISL_624853, EPI_ISL_624854, EPI_ISL_624855, EPI_ISL_624856, EPI_ISL_624857, EPI_ISL_624858, EPI_ISL_624859, EPI_ISL_624860, EPI_ISL_624861, EPI_ISL_624862, EPI_ISL_624863, EPI_ISL_624864, EPI_ISL_624865, EPI_ISL_624866, EPI_ISL_624867, EPI_ISL_624868, EPI_ISL_624869, EPI_ISL_624870, EPI_ISL_624871, EPI_ISL_624872, EPI_ISL_624873, EPI_ISL_624874, EPI_ISL_624875, EPI_ISL_624876, EPI_ISL_624877, EPI_ISL_624878, EPI_ISL_624879, EPI_ISL_624880, EPI_ISL_624881, EPI_ISL_624882, EPI_ISL_624883, EPI_ISL_624884, EPI_ISL_624885, EPI_ISL_624886, EPI_ISL_624887, EPI_ISL_624888, EPI_ISL_624889, EPI_ISL_624890, EPI_ISL_624891, EPI_ISL_624892, EPI_ISL_624893, EPI_ISL_624894, EPI_ISL_624895, EPI_ISL_624896, EPI_ISL_624897, EPI_ISL_624898, EPI_ISL_624899, EPI_ISL_624900, EPI_ISL_624901, EPI_ISL_624902, EPI_ISL_624903, EPI_ISL_624904, EPI_ISL_624905, EPI_ISL_624906, EPI_ISL_624907, EPI_ISL_624913, EPI_ISL_624923, EPI_ISL_624925, EPI_ISL_624934, | | | |

| | | | |
|---|---|---|---|
| EPI_ISL_624939, EPI_ISL_624941, EPI_ISL_624942, EPI_ISL_624947, EPI_ISL_624950, EPI_ISL_624953, EPI_ISL_624954, EPI_ISL_624965, EPI_ISL_624966, EPI_ISL_624967, EPI_ISL_624972, EPI_ISL_624975, EPI_ISL_624991, EPI_ISL_624992, EPI_ISL_625006, EPI_ISL_625007, EPI_ISL_625008, EPI_ISL_625012, EPI_ISL_625015, EPI_ISL_625021, EPI_ISL_625024, EPI_ISL_625026, EPI_ISL_625029, EPI_ISL_625033, EPI_ISL_625034, EPI_ISL_625044, EPI_ISL_625057, EPI_ISL_625058, EPI_ISL_625060, EPI_ISL_625063, EPI_ISL_625065, EPI_ISL_625066, EPI_ISL_625081, EPI_ISL_625085, EPI_ISL_625089, EPI_ISL_625090, EPI_ISL_625091, EPI_ISL_625095, EPI_ISL_625102, EPI_ISL_625110, EPI_ISL_625114, EPI_ISL_625116, EPI_ISL_625119, EPI_ISL_625124, EPI_ISL_625125, EPI_ISL_625128, EPI_ISL_625133, EPI_ISL_625137, EPI_ISL_625138, EPI_ISL_625143, EPI_ISL_625145, EPI_ISL_625146, EPI_ISL_625149, EPI_ISL_625153, EPI_ISL_625155, EPI_ISL_625157, EPI_ISL_625161, EPI_ISL_625163, EPI_ISL_625165, EPI_ISL_625168, EPI_ISL_625169, EPI_ISL_625172, EPI_ISL_625175, EPI_ISL_625180, EPI_ISL_625183, EPI_ISL_625185, EPI_ISL_625191, EPI_ISL_625197, EPI_ISL_625201, EPI_ISL_625210, EPI_ISL_625212, EPI_ISL_625215, EPI_ISL_625216, EPI_ISL_625217, EPI_ISL_625225, EPI_ISL_625230, EPI_ISL_625233, EPI_ISL_625234, EPI_ISL_625238, EPI_ISL_625245, EPI_ISL_625249, EPI_ISL_625252, EPI_ISL_625255, EPI_ISL_625257, EPI_ISL_625259, EPI_ISL_625266 | | | |
| see above | Lighthouse Lab in Alderley Park | Wellcome Sanger Institute for the COVID-19 Genomics UK (COG-UK) consortium | Jacquelyn Wynn, Mairead Hyland, The Lighthouse Lab in Alderley Park and Alex Alderton, Roberto Amato, Sonia Goncalves, Ewan Harrison, David K. Jackson, Ian Johnston, Dominic Kwiatkowski, Cordelia Langford, John Sillitoe on behalf of the Wellcome Sanger Institute COVID-19 Surveillance Team (http://www.sanger.ac.uk/covid-team) |
| EPI_ISL_625315, EPI_ISL_625316, EPI_ISL_625317, EPI_ISL_625318, EPI_ISL_625319, EPI_ISL_625320, EPI_ISL_625321, EPI_ISL_625322, EPI_ISL_625323, EPI_ISL_625324, EPI_ISL_625325, EPI_ISL_625326, EPI_ISL_625327, EPI_ISL_625328, EPI_ISL_625329, EPI_ISL_625330, EPI_ISL_625331, EPI_ISL_625332, EPI_ISL_625333, EPI_ISL_625334, EPI_ISL_625335, EPI_ISL_625336, EPI_ISL_625337, EPI_ISL_625338, EPI_ISL_625339, EPI_ISL_625340, EPI_ISL_625341, EPI_ISL_625342, EPI_ISL_625343, EPI_ISL_625344, EPI_ISL_625345, EPI_ISL_625346, EPI_ISL_625347, EPI_ISL_625348, EPI_ISL_625349, EPI_ISL_625350, EPI_ISL_625351, EPI_ISL_625352, EPI_ISL_625353, EPI_ISL_625354, EPI_ISL_625355, EPI_ISL_625356, EPI_ISL_625357, EPI_ISL_625358, EPI_ISL_625359, EPI_ISL_625360, EPI_ISL_625361, EPI_ISL_625362, EPI_ISL_625363, EPI_ISL_625364, EPI_ISL_625365, EPI_ISL_625366, EPI_ISL_625367, EPI_ISL_625368, EPI_ISL_625369, EPI_ISL_625370, EPI_ISL_625371, EPI_ISL_625372, EPI_ISL_625373, EPI_ISL_625374, EPI_ISL_625375, EPI_ISL_625376, EPI_ISL_625377, EPI_ISL_625378, EPI_ISL_625379, EPI_ISL_625380, EPI_ISL_625381, EPI_ISL_625382, EPI_ISL_625383, EPI_ISL_625384, EPI_ISL_625385, EPI_ISL_625386, EPI_ISL_625387, EPI_ISL_625388, EPI_ISL_625389, EPI_ISL_625390, EPI_ISL_625391, EPI_ISL_625392, EPI_ISL_625393, EPI_ISL_625394, EPI_ISL_625395 | | | |
| see above | Lighthouse Lab in Glasgow | Wellcome Sanger Institute for the COVID-19 Genomics UK (COG-UK) consortium | Harper VanSteenhouse, Yumi Kasai, David Gray, Carol Clugston, Anna Dominiczak and Alex Alderton, Roberto Amato, Sonia Goncalves, Ewan Harrison, David K. Jackson, Ian Johnston, Dominic Kwiatkowski, Cordelia Langford, John Sillitoe on behalf of the Wellcome Sanger Institute COVID-19 Surveillance Team (http://www.sanger.ac.uk/covid-team) |
| EPI_ISL_626624 | The National Institute of Public Health | State Veterinary Institute Prague | Nagy,A;Jirincova,H;Novakova,L;Trnka,D;Vecerova,J |
| EPI_ISL_626631 | National Public Health Laboratory, National Centre for Infectious Diseases | National Public Health Laboratory, National Centre for Infectious Diseases | Tze Minn Mak, Sophie Octavia, Zhenyang Zhou, Lin Cui, Raymond Tzer Pin Lin |
| EPI_ISL_626663 | Quadram Institute Bioscience | COVID-19 Genomics UK (COG-UK) Consortium | Dave J. Baker, Gemma L. Kay, Alp Aydin, Thanh Le-Viet, Steven Rudder, Ana P. Tedim, Anastasia Kolyva, Maria Diaz, Leonardo de Oliveira Martins, Nabil-Fareed Alikhan, Lizzie Meadows, Rachael Stanley, Ngozi Elumogo, Muhammed Yasir, Nicholas M. Thomson, Alexander J Trotter, Rachel Gilroy, Samuel Bloomfield, Claire Stuart, Andrew Bell, Reenesh Prakash, Samir Dervisevic, Alison E. Mather, John Wain, Mark Webber, Andrew J. Page, Justin O'Grady |
| EPI_ISL_626666 | University of Exeter | COVID-19 Genomics UK (COG-UK) Consortium | Ben Temperton,Aaron Jeffries,Michelle Michelsen,Joanna Warwick-Dugdale,Audrey Farbos,Robyn Manley,Stephen Michell,Jane Masoli |
| EPI_ISL_626667 | Quadram Institute Bioscience | COVID-19 Genomics UK (COG-UK) Consortium | Dave J. Baker, Gemma L. Kay, Alp Aydin, Thanh Le-Viet, Steven Rudder, Ana P. Tedim, Anastasia Kolyva, Maria Diaz, Leonardo de Oliveira Martins, Nabil-Fareed Alikhan, Lizzie Meadows, Rachael Stanley, Ngozi Elumogo, Muhammed Yasir, Nicholas M. Thomson, Alexander J Trotter, Rachel Gilroy, Samuel Bloomfield, Claire Stuart, Andrew Bell, Reenesh Prakash, Samir Dervisevic, Alison E. Mather, John Wain, Mark Webber, Andrew J. Page, Justin O'Grady |
| EPI_ISL_626672 | University of Exeter | COVID-19 Genomics UK (COG-UK) Consortium | Ben Temperton,Aaron Jeffries,Michelle Michelsen,Joanna Warwick-Dugdale,Audrey Farbos,Robyn Manley,Stephen Michell,Jane Masoli |
| EPI_ISL_626684 | Centre for Enzyme Innovation, University of Portsmouth / Translational Research Laboratory, Portsmouth Hospitals NHS Trust | COVID-19 Genomics UK (COG-UK) Consortium | Angela Beckett,Yann Bourgeois,Garry Scarlett,Sharon Glaysher,Scott Elliott,Kelly Bicknell,Robert Impey,Allyson Lloyd,Sarah Wyllie,Ethan Butcher,Anoop Chauhan,Samuel Robson |
| EPI_ISL_626691, EPI_ISL_626692 | West of Scotland Specialist Virology Centre, NHSGGC / MRC-University of Glasgow Centre for Virus Research | COVID-19 Genomics UK (COG-UK) Consortium | Ana da Silva Filipe, Natasha Johnson, Kathy Smollett, Daniel Mair, Stephen Carmichael, Lily Tong, Jenna Nichols, Elihu Aranday-Cortes, Kyriaki Nomikou; Sarah McDonald, Marc Niebel, Patawee Asamaphan; Richard Orton, Joseph Hughes, Sreenu Vattipally, David L Robertson; Alasdair MacLean, Rory Gunson; Kathy Li, Igor Starinskij, Natasha Jesudason, Rajiv Shah, James Shepherd, Antonia Ho, Emma Thomson |
| EPI_ISL_626693, EPI_ISL_626699, EPI_ISL_626702 | Quadram Institute Bioscience | COVID-19 Genomics UK (COG-UK) Consortium | Dave J. Baker, Gemma L. Kay, Alp Aydin, Thanh Le-Viet, Steven Rudder, Ana P. Tedim, Anastasia Kolyva, Maria Diaz, Leonardo de Oliveira Martins, Nabil-Fareed Alikhan, Lizzie Meadows, Rachael Stanley, Ngozi Elumogo, Muhammed Yasir, Nicholas M. Thomson, Alexander J Trotter, Rachel Gilroy, Samuel Bloomfield, Claire Stuart, Andrew Bell, Reenesh Prakash, Samir Dervisevic, Alison E. Mather, John Wain, Mark Webber, Andrew J. Page, Justin O'Grady |
| EPI_ISL_626703, EPI_ISL_626711 | Wales Specialist Virology Centre Sequencing lab: Pathogen Genomics Unit | COVID-19 Genomics UK (COG-UK) Consortium | Catherine Moore, Johnathan Evans, Laura Gifford, Malorie Perry, Simon Cottrell, Angela Marchbank, Alec Birchley, Alexander Adams, Amy Gaskin, Bree Gatica-Wilcox, Jason Coombes, Joel Southgate, Lauren Gilbert, Lee Graham, Nicole Pacchiarini, Sara Kumziene-Summerhayes, Sarah Taylor, Sophie Jones, Sara Rey, Matthew Bull, Joanne Watkins, Sally Corden, Tom Connor |
| EPI_ISL_626713, EPI_ISL_626730 | University of Exeter | COVID-19 Genomics UK (COG-UK) Consortium | Ben Temperton,Aaron Jeffries,Michelle Michelsen,Joanna Warwick-Dugdale,Audrey Farbos,Robyn Manley,Stephen Michell,Jane Masoli |
| EPI_ISL_626749 | West of Scotland Specialist Virology Centre, NHSGGC / MRC-University of Glasgow Centre for Virus Research | COVID-19 Genomics UK (COG-UK) Consortium | Ana da Silva Filipe, Natasha Johnson, Kathy Smollett, Daniel Mair, Stephen Carmichael, Lily Tong, Jenna Nichols, Elihu Aranday-Cortes, Kyriaki Nomikou; Sarah McDonald, Marc Niebel, Patawee Asamaphan; Richard Orton, Joseph Hughes, Sreenu Vattipally, David L Robertson; Alasdair MacLean, Rory Gunson; Kathy Li, Igor Starinskij, Natasha Jesudason, Rajiv Shah, James Shepherd, Antonia Ho, Emma Thomson |
| EPI_ISL_626764 | Quadram Institute Bioscience | COVID-19 Genomics UK (COG-UK) Consortium | Dave J. Baker, Gemma L. Kay, Alp Aydin, Thanh Le-Viet, Steven Rudder, Ana P. Tedim, Anastasia Kolyva, Maria Diaz, Leonardo de Oliveira Martins, Nabil-Fareed Alikhan, Lizzie Meadows, Rachael Stanley, Ngozi Elumogo, Muhammed Yasir, Nicholas M. Thomson, Alexander J Trotter, Rachel Gilroy, Samuel Bloomfield, Claire Stuart, Andrew Bell, Reenesh Prakash, Samir Dervisevic, Alison E. Mather, John Wain, Mark Webber, Andrew J. Page, Justin O'Grady |
| EPI_ISL_626777 | University of Exeter | COVID-19 Genomics UK (COG-UK) Consortium | Ben Temperton,Aaron Jeffries,Michelle Michelsen,Joanna Warwick-Dugdale,Audrey Farbos,Robyn Manley,Stephen Michell,Jane Masoli |
| EPI_ISL_626787, EPI_ISL_626796 | Wales Specialist Virology Centre Sequencing lab: Pathogen Genomics Unit | COVID-19 Genomics UK (COG-UK) Consortium | Catherine Moore, Johnathan Evans, Laura Gifford, Malorie Perry, Simon Cottrell, Angela Marchbank, Alec Birchley, Alexander Adams, Amy Gaskin, Bree Gatica-Wilcox, Jason Coombes, Joel Southgate, Lauren Gilbert, Lee Graham, Nicole Pacchiarini, Sara Kumziene-Summerhayes, Sarah Taylor, Sophie Jones, Sara Rey, Matthew Bull, Joanne Watkins, Sally Corden, Tom Connor |
| EPI_ISL_626802, EPI_ISL_626813 | University of Exeter | COVID-19 Genomics UK (COG-UK) Consortium | Ben Temperton,Aaron Jeffries,Michelle Michelsen,Joanna Warwick-Dugdale,Audrey Farbos,Robyn Manley,Stephen Michell,Jane Masoli |
| EPI_ISL_626829 | Wales Specialist Virology Centre Sequencing lab: Pathogen Genomics Unit | COVID-19 Genomics UK (COG-UK) Consortium | Catherine Moore, Johnathan Evans, Laura Gifford, Malorie Perry, Simon Cottrell, Angela Marchbank, Alec Birchley, Alexander Adams, Amy Gaskin, Bree Gatica-Wilcox, Jason Coombes, Joel Southgate, Lauren Gilbert, Lee Graham, Nicole Pacchiarini, Sara Kumziene-Summerhayes, Sarah Taylor, Sophie Jones, Sara Rey, Matthew Bull, Joanne Watkins, Sally Corden, Tom Connor |
| EPI_ISL_626831 | West of Scotland Specialist Virology Centre, NHSGGC / MRC-University of Glasgow Centre for Virus Research | COVID-19 Genomics UK (COG-UK) Consortium | Ana da Silva Filipe, Natasha Johnson, Kathy Smollett, Daniel Mair, Stephen Carmichael, Lily Tong, Jenna Nichols, Elihu Aranday-Cortes, Kyriaki Nomikou; Sarah McDonald, Marc Niebel, Patawee Asamaphan; Richard Orton, Joseph Hughes, Sreenu Vattipally, David L Robertson; Alasdair MacLean, Rory Gunson; Kathy Li, Igor Starinskij, Natasha Jesudason, Rajiv Shah, James Shepherd, Antonia Ho, Emma Thomson |
| EPI_ISL_626832 | Liverpool Clinical Laboratories | COVID-19 Genomics UK (COG-UK) Consortium | Sam Haldenby, Anita Lucaci, Steve Paterson, Julian Hiscox, Alistair Darby, M Almsaud, A Alrezaihi, Muhannad Alruwaili, Stuart D Armstrong, Jones Benjamin, Eleanor G Bentley, Anu Chawla, Jordan J Clark, Angela Cowell, Richard Eccles, Isabel García-Dorival, Matthew Gemmell, Alessandro Gerada, PKF Gilmore, Richard Gregory, Ximeng Han, Catherine Hartley, Margaret Hughes, Miren Iturriza-Gomara, James Johnson, L Luu, Jenifer Manson, Charlotte Nelson, Elaine O'Toole, Cassie Olateju, Rebekah Penrice-Randal , Lucille Rainbow, N.P Randle, Trevor Ian Robinson, Parul Sharma, Ghada T Shawli, James P Stewart, Neil Swainston, Ecaterina Vamos, Joanne Watts, Mark Whitehead |
| EPI_ISL_626835, EPI_ISL_626846, EPI_ISL_626849, EPI_ISL_626851 | West of Scotland Specialist Virology Centre, NHSGGC / MRC-University of Glasgow Centre for Virus Research | COVID-19 Genomics UK (COG-UK) Consortium | Ana da Silva Filipe, Natasha Johnson, Kathy Smollett, Daniel Mair, Stephen Carmichael, Lily Tong, Jenna Nichols, Elihu Aranday-Cortes, Kyriaki Nomikou; Sarah McDonald, Marc Niebel, Patawee Asamaphan; Richard Orton, Joseph Hughes, Sreenu Vattipally, David L Robertson; Alasdair MacLean, Rory Gunson; Kathy Li, Igor Starinskij, Natasha Jesudason, Rajiv Shah, James Shepherd, Antonia Ho, Emma Thomson |
| EPI_ISL_626854 | Quadram Institute Bioscience | COVID-19 Genomics UK (COG-UK) Consortium | Dave J. Baker, Gemma L. Kay, Alp Aydin, Thanh Le-Viet, Steven Rudder, Ana P. Tedim, Anastasia Kolyva, Maria Diaz, Leonardo de Oliveira Martins, Nabil-Fareed Alikhan, Lizzie Meadows, Rachael Stanley, Ngozi Elumogo, Muhammed Yasir, Nicholas M. Thomson, Alexander J Trotter, Rachel Gilroy, Samuel Bloomfield, Claire Stuart, Andrew Bell, Reenesh Prakash, Samir Dervisevic, Alison E. Mather, John Wain, Mark Webber, Andrew J. Page, Justin O'Grady |
| EPI_ISL_626855 | West of Scotland Specialist Virology Centre, NHSGGC / MRC-University of Glasgow Centre for Virus Research | COVID-19 Genomics UK (COG-UK) Consortium | Ana da Silva Filipe, Natasha Johnson, Kathy Smollett, Daniel Mair, Stephen Carmichael, Lily Tong, Jenna Nichols, Elihu Aranday-Cortes, Kyriaki Nomikou; Sarah McDonald, Marc Niebel, Patawee Asamaphan; Richard Orton, Joseph Hughes, Sreenu Vattipally, David L Robertson; Alasdair MacLean, Rory Gunson; |

| | | | |
|---|---|---|---|
| | | | Kathy Li, Igor Starinskij, Natasha Jesudason, Rajiv Shah, James Shepherd, Antonia Ho, Emma Thomson |
| EPI_ISL_626867 | Wales Specialist Virology Centre Sequencing lab: Pathogen Genomics Unit | COVID-19 Genomics UK (COG-UK) Consortium | Catherine Moore, Johnathan Evans, Laura Gifford, Malorie Perry, Simon Cottrell, Angela Marchbank, Alec Birchley, Alexander Adams, Amy Gaskin, Bree Gatica-Wilcox, Jason Coombes, Joel Southgate, Lauren Gilbert, Lee Graham, Nicole Pacchiarini, Sara Kumziene-Summerhayes, Sarah Taylor, Sophie Jones, Sara Rey, Matthew Bull, Joanne Watkins, Sally Corden, Tom Connor |
| EPI_ISL_626889 | University of Exeter | COVID-19 Genomics UK (COG-UK) Consortium | Ben Temperton,Aaron Jeffries,Michelle Michelsen,Joanna Warwick-Dugdale,Audrey Farbos,Robyn Manley,Stephen Michell,Jane Masoli |
| EPI_ISL_626895 | Centre for Enzyme Innovation, University of Portsmouth / Translational Research Laboratory, Portsmouth Hospitals NHS Trust | COVID-19 Genomics UK (COG-UK) Consortium | Angela Beckett,Yann Bourgeois,Garry Scarlett,Sharon Glaysher,Scott Elliott,Kelly Bicknell,Robert Impey,Allyson Lloyd,Sarah Wyllie,Ethan Butcher,Anoop Chauhan,Samuel Robson |
| EPI_ISL_626921 | Wales Specialist Virology Centre Sequencing lab: Pathogen Genomics Unit | COVID-19 Genomics UK (COG-UK) Consortium | Catherine Moore, Johnathan Evans, Laura Gifford, Malorie Perry, Simon Cottrell, Angela Marchbank, Alec Birchley, Alexander Adams, Amy Gaskin, Bree Gatica-Wilcox, Jason Coombes, Joel Southgate, Lauren Gilbert, Lee Graham, Nicole Pacchiarini, Sara Kumziene-Summerhayes, Sarah Taylor, Sophie Jones, Sara Rey, Matthew Bull, Joanne Watkins, Sally Corden, Tom Connor |
| EPI_ISL_626943, EPI_ISL_626945, EPI_ISL_626946 | University of Exeter | COVID-19 Genomics UK (COG-UK) Consortium | Ben Temperton,Aaron Jeffries,Michelle Michelsen,Joanna Warwick-Dugdale,Audrey Farbos,Robyn Manley,Stephen Michell,Jane Masoli |
| EPI_ISL_626957, EPI_ISL_626972, EPI_ISL_626978, EPI_ISL_626982, EPI_ISL_626983, EPI_ISL_626988, EPI_ISL_626991, EPI_ISL_626995 | West of Scotland Specialist Virology Centre, NHSGGC / MRC-University of Glasgow Centre for Virus Research | COVID-19 Genomics UK (COG-UK) Consortium | Ana da Silva Filipe, Natasha Johnson, Kathy Smollett, Daniel Mair, Stephen Carmichael, Lily Tong, Jenna Nichols, Elihu Aranday-Cortes, Kyriaki Nomikou; Sarah McDonald, Marc Niebel, Patawee Asamaphan; Richard Orton, Joseph Hughes, Sreenu Vattipally, David L Robertson; Alasdair MacLean, Rory Gunson; Kathy Li, Igor Starinskij, Natasha Jesudason, Rajiv Shah, James Shepherd, Antonia Ho, Emma Thomson |
| EPI_ISL_627016, EPI_ISL_627024 | Quadram Institute Bioscience | COVID-19 Genomics UK (COG-UK) Consortium | Dave J. Baker, Gemma L. Kay, Alp Aydin, Thanh Le-Viet, Steven Rudder, Ana P. Tedim, Anastasia Kolyva, Maria Diaz, Leonardo de Oliveira Martins, Nabil-Fareed Alikhan, Lizzie Meadows, Rachael Stanley, Ngozi Elumogo, Muhammed Yasir, Nicholas M. Thomson, Alexander J Trotter, Rachel Gilroy, Samuel Bloomfield, Claire Stuart, Andrew Bell, Reenesh Prakash, Samir Dervisevic, Alison E. Mather, John Wain, Mark Webber, Andrew J. Page, Justin O'Grady |
| EPI_ISL_627030 | West of Scotland Specialist Virology Centre, NHSGGC / MRC-University of Glasgow Centre for Virus Research | COVID-19 Genomics UK (COG-UK) Consortium | Ana da Silva Filipe, Natasha Johnson, Kathy Smollett, Daniel Mair, Stephen Carmichael, Lily Tong, Jenna Nichols, Elihu Aranday-Cortes, Kyriaki Nomikou; Sarah McDonald, Marc Niebel, Patawee Asamaphan; Richard Orton, Joseph Hughes, Sreenu Vattipally, David L Robertson; Alasdair MacLean, Rory Gunson; Kathy Li, Igor Starinskij, Natasha Jesudason, Rajiv Shah, James Shepherd, Antonia Ho, Emma Thomson |
| EPI_ISL_627033 | Wales Specialist Virology Centre Sequencing lab: Pathogen Genomics Unit | COVID-19 Genomics UK (COG-UK) Consortium | Catherine Moore, Johnathan Evans, Laura Gifford, Malorie Perry, Simon Cottrell, Angela Marchbank, Alec Birchley, Alexander Adams, Amy Gaskin, Bree Gatica-Wilcox, Jason Coombes, Joel Southgate, Lauren Gilbert, Lee Graham, Nicole Pacchiarini, Sara Kumziene-Summerhayes, Sarah Taylor, Sophie Jones, Sara Rey, Matthew Bull, Joanne Watkins, Sally Corden, Tom Connor |
| EPI_ISL_627041 | Quadram Institute Bioscience | COVID-19 Genomics UK (COG-UK) Consortium | Dave J. Baker, Gemma L. Kay, Alp Aydin, Thanh Le-Viet, Steven Rudder, Ana P. Tedim, Anastasia Kolyva, Maria Diaz, Leonardo de Oliveira Martins, Nabil-Fareed Alikhan, Lizzie Meadows, Rachael Stanley, Ngozi Elumogo, Muhammed Yasir, Nicholas M. Thomson, Alexander J Trotter, Rachel Gilroy, Samuel Bloomfield, Claire Stuart, Andrew Bell, Reenesh Prakash, Samir Dervisevic, Alison E. Mather, John Wain, Mark Webber, Andrew J. Page, Justin O'Grady |
| EPI_ISL_627042, EPI_ISL_627043 | Wales Specialist Virology Centre Sequencing lab: Pathogen Genomics Unit | COVID-19 Genomics UK (COG-UK) Consortium | Catherine Moore, Johnathan Evans, Laura Gifford, Malorie Perry, Simon Cottrell, Angela Marchbank, Alec Birchley, Alexander Adams, Amy Gaskin, Bree Gatica-Wilcox, Jason Coombes, Joel Southgate, Lauren Gilbert, Lee Graham, Nicole Pacchiarini, Sara Kumziene-Summerhayes, Sarah Taylor, Sophie Jones, Sara Rey, Matthew Bull, Joanne Watkins, Sally Corden, Tom Connor |
| EPI_ISL_627063 | West of Scotland Specialist Virology Centre, NHSGGC / MRC-University of Glasgow Centre for Virus Research | COVID-19 Genomics UK (COG-UK) Consortium | Ana da Silva Filipe, Natasha Johnson, Kathy Smollett, Daniel Mair, Stephen Carmichael, Lily Tong, Jenna Nichols, Elihu Aranday-Cortes, Kyriaki Nomikou; Sarah McDonald, Marc Niebel, Patawee Asamaphan; Richard Orton, Joseph Hughes, Sreenu Vattipally, David L Robertson; Alasdair MacLean, Rory Gunson; Kathy Li, Igor Starinskij, Natasha Jesudason, Rajiv Shah, James Shepherd, Antonia Ho, Emma Thomson |
| EPI_ISL_627072, EPI_ISL_627074, EPI_ISL_627075, EPI_ISL_627076, EPI_ISL_627077 | Quadram Institute Bioscience | COVID-19 Genomics UK (COG-UK) Consortium | Dave J. Baker, Gemma L. Kay, Alp Aydin, Thanh Le-Viet, Steven Rudder, Ana P. Tedim, Anastasia Kolyva, Maria Diaz, Leonardo de Oliveira Martins, Nabil-Fareed Alikhan, Lizzie Meadows, Rachael Stanley, Ngozi Elumogo, Muhammed Yasir, Nicholas M. Thomson, Alexander J Trotter, Rachel Gilroy, Samuel Bloomfield, Claire Stuart, Andrew Bell, Reenesh Prakash, Samir Dervisevic, Alison E. Mather, John Wain, Mark Webber, Andrew J. Page, Justin O'Grady |
| EPI_ISL_627091, EPI_ISL_627095 | Wales Specialist Virology Centre Sequencing lab: Pathogen Genomics Unit | COVID-19 Genomics UK (COG-UK) Consortium | Catherine Moore, Johnathan Evans, Laura Gifford, Malorie Perry, Simon Cottrell, Angela Marchbank, Alec Birchley, Alexander Adams, Amy Gaskin, Bree Gatica-Wilcox, Jason Coombes, Joel Southgate, Lauren Gilbert, Lee Graham, Nicole Pacchiarini, Sara Kumziene-Summerhayes, Sarah Taylor, Sophie Jones, Sara Rey, Matthew Bull, Joanne Watkins, Sally Corden, Tom Connor |
| EPI_ISL_627098 | Quadram Institute Bioscience | COVID-19 Genomics UK (COG-UK) Consortium | Dave J. Baker, Gemma L. Kay, Alp Aydin, Thanh Le-Viet, Steven Rudder, Ana P. Tedim, Anastasia Kolyva, Maria Diaz, Leonardo de Oliveira Martins, Nabil-Fareed Alikhan, Lizzie Meadows, Rachael Stanley, Ngozi Elumogo, Muhammed Yasir, Nicholas M. Thomson, Alexander J Trotter, Rachel Gilroy, Samuel Bloomfield, Claire Stuart, Andrew Bell, Reenesh Prakash, Samir Dervisevic, Alison E. Mather, John Wain, Mark Webber, Andrew J. Page, Justin O'Grady |
| EPI_ISL_627109 | Liverpool Clinical Laboratories | COVID-19 Genomics UK (COG-UK) Consortium | Sam Haldenby, Anita Lucaci, Steve Paterson, Julian Hiscox, Alistair Darby, M Almsaud, A Alrezaihi, Muhannad Alruwaili, Stuart D Armstrong, Jones Benjamin, Eleanor G Bentley, Anu Chawla, Jordan J Clark, Angela Cowell, Richard Eccles, Isabel García-Dorival, Matthew Gemmell, Alessandro Gerada, PKF Gilmore, Richard Gregory, Ximeng Han, Catherine Hartley, Margaret Hughes, Miren Iturriza-Gomara, James Johnson, L Luu, Jenifer Manson, Charlotte Nelson, Elaine O'Toole, Cassie Olateju, Rebekah Penrice-Randal , Lucille Rainbow, N.P Randle, Trevor Ian Robinson, Parul Sharma, Ghada T Shawli, James P Stewart, Neil Swainston, Ecaterina Vamos, Joanne Watts, Mark Whitehead |
| EPI_ISL_627120, EPI_ISL_627124 | Wales Specialist Virology Centre Sequencing lab: Pathogen Genomics Unit | COVID-19 Genomics UK (COG-UK) Consortium | Catherine Moore, Johnathan Evans, Laura Gifford, Malorie Perry, Simon Cottrell, Angela Marchbank, Alec Birchley, Alexander Adams, Amy Gaskin, Bree Gatica-Wilcox, Jason Coombes, Joel Southgate, Lauren Gilbert, Lee Graham, Nicole Pacchiarini, Sara Kumziene-Summerhayes, Sarah Taylor, Sophie Jones, Sara Rey, Matthew Bull, Joanne Watkins, Sally Corden, Tom Connor |
| EPI_ISL_627128 | Centre for Enzyme Innovation, University of Portsmouth / Translational Research Laboratory, Portsmouth Hospitals NHS Trust | COVID-19 Genomics UK (COG-UK) Consortium | Angela Beckett,Yann Bourgeois,Garry Scarlett,Sharon Glaysher,Scott Elliott,Kelly Bicknell,Robert Impey,Allyson Lloyd,Sarah Wyllie,Ethan Butcher,Anoop Chauhan,Samuel Robson |
| EPI_ISL_627145 | University of Exeter | COVID-19 Genomics UK (COG-UK) Consortium | Ben Temperton,Aaron Jeffries,Michelle Michelsen,Joanna Warwick-Dugdale,Audrey Farbos,Robyn Manley,Stephen Michell,Jane Masoli |
| EPI_ISL_627154 | Wales Specialist Virology Centre Sequencing lab: Pathogen Genomics Unit | COVID-19 Genomics UK (COG-UK) Consortium | Catherine Moore, Johnathan Evans, Laura Gifford, Malorie Perry, Simon Cottrell, Angela Marchbank, Alec Birchley, Alexander Adams, Amy Gaskin, Bree Gatica-Wilcox, Jason Coombes, Joel Southgate, Lauren Gilbert, Lee Graham, Nicole Pacchiarini, Sara Kumziene-Summerhayes, Sarah Taylor, Sophie Jones, Sara Rey, Matthew Bull, Joanne Watkins, Sally Corden, Tom Connor |
| EPI_ISL_627160 | Quadram Institute Bioscience | COVID-19 Genomics UK (COG-UK) Consortium | Dave J. Baker, Gemma L. Kay, Alp Aydin, Thanh Le-Viet, Steven Rudder, Ana P. Tedim, Anastasia Kolyva, Maria Diaz, Leonardo de Oliveira Martins, Nabil-Fareed Alikhan, Lizzie Meadows, Rachael Stanley, Ngozi Elumogo, Muhammed Yasir, Nicholas M. Thomson, Alexander J Trotter, Rachel Gilroy, Samuel Bloomfield, Claire Stuart, Andrew Bell, Reenesh Prakash, Samir Dervisevic, Alison E. Mather, John Wain, Mark Webber, Andrew J. Page, Justin O'Grady |
| EPI_ISL_627161 | Wales Specialist Virology Centre Sequencing lab: Pathogen Genomics Unit | COVID-19 Genomics UK (COG-UK) Consortium | Catherine Moore, Johnathan Evans, Laura Gifford, Malorie Perry, Simon Cottrell, Angela Marchbank, Alec Birchley, Alexander Adams, Amy Gaskin, Bree Gatica-Wilcox, Jason Coombes, Joel Southgate, Lauren Gilbert, Lee Graham, Nicole Pacchiarini, Sara Kumziene-Summerhayes, Sarah Taylor, Sophie Jones, Sara Rey, Matthew Bull, Joanne Watkins, Sally Corden, Tom Connor |
| EPI_ISL_627165 | West of Scotland Specialist Virology Centre, NHSGGC / MRC-University of Glasgow Centre for Virus Research | COVID-19 Genomics UK (COG-UK) Consortium | Ana da Silva Filipe, Natasha Johnson, Kathy Smollett, Daniel Mair, Stephen Carmichael, Lily Tong, Jenna Nichols, Elihu Aranday-Cortes, Kyriaki Nomikou; Sarah McDonald, Marc Niebel, Patawee Asamaphan; Richard Orton, Joseph Hughes, Sreenu Vattipally, David L Robertson; Alasdair MacLean, Rory Gunson; Kathy Li, Igor Starinskij, Natasha Jesudason, Rajiv Shah, James Shepherd, Antonia Ho, Emma Thomson |
| EPI_ISL_627182 | University of Exeter | COVID-19 Genomics UK (COG-UK) Consortium | Ben Temperton,Aaron Jeffries,Michelle Michelsen,Joanna Warwick-Dugdale,Audrey Farbos,Robyn Manley,Stephen Michell,Jane Masoli |
| EPI_ISL_627187 | Quadram Institute Bioscience | COVID-19 Genomics UK (COG-UK) Consortium | Dave J. Baker, Gemma L. Kay, Alp Aydin, Thanh Le-Viet, Steven Rudder, Ana P. Tedim, Anastasia Kolyva, Maria Diaz, Leonardo de Oliveira Martins, |

| | | | |
|---|---|---|---|
| | | | Nabil-Fareed Alikhan, Lizzie Meadows, Rachael Stanley, Ngozi Elumogo, Muhammed Yasir, Nicholas M. Thomson, Alexander J Trotter, Rachel Gilroy, Samuel Bloomfield, Claire Stuart, Andrew Bell, Reenesh Prakash, Samir Dervisevic, Alison E. Mather, John Wain, Mark Webber, Andrew J. Page, Justin O'Grady |
| EPI_ISL_627192 | Wales Specialist Virology Centre Sequencing lab: Pathogen Genomics Unit | COVID-19 Genomics UK (COG-UK) Consortium | Catherine Moore, Johnathan Evans, Laura Gifford, Malorie Perry, Simon Cottrell, Angela Marchbank, Alec Birchley, Alexander Adams, Amy Gaskin, Bree Gatica-Wilcox, Jason Coombes, Joel Southgate, Lauren Gilbert, Lee Graham, Nicole Pacchiarini, Sara Kumziene-Summerhayes, Sarah Taylor, Sophie Jones, Sara Rey, Matthew Bull, Joanne Watkins, Sally Corden, Tom Connor |
| EPI_ISL_627194 | University of Exeter | COVID-19 Genomics UK (COG-UK) Consortium | Ben Temperton,Aaron Jeffries,Michelle Michelsen,Joanna Warwick-Dugdale,Audrey Farbos,Robyn Manley,Stephen Michell,Jane Masoli |
| EPI_ISL_627209 | West of Scotland Specialist Virology Centre, NHSGGC / MRC-University of Glasgow Centre for Virus Research | COVID-19 Genomics UK (COG-UK) Consortium | Ana da Silva Filipe, Natasha Johnson, Kathy Smollett, Daniel Mair, Stephen Carmichael, Lily Tong, Jenna Nichols, Elihu Aranday-Cortes, Kyriaki Nomikou; Sarah McDonald, Marc Niebel, Patawee Asamaphan; Richard Orton, Joseph Hughes, Sreenu Vattipally, David L Robertson; Alasdair MacLean, Rory Gunson; Kathy Li, Igor Starinskij, Natasha Jesudason, Rajiv Shah, James Shepherd, Antonia Ho, Emma Thomson |
| EPI_ISL_627210 | University of Exeter | COVID-19 Genomics UK (COG-UK) Consortium | Ben Temperton,Aaron Jeffries,Michelle Michelsen,Joanna Warwick-Dugdale,Audrey Farbos,Robyn Manley,Stephen Michell,Jane Masoli |
| EPI_ISL_627213 | Quadram Institute Bioscience | COVID-19 Genomics UK (COG-UK) Consortium | Dave J. Baker, Gemma L. Kay, Alp Aydin, Thanh Le-Viet, Steven Rudder, Ana P. Tedim, Anastasia Kolyva, Maria Diaz, Leonardo de Oliveira Martins, Nabil-Fareed Alikhan, Lizzie Meadows, Rachael Stanley, Ngozi Elumogo, Muhammed Yasir, Nicholas M. Thomson, Alexander J Trotter, Rachel Gilroy, Samuel Bloomfield, Claire Stuart, Andrew Bell, Reenesh Prakash, Samir Dervisevic, Alison E. Mather, John Wain, Mark Webber, Andrew J. Page, Justin O'Grady |
| EPI_ISL_627218, EPI_ISL_627221, EPI_ISL_627222, EPI_ISL_627223, EPI_ISL_627225, EPI_ISL_627226, EPI_ISL_627238, EPI_ISL_627239, EPI_ISL_627241, EPI_ISL_627265, EPI_ISL_627268, EPI_ISL_627270, EPI_ISL_627271, EPI_ISL_627272, EPI_ISL_627274, EPI_ISL_627278 | | | |
| see above | Wales Specialist Virology Centre Sequencing lab: Pathogen Genomics Unit | COVID-19 Genomics UK (COG-UK) Consortium | Catherine Moore, Johnathan Evans, Laura Gifford, Malorie Perry, Simon Cottrell, Angela Marchbank, Alec Birchley, Alexander Adams, Amy Gaskin, Bree Gatica-Wilcox, Jason Coombes, Joel Southgate, Lauren Gilbert, Lee Graham, Nicole Pacchiarini, Sara Kumziene-Summerhayes, Sarah Taylor, Sophie Jones, Sara Rey, Matthew Bull, Joanne Watkins, Sally Corden, Tom Connor |
| EPI_ISL_627377, EPI_ISL_627379, EPI_ISL_627380, EPI_ISL_627384, EPI_ISL_627388, EPI_ISL_627391, EPI_ISL_627396, EPI_ISL_627397, EPI_ISL_627398, EPI_ISL_627404, EPI_ISL_627409, EPI_ISL_627410, EPI_ISL_627418, EPI_ISL_627419 | | | |
| see above | West of Scotland Specialist Virology Centre, NHSGGC / MRC-University of Glasgow Centre for Virus Research | COVID-19 Genomics UK (COG-UK) Consortium | Ana da Silva Filipe, Natasha Johnson, Kathy Smollett, Daniel Mair, Stephen Carmichael, Lily Tong, Jenna Nichols, Elihu Aranday-Cortes, Kyriaki Nomikou; Sarah McDonald, Marc Niebel, Patawee Asamaphan; Richard Orton, Joseph Hughes, Sreenu Vattipally, David L Robertson; Alasdair MacLean, Rory Gunson; Kathy Li, Igor Starinskij, Natasha Jesudason, Rajiv Shah, James Shepherd, Antonia Ho, Emma Thomson |
| EPI_ISL_627424, EPI_ISL_627428, EPI_ISL_627430, EPI_ISL_627431, EPI_ISL_627432, EPI_ISL_627433, EPI_ISL_627434 | University of Exeter | COVID-19 Genomics UK (COG-UK) Consortium | Ben Temperton,Aaron Jeffries,Michelle Michelsen,Joanna Warwick-Dugdale,Audrey Farbos,Robyn Manley,Stephen Michell,Jane Masoli |
| EPI_ISL_627597, EPI_ISL_627598, EPI_ISL_627599, EPI_ISL_627600, EPI_ISL_627604, EPI_ISL_627606, EPI_ISL_627611, EPI_ISL_627617, EPI_ISL_627618, EPI_ISL_627620, EPI_ISL_627625, EPI_ISL_627627, EPI_ISL_627628, EPI_ISL_627630, EPI_ISL_627632, EPI_ISL_627633, EPI_ISL_627634, EPI_ISL_627637, EPI_ISL_627638, EPI_ISL_627639, EPI_ISL_627642, EPI_ISL_627643, EPI_ISL_627644, EPI_ISL_627648, EPI_ISL_627652, EPI_ISL_627654, EPI_ISL_627656, EPI_ISL_627664, EPI_ISL_627665 | | | |
| see above | Quadram Institute Bioscience | COVID-19 Genomics UK (COG-UK) Consortium | Dave J. Baker, Gemma L. Kay, Alp Aydin, Thanh Le-Viet, Steven Rudder, Ana P. Tedim, Anastasia Kolyva, Maria Diaz, Leonardo de Oliveira Martins, Nabil-Fareed Alikhan, Lizzie Meadows, Rachael Stanley, Ngozi Elumogo, Muhammed Yasir, Nicholas M. Thomson, Alexander J Trotter, Rachel Gilroy, Samuel Bloomfield, Claire Stuart, Andrew Bell, Reenesh Prakash, Samir Dervisevic, Alison E. Mather, John Wain, Mark Webber, Andrew J. Page, Justin O'Grady |
| EPI_ISL_627667, EPI_ISL_627668, EPI_ISL_627669 | Queens Medical Centre, Clinical Microbiology Department / DeepSeq Nottingham | COVID-19 Genomics UK (COG-UK) Consortium | Gemma Clark, Wendy Smith, Manjinder Khakh, Vicki M Fleming, Michelle M Lister, Hannah Howson-Wells, Jonathan Ball, Patrick McClure, Joseph Chappell, Theocharis Tsoleridis, Nadine Holmes, Matthew Carlisle, Christopher Moore, Fei Sang, Johnny Debebe, Victoria Wright, Matthew Loose |
| EPI_ISL_627732, EPI_ISL_627733, EPI_ISL_627734, EPI_ISL_627736, EPI_ISL_627737, EPI_ISL_627742, EPI_ISL_627745, EPI_ISL_627747, EPI_ISL_627753, EPI_ISL_627756, EPI_ISL_627757, EPI_ISL_627758, EPI_ISL_627764, EPI_ISL_627765, EPI_ISL_627766, EPI_ISL_627768, EPI_ISL_627771, EPI_ISL_627773, EPI_ISL_627776, EPI_ISL_627779, EPI_ISL_627782, EPI_ISL_627787, EPI_ISL_627795, EPI_ISL_627796, EPI_ISL_627797, EPI_ISL_627800, EPI_ISL_627801, EPI_ISL_627804, EPI_ISL_627812, EPI_ISL_627815, EPI_ISL_627816, EPI_ISL_627819, EPI_ISL_627827, EPI_ISL_627831, EPI_ISL_627835, EPI_ISL_627844, EPI_ISL_627852, EPI_ISL_627853, EPI_ISL_627857, EPI_ISL_627860, EPI_ISL_627861, EPI_ISL_627862, EPI_ISL_627863, EPI_ISL_627866, EPI_ISL_627867, EPI_ISL_627870, EPI_ISL_627876, EPI_ISL_627881, EPI_ISL_627882, EPI_ISL_627883, EPI_ISL_627884, EPI_ISL_627885, EPI_ISL_627886, EPI_ISL_627889, EPI_ISL_627895, EPI_ISL_627900, EPI_ISL_627911, EPI_ISL_627912, EPI_ISL_627913, EPI_ISL_627915, EPI_ISL_627920, EPI_ISL_627923, EPI_ISL_627924, EPI_ISL_627927, EPI_ISL_627930, EPI_ISL_627935, EPI_ISL_627937, EPI_ISL_627952, EPI_ISL_627955, EPI_ISL_627959, EPI_ISL_627964, EPI_ISL_627968, EPI_ISL_627970, EPI_ISL_627975, EPI_ISL_627979, EPI_ISL_627980, EPI_ISL_627981, EPI_ISL_627986, EPI_ISL_627990, EPI_ISL_627993, EPI_ISL_627994, EPI_ISL_627995, EPI_ISL_627996, EPI_ISL_627997, EPI_ISL_628006, EPI_ISL_628010, EPI_ISL_628011, EPI_ISL_628013, EPI_ISL_628022, EPI_ISL_628038, EPI_ISL_628039, EPI_ISL_628044, EPI_ISL_628045, EPI_ISL_628050, EPI_ISL_628052, EPI_ISL_628057, EPI_ISL_628066, EPI_ISL_628067, EPI_ISL_628070, EPI_ISL_628073, EPI_ISL_628074, EPI_ISL_628081, EPI_ISL_628082, EPI_ISL_628083, EPI_ISL_628090, EPI_ISL_628094, EPI_ISL_628095, EPI_ISL_628098, EPI_ISL_628099, EPI_ISL_628102, EPI_ISL_628103, EPI_ISL_628106, EPI_ISL_628109, EPI_ISL_628111, EPI_ISL_628112, EPI_ISL_628113, EPI_ISL_628121, EPI_ISL_628129, EPI_ISL_628133, EPI_ISL_628134, EPI_ISL_628142, EPI_ISL_628143, EPI_ISL_628144, EPI_ISL_628148, EPI_ISL_628150, EPI_ISL_628171, EPI_ISL_628172, EPI_ISL_628175, EPI_ISL_628176, EPI_ISL_628184, EPI_ISL_628186, EPI_ISL_628191, EPI_ISL_628195, EPI_ISL_628197, EPI_ISL_628199, EPI_ISL_628210, EPI_ISL_628213, EPI_ISL_628222, EPI_ISL_628223, EPI_ISL_628228, EPI_ISL_628229, EPI_ISL_628235, EPI_ISL_628238, EPI_ISL_628239, EPI_ISL_628248, EPI_ISL_628249, EPI_ISL_628252, EPI_ISL_628257, EPI_ISL_628259, EPI_ISL_628261, EPI_ISL_628262, EPI_ISL_628267, EPI_ISL_628270, EPI_ISL_628274, EPI_ISL_628275, EPI_ISL_628276, EPI_ISL_628277, EPI_ISL_628280, EPI_ISL_628284, EPI_ISL_628285, EPI_ISL_628290, EPI_ISL_628293, EPI_ISL_628299, EPI_ISL_628300, EPI_ISL_628302, EPI_ISL_628305, EPI_ISL_628306 | | | |
| see above | Wales Specialist Virology Centre Sequencing lab: Pathogen Genomics Unit | COVID-19 Genomics UK (COG-UK) Consortium | Catherine Moore, Johnathan Evans, Laura Gifford, Malorie Perry, Simon Cottrell, Angela Marchbank, Alec Birchley, Alexander Adams, Amy Gaskin, Bree Gatica-Wilcox, Jason Coombes, Joel Southgate, Lauren Gilbert, Lee Graham, Nicole Pacchiarini, Sara Kumziene-Summerhayes, Sarah Taylor, Sophie Jones, Sara Rey, Matthew Bull, Joanne Watkins, Sally Corden, Tom Connor |
| EPI_ISL_628332, EPI_ISL_628333 | Centre for Enzyme Innovation, University of Portsmouth / Translational Research Laboratory, Portsmouth Hospitals NHS Trust | COVID-19 Genomics UK (COG-UK) Consortium | Angela Beckett,Yann Bourgeois,Garry Scarlett,Sharon Glaysher,Scott Elliott,Kelly Bicknell,Robert Impey,Allyson Lloyd,Sarah Wyllie,Ethan Butcher,Anoop Chauhan,Samuel Robson |
| EPI_ISL_628806, EPI_ISL_628858, EPI_ISL_628867, EPI_ISL_628868, EPI_ISL_628869, EPI_ISL_628870, EPI_ISL_628871, EPI_ISL_628872, EPI_ISL_628873, EPI_ISL_628874, EPI_ISL_628875, EPI_ISL_628876, EPI_ISL_628877, EPI_ISL_628878, EPI_ISL_628879, EPI_ISL_628880, EPI_ISL_628881, EPI_ISL_628882, EPI_ISL_628883, EPI_ISL_628884, EPI_ISL_628885, EPI_ISL_628886 | | | |
| see above | University of Michigan Clinical Microbiology Laboratory | Lauring Lab, University of Michigan, Department of Microbiology and Immunology | Valesano |
| EPI_ISL_629109, EPI_ISL_629110, EPI_ISL_629111, EPI_ISL_629112, EPI_ISL_629113, EPI_ISL_629114, EPI_ISL_629115, EPI_ISL_629116, EPI_ISL_629117, EPI_ISL_629118, EPI_ISL_629119, EPI_ISL_629120, EPI_ISL_629121, EPI_ISL_629122, EPI_ISL_629124, EPI_ISL_629125, EPI_ISL_629126, EPI_ISL_629127, EPI_ISL_629128, EPI_ISL_629129, EPI_ISL_629130, EPI_ISL_629131, EPI_ISL_629132, EPI_ISL_629133, EPI_ISL_629134, EPI_ISL_629136, EPI_ISL_629137 | | | |
| see above | Lighthouse Lab in Milton Keynes | Wellcome Sanger Institute for the COVID-19 Genomics UK (COG-UK) consortium | The Lighthouse Lab in Milton Keynes and Alex Alderton, Roberto Amato, Sonia Goncalves, Ewan Harrison, David K. Jackson, Ian Johnston, Dominic Kwiatkowski, Cordelia Langford, John Sillitoe on behalf of the Wellcome Sanger Institute COVID-19 Surveillance Team |
| EPI_ISL_629138 | Lighthouse Lab in Milton Keynes | Wellcome Sanger Institute for the COVID-19 Genomics UK (COG-UK) Consortium | The Lighthouse Lab in Milton Keynes and Alex Alderton, Roberto Amato, Sonia Goncalves, Ewan Harrison, David K. Jackson, Ian Johnston, Dominic Kwiatkowski, Cordelia Langford, John Sillitoe on behalf of the Wellcome Sanger Institute COVID-19 Surveillance Team |
| EPI_ISL_629139, EPI_ISL_629140, EPI_ISL_629141, EPI_ISL_629142, EPI_ISL_629143, EPI_ISL_629144, EPI_ISL_629145, EPI_ISL_629146, EPI_ISL_629147, EPI_ISL_629148, EPI_ISL_629149, EPI_ISL_629150, EPI_ISL_629151, EPI_ISL_629152, EPI_ISL_629153, EPI_ISL_629154, EPI_ISL_629155, EPI_ISL_629156, EPI_ISL_629157, EPI_ISL_629158, EPI_ISL_629159, EPI_ISL_629160, EPI_ISL_629161, EPI_ISL_629162, EPI_ISL_629163, EPI_ISL_629164, EPI_ISL_629165, EPI_ISL_629166, EPI_ISL_629167, EPI_ISL_629169, EPI_ISL_629170, EPI_ISL_629171, EPI_ISL_629172, EPI_ISL_629173, EPI_ISL_629174, EPI_ISL_629175, EPI_ISL_629176, EPI_ISL_629177, EPI_ISL_629178, EPI_ISL_629179, EPI_ISL_629180, EPI_ISL_629181, EPI_ISL_629182, EPI_ISL_629183, EPI_ISL_629184, EPI_ISL_629185, EPI_ISL_629186, EPI_ISL_629187, EPI_ISL_629188, EPI_ISL_629190, EPI_ISL_629191, EPI_ISL_629192, EPI_ISL_629193, EPI_ISL_629194, EPI_ISL_629196, EPI_ISL_629197, EPI_ISL_629198 | | | |
| see above | Lighthouse Lab in Milton Keynes | Wellcome Sanger Institute for the COVID-19 Genomics UK (COG-UK) consortium | The Lighthouse Lab in Milton Keynes and Alex Alderton, Roberto Amato, Sonia Goncalves, Ewan Harrison, David K. Jackson, Ian Johnston, Dominic Kwiatkowski, Cordelia Langford, John Sillitoe on behalf of the Wellcome Sanger Institute COVID-19 Surveillance Team |
| EPI_ISL_629199 | Lighthouse Lab in Milton Keynes | Wellcome Sanger Institute for the COVID-19 Genomics UK (COG-UK) Consortium | The Lighthouse Lab in Milton Keynes and Alex Alderton, Roberto Amato, Sonia Goncalves, Ewan Harrison, David K. Jackson, Ian Johnston, Dominic Kwiatkowski, Cordelia Langford, John Sillitoe on behalf of the Wellcome Sanger Institute COVID-19 Surveillance Team |
| EPI_ISL_629200, EPI_ISL_629201, EPI_ISL_629202, EPI_ISL_629203, EPI_ISL_629204, EPI_ISL_629205, EPI_ISL_629206, EPI_ISL_629207, EPI_ISL_629208, EPI_ISL_629209, EPI_ISL_629210, EPI_ISL_629211, EPI_ISL_629212, EPI_ISL_629213, EPI_ISL_629214, EPI_ISL_629215, EPI_ISL_629216, EPI_ISL_629217, EPI_ISL_629218 | | | |
| see above | Lighthouse Lab in Milton Keynes | Wellcome Sanger Institute for the COVID-19 Genomics UK (COG-UK) consortium | The Lighthouse Lab in Milton Keynes and Alex Alderton, Roberto Amato, Sonia Goncalves, Ewan Harrison, David K. Jackson, Ian Johnston, Dominic Kwiatkowski, Cordelia Langford, John Sillitoe on behalf of the Wellcome Sanger Institute COVID-19 Surveillance Team |
| EPI_ISL_629219 | Lighthouse Lab in Milton Keynes | Wellcome Sanger Institute for the COVID-19 Genomics UK | The Lighthouse Lab in Milton Keynes and Alex Alderton, Roberto Amato, Sonia Goncalves, Ewan Harrison, David K. Jackson, Ian Johnston, Dominic |

| | | | |
|---|---|---|---|
| | | (COG-UK) Consortium | Kwiatkowski, Cordelia Langford, John Sillitoe on behalf of the Wellcome Sanger Institute COVID-19 Surveillance Team |
| EPI_ISL_629220, EPI_ISL_629221, EPI_ISL_629222, EPI_ISL_629223, EPI_ISL_629224, EPI_ISL_629225, EPI_ISL_629226, EPI_ISL_629227, EPI_ISL_629228, EPI_ISL_629229, EPI_ISL_629230, EPI_ISL_629231, EPI_ISL_629232, EPI_ISL_629233, EPI_ISL_629234, EPI_ISL_629235, EPI_ISL_629236, EPI_ISL_629237, EPI_ISL_629238, EPI_ISL_629239, EPI_ISL_629240, EPI_ISL_629241, EPI_ISL_629242, EPI_ISL_629243, EPI_ISL_629244 | | | |
| see above | Lighthouse Lab in Milton Keynes | Wellcome Sanger Institute for the COVID-19 Genomics UK (COG-UK) consortium | The Lighthouse Lab in Milton Keynes and Alex Alderton, Roberto Amato, Sonia Goncalves, Ewan Harrison, David K. Jackson, Ian Johnston, Dominic Kwiatkowski, Cordelia Langford, John Sillitoe on behalf of the Wellcome Sanger Institute COVID-19 Surveillance Team |
| EPI_ISL_629245 | Lighthouse Lab in Milton Keynes | Wellcome Sanger Institute for the COVID-19 Genomics UK (COG-UK) Consortium | The Lighthouse Lab in Milton Keynes and Alex Alderton, Roberto Amato, Sonia Goncalves, Ewan Harrison, David K. Jackson, Ian Johnston, Dominic Kwiatkowski, Cordelia Langford, John Sillitoe on behalf of the Wellcome Sanger Institute COVID-19 Surveillance Team |
| EPI_ISL_629247, EPI_ISL_629248, EPI_ISL_629249, EPI_ISL_629250, EPI_ISL_629251, EPI_ISL_629252, EPI_ISL_629253, EPI_ISL_629254, EPI_ISL_629255, EPI_ISL_629256, EPI_ISL_629257, EPI_ISL_629258, EPI_ISL_629259, EPI_ISL_629260, EPI_ISL_629261, EPI_ISL_629262, EPI_ISL_629263, EPI_ISL_629264, EPI_ISL_629265, EPI_ISL_629266, EPI_ISL_629267, EPI_ISL_629268, EPI_ISL_629269, EPI_ISL_629270, EPI_ISL_629271, EPI_ISL_629272, EPI_ISL_629273, EPI_ISL_629274, EPI_ISL_629275, EPI_ISL_629276, EPI_ISL_629277, EPI_ISL_629278, EPI_ISL_629280, EPI_ISL_629281, EPI_ISL_629282, EPI_ISL_629283, EPI_ISL_629284, EPI_ISL_629285, EPI_ISL_629286, EPI_ISL_629287, EPI_ISL_629288, EPI_ISL_629289, EPI_ISL_629290, EPI_ISL_629292, EPI_ISL_629294, EPI_ISL_629295, EPI_ISL_629296, EPI_ISL_629297, EPI_ISL_629298, EPI_ISL_629299, EPI_ISL_629300, EPI_ISL_629301, EPI_ISL_629302, EPI_ISL_629303, EPI_ISL_629304, EPI_ISL_629305, EPI_ISL_629306, EPI_ISL_629307, EPI_ISL_629308, EPI_ISL_629309, EPI_ISL_629310, EPI_ISL_629311, EPI_ISL_629312, EPI_ISL_629313, EPI_ISL_629314, EPI_ISL_629315, EPI_ISL_629316, EPI_ISL_629317, EPI_ISL_629318, EPI_ISL_629319, EPI_ISL_629320, EPI_ISL_629321, EPI_ISL_629322, EPI_ISL_629323, EPI_ISL_629324, EPI_ISL_629325, EPI_ISL_629326, EPI_ISL_629327, EPI_ISL_629328, EPI_ISL_629329, EPI_ISL_629330, EPI_ISL_629331, EPI_ISL_629332, EPI_ISL_629333, EPI_ISL_629334, EPI_ISL_629335, EPI_ISL_629336, EPI_ISL_629337, EPI_ISL_629338, EPI_ISL_629339, EPI_ISL_629340, EPI_ISL_629342, EPI_ISL_629343, EPI_ISL_629344, EPI_ISL_629345, EPI_ISL_629346, EPI_ISL_629347, EPI_ISL_629348, EPI_ISL_629349, EPI_ISL_629350, EPI_ISL_629351, EPI_ISL_629352, EPI_ISL_629353, EPI_ISL_629354, EPI_ISL_629355, EPI_ISL_629356, EPI_ISL_629357, EPI_ISL_629358, EPI_ISL_629361, EPI_ISL_629363, EPI_ISL_629364, EPI_ISL_629365, EPI_ISL_629366, EPI_ISL_629367, EPI_ISL_629368, EPI_ISL_629369, EPI_ISL_629371, EPI_ISL_629372, EPI_ISL_629373, EPI_ISL_629374, EPI_ISL_629375, EPI_ISL_629376, EPI_ISL_629377, EPI_ISL_629378, EPI_ISL_629379, EPI_ISL_629381, EPI_ISL_629382, EPI_ISL_629383, EPI_ISL_629384, EPI_ISL_629385, EPI_ISL_629386, EPI_ISL_629387, EPI_ISL_629388, EPI_ISL_629389, EPI_ISL_629390, EPI_ISL_629391, EPI_ISL_629393, EPI_ISL_629394, EPI_ISL_629395, EPI_ISL_629396, EPI_ISL_629397, EPI_ISL_629398, EPI_ISL_629399, EPI_ISL_629400, EPI_ISL_629401, EPI_ISL_629402, EPI_ISL_629403, EPI_ISL_629404, EPI_ISL_629405, EPI_ISL_629406, EPI_ISL_629407, EPI_ISL_629408 | | | |
| see above | Lighthouse Lab in Milton Keynes | Wellcome Sanger Institute for the COVID-19 Genomics UK (COG-UK) consortium | The Lighthouse Lab in Milton Keynes and Alex Alderton, Roberto Amato, Sonia Goncalves, Ewan Harrison, David K. Jackson, Ian Johnston, Dominic Kwiatkowski, Cordelia Langford, John Sillitoe on behalf of the Wellcome Sanger Institute COVID-19 Surveillance Team |
| EPI_ISL_629409 | Lighthouse Lab in Milton Keynes | Wellcome Sanger Institute for the COVID-19 Genomics UK (COG-UK) Consortium | The Lighthouse Lab in Milton Keynes and Alex Alderton, Roberto Amato, Sonia Goncalves, Ewan Harrison, David K. Jackson, Ian Johnston, Dominic Kwiatkowski, Cordelia Langford, John Sillitoe on behalf of the Wellcome Sanger Institute COVID-19 Surveillance Team |
| EPI_ISL_629410, EPI_ISL_629411, EPI_ISL_629412, EPI_ISL_629413, EPI_ISL_629415, EPI_ISL_629416, EPI_ISL_629417, EPI_ISL_629418, EPI_ISL_629419, EPI_ISL_629420, EPI_ISL_629421, EPI_ISL_629422, EPI_ISL_629423, EPI_ISL_629424, EPI_ISL_629425, EPI_ISL_629426, EPI_ISL_629427, EPI_ISL_629428, EPI_ISL_629429, EPI_ISL_629430, EPI_ISL_629431, EPI_ISL_629432, EPI_ISL_629433, EPI_ISL_629434, EPI_ISL_629436, EPI_ISL_629439, EPI_ISL_629441, EPI_ISL_629444, EPI_ISL_629448, EPI_ISL_629450, EPI_ISL_629452, EPI_ISL_629454, EPI_ISL_629456, EPI_ISL_629458, EPI_ISL_629460, EPI_ISL_629462, EPI_ISL_629465, EPI_ISL_629466, EPI_ISL_629468, EPI_ISL_629470, EPI_ISL_629471, EPI_ISL_629473, EPI_ISL_629474, EPI_ISL_629475, EPI_ISL_629476, EPI_ISL_629482, EPI_ISL_629483, EPI_ISL_629485, EPI_ISL_629486, EPI_ISL_629487, EPI_ISL_629489 | | | |
| see above | Lighthouse Lab in Milton Keynes | Wellcome Sanger Institute for the COVID-19 Genomics UK (COG-UK) consortium | The Lighthouse Lab in Milton Keynes and Alex Alderton, Roberto Amato, Sonia Goncalves, Ewan Harrison, David K. Jackson, Ian Johnston, Dominic Kwiatkowski, Cordelia Langford, John Sillitoe on behalf of the Wellcome Sanger Institute COVID-19 Surveillance Team |
| EPI_ISL_629490 | Lighthouse Lab in Milton Keynes | Wellcome Sanger Institute for the COVID-19 Genomics UK (COG-UK) Consortium | The Lighthouse Lab in Milton Keynes and Alex Alderton, Roberto Amato, Sonia Goncalves, Ewan Harrison, David K. Jackson, Ian Johnston, Dominic Kwiatkowski, Cordelia Langford, John Sillitoe on behalf of the Wellcome Sanger Institute COVID-19 Surveillance Team |
| EPI_ISL_629494, EPI_ISL_629495, EPI_ISL_629499, EPI_ISL_629501, EPI_ISL_629502, EPI_ISL_629503, EPI_ISL_629506, EPI_ISL_629510, EPI_ISL_629514, EPI_ISL_629515, EPI_ISL_629517, EPI_ISL_629518, EPI_ISL_629519, EPI_ISL_629520, EPI_ISL_629522, EPI_ISL_629523, EPI_ISL_629524, EPI_ISL_629528, EPI_ISL_629531, EPI_ISL_629532, EPI_ISL_629534, EPI_ISL_629535, EPI_ISL_629537, EPI_ISL_629539, EPI_ISL_629541, EPI_ISL_629544, EPI_ISL_629545, EPI_ISL_629549, EPI_ISL_629555, EPI_ISL_629565, EPI_ISL_629569, EPI_ISL_629572, EPI_ISL_629577, EPI_ISL_629581, EPI_ISL_629587, EPI_ISL_629591 | | | |
| see above | Lighthouse Lab in Milton Keynes | Wellcome Sanger Institute for the COVID-19 Genomics UK (COG-UK) consortium | The Lighthouse Lab in Milton Keynes and Alex Alderton, Roberto Amato, Sonia Goncalves, Ewan Harrison, David K. Jackson, Ian Johnston, Dominic Kwiatkowski, Cordelia Langford, John Sillitoe on behalf of the Wellcome Sanger Institute COVID-19 Surveillance Team |
| EPI_ISL_629592 | Lighthouse Lab in Milton Keynes | Wellcome Sanger Institute for the COVID-19 Genomics UK (COG-UK) Consortium | The Lighthouse Lab in Milton Keynes and Alex Alderton, Roberto Amato, Sonia Goncalves, Ewan Harrison, David K. Jackson, Ian Johnston, Dominic Kwiatkowski, Cordelia Langford, John Sillitoe on behalf of the Wellcome Sanger Institute COVID-19 Surveillance Team |
| EPI_ISL_629594, EPI_ISL_629595, EPI_ISL_629596, EPI_ISL_629597, EPI_ISL_629598, EPI_ISL_629599, EPI_ISL_629603, EPI_ISL_629608, EPI_ISL_629614, EPI_ISL_629615, EPI_ISL_629617, EPI_ISL_629618, EPI_ISL_629619, EPI_ISL_629621, EPI_ISL_629622, EPI_ISL_629624, EPI_ISL_629625, EPI_ISL_629626, EPI_ISL_629628, EPI_ISL_629630, EPI_ISL_629631, EPI_ISL_629633, EPI_ISL_629635, EPI_ISL_629636, EPI_ISL_629637, EPI_ISL_629641, EPI_ISL_629644, EPI_ISL_629645, EPI_ISL_629648, EPI_ISL_629649, EPI_ISL_629651, EPI_ISL_629656, EPI_ISL_629657, EPI_ISL_629662, EPI_ISL_629663, EPI_ISL_629664, EPI_ISL_629665, EPI_ISL_629669, EPI_ISL_629671, EPI_ISL_629672, EPI_ISL_629675, EPI_ISL_629677, EPI_ISL_629678, EPI_ISL_629679, EPI_ISL_629683, EPI_ISL_629685, EPI_ISL_629689, EPI_ISL_629690, EPI_ISL_629692, EPI_ISL_629693, EPI_ISL_629696, EPI_ISL_629701, EPI_ISL_629702, EPI_ISL_629703, EPI_ISL_629709, EPI_ISL_629710, EPI_ISL_629711, EPI_ISL_629720, EPI_ISL_629721, EPI_ISL_629724, EPI_ISL_629725, EPI_ISL_629729, EPI_ISL_629730, EPI_ISL_629733, EPI_ISL_629734, EPI_ISL_629735, EPI_ISL_629736, EPI_ISL_629737, EPI_ISL_629738, EPI_ISL_629741, EPI_ISL_629745, EPI_ISL_629747, EPI_ISL_629748, EPI_ISL_629750 | | | |
| see above | Lighthouse Lab in Milton Keynes | Wellcome Sanger Institute for the COVID-19 Genomics UK (COG-UK) consortium | The Lighthouse Lab in Milton Keynes and Alex Alderton, Roberto Amato, Sonia Goncalves, Ewan Harrison, David K. Jackson, Ian Johnston, Dominic Kwiatkowski, Cordelia Langford, John Sillitoe on behalf of the Wellcome Sanger Institute COVID-19 Surveillance Team |
| EPI_ISL_629754 | Lighthouse Lab in Glasgow | Wellcome Sanger Institute for the COVID-19 Genomics UK (COG-UK) consortium | Harper VanSteenhouse, Yumi Kasai, David Gray, Carol Clugston, Anna Dominiczak and Alex Alderton, Roberto Amato, Sonia Goncalves, Ewan Harrison, David K. Jackson, Ian Johnston, Dominic Kwiatkowski, Cordelia Langford, John Sillitoe on behalf of the Wellcome Sanger Institute COVID-19 Surveillance Team |
| EPI_ISL_630007, EPI_ISL_630008, EPI_ISL_630009, EPI_ISL_630010, EPI_ISL_630012, EPI_ISL_630013, EPI_ISL_630014, EPI_ISL_630015, EPI_ISL_630016, EPI_ISL_630018, EPI_ISL_630019, EPI_ISL_630020, EPI_ISL_630021, EPI_ISL_630022, EPI_ISL_630025, EPI_ISL_630026, EPI_ISL_630027, EPI_ISL_630028, EPI_ISL_630029, EPI_ISL_630030, EPI_ISL_630033, EPI_ISL_630034, EPI_ISL_630035, EPI_ISL_630036, EPI_ISL_630039, EPI_ISL_630041, EPI_ISL_630042, EPI_ISL_630043, EPI_ISL_630045, EPI_ISL_630046, EPI_ISL_630048, EPI_ISL_630049, EPI_ISL_630051, EPI_ISL_630052, EPI_ISL_630053, EPI_ISL_630054, EPI_ISL_630056, EPI_ISL_630058, EPI_ISL_630059, EPI_ISL_630060, EPI_ISL_630061, EPI_ISL_630062, EPI_ISL_630063, EPI_ISL_630064, EPI_ISL_630065, EPI_ISL_630068, EPI_ISL_630069, EPI_ISL_630071, EPI_ISL_630072, EPI_ISL_630073, EPI_ISL_630074, EPI_ISL_630076, EPI_ISL_630077, EPI_ISL_630078, EPI_ISL_630079, EPI_ISL_630082, EPI_ISL_630083, EPI_ISL_630084, EPI_ISL_630085, EPI_ISL_630087, EPI_ISL_630088, EPI_ISL_630089, EPI_ISL_630090, EPI_ISL_630095, EPI_ISL_630097, EPI_ISL_630098, EPI_ISL_630099, EPI_ISL_630100, EPI_ISL_630101, EPI_ISL_630102, EPI_ISL_630103, EPI_ISL_630106, EPI_ISL_630107, EPI_ISL_630108, EPI_ISL_630110, EPI_ISL_630111, EPI_ISL_630112, EPI_ISL_630113, EPI_ISL_630114, EPI_ISL_630115, EPI_ISL_630116, EPI_ISL_630117, EPI_ISL_630118, EPI_ISL_630120, EPI_ISL_630121, EPI_ISL_630122, EPI_ISL_630123, EPI_ISL_630124, EPI_ISL_630125, EPI_ISL_630127, EPI_ISL_630128, EPI_ISL_630129, EPI_ISL_630130, EPI_ISL_630131, EPI_ISL_630133, EPI_ISL_630135, EPI_ISL_630136, EPI_ISL_630137, EPI_ISL_630138, EPI_ISL_630139, EPI_ISL_630140, EPI_ISL_630141, EPI_ISL_630142, EPI_ISL_630145, EPI_ISL_630146, EPI_ISL_630147, EPI_ISL_630148, EPI_ISL_630149, EPI_ISL_630150, EPI_ISL_630151, EPI_ISL_630152, EPI_ISL_630154, EPI_ISL_630155, EPI_ISL_630156, EPI_ISL_630157, EPI_ISL_630158, EPI_ISL_630159, EPI_ISL_630160, EPI_ISL_630161, EPI_ISL_630162, EPI_ISL_630163, EPI_ISL_630165, EPI_ISL_630166, EPI_ISL_630167, EPI_ISL_630168, EPI_ISL_630169, EPI_ISL_630170, EPI_ISL_630171, EPI_ISL_630172, EPI_ISL_630173, EPI_ISL_630175, EPI_ISL_630176, EPI_ISL_630178, EPI_ISL_630180, EPI_ISL_630181, EPI_ISL_630183, EPI_ISL_630184, EPI_ISL_630185, EPI_ISL_630186, EPI_ISL_630187, EPI_ISL_630188, EPI_ISL_630189, EPI_ISL_630190 | | | |
| see above | Lighthouse Lab in Milton Keynes | Wellcome Sanger Institute for the COVID-19 Genomics UK (COG-UK) consortium | The Lighthouse Lab in Milton Keynes and Alex Alderton, Roberto Amato, Sonia Goncalves, Ewan Harrison, David K. Jackson, Ian Johnston, Dominic Kwiatkowski, Cordelia Langford, John Sillitoe on behalf of the Wellcome Sanger Institute COVID-19 Surveillance Team |
| EPI_ISL_630191 | Lighthouse Lab in Milton Keynes | Wellcome Sanger Institute for the COVID-19 Genomics UK (COG-UK) Consortium | The Lighthouse Lab in Milton Keynes and Alex Alderton, Roberto Amato, Sonia Goncalves, Ewan Harrison, David K. Jackson, Ian Johnston, Dominic Kwiatkowski, Cordelia Langford, John Sillitoe on behalf of the Wellcome Sanger Institute COVID-19 Surveillance Team |
| EPI_ISL_630192, EPI_ISL_630193, EPI_ISL_630194, EPI_ISL_630195, EPI_ISL_630197, EPI_ISL_630198, EPI_ISL_630201, EPI_ISL_630202, EPI_ISL_630203, EPI_ISL_630204, EPI_ISL_630205, EPI_ISL_630207, EPI_ISL_630209, EPI_ISL_630210, EPI_ISL_630212, EPI_ISL_630213, EPI_ISL_630214, EPI_ISL_630215, EPI_ISL_630216, EPI_ISL_630218 | | | |
| see above | Lighthouse Lab in Milton Keynes | Wellcome Sanger Institute for the COVID-19 Genomics UK (COG-UK) consortium | The Lighthouse Lab in Milton Keynes and Alex Alderton, Roberto Amato, Sonia Goncalves, Ewan Harrison, David K. Jackson, Ian Johnston, Dominic Kwiatkowski, Cordelia Langford, John Sillitoe on behalf of the Wellcome Sanger Institute COVID-19 Surveillance Team |
| EPI_ISL_630219 | Lighthouse Lab in Milton Keynes | Wellcome Sanger Institute for the COVID-19 Genomics UK (COG-UK) Consortium | The Lighthouse Lab in Milton Keynes and Alex Alderton, Roberto Amato, Sonia Goncalves, Ewan Harrison, David K. Jackson, Ian Johnston, Dominic Kwiatkowski, Cordelia Langford, John Sillitoe on behalf of the Wellcome Sanger Institute COVID-19 Surveillance Team |
| EPI_ISL_630220, EPI_ISL_630221, EPI_ISL_630222, EPI_ISL_630225, EPI_ISL_630226, EPI_ISL_630228, EPI_ISL_630229, EPI_ISL_630230, EPI_ISL_630231, EPI_ISL_630232, EPI_ISL_630234, EPI_ISL_630235, EPI_ISL_630236, EPI_ISL_630237, EPI_ISL_630239, EPI_ISL_630240, EPI_ISL_630242, EPI_ISL_630243, EPI_ISL_630244, EPI_ISL_630245, EPI_ISL_630246, EPI_ISL_630247, EPI_ISL_630248 | | | |
| see above | Lighthouse Lab in Milton Keynes | Wellcome Sanger Institute for the COVID-19 Genomics UK (COG-UK) consortium | The Lighthouse Lab in Milton Keynes and Alex Alderton, Roberto Amato, Sonia Goncalves, Ewan Harrison, David K. Jackson, Ian Johnston, Dominic Kwiatkowski, Cordelia Langford, John Sillitoe on behalf of the Wellcome Sanger Institute COVID-19 Surveillance Team |
| EPI_ISL_630249 | Lighthouse Lab in Milton Keynes | Wellcome Sanger Institute for the COVID-19 Genomics UK (COG-UK) Consortium | The Lighthouse Lab in Milton Keynes and Alex Alderton, Roberto Amato, Sonia Goncalves, Ewan Harrison, David K. Jackson, Ian Johnston, Dominic Kwiatkowski, Cordelia Langford, John Sillitoe on behalf of the Wellcome Sanger Institute COVID-19 Surveillance Team |
| EPI_ISL_630250, EPI_ISL_630251, EPI_ISL_630252, EPI_ISL_630253, EPI_ISL_630254, EPI_ISL_630255, EPI_ISL_630256, EPI_ISL_630257, EPI_ISL_630258, EPI_ISL_630259, EPI_ISL_630260, EPI_ISL_630261, EPI_ISL_630263, EPI_ISL_630264, EPI_ISL_630265, EPI_ISL_630266, EPI_ISL_630267, EPI_ISL_630268, EPI_ISL_630270, EPI_ISL_630271, EPI_ISL_630273, EPI_ISL_630274, EPI_ISL_630275, EPI_ISL_630278, EPI_ISL_630280, EPI_ISL_630281, EPI_ISL_630282, EPI_ISL_630283, EPI_ISL_630284, EPI_ISL_630286, EPI_ISL_630287, EPI_ISL_630288, EPI_ISL_630289, EPI_ISL_630290, EPI_ISL_630291, EPI_ISL_630292, EPI_ISL_630293, EPI_ISL_630295, EPI_ISL_630296, EPI_ISL_630297, EPI_ISL_630298, EPI_ISL_630299, EPI_ISL_630300, EPI_ISL_630301, EPI_ISL_630302, EPI_ISL_630303, EPI_ISL_630305, EPI_ISL_630306, EPI_ISL_630309, EPI_ISL_630310, EPI_ISL_630311, EPI_ISL_630312, EPI_ISL_630314, EPI_ISL_630315, EPI_ISL_630317, EPI_ISL_630318, EPI_ISL_630319, EPI_ISL_630321, EPI_ISL_630322, EPI_ISL_630323, EPI_ISL_630324, EPI_ISL_630326, EPI_ISL_630327, EPI_ISL_630329, EPI_ISL_630330, EPI_ISL_630334, EPI_ISL_630335, EPI_ISL_630336, EPI_ISL_630337, EPI_ISL_630338, EPI_ISL_630339, EPI_ISL_630340, EPI_ISL_630341, EPI_ISL_630343, EPI_ISL_630344, EPI_ISL_630345 | | | |
| see above | Lighthouse Lab in Milton Keynes | Wellcome Sanger Institute for the COVID-19 Genomics UK (COG-UK) consortium | The Lighthouse Lab in Milton Keynes and Alex Alderton, Roberto Amato, Sonia Goncalves, Ewan Harrison, David K. Jackson, Ian Johnston, Dominic Kwiatkowski, Cordelia Langford, John Sillitoe on behalf of the Wellcome Sanger Institute COVID-19 Surveillance Team |

| | | | |
|---|---|---|---|
| EPI_ISL_630702, EPI_ISL_630704, EPI_ISL_630705 | Lighthouse Lab in Glasgow | Wellcome Sanger Institute for the COVID-19 Genomics UK (COG-UK) consortium | Harper VanSteenhouse, Yumi Kasai, David Gray, Carol Clugston, Anna Dominiczak and Alex Alderton, Roberto Amato, Sonia Goncalves, Ewan Harrison, David K. Jackson, Ian Johnston, Dominic Kwiatkowski, Cordelia Langford, John Sillitoe on behalf of the Wellcome Sanger Institute COVID-19 Surveillance Team |
| EPI_ISL_630708, EPI_ISL_630710, EPI_ISL_630711, EPI_ISL_630716, EPI_ISL_630717, EPI_ISL_630720, EPI_ISL_630722, EPI_ISL_630723, EPI_ISL_630724, EPI_ISL_630729 | Lighthouse Lab in Alderley Park | Wellcome Sanger Institute for the COVID-19 Genomics UK (COG-UK) consortium | Jacquelyn Wynn, Mairead Hyland, The Lighthouse Lab in Alderley Park and Alex Alderton, Roberto Amato, Sonia Goncalves, Ewan Harrison, David K. Jackson, Ian Johnston, Dominic Kwiatkowski, Cordelia Langford, John Sillitoe on behalf of the Wellcome Sanger Institute COVID-19 Surveillance Team |
| EPI_ISL_630731 | Lighthouse Lab in Glasgow | Wellcome Sanger Institute for the COVID-19 Genomics UK (COG-UK) consortium | Harper VanSteenhouse, Yumi Kasai, David Gray, Carol Clugston, Anna Dominiczak and Alex Alderton, Roberto Amato, Sonia Goncalves, Ewan Harrison, David K. Jackson, Ian Johnston, Dominic Kwiatkowski, Cordelia Langford, John Sillitoe on behalf of the Wellcome Sanger Institute COVID-19 Surveillance Team |
| EPI_ISL_630732, EPI_ISL_630736, EPI_ISL_630737, EPI_ISL_630746, EPI_ISL_630747, EPI_ISL_630748 | Lighthouse Lab in Alderley Park | Wellcome Sanger Institute for the COVID-19 Genomics UK (COG-UK) consortium | Jacquelyn Wynn, Mairead Hyland, The Lighthouse Lab in Alderley Park and Alex Alderton, Roberto Amato, Sonia Goncalves, Ewan Harrison, David K. Jackson, Ian Johnston, Dominic Kwiatkowski, Cordelia Langford, John Sillitoe on behalf of the Wellcome Sanger Institute COVID-19 Surveillance Team |
| EPI_ISL_630749 | Lighthouse Lab in Glasgow | Wellcome Sanger Institute for the COVID-19 Genomics UK (COG-UK) consortium | Harper VanSteenhouse, Yumi Kasai, David Gray, Carol Clugston, Anna Dominiczak and Alex Alderton, Roberto Amato, Sonia Goncalves, Ewan Harrison, David K. Jackson, Ian Johnston, Dominic Kwiatkowski, Cordelia Langford, John Sillitoe on behalf of the Wellcome Sanger Institute COVID-19 Surveillance Team |
| EPI_ISL_630753, EPI_ISL_630754, EPI_ISL_630755, EPI_ISL_630756, EPI_ISL_630758, EPI_ISL_630762 | Lighthouse Lab in Alderley Park | Wellcome Sanger Institute for the COVID-19 Genomics UK (COG-UK) consortium | Jacquelyn Wynn, Mairead Hyland, The Lighthouse Lab in Alderley Park and Alex Alderton, Roberto Amato, Sonia Goncalves, Ewan Harrison, David K. Jackson, Ian Johnston, Dominic Kwiatkowski, Cordelia Langford, John Sillitoe on behalf of the Wellcome Sanger Institute COVID-19 Surveillance Team |
| EPI_ISL_630764 | Lighthouse Lab in Glasgow | Wellcome Sanger Institute for the COVID-19 Genomics UK (COG-UK) consortium | Harper VanSteenhouse, Yumi Kasai, David Gray, Carol Clugston, Anna Dominiczak and Alex Alderton, Roberto Amato, Sonia Goncalves, Ewan Harrison, David K. Jackson, Ian Johnston, Dominic Kwiatkowski, Cordelia Langford, John Sillitoe on behalf of the Wellcome Sanger Institute COVID-19 Surveillance Team |
| EPI_ISL_630766, EPI_ISL_630767, EPI_ISL_630771 | Lighthouse Lab in Alderley Park | Wellcome Sanger Institute for the COVID-19 Genomics UK (COG-UK) consortium | Jacquelyn Wynn, Mairead Hyland, The Lighthouse Lab in Alderley Park and Alex Alderton, Roberto Amato, Sonia Goncalves, Ewan Harrison, David K. Jackson, Ian Johnston, Dominic Kwiatkowski, Cordelia Langford, John Sillitoe on behalf of the Wellcome Sanger Institute COVID-19 Surveillance Team |
| EPI_ISL_630775 | Lighthouse Lab in Glasgow | Wellcome Sanger Institute for the COVID-19 Genomics UK (COG-UK) consortium | Harper VanSteenhouse, Yumi Kasai, David Gray, Carol Clugston, Anna Dominiczak and Alex Alderton, Roberto Amato, Sonia Goncalves, Ewan Harrison, David K. Jackson, Ian Johnston, Dominic Kwiatkowski, Cordelia Langford, John Sillitoe on behalf of the Wellcome Sanger Institute COVID-19 Surveillance Team |
| EPI_ISL_630777, EPI_ISL_630781, EPI_ISL_630784, EPI_ISL_630786, EPI_ISL_630788, EPI_ISL_630790, EPI_ISL_630793, EPI_ISL_630803, EPI_ISL_630807, EPI_ISL_630808, EPI_ISL_630810, EPI_ISL_630812 | | | |
| see above | Lighthouse Lab in Alderley Park | Wellcome Sanger Institute for the COVID-19 Genomics UK (COG-UK) consortium | Jacquelyn Wynn, Mairead Hyland, The Lighthouse Lab in Alderley Park and Alex Alderton, Roberto Amato, Sonia Goncalves, Ewan Harrison, David K. Jackson, Ian Johnston, Dominic Kwiatkowski, Cordelia Langford, John Sillitoe on behalf of the Wellcome Sanger Institute COVID-19 Surveillance Team |
| EPI_ISL_630813 | Lighthouse Lab in Glasgow | Wellcome Sanger Institute for the COVID-19 Genomics UK (COG-UK) consortium | Harper VanSteenhouse, Yumi Kasai, David Gray, Carol Clugston, Anna Dominiczak and Alex Alderton, Roberto Amato, Sonia Goncalves, Ewan Harrison, David K. Jackson, Ian Johnston, Dominic Kwiatkowski, Cordelia Langford, John Sillitoe on behalf of the Wellcome Sanger Institute COVID-19 Surveillance Team |
| EPI_ISL_630822, EPI_ISL_630823, EPI_ISL_630830, EPI_ISL_630832, EPI_ISL_630834, EPI_ISL_630835, EPI_ISL_630839, EPI_ISL_630843, EPI_ISL_630844, EPI_ISL_630845, EPI_ISL_630855, EPI_ISL_630856, EPI_ISL_630858, EPI_ISL_630861, EPI_ISL_630863, EPI_ISL_630867, EPI_ISL_630868, EPI_ISL_630869 | | | |
| see above | Lighthouse Lab in Alderley Park | Wellcome Sanger Institute for the COVID-19 Genomics UK (COG-UK) consortium | Jacquelyn Wynn, Mairead Hyland, The Lighthouse Lab in Alderley Park and Alex Alderton, Roberto Amato, Sonia Goncalves, Ewan Harrison, David K. Jackson, Ian Johnston, Dominic Kwiatkowski, Cordelia Langford, John Sillitoe on behalf of the Wellcome Sanger Institute COVID-19 Surveillance Team |
| EPI_ISL_630876 | Lighthouse Lab in Glasgow | Wellcome Sanger Institute for the COVID-19 Genomics UK (COG-UK) consortium | Harper VanSteenhouse, Yumi Kasai, David Gray, Carol Clugston, Anna Dominiczak and Alex Alderton, Roberto Amato, Sonia Goncalves, Ewan Harrison, David K. Jackson, Ian Johnston, Dominic Kwiatkowski, Cordelia Langford, John Sillitoe on behalf of the Wellcome Sanger Institute COVID-19 Surveillance Team |
| EPI_ISL_630877, EPI_ISL_630879, EPI_ISL_630880, EPI_ISL_630882 | Lighthouse Lab in Alderley Park | Wellcome Sanger Institute for the COVID-19 Genomics UK (COG-UK) consortium | Jacquelyn Wynn, Mairead Hyland, The Lighthouse Lab in Alderley Park and Alex Alderton, Roberto Amato, Sonia Goncalves, Ewan Harrison, David K. Jackson, Ian Johnston, Dominic Kwiatkowski, Cordelia Langford, John Sillitoe on behalf of the Wellcome Sanger Institute COVID-19 Surveillance Team |
| EPI_ISL_630883 | Lighthouse Lab in Glasgow | Wellcome Sanger Institute for the COVID-19 Genomics UK (COG-UK) consortium | Harper VanSteenhouse, Yumi Kasai, David Gray, Carol Clugston, Anna Dominiczak and Alex Alderton, Roberto Amato, Sonia Goncalves, Ewan Harrison, David K. Jackson, Ian Johnston, Dominic Kwiatkowski, Cordelia Langford, John Sillitoe on behalf of the Wellcome Sanger Institute COVID-19 Surveillance Team |
| EPI_ISL_630884 | Lighthouse Lab in Alderley Park | Wellcome Sanger Institute for the COVID-19 Genomics UK (COG-UK) consortium | Jacquelyn Wynn, Mairead Hyland, The Lighthouse Lab in Alderley Park and Alex Alderton, Roberto Amato, Sonia Goncalves, Ewan Harrison, David K. Jackson, Ian Johnston, Dominic Kwiatkowski, Cordelia Langford, John Sillitoe on behalf of the Wellcome Sanger Institute COVID-19 Surveillance Team |
| EPI_ISL_630891 | Lighthouse Lab in Alderley Park | Wellcome Sanger Institute for the COVID-19 Genomics UK (COG-UK) Consortium | Jacquelyn Wynn, Mairead Hyland, The Lighthouse Lab in Alderley Park and Alex Alderton, Roberto Amato, Sonia Goncalves, Ewan Harrison, David K. Jackson, Ian Johnston, Dominic Kwiatkowski, Cordelia Langford, John Sillitoe on behalf of the Wellcome Sanger Institute COVID-19 Surveillance Team |
| EPI_ISL_630892, EPI_ISL_630901, EPI_ISL_630902, EPI_ISL_630904, EPI_ISL_630905, EPI_ISL_630912, EPI_ISL_630914 | Lighthouse Lab in Alderley Park | Wellcome Sanger Institute for the COVID-19 Genomics UK (COG-UK) consortium | Jacquelyn Wynn, Mairead Hyland, The Lighthouse Lab in Alderley Park and Alex Alderton, Roberto Amato, Sonia Goncalves, Ewan Harrison, David K. Jackson, Ian Johnston, Dominic Kwiatkowski, Cordelia Langford, John Sillitoe on behalf of the Wellcome Sanger Institute COVID-19 Surveillance Team |
| EPI_ISL_630915 | Lighthouse Lab in Glasgow | Wellcome Sanger Institute for the COVID-19 Genomics UK (COG-UK) consortium | Harper VanSteenhouse, Yumi Kasai, David Gray, Carol Clugston, Anna Dominiczak and Alex Alderton, Roberto Amato, Sonia Goncalves, Ewan Harrison, David K. Jackson, Ian Johnston, Dominic Kwiatkowski, Cordelia Langford, John Sillitoe on behalf of the Wellcome Sanger Institute COVID-19 Surveillance Team |
| EPI_ISL_630917, EPI_ISL_630918, EPI_ISL_630928, EPI_ISL_630929, EPI_ISL_630932, EPI_ISL_630934, EPI_ISL_630936, EPI_ISL_630939, EPI_ISL_630940, EPI_ISL_630943, EPI_ISL_630944, EPI_ISL_630945, EPI_ISL_630947, EPI_ISL_630950, EPI_ISL_630953, EPI_ISL_630956, EPI_ISL_630958, EPI_ISL_630963, EPI_ISL_630964 | | | |
| see above | Lighthouse Lab in Alderley Park | Wellcome Sanger Institute for the COVID-19 Genomics UK (COG-UK) consortium | Jacquelyn Wynn, Mairead Hyland, The Lighthouse Lab in Alderley Park and Alex Alderton, Roberto Amato, Sonia Goncalves, Ewan Harrison, David K. Jackson, Ian Johnston, Dominic Kwiatkowski, Cordelia Langford, John Sillitoe on behalf of the Wellcome Sanger Institute COVID-19 Surveillance Team |
| EPI_ISL_630965 | Lighthouse Lab in Glasgow | Wellcome Sanger Institute for the COVID-19 Genomics UK (COG-UK) consortium | Harper VanSteenhouse, Yumi Kasai, David Gray, Carol Clugston, Anna Dominiczak and Alex Alderton, Roberto Amato, Sonia Goncalves, Ewan Harrison, David K. Jackson, Ian Johnston, Dominic Kwiatkowski, Cordelia Langford, John Sillitoe on behalf of the Wellcome Sanger Institute COVID-19 Surveillance Team |
| EPI_ISL_630966, EPI_ISL_630967 | Lighthouse Lab in Alderley Park | Wellcome Sanger Institute for the COVID-19 Genomics UK (COG-UK) consortium | Jacquelyn Wynn, Mairead Hyland, The Lighthouse Lab in Alderley Park and Alex Alderton, Roberto Amato, Sonia Goncalves, Ewan Harrison, David K. Jackson, Ian Johnston, Dominic Kwiatkowski, Cordelia Langford, John Sillitoe on behalf of the Wellcome Sanger Institute COVID-19 Surveillance Team |
| EPI_ISL_630973, EPI_ISL_630980, EPI_ISL_630981, EPI_ISL_630986, EPI_ISL_630987, EPI_ISL_630993, EPI_ISL_630995, EPI_ISL_631002, EPI_ISL_631004, EPI_ISL_631007, EPI_ISL_631009, EPI_ISL_631010, EPI_ISL_631015 | | | |
| see above | Lighthouse Lab in Milton Keynes | Wellcome Sanger Institute for the COVID-19 Genomics UK (COG-UK) consortium | The Lighthouse Lab in Milton Keynes and Alex Alderton, Roberto Amato, Sonia Goncalves, Ewan Harrison, David K. Jackson, Ian Johnston, Dominic Kwiatkowski, Cordelia Langford, John Sillitoe on behalf of the Wellcome Sanger Institute COVID-19 Surveillance Team |
| EPI_ISL_631016 | Lighthouse Lab in Cambridge | Wellcome Sanger Institute for the COVID-19 Genomics UK (COG-UK) consortium | Rob Howes, The Lighthouse Lab in Cambridge and Alex Alderton, Roberto Amato, Sonia Goncalves, Ewan Harrison, David K. Jackson, Ian Johnston, Dominic Kwiatkowski, Cordelia Langford, John Sillitoe on behalf of the Wellcome Sanger Institute COVID-19 Surveillance Team |
| EPI_ISL_631019, EPI_ISL_631020, EPI_ISL_631021, EPI_ISL_631022, EPI_ISL_631026, EPI_ISL_631029, EPI_ISL_631030, EPI_ISL_631036, EPI_ISL_631037, EPI_ISL_631042, EPI_ISL_631047, EPI_ISL_631048, EPI_ISL_631051, EPI_ISL_631054, EPI_ISL_631061, EPI_ISL_631067, EPI_ISL_631070, EPI_ISL_631073, EPI_ISL_631075, EPI_ISL_631082, EPI_ISL_631083, EPI_ISL_631087, EPI_ISL_631088, EPI_ISL_631091, EPI_ISL_631097, EPI_ISL_631098, EPI_ISL_631104 | | | |
| see above | Lighthouse Lab in Milton Keynes | Wellcome Sanger Institute for the COVID-19 Genomics UK | The Lighthouse Lab in Milton Keynes and Alex Alderton, Roberto Amato, Sonia Goncalves, Ewan Harrison, David K. Jackson, Ian Johnston, Dominic |

| | | | |
|---|---|---|---|
| | | (COG-UK) consortium | Kwiatkowski, Cordelia Langford, John Sillitoe on behalf of the Wellcome Sanger Institute COVID-19 Surveillance Team |
| EPI_ISL_631105 | Lighthouse Lab in Cambridge | Wellcome Sanger Institute for the COVID-19 Genomics UK (COG-UK) consortium | Rob Howes, The Lighthouse Lab in Cambridge and Alex Alderton, Roberto Amato, Sonia Goncalves, Ewan Harrison, David K. Jackson, Ian Johnston, Dominic Kwiatkowski, Cordelia Langford, John Sillitoe on behalf of the Wellcome Sanger Institute COVID-19 Surveillance Team |
| EPI_ISL_631106, EPI_ISL_631108, EPI_ISL_631110, EPI_ISL_631112, EPI_ISL_631114, EPI_ISL_631115, EPI_ISL_631119, EPI_ISL_631122, EPI_ISL_631124, EPI_ISL_631125, EPI_ISL_631130, EPI_ISL_631134, EPI_ISL_631136, EPI_ISL_631138, EPI_ISL_631141, EPI_ISL_631143, EPI_ISL_631145, EPI_ISL_631146, EPI_ISL_631147, EPI_ISL_631152, EPI_ISL_631153, EPI_ISL_631156, EPI_ISL_631157, EPI_ISL_631158, EPI_ISL_631162, EPI_ISL_631166, EPI_ISL_631168, EPI_ISL_631172, EPI_ISL_631176, EPI_ISL_631178, EPI_ISL_631179 | | | |
| see above | Lighthouse Lab in Milton Keynes | Wellcome Sanger Institute for the COVID-19 Genomics UK (COG-UK) consortium | The Lighthouse Lab in Milton Keynes and Alex Alderton, Roberto Amato, Sonia Goncalves, Ewan Harrison, David K. Jackson, Ian Johnston, Dominic Kwiatkowski, Cordelia Langford, John Sillitoe on behalf of the Wellcome Sanger Institute COVID-19 Surveillance Team |
| EPI_ISL_631180, EPI_ISL_631181 | Lighthouse Lab in Cambridge | Wellcome Sanger Institute for the COVID-19 Genomics UK (COG-UK) consortium | Rob Howes, The Lighthouse Lab in Cambridge and Alex Alderton, Roberto Amato, Sonia Goncalves, Ewan Harrison, David K. Jackson, Ian Johnston, Dominic Kwiatkowski, Cordelia Langford, John Sillitoe on behalf of the Wellcome Sanger Institute COVID-19 Surveillance Team |
| EPI_ISL_631183, EPI_ISL_631188, EPI_ISL_631193, EPI_ISL_631198, EPI_ISL_631201, EPI_ISL_631213, EPI_ISL_631216, EPI_ISL_631218, EPI_ISL_631223, EPI_ISL_631226, EPI_ISL_631229, EPI_ISL_631230, EPI_ISL_631233, EPI_ISL_631234, EPI_ISL_631236, EPI_ISL_631239, EPI_ISL_631240, EPI_ISL_631242, EPI_ISL_631246, EPI_ISL_631247, EPI_ISL_631251, EPI_ISL_631252, EPI_ISL_631253, EPI_ISL_631255, EPI_ISL_631256, EPI_ISL_631258, EPI_ISL_631259, EPI_ISL_631263, EPI_ISL_631267 | | | |
| see above | Lighthouse Lab in Milton Keynes | Wellcome Sanger Institute for the COVID-19 Genomics UK (COG-UK) consortium | The Lighthouse Lab in Milton Keynes and Alex Alderton, Roberto Amato, Sonia Goncalves, Ewan Harrison, David K. Jackson, Ian Johnston, Dominic Kwiatkowski, Cordelia Langford, John Sillitoe on behalf of the Wellcome Sanger Institute COVID-19 Surveillance Team |
| EPI_ISL_632322, EPI_ISL_632339, EPI_ISL_632342, EPI_ISL_632402, EPI_ISL_632425, EPI_ISL_632447, EPI_ISL_632482, EPI_ISL_632710, EPI_ISL_632715, EPI_ISL_632721, EPI_ISL_632722, EPI_ISL_632732, EPI_ISL_632742, EPI_ISL_632743, EPI_ISL_632744, EPI_ISL_632745, EPI_ISL_632746, EPI_ISL_632761, EPI_ISL_632788 | | | |
| see above | Dutch COVID-19 response team | Erasmus Medical Center | Bas Oude Munnink, David Nieuwenhuijse, Reina Sikkema, Claudia Schapendonk, Irina Chestakova, Anne van der Linden, Theo Bestebroer, Stefan van Nieuwkoop, Mark Pronk, Pascal Lexmond, Corien Swaan, Manon Haverkate, Madelief Mollers, Mart Stein, Sandra Kengne Kamga Mobou, Jeroen van Kampen, Jolanda Voermans, Aura Timen, Corine GeurtsvanKessel, Annemiek van der Eijk, Richard Molenkamp, Marion Koopmans, on behalf of the Dutch national COVID-19 response team. |
| EPI_ISL_633091, EPI_ISL_633092, EPI_ISL_633093, EPI_ISL_633095, EPI_ISL_633096, EPI_ISL_633097, EPI_ISL_633098, EPI_ISL_633099 | MD PHL | MD PHL | Maryland Department of Health Laboratories Administration |
| EPI_ISL_633841, EPI_ISL_633918 | Lighthouse Lab in Glasgow | Wellcome Sanger Institute for the COVID-19 Genomics UK (COG-UK) consortium | Harper VanSteenhouse, Yumi Kasai, David Gray, Carol Clugston, Anna Dominiczak and Alex Alderton, Roberto Amato, Sonia Goncalves, Ewan Harrison, David K. Jackson, Ian Johnston, Dominic Kwiatkowski, Cordelia Langford, John Sillitoe on behalf of the Wellcome Sanger Institute COVID-19 Surveillance Team |
| EPI_ISL_634256, EPI_ISL_634257, EPI_ISL_634258, EPI_ISL_634259, EPI_ISL_634260, EPI_ISL_634261, EPI_ISL_634262, EPI_ISL_634263, EPI_ISL_634264, EPI_ISL_634265 | Lighthouse Lab in Milton Keynes | Wellcome Sanger Institute for the COVID-19 Genomics UK (COG-UK) consortium | The Lighthouse Lab in Milton Keynes and Alex Alderton, Roberto Amato, Sonia Goncalves, Ewan Harrison, David K. Jackson, Ian Johnston, Dominic Kwiatkowski, Cordelia Langford, John Sillitoe on behalf of the Wellcome Sanger Institute COVID-19 Surveillance Team |
| EPI_ISL_634266 | Lighthouse Lab in Milton Keynes | Wellcome Sanger Institute for the COVID-19 Genomics UK (COG-UK) Consortium | The Lighthouse Lab in Milton Keynes and Alex Alderton, Roberto Amato, Sonia Goncalves, Ewan Harrison, David K. Jackson, Ian Johnston, Dominic Kwiatkowski, Cordelia Langford, John Sillitoe on behalf of the Wellcome Sanger Institute COVID-19 Surveillance Team |
| EPI_ISL_634267, EPI_ISL_634269, EPI_ISL_634270, EPI_ISL_634271, EPI_ISL_634272, EPI_ISL_634273, EPI_ISL_634274, EPI_ISL_634276, EPI_ISL_634278, EPI_ISL_634279, EPI_ISL_634280, EPI_ISL_634281, EPI_ISL_634283, EPI_ISL_634284, EPI_ISL_634285, EPI_ISL_634286, EPI_ISL_634287, EPI_ISL_634288, EPI_ISL_634289, EPI_ISL_634291, EPI_ISL_634292, EPI_ISL_634293, EPI_ISL_634294, EPI_ISL_634297, EPI_ISL_634298, EPI_ISL_634299, EPI_ISL_634300, EPI_ISL_634301, EPI_ISL_634302, EPI_ISL_634303, EPI_ISL_634304, EPI_ISL_634306, EPI_ISL_634307, EPI_ISL_634308, EPI_ISL_634309, EPI_ISL_634310, EPI_ISL_634311, EPI_ISL_634312, EPI_ISL_634313, EPI_ISL_634315, EPI_ISL_634316, EPI_ISL_634317, EPI_ISL_634319, EPI_ISL_634321, EPI_ISL_634322, EPI_ISL_634323, EPI_ISL_634324, EPI_ISL_634325, EPI_ISL_634326, EPI_ISL_634327, EPI_ISL_634328, EPI_ISL_634329, EPI_ISL_634330, EPI_ISL_634331, EPI_ISL_634332, EPI_ISL_634333, EPI_ISL_634334, EPI_ISL_634335, EPI_ISL_634336, EPI_ISL_634337, EPI_ISL_634338, EPI_ISL_634340 | | | |
| see above | Lighthouse Lab in Milton Keynes | Wellcome Sanger Institute for the COVID-19 Genomics UK (COG-UK) consortium | The Lighthouse Lab in Milton Keynes and Alex Alderton, Roberto Amato, Sonia Goncalves, Ewan Harrison, David K. Jackson, Ian Johnston, Dominic Kwiatkowski, Cordelia Langford, John Sillitoe on behalf of the Wellcome Sanger Institute COVID-19 Surveillance Team |
| EPI_ISL_634341 | Lighthouse Lab in Milton Keynes | Wellcome Sanger Institute for the COVID-19 Genomics UK (COG-UK) Consortium | The Lighthouse Lab in Milton Keynes and Alex Alderton, Roberto Amato, Sonia Goncalves, Ewan Harrison, David K. Jackson, Ian Johnston, Dominic Kwiatkowski, Cordelia Langford, John Sillitoe on behalf of the Wellcome Sanger Institute COVID-19 Surveillance Team |
| EPI_ISL_634342, EPI_ISL_634343, EPI_ISL_634344, EPI_ISL_634345, EPI_ISL_634346, EPI_ISL_634347, EPI_ISL_634348, EPI_ISL_634349, EPI_ISL_634350, EPI_ISL_634351, EPI_ISL_634352, EPI_ISL_634353, EPI_ISL_634354, EPI_ISL_634355, EPI_ISL_634357, EPI_ISL_634360, EPI_ISL_634361, EPI_ISL_634362, EPI_ISL_634364, EPI_ISL_634365, EPI_ISL_634366, EPI_ISL_634367, EPI_ISL_634368, EPI_ISL_634369, EPI_ISL_634371, EPI_ISL_634372, EPI_ISL_634373, EPI_ISL_634374, EPI_ISL_634375 | | | |
| see above | Lighthouse Lab in Milton Keynes | Wellcome Sanger Institute for the COVID-19 Genomics UK (COG-UK) consortium | The Lighthouse Lab in Milton Keynes and Alex Alderton, Roberto Amato, Sonia Goncalves, Ewan Harrison, David K. Jackson, Ian Johnston, Dominic Kwiatkowski, Cordelia Langford, John Sillitoe on behalf of the Wellcome Sanger Institute COVID-19 Surveillance Team |
| EPI_ISL_634376 | Lighthouse Lab in Milton Keynes | Wellcome Sanger Institute for the COVID-19 Genomics UK (COG-UK) Consortium | The Lighthouse Lab in Milton Keynes and Alex Alderton, Roberto Amato, Sonia Goncalves, Ewan Harrison, David K. Jackson, Ian Johnston, Dominic Kwiatkowski, Cordelia Langford, John Sillitoe on behalf of the Wellcome Sanger Institute COVID-19 Surveillance Team |
| EPI_ISL_634377, EPI_ISL_634378, EPI_ISL_634379, EPI_ISL_634381, EPI_ISL_634382, EPI_ISL_634383 | Lighthouse Lab in Milton Keynes | Wellcome Sanger Institute for the COVID-19 Genomics UK (COG-UK) consortium | The Lighthouse Lab in Milton Keynes and Alex Alderton, Roberto Amato, Sonia Goncalves, Ewan Harrison, David K. Jackson, Ian Johnston, Dominic Kwiatkowski, Cordelia Langford, John Sillitoe on behalf of the Wellcome Sanger Institute COVID-19 Surveillance Team |
| EPI_ISL_634384 | Lighthouse Lab in Milton Keynes | Wellcome Sanger Institute for the COVID-19 Genomics UK (COG-UK) Consortium | The Lighthouse Lab in Milton Keynes and Alex Alderton, Roberto Amato, Sonia Goncalves, Ewan Harrison, David K. Jackson, Ian Johnston, Dominic Kwiatkowski, Cordelia Langford, John Sillitoe on behalf of the Wellcome Sanger Institute COVID-19 Surveillance Team |
| EPI_ISL_634386, EPI_ISL_634387, EPI_ISL_634388, EPI_ISL_634389, EPI_ISL_634390, EPI_ISL_634391, EPI_ISL_634392, EPI_ISL_634393, EPI_ISL_634394, EPI_ISL_634395, EPI_ISL_634396, EPI_ISL_634397, EPI_ISL_634398, EPI_ISL_634399, EPI_ISL_634401, EPI_ISL_634402, EPI_ISL_634403, EPI_ISL_634404, EPI_ISL_634405, EPI_ISL_634407, EPI_ISL_634408, EPI_ISL_634410, EPI_ISL_634411, EPI_ISL_634412, EPI_ISL_634414, EPI_ISL_634416, EPI_ISL_634417, EPI_ISL_634418, EPI_ISL_634419, EPI_ISL_634420, EPI_ISL_634421, EPI_ISL_634422, EPI_ISL_634423, EPI_ISL_634424, EPI_ISL_634425, EPI_ISL_634426, EPI_ISL_634427 | | | |
| see above | Lighthouse Lab in Milton Keynes | Wellcome Sanger Institute for the COVID-19 Genomics UK (COG-UK) consortium | The Lighthouse Lab in Milton Keynes and Alex Alderton, Roberto Amato, Sonia Goncalves, Ewan Harrison, David K. Jackson, Ian Johnston, Dominic Kwiatkowski, Cordelia Langford, John Sillitoe on behalf of the Wellcome Sanger Institute COVID-19 Surveillance Team |
| EPI_ISL_634428 | Lighthouse Lab in Milton Keynes | Wellcome Sanger Institute for the COVID-19 Genomics UK (COG-UK) Consortium | The Lighthouse Lab in Milton Keynes and Alex Alderton, Roberto Amato, Sonia Goncalves, Ewan Harrison, David K. Jackson, Ian Johnston, Dominic Kwiatkowski, Cordelia Langford, John Sillitoe on behalf of the Wellcome Sanger Institute COVID-19 Surveillance Team |
| EPI_ISL_634430, EPI_ISL_634431, EPI_ISL_634432, EPI_ISL_634434, EPI_ISL_634435, EPI_ISL_634436, EPI_ISL_634437, EPI_ISL_634438, EPI_ISL_634439, EPI_ISL_634440, EPI_ISL_634441, EPI_ISL_634442, EPI_ISL_634443, EPI_ISL_634444, EPI_ISL_634445, EPI_ISL_634446, EPI_ISL_634447, EPI_ISL_634448, EPI_ISL_634449, EPI_ISL_634450, EPI_ISL_634451, EPI_ISL_634452, EPI_ISL_634453, EPI_ISL_634454, EPI_ISL_634455, EPI_ISL_634457, EPI_ISL_634458, EPI_ISL_634459, EPI_ISL_634460, EPI_ISL_634461, EPI_ISL_634462, EPI_ISL_634463, EPI_ISL_634464 | | | |
| see above | Lighthouse Lab in Milton Keynes | Wellcome Sanger Institute for the COVID-19 Genomics UK (COG-UK) consortium | The Lighthouse Lab in Milton Keynes and Alex Alderton, Roberto Amato, Sonia Goncalves, Ewan Harrison, David K. Jackson, Ian Johnston, Dominic Kwiatkowski, Cordelia Langford, John Sillitoe on behalf of the Wellcome Sanger Institute COVID-19 Surveillance Team |
| EPI_ISL_634465 | Lighthouse Lab in Milton Keynes | Wellcome Sanger Institute for the COVID-19 Genomics UK (COG-UK) Consortium | The Lighthouse Lab in Milton Keynes and Alex Alderton, Roberto Amato, Sonia Goncalves, Ewan Harrison, David K. Jackson, Ian Johnston, Dominic Kwiatkowski, Cordelia Langford, John Sillitoe on behalf of the Wellcome Sanger Institute COVID-19 Surveillance Team |
| EPI_ISL_634466, EPI_ISL_634467, EPI_ISL_634468, EPI_ISL_634469, EPI_ISL_634470, EPI_ISL_634471, EPI_ISL_634474, EPI_ISL_634475, EPI_ISL_634476, EPI_ISL_634477, EPI_ISL_634478, EPI_ISL_634479, EPI_ISL_634480, EPI_ISL_634481 | | | |
| see above | Lighthouse Lab in Milton Keynes | Wellcome Sanger Institute for the COVID-19 Genomics UK (COG-UK) consortium | The Lighthouse Lab in Milton Keynes and Alex Alderton, Roberto Amato, Sonia Goncalves, Ewan Harrison, David K. Jackson, Ian Johnston, Dominic Kwiatkowski, Cordelia Langford, John Sillitoe on behalf of the Wellcome Sanger Institute COVID-19 Surveillance Team |
| EPI_ISL_634483 | Lighthouse Lab in Milton Keynes | Wellcome Sanger Institute for the COVID-19 Genomics UK (COG-UK) Consortium | The Lighthouse Lab in Milton Keynes and Alex Alderton, Roberto Amato, Sonia Goncalves, Ewan Harrison, David K. Jackson, Ian Johnston, Dominic Kwiatkowski, Cordelia Langford, John Sillitoe on behalf of the Wellcome Sanger Institute COVID-19 Surveillance Team |
| EPI_ISL_634484, EPI_ISL_634485, EPI_ISL_634487, EPI_ISL_634488, EPI_ISL_634489, EPI_ISL_634490, EPI_ISL_634491, EPI_ISL_634492, EPI_ISL_634493, EPI_ISL_634495, EPI_ISL_634496, EPI_ISL_634500, EPI_ISL_634501, EPI_ISL_634502, EPI_ISL_634503, EPI_ISL_634505, EPI_ISL_634506, EPI_ISL_634507, EPI_ISL_634511, EPI_ISL_634513, EPI_ISL_634514, EPI_ISL_634515, EPI_ISL_634516, EPI_ISL_634517, EPI_ISL_634518, EPI_ISL_634519 | | | |

| | | | |
|---|---|---|---|
| see above | Lighthouse Lab in Milton Keynes | Wellcome Sanger Institute for the COVID-19 Genomics UK (COG-UK) consortium | The Lighthouse Lab in Milton Keynes and Alex Alderton, Roberto Amato, Sonia Goncalves, Ewan Harrison, David K. Jackson, Ian Johnston, Dominic Kwiatkowski, Cordelia Langford, John Sillitoe on behalf of the Wellcome Sanger Institute COVID-19 Surveillance Team |
| EPI_ISL_634774, EPI_ISL_634786, EPI_ISL_634787 | Lighthouse Lab in Alderley Park | Wellcome Sanger Institute for the COVID-19 Genomics UK (COG-UK) consortium | Jacquelyn Wynn, Mairead Hyland, The Lighthouse Lab in Alderley Park and Alex Alderton, Roberto Amato, Sonia Goncalves, Ewan Harrison, David K. Jackson, Ian Johnston, Dominic Kwiatkowski, Cordelia Langford, John Sillitoe on behalf of the Wellcome Sanger Institute COVID-19 Surveillance Team (http://www.sanger.ac.uk/covid-team) |
| EPI_ISL_634790, EPI_ISL_634791, EPI_ISL_634792, EPI_ISL_634793, EPI_ISL_634794, EPI_ISL_634795, EPI_ISL_634796, EPI_ISL_634797, EPI_ISL_634799, EPI_ISL_634800, EPI_ISL_634802, EPI_ISL_634803, EPI_ISL_634804, EPI_ISL_634805, EPI_ISL_634806, EPI_ISL_634814 | | | |
| see above | Lighthouse Lab in Cambridge | Wellcome Sanger Institute for the COVID-19 Genomics UK (COG-UK) consortium | Rob Howes, The Lighthouse Lab in Cambridge and Alex Alderton, Roberto Amato, Sonia Goncalves, Ewan Harrison, David K. Jackson, Ian Johnston, Dominic Kwiatkowski, Cordelia Langford, John Sillitoe on behalf of the Wellcome Sanger Institute COVID-19 Surveillance Team (http://www.sanger.ac.uk/covid-team) |
| EPI_ISL_634857 | Minneapolis VA Health Care System | Minnesota Department of Health, Public Health Laboratory | Matt Plumb, Jacob Garfin, Alexandra Lorentz, and Xiong Wang |
| EPI_ISL_634881, EPI_ISL_634883 | Lab voor klinische biologie | Onderzoeksgroep Virologie | Laurens Lambrechts, Nick Vereecke, Marthe Pauwels, Bruno Verhasselt, Linos Vandekerckhove, Hans Nauwynck, Sebastiaan Theuns |
| EPI_ISL_634895 | Lab voor klinische biologie | Onderzoeksgroep Virologie | Nick Vereecke, Laurens Lambrechts, Marthe Pauwels, Bruno Verhasselt, Linos Vandekerckhove, Hans Nauwynck, Sebastiaan Theuns |
| EPI_ISL_635026, EPI_ISL_635027, EPI_ISL_635028, EPI_ISL_635029, EPI_ISL_635030, EPI_ISL_635031, EPI_ISL_635033, EPI_ISL_635034, EPI_ISL_635035, EPI_ISL_635039, EPI_ISL_635041, EPI_ISL_635042, EPI_ISL_635043, EPI_ISL_635044, EPI_ISL_635045, EPI_ISL_635046, EPI_ISL_635047, EPI_ISL_635048, EPI_ISL_635049, EPI_ISL_635050, EPI_ISL_635051, EPI_ISL_635054, EPI_ISL_635055, EPI_ISL_635056, EPI_ISL_635057, EPI_ISL_635058 | | | |
| see above | National Health Laboratory Service - Inkosi Albert Luthuli Central Hospital (NHLS-IALCH) | KRISP, KZN Research Innovation and Sequencing Platform | Giandhari J, Pillay S, Lessells R, Mdlalose K, York D, Khan S, Tegally H, Wilkinson E, de Oliveira T |
| EPI_ISL_635073 | University Hospital of Northern Norway, Department for Microbiology and Infectious Disease Control | Norwegian Institute of Public Health, Department of Virology | Kathrine Stene-Johansen, Kamilla Heddeland Instefjord, Hilde Elshaug, Marie Paulsen Madsen, Rasmus Riis Kopperud, Hilde Vollan, Karoline Bragstad, Olav Hungnes |
| EPI_ISL_635090 | Haukeland University Hospital, Department of Medical Microbiology | Norwegian Institute of Public Health, Department of Virology | Kathrine Stene-Johansen, Kamilla Heddeland Instefjord, Hilde Elshaug, Marie Paulsen Madsen, Rasmus Riis Kopperud, Hilde Vollan, Karoline Bragstad, Olav Hungnes |
| EPI_ISL_635102 | Akershus University Hospital, Department for Microbiology and Infectious Disease Control | Norwegian Institute of Public Health, Department of Virology | Kathrine Stene-Johansen, Kamilla Heddeland Instefjord, Hilde Elshaug, Marie Paulsen Madsen, Rasmus Riis Kopperud, Hilde Vollan, Karoline Bragstad, Olav Hungnes |
| EPI_ISL_635181 | University Hospital of Northern Norway, Department for Microbiology and Infectious Disease Control | Norwegian Institute of Public Health, Department of Virology | Kathrine Stene-Johansen, Kamilla Heddeland Instefjord, Hilde Elshaug, Marie Paulsen Madsen, Rasmus Riis Kopperud, Hilde Vollan, Karoline Bragstad, Olav Hungnes |
| EPI_ISL_635194 | Akershus University Hospital, Department for Microbiology and Infectious Disease Control | Norwegian Institute of Public Health, Department of Virology | Kathrine Stene-Johansen, Kamilla Heddeland Instefjord, Hilde Elshaug, Marie Paulsen Madsen, Rasmus Riis Kopperud, Hilde Vollan, Karoline Bragstad, Olav Hungnes |
| EPI_ISL_635198 | University Hospital of Northern Norway, Department for Microbiology and Infectious Disease Control | Norwegian Institute of Public Health, Department of Virology | Kathrine Stene-Johansen, Kamilla Heddeland Instefjord, Hilde Elshaug, Marie Paulsen Madsen, Rasmus Riis Kopperud, Hilde Vollan, Karoline Bragstad, Olav Hungnes |
| EPI_ISL_636537, EPI_ISL_636551, EPI_ISL_636559, EPI_ISL_636560, EPI_ISL_636591 | Dutch COVID-19 response team | National Institute for Public Health and the Environment (RIVM) | Adam Meijer, Harry Vennema, Jeroen Cremer, Sharon van den Brink, Bas van der Veer, AnneMarie van den Brandt, Florian Zwagemaker, Dennis Schmitz, Chantal Reusken, on behalf of the national COVID-19 response team |
| EPI_ISL_636716, EPI_ISL_636717, EPI_ISL_636719, EPI_ISL_636720, EPI_ISL_636721, EPI_ISL_636722, EPI_ISL_636723 | Respiratory Virus Unit, Microbiology Services Colindale, Public Health England | Respiratory Virus Unit, Microbiology Services Colindale, Public Health England | PHE Covid Sequencing Team |
| EPI_ISL_637118, EPI_ISL_637119 | Respiratory Virus Unit, Microbiology Services Colindale, Public Health England | COVID-19 Genomics UK (COG-UK) Consortium | PHE Covid Sequencing Team |
| EPI_ISL_637256 | Wales Specialist Virology Centre Sequencing lab: Pathogen Genomics Unit | COVID-19 Genomics UK (COG-UK) Consortium | Catherine Moore, Johnathan Evans, Laura Gifford, Malorie Perry, Simon Cottrell, Angela Marchbank, Alec Birchley, Alexander Adams, Amy Gaskin, Bree Gatica-Wilcox, Jason Coombes, Joel Southgate, Lauren Gilbert, Lee Graham, Nicole Pacchiarini, Sara Kumziene-Summerhayes, Sarah Taylor, Sophie Jones, Sara Rey, Matthew Bull, Joanne Watkins, Sally Corden, Tom Connor |
| EPI_ISL_637262, EPI_ISL_637265 | Liverpool Clinical Laboratories | COVID-19 Genomics UK (COG-UK) Consortium | Sam Haldenby, Anita Lucaci, Steve Paterson, Julian Hiscox, Alistair Darby, M Almsaud, A Alrezaihi, Muhannad Alruwaili, Stuart D Armstrong, Jones Benjamin, Eleanor G Bentley, Anu Chawla, Jordan J Clark, Angela Cowell, Richard Eccles, Isabel García-Dorival, Matthew Gemmell, Alessandro Gerada, PKF Gilmore, Richard Gregory, Ximeng Han, Catherine Hartley, Margaret Hughes, Miren Iturriza-Gomara, James Johnson, L Luu, Jenifer Manson, Charlotte Nelson, Elaine O'Toole, Cassie Olateju, Rebekah Penrice-Randal , Lucille Rainbow, N.P Randle, Trevor Ian Robinson, Parul Sharma, Ghada T Shawli, James P Stewart, Neil Swainston, Ecaterina Vamos, Joanne Watts, Mark Whitehead |
| EPI_ISL_637283 | Northumbria University / South Tees Hospitals NHS Foundation Trust / North Cumbria Integrated Care NHS Foundation Trust / North Tees and Hartlepool NHS Foundation Trust / Newcastle Hospitals NHS Foundation Trust | COVID-19 Genomics UK (COG-UK) Consortium | Darren L Smith,Andrew Nelson,Matthew Bashton,Greg R Young,Joshua Loh,John Allan,Mohammad A Tariq,Giles S Holt,Gary Black,Wen C Yew,Lynn Dover,Paul Baker,Steve Liggett,Sarah Essex,Jane Greenaway,Debra Padgett,Clive Graham,Garren Scott,Edward Barton,Emma Swindells,Brendan Payne,Jennifer Collins,Yusri Taha,Gary Eltringham |
| EPI_ISL_637285 | Virology Department, Sheffield Teaching Hospitals NHS Foundation Trust/Department of Infection, Immunity and Cardiovascular Disease, The Medical School, University of Sheffield | COVID-19 Genomics UK (COG-UK) Consortium | Thushan de Silva, Matthew Parker, Nikki Smith, Adri Angyal, Rebecca Brown, Luke Green, Rachel Tucker, Paul Parsons, Danielle Groves, Katie Johnson, Laura Carrilero, Alex Keeley, Dave Partridge, Matthew Wyles, Benjamin Lindsey, Mehmet Yavuz, Mohammad Raza, Cariad Evans |
| EPI_ISL_637294 | Department of Pathology, University of Cambridge | COVID-19 Genomics UK (COG-UK) Consortium | Aminu S. Jahun, Yasmin Chaudhry, Grant Hall, Iliana Georgana, Myra Hosmillo, Martin D. Curran, Malte Pinckert, Surendra Parmar, Ian Goodfellow |
| EPI_ISL_637295, EPI_ISL_637329, EPI_ISL_637350 | Liverpool Clinical Laboratories | COVID-19 Genomics UK (COG-UK) Consortium | Sam Haldenby, Anita Lucaci, Steve Paterson, Julian Hiscox, Alistair Darby, M Almsaud, A Alrezaihi, Muhannad Alruwaili, Stuart D Armstrong, Jones Benjamin, Eleanor G Bentley, Anu Chawla, Jordan J Clark, Angela Cowell, Richard Eccles, Isabel García-Dorival, Matthew Gemmell, Alessandro Gerada, PKF Gilmore, Richard Gregory, Ximeng Han, Catherine Hartley, Margaret Hughes, Miren Iturriza-Gomara, James Johnson, L Luu, Jenifer Manson, Charlotte Nelson, Elaine O'Toole, Cassie Olateju, Rebekah Penrice-Randal , Lucille Rainbow, N.P Randle, Trevor Ian Robinson, Parul Sharma, Ghada T Shawli, James P Stewart, Neil Swainston, Ecaterina Vamos, Joanne Watts, Mark Whitehead |
| EPI_ISL_637355 | Department of Pathology, University of Cambridge | COVID-19 Genomics UK (COG-UK) Consortium | Aminu S. Jahun, Yasmin Chaudhry, Grant Hall, Iliana Georgana, Myra Hosmillo, Martin D. Curran, Malte Pinckert, Surendra Parmar, Ian Goodfellow |
| EPI_ISL_637377 | Northumbria University / South Tees Hospitals NHS Foundation Trust / North Cumbria Integrated Care NHS Foundation Trust / North Tees and Hartlepool NHS Foundation Trust / Newcastle Hospitals NHS Foundation Trust | COVID-19 Genomics UK (COG-UK) Consortium | Darren L Smith,Andrew Nelson,Matthew Bashton,Greg R Young,Joshua Loh,John Allan,Mohammad A Tariq,Giles S Holt,Gary Black,Wen C Yew,Lynn Dover,Paul Baker,Steve Liggett,Sarah Essex,Jane Greenaway,Debra Padgett,Clive Graham,Garren Scott,Edward Barton,Emma Swindells,Brendan Payne,Jennifer Collins,Yusri Taha,Gary Eltringham |
| EPI_ISL_637394, EPI_ISL_637410, EPI_ISL_637418, EPI_ISL_637420, EPI_ISL_637433 | Department of Pathology, University of Cambridge | COVID-19 Genomics UK (COG-UK) Consortium | Aminu S. Jahun, Yasmin Chaudhry, Grant Hall, Iliana Georgana, Myra Hosmillo, Martin D. Curran, Malte Pinckert, Surendra Parmar, Ian Goodfellow |
| EPI_ISL_637444 | Liverpool Clinical Laboratories | COVID-19 Genomics UK (COG-UK) Consortium | Sam Haldenby, Anita Lucaci, Steve Paterson, Julian Hiscox, Alistair Darby, M Almsaud, A Alrezaihi, Muhannad Alruwaili, Stuart D Armstrong, Jones Benjamin, Eleanor G Bentley, Anu Chawla, Jordan J Clark, Angela Cowell, Richard Eccles, Isabel García-Dorival, Matthew Gemmell, Alessandro Gerada, PKF Gilmore, Richard Gregory, Ximeng Han, Catherine Hartley, Margaret Hughes, Miren Iturriza-Gomara, James Johnson, L Luu, Jenifer Manson, Charlotte Nelson, Elaine O'Toole, Cassie Olateju, Rebekah Penrice-Randal , Lucille Rainbow, N.P Randle, Trevor Ian Robinson, Parul Sharma, Ghada T Shawli, James P Stewart, Neil Swainston, Ecaterina Vamos, Joanne Watts, Mark Whitehead |

| | | | |
|---|---|---|---|
| EPI_ISL_637455 | Northumbria University / South Tees Hospitals NHS Foundation Trust / North Cumbria Integrated Care NHS Foundation Trust / North Tees and Hartlepool NHS Foundation Trust / Newcastle Hospitals NHS Foundation Trust | COVID-19 Genomics UK (COG-UK) Consortium | Darren L Smith,Andrew Nelson,Matthew Bashton,Greg R Young,Joshua Loh,John Allan,Mohammad A Tariq,Giles S Holt,Gary Black,Wen C Yew,Lynn Dover,Paul Baker,Steve Liggett,Sarah Essex,Jane Greenaway,Debra Padgett,Clive Graham,Garren Scott,Edward Barton,Emma Swindells,Brendan Payne,Jennifer Collins,Yusri Taha,Gary Eltringham |
| EPI_ISL_637468, EPI_ISL_637469, EPI_ISL_637509, EPI_ISL_637510, EPI_ISL_637511, EPI_ISL_637512, EPI_ISL_637529, EPI_ISL_637530 | Department of Pathology, University of Cambridge | COVID-19 Genomics UK (COG-UK) Consortium | Aminu S. Jahun, Yasmin Chaudhry, Grant Hall, Iliana Georgana, Myra Hosmillo, Martin D. Curran, Malte Pinckert, Surendra Parmar, Ian Goodfellow |
| EPI_ISL_637541 | Virology Department, Sheffield Teaching Hospitals NHS Foundation Trust/Department of Infection, Immunity and Cardiovascular Disease, The Medical School, University of Sheffield | COVID-19 Genomics UK (COG-UK) Consortium | Thushan de Silva, Matthew Parker, Nikki Smith, Adri Angyal, Rebecca Brown, Luke Green, Rachel Tucker, Paul Parsons, Danielle Groves, Katie Johnson, Laura Carrilero, Alex Keeley, Dave Partridge, Matthew Wyles, Benjamin Lindsey, Mehmet Yavuz, Mohammad Raza, Cariad Evans |
| EPI_ISL_637567 | Department of Pathology, University of Cambridge | COVID-19 Genomics UK (COG-UK) Consortium | Aminu S. Jahun, Yasmin Chaudhry, Grant Hall, Iliana Georgana, Myra Hosmillo, Martin D. Curran, Malte Pinckert, Surendra Parmar, Ian Goodfellow |
| EPI_ISL_637599 | Wales Specialist Virology Centre Sequencing lab: Pathogen Genomics Unit | COVID-19 Genomics UK (COG-UK) Consortium | Catherine Moore, Johnathan Evans, Laura Gifford, Malorie Perry, Simon Cottrell, Angela Marchbank, Alec Birchley, Alexander Adams, Amy Gaskin, Bree Gatica-Wilcox, Jason Coombes, Joel Southgate, Lauren Gilbert, Lee Graham, Nicole Pacchiarini, Sara Kumziene-Summerhayes, Sarah Taylor, Sophie Jones, Sara Rey, Matthew Bull, Joanne Watkins, Sally Corden, Tom Connor |
| EPI_ISL_637624, EPI_ISL_637640, EPI_ISL_637642, EPI_ISL_637695, EPI_ISL_637697, EPI_ISL_637700, EPI_ISL_637701, EPI_ISL_637702, EPI_ISL_637703, EPI_ISL_637705, EPI_ISL_637723, EPI_ISL_637724, EPI_ISL_637725, EPI_ISL_637737, EPI_ISL_637739, EPI_ISL_637740, EPI_ISL_637741, EPI_ISL_637742, EPI_ISL_637743, EPI_ISL_637744, EPI_ISL_637745, EPI_ISL_637746, EPI_ISL_637747 | | | |
| see above | Department of Pathology, University of Cambridge | COVID-19 Genomics UK (COG-UK) Consortium | Aminu S. Jahun, Yasmin Chaudhry, Grant Hall, Iliana Georgana, Myra Hosmillo, Martin D. Curran, Malte Pinckert, Surendra Parmar, Ian Goodfellow |
| EPI_ISL_637874 | Liverpool Clinical Laboratories | COVID-19 Genomics UK (COG-UK) Consortium | Sam Haldenby, Anita Lucaci, Steve Paterson, Julian Hiscox, Alistair Darby, M Almsaud, A Alrezaihi, Muhannad Alruwaili, Stuart D Armstrong, Jones Benjamin, Eleanor G Bentley, Anu Chawla, Jordan J Clark, Angela Cowell, Richard Eccles, Isabel García-Dorival, Matthew Gemmell, Alessandro Gerada, PKF Gilmore, Richard Gregory, Ximeng Han, Catherine Hartley, Margaret Hughes, Miren Iturriza-Gomara, James Johnson, L Luu, Jenifer Manson, Charlotte Nelson, Elaine O'Toole, Cassie Olateju, Rebekah Penrice-Randal , Lucille Rainbow, N.P Randle, Trevor Ian Robinson, Parul Sharma, Ghada T Shawli, James P Stewart, Neil Swainston, Ecaterina Vamos, Joanne Watts, Mark Whitehead |
| EPI_ISL_637876, EPI_ISL_637879, EPI_ISL_637881 | Virology Department, Sheffield Teaching Hospitals NHS Foundation Trust/Department of Infection, Immunity and Cardiovascular Disease, The Medical School, University of Sheffield | COVID-19 Genomics UK (COG-UK) Consortium | Thushan de Silva, Matthew Parker, Nikki Smith, Adri Angyal, Rebecca Brown, Luke Green, Rachel Tucker, Paul Parsons, Danielle Groves, Katie Johnson, Laura Carrilero, Alex Keeley, Dave Partridge, Matthew Wyles, Benjamin Lindsey, Mehmet Yavuz, Mohammad Raza, Cariad Evans |
| EPI_ISL_637893 | Liverpool Clinical Laboratories | COVID-19 Genomics UK (COG-UK) Consortium | Sam Haldenby, Anita Lucaci, Steve Paterson, Julian Hiscox, Alistair Darby, M Almsaud, A Alrezaihi, Muhannad Alruwaili, Stuart D Armstrong, Jones Benjamin, Eleanor G Bentley, Anu Chawla, Jordan J Clark, Angela Cowell, Richard Eccles, Isabel García-Dorival, Matthew Gemmell, Alessandro Gerada, PKF Gilmore, Richard Gregory, Ximeng Han, Catherine Hartley, Margaret Hughes, Miren Iturriza-Gomara, James Johnson, L Luu, Jenifer Manson, Charlotte Nelson, Elaine O'Toole, Cassie Olateju, Rebekah Penrice-Randal , Lucille Rainbow, N.P Randle, Trevor Ian Robinson, Parul Sharma, Ghada T Shawli, James P Stewart, Neil Swainston, Ecaterina Vamos, Joanne Watts, Mark Whitehead |
| EPI_ISL_637894 | Wales Specialist Virology Centre Sequencing lab: Pathogen Genomics Unit | COVID-19 Genomics UK (COG-UK) Consortium | Catherine Moore, Johnathan Evans, Laura Gifford, Malorie Perry, Simon Cottrell, Angela Marchbank, Alec Birchley, Alexander Adams, Amy Gaskin, Bree Gatica-Wilcox, Jason Coombes, Joel Southgate, Lauren Gilbert, Lee Graham, Nicole Pacchiarini, Sara Kumziene-Summerhayes, Sarah Taylor, Sophie Jones, Sara Rey, Matthew Bull, Joanne Watkins, Sally Corden, Tom Connor |
| EPI_ISL_637897, EPI_ISL_637905 | Virology Department, Sheffield Teaching Hospitals NHS Foundation Trust/Department of Infection, Immunity and Cardiovascular Disease, The Medical School, University of Sheffield | COVID-19 Genomics UK (COG-UK) Consortium | Thushan de Silva, Matthew Parker, Nikki Smith, Adri Angyal, Rebecca Brown, Luke Green, Rachel Tucker, Paul Parsons, Danielle Groves, Katie Johnson, Laura Carrilero, Alex Keeley, Dave Partridge, Matthew Wyles, Benjamin Lindsey, Mehmet Yavuz, Mohammad Raza, Cariad Evans |
| EPI_ISL_637914 | Liverpool Clinical Laboratories | COVID-19 Genomics UK (COG-UK) Consortium | Sam Haldenby, Anita Lucaci, Steve Paterson, Julian Hiscox, Alistair Darby, M Almsaud, A Alrezaihi, Muhannad Alruwaili, Stuart D Armstrong, Jones Benjamin, Eleanor G Bentley, Anu Chawla, Jordan J Clark, Angela Cowell, Richard Eccles, Isabel García-Dorival, Matthew Gemmell, Alessandro Gerada, PKF Gilmore, Richard Gregory, Ximeng Han, Catherine Hartley, Margaret Hughes, Miren Iturriza-Gomara, James Johnson, L Luu, Jenifer Manson, Charlotte Nelson, Elaine O'Toole, Cassie Olateju, Rebekah Penrice-Randal , Lucille Rainbow, N.P Randle, Trevor Ian Robinson, Parul Sharma, Ghada T Shawli, James P Stewart, Neil Swainston, Ecaterina Vamos, Joanne Watts, Mark Whitehead |
| EPI_ISL_637919 | Wales Specialist Virology Centre Sequencing lab: Pathogen Genomics Unit | COVID-19 Genomics UK (COG-UK) Consortium | Catherine Moore, Johnathan Evans, Laura Gifford, Malorie Perry, Simon Cottrell, Angela Marchbank, Alec Birchley, Alexander Adams, Amy Gaskin, Bree Gatica-Wilcox, Jason Coombes, Joel Southgate, Lauren Gilbert, Lee Graham, Nicole Pacchiarini, Sara Kumziene-Summerhayes, Sarah Taylor, Sophie Jones, Sara Rey, Matthew Bull, Joanne Watkins, Sally Corden, Tom Connor |
| EPI_ISL_637933 | Liverpool Clinical Laboratories | COVID-19 Genomics UK (COG-UK) Consortium | Sam Haldenby, Anita Lucaci, Steve Paterson, Julian Hiscox, Alistair Darby, M Almsaud, A Alrezaihi, Muhannad Alruwaili, Stuart D Armstrong, Jones Benjamin, Eleanor G Bentley, Anu Chawla, Jordan J Clark, Angela Cowell, Richard Eccles, Isabel García-Dorival, Matthew Gemmell, Alessandro Gerada, PKF Gilmore, Richard Gregory, Ximeng Han, Catherine Hartley, Margaret Hughes, Miren Iturriza-Gomara, James Johnson, L Luu, Jenifer Manson, Charlotte Nelson, Elaine O'Toole, Cassie Olateju, Rebekah Penrice-Randal , Lucille Rainbow, N.P Randle, Trevor Ian Robinson, Parul Sharma, Ghada T Shawli, James P Stewart, Neil Swainston, Ecaterina Vamos, Joanne Watts, Mark Whitehead |
| EPI_ISL_637943 | Northumbria University / South Tees Hospitals NHS Foundation Trust / North Cumbria Integrated Care NHS Foundation Trust / North Tees and Hartlepool NHS Foundation Trust / Newcastle Hospitals NHS Foundation Trust | COVID-19 Genomics UK (COG-UK) Consortium | Darren L Smith,Andrew Nelson,Matthew Bashton,Greg R Young,Joshua Loh,John Allan,Mohammad A Tariq,Giles S Holt,Gary Black,Wen C Yew,Lynn Dover,Paul Baker,Steve Liggett,Sarah Essex,Jane Greenaway,Debra Padgett,Clive Graham,Garren Scott,Edward Barton,Emma Swindells,Brendan Payne,Jennifer Collins,Yusri Taha,Gary Eltringham |
| EPI_ISL_637944 | Virology Department, Sheffield Teaching Hospitals NHS Foundation Trust/Department of Infection, Immunity and Cardiovascular Disease, The Medical School, University of Sheffield | COVID-19 Genomics UK (COG-UK) Consortium | Thushan de Silva, Matthew Parker, Nikki Smith, Adri Angyal, Rebecca Brown, Luke Green, Rachel Tucker, Paul Parsons, Danielle Groves, Katie Johnson, Laura Carrilero, Alex Keeley, Dave Partridge, Matthew Wyles, Benjamin Lindsey, Mehmet Yavuz, Mohammad Raza, Cariad Evans |
| EPI_ISL_637948 | Liverpool Clinical Laboratories | COVID-19 Genomics UK (COG-UK) Consortium | Sam Haldenby, Anita Lucaci, Steve Paterson, Julian Hiscox, Alistair Darby, M Almsaud, A Alrezaihi, Muhannad Alruwaili, Stuart D Armstrong, Jones Benjamin, Eleanor G Bentley, Anu Chawla, Jordan J Clark, Angela Cowell, Richard Eccles, Isabel García-Dorival, Matthew Gemmell, Alessandro Gerada, PKF Gilmore, Richard Gregory, Ximeng Han, Catherine Hartley, Margaret Hughes, Miren Iturriza-Gomara, James Johnson, L Luu, Jenifer Manson, Charlotte Nelson, Elaine O'Toole, Cassie Olateju, Rebekah Penrice-Randal , Lucille Rainbow, N.P Randle, Trevor Ian Robinson, Parul Sharma, Ghada T Shawli, James P Stewart, Neil Swainston, Ecaterina Vamos, Joanne Watts, Mark Whitehead |
| EPI_ISL_637957, EPI_ISL_637962, EPI_ISL_637964 | Wales Specialist Virology Centre Sequencing lab: Pathogen Genomics Unit | COVID-19 Genomics UK (COG-UK) Consortium | Catherine Moore, Johnathan Evans, Laura Gifford, Malorie Perry, Simon Cottrell, Angela Marchbank, Alec Birchley, Alexander Adams, Amy Gaskin, Bree Gatica-Wilcox, Jason Coombes, Joel Southgate, Lauren Gilbert, Lee Graham, Nicole Pacchiarini, Sara Kumziene-Summerhayes, Sarah Taylor, Sophie Jones, Sara Rey, Matthew Bull, Joanne Watkins, Sally Corden, Tom Connor |
| EPI_ISL_637968 | Liverpool Clinical Laboratories | COVID-19 Genomics UK (COG-UK) Consortium | Sam Haldenby, Anita Lucaci, Steve Paterson, Julian Hiscox, Alistair Darby, M Almsaud, A Alrezaihi, Muhannad Alruwaili, Stuart D Armstrong, Jones Benjamin, Eleanor G Bentley, Anu Chawla, Jordan J Clark, Angela Cowell, Richard Eccles, Isabel García-Dorival, Matthew Gemmell, Alessandro Gerada, PKF Gilmore, Richard Gregory, Ximeng Han, Catherine Hartley, Margaret Hughes, Miren Iturriza-Gomara, James Johnson, L Luu, Jenifer Manson, Charlotte Nelson, Elaine O'Toole, Cassie Olateju, Rebekah Penrice-Randal , Lucille Rainbow, N.P Randle, Trevor Ian Robinson, Parul Sharma, Ghada T Shawli, James P Stewart, Neil Swainston, Ecaterina Vamos, Joanne Watts, Mark Whitehead |
| EPI_ISL_637973 | Virology Department, Sheffield Teaching Hospitals NHS | COVID-19 Genomics UK (COG-UK) Consortium | Thushan de Silva, Matthew Parker, Nikki Smith, Adri Angyal, Rebecca Brown, Luke Green, Rachel Tucker, Paul Parsons, Danielle Groves, Katie Johnson, |

| | | | |
|---|---|---|---|
| | Foundation Trust/Department of Infection, Immunity and Cardiovascular Disease, The Medical School, University of Sheffield | | Laura Carrilero, Alex Keeley, Dave Partridge, Matthew Wyles, Benjamin Lindsey, Mehmet Yavuz, Mohammad Raza, Cariad Evans |
| EPI_ISL_637981, EPI_ISL_637984, EPI_ISL_637989 | Liverpool Clinical Laboratories | COVID-19 Genomics UK (COG-UK) Consortium | Sam Haldenby, Anita Lucaci, Steve Paterson, Julian Hiscox, Alistair Darby, M Almsaud, A Alrezaihi, Muhannad Alruwaili, Stuart D Armstrong, Jones Benjamin, Eleanor G Bentley, Anu Chawla, Jordan J Clark, Angela Cowell, Richard Eccles, Isabel García-Dorival, Matthew Gemmell, Alessandro Gerada, PKF Gilmore, Richard Gregory, Ximeng Han, Catherine Hartley, Margaret Hughes, Miren Iturriza-Gomara, James Johnson, L Luu, Jenifer Manson, Charlotte Nelson, Elaine O'Toole, Cassie Olateju, Rebekah Penrice-Randal , Lucille Rainbow, N.P Randle, Trevor Ian Robinson, Parul Sharma, Ghada T Shawli, James P Stewart, Neil Swainston, Ecaterina Vamos, Joanne Watts, Mark Whitehead |
| EPI_ISL_637992 | Virology Department, Sheffield Teaching Hospitals NHS Foundation Trust/Department of Infection, Immunity and Cardiovascular Disease, The Medical School, University of Sheffield | COVID-19 Genomics UK (COG-UK) Consortium | Thushan de Silva, Matthew Parker, Nikki Smith, Adri Angyal, Rebecca Brown, Luke Green, Rachel Tucker, Paul Parsons, Danielle Groves, Katie Johnson, Laura Carrilero, Alex Keeley, Dave Partridge, Matthew Wyles, Benjamin Lindsey, Mehmet Yavuz, Mohammad Raza, Cariad Evans |
| EPI_ISL_638001 | Department of Pathology, University of Cambridge | COVID-19 Genomics UK (COG-UK) Consortium | Aminu S. Jahun, Yasmin Chaudhry, Grant Hall, Iliana Georgana, Myra Hosmillo, Martin D. Curran, Malte Pinckert, Surendra Parmar, Ian Goodfellow |
| EPI_ISL_638009 | Northumbria University / South Tees Hospitals NHS Foundation Trust / North Cumbria Integrated Care NHS Foundation Trust / North Tees and Hartlepool NHS Foundation Trust / Newcastle Hospitals NHS Foundation Trust | COVID-19 Genomics UK (COG-UK) Consortium | Darren L Smith,Andrew Nelson,Matthew Bashton,Greg R Young,Joshua Loh,John Allan,Mohammad A Tariq,Giles S Holt,Gary Black,Wen C Yew,Lynn Dover,Paul Baker,Steve Liggett,Sarah Essex,Jane Greenaway,Debra Padgett,Clive Graham,Garren Scott,Edward Barton,Emma Swindells,Brendan Payne,Jennifer Collins,Yusri Taha,Gary Eltringham |
| EPI_ISL_638027 | Department of Pathology, University of Cambridge | COVID-19 Genomics UK (COG-UK) Consortium | Aminu S. Jahun, Yasmin Chaudhry, Grant Hall, Iliana Georgana, Myra Hosmillo, Martin D. Curran, Malte Pinckert, Surendra Parmar, Ian Goodfellow |
| EPI_ISL_638118, EPI_ISL_638122, EPI_ISL_638126, EPI_ISL_638130, EPI_ISL_638147, EPI_ISL_638151, EPI_ISL_638152, EPI_ISL_638154, EPI_ISL_638156 | Wales Specialist Virology Centre Sequencing lab: Pathogen Genomics Unit | COVID-19 Genomics UK (COG-UK) Consortium | Catherine Moore, Johnathan Evans, Laura Gifford, Malorie Perry, Simon Cottrell, Angela Marchbank, Alec Birchley, Alexander Adams, Amy Gaskin, Bree Gatica-Wilcox, Jason Coombes, Joel Southgate, Lauren Gilbert, Lee Graham, Nicole Pacchiarini, Sara Kumziene-Summerhayes, Sarah Taylor, Sophie Jones, Sara Rey, Matthew Bull, Joanne Watkins, Sally Corden, Tom Connor |
| EPI_ISL_638165, EPI_ISL_638195, EPI_ISL_638197, EPI_ISL_638198, EPI_ISL_638199, EPI_ISL_638200, EPI_ISL_638201, EPI_ISL_638203, EPI_ISL_638313, EPI_ISL_638314, EPI_ISL_638316, EPI_ISL_638320, EPI_ISL_638321, EPI_ISL_638322, EPI_ISL_638323, EPI_ISL_638324, EPI_ISL_638325, EPI_ISL_638327, EPI_ISL_638339, EPI_ISL_638340, EPI_ISL_638341, EPI_ISL_638342, EPI_ISL_638343, EPI_ISL_638344, EPI_ISL_638345, EPI_ISL_638346, EPI_ISL_638347, EPI_ISL_638348, EPI_ISL_638349, EPI_ISL_638350, EPI_ISL_638351, EPI_ISL_638352, EPI_ISL_638353, EPI_ISL_638354, EPI_ISL_638355, EPI_ISL_638356, EPI_ISL_638357, EPI_ISL_638358, EPI_ISL_638359, EPI_ISL_638360, EPI_ISL_638362, EPI_ISL_638363, EPI_ISL_638383, EPI_ISL_638384, EPI_ISL_638386, EPI_ISL_638387, EPI_ISL_638388, EPI_ISL_638390, EPI_ISL_638392, EPI_ISL_638394 | | | |
| see above | Department of Pathology, University of Cambridge | COVID-19 Genomics UK (COG-UK) Consortium | Aminu S. Jahun, Yasmin Chaudhry, Grant Hall, Iliana Georgana, Myra Hosmillo, Martin D. Curran, Malte Pinckert, Surendra Parmar, Ian Goodfellow |
| EPI_ISL_638531 | Oxford Viromics, NDM, University of Oxford; Oxford University Hospitals; Basingstoke and North Hampshire Hospital | COVID-19 Genomics UK (COG-UK) Consortium | Tanya Golubchik, David Bonsall, George Macintyre, Amy Trebes, Mariateresa de Cesare, Catrin Moore, Alex Mobbs, Anita Justice, Robert Shaw, Monique Andersson, Timothy Peto, Emma Wise, Nathan Moore, Jessica Lynch, Nick Cortes, Matilde Mori, Stephen Kidd, David Buck, John Todd, Christophe Fraser |
| EPI_ISL_638557 | Northumbria University / South Tees Hospitals NHS Foundation Trust / North Cumbria Integrated Care NHS Foundation Trust / North Tees and Hartlepool NHS Foundation Trust / Newcastle Hospitals NHS Foundation Trust | COVID-19 Genomics UK (COG-UK) Consortium | Darren L Smith,Andrew Nelson,Matthew Bashton,Greg R Young,Joshua Loh,John Allan,Mohammad A Tariq,Giles S Holt,Gary Black,Wen C Yew,Lynn Dover,Paul Baker,Steve Liggett,Sarah Essex,Jane Greenaway,Debra Padgett,Clive Graham,Garren Scott,Edward Barton,Emma Swindells,Brendan Payne,Jennifer Collins,Yusri Taha,Gary Eltringham |
| EPI_ISL_638585 | Queens Medical Centre, Clinical Microbiology Department / DeepSeq Nottingham | COVID-19 Genomics UK (COG-UK) Consortium | Gemma Clark, Wendy Smith, Manjinder Khakh, Vicki M Fleming, Michelle M Lister, Hannah Howson-Wells, Jonathan Ball, Patrick McClure, Joseph Chappell, Theocharis Tsoleridis, Nadine Holmes, Matthew Carlisle, Christopher Moore, Fei Sang, Johnny Debebe, Victoria Wright, Matthew Loose |
| EPI_ISL_638890 | Virology Department, Sheffield Teaching Hospitals NHS Foundation Trust/Department of Infection, Immunity and Cardiovascular Disease, The Medical School, University of Sheffield | COVID-19 Genomics UK (COG-UK) Consortium | Thushan de Silva, Matthew Parker, Nikki Smith, Adri Angyal, Rebecca Brown, Luke Green, Rachel Tucker, Paul Parsons, Danielle Groves, Katie Johnson, Laura Carrilero, Alex Keeley, Dave Partridge, Matthew Wyles, Benjamin Lindsey, Mehmet Yavuz, Mohammad Raza, Cariad Evans |
| EPI_ISL_639036, EPI_ISL_639048, EPI_ISL_639049, EPI_ISL_639059, EPI_ISL_639117, EPI_ISL_639124, EPI_ISL_639129, EPI_ISL_639180, EPI_ISL_639429, EPI_ISL_639446, EPI_ISL_639455, EPI_ISL_639458, EPI_ISL_639470, EPI_ISL_639478, EPI_ISL_639511, EPI_ISL_639522, EPI_ISL_639554, EPI_ISL_639589 | | | |
| see above | Wales Specialist Virology Centre Sequencing lab: Pathogen Genomics Unit | COVID-19 Genomics UK (COG-UK) Consortium | Catherine Moore, Johnathan Evans, Laura Gifford, Malorie Perry, Simon Cottrell, Angela Marchbank, Alec Birchley, Alexander Adams, Amy Gaskin, Bree Gatica-Wilcox, Jason Coombes, Joel Southgate, Lauren Gilbert, Lee Graham, Nicole Pacchiarini, Sara Kumziene-Summerhayes, Sarah Taylor, Sophie Jones, Sara Rey, Matthew Bull, Joanne Watkins, Sally Corden, Tom Connor |
| EPI_ISL_639736, EPI_ISL_639737 | South Eastern Area Laboratory Services (SEALS) | NSW Health Pathology - Institute of Clinical Pathology and Medical Research; Westmead Hospital; University of Sydney | CIDM-PH et al. |
| EPI_ISL_639750, EPI_ISL_639818 | unknown | Public Health Virology Laboratory, Forensic and Scientific Services (PHV-FSS) | Son Nguyen et al. |
| EPI_ISL_639853, EPI_ISL_639909 | National Virus Reference Laboratory | National Virus Reference Laboratory | Michael Carr, Gabriel Gonzalez, Jonathan Dean, Daniel Hare, Cillian F De Gascun |
| EPI_ISL_639997, EPI_ISL_640002 | CNR Virus des Infections Respiratoires - France SUD | CNR Virus des Infections Respiratoires - France SUD | Antonin Bal, Gregory Destras, Gwendolyne Burfin, Hadrien Règue, Alexandre Gaymard, Maude Bouscambert-Duchamp, Florence Morfin-Sherpa, Martine Valette, Bruno Lina, Laurence Josset |
| EPI_ISL_640142, EPI_ISL_640143, EPI_ISL_640144, EPI_ISL_640145, EPI_ISL_640146, EPI_ISL_640147, EPI_ISL_640148, EPI_ISL_640149, EPI_ISL_640150, EPI_ISL_640151, EPI_ISL_640152, EPI_ISL_640153, EPI_ISL_640154, EPI_ISL_640155, EPI_ISL_640156, EPI_ISL_640157, EPI_ISL_640159, EPI_ISL_640160, EPI_ISL_640162, EPI_ISL_640164, EPI_ISL_640165, EPI_ISL_640166, EPI_ISL_640168, EPI_ISL_640170, EPI_ISL_640171, EPI_ISL_640172, EPI_ISL_640173, EPI_ISL_640175, EPI_ISL_640176, EPI_ISL_640178, EPI_ISL_640179, EPI_ISL_640180, EPI_ISL_640181, EPI_ISL_640182, EPI_ISL_640184, EPI_ISL_640185, EPI_ISL_640187, EPI_ISL_640189, EPI_ISL_640191, EPI_ISL_640192, EPI_ISL_640193, EPI_ISL_640194, EPI_ISL_640195, EPI_ISL_640196, EPI_ISL_640199, EPI_ISL_640201, EPI_ISL_640202, EPI_ISL_640203, EPI_ISL_640204, EPI_ISL_640205, EPI_ISL_640206, EPI_ISL_640207, EPI_ISL_640209, EPI_ISL_640213 | | | |
| see above | University of Michigan Clinical Microbiology Laboratory | Lauring Lab, University of Michigan, Department of Microbiology and Immunology | Valesano |
| EPI_ISL_641304 | Microbiological Diagnostic Unit - Public Health Laboratory (MDU-PHL) | MDU-PHL | Seemann T., Schultz M.B., Sait, M.L., Sherry, N.L. |
| EPI_ISL_641404, EPI_ISL_641405, EPI_ISL_641406, EPI_ISL_641407, EPI_ISL_641408 | Department of Virus and Microbiological Special Diagnostics, Statens Serum Institut, Copenhagen, Denmark | Albertsen lab, Department of Chemistry and Bioscience, Aalborg University, Denmark | Thomas Bruun Rasmussen, Jannik Fonager, Morten Rasmussen |
| EPI_ISL_642851, EPI_ISL_642852, EPI_ISL_642882, EPI_ISL_642917, EPI_ISL_642926, EPI_ISL_642970 | Lighthouse Lab in Milton Keynes | Wellcome Sanger Institute for the COVID-19 Genomics UK (COG-UK) Consortium | The Lighthouse Lab in Milton Keynes and Alex Alderton, Roberto Amato, Sonia Goncalves, Ewan Harrison, David K. Jackson, Ian Johnston, Dominic Kwiatkowski, Cordelia Langford, John Sillitoe on behalf of the Wellcome Sanger Institute COVID-19 Surveillance Team |
| EPI_ISL_643117 | Lighthouse Lab in Glasgow | Wellcome Sanger Institute for the COVID-19 Genomics UK (COG-UK) Consortium | Harper VanSteenhouse, Yumi Kasai, David Gray, Carol Clugston, Anna Dominiczak and Alex Alderton, Roberto Amato, Sonia Goncalves, Ewan Harrison, David K. Jackson, Ian Johnston, Dominic Kwiatkowski, Cordelia Langford, John Sillitoe on behalf of the Wellcome Sanger Institute COVID-19 Surveillance Team |
| EPI_ISL_643985, EPI_ISL_643991, EPI_ISL_644052, EPI_ISL_644053, EPI_ISL_644054, EPI_ISL_644055, EPI_ISL_644056, EPI_ISL_644057, EPI_ISL_644058, EPI_ISL_644059, EPI_ISL_644060, EPI_ISL_644061, EPI_ISL_644063, EPI_ISL_644065, EPI_ISL_644067, EPI_ISL_644068, EPI_ISL_644069, EPI_ISL_644070, EPI_ISL_644071, EPI_ISL_644072, EPI_ISL_644073, EPI_ISL_644074, EPI_ISL_644076, EPI_ISL_644077, EPI_ISL_644078, EPI_ISL_644079, EPI_ISL_644080, EPI_ISL_644082, EPI_ISL_644083, EPI_ISL_644084, EPI_ISL_644085, EPI_ISL_644086, EPI_ISL_644087, EPI_ISL_644088, EPI_ISL_644089, EPI_ISL_644090, EPI_ISL_644091, EPI_ISL_644092 | | | |
| see above | Lighthouse Lab in Milton Keynes | Wellcome Sanger Institute for the COVID-19 Genomics UK (COG-UK) Consortium | The Lighthouse Lab in Milton Keynes and Alex Alderton, Roberto Amato, Sonia Goncalves, Ewan Harrison, David K. Jackson, Ian Johnston, Dominic Kwiatkowski, Cordelia Langford, John Sillitoe on behalf of the Wellcome Sanger Institute COVID-19 Surveillance Team |

| | | | |
|---|---|---|---|
| EPI_ISL_644096, EPI_ISL_644097, EPI_ISL_644100 | Lighthouse Lab in Alderley Park | Wellcome Sanger Institute for the COVID-19 Genomics UK (COG-UK) Consortium | Jacquelyn Wynn, Mairead Hyland, The Lighthouse Lab in Alderley Park and Alex Alderton, Roberto Amato, Sonia Goncalves, Ewan Harrison, David K. Jackson, Ian Johnston, Dominic Kwiatkowski, Cordelia Langford, John Sillitoe on behalf of the Wellcome Sanger Institute COVID-19 Surveillance Team |
| EPI_ISL_644102 | Lighthouse Lab in Glasgow | Wellcome Sanger Institute for the COVID-19 Genomics UK (COG-UK) Consortium | Harper VanSteenhouse, Yumi Kasai, David Gray, Carol Clugston, Anna Dominiczak and Alex Alderton, Roberto Amato, Sonia Goncalves, Ewan Harrison, David K. Jackson, Ian Johnston, Dominic Kwiatkowski, Cordelia Langford, John Sillitoe on behalf of the Wellcome Sanger Institute COVID-19 Surveillance Team |
| EPI_ISL_644105, EPI_ISL_644106 | Lighthouse Lab in Alderley Park | Wellcome Sanger Institute for the COVID-19 Genomics UK (COG-UK) Consortium | Jacquelyn Wynn, Mairead Hyland, The Lighthouse Lab in Alderley Park and Alex Alderton, Roberto Amato, Sonia Goncalves, Ewan Harrison, David K. Jackson, Ian Johnston, Dominic Kwiatkowski, Cordelia Langford, John Sillitoe on behalf of the Wellcome Sanger Institute COVID-19 Surveillance Team |
| EPI_ISL_644107 | Lighthouse Lab in Glasgow | Wellcome Sanger Institute for the COVID-19 Genomics UK (COG-UK) Consortium | Harper VanSteenhouse, Yumi Kasai, David Gray, Carol Clugston, Anna Dominiczak and Alex Alderton, Roberto Amato, Sonia Goncalves, Ewan Harrison, David K. Jackson, Ian Johnston, Dominic Kwiatkowski, Cordelia Langford, John Sillitoe on behalf of the Wellcome Sanger Institute COVID-19 Surveillance Team |
| EPI_ISL_644109, EPI_ISL_644110, EPI_ISL_644113 | Lighthouse Lab in Alderley Park | Wellcome Sanger Institute for the COVID-19 Genomics UK (COG-UK) Consortium | Jacquelyn Wynn, Mairead Hyland, The Lighthouse Lab in Alderley Park and Alex Alderton, Roberto Amato, Sonia Goncalves, Ewan Harrison, David K. Jackson, Ian Johnston, Dominic Kwiatkowski, Cordelia Langford, John Sillitoe on behalf of the Wellcome Sanger Institute COVID-19 Surveillance Team |
| EPI_ISL_644114 | Lighthouse Lab in Milton Keynes | Wellcome Sanger Institute for the COVID-19 Genomics UK (COG-UK) Consortium | The Lighthouse Lab in Milton Keynes and Alex Alderton, Roberto Amato, Sonia Goncalves, Ewan Harrison, David K. Jackson, Ian Johnston, Dominic Kwiatkowski, Cordelia Langford, John Sillitoe on behalf of the Wellcome Sanger Institute COVID-19 Surveillance Team |
| EPI_ISL_644115 | Lighthouse Lab in Cambridge | Wellcome Sanger Institute for the COVID-19 Genomics UK (COG-UK) Consortium | Rob Howes, The Lighthouse Lab in Cambridge and Alex Alderton, Roberto Amato, Sonia Goncalves, Ewan Harrison, David K. Jackson, Ian Johnston, Dominic Kwiatkowski, Cordelia Langford, John Sillitoe on behalf of the Wellcome Sanger Institute COVID-19 Surveillance Team |
| EPI_ISL_644116, EPI_ISL_644119 | Lighthouse Lab in Milton Keynes | Wellcome Sanger Institute for the COVID-19 Genomics UK (COG-UK) Consortium | The Lighthouse Lab in Milton Keynes and Alex Alderton, Roberto Amato, Sonia Goncalves, Ewan Harrison, David K. Jackson, Ian Johnston, Dominic Kwiatkowski, Cordelia Langford, John Sillitoe on behalf of the Wellcome Sanger Institute COVID-19 Surveillance Team |
| EPI_ISL_644346, EPI_ISL_644348, EPI_ISL_644352, EPI_ISL_644357, EPI_ISL_644372 | Michigan Department of Health and Human Services, Bureau of Laboratories | Michigan Department of Health and Human Services, Bureau of Laboratories | Blankenship HM, Riner D, Soehnlen MK |
| EPI_ISL_645297, EPI_ISL_645418, EPI_ISL_645452, EPI_ISL_645459, EPI_ISL_645464, EPI_ISL_645502, EPI_ISL_645503 | Lighthouse Lab in Alderley Park | Wellcome Sanger Institute for the COVID-19 Genomics UK (COG-UK) Consortium | Jacquelyn Wynn, Mairead Hyland, The Lighthouse Lab in Alderley Park and Alex Alderton, Roberto Amato, Sonia Goncalves, Ewan Harrison, David K. Jackson, Ian Johnston, Dominic Kwiatkowski, Cordelia Langford, John Sillitoe on behalf of the Wellcome Sanger Institute COVID-19 Surveillance Team |
| EPI_ISL_648153 | Stockholm | The Public Health Agency of Sweden | Anna-Malin Linde, Maria Lind Karlberg, Mattias Haukland, Reza Advani, Olov Svartstrom, Oskar Karlsson Lindsjo, Sandra Broddesson, Petra Edquist, Mia Brytting, Anna Risberg, Karin Tegmark-Wisell |
| EPI_ISL_648167 | Eskilstuna | The Public Health Agency of Sweden | Anna-Malin Linde, Maria Lind Karlberg, Mattias Haukland, Reza Advani, Olov Svartstrom, Oskar Karlsson Lindsjo, Sandra Broddesson, Petra Edquist, Mia Brytting, Anna Risberg, Karin Tegmark-Wisell |
| EPI_ISL_648176, EPI_ISL_648177, EPI_ISL_648178, EPI_ISL_648179 | The Public Health Agency of Sweden | The Public Health Agency of Sweden | Anna-Malin Linde, Maria Lind Karlberg, Mattias Haukland, Reza Advani, Olov Svartstrom, Oskar Karlsson Lindsjo, Sandra Broddesson, Petra Edquist, Mia Brytting, Anna Risberg, Karin Tegmark-Wisell |
| EPI_ISL_648192 | Sundsvall | The Public Health Agency of Sweden | Anna-Malin Linde, Maria Lind Karlberg, Mattias Haukland, Reza Advani, Olov Svartstrom, Oskar Karlsson Lindsjo, Sandra Broddesson, Petra Edquist, Mia Brytting, Anna Risberg, Karin Tegmark-Wisell |
| EPI_ISL_648287, EPI_ISL_648288, EPI_ISL_648289, EPI_ISL_648290, EPI_ISL_648291, EPI_ISL_648292, EPI_ISL_648293, EPI_ISL_648294, EPI_ISL_648295, EPI_ISL_648296 | MD PHL | MD PHL | Maryland Department of Health Laboratories Administration |
| EPI_ISL_648480, EPI_ISL_648481, EPI_ISL_648482, EPI_ISL_648483, EPI_ISL_648484, EPI_ISL_648485, EPI_ISL_648486, EPI_ISL_648487, EPI_ISL_648488, EPI_ISL_648489, EPI_ISL_648490, EPI_ISL_648491, EPI_ISL_648492, EPI_ISL_648493, EPI_ISL_648494, EPI_ISL_648495 | | | |
| see above | Santa Clara County Public Health Laboratory | Chan-Zuckerberg Biohub | CZB Cliahub Consortium |
| EPI_ISL_648923, EPI_ISL_648940, EPI_ISL_648947, EPI_ISL_648948, EPI_ISL_648961, EPI_ISL_649006, EPI_ISL_649010, EPI_ISL_649014, EPI_ISL_649030, EPI_ISL_649031, EPI_ISL_649038, EPI_ISL_649042, EPI_ISL_649044, EPI_ISL_649046, EPI_ISL_649049, EPI_ISL_649050, EPI_ISL_649053 | | | |
| see above | San Diego County Public Health Laboratory | Andersen lab at Scripps Research | SEARCH Alliance San Diego with Tracy Basler, Jovan Shephard, Brett Austin |
| EPI_ISL_649192, EPI_ISL_649205 | Lighthouse Lab in Milton Keynes | Wellcome Sanger Institute for the COVID-19 Genomics UK (COG-UK) Consortium | The Lighthouse Lab in Milton Keynes and Alex Alderton, Roberto Amato, Sonia Goncalves, Ewan Harrison, David K. Jackson, Ian Johnston, Dominic Kwiatkowski, Cordelia Langford, John Sillitoe on behalf of the Wellcome Sanger Institute COVID-19 Surveillance Team (http://www.sanger.ac.uk/covid-team) |
| EPI_ISL_649220 | Lighthouse Lab in Alderley Park | Wellcome Sanger Institute for the COVID-19 Genomics UK (COG-UK) Consortium | Jacquelyn Wynn, Mairead Hyland, The Lighthouse Lab in Alderley Park and Alex Alderton, Roberto Amato, Sonia Goncalves, Ewan Harrison, David K. Jackson, Ian Johnston, Dominic Kwiatkowski, Cordelia Langford, John Sillitoe on behalf of the Wellcome Sanger Institute COVID-19 Surveillance Team (http://www.sanger.ac.uk/covid-team) |
| EPI_ISL_649255, EPI_ISL_649283, EPI_ISL_649320, EPI_ISL_649362 | Lighthouse Lab in Milton Keynes | Wellcome Sanger Institute for the COVID-19 Genomics UK (COG-UK) Consortium | The Lighthouse Lab in Milton Keynes and Alex Alderton, Roberto Amato, Sonia Goncalves, Ewan Harrison, David K. Jackson, Ian Johnston, Dominic Kwiatkowski, Cordelia Langford, John Sillitoe on behalf of the Wellcome Sanger Institute COVID-19 Surveillance Team (http://www.sanger.ac.uk/covid-team) |
| EPI_ISL_649369, EPI_ISL_649370 | Lighthouse Lab in Alderley Park | Wellcome Sanger Institute for the COVID-19 Genomics UK (COG-UK) Consortium | Jacquelyn Wynn, Mairead Hyland, The Lighthouse Lab in Alderley Park and Alex Alderton, Roberto Amato, Sonia Goncalves, Ewan Harrison, David K. Jackson, Ian Johnston, Dominic Kwiatkowski, Cordelia Langford, John Sillitoe on behalf of the Wellcome Sanger Institute COVID-19 Surveillance Team (http://www.sanger.ac.uk/covid-team) |
| EPI_ISL_649386, EPI_ISL_649405, EPI_ISL_649424, EPI_ISL_649479, EPI_ISL_649552, EPI_ISL_649554 | Lighthouse Lab in Milton Keynes | Wellcome Sanger Institute for the COVID-19 Genomics UK (COG-UK) Consortium | The Lighthouse Lab in Milton Keynes and Alex Alderton, Roberto Amato, Sonia Goncalves, Ewan Harrison, David K. Jackson, Ian Johnston, Dominic Kwiatkowski, Cordelia Langford, John Sillitoe on behalf of the Wellcome Sanger Institute COVID-19 Surveillance Team (http://www.sanger.ac.uk/covid-team) |
| EPI_ISL_649813, EPI_ISL_649814, EPI_ISL_649815, EPI_ISL_649816, EPI_ISL_649818, EPI_ISL_649819, EPI_ISL_649820, EPI_ISL_649823, EPI_ISL_649824, EPI_ISL_649825, EPI_ISL_649826, EPI_ISL_649827, EPI_ISL_649828, EPI_ISL_649829, EPI_ISL_649830, EPI_ISL_649831, EPI_ISL_649832, EPI_ISL_649833, EPI_ISL_649834, EPI_ISL_649835, EPI_ISL_649836, EPI_ISL_649837, EPI_ISL_649838, EPI_ISL_649839, EPI_ISL_649840, EPI_ISL_649841, EPI_ISL_649842, EPI_ISL_649843, EPI_ISL_649847, EPI_ISL_649848, EPI_ISL_649850, EPI_ISL_649852, EPI_ISL_649854, EPI_ISL_649855, EPI_ISL_649859, EPI_ISL_649862, EPI_ISL_649865 | | | |
| see above | Respiratory Virus Unit, Microbiology Services Colindale, Public Health England | COVID-19 Genomics UK (COG-UK) Consortium | PHE Covid Sequencing Team |
| EPI_ISL_650114 | Liverpool Clinical Laboratories | COVID-19 Genomics UK (COG-UK) Consortium | Sam Haldenby, Anita Lucaci, Steve Paterson, Julian Hiscox, Alistair Darby, M Almsaud, A Alrezaihi, Muhannad Alruwaili, Stuart D Armstrong, Jones Benjamin, Eleanor G Bentley, Anu Chawla, Jordan J Clark, Angela Cowell, Richard Eccles, Isabel García-Dorival, Matthew Gemmell, Alessandro Gerada, PKF Gilmore, Richard Gregory, Ximeng Han, Catherine Hartley, Margaret Hughes, Miren Iturriza-Gomara, James Johnson, L Luu, Jenifer Manson, Charlotte Nelson, Elaine O'Toole, Cassie Olateju, Rebekah Penrice-Randal , Lucille Rainbow, N.P Randle, Trevor Ian Robinson, Parul Sharma, Ghada T Shawli, James P Stewart, Neil Swainston, Ecaterina Vamos, Joanne Watts, Mark Whitehead |
| EPI_ISL_650136 | Queens Medical Centre, Clinical Microbiology Department / DeepSeq Nottingham | COVID-19 Genomics UK (COG-UK) Consortium | Gemma Clark, Wendy Smith, Manjinder Khakh, Vicki M Fleming, Michelle M Lister, Hannah Howson-Wells, Jonathan Ball, Patrick McClure, Joseph Chappell, Theocharis Tsoleridis, Nadine Holmes, Matthew Carlisle, Christopher Moore, Fei Sang, Johnny Debebe, Victoria Wright, Matthew Loose |
| EPI_ISL_650138 | Virology Department, Royal Infirmary of Edinburgh, NHS Lothian / School of Biological Sciences, University of Edinburgh / Institute of Genetics and Molecular Medicine, University of Edinburgh | COVID-19 Genomics UK (COG-UK) Consortium | McHugh M, Dewar R, Rooke S, Gallagher M, Balcaza C, O'Toole Á, Scher E, Hill V, McCrone JT, Colquhoun R, Yu X, Jackson B, Rambaut A, Williams TC, Templeton K |
| EPI_ISL_650142 | Northumbria University / South Tees Hospitals NHS Foundation Trust / North Cumbria Integrated Care NHS | COVID-19 Genomics UK (COG-UK) Consortium | Darren L Smith,Andrew Nelson,Matthew Bashton,Greg R Young,Joshua Loh,John Allan,Mohammad A Tariq,Giles S Holt,Gary Black,Wen C Yew,Lynn Dover,Paul Baker,Steve Liggett,Sarah Essex,Jane Greenaway,Debra Padgett,Clive Graham,Garren Scott,Edward Barton,Emma Swindells,Brendan |

| | | | |
|---|---|---|---|
| | Foundation Trust / North Tees and Hartlepool NHS Foundation Trust / Newcastle Hospitals NHS Foundation Trust | | Payne,Jennifer Collins,Yusri Taha,Gary Eltringham |
| EPI_ISL_650150 | Queens Medical Centre, Clinical Microbiology Department / DeepSeq Nottingham | COVID-19 Genomics UK (COG-UK) Consortium | Gemma Clark, Wendy Smith, Manjinder Khakh, Vicki M Fleming, Michelle M Lister, Hannah Howson-Wells, Jonathan Ball, Patrick McClure, Joseph Chappell, Theocharis Tsoleridis, Nadine Holmes, Matthew Carlisle, Christopher Moore, Fei Sang, Johnny Debebe, Victoria Wright, Matthew Loose |
| EPI_ISL_650156 | Northumbria University / South Tees Hospitals NHS Foundation Trust / North Cumbria Integrated Care NHS Foundation Trust / North Tees and Hartlepool NHS Foundation Trust / Newcastle Hospitals NHS Foundation Trust | COVID-19 Genomics UK (COG-UK) Consortium | Darren L Smith,Andrew Nelson,Matthew Bashton,Greg R Young,Joshua Loh,John Allan,Mohammad A Tariq,Giles S Holt,Gary Black,Wen C Yew,Lynn Dover,Paul Baker,Steve Liggett,Sarah Essex,Jane Greenaway,Debra Padgett,Clive Graham,Garren Scott,Edward Barton,Emma Swindells,Brendan Payne,Jennifer Collins,Yusri Taha,Gary Eltringham |
| EPI_ISL_650166 | Liverpool Clinical Laboratories | COVID-19 Genomics UK (COG-UK) Consortium | Sam Haldenby, Anita Lucaci, Steve Paterson, Julian Hiscox, Alistair Darby, M Almsaud, A Alrezaihi, Muhannad Alruwaili, Stuart D Armstrong, Jones Benjamin, Eleanor G Bentley, Anu Chawla, Jordan J Clark, Angela Cowell, Richard Eccles, Isabel García-Dorival, Matthew Gemmell, Alessandro Gerada, PKF Gilmore, Richard Gregory, Ximeng Han, Catherine Hartley, Margaret Hughes, Miren Iturriza-Gomara, James Johnson, L Luu, Jenifer Manson, Charlotte Nelson, Elaine O'Toole, Cassie Olateju, Rebekah Penrice-Randal , Lucille Rainbow, N.P Randle, Trevor Ian Robinson, Parul Sharma, Ghada T Shawli, James P Stewart, Neil Swainston, Ecaterina Vamos, Joanne Watts, Mark Whitehead |
| EPI_ISL_650175 | West of Scotland Specialist Virology Centre, NHSGGC / MRC-University of Glasgow Centre for Virus Research | COVID-19 Genomics UK (COG-UK) Consortium | Ana da Silva Filipe, Natasha Johnson, Kathy Smollett, Daniel Mair, Stephen Carmichael, Alice Broos, Lily Tong, Jenna Nichols, Kyriaki Nomikou; Sarah McDonald; Richard Orton, Joseph Hughes, Sreenu Vattipally, David L Robertson; Alasdair MacLean, Rory Gunson; Sharif Shaaban, Matthew Holden; Rachel Blacow, Guy Mollett, Kathy Li, James Shepherd, Antonia Ho, Emma Thomson |
| EPI_ISL_650192, EPI_ISL_650194, EPI_ISL_650196, EPI_ISL_650198, EPI_ISL_650199, EPI_ISL_650202 | Liverpool Clinical Laboratories | COVID-19 Genomics UK (COG-UK) Consortium | Sam Haldenby, Anita Lucaci, Steve Paterson, Julian Hiscox, Alistair Darby, M Almsaud, A Alrezaihi, Muhannad Alruwaili, Stuart D Armstrong, Jones Benjamin, Eleanor G Bentley, Anu Chawla, Jordan J Clark, Angela Cowell, Richard Eccles, Isabel García-Dorival, Matthew Gemmell, Alessandro Gerada, PKF Gilmore, Richard Gregory, Ximeng Han, Catherine Hartley, Margaret Hughes, Miren Iturriza-Gomara, James Johnson, L Luu, Jenifer Manson, Charlotte Nelson, Elaine O'Toole, Cassie Olateju, Rebekah Penrice-Randal , Lucille Rainbow, N.P Randle, Trevor Ian Robinson, Parul Sharma, Ghada T Shawli, James P Stewart, Neil Swainston, Ecaterina Vamos, Joanne Watts, Mark Whitehead |
| EPI_ISL_650225 | University of Exeter | COVID-19 Genomics UK (COG-UK) Consortium | Ben Temperton,Aaron Jeffries,Michelle Michelsen,Joanna Warwick-Dugdale,Audrey Farbos,Robyn Manley,Stephen Michell,Jane Masoli |
| EPI_ISL_650255 | Virology Department, Royal Infirmary of Edinburgh, NHS Lothian / School of Biological Sciences, University of Edinburgh / Institute of Genetics and Molecular Medicine, University of Edinburgh | COVID-19 Genomics UK (COG-UK) Consortium | McHugh M, Dewar R, Rooke S, Gallagher M, Balcaza C, O'Toole Á, Scher E, Hill V, McCrone JT, Colquhoun R, Yu X, Jackson B, Rambaut A, Williams TC, Templeton K |
| EPI_ISL_650259, EPI_ISL_650260, EPI_ISL_650261 | University of Exeter | COVID-19 Genomics UK (COG-UK) Consortium | Ben Temperton,Aaron Jeffries,Michelle Michelsen,Joanna Warwick-Dugdale,Audrey Farbos,Robyn Manley,Stephen Michell,Jane Masoli |
| EPI_ISL_650270 | Regional Virus Laboratory, Belfast Health and Social Care Trust | COVID-19 Genomics UK (COG-UK) Consortium | Conall McCaughey, James McKenna, Tanya Curran, Susan Feeney, Alison Watt, Ciara Cox, Mairead Connor, Zoltan Molnar, David Simpson, Derek Fairley |
| EPI_ISL_650284 | Liverpool Clinical Laboratories | COVID-19 Genomics UK (COG-UK) Consortium | Sam Haldenby, Anita Lucaci, Steve Paterson, Julian Hiscox, Alistair Darby, M Almsaud, A Alrezaihi, Muhannad Alruwaili, Stuart D Armstrong, Jones Benjamin, Eleanor G Bentley, Anu Chawla, Jordan J Clark, Angela Cowell, Richard Eccles, Isabel García-Dorival, Matthew Gemmell, Alessandro Gerada, PKF Gilmore, Richard Gregory, Ximeng Han, Catherine Hartley, Margaret Hughes, Miren Iturriza-Gomara, James Johnson, L Luu, Jenifer Manson, Charlotte Nelson, Elaine O'Toole, Cassie Olateju, Rebekah Penrice-Randal , Lucille Rainbow, N.P Randle, Trevor Ian Robinson, Parul Sharma, Ghada T Shawli, James P Stewart, Neil Swainston, Ecaterina Vamos, Joanne Watts, Mark Whitehead |
| EPI_ISL_650296 | University of Exeter | COVID-19 Genomics UK (COG-UK) Consortium | Ben Temperton,Aaron Jeffries,Michelle Michelsen,Joanna Warwick-Dugdale,Audrey Farbos,Robyn Manley,Stephen Michell,Jane Masoli |
| EPI_ISL_650318, EPI_ISL_650324, EPI_ISL_650326 | Liverpool Clinical Laboratories | COVID-19 Genomics UK (COG-UK) Consortium | Sam Haldenby, Anita Lucaci, Steve Paterson, Julian Hiscox, Alistair Darby, M Almsaud, A Alrezaihi, Muhannad Alruwaili, Stuart D Armstrong, Jones Benjamin, Eleanor G Bentley, Anu Chawla, Jordan J Clark, Angela Cowell, Richard Eccles, Isabel García-Dorival, Matthew Gemmell, Alessandro Gerada, PKF Gilmore, Richard Gregory, Ximeng Han, Catherine Hartley, Margaret Hughes, Miren Iturriza-Gomara, James Johnson, L Luu, Jenifer Manson, Charlotte Nelson, Elaine O'Toole, Cassie Olateju, Rebekah Penrice-Randal , Lucille Rainbow, N.P Randle, Trevor Ian Robinson, Parul Sharma, Ghada T Shawli, James P Stewart, Neil Swainston, Ecaterina Vamos, Joanne Watts, Mark Whitehead |
| EPI_ISL_650329 | Regional Virus Laboratory, Belfast Health and Social Care Trust | COVID-19 Genomics UK (COG-UK) Consortium | Conall McCaughey, James McKenna, Tanya Curran, Susan Feeney, Alison Watt, Ciara Cox, Mairead Connor, Zoltan Molnar, David Simpson, Derek Fairley |
| EPI_ISL_650331 | Liverpool Clinical Laboratories | COVID-19 Genomics UK (COG-UK) Consortium | Sam Haldenby, Anita Lucaci, Steve Paterson, Julian Hiscox, Alistair Darby, M Almsaud, A Alrezaihi, Muhannad Alruwaili, Stuart D Armstrong, Jones Benjamin, Eleanor G Bentley, Anu Chawla, Jordan J Clark, Angela Cowell, Richard Eccles, Isabel García-Dorival, Matthew Gemmell, Alessandro Gerada, PKF Gilmore, Richard Gregory, Ximeng Han, Catherine Hartley, Margaret Hughes, Miren Iturriza-Gomara, James Johnson, L Luu, Jenifer Manson, Charlotte Nelson, Elaine O'Toole, Cassie Olateju, Rebekah Penrice-Randal , Lucille Rainbow, N.P Randle, Trevor Ian Robinson, Parul Sharma, Ghada T Shawli, James P Stewart, Neil Swainston, Ecaterina Vamos, Joanne Watts, Mark Whitehead |
| EPI_ISL_650347 | Queens Medical Centre, Clinical Microbiology Department / DeepSeq Nottingham | COVID-19 Genomics UK (COG-UK) Consortium | Gemma Clark, Wendy Smith, Manjinder Khakh, Vicki M Fleming, Michelle M Lister, Hannah Howson-Wells, Jonathan Ball, Patrick McClure, Joseph Chappell, Theocharis Tsoleridis, Nadine Holmes, Matthew Carlisle, Christopher Moore, Fei Sang, Johnny Debebe, Victoria Wright, Matthew Loose |
| EPI_ISL_650351, EPI_ISL_650359 | Northumbria University / South Tees Hospitals NHS Foundation Trust / North Cumbria Integrated Care NHS Foundation Trust / North Tees and Hartlepool NHS Foundation Trust / Newcastle Hospitals NHS Foundation Trust | COVID-19 Genomics UK (COG-UK) Consortium | Darren L Smith,Andrew Nelson,Matthew Bashton,Greg R Young,Joshua Loh,John Allan,Mohammad A Tariq,Giles S Holt,Gary Black,Wen C Yew,Lynn Dover,Paul Baker,Steve Liggett,Sarah Essex,Jane Greenaway,Debra Padgett,Clive Graham,Garren Scott,Edward Barton,Emma Swindells,Brendan Payne,Jennifer Collins,Yusri Taha,Gary Eltringham |
| EPI_ISL_650360, EPI_ISL_650361 | University of Exeter | COVID-19 Genomics UK (COG-UK) Consortium | Ben Temperton,Aaron Jeffries,Michelle Michelsen,Joanna Warwick-Dugdale,Audrey Farbos,Robyn Manley,Stephen Michell,Jane Masoli |
| EPI_ISL_650366 | Liverpool Clinical Laboratories | COVID-19 Genomics UK (COG-UK) Consortium | Sam Haldenby, Anita Lucaci, Steve Paterson, Julian Hiscox, Alistair Darby, M Almsaud, A Alrezaihi, Muhannad Alruwaili, Stuart D Armstrong, Jones Benjamin, Eleanor G Bentley, Anu Chawla, Jordan J Clark, Angela Cowell, Richard Eccles, Isabel García-Dorival, Matthew Gemmell, Alessandro Gerada, PKF Gilmore, Richard Gregory, Ximeng Han, Catherine Hartley, Margaret Hughes, Miren Iturriza-Gomara, James Johnson, L Luu, Jenifer Manson, Charlotte Nelson, Elaine O'Toole, Cassie Olateju, Rebekah Penrice-Randal , Lucille Rainbow, N.P Randle, Trevor Ian Robinson, Parul Sharma, Ghada T Shawli, James P Stewart, Neil Swainston, Ecaterina Vamos, Joanne Watts, Mark Whitehead |
| EPI_ISL_650378 | Regional Virus Laboratory, Belfast Health and Social Care Trust | COVID-19 Genomics UK (COG-UK) Consortium | Conall McCaughey, James McKenna, Tanya Curran, Susan Feeney, Alison Watt, Ciara Cox, Mairead Connor, Zoltan Molnar, David Simpson, Derek Fairley |
| EPI_ISL_650386 | Queens Medical Centre, Clinical Microbiology Department / DeepSeq Nottingham | COVID-19 Genomics UK (COG-UK) Consortium | Gemma Clark, Wendy Smith, Manjinder Khakh, Vicki M Fleming, Michelle M Lister, Hannah Howson-Wells, Jonathan Ball, Patrick McClure, Joseph Chappell, Theocharis Tsoleridis, Nadine Holmes, Matthew Carlisle, Christopher Moore, Fei Sang, Johnny Debebe, Victoria Wright, Matthew Loose |
| EPI_ISL_650393, EPI_ISL_650428 | Northumbria University / South Tees Hospitals NHS Foundation Trust / North Cumbria Integrated Care NHS Foundation Trust / North Tees and Hartlepool NHS Foundation Trust / Newcastle Hospitals NHS Foundation Trust | COVID-19 Genomics UK (COG-UK) Consortium | Darren L Smith,Andrew Nelson,Matthew Bashton,Greg R Young,Joshua Loh,John Allan,Mohammad A Tariq,Giles S Holt,Gary Black,Wen C Yew,Lynn Dover,Paul Baker,Steve Liggett,Sarah Essex,Jane Greenaway,Debra Padgett,Clive Graham,Garren Scott,Edward Barton,Emma Swindells,Brendan Payne,Jennifer Collins,Yusri Taha,Gary Eltringham |
| EPI_ISL_650444, EPI_ISL_650445 | University of Exeter | COVID-19 Genomics UK (COG-UK) Consortium | Ben Temperton,Aaron Jeffries,Michelle Michelsen,Joanna Warwick-Dugdale,Audrey Farbos,Robyn Manley,Stephen Michell,Jane Masoli |
| EPI_ISL_650479 | Liverpool Clinical Laboratories | COVID-19 Genomics UK (COG-UK) Consortium | Sam Haldenby, Anita Lucaci, Steve Paterson, Julian Hiscox, Alistair Darby, M Almsaud, A Alrezaihi, Muhannad Alruwaili, Stuart D Armstrong, Jones Benjamin, Eleanor G Bentley, Anu Chawla, Jordan J Clark, Angela Cowell, Richard Eccles, Isabel García-Dorival, Matthew Gemmell, Alessandro Gerada, PKF Gilmore, Richard Gregory, Ximeng Han, Catherine Hartley, Margaret Hughes, Miren Iturriza-Gomara, James Johnson, L Luu, Jenifer Manson, Charlotte |

| | | | |
|---|---|---|---|
| | | | Nelson, Elaine O'Toole, Cassie Olateju, Rebekah Penrice-Randal , Lucille Rainbow, N.P Randle, Trevor Ian Robinson, Parul Sharma, Ghada T Shawli, James P Stewart, Neil Swainston, Ecaterina Vamos, Joanne Watts, Mark Whitehead |
| EPI_ISL_650491, EPI_ISL_650492, EPI_ISL_650509 | West of Scotland Specialist Virology Centre, NHSGGC / MRC-University of Glasgow Centre for Virus Research | COVID-19 Genomics UK (COG-UK) Consortium | Ana da Silva Filipe, Natasha Johnson, Kathy Smollett, Daniel Mair, Stephen Carmichael, Alice Broos, Lily Tong, Jenna Nichols, Kyriaki Nomikou; Sarah McDonald; Richard Orton, Joseph Hughes, Sreenu Vattipally, David L Robertson; Alasdair MacLean, Rory Gunson; Sharif Shaaban, Matthew Holden; Rachel Blacow, Guy Mollett, Kathy Li, James Shepherd, Antonia Ho, Emma Thomson |
| EPI_ISL_650513, EPI_ISL_650524 | Regional Virus Laboratory, Belfast Health and Social Care Trust | COVID-19 Genomics UK (COG-UK) Consortium | Conall McCaughey, James McKenna, Tanya Curran, Susan Feeney, Alison Watt, Ciara Cox, Mairead Connor, Zoltan Molnar, David Simpson, Derek Fairley |
| EPI_ISL_650529 | Liverpool Clinical Laboratories | COVID-19 Genomics UK (COG-UK) Consortium | Sam Haldenby, Anita Lucaci, Steve Paterson, Julian Hiscox, Alistair Darby, M Almsaud, A Alrezaihi, Muhannad Alruwaili, Stuart D Armstrong, Jones Benjamin, Eleanor G Bentley, Anu Chawla, Jordan J Clark, Angela Cowell, Richard Eccles, Isabel García-Dorival, Matthew Gemmell, Alessandro Gerada, PKF Gilmore, Richard Gregory, Ximeng Han, Catherine Hartley, Margaret Hughes, Miren Iturriza-Gomara, James Johnson, L Luu, Jenifer Manson, Charlotte Nelson, Elaine O'Toole, Cassie Olateju, Rebekah Penrice-Randal , Lucille Rainbow, N.P Randle, Trevor Ian Robinson, Parul Sharma, Ghada T Shawli, James P Stewart, Neil Swainston, Ecaterina Vamos, Joanne Watts, Mark Whitehead |
| EPI_ISL_650531 | Virology Department, Royal Infirmary of Edinburgh, NHS Lothian / School of Biological Sciences, University of Edinburgh / Institute of Genetics and Molecular Medicine, University of Edinburgh | COVID-19 Genomics UK (COG-UK) Consortium | McHugh M, Dewar R, Rooke S, Gallagher M, Balcaza C, O'Toole Á, Scher E, Hill V, McCrone JT, Colquhoun R, Yu X, Jackson B, Rambaut A, Williams TC, Templeton K |
| EPI_ISL_650544, EPI_ISL_650547, EPI_ISL_650561 | Liverpool Clinical Laboratories | COVID-19 Genomics UK (COG-UK) Consortium | Sam Haldenby, Anita Lucaci, Steve Paterson, Julian Hiscox, Alistair Darby, M Almsaud, A Alrezaihi, Muhannad Alruwaili, Stuart D Armstrong, Jones Benjamin, Eleanor G Bentley, Anu Chawla, Jordan J Clark, Angela Cowell, Richard Eccles, Isabel García-Dorival, Matthew Gemmell, Alessandro Gerada, PKF Gilmore, Richard Gregory, Ximeng Han, Catherine Hartley, Margaret Hughes, Miren Iturriza-Gomara, James Johnson, L Luu, Jenifer Manson, Charlotte Nelson, Elaine O'Toole, Cassie Olateju, Rebekah Penrice-Randal , Lucille Rainbow, N.P Randle, Trevor Ian Robinson, Parul Sharma, Ghada T Shawli, James P Stewart, Neil Swainston, Ecaterina Vamos, Joanne Watts, Mark Whitehead |
| EPI_ISL_650586 | Northumbria University / South Tees Hospitals NHS Foundation Trust / North Cumbria Integrated Care NHS Foundation Trust / North Tees and Hartlepool NHS Foundation Trust / Newcastle Hospitals NHS Foundation Trust | COVID-19 Genomics UK (COG-UK) Consortium | Darren L Smith,Andrew Nelson,Matthew Bashton,Greg R Young,Joshua Loh,John Allan,Mohammad A Tariq,Giles S Holt,Gary Black,Wen C Yew,Lynn Dover,Paul Baker,Steve Liggett,Sarah Essex,Jane Greenaway,Debra Padgett,Clive Graham,Garren Scott,Edward Barton,Emma Swindells,Brendan Payne,Jennifer Collins,Yusri Taha,Gary Eltringham |
| EPI_ISL_650611 | Queens Medical Centre, Clinical Microbiology Department / DeepSeq Nottingham | COVID-19 Genomics UK (COG-UK) Consortium | Gemma Clark, Wendy Smith, Manjinder Khakh, Vicki M Fleming, Michelle M Lister, Hannah Howson-Wells, Jonathan Ball, Patrick McClure, Joseph Chappell, Theocharis Tsoleridis, Nadine Holmes, Matthew Carlisle, Christopher Moore, Fei Sang, Johnny Debebe, Victoria Wright, Matthew Loose |
| EPI_ISL_650619 | Liverpool Clinical Laboratories | COVID-19 Genomics UK (COG-UK) Consortium | Sam Haldenby, Anita Lucaci, Steve Paterson, Julian Hiscox, Alistair Darby, M Almsaud, A Alrezaihi, Muhannad Alruwaili, Stuart D Armstrong, Jones Benjamin, Eleanor G Bentley, Anu Chawla, Jordan J Clark, Angela Cowell, Richard Eccles, Isabel García-Dorival, Matthew Gemmell, Alessandro Gerada, PKF Gilmore, Richard Gregory, Ximeng Han, Catherine Hartley, Margaret Hughes, Miren Iturriza-Gomara, James Johnson, L Luu, Jenifer Manson, Charlotte Nelson, Elaine O'Toole, Cassie Olateju, Rebekah Penrice-Randal , Lucille Rainbow, N.P Randle, Trevor Ian Robinson, Parul Sharma, Ghada T Shawli, James P Stewart, Neil Swainston, Ecaterina Vamos, Joanne Watts, Mark Whitehead |
| EPI_ISL_650625 | Northumbria University / South Tees Hospitals NHS Foundation Trust / North Cumbria Integrated Care NHS Foundation Trust / North Tees and Hartlepool NHS Foundation Trust / Newcastle Hospitals NHS Foundation Trust | COVID-19 Genomics UK (COG-UK) Consortium | Darren L Smith,Andrew Nelson,Matthew Bashton,Greg R Young,Joshua Loh,John Allan,Mohammad A Tariq,Giles S Holt,Gary Black,Wen C Yew,Lynn Dover,Paul Baker,Steve Liggett,Sarah Essex,Jane Greenaway,Debra Padgett,Clive Graham,Garren Scott,Edward Barton,Emma Swindells,Brendan Payne,Jennifer Collins,Yusri Taha,Gary Eltringham |
| EPI_ISL_650641 | Queens Medical Centre, Clinical Microbiology Department / DeepSeq Nottingham | COVID-19 Genomics UK (COG-UK) Consortium | Gemma Clark, Wendy Smith, Manjinder Khakh, Vicki M Fleming, Michelle M Lister, Hannah Howson-Wells, Jonathan Ball, Patrick McClure, Joseph Chappell, Theocharis Tsoleridis, Nadine Holmes, Matthew Carlisle, Christopher Moore, Fei Sang, Johnny Debebe, Victoria Wright, Matthew Loose |
| EPI_ISL_650642, EPI_ISL_650643, EPI_ISL_650644, EPI_ISL_650645 | University of Exeter | COVID-19 Genomics UK (COG-UK) Consortium | Ben Temperton,Aaron Jeffries,Michelle Michelsen,Joanna Warwick-Dugdale,Audrey Farbos,Robyn Manley,Stephen Michell,Jane Masoli |
| EPI_ISL_650654, EPI_ISL_650656, EPI_ISL_650657, EPI_ISL_650658, EPI_ISL_650671, EPI_ISL_650688, EPI_ISL_650694, EPI_ISL_650710, EPI_ISL_650714, EPI_ISL_650755 | Liverpool Clinical Laboratories | COVID-19 Genomics UK (COG-UK) Consortium | Sam Haldenby, Anita Lucaci, Steve Paterson, Julian Hiscox, Alistair Darby, M Almsaud, A Alrezaihi, Muhannad Alruwaili, Stuart D Armstrong, Jones Benjamin, Eleanor G Bentley, Anu Chawla, Jordan J Clark, Angela Cowell, Richard Eccles, Isabel García-Dorival, Matthew Gemmell, Alessandro Gerada, PKF Gilmore, Richard Gregory, Ximeng Han, Catherine Hartley, Margaret Hughes, Miren Iturriza-Gomara, James Johnson, L Luu, Jenifer Manson, Charlotte Nelson, Elaine O'Toole, Cassie Olateju, Rebekah Penrice-Randal , Lucille Rainbow, N.P Randle, Trevor Ian Robinson, Parul Sharma, Ghada T Shawli, James P Stewart, Neil Swainston, Ecaterina Vamos, Joanne Watts, Mark Whitehead |
| EPI_ISL_650775 | Virology Department, Royal Infirmary of Edinburgh, NHS Lothian / School of Biological Sciences, University of Edinburgh / Institute of Genetics and Molecular Medicine, University of Edinburgh | COVID-19 Genomics UK (COG-UK) Consortium | McHugh M, Dewar R, Rooke S, Gallagher M, Balcaza C, O'Toole Á, Scher E, Hill V, McCrone JT, Colquhoun R, Yu X, Jackson B, Rambaut A, Williams TC, Templeton K |
| EPI_ISL_650782 | Liverpool Clinical Laboratories | COVID-19 Genomics UK (COG-UK) Consortium | Sam Haldenby, Anita Lucaci, Steve Paterson, Julian Hiscox, Alistair Darby, M Almsaud, A Alrezaihi, Muhannad Alruwaili, Stuart D Armstrong, Jones Benjamin, Eleanor G Bentley, Anu Chawla, Jordan J Clark, Angela Cowell, Richard Eccles, Isabel García-Dorival, Matthew Gemmell, Alessandro Gerada, PKF Gilmore, Richard Gregory, Ximeng Han, Catherine Hartley, Margaret Hughes, Miren Iturriza-Gomara, James Johnson, L Luu, Jenifer Manson, Charlotte Nelson, Elaine O'Toole, Cassie Olateju, Rebekah Penrice-Randal , Lucille Rainbow, N.P Randle, Trevor Ian Robinson, Parul Sharma, Ghada T Shawli, James P Stewart, Neil Swainston, Ecaterina Vamos, Joanne Watts, Mark Whitehead |
| EPI_ISL_650785 | Northumbria University / South Tees Hospitals NHS Foundation Trust / North Cumbria Integrated Care NHS Foundation Trust / North Tees and Hartlepool NHS Foundation Trust / Newcastle Hospitals NHS Foundation Trust | COVID-19 Genomics UK (COG-UK) Consortium | Darren L Smith,Andrew Nelson,Matthew Bashton,Greg R Young,Joshua Loh,John Allan,Mohammad A Tariq,Giles S Holt,Gary Black,Wen C Yew,Lynn Dover,Paul Baker,Steve Liggett,Sarah Essex,Jane Greenaway,Debra Padgett,Clive Graham,Garren Scott,Edward Barton,Emma Swindells,Brendan Payne,Jennifer Collins,Yusri Taha,Gary Eltringham |
| EPI_ISL_650788 | University of Exeter | COVID-19 Genomics UK (COG-UK) Consortium | Ben Temperton,Aaron Jeffries,Michelle Michelsen,Joanna Warwick-Dugdale,Audrey Farbos,Robyn Manley,Stephen Michell,Jane Masoli |
| EPI_ISL_650816 | West of Scotland Specialist Virology Centre, NHSGGC / MRC-University of Glasgow Centre for Virus Research | COVID-19 Genomics UK (COG-UK) Consortium | Ana da Silva Filipe, Natasha Johnson, Kathy Smollett, Daniel Mair, Stephen Carmichael, Alice Broos, Lily Tong, Jenna Nichols, Kyriaki Nomikou; Sarah McDonald; Richard Orton, Joseph Hughes, Sreenu Vattipally, David L Robertson; Alasdair MacLean, Rory Gunson; Sharif Shaaban, Matthew Holden; Rachel Blacow, Guy Mollett, Kathy Li, James Shepherd, Antonia Ho, Emma Thomson |
| EPI_ISL_650819, EPI_ISL_650833 | Virology Department, Royal Infirmary of Edinburgh, NHS Lothian / School of Biological Sciences, University of Edinburgh / Institute of Genetics and Molecular Medicine, University of Edinburgh | COVID-19 Genomics UK (COG-UK) Consortium | McHugh M, Dewar R, Rooke S, Gallagher M, Balcaza C, O'Toole Á, Scher E, Hill V, McCrone JT, Colquhoun R, Yu X, Jackson B, Rambaut A, Williams TC, Templeton K |
| EPI_ISL_650840 | West of Scotland Specialist Virology Centre, NHSGGC / MRC-University of Glasgow Centre for Virus Research | COVID-19 Genomics UK (COG-UK) Consortium | Ana da Silva Filipe, Natasha Johnson, Kathy Smollett, Daniel Mair, Stephen Carmichael, Alice Broos, Lily Tong, Jenna Nichols, Kyriaki Nomikou; Sarah McDonald; Richard Orton, Joseph Hughes, Sreenu Vattipally, David L Robertson; Alasdair MacLean, Rory Gunson; Sharif Shaaban, Matthew Holden; Rachel Blacow, Guy Mollett, Kathy Li, James Shepherd, Antonia Ho, Emma Thomson |
| EPI_ISL_650847 | Liverpool Clinical Laboratories | COVID-19 Genomics UK (COG-UK) Consortium | Sam Haldenby, Anita Lucaci, Steve Paterson, Julian Hiscox, Alistair Darby, M Almsaud, A Alrezaihi, Muhannad Alruwaili, Stuart D Armstrong, Jones Benjamin, Eleanor G Bentley, Anu Chawla, Jordan J Clark, Angela Cowell, Richard Eccles, Isabel García-Dorival, Matthew Gemmell, Alessandro Gerada, PKF Gilmore, Richard Gregory, Ximeng Han, Catherine Hartley, Margaret Hughes, Miren Iturriza-Gomara, James Johnson, L Luu, Jenifer Manson, Charlotte Nelson, Elaine O'Toole, Cassie Olateju, Rebekah Penrice-Randal , Lucille Rainbow, N.P Randle, Trevor Ian Robinson, Parul Sharma, Ghada T Shawli, James P Stewart, Neil Swainston, Ecaterina Vamos, Joanne Watts, Mark Whitehead |

| | | | |
|---|---|---|---|
| EPI_ISL_650854, EPI_ISL_650855, EPI_ISL_650856, EPI_ISL_650858 | West of Scotland Specialist Virology Centre, NHSGGC / MRC-University of Glasgow Centre for Virus Research | COVID-19 Genomics UK (COG-UK) Consortium | Ana da Silva Filipe, Natasha Johnson, Kathy Smollett, Daniel Mair, Stephen Carmichael, Alice Broos, Lily Tong, Jenna Nichols, Kyriaki Nomikou; Sarah McDonald; Richard Orton, Joseph Hughes, Sreenu Vattipally, David L Robertson; Alasdair MacLean, Rory Gunson; Sharif Shaaban, Matthew Holden; Rachel Blacow, Guy Mollett, Kathy Li, James Shepherd, Antonia Ho, Emma Thomson |
| EPI_ISL_650881, EPI_ISL_650886, EPI_ISL_650887, EPI_ISL_650889, EPI_ISL_650891, EPI_ISL_650892 | Liverpool Clinical Laboratories | COVID-19 Genomics UK (COG-UK) Consortium | Sam Haldenby, Anita Lucaci, Steve Paterson, Julian Hiscox, Alistair Darby, M Almsaud, A Alrezaihi, Muhannad Alruwaili, Stuart D Armstrong, Jones Benjamin, Eleanor G Bentley, Anu Chawla, Jordan J Clark, Angela Cowell, Richard Eccles, Isabel García-Dorival, Matthew Gemmell, Alessandro Gerada, PKF Gilmore, Richard Gregory, Ximeng Han, Catherine Hartley, Margaret Hughes, Miren Iturriza-Gomara, James Johnson, L Luu, Jenifer Manson, Charlotte Nelson, Elaine O'Toole, Cassie Olateju, Rebekah Penrice-Randal , Lucille Rainbow, N.P Randle, Trevor Ian Robinson, Parul Sharma, Ghada T Shawli, James P Stewart, Neil Swainston, Ecaterina Vamos, Joanne Watts, Mark Whitehead |
| EPI_ISL_650898 | Regional Virus Laboratory, Belfast Health and Social Care Trust | COVID-19 Genomics UK (COG-UK) Consortium | Conall McCaughey, James McKenna, Tanya Curran, Susan Feeney, Alison Watt, Ciara Cox, Mairead Connor, Zoltan Molnar, David Simpson, Derek Fairley |
| EPI_ISL_650904 | Liverpool Clinical Laboratories | COVID-19 Genomics UK (COG-UK) Consortium | Sam Haldenby, Anita Lucaci, Steve Paterson, Julian Hiscox, Alistair Darby, M Almsaud, A Alrezaihi, Muhannad Alruwaili, Stuart D Armstrong, Jones Benjamin, Eleanor G Bentley, Anu Chawla, Jordan J Clark, Angela Cowell, Richard Eccles, Isabel García-Dorival, Matthew Gemmell, Alessandro Gerada, PKF Gilmore, Richard Gregory, Ximeng Han, Catherine Hartley, Margaret Hughes, Miren Iturriza-Gomara, James Johnson, L Luu, Jenifer Manson, Charlotte Nelson, Elaine O'Toole, Cassie Olateju, Rebekah Penrice-Randal , Lucille Rainbow, N.P Randle, Trevor Ian Robinson, Parul Sharma, Ghada T Shawli, James P Stewart, Neil Swainston, Ecaterina Vamos, Joanne Watts, Mark Whitehead |
| EPI_ISL_650923 | Regional Virus Laboratory, Belfast Health and Social Care Trust | COVID-19 Genomics UK (COG-UK) Consortium | Conall McCaughey, James McKenna, Tanya Curran, Susan Feeney, Alison Watt, Ciara Cox, Mairead Connor, Zoltan Molnar, David Simpson, Derek Fairley |
| EPI_ISL_650926, EPI_ISL_650928 | Liverpool Clinical Laboratories | COVID-19 Genomics UK (COG-UK) Consortium | Sam Haldenby, Anita Lucaci, Steve Paterson, Julian Hiscox, Alistair Darby, M Almsaud, A Alrezaihi, Muhannad Alruwaili, Stuart D Armstrong, Jones Benjamin, Eleanor G Bentley, Anu Chawla, Jordan J Clark, Angela Cowell, Richard Eccles, Isabel García-Dorival, Matthew Gemmell, Alessandro Gerada, PKF Gilmore, Richard Gregory, Ximeng Han, Catherine Hartley, Margaret Hughes, Miren Iturriza-Gomara, James Johnson, L Luu, Jenifer Manson, Charlotte Nelson, Elaine O'Toole, Cassie Olateju, Rebekah Penrice-Randal , Lucille Rainbow, N.P Randle, Trevor Ian Robinson, Parul Sharma, Ghada T Shawli, James P Stewart, Neil Swainston, Ecaterina Vamos, Joanne Watts, Mark Whitehead |
| EPI_ISL_650930 | Regional Virus Laboratory, Belfast Health and Social Care Trust | COVID-19 Genomics UK (COG-UK) Consortium | Conall McCaughey, James McKenna, Tanya Curran, Susan Feeney, Alison Watt, Ciara Cox, Mairead Connor, Zoltan Molnar, David Simpson, Derek Fairley |
| EPI_ISL_650932, EPI_ISL_650933, EPI_ISL_650934, EPI_ISL_650935, EPI_ISL_650936, EPI_ISL_650939, EPI_ISL_650945, EPI_ISL_650946 | Liverpool Clinical Laboratories | COVID-19 Genomics UK (COG-UK) Consortium | Sam Haldenby, Anita Lucaci, Steve Paterson, Julian Hiscox, Alistair Darby, M Almsaud, A Alrezaihi, Muhannad Alruwaili, Stuart D Armstrong, Jones Benjamin, Eleanor G Bentley, Anu Chawla, Jordan J Clark, Angela Cowell, Richard Eccles, Isabel García-Dorival, Matthew Gemmell, Alessandro Gerada, PKF Gilmore, Richard Gregory, Ximeng Han, Catherine Hartley, Margaret Hughes, Miren Iturriza-Gomara, James Johnson, L Luu, Jenifer Manson, Charlotte Nelson, Elaine O'Toole, Cassie Olateju, Rebekah Penrice-Randal , Lucille Rainbow, N.P Randle, Trevor Ian Robinson, Parul Sharma, Ghada T Shawli, James P Stewart, Neil Swainston, Ecaterina Vamos, Joanne Watts, Mark Whitehead |
| EPI_ISL_651014, EPI_ISL_651015, EPI_ISL_651016, EPI_ISL_651017, EPI_ISL_651018 | Department of Pathology, University of Cambridge | COVID-19 Genomics UK (COG-UK) Consortium | Aminu S. Jahun, Yasmin Chaudhry, Grant Hall, Iliana Georgana, Myra Hosmillo, Martin D. Curran, Malte Pinckert, Surendra Parmar, Ian Goodfellow |
| EPI_ISL_651071, EPI_ISL_651072, EPI_ISL_651073 | Queens Medical Centre, Clinical Microbiology Department / DeepSeq Nottingham | COVID-19 Genomics UK (COG-UK) Consortium | Gemma Clark, Wendy Smith, Manjinder Khakh, Vicki M Fleming, Michelle M Lister, Hannah Howson-Wells, Jonathan Ball, Patrick McClure, Joseph Chappell, Theocharis Tsoleridis, Nadine Holmes, Matthew Carlisle, Christopher Moore, Fei Sang, Johnny Debebe, Victoria Wright, Matthew Loose |
| EPI_ISL_651074, EPI_ISL_651075 | University of Exeter | COVID-19 Genomics UK (COG-UK) Consortium | Ben Temperton,Aaron Jeffries,Michelle Michelsen,Joanna Warwick-Dugdale,Audrey Farbos,Robyn Manley,Stephen Michell,Jane Masoli |
| EPI_ISL_651086, EPI_ISL_651087, EPI_ISL_651088 | Oxford Viromics, NDM, University of Oxford; Oxford University Hospitals; Basingstoke and North Hampshire Hospital | COVID-19 Genomics UK (COG-UK) Consortium | Tanya Golubchik, David Bonsall, George Macintyre, Amy Trebes, Mariateresa de Cesare, Catrin Moore, Alex Mobbs, Anita Justice, Robert Shaw, Monique Andersson, Timothy Peto, Emma Wise, Nathan Moore, Jessica Lynch, Nick Cortes, Matilde Mori, Stephen Kidd, David Buck, John Todd, Christophe Fraser |
| EPI_ISL_651095 | Liverpool Clinical Laboratories | COVID-19 Genomics UK (COG-UK) Consortium | Sam Haldenby, Anita Lucaci, Steve Paterson, Julian Hiscox, Alistair Darby, M Almsaud, A Alrezaihi, Muhannad Alruwaili, Stuart D Armstrong, Jones Benjamin, Eleanor G Bentley, Anu Chawla, Jordan J Clark, Angela Cowell, Richard Eccles, Isabel García-Dorival, Matthew Gemmell, Alessandro Gerada, PKF Gilmore, Richard Gregory, Ximeng Han, Catherine Hartley, Margaret Hughes, Miren Iturriza-Gomara, James Johnson, L Luu, Jenifer Manson, Charlotte Nelson, Elaine O'Toole, Cassie Olateju, Rebekah Penrice-Randal , Lucille Rainbow, N.P Randle, Trevor Ian Robinson, Parul Sharma, Ghada T Shawli, James P Stewart, Neil Swainston, Ecaterina Vamos, Joanne Watts, Mark Whitehead |
| EPI_ISL_651097 | Regional Virus Laboratory, Belfast Health and Social Care Trust | COVID-19 Genomics UK (COG-UK) Consortium | Conall McCaughey, James McKenna, Tanya Curran, Susan Feeney, Alison Watt, Ciara Cox, Mairead Connor, Zoltan Molnar, David Simpson, Derek Fairley |
| EPI_ISL_651103, EPI_ISL_651104, EPI_ISL_651105, EPI_ISL_651106 | Oxford Viromics, NDM, University of Oxford; Oxford University Hospitals; Basingstoke and North Hampshire Hospital | COVID-19 Genomics UK (COG-UK) Consortium | Tanya Golubchik, David Bonsall, George Macintyre, Amy Trebes, Mariateresa de Cesare, Catrin Moore, Alex Mobbs, Anita Justice, Robert Shaw, Monique Andersson, Timothy Peto, Emma Wise, Nathan Moore, Jessica Lynch, Nick Cortes, Matilde Mori, Stephen Kidd, David Buck, John Todd, Christophe Fraser |
| EPI_ISL_651129, EPI_ISL_651131, EPI_ISL_651138 | Liverpool Clinical Laboratories | COVID-19 Genomics UK (COG-UK) Consortium | Sam Haldenby, Anita Lucaci, Steve Paterson, Julian Hiscox, Alistair Darby, M Almsaud, A Alrezaihi, Muhannad Alruwaili, Stuart D Armstrong, Jones Benjamin, Eleanor G Bentley, Anu Chawla, Jordan J Clark, Angela Cowell, Richard Eccles, Isabel García-Dorival, Matthew Gemmell, Alessandro Gerada, PKF Gilmore, Richard Gregory, Ximeng Han, Catherine Hartley, Margaret Hughes, Miren Iturriza-Gomara, James Johnson, L Luu, Jenifer Manson, Charlotte Nelson, Elaine O'Toole, Cassie Olateju, Rebekah Penrice-Randal , Lucille Rainbow, N.P Randle, Trevor Ian Robinson, Parul Sharma, Ghada T Shawli, James P Stewart, Neil Swainston, Ecaterina Vamos, Joanne Watts, Mark Whitehead |
| EPI_ISL_651143 | University of Exeter | COVID-19 Genomics UK (COG-UK) Consortium | Ben Temperton,Aaron Jeffries,Michelle Michelsen,Joanna Warwick-Dugdale,Audrey Farbos,Robyn Manley,Stephen Michell,Jane Masoli |
| EPI_ISL_651145, EPI_ISL_651161, EPI_ISL_651162, EPI_ISL_651192, EPI_ISL_651196, EPI_ISL_651199, EPI_ISL_651205, EPI_ISL_651225 | Liverpool Clinical Laboratories | COVID-19 Genomics UK (COG-UK) Consortium | Sam Haldenby, Anita Lucaci, Steve Paterson, Julian Hiscox, Alistair Darby, M Almsaud, A Alrezaihi, Muhannad Alruwaili, Stuart D Armstrong, Jones Benjamin, Eleanor G Bentley, Anu Chawla, Jordan J Clark, Angela Cowell, Richard Eccles, Isabel García-Dorival, Matthew Gemmell, Alessandro Gerada, PKF Gilmore, Richard Gregory, Ximeng Han, Catherine Hartley, Margaret Hughes, Miren Iturriza-Gomara, James Johnson, L Luu, Jenifer Manson, Charlotte Nelson, Elaine O'Toole, Cassie Olateju, Rebekah Penrice-Randal , Lucille Rainbow, N.P Randle, Trevor Ian Robinson, Parul Sharma, Ghada T Shawli, James P Stewart, Neil Swainston, Ecaterina Vamos, Joanne Watts, Mark Whitehead |
| EPI_ISL_651231 | University of Exeter | COVID-19 Genomics UK (COG-UK) Consortium | Ben Temperton,Aaron Jeffries,Michelle Michelsen,Joanna Warwick-Dugdale,Audrey Farbos,Robyn Manley,Stephen Michell,Jane Masoli |
| EPI_ISL_651236, EPI_ISL_651270 | Liverpool Clinical Laboratories | COVID-19 Genomics UK (COG-UK) Consortium | Sam Haldenby, Anita Lucaci, Steve Paterson, Julian Hiscox, Alistair Darby, M Almsaud, A Alrezaihi, Muhannad Alruwaili, Stuart D Armstrong, Jones Benjamin, Eleanor G Bentley, Anu Chawla, Jordan J Clark, Angela Cowell, Richard Eccles, Isabel García-Dorival, Matthew Gemmell, Alessandro Gerada, PKF Gilmore, Richard Gregory, Ximeng Han, Catherine Hartley, Margaret Hughes, Miren Iturriza-Gomara, James Johnson, L Luu, Jenifer Manson, Charlotte Nelson, Elaine O'Toole, Cassie Olateju, Rebekah Penrice-Randal , Lucille Rainbow, N.P Randle, Trevor Ian Robinson, Parul Sharma, Ghada T Shawli, James P Stewart, Neil Swainston, Ecaterina Vamos, Joanne Watts, Mark Whitehead |
| EPI_ISL_651285, EPI_ISL_651293 | Queens Medical Centre, Clinical Microbiology Department / DeepSeq Nottingham | COVID-19 Genomics UK (COG-UK) Consortium | Gemma Clark, Wendy Smith, Manjinder Khakh, Vicki M Fleming, Michelle M Lister, Hannah Howson-Wells, Jonathan Ball, Patrick McClure, Joseph Chappell, Theocharis Tsoleridis, Nadine Holmes, Matthew Carlisle, Christopher Moore, Fei Sang, Johnny Debebe, Victoria Wright, Matthew Loose |
| EPI_ISL_651300 | University of Exeter | COVID-19 Genomics UK (COG-UK) Consortium | Ben Temperton,Aaron Jeffries,Michelle Michelsen,Joanna Warwick-Dugdale,Audrey Farbos,Robyn Manley,Stephen Michell,Jane Masoli |
| EPI_ISL_651309 | Queens Medical Centre, Clinical Microbiology Department / DeepSeq Nottingham | COVID-19 Genomics UK (COG-UK) Consortium | Gemma Clark, Wendy Smith, Manjinder Khakh, Vicki M Fleming, Michelle M Lister, Hannah Howson-Wells, Jonathan Ball, Patrick McClure, Joseph Chappell, Theocharis Tsoleridis, Nadine Holmes, Matthew Carlisle, Christopher Moore, Fei Sang, Johnny Debebe, Victoria Wright, Matthew Loose |
| EPI_ISL_651311 | Liverpool Clinical Laboratories | COVID-19 Genomics UK (COG-UK) Consortium | Sam Haldenby, Anita Lucaci, Steve Paterson, Julian Hiscox, Alistair Darby, M Almsaud, A Alrezaihi, Muhannad Alruwaili, Stuart D Armstrong, Jones Benjamin, Eleanor G Bentley, Anu Chawla, Jordan J Clark, Angela Cowell, Richard Eccles, Isabel García-Dorival, Matthew Gemmell, Alessandro Gerada, PKF Gilmore, Richard Gregory, Ximeng Han, Catherine Hartley, Margaret Hughes, Miren Iturriza-Gomara, James Johnson, L Luu, Jenifer Manson, Charlotte Nelson, Elaine O'Toole, Cassie Olateju, Rebekah Penrice-Randal , Lucille Rainbow, N.P Randle, Trevor Ian Robinson, Parul Sharma, Ghada T Shawli, James P Stewart, Neil Swainston, Ecaterina Vamos, Joanne Watts, Mark Whitehead |

| | | | |
|---|---|---|---|
| EPI_ISL_651314 | Northumbria University / South Tees Hospitals NHS Foundation Trust / North Cumbria Integrated Care NHS Foundation Trust / North Tees and Hartlepool NHS Foundation Trust / Newcastle Hospitals NHS Foundation Trust | COVID-19 Genomics UK (COG-UK) Consortium | Darren L Smith,Andrew Nelson,Matthew Bashton,Greg R Young,Joshua Loh,John Allan,Mohammad A Tariq,Giles S Holt,Gary Black,Wen C Yew,Lynn Dover,Paul Baker,Steve Liggett,Sarah Essex,Jane Greenaway,Debra Padgett,Clive Graham,Garren Scott,Edward Barton,Emma Swindells,Brendan Payne,Jennifer Collins,Yusri Taha,Gary Eltringham |
| EPI_ISL_651318 | Regional Virus Laboratory, Belfast Health and Social Care Trust | COVID-19 Genomics UK (COG-UK) Consortium | Conall McCaughey, James McKenna, Tanya Curran, Susan Feeney, Alison Watt, Ciara Cox, Mairead Connor, Zoltan Molnar, David Simpson, Derek Fairley |
| EPI_ISL_651320 | Virology Department, Royal Infirmary of Edinburgh, NHS Lothian / School of Biological Sciences, University of Edinburgh / Institute of Genetics and Molecular Medicine, University of Edinburgh | COVID-19 Genomics UK (COG-UK) Consortium | McHugh M, Dewar R, Rooke S, Gallagher M, Balcaza C, O'Toole Á, Scher E, Hill V, McCrone JT, Colquhoun R, Yu X, Jackson B, Rambaut A, Williams TC, Templeton K |
| EPI_ISL_651344 | Liverpool Clinical Laboratories | COVID-19 Genomics UK (COG-UK) Consortium | Sam Haldenby, Anita Lucaci, Steve Paterson, Julian Hiscox, Alistair Darby, M Almsaud, A Alrezaihi, Muhannad Alruwaili, Stuart D Armstrong, Jones Benjamin, Eleanor G Bentley, Anu Chawla, Jordan J Clark, Angela Cowell, Richard Eccles, Isabel García-Dorival, Matthew Gemmell, Alessandro Gerada, PKF Gilmore, Richard Gregory, Ximeng Han, Catherine Hartley, Margaret Hughes, Miren Iturriza-Gomara, James Johnson, L Luu, Jenifer Manson, Charlotte Nelson, Elaine O'Toole, Cassie Olateju, Rebekah Penrice-Randal , Lucille Rainbow, N.P Randle, Trevor Ian Robinson, Parul Sharma, Ghada T Shawli, James P Stewart, Neil Swainston, Ecaterina Vamos, Joanne Watts, Mark Whitehead |
| EPI_ISL_651354 | Northumbria University / South Tees Hospitals NHS Foundation Trust / North Cumbria Integrated Care NHS Foundation Trust / North Tees and Hartlepool NHS Foundation Trust / Newcastle Hospitals NHS Foundation Trust | COVID-19 Genomics UK (COG-UK) Consortium | Darren L Smith,Andrew Nelson,Matthew Bashton,Greg R Young,Joshua Loh,John Allan,Mohammad A Tariq,Giles S Holt,Gary Black,Wen C Yew,Lynn Dover,Paul Baker,Steve Liggett,Sarah Essex,Jane Greenaway,Debra Padgett,Clive Graham,Garren Scott,Edward Barton,Emma Swindells,Brendan Payne,Jennifer Collins,Yusri Taha,Gary Eltringham |
| EPI_ISL_651358 | Liverpool Clinical Laboratories | COVID-19 Genomics UK (COG-UK) Consortium | Sam Haldenby, Anita Lucaci, Steve Paterson, Julian Hiscox, Alistair Darby, M Almsaud, A Alrezaihi, Muhannad Alruwaili, Stuart D Armstrong, Jones Benjamin, Eleanor G Bentley, Anu Chawla, Jordan J Clark, Angela Cowell, Richard Eccles, Isabel García-Dorival, Matthew Gemmell, Alessandro Gerada, PKF Gilmore, Richard Gregory, Ximeng Han, Catherine Hartley, Margaret Hughes, Miren Iturriza-Gomara, James Johnson, L Luu, Jenifer Manson, Charlotte Nelson, Elaine O'Toole, Cassie Olateju, Rebekah Penrice-Randal , Lucille Rainbow, N.P Randle, Trevor Ian Robinson, Parul Sharma, Ghada T Shawli, James P Stewart, Neil Swainston, Ecaterina Vamos, Joanne Watts, Mark Whitehead |
| EPI_ISL_651369 | Northumbria University / South Tees Hospitals NHS Foundation Trust / North Cumbria Integrated Care NHS Foundation Trust / North Tees and Hartlepool NHS Foundation Trust / Newcastle Hospitals NHS Foundation Trust | COVID-19 Genomics UK (COG-UK) Consortium | Darren L Smith,Andrew Nelson,Matthew Bashton,Greg R Young,Joshua Loh,John Allan,Mohammad A Tariq,Giles S Holt,Gary Black,Wen C Yew,Lynn Dover,Paul Baker,Steve Liggett,Sarah Essex,Jane Greenaway,Debra Padgett,Clive Graham,Garren Scott,Edward Barton,Emma Swindells,Brendan Payne,Jennifer Collins,Yusri Taha,Gary Eltringham |
| EPI_ISL_651372 | University of Exeter | COVID-19 Genomics UK (COG-UK) Consortium | Ben Temperton,Aaron Jeffries,Michelle Michelsen,Joanna Warwick-Dugdale,Audrey Farbos,Robyn Manley,Stephen Michell,Jane Masoli |
| EPI_ISL_651375 | Liverpool Clinical Laboratories | COVID-19 Genomics UK (COG-UK) Consortium | Sam Haldenby, Anita Lucaci, Steve Paterson, Julian Hiscox, Alistair Darby, M Almsaud, A Alrezaihi, Muhannad Alruwaili, Stuart D Armstrong, Jones Benjamin, Eleanor G Bentley, Anu Chawla, Jordan J Clark, Angela Cowell, Richard Eccles, Isabel García-Dorival, Matthew Gemmell, Alessandro Gerada, PKF Gilmore, Richard Gregory, Ximeng Han, Catherine Hartley, Margaret Hughes, Miren Iturriza-Gomara, James Johnson, L Luu, Jenifer Manson, Charlotte Nelson, Elaine O'Toole, Cassie Olateju, Rebekah Penrice-Randal , Lucille Rainbow, N.P Randle, Trevor Ian Robinson, Parul Sharma, Ghada T Shawli, James P Stewart, Neil Swainston, Ecaterina Vamos, Joanne Watts, Mark Whitehead |
| EPI_ISL_651384 | University of Exeter | COVID-19 Genomics UK (COG-UK) Consortium | Ben Temperton,Aaron Jeffries,Michelle Michelsen,Joanna Warwick-Dugdale,Audrey Farbos,Robyn Manley,Stephen Michell,Jane Masoli |
| EPI_ISL_651387 | Regional Virus Laboratory, Belfast Health and Social Care Trust | COVID-19 Genomics UK (COG-UK) Consortium | Conall McCaughey, James McKenna, Tanya Curran, Susan Feeney, Alison Watt, Ciara Cox, Mairead Connor, Zoltan Molnar, David Simpson, Derek Fairley |
| EPI_ISL_651396, EPI_ISL_651403 | Liverpool Clinical Laboratories | COVID-19 Genomics UK (COG-UK) Consortium | Sam Haldenby, Anita Lucaci, Steve Paterson, Julian Hiscox, Alistair Darby, M Almsaud, A Alrezaihi, Muhannad Alruwaili, Stuart D Armstrong, Jones Benjamin, Eleanor G Bentley, Anu Chawla, Jordan J Clark, Angela Cowell, Richard Eccles, Isabel García-Dorival, Matthew Gemmell, Alessandro Gerada, PKF Gilmore, Richard Gregory, Ximeng Han, Catherine Hartley, Margaret Hughes, Miren Iturriza-Gomara, James Johnson, L Luu, Jenifer Manson, Charlotte Nelson, Elaine O'Toole, Cassie Olateju, Rebekah Penrice-Randal , Lucille Rainbow, N.P Randle, Trevor Ian Robinson, Parul Sharma, Ghada T Shawli, James P Stewart, Neil Swainston, Ecaterina Vamos, Joanne Watts, Mark Whitehead |
| EPI_ISL_651404 | Regional Virus Laboratory, Belfast Health and Social Care Trust | COVID-19 Genomics UK (COG-UK) Consortium | Conall McCaughey, James McKenna, Tanya Curran, Susan Feeney, Alison Watt, Ciara Cox, Mairead Connor, Zoltan Molnar, David Simpson, Derek Fairley |
| EPI_ISL_651407 | Liverpool Clinical Laboratories | COVID-19 Genomics UK (COG-UK) Consortium | Sam Haldenby, Anita Lucaci, Steve Paterson, Julian Hiscox, Alistair Darby, M Almsaud, A Alrezaihi, Muhannad Alruwaili, Stuart D Armstrong, Jones Benjamin, Eleanor G Bentley, Anu Chawla, Jordan J Clark, Angela Cowell, Richard Eccles, Isabel García-Dorival, Matthew Gemmell, Alessandro Gerada, PKF Gilmore, Richard Gregory, Ximeng Han, Catherine Hartley, Margaret Hughes, Miren Iturriza-Gomara, James Johnson, L Luu, Jenifer Manson, Charlotte Nelson, Elaine O'Toole, Cassie Olateju, Rebekah Penrice-Randal , Lucille Rainbow, N.P Randle, Trevor Ian Robinson, Parul Sharma, Ghada T Shawli, James P Stewart, Neil Swainston, Ecaterina Vamos, Joanne Watts, Mark Whitehead |
| EPI_ISL_651410 | Regional Virus Laboratory, Belfast Health and Social Care Trust | COVID-19 Genomics UK (COG-UK) Consortium | Conall McCaughey, James McKenna, Tanya Curran, Susan Feeney, Alison Watt, Ciara Cox, Mairead Connor, Zoltan Molnar, David Simpson, Derek Fairley |
| EPI_ISL_651417, EPI_ISL_651421 | Oxford Viromics, NDM, University of Oxford; Oxford University Hospitals; Basingstoke and North Hampshire Hospital | COVID-19 Genomics UK (COG-UK) Consortium | Tanya Golubchik, David Bonsall, George Macintyre, Amy Trebes, Mariateresa de Cesare, Catrin Moore, Alex Mobbs, Anita Justice, Robert Shaw, Monique Andersson, Timothy Peto, Emma Wise, Nathan Moore, Jessica Lynch, Nick Cortes, Matilde Mori, Stephen Kidd, David Buck, John Todd, Christophe Fraser |
| EPI_ISL_651466 | Queens Medical Centre, Clinical Microbiology Department / DeepSeq Nottingham | COVID-19 Genomics UK (COG-UK) Consortium | Gemma Clark, Wendy Smith, Manjinder Khakh, Vicki M Fleming, Michelle M Lister, Hannah Howson-Wells, Jonathan Ball, Patrick McClure, Joseph Chappell, Theocharis Tsoleridis, Nadine Holmes, Matthew Carlisle, Christopher Moore, Fei Sang, Johnny Debebe, Victoria Wright, Matthew Loose |
| EPI_ISL_651467, EPI_ISL_651468 | University of Exeter | COVID-19 Genomics UK (COG-UK) Consortium | Ben Temperton,Aaron Jeffries,Michelle Michelsen,Joanna Warwick-Dugdale,Audrey Farbos,Robyn Manley,Stephen Michell,Jane Masoli |
| EPI_ISL_651469, EPI_ISL_651470, EPI_ISL_651471, EPI_ISL_651472, EPI_ISL_651473, EPI_ISL_651474, EPI_ISL_651475, EPI_ISL_651476, EPI_ISL_651477, EPI_ISL_651479, EPI_ISL_651480, EPI_ISL_651481, EPI_ISL_651485 | | | |
| see above | Oxford Viromics, NDM, University of Oxford; Oxford University Hospitals; Basingstoke and North Hampshire Hospital | COVID-19 Genomics UK (COG-UK) Consortium | Tanya Golubchik, David Bonsall, George Macintyre, Amy Trebes, Mariateresa de Cesare, Catrin Moore, Alex Mobbs, Anita Justice, Robert Shaw, Monique Andersson, Timothy Peto, Emma Wise, Nathan Moore, Jessica Lynch, Nick Cortes, Matilde Mori, Stephen Kidd, David Buck, John Todd, Christophe Fraser |
| EPI_ISL_651499 | Liverpool Clinical Laboratories | COVID-19 Genomics UK (COG-UK) Consortium | Sam Haldenby, Anita Lucaci, Steve Paterson, Julian Hiscox, Alistair Darby, M Almsaud, A Alrezaihi, Muhannad Alruwaili, Stuart D Armstrong, Jones Benjamin, Eleanor G Bentley, Anu Chawla, Jordan J Clark, Angela Cowell, Richard Eccles, Isabel García-Dorival, Matthew Gemmell, Alessandro Gerada, PKF Gilmore, Richard Gregory, Ximeng Han, Catherine Hartley, Margaret Hughes, Miren Iturriza-Gomara, James Johnson, L Luu, Jenifer Manson, Charlotte Nelson, Elaine O'Toole, Cassie Olateju, Rebekah Penrice-Randal , Lucille Rainbow, N.P Randle, Trevor Ian Robinson, Parul Sharma, Ghada T Shawli, James P Stewart, Neil Swainston, Ecaterina Vamos, Joanne Watts, Mark Whitehead |
| EPI_ISL_651504 | Queens Medical Centre, Clinical Microbiology Department / DeepSeq Nottingham | COVID-19 Genomics UK (COG-UK) Consortium | Gemma Clark, Wendy Smith, Manjinder Khakh, Vicki M Fleming, Michelle M Lister, Hannah Howson-Wells, Jonathan Ball, Patrick McClure, Joseph Chappell, Theocharis Tsoleridis, Nadine Holmes, Matthew Carlisle, Christopher Moore, Fei Sang, Johnny Debebe, Victoria Wright, Matthew Loose |
| EPI_ISL_651505, EPI_ISL_651506, EPI_ISL_651507, EPI_ISL_651509, EPI_ISL_651510, EPI_ISL_651513, EPI_ISL_651517 | Oxford Viromics, NDM, University of Oxford; Oxford University Hospitals; Basingstoke and North Hampshire Hospital | COVID-19 Genomics UK (COG-UK) Consortium | Tanya Golubchik, David Bonsall, George Macintyre, Amy Trebes, Mariateresa de Cesare, Catrin Moore, Alex Mobbs, Anita Justice, Robert Shaw, Monique Andersson, Timothy Peto, Emma Wise, Nathan Moore, Jessica Lynch, Nick Cortes, Matilde Mori, Stephen Kidd, David Buck, John Todd, Christophe Fraser |
| EPI_ISL_651521, EPI_ISL_651522 | Virology Department, Royal Infirmary of Edinburgh, NHS | COVID-19 Genomics UK (COG-UK) Consortium | McHugh M, Dewar R, Rooke S, Gallagher M, Balcaza C, O'Toole Á, Scher E, Hill V, McCrone JT, Colquhoun R, Yu X, Jackson B, Rambaut A, Williams TC, |

| | | | |
|---|---|---|---|
| | Lothian / School of Biological Sciences, University of Edinburgh / Institute of Genetics and Molecular Medicine, University of Edinburgh | | Templeton K |
| EPI_ISL_651528 | Queens Medical Centre, Clinical Microbiology Department / DeepSeq Nottingham | COVID-19 Genomics UK (COG-UK) Consortium | Gemma Clark, Wendy Smith, Manjinder Khakh, Vicki M Fleming, Michelle M Lister, Hannah Howson-Wells, Jonathan Ball, Patrick McClure, Joseph Chappell, Theocharis Tsoleridis, Nadine Holmes, Matthew Carlisle, Christopher Moore, Fei Sang, Johnny Debebe, Victoria Wright, Matthew Loose |
| EPI_ISL_651529, EPI_ISL_651541, EPI_ISL_651544 | Oxford Viromics, NDM, University of Oxford; Oxford University Hospitals; Basingstoke and North Hampshire Hospital | COVID-19 Genomics UK (COG-UK) Consortium | Tanya Golubchik, David Bonsall, George Macintyre, Amy Trebes, Mariateresa de Cesare, Catrin Moore, Alex Mobbs, Anita Justice, Robert Shaw, Monique Andersson, Timothy Peto, Emma Wise, Nathan Moore, Jessica Lynch, Nick Cortes, Matilde Mori, Stephen Kidd, David Buck, John Todd, Christophe Fraser |
| EPI_ISL_651547 | University of Exeter | COVID-19 Genomics UK (COG-UK) Consortium | Ben Temperton,Aaron Jeffries,Michelle Michelsen,Joanna Warwick-Dugdale,Audrey Farbos,Robyn Manley,Stephen Michell,Jane Masoli |
| EPI_ISL_651548, EPI_ISL_651549, EPI_ISL_651586, EPI_ISL_651587 | Oxford Viromics, NDM, University of Oxford; Oxford University Hospitals; Basingstoke and North Hampshire Hospital | COVID-19 Genomics UK (COG-UK) Consortium | Tanya Golubchik, David Bonsall, George Macintyre, Amy Trebes, Mariateresa de Cesare, Catrin Moore, Alex Mobbs, Anita Justice, Robert Shaw, Monique Andersson, Timothy Peto, Emma Wise, Nathan Moore, Jessica Lynch, Nick Cortes, Matilde Mori, Stephen Kidd, David Buck, John Todd, Christophe Fraser |
| EPI_ISL_651591 | University of Birmingham | COVID-19 Genomics UK (COG-UK) Consortium | Institute of Microbiology, University of Birmingham: Claire McMurray, Joanne Stockton, Samuel Nicholls, Radoslaw Poplawski, Will Rowe, Josh Quick, Nicholas Loman. University of Birmingham Testing Laboratory: Celina M Whalley, Andrew Bosworth, Charlotte Poxon, Kasun Wanigasooriya, Oliver Pickles, Mike Kidd, Alex Richter, Andrew D Beggs PHE Heartlands Lab: Husam Osman, Andrew Bosworth. Queen Elizabeth Hospital: Anna Casey |
| EPI_ISL_651741, EPI_ISL_651742, EPI_ISL_651817, EPI_ISL_651818, EPI_ISL_651819, EPI_ISL_651820, EPI_ISL_651884 | West of Scotland Specialist Virology Centre, NHSGGC / MRC-University of Glasgow Centre for Virus Research | COVID-19 Genomics UK (COG-UK) Consortium | Ana da Silva Filipe, Natasha Johnson, Kathy Smollett, Daniel Mair, Stephen Carmichael, Alice Broos, Lily Tong, Jenna Nichols, Kyriaki Nomikou; Sarah McDonald; Richard Orton, Joseph Hughes, Sreenu Vattipally, David L Robertson; Alasdair MacLean, Rory Gunson; Sharif Shaaban, Matthew Holden; Rachel Blacow, Guy Mollett, Kathy Li, James Shepherd, Antonia Ho, Emma Thomson |
| EPI_ISL_651960, EPI_ISL_651961, EPI_ISL_651962, EPI_ISL_651963, EPI_ISL_651964, EPI_ISL_651965, EPI_ISL_651966, EPI_ISL_651969, EPI_ISL_651970, EPI_ISL_651971, EPI_ISL_651972, EPI_ISL_651973, EPI_ISL_651974, EPI_ISL_651981, EPI_ISL_652062, EPI_ISL_652063 | | | |
| see above | Virology Department, Royal Infirmary of Edinburgh, NHS Lothian / School of Biological Sciences, University of Edinburgh / Institute of Genetics and Molecular Medicine, University of Edinburgh | COVID-19 Genomics UK (COG-UK) Consortium | McHugh M, Dewar R, Rooke S, Gallagher M, Balcaza C, O'Toole Á, Scher E, Hill V, McCrone JT, Colquhoun R, Yu X, Jackson B, Rambaut A, Williams TC, Templeton K |
| EPI_ISL_652073, EPI_ISL_652074, EPI_ISL_652075, EPI_ISL_652076, EPI_ISL_652077, EPI_ISL_652078, EPI_ISL_652079, EPI_ISL_652080, EPI_ISL_652081, EPI_ISL_652082, EPI_ISL_652083, EPI_ISL_652085, EPI_ISL_652086 | | | |
| see above | Liverpool Clinical Laboratories | COVID-19 Genomics UK (COG-UK) Consortium | Sam Haldenby, Anita Lucaci, Steve Paterson, Julian Hiscox, Alistair Darby, M Almsaud, A Alrezaihi, Muhannad Alruwaili, Stuart D Armstrong, Jones Benjamin, Eleanor G Bentley, Anu Chawla, Jordan J Clark, Angela Cowell, Richard Eccles, Isabel García-Dorival, Matthew Gemmell, Alessandro Gerada, PKF Gilmore, Richard Gregory, Ximeng Han, Catherine Hartley, Margaret Hughes, Miren Iturriza-Gomara, James Johnson, L Luu, Jenifer Manson, Charlotte Nelson, Elaine O'Toole, Cassie Olateju, Rebekah Penrice-Randal , Lucille Rainbow, N.P Randle, Trevor Ian Robinson, Parul Sharma, Ghada T Shawli, James P Stewart, Neil Swainston, Ecaterina Vamos, Joanne Watts, Mark Whitehead |
| EPI_ISL_652111, EPI_ISL_652113 | Northumbria University / South Tees Hospitals NHS Foundation Trust / North Cumbria Integrated Care NHS Foundation Trust / North Tees and Hartlepool NHS Foundation Trust / Newcastle Hospitals NHS Foundation Trust | COVID-19 Genomics UK (COG-UK) Consortium | Darren L Smith,Andrew Nelson,Matthew Bashton,Greg R Young,Joshua Loh,John Allan,Mohammad A Tariq,Giles S Holt,Gary Black,Wen C Yew,Lynn Dover,Paul Baker,Steve Liggett,Sarah Essex,Jane Greenaway,Debra Padgett,Clive Graham,Garren Scott,Edward Barton,Emma Swindells,Brendan Payne,Jennifer Collins,Yusri Taha,Gary Eltringham |
| EPI_ISL_652157, EPI_ISL_652158, EPI_ISL_652159, EPI_ISL_652160 | Department of Pathology, University of Cambridge | COVID-19 Genomics UK (COG-UK) Consortium | Aminu S. Jahun, Yasmin Chaudhry, Grant Hall, Iliana Georgana, Myra Hosmillo, Martin D. Curran, Malte Pinckert, Surendra Parmar, Ian Goodfellow |
| EPI_ISL_652304, EPI_ISL_652306, EPI_ISL_652308, EPI_ISL_652311, EPI_ISL_652312, EPI_ISL_652313, EPI_ISL_652314, EPI_ISL_652315, EPI_ISL_652316, EPI_ISL_652317, EPI_ISL_652318, EPI_ISL_652319, EPI_ISL_652320, EPI_ISL_652321, EPI_ISL_652322, EPI_ISL_652323, EPI_ISL_652324, EPI_ISL_652325, EPI_ISL_652326, EPI_ISL_652327, EPI_ISL_652328, EPI_ISL_652329 | | | |
| see above | Regional Virus Laboratory, Belfast Health and Social Care Trust | COVID-19 Genomics UK (COG-UK) Consortium | Conall McCaughey, James McKenna, Tanya Curran, Susan Feeney, Alison Watt, Ciara Cox, Mairead Connor, Zoltan Molnar, David Simpson, Derek Fairley |
| EPI_ISL_652342 | Northumbria University / South Tees Hospitals NHS Foundation Trust / North Cumbria Integrated Care NHS Foundation Trust / North Tees and Hartlepool NHS Foundation Trust / Newcastle Hospitals NHS Foundation Trust | COVID-19 Genomics UK (COG-UK) Consortium | Darren L Smith,Andrew Nelson,Matthew Bashton,Greg R Young,Joshua Loh,John Allan,Mohammad A Tariq,Giles S Holt,Gary Black,Wen C Yew,Lynn Dover,Paul Baker,Steve Liggett,Sarah Essex,Jane Greenaway,Debra Padgett,Clive Graham,Garren Scott,Edward Barton,Emma Swindells,Brendan Payne,Jennifer Collins,Yusri Taha,Gary Eltringham |
| EPI_ISL_652414 | Wales Specialist Virology Centre Sequencing lab: Pathogen Genomics Unit | COVID-19 Genomics UK (COG-UK) Consortium | Catherine Moore, Johnathan Evans, Laura Gifford, Malorie Perry, Simon Cottrell, Angela Marchbank, Alec Birchley, Alexander Adams, Amy Gaskin, Bree Gatica-Wilcox, Jason Coombes, Joel Southgate, Lauren Gilbert, Lee Graham, Nicole Pacchiarini, Sara Kumziene-Summerhayes, Sarah Taylor, Sophie Jones, Sara Rey, Matthew Bull, Joanne Watkins, Sally Corden, Tom Connor |
| EPI_ISL_652628, EPI_ISL_652629, EPI_ISL_652630, EPI_ISL_652631, EPI_ISL_652632, EPI_ISL_652633, EPI_ISL_652634, EPI_ISL_652635, EPI_ISL_652636, EPI_ISL_652637, EPI_ISL_652638, EPI_ISL_652639, EPI_ISL_652640, EPI_ISL_652641, EPI_ISL_652642, EPI_ISL_652643, EPI_ISL_652644, EPI_ISL_652645, EPI_ISL_652646, EPI_ISL_652647, EPI_ISL_652648, EPI_ISL_652649, EPI_ISL_652650, EPI_ISL_652651, EPI_ISL_652652, EPI_ISL_652653, EPI_ISL_652654, EPI_ISL_652655, EPI_ISL_652656, EPI_ISL_652657, EPI_ISL_652658, EPI_ISL_652659, EPI_ISL_652660, EPI_ISL_652661, EPI_ISL_652662, EPI_ISL_652663, EPI_ISL_652664, EPI_ISL_652665, EPI_ISL_652666, EPI_ISL_652667, EPI_ISL_652668, EPI_ISL_652669, EPI_ISL_652670, EPI_ISL_652671, EPI_ISL_652672, EPI_ISL_652673, EPI_ISL_652674 | | | |
| see above | Queens Medical Centre, Clinical Microbiology Department / DeepSeq Nottingham | COVID-19 Genomics UK (COG-UK) Consortium | Gemma Clark, Wendy Smith, Manjinder Khakh, Vicki M Fleming, Michelle M Lister, Hannah Howson-Wells, Jonathan Ball, Patrick McClure, Joseph Chappell, Theocharis Tsoleridis, Nadine Holmes, Matthew Carlisle, Christopher Moore, Fei Sang, Johnny Debebe, Victoria Wright, Matthew Loose |
| EPI_ISL_652675, EPI_ISL_652676, EPI_ISL_652677, EPI_ISL_652678, EPI_ISL_652679, EPI_ISL_652680, EPI_ISL_652681, EPI_ISL_652682, EPI_ISL_652683, EPI_ISL_652684, EPI_ISL_652685, EPI_ISL_652686, EPI_ISL_652687, EPI_ISL_652688, EPI_ISL_652689, EPI_ISL_652690, EPI_ISL_652691, EPI_ISL_652692, EPI_ISL_652693, EPI_ISL_652694, EPI_ISL_652695, EPI_ISL_652696, EPI_ISL_652697, EPI_ISL_652698, EPI_ISL_652699, EPI_ISL_652700, EPI_ISL_652702, EPI_ISL_652703, EPI_ISL_652704, EPI_ISL_652705, EPI_ISL_652706, EPI_ISL_652707, EPI_ISL_652708, EPI_ISL_652709, EPI_ISL_652710 | | | |
| see above | University of Exeter | COVID-19 Genomics UK (COG-UK) Consortium | Ben Temperton,Aaron Jeffries,Michelle Michelsen,Joanna Warwick-Dugdale,Audrey Farbos,Robyn Manley,Stephen Michell,Jane Masoli |
| EPI_ISL_652711, EPI_ISL_652712, EPI_ISL_652713, EPI_ISL_652714, EPI_ISL_652715, EPI_ISL_652716, EPI_ISL_652717, EPI_ISL_652718, EPI_ISL_652719, EPI_ISL_652720, EPI_ISL_652721, EPI_ISL_652722, EPI_ISL_652723, EPI_ISL_652724, EPI_ISL_652725, EPI_ISL_652726, EPI_ISL_652727, EPI_ISL_652728, EPI_ISL_652729, EPI_ISL_652730, EPI_ISL_652731, EPI_ISL_652732, EPI_ISL_652733, EPI_ISL_652734, EPI_ISL_652735, EPI_ISL_652736, EPI_ISL_652737, EPI_ISL_652738, EPI_ISL_652739, EPI_ISL_652740, EPI_ISL_652741, EPI_ISL_652742, EPI_ISL_652743, EPI_ISL_652744, EPI_ISL_652745, EPI_ISL_652746, EPI_ISL_652747, EPI_ISL_652748, EPI_ISL_652749, EPI_ISL_652750, EPI_ISL_652751, EPI_ISL_652752, EPI_ISL_652753, EPI_ISL_652754, EPI_ISL_652755, EPI_ISL_652756, EPI_ISL_652757, EPI_ISL_652758, EPI_ISL_652759, EPI_ISL_652760, EPI_ISL_652761, EPI_ISL_652762, EPI_ISL_652763, EPI_ISL_652764, EPI_ISL_652765, EPI_ISL_652766, EPI_ISL_652767, EPI_ISL_652768, EPI_ISL_652769, EPI_ISL_652770, EPI_ISL_652771, EPI_ISL_652772, EPI_ISL_652773, EPI_ISL_652774, EPI_ISL_652775, EPI_ISL_652776, EPI_ISL_652777, EPI_ISL_652778, EPI_ISL_652779, EPI_ISL_652780, EPI_ISL_652781, EPI_ISL_652782, EPI_ISL_652783, EPI_ISL_652784, EPI_ISL_652785, EPI_ISL_652786, EPI_ISL_652787, EPI_ISL_652788, EPI_ISL_652789, EPI_ISL_652790, EPI_ISL_652791, EPI_ISL_652792, EPI_ISL_652793, EPI_ISL_652794, EPI_ISL_652795, EPI_ISL_652796, EPI_ISL_652797, EPI_ISL_652798, EPI_ISL_652799, EPI_ISL_652800, EPI_ISL_652801, EPI_ISL_652802, EPI_ISL_652803, EPI_ISL_652804, EPI_ISL_652805, EPI_ISL_652806, EPI_ISL_652807, EPI_ISL_652808, EPI_ISL_652809, EPI_ISL_652810, EPI_ISL_652811, EPI_ISL_652812, EPI_ISL_652813, EPI_ISL_652814, EPI_ISL_652815, EPI_ISL_652816, EPI_ISL_652817, EPI_ISL_652818, EPI_ISL_652819, EPI_ISL_652820, EPI_ISL_652821, EPI_ISL_652822, EPI_ISL_652823, EPI_ISL_652824, EPI_ISL_652825, EPI_ISL_652826, EPI_ISL_652827, EPI_ISL_652828, EPI_ISL_652829, EPI_ISL_652830, EPI_ISL_652831, EPI_ISL_652832, EPI_ISL_652833, EPI_ISL_652834, EPI_ISL_652835, EPI_ISL_652836, EPI_ISL_652837, EPI_ISL_652838, EPI_ISL_652839, EPI_ISL_652840, EPI_ISL_652841, EPI_ISL_652842, EPI_ISL_652843, EPI_ISL_652844, EPI_ISL_652845, EPI_ISL_652846, EPI_ISL_652847, EPI_ISL_652848, EPI_ISL_652849, EPI_ISL_652850, EPI_ISL_652851, EPI_ISL_652852, EPI_ISL_652853, EPI_ISL_652854, EPI_ISL_652855, EPI_ISL_652856, EPI_ISL_652857, EPI_ISL_652858, EPI_ISL_652859, EPI_ISL_652860, EPI_ISL_652861, EPI_ISL_652862, EPI_ISL_652863, EPI_ISL_652864, EPI_ISL_652865, EPI_ISL_652866, EPI_ISL_652867, EPI_ISL_652868, EPI_ISL_652869, EPI_ISL_652870, EPI_ISL_652871, EPI_ISL_652872, EPI_ISL_652873, EPI_ISL_652874, EPI_ISL_652875, EPI_ISL_652876, EPI_ISL_652877, EPI_ISL_652878, EPI_ISL_652879, EPI_ISL_652880, EPI_ISL_652881, EPI_ISL_652882, EPI_ISL_652883, EPI_ISL_652884, EPI_ISL_652885, EPI_ISL_652886, EPI_ISL_652887, EPI_ISL_652888, EPI_ISL_652889, EPI_ISL_652890, EPI_ISL_652891, EPI_ISL_652892, EPI_ISL_652893, EPI_ISL_652894, EPI_ISL_652895, EPI_ISL_652896, EPI_ISL_652897, EPI_ISL_652898, EPI_ISL_652899, EPI_ISL_652900, EPI_ISL_652901, EPI_ISL_652902, EPI_ISL_652903, EPI_ISL_652904, EPI_ISL_652905, EPI_ISL_652906, EPI_ISL_652907, EPI_ISL_652908, EPI_ISL_652909, EPI_ISL_652910, EPI_ISL_652911, EPI_ISL_652912, EPI_ISL_652913, EPI_ISL_652914, EPI_ISL_652915, EPI_ISL_652916, EPI_ISL_652917, EPI_ISL_652918, EPI_ISL_652919, EPI_ISL_652920, EPI_ISL_652921, EPI_ISL_652922, EPI_ISL_652923, EPI_ISL_652924, EPI_ISL_652925, EPI_ISL_652926, EPI_ISL_652927, EPI_ISL_652928, EPI_ISL_652929, EPI_ISL_652930, EPI_ISL_652931, EPI_ISL_652932, EPI_ISL_652933, EPI_ISL_652934, EPI_ISL_652935, EPI_ISL_652936, EPI_ISL_652937, EPI_ISL_652938, EPI_ISL_652939, EPI_ISL_652940, EPI_ISL_652941, EPI_ISL_652942, EPI_ISL_652943, EPI_ISL_652944, EPI_ISL_652945, EPI_ISL_652946, EPI_ISL_652947, EPI_ISL_652948, EPI_ISL_652959, EPI_ISL_652965, EPI_ISL_652974, EPI_ISL_652977, EPI_ISL_652978, EPI_ISL_652979, EPI_ISL_652980, EPI_ISL_652981, EPI_ISL_652982, EPI_ISL_652983, EPI_ISL_652984, EPI_ISL_652985, EPI_ISL_652986, EPI_ISL_652987, EPI_ISL_652988, EPI_ISL_652989, EPI_ISL_652990, EPI_ISL_652991, EPI_ISL_652999, EPI_ISL_653000, EPI_ISL_653002, EPI_ISL_653004, EPI_ISL_653007, EPI_ISL_653010, EPI_ISL_653011, EPI_ISL_653012, EPI_ISL_653013, EPI_ISL_653022 | | | |
| see above | Oxford Viromics, NDM, University of Oxford; Oxford University Hospitals; Basingstoke and North Hampshire Hospital | COVID-19 Genomics UK (COG-UK) Consortium | Tanya Golubchik, David Bonsall, George Macintyre, Amy Trebes, Mariateresa de Cesare, Catrin Moore, Alex Mobbs, Anita Justice, Robert Shaw, Monique Andersson, Timothy Peto, Emma Wise, Nathan Moore, Jessica Lynch, Nick Cortes, Matilde Mori, Stephen Kidd, David Buck, John Todd, Christophe Fraser |
| EPI_ISL_654248, EPI_ISL_654249, EPI_ISL_654251, EPI_ISL_654252, EPI_ISL_654253, EPI_ISL_654256, EPI_ISL_654257, EPI_ISL_654260, EPI_ISL_654262, EPI_ISL_654267, EPI_ISL_654269, EPI_ISL_654270, EPI_ISL_654271, EPI_ISL_654272, EPI_ISL_654273, EPI_ISL_654274, EPI_ISL_654275, EPI_ISL_654312, | | | |

| | | | |
|---|---|---|---|
| EPI_ISL_654313, EPI_ISL_654314, EPI_ISL_654315, EPI_ISL_654316, EPI_ISL_654317, EPI_ISL_654318, EPI_ISL_654319, EPI_ISL_654320, EPI_ISL_654321, EPI_ISL_654322, EPI_ISL_654323, EPI_ISL_654324, EPI_ISL_654325, EPI_ISL_654326, EPI_ISL_654327, EPI_ISL_654342, EPI_ISL_654343, EPI_ISL_654359 | | | |
| see above | Hospital General Universitario Gregorio Marañón | SeqCOVID-SPAIN consortium/IBV(CSIC) | Darío García de Viedma, Laura Pérez-Lago, Marta Herranz, Jon Sicilia, Julia Suárez, Pilar Catalán, Patricia Muñoz and SeqCOVID-SPAIN consortium |
| EPI_ISL_654499, EPI_ISL_654500, EPI_ISL_654501 | The Public Health Agency of Sweden | The Public Health Agency of Sweden | Anna-Malin Linde, Maria Lind Karlberg, Mattias Haukland, Reza Advani, Olov Svartstrom, Oskar Karlsson Lindsjo, Sandra Broddesson, Petra Edquist, Mia Brytting, Anna Risberg, Karin Tegmark-Wisell |
| EPI_ISL_654592, EPI_ISL_654593, EPI_ISL_654594, EPI_ISL_654597, EPI_ISL_654598, EPI_ISL_654599, EPI_ISL_654600, EPI_ISL_654601, EPI_ISL_654602, EPI_ISL_654603, EPI_ISL_654604 | | | |
| see above | Servicio de Microbiología. Hospital Universitario Donostia. OSI Donostialdea. Área de Enfermedades Infecciosas, Grupo de Infección Respiratoria y Resistencia Antimicrobiana. Instituto de Investigación Sanitaria Biodonostia | SeqCOVID-SPAIN consortium/IBV(CSIC) | Gustavo Cilla Eguiluz, Milagrosa Montes Ros, Luis Piñeiro Vázquez, Ane Sorrarain, Jose Maria Marimón and SeqCOVID-SPAIN consortium |
| EPI_ISL_660252, EPI_ISL_660253, EPI_ISL_660254, EPI_ISL_660255 | NHLS-IALCH | KRISP, KZN Research Innovation and Sequencing Platform | Giandhari J, Pillay S, Lessells R, Mdlalose K, York D, Khan S, Tegally H, Wilkinson E, de Oliveira T |
| EPI_ISL_660259, EPI_ISL_660260, EPI_ISL_660261, EPI_ISL_660262, EPI_ISL_660263 | Molecualr Diagnostic Services (MDS) | KRISP, KZN Research Innovation and Sequencing Platform | Giandhari J, Pillay S, Lessells R, Mdlalose K, York D, Khan S, Tegally H, Wilkinson E, de Oliveira T |
| EPI_ISL_660489, EPI_ISL_660497 | Laboratoire de Microbiologie CHU Souro Sanou | Centre Muraz | Abdoul-Salam Ouedraogo, Yacouba Sawadogo, Essia Belarbi, Grit Schubert, Fabian Leendertz, Arsène Zongo, Soumeya Ouangraoua, Zekiba Tarnagda, Lassana Sangaré, Halidou Tinto |
| EPI_ISL_660546, EPI_ISL_660547 | Respiratory Virus Unit, Microbiology Services Colindale, Public Health England | COVID-19 Genomics UK (COG-UK) Consortium | PHE Covid Sequencing Team |
| EPI_ISL_660553, EPI_ISL_660566, EPI_ISL_660567, EPI_ISL_660569, EPI_ISL_660570, EPI_ISL_660571, EPI_ISL_660572, EPI_ISL_660573, EPI_ISL_660577, EPI_ISL_660580, EPI_ISL_660583, EPI_ISL_660584, EPI_ISL_660585, EPI_ISL_660586, EPI_ISL_660587, EPI_ISL_660588 | | | |
| see above | The National Institute of Public Health | State Veterinary Institute Prague | Nagy,A;Jirincova,H;Novakova,L;Trnka,D;Vecerova,J |
| EPI_ISL_660956, EPI_ISL_660957, EPI_ISL_660958, EPI_ISL_660959, EPI_ISL_660960, EPI_ISL_660961, EPI_ISL_660962, EPI_ISL_660963, EPI_ISL_660964, EPI_ISL_660965, EPI_ISL_660966, EPI_ISL_660967, EPI_ISL_660968 | | | |
| see above | Gundersen Molecular Diagnostics Laboratory | Kabara Cancer Research Institute | Craig S. Richmond, Paraic A. Kenny |
| EPI_ISL_661288 | Klinisk mikrobiologi | The Public Health Agency of Sweden | Department of Microbiology, The Public Health Agency of Sweden |
| EPI_ISL_661290 | Klinisk mikrobiologi, Viruslab | The Public Health Agency of Sweden | Department of Microbiology, The Public Health Agency of Sweden |
| EPI_ISL_661299 | Laboratoriemedicin, Klinisk mikrobiologi | The Public Health Agency of Sweden | Department of Microbiology, The Public Health Agency of Sweden |
| EPI_ISL_664342, EPI_ISL_664672, EPI_ISL_664724, EPI_ISL_664753, EPI_ISL_665202, EPI_ISL_665209, EPI_ISL_665210, EPI_ISL_665211, EPI_ISL_665212, EPI_ISL_665213, EPI_ISL_665214, EPI_ISL_665215, EPI_ISL_665216, EPI_ISL_665217, EPI_ISL_665224, EPI_ISL_665225, EPI_ISL_665226, EPI_ISL_665227 | | | |
| see above | University College London Hospital | COVID-19 Genomics UK (COG-UK) Consortium | Judith Heaney, Matthew Byott, Catherine Houlihan, Dan Frampton, Stuart Kirk, Moira Spyer and Eleni Nastouli |
| EPI_ISL_665275, EPI_ISL_665280, EPI_ISL_665300, EPI_ISL_665310, EPI_ISL_665317, EPI_ISL_665322, EPI_ISL_665329, EPI_ISL_665350, EPI_ISL_665368, EPI_ISL_665388, EPI_ISL_665389, EPI_ISL_665397, EPI_ISL_665398, EPI_ISL_665405, EPI_ISL_665407 | | | |
| see above | Northumbria University / South Tees Hospitals NHS Foundation Trust / North Cumbria Integrated Care NHS Foundation Trust / North Tees and Hartlepool NHS Foundation Trust / Newcastle Hospitals NHS Foundation Trust | COVID-19 Genomics UK (COG-UK) Consortium | Darren L Smith,Andrew Nelson,Matthew Bashton,Greg R Young,Joshua Loh,John Allan,Mohammad A Tariq,Giles S Holt,Gary Black,Wen C Yew,Lynn Dover,Paul Baker,Steve Liggett,Sarah Essex,Jane Greenaway,Debra Padgett,Clive Graham,Garren Scott,Edward Barton,Emma Swindells,Brendan Payne,Jennifer Collins,Yusri Taha,Gary Eltringham |
| EPI_ISL_665422 | University of Exeter | COVID-19 Genomics UK (COG-UK) Consortium | Ben Temperton,Aaron Jeffries,Michelle Michelsen,Joanna Warwick-Dugdale,Audrey Farbos,Robyn Manley,Stephen Michell,Jane Masoli |
| EPI_ISL_665489, EPI_ISL_665620, EPI_ISL_665627, EPI_ISL_665628, EPI_ISL_665631, EPI_ISL_665667, EPI_ISL_665681 | Northumbria University / South Tees Hospitals NHS Foundation Trust / North Cumbria Integrated Care NHS Foundation Trust / North Tees and Hartlepool NHS Foundation Trust / Newcastle Hospitals NHS Foundation Trust | COVID-19 Genomics UK (COG-UK) Consortium | Darren L Smith,Andrew Nelson,Matthew Bashton,Greg R Young,Joshua Loh,John Allan,Mohammad A Tariq,Giles S Holt,Gary Black,Wen C Yew,Lynn Dover,Paul Baker,Steve Liggett,Sarah Essex,Jane Greenaway,Debra Padgett,Clive Graham,Garren Scott,Edward Barton,Emma Swindells,Brendan Payne,Jennifer Collins,Yusri Taha,Gary Eltringham |
| EPI_ISL_665699 | Queens Medical Centre, Clinical Microbiology Department / DeepSeq Nottingham | COVID-19 Genomics UK (COG-UK) Consortium | Gemma Clark, Wendy Smith, Manjinder Khakh, Vicki M Fleming, Michelle M Lister, Hannah Howson-Wells, Jonathan Ball, Patrick McClure, Joseph Chappell, Theocharis Tsoleridis, Nadine Holmes, Matthew Carlisle, Christopher Moore, Fei Sang, Johnny Debebe, Victoria Wright, Matthew Loose |
| EPI_ISL_665706 | Northumbria University / South Tees Hospitals NHS Foundation Trust / North Cumbria Integrated Care NHS Foundation Trust / North Tees and Hartlepool NHS Foundation Trust / Newcastle Hospitals NHS Foundation Trust | COVID-19 Genomics UK (COG-UK) Consortium | Darren L Smith,Andrew Nelson,Matthew Bashton,Greg R Young,Joshua Loh,John Allan,Mohammad A Tariq,Giles S Holt,Gary Black,Wen C Yew,Lynn Dover,Paul Baker,Steve Liggett,Sarah Essex,Jane Greenaway,Debra Padgett,Clive Graham,Garren Scott,Edward Barton,Emma Swindells,Brendan Payne,Jennifer Collins,Yusri Taha,Gary Eltringham |
| EPI_ISL_665719 | University of Exeter | COVID-19 Genomics UK (COG-UK) Consortium | Ben Temperton,Aaron Jeffries,Michelle Michelsen,Joanna Warwick-Dugdale,Audrey Farbos,Robyn Manley,Stephen Michell,Jane Masoli |
| EPI_ISL_665739 | Northumbria University / South Tees Hospitals NHS Foundation Trust / North Cumbria Integrated Care NHS Foundation Trust / North Tees and Hartlepool NHS Foundation Trust / Newcastle Hospitals NHS Foundation Trust | COVID-19 Genomics UK (COG-UK) Consortium | Darren L Smith,Andrew Nelson,Matthew Bashton,Greg R Young,Joshua Loh,John Allan,Mohammad A Tariq,Giles S Holt,Gary Black,Wen C Yew,Lynn Dover,Paul Baker,Steve Liggett,Sarah Essex,Jane Greenaway,Debra Padgett,Clive Graham,Garren Scott,Edward Barton,Emma Swindells,Brendan Payne,Jennifer Collins,Yusri Taha,Gary Eltringham |
| EPI_ISL_665761 | Queens Medical Centre, Clinical Microbiology Department / DeepSeq Nottingham | COVID-19 Genomics UK (COG-UK) Consortium | Gemma Clark, Wendy Smith, Manjinder Khakh, Vicki M Fleming, Michelle M Lister, Hannah Howson-Wells, Jonathan Ball, Patrick McClure, Joseph Chappell, Theocharis Tsoleridis, Nadine Holmes, Matthew Carlisle, Christopher Moore, Fei Sang, Johnny Debebe, Victoria Wright, Matthew Loose |
| EPI_ISL_665784 | Northumbria University / South Tees Hospitals NHS Foundation Trust / North Cumbria Integrated Care NHS Foundation Trust / North Tees and Hartlepool NHS Foundation Trust / Newcastle Hospitals NHS Foundation Trust | COVID-19 Genomics UK (COG-UK) Consortium | Darren L Smith,Andrew Nelson,Matthew Bashton,Greg R Young,Joshua Loh,John Allan,Mohammad A Tariq,Giles S Holt,Gary Black,Wen C Yew,Lynn Dover,Paul Baker,Steve Liggett,Sarah Essex,Jane Greenaway,Debra Padgett,Clive Graham,Garren Scott,Edward Barton,Emma Swindells,Brendan Payne,Jennifer Collins,Yusri Taha,Gary Eltringham |
| EPI_ISL_665977, EPI_ISL_665978, EPI_ISL_665979, EPI_ISL_665980, EPI_ISL_665981, EPI_ISL_665982 | Queens Medical Centre, Clinical Microbiology Department / DeepSeq Nottingham | COVID-19 Genomics UK (COG-UK) Consortium | Gemma Clark, Wendy Smith, Manjinder Khakh, Vicki M Fleming, Michelle M Lister, Hannah Howson-Wells, Jonathan Ball, Patrick McClure, Joseph Chappell, Theocharis Tsoleridis, Nadine Holmes, Matthew Carlisle, Christopher Moore, Fei Sang, Johnny Debebe, Victoria Wright, Matthew Loose |
| EPI_ISL_665983, EPI_ISL_665984, EPI_ISL_665985, EPI_ISL_665986, EPI_ISL_665987, EPI_ISL_665988, EPI_ISL_665989, EPI_ISL_665990, EPI_ISL_665991, EPI_ISL_665992, EPI_ISL_665993, EPI_ISL_665994, EPI_ISL_665995, EPI_ISL_665996, EPI_ISL_665997, EPI_ISL_665998, EPI_ISL_665999, EPI_ISL_666000, EPI_ISL_666001, EPI_ISL_666002, EPI_ISL_666088, EPI_ISL_666089, EPI_ISL_666090, EPI_ISL_666091, EPI_ISL_666092, EPI_ISL_666093, EPI_ISL_666094, EPI_ISL_666095, EPI_ISL_666096, EPI_ISL_666097, EPI_ISL_666098, EPI_ISL_666099, EPI_ISL_666100 | | | |
| see above | Northumbria University / South Tees Hospitals NHS Foundation Trust / North Cumbria Integrated Care NHS Foundation Trust / North Tees and Hartlepool NHS Foundation Trust / Newcastle Hospitals NHS Foundation Trust | COVID-19 Genomics UK (COG-UK) Consortium | Darren L Smith,Andrew Nelson,Matthew Bashton,Greg R Young,Joshua Loh,John Allan,Mohammad A Tariq,Giles S Holt,Gary Black,Wen C Yew,Lynn Dover,Paul Baker,Steve Liggett,Sarah Essex,Jane Greenaway,Debra Padgett,Clive Graham,Garren Scott,Edward Barton,Emma Swindells,Brendan Payne,Jennifer Collins,Yusri Taha,Gary Eltringham |
| EPI_ISL_666140, EPI_ISL_666181, EPI_ISL_666364 | Wales Specialist Virology Centre Sequencing lab: Pathogen Genomics Unit | COVID-19 Genomics UK (COG-UK) Consortium | Catherine Moore, Johnathan Evans, Laura Gifford, Malorie Perry, Simon Cottrell, Angela Marchbank, Alec Birchley, Alexander Adams, Amy Gaskin, Bree Gatica-Wilcox, Jason Coombes, Joel Southgate, Lauren Gilbert, Lee Graham, Nicole Pacchiarini, Sara Kumziene-Summerhayes, Sarah Taylor, Sophie Jones, Sara Rey, Matthew Bull, Joanne Watkins, Sally Corden, Tom Connor |

| | | | |
|---|---|---|---|
| EPI_ISL_666891, EPI_ISL_666900, EPI_ISL_666905, EPI_ISL_666918, EPI_ISL_666935, EPI_ISL_666937, EPI_ISL_666943, EPI_ISL_666947 | Michigan Department of Health and Human Services, Bureau of Laboratories | Michigan Department of Health and Human Services, Bureau of Laboratories | Blankenship HM, Riner D, Soehnlen MK |
| EPI_ISL_666979, EPI_ISL_666980, EPI_ISL_666982, EPI_ISL_666984, EPI_ISL_666986, EPI_ISL_666988, EPI_ISL_666989, EPI_ISL_666990, EPI_ISL_667045, EPI_ISL_667048, EPI_ISL_667052, EPI_ISL_667053 | | | |
| see above | San Diego County Public Health Laboratory | Andersen lab at Scripps Research | SEARCH Alliance San Diego with Tracy Basler, Jovan Shephard, Brett Austin |
| EPI_ISL_667505, EPI_ISL_667506, EPI_ISL_667507, EPI_ISL_667509, EPI_ISL_667510, EPI_ISL_667511 | OHSU Lab Services Molecular Microbiology Lab | Oregon SARS-CoV-2 Genome Sequencing Center | Brendan L. O'Connell, Ruth V. Nichols, Sally Grindstaff, Alec J. Hirsch, Donna Hansel, Guang Fan, Daniel N. Streblow, William B. Messer, Andrew C. Adey, Benjamin N. Bimber, Brian J. O'Roak |
| EPI_ISL_667787, EPI_ISL_667792 | South Eastern Area Laboratory Services (SEALS) | NSW Health Pathology - Institute of Clinical Pathology and Medical Research; Westmead Hospital; University of Sydney | CIDM-PH et al. |
| EPI_ISL_668391 | Akershus University Hospital, Department for Microbiology and Infectious Disease Control | Norwegian Institute of Public Health, Department of Virology | Kathrine Stene-Johansen, Kamilla Heddeland Instefjord, Hilde Elshaug, Marie Paulsen Madsen, Rasmus Riis Kopperud, Hilde Vollan, Karoline Bragstad, Olav Hungnes |
| EPI_ISL_668394, EPI_ISL_668395 | Hospital of Southern Norway - Kristiansand, Department of Medical Microbiology | Norwegian Institute of Public Health, Department of Virology | Kathrine Stene-Johansen, Kamilla Heddeland Instefjord, Hilde Elshaug, Marie Paulsen Madsen, Rasmus Riis Kopperud, Hilde Vollan, Karoline Bragstad, Olav Hungnes |
| EPI_ISL_668399 | Nordland Hospital - Bodo, Laboratory Department, Molecular Biology Unit | Norwegian Institute of Public Health, Department of Virology | Kathrine Stene-Johansen, Kamilla Heddeland Instefjord, Hilde Elshaug, Marie Paulsen Madsen, Rasmus Riis Kopperud, Hilde Vollan, Karoline Bragstad, Olav Hungnes |
| EPI_ISL_668401 | Oslo University Hospital, Department of Medical Microbiology | Norwegian Institute of Public Health, Department of Virology | Kathrine Stene-Johansen, Kamilla Heddeland Instefjord, Hilde Elshaug, Marie Paulsen Madsen, Rasmus Riis Kopperud, Hilde Vollan, Karoline Bragstad, Olav Hungnes |
| EPI_ISL_668412, EPI_ISL_668414, EPI_ISL_668416 | Innlandet Hospital Trust, Division Lillehammer, Department for Medical Microbiology | Norwegian Institute of Public Health, Department of Virology | Kathrine Stene-Johansen, Kamilla Heddeland Instefjord, Hilde Elshaug, Marie Paulsen Madsen, Rasmus Riis Kopperud, Hilde Vollan, Karoline Bragstad, Olav Hungnes |
| EPI_ISL_668417, EPI_ISL_668420, EPI_ISL_668421 | Department of Medical Microbiology, St. Olavs hospital | Norwegian Institute of Public Health, Department of Virology | Kathrine Stene-Johansen, Kamilla Heddeland Instefjord, Hilde Elshaug, Marie Paulsen Madsen, Rasmus Riis Kopperud, Hilde Vollan, Karoline Bragstad, Olav Hungnes |
| EPI_ISL_668445 | Respiratory Virus Unit, Microbiology Services Colindale, Public Health England | COVID-19 Genomics UK (COG-UK) Consortium | PHE Covid Sequencing Team |
| EPI_ISL_670491, EPI_ISL_670501, EPI_ISL_670536, EPI_ISL_670537, EPI_ISL_670644, EPI_ISL_670645, EPI_ISL_670646, EPI_ISL_671219, EPI_ISL_671220, EPI_ISL_671221, EPI_ISL_671222, EPI_ISL_671223 | | | |
| see above | Department of Virus and Microbiological Special Diagnostics, Statens Serum Institut, Copenhagen, Denmark | Albertsen Lab, Department of Chemistry and Bioscience, Aalborg University, Denmark | Danish Covid-19 Genome Consortium |
| EPI_ISL_671415, EPI_ISL_671417 | National Laboratory of Virology, Szentágothai Research Centre | National Laboratory of Virology, Szentágothai Research Centre | Endre Gábor Tóth, Balázs Somogyi, Brigitta, Ferenc Jakab, Gábor Kemenesi |
| EPI_ISL_671673 | South Eastern Area Laboratory Services (SEALS) | NSW Health Pathology - Institute of Clinical Pathology and Medical Research; Westmead Hospital; University of Sydney | CIDM-PH et al. |
| EPI_ISL_671677 | DOHMH Corona | New York City Public Health Laboratory | Jade Wang, et al. |
| EPI_ISL_671678 | DOHMH Fort Greene | New York City Public Health Laboratory | Jade Wang, et al. |
| EPI_ISL_671679, EPI_ISL_671680 | DOHMH Morrisania | New York City Public Health Laboratory | Jade Wang, et al. |
| EPI_ISL_671681 | DOHMH Corona | New York City Public Health Laboratory | Jade Wang, et al. |
| EPI_ISL_671689 | DOHMH Central Harlem | New York City Public Health Laboratory | Jade Wang, et al. |
| EPI_ISL_671690 | DOHMH Crown Heights | New York City Public Health Laboratory | Jade Wang, et al. |
| EPI_ISL_671691 | DOHMH Corona | New York City Public Health Laboratory | Jade Wang, et al. |
| EPI_ISL_671692 | DOHMH Jamaica | New York City Public Health Laboratory | Jade Wang, et al. |
| EPI_ISL_671693 | DOHMH Chelsea | New York City Public Health Laboratory | Jade Wang, et al. |
| EPI_ISL_671694 | DOHMH Riverside | New York City Public Health Laboratory | Jade Wang, et al. |
| EPI_ISL_671695 | DOHMH Corona | New York City Public Health Laboratory | Jade Wang, et al. |
| EPI_ISL_671696 | DOHMH PHL | New York City Public Health Laboratory | Jade Wang, et al. |
| EPI_ISL_671697 | DOHMH Corona | New York City Public Health Laboratory | Jade Wang, et al. |
| EPI_ISL_671698 | DOHMH Jamaica | New York City Public Health Laboratory | Jade Wang, et al. |
| EPI_ISL_671699, EPI_ISL_671700 | DOHMH Corona | New York City Public Health Laboratory | Jade Wang, et al. |
| EPI_ISL_671701 | DOHMH Riverside | New York City Public Health Laboratory | Jade Wang, et al. |
| EPI_ISL_671702 | DOHMH Chelsea | New York City Public Health Laboratory | Jade Wang, et al. |
| EPI_ISL_671863 | Hospital Universitari i Politècnic la Fe, València, Spain | SeqCOVID-SPAIN consortium/IBV(CSIC) | Manuel Guerreiro, Marta Montero-Alonso, María Dolores Gómez and SeqCOVID-SPAIN consortium |
| EPI_ISL_672013, EPI_ISL_672035, EPI_ISL_672057, EPI_ISL_672069, EPI_ISL_672088, EPI_ISL_672292, EPI_ISL_672293, EPI_ISL_672294, EPI_ISL_672295, EPI_ISL_672296, EPI_ISL_672297 | | | |
| see above | Santa Clara County Public Health Laboratory | Chan-Zuckerberg Biohub | CZB Cliahub Consortium |
| EPI_ISL_672366, EPI_ISL_672367, EPI_ISL_672368, EPI_ISL_672369, EPI_ISL_672370 | Orange County Public Health Lab | Chan-Zuckerberg Biohub | CZB Cliahub Consortium |
| EPI_ISL_672525 | Madera County Department of Public Health | Chan-Zuckerberg Biohub | CZB Cliahub Consortium |
| EPI_ISL_677134, EPI_ISL_677135, EPI_ISL_677136, EPI_ISL_677137, EPI_ISL_677138, EPI_ISL_677139, EPI_ISL_677140, EPI_ISL_677141, EPI_ISL_677142, EPI_ISL_677144, EPI_ISL_677146, EPI_ISL_677179, EPI_ISL_677180, EPI_ISL_677181, EPI_ISL_677182, EPI_ISL_677183, EPI_ISL_677184, EPI_ISL_677185, EPI_ISL_677186, EPI_ISL_677187, EPI_ISL_677188, EPI_ISL_677189, EPI_ISL_677190, EPI_ISL_677191, EPI_ISL_677192, EPI_ISL_677193, EPI_ISL_677229, EPI_ISL_677230, EPI_ISL_677231, EPI_ISL_677232, EPI_ISL_677233, EPI_ISL_677236 | | | |
| see above | Virginia Division of Consolidated Laboratory Services | Virginia Division of Consolidated Laboratory Services | Virginia DCLS |
| EPI_ISL_677318 | Colorado Department of Public Health and Environment | Colorado Department of Puplic Health and Environment | Laura Bankers, Molly Hetherington-Rauth, Shannon Ely, Shannon R. Matzinger, Sarah Elizabeth Totten, Emily A. Travanty |
| EPI_ISL_677320, EPI_ISL_677477, EPI_ISL_677478, EPI_ISL_677484, EPI_ISL_677485, EPI_ISL_677586, EPI_ISL_677587, EPI_ISL_677588, EPI_ISL_677589, EPI_ISL_677590, EPI_ISL_677591, EPI_ISL_677592, EPI_ISL_677626, EPI_ISL_677663 | | | |
| see above | University of Wisconsin-Madison AIDS Vaccine Research Laboratories | University of Wisconsin-Madison AIDS Vaccine Research Laboratories | Gage Moreno, Katarina Braun, et al. AIDS Vaccine Research Laboratories |
| EPI_ISL_678380 | Area of Virology, Serology and Virology Division (SAViD), New South Wales Health Pathology Randwick | Virology Research Laboratory; Area of Virology, Serology and Virology Division (SAViD), New South Wales Health Pathology Randwick | Foster, C.; Au, J.; Ruiz Silva, M.; Deveson, I.; Bull, R.; Van Hal, S.; Rawlinson, W. |

| | | | |
|---|---|---|---|
| EPI_ISL_679924, EPI_ISL_679925 | Queens Medical Centre, Clinical Microbiology Department / DeepSeq Nottingham | COVID-19 Genomics UK (COG-UK) Consortium | Gemma Clark, Wendy Smith, Manjinder Khakh, Vicki M Fleming, Michelle M Lister, Hannah Howson-Wells, Jonathan Ball, Patrick McClure, Joseph Chappell, Theocharis Tsoleridis, Nadine Holmes, Matthew Carlisle, Christopher Moore, Fei Sang, Johnny Debebe, Victoria Wright, Matthew Loose |
| EPI_ISL_680016, EPI_ISL_680017, EPI_ISL_680018, EPI_ISL_680019, EPI_ISL_680020, EPI_ISL_680021, EPI_ISL_680022, EPI_ISL_680023, EPI_ISL_680024, EPI_ISL_680025, EPI_ISL_680026, EPI_ISL_680027, EPI_ISL_680028, EPI_ISL_680029, EPI_ISL_680030, EPI_ISL_680031, EPI_ISL_680032, EPI_ISL_680033, EPI_ISL_680034, EPI_ISL_680035, EPI_ISL_680036 | | | |
| see above | Centre for Enzyme Innovation, University of Portsmouth / Translational Research Laboratory, Portsmouth Hospitals NHS Trust | COVID-19 Genomics UK (COG-UK) Consortium | Angela Beckett,Yann Bourgeois,Garry Scarlett,Sharon Glaysher,Scott Elliott,Kelly Bicknell,Robert Impey,Allyson Lloyd,Sarah Wyllie,Ethan Butcher,Anoop Chauhan,Samuel Robson |
| EPI_ISL_680202, EPI_ISL_680203, EPI_ISL_680217, EPI_ISL_680330, EPI_ISL_680388, EPI_ISL_680389, EPI_ISL_680390, EPI_ISL_680391, EPI_ISL_680392, EPI_ISL_680393, EPI_ISL_680394, EPI_ISL_680395, EPI_ISL_680396, EPI_ISL_680397, EPI_ISL_680398, EPI_ISL_680399, EPI_ISL_680400, EPI_ISL_680401, EPI_ISL_680402, EPI_ISL_680403, EPI_ISL_680404, EPI_ISL_680405, EPI_ISL_680406, EPI_ISL_680407, EPI_ISL_680408 | | | |
| see above | Regional Virus Laboratory, Belfast Health and Social Care Trust | COVID-19 Genomics UK (COG-UK) Consortium | Conall McCaughey, James McKenna, Tanya Curran, Susan Feeney, Alison Watt, Ciara Cox, Mairead Connor, Zoltan Molnar, David Simpson, Derek Fairley |
| EPI_ISL_680485, EPI_ISL_680498, EPI_ISL_680499, EPI_ISL_680500, EPI_ISL_680501, EPI_ISL_680502 | Virology Department, Royal Infirmary of Edinburgh, NHS Lothian / School of Biological Sciences, University of Edinburgh / Institute of Genetics and Molecular Medicine, University of Edinburgh | COVID-19 Genomics UK (COG-UK) Consortium | McHugh M, Dewar R, Rooke S, Gallagher M, Balcaza C, O'Toole Á, Scher E, Hill V, McCrone JT, Colquhoun R, Yu X, Jackson B, Rambaut A, Williams TC, Templeton K |
| EPI_ISL_680562, EPI_ISL_680563, EPI_ISL_680564, EPI_ISL_680565, EPI_ISL_680566, EPI_ISL_680567, EPI_ISL_680568, EPI_ISL_680569, EPI_ISL_680570 | Centre for Enzyme Innovation, University of Portsmouth / Translational Research Laboratory, Portsmouth Hospitals NHS Trust | COVID-19 Genomics UK (COG-UK) Consortium | Angela Beckett,Yann Bourgeois,Garry Scarlett,Sharon Glaysher,Scott Elliott,Kelly Bicknell,Robert Impey,Allyson Lloyd,Sarah Wyllie,Ethan Butcher,Anoop Chauhan,Samuel Robson |
| EPI_ISL_682050, EPI_ISL_682052 | UPMC Clinical Microbiology Laboratory | Microbial Genomic Epidemiology Laboratory, University of Pittsburgh | Mustapha M. Mustapha, Jane W. Marsh, Dan Snyder, Marissa P. Griffith, Stephanie L. Mitchell, Vatsala R. Srinivasa, Kady D. Waggle, Chinelo Ezeonwuku, Vaughn S. Cooper, Lee H. Harrison |
| EPI_ISL_683002, EPI_ISL_683003, EPI_ISL_683204, EPI_ISL_683205, EPI_ISL_683206, EPI_ISL_683207, EPI_ISL_683208, EPI_ISL_683209, EPI_ISL_683210, EPI_ISL_683211, EPI_ISL_683235, EPI_ISL_683236, EPI_ISL_683237, EPI_ISL_683238, EPI_ISL_683239, EPI_ISL_683240, EPI_ISL_683241, EPI_ISL_683242, EPI_ISL_683243, EPI_ISL_683244, EPI_ISL_683245, EPI_ISL_683246, EPI_ISL_683247, EPI_ISL_683248, EPI_ISL_683249, EPI_ISL_683250, EPI_ISL_683251, EPI_ISL_683252, EPI_ISL_683253, EPI_ISL_683254, EPI_ISL_683261, EPI_ISL_683262, EPI_ISL_683263, EPI_ISL_683264, EPI_ISL_683265, EPI_ISL_683266, EPI_ISL_683272, EPI_ISL_683273, EPI_ISL_683274, EPI_ISL_683277, EPI_ISL_683278, EPI_ISL_683284, EPI_ISL_683285, EPI_ISL_683286, EPI_ISL_683287, EPI_ISL_683288, EPI_ISL_683289, EPI_ISL_683290, EPI_ISL_683291, EPI_ISL_683292, EPI_ISL_683293, EPI_ISL_683294, EPI_ISL_683295, EPI_ISL_683296, EPI_ISL_683297 | | | |
| see above | Department of Virus and Microbiological Special Diagnostics, Statens Serum Institut, Copenhagen, Denmark | Albertsen Lab, Department of Chemistry and Bioscience, Aalborg University, Denmark | Danish Covid-19 Genome Consortium |
| EPI_ISL_683726 | Mayo Clinic & Mayo Clinic Laboratories | Minnesota Department of Health, Public Health Laboratory | Alexandra Lorentz, Jacob Garfin, Matt Plumb, and Xiong Wang |
| EPI_ISL_692835, EPI_ISL_692836, EPI_ISL_692837, EPI_ISL_692838, EPI_ISL_692839, EPI_ISL_692840, EPI_ISL_692841, EPI_ISL_692842, EPI_ISL_692843, EPI_ISL_692846 | Massachusetts State Public Health Laboratory | Massachusetts State Public Health Laboratory | Andrew Lang, Timelia Fink, Glen Gallagher, Sandra Smole |
| EPI_ISL_700451, EPI_ISL_700452 | Pacaltsdorp Clinic wc PAC | NHLS/UCT | Houriiyah Tegally, Arash Iranzadeh, Deelan Doolabh, Lynn Tyers, Bruna Galvao, Innocent Mudau, Marvin Hsiao, Kruger Marais, Diana Hardie, Stephen Korsman, Carolyn Williamson |
| EPI_ISL_700469 | Knysna CDC wc WLC | NHLS/UCT | Houriiyah Tegally, Arash Iranzadeh, Deelan Doolabh, Lynn Tyers, Bruna Galvao, Innocent Mudau, Marvin Hsiao, Kruger Marais, Diana Hardie, Stephen Korsman, Carolyn Williamson |
| EPI_ISL_700476 | Alma CDC wc AHC | NHLS/UCT | Arash Iranzadeh, Deelan Doolabh, Lynn Tyers, Bruna Galvao, Innocent Mudau, Marvin Hsiao, Kruger Marais, Diana Hardie, Stephen Korsman, Carolyn Williamson |
| EPI_ISL_700478 | Pacaltsdorp Clinic wc PAC | NHLS/UCT | Houriiyah Tegally, Arash Iranzadeh, Deelan Doolabh, Lynn Tyers, Bruna Galvao, Innocent Mudau, Marvin Hsiao, Kruger Marais, Diana Hardie, Stephen Korsman, Carolyn Williamson |
| EPI_ISL_700484 | Knysna Hospital wc KNY | NHLS/UCT | Arash Iranzadeh, Deelan Doolabh, Lynn Tyers, Bruna Galvao, Innocent Mudau, Marvin Hsiao, Kruger Marais, Diana Hardie, Stephen Korsman, Carolyn Williamson |
| EPI_ISL_700510 | Knysna Hospital wc KNY | NHLS/UCT | Houriiyah Tegally, Arash Iranzadeh, Deelan Doolabh, Lynn Tyers, Bruna Galvao, Innocent Mudau, Marvin Hsiao, Kruger Marais, Diana Hardie, Stephen Korsman, Carolyn Williamson |
| EPI_ISL_700536, EPI_ISL_700556 | Conville CDC wc CVC | NHLS/UCT | Arash Iranzadeh, Deelan Doolabh, Lynn Tyers, Bruna Galvao, Innocent Mudau, Marvin Hsiao, Kruger Marais, Diana Hardie, Stephen Korsman, Carolyn Williamson |
| EPI_ISL_700703, EPI_ISL_700708 | Texas Department of State Health Services | Texas Department of State Health Services | Rashmi Tuladhar, Bonnie Oh, Jenny Zhang, Maliha Rahman, Anita Pokharel, Myong Koag, Chung Wang, Rachel Lee, Grace Kubin, Mayela Pedrueza, James Daniel Bonser |
| EPI_ISL_703144 | Virology Department, Royal Infirmary of Edinburgh, NHS Lothian / School of Biological Sciences, University of Edinburgh / Institute of Genetics and Molecular Medicine, University of Edinburgh | COVID-19 Genomics UK (COG-UK) Consortium | McHugh M, Dewar R, Rooke S, Gallagher M, Balcaza C, O'Toole Á, Scher E, Hill V, McCrone JT, Colquhoun R, Yu X, Jackson B, Rambaut A, Williams TC, Templeton K |
| EPI_ISL_703674 | Virology Department, Sheffield Teaching Hospitals NHS Foundation Trust/Department of Infection, Immunity and Cardiovascular Disease, The Medical School, University of Sheffield | COVID-19 Genomics UK (COG-UK) Consortium | Thushan de Silva, Matthew Parker, Nikki Smith, Adri Angyal, Rebecca Brown, Luke Green, Rachel Tucker, Paul Parsons, Danielle Groves, Katie Johnson, Laura Carrilero, Alex Keeley, Dave Partridge, Matthew Wyles, Benjamin Lindsey, Mehmet Yavuz, Mohammad Raza, Cariad Evans |
| EPI_ISL_704845, EPI_ISL_704866, EPI_ISL_704869 | Oxford Viromics, NDM, University of Oxford; Oxford University Hospitals; Basingstoke and North Hampshire Hospital | COVID-19 Genomics UK (COG-UK) Consortium | Tanya Golubchik, David Bonsall, George Macintyre, Amy Trebes, Mariateresa de Cesare, Catrin Moore, Alex Mobbs, Anita Justice, Robert Shaw, Monique Andersson, Timothy Peto, Emma Wise, Nathan Moore, Jessica Lynch, Nick Cortes, Matilde Mori, Stephen Kidd, David Buck, John Todd, Christophe Fraser |
| EPI_ISL_705225, EPI_ISL_705226, EPI_ISL_705302 | Northumbria University / South Tees Hospitals NHS Foundation Trust / North Cumbria Integrated Care NHS Foundation Trust / North Tees and Hartlepool NHS Foundation Trust / Newcastle Hospitals NHS Foundation Trust | COVID-19 Genomics UK (COG-UK) Consortium | Darren L Smith,Andrew Nelson,Matthew Bashton,Greg R Young,Joshua Loh,John Allan,Mohammad A Tariq,Giles S Holt,Gary Black,Wen C Yew,Lynn Dover,Paul Baker,Steve Liggett,Sarah Essex,Jane Greenaway,Debra Padgett,Clive Graham,Garren Scott,Edward Barton,Emma Swindells,Brendan Payne,Jennifer Collins,Yusri Taha,Gary Eltringham |
| EPI_ISL_705414, EPI_ISL_705417, EPI_ISL_705498 | Oxford Viromics, NDM, University of Oxford; Oxford University Hospitals; Basingstoke and North Hampshire Hospital | COVID-19 Genomics UK (COG-UK) Consortium | Tanya Golubchik, David Bonsall, George Macintyre, Amy Trebes, Mariateresa de Cesare, Catrin Moore, Alex Mobbs, Anita Justice, Robert Shaw, Monique Andersson, Timothy Peto, Emma Wise, Nathan Moore, Jessica Lynch, Nick Cortes, Matilde Mori, Stephen Kidd, David Buck, John Todd, Christophe Fraser |
| EPI_ISL_705537, EPI_ISL_705541 | Department of Pathology, University of Cambridge | COVID-19 Genomics UK (COG-UK) Consortium | Aminu S. Jahun, Yasmin Chaudhry, Grant Hall, Iliana Georgana, Myra Hosmillo, Martin D. Curran, Malte Pinckert, Surendra Parmar, Ian Goodfellow |
| EPI_ISL_705976, EPI_ISL_705977, EPI_ISL_705978, EPI_ISL_705979, EPI_ISL_705980, EPI_ISL_705981, EPI_ISL_705982, EPI_ISL_705983, EPI_ISL_705984, EPI_ISL_705985, EPI_ISL_705986, EPI_ISL_705987, EPI_ISL_705988, EPI_ISL_705989, EPI_ISL_705990, EPI_ISL_705991, EPI_ISL_705992, EPI_ISL_705993, EPI_ISL_705994, EPI_ISL_705995, EPI_ISL_705996, EPI_ISL_705997, EPI_ISL_705998, EPI_ISL_705999, EPI_ISL_706000, EPI_ISL_706001, EPI_ISL_706002, EPI_ISL_706003, EPI_ISL_706004, EPI_ISL_706005, EPI_ISL_706006, EPI_ISL_706007, EPI_ISL_706008, EPI_ISL_706009, EPI_ISL_706010, EPI_ISL_706011, EPI_ISL_706012 | | | |
| see above | Oxford Viromics, NDM, University of Oxford; Oxford | COVID-19 Genomics UK (COG-UK) Consortium | Tanya Golubchik, David Bonsall, George Macintyre, Amy Trebes, Mariateresa de Cesare, Catrin Moore, Alex Mobbs, Anita Justice, Robert Shaw, Monique |

| | | | |
|---|---|---|---|
| | University Hospitals; Basingstoke and North Hampshire Hospital | | Andersson, Timothy Peto, Emma Wise, Nathan Moore, Jessica Lynch, Nick Cortes, Matilde Mori, Stephen Kidd, David Buck, John Todd, Christophe Fraser |
| EPI_ISL_707789 | Rwanda National Reference Laboratory | Rwanda National Reference Laboratory | Enatha Mukantwari,Jeanne d'Arc |
| EPI_ISL_707895, EPI_ISL_707899 | Area of Virology, Serology and Virology Division (SAViD), New South Wales Health Pathology Randwick | Virology Research Laboratory; Area of Virology, Serology and Virology Division (SAViD), New South Wales Health Pathology Randwick | Foster, C.; Au, J.; Ruiz Silva, M.; Deveson, I.; Bull, R.; Van Hal, S.; Rawlinson, W. |
| EPI_ISL_707999, EPI_ISL_708000, EPI_ISL_708001, EPI_ISL_708002, EPI_ISL_708015 | Virology, Universitätsklinikum des Saarlandes | Epigenetics, Saarland University | Kathrin Kattler, Markus Vogelgesang, Stefan Lohse, Sascha Tierling, Sigrun Smola, Jörn Walter |
| EPI_ISL_708033 | Oslo University Hospital, Department of Medical Microbiology | Norwegian Institute of Public Health, Department of Virology | Kathrine Stene-Johansen, Kamilla Heddeland Instefjord, Hilde Elshaug, Marie Paulsen Madsen, Rasmus Riis Kopperud, Hilde Vollan, Karoline Bragstad, Olav Hungnes |
| EPI_ISL_708043 | Department of Medical Microbiology, St. Olavs hospital | Norwegian Institute of Public Health, Department of Virology | Kathrine Stene-Johansen, Kamilla Heddeland Instefjord, Hilde Elshaug, Marie Paulsen Madsen, Rasmus Riis Kopperud, Hilde Vollan, Karoline Bragstad, Olav Hungnes |
| EPI_ISL_708163 | Vestfold Hospital, Toensberg Department of Microbiology | Norwegian Institute of Public Health, Department of Virology | Kathrine Stene-Johansen, Kamilla Heddeland Instefjord, Hilde Elshaug, Marie Paulsen Madsen, Rasmus Riis Kopperud, Hilde Vollan, Karoline Bragstad, Olav Hungnes |
| EPI_ISL_708431 | Delaware Public Health Lab | Delaware Public Health Lab | Gregory Hovan |
| EPI_ISL_708804, EPI_ISL_708806 | Regional medical sciences center 6 chonburi | National Institute of Health, Department of Medical Sciences, Ministry of Public Health, Thailand | Pilailuk Okada; Siripaporn Phuygun; Thanutsapa Thanadachakul; Sittiporn Parnmen; Pakorn Piromtong; Warawan Wongboot; Sunthareeya Waicharoen; Malinee Chittaganpitch |
| EPI_ISL_709913, EPI_ISL_709914, EPI_ISL_709915, EPI_ISL_709919, EPI_ISL_709921, EPI_ISL_709924, EPI_ISL_709925, EPI_ISL_709926, EPI_ISL_709927, EPI_ISL_709928, EPI_ISL_709929, EPI_ISL_709930, EPI_ISL_709931 | | | |
| see above | Lighthouse Lab in Milton Keynes | Wellcome Sanger Institute for the COVID-19 Genomics UK (COG-UK) Consortium | The Lighthouse Lab in Milton Keynes and Alex Alderton, Roberto Amato, Sonia Goncalves, Ewan Harrison, David K. Jackson, Ian Johnston, Dominic Kwiatkowski, Cordelia Langford, John Sillitoe on behalf of the Wellcome Sanger Institute COVID-19 Surveillance Team |
| EPI_ISL_710155, EPI_ISL_710162, EPI_ISL_710174, EPI_ISL_710237, EPI_ISL_710238, EPI_ISL_710239, EPI_ISL_710269, EPI_ISL_710277, EPI_ISL_710355, EPI_ISL_710356, EPI_ISL_710357, EPI_ISL_710358, EPI_ISL_710359 | | | |
| see above | Colorado Department of Public Health and Environment | Colorado Department of Puplic Health and Environment | Laura Bankers, Molly C. Hetherington-Rauth, Shannon Ely, Shannon R. Matzinger, Sarah Elizabeth Totten, Emily A. Travanty |
| EPI_ISL_710503 | Laboratorio specialistico UOC Ematologia - Ospedale "San Francesco" - ATS-ASSL Nuoro | Laboratorio specialistico UOC Ematologia - Ospedale "San Francesco" - ATS-ASSL Nuoro | Giovanna Piras |
| EPI_ISL_710610 | Ektorps Vardcentral | The Public Health Agency of Sweden | Department of Microbiology, The Public Health Agency of Sweden |
| EPI_ISL_711057, EPI_ISL_712060 | Hôpital Fattouma-Bourguiba de Monastir | Laboratoire des Procédés de Criblage Moléculaire et Cellulaire-Centre de Biotechnologie de Sfax | Souissi,A., Abid,N., Ben Ayed,I., Gargouri,S., Abdelmoulah,F.,Elargoubi,A., Smeti,I., Bensaid,M., Stambouli,N., Kharat,N., Ajili,F., Fki-berrajah,L., Mhalla,S., Chtourou,A., Gaaloul,I., Nabli,A., Turki,M., Aouni,M., Hammami,A., Mastouri,M., Karray Hakim,H., Kamoun,S., Rebai,A. and Masmoudi,S. |
| EPI_ISL_714423, EPI_ISL_714424, EPI_ISL_714425, EPI_ISL_714426, EPI_ISL_714427, EPI_ISL_714428, EPI_ISL_714429, EPI_ISL_714430, EPI_ISL_714431, EPI_ISL_714432, EPI_ISL_714433, EPI_ISL_714434, EPI_ISL_714435, EPI_ISL_714436, EPI_ISL_714437, EPI_ISL_714438, EPI_ISL_714439, EPI_ISL_714440, EPI_ISL_714441, EPI_ISL_714442, EPI_ISL_714443, EPI_ISL_714928 | | | |
| see above | Department of Virus and Microbiological Special Diagnostics, Statens Serum Institut, Copenhagen, Denmark | Albertsen Lab, Department of Chemistry and Bioscience, Aalborg University, Denmark | Danish Covid-19 Genome Consortium |
| EPI_ISL_717574, EPI_ISL_717576 | Lab voor klinische biologie | Onderzoeksgroep Virologie | Laurens Lambrechts, Nick Vereecke, Marthe Pauwels, Bruno Verhasselt, Linos Vandekerckhove, Hans Nauwynck, Sebastiaan Theuns |
| EPI_ISL_717580 | Lab voor klinische biologie | Onderzoeksgroep Virologie | Nick Vereecke, Laurens Lambrechts, Marthe Pauwels, Bruno Verhasselt, Linos Vandekerckhove, Hans Nauwynck, Sebastiaan Theuns |
| EPI_ISL_717593, EPI_ISL_717595, EPI_ISL_717598 | Lab voor klinische biologie | Onderzoeksgroep Virologie | Laurens Lambrechts, Nick Vereecke, Marthe Pauwels, Bruno Verhasselt, Linos Vandekerckhove, Hans Nauwynck, Sebastiaan Theuns |
| EPI_ISL_717602, EPI_ISL_717604, EPI_ISL_717605 | Lab voor klinische biologie | Onderzoeksgroep Virologie | Nick Vereecke, Laurens Lambrechts, Marthe Pauwels, Bruno Verhasselt, Linos Vandekerckhove, Hans Nauwynck, Sebastiaan Theuns |
| EPI_ISL_717607, EPI_ISL_717608, EPI_ISL_717609, EPI_ISL_717616 | Lab voor klinische biologie | Onderzoeksgroep Virologie | Laurens Lambrechts, Nick Vereecke, Marthe Pauwels, Bruno Verhasselt, Linos Vandekerckhove, Hans Nauwynck, Sebastiaan Theuns |
| EPI_ISL_717617, EPI_ISL_717618, EPI_ISL_717619 | Lab voor klinische biologie | Onderzoeksgroep Virologie | Nick Vereecke, Laurens Lambrechts, Marthe Pauwels, Bruno Verhasselt, Linos Vandekerckhove, Hans Nauwynck, Sebastiaan Theuns |
| EPI_ISL_721542, EPI_ISL_721543, EPI_ISL_721545, EPI_ISL_721546, EPI_ISL_721547, EPI_ISL_721548, EPI_ISL_721549, EPI_ISL_721550, EPI_ISL_721551, EPI_ISL_721552, EPI_ISL_721553, EPI_ISL_721554, EPI_ISL_721555, EPI_ISL_721556, EPI_ISL_721557, EPI_ISL_721558, EPI_ISL_721559, EPI_ISL_721560, EPI_ISL_721561, EPI_ISL_721562, EPI_ISL_721563, EPI_ISL_721564, EPI_ISL_721565, EPI_ISL_721566, EPI_ISL_721567 | | | |
| see above | Lighthouse Lab in Glasgow | Wellcome Sanger Institute for the COVID-19 Genomics UK (COG-UK) Consortium | Harper VanSteenhouse, Yumi Kasai, David Gray, Carol Clugston, Anna Dominiczak and Alex Alderton, Roberto Amato, Sonia Goncalves, Ewan Harrison, David K. Jackson, Ian Johnston, Dominic Kwiatkowski, Cordelia Langford, John Sillitoe on behalf of the Wellcome Sanger Institute COVID-19 Surveillance Team |
| EPI_ISL_722273, EPI_ISL_722296, EPI_ISL_722308, EPI_ISL_722348, EPI_ISL_722373, EPI_ISL_722398 | Dutch COVID-19 response team | Erasmus Medical Center | Bas Oude Munnink, Reina Sikkema, David Nieuwenhuijse, Irina Chestakova, Anne van der Linden, Marjan Boter, Emmanuelle Munger, Corine GeurtsvanKessel, Annemiek van der Eijk, Richard Molenkamp, Marion Koopmans, on behalf of the Dutch national COVID-19 response team. |
| EPI_ISL_722414, EPI_ISL_722415, EPI_ISL_722416, EPI_ISL_722417, EPI_ISL_722418 | Dutch COVID-19 response team | Erasmus Medical Center | OH consortium |
| EPI_ISL_722514, EPI_ISL_722515, EPI_ISL_722516, EPI_ISL_722518, EPI_ISL_722519, EPI_ISL_722520, EPI_ISL_722521, EPI_ISL_722522, EPI_ISL_722523, EPI_ISL_722524, EPI_ISL_722525, EPI_ISL_722526, EPI_ISL_722527 | | | |
| see above | Dutch COVID-19 response team | Erasmus Medical Center | Bas Oude Munnink, Reina Sikkema, David Nieuwenhuijse, Irina Chestakova, Anne van der Linden, Marjan Boter, Emmanuelle Munger, Corine GeurtsvanKessel, Annemiek van der Eijk, Richard Molenkamp, Marion Koopmans, on behalf of the Dutch national COVID-19 response team. |
| EPI_ISL_722885, EPI_ISL_722886, EPI_ISL_722907, EPI_ISL_722908, EPI_ISL_722921, EPI_ISL_722924, EPI_ISL_722925 | Istituto Zooprofilattico Sperimentale della Puglia e della Basilicata | Istituto Zooprofilattico Sperimentale della Puglia e della Basilicata | Parisi A., Bianco A., Capozzi L., Del Sambro L., Manzulli V, Rondinone V., Pace L., Cipolletta D., Galante D. |
| EPI_ISL_724975 | Northumbria University / South Tees Hospitals NHS Foundation Trust / North Cumbria Integrated Care NHS Foundation Trust / North Tees and Hartlepool NHS Foundation Trust / Newcastle Hospitals NHS Foundation Trust | COVID-19 Genomics UK (COG-UK) Consortium | Darren L Smith,Andrew Nelson,Matthew Bashton,Greg R Young,Joshua Loh,John Allan,Mohammad A Tariq,Giles S Holt,Gary Black,Wen C Yew,Lynn Dover,Paul Baker,Steve Liggett,Sarah Essex,Jane Greenaway,Debra Padgett,Clive Graham,Garren Scott,Edward Barton,Emma Swindells,Brendan Payne,Jennifer Collins,Yusri Taha,Gary Eltringham |
| EPI_ISL_727710, EPI_ISL_727711, EPI_ISL_727774, EPI_ISL_727775, EPI_ISL_727776, EPI_ISL_727777, EPI_ISL_727778, EPI_ISL_727779, EPI_ISL_727780, EPI_ISL_727781, EPI_ISL_727782, EPI_ISL_727783, EPI_ISL_727784, EPI_ISL_727785, EPI_ISL_727786, EPI_ISL_727787, EPI_ISL_727788, EPI_ISL_727789, EPI_ISL_727790, EPI_ISL_727791, EPI_ISL_727792, EPI_ISL_727793, EPI_ISL_727794, EPI_ISL_727795, EPI_ISL_727796, EPI_ISL_727797, EPI_ISL_727798, EPI_ISL_727799, EPI_ISL_727800, EPI_ISL_727801, EPI_ISL_727802, EPI_ISL_727803, EPI_ISL_727804, EPI_ISL_727805, EPI_ISL_727806 | | | |
| see above | Centre for Enzyme Innovation, University of Portsmouth / Translational Research Laboratory, Portsmouth Hospitals NHS Trust | COVID-19 Genomics UK (COG-UK) Consortium | Angela Beckett,Yann Bourgeois,Garry Scarlett,Sharon Glaysher,Scott Elliott,Kelly Bicknell,Robert Impey,Allyson Lloyd,Sarah Wyllie,Ethan Butcher,Anoop Chauhan,Samuel Robson |
| EPI_ISL_728051 | University of Wisconsin-Madison AIDS Vaccine Research | University of Wisconsin-Madison AIDS Vaccine Research | Gage Moreno, Katarina Braun, et al. AIDS Vaccine Research Laboratories |

| | | | |
|---|---|---|---|
| | Laboratories | Laboratories | |
| EPI_ISL_730170 | San Diego County Public Health Laboratory | Andersen lab at Scripps Research | SEARCH Alliance San Diego with Tracy Basler, Jovan Shephard, Brett Austin |
| EPI_ISL_730376, EPI_ISL_730378, EPI_ISL_730389, EPI_ISL_730399, EPI_ISL_730403, EPI_ISL_730417, EPI_ISL_730422, EPI_ISL_730428, EPI_ISL_730431, EPI_ISL_730434, EPI_ISL_730442, EPI_ISL_730446, EPI_ISL_730452, EPI_ISL_730458, EPI_ISL_730471 | | | |
| see above | Biolab Diagnostic Laboratories | Andersen lab at Scripps Research | Issa Abu-Dayyeh, Ahmad Tibi, Lama Hussein, Lina Mohammad, Zein Naber, Amid Abdelnour with SEARCH Alliance San Diego |
| EPI_ISL_730616 | United Christian Hospital | Hong Kong Department of Health | Mak Gannon C.K., Lam Edman T.K., Chan Rickjason C.W., Tsang Dominic N.C. |
| EPI_ISL_732396, EPI_ISL_732489 | National Virus Reference Laboratory | National Virus Reference Laboratory | Michael Carr, Gabriel Gonzalez, Jonathan Dean, Daniel Hare, Cillian F De Gascun |
| EPI_ISL_732795 | Centro de Investigación Biomédica de La Rioja - Hospital San Pedro Logroño | SeqCOVID-SPAIN consortium/IBV(CSIC) | María de Toro, José Manuel Azcona Gutiérrez, María Pilar Bea Escudero, Miriam Blasco Alberdi and SeqCOVID-SPAIN consortium |
| EPI_ISL_733240 | UMMC-Health | WHO National Influenza Centre Russian Federation | Andrey Komissarov, Artem Fadeev, Anna Ivanova, Kseniya Komissarova, Dmitry Bazhenov, Tatiana Platonova, Daria Danilenko, Ksenia Safina, Elena Nabieva, Georgii Bazykin, Dmitry Lioznov |
| EPI_ISL_733254, EPI_ISL_733255, EPI_ISL_733256, EPI_ISL_733277, EPI_ISL_733278 | WHO National Influenza Centre Russian Federation | WHO National Influenza Centre Russian Federation | Andrey Komissarov, Artem Fadeev, Anna Ivanova, Kseniya Komissarova, Dmitry Bazhenov, Daria Danilenko, Ksenia Safina, Elena Nabieva, Georgii Bazykin, Dmitry Lioznov |
| EPI_ISL_734296, EPI_ISL_734297, EPI_ISL_734298, EPI_ISL_734299, EPI_ISL_734300, EPI_ISL_734301, EPI_ISL_734302, EPI_ISL_734369, EPI_ISL_734370, EPI_ISL_734371, EPI_ISL_734372, EPI_ISL_734373, EPI_ISL_734374, EPI_ISL_734375, EPI_ISL_734376, EPI_ISL_734377, EPI_ISL_734378, EPI_ISL_734379, EPI_ISL_734380, EPI_ISL_734381, EPI_ISL_734382, EPI_ISL_734383, EPI_ISL_734384, EPI_ISL_734385, EPI_ISL_734386, EPI_ISL_734387 | | | |
| see above | Wadsworth Center, New York State Department.of Health | Wadsworth Center, New York State Department.of Health | Kirsten St. George, Daryl M. Lamson, Alexis Russel, Jonathan Plitnick, Navjot Singh, John Kelly, Sara Griesemer, Erasmus Schneider, Erica Lasek-Nesselquist |
| EPI_ISL_734453, EPI_ISL_734454, EPI_ISL_734455, EPI_ISL_734456, EPI_ISL_734457, EPI_ISL_734458, EPI_ISL_734459, EPI_ISL_734461 | Masonic Medical Research Institute | Wadsworth Center, New York State Department.of Health | Kirsten St. George, Nathan Tucker, Ryan D. Pfeiffer, Daryl M. Lamson, Alexis Russel, Jonathan Plitnick, Navjot Singh, John Kelly, Sara Griesemer, Erasmus Schneider, Erica Lasek-Nesselquist |
| EPI_ISL_734482, EPI_ISL_734484, EPI_ISL_734490, EPI_ISL_734491, EPI_ISL_734496, EPI_ISL_734498, EPI_ISL_734499, EPI_ISL_734501, EPI_ISL_734502, EPI_ISL_735140, EPI_ISL_735141, EPI_ISL_735142, EPI_ISL_735143, EPI_ISL_735144, EPI_ISL_735145, EPI_ISL_735146, EPI_ISL_735147, EPI_ISL_735148, EPI_ISL_735149, EPI_ISL_735150, EPI_ISL_735151, EPI_ISL_735152, EPI_ISL_735153, EPI_ISL_735154, EPI_ISL_735155, EPI_ISL_735156, EPI_ISL_735157, EPI_ISL_735158, EPI_ISL_735159, EPI_ISL_735160, EPI_ISL_735162, EPI_ISL_735163, EPI_ISL_735164, EPI_ISL_735165, EPI_ISL_735166, EPI_ISL_735167, EPI_ISL_735168, EPI_ISL_735169, EPI_ISL_735170, EPI_ISL_735171, EPI_ISL_735172, EPI_ISL_735173, EPI_ISL_735174, EPI_ISL_735175, EPI_ISL_735176, EPI_ISL_735177, EPI_ISL_735178, EPI_ISL_735179, EPI_ISL_735180, EPI_ISL_735181, EPI_ISL_735182, EPI_ISL_735183, EPI_ISL_735184, EPI_ISL_735185, EPI_ISL_735186, EPI_ISL_735187, EPI_ISL_735188, EPI_ISL_735189, EPI_ISL_735190, EPI_ISL_735191, EPI_ISL_735192, EPI_ISL_735193, EPI_ISL_735194, EPI_ISL_735195, EPI_ISL_735196, EPI_ISL_735197, EPI_ISL_735198, EPI_ISL_735199, EPI_ISL_735200, EPI_ISL_735201, EPI_ISL_735202, EPI_ISL_735203, EPI_ISL_735204, EPI_ISL_735205, EPI_ISL_735206, EPI_ISL_735207, EPI_ISL_735208, EPI_ISL_735209, EPI_ISL_735210, EPI_ISL_735211, EPI_ISL_735212, EPI_ISL_735213, EPI_ISL_735214 | | | |
| see above | UZ Leuven, National Reference Laboratory for Coronaviruses, Laboratory Medicine, Leuven, Belgium | KU Leuven, Rega Institute, Clinical and Epidemiological Virology | Tony Wawina-Bokalanga, Joan Marti-Carerras, Bert Vanmechelen, Piet Maes |
| EPI_ISL_735444, EPI_ISL_735448 | Nucleic Acid Testing - Rwanda National Reference Laboratory | GIGA Medical Genomics | Yvan Butera, Keith Durkin, Maria Artesi, Bouchra Boujemla, Robert Rutayisire, Patrick Tuyisenge, Esperence Umumararungu, Sébastien Bontems, Marie-Pierre Hayette, Swaibu Gatare, Jacob Souopgui, Sabin Nsanzimana, Vincent Bours, Léon Mutesa |
| EPI_ISL_735488, EPI_ISL_737081, EPI_ISL_737082, EPI_ISL_737083, EPI_ISL_737084, EPI_ISL_737085, EPI_ISL_737086, EPI_ISL_737087 | UW Virology Lab | UW Virology Lab | Pavitra Roychoudhury, Hong Xie, Lasata Shrestha, Meei-Li Huang, Keith R Jerome, Alexander Greninger |
| EPI_ISL_738195 | UZ Leuven, National Reference Laboratory for Coronaviruses, Laboratory Medicine, Leuven, Belgium | KU Leuven, Rega Institute, Clinical and Epidemiological Virology | Tony Wawina-Bokalanga, Joan Marti-Carerras, Bert Vanmechelen, Piet Maes |
| EPI_ISL_738756 | Humboldt County Public Health Laboratory | Chan-Zuckerberg Biohub | CZB Cliahub Consortium |
| EPI_ISL_739674, EPI_ISL_739675, EPI_ISL_739676, EPI_ISL_739677 | Instituto Nacional de Salud, Bogotá, Colombia | Instituto Nacional de Salud, Bogotá, Colombia | Katherine Laiton-Donato, Diego A. Álvarez-Díaz, Carlos Franco-Muñoz, Mauricio Pacheco-Montealegre, Jonathan Reales, Diego Andrés Prada, Sheryl Corchuelo, Magdalena Weisner, Martha Lucia Ospina Martinez, Marcela Mercado-Reyes |
| EPI_ISL_739726, EPI_ISL_739729, EPI_ISL_739749, EPI_ISL_739766, EPI_ISL_739778, EPI_ISL_739781, EPI_ISL_739782, EPI_ISL_739789, EPI_ISL_739790, EPI_ISL_739804, EPI_ISL_739841, EPI_ISL_739856, EPI_ISL_739862, EPI_ISL_739867, EPI_ISL_739868, EPI_ISL_739877, EPI_ISL_739902, EPI_ISL_739919, EPI_ISL_739958, EPI_ISL_739987, EPI_ISL_740014, EPI_ISL_740049, EPI_ISL_740064, EPI_ISL_740070, EPI_ISL_740086, EPI_ISL_740090, EPI_ISL_740108, EPI_ISL_740138, EPI_ISL_740161, EPI_ISL_740191, EPI_ISL_740202, EPI_ISL_740210, EPI_ISL_740219, EPI_ISL_740220, EPI_ISL_740256, EPI_ISL_740279, EPI_ISL_740288, EPI_ISL_740316, EPI_ISL_740317, EPI_ISL_740327, EPI_ISL_740331, EPI_ISL_740333, EPI_ISL_740336, EPI_ISL_740342, EPI_ISL_740357, EPI_ISL_740359, EPI_ISL_740370, EPI_ISL_740406, EPI_ISL_740408, EPI_ISL_740409, EPI_ISL_740464, EPI_ISL_740473, EPI_ISL_740492, EPI_ISL_740500, EPI_ISL_740509, EPI_ISL_740521 | | | |
| see above | Laboratoire national de santé, Microbiology, Virology | Laboratoire national de santé, Microbiology, Microbial Genomics Platform | Anke Wienecke-Baldacchino, Catherine Ragimbeau,Jessica Tapp, Fatu Djabi, Lise Pignon, Raoul Salmon, Tamir Abdelrahman |
| EPI_ISL_741772 | Centre for Enzyme Innovation, University of Portsmouth / Translational Research Laboratory, Portsmouth Hospitals NHS Trust | COVID-19 Genomics UK (COG-UK) Consortium | Angela Beckett,Yann Bourgeois,Garry Scarlett,Sharon Glaysher,Scott Elliott,Kelly Bicknell,Robert Impey,Allyson Lloyd,Sarah Wyllie,Ethan Butcher,Anoop Chauhan,Samuel Robson |
| EPI_ISL_741773, EPI_ISL_741774, EPI_ISL_741775, EPI_ISL_741776 | Oxford Viromics, NDM, University of Oxford; Oxford University Hospitals; Basingstoke and North Hampshire Hospital | COVID-19 Genomics UK (COG-UK) Consortium | Tanya Golubchik, David Bonsall, George Macintyre, Amy Trebes, Mariateresa de Cesare, Catrin Moore, Alex Mobbs, Anita Justice, Robert Shaw, Monique Andersson, Timothy Peto, Emma Wise, Nathan Moore, Jessica Lynch, Nick Cortes, Matilde Mori, Stephen Kidd, David Buck, John Todd, Christophe Fraser |
| EPI_ISL_744135, EPI_ISL_744153, EPI_ISL_744157, EPI_ISL_744161, EPI_ISL_744174, EPI_ISL_744175, EPI_ISL_744181, EPI_ISL_744224, EPI_ISL_744246, EPI_ISL_744275, EPI_ISL_744296, EPI_ISL_744364, EPI_ISL_744380, EPI_ISL_744388, EPI_ISL_744391, EPI_ISL_744392, EPI_ISL_744411, EPI_ISL_744416, EPI_ISL_744423, EPI_ISL_744458, EPI_ISL_744463, EPI_ISL_744464, EPI_ISL_744474, EPI_ISL_744475, EPI_ISL_744481, EPI_ISL_744502, EPI_ISL_744506, EPI_ISL_744516, EPI_ISL_744538, EPI_ISL_744551, EPI_ISL_744555, EPI_ISL_744574, EPI_ISL_744578, EPI_ISL_744579, EPI_ISL_744591, EPI_ISL_744595, EPI_ISL_744601, EPI_ISL_744609, EPI_ISL_744612, EPI_ISL_744646, EPI_ISL_744660, EPI_ISL_744679, EPI_ISL_744682, EPI_ISL_744690, EPI_ISL_744693, EPI_ISL_744700, EPI_ISL_744703, EPI_ISL_744716, EPI_ISL_744738, EPI_ISL_744741, EPI_ISL_744754, EPI_ISL_744755, EPI_ISL_744774, EPI_ISL_744784, EPI_ISL_744785, EPI_ISL_744790, EPI_ISL_744814, EPI_ISL_744816, EPI_ISL_744824, EPI_ISL_744842, EPI_ISL_744859, EPI_ISL_744866, EPI_ISL_744870, EPI_ISL_744917, EPI_ISL_744925, EPI_ISL_744928, EPI_ISL_744942, EPI_ISL_744951, EPI_ISL_744971, EPI_ISL_744990, EPI_ISL_745002, EPI_ISL_745004, EPI_ISL_745015, EPI_ISL_745019 | | | |
| see above | Laboratoire national de santé, Microbiology, Virology | Laboratoire national de santé, Microbiology, Microbial Genomics Platform | Anke Wienecke-Baldacchino, Catherine Ragimbeau,Jessica Tapp, Fatu Djabi, Lise Pignon, Raoul Salmon, Tamir Abdelrahman |
| EPI_ISL_745520, EPI_ISL_745590, EPI_ISL_745648, EPI_ISL_746205, EPI_ISL_746209, EPI_ISL_746212, EPI_ISL_746216, EPI_ISL_746217, EPI_ISL_746230, EPI_ISL_746232, EPI_ISL_746236, EPI_ISL_746238, EPI_ISL_746240, EPI_ISL_746245, EPI_ISL_746252, EPI_ISL_746257, EPI_ISL_746260, EPI_ISL_746262, EPI_ISL_746271, EPI_ISL_746277, EPI_ISL_746279, EPI_ISL_746286, EPI_ISL_746291, EPI_ISL_746300, EPI_ISL_746301 | | | |
| see above | Ginkgo Bioworks Clinical Laboratory | Utah Public Health Laboratory | Erin L. Young, Kelly Oakeson, Tara Gallagher, Michael T. Pyne, E. Susan Slechta, Melanie A. Mallory, Jeffrey B. Stevenson, Salika M. Shakir, David R. Hillyard, Malaika McKenzie-Bennett, James McGann, Jim Griffin, Keith Robison, Alex Plocik, Becky Schilling, Martha Pierson, Rebecca Littlefield, Michelle Spencer, Birgitte Simen |
| EPI_ISL_747036, EPI_ISL_747037, EPI_ISL_747038, EPI_ISL_747115 | Respiratory Viruses Branch, Centers for Disease Control and Prevention | Respiratory Viruses Branch, Centers for Disease Control and Prevention | Queen,K., Li,Y., Tao,Y., Uehara,A., Montmayeur,A., Paden,C.R., Cook,P.W., Marine,R., Sheth,M., Wang,H., Lee,J., Tong,S. |
| EPI_ISL_747235 | Sukabumi Health Laboratory | West Java Health Laboratory; School of Life Sciences and Technology, Institut Teknologi Bandung | Azzania Fibriani, Ema Rahmawati, Ryan Bayusantika Ristandi, Rifky Waluyajati Rachman, Cut Nur Cinthia Alamanda, Isak Solihin, Rini Robiani, Miftahul Faridl, Karimatu Khoirunnisa |
| EPI_ISL_747379, EPI_ISL_747380, EPI_ISL_747381, EPI_ISL_747382 | Division of Emerging Infectious Diseases, Bureau of Infectious Diseases Diagnosis Control, Korea Disease Control and Prevention Agency | Division of Emerging Infectious Diseases, Bureau of Infectious Diseases Diagnosis Control, Korea Disease Control and Prevention Agency | Ae Kyung Park, Il-Hwan Kim, Heui Man Kim, Jeong-Min Kim, Namjoo Lee, Chaeyoung Lee, Sang Hee Woo, Eun-Jin Kim |
| EPI_ISL_748125, EPI_ISL_748126, EPI_ISL_748127, EPI_ISL_748128 | Department of Virus and Microbiological Special Diagnostics, Statens Serum Institut, Copenhagen, Denmark | Albertsen Lab, Department of Chemistry and Bioscience, Aalborg University, Denmark | Danish Covid-19 Genome Consortium |

| | | | |
|---|---|---|---|
| EPI_ISL_751488 | CHU Purpan - Laboratoire de Virologie - Institut Fédératif de Biologie | CHU Purpan - Laboratoire de Virologie - Institut Fédératif de Biologie | Latour J., Ranger N., Dubois M., Carcenac R., Harter A., Boyer P., Tremeaux P., Izopet J. |
| EPI_ISL_751543 | DE Public Health Laboratory | Genomics and Discovery, Respiratory Viruses Branch, Division of Viral Diseases, Centers for Disease Control and Prevention | Krista Queen, Yan Li, Ying Tao, Jing Zhang, Anna Uehara, Anna Montmayeur, Clinton R. Paden, Peter W. Cook,Rachel Marine, Mili Sheth, Haibin Wang, Justin Lee, Suxiang Tong |
| EPI_ISL_751622 | MI - Michigan Department of Health and Human Services - Bureau of Laboratories | Genomics and Discovery, Respiratory Viruses Branch, Division of Viral Diseases, Centers for Disease Control and Prevention | Krista Queen, Yan Li, Ying Tao, Jing Zhang, Anna Uehara, Anna Montmayeur, Clinton R. Paden, Peter W. Cook,Rachel Marine, Mili Sheth, Haibin Wang, Justin Lee, Suxiang Tong |
| EPI_ISL_751726 | DE Public Health Laboratory | Genomics and Discovery, Respiratory Viruses Branch, Division of Viral Diseases, Centers for Disease Control and Prevention | Krista Queen, Yan Li, Ying Tao, Jing Zhang, Anna Uehara, Anna Montmayeur, Clinton R. Paden, Peter W. Cook,Rachel Marine, Mili Sheth, Haibin Wang, Justin Lee, Suxiang Tong |
| EPI_ISL_753017, EPI_ISL_753018 | State Laboratories Division, Hawaii State Department of Health | State Laboratories Division, Hawaii State Department of Health | Pamela O'Brien, Sabrina Diemert, Drew Kuwazaki, Razvan Sultana, Edward Desmond |
| EPI_ISL_753790, EPI_ISL_753796, EPI_ISL_753804, EPI_ISL_753894, EPI_ISL_753902, EPI_ISL_754194 | Charité Universitätsmedizin Berlin, Institut für Virologie/Labor Berlin | Charité Universitätsmedizin Berlin, Institut für Virologie | Victor M Corman, Jörn Beheim-Schwarzbach, Barbara Mühlemann, Julia Schneider, Talitha Veith, Terry Jones, Christian Drosten |
| EPI_ISL_754691, EPI_ISL_754692, EPI_ISL_754693, EPI_ISL_754694, EPI_ISL_754713, EPI_ISL_754714, EPI_ISL_754715, EPI_ISL_754716, EPI_ISL_754717, EPI_ISL_754718, EPI_ISL_754719, EPI_ISL_754720, EPI_ISL_754721, EPI_ISL_754722, EPI_ISL_754723, EPI_ISL_754724, EPI_ISL_754725, EPI_ISL_754726, EPI_ISL_754727, EPI_ISL_754728, EPI_ISL_754729, EPI_ISL_754730, EPI_ISL_754731, EPI_ISL_754732, EPI_ISL_754734, EPI_ISL_754735, EPI_ISL_754736, EPI_ISL_754737, EPI_ISL_754738, EPI_ISL_754739, EPI_ISL_754740, EPI_ISL_754741, EPI_ISL_754742, EPI_ISL_754743, EPI_ISL_754779, EPI_ISL_754780, EPI_ISL_754801, EPI_ISL_754802, EPI_ISL_754803, EPI_ISL_754804, EPI_ISL_754805, EPI_ISL_754806, EPI_ISL_754807, EPI_ISL_754808, EPI_ISL_754809, EPI_ISL_754810, EPI_ISL_754811, EPI_ISL_754812, EPI_ISL_754813, EPI_ISL_754814, EPI_ISL_754815, EPI_ISL_754816, EPI_ISL_754817, EPI_ISL_754818, EPI_ISL_754819, EPI_ISL_754820, EPI_ISL_754821, EPI_ISL_754822, EPI_ISL_754823, EPI_ISL_754824, EPI_ISL_754825, EPI_ISL_754826, EPI_ISL_754827, EPI_ISL_754828, EPI_ISL_754829, EPI_ISL_754830, EPI_ISL_754831, EPI_ISL_754832, EPI_ISL_754833, EPI_ISL_754834, EPI_ISL_754835, EPI_ISL_754836, EPI_ISL_754837, EPI_ISL_754838, EPI_ISL_754839 | | | |
| see above | Wadsworth Center, New York State Department.of Health | Wadsworth Center, New York State Department.of Health | Kirsten St. George, Daryl M. Lamson, Alexis Russel, Matthew Shudt, Melissa A Leisner, Jonathan Plitnick, Navjot Singh, John Kelly, Sara Griesemer, Erasmus Schneider, Erica Lasek-Nesselquist |
| EPI_ISL_755137, EPI_ISL_755197 | UCSD EXCITE lab | Andersen lab at Scripps Research | SEARCH Alliance San Diego |
| EPI_ISL_755619 | Canterbury Health Laboratories | Institute of Environmental Science and Research (ESR) | Xiaoyun Ren, Matt Storey, Nikki Freed, Muhammad Faisal, Jing Wang, Hermes Perez, Anja Werno, Antje van der Linden, Arlo Upton, Chris Mansell, David Hammer, Dragana Drinkovic, Gary McAuliffe, Hana Sofia Andersson, James Ussher, Jill Sherwood, Josh Freeman, Julia Howard, Juliet Elvy, Mary DeAlmeida, Matt Blakiston, Matthew Rogers, Max Bloomfield, Michael Addidle, Michelle Balm, Sally Roberts, Sarah Jefferies, Sharmini Muttaiyah, Susan Morpeth, Susan Taylor, Timothy Blackmore, Vani Sathyendran, Veronica Playle, Virginia Hope, Erasmus Smit, Lauren Jelly, Olin Silander, Joep de Ligt |
| EPI_ISL_755742, EPI_ISL_755743, EPI_ISL_755744, EPI_ISL_755749 | Toronto Invasive Bacterial Diseases Network | McMaster University | Allison McGeer, Patryk Aftanas, Hooman Derakhshani, Angel Li, Kuganya Nirmalarajah, Emily Panousis, Ahmed Draia, Jalees Nasir, Michael Surette, Samira Mubareka, Andrew G. McArthur |
| EPI_ISL_756377 | Lighthouse Lab in Milton Keynes | Wellcome Sanger Institute for the COVID-19 Genomics UK (COG-UK) Consortium | The Lighthouse Lab in Milton Keynes and Alex Alderton, Roberto Amato, Sonia Goncalves, Ewan Harrison, David K. Jackson, Ian Johnston, Dominic Kwiatkowski, Cordelia Langford, John Sillitoe on behalf of the Wellcome Sanger Institute COVID-19 Surveillance Team |
| EPI_ISL_759703 | RSUD Bung Karno | Universitas Sebelas Maret (UNS), Rumah sakit Universitas Sebelas Maret (RS-UNS) Surakarta, National Institute of Health Research and Development, Indonesian Ministry of Health | Betty Suryawati, Yulia sari, hartono, maryani, Afif Avicena Gufron, Hana Apsari Pawestri, Kartika, Dewi Puspa, Hartanti Dian Ika, Arie Ardiansyah Nugraha, Vivi Setiawaty |
| EPI_ISL_759842 | Rumah Sakit Universitas Sebelas Maret (RS-UNS) | Universitas Sebelas Maret (UNS); Rumah Sakit Universitas Sebelas Maret (RS-UNS), Surakarta; National Institute of Health Research and Development, Indonesian Ministry of Health, Jakarta. | Betty Suryawati, Yulia Sari, Hartono, Maryani, Afif Avicena Ghufron, Hana Apsari Pawestri, Kartika, Dewi Puspa, Hartanti Dian Ika, Arie Ardiansyah Nugraha, Vivi Setiawaty. |
| EPI_ISL_760066, EPI_ISL_760106, EPI_ISL_760107, EPI_ISL_760108, EPI_ISL_760109, EPI_ISL_760110, EPI_ISL_760111, EPI_ISL_760137, EPI_ISL_760138, EPI_ISL_760139, EPI_ISL_760140, EPI_ISL_760141, EPI_ISL_760142, EPI_ISL_760143, EPI_ISL_760144, EPI_ISL_760189 | | | |
| see above | Division of Emerging Infectious Diseases, Bureau of Infectious Diseases Diagnosis Control, Korea Disease Control and Prevention Agency | Division of Emerging Infectious Diseases, Bureau of Infectious Diseases Diagnosis Control, Korea Disease Control and Prevention Agency | Ae Kyung Park, Il-Hwan Kim, Heui Man Kim, Jeong-Min Kim, Namjoo Lee, Chaeyoung Lee, Sang Hee Woo, Eun-Jin Kim |
| EPI_ISL_763068 | RSUD Kota Surakarta | Universitas Sebelas Maret (UNS); Rumah Sakit UNS (RS-UNS); National Institute of Health Research and Development, Indonesian Ministry of Health. | Maryani, Yulia Sari, Hartono, Afif A Ghufron, Revi G H Novika, Hana A Pawestri, Kartika, Dewi Puspa, Hartanti Dian Ika, Arie A Nugraha, Vivi Setiawaty, Betty Suryawati. |
| EPI_ISL_763070 | Puskesmas Jambu Kulon, Klaten | Universitas Sebelas Maret (UNS); Rumah Sakit UNS (RS-UNS); National Institute of Health Research and Development, Indonesian Ministry of Health. | Yulia Sari, Hartono, Maryani, Afif A Ghufron, Hana A Pawestri, Kartika, Dewi Puspa, Hartanti Dian Ika, Arie A Nugraha, Vivi Setiawaty, Betty Suryawati. |
| EPI_ISL_763095, EPI_ISL_763138, EPI_ISL_763323 | Istituto Zooprofilattico Sperimentale dell' Umbria e delle Marche -Togo Rosati | Istituto Superiore di Sanità | Massimo Biagetti , Monica Giammarioli, Luca De Sabato, Gabriele Vaccari, Ilaria Di Bartolo, Giovanni Ianiro |
| EPI_ISL_763395, EPI_ISL_763450, EPI_ISL_763524, EPI_ISL_763719, EPI_ISL_764127, EPI_ISL_764263, EPI_ISL_764395, EPI_ISL_764426, EPI_ISL_764427, EPI_ISL_764428, EPI_ISL_764429, EPI_ISL_764430 | | | |
| see above | Regional Virus Laboratory, Belfast Health and Social Care Trust | COVID-19 Genomics UK (COG-UK) Consortium | Conall McCaughey, James McKenna, Tanya Curran, Susan Feeney, Alison Watt, Ciara Cox, Mairead Connor, Zoltan Molnar, David Simpson, Derek Fairley |
| EPI_ISL_765652, EPI_ISL_765655, EPI_ISL_765657, EPI_ISL_765659, EPI_ISL_765762, EPI_ISL_765763, EPI_ISL_765764, EPI_ISL_765765, EPI_ISL_765766, EPI_ISL_765767 | Massachusetts General Hospital | Infectious Disease Program, Broad Institute of Harvard and MIT | Lemieux,J.E., Siddle,K.J., Shaw,B., Adams,G., Pierce,V., Turbett,S., Anahtar,M., Branda,J., Slater,D., Harris,J., Lin,A.E., Gladden-Young,A., Lagerborg,K., Rudy,M., DeRuff,K., Carter,A., Normandin,E., Bauer,M., Reilly,S., Tomkins-Tinch,C., Loreth,C., Chaluvadi,S., Neumann,A., Cusick,C., Chapman,S.B., Gnirke,A., Flowers,K., Cerrato,F., Birren,B.W., Gallagher,G., Smole,S., Park,D.J., MacInnis,B.L., Ryan,E., LaRocque,R., Rosenberg,E. and Sabeti,P.C. |
| EPI_ISL_766083 | Respiratory Virus Unit, National Infection Service, Public Health England | COVID-19 Genomics UK (COG-UK) Consortium | PHE Covid Sequencing Team |
| EPI_ISL_766619 | Klinisk mikrobiologi | The Public Health Agency of Sweden | Department of Microbiology, The Public Health Agency of Sweden |
| EPI_ISL_768780, EPI_ISL_768781, EPI_ISL_768782, EPI_ISL_768783 | AIID | Irish Coronavirus Sequencing Consortium - National Virus Reference Laboratory | Michael Carr, Gabriel Gonzalez, Alejandro Abner Garcia Leon, Patrick Mallon |
| EPI_ISL_770008 | Area De Salud Corredores | Inciensa, Instituto Costarricense de Investigación y Enseñanza en Nutrición y Salud | Francisco Duarte, Hebleen Porras, Claudio Soto-Garita, Estela Cordero, Adriana Godínez, Melany Calderón & Mariel López |
| EPI_ISL_770016 | Area De Salud Goicoechea 2 - Clinica Dr. Jimenez Nuñez | Inciensa, Instituto Costarricense de Investigación y Enseñanza en Nutrición y Salud | Francisco Duarte, Hebleen Porras, Claudio Soto-Garita, Estela Cordero, Adriana Godínez, Melany Calderón & Mariel López |
| EPI_ISL_770017 | Area De Salud Perez Zeledon | Inciensa, Instituto Costarricense de Investigación y Enseñanza en Nutrición y Salud | Francisco Duarte, Hebleen Porras, Claudio Soto-Garita, Estela Cordero, Adriana Godínez, Melany Calderón & Mariel López |
| EPI_ISL_770025 | Area De Salud San Juan-San Diego-Concepcion 2 | Inciensa, Instituto Costarricense de Investigación y Enseñanza en Nutrición y Salud | Francisco Duarte, Hebleen Porras, Claudio Soto-Garita, Estela Cordero, Adriana Godínez, Melany Calderón & Mariel López |

| | | | |
|---|---|---|---|
| EPI_ISL_775319, EPI_ISL_775320 | Nordland Hospital - Bodo, Laboratory Department, Molecular Biology Unit | Norwegian Institute of Public Health, Department of Virology | Kathrine Stene-Johansen, Kamilla Heddeland Instefjord, Hilde Elshaug, Atiya R Ali,Marie Paulsen Madsen, Rasmus Riis Kopperud, Hilde Vollan, Karoline Bragstad, Olav Hungnes |
| EPI_ISL_775325, EPI_ISL_775326 | Foerde Hospital, Department of Microbiology | Norwegian Institute of Public Health, Department of Virology | Kathrine Stene-Johansen, Kamilla Heddeland Instefjord, Hilde Elshaug, Atiya R Ali,Marie Paulsen Madsen, Rasmus Riis Kopperud, Hilde Vollan, Karoline Bragstad, Olav Hungnes |
| EPI_ISL_775441 | Innlandet Hospital Trust, Division Lillehammer, Department for Medical Microbiology | Norwegian Institute of Public Health, Department of Virology | Kathrine Stene-Johansen, Kamilla Heddeland Instefjord, Hilde Elshaug, Atiya R Ali,Marie Paulsen Madsen, Rasmus Riis Kopperud, Hilde Vollan, Karoline Bragstad, Olav Hungnes |
| EPI_ISL_775480 | Nordland Hospital - Bodo, Laboratory Department, Molecular Biology Unit | Norwegian Institute of Public Health, Department of Virology | Kathrine Stene-Johansen, Kamilla Heddeland Instefjord, Hilde Elshaug, Atiya R Ali,Marie Paulsen Madsen, Rasmus Riis Kopperud, Hilde Vollan, Karoline Bragstad, Olav Hungnes |
| EPI_ISL_779933 | Center of Medical Microbiology, Virology, and Hospital Hygiene, University of Duesseldorf | Center of Medical Microbiology, Virology, and Hospital Hygiene, University of Duesseldorf | Maximilian Damagnez, Alexander Dilthey, Ashley-Jane Duplessis, Torsten Houwaart, Lisanna Hülse, Malte Kohns Vasconcelos, Nadine Lübke, Jessica Nicolai, Klaus Pfeffer, Daniel Strelow, Teresa Tamayo, Jörg Timm, Andreas Walker, Tobias Wienemann |
| EPI_ISL_780036 | Hospital General Universitario Gregorio Marañón | SeqCOVID-SPAIN consortium/IBV(CSIC) | Darío García de Viedma, Laura Pérez-Lago, Marta Herranz, Jon Sicilia, Julia Suárez, Pilar Catalán, Patricia Muñoz and SeqCOVID-SPAIN consortium |
| EPI_ISL_785904, EPI_ISL_785934, EPI_ISL_785945, EPI_ISL_785994, EPI_ISL_786557, EPI_ISL_786558, EPI_ISL_786569, EPI_ISL_786570, EPI_ISL_786581, EPI_ISL_786582, EPI_ISL_786593, EPI_ISL_786594, EPI_ISL_786605, EPI_ISL_786606, EPI_ISL_786617, EPI_ISL_786618, EPI_ISL_786623, EPI_ISL_786629, EPI_ISL_786630, EPI_ISL_786640, EPI_ISL_786641, EPI_ISL_786642, EPI_ISL_786643, EPI_ISL_786644, EPI_ISL_786645, EPI_ISL_786646, EPI_ISL_786647, EPI_ISL_786648, EPI_ISL_786649, EPI_ISL_786650, EPI_ISL_786651, EPI_ISL_786652, EPI_ISL_786653, EPI_ISL_786654, EPI_ISL_786655, EPI_ISL_786656, EPI_ISL_786657, EPI_ISL_786658, EPI_ISL_786659, EPI_ISL_786660, EPI_ISL_786661, EPI_ISL_786662, EPI_ISL_786663, EPI_ISL_786664, EPI_ISL_786665, EPI_ISL_786666, EPI_ISL_786667, EPI_ISL_786668, EPI_ISL_786669, EPI_ISL_786670, EPI_ISL_786671, EPI_ISL_786672, EPI_ISL_786673, EPI_ISL_786674, EPI_ISL_786675, EPI_ISL_786676, EPI_ISL_786677, EPI_ISL_786678, EPI_ISL_786679, EPI_ISL_786680, EPI_ISL_786681, EPI_ISL_786682, EPI_ISL_786683, EPI_ISL_786684, EPI_ISL_786685, EPI_ISL_786686, EPI_ISL_786687, EPI_ISL_786688, EPI_ISL_786689, EPI_ISL_786690, EPI_ISL_786691, EPI_ISL_786692, EPI_ISL_786693, EPI_ISL_786694, EPI_ISL_786695, EPI_ISL_786696, EPI_ISL_786697, EPI_ISL_786698, EPI_ISL_786699, EPI_ISL_786700, EPI_ISL_786701, EPI_ISL_786702, EPI_ISL_786703, EPI_ISL_786704, EPI_ISL_786705, EPI_ISL_786706, EPI_ISL_786707, EPI_ISL_786708, EPI_ISL_786709, EPI_ISL_786710, EPI_ISL_786711, EPI_ISL_786712, EPI_ISL_786713, EPI_ISL_786714, EPI_ISL_786715, EPI_ISL_786716, EPI_ISL_786717, EPI_ISL_786718, EPI_ISL_786719, EPI_ISL_786720, EPI_ISL_786721, EPI_ISL_786722, EPI_ISL_786723, EPI_ISL_786724, EPI_ISL_786725, EPI_ISL_786726, EPI_ISL_786727, EPI_ISL_786728, EPI_ISL_786729, EPI_ISL_786730, EPI_ISL_786731, EPI_ISL_786732, EPI_ISL_786733, EPI_ISL_786734, EPI_ISL_786735, EPI_ISL_786736, EPI_ISL_786737, EPI_ISL_786738, EPI_ISL_786739, EPI_ISL_786740, EPI_ISL_786741, EPI_ISL_786742, EPI_ISL_786743, EPI_ISL_786744, EPI_ISL_786745, EPI_ISL_786746, EPI_ISL_786747, EPI_ISL_786749, EPI_ISL_786750, EPI_ISL_786751, EPI_ISL_786752, EPI_ISL_786753, EPI_ISL_786754, EPI_ISL_786755, EPI_ISL_786756, EPI_ISL_786757, EPI_ISL_786758, EPI_ISL_786759, EPI_ISL_786762, EPI_ISL_786763, EPI_ISL_786764, EPI_ISL_786765, EPI_ISL_786766, EPI_ISL_786767, EPI_ISL_786768, EPI_ISL_786770, EPI_ISL_786772, EPI_ISL_786773, EPI_ISL_786774, EPI_ISL_786775, EPI_ISL_786776, EPI_ISL_786777, EPI_ISL_786778, EPI_ISL_786779, EPI_ISL_786780, EPI_ISL_786781, EPI_ISL_786782, EPI_ISL_786784, EPI_ISL_786785, EPI_ISL_786786, EPI_ISL_786787, EPI_ISL_786788, EPI_ISL_786789, EPI_ISL_786790, EPI_ISL_786791, EPI_ISL_786792, EPI_ISL_786793, EPI_ISL_786794, EPI_ISL_786796, EPI_ISL_786797, EPI_ISL_786798, EPI_ISL_786799, EPI_ISL_786800, EPI_ISL_786801, EPI_ISL_786802, EPI_ISL_786803, EPI_ISL_786804, EPI_ISL_786805, EPI_ISL_786806, EPI_ISL_786807, EPI_ISL_786808, EPI_ISL_786809, EPI_ISL_786810, EPI_ISL_786811, EPI_ISL_786812, EPI_ISL_786813, EPI_ISL_786814, EPI_ISL_786815, EPI_ISL_786816, EPI_ISL_786817, EPI_ISL_786818, EPI_ISL_786819, EPI_ISL_786821, EPI_ISL_786822, EPI_ISL_786823, EPI_ISL_786824, EPI_ISL_786825, EPI_ISL_786826, EPI_ISL_786827, EPI_ISL_786828, EPI_ISL_786829, EPI_ISL_786830, EPI_ISL_786832, EPI_ISL_786834, EPI_ISL_786835, EPI_ISL_786836, EPI_ISL_786839, EPI_ISL_786841, EPI_ISL_786842, EPI_ISL_786848, EPI_ISL_786851, EPI_ISL_786852, EPI_ISL_786854, EPI_ISL_786855, EPI_ISL_786860, EPI_ISL_786861, EPI_ISL_786863, EPI_ISL_786864, EPI_ISL_786865, EPI_ISL_786869, EPI_ISL_786871, EPI_ISL_786873, EPI_ISL_786875, EPI_ISL_786877, EPI_ISL_786879, EPI_ISL_786883, EPI_ISL_786884, EPI_ISL_786887, EPI_ISL_786888, EPI_ISL_786889, EPI_ISL_786895, EPI_ISL_786896, EPI_ISL_786897, EPI_ISL_786899, EPI_ISL_786900, EPI_ISL_786901, EPI_ISL_786902, EPI_ISL_786903, EPI_ISL_786905, EPI_ISL_786910, EPI_ISL_786914, EPI_ISL_786919, EPI_ISL_786920, EPI_ISL_786921, EPI_ISL_786922, EPI_ISL_786923, EPI_ISL_786925 | | | |
| see above | Houston Methodist Hospital | Houston Methodist Hospital | S. Wesley Long, Randall J. Olsen, Paul A. Christensen, David W. Bernard, James J. Davis, Maulik Shukla, Marcus Nguyen, Matthew Ojeda Saavedra, Prasanti Yerramilli, Layne Pruitt, Sishir Subedi, Heather Hendrickson, and James M. Musser |
| EPI_ISL_791979 | RSUD Curup Bengkulu | National Institute of Health Research and Development | Nugraha,AA;Subangkit;Pawestri,HA;Ikawati,HD;Puspa,KD;Mardiani;Pangesti,KNA;Soekarso,T;Puspandari,N;Setiawaty,V |
| EPI_ISL_792018, EPI_ISL_792020 | Plateforme COVID IDF | National Reference Center for Viruses of Respiratory Infections, Institut Pasteur, Paris | Marion Barbet, Sylvie Behillil, Méline Bizard, Angela Brisebarre, Camille Capel, Etienne Simon-Lorière, Vincent Enouf, Maud Vanpeene, Sylvie van der Werf,Jacques Fourgeaud |
| EPI_ISL_792599, EPI_ISL_792625, EPI_ISL_792626 | LACEN-PB | Laboratory of Respiratory Viruses and Measles, Oswaldo Cruz Institute, FIOCRUZ | Paola Resende, Luciana Appolinario, Fernando Motta, Anna Carolina Paixao, Ana Carolina Mendonca, João Felipe Bezerra, Romero Henrique Teixeira de Vasconcelos, Dalane Loudal Florentino Teixeira, Thiago Franco de Oliveira Carneiro, Marilda Siqueira |
| EPI_ISL_794685 | PathWest Laboratory Medicine WA | PathWest Laboratory Medicine WA Microbial Surveillance Unit | PathWest Laboratory Medicine WA Microbial Surveillance Unit |
| EPI_ISL_796711 | Department of Medical Microbiology, St. Olavs hospital | Norwegian Institute of Public Health, Department of Virology | Kathrine Stene-Johansen, Kamilla Heddeland Instefjord, Hilde Elshaug, Atiya R Ali,Marie Paulsen Madsen, Rasmus Riis Kopperud, Hilde Vollan, Karoline Bragstad, Olav Hungnes |
| EPI_ISL_801406, EPI_ISL_801429, EPI_ISL_801486 | Dutch COVID-19 response team | Erasmus Medical Center | Bas Oude Munnink, Reina Sikkema, David Nieuwenhuijse, Irina Chestakova, Anne van der Linden, Marjan Boter, Emmanuelle Munger, Corine GeurtsvanKessel, Annemiek van der Eijk, Richard Molenkamp, Marion Koopmans, on behalf of the Dutch national COVID-19 response team. |
| EPI_ISL_802750, EPI_ISL_802751, EPI_ISL_802752, EPI_ISL_802753, EPI_ISL_802754, EPI_ISL_802755, EPI_ISL_802756, EPI_ISL_802757, EPI_ISL_802758, EPI_ISL_802759, EPI_ISL_802760, EPI_ISL_802761, EPI_ISL_802762, EPI_ISL_802763, EPI_ISL_802764, EPI_ISL_802766, EPI_ISL_802767, EPI_ISL_802768, EPI_ISL_802769, EPI_ISL_802770, EPI_ISL_802771, EPI_ISL_802772, EPI_ISL_802773, EPI_ISL_802774, EPI_ISL_802775, EPI_ISL_802776 | | | |
| see above | Hospital Santa María Nai | Instituto de Salud Carlos III | Iglesias-Caballero, M. Molinero Calamita, M. González-Esguevillas, M. Camarero, S. Pozo, F. Casas, I. Jiménez, P. Jiménez, M. Zaballos, A. Monzón, S. Varona, S. Juliá, M. Cuesta, I, J. García. |
| EPI_ISL_803105, EPI_ISL_803106 | Respiratory Viruses Branch, Centers for Disease Control and Prevention | Respiratory Viruses Branch, Centers for Disease Control and Prevention | Queen,K., Li,Y., Tao,Y., Uehara,A., Montmayeur,A., Paden,C.R., Cook,P.W., Marine,R., Sheth,M., Wang,H., Lee,J., Tong,S. |
| EPI_ISL_803401, EPI_ISL_803402, EPI_ISL_803403, EPI_ISL_803404, EPI_ISL_803405, EPI_ISL_803406, EPI_ISL_803407, EPI_ISL_803408, EPI_ISL_803409, EPI_ISL_803412, EPI_ISL_803440, EPI_ISL_803441, EPI_ISL_803442, EPI_ISL_803443, EPI_ISL_803444, EPI_ISL_803445, EPI_ISL_803446, EPI_ISL_803447, EPI_ISL_803448, EPI_ISL_803449, EPI_ISL_803450, EPI_ISL_803451, EPI_ISL_803452, EPI_ISL_803453, EPI_ISL_803454, EPI_ISL_803455, EPI_ISL_803456, EPI_ISL_803700, EPI_ISL_803701, EPI_ISL_803702, EPI_ISL_803703, EPI_ISL_803704, EPI_ISL_803705, EPI_ISL_803809, EPI_ISL_803810, EPI_ISL_803811, EPI_ISL_803812, EPI_ISL_803813, EPI_ISL_803814, EPI_ISL_803815, EPI_ISL_803816, EPI_ISL_803818, EPI_ISL_803819, EPI_ISL_803820, EPI_ISL_803821, EPI_ISL_803822, EPI_ISL_803823, EPI_ISL_803824, EPI_ISL_803826, EPI_ISL_803827, EPI_ISL_803831, EPI_ISL_803832, EPI_ISL_803833, EPI_ISL_803834, EPI_ISL_803835, EPI_ISL_803836, EPI_ISL_803837, EPI_ISL_803838, EPI_ISL_803839, EPI_ISL_803840, EPI_ISL_803841, EPI_ISL_803842, EPI_ISL_803843, EPI_ISL_803844, EPI_ISL_803845, EPI_ISL_803846, EPI_ISL_803847, EPI_ISL_803848, EPI_ISL_803849 | | | |
| see above | Wisconsin State Laboratory of Hygiene Communicable Disease Division | Wisconsin State Laboratory of Hygiene Communicable Disease Division | Kelsey R. Florek, Abigail C. Shockey |
| EPI_ISL_804371, EPI_ISL_804372 | CHU Purpan - Laboratoire de Virologie - Institut Fédératif de Biologie | CHU Purpan - Laboratoire de Virologie - Institut Fédératif de Biologie | Latour J., Ranger N., Dubois M., Carcenac R., Harter A., Boyer P., Tremeaux P., Izopet J. |
| EPI_ISL_804472, EPI_ISL_804480, EPI_ISL_804481, EPI_ISL_804514, EPI_ISL_804515, EPI_ISL_804516, EPI_ISL_804517, EPI_ISL_804518, EPI_ISL_804519, EPI_ISL_804520, EPI_ISL_804521, EPI_ISL_804522, EPI_ISL_804523, EPI_ISL_804524, EPI_ISL_804539, EPI_ISL_804540, EPI_ISL_804541, EPI_ISL_804542, EPI_ISL_804543, EPI_ISL_804544, EPI_ISL_804545, EPI_ISL_804546, EPI_ISL_804547, EPI_ISL_804548, EPI_ISL_804549, EPI_ISL_804550, EPI_ISL_804551, EPI_ISL_804552, EPI_ISL_804553, EPI_ISL_804554, EPI_ISL_804555, EPI_ISL_804556 | | | |
| see above | MEPHI, Aix Marseille University | MEPHI, Aix Marseille University | Anthony LEVASSEUR |
| EPI_ISL_804598 | Maryland Public Health Laboratory | Maryland Public Health Laboratory | Maryland Department of Health Laboratories Administration |
| EPI_ISL_819450 | Northumbria University / South Tees Hospitals NHS Foundation Trust / North Cumbria Integrated Care NHS Foundation Trust / North Tees and Hartlepool NHS Foundation Trust / Newcastle Hospitals NHS Foundation Trust | COVID-19 Genomics UK (COG-UK) Consortium | Darren L Smith,Andrew Nelson,Matthew Bashton,Greg R Young,Joshua Loh,John Allan,Mohammad A Tariq,Giles S Holt,Gary Black,Wen C Yew,Lynn Dover,Paul Baker,Steve Liggett,Sarah Essex,Jane Greenaway,Debra Padgett,Clive Graham,Garren Scott,Edward Barton,Emma Swindells,Brendan Payne,Jennifer Collins,Yusri Taha,Gary Eltringham |
| EPI_ISL_825548, EPI_ISL_825583, EPI_ISL_825584, EPI_ISL_825585, EPI_ISL_825587 | Respiratory Virus Unit, National Infection Service, Public Health England | COVID-19 Genomics UK (COG-UK) Consortium | PHE Covid Sequencing Team |
| EPI_ISL_826480, EPI_ISL_826481, EPI_ISL_826482, EPI_ISL_826483, EPI_ISL_826484, EPI_ISL_826485, EPI_ISL_826489, EPI_ISL_826490, EPI_ISL_826493, EPI_ISL_826497, EPI_ISL_826501, EPI_ISL_826502, EPI_ISL_826503, EPI_ISL_826505, EPI_ISL_826509, EPI_ISL_826510, EPI_ISL_826511, EPI_ISL_826512, EPI_ISL_826514, EPI_ISL_826515 | | | |
| see above | Lighthouse Lab in Alderley Park | Wellcome Sanger Institute for the COVID-19 Genomics UK (COG-UK) Consortium | Jacquelyn Wynn, Mairead Hyland, The Lighthouse Lab in Alderley Park and Alex Alderton, Roberto Amato, Sonia Goncalves, Ewan Harrison, David K. Jackson, Ian Johnston, Dominic Kwiatkowski, Cordelia Langford, John Sillitoe on behalf of the Wellcome Sanger Institute COVID-19 Surveillance Team |
| EPI_ISL_826516 | Lighthouse Lab in Glasgow | Wellcome Sanger Institute for the COVID-19 Genomics UK (COG-UK) Consortium | Harper VanSteenhouse, Yumi Kasai, David Gray, Carol Clugston, Anna Dominiczak and Alex Alderton, Roberto Amato, Sonia Goncalves, Ewan Harrison, David K. Jackson, Ian Johnston, Dominic Kwiatkowski, Cordelia Langford, John Sillitoe on behalf of the Wellcome Sanger Institute COVID-19 Surveillance Team |
| EPI_ISL_826674, EPI_ISL_826675, EPI_ISL_826676, EPI_ISL_826680, EPI_ISL_826682, EPI_ISL_826684, EPI_ISL_826696, EPI_ISL_826701, EPI_ISL_826702, EPI_ISL_826703, EPI_ISL_826705, EPI_ISL_826706, EPI_ISL_826707, EPI_ISL_826708, EPI_ISL_826709, EPI_ISL_826710, EPI_ISL_826713, EPI_ISL_826714, EPI_ISL_826743, EPI_ISL_826744, EPI_ISL_826766, EPI_ISL_826847, EPI_ISL_826848, EPI_ISL_826853, EPI_ISL_826857, EPI_ISL_826859, EPI_ISL_826860, EPI_ISL_826863, EPI_ISL_826876, EPI_ISL_827013, EPI_ISL_827057, EPI_ISL_827201 | | | |

| | | | |
|---|---|---|---|
| see above | deCODE genetics | deCODE genetics | Daniel F Gudbjartsson; Agnar Helgason; Hakon Jonsson; Olafur T Magnusson; Pall Melsted; Gudmundur L Norddahl; Jona Saemundsdottir; Asgeir Sigurdsson; Patrick Sulem; Arna B Agustsdottir; Hannes Eggertsson; Berglind Eiriksdottir; Run Fridriksdottir; Elisabet E Gardarsdottir; Gudmundur Georgsson; Olafia S Gretarsdottir; Kjartan R Gudmundsson; Thora R Gunnarsdottir; Arnaldur Gylfason; Hilma Holm; Brynjar O Jensson; Aslaug Jonasdottir; Kamilla S Josefsdottir; Thordur Kristjansson; Droplaug N Magnusdottir; Solvi Rognvaldsson; Louise le Roux; Gudrun Sigmundsdottir; Gardar Sveinbjornsson; Kristin E Sveinsdottir; Maney Sveinsdottir; Emil A Thorarensen; Bjarni Thorbjornsson; Gisli Masson; Ingileif Jonsdottir; Alma Moller; Thorolfur Gudnason; Karl G Kristinsson; Unnur Thorsteinsdottir; Kari Stefansson |
| EPI_ISL_827212, EPI_ISL_827229, EPI_ISL_827252, EPI_ISL_827253, EPI_ISL_827254, EPI_ISL_827256, EPI_ISL_827257, EPI_ISL_827266, EPI_ISL_827363, EPI_ISL_827419 | The National University Hospital of Iceland | deCODE genetics | Daniel F Gudbjartsson; Agnar Helgason; Hakon Jonsson; Olafur T Magnusson; Pall Melsted; Gudmundur L Norddahl; Jona Saemundsdottir; Asgeir Sigurdsson; Patrick Sulem; Arna B Agustsdottir; Hannes Eggertsson; Berglind Eiriksdottir; Run Fridriksdottir; Elisabet E Gardarsdottir; Gudmundur Georgsson; Olafia S Gretarsdottir; Kjartan R Gudmundsson; Thora R Gunnarsdottir; Arnaldur Gylfason; Hilma Holm; Brynjar O Jensson; Aslaug Jonasdottir; Kamilla S Josefsdottir; Thordur Kristjansson; Droplaug N Magnusdottir; Solvi Rognvaldsson; Louise le Roux; Gudrun Sigmundsdottir; Gardar Sveinbjornsson; Kristin E Sveinsdottir; Maney Sveinsdottir; Emil A Thorarensen; Bjarni Thorbjornsson; Gisli Masson; Ingileif Jonsdottir; Alma Moller; Thorolfur Gudnason; Karl G Kristinsson; Unnur Thorsteinsdottir; Kari Stefansson |
| EPI_ISL_827522, EPI_ISL_827526, EPI_ISL_827530 | deCODE genetics | deCODE genetics | Daniel F Gudbjartsson; Agnar Helgason; Hakon Jonsson; Olafur T Magnusson; Pall Melsted; Gudmundur L Norddahl; Jona Saemundsdottir; Asgeir Sigurdsson; Patrick Sulem; Arna B Agustsdottir; Hannes Eggertsson; Berglind Eiriksdottir; Run Fridriksdottir; Elisabet E Gardarsdottir; Gudmundur Georgsson; Olafia S Gretarsdottir; Kjartan R Gudmundsson; Thora R Gunnarsdottir; Arnaldur Gylfason; Hilma Holm; Brynjar O Jensson; Aslaug Jonasdottir; Kamilla S Josefsdottir; Thordur Kristjansson; Droplaug N Magnusdottir; Solvi Rognvaldsson; Louise le Roux; Gudrun Sigmundsdottir; Gardar Sveinbjornsson; Kristin E Sveinsdottir; Maney Sveinsdottir; Emil A Thorarensen; Bjarni Thorbjornsson; Gisli Masson; Ingileif Jonsdottir; Alma Moller; Thorolfur Gudnason; Karl G Kristinsson; Unnur Thorsteinsdottir; Kari Stefansson |
| EPI_ISL_827531, EPI_ISL_827534, EPI_ISL_827535 | The National University Hospital of Iceland | deCODE genetics | Daniel F Gudbjartsson; Agnar Helgason; Hakon Jonsson; Olafur T Magnusson; Pall Melsted; Gudmundur L Norddahl; Jona Saemundsdottir; Asgeir Sigurdsson; Patrick Sulem; Arna B Agustsdottir; Hannes Eggertsson; Berglind Eiriksdottir; Run Fridriksdottir; Elisabet E Gardarsdottir; Gudmundur Georgsson; Olafia S Gretarsdottir; Kjartan R Gudmundsson; Thora R Gunnarsdottir; Arnaldur Gylfason; Hilma Holm; Brynjar O Jensson; Aslaug Jonasdottir; Kamilla S Josefsdottir; Thordur Kristjansson; Droplaug N Magnusdottir; Solvi Rognvaldsson; Louise le Roux; Gudrun Sigmundsdottir; Gardar Sveinbjornsson; Kristin E Sveinsdottir; Maney Sveinsdottir; Emil A Thorarensen; Bjarni Thorbjornsson; Gisli Masson; Ingileif Jonsdottir; Alma Moller; Thorolfur Gudnason; Karl G Kristinsson; Unnur Thorsteinsdottir; Kari Stefansson |
| EPI_ISL_827538 | deCODE genetics | deCODE genetics | Daniel F Gudbjartsson; Agnar Helgason; Hakon Jonsson; Olafur T Magnusson; Pall Melsted; Gudmundur L Norddahl; Jona Saemundsdottir; Asgeir Sigurdsson; Patrick Sulem; Arna B Agustsdottir; Hannes Eggertsson; Berglind Eiriksdottir; Run Fridriksdottir; Elisabet E Gardarsdottir; Gudmundur Georgsson; Olafia S Gretarsdottir; Kjartan R Gudmundsson; Thora R Gunnarsdottir; Arnaldur Gylfason; Hilma Holm; Brynjar O Jensson; Aslaug Jonasdottir; Kamilla S Josefsdottir; Thordur Kristjansson; Droplaug N Magnusdottir; Solvi Rognvaldsson; Louise le Roux; Gudrun Sigmundsdottir; Gardar Sveinbjornsson; Kristin E Sveinsdottir; Maney Sveinsdottir; Emil A Thorarensen; Bjarni Thorbjornsson; Gisli Masson; Ingileif Jonsdottir; Alma Moller; Thorolfur Gudnason; Karl G Kristinsson; Unnur Thorsteinsdottir; Kari Stefansson |
| EPI_ISL_827548, EPI_ISL_827550, EPI_ISL_827553, EPI_ISL_827587, EPI_ISL_827589, EPI_ISL_827821 | The National University Hospital of Iceland | deCODE genetics | Daniel F Gudbjartsson; Agnar Helgason; Hakon Jonsson; Olafur T Magnusson; Pall Melsted; Gudmundur L Norddahl; Jona Saemundsdottir; Asgeir Sigurdsson; Patrick Sulem; Arna B Agustsdottir; Hannes Eggertsson; Berglind Eiriksdottir; Run Fridriksdottir; Elisabet E Gardarsdottir; Gudmundur Georgsson; Olafia S Gretarsdottir; Kjartan R Gudmundsson; Thora R Gunnarsdottir; Arnaldur Gylfason; Hilma Holm; Brynjar O Jensson; Aslaug Jonasdottir; Kamilla S Josefsdottir; Thordur Kristjansson; Droplaug N Magnusdottir; Solvi Rognvaldsson; Louise le Roux; Gudrun Sigmundsdottir; Gardar Sveinbjornsson; Kristin E Sveinsdottir; Maney Sveinsdottir; Emil A Thorarensen; Bjarni Thorbjornsson; Gisli Masson; Ingileif Jonsdottir; Alma Moller; Thorolfur Gudnason; Karl G Kristinsson; Unnur Thorsteinsdottir; Kari Stefansson |
| EPI_ISL_827953, EPI_ISL_827963 | deCODE genetics | deCODE genetics | Daniel F Gudbjartsson; Agnar Helgason; Hakon Jonsson; Olafur T Magnusson; Pall Melsted; Gudmundur L Norddahl; Jona Saemundsdottir; Asgeir Sigurdsson; Patrick Sulem; Arna B Agustsdottir; Hannes Eggertsson; Berglind Eiriksdottir; Run Fridriksdottir; Elisabet E Gardarsdottir; Gudmundur Georgsson; Olafia S Gretarsdottir; Kjartan R Gudmundsson; Thora R Gunnarsdottir; Arnaldur Gylfason; Hilma Holm; Brynjar O Jensson; Aslaug Jonasdottir; Kamilla S Josefsdottir; Thordur Kristjansson; Droplaug N Magnusdottir; Solvi Rognvaldsson; Louise le Roux; Gudrun Sigmundsdottir; Gardar Sveinbjornsson; Kristin E Sveinsdottir; Maney Sveinsdottir; Emil A Thorarensen; Bjarni Thorbjornsson; Gisli Masson; Ingileif Jonsdottir; Alma Moller; Thorolfur Gudnason; Karl G Kristinsson; Unnur Thorsteinsdottir; Kari Stefansson |
| EPI_ISL_828008, EPI_ISL_828205 | The National University Hospital of Iceland | deCODE genetics | Daniel F Gudbjartsson; Agnar Helgason; Hakon Jonsson; Olafur T Magnusson; Pall Melsted; Gudmundur L Norddahl; Jona Saemundsdottir; Asgeir Sigurdsson; Patrick Sulem; Arna B Agustsdottir; Hannes Eggertsson; Berglind Eiriksdottir; Run Fridriksdottir; Elisabet E Gardarsdottir; Gudmundur Georgsson; Olafia S Gretarsdottir; Kjartan R Gudmundsson; Thora R Gunnarsdottir; Arnaldur Gylfason; Hilma Holm; Brynjar O Jensson; Aslaug Jonasdottir; Kamilla S Josefsdottir; Thordur Kristjansson; Droplaug N Magnusdottir; Solvi Rognvaldsson; Louise le Roux; Gudrun Sigmundsdottir; Gardar Sveinbjornsson; Kristin E Sveinsdottir; Maney Sveinsdottir; Emil A Thorarensen; Bjarni Thorbjornsson; Gisli Masson; Ingileif Jonsdottir; Alma Moller; Thorolfur Gudnason; Karl G Kristinsson; Unnur Thorsteinsdottir; Kari Stefansson |
| EPI_ISL_828347, EPI_ISL_828382, EPI_ISL_828449, EPI_ISL_828456, EPI_ISL_828457, EPI_ISL_828458, EPI_ISL_828459, EPI_ISL_828460, EPI_ISL_828461, EPI_ISL_828462, EPI_ISL_828463 | | | |
| see above | deCODE genetics | deCODE genetics | Daniel F Gudbjartsson; Agnar Helgason; Hakon Jonsson; Olafur T Magnusson; Pall Melsted; Gudmundur L Norddahl; Jona Saemundsdottir; Asgeir Sigurdsson; Patrick Sulem; Arna B Agustsdottir; Hannes Eggertsson; Berglind Eiriksdottir; Run Fridriksdottir; Elisabet E Gardarsdottir; Gudmundur Georgsson; Olafia S Gretarsdottir; Kjartan R Gudmundsson; Thora R Gunnarsdottir; Arnaldur Gylfason; Hilma Holm; Brynjar O Jensson; Aslaug Jonasdottir; Kamilla S Josefsdottir; Thordur Kristjansson; Droplaug N Magnusdottir; Solvi Rognvaldsson; Louise le Roux; Gudrun Sigmundsdottir; Gardar Sveinbjornsson; Kristin E Sveinsdottir; Maney Sveinsdottir; Emil A Thorarensen; Bjarni Thorbjornsson; Gisli Masson; Ingileif Jonsdottir; Alma Moller; Thorolfur Gudnason; Karl G Kristinsson; Unnur Thorsteinsdottir; Kari Stefansson |
| EPI_ISL_828464, EPI_ISL_828465, EPI_ISL_828466, EPI_ISL_828467, EPI_ISL_828469 | The National University Hospital of Iceland | deCODE genetics | Daniel F Gudbjartsson; Agnar Helgason; Hakon Jonsson; Olafur T Magnusson; Pall Melsted; Gudmundur L Norddahl; Jona Saemundsdottir; Asgeir Sigurdsson; Patrick Sulem; Arna B Agustsdottir; Hannes Eggertsson; Berglind Eiriksdottir; Run Fridriksdottir; Elisabet E Gardarsdottir; Gudmundur Georgsson; Olafia S Gretarsdottir; Kjartan R Gudmundsson; Thora R Gunnarsdottir; Arnaldur Gylfason; Hilma Holm; Brynjar O Jensson; Aslaug Jonasdottir; Kamilla S Josefsdottir; Thordur Kristjansson; Droplaug N Magnusdottir; Solvi Rognvaldsson; Louise le Roux; Gudrun Sigmundsdottir; Gardar Sveinbjornsson; Kristin E Sveinsdottir; Maney Sveinsdottir; Emil A Thorarensen; Bjarni Thorbjornsson; Gisli Masson; Ingileif Jonsdottir; Alma Moller; Thorolfur Gudnason; Karl G Kristinsson; Unnur Thorsteinsdottir; Kari Stefansson |
| EPI_ISL_828481, EPI_ISL_828482 | deCODE genetics | deCODE genetics | Daniel F Gudbjartsson; Agnar Helgason; Hakon Jonsson; Olafur T Magnusson; Pall Melsted; Gudmundur L Norddahl; Jona Saemundsdottir; Asgeir Sigurdsson; Patrick Sulem; Arna B Agustsdottir; Hannes Eggertsson; Berglind Eiriksdottir; Run Fridriksdottir; Elisabet E Gardarsdottir; Gudmundur Georgsson; Olafia S Gretarsdottir; Kjartan R Gudmundsson; Thora R Gunnarsdottir; Arnaldur Gylfason; Hilma Holm; Brynjar O Jensson; Aslaug Jonasdottir; Kamilla S Josefsdottir; Thordur Kristjansson; Droplaug N Magnusdottir; Solvi Rognvaldsson; Louise le Roux; Gudrun Sigmundsdottir; Gardar Sveinbjornsson; Kristin E Sveinsdottir; Maney Sveinsdottir; Emil A Thorarensen; Bjarni Thorbjornsson; Gisli Masson; Ingileif Jonsdottir; Alma Moller; Thorolfur Gudnason; Karl G Kristinsson; Unnur Thorsteinsdottir; Kari Stefansson |
| EPI_ISL_828493, EPI_ISL_828495 | The National University Hospital of Iceland | deCODE genetics | Daniel F Gudbjartsson; Agnar Helgason; Hakon Jonsson; Olafur T Magnusson; Pall Melsted; Gudmundur L Norddahl; Jona Saemundsdottir; Asgeir Sigurdsson; Patrick Sulem; Arna B Agustsdottir; Hannes Eggertsson; Berglind Eiriksdottir; Run Fridriksdottir; Elisabet E Gardarsdottir; Gudmundur Georgsson; Olafia S Gretarsdottir; Kjartan R Gudmundsson; Thora R Gunnarsdottir; Arnaldur Gylfason; Hilma Holm; Brynjar O Jensson; Aslaug Jonasdottir; Kamilla S Josefsdottir; Thordur Kristjansson; Droplaug N Magnusdottir; Solvi Rognvaldsson; Louise le Roux; Gudrun Sigmundsdottir; Gardar Sveinbjornsson; Kristin E Sveinsdottir; Maney Sveinsdottir; Emil A Thorarensen; Bjarni Thorbjornsson; Gisli Masson; Ingileif Jonsdottir; Alma Moller; Thorolfur Gudnason; Karl G Kristinsson; Unnur Thorsteinsdottir; Kari Stefansson |
| EPI_ISL_828696, EPI_ISL_828698, EPI_ISL_828700, EPI_ISL_828701, EPI_ISL_828704, EPI_ISL_828718, EPI_ISL_828720, EPI_ISL_828721, EPI_ISL_828722, EPI_ISL_828723, EPI_ISL_828724, EPI_ISL_828727, EPI_ISL_828728, EPI_ISL_828927, EPI_ISL_828933, EPI_ISL_828935, EPI_ISL_828936, EPI_ISL_828937, EPI_ISL_829077, EPI_ISL_829078, EPI_ISL_829081, EPI_ISL_829091, EPI_ISL_829104, EPI_ISL_829117, EPI_ISL_829255, EPI_ISL_829256, EPI_ISL_829257, EPI_ISL_829258, EPI_ISL_829259, EPI_ISL_829260, EPI_ISL_829420, EPI_ISL_829458 | | | |
| see above | deCODE genetics | deCODE genetics | Daniel F Gudbjartsson; Agnar Helgason; Hakon Jonsson; Olafur T Magnusson; Pall Melsted; Gudmundur L Norddahl; Jona Saemundsdottir; Asgeir Sigurdsson; Patrick Sulem; Arna B Agustsdottir; Hannes Eggertsson; Berglind Eiriksdottir; Run Fridriksdottir; Elisabet E Gardarsdottir; Gudmundur Georgsson; Olafia S Gretarsdottir; Kjartan R Gudmundsson; Thora R Gunnarsdottir; Arnaldur Gylfason; Hilma Holm; Brynjar O Jensson; Aslaug Jonasdottir; |

| | | | |
|---|---|---|---|
| | | | Kamilla S Josefsdottir; Thordur Kristjansson; Droplaug N Magnusdottir; Solvi Rognvaldsson; Louise le Roux; Gudrun Sigmundsdottir; Gardar Sveinbjornsson; Kristin E Sveinsdottir; Maney Sveinsdottir; Emil A Thorarensen; Bjarni Thorbjornsson; Gisli Masson; Ingileif Jonsdottir; Alma Moller; Thorolfur Gudnason; Karl G Kristinsson; Unnur Thorsteinsdottir; Kari Stefansson |
| EPI_ISL_829476 | The National University Hospital of Iceland | deCODE genetics | Daniel F Gudbjartsson; Agnar Helgason; Hakon Jonsson; Olafur T Magnusson; Pall Melsted; Gudmundur L Norddahl; Jona Saemundsdottir; Asgeir Sigurdsson; Patrick Sulem; Arna B Agustsdottir; Hannes Eggertsson; Berglind Eiriksdottir; Run Fridriksdottir; Elisabet E Gardarsdottir; Gudmundur Georgsson; Olafia S Gretarsdottir; Kjartan R Gudmundsson; Thora R Gunnarsdottir; Arnaldur Gylfason; Hilma Holm; Brynjar O Jensson; Aslaug Jonasdottir; Kamilla S Josefsdottir; Thordur Kristjansson; Droplaug N Magnusdottir; Solvi Rognvaldsson; Louise le Roux; Gudrun Sigmundsdottir; Gardar Sveinbjornsson; Kristin E Sveinsdottir; Maney Sveinsdottir; Emil A Thorarensen; Bjarni Thorbjornsson; Gisli Masson; Ingileif Jonsdottir; Alma Moller; Thorolfur Gudnason; Karl G Kristinsson; Unnur Thorsteinsdottir; Kari Stefansson |
| EPI_ISL_829507, EPI_ISL_829508, EPI_ISL_829509, EPI_ISL_829510, EPI_ISL_829511, EPI_ISL_829513, EPI_ISL_829650, EPI_ISL_829736, EPI_ISL_829758 | deCODE genetics | deCODE genetics | Daniel F Gudbjartsson; Agnar Helgason; Hakon Jonsson; Olafur T Magnusson; Pall Melsted; Gudmundur L Norddahl; Jona Saemundsdottir; Asgeir Sigurdsson; Patrick Sulem; Arna B Agustsdottir; Hannes Eggertsson; Berglind Eiriksdottir; Run Fridriksdottir; Elisabet E Gardarsdottir; Gudmundur Georgsson; Olafia S Gretarsdottir; Kjartan R Gudmundsson; Thora R Gunnarsdottir; Arnaldur Gylfason; Hilma Holm; Brynjar O Jensson; Aslaug Jonasdottir; Kamilla S Josefsdottir; Thordur Kristjansson; Droplaug N Magnusdottir; Solvi Rognvaldsson; Louise le Roux; Gudrun Sigmundsdottir; Gardar Sveinbjornsson; Kristin E Sveinsdottir; Maney Sveinsdottir; Emil A Thorarensen; Bjarni Thorbjornsson; Gisli Masson; Ingileif Jonsdottir; Alma Moller; Thorolfur Gudnason; Karl G Kristinsson; Unnur Thorsteinsdottir; Kari Stefansson |
| EPI_ISL_829948, EPI_ISL_830105 | The National University Hospital of Iceland | deCODE genetics | Daniel F Gudbjartsson; Agnar Helgason; Hakon Jonsson; Olafur T Magnusson; Pall Melsted; Gudmundur L Norddahl; Jona Saemundsdottir; Asgeir Sigurdsson; Patrick Sulem; Arna B Agustsdottir; Hannes Eggertsson; Berglind Eiriksdottir; Run Fridriksdottir; Elisabet E Gardarsdottir; Gudmundur Georgsson; Olafia S Gretarsdottir; Kjartan R Gudmundsson; Thora R Gunnarsdottir; Arnaldur Gylfason; Hilma Holm; Brynjar O Jensson; Aslaug Jonasdottir; Kamilla S Josefsdottir; Thordur Kristjansson; Droplaug N Magnusdottir; Solvi Rognvaldsson; Louise le Roux; Gudrun Sigmundsdottir; Gardar Sveinbjornsson; Kristin E Sveinsdottir; Maney Sveinsdottir; Emil A Thorarensen; Bjarni Thorbjornsson; Gisli Masson; Ingileif Jonsdottir; Alma Moller; Thorolfur Gudnason; Karl G Kristinsson; Unnur Thorsteinsdottir; Kari Stefansson |
| EPI_ISL_830148, EPI_ISL_830149, EPI_ISL_830153, EPI_ISL_830154, EPI_ISL_830155, EPI_ISL_830156, EPI_ISL_830207, EPI_ISL_830271, EPI_ISL_830483, EPI_ISL_830484, EPI_ISL_830486, EPI_ISL_830488, EPI_ISL_830489, EPI_ISL_830490 | | | |
| see above | deCODE genetics | deCODE genetics | Daniel F Gudbjartsson; Agnar Helgason; Hakon Jonsson; Olafur T Magnusson; Pall Melsted; Gudmundur L Norddahl; Jona Saemundsdottir; Asgeir Sigurdsson; Patrick Sulem; Arna B Agustsdottir; Hannes Eggertsson; Berglind Eiriksdottir; Run Fridriksdottir; Elisabet E Gardarsdottir; Gudmundur Georgsson; Olafia S Gretarsdottir; Kjartan R Gudmundsson; Thora R Gunnarsdottir; Arnaldur Gylfason; Hilma Holm; Brynjar O Jensson; Aslaug Jonasdottir; Kamilla S Josefsdottir; Thordur Kristjansson; Droplaug N Magnusdottir; Solvi Rognvaldsson; Louise le Roux; Gudrun Sigmundsdottir; Gardar Sveinbjornsson; Kristin E Sveinsdottir; Maney Sveinsdottir; Emil A Thorarensen; Bjarni Thorbjornsson; Gisli Masson; Ingileif Jonsdottir; Alma Moller; Thorolfur Gudnason; Karl G Kristinsson; Unnur Thorsteinsdottir; Kari Stefansson |
| EPI_ISL_830753 | University Hospital Basel, Clinical Virology | University Hospital Basel, Clinical Bacteriology | Tim Roloff, Madlen Stange, Helena MB Seth-Smith, Alfredo Mari, Karoline Leuzinger, Julia Bielicki, Manuel Battegay, Hans Hirsch, Adrian Egli |
| EPI_ISL_831020 | INSPI-CRN DE INFLUENZA Y OTROS VIRUS RESPIRATORIOS | Instituto de Salud Publica de Chile | Javier Tognarelli, Barbara Parra, Loredana Arata, Jaime Lagos, Gisselle Barra, Alfredo Bruno, Domenica de Mora, Solon Narvaez, Jimmy Garcez, Michelle Paez, Martiza Olmedo, Manuel Gonzalez, Patricia Bustos, Rodrigo Fasce, Andres Castillo, Jorge Fernandez |
| EPI_ISL_831106 | Hospital Universitario La Paz (Madrid) | SeqCOVID-SPAIN consortium/IBV(CSIC) | María Rodríguez-Tejedor, Elias Dahdouh, Fernando Lázaro-Perona, Jesús Mingorance and SeqCOVID-SPAIN consortium |
| EPI_ISL_831778, EPI_ISL_831779, EPI_ISL_831780, EPI_ISL_831781 | United States Air Force School of Aerospace Medicine | United States Air Force School of Aerospace Medicine | Anthony Fries, Jennifer Meyer, William Gruner, Amanda Javorina, Sarah Purves, Clarise Starr, Elizabeth Macias |
| EPI_ISL_832144, EPI_ISL_832145 | Hospital | National Reference Center for Viruses of Respiratory Infections, Institut Pasteur, Paris | Marion Barbet, Sylvie Behillil, Méline Bizard, Angela Brisebarre, Camille Capel, Etienne Simon-Lorière, Vincent Enouf, Maud Vanpeene, Sylvie van der Werf, Léa Pilorge |
| EPI_ISL_832497, EPI_ISL_832498, EPI_ISL_832499, EPI_ISL_832500, EPI_ISL_832501, EPI_ISL_832502, EPI_ISL_832503, EPI_ISL_832504, EPI_ISL_832505, EPI_ISL_832506, EPI_ISL_832507, EPI_ISL_832508, EPI_ISL_832509, EPI_ISL_832510, EPI_ISL_832511, EPI_ISL_832512, EPI_ISL_832513, EPI_ISL_832514, EPI_ISL_832515, EPI_ISL_832516, EPI_ISL_832517, EPI_ISL_832518, EPI_ISL_832519, EPI_ISL_832539, EPI_ISL_832546, EPI_ISL_832550, EPI_ISL_832551, EPI_ISL_832552, EPI_ISL_832553, EPI_ISL_832554, EPI_ISL_832555, EPI_ISL_832556, EPI_ISL_832557, EPI_ISL_832558, EPI_ISL_832559, EPI_ISL_832560, EPI_ISL_832561, EPI_ISL_832562 | | | |
| see above | OHSU Lab Services Molecular Microbiology Lab | Oregon SARS-CoV-2 Genome Sequencing Center | Brendan L. O'Connell, Ruth V. Nichols, Sally Grindstaff, Alec J. Hirsch, Donna Hansel, Guang Fan, Daniel N. Streblow, William B. Messer, Andrew C. Adey, Benjamin N. Bimber, Brian J. O'Roak |
| EPI_ISL_833467 | CHU Purpan - Laboratoire de Virologie - Institut Fédératif de Biologie | CHU Purpan - Laboratoire de Virologie - Institut Fédératif de Biologie | Latour J., Ranger N., Dubois M., Carcenac R., Harter A., Boyer P., Tremeaux P., Izopet J. |
| EPI_ISL_842348, EPI_ISL_842350 | Virology Department, Sheffield Teaching Hospitals NHS Foundation Trust/Department of Infection, Immunity and Cardiovascular Disease, The Medical School, University of Sheffield | COVID-19 Genomics UK (COG-UK) Consortium | Thushan de Silva, Matthew Parker, Nikki Smith, Adri Angyal, Rebecca Brown, Luke Green, Rachel Tucker, Paul Parsons, Danielle Groves, Katie Johnson, Laura Carrilero, Alex Keeley, Dave Partridge, Matthew Wyles, Benjamin Lindsey, Mehmet Yavuz, Mohammad Raza, Cariad Evans |
| EPI_ISL_848083 | Laboratory of Clinical Research on Dermatozoonoses in Domestic Animals, Evandro Chagas National Institute of Infectious Diseases, Oswaldo Cruz Foundation (Fiocruz), Rio de Janeiro, RJ, Brazil. | Laboratory of Respiratory Viruses and Measles, Oswaldo Cruz Institute, FIOCRUZ | Guilherme Amaral Calvet, Michelle Fernanda Borges da Silva, Anielle de Pina Costa, Ezequias Batista Martins, Isabella Campos Vargas de Moraes, Lusiele Guaraldo, Patrícia Brasil, Sandro Antônio Pereira, Rodrigo Caldas Menezes, Isabella Dib Ferreira Gremião, Lucas Oliveira Keidel, Shanna Araujo dos Santos, Artur Augusto Velho Mendes Junior, Renato Orsini Ornellas, Maria Ogrzewalska, PaolaCristina Resende, Alex Pauvolid-Corrêa, Fernando do Couto Motta, Alice Sampaio Barreto da Rocha, Thiago C. Souza, Marilda Mendonça Siqueira |
| EPI_ISL_848270, EPI_ISL_848376, EPI_ISL_848522 | Illinois Department of Public Health | Gagnon Lab, Southern Illinois University | Keith Gagnon |
| EPI_ISL_848558, EPI_ISL_848559, EPI_ISL_848560 | Evandro Chagas Institute | Evandro Chagas Institute | Santos, M.C.; Silva, A.M.; Junior, W.D.C.; Barbagelata, L.S.; Ferreira, J.A.; Sousa, E.M.A.; da Silva, P.S.; Pinheiro, K.C.; L.C.; Sousa Junior, E.C. |
| EPI_ISL_848881, EPI_ISL_848882, EPI_ISL_848883, EPI_ISL_848884, EPI_ISL_848885, EPI_ISL_848886, EPI_ISL_848887, EPI_ISL_848888, EPI_ISL_848889, EPI_ISL_848890, EPI_ISL_848891, EPI_ISL_848892, EPI_ISL_848893, EPI_ISL_848894, EPI_ISL_848895, EPI_ISL_848896, EPI_ISL_848897, EPI_ISL_848898, EPI_ISL_848899, EPI_ISL_848900, EPI_ISL_848901, EPI_ISL_848902, EPI_ISL_848903, EPI_ISL_848904, EPI_ISL_848905, EPI_ISL_848906, EPI_ISL_848907, EPI_ISL_848908, EPI_ISL_848909, EPI_ISL_848910, EPI_ISL_848911, EPI_ISL_848912, EPI_ISL_848913, EPI_ISL_848914, EPI_ISL_848915, EPI_ISL_848916, EPI_ISL_848917, EPI_ISL_848918, EPI_ISL_848919, EPI_ISL_848920, EPI_ISL_848921, EPI_ISL_848922, EPI_ISL_848923, EPI_ISL_848924, EPI_ISL_848925, EPI_ISL_848926, EPI_ISL_848927, EPI_ISL_848928, EPI_ISL_848929, EPI_ISL_848930, EPI_ISL_848931 | | | |
| see above | Florida Bureau of Public Health Laboratories | Florida Bureau of Public Health Laboratories | Sarah Schmedes, Jason Blanton |
| EPI_ISL_849352, EPI_ISL_849355, EPI_ISL_849356, EPI_ISL_849358, EPI_ISL_849359, EPI_ISL_849365, EPI_ISL_849366 | Servicio Virosis Respiratorias-Departamento Virología-INEI | Instituto Nacional Enfermedades Infecciosas C.G.Malbran | Baumeister E., Avaro M., Benedetti E., Russo M., Dattero ME, Pontoriero A., Cisterna D., Molina V., Perandones C., Tuduri E., Lorenzo F., Poklepovich T., Campos J. |
| EPI_ISL_849492, EPI_ISL_849493, EPI_ISL_849494, EPI_ISL_849584, EPI_ISL_849603, EPI_ISL_849604 | Seattle Flu Study | Seattle Flu Study | Deborah A. Nickerson, Chris D. Frazar, Jover Lee, Benjamin Pelle, Matthew Richardson, Amanda Adler, Elisabeth Brandstetter, Peter D. Han, Kairsten Fay, Misja Ilcisin, Kirsten Lacombe, Thomas R. Sibley, Melissa Truong, Caitlin R. Wolf, Michael Boeckh, Janet A. Englund, Michael Famulare, Barry R. Lutz, Mark J. Rieder, Lea M. Starita, Matthew Thompson, Jay Shendure, Trevor Bedford, Helen Y. Chu |
| EPI_ISL_852599 | Max von Pettenkofer Institute, Virology, National Reference Center for Retroviruses, LMU München | Laboratory for Functional Genome Analysis, Dept. Genomics, Gene Center of the LMU Munich | Max Muenchhoff, Stefan Krebs, Alexander Graf, Oliver Keppler, Helmut Blum |
| EPI_ISL_852650, EPI_ISL_852651, EPI_ISL_852652 | Institute of Virology, Medical Center, University of Freiburg, Freiburg, Germany | Institute of Virology, Clinial Virus Genomics, Medical Center, University of Freiburg, Freiburg, Germany | Jonas Fuchs, Lisa Kern, Sandra Reuter, Hajo Grundmann, Marcus Panning |
| EPI_ISL_853718, EPI_ISL_853761 | Department of Microbiology, University Innsbruck | Bergthaler laboratory, CeMM Research Center for Molecular Medicine of the Austrian Academy of Sciences | Lukas Endler, Alexandra Popa, Benedikt Agerer, Jakob-Wendelin Genger, Alexander Lercher, Anna Schedl, Thomas Penz, Michael Schuster, Jan Laine, Martin Senekowitsch, Christoph Bock, Andreas Bergthaler |
| EPI_ISL_853870, EPI_ISL_853871, EPI_ISL_853872, EPI_ISL_853873 | Institute for Laboratory Diagnostics and Microbiology, Klinikum Klagenfurt am Worthersee | Bergthaler laboratory, CeMM Research Center for Molecular Medicine of the Austrian Academy of Sciences | Lukas Endler, Alexandra Popa, Benedikt Agerer, Jakob-Wendelin Genger, Alexander Lercher, Anna Schedl, Thomas Penz, Michael Schuster, Jan Laine, Martin Senekowitsch, Christoph Bock, Andreas Bergthaler |

| | | | |
|---|---|---|---|
| EPI_ISL_854222, EPI_ISL_854225 | Department of Microbiology, University Innsbruck | Bergthaler laboratory, CeMM Research Center for Molecular Medicine of the Austrian Academy of Sciences | Lukas Endler, Alexandra Popa, Benedikt Agerer, Jakob-Wendelin Genger, Alexander Lercher, Anna Schedl, Thomas Penz, Michael Schuster, Jan Laine, Martin Senekowitsch, Christoph Bock, Andreas Bergthaler |
| EPI_ISL_855495, EPI_ISL_855506, EPI_ISL_855510, EPI_ISL_855511 | KEMRI-Wellcome Trust Research Programme/KEMRI-CGMR-C Kilifi | KEMRI-Wellcome Trust Research Programme/KEMRI-CGMR-C Kilifi | Githinji et al |
| EPI_ISL_859768, EPI_ISL_859769, EPI_ISL_859770, EPI_ISL_859771, EPI_ISL_859772, EPI_ISL_859773, EPI_ISL_859774, EPI_ISL_859776, EPI_ISL_859777, EPI_ISL_859782, EPI_ISL_859783, EPI_ISL_859784, EPI_ISL_859785, EPI_ISL_859791, EPI_ISL_859808, EPI_ISL_859809, EPI_ISL_859811, EPI_ISL_859812, EPI_ISL_859813, EPI_ISL_859818, EPI_ISL_859820, EPI_ISL_859821, EPI_ISL_859822, EPI_ISL_859824, EPI_ISL_859826, EPI_ISL_859828, EPI_ISL_859829, EPI_ISL_859830, EPI_ISL_859832, EPI_ISL_859834, EPI_ISL_859835, EPI_ISL_859836, EPI_ISL_859837, EPI_ISL_859838, EPI_ISL_859839, EPI_ISL_859840, EPI_ISL_859841, EPI_ISL_859842 | | | |
| see above | BTC, Khalifa University | BTC, Khalifa University | Al Safar et al |
| EPI_ISL_860130 | Keio University School of Medicine | Keio University School of Medicine | Kenjiro Kosaki, Yuka Iwasaki, Hirotsugu Ishizu, Haruhiko Siomi, Kodai Abe |
| EPI_ISL_860733, EPI_ISL_860734, EPI_ISL_860739, EPI_ISL_860752, EPI_ISL_860753, EPI_ISL_860754, EPI_ISL_860755, EPI_ISL_860756, EPI_ISL_860757, EPI_ISL_860758, EPI_ISL_860759 | | | |
| see above | Swiss National Reference Centre for Influenza | Swiss National Reference Centre for Influenza | Ana Rita Gonçalves Cabecinhas, Samuel Cordey, Florian Laubscher,Christoph Grünig, Laurent Kaiser |
| EPI_ISL_861457 | WHO/Minsk | Charité Universitätsmedizin Berlin, Institut für Virologie | Victor M Corman, Barbara Mühlemann, Jörn Beheim-Schwarzbach, Talitha Veith, Julia Tesch, Tobias Bleicker, Julia Schneider, Shmialiova Natallia, Sivets Natallia, Terry Jones, Christian Drosten |
| EPI_ISL_861718, EPI_ISL_861719, EPI_ISL_861720 | Department of Medical Microbiology, Hospital Pengajar Universiti Putra Malaysia | Malaysia Genome Institute | Mohd Noor Mat Isa, Syafinaz Amin-Nordin, Irni Suhayu Sapian, Hui-Yee Chee, Yusuf Muhammad Noor, Nurhezreen Md Iqbal, Enizza Kasim, Siti Noraini Othman, Mohd Faizal Abu Bakar, Shamsidar Sopie, Azrin Ahmad, Narcisse Joseph, Muhammad MI, Avisha Richards, Nor Zahrin Hasran, Nor Azfa Johari |
| EPI_ISL_861876 | LATE - Laboratório de Técnicas Especiais - Hospital Israelita Albert Einstein | LATE - Laboratório de Técnicas Especiais - Hospital Israelita Albert Einstein | Deyvid Amgarten, Fernanda de Mello Malta, Raquel Riyuzo, Ana Paula Moreira Salles, Pedro Henrique Sebe Rodrigues, João Renato Rebello Pinho |
| EPI_ISL_867934 | Centre for Enzyme Innovation, University of Portsmouth / Translational Research Laboratory, Portsmouth Hospitals NHS Trust | COVID-19 Genomics UK (COG-UK) Consortium | Angela Beckett,Yann Bourgeois,Garry Scarlett,Sharon Glaysher,Scott Elliott,Kelly Bicknell,Robert Impey,Allyson Lloyd,Sarah Wyllie,Ethan Butcher,Anoop Chauhan,Samuel Robson |
| EPI_ISL_868938, EPI_ISL_868939, EPI_ISL_868940, EPI_ISL_868941, EPI_ISL_868942, EPI_ISL_868943, EPI_ISL_868944, EPI_ISL_868945, EPI_ISL_868946, EPI_ISL_868947, EPI_ISL_868948 | | | |
| see above | Bioinformatics and Biostatistics Lab, Advanced Sequencing Facility | COVID-19 Genomics UK (COG-UK) Consortium | Aengus Stewart,Jerome Nicod,Chelsea Sawyer,Laura Cubitt,Harshil Patel,Margaret Crawford |
| EPI_ISL_872469, EPI_ISL_872470, EPI_ISL_872471, EPI_ISL_872472, EPI_ISL_872473, EPI_ISL_872474, EPI_ISL_872475, EPI_ISL_872476, EPI_ISL_872477 | Eurofins Diatherix | Hudsonalpha Genome Sequencing Center | Jane Grimwood, Melissa Williams, Lori H. Handley, Joshua Stough, Leslie Malone, Stefan Brzezinski, Ada Stewart, Teresa Jones, Jenell Webber, John Lovell, Jennifer Cart, and Jeremy Schmutz |
| EPI_ISL_872903 | WHO National Influenza Centre Russian Federation | WHO National Influenza Centre Russian Federation | Andrey Komissarov, Artem Fadeev, Anna Ivanova, Kseniya Komissarova, Dmitry Bazhenov, Mikhail Bakaev, Daria Danilenko, Ksenia Safina, Elena Nabieva, Georgii Bazykin, Dmitry Lioznov |
| EPI_ISL_876524 | Florida Bureau of Public Health Laboratories | Florida Bureau of Public Health Laboratories | Sarah Schmedes, Jason Blanton |
| EPI_ISL_877411, EPI_ISL_877412, EPI_ISL_877413, EPI_ISL_877414, EPI_ISL_877415, EPI_ISL_877416, EPI_ISL_877417, EPI_ISL_877418 | Eurofins Diatherix | Hudsonalpha Genome Sequencing Center | Jane Grimwood, Melissa Williams, Lori H. Handley, Joshua Stough, Leslie Malone, Stefan Brzezinski, Ada Stewart, Teresa Jones, Jenell Webber, John Lovell, Jennifer Cart, and Jeremy Schmutz |
| EPI_ISL_878565 | Robert Garry lab | Andersen lab at Scripps Research | Allison Smither, Gilberto Sabino-Santos, Patricia Snarski, Lilia Melnik, Antoinette Bell, Kaylynn Genemaras, Arnaud Drouin, Dahlene Fusco, Robert Garry with SEARCH Alliance San Diego |
| EPI_ISL_878687, EPI_ISL_878693 | Rady's Childrens Hospital | Andersen lab at Scripps Research | SEARCH Alliance San Diego with Nanda Radamchar, David Dimmock, Linda Luo, Christina Clarke, Kathryn Bouic, Teresa Mueller, Denise Malicki |
| EPI_ISL_882682, EPI_ISL_882690, EPI_ISL_882725, EPI_ISL_882726, EPI_ISL_882752 | 1.AO Universitaria 'S. Giovanni di Dio e Ruggi D'Aragona, Scuola Medica Salernitana' Hospital / 2.UOC di Virologia e Microbiologia, Università della Campania 'L. Vanvitelli' / 3.AO Universitaria 'Federico II' Napoli Hospital / 4.AORN 'San Giuseppe Moscati' Avellino Hospital / 5.AO 'San Pio - presidio G. Rummo' Benevento Hospital / 6.AO 'Sant'Anna e San Sebastiano' Caserta Hospital / 7.PO 'Maria Santissima Addolorata' Eboli Hospital / 8.Biogem Istituto di Ricerche Genetiche | 1. Genome Research Center for Health (CRGS) / 2. Laboratory of Molecular Medicine and Genomics(LMMGe) / 3. Center for Research in Pure and Applied Mathematics (CRMPA) | Giorgio Giurato, Francesca Rizzo, Alessandro Weisz, Gianluigi Franci, Giovanni Nassa, Pasquale Pagliano, Roberta Tarallo, Elena Alexandrova, Ylenia D'Agostino, Carlo Ferravante, Jessica Lamberti, Viola Melone, Domenico Memoli, Valeria Mirici Cappa, Domenico Palumbo, Giovanni Pecoraro, Assunta Sellitto, Oriana Strianese, Ilaria Terenzi, Giuseppe Fenza, Aniello Gentile, Antonello Saccomanno, Sonia Amabile, Teresa Rocco, Annamaria Salvati, Emilia Vaccaro, Massimiliano Galdiero, Michele Cennamo, Giuseppe Portella, Maria Grazia Foti, Mariarosaria Ingino, Maria Landi, Maurizio Fumi, Vincenzo Rocco, Rita Greco, Vittoria Letizia, Arnolfo Petruzziello, Maddalena Schioppa, Gregorio Goffredi, Francesca Marciano, Michele Caraglia, Alessia Cossu, Marianna Scrima |
| EPI_ISL_882954 | State Veterinary Institute Prague | State Veterinary Institute Prague | Nagy,A;Cernikova,L;Stara,M |
| EPI_ISL_884342, EPI_ISL_884353, EPI_ISL_884368, EPI_ISL_884418, EPI_ISL_884419, EPI_ISL_884444 | Infectious Diseases, Quest Diagnostics | Infectious Diseases, Quest Diagnostics | Rosenthal,S.H., Gerasimova,A., Kagan,R.M., Anderson,B., Bernstein,L.E., Livingston,K.E., Hua,M., Liu,Y., Shalhout,D.F., Owen,R., Lacbawan,F. |
| EPI_ISL_888881, EPI_ISL_888882 | Michigan Department of Health and Human Services, Bureau of Laboratories | Michigan Department of Health and Human Services, Bureau of Laboratories | Blankenship HM, Riner D, Soehnlen MK |
| EPI_ISL_892221, EPI_ISL_892222, EPI_ISL_892223 | Lighthouse Lab in Alderley Park | Wellcome Sanger Institute for the COVID-19 Genomics UK (COG-UK) Consortium | Jacquelyn Wynn, Mairead Hyland, The Lighthouse Lab in Alderley Park and Alex Alderton, Roberto Amato, Sonia Goncalves, Ewan Harrison, David K. Jackson, Ian Johnston, Dominic Kwiatkowski, Cordelia Langford, John Sillitoe on behalf of the Wellcome Sanger Institute COVID-19 Surveillance Team |
| EPI_ISL_896118, EPI_ISL_896120, EPI_ISL_896171 | MEPHI, Aix Marseille University | MEPHI, Aix Marseille University | Anthony LEVASSEUR |
| EPI_ISL_896373 | University of Medicine and Pharmacy of Craiova | "Stefan cel Mare" University Metagenomics Lab | Lobiuc Andrei, Gheorghita Roxana |
| EPI_ISL_900097, EPI_ISL_900099, EPI_ISL_900122, EPI_ISL_900125, EPI_ISL_900154, EPI_ISL_900190, EPI_ISL_900231, EPI_ISL_900375, EPI_ISL_900388, EPI_ISL_900419, EPI_ISL_900435 | | | |
| see above | MEPHI, Aix Marseille University | MEPHI, Aix Marseille University | Anthony LEVASSEUR |
| EPI_ISL_904136, EPI_ISL_904154, EPI_ISL_904239, EPI_ISL_904437, EPI_ISL_904438, EPI_ISL_904439, EPI_ISL_904440, EPI_ISL_904441, EPI_ISL_904442 | Dutch COVID-19 response team | Erasmus Medical Center | Bas Oude Munnink, Reina Sikkema, David Nieuwenhuijse, Irina Chestakova, Anne van der Linden, Marjan Boter, Emmanuelle Munger, Corine GeurtsvanKessel, Annemiek van der Eijk, Richard Molenkamp, Marion Koopmans, on behalf of the Dutch national COVID-19 response team. |
| EPI_ISL_905383 | Dutch COVID-19 response team | National Institute for Public Health and the Environment (RIVM) | Adam Meijer, Harry Vennema, Dirk Eggink, Jeroen Cremer, Sharon van den Brink, Bas van der Veer, AnneMarie van den Brandt, Florian Zwagemaker, Dennis Schmitz, Chantal Reusken, on behalf of the national COVID-19 response team |
| EPI_ISL_906118 | Institute of Microbiology and Immunology, Faculty of Medicine, University of Ljubljana | Institute of Microbiology and Immunology, Faculty of Medicine, University of Ljubljana | Samo Zakotnik, Tomaž Mark Zorec, Matic Brvar, Miša Korva, Mario Poljak, Tatjana Avši - Županc |
| EPI_ISL_910330 | General Hospital - Veles | Research Center for Genetic Engineering and Biotechnology "Georgi D. Efremov" , Macedonian Academy of Sciences and Arts | RCGEB - MASA |

| | | | |
|---|---|---|---|
| EPI_ISL_910333 | General Hospital - Ohrid | Research Center for Genetic Engineering and Biotechnology "Georgi D. Efremov" , Macedoni | RCGEB - MASA |
| EPI_ISL_912456, EPI_ISL_912485, EPI_ISL_912488, EPI_ISL_912497, EPI_ISL_912504, EPI_ISL_912509, EPI_ISL_912513, EPI_ISL_912522 | NHLS Universitas Academic | UFS Virology | PA Bester, MM Nyaga, P Nthiga, MT Mogotsi, D Goedhals, T de Oliveira |
| EPI_ISL_913079, EPI_ISL_913087 | Center for Virology | Center for Virology | Jeremy V. Camp, Irene Goerzer, Monika Redlberger-Fritz, Stephan W. Aberle |
| EPI_ISL_913351 | Klinisk mikrobiologi | The Public Health Agency of Sweden | Anna-Malin Linde, Maria Lind Karlberg, Carlo Berg, Oskar Karlsson Lindsjo, Sofia Stamouli, Reza Advani, Mattias Haukland, Petra Holmstrom, Noura Walai, Petra Edquist, Mia Brytting, Anna Risberg, Karin Tegmark-Wisell |
| EPI_ISL_913925, EPI_ISL_913927, EPI_ISL_913928, EPI_ISL_913929, EPI_ISL_913930, EPI_ISL_913931, EPI_ISL_913932, EPI_ISL_913957 | Instituto de Diagnostico y Referencia Epidemiologicos INDRE_RNLSP | Instituto de Diagnostico y Referencia Epidemiologicos (INDRE) | Claudia Wong-Arambula, Abril Rodriguez-Maldonado, Fabiola Garces-Ayala, Adnan Araiza-Rodriguez, David Fragoso-Fonseca, Sergio Rangel-Guerrero, Mayra Jimenez-Morales, Nancy Munoz-Hernandez, Natividad Cruz-Ortiz, Tatiana Nunez-Garcia, Gisela Barrera-Badillo, Lucia Hernandez-Rivas, Irma Lopez-Martinez, Ernesto Ramirez-Gonzalez. |
| EPI_ISL_918484 | Gerontology Institute - Skopje | Research Center for Genetic Engineering and Biotechnology "Georgi D. Efremov" , Macedonian Academy of Sciences and Arts | RCGEB - MASA |
| EPI_ISL_918486 | General Hospital - Struga | Research Center for Genetic Engineering and Biotechnology "Georgi D. Efremov" , Macedonian Academy of Sciences and Arts | RCGEB - MASA |
| EPI_ISL_921025 | Regional Virus Laboratory, Belfast Health and Social Care Trust | COVID-19 Genomics UK (COG-UK) Consortium | Conall McCaughey, James McKenna, Tanya Curran, Susan Feeney, Alison Watt, Ciara Cox, Mairead Connor, Zoltan Molnar, David Simpson, Derek Fairley |
| EPI_ISL_924419, EPI_ISL_924423 | Virology Department, Sheffield Teaching Hospitals NHS Foundation Trust/Department of Infection, Immunity and Cardiovascular Disease, The Medical School, University of Sheffield | COVID-19 Genomics UK (COG-UK) Consortium | Thushan de Silva, Matthew Parker, Nikki Smith, Adri Angyal, Rebecca Brown, Luke Green, Rachel Tucker, Paul Parsons, Danielle Groves, Katie Johnson, Laura Carrilero, Alex Keeley, Dave Partridge, Matthew Wyles, Benjamin Lindsey, Mehmet Yavuz, Mohammad Raza, Cariad Evans |
| EPI_ISL_924613, EPI_ISL_925022 | Bioinformatics and Biostatistics Lab, Advanced Sequencing Facility | COVID-19 Genomics UK (COG-UK) Consortium | Aengus Stewart,Jerome Nicod,Chelsea Sawyer,Laura Cubitt,Harshil Patel,Margaret Crawford |
| EPI_ISL_925107, EPI_ISL_925108, EPI_ISL_925129, EPI_ISL_925130 | 1.AO Universitaria 'S. Giovanni di Dio e Ruggi D'Aragona, Scuola Medica Salernitana' Hospital / 2.UOC di Virologia e Microbiologia, Università della Campania 'L. Vanvitelli' / 3.AO Universitaria 'Federico II' Napoli Hospital / 4.AORN 'San Giuseppe Moscati' Avellino Hospital / 5.AO 'San Pio - presidio G. Rummo' Benevento Hospital / 6.AO 'Sant'Anna e San Sebastiano' Caserta Hospital / 7.PO 'Maria Santissima Addolorata' Eboli Hospital / 8.Biogem Istituto di Ricerche Genetiche | 1. Genome Research Center for Health (CRGS) / 2. Laboratory of Molecular Medicine and Genomics(LMMGe) / 3. Center for Research in Pure and Applied Mathematics (CRMPA) | Giorgio Giurato, Francesca Rizzo, Alessandro Weisz, Gianluigi Franci, Giovanni Nassa, Pasquale Pagliano, Roberta Tarallo, Elena Alexandrova, Ylenia D'Agostino, Carlo Ferravante, Jessica Lamberti, Viola Melone, Domenico Memoli, Valeria Mirici Cappa, Domenico Palumbo, Giovanni Pecoraro, Assunta Sellitto, Oriana Strianese, Ilaria Terenzi, Giuseppe Fenza, Aniello Gentile, Antonello Saccomanno, Sonia Amabile, Teresa Rocco, Annamaria Salvati, Emilia Vaccaro, Massimiliano Galdiero, Michele Cennamo, Giuseppe Portella, Maria Grazia Foti, Mariarosaria Ingino, Maria Landi, Maurizio Fumi, Vincenzo Rocco, Rita Greco, Vittoria Letizia, Arnolfo Petruzziello, Maddalena Schioppa, Gregorio Goffredi, Francesca Marciano, Michele Caraglia, Alessia Cossu, Marianna Scrima |
| EPI_ISL_925207 | Virginia DCLS | Virginia DCLS | Virginia DCLS |
| EPI_ISL_925442, EPI_ISL_925443, EPI_ISL_925444, EPI_ISL_925445, EPI_ISL_925479, EPI_ISL_925480, EPI_ISL_925481, EPI_ISL_925482 | Department of Clinical Microbiology | GIGA Medical Genomics | Keith Durkin, Maria Artesi, Sébastien Bontems, Raphaël Boreux, Bouchra Boujemla, Cécile Meex, Pierrette Melin, Marie-Pierre Hayette, Vincent Bours |
| EPI_ISL_925883 | Nucleic Acid Testing, National Reference Laboratory | GIGA Medical Genomics | Yvan Butera, Keith Durkin, Maria Artesi, Bouchra Boujemla, Robert Rutayisire, Patrick Tuyisenge, Esperence Umumararungu, Sébastien Bontems, Marie-Pierre Hayette, Nathalie Renotte, Swaibu Gatare, Jacob Souopgui, Sabin Nsanzimana, Vincent Bours, Léon Mutesa |
| EPI_ISL_930920, EPI_ISL_930921, EPI_ISL_930922, EPI_ISL_930923, EPI_ISL_930924, EPI_ISL_930925, EPI_ISL_930926, EPI_ISL_930927, EPI_ISL_930928, EPI_ISL_930929, EPI_ISL_930931, EPI_ISL_930934, EPI_ISL_930935, EPI_ISL_930936, EPI_ISL_930937, EPI_ISL_930938, EPI_ISL_930939, EPI_ISL_930940, EPI_ISL_930941, EPI_ISL_930942, EPI_ISL_930943, EPI_ISL_930944, EPI_ISL_930945, EPI_ISL_930946, EPI_ISL_930947, EPI_ISL_930948, EPI_ISL_930949, EPI_ISL_930950, EPI_ISL_930951, EPI_ISL_930952, EPI_ISL_930953, EPI_ISL_930954, EPI_ISL_930955, EPI_ISL_930956, EPI_ISL_930957, EPI_ISL_930958, EPI_ISL_930959, EPI_ISL_930960, EPI_ISL_930961, EPI_ISL_930962, EPI_ISL_930963, EPI_ISL_930966, EPI_ISL_930967, EPI_ISL_930968, EPI_ISL_930969, EPI_ISL_930970, EPI_ISL_930971, EPI_ISL_930972, EPI_ISL_930973, EPI_ISL_930974, EPI_ISL_930975, EPI_ISL_930976, EPI_ISL_930977, EPI_ISL_930978, EPI_ISL_930979, EPI_ISL_930980, EPI_ISL_930981, EPI_ISL_930982, EPI_ISL_930983, EPI_ISL_930984, EPI_ISL_930985, EPI_ISL_930986, EPI_ISL_930987, EPI_ISL_930988, EPI_ISL_930989, EPI_ISL_930990, EPI_ISL_930991, EPI_ISL_930992, EPI_ISL_930993, EPI_ISL_931396 | | | |
| see above | University Hospital Basel, Clinical Virology | University Hospital Basel, Clinical Bacteriology | Tim Roloff, Madlen Stange, Helena MB Seth-Smith, Alfredo Mari, Karoline Leuzinger, Julia Bielicki, Manuel Battegay, Hans Hirsch, Adrian Egli |
| EPI_ISL_933666, EPI_ISL_933667 | Instituto de Diagnostico y Referencia Epidemiologicos INDRE_RNLSP | Instituto de Diagnostico y Referencia Epidemiologicos (INDRE) | Claudia Wong-Arambula, Abril Rodriguez-Maldonado, Fabiola Garces-Ayala, Adnan Araiza-Rodriguez, David Fragoso-Fonseca, Sergio Rangel-Guerrero, Mayra Jimenez-Morales, Nancy Munoz-Hernandez, Natividad Cruz-Ortiz, Tatiana Nunez-Garcia, Gisela Barrera-Badillo, Lucia Hernandez-Rivas, Irma Lopez-Martinez, Ernesto Ramirez-Gonzalez. |
| EPI_ISL_935143, EPI_ISL_935144, EPI_ISL_935145, EPI_ISL_935146, EPI_ISL_935147, EPI_ISL_935148 | 1.AO Universitaria 'S. Giovanni di Dio e Ruggi D'Aragona, Scuola Medica Salernitana' Hospital / 2.UOC di Virologia e Microbiologia, Università della Campania 'L. Vanvitelli' / 3.AO Universitaria 'Federico II' Napoli Hospital / 4.AORN 'San Giuseppe Moscati' Avellino Hospital / 5.AO 'San Pio - presidio G. Rummo' Benevento Hospital / 6.AO 'Sant'Anna e San Sebastiano' Caserta Hospital / 7.PO 'Maria Santissima Addolorata' Eboli Hospital / 8.Biogem Istituto di Ricerche Genetiche | 1. Genome Research Center for Health (CRGS) / 2. Laboratory of Molecular Medicine and Genomics(LMMGe) / 3. Center for Research in Pure and Applied Mathematics (CRMPA) | Giorgio Giurato (Corresponding Author), Francesca Rizzo (Corresponding Author), Alessandro Weisz (Corresponding Author), Gianluigi Franci, Giovanni Nassa, Pasquale Pagliano, Roberta Tarallo, Elena Alexandrova, Ylenia D'Agostino, Carlo Ferravante, Jessica Lamberti, Viola Melone, Domenico Memoli, Valeria Mirici Cappa, Domenico Palumbo, Giovanni Pecoraro, Assunta Sellitto, Oriana Strianese, Ilaria Terenzi, Giuseppe Fenza, Aniello Gentile, Antonello Saccomanno, Sonia Amabile, Teresa Rocco, Annamaria Salvati, Emilia Vaccaro, Massimiliano Galdiero, Michele Cennamo, Giuseppe Portella, Maria Grazia Foti, Mariarosaria Ingino, Maria Landi, Maurizio Fumi, Vincenzo Rocco, Rita Greco, Vittoria Letizia, Arnolfo Petruzziello, Maddalena Schioppa, Gregorio Goffredi, Francesca Marciano, Michele Caraglia, Alessia Cossu, Marianna Scrima |
| EPI_ISL_937105, EPI_ISL_937119 | Quest Diagnostics | Quest Diagnostics | Rosenthal,S.H., Gerasimova,A., Kagan,R.M., Anderson, B., Livingston, K.E., Hua, M., Liu Y., Shalhout, D.F., Owen, R., Lacbawan, F. |
| EPI_ISL_940487 | Hôpital Bichat Claude Bernard, Laboratoire de Virologie | IAME UMR1137 Inserm, Université de Paris, Hôpital Bichat | Antoine Bridier-Nahmias, Amélie Recoing, Quentin Le Hingrat, Lena Daniel, Siham Hamri, Gilles Collin, Alexandre Storto, Mélanie Bertine, Charlotte Charpentier, Nadhira Houhou-Fidouh, Diane Descamps, Benoit Visseaux |
| EPI_ISL_940942, EPI_ISL_940943, EPI_ISL_940944, EPI_ISL_940945, EPI_ISL_940946, EPI_ISL_940947, EPI_ISL_940948, EPI_ISL_940949, EPI_ISL_940994 | Centers for Disease Control and Prevention, Dengue Branch | Centers for Disease Control and Prevention, Dengue Branch | Gilberto A. Santiago, Glenda Gonzalez, Betzabel Flores, Keyla Charriez, Gabriela Paz-Bailey, Jorge L. Munoz-Jordan |
| EPI_ISL_941199 | Servicio de Microbiología. Hospital Clínico Universitario de Valencia | SeqCOVID-SPAIN consortium/IBV(CSIC) | David Navarro Ortega, Eliseo Albert Vicent, Ignacio Torres and SeqCOVID-SPAIN consortium |
| EPI_ISL_942014, EPI_ISL_942015, EPI_ISL_942016, EPI_ISL_942017, EPI_ISL_942018, EPI_ISL_942019, EPI_ISL_942020, EPI_ISL_942021, EPI_ISL_942022, EPI_ISL_942023, EPI_ISL_942048, EPI_ISL_942049, EPI_ISL_942050, EPI_ISL_942051, EPI_ISL_942052, EPI_ISL_942053, EPI_ISL_942054, EPI_ISL_942055, EPI_ISL_942056, EPI_ISL_942057, EPI_ISL_942058, EPI_ISL_942059, EPI_ISL_942060, EPI_ISL_942061, EPI_ISL_942062, EPI_ISL_942063, EPI_ISL_942064, EPI_ISL_942065, EPI_ISL_942066, EPI_ISL_942067, EPI_ISL_942068 | | | |
| see above | Wisconsin State Laboratory of Hygiene Communicable Disease Division | Wisconsin State Laboratory of Hygiene Communicable Disease Division | Kelsey R. Florek, Abigail C. Shockey |
| EPI_ISL_949234 | Departamento de Microbiología, CDB, Hospital Clínic, | SeqCOVID-SPAIN consortium/IBV(CSIC) | Andrea Vergara, Mikel Martínez, Elisa Rubio, Jéssica Navero, Aida Peiró and SeqCOVID-SPAIN consortium |

| | | | |
|---|---|---|---|
| | Barcelona | | |
| EPI_ISL_952951, EPI_ISL_952952 | Liverpool Clinical Laboratories | COVID-19 Genomics UK (COG-UK) Consortium | Sam Haldenby, Anita Lucaci, Steve Paterson, Julian Hiscox, Alistair Darby, M Almsaud, A Alrezaihi, Muhannad Alruwaili, Stuart D Armstrong, Jones Benjamin, Eleanor G Bentley, Anu Chawla, Jordan J Clark, Angela Cowell, Richard Eccles, Isabel García-Dorival, Matthew Gemmell, Alessandro Gerada, PKF Gilmore, Richard Gregory, Ximeng Han, Catherine Hartley, Margaret Hughes, Miren Iturriza-Gomara, James Johnson, L Luu, Jenifer Manson, Charlotte Nelson, Elaine O'Toole, Cassie Olateju, Rebekah Penrice-Randal , Lucille Rainbow, N.P Randle, Trevor Ian Robinson, Parul Sharma, Ghada T Shawli, James P Stewart, Neil Swainston, Ecaterina Vamos, Joanne Watts, Mark Whitehead |
| EPI_ISL_954187, EPI_ISL_954188 | 1.AO Universitaria 'S. Giovanni di Dio e Ruggi D'Aragona, Scuola Medica Salernitana' Hospital / 2.UOC di Virologia e Microbiologia, Università della Campania 'L. Vanvitelli' / 3.AO Universitaria 'Federico II' Napoli Hospital / 4.AORN 'San Giuseppe Moscati' Avellino Hospital / 5.AO 'San Pio - presidio G. Rummo' Benevento Hospital / 6.AO 'Sant'Anna e San Sebastiano' Caserta Hospital / 7.PO 'Maria Santissima Addolorata' Eboli Hospital / 8.Biogem Istituto di Ricerche Genetiche | 1. Genome Research Center for Health (CRGS) / 2. Laboratory of Molecular Medicine and Genomics(LMMGe) / 3. Center for Research in Pure and Applied Mathematics (CRMPA) | Giorgio Giurato, Francesca Rizzo, Alessandro Weisz, Gianluigi Franci, Giovanni Nassa, Pasquale Pagliano, Roberta Tarallo, Elena Alexandrova, Ylenia D'Agostino, Carlo Ferravante, Jessica Lamberti, Viola Melone, Domenico Memoli, Valeria Mirici Cappa, Domenico Palumbo, Giovanni Pecoraro, Assunta Sellitto, Oriana Strianese, Ilaria Terenzi, Giuseppe Fenza, Aniello Gentile, Antonello Saccomanno, Sonia Amabile, Teresa Rocco, Annamaria Salvati, Emilia Vaccaro, Massimiliano Galdiero, Michele Cennamo, Giuseppe Portella, Maria Grazia Foti, Mariarosaria Ingino, Maria Landi, Maurizio Fumi, Vincenzo Rocco, Rita Greco, Vittoria Letizia, Arnolfo Petruzziello, Maddalena Schioppa, Gregorio Goffredi, Francesca Marciano, Michele Caraglia, Alessia Cossu, Marianna Scrima, Edmondo Adorisio, Morena D'Avenia, Michela Iacobellis, Rosanna Piluscio, Giorgio Dirani, Vittorio Sambri, Simona Semprini, Silvia Zanoli, Francesco Curcio, Stefania Marzinotto, Andreina Baj, Fausto Sessa. |
| EPI_ISL_960131 | Victoria Hospital wc VHW | National Health Laboratory Service/UCT | Arash Iranzadeh, Deelan Doolabh, Lynn Tyers, Bruna Galvao, Innocent Mudau, Marvin Hsiao, Kruger Marais, Diana Hardie, Stephen Korsman, Carolyn Williamson |
| EPI_ISL_960132, EPI_ISL_960133 | Groote Schuur Hospital wc GSH | National Health Laboratory Service/UCT | Arash Iranzadeh, Deelan Doolabh, Lynn Tyers, Bruna Galvao, Innocent Mudau, Marvin Hsiao, Kruger Marais, Diana Hardie, Stephen Korsman, Carolyn Williamson |
| EPI_ISL_960134 | Conville CDC wc CVC | National Health Laboratory Service/UCT | Arash Iranzadeh, Deelan Doolabh, Lynn Tyers, Bruna Galvao, Innocent Mudau, Marvin Hsiao, Kruger Marais, Diana Hardie, Stephen Korsman, Carolyn Williamson |
| EPI_ISL_960135 | Groote Schuur Hospital wc GSH | National Health Laboratory Service/UCT | Arash Iranzadeh, Deelan Doolabh, Lynn Tyers, Bruna Galvao, Innocent Mudau, Marvin Hsiao, Kruger Marais, Diana Hardie, Stephen Korsman, Carolyn Williamson |
| EPI_ISL_960237 | Nucleic Acid Testing, National Reference Laboratory | GIGA Medical Genomics | Yvan Butera, Keith Durkin, Maria Artesi, Bouchra Boujemla, Robert Rutayisire, Patrick Tuyisenge, Esperence Umumararungu, Sébastien Bontems, Marie-Pierre Hayette, Nathalie Renotte, Corinne Fasquelle, Swaibu Gatare, Jacob Souopgui, Sabin Nsanzimana, Vincent Bours, Léon Mutesa |
| EPI_ISL_965578, EPI_ISL_965619 | Dutch COVID-19 response team | Medical Microbiology, Maastricht University Medical Centre | Jozef Dingemans*, Brian van der Veer*, Erik Beuken, Carmen Reumkens, Lieke van Alphen, Christian Hoebe, Paul Savelkoul |
| EPI_ISL_968894, EPI_ISL_968898, EPI_ISL_968899, EPI_ISL_968961 | KEMRI-Wellcome Trust Research Programme/KEMRI-CGMR-C Kilifi | KEMRI-Wellcome Trust Research Programme/KEMRI-CGMR-C Kilifi | Githinji et al |
| EPI_ISL_969397 | Indian Council of Medical Research-National Institute of Virology, Pune | Indian Council of Medical Research-National Institute of Virology, Maximum Containment Laboratory | Pragya D. Yadav, Anita Shete-Aich, Dimpal A. Nyayanit, Abhinendra Kumar, Savita Patil, Triparna D. Majumdar |
| EPI_ISL_974465, EPI_ISL_974466, EPI_ISL_974467, EPI_ISL_974468, EPI_ISL_974469, EPI_ISL_974470, EPI_ISL_974471, EPI_ISL_974472, EPI_ISL_974473, EPI_ISL_974474, EPI_ISL_974475, EPI_ISL_974476, EPI_ISL_974477, EPI_ISL_974478, EPI_ISL_974479, EPI_ISL_974480, EPI_ISL_974481, EPI_ISL_974482, EPI_ISL_974483, EPI_ISL_974484, EPI_ISL_974485, EPI_ISL_974486, EPI_ISL_974487, EPI_ISL_974488, EPI_ISL_974489, EPI_ISL_974490, EPI_ISL_974491, EPI_ISL_974492, EPI_ISL_974493, EPI_ISL_974494, EPI_ISL_974495, EPI_ISL_974496, EPI_ISL_974497, EPI_ISL_974498, EPI_ISL_974499, EPI_ISL_974500, EPI_ISL_974501, EPI_ISL_974502, EPI_ISL_974503, EPI_ISL_974504, EPI_ISL_974505, EPI_ISL_974506, EPI_ISL_974507, EPI_ISL_974508, EPI_ISL_974509, EPI_ISL_974510, EPI_ISL_974511, EPI_ISL_974512, EPI_ISL_974513, EPI_ISL_974514, EPI_ISL_974515, EPI_ISL_974516, EPI_ISL_974517, EPI_ISL_974518, EPI_ISL_974519, EPI_ISL_974520, EPI_ISL_974521, EPI_ISL_974522, EPI_ISL_974523, EPI_ISL_974524, EPI_ISL_974525, EPI_ISL_974526, EPI_ISL_974527, EPI_ISL_974528, EPI_ISL_974529, EPI_ISL_974530, EPI_ISL_974531, EPI_ISL_974532, EPI_ISL_974533, EPI_ISL_974534, EPI_ISL_974535, EPI_ISL_974536, EPI_ISL_974537, EPI_ISL_974538, EPI_ISL_974539, EPI_ISL_974540, EPI_ISL_974541, EPI_ISL_974542, EPI_ISL_974543, EPI_ISL_974544, EPI_ISL_974545, EPI_ISL_974546, EPI_ISL_974547, EPI_ISL_974548, EPI_ISL_974549, EPI_ISL_974550, EPI_ISL_974551, EPI_ISL_974552, EPI_ISL_974553, EPI_ISL_974554, EPI_ISL_974555, EPI_ISL_974556 | | | |
| see above | BCCDC Public Health Laboratory | BCCDC Public Health Laboratory | Prystajecky Natalie, Linda Hoang, Dan Fornika, John Tyson, Shannon Russell, Kim Macdonald, Kimia Kamelian, Ana Pacagnella, Corrinne Ng, Loretta Janz, Robert Azana Terry Snutch, Mel Krajden |
| EPI_ISL_977586, EPI_ISL_977587, EPI_ISL_977588 | Department of Medical Microbiology, Hospital Pengajar Universiti Putra Malaysia | Malaysia Genome Institute | Mohd Noor Mat Isa, Syafinaz Amin-Nordin, Irni Suhayu Sapian, Hui-Yee Chee, Yusuf Muhammad Noor, Nurhezreen Md Iqbal, Enizza Kasim, Siti Noraini Othman, Mohd Faizal Abu Bakar, Shamsidar Sopie, Azrin Ahmad, Narcisse Joseph, Muhammad MI, Avisha Richards, Nor Zahrin Hasran, Nor Azfa Johari |
| EPI_ISL_977915, EPI_ISL_977916 | Chiu Laboratory, University of California, San Francisco | Chiu Laboratory, University of California, San Francisco | Charles Chiu, Xianding (Wayne) Deng, Candace Wang, Venice Servellita, Jill Hacker, Debra Wadford |
| EPI_ISL_982506 | Kentucky State Public Health Lab | Kentucky State Public Health Lab | Stephanie Lunn, Karim George, Joshua Tobias, William Grooms, Vaneet Arora, Matthew Johnson, Rachel Zinner, Rhonda Lucas |
| EPI_ISL_982930 | MEPHI Aix Marseille University (AMU) | MEPHI Aix Marseille University (AMU) | Anthony LEVASSEUR |
