## Supplementary material for "Detection and characterization of the SARS-CoV-2 lineage B.1.526 in New York": Supp. Table 3 GISAID Acknowledgment part 2: gisaid_hcov-19_acknowledgement_table_2021_02_13_01-2.pdf

We gratefully acknowledge the following Authors from the Originating laboratories responsible for obtaining the specimens, as well as the Submitting laboratories where the genome data were generated and shared via GISAID, on which this research is based.

All Submitters of data may be contacted directly via www.gisaid.org

Authors are sorted alphabetically.

| Accession ID | Originating Laboratory | Submitting Laboratory | Authors |
|---|---|---|---|
| EPI_ISL_906273, EPI_ISL_906274 | University of Wisconsin-Madison AIDS Vaccine Research Laboratories | University of Wisconsin-Madison AIDS Vaccine Research Laboratories | Gage Moreno, Katarina Braun, et al. AIDS Vaccine Research Laboratories |
| EPI_ISL_907237 | Lighthouse Lab in Glasgow | Wellcome Sanger Institute for the COVID-19 Genomics UK (COG-UK) Consortium | Harper VanSteenhouse, Yumi Kasai, David Gray, Carol Clugston, Anna Dominiczak and Alex Alderton, Roberto Amato, Sonia Goncalves, Ewan Harrison, David K. Jackson, Ian Johnston, Dominic Kwiatkowski, Cordelia Langford, John Sillitoe on behalf of the Wellcome Sanger Institute COVID-19 Surveillance Team |
| EPI_ISL_911372, EPI_ISL_911374, EPI_ISL_911375, EPI_ISL_911378, EPI_ISL_911379, EPI_ISL_911380 | University of Michigan Clinical Microbiology Laboratory | Lauring Lab, University of Michigan, Department of Microbiology and Immunology | Valesano |
| EPI_ISL_918430 | Bundeswehrkrankenhaus Westerstede | Bundeswehr Institute of Microbiology | Markus Antwerpen, Klaus Peter Ebert, Alexandra Rehn, Mathias Walter, Malena Bestehorn-Willmann, Sabine Zange, Enrico Georgi, Roman Wölfel |
| EPI_ISL_918431, EPI_ISL_918432, EPI_ISL_918433 | Bundeswehrkrankenhaus Berlin | Bundeswehr Institute of Microbiology | Markus Antwerpen, Martin Müller, Alexandra Rehn, Mathias Walter, Malena Bestehorn-Willmann, Sabine Zange, Enrico Georgi, Roman Wölfel |
| EPI_ISL_918483 | Laboratory of Microbiology, ASST Settelaghi, Varese, Italy | Laboratory of Microbiology, ASST Settelaghi, Varese, Italy | Novazzi,F., Genoni,A., Baj,A., Focosi,D., Spezia,P.G., Zago,C., Colombo,A., Cassani,G., Pasciuta,R., Tamborini,A., Rossi,A., Prestia,M., Capuano,R., Maggi,F. |
| EPI_ISL_924084, EPI_ISL_924101, EPI_ISL_924104, EPI_ISL_924108, EPI_ISL_924119, EPI_ISL_924122, EPI_ISL_924128, EPI_ISL_924148, EPI_ISL_924164, EPI_ISL_924166, EPI_ISL_924176, EPI_ISL_924196, EPI_ISL_924197, EPI_ISL_924203, EPI_ISL_924215, EPI_ISL_924225, EPI_ISL_924231, EPI_ISL_924240, EPI_ISL_924245, EPI_ISL_924252, EPI_ISL_924265, EPI_ISL_924274, EPI_ISL_924281, EPI_ISL_924290, EPI_ISL_924296, EPI_ISL_924302, EPI_ISL_924328, EPI_ISL_924332, EPI_ISL_924334, EPI_ISL_924351, EPI_ISL_924376, EPI_ISL_924384, EPI_ISL_924389, EPI_ISL_924392, EPI_ISL_924397, EPI_ISL_924406 | | | |
| see above | Virology Department, Sheffield Teaching Hospitals NHS Foundation Trust/Department of Infection, Immunity and Cardiovascular Disease, The Medical School, University of Sheffield | COVID-19 Genomics UK (COG-UK) Consortium | Thushan de Silva, Matthew Parker, Nikki Smith, Adri Angyal, Rebecca Brown, Luke Green, Rachel Tucker, Paul Parsons, Danielle Groves, Katie Johnson, Laura Carrilero, Alex Keeley, Dave Partridge, Matthew Wyles, Benjamin Lindsey, Mehmet Yavuz, Mohammad Raza, Cariad Evans |
| EPI_ISL_933798, EPI_ISL_933799, EPI_ISL_933800, EPI_ISL_933801 | PathWest Laboratory Medicine WA | PathWest Laboratory Medicine WA Microbial Surveillance Unit | PathWest Laboratory Medicine WA Microbial Surveillance Unit |
| EPI_ISL_934971, EPI_ISL_934972 | Thai Red Cross Emerging Infectious Diseases Health Science Centre, Chulalongkorn Hospital, Faculty of Medicine, Chulalongkorn University | Thai Red Cross Emerging Infectious Diseases Center and Faculty of Medicine, Chulalongkorn University | Rome Buathong, Sopon Iamsirithaworn, Sininat Petcharat, Yutthana Joyjinda, Weenassarin Ampoot, Apaporn Rodpan, Opass Putcharoen, Thiravat Hemachudha, Supaporn Wacharapluesadee |
| EPI_ISL_935017, EPI_ISL_935021 | Laboratory for Respiratory Viruses, Cantacuzino National Military-Medical Institute for Research and Development | Cantacuzino Institute Virology | Luiza Ustea, Nicoleta Paraschiv, Mihaela Lazar |
| EPI_ISL_935102 | Ministry of Health Turkey | Ministry of Health Turkey | Fatma Bayrakdar, Yasemin Cogun, Süleyman Yalcin, Aye Baak Alta, Gülay Korukluolu |
| EPI_ISL_935181, EPI_ISL_935182, EPI_ISL_935183, EPI_ISL_935184, EPI_ISL_935185, EPI_ISL_935186 | Wyoming Public Health Laboratory | Wyoming Public Health Laboratory | Noah Hull, Taylor Fearing, Lynette Gumbleton, Channing Weber, Ashley Norberg, Bailey Bowcutt, and Wanda Manley |
| EPI_ISL_937468, EPI_ISL_937469, EPI_ISL_937470, EPI_ISL_937471, EPI_ISL_937472, EPI_ISL_937473, EPI_ISL_937474, EPI_ISL_937475, EPI_ISL_937476, EPI_ISL_937477, EPI_ISL_937478, EPI_ISL_937479, EPI_ISL_937480, EPI_ISL_937481, EPI_ISL_937482, EPI_ISL_937483, EPI_ISL_937484, EPI_ISL_937485, EPI_ISL_937486, EPI_ISL_937487, EPI_ISL_937488, EPI_ISL_937489, EPI_ISL_937490, EPI_ISL_937506, EPI_ISL_937508, EPI_ISL_937509, EPI_ISL_937511 | | | |
| see above | Maine Health and Environmental Testing Laboratory (Maine HETL) | Tewhey Lab, The Jackson Laboratory | Matluk,N., Dewey,H., Iosue,F., Barter,M., Lynch,R., Munger,H. and Tewhey,R. |
| EPI_ISL_937515, EPI_ISL_937516, EPI_ISL_937517, EPI_ISL_937518, EPI_ISL_937519, EPI_ISL_937520, EPI_ISL_937521, EPI_ISL_937522, EPI_ISL_937523, EPI_ISL_937524, EPI_ISL_937525, EPI_ISL_937526, EPI_ISL_937527, EPI_ISL_937528, EPI_ISL_937529, EPI_ISL_937530 | | | |
| see above | National Public Health Laboratory, National Centre for Infectious Diseases | National Public Health Laboratory, National Centre for Infectious Diseases | Tze Minn Mak, Zhenyang Zhou, Lin Cui, Raymond Tzer Pin Lin |
| EPI_ISL_939647, EPI_ISL_940144 | Laboratory for Respiratory Viruses, Cantacuzino National Military-Medical Institute for Research and Development | Cantacuzino Institute Virology | Luiza Ustea, Nicoleta Paraschiv, Mihaela Lazar |
| EPI_ISL_940577, EPI_ISL_940578, EPI_ISL_940579 | Jessa | Jessa | Jessa_cmdLab |
| EPI_ISL_940741 | LHUB-ULB | UAntwerp, Laboratory of Medical Microbiology, Campus Drie Eiken S6.26, Universiteitsplein 1, 2610, Wilrijk, Belgium | Basil Britto Xavier, Jasmine Coppens, Marie Le Mercier, Christine Lammens, Veerle Matheeussen, Herman Goossens |
| EPI_ISL_941335 | New Mexico Department of Health Scientific Laboratory | New Mexico Department of Health Scientific Laboratory | Ellie Johnson, Anastacia Griego-Fisher, D'eldra Malone, Jennifer Benoit |
| EPI_ISL_942354, EPI_ISL_942366, EPI_ISL_942369, EPI_ISL_942370, EPI_ISL_942371, EPI_ISL_942372, EPI_ISL_942373 | MD PHL | MD PHL | Maryland Department of Health Laboratories Administration |
| EPI_ISL_942894 | Gundersen Molecular Diagnostics Laboratory | Kabara Cancer Research Institute | Craig S. Richmond, Paraic A. Kenny |
| EPI_ISL_942895 | South Eastern Area Laboratory Services (SEALS) | NSW Health Pathology - Institute of Clinical Pathology and Medical Research; Westmead Hospital; University of Sydney | CIDM-PH et al. |
| EPI_ISL_942935, EPI_ISL_942936, EPI_ISL_942937, EPI_ISL_942938, EPI_ISL_942939 | Wyoming Public Health Laboratory | Wyoming Public Health Laboratory | Noah Hull, Taylor Fearing, Lynette Gumbleton, Channing Weber, Ashley Norberg, Bailey Bowcutt, and Wanda Manley |
| EPI_ISL_943570 | Laboratorio de Referencia Nacional de Virus Respiratorio. Instituto Nacional de Salud Perú | Laboratorio de Referencia Nacional de Biotecnología y Biología Molecular. Instituto Nacional de Salud Perú | Carlos Padilla Rojas, Karolyn Vega Chozo, Luis Barcena, Priscila Lope Pari, Omar Caceres Rey, Marco Galarza Perez, Maribel Huaringa Nuñez, Johanna Balbuena Torrez, Henri Bailon Calderon, Nancy Rojas Serrano |
| EPI_ISL_944748 | unknown | Public Health Virology-Forensic and Scientific Services (PHV-FSS) | Son Nguyen et al. |
| EPI_ISL_944785 | Laboratory of Microbiology and Virology, Ospedale Amedeo di Savoia, ASL "Città di Torino" | U.O. Genomics, S.S. Genetics and Advanced Omics Techniques, Istituto Zooprofilattico Sperimentale del Piemonte Liguria e Valle d'Aosta | Chiara Beltramo, Francesco Cerutti, Valeria Ghisetti, Elena Bozzetta, Pier Luigi Acutis, Simone Peletto |

| | | | |
|---|---|---|---|
| EPI_ISL_944792, EPI_ISL_944793, EPI_ISL_944794, EPI_ISL_944795, EPI_ISL_944796, EPI_ISL_944797, EPI_ISL_944798, EPI_ISL_944800, EPI_ISL_944801, EPI_ISL_944802, EPI_ISL_944803, EPI_ISL_944804, EPI_ISL_944806, EPI_ISL_944807, EPI_ISL_944808, EPI_ISL_944809, EPI_ISL_944810, EPI_ISL_944811, EPI_ISL_944812, EPI_ISL_944813, EPI_ISL_944814, EPI_ISL_944815, EPI_ISL_944816, EPI_ISL_944817, EPI_ISL_944818, EPI_ISL_944819, EPI_ISL_944820, EPI_ISL_944821, EPI_ISL_944822, EPI_ISL_944823, EPI_ISL_944825, EPI_ISL_944826, EPI_ISL_944827, EPI_ISL_944828, EPI_ISL_944829, EPI_ISL_944830, EPI_ISL_944831, EPI_ISL_944832, EPI_ISL_944833, EPI_ISL_944834, EPI_ISL_944835, EPI_ISL_944836, EPI_ISL_944837, EPI_ISL_944838, EPI_ISL_944839, EPI_ISL_944840, EPI_ISL_944841, EPI_ISL_944842, EPI_ISL_944843, EPI_ISL_944845, EPI_ISL_944846, EPI_ISL_944847, EPI_ISL_944848, EPI_ISL_944849, EPI_ISL_944850, EPI_ISL_944851, EPI_ISL_944852, EPI_ISL_944853, EPI_ISL_944854, EPI_ISL_944855, EPI_ISL_944856, EPI_ISL_944857, EPI_ISL_944858, EPI_ISL_944859, EPI_ISL_944860, EPI_ISL_944861, EPI_ISL_944862, EPI_ISL_944864, EPI_ISL_944865, EPI_ISL_944866, EPI_ISL_944867, EPI_ISL_944868, EPI_ISL_944869, EPI_ISL_944870, EPI_ISL_944871, EPI_ISL_944872, EPI_ISL_944873, EPI_ISL_944874, EPI_ISL_944875, EPI_ISL_944876, EPI_ISL_944877, EPI_ISL_944878, EPI_ISL_944879, EPI_ISL_944880, EPI_ISL_944881, EPI_ISL_944882, EPI_ISL_944884, EPI_ISL_944885, EPI_ISL_944886, EPI_ISL_944887, EPI_ISL_944888, EPI_ISL_944889, EPI_ISL_944890, EPI_ISL_944891, EPI_ISL_944892, EPI_ISL_944893, EPI_ISL_944894, EPI_ISL_944895, EPI_ISL_944896, EPI_ISL_944897, EPI_ISL_944898, EPI_ISL_944899, EPI_ISL_944900, EPI_ISL_944901, EPI_ISL_944902, EPI_ISL_944903, EPI_ISL_944904, EPI_ISL_944905, EPI_ISL_944906, EPI_ISL_944907, EPI_ISL_944909, EPI_ISL_944910, EPI_ISL_944911, EPI_ISL_944912, EPI_ISL_944913, EPI_ISL_944915, EPI_ISL_944916, EPI_ISL_944917, EPI_ISL_944918, EPI_ISL_944919, EPI_ISL_944920, EPI_ISL_944921, EPI_ISL_944922, EPI_ISL_944924, EPI_ISL_944925, EPI_ISL_944926, EPI_ISL_944927, EPI_ISL_944928, EPI_ISL_944929, EPI_ISL_944930, EPI_ISL_944931, EPI_ISL_944932, EPI_ISL_944933, EPI_ISL_944934, EPI_ISL_944935, EPI_ISL_944936, EPI_ISL_944937, EPI_ISL_944938, EPI_ISL_944939, EPI_ISL_944940, EPI_ISL_944941, EPI_ISL_944942, EPI_ISL_944943, EPI_ISL_944944, EPI_ISL_944945, EPI_ISL_944946, EPI_ISL_944947, EPI_ISL_944948, EPI_ISL_944949, EPI_ISL_944950, EPI_ISL_944951, EPI_ISL_944952, EPI_ISL_944954, EPI_ISL_944955, EPI_ISL_944956, EPI_ISL_944957, EPI_ISL_944958, EPI_ISL_944959, EPI_ISL_944960, EPI_ISL_944961, EPI_ISL_944962, EPI_ISL_944963, EPI_ISL_944964, EPI_ISL_944965, EPI_ISL_944966, EPI_ISL_944967, EPI_ISL_944968, EPI_ISL_944969, EPI_ISL_944970, EPI_ISL_944971, EPI_ISL_944972, EPI_ISL_944973, EPI_ISL_944974, EPI_ISL_944975, EPI_ISL_944977, EPI_ISL_944979, EPI_ISL_944980, EPI_ISL_944981, EPI_ISL_944982, EPI_ISL_944983, EPI_ISL_944984, EPI_ISL_944985, EPI_ISL_944986, EPI_ISL_944987, EPI_ISL_944988, EPI_ISL_944989, EPI_ISL_944990, EPI_ISL_944991, EPI_ISL_944992, EPI_ISL_944993, EPI_ISL_944994, EPI_ISL_944995, EPI_ISL_944996, EPI_ISL_944997, EPI_ISL_944998, EPI_ISL_944999, EPI_ISL_945000, EPI_ISL_945001, EPI_ISL_945002, EPI_ISL_945003, EPI_ISL_945004, EPI_ISL_945005, EPI_ISL_945006, EPI_ISL_945007, EPI_ISL_945008, EPI_ISL_945009, EPI_ISL_945010, EPI_ISL_945011, EPI_ISL_945012, EPI_ISL_945013, EPI_ISL_945014, EPI_ISL_945015, EPI_ISL_945016, EPI_ISL_945017, EPI_ISL_945018, EPI_ISL_945019, EPI_ISL_945020, EPI_ISL_945021, EPI_ISL_945022, EPI_ISL_945024, EPI_ISL_945025, EPI_ISL_945026, EPI_ISL_945027, EPI_ISL_945028, EPI_ISL_945029, EPI_ISL_945030, EPI_ISL_945031, EPI_ISL_945033, EPI_ISL_945035, EPI_ISL_945036, EPI_ISL_945037, EPI_ISL_945038, EPI_ISL_945039, EPI_ISL_945040, EPI_ISL_945041, EPI_ISL_945042, EPI_ISL_945043, EPI_ISL_945044, EPI_ISL_945045, EPI_ISL_945046, EPI_ISL_945047, EPI_ISL_945048 | | | |
| see above | Lighthouse Lab in Glasgow | Wellcome Sanger Institute for the COVID-19 Genomics UK (COG-UK) Consortium | Harper VanSteenhouse, Yumi Kasai, David Gray, Carol Clugston, Anna Dominiczak and Alex Alderton, Roberto Amato, Sonia Goncalves, Ewan Harrison, David K. Jackson, Ian Johnston, Dominic Kwiatkowski, Cordelia Langford, John Sillitoe on behalf of the Wellcome Sanger Institute COVID-19 Surveillance Team |
| EPI_ISL_945374, EPI_ISL_945375, EPI_ISL_945376, EPI_ISL_945379, EPI_ISL_945386, EPI_ISL_945387, EPI_ISL_945389, EPI_ISL_945402, EPI_ISL_945408, EPI_ISL_945409, EPI_ISL_945419, EPI_ISL_945420, EPI_ISL_945422, EPI_ISL_945423, EPI_ISL_945433, EPI_ISL_945444, EPI_ISL_945446, EPI_ISL_945449, EPI_ISL_945458, EPI_ISL_945459, EPI_ISL_945466, EPI_ISL_945467, EPI_ISL_945468, EPI_ISL_945469, EPI_ISL_945475, EPI_ISL_945480, EPI_ISL_945485, EPI_ISL_945490, EPI_ISL_945494, EPI_ISL_945495, EPI_ISL_945498, EPI_ISL_945502, EPI_ISL_945506, EPI_ISL_945508, EPI_ISL_945514, EPI_ISL_945518, EPI_ISL_945522, EPI_ISL_945525, EPI_ISL_945529, EPI_ISL_945538, EPI_ISL_945539, EPI_ISL_945543, EPI_ISL_945545, EPI_ISL_945546, EPI_ISL_945548, EPI_ISL_945549, EPI_ISL_945555, EPI_ISL_945561, EPI_ISL_945563, EPI_ISL_945569, EPI_ISL_945579, EPI_ISL_945582, EPI_ISL_945587, EPI_ISL_945589, EPI_ISL_945591, EPI_ISL_945597, EPI_ISL_945619, EPI_ISL_945621, EPI_ISL_945627, EPI_ISL_945635, EPI_ISL_945636, EPI_ISL_945648, EPI_ISL_945650, EPI_ISL_945656, EPI_ISL_945664, EPI_ISL_945668, EPI_ISL_945672, EPI_ISL_945674, EPI_ISL_945677, EPI_ISL_945678, EPI_ISL_945681, EPI_ISL_945682, EPI_ISL_945692, EPI_ISL_945696, EPI_ISL_945697, EPI_ISL_945698, EPI_ISL_945701, EPI_ISL_945704 | | | |
| see above | Lighthouse Lab in Alderley Park | Wellcome Sanger Institute for the COVID-19 Genomics UK (COG-UK) Consortium | Jacquelyn Wynn, Mairead Hyland, The Lighthouse Lab in Alderley Park and Alex Alderton, Roberto Amato, Sonia Goncalves, Ewan Harrison, David K. Jackson, Ian Johnston, Dominic Kwiatkowski, Cordelia Langford, John Sillitoe on behalf of the Wellcome Sanger Institute COVID-19 Surveillance Team |
| EPI_ISL_945716, EPI_ISL_945718, EPI_ISL_945722, EPI_ISL_945725, EPI_ISL_945726, EPI_ISL_945728, EPI_ISL_945729, EPI_ISL_945730, EPI_ISL_945732, EPI_ISL_945733, EPI_ISL_945735, EPI_ISL_945736, EPI_ISL_945737, EPI_ISL_945738, EPI_ISL_945741, EPI_ISL_945742, EPI_ISL_945743, EPI_ISL_945744, EPI_ISL_945747, EPI_ISL_945753, EPI_ISL_945755, EPI_ISL_945757, EPI_ISL_945758, EPI_ISL_945759, EPI_ISL_945762, EPI_ISL_945763, EPI_ISL_945768, EPI_ISL_945771, EPI_ISL_945772, EPI_ISL_945776, EPI_ISL_945777, EPI_ISL_945781, EPI_ISL_945782, EPI_ISL_945788, EPI_ISL_945792, EPI_ISL_945795, EPI_ISL_945796, EPI_ISL_945799, EPI_ISL_945800, EPI_ISL_945802, EPI_ISL_945803, EPI_ISL_945804, EPI_ISL_945806, EPI_ISL_945807, EPI_ISL_945808, EPI_ISL_945809, EPI_ISL_945812, EPI_ISL_945813, EPI_ISL_945815, EPI_ISL_945816, EPI_ISL_945817, EPI_ISL_945818, EPI_ISL_945821, EPI_ISL_945822, EPI_ISL_945824, EPI_ISL_945826, EPI_ISL_945828, EPI_ISL_945831, EPI_ISL_945835, EPI_ISL_945836, EPI_ISL_945839, EPI_ISL_945841, EPI_ISL_945843, EPI_ISL_945844, EPI_ISL_945846, EPI_ISL_945848, EPI_ISL_945849, EPI_ISL_945850, EPI_ISL_945851, EPI_ISL_945853, EPI_ISL_945860, EPI_ISL_945862, EPI_ISL_945863, EPI_ISL_945865, EPI_ISL_945867, EPI_ISL_945868, EPI_ISL_945869, EPI_ISL_945870, EPI_ISL_945871, EPI_ISL_945872, EPI_ISL_945873, EPI_ISL_945874, EPI_ISL_945877, EPI_ISL_945878, EPI_ISL_945879, EPI_ISL_945882, EPI_ISL_945884, EPI_ISL_945885, EPI_ISL_945889, EPI_ISL_945895, EPI_ISL_945898, EPI_ISL_945899, EPI_ISL_945900, EPI_ISL_945901, EPI_ISL_945904, EPI_ISL_945906, EPI_ISL_945907, EPI_ISL_945908, EPI_ISL_945909, EPI_ISL_945910, EPI_ISL_945911, EPI_ISL_945914, EPI_ISL_945916, EPI_ISL_945917, EPI_ISL_945920, EPI_ISL_945921, EPI_ISL_945926, EPI_ISL_945933, EPI_ISL_945935, EPI_ISL_945938, EPI_ISL_945939, EPI_ISL_945940, EPI_ISL_945944, EPI_ISL_945945, EPI_ISL_945950, EPI_ISL_945951, EPI_ISL_945954, EPI_ISL_945956, EPI_ISL_945958, EPI_ISL_945960, EPI_ISL_945961, EPI_ISL_945962, EPI_ISL_945963, EPI_ISL_945965, EPI_ISL_945967, EPI_ISL_945969, EPI_ISL_945970, EPI_ISL_945971, EPI_ISL_945972, EPI_ISL_945976, EPI_ISL_945979, EPI_ISL_945980, EPI_ISL_945983, EPI_ISL_945984, EPI_ISL_945988, EPI_ISL_945990, EPI_ISL_945993, EPI_ISL_945994, EPI_ISL_945997, EPI_ISL_945998, EPI_ISL_945999, EPI_ISL_946000, EPI_ISL_946002, EPI_ISL_946005, EPI_ISL_946006, EPI_ISL_946007, EPI_ISL_946008, EPI_ISL_946009, EPI_ISL_946010, EPI_ISL_946011, EPI_ISL_946014, EPI_ISL_946021, EPI_ISL_946023, EPI_ISL_946026, EPI_ISL_946028, EPI_ISL_946030, EPI_ISL_946035, EPI_ISL_946037, EPI_ISL_946041, EPI_ISL_946048, EPI_ISL_946049, EPI_ISL_946052, EPI_ISL_946055, EPI_ISL_946057, EPI_ISL_946058, EPI_ISL_946059, EPI_ISL_946062, EPI_ISL_946066, EPI_ISL_946067, EPI_ISL_946068, EPI_ISL_946069, EPI_ISL_946070, EPI_ISL_946071, EPI_ISL_946072, EPI_ISL_946073, EPI_ISL_946076, EPI_ISL_946080, EPI_ISL_946086, EPI_ISL_946087, EPI_ISL_946088, EPI_ISL_946090, EPI_ISL_946093, EPI_ISL_946096, EPI_ISL_946106, EPI_ISL_946109, EPI_ISL_946110, EPI_ISL_946112, EPI_ISL_946113, EPI_ISL_946115, EPI_ISL_946123, EPI_ISL_946124, EPI_ISL_946126, EPI_ISL_946127, EPI_ISL_946129, EPI_ISL_946135, EPI_ISL_946137, EPI_ISL_946145, EPI_ISL_946147, EPI_ISL_946149, EPI_ISL_946150, EPI_ISL_946157, EPI_ISL_946160, EPI_ISL_946162, EPI_ISL_946169, EPI_ISL_946173, EPI_ISL_946174, EPI_ISL_946176, EPI_ISL_946177, EPI_ISL_946180, EPI_ISL_946181, EPI_ISL_946182, EPI_ISL_946188, EPI_ISL_946190, EPI_ISL_946192, EPI_ISL_946196, EPI_ISL_946199, EPI_ISL_946202, EPI_ISL_946206, EPI_ISL_946208, EPI_ISL_946210, EPI_ISL_946211, EPI_ISL_946213, EPI_ISL_946215, EPI_ISL_946218, EPI_ISL_946220, EPI_ISL_946221, EPI_ISL_946224, EPI_ISL_946225, EPI_ISL_946227, EPI_ISL_946229, EPI_ISL_946231, EPI_ISL_946232, EPI_ISL_946233, EPI_ISL_946234, EPI_ISL_946243, EPI_ISL_946250, EPI_ISL_946253, EPI_ISL_946254, EPI_ISL_946260, EPI_ISL_946264, EPI_ISL_946270, EPI_ISL_946271, EPI_ISL_946274, EPI_ISL_946281, EPI_ISL_946284, EPI_ISL_946285, EPI_ISL_946286, EPI_ISL_946292, EPI_ISL_946298, EPI_ISL_946301, EPI_ISL_946302, EPI_ISL_946304, EPI_ISL_946318, EPI_ISL_946319, EPI_ISL_946326, EPI_ISL_946328, EPI_ISL_946333, EPI_ISL_946335, EPI_ISL_946338, EPI_ISL_946339, EPI_ISL_946351, EPI_ISL_946356, EPI_ISL_946357, EPI_ISL_946360, EPI_ISL_946361, EPI_ISL_946365, EPI_ISL_946366, EPI_ISL_946367, EPI_ISL_946371, EPI_ISL_946374, EPI_ISL_946375, EPI_ISL_946378, EPI_ISL_946379, EPI_ISL_946380, EPI_ISL_946385, EPI_ISL_946391, EPI_ISL_946397, EPI_ISL_946400, EPI_ISL_946405, EPI_ISL_946407, EPI_ISL_946410, EPI_ISL_946411 | | | |
| see above | Lighthouse Lab in Glasgow | Wellcome Sanger Institute for the COVID-19 Genomics UK (COG-UK) Consortium | Harper VanSteenhouse, Yumi Kasai, David Gray, Carol Clugston, Anna Dominiczak and Alex Alderton, Roberto Amato, Sonia Goncalves, Ewan Harrison, David K. Jackson, Ian Johnston, Dominic Kwiatkowski, Cordelia Langford, John Sillitoe on behalf of the Wellcome Sanger Institute COVID-19 Surveillance Team |
| EPI_ISL_949405 | University of Birmingham | COVID-19 Genomics UK (COG-UK) Consortium | Institute of Microbiology, University of Birmingham: Claire McMurray, Joanne Stockton, Samuel Nicholls, Radoslaw Poplawski, Will Rowe, Josh Quick, Nicholas Loman. University of Birmingham Testing Laboratory: Celina M Whalley, Andrew Bosworth, Charlotte Poxon, Kasun Wanigasooriya, Oliver Pickles, Mike Kidd, Alex Richter, Andrew D Beggs PHE Heartlands Lab: Husam Osman, Andrew Bosworth. Queen Elizabeth Hospital: Anna Casey |
| EPI_ISL_949543, EPI_ISL_949545, EPI_ISL_949547, EPI_ISL_949551, EPI_ISL_949552, EPI_ISL_949557, EPI_ISL_949558, EPI_ISL_949559, EPI_ISL_949560, EPI_ISL_949561, EPI_ISL_949562, EPI_ISL_949563, EPI_ISL_949564, EPI_ISL_949565, EPI_ISL_949566, EPI_ISL_949567, EPI_ISL_949568, EPI_ISL_949569, EPI_ISL_949570, EPI_ISL_949571, EPI_ISL_949572, EPI_ISL_949573, EPI_ISL_949574, EPI_ISL_949575, EPI_ISL_949576, EPI_ISL_949577, EPI_ISL_949578, EPI_ISL_949579, EPI_ISL_949580, EPI_ISL_949582, EPI_ISL_949583, EPI_ISL_949584, EPI_ISL_949586, EPI_ISL_949587, EPI_ISL_949588, EPI_ISL_949589, EPI_ISL_949590, EPI_ISL_949591, EPI_ISL_949592, EPI_ISL_949593, EPI_ISL_949594, EPI_ISL_949595, EPI_ISL_949596, EPI_ISL_949597, EPI_ISL_949598, EPI_ISL_949599, EPI_ISL_949600, EPI_ISL_949601, EPI_ISL_949602, EPI_ISL_949603, EPI_ISL_949604, EPI_ISL_949605, EPI_ISL_949606, EPI_ISL_949607 | | | |
| see above | Department of Pathology, University of Cambridge | COVID-19 Genomics UK (COG-UK) Consortium | Aminu S. Jahun, Yasmin Chaudhry, Iliana Georgana, Myra Hosmillo, Rhys Izu, Martin D. Curran, Surendra Parmar, Ian Goodfellow |
| EPI_ISL_949613, EPI_ISL_949616, EPI_ISL_949617, EPI_ISL_949619, EPI_ISL_949621, EPI_ISL_949623, EPI_ISL_949625, EPI_ISL_949627, EPI_ISL_949629 | West of Scotland Specialist Virology Centre, NHSGGC / MRC-University of Glasgow Centre for Virus Research | COVID-19 Genomics UK (COG-UK) Consortium | Ana da Silva Filipe, Natasha Johnson, Kathy Smollett, Daniel Mair, Stephen Carmichael, Alice Broos, Lily Tong, Jenna Nichols, Kyriaki Nomikou; Sarah McDonald; Richard Orton, Joseph Hughes, Sreenu Vattipally, David L Robertson; Alasdair MacLean, Rory Gunson; Sharif Shaaban, Matthew Holden; Rachel Blacow, Guy Mollett, Kathy Li, James Shepherd, Antonia Ho, Emma Thomson |
| EPI_ISL_949662, EPI_ISL_949663, EPI_ISL_949664, EPI_ISL_949666, EPI_ISL_949668, EPI_ISL_949669, EPI_ISL_949670, EPI_ISL_949708, EPI_ISL_949709, EPI_ISL_949710, EPI_ISL_949711, EPI_ISL_949712, EPI_ISL_949713, EPI_ISL_949714, EPI_ISL_949715, EPI_ISL_949716, EPI_ISL_949717, EPI_ISL_949718, EPI_ISL_949719, EPI_ISL_949720, EPI_ISL_949721, EPI_ISL_949722, EPI_ISL_949723, EPI_ISL_949724, EPI_ISL_949725, EPI_ISL_949726, EPI_ISL_949727, EPI_ISL_949728, EPI_ISL_949729, EPI_ISL_949730, EPI_ISL_949731, EPI_ISL_949732, EPI_ISL_949733, EPI_ISL_949734, EPI_ISL_949735, EPI_ISL_949736, EPI_ISL_949737, EPI_ISL_949738, EPI_ISL_949739, EPI_ISL_949740, EPI_ISL_949741, EPI_ISL_949742, EPI_ISL_949743, EPI_ISL_949744, EPI_ISL_949745, EPI_ISL_949746 | | | |
| see above | Liverpool Clinical Laboratories | COVID-19 Genomics UK (COG-UK) Consortium | Sam Haldenby, Anita Lucaci, Steve Paterson, Julian Hiscox, Alistair Darby, M Almsaud, A Alrezaihi, Muhannad Alruwaili, Stuart D Armstrong, Jones Benjamin, Eleanor G Bentley, Anu Chawla, Jordan J Clark, Angela Cowell, Richard Eccles, Isabel García-Dorival, Matthew Gemmell, Alessandro Gerada, PKF Gilmore, Richard Gregory, Ximeng Han, Catherine Hartley, Margaret Hughes, Miren Iturriza-Gomara, James Johnson, L Luu, Jenifer Manson, Charlotte Nelson, Elaine O'Toole, Cassie Olateju, Rebekah Penrice-Randal , Lucille Rainbow, N.P Randle, Trevor Ian Robinson, Parul Sharma, Ghada T Shawli, James P Stewart, Neil Swainston, Ecaterina Vamos, Joanne Watts, Mark Whitehead |
| EPI_ISL_949795, EPI_ISL_949796, EPI_ISL_949797, EPI_ISL_949798, EPI_ISL_949799, EPI_ISL_949800, EPI_ISL_949801, EPI_ISL_949803, EPI_ISL_949804, EPI_ISL_949805, EPI_ISL_949806, EPI_ISL_949807, EPI_ISL_949808, EPI_ISL_949809, EPI_ISL_949810, EPI_ISL_949811, EPI_ISL_949812, EPI_ISL_949813, EPI_ISL_949814, EPI_ISL_949815, EPI_ISL_949816, EPI_ISL_949817, EPI_ISL_949818, EPI_ISL_949819, EPI_ISL_949820, EPI_ISL_949821, EPI_ISL_949822, EPI_ISL_949823, EPI_ISL_949824, EPI_ISL_949825, EPI_ISL_949826, EPI_ISL_949827, EPI_ISL_949828, EPI_ISL_949829, EPI_ISL_949830, EPI_ISL_949831, EPI_ISL_949832, EPI_ISL_949833, EPI_ISL_949834, EPI_ISL_949835, EPI_ISL_949836, EPI_ISL_949837, EPI_ISL_949838, EPI_ISL_949839, EPI_ISL_949840, EPI_ISL_949841, EPI_ISL_949842, EPI_ISL_949843, EPI_ISL_949844, EPI_ISL_949845, EPI_ISL_949846, EPI_ISL_949847, EPI_ISL_949848, EPI_ISL_949850, EPI_ISL_949851, EPI_ISL_949852, EPI_ISL_949853, EPI_ISL_949855, EPI_ISL_949856, EPI_ISL_949858, EPI_ISL_949859, EPI_ISL_949860, EPI_ISL_949861, EPI_ISL_949862, EPI_ISL_949863, EPI_ISL_949864, EPI_ISL_949865, EPI_ISL_949866, EPI_ISL_949867, EPI_ISL_949868, EPI_ISL_949869, EPI_ISL_949871, EPI_ISL_949872, EPI_ISL_949873, EPI_ISL_949874, EPI_ISL_949875, EPI_ISL_949876, EPI_ISL_949877, EPI_ISL_949878, EPI_ISL_949879, EPI_ISL_949880, EPI_ISL_949881, EPI_ISL_949883, EPI_ISL_949884, EPI_ISL_949885, EPI_ISL_949886, EPI_ISL_949887, EPI_ISL_949888, EPI_ISL_949889, EPI_ISL_949890, EPI_ISL_949891, EPI_ISL_949892, EPI_ISL_949893, EPI_ISL_949895, EPI_ISL_949896, EPI_ISL_949897, EPI_ISL_949898, EPI_ISL_949901, EPI_ISL_949903, EPI_ISL_949904, EPI_ISL_949915, EPI_ISL_949916, EPI_ISL_949918, EPI_ISL_949920, EPI_ISL_949921, EPI_ISL_949922, EPI_ISL_949923, EPI_ISL_949926, EPI_ISL_949928, EPI_ISL_949929, EPI_ISL_949930, EPI_ISL_949932, EPI_ISL_949933, EPI_ISL_949934, EPI_ISL_949935, EPI_ISL_949936, EPI_ISL_949937, EPI_ISL_949938, EPI_ISL_949939, EPI_ISL_949940, EPI_ISL_949941, EPI_ISL_949942, EPI_ISL_949943, EPI_ISL_949944, EPI_ISL_949945, EPI_ISL_949947, EPI_ISL_949949, EPI_ISL_949950, EPI_ISL_949951, EPI_ISL_949952, EPI_ISL_949953, EPI_ISL_949954, EPI_ISL_949955, EPI_ISL_949956, EPI_ISL_949957, EPI_ISL_949958, EPI_ISL_949961, EPI_ISL_949962, EPI_ISL_949964, EPI_ISL_949965, EPI_ISL_949966, EPI_ISL_949967, EPI_ISL_949968, EPI_ISL_949969, EPI_ISL_949970, EPI_ISL_949971, EPI_ISL_949972, EPI_ISL_949973, EPI_ISL_949974, EPI_ISL_950173, EPI_ISL_950177, EPI_ISL_950195, EPI_ISL_950198, EPI_ISL_950200, EPI_ISL_950201, EPI_ISL_950202, EPI_ISL_950203, EPI_ISL_950204, EPI_ISL_950205, EPI_ISL_950206, EPI_ISL_950207, EPI_ISL_950208, EPI_ISL_950209, EPI_ISL_950210 | | | |
| see above | University College London, Great Ormond Street Hospital for Children NHS Foundation Trust, Imperial College Healthcare NHS Trust | COVID-19 Genomics UK (COG-UK) Consortium | Sergi Castellano, Rachel Williams, Mark Kristiansen, Paola Resende Silva, Sunando Roy, Tony Brooks, Helena Tutill, Paola Niola, Patricia Dyal, Charlotte Williams, Leysa Forrest, Yasmin Panchbhaya, Jacqueline Findlay, Samuel Weeks, Julianne Brown, Kathryn Harris, Paul Randell, James Price, Alison Holmes, Judith Breuer |
| EPI_ISL_950402, EPI_ISL_950404, EPI_ISL_950405, EPI_ISL_950407, EPI_ISL_950408, EPI_ISL_950409, EPI_ISL_950411, EPI_ISL_950412, EPI_ISL_950413, EPI_ISL_950415, EPI_ISL_950424, EPI_ISL_950428, EPI_ISL_950431, EPI_ISL_950433, EPI_ISL_950434 | | | |
| see above | Northumbria University / South Tees Hospitals NHS Foundation Trust / North Cumbria Integrated Care NHS | COVID-19 Genomics UK (COG-UK) Consortium | Darren L Smith,Andrew Nelson,Matthew Bashton,Greg R Young,Joshua Loh,John Allan,Mohammad A Tariq,Giles S Holt,Gary Black,Wen C Yew,Lynn Dover,Paul Baker,Steve Liggett,Sarah Essex,Jane Greenaway,Debra Padgett,Clive Graham,Garren Scott,Edward Barton,Emma Swindells,Brendan |

| | | | |
|---|---|---|---|
| | Foundation Trust / North Tees and Hartlepool NHS Foundation Trust / Newcastle Hospitals NHS Foundation Trust | | Payne,Jennifer Collins,Yusri Taha,Gary Eltringham |
| EPI_ISL_950646, EPI_ISL_950647, EPI_ISL_950648, EPI_ISL_950649, EPI_ISL_950650, EPI_ISL_950651, EPI_ISL_950652, EPI_ISL_950653, EPI_ISL_950654, EPI_ISL_950655, EPI_ISL_950656, EPI_ISL_950657, EPI_ISL_950658, EPI_ISL_950659, EPI_ISL_950660, EPI_ISL_950661, EPI_ISL_950662, EPI_ISL_950663, EPI_ISL_950665, EPI_ISL_950666, EPI_ISL_950667, EPI_ISL_950668, EPI_ISL_950669, EPI_ISL_950670, EPI_ISL_950671, EPI_ISL_950672, EPI_ISL_950673, EPI_ISL_950674, EPI_ISL_950675, EPI_ISL_950676, EPI_ISL_950677, EPI_ISL_950678, EPI_ISL_950679, EPI_ISL_950680, EPI_ISL_950681, EPI_ISL_950682, EPI_ISL_950683, EPI_ISL_950684, EPI_ISL_950685, EPI_ISL_950686, EPI_ISL_950687, EPI_ISL_950688, EPI_ISL_950689, EPI_ISL_950690, EPI_ISL_950691, EPI_ISL_950692, EPI_ISL_950693, EPI_ISL_950694, EPI_ISL_950695, EPI_ISL_950696, EPI_ISL_950697, EPI_ISL_950717, EPI_ISL_950718, EPI_ISL_950719, EPI_ISL_950720, EPI_ISL_950721, EPI_ISL_950722, EPI_ISL_950762 | | | |
| see above | Queens Medical Centre, Clinical Microbiology Department / DeepSeq Nottingham | COVID-19 Genomics UK (COG-UK) Consortium | Gemma Clark, Wendy Smith, Manjinder Khakh, Vicki M Fleming, Michelle M Lister, Hannah Howson-Wells, Jonathan Ball, Patrick McClure, Joseph Chappell, Theocharis Tsoleridis, Nadine Holmes, Matthew Carlisle, Christopher Moore, Fei Sang, Johnny Debebe, Victoria Wright, Matthew Loose |
| EPI_ISL_951783, EPI_ISL_951971, EPI_ISL_951975, EPI_ISL_951976, EPI_ISL_951977, EPI_ISL_951978, EPI_ISL_951979, EPI_ISL_951980, EPI_ISL_951981, EPI_ISL_951982, EPI_ISL_951983, EPI_ISL_951984, EPI_ISL_951985, EPI_ISL_951986, EPI_ISL_952021, EPI_ISL_952023, EPI_ISL_952028, EPI_ISL_952044, EPI_ISL_952046, EPI_ISL_952047, EPI_ISL_952048, EPI_ISL_952049, EPI_ISL_952050, EPI_ISL_952052, EPI_ISL_952053, EPI_ISL_952054, EPI_ISL_952055, EPI_ISL_952056, EPI_ISL_952057, EPI_ISL_952058, EPI_ISL_952060, EPI_ISL_952251, EPI_ISL_952252, EPI_ISL_952253, EPI_ISL_952255, EPI_ISL_952256, EPI_ISL_952257, EPI_ISL_952258, EPI_ISL_952259, EPI_ISL_952261, EPI_ISL_952262, EPI_ISL_952263, EPI_ISL_952264, EPI_ISL_952266, EPI_ISL_952267, EPI_ISL_952268, EPI_ISL_952269, EPI_ISL_952270, EPI_ISL_952271, EPI_ISL_952272, EPI_ISL_952273, EPI_ISL_952274, EPI_ISL_952275, EPI_ISL_952276, EPI_ISL_952277, EPI_ISL_952278, EPI_ISL_952279, EPI_ISL_952280, EPI_ISL_952281, EPI_ISL_952282, EPI_ISL_952283, EPI_ISL_952284, EPI_ISL_952285, EPI_ISL_952286, EPI_ISL_952287, EPI_ISL_952288, EPI_ISL_952289, EPI_ISL_952290, EPI_ISL_952291, EPI_ISL_952292, EPI_ISL_952293, EPI_ISL_952294, EPI_ISL_952295, EPI_ISL_952296, EPI_ISL_952297, EPI_ISL_952299, EPI_ISL_952300, EPI_ISL_952301, EPI_ISL_952302, EPI_ISL_952303, EPI_ISL_952304, EPI_ISL_952305, EPI_ISL_952306, EPI_ISL_952307, EPI_ISL_952308, EPI_ISL_952309, EPI_ISL_952310, EPI_ISL_952311, EPI_ISL_952312, EPI_ISL_952313, EPI_ISL_952314, EPI_ISL_952315, EPI_ISL_952316, EPI_ISL_952317, EPI_ISL_952318, EPI_ISL_952319, EPI_ISL_952320, EPI_ISL_952321, EPI_ISL_952322, EPI_ISL_952323, EPI_ISL_952324, EPI_ISL_952325, EPI_ISL_952326, EPI_ISL_952327, EPI_ISL_952328, EPI_ISL_952329, EPI_ISL_952330, EPI_ISL_952331, EPI_ISL_952332, EPI_ISL_952333, EPI_ISL_952334, EPI_ISL_952335, EPI_ISL_952337, EPI_ISL_952338, EPI_ISL_952339, EPI_ISL_952340, EPI_ISL_952341, EPI_ISL_952342, EPI_ISL_952343, EPI_ISL_952344, EPI_ISL_952346, EPI_ISL_952347, EPI_ISL_952348, EPI_ISL_952349, EPI_ISL_952350, EPI_ISL_952351, EPI_ISL_952352, EPI_ISL_952353, EPI_ISL_952354, EPI_ISL_952355, EPI_ISL_952356, EPI_ISL_952357, EPI_ISL_952358, EPI_ISL_952359, EPI_ISL_952360, EPI_ISL_952361, EPI_ISL_952362, EPI_ISL_952363, EPI_ISL_952364, EPI_ISL_952366, EPI_ISL_952367, EPI_ISL_952368, EPI_ISL_952370, EPI_ISL_952371, EPI_ISL_952372, EPI_ISL_952373, EPI_ISL_952374, EPI_ISL_952375, EPI_ISL_952376 | | | |
| see above | Originating lab: Wales Specialist Virology Centre Sequencing lab: Pathogen Genomics Unit | Public Health Wales Microbiology Cardiff Wales Specialist Virology Centre | Catherine Moore, Johnathan Evans, Laura Gifford, Malorie Perry, Simon Cottrell, Angela Marchbank, Alec Birchley, Alexander Adams, Amy Gaskin, Bree Gatica-Wilcox, Jason Coombes, Joel Southgate, Lauren Gilbert, Lee Graham, Nicole Pacchiarini, Sara Kumziene-Summerhayes, Sarah Taylor, Sophie Jones, Sara Rey, Matthew Bull, Joanne Watkins, Sally Corden, Tom Connor |
| EPI_ISL_952486, EPI_ISL_952487, EPI_ISL_952488 | Centre for Enzyme Innovation, University of Portsmouth / Translational Research Laboratory, Portsmouth Hospitals NHS Trust | COVID-19 Genomics UK (COG-UK) Consortium | Angela Beckett,Salman Goudarzi,Christopher Fearn,Kate Cook,Katie Loveson,Sharon Glaysher,Scott Elliott,Samuel Robson |
| EPI_ISL_952954, EPI_ISL_952955, EPI_ISL_952957, EPI_ISL_952958, EPI_ISL_952959, EPI_ISL_952964, EPI_ISL_952965, EPI_ISL_952968, EPI_ISL_952970, EPI_ISL_952972, EPI_ISL_952975, EPI_ISL_952976, EPI_ISL_952977, EPI_ISL_952979, EPI_ISL_952980, EPI_ISL_952982, EPI_ISL_952984, EPI_ISL_952985, EPI_ISL_952986, EPI_ISL_952988, EPI_ISL_952990, EPI_ISL_952991, EPI_ISL_952994, EPI_ISL_952995, EPI_ISL_952997, EPI_ISL_952998, EPI_ISL_952999, EPI_ISL_953000, EPI_ISL_953002, EPI_ISL_953003, EPI_ISL_953004, EPI_ISL_953006, EPI_ISL_953009, EPI_ISL_953010, EPI_ISL_953012, EPI_ISL_953013, EPI_ISL_953015, EPI_ISL_953016, EPI_ISL_953017, EPI_ISL_953020, EPI_ISL_953021, EPI_ISL_953022 | | | |
| see above | Virology Department, Sheffield Teaching Hospitals NHS Foundation Trust/Department of Infection, Immunity and Cardiovascular Disease, The Medical School, University of Sheffield | COVID-19 Genomics UK (COG-UK) Consortium | Thushan de Silva, Matthew Parker, Nikki Smith, Adri Angyal, Rebecca Brown, Luke Green, Rachel Tucker, Paul Parsons, Danielle Groves, Katie Johnson, Laura Carrilero, Alex Keeley, Dave Partridge, Matthew Wyles, Benjamin Lindsey, Mehmet Yavuz, Mohammad Raza, Cariad Evans |
| EPI_ISL_953240, EPI_ISL_953245, EPI_ISL_953248, EPI_ISL_953249, EPI_ISL_953250, EPI_ISL_953252, EPI_ISL_953253, EPI_ISL_953255, EPI_ISL_953256, EPI_ISL_953257, EPI_ISL_953258, EPI_ISL_953264, EPI_ISL_953265, EPI_ISL_953266, EPI_ISL_953268, EPI_ISL_953269, EPI_ISL_953270, EPI_ISL_953271, EPI_ISL_953275, EPI_ISL_953276, EPI_ISL_953277, EPI_ISL_953279, EPI_ISL_953280, EPI_ISL_953282, EPI_ISL_953283, EPI_ISL_953284, EPI_ISL_953285 | | | |
| see above | Bioinformatics and Biostatistics Lab, Advanced Sequencing Facility | COVID-19 Genomics UK (COG-UK) Consortium | Aengus Stewart,Jerome Nicod,Chelsea Sawyer,Laura Cubitt,Harshil Patel,Margaret Crawford |
| EPI_ISL_953376 | Jessa | Jessa | Jessa_cmdLab |
| EPI_ISL_953387, EPI_ISL_953389, EPI_ISL_953391, EPI_ISL_953392, EPI_ISL_953393, EPI_ISL_953395, EPI_ISL_953396, EPI_ISL_953397, EPI_ISL_953398, EPI_ISL_953399, EPI_ISL_953400, EPI_ISL_953401 | | | |
| see above | National Public Health Laboratory, National Centre for Infectious Diseases | National Public Health Laboratory, National Centre for Infectious Diseases | Tze Minn Mak, Zhenyang Zhou, Lin Cui, Raymond Tzer Pin Lin |
| EPI_ISL_953428, EPI_ISL_953429, EPI_ISL_953430, EPI_ISL_953431, EPI_ISL_953432, EPI_ISL_953433, EPI_ISL_953434, EPI_ISL_953435, EPI_ISL_953436, EPI_ISL_953437, EPI_ISL_953438, EPI_ISL_953439, EPI_ISL_953440, EPI_ISL_953441, EPI_ISL_953442, EPI_ISL_953443, EPI_ISL_953444, EPI_ISL_953445, EPI_ISL_953446, EPI_ISL_953447, EPI_ISL_953448, EPI_ISL_953449, EPI_ISL_953450, EPI_ISL_953451, EPI_ISL_953453, EPI_ISL_953454, EPI_ISL_953455, EPI_ISL_953456, EPI_ISL_953457, EPI_ISL_953458, EPI_ISL_953459, EPI_ISL_953460, EPI_ISL_953461, EPI_ISL_953462, EPI_ISL_953463, EPI_ISL_953464, EPI_ISL_953465, EPI_ISL_953466, EPI_ISL_953467, EPI_ISL_953468, EPI_ISL_953469, EPI_ISL_953470, EPI_ISL_953471, EPI_ISL_953472, EPI_ISL_953473, EPI_ISL_953474, EPI_ISL_953475, EPI_ISL_953476, EPI_ISL_953477, EPI_ISL_953478, EPI_ISL_953479, EPI_ISL_953480, EPI_ISL_953481, EPI_ISL_953482, EPI_ISL_953483, EPI_ISL_953484, EPI_ISL_953485, EPI_ISL_953486, EPI_ISL_953487, EPI_ISL_953488 | | | |
| see above | Jessa | Jessa | Jessa_cmdLab |
| EPI_ISL_953499, EPI_ISL_953502, EPI_ISL_953503, EPI_ISL_953588, EPI_ISL_953589, EPI_ISL_953590, EPI_ISL_953591, EPI_ISL_953592, EPI_ISL_953593, EPI_ISL_953594, EPI_ISL_953595, EPI_ISL_953596, EPI_ISL_953597, EPI_ISL_953598, EPI_ISL_953621, EPI_ISL_953622, EPI_ISL_953623, EPI_ISL_953624, EPI_ISL_953625, EPI_ISL_953649, EPI_ISL_953650, EPI_ISL_953651, EPI_ISL_953922, EPI_ISL_953923, EPI_ISL_953924, EPI_ISL_953925, EPI_ISL_953926, EPI_ISL_953927, EPI_ISL_953928, EPI_ISL_953929, EPI_ISL_953930, EPI_ISL_953931, EPI_ISL_953932, EPI_ISL_953933, EPI_ISL_953934, EPI_ISL_953935, EPI_ISL_953936, EPI_ISL_953937 | | | |
| see above | University Hospitals of Geneva, Laboratory of Virology | HUG, Laboratory of Virology and the Health2030 Genome Center | Samuel Cordey, Ana Rita Goncalves, Laurent Kaiser, Lorenzo Cerutti, Henri Pegeot, Melyssa Elies, Deborah Penet, Keith Harshman, Ioannis Xenarios, Emmanouil Dermitzakis |
| EPI_ISL_954755, EPI_ISL_954756, EPI_ISL_954757, EPI_ISL_954758, EPI_ISL_954759, EPI_ISL_954760, EPI_ISL_954761, EPI_ISL_954762, EPI_ISL_954763, EPI_ISL_954764, EPI_ISL_954765, EPI_ISL_954766, EPI_ISL_954767, EPI_ISL_954768, EPI_ISL_954769, EPI_ISL_954770, EPI_ISL_954771, EPI_ISL_954772, EPI_ISL_954773, EPI_ISL_954774, EPI_ISL_954775, EPI_ISL_954777, EPI_ISL_954778, EPI_ISL_954779 | | | |
| see above | AZDelta | AZDelta | Geert Martens; Dieter De Smet |
| EPI_ISL_955104, EPI_ISL_955105, EPI_ISL_955106, EPI_ISL_955107, EPI_ISL_955108, EPI_ISL_955109, EPI_ISL_955110, EPI_ISL_955111 | Maryland Public Health Laboratory | Maryland Public Health Laboratory | Maryland Department of Health Laboratories Administration |
| EPI_ISL_955218, EPI_ISL_955220, EPI_ISL_955221 | Indiana Animal Disease Diagnostic Laboratory | Carpi Laboratory - Purdue University | Jack Dorman, Ilinca I Ciubotariu, Lev Gorenstein, Abebe A Fola, G Kenitra Hendrix, Rebecca P Wilkes, Giovanna Carpi |
| EPI_ISL_955290, EPI_ISL_955292 | TMC/Pathology | Pathogen Discovery, Respiratory Viruses Branch, Division of Viral Diseases, Centers for Disease Control and Prevention | Ying Tao, Jing Zhang, Yan Li, Krista Queen, Anna Uehara, Peter Cook, Clinton R. Paden, Haibin Wang, Suxiang Tong |
| EPI_ISL_955308 | OK Public Health Laboratory, Oklahoma State DOH | Pathogen Discovery, Respiratory Viruses Branch, Division of Viral Diseases, Centers for Disease Control and Prevention | Ying Tao, Jing Zhang, Yan Li, Krista Queen, Anna Uehara, Peter Cook, Clinton R. Paden, Haibin Wang, Suxiang Tong |
| EPI_ISL_955831, EPI_ISL_955832, EPI_ISL_955833, EPI_ISL_955834, EPI_ISL_955835, EPI_ISL_955836, EPI_ISL_955906, EPI_ISL_955907, EPI_ISL_955908, EPI_ISL_955909, EPI_ISL_955910, EPI_ISL_955911, EPI_ISL_955912, EPI_ISL_955913, EPI_ISL_955914, EPI_ISL_955915, EPI_ISL_955916, EPI_ISL_955917, EPI_ISL_955922, EPI_ISL_955923, EPI_ISL_955924, EPI_ISL_955925, EPI_ISL_955926, EPI_ISL_955927, EPI_ISL_955928, EPI_ISL_955929, EPI_ISL_955930, EPI_ISL_955931, EPI_ISL_955932, EPI_ISL_955933, EPI_ISL_955934, EPI_ISL_955935, EPI_ISL_955936, EPI_ISL_955937, EPI_ISL_955938 | | | |
| see above | University of Michigan Clinical Microbiology Laboratory | Lauring Lab, University of Michigan, Department of Microbiology and Immunology | Valesano |
| EPI_ISL_956272 | Thai Red Cross Emerging Infectious Diseases Health Science Centre, Chulalongkorn Hospital, Faculty of Medicine, Chulalongkorn University | Thai Red Cross Emerging Infectious Diseases Center and Faculty of Medicine, Chulalongkorn University | Rome Buathong, Sopon Iamsirithaworn, Sininat Petcharat, Yutthana Joyjinda, Weenassarin Ampoot, Apaporn Rodpan, Opass Putcharoen, Thiravat Hemachudha, Supaporn Wacharapluesadee |
| EPI_ISL_956296 | Instituto Nacional de Salud- Dirección de Redes de Laboratorios de Salud Pública | Instituto Nacional de Salud- Dirección de Investigación en Salud Pública | Katherine Laiton-Donato, Diego A. Álvarez-Díaz, Carlos Franco-Muñoz, Mauricio Pacheco-Montealegre, Hector Alejandro Ruiz-Moreno, Maria T. Herrera-Sepúlveda, Diego Andrés Prada, Jhonnatan Reales-González, Sheryll Corchuelo, Julian Naizaque, Gerardo Santamaría, Magdalena Wiesner, Martha Lucia Ospina Martinez, Marcela Mercado-Reyes |
| EPI_ISL_956422, EPI_ISL_956423, EPI_ISL_956424, EPI_ISL_956425, EPI_ISL_956426, EPI_ISL_956427, EPI_ISL_956428, EPI_ISL_956429, EPI_ISL_956430, EPI_ISL_956431, EPI_ISL_956432, EPI_ISL_956433, EPI_ISL_956434, EPI_ISL_956435, EPI_ISL_956436, EPI_ISL_956437, EPI_ISL_956438, EPI_ISL_956439, EPI_ISL_956440, EPI_ISL_956441, EPI_ISL_956442, EPI_ISL_956443, EPI_ISL_956444, EPI_ISL_956445, EPI_ISL_956447, EPI_ISL_956448, EPI_ISL_956449, EPI_ISL_956450, EPI_ISL_956451, EPI_ISL_956452, EPI_ISL_956453, EPI_ISL_956454, EPI_ISL_956455, EPI_ISL_956456, EPI_ISL_956457, EPI_ISL_956458, EPI_ISL_956459, EPI_ISL_956460, EPI_ISL_956461, EPI_ISL_956462, EPI_ISL_956463, EPI_ISL_956464, EPI_ISL_956465, EPI_ISL_956466, EPI_ISL_956467, EPI_ISL_956468, EPI_ISL_956469, EPI_ISL_956470, EPI_ISL_956471, EPI_ISL_956472, EPI_ISL_956473, EPI_ISL_956474, EPI_ISL_956475, EPI_ISL_956476, | | | |

| | | | |
|---|---|---|---|
| EPI_ISL_956477, EPI_ISL_956478, EPI_ISL_956479, EPI_ISL_956480, EPI_ISL_956481, EPI_ISL_956482, EPI_ISL_956483, EPI_ISL_956484, EPI_ISL_956485, EPI_ISL_956486, EPI_ISL_956488, EPI_ISL_956489, EPI_ISL_956490, EPI_ISL_956491, EPI_ISL_956492, EPI_ISL_956493, EPI_ISL_956494, EPI_ISL_956495, EPI_ISL_956496, EPI_ISL_956497, EPI_ISL_956498, EPI_ISL_956499, EPI_ISL_956500, EPI_ISL_956501, EPI_ISL_956502, EPI_ISL_956503, EPI_ISL_956504, EPI_ISL_956505, EPI_ISL_956506, EPI_ISL_956507, EPI_ISL_956509, EPI_ISL_956510, EPI_ISL_956511, EPI_ISL_956512, EPI_ISL_956513, EPI_ISL_956514, EPI_ISL_956515, EPI_ISL_956516, EPI_ISL_956517, EPI_ISL_956518, EPI_ISL_956519, EPI_ISL_956520, EPI_ISL_956521, EPI_ISL_956522, EPI_ISL_956523, EPI_ISL_956524, EPI_ISL_956525, EPI_ISL_956526, EPI_ISL_956527, EPI_ISL_956528, EPI_ISL_956529, EPI_ISL_956530, EPI_ISL_956531, EPI_ISL_956532, EPI_ISL_956533, EPI_ISL_956534, EPI_ISL_956535, EPI_ISL_956536, EPI_ISL_956537, EPI_ISL_956538, EPI_ISL_956539, EPI_ISL_956540, EPI_ISL_956541, EPI_ISL_956542, EPI_ISL_956543, EPI_ISL_956544, EPI_ISL_956545, EPI_ISL_956546, EPI_ISL_956547, EPI_ISL_956548, EPI_ISL_956549, EPI_ISL_956550, EPI_ISL_956551, EPI_ISL_956552, EPI_ISL_956553, EPI_ISL_956554, EPI_ISL_956555, EPI_ISL_956556, EPI_ISL_956557, EPI_ISL_956558, EPI_ISL_956559, EPI_ISL_956560, EPI_ISL_956561, EPI_ISL_956563, EPI_ISL_956564, EPI_ISL_956565, EPI_ISL_956566, EPI_ISL_956567, EPI_ISL_956568, EPI_ISL_956569, EPI_ISL_956570, EPI_ISL_956571, EPI_ISL_956572, EPI_ISL_956573, EPI_ISL_956574, EPI_ISL_956575, EPI_ISL_956577, EPI_ISL_956578, EPI_ISL_956579, EPI_ISL_956580, EPI_ISL_956581, EPI_ISL_956582, EPI_ISL_956583, EPI_ISL_956584, EPI_ISL_956586, EPI_ISL_956588, EPI_ISL_956589, EPI_ISL_956590, EPI_ISL_956591, EPI_ISL_956593, EPI_ISL_956594, EPI_ISL_956595, EPI_ISL_956596, EPI_ISL_956597, EPI_ISL_956598, EPI_ISL_956599, EPI_ISL_956600, EPI_ISL_956603, EPI_ISL_956604, EPI_ISL_956605, EPI_ISL_956607, EPI_ISL_956608, EPI_ISL_956609, EPI_ISL_956610, EPI_ISL_956611, EPI_ISL_956612, EPI_ISL_956613, EPI_ISL_956614, EPI_ISL_956615, EPI_ISL_956616, EPI_ISL_956617, EPI_ISL_956619, EPI_ISL_956620, EPI_ISL_956621, EPI_ISL_956622, EPI_ISL_956623, EPI_ISL_956624, EPI_ISL_956627, EPI_ISL_956628, EPI_ISL_956629, EPI_ISL_956630, EPI_ISL_956632, EPI_ISL_956633, EPI_ISL_956634, EPI_ISL_956635, EPI_ISL_956637, EPI_ISL_956638, EPI_ISL_956639, EPI_ISL_956640, EPI_ISL_956641, EPI_ISL_956642, EPI_ISL_956643, EPI_ISL_956644, EPI_ISL_956645, EPI_ISL_956646, EPI_ISL_956647, EPI_ISL_956648, EPI_ISL_956649, EPI_ISL_956650, EPI_ISL_956652, EPI_ISL_956653, EPI_ISL_956654, EPI_ISL_956655, EPI_ISL_956656, EPI_ISL_956657, EPI_ISL_956658, EPI_ISL_956659, EPI_ISL_956660, EPI_ISL_956661, EPI_ISL_956662, EPI_ISL_956663, EPI_ISL_956664, EPI_ISL_956665, EPI_ISL_956666, EPI_ISL_956667, EPI_ISL_956669, EPI_ISL_956670, EPI_ISL_956671, EPI_ISL_956672, EPI_ISL_956673, EPI_ISL_956674, EPI_ISL_956675, EPI_ISL_956676, EPI_ISL_956677, EPI_ISL_956678, EPI_ISL_956679, EPI_ISL_956680, EPI_ISL_956681, EPI_ISL_956682, EPI_ISL_956683, EPI_ISL_956684, EPI_ISL_956685, EPI_ISL_956686, EPI_ISL_956687, EPI_ISL_956688, EPI_ISL_956689, EPI_ISL_956692, EPI_ISL_956693, EPI_ISL_956695, EPI_ISL_956696, EPI_ISL_956697, EPI_ISL_956698, EPI_ISL_956700, EPI_ISL_956701, EPI_ISL_956702, EPI_ISL_956703, EPI_ISL_956704, EPI_ISL_956705, EPI_ISL_956706, EPI_ISL_956707, EPI_ISL_956709, EPI_ISL_956710, EPI_ISL_956711, EPI_ISL_956712, EPI_ISL_956713, EPI_ISL_956714, EPI_ISL_956715, EPI_ISL_956716, EPI_ISL_956717, EPI_ISL_956718, EPI_ISL_956719, EPI_ISL_956720, EPI_ISL_956721, EPI_ISL_956722, EPI_ISL_956723, EPI_ISL_956724, EPI_ISL_956725, EPI_ISL_956726, EPI_ISL_956727, EPI_ISL_956728, EPI_ISL_956730, EPI_ISL_956731, EPI_ISL_956732, EPI_ISL_956733, EPI_ISL_956734, EPI_ISL_956735, EPI_ISL_956736, EPI_ISL_956737, EPI_ISL_956738, EPI_ISL_956739, EPI_ISL_956740, EPI_ISL_956741, EPI_ISL_956742, EPI_ISL_956743, EPI_ISL_956744, EPI_ISL_956745, EPI_ISL_956746, EPI_ISL_956747, EPI_ISL_956748, EPI_ISL_956749, EPI_ISL_956750, EPI_ISL_956751, EPI_ISL_956752, EPI_ISL_956753, EPI_ISL_956754, EPI_ISL_956755, EPI_ISL_956756, EPI_ISL_956757, EPI_ISL_956758, EPI_ISL_956759, EPI_ISL_956760, EPI_ISL_956761, EPI_ISL_956762, EPI_ISL_956763, EPI_ISL_956764 | | | |
| see above | Lighthouse Lab in Alderley Park | Wellcome Sanger Institute for the COVID-19 Genomics UK (COG-UK) Consortium | Jacquelyn Wynn, Mairead Hyland, The Lighthouse Lab in Alderley Park and Alex Alderton, Roberto Amato, Sonia Goncalves, Ewan Harrison, David K. Jackson, Ian Johnston, Dominic Kwiatkowski, Cordelia Langford, John Sillitoe on behalf of the Wellcome Sanger Institute COVID-19 Surveillance Team |
| EPI_ISL_956765, EPI_ISL_956766, EPI_ISL_956767, EPI_ISL_956768, EPI_ISL_956769, EPI_ISL_956770, EPI_ISL_956771, EPI_ISL_956772, EPI_ISL_956773, EPI_ISL_956774, EPI_ISL_956775, EPI_ISL_956776, EPI_ISL_956777, EPI_ISL_956778, EPI_ISL_956779, EPI_ISL_956780, EPI_ISL_956781, EPI_ISL_956782, EPI_ISL_956783, EPI_ISL_956784, EPI_ISL_956785, EPI_ISL_956786, EPI_ISL_956787, EPI_ISL_956788, EPI_ISL_956789, EPI_ISL_956790, EPI_ISL_956791, EPI_ISL_956792, EPI_ISL_956793, EPI_ISL_956794, EPI_ISL_956795, EPI_ISL_956796, EPI_ISL_956797, EPI_ISL_956798, EPI_ISL_956799, EPI_ISL_956800, EPI_ISL_956801, EPI_ISL_956802, EPI_ISL_956803, EPI_ISL_956804, EPI_ISL_956805, EPI_ISL_956806, EPI_ISL_956807, EPI_ISL_956808, EPI_ISL_956809, EPI_ISL_956810, EPI_ISL_956811, EPI_ISL_956812, EPI_ISL_956813, EPI_ISL_956814, EPI_ISL_956815, EPI_ISL_956816, EPI_ISL_956817, EPI_ISL_956818, EPI_ISL_956819, EPI_ISL_956820, EPI_ISL_956821, EPI_ISL_956822, EPI_ISL_956823, EPI_ISL_956824, EPI_ISL_956825, EPI_ISL_956826, EPI_ISL_956827, EPI_ISL_956828, EPI_ISL_956829, EPI_ISL_956830, EPI_ISL_956831, EPI_ISL_956832, EPI_ISL_956833, EPI_ISL_956834, EPI_ISL_956835, EPI_ISL_956836, EPI_ISL_956837, EPI_ISL_956838, EPI_ISL_956839, EPI_ISL_956840, EPI_ISL_956841, EPI_ISL_956842, EPI_ISL_956843, EPI_ISL_956844, EPI_ISL_956845, EPI_ISL_956846, EPI_ISL_956847, EPI_ISL_956848, EPI_ISL_956849, EPI_ISL_956850, EPI_ISL_956851, EPI_ISL_956852, EPI_ISL_956853, EPI_ISL_956854, EPI_ISL_956855, EPI_ISL_956856, EPI_ISL_956857, EPI_ISL_956858, EPI_ISL_956859, EPI_ISL_956860, EPI_ISL_956861, EPI_ISL_956862, EPI_ISL_956863, EPI_ISL_956864, EPI_ISL_956865, EPI_ISL_956866, EPI_ISL_956867, EPI_ISL_956868, EPI_ISL_956869, EPI_ISL_956870, EPI_ISL_956871, EPI_ISL_956872, EPI_ISL_956873, EPI_ISL_956874, EPI_ISL_956875, EPI_ISL_956876, EPI_ISL_956877, EPI_ISL_956878, EPI_ISL_956879, EPI_ISL_956880, EPI_ISL_956881, EPI_ISL_956882, EPI_ISL_956883, EPI_ISL_956884, EPI_ISL_956885, EPI_ISL_956886, EPI_ISL_956887, EPI_ISL_956888, EPI_ISL_956889, EPI_ISL_956890, EPI_ISL_956891, EPI_ISL_956892, EPI_ISL_956893, EPI_ISL_956894, EPI_ISL_956895, EPI_ISL_956896, EPI_ISL_956897, EPI_ISL_956898, EPI_ISL_956899, EPI_ISL_956900, EPI_ISL_956901, EPI_ISL_956902, EPI_ISL_956903, EPI_ISL_956904, EPI_ISL_956905, EPI_ISL_956906, EPI_ISL_956907, EPI_ISL_956908, EPI_ISL_956909, EPI_ISL_956910, EPI_ISL_956911, EPI_ISL_956912, EPI_ISL_956913, EPI_ISL_956914, EPI_ISL_956915, EPI_ISL_956916, EPI_ISL_956917, EPI_ISL_956918, EPI_ISL_956919, EPI_ISL_956920, EPI_ISL_956921, EPI_ISL_956922, EPI_ISL_956923, EPI_ISL_956924, EPI_ISL_956925, EPI_ISL_956926, EPI_ISL_956927, EPI_ISL_956928, EPI_ISL_956929, EPI_ISL_956930, EPI_ISL_956931, EPI_ISL_956932, EPI_ISL_956933, EPI_ISL_956934, EPI_ISL_956935, EPI_ISL_956936, EPI_ISL_956937, EPI_ISL_956938, EPI_ISL_956939, EPI_ISL_956940, EPI_ISL_956941, EPI_ISL_956942, EPI_ISL_956943, EPI_ISL_956944, EPI_ISL_956945, EPI_ISL_956946, EPI_ISL_956947, EPI_ISL_956948, EPI_ISL_956949, EPI_ISL_956950, EPI_ISL_956951, EPI_ISL_956952, EPI_ISL_956953, EPI_ISL_956954, EPI_ISL_956955, EPI_ISL_956956, EPI_ISL_956957, EPI_ISL_956958, EPI_ISL_956959, EPI_ISL_956960, EPI_ISL_956961, EPI_ISL_956962, EPI_ISL_956963, EPI_ISL_956964, EPI_ISL_956965, EPI_ISL_956966, EPI_ISL_956967, EPI_ISL_956968, EPI_ISL_956969, EPI_ISL_956970, EPI_ISL_956971, EPI_ISL_956972, EPI_ISL_956973, EPI_ISL_956974, EPI_ISL_956975, EPI_ISL_956976, EPI_ISL_956977, EPI_ISL_956978, EPI_ISL_956979, EPI_ISL_956980, EPI_ISL_956981, EPI_ISL_956982, EPI_ISL_956983, EPI_ISL_956984, EPI_ISL_956985, EPI_ISL_956986, EPI_ISL_956987, EPI_ISL_956988, EPI_ISL_956989, EPI_ISL_956990, EPI_ISL_956991, EPI_ISL_956992, EPI_ISL_956993, EPI_ISL_956994, EPI_ISL_956995, EPI_ISL_956996, EPI_ISL_956997, EPI_ISL_956998, EPI_ISL_956999, EPI_ISL_957000, EPI_ISL_957001, EPI_ISL_957002, EPI_ISL_957003, EPI_ISL_957004, EPI_ISL_957005, EPI_ISL_957006, EPI_ISL_957007, EPI_ISL_957008, EPI_ISL_957009, EPI_ISL_957010, EPI_ISL_957011, EPI_ISL_957012, EPI_ISL_957013, EPI_ISL_957014, EPI_ISL_957015, EPI_ISL_957016, EPI_ISL_957017, EPI_ISL_957018, EPI_ISL_957019, EPI_ISL_957020, EPI_ISL_957021, EPI_ISL_957022, EPI_ISL_957023, EPI_ISL_957024, EPI_ISL_957025, EPI_ISL_957026, EPI_ISL_957027, EPI_ISL_957028, EPI_ISL_957029, EPI_ISL_957030, EPI_ISL_957031, EPI_ISL_957032, EPI_ISL_957033, EPI_ISL_957034, EPI_ISL_957035, EPI_ISL_957036, EPI_ISL_957037, EPI_ISL_957038, EPI_ISL_957039, EPI_ISL_957040, EPI_ISL_957041, EPI_ISL_957042, EPI_ISL_957043, EPI_ISL_957044, EPI_ISL_957045, EPI_ISL_957046, EPI_ISL_957047, EPI_ISL_957048, EPI_ISL_957049, EPI_ISL_957050, EPI_ISL_957051, EPI_ISL_957052, EPI_ISL_957053 | | | |
| see above | Lighthouse Lab in Milton Keynes | Wellcome Sanger Institute for the COVID-19 Genomics UK (COG-UK) Consortium | The Lighthouse Lab in Milton Keynes and Alex Alderton, Roberto Amato, Sonia Goncalves, Ewan Harrison, David K. Jackson, Ian Johnston, Dominic Kwiatkowski, Cordelia Langford, John Sillitoe on behalf of the Wellcome Sanger Institute COVID-19 Surveillance Team |
| EPI_ISL_957054, EPI_ISL_957055 | Lighthouse Lab in Glasgow | Wellcome Sanger Institute for the COVID-19 Genomics UK (COG-UK) Consortium | Harper VanSteenhouse, Yumi Kasai, David Gray, Carol Clugston, Anna Dominiczak and Alex Alderton, Roberto Amato, Sonia Goncalves, Ewan Harrison, David K. Jackson, Ian Johnston, Dominic Kwiatkowski, Cordelia Langford, John Sillitoe on behalf of the Wellcome Sanger Institute COVID-19 Surveillance Team |
| EPI_ISL_957056 | Lighthouse Lab in Milton Keynes | Wellcome Sanger Institute for the COVID-19 Genomics UK (COG-UK) Consortium | The Lighthouse Lab in Milton Keynes and Alex Alderton, Roberto Amato, Sonia Goncalves, Ewan Harrison, David K. Jackson, Ian Johnston, Dominic Kwiatkowski, Cordelia Langford, John Sillitoe on behalf of the Wellcome Sanger Institute COVID-19 Surveillance Team |
| EPI_ISL_957057 | Lighthouse Lab in Glasgow | Wellcome Sanger Institute for the COVID-19 Genomics UK (COG-UK) Consortium | Harper VanSteenhouse, Yumi Kasai, David Gray, Carol Clugston, Anna Dominiczak and Alex Alderton, Roberto Amato, Sonia Goncalves, Ewan Harrison, David K. Jackson, Ian Johnston, Dominic Kwiatkowski, Cordelia Langford, John Sillitoe on behalf of the Wellcome Sanger Institute COVID-19 Surveillance Team |
| EPI_ISL_957058, EPI_ISL_957059 | Lighthouse Lab in Milton Keynes | Wellcome Sanger Institute for the COVID-19 Genomics UK (COG-UK) Consortium | The Lighthouse Lab in Milton Keynes and Alex Alderton, Roberto Amato, Sonia Goncalves, Ewan Harrison, David K. Jackson, Ian Johnston, Dominic Kwiatkowski, Cordelia Langford, John Sillitoe on behalf of the Wellcome Sanger Institute COVID-19 Surveillance Team |
| EPI_ISL_957060, EPI_ISL_957061 | Lighthouse Lab in Glasgow | Wellcome Sanger Institute for the COVID-19 Genomics UK (COG-UK) Consortium | Harper VanSteenhouse, Yumi Kasai, David Gray, Carol Clugston, Anna Dominiczak and Alex Alderton, Roberto Amato, Sonia Goncalves, Ewan Harrison, David K. Jackson, Ian Johnston, Dominic Kwiatkowski, Cordelia Langford, John Sillitoe on behalf of the Wellcome Sanger Institute COVID-19 Surveillance Team |
| EPI_ISL_957062, EPI_ISL_957063, EPI_ISL_957064 | Lighthouse Lab in Milton Keynes | Wellcome Sanger Institute for the COVID-19 Genomics UK (COG-UK) Consortium | The Lighthouse Lab in Milton Keynes and Alex Alderton, Roberto Amato, Sonia Goncalves, Ewan Harrison, David K. Jackson, Ian Johnston, Dominic Kwiatkowski, Cordelia Langford, John Sillitoe on behalf of the Wellcome Sanger Institute COVID-19 Surveillance Team |
| EPI_ISL_957065 | Lighthouse Lab in Glasgow | Wellcome Sanger Institute for the COVID-19 Genomics UK (COG-UK) Consortium | Harper VanSteenhouse, Yumi Kasai, David Gray, Carol Clugston, Anna Dominiczak and Alex Alderton, Roberto Amato, Sonia Goncalves, Ewan Harrison, David K. Jackson, Ian Johnston, Dominic Kwiatkowski, Cordelia Langford, John Sillitoe on behalf of the Wellcome Sanger Institute COVID-19 Surveillance Team |
| EPI_ISL_957066 | Lighthouse Lab in Milton Keynes | Wellcome Sanger Institute for the COVID-19 Genomics UK (COG-UK) Consortium | The Lighthouse Lab in Milton Keynes and Alex Alderton, Roberto Amato, Sonia Goncalves, Ewan Harrison, David K. Jackson, Ian Johnston, Dominic Kwiatkowski, Cordelia Langford, John Sillitoe on behalf of the Wellcome Sanger Institute COVID-19 Surveillance Team |
| EPI_ISL_957067, EPI_ISL_957068 | Lighthouse Lab in Glasgow | Wellcome Sanger Institute for the COVID-19 Genomics UK (COG-UK) Consortium | Harper VanSteenhouse, Yumi Kasai, David Gray, Carol Clugston, Anna Dominiczak and Alex Alderton, Roberto Amato, Sonia Goncalves, Ewan Harrison, David K. Jackson, Ian Johnston, Dominic Kwiatkowski, Cordelia Langford, John Sillitoe on behalf of the Wellcome Sanger Institute COVID-19 Surveillance Team |
| EPI_ISL_957069, EPI_ISL_957070 | Lighthouse Lab in Milton Keynes | Wellcome Sanger Institute for the COVID-19 Genomics UK (COG-UK) Consortium | The Lighthouse Lab in Milton Keynes and Alex Alderton, Roberto Amato, Sonia Goncalves, Ewan Harrison, David K. Jackson, Ian Johnston, Dominic Kwiatkowski, Cordelia Langford, John Sillitoe on behalf of the Wellcome Sanger Institute COVID-19 Surveillance Team |
| EPI_ISL_957071, EPI_ISL_957072, EPI_ISL_957073 | Lighthouse Lab in Glasgow | Wellcome Sanger Institute for the COVID-19 Genomics UK (COG-UK) Consortium | Harper VanSteenhouse, Yumi Kasai, David Gray, Carol Clugston, Anna Dominiczak and Alex Alderton, Roberto Amato, Sonia Goncalves, Ewan Harrison, David K. Jackson, Ian Johnston, Dominic Kwiatkowski, Cordelia Langford, John Sillitoe on behalf of the Wellcome Sanger Institute COVID-19 Surveillance Team |
| EPI_ISL_957074 | Lighthouse Lab in Milton Keynes | Wellcome Sanger Institute for the COVID-19 Genomics UK (COG-UK) Consortium | The Lighthouse Lab in Milton Keynes and Alex Alderton, Roberto Amato, Sonia Goncalves, Ewan Harrison, David K. Jackson, Ian Johnston, Dominic Kwiatkowski, Cordelia Langford, John Sillitoe on behalf of the Wellcome Sanger Institute COVID-19 Surveillance Team |
| EPI_ISL_957075 | Lighthouse Lab in Glasgow | Wellcome Sanger Institute for the COVID-19 Genomics UK (COG-UK) Consortium | Harper VanSteenhouse, Yumi Kasai, David Gray, Carol Clugston, Anna Dominiczak and Alex Alderton, Roberto Amato, Sonia Goncalves, Ewan Harrison, David K. Jackson, Ian Johnston, Dominic Kwiatkowski, Cordelia Langford, John Sillitoe on behalf of the Wellcome Sanger Institute COVID-19 Surveillance Team |
| EPI_ISL_957076 | Lighthouse Lab in Milton Keynes | Wellcome Sanger Institute for the COVID-19 Genomics UK (COG-UK) Consortium | The Lighthouse Lab in Milton Keynes and Alex Alderton, Roberto Amato, Sonia Goncalves, Ewan Harrison, David K. Jackson, Ian Johnston, Dominic Kwiatkowski, Cordelia Langford, John Sillitoe on behalf of the Wellcome Sanger Institute COVID-19 Surveillance Team |
| EPI_ISL_957077, EPI_ISL_957078, EPI_ISL_957079, EPI_ISL_957080, EPI_ISL_957081, EPI_ISL_957082, EPI_ISL_957083, EPI_ISL_957084 | Lighthouse Lab in Glasgow | Wellcome Sanger Institute for the COVID-19 Genomics UK (COG-UK) Consortium | Harper VanSteenhouse, Yumi Kasai, David Gray, Carol Clugston, Anna Dominiczak and Alex Alderton, Roberto Amato, Sonia Goncalves, Ewan Harrison, David K. Jackson, Ian Johnston, Dominic Kwiatkowski, Cordelia Langford, John Sillitoe on behalf of the Wellcome Sanger Institute COVID-19 Surveillance Team |
| EPI_ISL_957085 | Lighthouse Lab in Milton Keynes | Wellcome Sanger Institute for the COVID-19 Genomics UK (COG-UK) Consortium | The Lighthouse Lab in Milton Keynes and Alex Alderton, Roberto Amato, Sonia Goncalves, Ewan Harrison, David K. Jackson, Ian Johnston, Dominic Kwiatkowski, Cordelia Langford, John Sillitoe on behalf of the Wellcome Sanger Institute COVID-19 Surveillance Team |

| | | | |
|---|---|---|---|
| EPI_ISL_957086 | Lighthouse Lab in Glasgow | Wellcome Sanger Institute for the COVID-19 Genomics UK (COG-UK) Consortium | Harper VanSteenhouse, Yumi Kasai, David Gray, Carol Clugston, Anna Dominiczak and Alex Alderton, Roberto Amato, Sonia Goncalves, Ewan Harrison, David K. Jackson, Ian Johnston, Dominic Kwiatkowski, Cordelia Langford, John Sillitoe on behalf of the Wellcome Sanger Institute COVID-19 Surveillance Team |
| EPI_ISL_957087, EPI_ISL_957088 | Lighthouse Lab in Milton Keynes | Wellcome Sanger Institute for the COVID-19 Genomics UK (COG-UK) Consortium | The Lighthouse Lab in Milton Keynes and Alex Alderton, Roberto Amato, Sonia Goncalves, Ewan Harrison, David K. Jackson, Ian Johnston, Dominic Kwiatkowski, Cordelia Langford, John Sillitoe on behalf of the Wellcome Sanger Institute COVID-19 Surveillance Team |
| EPI_ISL_957089, EPI_ISL_957090, EPI_ISL_957091, EPI_ISL_957092, EPI_ISL_957093, EPI_ISL_957094, EPI_ISL_957095 | Lighthouse Lab in Glasgow | Wellcome Sanger Institute for the COVID-19 Genomics UK (COG-UK) Consortium | Harper VanSteenhouse, Yumi Kasai, David Gray, Carol Clugston, Anna Dominiczak and Alex Alderton, Roberto Amato, Sonia Goncalves, Ewan Harrison, David K. Jackson, Ian Johnston, Dominic Kwiatkowski, Cordelia Langford, John Sillitoe on behalf of the Wellcome Sanger Institute COVID-19 Surveillance Team |
| EPI_ISL_957096 | Lighthouse Lab in Milton Keynes | Wellcome Sanger Institute for the COVID-19 Genomics UK (COG-UK) Consortium | The Lighthouse Lab in Milton Keynes and Alex Alderton, Roberto Amato, Sonia Goncalves, Ewan Harrison, David K. Jackson, Ian Johnston, Dominic Kwiatkowski, Cordelia Langford, John Sillitoe on behalf of the Wellcome Sanger Institute COVID-19 Surveillance Team |
| EPI_ISL_957097 | Lighthouse Lab in Glasgow | Wellcome Sanger Institute for the COVID-19 Genomics UK (COG-UK) Consortium | Harper VanSteenhouse, Yumi Kasai, David Gray, Carol Clugston, Anna Dominiczak and Alex Alderton, Roberto Amato, Sonia Goncalves, Ewan Harrison, David K. Jackson, Ian Johnston, Dominic Kwiatkowski, Cordelia Langford, John Sillitoe on behalf of the Wellcome Sanger Institute COVID-19 Surveillance Team |
| EPI_ISL_957098 | Lighthouse Lab in Milton Keynes | Wellcome Sanger Institute for the COVID-19 Genomics UK (COG-UK) Consortium | The Lighthouse Lab in Milton Keynes and Alex Alderton, Roberto Amato, Sonia Goncalves, Ewan Harrison, David K. Jackson, Ian Johnston, Dominic Kwiatkowski, Cordelia Langford, John Sillitoe on behalf of the Wellcome Sanger Institute COVID-19 Surveillance Team |
| EPI_ISL_957099 | Lighthouse Lab in Glasgow | Wellcome Sanger Institute for the COVID-19 Genomics UK (COG-UK) Consortium | Harper VanSteenhouse, Yumi Kasai, David Gray, Carol Clugston, Anna Dominiczak and Alex Alderton, Roberto Amato, Sonia Goncalves, Ewan Harrison, David K. Jackson, Ian Johnston, Dominic Kwiatkowski, Cordelia Langford, John Sillitoe on behalf of the Wellcome Sanger Institute COVID-19 Surveillance Team |
| EPI_ISL_957100 | Lighthouse Lab in Milton Keynes | Wellcome Sanger Institute for the COVID-19 Genomics UK (COG-UK) Consortium | The Lighthouse Lab in Milton Keynes and Alex Alderton, Roberto Amato, Sonia Goncalves, Ewan Harrison, David K. Jackson, Ian Johnston, Dominic Kwiatkowski, Cordelia Langford, John Sillitoe on behalf of the Wellcome Sanger Institute COVID-19 Surveillance Team |
| EPI_ISL_957102, EPI_ISL_957103, EPI_ISL_957104, EPI_ISL_957105 | Lighthouse Lab in Glasgow | Wellcome Sanger Institute for the COVID-19 Genomics UK (COG-UK) Consortium | Harper VanSteenhouse, Yumi Kasai, David Gray, Carol Clugston, Anna Dominiczak and Alex Alderton, Roberto Amato, Sonia Goncalves, Ewan Harrison, David K. Jackson, Ian Johnston, Dominic Kwiatkowski, Cordelia Langford, John Sillitoe on behalf of the Wellcome Sanger Institute COVID-19 Surveillance Team |
| EPI_ISL_957106 | Lighthouse Lab in Milton Keynes | Wellcome Sanger Institute for the COVID-19 Genomics UK (COG-UK) Consortium | The Lighthouse Lab in Milton Keynes and Alex Alderton, Roberto Amato, Sonia Goncalves, Ewan Harrison, David K. Jackson, Ian Johnston, Dominic Kwiatkowski, Cordelia Langford, John Sillitoe on behalf of the Wellcome Sanger Institute COVID-19 Surveillance Team |
| EPI_ISL_957107 | Lighthouse Lab in Glasgow | Wellcome Sanger Institute for the COVID-19 Genomics UK (COG-UK) Consortium | Harper VanSteenhouse, Yumi Kasai, David Gray, Carol Clugston, Anna Dominiczak and Alex Alderton, Roberto Amato, Sonia Goncalves, Ewan Harrison, David K. Jackson, Ian Johnston, Dominic Kwiatkowski, Cordelia Langford, John Sillitoe on behalf of the Wellcome Sanger Institute COVID-19 Surveillance Team |
| EPI_ISL_957108, EPI_ISL_957109 | Lighthouse Lab in Milton Keynes | Wellcome Sanger Institute for the COVID-19 Genomics UK (COG-UK) Consortium | The Lighthouse Lab in Milton Keynes and Alex Alderton, Roberto Amato, Sonia Goncalves, Ewan Harrison, David K. Jackson, Ian Johnston, Dominic Kwiatkowski, Cordelia Langford, John Sillitoe on behalf of the Wellcome Sanger Institute COVID-19 Surveillance Team |
| EPI_ISL_957110 | Lighthouse Lab in Glasgow | Wellcome Sanger Institute for the COVID-19 Genomics UK (COG-UK) Consortium | Harper VanSteenhouse, Yumi Kasai, David Gray, Carol Clugston, Anna Dominiczak and Alex Alderton, Roberto Amato, Sonia Goncalves, Ewan Harrison, David K. Jackson, Ian Johnston, Dominic Kwiatkowski, Cordelia Langford, John Sillitoe on behalf of the Wellcome Sanger Institute COVID-19 Surveillance Team |
| EPI_ISL_957111, EPI_ISL_957112, EPI_ISL_957113 | Lighthouse Lab in Milton Keynes | Wellcome Sanger Institute for the COVID-19 Genomics UK (COG-UK) Consortium | The Lighthouse Lab in Milton Keynes and Alex Alderton, Roberto Amato, Sonia Goncalves, Ewan Harrison, David K. Jackson, Ian Johnston, Dominic Kwiatkowski, Cordelia Langford, John Sillitoe on behalf of the Wellcome Sanger Institute COVID-19 Surveillance Team |
| EPI_ISL_957114, EPI_ISL_957115 | Lighthouse Lab in Glasgow | Wellcome Sanger Institute for the COVID-19 Genomics UK (COG-UK) Consortium | Harper VanSteenhouse, Yumi Kasai, David Gray, Carol Clugston, Anna Dominiczak and Alex Alderton, Roberto Amato, Sonia Goncalves, Ewan Harrison, David K. Jackson, Ian Johnston, Dominic Kwiatkowski, Cordelia Langford, John Sillitoe on behalf of the Wellcome Sanger Institute COVID-19 Surveillance Team |
| EPI_ISL_957116 | Lighthouse Lab in Milton Keynes | Wellcome Sanger Institute for the COVID-19 Genomics UK (COG-UK) Consortium | The Lighthouse Lab in Milton Keynes and Alex Alderton, Roberto Amato, Sonia Goncalves, Ewan Harrison, David K. Jackson, Ian Johnston, Dominic Kwiatkowski, Cordelia Langford, John Sillitoe on behalf of the Wellcome Sanger Institute COVID-19 Surveillance Team |
| EPI_ISL_957117, EPI_ISL_957118 | Lighthouse Lab in Glasgow | Wellcome Sanger Institute for the COVID-19 Genomics UK (COG-UK) Consortium | Harper VanSteenhouse, Yumi Kasai, David Gray, Carol Clugston, Anna Dominiczak and Alex Alderton, Roberto Amato, Sonia Goncalves, Ewan Harrison, David K. Jackson, Ian Johnston, Dominic Kwiatkowski, Cordelia Langford, John Sillitoe on behalf of the Wellcome Sanger Institute COVID-19 Surveillance Team |
| EPI_ISL_957119, EPI_ISL_957120 | Lighthouse Lab in Milton Keynes | Wellcome Sanger Institute for the COVID-19 Genomics UK (COG-UK) Consortium | The Lighthouse Lab in Milton Keynes and Alex Alderton, Roberto Amato, Sonia Goncalves, Ewan Harrison, David K. Jackson, Ian Johnston, Dominic Kwiatkowski, Cordelia Langford, John Sillitoe on behalf of the Wellcome Sanger Institute COVID-19 Surveillance Team |
| EPI_ISL_957121, EPI_ISL_957122 | Lighthouse Lab in Glasgow | Wellcome Sanger Institute for the COVID-19 Genomics UK (COG-UK) Consortium | Harper VanSteenhouse, Yumi Kasai, David Gray, Carol Clugston, Anna Dominiczak and Alex Alderton, Roberto Amato, Sonia Goncalves, Ewan Harrison, David K. Jackson, Ian Johnston, Dominic Kwiatkowski, Cordelia Langford, John Sillitoe on behalf of the Wellcome Sanger Institute COVID-19 Surveillance Team |
| EPI_ISL_957123 | Lighthouse Lab in Milton Keynes | Wellcome Sanger Institute for the COVID-19 Genomics UK (COG-UK) Consortium | The Lighthouse Lab in Milton Keynes and Alex Alderton, Roberto Amato, Sonia Goncalves, Ewan Harrison, David K. Jackson, Ian Johnston, Dominic Kwiatkowski, Cordelia Langford, John Sillitoe on behalf of the Wellcome Sanger Institute COVID-19 Surveillance Team |
| EPI_ISL_957124 | Lighthouse Lab in Glasgow | Wellcome Sanger Institute for the COVID-19 Genomics UK (COG-UK) Consortium | Harper VanSteenhouse, Yumi Kasai, David Gray, Carol Clugston, Anna Dominiczak and Alex Alderton, Roberto Amato, Sonia Goncalves, Ewan Harrison, David K. Jackson, Ian Johnston, Dominic Kwiatkowski, Cordelia Langford, John Sillitoe on behalf of the Wellcome Sanger Institute COVID-19 Surveillance Team |
| EPI_ISL_957125, EPI_ISL_957126, EPI_ISL_957127, EPI_ISL_957128, EPI_ISL_957129 | Lighthouse Lab in Milton Keynes | Wellcome Sanger Institute for the COVID-19 Genomics UK (COG-UK) Consortium | The Lighthouse Lab in Milton Keynes and Alex Alderton, Roberto Amato, Sonia Goncalves, Ewan Harrison, David K. Jackson, Ian Johnston, Dominic Kwiatkowski, Cordelia Langford, John Sillitoe on behalf of the Wellcome Sanger Institute COVID-19 Surveillance Team |
| EPI_ISL_957130 | Lighthouse Lab in Glasgow | Wellcome Sanger Institute for the COVID-19 Genomics UK (COG-UK) Consortium | Harper VanSteenhouse, Yumi Kasai, David Gray, Carol Clugston, Anna Dominiczak and Alex Alderton, Roberto Amato, Sonia Goncalves, Ewan Harrison, David K. Jackson, Ian Johnston, Dominic Kwiatkowski, Cordelia Langford, John Sillitoe on behalf of the Wellcome Sanger Institute COVID-19 Surveillance Team |
| EPI_ISL_957131, EPI_ISL_957132 | Lighthouse Lab in Milton Keynes | Wellcome Sanger Institute for the COVID-19 Genomics UK (COG-UK) Consortium | The Lighthouse Lab in Milton Keynes and Alex Alderton, Roberto Amato, Sonia Goncalves, Ewan Harrison, David K. Jackson, Ian Johnston, Dominic Kwiatkowski, Cordelia Langford, John Sillitoe on behalf of the Wellcome Sanger Institute COVID-19 Surveillance Team |
| EPI_ISL_957133 | Lighthouse Lab in Glasgow | Wellcome Sanger Institute for the COVID-19 Genomics UK (COG-UK) Consortium | Harper VanSteenhouse, Yumi Kasai, David Gray, Carol Clugston, Anna Dominiczak and Alex Alderton, Roberto Amato, Sonia Goncalves, Ewan Harrison, David K. Jackson, Ian Johnston, Dominic Kwiatkowski, Cordelia Langford, John Sillitoe on behalf of the Wellcome Sanger Institute COVID-19 Surveillance Team |
| EPI_ISL_957134, EPI_ISL_957135, EPI_ISL_957136, EPI_ISL_957137, EPI_ISL_957138, EPI_ISL_957139 | Lighthouse Lab in Milton Keynes | Wellcome Sanger Institute for the COVID-19 Genomics UK (COG-UK) Consortium | The Lighthouse Lab in Milton Keynes and Alex Alderton, Roberto Amato, Sonia Goncalves, Ewan Harrison, David K. Jackson, Ian Johnston, Dominic Kwiatkowski, Cordelia Langford, John Sillitoe on behalf of the Wellcome Sanger Institute COVID-19 Surveillance Team |
| EPI_ISL_957140, EPI_ISL_957141 | Lighthouse Lab in Glasgow | Wellcome Sanger Institute for the COVID-19 Genomics UK (COG-UK) Consortium | Harper VanSteenhouse, Yumi Kasai, David Gray, Carol Clugston, Anna Dominiczak and Alex Alderton, Roberto Amato, Sonia Goncalves, Ewan Harrison, David K. Jackson, Ian Johnston, Dominic Kwiatkowski, Cordelia Langford, John Sillitoe on behalf of the Wellcome Sanger Institute COVID-19 Surveillance Team |
| EPI_ISL_957142, EPI_ISL_957143, EPI_ISL_957144, EPI_ISL_957145, EPI_ISL_957146, EPI_ISL_957147, | Lighthouse Lab in Milton Keynes | Wellcome Sanger Institute for the COVID-19 Genomics UK (COG-UK) Consortium | The Lighthouse Lab in Milton Keynes and Alex Alderton, Roberto Amato, Sonia Goncalves, Ewan Harrison, David K. Jackson, Ian Johnston, Dominic Kwiatkowski, Cordelia Langford, John Sillitoe on behalf of the Wellcome Sanger Institute COVID-19 Surveillance Team |

| | | | |
|---|---|---|---|
| EPI_ISL_957148 | | | |
| EPI_ISL_957149, EPI_ISL_957150, EPI_ISL_957151 | Lighthouse Lab in Glasgow | Wellcome Sanger Institute for the COVID-19 Genomics UK (COG-UK) Consortium | Harper VanSteenhouse, Yumi Kasai, David Gray, Carol Clugston, Anna Dominiczak and Alex Alderton, Roberto Amato, Sonia Goncalves, Ewan Harrison, David K. Jackson, Ian Johnston, Dominic Kwiatkowski, Cordelia Langford, John Sillitoe on behalf of the Wellcome Sanger Institute COVID-19 Surveillance Team |
| EPI_ISL_957152, EPI_ISL_957153 | Lighthouse Lab in Milton Keynes | Wellcome Sanger Institute for the COVID-19 Genomics UK (COG-UK) Consortium | The Lighthouse Lab in Milton Keynes and Alex Alderton, Roberto Amato, Sonia Goncalves, Ewan Harrison, David K. Jackson, Ian Johnston, Dominic Kwiatkowski, Cordelia Langford, John Sillitoe on behalf of the Wellcome Sanger Institute COVID-19 Surveillance Team |
| EPI_ISL_957154, EPI_ISL_957155 | Lighthouse Lab in Glasgow | Wellcome Sanger Institute for the COVID-19 Genomics UK (COG-UK) Consortium | Harper VanSteenhouse, Yumi Kasai, David Gray, Carol Clugston, Anna Dominiczak and Alex Alderton, Roberto Amato, Sonia Goncalves, Ewan Harrison, David K. Jackson, Ian Johnston, Dominic Kwiatkowski, Cordelia Langford, John Sillitoe on behalf of the Wellcome Sanger Institute COVID-19 Surveillance Team |
| EPI_ISL_957156 | Lighthouse Lab in Milton Keynes | Wellcome Sanger Institute for the COVID-19 Genomics UK (COG-UK) Consortium | The Lighthouse Lab in Milton Keynes and Alex Alderton, Roberto Amato, Sonia Goncalves, Ewan Harrison, David K. Jackson, Ian Johnston, Dominic Kwiatkowski, Cordelia Langford, John Sillitoe on behalf of the Wellcome Sanger Institute COVID-19 Surveillance Team |
| EPI_ISL_957157, EPI_ISL_957158 | Lighthouse Lab in Glasgow | Wellcome Sanger Institute for the COVID-19 Genomics UK (COG-UK) Consortium | Harper VanSteenhouse, Yumi Kasai, David Gray, Carol Clugston, Anna Dominiczak and Alex Alderton, Roberto Amato, Sonia Goncalves, Ewan Harrison, David K. Jackson, Ian Johnston, Dominic Kwiatkowski, Cordelia Langford, John Sillitoe on behalf of the Wellcome Sanger Institute COVID-19 Surveillance Team |
| EPI_ISL_957159 | Lighthouse Lab in Milton Keynes | Wellcome Sanger Institute for the COVID-19 Genomics UK (COG-UK) Consortium | The Lighthouse Lab in Milton Keynes and Alex Alderton, Roberto Amato, Sonia Goncalves, Ewan Harrison, David K. Jackson, Ian Johnston, Dominic Kwiatkowski, Cordelia Langford, John Sillitoe on behalf of the Wellcome Sanger Institute COVID-19 Surveillance Team |
| EPI_ISL_957160 | Lighthouse Lab in Glasgow | Wellcome Sanger Institute for the COVID-19 Genomics UK (COG-UK) Consortium | Harper VanSteenhouse, Yumi Kasai, David Gray, Carol Clugston, Anna Dominiczak and Alex Alderton, Roberto Amato, Sonia Goncalves, Ewan Harrison, David K. Jackson, Ian Johnston, Dominic Kwiatkowski, Cordelia Langford, John Sillitoe on behalf of the Wellcome Sanger Institute COVID-19 Surveillance Team |
| EPI_ISL_957161 | Lighthouse Lab in Milton Keynes | Wellcome Sanger Institute for the COVID-19 Genomics UK (COG-UK) Consortium | The Lighthouse Lab in Milton Keynes and Alex Alderton, Roberto Amato, Sonia Goncalves, Ewan Harrison, David K. Jackson, Ian Johnston, Dominic Kwiatkowski, Cordelia Langford, John Sillitoe on behalf of the Wellcome Sanger Institute COVID-19 Surveillance Team |
| EPI_ISL_957162 | Lighthouse Lab in Glasgow | Wellcome Sanger Institute for the COVID-19 Genomics UK (COG-UK) Consortium | Harper VanSteenhouse, Yumi Kasai, David Gray, Carol Clugston, Anna Dominiczak and Alex Alderton, Roberto Amato, Sonia Goncalves, Ewan Harrison, David K. Jackson, Ian Johnston, Dominic Kwiatkowski, Cordelia Langford, John Sillitoe on behalf of the Wellcome Sanger Institute COVID-19 Surveillance Team |
| EPI_ISL_957163, EPI_ISL_957164 | Lighthouse Lab in Milton Keynes | Wellcome Sanger Institute for the COVID-19 Genomics UK (COG-UK) Consortium | The Lighthouse Lab in Milton Keynes and Alex Alderton, Roberto Amato, Sonia Goncalves, Ewan Harrison, David K. Jackson, Ian Johnston, Dominic Kwiatkowski, Cordelia Langford, John Sillitoe on behalf of the Wellcome Sanger Institute COVID-19 Surveillance Team |
| EPI_ISL_957165, EPI_ISL_957166 | Lighthouse Lab in Glasgow | Wellcome Sanger Institute for the COVID-19 Genomics UK (COG-UK) Consortium | Harper VanSteenhouse, Yumi Kasai, David Gray, Carol Clugston, Anna Dominiczak and Alex Alderton, Roberto Amato, Sonia Goncalves, Ewan Harrison, David K. Jackson, Ian Johnston, Dominic Kwiatkowski, Cordelia Langford, John Sillitoe on behalf of the Wellcome Sanger Institute COVID-19 Surveillance Team |
| EPI_ISL_957167 | Lighthouse Lab in Milton Keynes | Wellcome Sanger Institute for the COVID-19 Genomics UK (COG-UK) Consortium | The Lighthouse Lab in Milton Keynes and Alex Alderton, Roberto Amato, Sonia Goncalves, Ewan Harrison, David K. Jackson, Ian Johnston, Dominic Kwiatkowski, Cordelia Langford, John Sillitoe on behalf of the Wellcome Sanger Institute COVID-19 Surveillance Team |
| EPI_ISL_957168 | Lighthouse Lab in Glasgow | Wellcome Sanger Institute for the COVID-19 Genomics UK (COG-UK) Consortium | Harper VanSteenhouse, Yumi Kasai, David Gray, Carol Clugston, Anna Dominiczak and Alex Alderton, Roberto Amato, Sonia Goncalves, Ewan Harrison, David K. Jackson, Ian Johnston, Dominic Kwiatkowski, Cordelia Langford, John Sillitoe on behalf of the Wellcome Sanger Institute COVID-19 Surveillance Team |
| EPI_ISL_957169 | Lighthouse Lab in Milton Keynes | Wellcome Sanger Institute for the COVID-19 Genomics UK (COG-UK) Consortium | The Lighthouse Lab in Milton Keynes and Alex Alderton, Roberto Amato, Sonia Goncalves, Ewan Harrison, David K. Jackson, Ian Johnston, Dominic Kwiatkowski, Cordelia Langford, John Sillitoe on behalf of the Wellcome Sanger Institute COVID-19 Surveillance Team |
| EPI_ISL_957170, EPI_ISL_957171, EPI_ISL_957172 | Lighthouse Lab in Glasgow | Wellcome Sanger Institute for the COVID-19 Genomics UK (COG-UK) Consortium | Harper VanSteenhouse, Yumi Kasai, David Gray, Carol Clugston, Anna Dominiczak and Alex Alderton, Roberto Amato, Sonia Goncalves, Ewan Harrison, David K. Jackson, Ian Johnston, Dominic Kwiatkowski, Cordelia Langford, John Sillitoe on behalf of the Wellcome Sanger Institute COVID-19 Surveillance Team |
| EPI_ISL_957173, EPI_ISL_957174, EPI_ISL_957175 | Lighthouse Lab in Milton Keynes | Wellcome Sanger Institute for the COVID-19 Genomics UK (COG-UK) Consortium | The Lighthouse Lab in Milton Keynes and Alex Alderton, Roberto Amato, Sonia Goncalves, Ewan Harrison, David K. Jackson, Ian Johnston, Dominic Kwiatkowski, Cordelia Langford, John Sillitoe on behalf of the Wellcome Sanger Institute COVID-19 Surveillance Team |
| EPI_ISL_957176, EPI_ISL_957177 | Lighthouse Lab in Glasgow | Wellcome Sanger Institute for the COVID-19 Genomics UK (COG-UK) Consortium | Harper VanSteenhouse, Yumi Kasai, David Gray, Carol Clugston, Anna Dominiczak and Alex Alderton, Roberto Amato, Sonia Goncalves, Ewan Harrison, David K. Jackson, Ian Johnston, Dominic Kwiatkowski, Cordelia Langford, John Sillitoe on behalf of the Wellcome Sanger Institute COVID-19 Surveillance Team |
| EPI_ISL_957178, EPI_ISL_957179 | Lighthouse Lab in Milton Keynes | Wellcome Sanger Institute for the COVID-19 Genomics UK (COG-UK) Consortium | The Lighthouse Lab in Milton Keynes and Alex Alderton, Roberto Amato, Sonia Goncalves, Ewan Harrison, David K. Jackson, Ian Johnston, Dominic Kwiatkowski, Cordelia Langford, John Sillitoe on behalf of the Wellcome Sanger Institute COVID-19 Surveillance Team |
| EPI_ISL_957180 | Lighthouse Lab in Glasgow | Wellcome Sanger Institute for the COVID-19 Genomics UK (COG-UK) Consortium | Harper VanSteenhouse, Yumi Kasai, David Gray, Carol Clugston, Anna Dominiczak and Alex Alderton, Roberto Amato, Sonia Goncalves, Ewan Harrison, David K. Jackson, Ian Johnston, Dominic Kwiatkowski, Cordelia Langford, John Sillitoe on behalf of the Wellcome Sanger Institute COVID-19 Surveillance Team |
| EPI_ISL_957181, EPI_ISL_957182 | Lighthouse Lab in Milton Keynes | Wellcome Sanger Institute for the COVID-19 Genomics UK (COG-UK) Consortium | The Lighthouse Lab in Milton Keynes and Alex Alderton, Roberto Amato, Sonia Goncalves, Ewan Harrison, David K. Jackson, Ian Johnston, Dominic Kwiatkowski, Cordelia Langford, John Sillitoe on behalf of the Wellcome Sanger Institute COVID-19 Surveillance Team |
| EPI_ISL_957183 | Lighthouse Lab in Glasgow | Wellcome Sanger Institute for the COVID-19 Genomics UK (COG-UK) Consortium | Harper VanSteenhouse, Yumi Kasai, David Gray, Carol Clugston, Anna Dominiczak and Alex Alderton, Roberto Amato, Sonia Goncalves, Ewan Harrison, David K. Jackson, Ian Johnston, Dominic Kwiatkowski, Cordelia Langford, John Sillitoe on behalf of the Wellcome Sanger Institute COVID-19 Surveillance Team |
| EPI_ISL_957184, EPI_ISL_957185, EPI_ISL_957186 | Lighthouse Lab in Milton Keynes | Wellcome Sanger Institute for the COVID-19 Genomics UK (COG-UK) Consortium | The Lighthouse Lab in Milton Keynes and Alex Alderton, Roberto Amato, Sonia Goncalves, Ewan Harrison, David K. Jackson, Ian Johnston, Dominic Kwiatkowski, Cordelia Langford, John Sillitoe on behalf of the Wellcome Sanger Institute COVID-19 Surveillance Team |
| EPI_ISL_957187, EPI_ISL_957188 | Lighthouse Lab in Glasgow | Wellcome Sanger Institute for the COVID-19 Genomics UK (COG-UK) Consortium | Harper VanSteenhouse, Yumi Kasai, David Gray, Carol Clugston, Anna Dominiczak and Alex Alderton, Roberto Amato, Sonia Goncalves, Ewan Harrison, David K. Jackson, Ian Johnston, Dominic Kwiatkowski, Cordelia Langford, John Sillitoe on behalf of the Wellcome Sanger Institute COVID-19 Surveillance Team |
| EPI_ISL_957189 | Lighthouse Lab in Milton Keynes | Wellcome Sanger Institute for the COVID-19 Genomics UK (COG-UK) Consortium | The Lighthouse Lab in Milton Keynes and Alex Alderton, Roberto Amato, Sonia Goncalves, Ewan Harrison, David K. Jackson, Ian Johnston, Dominic Kwiatkowski, Cordelia Langford, John Sillitoe on behalf of the Wellcome Sanger Institute COVID-19 Surveillance Team |
| EPI_ISL_957190, EPI_ISL_957191 | Lighthouse Lab in Glasgow | Wellcome Sanger Institute for the COVID-19 Genomics UK (COG-UK) Consortium | Harper VanSteenhouse, Yumi Kasai, David Gray, Carol Clugston, Anna Dominiczak and Alex Alderton, Roberto Amato, Sonia Goncalves, Ewan Harrison, David K. Jackson, Ian Johnston, Dominic Kwiatkowski, Cordelia Langford, John Sillitoe on behalf of the Wellcome Sanger Institute COVID-19 Surveillance Team |
| EPI_ISL_957192, EPI_ISL_957193 | Lighthouse Lab in Milton Keynes | Wellcome Sanger Institute for the COVID-19 Genomics UK (COG-UK) Consortium | The Lighthouse Lab in Milton Keynes and Alex Alderton, Roberto Amato, Sonia Goncalves, Ewan Harrison, David K. Jackson, Ian Johnston, Dominic Kwiatkowski, Cordelia Langford, John Sillitoe on behalf of the Wellcome Sanger Institute COVID-19 Surveillance Team |
| EPI_ISL_957194 | Lighthouse Lab in Glasgow | Wellcome Sanger Institute for the COVID-19 Genomics UK (COG-UK) Consortium | Harper VanSteenhouse, Yumi Kasai, David Gray, Carol Clugston, Anna Dominiczak and Alex Alderton, Roberto Amato, Sonia Goncalves, Ewan Harrison, David K. Jackson, Ian Johnston, Dominic Kwiatkowski, Cordelia Langford, John Sillitoe on behalf of the Wellcome Sanger Institute COVID-19 Surveillance Team |
| EPI_ISL_957195, EPI_ISL_957196, EPI_ISL_957197 | Lighthouse Lab in Milton Keynes | Wellcome Sanger Institute for the COVID-19 Genomics UK (COG-UK) Consortium | The Lighthouse Lab in Milton Keynes and Alex Alderton, Roberto Amato, Sonia Goncalves, Ewan Harrison, David K. Jackson, Ian Johnston, Dominic Kwiatkowski, Cordelia Langford, John Sillitoe on behalf of the Wellcome Sanger Institute COVID-19 Surveillance Team |
| EPI_ISL_957198 | Lighthouse Lab in Glasgow | Wellcome Sanger Institute for the COVID-19 Genomics UK (COG-UK) Consortium | Harper VanSteenhouse, Yumi Kasai, David Gray, Carol Clugston, Anna Dominiczak and Alex Alderton, Roberto Amato, Sonia Goncalves, Ewan Harrison, David K. Jackson, Ian Johnston, Dominic Kwiatkowski, Cordelia Langford, John Sillitoe on behalf of the Wellcome Sanger Institute COVID-19 Surveillance |

| | | | |
|---|---|---|---|
| | | | Team |
| EPI_ISL_957199 | Lighthouse Lab in Milton Keynes | Wellcome Sanger Institute for the COVID-19 Genomics UK (COG-UK) Consortium | The Lighthouse Lab in Milton Keynes and Alex Alderton, Roberto Amato, Sonia Goncalves, Ewan Harrison, David K. Jackson, Ian Johnston, Dominic Kwiatkowski, Cordelia Langford, John Sillitoe on behalf of the Wellcome Sanger Institute COVID-19 Surveillance Team |
| EPI_ISL_957200, EPI_ISL_957201 | Lighthouse Lab in Glasgow | Wellcome Sanger Institute for the COVID-19 Genomics UK (COG-UK) Consortium | Harper VanSteenhouse, Yumi Kasai, David Gray, Carol Clugston, Anna Dominiczak and Alex Alderton, Roberto Amato, Sonia Goncalves, Ewan Harrison, David K. Jackson, Ian Johnston, Dominic Kwiatkowski, Cordelia Langford, John Sillitoe on behalf of the Wellcome Sanger Institute COVID-19 Surveillance Team |
| EPI_ISL_957202, EPI_ISL_957203, EPI_ISL_957204 | Lighthouse Lab in Milton Keynes | Wellcome Sanger Institute for the COVID-19 Genomics UK (COG-UK) Consortium | The Lighthouse Lab in Milton Keynes and Alex Alderton, Roberto Amato, Sonia Goncalves, Ewan Harrison, David K. Jackson, Ian Johnston, Dominic Kwiatkowski, Cordelia Langford, John Sillitoe on behalf of the Wellcome Sanger Institute COVID-19 Surveillance Team |
| EPI_ISL_957205, EPI_ISL_957206 | Lighthouse Lab in Glasgow | Wellcome Sanger Institute for the COVID-19 Genomics UK (COG-UK) Consortium | Harper VanSteenhouse, Yumi Kasai, David Gray, Carol Clugston, Anna Dominiczak and Alex Alderton, Roberto Amato, Sonia Goncalves, Ewan Harrison, David K. Jackson, Ian Johnston, Dominic Kwiatkowski, Cordelia Langford, John Sillitoe on behalf of the Wellcome Sanger Institute COVID-19 Surveillance Team |
| EPI_ISL_957207 | Lighthouse Lab in Milton Keynes | Wellcome Sanger Institute for the COVID-19 Genomics UK (COG-UK) Consortium | The Lighthouse Lab in Milton Keynes and Alex Alderton, Roberto Amato, Sonia Goncalves, Ewan Harrison, David K. Jackson, Ian Johnston, Dominic Kwiatkowski, Cordelia Langford, John Sillitoe on behalf of the Wellcome Sanger Institute COVID-19 Surveillance Team |
| EPI_ISL_957208 | Lighthouse Lab in Glasgow | Wellcome Sanger Institute for the COVID-19 Genomics UK (COG-UK) Consortium | Harper VanSteenhouse, Yumi Kasai, David Gray, Carol Clugston, Anna Dominiczak and Alex Alderton, Roberto Amato, Sonia Goncalves, Ewan Harrison, David K. Jackson, Ian Johnston, Dominic Kwiatkowski, Cordelia Langford, John Sillitoe on behalf of the Wellcome Sanger Institute COVID-19 Surveillance Team |
| EPI_ISL_957209 | Lighthouse Lab in Milton Keynes | Wellcome Sanger Institute for the COVID-19 Genomics UK (COG-UK) Consortium | The Lighthouse Lab in Milton Keynes and Alex Alderton, Roberto Amato, Sonia Goncalves, Ewan Harrison, David K. Jackson, Ian Johnston, Dominic Kwiatkowski, Cordelia Langford, John Sillitoe on behalf of the Wellcome Sanger Institute COVID-19 Surveillance Team |
| EPI_ISL_957210, EPI_ISL_957211, EPI_ISL_957212 | Lighthouse Lab in Glasgow | Wellcome Sanger Institute for the COVID-19 Genomics UK (COG-UK) Consortium | Harper VanSteenhouse, Yumi Kasai, David Gray, Carol Clugston, Anna Dominiczak and Alex Alderton, Roberto Amato, Sonia Goncalves, Ewan Harrison, David K. Jackson, Ian Johnston, Dominic Kwiatkowski, Cordelia Langford, John Sillitoe on behalf of the Wellcome Sanger Institute COVID-19 Surveillance Team |
| EPI_ISL_957213, EPI_ISL_957214 | Lighthouse Lab in Milton Keynes | Wellcome Sanger Institute for the COVID-19 Genomics UK (COG-UK) Consortium | The Lighthouse Lab in Milton Keynes and Alex Alderton, Roberto Amato, Sonia Goncalves, Ewan Harrison, David K. Jackson, Ian Johnston, Dominic Kwiatkowski, Cordelia Langford, John Sillitoe on behalf of the Wellcome Sanger Institute COVID-19 Surveillance Team |
| EPI_ISL_957215, EPI_ISL_957216 | Lighthouse Lab in Glasgow | Wellcome Sanger Institute for the COVID-19 Genomics UK (COG-UK) Consortium | Harper VanSteenhouse, Yumi Kasai, David Gray, Carol Clugston, Anna Dominiczak and Alex Alderton, Roberto Amato, Sonia Goncalves, Ewan Harrison, David K. Jackson, Ian Johnston, Dominic Kwiatkowski, Cordelia Langford, John Sillitoe on behalf of the Wellcome Sanger Institute COVID-19 Surveillance Team |
| EPI_ISL_957217, EPI_ISL_957218 | Lighthouse Lab in Milton Keynes | Wellcome Sanger Institute for the COVID-19 Genomics UK (COG-UK) Consortium | The Lighthouse Lab in Milton Keynes and Alex Alderton, Roberto Amato, Sonia Goncalves, Ewan Harrison, David K. Jackson, Ian Johnston, Dominic Kwiatkowski, Cordelia Langford, John Sillitoe on behalf of the Wellcome Sanger Institute COVID-19 Surveillance Team |
| EPI_ISL_957219 | Lighthouse Lab in Glasgow | Wellcome Sanger Institute for the COVID-19 Genomics UK (COG-UK) Consortium | Harper VanSteenhouse, Yumi Kasai, David Gray, Carol Clugston, Anna Dominiczak and Alex Alderton, Roberto Amato, Sonia Goncalves, Ewan Harrison, David K. Jackson, Ian Johnston, Dominic Kwiatkowski, Cordelia Langford, John Sillitoe on behalf of the Wellcome Sanger Institute COVID-19 Surveillance Team |
| EPI_ISL_957220, EPI_ISL_957221, EPI_ISL_957222 | Lighthouse Lab in Milton Keynes | Wellcome Sanger Institute for the COVID-19 Genomics UK (COG-UK) Consortium | The Lighthouse Lab in Milton Keynes and Alex Alderton, Roberto Amato, Sonia Goncalves, Ewan Harrison, David K. Jackson, Ian Johnston, Dominic Kwiatkowski, Cordelia Langford, John Sillitoe on behalf of the Wellcome Sanger Institute COVID-19 Surveillance Team |
| EPI_ISL_957223, EPI_ISL_957224, EPI_ISL_957225 | Lighthouse Lab in Glasgow | Wellcome Sanger Institute for the COVID-19 Genomics UK (COG-UK) Consortium | Harper VanSteenhouse, Yumi Kasai, David Gray, Carol Clugston, Anna Dominiczak and Alex Alderton, Roberto Amato, Sonia Goncalves, Ewan Harrison, David K. Jackson, Ian Johnston, Dominic Kwiatkowski, Cordelia Langford, John Sillitoe on behalf of the Wellcome Sanger Institute COVID-19 Surveillance Team |
| EPI_ISL_957226 | Lighthouse Lab in Milton Keynes | Wellcome Sanger Institute for the COVID-19 Genomics UK (COG-UK) Consortium | The Lighthouse Lab in Milton Keynes and Alex Alderton, Roberto Amato, Sonia Goncalves, Ewan Harrison, David K. Jackson, Ian Johnston, Dominic Kwiatkowski, Cordelia Langford, John Sillitoe on behalf of the Wellcome Sanger Institute COVID-19 Surveillance Team |
| EPI_ISL_957227 | Lighthouse Lab in Glasgow | Wellcome Sanger Institute for the COVID-19 Genomics UK (COG-UK) Consortium | Harper VanSteenhouse, Yumi Kasai, David Gray, Carol Clugston, Anna Dominiczak and Alex Alderton, Roberto Amato, Sonia Goncalves, Ewan Harrison, David K. Jackson, Ian Johnston, Dominic Kwiatkowski, Cordelia Langford, John Sillitoe on behalf of the Wellcome Sanger Institute COVID-19 Surveillance Team |
| EPI_ISL_957228 | Lighthouse Lab in Milton Keynes | Wellcome Sanger Institute for the COVID-19 Genomics UK (COG-UK) Consortium | The Lighthouse Lab in Milton Keynes and Alex Alderton, Roberto Amato, Sonia Goncalves, Ewan Harrison, David K. Jackson, Ian Johnston, Dominic Kwiatkowski, Cordelia Langford, John Sillitoe on behalf of the Wellcome Sanger Institute COVID-19 Surveillance Team |
| EPI_ISL_957229 | Lighthouse Lab in Glasgow | Wellcome Sanger Institute for the COVID-19 Genomics UK (COG-UK) Consortium | Harper VanSteenhouse, Yumi Kasai, David Gray, Carol Clugston, Anna Dominiczak and Alex Alderton, Roberto Amato, Sonia Goncalves, Ewan Harrison, David K. Jackson, Ian Johnston, Dominic Kwiatkowski, Cordelia Langford, John Sillitoe on behalf of the Wellcome Sanger Institute COVID-19 Surveillance Team |
| EPI_ISL_957230 | Lighthouse Lab in Milton Keynes | Wellcome Sanger Institute for the COVID-19 Genomics UK (COG-UK) Consortium | The Lighthouse Lab in Milton Keynes and Alex Alderton, Roberto Amato, Sonia Goncalves, Ewan Harrison, David K. Jackson, Ian Johnston, Dominic Kwiatkowski, Cordelia Langford, John Sillitoe on behalf of the Wellcome Sanger Institute COVID-19 Surveillance Team |
| EPI_ISL_957231 | Lighthouse Lab in Glasgow | Wellcome Sanger Institute for the COVID-19 Genomics UK (COG-UK) Consortium | Harper VanSteenhouse, Yumi Kasai, David Gray, Carol Clugston, Anna Dominiczak and Alex Alderton, Roberto Amato, Sonia Goncalves, Ewan Harrison, David K. Jackson, Ian Johnston, Dominic Kwiatkowski, Cordelia Langford, John Sillitoe on behalf of the Wellcome Sanger Institute COVID-19 Surveillance Team |
| EPI_ISL_957232, EPI_ISL_957233, EPI_ISL_957234, EPI_ISL_957235, EPI_ISL_957236 | Lighthouse Lab in Milton Keynes | Wellcome Sanger Institute for the COVID-19 Genomics UK (COG-UK) Consortium | The Lighthouse Lab in Milton Keynes and Alex Alderton, Roberto Amato, Sonia Goncalves, Ewan Harrison, David K. Jackson, Ian Johnston, Dominic Kwiatkowski, Cordelia Langford, John Sillitoe on behalf of the Wellcome Sanger Institute COVID-19 Surveillance Team |
| EPI_ISL_957237 | Lighthouse Lab in Glasgow | Wellcome Sanger Institute for the COVID-19 Genomics UK (COG-UK) Consortium | Harper VanSteenhouse, Yumi Kasai, David Gray, Carol Clugston, Anna Dominiczak and Alex Alderton, Roberto Amato, Sonia Goncalves, Ewan Harrison, David K. Jackson, Ian Johnston, Dominic Kwiatkowski, Cordelia Langford, John Sillitoe on behalf of the Wellcome Sanger Institute COVID-19 Surveillance Team |
| EPI_ISL_957238 | Lighthouse Lab in Milton Keynes | Wellcome Sanger Institute for the COVID-19 Genomics UK (COG-UK) Consortium | The Lighthouse Lab in Milton Keynes and Alex Alderton, Roberto Amato, Sonia Goncalves, Ewan Harrison, David K. Jackson, Ian Johnston, Dominic Kwiatkowski, Cordelia Langford, John Sillitoe on behalf of the Wellcome Sanger Institute COVID-19 Surveillance Team |
| EPI_ISL_957239 | Lighthouse Lab in Glasgow | Wellcome Sanger Institute for the COVID-19 Genomics UK (COG-UK) Consortium | Harper VanSteenhouse, Yumi Kasai, David Gray, Carol Clugston, Anna Dominiczak and Alex Alderton, Roberto Amato, Sonia Goncalves, Ewan Harrison, David K. Jackson, Ian Johnston, Dominic Kwiatkowski, Cordelia Langford, John Sillitoe on behalf of the Wellcome Sanger Institute COVID-19 Surveillance Team |
| EPI_ISL_957240, EPI_ISL_957241 | Lighthouse Lab in Milton Keynes | Wellcome Sanger Institute for the COVID-19 Genomics UK (COG-UK) Consortium | The Lighthouse Lab in Milton Keynes and Alex Alderton, Roberto Amato, Sonia Goncalves, Ewan Harrison, David K. Jackson, Ian Johnston, Dominic Kwiatkowski, Cordelia Langford, John Sillitoe on behalf of the Wellcome Sanger Institute COVID-19 Surveillance Team |
| EPI_ISL_957242, EPI_ISL_957243 | Lighthouse Lab in Glasgow | Wellcome Sanger Institute for the COVID-19 Genomics UK (COG-UK) Consortium | Harper VanSteenhouse, Yumi Kasai, David Gray, Carol Clugston, Anna Dominiczak and Alex Alderton, Roberto Amato, Sonia Goncalves, Ewan Harrison, David K. Jackson, Ian Johnston, Dominic Kwiatkowski, Cordelia Langford, John Sillitoe on behalf of the Wellcome Sanger Institute COVID-19 Surveillance Team |
| EPI_ISL_957244 | Lighthouse Lab in Milton Keynes | Wellcome Sanger Institute for the COVID-19 Genomics UK (COG-UK) Consortium | The Lighthouse Lab in Milton Keynes and Alex Alderton, Roberto Amato, Sonia Goncalves, Ewan Harrison, David K. Jackson, Ian Johnston, Dominic Kwiatkowski, Cordelia Langford, John Sillitoe on behalf of the Wellcome Sanger Institute COVID-19 Surveillance Team |
| EPI_ISL_957245, EPI_ISL_957246 | Lighthouse Lab in Glasgow | Wellcome Sanger Institute for the COVID-19 Genomics UK (COG-UK) Consortium | Harper VanSteenhouse, Yumi Kasai, David Gray, Carol Clugston, Anna Dominiczak and Alex Alderton, Roberto Amato, Sonia Goncalves, Ewan Harrison, David K. Jackson, Ian Johnston, Dominic Kwiatkowski, Cordelia Langford, John Sillitoe on behalf of the Wellcome Sanger Institute COVID-19 Surveillance Team |
| EPI_ISL_957247 | Lighthouse Lab in Milton Keynes | Wellcome Sanger Institute for the COVID-19 Genomics UK | The Lighthouse Lab in Milton Keynes and Alex Alderton, Roberto Amato, Sonia Goncalves, Ewan Harrison, David K. Jackson, Ian Johnston, Dominic |

| | | | |
|---|---|---|---|
| | | (COG-UK) Consortium | Kwiatkowski, Cordelia Langford, John Sillitoe on behalf of the Wellcome Sanger Institute COVID-19 Surveillance Team |
| EPI_ISL_957248 | Lighthouse Lab in Glasgow | Wellcome Sanger Institute for the COVID-19 Genomics UK (COG-UK) Consortium | Harper VanSteenhouse, Yumi Kasai, David Gray, Carol Clugston, Anna Dominiczak and Alex Alderton, Roberto Amato, Sonia Goncalves, Ewan Harrison, David K. Jackson, Ian Johnston, Dominic Kwiatkowski, Cordelia Langford, John Sillitoe on behalf of the Wellcome Sanger Institute COVID-19 Surveillance Team |
| EPI_ISL_957249 | Lighthouse Lab in Milton Keynes | Wellcome Sanger Institute for the COVID-19 Genomics UK (COG-UK) Consortium | The Lighthouse Lab in Milton Keynes and Alex Alderton, Roberto Amato, Sonia Goncalves, Ewan Harrison, David K. Jackson, Ian Johnston, Dominic Kwiatkowski, Cordelia Langford, John Sillitoe on behalf of the Wellcome Sanger Institute COVID-19 Surveillance Team |
| EPI_ISL_957250 | Lighthouse Lab in Glasgow | Wellcome Sanger Institute for the COVID-19 Genomics UK (COG-UK) Consortium | Harper VanSteenhouse, Yumi Kasai, David Gray, Carol Clugston, Anna Dominiczak and Alex Alderton, Roberto Amato, Sonia Goncalves, Ewan Harrison, David K. Jackson, Ian Johnston, Dominic Kwiatkowski, Cordelia Langford, John Sillitoe on behalf of the Wellcome Sanger Institute COVID-19 Surveillance Team |
| EPI_ISL_957251, EPI_ISL_957252 | Lighthouse Lab in Milton Keynes | Wellcome Sanger Institute for the COVID-19 Genomics UK (COG-UK) Consortium | The Lighthouse Lab in Milton Keynes and Alex Alderton, Roberto Amato, Sonia Goncalves, Ewan Harrison, David K. Jackson, Ian Johnston, Dominic Kwiatkowski, Cordelia Langford, John Sillitoe on behalf of the Wellcome Sanger Institute COVID-19 Surveillance Team |
| EPI_ISL_957253, EPI_ISL_957254, EPI_ISL_957255, EPI_ISL_957256, EPI_ISL_957257 | Lighthouse Lab in Glasgow | Wellcome Sanger Institute for the COVID-19 Genomics UK (COG-UK) Consortium | Harper VanSteenhouse, Yumi Kasai, David Gray, Carol Clugston, Anna Dominiczak and Alex Alderton, Roberto Amato, Sonia Goncalves, Ewan Harrison, David K. Jackson, Ian Johnston, Dominic Kwiatkowski, Cordelia Langford, John Sillitoe on behalf of the Wellcome Sanger Institute COVID-19 Surveillance Team |
| EPI_ISL_957258 | Lighthouse Lab in Milton Keynes | Wellcome Sanger Institute for the COVID-19 Genomics UK (COG-UK) Consortium | The Lighthouse Lab in Milton Keynes and Alex Alderton, Roberto Amato, Sonia Goncalves, Ewan Harrison, David K. Jackson, Ian Johnston, Dominic Kwiatkowski, Cordelia Langford, John Sillitoe on behalf of the Wellcome Sanger Institute COVID-19 Surveillance Team |
| EPI_ISL_957259, EPI_ISL_957260 | Lighthouse Lab in Glasgow | Wellcome Sanger Institute for the COVID-19 Genomics UK (COG-UK) Consortium | Harper VanSteenhouse, Yumi Kasai, David Gray, Carol Clugston, Anna Dominiczak and Alex Alderton, Roberto Amato, Sonia Goncalves, Ewan Harrison, David K. Jackson, Ian Johnston, Dominic Kwiatkowski, Cordelia Langford, John Sillitoe on behalf of the Wellcome Sanger Institute COVID-19 Surveillance Team |
| EPI_ISL_957261 | Lighthouse Lab in Milton Keynes | Wellcome Sanger Institute for the COVID-19 Genomics UK (COG-UK) Consortium | The Lighthouse Lab in Milton Keynes and Alex Alderton, Roberto Amato, Sonia Goncalves, Ewan Harrison, David K. Jackson, Ian Johnston, Dominic Kwiatkowski, Cordelia Langford, John Sillitoe on behalf of the Wellcome Sanger Institute COVID-19 Surveillance Team |
| EPI_ISL_957262, EPI_ISL_957263 | Lighthouse Lab in Glasgow | Wellcome Sanger Institute for the COVID-19 Genomics UK (COG-UK) Consortium | Harper VanSteenhouse, Yumi Kasai, David Gray, Carol Clugston, Anna Dominiczak and Alex Alderton, Roberto Amato, Sonia Goncalves, Ewan Harrison, David K. Jackson, Ian Johnston, Dominic Kwiatkowski, Cordelia Langford, John Sillitoe on behalf of the Wellcome Sanger Institute COVID-19 Surveillance Team |
| EPI_ISL_957264, EPI_ISL_957265 | Lighthouse Lab in Milton Keynes | Wellcome Sanger Institute for the COVID-19 Genomics UK (COG-UK) Consortium | The Lighthouse Lab in Milton Keynes and Alex Alderton, Roberto Amato, Sonia Goncalves, Ewan Harrison, David K. Jackson, Ian Johnston, Dominic Kwiatkowski, Cordelia Langford, John Sillitoe on behalf of the Wellcome Sanger Institute COVID-19 Surveillance Team |
| EPI_ISL_957266 | Lighthouse Lab in Glasgow | Wellcome Sanger Institute for the COVID-19 Genomics UK (COG-UK) Consortium | Harper VanSteenhouse, Yumi Kasai, David Gray, Carol Clugston, Anna Dominiczak and Alex Alderton, Roberto Amato, Sonia Goncalves, Ewan Harrison, David K. Jackson, Ian Johnston, Dominic Kwiatkowski, Cordelia Langford, John Sillitoe on behalf of the Wellcome Sanger Institute COVID-19 Surveillance Team |
| EPI_ISL_957267 | Lighthouse Lab in Milton Keynes | Wellcome Sanger Institute for the COVID-19 Genomics UK (COG-UK) Consortium | The Lighthouse Lab in Milton Keynes and Alex Alderton, Roberto Amato, Sonia Goncalves, Ewan Harrison, David K. Jackson, Ian Johnston, Dominic Kwiatkowski, Cordelia Langford, John Sillitoe on behalf of the Wellcome Sanger Institute COVID-19 Surveillance Team |
| EPI_ISL_957268 | Lighthouse Lab in Glasgow | Wellcome Sanger Institute for the COVID-19 Genomics UK (COG-UK) Consortium | Harper VanSteenhouse, Yumi Kasai, David Gray, Carol Clugston, Anna Dominiczak and Alex Alderton, Roberto Amato, Sonia Goncalves, Ewan Harrison, David K. Jackson, Ian Johnston, Dominic Kwiatkowski, Cordelia Langford, John Sillitoe on behalf of the Wellcome Sanger Institute COVID-19 Surveillance Team |
| EPI_ISL_957269 | Lighthouse Lab in Milton Keynes | Wellcome Sanger Institute for the COVID-19 Genomics UK (COG-UK) Consortium | The Lighthouse Lab in Milton Keynes and Alex Alderton, Roberto Amato, Sonia Goncalves, Ewan Harrison, David K. Jackson, Ian Johnston, Dominic Kwiatkowski, Cordelia Langford, John Sillitoe on behalf of the Wellcome Sanger Institute COVID-19 Surveillance Team |
| EPI_ISL_957270, EPI_ISL_957271, EPI_ISL_957272 | Lighthouse Lab in Glasgow | Wellcome Sanger Institute for the COVID-19 Genomics UK (COG-UK) Consortium | Harper VanSteenhouse, Yumi Kasai, David Gray, Carol Clugston, Anna Dominiczak and Alex Alderton, Roberto Amato, Sonia Goncalves, Ewan Harrison, David K. Jackson, Ian Johnston, Dominic Kwiatkowski, Cordelia Langford, John Sillitoe on behalf of the Wellcome Sanger Institute COVID-19 Surveillance Team |
| EPI_ISL_957273 | Lighthouse Lab in Milton Keynes | Wellcome Sanger Institute for the COVID-19 Genomics UK (COG-UK) Consortium | The Lighthouse Lab in Milton Keynes and Alex Alderton, Roberto Amato, Sonia Goncalves, Ewan Harrison, David K. Jackson, Ian Johnston, Dominic Kwiatkowski, Cordelia Langford, John Sillitoe on behalf of the Wellcome Sanger Institute COVID-19 Surveillance Team |
| EPI_ISL_957274, EPI_ISL_957275, EPI_ISL_957276 | Lighthouse Lab in Glasgow | Wellcome Sanger Institute for the COVID-19 Genomics UK (COG-UK) Consortium | Harper VanSteenhouse, Yumi Kasai, David Gray, Carol Clugston, Anna Dominiczak and Alex Alderton, Roberto Amato, Sonia Goncalves, Ewan Harrison, David K. Jackson, Ian Johnston, Dominic Kwiatkowski, Cordelia Langford, John Sillitoe on behalf of the Wellcome Sanger Institute COVID-19 Surveillance Team |
| EPI_ISL_957277, EPI_ISL_957278 | Lighthouse Lab in Milton Keynes | Wellcome Sanger Institute for the COVID-19 Genomics UK (COG-UK) Consortium | The Lighthouse Lab in Milton Keynes and Alex Alderton, Roberto Amato, Sonia Goncalves, Ewan Harrison, David K. Jackson, Ian Johnston, Dominic Kwiatkowski, Cordelia Langford, John Sillitoe on behalf of the Wellcome Sanger Institute COVID-19 Surveillance Team |
| EPI_ISL_957279 | Lighthouse Lab in Glasgow | Wellcome Sanger Institute for the COVID-19 Genomics UK (COG-UK) Consortium | Harper VanSteenhouse, Yumi Kasai, David Gray, Carol Clugston, Anna Dominiczak and Alex Alderton, Roberto Amato, Sonia Goncalves, Ewan Harrison, David K. Jackson, Ian Johnston, Dominic Kwiatkowski, Cordelia Langford, John Sillitoe on behalf of the Wellcome Sanger Institute COVID-19 Surveillance Team |
| EPI_ISL_957280, EPI_ISL_957281, EPI_ISL_957282 | Lighthouse Lab in Milton Keynes | Wellcome Sanger Institute for the COVID-19 Genomics UK (COG-UK) Consortium | The Lighthouse Lab in Milton Keynes and Alex Alderton, Roberto Amato, Sonia Goncalves, Ewan Harrison, David K. Jackson, Ian Johnston, Dominic Kwiatkowski, Cordelia Langford, John Sillitoe on behalf of the Wellcome Sanger Institute COVID-19 Surveillance Team |
| EPI_ISL_957283 | Lighthouse Lab in Glasgow | Wellcome Sanger Institute for the COVID-19 Genomics UK (COG-UK) Consortium | Harper VanSteenhouse, Yumi Kasai, David Gray, Carol Clugston, Anna Dominiczak and Alex Alderton, Roberto Amato, Sonia Goncalves, Ewan Harrison, David K. Jackson, Ian Johnston, Dominic Kwiatkowski, Cordelia Langford, John Sillitoe on behalf of the Wellcome Sanger Institute COVID-19 Surveillance Team |
| EPI_ISL_957284 | Lighthouse Lab in Milton Keynes | Wellcome Sanger Institute for the COVID-19 Genomics UK (COG-UK) Consortium | The Lighthouse Lab in Milton Keynes and Alex Alderton, Roberto Amato, Sonia Goncalves, Ewan Harrison, David K. Jackson, Ian Johnston, Dominic Kwiatkowski, Cordelia Langford, John Sillitoe on behalf of the Wellcome Sanger Institute COVID-19 Surveillance Team |
| EPI_ISL_957285, EPI_ISL_957286, EPI_ISL_957287 | Lighthouse Lab in Glasgow | Wellcome Sanger Institute for the COVID-19 Genomics UK (COG-UK) Consortium | Harper VanSteenhouse, Yumi Kasai, David Gray, Carol Clugston, Anna Dominiczak and Alex Alderton, Roberto Amato, Sonia Goncalves, Ewan Harrison, David K. Jackson, Ian Johnston, Dominic Kwiatkowski, Cordelia Langford, John Sillitoe on behalf of the Wellcome Sanger Institute COVID-19 Surveillance Team |
| EPI_ISL_957288 | Lighthouse Lab in Milton Keynes | Wellcome Sanger Institute for the COVID-19 Genomics UK (COG-UK) Consortium | The Lighthouse Lab in Milton Keynes and Alex Alderton, Roberto Amato, Sonia Goncalves, Ewan Harrison, David K. Jackson, Ian Johnston, Dominic Kwiatkowski, Cordelia Langford, John Sillitoe on behalf of the Wellcome Sanger Institute COVID-19 Surveillance Team |
| EPI_ISL_957289 | Lighthouse Lab in Glasgow | Wellcome Sanger Institute for the COVID-19 Genomics UK (COG-UK) Consortium | Harper VanSteenhouse, Yumi Kasai, David Gray, Carol Clugston, Anna Dominiczak and Alex Alderton, Roberto Amato, Sonia Goncalves, Ewan Harrison, David K. Jackson, Ian Johnston, Dominic Kwiatkowski, Cordelia Langford, John Sillitoe on behalf of the Wellcome Sanger Institute COVID-19 Surveillance Team |
| EPI_ISL_957290 | Lighthouse Lab in Milton Keynes | Wellcome Sanger Institute for the COVID-19 Genomics UK (COG-UK) Consortium | The Lighthouse Lab in Milton Keynes and Alex Alderton, Roberto Amato, Sonia Goncalves, Ewan Harrison, David K. Jackson, Ian Johnston, Dominic Kwiatkowski, Cordelia Langford, John Sillitoe on behalf of the Wellcome Sanger Institute COVID-19 Surveillance Team |
| EPI_ISL_957291, EPI_ISL_957292 | Lighthouse Lab in Glasgow | Wellcome Sanger Institute for the COVID-19 Genomics UK (COG-UK) Consortium | Harper VanSteenhouse, Yumi Kasai, David Gray, Carol Clugston, Anna Dominiczak and Alex Alderton, Roberto Amato, Sonia Goncalves, Ewan Harrison, David K. Jackson, Ian Johnston, Dominic Kwiatkowski, Cordelia Langford, John Sillitoe on behalf of the Wellcome Sanger Institute COVID-19 Surveillance Team |
| EPI_ISL_957293 | Lighthouse Lab in Milton Keynes | Wellcome Sanger Institute for the COVID-19 Genomics UK (COG-UK) Consortium | The Lighthouse Lab in Milton Keynes and Alex Alderton, Roberto Amato, Sonia Goncalves, Ewan Harrison, David K. Jackson, Ian Johnston, Dominic Kwiatkowski, Cordelia Langford, John Sillitoe on behalf of the Wellcome Sanger Institute COVID-19 Surveillance Team |
| EPI_ISL_957294 | Lighthouse Lab in Glasgow | Wellcome Sanger Institute for the COVID-19 Genomics UK (COG-UK) Consortium | Harper VanSteenhouse, Yumi Kasai, David Gray, Carol Clugston, Anna Dominiczak and Alex Alderton, Roberto Amato, Sonia Goncalves, Ewan Harrison, David K. Jackson, Ian Johnston, Dominic Kwiatkowski, Cordelia Langford, John Sillitoe on behalf of the Wellcome Sanger Institute COVID-19 Surveillance |

| | | | |
|---|---|---|---|
| | | | Team |
| EPI_ISL_957295, EPI_ISL_957296 | Lighthouse Lab in Milton Keynes | Wellcome Sanger Institute for the COVID-19 Genomics UK (COG-UK) Consortium | The Lighthouse Lab in Milton Keynes and Alex Alderton, Roberto Amato, Sonia Goncalves, Ewan Harrison, David K. Jackson, Ian Johnston, Dominic Kwiatkowski, Cordelia Langford, John Sillitoe on behalf of the Wellcome Sanger Institute COVID-19 Surveillance Team |
| EPI_ISL_957297 | Lighthouse Lab in Glasgow | Wellcome Sanger Institute for the COVID-19 Genomics UK (COG-UK) Consortium | Harper VanSteenhouse, Yumi Kasai, David Gray, Carol Clugston, Anna Dominiczak and Alex Alderton, Roberto Amato, Sonia Goncalves, Ewan Harrison, David K. Jackson, Ian Johnston, Dominic Kwiatkowski, Cordelia Langford, John Sillitoe on behalf of the Wellcome Sanger Institute COVID-19 Surveillance Team |
| EPI_ISL_957298 | Lighthouse Lab in Milton Keynes | Wellcome Sanger Institute for the COVID-19 Genomics UK (COG-UK) Consortium | The Lighthouse Lab in Milton Keynes and Alex Alderton, Roberto Amato, Sonia Goncalves, Ewan Harrison, David K. Jackson, Ian Johnston, Dominic Kwiatkowski, Cordelia Langford, John Sillitoe on behalf of the Wellcome Sanger Institute COVID-19 Surveillance Team |
| EPI_ISL_957299 | Lighthouse Lab in Glasgow | Wellcome Sanger Institute for the COVID-19 Genomics UK (COG-UK) Consortium | Harper VanSteenhouse, Yumi Kasai, David Gray, Carol Clugston, Anna Dominiczak and Alex Alderton, Roberto Amato, Sonia Goncalves, Ewan Harrison, David K. Jackson, Ian Johnston, Dominic Kwiatkowski, Cordelia Langford, John Sillitoe on behalf of the Wellcome Sanger Institute COVID-19 Surveillance Team |
| EPI_ISL_957300, EPI_ISL_957301 | Lighthouse Lab in Milton Keynes | Wellcome Sanger Institute for the COVID-19 Genomics UK (COG-UK) Consortium | The Lighthouse Lab in Milton Keynes and Alex Alderton, Roberto Amato, Sonia Goncalves, Ewan Harrison, David K. Jackson, Ian Johnston, Dominic Kwiatkowski, Cordelia Langford, John Sillitoe on behalf of the Wellcome Sanger Institute COVID-19 Surveillance Team |
| EPI_ISL_957302 | Lighthouse Lab in Glasgow | Wellcome Sanger Institute for the COVID-19 Genomics UK (COG-UK) Consortium | Harper VanSteenhouse, Yumi Kasai, David Gray, Carol Clugston, Anna Dominiczak and Alex Alderton, Roberto Amato, Sonia Goncalves, Ewan Harrison, David K. Jackson, Ian Johnston, Dominic Kwiatkowski, Cordelia Langford, John Sillitoe on behalf of the Wellcome Sanger Institute COVID-19 Surveillance Team |
| EPI_ISL_957303 | Lighthouse Lab in Milton Keynes | Wellcome Sanger Institute for the COVID-19 Genomics UK (COG-UK) Consortium | The Lighthouse Lab in Milton Keynes and Alex Alderton, Roberto Amato, Sonia Goncalves, Ewan Harrison, David K. Jackson, Ian Johnston, Dominic Kwiatkowski, Cordelia Langford, John Sillitoe on behalf of the Wellcome Sanger Institute COVID-19 Surveillance Team |
| EPI_ISL_957304 | Lighthouse Lab in Glasgow | Wellcome Sanger Institute for the COVID-19 Genomics UK (COG-UK) Consortium | Harper VanSteenhouse, Yumi Kasai, David Gray, Carol Clugston, Anna Dominiczak and Alex Alderton, Roberto Amato, Sonia Goncalves, Ewan Harrison, David K. Jackson, Ian Johnston, Dominic Kwiatkowski, Cordelia Langford, John Sillitoe on behalf of the Wellcome Sanger Institute COVID-19 Surveillance Team |
| EPI_ISL_957305 | Lighthouse Lab in Milton Keynes | Wellcome Sanger Institute for the COVID-19 Genomics UK (COG-UK) Consortium | The Lighthouse Lab in Milton Keynes and Alex Alderton, Roberto Amato, Sonia Goncalves, Ewan Harrison, David K. Jackson, Ian Johnston, Dominic Kwiatkowski, Cordelia Langford, John Sillitoe on behalf of the Wellcome Sanger Institute COVID-19 Surveillance Team |
| EPI_ISL_957306 | Lighthouse Lab in Glasgow | Wellcome Sanger Institute for the COVID-19 Genomics UK (COG-UK) Consortium | Harper VanSteenhouse, Yumi Kasai, David Gray, Carol Clugston, Anna Dominiczak and Alex Alderton, Roberto Amato, Sonia Goncalves, Ewan Harrison, David K. Jackson, Ian Johnston, Dominic Kwiatkowski, Cordelia Langford, John Sillitoe on behalf of the Wellcome Sanger Institute COVID-19 Surveillance Team |
| EPI_ISL_957307 | Lighthouse Lab in Milton Keynes | Wellcome Sanger Institute for the COVID-19 Genomics UK (COG-UK) Consortium | The Lighthouse Lab in Milton Keynes and Alex Alderton, Roberto Amato, Sonia Goncalves, Ewan Harrison, David K. Jackson, Ian Johnston, Dominic Kwiatkowski, Cordelia Langford, John Sillitoe on behalf of the Wellcome Sanger Institute COVID-19 Surveillance Team |
| EPI_ISL_957308 | Lighthouse Lab in Glasgow | Wellcome Sanger Institute for the COVID-19 Genomics UK (COG-UK) Consortium | Harper VanSteenhouse, Yumi Kasai, David Gray, Carol Clugston, Anna Dominiczak and Alex Alderton, Roberto Amato, Sonia Goncalves, Ewan Harrison, David K. Jackson, Ian Johnston, Dominic Kwiatkowski, Cordelia Langford, John Sillitoe on behalf of the Wellcome Sanger Institute COVID-19 Surveillance Team |
| EPI_ISL_957309 | Lighthouse Lab in Milton Keynes | Wellcome Sanger Institute for the COVID-19 Genomics UK (COG-UK) Consortium | The Lighthouse Lab in Milton Keynes and Alex Alderton, Roberto Amato, Sonia Goncalves, Ewan Harrison, David K. Jackson, Ian Johnston, Dominic Kwiatkowski, Cordelia Langford, John Sillitoe on behalf of the Wellcome Sanger Institute COVID-19 Surveillance Team |
| EPI_ISL_957310, EPI_ISL_957311, EPI_ISL_957312 | Lighthouse Lab in Glasgow | Wellcome Sanger Institute for the COVID-19 Genomics UK (COG-UK) Consortium | Harper VanSteenhouse, Yumi Kasai, David Gray, Carol Clugston, Anna Dominiczak and Alex Alderton, Roberto Amato, Sonia Goncalves, Ewan Harrison, David K. Jackson, Ian Johnston, Dominic Kwiatkowski, Cordelia Langford, John Sillitoe on behalf of the Wellcome Sanger Institute COVID-19 Surveillance Team |
| EPI_ISL_957313 | Lighthouse Lab in Milton Keynes | Wellcome Sanger Institute for the COVID-19 Genomics UK (COG-UK) Consortium | The Lighthouse Lab in Milton Keynes and Alex Alderton, Roberto Amato, Sonia Goncalves, Ewan Harrison, David K. Jackson, Ian Johnston, Dominic Kwiatkowski, Cordelia Langford, John Sillitoe on behalf of the Wellcome Sanger Institute COVID-19 Surveillance Team |
| EPI_ISL_957314, EPI_ISL_957315 | Lighthouse Lab in Glasgow | Wellcome Sanger Institute for the COVID-19 Genomics UK (COG-UK) Consortium | Harper VanSteenhouse, Yumi Kasai, David Gray, Carol Clugston, Anna Dominiczak and Alex Alderton, Roberto Amato, Sonia Goncalves, Ewan Harrison, David K. Jackson, Ian Johnston, Dominic Kwiatkowski, Cordelia Langford, John Sillitoe on behalf of the Wellcome Sanger Institute COVID-19 Surveillance Team |
| EPI_ISL_957316, EPI_ISL_957317, EPI_ISL_957318 | Lighthouse Lab in Milton Keynes | Wellcome Sanger Institute for the COVID-19 Genomics UK (COG-UK) Consortium | The Lighthouse Lab in Milton Keynes and Alex Alderton, Roberto Amato, Sonia Goncalves, Ewan Harrison, David K. Jackson, Ian Johnston, Dominic Kwiatkowski, Cordelia Langford, John Sillitoe on behalf of the Wellcome Sanger Institute COVID-19 Surveillance Team |
| EPI_ISL_957319, EPI_ISL_957320 | Lighthouse Lab in Glasgow | Wellcome Sanger Institute for the COVID-19 Genomics UK (COG-UK) Consortium | Harper VanSteenhouse, Yumi Kasai, David Gray, Carol Clugston, Anna Dominiczak and Alex Alderton, Roberto Amato, Sonia Goncalves, Ewan Harrison, David K. Jackson, Ian Johnston, Dominic Kwiatkowski, Cordelia Langford, John Sillitoe on behalf of the Wellcome Sanger Institute COVID-19 Surveillance Team |
| EPI_ISL_957321 | Lighthouse Lab in Milton Keynes | Wellcome Sanger Institute for the COVID-19 Genomics UK (COG-UK) Consortium | The Lighthouse Lab in Milton Keynes and Alex Alderton, Roberto Amato, Sonia Goncalves, Ewan Harrison, David K. Jackson, Ian Johnston, Dominic Kwiatkowski, Cordelia Langford, John Sillitoe on behalf of the Wellcome Sanger Institute COVID-19 Surveillance Team |
| EPI_ISL_957322, EPI_ISL_957323 | Lighthouse Lab in Glasgow | Wellcome Sanger Institute for the COVID-19 Genomics UK (COG-UK) Consortium | Harper VanSteenhouse, Yumi Kasai, David Gray, Carol Clugston, Anna Dominiczak and Alex Alderton, Roberto Amato, Sonia Goncalves, Ewan Harrison, David K. Jackson, Ian Johnston, Dominic Kwiatkowski, Cordelia Langford, John Sillitoe on behalf of the Wellcome Sanger Institute COVID-19 Surveillance Team |
| EPI_ISL_957324, EPI_ISL_957325, EPI_ISL_957326 | Lighthouse Lab in Milton Keynes | Wellcome Sanger Institute for the COVID-19 Genomics UK (COG-UK) Consortium | The Lighthouse Lab in Milton Keynes and Alex Alderton, Roberto Amato, Sonia Goncalves, Ewan Harrison, David K. Jackson, Ian Johnston, Dominic Kwiatkowski, Cordelia Langford, John Sillitoe on behalf of the Wellcome Sanger Institute COVID-19 Surveillance Team |
| EPI_ISL_957327, EPI_ISL_957328 | Lighthouse Lab in Glasgow | Wellcome Sanger Institute for the COVID-19 Genomics UK (COG-UK) Consortium | Harper VanSteenhouse, Yumi Kasai, David Gray, Carol Clugston, Anna Dominiczak and Alex Alderton, Roberto Amato, Sonia Goncalves, Ewan Harrison, David K. Jackson, Ian Johnston, Dominic Kwiatkowski, Cordelia Langford, John Sillitoe on behalf of the Wellcome Sanger Institute COVID-19 Surveillance Team |
| EPI_ISL_957329 | Lighthouse Lab in Milton Keynes | Wellcome Sanger Institute for the COVID-19 Genomics UK (COG-UK) Consortium | The Lighthouse Lab in Milton Keynes and Alex Alderton, Roberto Amato, Sonia Goncalves, Ewan Harrison, David K. Jackson, Ian Johnston, Dominic Kwiatkowski, Cordelia Langford, John Sillitoe on behalf of the Wellcome Sanger Institute COVID-19 Surveillance Team |
| EPI_ISL_957330, EPI_ISL_957331 | Lighthouse Lab in Glasgow | Wellcome Sanger Institute for the COVID-19 Genomics UK (COG-UK) Consortium | Harper VanSteenhouse, Yumi Kasai, David Gray, Carol Clugston, Anna Dominiczak and Alex Alderton, Roberto Amato, Sonia Goncalves, Ewan Harrison, David K. Jackson, Ian Johnston, Dominic Kwiatkowski, Cordelia Langford, John Sillitoe on behalf of the Wellcome Sanger Institute COVID-19 Surveillance Team |
| EPI_ISL_957332, EPI_ISL_957333 | Lighthouse Lab in Milton Keynes | Wellcome Sanger Institute for the COVID-19 Genomics UK (COG-UK) Consortium | The Lighthouse Lab in Milton Keynes and Alex Alderton, Roberto Amato, Sonia Goncalves, Ewan Harrison, David K. Jackson, Ian Johnston, Dominic Kwiatkowski, Cordelia Langford, John Sillitoe on behalf of the Wellcome Sanger Institute COVID-19 Surveillance Team |
| EPI_ISL_957334 | Lighthouse Lab in Glasgow | Wellcome Sanger Institute for the COVID-19 Genomics UK (COG-UK) Consortium | Harper VanSteenhouse, Yumi Kasai, David Gray, Carol Clugston, Anna Dominiczak and Alex Alderton, Roberto Amato, Sonia Goncalves, Ewan Harrison, David K. Jackson, Ian Johnston, Dominic Kwiatkowski, Cordelia Langford, John Sillitoe on behalf of the Wellcome Sanger Institute COVID-19 Surveillance Team |
| EPI_ISL_957335 | Lighthouse Lab in Milton Keynes | Wellcome Sanger Institute for the COVID-19 Genomics UK (COG-UK) Consortium | The Lighthouse Lab in Milton Keynes and Alex Alderton, Roberto Amato, Sonia Goncalves, Ewan Harrison, David K. Jackson, Ian Johnston, Dominic Kwiatkowski, Cordelia Langford, John Sillitoe on behalf of the Wellcome Sanger Institute COVID-19 Surveillance Team |
| EPI_ISL_957336, EPI_ISL_957337 | Lighthouse Lab in Glasgow | Wellcome Sanger Institute for the COVID-19 Genomics UK (COG-UK) Consortium | Harper VanSteenhouse, Yumi Kasai, David Gray, Carol Clugston, Anna Dominiczak and Alex Alderton, Roberto Amato, Sonia Goncalves, Ewan Harrison, David K. Jackson, Ian Johnston, Dominic Kwiatkowski, Cordelia Langford, John Sillitoe on behalf of the Wellcome Sanger Institute COVID-19 Surveillance Team |
| EPI_ISL_957338, EPI_ISL_957339, EPI_ISL_957340, EPI_ISL_957341, EPI_ISL_957342, EPI_ISL_957343, EPI_ISL_957344, EPI_ISL_957345, EPI_ISL_957346, EPI_ISL_957347, EPI_ISL_957348, EPI_ISL_957349, EPI_ISL_957350, EPI_ISL_957351, EPI_ISL_957352, EPI_ISL_957353, EPI_ISL_957354, EPI_ISL_957355, EPI_ISL_957356, EPI_ISL_957357, EPI_ISL_957358, EPI_ISL_957359, EPI_ISL_957360, EPI_ISL_957361, EPI_ISL_957362, EPI_ISL_957363, EPI_ISL_957364, EPI_ISL_957365, EPI_ISL_957366, EPI_ISL_957367, EPI_ISL_957368, EPI_ISL_957369, EPI_ISL_957370, EPI_ISL_957371, EPI_ISL_957372, EPI_ISL_957373, | | | |

EPI_ISL_957374, EPI_ISL_957375, EPI_ISL_957376, EPI_ISL_957377, EPI_ISL_957378, EPI_ISL_957379, EPI_ISL_957380, EPI_ISL_957381, EPI_ISL_957382, EPI_ISL_957383, EPI_ISL_957384, EPI_ISL_957385, EPI_ISL_957386, EPI_ISL_957387, EPI_ISL_957388, EPI_ISL_957389, EPI_ISL_957390, EPI_ISL_957391, EPI_ISL_957392, EPI_ISL_957393, EPI_ISL_957394, EPI_ISL_957395, EPI_ISL_957396, EPI_ISL_957397, EPI_ISL_957398, EPI_ISL_957399, EPI_ISL_957400, EPI_ISL_957401, EPI_ISL_957402, EPI_ISL_957403, EPI_ISL_957404, EPI_ISL_957405, EPI_ISL_957406, EPI_ISL_957407, EPI_ISL_957408, EPI_ISL_957409, EPI_ISL_957410, EPI_ISL_957411, EPI_ISL_957412, EPI_ISL_957413, EPI_ISL_957414, EPI_ISL_957415, EPI_ISL_957416, EPI_ISL_957417, EPI_ISL_957418, EPI_ISL_957419, EPI_ISL_957420, EPI_ISL_957421, EPI_ISL_957422, EPI_ISL_957424, EPI_ISL_957425, EPI_ISL_957426, EPI_ISL_957427, EPI_ISL_957428, EPI_ISL_957429, EPI_ISL_957430, EPI_ISL_957431, EPI_ISL_957432, EPI_ISL_957433, EPI_ISL_957434, EPI_ISL_957435, EPI_ISL_957436, EPI_ISL_957437, EPI_ISL_957438, EPI_ISL_957439, EPI_ISL_957440, EPI_ISL_957441, EPI_ISL_957442, EPI_ISL_957443, EPI_ISL_957444, EPI_ISL_957445, EPI_ISL_957446, EPI_ISL_957447, EPI_ISL_957448, EPI_ISL_957449, EPI_ISL_957450, EPI_ISL_957451, EPI_ISL_957452, EPI_ISL_957453, EPI_ISL_957454, EPI_ISL_957455, EPI_ISL_957456, EPI_ISL_957457, EPI_ISL_957458, EPI_ISL_957459, EPI_ISL_957460, EPI_ISL_957461, EPI_ISL_957462, EPI_ISL_957463, EPI_ISL_957464, EPI_ISL_957465, EPI_ISL_957466, EPI_ISL_957467, EPI_ISL_957468, EPI_ISL_957469, EPI_ISL_957470, EPI_ISL_957471, EPI_ISL_957472, EPI_ISL_957473, EPI_ISL_957474, EPI_ISL_957475, EPI_ISL_957476, EPI_ISL_957477, EPI_ISL_957478, EPI_ISL_957479, EPI_ISL_957480, EPI_ISL_957481, EPI_ISL_957482, EPI_ISL_957483, EPI_ISL_957484, EPI_ISL_957485, EPI_ISL_957486, EPI_ISL_957487, EPI_ISL_957488, EPI_ISL_957489, EPI_ISL_957490, EPI_ISL_957491, EPI_ISL_957492, EPI_ISL_957493, EPI_ISL_957494, EPI_ISL_957495, EPI_ISL_957496, EPI_ISL_957497, EPI_ISL_957498, EPI_ISL_957499, EPI_ISL_957500, EPI_ISL_957501, EPI_ISL_957502, EPI_ISL_957503, EPI_ISL_957504, EPI_ISL_957505, EPI_ISL_957506, EPI_ISL_957507, EPI_ISL_957508, EPI_ISL_957509, EPI_ISL_957510, EPI_ISL_957511, EPI_ISL_957512, EPI_ISL_957513, EPI_ISL_957514, EPI_ISL_957515, EPI_ISL_957516, EPI_ISL_957517, EPI_ISL_957518, EPI_ISL_957519, EPI_ISL_957520, EPI_ISL_957521, EPI_ISL_957522, EPI_ISL_957523, EPI_ISL_957524, EPI_ISL_957525, EPI_ISL_957526, EPI_ISL_957527, EPI_ISL_957528, EPI_ISL_957529, EPI_ISL_957530, EPI_ISL_957531, EPI_ISL_957532, EPI_ISL_957533, EPI_ISL_957534, EPI_ISL_957535, EPI_ISL_957536, EPI_ISL_957537, EPI_ISL_957538, EPI_ISL_957539, EPI_ISL_957540, EPI_ISL_957541, EPI_ISL_957542, EPI_ISL_957543, EPI_ISL_957544, EPI_ISL_957545, EPI_ISL_957546, EPI_ISL_957547, EPI_ISL_957548, EPI_ISL_957549, EPI_ISL_957550, EPI_ISL_957551, EPI_ISL_957552, EPI_ISL_957553, EPI_ISL_957554, EPI_ISL_957555, EPI_ISL_957556, EPI_ISL_957557, EPI_ISL_957558, EPI_ISL_957559, EPI_ISL_957560, EPI_ISL_957561, EPI_ISL_957562, EPI_ISL_957563, EPI_ISL_957564, EPI_ISL_957565, EPI_ISL_957566, EPI_ISL_957567, EPI_ISL_957568, EPI_ISL_957569, EPI_ISL_957570, EPI_ISL_957571, EPI_ISL_957572, EPI_ISL_957573, EPI_ISL_957574, EPI_ISL_957575, EPI_ISL_957576, EPI_ISL_957577, EPI_ISL_957578, EPI_ISL_957579, EPI_ISL_957580, EPI_ISL_957581, EPI_ISL_957582, EPI_ISL_957583, EPI_ISL_957584, EPI_ISL_957585, EPI_ISL_957586, EPI_ISL_957587, EPI_ISL_957588, EPI_ISL_957589, EPI_ISL_957590, EPI_ISL_957591, EPI_ISL_957592, EPI_ISL_957593, EPI_ISL_957594, EPI_ISL_957595, EPI_ISL_957596, EPI_ISL_957597, EPI_ISL_957598, EPI_ISL_957599, EPI_ISL_957600, EPI_ISL_957601, EPI_ISL_957602, EPI_ISL_957603, EPI_ISL_957604, EPI_ISL_957605, EPI_ISL_957606, EPI_ISL_957607, EPI_ISL_957608, EPI_ISL_957609, EPI_ISL_957610, EPI_ISL_957611

| | | | |
|---|---|---|---|
| see above | Lighthouse Lab in Milton Keynes | Wellcome Sanger Institute for the COVID-19 Genomics UK (COG-UK) Consortium | The Lighthouse Lab in Milton Keynes and Alex Alderton, Roberto Amato, Sonia Goncalves, Ewan Harrison, David K. Jackson, Ian Johnston, Dominic Kwiatkowski, Cordelia Langford, John Sillitoe on behalf of the Wellcome Sanger Institute COVID-19 Surveillance Team |
| EPI_ISL_957612 | Lighthouse Lab in Cambridge | Wellcome Sanger Institute for the COVID-19 Genomics UK (COG-UK) Consortium | Rob Howes, The Lighthouse Lab in Cambridge and Alex Alderton, Roberto Amato, Sonia Goncalves, Ewan Harrison, David K. Jackson, Ian Johnston, Dominic Kwiatkowski, Cordelia Langford, John Sillitoe on behalf of the Wellcome Sanger Institute COVID-19 Surveillance Team |
| EPI_ISL_957614 | Lighthouse Lab in Milton Keynes | Wellcome Sanger Institute for the COVID-19 Genomics UK (COG-UK) Consortium | The Lighthouse Lab in Milton Keynes and Alex Alderton, Roberto Amato, Sonia Goncalves, Ewan Harrison, David K. Jackson, Ian Johnston, Dominic Kwiatkowski, Cordelia Langford, John Sillitoe on behalf of the Wellcome Sanger Institute COVID-19 Surveillance Team |
| EPI_ISL_957615, EPI_ISL_957617, EPI_ISL_957618 | Lighthouse Lab in Cambridge | Wellcome Sanger Institute for the COVID-19 Genomics UK (COG-UK) Consortium | Rob Howes, The Lighthouse Lab in Cambridge and Alex Alderton, Roberto Amato, Sonia Goncalves, Ewan Harrison, David K. Jackson, Ian Johnston, Dominic Kwiatkowski, Cordelia Langford, John Sillitoe on behalf of the Wellcome Sanger Institute COVID-19 Surveillance Team |
| EPI_ISL_957619 | Lighthouse Lab in Milton Keynes | Wellcome Sanger Institute for the COVID-19 Genomics UK (COG-UK) Consortium | The Lighthouse Lab in Milton Keynes and Alex Alderton, Roberto Amato, Sonia Goncalves, Ewan Harrison, David K. Jackson, Ian Johnston, Dominic Kwiatkowski, Cordelia Langford, John Sillitoe on behalf of the Wellcome Sanger Institute COVID-19 Surveillance Team |
| EPI_ISL_957622 | Lighthouse Lab in Cambridge | Wellcome Sanger Institute for the COVID-19 Genomics UK (COG-UK) Consortium | Rob Howes, The Lighthouse Lab in Cambridge and Alex Alderton, Roberto Amato, Sonia Goncalves, Ewan Harrison, David K. Jackson, Ian Johnston, Dominic Kwiatkowski, Cordelia Langford, John Sillitoe on behalf of the Wellcome Sanger Institute COVID-19 Surveillance Team |
| EPI_ISL_957623 | Lighthouse Lab in Milton Keynes | Wellcome Sanger Institute for the COVID-19 Genomics UK (COG-UK) Consortium | The Lighthouse Lab in Milton Keynes and Alex Alderton, Roberto Amato, Sonia Goncalves, Ewan Harrison, David K. Jackson, Ian Johnston, Dominic Kwiatkowski, Cordelia Langford, John Sillitoe on behalf of the Wellcome Sanger Institute COVID-19 Surveillance Team |
| EPI_ISL_957625, EPI_ISL_957626 | Lighthouse Lab in Cambridge | Wellcome Sanger Institute for the COVID-19 Genomics UK (COG-UK) Consortium | Rob Howes, The Lighthouse Lab in Cambridge and Alex Alderton, Roberto Amato, Sonia Goncalves, Ewan Harrison, David K. Jackson, Ian Johnston, Dominic Kwiatkowski, Cordelia Langford, John Sillitoe on behalf of the Wellcome Sanger Institute COVID-19 Surveillance Team |
| EPI_ISL_957628, EPI_ISL_957629, EPI_ISL_957631, EPI_ISL_957632 | Lighthouse Lab in Milton Keynes | Wellcome Sanger Institute for the COVID-19 Genomics UK (COG-UK) Consortium | The Lighthouse Lab in Milton Keynes and Alex Alderton, Roberto Amato, Sonia Goncalves, Ewan Harrison, David K. Jackson, Ian Johnston, Dominic Kwiatkowski, Cordelia Langford, John Sillitoe on behalf of the Wellcome Sanger Institute COVID-19 Surveillance Team |
| EPI_ISL_957633 | Lighthouse Lab in Glasgow | Wellcome Sanger Institute for the COVID-19 Genomics UK (COG-UK) Consortium | Harper VanSteenhouse, Yumi Kasai, David Gray, Carol Clugston, Anna Dominiczak and Alex Alderton, Roberto Amato, Sonia Goncalves, Ewan Harrison, David K. Jackson, Ian Johnston, Dominic Kwiatkowski, Cordelia Langford, John Sillitoe on behalf of the Wellcome Sanger Institute COVID-19 Surveillance Team |
| EPI_ISL_957634, EPI_ISL_957636 | Lighthouse Lab in Cambridge | Wellcome Sanger Institute for the COVID-19 Genomics UK (COG-UK) Consortium | Rob Howes, The Lighthouse Lab in Cambridge and Alex Alderton, Roberto Amato, Sonia Goncalves, Ewan Harrison, David K. Jackson, Ian Johnston, Dominic Kwiatkowski, Cordelia Langford, John Sillitoe on behalf of the Wellcome Sanger Institute COVID-19 Surveillance Team |
| EPI_ISL_957637 | Lighthouse Lab in Glasgow | Wellcome Sanger Institute for the COVID-19 Genomics UK (COG-UK) Consortium | Harper VanSteenhouse, Yumi Kasai, David Gray, Carol Clugston, Anna Dominiczak and Alex Alderton, Roberto Amato, Sonia Goncalves, Ewan Harrison, David K. Jackson, Ian Johnston, Dominic Kwiatkowski, Cordelia Langford, John Sillitoe on behalf of the Wellcome Sanger Institute COVID-19 Surveillance Team |
| EPI_ISL_957638, EPI_ISL_957639, EPI_ISL_957640, EPI_ISL_957642, EPI_ISL_957644, EPI_ISL_957646, EPI_ISL_957647, EPI_ISL_957648 | Lighthouse Lab in Cambridge | Wellcome Sanger Institute for the COVID-19 Genomics UK (COG-UK) Consortium | Rob Howes, The Lighthouse Lab in Cambridge and Alex Alderton, Roberto Amato, Sonia Goncalves, Ewan Harrison, David K. Jackson, Ian Johnston, Dominic Kwiatkowski, Cordelia Langford, John Sillitoe on behalf of the Wellcome Sanger Institute COVID-19 Surveillance Team |
| EPI_ISL_957649, EPI_ISL_957651, EPI_ISL_957653 | Lighthouse Lab in Glasgow | Wellcome Sanger Institute for the COVID-19 Genomics UK (COG-UK) Consortium | Harper VanSteenhouse, Yumi Kasai, David Gray, Carol Clugston, Anna Dominiczak and Alex Alderton, Roberto Amato, Sonia Goncalves, Ewan Harrison, David K. Jackson, Ian Johnston, Dominic Kwiatkowski, Cordelia Langford, John Sillitoe on behalf of the Wellcome Sanger Institute COVID-19 Surveillance Team |
| EPI_ISL_957657, EPI_ISL_957658 | Lighthouse Lab in Cambridge | Wellcome Sanger Institute for the COVID-19 Genomics UK (COG-UK) Consortium | Rob Howes, The Lighthouse Lab in Cambridge and Alex Alderton, Roberto Amato, Sonia Goncalves, Ewan Harrison, David K. Jackson, Ian Johnston, Dominic Kwiatkowski, Cordelia Langford, John Sillitoe on behalf of the Wellcome Sanger Institute COVID-19 Surveillance Team |
| EPI_ISL_957659 | Lighthouse Lab in Milton Keynes | Wellcome Sanger Institute for the COVID-19 Genomics UK (COG-UK) Consortium | The Lighthouse Lab in Milton Keynes and Alex Alderton, Roberto Amato, Sonia Goncalves, Ewan Harrison, David K. Jackson, Ian Johnston, Dominic Kwiatkowski, Cordelia Langford, John Sillitoe on behalf of the Wellcome Sanger Institute COVID-19 Surveillance Team |
| EPI_ISL_957661 | Lighthouse Lab in Cambridge | Wellcome Sanger Institute for the COVID-19 Genomics UK (COG-UK) Consortium | Rob Howes, The Lighthouse Lab in Cambridge and Alex Alderton, Roberto Amato, Sonia Goncalves, Ewan Harrison, David K. Jackson, Ian Johnston, Dominic Kwiatkowski, Cordelia Langford, John Sillitoe on behalf of the Wellcome Sanger Institute COVID-19 Surveillance Team |
| EPI_ISL_957662 | Lighthouse Lab in Milton Keynes | Wellcome Sanger Institute for the COVID-19 Genomics UK (COG-UK) Consortium | The Lighthouse Lab in Milton Keynes and Alex Alderton, Roberto Amato, Sonia Goncalves, Ewan Harrison, David K. Jackson, Ian Johnston, Dominic Kwiatkowski, Cordelia Langford, John Sillitoe on behalf of the Wellcome Sanger Institute COVID-19 Surveillance Team |
| EPI_ISL_957663, EPI_ISL_957664 | Lighthouse Lab in Glasgow | Wellcome Sanger Institute for the COVID-19 Genomics UK (COG-UK) Consortium | Harper VanSteenhouse, Yumi Kasai, David Gray, Carol Clugston, Anna Dominiczak and Alex Alderton, Roberto Amato, Sonia Goncalves, Ewan Harrison, David K. Jackson, Ian Johnston, Dominic Kwiatkowski, Cordelia Langford, John Sillitoe on behalf of the Wellcome Sanger Institute COVID-19 Surveillance Team |
| EPI_ISL_957665 | Lighthouse Lab in Cambridge | Wellcome Sanger Institute for the COVID-19 Genomics UK (COG-UK) Consortium | Rob Howes, The Lighthouse Lab in Cambridge and Alex Alderton, Roberto Amato, Sonia Goncalves, Ewan Harrison, David K. Jackson, Ian Johnston, Dominic Kwiatkowski, Cordelia Langford, John Sillitoe on behalf of the Wellcome Sanger Institute COVID-19 Surveillance Team |
| EPI_ISL_957666 | Lighthouse Lab in Glasgow | Wellcome Sanger Institute for the COVID-19 Genomics UK (COG-UK) Consortium | Harper VanSteenhouse, Yumi Kasai, David Gray, Carol Clugston, Anna Dominiczak and Alex Alderton, Roberto Amato, Sonia Goncalves, Ewan Harrison, David K. Jackson, Ian Johnston, Dominic Kwiatkowski, Cordelia Langford, John Sillitoe on behalf of the Wellcome Sanger Institute COVID-19 Surveillance Team |
| EPI_ISL_957667, EPI_ISL_957668 | Lighthouse Lab in Milton Keynes | Wellcome Sanger Institute for the COVID-19 Genomics UK (COG-UK) Consortium | The Lighthouse Lab in Milton Keynes and Alex Alderton, Roberto Amato, Sonia Goncalves, Ewan Harrison, David K. Jackson, Ian Johnston, Dominic Kwiatkowski, Cordelia Langford, John Sillitoe on behalf of the Wellcome Sanger Institute COVID-19 Surveillance Team |
| EPI_ISL_957669 | Lighthouse Lab in Glasgow | Wellcome Sanger Institute for the COVID-19 Genomics UK (COG-UK) Consortium | Harper VanSteenhouse, Yumi Kasai, David Gray, Carol Clugston, Anna Dominiczak and Alex Alderton, Roberto Amato, Sonia Goncalves, Ewan Harrison, David K. Jackson, Ian Johnston, Dominic Kwiatkowski, Cordelia Langford, John Sillitoe on behalf of the Wellcome Sanger Institute COVID-19 Surveillance Team |
| EPI_ISL_957671 | Lighthouse Lab in Milton Keynes | Wellcome Sanger Institute for the COVID-19 Genomics UK (COG-UK) Consortium | The Lighthouse Lab in Milton Keynes and Alex Alderton, Roberto Amato, Sonia Goncalves, Ewan Harrison, David K. Jackson, Ian Johnston, Dominic Kwiatkowski, Cordelia Langford, John Sillitoe on behalf of the Wellcome Sanger Institute COVID-19 Surveillance Team |
| EPI_ISL_957673, EPI_ISL_957674 | Lighthouse Lab in Glasgow | Wellcome Sanger Institute for the COVID-19 Genomics UK (COG-UK) Consortium | Harper VanSteenhouse, Yumi Kasai, David Gray, Carol Clugston, Anna Dominiczak and Alex Alderton, Roberto Amato, Sonia Goncalves, Ewan Harrison, David K. Jackson, Ian Johnston, Dominic Kwiatkowski, Cordelia Langford, John Sillitoe on behalf of the Wellcome Sanger Institute COVID-19 Surveillance Team |
| EPI_ISL_957676 | Lighthouse Lab in Cambridge | Wellcome Sanger Institute for the COVID-19 Genomics UK | Rob Howes, The Lighthouse Lab in Cambridge and Alex Alderton, Roberto Amato, Sonia Goncalves, Ewan Harrison, David K. Jackson, Ian Johnston, |

| | | | |
|---|---|---|---|
| | | (COG-UK) Consortium | Dominic Kwiatkowski, Cordelia Langford, John Sillitoe on behalf of the Wellcome Sanger Institute COVID-19 Surveillance Team |
| EPI_ISL_957678, EPI_ISL_957679, EPI_ISL_957681, EPI_ISL_957682 | Lighthouse Lab in Glasgow | Wellcome Sanger Institute for the COVID-19 Genomics UK (COG-UK) Consortium | Harper VanSteenhouse, Yumi Kasai, David Gray, Carol Clugston, Anna Dominiczak and Alex Alderton, Roberto Amato, Sonia Goncalves, Ewan Harrison, David K. Jackson, Ian Johnston, Dominic Kwiatkowski, Cordelia Langford, John Sillitoe on behalf of the Wellcome Sanger Institute COVID-19 Surveillance Team |
| EPI_ISL_957684, EPI_ISL_957685 | Lighthouse Lab in Milton Keynes | Wellcome Sanger Institute for the COVID-19 Genomics UK (COG-UK) Consortium | The Lighthouse Lab in Milton Keynes and Alex Alderton, Roberto Amato, Sonia Goncalves, Ewan Harrison, David K. Jackson, Ian Johnston, Dominic Kwiatkowski, Cordelia Langford, John Sillitoe on behalf of the Wellcome Sanger Institute COVID-19 Surveillance Team |
| EPI_ISL_957688, EPI_ISL_957689 | Lighthouse Lab in Cambridge | Wellcome Sanger Institute for the COVID-19 Genomics UK (COG-UK) Consortium | Rob Howes, The Lighthouse Lab in Cambridge and Alex Alderton, Roberto Amato, Sonia Goncalves, Ewan Harrison, David K. Jackson, Ian Johnston, Dominic Kwiatkowski, Cordelia Langford, John Sillitoe on behalf of the Wellcome Sanger Institute COVID-19 Surveillance Team |
| EPI_ISL_957690, EPI_ISL_957692 | Lighthouse Lab in Milton Keynes | Wellcome Sanger Institute for the COVID-19 Genomics UK (COG-UK) Consortium | The Lighthouse Lab in Milton Keynes and Alex Alderton, Roberto Amato, Sonia Goncalves, Ewan Harrison, David K. Jackson, Ian Johnston, Dominic Kwiatkowski, Cordelia Langford, John Sillitoe on behalf of the Wellcome Sanger Institute COVID-19 Surveillance Team |
| EPI_ISL_957693, EPI_ISL_957694 | Lighthouse Lab in Glasgow | Wellcome Sanger Institute for the COVID-19 Genomics UK (COG-UK) Consortium | Harper VanSteenhouse, Yumi Kasai, David Gray, Carol Clugston, Anna Dominiczak and Alex Alderton, Roberto Amato, Sonia Goncalves, Ewan Harrison, David K. Jackson, Ian Johnston, Dominic Kwiatkowski, Cordelia Langford, John Sillitoe on behalf of the Wellcome Sanger Institute COVID-19 Surveillance Team |
| EPI_ISL_957695 | Lighthouse Lab in Milton Keynes | Wellcome Sanger Institute for the COVID-19 Genomics UK (COG-UK) Consortium | The Lighthouse Lab in Milton Keynes and Alex Alderton, Roberto Amato, Sonia Goncalves, Ewan Harrison, David K. Jackson, Ian Johnston, Dominic Kwiatkowski, Cordelia Langford, John Sillitoe on behalf of the Wellcome Sanger Institute COVID-19 Surveillance Team |
| EPI_ISL_957696 | Lighthouse Lab in Cambridge | Wellcome Sanger Institute for the COVID-19 Genomics UK (COG-UK) Consortium | Rob Howes, The Lighthouse Lab in Cambridge and Alex Alderton, Roberto Amato, Sonia Goncalves, Ewan Harrison, David K. Jackson, Ian Johnston, Dominic Kwiatkowski, Cordelia Langford, John Sillitoe on behalf of the Wellcome Sanger Institute COVID-19 Surveillance Team |
| EPI_ISL_957697 | Lighthouse Lab in Glasgow | Wellcome Sanger Institute for the COVID-19 Genomics UK (COG-UK) Consortium | Harper VanSteenhouse, Yumi Kasai, David Gray, Carol Clugston, Anna Dominiczak and Alex Alderton, Roberto Amato, Sonia Goncalves, Ewan Harrison, David K. Jackson, Ian Johnston, Dominic Kwiatkowski, Cordelia Langford, John Sillitoe on behalf of the Wellcome Sanger Institute COVID-19 Surveillance Team |
| EPI_ISL_957699 | Lighthouse Lab in Milton Keynes | Wellcome Sanger Institute for the COVID-19 Genomics UK (COG-UK) Consortium | The Lighthouse Lab in Milton Keynes and Alex Alderton, Roberto Amato, Sonia Goncalves, Ewan Harrison, David K. Jackson, Ian Johnston, Dominic Kwiatkowski, Cordelia Langford, John Sillitoe on behalf of the Wellcome Sanger Institute COVID-19 Surveillance Team |
| EPI_ISL_957702, EPI_ISL_957703, EPI_ISL_957704 | Lighthouse Lab in Cambridge | Wellcome Sanger Institute for the COVID-19 Genomics UK (COG-UK) Consortium | Rob Howes, The Lighthouse Lab in Cambridge and Alex Alderton, Roberto Amato, Sonia Goncalves, Ewan Harrison, David K. Jackson, Ian Johnston, Dominic Kwiatkowski, Cordelia Langford, John Sillitoe on behalf of the Wellcome Sanger Institute COVID-19 Surveillance Team |
| EPI_ISL_957705 | Lighthouse Lab in Milton Keynes | Wellcome Sanger Institute for the COVID-19 Genomics UK (COG-UK) Consortium | The Lighthouse Lab in Milton Keynes and Alex Alderton, Roberto Amato, Sonia Goncalves, Ewan Harrison, David K. Jackson, Ian Johnston, Dominic Kwiatkowski, Cordelia Langford, John Sillitoe on behalf of the Wellcome Sanger Institute COVID-19 Surveillance Team |
| EPI_ISL_957706 | Lighthouse Lab in Cambridge | Wellcome Sanger Institute for the COVID-19 Genomics UK (COG-UK) Consortium | Rob Howes, The Lighthouse Lab in Cambridge and Alex Alderton, Roberto Amato, Sonia Goncalves, Ewan Harrison, David K. Jackson, Ian Johnston, Dominic Kwiatkowski, Cordelia Langford, John Sillitoe on behalf of the Wellcome Sanger Institute COVID-19 Surveillance Team |
| EPI_ISL_957708 | Lighthouse Lab in Glasgow | Wellcome Sanger Institute for the COVID-19 Genomics UK (COG-UK) Consortium | Harper VanSteenhouse, Yumi Kasai, David Gray, Carol Clugston, Anna Dominiczak and Alex Alderton, Roberto Amato, Sonia Goncalves, Ewan Harrison, David K. Jackson, Ian Johnston, Dominic Kwiatkowski, Cordelia Langford, John Sillitoe on behalf of the Wellcome Sanger Institute COVID-19 Surveillance Team |
| EPI_ISL_957709 | Lighthouse Lab in Cambridge | Wellcome Sanger Institute for the COVID-19 Genomics UK (COG-UK) Consortium | Rob Howes, The Lighthouse Lab in Cambridge and Alex Alderton, Roberto Amato, Sonia Goncalves, Ewan Harrison, David K. Jackson, Ian Johnston, Dominic Kwiatkowski, Cordelia Langford, John Sillitoe on behalf of the Wellcome Sanger Institute COVID-19 Surveillance Team |
| EPI_ISL_957710, EPI_ISL_957711, EPI_ISL_957712 | Lighthouse Lab in Milton Keynes | Wellcome Sanger Institute for the COVID-19 Genomics UK (COG-UK) Consortium | The Lighthouse Lab in Milton Keynes and Alex Alderton, Roberto Amato, Sonia Goncalves, Ewan Harrison, David K. Jackson, Ian Johnston, Dominic Kwiatkowski, Cordelia Langford, John Sillitoe on behalf of the Wellcome Sanger Institute COVID-19 Surveillance Team |
| EPI_ISL_957713 | Lighthouse Lab in Glasgow | Wellcome Sanger Institute for the COVID-19 Genomics UK (COG-UK) Consortium | Harper VanSteenhouse, Yumi Kasai, David Gray, Carol Clugston, Anna Dominiczak and Alex Alderton, Roberto Amato, Sonia Goncalves, Ewan Harrison, David K. Jackson, Ian Johnston, Dominic Kwiatkowski, Cordelia Langford, John Sillitoe on behalf of the Wellcome Sanger Institute COVID-19 Surveillance Team |
| EPI_ISL_957714 | Lighthouse Lab in Cambridge | Wellcome Sanger Institute for the COVID-19 Genomics UK (COG-UK) Consortium | Rob Howes, The Lighthouse Lab in Cambridge and Alex Alderton, Roberto Amato, Sonia Goncalves, Ewan Harrison, David K. Jackson, Ian Johnston, Dominic Kwiatkowski, Cordelia Langford, John Sillitoe on behalf of the Wellcome Sanger Institute COVID-19 Surveillance Team |
| EPI_ISL_957715 | Lighthouse Lab in Glasgow | Wellcome Sanger Institute for the COVID-19 Genomics UK (COG-UK) Consortium | Harper VanSteenhouse, Yumi Kasai, David Gray, Carol Clugston, Anna Dominiczak and Alex Alderton, Roberto Amato, Sonia Goncalves, Ewan Harrison, David K. Jackson, Ian Johnston, Dominic Kwiatkowski, Cordelia Langford, John Sillitoe on behalf of the Wellcome Sanger Institute COVID-19 Surveillance Team |
| EPI_ISL_957716 | Lighthouse Lab in Cambridge | Wellcome Sanger Institute for the COVID-19 Genomics UK (COG-UK) Consortium | Rob Howes, The Lighthouse Lab in Cambridge and Alex Alderton, Roberto Amato, Sonia Goncalves, Ewan Harrison, David K. Jackson, Ian Johnston, Dominic Kwiatkowski, Cordelia Langford, John Sillitoe on behalf of the Wellcome Sanger Institute COVID-19 Surveillance Team |
| EPI_ISL_957717, EPI_ISL_957718 | Lighthouse Lab in Glasgow | Wellcome Sanger Institute for the COVID-19 Genomics UK (COG-UK) Consortium | Harper VanSteenhouse, Yumi Kasai, David Gray, Carol Clugston, Anna Dominiczak and Alex Alderton, Roberto Amato, Sonia Goncalves, Ewan Harrison, David K. Jackson, Ian Johnston, Dominic Kwiatkowski, Cordelia Langford, John Sillitoe on behalf of the Wellcome Sanger Institute COVID-19 Surveillance Team |
| EPI_ISL_957719 | Lighthouse Lab in Cambridge | Wellcome Sanger Institute for the COVID-19 Genomics UK (COG-UK) Consortium | Rob Howes, The Lighthouse Lab in Cambridge and Alex Alderton, Roberto Amato, Sonia Goncalves, Ewan Harrison, David K. Jackson, Ian Johnston, Dominic Kwiatkowski, Cordelia Langford, John Sillitoe on behalf of the Wellcome Sanger Institute COVID-19 Surveillance Team |
| EPI_ISL_957720 | Lighthouse Lab in Glasgow | Wellcome Sanger Institute for the COVID-19 Genomics UK (COG-UK) Consortium | Harper VanSteenhouse, Yumi Kasai, David Gray, Carol Clugston, Anna Dominiczak and Alex Alderton, Roberto Amato, Sonia Goncalves, Ewan Harrison, David K. Jackson, Ian Johnston, Dominic Kwiatkowski, Cordelia Langford, John Sillitoe on behalf of the Wellcome Sanger Institute COVID-19 Surveillance Team |
| EPI_ISL_957724 | Lighthouse Lab in Milton Keynes | Wellcome Sanger Institute for the COVID-19 Genomics UK (COG-UK) Consortium | The Lighthouse Lab in Milton Keynes and Alex Alderton, Roberto Amato, Sonia Goncalves, Ewan Harrison, David K. Jackson, Ian Johnston, Dominic Kwiatkowski, Cordelia Langford, John Sillitoe on behalf of the Wellcome Sanger Institute COVID-19 Surveillance Team |
| EPI_ISL_957725 | Lighthouse Lab in Glasgow | Wellcome Sanger Institute for the COVID-19 Genomics UK (COG-UK) Consortium | Harper VanSteenhouse, Yumi Kasai, David Gray, Carol Clugston, Anna Dominiczak and Alex Alderton, Roberto Amato, Sonia Goncalves, Ewan Harrison, David K. Jackson, Ian Johnston, Dominic Kwiatkowski, Cordelia Langford, John Sillitoe on behalf of the Wellcome Sanger Institute COVID-19 Surveillance Team |
| EPI_ISL_957727 | Lighthouse Lab in Milton Keynes | Wellcome Sanger Institute for the COVID-19 Genomics UK (COG-UK) Consortium | The Lighthouse Lab in Milton Keynes and Alex Alderton, Roberto Amato, Sonia Goncalves, Ewan Harrison, David K. Jackson, Ian Johnston, Dominic Kwiatkowski, Cordelia Langford, John Sillitoe on behalf of the Wellcome Sanger Institute COVID-19 Surveillance Team |
| EPI_ISL_957728 | Lighthouse Lab in Cambridge | Wellcome Sanger Institute for the COVID-19 Genomics UK (COG-UK) Consortium | Rob Howes, The Lighthouse Lab in Cambridge and Alex Alderton, Roberto Amato, Sonia Goncalves, Ewan Harrison, David K. Jackson, Ian Johnston, Dominic Kwiatkowski, Cordelia Langford, John Sillitoe on behalf of the Wellcome Sanger Institute COVID-19 Surveillance Team |
| EPI_ISL_957729, EPI_ISL_957730 | Lighthouse Lab in Glasgow | Wellcome Sanger Institute for the COVID-19 Genomics UK (COG-UK) Consortium | Harper VanSteenhouse, Yumi Kasai, David Gray, Carol Clugston, Anna Dominiczak and Alex Alderton, Roberto Amato, Sonia Goncalves, Ewan Harrison, David K. Jackson, Ian Johnston, Dominic Kwiatkowski, Cordelia Langford, John Sillitoe on behalf of the Wellcome Sanger Institute COVID-19 Surveillance Team |
| EPI_ISL_957731 | Lighthouse Lab in Cambridge | Wellcome Sanger Institute for the COVID-19 Genomics UK (COG-UK) Consortium | Rob Howes, The Lighthouse Lab in Cambridge and Alex Alderton, Roberto Amato, Sonia Goncalves, Ewan Harrison, David K. Jackson, Ian Johnston, Dominic Kwiatkowski, Cordelia Langford, John Sillitoe on behalf of the Wellcome Sanger Institute COVID-19 Surveillance Team |
| EPI_ISL_957732 | Lighthouse Lab in Glasgow | Wellcome Sanger Institute for the COVID-19 Genomics UK (COG-UK) Consortium | Harper VanSteenhouse, Yumi Kasai, David Gray, Carol Clugston, Anna Dominiczak and Alex Alderton, Roberto Amato, Sonia Goncalves, Ewan Harrison, David K. Jackson, Ian Johnston, Dominic Kwiatkowski, Cordelia Langford, John Sillitoe on behalf of the Wellcome Sanger Institute COVID-19 Surveillance Team |
| EPI_ISL_957733, EPI_ISL_957734 | Lighthouse Lab in Cambridge | Wellcome Sanger Institute for the COVID-19 Genomics UK (COG-UK) Consortium | Rob Howes, The Lighthouse Lab in Cambridge and Alex Alderton, Roberto Amato, Sonia Goncalves, Ewan Harrison, David K. Jackson, Ian Johnston, Dominic Kwiatkowski, Cordelia Langford, John Sillitoe on behalf of the Wellcome Sanger Institute COVID-19 Surveillance Team |

| | | | |
|---|---|---|---|
| EPI_ISL_957736 | Lighthouse Lab in Glasgow | Wellcome Sanger Institute for the COVID-19 Genomics UK (COG-UK) Consortium | Harper VanSteenhouse, Yumi Kasai, David Gray, Carol Clugston, Anna Dominiczak and Alex Alderton, Roberto Amato, Sonia Goncalves, Ewan Harrison, David K. Jackson, Ian Johnston, Dominic Kwiatkowski, Cordelia Langford, John Sillitoe on behalf of the Wellcome Sanger Institute COVID-19 Surveillance Team |
| EPI_ISL_957737, EPI_ISL_957738, EPI_ISL_957739, EPI_ISL_957742 | Lighthouse Lab in Cambridge | Wellcome Sanger Institute for the COVID-19 Genomics UK (COG-UK) Consortium | Rob Howes, The Lighthouse Lab in Cambridge and Alex Alderton, Roberto Amato, Sonia Goncalves, Ewan Harrison, David K. Jackson, Ian Johnston, Dominic Kwiatkowski, Cordelia Langford, John Sillitoe on behalf of the Wellcome Sanger Institute COVID-19 Surveillance Team |
| EPI_ISL_957743 | Lighthouse Lab in Glasgow | Wellcome Sanger Institute for the COVID-19 Genomics UK (COG-UK) Consortium | Harper VanSteenhouse, Yumi Kasai, David Gray, Carol Clugston, Anna Dominiczak and Alex Alderton, Roberto Amato, Sonia Goncalves, Ewan Harrison, David K. Jackson, Ian Johnston, Dominic Kwiatkowski, Cordelia Langford, John Sillitoe on behalf of the Wellcome Sanger Institute COVID-19 Surveillance Team |
| EPI_ISL_957744 | Lighthouse Lab in Milton Keynes | Wellcome Sanger Institute for the COVID-19 Genomics UK (COG-UK) Consortium | The Lighthouse Lab in Milton Keynes and Alex Alderton, Roberto Amato, Sonia Goncalves, Ewan Harrison, David K. Jackson, Ian Johnston, Dominic Kwiatkowski, Cordelia Langford, John Sillitoe on behalf of the Wellcome Sanger Institute COVID-19 Surveillance Team |
| EPI_ISL_957746 | Lighthouse Lab in Cambridge | Wellcome Sanger Institute for the COVID-19 Genomics UK (COG-UK) Consortium | Rob Howes, The Lighthouse Lab in Cambridge and Alex Alderton, Roberto Amato, Sonia Goncalves, Ewan Harrison, David K. Jackson, Ian Johnston, Dominic Kwiatkowski, Cordelia Langford, John Sillitoe on behalf of the Wellcome Sanger Institute COVID-19 Surveillance Team |
| EPI_ISL_957749, EPI_ISL_957750 | Lighthouse Lab in Glasgow | Wellcome Sanger Institute for the COVID-19 Genomics UK (COG-UK) Consortium | Harper VanSteenhouse, Yumi Kasai, David Gray, Carol Clugston, Anna Dominiczak and Alex Alderton, Roberto Amato, Sonia Goncalves, Ewan Harrison, David K. Jackson, Ian Johnston, Dominic Kwiatkowski, Cordelia Langford, John Sillitoe on behalf of the Wellcome Sanger Institute COVID-19 Surveillance Team |
| EPI_ISL_957751, EPI_ISL_957752, EPI_ISL_957754 | Lighthouse Lab in Cambridge | Wellcome Sanger Institute for the COVID-19 Genomics UK (COG-UK) Consortium | Rob Howes, The Lighthouse Lab in Cambridge and Alex Alderton, Roberto Amato, Sonia Goncalves, Ewan Harrison, David K. Jackson, Ian Johnston, Dominic Kwiatkowski, Cordelia Langford, John Sillitoe on behalf of the Wellcome Sanger Institute COVID-19 Surveillance Team |
| EPI_ISL_957756, EPI_ISL_957758 | Lighthouse Lab in Glasgow | Wellcome Sanger Institute for the COVID-19 Genomics UK (COG-UK) Consortium | Harper VanSteenhouse, Yumi Kasai, David Gray, Carol Clugston, Anna Dominiczak and Alex Alderton, Roberto Amato, Sonia Goncalves, Ewan Harrison, David K. Jackson, Ian Johnston, Dominic Kwiatkowski, Cordelia Langford, John Sillitoe on behalf of the Wellcome Sanger Institute COVID-19 Surveillance Team |
| EPI_ISL_957760, EPI_ISL_957761, EPI_ISL_957762 | Lighthouse Lab in Milton Keynes | Wellcome Sanger Institute for the COVID-19 Genomics UK (COG-UK) Consortium | The Lighthouse Lab in Milton Keynes and Alex Alderton, Roberto Amato, Sonia Goncalves, Ewan Harrison, David K. Jackson, Ian Johnston, Dominic Kwiatkowski, Cordelia Langford, John Sillitoe on behalf of the Wellcome Sanger Institute COVID-19 Surveillance Team |
| EPI_ISL_957763, EPI_ISL_957764 | Lighthouse Lab in Cambridge | Wellcome Sanger Institute for the COVID-19 Genomics UK (COG-UK) Consortium | Rob Howes, The Lighthouse Lab in Cambridge and Alex Alderton, Roberto Amato, Sonia Goncalves, Ewan Harrison, David K. Jackson, Ian Johnston, Dominic Kwiatkowski, Cordelia Langford, John Sillitoe on behalf of the Wellcome Sanger Institute COVID-19 Surveillance Team |
| EPI_ISL_957765 | Lighthouse Lab in Glasgow | Wellcome Sanger Institute for the COVID-19 Genomics UK (COG-UK) Consortium | Harper VanSteenhouse, Yumi Kasai, David Gray, Carol Clugston, Anna Dominiczak and Alex Alderton, Roberto Amato, Sonia Goncalves, Ewan Harrison, David K. Jackson, Ian Johnston, Dominic Kwiatkowski, Cordelia Langford, John Sillitoe on behalf of the Wellcome Sanger Institute COVID-19 Surveillance Team |
| EPI_ISL_957767 | Lighthouse Lab in Milton Keynes | Wellcome Sanger Institute for the COVID-19 Genomics UK (COG-UK) Consortium | The Lighthouse Lab in Milton Keynes and Alex Alderton, Roberto Amato, Sonia Goncalves, Ewan Harrison, David K. Jackson, Ian Johnston, Dominic Kwiatkowski, Cordelia Langford, John Sillitoe on behalf of the Wellcome Sanger Institute COVID-19 Surveillance Team |
| EPI_ISL_957768, EPI_ISL_957770, EPI_ISL_957773, EPI_ISL_957774 | Lighthouse Lab in Glasgow | Wellcome Sanger Institute for the COVID-19 Genomics UK (COG-UK) Consortium | Harper VanSteenhouse, Yumi Kasai, David Gray, Carol Clugston, Anna Dominiczak and Alex Alderton, Roberto Amato, Sonia Goncalves, Ewan Harrison, David K. Jackson, Ian Johnston, Dominic Kwiatkowski, Cordelia Langford, John Sillitoe on behalf of the Wellcome Sanger Institute COVID-19 Surveillance Team |
| EPI_ISL_957775 | Lighthouse Lab in Milton Keynes | Wellcome Sanger Institute for the COVID-19 Genomics UK (COG-UK) Consortium | The Lighthouse Lab in Milton Keynes and Alex Alderton, Roberto Amato, Sonia Goncalves, Ewan Harrison, David K. Jackson, Ian Johnston, Dominic Kwiatkowski, Cordelia Langford, John Sillitoe on behalf of the Wellcome Sanger Institute COVID-19 Surveillance Team |
| EPI_ISL_957776 | Lighthouse Lab in Cambridge | Wellcome Sanger Institute for the COVID-19 Genomics UK (COG-UK) Consortium | Rob Howes, The Lighthouse Lab in Cambridge and Alex Alderton, Roberto Amato, Sonia Goncalves, Ewan Harrison, David K. Jackson, Ian Johnston, Dominic Kwiatkowski, Cordelia Langford, John Sillitoe on behalf of the Wellcome Sanger Institute COVID-19 Surveillance Team |
| EPI_ISL_957777 | Lighthouse Lab in Milton Keynes | Wellcome Sanger Institute for the COVID-19 Genomics UK (COG-UK) Consortium | The Lighthouse Lab in Milton Keynes and Alex Alderton, Roberto Amato, Sonia Goncalves, Ewan Harrison, David K. Jackson, Ian Johnston, Dominic Kwiatkowski, Cordelia Langford, John Sillitoe on behalf of the Wellcome Sanger Institute COVID-19 Surveillance Team |
| EPI_ISL_957778 | Lighthouse Lab in Glasgow | Wellcome Sanger Institute for the COVID-19 Genomics UK (COG-UK) Consortium | Harper VanSteenhouse, Yumi Kasai, David Gray, Carol Clugston, Anna Dominiczak and Alex Alderton, Roberto Amato, Sonia Goncalves, Ewan Harrison, David K. Jackson, Ian Johnston, Dominic Kwiatkowski, Cordelia Langford, John Sillitoe on behalf of the Wellcome Sanger Institute COVID-19 Surveillance Team |
| EPI_ISL_957779, EPI_ISL_957781, EPI_ISL_957782, EPI_ISL_957784 | Lighthouse Lab in Cambridge | Wellcome Sanger Institute for the COVID-19 Genomics UK (COG-UK) Consortium | Rob Howes, The Lighthouse Lab in Cambridge and Alex Alderton, Roberto Amato, Sonia Goncalves, Ewan Harrison, David K. Jackson, Ian Johnston, Dominic Kwiatkowski, Cordelia Langford, John Sillitoe on behalf of the Wellcome Sanger Institute COVID-19 Surveillance Team |
| EPI_ISL_957785 | Lighthouse Lab in Glasgow | Wellcome Sanger Institute for the COVID-19 Genomics UK (COG-UK) Consortium | Harper VanSteenhouse, Yumi Kasai, David Gray, Carol Clugston, Anna Dominiczak and Alex Alderton, Roberto Amato, Sonia Goncalves, Ewan Harrison, David K. Jackson, Ian Johnston, Dominic Kwiatkowski, Cordelia Langford, John Sillitoe on behalf of the Wellcome Sanger Institute COVID-19 Surveillance Team |
| EPI_ISL_957786 | Lighthouse Lab in Milton Keynes | Wellcome Sanger Institute for the COVID-19 Genomics UK (COG-UK) Consortium | The Lighthouse Lab in Milton Keynes and Alex Alderton, Roberto Amato, Sonia Goncalves, Ewan Harrison, David K. Jackson, Ian Johnston, Dominic Kwiatkowski, Cordelia Langford, John Sillitoe on behalf of the Wellcome Sanger Institute COVID-19 Surveillance Team |
| EPI_ISL_957788, EPI_ISL_957789 | Lighthouse Lab in Cambridge | Wellcome Sanger Institute for the COVID-19 Genomics UK (COG-UK) Consortium | Rob Howes, The Lighthouse Lab in Cambridge and Alex Alderton, Roberto Amato, Sonia Goncalves, Ewan Harrison, David K. Jackson, Ian Johnston, Dominic Kwiatkowski, Cordelia Langford, John Sillitoe on behalf of the Wellcome Sanger Institute COVID-19 Surveillance Team |
| EPI_ISL_957790, EPI_ISL_957791 | Lighthouse Lab in Milton Keynes | Wellcome Sanger Institute for the COVID-19 Genomics UK (COG-UK) Consortium | The Lighthouse Lab in Milton Keynes and Alex Alderton, Roberto Amato, Sonia Goncalves, Ewan Harrison, David K. Jackson, Ian Johnston, Dominic Kwiatkowski, Cordelia Langford, John Sillitoe on behalf of the Wellcome Sanger Institute COVID-19 Surveillance Team |
| EPI_ISL_957792, EPI_ISL_957793, EPI_ISL_957794 | Lighthouse Lab in Glasgow | Wellcome Sanger Institute for the COVID-19 Genomics UK (COG-UK) Consortium | Harper VanSteenhouse, Yumi Kasai, David Gray, Carol Clugston, Anna Dominiczak and Alex Alderton, Roberto Amato, Sonia Goncalves, Ewan Harrison, David K. Jackson, Ian Johnston, Dominic Kwiatkowski, Cordelia Langford, John Sillitoe on behalf of the Wellcome Sanger Institute COVID-19 Surveillance Team |
| EPI_ISL_957795 | Lighthouse Lab in Milton Keynes | Wellcome Sanger Institute for the COVID-19 Genomics UK (COG-UK) Consortium | The Lighthouse Lab in Milton Keynes and Alex Alderton, Roberto Amato, Sonia Goncalves, Ewan Harrison, David K. Jackson, Ian Johnston, Dominic Kwiatkowski, Cordelia Langford, John Sillitoe on behalf of the Wellcome Sanger Institute COVID-19 Surveillance Team |
| EPI_ISL_957796 | Lighthouse Lab in Glasgow | Wellcome Sanger Institute for the COVID-19 Genomics UK (COG-UK) Consortium | Harper VanSteenhouse, Yumi Kasai, David Gray, Carol Clugston, Anna Dominiczak and Alex Alderton, Roberto Amato, Sonia Goncalves, Ewan Harrison, David K. Jackson, Ian Johnston, Dominic Kwiatkowski, Cordelia Langford, John Sillitoe on behalf of the Wellcome Sanger Institute COVID-19 Surveillance Team |
| EPI_ISL_957798 | Lighthouse Lab in Cambridge | Wellcome Sanger Institute for the COVID-19 Genomics UK (COG-UK) Consortium | Rob Howes, The Lighthouse Lab in Cambridge and Alex Alderton, Roberto Amato, Sonia Goncalves, Ewan Harrison, David K. Jackson, Ian Johnston, Dominic Kwiatkowski, Cordelia Langford, John Sillitoe on behalf of the Wellcome Sanger Institute COVID-19 Surveillance Team |
| EPI_ISL_957800, EPI_ISL_957801 | Lighthouse Lab in Milton Keynes | Wellcome Sanger Institute for the COVID-19 Genomics UK (COG-UK) Consortium | The Lighthouse Lab in Milton Keynes and Alex Alderton, Roberto Amato, Sonia Goncalves, Ewan Harrison, David K. Jackson, Ian Johnston, Dominic Kwiatkowski, Cordelia Langford, John Sillitoe on behalf of the Wellcome Sanger Institute COVID-19 Surveillance Team |
| EPI_ISL_957802 | Lighthouse Lab in Glasgow | Wellcome Sanger Institute for the COVID-19 Genomics UK (COG-UK) Consortium | Harper VanSteenhouse, Yumi Kasai, David Gray, Carol Clugston, Anna Dominiczak and Alex Alderton, Roberto Amato, Sonia Goncalves, Ewan Harrison, David K. Jackson, Ian Johnston, Dominic Kwiatkowski, Cordelia Langford, John Sillitoe on behalf of the Wellcome Sanger Institute COVID-19 Surveillance Team |
| EPI_ISL_957803, EPI_ISL_957804 | Lighthouse Lab in Milton Keynes | Wellcome Sanger Institute for the COVID-19 Genomics UK (COG-UK) Consortium | The Lighthouse Lab in Milton Keynes and Alex Alderton, Roberto Amato, Sonia Goncalves, Ewan Harrison, David K. Jackson, Ian Johnston, Dominic Kwiatkowski, Cordelia Langford, John Sillitoe on behalf of the Wellcome Sanger Institute COVID-19 Surveillance Team |
| EPI_ISL_957805 | Lighthouse Lab in Cambridge | Wellcome Sanger Institute for the COVID-19 Genomics UK (COG-UK) Consortium | Rob Howes, The Lighthouse Lab in Cambridge and Alex Alderton, Roberto Amato, Sonia Goncalves, Ewan Harrison, David K. Jackson, Ian Johnston, Dominic Kwiatkowski, Cordelia Langford, John Sillitoe on behalf of the Wellcome Sanger Institute COVID-19 Surveillance Team |
| EPI_ISL_957806 | Lighthouse Lab in Glasgow | Wellcome Sanger Institute for the COVID-19 Genomics UK (COG-UK) Consortium | Harper VanSteenhouse, Yumi Kasai, David Gray, Carol Clugston, Anna Dominiczak and Alex Alderton, Roberto Amato, Sonia Goncalves, Ewan Harrison, David K. Jackson, Ian Johnston, Dominic Kwiatkowski, Cordelia Langford, John Sillitoe on behalf of the Wellcome Sanger Institute COVID-19 Surveillance |

| | | | |
|---|---|---|---|
| | | | Team |
| EPI_ISL_957807, EPI_ISL_957808, EPI_ISL_957809 | Lighthouse Lab in Cambridge | Wellcome Sanger Institute for the COVID-19 Genomics UK (COG-UK) Consortium | Rob Howes, The Lighthouse Lab in Cambridge and Alex Alderton, Roberto Amato, Sonia Goncalves, Ewan Harrison, David K. Jackson, Ian Johnston, Dominic Kwiatkowski, Cordelia Langford, John Sillitoe on behalf of the Wellcome Sanger Institute COVID-19 Surveillance Team |
| EPI_ISL_957810 | Lighthouse Lab in Milton Keynes | Wellcome Sanger Institute for the COVID-19 Genomics UK (COG-UK) Consortium | The Lighthouse Lab in Milton Keynes and Alex Alderton, Roberto Amato, Sonia Goncalves, Ewan Harrison, David K. Jackson, Ian Johnston, Dominic Kwiatkowski, Cordelia Langford, John Sillitoe on behalf of the Wellcome Sanger Institute COVID-19 Surveillance Team |
| EPI_ISL_957811, EPI_ISL_957812 | Lighthouse Lab in Cambridge | Wellcome Sanger Institute for the COVID-19 Genomics UK (COG-UK) Consortium | Rob Howes, The Lighthouse Lab in Cambridge and Alex Alderton, Roberto Amato, Sonia Goncalves, Ewan Harrison, David K. Jackson, Ian Johnston, Dominic Kwiatkowski, Cordelia Langford, John Sillitoe on behalf of the Wellcome Sanger Institute COVID-19 Surveillance Team |
| EPI_ISL_957814 | Lighthouse Lab in Glasgow | Wellcome Sanger Institute for the COVID-19 Genomics UK (COG-UK) Consortium | Harper VanSteenhouse, Yumi Kasai, David Gray, Carol Clugston, Anna Dominiczak and Alex Alderton, Roberto Amato, Sonia Goncalves, Ewan Harrison, David K. Jackson, Ian Johnston, Dominic Kwiatkowski, Cordelia Langford, John Sillitoe on behalf of the Wellcome Sanger Institute COVID-19 Surveillance Team |
| EPI_ISL_957815 | Lighthouse Lab in Cambridge | Wellcome Sanger Institute for the COVID-19 Genomics UK (COG-UK) Consortium | Rob Howes, The Lighthouse Lab in Cambridge and Alex Alderton, Roberto Amato, Sonia Goncalves, Ewan Harrison, David K. Jackson, Ian Johnston, Dominic Kwiatkowski, Cordelia Langford, John Sillitoe on behalf of the Wellcome Sanger Institute COVID-19 Surveillance Team |
| EPI_ISL_957816 | Lighthouse Lab in Milton Keynes | Wellcome Sanger Institute for the COVID-19 Genomics UK (COG-UK) Consortium | The Lighthouse Lab in Milton Keynes and Alex Alderton, Roberto Amato, Sonia Goncalves, Ewan Harrison, David K. Jackson, Ian Johnston, Dominic Kwiatkowski, Cordelia Langford, John Sillitoe on behalf of the Wellcome Sanger Institute COVID-19 Surveillance Team |
| EPI_ISL_957818 | Lighthouse Lab in Cambridge | Wellcome Sanger Institute for the COVID-19 Genomics UK (COG-UK) Consortium | Rob Howes, The Lighthouse Lab in Cambridge and Alex Alderton, Roberto Amato, Sonia Goncalves, Ewan Harrison, David K. Jackson, Ian Johnston, Dominic Kwiatkowski, Cordelia Langford, John Sillitoe on behalf of the Wellcome Sanger Institute COVID-19 Surveillance Team |
| EPI_ISL_957819 | Lighthouse Lab in Glasgow | Wellcome Sanger Institute for the COVID-19 Genomics UK (COG-UK) Consortium | Harper VanSteenhouse, Yumi Kasai, David Gray, Carol Clugston, Anna Dominiczak and Alex Alderton, Roberto Amato, Sonia Goncalves, Ewan Harrison, David K. Jackson, Ian Johnston, Dominic Kwiatkowski, Cordelia Langford, John Sillitoe on behalf of the Wellcome Sanger Institute COVID-19 Surveillance Team |
| EPI_ISL_957820, EPI_ISL_957821 | Lighthouse Lab in Cambridge | Wellcome Sanger Institute for the COVID-19 Genomics UK (COG-UK) Consortium | Rob Howes, The Lighthouse Lab in Cambridge and Alex Alderton, Roberto Amato, Sonia Goncalves, Ewan Harrison, David K. Jackson, Ian Johnston, Dominic Kwiatkowski, Cordelia Langford, John Sillitoe on behalf of the Wellcome Sanger Institute COVID-19 Surveillance Team |
| EPI_ISL_957823 | Lighthouse Lab in Milton Keynes | Wellcome Sanger Institute for the COVID-19 Genomics UK (COG-UK) Consortium | The Lighthouse Lab in Milton Keynes and Alex Alderton, Roberto Amato, Sonia Goncalves, Ewan Harrison, David K. Jackson, Ian Johnston, Dominic Kwiatkowski, Cordelia Langford, John Sillitoe on behalf of the Wellcome Sanger Institute COVID-19 Surveillance Team |
| EPI_ISL_957824 | Lighthouse Lab in Glasgow | Wellcome Sanger Institute for the COVID-19 Genomics UK (COG-UK) Consortium | Harper VanSteenhouse, Yumi Kasai, David Gray, Carol Clugston, Anna Dominiczak and Alex Alderton, Roberto Amato, Sonia Goncalves, Ewan Harrison, David K. Jackson, Ian Johnston, Dominic Kwiatkowski, Cordelia Langford, John Sillitoe on behalf of the Wellcome Sanger Institute COVID-19 Surveillance Team |
| EPI_ISL_957825 | Lighthouse Lab in Cambridge | Wellcome Sanger Institute for the COVID-19 Genomics UK (COG-UK) Consortium | Rob Howes, The Lighthouse Lab in Cambridge and Alex Alderton, Roberto Amato, Sonia Goncalves, Ewan Harrison, David K. Jackson, Ian Johnston, Dominic Kwiatkowski, Cordelia Langford, John Sillitoe on behalf of the Wellcome Sanger Institute COVID-19 Surveillance Team |
| EPI_ISL_957827 | Lighthouse Lab in Milton Keynes | Wellcome Sanger Institute for the COVID-19 Genomics UK (COG-UK) Consortium | The Lighthouse Lab in Milton Keynes and Alex Alderton, Roberto Amato, Sonia Goncalves, Ewan Harrison, David K. Jackson, Ian Johnston, Dominic Kwiatkowski, Cordelia Langford, John Sillitoe on behalf of the Wellcome Sanger Institute COVID-19 Surveillance Team |
| EPI_ISL_957831, EPI_ISL_957833, EPI_ISL_957834 | Lighthouse Lab in Cambridge | Wellcome Sanger Institute for the COVID-19 Genomics UK (COG-UK) Consortium | Rob Howes, The Lighthouse Lab in Cambridge and Alex Alderton, Roberto Amato, Sonia Goncalves, Ewan Harrison, David K. Jackson, Ian Johnston, Dominic Kwiatkowski, Cordelia Langford, John Sillitoe on behalf of the Wellcome Sanger Institute COVID-19 Surveillance Team |
| EPI_ISL_957835 | Lighthouse Lab in Milton Keynes | Wellcome Sanger Institute for the COVID-19 Genomics UK (COG-UK) Consortium | The Lighthouse Lab in Milton Keynes and Alex Alderton, Roberto Amato, Sonia Goncalves, Ewan Harrison, David K. Jackson, Ian Johnston, Dominic Kwiatkowski, Cordelia Langford, John Sillitoe on behalf of the Wellcome Sanger Institute COVID-19 Surveillance Team |
| EPI_ISL_957836, EPI_ISL_957837, EPI_ISL_957839 | Lighthouse Lab in Glasgow | Wellcome Sanger Institute for the COVID-19 Genomics UK (COG-UK) Consortium | Harper VanSteenhouse, Yumi Kasai, David Gray, Carol Clugston, Anna Dominiczak and Alex Alderton, Roberto Amato, Sonia Goncalves, Ewan Harrison, David K. Jackson, Ian Johnston, Dominic Kwiatkowski, Cordelia Langford, John Sillitoe on behalf of the Wellcome Sanger Institute COVID-19 Surveillance Team |
| EPI_ISL_957840 | Lighthouse Lab in Cambridge | Wellcome Sanger Institute for the COVID-19 Genomics UK (COG-UK) Consortium | Rob Howes, The Lighthouse Lab in Cambridge and Alex Alderton, Roberto Amato, Sonia Goncalves, Ewan Harrison, David K. Jackson, Ian Johnston, Dominic Kwiatkowski, Cordelia Langford, John Sillitoe on behalf of the Wellcome Sanger Institute COVID-19 Surveillance Team |
| EPI_ISL_957842, EPI_ISL_957843 | Lighthouse Lab in Milton Keynes | Wellcome Sanger Institute for the COVID-19 Genomics UK (COG-UK) Consortium | The Lighthouse Lab in Milton Keynes and Alex Alderton, Roberto Amato, Sonia Goncalves, Ewan Harrison, David K. Jackson, Ian Johnston, Dominic Kwiatkowski, Cordelia Langford, John Sillitoe on behalf of the Wellcome Sanger Institute COVID-19 Surveillance Team |
| EPI_ISL_957845 | Lighthouse Lab in Glasgow | Wellcome Sanger Institute for the COVID-19 Genomics UK (COG-UK) Consortium | Harper VanSteenhouse, Yumi Kasai, David Gray, Carol Clugston, Anna Dominiczak and Alex Alderton, Roberto Amato, Sonia Goncalves, Ewan Harrison, David K. Jackson, Ian Johnston, Dominic Kwiatkowski, Cordelia Langford, John Sillitoe on behalf of the Wellcome Sanger Institute COVID-19 Surveillance Team |
| EPI_ISL_957846 | Lighthouse Lab in Cambridge | Wellcome Sanger Institute for the COVID-19 Genomics UK (COG-UK) Consortium | Rob Howes, The Lighthouse Lab in Cambridge and Alex Alderton, Roberto Amato, Sonia Goncalves, Ewan Harrison, David K. Jackson, Ian Johnston, Dominic Kwiatkowski, Cordelia Langford, John Sillitoe on behalf of the Wellcome Sanger Institute COVID-19 Surveillance Team |
| EPI_ISL_957847, EPI_ISL_957848 | Lighthouse Lab in Glasgow | Wellcome Sanger Institute for the COVID-19 Genomics UK (COG-UK) Consortium | Harper VanSteenhouse, Yumi Kasai, David Gray, Carol Clugston, Anna Dominiczak and Alex Alderton, Roberto Amato, Sonia Goncalves, Ewan Harrison, David K. Jackson, Ian Johnston, Dominic Kwiatkowski, Cordelia Langford, John Sillitoe on behalf of the Wellcome Sanger Institute COVID-19 Surveillance Team |
| EPI_ISL_957849, EPI_ISL_957850 | Lighthouse Lab in Milton Keynes | Wellcome Sanger Institute for the COVID-19 Genomics UK (COG-UK) Consortium | The Lighthouse Lab in Milton Keynes and Alex Alderton, Roberto Amato, Sonia Goncalves, Ewan Harrison, David K. Jackson, Ian Johnston, Dominic Kwiatkowski, Cordelia Langford, John Sillitoe on behalf of the Wellcome Sanger Institute COVID-19 Surveillance Team |
| EPI_ISL_957851 | Lighthouse Lab in Glasgow | Wellcome Sanger Institute for the COVID-19 Genomics UK (COG-UK) Consortium | Harper VanSteenhouse, Yumi Kasai, David Gray, Carol Clugston, Anna Dominiczak and Alex Alderton, Roberto Amato, Sonia Goncalves, Ewan Harrison, David K. Jackson, Ian Johnston, Dominic Kwiatkowski, Cordelia Langford, John Sillitoe on behalf of the Wellcome Sanger Institute COVID-19 Surveillance Team |
| EPI_ISL_957852 | Lighthouse Lab in Cambridge | Wellcome Sanger Institute for the COVID-19 Genomics UK (COG-UK) Consortium | Rob Howes, The Lighthouse Lab in Cambridge and Alex Alderton, Roberto Amato, Sonia Goncalves, Ewan Harrison, David K. Jackson, Ian Johnston, Dominic Kwiatkowski, Cordelia Langford, John Sillitoe on behalf of the Wellcome Sanger Institute COVID-19 Surveillance Team |
| EPI_ISL_957853, EPI_ISL_957854 | Lighthouse Lab in Glasgow | Wellcome Sanger Institute for the COVID-19 Genomics UK (COG-UK) Consortium | Harper VanSteenhouse, Yumi Kasai, David Gray, Carol Clugston, Anna Dominiczak and Alex Alderton, Roberto Amato, Sonia Goncalves, Ewan Harrison, David K. Jackson, Ian Johnston, Dominic Kwiatkowski, Cordelia Langford, John Sillitoe on behalf of the Wellcome Sanger Institute COVID-19 Surveillance Team |
| EPI_ISL_957855, EPI_ISL_957857 | Lighthouse Lab in Cambridge | Wellcome Sanger Institute for the COVID-19 Genomics UK (COG-UK) Consortium | Rob Howes, The Lighthouse Lab in Cambridge and Alex Alderton, Roberto Amato, Sonia Goncalves, Ewan Harrison, David K. Jackson, Ian Johnston, Dominic Kwiatkowski, Cordelia Langford, John Sillitoe on behalf of the Wellcome Sanger Institute COVID-19 Surveillance Team |
| EPI_ISL_957858 | Lighthouse Lab in Milton Keynes | Wellcome Sanger Institute for the COVID-19 Genomics UK (COG-UK) Consortium | The Lighthouse Lab in Milton Keynes and Alex Alderton, Roberto Amato, Sonia Goncalves, Ewan Harrison, David K. Jackson, Ian Johnston, Dominic Kwiatkowski, Cordelia Langford, John Sillitoe on behalf of the Wellcome Sanger Institute COVID-19 Surveillance Team |
| EPI_ISL_957860 | Lighthouse Lab in Glasgow | Wellcome Sanger Institute for the COVID-19 Genomics UK (COG-UK) Consortium | Harper VanSteenhouse, Yumi Kasai, David Gray, Carol Clugston, Anna Dominiczak and Alex Alderton, Roberto Amato, Sonia Goncalves, Ewan Harrison, David K. Jackson, Ian Johnston, Dominic Kwiatkowski, Cordelia Langford, John Sillitoe on behalf of the Wellcome Sanger Institute COVID-19 Surveillance Team |
| EPI_ISL_957861 | Lighthouse Lab in Cambridge | Wellcome Sanger Institute for the COVID-19 Genomics UK (COG-UK) Consortium | Rob Howes, The Lighthouse Lab in Cambridge and Alex Alderton, Roberto Amato, Sonia Goncalves, Ewan Harrison, David K. Jackson, Ian Johnston, Dominic Kwiatkowski, Cordelia Langford, John Sillitoe on behalf of the Wellcome Sanger Institute COVID-19 Surveillance Team |
| EPI_ISL_957864, EPI_ISL_957865 | Lighthouse Lab in Glasgow | Wellcome Sanger Institute for the COVID-19 Genomics UK (COG-UK) Consortium | Harper VanSteenhouse, Yumi Kasai, David Gray, Carol Clugston, Anna Dominiczak and Alex Alderton, Roberto Amato, Sonia Goncalves, Ewan Harrison, David K. Jackson, Ian Johnston, Dominic Kwiatkowski, Cordelia Langford, John Sillitoe on behalf of the Wellcome Sanger Institute COVID-19 Surveillance Team |
| EPI_ISL_957869, EPI_ISL_957870 | Lighthouse Lab in Cambridge | Wellcome Sanger Institute for the COVID-19 Genomics UK | Rob Howes, The Lighthouse Lab in Cambridge and Alex Alderton, Roberto Amato, Sonia Goncalves, Ewan Harrison, David K. Jackson, Ian Johnston, |

| | | | |
|---|---|---|---|
| | | (COG-UK) Consortium | Dominic Kwiatkowski, Cordelia Langford, John Sillitoe on behalf of the Wellcome Sanger Institute COVID-19 Surveillance Team |
| EPI_ISL_957871 | Lighthouse Lab in Glasgow | Wellcome Sanger Institute for the COVID-19 Genomics UK (COG-UK) Consortium | Harper VanSteenhouse, Yumi Kasai, David Gray, Carol Clugston, Anna Dominiczak and Alex Alderton, Roberto Amato, Sonia Goncalves, Ewan Harrison, David K. Jackson, Ian Johnston, Dominic Kwiatkowski, Cordelia Langford, John Sillitoe on behalf of the Wellcome Sanger Institute COVID-19 Surveillance Team |
| EPI_ISL_957873, EPI_ISL_957874 | Lighthouse Lab in Cambridge | Wellcome Sanger Institute for the COVID-19 Genomics UK (COG-UK) Consortium | Rob Howes, The Lighthouse Lab in Cambridge and Alex Alderton, Roberto Amato, Sonia Goncalves, Ewan Harrison, David K. Jackson, Ian Johnston, Dominic Kwiatkowski, Cordelia Langford, John Sillitoe on behalf of the Wellcome Sanger Institute COVID-19 Surveillance Team |
| EPI_ISL_957875 | Lighthouse Lab in Glasgow | Wellcome Sanger Institute for the COVID-19 Genomics UK (COG-UK) Consortium | Harper VanSteenhouse, Yumi Kasai, David Gray, Carol Clugston, Anna Dominiczak and Alex Alderton, Roberto Amato, Sonia Goncalves, Ewan Harrison, David K. Jackson, Ian Johnston, Dominic Kwiatkowski, Cordelia Langford, John Sillitoe on behalf of the Wellcome Sanger Institute COVID-19 Surveillance Team |
| EPI_ISL_957876 | Lighthouse Lab in Cambridge | Wellcome Sanger Institute for the COVID-19 Genomics UK (COG-UK) Consortium | Rob Howes, The Lighthouse Lab in Cambridge and Alex Alderton, Roberto Amato, Sonia Goncalves, Ewan Harrison, David K. Jackson, Ian Johnston, Dominic Kwiatkowski, Cordelia Langford, John Sillitoe on behalf of the Wellcome Sanger Institute COVID-19 Surveillance Team |
| EPI_ISL_957879 | Lighthouse Lab in Glasgow | Wellcome Sanger Institute for the COVID-19 Genomics UK (COG-UK) Consortium | Harper VanSteenhouse, Yumi Kasai, David Gray, Carol Clugston, Anna Dominiczak and Alex Alderton, Roberto Amato, Sonia Goncalves, Ewan Harrison, David K. Jackson, Ian Johnston, Dominic Kwiatkowski, Cordelia Langford, John Sillitoe on behalf of the Wellcome Sanger Institute COVID-19 Surveillance Team |
| EPI_ISL_957880, EPI_ISL_957881, EPI_ISL_957882, EPI_ISL_957883 | Lighthouse Lab in Cambridge | Wellcome Sanger Institute for the COVID-19 Genomics UK (COG-UK) Consortium | Rob Howes, The Lighthouse Lab in Cambridge and Alex Alderton, Roberto Amato, Sonia Goncalves, Ewan Harrison, David K. Jackson, Ian Johnston, Dominic Kwiatkowski, Cordelia Langford, John Sillitoe on behalf of the Wellcome Sanger Institute COVID-19 Surveillance Team |
| EPI_ISL_957884 | Lighthouse Lab in Glasgow | Wellcome Sanger Institute for the COVID-19 Genomics UK (COG-UK) Consortium | Harper VanSteenhouse, Yumi Kasai, David Gray, Carol Clugston, Anna Dominiczak and Alex Alderton, Roberto Amato, Sonia Goncalves, Ewan Harrison, David K. Jackson, Ian Johnston, Dominic Kwiatkowski, Cordelia Langford, John Sillitoe on behalf of the Wellcome Sanger Institute COVID-19 Surveillance Team |
| EPI_ISL_957885 | Lighthouse Lab in Milton Keynes | Wellcome Sanger Institute for the COVID-19 Genomics UK (COG-UK) Consortium | The Lighthouse Lab in Milton Keynes and Alex Alderton, Roberto Amato, Sonia Goncalves, Ewan Harrison, David K. Jackson, Ian Johnston, Dominic Kwiatkowski, Cordelia Langford, John Sillitoe on behalf of the Wellcome Sanger Institute COVID-19 Surveillance Team |
| EPI_ISL_957887 | Lighthouse Lab in Cambridge | Wellcome Sanger Institute for the COVID-19 Genomics UK (COG-UK) Consortium | Rob Howes, The Lighthouse Lab in Cambridge and Alex Alderton, Roberto Amato, Sonia Goncalves, Ewan Harrison, David K. Jackson, Ian Johnston, Dominic Kwiatkowski, Cordelia Langford, John Sillitoe on behalf of the Wellcome Sanger Institute COVID-19 Surveillance Team |
| EPI_ISL_957888 | Lighthouse Lab in Milton Keynes | Wellcome Sanger Institute for the COVID-19 Genomics UK (COG-UK) Consortium | The Lighthouse Lab in Milton Keynes and Alex Alderton, Roberto Amato, Sonia Goncalves, Ewan Harrison, David K. Jackson, Ian Johnston, Dominic Kwiatkowski, Cordelia Langford, John Sillitoe on behalf of the Wellcome Sanger Institute COVID-19 Surveillance Team |
| EPI_ISL_957889 | Lighthouse Lab in Glasgow | Wellcome Sanger Institute for the COVID-19 Genomics UK (COG-UK) Consortium | Harper VanSteenhouse, Yumi Kasai, David Gray, Carol Clugston, Anna Dominiczak and Alex Alderton, Roberto Amato, Sonia Goncalves, Ewan Harrison, David K. Jackson, Ian Johnston, Dominic Kwiatkowski, Cordelia Langford, John Sillitoe on behalf of the Wellcome Sanger Institute COVID-19 Surveillance Team |
| EPI_ISL_957890, EPI_ISL_957891 | Lighthouse Lab in Cambridge | Wellcome Sanger Institute for the COVID-19 Genomics UK (COG-UK) Consortium | Rob Howes, The Lighthouse Lab in Cambridge and Alex Alderton, Roberto Amato, Sonia Goncalves, Ewan Harrison, David K. Jackson, Ian Johnston, Dominic Kwiatkowski, Cordelia Langford, John Sillitoe on behalf of the Wellcome Sanger Institute COVID-19 Surveillance Team |
| EPI_ISL_957893 | Lighthouse Lab in Milton Keynes | Wellcome Sanger Institute for the COVID-19 Genomics UK (COG-UK) Consortium | The Lighthouse Lab in Milton Keynes and Alex Alderton, Roberto Amato, Sonia Goncalves, Ewan Harrison, David K. Jackson, Ian Johnston, Dominic Kwiatkowski, Cordelia Langford, John Sillitoe on behalf of the Wellcome Sanger Institute COVID-19 Surveillance Team |
| EPI_ISL_957894 | Lighthouse Lab in Glasgow | Wellcome Sanger Institute for the COVID-19 Genomics UK (COG-UK) Consortium | Harper VanSteenhouse, Yumi Kasai, David Gray, Carol Clugston, Anna Dominiczak and Alex Alderton, Roberto Amato, Sonia Goncalves, Ewan Harrison, David K. Jackson, Ian Johnston, Dominic Kwiatkowski, Cordelia Langford, John Sillitoe on behalf of the Wellcome Sanger Institute COVID-19 Surveillance Team |
| EPI_ISL_957896, EPI_ISL_957898 | Lighthouse Lab in Cambridge | Wellcome Sanger Institute for the COVID-19 Genomics UK (COG-UK) Consortium | Rob Howes, The Lighthouse Lab in Cambridge and Alex Alderton, Roberto Amato, Sonia Goncalves, Ewan Harrison, David K. Jackson, Ian Johnston, Dominic Kwiatkowski, Cordelia Langford, John Sillitoe on behalf of the Wellcome Sanger Institute COVID-19 Surveillance Team |
| EPI_ISL_957899, EPI_ISL_957900, EPI_ISL_957901 | Lighthouse Lab in Milton Keynes | Wellcome Sanger Institute for the COVID-19 Genomics UK (COG-UK) Consortium | The Lighthouse Lab in Milton Keynes and Alex Alderton, Roberto Amato, Sonia Goncalves, Ewan Harrison, David K. Jackson, Ian Johnston, Dominic Kwiatkowski, Cordelia Langford, John Sillitoe on behalf of the Wellcome Sanger Institute COVID-19 Surveillance Team |
| EPI_ISL_957902, EPI_ISL_957903, EPI_ISL_957905, EPI_ISL_957906, EPI_ISL_957908 | Lighthouse Lab in Cambridge | Wellcome Sanger Institute for the COVID-19 Genomics UK (COG-UK) Consortium | Rob Howes, The Lighthouse Lab in Cambridge and Alex Alderton, Roberto Amato, Sonia Goncalves, Ewan Harrison, David K. Jackson, Ian Johnston, Dominic Kwiatkowski, Cordelia Langford, John Sillitoe on behalf of the Wellcome Sanger Institute COVID-19 Surveillance Team |

EPI_ISL_957909, EPI_ISL_957910, EPI_ISL_957911, EPI_ISL_957912, EPI_ISL_957913, EPI_ISL_957914, EPI_ISL_957915, EPI_ISL_957916, EPI_ISL_957917, EPI_ISL_957918, EPI_ISL_957919, EPI_ISL_957920, EPI_ISL_957922, EPI_ISL_957923, EPI_ISL_957924, EPI_ISL_957925, EPI_ISL_957926, EPI_ISL_957927, EPI_ISL_957928, EPI_ISL_957929, EPI_ISL_957930, EPI_ISL_957931, EPI_ISL_957932, EPI_ISL_957933, EPI_ISL_957934, EPI_ISL_957935, EPI_ISL_957936, EPI_ISL_957937, EPI_ISL_957938, EPI_ISL_957939, EPI_ISL_957940, EPI_ISL_957941, EPI_ISL_957942, EPI_ISL_957943, EPI_ISL_957944, EPI_ISL_957945, EPI_ISL_957946, EPI_ISL_957947, EPI_ISL_957948, EPI_ISL_957949, EPI_ISL_957950, EPI_ISL_957951, EPI_ISL_957952, EPI_ISL_957953, EPI_ISL_957954, EPI_ISL_957955, EPI_ISL_957956, EPI_ISL_957957, EPI_ISL_957958, EPI_ISL_957959, EPI_ISL_957960, EPI_ISL_957963, EPI_ISL_957964, EPI_ISL_957965, EPI_ISL_957967, EPI_ISL_957968, EPI_ISL_957969, EPI_ISL_957970, EPI_ISL_957972, EPI_ISL_957973, EPI_ISL_957974, EPI_ISL_957975, EPI_ISL_957976, EPI_ISL_957977, EPI_ISL_957978, EPI_ISL_957979, EPI_ISL_957980, EPI_ISL_957981, EPI_ISL_957982, EPI_ISL_957983, EPI_ISL_957984, EPI_ISL_957985, EPI_ISL_957986, EPI_ISL_957987, EPI_ISL_957988, EPI_ISL_957989, EPI_ISL_957990, EPI_ISL_957991, EPI_ISL_957992, EPI_ISL_957993, EPI_ISL_957994, EPI_ISL_957995, EPI_ISL_957996, EPI_ISL_957997, EPI_ISL_957998, EPI_ISL_957999, EPI_ISL_958000, EPI_ISL_958001, EPI_ISL_958002, EPI_ISL_958003, EPI_ISL_958004, EPI_ISL_958005, EPI_ISL_958006, EPI_ISL_958007, EPI_ISL_958008, EPI_ISL_958009, EPI_ISL_958010, EPI_ISL_958011, EPI_ISL_958012, EPI_ISL_958013, EPI_ISL_958014, EPI_ISL_958015, EPI_ISL_958016, EPI_ISL_958017, EPI_ISL_958018, EPI_ISL_958019, EPI_ISL_958020, EPI_ISL_958021, EPI_ISL_958022, EPI_ISL_958023, EPI_ISL_958024, EPI_ISL_958025, EPI_ISL_958026, EPI_ISL_958027, EPI_ISL_958028, EPI_ISL_958029, EPI_ISL_958030, EPI_ISL_958031, EPI_ISL_958032, EPI_ISL_958033, EPI_ISL_958034, EPI_ISL_958035, EPI_ISL_958036, EPI_ISL_958037, EPI_ISL_958038, EPI_ISL_958039, EPI_ISL_958040, EPI_ISL_958041, EPI_ISL_958042, EPI_ISL_958043, EPI_ISL_958045, EPI_ISL_958046, EPI_ISL_958047, EPI_ISL_958048, EPI_ISL_958049, EPI_ISL_958050, EPI_ISL_958051, EPI_ISL_958052, EPI_ISL_958053, EPI_ISL_958054, EPI_ISL_958055, EPI_ISL_958056, EPI_ISL_958057, EPI_ISL_958058, EPI_ISL_958059, EPI_ISL_958060, EPI_ISL_958061, EPI_ISL_958062, EPI_ISL_958063, EPI_ISL_958064, EPI_ISL_958065, EPI_ISL_958066, EPI_ISL_958067, EPI_ISL_958068, EPI_ISL_958069, EPI_ISL_958070, EPI_ISL_958071, EPI_ISL_958072, EPI_ISL_958073, EPI_ISL_958074, EPI_ISL_958075, EPI_ISL_958076, EPI_ISL_958077, EPI_ISL_958078, EPI_ISL_958079, EPI_ISL_958080, EPI_ISL_958081, EPI_ISL_958082, EPI_ISL_958083, EPI_ISL_958084, EPI_ISL_958085, EPI_ISL_958086, EPI_ISL_958087, EPI_ISL_958088, EPI_ISL_958089, EPI_ISL_958090, EPI_ISL_958091, EPI_ISL_958092, EPI_ISL_958093, EPI_ISL_958094, EPI_ISL_958095, EPI_ISL_958096, EPI_ISL_958097, EPI_ISL_958098, EPI_ISL_958099, EPI_ISL_958100, EPI_ISL_958101, EPI_ISL_958102, EPI_ISL_958103, EPI_ISL_958104, EPI_ISL_958105, EPI_ISL_958106, EPI_ISL_958107, EPI_ISL_958108, EPI_ISL_958109, EPI_ISL_958110, EPI_ISL_958111, EPI_ISL_958112, EPI_ISL_958113, EPI_ISL_958114, EPI_ISL_958115, EPI_ISL_958116, EPI_ISL_958117, EPI_ISL_958118, EPI_ISL_958119, EPI_ISL_958120, EPI_ISL_958121, EPI_ISL_958122, EPI_ISL_958123, EPI_ISL_958124, EPI_ISL_958125, EPI_ISL_958126, EPI_ISL_958127, EPI_ISL_958128, EPI_ISL_958129, EPI_ISL_958130, EPI_ISL_958131, EPI_ISL_958132, EPI_ISL_958133, EPI_ISL_958134, EPI_ISL_958135, EPI_ISL_958136, EPI_ISL_958137, EPI_ISL_958138, EPI_ISL_958139, EPI_ISL_958140, EPI_ISL_958141, EPI_ISL_958142, EPI_ISL_958143, EPI_ISL_958144, EPI_ISL_958145, EPI_ISL_958146, EPI_ISL_958147, EPI_ISL_958148, EPI_ISL_958149, EPI_ISL_958150, EPI_ISL_958151, EPI_ISL_958152, EPI_ISL_958153, EPI_ISL_958154, EPI_ISL_958155, EPI_ISL_958156, EPI_ISL_958157, EPI_ISL_958158, EPI_ISL_958159, EPI_ISL_958160, EPI_ISL_958161, EPI_ISL_958162, EPI_ISL_958163, EPI_ISL_958164, EPI_ISL_958165, EPI_ISL_958166, EPI_ISL_958167, EPI_ISL_958168, EPI_ISL_958169, EPI_ISL_958170, EPI_ISL_958171, EPI_ISL_958172, EPI_ISL_958173, EPI_ISL_958174, EPI_ISL_958175, EPI_ISL_958176, EPI_ISL_958177, EPI_ISL_958178, EPI_ISL_958179, EPI_ISL_958180, EPI_ISL_958181, EPI_ISL_958182, EPI_ISL_958183, EPI_ISL_958184, EPI_ISL_958185, EPI_ISL_958186, EPI_ISL_958187, EPI_ISL_958188, EPI_ISL_958189, EPI_ISL_958190, EPI_ISL_958191, EPI_ISL_958192, EPI_ISL_958193, EPI_ISL_958195, EPI_ISL_958196, EPI_ISL_958197, EPI_ISL_958198, EPI_ISL_958199, EPI_ISL_958200, EPI_ISL_958201, EPI_ISL_958202, EPI_ISL_958203, EPI_ISL_958204, EPI_ISL_958205, EPI_ISL_958207, EPI_ISL_958208, EPI_ISL_958209, EPI_ISL_958210, EPI_ISL_958211, EPI_ISL_958212, EPI_ISL_958213, EPI_ISL_958214, EPI_ISL_958567, EPI_ISL_958568, EPI_ISL_958570, EPI_ISL_958571, EPI_ISL_958573, EPI_ISL_958574, EPI_ISL_958575, EPI_ISL_958576, EPI_ISL_958577, EPI_ISL_958580, EPI_ISL_958581, EPI_ISL_958582, EPI_ISL_958583, EPI_ISL_958584, EPI_ISL_958585, EPI_ISL_958588, EPI_ISL_958589, EPI_ISL_958592, EPI_ISL_958593, EPI_ISL_958594, EPI_ISL_958596, EPI_ISL_958597, EPI_ISL_958598, EPI_ISL_958599, EPI_ISL_958601, EPI_ISL_958603, EPI_ISL_958604, EPI_ISL_958605, EPI_ISL_958606, EPI_ISL_958607, EPI_ISL_958608, EPI_ISL_958609, EPI_ISL_958611, EPI_ISL_958612, EPI_ISL_958613, EPI_ISL_958614, EPI_ISL_958615, EPI_ISL_958616, EPI_ISL_958617, EPI_ISL_958618, EPI_ISL_958619, EPI_ISL_958620, EPI_ISL_958621, EPI_ISL_958622, EPI_ISL_958623, EPI_ISL_958625, EPI_ISL_958626, EPI_ISL_958627, EPI_ISL_958628, EPI_ISL_958629, EPI_ISL_958630, EPI_ISL_958631, EPI_ISL_958635, EPI_ISL_958637, EPI_ISL_958640, EPI_ISL_958642, EPI_ISL_958643, EPI_ISL_958644, EPI_ISL_958645, EPI_ISL_958647, EPI_ISL_958648, EPI_ISL_958649, EPI_ISL_958651, EPI_ISL_958652, EPI_ISL_958653, EPI_ISL_958654, EPI_ISL_958655, EPI_ISL_958656, EPI_ISL_958657, EPI_ISL_958658, EPI_ISL_958660, EPI_ISL_958661, EPI_ISL_958664, EPI_ISL_958665, EPI_ISL_958666, EPI_ISL_958667, EPI_ISL_958668, EPI_ISL_958669, EPI_ISL_958672, EPI_ISL_958674, EPI_ISL_958675, EPI_ISL_958676, EPI_ISL_958677, EPI_ISL_958678, EPI_ISL_958679, EPI_ISL_958681, EPI_ISL_958682, EPI_ISL_958683, EPI_ISL_958685, EPI_ISL_958686, EPI_ISL_958687, EPI_ISL_958688, EPI_ISL_958689, EPI_ISL_958692, EPI_ISL_958693, EPI_ISL_958694, EPI_ISL_958695, EPI_ISL_958697, EPI_ISL_958698, EPI_ISL_958699, EPI_ISL_958700, EPI_ISL_958701, EPI_ISL_958702, EPI_ISL_958703, EPI_ISL_958704, EPI_ISL_958707, EPI_ISL_958708, EPI_ISL_958709, EPI_ISL_958710, EPI_ISL_958711, EPI_ISL_958712, EPI_ISL_958714, EPI_ISL_958717, EPI_ISL_958718, EPI_ISL_958719, EPI_ISL_958720, EPI_ISL_958721, EPI_ISL_958722, EPI_ISL_958724, EPI_ISL_958727, EPI_ISL_958728, EPI_ISL_958729, EPI_ISL_958730, EPI_ISL_958732, EPI_ISL_958734, EPI_ISL_958736, EPI_ISL_958737, EPI_ISL_958738, EPI_ISL_958739, EPI_ISL_958740, EPI_ISL_958741, EPI_ISL_958742, EPI_ISL_958743, EPI_ISL_958744, EPI_ISL_958746, EPI_ISL_958749, EPI_ISL_958750, EPI_ISL_958752, EPI_ISL_958753, EPI_ISL_958755, EPI_ISL_958757, EPI_ISL_958758, EPI_ISL_958759, EPI_ISL_958761, EPI_ISL_958762, EPI_ISL_958764, EPI_ISL_958765, EPI_ISL_958766, EPI_ISL_958768, EPI_ISL_958770, EPI_ISL_958771, EPI_ISL_958772, EPI_ISL_958773, EPI_ISL_958774, EPI_ISL_958776, EPI_ISL_958777, EPI_ISL_958778, EPI_ISL_958780, EPI_ISL_958781, EPI_ISL_958782, EPI_ISL_958783, EPI_ISL_958785, EPI_ISL_958786, EPI_ISL_958788, EPI_ISL_958790, EPI_ISL_958794, EPI_ISL_958795, EPI_ISL_958796, EPI_ISL_958797, EPI_ISL_958798, EPI_ISL_958799, EPI_ISL_958800, EPI_ISL_958801, EPI_ISL_958802, EPI_ISL_958804, EPI_ISL_958805, EPI_ISL_958806, EPI_ISL_958809, EPI_ISL_958810, EPI_ISL_958812, EPI_ISL_958813, EPI_ISL_958814, EPI_ISL_958816, EPI_ISL_958817, EPI_ISL_958818, EPI_ISL_958819, EPI_ISL_958820, EPI_ISL_958821, EPI_ISL_958822, EPI_ISL_958824, EPI_ISL_958825, EPI_ISL_958827, EPI_ISL_958829, EPI_ISL_958830, EPI_ISL_958831, EPI_ISL_958832, EPI_ISL_958834, EPI_ISL_958835, EPI_ISL_958837, EPI_ISL_958838, EPI_ISL_958839, EPI_ISL_958841, EPI_ISL_958843, EPI_ISL_958844, EPI_ISL_958845, EPI_ISL_958846, EPI_ISL_958847, EPI_ISL_958848, EPI_ISL_958850, EPI_ISL_958852, EPI_ISL_958853, EPI_ISL_958854, EPI_ISL_958855, EPI_ISL_958856, EPI_ISL_958857, EPI_ISL_958858, EPI_ISL_958859, EPI_ISL_958860, EPI_ISL_958861, EPI_ISL_958862, EPI_ISL_958864, EPI_ISL_958865, EPI_ISL_958867, EPI_ISL_958868, EPI_ISL_958869, EPI_ISL_958870, EPI_ISL_958871, EPI_ISL_958872, EPI_ISL_958874, EPI_ISL_958875, EPI_ISL_958876, EPI_ISL_958877, EPI_ISL_958879, EPI_ISL_958880, EPI_ISL_958881, EPI_ISL_958882, EPI_ISL_958883, EPI_ISL_958886, EPI_ISL_958888, EPI_ISL_958890, EPI_ISL_958892, EPI_ISL_958894, EPI_ISL_958896, EPI_ISL_958897, EPI_ISL_958899, EPI_ISL_958900, EPI_ISL_958901, EPI_ISL_958905, EPI_ISL_958906, EPI_ISL_958907, EPI_ISL_958909, EPI_ISL_958910, EPI_ISL_958913, EPI_ISL_958914

| | | | |
|---|---|---|---|
| see above | Lighthouse Lab in Milton Keynes | Wellcome Sanger Institute for the COVID-19 Genomics UK (COG-UK) Consortium | The Lighthouse Lab in Milton Keynes and Alex Alderton, Roberto Amato, Sonia Goncalves, Ewan Harrison, David K. Jackson, Ian Johnston, Dominic Kwiatkowski, Cordelia Langford, John Sillitoe on behalf of the Wellcome Sanger Institute COVID-19 Surveillance Team |
| EPI_ISL_959419, EPI_ISL_959432, EPI_ISL_959433, EPI_ISL_959434, | Servicio de Microbiología, Hospital Universitario Son Espases | SeqCOVID-SPAIN consortium/IBV(CSIC) | Carla López-Causapé, Jordi Reina, Antonio Oliver and SeqCOVID-SPAIN consortium |

| | | | |
|---|---|---|---|
| EPI_ISL_959442, EPI_ISL_959447, EPI_ISL_959448, EPI_ISL_959449 | | | |
| EPI_ISL_959604, EPI_ISL_959605, EPI_ISL_959606 | Institute of Virology, Biomedical Research Center of the Slovak Academy of Sciences, Bratislava | Faculty of Natural Sciences, Comenius University, Bratislava | Broa Brejová, Viktória abanová, Kristína Boršová, Viktória Hodorová, Sabina Fumaová Havlíková, Juraj Kopáek, Martina Liková, ubomíra Lukáiková, Martina Neboháová, Monika Sláviková, Tomáš Vina, Jozef Nosek, Boris Klempa |
| EPI_ISL_959607 | Institute of Virology, Biomedical Research Center of the Slovak Academy of Sciences, Bratislava | Faculty of Natural Sciences, Comenius University, Bratislava | Viktória abanová, Kristína Boršová, Broa Brejová, Viktória Hodorová, Sabina Fumaová Havlíková, Juraj Kopáek, Martina Liková, ubomíra Lukáiková, Martina Neboháová, Monika Sláviková, Tomáš Vina, Jozef Nosek, Boris Klempa |
| EPI_ISL_959608 | Institute of Virology, Biomedical Research Center of the Slovak Academy of Sciences, Bratislava | Faculty of Natural Sciences, Comenius University, Bratislava | Kristína Boršová, Viktória abanová, Broa Brejová, Viktória Hodorová, Sabina Fumaová Havlíková, Juraj Kopáek, Martina Liková, ubomíra Lukáiková, Martina Neboháová, Monika Sláviková, Tomáš Vina, Boris Klempa, Jozef Nosek |
| EPI_ISL_959609 | Institute of Virology, Biomedical Research Center of the Slovak Academy of Sciences, Bratislava | Faculty of Natural Sciences, Comenius University, Bratislava | Viktória abanová, Kristína Boršová, Broa Brejová, Viktória Hodorová, Sabina Fumaová Havlíková, Juraj Kopáek, Martina Liková, ubomíra Lukáiková, Martina Neboháová, Monika Sláviková, Tomáš Vina, Jozef Nosek, Boris Klempa |
| EPI_ISL_959610 | Institute of Virology, Biomedical Research Center of the Slovak Academy of Sciences, Bratislava | Faculty of Natural Sciences, Comenius University, Bratislava | Kristína Boršová, Viktória abanová, Broa Brejová, Viktória Hodorová, Sabina Fumaová Havlíková, Juraj Kopáek, Martina Liková, ubomíra Lukáiková, Martina Neboháová, Monika Sláviková, Tomáš Vina, Boris Klempa, Jozef Nosek |
| EPI_ISL_959611 | Institute of Virology, Biomedical Research Center of the Slovak Academy of Sciences, Bratislava | Faculty of Natural Sciences, Comenius University, Bratislava | Viktória abanová, Kristína Boršová, Broa Brejová, Viktória Hodorová, Sabina Fumaová Havlíková, Juraj Kopáek, Martina Liková, ubomíra Lukáiková, Martina Neboháová, Monika Sláviková, Tomáš Vina, Jozef Nosek, Boris Klempa |
| EPI_ISL_959612 | Institute of Virology, Biomedical Research Center of the Slovak Academy of Sciences, Bratislava | Faculty of Natural Sciences, Comenius University, Bratislava | Kristína Boršová, Viktória abanová, Broa Brejová, Viktória Hodorová, Sabina Fumaová Havlíková, Juraj Kopáek, Martina Liková, ubomíra Lukáiková, Martina Neboháová, Monika Sláviková, Tomáš Vina, Boris Klempa, Jozef Nosek |
| EPI_ISL_959613 | Institute of Virology, Biomedical Research Center of the Slovak Academy of Sciences, Bratislava | Faculty of Natural Sciences, Comenius University, Bratislava | Viktória abanová, Kristína Boršová, Broa Brejová, Viktória Hodorová, Sabina Fumaová Havlíková, Juraj Kopáek, Martina Liková, ubomíra Lukáiková, Martina Neboháová, Monika Sláviková, Tomáš Vina, Jozef Nosek, Boris Klempa |
| EPI_ISL_959614 | Institute of Virology, Biomedical Research Center of the Slovak Academy of Sciences, Bratislava | Faculty of Natural Sciences, Comenius University, Bratislava | Kristína Boršová, Viktória abanová, Broa Brejová, Viktória Hodorová, Sabina Fumaová Havlíková, Juraj Kopáek, Martina Liková, ubomíra Lukáiková, Martina Neboháová, Monika Sláviková, Tomáš Vina, Boris Klempa, Jozef Nosek |
| EPI_ISL_959615, EPI_ISL_959616 | Institute of Virology, Biomedical Research Center of the Slovak Academy of Sciences, Bratislava | Faculty of Natural Sciences, Comenius University, Bratislava | Broa Brejová, Viktória abanová, Kristína Boršová, Viktória Hodorová, Sabina Fumaová Havlíková, Juraj Kopáek, Martina Liková, ubomíra Lukáiková, Martina Neboháová, Monika Sláviková, Tomáš Vina, Jozef Nosek, Boris Klempa |
| EPI_ISL_959617 | Institute of Virology, Biomedical Research Center of the Slovak Academy of Sciences, Bratislava | Faculty of Natural Sciences, Comenius University, Bratislava | Viktória abanová, Kristína Boršová, Broa Brejová, Viktória Hodorová, Sabina Fumaová Havlíková, Juraj Kopáek, Martina Liková, ubomíra Lukáiková, Martina Neboháová, Monika Sláviková, Tomáš Vina, Jozef Nosek, Boris Klempa |
| EPI_ISL_959618 | Institute of Virology, Biomedical Research Center of the Slovak Academy of Sciences, Bratislava | Faculty of Natural Sciences, Comenius University, Bratislava | Kristína Boršová, Viktória abanová, Broa Brejová, Viktória Hodorová, Sabina Fumaová Havlíková, Juraj Kopáek, Martina Liková, ubomíra Lukáiková, Martina Neboháová, Monika Sláviková, Tomáš Vina, Boris Klempa, Jozef Nosek |
| EPI_ISL_959619 | Institute of Virology, Biomedical Research Center of the Slovak Academy of Sciences, Bratislava | Faculty of Natural Sciences, Comenius University, Bratislava | Broa Brejová, Viktória abanová, Kristína Boršová, Viktória Hodorová, Sabina Fumaová Havlíková, Juraj Kopáek, Martina Liková, ubomíra Lukáiková, Martina Neboháová, Monika Sláviková, Tomáš Vina, Jozef Nosek, Boris Klempa |
| EPI_ISL_959620 | Institute of Virology, Biomedical Research Center of the Slovak Academy of Sciences, Bratislava | Faculty of Natural Sciences, Comenius University, Bratislava | Viktória abanová, Kristína Boršová, Broa Brejová, Viktória Hodorová, Sabina Fumaová Havlíková, Juraj Kopáek, Martina Liková, ubomíra Lukáiková, Martina Neboháová, Monika Sláviková, Tomáš Vina, Jozef Nosek, Boris Klempa |
| EPI_ISL_959626 | Institute of Virology, Biomedical Research Center of the Slovak Academy of Sciences, Bratislava | Faculty of Natural Sciences, Comenius University, Bratislava | Kristína Boršová, Viktória abanová, Broa Brejová, Viktória Hodorová, Sabina Fumaová Havlíková, Juraj Kopáek, Martina Liková, ubomíra Lukáiková, Martina Neboháová, Monika Sláviková, Tomáš Vina, Boris Klempa, Jozef Nosek |
| EPI_ISL_959627, EPI_ISL_959628 | Institute of Virology, Biomedical Research Center of the Slovak Academy of Sciences, Bratislava | Faculty of Natural Sciences, Comenius University, Bratislava | Viktória abanová, Kristína Boršová, Broa Brejová, Viktória Hodorová, Sabina Fumaová Havlíková, Juraj Kopáek, Martina Liková, ubomíra Lukáiková, Martina Neboháová, Monika Sláviková, Tomáš Vina, Jozef Nosek, Boris Klempa |
| EPI_ISL_959629 | Institute of Virology, Biomedical Research Center of the Slovak Academy of Sciences, Bratislava | Faculty of Natural Sciences, Comenius University, Bratislava | Broa Brejová, Viktória abanová, Kristína Boršová, Viktória Hodorová, Sabina Fumaová Havlíková, Juraj Kopáek, Martina Liková, ubomíra Lukáiková, Martina Neboháová, Monika Sláviková, Tomáš Vina, Jozef Nosek, Boris Klempa |
| EPI_ISL_959630 | Institute of Virology, Biomedical Research Center of the Slovak Academy of Sciences, Bratislava | Faculty of Natural Sciences, Comenius University, Bratislava | Kristína Boršová, Viktória abanová, Broa Brejová, Viktória Hodorová, Sabina Fumaová Havlíková, Juraj Kopáek, Martina Liková, ubomíra Lukáiková, Martina Neboháová, Monika Sláviková, Tomáš Vina, Boris Klempa, Jozef Nosek |
| EPI_ISL_959631 | Institute of Virology, Biomedical Research Center of the Slovak Academy of Sciences, Bratislava | Faculty of Natural Sciences, Comenius University, Bratislava | Broa Brejová, Viktória abanová, Kristína Boršová, Viktória Hodorová, Sabina Fumaová Havlíková, Juraj Kopáek, Martina Liková, ubomíra Lukáiková, Martina Neboháová, Monika Sláviková, Tomáš Vina, Jozef Nosek, Boris Klempa |
| EPI_ISL_959701, EPI_ISL_959702, EPI_ISL_959703, EPI_ISL_959704, EPI_ISL_959705, EPI_ISL_959706, EPI_ISL_959707, EPI_ISL_959708, EPI_ISL_959709, EPI_ISL_959710, EPI_ISL_959711, EPI_ISL_959712, EPI_ISL_959852, EPI_ISL_959853, EPI_ISL_959860, EPI_ISL_959861 | | | |
| see above | National Virus Reference Laboratory | National Virus Reference Laboratory | Michael Carr, Gabriel Gonzalez, Jonathan Dean, Cillian F De Gascun |
| EPI_ISL_960333, EPI_ISL_960385, EPI_ISL_960403 | University of Wisconsin-Madison AIDS Vaccine Research Laboratories | University of Wisconsin-Madison AIDS Vaccine Research Laboratories | Gage Moreno, Katarina Braun, et al. AIDS Vaccine Research Laboratories |
| EPI_ISL_960893 | Institute of Medical Microbiology and Hospital Hygiene | Institute of Medical Microbiology and Hospital Hygiene | Prof. Dr. Achim Kaasch, Aljoscha Tersteegen |
| EPI_ISL_961747 | Dipartimento Prevenzione Avezzano-Servizio Igiene epidemiologia Sanità Pubblica | Istituto Zooprofilattico Sperimentale dell'Abruzzo e Molise "G. Caporale" | Lorusso A, Marcacci M, Di Domenico M, Ancora M, Curini V, Mangone I, Rinaldi A, Scialabba S, Di Pasquale A, Cammà C, Puglia I, Calistri P, Savini G |
| EPI_ISL_962445 | Seattle Flu Study | Seattle Flu Study | Deborah A. Nickerson, Chris D. Frazar, Jover Lee, Benjamin Pelle, Erica Ryke, Matthew Richardson, Amanda Adler, Elisabeth Brandstetter, Peter D. Han, Kairsten Fay, Misja Ilcisin, Kirsten Lacombe, Thomas R. Sibley, Melissa Truong, Caitlin R. Wolf, Michael Boeckh, Janet A. Englund, Michael Famulare, Barry R. Lutz, Mark J. Rieder, Lea M. Starita, Matthew Thompson, Jay Shendure, Trevor Bedford, Helen Y. Chu |
| EPI_ISL_962449, EPI_ISL_962450, EPI_ISL_962451 | Seattle Flu Study | Seattle Flu Study | Deborah A. Nickerson, Chris D. Frazar, Jover Lee, Benjamin Pelle, Erica Ryke, Matthew Richardson, Amanda Adler, Elisabeth Brandstetter, Peter D. Han, Kairsten Fay, Misja Ilcisin, Kirsten Lacombe, Thomas R. Sibley, Melissa Truong, Caitlin R. Wolf, Karen Cowgill, Stephanie Schrag, Jeff Duchin, Michael Boeckh, Janet A. Englund, Michael Famulare, Barry R. Lutz, Mark J. Rieder, Lea M. Starita, Matthew Thompson, Helen Y. Chu, Trevor Bedford, Jay Shendure |
| EPI_ISL_962458 | Seattle Flu Study | Seattle Flu Study | Deborah A. Nickerson, Chris D. Frazar, Jover Lee, Benjamin Pelle, Erica Ryke, Matthew Richardson, Amanda Adler, Elisabeth Brandstetter, Peter D. Han, Kairsten Fay, Misja Ilcisin, Kirsten Lacombe, Thomas R. Sibley, Melissa Truong, Caitlin R. Wolf, Michael Boeckh, Janet A. Englund, Michael Famulare, Barry R. Lutz, Mark J. Rieder, Lea M. Starita, Matthew Thompson, Jay Shendure, Trevor Bedford, Helen Y. Chu |
| EPI_ISL_962474, EPI_ISL_962475, EPI_ISL_962476 | Seattle Flu Study | Seattle Flu Study | Deborah A. Nickerson, Chris D. Frazar, Jover Lee, Benjamin Pelle, Erica Ryke, Matthew Richardson, Amanda Adler, Elisabeth Brandstetter, Peter D. Han, Kairsten Fay, Misja Ilcisin, Kirsten Lacombe, Thomas R. Sibley, Melissa Truong, Caitlin R. Wolf, Karen Cowgill, Stephanie Schrag, Jeff Duchin, Michael Boeckh, Janet A. Englund, Michael Famulare, Barry R. Lutz, Mark J. Rieder, Lea M. Starita, Matthew Thompson, Helen Y. Chu, Trevor Bedford, Jay Shendure |
| EPI_ISL_962477 | Seattle Flu Study | Seattle Flu Study | Deborah A. Nickerson, Chris D. Frazar, Jover Lee, Benjamin Pelle, Erica Ryke, Matthew Richardson, Amanda Adler, Elisabeth Brandstetter, Peter D. Han, Kairsten Fay, Misja Ilcisin, Kirsten Lacombe, Thomas R. Sibley, Melissa Truong, Caitlin R. Wolf, Michael Boeckh, Janet A. Englund, Michael Famulare, Barry R. Lutz, Mark J. Rieder, Lea M. Starita, Matthew Thompson, Jay Shendure, Trevor Bedford, Helen Y. Chu |
| EPI_ISL_962478, EPI_ISL_962479, EPI_ISL_962480 | Seattle Flu Study | Seattle Flu Study | Deborah A. Nickerson, Chris D. Frazar, Jover Lee, Benjamin Pelle, Erica Ryke, Matthew Richardson, Amanda Adler, Elisabeth Brandstetter, Peter D. Han, Kairsten Fay, Misja Ilcisin, Kirsten Lacombe, Thomas R. Sibley, Melissa Truong, Caitlin R. Wolf, Karen Cowgill, Stephanie Schrag, Jeff Duchin, Michael Boeckh, Janet A. Englund, Michael Famulare, Barry R. Lutz, Mark J. Rieder, Lea M. Starita, Matthew Thompson, Helen Y. Chu, Trevor Bedford, Jay Shendure |
| EPI_ISL_962848, EPI_ISL_962849, EPI_ISL_962850, EPI_ISL_962851 | Los Angeles County PHL | Los Angeles County PHL | P. Hemarajata et al. |
| EPI_ISL_962975, EPI_ISL_962976, EPI_ISL_962977, EPI_ISL_962978, EPI_ISL_962979, EPI_ISL_962980, EPI_ISL_962981, EPI_ISL_962982, EPI_ISL_962983, EPI_ISL_962984, EPI_ISL_962985, EPI_ISL_962986, EPI_ISL_962987, EPI_ISL_962988, EPI_ISL_962989, EPI_ISL_962990, EPI_ISL_962991, EPI_ISL_962992, EPI_ISL_962993, EPI_ISL_962994, EPI_ISL_962995, EPI_ISL_962996, EPI_ISL_962997, EPI_ISL_962998, EPI_ISL_962999, EPI_ISL_963000, EPI_ISL_963001, EPI_ISL_963002, EPI_ISL_963003, EPI_ISL_963004, EPI_ISL_963005, EPI_ISL_963006, EPI_ISL_963007, EPI_ISL_963008, EPI_ISL_963009, EPI_ISL_963010, | | | |

EPI_ISL_963011, EPI_ISL_963012, EPI_ISL_963013, EPI_ISL_963014, EPI_ISL_963015, EPI_ISL_963016, EPI_ISL_963017, EPI_ISL_963018, EPI_ISL_963019, EPI_ISL_963020, EPI_ISL_963021, EPI_ISL_963022, EPI_ISL_963023, EPI_ISL_963024, EPI_ISL_963025, EPI_ISL_963026, EPI_ISL_963027, EPI_ISL_963028, EPI_ISL_963029, EPI_ISL_963030, EPI_ISL_963031, EPI_ISL_963032, EPI_ISL_963033, EPI_ISL_963034, EPI_ISL_963035, EPI_ISL_963036, EPI_ISL_963037, EPI_ISL_963038, EPI_ISL_963039, EPI_ISL_963040, EPI_ISL_963041, EPI_ISL_963042, EPI_ISL_963043, EPI_ISL_963044, EPI_ISL_963045, EPI_ISL_963046, EPI_ISL_963047, EPI_ISL_963048, EPI_ISL_963049, EPI_ISL_963050, EPI_ISL_963051, EPI_ISL_963052, EPI_ISL_963053, EPI_ISL_963054, EPI_ISL_963055, EPI_ISL_963056, EPI_ISL_963057, EPI_ISL_963058, EPI_ISL_963059, EPI_ISL_963060, EPI_ISL_963062, EPI_ISL_963063, EPI_ISL_963064, EPI_ISL_963065, EPI_ISL_963066, EPI_ISL_963067, EPI_ISL_963068, EPI_ISL_963069, EPI_ISL_963070, EPI_ISL_963071, EPI_ISL_963072, EPI_ISL_963073, EPI_ISL_963074, EPI_ISL_963075, EPI_ISL_963076, EPI_ISL_963077, EPI_ISL_963078, EPI_ISL_963079, EPI_ISL_963080, EPI_ISL_963081, EPI_ISL_963082, EPI_ISL_963083, EPI_ISL_963084, EPI_ISL_963085, EPI_ISL_963086, EPI_ISL_963087, EPI_ISL_963088, EPI_ISL_963089, EPI_ISL_963090, EPI_ISL_963091, EPI_ISL_963092, EPI_ISL_963093, EPI_ISL_963094, EPI_ISL_963095, EPI_ISL_963096, EPI_ISL_963097, EPI_ISL_963098, EPI_ISL_963099, EPI_ISL_963100, EPI_ISL_963101, EPI_ISL_963102, EPI_ISL_963103, EPI_ISL_963104, EPI_ISL_963105, EPI_ISL_963107, EPI_ISL_963108, EPI_ISL_963109, EPI_ISL_963110, EPI_ISL_963111, EPI_ISL_963112, EPI_ISL_963113, EPI_ISL_963114, EPI_ISL_963115, EPI_ISL_963116, EPI_ISL_963117, EPI_ISL_963118, EPI_ISL_963119, EPI_ISL_963120, EPI_ISL_963121, EPI_ISL_963122, EPI_ISL_963123, EPI_ISL_963124, EPI_ISL_963125, EPI_ISL_963126, EPI_ISL_963127, EPI_ISL_963128, EPI_ISL_963129, EPI_ISL_963130, EPI_ISL_963131, EPI_ISL_963132, EPI_ISL_963133, EPI_ISL_963134, EPI_ISL_963135, EPI_ISL_963136, EPI_ISL_963137, EPI_ISL_963138, EPI_ISL_963139, EPI_ISL_963140, EPI_ISL_963141, EPI_ISL_963142, EPI_ISL_963143, EPI_ISL_963144, EPI_ISL_963145, EPI_ISL_963146, EPI_ISL_963147, EPI_ISL_963148, EPI_ISL_963149, EPI_ISL_963150, EPI_ISL_963151, EPI_ISL_963152, EPI_ISL_963153, EPI_ISL_963154, EPI_ISL_963155, EPI_ISL_963156, EPI_ISL_963157, EPI_ISL_963158, EPI_ISL_963159, EPI_ISL_963160, EPI_ISL_963161, EPI_ISL_963162, EPI_ISL_963163, EPI_ISL_963164, EPI_ISL_963165, EPI_ISL_963166, EPI_ISL_963167, EPI_ISL_963168, EPI_ISL_963169, EPI_ISL_963170, EPI_ISL_963171, EPI_ISL_963172, EPI_ISL_963173, EPI_ISL_963174, EPI_ISL_963175, EPI_ISL_963176, EPI_ISL_963177, EPI_ISL_963178, EPI_ISL_963179, EPI_ISL_963180, EPI_ISL_963181, EPI_ISL_963182, EPI_ISL_963183, EPI_ISL_963184, EPI_ISL_963185, EPI_ISL_963186, EPI_ISL_963187, EPI_ISL_963188, EPI_ISL_963189, EPI_ISL_963190, EPI_ISL_963191, EPI_ISL_963192, EPI_ISL_963193, EPI_ISL_963194, EPI_ISL_963195, EPI_ISL_963196, EPI_ISL_963197, EPI_ISL_963198, EPI_ISL_963199, EPI_ISL_963200, EPI_ISL_963201, EPI_ISL_963202, EPI_ISL_963203, EPI_ISL_963204, EPI_ISL_963205, EPI_ISL_963206, EPI_ISL_963207, EPI_ISL_963208, EPI_ISL_963209, EPI_ISL_963210, EPI_ISL_963211, EPI_ISL_963212, EPI_ISL_963213, EPI_ISL_963214, EPI_ISL_963215, EPI_ISL_963216, EPI_ISL_963217, EPI_ISL_963218, EPI_ISL_963219, EPI_ISL_963220, EPI_ISL_963221, EPI_ISL_963222, EPI_ISL_963223, EPI_ISL_963224, EPI_ISL_963225, EPI_ISL_963226, EPI_ISL_963227, EPI_ISL_963228, EPI_ISL_963229, EPI_ISL_963230, EPI_ISL_963231, EPI_ISL_963232, EPI_ISL_963233, EPI_ISL_963234, EPI_ISL_963235, EPI_ISL_963236, EPI_ISL_963237, EPI_ISL_963238, EPI_ISL_963239, EPI_ISL_963240, EPI_ISL_963241, EPI_ISL_963242, EPI_ISL_963243, EPI_ISL_963244, EPI_ISL_963245, EPI_ISL_963246, EPI_ISL_963247, EPI_ISL_963248, EPI_ISL_963249, EPI_ISL_963250, EPI_ISL_963251, EPI_ISL_963252, EPI_ISL_963253, EPI_ISL_963254, EPI_ISL_963255, EPI_ISL_963256, EPI_ISL_963257, EPI_ISL_963258, EPI_ISL_963259, EPI_ISL_963260, EPI_ISL_963261, EPI_ISL_963262, EPI_ISL_963263, EPI_ISL_963264, EPI_ISL_963265, EPI_ISL_963266, EPI_ISL_963267, EPI_ISL_963268, EPI_ISL_963269, EPI_ISL_963270, EPI_ISL_963271, EPI_ISL_963272, EPI_ISL_963273, EPI_ISL_963274, EPI_ISL_963275, EPI_ISL_963276, EPI_ISL_963277, EPI_ISL_963278, EPI_ISL_963279, EPI_ISL_963280, EPI_ISL_963281, EPI_ISL_963282, EPI_ISL_963283, EPI_ISL_963284, EPI_ISL_963285, EPI_ISL_963286, EPI_ISL_963287, EPI_ISL_963288, EPI_ISL_963289, EPI_ISL_963290, EPI_ISL_963291, EPI_ISL_963292, EPI_ISL_963293, EPI_ISL_963294, EPI_ISL_963295, EPI_ISL_963297, EPI_ISL_963298, EPI_ISL_963299, EPI_ISL_963300, EPI_ISL_963301, EPI_ISL_963302, EPI_ISL_963303, EPI_ISL_963304, EPI_ISL_963305, EPI_ISL_963306, EPI_ISL_963307, EPI_ISL_963308, EPI_ISL_963309, EPI_ISL_963310, EPI_ISL_963311, EPI_ISL_963312, EPI_ISL_963313, EPI_ISL_963314, EPI_ISL_963315, EPI_ISL_963316, EPI_ISL_963317, EPI_ISL_963318, EPI_ISL_963319, EPI_ISL_963320, EPI_ISL_963322, EPI_ISL_963323

| | | | |
|---|---|---|---|
| see above | Lighthouse Lab in Alderley Park | Wellcome Sanger Institute for the COVID-19 Genomics UK (COG-UK) Consortium | Jacquelyn Wynn, Mairead Hyland, The Lighthouse Lab in Alderley Park and Alex Alderton, Roberto Amato, Sonia Goncalves, Ewan Harrison, David K. Jackson, Ian Johnston, Dominic Kwiatkowski, Cordelia Langford, John Sillitoe on behalf of the Wellcome Sanger Institute COVID-19 Surveillance Team |
| EPI_ISL_963324 | Lighthouse Lab in Milton Keynes | Wellcome Sanger Institute for the COVID-19 Genomics UK (COG-UK) Consortium | The Lighthouse Lab in Milton Keynes and Alex Alderton, Roberto Amato, Sonia Goncalves, Ewan Harrison, David K. Jackson, Ian Johnston, Dominic Kwiatkowski, Cordelia Langford, John Sillitoe on behalf of the Wellcome Sanger Institute COVID-19 Surveillance Team |
| EPI_ISL_963326 | Lighthouse Lab in Glasgow | Wellcome Sanger Institute for the COVID-19 Genomics UK (COG-UK) Consortium | Harper VanSteenhouse, Yumi Kasai, David Gray, Carol Clugston, Anna Dominiczak and Alex Alderton, Roberto Amato, Sonia Goncalves, Ewan Harrison, David K. Jackson, Ian Johnston, Dominic Kwiatkowski, Cordelia Langford, John Sillitoe on behalf of the Wellcome Sanger Institute COVID-19 Surveillance Team |
| EPI_ISL_963327, EPI_ISL_963328, EPI_ISL_963329 | Lighthouse Lab in Milton Keynes | Wellcome Sanger Institute for the COVID-19 Genomics UK (COG-UK) Consortium | The Lighthouse Lab in Milton Keynes and Alex Alderton, Roberto Amato, Sonia Goncalves, Ewan Harrison, David K. Jackson, Ian Johnston, Dominic Kwiatkowski, Cordelia Langford, John Sillitoe on behalf of the Wellcome Sanger Institute COVID-19 Surveillance Team |
| EPI_ISL_963330 | Lighthouse Lab in Glasgow | Wellcome Sanger Institute for the COVID-19 Genomics UK (COG-UK) Consortium | Harper VanSteenhouse, Yumi Kasai, David Gray, Carol Clugston, Anna Dominiczak and Alex Alderton, Roberto Amato, Sonia Goncalves, Ewan Harrison, David K. Jackson, Ian Johnston, Dominic Kwiatkowski, Cordelia Langford, John Sillitoe on behalf of the Wellcome Sanger Institute COVID-19 Surveillance Team |
| EPI_ISL_963331, EPI_ISL_963332, EPI_ISL_963333, EPI_ISL_963334, EPI_ISL_963335, EPI_ISL_963336, EPI_ISL_963337, EPI_ISL_963338, EPI_ISL_963339, EPI_ISL_963340 | Lighthouse Lab in Milton Keynes | Wellcome Sanger Institute for the COVID-19 Genomics UK (COG-UK) Consortium | The Lighthouse Lab in Milton Keynes and Alex Alderton, Roberto Amato, Sonia Goncalves, Ewan Harrison, David K. Jackson, Ian Johnston, Dominic Kwiatkowski, Cordelia Langford, John Sillitoe on behalf of the Wellcome Sanger Institute COVID-19 Surveillance Team |
| EPI_ISL_963341 | Lighthouse Lab in Glasgow | Wellcome Sanger Institute for the COVID-19 Genomics UK (COG-UK) Consortium | Harper VanSteenhouse, Yumi Kasai, David Gray, Carol Clugston, Anna Dominiczak and Alex Alderton, Roberto Amato, Sonia Goncalves, Ewan Harrison, David K. Jackson, Ian Johnston, Dominic Kwiatkowski, Cordelia Langford, John Sillitoe on behalf of the Wellcome Sanger Institute COVID-19 Surveillance Team |
| EPI_ISL_963342, EPI_ISL_963343 | Lighthouse Lab in Milton Keynes | Wellcome Sanger Institute for the COVID-19 Genomics UK (COG-UK) Consortium | The Lighthouse Lab in Milton Keynes and Alex Alderton, Roberto Amato, Sonia Goncalves, Ewan Harrison, David K. Jackson, Ian Johnston, Dominic Kwiatkowski, Cordelia Langford, John Sillitoe on behalf of the Wellcome Sanger Institute COVID-19 Surveillance Team |
| EPI_ISL_963344, EPI_ISL_963345, EPI_ISL_963346 | Lighthouse Lab in Glasgow | Wellcome Sanger Institute for the COVID-19 Genomics UK (COG-UK) Consortium | Harper VanSteenhouse, Yumi Kasai, David Gray, Carol Clugston, Anna Dominiczak and Alex Alderton, Roberto Amato, Sonia Goncalves, Ewan Harrison, David K. Jackson, Ian Johnston, Dominic Kwiatkowski, Cordelia Langford, John Sillitoe on behalf of the Wellcome Sanger Institute COVID-19 Surveillance Team |
| EPI_ISL_963347 | Lighthouse Lab in Milton Keynes | Wellcome Sanger Institute for the COVID-19 Genomics UK (COG-UK) Consortium | The Lighthouse Lab in Milton Keynes and Alex Alderton, Roberto Amato, Sonia Goncalves, Ewan Harrison, David K. Jackson, Ian Johnston, Dominic Kwiatkowski, Cordelia Langford, John Sillitoe on behalf of the Wellcome Sanger Institute COVID-19 Surveillance Team |
| EPI_ISL_963348 | Lighthouse Lab in Glasgow | Wellcome Sanger Institute for the COVID-19 Genomics UK (COG-UK) Consortium | Harper VanSteenhouse, Yumi Kasai, David Gray, Carol Clugston, Anna Dominiczak and Alex Alderton, Roberto Amato, Sonia Goncalves, Ewan Harrison, David K. Jackson, Ian Johnston, Dominic Kwiatkowski, Cordelia Langford, John Sillitoe on behalf of the Wellcome Sanger Institute COVID-19 Surveillance Team |
| EPI_ISL_963349, EPI_ISL_963350, EPI_ISL_963351, EPI_ISL_963352, EPI_ISL_963353, EPI_ISL_963354 | Lighthouse Lab in Milton Keynes | Wellcome Sanger Institute for the COVID-19 Genomics UK (COG-UK) Consortium | The Lighthouse Lab in Milton Keynes and Alex Alderton, Roberto Amato, Sonia Goncalves, Ewan Harrison, David K. Jackson, Ian Johnston, Dominic Kwiatkowski, Cordelia Langford, John Sillitoe on behalf of the Wellcome Sanger Institute COVID-19 Surveillance Team |
| EPI_ISL_963355, EPI_ISL_963356, EPI_ISL_963357, EPI_ISL_963358 | Lighthouse Lab in Glasgow | Wellcome Sanger Institute for the COVID-19 Genomics UK (COG-UK) Consortium | Harper VanSteenhouse, Yumi Kasai, David Gray, Carol Clugston, Anna Dominiczak and Alex Alderton, Roberto Amato, Sonia Goncalves, Ewan Harrison, David K. Jackson, Ian Johnston, Dominic Kwiatkowski, Cordelia Langford, John Sillitoe on behalf of the Wellcome Sanger Institute COVID-19 Surveillance Team |

EPI_ISL_963359, EPI_ISL_963360, EPI_ISL_963361, EPI_ISL_963362, EPI_ISL_963363, EPI_ISL_963364, EPI_ISL_963365, EPI_ISL_963366, EPI_ISL_963367, EPI_ISL_963368, EPI_ISL_963369

| | | | |
|---|---|---|---|
| see above | Lighthouse Lab in Milton Keynes | Wellcome Sanger Institute for the COVID-19 Genomics UK (COG-UK) Consortium | The Lighthouse Lab in Milton Keynes and Alex Alderton, Roberto Amato, Sonia Goncalves, Ewan Harrison, David K. Jackson, Ian Johnston, Dominic Kwiatkowski, Cordelia Langford, John Sillitoe on behalf of the Wellcome Sanger Institute COVID-19 Surveillance Team |
| EPI_ISL_963370, EPI_ISL_963371 | Lighthouse Lab in Glasgow | Wellcome Sanger Institute for the COVID-19 Genomics UK (COG-UK) Consortium | Harper VanSteenhouse, Yumi Kasai, David Gray, Carol Clugston, Anna Dominiczak and Alex Alderton, Roberto Amato, Sonia Goncalves, Ewan Harrison, David K. Jackson, Ian Johnston, Dominic Kwiatkowski, Cordelia Langford, John Sillitoe on behalf of the Wellcome Sanger Institute COVID-19 Surveillance Team |
| EPI_ISL_963372, EPI_ISL_963373, EPI_ISL_963374, EPI_ISL_963375 | Lighthouse Lab in Milton Keynes | Wellcome Sanger Institute for the COVID-19 Genomics UK (COG-UK) Consortium | The Lighthouse Lab in Milton Keynes and Alex Alderton, Roberto Amato, Sonia Goncalves, Ewan Harrison, David K. Jackson, Ian Johnston, Dominic Kwiatkowski, Cordelia Langford, John Sillitoe on behalf of the Wellcome Sanger Institute COVID-19 Surveillance Team |
| EPI_ISL_963376 | Lighthouse Lab in Glasgow | Wellcome Sanger Institute for the COVID-19 Genomics UK (COG-UK) Consortium | Harper VanSteenhouse, Yumi Kasai, David Gray, Carol Clugston, Anna Dominiczak and Alex Alderton, Roberto Amato, Sonia Goncalves, Ewan Harrison, David K. Jackson, Ian Johnston, Dominic Kwiatkowski, Cordelia Langford, John Sillitoe on behalf of the Wellcome Sanger Institute COVID-19 Surveillance Team |
| EPI_ISL_963377, EPI_ISL_963378, EPI_ISL_963379 | Lighthouse Lab in Milton Keynes | Wellcome Sanger Institute for the COVID-19 Genomics UK (COG-UK) Consortium | The Lighthouse Lab in Milton Keynes and Alex Alderton, Roberto Amato, Sonia Goncalves, Ewan Harrison, David K. Jackson, Ian Johnston, Dominic Kwiatkowski, Cordelia Langford, John Sillitoe on behalf of the Wellcome Sanger Institute COVID-19 Surveillance Team |
| EPI_ISL_963380 | Lighthouse Lab in Glasgow | Wellcome Sanger Institute for the COVID-19 Genomics UK (COG-UK) Consortium | Harper VanSteenhouse, Yumi Kasai, David Gray, Carol Clugston, Anna Dominiczak and Alex Alderton, Roberto Amato, Sonia Goncalves, Ewan Harrison, David K. Jackson, Ian Johnston, Dominic Kwiatkowski, Cordelia Langford, John Sillitoe on behalf of the Wellcome Sanger Institute COVID-19 Surveillance Team |
| EPI_ISL_963381, EPI_ISL_963382, EPI_ISL_963383, EPI_ISL_963384, EPI_ISL_963385 | Lighthouse Lab in Milton Keynes | Wellcome Sanger Institute for the COVID-19 Genomics UK (COG-UK) Consortium | The Lighthouse Lab in Milton Keynes and Alex Alderton, Roberto Amato, Sonia Goncalves, Ewan Harrison, David K. Jackson, Ian Johnston, Dominic Kwiatkowski, Cordelia Langford, John Sillitoe on behalf of the Wellcome Sanger Institute COVID-19 Surveillance Team |
| EPI_ISL_963386 | Lighthouse Lab in Glasgow | Wellcome Sanger Institute for the COVID-19 Genomics UK (COG-UK) Consortium | Harper VanSteenhouse, Yumi Kasai, David Gray, Carol Clugston, Anna Dominiczak and Alex Alderton, Roberto Amato, Sonia Goncalves, Ewan Harrison, David K. Jackson, Ian Johnston, Dominic Kwiatkowski, Cordelia Langford, John Sillitoe on behalf of the Wellcome Sanger Institute COVID-19 Surveillance Team |

| | | | |
|---|---|---|---|
| EPI_ISL_963387, EPI_ISL_963388, EPI_ISL_963389, EPI_ISL_963390, EPI_ISL_963391, EPI_ISL_963392, EPI_ISL_963393, EPI_ISL_963394, EPI_ISL_963395, EPI_ISL_963396 | Lighthouse Lab in Milton Keynes | Wellcome Sanger Institute for the COVID-19 Genomics UK (COG-UK) Consortium | The Lighthouse Lab in Milton Keynes and Alex Alderton, Roberto Amato, Sonia Goncalves, Ewan Harrison, David K. Jackson, Ian Johnston, Dominic Kwiatkowski, Cordelia Langford, John Sillitoe on behalf of the Wellcome Sanger Institute COVID-19 Surveillance Team |
| EPI_ISL_963397 | Lighthouse Lab in Glasgow | Wellcome Sanger Institute for the COVID-19 Genomics UK (COG-UK) Consortium | Harper VanSteenhouse, Yumi Kasai, David Gray, Carol Clugston, Anna Dominiczak and Alex Alderton, Roberto Amato, Sonia Goncalves, Ewan Harrison, David K. Jackson, Ian Johnston, Dominic Kwiatkowski, Cordelia Langford, John Sillitoe on behalf of the Wellcome Sanger Institute COVID-19 Surveillance Team |
| EPI_ISL_963398, EPI_ISL_963399, EPI_ISL_963400, EPI_ISL_963401, EPI_ISL_963402, EPI_ISL_963403, EPI_ISL_963404, EPI_ISL_963405, EPI_ISL_963406, EPI_ISL_963407, EPI_ISL_963409, EPI_ISL_963410, EPI_ISL_963411, EPI_ISL_963412, EPI_ISL_963413, EPI_ISL_963414, EPI_ISL_963415, EPI_ISL_963416, EPI_ISL_963417, EPI_ISL_963418 | | | |
| see above | Lighthouse Lab in Milton Keynes | Wellcome Sanger Institute for the COVID-19 Genomics UK (COG-UK) Consortium | The Lighthouse Lab in Milton Keynes and Alex Alderton, Roberto Amato, Sonia Goncalves, Ewan Harrison, David K. Jackson, Ian Johnston, Dominic Kwiatkowski, Cordelia Langford, John Sillitoe on behalf of the Wellcome Sanger Institute COVID-19 Surveillance Team |
| EPI_ISL_963419 | Lighthouse Lab in Glasgow | Wellcome Sanger Institute for the COVID-19 Genomics UK (COG-UK) Consortium | Harper VanSteenhouse, Yumi Kasai, David Gray, Carol Clugston, Anna Dominiczak and Alex Alderton, Roberto Amato, Sonia Goncalves, Ewan Harrison, David K. Jackson, Ian Johnston, Dominic Kwiatkowski, Cordelia Langford, John Sillitoe on behalf of the Wellcome Sanger Institute COVID-19 Surveillance Team |
| EPI_ISL_963420 | Lighthouse Lab in Milton Keynes | Wellcome Sanger Institute for the COVID-19 Genomics UK (COG-UK) Consortium | The Lighthouse Lab in Milton Keynes and Alex Alderton, Roberto Amato, Sonia Goncalves, Ewan Harrison, David K. Jackson, Ian Johnston, Dominic Kwiatkowski, Cordelia Langford, John Sillitoe on behalf of the Wellcome Sanger Institute COVID-19 Surveillance Team |
| EPI_ISL_963421 | Lighthouse Lab in Glasgow | Wellcome Sanger Institute for the COVID-19 Genomics UK (COG-UK) Consortium | Harper VanSteenhouse, Yumi Kasai, David Gray, Carol Clugston, Anna Dominiczak and Alex Alderton, Roberto Amato, Sonia Goncalves, Ewan Harrison, David K. Jackson, Ian Johnston, Dominic Kwiatkowski, Cordelia Langford, John Sillitoe on behalf of the Wellcome Sanger Institute COVID-19 Surveillance Team |
| EPI_ISL_963422, EPI_ISL_963423, EPI_ISL_963424 | Lighthouse Lab in Milton Keynes | Wellcome Sanger Institute for the COVID-19 Genomics UK (COG-UK) Consortium | The Lighthouse Lab in Milton Keynes and Alex Alderton, Roberto Amato, Sonia Goncalves, Ewan Harrison, David K. Jackson, Ian Johnston, Dominic Kwiatkowski, Cordelia Langford, John Sillitoe on behalf of the Wellcome Sanger Institute COVID-19 Surveillance Team |
| EPI_ISL_963425, EPI_ISL_963426 | Lighthouse Lab in Glasgow | Wellcome Sanger Institute for the COVID-19 Genomics UK (COG-UK) Consortium | Harper VanSteenhouse, Yumi Kasai, David Gray, Carol Clugston, Anna Dominiczak and Alex Alderton, Roberto Amato, Sonia Goncalves, Ewan Harrison, David K. Jackson, Ian Johnston, Dominic Kwiatkowski, Cordelia Langford, John Sillitoe on behalf of the Wellcome Sanger Institute COVID-19 Surveillance Team |
| EPI_ISL_963427, EPI_ISL_963429, EPI_ISL_963430 | Lighthouse Lab in Milton Keynes | Wellcome Sanger Institute for the COVID-19 Genomics UK (COG-UK) Consortium | The Lighthouse Lab in Milton Keynes and Alex Alderton, Roberto Amato, Sonia Goncalves, Ewan Harrison, David K. Jackson, Ian Johnston, Dominic Kwiatkowski, Cordelia Langford, John Sillitoe on behalf of the Wellcome Sanger Institute COVID-19 Surveillance Team |
| EPI_ISL_963431 | Lighthouse Lab in Glasgow | Wellcome Sanger Institute for the COVID-19 Genomics UK (COG-UK) Consortium | Harper VanSteenhouse, Yumi Kasai, David Gray, Carol Clugston, Anna Dominiczak and Alex Alderton, Roberto Amato, Sonia Goncalves, Ewan Harrison, David K. Jackson, Ian Johnston, Dominic Kwiatkowski, Cordelia Langford, John Sillitoe on behalf of the Wellcome Sanger Institute COVID-19 Surveillance Team |
| EPI_ISL_963432 | Lighthouse Lab in Milton Keynes | Wellcome Sanger Institute for the COVID-19 Genomics UK (COG-UK) Consortium | The Lighthouse Lab in Milton Keynes and Alex Alderton, Roberto Amato, Sonia Goncalves, Ewan Harrison, David K. Jackson, Ian Johnston, Dominic Kwiatkowski, Cordelia Langford, John Sillitoe on behalf of the Wellcome Sanger Institute COVID-19 Surveillance Team |
| EPI_ISL_963433, EPI_ISL_963434 | Lighthouse Lab in Glasgow | Wellcome Sanger Institute for the COVID-19 Genomics UK (COG-UK) Consortium | Harper VanSteenhouse, Yumi Kasai, David Gray, Carol Clugston, Anna Dominiczak and Alex Alderton, Roberto Amato, Sonia Goncalves, Ewan Harrison, David K. Jackson, Ian Johnston, Dominic Kwiatkowski, Cordelia Langford, John Sillitoe on behalf of the Wellcome Sanger Institute COVID-19 Surveillance Team |
| EPI_ISL_963435, EPI_ISL_963436, EPI_ISL_963437, EPI_ISL_963438, EPI_ISL_963439, EPI_ISL_963440, EPI_ISL_963441, EPI_ISL_963442, EPI_ISL_963443, EPI_ISL_963444, EPI_ISL_963445, EPI_ISL_963446, EPI_ISL_963447, EPI_ISL_963448 | | | |
| see above | Lighthouse Lab in Milton Keynes | Wellcome Sanger Institute for the COVID-19 Genomics UK (COG-UK) Consortium | The Lighthouse Lab in Milton Keynes and Alex Alderton, Roberto Amato, Sonia Goncalves, Ewan Harrison, David K. Jackson, Ian Johnston, Dominic Kwiatkowski, Cordelia Langford, John Sillitoe on behalf of the Wellcome Sanger Institute COVID-19 Surveillance Team |
| EPI_ISL_963449 | Lighthouse Lab in Glasgow | Wellcome Sanger Institute for the COVID-19 Genomics UK (COG-UK) Consortium | Harper VanSteenhouse, Yumi Kasai, David Gray, Carol Clugston, Anna Dominiczak and Alex Alderton, Roberto Amato, Sonia Goncalves, Ewan Harrison, David K. Jackson, Ian Johnston, Dominic Kwiatkowski, Cordelia Langford, John Sillitoe on behalf of the Wellcome Sanger Institute COVID-19 Surveillance Team |
| EPI_ISL_963450, EPI_ISL_963451 | Lighthouse Lab in Milton Keynes | Wellcome Sanger Institute for the COVID-19 Genomics UK (COG-UK) Consortium | The Lighthouse Lab in Milton Keynes and Alex Alderton, Roberto Amato, Sonia Goncalves, Ewan Harrison, David K. Jackson, Ian Johnston, Dominic Kwiatkowski, Cordelia Langford, John Sillitoe on behalf of the Wellcome Sanger Institute COVID-19 Surveillance Team |
| EPI_ISL_963452 | Lighthouse Lab in Glasgow | Wellcome Sanger Institute for the COVID-19 Genomics UK (COG-UK) Consortium | Harper VanSteenhouse, Yumi Kasai, David Gray, Carol Clugston, Anna Dominiczak and Alex Alderton, Roberto Amato, Sonia Goncalves, Ewan Harrison, David K. Jackson, Ian Johnston, Dominic Kwiatkowski, Cordelia Langford, John Sillitoe on behalf of the Wellcome Sanger Institute COVID-19 Surveillance Team |
| EPI_ISL_963453, EPI_ISL_963454, EPI_ISL_963455, EPI_ISL_963456, EPI_ISL_963457 | Lighthouse Lab in Milton Keynes | Wellcome Sanger Institute for the COVID-19 Genomics UK (COG-UK) Consortium | The Lighthouse Lab in Milton Keynes and Alex Alderton, Roberto Amato, Sonia Goncalves, Ewan Harrison, David K. Jackson, Ian Johnston, Dominic Kwiatkowski, Cordelia Langford, John Sillitoe on behalf of the Wellcome Sanger Institute COVID-19 Surveillance Team |
| EPI_ISL_963458 | Lighthouse Lab in Glasgow | Wellcome Sanger Institute for the COVID-19 Genomics UK (COG-UK) Consortium | Harper VanSteenhouse, Yumi Kasai, David Gray, Carol Clugston, Anna Dominiczak and Alex Alderton, Roberto Amato, Sonia Goncalves, Ewan Harrison, David K. Jackson, Ian Johnston, Dominic Kwiatkowski, Cordelia Langford, John Sillitoe on behalf of the Wellcome Sanger Institute COVID-19 Surveillance Team |
| EPI_ISL_963459, EPI_ISL_963460, EPI_ISL_963461, EPI_ISL_963462, EPI_ISL_963463, EPI_ISL_963464, EPI_ISL_963465, EPI_ISL_963466, EPI_ISL_963467, EPI_ISL_963468, EPI_ISL_963469, EPI_ISL_963470, EPI_ISL_963472, EPI_ISL_963473, EPI_ISL_963474 | | | |
| see above | Lighthouse Lab in Milton Keynes | Wellcome Sanger Institute for the COVID-19 Genomics UK (COG-UK) Consortium | The Lighthouse Lab in Milton Keynes and Alex Alderton, Roberto Amato, Sonia Goncalves, Ewan Harrison, David K. Jackson, Ian Johnston, Dominic Kwiatkowski, Cordelia Langford, John Sillitoe on behalf of the Wellcome Sanger Institute COVID-19 Surveillance Team |
| EPI_ISL_963475 | Lighthouse Lab in Glasgow | Wellcome Sanger Institute for the COVID-19 Genomics UK (COG-UK) Consortium | Harper VanSteenhouse, Yumi Kasai, David Gray, Carol Clugston, Anna Dominiczak and Alex Alderton, Roberto Amato, Sonia Goncalves, Ewan Harrison, David K. Jackson, Ian Johnston, Dominic Kwiatkowski, Cordelia Langford, John Sillitoe on behalf of the Wellcome Sanger Institute COVID-19 Surveillance Team |
| EPI_ISL_963476, EPI_ISL_963477, EPI_ISL_963478, EPI_ISL_963479, EPI_ISL_963480 | Lighthouse Lab in Milton Keynes | Wellcome Sanger Institute for the COVID-19 Genomics UK (COG-UK) Consortium | The Lighthouse Lab in Milton Keynes and Alex Alderton, Roberto Amato, Sonia Goncalves, Ewan Harrison, David K. Jackson, Ian Johnston, Dominic Kwiatkowski, Cordelia Langford, John Sillitoe on behalf of the Wellcome Sanger Institute COVID-19 Surveillance Team |
| EPI_ISL_963481 | Lighthouse Lab in Glasgow | Wellcome Sanger Institute for the COVID-19 Genomics UK (COG-UK) Consortium | Harper VanSteenhouse, Yumi Kasai, David Gray, Carol Clugston, Anna Dominiczak and Alex Alderton, Roberto Amato, Sonia Goncalves, Ewan Harrison, David K. Jackson, Ian Johnston, Dominic Kwiatkowski, Cordelia Langford, John Sillitoe on behalf of the Wellcome Sanger Institute COVID-19 Surveillance Team |
| EPI_ISL_963482, EPI_ISL_963483, EPI_ISL_963484, EPI_ISL_963485, EPI_ISL_963486, EPI_ISL_963487, EPI_ISL_963488 | Lighthouse Lab in Milton Keynes | Wellcome Sanger Institute for the COVID-19 Genomics UK (COG-UK) Consortium | The Lighthouse Lab in Milton Keynes and Alex Alderton, Roberto Amato, Sonia Goncalves, Ewan Harrison, David K. Jackson, Ian Johnston, Dominic Kwiatkowski, Cordelia Langford, John Sillitoe on behalf of the Wellcome Sanger Institute COVID-19 Surveillance Team |
| EPI_ISL_963489, EPI_ISL_963490 | Lighthouse Lab in Glasgow | Wellcome Sanger Institute for the COVID-19 Genomics UK (COG-UK) Consortium | Harper VanSteenhouse, Yumi Kasai, David Gray, Carol Clugston, Anna Dominiczak and Alex Alderton, Roberto Amato, Sonia Goncalves, Ewan Harrison, David K. Jackson, Ian Johnston, Dominic Kwiatkowski, Cordelia Langford, John Sillitoe on behalf of the Wellcome Sanger Institute COVID-19 Surveillance Team |
| EPI_ISL_963491, EPI_ISL_963492, EPI_ISL_963493, EPI_ISL_963494 | Lighthouse Lab in Milton Keynes | Wellcome Sanger Institute for the COVID-19 Genomics UK (COG-UK) Consortium | The Lighthouse Lab in Milton Keynes and Alex Alderton, Roberto Amato, Sonia Goncalves, Ewan Harrison, David K. Jackson, Ian Johnston, Dominic Kwiatkowski, Cordelia Langford, John Sillitoe on behalf of the Wellcome Sanger Institute COVID-19 Surveillance Team |

| | | | |
|---|---|---|---|
| EPI_ISL_963495 | Lighthouse Lab in Glasgow | Wellcome Sanger Institute for the COVID-19 Genomics UK (COG-UK) Consortium | Harper VanSteenhouse, Yumi Kasai, David Gray, Carol Clugston, Anna Dominiczak and Alex Alderton, Roberto Amato, Sonia Goncalves, Ewan Harrison, David K. Jackson, Ian Johnston, Dominic Kwiatkowski, Cordelia Langford, John Sillitoe on behalf of the Wellcome Sanger Institute COVID-19 Surveillance Team |
| EPI_ISL_963496, EPI_ISL_963497, EPI_ISL_963498, EPI_ISL_963499, EPI_ISL_963500, EPI_ISL_963501, EPI_ISL_963502, EPI_ISL_963503, EPI_ISL_963504, EPI_ISL_963505, EPI_ISL_963506, EPI_ISL_963507, EPI_ISL_963508 | | | |
| see above | Lighthouse Lab in Milton Keynes | Wellcome Sanger Institute for the COVID-19 Genomics UK (COG-UK) Consortium | The Lighthouse Lab in Milton Keynes and Alex Alderton, Roberto Amato, Sonia Goncalves, Ewan Harrison, David K. Jackson, Ian Johnston, Dominic Kwiatkowski, Cordelia Langford, John Sillitoe on behalf of the Wellcome Sanger Institute COVID-19 Surveillance Team |
| EPI_ISL_963509 | Lighthouse Lab in Glasgow | Wellcome Sanger Institute for the COVID-19 Genomics UK (COG-UK) Consortium | Harper VanSteenhouse, Yumi Kasai, David Gray, Carol Clugston, Anna Dominiczak and Alex Alderton, Roberto Amato, Sonia Goncalves, Ewan Harrison, David K. Jackson, Ian Johnston, Dominic Kwiatkowski, Cordelia Langford, John Sillitoe on behalf of the Wellcome Sanger Institute COVID-19 Surveillance Team |
| EPI_ISL_963510 | Lighthouse Lab in Milton Keynes | Wellcome Sanger Institute for the COVID-19 Genomics UK (COG-UK) Consortium | The Lighthouse Lab in Milton Keynes and Alex Alderton, Roberto Amato, Sonia Goncalves, Ewan Harrison, David K. Jackson, Ian Johnston, Dominic Kwiatkowski, Cordelia Langford, John Sillitoe on behalf of the Wellcome Sanger Institute COVID-19 Surveillance Team |
| EPI_ISL_963511 | Lighthouse Lab in Glasgow | Wellcome Sanger Institute for the COVID-19 Genomics UK (COG-UK) Consortium | Harper VanSteenhouse, Yumi Kasai, David Gray, Carol Clugston, Anna Dominiczak and Alex Alderton, Roberto Amato, Sonia Goncalves, Ewan Harrison, David K. Jackson, Ian Johnston, Dominic Kwiatkowski, Cordelia Langford, John Sillitoe on behalf of the Wellcome Sanger Institute COVID-19 Surveillance Team |
| EPI_ISL_963512, EPI_ISL_963513, EPI_ISL_963514, EPI_ISL_963515, EPI_ISL_963516, EPI_ISL_963517, EPI_ISL_963518, EPI_ISL_963519 | Lighthouse Lab in Milton Keynes | Wellcome Sanger Institute for the COVID-19 Genomics UK (COG-UK) Consortium | The Lighthouse Lab in Milton Keynes and Alex Alderton, Roberto Amato, Sonia Goncalves, Ewan Harrison, David K. Jackson, Ian Johnston, Dominic Kwiatkowski, Cordelia Langford, John Sillitoe on behalf of the Wellcome Sanger Institute COVID-19 Surveillance Team |
| EPI_ISL_963520, EPI_ISL_963521 | Lighthouse Lab in Glasgow | Wellcome Sanger Institute for the COVID-19 Genomics UK (COG-UK) Consortium | Harper VanSteenhouse, Yumi Kasai, David Gray, Carol Clugston, Anna Dominiczak and Alex Alderton, Roberto Amato, Sonia Goncalves, Ewan Harrison, David K. Jackson, Ian Johnston, Dominic Kwiatkowski, Cordelia Langford, John Sillitoe on behalf of the Wellcome Sanger Institute COVID-19 Surveillance Team |
| EPI_ISL_963522, EPI_ISL_963523, EPI_ISL_963525, EPI_ISL_963526, EPI_ISL_963527, EPI_ISL_963528 | Lighthouse Lab in Milton Keynes | Wellcome Sanger Institute for the COVID-19 Genomics UK (COG-UK) Consortium | The Lighthouse Lab in Milton Keynes and Alex Alderton, Roberto Amato, Sonia Goncalves, Ewan Harrison, David K. Jackson, Ian Johnston, Dominic Kwiatkowski, Cordelia Langford, John Sillitoe on behalf of the Wellcome Sanger Institute COVID-19 Surveillance Team |
| EPI_ISL_963529, EPI_ISL_963530 | Lighthouse Lab in Glasgow | Wellcome Sanger Institute for the COVID-19 Genomics UK (COG-UK) Consortium | Harper VanSteenhouse, Yumi Kasai, David Gray, Carol Clugston, Anna Dominiczak and Alex Alderton, Roberto Amato, Sonia Goncalves, Ewan Harrison, David K. Jackson, Ian Johnston, Dominic Kwiatkowski, Cordelia Langford, John Sillitoe on behalf of the Wellcome Sanger Institute COVID-19 Surveillance Team |
| EPI_ISL_963531, EPI_ISL_963532, EPI_ISL_963533, EPI_ISL_963534, EPI_ISL_963535, EPI_ISL_963536, EPI_ISL_963537 | Lighthouse Lab in Milton Keynes | Wellcome Sanger Institute for the COVID-19 Genomics UK (COG-UK) Consortium | The Lighthouse Lab in Milton Keynes and Alex Alderton, Roberto Amato, Sonia Goncalves, Ewan Harrison, David K. Jackson, Ian Johnston, Dominic Kwiatkowski, Cordelia Langford, John Sillitoe on behalf of the Wellcome Sanger Institute COVID-19 Surveillance Team |
| EPI_ISL_963538 | Lighthouse Lab in Glasgow | Wellcome Sanger Institute for the COVID-19 Genomics UK (COG-UK) Consortium | Harper VanSteenhouse, Yumi Kasai, David Gray, Carol Clugston, Anna Dominiczak and Alex Alderton, Roberto Amato, Sonia Goncalves, Ewan Harrison, David K. Jackson, Ian Johnston, Dominic Kwiatkowski, Cordelia Langford, John Sillitoe on behalf of the Wellcome Sanger Institute COVID-19 Surveillance Team |
| EPI_ISL_963539, EPI_ISL_963540, EPI_ISL_963541, EPI_ISL_963542, EPI_ISL_963543, EPI_ISL_963544, EPI_ISL_963545, EPI_ISL_963546, EPI_ISL_963547, EPI_ISL_963548, EPI_ISL_963549, EPI_ISL_963550, EPI_ISL_963551, EPI_ISL_963552, EPI_ISL_963553 | | | |
| see above | Lighthouse Lab in Milton Keynes | Wellcome Sanger Institute for the COVID-19 Genomics UK (COG-UK) Consortium | The Lighthouse Lab in Milton Keynes and Alex Alderton, Roberto Amato, Sonia Goncalves, Ewan Harrison, David K. Jackson, Ian Johnston, Dominic Kwiatkowski, Cordelia Langford, John Sillitoe on behalf of the Wellcome Sanger Institute COVID-19 Surveillance Team |
| EPI_ISL_963554 | Lighthouse Lab in Glasgow | Wellcome Sanger Institute for the COVID-19 Genomics UK (COG-UK) Consortium | Harper VanSteenhouse, Yumi Kasai, David Gray, Carol Clugston, Anna Dominiczak and Alex Alderton, Roberto Amato, Sonia Goncalves, Ewan Harrison, David K. Jackson, Ian Johnston, Dominic Kwiatkowski, Cordelia Langford, John Sillitoe on behalf of the Wellcome Sanger Institute COVID-19 Surveillance Team |
| EPI_ISL_963556, EPI_ISL_963557, EPI_ISL_963558 | Lighthouse Lab in Milton Keynes | Wellcome Sanger Institute for the COVID-19 Genomics UK (COG-UK) Consortium | The Lighthouse Lab in Milton Keynes and Alex Alderton, Roberto Amato, Sonia Goncalves, Ewan Harrison, David K. Jackson, Ian Johnston, Dominic Kwiatkowski, Cordelia Langford, John Sillitoe on behalf of the Wellcome Sanger Institute COVID-19 Surveillance Team |
| EPI_ISL_963559, EPI_ISL_963560 | Lighthouse Lab in Glasgow | Wellcome Sanger Institute for the COVID-19 Genomics UK (COG-UK) Consortium | Harper VanSteenhouse, Yumi Kasai, David Gray, Carol Clugston, Anna Dominiczak and Alex Alderton, Roberto Amato, Sonia Goncalves, Ewan Harrison, David K. Jackson, Ian Johnston, Dominic Kwiatkowski, Cordelia Langford, John Sillitoe on behalf of the Wellcome Sanger Institute COVID-19 Surveillance Team |
| EPI_ISL_963561 | Lighthouse Lab in Milton Keynes | Wellcome Sanger Institute for the COVID-19 Genomics UK (COG-UK) Consortium | The Lighthouse Lab in Milton Keynes and Alex Alderton, Roberto Amato, Sonia Goncalves, Ewan Harrison, David K. Jackson, Ian Johnston, Dominic Kwiatkowski, Cordelia Langford, John Sillitoe on behalf of the Wellcome Sanger Institute COVID-19 Surveillance Team |
| EPI_ISL_963562 | Lighthouse Lab in Glasgow | Wellcome Sanger Institute for the COVID-19 Genomics UK (COG-UK) Consortium | Harper VanSteenhouse, Yumi Kasai, David Gray, Carol Clugston, Anna Dominiczak and Alex Alderton, Roberto Amato, Sonia Goncalves, Ewan Harrison, David K. Jackson, Ian Johnston, Dominic Kwiatkowski, Cordelia Langford, John Sillitoe on behalf of the Wellcome Sanger Institute COVID-19 Surveillance Team |
| EPI_ISL_963563, EPI_ISL_963564, EPI_ISL_963565, EPI_ISL_963566, EPI_ISL_963567, EPI_ISL_963568, EPI_ISL_963569 | Lighthouse Lab in Milton Keynes | Wellcome Sanger Institute for the COVID-19 Genomics UK (COG-UK) Consortium | The Lighthouse Lab in Milton Keynes and Alex Alderton, Roberto Amato, Sonia Goncalves, Ewan Harrison, David K. Jackson, Ian Johnston, Dominic Kwiatkowski, Cordelia Langford, John Sillitoe on behalf of the Wellcome Sanger Institute COVID-19 Surveillance Team |
| EPI_ISL_963570 | Lighthouse Lab in Glasgow | Wellcome Sanger Institute for the COVID-19 Genomics UK (COG-UK) Consortium | Harper VanSteenhouse, Yumi Kasai, David Gray, Carol Clugston, Anna Dominiczak and Alex Alderton, Roberto Amato, Sonia Goncalves, Ewan Harrison, David K. Jackson, Ian Johnston, Dominic Kwiatkowski, Cordelia Langford, John Sillitoe on behalf of the Wellcome Sanger Institute COVID-19 Surveillance Team |
| EPI_ISL_963571, EPI_ISL_963572 | Lighthouse Lab in Milton Keynes | Wellcome Sanger Institute for the COVID-19 Genomics UK (COG-UK) Consortium | The Lighthouse Lab in Milton Keynes and Alex Alderton, Roberto Amato, Sonia Goncalves, Ewan Harrison, David K. Jackson, Ian Johnston, Dominic Kwiatkowski, Cordelia Langford, John Sillitoe on behalf of the Wellcome Sanger Institute COVID-19 Surveillance Team |
| EPI_ISL_963573 | Lighthouse Lab in Glasgow | Wellcome Sanger Institute for the COVID-19 Genomics UK (COG-UK) Consortium | Harper VanSteenhouse, Yumi Kasai, David Gray, Carol Clugston, Anna Dominiczak and Alex Alderton, Roberto Amato, Sonia Goncalves, Ewan Harrison, David K. Jackson, Ian Johnston, Dominic Kwiatkowski, Cordelia Langford, John Sillitoe on behalf of the Wellcome Sanger Institute COVID-19 Surveillance Team |
| EPI_ISL_963574, EPI_ISL_963575 | Lighthouse Lab in Milton Keynes | Wellcome Sanger Institute for the COVID-19 Genomics UK (COG-UK) Consortium | The Lighthouse Lab in Milton Keynes and Alex Alderton, Roberto Amato, Sonia Goncalves, Ewan Harrison, David K. Jackson, Ian Johnston, Dominic Kwiatkowski, Cordelia Langford, John Sillitoe on behalf of the Wellcome Sanger Institute COVID-19 Surveillance Team |
| EPI_ISL_963576 | Lighthouse Lab in Glasgow | Wellcome Sanger Institute for the COVID-19 Genomics UK (COG-UK) Consortium | Harper VanSteenhouse, Yumi Kasai, David Gray, Carol Clugston, Anna Dominiczak and Alex Alderton, Roberto Amato, Sonia Goncalves, Ewan Harrison, David K. Jackson, Ian Johnston, Dominic Kwiatkowski, Cordelia Langford, John Sillitoe on behalf of the Wellcome Sanger Institute COVID-19 Surveillance Team |
| EPI_ISL_963577, EPI_ISL_963578, EPI_ISL_963579, EPI_ISL_963580 | Lighthouse Lab in Milton Keynes | Wellcome Sanger Institute for the COVID-19 Genomics UK (COG-UK) Consortium | The Lighthouse Lab in Milton Keynes and Alex Alderton, Roberto Amato, Sonia Goncalves, Ewan Harrison, David K. Jackson, Ian Johnston, Dominic Kwiatkowski, Cordelia Langford, John Sillitoe on behalf of the Wellcome Sanger Institute COVID-19 Surveillance Team |
| EPI_ISL_963581 | Lighthouse Lab in Glasgow | Wellcome Sanger Institute for the COVID-19 Genomics UK (COG-UK) Consortium | Harper VanSteenhouse, Yumi Kasai, David Gray, Carol Clugston, Anna Dominiczak and Alex Alderton, Roberto Amato, Sonia Goncalves, Ewan Harrison, David K. Jackson, Ian Johnston, Dominic Kwiatkowski, Cordelia Langford, John Sillitoe on behalf of the Wellcome Sanger Institute COVID-19 Surveillance Team |
| EPI_ISL_963582, EPI_ISL_963583 | Lighthouse Lab in Milton Keynes | Wellcome Sanger Institute for the COVID-19 Genomics UK (COG-UK) Consortium | The Lighthouse Lab in Milton Keynes and Alex Alderton, Roberto Amato, Sonia Goncalves, Ewan Harrison, David K. Jackson, Ian Johnston, Dominic Kwiatkowski, Cordelia Langford, John Sillitoe on behalf of the Wellcome Sanger Institute COVID-19 Surveillance Team |

| | | | |
|---|---|---|---|
| EPI_ISL_963584 | Lighthouse Lab in Glasgow | Wellcome Sanger Institute for the COVID-19 Genomics UK (COG-UK) Consortium | Harper VanSteenhouse, Yumi Kasai, David Gray, Carol Clugston, Anna Dominiczak and Alex Alderton, Roberto Amato, Sonia Goncalves, Ewan Harrison, David K. Jackson, Ian Johnston, Dominic Kwiatkowski, Cordelia Langford, John Sillitoe on behalf of the Wellcome Sanger Institute COVID-19 Surveillance Team |
| EPI_ISL_963585, EPI_ISL_963586, EPI_ISL_963587, EPI_ISL_963588, EPI_ISL_963589, EPI_ISL_963590 | Lighthouse Lab in Milton Keynes | Wellcome Sanger Institute for the COVID-19 Genomics UK (COG-UK) Consortium | The Lighthouse Lab in Milton Keynes and Alex Alderton, Roberto Amato, Sonia Goncalves, Ewan Harrison, David K. Jackson, Ian Johnston, Dominic Kwiatkowski, Cordelia Langford, John Sillitoe on behalf of the Wellcome Sanger Institute COVID-19 Surveillance Team |
| EPI_ISL_963591 | Lighthouse Lab in Glasgow | Wellcome Sanger Institute for the COVID-19 Genomics UK (COG-UK) Consortium | Harper VanSteenhouse, Yumi Kasai, David Gray, Carol Clugston, Anna Dominiczak and Alex Alderton, Roberto Amato, Sonia Goncalves, Ewan Harrison, David K. Jackson, Ian Johnston, Dominic Kwiatkowski, Cordelia Langford, John Sillitoe on behalf of the Wellcome Sanger Institute COVID-19 Surveillance Team |
| EPI_ISL_963592, EPI_ISL_963593 | Lighthouse Lab in Milton Keynes | Wellcome Sanger Institute for the COVID-19 Genomics UK (COG-UK) Consortium | The Lighthouse Lab in Milton Keynes and Alex Alderton, Roberto Amato, Sonia Goncalves, Ewan Harrison, David K. Jackson, Ian Johnston, Dominic Kwiatkowski, Cordelia Langford, John Sillitoe on behalf of the Wellcome Sanger Institute COVID-19 Surveillance Team |
| EPI_ISL_963594 | Lighthouse Lab in Glasgow | Wellcome Sanger Institute for the COVID-19 Genomics UK (COG-UK) Consortium | Harper VanSteenhouse, Yumi Kasai, David Gray, Carol Clugston, Anna Dominiczak and Alex Alderton, Roberto Amato, Sonia Goncalves, Ewan Harrison, David K. Jackson, Ian Johnston, Dominic Kwiatkowski, Cordelia Langford, John Sillitoe on behalf of the Wellcome Sanger Institute COVID-19 Surveillance Team |
| EPI_ISL_963595, EPI_ISL_963596, EPI_ISL_963597, EPI_ISL_963598, EPI_ISL_963599, EPI_ISL_963600 | Lighthouse Lab in Milton Keynes | Wellcome Sanger Institute for the COVID-19 Genomics UK (COG-UK) Consortium | The Lighthouse Lab in Milton Keynes and Alex Alderton, Roberto Amato, Sonia Goncalves, Ewan Harrison, David K. Jackson, Ian Johnston, Dominic Kwiatkowski, Cordelia Langford, John Sillitoe on behalf of the Wellcome Sanger Institute COVID-19 Surveillance Team |
| EPI_ISL_963601 | Lighthouse Lab in Glasgow | Wellcome Sanger Institute for the COVID-19 Genomics UK (COG-UK) Consortium | Harper VanSteenhouse, Yumi Kasai, David Gray, Carol Clugston, Anna Dominiczak and Alex Alderton, Roberto Amato, Sonia Goncalves, Ewan Harrison, David K. Jackson, Ian Johnston, Dominic Kwiatkowski, Cordelia Langford, John Sillitoe on behalf of the Wellcome Sanger Institute COVID-19 Surveillance Team |
| EPI_ISL_963602 | Lighthouse Lab in Milton Keynes | Wellcome Sanger Institute for the COVID-19 Genomics UK (COG-UK) Consortium | The Lighthouse Lab in Milton Keynes and Alex Alderton, Roberto Amato, Sonia Goncalves, Ewan Harrison, David K. Jackson, Ian Johnston, Dominic Kwiatkowski, Cordelia Langford, John Sillitoe on behalf of the Wellcome Sanger Institute COVID-19 Surveillance Team |
| EPI_ISL_963603 | Lighthouse Lab in Glasgow | Wellcome Sanger Institute for the COVID-19 Genomics UK (COG-UK) Consortium | Harper VanSteenhouse, Yumi Kasai, David Gray, Carol Clugston, Anna Dominiczak and Alex Alderton, Roberto Amato, Sonia Goncalves, Ewan Harrison, David K. Jackson, Ian Johnston, Dominic Kwiatkowski, Cordelia Langford, John Sillitoe on behalf of the Wellcome Sanger Institute COVID-19 Surveillance Team |
| EPI_ISL_963604, EPI_ISL_963605, EPI_ISL_963606 | Lighthouse Lab in Milton Keynes | Wellcome Sanger Institute for the COVID-19 Genomics UK (COG-UK) Consortium | The Lighthouse Lab in Milton Keynes and Alex Alderton, Roberto Amato, Sonia Goncalves, Ewan Harrison, David K. Jackson, Ian Johnston, Dominic Kwiatkowski, Cordelia Langford, John Sillitoe on behalf of the Wellcome Sanger Institute COVID-19 Surveillance Team |
| EPI_ISL_963607 | Lighthouse Lab in Glasgow | Wellcome Sanger Institute for the COVID-19 Genomics UK (COG-UK) Consortium | Harper VanSteenhouse, Yumi Kasai, David Gray, Carol Clugston, Anna Dominiczak and Alex Alderton, Roberto Amato, Sonia Goncalves, Ewan Harrison, David K. Jackson, Ian Johnston, Dominic Kwiatkowski, Cordelia Langford, John Sillitoe on behalf of the Wellcome Sanger Institute COVID-19 Surveillance Team |
| EPI_ISL_963608, EPI_ISL_963609, EPI_ISL_963610, EPI_ISL_963611, EPI_ISL_963612, EPI_ISL_963613, EPI_ISL_963614, EPI_ISL_963615, EPI_ISL_963616, EPI_ISL_963617 | Lighthouse Lab in Milton Keynes | Wellcome Sanger Institute for the COVID-19 Genomics UK (COG-UK) Consortium | The Lighthouse Lab in Milton Keynes and Alex Alderton, Roberto Amato, Sonia Goncalves, Ewan Harrison, David K. Jackson, Ian Johnston, Dominic Kwiatkowski, Cordelia Langford, John Sillitoe on behalf of the Wellcome Sanger Institute COVID-19 Surveillance Team |
| EPI_ISL_963618 | Lighthouse Lab in Glasgow | Wellcome Sanger Institute for the COVID-19 Genomics UK (COG-UK) Consortium | Harper VanSteenhouse, Yumi Kasai, David Gray, Carol Clugston, Anna Dominiczak and Alex Alderton, Roberto Amato, Sonia Goncalves, Ewan Harrison, David K. Jackson, Ian Johnston, Dominic Kwiatkowski, Cordelia Langford, John Sillitoe on behalf of the Wellcome Sanger Institute COVID-19 Surveillance Team |
| EPI_ISL_963619, EPI_ISL_963620, EPI_ISL_963621, EPI_ISL_963622, EPI_ISL_963623, EPI_ISL_963624, EPI_ISL_963625 | Lighthouse Lab in Milton Keynes | Wellcome Sanger Institute for the COVID-19 Genomics UK (COG-UK) Consortium | The Lighthouse Lab in Milton Keynes and Alex Alderton, Roberto Amato, Sonia Goncalves, Ewan Harrison, David K. Jackson, Ian Johnston, Dominic Kwiatkowski, Cordelia Langford, John Sillitoe on behalf of the Wellcome Sanger Institute COVID-19 Surveillance Team |
| EPI_ISL_963626 | Lighthouse Lab in Glasgow | Wellcome Sanger Institute for the COVID-19 Genomics UK (COG-UK) Consortium | Harper VanSteenhouse, Yumi Kasai, David Gray, Carol Clugston, Anna Dominiczak and Alex Alderton, Roberto Amato, Sonia Goncalves, Ewan Harrison, David K. Jackson, Ian Johnston, Dominic Kwiatkowski, Cordelia Langford, John Sillitoe on behalf of the Wellcome Sanger Institute COVID-19 Surveillance Team |
| EPI_ISL_963627, EPI_ISL_963628, EPI_ISL_963629, EPI_ISL_963630, EPI_ISL_963631, EPI_ISL_963632 | Lighthouse Lab in Milton Keynes | Wellcome Sanger Institute for the COVID-19 Genomics UK (COG-UK) Consortium | The Lighthouse Lab in Milton Keynes and Alex Alderton, Roberto Amato, Sonia Goncalves, Ewan Harrison, David K. Jackson, Ian Johnston, Dominic Kwiatkowski, Cordelia Langford, John Sillitoe on behalf of the Wellcome Sanger Institute COVID-19 Surveillance Team |
| EPI_ISL_963633 | Lighthouse Lab in Glasgow | Wellcome Sanger Institute for the COVID-19 Genomics UK (COG-UK) Consortium | Harper VanSteenhouse, Yumi Kasai, David Gray, Carol Clugston, Anna Dominiczak and Alex Alderton, Roberto Amato, Sonia Goncalves, Ewan Harrison, David K. Jackson, Ian Johnston, Dominic Kwiatkowski, Cordelia Langford, John Sillitoe on behalf of the Wellcome Sanger Institute COVID-19 Surveillance Team |
| EPI_ISL_963634, EPI_ISL_963635, EPI_ISL_963636, EPI_ISL_963637, EPI_ISL_963638 | Lighthouse Lab in Milton Keynes | Wellcome Sanger Institute for the COVID-19 Genomics UK (COG-UK) Consortium | The Lighthouse Lab in Milton Keynes and Alex Alderton, Roberto Amato, Sonia Goncalves, Ewan Harrison, David K. Jackson, Ian Johnston, Dominic Kwiatkowski, Cordelia Langford, John Sillitoe on behalf of the Wellcome Sanger Institute COVID-19 Surveillance Team |
| EPI_ISL_963639 | Lighthouse Lab in Glasgow | Wellcome Sanger Institute for the COVID-19 Genomics UK (COG-UK) Consortium | Harper VanSteenhouse, Yumi Kasai, David Gray, Carol Clugston, Anna Dominiczak and Alex Alderton, Roberto Amato, Sonia Goncalves, Ewan Harrison, David K. Jackson, Ian Johnston, Dominic Kwiatkowski, Cordelia Langford, John Sillitoe on behalf of the Wellcome Sanger Institute COVID-19 Surveillance Team |
| EPI_ISL_963640 | Lighthouse Lab in Milton Keynes | Wellcome Sanger Institute for the COVID-19 Genomics UK (COG-UK) Consortium | The Lighthouse Lab in Milton Keynes and Alex Alderton, Roberto Amato, Sonia Goncalves, Ewan Harrison, David K. Jackson, Ian Johnston, Dominic Kwiatkowski, Cordelia Langford, John Sillitoe on behalf of the Wellcome Sanger Institute COVID-19 Surveillance Team |
| EPI_ISL_963641 | Lighthouse Lab in Glasgow | Wellcome Sanger Institute for the COVID-19 Genomics UK (COG-UK) Consortium | Harper VanSteenhouse, Yumi Kasai, David Gray, Carol Clugston, Anna Dominiczak and Alex Alderton, Roberto Amato, Sonia Goncalves, Ewan Harrison, David K. Jackson, Ian Johnston, Dominic Kwiatkowski, Cordelia Langford, John Sillitoe on behalf of the Wellcome Sanger Institute COVID-19 Surveillance Team |
| EPI_ISL_963642, EPI_ISL_963643, EPI_ISL_963644, EPI_ISL_963645, EPI_ISL_963646, EPI_ISL_963647, EPI_ISL_963648 | Lighthouse Lab in Milton Keynes | Wellcome Sanger Institute for the COVID-19 Genomics UK (COG-UK) Consortium | The Lighthouse Lab in Milton Keynes and Alex Alderton, Roberto Amato, Sonia Goncalves, Ewan Harrison, David K. Jackson, Ian Johnston, Dominic Kwiatkowski, Cordelia Langford, John Sillitoe on behalf of the Wellcome Sanger Institute COVID-19 Surveillance Team |
| EPI_ISL_963649 | Lighthouse Lab in Glasgow | Wellcome Sanger Institute for the COVID-19 Genomics UK (COG-UK) Consortium | Harper VanSteenhouse, Yumi Kasai, David Gray, Carol Clugston, Anna Dominiczak and Alex Alderton, Roberto Amato, Sonia Goncalves, Ewan Harrison, David K. Jackson, Ian Johnston, Dominic Kwiatkowski, Cordelia Langford, John Sillitoe on behalf of the Wellcome Sanger Institute COVID-19 Surveillance Team |
| EPI_ISL_963650 | Lighthouse Lab in Milton Keynes | Wellcome Sanger Institute for the COVID-19 Genomics UK (COG-UK) Consortium | The Lighthouse Lab in Milton Keynes and Alex Alderton, Roberto Amato, Sonia Goncalves, Ewan Harrison, David K. Jackson, Ian Johnston, Dominic Kwiatkowski, Cordelia Langford, John Sillitoe on behalf of the Wellcome Sanger Institute COVID-19 Surveillance Team |

EPI_ISL_963652, EPI_ISL_963653, EPI_ISL_963655, EPI_ISL_963656, EPI_ISL_963657, EPI_ISL_963660, EPI_ISL_963661, EPI_ISL_963662, EPI_ISL_963663, EPI_ISL_963664, EPI_ISL_963666, EPI_ISL_963667, EPI_ISL_963668, EPI_ISL_963669, EPI_ISL_963674, EPI_ISL_963675, EPI_ISL_963678, EPI_ISL_963680, EPI_ISL_963682, EPI_ISL_963683, EPI_ISL_963684, EPI_ISL_963688, EPI_ISL_963691, EPI_ISL_963692, EPI_ISL_963693, EPI_ISL_963696, EPI_ISL_963697, EPI_ISL_963698, EPI_ISL_963699, EPI_ISL_963702, EPI_ISL_963705, EPI_ISL_963708, EPI_ISL_963710, EPI_ISL_963711, EPI_ISL_963712, EPI_ISL_963713, EPI_ISL_963714, EPI_ISL_963715, EPI_ISL_963716, EPI_ISL_963717, EPI_ISL_963719, EPI_ISL_963722, EPI_ISL_963723, EPI_ISL_963725, EPI_ISL_963728, EPI_ISL_963729, EPI_ISL_963730, EPI_ISL_963739, EPI_ISL_963742, EPI_ISL_963743, EPI_ISL_963744, EPI_ISL_963745, EPI_ISL_963746, EPI_ISL_963748, EPI_ISL_963749, EPI_ISL_963751, EPI_ISL_963752, EPI_ISL_963755, EPI_ISL_963756, EPI_ISL_963757, EPI_ISL_963760, EPI_ISL_963762, EPI_ISL_963763, EPI_ISL_963767, EPI_ISL_963770, EPI_ISL_963771, EPI_ISL_963776, EPI_ISL_963777, EPI_ISL_963778, EPI_ISL_963782, EPI_ISL_963784, EPI_ISL_963785,

EPI_ISL_963787, EPI_ISL_963788, EPI_ISL_963791, EPI_ISL_963792, EPI_ISL_963793, EPI_ISL_963795, EPI_ISL_963796, EPI_ISL_963797, EPI_ISL_963799, EPI_ISL_963801, EPI_ISL_963802, EPI_ISL_963804, EPI_ISL_963809, EPI_ISL_963811, EPI_ISL_963812, EPI_ISL_963817, EPI_ISL_963818, EPI_ISL_963820, EPI_ISL_963827, EPI_ISL_963828, EPI_ISL_963829, EPI_ISL_963830, EPI_ISL_963834, EPI_ISL_963836, EPI_ISL_963837, EPI_ISL_963840, EPI_ISL_963842, EPI_ISL_963843, EPI_ISL_963847, EPI_ISL_963848, EPI_ISL_963854, EPI_ISL_963856, EPI_ISL_963858, EPI_ISL_963860, EPI_ISL_963862, EPI_ISL_963871, EPI_ISL_963876, EPI_ISL_963885, EPI_ISL_963886, EPI_ISL_963887, EPI_ISL_963888, EPI_ISL_963892, EPI_ISL_963894, EPI_ISL_963897, EPI_ISL_963898, EPI_ISL_963899, EPI_ISL_963905, EPI_ISL_963907, EPI_ISL_963910, EPI_ISL_963912, EPI_ISL_963913, EPI_ISL_963917, EPI_ISL_963918, EPI_ISL_963919, EPI_ISL_963920, EPI_ISL_963924, EPI_ISL_963926, EPI_ISL_963928, EPI_ISL_963929, EPI_ISL_963930, EPI_ISL_963931, EPI_ISL_963933, EPI_ISL_963935, EPI_ISL_963937, EPI_ISL_963939, EPI_ISL_963940, EPI_ISL_963943, EPI_ISL_963944, EPI_ISL_963946, EPI_ISL_963948, EPI_ISL_963950, EPI_ISL_963951, EPI_ISL_963954, EPI_ISL_963956, EPI_ISL_963957, EPI_ISL_963959, EPI_ISL_963961, EPI_ISL_963963, EPI_ISL_963965, EPI_ISL_963966, EPI_ISL_963967, EPI_ISL_963969, EPI_ISL_963972, EPI_ISL_963974, EPI_ISL_963975, EPI_ISL_963982, EPI_ISL_963983, EPI_ISL_963985, EPI_ISL_963986, EPI_ISL_963987, EPI_ISL_963988, EPI_ISL_963989, EPI_ISL_963990, EPI_ISL_963991, EPI_ISL_963992, EPI_ISL_963993, EPI_ISL_963994, EPI_ISL_963995, EPI_ISL_963996, EPI_ISL_963997, EPI_ISL_963998, EPI_ISL_963999, EPI_ISL_964000, EPI_ISL_964001, EPI_ISL_964002, EPI_ISL_964003, EPI_ISL_964004, EPI_ISL_964005, EPI_ISL_964006, EPI_ISL_964007, EPI_ISL_964008, EPI_ISL_964009, EPI_ISL_964010, EPI_ISL_964011, EPI_ISL_964012, EPI_ISL_964013, EPI_ISL_964014, EPI_ISL_964015, EPI_ISL_964016, EPI_ISL_964017, EPI_ISL_964018, EPI_ISL_964019, EPI_ISL_964020, EPI_ISL_964021, EPI_ISL_964022, EPI_ISL_964023, EPI_ISL_964024, EPI_ISL_964025, EPI_ISL_964026, EPI_ISL_964027, EPI_ISL_964028, EPI_ISL_964029, EPI_ISL_964030, EPI_ISL_964031, EPI_ISL_964032, EPI_ISL_964033, EPI_ISL_964034, EPI_ISL_964035, EPI_ISL_964036, EPI_ISL_964037, EPI_ISL_964038, EPI_ISL_964039, EPI_ISL_964040, EPI_ISL_964041, EPI_ISL_964042, EPI_ISL_964043, EPI_ISL_964044, EPI_ISL_964045, EPI_ISL_964046, EPI_ISL_964047, EPI_ISL_964048, EPI_ISL_964049, EPI_ISL_964050, EPI_ISL_964051, EPI_ISL_964052, EPI_ISL_964053, EPI_ISL_964054, EPI_ISL_964055, EPI_ISL_964056, EPI_ISL_964057, EPI_ISL_964058, EPI_ISL_964059, EPI_ISL_964060, EPI_ISL_964061, EPI_ISL_964062, EPI_ISL_964063, EPI_ISL_964064, EPI_ISL_964065, EPI_ISL_964066, EPI_ISL_964068, EPI_ISL_964069, EPI_ISL_964070, EPI_ISL_964071, EPI_ISL_964072, EPI_ISL_964073, EPI_ISL_964074, EPI_ISL_964075, EPI_ISL_964076, EPI_ISL_964077, EPI_ISL_964078, EPI_ISL_964079, EPI_ISL_964080, EPI_ISL_964081, EPI_ISL_964082, EPI_ISL_964083, EPI_ISL_964084, EPI_ISL_964085, EPI_ISL_964086, EPI_ISL_964088, EPI_ISL_964089, EPI_ISL_964090, EPI_ISL_964091, EPI_ISL_964092, EPI_ISL_964093, EPI_ISL_964094, EPI_ISL_964095, EPI_ISL_964096, EPI_ISL_964097, EPI_ISL_964098, EPI_ISL_964099, EPI_ISL_964100, EPI_ISL_964101, EPI_ISL_964102, EPI_ISL_964103, EPI_ISL_964104, EPI_ISL_964105, EPI_ISL_964106, EPI_ISL_964108, EPI_ISL_964109, EPI_ISL_964110, EPI_ISL_964111, EPI_ISL_964112, EPI_ISL_964113, EPI_ISL_964114, EPI_ISL_964115, EPI_ISL_964116, EPI_ISL_964117, EPI_ISL_964118, EPI_ISL_964119, EPI_ISL_964120, EPI_ISL_964121, EPI_ISL_964122, EPI_ISL_964123, EPI_ISL_964124, EPI_ISL_964125, EPI_ISL_964126, EPI_ISL_964127, EPI_ISL_964128, EPI_ISL_964129, EPI_ISL_964130, EPI_ISL_964131, EPI_ISL_964132, EPI_ISL_964133, EPI_ISL_964134, EPI_ISL_964135, EPI_ISL_964136, EPI_ISL_964137, EPI_ISL_964138, EPI_ISL_964139, EPI_ISL_964140, EPI_ISL_964141, EPI_ISL_964142, EPI_ISL_964143, EPI_ISL_964144, EPI_ISL_964145, EPI_ISL_964146, EPI_ISL_964147, EPI_ISL_964148, EPI_ISL_964149, EPI_ISL_964150, EPI_ISL_964151, EPI_ISL_964152, EPI_ISL_964153, EPI_ISL_964154, EPI_ISL_964155, EPI_ISL_964156, EPI_ISL_964157, EPI_ISL_964158, EPI_ISL_964159, EPI_ISL_964160, EPI_ISL_964161, EPI_ISL_964162, EPI_ISL_964163, EPI_ISL_964164, EPI_ISL_964165, EPI_ISL_964166, EPI_ISL_964167, EPI_ISL_964168, EPI_ISL_964169, EPI_ISL_964170, EPI_ISL_964171, EPI_ISL_964172, EPI_ISL_964173, EPI_ISL_964174, EPI_ISL_964175, EPI_ISL_964176, EPI_ISL_964177, EPI_ISL_964178, EPI_ISL_964179, EPI_ISL_964180, EPI_ISL_964181, EPI_ISL_964182, EPI_ISL_964183, EPI_ISL_964184, EPI_ISL_964185, EPI_ISL_964186, EPI_ISL_964187, EPI_ISL_964188, EPI_ISL_964189, EPI_ISL_964190, EPI_ISL_964191, EPI_ISL_964192, EPI_ISL_964193, EPI_ISL_964194, EPI_ISL_964195, EPI_ISL_964196, EPI_ISL_964197, EPI_ISL_964198, EPI_ISL_964199, EPI_ISL_964200, EPI_ISL_964201, EPI_ISL_964202, EPI_ISL_964203, EPI_ISL_964204, EPI_ISL_964205, EPI_ISL_964206, EPI_ISL_964207, EPI_ISL_964208, EPI_ISL_964209, EPI_ISL_964210, EPI_ISL_964211, EPI_ISL_964212, EPI_ISL_964213, EPI_ISL_964214, EPI_ISL_964215, EPI_ISL_964216, EPI_ISL_964217, EPI_ISL_964218, EPI_ISL_964220, EPI_ISL_964221, EPI_ISL_964222, EPI_ISL_964223, EPI_ISL_964224, EPI_ISL_964225, EPI_ISL_964226, EPI_ISL_964227, EPI_ISL_964228, EPI_ISL_964229, EPI_ISL_964230, EPI_ISL_964231, EPI_ISL_964232, EPI_ISL_964233, EPI_ISL_964235, EPI_ISL_964237, EPI_ISL_964238, EPI_ISL_964239, EPI_ISL_964240, EPI_ISL_964241, EPI_ISL_964242, EPI_ISL_964243, EPI_ISL_964244, EPI_ISL_964245, EPI_ISL_964246, EPI_ISL_964247, EPI_ISL_964248, EPI_ISL_964249, EPI_ISL_964250, EPI_ISL_964251, EPI_ISL_964252, EPI_ISL_964253, EPI_ISL_964255, EPI_ISL_964256, EPI_ISL_964258, EPI_ISL_964260, EPI_ISL_964261, EPI_ISL_964263, EPI_ISL_964264, EPI_ISL_964266, EPI_ISL_964268, EPI_ISL_964269, EPI_ISL_964271, EPI_ISL_964273, EPI_ISL_964274, EPI_ISL_964276, EPI_ISL_964277, EPI_ISL_964279, EPI_ISL_964280, EPI_ISL_964282, EPI_ISL_964283, EPI_ISL_964285, EPI_ISL_964286, EPI_ISL_964288, EPI_ISL_964290, EPI_ISL_964291, EPI_ISL_964293, EPI_ISL_964294, EPI_ISL_964296, EPI_ISL_964298, EPI_ISL_964299, EPI_ISL_964301, EPI_ISL_964302, EPI_ISL_964304, EPI_ISL_964306, EPI_ISL_964307, EPI_ISL_964309, EPI_ISL_964310, EPI_ISL_964311, EPI_ISL_964312, EPI_ISL_964313, EPI_ISL_964314, EPI_ISL_964315, EPI_ISL_964316, EPI_ISL_964317, EPI_ISL_964318, EPI_ISL_964319, EPI_ISL_964320, EPI_ISL_964321, EPI_ISL_964322, EPI_ISL_964323, EPI_ISL_964324, EPI_ISL_964325, EPI_ISL_964326, EPI_ISL_964327, EPI_ISL_964328, EPI_ISL_964329, EPI_ISL_964330, EPI_ISL_964331, EPI_ISL_964332, EPI_ISL_964333, EPI_ISL_964334, EPI_ISL_964335, EPI_ISL_964336, EPI_ISL_964337, EPI_ISL_964338, EPI_ISL_964339, EPI_ISL_964340, EPI_ISL_964341, EPI_ISL_964342, EPI_ISL_964343, EPI_ISL_964344, EPI_ISL_964345

| | | | |
|---|---|---|---|
| see above | Lighthouse Lab in Alderley Park | Wellcome Sanger Institute for the COVID-19 Genomics UK (COG-UK) Consortium | Jacquelyn Wynn, Mairead Hyland, The Lighthouse Lab in Alderley Park and Alex Alderton, Roberto Amato, Sonia Goncalves, Ewan Harrison, David K. Jackson, Ian Johnston, Dominic Kwiatkowski, Cordelia Langford, John Sillitoe on behalf of the Wellcome Sanger Institute COVID-19 Surveillance Team |
| EPI_ISL_964346, EPI_ISL_964347 | Lighthouse Lab in Milton Keynes | Wellcome Sanger Institute for the COVID-19 Genomics UK (COG-UK) Consortium | The Lighthouse Lab in Milton Keynes and Alex Alderton, Roberto Amato, Sonia Goncalves, Ewan Harrison, David K. Jackson, Ian Johnston, Dominic Kwiatkowski, Cordelia Langford, John Sillitoe on behalf of the Wellcome Sanger Institute COVID-19 Surveillance Team |
| EPI_ISL_964348, EPI_ISL_964349 | Lighthouse Lab in Glasgow | Wellcome Sanger Institute for the COVID-19 Genomics UK (COG-UK) Consortium | Harper VanSteenhouse, Yumi Kasai, David Gray, Carol Clugston, Anna Dominiczak and Alex Alderton, Roberto Amato, Sonia Goncalves, Ewan Harrison, David K. Jackson, Ian Johnston, Dominic Kwiatkowski, Cordelia Langford, John Sillitoe on behalf of the Wellcome Sanger Institute COVID-19 Surveillance Team |
| EPI_ISL_964350 | Lighthouse Lab in Milton Keynes | Wellcome Sanger Institute for the COVID-19 Genomics UK (COG-UK) Consortium | The Lighthouse Lab in Milton Keynes and Alex Alderton, Roberto Amato, Sonia Goncalves, Ewan Harrison, David K. Jackson, Ian Johnston, Dominic Kwiatkowski, Cordelia Langford, John Sillitoe on behalf of the Wellcome Sanger Institute COVID-19 Surveillance Team |
| EPI_ISL_964351, EPI_ISL_964352 | Lighthouse Lab in Glasgow | Wellcome Sanger Institute for the COVID-19 Genomics UK (COG-UK) Consortium | Harper VanSteenhouse, Yumi Kasai, David Gray, Carol Clugston, Anna Dominiczak and Alex Alderton, Roberto Amato, Sonia Goncalves, Ewan Harrison, David K. Jackson, Ian Johnston, Dominic Kwiatkowski, Cordelia Langford, John Sillitoe on behalf of the Wellcome Sanger Institute COVID-19 Surveillance Team |
| EPI_ISL_964353, EPI_ISL_964354 | Lighthouse Lab in Milton Keynes | Wellcome Sanger Institute for the COVID-19 Genomics UK (COG-UK) Consortium | The Lighthouse Lab in Milton Keynes and Alex Alderton, Roberto Amato, Sonia Goncalves, Ewan Harrison, David K. Jackson, Ian Johnston, Dominic Kwiatkowski, Cordelia Langford, John Sillitoe on behalf of the Wellcome Sanger Institute COVID-19 Surveillance Team |
| EPI_ISL_964355 | Lighthouse Lab in Glasgow | Wellcome Sanger Institute for the COVID-19 Genomics UK (COG-UK) Consortium | Harper VanSteenhouse, Yumi Kasai, David Gray, Carol Clugston, Anna Dominiczak and Alex Alderton, Roberto Amato, Sonia Goncalves, Ewan Harrison, David K. Jackson, Ian Johnston, Dominic Kwiatkowski, Cordelia Langford, John Sillitoe on behalf of the Wellcome Sanger Institute COVID-19 Surveillance Team |
| EPI_ISL_964356 | Lighthouse Lab in Milton Keynes | Wellcome Sanger Institute for the COVID-19 Genomics UK (COG-UK) Consortium | The Lighthouse Lab in Milton Keynes and Alex Alderton, Roberto Amato, Sonia Goncalves, Ewan Harrison, David K. Jackson, Ian Johnston, Dominic Kwiatkowski, Cordelia Langford, John Sillitoe on behalf of the Wellcome Sanger Institute COVID-19 Surveillance Team |
| EPI_ISL_964357, EPI_ISL_964358, EPI_ISL_964359, EPI_ISL_964360 | Lighthouse Lab in Glasgow | Wellcome Sanger Institute for the COVID-19 Genomics UK (COG-UK) Consortium | Harper VanSteenhouse, Yumi Kasai, David Gray, Carol Clugston, Anna Dominiczak and Alex Alderton, Roberto Amato, Sonia Goncalves, Ewan Harrison, David K. Jackson, Ian Johnston, Dominic Kwiatkowski, Cordelia Langford, John Sillitoe on behalf of the Wellcome Sanger Institute COVID-19 Surveillance Team |
| EPI_ISL_964361 | Lighthouse Lab in Milton Keynes | Wellcome Sanger Institute for the COVID-19 Genomics UK (COG-UK) Consortium | The Lighthouse Lab in Milton Keynes and Alex Alderton, Roberto Amato, Sonia Goncalves, Ewan Harrison, David K. Jackson, Ian Johnston, Dominic Kwiatkowski, Cordelia Langford, John Sillitoe on behalf of the Wellcome Sanger Institute COVID-19 Surveillance Team |
| EPI_ISL_964362 | Lighthouse Lab in Glasgow | Wellcome Sanger Institute for the COVID-19 Genomics UK (COG-UK) Consortium | Harper VanSteenhouse, Yumi Kasai, David Gray, Carol Clugston, Anna Dominiczak and Alex Alderton, Roberto Amato, Sonia Goncalves, Ewan Harrison, David K. Jackson, Ian Johnston, Dominic Kwiatkowski, Cordelia Langford, John Sillitoe on behalf of the Wellcome Sanger Institute COVID-19 Surveillance Team |
| EPI_ISL_964363 | Lighthouse Lab in Milton Keynes | Wellcome Sanger Institute for the COVID-19 Genomics UK (COG-UK) Consortium | The Lighthouse Lab in Milton Keynes and Alex Alderton, Roberto Amato, Sonia Goncalves, Ewan Harrison, David K. Jackson, Ian Johnston, Dominic Kwiatkowski, Cordelia Langford, John Sillitoe on behalf of the Wellcome Sanger Institute COVID-19 Surveillance Team |
| EPI_ISL_964364 | Lighthouse Lab in Glasgow | Wellcome Sanger Institute for the COVID-19 Genomics UK (COG-UK) Consortium | Harper VanSteenhouse, Yumi Kasai, David Gray, Carol Clugston, Anna Dominiczak and Alex Alderton, Roberto Amato, Sonia Goncalves, Ewan Harrison, David K. Jackson, Ian Johnston, Dominic Kwiatkowski, Cordelia Langford, John Sillitoe on behalf of the Wellcome Sanger Institute COVID-19 Surveillance Team |
| EPI_ISL_964365 | Lighthouse Lab in Milton Keynes | Wellcome Sanger Institute for the COVID-19 Genomics UK (COG-UK) Consortium | The Lighthouse Lab in Milton Keynes and Alex Alderton, Roberto Amato, Sonia Goncalves, Ewan Harrison, David K. Jackson, Ian Johnston, Dominic Kwiatkowski, Cordelia Langford, John Sillitoe on behalf of the Wellcome Sanger Institute COVID-19 Surveillance Team |
| EPI_ISL_964366, EPI_ISL_964367, EPI_ISL_964368, EPI_ISL_964369 | Lighthouse Lab in Glasgow | Wellcome Sanger Institute for the COVID-19 Genomics UK (COG-UK) Consortium | Harper VanSteenhouse, Yumi Kasai, David Gray, Carol Clugston, Anna Dominiczak and Alex Alderton, Roberto Amato, Sonia Goncalves, Ewan Harrison, David K. Jackson, Ian Johnston, Dominic Kwiatkowski, Cordelia Langford, John Sillitoe on behalf of the Wellcome Sanger Institute COVID-19 Surveillance Team |
| EPI_ISL_964370 | Lighthouse Lab in Milton Keynes | Wellcome Sanger Institute for the COVID-19 Genomics UK (COG-UK) Consortium | The Lighthouse Lab in Milton Keynes and Alex Alderton, Roberto Amato, Sonia Goncalves, Ewan Harrison, David K. Jackson, Ian Johnston, Dominic Kwiatkowski, Cordelia Langford, John Sillitoe on behalf of the Wellcome Sanger Institute COVID-19 Surveillance Team |
| EPI_ISL_964371, EPI_ISL_964372 | Lighthouse Lab in Glasgow | Wellcome Sanger Institute for the COVID-19 Genomics UK (COG-UK) Consortium | Harper VanSteenhouse, Yumi Kasai, David Gray, Carol Clugston, Anna Dominiczak and Alex Alderton, Roberto Amato, Sonia Goncalves, Ewan Harrison, David K. Jackson, Ian Johnston, Dominic Kwiatkowski, Cordelia Langford, John Sillitoe on behalf of the Wellcome Sanger Institute COVID-19 Surveillance Team |
| EPI_ISL_964373, EPI_ISL_964374 | Lighthouse Lab in Milton Keynes | Wellcome Sanger Institute for the COVID-19 Genomics UK (COG-UK) Consortium | The Lighthouse Lab in Milton Keynes and Alex Alderton, Roberto Amato, Sonia Goncalves, Ewan Harrison, David K. Jackson, Ian Johnston, Dominic Kwiatkowski, Cordelia Langford, John Sillitoe on behalf of the Wellcome Sanger Institute COVID-19 Surveillance Team |
| EPI_ISL_964375 | Lighthouse Lab in Glasgow | Wellcome Sanger Institute for the COVID-19 Genomics UK (COG-UK) Consortium | Harper VanSteenhouse, Yumi Kasai, David Gray, Carol Clugston, Anna Dominiczak and Alex Alderton, Roberto Amato, Sonia Goncalves, Ewan Harrison, David K. Jackson, Ian Johnston, Dominic Kwiatkowski, Cordelia Langford, John Sillitoe on behalf of the Wellcome Sanger Institute COVID-19 Surveillance Team |
| EPI_ISL_964376 | Lighthouse Lab in Milton Keynes | Wellcome Sanger Institute for the COVID-19 Genomics UK (COG-UK) Consortium | The Lighthouse Lab in Milton Keynes and Alex Alderton, Roberto Amato, Sonia Goncalves, Ewan Harrison, David K. Jackson, Ian Johnston, Dominic Kwiatkowski, Cordelia Langford, John Sillitoe on behalf of the Wellcome Sanger Institute COVID-19 Surveillance Team |
| EPI_ISL_964377, EPI_ISL_964378 | Lighthouse Lab in Glasgow | Wellcome Sanger Institute for the COVID-19 Genomics UK (COG-UK) Consortium | Harper VanSteenhouse, Yumi Kasai, David Gray, Carol Clugston, Anna Dominiczak and Alex Alderton, Roberto Amato, Sonia Goncalves, Ewan Harrison, David K. Jackson, Ian Johnston, Dominic Kwiatkowski, Cordelia Langford, John Sillitoe on behalf of the Wellcome Sanger Institute COVID-19 Surveillance Team |

| | | | |
|---|---|---|---|
| EPI_ISL_964379 | Lighthouse Lab in Milton Keynes | Wellcome Sanger Institute for the COVID-19 Genomics UK (COG-UK) Consortium | The Lighthouse Lab in Milton Keynes and Alex Alderton, Roberto Amato, Sonia Goncalves, Ewan Harrison, David K. Jackson, Ian Johnston, Dominic Kwiatkowski, Cordelia Langford, John Sillitoe on behalf of the Wellcome Sanger Institute COVID-19 Surveillance Team |
| EPI_ISL_964380 | Lighthouse Lab in Glasgow | Wellcome Sanger Institute for the COVID-19 Genomics UK (COG-UK) Consortium | Harper VanSteenhouse, Yumi Kasai, David Gray, Carol Clugston, Anna Dominiczak and Alex Alderton, Roberto Amato, Sonia Goncalves, Ewan Harrison, David K. Jackson, Ian Johnston, Dominic Kwiatkowski, Cordelia Langford, John Sillitoe on behalf of the Wellcome Sanger Institute COVID-19 Surveillance Team |
| EPI_ISL_964381 | Lighthouse Lab in Milton Keynes | Wellcome Sanger Institute for the COVID-19 Genomics UK (COG-UK) Consortium | The Lighthouse Lab in Milton Keynes and Alex Alderton, Roberto Amato, Sonia Goncalves, Ewan Harrison, David K. Jackson, Ian Johnston, Dominic Kwiatkowski, Cordelia Langford, John Sillitoe on behalf of the Wellcome Sanger Institute COVID-19 Surveillance Team |
| EPI_ISL_964382 | Lighthouse Lab in Glasgow | Wellcome Sanger Institute for the COVID-19 Genomics UK (COG-UK) Consortium | Harper VanSteenhouse, Yumi Kasai, David Gray, Carol Clugston, Anna Dominiczak and Alex Alderton, Roberto Amato, Sonia Goncalves, Ewan Harrison, David K. Jackson, Ian Johnston, Dominic Kwiatkowski, Cordelia Langford, John Sillitoe on behalf of the Wellcome Sanger Institute COVID-19 Surveillance Team |
| EPI_ISL_964383 | Lighthouse Lab in Milton Keynes | Wellcome Sanger Institute for the COVID-19 Genomics UK (COG-UK) Consortium | The Lighthouse Lab in Milton Keynes and Alex Alderton, Roberto Amato, Sonia Goncalves, Ewan Harrison, David K. Jackson, Ian Johnston, Dominic Kwiatkowski, Cordelia Langford, John Sillitoe on behalf of the Wellcome Sanger Institute COVID-19 Surveillance Team |
| EPI_ISL_964384 | Lighthouse Lab in Glasgow | Wellcome Sanger Institute for the COVID-19 Genomics UK (COG-UK) Consortium | Harper VanSteenhouse, Yumi Kasai, David Gray, Carol Clugston, Anna Dominiczak and Alex Alderton, Roberto Amato, Sonia Goncalves, Ewan Harrison, David K. Jackson, Ian Johnston, Dominic Kwiatkowski, Cordelia Langford, John Sillitoe on behalf of the Wellcome Sanger Institute COVID-19 Surveillance Team |
| EPI_ISL_964385, EPI_ISL_964386 | Lighthouse Lab in Milton Keynes | Wellcome Sanger Institute for the COVID-19 Genomics UK (COG-UK) Consortium | The Lighthouse Lab in Milton Keynes and Alex Alderton, Roberto Amato, Sonia Goncalves, Ewan Harrison, David K. Jackson, Ian Johnston, Dominic Kwiatkowski, Cordelia Langford, John Sillitoe on behalf of the Wellcome Sanger Institute COVID-19 Surveillance Team |
| EPI_ISL_964387, EPI_ISL_964388 | Lighthouse Lab in Glasgow | Wellcome Sanger Institute for the COVID-19 Genomics UK (COG-UK) Consortium | Harper VanSteenhouse, Yumi Kasai, David Gray, Carol Clugston, Anna Dominiczak and Alex Alderton, Roberto Amato, Sonia Goncalves, Ewan Harrison, David K. Jackson, Ian Johnston, Dominic Kwiatkowski, Cordelia Langford, John Sillitoe on behalf of the Wellcome Sanger Institute COVID-19 Surveillance Team |
| EPI_ISL_964389 | Lighthouse Lab in Milton Keynes | Wellcome Sanger Institute for the COVID-19 Genomics UK (COG-UK) Consortium | The Lighthouse Lab in Milton Keynes and Alex Alderton, Roberto Amato, Sonia Goncalves, Ewan Harrison, David K. Jackson, Ian Johnston, Dominic Kwiatkowski, Cordelia Langford, John Sillitoe on behalf of the Wellcome Sanger Institute COVID-19 Surveillance Team |
| EPI_ISL_964390, EPI_ISL_964391 | Lighthouse Lab in Glasgow | Wellcome Sanger Institute for the COVID-19 Genomics UK (COG-UK) Consortium | Harper VanSteenhouse, Yumi Kasai, David Gray, Carol Clugston, Anna Dominiczak and Alex Alderton, Roberto Amato, Sonia Goncalves, Ewan Harrison, David K. Jackson, Ian Johnston, Dominic Kwiatkowski, Cordelia Langford, John Sillitoe on behalf of the Wellcome Sanger Institute COVID-19 Surveillance Team |
| EPI_ISL_964392 | Lighthouse Lab in Milton Keynes | Wellcome Sanger Institute for the COVID-19 Genomics UK (COG-UK) Consortium | The Lighthouse Lab in Milton Keynes and Alex Alderton, Roberto Amato, Sonia Goncalves, Ewan Harrison, David K. Jackson, Ian Johnston, Dominic Kwiatkowski, Cordelia Langford, John Sillitoe on behalf of the Wellcome Sanger Institute COVID-19 Surveillance Team |
| EPI_ISL_964393 | Lighthouse Lab in Glasgow | Wellcome Sanger Institute for the COVID-19 Genomics UK (COG-UK) Consortium | Harper VanSteenhouse, Yumi Kasai, David Gray, Carol Clugston, Anna Dominiczak and Alex Alderton, Roberto Amato, Sonia Goncalves, Ewan Harrison, David K. Jackson, Ian Johnston, Dominic Kwiatkowski, Cordelia Langford, John Sillitoe on behalf of the Wellcome Sanger Institute COVID-19 Surveillance Team |
| EPI_ISL_964394, EPI_ISL_964395 | Lighthouse Lab in Milton Keynes | Wellcome Sanger Institute for the COVID-19 Genomics UK (COG-UK) Consortium | The Lighthouse Lab in Milton Keynes and Alex Alderton, Roberto Amato, Sonia Goncalves, Ewan Harrison, David K. Jackson, Ian Johnston, Dominic Kwiatkowski, Cordelia Langford, John Sillitoe on behalf of the Wellcome Sanger Institute COVID-19 Surveillance Team |
| EPI_ISL_964396, EPI_ISL_964397 | Lighthouse Lab in Glasgow | Wellcome Sanger Institute for the COVID-19 Genomics UK (COG-UK) Consortium | Harper VanSteenhouse, Yumi Kasai, David Gray, Carol Clugston, Anna Dominiczak and Alex Alderton, Roberto Amato, Sonia Goncalves, Ewan Harrison, David K. Jackson, Ian Johnston, Dominic Kwiatkowski, Cordelia Langford, John Sillitoe on behalf of the Wellcome Sanger Institute COVID-19 Surveillance Team |
| EPI_ISL_964398 | Lighthouse Lab in Milton Keynes | Wellcome Sanger Institute for the COVID-19 Genomics UK (COG-UK) Consortium | The Lighthouse Lab in Milton Keynes and Alex Alderton, Roberto Amato, Sonia Goncalves, Ewan Harrison, David K. Jackson, Ian Johnston, Dominic Kwiatkowski, Cordelia Langford, John Sillitoe on behalf of the Wellcome Sanger Institute COVID-19 Surveillance Team |
| EPI_ISL_964399 | Lighthouse Lab in Glasgow | Wellcome Sanger Institute for the COVID-19 Genomics UK (COG-UK) Consortium | Harper VanSteenhouse, Yumi Kasai, David Gray, Carol Clugston, Anna Dominiczak and Alex Alderton, Roberto Amato, Sonia Goncalves, Ewan Harrison, David K. Jackson, Ian Johnston, Dominic Kwiatkowski, Cordelia Langford, John Sillitoe on behalf of the Wellcome Sanger Institute COVID-19 Surveillance Team |
| EPI_ISL_964400 | Lighthouse Lab in Milton Keynes | Wellcome Sanger Institute for the COVID-19 Genomics UK (COG-UK) Consortium | The Lighthouse Lab in Milton Keynes and Alex Alderton, Roberto Amato, Sonia Goncalves, Ewan Harrison, David K. Jackson, Ian Johnston, Dominic Kwiatkowski, Cordelia Langford, John Sillitoe on behalf of the Wellcome Sanger Institute COVID-19 Surveillance Team |
| EPI_ISL_964401 | Lighthouse Lab in Glasgow | Wellcome Sanger Institute for the COVID-19 Genomics UK (COG-UK) Consortium | Harper VanSteenhouse, Yumi Kasai, David Gray, Carol Clugston, Anna Dominiczak and Alex Alderton, Roberto Amato, Sonia Goncalves, Ewan Harrison, David K. Jackson, Ian Johnston, Dominic Kwiatkowski, Cordelia Langford, John Sillitoe on behalf of the Wellcome Sanger Institute COVID-19 Surveillance Team |
| EPI_ISL_964402, EPI_ISL_964403 | Lighthouse Lab in Milton Keynes | Wellcome Sanger Institute for the COVID-19 Genomics UK (COG-UK) Consortium | The Lighthouse Lab in Milton Keynes and Alex Alderton, Roberto Amato, Sonia Goncalves, Ewan Harrison, David K. Jackson, Ian Johnston, Dominic Kwiatkowski, Cordelia Langford, John Sillitoe on behalf of the Wellcome Sanger Institute COVID-19 Surveillance Team |
| EPI_ISL_964404 | Lighthouse Lab in Glasgow | Wellcome Sanger Institute for the COVID-19 Genomics UK (COG-UK) Consortium | Harper VanSteenhouse, Yumi Kasai, David Gray, Carol Clugston, Anna Dominiczak and Alex Alderton, Roberto Amato, Sonia Goncalves, Ewan Harrison, David K. Jackson, Ian Johnston, Dominic Kwiatkowski, Cordelia Langford, John Sillitoe on behalf of the Wellcome Sanger Institute COVID-19 Surveillance Team |
| EPI_ISL_964405, EPI_ISL_964406 | Lighthouse Lab in Milton Keynes | Wellcome Sanger Institute for the COVID-19 Genomics UK (COG-UK) Consortium | The Lighthouse Lab in Milton Keynes and Alex Alderton, Roberto Amato, Sonia Goncalves, Ewan Harrison, David K. Jackson, Ian Johnston, Dominic Kwiatkowski, Cordelia Langford, John Sillitoe on behalf of the Wellcome Sanger Institute COVID-19 Surveillance Team |
| EPI_ISL_964407, EPI_ISL_964408, EPI_ISL_964409, EPI_ISL_964410, EPI_ISL_964411 | Lighthouse Lab in Glasgow | Wellcome Sanger Institute for the COVID-19 Genomics UK (COG-UK) Consortium | Harper VanSteenhouse, Yumi Kasai, David Gray, Carol Clugston, Anna Dominiczak and Alex Alderton, Roberto Amato, Sonia Goncalves, Ewan Harrison, David K. Jackson, Ian Johnston, Dominic Kwiatkowski, Cordelia Langford, John Sillitoe on behalf of the Wellcome Sanger Institute COVID-19 Surveillance Team |
| EPI_ISL_964412 | Lighthouse Lab in Milton Keynes | Wellcome Sanger Institute for the COVID-19 Genomics UK (COG-UK) Consortium | The Lighthouse Lab in Milton Keynes and Alex Alderton, Roberto Amato, Sonia Goncalves, Ewan Harrison, David K. Jackson, Ian Johnston, Dominic Kwiatkowski, Cordelia Langford, John Sillitoe on behalf of the Wellcome Sanger Institute COVID-19 Surveillance Team |
| EPI_ISL_964413 | Lighthouse Lab in Glasgow | Wellcome Sanger Institute for the COVID-19 Genomics UK (COG-UK) Consortium | Harper VanSteenhouse, Yumi Kasai, David Gray, Carol Clugston, Anna Dominiczak and Alex Alderton, Roberto Amato, Sonia Goncalves, Ewan Harrison, David K. Jackson, Ian Johnston, Dominic Kwiatkowski, Cordelia Langford, John Sillitoe on behalf of the Wellcome Sanger Institute COVID-19 Surveillance Team |
| EPI_ISL_964414 | Lighthouse Lab in Milton Keynes | Wellcome Sanger Institute for the COVID-19 Genomics UK (COG-UK) Consortium | The Lighthouse Lab in Milton Keynes and Alex Alderton, Roberto Amato, Sonia Goncalves, Ewan Harrison, David K. Jackson, Ian Johnston, Dominic Kwiatkowski, Cordelia Langford, John Sillitoe on behalf of the Wellcome Sanger Institute COVID-19 Surveillance Team |
| EPI_ISL_964415, EPI_ISL_964416, EPI_ISL_964417 | Lighthouse Lab in Glasgow | Wellcome Sanger Institute for the COVID-19 Genomics UK (COG-UK) Consortium | Harper VanSteenhouse, Yumi Kasai, David Gray, Carol Clugston, Anna Dominiczak and Alex Alderton, Roberto Amato, Sonia Goncalves, Ewan Harrison, David K. Jackson, Ian Johnston, Dominic Kwiatkowski, Cordelia Langford, John Sillitoe on behalf of the Wellcome Sanger Institute COVID-19 Surveillance Team |
| EPI_ISL_964418, EPI_ISL_964419, EPI_ISL_964420 | Lighthouse Lab in Milton Keynes | Wellcome Sanger Institute for the COVID-19 Genomics UK (COG-UK) Consortium | The Lighthouse Lab in Milton Keynes and Alex Alderton, Roberto Amato, Sonia Goncalves, Ewan Harrison, David K. Jackson, Ian Johnston, Dominic Kwiatkowski, Cordelia Langford, John Sillitoe on behalf of the Wellcome Sanger Institute COVID-19 Surveillance Team |
| EPI_ISL_964421, EPI_ISL_964422 | Lighthouse Lab in Glasgow | Wellcome Sanger Institute for the COVID-19 Genomics UK (COG-UK) Consortium | Harper VanSteenhouse, Yumi Kasai, David Gray, Carol Clugston, Anna Dominiczak and Alex Alderton, Roberto Amato, Sonia Goncalves, Ewan Harrison, David K. Jackson, Ian Johnston, Dominic Kwiatkowski, Cordelia Langford, John Sillitoe on behalf of the Wellcome Sanger Institute COVID-19 Surveillance Team |
| EPI_ISL_964423 | Lighthouse Lab in Milton Keynes | Wellcome Sanger Institute for the COVID-19 Genomics UK (COG-UK) Consortium | The Lighthouse Lab in Milton Keynes and Alex Alderton, Roberto Amato, Sonia Goncalves, Ewan Harrison, David K. Jackson, Ian Johnston, Dominic Kwiatkowski, Cordelia Langford, John Sillitoe on behalf of the Wellcome Sanger Institute COVID-19 Surveillance Team |
| EPI_ISL_964424, EPI_ISL_964425 | Lighthouse Lab in Glasgow | Wellcome Sanger Institute for the COVID-19 Genomics UK | Harper VanSteenhouse, Yumi Kasai, David Gray, Carol Clugston, Anna Dominiczak and Alex Alderton, Roberto Amato, Sonia Goncalves, Ewan Harrison, |

| | | | |
|---|---|---|---|
| | | (COG-UK) Consortium | David K. Jackson, Ian Johnston, Dominic Kwiatkowski, Cordelia Langford, John Sillitoe on behalf of the Wellcome Sanger Institute COVID-19 Surveillance Team |
| EPI_ISL_964426 | Lighthouse Lab in Milton Keynes | Wellcome Sanger Institute for the COVID-19 Genomics UK (COG-UK) Consortium | The Lighthouse Lab in Milton Keynes and Alex Alderton, Roberto Amato, Sonia Goncalves, Ewan Harrison, David K. Jackson, Ian Johnston, Dominic Kwiatkowski, Cordelia Langford, John Sillitoe on behalf of the Wellcome Sanger Institute COVID-19 Surveillance Team |
| EPI_ISL_964427 | Lighthouse Lab in Glasgow | Wellcome Sanger Institute for the COVID-19 Genomics UK (COG-UK) Consortium | Harper VanSteenhouse, Yumi Kasai, David Gray, Carol Clugston, Anna Dominiczak and Alex Alderton, Roberto Amato, Sonia Goncalves, Ewan Harrison, David K. Jackson, Ian Johnston, Dominic Kwiatkowski, Cordelia Langford, John Sillitoe on behalf of the Wellcome Sanger Institute COVID-19 Surveillance Team |
| EPI_ISL_964428, EPI_ISL_964429 | Lighthouse Lab in Milton Keynes | Wellcome Sanger Institute for the COVID-19 Genomics UK (COG-UK) Consortium | The Lighthouse Lab in Milton Keynes and Alex Alderton, Roberto Amato, Sonia Goncalves, Ewan Harrison, David K. Jackson, Ian Johnston, Dominic Kwiatkowski, Cordelia Langford, John Sillitoe on behalf of the Wellcome Sanger Institute COVID-19 Surveillance Team |
| EPI_ISL_964430 | Lighthouse Lab in Glasgow | Wellcome Sanger Institute for the COVID-19 Genomics UK (COG-UK) Consortium | Harper VanSteenhouse, Yumi Kasai, David Gray, Carol Clugston, Anna Dominiczak and Alex Alderton, Roberto Amato, Sonia Goncalves, Ewan Harrison, David K. Jackson, Ian Johnston, Dominic Kwiatkowski, Cordelia Langford, John Sillitoe on behalf of the Wellcome Sanger Institute COVID-19 Surveillance Team |
| EPI_ISL_964431, EPI_ISL_964432 | Lighthouse Lab in Milton Keynes | Wellcome Sanger Institute for the COVID-19 Genomics UK (COG-UK) Consortium | The Lighthouse Lab in Milton Keynes and Alex Alderton, Roberto Amato, Sonia Goncalves, Ewan Harrison, David K. Jackson, Ian Johnston, Dominic Kwiatkowski, Cordelia Langford, John Sillitoe on behalf of the Wellcome Sanger Institute COVID-19 Surveillance Team |
| EPI_ISL_964433, EPI_ISL_964434 | Lighthouse Lab in Glasgow | Wellcome Sanger Institute for the COVID-19 Genomics UK (COG-UK) Consortium | Harper VanSteenhouse, Yumi Kasai, David Gray, Carol Clugston, Anna Dominiczak and Alex Alderton, Roberto Amato, Sonia Goncalves, Ewan Harrison, David K. Jackson, Ian Johnston, Dominic Kwiatkowski, Cordelia Langford, John Sillitoe on behalf of the Wellcome Sanger Institute COVID-19 Surveillance Team |
| EPI_ISL_964435 | Lighthouse Lab in Milton Keynes | Wellcome Sanger Institute for the COVID-19 Genomics UK (COG-UK) Consortium | The Lighthouse Lab in Milton Keynes and Alex Alderton, Roberto Amato, Sonia Goncalves, Ewan Harrison, David K. Jackson, Ian Johnston, Dominic Kwiatkowski, Cordelia Langford, John Sillitoe on behalf of the Wellcome Sanger Institute COVID-19 Surveillance Team |
| EPI_ISL_964436, EPI_ISL_964437 | Lighthouse Lab in Glasgow | Wellcome Sanger Institute for the COVID-19 Genomics UK (COG-UK) Consortium | Harper VanSteenhouse, Yumi Kasai, David Gray, Carol Clugston, Anna Dominiczak and Alex Alderton, Roberto Amato, Sonia Goncalves, Ewan Harrison, David K. Jackson, Ian Johnston, Dominic Kwiatkowski, Cordelia Langford, John Sillitoe on behalf of the Wellcome Sanger Institute COVID-19 Surveillance Team |
| EPI_ISL_964438 | Lighthouse Lab in Milton Keynes | Wellcome Sanger Institute for the COVID-19 Genomics UK (COG-UK) Consortium | The Lighthouse Lab in Milton Keynes and Alex Alderton, Roberto Amato, Sonia Goncalves, Ewan Harrison, David K. Jackson, Ian Johnston, Dominic Kwiatkowski, Cordelia Langford, John Sillitoe on behalf of the Wellcome Sanger Institute COVID-19 Surveillance Team |
| EPI_ISL_964439, EPI_ISL_964440, EPI_ISL_964441, EPI_ISL_964442 | Lighthouse Lab in Glasgow | Wellcome Sanger Institute for the COVID-19 Genomics UK (COG-UK) Consortium | Harper VanSteenhouse, Yumi Kasai, David Gray, Carol Clugston, Anna Dominiczak and Alex Alderton, Roberto Amato, Sonia Goncalves, Ewan Harrison, David K. Jackson, Ian Johnston, Dominic Kwiatkowski, Cordelia Langford, John Sillitoe on behalf of the Wellcome Sanger Institute COVID-19 Surveillance Team |
| EPI_ISL_964443, EPI_ISL_964444, EPI_ISL_964445 | Lighthouse Lab in Milton Keynes | Wellcome Sanger Institute for the COVID-19 Genomics UK (COG-UK) Consortium | The Lighthouse Lab in Milton Keynes and Alex Alderton, Roberto Amato, Sonia Goncalves, Ewan Harrison, David K. Jackson, Ian Johnston, Dominic Kwiatkowski, Cordelia Langford, John Sillitoe on behalf of the Wellcome Sanger Institute COVID-19 Surveillance Team |
| EPI_ISL_964446 | Lighthouse Lab in Glasgow | Wellcome Sanger Institute for the COVID-19 Genomics UK (COG-UK) Consortium | Harper VanSteenhouse, Yumi Kasai, David Gray, Carol Clugston, Anna Dominiczak and Alex Alderton, Roberto Amato, Sonia Goncalves, Ewan Harrison, David K. Jackson, Ian Johnston, Dominic Kwiatkowski, Cordelia Langford, John Sillitoe on behalf of the Wellcome Sanger Institute COVID-19 Surveillance Team |
| EPI_ISL_964447, EPI_ISL_964448 | Lighthouse Lab in Milton Keynes | Wellcome Sanger Institute for the COVID-19 Genomics UK (COG-UK) Consortium | The Lighthouse Lab in Milton Keynes and Alex Alderton, Roberto Amato, Sonia Goncalves, Ewan Harrison, David K. Jackson, Ian Johnston, Dominic Kwiatkowski, Cordelia Langford, John Sillitoe on behalf of the Wellcome Sanger Institute COVID-19 Surveillance Team |
| EPI_ISL_964449 | Lighthouse Lab in Glasgow | Wellcome Sanger Institute for the COVID-19 Genomics UK (COG-UK) Consortium | Harper VanSteenhouse, Yumi Kasai, David Gray, Carol Clugston, Anna Dominiczak and Alex Alderton, Roberto Amato, Sonia Goncalves, Ewan Harrison, David K. Jackson, Ian Johnston, Dominic Kwiatkowski, Cordelia Langford, John Sillitoe on behalf of the Wellcome Sanger Institute COVID-19 Surveillance Team |
| EPI_ISL_964450 | Lighthouse Lab in Milton Keynes | Wellcome Sanger Institute for the COVID-19 Genomics UK (COG-UK) Consortium | The Lighthouse Lab in Milton Keynes and Alex Alderton, Roberto Amato, Sonia Goncalves, Ewan Harrison, David K. Jackson, Ian Johnston, Dominic Kwiatkowski, Cordelia Langford, John Sillitoe on behalf of the Wellcome Sanger Institute COVID-19 Surveillance Team |
| EPI_ISL_964451, EPI_ISL_964452 | Lighthouse Lab in Glasgow | Wellcome Sanger Institute for the COVID-19 Genomics UK (COG-UK) Consortium | Harper VanSteenhouse, Yumi Kasai, David Gray, Carol Clugston, Anna Dominiczak and Alex Alderton, Roberto Amato, Sonia Goncalves, Ewan Harrison, David K. Jackson, Ian Johnston, Dominic Kwiatkowski, Cordelia Langford, John Sillitoe on behalf of the Wellcome Sanger Institute COVID-19 Surveillance Team |
| EPI_ISL_964453, EPI_ISL_964454, EPI_ISL_964455 | Lighthouse Lab in Milton Keynes | Wellcome Sanger Institute for the COVID-19 Genomics UK (COG-UK) Consortium | The Lighthouse Lab in Milton Keynes and Alex Alderton, Roberto Amato, Sonia Goncalves, Ewan Harrison, David K. Jackson, Ian Johnston, Dominic Kwiatkowski, Cordelia Langford, John Sillitoe on behalf of the Wellcome Sanger Institute COVID-19 Surveillance Team |
| EPI_ISL_964456, EPI_ISL_964457, EPI_ISL_964458 | Lighthouse Lab in Glasgow | Wellcome Sanger Institute for the COVID-19 Genomics UK (COG-UK) Consortium | Harper VanSteenhouse, Yumi Kasai, David Gray, Carol Clugston, Anna Dominiczak and Alex Alderton, Roberto Amato, Sonia Goncalves, Ewan Harrison, David K. Jackson, Ian Johnston, Dominic Kwiatkowski, Cordelia Langford, John Sillitoe on behalf of the Wellcome Sanger Institute COVID-19 Surveillance Team |
| EPI_ISL_964459 | Lighthouse Lab in Milton Keynes | Wellcome Sanger Institute for the COVID-19 Genomics UK (COG-UK) Consortium | The Lighthouse Lab in Milton Keynes and Alex Alderton, Roberto Amato, Sonia Goncalves, Ewan Harrison, David K. Jackson, Ian Johnston, Dominic Kwiatkowski, Cordelia Langford, John Sillitoe on behalf of the Wellcome Sanger Institute COVID-19 Surveillance Team |
| EPI_ISL_964460 | Lighthouse Lab in Glasgow | Wellcome Sanger Institute for the COVID-19 Genomics UK (COG-UK) Consortium | Harper VanSteenhouse, Yumi Kasai, David Gray, Carol Clugston, Anna Dominiczak and Alex Alderton, Roberto Amato, Sonia Goncalves, Ewan Harrison, David K. Jackson, Ian Johnston, Dominic Kwiatkowski, Cordelia Langford, John Sillitoe on behalf of the Wellcome Sanger Institute COVID-19 Surveillance Team |
| EPI_ISL_964461, EPI_ISL_964462, EPI_ISL_964463, EPI_ISL_964464 | Lighthouse Lab in Milton Keynes | Wellcome Sanger Institute for the COVID-19 Genomics UK (COG-UK) Consortium | The Lighthouse Lab in Milton Keynes and Alex Alderton, Roberto Amato, Sonia Goncalves, Ewan Harrison, David K. Jackson, Ian Johnston, Dominic Kwiatkowski, Cordelia Langford, John Sillitoe on behalf of the Wellcome Sanger Institute COVID-19 Surveillance Team |
| EPI_ISL_964465 | Lighthouse Lab in Glasgow | Wellcome Sanger Institute for the COVID-19 Genomics UK (COG-UK) Consortium | Harper VanSteenhouse, Yumi Kasai, David Gray, Carol Clugston, Anna Dominiczak and Alex Alderton, Roberto Amato, Sonia Goncalves, Ewan Harrison, David K. Jackson, Ian Johnston, Dominic Kwiatkowski, Cordelia Langford, John Sillitoe on behalf of the Wellcome Sanger Institute COVID-19 Surveillance Team |
| EPI_ISL_964466, EPI_ISL_964467 | Lighthouse Lab in Milton Keynes | Wellcome Sanger Institute for the COVID-19 Genomics UK (COG-UK) Consortium | The Lighthouse Lab in Milton Keynes and Alex Alderton, Roberto Amato, Sonia Goncalves, Ewan Harrison, David K. Jackson, Ian Johnston, Dominic Kwiatkowski, Cordelia Langford, John Sillitoe on behalf of the Wellcome Sanger Institute COVID-19 Surveillance Team |
| EPI_ISL_964468, EPI_ISL_964469, EPI_ISL_964470, EPI_ISL_964471, EPI_ISL_964472, EPI_ISL_964473, EPI_ISL_964474, EPI_ISL_964475, EPI_ISL_964476, EPI_ISL_964477 | Lighthouse Lab in Glasgow | Wellcome Sanger Institute for the COVID-19 Genomics UK (COG-UK) Consortium | Harper VanSteenhouse, Yumi Kasai, David Gray, Carol Clugston, Anna Dominiczak and Alex Alderton, Roberto Amato, Sonia Goncalves, Ewan Harrison, David K. Jackson, Ian Johnston, Dominic Kwiatkowski, Cordelia Langford, John Sillitoe on behalf of the Wellcome Sanger Institute COVID-19 Surveillance Team |
| EPI_ISL_964478, EPI_ISL_964479 | Lighthouse Lab in Milton Keynes | Wellcome Sanger Institute for the COVID-19 Genomics UK (COG-UK) Consortium | The Lighthouse Lab in Milton Keynes and Alex Alderton, Roberto Amato, Sonia Goncalves, Ewan Harrison, David K. Jackson, Ian Johnston, Dominic Kwiatkowski, Cordelia Langford, John Sillitoe on behalf of the Wellcome Sanger Institute COVID-19 Surveillance Team |
| EPI_ISL_964480 | Lighthouse Lab in Glasgow | Wellcome Sanger Institute for the COVID-19 Genomics UK (COG-UK) Consortium | Harper VanSteenhouse, Yumi Kasai, David Gray, Carol Clugston, Anna Dominiczak and Alex Alderton, Roberto Amato, Sonia Goncalves, Ewan Harrison, David K. Jackson, Ian Johnston, Dominic Kwiatkowski, Cordelia Langford, John Sillitoe on behalf of the Wellcome Sanger Institute COVID-19 Surveillance Team |
| EPI_ISL_964481 | Lighthouse Lab in Milton Keynes | Wellcome Sanger Institute for the COVID-19 Genomics UK (COG-UK) Consortium | The Lighthouse Lab in Milton Keynes and Alex Alderton, Roberto Amato, Sonia Goncalves, Ewan Harrison, David K. Jackson, Ian Johnston, Dominic Kwiatkowski, Cordelia Langford, John Sillitoe on behalf of the Wellcome Sanger Institute COVID-19 Surveillance Team |
| EPI_ISL_964482, EPI_ISL_964483, EPI_ISL_964484 | Lighthouse Lab in Glasgow | Wellcome Sanger Institute for the COVID-19 Genomics UK (COG-UK) Consortium | Harper VanSteenhouse, Yumi Kasai, David Gray, Carol Clugston, Anna Dominiczak and Alex Alderton, Roberto Amato, Sonia Goncalves, Ewan Harrison, David K. Jackson, Ian Johnston, Dominic Kwiatkowski, Cordelia Langford, John Sillitoe on behalf of the Wellcome Sanger Institute COVID-19 Surveillance |

| | | | |
|---|---|---|---|
| | | | Team |
| EPI_ISL_964485, EPI_ISL_964486 | Lighthouse Lab in Milton Keynes | Wellcome Sanger Institute for the COVID-19 Genomics UK (COG-UK) Consortium | The Lighthouse Lab in Milton Keynes and Alex Alderton, Roberto Amato, Sonia Goncalves, Ewan Harrison, David K. Jackson, Ian Johnston, Dominic Kwiatkowski, Cordelia Langford, John Sillitoe on behalf of the Wellcome Sanger Institute COVID-19 Surveillance Team |
| EPI_ISL_964487, EPI_ISL_964488, EPI_ISL_964489, EPI_ISL_964490 | Lighthouse Lab in Glasgow | Wellcome Sanger Institute for the COVID-19 Genomics UK (COG-UK) Consortium | Harper VanSteenhouse, Yumi Kasai, David Gray, Carol Clugston, Anna Dominiczak and Alex Alderton, Roberto Amato, Sonia Goncalves, Ewan Harrison, David K. Jackson, Ian Johnston, Dominic Kwiatkowski, Cordelia Langford, John Sillitoe on behalf of the Wellcome Sanger Institute COVID-19 Surveillance Team |
| EPI_ISL_964491, EPI_ISL_964492 | Lighthouse Lab in Milton Keynes | Wellcome Sanger Institute for the COVID-19 Genomics UK (COG-UK) Consortium | The Lighthouse Lab in Milton Keynes and Alex Alderton, Roberto Amato, Sonia Goncalves, Ewan Harrison, David K. Jackson, Ian Johnston, Dominic Kwiatkowski, Cordelia Langford, John Sillitoe on behalf of the Wellcome Sanger Institute COVID-19 Surveillance Team |
| EPI_ISL_964493, EPI_ISL_964494 | Lighthouse Lab in Glasgow | Wellcome Sanger Institute for the COVID-19 Genomics UK (COG-UK) Consortium | Harper VanSteenhouse, Yumi Kasai, David Gray, Carol Clugston, Anna Dominiczak and Alex Alderton, Roberto Amato, Sonia Goncalves, Ewan Harrison, David K. Jackson, Ian Johnston, Dominic Kwiatkowski, Cordelia Langford, John Sillitoe on behalf of the Wellcome Sanger Institute COVID-19 Surveillance Team |
| EPI_ISL_964495, EPI_ISL_964496, EPI_ISL_964497, EPI_ISL_964498, EPI_ISL_964499 | Lighthouse Lab in Milton Keynes | Wellcome Sanger Institute for the COVID-19 Genomics UK (COG-UK) Consortium | The Lighthouse Lab in Milton Keynes and Alex Alderton, Roberto Amato, Sonia Goncalves, Ewan Harrison, David K. Jackson, Ian Johnston, Dominic Kwiatkowski, Cordelia Langford, John Sillitoe on behalf of the Wellcome Sanger Institute COVID-19 Surveillance Team |
| EPI_ISL_964500, EPI_ISL_964501 | Lighthouse Lab in Glasgow | Wellcome Sanger Institute for the COVID-19 Genomics UK (COG-UK) Consortium | Harper VanSteenhouse, Yumi Kasai, David Gray, Carol Clugston, Anna Dominiczak and Alex Alderton, Roberto Amato, Sonia Goncalves, Ewan Harrison, David K. Jackson, Ian Johnston, Dominic Kwiatkowski, Cordelia Langford, John Sillitoe on behalf of the Wellcome Sanger Institute COVID-19 Surveillance Team |
| EPI_ISL_964502 | Lighthouse Lab in Milton Keynes | Wellcome Sanger Institute for the COVID-19 Genomics UK (COG-UK) Consortium | The Lighthouse Lab in Milton Keynes and Alex Alderton, Roberto Amato, Sonia Goncalves, Ewan Harrison, David K. Jackson, Ian Johnston, Dominic Kwiatkowski, Cordelia Langford, John Sillitoe on behalf of the Wellcome Sanger Institute COVID-19 Surveillance Team |
| EPI_ISL_964503, EPI_ISL_964504 | Lighthouse Lab in Glasgow | Wellcome Sanger Institute for the COVID-19 Genomics UK (COG-UK) Consortium | Harper VanSteenhouse, Yumi Kasai, David Gray, Carol Clugston, Anna Dominiczak and Alex Alderton, Roberto Amato, Sonia Goncalves, Ewan Harrison, David K. Jackson, Ian Johnston, Dominic Kwiatkowski, Cordelia Langford, John Sillitoe on behalf of the Wellcome Sanger Institute COVID-19 Surveillance Team |
| EPI_ISL_964505, EPI_ISL_964506, EPI_ISL_964507 | Lighthouse Lab in Milton Keynes | Wellcome Sanger Institute for the COVID-19 Genomics UK (COG-UK) Consortium | The Lighthouse Lab in Milton Keynes and Alex Alderton, Roberto Amato, Sonia Goncalves, Ewan Harrison, David K. Jackson, Ian Johnston, Dominic Kwiatkowski, Cordelia Langford, John Sillitoe on behalf of the Wellcome Sanger Institute COVID-19 Surveillance Team |
| EPI_ISL_964508, EPI_ISL_964509 | Lighthouse Lab in Glasgow | Wellcome Sanger Institute for the COVID-19 Genomics UK (COG-UK) Consortium | Harper VanSteenhouse, Yumi Kasai, David Gray, Carol Clugston, Anna Dominiczak and Alex Alderton, Roberto Amato, Sonia Goncalves, Ewan Harrison, David K. Jackson, Ian Johnston, Dominic Kwiatkowski, Cordelia Langford, John Sillitoe on behalf of the Wellcome Sanger Institute COVID-19 Surveillance Team |
| EPI_ISL_964510 | Lighthouse Lab in Milton Keynes | Wellcome Sanger Institute for the COVID-19 Genomics UK (COG-UK) Consortium | The Lighthouse Lab in Milton Keynes and Alex Alderton, Roberto Amato, Sonia Goncalves, Ewan Harrison, David K. Jackson, Ian Johnston, Dominic Kwiatkowski, Cordelia Langford, John Sillitoe on behalf of the Wellcome Sanger Institute COVID-19 Surveillance Team |
| EPI_ISL_964511, EPI_ISL_964512 | Lighthouse Lab in Glasgow | Wellcome Sanger Institute for the COVID-19 Genomics UK (COG-UK) Consortium | Harper VanSteenhouse, Yumi Kasai, David Gray, Carol Clugston, Anna Dominiczak and Alex Alderton, Roberto Amato, Sonia Goncalves, Ewan Harrison, David K. Jackson, Ian Johnston, Dominic Kwiatkowski, Cordelia Langford, John Sillitoe on behalf of the Wellcome Sanger Institute COVID-19 Surveillance Team |
| EPI_ISL_964513 | Lighthouse Lab in Milton Keynes | Wellcome Sanger Institute for the COVID-19 Genomics UK (COG-UK) Consortium | The Lighthouse Lab in Milton Keynes and Alex Alderton, Roberto Amato, Sonia Goncalves, Ewan Harrison, David K. Jackson, Ian Johnston, Dominic Kwiatkowski, Cordelia Langford, John Sillitoe on behalf of the Wellcome Sanger Institute COVID-19 Surveillance Team |
| EPI_ISL_964514 | Lighthouse Lab in Glasgow | Wellcome Sanger Institute for the COVID-19 Genomics UK (COG-UK) Consortium | Harper VanSteenhouse, Yumi Kasai, David Gray, Carol Clugston, Anna Dominiczak and Alex Alderton, Roberto Amato, Sonia Goncalves, Ewan Harrison, David K. Jackson, Ian Johnston, Dominic Kwiatkowski, Cordelia Langford, John Sillitoe on behalf of the Wellcome Sanger Institute COVID-19 Surveillance Team |
| EPI_ISL_964515 | Lighthouse Lab in Milton Keynes | Wellcome Sanger Institute for the COVID-19 Genomics UK (COG-UK) Consortium | The Lighthouse Lab in Milton Keynes and Alex Alderton, Roberto Amato, Sonia Goncalves, Ewan Harrison, David K. Jackson, Ian Johnston, Dominic Kwiatkowski, Cordelia Langford, John Sillitoe on behalf of the Wellcome Sanger Institute COVID-19 Surveillance Team |
| EPI_ISL_964516 | Lighthouse Lab in Glasgow | Wellcome Sanger Institute for the COVID-19 Genomics UK (COG-UK) Consortium | Harper VanSteenhouse, Yumi Kasai, David Gray, Carol Clugston, Anna Dominiczak and Alex Alderton, Roberto Amato, Sonia Goncalves, Ewan Harrison, David K. Jackson, Ian Johnston, Dominic Kwiatkowski, Cordelia Langford, John Sillitoe on behalf of the Wellcome Sanger Institute COVID-19 Surveillance Team |
| EPI_ISL_964517 | Lighthouse Lab in Milton Keynes | Wellcome Sanger Institute for the COVID-19 Genomics UK (COG-UK) Consortium | The Lighthouse Lab in Milton Keynes and Alex Alderton, Roberto Amato, Sonia Goncalves, Ewan Harrison, David K. Jackson, Ian Johnston, Dominic Kwiatkowski, Cordelia Langford, John Sillitoe on behalf of the Wellcome Sanger Institute COVID-19 Surveillance Team |
| EPI_ISL_964518 | Lighthouse Lab in Glasgow | Wellcome Sanger Institute for the COVID-19 Genomics UK (COG-UK) Consortium | Harper VanSteenhouse, Yumi Kasai, David Gray, Carol Clugston, Anna Dominiczak and Alex Alderton, Roberto Amato, Sonia Goncalves, Ewan Harrison, David K. Jackson, Ian Johnston, Dominic Kwiatkowski, Cordelia Langford, John Sillitoe on behalf of the Wellcome Sanger Institute COVID-19 Surveillance Team |
| EPI_ISL_964519 | Lighthouse Lab in Milton Keynes | Wellcome Sanger Institute for the COVID-19 Genomics UK (COG-UK) Consortium | The Lighthouse Lab in Milton Keynes and Alex Alderton, Roberto Amato, Sonia Goncalves, Ewan Harrison, David K. Jackson, Ian Johnston, Dominic Kwiatkowski, Cordelia Langford, John Sillitoe on behalf of the Wellcome Sanger Institute COVID-19 Surveillance Team |
| EPI_ISL_964520, EPI_ISL_964521 | Lighthouse Lab in Glasgow | Wellcome Sanger Institute for the COVID-19 Genomics UK (COG-UK) Consortium | Harper VanSteenhouse, Yumi Kasai, David Gray, Carol Clugston, Anna Dominiczak and Alex Alderton, Roberto Amato, Sonia Goncalves, Ewan Harrison, David K. Jackson, Ian Johnston, Dominic Kwiatkowski, Cordelia Langford, John Sillitoe on behalf of the Wellcome Sanger Institute COVID-19 Surveillance Team |
| EPI_ISL_964522 | Lighthouse Lab in Milton Keynes | Wellcome Sanger Institute for the COVID-19 Genomics UK (COG-UK) Consortium | The Lighthouse Lab in Milton Keynes and Alex Alderton, Roberto Amato, Sonia Goncalves, Ewan Harrison, David K. Jackson, Ian Johnston, Dominic Kwiatkowski, Cordelia Langford, John Sillitoe on behalf of the Wellcome Sanger Institute COVID-19 Surveillance Team |
| EPI_ISL_964523 | Lighthouse Lab in Glasgow | Wellcome Sanger Institute for the COVID-19 Genomics UK (COG-UK) Consortium | Harper VanSteenhouse, Yumi Kasai, David Gray, Carol Clugston, Anna Dominiczak and Alex Alderton, Roberto Amato, Sonia Goncalves, Ewan Harrison, David K. Jackson, Ian Johnston, Dominic Kwiatkowski, Cordelia Langford, John Sillitoe on behalf of the Wellcome Sanger Institute COVID-19 Surveillance Team |
| EPI_ISL_964524 | Lighthouse Lab in Milton Keynes | Wellcome Sanger Institute for the COVID-19 Genomics UK (COG-UK) Consortium | The Lighthouse Lab in Milton Keynes and Alex Alderton, Roberto Amato, Sonia Goncalves, Ewan Harrison, David K. Jackson, Ian Johnston, Dominic Kwiatkowski, Cordelia Langford, John Sillitoe on behalf of the Wellcome Sanger Institute COVID-19 Surveillance Team |
| EPI_ISL_964525, EPI_ISL_964526, EPI_ISL_964527, EPI_ISL_964528 | Lighthouse Lab in Glasgow | Wellcome Sanger Institute for the COVID-19 Genomics UK (COG-UK) Consortium | Harper VanSteenhouse, Yumi Kasai, David Gray, Carol Clugston, Anna Dominiczak and Alex Alderton, Roberto Amato, Sonia Goncalves, Ewan Harrison, David K. Jackson, Ian Johnston, Dominic Kwiatkowski, Cordelia Langford, John Sillitoe on behalf of the Wellcome Sanger Institute COVID-19 Surveillance Team |
| EPI_ISL_964529, EPI_ISL_964530, EPI_ISL_964531 | Lighthouse Lab in Milton Keynes | Wellcome Sanger Institute for the COVID-19 Genomics UK (COG-UK) Consortium | The Lighthouse Lab in Milton Keynes and Alex Alderton, Roberto Amato, Sonia Goncalves, Ewan Harrison, David K. Jackson, Ian Johnston, Dominic Kwiatkowski, Cordelia Langford, John Sillitoe on behalf of the Wellcome Sanger Institute COVID-19 Surveillance Team |
| EPI_ISL_964532, EPI_ISL_964533 | Lighthouse Lab in Glasgow | Wellcome Sanger Institute for the COVID-19 Genomics UK (COG-UK) Consortium | Harper VanSteenhouse, Yumi Kasai, David Gray, Carol Clugston, Anna Dominiczak and Alex Alderton, Roberto Amato, Sonia Goncalves, Ewan Harrison, David K. Jackson, Ian Johnston, Dominic Kwiatkowski, Cordelia Langford, John Sillitoe on behalf of the Wellcome Sanger Institute COVID-19 Surveillance Team |
| EPI_ISL_964534, EPI_ISL_964535, EPI_ISL_964536 | Lighthouse Lab in Milton Keynes | Wellcome Sanger Institute for the COVID-19 Genomics UK (COG-UK) Consortium | The Lighthouse Lab in Milton Keynes and Alex Alderton, Roberto Amato, Sonia Goncalves, Ewan Harrison, David K. Jackson, Ian Johnston, Dominic Kwiatkowski, Cordelia Langford, John Sillitoe on behalf of the Wellcome Sanger Institute COVID-19 Surveillance Team |
| EPI_ISL_964537 | Lighthouse Lab in Glasgow | Wellcome Sanger Institute for the COVID-19 Genomics UK (COG-UK) Consortium | Harper VanSteenhouse, Yumi Kasai, David Gray, Carol Clugston, Anna Dominiczak and Alex Alderton, Roberto Amato, Sonia Goncalves, Ewan Harrison, David K. Jackson, Ian Johnston, Dominic Kwiatkowski, Cordelia Langford, John Sillitoe on behalf of the Wellcome Sanger Institute COVID-19 Surveillance Team |
| EPI_ISL_964538 | Lighthouse Lab in Milton Keynes | Wellcome Sanger Institute for the COVID-19 Genomics UK | The Lighthouse Lab in Milton Keynes and Alex Alderton, Roberto Amato, Sonia Goncalves, Ewan Harrison, David K. Jackson, Ian Johnston, Dominic |

| | | | |
|---|---|---|---|
| | | (COG-UK) Consortium | Kwiatkowski, Cordelia Langford, John Sillitoe on behalf of the Wellcome Sanger Institute COVID-19 Surveillance Team |
| EPI_ISL_964539, EPI_ISL_964540, EPI_ISL_964541, EPI_ISL_964542 | Lighthouse Lab in Glasgow | Wellcome Sanger Institute for the COVID-19 Genomics UK (COG-UK) Consortium | Harper VanSteenhouse, Yumi Kasai, David Gray, Carol Clugston, Anna Dominiczak and Alex Alderton, Roberto Amato, Sonia Goncalves, Ewan Harrison, David K. Jackson, Ian Johnston, Dominic Kwiatkowski, Cordelia Langford, John Sillitoe on behalf of the Wellcome Sanger Institute COVID-19 Surveillance Team |
| EPI_ISL_964543, EPI_ISL_964544, EPI_ISL_964545, EPI_ISL_964546, EPI_ISL_964547, EPI_ISL_964548, EPI_ISL_964549, EPI_ISL_964550, EPI_ISL_964551, EPI_ISL_964552, EPI_ISL_964553, EPI_ISL_964554, EPI_ISL_964555, EPI_ISL_964556, EPI_ISL_964557, EPI_ISL_964558, EPI_ISL_964559, EPI_ISL_964560, EPI_ISL_964561, EPI_ISL_964562, EPI_ISL_964563, EPI_ISL_964564, EPI_ISL_964565, EPI_ISL_964566, EPI_ISL_964567, EPI_ISL_964568, EPI_ISL_964569, EPI_ISL_964570, EPI_ISL_964571, EPI_ISL_964572, EPI_ISL_964573, EPI_ISL_964574, EPI_ISL_964575, EPI_ISL_964576, EPI_ISL_964577, EPI_ISL_964578, EPI_ISL_964579, EPI_ISL_964580, EPI_ISL_964581, EPI_ISL_964582, EPI_ISL_964583, EPI_ISL_964584, EPI_ISL_964585, EPI_ISL_964586, EPI_ISL_964587, EPI_ISL_964588, EPI_ISL_964589, EPI_ISL_964590, EPI_ISL_964591, EPI_ISL_964592, EPI_ISL_964593, EPI_ISL_964594, EPI_ISL_964595, EPI_ISL_964596, EPI_ISL_964597, EPI_ISL_964598, EPI_ISL_964599, EPI_ISL_964600, EPI_ISL_964601, EPI_ISL_964602, EPI_ISL_964603, EPI_ISL_964604, EPI_ISL_964605, EPI_ISL_964606, EPI_ISL_964607, EPI_ISL_964608, EPI_ISL_964609, EPI_ISL_964610, EPI_ISL_964611, EPI_ISL_964612, EPI_ISL_964613, EPI_ISL_964614, EPI_ISL_964615, EPI_ISL_964616, EPI_ISL_964617, EPI_ISL_964618, EPI_ISL_964619, EPI_ISL_964620, EPI_ISL_964621, EPI_ISL_964622, EPI_ISL_964623, EPI_ISL_964624, EPI_ISL_964625, EPI_ISL_964626, EPI_ISL_964627, EPI_ISL_964628, EPI_ISL_964629, EPI_ISL_964630, EPI_ISL_964631, EPI_ISL_964632, EPI_ISL_964633, EPI_ISL_964634, EPI_ISL_964635, EPI_ISL_964636, EPI_ISL_964637, EPI_ISL_964638, EPI_ISL_964639, EPI_ISL_964640, EPI_ISL_964641, EPI_ISL_964642, EPI_ISL_964643, EPI_ISL_964644, EPI_ISL_964645, EPI_ISL_964646, EPI_ISL_964647, EPI_ISL_964648, EPI_ISL_964649, EPI_ISL_964650, EPI_ISL_964651, EPI_ISL_964652, EPI_ISL_964653, EPI_ISL_964654, EPI_ISL_964655, EPI_ISL_964656, EPI_ISL_964657, EPI_ISL_964658, EPI_ISL_964659, EPI_ISL_964660, EPI_ISL_964661, EPI_ISL_964662, EPI_ISL_964663, EPI_ISL_964664, EPI_ISL_964665, EPI_ISL_964666, EPI_ISL_964667, EPI_ISL_964668, EPI_ISL_964669, EPI_ISL_964670, EPI_ISL_964671, EPI_ISL_964672, EPI_ISL_964673, EPI_ISL_964674, EPI_ISL_964675, EPI_ISL_964676, EPI_ISL_964677, EPI_ISL_964678, EPI_ISL_964679, EPI_ISL_964680, EPI_ISL_964681, EPI_ISL_964682, EPI_ISL_964683, EPI_ISL_964684, EPI_ISL_964685, EPI_ISL_964686, EPI_ISL_964687, EPI_ISL_964688, EPI_ISL_964689, EPI_ISL_964690, EPI_ISL_964691, EPI_ISL_964692, EPI_ISL_964693, EPI_ISL_964694, EPI_ISL_964695, EPI_ISL_964696, EPI_ISL_964697, EPI_ISL_964698, EPI_ISL_964699, EPI_ISL_964700, EPI_ISL_964701, EPI_ISL_964702, EPI_ISL_964703, EPI_ISL_964704, EPI_ISL_964705, EPI_ISL_964706, EPI_ISL_964707, EPI_ISL_964708, EPI_ISL_964709, EPI_ISL_964711, EPI_ISL_964712, EPI_ISL_964713, EPI_ISL_964714, EPI_ISL_964715, EPI_ISL_964716, EPI_ISL_964717, EPI_ISL_964718, EPI_ISL_964719, EPI_ISL_964720, EPI_ISL_964721, EPI_ISL_964722, EPI_ISL_964723, EPI_ISL_964724, EPI_ISL_964725, EPI_ISL_964727, EPI_ISL_964728, EPI_ISL_964729, EPI_ISL_964730, EPI_ISL_964731, EPI_ISL_964732, EPI_ISL_964733, EPI_ISL_964734, EPI_ISL_964735, EPI_ISL_964736, EPI_ISL_964737, EPI_ISL_964738, EPI_ISL_964739, EPI_ISL_964740, EPI_ISL_964741, EPI_ISL_964742, EPI_ISL_964743, EPI_ISL_964744, EPI_ISL_964745, EPI_ISL_964746, EPI_ISL_964747, EPI_ISL_964748, EPI_ISL_964750, EPI_ISL_964751, EPI_ISL_964752, EPI_ISL_964753, EPI_ISL_964754, EPI_ISL_964755, EPI_ISL_964756, EPI_ISL_964757, EPI_ISL_964758, EPI_ISL_964760, EPI_ISL_964761, EPI_ISL_964762, EPI_ISL_964763, EPI_ISL_964764, EPI_ISL_964765, EPI_ISL_964766, EPI_ISL_964767, EPI_ISL_964768, EPI_ISL_964769, EPI_ISL_964770, EPI_ISL_964771, EPI_ISL_964772, EPI_ISL_964773, EPI_ISL_964774, EPI_ISL_964775, EPI_ISL_964776, EPI_ISL_964777, EPI_ISL_964778, EPI_ISL_964779, EPI_ISL_964780, EPI_ISL_964781, EPI_ISL_964782, EPI_ISL_964783, EPI_ISL_964784, EPI_ISL_964785, EPI_ISL_964786, EPI_ISL_964787, EPI_ISL_964788, EPI_ISL_964789, EPI_ISL_964790, EPI_ISL_964791, EPI_ISL_964792, EPI_ISL_964793, EPI_ISL_964794, EPI_ISL_964795, EPI_ISL_964796, EPI_ISL_964797, EPI_ISL_964798, EPI_ISL_964799, EPI_ISL_964800, EPI_ISL_964801, EPI_ISL_964802, EPI_ISL_964803, EPI_ISL_964804, EPI_ISL_964805, EPI_ISL_964806, EPI_ISL_964807, EPI_ISL_964808, EPI_ISL_964809, EPI_ISL_964810, EPI_ISL_964811, EPI_ISL_964812, EPI_ISL_964813, EPI_ISL_964814, EPI_ISL_964815, EPI_ISL_964816, EPI_ISL_964818, EPI_ISL_964819, EPI_ISL_964820, EPI_ISL_964821, EPI_ISL_964822, EPI_ISL_964823, EPI_ISL_964824, EPI_ISL_964825, EPI_ISL_964826, EPI_ISL_964827, EPI_ISL_964828, EPI_ISL_964829, EPI_ISL_964830, EPI_ISL_964831, EPI_ISL_964832, EPI_ISL_964833, EPI_ISL_964834, EPI_ISL_964835, EPI_ISL_964836, EPI_ISL_964837, EPI_ISL_964838, EPI_ISL_964839, EPI_ISL_964840, EPI_ISL_964841, EPI_ISL_964842, EPI_ISL_964843, EPI_ISL_964844, EPI_ISL_964845, EPI_ISL_964846, EPI_ISL_964847, EPI_ISL_964848, EPI_ISL_964849, EPI_ISL_964850, EPI_ISL_964851, EPI_ISL_964852, EPI_ISL_964853, EPI_ISL_964854, EPI_ISL_964855, EPI_ISL_964856, EPI_ISL_964857, EPI_ISL_964858, EPI_ISL_964859, EPI_ISL_964860, EPI_ISL_964861, EPI_ISL_964862, EPI_ISL_964863, EPI_ISL_964864, EPI_ISL_964865, EPI_ISL_964866, EPI_ISL_964867, EPI_ISL_964868, EPI_ISL_964869, EPI_ISL_964870, EPI_ISL_964871, EPI_ISL_964872, EPI_ISL_964873, EPI_ISL_964874, EPI_ISL_964875 | | | |
| see above | Lighthouse Lab in Alderley Park | Wellcome Sanger Institute for the COVID-19 Genomics UK (COG-UK) Consortium | Jacquelyn Wynn, Mairead Hyland, The Lighthouse Lab in Alderley Park and Alex Alderton, Roberto Amato, Sonia Goncalves, Ewan Harrison, David K. Jackson, Ian Johnston, Dominic Kwiatkowski, Cordelia Langford, John Sillitoe on behalf of the Wellcome Sanger Institute COVID-19 Surveillance Team |
| EPI_ISL_964876 | Lighthouse Lab in Milton Keynes | Wellcome Sanger Institute for the COVID-19 Genomics UK (COG-UK) Consortium | The Lighthouse Lab in Milton Keynes and Alex Alderton, Roberto Amato, Sonia Goncalves, Ewan Harrison, David K. Jackson, Ian Johnston, Dominic Kwiatkowski, Cordelia Langford, John Sillitoe on behalf of the Wellcome Sanger Institute COVID-19 Surveillance Team |
| EPI_ISL_964988 | Furst Medical Laboratory | Norwegian Institute of Public Health, Department of Virology | Kathrine Stene-Johansen, Kamilla Heddeland Instefjord, Hilde Elshaug, Ignacio Garcia Llorente, Serina B Engebretsen, Atiya R Ali,Marie Paulsen Madsen, Rasmus Riis Kopperud, Hilde Vollan, Karoline Bragstad, Olav Hungnes |
| EPI_ISL_964994 | Akershus University Hospital, Department for Microbiology and Infectious Disease Control | Norwegian Institute of Public Health, Department of Virology | Kathrine Stene-Johansen, Kamilla Heddeland Instefjord, Hilde Elshaug, Ignacio Garcia Llorente, Serina B Engebretsen, Atiya R Ali,Marie Paulsen Madsen, Rasmus Riis Kopperud, Hilde Vollan, Karoline Bragstad, Olav Hungnes |
| EPI_ISL_965031 | Laboratorio Biologia Molecolare Sars Cov2 - UOC Laboratorio Analisi - Servizio Medicina di Laboratorio, Ospedale "San Francesco" - ATS-ASSL Nuoro | Laboratorio specialistico UOC Ematologia - Ospedale "San Francesco" - ATS-ASSL Nuoro | Piras Giovanna, Asproni Rosanna, Malune Paolo, Fiamma Maura, Monne Maria Itria, Palmas Angelo Domenico, Lo Maglio Iana, Mameli Giuseppe |
| EPI_ISL_965044 | Wyoming Public Health Laboratory | Wyoming Public Health Laboratory | Noah Hull, Taylor Fearing, Lynette Gumbleton, Channing Weber, Ashley Norberg, Bailey Bowcutt, and Wanda Manley |
| EPI_ISL_965122, EPI_ISL_965124 | Viral Respiratory Lab, National Institute for Biomedical Research (INRB) | Pathogen Sequencing Lab, National Institute for Biomedical Research (INRB) | Placide Mbala-Kingebeni, Edith Nkwembe, Eddy Kinganda-Lusamaki, Amuri Aziza, Francisca Muyembe Mawete, Emmanuel Lokilo Lofiko, Jean-Claude Makangara Cigolo, Catherine Pratt, Matthias Pauthner, Josh Quick, Allison Black, James Hadfield, Trevor Bedford, Ian Goodfellow, Andrew Rambaut, Nick Loman, Kristian Andersen, Michael Wiley, Steve Ahuka-Mundeke, Jean-Jacques Muyembe Tamfum |
| EPI_ISL_965155, EPI_ISL_965156 | Mary Eliza Mahoney HC | NJ_PHEL | Lindsey Bodnar, Shiv Verma, Dana Woell, Byeong Jeong |
| EPI_ISL_965224 | OSPEDALE SS. ANNUNZIATA CHIETI | Istituto Zooprofilattico Sperimentale dell'Abruzzo e Molise "G. Caporale" | Lorusso A, Marcacci M, Di Domenico M, Ancora M, Curini V, Mangone I, Rinaldi A, Scialabba S, Di Pasquale A, Cammà C, Puglia I, Calistri P, Savini G |
| EPI_ISL_965245 | OSPEDALE CIVILE TERAMO, DIV. PNEUMOLOGIA, PIA | Istituto Zooprofilattico Sperimentale dell'Abruzzo e Molise "G. Caporale" | Lorusso A, Marcacci M, Di Domenico M, Ancora M, Curini V, Mangone I, Rinaldi A, Scialabba S, Di Pasquale A, Cammà C, Puglia I, Calistri P, Savini G |
| EPI_ISL_965246 | OSP CIV GIULIANOVA, MED INT | Istituto Zooprofilattico Sperimentale dell'Abruzzo e Molise "G. Caporale" | Lorusso A, Marcacci M, Di Domenico M, Ancora M, Curini V, Mangone I, Rinaldi A, Scialabba S, Di Pasquale A, Cammà C, Puglia I, Calistri P, Savini G |
| EPI_ISL_965247, EPI_ISL_965248, EPI_ISL_965249, EPI_ISL_965250, EPI_ISL_965251, EPI_ISL_965252, EPI_ISL_965266 | SIESP CHIETI | Istituto Zooprofilattico Sperimentale dell'Abruzzo e Molise "G. Caporale" | Lorusso A, Marcacci M, Di Domenico M, Ancora M, Curini V, Mangone I, Rinaldi A, Scialabba S, Di Pasquale A, Cammà C, Puglia I, Calistri P, Savini G |
| EPI_ISL_965276 | OSP CIV ATRI | Istituto Zooprofilattico Sperimentale dell'Abruzzo e Molise "G. Caporale" | Lorusso A, Marcacci M, Di Domenico M, Ancora M, Curini V, Mangone I, Rinaldi A, Scialabba S, Di Pasquale A, Cammà C, Puglia I, Calistri P, Savini G |
| EPI_ISL_965314 | Synlab | GIGA Medical Genomics | Keith Durkin, Maria Artesi, Sébastien Bontems, Raphaël Boreux, Bouchra Boujemla, Cécile Meex, Pierrette Melin, Marie-Pierre Hayette, Vincent Bours |
| EPI_ISL_965397 | Clinique N.D de Grâce Gosselies | GIGA Medical Genomics | Keith Durkin, Maria Artesi, Sébastien Bontems, Raphaël Boreux, Bouchra Boujemla, Cécile Meex, Pierrette Melin, Marie-Pierre Hayette, Vincent Bours |
| EPI_ISL_965403, EPI_ISL_965404, EPI_ISL_965433 | Synlab | GIGA Medical Genomics | Keith Durkin, Maria Artesi, Sébastien Bontems, Raphaël Boreux, Bouchra Boujemla, Cécile Meex, Pierrette Melin, Marie-Pierre Hayette, Vincent Bours |
| EPI_ISL_965451 | Clinique N.D de Grâce Gosselies | GIGA Medical Genomics | Keith Durkin, Maria Artesi, Sébastien Bontems, Raphaël Boreux, Bouchra Boujemla, Cécile Meex, Pierrette Melin, Marie-Pierre Hayette, Vincent Bours |
| EPI_ISL_965470, EPI_ISL_965499, EPI_ISL_965508, EPI_ISL_965521, EPI_ISL_965524 | Synlab | GIGA Medical Genomics | Keith Durkin, Maria Artesi, Sébastien Bontems, Raphaël Boreux, Bouchra Boujemla, Cécile Meex, Pierrette Melin, Marie-Pierre Hayette, Vincent Bours |
| EPI_ISL_965855, EPI_ISL_965856, EPI_ISL_965857, EPI_ISL_965858, EPI_ISL_965859, EPI_ISL_965860, EPI_ISL_965861, EPI_ISL_965862, EPI_ISL_965864, EPI_ISL_965865, EPI_ISL_965866, EPI_ISL_965867, EPI_ISL_965868, EPI_ISL_965869, EPI_ISL_965870, EPI_ISL_965871, EPI_ISL_965872, EPI_ISL_965873, EPI_ISL_965874, EPI_ISL_965875, EPI_ISL_965876, EPI_ISL_965880, EPI_ISL_965881, EPI_ISL_965882, EPI_ISL_965883, EPI_ISL_965884, EPI_ISL_965886, EPI_ISL_965887, EPI_ISL_965888, EPI_ISL_965889, EPI_ISL_965904 | | | |
| see above | Massachusetts State Public Health Laboratory | Massachusetts State Public Health Laboratory | Andrew Lang, Timelia Fink, Glen Gallagher, Sandra Smole |
| EPI_ISL_965957 | Consejería de Sanidad y Asuntos Sociales de Castilla La Mancha | Instituto de Salud Carlos III | Iglesias-Caballero, M. Sandonís,V. Vázquez, S. Camarero, S. Pozo, F. Casas, I. Jiménez, P. Zaballos, A. Monzón, S. Varona, S. Cuesta, I. Gutiérrez, G. |
| EPI_ISL_966285 | Baylor Scott & White - Irving | Baylor Scott & White - Temple | Ari Rao, Linden Morales, Kimberly Walker, Marcus Volz, Shelby Hendrickson |
| EPI_ISL_966310, EPI_ISL_966313 | Baylor Scott & White - Temple | Baylor Scott & White - Temple | Ari Rao, Linden Morales, Kimberly Walker, Marcus Volz, Shelby Hendrickson |
| EPI_ISL_966334, EPI_ISL_966335, EPI_ISL_966336, EPI_ISL_966337, EPI_ISL_966339, EPI_ISL_966340, EPI_ISL_966341, EPI_ISL_966342, EPI_ISL_966343, EPI_ISL_966344, EPI_ISL_966345 | | | |
| see above | Kansas Health and Environmental Lab | Kansas Health and Environmental Lab | Mike Grose, Paige Drury, Carissa Robertson, Ben Olsen, and Phil Adam |
| EPI_ISL_966382, EPI_ISL_966384 | DOHMH Jamaica | New York City Public Health Laboratory | Jade Wang, et al. |
| EPI_ISL_966386 | DOHMH Corona | New York City Public Health Laboratory | Jade Wang, et al. |
| EPI_ISL_966387, EPI_ISL_966388 | DOHMH Jamaica | New York City Public Health Laboratory | Jade Wang, et al. |
| EPI_ISL_966391 | OCME Office Of Chief Medical Examiner | New York City Public Health Laboratory | Jade Wang, et al. |

| | | | |
|---|---|---|---|
| EPI_ISL_966395 | DOHMH Jamaica | New York City Public Health Laboratory | Jade Wang, et al. |
| EPI_ISL_966397, EPI_ISL_966400 | OCME Office Of Chief Medical Examiner | New York City Public Health Laboratory | Jade Wang, et al. |
| EPI_ISL_966401 | Department of Homeless Services | New York City Public Health Laboratory | Jade Wang, et al. |
| EPI_ISL_966402, EPI_ISL_966403 | OCME Office Of Chief Medical Examiner | New York City Public Health Laboratory | Jade Wang, et al. |
| EPI_ISL_966404 | DOHMH PHL | New York City Public Health Laboratory | Jade Wang, et al. |
| EPI_ISL_966405 | DOHMH Jamaica | New York City Public Health Laboratory | Jade Wang, et al. |
| EPI_ISL_966411 | OCME Office Of Chief Medical Examiner | New York City Public Health Laboratory | Jade Wang, et al. |
| EPI_ISL_966412 | DOHMH Jamaica | New York City Public Health Laboratory | Jade Wang, et al. |
| EPI_ISL_966413 | DOHMH Central Harlem | New York City Public Health Laboratory | Jade Wang, et al. |
| EPI_ISL_966414 | DOHMH Morrisania | New York City Public Health Laboratory | Jade Wang, et al. |
| EPI_ISL_966415 | DOHMH Corona | New York City Public Health Laboratory | Jade Wang, et al. |
| EPI_ISL_966416 | DOHMH Central Harlem | New York City Public Health Laboratory | Jade Wang, et al. |
| EPI_ISL_966417, EPI_ISL_966418, EPI_ISL_966419, EPI_ISL_966429, EPI_ISL_966430, EPI_ISL_966431 | OCME Office Of Chief Medical Examiner | New York City Public Health Laboratory | Jade Wang, et al. |
| EPI_ISL_966432, EPI_ISL_966433 | DOHMH Jamaica | New York City Public Health Laboratory | Jade Wang, et al. |
| EPI_ISL_966434, EPI_ISL_966435, EPI_ISL_966436, EPI_ISL_966437 | DOHMH Morrisania | New York City Public Health Laboratory | Jade Wang, et al. |
| EPI_ISL_966438, EPI_ISL_966439, EPI_ISL_966440, EPI_ISL_966441, EPI_ISL_966442 | OCME Office Of Chief Medical Examiner | New York City Public Health Laboratory | Jade Wang, et al. |
| EPI_ISL_966443 | DOHMH Fort Greene | New York City Public Health Laboratory | Jade Wang, et al. |
| EPI_ISL_966444 | DOHMH PHL | New York City Public Health Laboratory | Jade Wang, et al. |
| EPI_ISL_966445, EPI_ISL_966446 | DOHMH Corona | New York City Public Health Laboratory | Jade Wang, et al. |
| EPI_ISL_966447, EPI_ISL_966448, EPI_ISL_966449 | DOHMH Jamaica | New York City Public Health Laboratory | Jade Wang, et al. |
| EPI_ISL_966450 | DOHMH Central Harlem | New York City Public Health Laboratory | Jade Wang, et al. |
| EPI_ISL_966451 | DOHMH Morrisania | New York City Public Health Laboratory | Jade Wang, et al. |
| EPI_ISL_966453, EPI_ISL_966454, EPI_ISL_966455, EPI_ISL_966456, EPI_ISL_966457, EPI_ISL_966458 | OCME Office Of Chief Medical Examiner | New York City Public Health Laboratory | Jade Wang, et al. |
| EPI_ISL_966459 | DOHMH Corona | New York City Public Health Laboratory | Jade Wang, et al. |
| EPI_ISL_966460 | DOHMH Morrisania | New York City Public Health Laboratory | Jade Wang, et al. |
| EPI_ISL_966788, EPI_ISL_966789, EPI_ISL_966790, EPI_ISL_966791, EPI_ISL_966792, EPI_ISL_966793, EPI_ISL_966794, EPI_ISL_966795, EPI_ISL_966796, EPI_ISL_966797, EPI_ISL_966798, EPI_ISL_966799, EPI_ISL_966800, EPI_ISL_966801, EPI_ISL_966802, EPI_ISL_966803, EPI_ISL_966804, EPI_ISL_966805, EPI_ISL_966806 | | | |
| see above | Maine HETL | Tewhey Lab, The Jackson Laboratory | Matluk,N., Dewey,H., Iosue,F., Barter,M., Lynch,R., Munger,H. and Tewhey,R. |
| EPI_ISL_966852, EPI_ISL_966853, EPI_ISL_966854, EPI_ISL_966855, EPI_ISL_966856, EPI_ISL_966857, EPI_ISL_966858, EPI_ISL_966859, EPI_ISL_966860, EPI_ISL_966861, EPI_ISL_966862, EPI_ISL_966863, EPI_ISL_966864, EPI_ISL_966865, EPI_ISL_966866, EPI_ISL_966867, EPI_ISL_966868, EPI_ISL_966869, EPI_ISL_966870, EPI_ISL_966871, EPI_ISL_966872, EPI_ISL_966873, EPI_ISL_966874, EPI_ISL_966875, EPI_ISL_966876, EPI_ISL_966877, EPI_ISL_966878, EPI_ISL_966879, EPI_ISL_966880, EPI_ISL_966881, EPI_ISL_966882, EPI_ISL_966883, EPI_ISL_966884, EPI_ISL_966885, EPI_ISL_966886, EPI_ISL_966887, EPI_ISL_966888, EPI_ISL_966889, EPI_ISL_966890, EPI_ISL_966891, EPI_ISL_966892, EPI_ISL_966893, EPI_ISL_966894, EPI_ISL_966895, EPI_ISL_966896, EPI_ISL_966897, EPI_ISL_966898, EPI_ISL_966899, EPI_ISL_966900, EPI_ISL_966901, EPI_ISL_966902, EPI_ISL_966903, EPI_ISL_966904, EPI_ISL_966905, EPI_ISL_966906, EPI_ISL_966907, EPI_ISL_966908, EPI_ISL_966909, EPI_ISL_966910, EPI_ISL_966911, EPI_ISL_966912, EPI_ISL_966913, EPI_ISL_966914, EPI_ISL_966915, EPI_ISL_966916, EPI_ISL_966917, EPI_ISL_966918, EPI_ISL_966919, EPI_ISL_966920, EPI_ISL_966921, EPI_ISL_966922, EPI_ISL_966923, EPI_ISL_966924, EPI_ISL_966925, EPI_ISL_966926, EPI_ISL_966927, EPI_ISL_966928 | | | |
| see above | Helix/Illumina | Respiratory Viruses Branch, Division of Viral Diseases, Centers for Disease Control and Prevention | Peter W. Cook,Dakota Howard,Dhwani Batra,Ben L. Rambo-Martin,Eileen de Feo,Jan Antico,Christine Tran,Matthew Tolentino,Shannon Wickline,Kim Gietzen,Brad Sickler,Jingtao Liu,Eric Allen,Phil Febbo,Summer Galloway,Nicole L. Washington,Simon White,Geraint Levan,Kelly Schiabor Barrett,Elizabeth Cirulli,Alexandre Bolze,Ary Ascencio,Charlotte Rivera-Garcia,Ryan Cho,Jason Nguyen,Sherry Wang,Jimmy Ramirez,Tyler Cassens,Efren Sandoval,Magnus Isaksson,William Lee,David Becker,Marc Laurent,James Lu,Clinton R. Paden,Suxiang Tong,Duncan MacCannell, |
| EPI_ISL_967770 | State Laboratories Division, Hawaii State Department of Health | State Laboratories Division, Hawaii State Department of Health | Pamela O'Brien, Drew Kuwazaki, Ayana Garnet, Razvan Sultana, Edward Desmond |
| EPI_ISL_968078 | Monterey County Public Health Laboratory | Monterey County Public Health Laboratory | Monterey County Public Health Laboratory |
| EPI_ISL_968089 | Institute for Medical Research, Infectious Disease Research Centre, National Institutes of Health, Ministry of Health Malaysia | Institute for Medical Research, Infectious Disease Research Centre, National Institutes of Health, Ministry of Health Malaysia | Suppiah J, Kamel K, Azizan MA, Thayan R |
| EPI_ISL_969018, EPI_ISL_969047, EPI_ISL_969048, EPI_ISL_969049, EPI_ISL_969068, EPI_ISL_969069, EPI_ISL_969072, EPI_ISL_969073, EPI_ISL_969074 | KEMRI-Wellcome Trust Research Programme/KEMRI-CGMR-C Kilifi | KEMRI-Wellcome Trust Research Programme/KEMRI-CGMR-C Kilifi | Githinji et al |
| EPI_ISL_969455 | Lighthouse Lab in Glasgow | Wellcome Sanger Institute for the COVID-19 Genomics UK (COG-UK) Consortium | Harper VanSteenhouse, Yumi Kasai, David Gray, Carol Clugston, Anna Dominiczak and Alex Alderton, Roberto Amato, Sonia Goncalves, Ewan Harrison, David K. Jackson, Ian Johnston, Dominic Kwiatkowski, Cordelia Langford, John Sillitoe on behalf of the Wellcome Sanger Institute COVID-19 Surveillance Team |
| EPI_ISL_969457, EPI_ISL_969458, EPI_ISL_969459, EPI_ISL_969460, EPI_ISL_969461, EPI_ISL_969462, EPI_ISL_969463, EPI_ISL_969464, EPI_ISL_969465, EPI_ISL_969466, EPI_ISL_969467, EPI_ISL_969468, EPI_ISL_969469, EPI_ISL_969471, EPI_ISL_969472, EPI_ISL_969473, EPI_ISL_969474, EPI_ISL_969475, EPI_ISL_969476, EPI_ISL_969477, EPI_ISL_969478, EPI_ISL_969479, EPI_ISL_969480, EPI_ISL_969481, EPI_ISL_969482, EPI_ISL_969483, EPI_ISL_969484, EPI_ISL_969485, EPI_ISL_969486, EPI_ISL_969487, EPI_ISL_969488, EPI_ISL_969489, EPI_ISL_969490, EPI_ISL_969491, EPI_ISL_969492, EPI_ISL_969493, EPI_ISL_969494, EPI_ISL_969495, EPI_ISL_969496, EPI_ISL_969497, EPI_ISL_969498, EPI_ISL_969500, EPI_ISL_969501, EPI_ISL_969502, EPI_ISL_969503, EPI_ISL_969504, EPI_ISL_969505, EPI_ISL_969506, EPI_ISL_969507, EPI_ISL_969508, EPI_ISL_969510, EPI_ISL_969511, EPI_ISL_969512, EPI_ISL_969513, EPI_ISL_969514, EPI_ISL_969515, EPI_ISL_969516, EPI_ISL_969517, EPI_ISL_969518, EPI_ISL_969519, EPI_ISL_969520, EPI_ISL_969521, EPI_ISL_969522, EPI_ISL_969523, EPI_ISL_969524, EPI_ISL_969525, EPI_ISL_969526, EPI_ISL_969527, EPI_ISL_969528, EPI_ISL_969529, EPI_ISL_969530, EPI_ISL_969531, EPI_ISL_969532, EPI_ISL_969533, EPI_ISL_969534, EPI_ISL_969535, EPI_ISL_969536, EPI_ISL_969537, EPI_ISL_969538, EPI_ISL_969539, EPI_ISL_969540, EPI_ISL_969541, EPI_ISL_969542, EPI_ISL_969543, EPI_ISL_969544, EPI_ISL_969545, EPI_ISL_969546, EPI_ISL_969548, EPI_ISL_969549, EPI_ISL_969550, EPI_ISL_969551, EPI_ISL_969552, EPI_ISL_969553, EPI_ISL_969554, EPI_ISL_969555, EPI_ISL_969556, EPI_ISL_969557, EPI_ISL_969558, EPI_ISL_969559, EPI_ISL_969560, EPI_ISL_969561, EPI_ISL_969562, EPI_ISL_969563, EPI_ISL_969564, EPI_ISL_969565, EPI_ISL_969566, EPI_ISL_969567, EPI_ISL_969568, EPI_ISL_969569, EPI_ISL_969570, EPI_ISL_969571, EPI_ISL_969572, EPI_ISL_969573, EPI_ISL_969574, EPI_ISL_969575, EPI_ISL_969576, EPI_ISL_969577, EPI_ISL_969578, EPI_ISL_969579, EPI_ISL_969580, EPI_ISL_969581, EPI_ISL_969582, EPI_ISL_969583, EPI_ISL_969584, EPI_ISL_969585, EPI_ISL_969586, EPI_ISL_969587, EPI_ISL_969588, EPI_ISL_969589, EPI_ISL_969591, EPI_ISL_969592, EPI_ISL_969593, EPI_ISL_969594, EPI_ISL_969595, EPI_ISL_969596, EPI_ISL_969597, EPI_ISL_969598, EPI_ISL_969599, EPI_ISL_969600, EPI_ISL_969602, EPI_ISL_969603, EPI_ISL_969604, EPI_ISL_969605, EPI_ISL_969606, EPI_ISL_969607, EPI_ISL_969608, EPI_ISL_969609, EPI_ISL_969610, EPI_ISL_969611, EPI_ISL_969612, EPI_ISL_969613, EPI_ISL_969614, EPI_ISL_969615, EPI_ISL_969616, EPI_ISL_969617, EPI_ISL_969618, EPI_ISL_969619, EPI_ISL_969620, EPI_ISL_969621, EPI_ISL_969622, EPI_ISL_969623, EPI_ISL_969624, EPI_ISL_969625, EPI_ISL_969626, EPI_ISL_969627, EPI_ISL_969628, EPI_ISL_969629, EPI_ISL_969630, EPI_ISL_969631, EPI_ISL_969632, EPI_ISL_969633, EPI_ISL_969634, EPI_ISL_969635, EPI_ISL_969636, EPI_ISL_969637, EPI_ISL_969638, EPI_ISL_969639, EPI_ISL_969640, EPI_ISL_969641, EPI_ISL_969642, EPI_ISL_969643, EPI_ISL_969644, EPI_ISL_969645, EPI_ISL_969646, EPI_ISL_969647, EPI_ISL_969649, EPI_ISL_969650, EPI_ISL_969651, EPI_ISL_969652, EPI_ISL_969653, EPI_ISL_969654, EPI_ISL_969655, EPI_ISL_969656, EPI_ISL_969657, EPI_ISL_969658, EPI_ISL_969659, EPI_ISL_969660, EPI_ISL_969661, EPI_ISL_969662, EPI_ISL_969663, EPI_ISL_969664, EPI_ISL_969665, EPI_ISL_969666, EPI_ISL_969667, EPI_ISL_969668, EPI_ISL_969669, EPI_ISL_969670, EPI_ISL_969671, EPI_ISL_969672, EPI_ISL_969673, EPI_ISL_969674, EPI_ISL_969675, EPI_ISL_969677, EPI_ISL_969679, EPI_ISL_969682, EPI_ISL_969683, | | | |

| | | | |
|---|---|---|---|
| EPI_ISL_969685, EPI_ISL_969688, EPI_ISL_969690, EPI_ISL_969691, EPI_ISL_969693, EPI_ISL_969695, EPI_ISL_969697, EPI_ISL_969699, EPI_ISL_969701, EPI_ISL_969702, EPI_ISL_969704, EPI_ISL_969706, EPI_ISL_969708, EPI_ISL_969710, EPI_ISL_969712, EPI_ISL_969714, EPI_ISL_969716, EPI_ISL_969718, EPI_ISL_969720, EPI_ISL_969722, EPI_ISL_969724, EPI_ISL_969726, EPI_ISL_969728, EPI_ISL_969730, EPI_ISL_969732, EPI_ISL_969734, EPI_ISL_969736, EPI_ISL_969739, EPI_ISL_969742, EPI_ISL_969744, EPI_ISL_969746, EPI_ISL_969748, EPI_ISL_969750, EPI_ISL_969752, EPI_ISL_969754, EPI_ISL_969756, EPI_ISL_969759, EPI_ISL_969760, EPI_ISL_969762, EPI_ISL_969764, EPI_ISL_969766, EPI_ISL_969768, EPI_ISL_969769, EPI_ISL_969771, EPI_ISL_969774, EPI_ISL_969776, EPI_ISL_969779, EPI_ISL_969780, EPI_ISL_969782, EPI_ISL_969785, EPI_ISL_969786, EPI_ISL_969789, EPI_ISL_969791, EPI_ISL_969793, EPI_ISL_969795, EPI_ISL_969797, EPI_ISL_969799, EPI_ISL_969801, EPI_ISL_969802, EPI_ISL_969804, EPI_ISL_969806, EPI_ISL_969808, EPI_ISL_969809, EPI_ISL_969812, EPI_ISL_969813, EPI_ISL_969815, EPI_ISL_969817, EPI_ISL_969819, EPI_ISL_969821, EPI_ISL_969823, EPI_ISL_969824, EPI_ISL_969826, EPI_ISL_969828, EPI_ISL_969831, EPI_ISL_969833, EPI_ISL_969834, EPI_ISL_969836, EPI_ISL_969838, EPI_ISL_969840, EPI_ISL_969841, EPI_ISL_969843, EPI_ISL_969846, EPI_ISL_969848, EPI_ISL_969849, EPI_ISL_969852, EPI_ISL_969853, EPI_ISL_969856, EPI_ISL_969858, EPI_ISL_969859, EPI_ISL_969861, EPI_ISL_969863, EPI_ISL_969865, EPI_ISL_969867, EPI_ISL_969869, EPI_ISL_969871, EPI_ISL_969873, EPI_ISL_969875, EPI_ISL_969877, EPI_ISL_969879, EPI_ISL_969881, EPI_ISL_969883, EPI_ISL_969886, EPI_ISL_969887, EPI_ISL_969889, EPI_ISL_969891, EPI_ISL_969893, EPI_ISL_969895, EPI_ISL_969897, EPI_ISL_969899, EPI_ISL_969901, EPI_ISL_969903, EPI_ISL_969906, EPI_ISL_969907, EPI_ISL_969909, EPI_ISL_969911, EPI_ISL_969914, EPI_ISL_969916, EPI_ISL_969917, EPI_ISL_969919, EPI_ISL_969921, EPI_ISL_969923, EPI_ISL_969925, EPI_ISL_969928, EPI_ISL_969930, EPI_ISL_969932, EPI_ISL_969934, EPI_ISL_969936, EPI_ISL_969937, EPI_ISL_969939, EPI_ISL_969940, EPI_ISL_969943, EPI_ISL_969945, EPI_ISL_969947, EPI_ISL_969948, EPI_ISL_969951, EPI_ISL_969952, EPI_ISL_969955, EPI_ISL_969956, EPI_ISL_969958, EPI_ISL_969961, EPI_ISL_969963, EPI_ISL_969965, EPI_ISL_969967, EPI_ISL_969969, EPI_ISL_969971, EPI_ISL_969974, EPI_ISL_969975, EPI_ISL_969977, EPI_ISL_969979, EPI_ISL_969982, EPI_ISL_969983, EPI_ISL_969986, EPI_ISL_969988, EPI_ISL_969990, EPI_ISL_969992, EPI_ISL_969994, EPI_ISL_969996, EPI_ISL_969998, EPI_ISL_970000, EPI_ISL_970002, EPI_ISL_970004, EPI_ISL_970005, EPI_ISL_970007, EPI_ISL_970009, EPI_ISL_970011, EPI_ISL_970013, EPI_ISL_970015, EPI_ISL_970017, EPI_ISL_970019, EPI_ISL_970022, EPI_ISL_970024, EPI_ISL_970026, EPI_ISL_970028, EPI_ISL_970030, EPI_ISL_970033, EPI_ISL_970035, EPI_ISL_970037, EPI_ISL_970039, EPI_ISL_970041, EPI_ISL_970043, EPI_ISL_970045, EPI_ISL_970047, EPI_ISL_970049, EPI_ISL_970050, EPI_ISL_970052, EPI_ISL_970057, EPI_ISL_970059, EPI_ISL_970061, EPI_ISL_970064, EPI_ISL_970066, EPI_ISL_970068, EPI_ISL_970070, EPI_ISL_970073, EPI_ISL_970075, EPI_ISL_970076, EPI_ISL_970079, EPI_ISL_970080, EPI_ISL_970082, EPI_ISL_970084, EPI_ISL_970086, EPI_ISL_970087, EPI_ISL_970089, EPI_ISL_970091, EPI_ISL_970094, EPI_ISL_970096, EPI_ISL_970098, EPI_ISL_970099, EPI_ISL_970101, EPI_ISL_970103, EPI_ISL_970105, EPI_ISL_970107, EPI_ISL_970109, EPI_ISL_970111, EPI_ISL_970113, EPI_ISL_970115, EPI_ISL_970117, EPI_ISL_970119, EPI_ISL_970120, EPI_ISL_970122, EPI_ISL_970124, EPI_ISL_970126, EPI_ISL_970128, EPI_ISL_970130, EPI_ISL_970132, EPI_ISL_970134, EPI_ISL_970136, EPI_ISL_970138, EPI_ISL_970140, EPI_ISL_970145, EPI_ISL_970147, EPI_ISL_970148, EPI_ISL_970150, EPI_ISL_970152, EPI_ISL_970153, EPI_ISL_970155, EPI_ISL_970157, EPI_ISL_970160, EPI_ISL_970161, EPI_ISL_970163, EPI_ISL_970165, EPI_ISL_970166, EPI_ISL_970168, EPI_ISL_970170, EPI_ISL_970172, EPI_ISL_970173, EPI_ISL_970175, EPI_ISL_970177, EPI_ISL_970181, EPI_ISL_970183, EPI_ISL_970184, EPI_ISL_970186, EPI_ISL_970188, EPI_ISL_970190, EPI_ISL_970192, EPI_ISL_970195, EPI_ISL_970197, EPI_ISL_970198, EPI_ISL_970200, EPI_ISL_970202, EPI_ISL_970205, EPI_ISL_970206, EPI_ISL_970209, EPI_ISL_970211, EPI_ISL_970213, EPI_ISL_970215, EPI_ISL_970217, EPI_ISL_970219, EPI_ISL_970221, EPI_ISL_970224, EPI_ISL_970225, EPI_ISL_970227, EPI_ISL_970229, EPI_ISL_970231, EPI_ISL_970233, EPI_ISL_970235, EPI_ISL_970237, EPI_ISL_970239, EPI_ISL_970242, EPI_ISL_970247, EPI_ISL_970249, EPI_ISL_970251, EPI_ISL_970252, EPI_ISL_970254, EPI_ISL_970256, EPI_ISL_970258, EPI_ISL_970260, EPI_ISL_970262, EPI_ISL_970264, EPI_ISL_970266, EPI_ISL_970268, EPI_ISL_970270, EPI_ISL_970272, EPI_ISL_970274, EPI_ISL_970275, EPI_ISL_970278, EPI_ISL_970279, EPI_ISL_970281, EPI_ISL_970283, EPI_ISL_970285, EPI_ISL_970287, EPI_ISL_970288, EPI_ISL_970291, EPI_ISL_970292, EPI_ISL_970295, EPI_ISL_970297, EPI_ISL_970299, EPI_ISL_970301, EPI_ISL_970303, EPI_ISL_970306, EPI_ISL_970307, EPI_ISL_970309, EPI_ISL_970312, EPI_ISL_970314, EPI_ISL_970316, EPI_ISL_970319, EPI_ISL_970321, EPI_ISL_970323, EPI_ISL_970324, EPI_ISL_970326, EPI_ISL_970328, EPI_ISL_970330, EPI_ISL_970332, EPI_ISL_970334, EPI_ISL_970337, EPI_ISL_970338, EPI_ISL_970340, EPI_ISL_970342, EPI_ISL_970344, EPI_ISL_970346, EPI_ISL_970348, EPI_ISL_970350, EPI_ISL_970352, EPI_ISL_970354, EPI_ISL_970356, EPI_ISL_970358, EPI_ISL_970360, EPI_ISL_970362, EPI_ISL_970364, EPI_ISL_970365, EPI_ISL_970368, EPI_ISL_970370, EPI_ISL_970371, EPI_ISL_970373, EPI_ISL_970376, EPI_ISL_970377, EPI_ISL_970380, EPI_ISL_970382, EPI_ISL_970383, EPI_ISL_970386, EPI_ISL_970387, EPI_ISL_970390, EPI_ISL_970392, EPI_ISL_970394, EPI_ISL_970396, EPI_ISL_970398, EPI_ISL_970400, EPI_ISL_970402, EPI_ISL_970404, EPI_ISL_970406, EPI_ISL_970409, EPI_ISL_970411, EPI_ISL_970412, EPI_ISL_970415, EPI_ISL_970416, EPI_ISL_970419, EPI_ISL_970421, EPI_ISL_970422, EPI_ISL_970424, EPI_ISL_970426, EPI_ISL_970428, EPI_ISL_970430, EPI_ISL_970432, EPI_ISL_970435, EPI_ISL_970436, EPI_ISL_970439, EPI_ISL_970441, EPI_ISL_970443, EPI_ISL_970444, EPI_ISL_970446, EPI_ISL_970448, EPI_ISL_970450, EPI_ISL_970452, EPI_ISL_970454, EPI_ISL_970457, EPI_ISL_970460, EPI_ISL_970462, EPI_ISL_970464, EPI_ISL_970466, EPI_ISL_970468, EPI_ISL_970470, EPI_ISL_970473, EPI_ISL_970475, EPI_ISL_970477, EPI_ISL_970478, EPI_ISL_970481, EPI_ISL_970483, EPI_ISL_970486, EPI_ISL_970488, EPI_ISL_970490, EPI_ISL_970492, EPI_ISL_970493, EPI_ISL_970495, EPI_ISL_970497, EPI_ISL_970499, EPI_ISL_970500, EPI_ISL_970502, EPI_ISL_970504, EPI_ISL_970506, EPI_ISL_970508, EPI_ISL_970510, EPI_ISL_970512, EPI_ISL_970513, EPI_ISL_970515, EPI_ISL_970517, EPI_ISL_970519, EPI_ISL_970521, EPI_ISL_970523, EPI_ISL_970525, EPI_ISL_970527, EPI_ISL_970528, EPI_ISL_970530, EPI_ISL_970533, EPI_ISL_970535, EPI_ISL_970537, EPI_ISL_970540, EPI_ISL_970541, EPI_ISL_970543, EPI_ISL_970546, EPI_ISL_970548, EPI_ISL_970551, EPI_ISL_970553, EPI_ISL_970555, EPI_ISL_970558, EPI_ISL_970560, EPI_ISL_970562, EPI_ISL_970564, EPI_ISL_970566, EPI_ISL_970568, EPI_ISL_970569, EPI_ISL_970571, EPI_ISL_970573, EPI_ISL_970575, EPI_ISL_970577, EPI_ISL_970579, EPI_ISL_970581, EPI_ISL_970583, EPI_ISL_970585 | | | |
| see above | Lighthouse Lab in Alderley Park | Wellcome Sanger Institute for the COVID-19 Genomics UK (COG-UK) Consortium | Jacquelyn Wynn, Mairead Hyland, The Lighthouse Lab in Alderley Park and Alex Alderton, Roberto Amato, Sonia Goncalves, Ewan Harrison, David K. Jackson, Ian Johnston, Dominic Kwiatkowski, Cordelia Langford, John Sillitoe on behalf of the Wellcome Sanger Institute COVID-19 Surveillance Team |
| EPI_ISL_970587, EPI_ISL_970597, EPI_ISL_970616, EPI_ISL_970621, EPI_ISL_970628, EPI_ISL_970635, EPI_ISL_970639, EPI_ISL_970645, EPI_ISL_970664, EPI_ISL_970673, EPI_ISL_970680, EPI_ISL_970681, EPI_ISL_970687, EPI_ISL_970701, EPI_ISL_970708, EPI_ISL_970709, EPI_ISL_970715, EPI_ISL_970724, EPI_ISL_970732, EPI_ISL_970744, EPI_ISL_970747, EPI_ISL_970751, EPI_ISL_970753, EPI_ISL_970768, EPI_ISL_970775, EPI_ISL_970790 | | | |
| see above | Lighthouse Lab in Milton Keynes | Wellcome Sanger Institute for the COVID-19 Genomics UK (COG-UK) Consortium | The Lighthouse Lab in Milton Keynes and Alex Alderton, Roberto Amato, Sonia Goncalves, Ewan Harrison, David K. Jackson, Ian Johnston, Dominic Kwiatkowski, Cordelia Langford, John Sillitoe on behalf of the Wellcome Sanger Institute COVID-19 Surveillance Team |
| EPI_ISL_970812, EPI_ISL_970819, EPI_ISL_970820, EPI_ISL_970824 | Department of Virus and Microbiological Special Diagnostics, Statens Serum Institut, Copenhagen, Denmark | Aalborg University | Danish Covid-19 Genome Consortium |
| EPI_ISL_970826 | Lighthouse Lab in Milton Keynes | Wellcome Sanger Institute for the COVID-19 Genomics UK (COG-UK) Consortium | The Lighthouse Lab in Milton Keynes and Alex Alderton, Roberto Amato, Sonia Goncalves, Ewan Harrison, David K. Jackson, Ian Johnston, Dominic Kwiatkowski, Cordelia Langford, John Sillitoe on behalf of the Wellcome Sanger Institute COVID-19 Surveillance Team |
| EPI_ISL_970830, EPI_ISL_970831, EPI_ISL_970832, EPI_ISL_970837, EPI_ISL_970844, EPI_ISL_970850, EPI_ISL_970851, EPI_ISL_970859 | Department of Virus and Microbiological Special Diagnostics, Statens Serum Institut, Copenhagen, Denmark | Aalborg University | Danish Covid-19 Genome Consortium |
| EPI_ISL_970860 | Lighthouse Lab in Milton Keynes | Wellcome Sanger Institute for the COVID-19 Genomics UK (COG-UK) Consortium | The Lighthouse Lab in Milton Keynes and Alex Alderton, Roberto Amato, Sonia Goncalves, Ewan Harrison, David K. Jackson, Ian Johnston, Dominic Kwiatkowski, Cordelia Langford, John Sillitoe on behalf of the Wellcome Sanger Institute COVID-19 Surveillance Team |
| EPI_ISL_970862, EPI_ISL_970863, EPI_ISL_970874 | Department of Virus and Microbiological Special Diagnostics, Statens Serum Institut, Copenhagen, Denmark | Aalborg University | Danish Covid-19 Genome Consortium |
| EPI_ISL_970875 | Lighthouse Lab in Milton Keynes | Wellcome Sanger Institute for the COVID-19 Genomics UK (COG-UK) Consortium | The Lighthouse Lab in Milton Keynes and Alex Alderton, Roberto Amato, Sonia Goncalves, Ewan Harrison, David K. Jackson, Ian Johnston, Dominic Kwiatkowski, Cordelia Langford, John Sillitoe on behalf of the Wellcome Sanger Institute COVID-19 Surveillance Team |
| EPI_ISL_970880, EPI_ISL_970882, EPI_ISL_970885, EPI_ISL_970886, EPI_ISL_970891, EPI_ISL_970892, EPI_ISL_970895, EPI_ISL_970905, EPI_ISL_970908, EPI_ISL_970910, EPI_ISL_970927, EPI_ISL_970928, EPI_ISL_970929, EPI_ISL_970930, EPI_ISL_970932, EPI_ISL_970938, EPI_ISL_970939, EPI_ISL_970943, EPI_ISL_970946, EPI_ISL_970949, EPI_ISL_970950, EPI_ISL_970952, EPI_ISL_970953, EPI_ISL_970959, EPI_ISL_970960, EPI_ISL_970969, EPI_ISL_970970, EPI_ISL_970973, EPI_ISL_970975, EPI_ISL_970976, EPI_ISL_970978, EPI_ISL_970980 | | | |
| see above | Department of Virus and Microbiological Special Diagnostics, Statens Serum Institut, Copenhagen, Denmark | Aalborg University | Danish Covid-19 Genome Consortium |
| EPI_ISL_970986 | Lighthouse Lab in Milton Keynes | Wellcome Sanger Institute for the COVID-19 Genomics UK (COG-UK) Consortium | The Lighthouse Lab in Milton Keynes and Alex Alderton, Roberto Amato, Sonia Goncalves, Ewan Harrison, David K. Jackson, Ian Johnston, Dominic Kwiatkowski, Cordelia Langford, John Sillitoe on behalf of the Wellcome Sanger Institute COVID-19 Surveillance Team |
| EPI_ISL_970987, EPI_ISL_970989, EPI_ISL_970990, EPI_ISL_970991, EPI_ISL_970994, EPI_ISL_970996, EPI_ISL_970999, EPI_ISL_971000, EPI_ISL_971004, EPI_ISL_971009, EPI_ISL_971014, EPI_ISL_971021, EPI_ISL_971022, EPI_ISL_971032, EPI_ISL_971033, EPI_ISL_971035 | | | |
| see above | Department of Virus and Microbiological Special Diagnostics, Statens Serum Institut, Copenhagen, Denmark | Aalborg University | Danish Covid-19 Genome Consortium |
| EPI_ISL_971037, EPI_ISL_971042, EPI_ISL_971044 | Lighthouse Lab in Milton Keynes | Wellcome Sanger Institute for the COVID-19 Genomics UK (COG-UK) Consortium | The Lighthouse Lab in Milton Keynes and Alex Alderton, Roberto Amato, Sonia Goncalves, Ewan Harrison, David K. Jackson, Ian Johnston, Dominic Kwiatkowski, Cordelia Langford, John Sillitoe on behalf of the Wellcome Sanger Institute COVID-19 Surveillance Team |
| EPI_ISL_971045 | Department of Virus and Microbiological Special Diagnostics, Statens Serum Institut, Copenhagen, Denmark | Aalborg University | Danish Covid-19 Genome Consortium |
| EPI_ISL_971047, EPI_ISL_971053, EPI_ISL_971060, EPI_ISL_971071, EPI_ISL_971076 | Lighthouse Lab in Milton Keynes | Wellcome Sanger Institute for the COVID-19 Genomics UK (COG-UK) Consortium | The Lighthouse Lab in Milton Keynes and Alex Alderton, Roberto Amato, Sonia Goncalves, Ewan Harrison, David K. Jackson, Ian Johnston, Dominic Kwiatkowski, Cordelia Langford, John Sillitoe on behalf of the Wellcome Sanger Institute COVID-19 Surveillance Team |
| EPI_ISL_971081, EPI_ISL_971092 | Department of Virus and Microbiological Special Diagnostics, Statens Serum Institut, Copenhagen, Denmark | Aalborg University | Danish Covid-19 Genome Consortium |
| EPI_ISL_971093, EPI_ISL_971096 | Lighthouse Lab in Milton Keynes | Wellcome Sanger Institute for the COVID-19 Genomics UK (COG-UK) Consortium | The Lighthouse Lab in Milton Keynes and Alex Alderton, Roberto Amato, Sonia Goncalves, Ewan Harrison, David K. Jackson, Ian Johnston, Dominic Kwiatkowski, Cordelia Langford, John Sillitoe on behalf of the Wellcome Sanger Institute COVID-19 Surveillance Team |
| EPI_ISL_971100 | Department of Virus and Microbiological Special Diagnostics, Statens Serum Institut, Copenhagen, Denmark | Aalborg University | Danish Covid-19 Genome Consortium |
| EPI_ISL_971105 | Lighthouse Lab in Milton Keynes | Wellcome Sanger Institute for the COVID-19 Genomics UK (COG-UK) Consortium | The Lighthouse Lab in Milton Keynes and Alex Alderton, Roberto Amato, Sonia Goncalves, Ewan Harrison, David K. Jackson, Ian Johnston, Dominic Kwiatkowski, Cordelia Langford, John Sillitoe on behalf of the Wellcome Sanger Institute COVID-19 Surveillance Team |
| EPI_ISL_971112, EPI_ISL_971115, EPI_ISL_971121, EPI_ISL_971124 | Department of Virus and Microbiological Special Diagnostics, Statens Serum Institut, Copenhagen, Denmark | Aalborg University | Danish Covid-19 Genome Consortium |
| EPI_ISL_971125, EPI_ISL_971135, EPI_ISL_971142 | Lighthouse Lab in Milton Keynes | Wellcome Sanger Institute for the COVID-19 Genomics UK (COG-UK) Consortium | The Lighthouse Lab in Milton Keynes and Alex Alderton, Roberto Amato, Sonia Goncalves, Ewan Harrison, David K. Jackson, Ian Johnston, Dominic Kwiatkowski, Cordelia Langford, John Sillitoe on behalf of the Wellcome Sanger Institute COVID-19 Surveillance Team |
| EPI_ISL_971148, EPI_ISL_971158, EPI_ISL_971163, EPI_ISL_971166 | Department of Virus and Microbiological Special Diagnostics, Statens Serum Institut, Copenhagen, Denmark | Aalborg University | Danish Covid-19 Genome Consortium |

| | | | |
|---|---|---|---|
| EPI_ISL_971168, EPI_ISL_971171, EPI_ISL_971174, EPI_ISL_971176, EPI_ISL_971183 | Lighthouse Lab in Milton Keynes | Wellcome Sanger Institute for the COVID-19 Genomics UK (COG-UK) Consortium | The Lighthouse Lab in Milton Keynes and Alex Alderton, Roberto Amato, Sonia Goncalves, Ewan Harrison, David K. Jackson, Ian Johnston, Dominic Kwiatkowski, Cordelia Langford, John Sillitoe on behalf of the Wellcome Sanger Institute COVID-19 Surveillance Team |
| EPI_ISL_971184, EPI_ISL_971188 | Department of Virus and Microbiological Special Diagnostics, Statens Serum Institut, Copenhagen, Denmark | Aalborg University | Danish Covid-19 Genome Consortium |
| EPI_ISL_971191 | Lighthouse Lab in Milton Keynes | Wellcome Sanger Institute for the COVID-19 Genomics UK (COG-UK) Consortium | The Lighthouse Lab in Milton Keynes and Alex Alderton, Roberto Amato, Sonia Goncalves, Ewan Harrison, David K. Jackson, Ian Johnston, Dominic Kwiatkowski, Cordelia Langford, John Sillitoe on behalf of the Wellcome Sanger Institute COVID-19 Surveillance Team |
| EPI_ISL_971193 | Department of Virus and Microbiological Special Diagnostics, Statens Serum Institut, Copenhagen, Denmark | Aalborg University | Danish Covid-19 Genome Consortium |
| EPI_ISL_971194 | Lighthouse Lab in Milton Keynes | Wellcome Sanger Institute for the COVID-19 Genomics UK (COG-UK) Consortium | The Lighthouse Lab in Milton Keynes and Alex Alderton, Roberto Amato, Sonia Goncalves, Ewan Harrison, David K. Jackson, Ian Johnston, Dominic Kwiatkowski, Cordelia Langford, John Sillitoe on behalf of the Wellcome Sanger Institute COVID-19 Surveillance Team |
| EPI_ISL_971198 | Department of Virus and Microbiological Special Diagnostics, Statens Serum Institut, Copenhagen, Denmark | Aalborg University | Danish Covid-19 Genome Consortium |
| EPI_ISL_971206 | Lighthouse Lab in Milton Keynes | Wellcome Sanger Institute for the COVID-19 Genomics UK (COG-UK) Consortium | The Lighthouse Lab in Milton Keynes and Alex Alderton, Roberto Amato, Sonia Goncalves, Ewan Harrison, David K. Jackson, Ian Johnston, Dominic Kwiatkowski, Cordelia Langford, John Sillitoe on behalf of the Wellcome Sanger Institute COVID-19 Surveillance Team |
| EPI_ISL_971214, EPI_ISL_971220, EPI_ISL_971229 | Department of Virus and Microbiological Special Diagnostics, Statens Serum Institut, Copenhagen, Denmark | Aalborg University | Danish Covid-19 Genome Consortium |
| EPI_ISL_971231 | Lighthouse Lab in Milton Keynes | Wellcome Sanger Institute for the COVID-19 Genomics UK (COG-UK) Consortium | The Lighthouse Lab in Milton Keynes and Alex Alderton, Roberto Amato, Sonia Goncalves, Ewan Harrison, David K. Jackson, Ian Johnston, Dominic Kwiatkowski, Cordelia Langford, John Sillitoe on behalf of the Wellcome Sanger Institute COVID-19 Surveillance Team |
| EPI_ISL_971232 | Department of Virus and Microbiological Special Diagnostics, Statens Serum Institut, Copenhagen, Denmark | Aalborg University | Danish Covid-19 Genome Consortium |
| EPI_ISL_971239 | Lighthouse Lab in Milton Keynes | Wellcome Sanger Institute for the COVID-19 Genomics UK (COG-UK) Consortium | The Lighthouse Lab in Milton Keynes and Alex Alderton, Roberto Amato, Sonia Goncalves, Ewan Harrison, David K. Jackson, Ian Johnston, Dominic Kwiatkowski, Cordelia Langford, John Sillitoe on behalf of the Wellcome Sanger Institute COVID-19 Surveillance Team |
| EPI_ISL_971241 | Department of Virus and Microbiological Special Diagnostics, Statens Serum Institut, Copenhagen, Denmark | Aalborg University | Danish Covid-19 Genome Consortium |
| EPI_ISL_971242 | Lighthouse Lab in Milton Keynes | Wellcome Sanger Institute for the COVID-19 Genomics UK (COG-UK) Consortium | The Lighthouse Lab in Milton Keynes and Alex Alderton, Roberto Amato, Sonia Goncalves, Ewan Harrison, David K. Jackson, Ian Johnston, Dominic Kwiatkowski, Cordelia Langford, John Sillitoe on behalf of the Wellcome Sanger Institute COVID-19 Surveillance Team |
| EPI_ISL_971250 | Department of Virus and Microbiological Special Diagnostics, Statens Serum Institut, Copenhagen, Denmark | Aalborg University | Danish Covid-19 Genome Consortium |
| EPI_ISL_971255, EPI_ISL_971261, EPI_ISL_971269, EPI_ISL_971281 | Lighthouse Lab in Milton Keynes | Wellcome Sanger Institute for the COVID-19 Genomics UK (COG-UK) Consortium | The Lighthouse Lab in Milton Keynes and Alex Alderton, Roberto Amato, Sonia Goncalves, Ewan Harrison, David K. Jackson, Ian Johnston, Dominic Kwiatkowski, Cordelia Langford, John Sillitoe on behalf of the Wellcome Sanger Institute COVID-19 Surveillance Team |
| EPI_ISL_971290 | Department of Virus and Microbiological Special Diagnostics, Statens Serum Institut, Copenhagen, Denmark | Aalborg University | Danish Covid-19 Genome Consortium |
| EPI_ISL_971292 | Lighthouse Lab in Milton Keynes | Wellcome Sanger Institute for the COVID-19 Genomics UK (COG-UK) Consortium | The Lighthouse Lab in Milton Keynes and Alex Alderton, Roberto Amato, Sonia Goncalves, Ewan Harrison, David K. Jackson, Ian Johnston, Dominic Kwiatkowski, Cordelia Langford, John Sillitoe on behalf of the Wellcome Sanger Institute COVID-19 Surveillance Team |
| EPI_ISL_971293 | Department of Virus and Microbiological Special Diagnostics, Statens Serum Institut, Copenhagen, Denmark | Aalborg University | Danish Covid-19 Genome Consortium |
| EPI_ISL_971297 | Lighthouse Lab in Milton Keynes | Wellcome Sanger Institute for the COVID-19 Genomics UK (COG-UK) Consortium | The Lighthouse Lab in Milton Keynes and Alex Alderton, Roberto Amato, Sonia Goncalves, Ewan Harrison, David K. Jackson, Ian Johnston, Dominic Kwiatkowski, Cordelia Langford, John Sillitoe on behalf of the Wellcome Sanger Institute COVID-19 Surveillance Team |
| EPI_ISL_971299, EPI_ISL_971306, EPI_ISL_971315, EPI_ISL_971326 | Department of Virus and Microbiological Special Diagnostics, Statens Serum Institut, Copenhagen, Denmark | Aalborg University | Danish Covid-19 Genome Consortium |
| EPI_ISL_971327, EPI_ISL_971333 | Lighthouse Lab in Milton Keynes | Wellcome Sanger Institute for the COVID-19 Genomics UK (COG-UK) Consortium | The Lighthouse Lab in Milton Keynes and Alex Alderton, Roberto Amato, Sonia Goncalves, Ewan Harrison, David K. Jackson, Ian Johnston, Dominic Kwiatkowski, Cordelia Langford, John Sillitoe on behalf of the Wellcome Sanger Institute COVID-19 Surveillance Team |
| EPI_ISL_971335 | Department of Virus and Microbiological Special Diagnostics, Statens Serum Institut, Copenhagen, Denmark | Aalborg University | Danish Covid-19 Genome Consortium |
| EPI_ISL_971339, EPI_ISL_971347 | Lighthouse Lab in Milton Keynes | Wellcome Sanger Institute for the COVID-19 Genomics UK (COG-UK) Consortium | The Lighthouse Lab in Milton Keynes and Alex Alderton, Roberto Amato, Sonia Goncalves, Ewan Harrison, David K. Jackson, Ian Johnston, Dominic Kwiatkowski, Cordelia Langford, John Sillitoe on behalf of the Wellcome Sanger Institute COVID-19 Surveillance Team |
| EPI_ISL_971353, EPI_ISL_971355 | Department of Virus and Microbiological Special Diagnostics, Statens Serum Institut, Copenhagen, Denmark | Aalborg University | Danish Covid-19 Genome Consortium |
| EPI_ISL_971360 | Lighthouse Lab in Milton Keynes | Wellcome Sanger Institute for the COVID-19 Genomics UK (COG-UK) Consortium | The Lighthouse Lab in Milton Keynes and Alex Alderton, Roberto Amato, Sonia Goncalves, Ewan Harrison, David K. Jackson, Ian Johnston, Dominic Kwiatkowski, Cordelia Langford, John Sillitoe on behalf of the Wellcome Sanger Institute COVID-19 Surveillance Team |
| EPI_ISL_971361 | Department of Virus and Microbiological Special Diagnostics, Statens Serum Institut, Copenhagen, Denmark | Aalborg University | Danish Covid-19 Genome Consortium |
| EPI_ISL_971364, EPI_ISL_971365, EPI_ISL_971370 | Lighthouse Lab in Milton Keynes | Wellcome Sanger Institute for the COVID-19 Genomics UK (COG-UK) Consortium | The Lighthouse Lab in Milton Keynes and Alex Alderton, Roberto Amato, Sonia Goncalves, Ewan Harrison, David K. Jackson, Ian Johnston, Dominic Kwiatkowski, Cordelia Langford, John Sillitoe on behalf of the Wellcome Sanger Institute COVID-19 Surveillance Team |
| EPI_ISL_971375, EPI_ISL_971381 | Department of Virus and Microbiological Special Diagnostics, Statens Serum Institut, Copenhagen, Denmark | Aalborg University | Danish Covid-19 Genome Consortium |
| EPI_ISL_971385, EPI_ISL_971394 | Lighthouse Lab in Milton Keynes | Wellcome Sanger Institute for the COVID-19 Genomics UK (COG-UK) Consortium | The Lighthouse Lab in Milton Keynes and Alex Alderton, Roberto Amato, Sonia Goncalves, Ewan Harrison, David K. Jackson, Ian Johnston, Dominic Kwiatkowski, Cordelia Langford, John Sillitoe on behalf of the Wellcome Sanger Institute COVID-19 Surveillance Team |

EPI_ISL_971404, EPI_ISL_971418, EPI_ISL_971431, EPI_ISL_971437, EPI_ISL_971439, EPI_ISL_971447, EPI_ISL_971461, EPI_ISL_971467, EPI_ISL_971470, EPI_ISL_971480, EPI_ISL_971485, EPI_ISL_971494, EPI_ISL_971496, EPI_ISL_971497, EPI_ISL_971500, EPI_ISL_971502, EPI_ISL_971503, EPI_ISL_971504, EPI_ISL_971508, EPI_ISL_971509, EPI_ISL_971513, EPI_ISL_971515, EPI_ISL_971516, EPI_ISL_971529, EPI_ISL_971534, EPI_ISL_971539, EPI_ISL_971542, EPI_ISL_971547, EPI_ISL_971548, EPI_ISL_971549, EPI_ISL_971556, EPI_ISL_971557, EPI_ISL_971558, EPI_ISL_971565, EPI_ISL_971575, EPI_ISL_971578, EPI_ISL_971580, EPI_ISL_971581, EPI_ISL_971591, EPI_ISL_971615, EPI_ISL_971616, EPI_ISL_971623, EPI_ISL_971625, EPI_ISL_971628, EPI_ISL_971632, EPI_ISL_971633, EPI_ISL_971635, EPI_ISL_971638, EPI_ISL_971640, EPI_ISL_971648, EPI_ISL_971649, EPI_ISL_971650, EPI_ISL_971673, EPI_ISL_971674, EPI_ISL_971680, EPI_ISL_971684, EPI_ISL_971686, EPI_ISL_971697, EPI_ISL_971701, EPI_ISL_971705, EPI_ISL_971708, EPI_ISL_971710, EPI_ISL_971711, EPI_ISL_971714, EPI_ISL_971715, EPI_ISL_971716, EPI_ISL_971723, EPI_ISL_971724, EPI_ISL_971730, EPI_ISL_971733, EPI_ISL_971736, EPI_ISL_971737, EPI_ISL_971739, EPI_ISL_971742, EPI_ISL_971743, EPI_ISL_971745, EPI_ISL_971751, EPI_ISL_971755, EPI_ISL_971757, EPI_ISL_971761, EPI_ISL_971768, EPI_ISL_971769, EPI_ISL_971785, EPI_ISL_971808, EPI_ISL_971815, EPI_ISL_971822, EPI_ISL_971832, EPI_ISL_971849, EPI_ISL_971854, EPI_ISL_971856, EPI_ISL_971857, EPI_ISL_971859, EPI_ISL_971861, EPI_ISL_971865, EPI_ISL_971867, EPI_ISL_971876, EPI_ISL_971877, EPI_ISL_971880, EPI_ISL_971886, EPI_ISL_971899, EPI_ISL_971906, EPI_ISL_971908, EPI_ISL_971910, EPI_ISL_971915, EPI_ISL_971925, EPI_ISL_971931, EPI_ISL_971933, EPI_ISL_971934, EPI_ISL_971938, EPI_ISL_971942, EPI_ISL_971943, EPI_ISL_971946, EPI_ISL_971954, EPI_ISL_971960, EPI_ISL_971962, EPI_ISL_971963, EPI_ISL_971966, EPI_ISL_971968, EPI_ISL_971970, EPI_ISL_971971, EPI_ISL_971973, EPI_ISL_971975, EPI_ISL_971978, EPI_ISL_971981, EPI_ISL_971985, EPI_ISL_971986, EPI_ISL_971988, EPI_ISL_971990, EPI_ISL_971996, EPI_ISL_971999, EPI_ISL_972006, EPI_ISL_972008, EPI_ISL_972015, EPI_ISL_972016, EPI_ISL_972018, EPI_ISL_972023, EPI_ISL_972032, EPI_ISL_972034, EPI_ISL_972046, EPI_ISL_972049, EPI_ISL_972053, EPI_ISL_972054, EPI_ISL_972060, EPI_ISL_972068, EPI_ISL_972074, EPI_ISL_972080, EPI_ISL_972085, EPI_ISL_972088, EPI_ISL_972090, EPI_ISL_972092, EPI_ISL_972093, EPI_ISL_972101, EPI_ISL_972102, EPI_ISL_972105, EPI_ISL_972114, EPI_ISL_972119, EPI_ISL_972123, EPI_ISL_972130, EPI_ISL_972131, EPI_ISL_972132, EPI_ISL_972136, EPI_ISL_972138, EPI_ISL_972143, EPI_ISL_972150, EPI_ISL_972156, EPI_ISL_972164, EPI_ISL_972169, EPI_ISL_972170, EPI_ISL_972172, EPI_ISL_972184, EPI_ISL_972190, EPI_ISL_972193, EPI_ISL_972194, EPI_ISL_972195, EPI_ISL_972196, EPI_ISL_972197, EPI_ISL_972198, EPI_ISL_972201, EPI_ISL_972217, EPI_ISL_972222, EPI_ISL_972223, EPI_ISL_972241, EPI_ISL_972258, EPI_ISL_972261, EPI_ISL_972266, EPI_ISL_972267, EPI_ISL_972271, EPI_ISL_972274, EPI_ISL_972280, EPI_ISL_972288, EPI_ISL_972293, EPI_ISL_972309, EPI_ISL_972312, EPI_ISL_972313, EPI_ISL_972314, EPI_ISL_972324, EPI_ISL_972328, EPI_ISL_972332, EPI_ISL_972336, EPI_ISL_972338, EPI_ISL_972346, EPI_ISL_972348, EPI_ISL_972349, EPI_ISL_972351, EPI_ISL_972354, EPI_ISL_972358, EPI_ISL_972370, EPI_ISL_972375, EPI_ISL_972376, EPI_ISL_972377, EPI_ISL_972378, EPI_ISL_972381, EPI_ISL_972387, EPI_ISL_972391, EPI_ISL_972393, EPI_ISL_972394, EPI_ISL_972397, EPI_ISL_972406, EPI_ISL_972407, EPI_ISL_972423, EPI_ISL_972426, EPI_ISL_972431, EPI_ISL_972435, EPI_ISL_972436, EPI_ISL_972437, EPI_ISL_972438, EPI_ISL_972443, EPI_ISL_972448, EPI_ISL_972465, EPI_ISL_972466, EPI_ISL_972468, EPI_ISL_972471, EPI_ISL_972472, EPI_ISL_972474, EPI_ISL_972477, EPI_ISL_972478, EPI_ISL_972486, EPI_ISL_972488, EPI_ISL_972491, EPI_ISL_972493, EPI_ISL_972512, EPI_ISL_972513, EPI_ISL_972517, EPI_ISL_972528, EPI_ISL_972529, EPI_ISL_972530, EPI_ISL_972533, EPI_ISL_972536, EPI_ISL_972539, EPI_ISL_972545, EPI_ISL_972546, EPI_ISL_972547,

| | | | |
|---|---|---|---|
| EPI_ISL_972549, EPI_ISL_972552, EPI_ISL_972555, EPI_ISL_972556, EPI_ISL_972560, EPI_ISL_972565, EPI_ISL_972569, EPI_ISL_972574, EPI_ISL_972576, EPI_ISL_972577, EPI_ISL_972579, EPI_ISL_972580, EPI_ISL_972586, EPI_ISL_972590, EPI_ISL_972591, EPI_ISL_972592, EPI_ISL_972597, EPI_ISL_972601, EPI_ISL_972609, EPI_ISL_972610, EPI_ISL_972612, EPI_ISL_972614, EPI_ISL_972615, EPI_ISL_972625, EPI_ISL_972627, EPI_ISL_972639, EPI_ISL_972644, EPI_ISL_972661, EPI_ISL_972663, EPI_ISL_972673, EPI_ISL_972674, EPI_ISL_972680, EPI_ISL_972686, EPI_ISL_972687, EPI_ISL_972689, EPI_ISL_972690, EPI_ISL_972692, EPI_ISL_972707, EPI_ISL_972712, EPI_ISL_972713, EPI_ISL_972715, EPI_ISL_972718, EPI_ISL_972727, EPI_ISL_972728, EPI_ISL_972730, EPI_ISL_972737, EPI_ISL_972742, EPI_ISL_972743, EPI_ISL_972744, EPI_ISL_972751, EPI_ISL_972755, EPI_ISL_972758, EPI_ISL_972759, EPI_ISL_972763, EPI_ISL_972768, EPI_ISL_972769, EPI_ISL_972771, EPI_ISL_972772, EPI_ISL_972773, EPI_ISL_972774, EPI_ISL_972776, EPI_ISL_972777, EPI_ISL_972785, EPI_ISL_972787, EPI_ISL_972790, EPI_ISL_972791, EPI_ISL_972807, EPI_ISL_972809, EPI_ISL_972812, EPI_ISL_972813, EPI_ISL_972819, EPI_ISL_972821, EPI_ISL_972822, EPI_ISL_972825, EPI_ISL_972833, EPI_ISL_972834, EPI_ISL_972843, EPI_ISL_972848, EPI_ISL_972851, EPI_ISL_972853, EPI_ISL_972855, EPI_ISL_972858, EPI_ISL_972860, EPI_ISL_972872, EPI_ISL_972876, EPI_ISL_972881, EPI_ISL_972887, EPI_ISL_972888, EPI_ISL_972892, EPI_ISL_972895, EPI_ISL_972897, EPI_ISL_972898, EPI_ISL_972900, EPI_ISL_972903, EPI_ISL_972907, EPI_ISL_972911, EPI_ISL_972918, EPI_ISL_972922, EPI_ISL_972936, EPI_ISL_972938, EPI_ISL_972940, EPI_ISL_972941, EPI_ISL_972945, EPI_ISL_972948, EPI_ISL_972949, EPI_ISL_972957, EPI_ISL_972960, EPI_ISL_972962, EPI_ISL_972965, EPI_ISL_972975, EPI_ISL_972995, EPI_ISL_972998, EPI_ISL_973001, EPI_ISL_973014, EPI_ISL_973020, EPI_ISL_973024, EPI_ISL_973026, EPI_ISL_973027, EPI_ISL_973030, EPI_ISL_973038, EPI_ISL_973040, EPI_ISL_973047, EPI_ISL_973048, EPI_ISL_973052, EPI_ISL_973053, EPI_ISL_973055, EPI_ISL_973060, EPI_ISL_973063, EPI_ISL_973064, EPI_ISL_973069, EPI_ISL_973071, EPI_ISL_973073, EPI_ISL_973076, EPI_ISL_973082, EPI_ISL_973085, EPI_ISL_973086, EPI_ISL_973094, EPI_ISL_973096, EPI_ISL_973097, EPI_ISL_973111, EPI_ISL_973115, EPI_ISL_973120, EPI_ISL_973122, EPI_ISL_973123, EPI_ISL_973127, EPI_ISL_973130, EPI_ISL_973132, EPI_ISL_973135, EPI_ISL_973137, EPI_ISL_973143, EPI_ISL_973160, EPI_ISL_973167, EPI_ISL_973179, EPI_ISL_973182, EPI_ISL_973186, EPI_ISL_973189, EPI_ISL_973191, EPI_ISL_973195, EPI_ISL_973196, EPI_ISL_973199, EPI_ISL_973206, EPI_ISL_973208, EPI_ISL_973216, EPI_ISL_973224, EPI_ISL_973225, EPI_ISL_973229, EPI_ISL_973230, EPI_ISL_973232, EPI_ISL_973236, EPI_ISL_973238, EPI_ISL_973249, EPI_ISL_973250, EPI_ISL_973251, EPI_ISL_973256, EPI_ISL_973260, EPI_ISL_973264, EPI_ISL_973266, EPI_ISL_973273, EPI_ISL_973281, EPI_ISL_973288, EPI_ISL_973289, EPI_ISL_973296, EPI_ISL_973302, EPI_ISL_973304, EPI_ISL_973312, EPI_ISL_973317, EPI_ISL_973324, EPI_ISL_973328, EPI_ISL_973333, EPI_ISL_973338, EPI_ISL_973340, EPI_ISL_973341, EPI_ISL_973345, EPI_ISL_973349, EPI_ISL_973354, EPI_ISL_973358, EPI_ISL_973359, EPI_ISL_973363, EPI_ISL_973365, EPI_ISL_973368, EPI_ISL_973373, EPI_ISL_973383, EPI_ISL_973384, EPI_ISL_973396, EPI_ISL_973398, EPI_ISL_973399, EPI_ISL_973404, EPI_ISL_973406, EPI_ISL_973410, EPI_ISL_973412, EPI_ISL_973422, EPI_ISL_973423, EPI_ISL_973430, EPI_ISL_973432, EPI_ISL_973445, EPI_ISL_973447, EPI_ISL_973453, EPI_ISL_973456, EPI_ISL_973459, EPI_ISL_973461, EPI_ISL_973465, EPI_ISL_973467, EPI_ISL_973468, EPI_ISL_973471, EPI_ISL_973474, EPI_ISL_973484, EPI_ISL_973486, EPI_ISL_973487, EPI_ISL_973495, EPI_ISL_973499, EPI_ISL_973505, EPI_ISL_973508, EPI_ISL_973516, EPI_ISL_973518, EPI_ISL_973540, EPI_ISL_973553, EPI_ISL_973570, EPI_ISL_973580, EPI_ISL_973584, EPI_ISL_973586, EPI_ISL_973595, EPI_ISL_973599, EPI_ISL_973633, EPI_ISL_973636, EPI_ISL_973638, EPI_ISL_973641, EPI_ISL_973654, EPI_ISL_973664, EPI_ISL_973674, EPI_ISL_973677, EPI_ISL_973693, EPI_ISL_973694, EPI_ISL_973699, EPI_ISL_973706, EPI_ISL_973711, EPI_ISL_973713, EPI_ISL_973732, EPI_ISL_973741, EPI_ISL_973748, EPI_ISL_973762, EPI_ISL_973766, EPI_ISL_973772, EPI_ISL_973784, EPI_ISL_973796, EPI_ISL_973817, EPI_ISL_973820, EPI_ISL_973824, EPI_ISL_973829, EPI_ISL_973831, EPI_ISL_973833 | | | |
| see above | Department of Virus and Microbiological Special Diagnostics, Statens Serum Institut, Copenhagen, Denmark | Aalborg University | Danish Covid-19 Genome Consortium |
| EPI_ISL_974853 | Laboratorio de Virología / Hospital Universitario Central de Asturias (HUCA) | Laboratorio de Virología / Hospital Universitario Central de Asturias (HUCA) | Castelló C, Gómez de Oña J, Boga JA, Rojo S, Alvarez-Arguelles ME, Abreu F, Costales I, Sandoval M, Perez-Martínez Z, Martín-Rodríguez G, Coto E, Melón S |
| EPI_ISL_977417 | Hospital de Cabueñes | Laboratorio de Virología / Hospital Universitario Central de Asturias (HUCA) | Castelló C, Gómez de Oña J, Boga JA, Rojo S, Alvarez-Arguelles ME, Abreu F, Costales I, Sandoval M, Perez-Martínez Z, Martín-Rodríguez G, Coto E, Melón S |
| EPI_ISL_977489 | UPA Dr. Akira Tada | Instituto Adolfo Lutz, Interdiciplinary Procedures Center, Strategic Laboratory | Claudio Tavares Sacchi, Claudia Regina Gonçalves, Erica Valessa Ramos Gomes, Karoline Rodrigues Campos |
| EPI_ISL_977492 | Lab voor klinische biologie | Lab voor klinische biologie | Hannelore Hamerlinck, Marija Janevska, Bruno Verhasselt |
| EPI_ISL_977500 | LATE - Laboratório de Técnicas Especiais - Hospital Israelita Albert Einstein | LATE - Laboratório de Técnicas Especiais - Hospital Israelita Albert Einstein | Deyvid Amgarten, Fernanda de Mello Malta, Raquel Riyuzo, Ana Paula Moreira Salles, Pedro Henrique Sebe Rodrigues, João Renato Rebello Pinho |
| EPI_ISL_977501 | Hospital de Cabueñes | Laboratorio de Virología HUCA | Castelló C, Gómez de Oña J, Boga JA, Rojo S, Alvarez-Arguelles ME, Abreu F, Costales I, Sandoval M, Perez-Martínez Z, Martín-Rodríguez G, Coto E, Melón S |
| EPI_ISL_977502 | Hospital Oriente de Asturias | Laboratorio de Virología HUCA | Sandoval M, Castelló C, Gómez de Oña J, Boga JA, Rojo S, Alvarez-Arguelles ME, Abreu F, Costales I, Perez-Martínez Z, Martín-Rodríguez G, Coto E, Melón S |
| EPI_ISL_977505, EPI_ISL_977506 | Laboratorio de Virología HUCA | Laboratorio de Virología HUCA | Sandoval M, Castelló C, Gómez de Oña J, Boga JA, Rojo S, Alvarez-Arguelles ME, Abreu F, Costales I, Perez-Martínez Z, Martín-Rodríguez G, Coto E, Melón S |
| EPI_ISL_977507 | Hospital Valle del Nalon | Laboratorio de Virología HUCA | Sandoval M, Castelló C, Gómez de Oña J, Boga JA, Rojo S, Alvarez-Arguelles ME, Abreu F, Costales I, Perez-Martínez Z, Martín-Rodríguez G, Coto E, Melón S |
| EPI_ISL_977509 | Hospital Valle del Nalón | Laboratorio de Virología HUCA | Sandoval M, Castelló C, Gómez de Oña J, Boga JA, Rojo S, Alvarez-Arguelles ME, Abreu F, Costales I, Perez-Martínez Z, Martín-Rodríguez G, Coto E, Melón S |
| EPI_ISL_977510, EPI_ISL_977511, EPI_ISL_977512 | Laboratorio de Virología HUCA | Laboratorio de Virología HUCA | Sandoval M, Castelló C, Gómez de Oña J, Boga JA, Rojo S, Alvarez-Arguelles ME, Abreu F, Costales I, Perez-Martínez Z, Martín-Rodríguez G, Coto E, Melón S |
| EPI_ISL_977537 | Biolab Diagnostic Laboratories | Biolab Diagnostic Laboratories | Issa Abu-Dayyeh, Ahmad Tibi, Lama Hussein, Shayma Ali, Badia Saddedin, Amid Abdelnour |
| EPI_ISL_977566, EPI_ISL_977567, EPI_ISL_977575, EPI_ISL_977576, EPI_ISL_977577, EPI_ISL_977578, EPI_ISL_977579, EPI_ISL_977580 | University of Michigan Clinical Microbiology Laboratory | Lauring Lab, University of Michigan, Department of Microbiology and Immunology | Valesano |
| EPI_ISL_977613, EPI_ISL_977614, EPI_ISL_977615, EPI_ISL_977620 | SC (UCO) Igiene e Sanità Pubblica (funzione integrata con SC Microbiologia e Virologia) e Laboratory of Molecular Virology of the International Centre for Genetic Engineering and Biotechnology (ICGEB) | ARGO Laboratorio Genomica ed Epigenomica | Licastro D, Dal Monego S, Degasperi M, Marcello A, D'Agaro P, Pipan C |
| EPI_ISL_977660, EPI_ISL_977665 | Auburn Sports Medicine | Synergy Laboratories | Megan Cornwell |
| EPI_ISL_978373, EPI_ISL_978374, EPI_ISL_978375, EPI_ISL_978376, EPI_ISL_978377, EPI_ISL_978481, EPI_ISL_978482, EPI_ISL_978483, EPI_ISL_978484, EPI_ISL_978485, EPI_ISL_978486, EPI_ISL_978487 | | | |
| see above | Arizona State Public Health Laboratory | Arizona State Public Health Laboratory | Trung Huynh, Jessica Escobar, Katherine Fullerton, Nobuko Fukushima, Stacy White, Linda Getsinger, Victor Waddell |
| EPI_ISL_978555, EPI_ISL_978556, EPI_ISL_978557, EPI_ISL_978558, EPI_ISL_978559, EPI_ISL_978560, EPI_ISL_978561, EPI_ISL_978562, EPI_ISL_978563, EPI_ISL_978564, EPI_ISL_978565, EPI_ISL_978566, EPI_ISL_978567, EPI_ISL_978568, EPI_ISL_978569, EPI_ISL_978570, EPI_ISL_978571, EPI_ISL_978572, EPI_ISL_978573, EPI_ISL_978574, EPI_ISL_978575, EPI_ISL_978576, EPI_ISL_978577, EPI_ISL_978578, EPI_ISL_978579, EPI_ISL_978580, EPI_ISL_978581, EPI_ISL_978582, EPI_ISL_978583, EPI_ISL_978584, EPI_ISL_978585, EPI_ISL_978586, EPI_ISL_978587, EPI_ISL_978588, EPI_ISL_978589, EPI_ISL_978590, EPI_ISL_978591, EPI_ISL_978592, EPI_ISL_978593, EPI_ISL_978594, EPI_ISL_978595, EPI_ISL_978596, EPI_ISL_978597, EPI_ISL_978598, EPI_ISL_978599, EPI_ISL_978600, EPI_ISL_978601, EPI_ISL_978602, EPI_ISL_978603, EPI_ISL_978604, EPI_ISL_978605, EPI_ISL_978606, EPI_ISL_978607, EPI_ISL_978608, EPI_ISL_978609, EPI_ISL_978610, EPI_ISL_978611, EPI_ISL_978612, EPI_ISL_978613, EPI_ISL_978614, EPI_ISL_978615, EPI_ISL_978616, EPI_ISL_978617, EPI_ISL_978618, EPI_ISL_978619, EPI_ISL_978620 | | | |
| see above | Helix/Illumina | Respiratory Viruses Branch, Division of Viral Diseases, Centers for Disease Control and Prevention | Peter W. Cook,Dakota Howard,Dhwani Batra,Ben L. Rambo-Martin,Eileen de Feo,Jan Antico,Christine Tran,Matthew Tolentino,Shannon Wickline,Kim Gietzen,Brad Sickler,Jingtao Liu,Eric Allen,Phil Febbo,Summer Galloway,Nicole L. Washington,Simon White,Geraint Levan,Kelly Schiabor Barrett,Elizabeth Cirulli,Alexandre Bolze,Ary Ascencio,Charlotte Rivera-Garcia,Ryan Cho,Jason Nguyen,Sherry Wang,Jimmy Ramirez,Tyler Cassens,Efren Sandoval,Magnus Isaksson,William Lee,David Becker,Marc Laurent,James Lu,Clinton R. Paden,Suxiang Tong,Duncan MacCannell, |
| EPI_ISL_978889, EPI_ISL_978918, EPI_ISL_978919, EPI_ISL_978954, EPI_ISL_978984, EPI_ISL_978988 | Centre for Dengue Research and AICBU, Department of Immunology and Molecular Medicine | Centre for Dengue Research and AICBU, Department of Immunology and Molecular Medicine | Chandima Jeewandara, Deshni Jayathilaka, Dinuka Ariyaratne, Tibutius Thanesh Pramanayagam, Diyanath Ranasinghe, Laksiri Gomes, Gathsaurie Neelika Malavige |
| EPI_ISL_979317, EPI_ISL_979319, EPI_ISL_979320 | Cadham Provincial laboratory | National Microbiology Laboratory (NML) | Anna Majer, Shari Tyson, Grace Seo, Philip Mabon, Elsie Grudeski, Rhiannon Huzarewich, Russell Mandes, Anneliese Landgraff, Jennifer Tanner, Natalie Knox, Morag Graham, Gary Van Domselaar, Paul Van Caeseele, Jared Bullard, David Alexander, Kerry Dust, Nathalie Bastien, Yan Li, Timothy Booth, Darian Hole, Madison Chapel, Kirsten Biggar, CanCOGeN's metadata curation team, Public Health Agency of Canada CanCOGeN team |
| EPI_ISL_979356 | Victorian Infectious Diseases Reference Laboratory (VIDRL) | VIDRL and MDU-PHL | Caly L., Seemann T., Sait, M.L., Druce J., Sherry, N.L. |
| EPI_ISL_979357, EPI_ISL_979358, EPI_ISL_979359 | Microbiological Diagnostic Unit - Public Health Laboratory (MDU-PHL) | MDU-PHL | Seemann T., Sait, M.L., Sherry, N.L. |
| EPI_ISL_979429, EPI_ISL_979430, EPI_ISL_979431, EPI_ISL_979432, EPI_ISL_979433, EPI_ISL_979434, EPI_ISL_979435, EPI_ISL_979436, EPI_ISL_979437, EPI_ISL_979438, EPI_ISL_979439, EPI_ISL_979440, EPI_ISL_979441, EPI_ISL_979442, EPI_ISL_979443, EPI_ISL_979444, EPI_ISL_979445, EPI_ISL_979446, EPI_ISL_979447, EPI_ISL_979448, EPI_ISL_979449, EPI_ISL_979450, EPI_ISL_979451, EPI_ISL_979452 | | | |
| see above | New Mexico Department of Health Scientific Laboratory | New Mexico Department of Health Scientific Laboratory | Ellie Johnson, Anastacia Griego-Fisher, D'eldra Malone, Jennifer Benoit |
| EPI_ISL_979457 | State Hygienic Laboratory at the University of Iowa | State Hygienic Laboratory at the University of Iowa | Valerie Reeb, Erik Twait, Wes Hottel, Alankar Kampoowale |
| EPI_ISL_979803, EPI_ISL_979804, EPI_ISL_979893, EPI_ISL_979959, EPI_ISL_979960, EPI_ISL_979961, EPI_ISL_979963, EPI_ISL_979964, EPI_ISL_979965, EPI_ISL_979966, EPI_ISL_979967 | | | |
| see above | National Institute of Infectious Diseases-Prof. Dr. Matei Bals | National Institute of Infectious Diseases-Prof. Dr. Matei Bals | Leontina Banica, Marius Surleac, Corina Casangiu, Petre Milu, Andreea Tudor, Simona Paraschiv, Dan Otelea |

| | | | |
|---|---|---|---|
| | Molecular Diagnostics Laboratory | Molecular Diagnostics Laboratory | |
| EPI_ISL_979974, EPI_ISL_979975, EPI_ISL_979976, EPI_ISL_979977, EPI_ISL_979978, EPI_ISL_979979, EPI_ISL_979980, EPI_ISL_979981, EPI_ISL_979982, EPI_ISL_979983, EPI_ISL_979984, EPI_ISL_979985, EPI_ISL_979986, EPI_ISL_979987, EPI_ISL_979988, EPI_ISL_979989, EPI_ISL_979990, EPI_ISL_979991, EPI_ISL_979992, EPI_ISL_979993, EPI_ISL_979994, EPI_ISL_979995, EPI_ISL_979996, EPI_ISL_979997, EPI_ISL_979998, EPI_ISL_979999, EPI_ISL_980000, EPI_ISL_980001, EPI_ISL_980002, EPI_ISL_980003, EPI_ISL_980004, EPI_ISL_980005, EPI_ISL_980006, EPI_ISL_980007, EPI_ISL_980008, EPI_ISL_980009, EPI_ISL_980010, EPI_ISL_980011, EPI_ISL_980012, EPI_ISL_980013, EPI_ISL_980014, EPI_ISL_980015, EPI_ISL_980016, EPI_ISL_980017, EPI_ISL_980018, EPI_ISL_980019, EPI_ISL_980020, EPI_ISL_980021, EPI_ISL_980022, EPI_ISL_980023, EPI_ISL_980024, EPI_ISL_980025, EPI_ISL_980026, EPI_ISL_980027, EPI_ISL_980028, EPI_ISL_980029, EPI_ISL_980030, EPI_ISL_980031, EPI_ISL_980032, EPI_ISL_980033, EPI_ISL_980034, EPI_ISL_980035, EPI_ISL_980036, EPI_ISL_980037, EPI_ISL_980038, EPI_ISL_980039, EPI_ISL_980040, EPI_ISL_980041, EPI_ISL_980042, EPI_ISL_980043, EPI_ISL_980044, EPI_ISL_980045, EPI_ISL_980046, EPI_ISL_980047, EPI_ISL_980048, EPI_ISL_980049, EPI_ISL_980050, EPI_ISL_980051, EPI_ISL_980052, EPI_ISL_980053, EPI_ISL_980054, EPI_ISL_980055, EPI_ISL_980056, EPI_ISL_980057, EPI_ISL_980058, EPI_ISL_980059, EPI_ISL_980060, EPI_ISL_980061, EPI_ISL_980062, EPI_ISL_980063, EPI_ISL_980064, EPI_ISL_980065, EPI_ISL_980066, EPI_ISL_980067, EPI_ISL_980068, EPI_ISL_980069, EPI_ISL_980070, EPI_ISL_980071, EPI_ISL_980072, EPI_ISL_980073, EPI_ISL_980074, EPI_ISL_980075, EPI_ISL_980076, EPI_ISL_980077, EPI_ISL_980078, EPI_ISL_980079, EPI_ISL_980080, EPI_ISL_980081, EPI_ISL_980082, EPI_ISL_980083, EPI_ISL_980084, EPI_ISL_980085, EPI_ISL_980086, EPI_ISL_980087, EPI_ISL_980088, EPI_ISL_980089, EPI_ISL_980090, EPI_ISL_980091, EPI_ISL_980092, EPI_ISL_980093, EPI_ISL_980094, EPI_ISL_980095, EPI_ISL_980096, EPI_ISL_980097, EPI_ISL_980098, EPI_ISL_980099, EPI_ISL_980100, EPI_ISL_980101, EPI_ISL_980102, EPI_ISL_980103, EPI_ISL_980104, EPI_ISL_980105, EPI_ISL_980106, EPI_ISL_980107, EPI_ISL_980108, EPI_ISL_980109, EPI_ISL_980110, EPI_ISL_980111, EPI_ISL_980112, EPI_ISL_980113, EPI_ISL_980114, EPI_ISL_980115, EPI_ISL_980116, EPI_ISL_980117, EPI_ISL_980118, EPI_ISL_980119, EPI_ISL_980120, EPI_ISL_980121, EPI_ISL_980122, EPI_ISL_980123, EPI_ISL_980124, EPI_ISL_980125, EPI_ISL_980126, EPI_ISL_980127, EPI_ISL_980128, EPI_ISL_980129, EPI_ISL_980130, EPI_ISL_980131, EPI_ISL_980132, EPI_ISL_980133, EPI_ISL_980134, EPI_ISL_980135, EPI_ISL_980136, EPI_ISL_980137, EPI_ISL_980138, EPI_ISL_980139, EPI_ISL_980140, EPI_ISL_980141, EPI_ISL_980142, EPI_ISL_980143, EPI_ISL_980144, EPI_ISL_980145, EPI_ISL_980146, EPI_ISL_980147, EPI_ISL_980148, EPI_ISL_980149, EPI_ISL_980150, EPI_ISL_980151, EPI_ISL_980152, EPI_ISL_980153, EPI_ISL_980154, EPI_ISL_980155, EPI_ISL_980156, EPI_ISL_980157, EPI_ISL_980158, EPI_ISL_980159, EPI_ISL_980160, EPI_ISL_980161, EPI_ISL_980162, EPI_ISL_980163, EPI_ISL_980164, EPI_ISL_980165, EPI_ISL_980166, EPI_ISL_980167, EPI_ISL_980168, EPI_ISL_980169, EPI_ISL_980170, EPI_ISL_980171, EPI_ISL_980172, EPI_ISL_980173, EPI_ISL_980174, EPI_ISL_980175, EPI_ISL_980176, EPI_ISL_980177, EPI_ISL_980178, EPI_ISL_980179, EPI_ISL_980180, EPI_ISL_980181, EPI_ISL_980182, EPI_ISL_980183, EPI_ISL_980184, EPI_ISL_980185, EPI_ISL_980186, EPI_ISL_980187, EPI_ISL_980188, EPI_ISL_980189, EPI_ISL_980190, EPI_ISL_980191, EPI_ISL_980192, EPI_ISL_980193, EPI_ISL_980194, EPI_ISL_980195, EPI_ISL_980196, EPI_ISL_980197, EPI_ISL_980198, EPI_ISL_980199, EPI_ISL_980200, EPI_ISL_980201, EPI_ISL_980202, EPI_ISL_980203, EPI_ISL_980204, EPI_ISL_980205, EPI_ISL_980206, EPI_ISL_980207, EPI_ISL_980208, EPI_ISL_980209, EPI_ISL_980210, EPI_ISL_980211, EPI_ISL_980212, EPI_ISL_980213, EPI_ISL_980214, EPI_ISL_980215, EPI_ISL_980216, EPI_ISL_980217, EPI_ISL_980218, EPI_ISL_980219, EPI_ISL_980220, EPI_ISL_980221, EPI_ISL_980222, EPI_ISL_980223, EPI_ISL_980224, EPI_ISL_980225, EPI_ISL_980226, EPI_ISL_980227, EPI_ISL_980228, EPI_ISL_980229, EPI_ISL_980230, EPI_ISL_980231, EPI_ISL_980232, EPI_ISL_980233, EPI_ISL_980234 | | | |
| see above | Lighthouse Lab in Milton Keynes | Wellcome Sanger Institute for the COVID-19 Genomics UK (COG-UK) Consortium | The Lighthouse Lab in Milton Keynes and Alex Alderton, Roberto Amato, Sonia Goncalves, Ewan Harrison, David K. Jackson, Ian Johnston, Dominic Kwiatkowski, Cordelia Langford, John Sillitoe on behalf of the Wellcome Sanger Institute COVID-19 Surveillance Team |
| EPI_ISL_980280, EPI_ISL_980356, EPI_ISL_980416, EPI_ISL_980419, EPI_ISL_980433, EPI_ISL_980449, EPI_ISL_980512 | Lighthouse Lab in Cambridge | Wellcome Sanger Institute for the COVID-19 Genomics UK (COG-UK) Consortium | Rob Howes, The Lighthouse Lab in Cambridge and Alex Alderton, Roberto Amato, Sonia Goncalves, Ewan Harrison, David K. Jackson, Ian Johnston, Dominic Kwiatkowski, Cordelia Langford, John Sillitoe on behalf of the Wellcome Sanger Institute COVID-19 Surveillance Team |
| EPI_ISL_980551, EPI_ISL_980552, EPI_ISL_980553, EPI_ISL_980554, EPI_ISL_980555, EPI_ISL_980556, EPI_ISL_980557, EPI_ISL_980558, EPI_ISL_980559, EPI_ISL_980560, EPI_ISL_980561, EPI_ISL_980562, EPI_ISL_980563, EPI_ISL_980564, EPI_ISL_980565, EPI_ISL_980566, EPI_ISL_980567, EPI_ISL_980568, EPI_ISL_980569, EPI_ISL_980570, EPI_ISL_980571, EPI_ISL_980572, EPI_ISL_980573, EPI_ISL_980574, EPI_ISL_980575, EPI_ISL_980576, EPI_ISL_980577, EPI_ISL_980578, EPI_ISL_980579, EPI_ISL_980580, EPI_ISL_980581, EPI_ISL_980582, EPI_ISL_980583, EPI_ISL_980584, EPI_ISL_980585, EPI_ISL_980586, EPI_ISL_980587, EPI_ISL_980588, EPI_ISL_980589, EPI_ISL_980590, EPI_ISL_980591, EPI_ISL_980592, EPI_ISL_980593, EPI_ISL_980594, EPI_ISL_980595, EPI_ISL_980596, EPI_ISL_980597, EPI_ISL_980598, EPI_ISL_980599, EPI_ISL_980600, EPI_ISL_980601, EPI_ISL_980602, EPI_ISL_980603, EPI_ISL_980604, EPI_ISL_980605, EPI_ISL_980606, EPI_ISL_980607, EPI_ISL_980608, EPI_ISL_980609, EPI_ISL_980610, EPI_ISL_980611, EPI_ISL_980612, EPI_ISL_980613, EPI_ISL_980614, EPI_ISL_980615, EPI_ISL_980616, EPI_ISL_980617, EPI_ISL_980618, EPI_ISL_980619, EPI_ISL_980620, EPI_ISL_980621, EPI_ISL_980622, EPI_ISL_980623, EPI_ISL_980624, EPI_ISL_980625, EPI_ISL_980626, EPI_ISL_980627, EPI_ISL_980628, EPI_ISL_980629, EPI_ISL_980630, EPI_ISL_980631, EPI_ISL_980632, EPI_ISL_980633, EPI_ISL_980634, EPI_ISL_980635, EPI_ISL_980636, EPI_ISL_980637, EPI_ISL_980638, EPI_ISL_980639, EPI_ISL_980640, EPI_ISL_980641, EPI_ISL_980642, EPI_ISL_980643, EPI_ISL_980644, EPI_ISL_980645, EPI_ISL_980646, EPI_ISL_980647, EPI_ISL_980648, EPI_ISL_980649, EPI_ISL_980650, EPI_ISL_980651, EPI_ISL_980652, EPI_ISL_980653, EPI_ISL_980654, EPI_ISL_980655, EPI_ISL_980656, EPI_ISL_980657, EPI_ISL_980658, EPI_ISL_980659, EPI_ISL_980660, EPI_ISL_980661, EPI_ISL_980662, EPI_ISL_980663, EPI_ISL_980664, EPI_ISL_980665, EPI_ISL_980666, EPI_ISL_980667, EPI_ISL_980668, EPI_ISL_980669, EPI_ISL_980670, EPI_ISL_980671, EPI_ISL_980672, EPI_ISL_980673, EPI_ISL_980674, EPI_ISL_980675, EPI_ISL_980676, EPI_ISL_980677, EPI_ISL_980678, EPI_ISL_980679, EPI_ISL_980680, EPI_ISL_980681, EPI_ISL_980682, EPI_ISL_980683, EPI_ISL_980684, EPI_ISL_980685, EPI_ISL_980686, EPI_ISL_980687, EPI_ISL_980688, EPI_ISL_980689, EPI_ISL_980690, EPI_ISL_980691, EPI_ISL_980692, EPI_ISL_980693, EPI_ISL_980694, EPI_ISL_980695, EPI_ISL_980696, EPI_ISL_980697, EPI_ISL_980698, EPI_ISL_980699, EPI_ISL_980700, EPI_ISL_980701, EPI_ISL_980702, EPI_ISL_980703, EPI_ISL_980704, EPI_ISL_980705, EPI_ISL_980706, EPI_ISL_980707, EPI_ISL_980708, EPI_ISL_980709, EPI_ISL_980710, EPI_ISL_980711, EPI_ISL_980712, EPI_ISL_980713, EPI_ISL_980714, EPI_ISL_980715, EPI_ISL_980716, EPI_ISL_980717, EPI_ISL_980718, EPI_ISL_980719, EPI_ISL_980720, EPI_ISL_980721, EPI_ISL_980722, EPI_ISL_980723, EPI_ISL_980724, EPI_ISL_980725, EPI_ISL_980726, EPI_ISL_980727, EPI_ISL_980728, EPI_ISL_980729, EPI_ISL_980730, EPI_ISL_980731, EPI_ISL_980732, EPI_ISL_980733, EPI_ISL_980734, EPI_ISL_980735, EPI_ISL_980736, EPI_ISL_980737, EPI_ISL_980738, EPI_ISL_980739, EPI_ISL_980740, EPI_ISL_980741, EPI_ISL_980742, EPI_ISL_980743, EPI_ISL_980744, EPI_ISL_980745, EPI_ISL_980746, EPI_ISL_980747, EPI_ISL_980748, EPI_ISL_980749, EPI_ISL_980750, EPI_ISL_980751, EPI_ISL_980752, EPI_ISL_980753, EPI_ISL_980754, EPI_ISL_980755, EPI_ISL_980756, EPI_ISL_980757, EPI_ISL_980758, EPI_ISL_980759, EPI_ISL_980760, EPI_ISL_980761, EPI_ISL_980762, EPI_ISL_980763, EPI_ISL_980764, EPI_ISL_980765, EPI_ISL_980766, EPI_ISL_980767, EPI_ISL_980768, EPI_ISL_980769, EPI_ISL_980770, EPI_ISL_980771, EPI_ISL_980772, EPI_ISL_980773, EPI_ISL_980774, EPI_ISL_980775, EPI_ISL_980776, EPI_ISL_980777, EPI_ISL_980778, EPI_ISL_980779, EPI_ISL_980780, EPI_ISL_980781, EPI_ISL_980782, EPI_ISL_980783, EPI_ISL_980784, EPI_ISL_980785, EPI_ISL_980786, EPI_ISL_980787, EPI_ISL_980788, EPI_ISL_980789, EPI_ISL_980790, EPI_ISL_980791, EPI_ISL_980792, EPI_ISL_980793, EPI_ISL_980794, EPI_ISL_980795, EPI_ISL_980796, EPI_ISL_980797, EPI_ISL_980798, EPI_ISL_980799, EPI_ISL_980800, EPI_ISL_980801, EPI_ISL_980802, EPI_ISL_980803, EPI_ISL_980804, EPI_ISL_980805, EPI_ISL_980806 | | | |
| see above | Lighthouse Lab in Milton Keynes | Wellcome Sanger Institute for the COVID-19 Genomics UK (COG-UK) Consortium | The Lighthouse Lab in Milton Keynes and Alex Alderton, Roberto Amato, Sonia Goncalves, Ewan Harrison, David K. Jackson, Ian Johnston, Dominic Kwiatkowski, Cordelia Langford, John Sillitoe on behalf of the Wellcome Sanger Institute COVID-19 Surveillance Team |
| EPI_ISL_980843, EPI_ISL_980857, EPI_ISL_980858, EPI_ISL_980860, EPI_ISL_980861, EPI_ISL_980864, EPI_ISL_980868, EPI_ISL_980870, EPI_ISL_980885, EPI_ISL_980887, EPI_ISL_980888, EPI_ISL_980893, EPI_ISL_980900, EPI_ISL_980901, EPI_ISL_980903, EPI_ISL_980911, EPI_ISL_980917, EPI_ISL_980923, EPI_ISL_980930, EPI_ISL_980965, EPI_ISL_980966, EPI_ISL_980967, EPI_ISL_980968, EPI_ISL_980969, EPI_ISL_980970, EPI_ISL_980971, EPI_ISL_980972, EPI_ISL_980973, EPI_ISL_980975, EPI_ISL_980976, EPI_ISL_980989, EPI_ISL_980990, EPI_ISL_980991, EPI_ISL_980992, EPI_ISL_980993, EPI_ISL_980994, EPI_ISL_980995, EPI_ISL_980996, EPI_ISL_980997, EPI_ISL_980998, EPI_ISL_980999, EPI_ISL_981000, EPI_ISL_981001, EPI_ISL_981002, EPI_ISL_981003 | | | |
| see above | Innovative Genomics Institute, UC Berkeley | Innovative Genomics Institute, UC Berkeley | Stacia Wyman, Haridha Shivram, Phil Frankino, Liana Lareau |
| EPI_ISL_981060, EPI_ISL_981073, EPI_ISL_981074, EPI_ISL_981075, EPI_ISL_981078, EPI_ISL_981079, EPI_ISL_981080, EPI_ISL_981082, EPI_ISL_981085, EPI_ISL_981087, EPI_ISL_981089, EPI_ISL_981090, EPI_ISL_981091, EPI_ISL_981092, EPI_ISL_981096, EPI_ISL_981098, EPI_ISL_981099, EPI_ISL_981103, EPI_ISL_981104, EPI_ISL_981105, EPI_ISL_981106, EPI_ISL_981107, EPI_ISL_981110, EPI_ISL_981111, EPI_ISL_981112, EPI_ISL_981113, EPI_ISL_981119, EPI_ISL_981123, EPI_ISL_981124, EPI_ISL_981125, EPI_ISL_981126, EPI_ISL_981127, EPI_ISL_981147, EPI_ISL_981148, EPI_ISL_981149, EPI_ISL_981150, EPI_ISL_981151, EPI_ISL_981152, EPI_ISL_981153, EPI_ISL_981154, EPI_ISL_981155, EPI_ISL_981156, EPI_ISL_981157, EPI_ISL_981174, EPI_ISL_981179, EPI_ISL_981180, EPI_ISL_981181, EPI_ISL_981182, EPI_ISL_981183, EPI_ISL_981184, EPI_ISL_981185, EPI_ISL_981186, EPI_ISL_981187, EPI_ISL_981188, EPI_ISL_981189, EPI_ISL_981190 | | | |
| see above | Johns Hopkins Hospital Department of Pathology | Johns Hopkins Hospital Department of Pathology | C. Paul Morris, Chun Huai Luo, Adannaya Amadi, Matthew Schwartz, Nicholas Gallagher, Heba H. Mostafa |
| EPI_ISL_981317, EPI_ISL_981318, EPI_ISL_981322, EPI_ISL_981323, EPI_ISL_981345, EPI_ISL_981346, EPI_ISL_981347, EPI_ISL_981348, EPI_ISL_981349, EPI_ISL_981351, EPI_ISL_981353 | | | |
| see above | Hospital Universitari Vall d'Hebron - Vall d'Hebron Institut de Rercerca | Hospital Universitari Vall d'Hebron - Vall d'Hebron Institut de Rercerca | Cristina Andrés, Maria Piñana, Josep F Abril, Damir Garcia-Cehic, Ariadna Rando, Juliana Esperalba, Maria Gema Codina, Carla Castillo, Maria Carmen Martin, Tomàs Pumarola, Josep Quer, Andrés Antón |
| EPI_ISL_981375, EPI_ISL_981377 | Laboratory for Respiratory Viruses, Cantacuzino National Military-Medical Institute for Research and Development | Cantacuzino Institute Virology | Luiza Ustea, Nicoleta Paraschiv, Catalina Pascu, Mihaela Lazar |
| EPI_ISL_981378, EPI_ISL_981379 | AZ Klina | AZ Klina | Dr. C. Vael |
| EPI_ISL_981390, EPI_ISL_981391, EPI_ISL_981392, EPI_ISL_981393, EPI_ISL_981394, EPI_ISL_981396, EPI_ISL_981397, EPI_ISL_981398, EPI_ISL_981399, EPI_ISL_981400 | CH.INTERCOMMUNAL DE CRETEIL | Department of Virology, Henri Mondor University Hospital, Assistance Publique Hôpitaux de Paris, Université Paris-Est Créteil, INSERM U955 | Christophe Rodriguez, Slim Fourati, Vanessa Demontant, Guillaume Gricourt, Melissa N'Debi, Alexandre Soulier, Elisabeth Trawinski, Jean-Michel Pawlotsky |
| EPI_ISL_981401, EPI_ISL_981402, EPI_ISL_981403, EPI_ISL_981404, EPI_ISL_981405 | Groupe LCD | Department of Virology, Henri Mondor University Hospital, Assistance Publique Hôpitaux de Paris, Université Paris-Est Créteil, INSERM U955 | Christophe Rodriguez, Slim Fourati, Vanessa Demontant, Guillaume Gricourt, Melissa N'Debi, Alexandre Soulier, Elisabeth Trawinski, Jean-Michel Pawlotsky |
| EPI_ISL_981406, EPI_ISL_981407, EPI_ISL_981408 | CH.INTERCOMMUNAL DE CRETEIL | Department of Virology, Henri Mondor University Hospital, Assistance Publique Hôpitaux de Paris, Université Paris-Est Créteil, INSERM U955 | Christophe Rodriguez, Slim Fourati, Vanessa Demontant, Guillaume Gricourt, Melissa N'Debi, Alexandre Soulier, Elisabeth Trawinski, Jean-Michel Pawlotsky |
| EPI_ISL_981409, EPI_ISL_981410, EPI_ISL_981411, EPI_ISL_981412, EPI_ISL_981413, EPI_ISL_981414, EPI_ISL_981415, EPI_ISL_981416, EPI_ISL_981417, EPI_ISL_981418, EPI_ISL_981419, EPI_ISL_981420, EPI_ISL_981421, EPI_ISL_981422, EPI_ISL_981423, EPI_ISL_981424, EPI_ISL_981425, EPI_ISL_981426, EPI_ISL_981427, EPI_ISL_981428, EPI_ISL_981429, EPI_ISL_981430, EPI_ISL_981431, EPI_ISL_981432, EPI_ISL_981433, EPI_ISL_981434, EPI_ISL_981435, EPI_ISL_981436, EPI_ISL_981437, EPI_ISL_981438, EPI_ISL_981439, EPI_ISL_981440, EPI_ISL_981441, EPI_ISL_981442, EPI_ISL_981443, EPI_ISL_981444, EPI_ISL_981445, EPI_ISL_981446, EPI_ISL_981447, EPI_ISL_981448, EPI_ISL_981449, EPI_ISL_981450, EPI_ISL_981451 | | | |
| see above | Groupe LCD | Department of Virology, Henri Mondor University Hospital, Assistance Publique Hôpitaux de Paris, Université Paris-Est Créteil, INSERM U955 | Christophe Rodriguez, Slim Fourati, Vanessa Demontant, Guillaume Gricourt, Melissa N'Debi, Alexandre Soulier, Elisabeth Trawinski, Jean-Michel Pawlotsky |
| EPI_ISL_981452, EPI_ISL_981453 | CH.INTERCOMMUNAL DE CRETEIL | Department of Virology, Henri Mondor University Hospital, Assistance Publique Hôpitaux de Paris, Université Paris-Est Créteil, INSERM U955 | Christophe Rodriguez, Slim Fourati, Vanessa Demontant, Guillaume Gricourt, Melissa N'Debi, Alexandre Soulier, Elisabeth Trawinski, Jean-Michel Pawlotsky |
| EPI_ISL_981454, EPI_ISL_981455, EPI_ISL_981456, EPI_ISL_981457, EPI_ISL_981458, EPI_ISL_981459, EPI_ISL_981460, EPI_ISL_981461, EPI_ISL_981482, EPI_ISL_981483, EPI_ISL_981484 | | | |
| see above | CHI VILLENEUVE ST GEORGES | Department of Virology, Henri Mondor University Hospital, Assistance Publique Hôpitaux de Paris, Université Paris-Est | Christophe Rodriguez, Slim Fourati, Vanessa Demontant, Guillaume Gricourt, Melissa N'Debi, Alexandre Soulier, Elisabeth Trawinski, Jean-Michel Pawlotsky |

| | | | |
|---|---|---|---|
| | | Créteil, INSERM U955 | |
| EPI_ISL_981485 | INSTITUT GUSTAVE ROUSSY | Department of Virology, Henri Mondor University Hospital, Assistance Publique Hôpitaux de Paris, Université Paris-Est Créteil, INSERM U955 | Christophe Rodriguez, Slim Fourati, Vanessa Demontant, Guillaume Gricourt, Melissa N'Debi, Alexandre Soulier, Elisabeth Trawinski, Jean-Michel Pawlotsky |
| EPI_ISL_981487 | CH.INTERCOMMUNAL DE CRETEIL | Department of Virology, Henri Mondor University Hospital, Assistance Publique Hôpitaux de Paris, Université Paris-Est Créteil, INSERM U955 | Christophe Rodriguez, Slim Fourati, Vanessa Demontant, Guillaume Gricourt, Melissa N'Debi, Alexandre Soulier, Elisabeth Trawinski, Jean-Michel Pawlotsky |
| EPI_ISL_981489, EPI_ISL_981490, EPI_ISL_981491 | Groupe LCD | Department of Virology, Henri Mondor University Hospital, Assistance Publique Hôpitaux de Paris, Université Paris-Est Créteil, INSERM U955 | Christophe Rodriguez, Slim Fourati, Vanessa Demontant, Guillaume Gricourt, Melissa N'Debi, Alexandre Soulier, Elisabeth Trawinski, Jean-Michel Pawlotsky |
| EPI_ISL_981492 | LABM BIOLAM CLINIQUE ESTREE | Department of Virology, Henri Mondor University Hospital, Assistance Publique Hôpitaux de Paris, Université Paris-Est Créteil, INSERM U955 | Christophe Rodriguez, Slim Fourati, Vanessa Demontant, Guillaume Gricourt, Melissa N'Debi, Alexandre Soulier, Elisabeth Trawinski, Jean-Michel Pawlotsky |
| EPI_ISL_981493, EPI_ISL_981494, EPI_ISL_981495, EPI_ISL_981496 | Groupe LCD | Department of Virology, Henri Mondor University Hospital, Assistance Publique Hôpitaux de Paris, Université Paris-Est Créteil, INSERM U955 | Christophe Rodriguez, Slim Fourati, Vanessa Demontant, Guillaume Gricourt, Melissa N'Debi, Alexandre Soulier, Elisabeth Trawinski, Jean-Michel Pawlotsky |
| EPI_ISL_981497 | LABM BIOLAM CLINIQUE ESTREE | Department of Virology, Henri Mondor University Hospital, Assistance Publique Hôpitaux de Paris, Université Paris-Est Créteil, INSERM U955 | Christophe Rodriguez, Slim Fourati, Vanessa Demontant, Guillaume Gricourt, Melissa N'Debi, Alexandre Soulier, Elisabeth Trawinski, Jean-Michel Pawlotsky |
| EPI_ISL_981498, EPI_ISL_981499, EPI_ISL_981500, EPI_ISL_981501, EPI_ISL_981502, EPI_ISL_981503, EPI_ISL_981504, EPI_ISL_981505, EPI_ISL_981506, EPI_ISL_981507, EPI_ISL_981508, EPI_ISL_981509, EPI_ISL_981510, EPI_ISL_981511, EPI_ISL_981512, EPI_ISL_981513, EPI_ISL_981514, EPI_ISL_981515, EPI_ISL_981516, EPI_ISL_981517, EPI_ISL_981518, EPI_ISL_981519, EPI_ISL_981520, EPI_ISL_981521, EPI_ISL_981522, EPI_ISL_981523, EPI_ISL_981524, EPI_ISL_981525, EPI_ISL_981526, EPI_ISL_981527, EPI_ISL_981528, EPI_ISL_981529, EPI_ISL_981530, EPI_ISL_981531 | | | |
| see above | Groupe LCD | Department of Virology, Henri Mondor University Hospital, Assistance Publique Hôpitaux de Paris, Université Paris-Est Créteil, INSERM U955 | Christophe Rodriguez, Slim Fourati, Vanessa Demontant, Guillaume Gricourt, Melissa N'Debi, Alexandre Soulier, Elisabeth Trawinski, Jean-Michel Pawlotsky |
| EPI_ISL_981532 | Hôpital Pitié-Salpêtrière | Department of Virology, Henri Mondor University Hospital, Assistance Publique Hôpitaux de Paris, Université Paris-Est Créteil, INSERM U955 | Christophe Rodriguez, Slim Fourati, Vanessa Demontant, Guillaume Gricourt, Melissa N'Debi, Alexandre Soulier, Elisabeth Trawinski, Jean-Michel Pawlotsky |
| EPI_ISL_981533, EPI_ISL_981534 | Hôpital Avicenne | Department of Virology, Henri Mondor University Hospital, Assistance Publique Hôpitaux de Paris, Université Paris-Est Créteil, INSERM U955 | Christophe Rodriguez, Slim Fourati, Vanessa Demontant, Guillaume Gricourt, Melissa N'Debi, Alexandre Soulier, Elisabeth Trawinski, Jean-Michel Pawlotsky |
| EPI_ISL_981535, EPI_ISL_981536, EPI_ISL_981537 | Hôpital Pitié-Salpêtrière | Department of Virology, Henri Mondor University Hospital, Assistance Publique Hôpitaux de Paris, Université Paris-Est Créteil, INSERM U955 | Christophe Rodriguez, Slim Fourati, Vanessa Demontant, Guillaume Gricourt, Melissa N'Debi, Alexandre Soulier, Elisabeth Trawinski, Jean-Michel Pawlotsky |
| EPI_ISL_981549, EPI_ISL_981550, EPI_ISL_981552, EPI_ISL_981553, EPI_ISL_981554, EPI_ISL_981555, EPI_ISL_981556, EPI_ISL_981558, EPI_ISL_981559, EPI_ISL_981560, EPI_ISL_981561, EPI_ISL_981562, EPI_ISL_981563, EPI_ISL_981564, EPI_ISL_981565, EPI_ISL_981566, EPI_ISL_981567, EPI_ISL_981568, EPI_ISL_981569, EPI_ISL_981570, EPI_ISL_981571, EPI_ISL_981572, EPI_ISL_981573, EPI_ISL_981574, EPI_ISL_981575, EPI_ISL_981576, EPI_ISL_981577, EPI_ISL_981578, EPI_ISL_981579, EPI_ISL_981580, EPI_ISL_981581, EPI_ISL_981582, EPI_ISL_981583, EPI_ISL_981584, EPI_ISL_981585, EPI_ISL_981586, EPI_ISL_981587, EPI_ISL_981588, EPI_ISL_981589, EPI_ISL_981590, EPI_ISL_981591, EPI_ISL_981592, EPI_ISL_981593, EPI_ISL_981594, EPI_ISL_981595, EPI_ISL_981596, EPI_ISL_981597, EPI_ISL_981598, EPI_ISL_981599, EPI_ISL_981600, EPI_ISL_981601, EPI_ISL_981602, EPI_ISL_981603, EPI_ISL_981604, EPI_ISL_981605, EPI_ISL_981606, EPI_ISL_981607, EPI_ISL_981608, EPI_ISL_981609, EPI_ISL_981610, EPI_ISL_981642, EPI_ISL_981643, EPI_ISL_981651, EPI_ISL_981652, EPI_ISL_981653, EPI_ISL_981655, EPI_ISL_981656, EPI_ISL_981657, EPI_ISL_981658, EPI_ISL_981660, EPI_ISL_981661, EPI_ISL_981662, EPI_ISL_981663, EPI_ISL_981664, EPI_ISL_981665, EPI_ISL_981666, EPI_ISL_981667, EPI_ISL_981668, EPI_ISL_981669, EPI_ISL_981670, EPI_ISL_981671, EPI_ISL_981672, EPI_ISL_981673, EPI_ISL_981674, EPI_ISL_981675, EPI_ISL_981676, EPI_ISL_981677, EPI_ISL_981678, EPI_ISL_981679, EPI_ISL_981680, EPI_ISL_981681, EPI_ISL_981682, EPI_ISL_981683, EPI_ISL_981684, EPI_ISL_981685, EPI_ISL_981686, EPI_ISL_981687, EPI_ISL_981688, EPI_ISL_981689, EPI_ISL_981696, EPI_ISL_981697, EPI_ISL_981698, EPI_ISL_981699, EPI_ISL_981700, EPI_ISL_981701, EPI_ISL_981702, EPI_ISL_981703, EPI_ISL_981704, EPI_ISL_981706, EPI_ISL_981707, EPI_ISL_981708, EPI_ISL_981709, EPI_ISL_981713, EPI_ISL_981714, EPI_ISL_981715, EPI_ISL_981716, EPI_ISL_981717, EPI_ISL_981719, EPI_ISL_981721, EPI_ISL_981722, EPI_ISL_981723, EPI_ISL_981724, EPI_ISL_981725, EPI_ISL_981726, EPI_ISL_981727, EPI_ISL_981728, EPI_ISL_981729, EPI_ISL_981730, EPI_ISL_981731, EPI_ISL_981732, EPI_ISL_981733, EPI_ISL_981734, EPI_ISL_981735, EPI_ISL_981736, EPI_ISL_981737, EPI_ISL_981738, EPI_ISL_981739, EPI_ISL_981740, EPI_ISL_981741, EPI_ISL_981742, EPI_ISL_981743, EPI_ISL_981744, EPI_ISL_981745, EPI_ISL_981747, EPI_ISL_981748, EPI_ISL_981749, EPI_ISL_981750, EPI_ISL_981751, EPI_ISL_981752, EPI_ISL_981753, EPI_ISL_981754, EPI_ISL_981755, EPI_ISL_981756, EPI_ISL_981757, EPI_ISL_981758, EPI_ISL_981759, EPI_ISL_981760, EPI_ISL_981761, EPI_ISL_981762, EPI_ISL_981763, EPI_ISL_981764, EPI_ISL_981765, EPI_ISL_981766, EPI_ISL_981767, EPI_ISL_981768, EPI_ISL_981769, EPI_ISL_981770, EPI_ISL_981771, EPI_ISL_981772, EPI_ISL_981773, EPI_ISL_981774, EPI_ISL_981775, EPI_ISL_981776, EPI_ISL_981777, EPI_ISL_981778, EPI_ISL_981779, EPI_ISL_981780, EPI_ISL_981781, EPI_ISL_981782, EPI_ISL_981783, EPI_ISL_981784, EPI_ISL_981785, EPI_ISL_981786, EPI_ISL_981787, EPI_ISL_981788, EPI_ISL_981789, EPI_ISL_981790, EPI_ISL_981791, EPI_ISL_981792, EPI_ISL_981793, EPI_ISL_981794, EPI_ISL_981795, EPI_ISL_981796, EPI_ISL_981797, EPI_ISL_981798, EPI_ISL_981799, EPI_ISL_981800, EPI_ISL_981801, EPI_ISL_981802, EPI_ISL_981803, EPI_ISL_981804, EPI_ISL_981805, EPI_ISL_981806, EPI_ISL_981807, EPI_ISL_981827, EPI_ISL_981828, EPI_ISL_981829, EPI_ISL_981830, EPI_ISL_981831, EPI_ISL_981832, EPI_ISL_981833, EPI_ISL_981834, EPI_ISL_981835, EPI_ISL_981837, EPI_ISL_981838, EPI_ISL_981839, EPI_ISL_981840, EPI_ISL_981841, EPI_ISL_981842, EPI_ISL_981843, EPI_ISL_981844, EPI_ISL_981845, EPI_ISL_981846, EPI_ISL_981847, EPI_ISL_981848, EPI_ISL_981849 | | | |
| see above | University Hospitals of Geneva, Laboratory of Virology | HUG, Laboratory of Virology and the Health2030 Genome Center | Samuel Cordey, Ana Rita Goncalves, Laurent Kaiser, Lorenzo Cerutti, Henri Pegeot, Melyssa Elies, Deborah Penet, Keith Harshman, Ioannis Xenarios, Emmanouil Dermitzakis |
| EPI_ISL_982002, EPI_ISL_982019, EPI_ISL_982022, EPI_ISL_982029, EPI_ISL_982038, EPI_ISL_982056, EPI_ISL_982065, EPI_ISL_982106, EPI_ISL_982113 | TGen North | TGen North | "Jolene Bowers, Megan Folkerts, Chris French, Hayley Yaglom, Ashlyn Pfeiffer, Darrin Lemmer, Dave Engelthaler, The Arizona COVID Genomics Union (ACGU)" |
| EPI_ISL_982118 | CH.INTERCOMMUNAL DE CRETEIL | Department of Virology, Henri Mondor University Hospital, Assistance Publique Hôpitaux de Paris, Université Paris-Est Créteil, INSERM U955 | Christophe Rodriguez, Slim Fourati, Vanessa Demontant, Guillaume Gricourt, Melissa N'Debi, Alexandre Soulier, Elisabeth Trawinski, Jean-Michel Pawlotsky |
| EPI_ISL_982120, EPI_ISL_982121, EPI_ISL_982125, EPI_ISL_982127 | Hôpital Henri Mondor | Department of Virology, Henri Mondor University Hospital, Assistance Publique Hôpitaux de Paris, Université Paris-Est Créteil, INSERM U955 | Christophe Rodriguez, Slim Fourati, Vanessa Demontant, Guillaume Gricourt, Melissa N'Debi, Alexandre Soulier, Elisabeth Trawinski, Jean-Michel Pawlotsky |
| EPI_ISL_982128 | CHU SUD AMIENS | Department of Virology, Henri Mondor University Hospital, Assistance Publique Hôpitaux de Paris, Université Paris-Est Créteil, INSERM U955 | Christophe Rodriguez, Slim Fourati, Vanessa Demontant, Guillaume Gricourt, Melissa N'Debi, Alexandre Soulier, Elisabeth Trawinski, Jean-Michel Pawlotsky |
| EPI_ISL_982130, EPI_ISL_982131, EPI_ISL_982132, EPI_ISL_982134, EPI_ISL_982135, EPI_ISL_982136, EPI_ISL_982137, EPI_ISL_982138, EPI_ISL_982139, EPI_ISL_982140, EPI_ISL_982141 | | | |
| see above | Hôpital Henri Mondor | Department of Virology, Henri Mondor University Hospital, Assistance Publique Hôpitaux de Paris, Université Paris-Est Créteil, INSERM U955 | Christophe Rodriguez, Slim Fourati, Vanessa Demontant, Guillaume Gricourt, Melissa N'Debi, Alexandre Soulier, Elisabeth Trawinski, Jean-Michel Pawlotsky |
| EPI_ISL_982142, EPI_ISL_982143, EPI_ISL_982144 | CHU SUD AMIENS | Department of Virology, Henri Mondor University Hospital, Assistance Publique Hôpitaux de Paris, Université Paris-Est Créteil, INSERM U955 | Christophe Rodriguez, Slim Fourati, Vanessa Demontant, Guillaume Gricourt, Melissa N'Debi, Alexandre Soulier, Elisabeth Trawinski, Jean-Michel Pawlotsky |
| EPI_ISL_982145 | Groupe LCD | Department of Virology, Henri Mondor University Hospital, Assistance Publique Hôpitaux de Paris, Université Paris-Est Créteil, INSERM U955 | Christophe Rodriguez, Slim Fourati, Vanessa Demontant, Guillaume Gricourt, Melissa N'Debi, Alexandre Soulier, Elisabeth Trawinski, Jean-Michel Pawlotsky |
| EPI_ISL_982146 | CHU SUD AMIENS | Department of Virology, Henri Mondor University Hospital, Assistance Publique Hôpitaux de Paris, Université Paris-Est Créteil, INSERM U955 | Christophe Rodriguez, Slim Fourati, Vanessa Demontant, Guillaume Gricourt, Melissa N'Debi, Alexandre Soulier, Elisabeth Trawinski, Jean-Michel Pawlotsky |
| EPI_ISL_982147 | CH.INTERCOMMUNAL DE CRETEIL | Department of Virology, Henri Mondor University Hospital, | Christophe Rodriguez, Slim Fourati, Vanessa Demontant, Guillaume Gricourt, Melissa N'Debi, Alexandre Soulier, Elisabeth Trawinski, Jean-Michel |

| | | | |
|---|---|---|---|
| | | Assistance Publique Hôpitaux de Paris, Université Paris-Est Créteil, INSERM U955 | Pawlotsky |
| EPI_ISL_982148, EPI_ISL_982149 | Hôpital Henri Mondor | Department of Virology, Henri Mondor University Hospital, Assistance Publique Hôpitaux de Paris, Université Paris-Est Créteil, INSERM U955 | Christophe Rodriguez, Slim Fourati, Vanessa Demontant, Guillaume Gricourt, Melissa N'Debi, Alexandre Soulier, Elisabeth Trawinski, Jean-Michel Pawlotsky |
| EPI_ISL_982154 | CHU SUD AMIENS | Department of Virology, Henri Mondor University Hospital, Assistance Publique Hôpitaux de Paris, Université Paris-Est Créteil, INSERM U955 | Christophe Rodriguez, Slim Fourati, Vanessa Demontant, Guillaume Gricourt, Melissa N'Debi, Alexandre Soulier, Elisabeth Trawinski, Jean-Michel Pawlotsky |
| EPI_ISL_982155 | Hôpital Henri Mondor | Department of Virology, Henri Mondor University Hospital, Assistance Publique Hôpitaux de Paris, Université Paris-Est Créteil, INSERM U955 | Christophe Rodriguez, Slim Fourati, Vanessa Demontant, Guillaume Gricourt, Melissa N'Debi, Alexandre Soulier, Elisabeth Trawinski, Jean-Michel Pawlotsky |
| EPI_ISL_982158 | CHI VILLENEUVE ST GEORGES | Department of Virology, Henri Mondor University Hospital, Assistance Publique Hôpitaux de Paris, Université Paris-Est Créteil, INSERM U955 | Christophe Rodriguez, Slim Fourati, Vanessa Demontant, Guillaume Gricourt, Melissa N'Debi, Alexandre Soulier, Elisabeth Trawinski, Jean-Michel Pawlotsky |
| EPI_ISL_982159 | Hôpital Necker-Enfants malades | Department of Virology, Henri Mondor University Hospital, Assistance Publique Hôpitaux de Paris, Université Paris-Est Créteil, INSERM U955 | Christophe Rodriguez, Slim Fourati, Vanessa Demontant, Guillaume Gricourt, Melissa N'Debi, Alexandre Soulier, Elisabeth Trawinski, Jean-Michel Pawlotsky |
| EPI_ISL_982164 | Hôpital Henri Mondor | Department of Virology, Henri Mondor University Hospital, Assistance Publique Hôpitaux de Paris, Université Paris-Est Créteil, INSERM U955 | Christophe Rodriguez, Slim Fourati, Vanessa Demontant, Guillaume Gricourt, Melissa N'Debi, Alexandre Soulier, Elisabeth Trawinski, Jean-Michel Pawlotsky |
| EPI_ISL_982165 | CHU SUD AMIENS | Department of Virology, Henri Mondor University Hospital, Assistance Publique Hôpitaux de Paris, Université Paris-Est Créteil, INSERM U955 | Christophe Rodriguez, Slim Fourati, Vanessa Demontant, Guillaume Gricourt, Melissa N'Debi, Alexandre Soulier, Elisabeth Trawinski, Jean-Michel Pawlotsky |
| EPI_ISL_982166, EPI_ISL_982169, EPI_ISL_982170, EPI_ISL_982171 | Hôpital Henri Mondor | Department of Virology, Henri Mondor University Hospital, Assistance Publique Hôpitaux de Paris, Université Paris-Est Créteil, INSERM U955 | Christophe Rodriguez, Slim Fourati, Vanessa Demontant, Guillaume Gricourt, Melissa N'Debi, Alexandre Soulier, Elisabeth Trawinski, Jean-Michel Pawlotsky |
| EPI_ISL_982174, EPI_ISL_982175, EPI_ISL_982176 | CHU SUD AMIENS | Department of Virology, Henri Mondor University Hospital, Assistance Publique Hôpitaux de Paris, Université Paris-Est Créteil, INSERM U955 | Christophe Rodriguez, Slim Fourati, Vanessa Demontant, Guillaume Gricourt, Melissa N'Debi, Alexandre Soulier, Elisabeth Trawinski, Jean-Michel Pawlotsky |
| EPI_ISL_982177, EPI_ISL_982178 | Hôpital Henri Mondor | Department of Virology, Henri Mondor University Hospital, Assistance Publique Hôpitaux de Paris, Université Paris-Est Créteil, INSERM U955 | Christophe Rodriguez, Slim Fourati, Vanessa Demontant, Guillaume Gricourt, Melissa N'Debi, Alexandre Soulier, Elisabeth Trawinski, Jean-Michel Pawlotsky |
| EPI_ISL_982179 | CHI VILLENEUVE ST GEORGES | Department of Virology, Henri Mondor University Hospital, Assistance Publique Hôpitaux de Paris, Université Paris-Est Créteil, INSERM U955 | Christophe Rodriguez, Slim Fourati, Vanessa Demontant, Guillaume Gricourt, Melissa N'Debi, Alexandre Soulier, Elisabeth Trawinski, Jean-Michel Pawlotsky |
| EPI_ISL_982180 | Hôpital Henri Mondor | Department of Virology, Henri Mondor University Hospital, Assistance Publique Hôpitaux de Paris, Université Paris-Est Créteil, INSERM U955 | Christophe Rodriguez, Slim Fourati, Vanessa Demontant, Guillaume Gricourt, Melissa N'Debi, Alexandre Soulier, Elisabeth Trawinski, Jean-Michel Pawlotsky |
| EPI_ISL_982181 | CH.INTERCOMMUNAL DE CRETEIL | Department of Virology, Henri Mondor University Hospital, Assistance Publique Hôpitaux de Paris, Université Paris-Est Créteil, INSERM U955 | Christophe Rodriguez, Slim Fourati, Vanessa Demontant, Guillaume Gricourt, Melissa N'Debi, Alexandre Soulier, Elisabeth Trawinski, Jean-Michel Pawlotsky |
| EPI_ISL_982203, EPI_ISL_982204, EPI_ISL_982205, EPI_ISL_982206, EPI_ISL_982207, EPI_ISL_982208, EPI_ISL_982209, EPI_ISL_982210, EPI_ISL_982211, EPI_ISL_982212, EPI_ISL_982213, EPI_ISL_982214, EPI_ISL_982215, EPI_ISL_982216, EPI_ISL_982217, EPI_ISL_982218, EPI_ISL_982219, EPI_ISL_982220, EPI_ISL_982221, EPI_ISL_982222, EPI_ISL_982223, EPI_ISL_982224, EPI_ISL_982225 | | | |
| see above | Hôpital Henri Mondor | Department of Virology, Henri Mondor University Hospital, Assistance Publique Hôpitaux de Paris, Université Paris-Est Créteil, INSERM U955 | Christophe Rodriguez, Slim Fourati, Vanessa Demontant, Guillaume Gricourt, Melissa N'Debi, Alexandre Soulier, Elisabeth Trawinski, Jean-Michel Pawlotsky |
| EPI_ISL_982226, EPI_ISL_982227 | CH.INTERCOMMUNAL DE CRETEIL | Department of Virology, Henri Mondor University Hospital, Assistance Publique Hôpitaux de Paris, Université Paris-Est Créteil, INSERM U955 | Christophe Rodriguez, Slim Fourati, Vanessa Demontant, Guillaume Gricourt, Melissa N'Debi, Alexandre Soulier, Elisabeth Trawinski, Jean-Michel Pawlotsky |
| EPI_ISL_982274, EPI_ISL_982281, EPI_ISL_982282 | Lab voor klinische biologie | Lab voor klinische biologie | Hannelore Hamerlinck, Marija Janevska, Bruno Verhasselt |
| EPI_ISL_982299 | BCCDC Public Health Laboratory | BCCDC Public Health Laboratory | Prystajecky Natalie, John Tyson, Dan Fornika, Shannon Russell, Kim Macdonald, Kimia Kamelian, Ana Pacagnella, Corrinne Ng, Loretta Janz, Robert Azana, Mel Krajden |
| EPI_ISL_982862, EPI_ISL_982863, EPI_ISL_982864, EPI_ISL_982865, EPI_ISL_982870, EPI_ISL_982872, EPI_ISL_982873, EPI_ISL_982874, EPI_ISL_982875, EPI_ISL_982876, EPI_ISL_982887, EPI_ISL_982888, EPI_ISL_982895, EPI_ISL_982897, EPI_ISL_982900, EPI_ISL_982901, EPI_ISL_982904, EPI_ISL_982905, EPI_ISL_982906, EPI_ISL_982908, EPI_ISL_982911, EPI_ISL_982912, EPI_ISL_982913, EPI_ISL_982914, EPI_ISL_982915, EPI_ISL_982916, EPI_ISL_982917, EPI_ISL_982918, EPI_ISL_982919, EPI_ISL_982920, EPI_ISL_982921 | | | |
| see above | MEPHI Aix Marseille University (AMU) | MEPHI Aix Marseille University (AMU) | Anthony LEVASSEUR |
| EPI_ISL_983096, EPI_ISL_983098 | Microbiology and Virology Unit, Florence Careggi University Hospital | Microbiology and Virology Unit, Florence Careggi University Hospital | Vincenzo Di Pilato, Marco Coppi, Fabio Morecchiato, Noemi Aiezza, Ilaria Baccani, Alberto Antonelli, Emanuele Gori, Gian Maria Rossolini |
| EPI_ISL_983326 | Laboratorio di Genetica Medica Ospedale Belcolle | INMI Lazzaro Spallanzani IRCCS | F Messina, C.E.M Gruber, B Bartolini, E Giombini, M Rueca, O Butera, F Natoni, G Pessina, A Di Caro, MR Capobianchi |
| EPI_ISL_983327 | Dipartimento di Prevenzione ASL Roma 4 | INMI Lazzaro Spallanzani IRCCS | O Butera, F Messina, CEM Gruber, B Bartolini, E Giombini, M Rueca, S Ursino, MR Capobianchi, A Di Caro |
| EPI_ISL_983374 | Utah Public Health Laboratory | Utah Public Health Laboratory | Erin L. Young, Kelly F. Oakeson, Tara Gallagher |
| EPI_ISL_983439, EPI_ISL_983443, EPI_ISL_983447, EPI_ISL_983448, EPI_ISL_983449, EPI_ISL_983450, EPI_ISL_983451, EPI_ISL_983452, EPI_ISL_983453, EPI_ISL_983454, EPI_ISL_983455, EPI_ISL_983456, EPI_ISL_983457, EPI_ISL_983458, EPI_ISL_983459, EPI_ISL_983460 | | | |
| see above | URMC LABS | Wadsworth Center, New York State Department of Health | Kirsten St. George, Daryl M. Lamson, Alexis Russel, Matthew Shudt, Melissa A Leisner, Jonathan Plitnick, Navjot Singh, John Kelly, Erasmus Schneider, Erica Lasek-Nesselquist |
| EPI_ISL_983588, EPI_ISL_983589, EPI_ISL_983590, EPI_ISL_983591, EPI_ISL_983592, EPI_ISL_983593, EPI_ISL_983594, EPI_ISL_983595, EPI_ISL_983596, EPI_ISL_983597, EPI_ISL_983598 | | | |
| see above | Texas Department of State Health Services | Texas Department of State Health Services | Bonnie Oh, Anita Pokharel, James Daniel Bonser, Myong Koag, Chung Wang, Rachel Lee, Grace Kubin, Rashmi Tuladhar, Mayela Pedrueza, Maliha Rahman, Jenny Zhang |
| EPI_ISL_983640, EPI_ISL_983641, EPI_ISL_983642, EPI_ISL_983643, EPI_ISL_983644, EPI_ISL_983645, EPI_ISL_983646, EPI_ISL_983647 | University of Michigan Clinical Microbiology Laboratory | Lauring Lab, University of Michigan, Department of Microbiology and Immunology | Valesano |
| EPI_ISL_983649, EPI_ISL_983650, EPI_ISL_983651, EPI_ISL_983653, EPI_ISL_983654, EPI_ISL_983655, EPI_ISL_983656, EPI_ISL_983657, EPI_ISL_983658, EPI_ISL_983659, EPI_ISL_983660, EPI_ISL_983661, EPI_ISL_983662, EPI_ISL_983663, EPI_ISL_983664, EPI_ISL_983665, EPI_ISL_983666, EPI_ISL_983667, EPI_ISL_983668, EPI_ISL_983669, EPI_ISL_983670, EPI_ISL_983671, EPI_ISL_983672, EPI_ISL_983673, EPI_ISL_983674, EPI_ISL_983675, EPI_ISL_983676, EPI_ISL_983677, EPI_ISL_983678, EPI_ISL_983679, EPI_ISL_983680, EPI_ISL_983681, EPI_ISL_983682, EPI_ISL_983683, EPI_ISL_983684, EPI_ISL_983685, | | | |

| | | | |
|---|---|---|---|
| EPI_ISL_983686, EPI_ISL_983687, EPI_ISL_983688, EPI_ISL_983689, EPI_ISL_983690, EPI_ISL_983691 | | | |
| see above | Vault Health | Minnesota Department of Health, Public Health Laboratory | Alexandra Lorentz, Jacob Garfin, Matt Plumb, and Xiong Wang |
| EPI_ISL_983708, EPI_ISL_983709, EPI_ISL_983712, EPI_ISL_983714, EPI_ISL_983715, EPI_ISL_983716, EPI_ISL_983721, EPI_ISL_983722, EPI_ISL_983723, EPI_ISL_983724, EPI_ISL_983725, EPI_ISL_983732, EPI_ISL_983738, EPI_ISL_983739, EPI_ISL_983740, EPI_ISL_983741, EPI_ISL_983742, EPI_ISL_983743, EPI_ISL_983744, EPI_ISL_983745, EPI_ISL_983746 | | | |
| see above | Colorado Department of Public Health and Environment | Colorado Department of Puplic Health and Environment | Laura Bankers, Molly C. Hetherington-Rauth, Diana Ir, Shannon Ely, Shannon R. Matzinger, Sarah Elizabeth Totten, Emily A. Travanty |
| EPI_ISL_983863, EPI_ISL_983864, EPI_ISL_983865, EPI_ISL_983866, EPI_ISL_983867, EPI_ISL_983868, EPI_ISL_983869 | LACEN-RS/CEVS/SES-RS | State Center for Health Surveillance. Rio Grande do Sul State Secretary of Health (CEVS) | Aline Campos, Cynthia Molina, Lara Crescente, Leticia Garay, Ludmila Fiorenzano Baethgen, Richard Salvato, Tatiana Gregianini |
