## Supplementary material for "Detection and characterization of the SARS-CoV-2 lineage B.1.526 in New York": Supp. Table 3 GISAID Acknowledgment part 2: gisaid_hcov-19_acknowledgement_table_2021_02_13_01-3.pdf

We gratefully acknowledge the following Authors from the Originating laboratories responsible for obtaining the specimens, as well as the Submitting laboratories where the genome data were generated and shared via GISAID, on which this research is based.

All Submitters of data may be contacted directly via www.gisaid.org

Authors are sorted alphabetically.

| Accession ID | Originating Laboratory | Submitting Laboratory | Authors |
|---|---|---|---|
| EPI_ISL_936486 | Medicine and Surgery, University of Insubria | Medicine and Surgery, University of Insubria | Novazzi,F., Genoni,A., Baj,A., Spezia,P.G., Focosi,D., Zago,C.,Colombo,A., Cassani,G., Pasciuta,R., Tamborini,A., Rossi,A.,Prestia,M., Capuano,R. and Maggi,F. |
| EPI_ISL_949900, EPI_ISL_949902 | University College London, Great Ormond Street Hospital for Children NHS Foundation Trust, Imperial College Healthcare NHS Trust | COVID-19 Genomics UK (COG-UK) Consortium | Sergi Castellano, Rachel Williams, Mark Kristiansen, Paola Resende Silva, Sunando Roy, Tony Brooks, Helena Tutill, Paola Niola, Patricia Dyal, Charlotte Williams, Leysa Forrest, Yasmin Panchbhaya, Jacqueline Findlay, Samuel Weeks, Julianne Brown, Kathryn Harris, Paul Randell, James Price, Alison Holmes, Judith Breuer |
| EPI_ISL_950723, EPI_ISL_950724, EPI_ISL_950725, EPI_ISL_950726, EPI_ISL_950727, EPI_ISL_950728 | Queens Medical Centre, Clinical Microbiology Department / DeepSeq Nottingham | COVID-19 Genomics UK (COG-UK) Consortium | Gemma Clark, Wendy Smith, Manjinder Khakh, Vicki M Fleming, Michelle M Lister, Hannah Howson-Wells, Jonathan Ball, Patrick McClure, Joseph Chappell, Theocharis Tsoleridis, Nadine Holmes, Matthew Carlisle, Christopher Moore, Fei Sang, Johnny Debebe, Victoria Wright, Matthew Loose |
| EPI_ISL_952956, EPI_ISL_952960, EPI_ISL_952961, EPI_ISL_952962, EPI_ISL_952963, EPI_ISL_952966, EPI_ISL_952967, EPI_ISL_952971, EPI_ISL_952973, EPI_ISL_952974, EPI_ISL_952981, EPI_ISL_952987, EPI_ISL_952989, EPI_ISL_952992, EPI_ISL_952993, EPI_ISL_952996, EPI_ISL_953001, EPI_ISL_953005, EPI_ISL_953007, EPI_ISL_953008, EPI_ISL_953011, EPI_ISL_953018, EPI_ISL_953019, EPI_ISL_953023 | | | |
| see above | Virology Department, Sheffield Teaching Hospitals NHS Foundation Trust/Department of Infection, Immunity and Cardiovascular Disease, The Medical School, University of Sheffield | COVID-19 Genomics UK (COG-UK) Consortium | Thushan de Silva, Matthew Parker, Nikki Smith, Adri Angyal, Rebecca Brown, Luke Green, Rachel Tucker, Paul Parsons, Danielle Groves, Katie Johnson, Laura Carrilero, Alex Keeley, Dave Partridge, Matthew Wyles, Benjamin Lindsey, Mehmet Yavuz, Mohammad Raza, Cariad Evans |
| EPI_ISL_953377, EPI_ISL_953378, EPI_ISL_953379, EPI_ISL_953380, EPI_ISL_953381, EPI_ISL_953382, EPI_ISL_953383, EPI_ISL_953384 | Jessa | Jessa | Jessa_cmdLab |
| EPI_ISL_953385, EPI_ISL_953386, EPI_ISL_953388, EPI_ISL_953390, EPI_ISL_953394 | National Public Health Laboratory, National Centre for Infectious Diseases | National Public Health Laboratory, National Centre for Infectious Diseases | Tze Minn Mak, Zhenyang Zhou, Lin Cui, Raymond Tzer Pin Lin |
| EPI_ISL_960176, EPI_ISL_960177, EPI_ISL_960178, EPI_ISL_960179, EPI_ISL_960180, EPI_ISL_960181, EPI_ISL_960182, EPI_ISL_960183, EPI_ISL_960184, EPI_ISL_960185, EPI_ISL_960186, EPI_ISL_960187 | | | |
| see above | Jessa | Jessa | Jessa_cmdLab |
| EPI_ISL_960835, EPI_ISL_960843, EPI_ISL_960846, EPI_ISL_960849, EPI_ISL_960851, EPI_ISL_960852, EPI_ISL_960884, EPI_ISL_960885, EPI_ISL_960887, EPI_ISL_960891, EPI_ISL_960892, EPI_ISL_960894, EPI_ISL_960895, EPI_ISL_960896, EPI_ISL_960897, EPI_ISL_960898 | | | |
| see above | Landesamt für Verbraucherschutz Sachsen Anhalt, Magdeburg | Institute of Medical Microbiology and Hospital Hygiene | Prof. Dr. Achim Kaasch, Aljoscha Tersteegen |
| EPI_ISL_960900, EPI_ISL_960901 | Institute of Medical Microbiology and Hospital Hygiene | Institute of Medical Microbiology and Hospital Hygiene | Prof. Dr. Achim Kaasch, Aljoscha Tersteegen |
| EPI_ISL_960935, EPI_ISL_960936, EPI_ISL_960937, EPI_ISL_960938, EPI_ISL_960939, EPI_ISL_960940 | Landesamt für Verbraucherschutz Sachsen Anhalt, Magdeburg | Institute of Medical Microbiology and Hospital Hygiene | Prof. Dr. Achim Kaasch, Aljoscha Tersteegen |
| EPI_ISL_961181, EPI_ISL_961182, EPI_ISL_961183, EPI_ISL_961184, EPI_ISL_961185, EPI_ISL_961186, EPI_ISL_961187, EPI_ISL_961188, EPI_ISL_961189, EPI_ISL_961190, EPI_ISL_961191, EPI_ISL_961192, EPI_ISL_961193, EPI_ISL_961194, EPI_ISL_961195, EPI_ISL_961196, EPI_ISL_961197, EPI_ISL_961198, EPI_ISL_961199, EPI_ISL_961200, EPI_ISL_961201, EPI_ISL_961202, EPI_ISL_961203 | | | |
| see above | AZDelta | AZDelta | Geert Martens; Dieter De Smet |
| EPI_ISL_961673 | SIESP CHIETI DRIVE IN LANCIANO | Istituto Zooprofilattico Sperimentale dell'Abruzzo e Molise "G. Caporale" | Lorusso A, Marcacci M, Di Domenico M, Ancora M, Curini V, Mangone I, Rinaldi A, Scialabba S, Di Pasquale A, Cammà C, Puglia I, Calistri P, Savini G |
| EPI_ISL_961674, EPI_ISL_961675, EPI_ISL_961676, EPI_ISL_961677 | SIESP CHIETI - DRIVE IN CHIETI | Istituto Zooprofilattico Sperimentale dell'Abruzzo e Molise "G. Caporale" | Lorusso A, Marcacci M, Di Domenico M, Ancora M, Curini V, Mangone I, Rinaldi A, Scialabba S, Di Pasquale A, Cammà C, Puglia I, Calistri P, Savini G |
| EPI_ISL_961678 | SIESP TERAMO | Istituto Zooprofilattico Sperimentale dell'Abruzzo e Molise "G. Caporale" | Lorusso A, Marcacci M, Di Domenico M, Ancora M, Curini V, Mangone I, Rinaldi A, Scialabba S, Di Pasquale A, Cammà C, Puglia I, Calistri P, Savini G |
| EPI_ISL_961679, EPI_ISL_961680, EPI_ISL_961681, EPI_ISL_961682 | SIESP CHIETI - DRIVE IN CHIETI | Istituto Zooprofilattico Sperimentale dell'Abruzzo e Molise "G. Caporale" | Lorusso A, Marcacci M, Di Domenico M, Ancora M, Curini V, Mangone I, Rinaldi A, Scialabba S, Di Pasquale A, Cammà C, Puglia I, Calistri P, Savini G |
| EPI_ISL_961683 | SIESP TERAMO | Istituto Zooprofilattico Sperimentale dell'Abruzzo e Molise "G. Caporale" | Lorusso A, Marcacci M, Di Domenico M, Ancora M, Curini V, Mangone I, Rinaldi A, Scialabba S, Di Pasquale A, Cammà C, Puglia I, Calistri P, Savini G |
| EPI_ISL_961684 | SIESP CHIETI - DRIVE IN CHIETI | Istituto Zooprofilattico Sperimentale dell'Abruzzo e Molise "G. Caporale" | Lorusso A, Marcacci M, Di Domenico M, Ancora M, Curini V, Mangone I, Rinaldi A, Scialabba S, Di Pasquale A, Cammà C, Puglia I, Calistri P, Savini G |
| EPI_ISL_961685 | SIESP DIPARTIMENTO DI PREVENZIONE CHIETI | Istituto Zooprofilattico Sperimentale dell'Abruzzo e Molise "G. Caporale" | Lorusso A, Marcacci M, Di Domenico M, Ancora M, Curini V, Mangone I, Rinaldi A, Scialabba S, Di Pasquale A, Cammà C, Puglia I, Calistri P, Savini G |
| EPI_ISL_961686, EPI_ISL_961687 | SIESP CHIETI - DRIVE IN ORTONA | Istituto Zooprofilattico Sperimentale dell'Abruzzo e Molise "G. Caporale" | Lorusso A, Marcacci M, Di Domenico M, Ancora M, Curini V, Mangone I, Rinaldi A, Scialabba S, Di Pasquale A, Cammà C, Puglia I, Calistri P, Savini G |
| EPI_ISL_961688 | SIESP TERAMO | Istituto Zooprofilattico Sperimentale dell'Abruzzo e Molise "G. Caporale" | Lorusso A, Marcacci M, Di Domenico M, Ancora M, Curini V, Mangone I, Rinaldi A, Scialabba S, Di Pasquale A, Cammà C, Puglia I, Calistri P, Savini G |
| EPI_ISL_961689, EPI_ISL_961690 | SIESP CHIETI - DRIVE IN CHIETI | Istituto Zooprofilattico Sperimentale dell'Abruzzo e Molise "G. Caporale" | Lorusso A, Marcacci M, Di Domenico M, Ancora M, Curini V, Mangone I, Rinaldi A, Scialabba S, Di Pasquale A, Cammà C, Puglia I, Calistri P, Savini G |
| EPI_ISL_961691, EPI_ISL_961692, EPI_ISL_961693 | SIESP CHIETI - DRIVE IN ORTONA | Istituto Zooprofilattico Sperimentale dell'Abruzzo e Molise "G. Caporale" | Lorusso A, Marcacci M, Di Domenico M, Ancora M, Curini V, Mangone I, Rinaldi A, Scialabba S, Di Pasquale A, Cammà C, Puglia I, Calistri P, Savini G |
| EPI_ISL_961694 | SIESP CHIETI - DRIVE IN CHIETI | Istituto Zooprofilattico Sperimentale dell'Abruzzo e Molise "G. Caporale" | Lorusso A, Marcacci M, Di Domenico M, Ancora M, Curini V, Mangone I, Rinaldi A, Scialabba S, Di Pasquale A, Cammà C, Puglia I, Calistri P, Savini G |
| EPI_ISL_961695 | SIESP DIPARTIMENTO DI PREVENZIONE CHIETI | Istituto Zooprofilattico Sperimentale dell'Abruzzo e Molise "G. Caporale" | Lorusso A, Marcacci M, Di Domenico M, Ancora M, Curini V, Mangone I, Rinaldi A, Scialabba S, Di Pasquale A, Cammà C, Puglia I, Calistri P, Savini G |
| EPI_ISL_961696 | SIESP SULMONA | Istituto Zooprofilattico Sperimentale dell'Abruzzo e Molise "G. Caporale" | Lorusso A, Marcacci M, Di Domenico M, Ancora M, Curini V, Mangone I, Rinaldi A, Scialabba S, Di Pasquale A, Cammà C, Puglia I, Calistri P, Savini G |

| | | | |
|---|---|---|---|
| EPI_ISL_961697, EPI_ISL_961698 | SIESP CHIETI - DRIVE IN CHIETI | Istituto Zooprofilattico Sperimentale dell'Abruzzo e Molise "G. Caporale" | Lorusso A, Marcacci M, Di Domenico M, Ancora M, Curini V, Mangone I, Rinaldi A, Scialabba S, Di Pasquale A, Cammà C, Puglia I, Calistri P, Savini G |
| EPI_ISL_961699 | SIESP CHIETI - DRIVE IN ORTONA | Istituto Zooprofilattico Sperimentale dell'Abruzzo e Molise "G. Caporale" | Lorusso A, Marcacci M, Di Domenico M, Ancora M, Curini V, Mangone I, Rinaldi A, Scialabba S, Di Pasquale A, Cammà C, Puglia I, Calistri P, Savini G |
| EPI_ISL_961700 | SIESP CHIETI - DRIVE IN CHIETI | Istituto Zooprofilattico Sperimentale dell'Abruzzo e Molise "G. Caporale" | Lorusso A, Marcacci M, Di Domenico M, Ancora M, Curini V, Mangone I, Rinaldi A, Scialabba S, Di Pasquale A, Cammà C, Puglia I, Calistri P, Savini G |
| EPI_ISL_961701 | Ospedale Civile Teramo | Istituto Zooprofilattico Sperimentale dell'Abruzzo e Molise "G. Caporale" | Lorusso A, Marcacci M, Di Domenico M, Ancora M, Curini V, Mangone I, Rinaldi A, Scialabba S, Di Pasquale A, Cammà C, Puglia I, Calistri P, Savini G |
| EPI_ISL_961702 | SIESP CHIETI - DRIVE IN CHIETI | Istituto Zooprofilattico Sperimentale dell'Abruzzo e Molise "G. Caporale" | Lorusso A, Marcacci M, Di Domenico M, Ancora M, Curini V, Mangone I, Rinaldi A, Scialabba S, Di Pasquale A, Cammà C, Puglia I, Calistri P, Savini G |
| EPI_ISL_961703 | SIESP DIPARTIMENTO DI PREVENZIONE CHIETI | Istituto Zooprofilattico Sperimentale dell'Abruzzo e Molise "G. Caporale" | Lorusso A, Marcacci M, Di Domenico M, Ancora M, Curini V, Mangone I, Rinaldi A, Scialabba S, Di Pasquale A, Cammà C, Puglia I, Calistri P, Savini G |
| EPI_ISL_961704 | SIESP SULMONA | Istituto Zooprofilattico Sperimentale dell'Abruzzo e Molise "G. Caporale" | Lorusso A, Marcacci M, Di Domenico M, Ancora M, Curini V, Mangone I, Rinaldi A, Scialabba S, Di Pasquale A, Cammà C, Puglia I, Calistri P, Savini G |
| EPI_ISL_961705 | Ospedale San Salvatore UOC Malattie Infettive | Istituto Zooprofilattico Sperimentale dell'Abruzzo e Molise "G. Caporale" | Lorusso A, Marcacci M, Di Domenico M, Ancora M, Curini V, Mangone I, Rinaldi A, Scialabba S, Di Pasquale A, Cammà C, Puglia I, Calistri P, Savini G |
| EPI_ISL_961706 | SIESP CHIETI - DRIVE IN CHIETI | Istituto Zooprofilattico Sperimentale dell'Abruzzo e Molise "G. Caporale" | Lorusso A, Marcacci M, Di Domenico M, Ancora M, Curini V, Mangone I, Rinaldi A, Scialabba S, Di Pasquale A, Cammà C, Puglia I, Calistri P, Savini G |
| EPI_ISL_961707, EPI_ISL_961708 | SIESP DIPARTIMENTO DI PREVENZIONE CHIETI | Istituto Zooprofilattico Sperimentale dell'Abruzzo e Molise "G. Caporale" | Lorusso A, Marcacci M, Di Domenico M, Ancora M, Curini V, Mangone I, Rinaldi A, Scialabba S, Di Pasquale A, Cammà C, Puglia I, Calistri P, Savini G |
| EPI_ISL_961709, EPI_ISL_961710, EPI_ISL_961711 | SIESP CHIETI - DRIVE IN CHIETI | Istituto Zooprofilattico Sperimentale dell'Abruzzo e Molise "G. Caporale" | Lorusso A, Marcacci M, Di Domenico M, Ancora M, Curini V, Mangone I, Rinaldi A, Scialabba S, Di Pasquale A, Cammà C, Puglia I, Calistri P, Savini G |
| EPI_ISL_961712, EPI_ISL_961713 | SIESP CHIETI DRIVE IN LANCIANO | Istituto Zooprofilattico Sperimentale dell'Abruzzo e Molise "G. Caporale" | Lorusso A, Marcacci M, Di Domenico M, Ancora M, Curini V, Mangone I, Rinaldi A, Scialabba S, Di Pasquale A, Cammà C, Puglia I, Calistri P, Savini G |
| EPI_ISL_961714 | SIESP CHIETI - DRIVE IN CHIETI | Istituto Zooprofilattico Sperimentale dell'Abruzzo e Molise "G. Caporale" | Lorusso A, Marcacci M, Di Domenico M, Ancora M, Curini V, Mangone I, Rinaldi A, Scialabba S, Di Pasquale A, Cammà C, Puglia I, Calistri P, Savini G |
| EPI_ISL_961715, EPI_ISL_961716 | SIESP CHIETI - DRIVE IN ORTONA | Istituto Zooprofilattico Sperimentale dell'Abruzzo e Molise "G. Caporale" | Lorusso A, Marcacci M, Di Domenico M, Ancora M, Curini V, Mangone I, Rinaldi A, Scialabba S, Di Pasquale A, Cammà C, Puglia I, Calistri P, Savini G |
| EPI_ISL_961717 | SIESP CHIETI - DRIVE IN CHIETI | Istituto Zooprofilattico Sperimentale dell'Abruzzo e Molise "G. Caporale" | Lorusso A, Marcacci M, Di Domenico M, Ancora M, Curini V, Mangone I, Rinaldi A, Scialabba S, Di Pasquale A, Cammà C, Puglia I, Calistri P, Savini G |
| EPI_ISL_961718, EPI_ISL_961719, EPI_ISL_961720, EPI_ISL_961721, EPI_ISL_961722 | SIESP CHIETI - DRIVE IN ORTONA | Istituto Zooprofilattico Sperimentale dell'Abruzzo e Molise "G. Caporale" | Lorusso A, Marcacci M, Di Domenico M, Ancora M, Curini V, Mangone I, Rinaldi A, Scialabba S, Di Pasquale A, Cammà C, Puglia I, Calistri P, Savini G |
| EPI_ISL_961723 | SIESP CHIETI DRIVE IN LANCIANO | Istituto Zooprofilattico Sperimentale dell'Abruzzo e Molise "G. Caporale" | Lorusso A, Marcacci M, Di Domenico M, Ancora M, Curini V, Mangone I, Rinaldi A, Scialabba S, Di Pasquale A, Cammà C, Puglia I, Calistri P, Savini G |
| EPI_ISL_961724 | SIESP CHIETI - DRIVE IN ORTONA | Istituto Zooprofilattico Sperimentale dell'Abruzzo e Molise "G. Caporale" | Lorusso A, Marcacci M, Di Domenico M, Ancora M, Curini V, Mangone I, Rinaldi A, Scialabba S, Di Pasquale A, Cammà C, Puglia I, Calistri P, Savini G |
| EPI_ISL_961725 | SIESP CHIETI DRIVE IN GISSI | Istituto Zooprofilattico Sperimentale dell'Abruzzo e Molise "G. Caporale" | Lorusso A, Marcacci M, Di Domenico M, Ancora M, Curini V, Mangone I, Rinaldi A, Scialabba S, Di Pasquale A, Cammà C, Puglia I, Calistri P, Savini G |
| EPI_ISL_961726, EPI_ISL_961727 | SIESP CHIETI - DRIVE IN CHIETI | Istituto Zooprofilattico Sperimentale dell'Abruzzo e Molise "G. Caporale" | Lorusso A, Marcacci M, Di Domenico M, Ancora M, Curini V, Mangone I, Rinaldi A, Scialabba S, Di Pasquale A, Cammà C, Puglia I, Calistri P, Savini G |
| EPI_ISL_961728 | SIESP DIPARTIMENTO DI PREVENZIONE CHIETI | Istituto Zooprofilattico Sperimentale dell'Abruzzo e Molise "G. Caporale" | Lorusso A, Marcacci M, Di Domenico M, Ancora M, Curini V, Mangone I, Rinaldi A, Scialabba S, Di Pasquale A, Cammà C, Puglia I, Calistri P, Savini G |
| EPI_ISL_961729 | SIESP CHIETI - DRIVE IN CHIETI | Istituto Zooprofilattico Sperimentale dell'Abruzzo e Molise "G. Caporale" | Lorusso A, Marcacci M, Di Domenico M, Ancora M, Curini V, Mangone I, Rinaldi A, Scialabba S, Di Pasquale A, Cammà C, Puglia I, Calistri P, Savini G |
| EPI_ISL_961730 | Dipartimento Prevenzione Avezzano-Servizio Igiene epidemiologia Sanità Pubblica | Istituto Zooprofilattico Sperimentale dell'Abruzzo e Molise "G. Caporale" | Lorusso A, Marcacci M, Di Domenico M, Ancora M, Curini V, Mangone I, Rinaldi A, Scialabba S, Di Pasquale A, Cammà C, Puglia I, Calistri P, Savini G |
| EPI_ISL_961731, EPI_ISL_961732 | SIESP CHIETI - DRIVE IN CHIETI | Istituto Zooprofilattico Sperimentale dell'Abruzzo e Molise "G. Caporale" | Lorusso A, Marcacci M, Di Domenico M, Ancora M, Curini V, Mangone I, Rinaldi A, Scialabba S, Di Pasquale A, Cammà C, Puglia I, Calistri P, Savini G |
| EPI_ISL_961733 | SIESP CHIETI - DRIVE IN ORTONA | Istituto Zooprofilattico Sperimentale dell'Abruzzo e Molise "G. Caporale" | Lorusso A, Marcacci M, Di Domenico M, Ancora M, Curini V, Mangone I, Rinaldi A, Scialabba S, Di Pasquale A, Cammà C, Puglia I, Calistri P, Savini G |
| EPI_ISL_961734 | Dipartimento Prevenzione Avezzano-Servizio Igiene epidemiologia Sanità Pubblica | Istituto Zooprofilattico Sperimentale dell'Abruzzo e Molise "G. Caporale" | Lorusso A, Marcacci M, Di Domenico M, Ancora M, Curini V, Mangone I, Rinaldi A, Scialabba S, Di Pasquale A, Cammà C, Puglia I, Calistri P, Savini G |
| EPI_ISL_961735, EPI_ISL_961736 | SIESP CHIETI - DRIVE IN CHIETI | Istituto Zooprofilattico Sperimentale dell'Abruzzo e Molise "G. Caporale" | Lorusso A, Marcacci M, Di Domenico M, Ancora M, Curini V, Mangone I, Rinaldi A, Scialabba S, Di Pasquale A, Cammà C, Puglia I, Calistri P, Savini G |
| EPI_ISL_961737 | SIESP CHIETI- DRIVE IN VASTO | Istituto Zooprofilattico Sperimentale dell'Abruzzo e Molise "G. Caporale" | Lorusso A, Marcacci M, Di Domenico M, Ancora M, Curini V, Mangone I, Rinaldi A, Scialabba S, Di Pasquale A, Cammà C, Puglia I, Calistri P, Savini G |
| EPI_ISL_961738, EPI_ISL_961739 | SIESP CHIETI DRIVE IN LANCIANO | Istituto Zooprofilattico Sperimentale dell'Abruzzo e Molise "G. Caporale" | Lorusso A, Marcacci M, Di Domenico M, Ancora M, Curini V, Mangone I, Rinaldi A, Scialabba S, Di Pasquale A, Cammà C, Puglia I, Calistri P, Savini G |
| EPI_ISL_961740 | SIESP CHIETI - DRIVE IN CHIETI | Istituto Zooprofilattico Sperimentale dell'Abruzzo e Molise "G. Caporale" | Lorusso A, Marcacci M, Di Domenico M, Ancora M, Curini V, Mangone I, Rinaldi A, Scialabba S, Di Pasquale A, Cammà C, Puglia I, Calistri P, Savini G |
| EPI_ISL_961741 | SIESP CHIETI- DRIVE IN VASTO | Istituto Zooprofilattico Sperimentale dell'Abruzzo e Molise "G. Caporale" | Lorusso A, Marcacci M, Di Domenico M, Ancora M, Curini V, Mangone I, Rinaldi A, Scialabba S, Di Pasquale A, Cammà C, Puglia I, Calistri P, Savini G |
| EPI_ISL_961742 | SIESP CHIETI - DRIVE IN CHIETI | Istituto Zooprofilattico Sperimentale dell'Abruzzo e Molise "G. Caporale" | Lorusso A, Marcacci M, Di Domenico M, Ancora M, Curini V, Mangone I, Rinaldi A, Scialabba S, Di Pasquale A, Cammà C, Puglia I, Calistri P, Savini G |
| EPI_ISL_961743 | SIESP CHIETI DRIVE IN GISSI | Istituto Zooprofilattico Sperimentale dell'Abruzzo e Molise "G. Caporale" | Lorusso A, Marcacci M, Di Domenico M, Ancora M, Curini V, Mangone I, Rinaldi A, Scialabba S, Di Pasquale A, Cammà C, Puglia I, Calistri P, Savini G |
| EPI_ISL_961744 | SIESP CHIETI - DRIVE IN ORTONA | Istituto Zooprofilattico Sperimentale dell'Abruzzo e Molise "G. Caporale" | Lorusso A, Marcacci M, Di Domenico M, Ancora M, Curini V, Mangone I, Rinaldi A, Scialabba S, Di Pasquale A, Cammà C, Puglia I, Calistri P, Savini G |
| EPI_ISL_961745 | SIESP CHIETI DRIVE IN GISSI | Istituto Zooprofilattico Sperimentale dell'Abruzzo e Molise "G. Caporale" | Lorusso A, Marcacci M, Di Domenico M, Ancora M, Curini V, Mangone I, Rinaldi A, Scialabba S, Di Pasquale A, Cammà C, Puglia I, Calistri P, Savini G |

| | | | |
|---|---|---|---|
| EPI_ISL_961746 | SIESP CHIETI - DRIVE IN CHIETI | Istituto Zooprofilattico Sperimentale dell'Abruzzo e Molise "G. Caporale" | Lorusso A, Marcacci M, Di Domenico M, Ancora M, Curini V, Mangone I, Rinaldi A, Scialabba S, Di Pasquale A, Cammà C, Puglia I, Calistri P, Savini G |
| EPI_ISL_961748 | SIESP CHIETI DRIVE IN LANCIANO | Istituto Zooprofilattico Sperimentale dell'Abruzzo e Molise "G. Caporale" | Lorusso A, Marcacci M, Di Domenico M, Ancora M, Curini V, Mangone I, Rinaldi A, Scialabba S, Di Pasquale A, Cammà C, Puglia I, Calistri P, Savini G |
| EPI_ISL_961749 | SIESP DIPARTIMENTO DI PREVENZIONE CHIETI | Istituto Zooprofilattico Sperimentale dell'Abruzzo e Molise "G. Caporale" | Lorusso A, Marcacci M, Di Domenico M, Ancora M, Curini V, Mangone I, Rinaldi A, Scialabba S, Di Pasquale A, Cammà C, Puglia I, Calistri P, Savini G |
| EPI_ISL_961750 | Ospedale Cvile Atri Med. Interna | Istituto Zooprofilattico Sperimentale dell'Abruzzo e Molise "G. Caporale" | Lorusso A, Marcacci M, Di Domenico M, Ancora M, Curini V, Mangone I, Rinaldi A, Scialabba S, Di Pasquale A, Cammà C, Puglia I, Calistri P, Savini G |
| EPI_ISL_961751 | SIESP DIPARTIMENTO DI PREVENZIONE CHIETI | Istituto Zooprofilattico Sperimentale dell'Abruzzo e Molise "G. Caporale" | Lorusso A, Marcacci M, Di Domenico M, Ancora M, Curini V, Mangone I, Rinaldi A, Scialabba S, Di Pasquale A, Cammà C, Puglia I, Calistri P, Savini G |
| EPI_ISL_962180 | unknown | Public Health Virology-Forensic and Scientific Services (PHV-FSS) | Son Nguyen et al. |
| EPI_ISL_965025 | S.C. Microbiologia e Virologia Laboratorio Virologia -Speciale Centro Influenza - AOU di Sassari - Viale san Pietro 43/B Palazzo Infettivologia | Laboratorio specialistico UOC Ematologia - Ospedale "San Francesco" - ATS-ASSL Nuoro | Piras Giovanna, Malune Paolo, Asproni Rosanna, Monne Maria Itria, Palmas Angelo Domenico Serra Caterina, Rimini Elena, Rubino Salvatore |
| EPI_ISL_965028, EPI_ISL_965114 | Laboratorio Biologia Molecolare Sars Cov2 - UOC Laboratorio Analisi - Servizio Medicina di Laboratorio, Ospedale "San Francesco" - ATS-ASSL Nuoro | Laboratorio specialistico UOC Ematologia - Ospedale "San Francesco" - ATS-ASSL Nuoro | Piras Giovanna, Asproni Rosanna, Malune Paolo, Fiamma Maura, Monne Maria Itria, Palmas Angelo Domenico, Lo Maglio Iana, Mameli Giuseppe |
| EPI_ISL_965223 | SIESP SULMONA | Istituto Zooprofilattico Sperimentale dell'Abruzzo e Molise "G. Caporale" | Lorusso A, Marcacci M, Di Domenico M, Ancora M, Curini V, Mangone I, Rinaldi A, Scialabba S, Di Pasquale A, Cammà C, Puglia I, Calistri P, Savini G |
| EPI_ISL_965225, EPI_ISL_965226 | USCA SULMONA | Istituto Zooprofilattico Sperimentale dell'Abruzzo e Molise "G. Caporale" | Lorusso A, Marcacci M, Di Domenico M, Ancora M, Curini V, Mangone I, Rinaldi A, Scialabba S, Di Pasquale A, Cammà C, Puglia I, Calistri P, Savini G |
| EPI_ISL_965227, EPI_ISL_965228, EPI_ISL_965229 | SIESP CHIETI - DRIVE IN CHIETI | Istituto Zooprofilattico Sperimentale dell'Abruzzo e Molise "G. Caporale" | Lorusso A, Marcacci M, Di Domenico M, Ancora M, Curini V, Mangone I, Rinaldi A, Scialabba S, Di Pasquale A, Cammà C, Puglia I, Calistri P, Savini G |
| EPI_ISL_965230 | SIESP TERAMO DIP PREV | Istituto Zooprofilattico Sperimentale dell'Abruzzo e Molise "G. Caporale" | Lorusso A, Marcacci M, Di Domenico M, Ancora M, Curini V, Mangone I, Rinaldi A, Scialabba S, Di Pasquale A, Cammà C, Puglia I, Calistri P, Savini G |
| EPI_ISL_965231 | Ospedale San Salvatore UOC MED INT. | Istituto Zooprofilattico Sperimentale dell'Abruzzo e Molise "G. Caporale" | Lorusso A, Marcacci M, Di Domenico M, Ancora M, Curini V, Mangone I, Rinaldi A, Scialabba S, Di Pasquale A, Cammà C, Puglia I, Calistri P, Savini G |
| EPI_ISL_965232, EPI_ISL_965233 | SIESP CHIETI - DRIVE IN CHIETI | Istituto Zooprofilattico Sperimentale dell'Abruzzo e Molise "G. Caporale" | Lorusso A, Marcacci M, Di Domenico M, Ancora M, Curini V, Mangone I, Rinaldi A, Scialabba S, Di Pasquale A, Cammà C, Puglia I, Calistri P, Savini G |
| EPI_ISL_965234 | SIESP SULMONA | Istituto Zooprofilattico Sperimentale dell'Abruzzo e Molise "G. Caporale" | Lorusso A, Marcacci M, Di Domenico M, Ancora M, Curini V, Mangone I, Rinaldi A, Scialabba S, Di Pasquale A, Cammà C, Puglia I, Calistri P, Savini G |
| EPI_ISL_965235 | SIESP TERAMO DIP PREV | Istituto Zooprofilattico Sperimentale dell'Abruzzo e Molise "G. Caporale" | Lorusso A, Marcacci M, Di Domenico M, Ancora M, Curini V, Mangone I, Rinaldi A, Scialabba S, Di Pasquale A, Cammà C, Puglia I, Calistri P, Savini G |
| EPI_ISL_965236, EPI_ISL_965237, EPI_ISL_965238, EPI_ISL_965239 | SIESP CHIETI - DRIVE IN CHIETI | Istituto Zooprofilattico Sperimentale dell'Abruzzo e Molise "G. Caporale" | Lorusso A, Marcacci M, Di Domenico M, Ancora M, Curini V, Mangone I, Rinaldi A, Scialabba S, Di Pasquale A, Cammà C, Puglia I, Calistri P, Savini G |
| EPI_ISL_965240 | SIESP CHIETI | Istituto Zooprofilattico Sperimentale dell'Abruzzo e Molise "G. Caporale" | Lorusso A, Marcacci M, Di Domenico M, Ancora M, Curini V, Mangone I, Rinaldi A, Scialabba S, Di Pasquale A, Cammà C, Puglia I, Calistri P, Savini G |
| EPI_ISL_965241, EPI_ISL_965242 | USCA SULMONA | Istituto Zooprofilattico Sperimentale dell'Abruzzo e Molise "G. Caporale" | Lorusso A, Marcacci M, Di Domenico M, Ancora M, Curini V, Mangone I, Rinaldi A, Scialabba S, Di Pasquale A, Cammà C, Puglia I, Calistri P, Savini G |
| EPI_ISL_965243, EPI_ISL_965244 | SIESP CHIETI-DRIVE IN ORTONA | Istituto Zooprofilattico Sperimentale dell'Abruzzo e Molise "G. Caporale" | Lorusso A, Marcacci M, Di Domenico M, Ancora M, Curini V, Mangone I, Rinaldi A, Scialabba S, Di Pasquale A, Cammà C, Puglia I, Calistri P, Savini G |
| EPI_ISL_965253 | SIESP CHIETI | Istituto Zooprofilattico Sperimentale dell'Abruzzo e Molise "G. Caporale" | Lorusso A, Marcacci M, Di Domenico M, Ancora M, Curini V, Mangone I, Rinaldi A, Scialabba S, Di Pasquale A, Cammà C, Puglia I, Calistri P, Savini G |
| EPI_ISL_965254, EPI_ISL_965255 | SIESP CHIETI - DRIVE IN CHIETI | Istituto Zooprofilattico Sperimentale dell'Abruzzo e Molise "G. Caporale" | Lorusso A, Marcacci M, Di Domenico M, Ancora M, Curini V, Mangone I, Rinaldi A, Scialabba S, Di Pasquale A, Cammà C, Puglia I, Calistri P, Savini G |
| EPI_ISL_965256 | SIESP CHIETI-DRIVE IN VASTO | Istituto Zooprofilattico Sperimentale dell'Abruzzo e Molise "G. Caporale" | Lorusso A, Marcacci M, Di Domenico M, Ancora M, Curini V, Mangone I, Rinaldi A, Scialabba S, Di Pasquale A, Cammà C, Puglia I, Calistri P, Savini G |
| EPI_ISL_965257 | SIESP CHIETI-DRIVE IN GISSI | Istituto Zooprofilattico Sperimentale dell'Abruzzo e Molise "G. Caporale" | Lorusso A, Marcacci M, Di Domenico M, Ancora M, Curini V, Mangone I, Rinaldi A, Scialabba S, Di Pasquale A, Cammà C, Puglia I, Calistri P, Savini G |
| EPI_ISL_965258, EPI_ISL_965259 | SIESP CHIETI-DRIVE IN VASTO | Istituto Zooprofilattico Sperimentale dell'Abruzzo e Molise "G. Caporale" | Lorusso A, Marcacci M, Di Domenico M, Ancora M, Curini V, Mangone I, Rinaldi A, Scialabba S, Di Pasquale A, Cammà C, Puglia I, Calistri P, Savini G |
| EPI_ISL_965260 | SIESP CHIETI-DRIVE IN GISSI | Istituto Zooprofilattico Sperimentale dell'Abruzzo e Molise "G. Caporale" | Lorusso A, Marcacci M, Di Domenico M, Ancora M, Curini V, Mangone I, Rinaldi A, Scialabba S, Di Pasquale A, Cammà C, Puglia I, Calistri P, Savini G |
| EPI_ISL_965261 | SIESP CHIETI | Istituto Zooprofilattico Sperimentale dell'Abruzzo e Molise "G. Caporale" | Lorusso A, Marcacci M, Di Domenico M, Ancora M, Curini V, Mangone I, Rinaldi A, Scialabba S, Di Pasquale A, Cammà C, Puglia I, Calistri P, Savini G |
| EPI_ISL_965262 | SIESP CHIETI, DISTRETTO SAN CHIETI | Istituto Zooprofilattico Sperimentale dell'Abruzzo e Molise "G. Caporale" | Lorusso A, Marcacci M, Di Domenico M, Ancora M, Curini V, Mangone I, Rinaldi A, Scialabba S, Di Pasquale A, Cammà C, Puglia I, Calistri P, Savini G |
| EPI_ISL_965263 | SIESP CHIETI-DRIVE IN ORTONA | Istituto Zooprofilattico Sperimentale dell'Abruzzo e Molise "G. Caporale" | Lorusso A, Marcacci M, Di Domenico M, Ancora M, Curini V, Mangone I, Rinaldi A, Scialabba S, Di Pasquale A, Cammà C, Puglia I, Calistri P, Savini G |
| EPI_ISL_965265 | SIESP CHIETI - DRIVE IN CHIETI | Istituto Zooprofilattico Sperimentale dell'Abruzzo e Molise "G. Caporale" | Lorusso A, Marcacci M, Di Domenico M, Ancora M, Curini V, Mangone I, Rinaldi A, Scialabba S, Di Pasquale A, Cammà C, Puglia I, Calistri P, Savini G |
| EPI_ISL_965267 | SIESP CHIETI, DISTRETTON SAN CHIETI | Istituto Zooprofilattico Sperimentale dell'Abruzzo e Molise "G. Caporale" | Lorusso A, Marcacci M, Di Domenico M, Ancora M, Curini V, Mangone I, Rinaldi A, Scialabba S, Di Pasquale A, Cammà C, Puglia I, Calistri P, Savini G |
| EPI_ISL_965268, EPI_ISL_965269 | SIESP CHIETI, DISTRETTO SAN CHIETI | Istituto Zooprofilattico Sperimentale dell'Abruzzo e Molise "G. Caporale" | Lorusso A, Marcacci M, Di Domenico M, Ancora M, Curini V, Mangone I, Rinaldi A, Scialabba S, Di Pasquale A, Cammà C, Puglia I, Calistri P, Savini G |
| EPI_ISL_965270 | SIESP CHIETI | Istituto Zooprofilattico Sperimentale dell'Abruzzo e Molise "G. Caporale" | Lorusso A, Marcacci M, Di Domenico M, Ancora M, Curini V, Mangone I, Rinaldi A, Scialabba S, Di Pasquale A, Cammà C, Puglia I, Calistri P, Savini G |
| EPI_ISL_965271 | SIESP CHIETI, DISTRETTO SAN CHIETI | Istituto Zooprofilattico Sperimentale dell'Abruzzo e Molise "G. Caporale" | Lorusso A, Marcacci M, Di Domenico M, Ancora M, Curini V, Mangone I, Rinaldi A, Scialabba S, Di Pasquale A, Cammà C, Puglia I, Calistri P, Savini G |

| | | | |
|---|---|---|---|
| EPI_ISL_965272 | SIESP CHIETI | Istituto Zooprofilattico Sperimentale dell'Abruzzo e Molise "G. Caporale" | Lorusso A, Marcacci M, Di Domenico M, Ancora M, Curini V, Mangone I, Rinaldi A, Scialabba S, Di Pasquale A, Cammà C, Puglia I, Calistri P, Savini G |
| EPI_ISL_965273, EPI_ISL_965274, EPI_ISL_965275 | SIESP CHIETI, DISTRETTO SAN CHIETI | Istituto Zooprofilattico Sperimentale dell'Abruzzo e Molise "G. Caporale" | Lorusso A, Marcacci M, Di Domenico M, Ancora M, Curini V, Mangone I, Rinaldi A, Scialabba S, Di Pasquale A, Cammà C, Puglia I, Calistri P, Savini G |
| EPI_ISL_965278, EPI_ISL_965279, EPI_ISL_965280, EPI_ISL_965281, EPI_ISL_965282, EPI_ISL_965283, EPI_ISL_965284, EPI_ISL_965285, EPI_ISL_965286, EPI_ISL_965287, EPI_ISL_965288, EPI_ISL_965289, EPI_ISL_965290, EPI_ISL_965291, EPI_ISL_965292, EPI_ISL_965293, EPI_ISL_965294, EPI_ISL_965295, EPI_ISL_965296, EPI_ISL_965297, EPI_ISL_965298, EPI_ISL_965299, EPI_ISL_965300 | | | |
| see above | P.O.CARDARELLI | Istituto Zooprofilattico Sperimentale dell'Abruzzo e Molise "G. Caporale" | Scutellà M, Niro G, Lorusso A, Marcacci M, Di Domenico M, Ancora M, Curini V, Mangone I, Rinaldi A, Scialabba S, Di Pasquale A, Cammà C, Puglia I, Calistri P, Savini G |
| EPI_ISL_966245 | NYU Langone Health | Departments of Pathology and Medicine, New York University School of Medicine | Adriana Heguy, Dacia Dimartino, Emily Guzman, Christian Marier, Peter Meyn, Sitharam Ramaswami, Gael Westby, Paul Zappile, Yutong Zhang, Paolo Cotzia, Guiqing Wang |
| EPI_ISL_966346, EPI_ISL_966347, EPI_ISL_966348, EPI_ISL_966349, EPI_ISL_966350, EPI_ISL_966351, EPI_ISL_966352, EPI_ISL_966353, EPI_ISL_966354 | Kansas Health and Environmental Lab | Kansas Health and Environmental Lab | Mike Grose, Paige Drury, Carissa Robertson, Ben Olsen, and Phil Adam |
| EPI_ISL_966807, EPI_ISL_966808, EPI_ISL_966809, EPI_ISL_966810, EPI_ISL_966811, EPI_ISL_966812, EPI_ISL_966813, EPI_ISL_966814, EPI_ISL_966815, EPI_ISL_966816, EPI_ISL_966817, EPI_ISL_966818, EPI_ISL_966819, EPI_ISL_966820, EPI_ISL_966821, EPI_ISL_966822, EPI_ISL_966823, EPI_ISL_966824, EPI_ISL_966825, EPI_ISL_966826, EPI_ISL_966827, EPI_ISL_966828, EPI_ISL_966829, EPI_ISL_966830, EPI_ISL_966831, EPI_ISL_966832, EPI_ISL_966833, EPI_ISL_966834, EPI_ISL_966835, EPI_ISL_966836, EPI_ISL_966837, EPI_ISL_966838, EPI_ISL_966839, EPI_ISL_966840, EPI_ISL_966841, EPI_ISL_966842, EPI_ISL_966843, EPI_ISL_966844, EPI_ISL_966845, EPI_ISL_966846, EPI_ISL_966847, EPI_ISL_966848, EPI_ISL_966849, EPI_ISL_966850, EPI_ISL_966851 | | | |
| see above | Maine HETL | Tewhey Lab, The Jackson Laboratory | Matluk,N., Dewey,H., Iosue,F., Barter,M., Lynch,R., Munger,H. and Tewhey,R. |
| EPI_ISL_967882, EPI_ISL_967883 | Arizona State Public Health Laboratory | Arizona State Public Health Laboratory | Trung Huynh, Jessica Escobar, Katherine Fullerton, Nobuko Fukushima, Stacy White, Linda Getsinger, Victor Waddell |
| EPI_ISL_968212 | PHV-FSS | PHV-FSS | Son Nguyen et al. |
| EPI_ISL_969131 | Laboratorio Microbiologia e Virologia P.O. Cotugno A.O. dei Colli | Laboratorio Microbiologia e Virologia P.O. Cotugno A.O. dei Colli | Luigi Atripaldi, Claudia Tiberio, Anna Perfetti, |
| EPI_ISL_969227 | Laboratorio Microbiologia e Virologia P.O. Cotugno A.O. dei Colli | Laboratorio Microbiologia e Virologia P.O. Cotugno A.O. dei Colli | Luigi Atripaldi, Claudia Tiberio, Anna Perfetti |
| EPI_ISL_969297 | Laboratorio Microbiologia e Virologia, P.O. Cotugno, A.O. dei Colli | Laboratorio Microbiologia e Virologia, P.O. Cotugno, A.O. dei Colli | Luigi Atripaldi, Claudia Tiberio, Anna Perfetti |
| EPI_ISL_969456 | Lighthouse Lab in Milton Keynes | Wellcome Sanger Institute for the COVID-19 Genomics UK (COG-UK) Consortium | The Lighthouse Lab in Milton Keynes and Alex Alderton, Roberto Amato, Sonia Goncalves, Ewan Harrison, David K. Jackson, Ian Johnston, Dominic Kwiatkowski, Cordelia Langford, John Sillitoe on behalf of the Wellcome Sanger Institute COVID-19 Surveillance Team |
| EPI_ISL_969884 | Laboratorio Microbiologia e Virologia P.O. Cotugno A.O. dei Colli | Laboratorio Microbiologia e Virologia P.O. Cotugno A.O. dei Colli | Luigi Atripaldi,Claudia Tiberio, Anna Perfetti |
| EPI_ISL_970647 | Laboratorio Microbiologia e Virologia P.O. Cotugno A.O. dei Colli | Laboratorio Microbiologia e Virologia P.O. Cotugno A.O. dei Colli | Luigi Atripaldi, Claudia Tiberio,Anna Perfetti |
| EPI_ISL_974745 | Laboratorio Microbiologia e Virologia P.O. Cotugno A.O. dei Colli | Laboratorio Microbiologia e Virologia P.O. Cotugno A.O. dei Colli | Luigi Atripaldi, Claudia Tiberio, Anna Perfetti, |
| EPI_ISL_977495, EPI_ISL_977496, EPI_ISL_977497, EPI_ISL_977498 | University of Bari Biomedical Sciences and Human Oncology | University of Bari Biomedical Sciences and Human Oncology | Chironna M., Sallustio A., Loconsole D., Accogli M. |
| EPI_ISL_977568, EPI_ISL_977569, EPI_ISL_977570, EPI_ISL_977571, EPI_ISL_977572, EPI_ISL_977573, EPI_ISL_977574 | University of Michigan Clinical Microbiology Laboratory | Lauring Lab, University of Michigan, Department of Microbiology and Immunology | Valesano |
| EPI_ISL_977598 | Laboratorio Microbiologia e Virologia P.O. Cotugno A.O. dei Colli | Laboratorio Microbiologia e Virologia P.O. Cotugno A.O. dei Colli | Luigi Atripaldi, Claudia Tiberio, Anna Perfetti |
| EPI_ISL_977601, EPI_ISL_977603 | Laboratorio Microbiologia e Virologia P.O. Cotugno A.O. dei Colli | Laboratorio Microbiologia e Virologia P.O. Cotugno A.O. dei Colli | Luigi Atripaldi, Claudia Tiberio, Anna Perfetti, |
| EPI_ISL_977604, EPI_ISL_977605, EPI_ISL_977606, EPI_ISL_977607, EPI_ISL_977608, EPI_ISL_977609, EPI_ISL_977610, EPI_ISL_977611 | SC (UCO) Igiene e Sanità Pubblica (funzione integrata con SC Microbiologia e Virologia) e Laboratory of Molecular Virology of the International Centre for Genetic Engineering and Biotechnology (ICGEB) | ARGO Laboratorio Genomica ed Epigenomica | Licastro D, Dal Monego S, Degasperi M, Marcello A, D'Agaro P, De Rosa R |
| EPI_ISL_977612, EPI_ISL_977616, EPI_ISL_977617, EPI_ISL_977618, EPI_ISL_977619 | SC (UCO) Igiene e Sanità Pubblica (funzione integrata con SC Microbiologia e Virologia) e Laboratory of Molecular Virology of the International Centre for Genetic Engineering and Biotechnology (ICGEB) | ARGO Laboratorio Genomica ed Epigenomica | Licastro D, Dal Monego S, Degasperi M, Marcello A, D'Agaro P, Pipan C |
| EPI_ISL_977621 | SC (UCO) Igiene e Sanità Pubblica (funzione integrata con SC Microbiologia e Virologia) e Laboratory of Molecular Virology of the International Centre for Genetic Engineering and Biotechnology (ICGEB) | ARGO Laboratorio Genomica ed Epigenomica | Licastro D, Dal Monego S, Degasperi M, Marcello A, D'Agaro P |
| EPI_ISL_977622, EPI_ISL_977623, EPI_ISL_977624, EPI_ISL_977625, EPI_ISL_977626, EPI_ISL_977627, EPI_ISL_977628, EPI_ISL_977629, EPI_ISL_977630, EPI_ISL_977631, EPI_ISL_977632, EPI_ISL_977633, EPI_ISL_977634, EPI_ISL_977635, EPI_ISL_977636, EPI_ISL_977637, EPI_ISL_977638, EPI_ISL_977639, EPI_ISL_977640 | | | |
| see above | SC (UCO) Igiene e Sanità Pubblica (funzione integrata con SC Microbiologia e Virologia) e Laboratory of Molecular Virology of the International Centre for Genetic Engineering and Biotechnology (ICGEB) | ARGO Laboratorio Genomica ed Epigenomica | Licastro D, Dal Monego S, Degasperi M, Marcello A, Segat L, Piscianz E, D'Agaro P |
| EPI_ISL_977641, EPI_ISL_977642, EPI_ISL_977643, EPI_ISL_977644, EPI_ISL_977645, EPI_ISL_977646, EPI_ISL_977647, EPI_ISL_977648, EPI_ISL_977649, EPI_ISL_977650 | SC (UCO) Igiene e Sanità Pubblica (funzione integrata con SC Microbiologia e Virologia) e Laboratory of Molecular Virology of the International Centre for Genetic Engineering and Biotechnology (ICGEB) | ARGO Laboratorio Genomica ed Epigenomica | Licastro D, Dal Monego S, Degasperi M, Marcello A, D'Agaro P, Lombardo F |
| EPI_ISL_977651 | Laboratorio Microbiologia e Virologia P.O. Cotugno A.O. dei Colli | Laboratorio Microbiologia e Virologia P.O. Cotugno A.O. dei Colli | Luigi Atripaldi, Claudia Tiberio, Anna Perfetti, |
| EPI_ISL_978788, EPI_ISL_978789, EPI_ISL_978790, EPI_ISL_978791, EPI_ISL_978792, EPI_ISL_978793, EPI_ISL_978794, EPI_ISL_978795, EPI_ISL_978796, EPI_ISL_978797, EPI_ISL_978798, EPI_ISL_978799, EPI_ISL_978800, EPI_ISL_978801, EPI_ISL_978802, EPI_ISL_978803, EPI_ISL_978804, EPI_ISL_978805, EPI_ISL_978806, EPI_ISL_978807, EPI_ISL_978808, EPI_ISL_978809, EPI_ISL_978810, EPI_ISL_978812, EPI_ISL_978813, EPI_ISL_978814, EPI_ISL_978815, EPI_ISL_978816, EPI_ISL_978817, EPI_ISL_978818, EPI_ISL_978819, EPI_ISL_978820, EPI_ISL_978821, EPI_ISL_978822, EPI_ISL_978823, EPI_ISL_978824, EPI_ISL_978825, EPI_ISL_978826, EPI_ISL_978827, EPI_ISL_978828, EPI_ISL_978829, EPI_ISL_978830, EPI_ISL_978831, EPI_ISL_978832, EPI_ISL_978833, EPI_ISL_978834, EPI_ISL_978835, EPI_ISL_978836, EPI_ISL_978837, EPI_ISL_978838, EPI_ISL_978839, EPI_ISL_978840, EPI_ISL_978841, EPI_ISL_978842, | | | |

| | | | |
|---|---|---|---|
| EPI_ISL_978843, EPI_ISL_978844, EPI_ISL_978845, EPI_ISL_978846, EPI_ISL_978847, EPI_ISL_978848, EPI_ISL_978849, EPI_ISL_978850, EPI_ISL_978851, EPI_ISL_978852, EPI_ISL_978853, EPI_ISL_978854, EPI_ISL_978855, EPI_ISL_978856, EPI_ISL_978857, EPI_ISL_978858, EPI_ISL_978859, EPI_ISL_978860, EPI_ISL_978861, EPI_ISL_978862, EPI_ISL_978863, EPI_ISL_978864, EPI_ISL_978865, EPI_ISL_978866, EPI_ISL_978868, EPI_ISL_978869, EPI_ISL_978870, EPI_ISL_978871, EPI_ISL_978872, EPI_ISL_978873, EPI_ISL_978874, EPI_ISL_978875, EPI_ISL_978876, EPI_ISL_978877, EPI_ISL_978878, EPI_ISL_978879 | | | |
| see above | AZDelta | AZDelta | Geert Martens; Dieter De Smet |
| EPI_ISL_979360, EPI_ISL_979361 | Microbiological Diagnostic Unit - Public Health Laboratory (MDU-PHL) | MDU-PHL | Seemann T., Sait, M.L., Sherry, N.L. |
| EPI_ISL_979362, EPI_ISL_979363, EPI_ISL_979364 | Victorian Infectious Diseases Reference Laboratory (VIDRL) | VIDRL and MDU-PHL | Caly L., Seemann T., Sait, M.L., Druce J., Sherry, N.L. |
| EPI_ISL_979968, EPI_ISL_979969 | National Institute of Infectious Diseases-Prof. Dr. Matei Bals Molecular Diagnostics Laboratory | National Institute of Infectious Diseases-Prof. Dr. Matei Bals Molecular Diagnostics Laboratory | Leontina Banica, Marius Surleac, Corina Casangiu, Petre Milu, Andreea Tudor, Simona Paraschiv, Dan Otelea |
| EPI_ISL_980235, EPI_ISL_980236, EPI_ISL_980237, EPI_ISL_980238, EPI_ISL_980239, EPI_ISL_980240, EPI_ISL_980241, EPI_ISL_980242, EPI_ISL_980243, EPI_ISL_980244, EPI_ISL_980245, EPI_ISL_980246, EPI_ISL_980247, EPI_ISL_980248, EPI_ISL_980249, EPI_ISL_980250, EPI_ISL_980251, EPI_ISL_980252, EPI_ISL_980253, EPI_ISL_980254, EPI_ISL_980255, EPI_ISL_980256, EPI_ISL_980257, EPI_ISL_980258, EPI_ISL_980259, EPI_ISL_980260, EPI_ISL_980261, EPI_ISL_980262, EPI_ISL_980263, EPI_ISL_980264, EPI_ISL_980265, EPI_ISL_980266, EPI_ISL_980267, EPI_ISL_980268, EPI_ISL_980269, EPI_ISL_980270, EPI_ISL_980271, EPI_ISL_980272, EPI_ISL_980273, EPI_ISL_980274, EPI_ISL_980275, EPI_ISL_980276, EPI_ISL_980277, EPI_ISL_980278, EPI_ISL_980279, EPI_ISL_980281, EPI_ISL_980282, EPI_ISL_980283, EPI_ISL_980284, EPI_ISL_980285, EPI_ISL_980286, EPI_ISL_980287, EPI_ISL_980288, EPI_ISL_980289, EPI_ISL_980290, EPI_ISL_980291, EPI_ISL_980292, EPI_ISL_980293, EPI_ISL_980294, EPI_ISL_980295, EPI_ISL_980296, EPI_ISL_980297, EPI_ISL_980298, EPI_ISL_980299, EPI_ISL_980300, EPI_ISL_980301, EPI_ISL_980302, EPI_ISL_980303, EPI_ISL_980304, EPI_ISL_980305, EPI_ISL_980306, EPI_ISL_980307, EPI_ISL_980308, EPI_ISL_980309, EPI_ISL_980310, EPI_ISL_980311, EPI_ISL_980312, EPI_ISL_980313, EPI_ISL_980314, EPI_ISL_980315, EPI_ISL_980316, EPI_ISL_980317, EPI_ISL_980318, EPI_ISL_980319, EPI_ISL_980320, EPI_ISL_980321, EPI_ISL_980322, EPI_ISL_980323, EPI_ISL_980324, EPI_ISL_980325, EPI_ISL_980326, EPI_ISL_980327, EPI_ISL_980328, EPI_ISL_980329, EPI_ISL_980330, EPI_ISL_980331, EPI_ISL_980332, EPI_ISL_980333, EPI_ISL_980334, EPI_ISL_980335, EPI_ISL_980336, EPI_ISL_980337, EPI_ISL_980338, EPI_ISL_980339, EPI_ISL_980340, EPI_ISL_980341, EPI_ISL_980342, EPI_ISL_980343, EPI_ISL_980344, EPI_ISL_980345, EPI_ISL_980346, EPI_ISL_980347, EPI_ISL_980348, EPI_ISL_980349, EPI_ISL_980350, EPI_ISL_980351, EPI_ISL_980352, EPI_ISL_980353, EPI_ISL_980354, EPI_ISL_980355, EPI_ISL_980357, EPI_ISL_980358, EPI_ISL_980359, EPI_ISL_980360, EPI_ISL_980361, EPI_ISL_980362, EPI_ISL_980363, EPI_ISL_980364, EPI_ISL_980365, EPI_ISL_980366, EPI_ISL_980367, EPI_ISL_980368, EPI_ISL_980369, EPI_ISL_980370, EPI_ISL_980371, EPI_ISL_980372, EPI_ISL_980373, EPI_ISL_980374, EPI_ISL_980375, EPI_ISL_980376, EPI_ISL_980377, EPI_ISL_980378, EPI_ISL_980379, EPI_ISL_980380, EPI_ISL_980381, EPI_ISL_980382, EPI_ISL_980383, EPI_ISL_980384, EPI_ISL_980385, EPI_ISL_980386, EPI_ISL_980387, EPI_ISL_980388, EPI_ISL_980389, EPI_ISL_980390, EPI_ISL_980391, EPI_ISL_980392, EPI_ISL_980393, EPI_ISL_980394, EPI_ISL_980395, EPI_ISL_980396, EPI_ISL_980397, EPI_ISL_980398, EPI_ISL_980399, EPI_ISL_980400, EPI_ISL_980401, EPI_ISL_980402, EPI_ISL_980403, EPI_ISL_980404, EPI_ISL_980405, EPI_ISL_980406, EPI_ISL_980407, EPI_ISL_980408, EPI_ISL_980409, EPI_ISL_980410, EPI_ISL_980411, EPI_ISL_980412, EPI_ISL_980413, EPI_ISL_980414, EPI_ISL_980415, EPI_ISL_980417, EPI_ISL_980418, EPI_ISL_980420, EPI_ISL_980421, EPI_ISL_980422, EPI_ISL_980423, EPI_ISL_980424, EPI_ISL_980425, EPI_ISL_980426, EPI_ISL_980427, EPI_ISL_980428, EPI_ISL_980429, EPI_ISL_980430, EPI_ISL_980431, EPI_ISL_980432, EPI_ISL_980434, EPI_ISL_980435, EPI_ISL_980436, EPI_ISL_980437, EPI_ISL_980438, EPI_ISL_980439, EPI_ISL_980440, EPI_ISL_980441, EPI_ISL_980442, EPI_ISL_980443, EPI_ISL_980444, EPI_ISL_980445, EPI_ISL_980446, EPI_ISL_980447, EPI_ISL_980448, EPI_ISL_980450, EPI_ISL_980451, EPI_ISL_980452, EPI_ISL_980453, EPI_ISL_980454, EPI_ISL_980455, EPI_ISL_980456, EPI_ISL_980457, EPI_ISL_980458, EPI_ISL_980459, EPI_ISL_980460, EPI_ISL_980461, EPI_ISL_980462, EPI_ISL_980463, EPI_ISL_980464, EPI_ISL_980465, EPI_ISL_980466, EPI_ISL_980467, EPI_ISL_980468, EPI_ISL_980469, EPI_ISL_980470, EPI_ISL_980471, EPI_ISL_980472, EPI_ISL_980473, EPI_ISL_980474, EPI_ISL_980475, EPI_ISL_980476, EPI_ISL_980477, EPI_ISL_980478, EPI_ISL_980479, EPI_ISL_980480, EPI_ISL_980481, EPI_ISL_980482, EPI_ISL_980483, EPI_ISL_980484, EPI_ISL_980485, EPI_ISL_980486, EPI_ISL_980487, EPI_ISL_980488, EPI_ISL_980489, EPI_ISL_980490, EPI_ISL_980491, EPI_ISL_980492, EPI_ISL_980493, EPI_ISL_980494, EPI_ISL_980495, EPI_ISL_980496, EPI_ISL_980497, EPI_ISL_980498, EPI_ISL_980499, EPI_ISL_980500, EPI_ISL_980501, EPI_ISL_980502, EPI_ISL_980503, EPI_ISL_980504, EPI_ISL_980505, EPI_ISL_980506, EPI_ISL_980507, EPI_ISL_980508, EPI_ISL_980509, EPI_ISL_980510, EPI_ISL_980511, EPI_ISL_980513, EPI_ISL_980514, EPI_ISL_980515, EPI_ISL_980516, EPI_ISL_980517, EPI_ISL_980518, EPI_ISL_980519, EPI_ISL_980520, EPI_ISL_980521, EPI_ISL_980522, EPI_ISL_980523, EPI_ISL_980524, EPI_ISL_980525, EPI_ISL_980526, EPI_ISL_980527, EPI_ISL_980528, EPI_ISL_980529, EPI_ISL_980530, EPI_ISL_980531, EPI_ISL_980532, EPI_ISL_980533, EPI_ISL_980534, EPI_ISL_980535, EPI_ISL_980536, EPI_ISL_980537, EPI_ISL_980538, EPI_ISL_980539, EPI_ISL_980540, EPI_ISL_980541, EPI_ISL_980542, EPI_ISL_980543, EPI_ISL_980544, EPI_ISL_980545, EPI_ISL_980546, EPI_ISL_980547, EPI_ISL_980548, EPI_ISL_980549, EPI_ISL_980550 | | | |
| see above | Lighthouse Lab in Cambridge | Wellcome Sanger Institute for the COVID-19 Genomics UK (COG-UK) Consortium | Rob Howes, The Lighthouse Lab in Cambridge and Alex Alderton, Roberto Amato, Sonia Goncalves, Ewan Harrison, David K. Jackson, Ian Johnston, Dominic Kwiatkowski, Cordelia Langford, John Sillitoe on behalf of the Wellcome Sanger Institute COVID-19 Surveillance Team |
| EPI_ISL_981004 | Thai Red Cross Emerging Infectious Diseases Health Science Centre, Chulalongkorn Hospital, Faculty of Medicine, Chulalongkorn University | Thai Red Cross Emerging Infectious Diseases Center and Faculty of Medicine, Chulalongkorn University | Rome Buathong, Wichai Thanasopon, Sopon Iamsirithaworn, Opass Putcharoen,Sininat Petcharat, Yutthana Joyjinda, Weenassarin Ampoot, Apaporn Rodpan, Thiravat Hemachudha, Supaporn Wacharapluesadee |
| EPI_ISL_981007, EPI_ISL_981008, EPI_ISL_981009, EPI_ISL_981010, EPI_ISL_981011, EPI_ISL_981012, EPI_ISL_981013, EPI_ISL_981014, EPI_ISL_981015, EPI_ISL_981016, EPI_ISL_981017, EPI_ISL_981018, EPI_ISL_981019, EPI_ISL_981020, EPI_ISL_981021, EPI_ISL_981022, EPI_ISL_981023, EPI_ISL_981024, EPI_ISL_981025, EPI_ISL_981026, EPI_ISL_981027, EPI_ISL_981028, EPI_ISL_981029 | | | |
| see above | National Public Health Laboratory, National Centre for Infectious Diseases | National Public Health Laboratory, National Centre for Infectious Diseases | Tze Minn Mak, Zhenyang Zhou, Lin Cui, Raymond Tzer Pin Lin |
| EPI_ISL_981077, EPI_ISL_981088, EPI_ISL_981093, EPI_ISL_981094, EPI_ISL_981095, EPI_ISL_981097, EPI_ISL_981100, EPI_ISL_981114, EPI_ISL_981115, EPI_ISL_981116, EPI_ISL_981117, EPI_ISL_981120, EPI_ISL_981121, EPI_ISL_981128, EPI_ISL_981129, EPI_ISL_981130, EPI_ISL_981131, EPI_ISL_981132, EPI_ISL_981135, EPI_ISL_981158, EPI_ISL_981159, EPI_ISL_981160, EPI_ISL_981161, EPI_ISL_981162, EPI_ISL_981163, EPI_ISL_981164, EPI_ISL_981165, EPI_ISL_981166, EPI_ISL_981167, EPI_ISL_981168, EPI_ISL_981169, EPI_ISL_981170, EPI_ISL_981171, EPI_ISL_981172, EPI_ISL_981173, EPI_ISL_981175, EPI_ISL_981191, EPI_ISL_981192, EPI_ISL_981193, EPI_ISL_981194, EPI_ISL_981195, EPI_ISL_981196, EPI_ISL_981197, EPI_ISL_981198, EPI_ISL_981199, EPI_ISL_981200, EPI_ISL_981201 | | | |
| see above | Johns Hopkins Hospital Department of Pathology | Johns Hopkins Hospital Department of Pathology | C. Paul Morris, Chun Huai Luo, Adannaya Amadi, Matthew Schwartz, Nicholas Gallagher, Heba H. Mostafa |
| EPI_ISL_981372 | Laborarory for Respiratory Viruses, "Cantacuzino" National Military-Medical Institute for Resararch and Development | Cantacuzino Institute Virology | Luiza Ustea, Nicoleta Paraschiv, Catalina Pascu, Mihaela Lazar |
| EPI_ISL_981374 | Laboratory for Respiratory Viruses, Cantacuzino National Military-Medical Institute for Research and Development | Cantacuzino Institute Virology | Luiza Ustea, Nicoleta Paraschiv, Catalina Pascu, Mihaela Lazar |
| EPI_ISL_981551, EPI_ISL_981557, EPI_ISL_981611, EPI_ISL_981612, EPI_ISL_981613, EPI_ISL_981614, EPI_ISL_981615, EPI_ISL_981616, EPI_ISL_981617, EPI_ISL_981618, EPI_ISL_981619, EPI_ISL_981620, EPI_ISL_981621, EPI_ISL_981622, EPI_ISL_981623, EPI_ISL_981624, EPI_ISL_981625, EPI_ISL_981626, EPI_ISL_981627, EPI_ISL_981628, EPI_ISL_981629, EPI_ISL_981630, EPI_ISL_981631, EPI_ISL_981632, EPI_ISL_981633, EPI_ISL_981634, EPI_ISL_981635, EPI_ISL_981636, EPI_ISL_981637, EPI_ISL_981638, EPI_ISL_981639, EPI_ISL_981640, EPI_ISL_981641, EPI_ISL_981644, EPI_ISL_981645, EPI_ISL_981646, EPI_ISL_981647, EPI_ISL_981648, EPI_ISL_981649, EPI_ISL_981690, EPI_ISL_981691, EPI_ISL_981692, EPI_ISL_981693, EPI_ISL_981694, EPI_ISL_981695, EPI_ISL_981705, EPI_ISL_981710, EPI_ISL_981711, EPI_ISL_981712, EPI_ISL_981718, EPI_ISL_981720, EPI_ISL_981808, EPI_ISL_981809, EPI_ISL_981810, EPI_ISL_981811, EPI_ISL_981812, EPI_ISL_981814, EPI_ISL_981815, EPI_ISL_981816, EPI_ISL_981817, EPI_ISL_981818, EPI_ISL_981819, EPI_ISL_981820, EPI_ISL_981821, EPI_ISL_981822, EPI_ISL_981823, EPI_ISL_981824, EPI_ISL_981825, EPI_ISL_981826, EPI_ISL_981836 | | | |
| see above | University Hospitals of Geneva, Laboratory of Virology | HUG, Laboratory of Virology and the Health2030 Genome Center | Samuel Cordey, Ana Rita Goncalves, Laurent Kaiser, Lorenzo Cerutti, Henri Pegeot, Melyssa Elies, Deborah Penet, Keith Harshman, Ioannis Xenarios, Emmanouil Dermitzakis |
| EPI_ISL_983328 | Azienda Ospedaliera San Camillo Forlanini | INMI Lazzaro Spallanzani IRCCS | E Giombini, M. Rueca, B Bartolini, O Butera, C.E.M Gruber, F Messina, G Parisi, ML Guarino, A Di Caro, MR Capobianchi |
| EPI_ISL_983329 | Istituto Zooprofilattico Sperimentale Lazio e Toscana "M. Aleandri" | INMI Lazzaro Spallanzani IRCCS | CEM Gruber, B Bartolini, E Giombini, M Rueca, O Butera, F Messina, MT Scicluna, G Manna, A Cersini, A Di Caro, MR Capobianchi |
| EPI_ISL_983632, EPI_ISL_983633, EPI_ISL_983634, EPI_ISL_983635, EPI_ISL_983636, EPI_ISL_983637, EPI_ISL_983638, EPI_ISL_983639 | University of Michigan Clinical Microbiology Laboratory | Lauring Lab, University of Michigan, Department of Microbiology and Immunology | Valesano |
| EPI_ISL_983736, EPI_ISL_983737 | Colorado Department of Public Health and Environment | Colorado Department of Puplic Health and Environment | Laura Bankers, Molly C. Hetherington-Rauth, Diana Ir, Shannon Ely, Shannon R. Matzinger, Sarah Elizabeth Totten, Emily A. Travanty |
