## Supplementary material for "Detection and characterization of the SARS-CoV-2 lineage B.1.526 in New York": Supp. Table 3 GISAID Acknowledgment part 2: gisaid_hcov-19_acknowledgement_table_2021_02_13_01.pdf

We gratefully acknowledge the following Authors from the Originating laboratories responsible for obtaining the specimens, as well as the Submitting laboratories where the genome data were generated and shared via GISAID, on which this research is based.

All Submitters of data may be contacted directly via www.gisaid.org

Authors are sorted alphabetically.

| Accession ID | Originating Laboratory | Submitting Laboratory | Authors |
|---|---|---|---|
| EPI_ISL_888653, EPI_ISL_888654, EPI_ISL_888655, EPI_ISL_888656, EPI_ISL_888657, EPI_ISL_888658, EPI_ISL_888659, EPI_ISL_888660, EPI_ISL_888661, EPI_ISL_888662 | University of Michigan Clinical Microbiology Laboratory | Lauring Lab, University of Michigan, Department of Microbiology and Immunology | Valesano |
| EPI_ISL_892365, EPI_ISL_892367 | National Laboratory for Health, Environment and Food | National Laboratory for Health, Environment and Food | Aleksander Mahnic, Sandra Janezic, Maja Rupnik |
| EPI_ISL_903209, EPI_ISL_903210, EPI_ISL_903211, EPI_ISL_903212, EPI_ISL_903213, EPI_ISL_903214, EPI_ISL_903215, EPI_ISL_903216, EPI_ISL_903217 | University of Michigan Clinical Microbiology Laboratory | Lauring Lab, University of Michigan, Department of Microbiology and Immunology | Valesano |
| EPI_ISL_903349, EPI_ISL_903350, EPI_ISL_903351, EPI_ISL_903352, EPI_ISL_903354, EPI_ISL_903355, EPI_ISL_903356, EPI_ISL_903359, EPI_ISL_903360 | Wyoming Public Health Laboratory | Wyoming Public Health Laboratory | Noah Hull, Taylor Fearing, Lynette Gumbleton, Channing Weber, Ashley Norberg, Bailey Bowcutt, and Wanda Manley |
| EPI_ISL_905756, EPI_ISL_905757 | National Institute of Public Health - National Institute of Hygiene | National Institute of Public Health - National Institute of Hygiene | Wokowicz Tomasz, Zacharczuk Katarzyna |
| EPI_ISL_906146 | Gundersen Molecular Diagnostics Laboratory | Kabara Cancer Research Institute | Craig S. Richmond, Paraic A. Kenny |
| EPI_ISL_906189 | University of Wisconsin-Madison AIDS Vaccine Research Laboratories | University of Wisconsin-Madison AIDS Vaccine Research Laboratories | Gage Moreno, Katarina Braun, et al. AIDS Vaccine Research Laboratories |
| EPI_ISL_906833, EPI_ISL_906834, EPI_ISL_906835, EPI_ISL_906836, EPI_ISL_906837, EPI_ISL_906838, EPI_ISL_906839, EPI_ISL_906840, EPI_ISL_906841, EPI_ISL_906842, EPI_ISL_906843, EPI_ISL_906844, EPI_ISL_906845, EPI_ISL_906846 | | | |
| see above | National Public Health Laboratory, National Centre for Infectious Diseases | National Public Health Laboratory, National Centre for Infectious Diseases | Tze Minn Mak, Zhenyang Zhou, Lin Cui, Raymond Tzer Pin Lin |
| EPI_ISL_911373, EPI_ISL_911376, EPI_ISL_911377, EPI_ISL_911381, EPI_ISL_911382 | University of Michigan Clinical Microbiology Laboratory | Lauring Lab, University of Michigan, Department of Microbiology and Immunology | Valesano |
| EPI_ISL_911874, EPI_ISL_911875, EPI_ISL_911876, EPI_ISL_911877, EPI_ISL_911878, EPI_ISL_911879, EPI_ISL_911880, EPI_ISL_911881, EPI_ISL_911882, EPI_ISL_911883, EPI_ISL_911884, EPI_ISL_911886, EPI_ISL_911887, EPI_ISL_911888, EPI_ISL_911889, EPI_ISL_911893, EPI_ISL_911895, EPI_ISL_911896, EPI_ISL_911897, EPI_ISL_911898, EPI_ISL_911899, EPI_ISL_911912, EPI_ISL_911913, EPI_ISL_911914, EPI_ISL_911915, EPI_ISL_911916, EPI_ISL_911917 | | | |
| see above | Johns Hopkins Hospital Department of Pathology | Johns Hopkins Hospital Department of Pathology | C. Paul Morris, Chun Huai Luo, Adannaya Amadi, Matthew Schwartz, Nicholas Gallagher, Heba H. Mostafa |
| EPI_ISL_912178 | Tempus | Grubaugh Lab - Yale School of Public Health | Tara Alpert, Joseph Fauver, Anderson Brito, Mallery Breban, Anne Wyllie, Chantal Vogels, Mary Petrone, Annie Watkins, Chaney Kalinich, Isabel Ott, Nathan Grubaugh |
| EPI_ISL_912179, EPI_ISL_912180, EPI_ISL_912181, EPI_ISL_912182, EPI_ISL_912183, EPI_ISL_912184 | Yale Pathology Lab | Grubaugh Lab - Yale School of Public Health | Tara Alpert, Joseph Fauver, Chen Liu, Pei Hui, Jianhui Wang, Susan Bell and Han Zhou, Anderson Brito, Mallery Breban, Anne Wyllie, Chantal Vogels, Mary Petrone, Chaney Kalinich, Isabel Ott, Arnau Casanovas, Catherine Muenker, Adam Moore, Alice Lu, Maria Tokuyama, Patrick Wong, Peiwen Lu, Saad Omer, Richard Martinello, Allison Nelson, Shelli Farhadian, Akiko Iwasaki, Charlese Dela Cruz, Albert Ko, Nathan Grubaugh |
| EPI_ISL_912186 | Sydney South West Pathology Service (SSWPS) - Concord Repatriation General Hospital - NSW Health Pathology | NSW Health Pathology - Institute of Clinical Pathology and Medical Research; Westmead Hospital; University of Sydney | CIDM-PH et al. |
| EPI_ISL_912677, EPI_ISL_912811, EPI_ISL_912813, EPI_ISL_912815, EPI_ISL_912816, EPI_ISL_912817, EPI_ISL_912819, EPI_ISL_912820, EPI_ISL_912824, EPI_ISL_912825, EPI_ISL_912826, EPI_ISL_912829, EPI_ISL_912832, EPI_ISL_912833, EPI_ISL_912834, EPI_ISL_912835, EPI_ISL_912836, EPI_ISL_912839, EPI_ISL_912842, EPI_ISL_912843, EPI_ISL_912846, EPI_ISL_912847, EPI_ISL_912848, EPI_ISL_912849, EPI_ISL_912864, EPI_ISL_912865, EPI_ISL_912866, EPI_ISL_912867, EPI_ISL_912868, EPI_ISL_912869, EPI_ISL_912870, EPI_ISL_912871, EPI_ISL_912872, EPI_ISL_912873, EPI_ISL_912874, EPI_ISL_912875, EPI_ISL_912876, EPI_ISL_912877, EPI_ISL_912878, EPI_ISL_912879, EPI_ISL_912880, EPI_ISL_912881, EPI_ISL_912882, EPI_ISL_912883, EPI_ISL_912884, EPI_ISL_912885, EPI_ISL_912886, EPI_ISL_912887, EPI_ISL_912888, EPI_ISL_912889, EPI_ISL_912890 | | | |
| see above | Hôpital Henri Mondor | Department of Virology, Henri Mondor University Hospital, Assistance Publique Hôpitaux de Paris, Université Paris-Est Créteil, INSERM U955 | Christophe Rodriguez, Slim Fourati, Vanessa Demontant, Guillaume Gricourt, Melissa N'Debi, Alexandre Soulier, Elisabeth Trawinski, Jean-Michel Pawlotsky |
| EPI_ISL_913398, EPI_ISL_913408, EPI_ISL_913418 | Massachusetts State Public Health Laboratory | Massachusetts State Public Health Laboratory | Andrew Lang, Timelia Fink, Glen Gallagher, Sandra Smole |
| EPI_ISL_913440, EPI_ISL_913441, EPI_ISL_913442, EPI_ISL_913443, EPI_ISL_913444 | Institute for Infectious Diseases, University of Bern, Switzerland | Institute for Infectious Diseases, University of Bern, Switzerland | Michel C Koch, Christian Baumann, Miguel A Terrazos Miani, Cora Sägesser, Pascal Bittel, Stephen L Leib, Peter Keller, Franziska Suter-Riniker, Alban Ramette |
| EPI_ISL_915429 | Bundeswehr Institute of Microbiology | Bundeswehr Institute of Microbiology | Markus Antwerpen, Alexandra Rehn, Mathias Walter, Malena Bestehorn-Willmann, Sabine Zange, Enrico Georgi, Roman Wölfel |
| EPI_ISL_915430, EPI_ISL_915431, EPI_ISL_915432, EPI_ISL_915433 | Bundeswehrkrankenhaus Westerstede | Bundeswehr Institute of Microbiology | Markus Antwerpen, Klaus Peter Ebert, Alexandra Rehn, Mathias Walter, Malena Bestehorn-Willmann, Sabine Zange, Enrico Georgi, Roman Wölfel |
| EPI_ISL_918166 | Thai Red Cross Emerging Infectious Diseases Health Science Centre, Chulalongkorn Hospital, Faculty of Medicine, Chulalongkorn University | Thai Red Cross Emerging Infectious Diseases Center and Faculty of Medicine, Chulalongkorn University | Rome Buathong, Sopon Iamsirithaworn, Sininat Petcharat, Yutthana Joyjinda, Weenassarin Ampoot, Apaporn Rodpan, Opass Putcharoen, Thiravat Hemachudha, Supaporn Wacharapluesadee |
| EPI_ISL_918269 | SIESP DIPARTIMENTO DI PREVENZIONE TERAMO | Istituto Zooprofilattico Sperimentale dell'Abruzzo e Molise "G. Caporale" | Lorusso A, Marcacci M, Di Domenico M, Ancora M, Curini V, Mangone I, Rinaldi A, Scialabba S, Di Pasquale A, Cammà C, Puglia I, Calistri P, Savini G |
| EPI_ISL_918358 | Institute of Virology, Medical Center, University of Freiburg, Freiburg, Germany | Institute of Virology, Clinial Virus Genomics, Medical Center, University of Freiburg, Freiburg, Germany | Jonas Fuchs, Lisa Kern, Sandra Reuter, Hajo Grundmann, Marcus Panning |
| EPI_ISL_918423 | München Klinik GmbH | Bundeswehr Institute of Microbiology | Markus Antwerpen, Hans-Ulrich Schmidt, Alexandra Rehn, Mathias Walter, Malena Bestehorn-Willmann, Sabine Zange, Enrico Georgi, Roman Wölfel |
| EPI_ISL_918424, EPI_ISL_918425 | Bundeswehrzentralkrankenhaus Koblenz | Bundeswehr Institute of Microbiology | Markus Antwerpen, Ralf Hagen, Ingo Fengler, Alexandra Rehn, Mathias Walter, Malena Bestehorn-Willmann, Sabine Zange, Enrico Georgi, Roman Wölfel |
| EPI_ISL_918426, EPI_ISL_918427 | München Klinik GmbH | Bundeswehr Institute of Microbiology | Markus Antwerpen, Hans-Ulrich Schmidt, Alexandra Rehn, Mathias Walter, Malena Bestehorn-Willmann, Sabine Zange, Enrico Georgi, Roman Wölfel |
| EPI_ISL_918428 | Isar-Amper-Klinikum Haar | Bundeswehr Institute of Microbiology | Markus Antwerpen, Alexandra Rehn, Mathias Walter, Malena Bestehorn-Willmann, Sabine Zange, Enrico Georgi, Roman Wölfel |

| | | | |
|---|---|---|---|
| EPI_ISL_918429 | München Klinik GmbH | Bundeswehr Institute of Microbiology | Markus Antwerpen, Hans-Ulrich Schmidt, Alexandra Rehn, Mathias Walter, Malena Bestehorn-Willmann, Sabine Zange, Enrico Georgi, Roman Wölfel |
| EPI_ISL_918496 | BSWMC-Marble Falls | BSWMC-Temple Molecular | Ari Rao, Linden Morales, Kimberly Walker, Shelby Hendrickson |
| EPI_ISL_919149, EPI_ISL_919150, EPI_ISL_919151, EPI_ISL_919152, EPI_ISL_919153, EPI_ISL_919154, EPI_ISL_919155, EPI_ISL_919156, EPI_ISL_919157, EPI_ISL_919158, EPI_ISL_919159, EPI_ISL_919160, EPI_ISL_919161, EPI_ISL_919162, EPI_ISL_919163, EPI_ISL_919164, EPI_ISL_919165, EPI_ISL_919166, EPI_ISL_919167, EPI_ISL_919168, EPI_ISL_919169, EPI_ISL_919170, EPI_ISL_919171, EPI_ISL_919172, EPI_ISL_919173, EPI_ISL_919174, EPI_ISL_919175, EPI_ISL_919176, EPI_ISL_919177, EPI_ISL_919179, EPI_ISL_919180, EPI_ISL_919181, EPI_ISL_919182, EPI_ISL_919183, EPI_ISL_919184, EPI_ISL_919185, EPI_ISL_919186, EPI_ISL_919187, EPI_ISL_919188, EPI_ISL_919189 | | | |
| see above | Department of Pathology, University of Cambridge | COVID-19 Genomics UK (COG-UK) Consortium | Aminu S. Jahun, Yasmin Chaudhry, Iliana Georgana, Myra Hosmillo, Rhys Izu, Martin D. Curran, Surendra Parmar, Ian Goodfellow |
| EPI_ISL_919405 | Virology Department, Royal Infirmary of Edinburgh, NHS Lothian / School of Biological Sciences, University of Edinburgh / Institute of Genetics and Molecular Medicine, University of Edinburgh | COVID-19 Genomics UK (COG-UK) Consortium | McHugh M, Dewar R, Rooke S, Gallagher M, Balcaza C, O'Toole Á, Scher E, Hill V, McCrone JT, Colquhoun R, Yu X, Jackson B, Rambaut A, Williams TC, Templeton K |
| EPI_ISL_919629, EPI_ISL_919630, EPI_ISL_919663, EPI_ISL_919706, EPI_ISL_919707, EPI_ISL_919708, EPI_ISL_919709, EPI_ISL_919710, EPI_ISL_919712, EPI_ISL_919713, EPI_ISL_919714, EPI_ISL_919715, EPI_ISL_919716, EPI_ISL_919717, EPI_ISL_919718, EPI_ISL_919719, EPI_ISL_919720, EPI_ISL_919721, EPI_ISL_919722, EPI_ISL_919723, EPI_ISL_919724 | | | |
| see above | Liverpool Clinical Laboratories | COVID-19 Genomics UK (COG-UK) Consortium | Sam Haldenby, Anita Lucaci, Steve Paterson, Julian Hiscox, Alistair Darby, M Almsaud, A Alrezaihi, Muhannad Alruwaili, Stuart D Armstrong, Jones Benjamin, Eleanor G Bentley, Anu Chawla, Jordan J Clark, Angela Cowell, Richard Eccles, Isabel García-Dorival, Matthew Gemmell, Alessandro Gerada, PKF Gilmore, Richard Gregory, Ximeng Han, Catherine Hartley, Margaret Hughes, Miren Iturriza-Gomara, James Johnson, L Luu, Jenifer Manson, Charlotte Nelson, Elaine O'Toole, Cassie Olateju, Rebekah Penrice-Randal , Lucille Rainbow, N.P Randle, Trevor Ian Robinson, Parul Sharma, Ghada T Shawli, James P Stewart, Neil Swainston, Ecaterina Vamos, Joanne Watts, Mark Whitehead |
| EPI_ISL_919871, EPI_ISL_919872, EPI_ISL_919873, EPI_ISL_919881, EPI_ISL_919903, EPI_ISL_919904, EPI_ISL_919905, EPI_ISL_919906, EPI_ISL_919907, EPI_ISL_919908, EPI_ISL_919909, EPI_ISL_919910, EPI_ISL_919911, EPI_ISL_919912, EPI_ISL_919913, EPI_ISL_919914, EPI_ISL_919915, EPI_ISL_919916, EPI_ISL_919917, EPI_ISL_919918, EPI_ISL_919919, EPI_ISL_919927, EPI_ISL_919928, EPI_ISL_919929, EPI_ISL_919930, EPI_ISL_919931, EPI_ISL_919932, EPI_ISL_919934, EPI_ISL_919935, EPI_ISL_919936, EPI_ISL_919937, EPI_ISL_919938, EPI_ISL_919939, EPI_ISL_919940, EPI_ISL_919941, EPI_ISL_919942, EPI_ISL_919943, EPI_ISL_919944, EPI_ISL_919946, EPI_ISL_919947, EPI_ISL_919949, EPI_ISL_919950, EPI_ISL_919951, EPI_ISL_919953, EPI_ISL_919954, EPI_ISL_919956, EPI_ISL_919958, EPI_ISL_919959, EPI_ISL_919960, EPI_ISL_920092, EPI_ISL_920093, EPI_ISL_920094, EPI_ISL_920096, EPI_ISL_920097, EPI_ISL_920098, EPI_ISL_920099, EPI_ISL_920100, EPI_ISL_920101, EPI_ISL_920102, EPI_ISL_920103, EPI_ISL_920104, EPI_ISL_920105, EPI_ISL_920106, EPI_ISL_920107, EPI_ISL_920108, EPI_ISL_920109, EPI_ISL_920110, EPI_ISL_920111, EPI_ISL_920112, EPI_ISL_920113, EPI_ISL_920114, EPI_ISL_920115, EPI_ISL_920116, EPI_ISL_920152 | | | |
| see above | University College London, Great Ormond Street Hospital for Children NHS Foundation Trust, Imperial College Healthcare NHS Trust | COVID-19 Genomics UK (COG-UK) Consortium | Sergi Castellano, Rachel Williams, Mark Kristiansen, Paola Resende Silva, Sunando Roy, Tony Brooks, Helena Tutill, Paola Niola, Patricia Dyal, Charlotte Williams, Leysa Forrest, Yasmin Panchbhaya, Jacqueline Findlay, Samuel Weeks, Julianne Brown, Kathryn Harris, Paul Randell, James Price, Alison Holmes, Judith Breuer |
| EPI_ISL_920671, EPI_ISL_920685, EPI_ISL_920693, EPI_ISL_920701, EPI_ISL_920702, EPI_ISL_920723, EPI_ISL_920724, EPI_ISL_920725, EPI_ISL_920728, EPI_ISL_920731, EPI_ISL_920732, EPI_ISL_920733, EPI_ISL_920734, EPI_ISL_920752, EPI_ISL_920756, EPI_ISL_920757, EPI_ISL_920758, EPI_ISL_920761, EPI_ISL_920763, EPI_ISL_920771, EPI_ISL_920776, EPI_ISL_920777, EPI_ISL_920778, EPI_ISL_920779, EPI_ISL_920780 | | | |
| see above | University College London Hospital | COVID-19 Genomics UK (COG-UK) Consortium | Judith Heaney, Matthew Byott, Catherine Houlihan, Dan Frampton, Stuart Kirk, Moira Spyer and Eleni Nastouli |
| EPI_ISL_921659, EPI_ISL_921660, EPI_ISL_921661 | Northumbria University / South Tees Hospitals NHS Foundation Trust / North Cumbria Integrated Care NHS Foundation Trust / North Tees and Hartlepool NHS Foundation Trust / Newcastle Hospitals NHS Foundation Trust | COVID-19 Genomics UK (COG-UK) Consortium | Darren L Smith,Andrew Nelson,Matthew Bashton,Greg R Young,Joshua Loh,John Allan,Mohammad A Tariq,Giles S Holt,Gary Black,Wen C Yew,Lynn Dover,Paul Baker,Steve Liggett,Sarah Essex,Jane Greenaway,Debra Padgett,Clive Graham,Garren Scott,Edward Barton,Emma Swindells,Brendan Payne,Jennifer Collins,Yusri Taha,Gary Eltringham |
| EPI_ISL_922037, EPI_ISL_922038, EPI_ISL_922039, EPI_ISL_922040 | Queens Medical Centre, Clinical Microbiology Department / DeepSeq Nottingham | COVID-19 Genomics UK (COG-UK) Consortium | Gemma Clark, Wendy Smith, Manjinder Khakh, Vicki M Fleming, Michelle M Lister, Hannah Howson-Wells, Jonathan Ball, Patrick McClure, Joseph Chappell, Theocharis Tsoleridis, Nadine Holmes, Matthew Carlisle, Christopher Moore, Fei Sang, Johnny Debebe, Victoria Wright, Matthew Loose |
| EPI_ISL_923349 | Centre for Enzyme Innovation, University of Portsmouth / Translational Research Laboratory, Portsmouth Hospitals NHS Trust | COVID-19 Genomics UK (COG-UK) Consortium | Angela Beckett,Salman Goudarzi,Christopher Fearn,Kate Cook,Katie Loveson,Sharon Glaysher,Scott Elliott,Samuel Robson |
| EPI_ISL_924082, EPI_ISL_924085, EPI_ISL_924087, EPI_ISL_924089, EPI_ISL_924092, EPI_ISL_924097, EPI_ISL_924099, EPI_ISL_924103, EPI_ISL_924105, EPI_ISL_924109, EPI_ISL_924111, EPI_ISL_924112, EPI_ISL_924117, EPI_ISL_924118, EPI_ISL_924125, EPI_ISL_924126, EPI_ISL_924129, EPI_ISL_924131, EPI_ISL_924136, EPI_ISL_924142, EPI_ISL_924144, EPI_ISL_924146, EPI_ISL_924153, EPI_ISL_924160, EPI_ISL_924163, EPI_ISL_924165, EPI_ISL_924173, EPI_ISL_924183, EPI_ISL_924184, EPI_ISL_924185, EPI_ISL_924186, EPI_ISL_924191, EPI_ISL_924193, EPI_ISL_924198, EPI_ISL_924200, EPI_ISL_924201, EPI_ISL_924209, EPI_ISL_924212, EPI_ISL_924216, EPI_ISL_924217, EPI_ISL_924223, EPI_ISL_924227, EPI_ISL_924228, EPI_ISL_924229, EPI_ISL_924235, EPI_ISL_924238, EPI_ISL_924243, EPI_ISL_924249, EPI_ISL_924251, EPI_ISL_924258, EPI_ISL_924263, EPI_ISL_924269, EPI_ISL_924273, EPI_ISL_924276, EPI_ISL_924277, EPI_ISL_924282, EPI_ISL_924287, EPI_ISL_924293, EPI_ISL_924297, EPI_ISL_924298, EPI_ISL_924304, EPI_ISL_924305, EPI_ISL_924306, EPI_ISL_924308, EPI_ISL_924313, EPI_ISL_924330, EPI_ISL_924335, EPI_ISL_924339, EPI_ISL_924340, EPI_ISL_924350, EPI_ISL_924352, EPI_ISL_924356, EPI_ISL_924359, EPI_ISL_924361, EPI_ISL_924366, EPI_ISL_924368, EPI_ISL_924374, EPI_ISL_924377, EPI_ISL_924379, EPI_ISL_924382, EPI_ISL_924390, EPI_ISL_924410, EPI_ISL_924414 | | | |
| see above | Virology Department, Sheffield Teaching Hospitals NHS Foundation Trust/Department of Infection, Immunity and Cardiovascular Disease, The Medical School, University of Sheffield | COVID-19 Genomics UK (COG-UK) Consortium | Thushan de Silva, Matthew Parker, Nikki Smith, Adri Angyal, Rebecca Brown, Luke Green, Rachel Tucker, Paul Parsons, Danielle Groves, Katie Johnson, Laura Carrilero, Alex Keeley, Dave Partridge, Matthew Wyles, Benjamin Lindsey, Mehmet Yavuz, Mohammad Raza, Cariad Evans |
| EPI_ISL_925031, EPI_ISL_925032 | Mirialis | CNR Virus des Infections Respiratoires - France SUD | Antonin Bal, Gregory Destras, Gwendolyne Burfin, Hadrien Règue, Quentin Semanas, Martine Valette, Bruno Lina, Laurence Josset |
| EPI_ISL_925259, EPI_ISL_925260, EPI_ISL_925272, EPI_ISL_925289, EPI_ISL_925291, EPI_ISL_925300, EPI_ISL_925303 | Wyoming Public Health Laboratory | Wyoming Public Health Laboratory | Noah Hull, Taylor Fearing, Lynette Gumbleton, Channing Weber, Ashley Norberg, Bailey Bowcutt, and Wanda Manley |
| EPI_ISL_925918, EPI_ISL_925929, EPI_ISL_925939, EPI_ISL_925940, EPI_ISL_925944, EPI_ISL_925955, EPI_ISL_925958, EPI_ISL_925965, EPI_ISL_925968, EPI_ISL_925970, EPI_ISL_925985, EPI_ISL_925987, EPI_ISL_925990, EPI_ISL_926003, EPI_ISL_926011, EPI_ISL_926012, EPI_ISL_926017, EPI_ISL_926018, EPI_ISL_926029, EPI_ISL_926031, EPI_ISL_926037, EPI_ISL_926044, EPI_ISL_926045, EPI_ISL_926071, EPI_ISL_926079, EPI_ISL_926086, EPI_ISL_926088, EPI_ISL_926090, EPI_ISL_926099, EPI_ISL_926107, EPI_ISL_926113, EPI_ISL_926117, EPI_ISL_926118, EPI_ISL_926120, EPI_ISL_926126, EPI_ISL_926140, EPI_ISL_926142, EPI_ISL_926147, EPI_ISL_926156, EPI_ISL_926157, EPI_ISL_926158, EPI_ISL_926162, EPI_ISL_926163, EPI_ISL_926164, EPI_ISL_926174, EPI_ISL_926183, EPI_ISL_926199, EPI_ISL_926210, EPI_ISL_926211, EPI_ISL_926216, EPI_ISL_926222, EPI_ISL_926225, EPI_ISL_926227, EPI_ISL_926243, EPI_ISL_926245, EPI_ISL_926247, EPI_ISL_926257, EPI_ISL_926259, EPI_ISL_926261, EPI_ISL_926272, EPI_ISL_926274, EPI_ISL_926291, EPI_ISL_926295, EPI_ISL_926297, EPI_ISL_926303, EPI_ISL_926304, EPI_ISL_926308, EPI_ISL_926311, EPI_ISL_926320, EPI_ISL_926327, EPI_ISL_926328, EPI_ISL_926337, EPI_ISL_926342, EPI_ISL_926346, EPI_ISL_926347, EPI_ISL_926348, EPI_ISL_926360, EPI_ISL_926364, EPI_ISL_926367, EPI_ISL_926372, EPI_ISL_926374, EPI_ISL_926389, EPI_ISL_926390, EPI_ISL_926397, EPI_ISL_926398, EPI_ISL_926399, EPI_ISL_926406, EPI_ISL_926416, EPI_ISL_926422, EPI_ISL_926423, EPI_ISL_926435, EPI_ISL_926437, EPI_ISL_926440, EPI_ISL_926455, EPI_ISL_926461, EPI_ISL_926476, EPI_ISL_926478, EPI_ISL_926479, EPI_ISL_926483, EPI_ISL_926489, EPI_ISL_926492, EPI_ISL_926499, EPI_ISL_926512, EPI_ISL_926520, EPI_ISL_926521, EPI_ISL_926535, EPI_ISL_926549, EPI_ISL_926552, EPI_ISL_926558, EPI_ISL_926560, EPI_ISL_926567, EPI_ISL_926586, EPI_ISL_926601, EPI_ISL_926603, EPI_ISL_926616, EPI_ISL_926622, EPI_ISL_926628, EPI_ISL_926642, EPI_ISL_926643, EPI_ISL_926649, EPI_ISL_926655, EPI_ISL_926656, EPI_ISL_926666, EPI_ISL_926668, EPI_ISL_926674, EPI_ISL_926676, EPI_ISL_926687, EPI_ISL_926693, EPI_ISL_926710, EPI_ISL_926721, EPI_ISL_926722, EPI_ISL_926735, EPI_ISL_926752, EPI_ISL_926757, EPI_ISL_926758, EPI_ISL_926760, EPI_ISL_926763, EPI_ISL_926765, EPI_ISL_926778, EPI_ISL_926797, EPI_ISL_926799, EPI_ISL_926801, EPI_ISL_926805, EPI_ISL_926806, EPI_ISL_926810, EPI_ISL_926813, EPI_ISL_926817, EPI_ISL_926821, EPI_ISL_926825, EPI_ISL_926836, EPI_ISL_926853, EPI_ISL_926855, EPI_ISL_926872, EPI_ISL_926880, EPI_ISL_926899, EPI_ISL_926900, EPI_ISL_926909, EPI_ISL_926911, EPI_ISL_926915, EPI_ISL_926917, EPI_ISL_926923, EPI_ISL_926931, EPI_ISL_926934, EPI_ISL_926962, EPI_ISL_926967, EPI_ISL_926968, EPI_ISL_926973, EPI_ISL_926982, EPI_ISL_926989, EPI_ISL_927002, EPI_ISL_927004, EPI_ISL_927011, EPI_ISL_927014, EPI_ISL_927018, EPI_ISL_927020, EPI_ISL_927024, EPI_ISL_927032, EPI_ISL_927036, EPI_ISL_927066, EPI_ISL_927069, EPI_ISL_927070, EPI_ISL_927081, EPI_ISL_927082, EPI_ISL_927084, EPI_ISL_927097, EPI_ISL_927100, EPI_ISL_927106, EPI_ISL_927114, EPI_ISL_927122, EPI_ISL_927127, EPI_ISL_927132, EPI_ISL_927133, EPI_ISL_927134, EPI_ISL_927140, EPI_ISL_927141, EPI_ISL_927144, EPI_ISL_927148, EPI_ISL_927153, EPI_ISL_927167, EPI_ISL_927174, EPI_ISL_927177, EPI_ISL_927181, EPI_ISL_927182, EPI_ISL_927184, EPI_ISL_927189, EPI_ISL_927196, EPI_ISL_927199, EPI_ISL_927203, EPI_ISL_927213, EPI_ISL_927222, EPI_ISL_927236, EPI_ISL_927240, EPI_ISL_927241, EPI_ISL_927243, EPI_ISL_927248, EPI_ISL_927255, EPI_ISL_927263, EPI_ISL_927266, EPI_ISL_927278, EPI_ISL_927284, EPI_ISL_927288, EPI_ISL_927298, EPI_ISL_927300, EPI_ISL_927302, EPI_ISL_927319, EPI_ISL_927327, EPI_ISL_927341, EPI_ISL_927384, EPI_ISL_927385, EPI_ISL_927389, EPI_ISL_927397, EPI_ISL_927402, EPI_ISL_927408, EPI_ISL_927420, EPI_ISL_927424, EPI_ISL_927433, EPI_ISL_927441, EPI_ISL_927447, EPI_ISL_927455, EPI_ISL_927464, EPI_ISL_927470, EPI_ISL_927472, EPI_ISL_927473, EPI_ISL_927479, EPI_ISL_927484, EPI_ISL_927498, EPI_ISL_927522, EPI_ISL_927525, EPI_ISL_927534, EPI_ISL_927536, EPI_ISL_927550, EPI_ISL_927554, EPI_ISL_927555, EPI_ISL_927568, EPI_ISL_927577, EPI_ISL_927578, EPI_ISL_927587, EPI_ISL_927589, EPI_ISL_927598, EPI_ISL_927599, EPI_ISL_927600, EPI_ISL_927608, EPI_ISL_927610, EPI_ISL_927612, EPI_ISL_927615, EPI_ISL_927616, EPI_ISL_927617, EPI_ISL_927623, EPI_ISL_927628, EPI_ISL_927633, EPI_ISL_927634, EPI_ISL_927636, EPI_ISL_927641, EPI_ISL_927643, EPI_ISL_927644, EPI_ISL_927648, EPI_ISL_927653, EPI_ISL_927658, EPI_ISL_927669, EPI_ISL_927683, EPI_ISL_927711, EPI_ISL_927716, EPI_ISL_927723, EPI_ISL_927726, EPI_ISL_927733, EPI_ISL_927751, EPI_ISL_927758, EPI_ISL_927763, EPI_ISL_927769, EPI_ISL_927788, EPI_ISL_927794, EPI_ISL_927820, EPI_ISL_927822, EPI_ISL_927831, EPI_ISL_927833, EPI_ISL_927839, EPI_ISL_927849, EPI_ISL_927856, EPI_ISL_927861, EPI_ISL_927862, EPI_ISL_927864, EPI_ISL_927866, EPI_ISL_927868, EPI_ISL_927873, EPI_ISL_927876, EPI_ISL_927880, EPI_ISL_927905, EPI_ISL_927918, EPI_ISL_927919, EPI_ISL_927926, EPI_ISL_927927, EPI_ISL_927928, EPI_ISL_927943, EPI_ISL_927945, EPI_ISL_927952, EPI_ISL_927953, EPI_ISL_927956, EPI_ISL_927957, EPI_ISL_927964, EPI_ISL_927967, EPI_ISL_927972, EPI_ISL_927976, EPI_ISL_927980, EPI_ISL_927982, EPI_ISL_927984, EPI_ISL_927988, EPI_ISL_927998, EPI_ISL_928022, EPI_ISL_928025, EPI_ISL_928044, EPI_ISL_928049, EPI_ISL_928070, EPI_ISL_928072, EPI_ISL_928089, EPI_ISL_928096, EPI_ISL_928109, EPI_ISL_928125, EPI_ISL_928131, EPI_ISL_928135, EPI_ISL_928136, EPI_ISL_928141, EPI_ISL_928150, EPI_ISL_928154, EPI_ISL_928157, EPI_ISL_928163, EPI_ISL_928166, EPI_ISL_928167, EPI_ISL_928175, EPI_ISL_928191, EPI_ISL_928197, EPI_ISL_928200, EPI_ISL_928210, EPI_ISL_928212, EPI_ISL_928217, EPI_ISL_928225, EPI_ISL_928247, EPI_ISL_928249, EPI_ISL_928251, EPI_ISL_928259, EPI_ISL_928263, EPI_ISL_928273, EPI_ISL_928299, EPI_ISL_928315, EPI_ISL_928340, EPI_ISL_928362, EPI_ISL_928367, EPI_ISL_928383, EPI_ISL_928392, EPI_ISL_928417, EPI_ISL_928424, EPI_ISL_928425, EPI_ISL_928426, EPI_ISL_928428, EPI_ISL_928430, EPI_ISL_928431, EPI_ISL_928439, EPI_ISL_928441, EPI_ISL_928442, EPI_ISL_928444, EPI_ISL_928445, EPI_ISL_928454, EPI_ISL_928455, EPI_ISL_928456, EPI_ISL_928459, EPI_ISL_928462, EPI_ISL_928466, EPI_ISL_928470, EPI_ISL_928491, EPI_ISL_928494, EPI_ISL_928499, EPI_ISL_928504, EPI_ISL_928524, EPI_ISL_928525, EPI_ISL_928528, EPI_ISL_928548, EPI_ISL_928565, EPI_ISL_928568, EPI_ISL_928569, EPI_ISL_928586, EPI_ISL_928587, EPI_ISL_928595, EPI_ISL_928614, EPI_ISL_928621, EPI_ISL_928646, EPI_ISL_928648, EPI_ISL_928650, EPI_ISL_928652, EPI_ISL_928657, EPI_ISL_928660, EPI_ISL_928678, EPI_ISL_928688, EPI_ISL_928690, EPI_ISL_928705, EPI_ISL_928706, EPI_ISL_928711, EPI_ISL_928714, EPI_ISL_928716, EPI_ISL_928719, EPI_ISL_928723, EPI_ISL_928735, EPI_ISL_928743, EPI_ISL_928755, EPI_ISL_928757, EPI_ISL_928769, EPI_ISL_928780, EPI_ISL_928786, EPI_ISL_928788, EPI_ISL_928797, EPI_ISL_928808, EPI_ISL_928823, EPI_ISL_928833, EPI_ISL_928842, EPI_ISL_928853, EPI_ISL_928859, EPI_ISL_928860, EPI_ISL_928861, EPI_ISL_928863, EPI_ISL_928872, EPI_ISL_928878, EPI_ISL_928881, EPI_ISL_928885, EPI_ISL_928892, EPI_ISL_928895, EPI_ISL_928918, EPI_ISL_928926, EPI_ISL_928943, EPI_ISL_928944, EPI_ISL_928946, EPI_ISL_928965, EPI_ISL_928980, EPI_ISL_928987, EPI_ISL_928989, EPI_ISL_928994, EPI_ISL_928995, EPI_ISL_929010, EPI_ISL_929026, EPI_ISL_929039, EPI_ISL_929046, EPI_ISL_929055, EPI_ISL_929056, EPI_ISL_929058, EPI_ISL_929061, EPI_ISL_929062, EPI_ISL_929075, EPI_ISL_929076, EPI_ISL_929083, EPI_ISL_929088, EPI_ISL_929090, EPI_ISL_929097, EPI_ISL_929112, EPI_ISL_929120, EPI_ISL_929124, EPI_ISL_929140, EPI_ISL_929142, EPI_ISL_929145, EPI_ISL_929153, EPI_ISL_929159, EPI_ISL_929162, EPI_ISL_929165, EPI_ISL_929166, EPI_ISL_929168, EPI_ISL_929180, EPI_ISL_929182, EPI_ISL_929188, EPI_ISL_929194, EPI_ISL_929198, | | | |

EPI_ISL_929200, EPI_ISL_929203, EPI_ISL_929205, EPI_ISL_929207, EPI_ISL_929222, EPI_ISL_929223, EPI_ISL_929224, EPI_ISL_929226, EPI_ISL_929230, EPI_ISL_929235, EPI_ISL_929243, EPI_ISL_929250, EPI_ISL_929261, EPI_ISL_929268, EPI_ISL_929270, EPI_ISL_929308, EPI_ISL_929314, EPI_ISL_929323,
EPI_ISL_929326, EPI_ISL_929331, EPI_ISL_929332, EPI_ISL_929338, EPI_ISL_929340, EPI_ISL_929349, EPI_ISL_929360, EPI_ISL_929362, EPI_ISL_929372, EPI_ISL_929389, EPI_ISL_929418, EPI_ISL_929425, EPI_ISL_929435, EPI_ISL_929444, EPI_ISL_929451, EPI_ISL_929461, EPI_ISL_929462, EPI_ISL_929463,
EPI_ISL_929477, EPI_ISL_929486, EPI_ISL_929495, EPI_ISL_929515, EPI_ISL_929518, EPI_ISL_929524, EPI_ISL_929527, EPI_ISL_929538, EPI_ISL_929540, EPI_ISL_929546, EPI_ISL_929547, EPI_ISL_929560, EPI_ISL_929568, EPI_ISL_929575, EPI_ISL_929579, EPI_ISL_929600, EPI_ISL_929601, EPI_ISL_929607,
EPI_ISL_929609, EPI_ISL_929610, EPI_ISL_929614, EPI_ISL_929617, EPI_ISL_929618, EPI_ISL_929623, EPI_ISL_929631, EPI_ISL_929637, EPI_ISL_929643, EPI_ISL_929645, EPI_ISL_929653, EPI_ISL_929660, EPI_ISL_929661, EPI_ISL_929673, EPI_ISL_929686, EPI_ISL_929687, EPI_ISL_929688, EPI_ISL_929693,
EPI_ISL_929709, EPI_ISL_929711, EPI_ISL_929720, EPI_ISL_929723, EPI_ISL_929725, EPI_ISL_929741, EPI_ISL_929755, EPI_ISL_929756, EPI_ISL_929762, EPI_ISL_929783, EPI_ISL_929788, EPI_ISL_929807, EPI_ISL_929808, EPI_ISL_929810, EPI_ISL_929816, EPI_ISL_929823, EPI_ISL_929856, EPI_ISL_929860,
EPI_ISL_929866, EPI_ISL_929868, EPI_ISL_929872, EPI_ISL_929873, EPI_ISL_929890, EPI_ISL_929899, EPI_ISL_929907, EPI_ISL_929908, EPI_ISL_929910, EPI_ISL_929929, EPI_ISL_929931, EPI_ISL_929936, EPI_ISL_929945, EPI_ISL_929951, EPI_ISL_929956, EPI_ISL_929957, EPI_ISL_929958, EPI_ISL_929960,
EPI_ISL_929964, EPI_ISL_929976, EPI_ISL_929980, EPI_ISL_929987, EPI_ISL_929988, EPI_ISL_929993, EPI_ISL_929994, EPI_ISL_930008, EPI_ISL_930010, EPI_ISL_930011, EPI_ISL_930016, EPI_ISL_930034, EPI_ISL_930043, EPI_ISL_930049, EPI_ISL_930054, EPI_ISL_930074, EPI_ISL_930079, EPI_ISL_930080,
EPI_ISL_930083, EPI_ISL_930100, EPI_ISL_930107, EPI_ISL_930109, EPI_ISL_930115, EPI_ISL_930117, EPI_ISL_930128, EPI_ISL_930137, EPI_ISL_930138, EPI_ISL_930140, EPI_ISL_930153, EPI_ISL_930157, EPI_ISL_930160, EPI_ISL_930166, EPI_ISL_930172, EPI_ISL_930181, EPI_ISL_930184, EPI_ISL_930185,
EPI_ISL_930186, EPI_ISL_930188, EPI_ISL_930189, EPI_ISL_930210, EPI_ISL_930211, EPI_ISL_930215, EPI_ISL_930221, EPI_ISL_930224, EPI_ISL_930239, EPI_ISL_930247, EPI_ISL_930249, EPI_ISL_930254, EPI_ISL_930255, EPI_ISL_930258, EPI_ISL_930263, EPI_ISL_930268, EPI_ISL_930274, EPI_ISL_930286,
EPI_ISL_930288, EPI_ISL_930289, EPI_ISL_930292, EPI_ISL_930296, EPI_ISL_930314, EPI_ISL_930315, EPI_ISL_930319, EPI_ISL_930322, EPI_ISL_930333, EPI_ISL_930334, EPI_ISL_930335, EPI_ISL_930342, EPI_ISL_930360, EPI_ISL_930368, EPI_ISL_930372, EPI_ISL_930387, EPI_ISL_930396, EPI_ISL_930409,
EPI_ISL_930412, EPI_ISL_930419, EPI_ISL_930421, EPI_ISL_930422, EPI_ISL_930423, EPI_ISL_930442, EPI_ISL_930450, EPI_ISL_930452, EPI_ISL_930453, EPI_ISL_930454, EPI_ISL_930460, EPI_ISL_930478, EPI_ISL_930485, EPI_ISL_930488, EPI_ISL_930504, EPI_ISL_930511, EPI_ISL_930513, EPI_ISL_930519,
EPI_ISL_930522, EPI_ISL_930524, EPI_ISL_930526, EPI_ISL_930527, EPI_ISL_930546, EPI_ISL_930547, EPI_ISL_930552, EPI_ISL_930554, EPI_ISL_930563

| | | | |
|---|---|---|---|
| see above | Department of Virus and Microbiological Special Diagnostics, Statens Serum Institut, Copenhagen, Denmark | Aalborg University | Danish Covid-19 Genome Consortium |
| EPI_ISL_930568, EPI_ISL_930572, EPI_ISL_930573, EPI_ISL_930574, EPI_ISL_930576 | University of Liège COVID-19 testing center | GIGA Medical Genomics | Keith Durkin, Maria Artesi, Bouchra Boujemla, Emmanuel André, Marc Van Ranst, Fabrice Bureau, Laurent Gillet, Wouter Coppieters, Vincent Bours |
| EPI_ISL_930578 | St. Nikolaus-Hospital Eupen | GIGA Medical Genomics | Keith Durkin, Maria Artesi, Sébastien Bontems, Raphaël Boreux, Bouchra Boujemla, Cécile Meex, Pierrette Melin, Marie-Pierre Hayette, Vincent Bours |
| EPI_ISL_930580, EPI_ISL_930581, EPI_ISL_930582, EPI_ISL_930583, EPI_ISL_930584 | CHC | GIGA Medical Genomics | Keith Durkin, Maria Artesi, Sébastien Bontems, Raphaël Boreux, Bouchra Boujemla, Cécile Meex, Pierrette Melin, Marie-Pierre Hayette, Vincent Bours |
| EPI_ISL_930594, EPI_ISL_930595 | Labo Luc Olivier | GIGA Medical Genomics | Keith Durkin, Maria Artesi, Sébastien Bontems, Raphaël Boreux, Bouchra Boujemla, Cécile Meex, Pierrette Melin, Marie-Pierre Hayette, Vincent Bours |
| EPI_ISL_930596 | Synlab | GIGA Medical Genomics | Keith Durkin, Maria Artesi, Sébastien Bontems, Raphaël Boreux, Bouchra Boujemla, Cécile Meex, Pierrette Melin, Marie-Pierre Hayette, Vincent Bours |
| EPI_ISL_930604 | CHC | GIGA Medical Genomics | Keith Durkin, Maria Artesi, Sébastien Bontems, Raphaël Boreux, Bouchra Boujemla, Cécile Meex, Pierrette Melin, Marie-Pierre Hayette, Vincent Bours |
| EPI_ISL_930611, EPI_ISL_930614 | University of Liège COVID-19 testing center | GIGA Medical Genomics | Keith Durkin, Maria Artesi, Bouchra Boujemla, Emmanuel André, Marc Van Ranst, Fabrice Bureau, Laurent Gillet, Wouter Coppieters, Vincent Bours |
| EPI_ISL_930621 | St. Nikolaus-Hospital Eupen | GIGA Medical Genomics | Keith Durkin, Maria Artesi, Sébastien Bontems, Raphaël Boreux, Bouchra Boujemla, Cécile Meex, Pierrette Melin, Marie-Pierre Hayette, Vincent Bours |
| EPI_ISL_930622 | Synlab | GIGA Medical Genomics | Keith Durkin, Maria Artesi, Sébastien Bontems, Raphaël Boreux, Bouchra Boujemla, Cécile Meex, Pierrette Melin, Marie-Pierre Hayette, Vincent Bours |
| EPI_ISL_930623 | CHC | GIGA Medical Genomics | Keith Durkin, Maria Artesi, Sébastien Bontems, Raphaël Boreux, Bouchra Boujemla, Cécile Meex, Pierrette Melin, Marie-Pierre Hayette, Vincent Bours |
| EPI_ISL_930626, EPI_ISL_930632, EPI_ISL_930633 | University of Liège COVID-19 testing center | GIGA Medical Genomics | Keith Durkin, Maria Artesi, Bouchra Boujemla, Emmanuel André, Marc Van Ranst, Fabrice Bureau, Laurent Gillet, Wouter Coppieters, Vincent Bours |
| EPI_ISL_931490, EPI_ISL_931494, EPI_ISL_931495, EPI_ISL_931501, EPI_ISL_931502, EPI_ISL_931503, EPI_ISL_931504, EPI_ISL_931505, EPI_ISL_931506 | Maryland Public Health Laboratory (MD PHL) | Maryland Public Health Laboratory (MD PHL) | Maryland Department of Health Laboratories Administration |

EPI_ISL_931586, EPI_ISL_931587, EPI_ISL_931588, EPI_ISL_931590, EPI_ISL_931591, EPI_ISL_931592, EPI_ISL_931593, EPI_ISL_931594, EPI_ISL_931596, EPI_ISL_931597, EPI_ISL_931598, EPI_ISL_931599, EPI_ISL_931600, EPI_ISL_931601, EPI_ISL_931602, EPI_ISL_931603, EPI_ISL_931604, EPI_ISL_931605,
EPI_ISL_931606, EPI_ISL_931607, EPI_ISL_931608, EPI_ISL_931609, EPI_ISL_931610, EPI_ISL_931611, EPI_ISL_931612, EPI_ISL_931613, EPI_ISL_931614, EPI_ISL_931615, EPI_ISL_931616, EPI_ISL_931617, EPI_ISL_931618, EPI_ISL_931619, EPI_ISL_931620, EPI_ISL_931621, EPI_ISL_931622, EPI_ISL_931623,
EPI_ISL_931624, EPI_ISL_931625, EPI_ISL_931626, EPI_ISL_931627, EPI_ISL_931628, EPI_ISL_931629, EPI_ISL_931630, EPI_ISL_931631, EPI_ISL_931632, EPI_ISL_931633, EPI_ISL_931634, EPI_ISL_931635, EPI_ISL_931636, EPI_ISL_931637, EPI_ISL_931638, EPI_ISL_931639, EPI_ISL_931640, EPI_ISL_931641,
EPI_ISL_931642, EPI_ISL_931643, EPI_ISL_931644, EPI_ISL_931645, EPI_ISL_931646, EPI_ISL_931647, EPI_ISL_931648, EPI_ISL_931649, EPI_ISL_931650, EPI_ISL_931651, EPI_ISL_931652, EPI_ISL_931653, EPI_ISL_931654, EPI_ISL_931655, EPI_ISL_931656, EPI_ISL_931657, EPI_ISL_931658, EPI_ISL_931659,
EPI_ISL_931660, EPI_ISL_931661, EPI_ISL_931662, EPI_ISL_931663, EPI_ISL_931664, EPI_ISL_931665, EPI_ISL_931666, EPI_ISL_931667, EPI_ISL_931668, EPI_ISL_931669, EPI_ISL_931670, EPI_ISL_931671, EPI_ISL_931672, EPI_ISL_931673, EPI_ISL_931674, EPI_ISL_931675, EPI_ISL_931676, EPI_ISL_931677,
EPI_ISL_931678, EPI_ISL_931679, EPI_ISL_931680, EPI_ISL_931681, EPI_ISL_931682, EPI_ISL_931683, EPI_ISL_931684, EPI_ISL_931685, EPI_ISL_931686, EPI_ISL_931687, EPI_ISL_931688, EPI_ISL_931689, EPI_ISL_931691, EPI_ISL_931692, EPI_ISL_931693, EPI_ISL_931694, EPI_ISL_931695, EPI_ISL_931696,
EPI_ISL_931697, EPI_ISL_931698, EPI_ISL_931699, EPI_ISL_931700, EPI_ISL_931701, EPI_ISL_931702, EPI_ISL_931703, EPI_ISL_931704, EPI_ISL_931705, EPI_ISL_931706, EPI_ISL_931707, EPI_ISL_931708, EPI_ISL_931709, EPI_ISL_931710, EPI_ISL_931711, EPI_ISL_931712, EPI_ISL_931713, EPI_ISL_931714,
EPI_ISL_931715, EPI_ISL_931716, EPI_ISL_931717, EPI_ISL_931718, EPI_ISL_931719, EPI_ISL_931720, EPI_ISL_931721, EPI_ISL_931722, EPI_ISL_931723, EPI_ISL_931724, EPI_ISL_931725, EPI_ISL_931727, EPI_ISL_931728, EPI_ISL_931729, EPI_ISL_931730, EPI_ISL_931731, EPI_ISL_931732, EPI_ISL_931733,
EPI_ISL_931734, EPI_ISL_931735, EPI_ISL_931736, EPI_ISL_931737, EPI_ISL_931738, EPI_ISL_931739, EPI_ISL_931740, EPI_ISL_931741, EPI_ISL_931742, EPI_ISL_931743, EPI_ISL_931744, EPI_ISL_931745, EPI_ISL_931746, EPI_ISL_931747, EPI_ISL_931748, EPI_ISL_931749, EPI_ISL_931750, EPI_ISL_931751,
EPI_ISL_931752, EPI_ISL_931753, EPI_ISL_931754, EPI_ISL_931755, EPI_ISL_931756, EPI_ISL_931757, EPI_ISL_931758, EPI_ISL_931759, EPI_ISL_931760, EPI_ISL_931761, EPI_ISL_931762, EPI_ISL_931763, EPI_ISL_931764, EPI_ISL_931765, EPI_ISL_931766, EPI_ISL_931767, EPI_ISL_931768, EPI_ISL_931769,
EPI_ISL_931770, EPI_ISL_931771, EPI_ISL_931772, EPI_ISL_931773, EPI_ISL_931774, EPI_ISL_931775, EPI_ISL_931776, EPI_ISL_931777, EPI_ISL_931778, EPI_ISL_931779, EPI_ISL_931780, EPI_ISL_931781, EPI_ISL_931782, EPI_ISL_931783, EPI_ISL_931784, EPI_ISL_931785, EPI_ISL_931786, EPI_ISL_931787,
EPI_ISL_931788, EPI_ISL_931789, EPI_ISL_931790, EPI_ISL_931791, EPI_ISL_931792, EPI_ISL_931793, EPI_ISL_931794, EPI_ISL_931795, EPI_ISL_931796, EPI_ISL_931797, EPI_ISL_931798, EPI_ISL_931799, EPI_ISL_931800, EPI_ISL_931801, EPI_ISL_931802, EPI_ISL_931803, EPI_ISL_931804, EPI_ISL_931805,
EPI_ISL_931806, EPI_ISL_931807, EPI_ISL_931808, EPI_ISL_931809, EPI_ISL_931810, EPI_ISL_931811, EPI_ISL_931812, EPI_ISL_931813, EPI_ISL_931814, EPI_ISL_931815, EPI_ISL_931816, EPI_ISL_931817, EPI_ISL_931819, EPI_ISL_931820, EPI_ISL_931821, EPI_ISL_931823, EPI_ISL_931824, EPI_ISL_931825,
EPI_ISL_931826, EPI_ISL_931828, EPI_ISL_931829, EPI_ISL_931830, EPI_ISL_931831, EPI_ISL_931832, EPI_ISL_931833, EPI_ISL_931834, EPI_ISL_931835, EPI_ISL_931836, EPI_ISL_931837, EPI_ISL_931838, EPI_ISL_931839, EPI_ISL_931840, EPI_ISL_931841, EPI_ISL_931842, EPI_ISL_931843, EPI_ISL_931844,
EPI_ISL_931845, EPI_ISL_931846, EPI_ISL_931847, EPI_ISL_931848, EPI_ISL_931849, EPI_ISL_931850, EPI_ISL_931851, EPI_ISL_931852, EPI_ISL_931853, EPI_ISL_931854, EPI_ISL_931855, EPI_ISL_931856, EPI_ISL_931857, EPI_ISL_931858, EPI_ISL_931859, EPI_ISL_931860, EPI_ISL_931861, EPI_ISL_931862,
EPI_ISL_931863, EPI_ISL_931864, EPI_ISL_931865, EPI_ISL_931866, EPI_ISL_931867, EPI_ISL_931868, EPI_ISL_931869, EPI_ISL_931870, EPI_ISL_931871, EPI_ISL_931872, EPI_ISL_931873, EPI_ISL_931875, EPI_ISL_931876, EPI_ISL_931877, EPI_ISL_931878, EPI_ISL_931879, EPI_ISL_931880, EPI_ISL_931881,
EPI_ISL_931882, EPI_ISL_931883, EPI_ISL_931884, EPI_ISL_931885, EPI_ISL_931886, EPI_ISL_931887, EPI_ISL_931888, EPI_ISL_931889, EPI_ISL_931890, EPI_ISL_931891, EPI_ISL_931892, EPI_ISL_931893, EPI_ISL_931894, EPI_ISL_931895, EPI_ISL_931896, EPI_ISL_931897, EPI_ISL_931898, EPI_ISL_931899,
EPI_ISL_931900, EPI_ISL_931901, EPI_ISL_931902, EPI_ISL_931903, EPI_ISL_931904, EPI_ISL_931905, EPI_ISL_931906, EPI_ISL_931907, EPI_ISL_931908, EPI_ISL_931909, EPI_ISL_931910, EPI_ISL_931911, EPI_ISL_931912, EPI_ISL_931913, EPI_ISL_931914, EPI_ISL_931915, EPI_ISL_931916, EPI_ISL_931917,
EPI_ISL_931919, EPI_ISL_931920, EPI_ISL_931922, EPI_ISL_931923, EPI_ISL_931924, EPI_ISL_931925, EPI_ISL_931926, EPI_ISL_931927, EPI_ISL_931928, EPI_ISL_931929, EPI_ISL_931930, EPI_ISL_931931, EPI_ISL_931932, EPI_ISL_931933, EPI_ISL_931934, EPI_ISL_931935, EPI_ISL_931936, EPI_ISL_931938,
EPI_ISL_931940, EPI_ISL_931941, EPI_ISL_931942, EPI_ISL_931944, EPI_ISL_931945, EPI_ISL_931946, EPI_ISL_931947, EPI_ISL_931948, EPI_ISL_931949, EPI_ISL_931950, EPI_ISL_931951, EPI_ISL_931952, EPI_ISL_931954, EPI_ISL_931955, EPI_ISL_931956, EPI_ISL_931958, EPI_ISL_931960, EPI_ISL_931961,
EPI_ISL_931962, EPI_ISL_931963, EPI_ISL_931964, EPI_ISL_931965, EPI_ISL_931966, EPI_ISL_931967, EPI_ISL_931968, EPI_ISL_931969, EPI_ISL_931970, EPI_ISL_931971, EPI_ISL_931972, EPI_ISL_931973, EPI_ISL_931974, EPI_ISL_931975, EPI_ISL_931976, EPI_ISL_931977, EPI_ISL_931978, EPI_ISL_931980,
EPI_ISL_931981, EPI_ISL_931982, EPI_ISL_931983, EPI_ISL_931984, EPI_ISL_931985, EPI_ISL_931986, EPI_ISL_931987, EPI_ISL_931988, EPI_ISL_931989, EPI_ISL_931990, EPI_ISL_931991, EPI_ISL_931992, EPI_ISL_931993, EPI_ISL_931994, EPI_ISL_931995, EPI_ISL_931996, EPI_ISL_931997, EPI_ISL_931998,
EPI_ISL_931999, EPI_ISL_932000, EPI_ISL_932001, EPI_ISL_932002, EPI_ISL_932003, EPI_ISL_932004, EPI_ISL_932005, EPI_ISL_932006, EPI_ISL_932007, EPI_ISL_932008, EPI_ISL_932009, EPI_ISL_932010, EPI_ISL_932011, EPI_ISL_932012, EPI_ISL_932013, EPI_ISL_932014, EPI_ISL_932015, EPI_ISL_932016,
EPI_ISL_932018, EPI_ISL_932019, EPI_ISL_932020, EPI_ISL_932021, EPI_ISL_932022, EPI_ISL_932023, EPI_ISL_932024, EPI_ISL_932025, EPI_ISL_932026, EPI_ISL_932027, EPI_ISL_932028, EPI_ISL_932029, EPI_ISL_932030, EPI_ISL_932031, EPI_ISL_932032, EPI_ISL_932033, EPI_ISL_932034, EPI_ISL_932035,
EPI_ISL_932036, EPI_ISL_932037, EPI_ISL_932038, EPI_ISL_932040, EPI_ISL_932041, EPI_ISL_932042, EPI_ISL_932043, EPI_ISL_932044, EPI_ISL_932045, EPI_ISL_932047, EPI_ISL_932048, EPI_ISL_932049, EPI_ISL_932050, EPI_ISL_932051, EPI_ISL_932052, EPI_ISL_932053, EPI_ISL_932054, EPI_ISL_932055,
EPI_ISL_932056, EPI_ISL_932058, EPI_ISL_932059, EPI_ISL_932060, EPI_ISL_932062, EPI_ISL_932063, EPI_ISL_932064, EPI_ISL_932065, EPI_ISL_932066, EPI_ISL_932067, EPI_ISL_932069, EPI_ISL_932070, EPI_ISL_932071, EPI_ISL_932072, EPI_ISL_932073, EPI_ISL_932074, EPI_ISL_932075, EPI_ISL_932076,
EPI_ISL_932077, EPI_ISL_932078, EPI_ISL_932079, EPI_ISL_932080, EPI_ISL_932081, EPI_ISL_932082, EPI_ISL_932083, EPI_ISL_932084, EPI_ISL_932085, EPI_ISL_932087, EPI_ISL_932088, EPI_ISL_932089, EPI_ISL_932090, EPI_ISL_932091, EPI_ISL_932092, EPI_ISL_932093, EPI_ISL_932094, EPI_ISL_932095,
EPI_ISL_932096, EPI_ISL_932097, EPI_ISL_932098, EPI_ISL_932099, EPI_ISL_932100, EPI_ISL_932101, EPI_ISL_932102, EPI_ISL_932103, EPI_ISL_932104, EPI_ISL_932105, EPI_ISL_932106, EPI_ISL_932107, EPI_ISL_932109, EPI_ISL_932110, EPI_ISL_932111, EPI_ISL_932112, EPI_ISL_932113, EPI_ISL_932115,
EPI_ISL_932116, EPI_ISL_932117, EPI_ISL_932118, EPI_ISL_932119, EPI_ISL_932120, EPI_ISL_932121, EPI_ISL_932122, EPI_ISL_932123, EPI_ISL_932124, EPI_ISL_932125, EPI_ISL_932126, EPI_ISL_932127, EPI_ISL_932128, EPI_ISL_932129, EPI_ISL_932130, EPI_ISL_932131, EPI_ISL_932132, EPI_ISL_932133,
EPI_ISL_932134, EPI_ISL_932135, EPI_ISL_932136, EPI_ISL_932137, EPI_ISL_932138, EPI_ISL_932139, EPI_ISL_932140, EPI_ISL_932141, EPI_ISL_932142, EPI_ISL_932143, EPI_ISL_932144, EPI_ISL_932145, EPI_ISL_932146, EPI_ISL_932147, EPI_ISL_932148, EPI_ISL_932149, EPI_ISL_932150, EPI_ISL_932151,
EPI_ISL_932152, EPI_ISL_932153, EPI_ISL_932154, EPI_ISL_932155, EPI_ISL_932156, EPI_ISL_932158, EPI_ISL_932159, EPI_ISL_932160, EPI_ISL_932161, EPI_ISL_932162, EPI_ISL_932163, EPI_ISL_932164, EPI_ISL_932165, EPI_ISL_932166, EPI_ISL_932167, EPI_ISL_932168, EPI_ISL_932169, EPI_ISL_932170,
EPI_ISL_932171, EPI_ISL_932172, EPI_ISL_932173, EPI_ISL_932174, EPI_ISL_932175, EPI_ISL_932176, EPI_ISL_932177, EPI_ISL_932178, EPI_ISL_932179, EPI_ISL_932180, EPI_ISL_932181, EPI_ISL_932182, EPI_ISL_932183, EPI_ISL_932184, EPI_ISL_932185, EPI_ISL_932186, EPI_ISL_932187, EPI_ISL_932188,
EPI_ISL_932189, EPI_ISL_932190, EPI_ISL_932191, EPI_ISL_932192, EPI_ISL_932193, EPI_ISL_932194, EPI_ISL_932195, EPI_ISL_932196, EPI_ISL_932197, EPI_ISL_932198, EPI_ISL_932199, EPI_ISL_932200, EPI_ISL_932202, EPI_ISL_932203, EPI_ISL_932204, EPI_ISL_932205, EPI_ISL_932206, EPI_ISL_932207,
EPI_ISL_932208, EPI_ISL_932209, EPI_ISL_932210, EPI_ISL_932211, EPI_ISL_932212, EPI_ISL_932213, EPI_ISL_932214, EPI_ISL_932215, EPI_ISL_932216, EPI_ISL_932217, EPI_ISL_932218, EPI_ISL_932219, EPI_ISL_932220, EPI_ISL_932221, EPI_ISL_932223, EPI_ISL_932224, EPI_ISL_932225, EPI_ISL_932226,
EPI_ISL_932227, EPI_ISL_932228, EPI_ISL_932229, EPI_ISL_932230, EPI_ISL_932232, EPI_ISL_932233, EPI_ISL_932234, EPI_ISL_932235, EPI_ISL_932236, EPI_ISL_932237, EPI_ISL_932238, EPI_ISL_932239, EPI_ISL_932242, EPI_ISL_932243, EPI_ISL_932244, EPI_ISL_932245, EPI_ISL_932246, EPI_ISL_932247,
EPI_ISL_932248, EPI_ISL_932249, EPI_ISL_932250

| | | | |
|---|---|---|---|
| see above | Lighthouse Lab in Alderley Park | Wellcome Sanger Institute for the COVID-19 Genomics UK (COG-UK) Consortium | Jacquelyn Wynn, Mairead Hyland, The Lighthouse Lab in Alderley Park and Alex Alderton, Roberto Amato, Sonia Goncalves, Ewan Harrison, David K. Jackson, Ian Johnston, Dominic Kwiatkowski, Cordelia Langford, John Sillitoe on behalf of the Wellcome Sanger Institute COVID-19 Surveillance Team |

EPI_ISL_932850, EPI_ISL_932853, EPI_ISL_932855, EPI_ISL_932856, EPI_ISL_932857, EPI_ISL_932858, EPI_ISL_932859, EPI_ISL_932860, EPI_ISL_932863, EPI_ISL_932864, EPI_ISL_932865, EPI_ISL_932870, EPI_ISL_932871, EPI_ISL_932879, EPI_ISL_932884, EPI_ISL_932885, EPI_ISL_932894, EPI_ISL_932896,
EPI_ISL_932897, EPI_ISL_932899, EPI_ISL_932903, EPI_ISL_932905, EPI_ISL_932908, EPI_ISL_932914, EPI_ISL_932915, EPI_ISL_932917, EPI_ISL_932918, EPI_ISL_932919, EPI_ISL_932920, EPI_ISL_932921, EPI_ISL_932925, EPI_ISL_932929, EPI_ISL_932930, EPI_ISL_932931, EPI_ISL_932932, EPI_ISL_932936,
EPI_ISL_932939, EPI_ISL_932944, EPI_ISL_932948, EPI_ISL_932949, EPI_ISL_932955, EPI_ISL_932957, EPI_ISL_932958, EPI_ISL_932965, EPI_ISL_932966, EPI_ISL_932967, EPI_ISL_932970, EPI_ISL_932980, EPI_ISL_932986, EPI_ISL_932987, EPI_ISL_932991, EPI_ISL_932994, EPI_ISL_932995, EPI_ISL_932996,
EPI_ISL_933000, EPI_ISL_933002, EPI_ISL_933004, EPI_ISL_933006, EPI_ISL_933007, EPI_ISL_933009, EPI_ISL_933011, EPI_ISL_933018, EPI_ISL_933020, EPI_ISL_933021, EPI_ISL_933028, EPI_ISL_933033, EPI_ISL_933035, EPI_ISL_933036, EPI_ISL_933037, EPI_ISL_933038, EPI_ISL_933040, EPI_ISL_933042,
EPI_ISL_933044, EPI_ISL_933050, EPI_ISL_933051, EPI_ISL_933053, EPI_ISL_933054, EPI_ISL_933056, EPI_ISL_933058, EPI_ISL_933065, EPI_ISL_933067, EPI_ISL_933069, EPI_ISL_933070, EPI_ISL_933072, EPI_ISL_933073, EPI_ISL_933074, EPI_ISL_933079, EPI_ISL_933080, EPI_ISL_933096, EPI_ISL_933099,
EPI_ISL_933100, EPI_ISL_933109, EPI_ISL_933110

| | | | |
|---|---|---|---|
| see above | Lighthouse Lab in Glasgow | Wellcome Sanger Institute for the COVID-19 Genomics UK (COG-UK) Consortium | Harper VanSteenhouse, Yumi Kasai, David Gray, Carol Clugston, Anna Dominiczak and Alex Alderton, Roberto Amato, Sonia Goncalves, Ewan Harrison, David K. Jackson, Ian Johnston, Dominic Kwiatkowski, Cordelia Langford, John Sillitoe on behalf of the Wellcome Sanger Institute COVID-19 Surveillance Team |
| EPI_ISL_933777 | Victorian Infectious Diseases Reference Laboratory (VIDRL) | VIDRL and MDU-PHL | Caly L., Seemann T., Sait, M.L., Druce J., Sherry, N.L. |
| EPI_ISL_933778 | Microbiological Diagnostic Unit - Public Health Laboratory (MDU-PHL) | MDU-PHL | Seemann T., Sait, M.L., Sherry, N.L. |
| EPI_ISL_933779 | Victorian Infectious Diseases Reference Laboratory (VIDRL) | VIDRL and MDU-PHL | Caly L., Seemann T., Sait, M.L., Druce J., Sherry, N.L. |
| EPI_ISL_933794, EPI_ISL_933795, EPI_ISL_933796 | PathWest Laboratory Medicine WA | PathWest Laboratory Medicine WA Microbial Surveillance Unit | PathWest Laboratory Medicine WA Microbial Surveillance Unit |
| EPI_ISL_935012, EPI_ISL_935013, EPI_ISL_935014, EPI_ISL_935015, EPI_ISL_935016 | University of Michigan Clinical Microbiology Laboratory | Lauring Lab, University of Michigan, Department of Microbiology and Immunology | Valesano |
| EPI_ISL_935155 | Cytocheck Laboratory | Kansas Health and Environmental Lab | Mike Grose, Paige Drury, Carissa Robertson, Ben Olsen, and Phil Adam |
| EPI_ISL_935171, EPI_ISL_935172, EPI_ISL_935173, EPI_ISL_935174, EPI_ISL_935175, EPI_ISL_935176, EPI_ISL_935177, EPI_ISL_935178, EPI_ISL_935179, EPI_ISL_935180 | Wyoming Public Health Laboratory | Wyoming Public Health Laboratory | Noah Hull, Taylor Fearing, Lynette Gumbleton, Channing Weber, Ashley Norberg, Bailey Bowcutt, and Wanda Manley |
| EPI_ISL_935602, EPI_ISL_935603 | Labo Analyses med | National Reference Center for Viruses of Respiratory Infections, Institut Pasteur, Paris | Marion Barbet, Sylvie Behillil, Méline Bizard, Angela Brisebarre, Camille Capel, Etienne Simon-Lorière, Vincent Enouf, Maud Vanpeene, Sylvie van der Werf,Amzalag Jonas |
| EPI_ISL_935655, EPI_ISL_935656, EPI_ISL_935657, EPI_ISL_935658, EPI_ISL_935659, EPI_ISL_935660, EPI_ISL_935661, EPI_ISL_935662, EPI_ISL_935663, EPI_ISL_935664, EPI_ISL_935665, EPI_ISL_935666, EPI_ISL_935667, EPI_ISL_935668, EPI_ISL_935669, EPI_ISL_935670, EPI_ISL_935671, EPI_ISL_935672, EPI_ISL_935673 | | | |
| see above | Department of Clinical Microbiology | GIGA Medical Genomics | Keith Durkin, Maria Artesi, Sébastien Bontems, Raphaël Boreux, Bouchra Boujemla, Cécile Meex, Pierrette Melin, Marie-Pierre Hayette, Vincent Bours |
| EPI_ISL_935674, EPI_ISL_935675, EPI_ISL_935676, EPI_ISL_935677, EPI_ISL_935678, EPI_ISL_935679, EPI_ISL_935680, EPI_ISL_935681, EPI_ISL_935682, EPI_ISL_935683, EPI_ISL_935684, EPI_ISL_935685, EPI_ISL_935686, EPI_ISL_935687, EPI_ISL_935688, EPI_ISL_935689, EPI_ISL_935690, EPI_ISL_935691, EPI_ISL_935692, EPI_ISL_935693, EPI_ISL_935694, EPI_ISL_935695 | | | |
| see above | CHR de la Citadelle | GIGA Medical Genomics | Keith Durkin, Maria Artesi, Sébastien Bontems, Raphaël Boreux, Bouchra Boujemla, Cécile Meex, Pierrette Melin, Marie-Pierre Hayette, Vincent Bours |
| EPI_ISL_936485 | DPH, Massachusetts State Public Health Lab | DPH, Massachusetts State Public Health Lab | Lang,A.S., Fink,T., Gallagher,G.R., Smole,S.C. |
| EPI_ISL_937215 | DOHMH PHL | New York City Public Health Laboratory | Jade Wang, et al. |
| EPI_ISL_937216 | DOHMH Central Harlem | New York City Public Health Laboratory | Jade Wang, et al. |
| EPI_ISL_937217, EPI_ISL_937218 | DOHMH Chelsea | New York City Public Health Laboratory | Jade Wang, et al. |
| EPI_ISL_937219, EPI_ISL_937220, EPI_ISL_937221, EPI_ISL_937222, EPI_ISL_937223, EPI_ISL_937224, EPI_ISL_937225, EPI_ISL_937226 | DOHMH Jamaica | New York City Public Health Laboratory | Jade Wang, et al. |
| EPI_ISL_937227, EPI_ISL_937228, EPI_ISL_937229, EPI_ISL_937230, EPI_ISL_937231 | DOHMH Corona | New York City Public Health Laboratory | Jade Wang, et al. |
| EPI_ISL_937232, EPI_ISL_937233, EPI_ISL_937234, EPI_ISL_937235, EPI_ISL_937236 | DOHMH Morrisania | New York City Public Health Laboratory | Jade Wang, et al. |
| EPI_ISL_937237 | DOHMH Riverside | New York City Public Health Laboratory | Jade Wang, et al. |
| EPI_ISL_937238, EPI_ISL_937239 | DOHMH Morrisania | New York City Public Health Laboratory | Jade Wang, et al. |
| EPI_ISL_937251, EPI_ISL_937252, EPI_ISL_937253, EPI_ISL_937254, EPI_ISL_937255 | OCME Office Of Chief Medical Examiner | New York City Public Health Laboratory | Jade Wang, et al. |
| EPI_ISL_937265, EPI_ISL_937268 | NYC HH Elmhurst Hospital Medical Center | New York City Public Health Laboratory | Jade Wang, et al. |
| EPI_ISL_937459, EPI_ISL_937460, EPI_ISL_937461, EPI_ISL_937462, EPI_ISL_937463, EPI_ISL_937464, EPI_ISL_937465, EPI_ISL_937466, EPI_ISL_937467, EPI_ISL_937495, EPI_ISL_937496, EPI_ISL_937497, EPI_ISL_937498, EPI_ISL_937499, EPI_ISL_937500, EPI_ISL_937501, EPI_ISL_937502, EPI_ISL_937503, EPI_ISL_937504, EPI_ISL_937505, EPI_ISL_937507, EPI_ISL_937510, EPI_ISL_937512 | | | |
| see above | Maine Health and Environmental Testing Laboratory (Maine HETL) | Tewhey Lab, The Jackson Laboratory | Matluk,N., Dewey,H., Iosue,F., Barter,M., Lynch,R., Munger,H. and Tewhey,R. |
| EPI_ISL_937545, EPI_ISL_937546, EPI_ISL_937547, EPI_ISL_937548, EPI_ISL_937552, EPI_ISL_937555, EPI_ISL_937556, EPI_ISL_937560, EPI_ISL_937561, EPI_ISL_937563, EPI_ISL_937565, EPI_ISL_937566, EPI_ISL_937567, EPI_ISL_937572, EPI_ISL_937574, EPI_ISL_937575, EPI_ISL_937577, EPI_ISL_937582, EPI_ISL_937584, EPI_ISL_937585, EPI_ISL_937589, EPI_ISL_937591, EPI_ISL_937593, EPI_ISL_937597, EPI_ISL_937598, EPI_ISL_937600, EPI_ISL_937602, EPI_ISL_937603, EPI_ISL_937606, EPI_ISL_937607, EPI_ISL_937608, EPI_ISL_937611, EPI_ISL_937613, EPI_ISL_937615, EPI_ISL_937617, EPI_ISL_937619, EPI_ISL_937622, EPI_ISL_937624, EPI_ISL_937625, EPI_ISL_937626, EPI_ISL_937628, EPI_ISL_937629, EPI_ISL_937631, EPI_ISL_937632, EPI_ISL_937633, EPI_ISL_937635, EPI_ISL_937637, EPI_ISL_937642, EPI_ISL_937644, EPI_ISL_937645, EPI_ISL_937646, EPI_ISL_937647, EPI_ISL_937652, EPI_ISL_937654, EPI_ISL_937655, EPI_ISL_937656, EPI_ISL_937657, EPI_ISL_937659, EPI_ISL_937660, EPI_ISL_937661, EPI_ISL_937662, EPI_ISL_937667, EPI_ISL_937671, EPI_ISL_937673, EPI_ISL_937676, EPI_ISL_937677, EPI_ISL_937678, EPI_ISL_937679, EPI_ISL_937681, EPI_ISL_937683, EPI_ISL_937684, EPI_ISL_937686, EPI_ISL_937690, EPI_ISL_937691, EPI_ISL_937692, EPI_ISL_937694, EPI_ISL_937695, EPI_ISL_937696, EPI_ISL_937698, EPI_ISL_937701, EPI_ISL_937702, EPI_ISL_937705, EPI_ISL_937706, EPI_ISL_937708, EPI_ISL_937713, EPI_ISL_937718, EPI_ISL_937719, EPI_ISL_937722, EPI_ISL_937723, EPI_ISL_937725, EPI_ISL_937727, EPI_ISL_937728, EPI_ISL_937729, EPI_ISL_937731, EPI_ISL_937732, EPI_ISL_937733, EPI_ISL_937735, EPI_ISL_937736, EPI_ISL_937739, EPI_ISL_937740, EPI_ISL_937743, EPI_ISL_937747, EPI_ISL_937750, EPI_ISL_937754, EPI_ISL_937755, EPI_ISL_937756, EPI_ISL_937757, EPI_ISL_937758, EPI_ISL_937759, EPI_ISL_937760, EPI_ISL_937764, EPI_ISL_937771, EPI_ISL_937772, EPI_ISL_937773, EPI_ISL_937774, EPI_ISL_937775, EPI_ISL_937776, EPI_ISL_937778, EPI_ISL_937779, EPI_ISL_937782, EPI_ISL_937783, EPI_ISL_937785, EPI_ISL_937786, EPI_ISL_937788, EPI_ISL_937789, EPI_ISL_937791, EPI_ISL_937793, EPI_ISL_937795, EPI_ISL_937796, EPI_ISL_937797, EPI_ISL_937804, EPI_ISL_937807, EPI_ISL_937809, EPI_ISL_937810, EPI_ISL_937812, EPI_ISL_937813, EPI_ISL_937814, EPI_ISL_937816, EPI_ISL_937819, EPI_ISL_937821, EPI_ISL_937823, EPI_ISL_937825, EPI_ISL_937827, EPI_ISL_937828, EPI_ISL_937832, EPI_ISL_937833, EPI_ISL_937834, EPI_ISL_937836, EPI_ISL_937838, EPI_ISL_937841, EPI_ISL_937842, EPI_ISL_937844, EPI_ISL_937845, EPI_ISL_937846, EPI_ISL_937847, EPI_ISL_937848, EPI_ISL_937851, EPI_ISL_937855, EPI_ISL_937856, EPI_ISL_937857, EPI_ISL_937858, EPI_ISL_937859, EPI_ISL_937861, EPI_ISL_937866, EPI_ISL_937868, EPI_ISL_937869, EPI_ISL_937870, EPI_ISL_937871, EPI_ISL_937876, EPI_ISL_937878, EPI_ISL_937879, EPI_ISL_937880, EPI_ISL_937882, EPI_ISL_937884, EPI_ISL_937885 | | | |
| see above | Lighthouse Lab in Alderley Park | Wellcome Sanger Institute for the COVID-19 Genomics UK (COG-UK) Consortium | Jacquelyn Wynn, Mairead Hyland, The Lighthouse Lab in Alderley Park and Alex Alderton, Roberto Amato, Sonia Goncalves, Ewan Harrison, David K. Jackson, Ian Johnston, Dominic Kwiatkowski, Cordelia Langford, John Sillitoe on behalf of the Wellcome Sanger Institute COVID-19 Surveillance Team |
| EPI_ISL_938579, EPI_ISL_938580, EPI_ISL_938582, EPI_ISL_938583, EPI_ISL_938584, EPI_ISL_938585, EPI_ISL_938586, EPI_ISL_938588, EPI_ISL_938589, EPI_ISL_938590, EPI_ISL_938591, EPI_ISL_938593, EPI_ISL_938594, EPI_ISL_938595, EPI_ISL_938597, EPI_ISL_938598, EPI_ISL_938600, EPI_ISL_938601, EPI_ISL_938604, EPI_ISL_938605, EPI_ISL_938606, EPI_ISL_938608, EPI_ISL_938609, EPI_ISL_938611, EPI_ISL_938615, EPI_ISL_938616, EPI_ISL_938617, EPI_ISL_938618, EPI_ISL_938620, EPI_ISL_938621, EPI_ISL_938622, EPI_ISL_938623, EPI_ISL_938624, EPI_ISL_938625, EPI_ISL_938626, EPI_ISL_938627, EPI_ISL_938628, EPI_ISL_938629, EPI_ISL_938634, EPI_ISL_938635, EPI_ISL_938638, EPI_ISL_938639, EPI_ISL_938640, EPI_ISL_938641, EPI_ISL_938642, EPI_ISL_938644, EPI_ISL_938646, EPI_ISL_938647, EPI_ISL_938649, EPI_ISL_938651, EPI_ISL_938653, EPI_ISL_938654, EPI_ISL_938655, EPI_ISL_938656, EPI_ISL_938658, EPI_ISL_938659, EPI_ISL_938660, EPI_ISL_938661, EPI_ISL_938662, EPI_ISL_938663, EPI_ISL_938664, EPI_ISL_938665, EPI_ISL_938668, EPI_ISL_938669, EPI_ISL_938670, EPI_ISL_938671, EPI_ISL_938672, EPI_ISL_938673, EPI_ISL_938674, EPI_ISL_938675, EPI_ISL_938677, EPI_ISL_938678, EPI_ISL_938679, EPI_ISL_938680, EPI_ISL_938681, EPI_ISL_938682, EPI_ISL_938683, EPI_ISL_938685, EPI_ISL_938686, EPI_ISL_938687, EPI_ISL_938688, EPI_ISL_938689, EPI_ISL_938690, EPI_ISL_938691, EPI_ISL_938693, EPI_ISL_938695, EPI_ISL_938696, EPI_ISL_938697, EPI_ISL_938698, EPI_ISL_938699, EPI_ISL_938702, EPI_ISL_938703, EPI_ISL_938704, EPI_ISL_938705, EPI_ISL_938706, EPI_ISL_938708, EPI_ISL_938709, EPI_ISL_938710, EPI_ISL_938711, EPI_ISL_938712, EPI_ISL_938713, EPI_ISL_938714, EPI_ISL_938715, EPI_ISL_938716, EPI_ISL_938717, EPI_ISL_938718, EPI_ISL_938719, EPI_ISL_938720, EPI_ISL_938722, EPI_ISL_938723, EPI_ISL_938724, EPI_ISL_938725, EPI_ISL_938726, EPI_ISL_938727, EPI_ISL_938728, EPI_ISL_938730, EPI_ISL_938734, EPI_ISL_938735, EPI_ISL_938736, EPI_ISL_938738, EPI_ISL_938739, EPI_ISL_938740, EPI_ISL_938742, EPI_ISL_938745, EPI_ISL_938748, EPI_ISL_938749, EPI_ISL_938754, EPI_ISL_938755, EPI_ISL_938759, EPI_ISL_938760, EPI_ISL_938761, EPI_ISL_938763, EPI_ISL_938765, EPI_ISL_938766, EPI_ISL_938767, EPI_ISL_938768, EPI_ISL_938770, EPI_ISL_938772, EPI_ISL_938773, EPI_ISL_938774, EPI_ISL_938775, EPI_ISL_938776, EPI_ISL_938778, EPI_ISL_938779, EPI_ISL_938780, EPI_ISL_938781, EPI_ISL_938782, EPI_ISL_938784, EPI_ISL_938786, EPI_ISL_938788, EPI_ISL_938789, EPI_ISL_938790, EPI_ISL_938791, EPI_ISL_938792, EPI_ISL_938793, EPI_ISL_938795, EPI_ISL_938798, EPI_ISL_938799, EPI_ISL_938802, EPI_ISL_938805, EPI_ISL_938808, EPI_ISL_938809, EPI_ISL_938812, EPI_ISL_938813, EPI_ISL_938814, EPI_ISL_938816, EPI_ISL_938817, EPI_ISL_938818, EPI_ISL_938819, EPI_ISL_938820, EPI_ISL_938821, EPI_ISL_938822, EPI_ISL_938823, EPI_ISL_938824, EPI_ISL_938825, EPI_ISL_938826, EPI_ISL_938827, EPI_ISL_938828, EPI_ISL_938830, EPI_ISL_938831, EPI_ISL_938832, EPI_ISL_938833, EPI_ISL_938834, EPI_ISL_938835, EPI_ISL_938836, EPI_ISL_938838, EPI_ISL_938839, EPI_ISL_938841, EPI_ISL_938842, EPI_ISL_938843, EPI_ISL_938844, EPI_ISL_938847, EPI_ISL_938848, EPI_ISL_938849, EPI_ISL_938850, EPI_ISL_938851, EPI_ISL_938852, EPI_ISL_938853, | | | |

| | | | |
|---|---|---|---|
| EPI_ISL_938855, EPI_ISL_938856, EPI_ISL_938857, EPI_ISL_938858, EPI_ISL_938859, EPI_ISL_938860, EPI_ISL_938861, EPI_ISL_938862, EPI_ISL_938863, EPI_ISL_938864, EPI_ISL_938866, EPI_ISL_938867, EPI_ISL_938868, EPI_ISL_938869, EPI_ISL_938870, EPI_ISL_938871, EPI_ISL_938872, EPI_ISL_938873, EPI_ISL_938874, EPI_ISL_938877, EPI_ISL_938878, EPI_ISL_938879, EPI_ISL_938880, EPI_ISL_938881, EPI_ISL_938883, EPI_ISL_938884, EPI_ISL_938887, EPI_ISL_938888, EPI_ISL_938889, EPI_ISL_938890, EPI_ISL_938891, EPI_ISL_938892, EPI_ISL_938893, EPI_ISL_938894, EPI_ISL_938895, EPI_ISL_938896, EPI_ISL_938897, EPI_ISL_938898, EPI_ISL_938899, EPI_ISL_938902, EPI_ISL_938904, EPI_ISL_938905, EPI_ISL_938906, EPI_ISL_938907, EPI_ISL_938909, EPI_ISL_938910, EPI_ISL_938912, EPI_ISL_938914, EPI_ISL_938915, EPI_ISL_938917, EPI_ISL_938918, EPI_ISL_938919, EPI_ISL_938920, EPI_ISL_938922, EPI_ISL_938923, EPI_ISL_938924, EPI_ISL_938925, EPI_ISL_938926, EPI_ISL_938927, EPI_ISL_938928, EPI_ISL_938929, EPI_ISL_938930, EPI_ISL_938932 | | | |
| see above | Lighthouse Lab in Milton Keynes | Wellcome Sanger Institute for the COVID-19 Genomics UK (COG-UK) Consortium | The Lighthouse Lab in Milton Keynes and Alex Alderton, Roberto Amato, Sonia Goncalves, Ewan Harrison, David K. Jackson, Ian Johnston, Dominic Kwiatkowski, Cordelia Langford, John Sillitoe on behalf of the Wellcome Sanger Institute COVID-19 Surveillance Team |
| EPI_ISL_938935, EPI_ISL_938936, EPI_ISL_938937, EPI_ISL_938938, EPI_ISL_938939, EPI_ISL_938941, EPI_ISL_938942, EPI_ISL_938943 | Lighthouse Lab in Alderley Park | Wellcome Sanger Institute for the COVID-19 Genomics UK (COG-UK) Consortium | Jacquelyn Wynn, Mairead Hyland, The Lighthouse Lab in Alderley Park and Alex Alderton, Roberto Amato, Sonia Goncalves, Ewan Harrison, David K. Jackson, Ian Johnston, Dominic Kwiatkowski, Cordelia Langford, John Sillitoe on behalf of the Wellcome Sanger Institute COVID-19 Surveillance Team |
| EPI_ISL_939283, EPI_ISL_939284, EPI_ISL_939285, EPI_ISL_939287, EPI_ISL_939289, EPI_ISL_939295, EPI_ISL_939297, EPI_ISL_939298, EPI_ISL_939299, EPI_ISL_939303, EPI_ISL_939304, EPI_ISL_939305, EPI_ISL_939308, EPI_ISL_939309, EPI_ISL_939310, EPI_ISL_939311, EPI_ISL_939312, EPI_ISL_939316, EPI_ISL_939317, EPI_ISL_939322, EPI_ISL_939323, EPI_ISL_939324, EPI_ISL_939325, EPI_ISL_939326, EPI_ISL_939328, EPI_ISL_939330, EPI_ISL_939331, EPI_ISL_939333, EPI_ISL_939335, EPI_ISL_939336, EPI_ISL_939338, EPI_ISL_939341, EPI_ISL_939342, EPI_ISL_939344, EPI_ISL_939345, EPI_ISL_939347, EPI_ISL_939349, EPI_ISL_939353, EPI_ISL_939360, EPI_ISL_939362, EPI_ISL_939363, EPI_ISL_939366, EPI_ISL_939367, EPI_ISL_939371, EPI_ISL_939374, EPI_ISL_939375, EPI_ISL_939376, EPI_ISL_939377, EPI_ISL_939379, EPI_ISL_939381, EPI_ISL_939382, EPI_ISL_939383, EPI_ISL_939384, EPI_ISL_939385, EPI_ISL_939386, EPI_ISL_939387, EPI_ISL_939392, EPI_ISL_939394, EPI_ISL_939400, EPI_ISL_939408, EPI_ISL_939409, EPI_ISL_939410, EPI_ISL_939411, EPI_ISL_939412, EPI_ISL_939417, EPI_ISL_939419, EPI_ISL_939421, EPI_ISL_939423, EPI_ISL_939424, EPI_ISL_939426, EPI_ISL_939427, EPI_ISL_939430, EPI_ISL_939433, EPI_ISL_939434, EPI_ISL_939435, EPI_ISL_939436, EPI_ISL_939437, EPI_ISL_939440, EPI_ISL_939441, EPI_ISL_939442, EPI_ISL_939444, EPI_ISL_939445, EPI_ISL_939446, EPI_ISL_939447, EPI_ISL_939448, EPI_ISL_939449, EPI_ISL_939451, EPI_ISL_939452, EPI_ISL_939453, EPI_ISL_939455, EPI_ISL_939457, EPI_ISL_939463, EPI_ISL_939464, EPI_ISL_939465, EPI_ISL_939467, EPI_ISL_939470, EPI_ISL_939471, EPI_ISL_939474, EPI_ISL_939475, EPI_ISL_939476, EPI_ISL_939479, EPI_ISL_939480, EPI_ISL_939482, EPI_ISL_939484, EPI_ISL_939486, EPI_ISL_939488, EPI_ISL_939489, EPI_ISL_939490, EPI_ISL_939491, EPI_ISL_939495, EPI_ISL_939496, EPI_ISL_939499, EPI_ISL_939501, EPI_ISL_939502, EPI_ISL_939503, EPI_ISL_939511, EPI_ISL_939515, EPI_ISL_939516, EPI_ISL_939518, EPI_ISL_939525, EPI_ISL_939527, EPI_ISL_939529, EPI_ISL_939530, EPI_ISL_939531, EPI_ISL_939534, EPI_ISL_939538, EPI_ISL_939539, EPI_ISL_939540, EPI_ISL_939543, EPI_ISL_939544, EPI_ISL_939545, EPI_ISL_939546, EPI_ISL_939549, EPI_ISL_939553, EPI_ISL_939557, EPI_ISL_939558, EPI_ISL_939560, EPI_ISL_939564, EPI_ISL_939568, EPI_ISL_939569, EPI_ISL_939570, EPI_ISL_939575, EPI_ISL_939580, EPI_ISL_939583, EPI_ISL_939584, EPI_ISL_939586, EPI_ISL_939587, EPI_ISL_939589, EPI_ISL_939590, EPI_ISL_939591, EPI_ISL_939592, EPI_ISL_939593, EPI_ISL_939597, EPI_ISL_939602 | | | |
| see above | Lighthouse Lab in Glasgow | Wellcome Sanger Institute for the COVID-19 Genomics UK (COG-UK) Consortium | Harper VanSteenhouse, Yumi Kasai, David Gray, Carol Clugston, Anna Dominiczak and Alex Alderton, Roberto Amato, Sonia Goncalves, Ewan Harrison, David K. Jackson, Ian Johnston, Dominic Kwiatkowski, Cordelia Langford, John Sillitoe on behalf of the Wellcome Sanger Institute COVID-19 Surveillance Team |
| EPI_ISL_939618 | Los Angeles County PHL | Los Angeles County PHL | P. Hemarajata et al. |
| EPI_ISL_940554 | Laboratory for Respiratory Viruses, Cantacuzino National Military-Medical Institute for Research and Development | Cantacuzino Institute Virology | Luiza Ustea, Nicoleta Paraschiv, Mihaela Lazar |
| EPI_ISL_940566, EPI_ISL_940567, EPI_ISL_940568, EPI_ISL_940569, EPI_ISL_940570, EPI_ISL_940571, EPI_ISL_940572 | Jessa | Jessa | Jessa_cmdLab |
| EPI_ISL_940573, EPI_ISL_940574, EPI_ISL_940575 | AZT | Jessa | Jessa_cmdLab |
| EPI_ISL_940576, EPI_ISL_940587, EPI_ISL_940588, EPI_ISL_940589, EPI_ISL_940590, EPI_ISL_940591, EPI_ISL_940592, EPI_ISL_940593, EPI_ISL_940594, EPI_ISL_940595, EPI_ISL_940596, EPI_ISL_940597, EPI_ISL_940598, EPI_ISL_940599, EPI_ISL_940600, EPI_ISL_940601, EPI_ISL_940602, EPI_ISL_940603, EPI_ISL_940604, EPI_ISL_940605, EPI_ISL_940606 | | | |
| see above | Jessa | Jessa | Jessa_cmdLab |
| EPI_ISL_940629 | Hospital Municipal Josanias Castanha Braga | Instituto Adolfo Lutz, Interdiciplinary Procedures Center, Strategic Laboratory | Claudio Tavares Sacchi, Claudia Regina Gonçalves, Erica Valessa Ramos Gomes, Karoline Rodrigues Campos |
| EPI_ISL_940630 | Hospital Geral de Sao Paulo | Instituto Adolfo Lutz, Interdiciplinary Procedures Center, Strategic Laboratory | Claudio Tavares Sacchi, Claudia Regina Gonçalves, Erica Valessa Ramos Gomes, Karoline Rodrigues Campos |
| EPI_ISL_940739 | University of Bari Biomedical Sciences and Human Oncology | University of Bari Biomedical Sciences and Human Oncology | Chironna M., Sallustio A., Loconsole D., Accogli M. |
| EPI_ISL_941300, EPI_ISL_941301, EPI_ISL_941302, EPI_ISL_941303, EPI_ISL_941304, EPI_ISL_941305, EPI_ISL_941306, EPI_ISL_941307, EPI_ISL_941308, EPI_ISL_941309, EPI_ISL_941310, EPI_ISL_941311, EPI_ISL_941312, EPI_ISL_941313, EPI_ISL_941314, EPI_ISL_941315, EPI_ISL_941316, EPI_ISL_941317, EPI_ISL_941318, EPI_ISL_941319, EPI_ISL_941320, EPI_ISL_941321, EPI_ISL_941322, EPI_ISL_941323, EPI_ISL_941324, EPI_ISL_941325, EPI_ISL_941326, EPI_ISL_941327, EPI_ISL_941328, EPI_ISL_941329, EPI_ISL_941330, EPI_ISL_941331, EPI_ISL_941332, EPI_ISL_941333, EPI_ISL_941334 | | | |
| see above | New Mexico Department of Health Scientific Laboratory | New Mexico Department of Health Scientific Laboratory | Ellie Johnson, Anastacia Griego-Fisher, D'eldra Malone, Jennifer Benoit |
| EPI_ISL_941601, EPI_ISL_941620, EPI_ISL_941621, EPI_ISL_941622 | Instituto Nacional de Saude (INSA) | Instituto Nacional de Saude (INSA) | Borges et al |
| EPI_ISL_942326, EPI_ISL_942327, EPI_ISL_942328, EPI_ISL_942329, EPI_ISL_942330, EPI_ISL_942331, EPI_ISL_942332, EPI_ISL_942333, EPI_ISL_942334, EPI_ISL_942335, EPI_ISL_942336, EPI_ISL_942337, EPI_ISL_942338, EPI_ISL_942339, EPI_ISL_942340, EPI_ISL_942341, EPI_ISL_942342, EPI_ISL_942343, EPI_ISL_942344, EPI_ISL_942345, EPI_ISL_942346, EPI_ISL_942347, EPI_ISL_942348, EPI_ISL_942349, EPI_ISL_942350, EPI_ISL_942351, EPI_ISL_942352, EPI_ISL_942353, EPI_ISL_942355, EPI_ISL_942356, EPI_ISL_942357, EPI_ISL_942358, EPI_ISL_942359, EPI_ISL_942360, EPI_ISL_942361, EPI_ISL_942362, EPI_ISL_942363, EPI_ISL_942364, EPI_ISL_942365, EPI_ISL_942367, EPI_ISL_942368 | | | |
| see above | MD PHL | MD PHL | Maryland Department of Health Laboratories Administration |
| EPI_ISL_942880, EPI_ISL_942881, EPI_ISL_942882, EPI_ISL_942883, EPI_ISL_942884, EPI_ISL_942885, EPI_ISL_942886, EPI_ISL_942887, EPI_ISL_942888, EPI_ISL_942889, EPI_ISL_942890, EPI_ISL_942891, EPI_ISL_942892, EPI_ISL_942893, EPI_ISL_942949, EPI_ISL_942950 | | | |
| see above | Gundersen Molecular Diagnostics Laboratory | Kabara Cancer Research Institute | Craig S. Richmond, Paraic A. Kenny |
| EPI_ISL_943016, EPI_ISL_943023, EPI_ISL_943026, EPI_ISL_943029, EPI_ISL_943037, EPI_ISL_943038, EPI_ISL_943041, EPI_ISL_943045, EPI_ISL_943046, EPI_ISL_943047, EPI_ISL_943054, EPI_ISL_943126, EPI_ISL_943127, EPI_ISL_943128, EPI_ISL_943129, EPI_ISL_943130, EPI_ISL_943131, EPI_ISL_943132, EPI_ISL_943133, EPI_ISL_943134, EPI_ISL_943135, EPI_ISL_943136, EPI_ISL_943158, EPI_ISL_943159, EPI_ISL_943160, EPI_ISL_943161, EPI_ISL_943162, EPI_ISL_943221, EPI_ISL_943222, EPI_ISL_943223, EPI_ISL_943224, EPI_ISL_943225, EPI_ISL_943226, EPI_ISL_943227, EPI_ISL_943253, EPI_ISL_943254, EPI_ISL_943255, EPI_ISL_943297, EPI_ISL_943305, EPI_ISL_943306, EPI_ISL_943320, EPI_ISL_943326, EPI_ISL_943327, EPI_ISL_943332, EPI_ISL_943365, EPI_ISL_943366, EPI_ISL_943367, EPI_ISL_943368, EPI_ISL_943369, EPI_ISL_943370, EPI_ISL_943372, EPI_ISL_943373, EPI_ISL_943374, EPI_ISL_943375, EPI_ISL_943411, EPI_ISL_943412, EPI_ISL_943413, EPI_ISL_943414, EPI_ISL_943415, EPI_ISL_943416, EPI_ISL_943417, EPI_ISL_943418, EPI_ISL_943419, EPI_ISL_943436, EPI_ISL_943439, EPI_ISL_943445, EPI_ISL_943448, EPI_ISL_943486, EPI_ISL_943490, EPI_ISL_943498, EPI_ISL_943509, EPI_ISL_943525, EPI_ISL_943526, EPI_ISL_943542 | | | |
| see above | Dutch COVID-19 response team | National Institute for Public Health and the Environment (RIVM) | Adam Meijer, Harry Vennema, Dirk Eggink, Jeroen Cremer, Sharon van den Brink, Bas van der Veer, AnneMarie van den Brandt, Florian Zwagemaker, Dennis Schmitz, Chantal Reusken, on behalf of the national COVID-19 response team |
| EPI_ISL_943569 | Arizona State University | Arizona State University | Peter T. Skidmore, Emily A. Kaelin, LaRinda A. Holland, Nicholas J. Mellor, Kristina Buss, Joy M. Blain, Valerie Harris, Joshua LaBaer, Vel Murugan, Efrem S. Lim |
| EPI_ISL_943967, EPI_ISL_943968, EPI_ISL_943969, EPI_ISL_943970, EPI_ISL_943971, EPI_ISL_943972 | Hospital Geral de Sao Paulo | Instituto Adolfo Lutz, Interdiciplinary Procedures Center, Strategic Laboratory | Claudio Tavares Sacchi, Claudia Regina Gonçalves, Erica Valessa Ramos Gomes, Karoline Rodrigues Campos |
| EPI_ISL_944117 | Platform BIS UZA/UAntwerpen | UAntwerp, Laboratory of Medical Microbiology | Basil Britto Xavier, Jasmine Coppens, Marie Le Mercier, Christine Lammens, Veerle Matheeussen, Herman Goossens |
| EPI_ISL_944573 | National Institute of Infectious Diseases-Prof. Dr. Matei Bals Molecular Diagnostics Laboratory | National Institute of Infectious Diseases-Prof. Dr. Matei Bals Molecular Diagnostics Laboratory | Leontina Banica, Marius Surleac, Corina Casangiu, Petre Milu, Andreea Tudor, Simona Paraschiv, Dan Otelea |
| EPI_ISL_944746 | unknown | Public Health Virology-Forensic and Scientific Services (PHV-FSS) | Son Nguyen et al. |
| EPI_ISL_944770 | National Institute of Infectious Diseases-Prof. Dr. Matei Bals Molecular Diagnostics Laboratory | National Institute of Infectious Diseases-Prof. Dr. Matei Bals Molecular Diagnostics Laboratory | Leontina Banica, Marius Surleac, Corina Casangiu, Petre Milu, Andreea Tudor, Simona Paraschiv, Dan Otelea |
| EPI_ISL_944799, EPI_ISL_944805, EPI_ISL_944824, EPI_ISL_944863, EPI_ISL_944883, EPI_ISL_944908, EPI_ISL_944914, EPI_ISL_944923, EPI_ISL_944953, EPI_ISL_944976, EPI_ISL_944978, EPI_ISL_945023, EPI_ISL_945032, EPI_ISL_945034 | | | |
| see above | Lighthouse Lab in Glasgow | Wellcome Sanger Institute for the COVID-19 Genomics UK (COG-UK) Consortium | Harper VanSteenhouse, Yumi Kasai, David Gray, Carol Clugston, Anna Dominiczak and Alex Alderton, Roberto Amato, Sonia Goncalves, Ewan Harrison, David K. Jackson, Ian Johnston, Dominic Kwiatkowski, Cordelia Langford, John Sillitoe on behalf of the Wellcome Sanger Institute COVID-19 Surveillance Team |

| | | | |
|---|---|---|---|
| EPI_ISL_945377, EPI_ISL_945378, EPI_ISL_945380, EPI_ISL_945381, EPI_ISL_945382, EPI_ISL_945383, EPI_ISL_945384, EPI_ISL_945385, EPI_ISL_945388, EPI_ISL_945390, EPI_ISL_945391, EPI_ISL_945392, EPI_ISL_945393, EPI_ISL_945394, EPI_ISL_945395, EPI_ISL_945396, EPI_ISL_945397, EPI_ISL_945398, EPI_ISL_945399, EPI_ISL_945400, EPI_ISL_945401, EPI_ISL_945403, EPI_ISL_945404, EPI_ISL_945405, EPI_ISL_945406, EPI_ISL_945407, EPI_ISL_945410, EPI_ISL_945411, EPI_ISL_945412, EPI_ISL_945413, EPI_ISL_945414, EPI_ISL_945415, EPI_ISL_945416, EPI_ISL_945417, EPI_ISL_945418, EPI_ISL_945421, EPI_ISL_945424, EPI_ISL_945425, EPI_ISL_945426, EPI_ISL_945427, EPI_ISL_945428, EPI_ISL_945429, EPI_ISL_945430, EPI_ISL_945431, EPI_ISL_945432, EPI_ISL_945434, EPI_ISL_945435, EPI_ISL_945436, EPI_ISL_945437, EPI_ISL_945438, EPI_ISL_945439, EPI_ISL_945440, EPI_ISL_945441, EPI_ISL_945442, EPI_ISL_945443, EPI_ISL_945445, EPI_ISL_945447, EPI_ISL_945448, EPI_ISL_945450, EPI_ISL_945451, EPI_ISL_945452, EPI_ISL_945453, EPI_ISL_945454, EPI_ISL_945455, EPI_ISL_945456, EPI_ISL_945457, EPI_ISL_945460, EPI_ISL_945461, EPI_ISL_945462, EPI_ISL_945463, EPI_ISL_945464, EPI_ISL_945465, EPI_ISL_945470, EPI_ISL_945471, EPI_ISL_945472, EPI_ISL_945473, EPI_ISL_945474, EPI_ISL_945476, EPI_ISL_945477, EPI_ISL_945478, EPI_ISL_945479, EPI_ISL_945481, EPI_ISL_945482, EPI_ISL_945483, EPI_ISL_945484, EPI_ISL_945486, EPI_ISL_945487, EPI_ISL_945488, EPI_ISL_945489, EPI_ISL_945491, EPI_ISL_945492, EPI_ISL_945493, EPI_ISL_945496, EPI_ISL_945497, EPI_ISL_945499, EPI_ISL_945500, EPI_ISL_945501, EPI_ISL_945503, EPI_ISL_945504, EPI_ISL_945505, EPI_ISL_945507, EPI_ISL_945509, EPI_ISL_945510, EPI_ISL_945511, EPI_ISL_945512, EPI_ISL_945513, EPI_ISL_945515, EPI_ISL_945516, EPI_ISL_945517, EPI_ISL_945519, EPI_ISL_945520, EPI_ISL_945521, EPI_ISL_945523, EPI_ISL_945524, EPI_ISL_945526, EPI_ISL_945527, EPI_ISL_945528, EPI_ISL_945530, EPI_ISL_945531, EPI_ISL_945532, EPI_ISL_945533, EPI_ISL_945534, EPI_ISL_945535, EPI_ISL_945536, EPI_ISL_945537, EPI_ISL_945540, EPI_ISL_945541, EPI_ISL_945542, EPI_ISL_945544, EPI_ISL_945550, EPI_ISL_945551, EPI_ISL_945552, EPI_ISL_945553, EPI_ISL_945554, EPI_ISL_945556, EPI_ISL_945557, EPI_ISL_945558, EPI_ISL_945559, EPI_ISL_945560, EPI_ISL_945562, EPI_ISL_945564, EPI_ISL_945565, EPI_ISL_945566, EPI_ISL_945567, EPI_ISL_945568, EPI_ISL_945570, EPI_ISL_945571, EPI_ISL_945572, EPI_ISL_945573, EPI_ISL_945574, EPI_ISL_945575, EPI_ISL_945576, EPI_ISL_945577, EPI_ISL_945578, EPI_ISL_945580, EPI_ISL_945581, EPI_ISL_945583, EPI_ISL_945584, EPI_ISL_945585, EPI_ISL_945586, EPI_ISL_945588, EPI_ISL_945590, EPI_ISL_945592, EPI_ISL_945593, EPI_ISL_945594, EPI_ISL_945595, EPI_ISL_945596, EPI_ISL_945598, EPI_ISL_945599, EPI_ISL_945600, EPI_ISL_945601, EPI_ISL_945602, EPI_ISL_945603, EPI_ISL_945604, EPI_ISL_945605, EPI_ISL_945606, EPI_ISL_945607, EPI_ISL_945608, EPI_ISL_945609, EPI_ISL_945610, EPI_ISL_945611, EPI_ISL_945612, EPI_ISL_945613, EPI_ISL_945614, EPI_ISL_945615, EPI_ISL_945616, EPI_ISL_945617, EPI_ISL_945618, EPI_ISL_945620, EPI_ISL_945622, EPI_ISL_945623, EPI_ISL_945624, EPI_ISL_945625, EPI_ISL_945626, EPI_ISL_945628, EPI_ISL_945629, EPI_ISL_945630, EPI_ISL_945631, EPI_ISL_945632, EPI_ISL_945633, EPI_ISL_945634, EPI_ISL_945637, EPI_ISL_945638, EPI_ISL_945639, EPI_ISL_945640, EPI_ISL_945641, EPI_ISL_945642, EPI_ISL_945643, EPI_ISL_945644, EPI_ISL_945645, EPI_ISL_945646, EPI_ISL_945647, EPI_ISL_945649, EPI_ISL_945651, EPI_ISL_945652, EPI_ISL_945653, EPI_ISL_945654, EPI_ISL_945655, EPI_ISL_945657, EPI_ISL_945658, EPI_ISL_945659, EPI_ISL_945660, EPI_ISL_945661, EPI_ISL_945662, EPI_ISL_945663, EPI_ISL_945665, EPI_ISL_945666, EPI_ISL_945667, EPI_ISL_945669, EPI_ISL_945670, EPI_ISL_945671, EPI_ISL_945673, EPI_ISL_945675, EPI_ISL_945676, EPI_ISL_945679, EPI_ISL_945680, EPI_ISL_945683, EPI_ISL_945684, EPI_ISL_945685, EPI_ISL_945686, EPI_ISL_945687, EPI_ISL_945688, EPI_ISL_945689, EPI_ISL_945690, EPI_ISL_945691, EPI_ISL_945693, EPI_ISL_945694, EPI_ISL_945695, EPI_ISL_945699, EPI_ISL_945700, EPI_ISL_945702, EPI_ISL_945703, EPI_ISL_945705, EPI_ISL_945706, EPI_ISL_945707, EPI_ISL_945708, EPI_ISL_945709, EPI_ISL_945710, EPI_ISL_945711, EPI_ISL_945712, EPI_ISL_945713, EPI_ISL_945714 | | | |
| see above | Lighthouse Lab in Alderley Park | Wellcome Sanger Institute for the COVID-19 Genomics UK (COG-UK) Consortium | Jacquelyn Wynn, Mairead Hyland, The Lighthouse Lab in Alderley Park and Alex Alderton, Roberto Amato, Sonia Goncalves, Ewan Harrison, David K. Jackson, Ian Johnston, Dominic Kwiatkowski, Cordelia Langford, John Sillitoe on behalf of the Wellcome Sanger Institute COVID-19 Surveillance Team |
| EPI_ISL_945715, EPI_ISL_945717, EPI_ISL_945719, EPI_ISL_945720, EPI_ISL_945721, EPI_ISL_945723, EPI_ISL_945724, EPI_ISL_945727, EPI_ISL_945731, EPI_ISL_945734, EPI_ISL_945739, EPI_ISL_945740, EPI_ISL_945745, EPI_ISL_945746, EPI_ISL_945748, EPI_ISL_945749, EPI_ISL_945750, EPI_ISL_945751, EPI_ISL_945754, EPI_ISL_945756, EPI_ISL_945760 | | | |
| see above | Lighthouse Lab in Milton Keynes | Wellcome Sanger Institute for the COVID-19 Genomics UK (COG-UK) Consortium | The Lighthouse Lab in Milton Keynes and Alex Alderton, Roberto Amato, Sonia Goncalves, Ewan Harrison, David K. Jackson, Ian Johnston, Dominic Kwiatkowski, Cordelia Langford, John Sillitoe on behalf of the Wellcome Sanger Institute COVID-19 Surveillance Team |
| EPI_ISL_945761 | Lighthouse Lab in Glasgow | Wellcome Sanger Institute for the COVID-19 Genomics UK (COG-UK) Consortium | Harper VanSteenhouse, Yumi Kasai, David Gray, Carol Clugston, Anna Dominiczak and Alex Alderton, Roberto Amato, Sonia Goncalves, Ewan Harrison, David K. Jackson, Ian Johnston, Dominic Kwiatkowski, Cordelia Langford, John Sillitoe on behalf of the Wellcome Sanger Institute COVID-19 Surveillance Team |
| EPI_ISL_945764, EPI_ISL_945765, EPI_ISL_945766, EPI_ISL_945767, EPI_ISL_945769, EPI_ISL_945770, EPI_ISL_945773, EPI_ISL_945774, EPI_ISL_945775, EPI_ISL_945778, EPI_ISL_945779, EPI_ISL_945780, EPI_ISL_945783, EPI_ISL_945784, EPI_ISL_945785, EPI_ISL_945786, EPI_ISL_945787, EPI_ISL_945789, EPI_ISL_945790, EPI_ISL_945791, EPI_ISL_945793, EPI_ISL_945794, EPI_ISL_945797, EPI_ISL_945798, EPI_ISL_945801, EPI_ISL_945805, EPI_ISL_945810, EPI_ISL_945811, EPI_ISL_945814, EPI_ISL_945819, EPI_ISL_945820, EPI_ISL_945823, EPI_ISL_945825, EPI_ISL_945827, EPI_ISL_945829, EPI_ISL_945830, EPI_ISL_945832, EPI_ISL_945833, EPI_ISL_945834, EPI_ISL_945838, EPI_ISL_945840, EPI_ISL_945842, EPI_ISL_945845, EPI_ISL_945847, EPI_ISL_945852, EPI_ISL_945854, EPI_ISL_945855, EPI_ISL_945856, EPI_ISL_945857 | | | |
| see above | Lighthouse Lab in Milton Keynes | Wellcome Sanger Institute for the COVID-19 Genomics UK (COG-UK) Consortium | The Lighthouse Lab in Milton Keynes and Alex Alderton, Roberto Amato, Sonia Goncalves, Ewan Harrison, David K. Jackson, Ian Johnston, Dominic Kwiatkowski, Cordelia Langford, John Sillitoe on behalf of the Wellcome Sanger Institute COVID-19 Surveillance Team |
| EPI_ISL_945858 | Lighthouse Lab in Glasgow | Wellcome Sanger Institute for the COVID-19 Genomics UK (COG-UK) Consortium | Harper VanSteenhouse, Yumi Kasai, David Gray, Carol Clugston, Anna Dominiczak and Alex Alderton, Roberto Amato, Sonia Goncalves, Ewan Harrison, David K. Jackson, Ian Johnston, Dominic Kwiatkowski, Cordelia Langford, John Sillitoe on behalf of the Wellcome Sanger Institute COVID-19 Surveillance Team |
| EPI_ISL_945859, EPI_ISL_945861, EPI_ISL_945864, EPI_ISL_945866 | Lighthouse Lab in Milton Keynes | Wellcome Sanger Institute for the COVID-19 Genomics UK (COG-UK) Consortium | The Lighthouse Lab in Milton Keynes and Alex Alderton, Roberto Amato, Sonia Goncalves, Ewan Harrison, David K. Jackson, Ian Johnston, Dominic Kwiatkowski, Cordelia Langford, John Sillitoe on behalf of the Wellcome Sanger Institute COVID-19 Surveillance Team |
| EPI_ISL_945875 | Lighthouse Lab in Glasgow | Wellcome Sanger Institute for the COVID-19 Genomics UK (COG-UK) Consortium | Harper VanSteenhouse, Yumi Kasai, David Gray, Carol Clugston, Anna Dominiczak and Alex Alderton, Roberto Amato, Sonia Goncalves, Ewan Harrison, David K. Jackson, Ian Johnston, Dominic Kwiatkowski, Cordelia Langford, John Sillitoe on behalf of the Wellcome Sanger Institute COVID-19 Surveillance Team |
| EPI_ISL_945876, EPI_ISL_945880, EPI_ISL_945881, EPI_ISL_945883, EPI_ISL_945886, EPI_ISL_945887, EPI_ISL_945888, EPI_ISL_945890, EPI_ISL_945891, EPI_ISL_945892, EPI_ISL_945893, EPI_ISL_945894, EPI_ISL_945896 | | | |
| see above | Lighthouse Lab in Milton Keynes | Wellcome Sanger Institute for the COVID-19 Genomics UK (COG-UK) Consortium | The Lighthouse Lab in Milton Keynes and Alex Alderton, Roberto Amato, Sonia Goncalves, Ewan Harrison, David K. Jackson, Ian Johnston, Dominic Kwiatkowski, Cordelia Langford, John Sillitoe on behalf of the Wellcome Sanger Institute COVID-19 Surveillance Team |
| EPI_ISL_945897 | Lighthouse Lab in Glasgow | Wellcome Sanger Institute for the COVID-19 Genomics UK (COG-UK) Consortium | Harper VanSteenhouse, Yumi Kasai, David Gray, Carol Clugston, Anna Dominiczak and Alex Alderton, Roberto Amato, Sonia Goncalves, Ewan Harrison, David K. Jackson, Ian Johnston, Dominic Kwiatkowski, Cordelia Langford, John Sillitoe on behalf of the Wellcome Sanger Institute COVID-19 Surveillance Team |
| EPI_ISL_945902, EPI_ISL_945903, EPI_ISL_945905, EPI_ISL_945912, EPI_ISL_945913, EPI_ISL_945915, EPI_ISL_945918, EPI_ISL_945919, EPI_ISL_945922 | Lighthouse Lab in Milton Keynes | Wellcome Sanger Institute for the COVID-19 Genomics UK (COG-UK) Consortium | The Lighthouse Lab in Milton Keynes and Alex Alderton, Roberto Amato, Sonia Goncalves, Ewan Harrison, David K. Jackson, Ian Johnston, Dominic Kwiatkowski, Cordelia Langford, John Sillitoe on behalf of the Wellcome Sanger Institute COVID-19 Surveillance Team |
| EPI_ISL_945923 | Lighthouse Lab in Glasgow | Wellcome Sanger Institute for the COVID-19 Genomics UK (COG-UK) Consortium | Harper VanSteenhouse, Yumi Kasai, David Gray, Carol Clugston, Anna Dominiczak and Alex Alderton, Roberto Amato, Sonia Goncalves, Ewan Harrison, David K. Jackson, Ian Johnston, Dominic Kwiatkowski, Cordelia Langford, John Sillitoe on behalf of the Wellcome Sanger Institute COVID-19 Surveillance Team |
| EPI_ISL_945924, EPI_ISL_945925, EPI_ISL_945927, EPI_ISL_945928, EPI_ISL_945929, EPI_ISL_945930, EPI_ISL_945931, EPI_ISL_945932, EPI_ISL_945934, EPI_ISL_945936, EPI_ISL_945937, EPI_ISL_945941, EPI_ISL_945942, EPI_ISL_945943, EPI_ISL_945946, EPI_ISL_945947, EPI_ISL_945948, EPI_ISL_945949, EPI_ISL_945952, EPI_ISL_945953, EPI_ISL_945955, EPI_ISL_945957, EPI_ISL_945959, EPI_ISL_945964, EPI_ISL_945966, EPI_ISL_945968, EPI_ISL_945973, EPI_ISL_945974, EPI_ISL_945975 | | | |
| see above | Lighthouse Lab in Milton Keynes | Wellcome Sanger Institute for the COVID-19 Genomics UK (COG-UK) Consortium | The Lighthouse Lab in Milton Keynes and Alex Alderton, Roberto Amato, Sonia Goncalves, Ewan Harrison, David K. Jackson, Ian Johnston, Dominic Kwiatkowski, Cordelia Langford, John Sillitoe on behalf of the Wellcome Sanger Institute COVID-19 Surveillance Team |
| EPI_ISL_945977 | Lighthouse Lab in Glasgow | Wellcome Sanger Institute for the COVID-19 Genomics UK (COG-UK) Consortium | Harper VanSteenhouse, Yumi Kasai, David Gray, Carol Clugston, Anna Dominiczak and Alex Alderton, Roberto Amato, Sonia Goncalves, Ewan Harrison, David K. Jackson, Ian Johnston, Dominic Kwiatkowski, Cordelia Langford, John Sillitoe on behalf of the Wellcome Sanger Institute COVID-19 Surveillance Team |
| EPI_ISL_945978, EPI_ISL_945981, EPI_ISL_945982, EPI_ISL_945985, EPI_ISL_945986, EPI_ISL_945987, EPI_ISL_945989, EPI_ISL_945991, EPI_ISL_945992, EPI_ISL_945995, EPI_ISL_945996, EPI_ISL_946001 | | | |
| see above | Lighthouse Lab in Milton Keynes | Wellcome Sanger Institute for the COVID-19 Genomics UK (COG-UK) Consortium | The Lighthouse Lab in Milton Keynes and Alex Alderton, Roberto Amato, Sonia Goncalves, Ewan Harrison, David K. Jackson, Ian Johnston, Dominic Kwiatkowski, Cordelia Langford, John Sillitoe on behalf of the Wellcome Sanger Institute COVID-19 Surveillance Team |
| EPI_ISL_946003 | Lighthouse Lab in Glasgow | Wellcome Sanger Institute for the COVID-19 Genomics UK (COG-UK) Consortium | Harper VanSteenhouse, Yumi Kasai, David Gray, Carol Clugston, Anna Dominiczak and Alex Alderton, Roberto Amato, Sonia Goncalves, Ewan Harrison, David K. Jackson, Ian Johnston, Dominic Kwiatkowski, Cordelia Langford, John Sillitoe on behalf of the Wellcome Sanger Institute COVID-19 Surveillance Team |
| EPI_ISL_946004, EPI_ISL_946012, EPI_ISL_946013, EPI_ISL_946015, EPI_ISL_946016, EPI_ISL_946017, EPI_ISL_946018, EPI_ISL_946019, EPI_ISL_946020, EPI_ISL_946022, EPI_ISL_946024, EPI_ISL_946025, EPI_ISL_946027, EPI_ISL_946029, EPI_ISL_946031 | | | |
| see above | Lighthouse Lab in Milton Keynes | Wellcome Sanger Institute for the COVID-19 Genomics UK (COG-UK) Consortium | The Lighthouse Lab in Milton Keynes and Alex Alderton, Roberto Amato, Sonia Goncalves, Ewan Harrison, David K. Jackson, Ian Johnston, Dominic Kwiatkowski, Cordelia Langford, John Sillitoe on behalf of the Wellcome Sanger Institute COVID-19 Surveillance Team |
| EPI_ISL_946032 | Lighthouse Lab in Glasgow | Wellcome Sanger Institute for the COVID-19 Genomics UK (COG-UK) Consortium | Harper VanSteenhouse, Yumi Kasai, David Gray, Carol Clugston, Anna Dominiczak and Alex Alderton, Roberto Amato, Sonia Goncalves, Ewan Harrison, David K. Jackson, Ian Johnston, Dominic Kwiatkowski, Cordelia Langford, John Sillitoe on behalf of the Wellcome Sanger Institute COVID-19 Surveillance Team |
| EPI_ISL_946033, EPI_ISL_946034, EPI_ISL_946036, EPI_ISL_946038, EPI_ISL_946039, EPI_ISL_946040, EPI_ISL_946042, EPI_ISL_946043, EPI_ISL_946044, EPI_ISL_946045, EPI_ISL_946046 | | | |
| see above | Lighthouse Lab in Milton Keynes | Wellcome Sanger Institute for the COVID-19 Genomics UK (COG-UK) Consortium | The Lighthouse Lab in Milton Keynes and Alex Alderton, Roberto Amato, Sonia Goncalves, Ewan Harrison, David K. Jackson, Ian Johnston, Dominic Kwiatkowski, Cordelia Langford, John Sillitoe on behalf of the Wellcome Sanger Institute COVID-19 Surveillance Team |
| EPI_ISL_946047 | Lighthouse Lab in Glasgow | Wellcome Sanger Institute for the COVID-19 Genomics UK | Harper VanSteenhouse, Yumi Kasai, David Gray, Carol Clugston, Anna Dominiczak and Alex Alderton, Roberto Amato, Sonia Goncalves, Ewan Harrison, |

| | | | |
|---|---|---|---|
| | | (COG-UK) Consortium | David K. Jackson, Ian Johnston, Dominic Kwiatkowski, Cordelia Langford, John Sillitoe on behalf of the Wellcome Sanger Institute COVID-19 Surveillance Team |
| EPI_ISL_946050, EPI_ISL_946051, EPI_ISL_946053, EPI_ISL_946054, EPI_ISL_946056, EPI_ISL_946060, EPI_ISL_946061, EPI_ISL_946063, EPI_ISL_946064, EPI_ISL_946065, EPI_ISL_946074, EPI_ISL_946075, EPI_ISL_946077, EPI_ISL_946078, EPI_ISL_946079, EPI_ISL_946081, EPI_ISL_946082, EPI_ISL_946083, EPI_ISL_946084, EPI_ISL_946085, EPI_ISL_946089, EPI_ISL_946091, EPI_ISL_946092, EPI_ISL_946095, EPI_ISL_946097, EPI_ISL_946098, EPI_ISL_946099, EPI_ISL_946100, EPI_ISL_946101, EPI_ISL_946102, EPI_ISL_946103, EPI_ISL_946104, EPI_ISL_946105, EPI_ISL_946107, EPI_ISL_946108, EPI_ISL_946111, EPI_ISL_946114, EPI_ISL_946116 | | | |
| see above | Lighthouse Lab in Milton Keynes | Wellcome Sanger Institute for the COVID-19 Genomics UK (COG-UK) Consortium | The Lighthouse Lab in Milton Keynes and Alex Alderton, Roberto Amato, Sonia Goncalves, Ewan Harrison, David K. Jackson, Ian Johnston, Dominic Kwiatkowski, Cordelia Langford, John Sillitoe on behalf of the Wellcome Sanger Institute COVID-19 Surveillance Team |
| EPI_ISL_946117 | Lighthouse Lab in Glasgow | Wellcome Sanger Institute for the COVID-19 Genomics UK (COG-UK) Consortium | Harper VanSteenhouse, Yumi Kasai, David Gray, Carol Clugston, Anna Dominiczak and Alex Alderton, Roberto Amato, Sonia Goncalves, Ewan Harrison, David K. Jackson, Ian Johnston, Dominic Kwiatkowski, Cordelia Langford, John Sillitoe on behalf of the Wellcome Sanger Institute COVID-19 Surveillance Team |
| EPI_ISL_946118, EPI_ISL_946119, EPI_ISL_946120, EPI_ISL_946121, EPI_ISL_946122, EPI_ISL_946125, EPI_ISL_946128, EPI_ISL_946130, EPI_ISL_946131, EPI_ISL_946132, EPI_ISL_946133, EPI_ISL_946134, EPI_ISL_946136, EPI_ISL_946138, EPI_ISL_946139, EPI_ISL_946140, EPI_ISL_946141, EPI_ISL_946142, EPI_ISL_946143, EPI_ISL_946144, EPI_ISL_946146, EPI_ISL_946148, EPI_ISL_946151, EPI_ISL_946152, EPI_ISL_946154, EPI_ISL_946155, EPI_ISL_946156, EPI_ISL_946158, EPI_ISL_946159, EPI_ISL_946161, EPI_ISL_946163, EPI_ISL_946164, EPI_ISL_946165, EPI_ISL_946166, EPI_ISL_946167, EPI_ISL_946168, EPI_ISL_946170, EPI_ISL_946171, EPI_ISL_946172, EPI_ISL_946175, EPI_ISL_946178, EPI_ISL_946179, EPI_ISL_946183, EPI_ISL_946184, EPI_ISL_946185, EPI_ISL_946186, EPI_ISL_946187, EPI_ISL_946189, EPI_ISL_946191, EPI_ISL_946193, EPI_ISL_946194, EPI_ISL_946195, EPI_ISL_946197, EPI_ISL_946198, EPI_ISL_946200, EPI_ISL_946201, EPI_ISL_946203, EPI_ISL_946204, EPI_ISL_946205, EPI_ISL_946207, EPI_ISL_946209, EPI_ISL_946212, EPI_ISL_946214, EPI_ISL_946216, EPI_ISL_946217 | | | |
| see above | Lighthouse Lab in Milton Keynes | Wellcome Sanger Institute for the COVID-19 Genomics UK (COG-UK) Consortium | The Lighthouse Lab in Milton Keynes and Alex Alderton, Roberto Amato, Sonia Goncalves, Ewan Harrison, David K. Jackson, Ian Johnston, Dominic Kwiatkowski, Cordelia Langford, John Sillitoe on behalf of the Wellcome Sanger Institute COVID-19 Surveillance Team |
| EPI_ISL_946219 | Lighthouse Lab in Glasgow | Wellcome Sanger Institute for the COVID-19 Genomics UK (COG-UK) Consortium | Harper VanSteenhouse, Yumi Kasai, David Gray, Carol Clugston, Anna Dominiczak and Alex Alderton, Roberto Amato, Sonia Goncalves, Ewan Harrison, David K. Jackson, Ian Johnston, Dominic Kwiatkowski, Cordelia Langford, John Sillitoe on behalf of the Wellcome Sanger Institute COVID-19 Surveillance Team |
| EPI_ISL_946222, EPI_ISL_946223, EPI_ISL_946226, EPI_ISL_946228, EPI_ISL_946230, EPI_ISL_946235, EPI_ISL_946236, EPI_ISL_946237, EPI_ISL_946238, EPI_ISL_946239, EPI_ISL_946240, EPI_ISL_946241, EPI_ISL_946242, EPI_ISL_946244, EPI_ISL_946245, EPI_ISL_946246, EPI_ISL_946247, EPI_ISL_946248, EPI_ISL_946249, EPI_ISL_946251, EPI_ISL_946252, EPI_ISL_946255, EPI_ISL_946256, EPI_ISL_946257, EPI_ISL_946258 | | | |
| see above | Lighthouse Lab in Milton Keynes | Wellcome Sanger Institute for the COVID-19 Genomics UK (COG-UK) Consortium | The Lighthouse Lab in Milton Keynes and Alex Alderton, Roberto Amato, Sonia Goncalves, Ewan Harrison, David K. Jackson, Ian Johnston, Dominic Kwiatkowski, Cordelia Langford, John Sillitoe on behalf of the Wellcome Sanger Institute COVID-19 Surveillance Team |
| EPI_ISL_946259 | Lighthouse Lab in Glasgow | Wellcome Sanger Institute for the COVID-19 Genomics UK (COG-UK) Consortium | Harper VanSteenhouse, Yumi Kasai, David Gray, Carol Clugston, Anna Dominiczak and Alex Alderton, Roberto Amato, Sonia Goncalves, Ewan Harrison, David K. Jackson, Ian Johnston, Dominic Kwiatkowski, Cordelia Langford, John Sillitoe on behalf of the Wellcome Sanger Institute COVID-19 Surveillance Team |
| EPI_ISL_946261, EPI_ISL_946262, EPI_ISL_946263, EPI_ISL_946265, EPI_ISL_946266, EPI_ISL_946267, EPI_ISL_946268, EPI_ISL_946272, EPI_ISL_946273, EPI_ISL_946275, EPI_ISL_946276, EPI_ISL_946277, EPI_ISL_946278, EPI_ISL_946279, EPI_ISL_946280, EPI_ISL_946282, EPI_ISL_946283, EPI_ISL_946287, EPI_ISL_946288 | | | |
| see above | Lighthouse Lab in Milton Keynes | Wellcome Sanger Institute for the COVID-19 Genomics UK (COG-UK) Consortium | The Lighthouse Lab in Milton Keynes and Alex Alderton, Roberto Amato, Sonia Goncalves, Ewan Harrison, David K. Jackson, Ian Johnston, Dominic Kwiatkowski, Cordelia Langford, John Sillitoe on behalf of the Wellcome Sanger Institute COVID-19 Surveillance Team |
| EPI_ISL_946289, EPI_ISL_946290 | Lighthouse Lab in Glasgow | Wellcome Sanger Institute for the COVID-19 Genomics UK (COG-UK) Consortium | Harper VanSteenhouse, Yumi Kasai, David Gray, Carol Clugston, Anna Dominiczak and Alex Alderton, Roberto Amato, Sonia Goncalves, Ewan Harrison, David K. Jackson, Ian Johnston, Dominic Kwiatkowski, Cordelia Langford, John Sillitoe on behalf of the Wellcome Sanger Institute COVID-19 Surveillance Team |
| EPI_ISL_946291, EPI_ISL_946293, EPI_ISL_946294, EPI_ISL_946295, EPI_ISL_946296, EPI_ISL_946297, EPI_ISL_946299, EPI_ISL_946300 | Lighthouse Lab in Milton Keynes | Wellcome Sanger Institute for the COVID-19 Genomics UK (COG-UK) Consortium | The Lighthouse Lab in Milton Keynes and Alex Alderton, Roberto Amato, Sonia Goncalves, Ewan Harrison, David K. Jackson, Ian Johnston, Dominic Kwiatkowski, Cordelia Langford, John Sillitoe on behalf of the Wellcome Sanger Institute COVID-19 Surveillance Team |
| EPI_ISL_946303 | Lighthouse Lab in Glasgow | Wellcome Sanger Institute for the COVID-19 Genomics UK (COG-UK) Consortium | Harper VanSteenhouse, Yumi Kasai, David Gray, Carol Clugston, Anna Dominiczak and Alex Alderton, Roberto Amato, Sonia Goncalves, Ewan Harrison, David K. Jackson, Ian Johnston, Dominic Kwiatkowski, Cordelia Langford, John Sillitoe on behalf of the Wellcome Sanger Institute COVID-19 Surveillance Team |
| EPI_ISL_946305, EPI_ISL_946306, EPI_ISL_946307, EPI_ISL_946308, EPI_ISL_946309, EPI_ISL_946310, EPI_ISL_946311, EPI_ISL_946312, EPI_ISL_946313, EPI_ISL_946314, EPI_ISL_946315, EPI_ISL_946316, EPI_ISL_946317, EPI_ISL_946320, EPI_ISL_946321, EPI_ISL_946322, EPI_ISL_946324, EPI_ISL_946325, EPI_ISL_946327, EPI_ISL_946329, EPI_ISL_946330 | | | |
| see above | Lighthouse Lab in Milton Keynes | Wellcome Sanger Institute for the COVID-19 Genomics UK (COG-UK) Consortium | The Lighthouse Lab in Milton Keynes and Alex Alderton, Roberto Amato, Sonia Goncalves, Ewan Harrison, David K. Jackson, Ian Johnston, Dominic Kwiatkowski, Cordelia Langford, John Sillitoe on behalf of the Wellcome Sanger Institute COVID-19 Surveillance Team |
| EPI_ISL_946331 | Lighthouse Lab in Glasgow | Wellcome Sanger Institute for the COVID-19 Genomics UK (COG-UK) Consortium | Harper VanSteenhouse, Yumi Kasai, David Gray, Carol Clugston, Anna Dominiczak and Alex Alderton, Roberto Amato, Sonia Goncalves, Ewan Harrison, David K. Jackson, Ian Johnston, Dominic Kwiatkowski, Cordelia Langford, John Sillitoe on behalf of the Wellcome Sanger Institute COVID-19 Surveillance Team |
| EPI_ISL_946332, EPI_ISL_946334, EPI_ISL_946336, EPI_ISL_946337, EPI_ISL_946340, EPI_ISL_946341, EPI_ISL_946342, EPI_ISL_946343, EPI_ISL_946344, EPI_ISL_946345, EPI_ISL_946346, EPI_ISL_946347, EPI_ISL_946348, EPI_ISL_946349, EPI_ISL_946350, EPI_ISL_946352, EPI_ISL_946353, EPI_ISL_946354, EPI_ISL_946355, EPI_ISL_946358, EPI_ISL_946359, EPI_ISL_946362, EPI_ISL_946364, EPI_ISL_946368, EPI_ISL_946369, EPI_ISL_946370, EPI_ISL_946372 | | | |
| see above | Lighthouse Lab in Milton Keynes | Wellcome Sanger Institute for the COVID-19 Genomics UK (COG-UK) Consortium | The Lighthouse Lab in Milton Keynes and Alex Alderton, Roberto Amato, Sonia Goncalves, Ewan Harrison, David K. Jackson, Ian Johnston, Dominic Kwiatkowski, Cordelia Langford, John Sillitoe on behalf of the Wellcome Sanger Institute COVID-19 Surveillance Team |
| EPI_ISL_946373 | Lighthouse Lab in Glasgow | Wellcome Sanger Institute for the COVID-19 Genomics UK (COG-UK) Consortium | Harper VanSteenhouse, Yumi Kasai, David Gray, Carol Clugston, Anna Dominiczak and Alex Alderton, Roberto Amato, Sonia Goncalves, Ewan Harrison, David K. Jackson, Ian Johnston, Dominic Kwiatkowski, Cordelia Langford, John Sillitoe on behalf of the Wellcome Sanger Institute COVID-19 Surveillance Team |
| EPI_ISL_946376, EPI_ISL_946377, EPI_ISL_946382, EPI_ISL_946383, EPI_ISL_946384, EPI_ISL_946386 | Lighthouse Lab in Milton Keynes | Wellcome Sanger Institute for the COVID-19 Genomics UK (COG-UK) Consortium | The Lighthouse Lab in Milton Keynes and Alex Alderton, Roberto Amato, Sonia Goncalves, Ewan Harrison, David K. Jackson, Ian Johnston, Dominic Kwiatkowski, Cordelia Langford, John Sillitoe on behalf of the Wellcome Sanger Institute COVID-19 Surveillance Team |
| EPI_ISL_946387 | Lighthouse Lab in Glasgow | Wellcome Sanger Institute for the COVID-19 Genomics UK (COG-UK) Consortium | Harper VanSteenhouse, Yumi Kasai, David Gray, Carol Clugston, Anna Dominiczak and Alex Alderton, Roberto Amato, Sonia Goncalves, Ewan Harrison, David K. Jackson, Ian Johnston, Dominic Kwiatkowski, Cordelia Langford, John Sillitoe on behalf of the Wellcome Sanger Institute COVID-19 Surveillance Team |
| EPI_ISL_946388, EPI_ISL_946389, EPI_ISL_946390, EPI_ISL_946392, EPI_ISL_946393, EPI_ISL_946394 | Lighthouse Lab in Milton Keynes | Wellcome Sanger Institute for the COVID-19 Genomics UK (COG-UK) Consortium | The Lighthouse Lab in Milton Keynes and Alex Alderton, Roberto Amato, Sonia Goncalves, Ewan Harrison, David K. Jackson, Ian Johnston, Dominic Kwiatkowski, Cordelia Langford, John Sillitoe on behalf of the Wellcome Sanger Institute COVID-19 Surveillance Team |
| EPI_ISL_946395 | Lighthouse Lab in Glasgow | Wellcome Sanger Institute for the COVID-19 Genomics UK (COG-UK) Consortium | Harper VanSteenhouse, Yumi Kasai, David Gray, Carol Clugston, Anna Dominiczak and Alex Alderton, Roberto Amato, Sonia Goncalves, Ewan Harrison, David K. Jackson, Ian Johnston, Dominic Kwiatkowski, Cordelia Langford, John Sillitoe on behalf of the Wellcome Sanger Institute COVID-19 Surveillance Team |
| EPI_ISL_946396 | Lighthouse Lab in Milton Keynes | Wellcome Sanger Institute for the COVID-19 Genomics UK (COG-UK) Consortium | The Lighthouse Lab in Milton Keynes and Alex Alderton, Roberto Amato, Sonia Goncalves, Ewan Harrison, David K. Jackson, Ian Johnston, Dominic Kwiatkowski, Cordelia Langford, John Sillitoe on behalf of the Wellcome Sanger Institute COVID-19 Surveillance Team |
| EPI_ISL_946398 | Lighthouse Lab in Glasgow | Wellcome Sanger Institute for the COVID-19 Genomics UK (COG-UK) Consortium | Harper VanSteenhouse, Yumi Kasai, David Gray, Carol Clugston, Anna Dominiczak and Alex Alderton, Roberto Amato, Sonia Goncalves, Ewan Harrison, David K. Jackson, Ian Johnston, Dominic Kwiatkowski, Cordelia Langford, John Sillitoe on behalf of the Wellcome Sanger Institute COVID-19 Surveillance Team |
| EPI_ISL_946399, EPI_ISL_946401, EPI_ISL_946402, EPI_ISL_946404, EPI_ISL_946406 | Lighthouse Lab in Milton Keynes | Wellcome Sanger Institute for the COVID-19 Genomics UK (COG-UK) Consortium | The Lighthouse Lab in Milton Keynes and Alex Alderton, Roberto Amato, Sonia Goncalves, Ewan Harrison, David K. Jackson, Ian Johnston, Dominic Kwiatkowski, Cordelia Langford, John Sillitoe on behalf of the Wellcome Sanger Institute COVID-19 Surveillance Team |

| | | | |
|---|---|---|---|
| EPI_ISL_946408 | Lighthouse Lab in Glasgow | Wellcome Sanger Institute for the COVID-19 Genomics UK (COG-UK) Consortium | Harper VanSteenhouse, Yumi Kasai, David Gray, Carol Clugston, Anna Dominiczak and Alex Alderton, Roberto Amato, Sonia Goncalves, Ewan Harrison, David K. Jackson, Ian Johnston, Dominic Kwiatkowski, Cordelia Langford, John Sillitoe on behalf of the Wellcome Sanger Institute COVID-19 Surveillance Team |
| EPI_ISL_946409, EPI_ISL_946412, EPI_ISL_946413, EPI_ISL_946414, EPI_ISL_946415, EPI_ISL_946416, EPI_ISL_946417, EPI_ISL_946418 | Lighthouse Lab in Milton Keynes | Wellcome Sanger Institute for the COVID-19 Genomics UK (COG-UK) Consortium | The Lighthouse Lab in Milton Keynes and Alex Alderton, Roberto Amato, Sonia Goncalves, Ewan Harrison, David K. Jackson, Ian Johnston, Dominic Kwiatkowski, Cordelia Langford, John Sillitoe on behalf of the Wellcome Sanger Institute COVID-19 Surveillance Team |
| EPI_ISL_946419 | Lighthouse Lab in Glasgow | Wellcome Sanger Institute for the COVID-19 Genomics UK (COG-UK) Consortium | Harper VanSteenhouse, Yumi Kasai, David Gray, Carol Clugston, Anna Dominiczak and Alex Alderton, Roberto Amato, Sonia Goncalves, Ewan Harrison, David K. Jackson, Ian Johnston, Dominic Kwiatkowski, Cordelia Langford, John Sillitoe on behalf of the Wellcome Sanger Institute COVID-19 Surveillance Team |
| EPI_ISL_946420 | Lighthouse Lab in Milton Keynes | Wellcome Sanger Institute for the COVID-19 Genomics UK (COG-UK) Consortium | The Lighthouse Lab in Milton Keynes and Alex Alderton, Roberto Amato, Sonia Goncalves, Ewan Harrison, David K. Jackson, Ian Johnston, Dominic Kwiatkowski, Cordelia Langford, John Sillitoe on behalf of the Wellcome Sanger Institute COVID-19 Surveillance Team |
| EPI_ISL_946422, EPI_ISL_946423, EPI_ISL_946424, EPI_ISL_946425, EPI_ISL_946426, EPI_ISL_946428, EPI_ISL_946429, EPI_ISL_946430, EPI_ISL_946431, EPI_ISL_946432, EPI_ISL_946433, EPI_ISL_946434, EPI_ISL_946435, EPI_ISL_946436, EPI_ISL_946437, EPI_ISL_946438, EPI_ISL_946439, EPI_ISL_946440,<br>EPI_ISL_946441, EPI_ISL_946442, EPI_ISL_946443, EPI_ISL_946444, EPI_ISL_946446, EPI_ISL_946447, EPI_ISL_946448, EPI_ISL_946449, EPI_ISL_946450, EPI_ISL_946451, EPI_ISL_946452, EPI_ISL_946453, EPI_ISL_946455, EPI_ISL_946456, EPI_ISL_946457, EPI_ISL_946458, EPI_ISL_946459, EPI_ISL_946460,<br>EPI_ISL_946461, EPI_ISL_946462, EPI_ISL_946463, EPI_ISL_946464, EPI_ISL_946465, EPI_ISL_946466, EPI_ISL_946467, EPI_ISL_946468, EPI_ISL_946469, EPI_ISL_946470, EPI_ISL_946472, EPI_ISL_946473, EPI_ISL_946474, EPI_ISL_946475, EPI_ISL_946476, EPI_ISL_946477, EPI_ISL_946478, EPI_ISL_946479,<br>EPI_ISL_946480, EPI_ISL_946481, EPI_ISL_946482, EPI_ISL_946483, EPI_ISL_946484, EPI_ISL_946485, EPI_ISL_946486, EPI_ISL_946487, EPI_ISL_946488, EPI_ISL_946489, EPI_ISL_946490, EPI_ISL_946491, EPI_ISL_946492, EPI_ISL_946494, EPI_ISL_946495, EPI_ISL_946496, EPI_ISL_946497, EPI_ISL_946498,<br>EPI_ISL_946499, EPI_ISL_946500, EPI_ISL_946501, EPI_ISL_946502, EPI_ISL_946503, EPI_ISL_946504, EPI_ISL_946505, EPI_ISL_946506, EPI_ISL_946507, EPI_ISL_946508, EPI_ISL_946509, EPI_ISL_946510, EPI_ISL_946511, EPI_ISL_946512, EPI_ISL_946513, EPI_ISL_946514, EPI_ISL_946515, EPI_ISL_946516,<br>EPI_ISL_946517, EPI_ISL_946518, EPI_ISL_946519, EPI_ISL_946520, EPI_ISL_946521, EPI_ISL_946522, EPI_ISL_946523, EPI_ISL_946524, EPI_ISL_946525, EPI_ISL_946526, EPI_ISL_946527, EPI_ISL_946528, EPI_ISL_946529, EPI_ISL_946530, EPI_ISL_946531, EPI_ISL_946532, EPI_ISL_946533, EPI_ISL_946534,<br>EPI_ISL_946535, EPI_ISL_946536, EPI_ISL_946537, EPI_ISL_946538, EPI_ISL_946539, EPI_ISL_946540, EPI_ISL_946541, EPI_ISL_946542, EPI_ISL_946543, EPI_ISL_946544, EPI_ISL_946545, EPI_ISL_946546, EPI_ISL_946548, EPI_ISL_946549, EPI_ISL_946550, EPI_ISL_946551, EPI_ISL_946552, EPI_ISL_946553,<br>EPI_ISL_946554, EPI_ISL_946555, EPI_ISL_946556, EPI_ISL_946557, EPI_ISL_946558, EPI_ISL_946559, EPI_ISL_946560, EPI_ISL_946561, EPI_ISL_946562, EPI_ISL_946563, EPI_ISL_946564, EPI_ISL_946565, EPI_ISL_946566, EPI_ISL_946567, EPI_ISL_946568, EPI_ISL_946569, EPI_ISL_946570, EPI_ISL_946571,<br>EPI_ISL_946572, EPI_ISL_946573, EPI_ISL_946574, EPI_ISL_946575, EPI_ISL_946576, EPI_ISL_946577, EPI_ISL_946578, EPI_ISL_946579, EPI_ISL_946581, EPI_ISL_946582, EPI_ISL_946583, EPI_ISL_946584, EPI_ISL_946586, EPI_ISL_946587, EPI_ISL_946588, EPI_ISL_946589, EPI_ISL_946590, EPI_ISL_946591,<br>EPI_ISL_946592, EPI_ISL_946593, EPI_ISL_946594, EPI_ISL_946595, EPI_ISL_946596, EPI_ISL_946597, EPI_ISL_946598, EPI_ISL_946599, EPI_ISL_946600, EPI_ISL_946601, EPI_ISL_946602, EPI_ISL_946603, EPI_ISL_946604, EPI_ISL_946605, EPI_ISL_946606, EPI_ISL_946607, EPI_ISL_946608, EPI_ISL_946609,<br>EPI_ISL_946610, EPI_ISL_946611, EPI_ISL_946612, EPI_ISL_946613, EPI_ISL_946614, EPI_ISL_946615, EPI_ISL_946616, EPI_ISL_946617, EPI_ISL_946618, EPI_ISL_946619, EPI_ISL_946620, EPI_ISL_946621, EPI_ISL_946622, EPI_ISL_946623, EPI_ISL_946624, EPI_ISL_946625, EPI_ISL_946626, EPI_ISL_946627,<br>EPI_ISL_946628, EPI_ISL_946629, EPI_ISL_946630, EPI_ISL_946631, EPI_ISL_946632, EPI_ISL_946633, EPI_ISL_946634, EPI_ISL_946635, EPI_ISL_946636, EPI_ISL_946637, EPI_ISL_946638, EPI_ISL_946639, EPI_ISL_946640, EPI_ISL_946641, EPI_ISL_946642, EPI_ISL_946643, EPI_ISL_946644, EPI_ISL_946645,<br>EPI_ISL_946647, EPI_ISL_946648, EPI_ISL_946649, EPI_ISL_946650, EPI_ISL_946651, EPI_ISL_946652, EPI_ISL_946653, EPI_ISL_946654, EPI_ISL_946655, EPI_ISL_946656, EPI_ISL_946657, EPI_ISL_946658, EPI_ISL_946659, EPI_ISL_946660, EPI_ISL_946661, EPI_ISL_946662, EPI_ISL_946663, EPI_ISL_946664,<br>EPI_ISL_946665, EPI_ISL_946666, EPI_ISL_946667, EPI_ISL_946668, EPI_ISL_946669, EPI_ISL_946670, EPI_ISL_946671, EPI_ISL_946673, EPI_ISL_946674, EPI_ISL_946675, EPI_ISL_946676, EPI_ISL_946677, EPI_ISL_946678, EPI_ISL_946679, EPI_ISL_946680, EPI_ISL_946681, EPI_ISL_946682, EPI_ISL_946683,<br>EPI_ISL_946684, EPI_ISL_946685, EPI_ISL_946686, EPI_ISL_946687, EPI_ISL_946688, EPI_ISL_946689, EPI_ISL_946690, EPI_ISL_946691, EPI_ISL_946692, EPI_ISL_946693, EPI_ISL_946694, EPI_ISL_946695, EPI_ISL_946696, EPI_ISL_946697, EPI_ISL_946698, EPI_ISL_946699, EPI_ISL_946700, EPI_ISL_946702,<br>EPI_ISL_946703, EPI_ISL_946704, EPI_ISL_946705, EPI_ISL_946706, EPI_ISL_946707, EPI_ISL_946708, EPI_ISL_946709, EPI_ISL_946710, EPI_ISL_946711, EPI_ISL_946712, EPI_ISL_946713, EPI_ISL_946714, EPI_ISL_946715, EPI_ISL_946716, EPI_ISL_946717, EPI_ISL_946718, EPI_ISL_946719, EPI_ISL_946720,<br>EPI_ISL_946721, EPI_ISL_946722, EPI_ISL_946723, EPI_ISL_946724, EPI_ISL_946725, EPI_ISL_946726, EPI_ISL_946727, EPI_ISL_946728, EPI_ISL_946729, EPI_ISL_946730, EPI_ISL_946731, EPI_ISL_946732, EPI_ISL_946733, EPI_ISL_946734, EPI_ISL_946735, EPI_ISL_946736, EPI_ISL_946737, EPI_ISL_946738,<br>EPI_ISL_946739, EPI_ISL_946740, EPI_ISL_946741, EPI_ISL_946742, EPI_ISL_946743, EPI_ISL_946744, EPI_ISL_946745, EPI_ISL_946747, EPI_ISL_946748, EPI_ISL_946749, EPI_ISL_946750, EPI_ISL_946751, EPI_ISL_946752, EPI_ISL_946753, EPI_ISL_946754, EPI_ISL_946755, EPI_ISL_946756, EPI_ISL_946757,<br>EPI_ISL_946758, EPI_ISL_946759, EPI_ISL_946760, EPI_ISL_946761, EPI_ISL_946762, EPI_ISL_946763, EPI_ISL_946764, EPI_ISL_946765, EPI_ISL_946766, EPI_ISL_946767, EPI_ISL_946768, EPI_ISL_946769, EPI_ISL_946770, EPI_ISL_946771, EPI_ISL_946772, EPI_ISL_946773, EPI_ISL_946774, EPI_ISL_946775,<br>EPI_ISL_946776, EPI_ISL_946777, EPI_ISL_946778, EPI_ISL_946779, EPI_ISL_946781, EPI_ISL_946782, EPI_ISL_946783, EPI_ISL_946784, EPI_ISL_946785, EPI_ISL_946786, EPI_ISL_946787, EPI_ISL_946788, EPI_ISL_946789, EPI_ISL_946790, EPI_ISL_946791, EPI_ISL_946792, EPI_ISL_946793, EPI_ISL_946794,<br>EPI_ISL_946795, EPI_ISL_946796, EPI_ISL_946797, EPI_ISL_946798, EPI_ISL_946799, EPI_ISL_946800, EPI_ISL_946801, EPI_ISL_946802, EPI_ISL_946803, EPI_ISL_946804, EPI_ISL_946805, EPI_ISL_946806, EPI_ISL_946807, EPI_ISL_946808, EPI_ISL_946809, EPI_ISL_946810, EPI_ISL_946811, EPI_ISL_946812,<br>EPI_ISL_946813, EPI_ISL_946814, EPI_ISL_946815, EPI_ISL_946816, EPI_ISL_946817, EPI_ISL_946818, EPI_ISL_946819, EPI_ISL_946820, EPI_ISL_946821, EPI_ISL_946822, EPI_ISL_946823, EPI_ISL_946824, EPI_ISL_946825, EPI_ISL_946826, EPI_ISL_946827, EPI_ISL_946828, EPI_ISL_946829, EPI_ISL_946830,<br>EPI_ISL_946831, EPI_ISL_946832, EPI_ISL_946833, EPI_ISL_946834, EPI_ISL_946835, EPI_ISL_946836, EPI_ISL_946837, EPI_ISL_946838, EPI_ISL_946839, EPI_ISL_946840, EPI_ISL_946841, EPI_ISL_946842, EPI_ISL_946843, EPI_ISL_946844, EPI_ISL_946845, EPI_ISL_946846, EPI_ISL_946847, EPI_ISL_946848,<br>EPI_ISL_946849, EPI_ISL_946850, EPI_ISL_946851, EPI_ISL_946852, EPI_ISL_946853, EPI_ISL_946854, EPI_ISL_946855, EPI_ISL_946856, EPI_ISL_946857, EPI_ISL_946858, EPI_ISL_946859, EPI_ISL_946860, EPI_ISL_946861, EPI_ISL_946862, EPI_ISL_946863, EPI_ISL_946864, EPI_ISL_946865, EPI_ISL_946866,<br>EPI_ISL_946867, EPI_ISL_946868, EPI_ISL_946869, EPI_ISL_946870, EPI_ISL_946871, EPI_ISL_946872, EPI_ISL_946873, EPI_ISL_946874, EPI_ISL_946875, EPI_ISL_946876, EPI_ISL_946877, EPI_ISL_946878, EPI_ISL_946879, EPI_ISL_946880, EPI_ISL_946881, EPI_ISL_946882, EPI_ISL_946883, EPI_ISL_946884,<br>EPI_ISL_946885, EPI_ISL_946886, EPI_ISL_946887, EPI_ISL_946888, EPI_ISL_946889, EPI_ISL_946890, EPI_ISL_946891, EPI_ISL_946892, EPI_ISL_946893, EPI_ISL_946894, EPI_ISL_946895, EPI_ISL_946896, EPI_ISL_946897, EPI_ISL_946898, EPI_ISL_946899, EPI_ISL_946900, EPI_ISL_946901, EPI_ISL_946902,<br>EPI_ISL_946903, EPI_ISL_946904, EPI_ISL_946905, EPI_ISL_946906, EPI_ISL_946907, EPI_ISL_946908, EPI_ISL_946909, EPI_ISL_946910, EPI_ISL_946911, EPI_ISL_946912, EPI_ISL_946914, EPI_ISL_946915, EPI_ISL_946916, EPI_ISL_946917, EPI_ISL_946918, EPI_ISL_946919, EPI_ISL_946920, EPI_ISL_946921,<br>EPI_ISL_946922, EPI_ISL_946923, EPI_ISL_946924, EPI_ISL_946925, EPI_ISL_946926, EPI_ISL_946927, EPI_ISL_946928, EPI_ISL_946929, EPI_ISL_946930, EPI_ISL_946931, EPI_ISL_946932, EPI_ISL_946933, EPI_ISL_946934, EPI_ISL_946935, EPI_ISL_946936, EPI_ISL_946937, EPI_ISL_946938, EPI_ISL_946939,<br>EPI_ISL_946940, EPI_ISL_946941, EPI_ISL_946942, EPI_ISL_946943, EPI_ISL_946944, EPI_ISL_946945, EPI_ISL_946946, EPI_ISL_946947, EPI_ISL_946948, EPI_ISL_946949, EPI_ISL_946950, EPI_ISL_946951, EPI_ISL_946952, EPI_ISL_946953, EPI_ISL_946954, EPI_ISL_946955, EPI_ISL_946957, EPI_ISL_946958,<br>EPI_ISL_946959, EPI_ISL_946960, EPI_ISL_946961, EPI_ISL_946962, EPI_ISL_946964, EPI_ISL_946965, EPI_ISL_946966, EPI_ISL_946967, EPI_ISL_946968, EPI_ISL_946969, EPI_ISL_946970, EPI_ISL_946971, EPI_ISL_946972, EPI_ISL_946973, EPI_ISL_946974, EPI_ISL_946975, EPI_ISL_946976, EPI_ISL_946977,<br>EPI_ISL_946978, EPI_ISL_946979, EPI_ISL_946980, EPI_ISL_946981, EPI_ISL_946982, EPI_ISL_946983, EPI_ISL_946984, EPI_ISL_946985, EPI_ISL_946986, EPI_ISL_946987, EPI_ISL_946988, EPI_ISL_946989, EPI_ISL_946990, EPI_ISL_946991, EPI_ISL_946993, EPI_ISL_946994, EPI_ISL_946995, EPI_ISL_946996,<br>EPI_ISL_946997, EPI_ISL_946998, EPI_ISL_946999, EPI_ISL_947000, EPI_ISL_947001, EPI_ISL_947002, EPI_ISL_947003, EPI_ISL_947004, EPI_ISL_947005, EPI_ISL_947006, EPI_ISL_947007, EPI_ISL_947008, EPI_ISL_947009, EPI_ISL_947010, EPI_ISL_947011, EPI_ISL_947012, EPI_ISL_947013, EPI_ISL_947014,<br>EPI_ISL_947015, EPI_ISL_947016, EPI_ISL_947017, EPI_ISL_947018, EPI_ISL_947019, EPI_ISL_947020, EPI_ISL_947021, EPI_ISL_947022, EPI_ISL_947023, EPI_ISL_947024, EPI_ISL_947025, EPI_ISL_947026, EPI_ISL_947027, EPI_ISL_947028, EPI_ISL_947029, EPI_ISL_947030, EPI_ISL_947031, EPI_ISL_947032,<br>EPI_ISL_947033, EPI_ISL_947034, EPI_ISL_947035, EPI_ISL_947036, EPI_ISL_947037, EPI_ISL_947038, EPI_ISL_947039, EPI_ISL_947040, EPI_ISL_947041, EPI_ISL_947042, EPI_ISL_947043, EPI_ISL_947044, EPI_ISL_947045, EPI_ISL_947046, EPI_ISL_947047, EPI_ISL_947048, EPI_ISL_947049, EPI_ISL_947050,<br>EPI_ISL_947051, EPI_ISL_947052, EPI_ISL_947053, EPI_ISL_947054, EPI_ISL_947055, EPI_ISL_947056, EPI_ISL_947057, EPI_ISL_947058, EPI_ISL_947059, EPI_ISL_947060, EPI_ISL_947061, EPI_ISL_947062, EPI_ISL_947063, EPI_ISL_947064, EPI_ISL_947065, EPI_ISL_947066, EPI_ISL_947067, EPI_ISL_947068,<br>EPI_ISL_947069, EPI_ISL_947070, EPI_ISL_947071, EPI_ISL_947072, EPI_ISL_947073, EPI_ISL_947074, EPI_ISL_947075, EPI_ISL_947076, EPI_ISL_947077, EPI_ISL_947078, EPI_ISL_947079, EPI_ISL_947080, EPI_ISL_947081, EPI_ISL_947082, EPI_ISL_947083, EPI_ISL_947084, EPI_ISL_947085, EPI_ISL_947086,<br>EPI_ISL_947087, EPI_ISL_947088, EPI_ISL_947089, EPI_ISL_947090, EPI_ISL_947091, EPI_ISL_947092, EPI_ISL_947093, EPI_ISL_947094, EPI_ISL_947095, EPI_ISL_947096, EPI_ISL_947097, EPI_ISL_947098, EPI_ISL_947099, EPI_ISL_947100, EPI_ISL_947101, EPI_ISL_947102, EPI_ISL_947103, EPI_ISL_947104,<br>EPI_ISL_947105, EPI_ISL_947106 | | | |
| see above | Lighthouse Lab in Alderley Park | Wellcome Sanger Institute for the COVID-19 Genomics UK (COG-UK) Consortium | Jacquelyn Wynn, Mairead Hyland, The Lighthouse Lab in Alderley Park and Alex Alderton, Roberto Amato, Sonia Goncalves, Ewan Harrison, David K. Jackson, Ian Johnston, Dominic Kwiatkowski, Cordelia Langford, John Sillitoe on behalf of the Wellcome Sanger Institute COVID-19 Surveillance Team |
| EPI_ISL_947107, EPI_ISL_947108 | Lighthouse Lab in Glasgow | Wellcome Sanger Institute for the COVID-19 Genomics UK (COG-UK) Consortium | Harper VanSteenhouse, Yumi Kasai, David Gray, Carol Clugston, Anna Dominiczak and Alex Alderton, Roberto Amato, Sonia Goncalves, Ewan Harrison, David K. Jackson, Ian Johnston, Dominic Kwiatkowski, Cordelia Langford, John Sillitoe on behalf of the Wellcome Sanger Institute COVID-19 Surveillance Team |
| EPI_ISL_947109, EPI_ISL_947111, EPI_ISL_947112, EPI_ISL_947114, EPI_ISL_947115, EPI_ISL_947116, EPI_ISL_947117 | Lighthouse Lab in Milton Keynes | Wellcome Sanger Institute for the COVID-19 Genomics UK (COG-UK) Consortium | The Lighthouse Lab in Milton Keynes and Alex Alderton, Roberto Amato, Sonia Goncalves, Ewan Harrison, David K. Jackson, Ian Johnston, Dominic Kwiatkowski, Cordelia Langford, John Sillitoe on behalf of the Wellcome Sanger Institute COVID-19 Surveillance Team |
| EPI_ISL_947118, EPI_ISL_947119, EPI_ISL_947121 | Lighthouse Lab in Glasgow | Wellcome Sanger Institute for the COVID-19 Genomics UK (COG-UK) Consortium | Harper VanSteenhouse, Yumi Kasai, David Gray, Carol Clugston, Anna Dominiczak and Alex Alderton, Roberto Amato, Sonia Goncalves, Ewan Harrison, David K. Jackson, Ian Johnston, Dominic Kwiatkowski, Cordelia Langford, John Sillitoe on behalf of the Wellcome Sanger Institute COVID-19 Surveillance Team |
| EPI_ISL_947122, EPI_ISL_947123, EPI_ISL_947124, EPI_ISL_947125 | Lighthouse Lab in Milton Keynes | Wellcome Sanger Institute for the COVID-19 Genomics UK (COG-UK) Consortium | The Lighthouse Lab in Milton Keynes and Alex Alderton, Roberto Amato, Sonia Goncalves, Ewan Harrison, David K. Jackson, Ian Johnston, Dominic Kwiatkowski, Cordelia Langford, John Sillitoe on behalf of the Wellcome Sanger Institute COVID-19 Surveillance Team |
| EPI_ISL_947126, EPI_ISL_947127 | Lighthouse Lab in Glasgow | Wellcome Sanger Institute for the COVID-19 Genomics UK (COG-UK) Consortium | Harper VanSteenhouse, Yumi Kasai, David Gray, Carol Clugston, Anna Dominiczak and Alex Alderton, Roberto Amato, Sonia Goncalves, Ewan Harrison, David K. Jackson, Ian Johnston, Dominic Kwiatkowski, Cordelia Langford, John Sillitoe on behalf of the Wellcome Sanger Institute COVID-19 Surveillance Team |
| EPI_ISL_947128, EPI_ISL_947129, EPI_ISL_947130 | Lighthouse Lab in Milton Keynes | Wellcome Sanger Institute for the COVID-19 Genomics UK (COG-UK) Consortium | The Lighthouse Lab in Milton Keynes and Alex Alderton, Roberto Amato, Sonia Goncalves, Ewan Harrison, David K. Jackson, Ian Johnston, Dominic Kwiatkowski, Cordelia Langford, John Sillitoe on behalf of the Wellcome Sanger Institute COVID-19 Surveillance Team |
| EPI_ISL_947131 | Lighthouse Lab in Glasgow | Wellcome Sanger Institute for the COVID-19 Genomics UK (COG-UK) Consortium | Harper VanSteenhouse, Yumi Kasai, David Gray, Carol Clugston, Anna Dominiczak and Alex Alderton, Roberto Amato, Sonia Goncalves, Ewan Harrison, David K. Jackson, Ian Johnston, Dominic Kwiatkowski, Cordelia Langford, John Sillitoe on behalf of the Wellcome Sanger Institute COVID-19 Surveillance Team |
| EPI_ISL_947132, EPI_ISL_947133 | Lighthouse Lab in Milton Keynes | Wellcome Sanger Institute for the COVID-19 Genomics UK (COG-UK) Consortium | The Lighthouse Lab in Milton Keynes and Alex Alderton, Roberto Amato, Sonia Goncalves, Ewan Harrison, David K. Jackson, Ian Johnston, Dominic Kwiatkowski, Cordelia Langford, John Sillitoe on behalf of the Wellcome Sanger Institute COVID-19 Surveillance Team |
| EPI_ISL_947134 | Lighthouse Lab in Glasgow | Wellcome Sanger Institute for the COVID-19 Genomics UK (COG-UK) Consortium | Harper VanSteenhouse, Yumi Kasai, David Gray, Carol Clugston, Anna Dominiczak and Alex Alderton, Roberto Amato, Sonia Goncalves, Ewan Harrison, David K. Jackson, Ian Johnston, Dominic Kwiatkowski, Cordelia Langford, John Sillitoe on behalf of the Wellcome Sanger Institute COVID-19 Surveillance Team |
| EPI_ISL_947135, EPI_ISL_947136 | Lighthouse Lab in Milton Keynes | Wellcome Sanger Institute for the COVID-19 Genomics UK (COG-UK) Consortium | The Lighthouse Lab in Milton Keynes and Alex Alderton, Roberto Amato, Sonia Goncalves, Ewan Harrison, David K. Jackson, Ian Johnston, Dominic Kwiatkowski, Cordelia Langford, John Sillitoe on behalf of the Wellcome Sanger Institute COVID-19 Surveillance Team |

| | | | |
|---|---|---|---|
| EPI_ISL_947137 | Lighthouse Lab in Glasgow | Wellcome Sanger Institute for the COVID-19 Genomics UK (COG-UK) Consortium | Harper VanSteenhouse, Yumi Kasai, David Gray, Carol Clugston, Anna Dominiczak and Alex Alderton, Roberto Amato, Sonia Goncalves, Ewan Harrison, David K. Jackson, Ian Johnston, Dominic Kwiatkowski, Cordelia Langford, John Sillitoe on behalf of the Wellcome Sanger Institute COVID-19 Surveillance Team |
| EPI_ISL_947138 | Lighthouse Lab in Milton Keynes | Wellcome Sanger Institute for the COVID-19 Genomics UK (COG-UK) Consortium | The Lighthouse Lab in Milton Keynes and Alex Alderton, Roberto Amato, Sonia Goncalves, Ewan Harrison, David K. Jackson, Ian Johnston, Dominic Kwiatkowski, Cordelia Langford, John Sillitoe on behalf of the Wellcome Sanger Institute COVID-19 Surveillance Team |
| EPI_ISL_947139, EPI_ISL_947141 | Lighthouse Lab in Glasgow | Wellcome Sanger Institute for the COVID-19 Genomics UK (COG-UK) Consortium | Harper VanSteenhouse, Yumi Kasai, David Gray, Carol Clugston, Anna Dominiczak and Alex Alderton, Roberto Amato, Sonia Goncalves, Ewan Harrison, David K. Jackson, Ian Johnston, Dominic Kwiatkowski, Cordelia Langford, John Sillitoe on behalf of the Wellcome Sanger Institute COVID-19 Surveillance Team |
| EPI_ISL_947142, EPI_ISL_947143, EPI_ISL_947144, EPI_ISL_947145, EPI_ISL_947146 | Lighthouse Lab in Milton Keynes | Wellcome Sanger Institute for the COVID-19 Genomics UK (COG-UK) Consortium | The Lighthouse Lab in Milton Keynes and Alex Alderton, Roberto Amato, Sonia Goncalves, Ewan Harrison, David K. Jackson, Ian Johnston, Dominic Kwiatkowski, Cordelia Langford, John Sillitoe on behalf of the Wellcome Sanger Institute COVID-19 Surveillance Team |
| EPI_ISL_947147 | Lighthouse Lab in Glasgow | Wellcome Sanger Institute for the COVID-19 Genomics UK (COG-UK) Consortium | Harper VanSteenhouse, Yumi Kasai, David Gray, Carol Clugston, Anna Dominiczak and Alex Alderton, Roberto Amato, Sonia Goncalves, Ewan Harrison, David K. Jackson, Ian Johnston, Dominic Kwiatkowski, Cordelia Langford, John Sillitoe on behalf of the Wellcome Sanger Institute COVID-19 Surveillance Team |
| EPI_ISL_947148 | Lighthouse Lab in Milton Keynes | Wellcome Sanger Institute for the COVID-19 Genomics UK (COG-UK) Consortium | The Lighthouse Lab in Milton Keynes and Alex Alderton, Roberto Amato, Sonia Goncalves, Ewan Harrison, David K. Jackson, Ian Johnston, Dominic Kwiatkowski, Cordelia Langford, John Sillitoe on behalf of the Wellcome Sanger Institute COVID-19 Surveillance Team |
| EPI_ISL_947150 | Lighthouse Lab in Glasgow | Wellcome Sanger Institute for the COVID-19 Genomics UK (COG-UK) Consortium | Harper VanSteenhouse, Yumi Kasai, David Gray, Carol Clugston, Anna Dominiczak and Alex Alderton, Roberto Amato, Sonia Goncalves, Ewan Harrison, David K. Jackson, Ian Johnston, Dominic Kwiatkowski, Cordelia Langford, John Sillitoe on behalf of the Wellcome Sanger Institute COVID-19 Surveillance Team |
| EPI_ISL_947151, EPI_ISL_947152 | Lighthouse Lab in Milton Keynes | Wellcome Sanger Institute for the COVID-19 Genomics UK (COG-UK) Consortium | The Lighthouse Lab in Milton Keynes and Alex Alderton, Roberto Amato, Sonia Goncalves, Ewan Harrison, David K. Jackson, Ian Johnston, Dominic Kwiatkowski, Cordelia Langford, John Sillitoe on behalf of the Wellcome Sanger Institute COVID-19 Surveillance Team |
| EPI_ISL_947153 | Lighthouse Lab in Glasgow | Wellcome Sanger Institute for the COVID-19 Genomics UK (COG-UK) Consortium | Harper VanSteenhouse, Yumi Kasai, David Gray, Carol Clugston, Anna Dominiczak and Alex Alderton, Roberto Amato, Sonia Goncalves, Ewan Harrison, David K. Jackson, Ian Johnston, Dominic Kwiatkowski, Cordelia Langford, John Sillitoe on behalf of the Wellcome Sanger Institute COVID-19 Surveillance Team |
| EPI_ISL_947154, EPI_ISL_947155, EPI_ISL_947156, EPI_ISL_947157, EPI_ISL_947158, EPI_ISL_947159, EPI_ISL_947160, EPI_ISL_947161, EPI_ISL_947162 | Lighthouse Lab in Milton Keynes | Wellcome Sanger Institute for the COVID-19 Genomics UK (COG-UK) Consortium | The Lighthouse Lab in Milton Keynes and Alex Alderton, Roberto Amato, Sonia Goncalves, Ewan Harrison, David K. Jackson, Ian Johnston, Dominic Kwiatkowski, Cordelia Langford, John Sillitoe on behalf of the Wellcome Sanger Institute COVID-19 Surveillance Team |
| EPI_ISL_947163 | Lighthouse Lab in Glasgow | Wellcome Sanger Institute for the COVID-19 Genomics UK (COG-UK) Consortium | Harper VanSteenhouse, Yumi Kasai, David Gray, Carol Clugston, Anna Dominiczak and Alex Alderton, Roberto Amato, Sonia Goncalves, Ewan Harrison, David K. Jackson, Ian Johnston, Dominic Kwiatkowski, Cordelia Langford, John Sillitoe on behalf of the Wellcome Sanger Institute COVID-19 Surveillance Team |
| EPI_ISL_947164 | Lighthouse Lab in Milton Keynes | Wellcome Sanger Institute for the COVID-19 Genomics UK (COG-UK) Consortium | The Lighthouse Lab in Milton Keynes and Alex Alderton, Roberto Amato, Sonia Goncalves, Ewan Harrison, David K. Jackson, Ian Johnston, Dominic Kwiatkowski, Cordelia Langford, John Sillitoe on behalf of the Wellcome Sanger Institute COVID-19 Surveillance Team |
| EPI_ISL_947165 | Lighthouse Lab in Glasgow | Wellcome Sanger Institute for the COVID-19 Genomics UK (COG-UK) Consortium | Harper VanSteenhouse, Yumi Kasai, David Gray, Carol Clugston, Anna Dominiczak and Alex Alderton, Roberto Amato, Sonia Goncalves, Ewan Harrison, David K. Jackson, Ian Johnston, Dominic Kwiatkowski, Cordelia Langford, John Sillitoe on behalf of the Wellcome Sanger Institute COVID-19 Surveillance Team |
| EPI_ISL_947166, EPI_ISL_947167, EPI_ISL_947168 | Lighthouse Lab in Milton Keynes | Wellcome Sanger Institute for the COVID-19 Genomics UK (COG-UK) Consortium | The Lighthouse Lab in Milton Keynes and Alex Alderton, Roberto Amato, Sonia Goncalves, Ewan Harrison, David K. Jackson, Ian Johnston, Dominic Kwiatkowski, Cordelia Langford, John Sillitoe on behalf of the Wellcome Sanger Institute COVID-19 Surveillance Team |
| EPI_ISL_947169 | Lighthouse Lab in Glasgow | Wellcome Sanger Institute for the COVID-19 Genomics UK (COG-UK) Consortium | Harper VanSteenhouse, Yumi Kasai, David Gray, Carol Clugston, Anna Dominiczak and Alex Alderton, Roberto Amato, Sonia Goncalves, Ewan Harrison, David K. Jackson, Ian Johnston, Dominic Kwiatkowski, Cordelia Langford, John Sillitoe on behalf of the Wellcome Sanger Institute COVID-19 Surveillance Team |
| EPI_ISL_947170, EPI_ISL_947171, EPI_ISL_947173, EPI_ISL_947174, EPI_ISL_947175, EPI_ISL_947176, EPI_ISL_947177, EPI_ISL_947178 | Lighthouse Lab in Milton Keynes | Wellcome Sanger Institute for the COVID-19 Genomics UK (COG-UK) Consortium | The Lighthouse Lab in Milton Keynes and Alex Alderton, Roberto Amato, Sonia Goncalves, Ewan Harrison, David K. Jackson, Ian Johnston, Dominic Kwiatkowski, Cordelia Langford, John Sillitoe on behalf of the Wellcome Sanger Institute COVID-19 Surveillance Team |
| EPI_ISL_947179 | Lighthouse Lab in Glasgow | Wellcome Sanger Institute for the COVID-19 Genomics UK (COG-UK) Consortium | Harper VanSteenhouse, Yumi Kasai, David Gray, Carol Clugston, Anna Dominiczak and Alex Alderton, Roberto Amato, Sonia Goncalves, Ewan Harrison, David K. Jackson, Ian Johnston, Dominic Kwiatkowski, Cordelia Langford, John Sillitoe on behalf of the Wellcome Sanger Institute COVID-19 Surveillance Team |
| EPI_ISL_947180, EPI_ISL_947181 | Lighthouse Lab in Milton Keynes | Wellcome Sanger Institute for the COVID-19 Genomics UK (COG-UK) Consortium | The Lighthouse Lab in Milton Keynes and Alex Alderton, Roberto Amato, Sonia Goncalves, Ewan Harrison, David K. Jackson, Ian Johnston, Dominic Kwiatkowski, Cordelia Langford, John Sillitoe on behalf of the Wellcome Sanger Institute COVID-19 Surveillance Team |
| EPI_ISL_947182, EPI_ISL_947183 | Lighthouse Lab in Glasgow | Wellcome Sanger Institute for the COVID-19 Genomics UK (COG-UK) Consortium | Harper VanSteenhouse, Yumi Kasai, David Gray, Carol Clugston, Anna Dominiczak and Alex Alderton, Roberto Amato, Sonia Goncalves, Ewan Harrison, David K. Jackson, Ian Johnston, Dominic Kwiatkowski, Cordelia Langford, John Sillitoe on behalf of the Wellcome Sanger Institute COVID-19 Surveillance Team |
| EPI_ISL_947185 | Lighthouse Lab in Milton Keynes | Wellcome Sanger Institute for the COVID-19 Genomics UK (COG-UK) Consortium | The Lighthouse Lab in Milton Keynes and Alex Alderton, Roberto Amato, Sonia Goncalves, Ewan Harrison, David K. Jackson, Ian Johnston, Dominic Kwiatkowski, Cordelia Langford, John Sillitoe on behalf of the Wellcome Sanger Institute COVID-19 Surveillance Team |
| EPI_ISL_947186, EPI_ISL_947187 | Lighthouse Lab in Glasgow | Wellcome Sanger Institute for the COVID-19 Genomics UK (COG-UK) Consortium | Harper VanSteenhouse, Yumi Kasai, David Gray, Carol Clugston, Anna Dominiczak and Alex Alderton, Roberto Amato, Sonia Goncalves, Ewan Harrison, David K. Jackson, Ian Johnston, Dominic Kwiatkowski, Cordelia Langford, John Sillitoe on behalf of the Wellcome Sanger Institute COVID-19 Surveillance Team |
| EPI_ISL_947188, EPI_ISL_947190 | Lighthouse Lab in Milton Keynes | Wellcome Sanger Institute for the COVID-19 Genomics UK (COG-UK) Consortium | The Lighthouse Lab in Milton Keynes and Alex Alderton, Roberto Amato, Sonia Goncalves, Ewan Harrison, David K. Jackson, Ian Johnston, Dominic Kwiatkowski, Cordelia Langford, John Sillitoe on behalf of the Wellcome Sanger Institute COVID-19 Surveillance Team |
| EPI_ISL_947191 | Lighthouse Lab in Glasgow | Wellcome Sanger Institute for the COVID-19 Genomics UK (COG-UK) Consortium | Harper VanSteenhouse, Yumi Kasai, David Gray, Carol Clugston, Anna Dominiczak and Alex Alderton, Roberto Amato, Sonia Goncalves, Ewan Harrison, David K. Jackson, Ian Johnston, Dominic Kwiatkowski, Cordelia Langford, John Sillitoe on behalf of the Wellcome Sanger Institute COVID-19 Surveillance Team |
| EPI_ISL_947192, EPI_ISL_947193, EPI_ISL_947194, EPI_ISL_947195, EPI_ISL_947196 | Lighthouse Lab in Milton Keynes | Wellcome Sanger Institute for the COVID-19 Genomics UK (COG-UK) Consortium | The Lighthouse Lab in Milton Keynes and Alex Alderton, Roberto Amato, Sonia Goncalves, Ewan Harrison, David K. Jackson, Ian Johnston, Dominic Kwiatkowski, Cordelia Langford, John Sillitoe on behalf of the Wellcome Sanger Institute COVID-19 Surveillance Team |
| EPI_ISL_947197 | Lighthouse Lab in Glasgow | Wellcome Sanger Institute for the COVID-19 Genomics UK (COG-UK) Consortium | Harper VanSteenhouse, Yumi Kasai, David Gray, Carol Clugston, Anna Dominiczak and Alex Alderton, Roberto Amato, Sonia Goncalves, Ewan Harrison, David K. Jackson, Ian Johnston, Dominic Kwiatkowski, Cordelia Langford, John Sillitoe on behalf of the Wellcome Sanger Institute COVID-19 Surveillance Team |
| EPI_ISL_947198, EPI_ISL_947199, EPI_ISL_947200 | Lighthouse Lab in Milton Keynes | Wellcome Sanger Institute for the COVID-19 Genomics UK (COG-UK) Consortium | The Lighthouse Lab in Milton Keynes and Alex Alderton, Roberto Amato, Sonia Goncalves, Ewan Harrison, David K. Jackson, Ian Johnston, Dominic Kwiatkowski, Cordelia Langford, John Sillitoe on behalf of the Wellcome Sanger Institute COVID-19 Surveillance Team |
| EPI_ISL_947202 | Lighthouse Lab in Glasgow | Wellcome Sanger Institute for the COVID-19 Genomics UK (COG-UK) Consortium | Harper VanSteenhouse, Yumi Kasai, David Gray, Carol Clugston, Anna Dominiczak and Alex Alderton, Roberto Amato, Sonia Goncalves, Ewan Harrison, David K. Jackson, Ian Johnston, Dominic Kwiatkowski, Cordelia Langford, John Sillitoe on behalf of the Wellcome Sanger Institute COVID-19 Surveillance |

| | | | |
|---|---|---|---|
| | | | Team |
| EPI_ISL_947203 | Lighthouse Lab in Milton Keynes | Wellcome Sanger Institute for the COVID-19 Genomics UK (COG-UK) Consortium | The Lighthouse Lab in Milton Keynes and Alex Alderton, Roberto Amato, Sonia Goncalves, Ewan Harrison, David K. Jackson, Ian Johnston, Dominic Kwiatkowski, Cordelia Langford, John Sillitoe on behalf of the Wellcome Sanger Institute COVID-19 Surveillance Team |
| EPI_ISL_947204 | Lighthouse Lab in Glasgow | Wellcome Sanger Institute for the COVID-19 Genomics UK (COG-UK) Consortium | Harper VanSteenhouse, Yumi Kasai, David Gray, Carol Clugston, Anna Dominiczak and Alex Alderton, Roberto Amato, Sonia Goncalves, Ewan Harrison, David K. Jackson, Ian Johnston, Dominic Kwiatkowski, Cordelia Langford, John Sillitoe on behalf of the Wellcome Sanger Institute COVID-19 Surveillance Team |
| EPI_ISL_947205, EPI_ISL_947206, EPI_ISL_947207, EPI_ISL_947209, EPI_ISL_947210 | Lighthouse Lab in Milton Keynes | Wellcome Sanger Institute for the COVID-19 Genomics UK (COG-UK) Consortium | The Lighthouse Lab in Milton Keynes and Alex Alderton, Roberto Amato, Sonia Goncalves, Ewan Harrison, David K. Jackson, Ian Johnston, Dominic Kwiatkowski, Cordelia Langford, John Sillitoe on behalf of the Wellcome Sanger Institute COVID-19 Surveillance Team |
| EPI_ISL_947211, EPI_ISL_947212, EPI_ISL_947213, EPI_ISL_947214 | Lighthouse Lab in Glasgow | Wellcome Sanger Institute for the COVID-19 Genomics UK (COG-UK) Consortium | Harper VanSteenhouse, Yumi Kasai, David Gray, Carol Clugston, Anna Dominiczak and Alex Alderton, Roberto Amato, Sonia Goncalves, Ewan Harrison, David K. Jackson, Ian Johnston, Dominic Kwiatkowski, Cordelia Langford, John Sillitoe on behalf of the Wellcome Sanger Institute COVID-19 Surveillance Team |
| EPI_ISL_947215, EPI_ISL_947216, EPI_ISL_947217, EPI_ISL_947218, EPI_ISL_947219, EPI_ISL_947220, EPI_ISL_947221, EPI_ISL_947222, EPI_ISL_947223, EPI_ISL_947224, EPI_ISL_947225, EPI_ISL_947226, EPI_ISL_947227 | | | |
| see above | Lighthouse Lab in Milton Keynes | Wellcome Sanger Institute for the COVID-19 Genomics UK (COG-UK) Consortium | The Lighthouse Lab in Milton Keynes and Alex Alderton, Roberto Amato, Sonia Goncalves, Ewan Harrison, David K. Jackson, Ian Johnston, Dominic Kwiatkowski, Cordelia Langford, John Sillitoe on behalf of the Wellcome Sanger Institute COVID-19 Surveillance Team |
| EPI_ISL_947228, EPI_ISL_947229, EPI_ISL_947230 | Lighthouse Lab in Glasgow | Wellcome Sanger Institute for the COVID-19 Genomics UK (COG-UK) Consortium | Harper VanSteenhouse, Yumi Kasai, David Gray, Carol Clugston, Anna Dominiczak and Alex Alderton, Roberto Amato, Sonia Goncalves, Ewan Harrison, David K. Jackson, Ian Johnston, Dominic Kwiatkowski, Cordelia Langford, John Sillitoe on behalf of the Wellcome Sanger Institute COVID-19 Surveillance Team |
| EPI_ISL_947231, EPI_ISL_947232 | Lighthouse Lab in Milton Keynes | Wellcome Sanger Institute for the COVID-19 Genomics UK (COG-UK) Consortium | The Lighthouse Lab in Milton Keynes and Alex Alderton, Roberto Amato, Sonia Goncalves, Ewan Harrison, David K. Jackson, Ian Johnston, Dominic Kwiatkowski, Cordelia Langford, John Sillitoe on behalf of the Wellcome Sanger Institute COVID-19 Surveillance Team |
| EPI_ISL_947233 | Lighthouse Lab in Glasgow | Wellcome Sanger Institute for the COVID-19 Genomics UK (COG-UK) Consortium | Harper VanSteenhouse, Yumi Kasai, David Gray, Carol Clugston, Anna Dominiczak and Alex Alderton, Roberto Amato, Sonia Goncalves, Ewan Harrison, David K. Jackson, Ian Johnston, Dominic Kwiatkowski, Cordelia Langford, John Sillitoe on behalf of the Wellcome Sanger Institute COVID-19 Surveillance Team |
| EPI_ISL_947234, EPI_ISL_947235 | Lighthouse Lab in Milton Keynes | Wellcome Sanger Institute for the COVID-19 Genomics UK (COG-UK) Consortium | The Lighthouse Lab in Milton Keynes and Alex Alderton, Roberto Amato, Sonia Goncalves, Ewan Harrison, David K. Jackson, Ian Johnston, Dominic Kwiatkowski, Cordelia Langford, John Sillitoe on behalf of the Wellcome Sanger Institute COVID-19 Surveillance Team |
| EPI_ISL_947236 | Lighthouse Lab in Glasgow | Wellcome Sanger Institute for the COVID-19 Genomics UK (COG-UK) Consortium | Harper VanSteenhouse, Yumi Kasai, David Gray, Carol Clugston, Anna Dominiczak and Alex Alderton, Roberto Amato, Sonia Goncalves, Ewan Harrison, David K. Jackson, Ian Johnston, Dominic Kwiatkowski, Cordelia Langford, John Sillitoe on behalf of the Wellcome Sanger Institute COVID-19 Surveillance Team |
| EPI_ISL_947237 | Lighthouse Lab in Milton Keynes | Wellcome Sanger Institute for the COVID-19 Genomics UK (COG-UK) Consortium | The Lighthouse Lab in Milton Keynes and Alex Alderton, Roberto Amato, Sonia Goncalves, Ewan Harrison, David K. Jackson, Ian Johnston, Dominic Kwiatkowski, Cordelia Langford, John Sillitoe on behalf of the Wellcome Sanger Institute COVID-19 Surveillance Team |
| EPI_ISL_947238, EPI_ISL_947239 | Lighthouse Lab in Glasgow | Wellcome Sanger Institute for the COVID-19 Genomics UK (COG-UK) Consortium | Harper VanSteenhouse, Yumi Kasai, David Gray, Carol Clugston, Anna Dominiczak and Alex Alderton, Roberto Amato, Sonia Goncalves, Ewan Harrison, David K. Jackson, Ian Johnston, Dominic Kwiatkowski, Cordelia Langford, John Sillitoe on behalf of the Wellcome Sanger Institute COVID-19 Surveillance Team |
| EPI_ISL_947241, EPI_ISL_947242, EPI_ISL_947243, EPI_ISL_947244 | Lighthouse Lab in Milton Keynes | Wellcome Sanger Institute for the COVID-19 Genomics UK (COG-UK) Consortium | The Lighthouse Lab in Milton Keynes and Alex Alderton, Roberto Amato, Sonia Goncalves, Ewan Harrison, David K. Jackson, Ian Johnston, Dominic Kwiatkowski, Cordelia Langford, John Sillitoe on behalf of the Wellcome Sanger Institute COVID-19 Surveillance Team |
| EPI_ISL_947245 | Lighthouse Lab in Glasgow | Wellcome Sanger Institute for the COVID-19 Genomics UK (COG-UK) Consortium | Harper VanSteenhouse, Yumi Kasai, David Gray, Carol Clugston, Anna Dominiczak and Alex Alderton, Roberto Amato, Sonia Goncalves, Ewan Harrison, David K. Jackson, Ian Johnston, Dominic Kwiatkowski, Cordelia Langford, John Sillitoe on behalf of the Wellcome Sanger Institute COVID-19 Surveillance Team |
| EPI_ISL_947246, EPI_ISL_947248 | Lighthouse Lab in Milton Keynes | Wellcome Sanger Institute for the COVID-19 Genomics UK (COG-UK) Consortium | The Lighthouse Lab in Milton Keynes and Alex Alderton, Roberto Amato, Sonia Goncalves, Ewan Harrison, David K. Jackson, Ian Johnston, Dominic Kwiatkowski, Cordelia Langford, John Sillitoe on behalf of the Wellcome Sanger Institute COVID-19 Surveillance Team |
| EPI_ISL_947250 | Lighthouse Lab in Glasgow | Wellcome Sanger Institute for the COVID-19 Genomics UK (COG-UK) Consortium | Harper VanSteenhouse, Yumi Kasai, David Gray, Carol Clugston, Anna Dominiczak and Alex Alderton, Roberto Amato, Sonia Goncalves, Ewan Harrison, David K. Jackson, Ian Johnston, Dominic Kwiatkowski, Cordelia Langford, John Sillitoe on behalf of the Wellcome Sanger Institute COVID-19 Surveillance Team |
| EPI_ISL_947251 | Lighthouse Lab in Milton Keynes | Wellcome Sanger Institute for the COVID-19 Genomics UK (COG-UK) Consortium | The Lighthouse Lab in Milton Keynes and Alex Alderton, Roberto Amato, Sonia Goncalves, Ewan Harrison, David K. Jackson, Ian Johnston, Dominic Kwiatkowski, Cordelia Langford, John Sillitoe on behalf of the Wellcome Sanger Institute COVID-19 Surveillance Team |
| EPI_ISL_947253 | Lighthouse Lab in Glasgow | Wellcome Sanger Institute for the COVID-19 Genomics UK (COG-UK) Consortium | Harper VanSteenhouse, Yumi Kasai, David Gray, Carol Clugston, Anna Dominiczak and Alex Alderton, Roberto Amato, Sonia Goncalves, Ewan Harrison, David K. Jackson, Ian Johnston, Dominic Kwiatkowski, Cordelia Langford, John Sillitoe on behalf of the Wellcome Sanger Institute COVID-19 Surveillance Team |
| EPI_ISL_947254, EPI_ISL_947258, EPI_ISL_947259 | Lighthouse Lab in Milton Keynes | Wellcome Sanger Institute for the COVID-19 Genomics UK (COG-UK) Consortium | The Lighthouse Lab in Milton Keynes and Alex Alderton, Roberto Amato, Sonia Goncalves, Ewan Harrison, David K. Jackson, Ian Johnston, Dominic Kwiatkowski, Cordelia Langford, John Sillitoe on behalf of the Wellcome Sanger Institute COVID-19 Surveillance Team |
| EPI_ISL_947263 | Lighthouse Lab in Glasgow | Wellcome Sanger Institute for the COVID-19 Genomics UK (COG-UK) Consortium | Harper VanSteenhouse, Yumi Kasai, David Gray, Carol Clugston, Anna Dominiczak and Alex Alderton, Roberto Amato, Sonia Goncalves, Ewan Harrison, David K. Jackson, Ian Johnston, Dominic Kwiatkowski, Cordelia Langford, John Sillitoe on behalf of the Wellcome Sanger Institute COVID-19 Surveillance Team |
| EPI_ISL_947264, EPI_ISL_947266 | Lighthouse Lab in Milton Keynes | Wellcome Sanger Institute for the COVID-19 Genomics UK (COG-UK) Consortium | The Lighthouse Lab in Milton Keynes and Alex Alderton, Roberto Amato, Sonia Goncalves, Ewan Harrison, David K. Jackson, Ian Johnston, Dominic Kwiatkowski, Cordelia Langford, John Sillitoe on behalf of the Wellcome Sanger Institute COVID-19 Surveillance Team |
| EPI_ISL_947267, EPI_ISL_947269, EPI_ISL_947271, EPI_ISL_947272 | Lighthouse Lab in Glasgow | Wellcome Sanger Institute for the COVID-19 Genomics UK (COG-UK) Consortium | Harper VanSteenhouse, Yumi Kasai, David Gray, Carol Clugston, Anna Dominiczak and Alex Alderton, Roberto Amato, Sonia Goncalves, Ewan Harrison, David K. Jackson, Ian Johnston, Dominic Kwiatkowski, Cordelia Langford, John Sillitoe on behalf of the Wellcome Sanger Institute COVID-19 Surveillance Team |
| EPI_ISL_947276, EPI_ISL_947277, EPI_ISL_947279 | Lighthouse Lab in Milton Keynes | Wellcome Sanger Institute for the COVID-19 Genomics UK (COG-UK) Consortium | The Lighthouse Lab in Milton Keynes and Alex Alderton, Roberto Amato, Sonia Goncalves, Ewan Harrison, David K. Jackson, Ian Johnston, Dominic Kwiatkowski, Cordelia Langford, John Sillitoe on behalf of the Wellcome Sanger Institute COVID-19 Surveillance Team |
| EPI_ISL_947280 | Lighthouse Lab in Glasgow | Wellcome Sanger Institute for the COVID-19 Genomics UK (COG-UK) Consortium | Harper VanSteenhouse, Yumi Kasai, David Gray, Carol Clugston, Anna Dominiczak and Alex Alderton, Roberto Amato, Sonia Goncalves, Ewan Harrison, David K. Jackson, Ian Johnston, Dominic Kwiatkowski, Cordelia Langford, John Sillitoe on behalf of the Wellcome Sanger Institute COVID-19 Surveillance Team |
| EPI_ISL_947282 | Lighthouse Lab in Milton Keynes | Wellcome Sanger Institute for the COVID-19 Genomics UK (COG-UK) Consortium | The Lighthouse Lab in Milton Keynes and Alex Alderton, Roberto Amato, Sonia Goncalves, Ewan Harrison, David K. Jackson, Ian Johnston, Dominic Kwiatkowski, Cordelia Langford, John Sillitoe on behalf of the Wellcome Sanger Institute COVID-19 Surveillance Team |
| EPI_ISL_947283 | Lighthouse Lab in Glasgow | Wellcome Sanger Institute for the COVID-19 Genomics UK (COG-UK) Consortium | Harper VanSteenhouse, Yumi Kasai, David Gray, Carol Clugston, Anna Dominiczak and Alex Alderton, Roberto Amato, Sonia Goncalves, Ewan Harrison, David K. Jackson, Ian Johnston, Dominic Kwiatkowski, Cordelia Langford, John Sillitoe on behalf of the Wellcome Sanger Institute COVID-19 Surveillance Team |
| EPI_ISL_947285, EPI_ISL_947286, EPI_ISL_947288, EPI_ISL_947289, EPI_ISL_947291, EPI_ISL_947292 | Lighthouse Lab in Milton Keynes | Wellcome Sanger Institute for the COVID-19 Genomics UK (COG-UK) Consortium | The Lighthouse Lab in Milton Keynes and Alex Alderton, Roberto Amato, Sonia Goncalves, Ewan Harrison, David K. Jackson, Ian Johnston, Dominic Kwiatkowski, Cordelia Langford, John Sillitoe on behalf of the Wellcome Sanger Institute COVID-19 Surveillance Team |
| EPI_ISL_947294, EPI_ISL_947295 | Lighthouse Lab in Glasgow | Wellcome Sanger Institute for the COVID-19 Genomics UK (COG-UK) Consortium | Harper VanSteenhouse, Yumi Kasai, David Gray, Carol Clugston, Anna Dominiczak and Alex Alderton, Roberto Amato, Sonia Goncalves, Ewan Harrison, David K. Jackson, Ian Johnston, Dominic Kwiatkowski, Cordelia Langford, John Sillitoe on behalf of the Wellcome Sanger Institute COVID-19 Surveillance |

| | | | |
|---|---|---|---|
| | | | Team |
| EPI_ISL_947297, EPI_ISL_947299, EPI_ISL_947300, EPI_ISL_947302, EPI_ISL_947303 | Lighthouse Lab in Milton Keynes | Wellcome Sanger Institute for the COVID-19 Genomics UK (COG-UK) Consortium | The Lighthouse Lab in Milton Keynes and Alex Alderton, Roberto Amato, Sonia Goncalves, Ewan Harrison, David K. Jackson, Ian Johnston, Dominic Kwiatkowski, Cordelia Langford, John Sillitoe on behalf of the Wellcome Sanger Institute COVID-19 Surveillance Team |
| EPI_ISL_947305 | Lighthouse Lab in Glasgow | Wellcome Sanger Institute for the COVID-19 Genomics UK (COG-UK) Consortium | Harper VanSteenhouse, Yumi Kasai, David Gray, Carol Clugston, Anna Dominiczak and Alex Alderton, Roberto Amato, Sonia Goncalves, Ewan Harrison, David K. Jackson, Ian Johnston, Dominic Kwiatkowski, Cordelia Langford, John Sillitoe on behalf of the Wellcome Sanger Institute COVID-19 Surveillance Team |
| EPI_ISL_947306 | Lighthouse Lab in Milton Keynes | Wellcome Sanger Institute for the COVID-19 Genomics UK (COG-UK) Consortium | The Lighthouse Lab in Milton Keynes and Alex Alderton, Roberto Amato, Sonia Goncalves, Ewan Harrison, David K. Jackson, Ian Johnston, Dominic Kwiatkowski, Cordelia Langford, John Sillitoe on behalf of the Wellcome Sanger Institute COVID-19 Surveillance Team |
| EPI_ISL_947308 | Lighthouse Lab in Glasgow | Wellcome Sanger Institute for the COVID-19 Genomics UK (COG-UK) Consortium | Harper VanSteenhouse, Yumi Kasai, David Gray, Carol Clugston, Anna Dominiczak and Alex Alderton, Roberto Amato, Sonia Goncalves, Ewan Harrison, David K. Jackson, Ian Johnston, Dominic Kwiatkowski, Cordelia Langford, John Sillitoe on behalf of the Wellcome Sanger Institute COVID-19 Surveillance Team |
| EPI_ISL_947310, EPI_ISL_947311, EPI_ISL_947313 | Lighthouse Lab in Milton Keynes | Wellcome Sanger Institute for the COVID-19 Genomics UK (COG-UK) Consortium | The Lighthouse Lab in Milton Keynes and Alex Alderton, Roberto Amato, Sonia Goncalves, Ewan Harrison, David K. Jackson, Ian Johnston, Dominic Kwiatkowski, Cordelia Langford, John Sillitoe on behalf of the Wellcome Sanger Institute COVID-19 Surveillance Team |
| EPI_ISL_947314, EPI_ISL_947316 | Lighthouse Lab in Glasgow | Wellcome Sanger Institute for the COVID-19 Genomics UK (COG-UK) Consortium | Harper VanSteenhouse, Yumi Kasai, David Gray, Carol Clugston, Anna Dominiczak and Alex Alderton, Roberto Amato, Sonia Goncalves, Ewan Harrison, David K. Jackson, Ian Johnston, Dominic Kwiatkowski, Cordelia Langford, John Sillitoe on behalf of the Wellcome Sanger Institute COVID-19 Surveillance Team |
| EPI_ISL_947319, EPI_ISL_947321 | Lighthouse Lab in Milton Keynes | Wellcome Sanger Institute for the COVID-19 Genomics UK (COG-UK) Consortium | The Lighthouse Lab in Milton Keynes and Alex Alderton, Roberto Amato, Sonia Goncalves, Ewan Harrison, David K. Jackson, Ian Johnston, Dominic Kwiatkowski, Cordelia Langford, John Sillitoe on behalf of the Wellcome Sanger Institute COVID-19 Surveillance Team |
| EPI_ISL_947331, EPI_ISL_947332 | Lighthouse Lab in Glasgow | Wellcome Sanger Institute for the COVID-19 Genomics UK (COG-UK) Consortium | Harper VanSteenhouse, Yumi Kasai, David Gray, Carol Clugston, Anna Dominiczak and Alex Alderton, Roberto Amato, Sonia Goncalves, Ewan Harrison, David K. Jackson, Ian Johnston, Dominic Kwiatkowski, Cordelia Langford, John Sillitoe on behalf of the Wellcome Sanger Institute COVID-19 Surveillance Team |
| EPI_ISL_947333, EPI_ISL_947334 | Lighthouse Lab in Milton Keynes | Wellcome Sanger Institute for the COVID-19 Genomics UK (COG-UK) Consortium | The Lighthouse Lab in Milton Keynes and Alex Alderton, Roberto Amato, Sonia Goncalves, Ewan Harrison, David K. Jackson, Ian Johnston, Dominic Kwiatkowski, Cordelia Langford, John Sillitoe on behalf of the Wellcome Sanger Institute COVID-19 Surveillance Team |
| EPI_ISL_947335 | Lighthouse Lab in Glasgow | Wellcome Sanger Institute for the COVID-19 Genomics UK (COG-UK) Consortium | Harper VanSteenhouse, Yumi Kasai, David Gray, Carol Clugston, Anna Dominiczak and Alex Alderton, Roberto Amato, Sonia Goncalves, Ewan Harrison, David K. Jackson, Ian Johnston, Dominic Kwiatkowski, Cordelia Langford, John Sillitoe on behalf of the Wellcome Sanger Institute COVID-19 Surveillance Team |
| EPI_ISL_947336 | Lighthouse Lab in Milton Keynes | Wellcome Sanger Institute for the COVID-19 Genomics UK (COG-UK) Consortium | The Lighthouse Lab in Milton Keynes and Alex Alderton, Roberto Amato, Sonia Goncalves, Ewan Harrison, David K. Jackson, Ian Johnston, Dominic Kwiatkowski, Cordelia Langford, John Sillitoe on behalf of the Wellcome Sanger Institute COVID-19 Surveillance Team |
| EPI_ISL_947338 | Lighthouse Lab in Glasgow | Wellcome Sanger Institute for the COVID-19 Genomics UK (COG-UK) Consortium | Harper VanSteenhouse, Yumi Kasai, David Gray, Carol Clugston, Anna Dominiczak and Alex Alderton, Roberto Amato, Sonia Goncalves, Ewan Harrison, David K. Jackson, Ian Johnston, Dominic Kwiatkowski, Cordelia Langford, John Sillitoe on behalf of the Wellcome Sanger Institute COVID-19 Surveillance Team |
| EPI_ISL_947339, EPI_ISL_947340 | Lighthouse Lab in Milton Keynes | Wellcome Sanger Institute for the COVID-19 Genomics UK (COG-UK) Consortium | The Lighthouse Lab in Milton Keynes and Alex Alderton, Roberto Amato, Sonia Goncalves, Ewan Harrison, David K. Jackson, Ian Johnston, Dominic Kwiatkowski, Cordelia Langford, John Sillitoe on behalf of the Wellcome Sanger Institute COVID-19 Surveillance Team |
| EPI_ISL_947341 | Lighthouse Lab in Glasgow | Wellcome Sanger Institute for the COVID-19 Genomics UK (COG-UK) Consortium | Harper VanSteenhouse, Yumi Kasai, David Gray, Carol Clugston, Anna Dominiczak and Alex Alderton, Roberto Amato, Sonia Goncalves, Ewan Harrison, David K. Jackson, Ian Johnston, Dominic Kwiatkowski, Cordelia Langford, John Sillitoe on behalf of the Wellcome Sanger Institute COVID-19 Surveillance Team |
| EPI_ISL_947342 | Lighthouse Lab in Milton Keynes | Wellcome Sanger Institute for the COVID-19 Genomics UK (COG-UK) Consortium | The Lighthouse Lab in Milton Keynes and Alex Alderton, Roberto Amato, Sonia Goncalves, Ewan Harrison, David K. Jackson, Ian Johnston, Dominic Kwiatkowski, Cordelia Langford, John Sillitoe on behalf of the Wellcome Sanger Institute COVID-19 Surveillance Team |
| EPI_ISL_947343, EPI_ISL_947344 | Lighthouse Lab in Glasgow | Wellcome Sanger Institute for the COVID-19 Genomics UK (COG-UK) Consortium | Harper VanSteenhouse, Yumi Kasai, David Gray, Carol Clugston, Anna Dominiczak and Alex Alderton, Roberto Amato, Sonia Goncalves, Ewan Harrison, David K. Jackson, Ian Johnston, Dominic Kwiatkowski, Cordelia Langford, John Sillitoe on behalf of the Wellcome Sanger Institute COVID-19 Surveillance Team |
| EPI_ISL_947345, EPI_ISL_947346 | Lighthouse Lab in Milton Keynes | Wellcome Sanger Institute for the COVID-19 Genomics UK (COG-UK) Consortium | The Lighthouse Lab in Milton Keynes and Alex Alderton, Roberto Amato, Sonia Goncalves, Ewan Harrison, David K. Jackson, Ian Johnston, Dominic Kwiatkowski, Cordelia Langford, John Sillitoe on behalf of the Wellcome Sanger Institute COVID-19 Surveillance Team |
| EPI_ISL_947347 | Lighthouse Lab in Glasgow | Wellcome Sanger Institute for the COVID-19 Genomics UK (COG-UK) Consortium | Harper VanSteenhouse, Yumi Kasai, David Gray, Carol Clugston, Anna Dominiczak and Alex Alderton, Roberto Amato, Sonia Goncalves, Ewan Harrison, David K. Jackson, Ian Johnston, Dominic Kwiatkowski, Cordelia Langford, John Sillitoe on behalf of the Wellcome Sanger Institute COVID-19 Surveillance Team |
| EPI_ISL_947348, EPI_ISL_947349, EPI_ISL_947351 | Lighthouse Lab in Milton Keynes | Wellcome Sanger Institute for the COVID-19 Genomics UK (COG-UK) Consortium | The Lighthouse Lab in Milton Keynes and Alex Alderton, Roberto Amato, Sonia Goncalves, Ewan Harrison, David K. Jackson, Ian Johnston, Dominic Kwiatkowski, Cordelia Langford, John Sillitoe on behalf of the Wellcome Sanger Institute COVID-19 Surveillance Team |
| EPI_ISL_947353 | Lighthouse Lab in Glasgow | Wellcome Sanger Institute for the COVID-19 Genomics UK (COG-UK) Consortium | Harper VanSteenhouse, Yumi Kasai, David Gray, Carol Clugston, Anna Dominiczak and Alex Alderton, Roberto Amato, Sonia Goncalves, Ewan Harrison, David K. Jackson, Ian Johnston, Dominic Kwiatkowski, Cordelia Langford, John Sillitoe on behalf of the Wellcome Sanger Institute COVID-19 Surveillance Team |
| EPI_ISL_947355 | Lighthouse Lab in Milton Keynes | Wellcome Sanger Institute for the COVID-19 Genomics UK (COG-UK) Consortium | The Lighthouse Lab in Milton Keynes and Alex Alderton, Roberto Amato, Sonia Goncalves, Ewan Harrison, David K. Jackson, Ian Johnston, Dominic Kwiatkowski, Cordelia Langford, John Sillitoe on behalf of the Wellcome Sanger Institute COVID-19 Surveillance Team |
| EPI_ISL_947356, EPI_ISL_947357 | Lighthouse Lab in Glasgow | Wellcome Sanger Institute for the COVID-19 Genomics UK (COG-UK) Consortium | Harper VanSteenhouse, Yumi Kasai, David Gray, Carol Clugston, Anna Dominiczak and Alex Alderton, Roberto Amato, Sonia Goncalves, Ewan Harrison, David K. Jackson, Ian Johnston, Dominic Kwiatkowski, Cordelia Langford, John Sillitoe on behalf of the Wellcome Sanger Institute COVID-19 Surveillance Team |
| EPI_ISL_947358, EPI_ISL_947359 | Lighthouse Lab in Milton Keynes | Wellcome Sanger Institute for the COVID-19 Genomics UK (COG-UK) Consortium | The Lighthouse Lab in Milton Keynes and Alex Alderton, Roberto Amato, Sonia Goncalves, Ewan Harrison, David K. Jackson, Ian Johnston, Dominic Kwiatkowski, Cordelia Langford, John Sillitoe on behalf of the Wellcome Sanger Institute COVID-19 Surveillance Team |
| EPI_ISL_947360, EPI_ISL_947362, EPI_ISL_947363, EPI_ISL_947364, EPI_ISL_947365 | Lighthouse Lab in Glasgow | Wellcome Sanger Institute for the COVID-19 Genomics UK (COG-UK) Consortium | Harper VanSteenhouse, Yumi Kasai, David Gray, Carol Clugston, Anna Dominiczak and Alex Alderton, Roberto Amato, Sonia Goncalves, Ewan Harrison, David K. Jackson, Ian Johnston, Dominic Kwiatkowski, Cordelia Langford, John Sillitoe on behalf of the Wellcome Sanger Institute COVID-19 Surveillance Team |
| EPI_ISL_947366, EPI_ISL_947367, EPI_ISL_947369 | Lighthouse Lab in Milton Keynes | Wellcome Sanger Institute for the COVID-19 Genomics UK (COG-UK) Consortium | The Lighthouse Lab in Milton Keynes and Alex Alderton, Roberto Amato, Sonia Goncalves, Ewan Harrison, David K. Jackson, Ian Johnston, Dominic Kwiatkowski, Cordelia Langford, John Sillitoe on behalf of the Wellcome Sanger Institute COVID-19 Surveillance Team |
| EPI_ISL_947370 | Lighthouse Lab in Glasgow | Wellcome Sanger Institute for the COVID-19 Genomics UK (COG-UK) Consortium | Harper VanSteenhouse, Yumi Kasai, David Gray, Carol Clugston, Anna Dominiczak and Alex Alderton, Roberto Amato, Sonia Goncalves, Ewan Harrison, David K. Jackson, Ian Johnston, Dominic Kwiatkowski, Cordelia Langford, John Sillitoe on behalf of the Wellcome Sanger Institute COVID-19 Surveillance Team |
| EPI_ISL_947371, EPI_ISL_947372, EPI_ISL_947373, EPI_ISL_947375, EPI_ISL_947376 | Lighthouse Lab in Milton Keynes | Wellcome Sanger Institute for the COVID-19 Genomics UK (COG-UK) Consortium | The Lighthouse Lab in Milton Keynes and Alex Alderton, Roberto Amato, Sonia Goncalves, Ewan Harrison, David K. Jackson, Ian Johnston, Dominic Kwiatkowski, Cordelia Langford, John Sillitoe on behalf of the Wellcome Sanger Institute COVID-19 Surveillance Team |
| EPI_ISL_947377 | Lighthouse Lab in Glasgow | Wellcome Sanger Institute for the COVID-19 Genomics UK (COG-UK) Consortium | Harper VanSteenhouse, Yumi Kasai, David Gray, Carol Clugston, Anna Dominiczak and Alex Alderton, Roberto Amato, Sonia Goncalves, Ewan Harrison, David K. Jackson, Ian Johnston, Dominic Kwiatkowski, Cordelia Langford, John Sillitoe on behalf of the Wellcome Sanger Institute COVID-19 Surveillance Team |

| | | | |
|---|---|---|---|
| EPI_ISL_947378, EPI_ISL_947379 | Lighthouse Lab in Milton Keynes | Wellcome Sanger Institute for the COVID-19 Genomics UK (COG-UK) Consortium | The Lighthouse Lab in Milton Keynes and Alex Alderton, Roberto Amato, Sonia Goncalves, Ewan Harrison, David K. Jackson, Ian Johnston, Dominic Kwiatkowski, Cordelia Langford, John Sillitoe on behalf of the Wellcome Sanger Institute COVID-19 Surveillance Team |
| EPI_ISL_947380 | Lighthouse Lab in Glasgow | Wellcome Sanger Institute for the COVID-19 Genomics UK (COG-UK) Consortium | Harper VanSteenhouse, Yumi Kasai, David Gray, Carol Clugston, Anna Dominiczak and Alex Alderton, Roberto Amato, Sonia Goncalves, Ewan Harrison, David K. Jackson, Ian Johnston, Dominic Kwiatkowski, Cordelia Langford, John Sillitoe on behalf of the Wellcome Sanger Institute COVID-19 Surveillance Team |
| EPI_ISL_947381, EPI_ISL_947382 | Lighthouse Lab in Milton Keynes | Wellcome Sanger Institute for the COVID-19 Genomics UK (COG-UK) Consortium | The Lighthouse Lab in Milton Keynes and Alex Alderton, Roberto Amato, Sonia Goncalves, Ewan Harrison, David K. Jackson, Ian Johnston, Dominic Kwiatkowski, Cordelia Langford, John Sillitoe on behalf of the Wellcome Sanger Institute COVID-19 Surveillance Team |
| EPI_ISL_947383 | Lighthouse Lab in Glasgow | Wellcome Sanger Institute for the COVID-19 Genomics UK (COG-UK) Consortium | Harper VanSteenhouse, Yumi Kasai, David Gray, Carol Clugston, Anna Dominiczak and Alex Alderton, Roberto Amato, Sonia Goncalves, Ewan Harrison, David K. Jackson, Ian Johnston, Dominic Kwiatkowski, Cordelia Langford, John Sillitoe on behalf of the Wellcome Sanger Institute COVID-19 Surveillance Team |
| EPI_ISL_947384, EPI_ISL_947385 | Lighthouse Lab in Milton Keynes | Wellcome Sanger Institute for the COVID-19 Genomics UK (COG-UK) Consortium | The Lighthouse Lab in Milton Keynes and Alex Alderton, Roberto Amato, Sonia Goncalves, Ewan Harrison, David K. Jackson, Ian Johnston, Dominic Kwiatkowski, Cordelia Langford, John Sillitoe on behalf of the Wellcome Sanger Institute COVID-19 Surveillance Team |
| EPI_ISL_947386 | Lighthouse Lab in Glasgow | Wellcome Sanger Institute for the COVID-19 Genomics UK (COG-UK) Consortium | Harper VanSteenhouse, Yumi Kasai, David Gray, Carol Clugston, Anna Dominiczak and Alex Alderton, Roberto Amato, Sonia Goncalves, Ewan Harrison, David K. Jackson, Ian Johnston, Dominic Kwiatkowski, Cordelia Langford, John Sillitoe on behalf of the Wellcome Sanger Institute COVID-19 Surveillance Team |
| EPI_ISL_947387, EPI_ISL_947388 | Lighthouse Lab in Milton Keynes | Wellcome Sanger Institute for the COVID-19 Genomics UK (COG-UK) Consortium | The Lighthouse Lab in Milton Keynes and Alex Alderton, Roberto Amato, Sonia Goncalves, Ewan Harrison, David K. Jackson, Ian Johnston, Dominic Kwiatkowski, Cordelia Langford, John Sillitoe on behalf of the Wellcome Sanger Institute COVID-19 Surveillance Team |
| EPI_ISL_947389 | Lighthouse Lab in Glasgow | Wellcome Sanger Institute for the COVID-19 Genomics UK (COG-UK) Consortium | Harper VanSteenhouse, Yumi Kasai, David Gray, Carol Clugston, Anna Dominiczak and Alex Alderton, Roberto Amato, Sonia Goncalves, Ewan Harrison, David K. Jackson, Ian Johnston, Dominic Kwiatkowski, Cordelia Langford, John Sillitoe on behalf of the Wellcome Sanger Institute COVID-19 Surveillance Team |
| EPI_ISL_947390 | Lighthouse Lab in Milton Keynes | Wellcome Sanger Institute for the COVID-19 Genomics UK (COG-UK) Consortium | The Lighthouse Lab in Milton Keynes and Alex Alderton, Roberto Amato, Sonia Goncalves, Ewan Harrison, David K. Jackson, Ian Johnston, Dominic Kwiatkowski, Cordelia Langford, John Sillitoe on behalf of the Wellcome Sanger Institute COVID-19 Surveillance Team |
| EPI_ISL_947391 | Lighthouse Lab in Glasgow | Wellcome Sanger Institute for the COVID-19 Genomics UK (COG-UK) Consortium | Harper VanSteenhouse, Yumi Kasai, David Gray, Carol Clugston, Anna Dominiczak and Alex Alderton, Roberto Amato, Sonia Goncalves, Ewan Harrison, David K. Jackson, Ian Johnston, Dominic Kwiatkowski, Cordelia Langford, John Sillitoe on behalf of the Wellcome Sanger Institute COVID-19 Surveillance Team |
| EPI_ISL_947392, EPI_ISL_947393 | Lighthouse Lab in Milton Keynes | Wellcome Sanger Institute for the COVID-19 Genomics UK (COG-UK) Consortium | The Lighthouse Lab in Milton Keynes and Alex Alderton, Roberto Amato, Sonia Goncalves, Ewan Harrison, David K. Jackson, Ian Johnston, Dominic Kwiatkowski, Cordelia Langford, John Sillitoe on behalf of the Wellcome Sanger Institute COVID-19 Surveillance Team |
| EPI_ISL_947394 | Lighthouse Lab in Glasgow | Wellcome Sanger Institute for the COVID-19 Genomics UK (COG-UK) Consortium | Harper VanSteenhouse, Yumi Kasai, David Gray, Carol Clugston, Anna Dominiczak and Alex Alderton, Roberto Amato, Sonia Goncalves, Ewan Harrison, David K. Jackson, Ian Johnston, Dominic Kwiatkowski, Cordelia Langford, John Sillitoe on behalf of the Wellcome Sanger Institute COVID-19 Surveillance Team |
| EPI_ISL_947395 | Lighthouse Lab in Milton Keynes | Wellcome Sanger Institute for the COVID-19 Genomics UK (COG-UK) Consortium | The Lighthouse Lab in Milton Keynes and Alex Alderton, Roberto Amato, Sonia Goncalves, Ewan Harrison, David K. Jackson, Ian Johnston, Dominic Kwiatkowski, Cordelia Langford, John Sillitoe on behalf of the Wellcome Sanger Institute COVID-19 Surveillance Team |
| EPI_ISL_947396 | Lighthouse Lab in Glasgow | Wellcome Sanger Institute for the COVID-19 Genomics UK (COG-UK) Consortium | Harper VanSteenhouse, Yumi Kasai, David Gray, Carol Clugston, Anna Dominiczak and Alex Alderton, Roberto Amato, Sonia Goncalves, Ewan Harrison, David K. Jackson, Ian Johnston, Dominic Kwiatkowski, Cordelia Langford, John Sillitoe on behalf of the Wellcome Sanger Institute COVID-19 Surveillance Team |
| EPI_ISL_947397 | Lighthouse Lab in Milton Keynes | Wellcome Sanger Institute for the COVID-19 Genomics UK (COG-UK) Consortium | The Lighthouse Lab in Milton Keynes and Alex Alderton, Roberto Amato, Sonia Goncalves, Ewan Harrison, David K. Jackson, Ian Johnston, Dominic Kwiatkowski, Cordelia Langford, John Sillitoe on behalf of the Wellcome Sanger Institute COVID-19 Surveillance Team |
| EPI_ISL_947398, EPI_ISL_947399 | Lighthouse Lab in Glasgow | Wellcome Sanger Institute for the COVID-19 Genomics UK (COG-UK) Consortium | Harper VanSteenhouse, Yumi Kasai, David Gray, Carol Clugston, Anna Dominiczak and Alex Alderton, Roberto Amato, Sonia Goncalves, Ewan Harrison, David K. Jackson, Ian Johnston, Dominic Kwiatkowski, Cordelia Langford, John Sillitoe on behalf of the Wellcome Sanger Institute COVID-19 Surveillance Team |
| EPI_ISL_947401 | Lighthouse Lab in Milton Keynes | Wellcome Sanger Institute for the COVID-19 Genomics UK (COG-UK) Consortium | The Lighthouse Lab in Milton Keynes and Alex Alderton, Roberto Amato, Sonia Goncalves, Ewan Harrison, David K. Jackson, Ian Johnston, Dominic Kwiatkowski, Cordelia Langford, John Sillitoe on behalf of the Wellcome Sanger Institute COVID-19 Surveillance Team |
| EPI_ISL_947402 | Lighthouse Lab in Glasgow | Wellcome Sanger Institute for the COVID-19 Genomics UK (COG-UK) Consortium | Harper VanSteenhouse, Yumi Kasai, David Gray, Carol Clugston, Anna Dominiczak and Alex Alderton, Roberto Amato, Sonia Goncalves, Ewan Harrison, David K. Jackson, Ian Johnston, Dominic Kwiatkowski, Cordelia Langford, John Sillitoe on behalf of the Wellcome Sanger Institute COVID-19 Surveillance Team |
| EPI_ISL_947403 | Lighthouse Lab in Milton Keynes | Wellcome Sanger Institute for the COVID-19 Genomics UK (COG-UK) Consortium | The Lighthouse Lab in Milton Keynes and Alex Alderton, Roberto Amato, Sonia Goncalves, Ewan Harrison, David K. Jackson, Ian Johnston, Dominic Kwiatkowski, Cordelia Langford, John Sillitoe on behalf of the Wellcome Sanger Institute COVID-19 Surveillance Team |
| EPI_ISL_947404, EPI_ISL_947405, EPI_ISL_948100 | Lighthouse Lab in Glasgow | Wellcome Sanger Institute for the COVID-19 Genomics UK (COG-UK) Consortium | Harper VanSteenhouse, Yumi Kasai, David Gray, Carol Clugston, Anna Dominiczak and Alex Alderton, Roberto Amato, Sonia Goncalves, Ewan Harrison, David K. Jackson, Ian Johnston, Dominic Kwiatkowski, Cordelia Langford, John Sillitoe on behalf of the Wellcome Sanger Institute COVID-19 Surveillance Team |
| EPI_ISL_948102 | Lighthouse Lab in Cambridge | Wellcome Sanger Institute for the COVID-19 Genomics UK (COG-UK) Consortium | Rob Howes, The Lighthouse Lab in Cambridge and Alex Alderton, Roberto Amato, Sonia Goncalves, Ewan Harrison, David K. Jackson, Ian Johnston, Dominic Kwiatkowski, Cordelia Langford, John Sillitoe on behalf of the Wellcome Sanger Institute COVID-19 Surveillance Team |
| EPI_ISL_948103 | Lighthouse Lab in Glasgow | Wellcome Sanger Institute for the COVID-19 Genomics UK (COG-UK) Consortium | Harper VanSteenhouse, Yumi Kasai, David Gray, Carol Clugston, Anna Dominiczak and Alex Alderton, Roberto Amato, Sonia Goncalves, Ewan Harrison, David K. Jackson, Ian Johnston, Dominic Kwiatkowski, Cordelia Langford, John Sillitoe on behalf of the Wellcome Sanger Institute COVID-19 Surveillance Team |
| EPI_ISL_948104 | Lighthouse Lab in Cambridge | Wellcome Sanger Institute for the COVID-19 Genomics UK (COG-UK) Consortium | Rob Howes, The Lighthouse Lab in Cambridge and Alex Alderton, Roberto Amato, Sonia Goncalves, Ewan Harrison, David K. Jackson, Ian Johnston, Dominic Kwiatkowski, Cordelia Langford, John Sillitoe on behalf of the Wellcome Sanger Institute COVID-19 Surveillance Team |
| EPI_ISL_948105, EPI_ISL_948106 | Lighthouse Lab in Glasgow | Wellcome Sanger Institute for the COVID-19 Genomics UK (COG-UK) Consortium | Harper VanSteenhouse, Yumi Kasai, David Gray, Carol Clugston, Anna Dominiczak and Alex Alderton, Roberto Amato, Sonia Goncalves, Ewan Harrison, David K. Jackson, Ian Johnston, Dominic Kwiatkowski, Cordelia Langford, John Sillitoe on behalf of the Wellcome Sanger Institute COVID-19 Surveillance Team |
| EPI_ISL_948107 | Lighthouse Lab in Cambridge | Wellcome Sanger Institute for the COVID-19 Genomics UK (COG-UK) Consortium | Rob Howes, The Lighthouse Lab in Cambridge and Alex Alderton, Roberto Amato, Sonia Goncalves, Ewan Harrison, David K. Jackson, Ian Johnston, Dominic Kwiatkowski, Cordelia Langford, John Sillitoe on behalf of the Wellcome Sanger Institute COVID-19 Surveillance Team |
| EPI_ISL_948108, EPI_ISL_948109 | Lighthouse Lab in Glasgow | Wellcome Sanger Institute for the COVID-19 Genomics UK (COG-UK) Consortium | Harper VanSteenhouse, Yumi Kasai, David Gray, Carol Clugston, Anna Dominiczak and Alex Alderton, Roberto Amato, Sonia Goncalves, Ewan Harrison, David K. Jackson, Ian Johnston, Dominic Kwiatkowski, Cordelia Langford, John Sillitoe on behalf of the Wellcome Sanger Institute COVID-19 Surveillance Team |
| EPI_ISL_948110 | Lighthouse Lab in Cambridge | Wellcome Sanger Institute for the COVID-19 Genomics UK (COG-UK) Consortium | Rob Howes, The Lighthouse Lab in Cambridge and Alex Alderton, Roberto Amato, Sonia Goncalves, Ewan Harrison, David K. Jackson, Ian Johnston, Dominic Kwiatkowski, Cordelia Langford, John Sillitoe on behalf of the Wellcome Sanger Institute COVID-19 Surveillance Team |
| EPI_ISL_948111, EPI_ISL_948112, EPI_ISL_948113 | Lighthouse Lab in Glasgow | Wellcome Sanger Institute for the COVID-19 Genomics UK (COG-UK) Consortium | Harper VanSteenhouse, Yumi Kasai, David Gray, Carol Clugston, Anna Dominiczak and Alex Alderton, Roberto Amato, Sonia Goncalves, Ewan Harrison, David K. Jackson, Ian Johnston, Dominic Kwiatkowski, Cordelia Langford, John Sillitoe on behalf of the Wellcome Sanger Institute COVID-19 Surveillance Team |
| EPI_ISL_948114 | Lighthouse Lab in Cambridge | Wellcome Sanger Institute for the COVID-19 Genomics UK (COG-UK) Consortium | Rob Howes, The Lighthouse Lab in Cambridge and Alex Alderton, Roberto Amato, Sonia Goncalves, Ewan Harrison, David K. Jackson, Ian Johnston, Dominic Kwiatkowski, Cordelia Langford, John Sillitoe on behalf of the Wellcome Sanger Institute COVID-19 Surveillance Team |
| EPI_ISL_948115 | Lighthouse Lab in Glasgow | Wellcome Sanger Institute for the COVID-19 Genomics UK | Harper VanSteenhouse, Yumi Kasai, David Gray, Carol Clugston, Anna Dominiczak and Alex Alderton, Roberto Amato, Sonia Goncalves, Ewan Harrison, |

| | | | |
|---|---|---|---|
| | | (COG-UK) Consortium | David K. Jackson, Ian Johnston, Dominic Kwiatkowski, Cordelia Langford, John Sillitoe on behalf of the Wellcome Sanger Institute COVID-19 Surveillance Team |
| EPI_ISL_948117, EPI_ISL_948118 | Lighthouse Lab in Cambridge | Wellcome Sanger Institute for the COVID-19 Genomics UK (COG-UK) Consortium | Rob Howes, The Lighthouse Lab in Cambridge and Alex Alderton, Roberto Amato, Sonia Goncalves, Ewan Harrison, David K. Jackson, Ian Johnston, Dominic Kwiatkowski, Cordelia Langford, John Sillitoe on behalf of the Wellcome Sanger Institute COVID-19 Surveillance Team |
| EPI_ISL_948119 | Lighthouse Lab in Glasgow | Wellcome Sanger Institute for the COVID-19 Genomics UK (COG-UK) Consortium | Harper VanSteenhouse, Yumi Kasai, David Gray, Carol Clugston, Anna Dominiczak and Alex Alderton, Roberto Amato, Sonia Goncalves, Ewan Harrison, David K. Jackson, Ian Johnston, Dominic Kwiatkowski, Cordelia Langford, John Sillitoe on behalf of the Wellcome Sanger Institute COVID-19 Surveillance Team |
| EPI_ISL_948120 | Lighthouse Lab in Cambridge | Wellcome Sanger Institute for the COVID-19 Genomics UK (COG-UK) Consortium | Rob Howes, The Lighthouse Lab in Cambridge and Alex Alderton, Roberto Amato, Sonia Goncalves, Ewan Harrison, David K. Jackson, Ian Johnston, Dominic Kwiatkowski, Cordelia Langford, John Sillitoe on behalf of the Wellcome Sanger Institute COVID-19 Surveillance Team |
| EPI_ISL_948121 | Lighthouse Lab in Glasgow | Wellcome Sanger Institute for the COVID-19 Genomics UK (COG-UK) Consortium | Harper VanSteenhouse, Yumi Kasai, David Gray, Carol Clugston, Anna Dominiczak and Alex Alderton, Roberto Amato, Sonia Goncalves, Ewan Harrison, David K. Jackson, Ian Johnston, Dominic Kwiatkowski, Cordelia Langford, John Sillitoe on behalf of the Wellcome Sanger Institute COVID-19 Surveillance Team |
| EPI_ISL_948122, EPI_ISL_948123, EPI_ISL_948124 | Lighthouse Lab in Cambridge | Wellcome Sanger Institute for the COVID-19 Genomics UK (COG-UK) Consortium | Rob Howes, The Lighthouse Lab in Cambridge and Alex Alderton, Roberto Amato, Sonia Goncalves, Ewan Harrison, David K. Jackson, Ian Johnston, Dominic Kwiatkowski, Cordelia Langford, John Sillitoe on behalf of the Wellcome Sanger Institute COVID-19 Surveillance Team |
| EPI_ISL_948126, EPI_ISL_948127, EPI_ISL_948129, EPI_ISL_948130 | Lighthouse Lab in Glasgow | Wellcome Sanger Institute for the COVID-19 Genomics UK (COG-UK) Consortium | Harper VanSteenhouse, Yumi Kasai, David Gray, Carol Clugston, Anna Dominiczak and Alex Alderton, Roberto Amato, Sonia Goncalves, Ewan Harrison, David K. Jackson, Ian Johnston, Dominic Kwiatkowski, Cordelia Langford, John Sillitoe on behalf of the Wellcome Sanger Institute COVID-19 Surveillance Team |
| EPI_ISL_948131 | Lighthouse Lab in Cambridge | Wellcome Sanger Institute for the COVID-19 Genomics UK (COG-UK) Consortium | Rob Howes, The Lighthouse Lab in Cambridge and Alex Alderton, Roberto Amato, Sonia Goncalves, Ewan Harrison, David K. Jackson, Ian Johnston, Dominic Kwiatkowski, Cordelia Langford, John Sillitoe on behalf of the Wellcome Sanger Institute COVID-19 Surveillance Team |
| EPI_ISL_948132 | Lighthouse Lab in Glasgow | Wellcome Sanger Institute for the COVID-19 Genomics UK (COG-UK) Consortium | Harper VanSteenhouse, Yumi Kasai, David Gray, Carol Clugston, Anna Dominiczak and Alex Alderton, Roberto Amato, Sonia Goncalves, Ewan Harrison, David K. Jackson, Ian Johnston, Dominic Kwiatkowski, Cordelia Langford, John Sillitoe on behalf of the Wellcome Sanger Institute COVID-19 Surveillance Team |
| EPI_ISL_948134 | Lighthouse Lab in Cambridge | Wellcome Sanger Institute for the COVID-19 Genomics UK (COG-UK) Consortium | Rob Howes, The Lighthouse Lab in Cambridge and Alex Alderton, Roberto Amato, Sonia Goncalves, Ewan Harrison, David K. Jackson, Ian Johnston, Dominic Kwiatkowski, Cordelia Langford, John Sillitoe on behalf of the Wellcome Sanger Institute COVID-19 Surveillance Team |
| EPI_ISL_948135 | Lighthouse Lab in Milton Keynes | Wellcome Sanger Institute for the COVID-19 Genomics UK (COG-UK) Consortium | The Lighthouse Lab in Milton Keynes and Alex Alderton, Roberto Amato, Sonia Goncalves, Ewan Harrison, David K. Jackson, Ian Johnston, Dominic Kwiatkowski, Cordelia Langford, John Sillitoe on behalf of the Wellcome Sanger Institute COVID-19 Surveillance Team |
| EPI_ISL_948136, EPI_ISL_948137 | Lighthouse Lab in Cambridge | Wellcome Sanger Institute for the COVID-19 Genomics UK (COG-UK) Consortium | Rob Howes, The Lighthouse Lab in Cambridge and Alex Alderton, Roberto Amato, Sonia Goncalves, Ewan Harrison, David K. Jackson, Ian Johnston, Dominic Kwiatkowski, Cordelia Langford, John Sillitoe on behalf of the Wellcome Sanger Institute COVID-19 Surveillance Team |
| EPI_ISL_948138 | Lighthouse Lab in Glasgow | Wellcome Sanger Institute for the COVID-19 Genomics UK (COG-UK) Consortium | Harper VanSteenhouse, Yumi Kasai, David Gray, Carol Clugston, Anna Dominiczak and Alex Alderton, Roberto Amato, Sonia Goncalves, Ewan Harrison, David K. Jackson, Ian Johnston, Dominic Kwiatkowski, Cordelia Langford, John Sillitoe on behalf of the Wellcome Sanger Institute COVID-19 Surveillance Team |
| EPI_ISL_948139 | Lighthouse Lab in Cambridge | Wellcome Sanger Institute for the COVID-19 Genomics UK (COG-UK) Consortium | Rob Howes, The Lighthouse Lab in Cambridge and Alex Alderton, Roberto Amato, Sonia Goncalves, Ewan Harrison, David K. Jackson, Ian Johnston, Dominic Kwiatkowski, Cordelia Langford, John Sillitoe on behalf of the Wellcome Sanger Institute COVID-19 Surveillance Team |
| EPI_ISL_948140, EPI_ISL_948141, EPI_ISL_948142, EPI_ISL_948143, EPI_ISL_948144 | Lighthouse Lab in Glasgow | Wellcome Sanger Institute for the COVID-19 Genomics UK (COG-UK) Consortium | Harper VanSteenhouse, Yumi Kasai, David Gray, Carol Clugston, Anna Dominiczak and Alex Alderton, Roberto Amato, Sonia Goncalves, Ewan Harrison, David K. Jackson, Ian Johnston, Dominic Kwiatkowski, Cordelia Langford, John Sillitoe on behalf of the Wellcome Sanger Institute COVID-19 Surveillance Team |
| EPI_ISL_948145 | Lighthouse Lab in Cambridge | Wellcome Sanger Institute for the COVID-19 Genomics UK (COG-UK) Consortium | Rob Howes, The Lighthouse Lab in Cambridge and Alex Alderton, Roberto Amato, Sonia Goncalves, Ewan Harrison, David K. Jackson, Ian Johnston, Dominic Kwiatkowski, Cordelia Langford, John Sillitoe on behalf of the Wellcome Sanger Institute COVID-19 Surveillance Team |
| EPI_ISL_948146 | Lighthouse Lab in Glasgow | Wellcome Sanger Institute for the COVID-19 Genomics UK (COG-UK) Consortium | Harper VanSteenhouse, Yumi Kasai, David Gray, Carol Clugston, Anna Dominiczak and Alex Alderton, Roberto Amato, Sonia Goncalves, Ewan Harrison, David K. Jackson, Ian Johnston, Dominic Kwiatkowski, Cordelia Langford, John Sillitoe on behalf of the Wellcome Sanger Institute COVID-19 Surveillance Team |
| EPI_ISL_948147 | Lighthouse Lab in Cambridge | Wellcome Sanger Institute for the COVID-19 Genomics UK (COG-UK) Consortium | Rob Howes, The Lighthouse Lab in Cambridge and Alex Alderton, Roberto Amato, Sonia Goncalves, Ewan Harrison, David K. Jackson, Ian Johnston, Dominic Kwiatkowski, Cordelia Langford, John Sillitoe on behalf of the Wellcome Sanger Institute COVID-19 Surveillance Team |
| EPI_ISL_948148 | Lighthouse Lab in Glasgow | Wellcome Sanger Institute for the COVID-19 Genomics UK (COG-UK) Consortium | Harper VanSteenhouse, Yumi Kasai, David Gray, Carol Clugston, Anna Dominiczak and Alex Alderton, Roberto Amato, Sonia Goncalves, Ewan Harrison, David K. Jackson, Ian Johnston, Dominic Kwiatkowski, Cordelia Langford, John Sillitoe on behalf of the Wellcome Sanger Institute COVID-19 Surveillance Team |
| EPI_ISL_948149, EPI_ISL_948151 | Lighthouse Lab in Cambridge | Wellcome Sanger Institute for the COVID-19 Genomics UK (COG-UK) Consortium | Rob Howes, The Lighthouse Lab in Cambridge and Alex Alderton, Roberto Amato, Sonia Goncalves, Ewan Harrison, David K. Jackson, Ian Johnston, Dominic Kwiatkowski, Cordelia Langford, John Sillitoe on behalf of the Wellcome Sanger Institute COVID-19 Surveillance Team |
| EPI_ISL_948152 | Lighthouse Lab in Glasgow | Wellcome Sanger Institute for the COVID-19 Genomics UK (COG-UK) Consortium | Harper VanSteenhouse, Yumi Kasai, David Gray, Carol Clugston, Anna Dominiczak and Alex Alderton, Roberto Amato, Sonia Goncalves, Ewan Harrison, David K. Jackson, Ian Johnston, Dominic Kwiatkowski, Cordelia Langford, John Sillitoe on behalf of the Wellcome Sanger Institute COVID-19 Surveillance Team |
| EPI_ISL_948153 | Lighthouse Lab in Cambridge | Wellcome Sanger Institute for the COVID-19 Genomics UK (COG-UK) Consortium | Rob Howes, The Lighthouse Lab in Cambridge and Alex Alderton, Roberto Amato, Sonia Goncalves, Ewan Harrison, David K. Jackson, Ian Johnston, Dominic Kwiatkowski, Cordelia Langford, John Sillitoe on behalf of the Wellcome Sanger Institute COVID-19 Surveillance Team |
| EPI_ISL_948154, EPI_ISL_948156, EPI_ISL_948157, EPI_ISL_948158, EPI_ISL_948161, EPI_ISL_948162 | Lighthouse Lab in Glasgow | Wellcome Sanger Institute for the COVID-19 Genomics UK (COG-UK) Consortium | Harper VanSteenhouse, Yumi Kasai, David Gray, Carol Clugston, Anna Dominiczak and Alex Alderton, Roberto Amato, Sonia Goncalves, Ewan Harrison, David K. Jackson, Ian Johnston, Dominic Kwiatkowski, Cordelia Langford, John Sillitoe on behalf of the Wellcome Sanger Institute COVID-19 Surveillance Team |
| EPI_ISL_948163, EPI_ISL_948164, EPI_ISL_948165, EPI_ISL_948166 | Lighthouse Lab in Cambridge | Wellcome Sanger Institute for the COVID-19 Genomics UK (COG-UK) Consortium | Rob Howes, The Lighthouse Lab in Cambridge and Alex Alderton, Roberto Amato, Sonia Goncalves, Ewan Harrison, David K. Jackson, Ian Johnston, Dominic Kwiatkowski, Cordelia Langford, John Sillitoe on behalf of the Wellcome Sanger Institute COVID-19 Surveillance Team |
| EPI_ISL_948167, EPI_ISL_948168, EPI_ISL_948169, EPI_ISL_948170, EPI_ISL_948171 | Lighthouse Lab in Glasgow | Wellcome Sanger Institute for the COVID-19 Genomics UK (COG-UK) Consortium | Harper VanSteenhouse, Yumi Kasai, David Gray, Carol Clugston, Anna Dominiczak and Alex Alderton, Roberto Amato, Sonia Goncalves, Ewan Harrison, David K. Jackson, Ian Johnston, Dominic Kwiatkowski, Cordelia Langford, John Sillitoe on behalf of the Wellcome Sanger Institute COVID-19 Surveillance Team |
| EPI_ISL_948172 | Lighthouse Lab in Cambridge | Wellcome Sanger Institute for the COVID-19 Genomics UK (COG-UK) Consortium | Rob Howes, The Lighthouse Lab in Cambridge and Alex Alderton, Roberto Amato, Sonia Goncalves, Ewan Harrison, David K. Jackson, Ian Johnston, Dominic Kwiatkowski, Cordelia Langford, John Sillitoe on behalf of the Wellcome Sanger Institute COVID-19 Surveillance Team |
| EPI_ISL_948173 | Lighthouse Lab in Glasgow | Wellcome Sanger Institute for the COVID-19 Genomics UK (COG-UK) Consortium | Harper VanSteenhouse, Yumi Kasai, David Gray, Carol Clugston, Anna Dominiczak and Alex Alderton, Roberto Amato, Sonia Goncalves, Ewan Harrison, David K. Jackson, Ian Johnston, Dominic Kwiatkowski, Cordelia Langford, John Sillitoe on behalf of the Wellcome Sanger Institute COVID-19 Surveillance Team |
| EPI_ISL_948174 | Lighthouse Lab in Cambridge | Wellcome Sanger Institute for the COVID-19 Genomics UK (COG-UK) Consortium | Rob Howes, The Lighthouse Lab in Cambridge and Alex Alderton, Roberto Amato, Sonia Goncalves, Ewan Harrison, David K. Jackson, Ian Johnston, Dominic Kwiatkowski, Cordelia Langford, John Sillitoe on behalf of the Wellcome Sanger Institute COVID-19 Surveillance Team |
| EPI_ISL_948175 | Lighthouse Lab in Glasgow | Wellcome Sanger Institute for the COVID-19 Genomics UK (COG-UK) Consortium | Harper VanSteenhouse, Yumi Kasai, David Gray, Carol Clugston, Anna Dominiczak and Alex Alderton, Roberto Amato, Sonia Goncalves, Ewan Harrison, David K. Jackson, Ian Johnston, Dominic Kwiatkowski, Cordelia Langford, John Sillitoe on behalf of the Wellcome Sanger Institute COVID-19 Surveillance Team |
| EPI_ISL_948176 | Lighthouse Lab in Cambridge | Wellcome Sanger Institute for the COVID-19 Genomics UK (COG-UK) Consortium | Rob Howes, The Lighthouse Lab in Cambridge and Alex Alderton, Roberto Amato, Sonia Goncalves, Ewan Harrison, David K. Jackson, Ian Johnston, Dominic Kwiatkowski, Cordelia Langford, John Sillitoe on behalf of the Wellcome Sanger Institute COVID-19 Surveillance Team |

| | | | |
|---|---|---|---|
| EPI_ISL_948179 | Lighthouse Lab in Glasgow | Wellcome Sanger Institute for the COVID-19 Genomics UK (COG-UK) Consortium | Harper VanSteenhouse, Yumi Kasai, David Gray, Carol Clugston, Anna Dominiczak and Alex Alderton, Roberto Amato, Sonia Goncalves, Ewan Harrison, David K. Jackson, Ian Johnston, Dominic Kwiatkowski, Cordelia Langford, John Sillitoe on behalf of the Wellcome Sanger Institute COVID-19 Surveillance Team |
| EPI_ISL_948180, EPI_ISL_948181 | Lighthouse Lab in Cambridge | Wellcome Sanger Institute for the COVID-19 Genomics UK (COG-UK) Consortium | Rob Howes, The Lighthouse Lab in Cambridge and Alex Alderton, Roberto Amato, Sonia Goncalves, Ewan Harrison, David K. Jackson, Ian Johnston, Dominic Kwiatkowski, Cordelia Langford, John Sillitoe on behalf of the Wellcome Sanger Institute COVID-19 Surveillance Team |
| EPI_ISL_948182 | Lighthouse Lab in Glasgow | Wellcome Sanger Institute for the COVID-19 Genomics UK (COG-UK) Consortium | Harper VanSteenhouse, Yumi Kasai, David Gray, Carol Clugston, Anna Dominiczak and Alex Alderton, Roberto Amato, Sonia Goncalves, Ewan Harrison, David K. Jackson, Ian Johnston, Dominic Kwiatkowski, Cordelia Langford, John Sillitoe on behalf of the Wellcome Sanger Institute COVID-19 Surveillance Team |
| EPI_ISL_948183 | Lighthouse Lab in Cambridge | Wellcome Sanger Institute for the COVID-19 Genomics UK (COG-UK) Consortium | Rob Howes, The Lighthouse Lab in Cambridge and Alex Alderton, Roberto Amato, Sonia Goncalves, Ewan Harrison, David K. Jackson, Ian Johnston, Dominic Kwiatkowski, Cordelia Langford, John Sillitoe on behalf of the Wellcome Sanger Institute COVID-19 Surveillance Team |
| EPI_ISL_948184 | Lighthouse Lab in Glasgow | Wellcome Sanger Institute for the COVID-19 Genomics UK (COG-UK) Consortium | Harper VanSteenhouse, Yumi Kasai, David Gray, Carol Clugston, Anna Dominiczak and Alex Alderton, Roberto Amato, Sonia Goncalves, Ewan Harrison, David K. Jackson, Ian Johnston, Dominic Kwiatkowski, Cordelia Langford, John Sillitoe on behalf of the Wellcome Sanger Institute COVID-19 Surveillance Team |
| EPI_ISL_948187, EPI_ISL_948188, EPI_ISL_948189 | Lighthouse Lab in Cambridge | Wellcome Sanger Institute for the COVID-19 Genomics UK (COG-UK) Consortium | Rob Howes, The Lighthouse Lab in Cambridge and Alex Alderton, Roberto Amato, Sonia Goncalves, Ewan Harrison, David K. Jackson, Ian Johnston, Dominic Kwiatkowski, Cordelia Langford, John Sillitoe on behalf of the Wellcome Sanger Institute COVID-19 Surveillance Team |
| EPI_ISL_948190 | Lighthouse Lab in Glasgow | Wellcome Sanger Institute for the COVID-19 Genomics UK (COG-UK) Consortium | Harper VanSteenhouse, Yumi Kasai, David Gray, Carol Clugston, Anna Dominiczak and Alex Alderton, Roberto Amato, Sonia Goncalves, Ewan Harrison, David K. Jackson, Ian Johnston, Dominic Kwiatkowski, Cordelia Langford, John Sillitoe on behalf of the Wellcome Sanger Institute COVID-19 Surveillance Team |
| EPI_ISL_948191, EPI_ISL_948192, EPI_ISL_948193 | Lighthouse Lab in Cambridge | Wellcome Sanger Institute for the COVID-19 Genomics UK (COG-UK) Consortium | Rob Howes, The Lighthouse Lab in Cambridge and Alex Alderton, Roberto Amato, Sonia Goncalves, Ewan Harrison, David K. Jackson, Ian Johnston, Dominic Kwiatkowski, Cordelia Langford, John Sillitoe on behalf of the Wellcome Sanger Institute COVID-19 Surveillance Team |
| EPI_ISL_948195, EPI_ISL_948196 | Lighthouse Lab in Glasgow | Wellcome Sanger Institute for the COVID-19 Genomics UK (COG-UK) Consortium | Harper VanSteenhouse, Yumi Kasai, David Gray, Carol Clugston, Anna Dominiczak and Alex Alderton, Roberto Amato, Sonia Goncalves, Ewan Harrison, David K. Jackson, Ian Johnston, Dominic Kwiatkowski, Cordelia Langford, John Sillitoe on behalf of the Wellcome Sanger Institute COVID-19 Surveillance Team |
| EPI_ISL_948197, EPI_ISL_948198, EPI_ISL_948199, EPI_ISL_948200, EPI_ISL_948201 | Lighthouse Lab in Cambridge | Wellcome Sanger Institute for the COVID-19 Genomics UK (COG-UK) Consortium | Rob Howes, The Lighthouse Lab in Cambridge and Alex Alderton, Roberto Amato, Sonia Goncalves, Ewan Harrison, David K. Jackson, Ian Johnston, Dominic Kwiatkowski, Cordelia Langford, John Sillitoe on behalf of the Wellcome Sanger Institute COVID-19 Surveillance Team |
| EPI_ISL_948202 | Lighthouse Lab in Glasgow | Wellcome Sanger Institute for the COVID-19 Genomics UK (COG-UK) Consortium | Harper VanSteenhouse, Yumi Kasai, David Gray, Carol Clugston, Anna Dominiczak and Alex Alderton, Roberto Amato, Sonia Goncalves, Ewan Harrison, David K. Jackson, Ian Johnston, Dominic Kwiatkowski, Cordelia Langford, John Sillitoe on behalf of the Wellcome Sanger Institute COVID-19 Surveillance Team |
| EPI_ISL_948203, EPI_ISL_948205 | Lighthouse Lab in Cambridge | Wellcome Sanger Institute for the COVID-19 Genomics UK (COG-UK) Consortium | Rob Howes, The Lighthouse Lab in Cambridge and Alex Alderton, Roberto Amato, Sonia Goncalves, Ewan Harrison, David K. Jackson, Ian Johnston, Dominic Kwiatkowski, Cordelia Langford, John Sillitoe on behalf of the Wellcome Sanger Institute COVID-19 Surveillance Team |
| EPI_ISL_948206 | Lighthouse Lab in Glasgow | Wellcome Sanger Institute for the COVID-19 Genomics UK (COG-UK) Consortium | Harper VanSteenhouse, Yumi Kasai, David Gray, Carol Clugston, Anna Dominiczak and Alex Alderton, Roberto Amato, Sonia Goncalves, Ewan Harrison, David K. Jackson, Ian Johnston, Dominic Kwiatkowski, Cordelia Langford, John Sillitoe on behalf of the Wellcome Sanger Institute COVID-19 Surveillance Team |
| EPI_ISL_948207, EPI_ISL_948208, EPI_ISL_948209, EPI_ISL_948211, EPI_ISL_948212, EPI_ISL_948213, EPI_ISL_948214, EPI_ISL_948216 | Lighthouse Lab in Cambridge | Wellcome Sanger Institute for the COVID-19 Genomics UK (COG-UK) Consortium | Rob Howes, The Lighthouse Lab in Cambridge and Alex Alderton, Roberto Amato, Sonia Goncalves, Ewan Harrison, David K. Jackson, Ian Johnston, Dominic Kwiatkowski, Cordelia Langford, John Sillitoe on behalf of the Wellcome Sanger Institute COVID-19 Surveillance Team |
| EPI_ISL_948217, EPI_ISL_948218 | Lighthouse Lab in Glasgow | Wellcome Sanger Institute for the COVID-19 Genomics UK (COG-UK) Consortium | Harper VanSteenhouse, Yumi Kasai, David Gray, Carol Clugston, Anna Dominiczak and Alex Alderton, Roberto Amato, Sonia Goncalves, Ewan Harrison, David K. Jackson, Ian Johnston, Dominic Kwiatkowski, Cordelia Langford, John Sillitoe on behalf of the Wellcome Sanger Institute COVID-19 Surveillance Team |
| EPI_ISL_948219, EPI_ISL_948220 | Lighthouse Lab in Cambridge | Wellcome Sanger Institute for the COVID-19 Genomics UK (COG-UK) Consortium | Rob Howes, The Lighthouse Lab in Cambridge and Alex Alderton, Roberto Amato, Sonia Goncalves, Ewan Harrison, David K. Jackson, Ian Johnston, Dominic Kwiatkowski, Cordelia Langford, John Sillitoe on behalf of the Wellcome Sanger Institute COVID-19 Surveillance Team |
| EPI_ISL_948222 | Lighthouse Lab in Glasgow | Wellcome Sanger Institute for the COVID-19 Genomics UK (COG-UK) Consortium | Harper VanSteenhouse, Yumi Kasai, David Gray, Carol Clugston, Anna Dominiczak and Alex Alderton, Roberto Amato, Sonia Goncalves, Ewan Harrison, David K. Jackson, Ian Johnston, Dominic Kwiatkowski, Cordelia Langford, John Sillitoe on behalf of the Wellcome Sanger Institute COVID-19 Surveillance Team |
| EPI_ISL_948223, EPI_ISL_948224 | Lighthouse Lab in Cambridge | Wellcome Sanger Institute for the COVID-19 Genomics UK (COG-UK) Consortium | Rob Howes, The Lighthouse Lab in Cambridge and Alex Alderton, Roberto Amato, Sonia Goncalves, Ewan Harrison, David K. Jackson, Ian Johnston, Dominic Kwiatkowski, Cordelia Langford, John Sillitoe on behalf of the Wellcome Sanger Institute COVID-19 Surveillance Team |
| EPI_ISL_948225, EPI_ISL_948226 | Lighthouse Lab in Glasgow | Wellcome Sanger Institute for the COVID-19 Genomics UK (COG-UK) Consortium | Harper VanSteenhouse, Yumi Kasai, David Gray, Carol Clugston, Anna Dominiczak and Alex Alderton, Roberto Amato, Sonia Goncalves, Ewan Harrison, David K. Jackson, Ian Johnston, Dominic Kwiatkowski, Cordelia Langford, John Sillitoe on behalf of the Wellcome Sanger Institute COVID-19 Surveillance Team |
| EPI_ISL_948227, EPI_ISL_948228 | Lighthouse Lab in Cambridge | Wellcome Sanger Institute for the COVID-19 Genomics UK (COG-UK) Consortium | Rob Howes, The Lighthouse Lab in Cambridge and Alex Alderton, Roberto Amato, Sonia Goncalves, Ewan Harrison, David K. Jackson, Ian Johnston, Dominic Kwiatkowski, Cordelia Langford, John Sillitoe on behalf of the Wellcome Sanger Institute COVID-19 Surveillance Team |
| EPI_ISL_948229 | Lighthouse Lab in Glasgow | Wellcome Sanger Institute for the COVID-19 Genomics UK (COG-UK) Consortium | Harper VanSteenhouse, Yumi Kasai, David Gray, Carol Clugston, Anna Dominiczak and Alex Alderton, Roberto Amato, Sonia Goncalves, Ewan Harrison, David K. Jackson, Ian Johnston, Dominic Kwiatkowski, Cordelia Langford, John Sillitoe on behalf of the Wellcome Sanger Institute COVID-19 Surveillance Team |
| EPI_ISL_948230 | Lighthouse Lab in Cambridge | Wellcome Sanger Institute for the COVID-19 Genomics UK (COG-UK) Consortium | Rob Howes, The Lighthouse Lab in Cambridge and Alex Alderton, Roberto Amato, Sonia Goncalves, Ewan Harrison, David K. Jackson, Ian Johnston, Dominic Kwiatkowski, Cordelia Langford, John Sillitoe on behalf of the Wellcome Sanger Institute COVID-19 Surveillance Team |
| EPI_ISL_948232 | Lighthouse Lab in Glasgow | Wellcome Sanger Institute for the COVID-19 Genomics UK (COG-UK) Consortium | Harper VanSteenhouse, Yumi Kasai, David Gray, Carol Clugston, Anna Dominiczak and Alex Alderton, Roberto Amato, Sonia Goncalves, Ewan Harrison, David K. Jackson, Ian Johnston, Dominic Kwiatkowski, Cordelia Langford, John Sillitoe on behalf of the Wellcome Sanger Institute COVID-19 Surveillance Team |
| EPI_ISL_948233, EPI_ISL_948234, EPI_ISL_948235 | Lighthouse Lab in Cambridge | Wellcome Sanger Institute for the COVID-19 Genomics UK (COG-UK) Consortium | Rob Howes, The Lighthouse Lab in Cambridge and Alex Alderton, Roberto Amato, Sonia Goncalves, Ewan Harrison, David K. Jackson, Ian Johnston, Dominic Kwiatkowski, Cordelia Langford, John Sillitoe on behalf of the Wellcome Sanger Institute COVID-19 Surveillance Team |
| EPI_ISL_948236 | Lighthouse Lab in Glasgow | Wellcome Sanger Institute for the COVID-19 Genomics UK (COG-UK) Consortium | Harper VanSteenhouse, Yumi Kasai, David Gray, Carol Clugston, Anna Dominiczak and Alex Alderton, Roberto Amato, Sonia Goncalves, Ewan Harrison, David K. Jackson, Ian Johnston, Dominic Kwiatkowski, Cordelia Langford, John Sillitoe on behalf of the Wellcome Sanger Institute COVID-19 Surveillance Team |
| EPI_ISL_948237 | Lighthouse Lab in Cambridge | Wellcome Sanger Institute for the COVID-19 Genomics UK (COG-UK) Consortium | Rob Howes, The Lighthouse Lab in Cambridge and Alex Alderton, Roberto Amato, Sonia Goncalves, Ewan Harrison, David K. Jackson, Ian Johnston, Dominic Kwiatkowski, Cordelia Langford, John Sillitoe on behalf of the Wellcome Sanger Institute COVID-19 Surveillance Team |
| EPI_ISL_948238 | Lighthouse Lab in Glasgow | Wellcome Sanger Institute for the COVID-19 Genomics UK (COG-UK) Consortium | Harper VanSteenhouse, Yumi Kasai, David Gray, Carol Clugston, Anna Dominiczak and Alex Alderton, Roberto Amato, Sonia Goncalves, Ewan Harrison, David K. Jackson, Ian Johnston, Dominic Kwiatkowski, Cordelia Langford, John Sillitoe on behalf of the Wellcome Sanger Institute COVID-19 Surveillance Team |
| EPI_ISL_948239, EPI_ISL_948240, EPI_ISL_948241 | Lighthouse Lab in Cambridge | Wellcome Sanger Institute for the COVID-19 Genomics UK (COG-UK) Consortium | Rob Howes, The Lighthouse Lab in Cambridge and Alex Alderton, Roberto Amato, Sonia Goncalves, Ewan Harrison, David K. Jackson, Ian Johnston, Dominic Kwiatkowski, Cordelia Langford, John Sillitoe on behalf of the Wellcome Sanger Institute COVID-19 Surveillance Team |

| | | | |
|---|---|---|---|
| EPI_ISL_948242, EPI_ISL_948243 | Lighthouse Lab in Glasgow | Wellcome Sanger Institute for the COVID-19 Genomics UK (COG-UK) Consortium | Harper VanSteenhouse, Yumi Kasai, David Gray, Carol Clugston, Anna Dominiczak and Alex Alderton, Roberto Amato, Sonia Goncalves, Ewan Harrison, David K. Jackson, Ian Johnston, Dominic Kwiatkowski, Cordelia Langford, John Sillitoe on behalf of the Wellcome Sanger Institute COVID-19 Surveillance Team |
| EPI_ISL_948244 | Lighthouse Lab in Cambridge | Wellcome Sanger Institute for the COVID-19 Genomics UK (COG-UK) Consortium | Rob Howes, The Lighthouse Lab in Cambridge and Alex Alderton, Roberto Amato, Sonia Goncalves, Ewan Harrison, David K. Jackson, Ian Johnston, Dominic Kwiatkowski, Cordelia Langford, John Sillitoe on behalf of the Wellcome Sanger Institute COVID-19 Surveillance Team |
| EPI_ISL_948245, EPI_ISL_948246 | Lighthouse Lab in Glasgow | Wellcome Sanger Institute for the COVID-19 Genomics UK (COG-UK) Consortium | Harper VanSteenhouse, Yumi Kasai, David Gray, Carol Clugston, Anna Dominiczak and Alex Alderton, Roberto Amato, Sonia Goncalves, Ewan Harrison, David K. Jackson, Ian Johnston, Dominic Kwiatkowski, Cordelia Langford, John Sillitoe on behalf of the Wellcome Sanger Institute COVID-19 Surveillance Team |
| EPI_ISL_948247 | Lighthouse Lab in Cambridge | Wellcome Sanger Institute for the COVID-19 Genomics UK (COG-UK) Consortium | Rob Howes, The Lighthouse Lab in Cambridge and Alex Alderton, Roberto Amato, Sonia Goncalves, Ewan Harrison, David K. Jackson, Ian Johnston, Dominic Kwiatkowski, Cordelia Langford, John Sillitoe on behalf of the Wellcome Sanger Institute COVID-19 Surveillance Team |
| EPI_ISL_948248 | Lighthouse Lab in Glasgow | Wellcome Sanger Institute for the COVID-19 Genomics UK (COG-UK) Consortium | Harper VanSteenhouse, Yumi Kasai, David Gray, Carol Clugston, Anna Dominiczak and Alex Alderton, Roberto Amato, Sonia Goncalves, Ewan Harrison, David K. Jackson, Ian Johnston, Dominic Kwiatkowski, Cordelia Langford, John Sillitoe on behalf of the Wellcome Sanger Institute COVID-19 Surveillance Team |
| EPI_ISL_948249, EPI_ISL_948250, EPI_ISL_948251 | Lighthouse Lab in Cambridge | Wellcome Sanger Institute for the COVID-19 Genomics UK (COG-UK) Consortium | Rob Howes, The Lighthouse Lab in Cambridge and Alex Alderton, Roberto Amato, Sonia Goncalves, Ewan Harrison, David K. Jackson, Ian Johnston, Dominic Kwiatkowski, Cordelia Langford, John Sillitoe on behalf of the Wellcome Sanger Institute COVID-19 Surveillance Team |
| EPI_ISL_948252 | Lighthouse Lab in Milton Keynes | Wellcome Sanger Institute for the COVID-19 Genomics UK (COG-UK) Consortium | The Lighthouse Lab in Milton Keynes and Alex Alderton, Roberto Amato, Sonia Goncalves, Ewan Harrison, David K. Jackson, Ian Johnston, Dominic Kwiatkowski, Cordelia Langford, John Sillitoe on behalf of the Wellcome Sanger Institute COVID-19 Surveillance Team |
| EPI_ISL_948253 | Lighthouse Lab in Cambridge | Wellcome Sanger Institute for the COVID-19 Genomics UK (COG-UK) Consortium | Rob Howes, The Lighthouse Lab in Cambridge and Alex Alderton, Roberto Amato, Sonia Goncalves, Ewan Harrison, David K. Jackson, Ian Johnston, Dominic Kwiatkowski, Cordelia Langford, John Sillitoe on behalf of the Wellcome Sanger Institute COVID-19 Surveillance Team |
| EPI_ISL_948254 | Lighthouse Lab in Glasgow | Wellcome Sanger Institute for the COVID-19 Genomics UK (COG-UK) Consortium | Harper VanSteenhouse, Yumi Kasai, David Gray, Carol Clugston, Anna Dominiczak and Alex Alderton, Roberto Amato, Sonia Goncalves, Ewan Harrison, David K. Jackson, Ian Johnston, Dominic Kwiatkowski, Cordelia Langford, John Sillitoe on behalf of the Wellcome Sanger Institute COVID-19 Surveillance Team |
| EPI_ISL_948255 | Lighthouse Lab in Cambridge | Wellcome Sanger Institute for the COVID-19 Genomics UK (COG-UK) Consortium | Rob Howes, The Lighthouse Lab in Cambridge and Alex Alderton, Roberto Amato, Sonia Goncalves, Ewan Harrison, David K. Jackson, Ian Johnston, Dominic Kwiatkowski, Cordelia Langford, John Sillitoe on behalf of the Wellcome Sanger Institute COVID-19 Surveillance Team |
| EPI_ISL_948256 | Lighthouse Lab in Glasgow | Wellcome Sanger Institute for the COVID-19 Genomics UK (COG-UK) Consortium | Harper VanSteenhouse, Yumi Kasai, David Gray, Carol Clugston, Anna Dominiczak and Alex Alderton, Roberto Amato, Sonia Goncalves, Ewan Harrison, David K. Jackson, Ian Johnston, Dominic Kwiatkowski, Cordelia Langford, John Sillitoe on behalf of the Wellcome Sanger Institute COVID-19 Surveillance Team |
| EPI_ISL_948257, EPI_ISL_948258, EPI_ISL_948259 | Lighthouse Lab in Cambridge | Wellcome Sanger Institute for the COVID-19 Genomics UK (COG-UK) Consortium | Rob Howes, The Lighthouse Lab in Cambridge and Alex Alderton, Roberto Amato, Sonia Goncalves, Ewan Harrison, David K. Jackson, Ian Johnston, Dominic Kwiatkowski, Cordelia Langford, John Sillitoe on behalf of the Wellcome Sanger Institute COVID-19 Surveillance Team |
| EPI_ISL_948260 | Lighthouse Lab in Glasgow | Wellcome Sanger Institute for the COVID-19 Genomics UK (COG-UK) Consortium | Harper VanSteenhouse, Yumi Kasai, David Gray, Carol Clugston, Anna Dominiczak and Alex Alderton, Roberto Amato, Sonia Goncalves, Ewan Harrison, David K. Jackson, Ian Johnston, Dominic Kwiatkowski, Cordelia Langford, John Sillitoe on behalf of the Wellcome Sanger Institute COVID-19 Surveillance Team |
| EPI_ISL_948261 | Lighthouse Lab in Cambridge | Wellcome Sanger Institute for the COVID-19 Genomics UK (COG-UK) Consortium | Rob Howes, The Lighthouse Lab in Cambridge and Alex Alderton, Roberto Amato, Sonia Goncalves, Ewan Harrison, David K. Jackson, Ian Johnston, Dominic Kwiatkowski, Cordelia Langford, John Sillitoe on behalf of the Wellcome Sanger Institute COVID-19 Surveillance Team |
| EPI_ISL_948262, EPI_ISL_948263 | Lighthouse Lab in Glasgow | Wellcome Sanger Institute for the COVID-19 Genomics UK (COG-UK) Consortium | Harper VanSteenhouse, Yumi Kasai, David Gray, Carol Clugston, Anna Dominiczak and Alex Alderton, Roberto Amato, Sonia Goncalves, Ewan Harrison, David K. Jackson, Ian Johnston, Dominic Kwiatkowski, Cordelia Langford, John Sillitoe on behalf of the Wellcome Sanger Institute COVID-19 Surveillance Team |
| EPI_ISL_948264, EPI_ISL_948265 | Lighthouse Lab in Cambridge | Wellcome Sanger Institute for the COVID-19 Genomics UK (COG-UK) Consortium | Rob Howes, The Lighthouse Lab in Cambridge and Alex Alderton, Roberto Amato, Sonia Goncalves, Ewan Harrison, David K. Jackson, Ian Johnston, Dominic Kwiatkowski, Cordelia Langford, John Sillitoe on behalf of the Wellcome Sanger Institute COVID-19 Surveillance Team |
| EPI_ISL_948267, EPI_ISL_948268 | Lighthouse Lab in Glasgow | Wellcome Sanger Institute for the COVID-19 Genomics UK (COG-UK) Consortium | Harper VanSteenhouse, Yumi Kasai, David Gray, Carol Clugston, Anna Dominiczak and Alex Alderton, Roberto Amato, Sonia Goncalves, Ewan Harrison, David K. Jackson, Ian Johnston, Dominic Kwiatkowski, Cordelia Langford, John Sillitoe on behalf of the Wellcome Sanger Institute COVID-19 Surveillance Team |
| EPI_ISL_948269 | Lighthouse Lab in Cambridge | Wellcome Sanger Institute for the COVID-19 Genomics UK (COG-UK) Consortium | Rob Howes, The Lighthouse Lab in Cambridge and Alex Alderton, Roberto Amato, Sonia Goncalves, Ewan Harrison, David K. Jackson, Ian Johnston, Dominic Kwiatkowski, Cordelia Langford, John Sillitoe on behalf of the Wellcome Sanger Institute COVID-19 Surveillance Team |
| EPI_ISL_948270, EPI_ISL_948271, EPI_ISL_948272, EPI_ISL_948273 | Lighthouse Lab in Glasgow | Wellcome Sanger Institute for the COVID-19 Genomics UK (COG-UK) Consortium | Harper VanSteenhouse, Yumi Kasai, David Gray, Carol Clugston, Anna Dominiczak and Alex Alderton, Roberto Amato, Sonia Goncalves, Ewan Harrison, David K. Jackson, Ian Johnston, Dominic Kwiatkowski, Cordelia Langford, John Sillitoe on behalf of the Wellcome Sanger Institute COVID-19 Surveillance Team |
| EPI_ISL_948274 | Lighthouse Lab in Cambridge | Wellcome Sanger Institute for the COVID-19 Genomics UK (COG-UK) Consortium | Rob Howes, The Lighthouse Lab in Cambridge and Alex Alderton, Roberto Amato, Sonia Goncalves, Ewan Harrison, David K. Jackson, Ian Johnston, Dominic Kwiatkowski, Cordelia Langford, John Sillitoe on behalf of the Wellcome Sanger Institute COVID-19 Surveillance Team |
| EPI_ISL_948275, EPI_ISL_948276 | Lighthouse Lab in Glasgow | Wellcome Sanger Institute for the COVID-19 Genomics UK (COG-UK) Consortium | Harper VanSteenhouse, Yumi Kasai, David Gray, Carol Clugston, Anna Dominiczak and Alex Alderton, Roberto Amato, Sonia Goncalves, Ewan Harrison, David K. Jackson, Ian Johnston, Dominic Kwiatkowski, Cordelia Langford, John Sillitoe on behalf of the Wellcome Sanger Institute COVID-19 Surveillance Team |
| EPI_ISL_948277, EPI_ISL_948278, EPI_ISL_948279, EPI_ISL_948280, EPI_ISL_948281, EPI_ISL_948282, EPI_ISL_948283, EPI_ISL_948284 | Lighthouse Lab in Cambridge | Wellcome Sanger Institute for the COVID-19 Genomics UK (COG-UK) Consortium | Rob Howes, The Lighthouse Lab in Cambridge and Alex Alderton, Roberto Amato, Sonia Goncalves, Ewan Harrison, David K. Jackson, Ian Johnston, Dominic Kwiatkowski, Cordelia Langford, John Sillitoe on behalf of the Wellcome Sanger Institute COVID-19 Surveillance Team |
| EPI_ISL_948285 | Lighthouse Lab in Glasgow | Wellcome Sanger Institute for the COVID-19 Genomics UK (COG-UK) Consortium | Harper VanSteenhouse, Yumi Kasai, David Gray, Carol Clugston, Anna Dominiczak and Alex Alderton, Roberto Amato, Sonia Goncalves, Ewan Harrison, David K. Jackson, Ian Johnston, Dominic Kwiatkowski, Cordelia Langford, John Sillitoe on behalf of the Wellcome Sanger Institute COVID-19 Surveillance Team |
| EPI_ISL_948286 | Lighthouse Lab in Cambridge | Wellcome Sanger Institute for the COVID-19 Genomics UK (COG-UK) Consortium | Rob Howes, The Lighthouse Lab in Cambridge and Alex Alderton, Roberto Amato, Sonia Goncalves, Ewan Harrison, David K. Jackson, Ian Johnston, Dominic Kwiatkowski, Cordelia Langford, John Sillitoe on behalf of the Wellcome Sanger Institute COVID-19 Surveillance Team |
| EPI_ISL_948287 | Lighthouse Lab in Glasgow | Wellcome Sanger Institute for the COVID-19 Genomics UK (COG-UK) Consortium | Harper VanSteenhouse, Yumi Kasai, David Gray, Carol Clugston, Anna Dominiczak and Alex Alderton, Roberto Amato, Sonia Goncalves, Ewan Harrison, David K. Jackson, Ian Johnston, Dominic Kwiatkowski, Cordelia Langford, John Sillitoe on behalf of the Wellcome Sanger Institute COVID-19 Surveillance Team |
| EPI_ISL_948288 | Lighthouse Lab in Cambridge | Wellcome Sanger Institute for the COVID-19 Genomics UK (COG-UK) Consortium | Rob Howes, The Lighthouse Lab in Cambridge and Alex Alderton, Roberto Amato, Sonia Goncalves, Ewan Harrison, David K. Jackson, Ian Johnston, Dominic Kwiatkowski, Cordelia Langford, John Sillitoe on behalf of the Wellcome Sanger Institute COVID-19 Surveillance Team |
| EPI_ISL_948290, EPI_ISL_948291, EPI_ISL_948293, EPI_ISL_948295 | Lighthouse Lab in Glasgow | Wellcome Sanger Institute for the COVID-19 Genomics UK (COG-UK) Consortium | Harper VanSteenhouse, Yumi Kasai, David Gray, Carol Clugston, Anna Dominiczak and Alex Alderton, Roberto Amato, Sonia Goncalves, Ewan Harrison, David K. Jackson, Ian Johnston, Dominic Kwiatkowski, Cordelia Langford, John Sillitoe on behalf of the Wellcome Sanger Institute COVID-19 Surveillance Team |
| EPI_ISL_948296 | Lighthouse Lab in Cambridge | Wellcome Sanger Institute for the COVID-19 Genomics UK (COG-UK) Consortium | Rob Howes, The Lighthouse Lab in Cambridge and Alex Alderton, Roberto Amato, Sonia Goncalves, Ewan Harrison, David K. Jackson, Ian Johnston, Dominic Kwiatkowski, Cordelia Langford, John Sillitoe on behalf of the Wellcome Sanger Institute COVID-19 Surveillance Team |
| EPI_ISL_948297 | Lighthouse Lab in Glasgow | Wellcome Sanger Institute for the COVID-19 Genomics UK (COG-UK) Consortium | Harper VanSteenhouse, Yumi Kasai, David Gray, Carol Clugston, Anna Dominiczak and Alex Alderton, Roberto Amato, Sonia Goncalves, Ewan Harrison, David K. Jackson, Ian Johnston, Dominic Kwiatkowski, Cordelia Langford, John Sillitoe on behalf of the Wellcome Sanger Institute COVID-19 Surveillance |

| | | | |
|---|---|---|---|
| | | | Team |
| EPI_ISL_948298 | Lighthouse Lab in Cambridge | Wellcome Sanger Institute for the COVID-19 Genomics UK (COG-UK) Consortium | Rob Howes, The Lighthouse Lab in Cambridge and Alex Alderton, Roberto Amato, Sonia Goncalves, Ewan Harrison, David K. Jackson, Ian Johnston, Dominic Kwiatkowski, Cordelia Langford, John Sillitoe on behalf of the Wellcome Sanger Institute COVID-19 Surveillance Team |
| EPI_ISL_948299 | Lighthouse Lab in Glasgow | Wellcome Sanger Institute for the COVID-19 Genomics UK (COG-UK) Consortium | Harper VanSteenhouse, Yumi Kasai, David Gray, Carol Clugston, Anna Dominiczak and Alex Alderton, Roberto Amato, Sonia Goncalves, Ewan Harrison, David K. Jackson, Ian Johnston, Dominic Kwiatkowski, Cordelia Langford, John Sillitoe on behalf of the Wellcome Sanger Institute COVID-19 Surveillance Team |
| EPI_ISL_948300, EPI_ISL_948301, EPI_ISL_948302 | Lighthouse Lab in Cambridge | Wellcome Sanger Institute for the COVID-19 Genomics UK (COG-UK) Consortium | Rob Howes, The Lighthouse Lab in Cambridge and Alex Alderton, Roberto Amato, Sonia Goncalves, Ewan Harrison, David K. Jackson, Ian Johnston, Dominic Kwiatkowski, Cordelia Langford, John Sillitoe on behalf of the Wellcome Sanger Institute COVID-19 Surveillance Team |
| EPI_ISL_948303, EPI_ISL_948304 | Lighthouse Lab in Glasgow | Wellcome Sanger Institute for the COVID-19 Genomics UK (COG-UK) Consortium | Harper VanSteenhouse, Yumi Kasai, David Gray, Carol Clugston, Anna Dominiczak and Alex Alderton, Roberto Amato, Sonia Goncalves, Ewan Harrison, David K. Jackson, Ian Johnston, Dominic Kwiatkowski, Cordelia Langford, John Sillitoe on behalf of the Wellcome Sanger Institute COVID-19 Surveillance Team |
| EPI_ISL_948305, EPI_ISL_948306 | Lighthouse Lab in Cambridge | Wellcome Sanger Institute for the COVID-19 Genomics UK (COG-UK) Consortium | Rob Howes, The Lighthouse Lab in Cambridge and Alex Alderton, Roberto Amato, Sonia Goncalves, Ewan Harrison, David K. Jackson, Ian Johnston, Dominic Kwiatkowski, Cordelia Langford, John Sillitoe on behalf of the Wellcome Sanger Institute COVID-19 Surveillance Team |
| EPI_ISL_948307 | Lighthouse Lab in Glasgow | Wellcome Sanger Institute for the COVID-19 Genomics UK (COG-UK) Consortium | Harper VanSteenhouse, Yumi Kasai, David Gray, Carol Clugston, Anna Dominiczak and Alex Alderton, Roberto Amato, Sonia Goncalves, Ewan Harrison, David K. Jackson, Ian Johnston, Dominic Kwiatkowski, Cordelia Langford, John Sillitoe on behalf of the Wellcome Sanger Institute COVID-19 Surveillance Team |
| EPI_ISL_948308 | Lighthouse Lab in Cambridge | Wellcome Sanger Institute for the COVID-19 Genomics UK (COG-UK) Consortium | Rob Howes, The Lighthouse Lab in Cambridge and Alex Alderton, Roberto Amato, Sonia Goncalves, Ewan Harrison, David K. Jackson, Ian Johnston, Dominic Kwiatkowski, Cordelia Langford, John Sillitoe on behalf of the Wellcome Sanger Institute COVID-19 Surveillance Team |
| EPI_ISL_948309 | Lighthouse Lab in Glasgow | Wellcome Sanger Institute for the COVID-19 Genomics UK (COG-UK) Consortium | Harper VanSteenhouse, Yumi Kasai, David Gray, Carol Clugston, Anna Dominiczak and Alex Alderton, Roberto Amato, Sonia Goncalves, Ewan Harrison, David K. Jackson, Ian Johnston, Dominic Kwiatkowski, Cordelia Langford, John Sillitoe on behalf of the Wellcome Sanger Institute COVID-19 Surveillance Team |
| EPI_ISL_948310, EPI_ISL_948311, EPI_ISL_948312, EPI_ISL_948313, EPI_ISL_948314 | Lighthouse Lab in Cambridge | Wellcome Sanger Institute for the COVID-19 Genomics UK (COG-UK) Consortium | Rob Howes, The Lighthouse Lab in Cambridge and Alex Alderton, Roberto Amato, Sonia Goncalves, Ewan Harrison, David K. Jackson, Ian Johnston, Dominic Kwiatkowski, Cordelia Langford, John Sillitoe on behalf of the Wellcome Sanger Institute COVID-19 Surveillance Team |
| EPI_ISL_948315 | Lighthouse Lab in Glasgow | Wellcome Sanger Institute for the COVID-19 Genomics UK (COG-UK) Consortium | Harper VanSteenhouse, Yumi Kasai, David Gray, Carol Clugston, Anna Dominiczak and Alex Alderton, Roberto Amato, Sonia Goncalves, Ewan Harrison, David K. Jackson, Ian Johnston, Dominic Kwiatkowski, Cordelia Langford, John Sillitoe on behalf of the Wellcome Sanger Institute COVID-19 Surveillance Team |
| EPI_ISL_948316 | Lighthouse Lab in Cambridge | Wellcome Sanger Institute for the COVID-19 Genomics UK (COG-UK) Consortium | Rob Howes, The Lighthouse Lab in Cambridge and Alex Alderton, Roberto Amato, Sonia Goncalves, Ewan Harrison, David K. Jackson, Ian Johnston, Dominic Kwiatkowski, Cordelia Langford, John Sillitoe on behalf of the Wellcome Sanger Institute COVID-19 Surveillance Team |
| EPI_ISL_948317, EPI_ISL_948318, EPI_ISL_948319, EPI_ISL_948320 | Lighthouse Lab in Glasgow | Wellcome Sanger Institute for the COVID-19 Genomics UK (COG-UK) Consortium | Harper VanSteenhouse, Yumi Kasai, David Gray, Carol Clugston, Anna Dominiczak and Alex Alderton, Roberto Amato, Sonia Goncalves, Ewan Harrison, David K. Jackson, Ian Johnston, Dominic Kwiatkowski, Cordelia Langford, John Sillitoe on behalf of the Wellcome Sanger Institute COVID-19 Surveillance Team |
| EPI_ISL_948321, EPI_ISL_948322, EPI_ISL_948323 | Lighthouse Lab in Cambridge | Wellcome Sanger Institute for the COVID-19 Genomics UK (COG-UK) Consortium | Rob Howes, The Lighthouse Lab in Cambridge and Alex Alderton, Roberto Amato, Sonia Goncalves, Ewan Harrison, David K. Jackson, Ian Johnston, Dominic Kwiatkowski, Cordelia Langford, John Sillitoe on behalf of the Wellcome Sanger Institute COVID-19 Surveillance Team |
| EPI_ISL_948324, EPI_ISL_948325 | Lighthouse Lab in Glasgow | Wellcome Sanger Institute for the COVID-19 Genomics UK (COG-UK) Consortium | Harper VanSteenhouse, Yumi Kasai, David Gray, Carol Clugston, Anna Dominiczak and Alex Alderton, Roberto Amato, Sonia Goncalves, Ewan Harrison, David K. Jackson, Ian Johnston, Dominic Kwiatkowski, Cordelia Langford, John Sillitoe on behalf of the Wellcome Sanger Institute COVID-19 Surveillance Team |
| EPI_ISL_948326 | Lighthouse Lab in Cambridge | Wellcome Sanger Institute for the COVID-19 Genomics UK (COG-UK) Consortium | Rob Howes, The Lighthouse Lab in Cambridge and Alex Alderton, Roberto Amato, Sonia Goncalves, Ewan Harrison, David K. Jackson, Ian Johnston, Dominic Kwiatkowski, Cordelia Langford, John Sillitoe on behalf of the Wellcome Sanger Institute COVID-19 Surveillance Team |
| EPI_ISL_948327 | Lighthouse Lab in Milton Keynes | Wellcome Sanger Institute for the COVID-19 Genomics UK (COG-UK) Consortium | The Lighthouse Lab in Milton Keynes and Alex Alderton, Roberto Amato, Sonia Goncalves, Ewan Harrison, David K. Jackson, Ian Johnston, Dominic Kwiatkowski, Cordelia Langford, John Sillitoe on behalf of the Wellcome Sanger Institute COVID-19 Surveillance Team |
| EPI_ISL_948328 | Lighthouse Lab in Cambridge | Wellcome Sanger Institute for the COVID-19 Genomics UK (COG-UK) Consortium | Rob Howes, The Lighthouse Lab in Cambridge and Alex Alderton, Roberto Amato, Sonia Goncalves, Ewan Harrison, David K. Jackson, Ian Johnston, Dominic Kwiatkowski, Cordelia Langford, John Sillitoe on behalf of the Wellcome Sanger Institute COVID-19 Surveillance Team |
| EPI_ISL_948329 | Lighthouse Lab in Glasgow | Wellcome Sanger Institute for the COVID-19 Genomics UK (COG-UK) Consortium | Harper VanSteenhouse, Yumi Kasai, David Gray, Carol Clugston, Anna Dominiczak and Alex Alderton, Roberto Amato, Sonia Goncalves, Ewan Harrison, David K. Jackson, Ian Johnston, Dominic Kwiatkowski, Cordelia Langford, John Sillitoe on behalf of the Wellcome Sanger Institute COVID-19 Surveillance Team |
| EPI_ISL_948331, EPI_ISL_948332 | Lighthouse Lab in Cambridge | Wellcome Sanger Institute for the COVID-19 Genomics UK (COG-UK) Consortium | Rob Howes, The Lighthouse Lab in Cambridge and Alex Alderton, Roberto Amato, Sonia Goncalves, Ewan Harrison, David K. Jackson, Ian Johnston, Dominic Kwiatkowski, Cordelia Langford, John Sillitoe on behalf of the Wellcome Sanger Institute COVID-19 Surveillance Team |
| EPI_ISL_948333, EPI_ISL_948334, EPI_ISL_948335, EPI_ISL_948336 | Lighthouse Lab in Glasgow | Wellcome Sanger Institute for the COVID-19 Genomics UK (COG-UK) Consortium | Harper VanSteenhouse, Yumi Kasai, David Gray, Carol Clugston, Anna Dominiczak and Alex Alderton, Roberto Amato, Sonia Goncalves, Ewan Harrison, David K. Jackson, Ian Johnston, Dominic Kwiatkowski, Cordelia Langford, John Sillitoe on behalf of the Wellcome Sanger Institute COVID-19 Surveillance Team |
| EPI_ISL_948337, EPI_ISL_948338 | Lighthouse Lab in Cambridge | Wellcome Sanger Institute for the COVID-19 Genomics UK (COG-UK) Consortium | Rob Howes, The Lighthouse Lab in Cambridge and Alex Alderton, Roberto Amato, Sonia Goncalves, Ewan Harrison, David K. Jackson, Ian Johnston, Dominic Kwiatkowski, Cordelia Langford, John Sillitoe on behalf of the Wellcome Sanger Institute COVID-19 Surveillance Team |
| EPI_ISL_948339 | Lighthouse Lab in Glasgow | Wellcome Sanger Institute for the COVID-19 Genomics UK (COG-UK) Consortium | Harper VanSteenhouse, Yumi Kasai, David Gray, Carol Clugston, Anna Dominiczak and Alex Alderton, Roberto Amato, Sonia Goncalves, Ewan Harrison, David K. Jackson, Ian Johnston, Dominic Kwiatkowski, Cordelia Langford, John Sillitoe on behalf of the Wellcome Sanger Institute COVID-19 Surveillance Team |
| EPI_ISL_948340, EPI_ISL_948341, EPI_ISL_948342 | Lighthouse Lab in Cambridge | Wellcome Sanger Institute for the COVID-19 Genomics UK (COG-UK) Consortium | Rob Howes, The Lighthouse Lab in Cambridge and Alex Alderton, Roberto Amato, Sonia Goncalves, Ewan Harrison, David K. Jackson, Ian Johnston, Dominic Kwiatkowski, Cordelia Langford, John Sillitoe on behalf of the Wellcome Sanger Institute COVID-19 Surveillance Team |
| EPI_ISL_948343 | Lighthouse Lab in Glasgow | Wellcome Sanger Institute for the COVID-19 Genomics UK (COG-UK) Consortium | Harper VanSteenhouse, Yumi Kasai, David Gray, Carol Clugston, Anna Dominiczak and Alex Alderton, Roberto Amato, Sonia Goncalves, Ewan Harrison, David K. Jackson, Ian Johnston, Dominic Kwiatkowski, Cordelia Langford, John Sillitoe on behalf of the Wellcome Sanger Institute COVID-19 Surveillance Team |
| EPI_ISL_948344, EPI_ISL_948345, EPI_ISL_948346 | Lighthouse Lab in Cambridge | Wellcome Sanger Institute for the COVID-19 Genomics UK (COG-UK) Consortium | Rob Howes, The Lighthouse Lab in Cambridge and Alex Alderton, Roberto Amato, Sonia Goncalves, Ewan Harrison, David K. Jackson, Ian Johnston, Dominic Kwiatkowski, Cordelia Langford, John Sillitoe on behalf of the Wellcome Sanger Institute COVID-19 Surveillance Team |
| EPI_ISL_948347, EPI_ISL_948348, EPI_ISL_948349 | Lighthouse Lab in Glasgow | Wellcome Sanger Institute for the COVID-19 Genomics UK (COG-UK) Consortium | Harper VanSteenhouse, Yumi Kasai, David Gray, Carol Clugston, Anna Dominiczak and Alex Alderton, Roberto Amato, Sonia Goncalves, Ewan Harrison, David K. Jackson, Ian Johnston, Dominic Kwiatkowski, Cordelia Langford, John Sillitoe on behalf of the Wellcome Sanger Institute COVID-19 Surveillance Team |
| EPI_ISL_948350 | Lighthouse Lab in Cambridge | Wellcome Sanger Institute for the COVID-19 Genomics UK (COG-UK) Consortium | Rob Howes, The Lighthouse Lab in Cambridge and Alex Alderton, Roberto Amato, Sonia Goncalves, Ewan Harrison, David K. Jackson, Ian Johnston, Dominic Kwiatkowski, Cordelia Langford, John Sillitoe on behalf of the Wellcome Sanger Institute COVID-19 Surveillance Team |
| EPI_ISL_948351, EPI_ISL_948352, EPI_ISL_948353, EPI_ISL_948354 | Lighthouse Lab in Glasgow | Wellcome Sanger Institute for the COVID-19 Genomics UK (COG-UK) Consortium | Harper VanSteenhouse, Yumi Kasai, David Gray, Carol Clugston, Anna Dominiczak and Alex Alderton, Roberto Amato, Sonia Goncalves, Ewan Harrison, David K. Jackson, Ian Johnston, Dominic Kwiatkowski, Cordelia Langford, John Sillitoe on behalf of the Wellcome Sanger Institute COVID-19 Surveillance Team |
| EPI_ISL_948355 | Lighthouse Lab in Cambridge | Wellcome Sanger Institute for the COVID-19 Genomics UK (COG-UK) Consortium | Rob Howes, The Lighthouse Lab in Cambridge and Alex Alderton, Roberto Amato, Sonia Goncalves, Ewan Harrison, David K. Jackson, Ian Johnston, Dominic Kwiatkowski, Cordelia Langford, John Sillitoe on behalf of the Wellcome Sanger Institute COVID-19 Surveillance Team |

| | | | |
|---|---|---|---|
| EPI_ISL_948356, EPI_ISL_948357, EPI_ISL_948358 | Lighthouse Lab in Glasgow | Wellcome Sanger Institute for the COVID-19 Genomics UK (COG-UK) Consortium | Harper VanSteenhouse, Yumi Kasai, David Gray, Carol Clugston, Anna Dominiczak and Alex Alderton, Roberto Amato, Sonia Goncalves, Ewan Harrison, David K. Jackson, Ian Johnston, Dominic Kwiatkowski, Cordelia Langford, John Sillitoe on behalf of the Wellcome Sanger Institute COVID-19 Surveillance Team |
| EPI_ISL_948359 | Lighthouse Lab in Cambridge | Wellcome Sanger Institute for the COVID-19 Genomics UK (COG-UK) Consortium | Rob Howes, The Lighthouse Lab in Cambridge and Alex Alderton, Roberto Amato, Sonia Goncalves, Ewan Harrison, David K. Jackson, Ian Johnston, Dominic Kwiatkowski, Cordelia Langford, John Sillitoe on behalf of the Wellcome Sanger Institute COVID-19 Surveillance Team |
| EPI_ISL_948360 | Lighthouse Lab in Glasgow | Wellcome Sanger Institute for the COVID-19 Genomics UK (COG-UK) Consortium | Harper VanSteenhouse, Yumi Kasai, David Gray, Carol Clugston, Anna Dominiczak and Alex Alderton, Roberto Amato, Sonia Goncalves, Ewan Harrison, David K. Jackson, Ian Johnston, Dominic Kwiatkowski, Cordelia Langford, John Sillitoe on behalf of the Wellcome Sanger Institute COVID-19 Surveillance Team |
| EPI_ISL_948361 | Lighthouse Lab in Cambridge | Wellcome Sanger Institute for the COVID-19 Genomics UK (COG-UK) Consortium | Rob Howes, The Lighthouse Lab in Cambridge and Alex Alderton, Roberto Amato, Sonia Goncalves, Ewan Harrison, David K. Jackson, Ian Johnston, Dominic Kwiatkowski, Cordelia Langford, John Sillitoe on behalf of the Wellcome Sanger Institute COVID-19 Surveillance Team |
| EPI_ISL_948362, EPI_ISL_948363 | Lighthouse Lab in Glasgow | Wellcome Sanger Institute for the COVID-19 Genomics UK (COG-UK) Consortium | Harper VanSteenhouse, Yumi Kasai, David Gray, Carol Clugston, Anna Dominiczak and Alex Alderton, Roberto Amato, Sonia Goncalves, Ewan Harrison, David K. Jackson, Ian Johnston, Dominic Kwiatkowski, Cordelia Langford, John Sillitoe on behalf of the Wellcome Sanger Institute COVID-19 Surveillance Team |
| EPI_ISL_948364 | Lighthouse Lab in Cambridge | Wellcome Sanger Institute for the COVID-19 Genomics UK (COG-UK) Consortium | Rob Howes, The Lighthouse Lab in Cambridge and Alex Alderton, Roberto Amato, Sonia Goncalves, Ewan Harrison, David K. Jackson, Ian Johnston, Dominic Kwiatkowski, Cordelia Langford, John Sillitoe on behalf of the Wellcome Sanger Institute COVID-19 Surveillance Team |
| EPI_ISL_948365 | Lighthouse Lab in Glasgow | Wellcome Sanger Institute for the COVID-19 Genomics UK (COG-UK) Consortium | Harper VanSteenhouse, Yumi Kasai, David Gray, Carol Clugston, Anna Dominiczak and Alex Alderton, Roberto Amato, Sonia Goncalves, Ewan Harrison, David K. Jackson, Ian Johnston, Dominic Kwiatkowski, Cordelia Langford, John Sillitoe on behalf of the Wellcome Sanger Institute COVID-19 Surveillance Team |
| EPI_ISL_948366 | Lighthouse Lab in Cambridge | Wellcome Sanger Institute for the COVID-19 Genomics UK (COG-UK) Consortium | Rob Howes, The Lighthouse Lab in Cambridge and Alex Alderton, Roberto Amato, Sonia Goncalves, Ewan Harrison, David K. Jackson, Ian Johnston, Dominic Kwiatkowski, Cordelia Langford, John Sillitoe on behalf of the Wellcome Sanger Institute COVID-19 Surveillance Team |
| EPI_ISL_948367, EPI_ISL_948369, EPI_ISL_948370, EPI_ISL_948371 | Lighthouse Lab in Glasgow | Wellcome Sanger Institute for the COVID-19 Genomics UK (COG-UK) Consortium | Harper VanSteenhouse, Yumi Kasai, David Gray, Carol Clugston, Anna Dominiczak and Alex Alderton, Roberto Amato, Sonia Goncalves, Ewan Harrison, David K. Jackson, Ian Johnston, Dominic Kwiatkowski, Cordelia Langford, John Sillitoe on behalf of the Wellcome Sanger Institute COVID-19 Surveillance Team |
| EPI_ISL_948372 | Lighthouse Lab in Cambridge | Wellcome Sanger Institute for the COVID-19 Genomics UK (COG-UK) Consortium | Rob Howes, The Lighthouse Lab in Cambridge and Alex Alderton, Roberto Amato, Sonia Goncalves, Ewan Harrison, David K. Jackson, Ian Johnston, Dominic Kwiatkowski, Cordelia Langford, John Sillitoe on behalf of the Wellcome Sanger Institute COVID-19 Surveillance Team |
| EPI_ISL_948373, EPI_ISL_948375 | Lighthouse Lab in Glasgow | Wellcome Sanger Institute for the COVID-19 Genomics UK (COG-UK) Consortium | Harper VanSteenhouse, Yumi Kasai, David Gray, Carol Clugston, Anna Dominiczak and Alex Alderton, Roberto Amato, Sonia Goncalves, Ewan Harrison, David K. Jackson, Ian Johnston, Dominic Kwiatkowski, Cordelia Langford, John Sillitoe on behalf of the Wellcome Sanger Institute COVID-19 Surveillance Team |
| EPI_ISL_948378 | Lighthouse Lab in Cambridge | Wellcome Sanger Institute for the COVID-19 Genomics UK (COG-UK) Consortium | Rob Howes, The Lighthouse Lab in Cambridge and Alex Alderton, Roberto Amato, Sonia Goncalves, Ewan Harrison, David K. Jackson, Ian Johnston, Dominic Kwiatkowski, Cordelia Langford, John Sillitoe on behalf of the Wellcome Sanger Institute COVID-19 Surveillance Team |
| EPI_ISL_948379 | Lighthouse Lab in Glasgow | Wellcome Sanger Institute for the COVID-19 Genomics UK (COG-UK) Consortium | Harper VanSteenhouse, Yumi Kasai, David Gray, Carol Clugston, Anna Dominiczak and Alex Alderton, Roberto Amato, Sonia Goncalves, Ewan Harrison, David K. Jackson, Ian Johnston, Dominic Kwiatkowski, Cordelia Langford, John Sillitoe on behalf of the Wellcome Sanger Institute COVID-19 Surveillance Team |
| EPI_ISL_948380, EPI_ISL_948381, EPI_ISL_948382, EPI_ISL_948383 | Lighthouse Lab in Cambridge | Wellcome Sanger Institute for the COVID-19 Genomics UK (COG-UK) Consortium | Rob Howes, The Lighthouse Lab in Cambridge and Alex Alderton, Roberto Amato, Sonia Goncalves, Ewan Harrison, David K. Jackson, Ian Johnston, Dominic Kwiatkowski, Cordelia Langford, John Sillitoe on behalf of the Wellcome Sanger Institute COVID-19 Surveillance Team |
| EPI_ISL_948384, EPI_ISL_948385, EPI_ISL_948386, EPI_ISL_948387, EPI_ISL_948389, EPI_ISL_948390 | Lighthouse Lab in Glasgow | Wellcome Sanger Institute for the COVID-19 Genomics UK (COG-UK) Consortium | Harper VanSteenhouse, Yumi Kasai, David Gray, Carol Clugston, Anna Dominiczak and Alex Alderton, Roberto Amato, Sonia Goncalves, Ewan Harrison, David K. Jackson, Ian Johnston, Dominic Kwiatkowski, Cordelia Langford, John Sillitoe on behalf of the Wellcome Sanger Institute COVID-19 Surveillance Team |
| EPI_ISL_948391, EPI_ISL_948392, EPI_ISL_948393 | Lighthouse Lab in Cambridge | Wellcome Sanger Institute for the COVID-19 Genomics UK (COG-UK) Consortium | Rob Howes, The Lighthouse Lab in Cambridge and Alex Alderton, Roberto Amato, Sonia Goncalves, Ewan Harrison, David K. Jackson, Ian Johnston, Dominic Kwiatkowski, Cordelia Langford, John Sillitoe on behalf of the Wellcome Sanger Institute COVID-19 Surveillance Team |
| EPI_ISL_948394 | Lighthouse Lab in Glasgow | Wellcome Sanger Institute for the COVID-19 Genomics UK (COG-UK) Consortium | Harper VanSteenhouse, Yumi Kasai, David Gray, Carol Clugston, Anna Dominiczak and Alex Alderton, Roberto Amato, Sonia Goncalves, Ewan Harrison, David K. Jackson, Ian Johnston, Dominic Kwiatkowski, Cordelia Langford, John Sillitoe on behalf of the Wellcome Sanger Institute COVID-19 Surveillance Team |
| EPI_ISL_948395, EPI_ISL_948396, EPI_ISL_948397 | Lighthouse Lab in Cambridge | Wellcome Sanger Institute for the COVID-19 Genomics UK (COG-UK) Consortium | Rob Howes, The Lighthouse Lab in Cambridge and Alex Alderton, Roberto Amato, Sonia Goncalves, Ewan Harrison, David K. Jackson, Ian Johnston, Dominic Kwiatkowski, Cordelia Langford, John Sillitoe on behalf of the Wellcome Sanger Institute COVID-19 Surveillance Team |
| EPI_ISL_948398, EPI_ISL_948399, EPI_ISL_948400, EPI_ISL_948401 | Lighthouse Lab in Glasgow | Wellcome Sanger Institute for the COVID-19 Genomics UK (COG-UK) Consortium | Harper VanSteenhouse, Yumi Kasai, David Gray, Carol Clugston, Anna Dominiczak and Alex Alderton, Roberto Amato, Sonia Goncalves, Ewan Harrison, David K. Jackson, Ian Johnston, Dominic Kwiatkowski, Cordelia Langford, John Sillitoe on behalf of the Wellcome Sanger Institute COVID-19 Surveillance Team |
| EPI_ISL_948402 | Lighthouse Lab in Cambridge | Wellcome Sanger Institute for the COVID-19 Genomics UK (COG-UK) Consortium | Rob Howes, The Lighthouse Lab in Cambridge and Alex Alderton, Roberto Amato, Sonia Goncalves, Ewan Harrison, David K. Jackson, Ian Johnston, Dominic Kwiatkowski, Cordelia Langford, John Sillitoe on behalf of the Wellcome Sanger Institute COVID-19 Surveillance Team |
| EPI_ISL_948403, EPI_ISL_948404, EPI_ISL_948405, EPI_ISL_948406 | Lighthouse Lab in Glasgow | Wellcome Sanger Institute for the COVID-19 Genomics UK (COG-UK) Consortium | Harper VanSteenhouse, Yumi Kasai, David Gray, Carol Clugston, Anna Dominiczak and Alex Alderton, Roberto Amato, Sonia Goncalves, Ewan Harrison, David K. Jackson, Ian Johnston, Dominic Kwiatkowski, Cordelia Langford, John Sillitoe on behalf of the Wellcome Sanger Institute COVID-19 Surveillance Team |
| EPI_ISL_948408 | Lighthouse Lab in Cambridge | Wellcome Sanger Institute for the COVID-19 Genomics UK (COG-UK) Consortium | Rob Howes, The Lighthouse Lab in Cambridge and Alex Alderton, Roberto Amato, Sonia Goncalves, Ewan Harrison, David K. Jackson, Ian Johnston, Dominic Kwiatkowski, Cordelia Langford, John Sillitoe on behalf of the Wellcome Sanger Institute COVID-19 Surveillance Team |
| EPI_ISL_948409, EPI_ISL_948410, EPI_ISL_948411 | Lighthouse Lab in Glasgow | Wellcome Sanger Institute for the COVID-19 Genomics UK (COG-UK) Consortium | Harper VanSteenhouse, Yumi Kasai, David Gray, Carol Clugston, Anna Dominiczak and Alex Alderton, Roberto Amato, Sonia Goncalves, Ewan Harrison, David K. Jackson, Ian Johnston, Dominic Kwiatkowski, Cordelia Langford, John Sillitoe on behalf of the Wellcome Sanger Institute COVID-19 Surveillance Team |
| EPI_ISL_948412, EPI_ISL_948413, EPI_ISL_948414 | Lighthouse Lab in Cambridge | Wellcome Sanger Institute for the COVID-19 Genomics UK (COG-UK) Consortium | Rob Howes, The Lighthouse Lab in Cambridge and Alex Alderton, Roberto Amato, Sonia Goncalves, Ewan Harrison, David K. Jackson, Ian Johnston, Dominic Kwiatkowski, Cordelia Langford, John Sillitoe on behalf of the Wellcome Sanger Institute COVID-19 Surveillance Team |
| EPI_ISL_948415, EPI_ISL_948416, EPI_ISL_948417, EPI_ISL_948418, EPI_ISL_948419 | Lighthouse Lab in Glasgow | Wellcome Sanger Institute for the COVID-19 Genomics UK (COG-UK) Consortium | Harper VanSteenhouse, Yumi Kasai, David Gray, Carol Clugston, Anna Dominiczak and Alex Alderton, Roberto Amato, Sonia Goncalves, Ewan Harrison, David K. Jackson, Ian Johnston, Dominic Kwiatkowski, Cordelia Langford, John Sillitoe on behalf of the Wellcome Sanger Institute COVID-19 Surveillance Team |
| EPI_ISL_948420, EPI_ISL_948421, EPI_ISL_948422, EPI_ISL_948423, EPI_ISL_948424 | Lighthouse Lab in Cambridge | Wellcome Sanger Institute for the COVID-19 Genomics UK (COG-UK) Consortium | Rob Howes, The Lighthouse Lab in Cambridge and Alex Alderton, Roberto Amato, Sonia Goncalves, Ewan Harrison, David K. Jackson, Ian Johnston, Dominic Kwiatkowski, Cordelia Langford, John Sillitoe on behalf of the Wellcome Sanger Institute COVID-19 Surveillance Team |
| EPI_ISL_948425, EPI_ISL_948426, EPI_ISL_948427, EPI_ISL_948428, EPI_ISL_948429, EPI_ISL_948430 | Lighthouse Lab in Glasgow | Wellcome Sanger Institute for the COVID-19 Genomics UK (COG-UK) Consortium | Harper VanSteenhouse, Yumi Kasai, David Gray, Carol Clugston, Anna Dominiczak and Alex Alderton, Roberto Amato, Sonia Goncalves, Ewan Harrison, David K. Jackson, Ian Johnston, Dominic Kwiatkowski, Cordelia Langford, John Sillitoe on behalf of the Wellcome Sanger Institute COVID-19 Surveillance Team |
| EPI_ISL_948431, EPI_ISL_948433, EPI_ISL_948434, EPI_ISL_948435 | Lighthouse Lab in Cambridge | Wellcome Sanger Institute for the COVID-19 Genomics UK (COG-UK) Consortium | Rob Howes, The Lighthouse Lab in Cambridge and Alex Alderton, Roberto Amato, Sonia Goncalves, Ewan Harrison, David K. Jackson, Ian Johnston, Dominic Kwiatkowski, Cordelia Langford, John Sillitoe on behalf of the Wellcome Sanger Institute COVID-19 Surveillance Team |
| EPI_ISL_948436 | Lighthouse Lab in Glasgow | Wellcome Sanger Institute for the COVID-19 Genomics UK (COG-UK) Consortium | Harper VanSteenhouse, Yumi Kasai, David Gray, Carol Clugston, Anna Dominiczak and Alex Alderton, Roberto Amato, Sonia Goncalves, Ewan Harrison, David K. Jackson, Ian Johnston, Dominic Kwiatkowski, Cordelia Langford, John Sillitoe on behalf of the Wellcome Sanger Institute COVID-19 Surveillance |

| | | | |
|---|---|---|---|
| | | | Team |
| EPI_ISL_948437 | Lighthouse Lab in Cambridge | Wellcome Sanger Institute for the COVID-19 Genomics UK (COG-UK) Consortium | Rob Howes, The Lighthouse Lab in Cambridge and Alex Alderton, Roberto Amato, Sonia Goncalves, Ewan Harrison, David K. Jackson, Ian Johnston, Dominic Kwiatkowski, Cordelia Langford, John Sillitoe on behalf of the Wellcome Sanger Institute COVID-19 Surveillance Team |
| EPI_ISL_948438, EPI_ISL_948439, EPI_ISL_948440 | Lighthouse Lab in Glasgow | Wellcome Sanger Institute for the COVID-19 Genomics UK (COG-UK) Consortium | Harper VanSteenhouse, Yumi Kasai, David Gray, Carol Clugston, Anna Dominiczak and Alex Alderton, Roberto Amato, Sonia Goncalves, Ewan Harrison, David K. Jackson, Ian Johnston, Dominic Kwiatkowski, Cordelia Langford, John Sillitoe on behalf of the Wellcome Sanger Institute COVID-19 Surveillance Team |
| EPI_ISL_948441, EPI_ISL_948442 | Lighthouse Lab in Cambridge | Wellcome Sanger Institute for the COVID-19 Genomics UK (COG-UK) Consortium | Rob Howes, The Lighthouse Lab in Cambridge and Alex Alderton, Roberto Amato, Sonia Goncalves, Ewan Harrison, David K. Jackson, Ian Johnston, Dominic Kwiatkowski, Cordelia Langford, John Sillitoe on behalf of the Wellcome Sanger Institute COVID-19 Surveillance Team |
| EPI_ISL_948443, EPI_ISL_948444 | Lighthouse Lab in Glasgow | Wellcome Sanger Institute for the COVID-19 Genomics UK (COG-UK) Consortium | Harper VanSteenhouse, Yumi Kasai, David Gray, Carol Clugston, Anna Dominiczak and Alex Alderton, Roberto Amato, Sonia Goncalves, Ewan Harrison, David K. Jackson, Ian Johnston, Dominic Kwiatkowski, Cordelia Langford, John Sillitoe on behalf of the Wellcome Sanger Institute COVID-19 Surveillance Team |
| EPI_ISL_948445 | Lighthouse Lab in Cambridge | Wellcome Sanger Institute for the COVID-19 Genomics UK (COG-UK) Consortium | Rob Howes, The Lighthouse Lab in Cambridge and Alex Alderton, Roberto Amato, Sonia Goncalves, Ewan Harrison, David K. Jackson, Ian Johnston, Dominic Kwiatkowski, Cordelia Langford, John Sillitoe on behalf of the Wellcome Sanger Institute COVID-19 Surveillance Team |
| EPI_ISL_948446, EPI_ISL_948447, EPI_ISL_948448, EPI_ISL_948787, EPI_ISL_948788, EPI_ISL_948789, EPI_ISL_948790, EPI_ISL_948791, EPI_ISL_948792, EPI_ISL_948793, EPI_ISL_948794, EPI_ISL_948795, EPI_ISL_948796, EPI_ISL_948797, EPI_ISL_948798, EPI_ISL_948799, EPI_ISL_948800, EPI_ISL_948801, EPI_ISL_948802, EPI_ISL_948803, EPI_ISL_948804, EPI_ISL_948805, EPI_ISL_948806, EPI_ISL_948807, EPI_ISL_948808, EPI_ISL_948809, EPI_ISL_948810, EPI_ISL_948811, EPI_ISL_948812, EPI_ISL_948813, EPI_ISL_948814, EPI_ISL_948815, EPI_ISL_948816, EPI_ISL_948817, EPI_ISL_948818, EPI_ISL_948819, EPI_ISL_948820, EPI_ISL_948821, EPI_ISL_948822, EPI_ISL_948823, EPI_ISL_948824, EPI_ISL_948825, EPI_ISL_948826, EPI_ISL_948827, EPI_ISL_948828, EPI_ISL_948831, EPI_ISL_948832, EPI_ISL_948833, EPI_ISL_948835, EPI_ISL_948836, EPI_ISL_948837, EPI_ISL_948838, EPI_ISL_948839, EPI_ISL_948840, EPI_ISL_948841, EPI_ISL_948842, EPI_ISL_948843, EPI_ISL_948844, EPI_ISL_948846, EPI_ISL_948847, EPI_ISL_948849, EPI_ISL_948850, EPI_ISL_948851, EPI_ISL_948852, EPI_ISL_948853, EPI_ISL_948854, EPI_ISL_948855, EPI_ISL_948856, EPI_ISL_948857, EPI_ISL_948858, EPI_ISL_948859, EPI_ISL_948860, EPI_ISL_948861, EPI_ISL_948862, EPI_ISL_948863, EPI_ISL_948864, EPI_ISL_948865, EPI_ISL_948866, EPI_ISL_948867, EPI_ISL_948868, EPI_ISL_948869, EPI_ISL_948870, EPI_ISL_948871, EPI_ISL_948872, EPI_ISL_948873, EPI_ISL_948874, EPI_ISL_948875, EPI_ISL_948876, EPI_ISL_948877, EPI_ISL_948878, EPI_ISL_948879, EPI_ISL_948880, EPI_ISL_948881, EPI_ISL_948882, EPI_ISL_948883, EPI_ISL_948884, EPI_ISL_948885, EPI_ISL_948886, EPI_ISL_948887, EPI_ISL_948888, EPI_ISL_948889, EPI_ISL_948890, EPI_ISL_948891, EPI_ISL_948892, EPI_ISL_948893, EPI_ISL_948894, EPI_ISL_948895, EPI_ISL_948896, EPI_ISL_948897, EPI_ISL_948898, EPI_ISL_948899, EPI_ISL_948900, EPI_ISL_948901, EPI_ISL_948902, EPI_ISL_948903, EPI_ISL_948904, EPI_ISL_948905, EPI_ISL_948906, EPI_ISL_948907, EPI_ISL_948908, EPI_ISL_948909, EPI_ISL_948910, EPI_ISL_948911, EPI_ISL_948912, EPI_ISL_948913, EPI_ISL_948914, EPI_ISL_948915, EPI_ISL_948916, EPI_ISL_948917, EPI_ISL_948918, EPI_ISL_948919, EPI_ISL_948920, EPI_ISL_948921, EPI_ISL_948922, EPI_ISL_948923, EPI_ISL_948924, EPI_ISL_948925, EPI_ISL_948926, EPI_ISL_948927, EPI_ISL_948928, EPI_ISL_948929, EPI_ISL_948930, EPI_ISL_948931, EPI_ISL_948932, EPI_ISL_948933, EPI_ISL_948934, EPI_ISL_948935, EPI_ISL_948936, EPI_ISL_948937, EPI_ISL_948938, EPI_ISL_948939, EPI_ISL_948940, EPI_ISL_948941, EPI_ISL_948942, EPI_ISL_948943, EPI_ISL_948944 | | | |
| see above | Lighthouse Lab in Glasgow | Wellcome Sanger Institute for the COVID-19 Genomics UK (COG-UK) Consortium | Harper VanSteenhouse, Yumi Kasai, David Gray, Carol Clugston, Anna Dominiczak and Alex Alderton, Roberto Amato, Sonia Goncalves, Ewan Harrison, David K. Jackson, Ian Johnston, Dominic Kwiatkowski, Cordelia Langford, John Sillitoe on behalf of the Wellcome Sanger Institute COVID-19 Surveillance Team |
| EPI_ISL_949138, EPI_ISL_949139, EPI_ISL_949158, EPI_ISL_949159, EPI_ISL_949160, EPI_ISL_949161, EPI_ISL_949162, EPI_ISL_949163, EPI_ISL_949164, EPI_ISL_949166, EPI_ISL_949167, EPI_ISL_949168, EPI_ISL_949169, EPI_ISL_949170, EPI_ISL_949171, EPI_ISL_949172, EPI_ISL_949173, EPI_ISL_949174, EPI_ISL_949175 | | | |
| see above | Jessa | Jessa | Jessa_cmdLab |
| EPI_ISL_949181 | Laboratory of Virology and Molecular Diagnostics | Institute of Public Health of Republic of North Macedonia Laboratory of Virology and Molecular Diagnostics | M. Kuzmanovska, G. Boshevska |
| EPI_ISL_949184 | University of Bari Biomedical Sciences and Human Oncology | University of Bari Biomedical Sciences and Human Oncology | Chironna M., Sallustio A., Loconsole D., Accogli A. |
| EPI_ISL_949185, EPI_ISL_949191 | University of Bari Biomedical Sciences and Human Oncology | University of Bari Biomedical Sciences and Human Oncology | Chironna M., Sallustio A., Loconsole D., Accogli M. |
| EPI_ISL_949410 | University of Birmingham | COVID-19 Genomics UK (COG-UK) Consortium | Institute of Microbiology, University of Birmingham: Claire McMurray, Joanne Stockton, Samuel Nicholls, Radoslaw Poplawski, Will Rowe, Josh Quick, Nicholas Loman. University of Birmingham Testing Laboratory: Celina M Whalley, Andrew Bosworth, Charlotte Poxon, Kasun Wanigasooriya, Oliver Pickles, Mike Kidd, Alex Richter, Andrew D Beggs PHE Heartlands Lab: Husam Osman, Andrew Bosworth. Queen Elizabeth Hospital: Anna Casey |
| EPI_ISL_949462, EPI_ISL_949463, EPI_ISL_949464, EPI_ISL_949465, EPI_ISL_949467, EPI_ISL_949468, EPI_ISL_949469, EPI_ISL_949473, EPI_ISL_949475, EPI_ISL_949476, EPI_ISL_949477, EPI_ISL_949524, EPI_ISL_949525, EPI_ISL_949526, EPI_ISL_949527, EPI_ISL_949528, EPI_ISL_949529, EPI_ISL_949530, EPI_ISL_949531, EPI_ISL_949532, EPI_ISL_949533, EPI_ISL_949534, EPI_ISL_949535, EPI_ISL_949536, EPI_ISL_949537, EPI_ISL_949538, EPI_ISL_949539, EPI_ISL_949540, EPI_ISL_949542, EPI_ISL_949544, EPI_ISL_949546, EPI_ISL_949548, EPI_ISL_949549, EPI_ISL_949550, EPI_ISL_949553, EPI_ISL_949554, EPI_ISL_949555, EPI_ISL_949556 | | | |
| see above | Department of Pathology, University of Cambridge | COVID-19 Genomics UK (COG-UK) Consortium | Aminu S. Jahun, Yasmin Chaudhry, Iliana Georgana, Myra Hosmillo, Rhys Izu, Martin D. Curran, Surendra Parmar, Ian Goodfellow |
| EPI_ISL_949611, EPI_ISL_949615 | West of Scotland Specialist Virology Centre, NHSGGC / MRC-University of Glasgow Centre for Virus Research | COVID-19 Genomics UK (COG-UK) Consortium | Ana da Silva Filipe, Natasha Johnson, Kathy Smollett, Daniel Mair, Stephen Carmichael, Alice Broos, Lily Tong, Jenna Nichols, Kyriaki Nomikou; Sarah McDonald; Richard Orton, Joseph Hughes, Sreenu Vattipally, David L Robertson; Alasdair MacLean, Rory Gunson; Sharif Shaaban, Matthew Holden; Rachel Blacow, Guy Mollett, Kathy Li, James Shepherd, Antonia Ho, Emma Thomson |
| EPI_ISL_949645, EPI_ISL_949646 | Virology Department, Royal Infirmary of Edinburgh, NHS Lothian / School of Biological Sciences, University of Edinburgh / Institute of Genetics and Molecular Medicine, University of Edinburgh | COVID-19 Genomics UK (COG-UK) Consortium | McHugh M, Dewar R, Rooke S, Gallagher M, Balcaza C, O'Toole Á, Scher E, Hill V, McCrone JT, Colquhoun R, Yu X, Jackson B, Rambaut A, Williams TC, Templeton K |
| EPI_ISL_949655, EPI_ISL_949656, EPI_ISL_949657, EPI_ISL_949658, EPI_ISL_949659, EPI_ISL_949660, EPI_ISL_949661, EPI_ISL_949665, EPI_ISL_949667, EPI_ISL_949673, EPI_ISL_949674, EPI_ISL_949675, EPI_ISL_949676, EPI_ISL_949677, EPI_ISL_949678, EPI_ISL_949679, EPI_ISL_949680, EPI_ISL_949681, EPI_ISL_949682, EPI_ISL_949683, EPI_ISL_949688, EPI_ISL_949689, EPI_ISL_949690, EPI_ISL_949691, EPI_ISL_949692, EPI_ISL_949693, EPI_ISL_949694, EPI_ISL_949695, EPI_ISL_949696, EPI_ISL_949697, EPI_ISL_949698, EPI_ISL_949699, EPI_ISL_949700, EPI_ISL_949701, EPI_ISL_949702, EPI_ISL_949703, EPI_ISL_949704, EPI_ISL_949705, EPI_ISL_949706, EPI_ISL_949707 | | | |
| see above | Liverpool Clinical Laboratories | COVID-19 Genomics UK (COG-UK) Consortium | Sam Haldenby, Anita Lucaci, Steve Paterson, Julian Hiscox, Alistair Darby, M Almsaud, A Alrezaihi, Muhannad Alruwaili, Stuart D Armstrong, Jones Benjamin, Eleanor G Bentley, Anu Chawla, Jordan J Clark, Angela Cowell, Richard Eccles, Isabel García-Dorival, Matthew Gemmell, Alessandro Gerada, PKF Gilmore, Richard Gregory, Ximeng Han, Catherine Hartley, Margaret Hughes, Miren Iturriza-Gomara, James Johnson, L Luu, Jenifer Manson, Charlotte Nelson, Elaine O'Toole, Cassie Olateju, Rebekah Penrice-Randal , Lucille Rainbow, N.P Randle, Trevor Ian Robinson, Parul Sharma, Ghada T Shawli, James P Stewart, Neil Swainston, Ecaterina Vamos, Joanne Watts, Mark Whitehead |
| EPI_ISL_949793, EPI_ISL_949794, EPI_ISL_949802, EPI_ISL_949909, EPI_ISL_949910, EPI_ISL_949911, EPI_ISL_949912, EPI_ISL_949913, EPI_ISL_949914, EPI_ISL_949917, EPI_ISL_949919, EPI_ISL_949948, EPI_ISL_950019, EPI_ISL_950028, EPI_ISL_950034, EPI_ISL_950039, EPI_ISL_950040, EPI_ISL_950041, EPI_ISL_950042, EPI_ISL_950043, EPI_ISL_950044, EPI_ISL_950045, EPI_ISL_950046, EPI_ISL_950047, EPI_ISL_950049, EPI_ISL_950050, EPI_ISL_950051, EPI_ISL_950052, EPI_ISL_950053, EPI_ISL_950054, EPI_ISL_950055, EPI_ISL_950056, EPI_ISL_950057, EPI_ISL_950058, EPI_ISL_950059, EPI_ISL_950060, EPI_ISL_950061, EPI_ISL_950062, EPI_ISL_950063, EPI_ISL_950064, EPI_ISL_950065, EPI_ISL_950066, EPI_ISL_950067, EPI_ISL_950171, EPI_ISL_950172, EPI_ISL_950174, EPI_ISL_950175, EPI_ISL_950176, EPI_ISL_950178, EPI_ISL_950179, EPI_ISL_950180, EPI_ISL_950181, EPI_ISL_950182, EPI_ISL_950183, EPI_ISL_950184, EPI_ISL_950185, EPI_ISL_950186, EPI_ISL_950187, EPI_ISL_950188, EPI_ISL_950189, EPI_ISL_950190, EPI_ISL_950191, EPI_ISL_950192, EPI_ISL_950193, EPI_ISL_950194, EPI_ISL_950196, EPI_ISL_950199 | | | |
| see above | University College London, Great Ormond Street Hospital for Children NHS Foundation Trust, Imperial College Healthcare NHS Trust | COVID-19 Genomics UK (COG-UK) Consortium | Sergi Castellano, Rachel Williams, Mark Kristiansen, Paola Resende Silva, Sunando Roy, Tony Brooks, Helena Tutill, Paola Niola, Patricia Dyal, Charlotte Williams, Leysa Forrest, Yasmin Panchbhaya, Jacqueline Findlay, Samuel Weeks, Julianne Brown, Kathryn Harris, Paul Randell, James Price, Alison Holmes, Judith Breuer |
| EPI_ISL_950373, EPI_ISL_950376, EPI_ISL_950382, EPI_ISL_950391, EPI_ISL_950394, EPI_ISL_950396, EPI_ISL_950397, EPI_ISL_950399, EPI_ISL_950523, EPI_ISL_950525, EPI_ISL_950526, EPI_ISL_950527, EPI_ISL_950529, EPI_ISL_950531, EPI_ISL_950532, EPI_ISL_950533, EPI_ISL_950534, EPI_ISL_950535, EPI_ISL_950536, EPI_ISL_950537, EPI_ISL_950538, EPI_ISL_950539, EPI_ISL_950540, EPI_ISL_950541, EPI_ISL_950542, EPI_ISL_950543 | | | |
| see above | Northumbria University / South Tees Hospitals NHS Foundation Trust / North Cumbria Integrated Care NHS Foundation Trust / North Tees and Hartlepool NHS Foundation Trust / Newcastle Hospitals NHS Foundation Trust | COVID-19 Genomics UK (COG-UK) Consortium | Darren L Smith,Andrew Nelson,Matthew Bashton,Greg R Young,Joshua Loh,John Allan,Mohammad A Tariq,Giles S Holt,Gary Black,Wen C Yew,Lynn Dover,Paul Baker,Steve Liggett,Sarah Essex,Jane Greenaway,Debra Padgett,Clive Graham,Garren Scott,Edward Barton,Emma Swindells,Brendan Payne,Jennifer Collins,Yusri Taha,Gary Eltringham |
| EPI_ISL_950639, EPI_ISL_950640, EPI_ISL_950641, EPI_ISL_950642, EPI_ISL_950643, EPI_ISL_950644, EPI_ISL_950645 | Queens Medical Centre, Clinical Microbiology Department / DeepSeq Nottingham | COVID-19 Genomics UK (COG-UK) Consortium | Gemma Clark, Wendy Smith, Manjinder Khakh, Vicki M Fleming, Michelle M Lister, Hannah Howson-Wells, Jonathan Ball, Patrick McClure, Joseph Chappell, Theocharis Tsoleridis, Nadine Holmes, Matthew Carlisle, Christopher Moore, Fei Sang, Johnny Debebe, Victoria Wright, Matthew Loose |
| EPI_ISL_950755, EPI_ISL_950756, EPI_ISL_950757, EPI_ISL_950758, EPI_ISL_950759, EPI_ISL_950760, EPI_ISL_950761 | Lincolnshire Hospitals and DeepSeq Nottingham | COVID-19 Genomics UK (COG-UK) Consortium | Nichola Duckworth, Tim Sloan, Sarah Walsh, Jonathan Ball, Patrick McClure, Joeseph Chappell, Nadine Holmes, Matthew Carlisle, Christopher Moore, Fei Sang, Johnny Debebe, Victoria Wright, Matthew Loose |

| | | | |
|---|---|---|---|
| EPI_ISL_951492, EPI_ISL_951493, EPI_ISL_951494, EPI_ISL_951495, EPI_ISL_951496, EPI_ISL_951497, EPI_ISL_951498, EPI_ISL_951499, EPI_ISL_951501, EPI_ISL_951502, EPI_ISL_951503, EPI_ISL_951504, EPI_ISL_951505, EPI_ISL_951506, EPI_ISL_951508, EPI_ISL_951512, EPI_ISL_951513, EPI_ISL_951514, EPI_ISL_951515, EPI_ISL_951517, EPI_ISL_951518, EPI_ISL_951519, EPI_ISL_951521, EPI_ISL_951522, EPI_ISL_951523, EPI_ISL_951524, EPI_ISL_951525, EPI_ISL_951527, EPI_ISL_951528, EPI_ISL_951529, EPI_ISL_951530, EPI_ISL_951531, EPI_ISL_951532, EPI_ISL_951533, EPI_ISL_951534, EPI_ISL_951535, EPI_ISL_951536, EPI_ISL_951537, EPI_ISL_951538, EPI_ISL_951541, EPI_ISL_951542, EPI_ISL_951543, EPI_ISL_951544, EPI_ISL_951545, EPI_ISL_951546, EPI_ISL_951547, EPI_ISL_951548, EPI_ISL_951549, EPI_ISL_951550, EPI_ISL_951552, EPI_ISL_951553, EPI_ISL_951554, EPI_ISL_951556, EPI_ISL_951557, EPI_ISL_951558, EPI_ISL_951559, EPI_ISL_951560, EPI_ISL_951561, EPI_ISL_951562, EPI_ISL_951564, EPI_ISL_951567, EPI_ISL_951569, EPI_ISL_951570, EPI_ISL_951571, EPI_ISL_951572, EPI_ISL_951573, EPI_ISL_951574, EPI_ISL_951575, EPI_ISL_951576, EPI_ISL_951577, EPI_ISL_951578, EPI_ISL_951579, EPI_ISL_951580, EPI_ISL_951581, EPI_ISL_951582, EPI_ISL_951583, EPI_ISL_951585, EPI_ISL_951738, EPI_ISL_951739, EPI_ISL_951742, EPI_ISL_951744, EPI_ISL_951745, EPI_ISL_951753, EPI_ISL_951758, EPI_ISL_951759, EPI_ISL_951760, EPI_ISL_951761, EPI_ISL_951765, EPI_ISL_951766, EPI_ISL_951767, EPI_ISL_951769, EPI_ISL_951778, EPI_ISL_951779, EPI_ISL_951780, EPI_ISL_951781, EPI_ISL_951782, EPI_ISL_951785, EPI_ISL_951786, EPI_ISL_951787, EPI_ISL_951820, EPI_ISL_951821, EPI_ISL_951843, EPI_ISL_951844, EPI_ISL_951851, EPI_ISL_951852, EPI_ISL_951853, EPI_ISL_951855, EPI_ISL_951856, EPI_ISL_951857, EPI_ISL_951858, EPI_ISL_951859, EPI_ISL_951861, EPI_ISL_951862, EPI_ISL_951863, EPI_ISL_951866, EPI_ISL_951867, EPI_ISL_951869, EPI_ISL_951875, EPI_ISL_951876, EPI_ISL_951877, EPI_ISL_951878, EPI_ISL_951881, EPI_ISL_951882, EPI_ISL_951884, EPI_ISL_951886, EPI_ISL_951887, EPI_ISL_951888, EPI_ISL_951890, EPI_ISL_951893, EPI_ISL_951894, EPI_ISL_951895, EPI_ISL_951896, EPI_ISL_951897, EPI_ISL_951898, EPI_ISL_951900, EPI_ISL_951901, EPI_ISL_951902, EPI_ISL_951903, EPI_ISL_951904, EPI_ISL_951905, EPI_ISL_951906, EPI_ISL_951907, EPI_ISL_951908, EPI_ISL_951909, EPI_ISL_951910, EPI_ISL_951911, EPI_ISL_951912, EPI_ISL_951913, EPI_ISL_951916, EPI_ISL_951917, EPI_ISL_951918, EPI_ISL_951919, EPI_ISL_951920, EPI_ISL_951921, EPI_ISL_951922, EPI_ISL_951923, EPI_ISL_951924, EPI_ISL_951925, EPI_ISL_951926, EPI_ISL_951927, EPI_ISL_951928, EPI_ISL_951929, EPI_ISL_951930, EPI_ISL_951931, EPI_ISL_951932, EPI_ISL_951933, EPI_ISL_951934, EPI_ISL_951935, EPI_ISL_951936, EPI_ISL_951937, EPI_ISL_951938, EPI_ISL_951939, EPI_ISL_951940, EPI_ISL_951941, EPI_ISL_951942, EPI_ISL_951943, EPI_ISL_951944, EPI_ISL_951945, EPI_ISL_951946, EPI_ISL_951947, EPI_ISL_951948, EPI_ISL_951949, EPI_ISL_951950, EPI_ISL_951951, EPI_ISL_951952, EPI_ISL_951953, EPI_ISL_951954, EPI_ISL_951955, EPI_ISL_951956, EPI_ISL_951957, EPI_ISL_951958, EPI_ISL_951959, EPI_ISL_951960, EPI_ISL_951961, EPI_ISL_951962, EPI_ISL_951963, EPI_ISL_951964, EPI_ISL_951965, EPI_ISL_951966, EPI_ISL_951968, EPI_ISL_951969, EPI_ISL_951972, EPI_ISL_951973, EPI_ISL_952006, EPI_ISL_952007, EPI_ISL_952008, EPI_ISL_952009, EPI_ISL_952010, EPI_ISL_952011, EPI_ISL_952012, EPI_ISL_952013, EPI_ISL_952014, EPI_ISL_952015, EPI_ISL_952018, EPI_ISL_952019, EPI_ISL_952020, EPI_ISL_952022, EPI_ISL_952025, EPI_ISL_952027, EPI_ISL_952029, EPI_ISL_952030, EPI_ISL_952031, EPI_ISL_952032, EPI_ISL_952033, EPI_ISL_952035, EPI_ISL_952036, EPI_ISL_952037, EPI_ISL_952038, EPI_ISL_952039, EPI_ISL_952040, EPI_ISL_952041, EPI_ISL_952061, EPI_ISL_952062, EPI_ISL_952063, EPI_ISL_952064, EPI_ISL_952065, EPI_ISL_952066, EPI_ISL_952067, EPI_ISL_952069, EPI_ISL_952070, EPI_ISL_952071, EPI_ISL_952072, EPI_ISL_952073, EPI_ISL_952074, EPI_ISL_952075, EPI_ISL_952076, EPI_ISL_952077, EPI_ISL_952078, EPI_ISL_952079, EPI_ISL_952082, EPI_ISL_952083, EPI_ISL_952084, EPI_ISL_952085, EPI_ISL_952086, EPI_ISL_952087, EPI_ISL_952088, EPI_ISL_952089, EPI_ISL_952090, EPI_ISL_952091, EPI_ISL_952092, EPI_ISL_952093, EPI_ISL_952095, EPI_ISL_952096, EPI_ISL_952097, EPI_ISL_952100, EPI_ISL_952102, EPI_ISL_952104, EPI_ISL_952108, EPI_ISL_952109, EPI_ISL_952110, EPI_ISL_952111, EPI_ISL_952112, EPI_ISL_952113, EPI_ISL_952114, EPI_ISL_952119, EPI_ISL_952120, EPI_ISL_952122, EPI_ISL_952123, EPI_ISL_952124, EPI_ISL_952126, EPI_ISL_952128, EPI_ISL_952129, EPI_ISL_952130, EPI_ISL_952131, EPI_ISL_952132, EPI_ISL_952133, EPI_ISL_952134, EPI_ISL_952136, EPI_ISL_952137, EPI_ISL_952138, EPI_ISL_952139, EPI_ISL_952140, EPI_ISL_952141, EPI_ISL_952143, EPI_ISL_952144, EPI_ISL_952145, EPI_ISL_952146, EPI_ISL_952147, EPI_ISL_952148, EPI_ISL_952149, EPI_ISL_952150, EPI_ISL_952151, EPI_ISL_952152, EPI_ISL_952153, EPI_ISL_952154, EPI_ISL_952156, EPI_ISL_952157, EPI_ISL_952158, EPI_ISL_952159, EPI_ISL_952161, EPI_ISL_952162, EPI_ISL_952163, EPI_ISL_952164, EPI_ISL_952165, EPI_ISL_952166, EPI_ISL_952167 | | | |
| see above | Originating lab: Wales Specialist Virology Centre Sequencing lab: Pathogen Genomics Unit | Public Health Wales Microbiology Cardiff Wales Specialist Virology Centre | Catherine Moore, Johnathan Evans, Laura Gifford, Malorie Perry, Simon Cottrell, Angela Marchbank, Alec Birchley, Alexander Adams, Amy Gaskin, Bree Gatica-Wilcox, Jason Coombes, Joel Southgate, Lauren Gilbert, Lee Graham, Nicole Pacchiarini, Sara Kumziene-Summerhayes, Sarah Taylor, Sophie Jones, Sara Rey, Matthew Bull, Joanne Watkins, Sally Corden, Tom Connor |
| EPI_ISL_952453, EPI_ISL_952454, EPI_ISL_952456, EPI_ISL_952460, EPI_ISL_952462, EPI_ISL_952464, EPI_ISL_952465, EPI_ISL_952466, EPI_ISL_952467, EPI_ISL_952469, EPI_ISL_952470, EPI_ISL_952477, EPI_ISL_952478, EPI_ISL_952479, EPI_ISL_952480, EPI_ISL_952481, EPI_ISL_952482, EPI_ISL_952483, EPI_ISL_952484, EPI_ISL_952485, EPI_ISL_952558 | | | |
| see above | Centre for Enzyme Innovation, University of Portsmouth / Translational Research Laboratory, Portsmouth Hospitals NHS Trust | COVID-19 Genomics UK (COG-UK) Consortium | Angela Beckett,Salman Goudarzi,Christopher Fearn,Kate Cook,Katie Loveson,Sharon Glaysher,Scott Elliott,Samuel Robson |
| EPI_ISL_952953, EPI_ISL_952978 | Virology Department, Sheffield Teaching Hospitals NHS Foundation Trust/Department of Infection, Immunity and Cardiovascular Disease, The Medical School, University of Sheffield | COVID-19 Genomics UK (COG-UK) Consortium | Thushan de Silva, Matthew Parker, Nikki Smith, Adri Angyal, Rebecca Brown, Luke Green, Rachel Tucker, Paul Parsons, Danielle Groves, Katie Johnson, Laura Carrilero, Alex Keeley, Dave Partridge, Matthew Wyles, Benjamin Lindsey, Mehmet Yavuz, Mohammad Raza, Cariad Evans |
| EPI_ISL_953241, EPI_ISL_953243, EPI_ISL_953244, EPI_ISL_953246, EPI_ISL_953247, EPI_ISL_953251, EPI_ISL_953254, EPI_ISL_953259, EPI_ISL_953261, EPI_ISL_953262, EPI_ISL_953263, EPI_ISL_953267, EPI_ISL_953272, EPI_ISL_953281 | | | |
| see above | Bioinformatics and Biostatistics Lab, Advanced Sequencing Facility | COVID-19 Genomics UK (COG-UK) Consortium | Aengus Stewart,Jerome Nicod,Chelsea Sawyer,Laura Cubitt,Harshil Patel,Margaret Crawford |
| EPI_ISL_953426 | Laboratory of Virology and Molecular Diagnostics | Institute of Public Health of Republic of North Macedonia Laboratory of Virology and Molecular Diagnostics | M. Kuzmanovska, G. Boshevska |
| EPI_ISL_953452 | Jessa | Jessa | Jessa_cmdLab |
| EPI_ISL_953490, EPI_ISL_953491, EPI_ISL_953492, EPI_ISL_953494, EPI_ISL_953495, EPI_ISL_953496, EPI_ISL_953497, EPI_ISL_953500, EPI_ISL_953505, EPI_ISL_953506, EPI_ISL_953509, EPI_ISL_953510, EPI_ISL_953511, EPI_ISL_953514, EPI_ISL_953519, EPI_ISL_953520, EPI_ISL_953560, EPI_ISL_953561, EPI_ISL_953562, EPI_ISL_953563, EPI_ISL_953564, EPI_ISL_953565, EPI_ISL_953566, EPI_ISL_953567, EPI_ISL_953568, EPI_ISL_953569, EPI_ISL_953570, EPI_ISL_953571, EPI_ISL_953572, EPI_ISL_953573, EPI_ISL_953574, EPI_ISL_953575, EPI_ISL_953576, EPI_ISL_953577, EPI_ISL_953578, EPI_ISL_953579, EPI_ISL_953580, EPI_ISL_953581, EPI_ISL_953582, EPI_ISL_953583, EPI_ISL_953584, EPI_ISL_953585, EPI_ISL_953586, EPI_ISL_953587, EPI_ISL_953611, EPI_ISL_953612, EPI_ISL_953613, EPI_ISL_953614, EPI_ISL_953615, EPI_ISL_953616, EPI_ISL_953617, EPI_ISL_953618, EPI_ISL_953619, EPI_ISL_953620, EPI_ISL_953631, EPI_ISL_953634, EPI_ISL_953635, EPI_ISL_953642, EPI_ISL_953643, EPI_ISL_953644, EPI_ISL_953645, EPI_ISL_953646, EPI_ISL_953647, EPI_ISL_953648, EPI_ISL_953653, EPI_ISL_953658, EPI_ISL_953664, EPI_ISL_953854, EPI_ISL_953855, EPI_ISL_953856, EPI_ISL_953857, EPI_ISL_953858, EPI_ISL_953859, EPI_ISL_953860, EPI_ISL_953861, EPI_ISL_953862, EPI_ISL_953863, EPI_ISL_953864, EPI_ISL_953865, EPI_ISL_953866, EPI_ISL_953867, EPI_ISL_953868, EPI_ISL_953869, EPI_ISL_953870, EPI_ISL_953871, EPI_ISL_953872, EPI_ISL_953873, EPI_ISL_953874, EPI_ISL_953875, EPI_ISL_953876, EPI_ISL_953877, EPI_ISL_953878, EPI_ISL_953879, EPI_ISL_953880, EPI_ISL_953881, EPI_ISL_953882, EPI_ISL_953883, EPI_ISL_953884, EPI_ISL_953885, EPI_ISL_953886, EPI_ISL_953887, EPI_ISL_953888, EPI_ISL_953889, EPI_ISL_953890, EPI_ISL_953891, EPI_ISL_953892, EPI_ISL_953893, EPI_ISL_953894, EPI_ISL_953895, EPI_ISL_953896, EPI_ISL_953897, EPI_ISL_953898, EPI_ISL_953899, EPI_ISL_953900, EPI_ISL_953901, EPI_ISL_953902, EPI_ISL_953903, EPI_ISL_953904, EPI_ISL_953905, EPI_ISL_953906, EPI_ISL_953907, EPI_ISL_953908, EPI_ISL_953909, EPI_ISL_953910, EPI_ISL_953911, EPI_ISL_953912, EPI_ISL_953913, EPI_ISL_953914, EPI_ISL_953915, EPI_ISL_953916, EPI_ISL_953917, EPI_ISL_953918, EPI_ISL_953919, EPI_ISL_953920, EPI_ISL_953921 | | | |
| see above | University Hospitals of Geneva, Laboratory of Virology | HUG, Laboratory of Virology and the Health2030 Genome Center | Samuel Cordey, Ana Rita Goncalves, Laurent Kaiser, Lorenzo Cerutti, Henri Pegeot, Melyssa Elies, Deborah Penet, Keith Harshman, Ioannis Xenarios, Emmanouil Dermitzakis |
| EPI_ISL_953959 | Hopital | National Reference Center for Viruses of Respiratory Infections, Institut Pasteur, Paris | Marion Barbet, Sylvie Behillil, Méline Bizard, Angela Brisebarre, Camille Capel, Etienne Simon-Lorière, Vincent Enouf, Maud Vanpeene, Sylvie van der Werf,Ramirez José-Manuel |
| EPI_ISL_954009, EPI_ISL_954010 | Hopital | National Reference Center for Viruses of Respiratory Infections, Institut Pasteur, Paris | Marion Barbet, Sylvie Behillil, Méline Bizard, Angela Brisebarre, Camille Capel, Etienne Simon-Lorière, Vincent Enouf, Maud Vanpeene, Sylvie van der Werf,Fourgeaud Jacques |
| EPI_ISL_954030, EPI_ISL_954038 | Labo Analyses Med | National Reference Center for Viruses of Respiratory Infections, Institut Pasteur, Paris | Marion Barbet, Sylvie Behillil, Méline Bizard, Angela Brisebarre, Camille Capel, Etienne Simon-Lorière, Vincent Enouf, Maud Vanpeene, Sylvie van der Werf |
| EPI_ISL_954039 | Recherche | National Reference Center for Viruses of Respiratory Infections, Institut Pasteur, Paris | Marion Barbet, Sylvie Behillil, Méline Bizard, Angela Brisebarre, Camille Capel, Etienne Simon-Lorière, Vincent Enouf, Maud Vanpeene, Sylvie van der Werf,Leruez-Ville Marianne |
| EPI_ISL_954040 | Labo Analyses Med | National Reference Center for Viruses of Respiratory Infections, Institut Pasteur, Paris | Marion Barbet, Sylvie Behillil, Méline Bizard, Angela Brisebarre, Camille Capel, Etienne Simon-Lorière, Vincent Enouf, Maud Vanpeene, Sylvie van der Werf,Labm Hignard |
| EPI_ISL_954144, EPI_ISL_954180 | Hospital Universitari Vall d'Hebron - Vall d'Hebron Institut de Rercerca | Hospital Universitari Vall d'Hebron | Cristina Andrés, Maria Piñana, Josep F Abril, Damir Garcia-Cehic, Ariadna Rando, Juliana Esperalba, Maria Gema Codina, Carla Castillo, Maria Carmen Martin, Tomàs Pumarola, Josep Quer, Andrés Antón |
| EPI_ISL_954748 | National Institute of Laboratory Medicine and Referral Center | Genomic Research Lab, BCSIR | Md. Murshed Hasan Sarkar, Mohammad Samir Uzzaman, Eshrar Osman, Md. Ahashan Habib, Shahina Akter, Tanjina Akhtar Banu, Abu Sayeed Mohammad Mahmud, Barna Goswami, Iffat Jahan, Md. Saddam Hossain, Tasnim Nafisa, Md. Maruf Ahmed Molla, Mahmuda Yeasmin, Asish Kumar Ghosh, Arifa Akram, A.K.M.Shamsuzzaman, Md. Salim Khan |
| EPI_ISL_954749 | National Institute of Laboratory Medicine and Referral Center | Genomic Research Lab, BCSIR | Barna Goswami, Mohammad Samir Uzzaman, Eshrar Osman, Md. Ahashan Habib, Shahina Akter,Tanjina Akhtar Banu, Abu Sayeed Mohammad Mahmud, Md. Murshed Hasan Sarkar, Iffat Jahan, Md. Saddam Hossain, Tasnim Nafisa, Md. Maruf Ahmed Molla, Mahmuda Yeasmin, Asish Kumar Ghosh, Arifa Akram, A. K. M. Shamsuzzaman, Md. Salim Khan |
| EPI_ISL_954751 | National Institute of Laboratory Medicine and Referral Center | Genomic Research Lab, BCSIR | Iffat Jahan, Mohammad Samir Uzzaman, Eshrar Osman, Md. Ahashan Habib, Shahina Akter, Tanjina Akhtar Banu, Abu Sayeed Mohammad Mahmud, Md. Murshed Hasan Sarkar, Barna Goswami, Md. Saddam Hossain, Tasnim Nafisa, Md. Maruf Ahmed Molla, Mahmuda Yeasmin, Asish Kumar Ghosh, Arifa Akram, A. K. M. Shamsuzzaman, Md. Salim Khan |
| EPI_ISL_954814, EPI_ISL_954815, EPI_ISL_954816, EPI_ISL_954817 | AZDelta | AZDelta | Geert Martens; Dieter De Smet |
| EPI_ISL_955060 | Colorado Department of Public Health and Environment | Colorado Department of Puplic Health and Environment | Laura Bankers, Molly C. Hetherington-Rauth, Diana Ir, Shannon Ely, Shannon R. Matzinger, Sarah Elizabeth Totten, Emily A. Travanty |
| EPI_ISL_955061 | Hopital | National Reference Center for Viruses of Respiratory Infections, Institut Pasteur, Paris | Marion Barbet, Sylvie Behillil, Méline Bizard, Angela Brisebarre, Camille Capel, Etienne Simon-Lorière, Vincent Enouf, Maud Vanpeene, Sylvie van der Werf,Fourgeaud Jacques |
| EPI_ISL_955062, EPI_ISL_955063 | Labo analyses Med | National Reference Center for Viruses of Respiratory Infections, Institut Pasteur, Paris | Marion Barbet, Sylvie Behillil, Méline Bizard, Angela Brisebarre, Camille Capel, Etienne Simon-Lorière, Vincent Enouf, Maud Vanpeene, Sylvie van der Werf,Potiron GréGoire |
| EPI_ISL_955065 | Hopital | National Reference Center for Viruses of Respiratory | Marion Barbet, Sylvie Behillil, Méline Bizard, Angela Brisebarre, Camille Capel, Etienne Simon-Lorière, Vincent Enouf, Maud Vanpeene, Sylvie van der |

| | | | |
|---|---|---|---|
| | | Infections, Institut Pasteur, Paris | Werf,Fourgeaud Jacques |
| EPI_ISL_955066, EPI_ISL_955067 | Labo analyses med | National Reference Center for Viruses of Respiratory Infections, Institut Pasteur, Paris | Marion Barbet, Sylvie Behillil, Méline Bizard, Angela Brisebarre, Camille Capel, Etienne Simon-Lorière, Vincent Enouf, Maud Vanpeene, Sylvie van der Werf,Amzalag Jonas |
| EPI_ISL_955068, EPI_ISL_955069, EPI_ISL_955070, EPI_ISL_955071, EPI_ISL_955072 | Labo analyses Med | National Reference Center for Viruses of Respiratory Infections, Institut Pasteur, Paris | Marion Barbet, Sylvie Behillil, Méline Bizard, Angela Brisebarre, Camille Capel, Etienne Simon-Lorière, Vincent Enouf, Maud Vanpeene, Sylvie van der Werf,Potiron GréGoire |
| EPI_ISL_955073 | Labo Analyses Med | National Reference Center for Viruses of Respiratory Infections, Institut Pasteur, Paris | Marion Barbet, Sylvie Behillil, Méline Bizard, Angela Brisebarre, Camille Capel, Etienne Simon-Lorière, Vincent Enouf, Maud Vanpeene, Sylvie van der Werf,Jacques Alexandra |
| EPI_ISL_955076 | Labo analyses Med | National Reference Center for Viruses of Respiratory Infections, Institut Pasteur, Paris | Marion Barbet, Sylvie Behillil, Méline Bizard, Angela Brisebarre, Camille Capel, Etienne Simon-Lorière, Vincent Enouf, Maud Vanpeene, Sylvie van der Werf,Potiron GréGoire |
| EPI_ISL_955078 | Hopital | National Reference Center for Viruses of Respiratory Infections, Institut Pasteur, Paris | Marion Barbet, Sylvie Behillil, Méline Bizard, Angela Brisebarre, Camille Capel, Etienne Simon-Lorière, Vincent Enouf, Maud Vanpeene, Sylvie van der Werf,Fourgeaud Jacques |
| EPI_ISL_955079, EPI_ISL_955080, EPI_ISL_955082 | Labo analyses Med | National Reference Center for Viruses of Respiratory Infections, Institut Pasteur, Paris | Marion Barbet, Sylvie Behillil, Méline Bizard, Angela Brisebarre, Camille Capel, Etienne Simon-Lorière, Vincent Enouf, Maud Vanpeene, Sylvie van der Werf,Potiron GréGoire |
| EPI_ISL_955083, EPI_ISL_955084 | Hopital | National Reference Center for Viruses of Respiratory Infections, Institut Pasteur, Paris | Marion Barbet, Sylvie Behillil, Méline Bizard, Angela Brisebarre, Camille Capel, Etienne Simon-Lorière, Vincent Enouf, Maud Vanpeene, Sylvie van der Werf,Fourgeaud Jacques |
| EPI_ISL_955085 | Hopital | National Reference Center for Viruses of Respiratory Infections, Institut Pasteur, Paris | Marion Barbet, Sylvie Behillil, Méline Bizard, Angela Brisebarre, Camille Capel, Etienne Simon-Lorière, Vincent Enouf, Maud Vanpeene, Sylvie van der Werf,Joly Isabelle |
| EPI_ISL_955086, EPI_ISL_955087, EPI_ISL_955088, EPI_ISL_955089, EPI_ISL_955090, EPI_ISL_955091, EPI_ISL_955092, EPI_ISL_955093 | Labo analyses Med | National Reference Center for Viruses of Respiratory Infections, Institut Pasteur, Paris | Marion Barbet, Sylvie Behillil, Méline Bizard, Angela Brisebarre, Camille Capel, Etienne Simon-Lorière, Vincent Enouf, Maud Vanpeene, Sylvie van der Werf,Potiron GréGoire |
| EPI_ISL_955094 | Labo Analyses Med | National Reference Center for Viruses of Respiratory Infections, Institut Pasteur, Paris | Marion Barbet, Sylvie Behillil, Méline Bizard, Angela Brisebarre, Camille Capel, Etienne Simon-Lorière, Vincent Enouf, Maud Vanpeene, Sylvie van der Werf,Potiron GréGoire |
| EPI_ISL_955112, EPI_ISL_955113, EPI_ISL_955114, EPI_ISL_955115 | Maryland Public Health Laboratory | Maryland Public Health Laboratory | Maryland Department of Health Laboratories Administration |
| EPI_ISL_955172 | National Institute of Laboratory Medicine and Referral Center | Genomic Research Lab, BCSIR | Tasnim Nafisa, Mohammad Samir Uzzaman, Eshrar Osman, Md. Ahashan Habib, Shahina Akter,Tanjina Akhtar Banu, Abu Sayeed Mohammad Mahmud, Md. Murshed Hasan Sarkar, Barna Goswami, Iffat Jahan, Md. Saddam Hossain, Md. Maruf Ahmed Molla, ,Mahmuda Yeasmin, Asish Kumar Ghosh, Arifa Akram, A. K. M. Shamsuzzaman, Md. Salim Khan. |
| EPI_ISL_955215, EPI_ISL_955222 | Indiana Animal Disease Diagnostic Laboratory | Carpi Laboratory - Purdue University | Jack Dorman, Ilinca I Ciubotariu, Lev Gorenstein, Abebe A Fola, G Kenitra Hendrix, Rebecca P Wilkes, Giovanna Carpi |
| EPI_ISL_955267 | Clinical Pathology Lab | Pathogen Discovery, Respiratory Viruses Branch, Division of Viral Diseases, Centers for Disease Control and Prevention | Ying Tao, Jing Zhang, Yan Li, Krista Queen, Anna Uehara, Peter Cook, Clinton R. Paden, Haibin Wang, Suxiang Tong |
| EPI_ISL_955272 | AR Dept. of Health-PHL, Molecular Diagnostics | Pathogen Discovery, Respiratory Viruses Branch, Division of Viral Diseases, Centers for Disease Control and Prevention | Ying Tao, Jing Zhang, Yan Li, Krista Queen, Anna Uehara, Peter Cook, Clinton R. Paden, Haibin Wang, Suxiang Tong |
| EPI_ISL_955280, EPI_ISL_955282 | Clinical Pathology Lab | Pathogen Discovery, Respiratory Viruses Branch, Division of Viral Diseases, Centers for Disease Control and Prevention | Ying Tao, Jing Zhang, Yan Li, Krista Queen, Anna Uehara, Peter Cook, Clinton R. Paden, Haibin Wang, Suxiang Tong |
| EPI_ISL_955284 | NC State Laboratory of Public Health | Pathogen Discovery, Respiratory Viruses Branch, Division of Viral Diseases, Centers for Disease Control and Prevention | Ying Tao, Jing Zhang, Yan Li, Krista Queen, Anna Uehara, Peter Cook, Clinton R. Paden, Haibin Wang, Suxiang Tong |
| EPI_ISL_955287 | Clinical Pathology Lab | Pathogen Discovery, Respiratory Viruses Branch, Division of Viral Diseases, Centers for Disease Control and Prevention | Ying Tao, Jing Zhang, Yan Li, Krista Queen, Anna Uehara, Peter Cook, Clinton R. Paden, Haibin Wang, Suxiang Tong |
| EPI_ISL_955288 | GA Department of Public Health Laboratory | Pathogen Discovery, Respiratory Viruses Branch, Division of Viral Diseases, Centers for Disease Control and Prevention | Ying Tao, Jing Zhang, Yan Li, Krista Queen, Anna Uehara, Peter Cook, Clinton R. Paden, Haibin Wang, Suxiang Tong |
| EPI_ISL_955291 | IL Dept. of Public Health Springfield Laboratory | Pathogen Discovery, Respiratory Viruses Branch, Division of Viral Diseases, Centers for Disease Control and Prevention | Ying Tao, Jing Zhang, Yan Li, Krista Queen, Anna Uehara, Peter Cook, Clinton R. Paden, Haibin Wang, Suxiang Tong |
| EPI_ISL_955293 | TMC/Pathology | Pathogen Discovery, Respiratory Viruses Branch, Division of Viral Diseases, Centers for Disease Control and Prevention | Ying Tao, Jing Zhang, Yan Li, Krista Queen, Anna Uehara, Peter Cook, Clinton R. Paden, Haibin Wang, Suxiang Tong |
| EPI_ISL_955294 | Clinical Pathology Lab | Pathogen Discovery, Respiratory Viruses Branch, Division of Viral Diseases, Centers for Disease Control and Prevention | Ying Tao, Jing Zhang, Yan Li, Krista Queen, Anna Uehara, Peter Cook, Clinton R. Paden, Haibin Wang, Suxiang Tong |
| EPI_ISL_955298 | NC State Laboratory of Public Health | Pathogen Discovery, Respiratory Viruses Branch, Division of Viral Diseases, Centers for Disease Control and Prevention | Ying Tao, Jing Zhang, Yan Li, Krista Queen, Anna Uehara, Peter Cook, Clinton R. Paden, Haibin Wang, Suxiang Tong |
| EPI_ISL_955301 | Clinical Pathology Lab | Pathogen Discovery, Respiratory Viruses Branch, Division of Viral Diseases, Centers for Disease Control and Prevention | Ying Tao, Jing Zhang, Yan Li, Krista Queen, Anna Uehara, Peter Cook, Clinton R. Paden, Haibin Wang, Suxiang Tong |
| EPI_ISL_955307 | OK Public Health Laboratory, Oklahoma State DOH | Pathogen Discovery, Respiratory Viruses Branch, Division of Viral Diseases, Centers for Disease Control and Prevention | Ying Tao, Jing Zhang, Yan Li, Krista Queen, Anna Uehara, Peter Cook, Clinton R. Paden, Haibin Wang, Suxiang Tong |
| EPI_ISL_955309 | IL Dept. of Public Health Springfield Laboratory | Pathogen Discovery, Respiratory Viruses Branch, Division of Viral Diseases, Centers for Disease Control and Prevention | Ying Tao, Jing Zhang, Yan Li, Krista Queen, Anna Uehara, Peter Cook, Clinton R. Paden, Haibin Wang, Suxiang Tong |
| EPI_ISL_955313, EPI_ISL_955315, EPI_ISL_955331, EPI_ISL_955332 | AR Dept. of Health-PHL, Molecular Diagnostics | Pathogen Discovery, Respiratory Viruses Branch, Division of Viral Diseases, Centers for Disease Control and Prevention | Ying Tao, Jing Zhang, Yan Li, Krista Queen, Anna Uehara, Peter Cook, Clinton R. Paden, Haibin Wang, Suxiang Tong |
| EPI_ISL_955830, EPI_ISL_955862, EPI_ISL_955863, EPI_ISL_955864, EPI_ISL_955865, EPI_ISL_955866, EPI_ISL_955867, EPI_ISL_955869, EPI_ISL_955870, EPI_ISL_955871, EPI_ISL_955872, EPI_ISL_955873, EPI_ISL_955874, EPI_ISL_955876, EPI_ISL_955877, EPI_ISL_955878, EPI_ISL_955879, EPI_ISL_955881, EPI_ISL_955882, EPI_ISL_955883, EPI_ISL_955884, EPI_ISL_955885, EPI_ISL_955886, EPI_ISL_955887, EPI_ISL_955888, EPI_ISL_955889, EPI_ISL_955891, EPI_ISL_955892, EPI_ISL_955893, EPI_ISL_955894, EPI_ISL_955895, EPI_ISL_955896, EPI_ISL_955898, EPI_ISL_955899, EPI_ISL_955900, EPI_ISL_955901, EPI_ISL_955902, EPI_ISL_955903, EPI_ISL_955904, EPI_ISL_955918, EPI_ISL_955919, EPI_ISL_955920, EPI_ISL_955921 | | | |
| see above | University of Michigan Clinical Microbiology Laboratory | Lauring Lab, University of Michigan, Department of Microbiology and Immunology | Valesano |
| EPI_ISL_956275 | Thai Red Cross Emerging Infectious Diseases Health Science Centre, Chulalongkorn Hospital, Faculty of Medicine, Chulalongkorn University | Thai Red Cross Emerging Infectious Diseases Center and Faculty of Medicine, Chulalongkorn University | Ratima samorn, Panpit Suwangool, Sininat Petcharat, Yutthana Joyjinda, Weenassarin Ampoot, Apaporn Rodpan, Opass Putcharoen, Thiravat Hemachudha, Supaporn Wacharapluesadee |
| EPI_ISL_956331 | Laboratory Medicine | Department of Laboratory Medicine, Lin-Kou Chang Gung Memorial Hospital, Taoyuan, Taiwan | Kuo-Chien Tsao, Yu-Nong Gong, Shu-Li Yang, Yi-Chun Liu, Chung-Guei Huang, Mei-Jen Hsiao, Po-Wei Huang, Cheng-Ta Yang, Cheng-Hsun Chiu, Peng-Nien Huang, Kuo-Ming Lee, Guang-Wu Chen, Shin-Ru Shih |
| EPI_ISL_956446, EPI_ISL_956487, EPI_ISL_956508, EPI_ISL_956562, EPI_ISL_956576, EPI_ISL_956585, EPI_ISL_956587, EPI_ISL_956592, EPI_ISL_956601, EPI_ISL_956602, EPI_ISL_956606, EPI_ISL_956618, EPI_ISL_956625, EPI_ISL_956626, EPI_ISL_956631, EPI_ISL_956636, EPI_ISL_956651, EPI_ISL_956690, EPI_ISL_956691, EPI_ISL_956694, EPI_ISL_956699, EPI_ISL_956708, EPI_ISL_956729 | | | |
| see above | Lighthouse Lab in Alderley Park | Wellcome Sanger Institute for the COVID-19 Genomics UK | Jacquelyn Wynn, Mairead Hyland, The Lighthouse Lab in Alderley Park and Alex Alderton, Roberto Amato, Sonia Goncalves, Ewan Harrison, David K. |

| | | | |
|---|---|---|---|
| | | (COG-UK) Consortium | Jackson, Ian Johnston, Dominic Kwiatkowski, Cordelia Langford, John Sillitoe on behalf of the Wellcome Sanger Institute COVID-19 Surveillance Team |
| EPI_ISL_957613, EPI_ISL_957616, EPI_ISL_957620, EPI_ISL_957621, EPI_ISL_957624, EPI_ISL_957627, EPI_ISL_957630, EPI_ISL_957635, EPI_ISL_957641, EPI_ISL_957643, EPI_ISL_957645, EPI_ISL_957650, EPI_ISL_957652, EPI_ISL_957654, EPI_ISL_957655, EPI_ISL_957656, EPI_ISL_957660, EPI_ISL_957670, EPI_ISL_957672, EPI_ISL_957675, EPI_ISL_957677, EPI_ISL_957680, EPI_ISL_957683, EPI_ISL_957686, EPI_ISL_957687, EPI_ISL_957691, EPI_ISL_957698, EPI_ISL_957700, EPI_ISL_957701, EPI_ISL_957707, EPI_ISL_957721, EPI_ISL_957722, EPI_ISL_957723, EPI_ISL_957726, EPI_ISL_957735, EPI_ISL_957740, EPI_ISL_957741, EPI_ISL_957745, EPI_ISL_957747, EPI_ISL_957748, EPI_ISL_957753, EPI_ISL_957755, EPI_ISL_957757, EPI_ISL_957759, EPI_ISL_957766, EPI_ISL_957769, EPI_ISL_957771, EPI_ISL_957772, EPI_ISL_957780, EPI_ISL_957783, EPI_ISL_957787, EPI_ISL_957797, EPI_ISL_957799, EPI_ISL_957813, EPI_ISL_957817, EPI_ISL_957822, EPI_ISL_957826, EPI_ISL_957828, EPI_ISL_957829, EPI_ISL_957830, EPI_ISL_957832, EPI_ISL_957838, EPI_ISL_957841, EPI_ISL_957844, EPI_ISL_957856, EPI_ISL_957859, EPI_ISL_957862, EPI_ISL_957863, EPI_ISL_957866, EPI_ISL_957867, EPI_ISL_957868, EPI_ISL_957872, EPI_ISL_957877, EPI_ISL_957878, EPI_ISL_957886, EPI_ISL_957892, EPI_ISL_957895, EPI_ISL_957897, EPI_ISL_957904, EPI_ISL_957907, EPI_ISL_957921, EPI_ISL_957961, EPI_ISL_957962, EPI_ISL_957971, EPI_ISL_958044, EPI_ISL_958194, EPI_ISL_958206, EPI_ISL_958215, EPI_ISL_958216, EPI_ISL_958217, EPI_ISL_958218, EPI_ISL_958219, EPI_ISL_958220, EPI_ISL_958221, EPI_ISL_958222, EPI_ISL_958223, EPI_ISL_958224, EPI_ISL_958225, EPI_ISL_958226, EPI_ISL_958227, EPI_ISL_958228, EPI_ISL_958229, EPI_ISL_958230, EPI_ISL_958231, EPI_ISL_958232, EPI_ISL_958233, EPI_ISL_958234, EPI_ISL_958235, EPI_ISL_958236, EPI_ISL_958237, EPI_ISL_958238, EPI_ISL_958239, EPI_ISL_958240, EPI_ISL_958241, EPI_ISL_958242, EPI_ISL_958243, EPI_ISL_958244, EPI_ISL_958245, EPI_ISL_958246, EPI_ISL_958247, EPI_ISL_958248, EPI_ISL_958249, EPI_ISL_958250, EPI_ISL_958251, EPI_ISL_958252, EPI_ISL_958253, EPI_ISL_958254, EPI_ISL_958255, EPI_ISL_958256, EPI_ISL_958257, EPI_ISL_958258, EPI_ISL_958259, EPI_ISL_958260, EPI_ISL_958261, EPI_ISL_958262, EPI_ISL_958263, EPI_ISL_958264, EPI_ISL_958265, EPI_ISL_958266, EPI_ISL_958267, EPI_ISL_958268, EPI_ISL_958269, EPI_ISL_958270, EPI_ISL_958271, EPI_ISL_958272, EPI_ISL_958273, EPI_ISL_958274, EPI_ISL_958275, EPI_ISL_958277, EPI_ISL_958278, EPI_ISL_958279, EPI_ISL_958280, EPI_ISL_958281, EPI_ISL_958282, EPI_ISL_958283, EPI_ISL_958284, EPI_ISL_958285, EPI_ISL_958286, EPI_ISL_958287, EPI_ISL_958288, EPI_ISL_958289, EPI_ISL_958290, EPI_ISL_958291, EPI_ISL_958292, EPI_ISL_958293, EPI_ISL_958294, EPI_ISL_958295, EPI_ISL_958296, EPI_ISL_958297, EPI_ISL_958298, EPI_ISL_958299, EPI_ISL_958300, EPI_ISL_958301, EPI_ISL_958302, EPI_ISL_958303, EPI_ISL_958304, EPI_ISL_958305, EPI_ISL_958306, EPI_ISL_958307, EPI_ISL_958308, EPI_ISL_958309, EPI_ISL_958310, EPI_ISL_958311, EPI_ISL_958312, EPI_ISL_958313, EPI_ISL_958314, EPI_ISL_958315, EPI_ISL_958316, EPI_ISL_958317, EPI_ISL_958318, EPI_ISL_958319, EPI_ISL_958320, EPI_ISL_958321, EPI_ISL_958322, EPI_ISL_958323, EPI_ISL_958324, EPI_ISL_958325, EPI_ISL_958326, EPI_ISL_958327, EPI_ISL_958328, EPI_ISL_958329, EPI_ISL_958330, EPI_ISL_958331, EPI_ISL_958332, EPI_ISL_958333, EPI_ISL_958334, EPI_ISL_958335, EPI_ISL_958336, EPI_ISL_958337, EPI_ISL_958338, EPI_ISL_958339, EPI_ISL_958340, EPI_ISL_958341, EPI_ISL_958342, EPI_ISL_958343, EPI_ISL_958344, EPI_ISL_958345, EPI_ISL_958346, EPI_ISL_958347, EPI_ISL_958348, EPI_ISL_958349, EPI_ISL_958350, EPI_ISL_958351, EPI_ISL_958352, EPI_ISL_958353, EPI_ISL_958354, EPI_ISL_958355, EPI_ISL_958356, EPI_ISL_958357, EPI_ISL_958358, EPI_ISL_958359, EPI_ISL_958360, EPI_ISL_958361, EPI_ISL_958362, EPI_ISL_958363, EPI_ISL_958364, EPI_ISL_958365, EPI_ISL_958366, EPI_ISL_958367, EPI_ISL_958368, EPI_ISL_958369, EPI_ISL_958370, EPI_ISL_958371, EPI_ISL_958372, EPI_ISL_958373, EPI_ISL_958374, EPI_ISL_958375, EPI_ISL_958376, EPI_ISL_958377, EPI_ISL_958378, EPI_ISL_958379, EPI_ISL_958380, EPI_ISL_958381, EPI_ISL_958382, EPI_ISL_958383, EPI_ISL_958384, EPI_ISL_958385, EPI_ISL_958386, EPI_ISL_958387, EPI_ISL_958388, EPI_ISL_958389, EPI_ISL_958390, EPI_ISL_958391, EPI_ISL_958392, EPI_ISL_958393, EPI_ISL_958394, EPI_ISL_958395, EPI_ISL_958396, EPI_ISL_958397, EPI_ISL_958398, EPI_ISL_958399, EPI_ISL_958400, EPI_ISL_958401, EPI_ISL_958402, EPI_ISL_958403, EPI_ISL_958404, EPI_ISL_958405, EPI_ISL_958406, EPI_ISL_958407, EPI_ISL_958408, EPI_ISL_958409, EPI_ISL_958410, EPI_ISL_958411, EPI_ISL_958412, EPI_ISL_958413, EPI_ISL_958414, EPI_ISL_958415, EPI_ISL_958416, EPI_ISL_958417, EPI_ISL_958418, EPI_ISL_958419, EPI_ISL_958420, EPI_ISL_958421, EPI_ISL_958422, EPI_ISL_958423, EPI_ISL_958424, EPI_ISL_958425, EPI_ISL_958426, EPI_ISL_958427, EPI_ISL_958428, EPI_ISL_958429, EPI_ISL_958430, EPI_ISL_958431, EPI_ISL_958432, EPI_ISL_958433, EPI_ISL_958434, EPI_ISL_958435, EPI_ISL_958436, EPI_ISL_958437, EPI_ISL_958438, EPI_ISL_958439, EPI_ISL_958440, EPI_ISL_958441, EPI_ISL_958442, EPI_ISL_958443, EPI_ISL_958444, EPI_ISL_958445, EPI_ISL_958446, EPI_ISL_958447, EPI_ISL_958448, EPI_ISL_958449, EPI_ISL_958450, EPI_ISL_958451, EPI_ISL_958452, EPI_ISL_958453, EPI_ISL_958454, EPI_ISL_958455, EPI_ISL_958456, EPI_ISL_958457, EPI_ISL_958458, EPI_ISL_958459, EPI_ISL_958460, EPI_ISL_958461, EPI_ISL_958462, EPI_ISL_958463, EPI_ISL_958464, EPI_ISL_958465, EPI_ISL_958466, EPI_ISL_958467, EPI_ISL_958468, EPI_ISL_958469, EPI_ISL_958470, EPI_ISL_958471, EPI_ISL_958472, EPI_ISL_958473, EPI_ISL_958474, EPI_ISL_958475, EPI_ISL_958476, EPI_ISL_958477, EPI_ISL_958478, EPI_ISL_958479, EPI_ISL_958480, EPI_ISL_958481, EPI_ISL_958482, EPI_ISL_958483, EPI_ISL_958484, EPI_ISL_958485, EPI_ISL_958486, EPI_ISL_958487, EPI_ISL_958488, EPI_ISL_958489, EPI_ISL_958490, EPI_ISL_958491, EPI_ISL_958492, EPI_ISL_958493, EPI_ISL_958494, EPI_ISL_958495, EPI_ISL_958496, EPI_ISL_958497, EPI_ISL_958498, EPI_ISL_958499, EPI_ISL_958500, EPI_ISL_958501, EPI_ISL_958502, EPI_ISL_958503, EPI_ISL_958504, EPI_ISL_958505, EPI_ISL_958506, EPI_ISL_958507, EPI_ISL_958508, EPI_ISL_958509, EPI_ISL_958510, EPI_ISL_958511, EPI_ISL_958512, EPI_ISL_958513, EPI_ISL_958514, EPI_ISL_958515, EPI_ISL_958516, EPI_ISL_958517, EPI_ISL_958518, EPI_ISL_958519, EPI_ISL_958520, EPI_ISL_958521, EPI_ISL_958522, EPI_ISL_958523, EPI_ISL_958524, EPI_ISL_958525, EPI_ISL_958526, EPI_ISL_958527, EPI_ISL_958528, EPI_ISL_958529, EPI_ISL_958530, EPI_ISL_958531, EPI_ISL_958532, EPI_ISL_958533, EPI_ISL_958534, EPI_ISL_958535, EPI_ISL_958536, EPI_ISL_958537, EPI_ISL_958538, EPI_ISL_958539, EPI_ISL_958540, EPI_ISL_958541, EPI_ISL_958542, EPI_ISL_958543, EPI_ISL_958544, EPI_ISL_958545, EPI_ISL_958546, EPI_ISL_958547, EPI_ISL_958548, EPI_ISL_958549, EPI_ISL_958550, EPI_ISL_958551, EPI_ISL_958552, EPI_ISL_958553, EPI_ISL_958554, EPI_ISL_958555, EPI_ISL_958556, EPI_ISL_958557, EPI_ISL_958558, EPI_ISL_958559, EPI_ISL_958560, EPI_ISL_958561, EPI_ISL_958562, EPI_ISL_958563 | | | |
| see above | Lighthouse Lab in Milton Keynes | Wellcome Sanger Institute for the COVID-19 Genomics UK (COG-UK) Consortium | The Lighthouse Lab in Milton Keynes and Alex Alderton, Roberto Amato, Sonia Goncalves, Ewan Harrison, David K. Jackson, Ian Johnston, Dominic Kwiatkowski, Cordelia Langford, John Sillitoe on behalf of the Wellcome Sanger Institute COVID-19 Surveillance Team |
| EPI_ISL_958566 | Lighthouse Lab in Alderley Park | Wellcome Sanger Institute for the COVID-19 Genomics UK (COG-UK) Consortium | Jacquelyn Wynn, Mairead Hyland, The Lighthouse Lab in Alderley Park and Alex Alderton, Roberto Amato, Sonia Goncalves, Ewan Harrison, David K. Jackson, Ian Johnston, Dominic Kwiatkowski, Cordelia Langford, John Sillitoe on behalf of the Wellcome Sanger Institute COVID-19 Surveillance Team |
| EPI_ISL_958569, EPI_ISL_958572, EPI_ISL_958578, EPI_ISL_958579, EPI_ISL_958586, EPI_ISL_958587, EPI_ISL_958590, EPI_ISL_958591, EPI_ISL_958595, EPI_ISL_958600, EPI_ISL_958602, EPI_ISL_958610, EPI_ISL_958624, EPI_ISL_958632, EPI_ISL_958633, EPI_ISL_958634, EPI_ISL_958636, EPI_ISL_958638, EPI_ISL_958639, EPI_ISL_958641, EPI_ISL_958646, EPI_ISL_958650, EPI_ISL_958659, EPI_ISL_958662, EPI_ISL_958663, EPI_ISL_958670, EPI_ISL_958671, EPI_ISL_958673, EPI_ISL_958680, EPI_ISL_958684, EPI_ISL_958690, EPI_ISL_958691, EPI_ISL_958696, EPI_ISL_958705, EPI_ISL_958706, EPI_ISL_958713, EPI_ISL_958715, EPI_ISL_958716, EPI_ISL_958723, EPI_ISL_958725, EPI_ISL_958726, EPI_ISL_958731, EPI_ISL_958733, EPI_ISL_958735, EPI_ISL_958745, EPI_ISL_958747, EPI_ISL_958748, EPI_ISL_958754, EPI_ISL_958756, EPI_ISL_958760, EPI_ISL_958763, EPI_ISL_958767, EPI_ISL_958769, EPI_ISL_958775, EPI_ISL_958779, EPI_ISL_958784, EPI_ISL_958787, EPI_ISL_958789, EPI_ISL_958791, EPI_ISL_958792, EPI_ISL_958793, EPI_ISL_958803, EPI_ISL_958807, EPI_ISL_958808, EPI_ISL_958811, EPI_ISL_958815, EPI_ISL_958823, EPI_ISL_958826, EPI_ISL_958828, EPI_ISL_958833, EPI_ISL_958836, EPI_ISL_958840, EPI_ISL_958842, EPI_ISL_958849, EPI_ISL_958851, EPI_ISL_958863, EPI_ISL_958866, EPI_ISL_958873, EPI_ISL_958878, EPI_ISL_958884, EPI_ISL_958885, EPI_ISL_958887, EPI_ISL_958889, EPI_ISL_958891, EPI_ISL_958893, EPI_ISL_958895, EPI_ISL_958898, EPI_ISL_958902, EPI_ISL_958903, EPI_ISL_958904, EPI_ISL_958908, EPI_ISL_958911, EPI_ISL_958912, EPI_ISL_958915, EPI_ISL_958916 | | | |
| see above | Lighthouse Lab in Milton Keynes | Wellcome Sanger Institute for the COVID-19 Genomics UK (COG-UK) Consortium | The Lighthouse Lab in Milton Keynes and Alex Alderton, Roberto Amato, Sonia Goncalves, Ewan Harrison, David K. Jackson, Ian Johnston, Dominic Kwiatkowski, Cordelia Langford, John Sillitoe on behalf of the Wellcome Sanger Institute COVID-19 Surveillance Team |
| EPI_ISL_958917, EPI_ISL_958918, EPI_ISL_958919, EPI_ISL_958920, EPI_ISL_958921, EPI_ISL_958922, EPI_ISL_958923, EPI_ISL_958924, EPI_ISL_958925, EPI_ISL_958926, EPI_ISL_958927, EPI_ISL_958928, EPI_ISL_958929, EPI_ISL_958930, EPI_ISL_958931, EPI_ISL_958932, EPI_ISL_958933, EPI_ISL_958934, EPI_ISL_958935, EPI_ISL_958936, EPI_ISL_958937, EPI_ISL_958938, EPI_ISL_958939, EPI_ISL_958940, EPI_ISL_958941, EPI_ISL_958942, EPI_ISL_958943, EPI_ISL_958944, EPI_ISL_958945, EPI_ISL_958946, EPI_ISL_958947, EPI_ISL_958948, EPI_ISL_958949, EPI_ISL_958950, EPI_ISL_958951, EPI_ISL_958952, EPI_ISL_958953, EPI_ISL_958954, EPI_ISL_958955, EPI_ISL_958956, EPI_ISL_958957, EPI_ISL_958958, EPI_ISL_958959, EPI_ISL_958960, EPI_ISL_958961, EPI_ISL_958962, EPI_ISL_958963, EPI_ISL_958964, EPI_ISL_958965, EPI_ISL_958966, EPI_ISL_958967, EPI_ISL_958968, EPI_ISL_958969, EPI_ISL_958970, EPI_ISL_958971, EPI_ISL_958972, EPI_ISL_958973, EPI_ISL_958974, EPI_ISL_958975, EPI_ISL_958976, EPI_ISL_958977, EPI_ISL_958978, EPI_ISL_958979, EPI_ISL_958980, EPI_ISL_958981, EPI_ISL_958982, EPI_ISL_958983, EPI_ISL_958984, EPI_ISL_958985, EPI_ISL_958986, EPI_ISL_958987, EPI_ISL_958988, EPI_ISL_958989, EPI_ISL_958990, EPI_ISL_958991, EPI_ISL_958992, EPI_ISL_958993, EPI_ISL_958994, EPI_ISL_958995, EPI_ISL_958996, EPI_ISL_958997, EPI_ISL_958998, EPI_ISL_958999, EPI_ISL_959000, EPI_ISL_959001, EPI_ISL_959002, EPI_ISL_959003, EPI_ISL_959004, EPI_ISL_959005, EPI_ISL_959006, EPI_ISL_959007, EPI_ISL_959008, EPI_ISL_959009, EPI_ISL_959010, EPI_ISL_959011, EPI_ISL_959012, EPI_ISL_959013, EPI_ISL_959014, EPI_ISL_959015, EPI_ISL_959016, EPI_ISL_959017, EPI_ISL_959018, EPI_ISL_959019, EPI_ISL_959020, EPI_ISL_959021, EPI_ISL_959022, EPI_ISL_959023, EPI_ISL_959024, EPI_ISL_959025, EPI_ISL_959026, EPI_ISL_959027, EPI_ISL_959028, EPI_ISL_959029, EPI_ISL_959030, EPI_ISL_959031, EPI_ISL_959032, EPI_ISL_959033, EPI_ISL_959034, EPI_ISL_959035, EPI_ISL_959036, EPI_ISL_959037, EPI_ISL_959038, EPI_ISL_959039, EPI_ISL_959040, EPI_ISL_959041, EPI_ISL_959042, EPI_ISL_959043, EPI_ISL_959044, EPI_ISL_959045, EPI_ISL_959046, EPI_ISL_959047, EPI_ISL_959048, EPI_ISL_959049, EPI_ISL_959050, EPI_ISL_959051, EPI_ISL_959052, EPI_ISL_959053, EPI_ISL_959054, EPI_ISL_959055, EPI_ISL_959056, EPI_ISL_959057, EPI_ISL_959058, EPI_ISL_959059, EPI_ISL_959060, EPI_ISL_959061, EPI_ISL_959062, EPI_ISL_959063, EPI_ISL_959064, EPI_ISL_959065, EPI_ISL_959066, EPI_ISL_959067, EPI_ISL_959068, EPI_ISL_959069, EPI_ISL_959070, EPI_ISL_959071, EPI_ISL_959072, EPI_ISL_959073, EPI_ISL_959074, EPI_ISL_959075, EPI_ISL_959076, EPI_ISL_959077, EPI_ISL_959078, EPI_ISL_959079, EPI_ISL_959080, EPI_ISL_959081, EPI_ISL_959082, EPI_ISL_959083, EPI_ISL_959084, EPI_ISL_959085, EPI_ISL_959086, EPI_ISL_959087, EPI_ISL_959088, EPI_ISL_959089, EPI_ISL_959090, EPI_ISL_959091, EPI_ISL_959092, EPI_ISL_959093, EPI_ISL_959094, EPI_ISL_959095, EPI_ISL_959096, EPI_ISL_959097, EPI_ISL_959098, EPI_ISL_959099, EPI_ISL_959100, EPI_ISL_959101, EPI_ISL_959102, EPI_ISL_959103, EPI_ISL_959104, EPI_ISL_959105, EPI_ISL_959106, EPI_ISL_959107, EPI_ISL_959108, EPI_ISL_959109, EPI_ISL_959110, EPI_ISL_959111, EPI_ISL_959112, EPI_ISL_959113, EPI_ISL_959114, EPI_ISL_959115, EPI_ISL_959116, EPI_ISL_959117, EPI_ISL_959118, EPI_ISL_959119, EPI_ISL_959120, EPI_ISL_959121, EPI_ISL_959122, EPI_ISL_959123, EPI_ISL_959124, EPI_ISL_959125, EPI_ISL_959126, EPI_ISL_959127, EPI_ISL_959128, EPI_ISL_959129, EPI_ISL_959130, EPI_ISL_959131, EPI_ISL_959132, EPI_ISL_959133, EPI_ISL_959134, EPI_ISL_959135, EPI_ISL_959136, EPI_ISL_959137, EPI_ISL_959138, EPI_ISL_959139, EPI_ISL_959140, EPI_ISL_959141, EPI_ISL_959142, EPI_ISL_959143, EPI_ISL_959144, EPI_ISL_959145, EPI_ISL_959146, EPI_ISL_959147, EPI_ISL_959148, EPI_ISL_959149, EPI_ISL_959150, EPI_ISL_959151, EPI_ISL_959152, EPI_ISL_959153, EPI_ISL_959154, EPI_ISL_959155, EPI_ISL_959156, EPI_ISL_959157, EPI_ISL_959158, EPI_ISL_959159, EPI_ISL_959160, EPI_ISL_959161, EPI_ISL_959162, EPI_ISL_959163, EPI_ISL_959164, EPI_ISL_959165, EPI_ISL_959166, EPI_ISL_959167, EPI_ISL_959168, EPI_ISL_959169, EPI_ISL_959170, EPI_ISL_959171, EPI_ISL_959172, EPI_ISL_959173, EPI_ISL_959174, EPI_ISL_959175, EPI_ISL_959176, EPI_ISL_959177, EPI_ISL_959178, EPI_ISL_959179, EPI_ISL_959180, EPI_ISL_959181, EPI_ISL_959182, EPI_ISL_959183, EPI_ISL_959184, EPI_ISL_959185, EPI_ISL_959186, EPI_ISL_959187, EPI_ISL_959188, EPI_ISL_959189, EPI_ISL_959190, EPI_ISL_959191, EPI_ISL_959192, EPI_ISL_959193, EPI_ISL_959194, EPI_ISL_959195, EPI_ISL_959196, EPI_ISL_959197, EPI_ISL_959198, EPI_ISL_959199, EPI_ISL_959200, EPI_ISL_959201, EPI_ISL_959202, EPI_ISL_959203, EPI_ISL_959204, EPI_ISL_959205, EPI_ISL_959206, EPI_ISL_959207, EPI_ISL_959208, EPI_ISL_959209, EPI_ISL_959210, EPI_ISL_959211, EPI_ISL_959212, EPI_ISL_959213, EPI_ISL_959214, EPI_ISL_959215, EPI_ISL_959216, EPI_ISL_959217, EPI_ISL_959218, EPI_ISL_959219, EPI_ISL_959220, EPI_ISL_959221, EPI_ISL_959222, EPI_ISL_959223, EPI_ISL_959224, EPI_ISL_959225, EPI_ISL_959226, EPI_ISL_959227, EPI_ISL_959228, EPI_ISL_959229, EPI_ISL_959230, EPI_ISL_959231, EPI_ISL_959232, EPI_ISL_959233, EPI_ISL_959234, EPI_ISL_959235, EPI_ISL_959236, EPI_ISL_959237, EPI_ISL_959238, EPI_ISL_959239, EPI_ISL_959240, EPI_ISL_959241, EPI_ISL_959242, EPI_ISL_959243, EPI_ISL_959244, EPI_ISL_959245, EPI_ISL_959246, EPI_ISL_959247, EPI_ISL_959248, EPI_ISL_959249, EPI_ISL_959250, EPI_ISL_959251, EPI_ISL_959252, EPI_ISL_959253, EPI_ISL_959254, EPI_ISL_959255, EPI_ISL_959256, EPI_ISL_959257, EPI_ISL_959258, EPI_ISL_959259, EPI_ISL_959260, EPI_ISL_959261, EPI_ISL_959262, EPI_ISL_959263, EPI_ISL_959264, EPI_ISL_959265, EPI_ISL_959266 | | | |
| see above | Lighthouse Lab in Alderley Park | Wellcome Sanger Institute for the COVID-19 Genomics UK (COG-UK) Consortium | Jacquelyn Wynn, Mairead Hyland, The Lighthouse Lab in Alderley Park and Alex Alderton, Roberto Amato, Sonia Goncalves, Ewan Harrison, David K. Jackson, Ian Johnston, Dominic Kwiatkowski, Cordelia Langford, John Sillitoe on behalf of the Wellcome Sanger Institute COVID-19 Surveillance Team |
| EPI_ISL_959267 | Lighthouse Lab in Cambridge | Wellcome Sanger Institute for the COVID-19 Genomics UK (COG-UK) Consortium | Rob Howes, The Lighthouse Lab in Cambridge and Alex Alderton, Roberto Amato, Sonia Goncalves, Ewan Harrison, David K. Jackson, Ian Johnston, Dominic Kwiatkowski, Cordelia Langford, John Sillitoe on behalf of the Wellcome Sanger Institute COVID-19 Surveillance Team |
| EPI_ISL_959268, EPI_ISL_959269, EPI_ISL_959270 | Lighthouse Lab in Alderley Park | Wellcome Sanger Institute for the COVID-19 Genomics UK (COG-UK) Consortium | Jacquelyn Wynn, Mairead Hyland, The Lighthouse Lab in Alderley Park and Alex Alderton, Roberto Amato, Sonia Goncalves, Ewan Harrison, David K. Jackson, Ian Johnston, Dominic Kwiatkowski, Cordelia Langford, John Sillitoe on behalf of the Wellcome Sanger Institute COVID-19 Surveillance Team |
| EPI_ISL_959271, EPI_ISL_959272 | Lighthouse Lab in Milton Keynes | Wellcome Sanger Institute for the COVID-19 Genomics UK (COG-UK) Consortium | The Lighthouse Lab in Milton Keynes and Alex Alderton, Roberto Amato, Sonia Goncalves, Ewan Harrison, David K. Jackson, Ian Johnston, Dominic Kwiatkowski, Cordelia Langford, John Sillitoe on behalf of the Wellcome Sanger Institute COVID-19 Surveillance Team |
| EPI_ISL_959367 | National Institute of Laboratory Medicine and Referral Center | Genomic Research Lab, BCSIR | Md. Murshed Hasan Sarkar, Mohammad Samir Uzzaman, Eshrar Osman, Md. Ashahan Habib, Shahina Akter, Tanjina Akhtar Banu, Abu Sayeed Mohammad Mahmud, Barna Goswami, Iffat Jahan, Md. Saddam Hossain, Tasnim Nafisa, Md. Maruf Ahmed Molla, Mahmuda Yeasmin, Asish Kumar Ghosh, Arifa Akram, A. K. M.Shamsuzzaman, Md. Salim Khan |
| EPI_ISL_959394, EPI_ISL_959395, EPI_ISL_959396, EPI_ISL_959397, EPI_ISL_959398, EPI_ISL_959399, EPI_ISL_959400, EPI_ISL_959401, EPI_ISL_959402, EPI_ISL_959413 | genXone SA, Molecular Diagnostics Laboratory / NZOZ | genXone SA, Research & Development Laboratory | Maciej Sykulski, Grzegorz Nowicki, Jakub Grabowski, Natalia Drwska-Matelska, Anna Brylak-Baszków, Aleksandra Gidlewicz, Karol Szeszko, ukasz Krych, Micha Kaszuba |
| EPI_ISL_959418, EPI_ISL_959422, EPI_ISL_959435, EPI_ISL_959436, EPI_ISL_959437, EPI_ISL_959443, EPI_ISL_959445, EPI_ISL_959450, EPI_ISL_959451, EPI_ISL_959452, EPI_ISL_959453, EPI_ISL_959454 | | | |
| see above | Servicio de Microbiología, Hospital Universitario Son Espases | SeqCOVID-SPAIN consortium/IBV(CSIC) | Carla López-Causapé, Jordi Reina, Antonio Oliver and SeqCOVID-SPAIN consortium |
| EPI_ISL_959543 | University of Michigan Clinical Microbiology Laboratory | Lauring Lab, University of Michigan, Department of Microbiology and Immunology | Valesano |
| EPI_ISL_959598 | Vivalia - Clinique Saint-Joseph | GIGA Medical Genomics | Keith Durkin, Maria Artesi, Sébastien Bontems, Raphaël Boreux, Bouchra Boujemla, Cécile Meex, Pierrette Melin, Marie-Pierre Hayette, Vincent Bours |

| | | | |
|---|---|---|---|
| EPI_ISL_959599 | Synlab | GIGA Medical Genomics | Keith Durkin, Maria Artesi, Sébastien Bontems, Raphaël Boreux, Bouchra Boujemla, Cécile Meex, Pierrette Melin, Marie-Pierre Hayette, Vincent Bours |
| EPI_ISL_959600, EPI_ISL_959601, EPI_ISL_959602 | CHR de la Citadelle | GIGA Medical Genomics | Keith Durkin, Maria Artesi, Sébastien Bontems, Raphaël Boreux, Bouchra Boujemla, Cécile Meex, Pierrette Melin, Marie-Pierre Hayette, Vincent Bours |
| EPI_ISL_959603 | Synlab | GIGA Medical Genomics | Keith Durkin, Maria Artesi, Sébastien Bontems, Raphaël Boreux, Bouchra Boujemla, Cécile Meex, Pierrette Melin, Marie-Pierre Hayette, Vincent Bours |
| EPI_ISL_959621 | Institute of Virology, Biomedical Research Center of the Slovak Academy of Sciences, Bratislava | Faculty of Natural Sciences, Comenius University, Bratislava | Kristína Boršová, Viktória abanová, Broa Brejová, Viktória Hodorová, Sabina Fumaová Havlíková, Juraj Kopáek, Martina Liková, ubomíra Lukáiková, Martina Neboháová, Monika Sláviková, Tomáš Vina, Boris Klempa, Jozef Nosek |
| EPI_ISL_959632 | Institute of Virology, Biomedical Research Center of the Slovak Academy of Sciences, Bratislava | Faculty of Natural Sciences, Comenius University, Bratislava | Broa Brejová, Viktória abanová, Kristína Boršová, Viktória Hodorová, Sabina Fumaová Havlíková, Juraj Kopáek, Martina Liková, ubomíra Lukáiková, Martina Neboháová, Monika Sláviková, Tomáš Vina, Jozef Nosek, Boris Klempa |
| EPI_ISL_959633 | Institute of Virology, Biomedical Research Center of the Slovak Academy of Sciences, Bratislava | Faculty of Natural Sciences, Comenius University, Bratislava | Viktória abanová, Kristína Boršová, Broa Brejová, Viktória Hodorová, Sabina Fumaová Havlíková, Juraj Kopáek, Martina Liková, ubomíra Lukáiková, Martina Neboháová, Monika Sláviková, Tomáš Vina, Jozef Nosek, Boris Klempa |
| EPI_ISL_959634 | Institute of Virology, Biomedical Research Center of the Slovak Academy of Sciences, Bratislava | Faculty of Natural Sciences, Comenius University, Bratislava | Kristína Boršová, Viktória abanová, Broa Brejová, Viktória Hodorová, Sabina Fumaová Havlíková, Juraj Kopáek, Martina Liková, ubomíra Lukáiková, Martina Neboháová, Monika Sláviková, Tomáš Vina, Boris Klempa, Jozef Nosek |
| EPI_ISL_959635 | Institute of Virology, Biomedical Research Center of the Slovak Academy of Sciences, Bratislava | Faculty of Natural Sciences, Comenius University, Bratislava | Broa Brejová, Viktória abanová, Kristína Boršová, Viktória Hodorová, Sabina Fumaová Havlíková, Juraj Kopáek, Martina Liková, ubomíra Lukáiková, Martina Neboháová, Monika Sláviková, Tomáš Vina, Jozef Nosek, Boris Klempa |
| EPI_ISL_959636 | Institute of Virology, Biomedical Research Center of the Slovak Academy of Sciences, Bratislava | Faculty of Natural Sciences, Comenius University, Bratislava | Viktória abanová, Kristína Boršová, Broa Brejová, Viktória Hodorová, Sabina Fumaová Havlíková, Juraj Kopáek, Martina Liková, ubomíra Lukáiková, Martina Neboháová, Monika Sláviková, Tomáš Vina, Jozef Nosek, Boris Klempa |
| EPI_ISL_959713, EPI_ISL_959714, EPI_ISL_959715, EPI_ISL_959716, EPI_ISL_959717, EPI_ISL_959718, EPI_ISL_959719, EPI_ISL_959720, EPI_ISL_959721, EPI_ISL_959722, EPI_ISL_959723, EPI_ISL_959724, EPI_ISL_959725, EPI_ISL_959726, EPI_ISL_959727, EPI_ISL_959728, EPI_ISL_959729, EPI_ISL_959730, EPI_ISL_959731, EPI_ISL_959732, EPI_ISL_959733, EPI_ISL_959734, EPI_ISL_959735, EPI_ISL_959736, EPI_ISL_959737, EPI_ISL_959738, EPI_ISL_959739, EPI_ISL_959740, EPI_ISL_959741, EPI_ISL_959742, EPI_ISL_959743, EPI_ISL_959744, EPI_ISL_959745, EPI_ISL_959746, EPI_ISL_959747 | | | |
| see above | National Virus Reference Laboratory | National Virus Reference Laboratory | Michael Carr, Gabriel Gonzalez, Jonathan Dean, Cillian F De Gascun |
| EPI_ISL_960319, EPI_ISL_960323, EPI_ISL_960326, EPI_ISL_960328, EPI_ISL_960329, EPI_ISL_960331, EPI_ISL_960332, EPI_ISL_960337, EPI_ISL_960338, EPI_ISL_960345, EPI_ISL_960346, EPI_ISL_960350, EPI_ISL_960351, EPI_ISL_960371, EPI_ISL_960372, EPI_ISL_960373, EPI_ISL_960374, EPI_ISL_960377, EPI_ISL_960379, EPI_ISL_960384, EPI_ISL_960387, EPI_ISL_960400, EPI_ISL_960401, EPI_ISL_960402 | | | |
| see above | University of Wisconsin-Madison AIDS Vaccine Research Laboratories | University of Wisconsin-Madison AIDS Vaccine Research Laboratories | Gage Moreno, Katarina Braun, et al. AIDS Vaccine Research Laboratories |
| EPI_ISL_960838, EPI_ISL_960839, EPI_ISL_960840, EPI_ISL_960841 | Medizinisches Labor Prof. Dr. Schenk/ Dr. Ansorge & Kollegen, Magdeburg | Institute of Medical Microbiology and Hospital Hygiene | Prof. Dr. Achim Kaasch, Aljoscha Tersteegen |
| EPI_ISL_960842 | Institute of Medical Microbiology and Hospital Hygiene | Institute of Medical Microbiology and Hospital Hygiene | Prof. Dr. Achim Kaasch, Aljoscha Tersteegen |
| EPI_ISL_960844 | Medizinisches Labor Prof. Dr. Schenk/ Dr. Ansorge & Kollegen, Magdeburg | Institute of Medical Microbiology and Hospital Hygiene | Prof. Dr. Achim Kaasch, Aljoscha Tersteegen |
| EPI_ISL_960845, EPI_ISL_960847, EPI_ISL_960850, EPI_ISL_960858, EPI_ISL_960861, EPI_ISL_960864, EPI_ISL_960874, EPI_ISL_960875 | Institute of Medical Microbiology and Hospital Hygiene | Institute of Medical Microbiology and Hospital Hygiene | Prof. Dr. Achim Kaasch, Aljoscha Tersteegen |
| EPI_ISL_960883, EPI_ISL_960886, EPI_ISL_960904, EPI_ISL_960905, EPI_ISL_960906, EPI_ISL_960907, EPI_ISL_960908, EPI_ISL_960909, EPI_ISL_960910, EPI_ISL_960911, EPI_ISL_960912, EPI_ISL_960913, EPI_ISL_960914, EPI_ISL_960915, EPI_ISL_960916, EPI_ISL_960917, EPI_ISL_960918, EPI_ISL_960919, EPI_ISL_960920, EPI_ISL_960921, EPI_ISL_960922, EPI_ISL_960923, EPI_ISL_960924, EPI_ISL_960925, EPI_ISL_960926, EPI_ISL_960927, EPI_ISL_960928, EPI_ISL_960929, EPI_ISL_960930, EPI_ISL_960931, EPI_ISL_960932, EPI_ISL_960933, EPI_ISL_960934, EPI_ISL_960943, EPI_ISL_960944, EPI_ISL_960945, EPI_ISL_960946, EPI_ISL_960947, EPI_ISL_960948, EPI_ISL_960949, EPI_ISL_960950, EPI_ISL_960954, EPI_ISL_960955, EPI_ISL_960956, EPI_ISL_960958, EPI_ISL_960959, EPI_ISL_960960, EPI_ISL_960961, EPI_ISL_960962, EPI_ISL_960963, EPI_ISL_960964, EPI_ISL_960965, EPI_ISL_960966, EPI_ISL_960967, EPI_ISL_960968, EPI_ISL_960969, EPI_ISL_960970, EPI_ISL_960971, EPI_ISL_960972, EPI_ISL_960973, EPI_ISL_960974, EPI_ISL_960975, EPI_ISL_960976, EPI_ISL_960977, EPI_ISL_960978, EPI_ISL_960979, EPI_ISL_960980, EPI_ISL_960981 | | | |
| see above | Medizinisches Labor Prof. Dr. Schenk/ Dr. Ansorge & Kollegen, Magdeburg | Institute of Medical Microbiology and Hospital Hygiene | Prof. Dr. Achim Kaasch, Aljoscha Tersteegen |
| EPI_ISL_961204, EPI_ISL_961205, EPI_ISL_961206, EPI_ISL_961208, EPI_ISL_961209, EPI_ISL_961210, EPI_ISL_961211, EPI_ISL_961212, EPI_ISL_961213, EPI_ISL_961214, EPI_ISL_961216, EPI_ISL_961218, EPI_ISL_961226, EPI_ISL_961232, EPI_ISL_961233, EPI_ISL_961235, EPI_ISL_961236, EPI_ISL_961237, EPI_ISL_961239, EPI_ISL_961240, EPI_ISL_961241, EPI_ISL_961242, EPI_ISL_961243, EPI_ISL_961244, EPI_ISL_961245, EPI_ISL_961246, EPI_ISL_961247, EPI_ISL_961248, EPI_ISL_961249, EPI_ISL_961250, EPI_ISL_961251, EPI_ISL_961252, EPI_ISL_961254, EPI_ISL_961255, EPI_ISL_961256, EPI_ISL_961257, EPI_ISL_961258, EPI_ISL_961259, EPI_ISL_961260, EPI_ISL_961261, EPI_ISL_961262, EPI_ISL_961263, EPI_ISL_961264, EPI_ISL_961265, EPI_ISL_961266, EPI_ISL_961267, EPI_ISL_961268, EPI_ISL_961269, EPI_ISL_961272, EPI_ISL_961273, EPI_ISL_961274, EPI_ISL_961275, EPI_ISL_961276, EPI_ISL_961277, EPI_ISL_961278, EPI_ISL_961279, EPI_ISL_961280, EPI_ISL_961281, EPI_ISL_961282, EPI_ISL_961283, EPI_ISL_961284, EPI_ISL_961285 | | | |
| see above | Hospital General Universitario de Alicante - Instituto de Investigación Sanitaria y Biomédica de Alicante | SeqCOVID-SPAIN consortium/IBV(CSIC) | Maripaz Ventero Martín, Carmen Molina Pardines and SeqCOVID-SPAIN consortium |
| EPI_ISL_961355 | Centrum voor Medische Analyse | UAntwerp, Laboratory of Medical Microbiology | Basil Britto Xavier, Jasmine Coppens, Marie Le Mercier, Christine Lammens, Veerle Matheeussen, Herman Goossens |
| EPI_ISL_961359, EPI_ISL_961362 | Toronto Invasive Bacterial Diseases Network | McMaster University | Allison McGeer, Patryk Aftanas, Hooman Derakhshani, Angel Li, Kuganya Nirmalarajah, Emily Panousis, Ahmed Draia, Jalees Nasir, Michael Surette, Samira Mubareka, Andrew G. McArthur |
| EPI_ISL_961508, EPI_ISL_961522, EPI_ISL_961528 | Michigan Department of Health and Human Services, Bureau of Laboratories | Michigan Department of Health and Human Services, Bureau of Laboratories | Blankenship HM, Riner D, Soehnlen MK |
| EPI_ISL_961752, EPI_ISL_961753 | SIESP SULMONA | Istituto Zooprofilattico Sperimentale dell'Abruzzo e Molise "G. Caporale" | Lorusso A, Marcacci M, Di Domenico M, Ancora M, Curini V, Mangone I, Rinaldi A, Scialabba S, Di Pasquale A, Cammà C, Puglia I, Calistri P, Savini G |
| EPI_ISL_962189, EPI_ISL_962197, EPI_ISL_962198, EPI_ISL_962199, EPI_ISL_962200, EPI_ISL_962201, EPI_ISL_962202 | Toronto Invasive Bacterial Diseases Network | McMaster University | Allison McGeer, Patryk Aftanas, Hooman Derakhshani, Angel Li, Kuganya Nirmalarajah, Emily Panousis, Ahmed Draia, Jalees Nasir, Michael Surette, Samira Mubareka, Andrew G. McArthur |
| EPI_ISL_962303, EPI_ISL_962304 | Seattle Flu Study | Seattle Flu Study | Deborah A. Nickerson, Chris D. Frazar, Jover Lee, Benjamin Pelle, Erica Ryke, Matthew Richardson, Amanda Adler, Elisabeth Brandstetter, Peter D. Han, Kairsten Fay, Misja Ilcisin, Kirsten Lacombe, Thomas R. Sibley, Melissa Truong, Caitlin R. Wolf, Michael Boeckh, Janet A. Englund, Michael Famulare, Barry R. Lutz, Mark J. Rieder, Lea M. Starita, Matthew Thompson, Jay Shendure, Trevor Bedford, Helen Y. Chu |
| EPI_ISL_962372, EPI_ISL_962373, EPI_ISL_962374 | Seattle Flu Study | Seattle Flu Study | Deborah A. Nickerson, Chris D. Frazar, Jover Lee, Benjamin Pelle, Erica Ryke, Matthew Richardson, Amanda Adler, Elisabeth Brandstetter, Peter D. Han, Kairsten Fay, Misja Ilcisin, Kirsten Lacombe, Thomas R. Sibley, Melissa Truong, Caitlin R. Wolf, Karen Cowgill, Stephanie Schrag, Jeff Duchin, Michael Boeckh, Janet A. Englund, Michael Famulare, Barry R. Lutz, Mark J. Rieder, Lea M. Starita, Matthew Thompson, Helen Y. Chu, Trevor Bedford, Jay Shendure |
| EPI_ISL_962436, EPI_ISL_962437, EPI_ISL_962438, EPI_ISL_962439, EPI_ISL_962440, EPI_ISL_962441 | Washington State Department of Health | Seattle Flu Study | Deborah A. Nickerson, Chris D. Frazar, Jover Lee, Benjamin Pelle, Erica Ryke, Matthew Richardson, Amanda Adler, Elisabeth Brandstetter, Peter D. Han, Kairsten Fay, Misja Ilcisin, Kirsten Lacombe, Thomas R. Sibley, Melissa Truong, Caitlin R. Wolf, Romesh Gautom, Geoff Melly, Brian Hiatt, Philip Dykema, Scott Lindquist, Michael Boeckh, Janet A. Englund, Michael Famulare, Barry R. Lutz, Mark J. Rieder, Lea M. Starita, Matthew Thompson, Helen Y. Chu, Jay Shendure, Trevor Bedford |
| EPI_ISL_962443, EPI_ISL_962444, EPI_ISL_962465, EPI_ISL_962466, EPI_ISL_962468, EPI_ISL_962469, EPI_ISL_962470, EPI_ISL_962471, EPI_ISL_962472, EPI_ISL_962473 | Seattle Flu Study | Seattle Flu Study | Deborah A. Nickerson, Chris D. Frazar, Jover Lee, Benjamin Pelle, Erica Ryke, Matthew Richardson, Amanda Adler, Elisabeth Brandstetter, Peter D. Han, Kairsten Fay, Misja Ilcisin, Kirsten Lacombe, Thomas R. Sibley, Melissa Truong, Caitlin R. Wolf, Karen Cowgill, Stephanie Schrag, Jeff Duchin, Michael Boeckh, Janet A. Englund, Michael Famulare, Barry R. Lutz, Mark J. Rieder, Lea M. Starita, Matthew Thompson, Helen Y. Chu, Trevor Bedford, Jay Shendure |
| EPI_ISL_962517, EPI_ISL_962518, EPI_ISL_962519, EPI_ISL_962520 | VA Medical Center | Los Angeles County PHL | P. Hemarajata et al. |

| | | | |
|---|---|---|---|
| EPI_ISL_962523, EPI_ISL_962524 | Institute for Medical Research, Infectious Disease Research Centre, National Institutes of Health, Ministry of Health Malaysia | Institute for Medical Research, Infectious Disease Research Centre, National Institutes of Health, Ministry of Health Malaysia | Suppiah J, Kamel K, Azizan MA, Thayan R |
| EPI_ISL_962833, EPI_ISL_962834, EPI_ISL_962835, EPI_ISL_962836, EPI_ISL_962841, EPI_ISL_962842, EPI_ISL_962843, EPI_ISL_962844, EPI_ISL_962845 | UCLA Clinical Micro Lab | Los Angeles County PHL | P. Hemarajata et al. |
| EPI_ISL_962852, EPI_ISL_962853, EPI_ISL_962854, EPI_ISL_962855, EPI_ISL_962856, EPI_ISL_962857, EPI_ISL_962858, EPI_ISL_962859, EPI_ISL_962860, EPI_ISL_962861, EPI_ISL_962862, EPI_ISL_962863, EPI_ISL_962864, EPI_ISL_962865, EPI_ISL_962866, EPI_ISL_962867 | | | |
| see above | Los Angeles County PHL | Los Angeles County PHL | P. Hemarajata et al. |
| EPI_ISL_962879 | UCLA Clinical Micro Lab | Los Angeles County PHL | P. Hemarajata et al. |
| EPI_ISL_963325, EPI_ISL_963408, EPI_ISL_963471, EPI_ISL_963524, EPI_ISL_963555 | Lighthouse Lab in Milton Keynes | Wellcome Sanger Institute for the COVID-19 Genomics UK (COG-UK) Consortium | The Lighthouse Lab in Milton Keynes and Alex Alderton, Roberto Amato, Sonia Goncalves, Ewan Harrison, David K. Jackson, Ian Johnston, Dominic Kwiatkowski, Cordelia Langford, John Sillitoe on behalf of the Wellcome Sanger Institute COVID-19 Surveillance Team |
| EPI_ISL_963651, EPI_ISL_963654, EPI_ISL_963658, EPI_ISL_963659, EPI_ISL_963665, EPI_ISL_963670, EPI_ISL_963671, EPI_ISL_963672, EPI_ISL_963673, EPI_ISL_963676, EPI_ISL_963677, EPI_ISL_963679, EPI_ISL_963681, EPI_ISL_963685, EPI_ISL_963686, EPI_ISL_963687, EPI_ISL_963689, EPI_ISL_963690, EPI_ISL_963694, EPI_ISL_963695, EPI_ISL_963700, EPI_ISL_963701, EPI_ISL_963703, EPI_ISL_963704, EPI_ISL_963706, EPI_ISL_963707, EPI_ISL_963709, EPI_ISL_963718, EPI_ISL_963720, EPI_ISL_963721, EPI_ISL_963724, EPI_ISL_963726, EPI_ISL_963727, EPI_ISL_963731, EPI_ISL_963732, EPI_ISL_963733, EPI_ISL_963734, EPI_ISL_963735, EPI_ISL_963736, EPI_ISL_963737, EPI_ISL_963738, EPI_ISL_963740, EPI_ISL_963741, EPI_ISL_963747, EPI_ISL_963750, EPI_ISL_963753, EPI_ISL_963754, EPI_ISL_963758, EPI_ISL_963759, EPI_ISL_963761, EPI_ISL_963764, EPI_ISL_963765, EPI_ISL_963766, EPI_ISL_963768, EPI_ISL_963769, EPI_ISL_963772, EPI_ISL_963773, EPI_ISL_963774, EPI_ISL_963775, EPI_ISL_963779, EPI_ISL_963780, EPI_ISL_963781, EPI_ISL_963783, EPI_ISL_963786, EPI_ISL_963789, EPI_ISL_963790, EPI_ISL_963794, EPI_ISL_963798, EPI_ISL_963800, EPI_ISL_963803, EPI_ISL_963805, EPI_ISL_963806, EPI_ISL_963807, EPI_ISL_963808, EPI_ISL_963810, EPI_ISL_963813, EPI_ISL_963814, EPI_ISL_963815, EPI_ISL_963816, EPI_ISL_963819, EPI_ISL_963821, EPI_ISL_963822, EPI_ISL_963823, EPI_ISL_963824, EPI_ISL_963825, EPI_ISL_963826, EPI_ISL_963831, EPI_ISL_963832, EPI_ISL_963833, EPI_ISL_963835, EPI_ISL_963838, EPI_ISL_963839, EPI_ISL_963841, EPI_ISL_963844, EPI_ISL_963845, EPI_ISL_963846, EPI_ISL_963849, EPI_ISL_963850, EPI_ISL_963851, EPI_ISL_963852, EPI_ISL_963853, EPI_ISL_963855, EPI_ISL_963857, EPI_ISL_963859, EPI_ISL_963861, EPI_ISL_963863, EPI_ISL_963864, EPI_ISL_963865, EPI_ISL_963866, EPI_ISL_963867, EPI_ISL_963868, EPI_ISL_963869, EPI_ISL_963870, EPI_ISL_963872, EPI_ISL_963873, EPI_ISL_963874, EPI_ISL_963875, EPI_ISL_963877, EPI_ISL_963878, EPI_ISL_963879, EPI_ISL_963880, EPI_ISL_963881, EPI_ISL_963882, EPI_ISL_963883, EPI_ISL_963884, EPI_ISL_963889, EPI_ISL_963890, EPI_ISL_963891, EPI_ISL_963893, EPI_ISL_963895, EPI_ISL_963896, EPI_ISL_963900, EPI_ISL_963901, EPI_ISL_963902, EPI_ISL_963903, EPI_ISL_963904, EPI_ISL_963906, EPI_ISL_963908, EPI_ISL_963909, EPI_ISL_963911, EPI_ISL_963914, EPI_ISL_963915, EPI_ISL_963916, EPI_ISL_963921, EPI_ISL_963922, EPI_ISL_963923, EPI_ISL_963925, EPI_ISL_963927, EPI_ISL_963932, EPI_ISL_963934, EPI_ISL_963936, EPI_ISL_963938, EPI_ISL_963941, EPI_ISL_963942, EPI_ISL_963945, EPI_ISL_963947, EPI_ISL_963949, EPI_ISL_963952, EPI_ISL_963953, EPI_ISL_963958, EPI_ISL_963960, EPI_ISL_963962, EPI_ISL_963964, EPI_ISL_963968, EPI_ISL_963970, EPI_ISL_963971, EPI_ISL_963973, EPI_ISL_963976, EPI_ISL_963977, EPI_ISL_963978, EPI_ISL_963979, EPI_ISL_963981, EPI_ISL_963984, EPI_ISL_964067, EPI_ISL_964087, EPI_ISL_964236, EPI_ISL_964710, EPI_ISL_964726, EPI_ISL_964749, EPI_ISL_964759, EPI_ISL_964817, EPI_ISL_964877 | | | |
| see above | Lighthouse Lab in Alderley Park | Wellcome Sanger Institute for the COVID-19 Genomics UK (COG-UK) Consortium | Jacquelyn Wynn, Mairead Hyland, The Lighthouse Lab in Alderley Park and Alex Alderton, Roberto Amato, Sonia Goncalves, Ewan Harrison, David K. Jackson, Ian Johnston, Dominic Kwiatkowski, Cordelia Langford, John Sillitoe on behalf of the Wellcome Sanger Institute COVID-19 Surveillance Team |
| EPI_ISL_964917 | National Institute of Laboratory Medicine and Referral Center | Genomic Research Lab, BCSIR | Md. Maruf Ahmed Molla, Mohammad Samir Uzzaman, Eshrar Osman, Md. Ahashan Habib, Shahina Akter,Tanjina Akhtar Banu, Abu Sayeed Mohammad Mahmud, Md. Murshed Hasan Sarkar, Barna Goswami, Iffat Jahan, Md. Saddam Hossain, Tasnim Nafisa, Mahmuda Yeasmin, Asish Kumar Ghosh, Arifa Akram, A. K. M. Shamsuzzaman, Md. Salim Khan |
| EPI_ISL_964956, EPI_ISL_964957 | Nordland Hospital - Bodo, Laboratory Department, Molecular Biology Unit | Norwegian Institute of Public Health, Department of Virology | Kathrine Stene-Johansen, Kamilla Heddeland Instefjord, Hilde Elshaug, Ignacio Garcia Llorente, Serina B Engebretsen, Atiya R Ali,Marie Paulsen Madsen, Rasmus Riis Kopperud, Hilde Vollan, Karoline Bragstad, Olav Hungnes |
| EPI_ISL_964962 | Norwegian Institute of Public Health, Department of Virology | Norwegian Institute of Public Health, Department of Virology | Kathrine Stene-Johansen, Kamilla Heddeland Instefjord, Hilde Elshaug, Ignacio Garcia Llorente, Serina B Engebretsen, Atiya R Ali,Marie Paulsen Madsen, Rasmus Riis Kopperud, Hilde Vollan, Karoline Bragstad, Olav Hungnes |
| EPI_ISL_964965 | Akershus University Hospital, Department for Microbiology and Infectious Disease Control | Norwegian Institute of Public Health, Department of Virology | Kathrine Stene-Johansen, Kamilla Heddeland Instefjord, Hilde Elshaug, Ignacio Garcia Llorente, Serina B Engebretsen, Atiya R Ali,Marie Paulsen Madsen, Rasmus Riis Kopperud, Hilde Vollan, Karoline Bragstad, Olav Hungnes |
| EPI_ISL_964972 | Ostfold Hospital Trust - Kalnes, Centre for Laboratory Medicine, Section for gene technology and infection serology | Norwegian Institute of Public Health, Department of Virology | Kathrine Stene-Johansen, Kamilla Heddeland Instefjord, Hilde Elshaug, Ignacio Garcia Llorente, Serina B Engebretsen, Atiya R Ali,Marie Paulsen Madsen, Rasmus Riis Kopperud, Hilde Vollan, Karoline Bragstad, Olav Hungnes |
| EPI_ISL_964975 | Akershus University Hospital, Department for Microbiology and Infectious Disease Control | Norwegian Institute of Public Health, Department of Virology | Kathrine Stene-Johansen, Kamilla Heddeland Instefjord, Hilde Elshaug, Ignacio Garcia Llorente, Serina B Engebretsen, Atiya R Ali,Marie Paulsen Madsen, Rasmus Riis Kopperud, Hilde Vollan, Karoline Bragstad, Olav Hungnes |
| EPI_ISL_964977, EPI_ISL_964978 | Vestfold Hospital, Toensberg Department of Microbiology | Norwegian Institute of Public Health, Department of Virology | Kathrine Stene-Johansen, Kamilla Heddeland Instefjord, Hilde Elshaug, Ignacio Garcia Llorente, Serina B Engebretsen, Atiya R Ali,Marie Paulsen Madsen, Rasmus Riis Kopperud, Hilde Vollan, Karoline Bragstad, Olav Hungnes |
| EPI_ISL_964979 | Furst Medical Laboratory | Norwegian Institute of Public Health, Department of Virology | Kathrine Stene-Johansen, Kamilla Heddeland Instefjord, Hilde Elshaug, Ignacio Garcia Llorente, Serina B Engebretsen, Atiya R Ali,Marie Paulsen Madsen, Rasmus Riis Kopperud, Hilde Vollan, Karoline Bragstad, Olav Hungnes |
| EPI_ISL_964984, EPI_ISL_964985, EPI_ISL_964986 | Hospital of Southern Norway - Kristiansand, Department of Medical Microbiology | Norwegian Institute of Public Health, Department of Virology | Kathrine Stene-Johansen, Kamilla Heddeland Instefjord, Hilde Elshaug, Ignacio Garcia Llorente, Serina B Engebretsen, Atiya R Ali,Marie Paulsen Madsen, Rasmus Riis Kopperud, Hilde Vollan, Karoline Bragstad, Olav Hungnes |
| EPI_ISL_964989 | Furst Medical Laboratory | Norwegian Institute of Public Health, Department of Virology | Kathrine Stene-Johansen, Kamilla Heddeland Instefjord, Hilde Elshaug, Ignacio Garcia Llorente, Serina B Engebretsen, Atiya R Ali,Marie Paulsen Madsen, Rasmus Riis Kopperud, Hilde Vollan, Karoline Bragstad, Olav Hungnes |
| EPI_ISL_964991 | Nordland Hospital - Bodo, Laboratory Department, Molecular Biology Unit | Norwegian Institute of Public Health, Department of Virology | Kathrine Stene-Johansen, Kamilla Heddeland Instefjord, Hilde Elshaug, Ignacio Garcia Llorente, Serina B Engebretsen, Atiya R Ali,Marie Paulsen Madsen, Rasmus Riis Kopperud, Hilde Vollan, Karoline Bragstad, Olav Hungnes |
| EPI_ISL_964992, EPI_ISL_964993 | Akershus University Hospital, Department for Microbiology and Infectious Disease Control | Norwegian Institute of Public Health, Department of Virology | Kathrine Stene-Johansen, Kamilla Heddeland Instefjord, Hilde Elshaug, Ignacio Garcia Llorente, Serina B Engebretsen, Atiya R Ali,Marie Paulsen Madsen, Rasmus Riis Kopperud, Hilde Vollan, Karoline Bragstad, Olav Hungnes |
| EPI_ISL_964998, EPI_ISL_964999 | Oslo University Hospital, Department of Medical Microbiology | Norwegian Institute of Public Health, Department of Virology | Kathrine Stene-Johansen, Kamilla Heddeland Instefjord, Hilde Elshaug, Ignacio Garcia Llorente, Serina B Engebretsen, Atiya R Ali, Marie Paulsen Madsen, Rasmus Riis Kopperud, Hilde Vollan, Karoline Bragstad, Olav Hungnes |
| EPI_ISL_965017, EPI_ISL_965018 | Florida Bureau of Public Health Laboratories | Florida Bureau of Public Health Laboratories | Sarah Schmedes, Jason Blanton |
| EPI_ISL_965045, EPI_ISL_965046, EPI_ISL_965047, EPI_ISL_965048, EPI_ISL_965049, EPI_ISL_965050, EPI_ISL_965051, EPI_ISL_965052 | Wyoming Public Health Laboratory | Wyoming Public Health Laboratory | Noah Hull, Taylor Fearing, Lynette Gumbleton, Channing Weber, Ashley Norberg, Bailey Bowcutt, and Wanda Manley |
| EPI_ISL_965157, EPI_ISL_965158, EPI_ISL_965159, EPI_ISL_965160, EPI_ISL_965161, EPI_ISL_965162, EPI_ISL_965163, EPI_ISL_965164, EPI_ISL_965165, EPI_ISL_965166, EPI_ISL_965167, EPI_ISL_965169, EPI_ISL_965171, EPI_ISL_965172, EPI_ISL_965173, EPI_ISL_965174, EPI_ISL_965175, EPI_ISL_965176, EPI_ISL_965177 | | | |
| see above | Clinical Molecular Microbiology Laboratory, UNC Hospitals | Jeremy Wang | Jeremy Wang, Alexander Rubinsteyn, Colleen Rice, Jason Smedberg, Shawn Hawken, Melissa Miller, Corbin Jones, Robert Hagan |
| EPI_ISL_965306, EPI_ISL_965307, EPI_ISL_965308 | Labo Luc Olivier | GIGA Medical Genomics | Keith Durkin, Maria Artesi, Sébastien Bontems, Raphaël Boreux, Bouchra Boujemla, Cécile Meex, Pierrette Melin, Marie-Pierre Hayette, Vincent Bours |
| EPI_ISL_965311, EPI_ISL_965312, EPI_ISL_965313, EPI_ISL_965316, EPI_ISL_965317 | Synlab | GIGA Medical Genomics | Keith Durkin, Maria Artesi, Sébastien Bontems, Raphaël Boreux, Bouchra Boujemla, Cécile Meex, Pierrette Melin, Marie-Pierre Hayette, Vincent Bours |
| EPI_ISL_965318, EPI_ISL_965319 | Labo Luc Olivier | GIGA Medical Genomics | Keith Durkin, Maria Artesi, Sébastien Bontems, Raphaël Boreux, Bouchra Boujemla, Cécile Meex, Pierrette Melin, Marie-Pierre Hayette, Vincent Bours |
| EPI_ISL_965320, EPI_ISL_965325, EPI_ISL_965327 | Synlab | GIGA Medical Genomics | Keith Durkin, Maria Artesi, Sébastien Bontems, Raphaël Boreux, Bouchra Boujemla, Cécile Meex, Pierrette Melin, Marie-Pierre Hayette, Vincent Bours |
| EPI_ISL_965328, EPI_ISL_965330, EPI_ISL_965331 | Labo Luc Olivier | GIGA Medical Genomics | Keith Durkin, Maria Artesi, Sébastien Bontems, Raphaël Boreux, Bouchra Boujemla, Cécile Meex, Pierrette Melin, Marie-Pierre Hayette, Vincent Bours |

| | | | |
|---|---|---|---|
| EPI_ISL_965333, EPI_ISL_965334, EPI_ISL_965336, EPI_ISL_965337 | Synlab | GIGA Medical Genomics | Keith Durkin, Maria Artesi, Sébastien Bontems, Raphaël Boreux, Bouchra Boujemla, Cécile Meex, Pierrette Melin, Marie-Pierre Hayette, Vincent Bours |
| EPI_ISL_965343 | Labo Luc Olivier | GIGA Medical Genomics | Keith Durkin, Maria Artesi, Sébastien Bontems, Raphaël Boreux, Bouchra Boujemla, Cécile Meex, Pierrette Melin, Marie-Pierre Hayette, Vincent Bours |
| EPI_ISL_965393 | Clinique N.D de Grâce Gosselies | GIGA Medical Genomics | Keith Durkin, Maria Artesi, Sébastien Bontems, Raphaël Boreux, Bouchra Boujemla, Cécile Meex, Pierrette Melin, Marie-Pierre Hayette, Vincent Bours |
| EPI_ISL_965398 | Synlab | GIGA Medical Genomics | Keith Durkin, Maria Artesi, Sébastien Bontems, Raphaël Boreux, Bouchra Boujemla, Cécile Meex, Pierrette Melin, Marie-Pierre Hayette, Vincent Bours |
| EPI_ISL_965399 | Labo Luc Olivier | GIGA Medical Genomics | Keith Durkin, Maria Artesi, Sébastien Bontems, Raphaël Boreux, Bouchra Boujemla, Cécile Meex, Pierrette Melin, Marie-Pierre Hayette, Vincent Bours |
| EPI_ISL_965401 | Synlab | GIGA Medical Genomics | Keith Durkin, Maria Artesi, Sébastien Bontems, Raphaël Boreux, Bouchra Boujemla, Cécile Meex, Pierrette Melin, Marie-Pierre Hayette, Vincent Bours |
| EPI_ISL_965402 | Clinique N.D de Grâce Gosselies | GIGA Medical Genomics | Keith Durkin, Maria Artesi, Sébastien Bontems, Raphaël Boreux, Bouchra Boujemla, Cécile Meex, Pierrette Melin, Marie-Pierre Hayette, Vincent Bours |
| EPI_ISL_965405 | Synlab | GIGA Medical Genomics | Keith Durkin, Maria Artesi, Sébastien Bontems, Raphaël Boreux, Bouchra Boujemla, Cécile Meex, Pierrette Melin, Marie-Pierre Hayette, Vincent Bours |
| EPI_ISL_965406, EPI_ISL_965407 | Labo Luc Olivier | GIGA Medical Genomics | Keith Durkin, Maria Artesi, Sébastien Bontems, Raphaël Boreux, Bouchra Boujemla, Cécile Meex, Pierrette Melin, Marie-Pierre Hayette, Vincent Bours |
| EPI_ISL_965408 | Synlab | GIGA Medical Genomics | Keith Durkin, Maria Artesi, Sébastien Bontems, Raphaël Boreux, Bouchra Boujemla, Cécile Meex, Pierrette Melin, Marie-Pierre Hayette, Vincent Bours |
| EPI_ISL_965409 | Labo Luc Olivier | GIGA Medical Genomics | Keith Durkin, Maria Artesi, Sébastien Bontems, Raphaël Boreux, Bouchra Boujemla, Cécile Meex, Pierrette Melin, Marie-Pierre Hayette, Vincent Bours |
| EPI_ISL_965410 | Synlab | GIGA Medical Genomics | Keith Durkin, Maria Artesi, Sébastien Bontems, Raphaël Boreux, Bouchra Boujemla, Cécile Meex, Pierrette Melin, Marie-Pierre Hayette, Vincent Bours |
| EPI_ISL_965411, EPI_ISL_965412, EPI_ISL_965413 | Labo Luc Olivier | GIGA Medical Genomics | Keith Durkin, Maria Artesi, Sébastien Bontems, Raphaël Boreux, Bouchra Boujemla, Cécile Meex, Pierrette Melin, Marie-Pierre Hayette, Vincent Bours |
| EPI_ISL_965414, EPI_ISL_965415 | CHR de la Citadelle | GIGA Medical Genomics | Keith Durkin, Maria Artesi, Sébastien Bontems, Raphaël Boreux, Bouchra Boujemla, Cécile Meex, Pierrette Melin, Marie-Pierre Hayette, Vincent Bours |
| EPI_ISL_965416, EPI_ISL_965418 | Synlab | GIGA Medical Genomics | Keith Durkin, Maria Artesi, Sébastien Bontems, Raphaël Boreux, Bouchra Boujemla, Cécile Meex, Pierrette Melin, Marie-Pierre Hayette, Vincent Bours |
| EPI_ISL_965419, EPI_ISL_965420 | Labo Luc Olivier | GIGA Medical Genomics | Keith Durkin, Maria Artesi, Sébastien Bontems, Raphaël Boreux, Bouchra Boujemla, Cécile Meex, Pierrette Melin, Marie-Pierre Hayette, Vincent Bours |
| EPI_ISL_965421, EPI_ISL_965422, EPI_ISL_965423, EPI_ISL_965424, EPI_ISL_965425, EPI_ISL_965426, EPI_ISL_965427, EPI_ISL_965428, EPI_ISL_965429, EPI_ISL_965430 | CHR de la Citadelle | GIGA Medical Genomics | Keith Durkin, Maria Artesi, Sébastien Bontems, Raphaël Boreux, Bouchra Boujemla, Cécile Meex, Pierrette Melin, Marie-Pierre Hayette, Vincent Bours |
| EPI_ISL_965431 | Synlab | GIGA Medical Genomics | Keith Durkin, Maria Artesi, Sébastien Bontems, Raphaël Boreux, Bouchra Boujemla, Cécile Meex, Pierrette Melin, Marie-Pierre Hayette, Vincent Bours |
| EPI_ISL_965432 | CHR de la Citadelle | GIGA Medical Genomics | Keith Durkin, Maria Artesi, Sébastien Bontems, Raphaël Boreux, Bouchra Boujemla, Cécile Meex, Pierrette Melin, Marie-Pierre Hayette, Vincent Bours |
| EPI_ISL_965434, EPI_ISL_965435, EPI_ISL_965436 | Synlab | GIGA Medical Genomics | Keith Durkin, Maria Artesi, Sébastien Bontems, Raphaël Boreux, Bouchra Boujemla, Cécile Meex, Pierrette Melin, Marie-Pierre Hayette, Vincent Bours |
| EPI_ISL_965437, EPI_ISL_965438, EPI_ISL_965439, EPI_ISL_965440, EPI_ISL_965441, EPI_ISL_965442, EPI_ISL_965443, EPI_ISL_965444, EPI_ISL_965445, EPI_ISL_965446, EPI_ISL_965447, EPI_ISL_965448, EPI_ISL_965449, EPI_ISL_965450 | | | |
| see above | CHR de la Citadelle | GIGA Medical Genomics | Keith Durkin, Maria Artesi, Sébastien Bontems, Raphaël Boreux, Bouchra Boujemla, Cécile Meex, Pierrette Melin, Marie-Pierre Hayette, Vincent Bours |
| EPI_ISL_965452, EPI_ISL_965453, EPI_ISL_965454, EPI_ISL_965456 | Synlab | GIGA Medical Genomics | Keith Durkin, Maria Artesi, Sébastien Bontems, Raphaël Boreux, Bouchra Boujemla, Cécile Meex, Pierrette Melin, Marie-Pierre Hayette, Vincent Bours |
| EPI_ISL_965460 | St. Nikolaus-Hospital Eupen | GIGA Medical Genomics | Keith Durkin, Maria Artesi, Sébastien Bontems, Raphaël Boreux, Bouchra Boujemla, Cécile Meex, Pierrette Melin, Marie-Pierre Hayette, Vincent Bours |
| EPI_ISL_965467 | Synlab | GIGA Medical Genomics | Keith Durkin, Maria Artesi, Sébastien Bontems, Raphaël Boreux, Bouchra Boujemla, Cécile Meex, Pierrette Melin, Marie-Pierre Hayette, Vincent Bours |
| EPI_ISL_965469 | Labo Luc Olivier | GIGA Medical Genomics | Keith Durkin, Maria Artesi, Sébastien Bontems, Raphaël Boreux, Bouchra Boujemla, Cécile Meex, Pierrette Melin, Marie-Pierre Hayette, Vincent Bours |
| EPI_ISL_965472, EPI_ISL_965473 | Synlab | GIGA Medical Genomics | Keith Durkin, Maria Artesi, Sébastien Bontems, Raphaël Boreux, Bouchra Boujemla, Cécile Meex, Pierrette Melin, Marie-Pierre Hayette, Vincent Bours |
| EPI_ISL_965474, EPI_ISL_965476, EPI_ISL_965477, EPI_ISL_965478 | Labo Luc Olivier | GIGA Medical Genomics | Keith Durkin, Maria Artesi, Sébastien Bontems, Raphaël Boreux, Bouchra Boujemla, Cécile Meex, Pierrette Melin, Marie-Pierre Hayette, Vincent Bours |
| EPI_ISL_965479, EPI_ISL_965480, EPI_ISL_965482, EPI_ISL_965483 | Synlab | GIGA Medical Genomics | Keith Durkin, Maria Artesi, Sébastien Bontems, Raphaël Boreux, Bouchra Boujemla, Cécile Meex, Pierrette Melin, Marie-Pierre Hayette, Vincent Bours |
| EPI_ISL_965487, EPI_ISL_965488, EPI_ISL_965489, EPI_ISL_965490, EPI_ISL_965491, EPI_ISL_965492 | St. Nikolaus-Hospital Eupen | GIGA Medical Genomics | Keith Durkin, Maria Artesi, Sébastien Bontems, Raphaël Boreux, Bouchra Boujemla, Cécile Meex, Pierrette Melin, Marie-Pierre Hayette, Vincent Bours |
| EPI_ISL_965494, EPI_ISL_965497 | Synlab | GIGA Medical Genomics | Keith Durkin, Maria Artesi, Sébastien Bontems, Raphaël Boreux, Bouchra Boujemla, Cécile Meex, Pierrette Melin, Marie-Pierre Hayette, Vincent Bours |
| EPI_ISL_965498, EPI_ISL_965500 | St. Nikolaus-Hospital Eupen | GIGA Medical Genomics | Keith Durkin, Maria Artesi, Sébastien Bontems, Raphaël Boreux, Bouchra Boujemla, Cécile Meex, Pierrette Melin, Marie-Pierre Hayette, Vincent Bours |
| EPI_ISL_965501, EPI_ISL_965502 | Synlab | GIGA Medical Genomics | Keith Durkin, Maria Artesi, Sébastien Bontems, Raphaël Boreux, Bouchra Boujemla, Cécile Meex, Pierrette Melin, Marie-Pierre Hayette, Vincent Bours |
| EPI_ISL_965504 | Labo Luc Olivier | GIGA Medical Genomics | Keith Durkin, Maria Artesi, Sébastien Bontems, Raphaël Boreux, Bouchra Boujemla, Cécile Meex, Pierrette Melin, Marie-Pierre Hayette, Vincent Bours |
| EPI_ISL_965507, EPI_ISL_965514, EPI_ISL_965516, EPI_ISL_965519 | Synlab | GIGA Medical Genomics | Keith Durkin, Maria Artesi, Sébastien Bontems, Raphaël Boreux, Bouchra Boujemla, Cécile Meex, Pierrette Melin, Marie-Pierre Hayette, Vincent Bours |
| EPI_ISL_965520, EPI_ISL_965522, EPI_ISL_965523 | St. Nikolaus-Hospital Eupen | GIGA Medical Genomics | Keith Durkin, Maria Artesi, Sébastien Bontems, Raphaël Boreux, Bouchra Boujemla, Cécile Meex, Pierrette Melin, Marie-Pierre Hayette, Vincent Bours |
| EPI_ISL_965526 | Synlab | GIGA Medical Genomics | Keith Durkin, Maria Artesi, Sébastien Bontems, Raphaël Boreux, Bouchra Boujemla, Cécile Meex, Pierrette Melin, Marie-Pierre Hayette, Vincent Bours |
| EPI_ISL_965643, EPI_ISL_965711 | Dutch COVID-19 response team | Medical Microbiology, Maastricht University Medical Centre | Jozef Dingemans*, Brian van der Veer*, Erik Beuken, Carmen Reumkens, Lieke van Alphen, Christian Hoebe, Paul Savelkoul |
| EPI_ISL_965877, EPI_ISL_965878, EPI_ISL_965885, EPI_ISL_965895, EPI_ISL_965896, EPI_ISL_965898, EPI_ISL_965899, EPI_ISL_965901, EPI_ISL_965902 | Massachusetts State Public Health Laboratory | Massachusetts State Public Health Laboratory | Andrew Lang, Timelia Fink, Glen Gallagher, Sandra Smole |
| EPI_ISL_965907, EPI_ISL_965908 | Clinical Molecular Microbiology Laboratory, UNC Hospitals | Jeremy Wang | Jeremy Wang, Alexander Rubinsteyn, Colleen Rice, Jason Smedberg, Shawn Hawken, Melissa Miller, Corbin Jones, Robert Hagan |
| EPI_ISL_965917, EPI_ISL_965918, EPI_ISL_965919, EPI_ISL_965923 | Hospital Los Arcos del Mar Menor | Instituto de Salud Carlos III | Vázquez, S. Iglesias-Caballero, M. Sandonís,V. Camarero, S. Pozo, F. Casas, I. Jiménez, P. Zaballos, A. Monzón, S. Varona, S. Cuesta, I. Cámara, M. |
| EPI_ISL_965930 | Hospital Comarcal de Melilla | Instituto de Salud Carlos III | Sandonís,V. Vázquez, S. Iglesias-Caballero, M. Camarero, S. Pozo, F. Casas, I. Jiménez, P. Zaballos, A. Monzón, S. Varona, S. Cuesta, I. Román, S. |
| EPI_ISL_965935, EPI_ISL_965939, EPI_ISL_965941 | Hospital Los Arcos del Mar Menor | Instituto de Salud Carlos III | Sandonís,V. Vázquez, S. Iglesias-Caballero, M. Camarero, S. Pozo, F. Casas, I. Jiménez, P. Zaballos, A. Monzón, S. Varona, S. Cuesta, I. Cámara, M. |
| EPI_ISL_965945, EPI_ISL_965947, EPI_ISL_965951, EPI_ISL_965958 | Hospital Los Arcos del Mar Menor | Instituto de Salud Carlos III | Iglesias-Caballero, M. Sandonís,V. Vázquez, S. Camarero, S. Pozo, F. Casas, I. Jiménez, P. Zaballos, A. Monzón, S. Varona, S. Cuesta, I. Cámara, M. |
| EPI_ISL_966379, EPI_ISL_966381, | OCME Office Of Chief Medical Examiner | New York City Public Health Laboratory | Jade Wang, et al. |

| | | | |
|---|---|---|---|
| EPI_ISL_966383 | | | |
| EPI_ISL_966385 | Department of Homeless Services | New York City Public Health Laboratory | Jade Wang, et al. |
| EPI_ISL_966389 | DOHMH Central Harlem | New York City Public Health Laboratory | Jade Wang, et al. |
| EPI_ISL_966390 | OCME Office Of Chief Medical Examiner | New York City Public Health Laboratory | Jade Wang, et al. |
| EPI_ISL_966392 | DOHMH Corona | New York City Public Health Laboratory | Jade Wang, et al. |
| EPI_ISL_966393 | OCME Office Of Chief Medical Examiner | New York City Public Health Laboratory | Jade Wang, et al. |
| EPI_ISL_966394 | DOHMH Corona | New York City Public Health Laboratory | Jade Wang, et al. |
| EPI_ISL_966396 | OCME Office Of Chief Medical Examiner | New York City Public Health Laboratory | Jade Wang, et al. |
| EPI_ISL_966398 | DOHMH Jamaica | New York City Public Health Laboratory | Jade Wang, et al. |
| EPI_ISL_966399 | DOHMH Corona | New York City Public Health Laboratory | Jade Wang, et al. |
| EPI_ISL_966407, EPI_ISL_966408 | DOHMH Morrisania | New York City Public Health Laboratory | Jade Wang, et al. |
| EPI_ISL_966409, EPI_ISL_966410 | DOHMH Corona | New York City Public Health Laboratory | Jade Wang, et al. |
| EPI_ISL_966420 | DOHMH Morrisania | New York City Public Health Laboratory | Jade Wang, et al. |
| EPI_ISL_966421, EPI_ISL_966422 | DOHMH Jamaica | New York City Public Health Laboratory | Jade Wang, et al. |
| EPI_ISL_966423 | DOHMH Morrisania | New York City Public Health Laboratory | Jade Wang, et al. |
| EPI_ISL_966424 | DOHMH Jamaica | New York City Public Health Laboratory | Jade Wang, et al. |
| EPI_ISL_966425 | DOHMH Corona | New York City Public Health Laboratory | Jade Wang, et al. |
| EPI_ISL_966426 | DOHMH PHL | New York City Public Health Laboratory | Jade Wang, et al. |
| EPI_ISL_966461, EPI_ISL_966462, EPI_ISL_966463 | DOHMH Jamaica | New York City Public Health Laboratory | Jade Wang, et al. |
| EPI_ISL_966464, EPI_ISL_966465, EPI_ISL_966466 | DOHMH PHL | New York City Public Health Laboratory | Jade Wang, et al. |
| EPI_ISL_966467 | DOHMH Corona | New York City Public Health Laboratory | Jade Wang, et al. |
| EPI_ISL_966468 | DOHMH Chelsea | New York City Public Health Laboratory | Jade Wang, et al. |
| EPI_ISL_966469 | Department of Homeless Services | New York City Public Health Laboratory | Jade Wang, et al. |
| EPI_ISL_966470, EPI_ISL_966471, EPI_ISL_966473 | OCME Office Of Chief Medical Examiner | New York City Public Health Laboratory | Jade Wang, et al. |
| EPI_ISL_966474 | DOHMH Corona | New York City Public Health Laboratory | Jade Wang, et al. |
| EPI_ISL_966475 | DOHMH Central Harlem | New York City Public Health Laboratory | Jade Wang, et al. |
| EPI_ISL_966476, EPI_ISL_966477, EPI_ISL_966478 | DOHMH Corona | New York City Public Health Laboratory | Jade Wang, et al. |
| EPI_ISL_966479 | DOHMH Jamaica | New York City Public Health Laboratory | Jade Wang, et al. |
| EPI_ISL_966480 | DOHMH Corona | New York City Public Health Laboratory | Jade Wang, et al. |
| EPI_ISL_966481 | DOHMH Jamaica | New York City Public Health Laboratory | Jade Wang, et al. |
| EPI_ISL_966482 | DOHMH Corona | New York City Public Health Laboratory | Jade Wang, et al. |
| EPI_ISL_966483 | DOHMH Chelsea | New York City Public Health Laboratory | Jade Wang, et al. |
| EPI_ISL_966484 | DOHMH Crown Heights | New York City Public Health Laboratory | Jade Wang, et al. |
| EPI_ISL_966485, EPI_ISL_966486, EPI_ISL_966487, EPI_ISL_966488, EPI_ISL_966490 | OCME Office Of Chief Medical Examiner | New York City Public Health Laboratory | Jade Wang, et al. |
| EPI_ISL_966491 | DOHMH Corona | New York City Public Health Laboratory | Jade Wang, et al. |
| EPI_ISL_966492, EPI_ISL_966493, EPI_ISL_966494 | OCME Office Of Chief Medical Examiner | New York City Public Health Laboratory | Jade Wang, et al. |
| EPI_ISL_966786, EPI_ISL_966787 | Maine HETL | Tewhey Lab, The Jackson Laboratory | Matluk,N., Dewey,H., Iosue,F., Barter,M., Lynch,R., Munger,H. and Tewhey,R. |
| EPI_ISL_966929, EPI_ISL_966930, EPI_ISL_966931, EPI_ISL_966932, EPI_ISL_966933, EPI_ISL_966934, EPI_ISL_966935, EPI_ISL_966936, EPI_ISL_966937, EPI_ISL_966938, EPI_ISL_966939, EPI_ISL_966941, EPI_ISL_966942, EPI_ISL_966943, EPI_ISL_966944, EPI_ISL_966945, EPI_ISL_966946, EPI_ISL_966947, EPI_ISL_966948, EPI_ISL_966949, EPI_ISL_966950, EPI_ISL_966951, EPI_ISL_966952, EPI_ISL_966953, EPI_ISL_966954, EPI_ISL_966955, EPI_ISL_966956, EPI_ISL_966957, EPI_ISL_966958, EPI_ISL_966959, EPI_ISL_966960, EPI_ISL_966961, EPI_ISL_966962, EPI_ISL_966963, EPI_ISL_966964, EPI_ISL_966965, EPI_ISL_966966, EPI_ISL_966967, EPI_ISL_966968, EPI_ISL_966969, EPI_ISL_966970, EPI_ISL_966971, EPI_ISL_966972, EPI_ISL_966973, EPI_ISL_966974, EPI_ISL_966975, EPI_ISL_966976, EPI_ISL_966977, EPI_ISL_966978, EPI_ISL_966979, EPI_ISL_966980, EPI_ISL_966981, EPI_ISL_966982, EPI_ISL_966983, EPI_ISL_966984, EPI_ISL_966985, EPI_ISL_966986, EPI_ISL_966987, EPI_ISL_966988, EPI_ISL_966989, EPI_ISL_966990, EPI_ISL_966991, EPI_ISL_966992, EPI_ISL_966993, EPI_ISL_966994, EPI_ISL_966995, EPI_ISL_966996, EPI_ISL_966997, EPI_ISL_966998, EPI_ISL_966999, EPI_ISL_967000, EPI_ISL_967001, EPI_ISL_967002, EPI_ISL_967003, EPI_ISL_967004, EPI_ISL_967005, EPI_ISL_967006, EPI_ISL_967007, EPI_ISL_967008, EPI_ISL_967009, EPI_ISL_967010, EPI_ISL_967011, EPI_ISL_967012, EPI_ISL_967013, EPI_ISL_967014, EPI_ISL_967015, EPI_ISL_967016, EPI_ISL_967017, EPI_ISL_967018, EPI_ISL_967019, EPI_ISL_967020, EPI_ISL_967021, EPI_ISL_967022, EPI_ISL_967023, EPI_ISL_967024, EPI_ISL_967025, EPI_ISL_967026, EPI_ISL_967027, EPI_ISL_967028, EPI_ISL_967029, EPI_ISL_967030, EPI_ISL_967031, EPI_ISL_967032, EPI_ISL_967033, EPI_ISL_967034, EPI_ISL_967035, EPI_ISL_967036, EPI_ISL_967037, EPI_ISL_967038, EPI_ISL_967039, EPI_ISL_967040, EPI_ISL_967041, EPI_ISL_967042, EPI_ISL_967043, EPI_ISL_967044, EPI_ISL_967045, EPI_ISL_967046, EPI_ISL_967047, EPI_ISL_967048, EPI_ISL_967049, EPI_ISL_967050, EPI_ISL_967051, EPI_ISL_967052, EPI_ISL_967053, EPI_ISL_967054, EPI_ISL_967055, EPI_ISL_967056, EPI_ISL_967057 | | | |
| see above | Helix/Illumina | Respiratory Viruses Branch, Division of Viral Diseases, Centers for Disease Control and Prevention | Peter W. Cook,Dakota Howard,Dhwani Batra,Ben L. Rambo-Martin,Eileen de Feo,Jan Antico,Christine Tran,Matthew Tolentino,Shannon Wickline,Kim Gietzen,Brad Sickler,Jingtao Liu,Eric Allen,Phil Febbo,Summer Galloway,Nicole L. Washington,Simon White,Geraint Levan,Kelly Schiabor Barrett,Elizabeth Cirulli,Alexandre Bolze,Ary Ascencio,Charlotte Rivera-Garcia,Ryan Cho,Jason Nguyen,Sherry Wang,Jimmy Ramirez,Tyler Cassens,Efren Sandoval,Magnus Isaksson,William Lee,David Becker,Marc Laurent,James Lu,Clinton R. Paden,Suxiang Tong,Duncan MacCannell, |
| EPI_ISL_967764 | State Laboratories Division, Hawaii State Department of Health | State Laboratories Division, Hawaii State Department of Health | Pamela O'Brien, Drew Kuwazaki, Ayana Garnet, Razvan Sultana, Edward Desmond |
| EPI_ISL_969007, EPI_ISL_969008, EPI_ISL_969012, EPI_ISL_969013, EPI_ISL_969014, EPI_ISL_969016, EPI_ISL_969017, EPI_ISL_969025, EPI_ISL_969026, EPI_ISL_969030, EPI_ISL_969031, EPI_ISL_969043, EPI_ISL_969044, EPI_ISL_969046, EPI_ISL_969053, EPI_ISL_969056, EPI_ISL_969061, EPI_ISL_969063, EPI_ISL_969064, EPI_ISL_969065, EPI_ISL_969066, EPI_ISL_969067, EPI_ISL_969071, EPI_ISL_969245 | | | |
| see above | KEMRI-Wellcome Trust Research Programme/KEMRI-CGMR-C Kilifi | KEMRI-Wellcome Trust Research Programme/KEMRI-CGMR-C Kilifi | Githinji et al |
| EPI_ISL_969454, EPI_ISL_970032, EPI_ISL_970055, EPI_ISL_970142 | Lighthouse Lab in Alderley Park | Wellcome Sanger Institute for the COVID-19 Genomics UK (COG-UK) Consortium | Jacquelyn Wynn, Mairead Hyland, The Lighthouse Lab in Alderley Park and Alex Alderton, Roberto Amato, Sonia Goncalves, Ewan Harrison, David K. Jackson, Ian Johnston, Dominic Kwiatkowski, Cordelia Langford, John Sillitoe on behalf of the Wellcome Sanger Institute COVID-19 Surveillance Team |
| EPI_ISL_970469 | EHNV | Laboratory of genomics and metagenomics, Institute of Microbiology, University Hospital Centre and University of | Trestan Pillonel, Damien Jacot, Sébastien Aeby, Gilbert Greub, Claire Bertelli |

| | | | |
|---|---|---|---|
| | | Lausanne, Switzerland | |
| EPI_ISL_970589, EPI_ISL_970591, EPI_ISL_970593, EPI_ISL_970595, EPI_ISL_970599, EPI_ISL_970601, EPI_ISL_970602, EPI_ISL_970605, EPI_ISL_970607, EPI_ISL_970609, EPI_ISL_970612, EPI_ISL_970614, EPI_ISL_970618, EPI_ISL_970619, EPI_ISL_970623, EPI_ISL_970625, EPI_ISL_970630, EPI_ISL_970632, EPI_ISL_970633, EPI_ISL_970637, EPI_ISL_970641, EPI_ISL_970643, EPI_ISL_970648, EPI_ISL_970650, EPI_ISL_970652, EPI_ISL_970654, EPI_ISL_970656, EPI_ISL_970658, EPI_ISL_970660, EPI_ISL_970661, EPI_ISL_970666, EPI_ISL_970668, EPI_ISL_970670, EPI_ISL_970675, EPI_ISL_970677, EPI_ISL_970683, EPI_ISL_970685, EPI_ISL_970689, EPI_ISL_970690, EPI_ISL_970692, EPI_ISL_970694, EPI_ISL_970697, EPI_ISL_970699, EPI_ISL_970703, EPI_ISL_970704, EPI_ISL_970706, EPI_ISL_970711, EPI_ISL_970713, EPI_ISL_970717, EPI_ISL_970720, EPI_ISL_970722, EPI_ISL_970726, EPI_ISL_970727, EPI_ISL_970729, EPI_ISL_970734, EPI_ISL_970736, EPI_ISL_970738, EPI_ISL_970739, EPI_ISL_970741, EPI_ISL_970746, EPI_ISL_970749, EPI_ISL_970755, EPI_ISL_970757, EPI_ISL_970758, EPI_ISL_970760, EPI_ISL_970762, EPI_ISL_970764, EPI_ISL_970766, EPI_ISL_970770, EPI_ISL_970772, EPI_ISL_970774, EPI_ISL_970777, EPI_ISL_970779, EPI_ISL_970781, EPI_ISL_970782, EPI_ISL_970784, EPI_ISL_970786, EPI_ISL_970788, EPI_ISL_970792, EPI_ISL_970794, EPI_ISL_970796, EPI_ISL_970797, EPI_ISL_970800, EPI_ISL_970802, EPI_ISL_970803, EPI_ISL_970805, EPI_ISL_970807, EPI_ISL_970808 | | | |
| see above | Lighthouse Lab in Milton Keynes | Wellcome Sanger Institute for the COVID-19 Genomics UK (COG-UK) Consortium | The Lighthouse Lab in Milton Keynes and Alex Alderton, Roberto Amato, Sonia Goncalves, Ewan Harrison, David K. Jackson, Ian Johnston, Dominic Kwiatkowski, Cordelia Langford, John Sillitoe on behalf of the Wellcome Sanger Institute COVID-19 Surveillance Team |
| EPI_ISL_970810, EPI_ISL_970813, EPI_ISL_970814, EPI_ISL_970815, EPI_ISL_970816, EPI_ISL_970817, EPI_ISL_970822, EPI_ISL_970823, EPI_ISL_970825, EPI_ISL_970828, EPI_ISL_970833, EPI_ISL_970835, EPI_ISL_970836, EPI_ISL_970838, EPI_ISL_970839, EPI_ISL_970841, EPI_ISL_970842, EPI_ISL_970843 | | | |
| see above | Department of Virus and Microbiological Special Diagnostics, Statens Serum Institut, Copenhagen, Denmark | Aalborg University | Danish Covid-19 Genome Consortium |
| EPI_ISL_970845 | Lighthouse Lab in Milton Keynes | Wellcome Sanger Institute for the COVID-19 Genomics UK (COG-UK) Consortium | The Lighthouse Lab in Milton Keynes and Alex Alderton, Roberto Amato, Sonia Goncalves, Ewan Harrison, David K. Jackson, Ian Johnston, Dominic Kwiatkowski, Cordelia Langford, John Sillitoe on behalf of the Wellcome Sanger Institute COVID-19 Surveillance Team |
| EPI_ISL_970847, EPI_ISL_970848, EPI_ISL_970852, EPI_ISL_970853, EPI_ISL_970854, EPI_ISL_970855, EPI_ISL_970857, EPI_ISL_970858, EPI_ISL_970861, EPI_ISL_970864, EPI_ISL_970865, EPI_ISL_970866, EPI_ISL_970869, EPI_ISL_970870, EPI_ISL_970871, EPI_ISL_970872, EPI_ISL_970873, EPI_ISL_970876, EPI_ISL_970877, EPI_ISL_970881, EPI_ISL_970883, EPI_ISL_970884, EPI_ISL_970887, EPI_ISL_970888, EPI_ISL_970889, EPI_ISL_970890, EPI_ISL_970893, EPI_ISL_970894, EPI_ISL_970896, EPI_ISL_970897, EPI_ISL_970898 | | | |
| see above | Department of Virus and Microbiological Special Diagnostics, Statens Serum Institut, Copenhagen, Denmark | Aalborg University | Danish Covid-19 Genome Consortium |
| EPI_ISL_970899 | Lighthouse Lab in Milton Keynes | Wellcome Sanger Institute for the COVID-19 Genomics UK (COG-UK) Consortium | The Lighthouse Lab in Milton Keynes and Alex Alderton, Roberto Amato, Sonia Goncalves, Ewan Harrison, David K. Jackson, Ian Johnston, Dominic Kwiatkowski, Cordelia Langford, John Sillitoe on behalf of the Wellcome Sanger Institute COVID-19 Surveillance Team |
| EPI_ISL_970900, EPI_ISL_970903, EPI_ISL_970904, EPI_ISL_970906, EPI_ISL_970909, EPI_ISL_970911, EPI_ISL_970912, EPI_ISL_970913, EPI_ISL_970914, EPI_ISL_970915, EPI_ISL_970916, EPI_ISL_970917, EPI_ISL_970918, EPI_ISL_970919, EPI_ISL_970920, EPI_ISL_970921, EPI_ISL_970922, EPI_ISL_970923, EPI_ISL_970924 | | | |
| see above | Department of Virus and Microbiological Special Diagnostics, Statens Serum Institut, Copenhagen, Denmark | Aalborg University | Danish Covid-19 Genome Consortium |
| EPI_ISL_970925 | Lighthouse Lab in Milton Keynes | Wellcome Sanger Institute for the COVID-19 Genomics UK (COG-UK) Consortium | The Lighthouse Lab in Milton Keynes and Alex Alderton, Roberto Amato, Sonia Goncalves, Ewan Harrison, David K. Jackson, Ian Johnston, Dominic Kwiatkowski, Cordelia Langford, John Sillitoe on behalf of the Wellcome Sanger Institute COVID-19 Surveillance Team |
| EPI_ISL_970926, EPI_ISL_970933, EPI_ISL_970934, EPI_ISL_970935, EPI_ISL_970937, EPI_ISL_970940, EPI_ISL_970941 | Department of Virus and Microbiological Special Diagnostics, Statens Serum Institut, Copenhagen, Denmark | Aalborg University | Danish Covid-19 Genome Consortium |
| EPI_ISL_970944 | Lighthouse Lab in Milton Keynes | Wellcome Sanger Institute for the COVID-19 Genomics UK (COG-UK) Consortium | The Lighthouse Lab in Milton Keynes and Alex Alderton, Roberto Amato, Sonia Goncalves, Ewan Harrison, David K. Jackson, Ian Johnston, Dominic Kwiatkowski, Cordelia Langford, John Sillitoe on behalf of the Wellcome Sanger Institute COVID-19 Surveillance Team |
| EPI_ISL_970945, EPI_ISL_970948, EPI_ISL_970951, EPI_ISL_970954, EPI_ISL_970955, EPI_ISL_970956, EPI_ISL_970957, EPI_ISL_970958, EPI_ISL_970962, EPI_ISL_970963, EPI_ISL_970964, EPI_ISL_970967 | | | |
| see above | Department of Virus and Microbiological Special Diagnostics, Statens Serum Institut, Copenhagen, Denmark | Aalborg University | Danish Covid-19 Genome Consortium |
| EPI_ISL_970968 | Lighthouse Lab in Milton Keynes | Wellcome Sanger Institute for the COVID-19 Genomics UK (COG-UK) Consortium | The Lighthouse Lab in Milton Keynes and Alex Alderton, Roberto Amato, Sonia Goncalves, Ewan Harrison, David K. Jackson, Ian Johnston, Dominic Kwiatkowski, Cordelia Langford, John Sillitoe on behalf of the Wellcome Sanger Institute COVID-19 Surveillance Team |
| EPI_ISL_970971, EPI_ISL_970972, EPI_ISL_970974, EPI_ISL_970977, EPI_ISL_970979, EPI_ISL_970981, EPI_ISL_970983, EPI_ISL_970985, EPI_ISL_970988, EPI_ISL_970993, EPI_ISL_970997, EPI_ISL_970998, EPI_ISL_971001, EPI_ISL_971002, EPI_ISL_971003, EPI_ISL_971005, EPI_ISL_971007 | | | |
| see above | Department of Virus and Microbiological Special Diagnostics, Statens Serum Institut, Copenhagen, Denmark | Aalborg University | Danish Covid-19 Genome Consortium |
| EPI_ISL_971008 | Lighthouse Lab in Milton Keynes | Wellcome Sanger Institute for the COVID-19 Genomics UK (COG-UK) Consortium | The Lighthouse Lab in Milton Keynes and Alex Alderton, Roberto Amato, Sonia Goncalves, Ewan Harrison, David K. Jackson, Ian Johnston, Dominic Kwiatkowski, Cordelia Langford, John Sillitoe on behalf of the Wellcome Sanger Institute COVID-19 Surveillance Team |
| EPI_ISL_971010, EPI_ISL_971011, EPI_ISL_971013, EPI_ISL_971015, EPI_ISL_971016, EPI_ISL_971017, EPI_ISL_971018, EPI_ISL_971019, EPI_ISL_971020 | Department of Virus and Microbiological Special Diagnostics, Statens Serum Institut, Copenhagen, Denmark | Aalborg University | Danish Covid-19 Genome Consortium |
| EPI_ISL_971023 | Lighthouse Lab in Milton Keynes | Wellcome Sanger Institute for the COVID-19 Genomics UK (COG-UK) Consortium | The Lighthouse Lab in Milton Keynes and Alex Alderton, Roberto Amato, Sonia Goncalves, Ewan Harrison, David K. Jackson, Ian Johnston, Dominic Kwiatkowski, Cordelia Langford, John Sillitoe on behalf of the Wellcome Sanger Institute COVID-19 Surveillance Team |
| EPI_ISL_971024, EPI_ISL_971025, EPI_ISL_971026, EPI_ISL_971027, EPI_ISL_971029, EPI_ISL_971031, EPI_ISL_971034, EPI_ISL_971038 | Department of Virus and Microbiological Special Diagnostics, Statens Serum Institut, Copenhagen, Denmark | Aalborg University | Danish Covid-19 Genome Consortium |
| EPI_ISL_971039 | Lighthouse Lab in Milton Keynes | Wellcome Sanger Institute for the COVID-19 Genomics UK (COG-UK) Consortium | The Lighthouse Lab in Milton Keynes and Alex Alderton, Roberto Amato, Sonia Goncalves, Ewan Harrison, David K. Jackson, Ian Johnston, Dominic Kwiatkowski, Cordelia Langford, John Sillitoe on behalf of the Wellcome Sanger Institute COVID-19 Surveillance Team |
| EPI_ISL_971041, EPI_ISL_971048 | Department of Virus and Microbiological Special Diagnostics, Statens Serum Institut, Copenhagen, Denmark | Aalborg University | Danish Covid-19 Genome Consortium |
| EPI_ISL_971050, EPI_ISL_971055 | Lighthouse Lab in Milton Keynes | Wellcome Sanger Institute for the COVID-19 Genomics UK (COG-UK) Consortium | The Lighthouse Lab in Milton Keynes and Alex Alderton, Roberto Amato, Sonia Goncalves, Ewan Harrison, David K. Jackson, Ian Johnston, Dominic Kwiatkowski, Cordelia Langford, John Sillitoe on behalf of the Wellcome Sanger Institute COVID-19 Surveillance Team |
| EPI_ISL_971056, EPI_ISL_971058 | Department of Virus and Microbiological Special Diagnostics, Statens Serum Institut, Copenhagen, Denmark | Aalborg University | Danish Covid-19 Genome Consortium |
| EPI_ISL_971059 | Lighthouse Lab in Milton Keynes | Wellcome Sanger Institute for the COVID-19 Genomics UK (COG-UK) Consortium | The Lighthouse Lab in Milton Keynes and Alex Alderton, Roberto Amato, Sonia Goncalves, Ewan Harrison, David K. Jackson, Ian Johnston, Dominic Kwiatkowski, Cordelia Langford, John Sillitoe on behalf of the Wellcome Sanger Institute COVID-19 Surveillance Team |
| EPI_ISL_971062 | Department of Virus and Microbiological Special Diagnostics, Statens Serum Institut, Copenhagen, Denmark | Aalborg University | Danish Covid-19 Genome Consortium |
| EPI_ISL_971063, EPI_ISL_971066 | Lighthouse Lab in Milton Keynes | Wellcome Sanger Institute for the COVID-19 Genomics UK (COG-UK) Consortium | The Lighthouse Lab in Milton Keynes and Alex Alderton, Roberto Amato, Sonia Goncalves, Ewan Harrison, David K. Jackson, Ian Johnston, Dominic Kwiatkowski, Cordelia Langford, John Sillitoe on behalf of the Wellcome Sanger Institute COVID-19 Surveillance Team |
| EPI_ISL_971067 | Department of Virus and Microbiological Special Diagnostics, Statens Serum Institut, Copenhagen, Denmark | Aalborg University | Danish Covid-19 Genome Consortium |
| EPI_ISL_971069 | Lighthouse Lab in Milton Keynes | Wellcome Sanger Institute for the COVID-19 Genomics UK (COG-UK) Consortium | The Lighthouse Lab in Milton Keynes and Alex Alderton, Roberto Amato, Sonia Goncalves, Ewan Harrison, David K. Jackson, Ian Johnston, Dominic Kwiatkowski, Cordelia Langford, John Sillitoe on behalf of the Wellcome Sanger Institute COVID-19 Surveillance Team |
| EPI_ISL_971070, EPI_ISL_971073 | Department of Virus and Microbiological Special Diagnostics, Statens Serum Institut, Copenhagen, Denmark | Aalborg University | Danish Covid-19 Genome Consortium |

| | | | |
|---|---|---|---|
| EPI_ISL_971074 | Lighthouse Lab in Milton Keynes | Wellcome Sanger Institute for the COVID-19 Genomics UK (COG-UK) Consortium | The Lighthouse Lab in Milton Keynes and Alex Alderton, Roberto Amato, Sonia Goncalves, Ewan Harrison, David K. Jackson, Ian Johnston, Dominic Kwiatkowski, Cordelia Langford, John Sillitoe on behalf of the Wellcome Sanger Institute COVID-19 Surveillance Team |
| EPI_ISL_971078 | Department of Virus and Microbiological Special Diagnostics, Statens Serum Institut, Copenhagen, Denmark | Aalborg University | Danish Covid-19 Genome Consortium |
| EPI_ISL_971079 | Lighthouse Lab in Milton Keynes | Wellcome Sanger Institute for the COVID-19 Genomics UK (COG-UK) Consortium | The Lighthouse Lab in Milton Keynes and Alex Alderton, Roberto Amato, Sonia Goncalves, Ewan Harrison, David K. Jackson, Ian Johnston, Dominic Kwiatkowski, Cordelia Langford, John Sillitoe on behalf of the Wellcome Sanger Institute COVID-19 Surveillance Team |
| EPI_ISL_971082 | Department of Virus and Microbiological Special Diagnostics, Statens Serum Institut, Copenhagen, Denmark | Aalborg University | Danish Covid-19 Genome Consortium |
| EPI_ISL_971083, EPI_ISL_971087 | Lighthouse Lab in Milton Keynes | Wellcome Sanger Institute for the COVID-19 Genomics UK (COG-UK) Consortium | The Lighthouse Lab in Milton Keynes and Alex Alderton, Roberto Amato, Sonia Goncalves, Ewan Harrison, David K. Jackson, Ian Johnston, Dominic Kwiatkowski, Cordelia Langford, John Sillitoe on behalf of the Wellcome Sanger Institute COVID-19 Surveillance Team |
| EPI_ISL_971088 | Department of Virus and Microbiological Special Diagnostics, Statens Serum Institut, Copenhagen, Denmark | Aalborg University | Danish Covid-19 Genome Consortium |
| EPI_ISL_971090 | Lighthouse Lab in Milton Keynes | Wellcome Sanger Institute for the COVID-19 Genomics UK (COG-UK) Consortium | The Lighthouse Lab in Milton Keynes and Alex Alderton, Roberto Amato, Sonia Goncalves, Ewan Harrison, David K. Jackson, Ian Johnston, Dominic Kwiatkowski, Cordelia Langford, John Sillitoe on behalf of the Wellcome Sanger Institute COVID-19 Surveillance Team |
| EPI_ISL_971095, EPI_ISL_971098 | Department of Virus and Microbiological Special Diagnostics, Statens Serum Institut, Copenhagen, Denmark | Aalborg University | Danish Covid-19 Genome Consortium |
| EPI_ISL_971099, EPI_ISL_971102 | Lighthouse Lab in Milton Keynes | Wellcome Sanger Institute for the COVID-19 Genomics UK (COG-UK) Consortium | The Lighthouse Lab in Milton Keynes and Alex Alderton, Roberto Amato, Sonia Goncalves, Ewan Harrison, David K. Jackson, Ian Johnston, Dominic Kwiatkowski, Cordelia Langford, John Sillitoe on behalf of the Wellcome Sanger Institute COVID-19 Surveillance Team |
| EPI_ISL_971103, EPI_ISL_971106 | Department of Virus and Microbiological Special Diagnostics, Statens Serum Institut, Copenhagen, Denmark | Aalborg University | Danish Covid-19 Genome Consortium |
| EPI_ISL_971108, EPI_ISL_971109 | Lighthouse Lab in Milton Keynes | Wellcome Sanger Institute for the COVID-19 Genomics UK (COG-UK) Consortium | The Lighthouse Lab in Milton Keynes and Alex Alderton, Roberto Amato, Sonia Goncalves, Ewan Harrison, David K. Jackson, Ian Johnston, Dominic Kwiatkowski, Cordelia Langford, John Sillitoe on behalf of the Wellcome Sanger Institute COVID-19 Surveillance Team |
| EPI_ISL_971110 | Department of Virus and Microbiological Special Diagnostics, Statens Serum Institut, Copenhagen, Denmark | Aalborg University | Danish Covid-19 Genome Consortium |
| EPI_ISL_971113, EPI_ISL_971117 | Lighthouse Lab in Milton Keynes | Wellcome Sanger Institute for the COVID-19 Genomics UK (COG-UK) Consortium | The Lighthouse Lab in Milton Keynes and Alex Alderton, Roberto Amato, Sonia Goncalves, Ewan Harrison, David K. Jackson, Ian Johnston, Dominic Kwiatkowski, Cordelia Langford, John Sillitoe on behalf of the Wellcome Sanger Institute COVID-19 Surveillance Team |
| EPI_ISL_971118 | Department of Virus and Microbiological Special Diagnostics, Statens Serum Institut, Copenhagen, Denmark | Aalborg University | Danish Covid-19 Genome Consortium |
| EPI_ISL_971119, EPI_ISL_971123 | Lighthouse Lab in Milton Keynes | Wellcome Sanger Institute for the COVID-19 Genomics UK (COG-UK) Consortium | The Lighthouse Lab in Milton Keynes and Alex Alderton, Roberto Amato, Sonia Goncalves, Ewan Harrison, David K. Jackson, Ian Johnston, Dominic Kwiatkowski, Cordelia Langford, John Sillitoe on behalf of the Wellcome Sanger Institute COVID-19 Surveillance Team |
| EPI_ISL_971127 | Department of Virus and Microbiological Special Diagnostics, Statens Serum Institut, Copenhagen, Denmark | Aalborg University | Danish Covid-19 Genome Consortium |
| EPI_ISL_971128, EPI_ISL_971130 | Lighthouse Lab in Milton Keynes | Wellcome Sanger Institute for the COVID-19 Genomics UK (COG-UK) Consortium | The Lighthouse Lab in Milton Keynes and Alex Alderton, Roberto Amato, Sonia Goncalves, Ewan Harrison, David K. Jackson, Ian Johnston, Dominic Kwiatkowski, Cordelia Langford, John Sillitoe on behalf of the Wellcome Sanger Institute COVID-19 Surveillance Team |
| EPI_ISL_971131 | Department of Virus and Microbiological Special Diagnostics, Statens Serum Institut, Copenhagen, Denmark | Aalborg University | Danish Covid-19 Genome Consortium |
| EPI_ISL_971134 | Lighthouse Lab in Milton Keynes | Wellcome Sanger Institute for the COVID-19 Genomics UK (COG-UK) Consortium | The Lighthouse Lab in Milton Keynes and Alex Alderton, Roberto Amato, Sonia Goncalves, Ewan Harrison, David K. Jackson, Ian Johnston, Dominic Kwiatkowski, Cordelia Langford, John Sillitoe on behalf of the Wellcome Sanger Institute COVID-19 Surveillance Team |
| EPI_ISL_971136 | Department of Virus and Microbiological Special Diagnostics, Statens Serum Institut, Copenhagen, Denmark | Aalborg University | Danish Covid-19 Genome Consortium |
| EPI_ISL_971138, EPI_ISL_971146, EPI_ISL_971149 | Lighthouse Lab in Milton Keynes | Wellcome Sanger Institute for the COVID-19 Genomics UK (COG-UK) Consortium | The Lighthouse Lab in Milton Keynes and Alex Alderton, Roberto Amato, Sonia Goncalves, Ewan Harrison, David K. Jackson, Ian Johnston, Dominic Kwiatkowski, Cordelia Langford, John Sillitoe on behalf of the Wellcome Sanger Institute COVID-19 Surveillance Team |
| EPI_ISL_971152 | Department of Virus and Microbiological Special Diagnostics, Statens Serum Institut, Copenhagen, Denmark | Aalborg University | Danish Covid-19 Genome Consortium |
| EPI_ISL_971153, EPI_ISL_971156 | Lighthouse Lab in Milton Keynes | Wellcome Sanger Institute for the COVID-19 Genomics UK (COG-UK) Consortium | The Lighthouse Lab in Milton Keynes and Alex Alderton, Roberto Amato, Sonia Goncalves, Ewan Harrison, David K. Jackson, Ian Johnston, Dominic Kwiatkowski, Cordelia Langford, John Sillitoe on behalf of the Wellcome Sanger Institute COVID-19 Surveillance Team |
| EPI_ISL_971159 | Department of Virus and Microbiological Special Diagnostics, Statens Serum Institut, Copenhagen, Denmark | Aalborg University | Danish Covid-19 Genome Consortium |
| EPI_ISL_971161, EPI_ISL_971164, EPI_ISL_971165 | Lighthouse Lab in Milton Keynes | Wellcome Sanger Institute for the COVID-19 Genomics UK (COG-UK) Consortium | The Lighthouse Lab in Milton Keynes and Alex Alderton, Roberto Amato, Sonia Goncalves, Ewan Harrison, David K. Jackson, Ian Johnston, Dominic Kwiatkowski, Cordelia Langford, John Sillitoe on behalf of the Wellcome Sanger Institute COVID-19 Surveillance Team |
| EPI_ISL_971169, EPI_ISL_971172, EPI_ISL_971175, EPI_ISL_971178 | Department of Virus and Microbiological Special Diagnostics, Statens Serum Institut, Copenhagen, Denmark | Aalborg University | Danish Covid-19 Genome Consortium |
| EPI_ISL_971180 | Lighthouse Lab in Milton Keynes | Wellcome Sanger Institute for the COVID-19 Genomics UK (COG-UK) Consortium | The Lighthouse Lab in Milton Keynes and Alex Alderton, Roberto Amato, Sonia Goncalves, Ewan Harrison, David K. Jackson, Ian Johnston, Dominic Kwiatkowski, Cordelia Langford, John Sillitoe on behalf of the Wellcome Sanger Institute COVID-19 Surveillance Team |
| EPI_ISL_971181 | Department of Virus and Microbiological Special Diagnostics, Statens Serum Institut, Copenhagen, Denmark | Aalborg University | Danish Covid-19 Genome Consortium |
| EPI_ISL_971185 | Lighthouse Lab in Milton Keynes | Wellcome Sanger Institute for the COVID-19 Genomics UK (COG-UK) Consortium | The Lighthouse Lab in Milton Keynes and Alex Alderton, Roberto Amato, Sonia Goncalves, Ewan Harrison, David K. Jackson, Ian Johnston, Dominic Kwiatkowski, Cordelia Langford, John Sillitoe on behalf of the Wellcome Sanger Institute COVID-19 Surveillance Team |
| EPI_ISL_971186 | Department of Virus and Microbiological Special Diagnostics, Statens Serum Institut, Copenhagen, Denmark | Aalborg University | Danish Covid-19 Genome Consortium |
| EPI_ISL_971190 | Lighthouse Lab in Milton Keynes | Wellcome Sanger Institute for the COVID-19 Genomics UK (COG-UK) Consortium | The Lighthouse Lab in Milton Keynes and Alex Alderton, Roberto Amato, Sonia Goncalves, Ewan Harrison, David K. Jackson, Ian Johnston, Dominic Kwiatkowski, Cordelia Langford, John Sillitoe on behalf of the Wellcome Sanger Institute COVID-19 Surveillance Team |
| EPI_ISL_971195 | Department of Virus and Microbiological Special Diagnostics, Statens Serum Institut, Copenhagen, Denmark | Aalborg University | Danish Covid-19 Genome Consortium |
| EPI_ISL_971197, EPI_ISL_971200 | Lighthouse Lab in Milton Keynes | Wellcome Sanger Institute for the COVID-19 Genomics UK (COG-UK) Consortium | The Lighthouse Lab in Milton Keynes and Alex Alderton, Roberto Amato, Sonia Goncalves, Ewan Harrison, David K. Jackson, Ian Johnston, Dominic Kwiatkowski, Cordelia Langford, John Sillitoe on behalf of the Wellcome Sanger Institute COVID-19 Surveillance Team |
| EPI_ISL_971201, EPI_ISL_971203 | Department of Virus and Microbiological Special Diagnostics, Statens Serum Institut, Copenhagen, Denmark | Aalborg University | Danish Covid-19 Genome Consortium |
| EPI_ISL_971204 | Lighthouse Lab in Milton Keynes | Wellcome Sanger Institute for the COVID-19 Genomics UK (COG-UK) Consortium | The Lighthouse Lab in Milton Keynes and Alex Alderton, Roberto Amato, Sonia Goncalves, Ewan Harrison, David K. Jackson, Ian Johnston, Dominic Kwiatkowski, Cordelia Langford, John Sillitoe on behalf of the Wellcome Sanger Institute COVID-19 Surveillance Team |

| | | | |
|---|---|---|---|
| EPI_ISL_971207 | Department of Virus and Microbiological Special Diagnostics, Statens Serum Institut, Copenhagen, Denmark | Aalborg University | Danish Covid-19 Genome Consortium |
| EPI_ISL_971209, EPI_ISL_971211 | Lighthouse Lab in Milton Keynes | Wellcome Sanger Institute for the COVID-19 Genomics UK (COG-UK) Consortium | The Lighthouse Lab in Milton Keynes and Alex Alderton, Roberto Amato, Sonia Goncalves, Ewan Harrison, David K. Jackson, Ian Johnston, Dominic Kwiatkowski, Cordelia Langford, John Sillitoe on behalf of the Wellcome Sanger Institute COVID-19 Surveillance Team |
| EPI_ISL_971213 | Department of Virus and Microbiological Special Diagnostics, Statens Serum Institut, Copenhagen, Denmark | Aalborg University | Danish Covid-19 Genome Consortium |
| EPI_ISL_971215, EPI_ISL_971218, EPI_ISL_971221 | Lighthouse Lab in Milton Keynes | Wellcome Sanger Institute for the COVID-19 Genomics UK (COG-UK) Consortium | The Lighthouse Lab in Milton Keynes and Alex Alderton, Roberto Amato, Sonia Goncalves, Ewan Harrison, David K. Jackson, Ian Johnston, Dominic Kwiatkowski, Cordelia Langford, John Sillitoe on behalf of the Wellcome Sanger Institute COVID-19 Surveillance Team |
| EPI_ISL_971223 | Department of Virus and Microbiological Special Diagnostics, Statens Serum Institut, Copenhagen, Denmark | Aalborg University | Danish Covid-19 Genome Consortium |
| EPI_ISL_971224, EPI_ISL_971228, EPI_ISL_971234, EPI_ISL_971236 | Lighthouse Lab in Milton Keynes | Wellcome Sanger Institute for the COVID-19 Genomics UK (COG-UK) Consortium | The Lighthouse Lab in Milton Keynes and Alex Alderton, Roberto Amato, Sonia Goncalves, Ewan Harrison, David K. Jackson, Ian Johnston, Dominic Kwiatkowski, Cordelia Langford, John Sillitoe on behalf of the Wellcome Sanger Institute COVID-19 Surveillance Team |
| EPI_ISL_971237, EPI_ISL_971244 | Department of Virus and Microbiological Special Diagnostics, Statens Serum Institut, Copenhagen, Denmark | Aalborg University | Danish Covid-19 Genome Consortium |
| EPI_ISL_971246 | Lighthouse Lab in Milton Keynes | Wellcome Sanger Institute for the COVID-19 Genomics UK (COG-UK) Consortium | The Lighthouse Lab in Milton Keynes and Alex Alderton, Roberto Amato, Sonia Goncalves, Ewan Harrison, David K. Jackson, Ian Johnston, Dominic Kwiatkowski, Cordelia Langford, John Sillitoe on behalf of the Wellcome Sanger Institute COVID-19 Surveillance Team |
| EPI_ISL_971247 | Department of Virus and Microbiological Special Diagnostics, Statens Serum Institut, Copenhagen, Denmark | Aalborg University | Danish Covid-19 Genome Consortium |
| EPI_ISL_971248, EPI_ISL_971251 | Lighthouse Lab in Milton Keynes | Wellcome Sanger Institute for the COVID-19 Genomics UK (COG-UK) Consortium | The Lighthouse Lab in Milton Keynes and Alex Alderton, Roberto Amato, Sonia Goncalves, Ewan Harrison, David K. Jackson, Ian Johnston, Dominic Kwiatkowski, Cordelia Langford, John Sillitoe on behalf of the Wellcome Sanger Institute COVID-19 Surveillance Team |
| EPI_ISL_971252 | Department of Virus and Microbiological Special Diagnostics, Statens Serum Institut, Copenhagen, Denmark | Aalborg University | Danish Covid-19 Genome Consortium |
| EPI_ISL_971254 | Lighthouse Lab in Milton Keynes | Wellcome Sanger Institute for the COVID-19 Genomics UK (COG-UK) Consortium | The Lighthouse Lab in Milton Keynes and Alex Alderton, Roberto Amato, Sonia Goncalves, Ewan Harrison, David K. Jackson, Ian Johnston, Dominic Kwiatkowski, Cordelia Langford, John Sillitoe on behalf of the Wellcome Sanger Institute COVID-19 Surveillance Team |
| EPI_ISL_971257 | Department of Virus and Microbiological Special Diagnostics, Statens Serum Institut, Copenhagen, Denmark | Aalborg University | Danish Covid-19 Genome Consortium |
| EPI_ISL_971258 | Lighthouse Lab in Milton Keynes | Wellcome Sanger Institute for the COVID-19 Genomics UK (COG-UK) Consortium | The Lighthouse Lab in Milton Keynes and Alex Alderton, Roberto Amato, Sonia Goncalves, Ewan Harrison, David K. Jackson, Ian Johnston, Dominic Kwiatkowski, Cordelia Langford, John Sillitoe on behalf of the Wellcome Sanger Institute COVID-19 Surveillance Team |
| EPI_ISL_971263 | Department of Virus and Microbiological Special Diagnostics, Statens Serum Institut, Copenhagen, Denmark | Aalborg University | Danish Covid-19 Genome Consortium |
| EPI_ISL_971264, EPI_ISL_971267 | Lighthouse Lab in Milton Keynes | Wellcome Sanger Institute for the COVID-19 Genomics UK (COG-UK) Consortium | The Lighthouse Lab in Milton Keynes and Alex Alderton, Roberto Amato, Sonia Goncalves, Ewan Harrison, David K. Jackson, Ian Johnston, Dominic Kwiatkowski, Cordelia Langford, John Sillitoe on behalf of the Wellcome Sanger Institute COVID-19 Surveillance Team |
| EPI_ISL_971270, EPI_ISL_971272 | Department of Virus and Microbiological Special Diagnostics, Statens Serum Institut, Copenhagen, Denmark | Aalborg University | Danish Covid-19 Genome Consortium |
| EPI_ISL_971273 | Lighthouse Lab in Milton Keynes | Wellcome Sanger Institute for the COVID-19 Genomics UK (COG-UK) Consortium | The Lighthouse Lab in Milton Keynes and Alex Alderton, Roberto Amato, Sonia Goncalves, Ewan Harrison, David K. Jackson, Ian Johnston, Dominic Kwiatkowski, Cordelia Langford, John Sillitoe on behalf of the Wellcome Sanger Institute COVID-19 Surveillance Team |
| EPI_ISL_971275 | Department of Virus and Microbiological Special Diagnostics, Statens Serum Institut, Copenhagen, Denmark | Aalborg University | Danish Covid-19 Genome Consortium |
| EPI_ISL_971277 | Lighthouse Lab in Milton Keynes | Wellcome Sanger Institute for the COVID-19 Genomics UK (COG-UK) Consortium | The Lighthouse Lab in Milton Keynes and Alex Alderton, Roberto Amato, Sonia Goncalves, Ewan Harrison, David K. Jackson, Ian Johnston, Dominic Kwiatkowski, Cordelia Langford, John Sillitoe on behalf of the Wellcome Sanger Institute COVID-19 Surveillance Team |
| EPI_ISL_971283 | Department of Virus and Microbiological Special Diagnostics, Statens Serum Institut, Copenhagen, Denmark | Aalborg University | Danish Covid-19 Genome Consortium |
| EPI_ISL_971284, EPI_ISL_971286 | Lighthouse Lab in Milton Keynes | Wellcome Sanger Institute for the COVID-19 Genomics UK (COG-UK) Consortium | The Lighthouse Lab in Milton Keynes and Alex Alderton, Roberto Amato, Sonia Goncalves, Ewan Harrison, David K. Jackson, Ian Johnston, Dominic Kwiatkowski, Cordelia Langford, John Sillitoe on behalf of the Wellcome Sanger Institute COVID-19 Surveillance Team |
| EPI_ISL_971287 | Department of Virus and Microbiological Special Diagnostics, Statens Serum Institut, Copenhagen, Denmark | Aalborg University | Danish Covid-19 Genome Consortium |
| EPI_ISL_971289, EPI_ISL_971295 | Lighthouse Lab in Milton Keynes | Wellcome Sanger Institute for the COVID-19 Genomics UK (COG-UK) Consortium | The Lighthouse Lab in Milton Keynes and Alex Alderton, Roberto Amato, Sonia Goncalves, Ewan Harrison, David K. Jackson, Ian Johnston, Dominic Kwiatkowski, Cordelia Langford, John Sillitoe on behalf of the Wellcome Sanger Institute COVID-19 Surveillance Team |
| EPI_ISL_971296 | Department of Virus and Microbiological Special Diagnostics, Statens Serum Institut, Copenhagen, Denmark | Aalborg University | Danish Covid-19 Genome Consortium |
| EPI_ISL_971301 | Lighthouse Lab in Milton Keynes | Wellcome Sanger Institute for the COVID-19 Genomics UK (COG-UK) Consortium | The Lighthouse Lab in Milton Keynes and Alex Alderton, Roberto Amato, Sonia Goncalves, Ewan Harrison, David K. Jackson, Ian Johnston, Dominic Kwiatkowski, Cordelia Langford, John Sillitoe on behalf of the Wellcome Sanger Institute COVID-19 Surveillance Team |
| EPI_ISL_971302 | Department of Virus and Microbiological Special Diagnostics, Statens Serum Institut, Copenhagen, Denmark | Aalborg University | Danish Covid-19 Genome Consortium |
| EPI_ISL_971304, EPI_ISL_971307 | Lighthouse Lab in Milton Keynes | Wellcome Sanger Institute for the COVID-19 Genomics UK (COG-UK) Consortium | The Lighthouse Lab in Milton Keynes and Alex Alderton, Roberto Amato, Sonia Goncalves, Ewan Harrison, David K. Jackson, Ian Johnston, Dominic Kwiatkowski, Cordelia Langford, John Sillitoe on behalf of the Wellcome Sanger Institute COVID-19 Surveillance Team |
| EPI_ISL_971309 | Department of Virus and Microbiological Special Diagnostics, Statens Serum Institut, Copenhagen, Denmark | Aalborg University | Danish Covid-19 Genome Consortium |
| EPI_ISL_971310 | Lighthouse Lab in Milton Keynes | Wellcome Sanger Institute for the COVID-19 Genomics UK (COG-UK) Consortium | The Lighthouse Lab in Milton Keynes and Alex Alderton, Roberto Amato, Sonia Goncalves, Ewan Harrison, David K. Jackson, Ian Johnston, Dominic Kwiatkowski, Cordelia Langford, John Sillitoe on behalf of the Wellcome Sanger Institute COVID-19 Surveillance Team |
| EPI_ISL_971312 | Department of Virus and Microbiological Special Diagnostics, Statens Serum Institut, Copenhagen, Denmark | Aalborg University | Danish Covid-19 Genome Consortium |
| EPI_ISL_971313, EPI_ISL_971316 | Lighthouse Lab in Milton Keynes | Wellcome Sanger Institute for the COVID-19 Genomics UK (COG-UK) Consortium | The Lighthouse Lab in Milton Keynes and Alex Alderton, Roberto Amato, Sonia Goncalves, Ewan Harrison, David K. Jackson, Ian Johnston, Dominic Kwiatkowski, Cordelia Langford, John Sillitoe on behalf of the Wellcome Sanger Institute COVID-19 Surveillance Team |
| EPI_ISL_971318 | Department of Virus and Microbiological Special Diagnostics, Statens Serum Institut, Copenhagen, Denmark | Aalborg University | Danish Covid-19 Genome Consortium |
| EPI_ISL_971319, EPI_ISL_971320 | Lighthouse Lab in Milton Keynes | Wellcome Sanger Institute for the COVID-19 Genomics UK (COG-UK) Consortium | The Lighthouse Lab in Milton Keynes and Alex Alderton, Roberto Amato, Sonia Goncalves, Ewan Harrison, David K. Jackson, Ian Johnston, Dominic Kwiatkowski, Cordelia Langford, John Sillitoe on behalf of the Wellcome Sanger Institute COVID-19 Surveillance Team |
| EPI_ISL_971322, EPI_ISL_971323 | Department of Virus and Microbiological Special Diagnostics, Statens Serum Institut, Copenhagen, Denmark | Aalborg University | Danish Covid-19 Genome Consortium |

| | | | |
|---|---|---|---|
| EPI_ISL_971324 | Lighthouse Lab in Milton Keynes | Wellcome Sanger Institute for the COVID-19 Genomics UK (COG-UK) Consortium | The Lighthouse Lab in Milton Keynes and Alex Alderton, Roberto Amato, Sonia Goncalves, Ewan Harrison, David K. Jackson, Ian Johnston, Dominic Kwiatkowski, Cordelia Langford, John Sillitoe on behalf of the Wellcome Sanger Institute COVID-19 Surveillance Team |
| EPI_ISL_971329 | Department of Virus and Microbiological Special Diagnostics, Statens Serum Institut, Copenhagen, Denmark | Aalborg University | Danish Covid-19 Genome Consortium |
| EPI_ISL_971330 | Lighthouse Lab in Milton Keynes | Wellcome Sanger Institute for the COVID-19 Genomics UK (COG-UK) Consortium | The Lighthouse Lab in Milton Keynes and Alex Alderton, Roberto Amato, Sonia Goncalves, Ewan Harrison, David K. Jackson, Ian Johnston, Dominic Kwiatkowski, Cordelia Langford, John Sillitoe on behalf of the Wellcome Sanger Institute COVID-19 Surveillance Team |
| EPI_ISL_971332 | Department of Virus and Microbiological Special Diagnostics, Statens Serum Institut, Copenhagen, Denmark | Aalborg University | Danish Covid-19 Genome Consortium |
| EPI_ISL_971336 | Lighthouse Lab in Milton Keynes | Wellcome Sanger Institute for the COVID-19 Genomics UK (COG-UK) Consortium | The Lighthouse Lab in Milton Keynes and Alex Alderton, Roberto Amato, Sonia Goncalves, Ewan Harrison, David K. Jackson, Ian Johnston, Dominic Kwiatkowski, Cordelia Langford, John Sillitoe on behalf of the Wellcome Sanger Institute COVID-19 Surveillance Team |
| EPI_ISL_971338 | Department of Virus and Microbiological Special Diagnostics, Statens Serum Institut, Copenhagen, Denmark | Aalborg University | Danish Covid-19 Genome Consortium |
| EPI_ISL_971343, EPI_ISL_971350, EPI_ISL_971352, EPI_ISL_971357 | Lighthouse Lab in Milton Keynes | Wellcome Sanger Institute for the COVID-19 Genomics UK (COG-UK) Consortium | The Lighthouse Lab in Milton Keynes and Alex Alderton, Roberto Amato, Sonia Goncalves, Ewan Harrison, David K. Jackson, Ian Johnston, Dominic Kwiatkowski, Cordelia Langford, John Sillitoe on behalf of the Wellcome Sanger Institute COVID-19 Surveillance Team |
| EPI_ISL_971358, EPI_ISL_971363, EPI_ISL_971368, EPI_ISL_971371 | Department of Virus and Microbiological Special Diagnostics, Statens Serum Institut, Copenhagen, Denmark | Aalborg University | Danish Covid-19 Genome Consortium |
| EPI_ISL_971373, EPI_ISL_971376, EPI_ISL_971378 | Lighthouse Lab in Milton Keynes | Wellcome Sanger Institute for the COVID-19 Genomics UK (COG-UK) Consortium | The Lighthouse Lab in Milton Keynes and Alex Alderton, Roberto Amato, Sonia Goncalves, Ewan Harrison, David K. Jackson, Ian Johnston, Dominic Kwiatkowski, Cordelia Langford, John Sillitoe on behalf of the Wellcome Sanger Institute COVID-19 Surveillance Team |
| EPI_ISL_971379 | Department of Virus and Microbiological Special Diagnostics, Statens Serum Institut, Copenhagen, Denmark | Aalborg University | Danish Covid-19 Genome Consortium |
| EPI_ISL_971382 | Lighthouse Lab in Milton Keynes | Wellcome Sanger Institute for the COVID-19 Genomics UK (COG-UK) Consortium | The Lighthouse Lab in Milton Keynes and Alex Alderton, Roberto Amato, Sonia Goncalves, Ewan Harrison, David K. Jackson, Ian Johnston, Dominic Kwiatkowski, Cordelia Langford, John Sillitoe on behalf of the Wellcome Sanger Institute COVID-19 Surveillance Team |
| EPI_ISL_971384 | Department of Virus and Microbiological Special Diagnostics, Statens Serum Institut, Copenhagen, Denmark | Aalborg University | Danish Covid-19 Genome Consortium |
| EPI_ISL_971388 | Lighthouse Lab in Milton Keynes | Wellcome Sanger Institute for the COVID-19 Genomics UK (COG-UK) Consortium | The Lighthouse Lab in Milton Keynes and Alex Alderton, Roberto Amato, Sonia Goncalves, Ewan Harrison, David K. Jackson, Ian Johnston, Dominic Kwiatkowski, Cordelia Langford, John Sillitoe on behalf of the Wellcome Sanger Institute COVID-19 Surveillance Team |
| EPI_ISL_971389 | Department of Virus and Microbiological Special Diagnostics, Statens Serum Institut, Copenhagen, Denmark | Aalborg University | Danish Covid-19 Genome Consortium |
| EPI_ISL_971392, EPI_ISL_971397 | Lighthouse Lab in Milton Keynes | Wellcome Sanger Institute for the COVID-19 Genomics UK (COG-UK) Consortium | The Lighthouse Lab in Milton Keynes and Alex Alderton, Roberto Amato, Sonia Goncalves, Ewan Harrison, David K. Jackson, Ian Johnston, Dominic Kwiatkowski, Cordelia Langford, John Sillitoe on behalf of the Wellcome Sanger Institute COVID-19 Surveillance Team |
| EPI_ISL_971398 | Department of Virus and Microbiological Special Diagnostics, Statens Serum Institut, Copenhagen, Denmark | Aalborg University | Danish Covid-19 Genome Consortium |
| EPI_ISL_971400 | Lighthouse Lab in Milton Keynes | Wellcome Sanger Institute for the COVID-19 Genomics UK (COG-UK) Consortium | The Lighthouse Lab in Milton Keynes and Alex Alderton, Roberto Amato, Sonia Goncalves, Ewan Harrison, David K. Jackson, Ian Johnston, Dominic Kwiatkowski, Cordelia Langford, John Sillitoe on behalf of the Wellcome Sanger Institute COVID-19 Surveillance Team |
| EPI_ISL_971401 | Department of Virus and Microbiological Special Diagnostics, Statens Serum Institut, Copenhagen, Denmark | Aalborg University | Danish Covid-19 Genome Consortium |
| EPI_ISL_971402, EPI_ISL_971405, EPI_ISL_971407 | Lighthouse Lab in Milton Keynes | Wellcome Sanger Institute for the COVID-19 Genomics UK (COG-UK) Consortium | The Lighthouse Lab in Milton Keynes and Alex Alderton, Roberto Amato, Sonia Goncalves, Ewan Harrison, David K. Jackson, Ian Johnston, Dominic Kwiatkowski, Cordelia Langford, John Sillitoe on behalf of the Wellcome Sanger Institute COVID-19 Surveillance Team |
| EPI_ISL_971408 | Department of Virus and Microbiological Special Diagnostics, Statens Serum Institut, Copenhagen, Denmark | Aalborg University | Danish Covid-19 Genome Consortium |
| EPI_ISL_971410 | Lighthouse Lab in Milton Keynes | Wellcome Sanger Institute for the COVID-19 Genomics UK (COG-UK) Consortium | The Lighthouse Lab in Milton Keynes and Alex Alderton, Roberto Amato, Sonia Goncalves, Ewan Harrison, David K. Jackson, Ian Johnston, Dominic Kwiatkowski, Cordelia Langford, John Sillitoe on behalf of the Wellcome Sanger Institute COVID-19 Surveillance Team |
| EPI_ISL_971411 | Department of Virus and Microbiological Special Diagnostics, Statens Serum Institut, Copenhagen, Denmark | Aalborg University | Danish Covid-19 Genome Consortium |
| EPI_ISL_971413 | Lighthouse Lab in Milton Keynes | Wellcome Sanger Institute for the COVID-19 Genomics UK (COG-UK) Consortium | The Lighthouse Lab in Milton Keynes and Alex Alderton, Roberto Amato, Sonia Goncalves, Ewan Harrison, David K. Jackson, Ian Johnston, Dominic Kwiatkowski, Cordelia Langford, John Sillitoe on behalf of the Wellcome Sanger Institute COVID-19 Surveillance Team |
| EPI_ISL_971415 | Department of Virus and Microbiological Special Diagnostics, Statens Serum Institut, Copenhagen, Denmark | Aalborg University | Danish Covid-19 Genome Consortium |
| EPI_ISL_971417 | Lighthouse Lab in Milton Keynes | Wellcome Sanger Institute for the COVID-19 Genomics UK (COG-UK) Consortium | The Lighthouse Lab in Milton Keynes and Alex Alderton, Roberto Amato, Sonia Goncalves, Ewan Harrison, David K. Jackson, Ian Johnston, Dominic Kwiatkowski, Cordelia Langford, John Sillitoe on behalf of the Wellcome Sanger Institute COVID-19 Surveillance Team |
| EPI_ISL_971419 | Department of Virus and Microbiological Special Diagnostics, Statens Serum Institut, Copenhagen, Denmark | Aalborg University | Danish Covid-19 Genome Consortium |
| EPI_ISL_971420 | Lighthouse Lab in Milton Keynes | Wellcome Sanger Institute for the COVID-19 Genomics UK (COG-UK) Consortium | The Lighthouse Lab in Milton Keynes and Alex Alderton, Roberto Amato, Sonia Goncalves, Ewan Harrison, David K. Jackson, Ian Johnston, Dominic Kwiatkowski, Cordelia Langford, John Sillitoe on behalf of the Wellcome Sanger Institute COVID-19 Surveillance Team |
| EPI_ISL_971422 | Department of Virus and Microbiological Special Diagnostics, Statens Serum Institut, Copenhagen, Denmark | Aalborg University | Danish Covid-19 Genome Consortium |
| EPI_ISL_971423 | Lighthouse Lab in Milton Keynes | Wellcome Sanger Institute for the COVID-19 Genomics UK (COG-UK) Consortium | The Lighthouse Lab in Milton Keynes and Alex Alderton, Roberto Amato, Sonia Goncalves, Ewan Harrison, David K. Jackson, Ian Johnston, Dominic Kwiatkowski, Cordelia Langford, John Sillitoe on behalf of the Wellcome Sanger Institute COVID-19 Surveillance Team |
| EPI_ISL_971425 | Lighthouse Lab in Cambridge | Wellcome Sanger Institute for the COVID-19 Genomics UK (COG-UK) Consortium | Rob Howes, The Lighthouse Lab in Cambridge and Alex Alderton, Roberto Amato, Sonia Goncalves, Ewan Harrison, David K. Jackson, Ian Johnston, Dominic Kwiatkowski, Cordelia Langford, John Sillitoe on behalf of the Wellcome Sanger Institute COVID-19 Surveillance Team |
| EPI_ISL_971426 | Department of Virus and Microbiological Special Diagnostics, Statens Serum Institut, Copenhagen, Denmark | Aalborg University | Danish Covid-19 Genome Consortium |
| EPI_ISL_971428 | Lighthouse Lab in Milton Keynes | Wellcome Sanger Institute for the COVID-19 Genomics UK (COG-UK) Consortium | The Lighthouse Lab in Milton Keynes and Alex Alderton, Roberto Amato, Sonia Goncalves, Ewan Harrison, David K. Jackson, Ian Johnston, Dominic Kwiatkowski, Cordelia Langford, John Sillitoe on behalf of the Wellcome Sanger Institute COVID-19 Surveillance Team |

EPI_ISL_971429, EPI_ISL_971434, EPI_ISL_971441, EPI_ISL_971443, EPI_ISL_971445, EPI_ISL_971450, EPI_ISL_971452, EPI_ISL_971455, EPI_ISL_971456, EPI_ISL_971458, EPI_ISL_971460, EPI_ISL_971462, EPI_ISL_971463, EPI_ISL_971464, EPI_ISL_971465, EPI_ISL_971466, EPI_ISL_971468, EPI_ISL_971469, EPI_ISL_971471, EPI_ISL_971472, EPI_ISL_971474, EPI_ISL_971475, EPI_ISL_971476, EPI_ISL_971479, EPI_ISL_971481, EPI_ISL_971482, EPI_ISL_971483, EPI_ISL_971484, EPI_ISL_971486, EPI_ISL_971487, EPI_ISL_971488, EPI_ISL_971489, EPI_ISL_971490, EPI_ISL_971491, EPI_ISL_971492, EPI_ISL_971493, EPI_ISL_971495, EPI_ISL_971498, EPI_ISL_971499, EPI_ISL_971501, EPI_ISL_971505, EPI_ISL_971506, EPI_ISL_971507, EPI_ISL_971510, EPI_ISL_971511, EPI_ISL_971512, EPI_ISL_971514, EPI_ISL_971517, EPI_ISL_971520, EPI_ISL_971521, EPI_ISL_971522, EPI_ISL_971523, EPI_ISL_971524, EPI_ISL_971525, EPI_ISL_971526, EPI_ISL_971528, EPI_ISL_971530, EPI_ISL_971531, EPI_ISL_971533, EPI_ISL_971535, EPI_ISL_971536, EPI_ISL_971537, EPI_ISL_971538, EPI_ISL_971540, EPI_ISL_971541, EPI_ISL_971543, EPI_ISL_971544, EPI_ISL_971545, EPI_ISL_971546, EPI_ISL_971551, EPI_ISL_971552, EPI_ISL_971554, EPI_ISL_971555, EPI_ISL_971559, EPI_ISL_971561, EPI_ISL_971562, EPI_ISL_971563, EPI_ISL_971564, EPI_ISL_971567, EPI_ISL_971568, EPI_ISL_971569, EPI_ISL_971570, EPI_ISL_971571, EPI_ISL_971572, EPI_ISL_971573, EPI_ISL_971574, EPI_ISL_971576, EPI_ISL_971577, EPI_ISL_971582, EPI_ISL_971583, EPI_ISL_971584, EPI_ISL_971585, EPI_ISL_971587, EPI_ISL_971588, EPI_ISL_971589, EPI_ISL_971590, EPI_ISL_971592, EPI_ISL_971593, EPI_ISL_971594, EPI_ISL_971595, EPI_ISL_971596, EPI_ISL_971597, EPI_ISL_971598, EPI_ISL_971599, EPI_ISL_971600, EPI_ISL_971601, EPI_ISL_971602, EPI_ISL_971603, EPI_ISL_971604, EPI_ISL_971605, EPI_ISL_971606, EPI_ISL_971607, EPI_ISL_971609, EPI_ISL_971611, EPI_ISL_971612, EPI_ISL_971618, EPI_ISL_971619, EPI_ISL_971620, EPI_ISL_971621, EPI_ISL_971622, EPI_ISL_971624, EPI_ISL_971626, EPI_ISL_971627, EPI_ISL_971629, EPI_ISL_971630, EPI_ISL_971631, EPI_ISL_971634, EPI_ISL_971636, EPI_ISL_971637, EPI_ISL_971639, EPI_ISL_971641, EPI_ISL_971644, EPI_ISL_971645, EPI_ISL_971646, EPI_ISL_971647, EPI_ISL_971651, EPI_ISL_971652, EPI_ISL_971653, EPI_ISL_971654, EPI_ISL_971656, EPI_ISL_971657, EPI_ISL_971658, EPI_ISL_971659, EPI_ISL_971660,

EPI_ISL_971661, EPI_ISL_971662, EPI_ISL_971663, EPI_ISL_971664, EPI_ISL_971665, EPI_ISL_971666, EPI_ISL_971667, EPI_ISL_971668, EPI_ISL_971669, EPI_ISL_971670, EPI_ISL_971671, EPI_ISL_971672, EPI_ISL_971676, EPI_ISL_971677, EPI_ISL_971678, EPI_ISL_971679, EPI_ISL_971681, EPI_ISL_971682, EPI_ISL_971683, EPI_ISL_971685, EPI_ISL_971687, EPI_ISL_971688, EPI_ISL_971689, EPI_ISL_971691, EPI_ISL_971692, EPI_ISL_971693, EPI_ISL_971694, EPI_ISL_971696, EPI_ISL_971698, EPI_ISL_971700, EPI_ISL_971702, EPI_ISL_971703, EPI_ISL_971704, EPI_ISL_971707, EPI_ISL_971709, EPI_ISL_971712, EPI_ISL_971713, EPI_ISL_971717, EPI_ISL_971718, EPI_ISL_971720, EPI_ISL_971721, EPI_ISL_971722, EPI_ISL_971725, EPI_ISL_971726, EPI_ISL_971727, EPI_ISL_971728, EPI_ISL_971729, EPI_ISL_971731, EPI_ISL_971732, EPI_ISL_971734, EPI_ISL_971735, EPI_ISL_971740, EPI_ISL_971741, EPI_ISL_971744, EPI_ISL_971746, EPI_ISL_971747, EPI_ISL_971748, EPI_ISL_971750, EPI_ISL_971752, EPI_ISL_971753, EPI_ISL_971754, EPI_ISL_971756, EPI_ISL_971758, EPI_ISL_971759, EPI_ISL_971760, EPI_ISL_971762, EPI_ISL_971763, EPI_ISL_971764, EPI_ISL_971765, EPI_ISL_971766, EPI_ISL_971770, EPI_ISL_971771, EPI_ISL_971772, EPI_ISL_971774, EPI_ISL_971775, EPI_ISL_971776, EPI_ISL_971777, EPI_ISL_971778, EPI_ISL_971779, EPI_ISL_971780, EPI_ISL_971781, EPI_ISL_971782, EPI_ISL_971783, EPI_ISL_971786, EPI_ISL_971787, EPI_ISL_971788, EPI_ISL_971789, EPI_ISL_971792, EPI_ISL_971794, EPI_ISL_971795, EPI_ISL_971796, EPI_ISL_971797, EPI_ISL_971799, EPI_ISL_971800, EPI_ISL_971801, EPI_ISL_971802, EPI_ISL_971803, EPI_ISL_971804, EPI_ISL_971806, EPI_ISL_971807, EPI_ISL_971809, EPI_ISL_971810, EPI_ISL_971812, EPI_ISL_971813, EPI_ISL_971814, EPI_ISL_971816, EPI_ISL_971817, EPI_ISL_971818, EPI_ISL_971820, EPI_ISL_971821, EPI_ISL_971823, EPI_ISL_971824, EPI_ISL_971825, EPI_ISL_971826, EPI_ISL_971827, EPI_ISL_971828, EPI_ISL_971829, EPI_ISL_971830, EPI_ISL_971831, EPI_ISL_971833, EPI_ISL_971835, EPI_ISL_971836, EPI_ISL_971837, EPI_ISL_971838, EPI_ISL_971840, EPI_ISL_971841, EPI_ISL_971842, EPI_ISL_971843, EPI_ISL_971845, EPI_ISL_971847, EPI_ISL_971848, EPI_ISL_971850, EPI_ISL_971851, EPI_ISL_971852, EPI_ISL_971853, EPI_ISL_971855, EPI_ISL_971858, EPI_ISL_971860, EPI_ISL_971863, EPI_ISL_971864, EPI_ISL_971866, EPI_ISL_971868, EPI_ISL_971869, EPI_ISL_971870, EPI_ISL_971871, EPI_ISL_971873, EPI_ISL_971874, EPI_ISL_971878, EPI_ISL_971879, EPI_ISL_971881, EPI_ISL_971882, EPI_ISL_971883, EPI_ISL_971884, EPI_ISL_971887, EPI_ISL_971888, EPI_ISL_971889, EPI_ISL_971890, EPI_ISL_971891, EPI_ISL_971892, EPI_ISL_971893, EPI_ISL_971894, EPI_ISL_971895, EPI_ISL_971896, EPI_ISL_971897, EPI_ISL_971898, EPI_ISL_971900, EPI_ISL_971901, EPI_ISL_971902, EPI_ISL_971903, EPI_ISL_971904, EPI_ISL_971905, EPI_ISL_971907, EPI_ISL_971909, EPI_ISL_971912, EPI_ISL_971913, EPI_ISL_971916, EPI_ISL_971917, EPI_ISL_971918, EPI_ISL_971922, EPI_ISL_971923, EPI_ISL_971924, EPI_ISL_971926, EPI_ISL_971927, EPI_ISL_971928, EPI_ISL_971929, EPI_ISL_971930, EPI_ISL_971932, EPI_ISL_971935, EPI_ISL_971937, EPI_ISL_971940, EPI_ISL_971941, EPI_ISL_971944, EPI_ISL_971947, EPI_ISL_971948, EPI_ISL_971950, EPI_ISL_971952, EPI_ISL_971953, EPI_ISL_971955, EPI_ISL_971956, EPI_ISL_971957, EPI_ISL_971958, EPI_ISL_971959, EPI_ISL_971961, EPI_ISL_971964, EPI_ISL_971965, EPI_ISL_971967, EPI_ISL_971969, EPI_ISL_971972, EPI_ISL_971974, EPI_ISL_971976, EPI_ISL_971979, EPI_ISL_971980, EPI_ISL_971982, EPI_ISL_971983, EPI_ISL_971984, EPI_ISL_971989, EPI_ISL_971992, EPI_ISL_971993, EPI_ISL_971994, EPI_ISL_971998, EPI_ISL_972000, EPI_ISL_972001, EPI_ISL_972002, EPI_ISL_972003, EPI_ISL_972004, EPI_ISL_972005, EPI_ISL_972007, EPI_ISL_972009, EPI_ISL_972010, EPI_ISL_972011, EPI_ISL_972012, EPI_ISL_972013, EPI_ISL_972014, EPI_ISL_972017, EPI_ISL_972019, EPI_ISL_972020, EPI_ISL_972021, EPI_ISL_972022, EPI_ISL_972024, EPI_ISL_972026, EPI_ISL_972028, EPI_ISL_972029, EPI_ISL_972030, EPI_ISL_972031, EPI_ISL_972033, EPI_ISL_972035, EPI_ISL_972036, EPI_ISL_972037, EPI_ISL_972038, EPI_ISL_972039, EPI_ISL_972040, EPI_ISL_972041, EPI_ISL_972042, EPI_ISL_972043, EPI_ISL_972045, EPI_ISL_972047, EPI_ISL_972048, EPI_ISL_972052, EPI_ISL_972056, EPI_ISL_972057, EPI_ISL_972059, EPI_ISL_972061, EPI_ISL_972062, EPI_ISL_972063, EPI_ISL_972064, EPI_ISL_972065, EPI_ISL_972066, EPI_ISL_972067, EPI_ISL_972069, EPI_ISL_972071, EPI_ISL_972073, EPI_ISL_972075, EPI_ISL_972076, EPI_ISL_972077, EPI_ISL_972078, EPI_ISL_972079, EPI_ISL_972081, EPI_ISL_972083, EPI_ISL_972084, EPI_ISL_972086, EPI_ISL_972087, EPI_ISL_972095, EPI_ISL_972096, EPI_ISL_972097, EPI_ISL_972099, EPI_ISL_972100, EPI_ISL_972103, EPI_ISL_972104, EPI_ISL_972106, EPI_ISL_972107, EPI_ISL_972108, EPI_ISL_972110, EPI_ISL_972112, EPI_ISL_972113, EPI_ISL_972115, EPI_ISL_972120, EPI_ISL_972121, EPI_ISL_972122, EPI_ISL_972124, EPI_ISL_972125, EPI_ISL_972126, EPI_ISL_972127, EPI_ISL_972128, EPI_ISL_972129, EPI_ISL_972133, EPI_ISL_972134, EPI_ISL_972140, EPI_ISL_972141, EPI_ISL_972142, EPI_ISL_972144, EPI_ISL_972145, EPI_ISL_972146, EPI_ISL_972147, EPI_ISL_972148, EPI_ISL_972149, EPI_ISL_972151, EPI_ISL_972152, EPI_ISL_972154, EPI_ISL_972155, EPI_ISL_972157, EPI_ISL_972158, EPI_ISL_972159, EPI_ISL_972160, EPI_ISL_972161, EPI_ISL_972162, EPI_ISL_972163, EPI_ISL_972165, EPI_ISL_972166, EPI_ISL_972167, EPI_ISL_972168, EPI_ISL_972171, EPI_ISL_972173, EPI_ISL_972174, EPI_ISL_972175, EPI_ISL_972177, EPI_ISL_972179, EPI_ISL_972180, EPI_ISL_972181, EPI_ISL_972182, EPI_ISL_972183, EPI_ISL_972185, EPI_ISL_972186, EPI_ISL_972187, EPI_ISL_972188, EPI_ISL_972189, EPI_ISL_972191, EPI_ISL_972192, EPI_ISL_972199, EPI_ISL_972200, EPI_ISL_972203, EPI_ISL_972204, EPI_ISL_972205, EPI_ISL_972206, EPI_ISL_972208, EPI_ISL_972209, EPI_ISL_972210, EPI_ISL_972211, EPI_ISL_972214, EPI_ISL_972215, EPI_ISL_972219, EPI_ISL_972220, EPI_ISL_972224, EPI_ISL_972227, EPI_ISL_972228, EPI_ISL_972229, EPI_ISL_972230, EPI_ISL_972231, EPI_ISL_972233, EPI_ISL_972234, EPI_ISL_972235, EPI_ISL_972237, EPI_ISL_972239, EPI_ISL_972242, EPI_ISL_972243, EPI_ISL_972245, EPI_ISL_972246, EPI_ISL_972247, EPI_ISL_972248, EPI_ISL_972249, EPI_ISL_972250, EPI_ISL_972252, EPI_ISL_972253, EPI_ISL_972254, EPI_ISL_972255, EPI_ISL_972256, EPI_ISL_972257, EPI_ISL_972259, EPI_ISL_972260, EPI_ISL_972262, EPI_ISL_972264, EPI_ISL_972265, EPI_ISL_972268, EPI_ISL_972269, EPI_ISL_972272, EPI_ISL_972276, EPI_ISL_972278, EPI_ISL_972279, EPI_ISL_972281, EPI_ISL_972283, EPI_ISL_972284, EPI_ISL_972285, EPI_ISL_972286, EPI_ISL_972287, EPI_ISL_972289, EPI_ISL_972292, EPI_ISL_972294, EPI_ISL_972295, EPI_ISL_972296, EPI_ISL_972298, EPI_ISL_972299, EPI_ISL_972300, EPI_ISL_972301, EPI_ISL_972303, EPI_ISL_972304, EPI_ISL_972305, EPI_ISL_972306, EPI_ISL_972308, EPI_ISL_972310, EPI_ISL_972311, EPI_ISL_972315, EPI_ISL_972316, EPI_ISL_972317, EPI_ISL_972318, EPI_ISL_972319, EPI_ISL_972320, EPI_ISL_972321, EPI_ISL_972322, EPI_ISL_972323, EPI_ISL_972325, EPI_ISL_972326, EPI_ISL_972327, EPI_ISL_972329, EPI_ISL_972330, EPI_ISL_972331, EPI_ISL_972333, EPI_ISL_972334, EPI_ISL_972335, EPI_ISL_972337, EPI_ISL_972339, EPI_ISL_972340, EPI_ISL_972343, EPI_ISL_972344, EPI_ISL_972345, EPI_ISL_972347, EPI_ISL_972350, EPI_ISL_972355, EPI_ISL_972356, EPI_ISL_972357, EPI_ISL_972359, EPI_ISL_972360, EPI_ISL_972361, EPI_ISL_972362, EPI_ISL_972364, EPI_ISL_972365, EPI_ISL_972366, EPI_ISL_972367, EPI_ISL_972368, EPI_ISL_972372, EPI_ISL_972373, EPI_ISL_972374, EPI_ISL_972379, EPI_ISL_972380, EPI_ISL_972383, EPI_ISL_972384, EPI_ISL_972385, EPI_ISL_972386, EPI_ISL_972388, EPI_ISL_972389, EPI_ISL_972390, EPI_ISL_972395, EPI_ISL_972396, EPI_ISL_972400, EPI_ISL_972401, EPI_ISL_972402, EPI_ISL_972403, EPI_ISL_972404, EPI_ISL_972405, EPI_ISL_972408, EPI_ISL_972409, EPI_ISL_972411, EPI_ISL_972412, EPI_ISL_972414, EPI_ISL_972415, EPI_ISL_972416, EPI_ISL_972417, EPI_ISL_972419, EPI_ISL_972420, EPI_ISL_972421, EPI_ISL_972422, EPI_ISL_972424, EPI_ISL_972425, EPI_ISL_972427, EPI_ISL_972428, EPI_ISL_972429, EPI_ISL_972430, EPI_ISL_972432, EPI_ISL_972433, EPI_ISL_972434, EPI_ISL_972439, EPI_ISL_972440, EPI_ISL_972441, EPI_ISL_972444, EPI_ISL_972445, EPI_ISL_972446, EPI_ISL_972447, EPI_ISL_972449, EPI_ISL_972450, EPI_ISL_972451, EPI_ISL_972453, EPI_ISL_972454, EPI_ISL_972455, EPI_ISL_972457, EPI_ISL_972458, EPI_ISL_972460, EPI_ISL_972461, EPI_ISL_972462, EPI_ISL_972463, EPI_ISL_972464, EPI_ISL_972467, EPI_ISL_972469, EPI_ISL_972470, EPI_ISL_972473, EPI_ISL_972475, EPI_ISL_972476, EPI_ISL_972480, EPI_ISL_972481, EPI_ISL_972482, EPI_ISL_972483, EPI_ISL_972485, EPI_ISL_972487, EPI_ISL_972489, EPI_ISL_972490, EPI_ISL_972492, EPI_ISL_972494, EPI_ISL_972495, EPI_ISL_972497, EPI_ISL_972498, EPI_ISL_972499, EPI_ISL_972500, EPI_ISL_972503, EPI_ISL_972504, EPI_ISL_972505, EPI_ISL_972506, EPI_ISL_972507, EPI_ISL_972508, EPI_ISL_972509, EPI_ISL_972510, EPI_ISL_972511, EPI_ISL_972515, EPI_ISL_972516, EPI_ISL_972518, EPI_ISL_972519, EPI_ISL_972520, EPI_ISL_972521, EPI_ISL_972523, EPI_ISL_972524, EPI_ISL_972526, EPI_ISL_972527, EPI_ISL_972531, EPI_ISL_972532, EPI_ISL_972534, EPI_ISL_972535, EPI_ISL_972537, EPI_ISL_972538, EPI_ISL_972540, EPI_ISL_972542, EPI_ISL_972543, EPI_ISL_972548, EPI_ISL_972550, EPI_ISL_972551, EPI_ISL_972553, EPI_ISL_972558, EPI_ISL_972561, EPI_ISL_972562, EPI_ISL_972563, EPI_ISL_972564, EPI_ISL_972567, EPI_ISL_972568, EPI_ISL_972570, EPI_ISL_972571, EPI_ISL_972572, EPI_ISL_972573, EPI_ISL_972578, EPI_ISL_972581, EPI_ISL_972582, EPI_ISL_972583, EPI_ISL_972584, EPI_ISL_972588, EPI_ISL_972589, EPI_ISL_972593, EPI_ISL_972594, EPI_ISL_972595, EPI_ISL_972596, EPI_ISL_972598, EPI_ISL_972599, EPI_ISL_972600, EPI_ISL_972603, EPI_ISL_972604, EPI_ISL_972605, EPI_ISL_972606, EPI_ISL_972611, EPI_ISL_972613, EPI_ISL_972616, EPI_ISL_972617, EPI_ISL_972619, EPI_ISL_972620, EPI_ISL_972621, EPI_ISL_972622, EPI_ISL_972623, EPI_ISL_972626, EPI_ISL_972628, EPI_ISL_972629, EPI_ISL_972630, EPI_ISL_972631, EPI_ISL_972632, EPI_ISL_972636, EPI_ISL_972637, EPI_ISL_972638, EPI_ISL_972640, EPI_ISL_972641, EPI_ISL_972642, EPI_ISL_972643, EPI_ISL_972645, EPI_ISL_972646, EPI_ISL_972647, EPI_ISL_972648, EPI_ISL_972649, EPI_ISL_972650, EPI_ISL_972651, EPI_ISL_972652, EPI_ISL_972654, EPI_ISL_972655, EPI_ISL_972656, EPI_ISL_972657, EPI_ISL_972658, EPI_ISL_972659, EPI_ISL_972660, EPI_ISL_972664, EPI_ISL_972665, EPI_ISL_972666, EPI_ISL_972667, EPI_ISL_972669, EPI_ISL_972670, EPI_ISL_972672, EPI_ISL_972675, EPI_ISL_972676, EPI_ISL_972677, EPI_ISL_972678, EPI_ISL_972679, EPI_ISL_972681, EPI_ISL_972682, EPI_ISL_972683, EPI_ISL_972684, EPI_ISL_972685, EPI_ISL_972691, EPI_ISL_972693, EPI_ISL_972694, EPI_ISL_972695, EPI_ISL_972696, EPI_ISL_972698, EPI_ISL_972699, EPI_ISL_972700, EPI_ISL_972701, EPI_ISL_972703, EPI_ISL_972704, EPI_ISL_972705, EPI_ISL_972706, EPI_ISL_972708, EPI_ISL_972710, EPI_ISL_972711, EPI_ISL_972716, EPI_ISL_972717, EPI_ISL_972719, EPI_ISL_972720, EPI_ISL_972721, EPI_ISL_972723, EPI_ISL_972724, EPI_ISL_972725, EPI_ISL_972726, EPI_ISL_972729, EPI_ISL_972731, EPI_ISL_972732, EPI_ISL_972733, EPI_ISL_972735, EPI_ISL_972736, EPI_ISL_972739, EPI_ISL_972740, EPI_ISL_972745, EPI_ISL_972746, EPI_ISL_972747, EPI_ISL_972748, EPI_ISL_972749, EPI_ISL_972753, EPI_ISL_972754, EPI_ISL_972756, EPI_ISL_972757, EPI_ISL_972760, EPI_ISL_972762, EPI_ISL_972764, EPI_ISL_972765, EPI_ISL_972767, EPI_ISL_972770, EPI_ISL_972775, EPI_ISL_972780, EPI_ISL_972781, EPI_ISL_972782, EPI_ISL_972783, EPI_ISL_972784, EPI_ISL_972786, EPI_ISL_972788, EPI_ISL_972789, EPI_ISL_972792, EPI_ISL_972794, EPI_ISL_972795, EPI_ISL_972796, EPI_ISL_972797, EPI_ISL_972798, EPI_ISL_972799, EPI_ISL_972800, EPI_ISL_972801, EPI_ISL_972802, EPI_ISL_972803, EPI_ISL_972804, EPI_ISL_972806, EPI_ISL_972808, EPI_ISL_972814, EPI_ISL_972815, EPI_ISL_972817, EPI_ISL_972818, EPI_ISL_972820, EPI_ISL_972823, EPI_ISL_972824, EPI_ISL_972826, EPI_ISL_972827, EPI_ISL_972828, EPI_ISL_972829, EPI_ISL_972830, EPI_ISL_972831, EPI_ISL_972832, EPI_ISL_972836, EPI_ISL_972837, EPI_ISL_972838, EPI_ISL_972839, EPI_ISL_972840, EPI_ISL_972841, EPI_ISL_972842, EPI_ISL_972844, EPI_ISL_972845, EPI_ISL_972847, EPI_ISL_972849, EPI_ISL_972850, EPI_ISL_972852, EPI_ISL_972854, EPI_ISL_972856, EPI_ISL_972857, EPI_ISL_972859, EPI_ISL_972862, EPI_ISL_972863, EPI_ISL_972864, EPI_ISL_972865, EPI_ISL_972866, EPI_ISL_972867, EPI_ISL_972868, EPI_ISL_972869, EPI_ISL_972870, EPI_ISL_972873, EPI_ISL_972878, EPI_ISL_972879, EPI_ISL_972880, EPI_ISL_972883, EPI_ISL_972886, EPI_ISL_972889, EPI_ISL_972890, EPI_ISL_972893, EPI_ISL_972894, EPI_ISL_972896, EPI_ISL_972901, EPI_ISL_972902, EPI_ISL_972904, EPI_ISL_972906, EPI_ISL_972909, EPI_ISL_972910, EPI_ISL_972912, EPI_ISL_972914, EPI_ISL_972915, EPI_ISL_972919, EPI_ISL_972920, EPI_ISL_972921, EPI_ISL_972923, EPI_ISL_972925, EPI_ISL_972926, EPI_ISL_972927, EPI_ISL_972928, EPI_ISL_972930, EPI_ISL_972931, EPI_ISL_972932, EPI_ISL_972934, EPI_ISL_972935, EPI_ISL_972937, EPI_ISL_972939, EPI_ISL_972942, EPI_ISL_972943, EPI_ISL_972946, EPI_ISL_972947, EPI_ISL_972951, EPI_ISL_972952, EPI_ISL_972953, EPI_ISL_972955, EPI_ISL_972956, EPI_ISL_972958, EPI_ISL_972959, EPI_ISL_972961, EPI_ISL_972964, EPI_ISL_972966, EPI_ISL_972967, EPI_ISL_972968, EPI_ISL_972971, EPI_ISL_972974, EPI_ISL_972976, EPI_ISL_972977, EPI_ISL_972978, EPI_ISL_972979, EPI_ISL_972980, EPI_ISL_972981, EPI_ISL_972982, EPI_ISL_972983, EPI_ISL_972984, EPI_ISL_972985, EPI_ISL_972986, EPI_ISL_972987, EPI_ISL_972988, EPI_ISL_972989, EPI_ISL_972990, EPI_ISL_972991, EPI_ISL_972992, EPI_ISL_972993, EPI_ISL_972994, EPI_ISL_972997, EPI_ISL_972999, EPI_ISL_973000, EPI_ISL_973003, EPI_ISL_973004, EPI_ISL_973005, EPI_ISL_973006, EPI_ISL_973007, EPI_ISL_973008, EPI_ISL_973010, EPI_ISL_973011, EPI_ISL_973012, EPI_ISL_973013, EPI_ISL_973015, EPI_ISL_973016, EPI_ISL_973017, EPI_ISL_973018, EPI_ISL_973021, EPI_ISL_973023, EPI_ISL_973025, EPI_ISL_973029, EPI_ISL_973031, EPI_ISL_973032, EPI_ISL_973033, EPI_ISL_973034, EPI_ISL_973036, EPI_ISL_973037, EPI_ISL_973039, EPI_ISL_973041, EPI_ISL_973042, EPI_ISL_973043, EPI_ISL_973044, EPI_ISL_973045, EPI_ISL_973046, EPI_ISL_973049, EPI_ISL_973050, EPI_ISL_973051, EPI_ISL_973054, EPI_ISL_973056, EPI_ISL_973057, EPI_ISL_973058, EPI_ISL_973059, EPI_ISL_973061, EPI_ISL_973062, EPI_ISL_973065, EPI_ISL_973066, EPI_ISL_973067, EPI_ISL_973068, EPI_ISL_973070, EPI_ISL_973072, EPI_ISL_973074, EPI_ISL_973075, EPI_ISL_973079, EPI_ISL_973081, EPI_ISL_973083, EPI_ISL_973084, EPI_ISL_973087, EPI_ISL_973089, EPI_ISL_973090, EPI_ISL_973091, EPI_ISL_973092, EPI_ISL_973095, EPI_ISL_973098, EPI_ISL_973099, EPI_ISL_973100, EPI_ISL_973102, EPI_ISL_973103, EPI_ISL_973104, EPI_ISL_973106, EPI_ISL_973107, EPI_ISL_973108, EPI_ISL_973109, EPI_ISL_973110, EPI_ISL_973112, EPI_ISL_973114, EPI_ISL_973117, EPI_ISL_973118, EPI_ISL_973119, EPI_ISL_973121, EPI_ISL_973125, EPI_ISL_973126, EPI_ISL_973129, EPI_ISL_973133, EPI_ISL_973134, EPI_ISL_973136, EPI_ISL_973139, EPI_ISL_973140, EPI_ISL_973141, EPI_ISL_973142, EPI_ISL_973144, EPI_ISL_973146, EPI_ISL_973147, EPI_ISL_973149, EPI_ISL_973150, EPI_ISL_973151, EPI_ISL_973152, EPI_ISL_973153, EPI_ISL_973154, EPI_ISL_973155, EPI_ISL_973156, EPI_ISL_973158, EPI_ISL_973159, EPI_ISL_973161, EPI_ISL_973162, EPI_ISL_973163, EPI_ISL_973164, EPI_ISL_973166, EPI_ISL_973168, EPI_ISL_973169, EPI_ISL_973170, EPI_ISL_973171, EPI_ISL_973172, EPI_ISL_973173, EPI_ISL_973174, EPI_ISL_973176, EPI_ISL_973177, EPI_ISL_973181, EPI_ISL_973183, EPI_ISL_973184, EPI_ISL_973185, EPI_ISL_973187, EPI_ISL_973188, EPI_ISL_973190, EPI_ISL_973192, EPI_ISL_973193, EPI_ISL_973198, EPI_ISL_973200, EPI_ISL_973201, EPI_ISL_973202, EPI_ISL_973203, EPI_ISL_973204, EPI_ISL_973205, EPI_ISL_973209, EPI_ISL_973210, EPI_ISL_973211, EPI_ISL_973212, EPI_ISL_973214, EPI_ISL_973215, EPI_ISL_973217, EPI_ISL_973218, EPI_ISL_973219, EPI_ISL_973220, EPI_ISL_973221, EPI_ISL_973222, EPI_ISL_973223, EPI_ISL_973226, EPI_ISL_973228, EPI_ISL_973231, EPI_ISL_973235, EPI_ISL_973237, EPI_ISL_973240, EPI_ISL_973241, EPI_ISL_973242, EPI_ISL_973243, EPI_ISL_973244, EPI_ISL_973245, EPI_ISL_973246, EPI_ISL_973247, EPI_ISL_973248, EPI_ISL_973253, EPI_ISL_973254, EPI_ISL_973255, EPI_ISL_973257, EPI_ISL_973258, EPI_ISL_973259, EPI_ISL_973261, EPI_ISL_973262, EPI_ISL_973267, EPI_ISL_973268, EPI_ISL_973269, EPI_ISL_973270, EPI_ISL_973272, EPI_ISL_973274, EPI_ISL_973275, EPI_ISL_973279, EPI_ISL_973280, EPI_ISL_973282, EPI_ISL_973283, EPI_ISL_973284, EPI_ISL_973285, EPI_ISL_973286, EPI_ISL_973287, EPI_ISL_973290, EPI_ISL_973291, EPI_ISL_973292, EPI_ISL_973293, EPI_ISL_973294, EPI_ISL_973295, EPI_ISL_973297, EPI_ISL_973299, EPI_ISL_973300, EPI_ISL_973301, EPI_ISL_973303, EPI_ISL_973305, EPI_ISL_973306, EPI_ISL_973307, EPI_ISL_973308, EPI_ISL_973309, EPI_ISL_973311, EPI_ISL_973313, EPI_ISL_973314, EPI_ISL_973315, EPI_ISL_973316, EPI_ISL_973318, EPI_ISL_973319, EPI_ISL_973320, EPI_ISL_973321, EPI_ISL_973322, EPI_ISL_973323, EPI_ISL_973326, EPI_ISL_973327, EPI_ISL_973329, EPI_ISL_973330, EPI_ISL_973331, EPI_ISL_973334, EPI_ISL_973335, EPI_ISL_973336, EPI_ISL_973337, EPI_ISL_973343, EPI_ISL_973344, EPI_ISL_973347, EPI_ISL_973348, EPI_ISL_973352, EPI_ISL_973353, EPI_ISL_973355, EPI_ISL_973356, EPI_ISL_973360, EPI_ISL_973361, EPI_ISL_973362, EPI_ISL_973364, EPI_ISL_973366, EPI_ISL_973367, EPI_ISL_973370, EPI_ISL_973371, EPI_ISL_973372, EPI_ISL_973374, EPI_ISL_973375, EPI_ISL_973376, EPI_ISL_973377, EPI_ISL_973378, EPI_ISL_973380, EPI_ISL_973381, EPI_ISL_973382, EPI_ISL_973385, EPI_ISL_973386, EPI_ISL_973387, EPI_ISL_973388, EPI_ISL_973391, EPI_ISL_973392, EPI_ISL_973393, EPI_ISL_973394, EPI_ISL_973395, EPI_ISL_973400, EPI_ISL_973401, EPI_ISL_973402, EPI_ISL_973403, EPI_ISL_973405, EPI_ISL_973408, EPI_ISL_973409, EPI_ISL_973411, EPI_ISL_973413, EPI_ISL_973414, EPI_ISL_973416, EPI_ISL_973418, EPI_ISL_973419, EPI_ISL_973421, EPI_ISL_973424, EPI_ISL_973425, EPI_ISL_973426, EPI_ISL_973427, EPI_ISL_973428, EPI_ISL_973429, EPI_ISL_973431, EPI_ISL_973433, EPI_ISL_973434, EPI_ISL_973435, EPI_ISL_973436, EPI_ISL_973437, EPI_ISL_973439, EPI_ISL_973440, EPI_ISL_973441, EPI_ISL_973442, EPI_ISL_973444, EPI_ISL_973446, EPI_ISL_973448, EPI_ISL_973449, EPI_ISL_973450, EPI_ISL_973451, EPI_ISL_973452, EPI_ISL_973454, EPI_ISL_973455, EPI_ISL_973457, EPI_ISL_973458, EPI_ISL_973460, EPI_ISL_973462, EPI_ISL_973464, EPI_ISL_973466, EPI_ISL_973469, EPI_ISL_973470, EPI_ISL_973472, EPI_ISL_973473, EPI_ISL_973475, EPI_ISL_973476, EPI_ISL_973480, EPI_ISL_973482, EPI_ISL_973489, EPI_ISL_973493, EPI_ISL_973497, EPI_ISL_973501, EPI_ISL_973503, EPI_ISL_973510, EPI_ISL_973512, EPI_ISL_973519, EPI_ISL_973521, EPI_ISL_973525, EPI_ISL_973527, EPI_ISL_973529, EPI_ISL_973532, EPI_ISL_973533, EPI_ISL_973538, EPI_ISL_973542, EPI_ISL_973546, EPI_ISL_973548, EPI_ISL_973550, EPI_ISL_973555, EPI_ISL_973557, EPI_ISL_973559, EPI_ISL_973561, EPI_ISL_973563, EPI_ISL_973566, EPI_ISL_973572, EPI_ISL_973574, EPI_ISL_973576, EPI_ISL_973589, EPI_ISL_973593, EPI_ISL_973597, EPI_ISL_973603, EPI_ISL_973605, EPI_ISL_973609, EPI_ISL_973611, EPI_ISL_973616, EPI_ISL_973618, EPI_ISL_973621, EPI_ISL_973623, EPI_ISL_973624, EPI_ISL_973626, EPI_ISL_973628, EPI_ISL_973629, EPI_ISL_973631, EPI_ISL_973640, EPI_ISL_973645, EPI_ISL_973647, EPI_ISL_973648, EPI_ISL_973651, EPI_ISL_973652, EPI_ISL_973658, EPI_ISL_973660, EPI_ISL_973662, EPI_ISL_973668, EPI_ISL_973670, EPI_ISL_973680, EPI_ISL_973682, EPI_ISL_973684, EPI_ISL_973686, EPI_ISL_973688, EPI_ISL_973689, EPI_ISL_973691, EPI_ISL_973696, EPI_ISL_973698, EPI_ISL_973703, EPI_ISL_973708, EPI_ISL_973710, EPI_ISL_973715, EPI_ISL_973717, EPI_ISL_973720, EPI_ISL_973726, EPI_ISL_973728, EPI_ISL_973730, EPI_ISL_973734, EPI_ISL_973735, EPI_ISL_973738, EPI_ISL_973739, EPI_ISL_973743, EPI_ISL_973744, EPI_ISL_973746, EPI_ISL_973750, EPI_ISL_973754, EPI_ISL_973756, EPI_ISL_973757, EPI_ISL_973759, EPI_ISL_973761, EPI_ISL_973763, EPI_ISL_973765, EPI_ISL_973768, EPI_ISL_973770, EPI_ISL_973773, EPI_ISL_973777, EPI_ISL_973778, EPI_ISL_973782, EPI_ISL_973786, EPI_ISL_973787, EPI_ISL_973789, EPI_ISL_973790, EPI_ISL_973792, EPI_ISL_973793, EPI_ISL_973798, EPI_ISL_973801, EPI_ISL_973803, EPI_ISL_973806, EPI_ISL_973808, EPI_ISL_973809, EPI_ISL_973811, EPI_ISL_973812, EPI_ISL_973814, EPI_ISL_973816, EPI_ISL_973819, EPI_ISL_973822, EPI_ISL_973825, EPI_ISL_973835, EPI_ISL_973836

| | | | |
|---|---|---|---|
| see above | Department of Virus and Microbiological Special Diagnostics, Statens Serum Institut, Copenhagen, Denmark | Aalborg University | Danish Covid-19 Genome Consortium |
| EPI_ISL_977490 | Hospital Municipal Guido Guida | Instituto Adolfo Lutz, Interdiciplinary Procedures Center, Strategic Laboratory | Claudio Tavares Sacchi, Claudia Regina Gonçalves, Erica Valessa Ramos Gomes, Karoline Rodrigues Campos |
| EPI_ISL_977493 | Lab voor klinische biologie | Lab voor klinische biologie | Hannelore Hamerlinck, Marija Janevska, Bruno Verhasselt |
| EPI_ISL_977503 | Szpital Miejski im. Franciszka Raszei w Poznaniu, Oddzia Chirurgii Ogólnej i Minimalnie Inwazyjnej | genXone SA, Research & Development Laboratory | Maciej Sykulski, Grzegorz Nowicki, Jakub Grabowski, Natalia Drwska-Matelska, Anna Brylak-Baszków, Aleksandra Gidlewicz, Karol Szeszko, ukasz Krych, Micha Kaszuba |
| EPI_ISL_977600 | Biolab Diagnostic Laboratories | Biolab Diagnostic Laboratories | Issa Abu-Dayyeh, Ahmad Tibi, Lama Hussein, Shayma Ali, Badia Saddedin, Amid Abdelnour |

EPI_ISL_978169, EPI_ISL_978170, EPI_ISL_978171, EPI_ISL_978172, EPI_ISL_978173, EPI_ISL_978174, EPI_ISL_978175, EPI_ISL_978176, EPI_ISL_978177, EPI_ISL_978178, EPI_ISL_978179, EPI_ISL_978180

| | | | |
|---|---|---|---|
| see above | Virginia Division of Consolidated Laboratory Services | Virginia Division of Consolidated Laboratory Services | Virginia DCLS |
| EPI_ISL_978310, EPI_ISL_978319, EPI_ISL_978320, EPI_ISL_978321, EPI_ISL_978322, EPI_ISL_978323, | Texas Department of State Health Services | Texas Department of State Health Services | Bonnie Oh, Anita Pokharel, James Daniel Bonser, Myong Koag, Chung Wang, Rachel Lee, Grace Kubin, Rashmi Tuladhar, Mayela Pedrueza, Maliha Rahman, Jenny Zhang |

| | | | |
|---|---|---|---|
| EPI_ISL_978324, EPI_ISL_978325 | | | |
| EPI_ISL_978429, EPI_ISL_978430, EPI_ISL_978431, EPI_ISL_978477, EPI_ISL_978478, EPI_ISL_978479, EPI_ISL_978480 | Arizona State Public Health Laboratory | Arizona State Public Health Laboratory | Trung Huynh, Jessica Escobar, Katherine Fullerton, Nobuko Fukushima, Stacy White, Linda Getsinger, Victor Waddell |
| EPI_ISL_978553 | Kansas Health and Environmental Lab | Kansas Health and Environmental Lab | Mike Grose, Paige Drury, Carissa Robertson, Ben Olsen, and Phil Adam |
| EPI_ISL_978621 | Helix/Illumina | Respiratory Viruses Branch, Division of Viral Diseases, Centers for Disease Control and Prevention | Peter W. Cook,Dakota Howard,Dhwani Batra,Ben L. Rambo-Martin,Eileen de Feo,Jan Antico,Christine Tran,Matthew Tolentino,Shannon Wickline,Kim Gietzen,Brad Sickler,Jingtao Liu,Eric Allen,Phil Febbo,Summer Galloway,Nicole L. Washington,Simon White,Geraint Levan,Kelly Schiabor Barrett,Elizabeth Cirulli,Alexandre Bolze,Ary Ascencio,Charlotte Rivera-Garcia,Ryan Cho,Jason Nguyen,Sherry Wang,Jimmy Ramirez,Tyler Cassens,Efren Sandoval,Magnus Isaksson,William Lee,David Becker,Marc Laurent,James Lu,Clinton R. Paden,Suxiang Tong,Duncan MacCannell, |
| EPI_ISL_978887, EPI_ISL_978888, EPI_ISL_978952, EPI_ISL_978953, EPI_ISL_978985, EPI_ISL_978986, EPI_ISL_978987 | Centre for Dengue Research and AICBU, Department of Immunology and Molecular Medicine | Centre for Dengue Research and AICBU, Department of Immunology and Molecular Medicine | Chandima Jeewandara, Deshni Jayathilaka, Dinuka Ariyaratne, Tibutius Thanesh Pramanayagam, Diyanath Ranasinghe, Laksiri Gomes, Gathsaurie Neelika Malavige |
| EPI_ISL_979253 | Institute of Microbiology and Immunology, Faculty of Medicine, University of Ljubljana | Institute of Microbiology and Immunology, Faculty of Medicine, University of Ljubljana | Samo Zakotnik, Tomaž Mark Zorec, Matic Brvar, Doroteja Vlaj, Patricija Pozvek,Špela Pleh, Miša Korva, Mario Poljak, Tatjana Avši - Županc |
| EPI_ISL_979313, EPI_ISL_979318, EPI_ISL_979326 | Cadham Provincial laboratory | National Microbiology Laboratory (NML) | Anna Majer, Shari Tyson, Grace Seo, Philip Mabon, Elsie Grudeski, Rhiannon Huzarewich, Russell Mandes, Anneliese Landgraff, Jennifer Tanner, Natalie Knox, Morag Graham, Gary Van Domselaar, Paul Van Caeseele, Jared Bullard, David Alexander, Kerry Dust, Nathalie Bastien, Yan Li, Timothy Booth, Darian Hole, Madison Chapel, Kirsten Biggar, CanCOGeN's metadata curation team, Public Health Agency of Canada CanCOGeN team |
| EPI_ISL_979365, EPI_ISL_979366, EPI_ISL_979367, EPI_ISL_979369 | The Jackson Laboratory | The Jackson Laboratory | Lloyd M, Sanderson B, Srivastava A, Maurya R, Renzette N, Omerza G, Kelly K, Li L, Wei C L, Adams M |
| EPI_ISL_979410, EPI_ISL_979411, EPI_ISL_979412, EPI_ISL_979413, EPI_ISL_979414, EPI_ISL_979415, EPI_ISL_979416, EPI_ISL_979417, EPI_ISL_979418, EPI_ISL_979419, EPI_ISL_979420, EPI_ISL_979421, EPI_ISL_979422, EPI_ISL_979423, EPI_ISL_979424, EPI_ISL_979425, EPI_ISL_979426, EPI_ISL_979427, EPI_ISL_979428 | | | |
| see above | New Mexico Department of Health Scientific Laboratory | New Mexico Department of Health Scientific Laboratory | Ellie Johnson, Anastacia Griego-Fisher, D'eldra Malone, Jennifer Benoit |
| EPI_ISL_979458 | Test Iowa | State Hygienic Laboratory at the University of Iowa | Valerie Reeb, Erik Twait, Wes Hottel, Alankar Kampoowale |
| EPI_ISL_979469, EPI_ISL_979470, EPI_ISL_979471, EPI_ISL_979472, EPI_ISL_979473, EPI_ISL_979474, EPI_ISL_979475, EPI_ISL_979476, EPI_ISL_979477, EPI_ISL_979478, EPI_ISL_979479, EPI_ISL_979480, EPI_ISL_979481, EPI_ISL_979482, EPI_ISL_979483, EPI_ISL_979484, EPI_ISL_979485, EPI_ISL_979486, EPI_ISL_979487, EPI_ISL_979488, EPI_ISL_979489, EPI_ISL_979490, EPI_ISL_979491, EPI_ISL_979492, EPI_ISL_979493, EPI_ISL_979494, EPI_ISL_979495, EPI_ISL_979496, EPI_ISL_979497, EPI_ISL_979498, EPI_ISL_979499, EPI_ISL_979500, EPI_ISL_979501, EPI_ISL_979502, EPI_ISL_979503, EPI_ISL_979504, EPI_ISL_979505, EPI_ISL_979506, EPI_ISL_979507, EPI_ISL_979508, EPI_ISL_979509, EPI_ISL_979510, EPI_ISL_979511, EPI_ISL_979512, EPI_ISL_979513, EPI_ISL_979514, EPI_ISL_979515, EPI_ISL_979516, EPI_ISL_979517, EPI_ISL_979518, EPI_ISL_979519, EPI_ISL_979520, EPI_ISL_979521, EPI_ISL_979522, EPI_ISL_979523, EPI_ISL_979524, EPI_ISL_979525, EPI_ISL_979526, EPI_ISL_979527, EPI_ISL_979528, EPI_ISL_979529, EPI_ISL_979530, EPI_ISL_979531, EPI_ISL_979532, EPI_ISL_979533, EPI_ISL_979534, EPI_ISL_979535, EPI_ISL_979536, EPI_ISL_979537, EPI_ISL_979538, EPI_ISL_979539, EPI_ISL_979540, EPI_ISL_979541, EPI_ISL_979542, EPI_ISL_979543 | | | |
| see above | Eurofins Diatherix | Hudsonalpha Genome Sequencing Center | Jane Grimwood, Melissa Williams, Lori H. Handley, Joshua Stough, Leslie Malone, Stefan Brzezinski, Ada Stewart, Teresa Jones, Jenell Webber, John Lovell, Jennifer Cart, and Jeremy Schmutz |
| EPI_ISL_979799, EPI_ISL_979800, EPI_ISL_979801, EPI_ISL_979802 | National Institute of Infectious Diseases-Prof. Dr. Matei Bals Molecular Diagnostics Laboratory | National Institute of Infectious Diseases-Prof. Dr. Matei Bals Molecular Diagnostics Laboratory | Leontina Banica, Marius Surleac, Corina Casangiu, Petre Milu, Andreea Tudor, Simona Paraschiv, Dan Otelea |
| EPI_ISL_980807 | genXone SA, Molecular Diagnostics Laboratory / NZOZ | genXone SA, Research & Development Laboratory | Maciej Sykulski, Grzegorz Nowicki, Jakub Grabowski, Natalia Drwska-Matelska, Anna Brylak-Baszków, Aleksandra Gidlewicz, Karol Szeszko, ukasz Krych, Micha Kaszuba |
| EPI_ISL_980841, EPI_ISL_980842, EPI_ISL_980844, EPI_ISL_980846, EPI_ISL_980847, EPI_ISL_980848, EPI_ISL_980849, EPI_ISL_980850, EPI_ISL_980853, EPI_ISL_980856, EPI_ISL_980862, EPI_ISL_980865, EPI_ISL_980866, EPI_ISL_980867, EPI_ISL_980872, EPI_ISL_980874, EPI_ISL_980875, EPI_ISL_980877, EPI_ISL_980878, EPI_ISL_980879, EPI_ISL_980882, EPI_ISL_980883, EPI_ISL_980886, EPI_ISL_980889, EPI_ISL_980890, EPI_ISL_980891, EPI_ISL_980892, EPI_ISL_980894, EPI_ISL_980895, EPI_ISL_980906, EPI_ISL_980909, EPI_ISL_980912, EPI_ISL_980913, EPI_ISL_980915, EPI_ISL_980918, EPI_ISL_980919, EPI_ISL_980946, EPI_ISL_980947, EPI_ISL_980948, EPI_ISL_980949, EPI_ISL_980950, EPI_ISL_980951, EPI_ISL_980952, EPI_ISL_980953, EPI_ISL_980954, EPI_ISL_980955, EPI_ISL_980956, EPI_ISL_980957, EPI_ISL_980958, EPI_ISL_980959, EPI_ISL_980960, EPI_ISL_980961, EPI_ISL_980962, EPI_ISL_980963, EPI_ISL_980964, EPI_ISL_980984, EPI_ISL_980985, EPI_ISL_980986, EPI_ISL_980987, EPI_ISL_980988 | | | |
| see above | Innovative Genomics Institute, UC Berkeley | Innovative Genomics Institute, UC Berkeley | Stacia Wyman, Haridha Shivram, Phil Frankino, Liana Lareau |
| EPI_ISL_981070, EPI_ISL_981076, EPI_ISL_981081, EPI_ISL_981083, EPI_ISL_981084, EPI_ISL_981086, EPI_ISL_981101, EPI_ISL_981102, EPI_ISL_981108, EPI_ISL_981109, EPI_ISL_981118, EPI_ISL_981122, EPI_ISL_981133, EPI_ISL_981134, EPI_ISL_981139, EPI_ISL_981140, EPI_ISL_981141, EPI_ISL_981142, EPI_ISL_981143, EPI_ISL_981144, EPI_ISL_981145, EPI_ISL_981146 | | | |
| see above | Johns Hopkins Hospital Department of Pathology | Johns Hopkins Hospital Department of Pathology | C. Paul Morris, Chun Huai Luo, Adannaya Amadi, Matthew Schwartz, Nicholas Gallagher, Heba H. Mostafa |
| EPI_ISL_981308, EPI_ISL_981309, EPI_ISL_981310, EPI_ISL_981311, EPI_ISL_981312, EPI_ISL_981313, EPI_ISL_981314, EPI_ISL_981315, EPI_ISL_981316, EPI_ISL_981319, EPI_ISL_981320, EPI_ISL_981321, EPI_ISL_981324, EPI_ISL_981326, EPI_ISL_981327, EPI_ISL_981328, EPI_ISL_981329, EPI_ISL_981330, EPI_ISL_981331, EPI_ISL_981332, EPI_ISL_981333, EPI_ISL_981334, EPI_ISL_981335, EPI_ISL_981336, EPI_ISL_981337, EPI_ISL_981338, EPI_ISL_981339, EPI_ISL_981340, EPI_ISL_981341, EPI_ISL_981342, EPI_ISL_981343, EPI_ISL_981344, EPI_ISL_981350, EPI_ISL_981352, EPI_ISL_981354, EPI_ISL_981355, EPI_ISL_981356, EPI_ISL_981357, EPI_ISL_981358, EPI_ISL_981359, EPI_ISL_981360, EPI_ISL_981361, EPI_ISL_981362, EPI_ISL_981363, EPI_ISL_981364, EPI_ISL_981365, EPI_ISL_981366, EPI_ISL_981367 | | | |
| see above | Hospital Universitari Vall d'Hebron - Vall d'Hebron Institut de Rercerca | Hospital Universitari Vall d'Hebron - Vall d'Hebron Institut de Rercerca | Cristina Andrés, Maria Piñana, Josep F Abril, Damir Garcia-Cehic, Ariadna Rando, Juliana Esperalba, Maria Gema Codina, Carla Castillo, Maria Carmen Martin, Tomàs Pumarola, Josep Quer, Andrés Antón |
| EPI_ISL_981376 | AZ Klina | AZ Klina | Dr. C. Vael |
| EPI_ISL_981385 | IAL Regional de Bauru | Instituto Adolfo Lutz, Interdiciplinary Procedures Center, Strategic Laboratory | Claudio Tavares Sacchi, Claudia Regina Gonçalves, Erica Valessa Ramos Gomes, Karoline Rodrigues Campos |
| EPI_ISL_981389 | Hospital Geral de Sao Paulo | Instituto Adolfo Lutz, Interdiciplinary Procedures Center, Strategic Laboratory | Claudio Tavares Sacchi, Claudia Regina Gonçalves, Erica Valessa Ramos Gomes, Karoline Rodrigues Campos |
| EPI_ISL_981395, EPI_ISL_981462, EPI_ISL_981463, EPI_ISL_981464, EPI_ISL_981465 | CH.INTERCOMMUNAL DE CRETEIL | Department of Virology, Henri Mondor University Hospital, Assistance Publique Hôpitaux de Paris, Université Paris-Est Créteil, INSERM U955 | Christophe Rodriguez, Slim Fourati, Vanessa Demontant, Guillaume Gricourt, Melissa N'Debi, Alexandre Soulier, Elisabeth Trawinski, Jean-Michel Pawlotsky |
| EPI_ISL_981466, EPI_ISL_981467, EPI_ISL_981468, EPI_ISL_981469, EPI_ISL_981470, EPI_ISL_981471, EPI_ISL_981472, EPI_ISL_981473, EPI_ISL_981474, EPI_ISL_981475, EPI_ISL_981478, EPI_ISL_981479, EPI_ISL_981480, EPI_ISL_981481 | | | |
| see above | CHI VILLENEUVE ST GEORGES | Department of Virology, Henri Mondor University Hospital, Assistance Publique Hôpitaux de Paris, Université Paris-Est Créteil, INSERM U955 | Christophe Rodriguez, Slim Fourati, Vanessa Demontant, Guillaume Gricourt, Melissa N'Debi, Alexandre Soulier, Elisabeth Trawinski, Jean-Michel Pawlotsky |
| EPI_ISL_981486, EPI_ISL_981488, EPI_ISL_981538, EPI_ISL_981539, EPI_ISL_981540, EPI_ISL_981541, EPI_ISL_981542, EPI_ISL_981543, EPI_ISL_981544, EPI_ISL_981545, EPI_ISL_981546 | | | |
| see above | CH.INTERCOMMUNAL DE CRETEIL | Department of Virology, Henri Mondor University Hospital, Assistance Publique Hôpitaux de Paris, Université Paris-Est Créteil, INSERM U955 | Christophe Rodriguez, Slim Fourati, Vanessa Demontant, Guillaume Gricourt, Melissa N'Debi, Alexandre Soulier, Elisabeth Trawinski, Jean-Michel Pawlotsky |
| EPI_ISL_981547 | G.H.E.F.Grand Hôpital EST Francilien | Department of Virology, Henri Mondor University Hospital, Assistance Publique Hôpitaux de Paris, Université Paris-Est Créteil, INSERM U955 | Christophe Rodriguez, Slim Fourati, Vanessa Demontant, Guillaume Gricourt, Melissa N'Debi, Alexandre Soulier, Elisabeth Trawinski, Jean-Michel Pawlotsky |
| EPI_ISL_981650, EPI_ISL_981654, EPI_ISL_981659, EPI_ISL_981746, EPI_ISL_981813 | University Hospitals of Geneva, Laboratory of Virology | HUG, Laboratory of Virology and the Health2030 Genome Center | Samuel Cordey, Ana Rita Goncalves, Laurent Kaiser, Lorenzo Cerutti, Henri Pegeot, Melyssa Elies, Deborah Penet, Keith Harshman, Ioannis Xenarios, Emmanouil Dermitzakis |
| EPI_ISL_981977 | National Institute of Laboratory Medicine and Referral Center | Genomic Research Lab, BCSIR | Md. Murshed Hasan Sarkar, Mohammad Samir Uzzaman, Eshrar Osman, Md. Ahashan Habib, Shahina Akter, Tanjina Akhtar Banu, Abu Sayeed |

| | | | |
|---|---|---|---|
| | | | Mohammad Mahmud, Barna Goswami, Iffat Jahan, Md. Saddam Hossain, Tasnim Nafisa, Md. Maruf Ahmed Molla, Mahmuda Yeasmin, Asish Kumar Ghosh, Arifa Akram, A. K. M.Shamsuzzaman, Md. Salim Khan |
| EPI_ISL_982021, EPI_ISL_982026, EPI_ISL_982027, EPI_ISL_982043, EPI_ISL_982044, EPI_ISL_982063, EPI_ISL_982086 | TGen North | TGen North | "Jolene Bowers, Megan Folkerts, Chris French, Hayley Yaglom, Ashlyn Pfeiffer, Darrin Lemmer, Dave Engelthaler, The Arizona COVID Genomics Union (ACGU)" |
| EPI_ISL_982123 | EFS ILE DE FRANCE BANQUE DE TISSUS | Department of Virology, Henri Mondor University Hospital, Assistance Publique Hôpitaux de Paris, Université Paris-Est Créteil, INSERM U955 | Christophe Rodriguez, Slim Fourati, Vanessa Demontant, Guillaume Gricourt, Melissa N'Debi, Alexandre Soulier, Elisabeth Trawinski, Jean-Michel Pawlotsky |
| EPI_ISL_982153 | Hôpital Henri Mondor | Department of Virology, Henri Mondor University Hospital, Assistance Publique Hôpitaux de Paris, Université Paris-Est Créteil, INSERM U955 | Christophe Rodriguez, Slim Fourati, Vanessa Demontant, Guillaume Gricourt, Melissa N'Debi, Alexandre Soulier, Elisabeth Trawinski, Jean-Michel Pawlotsky |
| EPI_ISL_982156 | Hôpital Pitié-Salpêtrière | Department of Virology, Henri Mondor University Hospital, Assistance Publique Hôpitaux de Paris, Université Paris-Est Créteil, INSERM U955 | Christophe Rodriguez, Slim Fourati, Vanessa Demontant, Guillaume Gricourt, Melissa N'Debi, Alexandre Soulier, Elisabeth Trawinski, Jean-Michel Pawlotsky |
| EPI_ISL_982161 | Hôpital Henri Mondor | Department of Virology, Henri Mondor University Hospital, Assistance Publique Hôpitaux de Paris, Université Paris-Est Créteil, INSERM U955 | Christophe Rodriguez, Slim Fourati, Vanessa Demontant, Guillaume Gricourt, Melissa N'Debi, Alexandre Soulier, Elisabeth Trawinski, Jean-Michel Pawlotsky |
| EPI_ISL_982162 | G.H.E.F.Grand Hôpital EST Francilien | Department of Virology, Henri Mondor University Hospital, Assistance Publique Hôpitaux de Paris, Université Paris-Est Créteil, INSERM U955 | Christophe Rodriguez, Slim Fourati, Vanessa Demontant, Guillaume Gricourt, Melissa N'Debi, Alexandre Soulier, Elisabeth Trawinski, Jean-Michel Pawlotsky |
| EPI_ISL_982234, EPI_ISL_982235 | MEPHI, Aix Marseille University | MEPHI, Aix Marseille University | Anthony LEVASSEUR |
| EPI_ISL_982237, EPI_ISL_982238, EPI_ISL_982270, EPI_ISL_982275, EPI_ISL_982277, EPI_ISL_982278, EPI_ISL_982295, EPI_ISL_982296 | Lab voor klinische biologie | Lab voor klinische biologie | Hannelore Hamerlinck, Marija Janevska, Bruno Verhasselt |
| EPI_ISL_982303, EPI_ISL_982305, EPI_ISL_982306 | CH.INTERCOMMUNAL DE CRETEIL | Department of Virology, Henri Mondor University Hospital, Assistance Publique Hôpitaux de Paris, Université Paris-Est Créteil, INSERM U955 | Christophe Rodriguez, Slim Fourati, Vanessa Demontant, Guillaume Gricourt, Melissa N'Debi, Alexandre Soulier, Elisabeth Trawinski, Jean-Michel Pawlotsky |
| EPI_ISL_982310 | G.H.E.F.Grand Hôpital EST Francilien | Department of Virology, Henri Mondor University Hospital, Assistance Publique Hôpitaux de Paris, Université Paris-Est Créteil, INSERM U955 | Christophe Rodriguez, Slim Fourati, Vanessa Demontant, Guillaume Gricourt, Melissa N'Debi, Alexandre Soulier, Elisabeth Trawinski, Jean-Michel Pawlotsky |
| EPI_ISL_982312, EPI_ISL_982314, EPI_ISL_982315, EPI_ISL_982316, EPI_ISL_982317, EPI_ISL_982318, EPI_ISL_982319, EPI_ISL_982320 | CH.INTERCOMMUNAL DE CRETEIL | Department of Virology, Henri Mondor University Hospital, Assistance Publique Hôpitaux de Paris, Université Paris-Est Créteil, INSERM U955 | Christophe Rodriguez, Slim Fourati, Vanessa Demontant, Guillaume Gricourt, Melissa N'Debi, Alexandre Soulier, Elisabeth Trawinski, Jean-Michel Pawlotsky |
| EPI_ISL_982861, EPI_ISL_982866, EPI_ISL_982867, EPI_ISL_982869, EPI_ISL_982871, EPI_ISL_982877, EPI_ISL_982878, EPI_ISL_982879, EPI_ISL_982880, EPI_ISL_982881, EPI_ISL_982882, EPI_ISL_982889, EPI_ISL_982894, EPI_ISL_982896, EPI_ISL_982898, EPI_ISL_982909, EPI_ISL_982922, EPI_ISL_982923 | | | |
| see above | MEPHI Aix Marseille University (AMU) | MEPHI Aix Marseille University (AMU) | Anthony LEVASSEUR |
| EPI_ISL_983001 | National Institute of Laboratory Medicine and Referral Center | Genomic Research Lab, BCSIR | Tanjina Akhtar Banu, Mohammad Samir Uzzaman, Eshrar Osman, Md. Ahashan Habib, Shahina Akter, Abu Sayeed Mohammad Mahmud, Md. Murshed Hasan Sarkar, Barna Goswami, Iffat Jahan, Md. Saddam Hossain, Tasnim Nafisa, Md. Maruf Ahmed Molla, Mahmuda Yeasmin, Asish Kumar Ghosh, Arifa Akram, A. K. M.Shamsuzzaman, Md. Salim Khan |
| EPI_ISL_983097 | Microbiology and Virology Unit, Florence Careggi University Hospital | Microbiology and Virology Unit, Florence Careggi University Hospital | Vincenzo Di Pilato, Marco Coppi, Fabio Morecchiato, Noemi Aiezza, Ilaria Baccani, Alberto Antonelli, Emanuele Gori, Gian Maria Rossolini |
| EPI_ISL_983344 | National Institute of Laboratory Medicine and Referral Center | Genomic Research Lab, BCSIR | Tasnim Nafisa, Mohammad Samir Uzzaman, Eshrar Osman, Md. Ahashan Habib, Shahina Akter, Tanjina Akhtar Banu, Abu Sayeed Mohammad Mahmud, Md. Murshed Hasan Sarkar, Barna Goswami, Iffat Jahan, Md. Saddam Hossain, Md. Maruf Ahmed Molla, Mahmuda Yeasmin, Asish Kumar Ghosh, Arifa Akram, A. K.M.Shamsuzzaman, Md. Salim Khan |
| EPI_ISL_983371 | Utah Public Health Laboratory | Utah Public Health Laboratory | Erin L. Young, Kelly F. Oakeson, Tara Gallagher |
| EPI_ISL_983438, EPI_ISL_983440, EPI_ISL_983441, EPI_ISL_983442, EPI_ISL_983444 | URMC LABS | Wadsworth Center, New York State Department of Health | Kirsten St. George, Daryl M. Lamson, Alexis Russel, Matthew Shudt, Melissa A Leisner, Jonathan Plitnick, Navjot Singh, John Kelly, Erasmus Schneider, Erica Lasek-Nesselquist |
| EPI_ISL_983446 | THE MARY IMOGENE BASSETT HOSPITAL | Wadsworth Center, New York State Department of Health | Kirsten St. George, Daryl M. Lamson, Alexis Russel, Matthew Shudt, Melissa A Leisner, Jonathan Plitnick, Navjot Singh, John Kelly, Erasmus Schneider, Erica Lasek-Nesselquist |
| EPI_ISL_983461, EPI_ISL_983462, EPI_ISL_983463, EPI_ISL_983464, EPI_ISL_983465, EPI_ISL_983466, EPI_ISL_983467, EPI_ISL_983468, EPI_ISL_983469, EPI_ISL_983470, EPI_ISL_983471, EPI_ISL_983472, EPI_ISL_983473, EPI_ISL_983474, EPI_ISL_983475 | | | |
| see above | URMC LABS | Wadsworth Center, New York State Department of Health | Kirsten St. George, Daryl M. Lamson, Alexis Russel, Matthew Shudt, Melissa A Leisner, Jonathan Plitnick, Navjot Singh, John Kelly, Erasmus Schneider, Erica Lasek-Nesselquist |
| EPI_ISL_983476 | Wadsworth Center, New York State Department of Health | Wadsworth Center, New York State Department of Health | Kirsten St. George, Daryl M. Lamson, Alexis Russel, Matthew Shudt, Melissa A Leisner, Jonathan Plitnick, Navjot Singh, John Kelly, Erasmus Schneider, Erica Lasek-Nesselquist |
| EPI_ISL_983477 | THE MARY IMOGENE BASSETT HOSPITAL | Wadsworth Center, New York State Department of Health | Kirsten St. George, Daryl M. Lamson, Alexis Russel, Matthew Shudt, Melissa A Leisner, Jonathan Plitnick, Navjot Singh, John Kelly, Erasmus Schneider, Erica Lasek-Nesselquist |
| EPI_ISL_983478, EPI_ISL_983479, EPI_ISL_983480, EPI_ISL_983481, EPI_ISL_983482, EPI_ISL_983483, EPI_ISL_983484 | URMC LABS | Wadsworth Center, New York State Department of Health | Kirsten St. George, Daryl M. Lamson, Alexis Russel, Matthew Shudt, Melissa A Leisner, Jonathan Plitnick, Navjot Singh, John Kelly, Erasmus Schneider, Erica Lasek-Nesselquist |
| EPI_ISL_983485, EPI_ISL_983486, EPI_ISL_983487 | THE MARY IMOGENE BASSETT HOSPITAL | Wadsworth Center, New York State Department of Health | Kirsten St. George, Daryl M. Lamson, Alexis Russel, Matthew Shudt, Melissa A Leisner, Jonathan Plitnick, Navjot Singh, John Kelly, Erasmus Schneider, Erica Lasek-Nesselquist |
| EPI_ISL_983494, EPI_ISL_983495, EPI_ISL_983496, EPI_ISL_983497, EPI_ISL_983498, EPI_ISL_983499 | Kansas Health and Environmental Lab | Kansas Health and Environmental Lab | Mike Grose, Paige Drury, Carissa Robertson, Ben Olsen, and Phil Adam |
| EPI_ISL_983599, EPI_ISL_983600, EPI_ISL_983601, EPI_ISL_983602, EPI_ISL_983603, EPI_ISL_983604 | Texas Department of State Health Services | Texas Department of State Health Services | Bonnie Oh, Anita Pokharel, James Daniel Bonser, Myong Koag, Chung Wang, Rachel Lee, Grace Kubin, Rashmi Tuladhar, Mayela Pedrueza, Maliha Rahman, Jenny Zhang |
| EPI_ISL_983652, EPI_ISL_983692, EPI_ISL_983693, EPI_ISL_983694, | Vault Health | Minnesota Department of Health, Public Health Laboratory | Alexandra Lorentz, Jacob Garfin, Matt Plumb, and Xiong Wang |

| | | | |
|---|---|---|---|
| EPI_ISL_983695, EPI_ISL_983696, EPI_ISL_983697 | | | |
| EPI_ISL_983705, EPI_ISL_983706, EPI_ISL_983718, EPI_ISL_983719, EPI_ISL_983726, EPI_ISL_983727, EPI_ISL_983731, EPI_ISL_983733, EPI_ISL_983734, EPI_ISL_983747, EPI_ISL_983748, EPI_ISL_983749, EPI_ISL_983750, EPI_ISL_983751, EPI_ISL_983752, EPI_ISL_983753, EPI_ISL_983754, EPI_ISL_983755, EPI_ISL_983756, EPI_ISL_983757, EPI_ISL_983758, EPI_ISL_983759, EPI_ISL_983760, EPI_ISL_983761, EPI_ISL_983762, EPI_ISL_983763, EPI_ISL_983764, EPI_ISL_983765, EPI_ISL_983766, EPI_ISL_983767, EPI_ISL_983768, EPI_ISL_983769, EPI_ISL_983770, EPI_ISL_983771, EPI_ISL_983772, EPI_ISL_983773, EPI_ISL_983774, EPI_ISL_983775, EPI_ISL_983776, EPI_ISL_983777, EPI_ISL_983778, EPI_ISL_983779, EPI_ISL_983780, EPI_ISL_983781, EPI_ISL_983782 | | | |
| see above | Colorado Department of Public Health and Environment | Colorado Department of Puplic Health and Environment | Laura Bankers, Molly C. Hetherington-Rauth, Diana Ir, Shannon Ely, Shannon R. Matzinger, Sarah Elizabeth Totten, Emily A. Travanty |
