## Supplementary material for "Detection and characterization of the SARS-CoV-2 lineage B.1.526 in New York": Supp. Table 3 GISAID Acknowledgment part 2: gisaid_hcov-19_acknowledgement_table_2021_02_13_010-3.pdf

We gratefully acknowledge the following Authors from the Originating laboratories responsible for obtaining the specimens, as well as the Submitting laboratories where the genome data were generated and shared via GISAID, on which this research is based.

All Submitters of data may be contacted directly via www.gisaid.org

Authors are sorted alphabetically.

| Accession ID | Originating Laboratory | Submitting Laboratory | Authors |
|---|---|---|---|
| EPI_ISL_728587 | Dutch COVID-19 response team | National Institute for Public Health and the Environment (RIVM) | Adam Meijer, Harry Vennema, Jeroen Cremer, Sharon van den Brink, Bas van der Veer, AnneMarie van den Brandt, Florian Zwagemaker, Dennis Schmitz, Chantal Reusken, on behalf of the national COVID-19 response team |
| EPI_ISL_737930 | Hospital Universitario 12 de Octubre | Hospital Universitario 12 de Octubre | Raul Recio, Esther Viedma, Sara González, Irene Múñoz-Gallego, Mª Dolores Folgueira, Rafael Delgado |
| EPI_ISL_738137 | Area of Virology, Serology and Virology Division (SAViD), New South Wales Health Pathology Randwick | Area of Virology, Serology and Virology Division (SAViD), New South Wales Health Pathology Randwick | Rawlinson, W., Deveson, I., Van Haal, S., Foster, C. |
| EPI_ISL_738139, EPI_ISL_738141, EPI_ISL_738142 | Division of Emerging Infectious Diseases, Bureau of Infectious Diseases Diagnosis Control, Korea Disease Control and Prevention Agency | Division of Emerging Infectious Diseases, Bureau of Infectious Diseases Diagnosis Control, Korea Disease Control and Prevention Agency | Jeong-Min Kim, Il-Hwan Kim, Ae Kyung Park, Namjoo Lee, Sang Hee Woo, Heui Man Kim, Eun-Jin Kim |
| EPI_ISL_740896 | Histopath | NSW Health Pathology - Institute of Clinical Pathology and Medical Research; Westmead Hospital; University of Sydney | CIDM-PH et al. |
| EPI_ISL_745194, EPI_ISL_745195, EPI_ISL_745196 | Osmania Medical College | CSIR-Centre for Cellular and Molecular Biology | Dr.V.Sudha Rani,Dr.S.Pavani,Dr.Satyaprasad,Dr.P.Shashikala Reddy,Namami Gaur,Sakshi Shambhavi,Lamuk Zaveri,Shagufta Khan,Nikhil Hajirnis,M Soujanya Reddy,Pratheusa Maccha,Tulasi Nagabandi,Purushotham Vodnala,Blessy B John,Viswagithe S L,B Himasri,Onkar Kulkarni,Payel Mukherjee,Sofia Banu,Priya Singh,Archana Bharadwaj Siva,Karthik Bharadwaj Tallapaka,Rakesh K Mishra,Divya Tej Sowpati |
| EPI_ISL_745197 | Kakatiya Medical College | CSIR-Centre for Cellular and Molecular Biology | Dr.V.Sudha Rani,Dr.S.Pavani,Dr.Satyaprasad,Dr.P.Shashikala Reddy,Namami Gaur,Sakshi Shambhavi,Lamuk Zaveri,Shagufta Khan,Nikhil Hajirnis,M Soujanya Reddy,Pratheusa Maccha,Tulasi Nagabandi,Purushotham Vodnala,Blessy B John,Viswagithe S L,B Himasri,Onkar Kulkarni,Payel Mukherjee,Sofia Banu,Priya Singh,Archana Bharadwaj Siva,Karthik Bharadwaj Tallapaka,Rakesh K Mishra,Divya Tej Sowpati |
| EPI_ISL_745198 | ESI Hospital | CSIR-Centre for Cellular and Molecular Biology | Dr.V.Sudha Rani,Dr.S.Pavani,Dr.Satyaprasad,Dr.P.Shashikala Reddy,Namami Gaur,Sakshi Shambhavi,Lamuk Zaveri,Shagufta Khan,Nikhil Hajirnis,M Soujanya Reddy,Pratheusa Maccha,Tulasi Nagabandi,Purushotham Vodnala,Blessy B John,Viswagithe S L,B Himasri,Onkar Kulkarni,Payel Mukherjee,Sofia Banu,Priya Singh,Archana Bharadwaj Siva,Karthik Bharadwaj Tallapaka,Rakesh K Mishra,Divya Tej Sowpati |
| EPI_ISL_745199 | Rangaraya Medical College | CSIR-Centre for Cellular and Molecular Biology | Dr.V.Sudha Rani,Dr.S.Pavani,Dr.Satyaprasad,Dr.P.Shashikala Reddy,Namami Gaur,Sakshi Shambhavi,Lamuk Zaveri,Shagufta Khan,Nikhil Hajirnis,M Soujanya Reddy,Pratheusa Maccha,Tulasi Nagabandi,Purushotham Vodnala,Blessy B John,Viswagithe S L,B Himasri,Onkar Kulkarni,Payel Mukherjee,Sofia Banu,Priya Singh,Archana Bharadwaj Siva,Karthik Bharadwaj Tallapaka,Rakesh K Mishra,Divya Tej Sowpati |
| EPI_ISL_745200 | Nizams Institute of Medical Sciences | CSIR-Centre for Cellular and Molecular Biology | Dr.V.Sudha Rani,Dr.S.Pavani,Dr.Satyaprasad,Dr.P.Shashikala Reddy,Namami Gaur,Sakshi Shambhavi,Lamuk Zaveri,Shagufta Khan,Nikhil Hajirnis,M Soujanya Reddy,Pratheusa Maccha,Tulasi Nagabandi,Purushotham Vodnala,Blessy B John,Viswagithe S L,B Himasri,Onkar Kulkarni,Payel Mukherjee,Sofia Banu,Priya Singh,Archana Bharadwaj Siva,Karthik Bharadwaj Tallapaka,Rakesh K Mishra,Divya Tej Sowpati |
| EPI_ISL_745201 | Guntur General Hospital | CSIR-Centre for Cellular and Molecular Biology | Dr.V.Sudha Rani,Dr.S.Pavani,Dr.Satyaprasad,Dr.P.Shashikala Reddy,Namami Gaur,Sakshi Shambhavi,Lamuk Zaveri,Shagufta Khan,Nikhil Hajirnis,M Soujanya Reddy,Pratheusa Maccha,Tulasi Nagabandi,Purushotham Vodnala,Blessy B John,Viswagithe S L,B Himasri,Onkar Kulkarni,Payel Mukherjee,Sofia Banu,Priya Singh,Archana Bharadwaj Siva,Karthik Bharadwaj Tallapaka,Rakesh K Mishra,Divya Tej Sowpati |
| EPI_ISL_746835 | National Institute for Infectious Diseases, INMI, "L. Spallanzani" IRCCS | National Institute for Infectious Diseases, INMI, "L. Spallanzani" IRCCS | M Rueca, E Giombini, B Bartolini, C.E.M Gruber, F Messina, A Di Caro, MR Capobianchi |
| EPI_ISL_747244 | BBMP Urban PHC | Department of Neurovirology, National Institute of Mental Health and Neurosciences (NIMHANS) | Chitra Pattabiraman, Pramada Prasad, Anita S Desai, V Ravi |
| EPI_ISL_747523, EPI_ISL_747525 | Dutch COVID-19 response team | National Institute for Public Health and the Environment (RIVM) | Adam Meijer, Harry Vennema, Jeroen Cremer, Sharon van den Brink, Bas van der Veer, AnneMarie van den Brandt, Florian Zwagemaker, Dennis Schmitz, Chantal Reusken, on behalf of the national COVID-19 response team |
| EPI_ISL_751194, EPI_ISL_751195, EPI_ISL_751196, EPI_ISL_751197, EPI_ISL_751198, EPI_ISL_751199, EPI_ISL_751200 | Dienststelle Gesundheit und Sport Kanton Luzern | Institute of Medical Virology, University of Zurich | Stefan Schmutz, Kevin Steiner, Verena Kufner, Maryam Zaheri, Gabriela Ziltener, Jürg Böni, Michael Huber, Alexandra Trkola, Claudia Schmutz, Eva Spieler |
| EPI_ISL_751800 | Colorado Department of Public Health and Environment | Colorado Department of Public Health & Environment | Laura Bankers, Molly C. Hetherington-Rauth, Diana Ir, Shannon Ely, Shannon R. Matzinger, Sarah Elizabeth Totten, Emily A. Travanty |
| EPI_ISL_752598 | SA Pathology | SA Pathology | Lex Leong, Julien Soubrier, Chuan Kok Lim, Song Gao, Mark Turra, Karin Kassahn, Ivan Bastian, Geoff Higgins |
| EPI_ISL_754076, EPI_ISL_754100, EPI_ISL_754101, EPI_ISL_754104, EPI_ISL_754106, EPI_ISL_754107, EPI_ISL_754108 | National Public Health Laboratory, National Centre for Infectious Diseases | National Public Health Laboratory, National Centre for Infectious Diseases | Tze Minn Mak, Sophie Octavia, Zhenyang Zhou, Lin Cui, Raymond Tzer Pin Lin |
| EPI_ISL_754123 | UCLA Clinical Micro Lab | Los Angeles County PHL | P. Hemarajata et al. |
| EPI_ISL_754236 | Diagnósticos da América - DASA | Instituto de Medicina Tropical Universidade de São Paulo | Brazil-UK Centre for Arbovirus Discovery Diagnosis Genomics and Epidemiology (CADDE) Genomic Network - Instituto de Medicina Tropical |
| EPI_ISL_754246, EPI_ISL_754255, EPI_ISL_754279, EPI_ISL_754285, EPI_ISL_754330 | Respiratory Virus Unit, National Infection Service, Public Health England | COVID-19 Genomics UK (COG-UK) Consortium | PHE Covid Sequencing Team |
| EPI_ISL_755572 | Center of Advanced Studies and Technology, CAST | Center of Advanced Studies and Technology, CAST | Ferrante,R., Mandatori,D., De Fabritiis,S. |
| EPI_ISL_755588 | Regional medical sciences center 6 Chonburi | National Institute of Health, Department of Medical Sciences, Ministry of Public Health, Thailand | Pilailuk Okada; Siripaporn Phuygun; Thanutsapa Thanadachakul; Sittiporn Parnmen; Pakorn Piromtong; Warawan Wongboot; Sunthareeya Waicharoen; Malinee Chittaganpitch |
| EPI_ISL_755589 | Laboratoire national de santé, Microbiology, Virology | Laboratoire national de santé, Microbiology, Microbial Genomics Platform | Anke Wienecke-Baldacchino, Catherine Ragimbeau,Jessica Tapp, Fatu Djabi, Lise Pignon, Raoul Salmon, Tamir Abdelrahman |
| EPI_ISL_755630, EPI_ISL_755631 | LabPLUS | Institute of Environmental Science and Research (ESR) | Xiaoyun Ren, Matt Storey, Nikki Freed, Muhammad Faisal, Jing Wang, Hermes Perez, Anja Werno, Antje van der Linden, Arlo Upton, Chris Mansell, David Hammer, Dragana Drinkovic, Gary McAuliffe, Hana Sofia Andersson, James Ussher, Jill Sherwood, Josh Freeman, Julia Howard, Juliet Elvy, Mary DeAlmeida, Matt Blakiston, Matthew Rogers, Max Bloomfield, Michael Addidle, Michelle Balm, Sally Roberts, Sarah Jefferies, Sharmini Muttaiyah, Susan Morpeth, Susan Taylor, Timothy Blackmore, Vani Sathyendran, Veronica Playle, Virginia Hope, Erasmus Smit, Lauren Jelly, Olin Silander, Joep de Ligt |
| EPI_ISL_755634 | Middlemore Hospital | Institute of Environmental Science and Research (ESR) | Xiaoyun Ren, Matt Storey, Nikki Freed, Muhammad Faisal, Jing Wang, Hermes Perez, Anja Werno, Antje van der Linden, Arlo Upton, Chris Mansell, David Hammer, Dragana Drinkovic, Gary McAuliffe, Hana Sofia Andersson, James Ussher, Jill Sherwood, Josh Freeman, Julia Howard, Juliet Elvy, Mary DeAlmeida, Matt Blakiston, Matthew Rogers, Max Bloomfield, Michael Addidle, Michelle Balm, Sally Roberts, Sarah Jefferies, Sharmini Muttaiyah, Susan Morpeth, Susan Taylor, Timothy Blackmore, Vani Sathyendran, Veronica Playle, Virginia Hope, Erasmus Smit, Lauren Jelly, Olin Silander, Joep de Ligt |

| | | | |
|---|---|---|---|
| EPI_ISL_755637 | LabPLUS | Institute of Environmental Science and Research (ESR) | Xiaoyun Ren, Matt Storey, Nikki Freed, Muhammad Faisal, Jing Wang, Hermes Perez, Anja Werno, Antje van der Linden, Arlo Upton, Chris Mansell, David Hammer, Dragana Drinkovic, Gary McAuliffe, Hana Sofia Andersson, James Ussher, Jill Sherwood, Josh Freeman, Julia Howard, Juliet Elvy, Mary DeAlmeida, Matt Blakiston, Matthew Rogers, Max Bloomfield, Michael Addidle, Michelle Balm, Sally Roberts, Sarah Jefferies, Sharmini Muttaiyah, Susan Morpeth, Susan Taylor, Timothy Blackmore, Vani Sathyendran, Veronica Playle, Virginia Hope, Erasmus Smit, Lauren Jelly, Olin Silander, Joep de Ligt |
| EPI_ISL_755638 | Canterbury Health Laboratories | Institute of Environmental Science and Research (ESR) | Xiaoyun Ren, Matt Storey, Nikki Freed, Muhammad Faisal, Jing Wang, Hermes Perez, Anja Werno, Antje van der Linden, Arlo Upton, Chris Mansell, David Hammer, Dragana Drinkovic, Gary McAuliffe, Hana Sofia Andersson, James Ussher, Jill Sherwood, Josh Freeman, Julia Howard, Juliet Elvy, Mary DeAlmeida, Matt Blakiston, Matthew Rogers, Max Bloomfield, Michael Addidle, Michelle Balm, Sally Roberts, Sarah Jefferies, Sharmini Muttaiyah, Susan Morpeth, Susan Taylor, Timothy Blackmore, Vani Sathyendran, Veronica Playle, Virginia Hope, Erasmus Smit, Lauren Jelly, Olin Silander, Joep de Ligt |
| EPI_ISL_756260, EPI_ISL_756261, EPI_ISL_756262, EPI_ISL_756263, EPI_ISL_756264, EPI_ISL_756265, EPI_ISL_756266, EPI_ISL_756267, EPI_ISL_756268, EPI_ISL_756269, EPI_ISL_756270 | | | |
| see above | Wyoming Public Health Laboratory | Wyoming Public Health Laboratory | Noah Hull, Taylor Fearing, Channing Weber, Ashley Norberg, Bailey Bowcutt, and Wanda Manley |
| EPI_ISL_756271, EPI_ISL_756273 | Hospital General Universitario Gregorio Marañón | Hospital General Universitario Gregorio Marañón | Sergio Buenestado Serrano, Pedro Sola Campoy, Pilar Catalán, Patricia Muñoz, Darío García de Viedma. |
| EPI_ISL_757285 | Department of Virology, Public Health Laboratories Division | Department of Virology, Public Health Laboratories Division | Massab Umair, Aamer Ikram, Muhammad Salman |
| EPI_ISL_759706 | University of Wisconsin-Madison AIDS Vaccine Research Laboratories | University of Wisconsin-Madison AIDS Vaccine Research Laboratories | Gage Moreno, Katarina Braun, et al. AIDS Vaccine Research Laboratories |
| EPI_ISL_759707, EPI_ISL_759708, EPI_ISL_759709, EPI_ISL_759713, EPI_ISL_759714, EPI_ISL_759715, EPI_ISL_759716, EPI_ISL_759717, EPI_ISL_759719, EPI_ISL_759720 | Instituto Nacional de Saude (INSA) | Instituto Nacional de Saude (INSA) | Borges et al |
| EPI_ISL_759725, EPI_ISL_759730, EPI_ISL_759732, EPI_ISL_759733, EPI_ISL_759744, EPI_ISL_759752, EPI_ISL_759753, EPI_ISL_759754, EPI_ISL_759755, EPI_ISL_759756, EPI_ISL_759757 | | | |
| see above | University of Wisconsin-Madison AIDS Vaccine Research Laboratories | University of Wisconsin-Madison AIDS Vaccine Research Laboratories | Gage Moreno, Katarina Braun, et al. AIDS Vaccine Research Laboratories |
| EPI_ISL_759761, EPI_ISL_759762 | Instituto Nacional de Saude (INSA) | Instituto Nacional de Saude (INSA) | Borges et al |
| EPI_ISL_759893, EPI_ISL_759894 | Temporary Specimen Collection Centre | Hong Kong Department of Health | Alan K.L. Tsang, Peter C.W. Yip, Edman T.K. Lam, Rickjason C.W. Chan, Dominic N.C. Tsang |
| EPI_ISL_759895, EPI_ISL_759897, EPI_ISL_759898 | United Christian Hospital | Hong Kong Department of Health | Alan K.L. Tsang, Peter C.W. Yip, Edman T.K. Lam, Rickjason C.W. Chan, Dominic N.C. Tsang |
| EPI_ISL_760251, EPI_ISL_760258, EPI_ISL_760260, EPI_ISL_760269, EPI_ISL_760270, EPI_ISL_760271, EPI_ISL_760275, EPI_ISL_760290, EPI_ISL_760310, EPI_ISL_760319, EPI_ISL_760325, EPI_ISL_760330, EPI_ISL_760332, EPI_ISL_760334, EPI_ISL_760336, EPI_ISL_760338, EPI_ISL_760339, EPI_ISL_760343, EPI_ISL_760347, EPI_ISL_760348, EPI_ISL_760361, EPI_ISL_760366, EPI_ISL_760367, EPI_ISL_760376, EPI_ISL_760388, EPI_ISL_760394, EPI_ISL_760398, EPI_ISL_760399, EPI_ISL_760400, EPI_ISL_760401, EPI_ISL_760407, EPI_ISL_760410, EPI_ISL_760412, EPI_ISL_760420, EPI_ISL_760423, EPI_ISL_760426, EPI_ISL_760429, EPI_ISL_760437, EPI_ISL_760438, EPI_ISL_760444, EPI_ISL_760451, EPI_ISL_760454, EPI_ISL_760456, EPI_ISL_760458, EPI_ISL_760459, EPI_ISL_760464, EPI_ISL_760467, EPI_ISL_760472, EPI_ISL_760480, EPI_ISL_760481, EPI_ISL_760493, EPI_ISL_760502, EPI_ISL_760503, EPI_ISL_760506, EPI_ISL_760513, EPI_ISL_760514, EPI_ISL_760517, EPI_ISL_760526, EPI_ISL_760533 | | | |
| see above | Lighthouse Lab in Glasgow | Wellcome Sanger Institute for the COVID-19 Genomics UK (COG-UK) Consortium | Harper VanSteenhouse, Yumi Kasai, David Gray, Carol Clugston, Anna Dominiczak and Alex Alderton, Roberto Amato, Sonia Goncalves, Ewan Harrison, David K. Jackson, Ian Johnston, Dominic Kwiatkowski, Cordelia Langford, John Sillitoe on behalf of the Wellcome Sanger Institute COVID-19 Surveillance Team |
| EPI_ISL_761185 | Lighthouse Lab in Alderley Park | Wellcome Sanger Institute for the COVID-19 Genomics UK (COG-UK) Consortium | Jacquelyn Wynn, Mairead Hyland, The Lighthouse Lab in Alderley Park and Alex Alderton, Roberto Amato, Sonia Goncalves, Ewan Harrison, David K. Jackson, Ian Johnston, Dominic Kwiatkowski, Cordelia Langford, John Sillitoe on behalf of the Wellcome Sanger Institute COVID-19 Surveillance Team |
| EPI_ISL_762992 | Division of Emerging Infectious Diseases, Bureau of Infectious Diseases Diagnosis Control, Korea Disease Control and Prevention Agency | Division of Emerging Infectious Diseases, Bureau of Infectious Diseases Diagnosis Control, Korea Disease Control and Prevention Agency | Ae Kyung Park, Il-Hwan Kim, Heui Man Kim, Jeong-Min Kim, Namjoo Lee, Chaeyoung Lee, Sang Hee Woo, Eun-Jin Kim |
| EPI_ISL_763075 | Diagnosticos da America - DASA | Instituto Adolfo Lutz, Interdiciplinary Procedures Center, Strategic Laboratory | Claudio Tavares Sacchi, Claudia Regina Gonçalves, Erica Valessa Ramos Gomes, Karoline Rodrigues Campos |
| EPI_ISL_763294, EPI_ISL_763295, EPI_ISL_763296, EPI_ISL_763297, EPI_ISL_763299, EPI_ISL_763300, EPI_ISL_763301, EPI_ISL_763302, EPI_ISL_763303, EPI_ISL_763304, EPI_ISL_763309, EPI_ISL_763311, EPI_ISL_763312, EPI_ISL_763314, EPI_ISL_763357 | | | |
| see above | Dutch COVID-19 response team | Erasmus Medical Center | Bas Oude Munnink, Reina Sikkema, David Nieuwenhuijse, Irina Chestakova, Anne van der Linden, Marjan Boter, Emmanuelle Munger, Corine GeurtsvanKessel, Annemiek van der Eijk, Richard Molenkamp, Marion Koopmans, on behalf of the Dutch national COVID-19 response team. |
| EPI_ISL_763369 | Virology Department, Sheffield Teaching Hospitals NHS Foundation Trust/Department of Infection, Immunity and Cardiovascular Disease, The Medical School, University of Sheffield | COVID-19 Genomics UK (COG-UK) Consortium | Thushan de Silva, Matthew Parker, Nikki Smith, Adri Angyal, Rebecca Brown, Luke Green, Rachel Tucker, Paul Parsons, Danielle Groves, Katie Johnson, Laura Carrilero, Alex Keeley, Dave Partridge, Matthew Wyles, Benjamin Lindsey, Mehmet Yavuz, Mohammad Raza, Cariad Evans |
| EPI_ISL_763379, EPI_ISL_763380 | Department of Pathology, University of Cambridge | COVID-19 Genomics UK (COG-UK) Consortium | Aminu S. Jahun, Yasmin Chaudhry, Grant Hall, Iliana Georgana, Myra Hosmillo, Martin D. Curran, Malte Pinckert, Surendra Parmar, Ian Goodfellow |
| EPI_ISL_763389 | Queens Medical Centre, Clinical Microbiology Department / DeepSeq Nottingham | COVID-19 Genomics UK (COG-UK) Consortium | Gemma Clark, Wendy Smith, Manjinder Khakh, Vicki M Fleming, Michelle M Lister, Hannah Howson-Wells, Jonathan Ball, Patrick McClure, Joseph Chappell, Theocharis Tsoleridis, Nadine Holmes, Matthew Carlisle, Christopher Moore, Fei Sang, Johnny Debebe, Victoria Wright, Matthew Loose |
| EPI_ISL_763394 | Virology Department, Sheffield Teaching Hospitals NHS Foundation Trust/Department of Infection, Immunity and Cardiovascular Disease, The Medical School, University of Sheffield | COVID-19 Genomics UK (COG-UK) Consortium | Thushan de Silva, Matthew Parker, Nikki Smith, Adri Angyal, Rebecca Brown, Luke Green, Rachel Tucker, Paul Parsons, Danielle Groves, Katie Johnson, Laura Carrilero, Alex Keeley, Dave Partridge, Matthew Wyles, Benjamin Lindsey, Mehmet Yavuz, Mohammad Raza, Cariad Evans |
| EPI_ISL_763406 | Quadram Institute Bioscience | COVID-19 Genomics UK (COG-UK) Consortium | Dave J. Baker, Gemma L. Kay, Alp Aydin, Thanh Le-Viet, Steven Rudder, Ana P. Tedim, Anastasia Kolyva, Maria Diaz, Leonardo de Oliveira Martins, Nabil-Fareed Alikhan, Lizzie Meadows, Rachael Stanley, Ngozi Elumogo, Muhammed Yasir, Nicholas M. Thomson, Alexander J Trotter, Rachel Gilroy, Samuel Bloomfield, Claire Stuart, Andrew Bell, Reenesh Prakash, Samir Dervisevic, Alison E. Mather, John Wain, Mark Webber, Andrew J. Page, Justin O'Grady |
| EPI_ISL_763415, EPI_ISL_763424, EPI_ISL_763439 | Virology Department, Sheffield Teaching Hospitals NHS Foundation Trust/Department of Infection, Immunity and Cardiovascular Disease, The Medical School, University of Sheffield | COVID-19 Genomics UK (COG-UK) Consortium | Thushan de Silva, Matthew Parker, Nikki Smith, Adri Angyal, Rebecca Brown, Luke Green, Rachel Tucker, Paul Parsons, Danielle Groves, Katie Johnson, Laura Carrilero, Alex Keeley, Dave Partridge, Matthew Wyles, Benjamin Lindsey, Mehmet Yavuz, Mohammad Raza, Cariad Evans |
| EPI_ISL_763447 | Virology Department, Royal Infirmary of Edinburgh, NHS Lothian / School of Biological Sciences, University of Edinburgh / Institute of Genetics and Molecular Medicine, University of Edinburgh | COVID-19 Genomics UK (COG-UK) Consortium | McHugh M, Dewar R, Rooke S, Gallagher M, Balcaza C, O'Toole Á, Scher E, Hill V, McCrone JT, Colquhoun R, Yu X, Jackson B, Rambaut A, Williams TC, Templeton K |
| EPI_ISL_763449 | Queens Medical Centre, Clinical Microbiology Department / DeepSeq Nottingham | COVID-19 Genomics UK (COG-UK) Consortium | Gemma Clark, Wendy Smith, Manjinder Khakh, Vicki M Fleming, Michelle M Lister, Hannah Howson-Wells, Jonathan Ball, Patrick McClure, Joseph Chappell, Theocharis Tsoleridis, Nadine Holmes, Matthew Carlisle, Christopher Moore, Fei Sang, Johnny Debebe, Victoria Wright, Matthew Loose |
| EPI_ISL_763464, EPI_ISL_763469 | Department of Pathology, University of Cambridge | COVID-19 Genomics UK (COG-UK) Consortium | Aminu S. Jahun, Yasmin Chaudhry, Grant Hall, Iliana Georgana, Myra Hosmillo, Martin D. Curran, Malte Pinckert, Surendra Parmar, Ian Goodfellow |
| EPI_ISL_763478 | Quadram Institute Bioscience | COVID-19 Genomics UK (COG-UK) Consortium | Dave J. Baker, Gemma L. Kay, Alp Aydin, Thanh Le-Viet, Steven Rudder, Ana P. Tedim, Anastasia Kolyva, Maria Diaz, Leonardo de Oliveira Martins, Nabil-Fareed Alikhan, Lizzie Meadows, Rachael Stanley, Ngozi Elumogo, Muhammed Yasir, Nicholas M. Thomson, Alexander J Trotter, Rachel Gilroy, Samuel Bloomfield, Claire Stuart, Andrew Bell, Reenesh Prakash, Samir Dervisevic, Alison E. Mather, John Wain, Mark Webber, Andrew J. Page, Justin O'Grady |

| | | | |
|---|---|---|---|
| EPI_ISL_763499, EPI_ISL_763504 | Virology Department, Sheffield Teaching Hospitals NHS Foundation Trust/Department of Infection, Immunity and Cardiovascular Disease, The Medical School, University of Sheffield | COVID-19 Genomics UK (COG-UK) Consortium | Thushan de Silva, Matthew Parker, Nikki Smith, Adri Angyal, Rebecca Brown, Luke Green, Rachel Tucker, Paul Parsons, Danielle Groves, Katie Johnson, Laura Carrilero, Alex Keeley, Dave Partridge, Matthew Wyles, Benjamin Lindsey, Mehmet Yavuz, Mohammad Raza, Cariad Evans |
| EPI_ISL_763508, EPI_ISL_763518 | Department of Pathology, University of Cambridge | COVID-19 Genomics UK (COG-UK) Consortium | Aminu S. Jahun, Yasmin Chaudhry, Grant Hall, Iliana Georgana, Myra Hosmillo, Martin D. Curran, Malte Pinckert, Surendra Parmar, Ian Goodfellow |
| EPI_ISL_763547 | Queens Medical Centre, Clinical Microbiology Department / DeepSeq Nottingham | COVID-19 Genomics UK (COG-UK) Consortium | Gemma Clark, Wendy Smith, Manjinder Khakh, Vicki M Fleming, Michelle M Lister, Hannah Howson-Wells, Jonathan Ball, Patrick McClure, Joseph Chappell, Theocharis Tsoleridis, Nadine Holmes, Matthew Carlisle, Christopher Moore, Fei Sang, Johnny Debebe, Victoria Wright, Matthew Loose |
| EPI_ISL_763548 | Virology Department, Royal Infirmary of Edinburgh, NHS Lothian / School of Biological Sciences, University of Edinburgh / Institute of Genetics and Molecular Medicine, University of Edinburgh | COVID-19 Genomics UK (COG-UK) Consortium | McHugh M, Dewar R, Rooke S, Gallagher M, Balcaza C, O'Toole Á, Scher E, Hill V, McCrone JT, Colquhoun R, Yu X, Jackson B, Rambaut A, Williams TC, Templeton K |
| EPI_ISL_763580 | Queens Medical Centre, Clinical Microbiology Department / DeepSeq Nottingham | COVID-19 Genomics UK (COG-UK) Consortium | Gemma Clark, Wendy Smith, Manjinder Khakh, Vicki M Fleming, Michelle M Lister, Hannah Howson-Wells, Jonathan Ball, Patrick McClure, Joseph Chappell, Theocharis Tsoleridis, Nadine Holmes, Matthew Carlisle, Christopher Moore, Fei Sang, Johnny Debebe, Victoria Wright, Matthew Loose |
| EPI_ISL_763592 | Department of Pathology, University of Cambridge | COVID-19 Genomics UK (COG-UK) Consortium | Aminu S. Jahun, Yasmin Chaudhry, Grant Hall, Iliana Georgana, Myra Hosmillo, Martin D. Curran, Malte Pinckert, Surendra Parmar, Ian Goodfellow |
| EPI_ISL_763632 | Virology Department, Sheffield Teaching Hospitals NHS Foundation Trust/Department of Infection, Immunity and Cardiovascular Disease, The Medical School, University of Sheffield | COVID-19 Genomics UK (COG-UK) Consortium | Thushan de Silva, Matthew Parker, Nikki Smith, Adri Angyal, Rebecca Brown, Luke Green, Rachel Tucker, Paul Parsons, Danielle Groves, Katie Johnson, Laura Carrilero, Alex Keeley, Dave Partridge, Matthew Wyles, Benjamin Lindsey, Mehmet Yavuz, Mohammad Raza, Cariad Evans |
| EPI_ISL_763650, EPI_ISL_763663 | Queens Medical Centre, Clinical Microbiology Department / DeepSeq Nottingham | COVID-19 Genomics UK (COG-UK) Consortium | Gemma Clark, Wendy Smith, Manjinder Khakh, Vicki M Fleming, Michelle M Lister, Hannah Howson-Wells, Jonathan Ball, Patrick McClure, Joseph Chappell, Theocharis Tsoleridis, Nadine Holmes, Matthew Carlisle, Christopher Moore, Fei Sang, Johnny Debebe, Victoria Wright, Matthew Loose |
| EPI_ISL_763666 | Department of Pathology, University of Cambridge | COVID-19 Genomics UK (COG-UK) Consortium | Aminu S. Jahun, Yasmin Chaudhry, Grant Hall, Iliana Georgana, Myra Hosmillo, Martin D. Curran, Malte Pinckert, Surendra Parmar, Ian Goodfellow |
| EPI_ISL_763673, EPI_ISL_763678 | Queens Medical Centre, Clinical Microbiology Department / DeepSeq Nottingham | COVID-19 Genomics UK (COG-UK) Consortium | Gemma Clark, Wendy Smith, Manjinder Khakh, Vicki M Fleming, Michelle M Lister, Hannah Howson-Wells, Jonathan Ball, Patrick McClure, Joseph Chappell, Theocharis Tsoleridis, Nadine Holmes, Matthew Carlisle, Christopher Moore, Fei Sang, Johnny Debebe, Victoria Wright, Matthew Loose |
| EPI_ISL_763681 | Quadram Institute Bioscience | COVID-19 Genomics UK (COG-UK) Consortium | Dave J. Baker, Gemma L. Kay, Alp Aydin, Thanh Le-Viet, Steven Rudder, Ana P. Tedim, Anastasia Kolyva, Maria Diaz, Leonardo de Oliveira Martins, Nabil-Fareed Alikhan, Lizzie Meadows, Rachael Stanley, Ngozi Elumogo, Muhammed Yasir, Nicholas M. Thomson, Alexander J Trotter, Rachel Gilroy, Samuel Bloomfield, Claire Stuart, Andrew Bell, Reenesh Prakash, Samir Dervisevic, Alison E. Mather, John Wain, Mark Webber, Andrew J. Page, Justin O'Grady |
| EPI_ISL_763688 | Department of Pathology, University of Cambridge | COVID-19 Genomics UK (COG-UK) Consortium | Aminu S. Jahun, Yasmin Chaudhry, Grant Hall, Iliana Georgana, Myra Hosmillo, Martin D. Curran, Malte Pinckert, Surendra Parmar, Ian Goodfellow |
| EPI_ISL_763694, EPI_ISL_763695 | Virology Department, Royal Infirmary of Edinburgh, NHS Lothian / School of Biological Sciences, University of Edinburgh / Institute of Genetics and Molecular Medicine, University of Edinburgh | COVID-19 Genomics UK (COG-UK) Consortium | McHugh M, Dewar R, Rooke S, Gallagher M, Balcaza C, O'Toole Á, Scher E, Hill V, McCrone JT, Colquhoun R, Yu X, Jackson B, Rambaut A, Williams TC, Templeton K |
| EPI_ISL_763702 | Department of Pathology, University of Cambridge | COVID-19 Genomics UK (COG-UK) Consortium | Aminu S. Jahun, Yasmin Chaudhry, Grant Hall, Iliana Georgana, Myra Hosmillo, Martin D. Curran, Malte Pinckert, Surendra Parmar, Ian Goodfellow |
| EPI_ISL_763737, EPI_ISL_763797 | Quadram Institute Bioscience | COVID-19 Genomics UK (COG-UK) Consortium | Dave J. Baker, Gemma L. Kay, Alp Aydin, Thanh Le-Viet, Steven Rudder, Ana P. Tedim, Anastasia Kolyva, Maria Diaz, Leonardo de Oliveira Martins, Nabil-Fareed Alikhan, Lizzie Meadows, Rachael Stanley, Ngozi Elumogo, Muhammed Yasir, Nicholas M. Thomson, Alexander J Trotter, Rachel Gilroy, Samuel Bloomfield, Claire Stuart, Andrew Bell, Reenesh Prakash, Samir Dervisevic, Alison E. Mather, John Wain, Mark Webber, Andrew J. Page, Justin O'Grady |
| EPI_ISL_763816 | Department of Pathology, University of Cambridge | COVID-19 Genomics UK (COG-UK) Consortium | Aminu S. Jahun, Yasmin Chaudhry, Grant Hall, Iliana Georgana, Myra Hosmillo, Martin D. Curran, Malte Pinckert, Surendra Parmar, Ian Goodfellow |
| EPI_ISL_763817 | Virology Department, Royal Infirmary of Edinburgh, NHS Lothian / School of Biological Sciences, University of Edinburgh / Institute of Genetics and Molecular Medicine, University of Edinburgh | COVID-19 Genomics UK (COG-UK) Consortium | McHugh M, Dewar R, Rooke S, Gallagher M, Balcaza C, O'Toole Á, Scher E, Hill V, McCrone JT, Colquhoun R, Yu X, Jackson B, Rambaut A, Williams TC, Templeton K |
| EPI_ISL_763839 | Department of Pathology, University of Cambridge | COVID-19 Genomics UK (COG-UK) Consortium | Aminu S. Jahun, Yasmin Chaudhry, Grant Hall, Iliana Georgana, Myra Hosmillo, Martin D. Curran, Malte Pinckert, Surendra Parmar, Ian Goodfellow |
| EPI_ISL_763842 | Virology Department, Sheffield Teaching Hospitals NHS Foundation Trust/Department of Infection, Immunity and Cardiovascular Disease, The Medical School, University of Sheffield | COVID-19 Genomics UK (COG-UK) Consortium | Thushan de Silva, Matthew Parker, Nikki Smith, Adri Angyal, Rebecca Brown, Luke Green, Rachel Tucker, Paul Parsons, Danielle Groves, Katie Johnson, Laura Carrilero, Alex Keeley, Dave Partridge, Matthew Wyles, Benjamin Lindsey, Mehmet Yavuz, Mohammad Raza, Cariad Evans |
| EPI_ISL_763843, EPI_ISL_763850 | Department of Pathology, University of Cambridge | COVID-19 Genomics UK (COG-UK) Consortium | Aminu S. Jahun, Yasmin Chaudhry, Grant Hall, Iliana Georgana, Myra Hosmillo, Martin D. Curran, Malte Pinckert, Surendra Parmar, Ian Goodfellow |
| EPI_ISL_763853 | Quadram Institute Bioscience | COVID-19 Genomics UK (COG-UK) Consortium | Dave J. Baker, Gemma L. Kay, Alp Aydin, Thanh Le-Viet, Steven Rudder, Ana P. Tedim, Anastasia Kolyva, Maria Diaz, Leonardo de Oliveira Martins, Nabil-Fareed Alikhan, Lizzie Meadows, Rachael Stanley, Ngozi Elumogo, Muhammed Yasir, Nicholas M. Thomson, Alexander J Trotter, Rachel Gilroy, Samuel Bloomfield, Claire Stuart, Andrew Bell, Reenesh Prakash, Samir Dervisevic, Alison E. Mather, John Wain, Mark Webber, Andrew J. Page, Justin O'Grady |
| EPI_ISL_763855, EPI_ISL_763856, EPI_ISL_763857, EPI_ISL_763858 | Queens Medical Centre, Clinical Microbiology Department / DeepSeq Nottingham | COVID-19 Genomics UK (COG-UK) Consortium | Gemma Clark, Wendy Smith, Manjinder Khakh, Vicki M Fleming, Michelle M Lister, Hannah Howson-Wells, Jonathan Ball, Patrick McClure, Joseph Chappell, Theocharis Tsoleridis, Nadine Holmes, Matthew Carlisle, Christopher Moore, Fei Sang, Johnny Debebe, Victoria Wright, Matthew Loose |
| EPI_ISL_763866 | Virology Department, Sheffield Teaching Hospitals NHS Foundation Trust/Department of Infection, Immunity and Cardiovascular Disease, The Medical School, University of Sheffield | COVID-19 Genomics UK (COG-UK) Consortium | Thushan de Silva, Matthew Parker, Nikki Smith, Adri Angyal, Rebecca Brown, Luke Green, Rachel Tucker, Paul Parsons, Danielle Groves, Katie Johnson, Laura Carrilero, Alex Keeley, Dave Partridge, Matthew Wyles, Benjamin Lindsey, Mehmet Yavuz, Mohammad Raza, Cariad Evans |
| EPI_ISL_763883, EPI_ISL_763884, EPI_ISL_763885, EPI_ISL_763886, EPI_ISL_763887, EPI_ISL_763889, EPI_ISL_763890, EPI_ISL_763891 | Department of Pathology, University of Cambridge | COVID-19 Genomics UK (COG-UK) Consortium | Aminu S. Jahun, Yasmin Chaudhry, Grant Hall, Iliana Georgana, Myra Hosmillo, Martin D. Curran, Malte Pinckert, Surendra Parmar, Ian Goodfellow |
| EPI_ISL_763926, EPI_ISL_763927 | Queens Medical Centre, Clinical Microbiology Department / DeepSeq Nottingham | COVID-19 Genomics UK (COG-UK) Consortium | Gemma Clark, Wendy Smith, Manjinder Khakh, Vicki M Fleming, Michelle M Lister, Hannah Howson-Wells, Jonathan Ball, Patrick McClure, Joseph Chappell, Theocharis Tsoleridis, Nadine Holmes, Matthew Carlisle, Christopher Moore, Fei Sang, Johnny Debebe, Victoria Wright, Matthew Loose |
| EPI_ISL_764105, EPI_ISL_764106 | Virology Department, Sheffield Teaching Hospitals NHS Foundation Trust/Department of Infection, Immunity and Cardiovascular Disease, The Medical School, University of Sheffield | COVID-19 Genomics UK (COG-UK) Consortium | Thushan de Silva, Matthew Parker, Nikki Smith, Adri Angyal, Rebecca Brown, Luke Green, Rachel Tucker, Paul Parsons, Danielle Groves, Katie Johnson, Laura Carrilero, Alex Keeley, Dave Partridge, Matthew Wyles, Benjamin Lindsey, Mehmet Yavuz, Mohammad Raza, Cariad Evans |
| EPI_ISL_764113, EPI_ISL_764114 | Quadram Institute Bioscience | COVID-19 Genomics UK (COG-UK) Consortium | Dave J. Baker, Gemma L. Kay, Alp Aydin, Thanh Le-Viet, Steven Rudder, Ana P. Tedim, Anastasia Kolyva, Maria Diaz, Leonardo de Oliveira Martins, Nabil-Fareed Alikhan, Lizzie Meadows, Rachael Stanley, Ngozi Elumogo, Muhammed Yasir, Nicholas M. Thomson, Alexander J Trotter, Rachel Gilroy, Samuel Bloomfield, Claire Stuart, Andrew Bell, Reenesh Prakash, Samir Dervisevic, Alison E. Mather, John Wain, Mark Webber, Andrew J. Page, Justin O'Grady |
| EPI_ISL_764144, EPI_ISL_764149, EPI_ISL_764153 | Virology Department, Sheffield Teaching Hospitals NHS Foundation Trust/Department of Infection, Immunity and | COVID-19 Genomics UK (COG-UK) Consortium | Thushan de Silva, Matthew Parker, Nikki Smith, Adri Angyal, Rebecca Brown, Luke Green, Rachel Tucker, Paul Parsons, Danielle Groves, Katie Johnson, Laura Carrilero, Alex Keeley, Dave Partridge, Matthew Wyles, Benjamin Lindsey, Mehmet Yavuz, Mohammad Raza, Cariad Evans |

| | | | |
|---|---|---|---|
| | Cardiovascular Disease, The Medical School, University of Sheffield | | |
| EPI_ISL_764164 | Department of Pathology, University of Cambridge | COVID-19 Genomics UK (COG-UK) Consortium | Aminu S. Jahun, Yasmin Chaudhry, Grant Hall, Iliana Georgana, Myra Hosmillo, Martin D. Curran, Malte Pinckert, Surendra Parmar, Ian Goodfellow |
| EPI_ISL_764167 | Virology Department, Sheffield Teaching Hospitals NHS Foundation Trust/Department of Infection, Immunity and Cardiovascular Disease, The Medical School, University of Sheffield | COVID-19 Genomics UK (COG-UK) Consortium | Thushan de Silva, Matthew Parker, Nikki Smith, Adri Angyal, Rebecca Brown, Luke Green, Rachel Tucker, Paul Parsons, Danielle Groves, Katie Johnson, Laura Carrilero, Alex Keeley, Dave Partridge, Matthew Wyles, Benjamin Lindsey, Mehmet Yavuz, Mohammad Raza, Cariad Evans |
| EPI_ISL_764188 | University College London, Great Ormond Street Hospital for Children NHS Foundation Trust, Imperial College Healthcare NHS Trust | COVID-19 Genomics UK (COG-UK) Consortium | Sergi Castellano, Rachel Williams, Mark Kristiansen, Paola Resende Silva, Sunando Roy, Tony Brooks, Helena Tutill, Paola Niola, Patricia Dyal, Charlotte Williams, Leysa Forrest, Yasmin Panchbhaya, Jacqueline Findlay, Samuel Weeks, Julianne Brown, Kathryn Harris, Paul Randell, James Price, Alison Holmes, Judith Breuer |
| EPI_ISL_764189, EPI_ISL_764190 | Queens Medical Centre, Clinical Microbiology Department / DeepSeq Nottingham | COVID-19 Genomics UK (COG-UK) Consortium | Gemma Clark, Wendy Smith, Manjinder Khakh, Vicki M Fleming, Michelle M Lister, Hannah Howson-Wells, Jonathan Ball, Patrick McClure, Joseph Chappell, Theocharis Tsoleridis, Nadine Holmes, Matthew Carlisle, Christopher Moore, Fei Sang, Johnny Debebe, Victoria Wright, Matthew Loose |
| EPI_ISL_764195 | Virology Department, Royal Infirmary of Edinburgh, NHS Lothian / School of Biological Sciences, University of Edinburgh / Institute of Genetics and Molecular Medicine, University of Edinburgh | COVID-19 Genomics UK (COG-UK) Consortium | McHugh M, Dewar R, Rooke S, Gallagher M, Balcaza C, O'Toole Á, Scher E, Hill V, McCrone JT, Colquhoun R, Yu X, Jackson B, Rambaut A, Williams TC, Templeton K |
| EPI_ISL_764213, EPI_ISL_764216 | Virology Department, Sheffield Teaching Hospitals NHS Foundation Trust/Department of Infection, Immunity and Cardiovascular Disease, The Medical School, University of Sheffield | COVID-19 Genomics UK (COG-UK) Consortium | Thushan de Silva, Matthew Parker, Nikki Smith, Adri Angyal, Rebecca Brown, Luke Green, Rachel Tucker, Paul Parsons, Danielle Groves, Katie Johnson, Laura Carrilero, Alex Keeley, Dave Partridge, Matthew Wyles, Benjamin Lindsey, Mehmet Yavuz, Mohammad Raza, Cariad Evans |
| EPI_ISL_764221 | Virology Department, Royal Infirmary of Edinburgh, NHS Lothian / School of Biological Sciences, University of Edinburgh / Institute of Genetics and Molecular Medicine, University of Edinburgh | COVID-19 Genomics UK (COG-UK) Consortium | McHugh M, Dewar R, Rooke S, Gallagher M, Balcaza C, O'Toole Á, Scher E, Hill V, McCrone JT, Colquhoun R, Yu X, Jackson B, Rambaut A, Williams TC, Templeton K |
| EPI_ISL_764233 | Queens Medical Centre, Clinical Microbiology Department / DeepSeq Nottingham | COVID-19 Genomics UK (COG-UK) Consortium | Gemma Clark, Wendy Smith, Manjinder Khakh, Vicki M Fleming, Michelle M Lister, Hannah Howson-Wells, Jonathan Ball, Patrick McClure, Joseph Chappell, Theocharis Tsoleridis, Nadine Holmes, Matthew Carlisle, Christopher Moore, Fei Sang, Johnny Debebe, Victoria Wright, Matthew Loose |
| EPI_ISL_764236 | Department of Pathology, University of Cambridge | COVID-19 Genomics UK (COG-UK) Consortium | Aminu S. Jahun, Yasmin Chaudhry, Grant Hall, Iliana Georgana, Myra Hosmillo, Martin D. Curran, Malte Pinckert, Surendra Parmar, Ian Goodfellow |
| EPI_ISL_764239 | Virology Department, Sheffield Teaching Hospitals NHS Foundation Trust/Department of Infection, Immunity and Cardiovascular Disease, The Medical School, University of Sheffield | COVID-19 Genomics UK (COG-UK) Consortium | Thushan de Silva, Matthew Parker, Nikki Smith, Adri Angyal, Rebecca Brown, Luke Green, Rachel Tucker, Paul Parsons, Danielle Groves, Katie Johnson, Laura Carrilero, Alex Keeley, Dave Partridge, Matthew Wyles, Benjamin Lindsey, Mehmet Yavuz, Mohammad Raza, Cariad Evans |
| EPI_ISL_764278, EPI_ISL_764279 | Virology Department, Royal Infirmary of Edinburgh, NHS Lothian / School of Biological Sciences, University of Edinburgh / Institute of Genetics and Molecular Medicine, University of Edinburgh | COVID-19 Genomics UK (COG-UK) Consortium | McHugh M, Dewar R, Rooke S, Gallagher M, Balcaza C, O'Toole Á, Scher E, Hill V, McCrone JT, Colquhoun R, Yu X, Jackson B, Rambaut A, Williams TC, Templeton K |
| EPI_ISL_764343, EPI_ISL_764344, EPI_ISL_764345, EPI_ISL_764346, EPI_ISL_764347, EPI_ISL_764348, EPI_ISL_764349, EPI_ISL_764350, EPI_ISL_764351, EPI_ISL_764352, EPI_ISL_764353, EPI_ISL_764355 | | | |
| see above | Department of Pathology, University of Cambridge | COVID-19 Genomics UK (COG-UK) Consortium | Aminu S. Jahun, Yasmin Chaudhry, Grant Hall, Iliana Georgana, Myra Hosmillo, Martin D. Curran, Malte Pinckert, Surendra Parmar, Ian Goodfellow |
| EPI_ISL_764361, EPI_ISL_764362, EPI_ISL_764363, EPI_ISL_764364, EPI_ISL_764365, EPI_ISL_764366, EPI_ISL_764367, EPI_ISL_764368, EPI_ISL_764369, EPI_ISL_764370, EPI_ISL_764371, EPI_ISL_764372 | | | |
| see above | Virology Department, Royal Infirmary of Edinburgh, NHS Lothian / School of Biological Sciences, University of Edinburgh / Institute of Genetics and Molecular Medicine, University of Edinburgh | COVID-19 Genomics UK (COG-UK) Consortium | McHugh M, Dewar R, Rooke S, Gallagher M, Balcaza C, O'Toole Á, Scher E, Hill V, McCrone JT, Colquhoun R, Yu X, Jackson B, Rambaut A, Williams TC, Templeton K |
| EPI_ISL_764485, EPI_ISL_764486, EPI_ISL_764488 | Quadram Institute Bioscience | COVID-19 Genomics UK (COG-UK) Consortium | Dave J. Baker, Gemma L. Kay, Alp Aydin, Thanh Le-Viet, Steven Rudder, Ana P. Tedim, Anastasia Kolyva, Maria Diaz, Leonardo de Oliveira Martins, Nabil-Fareed Alikhan, Lizzie Meadows, Rachael Stanley, Ngozi Elumogo, Muhammed Yasir, Nicholas M. Thomson, Alexander J Trotter, Rachel Gilroy, Samuel Bloomfield, Claire Stuart, Andrew Bell, Reenesh Prakash, Samir Dervisevic, Alison E. Mather, John Wain, Mark Webber, Andrew J. Page, Justin O'Grady |
| EPI_ISL_764498, EPI_ISL_764499, EPI_ISL_764500, EPI_ISL_764505, EPI_ISL_764506, EPI_ISL_764507, EPI_ISL_764508, EPI_ISL_764509, EPI_ISL_764510, EPI_ISL_764511, EPI_ISL_764512, EPI_ISL_764513, EPI_ISL_764514, EPI_ISL_764515, EPI_ISL_764516, EPI_ISL_764517, EPI_ISL_764518, EPI_ISL_764519, EPI_ISL_764520, EPI_ISL_764521, EPI_ISL_764522, EPI_ISL_764523, EPI_ISL_764524 | | | |
| see above | Queens Medical Centre, Clinical Microbiology Department / DeepSeq Nottingham | COVID-19 Genomics UK (COG-UK) Consortium | Gemma Clark, Wendy Smith, Manjinder Khakh, Vicki M Fleming, Michelle M Lister, Hannah Howson-Wells, Jonathan Ball, Patrick McClure, Joseph Chappell, Theocharis Tsoleridis, Nadine Holmes, Matthew Carlisle, Christopher Moore, Fei Sang, Johnny Debebe, Victoria Wright, Matthew Loose |
| EPI_ISL_765150, EPI_ISL_765169, EPI_ISL_765170, EPI_ISL_765171, EPI_ISL_765186, EPI_ISL_765187 | Virology Department, Sheffield Teaching Hospitals NHS Foundation Trust/Department of Infection, Immunity and Cardiovascular Disease, The Medical School, University of Sheffield | COVID-19 Genomics UK (COG-UK) Consortium | Thushan de Silva, Matthew Parker, Nikki Smith, Adri Angyal, Rebecca Brown, Luke Green, Rachel Tucker, Paul Parsons, Danielle Groves, Katie Johnson, Laura Carrilero, Alex Keeley, Dave Partridge, Matthew Wyles, Benjamin Lindsey, Mehmet Yavuz, Mohammad Raza, Cariad Evans |
| EPI_ISL_765567 | National Institute for Infectious Diseases, INMI, "L. Spallanzani" IRCCS | National Institute for Infectious Diseases, INMI, "L. Spallanzani" IRCCS | E Giombini, C.E.M Gruber, M Rueca, B Bartolini, O Butera, F Messina, A Di Caro, G Parisi, MR Capobianchi |
| EPI_ISL_765568 | National Institute for Infectious Diseases, INMI, "L. Spallanzani" IRCCS | National Institute for Infectious Diseases, INMI, "L. Spallanzani" IRCCS | O Butera, C.E.M Gruber, F Messina, M Rueca, B Bartolini, E Giombini, MR Capobianchi, A Di Caro |
| EPI_ISL_765570 | National Institute for Infectious Diseases, INMI, "L. Spallanzani" IRCCS | National Institute for Infectious Diseases, INMI, "L. Spallanzani" IRCCS | M. Rueca, C.E.M Gruber, B Bartolini, O Butera, F Messina, E Giombini, A Di Caro, MR Capobianchi |
| EPI_ISL_765572 | National Institute for Infectious Diseases, INMI, "L. Spallanzani" IRCCS | National Institute for Infectious Diseases, INMI, "L. Spallanzani" IRCCS | C.E.M Gruber, M Rueca, B Bartolini, F Messina, E Giombini,O Butera, A Di Caro, MR Capobianchi |
| EPI_ISL_765573 | National Institute for Infectious Diseases, INMI, "L. Spallanzani" IRCCS | National Institute for Infectious Diseases, INMI, "L. Spallanzani" IRCCS | F Messina, E Giombini, M Rueca, B Bartolini, C.E.M Gruber, MR Capobianchi, O Butera, A Di Caro |
| EPI_ISL_766016, EPI_ISL_766017, EPI_ISL_766018, EPI_ISL_766019, EPI_ISL_766020, EPI_ISL_766021, EPI_ISL_766024, EPI_ISL_766026 | Department of Virology and Immunology, University of Helsinki and Helsinki University Hospital, Huslab Finland | Department of Virology, Faculty of Medicine, University of Helsinki, Helsinki, Finland | Teemu Smura, Olli Vapalahti, Maija Lappalainen, Satu Kurkela |
| EPI_ISL_766049 | Nigeria Centre for Disease Control (NCDC), National Reference Laboratory (NRL) | National Reference Laboratory, NCDC, Abuja | Olusola Akanbi, Chimaobi Chukwu, Dr Omoare Adesuyi, Mrs Nwando Mba, Dr Ndodo Nnaemeka, Dr Chikwe Ihekweazu |
| EPI_ISL_766051 | Nigeria Centre For Disease Control, | National reference Laboratory, NCDC, Gaduwa, Abuja | Dr Ndodo Nnaemeka, Olusola Akanbi, Chimaobi Chukwu, Dr Omoare Adesuyi, Kingsley Madubuike, Anthony Ahumibe, Naidoo Dhamari, Nwando Mba, Dr Chikwe Ihekweazu |
| EPI_ISL_766059, EPI_ISL_766063, EPI_ISL_766068, EPI_ISL_766092, EPI_ISL_766113, EPI_ISL_766114, EPI_ISL_766206, EPI_ISL_766207, EPI_ISL_766208, EPI_ISL_766209, EPI_ISL_766210, EPI_ISL_766211, EPI_ISL_766212, EPI_ISL_766220, EPI_ISL_766232, EPI_ISL_766241, EPI_ISL_766245, EPI_ISL_766505 | | | |

| | | | |
|---|---|---|---|
| see above | Respiratory Virus Unit, National Infection Service, Public Health England | COVID-19 Genomics UK (COG-UK) Consortium | PHE Covid Sequencing Team |
| EPI_ISL_766569, EPI_ISL_766570 | Oman-National Influenza Center | Oman-National Influenza Center | Samiha Al-Kharusi, Laila Al-Balushi, Hamida Al-Barwani, Aisha Al-Busaidi, Intisar Al-Shukri, Samira Al-Mahruqi, Hanan Al-Kindi, Amina Al-Jardani |
| EPI_ISL_766586 | Limmattal Hospital | Institute of Medical Virology, University of Zurich | Stefan Schmutz, Maryam Zaheri, Verena Kufner, Annette Audigé, Maria Grünberg, Kevin Steiner, Jon Huder, Cyril Shah, Riccarda Capaul, Jürg Böni, Michael Huber, Alexandra Trkola |
| EPI_ISL_766588 | Xundheitszentrum Wengen | Institute of Medical Virology, University of Zurich | Stefan Schmutz, Maryam Zaheri, Verena Kufner, Annette Audigé, Maria Grünberg, Kevin Steiner, Jon Huder, Cyril Shah, Riccarda Capaul, Jürg Böni, Michael Huber, Alexandra Trkola |
| EPI_ISL_766589 | Dr. Boubaker Karim laboratory | Institute of Medical Virology, University of Zurich | Stefan Schmutz, Maryam Zaheri, Verena Kufner, Annette Audigé, Maria Grünberg, Kevin Steiner, Jon Huder, Cyril Shah, Riccarda Capaul, Jürg Böni, Michael Huber, Alexandra Trkola |
| EPI_ISL_766594, EPI_ISL_766595, EPI_ISL_766596, EPI_ISL_766597, EPI_ISL_766598, EPI_ISL_766599, EPI_ISL_766600, EPI_ISL_766601, EPI_ISL_766602, EPI_ISL_766603, EPI_ISL_766604, EPI_ISL_766605, EPI_ISL_766606, EPI_ISL_766607, EPI_ISL_766612 | | | |
| see above | The Public Health Agency of Sweden | The Public Health Agency of Sweden | Department of Microbiology, The Public Health Agency of Sweden |
| EPI_ISL_766692, EPI_ISL_766693 | Klinisk mikrobiologi, Laboratoriemedicin | The Public Health Agency of Sweden | Department of Microbiology, The Public Health Agency of Sweden |
| EPI_ISL_766695 | A05 Biomedicum | The Public Health Agency of Sweden | Department of Microbiology, The Public Health Agency of Sweden |
| EPI_ISL_766696 | Klinisk Mikrobiologi | The Public Health Agency of Sweden | Department of Microbiology, The Public Health Agency of Sweden |
| EPI_ISL_766698 | A05 Biomedicum | The Public Health Agency of Sweden | Department of Microbiology, The Public Health Agency of Sweden |
| EPI_ISL_766700, EPI_ISL_766701, EPI_ISL_766702, EPI_ISL_766703, EPI_ISL_766704, EPI_ISL_766705, EPI_ISL_766706 | The Public Health Agency of Sweden | The Public Health Agency of Sweden | Department of Microbiology, The Public Health Agency of Sweden |
| EPI_ISL_766708, EPI_ISL_766709, EPI_ISL_766715, EPI_ISL_766716, EPI_ISL_766717, EPI_ISL_766720, EPI_ISL_766721, EPI_ISL_766722, EPI_ISL_766723 | A05 Biomedicum | The Public Health Agency of Sweden | Department of Microbiology, The Public Health Agency of Sweden |
| EPI_ISL_766724 | TATAA Biocenter | The Public Health Agency of Sweden | Department of Microbiology, The Public Health Agency of Sweden |
| EPI_ISL_766730 | Klinisk mikrobiologi | The Public Health Agency of Sweden | Department of Microbiology, The Public Health Agency of Sweden |
| EPI_ISL_767045, EPI_ISL_767046, EPI_ISL_767049, EPI_ISL_767051, EPI_ISL_767052, EPI_ISL_767053, EPI_ISL_767054, EPI_ISL_767055, EPI_ISL_767056, EPI_ISL_767057, EPI_ISL_767058, EPI_ISL_767059, EPI_ISL_767060, EPI_ISL_767061, EPI_ISL_767062, EPI_ISL_767063, EPI_ISL_767067, EPI_ISL_767070, EPI_ISL_767071, EPI_ISL_767072 | | | |
| see above | New Mexico Department of Health Scientific Laboratory | New Mexico Department of Health Scientific Laboratory | D'eldra Malone, Ellie Johnson, Anastacia Griego-Fisher |
| EPI_ISL_767096 | Lighthouse Lab in Glasgow | Wellcome Sanger Institute for the COVID-19 Genomics UK (COG-UK) Consortium | Harper VanSteenhouse, Yumi Kasai, David Gray, Carol Clugston, Anna Dominiczak and Alex Alderton, Roberto Amato, Sonia Goncalves, Ewan Harrison, David K. Jackson, Ian Johnston, Dominic Kwiatkowski, Cordelia Langford, John Sillitoe on behalf of the Wellcome Sanger Institute COVID-19 Surveillance Team |
| EPI_ISL_767418, EPI_ISL_767422, EPI_ISL_767509, EPI_ISL_767510, EPI_ISL_767511, EPI_ISL_767512, EPI_ISL_767513, EPI_ISL_767514, EPI_ISL_767515, EPI_ISL_767516, EPI_ISL_767524, EPI_ISL_767525, EPI_ISL_767526, EPI_ISL_767527, EPI_ISL_767528, EPI_ISL_767529, EPI_ISL_767530, EPI_ISL_767531, EPI_ISL_767532, EPI_ISL_767533, EPI_ISL_767534, EPI_ISL_767535, EPI_ISL_767536, EPI_ISL_767537 | | | |
| see above | Wadsworth Center, New York State Department.of Health | Wadsworth Center, New York State Department.of Health | Kirsten St. George, Daryl M. Lamson, Alexis Russel, Matthew Shudt, Melissa A Leisner, Jonathan Plitnick, Navjot Singh, John Kelly, Sara Griesemer, Erasmus Schneider, Erica Lasek-Nesselquist |
| EPI_ISL_767557, EPI_ISL_767558, EPI_ISL_767561, EPI_ISL_767566 | URMC LABS | Wadsworth Center, New York State Department.of Health | Kirsten St. George, Daryl M. Lamson, Alexis Russel, Matthew Shudt, Melissa A Leisner, Jonathan Plitnick, Navjot Singh, John Kelly, Sara Griesemer, Erasmus Schneider, Erica Lasek-Nesselquist |
| EPI_ISL_767580, EPI_ISL_767584 | WHITE PLAINS HOSPITAL CENTER LABORATORY | Wadsworth Center, New York State Department.of Health | Kirsten St. George, Daryl M. Lamson, Alexis Russel, Matthew Shudt, Melissa A Leisner, Jonathan Plitnick, Navjot Singh, John Kelly, Sara Griesemer, Erasmus Schneider, Erica Lasek-Nesselquist |
| EPI_ISL_767589, EPI_ISL_767594, EPI_ISL_767599, EPI_ISL_767600, EPI_ISL_767601, EPI_ISL_767602, EPI_ISL_767603, EPI_ISL_767604, EPI_ISL_767605, EPI_ISL_767606, EPI_ISL_767607, EPI_ISL_767608, EPI_ISL_767609, EPI_ISL_767610, EPI_ISL_767611, EPI_ISL_767612, EPI_ISL_767614 | | | |
| see above | SARATOGA HOSPITAL LABORATORY | Wadsworth Center, New York State Department.of Health | Kirsten St. George, Daryl M. Lamson, Alexis Russel, Matthew Shudt, Melissa A Leisner, Jonathan Plitnick, Navjot Singh, John Kelly, Sara Griesemer, Erasmus Schneider, Erica Lasek-Nesselquist |
| EPI_ISL_767615 | WHITE PLAINS HOSPITAL CENTER LABORATORY | Wadsworth Center, New York State Department.of Health | Kirsten St. George, Daryl M. Lamson, Alexis Russel, Matthew Shudt, Melissa A Leisner, Jonathan Plitnick, Navjot Singh, John Kelly, Sara Griesemer, Erasmus Schneider, Erica Lasek-Nesselquist |
| EPI_ISL_767893 | Laverty Pathology | NSW Health Pathology - Institute of Clinical Pathology and Medical Research; Westmead Hospital; University of Sydney | CIDM-PH et al. |
| EPI_ISL_767894, EPI_ISL_767895 | St Vincent's Pathology (SydPath) | NSW Health Pathology - Institute of Clinical Pathology and Medical Research; Westmead Hospital; University of Sydney | CIDM-PH et al. |
| EPI_ISL_767896 | Australian Clinical Labs | NSW Health Pathology - Institute of Clinical Pathology and Medical Research; Westmead Hospital; University of Sydney | CIDM-PH et al. |
| EPI_ISL_767897 | PALMS | NSW Health Pathology - Institute of Clinical Pathology and Medical Research; Westmead Hospital; University of Sydney | CIDM-PH et al. |
| EPI_ISL_767898 | Pathology West - NSW Health Pathology | NSW Health Pathology - Institute of Clinical Pathology and Medical Research; Westmead Hospital; University of Sydney | CIDM-PH et al. |
| EPI_ISL_767899 | Laverty Pathology | NSW Health Pathology - Institute of Clinical Pathology and Medical Research; Westmead Hospital; University of Sydney | CIDM-PH et al. |
| EPI_ISL_767900, EPI_ISL_767901, EPI_ISL_767904 | Sydney South West Pathology Service (SSWPS) - Liverpool Hospital - NSW Health Pathology | NSW Health Pathology - Institute of Clinical Pathology and Medical Research; Westmead Hospital; University of Sydney | CIDM-PH et al. |
| EPI_ISL_767915 | St Vincent's Pathology (SydPath) | NSW Health Pathology - Institute of Clinical Pathology and Medical Research; Westmead Hospital; University of Sydney | CIDM-PH et al. |
| EPI_ISL_767922 | South Eastern Area Laboratory Services (SEALS) | NSW Health Pathology - Institute of Clinical Pathology and Medical Research; Westmead Hospital; University of Sydney | CIDM-PH et al. |
| EPI_ISL_767931 | 4Cyte Pathology | NSW Health Pathology - Institute of Clinical Pathology and Medical Research; Westmead Hospital; University of Sydney | CIDM-PH et al. |
| EPI_ISL_767932 | Southern. IML Pathology | NSW Health Pathology - Institute of Clinical Pathology and Medical Research; Westmead Hospital; University of Sydney | CIDM-PH et al. |
| EPI_ISL_767941, EPI_ISL_767943, EPI_ISL_767945, EPI_ISL_767946, EPI_ISL_767947, EPI_ISL_767948, EPI_ISL_767951, EPI_ISL_767954, EPI_ISL_767956, EPI_ISL_767963, EPI_ISL_767971, EPI_ISL_767972, EPI_ISL_767975, EPI_ISL_767981, EPI_ISL_767982, EPI_ISL_767984, EPI_ISL_767985, EPI_ISL_767987, EPI_ISL_767988, EPI_ISL_767993, EPI_ISL_767998, EPI_ISL_768000, EPI_ISL_768001, EPI_ISL_768009, EPI_ISL_768011, EPI_ISL_768013, EPI_ISL_768014, EPI_ISL_768016, EPI_ISL_768017, EPI_ISL_768019, EPI_ISL_768022, EPI_ISL_768024, EPI_ISL_768025, EPI_ISL_768028, EPI_ISL_768029, EPI_ISL_768031, EPI_ISL_768032, EPI_ISL_768035, EPI_ISL_768036, EPI_ISL_768038, EPI_ISL_768040, EPI_ISL_768042, EPI_ISL_768043, EPI_ISL_768045, EPI_ISL_768047, EPI_ISL_768048, EPI_ISL_768049, EPI_ISL_768051, EPI_ISL_768052, EPI_ISL_768057, EPI_ISL_768059, EPI_ISL_768066, EPI_ISL_768068, EPI_ISL_768071, | | | |

| | | | |
|---|---|---|---|
| EPI_ISL_768073, EPI_ISL_768074, EPI_ISL_768075, EPI_ISL_768076, EPI_ISL_768083, EPI_ISL_768085, EPI_ISL_768093, EPI_ISL_768095, EPI_ISL_768097, EPI_ISL_768099, EPI_ISL_768100, EPI_ISL_768102, EPI_ISL_768103, EPI_ISL_768104, EPI_ISL_768105, EPI_ISL_768107, EPI_ISL_768108, EPI_ISL_768109, EPI_ISL_768114, EPI_ISL_768116, EPI_ISL_768117, EPI_ISL_768122, EPI_ISL_768124, EPI_ISL_768125, EPI_ISL_768127, EPI_ISL_768128, EPI_ISL_768129, EPI_ISL_768130, EPI_ISL_768131, EPI_ISL_768134, EPI_ISL_768135, EPI_ISL_768136, EPI_ISL_768137, EPI_ISL_768138, EPI_ISL_768139, EPI_ISL_768141, EPI_ISL_768143, EPI_ISL_768145, EPI_ISL_768147, EPI_ISL_768148, EPI_ISL_768150, EPI_ISL_768151, EPI_ISL_768153, EPI_ISL_768155, EPI_ISL_768156, EPI_ISL_768158, EPI_ISL_768159, EPI_ISL_768170, EPI_ISL_768172, EPI_ISL_768175, EPI_ISL_768178, EPI_ISL_768184, EPI_ISL_768185, EPI_ISL_768191, EPI_ISL_768194, EPI_ISL_768195, EPI_ISL_768209, EPI_ISL_768211, EPI_ISL_768212, EPI_ISL_768213, EPI_ISL_768217, EPI_ISL_768223, EPI_ISL_768226, EPI_ISL_768228, EPI_ISL_768230, EPI_ISL_768232, EPI_ISL_768233, EPI_ISL_768234, EPI_ISL_768237, EPI_ISL_768238, EPI_ISL_768240, EPI_ISL_768241, EPI_ISL_768242, EPI_ISL_768243, EPI_ISL_768245, EPI_ISL_768249, EPI_ISL_768255, EPI_ISL_768259, EPI_ISL_768262, EPI_ISL_768263, EPI_ISL_768264, EPI_ISL_768266, EPI_ISL_768267, EPI_ISL_768268, EPI_ISL_768269, EPI_ISL_768271, EPI_ISL_768272, EPI_ISL_768273, EPI_ISL_768274, EPI_ISL_768275, EPI_ISL_768278, EPI_ISL_768284, EPI_ISL_768286, EPI_ISL_768287, EPI_ISL_768289, EPI_ISL_768290, EPI_ISL_768295, EPI_ISL_768298, EPI_ISL_768299, EPI_ISL_768300, EPI_ISL_768301, EPI_ISL_768303, EPI_ISL_768304, EPI_ISL_768305, EPI_ISL_768311, EPI_ISL_768314, EPI_ISL_768315, EPI_ISL_768320, EPI_ISL_768322, EPI_ISL_768323 | | | |
| see above | Viollier AG | Department of Biosystems Science and Engineering, ETH Zürich | Chaoran Chen, Sarah Nadeau, Catharine Aquino, Ivan Topolsky, Philipp Jablonski, Lara Fuhrmann, David Dreifuss, Katharina Jahn, Andreia Cabral de Gouvea, Maria Domenica Moccia, Simon Grüter, Timothy Sykes, Lennart Opitz, Griffin White, Laura Neff, Doris Popovic, Andrea Patrignani, Jay Tracy, Ralph Schlapbach, Christiane Beckmann, Maurice Redondo, Olivier Kobel, Christoph Noppen, Sophie Seidel, Noemie Santamaria de Souza, Niko Beerenwinkel, Tanja Stadler |
| EPI_ISL_768603, EPI_ISL_768604, EPI_ISL_768605, EPI_ISL_768606, EPI_ISL_768607 | South Eastern Area Laboratory Services (SEALS) | NSW Health Pathology - Institute of Clinical Pathology and Medical Research; Westmead Hospital; University of Sydney | CIDM-PH et al. |
| EPI_ISL_768620, EPI_ISL_768623, EPI_ISL_768624, EPI_ISL_768628, EPI_ISL_768629, EPI_ISL_768630, EPI_ISL_768631, EPI_ISL_768632 | National Public Health Laboratory, National Centre for Infectious Diseases | National Public Health Laboratory, National Centre for Infectious Diseases | Tze Minn Mak, Sophie Octavia, Zhenyang Zhou, Lin Cui, Raymond Tzer Pin Lin |
| EPI_ISL_768743 | MoPH (Lebanese ministry of public health) | Microbial Genomics Lab LAU Byblos | Hamad Hassan, Sima Tokajian, Georgi Merhi, Tamara Salloum, Jad Koweyes |
| EPI_ISL_768817 | Laboratoire Mirialis | CNR Virus des Infections Respiratoires - France SUD | Antonin Bal, Gregory Destras, Gwendolyne Burfin, Hadrien Règue, Quentin Semanas, Martine Valette, Bruno Lina, Laurence Josset |
| EPI_ISL_768819, EPI_ISL_768820 | Laboratoire BPO-BIOEPINE | CNR Virus des Infections Respiratoires - France SUD | Antonin Bal, Gregory Destras, Gwendolyne Burfin, Hadrien Règue, Quentin Semanas, Martine Valette, Bruno Lina, Laurence Josset |
| EPI_ISL_768821 | Laboratoire Novabio | CNR Virus des Infections Respiratoires - France SUD | Antonin Bal, Gregory Destras, Gwendolyne Burfin, Hadrien Règue, Quentin Semanas, Martine Valette, Bruno Lina, Laurence Josset |
| EPI_ISL_768822 | CHU Poitiers | CNR Virus des Infections Respiratoires - France SUD | Antonin Bal, Gregory Destras, Gwendolyne Burfin, Hadrien Règue, Quentin Semanas, Martine Valette, Bruno Lina, Agnès Beby-Defaux, Magali Garcia, Clément Jousselin, Nicolas Lévêque, Laurence Josset |
| EPI_ISL_768823 | CHU Bordeaux | CNR Virus des Infections Respiratoires - France SUD | Antonin Bal, Gregory Destras, Gwendolyne Burfin, Hadrien Règue, Quentin Semanas, Martine Valette, Bruno Lina, Pantxika Bellecave, Camille Ciccone, Isabelle Garrigue, Marie-Edith Lafon, Pascale Trimoulet, Laurence Josset |
| EPI_ISL_768825, EPI_ISL_768826 | Laboratoire AX Bio Ocean | CNR Virus des Infections Respiratoires - France SUD | Antonin Bal, Gregory Destras, Gwendolyne Burfin, Hadrien Règue, Quentin Semanas, Martine Valette, Bruno Lina, Laurence Josset |
| EPI_ISL_768827, EPI_ISL_768828 | CNR Virus des Infections Respiratoires - France SUD | CNR Virus des Infections Respiratoires - France SUD | Antonin Bal, Gregory Destras, Gwendolyne Burfin, Hadrien Règue, Quentin Semanas, Martine Valette, Bruno Lina, Laurence Josset |
| EPI_ISL_769874, EPI_ISL_769875, EPI_ISL_769876 | Respiratory Virus Unit, National Infection Service, Public Health England | COVID-19 Genomics UK (COG-UK) Consortium | PHE Covid Sequencing Team |
| EPI_ISL_769916, EPI_ISL_769919, EPI_ISL_769921, EPI_ISL_769936, EPI_ISL_769937, EPI_ISL_769938, EPI_ISL_769939, EPI_ISL_769940, EPI_ISL_769942, EPI_ISL_769943, EPI_ISL_769944, EPI_ISL_769945, EPI_ISL_769947, EPI_ISL_769948, EPI_ISL_769949, EPI_ISL_769959, EPI_ISL_769960, EPI_ISL_769961, EPI_ISL_769962, EPI_ISL_769963, EPI_ISL_769964, EPI_ISL_769965, EPI_ISL_769966, EPI_ISL_769967, EPI_ISL_769968, EPI_ISL_769969, EPI_ISL_769970 | | | |
| see above | Albany Medical Center Hospital Clinical Laboratories | Wadsworth Center, New York State Department.of Health | Kirsten St. George, Daryl M. Lamson, Alexis Russel, Matthew Shudt, Melissa A Leisner, Jonathan Plitnick, Navjot Singh, John Kelly, Sara Griesemer, Erasmus Schneider, Erica Lasek-Nesselquist |
| EPI_ISL_770439, EPI_ISL_770440, EPI_ISL_770441 | Respiratory Virus Unit, National Infection Service, Public Health England | COVID-19 Genomics UK (COG-UK) Consortium | PHE Covid Sequencing Team |
| EPI_ISL_770470, EPI_ISL_770473 | National Health laboratory | Botswana Institute for Technology Research and Innovation | Kefentse Arnold Tumedi, Madisa Mine, Dineo Emang Tshiamo. Gape Nyepetsi, Thongbotho Mphoyakgosi, Malebogo Kebabonye, Maitshwarelo Ignatius Matsheka |
| EPI_ISL_770475 | National health Laboratory | Botswana Institute for Technology Research and Innovation | Kefentse Arnold Tumedi, Madisa Mine, Dineo Emang Tshiamo. Gape Nyepetsi, Thongbotho Mphoyakgosi, Malebogo Kebabonye, Maitshwarelo Ignatius Matsheka |
| EPI_ISL_770498, EPI_ISL_770516, EPI_ISL_770517, EPI_ISL_770518, EPI_ISL_770519, EPI_ISL_770520, EPI_ISL_770521, EPI_ISL_770522, EPI_ISL_770523, EPI_ISL_770524, EPI_ISL_770525, EPI_ISL_770526, EPI_ISL_770527, EPI_ISL_770528, EPI_ISL_770529, EPI_ISL_770530, EPI_ISL_770531, EPI_ISL_770532 | | | |
| see above | Lithuanian University of Health Sciences Hospital, Department of Laboratory Medicine | Lithuanian University of Health Sciences, Molecular cardiology lab. | Lukas Zemaitis, Ingrida Olendrait, Arnoldas Pautienius, Kamile Tamusauskaite, Dovydas Gecys, Laura Pareckaite, Vaiva Lesauskaite, Astra Vitkauskiene |
| EPI_ISL_770816, EPI_ISL_770817, EPI_ISL_770819 | Minnesota Department of Health, Public Health Laboratory | Minnesota Department of Health, Public Health Laboratory | Alexandra Lorentz, Jacob Garfin, Matt Plumb, and Xiong Wang |
| EPI_ISL_770846, EPI_ISL_770849, EPI_ISL_770853, EPI_ISL_770857, EPI_ISL_770859, EPI_ISL_770861, EPI_ISL_770863, EPI_ISL_770864, EPI_ISL_770865, EPI_ISL_770877, EPI_ISL_770884, EPI_ISL_770901, EPI_ISL_770902, EPI_ISL_771039, EPI_ISL_771040, EPI_ISL_771041, EPI_ISL_771042, EPI_ISL_771043, EPI_ISL_771044, EPI_ISL_771045, EPI_ISL_771046, EPI_ISL_771047, EPI_ISL_771048, EPI_ISL_771049, EPI_ISL_771050, EPI_ISL_771053, EPI_ISL_771054, EPI_ISL_771055, EPI_ISL_771056, EPI_ISL_771057, EPI_ISL_771058, EPI_ISL_771059, EPI_ISL_771061, EPI_ISL_771062, EPI_ISL_771063, EPI_ISL_771064, EPI_ISL_771065, EPI_ISL_771066, EPI_ISL_771067, EPI_ISL_771068, EPI_ISL_771069, EPI_ISL_771070, EPI_ISL_771071, EPI_ISL_771072, EPI_ISL_771073, EPI_ISL_771074, EPI_ISL_771075, EPI_ISL_771076, EPI_ISL_771077, EPI_ISL_771078, EPI_ISL_771079, EPI_ISL_771080, EPI_ISL_771081, EPI_ISL_771082, EPI_ISL_771083, EPI_ISL_771084, EPI_ISL_771085, EPI_ISL_771086, EPI_ISL_771087, EPI_ISL_771088, EPI_ISL_771089, EPI_ISL_771091, EPI_ISL_771094 | | | |
| see above | Laboratoire national de santé, Microbiology, Virology | Laboratoire national de santé, Microbiology, Microbial Genomics Platform | Anke Wienecke-Baldacchino, Catherine Ragimbeau,Jessica Tapp, Fatu Djabi, Lise Pignon, Raoul Salmon, Tamir Abdelrahman |
| EPI_ISL_775273, EPI_ISL_775274 | Furst Medical Laboratory | Norwegian Institute of Public Health, Department of Virology | Kathrine Stene-Johansen, Kamilla Heddeland Instefjord, Hilde Elshaug, Atiya R Ali,Marie Paulsen Madsen, Rasmus Riis Kopperud, Hilde Vollan, Karoline Bragstad, Olav Hungnes |
| EPI_ISL_775434, EPI_ISL_775435, EPI_ISL_775437 | Department of Medical Microbiology - section Molde, Molde Hospital | Norwegian Institute of Public Health, Department of Virology | Kathrine Stene-Johansen, Kamilla Heddeland Instefjord, Hilde Elshaug, Atiya R Ali,Marie Paulsen Madsen, Rasmus Riis Kopperud, Hilde Vollan, Karoline Bragstad, Olav Hungnes |
| EPI_ISL_775533, EPI_ISL_775534, EPI_ISL_775535, EPI_ISL_775536, EPI_ISL_775537, EPI_ISL_775538, EPI_ISL_775539, EPI_ISL_775540, EPI_ISL_775541, EPI_ISL_775576, EPI_ISL_775577, EPI_ISL_775578, EPI_ISL_775579, EPI_ISL_775580, EPI_ISL_775581, EPI_ISL_775582 | | | |
| see above | ABC | The Public Health Agency of Sweden | Department of Microbiology, The Public Health Agency of Sweden |
| EPI_ISL_775586 | Unilabs AB | The Public Health Agency of Sweden | Department of Microbiology, The Public Health Agency of Sweden |
| EPI_ISL_776570, EPI_ISL_776571, EPI_ISL_776572, EPI_ISL_776573, EPI_ISL_776574, EPI_ISL_776576, EPI_ISL_776577, EPI_ISL_776578, EPI_ISL_776579, EPI_ISL_776580, EPI_ISL_776581, EPI_ISL_776582, EPI_ISL_776583, EPI_ISL_776584, EPI_ISL_776585, EPI_ISL_776586, EPI_ISL_776587, EPI_ISL_776588, EPI_ISL_776589, EPI_ISL_776591, EPI_ISL_776592, EPI_ISL_776593, EPI_ISL_776594, EPI_ISL_776595, EPI_ISL_776596, EPI_ISL_776597, EPI_ISL_776598, EPI_ISL_776599, EPI_ISL_776600, EPI_ISL_776601, EPI_ISL_776602, EPI_ISL_776603, EPI_ISL_776605, EPI_ISL_776606, EPI_ISL_776607, EPI_ISL_776608, EPI_ISL_776610, EPI_ISL_776611, EPI_ISL_776612, EPI_ISL_776613, EPI_ISL_776614, EPI_ISL_776615, EPI_ISL_776616, EPI_ISL_776617, EPI_ISL_776618, EPI_ISL_776619, EPI_ISL_776620, EPI_ISL_776621, EPI_ISL_776622, EPI_ISL_776623, EPI_ISL_776624, EPI_ISL_776625, EPI_ISL_776626, EPI_ISL_776627, EPI_ISL_776628, EPI_ISL_776629, EPI_ISL_776630, EPI_ISL_776631, EPI_ISL_776632, EPI_ISL_776633, EPI_ISL_776634, EPI_ISL_776635, EPI_ISL_776636, EPI_ISL_776637, EPI_ISL_776638, EPI_ISL_776639, EPI_ISL_776640, EPI_ISL_776641, EPI_ISL_776642, EPI_ISL_776643, EPI_ISL_776644, EPI_ISL_776645, EPI_ISL_776646, EPI_ISL_776647, EPI_ISL_776648, EPI_ISL_776649, EPI_ISL_776650, EPI_ISL_776651, EPI_ISL_776652 | | | |
| see above | Israel Central Virology laboratory | Israel Central Virology laboratory | Neta Zuckerman, Efrat Dahan Bucris, Oran Erster, Michal Mandelboim, Orna Mor, Ella Mendelson |
| EPI_ISL_778865 | Clinical Pathology Labs | Centers for Disease Control and Prevention Division of Viral Diseases, Pathogen Discovery | Ying Tao Yan Li Jing Zhang Krista Queen Anna Uehara Peter Cook Clinton R. Paden Haibin Wang Suxiang Tong |
| EPI_ISL_778878, EPI_ISL_778879, EPI_ISL_778880, EPI_ISL_778881 | Maryland Public Health Laboratory | Maryland Public Health Laboratory | Maryland Department of Health Laboratories Administration |
| EPI_ISL_779622, EPI_ISL_779623 | Victorian Infectious Diseases Reference Laboratory (VIDRL) | VIDRL and MDU-PHL | Caly L., Seemann T., Sait, M.L., Druce J., Sherry, N.L. |
| EPI_ISL_779730 | Servicio de Microbiología Clínica (Complejo Hospitalario de Navarra, Pamplona), Instituto de Investigación Sanitaria de | SeqCOVID-SPAIN consortium/IBV(CSIC) | Carmen Ezpeleta Baquedano, Ana Navascués, Ana Miqueleiz and SeqCOVID-SPAIN consortium |

| | | | |
|---|---|---|---|
| | Navarra (IdiSNA) | | |
| EPI_ISL_779731 | Servicio de Microbiología, Laboratori Clínic Metropolitana Nord. Hospital Universitari Germans Trias i Pujol. Institut d'Investigació en Ciències de la Salut Germans Trias i Pujol (IGTP) | SeqCOVID-SPAIN consortium/IBV(CSIC) | Elisa Martró, Antoni E. Bordoy, Anna Not, Adrián Antuori, Anabel Fernández, Nona Romaní, Verónica Saludes, Cristina Casañ and SeqCOVID-SPAIN consortium |
| EPI_ISL_779780, EPI_ISL_779783, EPI_ISL_779801, EPI_ISL_779802, EPI_ISL_779803, EPI_ISL_779804, EPI_ISL_779805, EPI_ISL_779806, EPI_ISL_779807, EPI_ISL_779808, EPI_ISL_779809 | | | |
| see above | CNR Virus des Infections Respiratoires - France SUD | CNR Virus des Infections Respiratoires - France SUD | Antonin Bal, Gregory Destras, Gwendolyne Burfin, Hadrien Règue, Quentin Semanas, Martine Valette, Bruno Lina, Laurence Josset |
| EPI_ISL_779846 | CHU de Nice - Hôpital Archet 2 | CNR Virus des Infections Respiratoires - France SUD | Antonin Bal, Géraldine Gonfrier, Gregory Destras, Gwendolyne Burfin, Hadrien Règue, Quentin Semanas, Martine Valette, Bruno Lina, Valérie Giordanengo, Laurence Josset |
| EPI_ISL_779865, EPI_ISL_779883, EPI_ISL_779902, EPI_ISL_779914, EPI_ISL_779915 | Servicio de Microbiología, Hospital Universitario Son Espases | SeqCOVID-SPAIN consortium/IBV(CSIC) | Carla López-Causapé, Jordi Reina, Antonio Oliver and SeqCOVID-SPAIN consortium |
| EPI_ISL_781517, EPI_ISL_781518, EPI_ISL_781525, EPI_ISL_781526, EPI_ISL_781559, EPI_ISL_781563, EPI_ISL_781573, EPI_ISL_781574, EPI_ISL_781589, EPI_ISL_781596, EPI_ISL_781614, EPI_ISL_781628, EPI_ISL_781630, EPI_ISL_781631, EPI_ISL_781644, EPI_ISL_781648, EPI_ISL_781664, EPI_ISL_781672, EPI_ISL_781675, EPI_ISL_781683, EPI_ISL_781693, EPI_ISL_781727, EPI_ISL_781751, EPI_ISL_781761, EPI_ISL_781769, EPI_ISL_781778, EPI_ISL_781781, EPI_ISL_781787, EPI_ISL_781790, EPI_ISL_781802, EPI_ISL_781820, EPI_ISL_781821, EPI_ISL_781825, EPI_ISL_781829, EPI_ISL_781835, EPI_ISL_781847, EPI_ISL_781865, EPI_ISL_781868 | | | |
| see above | Lighthouse Lab in Cambridge | Wellcome Sanger Institute for the COVID-19 Genomics UK (COG-UK) Consortium | Rob Howes, The Lighthouse Lab in Cambridge and Alex Alderton, Roberto Amato, Sonia Goncalves, Ewan Harrison, David K. Jackson, Ian Johnston, Dominic Kwiatkowski, Cordelia Langford, John Sillitoe on behalf of the Wellcome Sanger Institute COVID-19 Surveillance Team |
| EPI_ISL_781870, EPI_ISL_781871, EPI_ISL_781875, EPI_ISL_781882, EPI_ISL_781886, EPI_ISL_781890, EPI_ISL_781895, EPI_ISL_781910, EPI_ISL_781918, EPI_ISL_781922, EPI_ISL_781928, EPI_ISL_781930, EPI_ISL_781953, EPI_ISL_781956, EPI_ISL_781960, EPI_ISL_781967, EPI_ISL_781971, EPI_ISL_781980, EPI_ISL_781990, EPI_ISL_781992, EPI_ISL_781997, EPI_ISL_782001, EPI_ISL_782003, EPI_ISL_782004, EPI_ISL_782008, EPI_ISL_782014, EPI_ISL_782018, EPI_ISL_782027, EPI_ISL_782036, EPI_ISL_782038, EPI_ISL_782039, EPI_ISL_782056, EPI_ISL_782059, EPI_ISL_782062, EPI_ISL_782064, EPI_ISL_782065, EPI_ISL_782066, EPI_ISL_782074, EPI_ISL_782075, EPI_ISL_782079, EPI_ISL_782084, EPI_ISL_782101, EPI_ISL_782107, EPI_ISL_782112, EPI_ISL_782114, EPI_ISL_782115, EPI_ISL_782116, EPI_ISL_782121, EPI_ISL_782127, EPI_ISL_782128, EPI_ISL_782145, EPI_ISL_782146, EPI_ISL_782151, EPI_ISL_782152, EPI_ISL_782153, EPI_ISL_782154, EPI_ISL_782155, EPI_ISL_782156, EPI_ISL_782157, EPI_ISL_782158, EPI_ISL_782159, EPI_ISL_782160, EPI_ISL_782162, EPI_ISL_782163, EPI_ISL_782164, EPI_ISL_782166, EPI_ISL_782167, EPI_ISL_782168, EPI_ISL_782169, EPI_ISL_782170, EPI_ISL_782172, EPI_ISL_782173, EPI_ISL_782174, EPI_ISL_782175, EPI_ISL_782176, EPI_ISL_782177, EPI_ISL_782178, EPI_ISL_782179, EPI_ISL_782180, EPI_ISL_782181, EPI_ISL_782182, EPI_ISL_782183, EPI_ISL_782185, EPI_ISL_782186, EPI_ISL_782187, EPI_ISL_782188, EPI_ISL_782189, EPI_ISL_782190, EPI_ISL_782191, EPI_ISL_782192, EPI_ISL_782193, EPI_ISL_782195, EPI_ISL_782196, EPI_ISL_782198, EPI_ISL_782199, EPI_ISL_782200, EPI_ISL_782203, EPI_ISL_782204, EPI_ISL_782205, EPI_ISL_782206, EPI_ISL_782208, EPI_ISL_782209, EPI_ISL_782210, EPI_ISL_782213, EPI_ISL_782214, EPI_ISL_782215, EPI_ISL_782216, EPI_ISL_782217, EPI_ISL_782218, EPI_ISL_782219, EPI_ISL_782221, EPI_ISL_782222, EPI_ISL_782223, EPI_ISL_782225, EPI_ISL_782227, EPI_ISL_782229, EPI_ISL_782230, EPI_ISL_782231, EPI_ISL_782232, EPI_ISL_782233, EPI_ISL_782234, EPI_ISL_782236, EPI_ISL_782237, EPI_ISL_782238, EPI_ISL_782239, EPI_ISL_782240, EPI_ISL_782241, EPI_ISL_782244, EPI_ISL_782246, EPI_ISL_782247, EPI_ISL_782248, EPI_ISL_782251, EPI_ISL_782253, EPI_ISL_782254, EPI_ISL_782255, EPI_ISL_782257, EPI_ISL_782258, EPI_ISL_782260, EPI_ISL_782261, EPI_ISL_782262, EPI_ISL_782263, EPI_ISL_782265, EPI_ISL_782266, EPI_ISL_782269, EPI_ISL_782270, EPI_ISL_782272, EPI_ISL_782273, EPI_ISL_782275, EPI_ISL_782276, EPI_ISL_782277, EPI_ISL_782279, EPI_ISL_782280, EPI_ISL_782281, EPI_ISL_782283, EPI_ISL_782284, EPI_ISL_782285, EPI_ISL_782286, EPI_ISL_782289, EPI_ISL_782290, EPI_ISL_782291, EPI_ISL_782293, EPI_ISL_782294, EPI_ISL_782295, EPI_ISL_782296, EPI_ISL_782297, EPI_ISL_782298, EPI_ISL_782299, EPI_ISL_782300, EPI_ISL_782301, EPI_ISL_782302, EPI_ISL_782303, EPI_ISL_782304, EPI_ISL_782305, EPI_ISL_782306, EPI_ISL_782308, EPI_ISL_782310, EPI_ISL_782311, EPI_ISL_782312, EPI_ISL_782318, EPI_ISL_782319, EPI_ISL_782320, EPI_ISL_782321, EPI_ISL_782322, EPI_ISL_782323, EPI_ISL_782324, EPI_ISL_782329, EPI_ISL_782330, EPI_ISL_782331, EPI_ISL_782332, EPI_ISL_782333, EPI_ISL_782334, EPI_ISL_782336, EPI_ISL_782337, EPI_ISL_782338, EPI_ISL_782339, EPI_ISL_782340, EPI_ISL_782341, EPI_ISL_782342, EPI_ISL_782343, EPI_ISL_782344, EPI_ISL_782346, EPI_ISL_782347, EPI_ISL_782348, EPI_ISL_782349, EPI_ISL_782350, EPI_ISL_782351, EPI_ISL_782354, EPI_ISL_782356, EPI_ISL_782357, EPI_ISL_782359, EPI_ISL_782360, EPI_ISL_782363, EPI_ISL_782364, EPI_ISL_782365, EPI_ISL_782366, EPI_ISL_782367, EPI_ISL_782368, EPI_ISL_782370, EPI_ISL_782373, EPI_ISL_782374, EPI_ISL_782375, EPI_ISL_782376, EPI_ISL_782378, EPI_ISL_782379, EPI_ISL_782381, EPI_ISL_782382, EPI_ISL_782383, EPI_ISL_782384, EPI_ISL_782385, EPI_ISL_782386, EPI_ISL_782387, EPI_ISL_782389, EPI_ISL_782392, EPI_ISL_782394, EPI_ISL_782395, EPI_ISL_782396, EPI_ISL_782397, EPI_ISL_782398, EPI_ISL_782399, EPI_ISL_782400, EPI_ISL_782401, EPI_ISL_782403, EPI_ISL_782404, EPI_ISL_782405, EPI_ISL_782406, EPI_ISL_782407, EPI_ISL_782408, EPI_ISL_782410, EPI_ISL_782411, EPI_ISL_782413, EPI_ISL_782414, EPI_ISL_782415, EPI_ISL_782416, EPI_ISL_782417, EPI_ISL_782418, EPI_ISL_782421, EPI_ISL_782422, EPI_ISL_782424, EPI_ISL_782427, EPI_ISL_782428, EPI_ISL_782429, EPI_ISL_782430, EPI_ISL_782431, EPI_ISL_782433, EPI_ISL_782434, EPI_ISL_782436, EPI_ISL_782437, EPI_ISL_782438, EPI_ISL_782440, EPI_ISL_782442, EPI_ISL_782443, EPI_ISL_782444, EPI_ISL_782445, EPI_ISL_782446, EPI_ISL_782447, EPI_ISL_782448, EPI_ISL_782449, EPI_ISL_782450, EPI_ISL_782452, EPI_ISL_782453, EPI_ISL_782455, EPI_ISL_782456, EPI_ISL_782457, EPI_ISL_782458, EPI_ISL_782459, EPI_ISL_782460, EPI_ISL_782462, EPI_ISL_782463, EPI_ISL_782464, EPI_ISL_782465, EPI_ISL_782467, EPI_ISL_782468, EPI_ISL_782469, EPI_ISL_782470, EPI_ISL_782472, EPI_ISL_782473, EPI_ISL_782475, EPI_ISL_782476, EPI_ISL_782477, EPI_ISL_782479, EPI_ISL_782480, EPI_ISL_782481 | | | |
| see above | Lighthouse Lab in Glasgow | Wellcome Sanger Institute for the COVID-19 Genomics UK (COG-UK) Consortium | Harper VanSteenhouse, Yumi Kasai, David Gray, Carol Clugston, Anna Dominiczak and Alex Alderton, Roberto Amato, Sonia Goncalves, Ewan Harrison, David K. Jackson, Ian Johnston, Dominic Kwiatkowski, Cordelia Langford, John Sillitoe on behalf of the Wellcome Sanger Institute COVID-19 Surveillance Team |
| EPI_ISL_788989, EPI_ISL_788990 | Klinisk mikrobiologi | The Public Health Agency of Sweden | Department of Microbiology, The Public Health Agency of Sweden |
| EPI_ISL_788991, EPI_ISL_788992, EPI_ISL_788994, EPI_ISL_788995, EPI_ISL_788996, EPI_ISL_788997, EPI_ISL_788998, EPI_ISL_789014, EPI_ISL_789015, EPI_ISL_789016, EPI_ISL_789025, EPI_ISL_789026, EPI_ISL_789027, EPI_ISL_789028 | | | |
| see above | ABC | The Public Health Agency of Sweden | Department of Microbiology, The Public Health Agency of Sweden |
| EPI_ISL_789058, EPI_ISL_789059 | Klinisk mikrobiologi, Viruslab | The Public Health Agency of Sweden | Department of Microbiology, The Public Health Agency of Sweden |
| EPI_ISL_789060, EPI_ISL_789068, EPI_ISL_789071, EPI_ISL_789072, EPI_ISL_789084, EPI_ISL_789085, EPI_ISL_789087, EPI_ISL_789088, EPI_ISL_789089, EPI_ISL_789090, EPI_ISL_789093 | | | |
| see above | National Virus Reference Laboratory | National Virus Reference Laboratory | Michael Carr, Gabriel Gonzalez, Dana Alalwan, Jonathan Dean, Daniel Hare, Cillian F De Gascun |
| EPI_ISL_790571, EPI_ISL_790579, EPI_ISL_790580, EPI_ISL_790581, EPI_ISL_790584, EPI_ISL_790591, EPI_ISL_790604, EPI_ISL_790605, EPI_ISL_790606, EPI_ISL_790607, EPI_ISL_790608, EPI_ISL_790609, EPI_ISL_790610, EPI_ISL_790612, EPI_ISL_790613, EPI_ISL_790614, EPI_ISL_790616, EPI_ISL_790619, EPI_ISL_790620, EPI_ISL_790621, EPI_ISL_790630, EPI_ISL_790711, EPI_ISL_790712, EPI_ISL_790713, EPI_ISL_790728, EPI_ISL_790740, EPI_ISL_790749, EPI_ISL_790750, EPI_ISL_790751, EPI_ISL_790752, EPI_ISL_790753, EPI_ISL_790754, EPI_ISL_790755, EPI_ISL_790756, EPI_ISL_790757, EPI_ISL_790758, EPI_ISL_790759, EPI_ISL_790760, EPI_ISL_790774, EPI_ISL_790775, EPI_ISL_790776, EPI_ISL_790778, EPI_ISL_790802, EPI_ISL_790803, EPI_ISL_790804, EPI_ISL_790805, EPI_ISL_790806, EPI_ISL_790808, EPI_ISL_790817, EPI_ISL_790824, EPI_ISL_790825, EPI_ISL_790839, EPI_ISL_790840, EPI_ISL_790848, EPI_ISL_790862, EPI_ISL_790863, EPI_ISL_790864, EPI_ISL_790865, EPI_ISL_790873, EPI_ISL_790879, EPI_ISL_790880, EPI_ISL_790881, EPI_ISL_790882, EPI_ISL_790883, EPI_ISL_790888, EPI_ISL_790907, EPI_ISL_790908, EPI_ISL_790909, EPI_ISL_790910, EPI_ISL_790914, EPI_ISL_790923, EPI_ISL_790924, EPI_ISL_790925, EPI_ISL_790940, EPI_ISL_790946, EPI_ISL_790947, EPI_ISL_790948, EPI_ISL_790949, EPI_ISL_790950, EPI_ISL_790951, EPI_ISL_790961, EPI_ISL_790962, EPI_ISL_790963, EPI_ISL_790964, EPI_ISL_790966, EPI_ISL_790967, EPI_ISL_790972, EPI_ISL_790973, EPI_ISL_790978, EPI_ISL_790979, EPI_ISL_790980, EPI_ISL_790983, EPI_ISL_790999, EPI_ISL_791000, EPI_ISL_791001, EPI_ISL_791006, EPI_ISL_791007, EPI_ISL_791008, EPI_ISL_791010, EPI_ISL_791011, EPI_ISL_791012, EPI_ISL_791013, EPI_ISL_791014, EPI_ISL_791015, EPI_ISL_791016, EPI_ISL_791017, EPI_ISL_791018, EPI_ISL_791021, EPI_ISL_791022, EPI_ISL_791023, EPI_ISL_791024, EPI_ISL_791025, EPI_ISL_791028, EPI_ISL_791029, EPI_ISL_791030, EPI_ISL_791031, EPI_ISL_791046, EPI_ISL_791047, EPI_ISL_791048, EPI_ISL_791053, EPI_ISL_791064, EPI_ISL_791076, EPI_ISL_791078, EPI_ISL_791080 | | | |
| see above | Dutch COVID-19 response team | National Institute for Public Health and the Environment (RIVM) | Adam Meijer, Harry Vennema, Jeroen Cremer, Sharon van den Brink, Bas van der Veer, AnneMarie van den Brandt, Florian Zwagemaker, Dennis Schmitz, Chantal Reusken, on behalf of the national COVID-19 response team |
| EPI_ISL_791211 | Respiratory Virus Unit, National Infection Service, Public Health England | COVID-19 Genomics UK (COG-UK) Consortium | PHE Covid Sequencing Team |
| EPI_ISL_791332 | UZ Leuven, National Reference Laboratory for Coronaviruses, Laboratory Medicine, Leuven, Belgium | KU Leuven, Rega Institute, Clinical and Epidemiological Virology | Tony Wawina-Bokalanga, Joan Marti-Carerras, Bert Vanmechelen, Piet Maes |
| EPI_ISL_791344, EPI_ISL_791418, EPI_ISL_791419, EPI_ISL_791420, EPI_ISL_791428, EPI_ISL_791431, EPI_ISL_791432, EPI_ISL_791433, EPI_ISL_791434, EPI_ISL_791440, EPI_ISL_791447, EPI_ISL_791448, EPI_ISL_791449, EPI_ISL_791450, EPI_ISL_791454, EPI_ISL_791459, EPI_ISL_791460, EPI_ISL_791462, EPI_ISL_791464, EPI_ISL_791467, EPI_ISL_791470, EPI_ISL_791473, EPI_ISL_791475, EPI_ISL_791476, EPI_ISL_791478, EPI_ISL_791481, EPI_ISL_791482 | | | |
| see above | Johns Hopkins Hospital Department of Pathology | Johns Hopkins Hospital Department of Pathology | C. Paul Morris, Chun Huai Luo, Adannaya Amadi, Nicholas Gallagher, Heba H. Mostafa |
| EPI_ISL_792032 | Pathology West - NSW Health Pathology | NSW Health Pathology - Institute of Clinical Pathology and Medical Research; Westmead Hospital; University of Sydney | CIDM-PH et al. |
| EPI_ISL_792033 | Laverty Pathology | NSW Health Pathology - Institute of Clinical Pathology and Medical Research; Westmead Hospital; University of Sydney | CIDM-PH et al. |
| EPI_ISL_792034 | Douglass Hanly Moir Pathology | NSW Health Pathology - Institute of Clinical Pathology and Medical Research; Westmead Hospital; University of Sydney | CIDM-PH et al. |
| EPI_ISL_792041 | hospital | National Reference Center for Viruses of Respiratory Infections, Institut Pasteur, Paris | Marion Barbet, Sylvie Behillil, Méline Bizard, Angela Brisebarre, Camille Capel, Etienne Simon-Lorière, Vincent Enouf, Maud Vanpeene, Sylvie van der Werf,Laurent Roudière |
| EPI_ISL_792042 | General practitioner | National Reference Center for Viruses of Respiratory Infections, Institut Pasteur, Paris | Marion Barbet, Sylvie Behillil, Méline Bizard, Angela Brisebarre, Camille Capel, Etienne Simon-Lorière, Vincent Enouf, Maud Vanpeene, Sylvie van der Werf, |
| EPI_ISL_792043, EPI_ISL_792044, EPI_ISL_792045, EPI_ISL_792046, EPI_ISL_792047, EPI_ISL_792048, EPI_ISL_792049, EPI_ISL_792052 | General practitioner | National Reference Center for Viruses of Respiratory Infections, Institut Pasteur, Paris | Marion Barbet, Sylvie Behillil, Méline Bizard, Angela Brisebarre, Camille Capel, Etienne Simon-Lorière, Vincent Enouf, Maud Vanpeene, Sylvie van der Werf,Brieuc Lefaure |

| | | | |
|---|---|---|---|
| EPI_ISL_792053 | General practitioner | National Reference Center for Viruses of Respiratory Infections, Institut Pasteur, Paris | Marion Barbet, Sylvie Behillil, Méline Bizard, Angela Brisebarre, Camille Capel, Etienne Simon-Lorière, Vincent Enouf, Maud Vanpeene, Sylvie van der Werf |
| EPI_ISL_792056 | Hopital | National Reference Center for Viruses of Respiratory Infections, Institut Pasteur, Paris | Marion Barbet, Sylvie Behillil, Méline Bizard, Angela Brisebarre, Camille Capel, Etienne Simon-Lorière, Vincent Enouf, Maud Vanpeene, Sylvie van der Werf,Patricia Stoessel |
| EPI_ISL_792086, EPI_ISL_792087 | Toronto Invasive Bacterial Diseases Network | McMaster University | Allison McGeer, Patryk Aftanas, Hooman Derakhshani, Angel Li, Kuganya Nirmalarajah, Emily Panousis, Ahmed Draia, Jalees Nasir, Michael Surette, Samira Mubareka, Andrew G. McArthur |
| EPI_ISL_792525 | Laboratorio del Hospital Interzonal General de Agudos Evita | Grupo de Genómica y Bioinformática del Instituto de Investigación de la Cadena Láctea CONICET-INTA on behalf of 'Proyecto Argentino Interinstitucional de genomica de SARS-CoV-2' (PAIS Consortium) | Amadio, AF, Eberhardt, MF; Irazoqui, M; Torres, C; Aulicino, P; König, G; Desimone, I; Luczac, E; Serrano, L; Grossi, O; Musto; Alexay, S; Goya, S; Nabaes Jodar, MS; Viegas, M. |
| EPI_ISL_792559 | Grubaugh Lab - Yale School of Public Health | Grubaugh Lab - Yale School of Public Health | Joseph Fauver, Tara Alpert, Anderson Brito, Nathan Grubaugh |
| EPI_ISL_792655, EPI_ISL_792656, EPI_ISL_792657, EPI_ISL_792658, EPI_ISL_792661, EPI_ISL_792662, EPI_ISL_792663, EPI_ISL_792664, EPI_ISL_792665, EPI_ISL_792666, EPI_ISL_792667, EPI_ISL_792676, EPI_ISL_792677, EPI_ISL_792678 | | | |
| see above | Los Angeles County PHL | Los Angeles County PHL | P. Hemarajata et al. |
| EPI_ISL_792679 | Olive View Medical Center | Los Angeles County PHL | P. Hemarajata et al. |
| EPI_ISL_794104 | URMC LABS | Wadsworth Center, New York State Department.of Health | Kirsten St. George, Daryl M. Lamson, Alexis Russel, Matthew Shudt, Melissa A Leisner, Jonathan Plitnick, Navjot Singh, John Kelly, Sara Griesemer, Erasmus Schneider, Erica Lasek-Nesselquist |
| EPI_ISL_794107, EPI_ISL_794108, EPI_ISL_794109, EPI_ISL_794110, EPI_ISL_794111, EPI_ISL_794114 | GLENS FALLS HOSPITAL LABORATORY | Wadsworth Center, New York State Department.of Health | Kirsten St. George, Daryl M. Lamson, Alexis Russel, Matthew Shudt, Melissa A Leisner, Jonathan Plitnick, Navjot Singh, John Kelly, Sara Griesemer, Erasmus Schneider, Erica Lasek-Nesselquist |
| EPI_ISL_794119, EPI_ISL_794120, EPI_ISL_794121, EPI_ISL_794122, EPI_ISL_794123, EPI_ISL_794124, EPI_ISL_794125, EPI_ISL_794126, EPI_ISL_794127, EPI_ISL_794128, EPI_ISL_794129, EPI_ISL_794130, EPI_ISL_794131, EPI_ISL_794132, EPI_ISL_794133, EPI_ISL_794134, EPI_ISL_794135, EPI_ISL_794136, EPI_ISL_794137, EPI_ISL_794140, EPI_ISL_794142, EPI_ISL_794146, EPI_ISL_794147 | | | |
| see above | NORTHWELL HEALTH LABORATORIES | Wadsworth Center, New York State Department.of Health | Kirsten St. George, Daryl M. Lamson, Alexis Russel, Matthew Shudt, Melissa A Leisner, Jonathan Plitnick, Navjot Singh, John Kelly, Sara Griesemer, Erasmus Schneider, Erica Lasek-Nesselquist |
| EPI_ISL_794208, EPI_ISL_794209, EPI_ISL_794210, EPI_ISL_794211, EPI_ISL_794212, EPI_ISL_794213, EPI_ISL_794214, EPI_ISL_794215, EPI_ISL_794216, EPI_ISL_794217, EPI_ISL_794218, EPI_ISL_794239, EPI_ISL_794240 | | | |
| see above | WESTCHESTER MEDICAL CENTER | Wadsworth Center, New York State Department.of Health | Kirsten St. George, Daryl M. Lamson, Alexis Russel, Matthew Shudt, Melissa A Leisner, Jonathan Plitnick, Navjot Singh, John Kelly, Sara Griesemer, Erasmus Schneider, Erica Lasek-Nesselquist |
| EPI_ISL_794746 | SISP ASL BA AREA SUD | Istituto Zooprofilattico Sperimentale della Puglia e della Basilicata | Parisi A., Bianco A., Capozzi L., Del Sambro L., Manzulli V, Rondinone V., Pace L., Cipolletta D., Galante D. |
| EPI_ISL_794747 | Ospedale Vito Fazzi | Istituto Zooprofilattico Sperimentale della Puglia e della Basilicata | Parisi A., Bianco A., Capozzi L., Del Sambro L., Manzulli V, Rondinone V., Pace L., Cipolletta D., Galante D. |
| EPI_ISL_794748 | SISP ASL BA AREA SUD | Istituto Zooprofilattico Sperimentale della Puglia e della Basilicata | Parisi A., Bianco A., Capozzi L., Del Sambro L., Manzulli V, Rondinone V., Pace L., Cipolletta D., Galante D. |
| EPI_ISL_794750 | Ospedale Santa Caterina Novella | Istituto Zooprofilattico Sperimentale della Puglia e della Basilicata | Parisi A., Bianco A., Capozzi L., Del Sambro L., Manzulli V, Rondinone V., Pace L., Cipolletta D., Galante D. |
| EPI_ISL_794751 | Ospedale Vito Fazzi | Istituto Zooprofilattico Sperimentale della Puglia e della Basilicata | Parisi A., Bianco A., Capozzi L., Del Sambro L., Manzulli V, Rondinone V., Pace L., Cipolletta D., Galante D. |
| EPI_ISL_794752 | Istituto Zooprofilattico Sperimentale della Puglia e della Basilicata | Istituto Zooprofilattico Sperimentale della Puglia e della Basilicata | Parisi A., Bianco A., Capozzi L., Del Sambro L., Manzulli V, Rondinone V., Pace L., Cipolletta D., Galante D. |
| EPI_ISL_794753 | Ospedale Cardinale G. Panico | Istituto Zooprofilattico Sperimentale della Puglia e della Basilicata | Parisi A., Bianco A., Capozzi L., Del Sambro L., Manzulli V, Rondinone V., Pace L., Cipolletta D., Galante D. |
| EPI_ISL_794756, EPI_ISL_794757, EPI_ISL_794758 | Presidio di Brindisi Di Summa - Perrino | Istituto Zooprofilattico Sperimentale della Puglia e della Basilicata | Parisi A., Bianco A., Capozzi L., Del Sambro L., Manzulli V, Rondinone V., Pace L., Cipolletta D., Galante D. |
| EPI_ISL_794759, EPI_ISL_794779, EPI_ISL_794780, EPI_ISL_794781, EPI_ISL_794782, EPI_ISL_794783, EPI_ISL_794785, EPI_ISL_794786, EPI_ISL_794787 | Istituto Zooprofilattico Sperimentale della Puglia e della Basilicata | Istituto Zooprofilattico Sperimentale della Puglia e della Basilicata | Parisi A., Bianco A., Capozzi L., Del Sambro L., Manzulli V, Rondinone V., Pace L., Cipolletta D., Galante D. |
| EPI_ISL_794825 | Lab voor klinische biologie | Onderzoeksgroep Virologie | Laurens Lambrechts, Nick Vereecke, Marthe Pauwels, Bruno Verhasselt, Linos Vandekerckhove, Hans Nauwynck, Sebastiaan Theuns |
| EPI_ISL_796163, EPI_ISL_796164, EPI_ISL_796165, EPI_ISL_796170, EPI_ISL_796178, EPI_ISL_796181, EPI_ISL_796186, EPI_ISL_796187, EPI_ISL_796193, EPI_ISL_796200, EPI_ISL_796205, EPI_ISL_796207, EPI_ISL_796213, EPI_ISL_796221, EPI_ISL_796230, EPI_ISL_796231, EPI_ISL_796232, EPI_ISL_796238, EPI_ISL_796239, EPI_ISL_796241, EPI_ISL_796244, EPI_ISL_796248, EPI_ISL_796254, EPI_ISL_796257, EPI_ISL_796258, EPI_ISL_796260, EPI_ISL_796261, EPI_ISL_796262, EPI_ISL_796263, EPI_ISL_796272, EPI_ISL_796274, EPI_ISL_796277, EPI_ISL_796278, EPI_ISL_796281, EPI_ISL_796292, EPI_ISL_796296, EPI_ISL_796299, EPI_ISL_796308, EPI_ISL_796311, EPI_ISL_796313, EPI_ISL_796314, EPI_ISL_796315, EPI_ISL_796323, EPI_ISL_796328, EPI_ISL_796329, EPI_ISL_796330, EPI_ISL_796331, EPI_ISL_796332, EPI_ISL_796335, EPI_ISL_796336, EPI_ISL_796338, EPI_ISL_796339, EPI_ISL_796341, EPI_ISL_796343, EPI_ISL_796344, EPI_ISL_796345, EPI_ISL_796346, EPI_ISL_796347, EPI_ISL_796349, EPI_ISL_796350, EPI_ISL_796352, EPI_ISL_796354, EPI_ISL_796357, EPI_ISL_796358, EPI_ISL_796360, EPI_ISL_796365, EPI_ISL_796366, EPI_ISL_796367, EPI_ISL_796370, EPI_ISL_796375, EPI_ISL_796379, EPI_ISL_796380, EPI_ISL_796381, EPI_ISL_796383, EPI_ISL_796387, EPI_ISL_796388, EPI_ISL_796389, EPI_ISL_796392, EPI_ISL_796394, EPI_ISL_796397, EPI_ISL_796402, EPI_ISL_796404, EPI_ISL_796405, EPI_ISL_796406, EPI_ISL_796408, EPI_ISL_796409, EPI_ISL_796410, EPI_ISL_796412, EPI_ISL_796414, EPI_ISL_796415, EPI_ISL_796419, EPI_ISL_796424, EPI_ISL_796425, EPI_ISL_796429, EPI_ISL_796430, EPI_ISL_796432, EPI_ISL_796433, EPI_ISL_796434, EPI_ISL_796435, EPI_ISL_796442, EPI_ISL_796445, EPI_ISL_796460, EPI_ISL_796476, EPI_ISL_796477, EPI_ISL_796479, EPI_ISL_796483, EPI_ISL_796485, EPI_ISL_796487, EPI_ISL_796490, EPI_ISL_796492, EPI_ISL_796494, EPI_ISL_796495, EPI_ISL_796496, EPI_ISL_796499, EPI_ISL_796500, EPI_ISL_796502, EPI_ISL_796503, EPI_ISL_796507, EPI_ISL_796508, EPI_ISL_796509, EPI_ISL_796510, EPI_ISL_796512, EPI_ISL_796513, EPI_ISL_796519, EPI_ISL_796520, EPI_ISL_796521, EPI_ISL_796522, EPI_ISL_796524, EPI_ISL_796525, EPI_ISL_796526, EPI_ISL_796527, EPI_ISL_796529, EPI_ISL_796530, EPI_ISL_796532, EPI_ISL_796533, EPI_ISL_796534, EPI_ISL_796552, EPI_ISL_796554, EPI_ISL_796555, EPI_ISL_796556, EPI_ISL_796557, EPI_ISL_796559, EPI_ISL_796560, EPI_ISL_796562, EPI_ISL_796563, EPI_ISL_796566, EPI_ISL_796568, EPI_ISL_796569, EPI_ISL_796570, EPI_ISL_796580, EPI_ISL_796583, EPI_ISL_796588, EPI_ISL_796590, EPI_ISL_796593, EPI_ISL_796594, EPI_ISL_796596, EPI_ISL_796597, EPI_ISL_796603, EPI_ISL_796605, EPI_ISL_796618, EPI_ISL_796641, EPI_ISL_796642, EPI_ISL_796644, EPI_ISL_796645, EPI_ISL_796646 | | | |
| see above | Viollier AG | Department of Biosystems Science and Engineering, ETH Zürich | Chaoran Chen, Sarah Nadeau, Catharine Aquino, Ivan Topolsky, Philipp Jablonski, Lara Fuhrmann, David Dreifuss, Katharina Jahn, Andreia Cabral de Gouvea, Maria Domenica Moccia, Simon Grüter, Timothy Sykes, Lennart Opitz, Griffin White, Laura Neff, Doris Popovic, Andrea Patrignani, Jay Tracy, Ralph Schlapbach, Christiane Beckmann, Maurice Redondo, Olivier Kobel, Christoph Noppen, Sophie Seidel, Noemie Santamaria de Souza, Niko Beerenwinkel, Tanja Stadler |
| EPI_ISL_796652 | Haukeland University Hospital, Dept. of Microbiology | Norwegian Institute of Public Health, Department of Virology | Kathrine Stene-Johansen, Kamilla Heddeland Instefjord, Hilde Elshaug, Atiya R Ali,Marie Paulsen Madsen, Rasmus Riis Kopperud, Hilde Vollan, Karoline Bragstad, Olav Hungnes |
| EPI_ISL_796662 | Norwegian Institute of Public Health, Department of Virology | Norwegian Institute of Public Health, Department of Virology | Kathrine Stene-Johansen, Kamilla Heddeland Instefjord, Hilde Elshaug, Atiya R Ali,Marie Paulsen Madsen, Rasmus Riis Kopperud, Hilde Vollan, Karoline Bragstad, Olav Hungnes |
| EPI_ISL_796664 | Hospital of Southern Norway - Kristiansand, Department of Medical Microbiology | Norwegian Institute of Public Health, Department of Virology | Kathrine Stene-Johansen, Kamilla Heddeland Instefjord, Hilde Elshaug, Atiya R Ali,Marie Paulsen Madsen, Rasmus Riis Kopperud, Hilde Vollan, Karoline Bragstad, Olav Hungnes |
| EPI_ISL_796670 | University Hospital of Northern Norway, Department for Microbiology and Infectious Disease Control | Norwegian Institute of Public Health, Department of Virology | Kathrine Stene-Johansen, Kamilla Heddeland Instefjord, Hilde Elshaug, Atiya R Ali,Marie Paulsen Madsen, Rasmus Riis Kopperud, Hilde Vollan, Karoline Bragstad, Olav Hungnes |
| EPI_ISL_796674 | Ostfold Hospital Trust - Kalnes, Centre for Laboratory Medicine, Section for gene technology and infection serology | Norwegian Institute of Public Health, Department of Virology | Kathrine Stene-Johansen, Kamilla Heddeland Instefjord, Hilde Elshaug, Atiya R Ali,Marie Paulsen Madsen, Rasmus Riis Kopperud, Hilde Vollan, Karoline Bragstad, Olav Hungnes |

| | | | |
|---|---|---|---|
| EPI_ISL_796679, EPI_ISL_796680 | Unilabs Laboratory Medicine | Norwegian Institute of Public Health, Department of Virology | Kathrine Stene-Johansen, Kamilla Heddeland Instefjord, Hilde Elshaug, Atiya R Ali,Marie Paulsen Madsen, Rasmus Riis Kopperud, Hilde Vollan, Karoline Bragstad, Olav Hungnes |
| EPI_ISL_796703 | Oslo University Hospital, Department of Medical Microbiology | Norwegian Institute of Public Health, Department of Virology | Kathrine Stene-Johansen, Kamilla Heddeland Instefjord, Hilde Elshaug, Atiya R Ali,Marie Paulsen Madsen, Rasmus Riis Kopperud, Hilde Vollan, Karoline Bragstad, Olav Hungnes |
| EPI_ISL_796704 | University Hospital of Northern Norway, Department for Microbiology and Infectious Disease Control | Norwegian Institute of Public Health, Department of Virology | Kathrine Stene-Johansen, Kamilla Heddeland Instefjord, Hilde Elshaug, Atiya R Ali,Marie Paulsen Madsen, Rasmus Riis Kopperud, Hilde Vollan, Karoline Bragstad, Olav Hungnes |
| EPI_ISL_796712 | Akershus University Hospital, Department for Microbiology and Infectious Disease Control | Norwegian Institute of Public Health, Department of Virology | Kathrine Stene-Johansen, Kamilla Heddeland Instefjord, Hilde Elshaug, Atiya R Ali,Marie Paulsen Madsen, Rasmus Riis Kopperud, Hilde Vollan, Karoline Bragstad, Olav Hungnes |
| EPI_ISL_796735, EPI_ISL_796740, EPI_ISL_796744, EPI_ISL_796745, EPI_ISL_796746, EPI_ISL_796747, EPI_ISL_796748, EPI_ISL_796749, EPI_ISL_796750, EPI_ISL_796751, EPI_ISL_796760, EPI_ISL_796769, EPI_ISL_796772, EPI_ISL_796781 | | | |
| see above | Instituto Nacional de Saude (INSA) | Instituto Nacional de Saude (INSA) | Borges et al |
| EPI_ISL_796782 | Laboratory of Microbiology, National Reference Lab, Charles Nicolle Hospital; 2-University of Tunis ElManar, Faculty of Medicine of Tunis, LR99ES09, Tunis, Tunisia | Clinical and Experimental Pharmacology Lab, LR16SP02, National Center of Pharmacovigilance, University of Tunis El Manar, Tunis, Tunisia. 2-Neurodegenerative diseases and psychiatric troubles, LR18SP03, Razi Hospital, University of Tunis El Manar, Tunis, Tunisia. 3- Ministry of Health, National Observatory of New and Emerging Diseases, 1006, Tunis, Tunisia | Ilhem Boutiba-Ben Boubaker, Sameh Trabelsi, Nissaf Ben Alaya, Maher Kharrat, Alia BenKahla, Jalila Ben Khelil, Salma Abid, Sana Ferjani, Mouna Ben Sassi, Mouna Safer, Zaineb Hamzaoui, Guedi Ali Barreh, Habiba Ben Romdhane, Souissi Amira, Sarra Chamman, Hanen El Jebari, Ahmed Fakhfakh, Gaies Emna, Riadh Daghfous, Riadh Gouider. |
| EPI_ISL_797087 | Lighthouse Lab in Alderley Park | Wellcome Sanger Institute for the COVID-19 Genomics UK (COG-UK) Consortium | Jacquelyn Wynn, Mairead Hyland, The Lighthouse Lab in Alderley Park and Alex Alderton, Roberto Amato, Sonia Goncalves, Ewan Harrison, David K. Jackson, Ian Johnston, Dominic Kwiatkowski, Cordelia Langford, John Sillitoe on behalf of the Wellcome Sanger Institute COVID-19 Surveillance Team |
| EPI_ISL_797766, EPI_ISL_797825, EPI_ISL_797829, EPI_ISL_797830, EPI_ISL_797850, EPI_ISL_797853, EPI_ISL_797885, EPI_ISL_797994 | Lighthouse Lab in Glasgow | Wellcome Sanger Institute for the COVID-19 Genomics UK (COG-UK) Consortium | Harper VanSteenhouse, Yumi Kasai, David Gray, Carol Clugston, Anna Dominiczak and Alex Alderton, Roberto Amato, Sonia Goncalves, Ewan Harrison, David K. Jackson, Ian Johnston, Dominic Kwiatkowski, Cordelia Langford, John Sillitoe on behalf of the Wellcome Sanger Institute COVID-19 Surveillance Team |
| EPI_ISL_798000, EPI_ISL_798003, EPI_ISL_798018, EPI_ISL_798032, EPI_ISL_798038, EPI_ISL_798040, EPI_ISL_798046, EPI_ISL_798048, EPI_ISL_798053, EPI_ISL_798055, EPI_ISL_798058, EPI_ISL_798060, EPI_ISL_798069, EPI_ISL_798072, EPI_ISL_798076, EPI_ISL_798077, EPI_ISL_798078, EPI_ISL_798079, EPI_ISL_798086, EPI_ISL_798088, EPI_ISL_798093, EPI_ISL_798094, EPI_ISL_798095, EPI_ISL_798097, EPI_ISL_798105, EPI_ISL_798110, EPI_ISL_798111, EPI_ISL_798112, EPI_ISL_798113, EPI_ISL_798115, EPI_ISL_798117, EPI_ISL_798121, EPI_ISL_798126, EPI_ISL_798128, EPI_ISL_798130, EPI_ISL_798132, EPI_ISL_798137, EPI_ISL_798140, EPI_ISL_798143, EPI_ISL_798145, EPI_ISL_798148, EPI_ISL_798149, EPI_ISL_798151, EPI_ISL_798153, EPI_ISL_798160, EPI_ISL_798161, EPI_ISL_798166, EPI_ISL_798167, EPI_ISL_798168, EPI_ISL_798183, EPI_ISL_798185, EPI_ISL_798186, EPI_ISL_798193, EPI_ISL_798197, EPI_ISL_798199, EPI_ISL_798201, EPI_ISL_798203, EPI_ISL_798211, EPI_ISL_798213, EPI_ISL_798214, EPI_ISL_798216, EPI_ISL_798217, EPI_ISL_798218, EPI_ISL_798224, EPI_ISL_798230, EPI_ISL_798243, EPI_ISL_798244, EPI_ISL_798251, EPI_ISL_798252, EPI_ISL_798253, EPI_ISL_798254, EPI_ISL_798256, EPI_ISL_798258, EPI_ISL_798266, EPI_ISL_798270, EPI_ISL_798272, EPI_ISL_798273, EPI_ISL_798281, EPI_ISL_798288, EPI_ISL_798289, EPI_ISL_798294, EPI_ISL_798298, EPI_ISL_798300, EPI_ISL_798307, EPI_ISL_798311, EPI_ISL_798312, EPI_ISL_798315, EPI_ISL_798316, EPI_ISL_798319, EPI_ISL_798322, EPI_ISL_798326, EPI_ISL_798327 | | | |
| see above | Lighthouse Lab in Cambridge | Wellcome Sanger Institute for the COVID-19 Genomics UK (COG-UK) Consortium | Rob Howes, The Lighthouse Lab in Cambridge and Alex Alderton, Roberto Amato, Sonia Goncalves, Ewan Harrison, David K. Jackson, Ian Johnston, Dominic Kwiatkowski, Cordelia Langford, John Sillitoe on behalf of the Wellcome Sanger Institute COVID-19 Surveillance Team |
| EPI_ISL_798333, EPI_ISL_798335, EPI_ISL_798337, EPI_ISL_798338, EPI_ISL_798339, EPI_ISL_798340, EPI_ISL_798342, EPI_ISL_798343, EPI_ISL_798344, EPI_ISL_798347, EPI_ISL_798348, EPI_ISL_798350, EPI_ISL_798351, EPI_ISL_798352, EPI_ISL_798353, EPI_ISL_798354, EPI_ISL_798355, EPI_ISL_798357, EPI_ISL_798358, EPI_ISL_798359, EPI_ISL_798360, EPI_ISL_798361, EPI_ISL_798363, EPI_ISL_798364, EPI_ISL_798365, EPI_ISL_798366, EPI_ISL_798367, EPI_ISL_798371, EPI_ISL_798372, EPI_ISL_798373, EPI_ISL_798374, EPI_ISL_798375, EPI_ISL_798376, EPI_ISL_798377, EPI_ISL_798380, EPI_ISL_798381, EPI_ISL_798382, EPI_ISL_798383, EPI_ISL_798385, EPI_ISL_798387, EPI_ISL_798390, EPI_ISL_798391, EPI_ISL_798392, EPI_ISL_798393, EPI_ISL_798394, EPI_ISL_798395, EPI_ISL_798397, EPI_ISL_798398, EPI_ISL_798400 | | | |
| see above | Lighthouse Lab in Milton Keynes | Wellcome Sanger Institute for the COVID-19 Genomics UK (COG-UK) Consortium | The Lighthouse Lab in Milton Keynes and Alex Alderton, Roberto Amato, Sonia Goncalves, Ewan Harrison, David K. Jackson, Ian Johnston, Dominic Kwiatkowski, Cordelia Langford, John Sillitoe on behalf of the Wellcome Sanger Institute COVID-19 Surveillance Team |
| EPI_ISL_798401 | Lighthouse Lab in Alderley Park | Wellcome Sanger Institute for the COVID-19 Genomics UK (COG-UK) Consortium | Jacquelyn Wynn, Mairead Hyland, The Lighthouse Lab in Alderley Park and Alex Alderton, Roberto Amato, Sonia Goncalves, Ewan Harrison, David K. Jackson, Ian Johnston, Dominic Kwiatkowski, Cordelia Langford, John Sillitoe on behalf of the Wellcome Sanger Institute COVID-19 Surveillance Team |
| EPI_ISL_798402, EPI_ISL_798403, EPI_ISL_798406, EPI_ISL_798407, EPI_ISL_798408, EPI_ISL_798409, EPI_ISL_798410, EPI_ISL_798411, EPI_ISL_798412, EPI_ISL_798414, EPI_ISL_798415, EPI_ISL_798418, EPI_ISL_798419, EPI_ISL_798420, EPI_ISL_798421, EPI_ISL_798422, EPI_ISL_798423, EPI_ISL_798424, EPI_ISL_798425, EPI_ISL_798426, EPI_ISL_798427, EPI_ISL_798428, EPI_ISL_798429, EPI_ISL_798430, EPI_ISL_798432, EPI_ISL_798433, EPI_ISL_798434, EPI_ISL_798435, EPI_ISL_798437, EPI_ISL_798438, EPI_ISL_798439, EPI_ISL_798440, EPI_ISL_798441, EPI_ISL_798442, EPI_ISL_798443, EPI_ISL_798445, EPI_ISL_798446, EPI_ISL_798447, EPI_ISL_798448, EPI_ISL_798449, EPI_ISL_798450, EPI_ISL_798451, EPI_ISL_798452, EPI_ISL_798453, EPI_ISL_798454, EPI_ISL_798456, EPI_ISL_798457, EPI_ISL_798458, EPI_ISL_798459, EPI_ISL_798460, EPI_ISL_798461, EPI_ISL_798462, EPI_ISL_798463, EPI_ISL_798464, EPI_ISL_798466, EPI_ISL_798467, EPI_ISL_798469, EPI_ISL_798470, EPI_ISL_798471, EPI_ISL_798472, EPI_ISL_798473, EPI_ISL_798474, EPI_ISL_798476, EPI_ISL_798477, EPI_ISL_798478, EPI_ISL_798480, EPI_ISL_798481, EPI_ISL_798482, EPI_ISL_798484, EPI_ISL_798485, EPI_ISL_798486, EPI_ISL_798488, EPI_ISL_798489, EPI_ISL_798490, EPI_ISL_798491, EPI_ISL_798494, EPI_ISL_798495, EPI_ISL_798496, EPI_ISL_798497, EPI_ISL_798498, EPI_ISL_798499, EPI_ISL_798501, EPI_ISL_798503, EPI_ISL_798504, EPI_ISL_798505, EPI_ISL_798506, EPI_ISL_798507, EPI_ISL_798509, EPI_ISL_798511, EPI_ISL_798512, EPI_ISL_798514, EPI_ISL_798515, EPI_ISL_798518, EPI_ISL_798520, EPI_ISL_798521, EPI_ISL_798522, EPI_ISL_798523, EPI_ISL_798525, EPI_ISL_798526, EPI_ISL_798527, EPI_ISL_798528, EPI_ISL_798529, EPI_ISL_798530, EPI_ISL_798532, EPI_ISL_798533, EPI_ISL_798535, EPI_ISL_798536, EPI_ISL_798537, EPI_ISL_798538, EPI_ISL_798540, EPI_ISL_798543 | | | |
| see above | Lighthouse Lab in Milton Keynes | Wellcome Sanger Institute for the COVID-19 Genomics UK (COG-UK) Consortium | The Lighthouse Lab in Milton Keynes and Alex Alderton, Roberto Amato, Sonia Goncalves, Ewan Harrison, David K. Jackson, Ian Johnston, Dominic Kwiatkowski, Cordelia Langford, John Sillitoe on behalf of the Wellcome Sanger Institute COVID-19 Surveillance Team |
| EPI_ISL_798544 | Lighthouse Lab in Alderley Park | Wellcome Sanger Institute for the COVID-19 Genomics UK (COG-UK) Consortium | Jacquelyn Wynn, Mairead Hyland, The Lighthouse Lab in Alderley Park and Alex Alderton, Roberto Amato, Sonia Goncalves, Ewan Harrison, David K. Jackson, Ian Johnston, Dominic Kwiatkowski, Cordelia Langford, John Sillitoe on behalf of the Wellcome Sanger Institute COVID-19 Surveillance Team |
| EPI_ISL_798545, EPI_ISL_798546, EPI_ISL_798547, EPI_ISL_798549, EPI_ISL_798551, EPI_ISL_798552, EPI_ISL_798553, EPI_ISL_798554, EPI_ISL_798556, EPI_ISL_798557, EPI_ISL_798558, EPI_ISL_798559, EPI_ISL_798560, EPI_ISL_798562, EPI_ISL_798563, EPI_ISL_798565, EPI_ISL_798566, EPI_ISL_798567, EPI_ISL_798568, EPI_ISL_798569, EPI_ISL_798570, EPI_ISL_798571, EPI_ISL_798572, EPI_ISL_798573, EPI_ISL_798574, EPI_ISL_798575, EPI_ISL_798576, EPI_ISL_798577, EPI_ISL_798578, EPI_ISL_798580, EPI_ISL_798581, EPI_ISL_798582, EPI_ISL_798583, EPI_ISL_798584, EPI_ISL_798585, EPI_ISL_798586, EPI_ISL_798587, EPI_ISL_798588, EPI_ISL_798589, EPI_ISL_798590, EPI_ISL_798592, EPI_ISL_798594, EPI_ISL_798595, EPI_ISL_798596, EPI_ISL_798597, EPI_ISL_798599, EPI_ISL_798600, EPI_ISL_798602, EPI_ISL_798603, EPI_ISL_798604, EPI_ISL_798605, EPI_ISL_798606, EPI_ISL_798608, EPI_ISL_798610, EPI_ISL_798611, EPI_ISL_798613, EPI_ISL_798614 | | | |
| see above | Lighthouse Lab in Milton Keynes | Wellcome Sanger Institute for the COVID-19 Genomics UK (COG-UK) Consortium | The Lighthouse Lab in Milton Keynes and Alex Alderton, Roberto Amato, Sonia Goncalves, Ewan Harrison, David K. Jackson, Ian Johnston, Dominic Kwiatkowski, Cordelia Langford, John Sillitoe on behalf of the Wellcome Sanger Institute COVID-19 Surveillance Team |
| EPI_ISL_798615 | Lighthouse Lab in Alderley Park | Wellcome Sanger Institute for the COVID-19 Genomics UK (COG-UK) Consortium | Jacquelyn Wynn, Mairead Hyland, The Lighthouse Lab in Alderley Park and Alex Alderton, Roberto Amato, Sonia Goncalves, Ewan Harrison, David K. Jackson, Ian Johnston, Dominic Kwiatkowski, Cordelia Langford, John Sillitoe on behalf of the Wellcome Sanger Institute COVID-19 Surveillance Team |
| EPI_ISL_798616, EPI_ISL_798617, EPI_ISL_798618, EPI_ISL_798619, EPI_ISL_798620, EPI_ISL_798621, EPI_ISL_798623, EPI_ISL_798624, EPI_ISL_798625, EPI_ISL_798626, EPI_ISL_798627, EPI_ISL_798628, EPI_ISL_798629, EPI_ISL_798630, EPI_ISL_798634, EPI_ISL_798636, EPI_ISL_798638, EPI_ISL_798642, EPI_ISL_798644, EPI_ISL_798645, EPI_ISL_798646, EPI_ISL_798647, EPI_ISL_798648, EPI_ISL_798649, EPI_ISL_798650, EPI_ISL_798651, EPI_ISL_798653, EPI_ISL_798654, EPI_ISL_798655, EPI_ISL_798656, EPI_ISL_798657, EPI_ISL_798658, EPI_ISL_798660, EPI_ISL_798662, EPI_ISL_798663 | | | |
| see above | Lighthouse Lab in Milton Keynes | Wellcome Sanger Institute for the COVID-19 Genomics UK (COG-UK) Consortium | The Lighthouse Lab in Milton Keynes and Alex Alderton, Roberto Amato, Sonia Goncalves, Ewan Harrison, David K. Jackson, Ian Johnston, Dominic Kwiatkowski, Cordelia Langford, John Sillitoe on behalf of the Wellcome Sanger Institute COVID-19 Surveillance Team |
| EPI_ISL_798664 | Lighthouse Lab in Cambridge | Wellcome Sanger Institute for the COVID-19 Genomics UK (COG-UK) Consortium | Rob Howes, The Lighthouse Lab in Cambridge and Alex Alderton, Roberto Amato, Sonia Goncalves, Ewan Harrison, David K. Jackson, Ian Johnston, Dominic Kwiatkowski, Cordelia Langford, John Sillitoe on behalf of the Wellcome Sanger Institute COVID-19 Surveillance Team |
| EPI_ISL_798666, EPI_ISL_798670, EPI_ISL_798672 | Lighthouse Lab in Milton Keynes | Wellcome Sanger Institute for the COVID-19 Genomics UK (COG-UK) Consortium | The Lighthouse Lab in Milton Keynes and Alex Alderton, Roberto Amato, Sonia Goncalves, Ewan Harrison, David K. Jackson, Ian Johnston, Dominic Kwiatkowski, Cordelia Langford, John Sillitoe on behalf of the Wellcome Sanger Institute COVID-19 Surveillance Team |
| EPI_ISL_798673 | Lighthouse Lab in Cambridge | Wellcome Sanger Institute for the COVID-19 Genomics UK (COG-UK) Consortium | Rob Howes, The Lighthouse Lab in Cambridge and Alex Alderton, Roberto Amato, Sonia Goncalves, Ewan Harrison, David K. Jackson, Ian Johnston, Dominic Kwiatkowski, Cordelia Langford, John Sillitoe on behalf of the Wellcome Sanger Institute COVID-19 Surveillance Team |
| EPI_ISL_798675 | Lighthouse Lab in Milton Keynes | Wellcome Sanger Institute for the COVID-19 Genomics UK (COG-UK) Consortium | The Lighthouse Lab in Milton Keynes and Alex Alderton, Roberto Amato, Sonia Goncalves, Ewan Harrison, David K. Jackson, Ian Johnston, Dominic Kwiatkowski, Cordelia Langford, John Sillitoe on behalf of the Wellcome Sanger Institute COVID-19 Surveillance Team |
| EPI_ISL_798677, EPI_ISL_798679 | Lighthouse Lab in Cambridge | Wellcome Sanger Institute for the COVID-19 Genomics UK (COG-UK) Consortium | Rob Howes, The Lighthouse Lab in Cambridge and Alex Alderton, Roberto Amato, Sonia Goncalves, Ewan Harrison, David K. Jackson, Ian Johnston, Dominic Kwiatkowski, Cordelia Langford, John Sillitoe on behalf of the Wellcome Sanger Institute COVID-19 Surveillance Team |
| EPI_ISL_798680 | Lighthouse Lab in Milton Keynes | Wellcome Sanger Institute for the COVID-19 Genomics UK (COG-UK) Consortium | The Lighthouse Lab in Milton Keynes and Alex Alderton, Roberto Amato, Sonia Goncalves, Ewan Harrison, David K. Jackson, Ian Johnston, Dominic Kwiatkowski, Cordelia Langford, John Sillitoe on behalf of the Wellcome Sanger Institute COVID-19 Surveillance Team |

| | | | |
|---|---|---|---|
| EPI_ISL_798681, EPI_ISL_798682, EPI_ISL_798683 | Lighthouse Lab in Cambridge | Wellcome Sanger Institute for the COVID-19 Genomics UK (COG-UK) Consortium | Rob Howes, The Lighthouse Lab in Cambridge and Alex Alderton, Roberto Amato, Sonia Goncalves, Ewan Harrison, David K. Jackson, Ian Johnston, Dominic Kwiatkowski, Cordelia Langford, John Sillitoe on behalf of the Wellcome Sanger Institute COVID-19 Surveillance Team |
| EPI_ISL_798684 | Lighthouse Lab in Alderley Park | Wellcome Sanger Institute for the COVID-19 Genomics UK (COG-UK) Consortium | Jacquelyn Wynn, Mairead Hyland, The Lighthouse Lab in Alderley Park and Alex Alderton, Roberto Amato, Sonia Goncalves, Ewan Harrison, David K. Jackson, Ian Johnston, Dominic Kwiatkowski, Cordelia Langford, John Sillitoe on behalf of the Wellcome Sanger Institute COVID-19 Surveillance Team |
| EPI_ISL_798685, EPI_ISL_798686 | Lighthouse Lab in Cambridge | Wellcome Sanger Institute for the COVID-19 Genomics UK (COG-UK) Consortium | Rob Howes, The Lighthouse Lab in Cambridge and Alex Alderton, Roberto Amato, Sonia Goncalves, Ewan Harrison, David K. Jackson, Ian Johnston, Dominic Kwiatkowski, Cordelia Langford, John Sillitoe on behalf of the Wellcome Sanger Institute COVID-19 Surveillance Team |
| EPI_ISL_798691, EPI_ISL_798693 | Lighthouse Lab in Milton Keynes | Wellcome Sanger Institute for the COVID-19 Genomics UK (COG-UK) Consortium | The Lighthouse Lab in Milton Keynes and Alex Alderton, Roberto Amato, Sonia Goncalves, Ewan Harrison, David K. Jackson, Ian Johnston, Dominic Kwiatkowski, Cordelia Langford, John Sillitoe on behalf of the Wellcome Sanger Institute COVID-19 Surveillance Team |
| EPI_ISL_798694 | Lighthouse Lab in Cambridge | Wellcome Sanger Institute for the COVID-19 Genomics UK (COG-UK) Consortium | Rob Howes, The Lighthouse Lab in Cambridge and Alex Alderton, Roberto Amato, Sonia Goncalves, Ewan Harrison, David K. Jackson, Ian Johnston, Dominic Kwiatkowski, Cordelia Langford, John Sillitoe on behalf of the Wellcome Sanger Institute COVID-19 Surveillance Team |
| EPI_ISL_798696 | Lighthouse Lab in Milton Keynes | Wellcome Sanger Institute for the COVID-19 Genomics UK (COG-UK) Consortium | The Lighthouse Lab in Milton Keynes and Alex Alderton, Roberto Amato, Sonia Goncalves, Ewan Harrison, David K. Jackson, Ian Johnston, Dominic Kwiatkowski, Cordelia Langford, John Sillitoe on behalf of the Wellcome Sanger Institute COVID-19 Surveillance Team |
| EPI_ISL_798697, EPI_ISL_798698, EPI_ISL_798699 | Lighthouse Lab in Cambridge | Wellcome Sanger Institute for the COVID-19 Genomics UK (COG-UK) Consortium | Rob Howes, The Lighthouse Lab in Cambridge and Alex Alderton, Roberto Amato, Sonia Goncalves, Ewan Harrison, David K. Jackson, Ian Johnston, Dominic Kwiatkowski, Cordelia Langford, John Sillitoe on behalf of the Wellcome Sanger Institute COVID-19 Surveillance Team |
| EPI_ISL_798701, EPI_ISL_798702 | Lighthouse Lab in Milton Keynes | Wellcome Sanger Institute for the COVID-19 Genomics UK (COG-UK) Consortium | The Lighthouse Lab in Milton Keynes and Alex Alderton, Roberto Amato, Sonia Goncalves, Ewan Harrison, David K. Jackson, Ian Johnston, Dominic Kwiatkowski, Cordelia Langford, John Sillitoe on behalf of the Wellcome Sanger Institute COVID-19 Surveillance Team |
| EPI_ISL_798703, EPI_ISL_798706 | Lighthouse Lab in Cambridge | Wellcome Sanger Institute for the COVID-19 Genomics UK (COG-UK) Consortium | Rob Howes, The Lighthouse Lab in Cambridge and Alex Alderton, Roberto Amato, Sonia Goncalves, Ewan Harrison, David K. Jackson, Ian Johnston, Dominic Kwiatkowski, Cordelia Langford, John Sillitoe on behalf of the Wellcome Sanger Institute COVID-19 Surveillance Team |
| EPI_ISL_798707 | Lighthouse Lab in Milton Keynes | Wellcome Sanger Institute for the COVID-19 Genomics UK (COG-UK) Consortium | The Lighthouse Lab in Milton Keynes and Alex Alderton, Roberto Amato, Sonia Goncalves, Ewan Harrison, David K. Jackson, Ian Johnston, Dominic Kwiatkowski, Cordelia Langford, John Sillitoe on behalf of the Wellcome Sanger Institute COVID-19 Surveillance Team |
| EPI_ISL_798708 | Lighthouse Lab in Cambridge | Wellcome Sanger Institute for the COVID-19 Genomics UK (COG-UK) Consortium | Rob Howes, The Lighthouse Lab in Cambridge and Alex Alderton, Roberto Amato, Sonia Goncalves, Ewan Harrison, David K. Jackson, Ian Johnston, Dominic Kwiatkowski, Cordelia Langford, John Sillitoe on behalf of the Wellcome Sanger Institute COVID-19 Surveillance Team |
| EPI_ISL_798709 | Lighthouse Lab in Alderley Park | Wellcome Sanger Institute for the COVID-19 Genomics UK (COG-UK) Consortium | Jacquelyn Wynn, Mairead Hyland, The Lighthouse Lab in Alderley Park and Alex Alderton, Roberto Amato, Sonia Goncalves, Ewan Harrison, David K. Jackson, Ian Johnston, Dominic Kwiatkowski, Cordelia Langford, John Sillitoe on behalf of the Wellcome Sanger Institute COVID-19 Surveillance Team |
| EPI_ISL_798710 | Lighthouse Lab in Milton Keynes | Wellcome Sanger Institute for the COVID-19 Genomics UK (COG-UK) Consortium | The Lighthouse Lab in Milton Keynes and Alex Alderton, Roberto Amato, Sonia Goncalves, Ewan Harrison, David K. Jackson, Ian Johnston, Dominic Kwiatkowski, Cordelia Langford, John Sillitoe on behalf of the Wellcome Sanger Institute COVID-19 Surveillance Team |
| EPI_ISL_798711 | Lighthouse Lab in Cambridge | Wellcome Sanger Institute for the COVID-19 Genomics UK (COG-UK) Consortium | Rob Howes, The Lighthouse Lab in Cambridge and Alex Alderton, Roberto Amato, Sonia Goncalves, Ewan Harrison, David K. Jackson, Ian Johnston, Dominic Kwiatkowski, Cordelia Langford, John Sillitoe on behalf of the Wellcome Sanger Institute COVID-19 Surveillance Team |
| EPI_ISL_798712 | Lighthouse Lab in Milton Keynes | Wellcome Sanger Institute for the COVID-19 Genomics UK (COG-UK) Consortium | The Lighthouse Lab in Milton Keynes and Alex Alderton, Roberto Amato, Sonia Goncalves, Ewan Harrison, David K. Jackson, Ian Johnston, Dominic Kwiatkowski, Cordelia Langford, John Sillitoe on behalf of the Wellcome Sanger Institute COVID-19 Surveillance Team |
| EPI_ISL_798713, EPI_ISL_798714, EPI_ISL_798715 | Lighthouse Lab in Cambridge | Wellcome Sanger Institute for the COVID-19 Genomics UK (COG-UK) Consortium | Rob Howes, The Lighthouse Lab in Cambridge and Alex Alderton, Roberto Amato, Sonia Goncalves, Ewan Harrison, David K. Jackson, Ian Johnston, Dominic Kwiatkowski, Cordelia Langford, John Sillitoe on behalf of the Wellcome Sanger Institute COVID-19 Surveillance Team |
| EPI_ISL_798716 | Lighthouse Lab in Milton Keynes | Wellcome Sanger Institute for the COVID-19 Genomics UK (COG-UK) Consortium | The Lighthouse Lab in Milton Keynes and Alex Alderton, Roberto Amato, Sonia Goncalves, Ewan Harrison, David K. Jackson, Ian Johnston, Dominic Kwiatkowski, Cordelia Langford, John Sillitoe on behalf of the Wellcome Sanger Institute COVID-19 Surveillance Team |
| EPI_ISL_798717, EPI_ISL_798718 | Lighthouse Lab in Cambridge | Wellcome Sanger Institute for the COVID-19 Genomics UK (COG-UK) Consortium | Rob Howes, The Lighthouse Lab in Cambridge and Alex Alderton, Roberto Amato, Sonia Goncalves, Ewan Harrison, David K. Jackson, Ian Johnston, Dominic Kwiatkowski, Cordelia Langford, John Sillitoe on behalf of the Wellcome Sanger Institute COVID-19 Surveillance Team |
| EPI_ISL_798719 | Lighthouse Lab in Alderley Park | Wellcome Sanger Institute for the COVID-19 Genomics UK (COG-UK) Consortium | Jacquelyn Wynn, Mairead Hyland, The Lighthouse Lab in Alderley Park and Alex Alderton, Roberto Amato, Sonia Goncalves, Ewan Harrison, David K. Jackson, Ian Johnston, Dominic Kwiatkowski, Cordelia Langford, John Sillitoe on behalf of the Wellcome Sanger Institute COVID-19 Surveillance Team |
| EPI_ISL_798720, EPI_ISL_798721 | Lighthouse Lab in Milton Keynes | Wellcome Sanger Institute for the COVID-19 Genomics UK (COG-UK) Consortium | The Lighthouse Lab in Milton Keynes and Alex Alderton, Roberto Amato, Sonia Goncalves, Ewan Harrison, David K. Jackson, Ian Johnston, Dominic Kwiatkowski, Cordelia Langford, John Sillitoe on behalf of the Wellcome Sanger Institute COVID-19 Surveillance Team |
| EPI_ISL_798723 | Lighthouse Lab in Cambridge | Wellcome Sanger Institute for the COVID-19 Genomics UK (COG-UK) Consortium | Rob Howes, The Lighthouse Lab in Cambridge and Alex Alderton, Roberto Amato, Sonia Goncalves, Ewan Harrison, David K. Jackson, Ian Johnston, Dominic Kwiatkowski, Cordelia Langford, John Sillitoe on behalf of the Wellcome Sanger Institute COVID-19 Surveillance Team |
| EPI_ISL_798724, EPI_ISL_798725 | Lighthouse Lab in Milton Keynes | Wellcome Sanger Institute for the COVID-19 Genomics UK (COG-UK) Consortium | The Lighthouse Lab in Milton Keynes and Alex Alderton, Roberto Amato, Sonia Goncalves, Ewan Harrison, David K. Jackson, Ian Johnston, Dominic Kwiatkowski, Cordelia Langford, John Sillitoe on behalf of the Wellcome Sanger Institute COVID-19 Surveillance Team |
| EPI_ISL_798726 | Lighthouse Lab in Cambridge | Wellcome Sanger Institute for the COVID-19 Genomics UK (COG-UK) Consortium | Rob Howes, The Lighthouse Lab in Cambridge and Alex Alderton, Roberto Amato, Sonia Goncalves, Ewan Harrison, David K. Jackson, Ian Johnston, Dominic Kwiatkowski, Cordelia Langford, John Sillitoe on behalf of the Wellcome Sanger Institute COVID-19 Surveillance Team |
| EPI_ISL_798728 | Lighthouse Lab in Milton Keynes | Wellcome Sanger Institute for the COVID-19 Genomics UK (COG-UK) Consortium | The Lighthouse Lab in Milton Keynes and Alex Alderton, Roberto Amato, Sonia Goncalves, Ewan Harrison, David K. Jackson, Ian Johnston, Dominic Kwiatkowski, Cordelia Langford, John Sillitoe on behalf of the Wellcome Sanger Institute COVID-19 Surveillance Team |
| EPI_ISL_798729 | Lighthouse Lab in Alderley Park | Wellcome Sanger Institute for the COVID-19 Genomics UK (COG-UK) Consortium | Jacquelyn Wynn, Mairead Hyland, The Lighthouse Lab in Alderley Park and Alex Alderton, Roberto Amato, Sonia Goncalves, Ewan Harrison, David K. Jackson, Ian Johnston, Dominic Kwiatkowski, Cordelia Langford, John Sillitoe on behalf of the Wellcome Sanger Institute COVID-19 Surveillance Team |
| EPI_ISL_798730, EPI_ISL_798731, EPI_ISL_798732, EPI_ISL_798733, EPI_ISL_798734, EPI_ISL_798736, EPI_ISL_798737 | Lighthouse Lab in Cambridge | Wellcome Sanger Institute for the COVID-19 Genomics UK (COG-UK) Consortium | Rob Howes, The Lighthouse Lab in Cambridge and Alex Alderton, Roberto Amato, Sonia Goncalves, Ewan Harrison, David K. Jackson, Ian Johnston, Dominic Kwiatkowski, Cordelia Langford, John Sillitoe on behalf of the Wellcome Sanger Institute COVID-19 Surveillance Team |
| EPI_ISL_798738, EPI_ISL_798739, EPI_ISL_798740, EPI_ISL_798741 | Lighthouse Lab in Milton Keynes | Wellcome Sanger Institute for the COVID-19 Genomics UK (COG-UK) Consortium | The Lighthouse Lab in Milton Keynes and Alex Alderton, Roberto Amato, Sonia Goncalves, Ewan Harrison, David K. Jackson, Ian Johnston, Dominic Kwiatkowski, Cordelia Langford, John Sillitoe on behalf of the Wellcome Sanger Institute COVID-19 Surveillance Team |
| EPI_ISL_798742 | Lighthouse Lab in Alderley Park | Wellcome Sanger Institute for the COVID-19 Genomics UK (COG-UK) Consortium | Jacquelyn Wynn, Mairead Hyland, The Lighthouse Lab in Alderley Park and Alex Alderton, Roberto Amato, Sonia Goncalves, Ewan Harrison, David K. Jackson, Ian Johnston, Dominic Kwiatkowski, Cordelia Langford, John Sillitoe on behalf of the Wellcome Sanger Institute COVID-19 Surveillance Team |
| EPI_ISL_798743, EPI_ISL_798745, EPI_ISL_798746, EPI_ISL_798747, EPI_ISL_798748 | Lighthouse Lab in Cambridge | Wellcome Sanger Institute for the COVID-19 Genomics UK (COG-UK) Consortium | Rob Howes, The Lighthouse Lab in Cambridge and Alex Alderton, Roberto Amato, Sonia Goncalves, Ewan Harrison, David K. Jackson, Ian Johnston, Dominic Kwiatkowski, Cordelia Langford, John Sillitoe on behalf of the Wellcome Sanger Institute COVID-19 Surveillance Team |
| EPI_ISL_798750 | Lighthouse Lab in Milton Keynes | Wellcome Sanger Institute for the COVID-19 Genomics UK (COG-UK) Consortium | The Lighthouse Lab in Milton Keynes and Alex Alderton, Roberto Amato, Sonia Goncalves, Ewan Harrison, David K. Jackson, Ian Johnston, Dominic Kwiatkowski, Cordelia Langford, John Sillitoe on behalf of the Wellcome Sanger Institute COVID-19 Surveillance Team |
| EPI_ISL_798751 | Lighthouse Lab in Cambridge | Wellcome Sanger Institute for the COVID-19 Genomics UK (COG-UK) Consortium | Rob Howes, The Lighthouse Lab in Cambridge and Alex Alderton, Roberto Amato, Sonia Goncalves, Ewan Harrison, David K. Jackson, Ian Johnston, Dominic Kwiatkowski, Cordelia Langford, John Sillitoe on behalf of the Wellcome Sanger Institute COVID-19 Surveillance Team |
| EPI_ISL_798752 | Lighthouse Lab in Milton Keynes | Wellcome Sanger Institute for the COVID-19 Genomics UK (COG-UK) Consortium | The Lighthouse Lab in Milton Keynes and Alex Alderton, Roberto Amato, Sonia Goncalves, Ewan Harrison, David K. Jackson, Ian Johnston, Dominic Kwiatkowski, Cordelia Langford, John Sillitoe on behalf of the Wellcome Sanger Institute COVID-19 Surveillance Team |
| EPI_ISL_798753, EPI_ISL_798754 | Lighthouse Lab in Cambridge | Wellcome Sanger Institute for the COVID-19 Genomics UK (COG-UK) Consortium | Rob Howes, The Lighthouse Lab in Cambridge and Alex Alderton, Roberto Amato, Sonia Goncalves, Ewan Harrison, David K. Jackson, Ian Johnston, Dominic Kwiatkowski, Cordelia Langford, John Sillitoe on behalf of the Wellcome Sanger Institute COVID-19 Surveillance Team |
| EPI_ISL_798755 | Lighthouse Lab in Milton Keynes | Wellcome Sanger Institute for the COVID-19 Genomics UK (COG-UK) Consortium | The Lighthouse Lab in Milton Keynes and Alex Alderton, Roberto Amato, Sonia Goncalves, Ewan Harrison, David K. Jackson, Ian Johnston, Dominic Kwiatkowski, Cordelia Langford, John Sillitoe on behalf of the Wellcome Sanger Institute COVID-19 Surveillance Team |

| | | | |
|---|---|---|---|
| EPI_ISL_798756, EPI_ISL_798757, EPI_ISL_798758, EPI_ISL_798759, EPI_ISL_798760, EPI_ISL_798761, EPI_ISL_798762, EPI_ISL_798763 | Lighthouse Lab in Cambridge | Wellcome Sanger Institute for the COVID-19 Genomics UK (COG-UK) Consortium | Rob Howes, The Lighthouse Lab in Cambridge and Alex Alderton, Roberto Amato, Sonia Goncalves, Ewan Harrison, David K. Jackson, Ian Johnston, Dominic Kwiatkowski, Cordelia Langford, John Sillitoe on behalf of the Wellcome Sanger Institute COVID-19 Surveillance Team |
| EPI_ISL_798764 | Lighthouse Lab in Milton Keynes | Wellcome Sanger Institute for the COVID-19 Genomics UK (COG-UK) Consortium | The Lighthouse Lab in Milton Keynes and Alex Alderton, Roberto Amato, Sonia Goncalves, Ewan Harrison, David K. Jackson, Ian Johnston, Dominic Kwiatkowski, Cordelia Langford, John Sillitoe on behalf of the Wellcome Sanger Institute COVID-19 Surveillance Team |
| EPI_ISL_798765, EPI_ISL_798766 | Lighthouse Lab in Cambridge | Wellcome Sanger Institute for the COVID-19 Genomics UK (COG-UK) Consortium | Rob Howes, The Lighthouse Lab in Cambridge and Alex Alderton, Roberto Amato, Sonia Goncalves, Ewan Harrison, David K. Jackson, Ian Johnston, Dominic Kwiatkowski, Cordelia Langford, John Sillitoe on behalf of the Wellcome Sanger Institute COVID-19 Surveillance Team |
| EPI_ISL_798767, EPI_ISL_798769, EPI_ISL_798770 | Lighthouse Lab in Milton Keynes | Wellcome Sanger Institute for the COVID-19 Genomics UK (COG-UK) Consortium | The Lighthouse Lab in Milton Keynes and Alex Alderton, Roberto Amato, Sonia Goncalves, Ewan Harrison, David K. Jackson, Ian Johnston, Dominic Kwiatkowski, Cordelia Langford, John Sillitoe on behalf of the Wellcome Sanger Institute COVID-19 Surveillance Team |
| EPI_ISL_798772, EPI_ISL_798773 | Lighthouse Lab in Cambridge | Wellcome Sanger Institute for the COVID-19 Genomics UK (COG-UK) Consortium | Rob Howes, The Lighthouse Lab in Cambridge and Alex Alderton, Roberto Amato, Sonia Goncalves, Ewan Harrison, David K. Jackson, Ian Johnston, Dominic Kwiatkowski, Cordelia Langford, John Sillitoe on behalf of the Wellcome Sanger Institute COVID-19 Surveillance Team |
| EPI_ISL_798774 | Lighthouse Lab in Milton Keynes | Wellcome Sanger Institute for the COVID-19 Genomics UK (COG-UK) Consortium | The Lighthouse Lab in Milton Keynes and Alex Alderton, Roberto Amato, Sonia Goncalves, Ewan Harrison, David K. Jackson, Ian Johnston, Dominic Kwiatkowski, Cordelia Langford, John Sillitoe on behalf of the Wellcome Sanger Institute COVID-19 Surveillance Team |
| EPI_ISL_798775 | Lighthouse Lab in Cambridge | Wellcome Sanger Institute for the COVID-19 Genomics UK (COG-UK) Consortium | Rob Howes, The Lighthouse Lab in Cambridge and Alex Alderton, Roberto Amato, Sonia Goncalves, Ewan Harrison, David K. Jackson, Ian Johnston, Dominic Kwiatkowski, Cordelia Langford, John Sillitoe on behalf of the Wellcome Sanger Institute COVID-19 Surveillance Team |
| EPI_ISL_798776, EPI_ISL_798777, EPI_ISL_798779 | Lighthouse Lab in Milton Keynes | Wellcome Sanger Institute for the COVID-19 Genomics UK (COG-UK) Consortium | The Lighthouse Lab in Milton Keynes and Alex Alderton, Roberto Amato, Sonia Goncalves, Ewan Harrison, David K. Jackson, Ian Johnston, Dominic Kwiatkowski, Cordelia Langford, John Sillitoe on behalf of the Wellcome Sanger Institute COVID-19 Surveillance Team |
| EPI_ISL_798780 | Lighthouse Lab in Cambridge | Wellcome Sanger Institute for the COVID-19 Genomics UK (COG-UK) Consortium | Rob Howes, The Lighthouse Lab in Cambridge and Alex Alderton, Roberto Amato, Sonia Goncalves, Ewan Harrison, David K. Jackson, Ian Johnston, Dominic Kwiatkowski, Cordelia Langford, John Sillitoe on behalf of the Wellcome Sanger Institute COVID-19 Surveillance Team |
| EPI_ISL_798781 | Lighthouse Lab in Milton Keynes | Wellcome Sanger Institute for the COVID-19 Genomics UK (COG-UK) Consortium | The Lighthouse Lab in Milton Keynes and Alex Alderton, Roberto Amato, Sonia Goncalves, Ewan Harrison, David K. Jackson, Ian Johnston, Dominic Kwiatkowski, Cordelia Langford, John Sillitoe on behalf of the Wellcome Sanger Institute COVID-19 Surveillance Team |
| EPI_ISL_798782, EPI_ISL_798783, EPI_ISL_798785, EPI_ISL_798787, EPI_ISL_798788, EPI_ISL_798789, EPI_ISL_798792, EPI_ISL_798794, EPI_ISL_798795, EPI_ISL_798796, EPI_ISL_798797, EPI_ISL_798799, EPI_ISL_798800 | | | |
| see above | Lighthouse Lab in Cambridge | Wellcome Sanger Institute for the COVID-19 Genomics UK (COG-UK) Consortium | Rob Howes, The Lighthouse Lab in Cambridge and Alex Alderton, Roberto Amato, Sonia Goncalves, Ewan Harrison, David K. Jackson, Ian Johnston, Dominic Kwiatkowski, Cordelia Langford, John Sillitoe on behalf of the Wellcome Sanger Institute COVID-19 Surveillance Team |
| EPI_ISL_798801 | Lighthouse Lab in Milton Keynes | Wellcome Sanger Institute for the COVID-19 Genomics UK (COG-UK) Consortium | The Lighthouse Lab in Milton Keynes and Alex Alderton, Roberto Amato, Sonia Goncalves, Ewan Harrison, David K. Jackson, Ian Johnston, Dominic Kwiatkowski, Cordelia Langford, John Sillitoe on behalf of the Wellcome Sanger Institute COVID-19 Surveillance Team |
| EPI_ISL_798802 | Lighthouse Lab in Cambridge | Wellcome Sanger Institute for the COVID-19 Genomics UK (COG-UK) Consortium | Rob Howes, The Lighthouse Lab in Cambridge and Alex Alderton, Roberto Amato, Sonia Goncalves, Ewan Harrison, David K. Jackson, Ian Johnston, Dominic Kwiatkowski, Cordelia Langford, John Sillitoe on behalf of the Wellcome Sanger Institute COVID-19 Surveillance Team |
| EPI_ISL_798803 | Lighthouse Lab in Milton Keynes | Wellcome Sanger Institute for the COVID-19 Genomics UK (COG-UK) Consortium | The Lighthouse Lab in Milton Keynes and Alex Alderton, Roberto Amato, Sonia Goncalves, Ewan Harrison, David K. Jackson, Ian Johnston, Dominic Kwiatkowski, Cordelia Langford, John Sillitoe on behalf of the Wellcome Sanger Institute COVID-19 Surveillance Team |
| EPI_ISL_798805, EPI_ISL_798806, EPI_ISL_798807 | Lighthouse Lab in Cambridge | Wellcome Sanger Institute for the COVID-19 Genomics UK (COG-UK) Consortium | Rob Howes, The Lighthouse Lab in Cambridge and Alex Alderton, Roberto Amato, Sonia Goncalves, Ewan Harrison, David K. Jackson, Ian Johnston, Dominic Kwiatkowski, Cordelia Langford, John Sillitoe on behalf of the Wellcome Sanger Institute COVID-19 Surveillance Team |
| EPI_ISL_798808, EPI_ISL_798809 | Lighthouse Lab in Milton Keynes | Wellcome Sanger Institute for the COVID-19 Genomics UK (COG-UK) Consortium | The Lighthouse Lab in Milton Keynes and Alex Alderton, Roberto Amato, Sonia Goncalves, Ewan Harrison, David K. Jackson, Ian Johnston, Dominic Kwiatkowski, Cordelia Langford, John Sillitoe on behalf of the Wellcome Sanger Institute COVID-19 Surveillance Team |
| EPI_ISL_798810 | Lighthouse Lab in Cambridge | Wellcome Sanger Institute for the COVID-19 Genomics UK (COG-UK) Consortium | Rob Howes, The Lighthouse Lab in Cambridge and Alex Alderton, Roberto Amato, Sonia Goncalves, Ewan Harrison, David K. Jackson, Ian Johnston, Dominic Kwiatkowski, Cordelia Langford, John Sillitoe on behalf of the Wellcome Sanger Institute COVID-19 Surveillance Team |
| EPI_ISL_798812 | Lighthouse Lab in Milton Keynes | Wellcome Sanger Institute for the COVID-19 Genomics UK (COG-UK) Consortium | The Lighthouse Lab in Milton Keynes and Alex Alderton, Roberto Amato, Sonia Goncalves, Ewan Harrison, David K. Jackson, Ian Johnston, Dominic Kwiatkowski, Cordelia Langford, John Sillitoe on behalf of the Wellcome Sanger Institute COVID-19 Surveillance Team |
| EPI_ISL_798813, EPI_ISL_798815, EPI_ISL_798816 | Lighthouse Lab in Cambridge | Wellcome Sanger Institute for the COVID-19 Genomics UK (COG-UK) Consortium | Rob Howes, The Lighthouse Lab in Cambridge and Alex Alderton, Roberto Amato, Sonia Goncalves, Ewan Harrison, David K. Jackson, Ian Johnston, Dominic Kwiatkowski, Cordelia Langford, John Sillitoe on behalf of the Wellcome Sanger Institute COVID-19 Surveillance Team |
| EPI_ISL_798817 | Lighthouse Lab in Milton Keynes | Wellcome Sanger Institute for the COVID-19 Genomics UK (COG-UK) Consortium | The Lighthouse Lab in Milton Keynes and Alex Alderton, Roberto Amato, Sonia Goncalves, Ewan Harrison, David K. Jackson, Ian Johnston, Dominic Kwiatkowski, Cordelia Langford, John Sillitoe on behalf of the Wellcome Sanger Institute COVID-19 Surveillance Team |
| EPI_ISL_798818, EPI_ISL_798820 | Lighthouse Lab in Cambridge | Wellcome Sanger Institute for the COVID-19 Genomics UK (COG-UK) Consortium | Rob Howes, The Lighthouse Lab in Cambridge and Alex Alderton, Roberto Amato, Sonia Goncalves, Ewan Harrison, David K. Jackson, Ian Johnston, Dominic Kwiatkowski, Cordelia Langford, John Sillitoe on behalf of the Wellcome Sanger Institute COVID-19 Surveillance Team |
| EPI_ISL_798821, EPI_ISL_798822, EPI_ISL_798823 | Lighthouse Lab in Milton Keynes | Wellcome Sanger Institute for the COVID-19 Genomics UK (COG-UK) Consortium | The Lighthouse Lab in Milton Keynes and Alex Alderton, Roberto Amato, Sonia Goncalves, Ewan Harrison, David K. Jackson, Ian Johnston, Dominic Kwiatkowski, Cordelia Langford, John Sillitoe on behalf of the Wellcome Sanger Institute COVID-19 Surveillance Team |
| EPI_ISL_798827, EPI_ISL_798828, EPI_ISL_798829 | Lighthouse Lab in Cambridge | Wellcome Sanger Institute for the COVID-19 Genomics UK (COG-UK) Consortium | Rob Howes, The Lighthouse Lab in Cambridge and Alex Alderton, Roberto Amato, Sonia Goncalves, Ewan Harrison, David K. Jackson, Ian Johnston, Dominic Kwiatkowski, Cordelia Langford, John Sillitoe on behalf of the Wellcome Sanger Institute COVID-19 Surveillance Team |
| EPI_ISL_798830 | Lighthouse Lab in Milton Keynes | Wellcome Sanger Institute for the COVID-19 Genomics UK (COG-UK) Consortium | The Lighthouse Lab in Milton Keynes and Alex Alderton, Roberto Amato, Sonia Goncalves, Ewan Harrison, David K. Jackson, Ian Johnston, Dominic Kwiatkowski, Cordelia Langford, John Sillitoe on behalf of the Wellcome Sanger Institute COVID-19 Surveillance Team |
| EPI_ISL_798831 | Lighthouse Lab in Cambridge | Wellcome Sanger Institute for the COVID-19 Genomics UK (COG-UK) Consortium | Rob Howes, The Lighthouse Lab in Cambridge and Alex Alderton, Roberto Amato, Sonia Goncalves, Ewan Harrison, David K. Jackson, Ian Johnston, Dominic Kwiatkowski, Cordelia Langford, John Sillitoe on behalf of the Wellcome Sanger Institute COVID-19 Surveillance Team |
| EPI_ISL_798832 | Lighthouse Lab in Milton Keynes | Wellcome Sanger Institute for the COVID-19 Genomics UK (COG-UK) Consortium | The Lighthouse Lab in Milton Keynes and Alex Alderton, Roberto Amato, Sonia Goncalves, Ewan Harrison, David K. Jackson, Ian Johnston, Dominic Kwiatkowski, Cordelia Langford, John Sillitoe on behalf of the Wellcome Sanger Institute COVID-19 Surveillance Team |
| EPI_ISL_798835, EPI_ISL_798836 | Lighthouse Lab in Cambridge | Wellcome Sanger Institute for the COVID-19 Genomics UK (COG-UK) Consortium | Rob Howes, The Lighthouse Lab in Cambridge and Alex Alderton, Roberto Amato, Sonia Goncalves, Ewan Harrison, David K. Jackson, Ian Johnston, Dominic Kwiatkowski, Cordelia Langford, John Sillitoe on behalf of the Wellcome Sanger Institute COVID-19 Surveillance Team |
| EPI_ISL_798837, EPI_ISL_798838 | Lighthouse Lab in Milton Keynes | Wellcome Sanger Institute for the COVID-19 Genomics UK (COG-UK) Consortium | The Lighthouse Lab in Milton Keynes and Alex Alderton, Roberto Amato, Sonia Goncalves, Ewan Harrison, David K. Jackson, Ian Johnston, Dominic Kwiatkowski, Cordelia Langford, John Sillitoe on behalf of the Wellcome Sanger Institute COVID-19 Surveillance Team |
| EPI_ISL_798839, EPI_ISL_798840, EPI_ISL_798842, EPI_ISL_798843 | Lighthouse Lab in Cambridge | Wellcome Sanger Institute for the COVID-19 Genomics UK (COG-UK) Consortium | Rob Howes, The Lighthouse Lab in Cambridge and Alex Alderton, Roberto Amato, Sonia Goncalves, Ewan Harrison, David K. Jackson, Ian Johnston, Dominic Kwiatkowski, Cordelia Langford, John Sillitoe on behalf of the Wellcome Sanger Institute COVID-19 Surveillance Team |
| EPI_ISL_798844 | Lighthouse Lab in Milton Keynes | Wellcome Sanger Institute for the COVID-19 Genomics UK (COG-UK) Consortium | The Lighthouse Lab in Milton Keynes and Alex Alderton, Roberto Amato, Sonia Goncalves, Ewan Harrison, David K. Jackson, Ian Johnston, Dominic Kwiatkowski, Cordelia Langford, John Sillitoe on behalf of the Wellcome Sanger Institute COVID-19 Surveillance Team |
| EPI_ISL_798846, EPI_ISL_798847, EPI_ISL_798849 | Lighthouse Lab in Cambridge | Wellcome Sanger Institute for the COVID-19 Genomics UK (COG-UK) Consortium | Rob Howes, The Lighthouse Lab in Cambridge and Alex Alderton, Roberto Amato, Sonia Goncalves, Ewan Harrison, David K. Jackson, Ian Johnston, Dominic Kwiatkowski, Cordelia Langford, John Sillitoe on behalf of the Wellcome Sanger Institute COVID-19 Surveillance Team |
| EPI_ISL_798851 | Lighthouse Lab in Milton Keynes | Wellcome Sanger Institute for the COVID-19 Genomics UK (COG-UK) Consortium | The Lighthouse Lab in Milton Keynes and Alex Alderton, Roberto Amato, Sonia Goncalves, Ewan Harrison, David K. Jackson, Ian Johnston, Dominic Kwiatkowski, Cordelia Langford, John Sillitoe on behalf of the Wellcome Sanger Institute COVID-19 Surveillance Team |
| EPI_ISL_798853 | Lighthouse Lab in Alderley Park | Wellcome Sanger Institute for the COVID-19 Genomics UK (COG-UK) Consortium | Jacquelyn Wynn, Mairead Hyland, The Lighthouse Lab in Alderley Park and Alex Alderton, Roberto Amato, Sonia Goncalves, Ewan Harrison, David K. Jackson, Ian Johnston, Dominic Kwiatkowski, Cordelia Langford, John Sillitoe on behalf of the Wellcome Sanger Institute COVID-19 Surveillance Team |
| EPI_ISL_798854 | Lighthouse Lab in Cambridge | Wellcome Sanger Institute for the COVID-19 Genomics UK (COG-UK) Consortium | Rob Howes, The Lighthouse Lab in Cambridge and Alex Alderton, Roberto Amato, Sonia Goncalves, Ewan Harrison, David K. Jackson, Ian Johnston, Dominic Kwiatkowski, Cordelia Langford, John Sillitoe on behalf of the Wellcome Sanger Institute COVID-19 Surveillance Team |

| | | | |
|---|---|---|---|
| EPI_ISL_798855, EPI_ISL_798856 | Lighthouse Lab in Milton Keynes | Wellcome Sanger Institute for the COVID-19 Genomics UK (COG-UK) Consortium | The Lighthouse Lab in Milton Keynes and Alex Alderton, Roberto Amato, Sonia Goncalves, Ewan Harrison, David K. Jackson, Ian Johnston, Dominic Kwiatkowski, Cordelia Langford, John Sillitoe on behalf of the Wellcome Sanger Institute COVID-19 Surveillance Team |
| EPI_ISL_798857, EPI_ISL_798858, EPI_ISL_798859, EPI_ISL_798860, EPI_ISL_798861, EPI_ISL_798862 | Lighthouse Lab in Cambridge | Wellcome Sanger Institute for the COVID-19 Genomics UK (COG-UK) Consortium | Rob Howes, The Lighthouse Lab in Cambridge and Alex Alderton, Roberto Amato, Sonia Goncalves, Ewan Harrison, David K. Jackson, Ian Johnston, Dominic Kwiatkowski, Cordelia Langford, John Sillitoe on behalf of the Wellcome Sanger Institute COVID-19 Surveillance Team |
| EPI_ISL_798863, EPI_ISL_798864 | Lighthouse Lab in Milton Keynes | Wellcome Sanger Institute for the COVID-19 Genomics UK (COG-UK) Consortium | The Lighthouse Lab in Milton Keynes and Alex Alderton, Roberto Amato, Sonia Goncalves, Ewan Harrison, David K. Jackson, Ian Johnston, Dominic Kwiatkowski, Cordelia Langford, John Sillitoe on behalf of the Wellcome Sanger Institute COVID-19 Surveillance Team |
| EPI_ISL_798865, EPI_ISL_798866, EPI_ISL_798867, EPI_ISL_798868 | Lighthouse Lab in Cambridge | Wellcome Sanger Institute for the COVID-19 Genomics UK (COG-UK) Consortium | Rob Howes, The Lighthouse Lab in Cambridge and Alex Alderton, Roberto Amato, Sonia Goncalves, Ewan Harrison, David K. Jackson, Ian Johnston, Dominic Kwiatkowski, Cordelia Langford, John Sillitoe on behalf of the Wellcome Sanger Institute COVID-19 Surveillance Team |
| EPI_ISL_798870 | Lighthouse Lab in Milton Keynes | Wellcome Sanger Institute for the COVID-19 Genomics UK (COG-UK) Consortium | The Lighthouse Lab in Milton Keynes and Alex Alderton, Roberto Amato, Sonia Goncalves, Ewan Harrison, David K. Jackson, Ian Johnston, Dominic Kwiatkowski, Cordelia Langford, John Sillitoe on behalf of the Wellcome Sanger Institute COVID-19 Surveillance Team |
| EPI_ISL_798871, EPI_ISL_798873 | Lighthouse Lab in Cambridge | Wellcome Sanger Institute for the COVID-19 Genomics UK (COG-UK) Consortium | Rob Howes, The Lighthouse Lab in Cambridge and Alex Alderton, Roberto Amato, Sonia Goncalves, Ewan Harrison, David K. Jackson, Ian Johnston, Dominic Kwiatkowski, Cordelia Langford, John Sillitoe on behalf of the Wellcome Sanger Institute COVID-19 Surveillance Team |
| EPI_ISL_798874, EPI_ISL_798875, EPI_ISL_798876 | Lighthouse Lab in Milton Keynes | Wellcome Sanger Institute for the COVID-19 Genomics UK (COG-UK) Consortium | The Lighthouse Lab in Milton Keynes and Alex Alderton, Roberto Amato, Sonia Goncalves, Ewan Harrison, David K. Jackson, Ian Johnston, Dominic Kwiatkowski, Cordelia Langford, John Sillitoe on behalf of the Wellcome Sanger Institute COVID-19 Surveillance Team |
| EPI_ISL_798877 | Lighthouse Lab in Cambridge | Wellcome Sanger Institute for the COVID-19 Genomics UK (COG-UK) Consortium | Rob Howes, The Lighthouse Lab in Cambridge and Alex Alderton, Roberto Amato, Sonia Goncalves, Ewan Harrison, David K. Jackson, Ian Johnston, Dominic Kwiatkowski, Cordelia Langford, John Sillitoe on behalf of the Wellcome Sanger Institute COVID-19 Surveillance Team |
| EPI_ISL_798879 | Lighthouse Lab in Milton Keynes | Wellcome Sanger Institute for the COVID-19 Genomics UK (COG-UK) Consortium | The Lighthouse Lab in Milton Keynes and Alex Alderton, Roberto Amato, Sonia Goncalves, Ewan Harrison, David K. Jackson, Ian Johnston, Dominic Kwiatkowski, Cordelia Langford, John Sillitoe on behalf of the Wellcome Sanger Institute COVID-19 Surveillance Team |
| EPI_ISL_798880 | Lighthouse Lab in Alderley Park | Wellcome Sanger Institute for the COVID-19 Genomics UK (COG-UK) Consortium | Jacquelyn Wynn, Mairead Hyland, The Lighthouse Lab in Alderley Park and Alex Alderton, Roberto Amato, Sonia Goncalves, Ewan Harrison, David K. Jackson, Ian Johnston, Dominic Kwiatkowski, Cordelia Langford, John Sillitoe on behalf of the Wellcome Sanger Institute COVID-19 Surveillance Team |
| EPI_ISL_798881 | Lighthouse Lab in Cambridge | Wellcome Sanger Institute for the COVID-19 Genomics UK (COG-UK) Consortium | Rob Howes, The Lighthouse Lab in Cambridge and Alex Alderton, Roberto Amato, Sonia Goncalves, Ewan Harrison, David K. Jackson, Ian Johnston, Dominic Kwiatkowski, Cordelia Langford, John Sillitoe on behalf of the Wellcome Sanger Institute COVID-19 Surveillance Team |
| EPI_ISL_798882 | Lighthouse Lab in Milton Keynes | Wellcome Sanger Institute for the COVID-19 Genomics UK (COG-UK) Consortium | The Lighthouse Lab in Milton Keynes and Alex Alderton, Roberto Amato, Sonia Goncalves, Ewan Harrison, David K. Jackson, Ian Johnston, Dominic Kwiatkowski, Cordelia Langford, John Sillitoe on behalf of the Wellcome Sanger Institute COVID-19 Surveillance Team |
| EPI_ISL_798883 | Lighthouse Lab in Cambridge | Wellcome Sanger Institute for the COVID-19 Genomics UK (COG-UK) Consortium | Rob Howes, The Lighthouse Lab in Cambridge and Alex Alderton, Roberto Amato, Sonia Goncalves, Ewan Harrison, David K. Jackson, Ian Johnston, Dominic Kwiatkowski, Cordelia Langford, John Sillitoe on behalf of the Wellcome Sanger Institute COVID-19 Surveillance Team |
| EPI_ISL_798885 | Lighthouse Lab in Alderley Park | Wellcome Sanger Institute for the COVID-19 Genomics UK (COG-UK) Consortium | Jacquelyn Wynn, Mairead Hyland, The Lighthouse Lab in Alderley Park and Alex Alderton, Roberto Amato, Sonia Goncalves, Ewan Harrison, David K. Jackson, Ian Johnston, Dominic Kwiatkowski, Cordelia Langford, John Sillitoe on behalf of the Wellcome Sanger Institute COVID-19 Surveillance Team |
| EPI_ISL_798886, EPI_ISL_798887 | Lighthouse Lab in Cambridge | Wellcome Sanger Institute for the COVID-19 Genomics UK (COG-UK) Consortium | Rob Howes, The Lighthouse Lab in Cambridge and Alex Alderton, Roberto Amato, Sonia Goncalves, Ewan Harrison, David K. Jackson, Ian Johnston, Dominic Kwiatkowski, Cordelia Langford, John Sillitoe on behalf of the Wellcome Sanger Institute COVID-19 Surveillance Team |
| EPI_ISL_798889 | Lighthouse Lab in Milton Keynes | Wellcome Sanger Institute for the COVID-19 Genomics UK (COG-UK) Consortium | The Lighthouse Lab in Milton Keynes and Alex Alderton, Roberto Amato, Sonia Goncalves, Ewan Harrison, David K. Jackson, Ian Johnston, Dominic Kwiatkowski, Cordelia Langford, John Sillitoe on behalf of the Wellcome Sanger Institute COVID-19 Surveillance Team |
| EPI_ISL_798890 | Lighthouse Lab in Cambridge | Wellcome Sanger Institute for the COVID-19 Genomics UK (COG-UK) Consortium | Rob Howes, The Lighthouse Lab in Cambridge and Alex Alderton, Roberto Amato, Sonia Goncalves, Ewan Harrison, David K. Jackson, Ian Johnston, Dominic Kwiatkowski, Cordelia Langford, John Sillitoe on behalf of the Wellcome Sanger Institute COVID-19 Surveillance Team |
| EPI_ISL_798892 | Lighthouse Lab in Milton Keynes | Wellcome Sanger Institute for the COVID-19 Genomics UK (COG-UK) Consortium | The Lighthouse Lab in Milton Keynes and Alex Alderton, Roberto Amato, Sonia Goncalves, Ewan Harrison, David K. Jackson, Ian Johnston, Dominic Kwiatkowski, Cordelia Langford, John Sillitoe on behalf of the Wellcome Sanger Institute COVID-19 Surveillance Team |
| EPI_ISL_798893, EPI_ISL_798894, EPI_ISL_798895, EPI_ISL_798896, EPI_ISL_798897, EPI_ISL_798899, EPI_ISL_798900, EPI_ISL_798901, EPI_ISL_798903, EPI_ISL_798904 | Lighthouse Lab in Cambridge | Wellcome Sanger Institute for the COVID-19 Genomics UK (COG-UK) Consortium | Rob Howes, The Lighthouse Lab in Cambridge and Alex Alderton, Roberto Amato, Sonia Goncalves, Ewan Harrison, David K. Jackson, Ian Johnston, Dominic Kwiatkowski, Cordelia Langford, John Sillitoe on behalf of the Wellcome Sanger Institute COVID-19 Surveillance Team |
| EPI_ISL_798905 | Lighthouse Lab in Alderley Park | Wellcome Sanger Institute for the COVID-19 Genomics UK (COG-UK) Consortium | Jacquelyn Wynn, Mairead Hyland, The Lighthouse Lab in Alderley Park and Alex Alderton, Roberto Amato, Sonia Goncalves, Ewan Harrison, David K. Jackson, Ian Johnston, Dominic Kwiatkowski, Cordelia Langford, John Sillitoe on behalf of the Wellcome Sanger Institute COVID-19 Surveillance Team |
| EPI_ISL_798908, EPI_ISL_798909 | Lighthouse Lab in Cambridge | Wellcome Sanger Institute for the COVID-19 Genomics UK (COG-UK) Consortium | Rob Howes, The Lighthouse Lab in Cambridge and Alex Alderton, Roberto Amato, Sonia Goncalves, Ewan Harrison, David K. Jackson, Ian Johnston, Dominic Kwiatkowski, Cordelia Langford, John Sillitoe on behalf of the Wellcome Sanger Institute COVID-19 Surveillance Team |
| EPI_ISL_798910, EPI_ISL_798911 | Lighthouse Lab in Milton Keynes | Wellcome Sanger Institute for the COVID-19 Genomics UK (COG-UK) Consortium | The Lighthouse Lab in Milton Keynes and Alex Alderton, Roberto Amato, Sonia Goncalves, Ewan Harrison, David K. Jackson, Ian Johnston, Dominic Kwiatkowski, Cordelia Langford, John Sillitoe on behalf of the Wellcome Sanger Institute COVID-19 Surveillance Team |
| EPI_ISL_798912, EPI_ISL_798913 | Lighthouse Lab in Cambridge | Wellcome Sanger Institute for the COVID-19 Genomics UK (COG-UK) Consortium | Rob Howes, The Lighthouse Lab in Cambridge and Alex Alderton, Roberto Amato, Sonia Goncalves, Ewan Harrison, David K. Jackson, Ian Johnston, Dominic Kwiatkowski, Cordelia Langford, John Sillitoe on behalf of the Wellcome Sanger Institute COVID-19 Surveillance Team |
| EPI_ISL_798915 | Lighthouse Lab in Milton Keynes | Wellcome Sanger Institute for the COVID-19 Genomics UK (COG-UK) Consortium | The Lighthouse Lab in Milton Keynes and Alex Alderton, Roberto Amato, Sonia Goncalves, Ewan Harrison, David K. Jackson, Ian Johnston, Dominic Kwiatkowski, Cordelia Langford, John Sillitoe on behalf of the Wellcome Sanger Institute COVID-19 Surveillance Team |
| EPI_ISL_798916, EPI_ISL_798917, EPI_ISL_798918 | Lighthouse Lab in Cambridge | Wellcome Sanger Institute for the COVID-19 Genomics UK (COG-UK) Consortium | Rob Howes, The Lighthouse Lab in Cambridge and Alex Alderton, Roberto Amato, Sonia Goncalves, Ewan Harrison, David K. Jackson, Ian Johnston, Dominic Kwiatkowski, Cordelia Langford, John Sillitoe on behalf of the Wellcome Sanger Institute COVID-19 Surveillance Team |
| EPI_ISL_798919 | Lighthouse Lab in Milton Keynes | Wellcome Sanger Institute for the COVID-19 Genomics UK (COG-UK) Consortium | The Lighthouse Lab in Milton Keynes and Alex Alderton, Roberto Amato, Sonia Goncalves, Ewan Harrison, David K. Jackson, Ian Johnston, Dominic Kwiatkowski, Cordelia Langford, John Sillitoe on behalf of the Wellcome Sanger Institute COVID-19 Surveillance Team |
| EPI_ISL_798920 | Lighthouse Lab in Cambridge | Wellcome Sanger Institute for the COVID-19 Genomics UK (COG-UK) Consortium | Rob Howes, The Lighthouse Lab in Cambridge and Alex Alderton, Roberto Amato, Sonia Goncalves, Ewan Harrison, David K. Jackson, Ian Johnston, Dominic Kwiatkowski, Cordelia Langford, John Sillitoe on behalf of the Wellcome Sanger Institute COVID-19 Surveillance Team |
| EPI_ISL_798922 | Lighthouse Lab in Milton Keynes | Wellcome Sanger Institute for the COVID-19 Genomics UK (COG-UK) Consortium | The Lighthouse Lab in Milton Keynes and Alex Alderton, Roberto Amato, Sonia Goncalves, Ewan Harrison, David K. Jackson, Ian Johnston, Dominic Kwiatkowski, Cordelia Langford, John Sillitoe on behalf of the Wellcome Sanger Institute COVID-19 Surveillance Team |
| EPI_ISL_798923 | Lighthouse Lab in Cambridge | Wellcome Sanger Institute for the COVID-19 Genomics UK (COG-UK) Consortium | Rob Howes, The Lighthouse Lab in Cambridge and Alex Alderton, Roberto Amato, Sonia Goncalves, Ewan Harrison, David K. Jackson, Ian Johnston, Dominic Kwiatkowski, Cordelia Langford, John Sillitoe on behalf of the Wellcome Sanger Institute COVID-19 Surveillance Team |
| EPI_ISL_798925 | Lighthouse Lab in Milton Keynes | Wellcome Sanger Institute for the COVID-19 Genomics UK (COG-UK) Consortium | The Lighthouse Lab in Milton Keynes and Alex Alderton, Roberto Amato, Sonia Goncalves, Ewan Harrison, David K. Jackson, Ian Johnston, Dominic Kwiatkowski, Cordelia Langford, John Sillitoe on behalf of the Wellcome Sanger Institute COVID-19 Surveillance Team |
| EPI_ISL_798927, EPI_ISL_798929 | Lighthouse Lab in Cambridge | Wellcome Sanger Institute for the COVID-19 Genomics UK (COG-UK) Consortium | Rob Howes, The Lighthouse Lab in Cambridge and Alex Alderton, Roberto Amato, Sonia Goncalves, Ewan Harrison, David K. Jackson, Ian Johnston, Dominic Kwiatkowski, Cordelia Langford, John Sillitoe on behalf of the Wellcome Sanger Institute COVID-19 Surveillance Team |
| EPI_ISL_798930 | Lighthouse Lab in Milton Keynes | Wellcome Sanger Institute for the COVID-19 Genomics UK (COG-UK) Consortium | The Lighthouse Lab in Milton Keynes and Alex Alderton, Roberto Amato, Sonia Goncalves, Ewan Harrison, David K. Jackson, Ian Johnston, Dominic Kwiatkowski, Cordelia Langford, John Sillitoe on behalf of the Wellcome Sanger Institute COVID-19 Surveillance Team |
| EPI_ISL_798932 | Lighthouse Lab in Cambridge | Wellcome Sanger Institute for the COVID-19 Genomics UK (COG-UK) Consortium | Rob Howes, The Lighthouse Lab in Cambridge and Alex Alderton, Roberto Amato, Sonia Goncalves, Ewan Harrison, David K. Jackson, Ian Johnston, Dominic Kwiatkowski, Cordelia Langford, John Sillitoe on behalf of the Wellcome Sanger Institute COVID-19 Surveillance Team |
| EPI_ISL_798933 | Lighthouse Lab in Milton Keynes | Wellcome Sanger Institute for the COVID-19 Genomics UK | The Lighthouse Lab in Milton Keynes and Alex Alderton, Roberto Amato, Sonia Goncalves, Ewan Harrison, David K. Jackson, Ian Johnston, Dominic |

| | | | |
|---|---|---|---|
| | | (COG-UK) Consortium | Kwiatkowski, Cordelia Langford, John Sillitoe on behalf of the Wellcome Sanger Institute COVID-19 Surveillance Team |
| EPI_ISL_798935 | Lighthouse Lab in Alderley Park | Wellcome Sanger Institute for the COVID-19 Genomics UK (COG-UK) Consortium | Jacquelyn Wynn, Mairead Hyland, The Lighthouse Lab in Alderley Park and Alex Alderton, Roberto Amato, Sonia Goncalves, Ewan Harrison, David K. Jackson, Ian Johnston, Dominic Kwiatkowski, Cordelia Langford, John Sillitoe on behalf of the Wellcome Sanger Institute COVID-19 Surveillance Team |
| EPI_ISL_798936 | Lighthouse Lab in Milton Keynes | Wellcome Sanger Institute for the COVID-19 Genomics UK (COG-UK) Consortium | The Lighthouse Lab in Milton Keynes and Alex Alderton, Roberto Amato, Sonia Goncalves, Ewan Harrison, David K. Jackson, Ian Johnston, Dominic Kwiatkowski, Cordelia Langford, John Sillitoe on behalf of the Wellcome Sanger Institute COVID-19 Surveillance Team |
| EPI_ISL_798937, EPI_ISL_798938, EPI_ISL_798940 | Lighthouse Lab in Cambridge | Wellcome Sanger Institute for the COVID-19 Genomics UK (COG-UK) Consortium | Rob Howes, The Lighthouse Lab in Cambridge and Alex Alderton, Roberto Amato, Sonia Goncalves, Ewan Harrison, David K. Jackson, Ian Johnston, Dominic Kwiatkowski, Cordelia Langford, John Sillitoe on behalf of the Wellcome Sanger Institute COVID-19 Surveillance Team |
| EPI_ISL_798941 | Lighthouse Lab in Alderley Park | Wellcome Sanger Institute for the COVID-19 Genomics UK (COG-UK) Consortium | Jacquelyn Wynn, Mairead Hyland, The Lighthouse Lab in Alderley Park and Alex Alderton, Roberto Amato, Sonia Goncalves, Ewan Harrison, David K. Jackson, Ian Johnston, Dominic Kwiatkowski, Cordelia Langford, John Sillitoe on behalf of the Wellcome Sanger Institute COVID-19 Surveillance Team |
| EPI_ISL_798942 | Lighthouse Lab in Milton Keynes | Wellcome Sanger Institute for the COVID-19 Genomics UK (COG-UK) Consortium | The Lighthouse Lab in Milton Keynes and Alex Alderton, Roberto Amato, Sonia Goncalves, Ewan Harrison, David K. Jackson, Ian Johnston, Dominic Kwiatkowski, Cordelia Langford, John Sillitoe on behalf of the Wellcome Sanger Institute COVID-19 Surveillance Team |
| EPI_ISL_798943, EPI_ISL_798945, EPI_ISL_798946, EPI_ISL_798947 | Lighthouse Lab in Cambridge | Wellcome Sanger Institute for the COVID-19 Genomics UK (COG-UK) Consortium | Rob Howes, The Lighthouse Lab in Cambridge and Alex Alderton, Roberto Amato, Sonia Goncalves, Ewan Harrison, David K. Jackson, Ian Johnston, Dominic Kwiatkowski, Cordelia Langford, John Sillitoe on behalf of the Wellcome Sanger Institute COVID-19 Surveillance Team |
| EPI_ISL_798948 | Lighthouse Lab in Milton Keynes | Wellcome Sanger Institute for the COVID-19 Genomics UK (COG-UK) Consortium | The Lighthouse Lab in Milton Keynes and Alex Alderton, Roberto Amato, Sonia Goncalves, Ewan Harrison, David K. Jackson, Ian Johnston, Dominic Kwiatkowski, Cordelia Langford, John Sillitoe on behalf of the Wellcome Sanger Institute COVID-19 Surveillance Team |
| EPI_ISL_798949 | Lighthouse Lab in Cambridge | Wellcome Sanger Institute for the COVID-19 Genomics UK (COG-UK) Consortium | Rob Howes, The Lighthouse Lab in Cambridge and Alex Alderton, Roberto Amato, Sonia Goncalves, Ewan Harrison, David K. Jackson, Ian Johnston, Dominic Kwiatkowski, Cordelia Langford, John Sillitoe on behalf of the Wellcome Sanger Institute COVID-19 Surveillance Team |
| EPI_ISL_798950 | Lighthouse Lab in Milton Keynes | Wellcome Sanger Institute for the COVID-19 Genomics UK (COG-UK) Consortium | The Lighthouse Lab in Milton Keynes and Alex Alderton, Roberto Amato, Sonia Goncalves, Ewan Harrison, David K. Jackson, Ian Johnston, Dominic Kwiatkowski, Cordelia Langford, John Sillitoe on behalf of the Wellcome Sanger Institute COVID-19 Surveillance Team |
| EPI_ISL_798953, EPI_ISL_798954 | Lighthouse Lab in Cambridge | Wellcome Sanger Institute for the COVID-19 Genomics UK (COG-UK) Consortium | Rob Howes, The Lighthouse Lab in Cambridge and Alex Alderton, Roberto Amato, Sonia Goncalves, Ewan Harrison, David K. Jackson, Ian Johnston, Dominic Kwiatkowski, Cordelia Langford, John Sillitoe on behalf of the Wellcome Sanger Institute COVID-19 Surveillance Team |
| EPI_ISL_798955 | Lighthouse Lab in Milton Keynes | Wellcome Sanger Institute for the COVID-19 Genomics UK (COG-UK) Consortium | The Lighthouse Lab in Milton Keynes and Alex Alderton, Roberto Amato, Sonia Goncalves, Ewan Harrison, David K. Jackson, Ian Johnston, Dominic Kwiatkowski, Cordelia Langford, John Sillitoe on behalf of the Wellcome Sanger Institute COVID-19 Surveillance Team |
| EPI_ISL_798956 | Lighthouse Lab in Cambridge | Wellcome Sanger Institute for the COVID-19 Genomics UK (COG-UK) Consortium | Rob Howes, The Lighthouse Lab in Cambridge and Alex Alderton, Roberto Amato, Sonia Goncalves, Ewan Harrison, David K. Jackson, Ian Johnston, Dominic Kwiatkowski, Cordelia Langford, John Sillitoe on behalf of the Wellcome Sanger Institute COVID-19 Surveillance Team |
| EPI_ISL_798957 | Lighthouse Lab in Alderley Park | Wellcome Sanger Institute for the COVID-19 Genomics UK (COG-UK) Consortium | Jacquelyn Wynn, Mairead Hyland, The Lighthouse Lab in Alderley Park and Alex Alderton, Roberto Amato, Sonia Goncalves, Ewan Harrison, David K. Jackson, Ian Johnston, Dominic Kwiatkowski, Cordelia Langford, John Sillitoe on behalf of the Wellcome Sanger Institute COVID-19 Surveillance Team |
| EPI_ISL_798958, EPI_ISL_798959, EPI_ISL_798960 | Lighthouse Lab in Cambridge | Wellcome Sanger Institute for the COVID-19 Genomics UK (COG-UK) Consortium | Rob Howes, The Lighthouse Lab in Cambridge and Alex Alderton, Roberto Amato, Sonia Goncalves, Ewan Harrison, David K. Jackson, Ian Johnston, Dominic Kwiatkowski, Cordelia Langford, John Sillitoe on behalf of the Wellcome Sanger Institute COVID-19 Surveillance Team |
| EPI_ISL_798961 | Lighthouse Lab in Milton Keynes | Wellcome Sanger Institute for the COVID-19 Genomics UK (COG-UK) Consortium | The Lighthouse Lab in Milton Keynes and Alex Alderton, Roberto Amato, Sonia Goncalves, Ewan Harrison, David K. Jackson, Ian Johnston, Dominic Kwiatkowski, Cordelia Langford, John Sillitoe on behalf of the Wellcome Sanger Institute COVID-19 Surveillance Team |
| EPI_ISL_798965 | Lighthouse Lab in Cambridge | Wellcome Sanger Institute for the COVID-19 Genomics UK (COG-UK) Consortium | Rob Howes, The Lighthouse Lab in Cambridge and Alex Alderton, Roberto Amato, Sonia Goncalves, Ewan Harrison, David K. Jackson, Ian Johnston, Dominic Kwiatkowski, Cordelia Langford, John Sillitoe on behalf of the Wellcome Sanger Institute COVID-19 Surveillance Team |
| EPI_ISL_798966, EPI_ISL_798967, EPI_ISL_798970 | Lighthouse Lab in Milton Keynes | Wellcome Sanger Institute for the COVID-19 Genomics UK (COG-UK) Consortium | The Lighthouse Lab in Milton Keynes and Alex Alderton, Roberto Amato, Sonia Goncalves, Ewan Harrison, David K. Jackson, Ian Johnston, Dominic Kwiatkowski, Cordelia Langford, John Sillitoe on behalf of the Wellcome Sanger Institute COVID-19 Surveillance Team |
| EPI_ISL_798971 | Lighthouse Lab in Cambridge | Wellcome Sanger Institute for the COVID-19 Genomics UK (COG-UK) Consortium | Rob Howes, The Lighthouse Lab in Cambridge and Alex Alderton, Roberto Amato, Sonia Goncalves, Ewan Harrison, David K. Jackson, Ian Johnston, Dominic Kwiatkowski, Cordelia Langford, John Sillitoe on behalf of the Wellcome Sanger Institute COVID-19 Surveillance Team |
| EPI_ISL_798972, EPI_ISL_798973 | Lighthouse Lab in Milton Keynes | Wellcome Sanger Institute for the COVID-19 Genomics UK (COG-UK) Consortium | The Lighthouse Lab in Milton Keynes and Alex Alderton, Roberto Amato, Sonia Goncalves, Ewan Harrison, David K. Jackson, Ian Johnston, Dominic Kwiatkowski, Cordelia Langford, John Sillitoe on behalf of the Wellcome Sanger Institute COVID-19 Surveillance Team |
| EPI_ISL_798974 | Lighthouse Lab in Cambridge | Wellcome Sanger Institute for the COVID-19 Genomics UK (COG-UK) Consortium | Rob Howes, The Lighthouse Lab in Cambridge and Alex Alderton, Roberto Amato, Sonia Goncalves, Ewan Harrison, David K. Jackson, Ian Johnston, Dominic Kwiatkowski, Cordelia Langford, John Sillitoe on behalf of the Wellcome Sanger Institute COVID-19 Surveillance Team |
| EPI_ISL_798975 | Lighthouse Lab in Milton Keynes | Wellcome Sanger Institute for the COVID-19 Genomics UK (COG-UK) Consortium | The Lighthouse Lab in Milton Keynes and Alex Alderton, Roberto Amato, Sonia Goncalves, Ewan Harrison, David K. Jackson, Ian Johnston, Dominic Kwiatkowski, Cordelia Langford, John Sillitoe on behalf of the Wellcome Sanger Institute COVID-19 Surveillance Team |
| EPI_ISL_798976 | Lighthouse Lab in Cambridge | Wellcome Sanger Institute for the COVID-19 Genomics UK (COG-UK) Consortium | Rob Howes, The Lighthouse Lab in Cambridge and Alex Alderton, Roberto Amato, Sonia Goncalves, Ewan Harrison, David K. Jackson, Ian Johnston, Dominic Kwiatkowski, Cordelia Langford, John Sillitoe on behalf of the Wellcome Sanger Institute COVID-19 Surveillance Team |
| EPI_ISL_798977 | Lighthouse Lab in Milton Keynes | Wellcome Sanger Institute for the COVID-19 Genomics UK (COG-UK) Consortium | The Lighthouse Lab in Milton Keynes and Alex Alderton, Roberto Amato, Sonia Goncalves, Ewan Harrison, David K. Jackson, Ian Johnston, Dominic Kwiatkowski, Cordelia Langford, John Sillitoe on behalf of the Wellcome Sanger Institute COVID-19 Surveillance Team |
| EPI_ISL_798978 | Lighthouse Lab in Cambridge | Wellcome Sanger Institute for the COVID-19 Genomics UK (COG-UK) Consortium | Rob Howes, The Lighthouse Lab in Cambridge and Alex Alderton, Roberto Amato, Sonia Goncalves, Ewan Harrison, David K. Jackson, Ian Johnston, Dominic Kwiatkowski, Cordelia Langford, John Sillitoe on behalf of the Wellcome Sanger Institute COVID-19 Surveillance Team |
| EPI_ISL_798979 | Lighthouse Lab in Milton Keynes | Wellcome Sanger Institute for the COVID-19 Genomics UK (COG-UK) Consortium | The Lighthouse Lab in Milton Keynes and Alex Alderton, Roberto Amato, Sonia Goncalves, Ewan Harrison, David K. Jackson, Ian Johnston, Dominic Kwiatkowski, Cordelia Langford, John Sillitoe on behalf of the Wellcome Sanger Institute COVID-19 Surveillance Team |
| EPI_ISL_798980 | Lighthouse Lab in Cambridge | Wellcome Sanger Institute for the COVID-19 Genomics UK (COG-UK) Consortium | Rob Howes, The Lighthouse Lab in Cambridge and Alex Alderton, Roberto Amato, Sonia Goncalves, Ewan Harrison, David K. Jackson, Ian Johnston, Dominic Kwiatkowski, Cordelia Langford, John Sillitoe on behalf of the Wellcome Sanger Institute COVID-19 Surveillance Team |
| EPI_ISL_798981 | Lighthouse Lab in Milton Keynes | Wellcome Sanger Institute for the COVID-19 Genomics UK (COG-UK) Consortium | The Lighthouse Lab in Milton Keynes and Alex Alderton, Roberto Amato, Sonia Goncalves, Ewan Harrison, David K. Jackson, Ian Johnston, Dominic Kwiatkowski, Cordelia Langford, John Sillitoe on behalf of the Wellcome Sanger Institute COVID-19 Surveillance Team |
| EPI_ISL_798982 | Lighthouse Lab in Cambridge | Wellcome Sanger Institute for the COVID-19 Genomics UK (COG-UK) Consortium | Rob Howes, The Lighthouse Lab in Cambridge and Alex Alderton, Roberto Amato, Sonia Goncalves, Ewan Harrison, David K. Jackson, Ian Johnston, Dominic Kwiatkowski, Cordelia Langford, John Sillitoe on behalf of the Wellcome Sanger Institute COVID-19 Surveillance Team |
| EPI_ISL_798983, EPI_ISL_798987 | Lighthouse Lab in Milton Keynes | Wellcome Sanger Institute for the COVID-19 Genomics UK (COG-UK) Consortium | The Lighthouse Lab in Milton Keynes and Alex Alderton, Roberto Amato, Sonia Goncalves, Ewan Harrison, David K. Jackson, Ian Johnston, Dominic Kwiatkowski, Cordelia Langford, John Sillitoe on behalf of the Wellcome Sanger Institute COVID-19 Surveillance Team |
| EPI_ISL_798989, EPI_ISL_798990, EPI_ISL_798992, EPI_ISL_798993 | Lighthouse Lab in Cambridge | Wellcome Sanger Institute for the COVID-19 Genomics UK (COG-UK) Consortium | Rob Howes, The Lighthouse Lab in Cambridge and Alex Alderton, Roberto Amato, Sonia Goncalves, Ewan Harrison, David K. Jackson, Ian Johnston, Dominic Kwiatkowski, Cordelia Langford, John Sillitoe on behalf of the Wellcome Sanger Institute COVID-19 Surveillance Team |
| EPI_ISL_798994, EPI_ISL_798995, EPI_ISL_798996 | Lighthouse Lab in Milton Keynes | Wellcome Sanger Institute for the COVID-19 Genomics UK (COG-UK) Consortium | The Lighthouse Lab in Milton Keynes and Alex Alderton, Roberto Amato, Sonia Goncalves, Ewan Harrison, David K. Jackson, Ian Johnston, Dominic Kwiatkowski, Cordelia Langford, John Sillitoe on behalf of the Wellcome Sanger Institute COVID-19 Surveillance Team |
| EPI_ISL_798997, EPI_ISL_798998, EPI_ISL_798999, EPI_ISL_799001, EPI_ISL_799002 | Lighthouse Lab in Cambridge | Wellcome Sanger Institute for the COVID-19 Genomics UK (COG-UK) Consortium | Rob Howes, The Lighthouse Lab in Cambridge and Alex Alderton, Roberto Amato, Sonia Goncalves, Ewan Harrison, David K. Jackson, Ian Johnston, Dominic Kwiatkowski, Cordelia Langford, John Sillitoe on behalf of the Wellcome Sanger Institute COVID-19 Surveillance Team |
| EPI_ISL_799004 | Lighthouse Lab in Alderley Park | Wellcome Sanger Institute for the COVID-19 Genomics UK (COG-UK) Consortium | Jacquelyn Wynn, Mairead Hyland, The Lighthouse Lab in Alderley Park and Alex Alderton, Roberto Amato, Sonia Goncalves, Ewan Harrison, David K. Jackson, Ian Johnston, Dominic Kwiatkowski, Cordelia Langford, John Sillitoe on behalf of the Wellcome Sanger Institute COVID-19 Surveillance Team |
| EPI_ISL_799007, EPI_ISL_799008, EPI_ISL_799013, EPI_ISL_799017, EPI_ISL_799018, EPI_ISL_799032, EPI_ISL_799035, EPI_ISL_799036, EPI_ISL_799037, EPI_ISL_799039, EPI_ISL_799048, EPI_ISL_799052, EPI_ISL_799058, EPI_ISL_799061, EPI_ISL_799076, EPI_ISL_799078, EPI_ISL_799088, EPI_ISL_799091, EPI_ISL_799092, EPI_ISL_799093, EPI_ISL_799094, EPI_ISL_799095, EPI_ISL_799097, EPI_ISL_799109, EPI_ISL_799110, EPI_ISL_799116, EPI_ISL_799117, EPI_ISL_799119, EPI_ISL_799121, EPI_ISL_799128, EPI_ISL_799133, EPI_ISL_799135, EPI_ISL_799136, EPI_ISL_799141, EPI_ISL_799142, EPI_ISL_799146, EPI_ISL_799148, EPI_ISL_799159, EPI_ISL_799168, EPI_ISL_799173, EPI_ISL_799174, EPI_ISL_799181, EPI_ISL_799184, EPI_ISL_799185 | | | |

| | | | |
|---|---|---|---|
| see above | Lighthouse Lab in Cambridge | Wellcome Sanger Institute for the COVID-19 Genomics UK (COG-UK) Consortium | Rob Howes, The Lighthouse Lab in Cambridge and Alex Alderton, Roberto Amato, Sonia Goncalves, Ewan Harrison, David K. Jackson, Ian Johnston, Dominic Kwiatkowski, Cordelia Langford, John Sillitoe on behalf of the Wellcome Sanger Institute COVID-19 Surveillance Team |
| EPI_ISL_799187 | Lighthouse Lab in Alderley Park | Wellcome Sanger Institute for the COVID-19 Genomics UK (COG-UK) Consortium | Jacquelyn Wynn, Mairead Hyland, The Lighthouse Lab in Alderley Park and Alex Alderton, Roberto Amato, Sonia Goncalves, Ewan Harrison, David K. Jackson, Ian Johnston, Dominic Kwiatkowski, Cordelia Langford, John Sillitoe on behalf of the Wellcome Sanger Institute COVID-19 Surveillance Team |
| EPI_ISL_799195, EPI_ISL_799199, EPI_ISL_799206, EPI_ISL_799211, EPI_ISL_799217, EPI_ISL_799220, EPI_ISL_799223, EPI_ISL_799225, EPI_ISL_799238, EPI_ISL_799239, EPI_ISL_799241, EPI_ISL_799242, EPI_ISL_799247, EPI_ISL_799251, EPI_ISL_799253, EPI_ISL_799254, EPI_ISL_799259, EPI_ISL_799266, EPI_ISL_799284, EPI_ISL_799293, EPI_ISL_799295 | | | |
| see above | Lighthouse Lab in Cambridge | Wellcome Sanger Institute for the COVID-19 Genomics UK (COG-UK) Consortium | Rob Howes, The Lighthouse Lab in Cambridge and Alex Alderton, Roberto Amato, Sonia Goncalves, Ewan Harrison, David K. Jackson, Ian Johnston, Dominic Kwiatkowski, Cordelia Langford, John Sillitoe on behalf of the Wellcome Sanger Institute COVID-19 Surveillance Team |
| EPI_ISL_799296 | Lighthouse Lab in Milton Keynes | Wellcome Sanger Institute for the COVID-19 Genomics UK (COG-UK) Consortium | The Lighthouse Lab in Milton Keynes and Alex Alderton, Roberto Amato, Sonia Goncalves, Ewan Harrison, David K. Jackson, Ian Johnston, Dominic Kwiatkowski, Cordelia Langford, John Sillitoe on behalf of the Wellcome Sanger Institute COVID-19 Surveillance Team |
| EPI_ISL_799298 | Lighthouse Lab in Glasgow | Wellcome Sanger Institute for the COVID-19 Genomics UK (COG-UK) Consortium | Harper VanSteenhouse, Yumi Kasai, David Gray, Carol Clugston, Anna Dominiczak and Alex Alderton, Roberto Amato, Sonia Goncalves, Ewan Harrison, David K. Jackson, Ian Johnston, Dominic Kwiatkowski, Cordelia Langford, John Sillitoe on behalf of the Wellcome Sanger Institute COVID-19 Surveillance Team |
| EPI_ISL_799299 | Lighthouse Lab in Milton Keynes | Wellcome Sanger Institute for the COVID-19 Genomics UK (COG-UK) Consortium | The Lighthouse Lab in Milton Keynes and Alex Alderton, Roberto Amato, Sonia Goncalves, Ewan Harrison, David K. Jackson, Ian Johnston, Dominic Kwiatkowski, Cordelia Langford, John Sillitoe on behalf of the Wellcome Sanger Institute COVID-19 Surveillance Team |
| EPI_ISL_799302 | Lighthouse Lab in Glasgow | Wellcome Sanger Institute for the COVID-19 Genomics UK (COG-UK) Consortium | Harper VanSteenhouse, Yumi Kasai, David Gray, Carol Clugston, Anna Dominiczak and Alex Alderton, Roberto Amato, Sonia Goncalves, Ewan Harrison, David K. Jackson, Ian Johnston, Dominic Kwiatkowski, Cordelia Langford, John Sillitoe on behalf of the Wellcome Sanger Institute COVID-19 Surveillance Team |
| EPI_ISL_799303, EPI_ISL_799305, EPI_ISL_799306, EPI_ISL_799308 | Lighthouse Lab in Milton Keynes | Wellcome Sanger Institute for the COVID-19 Genomics UK (COG-UK) Consortium | The Lighthouse Lab in Milton Keynes and Alex Alderton, Roberto Amato, Sonia Goncalves, Ewan Harrison, David K. Jackson, Ian Johnston, Dominic Kwiatkowski, Cordelia Langford, John Sillitoe on behalf of the Wellcome Sanger Institute COVID-19 Surveillance Team |
| EPI_ISL_799310 | Lighthouse Lab in Glasgow | Wellcome Sanger Institute for the COVID-19 Genomics UK (COG-UK) Consortium | Harper VanSteenhouse, Yumi Kasai, David Gray, Carol Clugston, Anna Dominiczak and Alex Alderton, Roberto Amato, Sonia Goncalves, Ewan Harrison, David K. Jackson, Ian Johnston, Dominic Kwiatkowski, Cordelia Langford, John Sillitoe on behalf of the Wellcome Sanger Institute COVID-19 Surveillance Team |
| EPI_ISL_799311, EPI_ISL_799314 | Lighthouse Lab in Milton Keynes | Wellcome Sanger Institute for the COVID-19 Genomics UK (COG-UK) Consortium | The Lighthouse Lab in Milton Keynes and Alex Alderton, Roberto Amato, Sonia Goncalves, Ewan Harrison, David K. Jackson, Ian Johnston, Dominic Kwiatkowski, Cordelia Langford, John Sillitoe on behalf of the Wellcome Sanger Institute COVID-19 Surveillance Team |
| EPI_ISL_799315 | Lighthouse Lab in Glasgow | Wellcome Sanger Institute for the COVID-19 Genomics UK (COG-UK) Consortium | Harper VanSteenhouse, Yumi Kasai, David Gray, Carol Clugston, Anna Dominiczak and Alex Alderton, Roberto Amato, Sonia Goncalves, Ewan Harrison, David K. Jackson, Ian Johnston, Dominic Kwiatkowski, Cordelia Langford, John Sillitoe on behalf of the Wellcome Sanger Institute COVID-19 Surveillance Team |
| EPI_ISL_799317 | Lighthouse Lab in Milton Keynes | Wellcome Sanger Institute for the COVID-19 Genomics UK (COG-UK) Consortium | The Lighthouse Lab in Milton Keynes and Alex Alderton, Roberto Amato, Sonia Goncalves, Ewan Harrison, David K. Jackson, Ian Johnston, Dominic Kwiatkowski, Cordelia Langford, John Sillitoe on behalf of the Wellcome Sanger Institute COVID-19 Surveillance Team |
| EPI_ISL_799318 | Lighthouse Lab in Glasgow | Wellcome Sanger Institute for the COVID-19 Genomics UK (COG-UK) Consortium | Harper VanSteenhouse, Yumi Kasai, David Gray, Carol Clugston, Anna Dominiczak and Alex Alderton, Roberto Amato, Sonia Goncalves, Ewan Harrison, David K. Jackson, Ian Johnston, Dominic Kwiatkowski, Cordelia Langford, John Sillitoe on behalf of the Wellcome Sanger Institute COVID-19 Surveillance Team |
| EPI_ISL_799319, EPI_ISL_799320 | Lighthouse Lab in Milton Keynes | Wellcome Sanger Institute for the COVID-19 Genomics UK (COG-UK) Consortium | The Lighthouse Lab in Milton Keynes and Alex Alderton, Roberto Amato, Sonia Goncalves, Ewan Harrison, David K. Jackson, Ian Johnston, Dominic Kwiatkowski, Cordelia Langford, John Sillitoe on behalf of the Wellcome Sanger Institute COVID-19 Surveillance Team |
| EPI_ISL_799322 | Lighthouse Lab in Glasgow | Wellcome Sanger Institute for the COVID-19 Genomics UK (COG-UK) Consortium | Harper VanSteenhouse, Yumi Kasai, David Gray, Carol Clugston, Anna Dominiczak and Alex Alderton, Roberto Amato, Sonia Goncalves, Ewan Harrison, David K. Jackson, Ian Johnston, Dominic Kwiatkowski, Cordelia Langford, John Sillitoe on behalf of the Wellcome Sanger Institute COVID-19 Surveillance Team |
| EPI_ISL_799323, EPI_ISL_799325 | Lighthouse Lab in Milton Keynes | Wellcome Sanger Institute for the COVID-19 Genomics UK (COG-UK) Consortium | The Lighthouse Lab in Milton Keynes and Alex Alderton, Roberto Amato, Sonia Goncalves, Ewan Harrison, David K. Jackson, Ian Johnston, Dominic Kwiatkowski, Cordelia Langford, John Sillitoe on behalf of the Wellcome Sanger Institute COVID-19 Surveillance Team |
| EPI_ISL_799326 | Lighthouse Lab in Glasgow | Wellcome Sanger Institute for the COVID-19 Genomics UK (COG-UK) Consortium | Harper VanSteenhouse, Yumi Kasai, David Gray, Carol Clugston, Anna Dominiczak and Alex Alderton, Roberto Amato, Sonia Goncalves, Ewan Harrison, David K. Jackson, Ian Johnston, Dominic Kwiatkowski, Cordelia Langford, John Sillitoe on behalf of the Wellcome Sanger Institute COVID-19 Surveillance Team |
| EPI_ISL_799327, EPI_ISL_799333, EPI_ISL_799334 | Lighthouse Lab in Milton Keynes | Wellcome Sanger Institute for the COVID-19 Genomics UK (COG-UK) Consortium | The Lighthouse Lab in Milton Keynes and Alex Alderton, Roberto Amato, Sonia Goncalves, Ewan Harrison, David K. Jackson, Ian Johnston, Dominic Kwiatkowski, Cordelia Langford, John Sillitoe on behalf of the Wellcome Sanger Institute COVID-19 Surveillance Team |
| EPI_ISL_799345 | Lighthouse Lab in Glasgow | Wellcome Sanger Institute for the COVID-19 Genomics UK (COG-UK) Consortium | Harper VanSteenhouse, Yumi Kasai, David Gray, Carol Clugston, Anna Dominiczak and Alex Alderton, Roberto Amato, Sonia Goncalves, Ewan Harrison, David K. Jackson, Ian Johnston, Dominic Kwiatkowski, Cordelia Langford, John Sillitoe on behalf of the Wellcome Sanger Institute COVID-19 Surveillance Team |
| EPI_ISL_799346, EPI_ISL_799347, EPI_ISL_799348, EPI_ISL_799350, EPI_ISL_799351, EPI_ISL_799352, EPI_ISL_799353, EPI_ISL_799354 | Lighthouse Lab in Milton Keynes | Wellcome Sanger Institute for the COVID-19 Genomics UK (COG-UK) Consortium | The Lighthouse Lab in Milton Keynes and Alex Alderton, Roberto Amato, Sonia Goncalves, Ewan Harrison, David K. Jackson, Ian Johnston, Dominic Kwiatkowski, Cordelia Langford, John Sillitoe on behalf of the Wellcome Sanger Institute COVID-19 Surveillance Team |
| EPI_ISL_799355 | Lighthouse Lab in Glasgow | Wellcome Sanger Institute for the COVID-19 Genomics UK (COG-UK) Consortium | Harper VanSteenhouse, Yumi Kasai, David Gray, Carol Clugston, Anna Dominiczak and Alex Alderton, Roberto Amato, Sonia Goncalves, Ewan Harrison, David K. Jackson, Ian Johnston, Dominic Kwiatkowski, Cordelia Langford, John Sillitoe on behalf of the Wellcome Sanger Institute COVID-19 Surveillance Team |
| EPI_ISL_799356, EPI_ISL_799357, EPI_ISL_799362, EPI_ISL_799363, EPI_ISL_799364, EPI_ISL_799366, EPI_ISL_799368 | Lighthouse Lab in Milton Keynes | Wellcome Sanger Institute for the COVID-19 Genomics UK (COG-UK) Consortium | The Lighthouse Lab in Milton Keynes and Alex Alderton, Roberto Amato, Sonia Goncalves, Ewan Harrison, David K. Jackson, Ian Johnston, Dominic Kwiatkowski, Cordelia Langford, John Sillitoe on behalf of the Wellcome Sanger Institute COVID-19 Surveillance Team |
| EPI_ISL_799369 | Lighthouse Lab in Glasgow | Wellcome Sanger Institute for the COVID-19 Genomics UK (COG-UK) Consortium | Harper VanSteenhouse, Yumi Kasai, David Gray, Carol Clugston, Anna Dominiczak and Alex Alderton, Roberto Amato, Sonia Goncalves, Ewan Harrison, David K. Jackson, Ian Johnston, Dominic Kwiatkowski, Cordelia Langford, John Sillitoe on behalf of the Wellcome Sanger Institute COVID-19 Surveillance Team |
| EPI_ISL_799370, EPI_ISL_799372, EPI_ISL_799375, EPI_ISL_799376 | Lighthouse Lab in Milton Keynes | Wellcome Sanger Institute for the COVID-19 Genomics UK (COG-UK) Consortium | The Lighthouse Lab in Milton Keynes and Alex Alderton, Roberto Amato, Sonia Goncalves, Ewan Harrison, David K. Jackson, Ian Johnston, Dominic Kwiatkowski, Cordelia Langford, John Sillitoe on behalf of the Wellcome Sanger Institute COVID-19 Surveillance Team |
| EPI_ISL_799377 | Lighthouse Lab in Glasgow | Wellcome Sanger Institute for the COVID-19 Genomics UK (COG-UK) Consortium | Harper VanSteenhouse, Yumi Kasai, David Gray, Carol Clugston, Anna Dominiczak and Alex Alderton, Roberto Amato, Sonia Goncalves, Ewan Harrison, David K. Jackson, Ian Johnston, Dominic Kwiatkowski, Cordelia Langford, John Sillitoe on behalf of the Wellcome Sanger Institute COVID-19 Surveillance Team |
| EPI_ISL_799378, EPI_ISL_799380, EPI_ISL_799384, EPI_ISL_799386, EPI_ISL_799387, EPI_ISL_799388, EPI_ISL_799392, EPI_ISL_799394, EPI_ISL_799395, EPI_ISL_799397 | Lighthouse Lab in Milton Keynes | Wellcome Sanger Institute for the COVID-19 Genomics UK (COG-UK) Consortium | The Lighthouse Lab in Milton Keynes and Alex Alderton, Roberto Amato, Sonia Goncalves, Ewan Harrison, David K. Jackson, Ian Johnston, Dominic Kwiatkowski, Cordelia Langford, John Sillitoe on behalf of the Wellcome Sanger Institute COVID-19 Surveillance Team |
| EPI_ISL_799398, EPI_ISL_799399 | Lighthouse Lab in Glasgow | Wellcome Sanger Institute for the COVID-19 Genomics UK (COG-UK) Consortium | Harper VanSteenhouse, Yumi Kasai, David Gray, Carol Clugston, Anna Dominiczak and Alex Alderton, Roberto Amato, Sonia Goncalves, Ewan Harrison, David K. Jackson, Ian Johnston, Dominic Kwiatkowski, Cordelia Langford, John Sillitoe on behalf of the Wellcome Sanger Institute COVID-19 Surveillance |

| | | | |
|---|---|---|---|
| | | | Team |
| EPI_ISL_799400 | Lighthouse Lab in Milton Keynes | Wellcome Sanger Institute for the COVID-19 Genomics UK (COG-UK) Consortium | The Lighthouse Lab in Milton Keynes and Alex Alderton, Roberto Amato, Sonia Goncalves, Ewan Harrison, David K. Jackson, Ian Johnston, Dominic Kwiatkowski, Cordelia Langford, John Sillitoe on behalf of the Wellcome Sanger Institute COVID-19 Surveillance Team |
| EPI_ISL_799401 | Lighthouse Lab in Glasgow | Wellcome Sanger Institute for the COVID-19 Genomics UK (COG-UK) Consortium | Harper VanSteenhouse, Yumi Kasai, David Gray, Carol Clugston, Anna Dominiczak and Alex Alderton, Roberto Amato, Sonia Goncalves, Ewan Harrison, David K. Jackson, Ian Johnston, Dominic Kwiatkowski, Cordelia Langford, John Sillitoe on behalf of the Wellcome Sanger Institute COVID-19 Surveillance Team |
| EPI_ISL_799403, EPI_ISL_799404 | Lighthouse Lab in Milton Keynes | Wellcome Sanger Institute for the COVID-19 Genomics UK (COG-UK) Consortium | The Lighthouse Lab in Milton Keynes and Alex Alderton, Roberto Amato, Sonia Goncalves, Ewan Harrison, David K. Jackson, Ian Johnston, Dominic Kwiatkowski, Cordelia Langford, John Sillitoe on behalf of the Wellcome Sanger Institute COVID-19 Surveillance Team |
| EPI_ISL_799405 | Lighthouse Lab in Glasgow | Wellcome Sanger Institute for the COVID-19 Genomics UK (COG-UK) Consortium | Harper VanSteenhouse, Yumi Kasai, David Gray, Carol Clugston, Anna Dominiczak and Alex Alderton, Roberto Amato, Sonia Goncalves, Ewan Harrison, David K. Jackson, Ian Johnston, Dominic Kwiatkowski, Cordelia Langford, John Sillitoe on behalf of the Wellcome Sanger Institute COVID-19 Surveillance Team |
| EPI_ISL_799406, EPI_ISL_799410, EPI_ISL_799411, EPI_ISL_799412, EPI_ISL_799413, EPI_ISL_799415, EPI_ISL_799416 | Lighthouse Lab in Milton Keynes | Wellcome Sanger Institute for the COVID-19 Genomics UK (COG-UK) Consortium | The Lighthouse Lab in Milton Keynes and Alex Alderton, Roberto Amato, Sonia Goncalves, Ewan Harrison, David K. Jackson, Ian Johnston, Dominic Kwiatkowski, Cordelia Langford, John Sillitoe on behalf of the Wellcome Sanger Institute COVID-19 Surveillance Team |
| EPI_ISL_799417, EPI_ISL_799418, EPI_ISL_799419, EPI_ISL_799422, EPI_ISL_799424, EPI_ISL_799425 | Lighthouse Lab in Glasgow | Wellcome Sanger Institute for the COVID-19 Genomics UK (COG-UK) Consortium | Harper VanSteenhouse, Yumi Kasai, David Gray, Carol Clugston, Anna Dominiczak and Alex Alderton, Roberto Amato, Sonia Goncalves, Ewan Harrison, David K. Jackson, Ian Johnston, Dominic Kwiatkowski, Cordelia Langford, John Sillitoe on behalf of the Wellcome Sanger Institute COVID-19 Surveillance Team |
| EPI_ISL_799427, EPI_ISL_799432 | Lighthouse Lab in Milton Keynes | Wellcome Sanger Institute for the COVID-19 Genomics UK (COG-UK) Consortium | The Lighthouse Lab in Milton Keynes and Alex Alderton, Roberto Amato, Sonia Goncalves, Ewan Harrison, David K. Jackson, Ian Johnston, Dominic Kwiatkowski, Cordelia Langford, John Sillitoe on behalf of the Wellcome Sanger Institute COVID-19 Surveillance Team |
| EPI_ISL_799433 | Lighthouse Lab in Glasgow | Wellcome Sanger Institute for the COVID-19 Genomics UK (COG-UK) Consortium | Harper VanSteenhouse, Yumi Kasai, David Gray, Carol Clugston, Anna Dominiczak and Alex Alderton, Roberto Amato, Sonia Goncalves, Ewan Harrison, David K. Jackson, Ian Johnston, Dominic Kwiatkowski, Cordelia Langford, John Sillitoe on behalf of the Wellcome Sanger Institute COVID-19 Surveillance Team |
| EPI_ISL_799434, EPI_ISL_799436, EPI_ISL_799437 | Lighthouse Lab in Milton Keynes | Wellcome Sanger Institute for the COVID-19 Genomics UK (COG-UK) Consortium | The Lighthouse Lab in Milton Keynes and Alex Alderton, Roberto Amato, Sonia Goncalves, Ewan Harrison, David K. Jackson, Ian Johnston, Dominic Kwiatkowski, Cordelia Langford, John Sillitoe on behalf of the Wellcome Sanger Institute COVID-19 Surveillance Team |
| EPI_ISL_799438 | Lighthouse Lab in Glasgow | Wellcome Sanger Institute for the COVID-19 Genomics UK (COG-UK) Consortium | Harper VanSteenhouse, Yumi Kasai, David Gray, Carol Clugston, Anna Dominiczak and Alex Alderton, Roberto Amato, Sonia Goncalves, Ewan Harrison, David K. Jackson, Ian Johnston, Dominic Kwiatkowski, Cordelia Langford, John Sillitoe on behalf of the Wellcome Sanger Institute COVID-19 Surveillance Team |
| EPI_ISL_799439, EPI_ISL_799440, EPI_ISL_799445, EPI_ISL_799446, EPI_ISL_799447, EPI_ISL_799448 | Lighthouse Lab in Milton Keynes | Wellcome Sanger Institute for the COVID-19 Genomics UK (COG-UK) Consortium | The Lighthouse Lab in Milton Keynes and Alex Alderton, Roberto Amato, Sonia Goncalves, Ewan Harrison, David K. Jackson, Ian Johnston, Dominic Kwiatkowski, Cordelia Langford, John Sillitoe on behalf of the Wellcome Sanger Institute COVID-19 Surveillance Team |
| EPI_ISL_799452 | Lighthouse Lab in Glasgow | Wellcome Sanger Institute for the COVID-19 Genomics UK (COG-UK) Consortium | Harper VanSteenhouse, Yumi Kasai, David Gray, Carol Clugston, Anna Dominiczak and Alex Alderton, Roberto Amato, Sonia Goncalves, Ewan Harrison, David K. Jackson, Ian Johnston, Dominic Kwiatkowski, Cordelia Langford, John Sillitoe on behalf of the Wellcome Sanger Institute COVID-19 Surveillance Team |
| EPI_ISL_799454 | Lighthouse Lab in Milton Keynes | Wellcome Sanger Institute for the COVID-19 Genomics UK (COG-UK) Consortium | The Lighthouse Lab in Milton Keynes and Alex Alderton, Roberto Amato, Sonia Goncalves, Ewan Harrison, David K. Jackson, Ian Johnston, Dominic Kwiatkowski, Cordelia Langford, John Sillitoe on behalf of the Wellcome Sanger Institute COVID-19 Surveillance Team |
| EPI_ISL_799456, EPI_ISL_799458, EPI_ISL_799461 | Lighthouse Lab in Glasgow | Wellcome Sanger Institute for the COVID-19 Genomics UK (COG-UK) Consortium | Harper VanSteenhouse, Yumi Kasai, David Gray, Carol Clugston, Anna Dominiczak and Alex Alderton, Roberto Amato, Sonia Goncalves, Ewan Harrison, David K. Jackson, Ian Johnston, Dominic Kwiatkowski, Cordelia Langford, John Sillitoe on behalf of the Wellcome Sanger Institute COVID-19 Surveillance Team |
| EPI_ISL_799467, EPI_ISL_799468, EPI_ISL_799469 | Lighthouse Lab in Milton Keynes | Wellcome Sanger Institute for the COVID-19 Genomics UK (COG-UK) Consortium | The Lighthouse Lab in Milton Keynes and Alex Alderton, Roberto Amato, Sonia Goncalves, Ewan Harrison, David K. Jackson, Ian Johnston, Dominic Kwiatkowski, Cordelia Langford, John Sillitoe on behalf of the Wellcome Sanger Institute COVID-19 Surveillance Team |
| EPI_ISL_799474 | Lighthouse Lab in Glasgow | Wellcome Sanger Institute for the COVID-19 Genomics UK (COG-UK) Consortium | Harper VanSteenhouse, Yumi Kasai, David Gray, Carol Clugston, Anna Dominiczak and Alex Alderton, Roberto Amato, Sonia Goncalves, Ewan Harrison, David K. Jackson, Ian Johnston, Dominic Kwiatkowski, Cordelia Langford, John Sillitoe on behalf of the Wellcome Sanger Institute COVID-19 Surveillance Team |

EPI_ISL_799476, EPI_ISL_799477, EPI_ISL_799479, EPI_ISL_799482, EPI_ISL_799485, EPI_ISL_799487, EPI_ISL_799488, EPI_ISL_799490, EPI_ISL_799492, EPI_ISL_799494, EPI_ISL_799496, EPI_ISL_799497, EPI_ISL_799498, EPI_ISL_799500, EPI_ISL_799501, EPI_ISL_799505, EPI_ISL_799506, EPI_ISL_799508, EPI_ISL_799509

| | | | |
|---|---|---|---|
| see above | Lighthouse Lab in Milton Keynes | Wellcome Sanger Institute for the COVID-19 Genomics UK (COG-UK) Consortium | The Lighthouse Lab in Milton Keynes and Alex Alderton, Roberto Amato, Sonia Goncalves, Ewan Harrison, David K. Jackson, Ian Johnston, Dominic Kwiatkowski, Cordelia Langford, John Sillitoe on behalf of the Wellcome Sanger Institute COVID-19 Surveillance Team |
| EPI_ISL_799510 | Lighthouse Lab in Glasgow | Wellcome Sanger Institute for the COVID-19 Genomics UK (COG-UK) Consortium | Harper VanSteenhouse, Yumi Kasai, David Gray, Carol Clugston, Anna Dominiczak and Alex Alderton, Roberto Amato, Sonia Goncalves, Ewan Harrison, David K. Jackson, Ian Johnston, Dominic Kwiatkowski, Cordelia Langford, John Sillitoe on behalf of the Wellcome Sanger Institute COVID-19 Surveillance Team |
| EPI_ISL_799512, EPI_ISL_799514, EPI_ISL_799515, EPI_ISL_799519, EPI_ISL_799522, EPI_ISL_799523 | Lighthouse Lab in Milton Keynes | Wellcome Sanger Institute for the COVID-19 Genomics UK (COG-UK) Consortium | The Lighthouse Lab in Milton Keynes and Alex Alderton, Roberto Amato, Sonia Goncalves, Ewan Harrison, David K. Jackson, Ian Johnston, Dominic Kwiatkowski, Cordelia Langford, John Sillitoe on behalf of the Wellcome Sanger Institute COVID-19 Surveillance Team |
| EPI_ISL_799525, EPI_ISL_799526 | Lighthouse Lab in Glasgow | Wellcome Sanger Institute for the COVID-19 Genomics UK (COG-UK) Consortium | Harper VanSteenhouse, Yumi Kasai, David Gray, Carol Clugston, Anna Dominiczak and Alex Alderton, Roberto Amato, Sonia Goncalves, Ewan Harrison, David K. Jackson, Ian Johnston, Dominic Kwiatkowski, Cordelia Langford, John Sillitoe on behalf of the Wellcome Sanger Institute COVID-19 Surveillance Team |
| EPI_ISL_799528, EPI_ISL_799529 | Lighthouse Lab in Milton Keynes | Wellcome Sanger Institute for the COVID-19 Genomics UK (COG-UK) Consortium | The Lighthouse Lab in Milton Keynes and Alex Alderton, Roberto Amato, Sonia Goncalves, Ewan Harrison, David K. Jackson, Ian Johnston, Dominic Kwiatkowski, Cordelia Langford, John Sillitoe on behalf of the Wellcome Sanger Institute COVID-19 Surveillance Team |
| EPI_ISL_799530 | Lighthouse Lab in Glasgow | Wellcome Sanger Institute for the COVID-19 Genomics UK (COG-UK) Consortium | Harper VanSteenhouse, Yumi Kasai, David Gray, Carol Clugston, Anna Dominiczak and Alex Alderton, Roberto Amato, Sonia Goncalves, Ewan Harrison, David K. Jackson, Ian Johnston, Dominic Kwiatkowski, Cordelia Langford, John Sillitoe on behalf of the Wellcome Sanger Institute COVID-19 Surveillance Team |
| EPI_ISL_799533, EPI_ISL_799535, EPI_ISL_799536 | Lighthouse Lab in Milton Keynes | Wellcome Sanger Institute for the COVID-19 Genomics UK (COG-UK) Consortium | The Lighthouse Lab in Milton Keynes and Alex Alderton, Roberto Amato, Sonia Goncalves, Ewan Harrison, David K. Jackson, Ian Johnston, Dominic Kwiatkowski, Cordelia Langford, John Sillitoe on behalf of the Wellcome Sanger Institute COVID-19 Surveillance Team |
| EPI_ISL_799537 | Lighthouse Lab in Glasgow | Wellcome Sanger Institute for the COVID-19 Genomics UK (COG-UK) Consortium | Harper VanSteenhouse, Yumi Kasai, David Gray, Carol Clugston, Anna Dominiczak and Alex Alderton, Roberto Amato, Sonia Goncalves, Ewan Harrison, David K. Jackson, Ian Johnston, Dominic Kwiatkowski, Cordelia Langford, John Sillitoe on behalf of the Wellcome Sanger Institute COVID-19 Surveillance Team |
| EPI_ISL_799538, EPI_ISL_799540 | Lighthouse Lab in Milton Keynes | Wellcome Sanger Institute for the COVID-19 Genomics UK (COG-UK) Consortium | The Lighthouse Lab in Milton Keynes and Alex Alderton, Roberto Amato, Sonia Goncalves, Ewan Harrison, David K. Jackson, Ian Johnston, Dominic Kwiatkowski, Cordelia Langford, John Sillitoe on behalf of the Wellcome Sanger Institute COVID-19 Surveillance Team |
| EPI_ISL_799541, EPI_ISL_799542, EPI_ISL_799543 | Lighthouse Lab in Glasgow | Wellcome Sanger Institute for the COVID-19 Genomics UK (COG-UK) Consortium | Harper VanSteenhouse, Yumi Kasai, David Gray, Carol Clugston, Anna Dominiczak and Alex Alderton, Roberto Amato, Sonia Goncalves, Ewan Harrison, David K. Jackson, Ian Johnston, Dominic Kwiatkowski, Cordelia Langford, John Sillitoe on behalf of the Wellcome Sanger Institute COVID-19 Surveillance Team |
| EPI_ISL_799546, EPI_ISL_799547 | Lighthouse Lab in Milton Keynes | Wellcome Sanger Institute for the COVID-19 Genomics UK | The Lighthouse Lab in Milton Keynes and Alex Alderton, Roberto Amato, Sonia Goncalves, Ewan Harrison, David K. Jackson, Ian Johnston, Dominic |

| | | | |
|---|---|---|---|
| | | (COG-UK) Consortium | Kwiatkowski, Cordelia Langford, John Sillitoe on behalf of the Wellcome Sanger Institute COVID-19 Surveillance Team |
| EPI_ISL_799549 | Lighthouse Lab in Glasgow | Wellcome Sanger Institute for the COVID-19 Genomics UK (COG-UK) Consortium | Harper VanSteenhouse, Yumi Kasai, David Gray, Carol Clugston, Anna Dominiczak and Alex Alderton, Roberto Amato, Sonia Goncalves, Ewan Harrison, David K. Jackson, Ian Johnston, Dominic Kwiatkowski, Cordelia Langford, John Sillitoe on behalf of the Wellcome Sanger Institute COVID-19 Surveillance Team |
| EPI_ISL_799550, EPI_ISL_799551, EPI_ISL_799555 | Lighthouse Lab in Milton Keynes | Wellcome Sanger Institute for the COVID-19 Genomics UK (COG-UK) Consortium | The Lighthouse Lab in Milton Keynes and Alex Alderton, Roberto Amato, Sonia Goncalves, Ewan Harrison, David K. Jackson, Ian Johnston, Dominic Kwiatkowski, Cordelia Langford, John Sillitoe on behalf of the Wellcome Sanger Institute COVID-19 Surveillance Team |
| EPI_ISL_799556, EPI_ISL_799557, EPI_ISL_799559, EPI_ISL_799560 | Lighthouse Lab in Glasgow | Wellcome Sanger Institute for the COVID-19 Genomics UK (COG-UK) Consortium | Harper VanSteenhouse, Yumi Kasai, David Gray, Carol Clugston, Anna Dominiczak and Alex Alderton, Roberto Amato, Sonia Goncalves, Ewan Harrison, David K. Jackson, Ian Johnston, Dominic Kwiatkowski, Cordelia Langford, John Sillitoe on behalf of the Wellcome Sanger Institute COVID-19 Surveillance Team |
| EPI_ISL_799562, EPI_ISL_799563, EPI_ISL_799565 | Lighthouse Lab in Milton Keynes | Wellcome Sanger Institute for the COVID-19 Genomics UK (COG-UK) Consortium | The Lighthouse Lab in Milton Keynes and Alex Alderton, Roberto Amato, Sonia Goncalves, Ewan Harrison, David K. Jackson, Ian Johnston, Dominic Kwiatkowski, Cordelia Langford, John Sillitoe on behalf of the Wellcome Sanger Institute COVID-19 Surveillance Team |
| EPI_ISL_799566 | Lighthouse Lab in Glasgow | Wellcome Sanger Institute for the COVID-19 Genomics UK (COG-UK) Consortium | Harper VanSteenhouse, Yumi Kasai, David Gray, Carol Clugston, Anna Dominiczak and Alex Alderton, Roberto Amato, Sonia Goncalves, Ewan Harrison, David K. Jackson, Ian Johnston, Dominic Kwiatkowski, Cordelia Langford, John Sillitoe on behalf of the Wellcome Sanger Institute COVID-19 Surveillance Team |
| EPI_ISL_799567, EPI_ISL_799570, EPI_ISL_799572, EPI_ISL_799573, EPI_ISL_799575, EPI_ISL_799577, EPI_ISL_799578, EPI_ISL_799579, EPI_ISL_799580, EPI_ISL_799582, EPI_ISL_799583 | | | |
| see above | Lighthouse Lab in Milton Keynes | Wellcome Sanger Institute for the COVID-19 Genomics UK (COG-UK) Consortium | The Lighthouse Lab in Milton Keynes and Alex Alderton, Roberto Amato, Sonia Goncalves, Ewan Harrison, David K. Jackson, Ian Johnston, Dominic Kwiatkowski, Cordelia Langford, John Sillitoe on behalf of the Wellcome Sanger Institute COVID-19 Surveillance Team |
| EPI_ISL_799586 | Lighthouse Lab in Glasgow | Wellcome Sanger Institute for the COVID-19 Genomics UK (COG-UK) Consortium | Harper VanSteenhouse, Yumi Kasai, David Gray, Carol Clugston, Anna Dominiczak and Alex Alderton, Roberto Amato, Sonia Goncalves, Ewan Harrison, David K. Jackson, Ian Johnston, Dominic Kwiatkowski, Cordelia Langford, John Sillitoe on behalf of the Wellcome Sanger Institute COVID-19 Surveillance Team |
| EPI_ISL_799587, EPI_ISL_799588, EPI_ISL_799590, EPI_ISL_799595, EPI_ISL_799597, EPI_ISL_799598, EPI_ISL_799602, EPI_ISL_799603, EPI_ISL_799604, EPI_ISL_799608, EPI_ISL_799610, EPI_ISL_799611, EPI_ISL_799614, EPI_ISL_799616, EPI_ISL_799617, EPI_ISL_799618 | | | |
| see above | Lighthouse Lab in Milton Keynes | Wellcome Sanger Institute for the COVID-19 Genomics UK (COG-UK) Consortium | The Lighthouse Lab in Milton Keynes and Alex Alderton, Roberto Amato, Sonia Goncalves, Ewan Harrison, David K. Jackson, Ian Johnston, Dominic Kwiatkowski, Cordelia Langford, John Sillitoe on behalf of the Wellcome Sanger Institute COVID-19 Surveillance Team |
| EPI_ISL_799619, EPI_ISL_799622 | Lighthouse Lab in Glasgow | Wellcome Sanger Institute for the COVID-19 Genomics UK (COG-UK) Consortium | Harper VanSteenhouse, Yumi Kasai, David Gray, Carol Clugston, Anna Dominiczak and Alex Alderton, Roberto Amato, Sonia Goncalves, Ewan Harrison, David K. Jackson, Ian Johnston, Dominic Kwiatkowski, Cordelia Langford, John Sillitoe on behalf of the Wellcome Sanger Institute COVID-19 Surveillance Team |
| EPI_ISL_799624, EPI_ISL_799625, EPI_ISL_799626, EPI_ISL_799627, EPI_ISL_799629 | Lighthouse Lab in Milton Keynes | Wellcome Sanger Institute for the COVID-19 Genomics UK (COG-UK) Consortium | The Lighthouse Lab in Milton Keynes and Alex Alderton, Roberto Amato, Sonia Goncalves, Ewan Harrison, David K. Jackson, Ian Johnston, Dominic Kwiatkowski, Cordelia Langford, John Sillitoe on behalf of the Wellcome Sanger Institute COVID-19 Surveillance Team |
| EPI_ISL_799630 | Lighthouse Lab in Glasgow | Wellcome Sanger Institute for the COVID-19 Genomics UK (COG-UK) Consortium | Harper VanSteenhouse, Yumi Kasai, David Gray, Carol Clugston, Anna Dominiczak and Alex Alderton, Roberto Amato, Sonia Goncalves, Ewan Harrison, David K. Jackson, Ian Johnston, Dominic Kwiatkowski, Cordelia Langford, John Sillitoe on behalf of the Wellcome Sanger Institute COVID-19 Surveillance Team |
| EPI_ISL_799631, EPI_ISL_799632, EPI_ISL_799633, EPI_ISL_799636 | Lighthouse Lab in Milton Keynes | Wellcome Sanger Institute for the COVID-19 Genomics UK (COG-UK) Consortium | The Lighthouse Lab in Milton Keynes and Alex Alderton, Roberto Amato, Sonia Goncalves, Ewan Harrison, David K. Jackson, Ian Johnston, Dominic Kwiatkowski, Cordelia Langford, John Sillitoe on behalf of the Wellcome Sanger Institute COVID-19 Surveillance Team |
| EPI_ISL_799641 | Lighthouse Lab in Glasgow | Wellcome Sanger Institute for the COVID-19 Genomics UK (COG-UK) Consortium | Harper VanSteenhouse, Yumi Kasai, David Gray, Carol Clugston, Anna Dominiczak and Alex Alderton, Roberto Amato, Sonia Goncalves, Ewan Harrison, David K. Jackson, Ian Johnston, Dominic Kwiatkowski, Cordelia Langford, John Sillitoe on behalf of the Wellcome Sanger Institute COVID-19 Surveillance Team |
| EPI_ISL_799642, EPI_ISL_799643 | Lighthouse Lab in Milton Keynes | Wellcome Sanger Institute for the COVID-19 Genomics UK (COG-UK) Consortium | The Lighthouse Lab in Milton Keynes and Alex Alderton, Roberto Amato, Sonia Goncalves, Ewan Harrison, David K. Jackson, Ian Johnston, Dominic Kwiatkowski, Cordelia Langford, John Sillitoe on behalf of the Wellcome Sanger Institute COVID-19 Surveillance Team |
| EPI_ISL_799644 | Lighthouse Lab in Alderley Park | Wellcome Sanger Institute for the COVID-19 Genomics UK (COG-UK) Consortium | Jacquelyn Wynn, Mairead Hyland, The Lighthouse Lab in Alderley Park and Alex Alderton, Roberto Amato, Sonia Goncalves, Ewan Harrison, David K. Jackson, Ian Johnston, Dominic Kwiatkowski, Cordelia Langford, John Sillitoe on behalf of the Wellcome Sanger Institute COVID-19 Surveillance Team |
| EPI_ISL_799648, EPI_ISL_799649, EPI_ISL_799650 | Lighthouse Lab in Glasgow | Wellcome Sanger Institute for the COVID-19 Genomics UK (COG-UK) Consortium | Harper VanSteenhouse, Yumi Kasai, David Gray, Carol Clugston, Anna Dominiczak and Alex Alderton, Roberto Amato, Sonia Goncalves, Ewan Harrison, David K. Jackson, Ian Johnston, Dominic Kwiatkowski, Cordelia Langford, John Sillitoe on behalf of the Wellcome Sanger Institute COVID-19 Surveillance Team |
| EPI_ISL_799652, EPI_ISL_799655 | Lighthouse Lab in Alderley Park | Wellcome Sanger Institute for the COVID-19 Genomics UK (COG-UK) Consortium | Jacquelyn Wynn, Mairead Hyland, The Lighthouse Lab in Alderley Park and Alex Alderton, Roberto Amato, Sonia Goncalves, Ewan Harrison, David K. Jackson, Ian Johnston, Dominic Kwiatkowski, Cordelia Langford, John Sillitoe on behalf of the Wellcome Sanger Institute COVID-19 Surveillance Team |
| EPI_ISL_799660 | Lighthouse Lab in Milton Keynes | Wellcome Sanger Institute for the COVID-19 Genomics UK (COG-UK) Consortium | The Lighthouse Lab in Milton Keynes and Alex Alderton, Roberto Amato, Sonia Goncalves, Ewan Harrison, David K. Jackson, Ian Johnston, Dominic Kwiatkowski, Cordelia Langford, John Sillitoe on behalf of the Wellcome Sanger Institute COVID-19 Surveillance Team |
| EPI_ISL_799661 | Lighthouse Lab in Glasgow | Wellcome Sanger Institute for the COVID-19 Genomics UK (COG-UK) Consortium | Harper VanSteenhouse, Yumi Kasai, David Gray, Carol Clugston, Anna Dominiczak and Alex Alderton, Roberto Amato, Sonia Goncalves, Ewan Harrison, David K. Jackson, Ian Johnston, Dominic Kwiatkowski, Cordelia Langford, John Sillitoe on behalf of the Wellcome Sanger Institute COVID-19 Surveillance Team |
| EPI_ISL_799662 | Lighthouse Lab in Alderley Park | Wellcome Sanger Institute for the COVID-19 Genomics UK (COG-UK) Consortium | Jacquelyn Wynn, Mairead Hyland, The Lighthouse Lab in Alderley Park and Alex Alderton, Roberto Amato, Sonia Goncalves, Ewan Harrison, David K. Jackson, Ian Johnston, Dominic Kwiatkowski, Cordelia Langford, John Sillitoe on behalf of the Wellcome Sanger Institute COVID-19 Surveillance Team |
| EPI_ISL_799663 | Lighthouse Lab in Milton Keynes | Wellcome Sanger Institute for the COVID-19 Genomics UK (COG-UK) Consortium | The Lighthouse Lab in Milton Keynes and Alex Alderton, Roberto Amato, Sonia Goncalves, Ewan Harrison, David K. Jackson, Ian Johnston, Dominic Kwiatkowski, Cordelia Langford, John Sillitoe on behalf of the Wellcome Sanger Institute COVID-19 Surveillance Team |
| EPI_ISL_799664 | Lighthouse Lab in Glasgow | Wellcome Sanger Institute for the COVID-19 Genomics UK (COG-UK) Consortium | Harper VanSteenhouse, Yumi Kasai, David Gray, Carol Clugston, Anna Dominiczak and Alex Alderton, Roberto Amato, Sonia Goncalves, Ewan Harrison, David K. Jackson, Ian Johnston, Dominic Kwiatkowski, Cordelia Langford, John Sillitoe on behalf of the Wellcome Sanger Institute COVID-19 Surveillance Team |
| EPI_ISL_799666, EPI_ISL_799671, EPI_ISL_799674, EPI_ISL_799675 | Lighthouse Lab in Milton Keynes | Wellcome Sanger Institute for the COVID-19 Genomics UK (COG-UK) Consortium | The Lighthouse Lab in Milton Keynes and Alex Alderton, Roberto Amato, Sonia Goncalves, Ewan Harrison, David K. Jackson, Ian Johnston, Dominic Kwiatkowski, Cordelia Langford, John Sillitoe on behalf of the Wellcome Sanger Institute COVID-19 Surveillance Team |
| EPI_ISL_799676 | Lighthouse Lab in Glasgow | Wellcome Sanger Institute for the COVID-19 Genomics UK (COG-UK) Consortium | Harper VanSteenhouse, Yumi Kasai, David Gray, Carol Clugston, Anna Dominiczak and Alex Alderton, Roberto Amato, Sonia Goncalves, Ewan Harrison, David K. Jackson, Ian Johnston, Dominic Kwiatkowski, Cordelia Langford, John Sillitoe on behalf of the Wellcome Sanger Institute COVID-19 Surveillance Team |
| EPI_ISL_799677 | Lighthouse Lab in Milton Keynes | Wellcome Sanger Institute for the COVID-19 Genomics UK (COG-UK) Consortium | The Lighthouse Lab in Milton Keynes and Alex Alderton, Roberto Amato, Sonia Goncalves, Ewan Harrison, David K. Jackson, Ian Johnston, Dominic Kwiatkowski, Cordelia Langford, John Sillitoe on behalf of the Wellcome Sanger Institute COVID-19 Surveillance Team |
| EPI_ISL_799678 | Lighthouse Lab in Alderley Park | Wellcome Sanger Institute for the COVID-19 Genomics UK (COG-UK) Consortium | Jacquelyn Wynn, Mairead Hyland, The Lighthouse Lab in Alderley Park and Alex Alderton, Roberto Amato, Sonia Goncalves, Ewan Harrison, David K. Jackson, Ian Johnston, Dominic Kwiatkowski, Cordelia Langford, John Sillitoe on behalf of the Wellcome Sanger Institute COVID-19 Surveillance Team |
| EPI_ISL_799679 | Lighthouse Lab in Glasgow | Wellcome Sanger Institute for the COVID-19 Genomics UK (COG-UK) Consortium | Harper VanSteenhouse, Yumi Kasai, David Gray, Carol Clugston, Anna Dominiczak and Alex Alderton, Roberto Amato, Sonia Goncalves, Ewan Harrison, David K. Jackson, Ian Johnston, Dominic Kwiatkowski, Cordelia Langford, John Sillitoe on behalf of the Wellcome Sanger Institute COVID-19 Surveillance Team |
| EPI_ISL_799688 | Lighthouse Lab in Milton Keynes | Wellcome Sanger Institute for the COVID-19 Genomics UK (COG-UK) Consortium | The Lighthouse Lab in Milton Keynes and Alex Alderton, Roberto Amato, Sonia Goncalves, Ewan Harrison, David K. Jackson, Ian Johnston, Dominic Kwiatkowski, Cordelia Langford, John Sillitoe on behalf of the Wellcome Sanger Institute COVID-19 Surveillance Team |

| | | | |
|---|---|---|---|
| EPI_ISL_799689 | Lighthouse Lab in Glasgow | Wellcome Sanger Institute for the COVID-19 Genomics UK (COG-UK) Consortium | Harper VanSteenhouse, Yumi Kasai, David Gray, Carol Clugston, Anna Dominiczak and Alex Alderton, Roberto Amato, Sonia Goncalves, Ewan Harrison, David K. Jackson, Ian Johnston, Dominic Kwiatkowski, Cordelia Langford, John Sillitoe on behalf of the Wellcome Sanger Institute COVID-19 Surveillance Team |
| EPI_ISL_799690 | Lighthouse Lab in Milton Keynes | Wellcome Sanger Institute for the COVID-19 Genomics UK (COG-UK) Consortium | The Lighthouse Lab in Milton Keynes and Alex Alderton, Roberto Amato, Sonia Goncalves, Ewan Harrison, David K. Jackson, Ian Johnston, Dominic Kwiatkowski, Cordelia Langford, John Sillitoe on behalf of the Wellcome Sanger Institute COVID-19 Surveillance Team |
| EPI_ISL_799691 | Lighthouse Lab in Alderley Park | Wellcome Sanger Institute for the COVID-19 Genomics UK (COG-UK) Consortium | Jacquelyn Wynn, Mairead Hyland, The Lighthouse Lab in Alderley Park and Alex Alderton, Roberto Amato, Sonia Goncalves, Ewan Harrison, David K. Jackson, Ian Johnston, Dominic Kwiatkowski, Cordelia Langford, John Sillitoe on behalf of the Wellcome Sanger Institute COVID-19 Surveillance Team |
| EPI_ISL_799694, EPI_ISL_799696, EPI_ISL_799698, EPI_ISL_799700, EPI_ISL_799703 | Lighthouse Lab in Milton Keynes | Wellcome Sanger Institute for the COVID-19 Genomics UK (COG-UK) Consortium | The Lighthouse Lab in Milton Keynes and Alex Alderton, Roberto Amato, Sonia Goncalves, Ewan Harrison, David K. Jackson, Ian Johnston, Dominic Kwiatkowski, Cordelia Langford, John Sillitoe on behalf of the Wellcome Sanger Institute COVID-19 Surveillance Team |
| EPI_ISL_799708 | Lighthouse Lab in Glasgow | Wellcome Sanger Institute for the COVID-19 Genomics UK (COG-UK) Consortium | Harper VanSteenhouse, Yumi Kasai, David Gray, Carol Clugston, Anna Dominiczak and Alex Alderton, Roberto Amato, Sonia Goncalves, Ewan Harrison, David K. Jackson, Ian Johnston, Dominic Kwiatkowski, Cordelia Langford, John Sillitoe on behalf of the Wellcome Sanger Institute COVID-19 Surveillance Team |
| EPI_ISL_799711 | Lighthouse Lab in Alderley Park | Wellcome Sanger Institute for the COVID-19 Genomics UK (COG-UK) Consortium | Jacquelyn Wynn, Mairead Hyland, The Lighthouse Lab in Alderley Park and Alex Alderton, Roberto Amato, Sonia Goncalves, Ewan Harrison, David K. Jackson, Ian Johnston, Dominic Kwiatkowski, Cordelia Langford, John Sillitoe on behalf of the Wellcome Sanger Institute COVID-19 Surveillance Team |
| EPI_ISL_799713 | Lighthouse Lab in Glasgow | Wellcome Sanger Institute for the COVID-19 Genomics UK (COG-UK) Consortium | Harper VanSteenhouse, Yumi Kasai, David Gray, Carol Clugston, Anna Dominiczak and Alex Alderton, Roberto Amato, Sonia Goncalves, Ewan Harrison, David K. Jackson, Ian Johnston, Dominic Kwiatkowski, Cordelia Langford, John Sillitoe on behalf of the Wellcome Sanger Institute COVID-19 Surveillance Team |
| EPI_ISL_799714, EPI_ISL_799715, EPI_ISL_799716, EPI_ISL_799717 | Lighthouse Lab in Milton Keynes | Wellcome Sanger Institute for the COVID-19 Genomics UK (COG-UK) Consortium | The Lighthouse Lab in Milton Keynes and Alex Alderton, Roberto Amato, Sonia Goncalves, Ewan Harrison, David K. Jackson, Ian Johnston, Dominic Kwiatkowski, Cordelia Langford, John Sillitoe on behalf of the Wellcome Sanger Institute COVID-19 Surveillance Team |
| EPI_ISL_799718 | Lighthouse Lab in Glasgow | Wellcome Sanger Institute for the COVID-19 Genomics UK (COG-UK) Consortium | Harper VanSteenhouse, Yumi Kasai, David Gray, Carol Clugston, Anna Dominiczak and Alex Alderton, Roberto Amato, Sonia Goncalves, Ewan Harrison, David K. Jackson, Ian Johnston, Dominic Kwiatkowski, Cordelia Langford, John Sillitoe on behalf of the Wellcome Sanger Institute COVID-19 Surveillance Team |
| EPI_ISL_799720, EPI_ISL_799721 | Lighthouse Lab in Milton Keynes | Wellcome Sanger Institute for the COVID-19 Genomics UK (COG-UK) Consortium | The Lighthouse Lab in Milton Keynes and Alex Alderton, Roberto Amato, Sonia Goncalves, Ewan Harrison, David K. Jackson, Ian Johnston, Dominic Kwiatkowski, Cordelia Langford, John Sillitoe on behalf of the Wellcome Sanger Institute COVID-19 Surveillance Team |
| EPI_ISL_799722 | Lighthouse Lab in Glasgow | Wellcome Sanger Institute for the COVID-19 Genomics UK (COG-UK) Consortium | Harper VanSteenhouse, Yumi Kasai, David Gray, Carol Clugston, Anna Dominiczak and Alex Alderton, Roberto Amato, Sonia Goncalves, Ewan Harrison, David K. Jackson, Ian Johnston, Dominic Kwiatkowski, Cordelia Langford, John Sillitoe on behalf of the Wellcome Sanger Institute COVID-19 Surveillance Team |
| EPI_ISL_799723 | Lighthouse Lab in Milton Keynes | Wellcome Sanger Institute for the COVID-19 Genomics UK (COG-UK) Consortium | The Lighthouse Lab in Milton Keynes and Alex Alderton, Roberto Amato, Sonia Goncalves, Ewan Harrison, David K. Jackson, Ian Johnston, Dominic Kwiatkowski, Cordelia Langford, John Sillitoe on behalf of the Wellcome Sanger Institute COVID-19 Surveillance Team |
| EPI_ISL_799729, EPI_ISL_799730, EPI_ISL_799731 | Lighthouse Lab in Glasgow | Wellcome Sanger Institute for the COVID-19 Genomics UK (COG-UK) Consortium | Harper VanSteenhouse, Yumi Kasai, David Gray, Carol Clugston, Anna Dominiczak and Alex Alderton, Roberto Amato, Sonia Goncalves, Ewan Harrison, David K. Jackson, Ian Johnston, Dominic Kwiatkowski, Cordelia Langford, John Sillitoe on behalf of the Wellcome Sanger Institute COVID-19 Surveillance Team |
| EPI_ISL_799732 | Lighthouse Lab in Alderley Park | Wellcome Sanger Institute for the COVID-19 Genomics UK (COG-UK) Consortium | Jacquelyn Wynn, Mairead Hyland, The Lighthouse Lab in Alderley Park and Alex Alderton, Roberto Amato, Sonia Goncalves, Ewan Harrison, David K. Jackson, Ian Johnston, Dominic Kwiatkowski, Cordelia Langford, John Sillitoe on behalf of the Wellcome Sanger Institute COVID-19 Surveillance Team |
| EPI_ISL_799733 | Lighthouse Lab in Milton Keynes | Wellcome Sanger Institute for the COVID-19 Genomics UK (COG-UK) Consortium | The Lighthouse Lab in Milton Keynes and Alex Alderton, Roberto Amato, Sonia Goncalves, Ewan Harrison, David K. Jackson, Ian Johnston, Dominic Kwiatkowski, Cordelia Langford, John Sillitoe on behalf of the Wellcome Sanger Institute COVID-19 Surveillance Team |
| EPI_ISL_799735 | Lighthouse Lab in Glasgow | Wellcome Sanger Institute for the COVID-19 Genomics UK (COG-UK) Consortium | Harper VanSteenhouse, Yumi Kasai, David Gray, Carol Clugston, Anna Dominiczak and Alex Alderton, Roberto Amato, Sonia Goncalves, Ewan Harrison, David K. Jackson, Ian Johnston, Dominic Kwiatkowski, Cordelia Langford, John Sillitoe on behalf of the Wellcome Sanger Institute COVID-19 Surveillance Team |
| EPI_ISL_799736 | Lighthouse Lab in Milton Keynes | Wellcome Sanger Institute for the COVID-19 Genomics UK (COG-UK) Consortium | The Lighthouse Lab in Milton Keynes and Alex Alderton, Roberto Amato, Sonia Goncalves, Ewan Harrison, David K. Jackson, Ian Johnston, Dominic Kwiatkowski, Cordelia Langford, John Sillitoe on behalf of the Wellcome Sanger Institute COVID-19 Surveillance Team |
| EPI_ISL_799737, EPI_ISL_799739 | Lighthouse Lab in Glasgow | Wellcome Sanger Institute for the COVID-19 Genomics UK (COG-UK) Consortium | Harper VanSteenhouse, Yumi Kasai, David Gray, Carol Clugston, Anna Dominiczak and Alex Alderton, Roberto Amato, Sonia Goncalves, Ewan Harrison, David K. Jackson, Ian Johnston, Dominic Kwiatkowski, Cordelia Langford, John Sillitoe on behalf of the Wellcome Sanger Institute COVID-19 Surveillance Team |
| EPI_ISL_799741, EPI_ISL_799742, EPI_ISL_799745, EPI_ISL_799746 | Lighthouse Lab in Milton Keynes | Wellcome Sanger Institute for the COVID-19 Genomics UK (COG-UK) Consortium | The Lighthouse Lab in Milton Keynes and Alex Alderton, Roberto Amato, Sonia Goncalves, Ewan Harrison, David K. Jackson, Ian Johnston, Dominic Kwiatkowski, Cordelia Langford, John Sillitoe on behalf of the Wellcome Sanger Institute COVID-19 Surveillance Team |
| EPI_ISL_799751 | Lighthouse Lab in Alderley Park | Wellcome Sanger Institute for the COVID-19 Genomics UK (COG-UK) Consortium | Jacquelyn Wynn, Mairead Hyland, The Lighthouse Lab in Alderley Park and Alex Alderton, Roberto Amato, Sonia Goncalves, Ewan Harrison, David K. Jackson, Ian Johnston, Dominic Kwiatkowski, Cordelia Langford, John Sillitoe on behalf of the Wellcome Sanger Institute COVID-19 Surveillance Team |
| EPI_ISL_799753 | Lighthouse Lab in Milton Keynes | Wellcome Sanger Institute for the COVID-19 Genomics UK (COG-UK) Consortium | The Lighthouse Lab in Milton Keynes and Alex Alderton, Roberto Amato, Sonia Goncalves, Ewan Harrison, David K. Jackson, Ian Johnston, Dominic Kwiatkowski, Cordelia Langford, John Sillitoe on behalf of the Wellcome Sanger Institute COVID-19 Surveillance Team |
| EPI_ISL_799754 | Lighthouse Lab in Alderley Park | Wellcome Sanger Institute for the COVID-19 Genomics UK (COG-UK) Consortium | Jacquelyn Wynn, Mairead Hyland, The Lighthouse Lab in Alderley Park and Alex Alderton, Roberto Amato, Sonia Goncalves, Ewan Harrison, David K. Jackson, Ian Johnston, Dominic Kwiatkowski, Cordelia Langford, John Sillitoe on behalf of the Wellcome Sanger Institute COVID-19 Surveillance Team |
| EPI_ISL_799755, EPI_ISL_799756, EPI_ISL_799757 | Lighthouse Lab in Milton Keynes | Wellcome Sanger Institute for the COVID-19 Genomics UK (COG-UK) Consortium | The Lighthouse Lab in Milton Keynes and Alex Alderton, Roberto Amato, Sonia Goncalves, Ewan Harrison, David K. Jackson, Ian Johnston, Dominic Kwiatkowski, Cordelia Langford, John Sillitoe on behalf of the Wellcome Sanger Institute COVID-19 Surveillance Team |
| EPI_ISL_799759 | Lighthouse Lab in Glasgow | Wellcome Sanger Institute for the COVID-19 Genomics UK (COG-UK) Consortium | Harper VanSteenhouse, Yumi Kasai, David Gray, Carol Clugston, Anna Dominiczak and Alex Alderton, Roberto Amato, Sonia Goncalves, Ewan Harrison, David K. Jackson, Ian Johnston, Dominic Kwiatkowski, Cordelia Langford, John Sillitoe on behalf of the Wellcome Sanger Institute COVID-19 Surveillance Team |
| EPI_ISL_799760 | Lighthouse Lab in Alderley Park | Wellcome Sanger Institute for the COVID-19 Genomics UK (COG-UK) Consortium | Jacquelyn Wynn, Mairead Hyland, The Lighthouse Lab in Alderley Park and Alex Alderton, Roberto Amato, Sonia Goncalves, Ewan Harrison, David K. Jackson, Ian Johnston, Dominic Kwiatkowski, Cordelia Langford, John Sillitoe on behalf of the Wellcome Sanger Institute COVID-19 Surveillance Team |
| EPI_ISL_799761 | Lighthouse Lab in Glasgow | Wellcome Sanger Institute for the COVID-19 Genomics UK (COG-UK) Consortium | Harper VanSteenhouse, Yumi Kasai, David Gray, Carol Clugston, Anna Dominiczak and Alex Alderton, Roberto Amato, Sonia Goncalves, Ewan Harrison, David K. Jackson, Ian Johnston, Dominic Kwiatkowski, Cordelia Langford, John Sillitoe on behalf of the Wellcome Sanger Institute COVID-19 Surveillance Team |
| EPI_ISL_799762, EPI_ISL_799764, EPI_ISL_799771, EPI_ISL_799772, EPI_ISL_799774 | Lighthouse Lab in Milton Keynes | Wellcome Sanger Institute for the COVID-19 Genomics UK (COG-UK) Consortium | The Lighthouse Lab in Milton Keynes and Alex Alderton, Roberto Amato, Sonia Goncalves, Ewan Harrison, David K. Jackson, Ian Johnston, Dominic Kwiatkowski, Cordelia Langford, John Sillitoe on behalf of the Wellcome Sanger Institute COVID-19 Surveillance Team |
| EPI_ISL_799775, EPI_ISL_799776 | Lighthouse Lab in Glasgow | Wellcome Sanger Institute for the COVID-19 Genomics UK (COG-UK) Consortium | Harper VanSteenhouse, Yumi Kasai, David Gray, Carol Clugston, Anna Dominiczak and Alex Alderton, Roberto Amato, Sonia Goncalves, Ewan Harrison, David K. Jackson, Ian Johnston, Dominic Kwiatkowski, Cordelia Langford, John Sillitoe on behalf of the Wellcome Sanger Institute COVID-19 Surveillance Team |
| EPI_ISL_799779 | Lighthouse Lab in Milton Keynes | Wellcome Sanger Institute for the COVID-19 Genomics UK (COG-UK) Consortium | The Lighthouse Lab in Milton Keynes and Alex Alderton, Roberto Amato, Sonia Goncalves, Ewan Harrison, David K. Jackson, Ian Johnston, Dominic Kwiatkowski, Cordelia Langford, John Sillitoe on behalf of the Wellcome Sanger Institute COVID-19 Surveillance Team |
| EPI_ISL_799780, EPI_ISL_799781, EPI_ISL_799783, EPI_ISL_799786 | Lighthouse Lab in Glasgow | Wellcome Sanger Institute for the COVID-19 Genomics UK (COG-UK) Consortium | Harper VanSteenhouse, Yumi Kasai, David Gray, Carol Clugston, Anna Dominiczak and Alex Alderton, Roberto Amato, Sonia Goncalves, Ewan Harrison, David K. Jackson, Ian Johnston, Dominic Kwiatkowski, Cordelia Langford, John Sillitoe on behalf of the Wellcome Sanger Institute COVID-19 Surveillance |

| | | | |
|---|---|---|---|
| | | | Team |
| EPI_ISL_799790 | Lighthouse Lab in Milton Keynes | Wellcome Sanger Institute for the COVID-19 Genomics UK (COG-UK) Consortium | The Lighthouse Lab in Milton Keynes and Alex Alderton, Roberto Amato, Sonia Goncalves, Ewan Harrison, David K. Jackson, Ian Johnston, Dominic Kwiatkowski, Cordelia Langford, John Sillitoe on behalf of the Wellcome Sanger Institute COVID-19 Surveillance Team |
| EPI_ISL_799791, EPI_ISL_799792 | Lighthouse Lab in Glasgow | Wellcome Sanger Institute for the COVID-19 Genomics UK (COG-UK) Consortium | Harper VanSteenhouse, Yumi Kasai, David Gray, Carol Clugston, Anna Dominiczak and Alex Alderton, Roberto Amato, Sonia Goncalves, Ewan Harrison, David K. Jackson, Ian Johnston, Dominic Kwiatkowski, Cordelia Langford, John Sillitoe on behalf of the Wellcome Sanger Institute COVID-19 Surveillance Team |
| EPI_ISL_799793, EPI_ISL_799794 | Lighthouse Lab in Milton Keynes | Wellcome Sanger Institute for the COVID-19 Genomics UK (COG-UK) Consortium | The Lighthouse Lab in Milton Keynes and Alex Alderton, Roberto Amato, Sonia Goncalves, Ewan Harrison, David K. Jackson, Ian Johnston, Dominic Kwiatkowski, Cordelia Langford, John Sillitoe on behalf of the Wellcome Sanger Institute COVID-19 Surveillance Team |
| EPI_ISL_799797 | Lighthouse Lab in Alderley Park | Wellcome Sanger Institute for the COVID-19 Genomics UK (COG-UK) Consortium | Jacquelyn Wynn, Mairead Hyland, The Lighthouse Lab in Alderley Park and Alex Alderton, Roberto Amato, Sonia Goncalves, Ewan Harrison, David K. Jackson, Ian Johnston, Dominic Kwiatkowski, Cordelia Langford, John Sillitoe on behalf of the Wellcome Sanger Institute COVID-19 Surveillance Team |
| EPI_ISL_799798, EPI_ISL_799802, EPI_ISL_799803 | Lighthouse Lab in Milton Keynes | Wellcome Sanger Institute for the COVID-19 Genomics UK (COG-UK) Consortium | The Lighthouse Lab in Milton Keynes and Alex Alderton, Roberto Amato, Sonia Goncalves, Ewan Harrison, David K. Jackson, Ian Johnston, Dominic Kwiatkowski, Cordelia Langford, John Sillitoe on behalf of the Wellcome Sanger Institute COVID-19 Surveillance Team |
| EPI_ISL_799809 | Lighthouse Lab in Alderley Park | Wellcome Sanger Institute for the COVID-19 Genomics UK (COG-UK) Consortium | Jacquelyn Wynn, Mairead Hyland, The Lighthouse Lab in Alderley Park and Alex Alderton, Roberto Amato, Sonia Goncalves, Ewan Harrison, David K. Jackson, Ian Johnston, Dominic Kwiatkowski, Cordelia Langford, John Sillitoe on behalf of the Wellcome Sanger Institute COVID-19 Surveillance Team |
| EPI_ISL_799810 | Lighthouse Lab in Milton Keynes | Wellcome Sanger Institute for the COVID-19 Genomics UK (COG-UK) Consortium | The Lighthouse Lab in Milton Keynes and Alex Alderton, Roberto Amato, Sonia Goncalves, Ewan Harrison, David K. Jackson, Ian Johnston, Dominic Kwiatkowski, Cordelia Langford, John Sillitoe on behalf of the Wellcome Sanger Institute COVID-19 Surveillance Team |
| EPI_ISL_799811 | Lighthouse Lab in Glasgow | Wellcome Sanger Institute for the COVID-19 Genomics UK (COG-UK) Consortium | Harper VanSteenhouse, Yumi Kasai, David Gray, Carol Clugston, Anna Dominiczak and Alex Alderton, Roberto Amato, Sonia Goncalves, Ewan Harrison, David K. Jackson, Ian Johnston, Dominic Kwiatkowski, Cordelia Langford, John Sillitoe on behalf of the Wellcome Sanger Institute COVID-19 Surveillance Team |
| EPI_ISL_799813 | Lighthouse Lab in Milton Keynes | Wellcome Sanger Institute for the COVID-19 Genomics UK (COG-UK) Consortium | The Lighthouse Lab in Milton Keynes and Alex Alderton, Roberto Amato, Sonia Goncalves, Ewan Harrison, David K. Jackson, Ian Johnston, Dominic Kwiatkowski, Cordelia Langford, John Sillitoe on behalf of the Wellcome Sanger Institute COVID-19 Surveillance Team |
| EPI_ISL_799814 | Lighthouse Lab in Alderley Park | Wellcome Sanger Institute for the COVID-19 Genomics UK (COG-UK) Consortium | Jacquelyn Wynn, Mairead Hyland, The Lighthouse Lab in Alderley Park and Alex Alderton, Roberto Amato, Sonia Goncalves, Ewan Harrison, David K. Jackson, Ian Johnston, Dominic Kwiatkowski, Cordelia Langford, John Sillitoe on behalf of the Wellcome Sanger Institute COVID-19 Surveillance Team |
| EPI_ISL_799815 | Lighthouse Lab in Milton Keynes | Wellcome Sanger Institute for the COVID-19 Genomics UK (COG-UK) Consortium | The Lighthouse Lab in Milton Keynes and Alex Alderton, Roberto Amato, Sonia Goncalves, Ewan Harrison, David K. Jackson, Ian Johnston, Dominic Kwiatkowski, Cordelia Langford, John Sillitoe on behalf of the Wellcome Sanger Institute COVID-19 Surveillance Team |
| EPI_ISL_799825 | Lighthouse Lab in Glasgow | Wellcome Sanger Institute for the COVID-19 Genomics UK (COG-UK) Consortium | Harper VanSteenhouse, Yumi Kasai, David Gray, Carol Clugston, Anna Dominiczak and Alex Alderton, Roberto Amato, Sonia Goncalves, Ewan Harrison, David K. Jackson, Ian Johnston, Dominic Kwiatkowski, Cordelia Langford, John Sillitoe on behalf of the Wellcome Sanger Institute COVID-19 Surveillance Team |
| EPI_ISL_799826, EPI_ISL_799829 | Lighthouse Lab in Milton Keynes | Wellcome Sanger Institute for the COVID-19 Genomics UK (COG-UK) Consortium | The Lighthouse Lab in Milton Keynes and Alex Alderton, Roberto Amato, Sonia Goncalves, Ewan Harrison, David K. Jackson, Ian Johnston, Dominic Kwiatkowski, Cordelia Langford, John Sillitoe on behalf of the Wellcome Sanger Institute COVID-19 Surveillance Team |
| EPI_ISL_799830 | Lighthouse Lab in Glasgow | Wellcome Sanger Institute for the COVID-19 Genomics UK (COG-UK) Consortium | Harper VanSteenhouse, Yumi Kasai, David Gray, Carol Clugston, Anna Dominiczak and Alex Alderton, Roberto Amato, Sonia Goncalves, Ewan Harrison, David K. Jackson, Ian Johnston, Dominic Kwiatkowski, Cordelia Langford, John Sillitoe on behalf of the Wellcome Sanger Institute COVID-19 Surveillance Team |
| EPI_ISL_799833, EPI_ISL_799834, EPI_ISL_799837 | Lighthouse Lab in Milton Keynes | Wellcome Sanger Institute for the COVID-19 Genomics UK (COG-UK) Consortium | The Lighthouse Lab in Milton Keynes and Alex Alderton, Roberto Amato, Sonia Goncalves, Ewan Harrison, David K. Jackson, Ian Johnston, Dominic Kwiatkowski, Cordelia Langford, John Sillitoe on behalf of the Wellcome Sanger Institute COVID-19 Surveillance Team |
| EPI_ISL_799843 | Lighthouse Lab in Glasgow | Wellcome Sanger Institute for the COVID-19 Genomics UK (COG-UK) Consortium | Harper VanSteenhouse, Yumi Kasai, David Gray, Carol Clugston, Anna Dominiczak and Alex Alderton, Roberto Amato, Sonia Goncalves, Ewan Harrison, David K. Jackson, Ian Johnston, Dominic Kwiatkowski, Cordelia Langford, John Sillitoe on behalf of the Wellcome Sanger Institute COVID-19 Surveillance Team |
| EPI_ISL_799848, EPI_ISL_799850, EPI_ISL_799852, EPI_ISL_799853, EPI_ISL_799854, EPI_ISL_799856, EPI_ISL_799857 | Lighthouse Lab in Milton Keynes | Wellcome Sanger Institute for the COVID-19 Genomics UK (COG-UK) Consortium | The Lighthouse Lab in Milton Keynes and Alex Alderton, Roberto Amato, Sonia Goncalves, Ewan Harrison, David K. Jackson, Ian Johnston, Dominic Kwiatkowski, Cordelia Langford, John Sillitoe on behalf of the Wellcome Sanger Institute COVID-19 Surveillance Team |
| EPI_ISL_799861 | Lighthouse Lab in Glasgow | Wellcome Sanger Institute for the COVID-19 Genomics UK (COG-UK) Consortium | Harper VanSteenhouse, Yumi Kasai, David Gray, Carol Clugston, Anna Dominiczak and Alex Alderton, Roberto Amato, Sonia Goncalves, Ewan Harrison, David K. Jackson, Ian Johnston, Dominic Kwiatkowski, Cordelia Langford, John Sillitoe on behalf of the Wellcome Sanger Institute COVID-19 Surveillance Team |
| EPI_ISL_799862 | Lighthouse Lab in Milton Keynes | Wellcome Sanger Institute for the COVID-19 Genomics UK (COG-UK) Consortium | The Lighthouse Lab in Milton Keynes and Alex Alderton, Roberto Amato, Sonia Goncalves, Ewan Harrison, David K. Jackson, Ian Johnston, Dominic Kwiatkowski, Cordelia Langford, John Sillitoe on behalf of the Wellcome Sanger Institute COVID-19 Surveillance Team |
| EPI_ISL_799864, EPI_ISL_799866, EPI_ISL_799869 | Lighthouse Lab in Glasgow | Wellcome Sanger Institute for the COVID-19 Genomics UK (COG-UK) Consortium | Harper VanSteenhouse, Yumi Kasai, David Gray, Carol Clugston, Anna Dominiczak and Alex Alderton, Roberto Amato, Sonia Goncalves, Ewan Harrison, David K. Jackson, Ian Johnston, Dominic Kwiatkowski, Cordelia Langford, John Sillitoe on behalf of the Wellcome Sanger Institute COVID-19 Surveillance Team |
| EPI_ISL_799870 | Lighthouse Lab in Milton Keynes | Wellcome Sanger Institute for the COVID-19 Genomics UK (COG-UK) Consortium | The Lighthouse Lab in Milton Keynes and Alex Alderton, Roberto Amato, Sonia Goncalves, Ewan Harrison, David K. Jackson, Ian Johnston, Dominic Kwiatkowski, Cordelia Langford, John Sillitoe on behalf of the Wellcome Sanger Institute COVID-19 Surveillance Team |
| EPI_ISL_799871 | Lighthouse Lab in Glasgow | Wellcome Sanger Institute for the COVID-19 Genomics UK (COG-UK) Consortium | Harper VanSteenhouse, Yumi Kasai, David Gray, Carol Clugston, Anna Dominiczak and Alex Alderton, Roberto Amato, Sonia Goncalves, Ewan Harrison, David K. Jackson, Ian Johnston, Dominic Kwiatkowski, Cordelia Langford, John Sillitoe on behalf of the Wellcome Sanger Institute COVID-19 Surveillance Team |
| EPI_ISL_799873, EPI_ISL_799874 | Lighthouse Lab in Milton Keynes | Wellcome Sanger Institute for the COVID-19 Genomics UK (COG-UK) Consortium | The Lighthouse Lab in Milton Keynes and Alex Alderton, Roberto Amato, Sonia Goncalves, Ewan Harrison, David K. Jackson, Ian Johnston, Dominic Kwiatkowski, Cordelia Langford, John Sillitoe on behalf of the Wellcome Sanger Institute COVID-19 Surveillance Team |
| EPI_ISL_799875 | Lighthouse Lab in Glasgow | Wellcome Sanger Institute for the COVID-19 Genomics UK (COG-UK) Consortium | Harper VanSteenhouse, Yumi Kasai, David Gray, Carol Clugston, Anna Dominiczak and Alex Alderton, Roberto Amato, Sonia Goncalves, Ewan Harrison, David K. Jackson, Ian Johnston, Dominic Kwiatkowski, Cordelia Langford, John Sillitoe on behalf of the Wellcome Sanger Institute COVID-19 Surveillance Team |
| EPI_ISL_799879 | Lighthouse Lab in Milton Keynes | Wellcome Sanger Institute for the COVID-19 Genomics UK (COG-UK) Consortium | The Lighthouse Lab in Milton Keynes and Alex Alderton, Roberto Amato, Sonia Goncalves, Ewan Harrison, David K. Jackson, Ian Johnston, Dominic Kwiatkowski, Cordelia Langford, John Sillitoe on behalf of the Wellcome Sanger Institute COVID-19 Surveillance Team |
| EPI_ISL_799881 | Lighthouse Lab in Alderley Park | Wellcome Sanger Institute for the COVID-19 Genomics UK (COG-UK) Consortium | Jacquelyn Wynn, Mairead Hyland, The Lighthouse Lab in Alderley Park and Alex Alderton, Roberto Amato, Sonia Goncalves, Ewan Harrison, David K. Jackson, Ian Johnston, Dominic Kwiatkowski, Cordelia Langford, John Sillitoe on behalf of the Wellcome Sanger Institute COVID-19 Surveillance Team |
| EPI_ISL_799883 | Lighthouse Lab in Glasgow | Wellcome Sanger Institute for the COVID-19 Genomics UK (COG-UK) Consortium | Harper VanSteenhouse, Yumi Kasai, David Gray, Carol Clugston, Anna Dominiczak and Alex Alderton, Roberto Amato, Sonia Goncalves, Ewan Harrison, David K. Jackson, Ian Johnston, Dominic Kwiatkowski, Cordelia Langford, John Sillitoe on behalf of the Wellcome Sanger Institute COVID-19 Surveillance Team |
| EPI_ISL_799886 | Lighthouse Lab in Milton Keynes | Wellcome Sanger Institute for the COVID-19 Genomics UK (COG-UK) Consortium | The Lighthouse Lab in Milton Keynes and Alex Alderton, Roberto Amato, Sonia Goncalves, Ewan Harrison, David K. Jackson, Ian Johnston, Dominic Kwiatkowski, Cordelia Langford, John Sillitoe on behalf of the Wellcome Sanger Institute COVID-19 Surveillance Team |
| EPI_ISL_799889, EPI_ISL_799893 | Lighthouse Lab in Glasgow | Wellcome Sanger Institute for the COVID-19 Genomics UK (COG-UK) Consortium | Harper VanSteenhouse, Yumi Kasai, David Gray, Carol Clugston, Anna Dominiczak and Alex Alderton, Roberto Amato, Sonia Goncalves, Ewan Harrison, David K. Jackson, Ian Johnston, Dominic Kwiatkowski, Cordelia Langford, John Sillitoe on behalf of the Wellcome Sanger Institute COVID-19 Surveillance Team |
| EPI_ISL_799896 | Lighthouse Lab in Milton Keynes | Wellcome Sanger Institute for the COVID-19 Genomics UK | The Lighthouse Lab in Milton Keynes and Alex Alderton, Roberto Amato, Sonia Goncalves, Ewan Harrison, David K. Jackson, Ian Johnston, Dominic |

| | | | |
|---|---|---|---|
| | | (COG-UK) Consortium | Kwiatkowski, Cordelia Langford, John Sillitoe on behalf of the Wellcome Sanger Institute COVID-19 Surveillance Team |
| EPI_ISL_799897, EPI_ISL_799899 | Lighthouse Lab in Alderley Park | Wellcome Sanger Institute for the COVID-19 Genomics UK (COG-UK) Consortium | Jacquelyn Wynn, Mairead Hyland, The Lighthouse Lab in Alderley Park and Alex Alderton, Roberto Amato, Sonia Goncalves, Ewan Harrison, David K. Jackson, Ian Johnston, Dominic Kwiatkowski, Cordelia Langford, John Sillitoe on behalf of the Wellcome Sanger Institute COVID-19 Surveillance Team |
| EPI_ISL_799901 | Lighthouse Lab in Glasgow | Wellcome Sanger Institute for the COVID-19 Genomics UK (COG-UK) Consortium | Harper VanSteenhouse, Yumi Kasai, David Gray, Carol Clugston, Anna Dominiczak and Alex Alderton, Roberto Amato, Sonia Goncalves, Ewan Harrison, David K. Jackson, Ian Johnston, Dominic Kwiatkowski, Cordelia Langford, John Sillitoe on behalf of the Wellcome Sanger Institute COVID-19 Surveillance Team |
| EPI_ISL_799902 | Lighthouse Lab in Alderley Park | Wellcome Sanger Institute for the COVID-19 Genomics UK (COG-UK) Consortium | Jacquelyn Wynn, Mairead Hyland, The Lighthouse Lab in Alderley Park and Alex Alderton, Roberto Amato, Sonia Goncalves, Ewan Harrison, David K. Jackson, Ian Johnston, Dominic Kwiatkowski, Cordelia Langford, John Sillitoe on behalf of the Wellcome Sanger Institute COVID-19 Surveillance Team |
| EPI_ISL_799904, EPI_ISL_799907, EPI_ISL_799909 | Lighthouse Lab in Milton Keynes | Wellcome Sanger Institute for the COVID-19 Genomics UK (COG-UK) Consortium | The Lighthouse Lab in Milton Keynes and Alex Alderton, Roberto Amato, Sonia Goncalves, Ewan Harrison, David K. Jackson, Ian Johnston, Dominic Kwiatkowski, Cordelia Langford, John Sillitoe on behalf of the Wellcome Sanger Institute COVID-19 Surveillance Team |
| EPI_ISL_799910, EPI_ISL_799912 | Lighthouse Lab in Glasgow | Wellcome Sanger Institute for the COVID-19 Genomics UK (COG-UK) Consortium | Harper VanSteenhouse, Yumi Kasai, David Gray, Carol Clugston, Anna Dominiczak and Alex Alderton, Roberto Amato, Sonia Goncalves, Ewan Harrison, David K. Jackson, Ian Johnston, Dominic Kwiatkowski, Cordelia Langford, John Sillitoe on behalf of the Wellcome Sanger Institute COVID-19 Surveillance Team |
| EPI_ISL_799913 | Lighthouse Lab in Milton Keynes | Wellcome Sanger Institute for the COVID-19 Genomics UK (COG-UK) Consortium | The Lighthouse Lab in Milton Keynes and Alex Alderton, Roberto Amato, Sonia Goncalves, Ewan Harrison, David K. Jackson, Ian Johnston, Dominic Kwiatkowski, Cordelia Langford, John Sillitoe on behalf of the Wellcome Sanger Institute COVID-19 Surveillance Team |
| EPI_ISL_799914 | Lighthouse Lab in Alderley Park | Wellcome Sanger Institute for the COVID-19 Genomics UK (COG-UK) Consortium | Jacquelyn Wynn, Mairead Hyland, The Lighthouse Lab in Alderley Park and Alex Alderton, Roberto Amato, Sonia Goncalves, Ewan Harrison, David K. Jackson, Ian Johnston, Dominic Kwiatkowski, Cordelia Langford, John Sillitoe on behalf of the Wellcome Sanger Institute COVID-19 Surveillance Team |
| EPI_ISL_799916, EPI_ISL_799917, EPI_ISL_799918, EPI_ISL_799919 | Lighthouse Lab in Glasgow | Wellcome Sanger Institute for the COVID-19 Genomics UK (COG-UK) Consortium | Harper VanSteenhouse, Yumi Kasai, David Gray, Carol Clugston, Anna Dominiczak and Alex Alderton, Roberto Amato, Sonia Goncalves, Ewan Harrison, David K. Jackson, Ian Johnston, Dominic Kwiatkowski, Cordelia Langford, John Sillitoe on behalf of the Wellcome Sanger Institute COVID-19 Surveillance Team |
| EPI_ISL_799921, EPI_ISL_799926 | Lighthouse Lab in Milton Keynes | Wellcome Sanger Institute for the COVID-19 Genomics UK (COG-UK) Consortium | The Lighthouse Lab in Milton Keynes and Alex Alderton, Roberto Amato, Sonia Goncalves, Ewan Harrison, David K. Jackson, Ian Johnston, Dominic Kwiatkowski, Cordelia Langford, John Sillitoe on behalf of the Wellcome Sanger Institute COVID-19 Surveillance Team |
| EPI_ISL_799935 | Lighthouse Lab in Cambridge | Wellcome Sanger Institute for the COVID-19 Genomics UK (COG-UK) Consortium | Rob Howes, The Lighthouse Lab in Cambridge and Alex Alderton, Roberto Amato, Sonia Goncalves, Ewan Harrison, David K. Jackson, Ian Johnston, Dominic Kwiatkowski, Cordelia Langford, John Sillitoe on behalf of the Wellcome Sanger Institute COVID-19 Surveillance Team |
| EPI_ISL_799938 | Lighthouse Lab in Alderley Park | Wellcome Sanger Institute for the COVID-19 Genomics UK (COG-UK) Consortium | Jacquelyn Wynn, Mairead Hyland, The Lighthouse Lab in Alderley Park and Alex Alderton, Roberto Amato, Sonia Goncalves, Ewan Harrison, David K. Jackson, Ian Johnston, Dominic Kwiatkowski, Cordelia Langford, John Sillitoe on behalf of the Wellcome Sanger Institute COVID-19 Surveillance Team |
| EPI_ISL_799939 | Lighthouse Lab in Cambridge | Wellcome Sanger Institute for the COVID-19 Genomics UK (COG-UK) Consortium | Rob Howes, The Lighthouse Lab in Cambridge and Alex Alderton, Roberto Amato, Sonia Goncalves, Ewan Harrison, David K. Jackson, Ian Johnston, Dominic Kwiatkowski, Cordelia Langford, John Sillitoe on behalf of the Wellcome Sanger Institute COVID-19 Surveillance Team |
| EPI_ISL_799942 | Lighthouse Lab in Alderley Park | Wellcome Sanger Institute for the COVID-19 Genomics UK (COG-UK) Consortium | Jacquelyn Wynn, Mairead Hyland, The Lighthouse Lab in Alderley Park and Alex Alderton, Roberto Amato, Sonia Goncalves, Ewan Harrison, David K. Jackson, Ian Johnston, Dominic Kwiatkowski, Cordelia Langford, John Sillitoe on behalf of the Wellcome Sanger Institute COVID-19 Surveillance Team |
| EPI_ISL_799944 | Lighthouse Lab in Cambridge | Wellcome Sanger Institute for the COVID-19 Genomics UK (COG-UK) Consortium | Rob Howes, The Lighthouse Lab in Cambridge and Alex Alderton, Roberto Amato, Sonia Goncalves, Ewan Harrison, David K. Jackson, Ian Johnston, Dominic Kwiatkowski, Cordelia Langford, John Sillitoe on behalf of the Wellcome Sanger Institute COVID-19 Surveillance Team |
| EPI_ISL_799945 | Lighthouse Lab in Glasgow | Wellcome Sanger Institute for the COVID-19 Genomics UK (COG-UK) Consortium | Harper VanSteenhouse, Yumi Kasai, David Gray, Carol Clugston, Anna Dominiczak and Alex Alderton, Roberto Amato, Sonia Goncalves, Ewan Harrison, David K. Jackson, Ian Johnston, Dominic Kwiatkowski, Cordelia Langford, John Sillitoe on behalf of the Wellcome Sanger Institute COVID-19 Surveillance Team |
| EPI_ISL_799947, EPI_ISL_799950 | Lighthouse Lab in Cambridge | Wellcome Sanger Institute for the COVID-19 Genomics UK (COG-UK) Consortium | Rob Howes, The Lighthouse Lab in Cambridge and Alex Alderton, Roberto Amato, Sonia Goncalves, Ewan Harrison, David K. Jackson, Ian Johnston, Dominic Kwiatkowski, Cordelia Langford, John Sillitoe on behalf of the Wellcome Sanger Institute COVID-19 Surveillance Team |
| EPI_ISL_799951 | Lighthouse Lab in Glasgow | Wellcome Sanger Institute for the COVID-19 Genomics UK (COG-UK) Consortium | Harper VanSteenhouse, Yumi Kasai, David Gray, Carol Clugston, Anna Dominiczak and Alex Alderton, Roberto Amato, Sonia Goncalves, Ewan Harrison, David K. Jackson, Ian Johnston, Dominic Kwiatkowski, Cordelia Langford, John Sillitoe on behalf of the Wellcome Sanger Institute COVID-19 Surveillance Team |
| EPI_ISL_799952, EPI_ISL_799953, EPI_ISL_799955, EPI_ISL_799957, EPI_ISL_799958, EPI_ISL_799959, EPI_ISL_799960, EPI_ISL_799962, EPI_ISL_799963 | Lighthouse Lab in Cambridge | Wellcome Sanger Institute for the COVID-19 Genomics UK (COG-UK) Consortium | Rob Howes, The Lighthouse Lab in Cambridge and Alex Alderton, Roberto Amato, Sonia Goncalves, Ewan Harrison, David K. Jackson, Ian Johnston, Dominic Kwiatkowski, Cordelia Langford, John Sillitoe on behalf of the Wellcome Sanger Institute COVID-19 Surveillance Team |
| EPI_ISL_799966 | Lighthouse Lab in Alderley Park | Wellcome Sanger Institute for the COVID-19 Genomics UK (COG-UK) Consortium | Jacquelyn Wynn, Mairead Hyland, The Lighthouse Lab in Alderley Park and Alex Alderton, Roberto Amato, Sonia Goncalves, Ewan Harrison, David K. Jackson, Ian Johnston, Dominic Kwiatkowski, Cordelia Langford, John Sillitoe on behalf of the Wellcome Sanger Institute COVID-19 Surveillance Team |
| EPI_ISL_799973 | Lighthouse Lab in Glasgow | Wellcome Sanger Institute for the COVID-19 Genomics UK (COG-UK) Consortium | Harper VanSteenhouse, Yumi Kasai, David Gray, Carol Clugston, Anna Dominiczak and Alex Alderton, Roberto Amato, Sonia Goncalves, Ewan Harrison, David K. Jackson, Ian Johnston, Dominic Kwiatkowski, Cordelia Langford, John Sillitoe on behalf of the Wellcome Sanger Institute COVID-19 Surveillance Team |
| EPI_ISL_799974 | Lighthouse Lab in Cambridge | Wellcome Sanger Institute for the COVID-19 Genomics UK (COG-UK) Consortium | Rob Howes, The Lighthouse Lab in Cambridge and Alex Alderton, Roberto Amato, Sonia Goncalves, Ewan Harrison, David K. Jackson, Ian Johnston, Dominic Kwiatkowski, Cordelia Langford, John Sillitoe on behalf of the Wellcome Sanger Institute COVID-19 Surveillance Team |
| EPI_ISL_799976 | Lighthouse Lab in Glasgow | Wellcome Sanger Institute for the COVID-19 Genomics UK (COG-UK) Consortium | Harper VanSteenhouse, Yumi Kasai, David Gray, Carol Clugston, Anna Dominiczak and Alex Alderton, Roberto Amato, Sonia Goncalves, Ewan Harrison, David K. Jackson, Ian Johnston, Dominic Kwiatkowski, Cordelia Langford, John Sillitoe on behalf of the Wellcome Sanger Institute COVID-19 Surveillance Team |
| EPI_ISL_799978 | Lighthouse Lab in Alderley Park | Wellcome Sanger Institute for the COVID-19 Genomics UK (COG-UK) Consortium | Jacquelyn Wynn, Mairead Hyland, The Lighthouse Lab in Alderley Park and Alex Alderton, Roberto Amato, Sonia Goncalves, Ewan Harrison, David K. Jackson, Ian Johnston, Dominic Kwiatkowski, Cordelia Langford, John Sillitoe on behalf of the Wellcome Sanger Institute COVID-19 Surveillance Team |
| EPI_ISL_799981 | Lighthouse Lab in Glasgow | Wellcome Sanger Institute for the COVID-19 Genomics UK (COG-UK) Consortium | Harper VanSteenhouse, Yumi Kasai, David Gray, Carol Clugston, Anna Dominiczak and Alex Alderton, Roberto Amato, Sonia Goncalves, Ewan Harrison, David K. Jackson, Ian Johnston, Dominic Kwiatkowski, Cordelia Langford, John Sillitoe on behalf of the Wellcome Sanger Institute COVID-19 Surveillance Team |
| EPI_ISL_799982, EPI_ISL_799983 | Lighthouse Lab in Alderley Park | Wellcome Sanger Institute for the COVID-19 Genomics UK (COG-UK) Consortium | Jacquelyn Wynn, Mairead Hyland, The Lighthouse Lab in Alderley Park and Alex Alderton, Roberto Amato, Sonia Goncalves, Ewan Harrison, David K. Jackson, Ian Johnston, Dominic Kwiatkowski, Cordelia Langford, John Sillitoe on behalf of the Wellcome Sanger Institute COVID-19 Surveillance Team |
| EPI_ISL_799985 | Lighthouse Lab in Glasgow | Wellcome Sanger Institute for the COVID-19 Genomics UK (COG-UK) Consortium | Harper VanSteenhouse, Yumi Kasai, David Gray, Carol Clugston, Anna Dominiczak and Alex Alderton, Roberto Amato, Sonia Goncalves, Ewan Harrison, David K. Jackson, Ian Johnston, Dominic Kwiatkowski, Cordelia Langford, John Sillitoe on behalf of the Wellcome Sanger Institute COVID-19 Surveillance Team |
| EPI_ISL_799988 | Lighthouse Lab in Cambridge | Wellcome Sanger Institute for the COVID-19 Genomics UK (COG-UK) Consortium | Rob Howes, The Lighthouse Lab in Cambridge and Alex Alderton, Roberto Amato, Sonia Goncalves, Ewan Harrison, David K. Jackson, Ian Johnston, Dominic Kwiatkowski, Cordelia Langford, John Sillitoe on behalf of the Wellcome Sanger Institute COVID-19 Surveillance Team |
| EPI_ISL_799989 | Lighthouse Lab in Alderley Park | Wellcome Sanger Institute for the COVID-19 Genomics UK (COG-UK) Consortium | Jacquelyn Wynn, Mairead Hyland, The Lighthouse Lab in Alderley Park and Alex Alderton, Roberto Amato, Sonia Goncalves, Ewan Harrison, David K. Jackson, Ian Johnston, Dominic Kwiatkowski, Cordelia Langford, John Sillitoe on behalf of the Wellcome Sanger Institute COVID-19 Surveillance Team |
| EPI_ISL_799990 | Lighthouse Lab in Glasgow | Wellcome Sanger Institute for the COVID-19 Genomics UK (COG-UK) Consortium | Harper VanSteenhouse, Yumi Kasai, David Gray, Carol Clugston, Anna Dominiczak and Alex Alderton, Roberto Amato, Sonia Goncalves, Ewan Harrison, David K. Jackson, Ian Johnston, Dominic Kwiatkowski, Cordelia Langford, John Sillitoe on behalf of the Wellcome Sanger Institute COVID-19 Surveillance Team |
| EPI_ISL_799991 | Lighthouse Lab in Alderley Park | Wellcome Sanger Institute for the COVID-19 Genomics UK | Jacquelyn Wynn, Mairead Hyland, The Lighthouse Lab in Alderley Park and Alex Alderton, Roberto Amato, Sonia Goncalves, Ewan Harrison, David K. |

| | | | |
|---|---|---|---|
| | | (COG-UK) Consortium | Jackson, Ian Johnston, Dominic Kwiatkowski, Cordelia Langford, John Sillitoe on behalf of the Wellcome Sanger Institute COVID-19 Surveillance Team |
| EPI_ISL_799996 | Lighthouse Lab in Cambridge | Wellcome Sanger Institute for the COVID-19 Genomics UK (COG-UK) Consortium | Rob Howes, The Lighthouse Lab in Cambridge and Alex Alderton, Roberto Amato, Sonia Goncalves, Ewan Harrison, David K. Jackson, Ian Johnston, Dominic Kwiatkowski, Cordelia Langford, John Sillitoe on behalf of the Wellcome Sanger Institute COVID-19 Surveillance Team |
| EPI_ISL_799997 | Lighthouse Lab in Glasgow | Wellcome Sanger Institute for the COVID-19 Genomics UK (COG-UK) Consortium | Harper VanSteenhouse, Yumi Kasai, David Gray, Carol Clugston, Anna Dominiczak and Alex Alderton, Roberto Amato, Sonia Goncalves, Ewan Harrison, David K. Jackson, Ian Johnston, Dominic Kwiatkowski, Cordelia Langford, John Sillitoe on behalf of the Wellcome Sanger Institute COVID-19 Surveillance Team |
| EPI_ISL_799998 | Lighthouse Lab in Alderley Park | Wellcome Sanger Institute for the COVID-19 Genomics UK (COG-UK) Consortium | Jacquelyn Wynn, Mairead Hyland, The Lighthouse Lab in Alderley Park and Alex Alderton, Roberto Amato, Sonia Goncalves, Ewan Harrison, David K. Jackson, Ian Johnston, Dominic Kwiatkowski, Cordelia Langford, John Sillitoe on behalf of the Wellcome Sanger Institute COVID-19 Surveillance Team |
| EPI_ISL_800001 | Lighthouse Lab in Glasgow | Wellcome Sanger Institute for the COVID-19 Genomics UK (COG-UK) Consortium | Harper VanSteenhouse, Yumi Kasai, David Gray, Carol Clugston, Anna Dominiczak and Alex Alderton, Roberto Amato, Sonia Goncalves, Ewan Harrison, David K. Jackson, Ian Johnston, Dominic Kwiatkowski, Cordelia Langford, John Sillitoe on behalf of the Wellcome Sanger Institute COVID-19 Surveillance Team |
| EPI_ISL_800004 | Lighthouse Lab in Cambridge | Wellcome Sanger Institute for the COVID-19 Genomics UK (COG-UK) Consortium | Rob Howes, The Lighthouse Lab in Cambridge and Alex Alderton, Roberto Amato, Sonia Goncalves, Ewan Harrison, David K. Jackson, Ian Johnston, Dominic Kwiatkowski, Cordelia Langford, John Sillitoe on behalf of the Wellcome Sanger Institute COVID-19 Surveillance Team |
| EPI_ISL_800005 | Lighthouse Lab in Glasgow | Wellcome Sanger Institute for the COVID-19 Genomics UK (COG-UK) Consortium | Harper VanSteenhouse, Yumi Kasai, David Gray, Carol Clugston, Anna Dominiczak and Alex Alderton, Roberto Amato, Sonia Goncalves, Ewan Harrison, David K. Jackson, Ian Johnston, Dominic Kwiatkowski, Cordelia Langford, John Sillitoe on behalf of the Wellcome Sanger Institute COVID-19 Surveillance Team |
| EPI_ISL_800006, EPI_ISL_800007, EPI_ISL_800008 | Lighthouse Lab in Cambridge | Wellcome Sanger Institute for the COVID-19 Genomics UK (COG-UK) Consortium | Rob Howes, The Lighthouse Lab in Cambridge and Alex Alderton, Roberto Amato, Sonia Goncalves, Ewan Harrison, David K. Jackson, Ian Johnston, Dominic Kwiatkowski, Cordelia Langford, John Sillitoe on behalf of the Wellcome Sanger Institute COVID-19 Surveillance Team |
| EPI_ISL_800010 | Lighthouse Lab in Glasgow | Wellcome Sanger Institute for the COVID-19 Genomics UK (COG-UK) Consortium | Harper VanSteenhouse, Yumi Kasai, David Gray, Carol Clugston, Anna Dominiczak and Alex Alderton, Roberto Amato, Sonia Goncalves, Ewan Harrison, David K. Jackson, Ian Johnston, Dominic Kwiatkowski, Cordelia Langford, John Sillitoe on behalf of the Wellcome Sanger Institute COVID-19 Surveillance Team |
| EPI_ISL_800012 | Lighthouse Lab in Cambridge | Wellcome Sanger Institute for the COVID-19 Genomics UK (COG-UK) Consortium | Rob Howes, The Lighthouse Lab in Cambridge and Alex Alderton, Roberto Amato, Sonia Goncalves, Ewan Harrison, David K. Jackson, Ian Johnston, Dominic Kwiatkowski, Cordelia Langford, John Sillitoe on behalf of the Wellcome Sanger Institute COVID-19 Surveillance Team |
| EPI_ISL_800017 | Lighthouse Lab in Alderley Park | Wellcome Sanger Institute for the COVID-19 Genomics UK (COG-UK) Consortium | Jacquelyn Wynn, Mairead Hyland, The Lighthouse Lab in Alderley Park and Alex Alderton, Roberto Amato, Sonia Goncalves, Ewan Harrison, David K. Jackson, Ian Johnston, Dominic Kwiatkowski, Cordelia Langford, John Sillitoe on behalf of the Wellcome Sanger Institute COVID-19 Surveillance Team |
| EPI_ISL_800018 | Lighthouse Lab in Cambridge | Wellcome Sanger Institute for the COVID-19 Genomics UK (COG-UK) Consortium | Rob Howes, The Lighthouse Lab in Cambridge and Alex Alderton, Roberto Amato, Sonia Goncalves, Ewan Harrison, David K. Jackson, Ian Johnston, Dominic Kwiatkowski, Cordelia Langford, John Sillitoe on behalf of the Wellcome Sanger Institute COVID-19 Surveillance Team |
| EPI_ISL_800022 | Lighthouse Lab in Alderley Park | Wellcome Sanger Institute for the COVID-19 Genomics UK (COG-UK) Consortium | Jacquelyn Wynn, Mairead Hyland, The Lighthouse Lab in Alderley Park and Alex Alderton, Roberto Amato, Sonia Goncalves, Ewan Harrison, David K. Jackson, Ian Johnston, Dominic Kwiatkowski, Cordelia Langford, John Sillitoe on behalf of the Wellcome Sanger Institute COVID-19 Surveillance Team |
| EPI_ISL_800024 | Lighthouse Lab in Cambridge | Wellcome Sanger Institute for the COVID-19 Genomics UK (COG-UK) Consortium | Rob Howes, The Lighthouse Lab in Cambridge and Alex Alderton, Roberto Amato, Sonia Goncalves, Ewan Harrison, David K. Jackson, Ian Johnston, Dominic Kwiatkowski, Cordelia Langford, John Sillitoe on behalf of the Wellcome Sanger Institute COVID-19 Surveillance Team |
| EPI_ISL_800025 | Lighthouse Lab in Alderley Park | Wellcome Sanger Institute for the COVID-19 Genomics UK (COG-UK) Consortium | Jacquelyn Wynn, Mairead Hyland, The Lighthouse Lab in Alderley Park and Alex Alderton, Roberto Amato, Sonia Goncalves, Ewan Harrison, David K. Jackson, Ian Johnston, Dominic Kwiatkowski, Cordelia Langford, John Sillitoe on behalf of the Wellcome Sanger Institute COVID-19 Surveillance Team |
| EPI_ISL_800026 | Lighthouse Lab in Cambridge | Wellcome Sanger Institute for the COVID-19 Genomics UK (COG-UK) Consortium | Rob Howes, The Lighthouse Lab in Cambridge and Alex Alderton, Roberto Amato, Sonia Goncalves, Ewan Harrison, David K. Jackson, Ian Johnston, Dominic Kwiatkowski, Cordelia Langford, John Sillitoe on behalf of the Wellcome Sanger Institute COVID-19 Surveillance Team |
| EPI_ISL_800027 | Lighthouse Lab in Glasgow | Wellcome Sanger Institute for the COVID-19 Genomics UK (COG-UK) Consortium | Harper VanSteenhouse, Yumi Kasai, David Gray, Carol Clugston, Anna Dominiczak and Alex Alderton, Roberto Amato, Sonia Goncalves, Ewan Harrison, David K. Jackson, Ian Johnston, Dominic Kwiatkowski, Cordelia Langford, John Sillitoe on behalf of the Wellcome Sanger Institute COVID-19 Surveillance Team |
| EPI_ISL_800028 | Lighthouse Lab in Cambridge | Wellcome Sanger Institute for the COVID-19 Genomics UK (COG-UK) Consortium | Rob Howes, The Lighthouse Lab in Cambridge and Alex Alderton, Roberto Amato, Sonia Goncalves, Ewan Harrison, David K. Jackson, Ian Johnston, Dominic Kwiatkowski, Cordelia Langford, John Sillitoe on behalf of the Wellcome Sanger Institute COVID-19 Surveillance Team |
| EPI_ISL_800029, EPI_ISL_800030, EPI_ISL_800034 | Lighthouse Lab in Glasgow | Wellcome Sanger Institute for the COVID-19 Genomics UK (COG-UK) Consortium | Harper VanSteenhouse, Yumi Kasai, David Gray, Carol Clugston, Anna Dominiczak and Alex Alderton, Roberto Amato, Sonia Goncalves, Ewan Harrison, David K. Jackson, Ian Johnston, Dominic Kwiatkowski, Cordelia Langford, John Sillitoe on behalf of the Wellcome Sanger Institute COVID-19 Surveillance Team |
| EPI_ISL_800036 | Lighthouse Lab in Alderley Park | Wellcome Sanger Institute for the COVID-19 Genomics UK (COG-UK) Consortium | Jacquelyn Wynn, Mairead Hyland, The Lighthouse Lab in Alderley Park and Alex Alderton, Roberto Amato, Sonia Goncalves, Ewan Harrison, David K. Jackson, Ian Johnston, Dominic Kwiatkowski, Cordelia Langford, John Sillitoe on behalf of the Wellcome Sanger Institute COVID-19 Surveillance Team |
| EPI_ISL_800037 | Lighthouse Lab in Cambridge | Wellcome Sanger Institute for the COVID-19 Genomics UK (COG-UK) Consortium | Rob Howes, The Lighthouse Lab in Cambridge and Alex Alderton, Roberto Amato, Sonia Goncalves, Ewan Harrison, David K. Jackson, Ian Johnston, Dominic Kwiatkowski, Cordelia Langford, John Sillitoe on behalf of the Wellcome Sanger Institute COVID-19 Surveillance Team |
| EPI_ISL_800038 | Lighthouse Lab in Glasgow | Wellcome Sanger Institute for the COVID-19 Genomics UK (COG-UK) Consortium | Harper VanSteenhouse, Yumi Kasai, David Gray, Carol Clugston, Anna Dominiczak and Alex Alderton, Roberto Amato, Sonia Goncalves, Ewan Harrison, David K. Jackson, Ian Johnston, Dominic Kwiatkowski, Cordelia Langford, John Sillitoe on behalf of the Wellcome Sanger Institute COVID-19 Surveillance Team |
| EPI_ISL_800039, EPI_ISL_800040, EPI_ISL_800043, EPI_ISL_800044 | Lighthouse Lab in Alderley Park | Wellcome Sanger Institute for the COVID-19 Genomics UK (COG-UK) Consortium | Jacquelyn Wynn, Mairead Hyland, The Lighthouse Lab in Alderley Park and Alex Alderton, Roberto Amato, Sonia Goncalves, Ewan Harrison, David K. Jackson, Ian Johnston, Dominic Kwiatkowski, Cordelia Langford, John Sillitoe on behalf of the Wellcome Sanger Institute COVID-19 Surveillance Team |
| EPI_ISL_800045, EPI_ISL_800047 | Lighthouse Lab in Cambridge | Wellcome Sanger Institute for the COVID-19 Genomics UK (COG-UK) Consortium | Rob Howes, The Lighthouse Lab in Cambridge and Alex Alderton, Roberto Amato, Sonia Goncalves, Ewan Harrison, David K. Jackson, Ian Johnston, Dominic Kwiatkowski, Cordelia Langford, John Sillitoe on behalf of the Wellcome Sanger Institute COVID-19 Surveillance Team |
| EPI_ISL_800050, EPI_ISL_800051 | Lighthouse Lab in Alderley Park | Wellcome Sanger Institute for the COVID-19 Genomics UK (COG-UK) Consortium | Jacquelyn Wynn, Mairead Hyland, The Lighthouse Lab in Alderley Park and Alex Alderton, Roberto Amato, Sonia Goncalves, Ewan Harrison, David K. Jackson, Ian Johnston, Dominic Kwiatkowski, Cordelia Langford, John Sillitoe on behalf of the Wellcome Sanger Institute COVID-19 Surveillance Team |
| EPI_ISL_800055 | Lighthouse Lab in Cambridge | Wellcome Sanger Institute for the COVID-19 Genomics UK (COG-UK) Consortium | Rob Howes, The Lighthouse Lab in Cambridge and Alex Alderton, Roberto Amato, Sonia Goncalves, Ewan Harrison, David K. Jackson, Ian Johnston, Dominic Kwiatkowski, Cordelia Langford, John Sillitoe on behalf of the Wellcome Sanger Institute COVID-19 Surveillance Team |
| EPI_ISL_800056 | Lighthouse Lab in Glasgow | Wellcome Sanger Institute for the COVID-19 Genomics UK (COG-UK) Consortium | Harper VanSteenhouse, Yumi Kasai, David Gray, Carol Clugston, Anna Dominiczak and Alex Alderton, Roberto Amato, Sonia Goncalves, Ewan Harrison, David K. Jackson, Ian Johnston, Dominic Kwiatkowski, Cordelia Langford, John Sillitoe on behalf of the Wellcome Sanger Institute COVID-19 Surveillance Team |
| EPI_ISL_800059, EPI_ISL_800061 | Lighthouse Lab in Alderley Park | Wellcome Sanger Institute for the COVID-19 Genomics UK (COG-UK) Consortium | Jacquelyn Wynn, Mairead Hyland, The Lighthouse Lab in Alderley Park and Alex Alderton, Roberto Amato, Sonia Goncalves, Ewan Harrison, David K. Jackson, Ian Johnston, Dominic Kwiatkowski, Cordelia Langford, John Sillitoe on behalf of the Wellcome Sanger Institute COVID-19 Surveillance Team |
| EPI_ISL_800065 | Lighthouse Lab in Cambridge | Wellcome Sanger Institute for the COVID-19 Genomics UK (COG-UK) Consortium | Rob Howes, The Lighthouse Lab in Cambridge and Alex Alderton, Roberto Amato, Sonia Goncalves, Ewan Harrison, David K. Jackson, Ian Johnston, Dominic Kwiatkowski, Cordelia Langford, John Sillitoe on behalf of the Wellcome Sanger Institute COVID-19 Surveillance Team |
| EPI_ISL_800067 | Lighthouse Lab in Alderley Park | Wellcome Sanger Institute for the COVID-19 Genomics UK (COG-UK) Consortium | Jacquelyn Wynn, Mairead Hyland, The Lighthouse Lab in Alderley Park and Alex Alderton, Roberto Amato, Sonia Goncalves, Ewan Harrison, David K. Jackson, Ian Johnston, Dominic Kwiatkowski, Cordelia Langford, John Sillitoe on behalf of the Wellcome Sanger Institute COVID-19 Surveillance Team |
| EPI_ISL_800068, EPI_ISL_800070 | Lighthouse Lab in Glasgow | Wellcome Sanger Institute for the COVID-19 Genomics UK (COG-UK) Consortium | Harper VanSteenhouse, Yumi Kasai, David Gray, Carol Clugston, Anna Dominiczak and Alex Alderton, Roberto Amato, Sonia Goncalves, Ewan Harrison, David K. Jackson, Ian Johnston, Dominic Kwiatkowski, Cordelia Langford, John Sillitoe on behalf of the Wellcome Sanger Institute COVID-19 Surveillance Team |
| EPI_ISL_800072 | Lighthouse Lab in Cambridge | Wellcome Sanger Institute for the COVID-19 Genomics UK (COG-UK) Consortium | Rob Howes, The Lighthouse Lab in Cambridge and Alex Alderton, Roberto Amato, Sonia Goncalves, Ewan Harrison, David K. Jackson, Ian Johnston, Dominic Kwiatkowski, Cordelia Langford, John Sillitoe on behalf of the Wellcome Sanger Institute COVID-19 Surveillance Team |

| | | | |
|---|---|---|---|
| EPI_ISL_800074 | Lighthouse Lab in Alderley Park | Wellcome Sanger Institute for the COVID-19 Genomics UK (COG-UK) Consortium | Jacquelyn Wynn, Mairead Hyland, The Lighthouse Lab in Alderley Park and Alex Alderton, Roberto Amato, Sonia Goncalves, Ewan Harrison, David K. Jackson, Ian Johnston, Dominic Kwiatkowski, Cordelia Langford, John Sillitoe on behalf of the Wellcome Sanger Institute COVID-19 Surveillance Team |
| EPI_ISL_800077 | Lighthouse Lab in Cambridge | Wellcome Sanger Institute for the COVID-19 Genomics UK (COG-UK) Consortium | Rob Howes, The Lighthouse Lab in Cambridge and Alex Alderton, Roberto Amato, Sonia Goncalves, Ewan Harrison, David K. Jackson, Ian Johnston, Dominic Kwiatkowski, Cordelia Langford, John Sillitoe on behalf of the Wellcome Sanger Institute COVID-19 Surveillance Team |
| EPI_ISL_800078, EPI_ISL_800079 | Lighthouse Lab in Glasgow | Wellcome Sanger Institute for the COVID-19 Genomics UK (COG-UK) Consortium | Harper VanSteenhouse, Yumi Kasai, David Gray, Carol Clugston, Anna Dominiczak and Alex Alderton, Roberto Amato, Sonia Goncalves, Ewan Harrison, David K. Jackson, Ian Johnston, Dominic Kwiatkowski, Cordelia Langford, John Sillitoe on behalf of the Wellcome Sanger Institute COVID-19 Surveillance Team |
| EPI_ISL_800082, EPI_ISL_800083 | Lighthouse Lab in Cambridge | Wellcome Sanger Institute for the COVID-19 Genomics UK (COG-UK) Consortium | Rob Howes, The Lighthouse Lab in Cambridge and Alex Alderton, Roberto Amato, Sonia Goncalves, Ewan Harrison, David K. Jackson, Ian Johnston, Dominic Kwiatkowski, Cordelia Langford, John Sillitoe on behalf of the Wellcome Sanger Institute COVID-19 Surveillance Team |
| EPI_ISL_800084 | Lighthouse Lab in Glasgow | Wellcome Sanger Institute for the COVID-19 Genomics UK (COG-UK) Consortium | Harper VanSteenhouse, Yumi Kasai, David Gray, Carol Clugston, Anna Dominiczak and Alex Alderton, Roberto Amato, Sonia Goncalves, Ewan Harrison, David K. Jackson, Ian Johnston, Dominic Kwiatkowski, Cordelia Langford, John Sillitoe on behalf of the Wellcome Sanger Institute COVID-19 Surveillance Team |
| EPI_ISL_800085 | Lighthouse Lab in Cambridge | Wellcome Sanger Institute for the COVID-19 Genomics UK (COG-UK) Consortium | Rob Howes, The Lighthouse Lab in Cambridge and Alex Alderton, Roberto Amato, Sonia Goncalves, Ewan Harrison, David K. Jackson, Ian Johnston, Dominic Kwiatkowski, Cordelia Langford, John Sillitoe on behalf of the Wellcome Sanger Institute COVID-19 Surveillance Team |
| EPI_ISL_800086 | Lighthouse Lab in Glasgow | Wellcome Sanger Institute for the COVID-19 Genomics UK (COG-UK) Consortium | Harper VanSteenhouse, Yumi Kasai, David Gray, Carol Clugston, Anna Dominiczak and Alex Alderton, Roberto Amato, Sonia Goncalves, Ewan Harrison, David K. Jackson, Ian Johnston, Dominic Kwiatkowski, Cordelia Langford, John Sillitoe on behalf of the Wellcome Sanger Institute COVID-19 Surveillance Team |
| EPI_ISL_800088, EPI_ISL_800092, EPI_ISL_800093 | Lighthouse Lab in Cambridge | Wellcome Sanger Institute for the COVID-19 Genomics UK (COG-UK) Consortium | Rob Howes, The Lighthouse Lab in Cambridge and Alex Alderton, Roberto Amato, Sonia Goncalves, Ewan Harrison, David K. Jackson, Ian Johnston, Dominic Kwiatkowski, Cordelia Langford, John Sillitoe on behalf of the Wellcome Sanger Institute COVID-19 Surveillance Team |
| EPI_ISL_800094 | Lighthouse Lab in Alderley Park | Wellcome Sanger Institute for the COVID-19 Genomics UK (COG-UK) Consortium | Jacquelyn Wynn, Mairead Hyland, The Lighthouse Lab in Alderley Park and Alex Alderton, Roberto Amato, Sonia Goncalves, Ewan Harrison, David K. Jackson, Ian Johnston, Dominic Kwiatkowski, Cordelia Langford, John Sillitoe on behalf of the Wellcome Sanger Institute COVID-19 Surveillance Team |
| EPI_ISL_800095, EPI_ISL_800096, EPI_ISL_800098 | Lighthouse Lab in Cambridge | Wellcome Sanger Institute for the COVID-19 Genomics UK (COG-UK) Consortium | Rob Howes, The Lighthouse Lab in Cambridge and Alex Alderton, Roberto Amato, Sonia Goncalves, Ewan Harrison, David K. Jackson, Ian Johnston, Dominic Kwiatkowski, Cordelia Langford, John Sillitoe on behalf of the Wellcome Sanger Institute COVID-19 Surveillance Team |
| EPI_ISL_800101 | Lighthouse Lab in Alderley Park | Wellcome Sanger Institute for the COVID-19 Genomics UK (COG-UK) Consortium | Jacquelyn Wynn, Mairead Hyland, The Lighthouse Lab in Alderley Park and Alex Alderton, Roberto Amato, Sonia Goncalves, Ewan Harrison, David K. Jackson, Ian Johnston, Dominic Kwiatkowski, Cordelia Langford, John Sillitoe on behalf of the Wellcome Sanger Institute COVID-19 Surveillance Team |
| EPI_ISL_800102, EPI_ISL_800103 | Lighthouse Lab in Cambridge | Wellcome Sanger Institute for the COVID-19 Genomics UK (COG-UK) Consortium | Rob Howes, The Lighthouse Lab in Cambridge and Alex Alderton, Roberto Amato, Sonia Goncalves, Ewan Harrison, David K. Jackson, Ian Johnston, Dominic Kwiatkowski, Cordelia Langford, John Sillitoe on behalf of the Wellcome Sanger Institute COVID-19 Surveillance Team |
| EPI_ISL_800105, EPI_ISL_800110, EPI_ISL_800112 | Lighthouse Lab in Alderley Park | Wellcome Sanger Institute for the COVID-19 Genomics UK (COG-UK) Consortium | Jacquelyn Wynn, Mairead Hyland, The Lighthouse Lab in Alderley Park and Alex Alderton, Roberto Amato, Sonia Goncalves, Ewan Harrison, David K. Jackson, Ian Johnston, Dominic Kwiatkowski, Cordelia Langford, John Sillitoe on behalf of the Wellcome Sanger Institute COVID-19 Surveillance Team |
| EPI_ISL_800114, EPI_ISL_800115, EPI_ISL_800116 | Lighthouse Lab in Cambridge | Wellcome Sanger Institute for the COVID-19 Genomics UK (COG-UK) Consortium | Rob Howes, The Lighthouse Lab in Cambridge and Alex Alderton, Roberto Amato, Sonia Goncalves, Ewan Harrison, David K. Jackson, Ian Johnston, Dominic Kwiatkowski, Cordelia Langford, John Sillitoe on behalf of the Wellcome Sanger Institute COVID-19 Surveillance Team |
| EPI_ISL_800118 | Lighthouse Lab in Alderley Park | Wellcome Sanger Institute for the COVID-19 Genomics UK (COG-UK) Consortium | Jacquelyn Wynn, Mairead Hyland, The Lighthouse Lab in Alderley Park and Alex Alderton, Roberto Amato, Sonia Goncalves, Ewan Harrison, David K. Jackson, Ian Johnston, Dominic Kwiatkowski, Cordelia Langford, John Sillitoe on behalf of the Wellcome Sanger Institute COVID-19 Surveillance Team |
| EPI_ISL_800119, EPI_ISL_800121, EPI_ISL_800123 | Lighthouse Lab in Cambridge | Wellcome Sanger Institute for the COVID-19 Genomics UK (COG-UK) Consortium | Rob Howes, The Lighthouse Lab in Cambridge and Alex Alderton, Roberto Amato, Sonia Goncalves, Ewan Harrison, David K. Jackson, Ian Johnston, Dominic Kwiatkowski, Cordelia Langford, John Sillitoe on behalf of the Wellcome Sanger Institute COVID-19 Surveillance Team |
| EPI_ISL_800124 | Lighthouse Lab in Glasgow | Wellcome Sanger Institute for the COVID-19 Genomics UK (COG-UK) Consortium | Harper VanSteenhouse, Yumi Kasai, David Gray, Carol Clugston, Anna Dominiczak and Alex Alderton, Roberto Amato, Sonia Goncalves, Ewan Harrison, David K. Jackson, Ian Johnston, Dominic Kwiatkowski, Cordelia Langford, John Sillitoe on behalf of the Wellcome Sanger Institute COVID-19 Surveillance Team |
| EPI_ISL_800125 | Lighthouse Lab in Cambridge | Wellcome Sanger Institute for the COVID-19 Genomics UK (COG-UK) Consortium | Rob Howes, The Lighthouse Lab in Cambridge and Alex Alderton, Roberto Amato, Sonia Goncalves, Ewan Harrison, David K. Jackson, Ian Johnston, Dominic Kwiatkowski, Cordelia Langford, John Sillitoe on behalf of the Wellcome Sanger Institute COVID-19 Surveillance Team |
| EPI_ISL_800127, EPI_ISL_800128, EPI_ISL_800130, EPI_ISL_800132 | Lighthouse Lab in Alderley Park | Wellcome Sanger Institute for the COVID-19 Genomics UK (COG-UK) Consortium | Jacquelyn Wynn, Mairead Hyland, The Lighthouse Lab in Alderley Park and Alex Alderton, Roberto Amato, Sonia Goncalves, Ewan Harrison, David K. Jackson, Ian Johnston, Dominic Kwiatkowski, Cordelia Langford, John Sillitoe on behalf of the Wellcome Sanger Institute COVID-19 Surveillance Team |
| EPI_ISL_800133 | Lighthouse Lab in Cambridge | Wellcome Sanger Institute for the COVID-19 Genomics UK (COG-UK) Consortium | Rob Howes, The Lighthouse Lab in Cambridge and Alex Alderton, Roberto Amato, Sonia Goncalves, Ewan Harrison, David K. Jackson, Ian Johnston, Dominic Kwiatkowski, Cordelia Langford, John Sillitoe on behalf of the Wellcome Sanger Institute COVID-19 Surveillance Team |
| EPI_ISL_800134 | Lighthouse Lab in Alderley Park | Wellcome Sanger Institute for the COVID-19 Genomics UK (COG-UK) Consortium | Jacquelyn Wynn, Mairead Hyland, The Lighthouse Lab in Alderley Park and Alex Alderton, Roberto Amato, Sonia Goncalves, Ewan Harrison, David K. Jackson, Ian Johnston, Dominic Kwiatkowski, Cordelia Langford, John Sillitoe on behalf of the Wellcome Sanger Institute COVID-19 Surveillance Team |
| EPI_ISL_800137 | Lighthouse Lab in Cambridge | Wellcome Sanger Institute for the COVID-19 Genomics UK (COG-UK) Consortium | Rob Howes, The Lighthouse Lab in Cambridge and Alex Alderton, Roberto Amato, Sonia Goncalves, Ewan Harrison, David K. Jackson, Ian Johnston, Dominic Kwiatkowski, Cordelia Langford, John Sillitoe on behalf of the Wellcome Sanger Institute COVID-19 Surveillance Team |
| EPI_ISL_800141 | Lighthouse Lab in Glasgow | Wellcome Sanger Institute for the COVID-19 Genomics UK (COG-UK) Consortium | Harper VanSteenhouse, Yumi Kasai, David Gray, Carol Clugston, Anna Dominiczak and Alex Alderton, Roberto Amato, Sonia Goncalves, Ewan Harrison, David K. Jackson, Ian Johnston, Dominic Kwiatkowski, Cordelia Langford, John Sillitoe on behalf of the Wellcome Sanger Institute COVID-19 Surveillance Team |
| EPI_ISL_800143, EPI_ISL_800148 | Lighthouse Lab in Alderley Park | Wellcome Sanger Institute for the COVID-19 Genomics UK (COG-UK) Consortium | Jacquelyn Wynn, Mairead Hyland, The Lighthouse Lab in Alderley Park and Alex Alderton, Roberto Amato, Sonia Goncalves, Ewan Harrison, David K. Jackson, Ian Johnston, Dominic Kwiatkowski, Cordelia Langford, John Sillitoe on behalf of the Wellcome Sanger Institute COVID-19 Surveillance Team |
| EPI_ISL_800149 | Lighthouse Lab in Cambridge | Wellcome Sanger Institute for the COVID-19 Genomics UK (COG-UK) Consortium | Rob Howes, The Lighthouse Lab in Cambridge and Alex Alderton, Roberto Amato, Sonia Goncalves, Ewan Harrison, David K. Jackson, Ian Johnston, Dominic Kwiatkowski, Cordelia Langford, John Sillitoe on behalf of the Wellcome Sanger Institute COVID-19 Surveillance Team |
| EPI_ISL_800150 | Lighthouse Lab in Alderley Park | Wellcome Sanger Institute for the COVID-19 Genomics UK (COG-UK) Consortium | Jacquelyn Wynn, Mairead Hyland, The Lighthouse Lab in Alderley Park and Alex Alderton, Roberto Amato, Sonia Goncalves, Ewan Harrison, David K. Jackson, Ian Johnston, Dominic Kwiatkowski, Cordelia Langford, John Sillitoe on behalf of the Wellcome Sanger Institute COVID-19 Surveillance Team |
| EPI_ISL_800151 | Lighthouse Lab in Glasgow | Wellcome Sanger Institute for the COVID-19 Genomics UK (COG-UK) Consortium | Harper VanSteenhouse, Yumi Kasai, David Gray, Carol Clugston, Anna Dominiczak and Alex Alderton, Roberto Amato, Sonia Goncalves, Ewan Harrison, David K. Jackson, Ian Johnston, Dominic Kwiatkowski, Cordelia Langford, John Sillitoe on behalf of the Wellcome Sanger Institute COVID-19 Surveillance Team |
| EPI_ISL_800152 | Lighthouse Lab in Alderley Park | Wellcome Sanger Institute for the COVID-19 Genomics UK (COG-UK) Consortium | Jacquelyn Wynn, Mairead Hyland, The Lighthouse Lab in Alderley Park and Alex Alderton, Roberto Amato, Sonia Goncalves, Ewan Harrison, David K. Jackson, Ian Johnston, Dominic Kwiatkowski, Cordelia Langford, John Sillitoe on behalf of the Wellcome Sanger Institute COVID-19 Surveillance Team |
| EPI_ISL_800153 | Lighthouse Lab in Cambridge | Wellcome Sanger Institute for the COVID-19 Genomics UK (COG-UK) Consortium | Rob Howes, The Lighthouse Lab in Cambridge and Alex Alderton, Roberto Amato, Sonia Goncalves, Ewan Harrison, David K. Jackson, Ian Johnston, Dominic Kwiatkowski, Cordelia Langford, John Sillitoe on behalf of the Wellcome Sanger Institute COVID-19 Surveillance Team |
| EPI_ISL_800154, EPI_ISL_800155 | Lighthouse Lab in Alderley Park | Wellcome Sanger Institute for the COVID-19 Genomics UK (COG-UK) Consortium | Jacquelyn Wynn, Mairead Hyland, The Lighthouse Lab in Alderley Park and Alex Alderton, Roberto Amato, Sonia Goncalves, Ewan Harrison, David K. Jackson, Ian Johnston, Dominic Kwiatkowski, Cordelia Langford, John Sillitoe on behalf of the Wellcome Sanger Institute COVID-19 Surveillance Team |
| EPI_ISL_800156 | Lighthouse Lab in Cambridge | Wellcome Sanger Institute for the COVID-19 Genomics UK (COG-UK) Consortium | Rob Howes, The Lighthouse Lab in Cambridge and Alex Alderton, Roberto Amato, Sonia Goncalves, Ewan Harrison, David K. Jackson, Ian Johnston, Dominic Kwiatkowski, Cordelia Langford, John Sillitoe on behalf of the Wellcome Sanger Institute COVID-19 Surveillance Team |
| EPI_ISL_800161 | Lighthouse Lab in Glasgow | Wellcome Sanger Institute for the COVID-19 Genomics UK (COG-UK) Consortium | Harper VanSteenhouse, Yumi Kasai, David Gray, Carol Clugston, Anna Dominiczak and Alex Alderton, Roberto Amato, Sonia Goncalves, Ewan Harrison, David K. Jackson, Ian Johnston, Dominic Kwiatkowski, Cordelia Langford, John Sillitoe on behalf of the Wellcome Sanger Institute COVID-19 Surveillance Team |
| EPI_ISL_800163 | Lighthouse Lab in Alderley Park | Wellcome Sanger Institute for the COVID-19 Genomics UK | Jacquelyn Wynn, Mairead Hyland, The Lighthouse Lab in Alderley Park and Alex Alderton, Roberto Amato, Sonia Goncalves, Ewan Harrison, David K. |

| | | | |
|---|---|---|---|
| | | (COG-UK) Consortium | Jackson, Ian Johnston, Dominic Kwiatkowski, Cordelia Langford, John Sillitoe on behalf of the Wellcome Sanger Institute COVID-19 Surveillance Team |
| EPI_ISL_800164 | Lighthouse Lab in Glasgow | Wellcome Sanger Institute for the COVID-19 Genomics UK (COG-UK) Consortium | Harper VanSteenhouse, Yumi Kasai, David Gray, Carol Clugston, Anna Dominiczak and Alex Alderton, Roberto Amato, Sonia Goncalves, Ewan Harrison, David K. Jackson, Ian Johnston, Dominic Kwiatkowski, Cordelia Langford, John Sillitoe on behalf of the Wellcome Sanger Institute COVID-19 Surveillance Team |
| EPI_ISL_800166, EPI_ISL_800167, EPI_ISL_800168 | Lighthouse Lab in Cambridge | Wellcome Sanger Institute for the COVID-19 Genomics UK (COG-UK) Consortium | Rob Howes, The Lighthouse Lab in Cambridge and Alex Alderton, Roberto Amato, Sonia Goncalves, Ewan Harrison, David K. Jackson, Ian Johnston, Dominic Kwiatkowski, Cordelia Langford, John Sillitoe on behalf of the Wellcome Sanger Institute COVID-19 Surveillance Team |
| EPI_ISL_800169, EPI_ISL_800170 | Lighthouse Lab in Alderley Park | Wellcome Sanger Institute for the COVID-19 Genomics UK (COG-UK) Consortium | Jacquelyn Wynn, Mairead Hyland, The Lighthouse Lab in Alderley Park and Alex Alderton, Roberto Amato, Sonia Goncalves, Ewan Harrison, David K. Jackson, Ian Johnston, Dominic Kwiatkowski, Cordelia Langford, John Sillitoe on behalf of the Wellcome Sanger Institute COVID-19 Surveillance Team |
| EPI_ISL_800172, EPI_ISL_800173 | Lighthouse Lab in Cambridge | Wellcome Sanger Institute for the COVID-19 Genomics UK (COG-UK) Consortium | Rob Howes, The Lighthouse Lab in Cambridge and Alex Alderton, Roberto Amato, Sonia Goncalves, Ewan Harrison, David K. Jackson, Ian Johnston, Dominic Kwiatkowski, Cordelia Langford, John Sillitoe on behalf of the Wellcome Sanger Institute COVID-19 Surveillance Team |
| EPI_ISL_800175, EPI_ISL_800176 | Lighthouse Lab in Glasgow | Wellcome Sanger Institute for the COVID-19 Genomics UK (COG-UK) Consortium | Harper VanSteenhouse, Yumi Kasai, David Gray, Carol Clugston, Anna Dominiczak and Alex Alderton, Roberto Amato, Sonia Goncalves, Ewan Harrison, David K. Jackson, Ian Johnston, Dominic Kwiatkowski, Cordelia Langford, John Sillitoe on behalf of the Wellcome Sanger Institute COVID-19 Surveillance Team |
| EPI_ISL_800177, EPI_ISL_800178 | Lighthouse Lab in Cambridge | Wellcome Sanger Institute for the COVID-19 Genomics UK (COG-UK) Consortium | Rob Howes, The Lighthouse Lab in Cambridge and Alex Alderton, Roberto Amato, Sonia Goncalves, Ewan Harrison, David K. Jackson, Ian Johnston, Dominic Kwiatkowski, Cordelia Langford, John Sillitoe on behalf of the Wellcome Sanger Institute COVID-19 Surveillance Team |
| EPI_ISL_800179 | Lighthouse Lab in Alderley Park | Wellcome Sanger Institute for the COVID-19 Genomics UK (COG-UK) Consortium | Jacquelyn Wynn, Mairead Hyland, The Lighthouse Lab in Alderley Park and Alex Alderton, Roberto Amato, Sonia Goncalves, Ewan Harrison, David K. Jackson, Ian Johnston, Dominic Kwiatkowski, Cordelia Langford, John Sillitoe on behalf of the Wellcome Sanger Institute COVID-19 Surveillance Team |
| EPI_ISL_800180 | Lighthouse Lab in Glasgow | Wellcome Sanger Institute for the COVID-19 Genomics UK (COG-UK) Consortium | Harper VanSteenhouse, Yumi Kasai, David Gray, Carol Clugston, Anna Dominiczak and Alex Alderton, Roberto Amato, Sonia Goncalves, Ewan Harrison, David K. Jackson, Ian Johnston, Dominic Kwiatkowski, Cordelia Langford, John Sillitoe on behalf of the Wellcome Sanger Institute COVID-19 Surveillance Team |
| EPI_ISL_800181 | Lighthouse Lab in Alderley Park | Wellcome Sanger Institute for the COVID-19 Genomics UK (COG-UK) Consortium | Jacquelyn Wynn, Mairead Hyland, The Lighthouse Lab in Alderley Park and Alex Alderton, Roberto Amato, Sonia Goncalves, Ewan Harrison, David K. Jackson, Ian Johnston, Dominic Kwiatkowski, Cordelia Langford, John Sillitoe on behalf of the Wellcome Sanger Institute COVID-19 Surveillance Team |
| EPI_ISL_800182 | Lighthouse Lab in Cambridge | Wellcome Sanger Institute for the COVID-19 Genomics UK (COG-UK) Consortium | Rob Howes, The Lighthouse Lab in Cambridge and Alex Alderton, Roberto Amato, Sonia Goncalves, Ewan Harrison, David K. Jackson, Ian Johnston, Dominic Kwiatkowski, Cordelia Langford, John Sillitoe on behalf of the Wellcome Sanger Institute COVID-19 Surveillance Team |
| EPI_ISL_800184, EPI_ISL_800187 | Lighthouse Lab in Alderley Park | Wellcome Sanger Institute for the COVID-19 Genomics UK (COG-UK) Consortium | Jacquelyn Wynn, Mairead Hyland, The Lighthouse Lab in Alderley Park and Alex Alderton, Roberto Amato, Sonia Goncalves, Ewan Harrison, David K. Jackson, Ian Johnston, Dominic Kwiatkowski, Cordelia Langford, John Sillitoe on behalf of the Wellcome Sanger Institute COVID-19 Surveillance Team |
| EPI_ISL_800188, EPI_ISL_800190, EPI_ISL_800191 | Lighthouse Lab in Glasgow | Wellcome Sanger Institute for the COVID-19 Genomics UK (COG-UK) Consortium | Harper VanSteenhouse, Yumi Kasai, David Gray, Carol Clugston, Anna Dominiczak and Alex Alderton, Roberto Amato, Sonia Goncalves, Ewan Harrison, David K. Jackson, Ian Johnston, Dominic Kwiatkowski, Cordelia Langford, John Sillitoe on behalf of the Wellcome Sanger Institute COVID-19 Surveillance Team |
| EPI_ISL_800195, EPI_ISL_800196 | Lighthouse Lab in Cambridge | Wellcome Sanger Institute for the COVID-19 Genomics UK (COG-UK) Consortium | Rob Howes, The Lighthouse Lab in Cambridge and Alex Alderton, Roberto Amato, Sonia Goncalves, Ewan Harrison, David K. Jackson, Ian Johnston, Dominic Kwiatkowski, Cordelia Langford, John Sillitoe on behalf of the Wellcome Sanger Institute COVID-19 Surveillance Team |
| EPI_ISL_800200, EPI_ISL_800201 | Lighthouse Lab in Glasgow | Wellcome Sanger Institute for the COVID-19 Genomics UK (COG-UK) Consortium | Harper VanSteenhouse, Yumi Kasai, David Gray, Carol Clugston, Anna Dominiczak and Alex Alderton, Roberto Amato, Sonia Goncalves, Ewan Harrison, David K. Jackson, Ian Johnston, Dominic Kwiatkowski, Cordelia Langford, John Sillitoe on behalf of the Wellcome Sanger Institute COVID-19 Surveillance Team |
| EPI_ISL_800211, EPI_ISL_800214, EPI_ISL_800218 | Lighthouse Lab in Cambridge | Wellcome Sanger Institute for the COVID-19 Genomics UK (COG-UK) Consortium | Rob Howes, The Lighthouse Lab in Cambridge and Alex Alderton, Roberto Amato, Sonia Goncalves, Ewan Harrison, David K. Jackson, Ian Johnston, Dominic Kwiatkowski, Cordelia Langford, John Sillitoe on behalf of the Wellcome Sanger Institute COVID-19 Surveillance Team |
| EPI_ISL_800222, EPI_ISL_800224, EPI_ISL_800227 | Lighthouse Lab in Alderley Park | Wellcome Sanger Institute for the COVID-19 Genomics UK (COG-UK) Consortium | Jacquelyn Wynn, Mairead Hyland, The Lighthouse Lab in Alderley Park and Alex Alderton, Roberto Amato, Sonia Goncalves, Ewan Harrison, David K. Jackson, Ian Johnston, Dominic Kwiatkowski, Cordelia Langford, John Sillitoe on behalf of the Wellcome Sanger Institute COVID-19 Surveillance Team |
| EPI_ISL_800228, EPI_ISL_800233 | Lighthouse Lab in Cambridge | Wellcome Sanger Institute for the COVID-19 Genomics UK (COG-UK) Consortium | Rob Howes, The Lighthouse Lab in Cambridge and Alex Alderton, Roberto Amato, Sonia Goncalves, Ewan Harrison, David K. Jackson, Ian Johnston, Dominic Kwiatkowski, Cordelia Langford, John Sillitoe on behalf of the Wellcome Sanger Institute COVID-19 Surveillance Team |
| EPI_ISL_800234, EPI_ISL_800235, EPI_ISL_800236 | Lighthouse Lab in Glasgow | Wellcome Sanger Institute for the COVID-19 Genomics UK (COG-UK) Consortium | Harper VanSteenhouse, Yumi Kasai, David Gray, Carol Clugston, Anna Dominiczak and Alex Alderton, Roberto Amato, Sonia Goncalves, Ewan Harrison, David K. Jackson, Ian Johnston, Dominic Kwiatkowski, Cordelia Langford, John Sillitoe on behalf of the Wellcome Sanger Institute COVID-19 Surveillance Team |
| EPI_ISL_800238 | Lighthouse Lab in Cambridge | Wellcome Sanger Institute for the COVID-19 Genomics UK (COG-UK) Consortium | Rob Howes, The Lighthouse Lab in Cambridge and Alex Alderton, Roberto Amato, Sonia Goncalves, Ewan Harrison, David K. Jackson, Ian Johnston, Dominic Kwiatkowski, Cordelia Langford, John Sillitoe on behalf of the Wellcome Sanger Institute COVID-19 Surveillance Team |
| EPI_ISL_800240 | Lighthouse Lab in Alderley Park | Wellcome Sanger Institute for the COVID-19 Genomics UK (COG-UK) Consortium | Jacquelyn Wynn, Mairead Hyland, The Lighthouse Lab in Alderley Park and Alex Alderton, Roberto Amato, Sonia Goncalves, Ewan Harrison, David K. Jackson, Ian Johnston, Dominic Kwiatkowski, Cordelia Langford, John Sillitoe on behalf of the Wellcome Sanger Institute COVID-19 Surveillance Team |
| EPI_ISL_800241 | Lighthouse Lab in Glasgow | Wellcome Sanger Institute for the COVID-19 Genomics UK (COG-UK) Consortium | Harper VanSteenhouse, Yumi Kasai, David Gray, Carol Clugston, Anna Dominiczak and Alex Alderton, Roberto Amato, Sonia Goncalves, Ewan Harrison, David K. Jackson, Ian Johnston, Dominic Kwiatkowski, Cordelia Langford, John Sillitoe on behalf of the Wellcome Sanger Institute COVID-19 Surveillance Team |
| EPI_ISL_800243, EPI_ISL_800246, EPI_ISL_800248 | Lighthouse Lab in Alderley Park | Wellcome Sanger Institute for the COVID-19 Genomics UK (COG-UK) Consortium | Jacquelyn Wynn, Mairead Hyland, The Lighthouse Lab in Alderley Park and Alex Alderton, Roberto Amato, Sonia Goncalves, Ewan Harrison, David K. Jackson, Ian Johnston, Dominic Kwiatkowski, Cordelia Langford, John Sillitoe on behalf of the Wellcome Sanger Institute COVID-19 Surveillance Team |
| EPI_ISL_800250, EPI_ISL_800252, EPI_ISL_800254, EPI_ISL_800255, EPI_ISL_800256, EPI_ISL_800257, EPI_ISL_800258, EPI_ISL_800259, EPI_ISL_800260, EPI_ISL_800261 | Lighthouse Lab in Cambridge | Wellcome Sanger Institute for the COVID-19 Genomics UK (COG-UK) Consortium | Rob Howes, The Lighthouse Lab in Cambridge and Alex Alderton, Roberto Amato, Sonia Goncalves, Ewan Harrison, David K. Jackson, Ian Johnston, Dominic Kwiatkowski, Cordelia Langford, John Sillitoe on behalf of the Wellcome Sanger Institute COVID-19 Surveillance Team |
| EPI_ISL_800262 | Lighthouse Lab in Alderley Park | Wellcome Sanger Institute for the COVID-19 Genomics UK (COG-UK) Consortium | Jacquelyn Wynn, Mairead Hyland, The Lighthouse Lab in Alderley Park and Alex Alderton, Roberto Amato, Sonia Goncalves, Ewan Harrison, David K. Jackson, Ian Johnston, Dominic Kwiatkowski, Cordelia Langford, John Sillitoe on behalf of the Wellcome Sanger Institute COVID-19 Surveillance Team |
| EPI_ISL_800263, EPI_ISL_800264, EPI_ISL_800265, EPI_ISL_800266, EPI_ISL_800268, EPI_ISL_800269, EPI_ISL_800270, EPI_ISL_800272, EPI_ISL_800273, EPI_ISL_800275, EPI_ISL_800276, EPI_ISL_800277, EPI_ISL_800278, EPI_ISL_800279, EPI_ISL_800283, EPI_ISL_800284, EPI_ISL_800285, EPI_ISL_800287, EPI_ISL_800288, EPI_ISL_800289, EPI_ISL_800290, EPI_ISL_800291, EPI_ISL_800292, EPI_ISL_800293, EPI_ISL_800294, EPI_ISL_800295, EPI_ISL_800296, EPI_ISL_800298, EPI_ISL_800300, EPI_ISL_800302, EPI_ISL_800304, EPI_ISL_800305, EPI_ISL_800306, EPI_ISL_800307, EPI_ISL_800309, EPI_ISL_800310, EPI_ISL_800311, EPI_ISL_800312, EPI_ISL_800313, EPI_ISL_800314, EPI_ISL_800315, EPI_ISL_800316, EPI_ISL_800318, EPI_ISL_800319, EPI_ISL_800320, EPI_ISL_800321, EPI_ISL_800322, EPI_ISL_800323, EPI_ISL_800324, EPI_ISL_800325, EPI_ISL_800326, EPI_ISL_800327, EPI_ISL_800328, EPI_ISL_800329, EPI_ISL_800330, EPI_ISL_800331, EPI_ISL_800332, EPI_ISL_800333, EPI_ISL_800334, EPI_ISL_800335, EPI_ISL_800336, EPI_ISL_800337, EPI_ISL_800339, EPI_ISL_800341, EPI_ISL_800343, EPI_ISL_800344, EPI_ISL_800345, EPI_ISL_800346, EPI_ISL_800349, EPI_ISL_800352, EPI_ISL_800353 | | | |
| see above | Lighthouse Lab in Cambridge | Wellcome Sanger Institute for the COVID-19 Genomics UK (COG-UK) Consortium | Rob Howes, The Lighthouse Lab in Cambridge and Alex Alderton, Roberto Amato, Sonia Goncalves, Ewan Harrison, David K. Jackson, Ian Johnston, Dominic Kwiatkowski, Cordelia Langford, John Sillitoe on behalf of the Wellcome Sanger Institute COVID-19 Surveillance Team |
| EPI_ISL_800355 | Lighthouse Lab in Alderley Park | Wellcome Sanger Institute for the COVID-19 Genomics UK (COG-UK) Consortium | Jacquelyn Wynn, Mairead Hyland, The Lighthouse Lab in Alderley Park and Alex Alderton, Roberto Amato, Sonia Goncalves, Ewan Harrison, David K. Jackson, Ian Johnston, Dominic Kwiatkowski, Cordelia Langford, John Sillitoe on behalf of the Wellcome Sanger Institute COVID-19 Surveillance Team |
| EPI_ISL_800356, EPI_ISL_800357, EPI_ISL_800358, EPI_ISL_800361 | Lighthouse Lab in Cambridge | Wellcome Sanger Institute for the COVID-19 Genomics UK (COG-UK) Consortium | Rob Howes, The Lighthouse Lab in Cambridge and Alex Alderton, Roberto Amato, Sonia Goncalves, Ewan Harrison, David K. Jackson, Ian Johnston, Dominic Kwiatkowski, Cordelia Langford, John Sillitoe on behalf of the Wellcome Sanger Institute COVID-19 Surveillance Team |
| EPI_ISL_800362 | Lighthouse Lab in Alderley Park | Wellcome Sanger Institute for the COVID-19 Genomics UK (COG-UK) Consortium | Jacquelyn Wynn, Mairead Hyland, The Lighthouse Lab in Alderley Park and Alex Alderton, Roberto Amato, Sonia Goncalves, Ewan Harrison, David K. Jackson, Ian Johnston, Dominic Kwiatkowski, Cordelia Langford, John Sillitoe on behalf of the Wellcome Sanger Institute COVID-19 Surveillance Team |
| EPI_ISL_800363, EPI_ISL_800364, EPI_ISL_800366, EPI_ISL_800368, EPI_ISL_800369, EPI_ISL_800371, EPI_ISL_800373, EPI_ISL_800374, EPI_ISL_800375, EPI_ISL_800377, EPI_ISL_800378, EPI_ISL_800379, EPI_ISL_800380, EPI_ISL_800381, EPI_ISL_800382, EPI_ISL_800383, EPI_ISL_800385, EPI_ISL_800386, EPI_ISL_800387, EPI_ISL_800388, EPI_ISL_800390, EPI_ISL_800391, EPI_ISL_800393, EPI_ISL_800394 | | | |

| | | | |
|---|---|---|---|
| see above | Lighthouse Lab in Cambridge | Wellcome Sanger Institute for the COVID-19 Genomics UK (COG-UK) Consortium | Rob Howes, The Lighthouse Lab in Cambridge and Alex Alderton, Roberto Amato, Sonia Goncalves, Ewan Harrison, David K. Jackson, Ian Johnston, Dominic Kwiatkowski, Cordelia Langford, John Sillitoe on behalf of the Wellcome Sanger Institute COVID-19 Surveillance Team |
| EPI_ISL_800395 | Lighthouse Lab in Alderley Park | Wellcome Sanger Institute for the COVID-19 Genomics UK (COG-UK) Consortium | Jacquelyn Wynn, Mairead Hyland, The Lighthouse Lab in Alderley Park and Alex Alderton, Roberto Amato, Sonia Goncalves, Ewan Harrison, David K. Jackson, Ian Johnston, Dominic Kwiatkowski, Cordelia Langford, John Sillitoe on behalf of the Wellcome Sanger Institute COVID-19 Surveillance Team |
| EPI_ISL_800396, EPI_ISL_800397, EPI_ISL_800398, EPI_ISL_800399, EPI_ISL_800400, EPI_ISL_800401, EPI_ISL_800402, EPI_ISL_800403, EPI_ISL_800404, EPI_ISL_800405, EPI_ISL_800406, EPI_ISL_800408, EPI_ISL_800409, EPI_ISL_800410, EPI_ISL_800411, EPI_ISL_800412, EPI_ISL_800413, EPI_ISL_800414, EPI_ISL_800415, EPI_ISL_800416, EPI_ISL_800417, EPI_ISL_800418, EPI_ISL_800419, EPI_ISL_800421, EPI_ISL_800422, EPI_ISL_800423, EPI_ISL_800424, EPI_ISL_800425, EPI_ISL_800426, EPI_ISL_800427, EPI_ISL_800429, EPI_ISL_800430, EPI_ISL_800431, EPI_ISL_800432, EPI_ISL_800433, EPI_ISL_800434, EPI_ISL_800435, EPI_ISL_800436, EPI_ISL_800437, EPI_ISL_800438, EPI_ISL_800439, EPI_ISL_800440, EPI_ISL_800443, EPI_ISL_800445, EPI_ISL_800447, EPI_ISL_800448, EPI_ISL_800449, EPI_ISL_800450, EPI_ISL_800451, EPI_ISL_800455, EPI_ISL_800458, EPI_ISL_800459 | | | |
| see above | Lighthouse Lab in Cambridge | Wellcome Sanger Institute for the COVID-19 Genomics UK (COG-UK) Consortium | Rob Howes, The Lighthouse Lab in Cambridge and Alex Alderton, Roberto Amato, Sonia Goncalves, Ewan Harrison, David K. Jackson, Ian Johnston, Dominic Kwiatkowski, Cordelia Langford, John Sillitoe on behalf of the Wellcome Sanger Institute COVID-19 Surveillance Team |
| EPI_ISL_800460 | Lighthouse Lab in Alderley Park | Wellcome Sanger Institute for the COVID-19 Genomics UK (COG-UK) Consortium | Jacquelyn Wynn, Mairead Hyland, The Lighthouse Lab in Alderley Park and Alex Alderton, Roberto Amato, Sonia Goncalves, Ewan Harrison, David K. Jackson, Ian Johnston, Dominic Kwiatkowski, Cordelia Langford, John Sillitoe on behalf of the Wellcome Sanger Institute COVID-19 Surveillance Team |
| EPI_ISL_800461, EPI_ISL_800462, EPI_ISL_800463, EPI_ISL_800464, EPI_ISL_800465, EPI_ISL_800466, EPI_ISL_800467, EPI_ISL_800469, EPI_ISL_800470, EPI_ISL_800471, EPI_ISL_800472, EPI_ISL_800473, EPI_ISL_800475, EPI_ISL_800476, EPI_ISL_800477, EPI_ISL_800478, EPI_ISL_800479, EPI_ISL_800480, EPI_ISL_800482, EPI_ISL_800483, EPI_ISL_800484, EPI_ISL_800486, EPI_ISL_800487, EPI_ISL_800488, EPI_ISL_800489, EPI_ISL_800490, EPI_ISL_800491, EPI_ISL_800492, EPI_ISL_800493, EPI_ISL_800496, EPI_ISL_800497, EPI_ISL_800498, EPI_ISL_800499, EPI_ISL_800500, EPI_ISL_800503, EPI_ISL_800504, EPI_ISL_800505 | | | |
| see above | Lighthouse Lab in Cambridge | Wellcome Sanger Institute for the COVID-19 Genomics UK (COG-UK) Consortium | Rob Howes, The Lighthouse Lab in Cambridge and Alex Alderton, Roberto Amato, Sonia Goncalves, Ewan Harrison, David K. Jackson, Ian Johnston, Dominic Kwiatkowski, Cordelia Langford, John Sillitoe on behalf of the Wellcome Sanger Institute COVID-19 Surveillance Team |
| EPI_ISL_800506 | Lighthouse Lab in Alderley Park | Wellcome Sanger Institute for the COVID-19 Genomics UK (COG-UK) Consortium | Jacquelyn Wynn, Mairead Hyland, The Lighthouse Lab in Alderley Park and Alex Alderton, Roberto Amato, Sonia Goncalves, Ewan Harrison, David K. Jackson, Ian Johnston, Dominic Kwiatkowski, Cordelia Langford, John Sillitoe on behalf of the Wellcome Sanger Institute COVID-19 Surveillance Team |
| EPI_ISL_800507, EPI_ISL_800508, EPI_ISL_800510, EPI_ISL_800511, EPI_ISL_800512, EPI_ISL_800513, EPI_ISL_800514, EPI_ISL_800515, EPI_ISL_800517, EPI_ISL_800519, EPI_ISL_800520, EPI_ISL_800521, EPI_ISL_800522, EPI_ISL_800523 | | | |
| see above | Lighthouse Lab in Cambridge | Wellcome Sanger Institute for the COVID-19 Genomics UK (COG-UK) Consortium | Rob Howes, The Lighthouse Lab in Cambridge and Alex Alderton, Roberto Amato, Sonia Goncalves, Ewan Harrison, David K. Jackson, Ian Johnston, Dominic Kwiatkowski, Cordelia Langford, John Sillitoe on behalf of the Wellcome Sanger Institute COVID-19 Surveillance Team |
| EPI_ISL_800526 | Lighthouse Lab in Alderley Park | Wellcome Sanger Institute for the COVID-19 Genomics UK (COG-UK) Consortium | Jacquelyn Wynn, Mairead Hyland, The Lighthouse Lab in Alderley Park and Alex Alderton, Roberto Amato, Sonia Goncalves, Ewan Harrison, David K. Jackson, Ian Johnston, Dominic Kwiatkowski, Cordelia Langford, John Sillitoe on behalf of the Wellcome Sanger Institute COVID-19 Surveillance Team |
| EPI_ISL_800527, EPI_ISL_800529, EPI_ISL_800530, EPI_ISL_800532, EPI_ISL_800533, EPI_ISL_800534, EPI_ISL_800535, EPI_ISL_800536, EPI_ISL_800537, EPI_ISL_800538, EPI_ISL_800539, EPI_ISL_800540, EPI_ISL_800541 | | | |
| see above | Lighthouse Lab in Cambridge | Wellcome Sanger Institute for the COVID-19 Genomics UK (COG-UK) Consortium | Rob Howes, The Lighthouse Lab in Cambridge and Alex Alderton, Roberto Amato, Sonia Goncalves, Ewan Harrison, David K. Jackson, Ian Johnston, Dominic Kwiatkowski, Cordelia Langford, John Sillitoe on behalf of the Wellcome Sanger Institute COVID-19 Surveillance Team |
| EPI_ISL_800542 | Lighthouse Lab in Alderley Park | Wellcome Sanger Institute for the COVID-19 Genomics UK (COG-UK) Consortium | Jacquelyn Wynn, Mairead Hyland, The Lighthouse Lab in Alderley Park and Alex Alderton, Roberto Amato, Sonia Goncalves, Ewan Harrison, David K. Jackson, Ian Johnston, Dominic Kwiatkowski, Cordelia Langford, John Sillitoe on behalf of the Wellcome Sanger Institute COVID-19 Surveillance Team |
| EPI_ISL_800543, EPI_ISL_800544, EPI_ISL_800545, EPI_ISL_800546, EPI_ISL_800547, EPI_ISL_800548, EPI_ISL_800549, EPI_ISL_800550, EPI_ISL_800551, EPI_ISL_800552, EPI_ISL_800553, EPI_ISL_800554 | | | |
| see above | Lighthouse Lab in Cambridge | Wellcome Sanger Institute for the COVID-19 Genomics UK (COG-UK) Consortium | Rob Howes, The Lighthouse Lab in Cambridge and Alex Alderton, Roberto Amato, Sonia Goncalves, Ewan Harrison, David K. Jackson, Ian Johnston, Dominic Kwiatkowski, Cordelia Langford, John Sillitoe on behalf of the Wellcome Sanger Institute COVID-19 Surveillance Team |
| EPI_ISL_800555 | Lighthouse Lab in Alderley Park | Wellcome Sanger Institute for the COVID-19 Genomics UK (COG-UK) Consortium | Jacquelyn Wynn, Mairead Hyland, The Lighthouse Lab in Alderley Park and Alex Alderton, Roberto Amato, Sonia Goncalves, Ewan Harrison, David K. Jackson, Ian Johnston, Dominic Kwiatkowski, Cordelia Langford, John Sillitoe on behalf of the Wellcome Sanger Institute COVID-19 Surveillance Team |
| EPI_ISL_800556, EPI_ISL_800557, EPI_ISL_800559, EPI_ISL_800560, EPI_ISL_800561, EPI_ISL_800562, EPI_ISL_800563, EPI_ISL_800564, EPI_ISL_800565, EPI_ISL_800567, EPI_ISL_800568, EPI_ISL_800569, EPI_ISL_800570, EPI_ISL_800571, EPI_ISL_800572, EPI_ISL_800573, EPI_ISL_800574 | | | |
| see above | Lighthouse Lab in Cambridge | Wellcome Sanger Institute for the COVID-19 Genomics UK (COG-UK) Consortium | Rob Howes, The Lighthouse Lab in Cambridge and Alex Alderton, Roberto Amato, Sonia Goncalves, Ewan Harrison, David K. Jackson, Ian Johnston, Dominic Kwiatkowski, Cordelia Langford, John Sillitoe on behalf of the Wellcome Sanger Institute COVID-19 Surveillance Team |
| EPI_ISL_800576 | Lighthouse Lab in Glasgow | Wellcome Sanger Institute for the COVID-19 Genomics UK (COG-UK) Consortium | Harper VanSteenhouse, Yumi Kasai, David Gray, Carol Clugston, Anna Dominiczak and Alex Alderton, Roberto Amato, Sonia Goncalves, Ewan Harrison, David K. Jackson, Ian Johnston, Dominic Kwiatkowski, Cordelia Langford, John Sillitoe on behalf of the Wellcome Sanger Institute COVID-19 Surveillance Team |
| EPI_ISL_801411, EPI_ISL_801415, EPI_ISL_801419, EPI_ISL_801422, EPI_ISL_801438, EPI_ISL_801441, EPI_ISL_801454, EPI_ISL_801455, EPI_ISL_801456, EPI_ISL_801457, EPI_ISL_801459, EPI_ISL_801491, EPI_ISL_801492, EPI_ISL_801493, EPI_ISL_801494, EPI_ISL_801495, EPI_ISL_801497, EPI_ISL_801510, EPI_ISL_801512, EPI_ISL_801513 | | | |
| see above | Dutch COVID-19 response team | Erasmus Medical Center | Bas Oude Munnink, Reina Sikkema, David Nieuwenhuijse, Irina Chestakova, Anne van der Linden, Marjan Boter, Emmanuelle Munger, Corine GeurtsvanKessel, Annemiek van der Eijk, Richard Molenkamp, Marion Koopmans, on behalf of the Dutch national COVID-19 response team. |
| EPI_ISL_801516 | Instituto Nacional de Saude (INSA) | Instituto Nacional de Saude (INSA) | Borges et al |
| EPI_ISL_801537 | Sunrise Medical Labs | Pathogen Discovery, Respiratory Viruses Branch, Division of Viral Diseases, Centers for Disease Control and Prevention | Ying Tao, Yan Li, Jing Zhang, Krista Queen, Anna Uehara, Peter Cook, Clinton R. Paden, Haibin Wang, Suxiang Tong |
| EPI_ISL_802433 | Wadsworth Center, New York State Department.of Health | Wadsworth Center, New York State Department.of Health | Kirsten St. George, Daryl M. Lamson, Alexis Russel, Matthew Shudt, Melissa A Leisner, Jonathan Plitnick, Navjot Singh, John Kelly, Sara Griesemer, Erasmus Schneider, Erica Lasek-Nesselquist |
| EPI_ISL_802436 | TEMPUS LABS INC | Wadsworth Center, New York State Department.of Health | Kirsten St. George, Daryl M. Lamson, Alexis Russel, Matthew Shudt, Melissa A Leisner, Jonathan Plitnick, Navjot Singh, John Kelly, Sara Griesemer, Erasmus Schneider, Erica Lasek-Nesselquist |
| EPI_ISL_802444 | SUNRISE MEDICAL LABORATORIES | Wadsworth Center, New York State Department.of Health | Kirsten St. George, Daryl M. Lamson, Alexis Russel, Matthew Shudt, Melissa A Leisner, Jonathan Plitnick, Navjot Singh, John Kelly, Sara Griesemer, Erasmus Schneider, Erica Lasek-Nesselquist |
| EPI_ISL_802507, EPI_ISL_802521, EPI_ISL_802537, EPI_ISL_802540, EPI_ISL_802541, EPI_ISL_802543 | Dutch COVID-19 response team | Erasmus Medical Center | Bas Oude Munnink, Reina Sikkema, David Nieuwenhuijse, Irina Chestakova, Anne van der Linden, Marjan Boter, Emmanuelle Munger, Corine GeurtsvanKessel, Annemiek van der Eijk, Richard Molenkamp, Marion Koopmans, on behalf of the Dutch national COVID-19 response team. |
| EPI_ISL_802574, EPI_ISL_802576, EPI_ISL_802578, EPI_ISL_802581, EPI_ISL_802582, EPI_ISL_802583, EPI_ISL_802584, EPI_ISL_802588, EPI_ISL_802595, EPI_ISL_802596, EPI_ISL_802598 | | | |
| see above | Vault Health | Minnesota Department of Health, Public Health Laboratory | Alexandra Lorentz, Jacob Garfin, Matt Plumb, and Xiong Wang |
| EPI_ISL_802602, EPI_ISL_802603, EPI_ISL_802612, EPI_ISL_802613, EPI_ISL_802614, EPI_ISL_802615, EPI_ISL_802616, EPI_ISL_802617, EPI_ISL_802632, EPI_ISL_802633, EPI_ISL_802634, EPI_ISL_802635, EPI_ISL_802636, EPI_ISL_802647, EPI_ISL_802651, EPI_ISL_802653, EPI_ISL_802655, EPI_ISL_802661, EPI_ISL_802675, EPI_ISL_802676, EPI_ISL_802677, EPI_ISL_802678, EPI_ISL_802697, EPI_ISL_802698, EPI_ISL_802701, EPI_ISL_802702, EPI_ISL_802703, EPI_ISL_802704, EPI_ISL_802705, EPI_ISL_802714, EPI_ISL_802718, EPI_ISL_802720, EPI_ISL_802721, EPI_ISL_802723 | | | |
| see above | Helix/Illumina | Genomics and Discovery, Respiratory Viruses Branch, Division of Viral Diseases, Centers for Disease Control and Prevention | Peter W. Cook, Dhwani Batra, Ben L. Rambo-Martin Eileen de Feo, Jan Antico, Christine Tran, Matthew Tolentino, Shannon Wickline, Kim Gietzen, Brad Sickler, Jingtao Liu, Eric Allen, Phil Febbo, Summer Galloway, Nicole L. Washington, Simon White, Geraint Levan, Kelly Schiabor Barrett, Elizabeth Cirulli, Alexandre Bolze, Ary Ascencio, Charlotte Rivera-Garcia, Ryan Cho, Jason Nguyen, Sherry Wang, Jimmy Ramirez, Tyler Cassens, Efren Sandoval, Magnus Isaksson, William Lee, David Becker, Marc Laurent, James Lu, Clinton R. Paden, Suxiang Tong, Duncan MacCannell |
| EPI_ISL_802777, EPI_ISL_802778, EPI_ISL_802779, EPI_ISL_802780, EPI_ISL_802781, EPI_ISL_802782, EPI_ISL_802783, EPI_ISL_802784, EPI_ISL_802785, EPI_ISL_802786, EPI_ISL_802787 | | | |
| see above | Columbia University Irving Medical Center | Wadsworth Center, New York State Department.of Health | Kirsten St. George, Daryl M. Lamson, Alexis Russel, Matthew Shudt, Melissa A Leisner, Jonathan Plitnick, Navjot Singh, John Kelly, Sara Griesemer, Erasmus Schneider, Erica Lasek-Nesselquist |
| EPI_ISL_802994, EPI_ISL_802995 | Charite Universitatsmedizin Berlin, Institut fur Virologie, Charitéplatz 1, 10117 Berlin, Germany | Charite Universitatsmedizin Berlin, Institut fur Virologie, Charitéplatz 1, 10117 Berlin, Germany | Victor M Corman, Tobias Bleicker, Julia Tesch, Jörn Beheim-Schwarzbach, Barbara Mühlemann, Talitha Veith, Julia Schneider, Terry Jones, Christian Drosten |
| EPI_ISL_802996, EPI_ISL_802997, EPI_ISL_802998, EPI_ISL_802999, EPI_ISL_803000, EPI_ISL_803001, EPI_ISL_803002, EPI_ISL_803003, EPI_ISL_803004, EPI_ISL_803005, EPI_ISL_803006, EPI_ISL_803007, EPI_ISL_803008, EPI_ISL_803009, EPI_ISL_803010, EPI_ISL_803011, EPI_ISL_803012, EPI_ISL_803013, EPI_ISL_803014, EPI_ISL_803015, EPI_ISL_803016, EPI_ISL_803017, EPI_ISL_803018, EPI_ISL_803019, EPI_ISL_803020, EPI_ISL_803021, EPI_ISL_803022, EPI_ISL_803023, EPI_ISL_803024, EPI_ISL_803025, EPI_ISL_803026, EPI_ISL_803027, EPI_ISL_803028, EPI_ISL_803029, EPI_ISL_803030, EPI_ISL_803031, | | | |

| | | | |
|---|---|---|---|
| EPI_ISL_803032, EPI_ISL_803033, EPI_ISL_803034, EPI_ISL_803035, EPI_ISL_803036, EPI_ISL_803037, EPI_ISL_803038, EPI_ISL_803039, EPI_ISL_803040, EPI_ISL_803041, EPI_ISL_803042, EPI_ISL_803043, EPI_ISL_803044, EPI_ISL_803045, EPI_ISL_803046, EPI_ISL_803047, EPI_ISL_803048, EPI_ISL_803049, EPI_ISL_803050, EPI_ISL_803051, EPI_ISL_803052, EPI_ISL_803053, EPI_ISL_803054, EPI_ISL_803055, EPI_ISL_803056, EPI_ISL_803057, EPI_ISL_803058, EPI_ISL_803059, EPI_ISL_803060, EPI_ISL_803061, EPI_ISL_803062, EPI_ISL_803063, EPI_ISL_803064, EPI_ISL_803065, EPI_ISL_803066, EPI_ISL_803067, EPI_ISL_803068, EPI_ISL_803069, EPI_ISL_803070, EPI_ISL_803071, EPI_ISL_803072, EPI_ISL_803073, EPI_ISL_803074, EPI_ISL_803075, EPI_ISL_803076, EPI_ISL_803077, EPI_ISL_803078, EPI_ISL_803080, EPI_ISL_803081, EPI_ISL_803082, EPI_ISL_803083, EPI_ISL_803084, EPI_ISL_803085, EPI_ISL_803086, EPI_ISL_803087, EPI_ISL_803088, EPI_ISL_803089, EPI_ISL_803090, EPI_ISL_803091, EPI_ISL_803092, EPI_ISL_803093, EPI_ISL_803094, EPI_ISL_803095, EPI_ISL_803096, EPI_ISL_803097 | | | |
| see above | Quest Diagnostics | Quest Diagnostics | Rosenthal,S.H., Gerasimova,A., Kagan,R.M., Anderson, B., Livingston, K.E., Hua, M., Liu Y., Shalhout, D.F., Owen, R., Lacbawan, F. |
| EPI_ISL_803854 | Hospital General Universitario Gregorio Marañón | Hospital General Universitario Gregorio Marañón | Sergio Buenestado Serrano, Pedro Sola Campoy, Laura Pérez-Lago, Pilar Catalán, Patricia Muñoz, Darío García de Viedma |
| EPI_ISL_803898 | National Institute for Infectious Diseases, INMI, "L. Spallanzani" IRCCS | National Institute for Infectious Diseases, INMI, "L. Spallanzani" IRCCS | C.E.M Gruber, B Bartolini, E Giombini, M Rueca, O Butera, F Messina, A Di Caro, MR Capobianchi |
| EPI_ISL_803903 | Shimoga Institute of Medical Sciences | Department of Neurovirology, National Institute of Mental Health and Neurosciences (NIMHANS) | Chitra Pattabiraman, Pramada Prasad, Anita S Desai, V Ravi |
| EPI_ISL_803906, EPI_ISL_803907 | Kidwai Memorial Institute of Oncology | Department of Neurovirology, National Institute of Mental Health and Neurosciences (NIMHANS) | Chitra Pattabiraman, Pramada Prasad, Anita S Desai, V Ravi |
| EPI_ISL_803908 | BBMP Urban PHC | Department of Neurovirology, National Institute of Mental Health and Neurosciences (NIMHANS) | Chitra Pattabiraman, Pramada Prasad, Risha Rasheed, Anita S Desai, V Ravi |
| EPI_ISL_803909 | Shimoga Institute of Medical Sciences | Department of Neurovirology, National Institute of Mental Health and Neurosciences (NIMHANS) | Chitra Pattabiraman, Pramada Prasad, Anita S Desai, V Ravi |
| EPI_ISL_803910 | Kidwai Memorial Institute of Oncology | Department of Neurovirology, National Institute of Mental Health and Neurosciences (NIMHANS) | Chitra Pattabiraman, Pramada Prasad, Anita S Desai, V Ravi |
| EPI_ISL_803912 | BBMP Urban PHC | Department of Neurovirology, National Institute of Mental Health and Neurosciences (NIMHANS) | Chitra Pattabiraman, Pramada Prasad, Anita S Desai, V Ravi |
| EPI_ISL_803913 | Shimoga Institute of Medical Sciences | Department of Neurovirology, National Institute of Mental Health and Neurosciences (NIMHANS) | Chitra Pattabiraman, Pramada Prasad, Risha Rasheed, Anita S Desai, V Ravi |
| EPI_ISL_803914 | Kidwai Memorial Institute of Oncology | Department of Neurovirology, National Institute of Mental Health and Neurosciences (NIMHANS) | Chitra Pattabiraman, Pramada Prasad, Risha Rasheed, Anita S Desai, V Ravi |
| EPI_ISL_803915 | Kidwai Memorial Institute of Oncology | Department of Neurovirology, National Institute of Mental Health and Neurosciences (NIMHANS) | Chitra Pattabiraman, Pramada Prasad, Anita S Desai, V Ravi |
| EPI_ISL_803916 | Shimoga Institute of Medical Sciences | Department of Neurovirology, National Institute of Mental Health and Neurosciences (NIMHANS) | Chitra Pattabiraman, Pramada Prasad, Anita S Desai, V Ravi |
| EPI_ISL_804019 | Maryland Public Health Laboratory | Maryland Public Health Laboratory | Maryland Department of Health Laboratories Administration |
| EPI_ISL_804054 | SC (UCO) Igiene e Sanità Pubblica (funzione integrata con SC Microbiologia e Virologia) e Laboratory of Molecular Virology of the International Centre for Genetic Engineering and Biotechnology (ICGEB) | ARGO Laboratorio Genomica ed Epigenomica | Licastro D, Dal Monego S, Degasperi M, Marcello A, D'Agaro P |
| EPI_ISL_804218, EPI_ISL_804219, EPI_ISL_804220, EPI_ISL_804221, EPI_ISL_804240, EPI_ISL_804241, EPI_ISL_804250, EPI_ISL_804273, EPI_ISL_804294, EPI_ISL_804295, EPI_ISL_804296, EPI_ISL_804297, EPI_ISL_804298, EPI_ISL_804299, EPI_ISL_804300, EPI_ISL_804301, EPI_ISL_804302, EPI_ISL_804303, EPI_ISL_804304, EPI_ISL_804305, EPI_ISL_804306, EPI_ISL_804307, EPI_ISL_804308, EPI_ISL_804309, EPI_ISL_804310, EPI_ISL_804311, EPI_ISL_804312, EPI_ISL_804313, EPI_ISL_804314, EPI_ISL_804315, EPI_ISL_804316, EPI_ISL_804317, EPI_ISL_804318, EPI_ISL_804319, EPI_ISL_804320, EPI_ISL_804321, EPI_ISL_804322, EPI_ISL_804324, EPI_ISL_804325, EPI_ISL_804326, EPI_ISL_804327, EPI_ISL_804328, EPI_ISL_804329, EPI_ISL_804336, EPI_ISL_804339, EPI_ISL_804340, EPI_ISL_804341, EPI_ISL_804342, EPI_ISL_804343, EPI_ISL_804344, EPI_ISL_804345, EPI_ISL_804346, EPI_ISL_804347, EPI_ISL_804348, EPI_ISL_804349, EPI_ISL_804350, EPI_ISL_804351, EPI_ISL_804352, EPI_ISL_804353, EPI_ISL_804354, EPI_ISL_804355, EPI_ISL_804356, EPI_ISL_804361, EPI_ISL_804362, EPI_ISL_804363, EPI_ISL_804364, EPI_ISL_804365, EPI_ISL_804366, EPI_ISL_804368, EPI_ISL_804812 | | | |
| see above | Respiratory Virus Unit, National Infection Service, Public Health England | COVID-19 Genomics UK (COG-UK) Consortium | PHE Covid Sequencing Team |
| EPI_ISL_804816, EPI_ISL_804833, EPI_ISL_804834, EPI_ISL_804836, EPI_ISL_804837, EPI_ISL_804841 | DB Diagnosticos do Brasil | Laboratório de Parasitologia Médica - Instituto de Medicina Tropical - Universidade de São Paulo | Nuno Faria, Ingra Morales Claro, Darlan Candido, Lucas A. Moyses Franco, Pamela dos Santos Andrade, Thais de Moura Coletti, Camila A. Maia da Silva, Flavia Cristina Sales, Erika Regina Manuli, Renato A. Santana, Nelson Gaburo, Cecília da Cunha Camilo, Nelson Abrahim Fraiji, Myuki Alfaia Esashika Crispim, Maria do Perpétuo Socorro Sampaio Carvalho, Andrew Rambaut, Nick Loman, Oliver G. Pybus, Ester C. Sabino; DB; HEMOAM; CDL; CADDE Genomic Network. |
| EPI_ISL_804859, EPI_ISL_804941 | DC Public Health Lab/ Dept. of Forensic Sciences | DC Public Health Lab/ Dept. of Forensic Sciences | Scott Nguyen, Elizabeth Zelaya, Connie Maza, Monica Mann, Brittany Hamilton, David Payne, Jocelyn Hauser |
| EPI_ISL_804980 | Columbia University Irving Medical Center | Wadsworth Center, New York State Department of Health | Kirsten St. George, Daryl M. Lamson, Alexis Russel, Matthew Shudt, Melissa A Leisner, Jonathan Plitnick, Navjot Singh, John Kelly, Erasmus Schneider, Erica Lasek-Nesselquist |
| EPI_ISL_806719 | Pathology North - Royal North Shore Hospital - NSW Health Pathology | NSW Health Pathology - Institute of Clinical Pathology and Medical Research; Westmead Hospital; University of Sydney | CIDM-PH et al. |
| EPI_ISL_806720 | Laverty Pathology | NSW Health Pathology - Institute of Clinical Pathology and Medical Research; Westmead Hospital; University of Sydney | CIDM-PH et al. |
| EPI_ISL_807154, EPI_ISL_807155 | Deva County Emergency Hospital | National Institute of Infectious Diseases-Prof. Dr. Matei Bals Molecular Diagnostics Laboratory | Leontina Banica, Marius Surleac, Corina Casangiu, Petre Milu, Andreea Tudor, Simona Paraschiv, Dan Otelea |
| EPI_ISL_810793, EPI_ISL_810794, EPI_ISL_810796, EPI_ISL_810798, EPI_ISL_810799, EPI_ISL_810801, EPI_ISL_810803 | PathWest Laboratory Medicine WA | PathWest Laboratory Medicine WA Microbial Surveillance Unit | PathWest Laboratory Medicine WA Microbial Surveillance Unit |
| EPI_ISL_811120 | Respiratory Virus Unit, National Infection Service, Public Health England | COVID-19 Genomics UK (COG-UK) Consortium | PHE Covid Sequencing Team |
| EPI_ISL_811132 | Viollier AG | Department of Biosystems Science and Engineering, ETH Zürich | Chaoran Chen, Sarah Nadeau, Catharine Aquino, Ivan Topolsky, Philipp Jablonski, Lara Fuhrmann, David Dreifuss, Katharina Jahn, Andreia Cabral de Gouvea, Maria Domenica Moccia, Simon Grüter, Timothy Sykes, Lennart Opitz, Griffin White, Laura Neff, Doris Popovic, Andrea Patrignani, Jay Tracy, Ralph Schlapbach, Christiane Beckmann, Maurice Redondo, Olivier Kobel, Christoph Noppen, Sophie Seidel, Noemie Santamaria de Souza, Niko Beerenwinkel, Tanja Stadler |
| EPI_ISL_811136 | Ministry of Health Turkey | Ministry of Health Turkey | Fatma Bayrakdar, Yasemin Cogun, Süleyman Yalcin, Aye Baak Alta, Gülay Korukluolu |
| EPI_ISL_812290, EPI_ISL_812291, EPI_ISL_812292, EPI_ISL_812293, EPI_ISL_812294, EPI_ISL_812295, EPI_ISL_812296 | Delaware Public Health Lab | Delaware Public Health Lab | Gregory Hovan |
| EPI_ISL_812482 | Laboratorio de Referencia Nacional de Virus Respiratorios, Instituto Nacional de Salud Peru | Laboratorio de Genómica Microbiana, Universidad Peruana Cayetano Heredia | Pablo Tsukayama, Alejandra Dávila-Barclay, Guillermo Salvatierra, Luis González, Pedro E. Romero, Brenda Ayzanoa, Janet Huancachoque, Pool Marcos, Camila Castillo-Vilcahuamán, Oscar Escalante, Priscila Lope, Nancy Rojas |
| EPI_ISL_812761, EPI_ISL_812762, EPI_ISL_812774, EPI_ISL_812775, EPI_ISL_812776, EPI_ISL_812873 | Ministry of Health Turkey | Ministry of Health Turkey | Fatma Bayrakdar, Yasemin Cogun, Süleyman Yalcin, Aye Baak Alta, Gülay Korukluolu |

| | | | |
|---|---|---|---|
| EPI_ISL_812922 | Pasteur Institute in Ho Chi Minh city | Department of Microbiology and Immunology - Pasteur Institute in Ho Chi Minh city | Manh Huy Dao, Hang Thi Thu Pham, Nhung Pham Hong Vu, Hieu Minh Le, Thang Minh Cao, Loan Thi Kim Huynh, Long Thanh Nguyen, Anh Hoang Nguyen, Hieu Trung Nguyen, Thao Thi Ngoc Nguyen, Thinh Viet Nguyen, Quang Duy Pham, Quang Chan Luong, Lan Trong Phan, Thuong Vu Nguyen. |
| EPI_ISL_812924, EPI_ISL_812925, EPI_ISL_812926, EPI_ISL_812927, EPI_ISL_812928 | Ramathibodi Hospital | COVID-19 Network Investigations (CONI) Alliance | Elizabeth Batty, Wasun Chantratita, Thanat Chookajorn, Stefan Fernandez, Angkana Huang, Anthony R. Jones, Khajohn Joonsalak, Chonticha Klungtong, Theerarat Kochakarn, Namfon Kotanan, Krittikorn Kumpornsin, Duangkamon Loesbanluechai, Wudtichai Manasatienkij, Bhakbhoom Panthan, Ekawat Pasomsub, Kingkan Rakmanee, Insee Sensorn, Janjira Thaipadungpanit, Arporn Wangwiwatsin,Treewat Watthanachockchai |
| EPI_ISL_812929, EPI_ISL_812930, EPI_ISL_812931, EPI_ISL_812933, EPI_ISL_812934, EPI_ISL_812935, EPI_ISL_812936, EPI_ISL_812937, EPI_ISL_812938, EPI_ISL_812939, EPI_ISL_812940, EPI_ISL_812941, EPI_ISL_812942, EPI_ISL_812943 | | | |
| see above | Institute for Urban Disease Control and Prevention | COVID-19 Network Investigations (CONI) Alliance | Kamolthip Atsawawaranunt, Elizabeth Batty, Wasun Chantratita, Thanat Chookajorn, Stefan Fernandez, Angkana Huang, Anthony R. Jones, Khajohn Joonsalak, Chonticha Klungtong, Theerarat Kochakarn, Prayuth Kaewmalang, Amornmas Kongklieng, Namfon Kotanan, Krittikorn Kumpornsin, Duangkamon Loesbanluechai, Wudtichai Manasatienkij, Anek Mungaomklang, Bhakbhoom Panthan, Pukkaporn Parnwijitkul, Ekawat Pasomsub, Kingkan Rakmanee, Insee Sensorn, Janjira Thaipadungpanit, Arporn Wangwiwatsin, Treewat Watthanachockchai |
| EPI_ISL_813102, EPI_ISL_813103, EPI_ISL_813104, EPI_ISL_813105, EPI_ISL_813106, EPI_ISL_813107, EPI_ISL_813108, EPI_ISL_813109, EPI_ISL_813110, EPI_ISL_813111, EPI_ISL_813112, EPI_ISL_813113, EPI_ISL_813114, EPI_ISL_813115, EPI_ISL_813117, EPI_ISL_813118, EPI_ISL_813119, EPI_ISL_813120, EPI_ISL_813121, EPI_ISL_813122, EPI_ISL_813123, EPI_ISL_813124, EPI_ISL_813125, EPI_ISL_813126, EPI_ISL_813127, EPI_ISL_813128, EPI_ISL_813129, EPI_ISL_813130, EPI_ISL_813131, EPI_ISL_813132, EPI_ISL_813137 | | | |
| see above | University of Birmingham | COVID-19 Genomics UK (COG-UK) Consortium | Institute of Microbiology, University of Birmingham: Claire McMurray, Joanne Stockton, Samuel Nicholls, Radoslaw Poplawski, Will Rowe, Josh Quick, Nicholas Loman. University of Birmingham Testing Laboratory: Celina M Whalley, Andrew Bosworth, Charlotte Poxon, Kasun Wanigasooriya, Oliver Pickles, Mike Kidd, Alex Richter, Andrew D Beggs PHE Heartlands Lab: Husam Osman, Andrew Bosworth. Queen Elizabeth Hospital: Anna Casey |
| EPI_ISL_813166, EPI_ISL_813168, EPI_ISL_813169, EPI_ISL_813170, EPI_ISL_813176, EPI_ISL_813177, EPI_ISL_813178, EPI_ISL_813179, EPI_ISL_813180, EPI_ISL_813181, EPI_ISL_813184, EPI_ISL_813185, EPI_ISL_813186, EPI_ISL_813191, EPI_ISL_813192, EPI_ISL_813193, EPI_ISL_813194, EPI_ISL_813231, EPI_ISL_813293, EPI_ISL_813317 | | | |
| see above | Department of Pathology, University of Cambridge | COVID-19 Genomics UK (COG-UK) Consortium | Aminu S. Jahun, Yasmin Chaudhry, Grant Hall, Iliana Georgana, Myra Hosmillo, Martin D. Curran, Malte Pinckert, Surendra Parmar, Ian Goodfellow |
| EPI_ISL_813487, EPI_ISL_813488, EPI_ISL_813489, EPI_ISL_813500, EPI_ISL_813501, EPI_ISL_813502, EPI_ISL_813503, EPI_ISL_813504, EPI_ISL_813505, EPI_ISL_813507, EPI_ISL_813508, EPI_ISL_813509, EPI_ISL_813511, EPI_ISL_813512, EPI_ISL_813513, EPI_ISL_813514, EPI_ISL_813515, EPI_ISL_813516, EPI_ISL_813517, EPI_ISL_813518, EPI_ISL_813519, EPI_ISL_813520, EPI_ISL_813521, EPI_ISL_813522, EPI_ISL_813523, EPI_ISL_813524, EPI_ISL_813525, EPI_ISL_813526, EPI_ISL_813527, EPI_ISL_813528, EPI_ISL_813529, EPI_ISL_813530, EPI_ISL_813531, EPI_ISL_813532, EPI_ISL_813533, EPI_ISL_813534, EPI_ISL_813535, EPI_ISL_813536, EPI_ISL_813537, EPI_ISL_813538, EPI_ISL_813539, EPI_ISL_813540, EPI_ISL_813541, EPI_ISL_813542, EPI_ISL_813543, EPI_ISL_813544, EPI_ISL_813545, EPI_ISL_813546, EPI_ISL_813547, EPI_ISL_813548, EPI_ISL_813549, EPI_ISL_813550, EPI_ISL_813551, EPI_ISL_813552, EPI_ISL_813553, EPI_ISL_813554, EPI_ISL_813555, EPI_ISL_813556, EPI_ISL_813557, EPI_ISL_813558, EPI_ISL_813559, EPI_ISL_813560, EPI_ISL_813561, EPI_ISL_813562, EPI_ISL_813563, EPI_ISL_813564, EPI_ISL_813565, EPI_ISL_813566, EPI_ISL_813567, EPI_ISL_813568, EPI_ISL_813569, EPI_ISL_813570, EPI_ISL_813571, EPI_ISL_813572, EPI_ISL_813573, EPI_ISL_813574, EPI_ISL_813575, EPI_ISL_813576, EPI_ISL_813577, EPI_ISL_813578, EPI_ISL_813579, EPI_ISL_813580, EPI_ISL_813584, EPI_ISL_813585, EPI_ISL_813586 | | | |
| see above | University of Exeter | COVID-19 Genomics UK (COG-UK) Consortium | Ben Temperton,Aaron Jeffries,Michelle Michelsen,Joanna Warwick-Dugdale,Audrey Farbos,Robyn Manley,Stephen Michell,Jane Masoli |
| EPI_ISL_813783, EPI_ISL_813785, EPI_ISL_813786, EPI_ISL_813787, EPI_ISL_813788, EPI_ISL_813789, EPI_ISL_813790, EPI_ISL_813791, EPI_ISL_813792, EPI_ISL_813793, EPI_ISL_813794, EPI_ISL_813795, EPI_ISL_813796, EPI_ISL_813797, EPI_ISL_813798, EPI_ISL_813799, EPI_ISL_813800, EPI_ISL_813801, EPI_ISL_813802, EPI_ISL_813804, EPI_ISL_813806 | | | |
| see above | Liverpool Clinical Laboratories | COVID-19 Genomics UK (COG-UK) Consortium | Sam Haldenby, Anita Lucaci, Steve Paterson, Julian Hiscox, Alistair Darby, M Almsaud, A Alrezaihi, Muhannad Alruwaili, Stuart D Armstrong, Jones Benjamin, Eleanor G Bentley, Anu Chawla, Jordan J Clark, Angela Cowell, Richard Eccles, Isabel García-Dorival, Matthew Gemmell, Alessandro Gerada, PKF Gilmore, Richard Gregory, Ximeng Han, Catherine Hartley, Margaret Hughes, Miren Iturriza-Gomara, James Johnson, L Luu, Jenifer Manson, Charlotte Nelson, Elaine O'Toole, Cassie Olateju, Rebekah Penrice-Randal , Lucille Rainbow, N.P Randle, Trevor Ian Robinson, Parul Sharma, Ghada T Shawli, James P Stewart, Neil Swainston, Ecaterina Vamos, Joanne Watts, Mark Whitehead |
| EPI_ISL_813845, EPI_ISL_813846, EPI_ISL_813848, EPI_ISL_813849, EPI_ISL_813850, EPI_ISL_813851, EPI_ISL_813852, EPI_ISL_813853, EPI_ISL_813854, EPI_ISL_813855, EPI_ISL_813856, EPI_ISL_813857, EPI_ISL_813858, EPI_ISL_813859, EPI_ISL_813860, EPI_ISL_813861, EPI_ISL_813862, EPI_ISL_813863, EPI_ISL_813864, EPI_ISL_813865, EPI_ISL_813867, EPI_ISL_813868, EPI_ISL_813869, EPI_ISL_813870, EPI_ISL_813871, EPI_ISL_813872, EPI_ISL_813873, EPI_ISL_813874, EPI_ISL_813875, EPI_ISL_813876, EPI_ISL_813877, EPI_ISL_813878, EPI_ISL_813879, EPI_ISL_813896, EPI_ISL_813897, EPI_ISL_813898, EPI_ISL_813899, EPI_ISL_813936, EPI_ISL_813946, EPI_ISL_813948, EPI_ISL_813949, EPI_ISL_813950, EPI_ISL_813951, EPI_ISL_813952, EPI_ISL_813953, EPI_ISL_813954, EPI_ISL_813955, EPI_ISL_813957, EPI_ISL_813960, EPI_ISL_813961, EPI_ISL_813965, EPI_ISL_813967, EPI_ISL_813968, EPI_ISL_813969, EPI_ISL_813970 | | | |
| see above | University College London, Great Ormond Street Hospital for Children NHS Foundation Trust, Imperial College Healthcare NHS Trust | COVID-19 Genomics UK (COG-UK) Consortium | Sergi Castellano, Rachel Williams, Mark Kristiansen, Paola Resende Silva, Sunando Roy, Tony Brooks, Helena Tutill, Paola Niola, Patricia Dyal, Charlotte Williams, Leysa Forrest, Yasmin Panchbhaya, Jacqueline Findlay, Samuel Weeks, Julianne Brown, Kathryn Harris, Paul Randell, James Price, Alison Holmes, Judith Breuer |
| EPI_ISL_814097, EPI_ISL_814102, EPI_ISL_814103, EPI_ISL_814104, EPI_ISL_814105, EPI_ISL_814106, EPI_ISL_814107, EPI_ISL_814108, EPI_ISL_814109, EPI_ISL_814110, EPI_ISL_814111, EPI_ISL_814112, EPI_ISL_814113, EPI_ISL_814114, EPI_ISL_814115, EPI_ISL_814116, EPI_ISL_814117, EPI_ISL_814118, EPI_ISL_814119, EPI_ISL_814120, EPI_ISL_814121, EPI_ISL_814122, EPI_ISL_814123, EPI_ISL_814124, EPI_ISL_814125, EPI_ISL_814126, EPI_ISL_814127, EPI_ISL_814128, EPI_ISL_814129, EPI_ISL_814130, EPI_ISL_814131, EPI_ISL_814132, EPI_ISL_814133, EPI_ISL_814134, EPI_ISL_814135, EPI_ISL_814136, EPI_ISL_814137, EPI_ISL_814138, EPI_ISL_814139, EPI_ISL_814140, EPI_ISL_814141, EPI_ISL_814142, EPI_ISL_814143, EPI_ISL_814144, EPI_ISL_814145, EPI_ISL_814146, EPI_ISL_814147, EPI_ISL_814148, EPI_ISL_814149, EPI_ISL_814150, EPI_ISL_814151, EPI_ISL_814152, EPI_ISL_814153, EPI_ISL_814154, EPI_ISL_814155, EPI_ISL_814156, EPI_ISL_814157, EPI_ISL_814158, EPI_ISL_814159, EPI_ISL_814160, EPI_ISL_814161, EPI_ISL_814162, EPI_ISL_814163, EPI_ISL_814164, EPI_ISL_814165, EPI_ISL_814166, EPI_ISL_814167, EPI_ISL_814168, EPI_ISL_814169, EPI_ISL_814170, EPI_ISL_814171, EPI_ISL_814172, EPI_ISL_814173, EPI_ISL_814174, EPI_ISL_814175, EPI_ISL_814176, EPI_ISL_814177, EPI_ISL_814178, EPI_ISL_814179 | | | |
| see above | Israel Central Virology laboratory | Israel National Consortium for SARS-CoV-2 sequencing | Neta Zuckerman, Efrat Dahan Bucris, Michal Mandelboim, Dana Bar-Ilan, Oran Erster, Tzvia Mann, Omer Murik, David A. Zeevi, Assaf Rokney, Joseph Jaffe, Eva Nachum, Maya Davidovich Cohen, Ephraim Fass, Gal Zizelski Valenci, Mor Rubinstein, Efrat Rorman, Israel Nissan, Efrat Glick-Saar, Omri Nayshool, Gideon Rechavi, Ella Mendelson, Orna Mor |
| EPI_ISL_814275, EPI_ISL_814280, EPI_ISL_814283, EPI_ISL_814301, EPI_ISL_814305, EPI_ISL_814314, EPI_ISL_814318, EPI_ISL_814325, EPI_ISL_814330, EPI_ISL_814332, EPI_ISL_814336, EPI_ISL_814338, EPI_ISL_814342, EPI_ISL_814347, EPI_ISL_814374, EPI_ISL_814376, EPI_ISL_814379, EPI_ISL_814381, EPI_ISL_814382, EPI_ISL_814385, EPI_ISL_814388, EPI_ISL_814398, EPI_ISL_814470, EPI_ISL_814471, EPI_ISL_814488, EPI_ISL_814489, EPI_ISL_814490, EPI_ISL_814491, EPI_ISL_814503, EPI_ISL_814507, EPI_ISL_814527, EPI_ISL_814554 | | | |
| see above | Wales Specialist Virology Centre Sequencing lab: Pathogen Genomics Unit | COVID-19 Genomics UK (COG-UK) Consortium | Catherine Moore, Johnathan Evans, Laura Gifford, Malorie Perry, Simon Cottrell, Angela Marchbank, Alec Birchley, Alexander Adams, Amy Gaskin, Bree Gatica-Wilcox, Jason Coombes, Joel Southgate, Lauren Gilbert, Lee Graham, Nicole Pacchiarini, Sara Kumziene-Summerhayes, Sarah Taylor, Sophie Jones, Sara Rey, Matthew Bull, Joanne Watkins, Sally Corden, Tom Connor |
| EPI_ISL_814557 | Liverpool Clinical Laboratories | COVID-19 Genomics UK (COG-UK) Consortium | Sam Haldenby, Anita Lucaci, Steve Paterson, Julian Hiscox, Alistair Darby, M Almsaud, A Alrezaihi, Muhannad Alruwaili, Stuart D Armstrong, Jones Benjamin, Eleanor G Bentley, Anu Chawla, Jordan J Clark, Angela Cowell, Richard Eccles, Isabel García-Dorival, Matthew Gemmell, Alessandro Gerada, PKF Gilmore, Richard Gregory, Ximeng Han, Catherine Hartley, Margaret Hughes, Miren Iturriza-Gomara, James Johnson, L Luu, Jenifer Manson, Charlotte Nelson, Elaine O'Toole, Cassie Olateju, Rebekah Penrice-Randal , Lucille Rainbow, N.P Randle, Trevor Ian Robinson, Parul Sharma, Ghada T Shawli, James P Stewart, Neil Swainston, Ecaterina Vamos, Joanne Watts, Mark Whitehead |
| EPI_ISL_814567, EPI_ISL_814585, EPI_ISL_814586, EPI_ISL_814601, EPI_ISL_814762, EPI_ISL_814763, EPI_ISL_815047, EPI_ISL_815051, EPI_ISL_815063, EPI_ISL_815064, EPI_ISL_815065, EPI_ISL_815066, EPI_ISL_815067, EPI_ISL_815076, EPI_ISL_815077, EPI_ISL_815078, EPI_ISL_815079, EPI_ISL_815080, EPI_ISL_815081, EPI_ISL_815082, EPI_ISL_815083, EPI_ISL_815084, EPI_ISL_815085, EPI_ISL_815086, EPI_ISL_815087, EPI_ISL_815137, EPI_ISL_815150, EPI_ISL_815205, EPI_ISL_815206, EPI_ISL_815208, EPI_ISL_815209, EPI_ISL_815210, EPI_ISL_815211, EPI_ISL_815212, EPI_ISL_815213, EPI_ISL_815214, EPI_ISL_815215, EPI_ISL_815216, EPI_ISL_815217, EPI_ISL_815218, EPI_ISL_815219, EPI_ISL_815220, EPI_ISL_815221, EPI_ISL_815222, EPI_ISL_815223, EPI_ISL_815224, EPI_ISL_815225, EPI_ISL_815226, EPI_ISL_815227, EPI_ISL_815228, EPI_ISL_815229, EPI_ISL_815230, EPI_ISL_815231, EPI_ISL_815232 | | | |
| see above | Wales Specialist Virology Centre Sequencing lab: Pathogen Genomics Unit | COVID-19 Genomics UK (COG-UK) Consortium | Catherine Moore, Johnathan Evans, Laura Gifford, Malorie Perry, Simon Cottrell, Angela Marchbank, Alec Birchley, Alexander Adams, Amy Gaskin, Bree Gatica-Wilcox, Jason Coombes, Joel Southgate, Lauren Gilbert, Lee Graham, Nicole Pacchiarini, Sara Kumziene-Summerhayes, Sarah Taylor, Sophie Jones, Sara Rey, Matthew Bull, Joanne Watkins, Sally Corden, Tom Connor |
| EPI_ISL_816153, EPI_ISL_816172, EPI_ISL_816174, EPI_ISL_816197, EPI_ISL_816198, EPI_ISL_816199, EPI_ISL_816200, EPI_ISL_816201, EPI_ISL_816204, EPI_ISL_816205, EPI_ISL_816206, EPI_ISL_816207 | | | |
| see above | Centre for Enzyme Innovation, University of Portsmouth / Translational Research Laboratory, Portsmouth Hospitals NHS Trust | COVID-19 Genomics UK (COG-UK) Consortium | Angela Beckett,Yann Bourgeois,Garry Scarlett,Sharon Glaysher,Scott Elliott,Kelly Bicknell,Robert Impey,Allyson Lloyd,Sarah Wyllie,Ethan Butcher,Anoop Chauhan,Samuel Robson |
| EPI_ISL_816231, EPI_ISL_816242, EPI_ISL_816247, EPI_ISL_816271, EPI_ISL_816273, EPI_ISL_816280, EPI_ISL_816283, EPI_ISL_816288, EPI_ISL_816289, EPI_ISL_816308, EPI_ISL_816332, EPI_ISL_816335, EPI_ISL_816339, EPI_ISL_816340, EPI_ISL_816345, EPI_ISL_816351, EPI_ISL_816352, EPI_ISL_816359, EPI_ISL_816372, EPI_ISL_816374, EPI_ISL_816379, EPI_ISL_816381, EPI_ISL_816384, EPI_ISL_816401, EPI_ISL_816412, EPI_ISL_816414, EPI_ISL_816429, EPI_ISL_816456, EPI_ISL_816457, EPI_ISL_816461, EPI_ISL_816467, EPI_ISL_816478, EPI_ISL_816491, EPI_ISL_816497, EPI_ISL_816534, EPI_ISL_816540, EPI_ISL_816549, EPI_ISL_816550, EPI_ISL_816557, EPI_ISL_816558, EPI_ISL_816575, EPI_ISL_816578, EPI_ISL_816586, EPI_ISL_816609, EPI_ISL_816631, EPI_ISL_816637, EPI_ISL_816638, EPI_ISL_816647, EPI_ISL_816651 | | | |
| see above | Virology Department, Sheffield Teaching Hospitals NHS Foundation Trust/Department of Infection, Immunity and Cardiovascular Disease, The Medical School, University of Sheffield | COVID-19 Genomics UK (COG-UK) Consortium | Thushan de Silva, Matthew Parker, Nikki Smith, Adri Angyal, Rebecca Brown, Luke Green, Rachel Tucker, Paul Parsons, Danielle Groves, Katie Johnson, Laura Carrilero, Alex Keeley, Dave Partridge, Matthew Wyles, Benjamin Lindsey, Mehmet Yavuz, Mohammad Raza, Cariad Evans |
| EPI_ISL_819130, EPI_ISL_819131, EPI_ISL_819132, EPI_ISL_819136, EPI_ISL_819137, EPI_ISL_819143, EPI_ISL_819148, EPI_ISL_819150, EPI_ISL_819167, EPI_ISL_819168, EPI_ISL_819172, EPI_ISL_819173, EPI_ISL_819174, EPI_ISL_819176, EPI_ISL_819178, EPI_ISL_819179, EPI_ISL_819181, EPI_ISL_819182, EPI_ISL_819183, EPI_ISL_819184, EPI_ISL_819185, EPI_ISL_819193, EPI_ISL_819196, EPI_ISL_819199, EPI_ISL_819202 | | | |

| | | | |
|---|---|---|---|
| see above | Servicio de Microbiología, Hospital Universitario Son Espases | SeqCOVID-SPAIN consortium/IBV(CSIC) | Carla López-Causapé, Jordi Reina, Antonio Oliver and SeqCOVID-SPAIN consortium |
| EPI_ISL_819430 | Quadram Institute Bioscience | COVID-19 Genomics UK (COG-UK) Consortium | Dave J. Baker, Gemma L. Kay, Alp Aydin, Thanh Le-Viet, Steven Rudder, Ana P. Tedim, Anastasia Kolyva, Maria Diaz, Leonardo de Oliveira Martins, Nabil-Fareed Alikhan, Lizzie Meadows, Rachael Stanley, Ngozi Elumogo, Muhammed Yasir, Nicholas M. Thomson, Alexander J Trotter, Rachel Gilroy, Samuel Bloomfield, Claire Stuart, Andrew Bell, Reenesh Prakash, Samir Dervisevic, Alison E. Mather, John Wain, Mark Webber, Andrew J. Page, Justin O'Grady |
| EPI_ISL_819882, EPI_ISL_819883, EPI_ISL_819884 | University College London, Great Ormond Street Hospital for Children NHS Foundation Trust, Imperial College Healthcare NHS Trust | COVID-19 Genomics UK (COG-UK) Consortium | Sergi Castellano, Rachel Williams, Mark Kristiansen, Paola Resende Silva, Sunando Roy, Tony Brooks, Helena Tutill, Paola Niola, Patricia Dyal, Charlotte Williams, Leysa Forrest, Yasmin Panchbhaya, Jacqueline Findlay, Samuel Weeks, Julianne Brown, Kathryn Harris, Paul Randell, James Price, Alison Holmes, Judith Breuer |
| EPI_ISL_819885, EPI_ISL_819886, EPI_ISL_819887 | Quadram Institute Bioscience | COVID-19 Genomics UK (COG-UK) Consortium | Dave J. Baker, Gemma L. Kay, Alp Aydin, Thanh Le-Viet, Steven Rudder, Ana P. Tedim, Anastasia Kolyva, Maria Diaz, Leonardo de Oliveira Martins, Nabil-Fareed Alikhan, Lizzie Meadows, Rachael Stanley, Ngozi Elumogo, Muhammed Yasir, Nicholas M. Thomson, Alexander J Trotter, Rachel Gilroy, Samuel Bloomfield, Claire Stuart, Andrew Bell, Reenesh Prakash, Samir Dervisevic, Alison E. Mather, John Wain, Mark Webber, Andrew J. Page, Justin O'Grady |
| EPI_ISL_819888, EPI_ISL_819889 | University College London, Great Ormond Street Hospital for Children NHS Foundation Trust, Imperial College Healthcare NHS Trust | COVID-19 Genomics UK (COG-UK) Consortium | Sergi Castellano, Rachel Williams, Mark Kristiansen, Paola Resende Silva, Sunando Roy, Tony Brooks, Helena Tutill, Paola Niola, Patricia Dyal, Charlotte Williams, Leysa Forrest, Yasmin Panchbhaya, Jacqueline Findlay, Samuel Weeks, Julianne Brown, Kathryn Harris, Paul Randell, James Price, Alison Holmes, Judith Breuer |
| EPI_ISL_819890, EPI_ISL_819891 | Quadram Institute Bioscience | COVID-19 Genomics UK (COG-UK) Consortium | Dave J. Baker, Gemma L. Kay, Alp Aydin, Thanh Le-Viet, Steven Rudder, Ana P. Tedim, Anastasia Kolyva, Maria Diaz, Leonardo de Oliveira Martins, Nabil-Fareed Alikhan, Lizzie Meadows, Rachael Stanley, Ngozi Elumogo, Muhammed Yasir, Nicholas M. Thomson, Alexander J Trotter, Rachel Gilroy, Samuel Bloomfield, Claire Stuart, Andrew Bell, Reenesh Prakash, Samir Dervisevic, Alison E. Mather, John Wain, Mark Webber, Andrew J. Page, Justin O'Grady |
| EPI_ISL_819892, EPI_ISL_819893 | University College London, Great Ormond Street Hospital for Children NHS Foundation Trust, Imperial College Healthcare NHS Trust | COVID-19 Genomics UK (COG-UK) Consortium | Sergi Castellano, Rachel Williams, Mark Kristiansen, Paola Resende Silva, Sunando Roy, Tony Brooks, Helena Tutill, Paola Niola, Patricia Dyal, Charlotte Williams, Leysa Forrest, Yasmin Panchbhaya, Jacqueline Findlay, Samuel Weeks, Julianne Brown, Kathryn Harris, Paul Randell, James Price, Alison Holmes, Judith Breuer |
| EPI_ISL_819894, EPI_ISL_819895, EPI_ISL_819896, EPI_ISL_819897, EPI_ISL_819898, EPI_ISL_819899, EPI_ISL_819900, EPI_ISL_819901, EPI_ISL_819902, EPI_ISL_819903, EPI_ISL_819904, EPI_ISL_819905, EPI_ISL_819906, EPI_ISL_819907, EPI_ISL_819908, EPI_ISL_819909, EPI_ISL_819910, EPI_ISL_819911, EPI_ISL_819912, EPI_ISL_819913, EPI_ISL_819914, EPI_ISL_819915, EPI_ISL_819916, EPI_ISL_819917, EPI_ISL_819918 | | | |
| see above | Quadram Institute Bioscience | COVID-19 Genomics UK (COG-UK) Consortium | Dave J. Baker, Gemma L. Kay, Alp Aydin, Thanh Le-Viet, Steven Rudder, Ana P. Tedim, Anastasia Kolyva, Maria Diaz, Leonardo de Oliveira Martins, Nabil-Fareed Alikhan, Lizzie Meadows, Rachael Stanley, Ngozi Elumogo, Muhammed Yasir, Nicholas M. Thomson, Alexander J Trotter, Rachel Gilroy, Samuel Bloomfield, Claire Stuart, Andrew Bell, Reenesh Prakash, Samir Dervisevic, Alison E. Mather, John Wain, Mark Webber, Andrew J. Page, Justin O'Grady |
| EPI_ISL_819919, EPI_ISL_819920, EPI_ISL_819921, EPI_ISL_819922, EPI_ISL_819923, EPI_ISL_819924, EPI_ISL_819925, EPI_ISL_819926, EPI_ISL_819927 | University College London, Great Ormond Street Hospital for Children NHS Foundation Trust, Imperial College Healthcare NHS Trust | COVID-19 Genomics UK (COG-UK) Consortium | Sergi Castellano, Rachel Williams, Mark Kristiansen, Paola Resende Silva, Sunando Roy, Tony Brooks, Helena Tutill, Paola Niola, Patricia Dyal, Charlotte Williams, Leysa Forrest, Yasmin Panchbhaya, Jacqueline Findlay, Samuel Weeks, Julianne Brown, Kathryn Harris, Paul Randell, James Price, Alison Holmes, Judith Breuer |
| EPI_ISL_819928 | Quadram Institute Bioscience | COVID-19 Genomics UK (COG-UK) Consortium | Dave J. Baker, Gemma L. Kay, Alp Aydin, Thanh Le-Viet, Steven Rudder, Ana P. Tedim, Anastasia Kolyva, Maria Diaz, Leonardo de Oliveira Martins, Nabil-Fareed Alikhan, Lizzie Meadows, Rachael Stanley, Ngozi Elumogo, Muhammed Yasir, Nicholas M. Thomson, Alexander J Trotter, Rachel Gilroy, Samuel Bloomfield, Claire Stuart, Andrew Bell, Reenesh Prakash, Samir Dervisevic, Alison E. Mather, John Wain, Mark Webber, Andrew J. Page, Justin O'Grady |
| EPI_ISL_819929, EPI_ISL_819930 | University College London, Great Ormond Street Hospital for Children NHS Foundation Trust, Imperial College Healthcare NHS Trust | COVID-19 Genomics UK (COG-UK) Consortium | Sergi Castellano, Rachel Williams, Mark Kristiansen, Paola Resende Silva, Sunando Roy, Tony Brooks, Helena Tutill, Paola Niola, Patricia Dyal, Charlotte Williams, Leysa Forrest, Yasmin Panchbhaya, Jacqueline Findlay, Samuel Weeks, Julianne Brown, Kathryn Harris, Paul Randell, James Price, Alison Holmes, Judith Breuer |
| EPI_ISL_819931, EPI_ISL_819932, EPI_ISL_819933, EPI_ISL_819934, EPI_ISL_819935 | Quadram Institute Bioscience | COVID-19 Genomics UK (COG-UK) Consortium | Dave J. Baker, Gemma L. Kay, Alp Aydin, Thanh Le-Viet, Steven Rudder, Ana P. Tedim, Anastasia Kolyva, Maria Diaz, Leonardo de Oliveira Martins, Nabil-Fareed Alikhan, Lizzie Meadows, Rachael Stanley, Ngozi Elumogo, Muhammed Yasir, Nicholas M. Thomson, Alexander J Trotter, Rachel Gilroy, Samuel Bloomfield, Claire Stuart, Andrew Bell, Reenesh Prakash, Samir Dervisevic, Alison E. Mather, John Wain, Mark Webber, Andrew J. Page, Justin O'Grady |
| EPI_ISL_819936, EPI_ISL_819937, EPI_ISL_819938, EPI_ISL_819939, EPI_ISL_819940, EPI_ISL_819941 | University College London, Great Ormond Street Hospital for Children NHS Foundation Trust, Imperial College Healthcare NHS Trust | COVID-19 Genomics UK (COG-UK) Consortium | Sergi Castellano, Rachel Williams, Mark Kristiansen, Paola Resende Silva, Sunando Roy, Tony Brooks, Helena Tutill, Paola Niola, Patricia Dyal, Charlotte Williams, Leysa Forrest, Yasmin Panchbhaya, Jacqueline Findlay, Samuel Weeks, Julianne Brown, Kathryn Harris, Paul Randell, James Price, Alison Holmes, Judith Breuer |
| EPI_ISL_819942 | Queens Medical Centre, Clinical Microbiology Department / DeepSeq Nottingham | COVID-19 Genomics UK (COG-UK) Consortium | Gemma Clark, Wendy Smith, Manjinder Khakh, Vicki M Fleming, Michelle M Lister, Hannah Howson-Wells, Jonathan Ball, Patrick McClure, Joseph Chappell, Theocharis Tsoleridis, Nadine Holmes, Matthew Carlisle, Christopher Moore, Fei Sang, Johnny Debebe, Victoria Wright, Matthew Loose |
| EPI_ISL_819943, EPI_ISL_819944 | University College London, Great Ormond Street Hospital for Children NHS Foundation Trust, Imperial College Healthcare NHS Trust | COVID-19 Genomics UK (COG-UK) Consortium | Sergi Castellano, Rachel Williams, Mark Kristiansen, Paola Resende Silva, Sunando Roy, Tony Brooks, Helena Tutill, Paola Niola, Patricia Dyal, Charlotte Williams, Leysa Forrest, Yasmin Panchbhaya, Jacqueline Findlay, Samuel Weeks, Julianne Brown, Kathryn Harris, Paul Randell, James Price, Alison Holmes, Judith Breuer |
| EPI_ISL_819945 | Quadram Institute Bioscience | COVID-19 Genomics UK (COG-UK) Consortium | Dave J. Baker, Gemma L. Kay, Alp Aydin, Thanh Le-Viet, Steven Rudder, Ana P. Tedim, Anastasia Kolyva, Maria Diaz, Leonardo de Oliveira Martins, Nabil-Fareed Alikhan, Lizzie Meadows, Rachael Stanley, Ngozi Elumogo, Muhammed Yasir, Nicholas M. Thomson, Alexander J Trotter, Rachel Gilroy, Samuel Bloomfield, Claire Stuart, Andrew Bell, Reenesh Prakash, Samir Dervisevic, Alison E. Mather, John Wain, Mark Webber, Andrew J. Page, Justin O'Grady |
| EPI_ISL_819946, EPI_ISL_819947, EPI_ISL_819948, EPI_ISL_819949, EPI_ISL_819950, EPI_ISL_819951, EPI_ISL_819952 | University College London, Great Ormond Street Hospital for Children NHS Foundation Trust, Imperial College Healthcare NHS Trust | COVID-19 Genomics UK (COG-UK) Consortium | Sergi Castellano, Rachel Williams, Mark Kristiansen, Paola Resende Silva, Sunando Roy, Tony Brooks, Helena Tutill, Paola Niola, Patricia Dyal, Charlotte Williams, Leysa Forrest, Yasmin Panchbhaya, Jacqueline Findlay, Samuel Weeks, Julianne Brown, Kathryn Harris, Paul Randell, James Price, Alison Holmes, Judith Breuer |
| EPI_ISL_819953 | Quadram Institute Bioscience | COVID-19 Genomics UK (COG-UK) Consortium | Dave J. Baker, Gemma L. Kay, Alp Aydin, Thanh Le-Viet, Steven Rudder, Ana P. Tedim, Anastasia Kolyva, Maria Diaz, Leonardo de Oliveira Martins, Nabil-Fareed Alikhan, Lizzie Meadows, Rachael Stanley, Ngozi Elumogo, Muhammed Yasir, Nicholas M. Thomson, Alexander J Trotter, Rachel Gilroy, Samuel Bloomfield, Claire Stuart, Andrew Bell, Reenesh Prakash, Samir Dervisevic, Alison E. Mather, John Wain, Mark Webber, Andrew J. Page, Justin O'Grady |
| EPI_ISL_819954, EPI_ISL_819955 | University College London, Great Ormond Street Hospital for Children NHS Foundation Trust, Imperial College Healthcare NHS Trust | COVID-19 Genomics UK (COG-UK) Consortium | Sergi Castellano, Rachel Williams, Mark Kristiansen, Paola Resende Silva, Sunando Roy, Tony Brooks, Helena Tutill, Paola Niola, Patricia Dyal, Charlotte Williams, Leysa Forrest, Yasmin Panchbhaya, Jacqueline Findlay, Samuel Weeks, Julianne Brown, Kathryn Harris, Paul Randell, James Price, Alison Holmes, Judith Breuer |
| EPI_ISL_819956 | Quadram Institute Bioscience | COVID-19 Genomics UK (COG-UK) Consortium | Dave J. Baker, Gemma L. Kay, Alp Aydin, Thanh Le-Viet, Steven Rudder, Ana P. Tedim, Anastasia Kolyva, Maria Diaz, Leonardo de Oliveira Martins, Nabil-Fareed Alikhan, Lizzie Meadows, Rachael Stanley, Ngozi Elumogo, Muhammed Yasir, Nicholas M. Thomson, Alexander J Trotter, Rachel Gilroy, Samuel Bloomfield, Claire Stuart, Andrew Bell, Reenesh Prakash, Samir Dervisevic, Alison E. Mather, John Wain, Mark Webber, Andrew J. Page, Justin O'Grady |
| EPI_ISL_819957, EPI_ISL_819958 | University College London, Great Ormond Street Hospital for Children NHS Foundation Trust, Imperial College Healthcare NHS Trust | COVID-19 Genomics UK (COG-UK) Consortium | Sergi Castellano, Rachel Williams, Mark Kristiansen, Paola Resende Silva, Sunando Roy, Tony Brooks, Helena Tutill, Paola Niola, Patricia Dyal, Charlotte Williams, Leysa Forrest, Yasmin Panchbhaya, Jacqueline Findlay, Samuel Weeks, Julianne Brown, Kathryn Harris, Paul Randell, James Price, Alison Holmes, Judith Breuer |
| EPI_ISL_819959, EPI_ISL_819960 | Quadram Institute Bioscience | COVID-19 Genomics UK (COG-UK) Consortium | Dave J. Baker, Gemma L. Kay, Alp Aydin, Thanh Le-Viet, Steven Rudder, Ana P. Tedim, Anastasia Kolyva, Maria Diaz, Leonardo de Oliveira Martins, Nabil-Fareed Alikhan, Lizzie Meadows, Rachael Stanley, Ngozi Elumogo, Muhammed Yasir, Nicholas M. Thomson, Alexander J Trotter, Rachel Gilroy, |

| | | | |
|---|---|---|---|
| | | | Samuel Bloomfield, Claire Stuart, Andrew Bell, Reenesh Prakash, Samir Dervisevic, Alison E. Mather, John Wain, Mark Webber, Andrew J. Page, Justin O'Grady |
| EPI_ISL_819961, EPI_ISL_819962 | University College London, Great Ormond Street Hospital for Children NHS Foundation Trust, Imperial College Healthcare NHS Trust | COVID-19 Genomics UK (COG-UK) Consortium | Sergi Castellano, Rachel Williams, Mark Kristiansen, Paola Resende Silva, Sunando Roy, Tony Brooks, Helena Tutill, Paola Niola, Patricia Dyal, Charlotte Williams, Leysa Forrest, Yasmin Panchbhaya, Jacqueline Findlay, Samuel Weeks, Julianne Brown, Kathryn Harris, Paul Randell, James Price, Alison Holmes, Judith Breuer |
| EPI_ISL_819963 | Quadram Institute Bioscience | COVID-19 Genomics UK (COG-UK) Consortium | Dave J. Baker, Gemma L. Kay, Alp Aydin, Thanh Le-Viet, Steven Rudder, Ana P. Tedim, Anastasia Kolyva, Maria Diaz, Leonardo de Oliveira Martins, Nabil-Fareed Alikhan, Lizzie Meadows, Rachael Stanley, Ngozi Elumogo, Muhammed Yasir, Nicholas M. Thomson, Alexander J Trotter, Rachel Gilroy, Samuel Bloomfield, Claire Stuart, Andrew Bell, Reenesh Prakash, Samir Dervisevic, Alison E. Mather, John Wain, Mark Webber, Andrew J. Page, Justin O'Grady |
| EPI_ISL_819964, EPI_ISL_819965, EPI_ISL_819966, EPI_ISL_819967, EPI_ISL_819968, EPI_ISL_819969 | University College London, Great Ormond Street Hospital for Children NHS Foundation Trust, Imperial College Healthcare NHS Trust | COVID-19 Genomics UK (COG-UK) Consortium | Sergi Castellano, Rachel Williams, Mark Kristiansen, Paola Resende Silva, Sunando Roy, Tony Brooks, Helena Tutill, Paola Niola, Patricia Dyal, Charlotte Williams, Leysa Forrest, Yasmin Panchbhaya, Jacqueline Findlay, Samuel Weeks, Julianne Brown, Kathryn Harris, Paul Randell, James Price, Alison Holmes, Judith Breuer |
| EPI_ISL_819970, EPI_ISL_819971, EPI_ISL_819972, EPI_ISL_819973, EPI_ISL_819974, EPI_ISL_819975, EPI_ISL_819976, EPI_ISL_819977 | Quadram Institute Bioscience | COVID-19 Genomics UK (COG-UK) Consortium | Dave J. Baker, Gemma L. Kay, Alp Aydin, Thanh Le-Viet, Steven Rudder, Ana P. Tedim, Anastasia Kolyva, Maria Diaz, Leonardo de Oliveira Martins, Nabil-Fareed Alikhan, Lizzie Meadows, Rachael Stanley, Ngozi Elumogo, Muhammed Yasir, Nicholas M. Thomson, Alexander J Trotter, Rachel Gilroy, Samuel Bloomfield, Claire Stuart, Andrew Bell, Reenesh Prakash, Samir Dervisevic, Alison E. Mather, John Wain, Mark Webber, Andrew J. Page, Justin O'Grady |
| EPI_ISL_819978, EPI_ISL_819979, EPI_ISL_819980, EPI_ISL_819981, EPI_ISL_819982, EPI_ISL_819983, EPI_ISL_819984, EPI_ISL_819985, EPI_ISL_819986, EPI_ISL_819987, EPI_ISL_819988, EPI_ISL_819989 | | | |
| see above | University College London, Great Ormond Street Hospital for Children NHS Foundation Trust, Imperial College Healthcare NHS Trust | COVID-19 Genomics UK (COG-UK) Consortium | Sergi Castellano, Rachel Williams, Mark Kristiansen, Paola Resende Silva, Sunando Roy, Tony Brooks, Helena Tutill, Paola Niola, Patricia Dyal, Charlotte Williams, Leysa Forrest, Yasmin Panchbhaya, Jacqueline Findlay, Samuel Weeks, Julianne Brown, Kathryn Harris, Paul Randell, James Price, Alison Holmes, Judith Breuer |
| EPI_ISL_819990 | Quadram Institute Bioscience | COVID-19 Genomics UK (COG-UK) Consortium | Dave J. Baker, Gemma L. Kay, Alp Aydin, Thanh Le-Viet, Steven Rudder, Ana P. Tedim, Anastasia Kolyva, Maria Diaz, Leonardo de Oliveira Martins, Nabil-Fareed Alikhan, Lizzie Meadows, Rachael Stanley, Ngozi Elumogo, Muhammed Yasir, Nicholas M. Thomson, Alexander J Trotter, Rachel Gilroy, Samuel Bloomfield, Claire Stuart, Andrew Bell, Reenesh Prakash, Samir Dervisevic, Alison E. Mather, John Wain, Mark Webber, Andrew J. Page, Justin O'Grady |
| EPI_ISL_819991, EPI_ISL_819992, EPI_ISL_819993, EPI_ISL_819994, EPI_ISL_819995 | University College London, Great Ormond Street Hospital for Children NHS Foundation Trust, Imperial College Healthcare NHS Trust | COVID-19 Genomics UK (COG-UK) Consortium | Sergi Castellano, Rachel Williams, Mark Kristiansen, Paola Resende Silva, Sunando Roy, Tony Brooks, Helena Tutill, Paola Niola, Patricia Dyal, Charlotte Williams, Leysa Forrest, Yasmin Panchbhaya, Jacqueline Findlay, Samuel Weeks, Julianne Brown, Kathryn Harris, Paul Randell, James Price, Alison Holmes, Judith Breuer |
| EPI_ISL_819996 | Quadram Institute Bioscience | COVID-19 Genomics UK (COG-UK) Consortium | Dave J. Baker, Gemma L. Kay, Alp Aydin, Thanh Le-Viet, Steven Rudder, Ana P. Tedim, Anastasia Kolyva, Maria Diaz, Leonardo de Oliveira Martins, Nabil-Fareed Alikhan, Lizzie Meadows, Rachael Stanley, Ngozi Elumogo, Muhammed Yasir, Nicholas M. Thomson, Alexander J Trotter, Rachel Gilroy, Samuel Bloomfield, Claire Stuart, Andrew Bell, Reenesh Prakash, Samir Dervisevic, Alison E. Mather, John Wain, Mark Webber, Andrew J. Page, Justin O'Grady |
| EPI_ISL_819997, EPI_ISL_819998, EPI_ISL_819999, EPI_ISL_820000 | University College London, Great Ormond Street Hospital for Children NHS Foundation Trust, Imperial College Healthcare NHS Trust | COVID-19 Genomics UK (COG-UK) Consortium | Sergi Castellano, Rachel Williams, Mark Kristiansen, Paola Resende Silva, Sunando Roy, Tony Brooks, Helena Tutill, Paola Niola, Patricia Dyal, Charlotte Williams, Leysa Forrest, Yasmin Panchbhaya, Jacqueline Findlay, Samuel Weeks, Julianne Brown, Kathryn Harris, Paul Randell, James Price, Alison Holmes, Judith Breuer |
| EPI_ISL_820001, EPI_ISL_820002, EPI_ISL_820003, EPI_ISL_820004, EPI_ISL_820005 | Quadram Institute Bioscience | COVID-19 Genomics UK (COG-UK) Consortium | Dave J. Baker, Gemma L. Kay, Alp Aydin, Thanh Le-Viet, Steven Rudder, Ana P. Tedim, Anastasia Kolyva, Maria Diaz, Leonardo de Oliveira Martins, Nabil-Fareed Alikhan, Lizzie Meadows, Rachael Stanley, Ngozi Elumogo, Muhammed Yasir, Nicholas M. Thomson, Alexander J Trotter, Rachel Gilroy, Samuel Bloomfield, Claire Stuart, Andrew Bell, Reenesh Prakash, Samir Dervisevic, Alison E. Mather, John Wain, Mark Webber, Andrew J. Page, Justin O'Grady |
| EPI_ISL_820006 | Queens Medical Centre, Clinical Microbiology Department / DeepSeq Nottingham | COVID-19 Genomics UK (COG-UK) Consortium | Gemma Clark, Wendy Smith, Manjinder Khakh, Vicki M Fleming, Michelle M Lister, Hannah Howson-Wells, Jonathan Ball, Patrick McClure, Joseph Chappell, Theocharis Tsoleridis, Nadine Holmes, Matthew Carlisle, Christopher Moore, Fei Sang, Johnny Debebe, Victoria Wright, Matthew Loose |
| EPI_ISL_820007, EPI_ISL_820008, EPI_ISL_820009, EPI_ISL_820010 | University College London, Great Ormond Street Hospital for Children NHS Foundation Trust, Imperial College Healthcare NHS Trust | COVID-19 Genomics UK (COG-UK) Consortium | Sergi Castellano, Rachel Williams, Mark Kristiansen, Paola Resende Silva, Sunando Roy, Tony Brooks, Helena Tutill, Paola Niola, Patricia Dyal, Charlotte Williams, Leysa Forrest, Yasmin Panchbhaya, Jacqueline Findlay, Samuel Weeks, Julianne Brown, Kathryn Harris, Paul Randell, James Price, Alison Holmes, Judith Breuer |
| EPI_ISL_820011 | Quadram Institute Bioscience | COVID-19 Genomics UK (COG-UK) Consortium | Dave J. Baker, Gemma L. Kay, Alp Aydin, Thanh Le-Viet, Steven Rudder, Ana P. Tedim, Anastasia Kolyva, Maria Diaz, Leonardo de Oliveira Martins, Nabil-Fareed Alikhan, Lizzie Meadows, Rachael Stanley, Ngozi Elumogo, Muhammed Yasir, Nicholas M. Thomson, Alexander J Trotter, Rachel Gilroy, Samuel Bloomfield, Claire Stuart, Andrew Bell, Reenesh Prakash, Samir Dervisevic, Alison E. Mather, John Wain, Mark Webber, Andrew J. Page, Justin O'Grady |
| EPI_ISL_820012, EPI_ISL_820013, EPI_ISL_820014 | University College London, Great Ormond Street Hospital for Children NHS Foundation Trust, Imperial College Healthcare NHS Trust | COVID-19 Genomics UK (COG-UK) Consortium | Sergi Castellano, Rachel Williams, Mark Kristiansen, Paola Resende Silva, Sunando Roy, Tony Brooks, Helena Tutill, Paola Niola, Patricia Dyal, Charlotte Williams, Leysa Forrest, Yasmin Panchbhaya, Jacqueline Findlay, Samuel Weeks, Julianne Brown, Kathryn Harris, Paul Randell, James Price, Alison Holmes, Judith Breuer |
| EPI_ISL_820015 | Quadram Institute Bioscience | COVID-19 Genomics UK (COG-UK) Consortium | Dave J. Baker, Gemma L. Kay, Alp Aydin, Thanh Le-Viet, Steven Rudder, Ana P. Tedim, Anastasia Kolyva, Maria Diaz, Leonardo de Oliveira Martins, Nabil-Fareed Alikhan, Lizzie Meadows, Rachael Stanley, Ngozi Elumogo, Muhammed Yasir, Nicholas M. Thomson, Alexander J Trotter, Rachel Gilroy, Samuel Bloomfield, Claire Stuart, Andrew Bell, Reenesh Prakash, Samir Dervisevic, Alison E. Mather, John Wain, Mark Webber, Andrew J. Page, Justin O'Grady |
| EPI_ISL_820016, EPI_ISL_820017, EPI_ISL_820018, EPI_ISL_820019, EPI_ISL_820020 | University College London, Great Ormond Street Hospital for Children NHS Foundation Trust, Imperial College Healthcare NHS Trust | COVID-19 Genomics UK (COG-UK) Consortium | Sergi Castellano, Rachel Williams, Mark Kristiansen, Paola Resende Silva, Sunando Roy, Tony Brooks, Helena Tutill, Paola Niola, Patricia Dyal, Charlotte Williams, Leysa Forrest, Yasmin Panchbhaya, Jacqueline Findlay, Samuel Weeks, Julianne Brown, Kathryn Harris, Paul Randell, James Price, Alison Holmes, Judith Breuer |
| EPI_ISL_820021, EPI_ISL_820022, EPI_ISL_820023 | Quadram Institute Bioscience | COVID-19 Genomics UK (COG-UK) Consortium | Dave J. Baker, Gemma L. Kay, Alp Aydin, Thanh Le-Viet, Steven Rudder, Ana P. Tedim, Anastasia Kolyva, Maria Diaz, Leonardo de Oliveira Martins, Nabil-Fareed Alikhan, Lizzie Meadows, Rachael Stanley, Ngozi Elumogo, Muhammed Yasir, Nicholas M. Thomson, Alexander J Trotter, Rachel Gilroy, Samuel Bloomfield, Claire Stuart, Andrew Bell, Reenesh Prakash, Samir Dervisevic, Alison E. Mather, John Wain, Mark Webber, Andrew J. Page, Justin O'Grady |
| EPI_ISL_820024 | University College London, Great Ormond Street Hospital for Children NHS Foundation Trust, Imperial College Healthcare NHS Trust | COVID-19 Genomics UK (COG-UK) Consortium | Sergi Castellano, Rachel Williams, Mark Kristiansen, Paola Resende Silva, Sunando Roy, Tony Brooks, Helena Tutill, Paola Niola, Patricia Dyal, Charlotte Williams, Leysa Forrest, Yasmin Panchbhaya, Jacqueline Findlay, Samuel Weeks, Julianne Brown, Kathryn Harris, Paul Randell, James Price, Alison Holmes, Judith Breuer |
| EPI_ISL_820025, EPI_ISL_820026, EPI_ISL_820027, EPI_ISL_820028, EPI_ISL_820029, EPI_ISL_820030 | Quadram Institute Bioscience | COVID-19 Genomics UK (COG-UK) Consortium | Dave J. Baker, Gemma L. Kay, Alp Aydin, Thanh Le-Viet, Steven Rudder, Ana P. Tedim, Anastasia Kolyva, Maria Diaz, Leonardo de Oliveira Martins, Nabil-Fareed Alikhan, Lizzie Meadows, Rachael Stanley, Ngozi Elumogo, Muhammed Yasir, Nicholas M. Thomson, Alexander J Trotter, Rachel Gilroy, Samuel Bloomfield, Claire Stuart, Andrew Bell, Reenesh Prakash, Samir Dervisevic, Alison E. Mather, John Wain, Mark Webber, Andrew J. Page, Justin O'Grady |
| EPI_ISL_820031, EPI_ISL_820032, EPI_ISL_820033, EPI_ISL_820034, EPI_ISL_820035, EPI_ISL_820036, EPI_ISL_820037, EPI_ISL_820038, EPI_ISL_820039, EPI_ISL_820040 | University College London, Great Ormond Street Hospital for Children NHS Foundation Trust, Imperial College Healthcare NHS Trust | COVID-19 Genomics UK (COG-UK) Consortium | Sergi Castellano, Rachel Williams, Mark Kristiansen, Paola Resende Silva, Sunando Roy, Tony Brooks, Helena Tutill, Paola Niola, Patricia Dyal, Charlotte Williams, Leysa Forrest, Yasmin Panchbhaya, Jacqueline Findlay, Samuel Weeks, Julianne Brown, Kathryn Harris, Paul Randell, James Price, Alison Holmes, Judith Breuer |
| EPI_ISL_820041, EPI_ISL_820042, EPI_ISL_820043, EPI_ISL_820044 | Quadram Institute Bioscience | COVID-19 Genomics UK (COG-UK) Consortium | Dave J. Baker, Gemma L. Kay, Alp Aydin, Thanh Le-Viet, Steven Rudder, Ana P. Tedim, Anastasia Kolyva, Maria Diaz, Leonardo de Oliveira Martins, Nabil-Fareed Alikhan, Lizzie Meadows, Rachael Stanley, Ngozi Elumogo, Muhammed Yasir, Nicholas M. Thomson, Alexander J Trotter, Rachel Gilroy, Samuel Bloomfield, Claire Stuart, Andrew Bell, Reenesh Prakash, Samir Dervisevic, Alison E. Mather, John Wain, Mark Webber, Andrew J. Page, Justin |

| | | | |
|---|---|---|---|
| | | | O'Grady |
| EPI_ISL_820045, EPI_ISL_820046, EPI_ISL_820047, EPI_ISL_820048, EPI_ISL_820049, EPI_ISL_820050, EPI_ISL_820051, EPI_ISL_820052, EPI_ISL_820053, EPI_ISL_820054 | University College London, Great Ormond Street Hospital for Children NHS Foundation Trust, Imperial College Healthcare NHS Trust | COVID-19 Genomics UK (COG-UK) Consortium | Sergi Castellano, Rachel Williams, Mark Kristiansen, Paola Resende Silva, Sunando Roy, Tony Brooks, Helena Tutill, Paola Niola, Patricia Dyal, Charlotte Williams, Leysa Forrest, Yasmin Panchbhaya, Jacqueline Findlay, Samuel Weeks, Julianne Brown, Kathryn Harris, Paul Randell, James Price, Alison Holmes, Judith Breuer |
| EPI_ISL_820055, EPI_ISL_820056 | Quadram Institute Bioscience | COVID-19 Genomics UK (COG-UK) Consortium | Dave J. Baker, Gemma L. Kay, Alp Aydin, Thanh Le-Viet, Steven Rudder, Ana P. Tedim, Anastasia Kolyva, Maria Diaz, Leonardo de Oliveira Martins, Nabil-Fareed Alikhan, Lizzie Meadows, Rachael Stanley, Ngozi Elumogo, Muhammed Yasir, Nicholas M. Thomson, Alexander J Trotter, Rachel Gilroy, Samuel Bloomfield, Claire Stuart, Andrew Bell, Reenesh Prakash, Samir Dervisevic, Alison E. Mather, John Wain, Mark Webber, Andrew J. Page, Justin O'Grady |
| EPI_ISL_820057, EPI_ISL_820058, EPI_ISL_820059, EPI_ISL_820060, EPI_ISL_820061 | University College London, Great Ormond Street Hospital for Children NHS Foundation Trust, Imperial College Healthcare NHS Trust | COVID-19 Genomics UK (COG-UK) Consortium | Sergi Castellano, Rachel Williams, Mark Kristiansen, Paola Resende Silva, Sunando Roy, Tony Brooks, Helena Tutill, Paola Niola, Patricia Dyal, Charlotte Williams, Leysa Forrest, Yasmin Panchbhaya, Jacqueline Findlay, Samuel Weeks, Julianne Brown, Kathryn Harris, Paul Randell, James Price, Alison Holmes, Judith Breuer |
| EPI_ISL_820062 | Quadram Institute Bioscience | COVID-19 Genomics UK (COG-UK) Consortium | Dave J. Baker, Gemma L. Kay, Alp Aydin, Thanh Le-Viet, Steven Rudder, Ana P. Tedim, Anastasia Kolyva, Maria Diaz, Leonardo de Oliveira Martins, Nabil-Fareed Alikhan, Lizzie Meadows, Rachael Stanley, Ngozi Elumogo, Muhammed Yasir, Nicholas M. Thomson, Alexander J Trotter, Rachel Gilroy, Samuel Bloomfield, Claire Stuart, Andrew Bell, Reenesh Prakash, Samir Dervisevic, Alison E. Mather, John Wain, Mark Webber, Andrew J. Page, Justin O'Grady |
| EPI_ISL_820063, EPI_ISL_820064, EPI_ISL_820065, EPI_ISL_820066 | University College London, Great Ormond Street Hospital for Children NHS Foundation Trust, Imperial College Healthcare NHS Trust | COVID-19 Genomics UK (COG-UK) Consortium | Sergi Castellano, Rachel Williams, Mark Kristiansen, Paola Resende Silva, Sunando Roy, Tony Brooks, Helena Tutill, Paola Niola, Patricia Dyal, Charlotte Williams, Leysa Forrest, Yasmin Panchbhaya, Jacqueline Findlay, Samuel Weeks, Julianne Brown, Kathryn Harris, Paul Randell, James Price, Alison Holmes, Judith Breuer |
| EPI_ISL_820067, EPI_ISL_820068, EPI_ISL_820069, EPI_ISL_820070, EPI_ISL_820071, EPI_ISL_820072, EPI_ISL_820073, EPI_ISL_820074, EPI_ISL_820075, EPI_ISL_820076, EPI_ISL_820077, EPI_ISL_820078, EPI_ISL_820079, EPI_ISL_820080 | | | |
| see above | Quadram Institute Bioscience | COVID-19 Genomics UK (COG-UK) Consortium | Dave J. Baker, Gemma L. Kay, Alp Aydin, Thanh Le-Viet, Steven Rudder, Ana P. Tedim, Anastasia Kolyva, Maria Diaz, Leonardo de Oliveira Martins, Nabil-Fareed Alikhan, Lizzie Meadows, Rachael Stanley, Ngozi Elumogo, Muhammed Yasir, Nicholas M. Thomson, Alexander J Trotter, Rachel Gilroy, Samuel Bloomfield, Claire Stuart, Andrew Bell, Reenesh Prakash, Samir Dervisevic, Alison E. Mather, John Wain, Mark Webber, Andrew J. Page, Justin O'Grady |
| EPI_ISL_820081 | University College London, Great Ormond Street Hospital for Children NHS Foundation Trust, Imperial College Healthcare NHS Trust | COVID-19 Genomics UK (COG-UK) Consortium | Sergi Castellano, Rachel Williams, Mark Kristiansen, Paola Resende Silva, Sunando Roy, Tony Brooks, Helena Tutill, Paola Niola, Patricia Dyal, Charlotte Williams, Leysa Forrest, Yasmin Panchbhaya, Jacqueline Findlay, Samuel Weeks, Julianne Brown, Kathryn Harris, Paul Randell, James Price, Alison Holmes, Judith Breuer |
| EPI_ISL_820082 | Quadram Institute Bioscience | COVID-19 Genomics UK (COG-UK) Consortium | Dave J. Baker, Gemma L. Kay, Alp Aydin, Thanh Le-Viet, Steven Rudder, Ana P. Tedim, Anastasia Kolyva, Maria Diaz, Leonardo de Oliveira Martins, Nabil-Fareed Alikhan, Lizzie Meadows, Rachael Stanley, Ngozi Elumogo, Muhammed Yasir, Nicholas M. Thomson, Alexander J Trotter, Rachel Gilroy, Samuel Bloomfield, Claire Stuart, Andrew Bell, Reenesh Prakash, Samir Dervisevic, Alison E. Mather, John Wain, Mark Webber, Andrew J. Page, Justin O'Grady |
| EPI_ISL_820083, EPI_ISL_820084, EPI_ISL_820085, EPI_ISL_820086, EPI_ISL_820087 | University College London, Great Ormond Street Hospital for Children NHS Foundation Trust, Imperial College Healthcare NHS Trust | COVID-19 Genomics UK (COG-UK) Consortium | Sergi Castellano, Rachel Williams, Mark Kristiansen, Paola Resende Silva, Sunando Roy, Tony Brooks, Helena Tutill, Paola Niola, Patricia Dyal, Charlotte Williams, Leysa Forrest, Yasmin Panchbhaya, Jacqueline Findlay, Samuel Weeks, Julianne Brown, Kathryn Harris, Paul Randell, James Price, Alison Holmes, Judith Breuer |
| EPI_ISL_820088 | Quadram Institute Bioscience | COVID-19 Genomics UK (COG-UK) Consortium | Dave J. Baker, Gemma L. Kay, Alp Aydin, Thanh Le-Viet, Steven Rudder, Ana P. Tedim, Anastasia Kolyva, Maria Diaz, Leonardo de Oliveira Martins, Nabil-Fareed Alikhan, Lizzie Meadows, Rachael Stanley, Ngozi Elumogo, Muhammed Yasir, Nicholas M. Thomson, Alexander J Trotter, Rachel Gilroy, Samuel Bloomfield, Claire Stuart, Andrew Bell, Reenesh Prakash, Samir Dervisevic, Alison E. Mather, John Wain, Mark Webber, Andrew J. Page, Justin O'Grady |
| EPI_ISL_820089 | University College London, Great Ormond Street Hospital for Children NHS Foundation Trust, Imperial College Healthcare NHS Trust | COVID-19 Genomics UK (COG-UK) Consortium | Sergi Castellano, Rachel Williams, Mark Kristiansen, Paola Resende Silva, Sunando Roy, Tony Brooks, Helena Tutill, Paola Niola, Patricia Dyal, Charlotte Williams, Leysa Forrest, Yasmin Panchbhaya, Jacqueline Findlay, Samuel Weeks, Julianne Brown, Kathryn Harris, Paul Randell, James Price, Alison Holmes, Judith Breuer |
| EPI_ISL_820090, EPI_ISL_820091, EPI_ISL_820092, EPI_ISL_820093 | Quadram Institute Bioscience | COVID-19 Genomics UK (COG-UK) Consortium | Dave J. Baker, Gemma L. Kay, Alp Aydin, Thanh Le-Viet, Steven Rudder, Ana P. Tedim, Anastasia Kolyva, Maria Diaz, Leonardo de Oliveira Martins, Nabil-Fareed Alikhan, Lizzie Meadows, Rachael Stanley, Ngozi Elumogo, Muhammed Yasir, Nicholas M. Thomson, Alexander J Trotter, Rachel Gilroy, Samuel Bloomfield, Claire Stuart, Andrew Bell, Reenesh Prakash, Samir Dervisevic, Alison E. Mather, John Wain, Mark Webber, Andrew J. Page, Justin O'Grady |
| EPI_ISL_820094 | University College London, Great Ormond Street Hospital for Children NHS Foundation Trust, Imperial College Healthcare NHS Trust | COVID-19 Genomics UK (COG-UK) Consortium | Sergi Castellano, Rachel Williams, Mark Kristiansen, Paola Resende Silva, Sunando Roy, Tony Brooks, Helena Tutill, Paola Niola, Patricia Dyal, Charlotte Williams, Leysa Forrest, Yasmin Panchbhaya, Jacqueline Findlay, Samuel Weeks, Julianne Brown, Kathryn Harris, Paul Randell, James Price, Alison Holmes, Judith Breuer |
| EPI_ISL_820095, EPI_ISL_820096, EPI_ISL_820097, EPI_ISL_820098 | Quadram Institute Bioscience | COVID-19 Genomics UK (COG-UK) Consortium | Dave J. Baker, Gemma L. Kay, Alp Aydin, Thanh Le-Viet, Steven Rudder, Ana P. Tedim, Anastasia Kolyva, Maria Diaz, Leonardo de Oliveira Martins, Nabil-Fareed Alikhan, Lizzie Meadows, Rachael Stanley, Ngozi Elumogo, Muhammed Yasir, Nicholas M. Thomson, Alexander J Trotter, Rachel Gilroy, Samuel Bloomfield, Claire Stuart, Andrew Bell, Reenesh Prakash, Samir Dervisevic, Alison E. Mather, John Wain, Mark Webber, Andrew J. Page, Justin O'Grady |
| EPI_ISL_820099, EPI_ISL_820100, EPI_ISL_820101 | University College London, Great Ormond Street Hospital for Children NHS Foundation Trust, Imperial College Healthcare NHS Trust | COVID-19 Genomics UK (COG-UK) Consortium | Sergi Castellano, Rachel Williams, Mark Kristiansen, Paola Resende Silva, Sunando Roy, Tony Brooks, Helena Tutill, Paola Niola, Patricia Dyal, Charlotte Williams, Leysa Forrest, Yasmin Panchbhaya, Jacqueline Findlay, Samuel Weeks, Julianne Brown, Kathryn Harris, Paul Randell, James Price, Alison Holmes, Judith Breuer |
| EPI_ISL_820102, EPI_ISL_820103, EPI_ISL_820104, EPI_ISL_820105, EPI_ISL_820106, EPI_ISL_820107, EPI_ISL_820108 | Quadram Institute Bioscience | COVID-19 Genomics UK (COG-UK) Consortium | Dave J. Baker, Gemma L. Kay, Alp Aydin, Thanh Le-Viet, Steven Rudder, Ana P. Tedim, Anastasia Kolyva, Maria Diaz, Leonardo de Oliveira Martins, Nabil-Fareed Alikhan, Lizzie Meadows, Rachael Stanley, Ngozi Elumogo, Muhammed Yasir, Nicholas M. Thomson, Alexander J Trotter, Rachel Gilroy, Samuel Bloomfield, Claire Stuart, Andrew Bell, Reenesh Prakash, Samir Dervisevic, Alison E. Mather, John Wain, Mark Webber, Andrew J. Page, Justin O'Grady |
| EPI_ISL_822299, EPI_ISL_822301 | Lighthouse Lab in Cambridge | Wellcome Sanger Institute for the COVID-19 Genomics UK (COG-UK) Consortium | Rob Howes, The Lighthouse Lab in Cambridge and Alex Alderton, Roberto Amato, Sonia Goncalves, Ewan Harrison, David K. Jackson, Ian Johnston, Dominic Kwiatkowski, Cordelia Langford, John Sillitoe on behalf of the Wellcome Sanger Institute COVID-19 Surveillance Team |
| EPI_ISL_822305, EPI_ISL_822306 | Lighthouse Lab in Milton Keynes | Wellcome Sanger Institute for the COVID-19 Genomics UK (COG-UK) Consortium | The Lighthouse Lab in Milton Keynes and Alex Alderton, Roberto Amato, Sonia Goncalves, Ewan Harrison, David K. Jackson, Ian Johnston, Dominic Kwiatkowski, Cordelia Langford, John Sillitoe on behalf of the Wellcome Sanger Institute COVID-19 Surveillance Team |
| EPI_ISL_822308 | Lighthouse Lab in Alderley Park | Wellcome Sanger Institute for the COVID-19 Genomics UK (COG-UK) Consortium | Jacquelyn Wynn, Mairead Hyland, The Lighthouse Lab in Alderley Park and Alex Alderton, Roberto Amato, Sonia Goncalves, Ewan Harrison, David K. Jackson, Ian Johnston, Dominic Kwiatkowski, Cordelia Langford, John Sillitoe on behalf of the Wellcome Sanger Institute COVID-19 Surveillance Team |
| EPI_ISL_822372, EPI_ISL_822373, EPI_ISL_822374, EPI_ISL_822375, EPI_ISL_822376, EPI_ISL_822377, EPI_ISL_822378, EPI_ISL_822379, EPI_ISL_822380, EPI_ISL_822381, EPI_ISL_822382, EPI_ISL_822383, EPI_ISL_822384, EPI_ISL_822385, EPI_ISL_822386, EPI_ISL_822387, EPI_ISL_822388, EPI_ISL_822389, EPI_ISL_822390, EPI_ISL_822391, EPI_ISL_822392, EPI_ISL_822393, EPI_ISL_822394, EPI_ISL_822395, EPI_ISL_822396, EPI_ISL_822397, EPI_ISL_822398, EPI_ISL_822399, EPI_ISL_822400, EPI_ISL_822401, EPI_ISL_822402, EPI_ISL_822403, EPI_ISL_822404, EPI_ISL_822405, EPI_ISL_822406, EPI_ISL_822407, EPI_ISL_822408, EPI_ISL_822409, EPI_ISL_822410, EPI_ISL_822411, EPI_ISL_822412, EPI_ISL_822413, EPI_ISL_822414, EPI_ISL_822415, EPI_ISL_822416, EPI_ISL_822417, EPI_ISL_822418, EPI_ISL_822419, EPI_ISL_822420, EPI_ISL_822421, EPI_ISL_822422, EPI_ISL_822423, EPI_ISL_822424, EPI_ISL_822425, EPI_ISL_822426, EPI_ISL_822427, EPI_ISL_822428, EPI_ISL_822429, EPI_ISL_822430, EPI_ISL_822431, EPI_ISL_822432, EPI_ISL_822433, EPI_ISL_822434, EPI_ISL_822435, EPI_ISL_822436, EPI_ISL_822437, EPI_ISL_822438, EPI_ISL_822439, EPI_ISL_822440, EPI_ISL_822441, EPI_ISL_822442, EPI_ISL_822443, EPI_ISL_822444, EPI_ISL_822445, EPI_ISL_822446, EPI_ISL_822447, EPI_ISL_822448, EPI_ISL_822449, EPI_ISL_822450, EPI_ISL_822451, EPI_ISL_822452, EPI_ISL_822453, EPI_ISL_822454, EPI_ISL_822455, EPI_ISL_822456, EPI_ISL_822457, EPI_ISL_822458, EPI_ISL_822459, EPI_ISL_822460, EPI_ISL_822461, EPI_ISL_822462, EPI_ISL_822463, EPI_ISL_822464, EPI_ISL_822465, EPI_ISL_822466, EPI_ISL_822467, EPI_ISL_822468, EPI_ISL_822469, EPI_ISL_822470, EPI_ISL_822471, EPI_ISL_822472, EPI_ISL_822473, EPI_ISL_822474, EPI_ISL_822475, EPI_ISL_822476, EPI_ISL_822477, EPI_ISL_822478, EPI_ISL_822479, EPI_ISL_822480, EPI_ISL_822481, EPI_ISL_822482, EPI_ISL_822483, EPI_ISL_822484, EPI_ISL_822485, EPI_ISL_822486, EPI_ISL_822487, EPI_ISL_822488, EPI_ISL_822489, EPI_ISL_822490, EPI_ISL_822491, EPI_ISL_822492, EPI_ISL_822493, EPI_ISL_822494, EPI_ISL_822495, EPI_ISL_822496, EPI_ISL_822497, EPI_ISL_822498, EPI_ISL_822499, EPI_ISL_822500, EPI_ISL_822501, EPI_ISL_822502, EPI_ISL_822503, EPI_ISL_822504, EPI_ISL_822505, EPI_ISL_822506, EPI_ISL_822507, EPI_ISL_822508, EPI_ISL_822509, EPI_ISL_822510, EPI_ISL_822511, EPI_ISL_822512, EPI_ISL_822513, EPI_ISL_822514, EPI_ISL_822515, EPI_ISL_822516, EPI_ISL_822517, EPI_ISL_822518, EPI_ISL_822519, EPI_ISL_822520, EPI_ISL_822521, EPI_ISL_822522, EPI_ISL_822523, EPI_ISL_822524, EPI_ISL_822525, EPI_ISL_822526, EPI_ISL_822527, EPI_ISL_822528, EPI_ISL_822838, EPI_ISL_822841, EPI_ISL_822843, EPI_ISL_822845, EPI_ISL_822867, EPI_ISL_822874, EPI_ISL_822875, EPI_ISL_822879, EPI_ISL_822881, EPI_ISL_822882, EPI_ISL_822884, EPI_ISL_822889, EPI_ISL_822892, EPI_ISL_822895, EPI_ISL_822898, EPI_ISL_822905, EPI_ISL_822914, EPI_ISL_822920, EPI_ISL_822923, EPI_ISL_822931, EPI_ISL_822944, EPI_ISL_822950, EPI_ISL_822953, EPI_ISL_822961, EPI_ISL_822962, EPI_ISL_822974, EPI_ISL_822975, EPI_ISL_823005, EPI_ISL_823013, EPI_ISL_823017, EPI_ISL_823018, EPI_ISL_823069, EPI_ISL_823073, EPI_ISL_823074, EPI_ISL_823075, EPI_ISL_823076, EPI_ISL_823077, EPI_ISL_823078, EPI_ISL_823079, EPI_ISL_823080, EPI_ISL_823081, | | | |

| | | | |
|---|---|---|---|
| EPI_ISL_823082, EPI_ISL_823083, EPI_ISL_823084, EPI_ISL_823085, EPI_ISL_823086, EPI_ISL_823087, EPI_ISL_823088, EPI_ISL_823089, EPI_ISL_823090, EPI_ISL_823091, EPI_ISL_823092, EPI_ISL_823105, EPI_ISL_823117, EPI_ISL_823119, EPI_ISL_823121, EPI_ISL_823132, EPI_ISL_823135, EPI_ISL_823136, EPI_ISL_823148, EPI_ISL_823150, EPI_ISL_823151, EPI_ISL_823160, EPI_ISL_823161, EPI_ISL_823172, EPI_ISL_823173, EPI_ISL_823378, EPI_ISL_823390, EPI_ISL_823391, EPI_ISL_823392, EPI_ISL_823393, EPI_ISL_823394, EPI_ISL_823395, EPI_ISL_823396, EPI_ISL_823458, EPI_ISL_823552, EPI_ISL_823553, EPI_ISL_823554, EPI_ISL_823555, EPI_ISL_823556, EPI_ISL_823557, EPI_ISL_823558, EPI_ISL_823559, EPI_ISL_823560, EPI_ISL_823561, EPI_ISL_823562, EPI_ISL_823563, EPI_ISL_823564, EPI_ISL_823565, EPI_ISL_823566, EPI_ISL_823567, EPI_ISL_823568, EPI_ISL_823634, EPI_ISL_823636, EPI_ISL_823637, EPI_ISL_823640, EPI_ISL_823641, EPI_ISL_823642, EPI_ISL_823650, EPI_ISL_823651, EPI_ISL_823652, EPI_ISL_823659, EPI_ISL_823663, EPI_ISL_823674, EPI_ISL_823677, EPI_ISL_823683, EPI_ISL_823690, EPI_ISL_823694, EPI_ISL_823696, EPI_ISL_823697, EPI_ISL_823698, EPI_ISL_823699, EPI_ISL_823700, EPI_ISL_823701, EPI_ISL_823702, EPI_ISL_823703, EPI_ISL_823704, EPI_ISL_823705, EPI_ISL_823706, EPI_ISL_823707, EPI_ISL_823708, EPI_ISL_823709, EPI_ISL_823710, EPI_ISL_823711, EPI_ISL_823712, EPI_ISL_823713, EPI_ISL_823714, EPI_ISL_823715, EPI_ISL_823716, EPI_ISL_823717, EPI_ISL_823718, EPI_ISL_823719, EPI_ISL_823720, EPI_ISL_823721, EPI_ISL_823722, EPI_ISL_823723, EPI_ISL_823724, EPI_ISL_823725, EPI_ISL_823726, EPI_ISL_823727, EPI_ISL_823728, EPI_ISL_823729, EPI_ISL_823730, EPI_ISL_823731, EPI_ISL_823732, EPI_ISL_823733, EPI_ISL_823734, EPI_ISL_823735, EPI_ISL_823736, EPI_ISL_823737, EPI_ISL_823738, EPI_ISL_823739, EPI_ISL_823740, EPI_ISL_823741, EPI_ISL_823742, EPI_ISL_823743, EPI_ISL_823744, EPI_ISL_823745, EPI_ISL_823746, EPI_ISL_823747, EPI_ISL_823748, EPI_ISL_823749, EPI_ISL_823750, EPI_ISL_823751, EPI_ISL_823752, EPI_ISL_823753, EPI_ISL_823754, EPI_ISL_823755, EPI_ISL_823756, EPI_ISL_823757, EPI_ISL_823758, EPI_ISL_823759, EPI_ISL_823760, EPI_ISL_823761, EPI_ISL_823762, EPI_ISL_823763, EPI_ISL_823764, EPI_ISL_823765, EPI_ISL_823766, EPI_ISL_823767, EPI_ISL_823768, EPI_ISL_823796, EPI_ISL_823797, EPI_ISL_823798, EPI_ISL_823799, EPI_ISL_823802, EPI_ISL_823805, EPI_ISL_823807, EPI_ISL_823811, EPI_ISL_823812 | | | |
| see above | Wales Specialist Virology Centre Sequencing lab: Pathogen Genomics Unit | COVID-19 Genomics UK (COG-UK) Consortium | Catherine Moore, Johnathan Evans, Laura Gifford, Malorie Perry, Simon Cottrell, Angela Marchbank, Alec Birchley, Alexander Adams, Amy Gaskin, Bree Gatica-Wilcox, Jason Coombes, Joel Southgate, Lauren Gilbert, Lee Graham, Nicole Pacchiarini, Sara Kumziene-Summerhayes, Sarah Taylor, Sophie Jones, Sara Rey, Matthew Bull, Joanne Watkins, Sally Corden, Tom Connor |
| EPI_ISL_823977, EPI_ISL_823978, EPI_ISL_823982, EPI_ISL_823997, EPI_ISL_823998, EPI_ISL_823999, EPI_ISL_824023, EPI_ISL_824035, EPI_ISL_824054, EPI_ISL_824078, EPI_ISL_824080, EPI_ISL_824082, EPI_ISL_824083, EPI_ISL_824084, EPI_ISL_824085, EPI_ISL_824086, EPI_ISL_824087, EPI_ISL_824089, EPI_ISL_824090, EPI_ISL_824091, EPI_ISL_824092, EPI_ISL_824093, EPI_ISL_824095, EPI_ISL_824104, EPI_ISL_824105, EPI_ISL_824106, EPI_ISL_824107, EPI_ISL_824108, EPI_ISL_824130, EPI_ISL_824132, EPI_ISL_824133, EPI_ISL_824135, EPI_ISL_824138, EPI_ISL_824139, EPI_ISL_824142, EPI_ISL_824143, EPI_ISL_824144, EPI_ISL_824151, EPI_ISL_824155, EPI_ISL_824156, EPI_ISL_824188, EPI_ISL_824189, EPI_ISL_824190, EPI_ISL_824191, EPI_ISL_824192, EPI_ISL_824193, EPI_ISL_824194, EPI_ISL_824224, EPI_ISL_824225, EPI_ISL_824226, EPI_ISL_824227, EPI_ISL_824228, EPI_ISL_824229, EPI_ISL_824255, EPI_ISL_824256 | | | |
| see above | Dutch COVID-19 response team | National Institute for Public Health and the Environment (RIVM) | Adam Meijer, Harry Vennema, Jeroen Cremer, Sharon van den Brink, Bas van der Veer, AnneMarie van den Brandt, Florian Zwagemaker, Dennis Schmitz, Chantal Reusken, on behalf of the national COVID-19 response team |
| EPI_ISL_824558, EPI_ISL_824565, EPI_ISL_824570, EPI_ISL_824580, EPI_ISL_824590, EPI_ISL_824591, EPI_ISL_824592, EPI_ISL_824602, EPI_ISL_824603, EPI_ISL_824613, EPI_ISL_824624, EPI_ISL_824636, EPI_ISL_824648, EPI_ISL_824649, EPI_ISL_824653, EPI_ISL_824654, EPI_ISL_824655, EPI_ISL_824660, EPI_ISL_824664, EPI_ISL_824665, EPI_ISL_824671, EPI_ISL_824676, EPI_ISL_824677, EPI_ISL_824683, EPI_ISL_824687, EPI_ISL_824688, EPI_ISL_824689, EPI_ISL_824699, EPI_ISL_824700, EPI_ISL_824701, EPI_ISL_824707, EPI_ISL_824711, EPI_ISL_824712, EPI_ISL_824713, EPI_ISL_824723, EPI_ISL_824724, EPI_ISL_824731, EPI_ISL_824734, EPI_ISL_824735 | | | |
| see above | Cedars-Sinai Medical Center, Department of Pathology & Laboratory Medicine, Molecular Pathology Laboratory | Cedars-Sinai Medical Center, Molecular Pathology Laboratory of Department of Pathology & Laboratory Medicine and Genomic Core | Wenjuan Zhang, Brian Davis, Stephanie Chen, Jorge Mario Sincuir Martinez, Jasmine T Plummer, Eric Vail |
| EPI_ISL_824799 | Department of Clinical Microbiology | GIGA Medical Genomics | Keith Durkin, Maria Artesi, Sébastien Bontems, Raphaël Boreux, Bouchra Boujemla, Cécile Meex, Pierrette Melin, Marie-Pierre Hayette, Vincent Bours |
| EPI_ISL_824920, EPI_ISL_824942 | Charité Universitätsmedizin Berlin, Institute of Virology, Charitéplatz 1, 10117 Berlin, Germany | Charité Universitätsmedizin Berlin, Institute of Virology, Charitéplatz 1, 10117 Berlin, Germany | Victor M Corman, Jörn Beheim-Schwarzbach, Tobias Bleicker, Julia Tesch, Barbara Mühlemann, Talitha Veith, Julia Schneider, Terry Jones, Christian Drosten |
| EPI_ISL_825043 | NHL Municipal Medical College, Ahmedbad | Gujarat Biotechnology Research Centre | Jayshri Pethani, Ramesh Pandit, Janvi Raval, Zarna Patel, Nitin Savaliya, Dinesh Kumar, Zuber Saiyed, Labdhi Pandya, Afzal Ansari, Nikha Trivedi, Apurvasinh Puvar, Monila Patel, Atit Shah, NM Shaikh, Bimal Chauhan, Tanmay Mehta, Bhavin Prajapati, Chaitanya Joshi, Madhvi Joshi |
| EPI_ISL_825044 | NHL Municipal Medical College, Ahmedbad | Gujarat Biotechnology Research Centre | Monila Patel, Janvi Raval, Zarna Patel, Nitin Savaliya, Dinesh Kumar, Zuber Saiyed, Labdhi Pandya, Afzal Ansari, Nikha Trivedi, Apurvasinh Puvar, Ramesh Pandit, Jayshri Pethani, Atit Shah, NM Shaikh, Bimal Chauhan, Tanmay Mehta, Bhavin Prajapati, Chaitanya Joshi, Madhvi Joshi |
| EPI_ISL_825045 | NHL Municipal Medical College, Ahmedbad | Gujarat Biotechnology Research Centre | Atit Shah, Zarna Patel, Nitin Savaliya, Dinesh Kumar, Zuber Saiyed, Labdhi Pandya, Afzal Ansari, Nikha Trivedi, Apurvasinh Puvar, Ramesh Pandit, Janvi Raval, Jayshri Pethani, Monila Patel, NM Shaikh, Bimal Chauhan, Tanmay Mehta, Bhavin Prajapati, Chaitanya Joshi, Madhvi Joshi |
| EPI_ISL_825046 | NHL Municipal Medical College, Ahmedbad | Gujarat Biotechnology Research Centre | NM Shaikh, Nitin Savaliya, Dinesh Kumar, Zuber Saiyed, Labdhi Pandya, Afzal Ansari, Nikha Trivedi, Apurvasinh Puvar, Ramesh Pandit, Janvi Raval, Zarna Patel, Jayshri Pethani, Monila Patel, Atit Shah, Bimal Chauhan, Tanmay Mehta, Bhavin Prajapati, Chaitanya Joshi, Madhvi Joshi |
| EPI_ISL_825047 | NHL Municipal Medical College, Ahmedbad | Gujarat Biotechnology Research Centre | Bimal Chauhan, Dinesh Kumar, Zuber Saiyed, Labdhi Pandya, Afzal Ansari, Nikha Trivedi, Apurvasinh Puvar, Ramesh Pandit, Janvi Raval, Zarna Patel, Nitin Savaliya, Jayshri Pethani, Monila Patel, Atit Shah, NM Shaikh, Tanmay Mehta, Bhavin Prajapati, Chaitanya Joshi, Madhvi Joshi |
| EPI_ISL_825048 | GMERS Medical College and Hospital, Gotri, Vadodara | Gujarat Biotechnology Research Centre | Mitesh Kamothi, Zuber Saiyed, Labdhi Pandya, Afzal Ansari, Nikha Trivedi, Apurvasinh Puvar, Ramesh Pandit, Janvi Raval, Zarna Patel, Nitin Savaliya, Dinesh Kumar, Bithika Duttaroy, Kalpesh Mistry, Chaitanya Joshi, Madhvi Joshi |
| EPI_ISL_825049 | GMERS Medical College and Hospital, Gotri, Vadodara | Gujarat Biotechnology Research Centre | Bithika Duttaroy, Labdhi Pandya, Afzal Ansari, Nikha Trivedi, Apurvasinh Puvar, Ramesh Pandit, Janvi Raval, Zarna Patel, Nitin Savaliya, Dinesh Kumar, Zuber Saiyed, Mitesh Kamothi, Kalpesh Mistry, Chaitanya Joshi, Madhvi Joshi |
| EPI_ISL_825050 | NHL Municipal Medical College, Ahmedbad | Gujarat Biotechnology Research Centre | Tanmay Mehta, Afzal Ansari, Nikha Trivedi, Apurvasinh Puvar, Ramesh Pandit, Janvi Raval, Zarna Patel, Nitin Savaliya, Dinesh Kumar, Zuber Saiyed, Labdhi Pandya, Jayshri Pethani, Monila Patel, Atit Shah, NM Shaikh, Bimal Chauhan, Bhavin Prajapati, Chaitanya Joshi, Madhvi Joshi |
| EPI_ISL_825051 | NHL Municipal Medical College, Ahmedbad | Gujarat Biotechnology Research Centre | Bhavin Prajapati, Nikha Trivedi, Apurvasinh Puvar, Ramesh Pandit, Janvi Raval, Zarna Patel, Nitin Savaliya, Dinesh Kumar, Zuber Saiyed, Labdhi Pandya, Afzal Ansari, Jayshri Pethani, Monila Patel, Atit Shah, NM Shaikh, Bimal Chauhan, Tanmay Mehta, Chaitanya Joshi, Madhvi Joshi |
| EPI_ISL_825052 | NHL Municipal Medical College, Ahmedbad | Gujarat Biotechnology Research Centre | Apurvasinh Puvar, Ramesh Pandit, Janvi Raval, Zarna Patel, Nitin Savaliya, Dinesh Kumar, Zuber Saiyed, Labdhi Pandya, Afzal Ansari, Nikha Trivedi, Jayshri Pethani, Monila Patel, Atit Shah, NM Shaikh, Bimal Chauhan, Tanmay Mehta, Bhavin Prajapati, Chaitanya Joshi, Madhvi Joshi |
| EPI_ISL_825053 | NHL Municipal Medical College, Ahmedbad | Gujarat Biotechnology Research Centre | Ramesh Pandit, Janvi Raval, Zarna Patel, Nitin Savaliya, Dinesh Kumar, Zuber Saiyed, Labdhi Pandya, Afzal Ansari, Nikha Trivedi, Apurvasinh Puvar, Jayshri Pethani, Monila Patel, Atit Shah, NM Shaikh, Bimal Chauhan, Tanmay Mehta, Bhavin Prajapati, Chaitanya Joshi, Madhvi Joshi |
| EPI_ISL_825054 | GMERS Medical College and Hospital, Gotri, Vadodara | Gujarat Biotechnology Research Centre | Kalpesh Mistry, Janvi Raval, Zarna Patel, Nitin Savaliya, Dinesh Kumar, Zuber Saiyed, Afzal Ansari, Nikha Trivedi, Apurvasinh Puvar, Ramesh Pandit, Mitesh Kamothi, Bithika Duttaroy, Chaitanya Joshi, Madhvi Joshi |
| EPI_ISL_825055 | GMERS Medical College and Hospital, Gotri, Vadodara | Gujarat Biotechnology Research Centre | Zarna Patel, Nitin Savaliya, Dinesh Kumar, Zuber Saiyed, Afzal Ansari, Nikha Trivedi, Apurvasinh Puvar, Ramesh Pandit, Janvi Raval, Mitesh Kamothi, Bithika Duttaroy, Kalpesh Mistry, Chaitanya Joshi, Madhvi Joshi |
| EPI_ISL_825056 | GMERS Medical College and Hospital, Gotri, Vadodara | Gujarat Biotechnology Research Centre | Nitin Savaliya, Dinesh Kumar, Zuber Saiyed, Afzal Ansari, Nikha Trivedi, Apurvasinh Puvar, Ramesh Pandit, Janvi Raval, Zarna Patel, Mitesh Kamothi, Bithika Duttaroy, Kalpesh Mistry, Chaitanya Joshi, Madhvi Joshi |
| EPI_ISL_825057 | GMERS Medical College and Hospital, Gotri, Vadodara | Gujarat Biotechnology Research Centre | Dinesh Kumar, Zuber Saiyed, Afzal Ansari, Nikha Trivedi, Apurvasinh Puvar, Ramesh Pandit, Janvi Raval, Zarna Patel, Nitin Savaliya, Mitesh Kamothi, Bithika Duttaroy, Kalpesh Mistry, Chaitanya Joshi, Madhvi Joshi |
| EPI_ISL_825058 | GMERS Medical College and Hospital, Gotri, Vadodara | Gujarat Biotechnology Research Centre | Zuber Saiyed, Afzal Ansari, Nikha Trivedi, Apurvasinh Puvar, Ramesh Pandit, Janvi Raval, Zarna Patel, Nitin Savaliya, Dinesh Kumar, Mitesh Kamothi, Bithika Duttaroy, Kalpesh Mistry, Chaitanya Joshi, Madhvi Joshi |
| EPI_ISL_825059 | NHL Municipal Medical College, Ahmedbad | Gujarat Biotechnology Research Centre | Afzal Ansari, Nikha Trivedi, Apurvasinh Puvar, Ramesh Pandit, Janvi Raval, Zarna Patel, Nitin Savaliya, Dinesh Kumar, Zuber Saiyed, Jayshri Pethani, Monila Patel, Atit Shah, NM Shaikh, Bimal Chauhan, Tanmay Mehta, Bhavin Prajapati, Chaitanya Joshi, Madhvi Joshi |
| EPI_ISL_825060 | NHL Municipal Medical College, Ahmedbad | Gujarat Biotechnology Research Centre | Nikha Trivedi, Apurvasinh Puvar, Ramesh Pandit, Janvi Raval, Zarna Patel, Nitin Savaliya, Dinesh Kumar, Zuber Saiyed, Afzal Ansari, Jayshri Pethani, Monila Patel, Atit Shah, NM Shaikh, Bimal Chauhan, Tanmay Mehta, Bhavin Prajapati, Chaitanya Joshi, Madhvi Joshi |
| EPI_ISL_825140 | Sanjay Gandhi Institute of Trauma & Orthopaedics | Department of Neurovirology, National Institute of Mental Health and Neurosciences (NIMHANS) | Chitra Pattabiraman, Pramada Prasad, Anita S Desai, V Ravi |
| EPI_ISL_825141, EPI_ISL_825142, EPI_ISL_825143 | BBMP Urban PHC | Department of Neurovirology, National Institute of Mental Health and Neurosciences (NIMHANS) | Chitra Pattabiraman, Pramada Prasad, Anita S Desai, V Ravi |
| EPI_ISL_825144 | Bangalore Medical College and Research Institute | Department of Neurovirology, National Institute of Mental Health and Neurosciences (NIMHANS) | Chitra Pattabiraman, Pramada Prasad, Anita S Desai, V Ravi |
| EPI_ISL_825145 | Virus Research and Diagnostic Laboratory (DH-VRDL) | Department of Neurovirology, National Institute of Mental Health and Neurosciences (NIMHANS) | Chitra Pattabiraman, Pramada Prasad, Anita S Desai, V Ravi |
| EPI_ISL_825146 | Kidwai Memorial Institute of Oncology | Department of Neurovirology, National Institute of Mental Health and Neurosciences (NIMHANS) | Chitra Pattabiraman, Pramada Prasad, Anita S Desai, V Ravi |

| | | | |
|---|---|---|---|
| EPI_ISL_825147 | Narayana Nethralaya lab | Department of Neurovirology, National Institute of Mental Health and Neurosciences (NIMHANS) | Chitra Pattabiraman, Pramada Prasad, Anita S Desai, V Ravi |
| EPI_ISL_825148 | BBMP Urban PHC | Department of Neurovirology, National Institute of Mental Health and Neurosciences (NIMHANS) | Chitra Pattabiraman, Pramada Prasad, Anita S Desai, V Ravi |
| EPI_ISL_825149 | National Institute of Virology Bangalore Unit (NIV BU) | Department of Neurovirology, National Institute of Mental Health and Neurosciences (NIMHANS) | Chitra Pattabiraman, Pramada Prasad, Anita S Desai, V Ravi |
| EPI_ISL_825150, EPI_ISL_825151 | Kidwai Memorial Institute of Oncology | Department of Neurovirology, National Institute of Mental Health and Neurosciences (NIMHANS) | Chitra Pattabiraman, Pramada Prasad, Anita S Desai, V Ravi |
| EPI_ISL_825153, EPI_ISL_825154 | Virus Research and Diagnostic Laboratory (DH-VRDL) | Department of Neurovirology, National Institute of Mental Health and Neurosciences (NIMHANS) | Chitra Pattabiraman, Pramada Prasad, Anita S Desai, V Ravi |
| EPI_ISL_825155 | Microbiology and Virology Unit, Florence Careggi University Hospital | Microbiology and Virology Unit, Florence Careggi University Hospital | Vincenzo Di Pilato, Marco Coppi, Fabio Morecchiato, Alberto Antonelli, Emanuele Gori, Gian Maria Rossolini |
| EPI_ISL_825161, EPI_ISL_825162, EPI_ISL_825163, EPI_ISL_825164, EPI_ISL_825167, EPI_ISL_825168 | Robert Koch Institute, ZBS1 Highly Pathogenic Viruses, Berlin, Germany | Robert Koch Institute, ZBS1 Highly Pathogenic Viruses & Bioinformatics MF1, Berlin, Germany | Annika Brinkmann, Janine Michel, Livia Schrick, Steven Uddin, Dominique Seifert, Alexander Dalpke, Leo Büttner, Kristina Hochauf-Stange, Dirk Lindemann, Lars Schaade, Andreas Nitsche |
| EPI_ISL_825498, EPI_ISL_825501, EPI_ISL_825502, EPI_ISL_825503, EPI_ISL_825505, EPI_ISL_825506, EPI_ISL_825507, EPI_ISL_825508, EPI_ISL_825509, EPI_ISL_825510, EPI_ISL_825511, EPI_ISL_825512, EPI_ISL_825513, EPI_ISL_825514, EPI_ISL_825515, EPI_ISL_825516, EPI_ISL_825517, EPI_ISL_825518, EPI_ISL_825519, EPI_ISL_825520, EPI_ISL_825521, EPI_ISL_825522, EPI_ISL_825523, EPI_ISL_825524, EPI_ISL_825525, EPI_ISL_825526, EPI_ISL_825527, EPI_ISL_825528, EPI_ISL_825529, EPI_ISL_825530, EPI_ISL_825531, EPI_ISL_825546, EPI_ISL_825547, EPI_ISL_825554, EPI_ISL_825555, EPI_ISL_825557, EPI_ISL_825558, EPI_ISL_825559, EPI_ISL_825560, EPI_ISL_825561, EPI_ISL_825562, EPI_ISL_825563, EPI_ISL_825564, EPI_ISL_825565, EPI_ISL_825566, EPI_ISL_825567, EPI_ISL_825572, EPI_ISL_825605, EPI_ISL_825606, EPI_ISL_825607, EPI_ISL_825608, EPI_ISL_825609 | | | |
| see above | Respiratory Virus Unit, National Infection Service, Public Health England | COVID-19 Genomics UK (COG-UK) Consortium | PHE Covid Sequencing Team |
| EPI_ISL_825619, EPI_ISL_825620, EPI_ISL_825621, EPI_ISL_825624 | Hospital Universitari Vall d'Hebron - Vall d'Hebron Institut de Rercerca | Hospital Universitari Vall d'Hebron | Cristina Andrés, Maria Piñana, Josep F Abril, Damir Garcia-Cehic, Ariadna Rando, Juliana Esperalba, Maria Gema Codina, Carla Castillo, Maria Carmen Martin, Tomàs Pumarola, Josep Quer, Andrés Antón |
| EPI_ISL_826284 | University of Bari Biomedical Sciences and Human Oncology | University of Bari Biomedical Sciences and Human Oncology | Chironna Maria, Sallustio Anna, Loconsole Daniela, Accogli Marisa |
| EPI_ISL_826287 | The Jackson Laboratory | The Jackson Laboratory | Lloyd M, Maurya R, Renzette N, Omerza G, Kelly K, Li L, Wei C L, Adams M |
| EPI_ISL_826671, EPI_ISL_826695, EPI_ISL_826698, EPI_ISL_826699, EPI_ISL_827166, EPI_ISL_827167, EPI_ISL_827168, EPI_ISL_827430, EPI_ISL_827444, EPI_ISL_827459, EPI_ISL_827460, EPI_ISL_827464, EPI_ISL_827479, EPI_ISL_827481, EPI_ISL_827499, EPI_ISL_827606, EPI_ISL_827785, EPI_ISL_827787 | | | |
| see above | deCODE genetics | deCODE genetics | Daniel F Gudbjartsson; Agnar Helgason; Hakon Jonsson; Olafur T Magnusson; Pall Melsted; Gudmundur L Norddahl; Jona Saemundsdottir; Asgeir Sigurdsson; Patrick Sulem; Arna B Agustsdottir; Hannes Eggertsson; Berglind Eiriksdottir; Run Fridriksdottir; Elisabet E Gardarsdottir; Gudmundur Georgsson; Olafia S Gretarsdottir; Kjartan R Gudmundsson; Thora R Gunnarsdottir; Arnaldur Gylfason; Hilma Holm; Brynjar O Jensson; Aslaug Jonasdottir; Kamilla S Josefsdottir; Thordur Kristjansson; Droplaug N Magnusdottir; Solvi Rognvaldsson; Louise le Roux; Gudrun Sigmundsdottir; Gardar Sveinbjornsson; Kristin E Sveinsdottir; Maney Sveinsdottir; Emil A Thorarensen; Bjarni Thorbjornsson; Gisli Masson; Ingileif Jonsdottir; Alma Moller; Thorolfur Gudnason; Karl G Kristinsson; Unnur Thorsteinsdottir; Kari Stefansson |
| EPI_ISL_828087 | The National University Hospital of Iceland | deCODE genetics | Daniel F Gudbjartsson; Agnar Helgason; Hakon Jonsson; Olafur T Magnusson; Pall Melsted; Gudmundur L Norddahl; Jona Saemundsdottir; Asgeir Sigurdsson; Patrick Sulem; Arna B Agustsdottir; Hannes Eggertsson; Berglind Eiriksdottir; Run Fridriksdottir; Elisabet E Gardarsdottir; Gudmundur Georgsson; Olafia S Gretarsdottir; Kjartan R Gudmundsson; Thora R Gunnarsdottir; Arnaldur Gylfason; Hilma Holm; Brynjar O Jensson; Aslaug Jonasdottir; Kamilla S Josefsdottir; Thordur Kristjansson; Droplaug N Magnusdottir; Solvi Rognvaldsson; Louise le Roux; Gudrun Sigmundsdottir; Gardar Sveinbjornsson; Kristin E Sveinsdottir; Maney Sveinsdottir; Emil A Thorarensen; Bjarni Thorbjornsson; Gisli Masson; Ingileif Jonsdottir; Alma Moller; Thorolfur Gudnason; Karl G Kristinsson; Unnur Thorsteinsdottir; Kari Stefansson |
| EPI_ISL_828492 | deCODE genetics | deCODE genetics | Daniel F Gudbjartsson; Agnar Helgason; Hakon Jonsson; Olafur T Magnusson; Pall Melsted; Gudmundur L Norddahl; Jona Saemundsdottir; Asgeir Sigurdsson; Patrick Sulem; Arna B Agustsdottir; Hannes Eggertsson; Berglind Eiriksdottir; Run Fridriksdottir; Elisabet E Gardarsdottir; Gudmundur Georgsson; Olafia S Gretarsdottir; Kjartan R Gudmundsson; Thora R Gunnarsdottir; Arnaldur Gylfason; Hilma Holm; Brynjar O Jensson; Aslaug Jonasdottir; Kamilla S Josefsdottir; Thordur Kristjansson; Droplaug N Magnusdottir; Solvi Rognvaldsson; Louise le Roux; Gudrun Sigmundsdottir; Gardar Sveinbjornsson; Kristin E Sveinsdottir; Maney Sveinsdottir; Emil A Thorarensen; Bjarni Thorbjornsson; Gisli Masson; Ingileif Jonsdottir; Alma Moller; Thorolfur Gudnason; Karl G Kristinsson; Unnur Thorsteinsdottir; Kari Stefansson |
| EPI_ISL_828653 | The National University Hospital of Iceland | deCODE genetics | Daniel F Gudbjartsson; Agnar Helgason; Hakon Jonsson; Olafur T Magnusson; Pall Melsted; Gudmundur L Norddahl; Jona Saemundsdottir; Asgeir Sigurdsson; Patrick Sulem; Arna B Agustsdottir; Hannes Eggertsson; Berglind Eiriksdottir; Run Fridriksdottir; Elisabet E Gardarsdottir; Gudmundur Georgsson; Olafia S Gretarsdottir; Kjartan R Gudmundsson; Thora R Gunnarsdottir; Arnaldur Gylfason; Hilma Holm; Brynjar O Jensson; Aslaug Jonasdottir; Kamilla S Josefsdottir; Thordur Kristjansson; Droplaug N Magnusdottir; Solvi Rognvaldsson; Louise le Roux; Gudrun Sigmundsdottir; Gardar Sveinbjornsson; Kristin E Sveinsdottir; Maney Sveinsdottir; Emil A Thorarensen; Bjarni Thorbjornsson; Gisli Masson; Ingileif Jonsdottir; Alma Moller; Thorolfur Gudnason; Karl G Kristinsson; Unnur Thorsteinsdottir; Kari Stefansson |
| EPI_ISL_828694, EPI_ISL_828702, EPI_ISL_828715, EPI_ISL_828801, EPI_ISL_828824, EPI_ISL_828842, EPI_ISL_828844, EPI_ISL_828845, EPI_ISL_828846, EPI_ISL_829090, EPI_ISL_829094, EPI_ISL_829100 | | | |
| see above | deCODE genetics | deCODE genetics | Daniel F Gudbjartsson; Agnar Helgason; Hakon Jonsson; Olafur T Magnusson; Pall Melsted; Gudmundur L Norddahl; Jona Saemundsdottir; Asgeir Sigurdsson; Patrick Sulem; Arna B Agustsdottir; Hannes Eggertsson; Berglind Eiriksdottir; Run Fridriksdottir; Elisabet E Gardarsdottir; Gudmundur Georgsson; Olafia S Gretarsdottir; Kjartan R Gudmundsson; Thora R Gunnarsdottir; Arnaldur Gylfason; Hilma Holm; Brynjar O Jensson; Aslaug Jonasdottir; Kamilla S Josefsdottir; Thordur Kristjansson; Droplaug N Magnusdottir; Solvi Rognvaldsson; Louise le Roux; Gudrun Sigmundsdottir; Gardar Sveinbjornsson; Kristin E Sveinsdottir; Maney Sveinsdottir; Emil A Thorarensen; Bjarni Thorbjornsson; Gisli Masson; Ingileif Jonsdottir; Alma Moller; Thorolfur Gudnason; Karl G Kristinsson; Unnur Thorsteinsdottir; Kari Stefansson |
| EPI_ISL_829132 | The National University Hospital of Iceland | deCODE genetics | Daniel F Gudbjartsson; Agnar Helgason; Hakon Jonsson; Olafur T Magnusson; Pall Melsted; Gudmundur L Norddahl; Jona Saemundsdottir; Asgeir Sigurdsson; Patrick Sulem; Arna B Agustsdottir; Hannes Eggertsson; Berglind Eiriksdottir; Run Fridriksdottir; Elisabet E Gardarsdottir; Gudmundur Georgsson; Olafia S Gretarsdottir; Kjartan R Gudmundsson; Thora R Gunnarsdottir; Arnaldur Gylfason; Hilma Holm; Brynjar O Jensson; Aslaug Jonasdottir; Kamilla S Josefsdottir; Thordur Kristjansson; Droplaug N Magnusdottir; Solvi Rognvaldsson; Louise le Roux; Gudrun Sigmundsdottir; Gardar Sveinbjornsson; Kristin E Sveinsdottir; Maney Sveinsdottir; Emil A Thorarensen; Bjarni Thorbjornsson; Gisli Masson; Ingileif Jonsdottir; Alma Moller; Thorolfur Gudnason; Karl G Kristinsson; Unnur Thorsteinsdottir; Kari Stefansson |
| EPI_ISL_829140, EPI_ISL_829365, EPI_ISL_829367, EPI_ISL_829382, EPI_ISL_829391, EPI_ISL_829416, EPI_ISL_829417, EPI_ISL_829418, EPI_ISL_829419, EPI_ISL_829422, EPI_ISL_829423, EPI_ISL_829424, EPI_ISL_829425 | | | |
| see above | deCODE genetics | deCODE genetics | Daniel F Gudbjartsson; Agnar Helgason; Hakon Jonsson; Olafur T Magnusson; Pall Melsted; Gudmundur L Norddahl; Jona Saemundsdottir; Asgeir Sigurdsson; Patrick Sulem; Arna B Agustsdottir; Hannes Eggertsson; Berglind Eiriksdottir; Run Fridriksdottir; Elisabet E Gardarsdottir; Gudmundur Georgsson; Olafia S Gretarsdottir; Kjartan R Gudmundsson; Thora R Gunnarsdottir; Arnaldur Gylfason; Hilma Holm; Brynjar O Jensson; Aslaug Jonasdottir; Kamilla S Josefsdottir; Thordur Kristjansson; Droplaug N Magnusdottir; Solvi Rognvaldsson; Louise le Roux; Gudrun Sigmundsdottir; Gardar Sveinbjornsson; Kristin E Sveinsdottir; Maney Sveinsdottir; Emil A Thorarensen; Bjarni Thorbjornsson; Gisli Masson; Ingileif Jonsdottir; Alma Moller; Thorolfur Gudnason; Karl G Kristinsson; Unnur Thorsteinsdottir; Kari Stefansson |
| EPI_ISL_829544 | The National University Hospital of Iceland | deCODE genetics | Daniel F Gudbjartsson; Agnar Helgason; Hakon Jonsson; Olafur T Magnusson; Pall Melsted; Gudmundur L Norddahl; Jona Saemundsdottir; Asgeir Sigurdsson; Patrick Sulem; Arna B Agustsdottir; Hannes Eggertsson; Berglind Eiriksdottir; Run Fridriksdottir; Elisabet E Gardarsdottir; Gudmundur Georgsson; Olafia S Gretarsdottir; Kjartan R Gudmundsson; Thora R Gunnarsdottir; Arnaldur Gylfason; Hilma Holm; Brynjar O Jensson; Aslaug Jonasdottir; Kamilla S Josefsdottir; Thordur Kristjansson; Droplaug N Magnusdottir; Solvi Rognvaldsson; Louise le Roux; Gudrun Sigmundsdottir; Gardar Sveinbjornsson; Kristin E Sveinsdottir; Maney Sveinsdottir; Emil A Thorarensen; Bjarni Thorbjornsson; Gisli Masson; Ingileif Jonsdottir; Alma Moller; Thorolfur Gudnason; Karl G Kristinsson; Unnur Thorsteinsdottir; Kari Stefansson |

| | | | |
|---|---|---|---|
| EPI_ISL_829604, EPI_ISL_829674 | deCODE genetics | deCODE genetics | Daniel F Gudbjartsson; Agnar Helgason; Hakon Jonsson; Olafur T Magnusson; Pall Melsted; Gudmundur L Norddahl; Jona Saemundsdottir; Asgeir Sigurdsson; Patrick Sulem; Arna B Agustsdottir; Hannes Eggertsson; Berglind Eiriksdottir; Run Fridriksdottir; Elisabet E Gardarsdottir; Gudmundur Georgsson; Olafia S Gretarsdottir; Kjartan R Gudmundsson; Thora R Gunnarsdottir; Arnaldur Gylfason; Hilma Holm; Brynjar O Jensson; Aslaug Jonasdottir; Kamilla S Josefsdottir; Thordur Kristjansson; Droplaug N Magnusdottir; Solvi Rognvaldsson; Louise le Roux; Gudrun Sigmundsdottir; Gardar Sveinbjornsson; Kristin E Sveinsdottir; Maney Sveinsdottir; Emil A Thorarensen; Bjarni Thorbjornsson; Gisli Masson; Ingileif Jonsdottir; Alma Moller; Thorolfur Gudnason; Karl G Kristinsson; Unnur Thorsteinsdottir; Kari Stefansson |
| EPI_ISL_830208, EPI_ISL_830210, EPI_ISL_830223, EPI_ISL_830225, EPI_ISL_830227, EPI_ISL_830229, EPI_ISL_830230, EPI_ISL_830232, EPI_ISL_830234, EPI_ISL_830236, EPI_ISL_830238, EPI_ISL_830239, EPI_ISL_830241 | | | |
| see above | KALEIDA CENTER FOR LABORATORY MEDICINE | Wadsworth Center, New York State Department of Health | Kirsten St. George, Daryl M. Lamson, Alexis Russel, Matthew Shudt, Melissa A Leisner, Jonathan Plitnick, Navjot Singh, John Kelly, Erasmus Schneider, Erica Lasek-Nesselquist |
| EPI_ISL_830299, EPI_ISL_830532 | deCODE genetics | deCODE genetics | Daniel F Gudbjartsson; Agnar Helgason; Hakon Jonsson; Olafur T Magnusson; Pall Melsted; Gudmundur L Norddahl; Jona Saemundsdottir; Asgeir Sigurdsson; Patrick Sulem; Arna B Agustsdottir; Hannes Eggertsson; Berglind Eiriksdottir; Run Fridriksdottir; Elisabet E Gardarsdottir; Gudmundur Georgsson; Olafia S Gretarsdottir; Kjartan R Gudmundsson; Thora R Gunnarsdottir; Arnaldur Gylfason; Hilma Holm; Brynjar O Jensson; Aslaug Jonasdottir; Kamilla S Josefsdottir; Thordur Kristjansson; Droplaug N Magnusdottir; Solvi Rognvaldsson; Louise le Roux; Gudrun Sigmundsdottir; Gardar Sveinbjornsson; Kristin E Sveinsdottir; Maney Sveinsdottir; Emil A Thorarensen; Bjarni Thorbjornsson; Gisli Masson; Ingileif Jonsdottir; Alma Moller; Thorolfur Gudnason; Karl G Kristinsson; Unnur Thorsteinsdottir; Kari Stefansson |
| EPI_ISL_830575, EPI_ISL_830579, EPI_ISL_830580, EPI_ISL_830581, EPI_ISL_830582 | KALEIDA CENTER FOR LABORATORY MEDICINE | Wadsworth Center, New York State Department of Health | Kirsten St. George, Daryl M. Lamson, Alexis Russel, Matthew Shudt, Melissa A Leisner, Jonathan Plitnick, Navjot Singh, John Kelly, Erasmus Schneider, Erica Lasek-Nesselquist |
| EPI_ISL_831500, EPI_ISL_831501, EPI_ISL_831502 | University of Wisconsin-Madison AIDS Vaccine Research Laboratories | University of Wisconsin-Madison AIDS Vaccine Research Laboratories | Gage Moreno, Katarina Braun, et al. AIDS Vaccine Research Laboratories |
| EPI_ISL_831752, EPI_ISL_831891 | United States Air Force School of Aerospace Medicine | United States Air Force School of Aerospace Medicine | Anthony Fries, Jennifer Meyer, William Gruner, Amanda Javorina, Sarah Purves, Clarise Starr, Elizabeth Macias |
| EPI_ISL_831953, EPI_ISL_831954, EPI_ISL_831955, EPI_ISL_831956, EPI_ISL_831957 | Laboratoriemedicin, Klinisk mikrobiologi | The Public Health Agency of Sweden | Department of Microbiology, The Public Health Agency of Sweden |
| EPI_ISL_831959, EPI_ISL_831982, EPI_ISL_831983, EPI_ISL_831984, EPI_ISL_831985 | ABC | The Public Health Agency of Sweden | Department of Microbiology, The Public Health Agency of Sweden |
| EPI_ISL_831998, EPI_ISL_831999 | Laboratoriemedicin, Klinisk mikrobiologi | The Public Health Agency of Sweden | Department of Microbiology, The Public Health Agency of Sweden |
| EPI_ISL_832000 | Unilabs, Mikrobiologiska laboratoriet | The Public Health Agency of Sweden | Department of Microbiology, The Public Health Agency of Sweden |
| EPI_ISL_832044, EPI_ISL_832045, EPI_ISL_832046 | Wyoming Public Health Laboratory | Wyoming Public Health Laboratory | Noah Hull, Taylor Fearing, Lynette Gumbleton, Channing Weber, Ashley Norberg, Bailey Bowcutt, and Wanda Manley |
| EPI_ISL_832107, EPI_ISL_832108, EPI_ISL_832109 | MD Laboratories | Los Angeles County PHL | P. Hemarajata et al. |
| EPI_ISL_832169 | Hospital | National Reference Center for Viruses of Respiratory Infections, Institut Pasteur, Paris | Marion Barbet, Sylvie Behillil, Méline Bizard, Angela Brisebarre, Camille Capel, Etienne Simon-Lorière, Vincent Enouf, Maud Vanpeene, Sylvie van der Werf, Clémence Guillaume |
| EPI_ISL_832171, EPI_ISL_832172, EPI_ISL_832173, EPI_ISL_832174, EPI_ISL_832175 | Hospital | National Reference Center for Viruses of Respiratory Infections, Institut Pasteur, Paris | Marion Barbet, Sylvie Behillil, Méline Bizard, Angela Brisebarre, Camille Capel, Etienne Simon-Lorière, Vincent Enouf, Maud Vanpeene, Sylvie van der Werf, Patricia Stoessel-Thouvenin |
| EPI_ISL_832176, EPI_ISL_832177 | Hospital | National Reference Center for Viruses of Respiratory Infections, Institut Pasteur, Paris | Marion Barbet, Sylvie Behillil, Méline Bizard, Angela Brisebarre, Camille Capel, Etienne Simon-Lorière, Vincent Enouf, Maud Vanpeene, Sylvie van der Werf, Aurélie Guigon |
| EPI_ISL_832180 | Hospital | National Reference Center for Viruses of Respiratory Infections, Institut Pasteur, Paris | Marion Barbet, Sylvie Behillil, Méline Bizard, Angela Brisebarre, Camille Capel, Etienne Simon-Lorière, Vincent Enouf, Maud Vanpeene, Sylvie van der Werf, Laurent Roudiere |
| EPI_ISL_832181 | Sentinelles IDF | National Reference Center for Viruses of Respiratory Infections, Institut Pasteur, Paris | Marion Barbet, Sylvie Behillil, Méline Bizard, Angela Brisebarre, Camille Capel, Etienne Simon-Lorière, Vincent Enouf, Maud Vanpeene, Sylvie van der Werf, Ghatfan Hassan |
| EPI_ISL_832182, EPI_ISL_832183, EPI_ISL_832184, EPI_ISL_832185, EPI_ISL_832186, EPI_ISL_832187, EPI_ISL_832188, EPI_ISL_832191, EPI_ISL_832192, EPI_ISL_832193 | Laboratory Analyses Med | National Reference Center for Viruses of Respiratory Infections, Institut Pasteur, Paris | Marion Barbet, Sylvie Behillil, Méline Bizard, Angela Brisebarre, Camille Capel, Etienne Simon-Lorière, Vincent Enouf, Maud Vanpeene, Sylvie van der Werf, Brieuc Lefaure |
| EPI_ISL_832194, EPI_ISL_832195 | Laboratory Analyses Med | National Reference Center for Viruses of Respiratory Infections, Institut Pasteur, Paris | Marion Barbet, Sylvie Behillil, Méline Bizard, Angela Brisebarre, Camille Capel, Etienne Simon-Lorière, Vincent Enouf, Maud Vanpeene, Sylvie van der Werf, Brieuc Gestin |
| EPI_ISL_832196 | Laboratory Analyses Med | National Reference Center for Viruses of Respiratory Infections, Institut Pasteur, Paris | Marion Barbet, Sylvie Behillil, Méline Bizard, Angela Brisebarre, Camille Capel, Etienne Simon-Lorière, Vincent Enouf, Maud Vanpeene, Sylvie van der Werf, Heugas |
| EPI_ISL_832200 | Hospital | National Reference Center for Viruses of Respiratory Infections, Institut Pasteur, Paris | Marion Barbet, Sylvie Behillil, Méline Bizard, Angela Brisebarre, Camille Capel, Etienne Simon-Lorière, Vincent Enouf, Maud Vanpeene, Sylvie van der Werf, Patricia Stoessel-Thouvenin |
| EPI_ISL_832243 | DOHMH Corona | New York City Public Health Laboratory | Jade Wang, et al. |
| EPI_ISL_832246 | DOHMH Jamaica | New York City Public Health Laboratory | Jade Wang, et al. |
| EPI_ISL_832247, EPI_ISL_832248 | DOHMH Morrisania | New York City Public Health Laboratory | Jade Wang, et al. |
| EPI_ISL_832249 | DOHMH PHL | New York City Public Health Laboratory | Jade Wang, et al. |
| EPI_ISL_832252 | DOHMH Corona | New York City Public Health Laboratory | Jade Wang, et al. |
| EPI_ISL_832257 | DOHMH Morrisania | New York City Public Health Laboratory | Jade Wang, et al. |
| EPI_ISL_832263 | DOHMH Jamaica | New York City Public Health Laboratory | Jade Wang, et al. |
| EPI_ISL_832266, EPI_ISL_832267 | DOHMH Central Harlem | New York City Public Health Laboratory | Jade Wang, et al. |
| EPI_ISL_832269 | DOHMH Fort Greene | New York City Public Health Laboratory | Jade Wang, et al. |
| EPI_ISL_832270, EPI_ISL_832271 | DOHMH Riverside | New York City Public Health Laboratory | Jade Wang, et al. |
| EPI_ISL_832312 | DOHMH Central Harlem | New York City Public Health Laboratory | Jade Wang, et al. |
| EPI_ISL_832313, EPI_ISL_832314 | DOHMH Chelsea | New York City Public Health Laboratory | Jade Wang, et al. |
| EPI_ISL_832315, EPI_ISL_832316 | DOHMH Morrisania | New York City Public Health Laboratory | Jade Wang, et al. |
| EPI_ISL_832317, EPI_ISL_832318, EPI_ISL_832319 | DOHMH Jamaica | New York City Public Health Laboratory | Jade Wang, et al. |

| | | | |
|---|---|---|---|
| EPI_ISL_832320, EPI_ISL_832321, EPI_ISL_832322 | DOHMH Corona | New York City Public Health Laboratory | Jade Wang, et al. |
| EPI_ISL_832323, EPI_ISL_832324 | DOHMH Jamaica | New York City Public Health Laboratory | Jade Wang, et al. |
| EPI_ISL_832325 | DOHMH Corona | New York City Public Health Laboratory | Jade Wang, et al. |
| EPI_ISL_832326 | DOHMH Riverside | New York City Public Health Laboratory | Jade Wang, et al. |
| EPI_ISL_832327 | DOHMH Morrisania | New York City Public Health Laboratory | Jade Wang, et al. |
| EPI_ISL_832328, EPI_ISL_832329, EPI_ISL_832330 | DOHMH Jamaica | New York City Public Health Laboratory | Jade Wang, et al. |
| EPI_ISL_832331, EPI_ISL_832332, EPI_ISL_832333, EPI_ISL_832334, EPI_ISL_832335 | DOHMH Corona | New York City Public Health Laboratory | Jade Wang, et al. |
| EPI_ISL_832336, EPI_ISL_832337 | DOHMH Central Harlem | New York City Public Health Laboratory | Jade Wang, et al. |
| EPI_ISL_832338 | DOHMH Crown Heights | New York City Public Health Laboratory | Jade Wang, et al. |
| EPI_ISL_832339 | DOHMH Riverside | New York City Public Health Laboratory | Jade Wang, et al. |
| EPI_ISL_832340 | DOHMH Crown Heights | New York City Public Health Laboratory | Jade Wang, et al. |
| EPI_ISL_832341 | DOHMH Chelsea | New York City Public Health Laboratory | Jade Wang, et al. |
| EPI_ISL_832342, EPI_ISL_832343, EPI_ISL_832344, EPI_ISL_832345 | DOHMH Morrisania | New York City Public Health Laboratory | Jade Wang, et al. |
| EPI_ISL_832346, EPI_ISL_832347, EPI_ISL_832348, EPI_ISL_832349, EPI_ISL_832350 | DOHMH Central Harlem | New York City Public Health Laboratory | Jade Wang, et al. |
| EPI_ISL_832351 | DOHMH Corona | New York City Public Health Laboratory | Jade Wang, et al. |
| EPI_ISL_832352, EPI_ISL_832353 | DOHMH PHL | New York City Public Health Laboratory | Jade Wang, et al. |
| EPI_ISL_832354 | DOHMH Jamaica | New York City Public Health Laboratory | Jade Wang, et al. |
| EPI_ISL_832364, EPI_ISL_832365, EPI_ISL_832366, EPI_ISL_832367 | Center of Medical Microbiology, Virology, and Hospital Hygiene, University of Duesseldorf | Center of Medical Microbiology, Virology, and Hospital Hygiene, University of Duesseldorf | Maximilian Damagnez, Alexander Dilthey, Ashley-Jane Duplessis, Torsten Houwaart, Lisanna Hülse, Malte Kohns Vasconcelos, Nadine Lübke, Jessica Nicolai, Klaus Pfeffer, Daniel Strelow, Teresa Tamayo, Jörg Timm, Andreas Walker, Tobias Wienemann |
| EPI_ISL_832375, EPI_ISL_832376, EPI_ISL_832377, EPI_ISL_832379, EPI_ISL_832380, EPI_ISL_832381, EPI_ISL_832382 | Santa Clara County Public Health Laboratory | Santa Clara County Public Health Laboratory | Santa Clara County Public Health Department |
| EPI_ISL_832898, EPI_ISL_832899, EPI_ISL_832900, EPI_ISL_832901, EPI_ISL_832902, EPI_ISL_832903, EPI_ISL_832904, EPI_ISL_832905, EPI_ISL_832906, EPI_ISL_832907, EPI_ISL_832908, EPI_ISL_832909, EPI_ISL_832910, EPI_ISL_832919, EPI_ISL_832920, EPI_ISL_832921, EPI_ISL_832922, EPI_ISL_832923, EPI_ISL_832924, EPI_ISL_832925, EPI_ISL_832930, EPI_ISL_832931, EPI_ISL_832932 | | | |
| see above | Maine HETL | Tewhey Lab, The Jackson Laboratory | Matluk,N., Dewey,H., Iosue,F., Barter,M., Lynch,R., Munger,H. and Tewhey,R. |
| EPI_ISL_833043 | SIESP DIPARTIMENTO DI PREVENZIONE CHIETI | Istituto Zooprofilattico Sperimentale dell'Abruzzo e Molise "G. Caporale" | Lorusso A, Marcacci M, Di Domenico M, Ancora M, Curini V, Mangone I, Rinaldi A, Delli Compagni E, Di Pasquale A, Cammà C, Puglia I, Calistri P, Savini G |
| EPI_ISL_833044 | SIESP CHIETI - DRIVE IN CHIETI | Istituto Zooprofilattico Sperimentale dell'Abruzzo e Molise "G. Caporale" | Lorusso A, Marcacci M, Di Domenico M, Ancora M, Curini V, Mangone I, Rinaldi A, Delli Compagni E, Di Pasquale A, Cammà C, Puglia I, Calistri P, Savini G |
| EPI_ISL_833046 | FPAM | Istituto Zooprofilattico Sperimentale dell'Abruzzo e Molise "G. Caporale" | Lorusso A, Marcacci M, Di Domenico M, Ancora M, Curini V, Mangone I, Rinaldi A, Delli Compagni E, Di Pasquale A, Cammà C, Puglia I, Calistri P, Savini G |
| EPI_ISL_833047, EPI_ISL_833048 | OSPEDALE CIVILE ATRI | Istituto Zooprofilattico Sperimentale dell'Abruzzo e Molise "G. Caporale" | Lorusso A, Marcacci M, Di Domenico M, Ancora M, Curini V, Mangone I, Rinaldi A, Delli Compagni E, Di Pasquale A, Cammà C, Puglia I, Calistri P, Savini G |
| EPI_ISL_833052, EPI_ISL_833053, EPI_ISL_833054 | "Presidio Ospedaliero "San Liberatore" Atri | Istituto Zooprofilattico Sperimentale dell'Abruzzo e Molise "G. Caporale" | Lorusso A, Marcacci M, Di Domenico M, Ancora M, Curini V, Mangone I, Rinaldi A, Delli Compagni E, Di Pasquale A, Cammà C, Puglia I, Calistri P, Savini G |
| EPI_ISL_833061 | DIP. PREV. AVEZZANO SERVIZIO DI IGIENE EPIDEMIOLOGIAE SANITA' PUBBLICA | Istituto Zooprofilattico Sperimentale dell'Abruzzo e Molise "G. Caporale" | Lorusso A, Marcacci M, Di Domenico M, Ancora M, Curini V, Mangone I, Rinaldi A, Delli Compagni E, Di Pasquale A, Cammà C, Puglia I, Calistri P, Savini G |
| EPI_ISL_833068, EPI_ISL_833069, EPI_ISL_833070, EPI_ISL_833071, EPI_ISL_833072, EPI_ISL_833073, EPI_ISL_833074, EPI_ISL_833075 | FPAM | Istituto Zooprofilattico Sperimentale dell'Abruzzo e Molise "G. Caporale" | Lorusso A, Marcacci M, Di Domenico M, Ancora M, Curini V, Mangone I, Rinaldi A, Delli Compagni E, Di Pasquale A, Cammà C, Puglia I, Calistri P, Savini G |
| EPI_ISL_833076 | IZSAM | Istituto Zooprofilattico Sperimentale dell'Abruzzo e Molise "G. Caporale" | Lorusso A, Marcacci M, Di Domenico M, Ancora M, Curini V, Mangone I, Rinaldi A, Delli Compagni E, Di Pasquale A, Cammà C, Puglia I, Calistri P, Savini G |
| EPI_ISL_833077, EPI_ISL_833078 | OSPEDALE CIVILE ATRI | Istituto Zooprofilattico Sperimentale dell'Abruzzo e Molise "G. Caporale" | Lorusso A, Marcacci M, Di Domenico M, Ancora M, Curini V, Mangone I, Rinaldi A, Delli Compagni E, Di Pasquale A, Cammà C, Puglia I, Calistri P, Savini G |
| EPI_ISL_833082 | OSPEDALE SAN SALVATORE | Istituto Zooprofilattico Sperimentale dell'Abruzzo e Molise "G. Caporale" | Lorusso A, Marcacci M, Di Domenico M, Ancora M, Curini V, Mangone I, Rinaldi A, Delli Compagni E, Di Pasquale A, Cammà C, Puglia I, Calistri P, Savini G |
| EPI_ISL_833083, EPI_ISL_833084 | RPS | Istituto Zooprofilattico Sperimentale dell'Abruzzo e Molise "G. Caporale" | Lorusso A, Marcacci M, Di Domenico M, Ancora M, Curini V, Mangone I, Rinaldi A, Delli Compagni E, Di Pasquale A, Cammà C, Puglia I, Calistri P, Savini G |
| EPI_ISL_833085, EPI_ISL_833086 | Servizio di Igiene Epidemiologia e Sanità Pubblica (SIESP) CHIETI - DRIVE IN GISSI | Istituto Zooprofilattico Sperimentale dell'Abruzzo e Molise "G. Caporale" | Lorusso A, Marcacci M, Di Domenico M, Ancora M, Curini V, Mangone I, Rinaldi A, Delli Compagni E, Di Pasquale A, Cammà C, Puglia I, Calistri P, Savini G |
| EPI_ISL_833088, EPI_ISL_833089 | Servizio di Igiene Epidemiologia e Sanità Pubblica (SIESP) SULMONA | Istituto Zooprofilattico Sperimentale dell'Abruzzo e Molise "G. Caporale" | Lorusso A, Marcacci M, Di Domenico M, Ancora M, Curini V, Mangone I, Rinaldi A, Delli Compagni E, Di Pasquale A, Cammà C, Puglia I, Calistri P, Savini G |
| EPI_ISL_833090, EPI_ISL_833091, EPI_ISL_833092, EPI_ISL_833093, EPI_ISL_833094, EPI_ISL_833096 | SIESP CHIETI - DRIVE IN CHIETI | Istituto Zooprofilattico Sperimentale dell'Abruzzo e Molise "G. Caporale" | Lorusso A, Marcacci M, Di Domenico M, Ancora M, Curini V, Mangone I, Rinaldi A, Delli Compagni E, Di Pasquale A, Cammà C, Puglia I, Calistri P, Savini G |
| EPI_ISL_833097 | SIESP CHIETI - DRIVE IN LANCIANO | Istituto Zooprofilattico Sperimentale dell'Abruzzo e Molise "G. Caporale" | Lorusso A, Marcacci M, Di Domenico M, Ancora M, Curini V, Mangone I, Rinaldi A, Delli Compagni E, Di Pasquale A, Cammà C, Puglia I, Calistri P, Savini G |
| EPI_ISL_833101, EPI_ISL_833102 | SIESP DIPARTIMENTO DI PREVENZIONE CHIETI | Istituto Zooprofilattico Sperimentale dell'Abruzzo e Molise "G. Caporale" | Lorusso A, Marcacci M, Di Domenico M, Ancora M, Curini V, Mangone I, Rinaldi A, Delli Compagni E, Di Pasquale A, Cammà C, Puglia I, Calistri P, Savini G |
| EPI_ISL_833103, EPI_ISL_833104, EPI_ISL_833105, EPI_ISL_833106, EPI_ISL_833107, EPI_ISL_833108, EPI_ISL_833109, EPI_ISL_833110, EPI_ISL_833111, EPI_ISL_833112, EPI_ISL_833113, EPI_ISL_833115, EPI_ISL_833116, EPI_ISL_833117, EPI_ISL_833118 | | | |

| | | | |
|---|---|---|---|
| see above | SIESP DIPARTIMENTO DI PREVENZIONE TERAMO C.DA CASALENA | Istituto Zooprofilattico Sperimentale dell'Abruzzo e Molise "G. Caporale" | Lorusso A, Marcacci M, Di Domenico M, Ancora M, Curini V, Mangone I, Rinaldi A, Delli Compagni E, Di Pasquale A, Cammà C, Puglia I, Calistri P, Savini G |
| EPI_ISL_833119 | SIESP L'AQUILA | Istituto Zooprofilattico Sperimentale dell'Abruzzo e Molise "G. Caporale" | Lorusso A, Marcacci M, Di Domenico M, Ancora M, Curini V, Mangone I, Rinaldi A, Delli Compagni E, Di Pasquale A, Cammà C, Puglia I, Calistri P, Savini G |
| EPI_ISL_833120, EPI_ISL_833121 | SIESP SULMONA | Istituto Zooprofilattico Sperimentale dell'Abruzzo e Molise "G. Caporale" | Lorusso A, Marcacci M, Di Domenico M, Ancora M, Curini V, Mangone I, Rinaldi A, Delli Compagni E, Di Pasquale A, Cammà C, Puglia I, Calistri P, Savini G |
| EPI_ISL_833123 | Usca Pizzoli | Istituto Zooprofilattico Sperimentale dell'Abruzzo e Molise "G. Caporale" | Lorusso A, Marcacci M, Di Domenico M, Ancora M, Curini V, Mangone I, Rinaldi A, Delli Compagni E, Di Pasquale A, Cammà C, Puglia I, Calistri P, Savini G |
| EPI_ISL_833125, EPI_ISL_833126, EPI_ISL_833127, EPI_ISL_833128 | USCA Sulmona | Istituto Zooprofilattico Sperimentale dell'Abruzzo e Molise "G. Caporale" | Lorusso A, Marcacci M, Di Domenico M, Ancora M, Curini V, Mangone I, Rinaldi A, Delli Compagni E, Di Pasquale A, Cammà C, Puglia I, Calistri P, Savini G |
| EPI_ISL_833129 | USCA Tagliacozzo | Istituto Zooprofilattico Sperimentale dell'Abruzzo e Molise "G. Caporale" | Lorusso A, Marcacci M, Di Domenico M, Ancora M, Curini V, Mangone I, Rinaldi A, Delli Compagni E, Di Pasquale A, Cammà C, Puglia I, Calistri P, Savini G |
| EPI_ISL_833139, EPI_ISL_833140 | Laboratorio de Ecologia de Doencas Transmissiveis na Amazonia, Instituto Leonidas e Maria Deane - Fiocruz Amazonia | Laboratorio de Ecologia de Doencas Transmissiveis na Amazonia, Instituto Leonidas e Maria Deane - Fiocruz Amazonia | Valdinete Nascimento, Victor Souza, André Corado, Fernanda Nascimento, George Silva, Ágatha Costa, Debora Duarte, Karina Pessoa, Matilde Mejía, Luciana Gonçalves, Maria Júlia Brandão, Michele Jesus, Felipe Naveca |
| EPI_ISL_833168, EPI_ISL_833169, EPI_ISL_833170, EPI_ISL_833174, EPI_ISL_833175 | DB Diagnosticos do Brasil | Instituto Adolfo Lutz, Interdiciplinary Procedures Center, Strategic Laboratory | Claudio Tavares Sacchi, Claudia Regina Gonçalves, Erica Valessa Ramos Gomes, Karoline Rodrigues Campos |
| EPI_ISL_833184 | Microbiology Department. Hospital Clinico Universitario de Santiago | Microbiology Department. Complexo Hospitalario Universitario de Vigo | Microbiology Departments from Complexo Hospitalario Universitario de Vigo and Hospital Clinico Universitario de Santiago. EPICOVIGAL. |
| EPI_ISL_833201 | Department of Virology and Immunology, University of Helsinki and Helsinki University Hospital, Huslab Finland | Department of Virology, Faculty of Medicine, University of Helsinki, Helsinki, Finland | Teemu Smura, Ravi Kant, Phuoc Truong, Hussein Alburkat, Hannimari Kallio-Kokko, Jenni Virtanen, Maija Suvanto, Fathiah Zakham, Essi Korhonen, Sari Hannula, Harri Kangas, Pekka Ellonen, Olli Vapalahti |
| EPI_ISL_833271 | SIESP CHIETI - DRIVE IN GISSI | Istituto Zooprofilattico Sperimentale dell'Abruzzo e Molise "G. Caporale" | Lorusso A, Marcacci M, Di Domenico M, Ancora M, Curini V, Mangone I, Rinaldi A, Di Pasquale A, Cammà C, Puglia I, Calistri P, Savini G |
| EPI_ISL_833277 | USCA-Sulmona | Istituto Zooprofilattico Sperimentale dell'Abruzzo e Molise "G. Caporale" | Lorusso A, Marcacci M, Di Domenico M, Ancora M, Curini V, Mangone I, Rinaldi A, Di Pasquale A, Cammà C, Puglia I, Calistri P, Savini G |
| EPI_ISL_833289 | Ospedale SS Annunziata Chieti | Istituto Zooprofilattico Sperimentale dell'Abruzzo e Molise "G. Caporale" | Lorusso A, Marcacci M, Di Domenico M, Ancora M, Curini V, Mangone I, Rinaldi A, Di Pasquale A, Cammà C, Puglia I, Calistri P, Savini G |
| EPI_ISL_833433 | UZA/UAntwerpen (BIS lab), University Hospital Antwerp, Edegem, Belgium | Laboratory of Medical Microbiology, University of Antwerp, Campus Drie Eiken, S6.26, Universiteitsplein 1, 2610, Wilrijk, Antwerp, Belgium | Basil Britto Xavier, Jasmine Coppens, Sahaya Glingston Rajakani, Christine Lammens, Veerle Matheeussen, Herman Goossens |
| EPI_ISL_834343 | Lighthouse Lab in Alderley Park | Wellcome Sanger Institute for the COVID-19 Genomics UK (COG-UK) Consortium | Jacquelyn Wynn, Mairead Hyland, The Lighthouse Lab in Alderley Park and Alex Alderton, Roberto Amato, Sonia Goncalves, Ewan Harrison, David K. Jackson, Ian Johnston, Dominic Kwiatkowski, Cordelia Langford, John Sillitoe on behalf of the Wellcome Sanger Institute COVID-19 Surveillance Team |
| EPI_ISL_837005, EPI_ISL_837006, EPI_ISL_837007, EPI_ISL_837008, EPI_ISL_837009, EPI_ISL_837010, EPI_ISL_837011, EPI_ISL_837012, EPI_ISL_837013, EPI_ISL_837014, EPI_ISL_837015, EPI_ISL_837016, EPI_ISL_837017, EPI_ISL_837018, EPI_ISL_837019, EPI_ISL_837020, EPI_ISL_837021, EPI_ISL_837022, EPI_ISL_837023, EPI_ISL_837024, EPI_ISL_837025, EPI_ISL_837026, EPI_ISL_837027, EPI_ISL_837028, EPI_ISL_837029, EPI_ISL_837030, EPI_ISL_837031, EPI_ISL_837032, EPI_ISL_837033, EPI_ISL_837034, EPI_ISL_837035, EPI_ISL_837036, EPI_ISL_837037, EPI_ISL_837038, EPI_ISL_837039, EPI_ISL_837040, EPI_ISL_837061, EPI_ISL_837062, EPI_ISL_837072, EPI_ISL_837075, EPI_ISL_837079, EPI_ISL_837081, EPI_ISL_837088, EPI_ISL_837089, EPI_ISL_837091, EPI_ISL_837095, EPI_ISL_837103, EPI_ISL_837104, EPI_ISL_837105, EPI_ISL_837123, EPI_ISL_837124, EPI_ISL_837125, EPI_ISL_837126, EPI_ISL_837127, EPI_ISL_837128, EPI_ISL_837129, EPI_ISL_837130, EPI_ISL_837131, EPI_ISL_837132, EPI_ISL_837133, EPI_ISL_837134, EPI_ISL_837210, EPI_ISL_837211, EPI_ISL_837212, EPI_ISL_837213, EPI_ISL_837214, EPI_ISL_837215, EPI_ISL_837216, EPI_ISL_837217, EPI_ISL_837218, EPI_ISL_837219, EPI_ISL_837220, EPI_ISL_837221, EPI_ISL_837222, EPI_ISL_837223, EPI_ISL_837224 | | | |
| see above | Respiratory Virus Unit, National Infection Service, Public Health England | COVID-19 Genomics UK (COG-UK) Consortium | PHE Covid Sequencing Team |
| EPI_ISL_837433, EPI_ISL_837435, EPI_ISL_837438, EPI_ISL_837439, EPI_ISL_837440, EPI_ISL_837442, EPI_ISL_837446, EPI_ISL_837473, EPI_ISL_837474, EPI_ISL_837475, EPI_ISL_837476, EPI_ISL_837477, EPI_ISL_837478, EPI_ISL_837489, EPI_ISL_837490, EPI_ISL_837491, EPI_ISL_837492, EPI_ISL_837493, EPI_ISL_837494, EPI_ISL_837495, EPI_ISL_837496, EPI_ISL_837497, EPI_ISL_837498, EPI_ISL_837499, EPI_ISL_837500, EPI_ISL_837501, EPI_ISL_837502, EPI_ISL_837503, EPI_ISL_837504 | | | |
| see above | Istituto Zooprofilattico Sperimentale del Mezzogiorno | TIGEM | Patrizia Annunziata, Andrea Ballabio, Valentina Bouche, Davide Cacchiarelli (CorrespAuthor), Pellegrino Cerino, Chiara Colantuono, Lucio Di Filippo, Antonio Grimaldi, Antonio Limone, Gabriella Loconte, Anna Manfredi, Francesco Panariello, Biancamaria Pierri, Marcello Salvi, Lucia Vassallo |
| EPI_ISL_837509, EPI_ISL_837519, EPI_ISL_837520, EPI_ISL_837523, EPI_ISL_837524, EPI_ISL_837528, EPI_ISL_837544, EPI_ISL_837545, EPI_ISL_837547, EPI_ISL_837548, EPI_ISL_837549 | | | |
| see above | UW Virology Lab | UW Virology Lab | Pavitra Roychoudhury, Hong Xie, Lasata Shrestha, Meei-Li Huang, Keith R Jerome, Alexander Greninger |
| EPI_ISL_837815, EPI_ISL_837816, EPI_ISL_837818, EPI_ISL_837819, EPI_ISL_837822, EPI_ISL_837823, EPI_ISL_837824, EPI_ISL_837825, EPI_ISL_837827, EPI_ISL_837833 | Wyoming Public Health Laboratory | Wyoming Public Health Laboratory | Noah Hull, Taylor Fearing, Lynette Gumbleton, Channing Weber, Ashley Norberg, Bailey Bowcutt, and Wanda Manley |
| EPI_ISL_837882, EPI_ISL_837902 | Department of Pathology, University of Cambridge | COVID-19 Genomics UK (COG-UK) Consortium | Aminu S. Jahun, Yasmin Chaudhry, Grant Hall, Iliana Georgana, Myra Hosmillo, Martin D. Curran, Malte Pinckert, Surendra Parmar, Ian Goodfellow |
| EPI_ISL_838142, EPI_ISL_838143, EPI_ISL_838144, EPI_ISL_838145, EPI_ISL_838146, EPI_ISL_838147, EPI_ISL_838148, EPI_ISL_838149, EPI_ISL_838150, EPI_ISL_838151, EPI_ISL_838152, EPI_ISL_838153, EPI_ISL_838154, EPI_ISL_838155, EPI_ISL_838156, EPI_ISL_838157, EPI_ISL_838158, EPI_ISL_838159, EPI_ISL_838160, EPI_ISL_838161, EPI_ISL_838162, EPI_ISL_838163, EPI_ISL_838164, EPI_ISL_838165, EPI_ISL_838166, EPI_ISL_838167, EPI_ISL_838168, EPI_ISL_838169, EPI_ISL_838170, EPI_ISL_838171, EPI_ISL_838172, EPI_ISL_838173, EPI_ISL_838174, EPI_ISL_838175, EPI_ISL_838176, EPI_ISL_838177, EPI_ISL_838178, EPI_ISL_838179, EPI_ISL_838180, EPI_ISL_838181, EPI_ISL_838182, EPI_ISL_838183, EPI_ISL_838184, EPI_ISL_838186, EPI_ISL_838187, EPI_ISL_838198, EPI_ISL_838206, EPI_ISL_838207, EPI_ISL_838213 | | | |
| see above | West of Scotland Specialist Virology Centre, NHSGGC / MRC-University of Glasgow Centre for Virus Research | COVID-19 Genomics UK (COG-UK) Consortium | Ana da Silva Filipe, Natasha Johnson, Kathy Smollett, Daniel Mair, Stephen Carmichael, Alice Broos, Lily Tong, Jenna Nichols, Kyriaki Nomikou; Sarah McDonald; Richard Orton, Joseph Hughes, Sreenu Vattipally, David L Robertson; Alasdair MacLean, Rory Gunson; Sharif Shaaban, Matthew Holden; Rachel Blacow, Guy Mollett, Kathy Li, James Shepherd, Antonia Ho, Emma Thomson |
| EPI_ISL_838241, EPI_ISL_838242, EPI_ISL_838244, EPI_ISL_838255, EPI_ISL_838262, EPI_ISL_838263 | Virology Department, Royal Infirmary of Edinburgh, NHS Lothian / School of Biological Sciences, University of Edinburgh / Institute of Genetics and Molecular Medicine, University of Edinburgh | COVID-19 Genomics UK (COG-UK) Consortium | McHugh M, Dewar R, Rooke S, Gallagher M, Balcaza C, O'Toole Á, Scher E, Hill V, McCrone JT, Colquhoun R, Yu X, Jackson B, Rambaut A, Williams TC, Templeton K |
| EPI_ISL_838571, EPI_ISL_838572, EPI_ISL_838573, EPI_ISL_838574, EPI_ISL_838575 | Liverpool Clinical Laboratories | COVID-19 Genomics UK (COG-UK) Consortium | Sam Haldenby, Anita Lucaci, Steve Paterson, Julian Hiscox, Alistair Darby, M Almsaud, A Alrezaihi, Muhannad Alruwaili, Stuart D Armstrong, Jones Benjamin, Eleanor G Bentley, Anu Chawla, Jordan J Clark, Angela Cowell, Richard Eccles, Isabel García-Dorival, Matthew Gemmell, Alessandro Gerada, PKF Gilmore, Richard Gregory, Ximeng Han, Catherine Hartley, Margaret Hughes, Miren Iturriza-Gomara, James Johnson, L Luu, Jenifer Manson, Charlotte Nelson, Elaine O'Toole, Cassie Olateju, Rebekah Penrice-Randal , Lucille Rainbow, N.P Randle, Trevor Ian Robinson, Parul Sharma, Ghada T Shawli, James P Stewart, Neil Swainston, Ecaterina Vamos, Joanne Watts, Mark Whitehead |
| EPI_ISL_838734, EPI_ISL_838735, EPI_ISL_838736, EPI_ISL_838737, EPI_ISL_838738, EPI_ISL_838740, EPI_ISL_838741, EPI_ISL_838742, EPI_ISL_838743, EPI_ISL_838745, EPI_ISL_838746, EPI_ISL_838747, EPI_ISL_838748, EPI_ISL_838749, EPI_ISL_838750, EPI_ISL_838751, EPI_ISL_838752, EPI_ISL_838753, EPI_ISL_838754, EPI_ISL_838755, EPI_ISL_838756, EPI_ISL_838757, EPI_ISL_838758, EPI_ISL_838759, EPI_ISL_838760, EPI_ISL_838762, EPI_ISL_838763, EPI_ISL_838764, EPI_ISL_839312, EPI_ISL_839315, EPI_ISL_839318, EPI_ISL_839319, EPI_ISL_839320, EPI_ISL_839321, EPI_ISL_839322, EPI_ISL_839323 | | | |
| see above | University College London, Great Ormond Street Hospital for Children NHS Foundation Trust, Imperial College Healthcare NHS Trust | COVID-19 Genomics UK (COG-UK) Consortium | Sergi Castellano, Rachel Williams, Mark Kristiansen, Paola Resende Silva, Sunando Roy, Tony Brooks, Helena Tutill, Paola Niola, Patricia Dyal, Charlotte Williams, Leysa Forrest, Yasmin Panchbhaya, Jacqueline Findlay, Samuel Weeks, Julianne Brown, Kathryn Harris, Paul Randell, James Price, Alison Holmes, Judith Breuer |
| EPI_ISL_839607, EPI_ISL_839608, EPI_ISL_839609, EPI_ISL_839610, EPI_ISL_839611, EPI_ISL_839612, EPI_ISL_839613, EPI_ISL_839614, EPI_ISL_839616, EPI_ISL_839617, EPI_ISL_839620, EPI_ISL_839623, EPI_ISL_839625, EPI_ISL_839627, EPI_ISL_839628, EPI_ISL_839629, EPI_ISL_839630, EPI_ISL_839641, | | | |

| | | | |
|---|---|---|---|
| EPI_ISL_839649, EPI_ISL_839650, EPI_ISL_839654, EPI_ISL_839655, EPI_ISL_839656, EPI_ISL_839667, EPI_ISL_839816, EPI_ISL_839818, EPI_ISL_839819, EPI_ISL_839820, EPI_ISL_839821, EPI_ISL_839822 | | | |
| see above | Northumbria University / South Tees Hospitals NHS Foundation Trust / North Cumbria Integrated Care NHS Foundation Trust / North Tees and Hartlepool NHS Foundation Trust / Newcastle Hospitals NHS Foundation Trust | COVID-19 Genomics UK (COG-UK) Consortium | Darren L Smith,Andrew Nelson,Matthew Bashton,Greg R Young,Joshua Loh,John Allan,Mohammad A Tariq,Giles S Holt,Gary Black,Wen C Yew,Lynn Dover,Paul Baker,Steve Liggett,Sarah Essex,Jane Greenaway,Debra Padgett,Clive Graham,Garren Scott,Edward Barton,Emma Swindells,Brendan Payne,Jennifer Collins,Yusri Taha,Gary Eltringham |
| EPI_ISL_839991, EPI_ISL_839993 | Queens Medical Centre, Clinical Microbiology Department / DeepSeq Nottingham | COVID-19 Genomics UK (COG-UK) Consortium | Gemma Clark, Wendy Smith, Manjinder Khakh, Vicki M Fleming, Michelle M Lister, Hannah Howson-Wells, Jonathan Ball, Patrick McClure, Joseph Chappell, Theocharis Tsoleridis, Nadine Holmes, Matthew Carlisle, Christopher Moore, Fei Sang, Johnny Debebe, Victoria Wright, Matthew Loose |
| EPI_ISL_840088, EPI_ISL_840089, EPI_ISL_840090, EPI_ISL_840091, EPI_ISL_840092, EPI_ISL_840093, EPI_ISL_840094, EPI_ISL_840095, EPI_ISL_840096, EPI_ISL_840097, EPI_ISL_840098, EPI_ISL_840099, EPI_ISL_840100, EPI_ISL_840101, EPI_ISL_840102, EPI_ISL_840103 | | | |
| see above | Lincolnshire Hospitals and DeepSeq Nottingham | COVID-19 Genomics UK (COG-UK) Consortium | Nichola Duckworth, Tim Sloan, Sarah Walsh, Jonathan Ball, Patrick McClure, Joeseph Chappell, Nadine Holmes, Matthew Carlisle, Christopher Moore, Fei Sang, Johnny Debebe, Victoria Wright, Matthew Loose |
| EPI_ISL_840288, EPI_ISL_840289, EPI_ISL_840290, EPI_ISL_840291, EPI_ISL_840292, EPI_ISL_840294, EPI_ISL_840295, EPI_ISL_840296, EPI_ISL_840297, EPI_ISL_840298, EPI_ISL_840314, EPI_ISL_840315, EPI_ISL_840316, EPI_ISL_840317, EPI_ISL_840318, EPI_ISL_840319, EPI_ISL_840320, EPI_ISL_840321, EPI_ISL_840322, EPI_ISL_840323, EPI_ISL_840324, EPI_ISL_840325, EPI_ISL_840326, EPI_ISL_840327, EPI_ISL_840328, EPI_ISL_840329, EPI_ISL_840330, EPI_ISL_840331, EPI_ISL_840339, EPI_ISL_840340, EPI_ISL_840341, EPI_ISL_840342, EPI_ISL_840365, EPI_ISL_840366 | | | |
| see above | Oxford Viromics, NDM, University of Oxford; Oxford University Hospitals; Basingstoke and North Hampshire Hospital | COVID-19 Genomics UK (COG-UK) Consortium | Tanya Golubchik, David Bonsall, George Macintyre, Amy Trebes, Mariateresa de Cesare, Catrin Moore, Alex Mobbs, Anita Justice, Robert Shaw, Monique Andersson, Timothy Peto, Emma Wise, Nathan Moore, Jessica Lynch, Nick Cortes, Matilde Mori, Stephen Kidd, David Buck, John Todd, Christophe Fraser |
| EPI_ISL_840374, EPI_ISL_840375, EPI_ISL_840376, EPI_ISL_840377, EPI_ISL_840378, EPI_ISL_840380, EPI_ISL_840381, EPI_ISL_840382, EPI_ISL_840383, EPI_ISL_840384, EPI_ISL_840385, EPI_ISL_840386, EPI_ISL_840398, EPI_ISL_840400, EPI_ISL_840401, EPI_ISL_840402, EPI_ISL_840403, EPI_ISL_840404, EPI_ISL_840405, EPI_ISL_840406, EPI_ISL_840407, EPI_ISL_840408, EPI_ISL_840409, EPI_ISL_840410, EPI_ISL_840411, EPI_ISL_840412, EPI_ISL_840413, EPI_ISL_840414, EPI_ISL_840415, EPI_ISL_840416, EPI_ISL_840417, EPI_ISL_840418, EPI_ISL_840419, EPI_ISL_840420, EPI_ISL_840421, EPI_ISL_840422, EPI_ISL_840423, EPI_ISL_840424, EPI_ISL_840425, EPI_ISL_840426, EPI_ISL_840427, EPI_ISL_840428, EPI_ISL_840429, EPI_ISL_840430, EPI_ISL_840431, EPI_ISL_840432, EPI_ISL_840433, EPI_ISL_840434, EPI_ISL_840435, EPI_ISL_840436, EPI_ISL_840437, EPI_ISL_840438, EPI_ISL_840439, EPI_ISL_840440, EPI_ISL_840441, EPI_ISL_840442, EPI_ISL_840443, EPI_ISL_840444, EPI_ISL_840445, EPI_ISL_840446, EPI_ISL_840447, EPI_ISL_840448, EPI_ISL_840449, EPI_ISL_840450, EPI_ISL_840451, EPI_ISL_840452, EPI_ISL_840453, EPI_ISL_840454, EPI_ISL_840455, EPI_ISL_840456, EPI_ISL_840457, EPI_ISL_840458, EPI_ISL_840459, EPI_ISL_840460, EPI_ISL_840461, EPI_ISL_840462, EPI_ISL_840463, EPI_ISL_840464, EPI_ISL_840465, EPI_ISL_840466, EPI_ISL_840467, EPI_ISL_840468, EPI_ISL_840469, EPI_ISL_840470, EPI_ISL_840471, EPI_ISL_840472, EPI_ISL_840473, EPI_ISL_840474, EPI_ISL_840475, EPI_ISL_840476, EPI_ISL_840477, EPI_ISL_840478, EPI_ISL_840479, EPI_ISL_840480, EPI_ISL_840481, EPI_ISL_840482, EPI_ISL_840483, EPI_ISL_840484, EPI_ISL_840485, EPI_ISL_840486, EPI_ISL_840487, EPI_ISL_840488, EPI_ISL_840489, EPI_ISL_840490, EPI_ISL_840491, EPI_ISL_840492, EPI_ISL_840493, EPI_ISL_840494, EPI_ISL_840495, EPI_ISL_840496, EPI_ISL_840497, EPI_ISL_840498, EPI_ISL_840499, EPI_ISL_840500, EPI_ISL_840501, EPI_ISL_840502, EPI_ISL_840503, EPI_ISL_840505, EPI_ISL_840506, EPI_ISL_840507, EPI_ISL_840509, EPI_ISL_840510, EPI_ISL_840516, EPI_ISL_840517, EPI_ISL_840518, EPI_ISL_840662, EPI_ISL_840667, EPI_ISL_840668, EPI_ISL_840669, EPI_ISL_840670, EPI_ISL_840705, EPI_ISL_840706, EPI_ISL_840707, EPI_ISL_840708, EPI_ISL_840709, EPI_ISL_840710, EPI_ISL_840726, EPI_ISL_840728, EPI_ISL_840729, EPI_ISL_840730, EPI_ISL_840737, EPI_ISL_840738, EPI_ISL_840739, EPI_ISL_840740, EPI_ISL_840742, EPI_ISL_840743, EPI_ISL_840744, EPI_ISL_840745, EPI_ISL_840746, EPI_ISL_840749, EPI_ISL_840750, EPI_ISL_840751, EPI_ISL_840752, EPI_ISL_840754, EPI_ISL_840756, EPI_ISL_840757, EPI_ISL_840758, EPI_ISL_840759, EPI_ISL_840760, EPI_ISL_840761, EPI_ISL_840762, EPI_ISL_840763, EPI_ISL_840764, EPI_ISL_840765, EPI_ISL_840766, EPI_ISL_840767, EPI_ISL_840769 | | | |
| see above | Originating lab: Wales Specialist Virology Centre Sequencing lab: Pathogen Genomics Unit | Public Health Wales Microbiology Cardiff Wales Specialist Virology Centre | Catherine Moore, Johnathan Evans, Laura Gifford, Malorie Perry, Simon Cottrell, Angela Marchbank, Alec Birchley, Alexander Adams, Amy Gaskin, Bree Gatica-Wilcox, Jason Coombes, Joel Southgate, Lauren Gilbert, Lee Graham, Nicole Pacchiarini, Sara Kumziene-Summerhayes, Sarah Taylor, Sophie Jones, Sara Rey, Matthew Bull, Joanne Watkins, Sally Corden, Tom Connor |
| EPI_ISL_840844, EPI_ISL_840893, EPI_ISL_840894, EPI_ISL_841065, EPI_ISL_841066 | Wales Specialist Virology Centre Sequencing lab: Pathogen Genomics Unit | Public Health Wales Microbiology Cardiff Wales Specialist Virology Centre | Catherine Moore, Johnathan Evans, Laura Gifford, Malorie Perry, Simon Cottrell, Angela Marchbank, Alec Birchley, Alexander Adams, Amy Gaskin, Bree Gatica-Wilcox, Jason Coombes, Joel Southgate, Lauren Gilbert, Lee Graham, Nicole Pacchiarini, Sara Kumziene-Summerhayes, Sarah Taylor, Sophie Jones, Sara Rey, Matthew Bull, Joanne Watkins, Sally Corden, Tom Connor |
| EPI_ISL_841846, EPI_ISL_841847 | Centre for Enzyme Innovation, University of Portsmouth / Translational Research Laboratory, Portsmouth Hospitals NHS Trust | COVID-19 Genomics UK (COG-UK) Consortium | Angela Beckett,Yann Bourgeois,Garry Scarlett,Sharon Glaysher,Scott Elliott,Kelly Bicknell,Robert Impey,Allyson Lloyd,Sarah Wyllie,Ethan Butcher,Anoop Chauhan,Samuel Robson |
| EPI_ISL_842214, EPI_ISL_842217, EPI_ISL_842218, EPI_ISL_842233, EPI_ISL_842235, EPI_ISL_842254, EPI_ISL_842255, EPI_ISL_842256, EPI_ISL_842263, EPI_ISL_842264, EPI_ISL_842266, EPI_ISL_842272, EPI_ISL_842281, EPI_ISL_842282, EPI_ISL_842291, EPI_ISL_842295, EPI_ISL_842306, EPI_ISL_842316, EPI_ISL_842318, EPI_ISL_842319, EPI_ISL_842335 | | | |
| see above | Virology Department, Sheffield Teaching Hospitals NHS Foundation Trust/Department of Infection, Immunity and Cardiovascular Disease, The Medical School, University of Sheffield | COVID-19 Genomics UK (COG-UK) Consortium | Thushan de Silva, Matthew Parker, Nikki Smith, Adri Angyal, Rebecca Brown, Luke Green, Rachel Tucker, Paul Parsons, Danielle Groves, Katie Johnson, Laura Carrilero, Alex Keeley, Dave Partridge, Matthew Wyles, Benjamin Lindsey, Mehmet Yavuz, Mohammad Raza, Cariad Evans |
| EPI_ISL_842514, EPI_ISL_842516, EPI_ISL_842517, EPI_ISL_842518, EPI_ISL_842519, EPI_ISL_842520, EPI_ISL_842521, EPI_ISL_842522, EPI_ISL_842523, EPI_ISL_842524, EPI_ISL_842525, EPI_ISL_842526, EPI_ISL_842527, EPI_ISL_842528, EPI_ISL_842529, EPI_ISL_842530, EPI_ISL_842531, EPI_ISL_842532, EPI_ISL_842533, EPI_ISL_842534, EPI_ISL_842535, EPI_ISL_842536, EPI_ISL_842537, EPI_ISL_842538, EPI_ISL_842539, EPI_ISL_842540, EPI_ISL_842541, EPI_ISL_842542, EPI_ISL_842543, EPI_ISL_842544, EPI_ISL_842545, EPI_ISL_842546, EPI_ISL_842547, EPI_ISL_842548, EPI_ISL_842549, EPI_ISL_842550, EPI_ISL_842551, EPI_ISL_842552, EPI_ISL_842553, EPI_ISL_842554, EPI_ISL_842555, EPI_ISL_842556, EPI_ISL_842557, EPI_ISL_842558, EPI_ISL_842559, EPI_ISL_842560, EPI_ISL_842561, EPI_ISL_842562, EPI_ISL_842563, EPI_ISL_842564, EPI_ISL_842565, EPI_ISL_842566, EPI_ISL_842567, EPI_ISL_842568, EPI_ISL_842569, EPI_ISL_842570, EPI_ISL_842571, EPI_ISL_842572, EPI_ISL_842573, EPI_ISL_842574, EPI_ISL_842575, EPI_ISL_842576, EPI_ISL_842577, EPI_ISL_842578, EPI_ISL_842579, EPI_ISL_842580, EPI_ISL_842581, EPI_ISL_842582, EPI_ISL_842584, EPI_ISL_842595 | | | |
| see above | Bioinformatics and Biostatistics Lab, Advanced Sequencing Facility | COVID-19 Genomics UK (COG-UK) Consortium | Aengus Stewart,Jerome Nicod,Chelsea Sawyer,Laura Cubitt,Harshil Patel,Margaret Crawford |
| EPI_ISL_843077, EPI_ISL_843080, EPI_ISL_843084, EPI_ISL_843085, EPI_ISL_843086, EPI_ISL_843087, EPI_ISL_843093, EPI_ISL_843094, EPI_ISL_843095, EPI_ISL_843096, EPI_ISL_843097, EPI_ISL_843098, EPI_ISL_843099, EPI_ISL_843100, EPI_ISL_843101, EPI_ISL_843102, EPI_ISL_843103, EPI_ISL_843104, EPI_ISL_843115, EPI_ISL_843126, EPI_ISL_843127, EPI_ISL_843128, EPI_ISL_843129, EPI_ISL_843130, EPI_ISL_843131, EPI_ISL_843132, EPI_ISL_843133, EPI_ISL_843134, EPI_ISL_843135, EPI_ISL_843136, EPI_ISL_843137, EPI_ISL_843138, EPI_ISL_843139, EPI_ISL_843140, EPI_ISL_843141, EPI_ISL_843142, EPI_ISL_843143 | | | |
| see above | Barts Health NHS Trust | COVID-19 Genomics UK (COG-UK) Consortium | CUTINO-MOGUEL, Maria-Teresa; HARRINGTON, David; OWOYEMI, Dola; SHYLINI, Raghavendran; BROAD, Claire; KELE, Beatrix |
| EPI_ISL_845578, EPI_ISL_845580, EPI_ISL_845583, EPI_ISL_845585, EPI_ISL_845587, EPI_ISL_845589, EPI_ISL_845602 | KU Leuven, Rega Institute, Clinical and Epidemiological Virology | KU Leuven, Rega Institute, Clinical and Epidemiological Virology | Tony Wawina-Bokalanga, Bert Vanmechelen, Joan Marti-Carerras, Piet Maes |
| EPI_ISL_845618, EPI_ISL_845619 | Ayudas diagnósticas SURA | Instituto Nacional de Salud - Dirección de Investigación en Salud Pública | Katherine Laiton-Donato, Diego A. Álvarez-Díaz, Carlos Franco-Muñoz, Mauricio Pacheco-Montealegre, Maria T. Herrera-Sepúlveda, Jonathan Reales, Sheryll Corchuelo, Julian Naizaque, Gerardo Santamaría, Paola Muñoz-Laiton, Diego Andrés Prada, Magdalena Wiesner, Martha Lucia Ospina Martinez, Marcela Mercado-Reyes |
| EPI_ISL_845627 | SYNLAB COLOMBIA S.A.S | Instituto Nacional de Salud - Dirección de Investigación en Salud Pública | Katherine Laiton-Donato, Diego A. Álvarez-Díaz, Carlos Franco-Muñoz, Mauricio Pacheco-Montealegre, Maria T. Herrera-Sepúlveda, Jonathan Reales, Sheryll Corchuelo, Julian Naizaque, Gerardo Santamaría, Paola Muñoz-Laiton, Diego Andrés Prada, Magdalena Wiesner, Martha Lucia Ospina Martinez, Marcela Mercado-Reyes |
| EPI_ISL_845635 | Laboratorio de Salud Pública - Secretaría Distrital de Salud | Instituto Nacional de Salud - Dirección de Investigación en Salud Pública | Katherine Laiton-Donato, Diego A. Álvarez-Díaz, Carlos Franco-Muñoz, Mauricio Pacheco-Montealegre, Maria T. Herrera-Sepúlveda, Jonathan Reales, Sheryll Corchuelo, Julian Naizaque, Gerardo Santamaría, Paola Muñoz-Laiton, Diego Andrés Prada, Magdalena Wiesner, Martha Lucia Ospina Martinez, Marcela Mercado-Reyes |
| EPI_ISL_845814, EPI_ISL_845815, EPI_ISL_845816, EPI_ISL_845817, EPI_ISL_845818, EPI_ISL_845819, EPI_ISL_845820, EPI_ISL_845821, EPI_ISL_845822, EPI_ISL_845823, EPI_ISL_845824 | | | |
| see above | TGen North | TGen North | Jolene Bowers, Megan Folkerts, Chris French, Hayley Yaglom, Ashlyn Pfeiffer, Darrin Lemmer, Dave Engelthaler, The Arizona COVID Genomics Union (ACGU) |
| EPI_ISL_846592, EPI_ISL_846594 | Respiratory Virus Unit, National Infection Service, Public Health England | COVID-19 Genomics UK (COG-UK) Consortium | PHE Covid Sequencing Team |
| EPI_ISL_846653, EPI_ISL_846654 | Servicio de Microbiología Clínica (Complejo Hospitalario de Navarra, Pamplona), Instituto de Investigación Sanitaria de Navarra (IdiSNA) | SeqCOVID-SPAIN consortium/IBV(CSIC) | Carmen Ezpeleta Baquedano, Ana Navascués, Ana Miqueleiz and SeqCOVID-SPAIN consortium |
| EPI_ISL_847530, EPI_ISL_847538, EPI_ISL_847541, EPI_ISL_847578, EPI_ISL_847580, EPI_ISL_847608, EPI_ISL_847609, EPI_ISL_847610, EPI_ISL_847611, EPI_ISL_847612, EPI_ISL_847714, EPI_ISL_847723, EPI_ISL_847725 | | | |
| see above | California Department of Public Health | Chiu Laboratory, University of California, San Francisco | Charles Chiu, Xianding (Wayne) Deng, Candace Wang, Brian Bushnell, Scot Federman, Jill Hacker, Debra Wadford |
| EPI_ISL_847747 | Chiu Laboratory, University of California, San Francisco | Chiu Laboratory, University of California, San Francisco | Charles Chiu, Xianding (Wayne) Deng, Candace Wang, Brian Bushnell, Scot Federman, Jill Hacker, Debra Wadford |

| | | | |
|---|---|---|---|
| EPI_ISL_847751, EPI_ISL_847753, EPI_ISL_847772 | California Department of Public Health | Chiu Laboratory, University of California, San Francisco | Charles Chiu, Xianding (Wayne) Deng, Candace Wang, Brian Bushnell, Scot Federman, Jill Hacker, Debra Wadford |
| EPI_ISL_847840 | Tempus | Grubaugh Lab - Yale School of Public Health | Tara Alpert, Joseph Fauver, Anderson Brito, Mallery Breban, Anne Wyllie, Chantal Vogels, Mary Petrone, Chaney Kalinich, Isabel Ott, Nathan Grubaugh |
| EPI_ISL_847843, EPI_ISL_847846, EPI_ISL_847850, EPI_ISL_847851, EPI_ISL_847854, EPI_ISL_847902, EPI_ISL_847903, EPI_ISL_847904, EPI_ISL_847905, EPI_ISL_847906, EPI_ISL_847907, EPI_ISL_847908, EPI_ISL_847909, EPI_ISL_847910, EPI_ISL_847911, EPI_ISL_847912, EPI_ISL_847913, EPI_ISL_847914, EPI_ISL_847915, EPI_ISL_847916, EPI_ISL_847917, EPI_ISL_847918, EPI_ISL_847919, EPI_ISL_847922, EPI_ISL_847923, EPI_ISL_847926, EPI_ISL_847939, EPI_ISL_847968, EPI_ISL_847969, EPI_ISL_847970, EPI_ISL_847971, EPI_ISL_847972, EPI_ISL_847973, EPI_ISL_847974, EPI_ISL_847975, EPI_ISL_847976, EPI_ISL_847977 | | | |
| see above | University Hospitals of Geneva, Laboratory of Virology | HUG, Laboratory of Virology and the Health2030 Genome Center | Samuel Cordey, Ana Rita Goncalves, Laurent Kaiser, Lorenzo Cerutti, Henri Pegeot, Melyssa Elies, Keith Harshman, Ioannis Xenarios, Emmanouil Dermitzakis |
| EPI_ISL_847983, EPI_ISL_847992, EPI_ISL_848014, EPI_ISL_848022 | Michigan Department of Health and Human Services, Bureau of Laboratories | Michigan Department of Health and Human Services, Bureau of Laboratories | Blankenship HM, Riner D, Soehnlen MK |
| EPI_ISL_848188, EPI_ISL_848189, EPI_ISL_848190, EPI_ISL_848191, EPI_ISL_848193, EPI_ISL_848194, EPI_ISL_848195, EPI_ISL_848196, EPI_ISL_848197, EPI_ISL_848198 | Microbiology Lab | NBCC Sequencing Facility | Dr. Jeff Wrana |
| EPI_ISL_848199 | Microbiology Department. Complexo Hospitalario Universitario de Vigo | Microbiology Department. Complexo Hospitalario Universitario de Vigo | Microbiology Department, Complexo Hospitalario Universitario de Vigo (CHUVI). EPICOVIGAL. |
| EPI_ISL_849745, EPI_ISL_849746, EPI_ISL_849747, EPI_ISL_849748, EPI_ISL_849749 | unknown | PHV-FSS | Son Nguyen et al. |
| EPI_ISL_849794, EPI_ISL_849796, EPI_ISL_849802, EPI_ISL_849803, EPI_ISL_849805, EPI_ISL_849806, EPI_ISL_849807, EPI_ISL_849808, EPI_ISL_849809, EPI_ISL_849810, EPI_ISL_849811, EPI_ISL_849812, EPI_ISL_849813, EPI_ISL_849814, EPI_ISL_849815, EPI_ISL_849816, EPI_ISL_849817, EPI_ISL_849818, EPI_ISL_849819, EPI_ISL_849820, EPI_ISL_849821, EPI_ISL_849822, EPI_ISL_849823, EPI_ISL_849824, EPI_ISL_849825, EPI_ISL_849826, EPI_ISL_849827, EPI_ISL_849828, EPI_ISL_849829, EPI_ISL_849830, EPI_ISL_849831, EPI_ISL_849832, EPI_ISL_849833, EPI_ISL_849834, EPI_ISL_849835, EPI_ISL_849836, EPI_ISL_849837, EPI_ISL_849838, EPI_ISL_849839, EPI_ISL_849840, EPI_ISL_849841, EPI_ISL_849842, EPI_ISL_849843, EPI_ISL_849844, EPI_ISL_849845, EPI_ISL_849846, EPI_ISL_849847, EPI_ISL_849848, EPI_ISL_849849, EPI_ISL_849850, EPI_ISL_849851, EPI_ISL_849852, EPI_ISL_849853, EPI_ISL_849854, EPI_ISL_849855, EPI_ISL_849856, EPI_ISL_849857, EPI_ISL_849858, EPI_ISL_849859, EPI_ISL_849860, EPI_ISL_849861, EPI_ISL_849862, EPI_ISL_849863, EPI_ISL_849864 | | | |
| see above | A. Krumbholz, Labor Dr. Krause und Kollegen MVZ GmbH, Kiel | Charité Universitätsmedizin Berlin, Institut für Virologie | Victor M Corman, Jörn Beheim-Schwarzbach, Talitha Veith, Julia Schneider, Tobias Bleicker, Julia Tesch, Barbara Mühlemann, Terry Jones, Christian Drosten |
| EPI_ISL_850654, EPI_ISL_850655, EPI_ISL_850656, EPI_ISL_850659 | Division of Emerging Infectious Diseases, Bureau of Infectious Diseases Diagnosis Control, Korea Disease Control and Prevention Agency | Division of Emerging Infectious Diseases, Bureau of Infectious Diseases Diagnosis Control, Korea Disease Control and Prevention Agency | Ae Kyung Park, Il-Hwan Kim, Heui Man Kim, Jeong-Min Kim, Namjoo Lee, Chaeyoung Lee, Sang Hee Woo, Eun-Jin Kim |
| EPI_ISL_852814, EPI_ISL_852815, EPI_ISL_852816 | CHU Purpan - Laboratoire de Virologie - Institut Fédératif de Biologie | CHU Purpan - Laboratoire de Virologie - Institut Fédératif de Biologie | Latour J., Ranger N., Dubois M., Carcenac R., Harter A., Boyer P., Tremeaux P., Izopet J. |
| EPI_ISL_852820, EPI_ISL_852822, EPI_ISL_852824, EPI_ISL_852826, EPI_ISL_852827, EPI_ISL_852829, EPI_ISL_852830, EPI_ISL_852831, EPI_ISL_852832, EPI_ISL_852833 | Microbiology Lab | NBCC Sequencing Facility | Dr. Jeff Wrana |
| EPI_ISL_852842 | Laboratory of Molecular Biology, Diagnostyka sp. z o.o. | genXone SA, Research & Development Laboratory | Maciej Sykulski, Grzegorz Nowicki, Jakub Grabowski, Natalia Drwska-Matelska, ukasz Krych, Micha Kaszuba |
| EPI_ISL_852988, EPI_ISL_852998, EPI_ISL_853000, EPI_ISL_853016 | Hospital General Universitario Gregorio Marañón | SeqCOVID-SPAIN consortium/IBV(CSIC) | Darío García de Viedma, Laura Pérez-Lago, Pedro J Sola-Campoy, Sergio Buenestado-Serrano, Marta Herranz, Victor Manuel de la Cueva, Julia Suárez, Pilar Catalán, Patricia Muñoz and SeqCOVID-SPAIN consortium |
| EPI_ISL_853274, EPI_ISL_853315 | Microbiology Lab | NBCC Sequencing Facility | Dr. Jeff Wrana |
| EPI_ISL_853357, EPI_ISL_853358, EPI_ISL_853359, EPI_ISL_853362, EPI_ISL_853365, EPI_ISL_853366, EPI_ISL_853368, EPI_ISL_853371 | UPMC Clinical Microbiology Laboratory | Microbial Genome Sequencing Center; Microbial Genomic Epidemiology Laboratory | Mustapha M. Mustapha, Jane W. Marsh, Dan Snyder, Marissa P. Griffith, Stephanie L. Mitchell, Vatsala R. Srinivasa, Kady D. Waggle, Chinelo Ezeonwuku, Vaughn S. Cooper, Lee H. Harrison |
| EPI_ISL_853404 | Charité Universitätsmedizin Berlin, Institut für Virologie/Labor Berlin | Charité Universitätsmedizin Berlin, Institut für Virologie | Victor M Corman, Julia Schneider, Barbara Mühlemann, Jörn Beheim-Schwarzbach, Talitha Veith, Julia Tesch, Tobias Bleicker, Terry Jones, Christian Drosten |
| EPI_ISL_853788, EPI_ISL_853806 | Department of Microbiology, University Innsbruck | Bergthaler laboratory, CeMM Research Center for Molecular Medicine of the Austrian Academy of Sciences | Lukas Endler, Alexandra Popa, Benedikt Agerer, Jakob-Wendelin Genger, Alexander Lercher, Anna Schedl, Thomas Penz, Michael Schuster, Jan Laine, Martin Senekowitsch, Christoph Bock, Andreas Bergthaler |
| EPI_ISL_853926, EPI_ISL_853927, EPI_ISL_853928 | ILV Kärnten | Bergthaler laboratory, CeMM Research Center for Molecular Medicine of the Austrian Academy of Sciences | Lukas Endler, Alexandra Popa, Benedikt Agerer, Jakob-Wendelin Genger, Alexander Lercher, Anna Schedl, Thomas Penz, Michael Schuster, Jan Laine, Martin Senekowitsch, Christoph Bock, Andreas Bergthaler |
| EPI_ISL_854373, EPI_ISL_854427, EPI_ISL_854428, EPI_ISL_854429, EPI_ISL_854430, EPI_ISL_854431, EPI_ISL_854432, EPI_ISL_854433, EPI_ISL_854434, EPI_ISL_854435 | MONTEFIORE MEDICAL CENTER LABORATORIES | Wadsworth Center, New York State Department of Health | Kirsten St. George, Daryl M. Lamson, Alexis Russel, Matthew Shudt, Melissa A Leisner, Jonathan Plitnick, Navjot Singh, John Kelly, Erasmus Schneider, Erica Lasek-Nesselquist |
| EPI_ISL_855382 | Hospital | National Reference Center for Viruses of Respiratory Infections, Institut Pasteur, Paris | Marion Barbet, Sylvie Behillil, Méline Bizard, Angela Brisebarre, Camille Capel, Etienne Simon-Lorière, Vincent Enouf, Maud Vanpeene, Sylvie van der Werf |
| EPI_ISL_855576, EPI_ISL_855580, EPI_ISL_855583, EPI_ISL_855586, EPI_ISL_855587 | Respiratory Virus Unit, National Infection Service, Public Health England | COVID-19 Genomics UK (COG-UK) Consortium | PHE Covid Sequencing Team |
| EPI_ISL_855921 | Lab voor klinische biologie | Onderzoeksgroep Virologie | Laurens Lambrechts, Nick Vereecke, Marthe Pauwels, Bruno Verhasselt, Linos Vandekerckhove, Hans Nauwynck, Sebastiaan Theuns |
| EPI_ISL_856681 | Charité Universitätsmedizin Berlin, Institute of Virology, Charitéplatz 1, 10117 Berlin, Germany | Charité Universitätsmedizin Berlin, Institute of Virology, Charitéplatz 1, 10117 Berlin, Germany | Victor M Corman, Julia Schneider, Jörn Beheim-Schwarzbach, Tobias Bleicker, Julia Tesch, Barbara Mühlemann, Talitha Veith, Terry Jones, Christian Drosten |
| EPI_ISL_856757, EPI_ISL_856758, EPI_ISL_856759 | Servicio Virosis Respiratorias-Departamento Virología-INEI | Instituto Nacional Enfermedades Infecciosas C.G.Malbran | Baumeister E., Avaro M., Benedetti E., Russo M., Dattero ME, Pontoriero A., Cisterna D., Molina V., Perandones C., Tuduri E., Lorenzo F., Poklepovich T., Campos J. |
| EPI_ISL_857186 | OCME Office Of Chief Medical Examiner | New York City Public Health Laboratory | Jade Wang, et al. |
| EPI_ISL_857188 | Department of Homeless Services | New York City Public Health Laboratory | Jade Wang, et al. |
| EPI_ISL_857193, EPI_ISL_857199, EPI_ISL_857291, EPI_ISL_857292 | OCME Office Of Chief Medical Examiner | New York City Public Health Laboratory | Jade Wang, et al. |
| EPI_ISL_857293 | Department of Homeless Services | New York City Public Health Laboratory | Jade Wang, et al. |
| EPI_ISL_857294, EPI_ISL_857295, EPI_ISL_857296, EPI_ISL_857297 | OCME Office Of Chief Medical Examiner | New York City Public Health Laboratory | Jade Wang, et al. |

| | | | |
|---|---|---|---|
| EPI_ISL_857298 | Department of Homeless Services | New York City Public Health Laboratory | Jade Wang, et al. |
| EPI_ISL_857299, EPI_ISL_857300 | OCME Office Of Chief Medical Examiner | New York City Public Health Laboratory | Jade Wang, et al. |
| EPI_ISL_857301 | Department of Homeless Services | New York City Public Health Laboratory | Jade Wang, et al. |
| EPI_ISL_857302, EPI_ISL_857303, EPI_ISL_857304, EPI_ISL_857305 | OCME Office Of Chief Medical Examiner | New York City Public Health Laboratory | Jade Wang, et al. |
| EPI_ISL_857306 | Department of Homeless Services | New York City Public Health Laboratory | Jade Wang, et al. |
| EPI_ISL_857315, EPI_ISL_857316, EPI_ISL_857317, EPI_ISL_857318, EPI_ISL_857319, EPI_ISL_857320, EPI_ISL_857321, EPI_ISL_857322, EPI_ISL_857323, EPI_ISL_857324, EPI_ISL_857325, EPI_ISL_857326, EPI_ISL_857327, EPI_ISL_857328, EPI_ISL_857329, EPI_ISL_857330, EPI_ISL_857331, EPI_ISL_857332, EPI_ISL_857333, EPI_ISL_857334, EPI_ISL_857335, EPI_ISL_857336, EPI_ISL_857337, EPI_ISL_857338, EPI_ISL_857339, EPI_ISL_857340, EPI_ISL_857341, EPI_ISL_857342, EPI_ISL_857343, EPI_ISL_857344, EPI_ISL_857345, EPI_ISL_857346, EPI_ISL_857347 | | | |
| see above | Cancer Biology Department, National Cancer Institute | Cancer Biology Department, National Cancer Institute | Zekri,A.N., Sedawy,M.G., Ahmed,O.S., Hafez,M.M., Soliman,H.K., Bahnassy,A.A., Elhosieny,F.W., Gad,A.E., Hamdy,M.S., Soliman,M.S., Soliman,L., Abouelhoda,M. |
| EPI_ISL_858703, EPI_ISL_858705, EPI_ISL_858710, EPI_ISL_858787 | Lighthouse Lab in Alderley Park | Wellcome Sanger Institute for the COVID-19 Genomics UK (COG-UK) Consortium | Jacquelyn Wynn, Mairead Hyland, The Lighthouse Lab in Alderley Park and Alex Alderton, Roberto Amato, Sonia Goncalves, Ewan Harrison, David K. Jackson, Ian Johnston, Dominic Kwiatkowski, Cordelia Langford, John Sillitoe on behalf of the Wellcome Sanger Institute COVID-19 Surveillance Team |
| EPI_ISL_858792, EPI_ISL_858795, EPI_ISL_858798, EPI_ISL_858799, EPI_ISL_858800, EPI_ISL_858803, EPI_ISL_858804, EPI_ISL_858805, EPI_ISL_858807, EPI_ISL_858808 | Lighthouse Lab in Milton Keynes | Wellcome Sanger Institute for the COVID-19 Genomics UK (COG-UK) Consortium | The Lighthouse Lab in Milton Keynes and Alex Alderton, Roberto Amato, Sonia Goncalves, Ewan Harrison, David K. Jackson, Ian Johnston, Dominic Kwiatkowski, Cordelia Langford, John Sillitoe on behalf of the Wellcome Sanger Institute COVID-19 Surveillance Team |
| EPI_ISL_858812, EPI_ISL_858818, EPI_ISL_858819, EPI_ISL_858820, EPI_ISL_858823, EPI_ISL_858826, EPI_ISL_858829, EPI_ISL_858833, EPI_ISL_858836, EPI_ISL_858837, EPI_ISL_858838, EPI_ISL_858839, EPI_ISL_858840, EPI_ISL_858841, EPI_ISL_858845, EPI_ISL_858846, EPI_ISL_858848, EPI_ISL_858849, EPI_ISL_858850, EPI_ISL_858852, EPI_ISL_858853, EPI_ISL_858855, EPI_ISL_858857, EPI_ISL_858858, EPI_ISL_858861, EPI_ISL_858862, EPI_ISL_858864, EPI_ISL_858866, EPI_ISL_858868, EPI_ISL_858869, EPI_ISL_858870, EPI_ISL_858871, EPI_ISL_858872, EPI_ISL_858875, EPI_ISL_858876, EPI_ISL_858877, EPI_ISL_858878, EPI_ISL_858880, EPI_ISL_858881, EPI_ISL_858882, EPI_ISL_858883, EPI_ISL_858884, EPI_ISL_858885, EPI_ISL_858886, EPI_ISL_858888, EPI_ISL_858889, EPI_ISL_858890, EPI_ISL_858892, EPI_ISL_858893, EPI_ISL_858894, EPI_ISL_858896, EPI_ISL_858897, EPI_ISL_858898 | | | |
| see above | Lighthouse Lab in Alderley Park | Wellcome Sanger Institute for the COVID-19 Genomics UK (COG-UK) Consortium | Jacquelyn Wynn, Mairead Hyland, The Lighthouse Lab in Alderley Park and Alex Alderton, Roberto Amato, Sonia Goncalves, Ewan Harrison, David K. Jackson, Ian Johnston, Dominic Kwiatkowski, Cordelia Langford, John Sillitoe on behalf of the Wellcome Sanger Institute COVID-19 Surveillance Team |
| EPI_ISL_858900, EPI_ISL_858901, EPI_ISL_858903, EPI_ISL_858904, EPI_ISL_858906, EPI_ISL_858908, EPI_ISL_858910, EPI_ISL_858912, EPI_ISL_858916, EPI_ISL_858919, EPI_ISL_858921, EPI_ISL_858922, EPI_ISL_858927, EPI_ISL_858928, EPI_ISL_858930, EPI_ISL_858935, EPI_ISL_858936, EPI_ISL_858937, EPI_ISL_858943, EPI_ISL_858945, EPI_ISL_858953, EPI_ISL_858958, EPI_ISL_858969, EPI_ISL_858970, EPI_ISL_858974, EPI_ISL_858975, EPI_ISL_858990, EPI_ISL_859149, EPI_ISL_859150, EPI_ISL_859151, EPI_ISL_859153, EPI_ISL_859155, EPI_ISL_859156, EPI_ISL_859157, EPI_ISL_859158, EPI_ISL_859159, EPI_ISL_859160, EPI_ISL_859162, EPI_ISL_859163, EPI_ISL_859166, EPI_ISL_859167, EPI_ISL_859168, EPI_ISL_859169, EPI_ISL_859170, EPI_ISL_859172, EPI_ISL_859174, EPI_ISL_859175, EPI_ISL_859176, EPI_ISL_859177, EPI_ISL_859178, EPI_ISL_859180, EPI_ISL_859182, EPI_ISL_859184, EPI_ISL_859185, EPI_ISL_859191, EPI_ISL_859192, EPI_ISL_859195, EPI_ISL_859196, EPI_ISL_859197, EPI_ISL_859200, EPI_ISL_859294, EPI_ISL_859309, EPI_ISL_859312, EPI_ISL_859316, EPI_ISL_859317, EPI_ISL_859319 | | | |
| see above | Lighthouse Lab in Glasgow | Wellcome Sanger Institute for the COVID-19 Genomics UK (COG-UK) Consortium | Harper VanSteenhouse, Yumi Kasai, David Gray, Carol Clugston, Anna Dominiczak and Alex Alderton, Roberto Amato, Sonia Goncalves, Ewan Harrison, David K. Jackson, Ian Johnston, Dominic Kwiatkowski, Cordelia Langford, John Sillitoe on behalf of the Wellcome Sanger Institute COVID-19 Surveillance Team |
| EPI_ISL_860050, EPI_ISL_860058, EPI_ISL_860059, EPI_ISL_860060, EPI_ISL_860061, EPI_ISL_860062, EPI_ISL_860065, EPI_ISL_860077, EPI_ISL_860081, EPI_ISL_860082, EPI_ISL_860083, EPI_ISL_860084, EPI_ISL_860085, EPI_ISL_860086, EPI_ISL_860087, EPI_ISL_860088 | | | |
| see above | BTC, Khalifa University | BTC, Khalifa University | Al Safar et al |
| EPI_ISL_860214 | Hospital of Southern Norway - Kristiansand, Department of Medical Microbiology | Norwegian Institute of Public Health, Department of Virology | Kathrine Stene-Johansen, Kamilla Heddeland Instefjord, Hilde Elshaug, Atiya R Ali,Marie Paulsen Madsen, Rasmus Riis Kopperud, Hilde Vollan, Karoline Bragstad, Olav Hungnes |
| EPI_ISL_860220, EPI_ISL_860221 | Oslo University Hospital, Department of Medical Microbiology | Norwegian Institute of Public Health, Department of Virology | Kathrine Stene-Johansen, Kamilla Heddeland Instefjord, Hilde Elshaug, Atiya R Ali,Marie Paulsen Madsen, Rasmus Riis Kopperud, Hilde Vollan, Karoline Bragstad, Olav Hungnes |
| EPI_ISL_860222 | Hospital of Southern Norway - Kristiansand, Department of Medical Microbiology | Norwegian Institute of Public Health, Department of Virology | Kathrine Stene-Johansen, Kamilla Heddeland Instefjord, Hilde Elshaug, Atiya R Ali,Marie Paulsen Madsen, Rasmus Riis Kopperud, Hilde Vollan, Karoline Bragstad, Olav Hungnes |
| EPI_ISL_860223 | Akershus University Hospital, Department for Microbiology and Infectious Disease Control | Norwegian Institute of Public Health, Department of Virology | Kathrine Stene-Johansen, Kamilla Heddeland Instefjord, Hilde Elshaug, Atiya R Ali,Marie Paulsen Madsen, Rasmus Riis Kopperud, Hilde Vollan, Karoline Bragstad, Olav Hungnes |
| EPI_ISL_860224, EPI_ISL_860225 | Medical Microbiology Unit, Department for Laboratory Medicine, Drammen Hospital, Vestre Viken Health Trust, | Norwegian Institute of Public Health, Department of Virology | Kathrine Stene-Johansen, Kamilla Heddeland Instefjord, Hilde Elshaug, Atiya R Ali,Marie Paulsen Madsen, Rasmus Riis Kopperud, Hilde Vollan, Karoline Bragstad, Olav Hungnes |
| EPI_ISL_860233, EPI_ISL_860234 | Norwegian Institute of Public Health, Department of Virology | Norwegian Institute of Public Health, Department of Virology | Kathrine Stene-Johansen, Kamilla Heddeland Instefjord, Hilde Elshaug, Atiya R Ali,Marie Paulsen Madsen, Rasmus Riis Kopperud, Hilde Vollan, Karoline Bragstad, Olav Hungnes |
| EPI_ISL_860235 | Oslo University Hospital, Department of Medical Microbiology | Norwegian Institute of Public Health, Department of Virology | Kathrine Stene-Johansen, Kamilla Heddeland Instefjord, Hilde Elshaug, Atiya R Ali,Marie Paulsen Madsen, Rasmus Riis Kopperud, Hilde Vollan, Karoline Bragstad, Olav Hungnes |
| EPI_ISL_860293 | Ostfold Hospital Trust - Kalnes, Centre for Laboratory Medicine, Section for gene technology and infection serology | Norwegian Institute of Public Health, Department of Virology | Kathrine Stene-Johansen, Kamilla Heddeland Instefjord, Hilde Elshaug, Atiya R Ali,Marie Paulsen Madsen, Rasmus Riis Kopperud, Hilde Vollan, Karoline Bragstad, Olav Hungnes |
| EPI_ISL_860300 | Unit 17: Influenza & Other Respiratory Viruses, German National Influenza Center | Project group Epidemiology of Highly Pathogenic Microorganisms, Robert Koch-Institute | Andreas Sachse, Grit Schubert, Essia Belarbi, Sébastien Calvignac-Spencer, Thorsten Wolff, Ralf Dürrwald, Djin-Ye Oh, Marianne Wedde |
| EPI_ISL_860566, EPI_ISL_860603, EPI_ISL_860605, EPI_ISL_860606, EPI_ISL_860607, EPI_ISL_860608, EPI_ISL_860609, EPI_ISL_860610, EPI_ISL_860613, EPI_ISL_860614, EPI_ISL_860615, EPI_ISL_860616, EPI_ISL_860617, EPI_ISL_860618, EPI_ISL_860620, EPI_ISL_860621, EPI_ISL_860622, EPI_ISL_860623, EPI_ISL_860624, EPI_ISL_860625 | | | |
| see above | NHLS-IALCH | KRISP, KZn Research Innovation and Sequencing Platform | Giandhari J, Pillay S, Lessells R, Mdlalose K, York D, Khan S, Tegally H, Wilkinson E, de Oliveira T |
| EPI_ISL_860635, EPI_ISL_860636, EPI_ISL_860637, EPI_ISL_860638, EPI_ISL_860639, EPI_ISL_860640 | Respiratory Virus Unit, National Infection Service, Public Health England | COVID-19 Genomics UK (COG-UK) Consortium | PHE Covid Sequencing Team |
| EPI_ISL_860939, EPI_ISL_860941, EPI_ISL_860944, EPI_ISL_860958, EPI_ISL_861026 | Johns Hopkins Hospital Department of Pathology | Johns Hopkins Hospital Department of Pathology | C. Paul Morris, Chun Huai Luo, Adannaya Amadi, Nicholas Gallagher, Heba H. Mostafa |
| EPI_ISL_861106, EPI_ISL_861107, EPI_ISL_861108, EPI_ISL_861134, EPI_ISL_861144, EPI_ISL_861145, EPI_ISL_861147, EPI_ISL_861148, EPI_ISL_861149, EPI_ISL_861150, EPI_ISL_861151, EPI_ISL_861152, EPI_ISL_861153, EPI_ISL_861154, EPI_ISL_861155, EPI_ISL_861156, EPI_ISL_861157, EPI_ISL_861158, EPI_ISL_861159, EPI_ISL_861160, EPI_ISL_861161, EPI_ISL_861162, EPI_ISL_861163, EPI_ISL_861164, EPI_ISL_861165, EPI_ISL_861166, EPI_ISL_861167, EPI_ISL_861168, EPI_ISL_861169, EPI_ISL_861170, EPI_ISL_861171, EPI_ISL_861172, EPI_ISL_861173, EPI_ISL_861174 | | | |
| see above | KALEIDA CENTER FOR LABORATORY MEDICINE | Wadsworth Center, New York State Department of Health | Kirsten St. George, Daryl M. Lamson, Alexis Russel, Matthew Shudt, Melissa A Leisner, Jonathan Plitnick, Navjot Singh, John Kelly, Erasmus Schneider, Erica Lasek-Nesselquist |
| EPI_ISL_861233, EPI_ISL_861240, EPI_ISL_861241 | Ohio Department of Health Laboratory | Ohio Department of Health Laboratory | Holmes, Jennifer; Eric Brandt, Keoni Omura, Glen McGillivary, Caitlin McDonnell, Kirtana Ramadugu, Erica Leasure, Kelsey Florek, Heather Blankenship, Quanta Brown, and Tammy Bannerman |
| EPI_ISL_861244, EPI_ISL_861248, EPI_ISL_861250, EPI_ISL_861251, EPI_ISL_861252, EPI_ISL_861254, EPI_ISL_861255, EPI_ISL_861257, EPI_ISL_861258, EPI_ISL_861259, EPI_ISL_861260, EPI_ISL_861262, EPI_ISL_861263, EPI_ISL_861264, EPI_ISL_861265, EPI_ISL_861266, EPI_ISL_861268, EPI_ISL_861269, EPI_ISL_861270, EPI_ISL_861271, EPI_ISL_861272, EPI_ISL_861273, EPI_ISL_861274, EPI_ISL_861275, EPI_ISL_861276, EPI_ISL_861277, EPI_ISL_861278, EPI_ISL_861279, EPI_ISL_861280, EPI_ISL_861281, EPI_ISL_861282, EPI_ISL_861283, EPI_ISL_861284, EPI_ISL_861285, EPI_ISL_861286, EPI_ISL_861287, EPI_ISL_861288, EPI_ISL_861303, EPI_ISL_861304, EPI_ISL_861307, EPI_ISL_861316, EPI_ISL_861322, EPI_ISL_861328, EPI_ISL_861330, EPI_ISL_861331 | | | |
| see above | MONTEFIORE MEDICAL CENTER LABORATORIES | Wadsworth Center, New York State Department of Health | Kirsten St. George, Daryl M. Lamson, Alexis Russel, Matthew Shudt, Melissa A Leisner, Jonathan Plitnick, Navjot Singh, John Kelly, Erasmus Schneider, Erica Lasek-Nesselquist |
| EPI_ISL_861466 | UHAS COVID-19 Lab | UHAS COVID-19 Lab | Kwabena O. Duedu, Jones Gyamfi, Reuben Ayivor-Djanie, John O. Gyapong and the UHAS COVID-19 Lab Team |
| EPI_ISL_861528 | ZOTZ KLIMAS MVZ Düsseldorf-Centrum GbR ÜBAG für | Center of Medical Microbiology, Virology, and Hospital | Maximilian Damagnez, Alexander Dilthey, Ashley-Jane Duplessis, Patrick Finzer, Katrin Hoffmann, Torsten Houwaart, Lisanna Hülse, Malte Kohns |

| | | | |
|---|---|---|---|
| | Labormedizin, Genetik, Zytologie, Pathologie | Hygiene, University of Duesseldorf | Vasconcelos, Marek Korencak, Nadine Lübke, Jessica Nicolai, Klaus Pfeffer, Daniel Strelow, Jörg Timm, Andreas Walker, Tobias Wienemann, Rainer Zotz |
| EPI_ISL_861556, EPI_ISL_861567, EPI_ISL_861575, EPI_ISL_861587 | Instituto Nacional de Saude (INSA) | Instituto Nacional de Saude (INSA) | Borges et al |
| EPI_ISL_861676 | UPA Vila Santa Catarina | Instituto Adolfo Lutz, Interdiciplinary Procedures Center, Strategic Laboratory | Claudio Tavares Sacchi, Claudia Regina Gonçalves, Erica Valessa Ramos Gomes, Karoline Rodrigues Campos |
| EPI_ISL_861677 | Instituto Adolfo Lutz - Central | Instituto Adolfo Lutz, Interdiciplinary Procedures Center, Strategic Laboratory | Claudio Tavares Sacchi, Claudia Regina Gonçalves, Erica Valessa Ramos Gomes, Karoline Rodrigues Campos |
| EPI_ISL_861678 | Laboratorio Municipal de Guarulhos | Instituto Adolfo Lutz, Interdiciplinary Procedures Center, Strategic Laboratory | Claudio Tavares Sacchi, Claudia Regina Gonçalves, Erica Valessa Ramos Gomes, Karoline Rodrigues Campos |
| EPI_ISL_861680 | Hospital e Pronto Socorro Portinari | Instituto Adolfo Lutz, Interdiciplinary Procedures Center, Strategic Laboratory | Claudio Tavares Sacchi, Claudia Regina Gonçalves, Erica Valessa Ramos Gomes, Karoline Rodrigues Campos |
| EPI_ISL_861698 | Los Angeles County PHL | Los Angeles County PHL | P. Hemarajata et al. |
| EPI_ISL_861859, EPI_ISL_861860, EPI_ISL_861861, EPI_ISL_861863, EPI_ISL_861864 | Labormedizinisches Zentrum Dr Risch | University Hospital Basel, Clinical Bacteriology | Tim Roloff, Madlen Stange, Helena MB Seth-Smith, Alfredo Mari, Karoline Leuzinger, Julia Bielicki, Nadia Wohlwend,Martin Risch, Lorenz Risch, Manuel Battegay, Hans Hirsch, Adrian Egli |
| EPI_ISL_861973, EPI_ISL_861974, EPI_ISL_861975, EPI_ISL_861976, EPI_ISL_861977, EPI_ISL_861978, EPI_ISL_861979, EPI_ISL_861980, EPI_ISL_861981, EPI_ISL_861982, EPI_ISL_861983, EPI_ISL_861984, EPI_ISL_861985, EPI_ISL_861986, EPI_ISL_861987, EPI_ISL_861988, EPI_ISL_861989, EPI_ISL_861990, EPI_ISL_861991, EPI_ISL_861992, EPI_ISL_861993, EPI_ISL_861994, EPI_ISL_861995, EPI_ISL_861996 | | | |
| see above | OHSU Lab Services Molecular Microbiology Lab | Oregon SARS-CoV-2 Genome Sequencing Center | Brendan L. O'Connell, Sally Grindstaff, Kayla Carter, Ruth V. Nichols, Alec J. Hirsch, Donna Hansel, Guang Fan, Xuan Qin, Daniel N. Streblow, William B. Messer, Andrew C. Adey, Benjamin N. Bimber, Brian J. O'Roak |
| EPI_ISL_862138, EPI_ISL_862145, EPI_ISL_862159 | Charité Universitätsmedizin Berlin, Institut fur Virologie/Labor Berlin | Charité Universitätsmedizin Berlin, Institut für Virologie | Victor M Corman, Barbara Mühlemann, Jörn Beheim-Schwarzbach, Tobias Bleicker, Julia Tesch, Talitha Veith, Julia Schneider, Terry Jones, Christian Drosten |
| EPI_ISL_862544 | Compejo Hospitalario Xeral-Calde | Instituto de Salud Carlos III | Iglesias-Caballero, M.Camarero, S. Molinero Calamita, M. González-Esguevillas, M. Pozo, F. Casas, I. Jiménez, P. Jiménez, M. Zaballos, A. Monzón, S. Varona, S. Juliá, M. Cuesta, I. Alonso, P. |
| EPI_ISL_862545 | Hospital Universitario Virgen de la Arrixaca | Instituto de Salud Carlos III | Iglesias-Caballero, M. Camarero, S. Molinero Calamita, M. González-Esguevillas, M. Pozo, F. Casas, I. Jiménez, P. Jiménez, M. Zaballos, A. Monzón, S. Varona, S. Juliá, M. Cuesta, I, Moreno, A. |
| EPI_ISL_862547 | Consejería de Sanidad y Asuntos Sociales | Instituto de Salud Carlos III | Iglesias-Caballero, M. Camarero, S. Molinero Calamita, M. González-Esguevillas, M. Pozo, F. Casas, I. Jiménez, P. Jiménez, M. Zaballos, A. Monzón, S. Varona, S. Juliá, M. Cuesta, I. Gutiérrez, G. |
| EPI_ISL_862549 | Hospital Universitario Virgen de la Arrixaca | Instituto de Salud Carlos III | Iglesias-Caballero, M. Camarero, S. Molinero Calamita, M. González-Esguevillas, M. Pozo, F. Casas, I. Jiménez, P. Jiménez, M. Zaballos, A. Monzón, S. Varona, S. Juliá, M. Cuesta, I, Moreno, A. |
| EPI_ISL_862550 | Complejo Hospitalario de Navarra | Instituto de Salud Carlos III | Iglesias-Caballero, M.Camarero, S. Molinero Calamita, M. González-Esguevillas, M. Pozo, F. Casas, I. Jiménez, P. Jiménez, M. Zaballos, A. Monzón, S. Varona, S. Juliá, M. Cuesta, I. Ezpeleta, C. |
| EPI_ISL_862553, EPI_ISL_862554, EPI_ISL_862555 | Compejo Hospitalario Xeral-Calde | Instituto de Salud Carlos III | Iglesias-Caballero, M.Camarero, S. Molinero Calamita, M. González-Esguevillas, M. Pozo, F. Casas, I. Jiménez, P. Jiménez, M. Zaballos, A. Monzón, S. Varona, S. Juliá, M. Cuesta, I. Alonso, P. |
| EPI_ISL_862558 | Consejería de Sanidad y Asuntos Sociales | Instituto de Salud Carlos III | Iglesias-Caballero, M. Camarero, S. Molinero Calamita, M. González-Esguevillas, M. Pozo, F. Casas, I. Jiménez, P. Jiménez, M. Zaballos, A. Monzón, S. Varona, S. Juliá, M. Cuesta, I. Gutiérrez, G. |
| EPI_ISL_862564, EPI_ISL_862565 | Gerencia de Asistencia Sanitaria de Soria | Instituto de Salud Carlos III | Iglesias-Caballero, M.Camarero, S. Molinero Calamita, M. González-Esguevillas, M. Pozo, F. Casas, I. Jiménez, P. Jiménez, M. Zaballos, A. Monzón, S. Varona, S. Juliá, M. Cuesta, I. Aldea, C. |
| EPI_ISL_862568 | Hospital Universitario Virgen de la Arrixaca | Instituto de Salud Carlos III | Iglesias-Caballero, M. Camarero, S. Molinero Calamita, M. González-Esguevillas, M. Pozo, F. Casas, I. Jiménez, P. Jiménez, M. Zaballos, A. Monzón, S. Varona, S. Juliá, M. Cuesta, I, Moreno, A. |
| EPI_ISL_862570, EPI_ISL_862571, EPI_ISL_862572 | Complejo Hospitalario de Navarra | Instituto de Salud Carlos III | Iglesias-Caballero, M.Camarero, S. Molinero Calamita, M. González-Esguevillas, M. Pozo, F. Casas, I. Jiménez, P. Jiménez, M. Zaballos, A. Monzón, S. Varona, S. Juliá, M. Cuesta, I. Ezpeleta, C. |
| EPI_ISL_862586 | Hospital Clínic | Instituto de Salud Carlos III | Iglesias-Caballero, M.Camarero, S. Molinero Calamita, M. González-Esguevillas, M. Pozo, F. Casas, I. Jiménez, P. Jiménez, M. Zaballos, A. Monzón, S. Varona, S. Juliá, M. Cuesta, I. Marcos, M.A. |
| EPI_ISL_862736, EPI_ISL_862757 | Utah Public Health Laboratory, Utah Public Health Laboratory Infectious Disease submission group | Utah Public Health Laboratory, Utah Public Health Laboratory Infectious Disease submission group | Young,E.L., Oakeson,K.F., Gallagher,T. |
| EPI_ISL_862783, EPI_ISL_862784, EPI_ISL_862785 | Cancer Biology Department, National Cancer Institute | Cancer Biology Department, National Cancer Institute | Zekri,A.N., Sedawy,M.G., Ahmed,O.S., Hafez,M.M., Soliman,H.K., Bahnassy,A.A., Elhosieny,F.W., Gad,A.E., Hamdy,M.S., Soliman,M.S., Soliman,L., Abouelhoda,M. |
| EPI_ISL_865090, EPI_ISL_865091, EPI_ISL_865146, EPI_ISL_865147, EPI_ISL_865148 | Virology Department, Royal Infirmary of Edinburgh, NHS Lothian / School of Biological Sciences, University of Edinburgh / Institute of Genetics and Molecular Medicine, University of Edinburgh | COVID-19 Genomics UK (COG-UK) Consortium | McHugh M, Dewar R, Rooke S, Gallagher M, Balcaza C, O'Toole Á, Scher E, Hill V, McCrone JT, Colquhoun R, Yu X, Jackson B, Rambaut A, Williams TC, Templeton K |
| EPI_ISL_865217, EPI_ISL_865218, EPI_ISL_865219, EPI_ISL_865220, EPI_ISL_865221, EPI_ISL_865222, EPI_ISL_865223, EPI_ISL_865224, EPI_ISL_865225, EPI_ISL_865226, EPI_ISL_865227, EPI_ISL_865228, EPI_ISL_865229, EPI_ISL_865230, EPI_ISL_865231, EPI_ISL_865232, EPI_ISL_865233, EPI_ISL_865234, EPI_ISL_865235, EPI_ISL_865236, EPI_ISL_865237, EPI_ISL_865391, EPI_ISL_865392, EPI_ISL_865393, EPI_ISL_865394, EPI_ISL_865395, EPI_ISL_865396, EPI_ISL_865397, EPI_ISL_865399, EPI_ISL_865401, EPI_ISL_865402, EPI_ISL_865403, EPI_ISL_865404, EPI_ISL_865406, EPI_ISL_865407, EPI_ISL_865408, EPI_ISL_865409, EPI_ISL_865410, EPI_ISL_865423, EPI_ISL_865424, EPI_ISL_865425, EPI_ISL_865426, EPI_ISL_865429, EPI_ISL_865431, EPI_ISL_865434, EPI_ISL_865438, EPI_ISL_865439, EPI_ISL_865440, EPI_ISL_865441, EPI_ISL_865443, EPI_ISL_865444, EPI_ISL_865445, EPI_ISL_865446, EPI_ISL_865447, EPI_ISL_865451, EPI_ISL_865452, EPI_ISL_865453, EPI_ISL_865454, EPI_ISL_865455, EPI_ISL_865457, EPI_ISL_865458 | | | |
| see above | Liverpool Clinical Laboratories | COVID-19 Genomics UK (COG-UK) Consortium | Sam Haldenby, Anita Lucaci, Steve Paterson, Julian Hiscox, Alistair Darby, M Almsaud, A Alrezaihi, Muhannad Alruwaili, Stuart D Armstrong, Jones Benjamin, Eleanor G Bentley, Anu Chawla, Jordan J Clark, Angela Cowell, Richard Eccles, Isabel García-Dorival, Matthew Gemmell, Alessandro Gerada, PKF Gilmore, Richard Gregory, Ximeng Han, Catherine Hartley, Margaret Hughes, Miren Iturriza-Gomara, James Johnson, L Luu, Jenifer Manson, Charlotte Nelson, Elaine O'Toole, Cassie Olateju, Rebekah Penrice-Randal , Lucille Rainbow, N.P Randle, Trevor Ian Robinson, Parul Sharma, Ghada T Shawli, James P Stewart, Neil Swainston, Ecaterina Vamos, Joanne Watts, Mark Whitehead |
| EPI_ISL_865490, EPI_ISL_865491, EPI_ISL_865493, EPI_ISL_865503, EPI_ISL_865504, EPI_ISL_865505, EPI_ISL_865506, EPI_ISL_865507, EPI_ISL_865508, EPI_ISL_865509, EPI_ISL_865510, EPI_ISL_865511, EPI_ISL_865512, EPI_ISL_865513, EPI_ISL_865543, EPI_ISL_865544, EPI_ISL_865545, EPI_ISL_865559, EPI_ISL_865560 | | | |
| see above | Barts Health NHS Trust | COVID-19 Genomics UK (COG-UK) Consortium | CUTINO-MOGUEL, Maria-Teresa; HARRINGTON, David; OWOYEMI, Dola; KULASEGARAN-SHYLINI, Raghavendran; BROAD, Claire; KELE, Beatrix |
| EPI_ISL_865754, EPI_ISL_865758, EPI_ISL_865759, EPI_ISL_865760, EPI_ISL_865762, EPI_ISL_865763, EPI_ISL_865764, EPI_ISL_865765, EPI_ISL_865766, EPI_ISL_865767, EPI_ISL_865768, EPI_ISL_865769, EPI_ISL_865770, EPI_ISL_865771, EPI_ISL_865772, EPI_ISL_865773, EPI_ISL_865775, EPI_ISL_865776, EPI_ISL_865777, EPI_ISL_865778, EPI_ISL_865779, EPI_ISL_865780, EPI_ISL_865781, EPI_ISL_865782, EPI_ISL_865783, EPI_ISL_865784, EPI_ISL_865785, EPI_ISL_865786, EPI_ISL_865787, EPI_ISL_865788, EPI_ISL_865789, EPI_ISL_865790, EPI_ISL_865791, EPI_ISL_865792, EPI_ISL_865793, EPI_ISL_865794, EPI_ISL_865795, EPI_ISL_865796 | | | |
| see above | University College London, Great Ormond Street Hospital for Children NHS Foundation Trust, Imperial College Healthcare NHS Trust | COVID-19 Genomics UK (COG-UK) Consortium | Sergi Castellano, Rachel Williams, Mark Kristiansen, Paola Resende Silva, Sunando Roy, Tony Brooks, Helena Tutill, Paola Niola, Patricia Dyal, Charlotte Williams, Leysa Forrest, Yasmin Panchbhaya, Jacqueline Findlay, Samuel Weeks, Julianne Brown, Kathryn Harris, Paul Randell, James Price, Alison Holmes, Judith Breuer |
| EPI_ISL_866123, EPI_ISL_866128, EPI_ISL_866130, EPI_ISL_866137, EPI_ISL_866139, EPI_ISL_866141, | University College London Hospital | COVID-19 Genomics UK (COG-UK) Consortium | Judith Heaney, Matthew Byott, Catherine Houlihan, Dan Frampton, Stuart Kirk, Moira Spyer and Eleni Nastouli |

| | | | |
|---|---|---|---|
| EPI_ISL_866145 | | | |
| EPI_ISL_866582, EPI_ISL_866824 | Quadram Institute Bioscience | COVID-19 Genomics UK (COG-UK) Consortium | Dave J. Baker, Gemma L. Kay, Alp Aydin, Thanh Le-Viet, Steven Rudder, Ana P. Tedim, Anastasia Kolyva, Maria Diaz, Leonardo de Oliveira Martins, Nabil-Fareed Alikhan, Lizzie Meadows, Rachael Stanley, Ngozi Elumogo, Muhammed Yasir, Nicholas M. Thomson, Alexander J Trotter, Rachel Gilroy, Samuel Bloomfield, Claire Stuart, Andrew Bell, Reenesh Prakash, Samir Dervisevic, Alison E. Mather, John Wain, Mark Webber, Andrew J. Page, Justin O'Grady |
| EPI_ISL_867172, EPI_ISL_867173, EPI_ISL_867174, EPI_ISL_867177, EPI_ISL_867183 | Oxford Viromics, NDM, University of Oxford; Oxford University Hospitals; Basingstoke and North Hampshire Hospital | COVID-19 Genomics UK (COG-UK) Consortium | Tanya Golubchik, David Bonsall, George Macintyre, Amy Trebes, Mariateresa de Cesare, Catrin Moore, Alex Mobbs, Anita Justice, Robert Shaw, Monique Andersson, Timothy Peto, Emma Wise, Nathan Moore, Jessica Lynch, Nick Cortes, Matilde Mori, Stephen Kidd, David Buck, John Todd, Christophe Fraser |
| EPI_ISL_868411, EPI_ISL_868414, EPI_ISL_868419, EPI_ISL_868422, EPI_ISL_868436, EPI_ISL_868452, EPI_ISL_868475, EPI_ISL_868483, EPI_ISL_868512, EPI_ISL_868577, EPI_ISL_868614, EPI_ISL_868628, EPI_ISL_868635, EPI_ISL_868645, EPI_ISL_868649, EPI_ISL_868669, EPI_ISL_868673 | | | |
| see above | Virology Department, Sheffield Teaching Hospitals NHS Foundation Trust/Department of Infection, Immunity and Cardiovascular Disease, The Medical School, University of Sheffield | COVID-19 Genomics UK (COG-UK) Consortium | Thushan de Silva, Matthew Parker, Nikki Smith, Adri Angyal, Rebecca Brown, Luke Green, Rachel Tucker, Paul Parsons, Danielle Groves, Katie Johnson, Laura Carrilero, Alex Keeley, Dave Partridge, Matthew Wyles, Benjamin Lindsey, Mehmet Yavuz, Mohammad Raza, Cariad Evans |
| EPI_ISL_871793, EPI_ISL_871796, EPI_ISL_871798, EPI_ISL_871799, EPI_ISL_871805, EPI_ISL_871809, EPI_ISL_871813, EPI_ISL_871814 | AIID | Irish Coronavirus Sequencing Consortium-Teagasc Grange | Matthew McCabe, Aljandro Abner Garcia Leon, Fiona Crispie, Calum Walsh, Michael Carr, John Kenny, Paul Cotter, Patrick Mallon, Gabriel Gonzalez |
| EPI_ISL_871896 | Hospital Universitario Marqués de Valdecilla - IDIVAL (Santander, Cantabria) | SeqCOVID-SPAIN consortium/IBV(CSIC) | Mónica Gozalo Margüello, María Eliecer Cano García, Jose Manuel Méndez Legaza, Daniel Pablo Marcos, Jesús Rodríguez Rodríguez, María Siller Ruiz and SeqCOVID-SPAIN consortium |
| EPI_ISL_871912, EPI_ISL_871919, EPI_ISL_871920, EPI_ISL_871923, EPI_ISL_871925, EPI_ISL_871926, EPI_ISL_871934 | Servicio de Microbiología, Laboratori Clínic Metropolitana Nord. Hospital Universitari Germans Trias i Pujol. Institut d'Investigació en Ciències de la Salut Germans Trias i Pujol (IGTP) | SeqCOVID-SPAIN consortium/IBV(CSIC) | Elisa Martró, Antoni E. Bordoy, Anna Not, Adrián Antuori, Anabel Fernández, Nona Romaní, Verónica Saludes, Cristina Casañ and SeqCOVID-SPAIN consortium |
| EPI_ISL_871939, EPI_ISL_871940, EPI_ISL_871954, EPI_ISL_871960, EPI_ISL_871962, EPI_ISL_871963, EPI_ISL_871964 | Servicio de Microbiología Clínica (Complejo Hospitalario de Navarra, Pamplona), Instituto de Investigación Sanitaria de Navarra (IdiSNA) | SeqCOVID-SPAIN consortium/IBV(CSIC) | Carmen Ezpeleta Baquedano, Ana Navascués, Ana Miqueleiz and SeqCOVID-SPAIN consortium |
| EPI_ISL_872060 | Department of Clinical Microbiology | GIGA Medical Genomics | Keith Durkin, Maria Artesi, Sébastien Bontems, Raphaël Boreux, Bouchra Boujemla, Cécile Meex, Pierrette Melin, Marie-Pierre Hayette, Vincent Bours |
| EPI_ISL_872100, EPI_ISL_872103 | Chu Tivoli | GIGA Medical Genomics | Keith Durkin, Maria Artesi, Sébastien Bontems, Raphaël Boreux, Bouchra Boujemla, Cécile Meex, Pierrette Melin, Marie-Pierre Hayette, Vincent Bours |
| EPI_ISL_872109 | Department of Clinical Microbiology | GIGA Medical Genomics | Keith Durkin, Maria Artesi, Sébastien Bontems, Raphaël Boreux, Bouchra Boujemla, Cécile Meex, Pierrette Melin, Marie-Pierre Hayette, Vincent Bours |
| EPI_ISL_872193 | Caribbean Public Health Agency | Carrington Lab, Department ofBuilding 36, First Floor Biochemistry Unit, Faculty of Medical Sciences, The University of the West Indies | Nikita S. D. Sahadeo, Arianne Brown-Jordan, Vernie Ramkissoon, Sarah Hill, Naresh Nandram, Avery Hinds, Jerome Foster, Stanley Giddings, Karla Georges, Marsha Ivey, Rahul Naidu, Risha Singh, SueMin Nathaniel, Rajini Haraksingh, Jaya Jayaraman, Chinna Chinnadurai, Adesh Ramsubhag, Nuno Faria, Oliver Pybus, Christopher Oura, Gabriel Escobar, Christine V. F. Carrington |
| EPI_ISL_872194, EPI_ISL_872195 | Caribbean Public Health Agency | Carrington Lab, Department of PreClinical Sciences, Faculty of Medical Sciences, The University of the West Indies | Nikita S. D. Sahadeo, Arianne Brown-Jordan, Vernie Ramkissoon, Sarah Hill, Naresh Nandram, Avery Hinds, Jerome Foster, Stanley Giddings, Karla Georges, Marsha Ivey, Rahul Naidu, Risha Singh, SueMin Nathaniel, Rajini Haraksingh, Jaya Jayaraman, Chinna Chinnadurai, Adesh Ramsubhag, Nuno Faria, Oliver Pybus, Christopher Oura, Gabriel Escobar, Christine V. F. Carrington |
| EPI_ISL_872203 | Labo Analyses Med | National Reference Center for Viruses of Respiratory Infections, Institut Pasteur, Paris | Marion Barbet, Sylvie Behillil, Méline Bizard, Angela Brisebarre, Camille Capel, Etienne Simon-Lorière, Vincent Enouf, Maud Vanpeene, Sylvie van der Werf |
| EPI_ISL_872574 | South Eastern Area Laboratory Services (SEALS) | NSW Health Pathology - Institute of Clinical Pathology and Medical Research; Westmead Hospital; University of Sydney | CIDM-PH et al. |
| EPI_ISL_872620, EPI_ISL_872621, EPI_ISL_872622 | Nigeria Centre for Disease Control (NCDC) | African Centre of Excellence for Genomics of Infectious Diseases (ACEGID), Redeemer's University | Oluniyi P.E. et al |
| EPI_ISL_872985, EPI_ISL_872986, EPI_ISL_872987, EPI_ISL_872988, EPI_ISL_872989, EPI_ISL_872990, EPI_ISL_872991, EPI_ISL_872992, EPI_ISL_872993, EPI_ISL_872994, EPI_ISL_872995, EPI_ISL_872996, EPI_ISL_872997, EPI_ISL_872998, EPI_ISL_872999, EPI_ISL_873000, EPI_ISL_873001, EPI_ISL_873002, EPI_ISL_873003, EPI_ISL_873004, EPI_ISL_873005, EPI_ISL_873006, EPI_ISL_873007, EPI_ISL_873008, EPI_ISL_873009, EPI_ISL_873010, EPI_ISL_873011, EPI_ISL_873012, EPI_ISL_873013, EPI_ISL_873014, EPI_ISL_873015, EPI_ISL_873016, EPI_ISL_873017, EPI_ISL_873018, EPI_ISL_873019, EPI_ISL_873020 | | | |
| see above | HELIX LLC | WHO National Influenza Centre Russian Federation | Andrey Komissarov, Artem Fadeev, Anna Ivanova, Kseniya Komissarova, Dmitry Bazhenov, Mikhail Bakaev, Daria Danilenko, Ksenia Safina, Elena Nabieva, Georgii Bazykin, Dmitry Lioznov |
| EPI_ISL_873222, EPI_ISL_873227, EPI_ISL_873228, EPI_ISL_873229, EPI_ISL_873230, EPI_ISL_873231, EPI_ISL_873232, EPI_ISL_873233 | M Health Fairview | Minnesota Department of Health, Public Health Laboratory | Alexandra Lorentz, Jacob Garfin, Matt Plumb, and Xiong Wang |
| EPI_ISL_875551 | ULSS 1 Dolomiti | Istituto Zooprofilattico Sperimentale delle Venezie | Adelaide Milani, Alessia Schivo, Annalisa Salviato, Erika Giorgia Quaranta, Ambra Pastori, Bianca Zecchin, Alice Fusaro, Isabella Monne, Calogero Terregino, Antonia Ricci |
| EPI_ISL_875959, EPI_ISL_875960, EPI_ISL_875961, EPI_ISL_875962, EPI_ISL_875963, EPI_ISL_875964, EPI_ISL_875965, EPI_ISL_875966, EPI_ISL_875967, EPI_ISL_875968, EPI_ISL_875969, EPI_ISL_875970, EPI_ISL_875971, EPI_ISL_875972, EPI_ISL_875973, EPI_ISL_875974, EPI_ISL_875975, EPI_ISL_875976, EPI_ISL_875977, EPI_ISL_875978, EPI_ISL_875979, EPI_ISL_875980, EPI_ISL_875981, EPI_ISL_875982, EPI_ISL_875983, EPI_ISL_875984, EPI_ISL_875985, EPI_ISL_875986, EPI_ISL_875987, EPI_ISL_875988, EPI_ISL_875989, EPI_ISL_875990, EPI_ISL_875991, EPI_ISL_875992, EPI_ISL_875993, EPI_ISL_875994, EPI_ISL_875995, EPI_ISL_875996, EPI_ISL_875997, EPI_ISL_875998, EPI_ISL_875999, EPI_ISL_876000, EPI_ISL_876001, EPI_ISL_876002, EPI_ISL_876003, EPI_ISL_876004, EPI_ISL_876005, EPI_ISL_876006, EPI_ISL_876007, EPI_ISL_876008, EPI_ISL_876009, EPI_ISL_876010, EPI_ISL_876011, EPI_ISL_876012, EPI_ISL_876013, EPI_ISL_876014, EPI_ISL_876015, EPI_ISL_876016, EPI_ISL_876018, EPI_ISL_876019, EPI_ISL_876020, EPI_ISL_876021, EPI_ISL_876022, EPI_ISL_876023, EPI_ISL_876024, EPI_ISL_876025, EPI_ISL_876026, EPI_ISL_876027, EPI_ISL_876028, EPI_ISL_876029, EPI_ISL_876030, EPI_ISL_876031 | | | |
| see above | Laboratory of Molecular Biology, Diagnostyka sp. z o.o. | genXone SA, Research & Development Laboratory | Maciej Sykulski, Grzegorz Nowicki, Monika Makowska-Woniak, Jakub Grabowski, Natalia Drwska-Matelska, ukasz Krych, Micha Kaszuba |
| EPI_ISL_876044 | M Health Fairview | Minnesota Department of Health, Public Health Laboratory | Alexandra Lorentz, Jacob Garfin, Matt Plumb, and Xiong Wang |
| EPI_ISL_876057, EPI_ISL_876064, EPI_ISL_876082, EPI_ISL_876091, EPI_ISL_876094, EPI_ISL_876103, EPI_ISL_876115, EPI_ISL_876125, EPI_ISL_876140, EPI_ISL_876158, EPI_ISL_876209, EPI_ISL_876210, EPI_ISL_876216, EPI_ISL_876273, EPI_ISL_876274, EPI_ISL_876275, EPI_ISL_876276, EPI_ISL_876277, EPI_ISL_876278, EPI_ISL_876279 | | | |
| see above | Massachusetts State Public Health Laboratory | Massachusetts State Public Health Laboratory | Andrew Lang, Timelia Fink, Glen Gallagher, Sandra Smole |
| EPI_ISL_876748, EPI_ISL_876749, EPI_ISL_876750, EPI_ISL_876751 | Istituto Zooprofilattico Sperimentale della Puglia e della Basilicata | Istituto Zooprofilattico Sperimentale della Puglia e della Basilicata | Parisi A., Bianco A., Capozzi L., Del Sambro L., Manzulli V, Rondinone V., Pace L., Cipolletta D., Galante D. |
| EPI_ISL_877460 | Siti Khodijah Hospital | Institute of Tropical Disease, Universitas Airlangga | Maria I Lusida, Krisnoadi Rahardjo, Aldise M Nastri, Jezzy R Dewantari, Rima R Prasetya, Muhammad Hamdan, Gatot Soegiarto, Laksmi Wulandari, Resti Yudhawati, Soetjipto, Yasuko Mori, Kazufumi Shimizu |
| EPI_ISL_878571, EPI_ISL_878574 | Biolab Diagnostic Laboratories | Andersen lab at Scripps Research | Issa Abu-Dayyeh, Ahmad Tibi, Lama Hussein, Lina Mohammad, Zein Naber, Amid Abdelnour with SEARCH Alliance San Diego |
| EPI_ISL_878700, EPI_ISL_878739, EPI_ISL_880188, EPI_ISL_880193 | Rady's Childrens Hospital | Andersen lab at Scripps Research | SEARCH Alliance San Diego with Nanda Radamchar, David Dimmock, Linda Luo, Christina Clarke, Kathryn Bouic, Teresa Mueller, Denise Malicki |
| EPI_ISL_882590 | Lighthouse Lab in Alderley Park | Wellcome Sanger Institute for the COVID-19 Genomics UK (COG-UK) Consortium | Jacquelyn Wynn, Mairead Hyland, The Lighthouse Lab in Alderley Park and Alex Alderton, Roberto Amato, Sonia Goncalves, Ewan Harrison, David K. Jackson, Ian Johnston, Dominic Kwiatkowski, Cordelia Langford, John Sillitoe on behalf of the Wellcome Sanger Institute COVID-19 Surveillance Team |
| EPI_ISL_882661, EPI_ISL_882662 | Hospital de Santa Barbara de Goias | Instituto Adolfo Lutz, Interdiciplinary Procedures Center, Strategic Laboratory | Claudio Tavares Sacchi, Claudia Regina Gonçalves, Erica Valessa Ramos Gomes, Karoline Rodrigues Campos |

| | | | |
|---|---|---|---|
| EPI_ISL_882770 | Darus Syifa' Islamic Hospital | Institute of Tropical Disease, Universitas Airlangga | Aldise M Nastri, Jezzy R Dewantari, Rima R Prasetya, Krisnoadi Rahardjo, HM Faiz, Gatot Soegiarto, Laksmi Wulandari, Resti Yudhawati, Soetjipto, Yasuko Mori, Kazufumi Shimizu, Maria I Lusida |
| EPI_ISL_883999 | Labo Analyses Med | National Reference Center for Viruses of Respiratory Infections, Institut Pasteur, Paris | Marion Barbet, Sylvie Behillil, Méline Bizard, Angela Brisebarre, Camille Capel, Etienne Simon-Lorière, Vincent Enouf, Maud Vanpeene, Sylvie van der Werf,Rousset Dominique |
| EPI_ISL_884262, EPI_ISL_884263, EPI_ISL_884267, EPI_ISL_884271, EPI_ISL_884283, EPI_ISL_884284, EPI_ISL_884285 | Institute of Medical Microbiology and Hospital Hygiene | Institute of Medical Microbiology and Hospital Hygiene | Prof. Dr. Achim Kaasch, Aljoscha Tersteegen |
| EPI_ISL_884857, EPI_ISL_884858, EPI_ISL_884859, EPI_ISL_884861, EPI_ISL_884862, EPI_ISL_884863 | Bacteriology, Georgia Public Health Laboratory (GPHL) | Bacteriology, Georgia Public Health Laboratory (GPHL) | Reeves,S., Edwards,J., Dixey,C., Parrott,T. |
| EPI_ISL_884956, EPI_ISL_884981, EPI_ISL_884997, EPI_ISL_884998, EPI_ISL_884999, EPI_ISL_885000, EPI_ISL_885001, EPI_ISL_885002, EPI_ISL_885003, EPI_ISL_885004, EPI_ISL_885005, EPI_ISL_885006, EPI_ISL_885007, EPI_ISL_885008, EPI_ISL_885012 | | | |
| see above | Santa Clara County Public Health Laboratory | Chan-Zuckerberg Biohub | CZB Cliahub Consortium |
| EPI_ISL_885068, EPI_ISL_885070, EPI_ISL_885071, EPI_ISL_885072, EPI_ISL_885073, EPI_ISL_885074, EPI_ISL_885077, EPI_ISL_885078, EPI_ISL_885079, EPI_ISL_885080, EPI_ISL_885081, EPI_ISL_885082, EPI_ISL_885083, EPI_ISL_885084, EPI_ISL_885085, EPI_ISL_885086, EPI_ISL_885087, EPI_ISL_885090, EPI_ISL_885091, EPI_ISL_885092, EPI_ISL_885093, EPI_ISL_885094, EPI_ISL_885095, EPI_ISL_885096, EPI_ISL_885097, EPI_ISL_885098, EPI_ISL_885099, EPI_ISL_885100, EPI_ISL_885101, EPI_ISL_885102, EPI_ISL_885103, EPI_ISL_885104, EPI_ISL_885105, EPI_ISL_885106, EPI_ISL_885107 | | | |
| see above | Orange County Public Health Lab | Chan-Zuckerberg Biohub | CZB Cliahub Consortium |
| EPI_ISL_885140 | PathWest Laboratory Medicine WA | PathWest Laboratory Medicine WA Microbial Surveillance Unit | PathWest Laboratory Medicine WA Microbial Surveillance Unit |
| EPI_ISL_887424, EPI_ISL_887428, EPI_ISL_887439, EPI_ISL_887441, EPI_ISL_887442, EPI_ISL_887443, EPI_ISL_887446, EPI_ISL_887447, EPI_ISL_887482, EPI_ISL_887483, EPI_ISL_887484, EPI_ISL_887485, EPI_ISL_887494, EPI_ISL_887495, EPI_ISL_887496, EPI_ISL_887497, EPI_ISL_887498, EPI_ISL_887499 | | | |
| see above | Instituto Nacional de Saude (INS), Mozambique | KRISP, KZN Research Innovation and Sequencing Platform | Nalia Ismael, Nadia Sitoe, Paulo Arnaldo, Nedio Mabunda, Giandhari J, Pillay S, Tegally H, Wilkinson E, de Oliveira T |
| EPI_ISL_887592, EPI_ISL_887598, EPI_ISL_887599, EPI_ISL_887600, EPI_ISL_887604, EPI_ISL_887610, EPI_ISL_887611, EPI_ISL_887612, EPI_ISL_887614, EPI_ISL_887616, EPI_ISL_887621, EPI_ISL_887623, EPI_ISL_887624, EPI_ISL_887625, EPI_ISL_887626, EPI_ISL_887629, EPI_ISL_887632, EPI_ISL_887633, EPI_ISL_887636, EPI_ISL_887637, EPI_ISL_887638, EPI_ISL_887639, EPI_ISL_887642, EPI_ISL_887643, EPI_ISL_887644, EPI_ISL_887645, EPI_ISL_887650, EPI_ISL_887653, EPI_ISL_887654, EPI_ISL_887655, EPI_ISL_887656, EPI_ISL_887657, EPI_ISL_887658, EPI_ISL_887659, EPI_ISL_887662, EPI_ISL_887663, EPI_ISL_887665, EPI_ISL_887668, EPI_ISL_887669, EPI_ISL_887670, EPI_ISL_887673, EPI_ISL_887678, EPI_ISL_887680, EPI_ISL_887681, EPI_ISL_887683, EPI_ISL_887684, EPI_ISL_887686, EPI_ISL_887687, EPI_ISL_887692, EPI_ISL_887694, EPI_ISL_887696, EPI_ISL_887697, EPI_ISL_887698, EPI_ISL_887699, EPI_ISL_887700, EPI_ISL_887701, EPI_ISL_887703, EPI_ISL_887706, EPI_ISL_887708, EPI_ISL_887709, EPI_ISL_887711, EPI_ISL_887715, EPI_ISL_887721, EPI_ISL_887726, EPI_ISL_887728, EPI_ISL_887731, EPI_ISL_887732, EPI_ISL_887738, EPI_ISL_887740, EPI_ISL_887742, EPI_ISL_887743, EPI_ISL_887744, EPI_ISL_887745, EPI_ISL_887747, EPI_ISL_887751, EPI_ISL_887752, EPI_ISL_887755, EPI_ISL_887759, EPI_ISL_887762, EPI_ISL_887766, EPI_ISL_887767, EPI_ISL_887768, EPI_ISL_887770, EPI_ISL_887771, EPI_ISL_887776, EPI_ISL_887777, EPI_ISL_887778, EPI_ISL_887779, EPI_ISL_887780, EPI_ISL_887782, EPI_ISL_887785, EPI_ISL_887786, EPI_ISL_887787, EPI_ISL_887788, EPI_ISL_887789, EPI_ISL_887791, EPI_ISL_887793, EPI_ISL_887797, EPI_ISL_887808, EPI_ISL_887811, EPI_ISL_887812, EPI_ISL_887813, EPI_ISL_887816, EPI_ISL_887821, EPI_ISL_887822, EPI_ISL_887823, EPI_ISL_887826, EPI_ISL_887828, EPI_ISL_887830, EPI_ISL_887831, EPI_ISL_887832, EPI_ISL_887834, EPI_ISL_887837, EPI_ISL_887838, EPI_ISL_887849, EPI_ISL_887851, EPI_ISL_887856, EPI_ISL_887859, EPI_ISL_887861, EPI_ISL_887862, EPI_ISL_887864, EPI_ISL_887867, EPI_ISL_887868, EPI_ISL_887869, EPI_ISL_887870, EPI_ISL_887871, EPI_ISL_887872, EPI_ISL_887873, EPI_ISL_887874, EPI_ISL_887877, EPI_ISL_887880, EPI_ISL_887885, EPI_ISL_887886, EPI_ISL_887887, EPI_ISL_887890, EPI_ISL_887891, EPI_ISL_887892, EPI_ISL_887893, EPI_ISL_887898, EPI_ISL_887901, EPI_ISL_887902, EPI_ISL_887903, EPI_ISL_887905, EPI_ISL_887908, EPI_ISL_887915, EPI_ISL_887916, EPI_ISL_887918, EPI_ISL_887920, EPI_ISL_887924, EPI_ISL_887927, EPI_ISL_887930, EPI_ISL_887931, EPI_ISL_887932, EPI_ISL_887935, EPI_ISL_887938, EPI_ISL_887940, EPI_ISL_887944, EPI_ISL_887946, EPI_ISL_887951, EPI_ISL_887953, EPI_ISL_887957, EPI_ISL_887959, EPI_ISL_887960, EPI_ISL_887961, EPI_ISL_887964, EPI_ISL_887968, EPI_ISL_887971, EPI_ISL_887972, EPI_ISL_887973, EPI_ISL_887976, EPI_ISL_887979, EPI_ISL_887986, EPI_ISL_887987, EPI_ISL_887988, EPI_ISL_887989, EPI_ISL_887991, EPI_ISL_887992, EPI_ISL_887994, EPI_ISL_887997, EPI_ISL_887998, EPI_ISL_888000, EPI_ISL_888001, EPI_ISL_888002, EPI_ISL_888003, EPI_ISL_888008, EPI_ISL_888010, EPI_ISL_888013, EPI_ISL_888014, EPI_ISL_888015, EPI_ISL_888018, EPI_ISL_888022, EPI_ISL_888024, EPI_ISL_888026, EPI_ISL_888027, EPI_ISL_888029, EPI_ISL_888030, EPI_ISL_888036, EPI_ISL_888037, EPI_ISL_888042, EPI_ISL_888043, EPI_ISL_888044, EPI_ISL_888050, EPI_ISL_888051, EPI_ISL_888057, EPI_ISL_888058, EPI_ISL_888063, EPI_ISL_888064, EPI_ISL_888066, EPI_ISL_888068, EPI_ISL_888069, EPI_ISL_888074, EPI_ISL_888075, EPI_ISL_888076, EPI_ISL_888080, EPI_ISL_888081, EPI_ISL_888083, EPI_ISL_888084, EPI_ISL_888085, EPI_ISL_888088, EPI_ISL_888095, EPI_ISL_888101, EPI_ISL_888102, EPI_ISL_888103, EPI_ISL_888105, EPI_ISL_888107, EPI_ISL_888109, EPI_ISL_888110, EPI_ISL_888111, EPI_ISL_888113, EPI_ISL_888114, EPI_ISL_888116, EPI_ISL_888117, EPI_ISL_888122, EPI_ISL_888128, EPI_ISL_888132, EPI_ISL_888133, EPI_ISL_888139, EPI_ISL_888140, EPI_ISL_888143, EPI_ISL_888144, EPI_ISL_888147, EPI_ISL_888149, EPI_ISL_888150, EPI_ISL_888151, EPI_ISL_888152, EPI_ISL_888153, EPI_ISL_888155, EPI_ISL_888156, EPI_ISL_888158, EPI_ISL_888162, EPI_ISL_888164, EPI_ISL_888165, EPI_ISL_888166, EPI_ISL_888174, EPI_ISL_888175, EPI_ISL_888179, EPI_ISL_888183, EPI_ISL_888184, EPI_ISL_888185, EPI_ISL_888187, EPI_ISL_888189, EPI_ISL_888191, EPI_ISL_888192, EPI_ISL_888195, EPI_ISL_888199, EPI_ISL_888201, EPI_ISL_888202, EPI_ISL_888204, EPI_ISL_888205, EPI_ISL_888208, EPI_ISL_888209, EPI_ISL_888210, EPI_ISL_888215, EPI_ISL_888217, EPI_ISL_888218, EPI_ISL_888220, EPI_ISL_888221, EPI_ISL_888222, EPI_ISL_888225, EPI_ISL_888226, EPI_ISL_888228, EPI_ISL_888229, EPI_ISL_888232, EPI_ISL_888235, EPI_ISL_888237, EPI_ISL_888241, EPI_ISL_888242, EPI_ISL_888243, EPI_ISL_888245, EPI_ISL_888247, EPI_ISL_888248, EPI_ISL_888249, EPI_ISL_888250, EPI_ISL_888251, EPI_ISL_888252, EPI_ISL_888253, EPI_ISL_888255, EPI_ISL_888256, EPI_ISL_888257, EPI_ISL_888258, EPI_ISL_888259, EPI_ISL_888260, EPI_ISL_888261, EPI_ISL_888262, EPI_ISL_888265, EPI_ISL_888266, EPI_ISL_888268, EPI_ISL_888271, EPI_ISL_888272, EPI_ISL_888273, EPI_ISL_888274, EPI_ISL_888275, EPI_ISL_888276, EPI_ISL_888277, EPI_ISL_888280, EPI_ISL_888281, EPI_ISL_888283, EPI_ISL_888284, EPI_ISL_888287, EPI_ISL_888291, EPI_ISL_888292, EPI_ISL_888297, EPI_ISL_888299, EPI_ISL_888300, EPI_ISL_888302, EPI_ISL_888310, EPI_ISL_888312, EPI_ISL_888314, EPI_ISL_888315, EPI_ISL_888316, EPI_ISL_888320, EPI_ISL_888323, EPI_ISL_888325, EPI_ISL_888326, EPI_ISL_888327, EPI_ISL_888328, EPI_ISL_888329, EPI_ISL_888330, EPI_ISL_888332, EPI_ISL_888333, EPI_ISL_888334, EPI_ISL_888337, EPI_ISL_888339, EPI_ISL_888341, EPI_ISL_888342, EPI_ISL_888344, EPI_ISL_888347, EPI_ISL_888348, EPI_ISL_888350, EPI_ISL_888351, EPI_ISL_888352, EPI_ISL_888353, EPI_ISL_888354, EPI_ISL_888356, EPI_ISL_888360, EPI_ISL_888363, EPI_ISL_888365, EPI_ISL_888366, EPI_ISL_888367, EPI_ISL_888368, EPI_ISL_888369, EPI_ISL_888370, EPI_ISL_888372, EPI_ISL_888373, EPI_ISL_888375, EPI_ISL_888377, EPI_ISL_888378, EPI_ISL_888379, EPI_ISL_888380, EPI_ISL_888381, EPI_ISL_888382, EPI_ISL_888384, EPI_ISL_888385, EPI_ISL_888386, EPI_ISL_888387, EPI_ISL_888388, EPI_ISL_888389, EPI_ISL_888391, EPI_ISL_888392, EPI_ISL_888393, EPI_ISL_888394, EPI_ISL_888395, EPI_ISL_888396, EPI_ISL_888397, EPI_ISL_888399, EPI_ISL_888400, EPI_ISL_888402, EPI_ISL_888404, EPI_ISL_888405, EPI_ISL_888406, EPI_ISL_888407, EPI_ISL_888408, EPI_ISL_888409, EPI_ISL_888410, EPI_ISL_888411, EPI_ISL_888413, EPI_ISL_888414, EPI_ISL_888415, EPI_ISL_888417, EPI_ISL_888419, EPI_ISL_888420, EPI_ISL_888421, EPI_ISL_888422, EPI_ISL_888424, EPI_ISL_888426, EPI_ISL_888427, EPI_ISL_888429, EPI_ISL_888432, EPI_ISL_888435, EPI_ISL_888439, EPI_ISL_888443, EPI_ISL_888448, EPI_ISL_888449, EPI_ISL_888451, EPI_ISL_888452, EPI_ISL_888453, EPI_ISL_888454, EPI_ISL_888458, EPI_ISL_888460, EPI_ISL_888462, EPI_ISL_888463, EPI_ISL_888464, EPI_ISL_888466, EPI_ISL_888467, EPI_ISL_888468, EPI_ISL_888471, EPI_ISL_888475, EPI_ISL_888476, EPI_ISL_888478, EPI_ISL_888479, EPI_ISL_888480, EPI_ISL_888481, EPI_ISL_888483, EPI_ISL_888486, EPI_ISL_888488, EPI_ISL_888490, EPI_ISL_888491, EPI_ISL_888492, EPI_ISL_888493, EPI_ISL_888496, EPI_ISL_888498, EPI_ISL_888510, EPI_ISL_888512, EPI_ISL_888514, EPI_ISL_888527, EPI_ISL_888528, EPI_ISL_888538, EPI_ISL_888541, EPI_ISL_888544, EPI_ISL_888547, EPI_ISL_888562, EPI_ISL_888566, EPI_ISL_888567, EPI_ISL_888568, EPI_ISL_888571, EPI_ISL_888582 | | | |
| see above | Labcorp | Genomics and Discovery, Respiratory Viruses Branch, Division of Viral Diseases, Centers for Disease Control and Prevention | Peter W. Cook,Dhwani Batra,Ben L. Rambo-Martin,Summer Galloway,Brian Krueger,Minoo Agarwal,Eyad Almasri,Debbie Boles,Ayla Burns,Nuthawin Charoensri,Oren Cohen,Susan Countryman,Mary Ann Cristobal,Bobbi Croy,Suzanne Dale,Hrushikesh Deshmukh,Amanda Douglas,Vincent Drouillon,Marcia Eisenberg,Howard Engler,Rama Ghatti,Prashant Gupta,Susan Hicks,Jake Humphrey,Lax Iyer,Manoj Jain,Mohan Kolli,Tim Kuphal,Stanley Letovsky,Michael Levandoski,Craig Lukasik,Jonathan Meltzer,Brian Norvell,Mindy Nye,Scott Parker,Christos Petropoulos,John Pruitt,Steven Ragan,Scott Ryan,Mike Sapeta,Jana Schroth,Suresh Babu Selvaraju,Goran Stevovic,Amanda Suchanek,Andrea Throop,Lyndon Tilson,Thomas Urban,Joe Voshell,Kimberly Wagner,Jonathan Williams,Mary Williamson,Qian Zeng,Tricia Zwiefelhofer,Clinton R. Paden,Suxiang Tong,Duncan MacCannell, |
| EPI_ISL_888591 | Siti Khodijah Hospital | Institute of Tropical Disease, Universitas Airlangga | Krisnoadi Rahardjo, Aldise M Nastri, Jezzy R Dewantari, Rima R Prasetya, Muhammad Hamdan, Gatot Soegiarto, Laksmi Wulandari, Resti Yudhawati, Yasuko Mori, Soetjipto, Kazufumi Shimizu, Maria I Lusida |
| EPI_ISL_888593 | Siti Khodijah Hospital | Institute of Tropical Disease, Universitas Airlangga | Rima R Prasetya, Krisnoadi Rahardjo, Aldise M Nastri, Jezzy R Dewantari, Muhammad Hamdan, Gatot Soegiarto, Laksmi Wulandari, Resti Yudhawati, Soetjipto, Yasuko Mori, Kazufumi Shimizu, Maria I Lusida |
| EPI_ISL_888674 | Center of Advanced Studies and Technology, Molecular Genetics Laboratory | Center of Advanced Studies and Technology, Molecular Genetics Laboratory | De Fabritiis Simone, Mandatori Domitilla, Ferrante Rossella |
| EPI_ISL_888833 | National Virus Reference Laboratory | National Virus Reference Laboratory | Michael Carr, Gabriel Gonzalez, Jonathan Dean, Cillian F De Gascun |
| EPI_ISL_888862, EPI_ISL_888863, EPI_ISL_888864, EPI_ISL_888866, EPI_ISL_888867, EPI_ISL_888868, EPI_ISL_888873 | Michigan Department of Health and Human Services, Bureau of Laboratories | Michigan Department of Health and Human Services, Bureau of Laboratories | Blankenship HM, Riner D, Soehnlen MK |
| EPI_ISL_889444, EPI_ISL_889445, EPI_ISL_889446, EPI_ISL_889447, EPI_ISL_889448, EPI_ISL_889449, EPI_ISL_889450, EPI_ISL_889451, EPI_ISL_889452, EPI_ISL_889453, EPI_ISL_889454, EPI_ISL_889455, EPI_ISL_889456, EPI_ISL_889457, EPI_ISL_889458, EPI_ISL_889459, EPI_ISL_889460 | | | |
| see above | LSUHS Emerging Viral Threat Laboratory | Microbial Genome Sequencing Center | Jeremy P. Kamil, Jennifer L. Carroll, Camille F. Abshire, Maarten Van Diest, Mohammed N.A. Siddiquey, Andrew D. Yurochko, Martin J. Sapp, Rona S. Scott, Christopher G. Kevil, Daniel J. Snyder, Vaughn S. Cooper, John A. Vanchiere |
| EPI_ISL_890242, EPI_ISL_890247, EPI_ISL_890251, EPI_ISL_890256, EPI_ISL_890262, EPI_ISL_890267, EPI_ISL_890268, EPI_ISL_890273, EPI_ISL_890312, EPI_ISL_890315, EPI_ISL_890318, EPI_ISL_890325, EPI_ISL_890335, EPI_ISL_890339, EPI_ISL_890342, EPI_ISL_890345, EPI_ISL_890349 | | | |
| see above | KU Leuven, Rega Institute, Clinical and Epidemiological Virology | KU Leuven, Rega Institute, Clinical and Epidemiological Virology | Tony Wawina-Bokalanga, Bert Vanmechelen, Joan Marti-Carerras, Piet Maes |
| EPI_ISL_890890, EPI_ISL_890891, EPI_ISL_890893 | Hospital | National Reference Center for Viruses of Respiratory Infections, Institut Pasteur, Paris | Marion Barbet, Sylvie Behillil, Méline Bizard, Angela Brisebarre, Camille Capel, Etienne Simon-Lorière, Vincent Enouf, Maud Vanpeene, Sylvie van der Werf, Patricia Stoessel-Thouvenin |
| EPI_ISL_890895 | Laboratory Analyses Med | National Reference Center for Viruses of Respiratory Infections, Institut Pasteur, Paris | Marion Barbet, Sylvie Behillil, Méline Bizard, Angela Brisebarre, Camille Capel, Etienne Simon-Lorière, Vincent Enouf, Maud Vanpeene, Sylvie van der Werf, Céline Menaut |
| EPI_ISL_891012, EPI_ISL_891013, EPI_ISL_891014, EPI_ISL_891015, | Washington State Department of Health | Seattle Flu Study | Deborah A. Nickerson, Chris D. Frazar, Jover Lee, Benjamin Pelle, Erica Ryke, Matthew Richardson, Amanda Adler, Elisabeth Brandstetter, Peter D. Han, Kairsten Fay, Misja Ilcisin, Kirsten Lacombe, Thomas R. Sibley, Melissa Truong, Caitlin R. Wolf, Romesh Gautom, Geoff Melly, Brian Hiatt, Philip Dykema, |

| | | | |
|---|---|---|---|
| EPI_ISL_891016, EPI_ISL_891017, EPI_ISL_891018, EPI_ISL_891019 | | | Scott Lindquist, Michael Boeckh, Janet A. Englund, Michael Famulare, Barry R. Lutz, Mark J. Rieder, Lea M. Starita, Matthew Thompson, Helen Y. Chu, Jay Shendure, Trevor Bedford |
| EPI_ISL_891142, EPI_ISL_891143 | Indiana Animal Disease Diagnostic Laboratory | Carpi Laboratory - Purdue University | Jack Dorman, Ilinca I Ciubotariu, Lev Gorenstein, Abebe A Fola, G Kenitra Hendrix, Rebecca P Wilkes, Giovanna Carpi |
| EPI_ISL_891151 | Academic Hospital of Gadjah Mada University (RSA UGM) | Genetics Working Group (Pokja Genetik) Faculty of Medicine, Public Health and Nursing Universitas Gadjah Mada (FK-KMK UGM); Disease Investigation Center Wates Ministry of Agriculture Indonesia; Department of Microbiology FK-KMK UGM; Laboratorium Diagnostik Yayasan Tahija World Mosquito Program (WMP) Yogyakarta Center for Tropical Medicine FK-KMK UGM; Integrated Research center FK-KMK UGM; Department of Computer Science and Electronics FMIPA UGM | Gunadi, Hendra Wibawa, Marcellus, Mohamad S. Hakim, Edwin W. Daniwijaya, Ludhang P. Rizki, Endah Supriyati, Eggi Arguni, Titik Nuryastuti, Tri Wibawa, Dwi AA Nugrahaningsih, Afiahayati, Siswanto, Kristy Iskandar, Nungki Anggorowati, Alvin Santoso Kalim, Susan Simanjaya |
| EPI_ISL_891152 | SMS Medical College Jaipur | SMS Medical College Jaipur | Bharti Malhotra, Swati Gautam,Himanshu Sharma, Pratibha Sharma, Neha Bhomia, Nivedita Gupta, Pragya Yadav, Varsha Potdar |
| EPI_ISL_891197 | DPH, Massachusetts State Public Health Lab | DPH, Massachusetts State Public Health Lab | Lang,A.S., Fink,T., Gallagher,G.R., Smole,S.C. |
| EPI_ISL_891207 | SMS Medical College Jaipur | SMS Medical College jaipur | Bharti Malhotra, Swati Gautam, Himanshu Sharma,Pratibha Sharma, Neha Bhomia, Nivedita Gupta, Pragya Yadav, Varsha Potdar |
| EPI_ISL_891210 | SMS Medical College jaipur | SMS Medical College Jaipur | Bharti Malhotra, Swati Gautam, Himanshu Sharma, Pratibha Sharma, Neha Bhomia, Nivedita Gupta , Pragya Yadav,Varsha Potdar |
| EPI_ISL_893743, EPI_ISL_893783 | Institute of Virology, Medical Center, University of Freiburg, Freiburg, Germany | Institute of Virology, Clinial Virus Genomics, Medical Center, University of Freiburg, Freiburg, Germany | Jonas Fuchs, Lisa Kern, Sandra Reuter, Hajo Grundmann, Marcus Panning |
| EPI_ISL_894163, EPI_ISL_894167 | Institute of Medical Microbiology and Hospital Hygiene | Institute of Medical Microbiology and Hospital Hygiene | Prof. Dr. Achim Kaasch, Aljoscha Tersteegen |
| EPI_ISL_896116 | Kantonsspital Aarau, Institut für Labormedizin | University Hospital Basel, Clinical Bacteriology | Tim Roloff, Madlen Stange, Helena MB Seth-Smith, Alfredo Mari, Karoline Leuzinger, Julia Bielicki, Michael Oberle, Manuel Battegay, Hans Hirsch, Adrian Egli |
| EPI_ISL_896143, EPI_ISL_896163, EPI_ISL_896164, EPI_ISL_896165, EPI_ISL_896166, EPI_ISL_896197, EPI_ISL_896198, EPI_ISL_896199, EPI_ISL_896200, EPI_ISL_896201, EPI_ISL_896202, EPI_ISL_896203 | | | |
| see above | MEPHI, Aix Marseille University | MEPHI, Aix Marseille University | Anthony LEVASSEUR |
| EPI_ISL_898012, EPI_ISL_898013, EPI_ISL_898014 | KU Leuven, Rega Institute, Clinical and Epidemiological Virology | KU Leuven, Rega Institute, Clinical and Epidemiological Virology | Tony Wawina-Bokalanga, Bert Vanmechelen, Joan Marti-Carerras, Piet Maes |
| EPI_ISL_899003, EPI_ISL_899088, EPI_ISL_899105, EPI_ISL_899110, EPI_ISL_899116, EPI_ISL_899130, EPI_ISL_899140, EPI_ISL_899141, EPI_ISL_899142, EPI_ISL_899165, EPI_ISL_899166, EPI_ISL_899171, EPI_ISL_899177, EPI_ISL_899182, EPI_ISL_899183, EPI_ISL_899184, EPI_ISL_899278, EPI_ISL_899279, EPI_ISL_899280, EPI_ISL_899281, EPI_ISL_899282, EPI_ISL_899283, EPI_ISL_899284, EPI_ISL_899385, EPI_ISL_899386, EPI_ISL_899387, EPI_ISL_899388, EPI_ISL_899389, EPI_ISL_899390, EPI_ISL_899391, EPI_ISL_899392, EPI_ISL_899393, EPI_ISL_899394, EPI_ISL_899395, EPI_ISL_899396, EPI_ISL_899397, EPI_ISL_899398, EPI_ISL_899399, EPI_ISL_899400, EPI_ISL_899401, EPI_ISL_899402, EPI_ISL_899403, EPI_ISL_899404, EPI_ISL_899516, EPI_ISL_899562, EPI_ISL_899576, EPI_ISL_899577, EPI_ISL_899614, EPI_ISL_899615, EPI_ISL_899616, EPI_ISL_899617, EPI_ISL_899618, EPI_ISL_899619, EPI_ISL_899620, EPI_ISL_899647, EPI_ISL_899648, EPI_ISL_899649, EPI_ISL_899650, EPI_ISL_899651, EPI_ISL_899652, EPI_ISL_899653, EPI_ISL_899654, EPI_ISL_899655, EPI_ISL_899656, EPI_ISL_899657, EPI_ISL_899658, EPI_ISL_899659, EPI_ISL_899660, EPI_ISL_899661, EPI_ISL_899662, EPI_ISL_899663, EPI_ISL_899759, EPI_ISL_899760, EPI_ISL_899761, EPI_ISL_899762, EPI_ISL_899763, EPI_ISL_899764, EPI_ISL_899765, EPI_ISL_899766, EPI_ISL_899767, EPI_ISL_899810, EPI_ISL_899862, EPI_ISL_899923, EPI_ISL_899924, EPI_ISL_899925, EPI_ISL_899926, EPI_ISL_899927, EPI_ISL_899928, EPI_ISL_899929, EPI_ISL_899930, EPI_ISL_899931, EPI_ISL_899932, EPI_ISL_899934, EPI_ISL_899935, EPI_ISL_899936 | | | |
| see above | Viollier AG | Department of Biosystems Science and Engineering, ETH Zürich | Christian Beisel, Sarah Nadeau, Chaoran Chen, Ivan Topolsky, Philipp Jablonski, Lara Fuhrmann, David Dreifuss, Katharina Jahn, Tobias Schär, Ina Nissen, Natascha Santacroce, Elodie Burcklen, Christiane Beckmann, Maurice Redondo, Olivier Kobel, Christoph Noppen, Sophie Seidel, Noemie Santamaria de Souza, Niko Beerenwinkel, Tanja Stadler |
| EPI_ISL_900470 | MEPHI, Aix Marseille University | MEPHI, Aix Marseille University | Anthony LEVASSEUR |
| EPI_ISL_900505 | Mirialis | CNR Virus des Infections Respiratoires - France SUD | Antonin Bal, Gregory Destras, Gwendolyne Burfin, Hadrien Règue, Quentin Semanas, Martine Valette, Bruno Lina, Laurence Josset |
| EPI_ISL_900506 | AX BIO OCEAN | CNR Virus des Infections Respiratoires - France SUD | Antonin Bal, Gregory Destras, Gwendolyne Burfin, Hadrien Règue, Quentin Semanas, Martine Valette, Bruno Lina, Laurence Josset |
| EPI_ISL_900528 | CHU Clermont-Ferrand | CNR Virus des Infections Respiratoires - France SUD | Antonin Bal, Gregory Destras, Gwendolyne Burfin, Hadrien Règue, Quentin Semanas, Martine Valette, Bruno Lina, Christine Archimbaud, Amélie Brebion, Hélène Chabrolles, Martine Chambon, Audrey Mirand, Christel Regagnon, Maxime Bisseux, Patricia Combes, Cécile Henquell, Laurence Josset |
| EPI_ISL_900534 | CH VILLEFRANCHE | CNR Virus des Infections Respiratoires - France SUD | Antonin Bal, Gregory Destras, Gwendolyne Burfin, Hadrien Règue, Quentin Semanas, Martine Valette, Bruno Lina, Laurence Josset |
| EPI_ISL_900535 | Hôpital Paris Saint-Joseph | CNR Virus des Infections Respiratoires - France SUD | Antonin Bal, Gregory Destras, Gwendolyne Burfin, Hadrien Règue, Quentin Semanas, Martine Valette, Bruno Lina, Sylvie Larrat, Laurence Josset |
| EPI_ISL_900543 | Centre hospitalier Emile Roux | CNR Virus des Infections Respiratoires - France SUD | Antonin Bal, Gregory Destras, Gwendolyne Burfin, Hadrien Règue, Quentin Semanas, Martine Valette, Bruno Lina, Laurence Josset |
| EPI_ISL_900575, EPI_ISL_900587, EPI_ISL_900588, EPI_ISL_900589, EPI_ISL_900590, EPI_ISL_900591, EPI_ISL_900647, EPI_ISL_900651, EPI_ISL_900652 | IZSM | TIGEM | Patrizia Annunziata, Andrea Ballabio, Valentina Bouche, Davide Cacchiarelli (CorrespAuthor), Pellegrino Cerino, Chiara Colantuono, Maria Concetta Cuomo, Denise Di Concilio, Lucio Di Filippo, Antonio Grimaldi, Antonio Limone, Anna Manfredi, Francesco Panariello, Biancamaria Pierri, Marcello Salvi |
| EPI_ISL_902737 | RSA UGM | Genetics Working Group (Pokja Genetik) Faculty of Medicine, Public Health and Nursing Universitas Gadjah Mada (FK-KMK UGM); Disease Investigation Center Wates Ministry of Agriculture Indonesia; Department of Microbiology FK-KMK UGM; Laboratorium Diagnostik Yayasan Tahija World Mosquito Program (WMP) Yogyakarta Center for Tropical Medicine FK-KMK UGM; Integrated Research center FK-KMK UGM; Department of Computer Science and Electronics FMIPA UGM | Gunadi, Hendra Wibawa, Marcellus, Mohamad S. Hakim, Edwin W. Daniwijaya, Ludhang P. Rizki, Endah Supriyati, Eggi Arguni, Titik Nuryastuti, Tri Wibawa, Dwi AA Nugrahaningsih, Afiahayati, Siswanto, Kristy Iskandar, Nungki Anggorowati, Aditya Rifqi Fauzi, Fadil Fahri |
| EPI_ISL_902749 | RSA UGM | Genetics Working Group (Pokja Genetik) Faculty of Medicine, Public Health and Nursing Universitas Gadjah Mada (FK-KMK UGM); Disease Investigation Center Wates Ministry of Agriculture Indonesia; Department of Microbiology FK-KMK UGM; Laboratorium Diagnostik Yayasan Tahija World Mosquito Program (WMP) Yogyakarta Center for Tropical Medicine FK-KMK UGM; Integrated Research center FK-KMK UGM; Department of Computer Science and Electronics FMIPA UGM | Gunadi, Hendra Wibawa, Marcellus, Mohamad S. Hakim, Edwin W. Daniwijaya, Ludhang P. Rizki, Endah Supriyati, Eggi Arguni, Titik Nuryastuti, Tri Wibawa, Dwi AA Nugrahaningsih, Afiahayati, Siswanto, Kristy Iskandar, Nungki Anggorowati, Dyah Ayu Puspitarani, Dwiki afandy |
| EPI_ISL_903026 | Seattle Flu Study | Seattle Flu Study | Deborah A. Nickerson, Chris D. Frazar, Jover Lee, Benjamin Pelle, Erica Ryke, Matthew Richardson, Amanda Adler, Elisabeth Brandstetter, Peter D. Han, Kairsten Fay, Misja Ilcisin, Kirsten Lacombe, Thomas R. Sibley, Melissa Truong, Caitlin R. Wolf, Karen Cowgill, Stephanie Schrag, Jeff Duchin, Michael Boeckh, Janet A. Englund, Michael Famulare, Barry R. Lutz, Mark J. Rieder, Lea M. Starita, Matthew Thompson, Helen Y. Chu, Trevor Bedford, Jay Shendure |
| EPI_ISL_903027 | Seattle Flu Study | Seattle Flu Study | Deborah A. Nickerson, Chris D. Frazar, Jover Lee, Benjamin Pelle, Erica Ryke, Matthew Richardson, Amanda Adler, Elisabeth Brandstetter, Peter D. Han, Kairsten Fay, Misja Ilcisin, Kirsten Lacombe, Thomas R. Sibley, Melissa Truong, Caitlin R. Wolf, Michael Boeckh, Janet A. Englund, Michael Famulare, Barry R. Lutz, Mark J. Rieder, Lea M. Starita, Matthew Thompson, Jay Shendure, Trevor Bedford, Helen Y. Chu |
| EPI_ISL_903028, EPI_ISL_903029, EPI_ISL_903030, EPI_ISL_903031, | Seattle Flu Study | Seattle Flu Study | Deborah A. Nickerson, Chris D. Frazar, Jover Lee, Benjamin Pelle, Erica Ryke, Matthew Richardson, Amanda Adler, Elisabeth Brandstetter, Peter D. Han, Kairsten Fay, Misja Ilcisin, Kirsten Lacombe, Thomas R. Sibley, Melissa Truong, Caitlin R. Wolf, Karen Cowgill, Stephanie Schrag, Jeff Duchin, Michael |

| | | | |
|---|---|---|---|
| EPI_ISL_903032, EPI_ISL_903033, EPI_ISL_903034, EPI_ISL_903035, EPI_ISL_903036 | | | Boeckh, Janet A. Englund, Michael Famulare, Barry R. Lutz, Mark J. Rieder, Lea M. Starita, Matthew Thompson, Helen Y. Chu, Trevor Bedford, Jay Shendure |
| EPI_ISL_903229, EPI_ISL_903233, EPI_ISL_903234, EPI_ISL_903236, EPI_ISL_903240, EPI_ISL_903258, EPI_ISL_903259, EPI_ISL_903260, EPI_ISL_903261, EPI_ISL_903262, EPI_ISL_903263, EPI_ISL_903265, EPI_ISL_903266, EPI_ISL_903267, EPI_ISL_903268, EPI_ISL_903269, EPI_ISL_903270, EPI_ISL_903271, EPI_ISL_903272 | | | |
| see above | M Health Fairview | Minnesota Department of Health, Public Health Laboratory | Alexandra Lorentz, Jacob Garfin, Matt Plumb, and Xiong Wang |
| EPI_ISL_903367, EPI_ISL_903368, EPI_ISL_903369, EPI_ISL_903370 | Los Angeles County PHL | Los Angeles County PHL | P. Hemarajata et al. |
| EPI_ISL_903376, EPI_ISL_903382, EPI_ISL_903384, EPI_ISL_903385, EPI_ISL_903386, EPI_ISL_903387 | MOH - Jaber Al-Ahmad Hospital (Innovation Research Laboratory) | MOH - Jaber Al-Ahmad Hospital (Innovation Research Laboratory) | Salman Al-Sabah , Mohammad Alghounaim |
| EPI_ISL_903565 | AR Dept. of Health-Public Health Lab | Genomics and Discovery, Respiratory Viruses Branch, Division of Viral Diseases, Centers for Disease Control and Prevention | Krista Queen, Yan Li, Ying Tao, Jing Zhang, Anna Uehara, Anna Montmayeur, Clinton R. Paden, Peter W. Cook, Rachel Marine, Mili Sheth, Jasmine Padilla, Sarah Nobles, Mark Burroughs, Lori Rowe, Haibin Wang, Ben L. Rambo-Martin, Dhwani Batra, Justin Lee, Suxiang Tong |
| EPI_ISL_903582 | TN Division of Laboratory Services | Genomics and Discovery, Respiratory Viruses Branch, Division of Viral Diseases, Centers for Disease Control and Prevention | Krista Queen, Yan Li, Ying Tao, Jing Zhang, Anna Uehara, Anna Montmayeur, Clinton R. Paden, Peter W. Cook, Rachel Marine, Mili Sheth, Jasmine Padilla, Sarah Nobles, Mark Burroughs, Lori Rowe, Haibin Wang, Ben L. Rambo-Martin, Dhwani Batra, Justin Lee, Suxiang Tong |
| EPI_ISL_903583 | HI Dept. of Health, State Laboratories Division | Genomics and Discovery, Respiratory Viruses Branch, Division of Viral Diseases, Centers for Disease Control and Prevention | Krista Queen, Yan Li, Ying Tao, Jing Zhang, Anna Uehara, Anna Montmayeur, Clinton R. Paden, Peter W. Cook, Rachel Marine, Mili Sheth, Jasmine Padilla, Sarah Nobles, Mark Burroughs, Lori Rowe, Haibin Wang, Ben L. Rambo-Martin, Dhwani Batra, Justin Lee, Suxiang Tong |
| EPI_ISL_903599 | WY Public Health Laboratory | Genomics and Discovery, Respiratory Viruses Branch, Division of Viral Diseases, Centers for Disease Control and Prevention | Krista Queen, Yan Li, Ying Tao, Jing Zhang, Anna Uehara, Anna Montmayeur, Clinton R. Paden, Peter W. Cook, Rachel Marine, Mili Sheth, Jasmine Padilla, Sarah Nobles, Mark Burroughs, Lori Rowe, Haibin Wang, Ben L. Rambo-Martin, Dhwani Batra, Justin Lee, Suxiang Tong |
| EPI_ISL_903603 | ID Bureau of Laboratories | Genomics and Discovery, Respiratory Viruses Branch, Division of Viral Diseases, Centers for Disease Control and Prevention | Krista Queen, Yan Li, Ying Tao, Jing Zhang, Anna Uehara, Anna Montmayeur, Clinton R. Paden, Peter W. Cook, Rachel Marine, Mili Sheth, Jasmine Padilla, Sarah Nobles, Mark Burroughs, Lori Rowe, Haibin Wang, Ben L. Rambo-Martin, Dhwani Batra, Justin Lee, Suxiang Tong |
| EPI_ISL_903639 | AR Dept. of Health-Public Health Lab | Genomics and Discovery, Respiratory Viruses Branch, Division of Viral Diseases, Centers for Disease Control and Prevention | Krista Queen, Yan Li, Ying Tao, Jing Zhang, Anna Uehara, Anna Montmayeur, Clinton R. Paden, Peter W. Cook, Rachel Marine, Mili Sheth, Jasmine Padilla, Sarah Nobles, Mark Burroughs, Lori Rowe, Haibin Wang, Ben L. Rambo-Martin, Dhwani Batra, Justin Lee, Suxiang Tong |
| EPI_ISL_903657 | VT Dept. of Health Laboratory | Genomics and Discovery, Respiratory Viruses Branch, Division of Viral Diseases, Centers for Disease Control and Prevention | Krista Queen, Yan Li, Ying Tao, Jing Zhang, Anna Uehara, Anna Montmayeur, Clinton R. Paden, Peter W. Cook, Rachel Marine, Mili Sheth, Jasmine Padilla, Sarah Nobles, Mark Burroughs, Lori Rowe, Haibin Wang, Ben L. Rambo-Martin, Dhwani Batra, Justin Lee, Suxiang Tong |
| EPI_ISL_903659 | MN PHL Division, Minnesota Department of Health | Genomics and Discovery, Respiratory Viruses Branch, Division of Viral Diseases, Centers for Disease Control and Prevention | Krista Queen, Yan Li, Ying Tao, Jing Zhang, Anna Uehara, Anna Montmayeur, Clinton R. Paden, Peter W. Cook, Rachel Marine, Mili Sheth, Jasmine Padilla, Sarah Nobles, Mark Burroughs, Lori Rowe, Haibin Wang, Ben L. Rambo-Martin, Dhwani Batra, Justin Lee, Suxiang Tong |
| EPI_ISL_903665, EPI_ISL_903666 | OR State PHL-Virology/Immunology Section | Genomics and Discovery, Respiratory Viruses Branch, Division of Viral Diseases, Centers for Disease Control and Prevention | Krista Queen, Yan Li, Ying Tao, Jing Zhang, Anna Uehara, Anna Montmayeur, Clinton R. Paden, Peter W. Cook, Rachel Marine, Mili Sheth, Jasmine Padilla, Sarah Nobles, Mark Burroughs, Lori Rowe, Haibin Wang, Ben L. Rambo-Martin, Dhwani Batra, Justin Lee, Suxiang Tong |
| EPI_ISL_903669 | IA State Hygienic Laboratory | Genomics and Discovery, Respiratory Viruses Branch, Division of Viral Diseases, Centers for Disease Control and Prevention | Krista Queen, Yan Li, Ying Tao, Jing Zhang, Anna Uehara, Anna Montmayeur, Clinton R. Paden, Peter W. Cook, Rachel Marine, Mili Sheth, Jasmine Padilla, Sarah Nobles, Mark Burroughs, Lori Rowe, Haibin Wang, Ben L. Rambo-Martin, Dhwani Batra, Justin Lee, Suxiang Tong |
| EPI_ISL_903676 | OR State PHL-Virology/Immunology Section | Genomics and Discovery, Respiratory Viruses Branch, Division of Viral Diseases, Centers for Disease Control and Prevention | Krista Queen, Yan Li, Ying Tao, Jing Zhang, Anna Uehara, Anna Montmayeur, Clinton R. Paden, Peter W. Cook, Rachel Marine, Mili Sheth, Jasmine Padilla, Sarah Nobles, Mark Burroughs, Lori Rowe, Haibin Wang, Ben L. Rambo-Martin, Dhwani Batra, Justin Lee, Suxiang Tong |
| EPI_ISL_903679 | AR Dept. of Health-Public Health Lab | Genomics and Discovery, Respiratory Viruses Branch, Division of Viral Diseases, Centers for Disease Control and Prevention | Krista Queen, Yan Li, Ying Tao, Jing Zhang, Anna Uehara, Anna Montmayeur, Clinton R. Paden, Peter W. Cook, Rachel Marine, Mili Sheth, Jasmine Padilla, Sarah Nobles, Mark Burroughs, Lori Rowe, Haibin Wang, Ben L. Rambo-Martin, Dhwani Batra, Justin Lee, Suxiang Tong |
| EPI_ISL_903694, EPI_ISL_903696 | VT Dept. of Health Laboratory | Genomics and Discovery, Respiratory Viruses Branch, Division of Viral Diseases, Centers for Disease Control and Prevention | Krista Queen, Yan Li, Ying Tao, Jing Zhang, Anna Uehara, Anna Montmayeur, Clinton R. Paden, Peter W. Cook, Rachel Marine, Mili Sheth, Jasmine Padilla, Sarah Nobles, Mark Burroughs, Lori Rowe, Haibin Wang, Ben L. Rambo-Martin, Dhwani Batra, Justin Lee, Suxiang Tong |
| EPI_ISL_903697 | AR Dept. of Health-Public Health Lab | Genomics and Discovery, Respiratory Viruses Branch, Division of Viral Diseases, Centers for Disease Control and Prevention | Krista Queen, Yan Li, Ying Tao, Jing Zhang, Anna Uehara, Anna Montmayeur, Clinton R. Paden, Peter W. Cook, Rachel Marine, Mili Sheth, Jasmine Padilla, Sarah Nobles, Mark Burroughs, Lori Rowe, Haibin Wang, Ben L. Rambo-Martin, Dhwani Batra, Justin Lee, Suxiang Tong |
| EPI_ISL_903729 | AL Dept. of Public Health Bureau of Clinical Laboratories | Genomics and Discovery, Respiratory Viruses Branch, Division of Viral Diseases, Centers for Disease Control and Prevention | Krista Queen, Yan Li, Ying Tao, Jing Zhang, Anna Uehara, Anna Montmayeur, Clinton R. Paden, Peter W. Cook, Rachel Marine, Mili Sheth, Jasmine Padilla, Sarah Nobles, Mark Burroughs, Lori Rowe, Haibin Wang, Ben L. Rambo-Martin, Dhwani Batra, Justin Lee, Suxiang Tong |
| EPI_ISL_903730 | VT Dept. of Health Laboratory | Genomics and Discovery, Respiratory Viruses Branch, Division of Viral Diseases, Centers for Disease Control and Prevention | Krista Queen, Yan Li, Ying Tao, Jing Zhang, Anna Uehara, Anna Montmayeur, Clinton R. Paden, Peter W. Cook, Rachel Marine, Mili Sheth, Jasmine Padilla, Sarah Nobles, Mark Burroughs, Lori Rowe, Haibin Wang, Ben L. Rambo-Martin, Dhwani Batra, Justin Lee, Suxiang Tong |
| EPI_ISL_903743 | DC Public Health Lab/ Dept. of Forensic Sciences | Genomics and Discovery, Respiratory Viruses Branch, Division of Viral Diseases, Centers for Disease Control and Prevention | Krista Queen, Yan Li, Ying Tao, Jing Zhang, Anna Uehara, Anna Montmayeur, Clinton R. Paden, Peter W. Cook, Rachel Marine, Mili Sheth, Jasmine Padilla, Sarah Nobles, Mark Burroughs, Lori Rowe, Haibin Wang, Ben L. Rambo-Martin, Dhwani Batra, Justin Lee, Suxiang Tong |
| EPI_ISL_903749 | WY Public Health Laboratory | Genomics and Discovery, Respiratory Viruses Branch, Division of Viral Diseases, Centers for Disease Control and Prevention | Krista Queen, Yan Li, Ying Tao, Jing Zhang, Anna Uehara, Anna Montmayeur, Clinton R. Paden, Peter W. Cook, Rachel Marine, Mili Sheth, Jasmine Padilla, Sarah Nobles, Mark Burroughs, Lori Rowe, Haibin Wang, Ben L. Rambo-Martin, Dhwani Batra, Justin Lee, Suxiang Tong |
| EPI_ISL_903757 | TN Division of Laboratory Services | Genomics and Discovery, Respiratory Viruses Branch, Division of Viral Diseases, Centers for Disease Control and Prevention | Krista Queen, Yan Li, Ying Tao, Jing Zhang, Anna Uehara, Anna Montmayeur, Clinton R. Paden, Peter W. Cook, Rachel Marine, Mili Sheth, Jasmine Padilla, Sarah Nobles, Mark Burroughs, Lori Rowe, Haibin Wang, Ben L. Rambo-Martin, Dhwani Batra, Justin Lee, Suxiang Tong |
| EPI_ISL_903763, EPI_ISL_903764 | WY Public Health Laboratory | Genomics and Discovery, Respiratory Viruses Branch, Division of Viral Diseases, Centers for Disease Control and Prevention | Krista Queen, Yan Li, Ying Tao, Jing Zhang, Anna Uehara, Anna Montmayeur, Clinton R. Paden, Peter W. Cook, Rachel Marine, Mili Sheth, Jasmine Padilla, Sarah Nobles, Mark Burroughs, Lori Rowe, Haibin Wang, Ben L. Rambo-Martin, Dhwani Batra, Justin Lee, Suxiang Tong |
| EPI_ISL_903774 | NE Public Health Laboratory | Genomics and Discovery, Respiratory Viruses Branch, Division of Viral Diseases, Centers for Disease Control and Prevention | Krista Queen, Yan Li, Ying Tao, Jing Zhang, Anna Uehara, Anna Montmayeur, Clinton R. Paden, Peter W. Cook, Rachel Marine, Mili Sheth, Jasmine Padilla, Sarah Nobles, Mark Burroughs, Lori Rowe, Haibin Wang, Ben L. Rambo-Martin, Dhwani Batra, Justin Lee, Suxiang Tong |

| | | | |
|---|---|---|---|
| EPI_ISL_903793 | DC Public Health Lab/ Dept. of Forensic Sciences | Genomics and Discovery, Respiratory Viruses Branch, Division of Viral Diseases, Centers for Disease Control and Prevention | Krista Queen, Yan Li, Ying Tao, Jing Zhang, Anna Uehara, Anna Montmayeur, Clinton R. Paden, Peter W. Cook, Rachel Marine, Mili Sheth, Jasmine Padilla, Sarah Nobles, Mark Burroughs, Lori Rowe, Haibin Wang, Ben L. Rambo-Martin, Dhwani Batra, Justin Lee, Suxiang Tong |
| EPI_ISL_903797, EPI_ISL_903802 | TN Division of Laboratory Services | Genomics and Discovery, Respiratory Viruses Branch, Division of Viral Diseases, Centers for Disease Control and Prevention | Krista Queen, Yan Li, Ying Tao, Jing Zhang, Anna Uehara, Anna Montmayeur, Clinton R. Paden, Peter W. Cook, Rachel Marine, Mili Sheth, Jasmine Padilla, Sarah Nobles, Mark Burroughs, Lori Rowe, Haibin Wang, Ben L. Rambo-Martin, Dhwani Batra, Justin Lee, Suxiang Tong |
| EPI_ISL_903803 | WY Public Health Laboratory | Genomics and Discovery, Respiratory Viruses Branch, Division of Viral Diseases, Centers for Disease Control and Prevention | Krista Queen, Yan Li, Ying Tao, Jing Zhang, Anna Uehara, Anna Montmayeur, Clinton R. Paden, Peter W. Cook, Rachel Marine, Mili Sheth, Jasmine Padilla, Sarah Nobles, Mark Burroughs, Lori Rowe, Haibin Wang, Ben L. Rambo-Martin, Dhwani Batra, Justin Lee, Suxiang Tong |
| EPI_ISL_903805 | TN Division of Laboratory Services | Genomics and Discovery, Respiratory Viruses Branch, Division of Viral Diseases, Centers for Disease Control and Prevention | Krista Queen, Yan Li, Ying Tao, Jing Zhang, Anna Uehara, Anna Montmayeur, Clinton R. Paden, Peter W. Cook, Rachel Marine, Mili Sheth, Jasmine Padilla, Sarah Nobles, Mark Burroughs, Lori Rowe, Haibin Wang, Ben L. Rambo-Martin, Dhwani Batra, Justin Lee, Suxiang Tong |
| EPI_ISL_903809 | WY Public Health Laboratory | Genomics and Discovery, Respiratory Viruses Branch, Division of Viral Diseases, Centers for Disease Control and Prevention | Krista Queen, Yan Li, Ying Tao, Jing Zhang, Anna Uehara, Anna Montmayeur, Clinton R. Paden, Peter W. Cook, Rachel Marine, Mili Sheth, Jasmine Padilla, Sarah Nobles, Mark Burroughs, Lori Rowe, Haibin Wang, Ben L. Rambo-Martin, Dhwani Batra, Justin Lee, Suxiang Tong |
| EPI_ISL_903816 | VT Dept. of Health Laboratory | Genomics and Discovery, Respiratory Viruses Branch, Division of Viral Diseases, Centers for Disease Control and Prevention | Krista Queen, Yan Li, Ying Tao, Jing Zhang, Anna Uehara, Anna Montmayeur, Clinton R. Paden, Peter W. Cook, Rachel Marine, Mili Sheth, Jasmine Padilla, Sarah Nobles, Mark Burroughs, Lori Rowe, Haibin Wang, Ben L. Rambo-Martin, Dhwani Batra, Justin Lee, Suxiang Tong |
| EPI_ISL_903835 | NM Dept. Health, Scientific Laboratory Division | Genomics and Discovery, Respiratory Viruses Branch, Division of Viral Diseases, Centers for Disease Control and Prevention | Krista Queen, Yan Li, Ying Tao, Jing Zhang, Anna Uehara, Anna Montmayeur, Clinton R. Paden, Peter W. Cook, Rachel Marine, Mili Sheth, Jasmine Padilla, Sarah Nobles, Mark Burroughs, Lori Rowe, Haibin Wang, Ben L. Rambo-Martin, Dhwani Batra, Justin Lee, Suxiang Tong |
| EPI_ISL_903840 | IA State Hygienic Laboratory | Genomics and Discovery, Respiratory Viruses Branch, Division of Viral Diseases, Centers for Disease Control and Prevention | Krista Queen, Yan Li, Ying Tao, Jing Zhang, Anna Uehara, Anna Montmayeur, Clinton R. Paden, Peter W. Cook, Rachel Marine, Mili Sheth, Jasmine Padilla, Sarah Nobles, Mark Burroughs, Lori Rowe, Haibin Wang, Ben L. Rambo-Martin, Dhwani Batra, Justin Lee, Suxiang Tong |
| EPI_ISL_903849, EPI_ISL_903853 | MT Public Health Laboratory | Genomics and Discovery, Respiratory Viruses Branch, Division of Viral Diseases, Centers for Disease Control and Prevention | Krista Queen, Yan Li, Ying Tao, Jing Zhang, Anna Uehara, Anna Montmayeur, Clinton R. Paden, Peter W. Cook, Rachel Marine, Mili Sheth, Jasmine Padilla, Sarah Nobles, Mark Burroughs, Lori Rowe, Haibin Wang, Ben L. Rambo-Martin, Dhwani Batra, Justin Lee, Suxiang Tong |
| EPI_ISL_903857 | MN PHL Division, Minnesota Department of Health | Genomics and Discovery, Respiratory Viruses Branch, Division of Viral Diseases, Centers for Disease Control and Prevention | Krista Queen, Yan Li, Ying Tao, Jing Zhang, Anna Uehara, Anna Montmayeur, Clinton R. Paden, Peter W. Cook, Rachel Marine, Mili Sheth, Jasmine Padilla, Sarah Nobles, Mark Burroughs, Lori Rowe, Haibin Wang, Ben L. Rambo-Martin, Dhwani Batra, Justin Lee, Suxiang Tong |
| EPI_ISL_903860 | NH Dept. of Health and Human Services Public Health Labs | Genomics and Discovery, Respiratory Viruses Branch, Division of Viral Diseases, Centers for Disease Control and Prevention | Krista Queen, Yan Li, Ying Tao, Jing Zhang, Anna Uehara, Anna Montmayeur, Clinton R. Paden, Peter W. Cook, Rachel Marine, Mili Sheth, Jasmine Padilla, Sarah Nobles, Mark Burroughs, Lori Rowe, Haibin Wang, Ben L. Rambo-Martin, Dhwani Batra, Justin Lee, Suxiang Tong |
| EPI_ISL_903863 | WY Public Health Laboratory | Genomics and Discovery, Respiratory Viruses Branch, Division of Viral Diseases, Centers for Disease Control and Prevention | Krista Queen, Yan Li, Ying Tao, Jing Zhang, Anna Uehara, Anna Montmayeur, Clinton R. Paden, Peter W. Cook, Rachel Marine, Mili Sheth, Jasmine Padilla, Sarah Nobles, Mark Burroughs, Lori Rowe, Haibin Wang, Ben L. Rambo-Martin, Dhwani Batra, Justin Lee, Suxiang Tong |
| EPI_ISL_903871 | NC State Laboratory of Public Health | Genomics and Discovery, Respiratory Viruses Branch, Division of Viral Diseases, Centers for Disease Control and Prevention | Krista Queen, Yan Li, Ying Tao, Jing Zhang, Anna Uehara, Anna Montmayeur, Clinton R. Paden, Peter W. Cook, Rachel Marine, Mili Sheth, Jasmine Padilla, Sarah Nobles, Mark Burroughs, Lori Rowe, Haibin Wang, Ben L. Rambo-Martin, Dhwani Batra, Justin Lee, Suxiang Tong |
| EPI_ISL_903880 | NH Dept. of Health and Human Services Public Health Labs | Genomics and Discovery, Respiratory Viruses Branch, Division of Viral Diseases, Centers for Disease Control and Prevention | Krista Queen, Yan Li, Ying Tao, Jing Zhang, Anna Uehara, Anna Montmayeur, Clinton R. Paden, Peter W. Cook, Rachel Marine, Mili Sheth, Jasmine Padilla, Sarah Nobles, Mark Burroughs, Lori Rowe, Haibin Wang, Ben L. Rambo-Martin, Dhwani Batra, Justin Lee, Suxiang Tong |
| EPI_ISL_903882 | TN Division of Laboratory Services | Genomics and Discovery, Respiratory Viruses Branch, Division of Viral Diseases, Centers for Disease Control and Prevention | Krista Queen, Yan Li, Ying Tao, Jing Zhang, Anna Uehara, Anna Montmayeur, Clinton R. Paden, Peter W. Cook, Rachel Marine, Mili Sheth, Jasmine Padilla, Sarah Nobles, Mark Burroughs, Lori Rowe, Haibin Wang, Ben L. Rambo-Martin, Dhwani Batra, Justin Lee, Suxiang Tong |
| EPI_ISL_903895 | WY Public Health Laboratory | Genomics and Discovery, Respiratory Viruses Branch, Division of Viral Diseases, Centers for Disease Control and Prevention | Krista Queen, Yan Li, Ying Tao, Jing Zhang, Anna Uehara, Anna Montmayeur, Clinton R. Paden, Peter W. Cook, Rachel Marine, Mili Sheth, Jasmine Padilla, Sarah Nobles, Mark Burroughs, Lori Rowe, Haibin Wang, Ben L. Rambo-Martin, Dhwani Batra, Justin Lee, Suxiang Tong |
| EPI_ISL_903899 | AL Dept. of Public Health Bureau of Clinical Laboratories | Genomics and Discovery, Respiratory Viruses Branch, Division of Viral Diseases, Centers for Disease Control and Prevention | Krista Queen, Yan Li, Ying Tao, Jing Zhang, Anna Uehara, Anna Montmayeur, Clinton R. Paden, Peter W. Cook, Rachel Marine, Mili Sheth, Jasmine Padilla, Sarah Nobles, Mark Burroughs, Lori Rowe, Haibin Wang, Ben L. Rambo-Martin, Dhwani Batra, Justin Lee, Suxiang Tong |
| EPI_ISL_903900 | IA State Hygienic Laboratory | Genomics and Discovery, Respiratory Viruses Branch, Division of Viral Diseases, Centers for Disease Control and Prevention | Krista Queen, Yan Li, Ying Tao, Jing Zhang, Anna Uehara, Anna Montmayeur, Clinton R. Paden, Peter W. Cook, Rachel Marine, Mili Sheth, Jasmine Padilla, Sarah Nobles, Mark Burroughs, Lori Rowe, Haibin Wang, Ben L. Rambo-Martin, Dhwani Batra, Justin Lee, Suxiang Tong |
| EPI_ISL_903901 | NE Public Health Laboratory | Genomics and Discovery, Respiratory Viruses Branch, Division of Viral Diseases, Centers for Disease Control and Prevention | Krista Queen, Yan Li, Ying Tao, Jing Zhang, Anna Uehara, Anna Montmayeur, Clinton R. Paden, Peter W. Cook, Rachel Marine, Mili Sheth, Jasmine Padilla, Sarah Nobles, Mark Burroughs, Lori Rowe, Haibin Wang, Ben L. Rambo-Martin, Dhwani Batra, Justin Lee, Suxiang Tong |
| EPI_ISL_903912 | TN Division of Laboratory Services | Genomics and Discovery, Respiratory Viruses Branch, Division of Viral Diseases, Centers for Disease Control and Prevention | Krista Queen, Yan Li, Ying Tao, Jing Zhang, Anna Uehara, Anna Montmayeur, Clinton R. Paden, Peter W. Cook, Rachel Marine, Mili Sheth, Jasmine Padilla, Sarah Nobles, Mark Burroughs, Lori Rowe, Haibin Wang, Ben L. Rambo-Martin, Dhwani Batra, Justin Lee, Suxiang Tong |
| EPI_ISL_903920 | IA State Hygienic Laboratory | Genomics and Discovery, Respiratory Viruses Branch, Division of Viral Diseases, Centers for Disease Control and Prevention | Krista Queen, Yan Li, Ying Tao, Jing Zhang, Anna Uehara, Anna Montmayeur, Clinton R. Paden, Peter W. Cook, Rachel Marine, Mili Sheth, Jasmine Padilla, Sarah Nobles, Mark Burroughs, Lori Rowe, Haibin Wang, Ben L. Rambo-Martin, Dhwani Batra, Justin Lee, Suxiang Tong |
| EPI_ISL_903922 | DC Public Health Lab/ Dept. of Forensic Sciences | Genomics and Discovery, Respiratory Viruses Branch, Division of Viral Diseases, Centers for Disease Control and Prevention | Krista Queen, Yan Li, Ying Tao, Jing Zhang, Anna Uehara, Anna Montmayeur, Clinton R. Paden, Peter W. Cook, Rachel Marine, Mili Sheth, Jasmine Padilla, Sarah Nobles, Mark Burroughs, Lori Rowe, Haibin Wang, Ben L. Rambo-Martin, Dhwani Batra, Justin Lee, Suxiang Tong |
| EPI_ISL_903924 | TN Division of Laboratory Services | Genomics and Discovery, Respiratory Viruses Branch, Division of Viral Diseases, Centers for Disease Control and Prevention | Krista Queen, Yan Li, Ying Tao, Jing Zhang, Anna Uehara, Anna Montmayeur, Clinton R. Paden, Peter W. Cook, Rachel Marine, Mili Sheth, Jasmine Padilla, Sarah Nobles, Mark Burroughs, Lori Rowe, Haibin Wang, Ben L. Rambo-Martin, Dhwani Batra, Justin Lee, Suxiang Tong |
| EPI_ISL_903927 | IA State Hygienic Laboratory | Genomics and Discovery, Respiratory Viruses Branch, Division of Viral Diseases, Centers for Disease Control and Prevention | Krista Queen, Yan Li, Ying Tao, Jing Zhang, Anna Uehara, Anna Montmayeur, Clinton R. Paden, Peter W. Cook, Rachel Marine, Mili Sheth, Jasmine Padilla, Sarah Nobles, Mark Burroughs, Lori Rowe, Haibin Wang, Ben L. Rambo-Martin, Dhwani Batra, Justin Lee, Suxiang Tong |
| EPI_ISL_903939 | TN Division of Laboratory Services | Genomics and Discovery, Respiratory Viruses Branch, Division of Viral Diseases, Centers for Disease Control and Prevention | Krista Queen, Yan Li, Ying Tao, Jing Zhang, Anna Uehara, Anna Montmayeur, Clinton R. Paden, Peter W. Cook, Rachel Marine, Mili Sheth, Jasmine Padilla, Sarah Nobles, Mark Burroughs, Lori Rowe, Haibin Wang, Ben L. Rambo-Martin, Dhwani Batra, Justin Lee, Suxiang Tong |

| | | | |
|---|---|---|---|
| EPI_ISL_903969, EPI_ISL_903973 | DC Public Health Lab/ Dept. of Forensic Sciences | Genomics and Discovery, Respiratory Viruses Branch, Division of Viral Diseases, Centers for Disease Control and Prevention | Krista Queen, Yan Li, Ying Tao, Jing Zhang, Anna Uehara, Anna Montmayeur, Clinton R. Paden, Peter W. Cook, Rachel Marine, Mili Sheth, Jasmine Padilla, Sarah Nobles, Mark Burroughs, Lori Rowe, Haibin Wang, Ben L. Rambo-Martin, Dhwani Batra, Justin Lee, Suxiang Tong |
| EPI_ISL_903980 | Institute of Virology, Biomedical Research Center of the Slovak Academy of Sciences, Bratislava | Faculty of Natural Sciences, Comenius University, Bratislava | Viktória abanová, Kristína Boršová, Broa Brejová, Viktória Hodorová, Sabina Fumaová Havlíková, Juraj Kopáek, Martina Liková, ubomíra Lukáiková, Martina Neboháová, Monika Sláviková, Tomáš Vina, Jozef Nosek, Boris Klempa |
| EPI_ISL_903981 | Institute of Virology, Biomedical Research Center of the Slovak Academy of Sciences, Bratislava | Faculty of Natural Sciences, Comenius University, Bratislava | Kristína Boršová, Viktória abanová, Broa Brejová, Viktória Hodorová, Sabina Fumaová Havlíková, Juraj Kopáek, Martina Liková, ubomíra Lukáiková, Martina Neboháová, Monika Sláviková, Tomáš Vina, Boris Klempa, Jozef Nosek |
| EPI_ISL_903982 | Institute of Virology, Biomedical Research Center of the Slovak Academy of Sciences, Bratislava | Faculty of Natural Sciences, Comenius University, Bratislava | Viktória abanová, Kristína Boršová, Broa Brejová, Viktória Hodorová, Sabina Fumaová Havlíková, Juraj Kopáek, Martina Liková, ubomíra Lukáiková, Martina Neboháová, Monika Sláviková, Tomáš Vina, Jozef Nosek, Boris Klempa |
| EPI_ISL_903983 | Institute of Virology, Biomedical Research Center of the Slovak Academy of Sciences, Bratislava | Faculty of Natural Sciences, Comenius University, Bratislava | Kristína Boršová, Viktória abanová, Broa Brejová, Viktória Hodorová, Sabina Fumaová Havlíková, Juraj Kopáek, Martina Liková, ubomíra Lukáiková, Martina Neboháová, Monika Sláviková, Tomáš Vina, Boris Klempa, Jozef Nosek |
| EPI_ISL_903984 | Institute of Virology, Biomedical Research Center of the Slovak Academy of Sciences, Bratislava | Faculty of Natural Sciences, Comenius University, Bratislava | Viktória abanová, Kristína Boršová, Broa Brejová, Viktória Hodorová, Sabina Fumaová Havlíková, Juraj Kopáek, Martina Liková, ubomíra Lukáiková, Martina Neboháová, Monika Sláviková, Tomáš Vina, Jozef Nosek, Boris Klempa |
| EPI_ISL_903985 | Institute of Virology, Biomedical Research Center of the Slovak Academy of Sciences, Bratislava | Faculty of Natural Sciences, Comenius University, Bratislava | Kristína Boršová, Viktória abanová, Broa Brejová, Viktória Hodorová, Sabina Fumaová Havlíková, Juraj Kopáek, Martina Liková, ubomíra Lukáiková, Martina Neboháová, Monika Sláviková, Tomáš Vina, Boris Klempa, Jozef Nosek |
| EPI_ISL_903986, EPI_ISL_903987, EPI_ISL_903988 | Institute of Virology, Biomedical Research Center of the Slovak Academy of Sciences, Bratislava | Faculty of Natural Sciences, Comenius University, Bratislava | Broa Brejová, Viktória abanová, Kristína Boršová, Viktória Hodorová, Sabina Fumaová Havlíková, Juraj Kopáek, Martina Liková, ubomíra Lukáiková, Martina Neboháová, Monika Sláviková, Tomáš Vina, Jozef Nosek, Boris Klempa |
| EPI_ISL_903989 | Institute of Virology, Biomedical Research Center of the Slovak Academy of Sciences, Bratislava | Faculty of Natural Sciences, Comenius University, Bratislava | Viktória abanová, Kristína Boršová, Broa Brejová, Viktória Hodorová, Sabina Fumaová Havlíková, Juraj Kopáek, Martina Liková, ubomíra Lukáiková, Martina Neboháová, Monika Sláviková, Tomáš Vina, Jozef Nosek, Boris Klempa |
| EPI_ISL_903990 | Institute of Virology, Biomedical Research Center of the Slovak Academy of Sciences, Bratislava | Faculty of Natural Sciences, Comenius University, Bratislava | Kristína Boršová, Viktória abanová, Broa Brejová, Viktória Hodorová, Sabina Fumaová Havlíková, Juraj Kopáek, Martina Liková, ubomíra Lukáiková, Martina Neboháová, Monika Sláviková, Tomáš Vina, Boris Klempa, Jozef Nosek |
| EPI_ISL_903991 | Institute of Virology, Biomedical Research Center of the Slovak Academy of Sciences, Bratislava | Faculty of Natural Sciences, Comenius University, Bratislava | Viktória abanová, Kristína Boršová, Broa Brejová, Viktória Hodorová, Sabina Fumaová Havlíková, Juraj Kopáek, Martina Liková, ubomíra Lukáiková, Martina Neboháová, Monika Sláviková, Tomáš Vina, Jozef Nosek, Boris Klempa |
| EPI_ISL_903992 | Institute of Virology, Biomedical Research Center of the Slovak Academy of Sciences, Bratislava | Faculty of Natural Sciences, Comenius University, Bratislava | Kristína Boršová, Viktória abanová, Broa Brejová, Viktória Hodorová, Sabina Fumaová Havlíková, Juraj Kopáek, Martina Liková, ubomíra Lukáiková, Martina Neboháová, Monika Sláviková, Tomáš Vina, Boris Klempa, Jozef Nosek |
| EPI_ISL_903993 | Institute of Virology, Biomedical Research Center of the Slovak Academy of Sciences, Bratislava | Faculty of Natural Sciences, Comenius University, Bratislava | Viktória abanová, Kristína Boršová, Broa Brejová, Viktória Hodorová, Sabina Fumaová Havlíková, Juraj Kopáek, Martina Liková, ubomíra Lukáiková, Martina Neboháová, Monika Sláviková, Tomáš Vina, Jozef Nosek, Boris Klempa |
| EPI_ISL_903994 | Institute of Virology, Biomedical Research Center of the Slovak Academy of Sciences, Bratislava | Faculty of Natural Sciences, Comenius University, Bratislava | Kristína Boršová, Viktória abanová, Broa Brejová, Viktória Hodorová, Sabina Fumaová Havlíková, Juraj Kopáek, Martina Liková, ubomíra Lukáiková, Martina Neboháová, Monika Sláviková, Tomáš Vina, Boris Klempa, Jozef Nosek |
| EPI_ISL_903995 | Institute of Virology, Biomedical Research Center of the Slovak Academy of Sciences, Bratislava | Faculty of Natural Sciences, Comenius University, Bratislava | Viktória abanová, Kristína Boršová, Broa Brejová, Viktória Hodorová, Sabina Fumaová Havlíková, Juraj Kopáek, Martina Liková, ubomíra Lukáiková, Martina Neboháová, Monika Sláviková, Tomáš Vina, Jozef Nosek, Boris Klempa |
| EPI_ISL_903996 | Institute of Virology, Biomedical Research Center of the Slovak Academy of Sciences, Bratislava | Faculty of Natural Sciences, Comenius University, Bratislava | Kristína Boršová, Viktória abanová, Broa Brejová, Viktória Hodorová, Sabina Fumaová Havlíková, Juraj Kopáek, Martina Liková, ubomíra Lukáiková, Martina Neboháová, Monika Sláviková, Tomáš Vina, Boris Klempa, Jozef Nosek |
| EPI_ISL_903997, EPI_ISL_903998, EPI_ISL_903999 | Institute of Virology, Biomedical Research Center of the Slovak Academy of Sciences, Bratislava | Faculty of Natural Sciences, Comenius University, Bratislava | Broa Brejová, Viktória abanová, Kristína Boršová, Viktória Hodorová, Sabina Fumaová Havlíková, Juraj Kopáek, Martina Liková, ubomíra Lukáiková, Martina Neboháová, Monika Sláviková, Tomáš Vina, Jozef Nosek, Boris Klempa |
| EPI_ISL_904000 | Institute of Virology, Biomedical Research Center of the Slovak Academy of Sciences, Bratislava | Faculty of Natural Sciences, Comenius University, Bratislava | Viktória abanová, Kristína Boršová, Broa Brejová, Viktória Hodorová, Sabina Fumaová Havlíková, Juraj Kopáek, Martina Liková, ubomíra Lukáiková, Martina Neboháová, Monika Sláviková, Tomáš Vina, Jozef Nosek, Boris Klempa |
| EPI_ISL_904001 | Institute of Virology, Biomedical Research Center of the Slovak Academy of Sciences, Bratislava | Faculty of Natural Sciences, Comenius University, Bratislava | Kristína Boršová, Viktória abanová, Broa Brejová, Viktória Hodorová, Sabina Fumaová Havlíková, Juraj Kopáek, Martina Liková, ubomíra Lukáiková, Martina Neboháová, Monika Sláviková, Tomáš Vina, Boris Klempa, Jozef Nosek |
| EPI_ISL_904002 | Institute of Virology, Biomedical Research Center of the Slovak Academy of Sciences, Bratislava | Faculty of Natural Sciences, Comenius University, Bratislava | Broa Brejová, Viktória abanová, Kristína Boršová, Viktória Hodorová, Sabina Fumaová Havlíková, Juraj Kopáek, Martina Liková, ubomíra Lukáiková, Martina Neboháová, Monika Sláviková, Tomáš Vina, Jozef Nosek, Boris Klempa |
| EPI_ISL_904003 | Institute of Virology, Biomedical Research Center of the Slovak Academy of Sciences, Bratislava | Faculty of Natural Sciences, Comenius University, Bratislava | Viktória abanová, Kristína Boršová, Broa Brejová, Viktória Hodorová, Sabina Fumaová Havlíková, Juraj Kopáek, Martina Liková, ubomíra Lukáiková, Martina Neboháová, Monika Sláviková, Tomáš Vina, Jozef Nosek, Boris Klempa |
| EPI_ISL_904151, EPI_ISL_904188, EPI_ISL_904189, EPI_ISL_904231, EPI_ISL_904235, EPI_ISL_904248, EPI_ISL_904264, EPI_ISL_904278, EPI_ISL_904279, EPI_ISL_904291, EPI_ISL_904292, EPI_ISL_904293, EPI_ISL_904294, EPI_ISL_904295, EPI_ISL_904296, EPI_ISL_904297, EPI_ISL_904298, EPI_ISL_904299, EPI_ISL_904315, EPI_ISL_904370, EPI_ISL_904375, EPI_ISL_904497, EPI_ISL_904498, EPI_ISL_904499, EPI_ISL_904500, EPI_ISL_904501, EPI_ISL_904502, EPI_ISL_904503, EPI_ISL_904504, EPI_ISL_904505, EPI_ISL_904616, EPI_ISL_904617 | | | |
| see above | Dutch COVID-19 response team | Erasmus Medical Center | Bas Oude Munnink, Reina Sikkema, David Nieuwenhuijse, Irina Chestakova, Anne van der Linden, Marjan Boter, Emmanuelle Munger, Corine GeurtsvanKessel, Annemiek van der Eijk, Richard Molenkamp, Marion Koopmans, on behalf of the Dutch national COVID-19 response team. |
| EPI_ISL_904639 | Servicio de Microbiología, Laboratori Clínic Metropolitana Nord. Hospital Universitari Germans Trias i Pujol. Institut d'Investigació en Ciències de la Salut Germans Trias i Pujol (IGTP) | IrsiCaixa - Can Ruti CovidSeq | Marc Noguera-Julian, Mariona Parera, Maria Casadellà, Pilar Armengol, Francesc Catala-Moll, Roger Paredes, Bonaventura Clotet Elisa Martró, Verónica Saludes, Anna Not, Ana Pérez, Montserrat Giménez, Ignacio Blanco, Cristina Casañ, Antoni E. Bordoy, Adrián Antuori |
| EPI_ISL_904748, EPI_ISL_904749, EPI_ISL_904750, EPI_ISL_904766, EPI_ISL_904796, EPI_ISL_904889, EPI_ISL_904896, EPI_ISL_904897, EPI_ISL_904898, EPI_ISL_904900, EPI_ISL_904902, EPI_ISL_904905, EPI_ISL_905091, EPI_ISL_905126, EPI_ISL_905135, EPI_ISL_905140, EPI_ISL_905144, EPI_ISL_905145, EPI_ISL_905146, EPI_ISL_905147, EPI_ISL_905148, EPI_ISL_905152, EPI_ISL_905155, EPI_ISL_905158, EPI_ISL_905165, EPI_ISL_905166, EPI_ISL_905168, EPI_ISL_905170, EPI_ISL_905172, EPI_ISL_905173, EPI_ISL_905181, EPI_ISL_905182, EPI_ISL_905183, EPI_ISL_905200, EPI_ISL_905211, EPI_ISL_905212, EPI_ISL_905216, EPI_ISL_905217, EPI_ISL_905220, EPI_ISL_905221, EPI_ISL_905222, EPI_ISL_905223, EPI_ISL_905228, EPI_ISL_905230, EPI_ISL_905231, EPI_ISL_905295, EPI_ISL_905297, EPI_ISL_905414, EPI_ISL_905484, EPI_ISL_905537, EPI_ISL_905552, EPI_ISL_905606, EPI_ISL_905650, EPI_ISL_905658 | | | |
| see above | Dutch COVID-19 response team | National Institute for Public Health and the Environment (RIVM) | Adam Meijer, Harry Vennema, Dirk Eggink, Jeroen Cremer, Sharon van den Brink, Bas van der Veer, AnneMarie van den Brandt, Florian Zwagemaker, Dennis Schmitz, Chantal Reusken, on behalf of the national COVID-19 response team |
| EPI_ISL_905731 | Academic Hospital of Gadjah Mada University (RSA UGM) | Genetics Working Group (Pokja Genetik) Faculty of Medicine, Public Health and Nursing Universitas Gadjah Mada (FK-KMK UGM); Disease Investigation Center Wates Ministry of Agriculture Indonesia; Department of Microbiology FK-KMK UGM; Laboratorium Diagnostik Yayasan Tahija World Mosquito Program (WMP) Yogyakarta Center for Tropical Medicine FK-KMK UGM; Integrated Research center FK-KMK UGM; Department of Computer Science and Electronics FMIPA UGM | Gunadi, Hendra Wibawa, Marcellus, Mohamad S. Hakim, Edwin W. Daniwijaya, Ludhang P. Rizki, Endah Supriyati, Eggi Arguni, Titik Nuryastuti, Tri Wibawa, Dwi AA Nugrahaningsih, Afiahayati, Siswanto, Kristy Iskandar,Nungki Anggorowati, Dyah Ayu Puspitarani, Dwiki afandy |
| EPI_ISL_905752, EPI_ISL_905753 | Laboratorium Mikrobiologiczne SYNEVO | National Institute of Public Health - National Institute of Hygiene | Wokowicz Tomasz, Zacharczuk Katarzyna |
| EPI_ISL_905902, EPI_ISL_905903, EPI_ISL_905904, EPI_ISL_905905, EPI_ISL_905906 | OHSU Lab Services Molecular Microbiology Lab | Oregon SARS-CoV-2 Genome Sequencing Center | Brendan L. O'Connell, Sally Grindstaff, Kayla Carter, Ruth V. Nichols, Alec J. Hirsch, Donna Hansel, Guang Fan, Xuan, Qin, Daniel N. Streblow, William B. Messer, Andrew C. Adey, Benjamin N. Bimber, Brian J. O'Roak |
| EPI_ISL_906050 | RSA UGM | Genetics Working Group (Pokja Genetik) Faculty of Medicine, Public Health and Nursing Universitas Gadjah Mada (FK-KMK UGM); Disease Investigation Center Wates Ministry of | Gunadi, Hendra Wibawa, Marcellus, Mohamad S. Hakim, Edwin W. Daniwijaya, Ludhang P. Rizki, Endah Supriyati, Eggi Arguni, Titik Nuryastuti, Tri Wibawa, Dwi AA Nugrahaningsih, Afiahayati, Siswanto,Kristy Iskandar,Nungki Anggorowati, Susan Simanjaya, Kemala Athollah |

| | | | |
|---|---|---|---|
| | | Agriculture Indonesia; Department of Microbiology FK-KMK UGM; Laboratorium Diagnostik Yayasan Tahija World Mosquito Program (WMP) Yogyakarta Center for Tropical Medicine FK-KMK UGM; Integrated Research center FK-KMK UGM; Department of Computer Science and Electronics FMIPA UGM | |
| EPI_ISL_906051 | RSA UGM | Genetics Working Group (Pokja Genetik) Faculty of Medicine, Public Health and Nursing Universitas Gadjah Mada (FK-KMK UGM); Disease Investigation Center Wates Ministry of Agriculture Indonesia; Department of Microbiology FK-KMK UGM; Laboratorium Diagnostik Yayasan Tahija World Mosquito Program (WMP) Yogyakarta Center for Tropical Medicine FK-KMK UGM; Integrated Research center FK-KMK UGM; Department of Computer Science and Electronics FMIPA UGM | Gunadi, Hendra Wibawa, Marcellus, Mohamad S. Hakim, Edwin W. Daniwijaya, Ludhang P. Rizki, Endah Supriyati, Eggi Arguni, Titik Nuryastuti, Tri Wibawa, Dwi AA Nugrahaningsih, Afiahayati, Siswanto, Kristy Iskandar, Nungki Anggorowati, Alvin Santoso Kalim, Dwiki Afandy |
| EPI_ISL_906052 | RSA UGM | Genetics Working Group (Pokja Genetik) Faculty of Medicine, Public Health and Nursing Universitas Gadjah Mada (FK-KMK UGM); Disease Investigation Center Wates Ministry of Agriculture Indonesia; Department of Microbiology FK-KMK UGM; Laboratorium Diagnostik Yayasan Tahija World Mosquito Program (WMP) Yogyakarta Center for Tropical Medicine FK-KMK UGM; Integrated Research center FK-KMK UGM; Department of Computer Science and Electronics FMIPA UGM | Gunadi, Hendra Wibawa, Marcellus, Mohamad S. Hakim, Edwin W. Daniwijaya, Ludhang P. Rizki, Endah Supriyati, Eggi Arguni, Titik Nuryastuti, Tri Wibawa, Dwi AA Nugrahaningsih, Afiahayati, Siswanto, Kristy Iskandar, Nungki Anggorowati, William Widitjiarso, Untung Riawan |
| EPI_ISL_906061 | Tilia Laboratories s.r.o. | Tilia Laboratories s.r.o. | Sona Pekova, MD, PhD. |
| EPI_ISL_906083 | Child Health Research Foundation | Child Health Research Foundation | Senjuti Saha, Arif Mohammad Tanmoy, Sharmistha Goswami, Afroza Akter Tanni, Syed Muktadir Al Sium, Roly Malaker, Md Hafizur Rahman, Samir K Saha |
| EPI_ISL_906548 | Laboratorio de Salud Pública Fronterizo de Arauca | Instituto Nacional de Salud- Dirección de Investigación en Salud Pública, Universidad de los Andes- Applied genomics research group, Vicerrectoria de Investigación y Creación, Universidad de los Andes- Systems and Computing Engineering Department | Katherine Laiton-Donato, Diego A. Álvarez-Díaz, Carlos Franco-Muñoz, Mauricio Pacheco-Montealegre, Héctor Alejandro Ruiz-Moreno, Maria T. Herrera-Sepúlveda, Diego Andrés Prada, Jhonnatan Reales-González, Sheryll Corchuelo, Julian Naizaque, Gerardo Santamaría Jorge Duitama, Laura Natalia Gonzalez, Jorge Ivan Diaz, Silvia Restrepo-Restrepo, Magdalena Wiesner, Martha Lucia Ospina Martinez, Marcela Mercado-Reyes |
| EPI_ISL_906554 | Laboratorio Bienestar | Instituto Nacional de Salud- Dirección de Investigación en Salud Pública, Universidad de los Andes- Applied genomics research group, Vicerrectoria de Investigación y Creación, Universidad de los Andes- Systems and Computing Engineering Department | Katherine Laiton-Donato, Diego A. Álvarez-Díaz, Carlos Franco-Muñoz, Mauricio Pacheco-Montealegre, Héctor Alejandro Ruiz-Moreno, Maria T. Herrera-Sepúlveda, Diego Andrés Prada, Jhonnatan Reales-González, Sheryll Corchuelo, Julian Naizaque, Gerardo Santamaría Jorge Duitama, Laura Natalia Gonzalez, Jorge Ivan Diaz, Silvia Restrepo-Restrepo, Magdalena Wiesner, Martha Lucia Ospina Martinez, Marcela Mercado-Reyes |
| EPI_ISL_906569, EPI_ISL_906570 | Maine Health and Environmental Testing Laboratory (Maine HETL) | Tewhey Lab, The Jackson Laboratory | Matluk,N., Dewey,H., Iosue,F., Barter,M., Lynch,R., Munger,H. and Tewhey,R. |
| EPI_ISL_906726 | Hematology Laboratory, Section of Molecular Diagnostics, University Clinical Centre, Medical University of Gdansk | Laboratory of Recombinant Vaccines | Lukasz Rabalski, Maciej Kosinski, Maciej Grzybek, Adam Sodol, Aneta Szulc, Krzysztof Lewandowski, Ewa Milosz, Marlena Robakowska, Boguslaw Szewczyk, Krystyna Bienkowska-Szewczyk |
| EPI_ISL_906849 | Respiratory Viruses Branch, Centers for Disease Control and Prevention | Respiratory Viruses Branch, Centers for Disease Control and Prevention | Tao,Y., Li,Y., Zhang,J., Queen,K., Uehara,A., Cook,P., Paden,C.R., Wang,H., Tong,S. |
| EPI_ISL_907086, EPI_ISL_907087, EPI_ISL_907088, EPI_ISL_907089, EPI_ISL_907090, EPI_ISL_907091, EPI_ISL_907092, EPI_ISL_907094, EPI_ISL_907095, EPI_ISL_907096, EPI_ISL_907097, EPI_ISL_907098, EPI_ISL_907099, EPI_ISL_907100 | | | |
| see above | Cancer Biology Department, National Cancer Institute | Cancer Biology Department, National Cancer Institute | Zekri,A.N., Sedawy,M.G., Ahmed,O.S., Hafez,M.M., Soliman,H.K., Bahnassy,A.A., Elhosieny,F.W., Gad,A.E., Hamdy,M.S., Soliman,M.S., Soliman,L., Abouelhoda,M. |
| EPI_ISL_909966, EPI_ISL_909968, EPI_ISL_909975, EPI_ISL_909977, EPI_ISL_910004, EPI_ISL_910006 | A. Krumbholz, Labor Dr. Krause und Kollegen MVZ GmbH, Kiel | Charité Universitätsmedizin Berlin, Institut für Virologie | Victor M Corman, Tobias Bleicker, Julia Tesch, Barbara Mühlemann, Jörn Beheim-Schwarzbach, Talitha Veith, Julia Schneider, Cornelia Schlee, Tomasz Zemojtel, Terry Jones, Christian Drosten |
| EPI_ISL_910634 | Laboratoire national de sante, Microbiology, Virology | Laboratoire national de sante, Microbiology, Microbial Genomics Platform | Anke Wienecke-Baldacchino, Catherine Ragimbeau,Jessica Tapp, Fatu Djabi, Lise Pignon, Raoul Salmon, Tamir Abdelrahman |
| EPI_ISL_911529 | M Health Fairview | Minnesota Department of Health, Public Health Laboratory | Alexandra Lorentz, Jacob Garfin, Matt Plumb, and Xiong Wang |
| EPI_ISL_911702 | Alaska State Virology Laboratory | Alaska State Virology Laboratory | Stephanie DeRonde, Lisa Smith, Ph.D., Jack Chen, Ph.D. |
| EPI_ISL_911707 | Academic Hospital of Gadjah Mada University (RSA UGM) | Genetics Working Group (Pokja Genetik) Faculty of Medicine, Public Health and Nursing Universitas Gadjah Mada (FK-KMK UGM); Disease Investigation Center Wates Ministry of Agriculture Indonesia; Department of Microbiology FK-KMK UGM; Laboratorium Diagnostik Yayasan Tahija World Mosquito Program (WMP) Yogyakarta Center for Tropical Medicine FK-KMK UGM; Integrated Research center FK-KMK UGM; Department of Computer Science and Electronics FMIPA UGM | Gunadi, Hendra Wibawa, Marcellus, Mohamad S. Hakim, Edwin W. Daniwijaya, Ludhang P. Rizki, Endah Supriyati, Eggi Arguni, Titik Nuryastuti, Tri Wibawa, Dwi AA Nugrahaningsih, Afiahayati, Siswanto, Kristy Iskandar, Nungki Anggorowati, Untung Riawan, Kemala Athollah |
| EPI_ISL_911709 | Academic Hospital of Gadjah Mada University (RSA UGM) | Genetics Working Group (Pokja Genetik) Faculty of Medicine, Public Health and Nursing Universitas Gadjah Mada (FK-KMK UGM); Disease Investigation Center Wates Ministry of Agriculture Indonesia; Department of Microbiology FK-KMK UGM; Laboratorium Diagnostik Yayasan Tahija World Mosquito Program (WMP) Yogyakarta Center for Tropical Medicine FK-KMK UGM; Integrated Research center FK-KMK UGM; Department of Computer Science and Electronics FMIPA UGM | Gunadi, Hendra Wibawa, Marcellus, Mohamad S. Hakim, Edwin W. Daniwijaya, Ludhang P. Rizki, Endah Supriyati, Eggi Arguni, Titik Nuryastuti, Tri Wibawa, Dwi AA Nugrahaningsih, Afiahayati, Siswanto, Kristy Iskandar, Nungki Anggorowati, Dyah Ayu Puspitarani, Dwiki afandy |
| EPI_ISL_911827 | Johns Hopkins Hospital Department of Pathology | Johns Hopkins Hospital Department of Pathology | C. Paul Morris, Chun Huai Luo, Adannaya Amadi, Matthew Schwartz, Nicholas Gallagher, Heba H. Mostafa |
| EPI_ISL_911971 | Seattle Flu Study | Seattle Flu Study | Deborah A. Nickerson, Chris D. Frazar, Jover Lee, Benjamin Pelle, Erica Ryke, Matthew Richardson, Amanda Adler, Elisabeth Brandstetter, Peter D. Han, Kairsten Fay, Misja Ilcisin, Kirsten Lacombe, Thomas R. Sibley, Melissa Truong, Caitlin R. Wolf, Michael Boeckh, Janet A. Englund, Michael Famulare, Barry R. Lutz, Mark J. Rieder, Lea M. Starita, Matthew Thompson, Jay Shendure, Trevor Bedford, Helen Y. Chu |
| EPI_ISL_912288 | Hospital General Universitario Gregorio Marañón | SeqCOVID-SPAIN consortium / IBV (CSIC) | Darío García de Viedma, Laura Pérez-Lago, Pedro J Sola-Campoy, Sergio Buenestado-Serrano, Marta Herranz, Victor Manuel de la Cueva, Julia Suárez, Pilar Catalán, Patricia Muñoz and SeqCOVID-SPAIN consortium |

| | | | |
|---|---|---|---|
| EPI_ISL_912364, EPI_ISL_912365, EPI_ISL_912367, EPI_ISL_912370, EPI_ISL_912371, EPI_ISL_912372, EPI_ISL_912373 | Fondation Congolaise pour la recherche medicale (FCRM), Francine Ntoumi | NGS Competence Center Tuebingen, Institut für Medizinische Mikrobiologie und Hygiene, Universitaetsklinikum Tübingen | Angel Angelov |
| EPI_ISL_912408 | KU Leuven, Rega Institute, Clinical and Epidemiological Virology | KU Leuven, Rega Institute, Clinical and Epidemiological Virology | Tony Wawina-Bokalanga, Bert Vanmechelen, Joan Marti-Carerras, Piet Maes |
| EPI_ISL_912483, EPI_ISL_912507 | NHLS Universitas Academic | UFS Virology | PA Bester, MM Nyaga, P Nthiga, MT Mogotsi, D Goedhals, T de Oliveira |
| EPI_ISL_912893, EPI_ISL_912894, EPI_ISL_912895, EPI_ISL_912902, EPI_ISL_912903, EPI_ISL_912904, EPI_ISL_912905, EPI_ISL_912927, EPI_ISL_912928, EPI_ISL_912930, EPI_ISL_912935, EPI_ISL_912938, EPI_ISL_912939, EPI_ISL_912941, EPI_ISL_912982, EPI_ISL_912983, EPI_ISL_912984, EPI_ISL_912985, EPI_ISL_912986, EPI_ISL_912987, EPI_ISL_912988, EPI_ISL_912989, EPI_ISL_912990, EPI_ISL_912991, EPI_ISL_912992, EPI_ISL_912993, EPI_ISL_912994, EPI_ISL_912995, EPI_ISL_912996, EPI_ISL_912997, EPI_ISL_912998, EPI_ISL_912999, EPI_ISL_913000, EPI_ISL_913001, EPI_ISL_913002, EPI_ISL_913003, EPI_ISL_913004, EPI_ISL_913005, EPI_ISL_913006, EPI_ISL_913007, EPI_ISL_913008, EPI_ISL_913009, EPI_ISL_913010, EPI_ISL_913011 | | | |
| see above | Hôpital Henri Mondor | Department of Virology, Henri Mondor University Hospital, Assistance Publique Hôpitaux de Paris, Université Paris-Est Créteil, INSERM U955 | Christophe Rodriguez, Slim Fourati, Vanessa Demontant, Guillaume Gricourt, Melissa N'Debi, Alexandre Soulier, Elisabeth Trawinski, Jean-Michel Pawlotsky |
| EPI_ISL_913282, EPI_ISL_913287, EPI_ISL_913288, EPI_ISL_913289, EPI_ISL_913290 | ABC Labs | The Public Health Agency of Sweden | Anna-Malin Linde, Maria Lind Karlberg, Carlo Berg, Oskar Karlsson Lindsjo, Sofia Stamouli, Reza Advani, Mattias Haukland, Petra Holmstrom, Noura Walai, Petra Edquist, Mia Brytting, Anna Risberg, Karin Tegmark-Wisell |
| EPI_ISL_913294 | Laboratoriemedicin, Klinisk mikrobiologi | The Public Health Agency of Sweden | Anna-Malin Linde, Maria Lind Karlberg, Carlo Berg, Oskar Karlsson Lindsjo, Sofia Stamouli, Reza Advani, Mattias Haukland, Petra Holmstrom, Noura Walai, Petra Edquist, Mia Brytting, Anna Risberg, Karin Tegmark-Wisell |
| EPI_ISL_913302 | Synlab Medilab, Mikrobiologi | The Public Health Agency of Sweden | Anna-Malin Linde, Maria Lind Karlberg, Carlo Berg, Oskar Karlsson Lindsjo, Sofia Stamouli, Reza Advani, Mattias Haukland, Petra Holmstrom, Noura Walai, Petra Edquist, Mia Brytting, Anna Risberg, Karin Tegmark-Wisell |
| EPI_ISL_913316 | ABC Labs | The Public Health Agency of Sweden | Anna-Malin Linde, Maria Lind Karlberg, Carlo Berg, Oskar Karlsson Lindsjo, Sofia Stamouli, Reza Advani, Mattias Haukland, Petra Holmstrom, Noura Walai, Petra Edquist, Mia Brytting, Anna Risberg, Karin Tegmark-Wisell |
| EPI_ISL_913460 | Klinisk mikrobiologi | The Public Health Agency of Sweden | Anna-Malin Linde, Maria Lind Karlberg, Carlo Berg, Oskar Karlsson Lindsjo, Sofia Stamouli, Reza Advani, Mattias Haukland, Petra Holmstrom, Noura Walai, Petra Edquist, Mia Brytting, Anna Risberg, Karin Tegmark-Wisell |
| EPI_ISL_913630, EPI_ISL_913631, EPI_ISL_913632, EPI_ISL_913633, EPI_ISL_913636 | Michigan Department of Health and Human Services, Bureau of Laboratories | Michigan Department of Health and Human Services, Bureau of Laboratories | Blankenship HM, Riner D, Soehnlen MK |
| EPI_ISL_913740, EPI_ISL_913741, EPI_ISL_913742, EPI_ISL_913743, EPI_ISL_913744, EPI_ISL_913745 | DC Public Health Lab/ Dept. of Forensic Sciences | DC Public Health Lab/ Dept. of Forensic Sciences | Scott Nguyen, Elizabeth Zelaya, Connie Maza, Monica Mann, Brittany Hamilton, David Payne, Jocelyn Hauser |
| EPI_ISL_913802, EPI_ISL_913825, EPI_ISL_913887 | TGen North | TGen North | "Jolene Bowers, Megan Folkerts, Chris French, Hayley Yaglom, Ashlyn Pfeiffer, Darrin Lemmer, Dave Engelthaler, The Arizona COVID Genomics Union (ACGU)" |
| EPI_ISL_914646 | Santa Clara County Public Health Laboratory | Santa Clara County Public Health Laboratory | Santa Clara County Public Health Department |
| EPI_ISL_914647, EPI_ISL_914662, EPI_ISL_914667, EPI_ISL_914668, EPI_ISL_914669, EPI_ISL_914670, EPI_ISL_914671, EPI_ISL_914672, EPI_ISL_914673, EPI_ISL_914674, EPI_ISL_914675, EPI_ISL_914676, EPI_ISL_914677, EPI_ISL_914678, EPI_ISL_914679, EPI_ISL_914680, EPI_ISL_914681, EPI_ISL_914682, EPI_ISL_914683, EPI_ISL_914684, EPI_ISL_914685, EPI_ISL_914686, EPI_ISL_914687, EPI_ISL_914688, EPI_ISL_914689, EPI_ISL_914690, EPI_ISL_914691, EPI_ISL_914692, EPI_ISL_914693, EPI_ISL_914694, EPI_ISL_914695 | | | |
| see above | Utah Public Health Laboratory | Utah Public Health Laboratory | Erin L. Young, Kelly F. Oakeson, Tara Gallagher |
| EPI_ISL_914808 | HLE - ASOCIACION HOGAR PARA ANCIANOS PRESBITERO JAFET JIMENEZ MORALES DE GRECIA | Inciensa, Instituto Costarricense de Investigación y Enseñanza en Nutrición y Salud | Francisco Duarte, Hebleen Porras, Claudio Soto-Garita, Estela Cordero, Adriana Godínez, Melany Calderón & Mariel López |
| EPI_ISL_914810 | AREA DE SALUD SAN JUAN-SAN DIEGO-CONCEPCION 2 | Inciensa, Instituto Costarricense de Investigación y Enseñanza en Nutrición y Salud | Francisco Duarte, Hebleen Porras, Claudio Soto-Garita, Estela Cordero, Adriana Godínez, Melany Calderón & Mariel López |
| EPI_ISL_914811, EPI_ISL_914812 | AREA DE SALUD GOICOECHEA 2 - CLINICA DR. JIMENEZ NUÑEZ | Inciensa, Instituto Costarricense de Investigación y Enseñanza en Nutrición y Salud | Francisco Duarte, Hebleen Porras, Claudio Soto-Garita, Estela Cordero, Adriana Godínez, Melany Calderón & Mariel López |
| EPI_ISL_914813 | TAMIZAJE COMUNITARIO - PASO CANOAS | Inciensa, Instituto Costarricense de Investigación y Enseñanza en Nutrición y Salud | Francisco Duarte, Hebleen Porras, Claudio Soto-Garita, Estela Cordero, Adriana Godínez, Melany Calderón & Mariel López |
| EPI_ISL_914821, EPI_ISL_914822 | HOSPITAL DR. ENRIQUE BALTODANO BRICEÑO | Inciensa, Instituto Costarricense de Investigación y Enseñanza en Nutrición y Salud | Francisco Duarte, Hebleen Porras, Claudio Soto-Garita, Estela Cordero, Adriana Godínez, Melany Calderón & Mariel López |
| EPI_ISL_914888, EPI_ISL_914889 | Vilnius university hospital Santaros Klinikos, Center of Laboratory Medicine | Vilnius University Hospital Santaros Klinikos | Ingrida Olendraite, Daniel Naumovas, Rimvydas Norvilas, Dovil Ežerskyt, Justinas Šlikas |
| EPI_ISL_915438 | Los Angeles County PHL | Los Angeles County PHL | P. Hemarajata et al. |
| EPI_ISL_918171, EPI_ISL_918175 | Department of Infectious Diseases and Immunology, National Hospital Organization Nagoya Medical Center | Clinical Research Center, National Hospital Organization Nagoya Medical Center | Yoshihiro Nakata, Hirotaka Ode, Mai Kubota, Masakazu Matsuda, Kazuhiro Matsuoka, Miho Nakasuji, Mikiko Mori, Mayumi Imahashi, Yoshiyuki Yokomaku, Yasumasa Iwatani |
| EPI_ISL_918344, EPI_ISL_918345, EPI_ISL_918346, EPI_ISL_918347, EPI_ISL_918354, EPI_ISL_918355 | Institute of Virology, Medical Center, University of Freiburg, Freiburg, Germany | Institute of Virology, Clinial Virus Genomics, Medical Center, University of Freiburg, Freiburg, Germany | Jonas Fuchs, Lisa Kern, Sandra Reuter, Hajo Grundmann, Marcus Panning |
| EPI_ISL_918522 | LACEN - Laboratório Central de Saúde Pública do Para | Evandro Chagas Institute | Santos, M.C.; Silva, A.M.; Junior, W.D.C.; Barbagelata, L.S.; Ferreira, J.A.; Sousa, E.M.A.; da Silva, P.S.; Pinheiro, K.C.; L.C.; Sousa Junior, E.C. |
| EPI_ISL_918555, EPI_ISL_918556 | LACEN - Laboratório Central de Saúde Pública do Amapa | Evandro Chagas Institute | Santos, M.C.; Silva, A.M.; Junior, W.D.C.; Barbagelata, L.S.; Ferreira, J.A.; Sousa, E.M.A.; da Silva, P.S.; Pinheiro, K.C.; L.C.; Sousa Junior, E.C. |
| EPI_ISL_918758, EPI_ISL_918759, EPI_ISL_918858 | University of Birmingham | COVID-19 Genomics UK (COG-UK) Consortium | Institute of Microbiology, University of Birmingham: Claire McMurray, Joanne Stockton, Samuel Nicholls, Radoslaw Poplawski, Will Rowe, Josh Quick, Nicholas Loman. University of Birmingham Testing Laboratory: Celina M Whalley, Andrew Bosworth, Charlotte Poxon, Kasun Wanigasooriya, Oliver Pickles, Mike Kidd, Alex Richter, Andrew D Beggs PHE Heartlands Lab: Husam Osman, Andrew Bosworth. Queen Elizabeth Hospital: Anna Casey |
| EPI_ISL_919243, EPI_ISL_919245, EPI_ISL_919247, EPI_ISL_919248, EPI_ISL_919250, EPI_ISL_919255, EPI_ISL_919256, EPI_ISL_919260, EPI_ISL_919276 | West of Scotland Specialist Virology Centre, NHSGGC / MRC-University of Glasgow Centre for Virus Research | COVID-19 Genomics UK (COG-UK) Consortium | Ana da Silva Filipe, Natasha Johnson, Kathy Smollett, Daniel Mair, Stephen Carmichael, Alice Broos, Lily Tong, Jenna Nichols, Kyriaki Nomikou; Sarah McDonald; Richard Orton, Joseph Hughes, Sreenu Vattipally, David L Robertson; Alasdair MacLean, Rory Gunson; Sharif Shaaban, Matthew Holden; Rachel Blacow, Guy Mollett, Kathy Li, James Shepherd, Antonia Ho, Emma Thomson |
| EPI_ISL_919343, EPI_ISL_919344, EPI_ISL_919379 | Virology Department, Royal Infirmary of Edinburgh, NHS Lothian / School of Biological Sciences, University of Edinburgh / Institute of Genetics and Molecular Medicine, University of Edinburgh | COVID-19 Genomics UK (COG-UK) Consortium | McHugh M, Dewar R, Rooke S, Gallagher M, Balcaza C, O'Toole Á, Scher E, Hill V, McCrone JT, Colquhoun R, Yu X, Jackson B, Rambaut A, Williams TC, Templeton K |
| EPI_ISL_919728, EPI_ISL_919729, EPI_ISL_919730, EPI_ISL_919731, EPI_ISL_919732, EPI_ISL_919733, EPI_ISL_919734, EPI_ISL_919735, EPI_ISL_919736, EPI_ISL_919738, EPI_ISL_919739, EPI_ISL_919740, EPI_ISL_919741, EPI_ISL_919742, EPI_ISL_919743, EPI_ISL_919745, EPI_ISL_919746, EPI_ISL_919747, EPI_ISL_919755, EPI_ISL_919756, EPI_ISL_919757, EPI_ISL_919758, EPI_ISL_919759, EPI_ISL_919768, EPI_ISL_919769, EPI_ISL_919770, EPI_ISL_919771, EPI_ISL_919772, EPI_ISL_919773, EPI_ISL_919774, EPI_ISL_919775 | | | |
| see above | University College London, Great Ormond Street Hospital for Children NHS Foundation Trust, Imperial College Healthcare | COVID-19 Genomics UK (COG-UK) Consortium | Sergi Castellano, Rachel Williams, Mark Kristiansen, Paola Resende Silva, Sunando Roy, Tony Brooks, Helena Tutill, Paola Niola, Patricia Dyal, Charlotte Williams, Leysa Forrest, Yasmin Panchbhaya, Jacqueline Findlay, Samuel Weeks, Julianne Brown, Kathryn Harris, Paul Randell, James Price, Alison |

| | | | |
|---|---|---|---|
| | NHS Trust | | Holmes, Judith Breuer |
| EPI_ISL_920192, EPI_ISL_920475, EPI_ISL_920484, EPI_ISL_920493, EPI_ISL_920512, EPI_ISL_920515, EPI_ISL_920518, EPI_ISL_920519, EPI_ISL_920537, EPI_ISL_920680 | University College London Hospital | COVID-19 Genomics UK (COG-UK) Consortium | Judith Heaney, Matthew Byott, Catherine Houlihan, Dan Frampton, Stuart Kirk, Moira Spyer and Eleni Nastouli |
| EPI_ISL_921198, EPI_ISL_921199, EPI_ISL_921201, EPI_ISL_921205, EPI_ISL_921209, EPI_ISL_921210, EPI_ISL_921211, EPI_ISL_921213, EPI_ISL_921216, EPI_ISL_921217, EPI_ISL_921218, EPI_ISL_921219, EPI_ISL_921220, EPI_ISL_921221, EPI_ISL_921222, EPI_ISL_921223, EPI_ISL_921224, EPI_ISL_921226, EPI_ISL_921322, EPI_ISL_921323, EPI_ISL_921324, EPI_ISL_921325, EPI_ISL_921326, EPI_ISL_921327, EPI_ISL_921328, EPI_ISL_921330, EPI_ISL_921331, EPI_ISL_921332, EPI_ISL_921333, EPI_ISL_921334, EPI_ISL_921336, EPI_ISL_921337, EPI_ISL_921338, EPI_ISL_921339, EPI_ISL_921340, EPI_ISL_921342, EPI_ISL_921343, EPI_ISL_921350, EPI_ISL_921360, EPI_ISL_921362, EPI_ISL_921363, EPI_ISL_921664, EPI_ISL_921727, EPI_ISL_921728, EPI_ISL_921730, EPI_ISL_921731, EPI_ISL_921734, EPI_ISL_921737, EPI_ISL_921738, EPI_ISL_921739, EPI_ISL_921740, EPI_ISL_921744, EPI_ISL_921748, EPI_ISL_921749, EPI_ISL_921750, EPI_ISL_921760, EPI_ISL_921761, EPI_ISL_921762, EPI_ISL_921768, EPI_ISL_921775, EPI_ISL_921780, EPI_ISL_921781, EPI_ISL_921782 | | | |
| see above | Northumbria University / South Tees Hospitals NHS Foundation Trust / North Cumbria Integrated Care NHS Foundation Trust / North Tees and Hartlepool NHS Foundation Trust / Newcastle Hospitals NHS Foundation Trust | COVID-19 Genomics UK (COG-UK) Consortium | Darren L Smith,Andrew Nelson,Matthew Bashton,Greg R Young,Joshua Loh,John Allan,Mohammad A Tariq,Giles S Holt,Gary Black,Wen C Yew,Lynn Dover,Paul Baker,Steve Liggett,Sarah Essex,Jane Greenaway,Debra Padgett,Clive Graham,Garren Scott,Edward Barton,Emma Swindells,Brendan Payne,Jennifer Collins,Yusri Taha,Gary Eltringham |
| EPI_ISL_921783, EPI_ISL_921784, EPI_ISL_921785, EPI_ISL_921794, EPI_ISL_921795, EPI_ISL_921797, EPI_ISL_921803, EPI_ISL_921804, EPI_ISL_921806, EPI_ISL_921807, EPI_ISL_921814, EPI_ISL_921816, EPI_ISL_921817, EPI_ISL_921818, EPI_ISL_921822, EPI_ISL_921823, EPI_ISL_921827, EPI_ISL_921829, EPI_ISL_921830, EPI_ISL_921831, EPI_ISL_921832, EPI_ISL_921835, EPI_ISL_921836, EPI_ISL_921837, EPI_ISL_921838, EPI_ISL_921840, EPI_ISL_921843, EPI_ISL_921844, EPI_ISL_921850 | | | |
| see above | Quadram Institute Bioscience | COVID-19 Genomics UK (COG-UK) Consortium | Dave J. Baker, Gemma L. Kay, Alp Aydin, Thanh Le-Viet, Steven Rudder, Ana P. Tedim, Anastasia Kolyva, Maria Diaz, Leonardo de Oliveira Martins, Nabil-Fareed Alikhan, Lizzie Meadows, Rachael Stanley, Ngozi Elumogo, Muhammed Yasir, Nicholas M. Thomson, Alexander J Trotter, Rachel Gilroy, Samuel Bloomfield, Claire Stuart, Andrew Bell, Reenesh Prakash, Samir Dervisevic, Alison E. Mather, John Wain, Mark Webber, Andrew J. Page, Justin O'Grady |
| EPI_ISL_922172, EPI_ISL_922173, EPI_ISL_922253, EPI_ISL_922269, EPI_ISL_922273, EPI_ISL_922312, EPI_ISL_922315, EPI_ISL_922316, EPI_ISL_922319 | Oxford Viromics, NDM, University of Oxford; Oxford University Hospitals; Basingstoke and North Hampshire Hospital | COVID-19 Genomics UK (COG-UK) Consortium | Tanya Golubchik, David Bonsall, George Macintyre, Amy Trebes, Mariateresa de Cesare, Catrin Moore, Alex Mobbs, Anita Justice, Robert Shaw, Monique Andersson, Timothy Peto, Emma Wise, Nathan Moore, Jessica Lynch, Nick Cortes, Matilde Mori, Stephen Kidd, David Buck, John Todd, Christophe Fraser |
| EPI_ISL_923227, EPI_ISL_923229, EPI_ISL_923230, EPI_ISL_923231, EPI_ISL_923237, EPI_ISL_923240, EPI_ISL_923243, EPI_ISL_923245, EPI_ISL_923246, EPI_ISL_923247, EPI_ISL_923248, EPI_ISL_923249, EPI_ISL_923250, EPI_ISL_923251 | | | |
| see above | Centre for Enzyme Innovation, University of Portsmouth / Translational Research Laboratory, Portsmouth Hospitals NHS Trust | COVID-19 Genomics UK (COG-UK) Consortium | Angela Beckett,Salman Goudarzi,Christopher Fearn,Kate Cook,Katie Loveson,Sharon Glaysher,Scott Elliott,Samuel Robson |
| EPI_ISL_925033, EPI_ISL_925034, EPI_ISL_925035, EPI_ISL_925036, EPI_ISL_925037, EPI_ISL_925038, EPI_ISL_925039, EPI_ISL_925040, EPI_ISL_925041, EPI_ISL_925042, EPI_ISL_925043, EPI_ISL_925044, EPI_ISL_925045, EPI_ISL_925046, EPI_ISL_925047 | | | |
| see above | Laboratory of Molecular Biology, Diagnostyka sp. z o.o. | genXone SA, Research & Development Laboratory | Maciej Sykulski, Grzegorz Nowicki, Monika Makowska-Woniak, Jakub Grabowski, Natalia Drwska-Matelska, ukasz Krych, Micha Kaszuba |
| EPI_ISL_930734 | Utah Public Health Laboratory | Utah Public Health Laboratory | Erin L. Young, Kelly F. Oakeson, Tara Gallagher |
| EPI_ISL_933430, EPI_ISL_933435, EPI_ISL_933479, EPI_ISL_933480 | Lighthouse Lab in Milton Keynes | Wellcome Sanger Institute for the COVID-19 Genomics UK (COG-UK) Consortium | The Lighthouse Lab in Milton Keynes and Alex Alderton, Roberto Amato, Sonia Goncalves, Ewan Harrison, David K. Jackson, Ian Johnston, Dominic Kwiatkowski, Cordelia Langford, John Sillitoe on behalf of the Wellcome Sanger Institute COVID-19 Surveillance Team |
| EPI_ISL_934428, EPI_ISL_934429, EPI_ISL_934430 | ILV Kärnten | Bergthaler laboratory, CeMM Research Center for Molecular Medicine of the Austrian Academy of Sciences | Lukas Endler, Anna Schedl, Thomas Penz, Benedikt Agerer, Maelle Le Moing, Michael Schuster, Bekir Erguner, Jan Laine, Martin Senekowitsch, Christoph Bock, Andreas Bergthaler |
| EPI_ISL_934434, EPI_ISL_934435 | Austrian Agency for Health and Food Safety (AGES) | Bergthaler laboratory, CeMM Research Center for Molecular Medicine of the Austrian Academy of Sciences | Lukas Endler, Anna Schedl, Thomas Penz, Benedikt Agerer, Maelle Le Moing, Michael Schuster, Bekir Erguner, Jan Laine, Martin Senekowitsch, Christoph Bock, Andreas Bergthaler |
| EPI_ISL_934579, EPI_ISL_934580, EPI_ISL_934581, EPI_ISL_934582, EPI_ISL_934583, EPI_ISL_934584, EPI_ISL_934585, EPI_ISL_934586, EPI_ISL_934587, EPI_ISL_934588 | Department of Laboratory Medicine, Division of Clinical Virology, University of Medicine, Vienna | Bergthaler laboratory, CeMM Research Center for Molecular Medicine of the Austrian Academy of Sciences | Lukas Endler, Anna Schedl, Thomas Penz, Benedikt Agerer, Maelle Le Moing, Michael Schuster, Bekir Erguner, Jan Laine, Martin Senekowitsch, Christoph Bock, Andreas Bergthaler |
| EPI_ISL_934644, EPI_ISL_934647 | Department of Microbiology, University Innsbruck | Bergthaler laboratory, CeMM Research Center for Molecular Medicine of the Austrian Academy of Sciences | Lukas Endler, Anna Schedl, Thomas Penz, Benedikt Agerer, Maelle Le Moing, Michael Schuster, Bekir Erguner, Jan Laine, Martin Senekowitsch, Christoph Bock, Andreas Bergthaler |
| EPI_ISL_935033, EPI_ISL_935035 | Laboratory of Molecular Biology, Diagnostyka sp. z o.o. | genXone SA, Research & Development Laboratory | Maciej Sykulski, Grzegorz Nowicki, Monika Makowska-Woniak, Jakub Grabowski, Natalia Drwska-Matelska, ukasz Krych, Micha Kaszuba |
| EPI_ISL_935696, EPI_ISL_935697, EPI_ISL_935698, EPI_ISL_935699, EPI_ISL_935700, EPI_ISL_935701, EPI_ISL_935702, EPI_ISL_935703, EPI_ISL_935704, EPI_ISL_935705, EPI_ISL_935706, EPI_ISL_935707, EPI_ISL_935723, EPI_ISL_935724, EPI_ISL_935725, EPI_ISL_935726, EPI_ISL_935727, EPI_ISL_935728, EPI_ISL_935729, EPI_ISL_935730, EPI_ISL_935731, EPI_ISL_935732, EPI_ISL_935733, EPI_ISL_935734, EPI_ISL_935735, EPI_ISL_935736, EPI_ISL_935737, EPI_ISL_935738, EPI_ISL_935739, EPI_ISL_935740, EPI_ISL_935741, EPI_ISL_935742, EPI_ISL_935743, EPI_ISL_935744, EPI_ISL_935745, EPI_ISL_935746, EPI_ISL_935747, EPI_ISL_935748, EPI_ISL_935749 | | | |
| see above | Houston Health Dept. | Houston Health Dept. | Ryker Penn, Pamela Brown, Adolpho Lara |
| EPI_ISL_936133 | New York Presbyterian Hospital | Wadsworth Center, New York State Department of Health | Kirsten St. George, Daryl M. Lamson, Alexis Russel, Matthew Shudt, Melissa A Leisner, Jonathan Plitnick, Navjot Singh, John Kelly, Erasmus Schneider, Erica Lasek-Nesselquist |
| EPI_ISL_936149, EPI_ISL_936150, EPI_ISL_936151, EPI_ISL_936152, EPI_ISL_936154, EPI_ISL_936155, EPI_ISL_936156, EPI_ISL_936157, EPI_ISL_936158, EPI_ISL_936159, EPI_ISL_936160, EPI_ISL_936161, EPI_ISL_936162, EPI_ISL_936163, EPI_ISL_936164, EPI_ISL_936165, EPI_ISL_936166, EPI_ISL_936167, EPI_ISL_936168, EPI_ISL_936169, EPI_ISL_936170, EPI_ISL_936171, EPI_ISL_936172, EPI_ISL_936174, EPI_ISL_936175, EPI_ISL_936176, EPI_ISL_936177, EPI_ISL_936178, EPI_ISL_936179 | | | |
| see above | MEMORIAL SLOAN KETTERING CANCER CENTER | Wadsworth Center, New York State Department of Health | Kirsten St. George, Daryl M. Lamson, Alexis Russel, Matthew Shudt, Melissa A Leisner, Jonathan Plitnick, Navjot Singh, John Kelly, Erasmus Schneider, Erica Lasek-Nesselquist |
| EPI_ISL_936181 | ALBANY MEDICAL CENTER | Wadsworth Center, New York State Department of Health | Kirsten St. George, Daryl M. Lamson, Alexis Russel, Matthew Shudt, Melissa A Leisner, Jonathan Plitnick, Navjot Singh, John Kelly, Erasmus Schneider, Erica Lasek-Nesselquist |
| EPI_ISL_936697, EPI_ISL_936698, EPI_ISL_936702, EPI_ISL_936704, EPI_ISL_936708, EPI_ISL_936709, EPI_ISL_936710, EPI_ISL_936711, EPI_ISL_936712, EPI_ISL_936713, EPI_ISL_936714, EPI_ISL_936715, EPI_ISL_936716, EPI_ISL_936717, EPI_ISL_936718, EPI_ISL_936719, EPI_ISL_936720, EPI_ISL_936721, EPI_ISL_936722, EPI_ISL_936723, EPI_ISL_936724, EPI_ISL_936725, EPI_ISL_936726, EPI_ISL_936727, EPI_ISL_936728, EPI_ISL_936729, EPI_ISL_936730, EPI_ISL_936731, EPI_ISL_936732, EPI_ISL_936733, EPI_ISL_936734, EPI_ISL_936735, EPI_ISL_936736, EPI_ISL_936737, EPI_ISL_936738, EPI_ISL_936739, EPI_ISL_936740, EPI_ISL_936741, EPI_ISL_936742, EPI_ISL_936743, EPI_ISL_936744, EPI_ISL_936745, EPI_ISL_936746, EPI_ISL_936747, EPI_ISL_936748, EPI_ISL_936749, EPI_ISL_936750, EPI_ISL_936751, EPI_ISL_936752, EPI_ISL_936753, EPI_ISL_936754, EPI_ISL_936755, EPI_ISL_936756, EPI_ISL_936757, EPI_ISL_936758, EPI_ISL_936759, EPI_ISL_936760, EPI_ISL_936761, EPI_ISL_936762, EPI_ISL_936763, EPI_ISL_936764, EPI_ISL_936765, EPI_ISL_936766, EPI_ISL_936767, EPI_ISL_936768, EPI_ISL_936769, EPI_ISL_936770, EPI_ISL_936771, EPI_ISL_936772, EPI_ISL_936773, EPI_ISL_936774, EPI_ISL_936775, EPI_ISL_936776, EPI_ISL_936777, EPI_ISL_936778, EPI_ISL_936779, EPI_ISL_936780, EPI_ISL_936781, EPI_ISL_936782, EPI_ISL_936783, EPI_ISL_936784, EPI_ISL_936785, EPI_ISL_936786, EPI_ISL_936787, EPI_ISL_936788, EPI_ISL_936789, EPI_ISL_936790, EPI_ISL_936791, EPI_ISL_936792, EPI_ISL_936793, EPI_ISL_936794, EPI_ISL_936795, EPI_ISL_936796, EPI_ISL_936797, EPI_ISL_936798, EPI_ISL_936799, EPI_ISL_936800, EPI_ISL_936801, EPI_ISL_936802, EPI_ISL_936803, EPI_ISL_936804, EPI_ISL_936805, EPI_ISL_936806, EPI_ISL_936807, EPI_ISL_936808, EPI_ISL_936809, EPI_ISL_936810, EPI_ISL_936811, EPI_ISL_936812 | | | |
| see above | Northwestern Memorial Hospital | Ozer Lab | Ramon Lorenzo-Redondo, Lacy M. Simons, Chad J. Achenbach, Lawrence J. Jennings, Michael G. Ison, Judd F. Hultquist, Egon A. Ozer |
| EPI_ISL_937260, EPI_ISL_937261 | NYC HH Elmhurst Hospital Medical Center | New York City Public Health Laboratory | Jade Wang, et al. |
| EPI_ISL_940793, EPI_ISL_940794, EPI_ISL_940795, EPI_ISL_940796, EPI_ISL_940797, EPI_ISL_940798, EPI_ISL_940799, EPI_ISL_940800, EPI_ISL_940801, EPI_ISL_940803, EPI_ISL_940804, EPI_ISL_940805, EPI_ISL_940806, EPI_ISL_940807, EPI_ISL_940808, EPI_ISL_940809, EPI_ISL_940810, EPI_ISL_940811, EPI_ISL_940812, EPI_ISL_940813, EPI_ISL_940814, EPI_ISL_940815, EPI_ISL_940816, EPI_ISL_940817, EPI_ISL_940818 | | | |
| see above | Houston Health Dept. | Houston Health Dept. | Ryker Penn, Pamela Brown, Adolpho Lara |
| EPI_ISL_940850, EPI_ISL_940854, EPI_ISL_940857 | Vaccines and Infectious Diseases Analytics Research Unit (VIDA) | KRISP, KZN Research Innovation and Sequencing Platform | Baillie Vicky, du Plessis Jeanine, Giandhari Jennifer, Pillay Sureshnee, Naidoo Yeshnee, Tegally Houriiyah, de Oliveira Tulio, Madhi Shabir |
| EPI_ISL_941270 | HIA PERCY - lab biologie | National Reference Center for Viruses of Respiratory | Marion Barbet, Sylvie Behillil, Méline Bizard, Angela Brisebarre, Camille Capel, Etienne Simon-Lorière, Vincent Enouf, Maud Vanpeene, Sylvie van der |

| | | | |
|---|---|---|---|
| | | Infections, Institut Pasteur, Paris | Werf, Vincent Foissaud |
| EPI_ISL_941369, EPI_ISL_941600 | Instituto Nacional de Saude (INSA) | Instituto Nacional de Saude (INSA) | Borges et al |
| EPI_ISL_941902, EPI_ISL_941922, EPI_ISL_941923, EPI_ISL_941924 | Virginia DCLS | Virginia DCLS | Virginia DCLS |
| EPI_ISL_941925 | Nigeria Centre for Disease Control (NCDC) | African Centre of Excellence for Genomics of Infectious Diseases (ACEGID), Redeemer's University | Oluniyi P.E. et al |
| EPI_ISL_942033, EPI_ISL_942034, EPI_ISL_942035, EPI_ISL_942036, EPI_ISL_942037, EPI_ISL_942039, EPI_ISL_942040, EPI_ISL_942041, EPI_ISL_942042, EPI_ISL_942043, EPI_ISL_942044, EPI_ISL_942045, EPI_ISL_942046, EPI_ISL_942047, EPI_ISL_942076, EPI_ISL_942077, EPI_ISL_942078, EPI_ISL_942079, EPI_ISL_942080, EPI_ISL_942081, EPI_ISL_942082, EPI_ISL_942083, EPI_ISL_942084, EPI_ISL_942085, EPI_ISL_942086, EPI_ISL_942087, EPI_ISL_942088, EPI_ISL_942089, EPI_ISL_942090, EPI_ISL_942091, EPI_ISL_942092, EPI_ISL_942093, EPI_ISL_942094, EPI_ISL_942095, EPI_ISL_942140, EPI_ISL_942141 | | | |
| see above | Wisconsin State Laboratory of Hygiene Communicable Disease Division | Wisconsin State Laboratory of Hygiene Communicable Disease Division | Kelsey R. Florek, Abigail C. Shockey |
| EPI_ISL_942734, EPI_ISL_942735, EPI_ISL_942736, EPI_ISL_942737, EPI_ISL_942738, EPI_ISL_942739, EPI_ISL_942740, EPI_ISL_942741, EPI_ISL_942742, EPI_ISL_942743, EPI_ISL_942744, EPI_ISL_942745, EPI_ISL_942746, EPI_ISL_942747, EPI_ISL_942748, EPI_ISL_942749, EPI_ISL_942750, EPI_ISL_942751, EPI_ISL_942752 | | | |
| see above | Gundersen Molecular Diagnostics Laboratory | Kabara Cancer Research Institute | Craig S. Richmond, Paraic A. Kenny |
| EPI_ISL_943986 | LACEN do Estado de Tocantins | Instituto Adolfo Lutz, Interdiciplinary Procedures Center, Strategic Laboratory | Claudio Tavares Sacchi, Claudia Regina Gonçalves, Erica Valessa Ramos Gomes, Karoline Rodrigues Campos |
| EPI_ISL_944100 | Institute for Medical Research, Infectious Disease Research Centre, National Institutes of Health, Ministry of Health Malaysia | Institute for Medical Research, Infectious Disease Research Centre, National Institutes of Health, Ministry of Health Malaysia | Suppiah J, Kamel K, Azizan MA, Thayan R |
| EPI_ISL_944188, EPI_ISL_944193, EPI_ISL_944196, EPI_ISL_944204, EPI_ISL_944216, EPI_ISL_944226, EPI_ISL_944232, EPI_ISL_944233, EPI_ISL_944236, EPI_ISL_944237, EPI_ISL_944238, EPI_ISL_944239, EPI_ISL_944240, EPI_ISL_944243, EPI_ISL_944244, EPI_ISL_944245, EPI_ISL_944256, EPI_ISL_944257, EPI_ISL_944258, EPI_ISL_944259, EPI_ISL_944260, EPI_ISL_944261, EPI_ISL_944262, EPI_ISL_944263, EPI_ISL_944264, EPI_ISL_944265, EPI_ISL_944266, EPI_ISL_944267, EPI_ISL_944268, EPI_ISL_944269, EPI_ISL_944270, EPI_ISL_944271, EPI_ISL_944272, EPI_ISL_944273, EPI_ISL_944274, EPI_ISL_944275, EPI_ISL_944371, EPI_ISL_944373, EPI_ISL_944381, EPI_ISL_944383, EPI_ISL_944388, EPI_ISL_944389, EPI_ISL_944403, EPI_ISL_944413, EPI_ISL_944414, EPI_ISL_944415, EPI_ISL_944429, EPI_ISL_944431, EPI_ISL_944432, EPI_ISL_944433, EPI_ISL_944434, EPI_ISL_944435, EPI_ISL_944436, EPI_ISL_944437, EPI_ISL_944438, EPI_ISL_944439, EPI_ISL_944440, EPI_ISL_944441, EPI_ISL_944442, EPI_ISL_944443, EPI_ISL_944444, EPI_ISL_944445, EPI_ISL_944446, EPI_ISL_944447, EPI_ISL_944448, EPI_ISL_944449, EPI_ISL_944450, EPI_ISL_944451, EPI_ISL_944452, EPI_ISL_944453, EPI_ISL_944454, EPI_ISL_944455, EPI_ISL_944456, EPI_ISL_944457 | | | |
| see above | Israel Central Virology laboratory | Israel National Consortium for SARS-CoV-2 sequencing | Neta Zuckerman, Efrat Dahan Bucris, Michal Mandelboim, Dana Bar-Ilan, Oran Erster, Tzvia Mann, Omer Murik, David A. Zeevi, Assaf Rokney, Joseph Jaffe, Eva Nachum, Maya Davidovich Cohen, Ephraim Fass, Gal Zizelski Valenci, Mor Rubinstein, Efrat Rorman, Israel Nissan, Efrat Glick-Saar, Omri Nayshool, Gideon Rechavi, Ella Mendelson, Orna Mor |
| EPI_ISL_944736 | unknown | Public Health Virology-Forensic and Scientific Services (PHV-FSS) | Son Nguyen et al. |
| EPI_ISL_944749, EPI_ISL_944750, EPI_ISL_944751, EPI_ISL_944752, EPI_ISL_944753, EPI_ISL_944754, EPI_ISL_944755 | State Virus Research and Diagnostic Laboratory (VRDL), AIIMS Raipur | State Virus Research and Diagnostic Laboratory (VRDL), AIIMS Raipur | Pushpendra Singh, Kuldeep Sharma, Pragya Agarwala, Priyanka Singh, Somya Sharma, Sanjay Singh Negi, Anudita Bhargava |
| EPI_ISL_944779 | General Hospital - Tetovo | Research Center for Genetic Engineering and Biotechnology "Georgi D. Efremov" , Macedonian Academy of Sciences and Arts | Aleksandar J. Dimovski, Dijana Plasheska-Karanfilska, Predrag Noveski, Gjorgji Bozinovski, Milena Jakimovska |
| EPI_ISL_944786 | General Hospital - Prilep | Research Center for Genetic Engineering and Biotechnology "Georgi D. Efremov" , Macedonian Academy of Sciences and Arts | Aleksandar J. Dimovski, Dijana Plasheska-Karanfilska, Predrag Noveski, Gjorgji Bozinovski, Milena Jakimovska |
| EPI_ISL_945051 | Lighthouse Lab in Cambridge | Wellcome Sanger Institute for the COVID-19 Genomics UK (COG-UK) Consortium | Rob Howes, The Lighthouse Lab in Cambridge and Alex Alderton, Roberto Amato, Sonia Goncalves, Ewan Harrison, David K. Jackson, Ian Johnston, Dominic Kwiatkowski, Cordelia Langford, John Sillitoe on behalf of the Wellcome Sanger Institute COVID-19 Surveillance Team |
| EPI_ISL_945052, EPI_ISL_945138, EPI_ISL_945195, EPI_ISL_945224 | Lighthouse Lab in Milton Keynes | Wellcome Sanger Institute for the COVID-19 Genomics UK (COG-UK) Consortium | The Lighthouse Lab in Milton Keynes and Alex Alderton, Roberto Amato, Sonia Goncalves, Ewan Harrison, David K. Jackson, Ian Johnston, Dominic Kwiatkowski, Cordelia Langford, John Sillitoe on behalf of the Wellcome Sanger Institute COVID-19 Surveillance Team |
| EPI_ISL_945226 | Lighthouse Lab in Cambridge | Wellcome Sanger Institute for the COVID-19 Genomics UK (COG-UK) Consortium | Rob Howes, The Lighthouse Lab in Cambridge and Alex Alderton, Roberto Amato, Sonia Goncalves, Ewan Harrison, David K. Jackson, Ian Johnston, Dominic Kwiatkowski, Cordelia Langford, John Sillitoe on behalf of the Wellcome Sanger Institute COVID-19 Surveillance Team |
| EPI_ISL_945234 | Lighthouse Lab in Glasgow | Wellcome Sanger Institute for the COVID-19 Genomics UK (COG-UK) Consortium | Harper VanSteenhouse, Yumi Kasai, David Gray, Carol Clugston, Anna Dominiczak and Alex Alderton, Roberto Amato, Sonia Goncalves, Ewan Harrison, David K. Jackson, Ian Johnston, Dominic Kwiatkowski, Cordelia Langford, John Sillitoe on behalf of the Wellcome Sanger Institute COVID-19 Surveillance Team |
| EPI_ISL_945235 | Lighthouse Lab in Alderley Park | Wellcome Sanger Institute for the COVID-19 Genomics UK (COG-UK) Consortium | Jacquelyn Wynn, Mairead Hyland, The Lighthouse Lab in Alderley Park and Alex Alderton, Roberto Amato, Sonia Goncalves, Ewan Harrison, David K. Jackson, Ian Johnston, Dominic Kwiatkowski, Cordelia Langford, John Sillitoe on behalf of the Wellcome Sanger Institute COVID-19 Surveillance Team |
| EPI_ISL_945358, EPI_ISL_945360 | Lighthouse Lab in Milton Keynes | Wellcome Sanger Institute for the COVID-19 Genomics UK (COG-UK) Consortium | The Lighthouse Lab in Milton Keynes and Alex Alderton, Roberto Amato, Sonia Goncalves, Ewan Harrison, David K. Jackson, Ian Johnston, Dominic Kwiatkowski, Cordelia Langford, John Sillitoe on behalf of the Wellcome Sanger Institute COVID-19 Surveillance Team |
| EPI_ISL_947678 | General Hospital - Tetovo | Research Center for Genetic Engineering and Biotechnology "Georgi D. Efremov" , Macedonian Academy of Sciences and Arts | Aleksandar J. Dimovski, Dijana Plasheska-Karanfilska, Predrag Noveski, Gjorgji Bozinovski, Milena Jakimovska |
| EPI_ISL_949067 | PHI Institute of Nephrology - Struga | Research Center for Genetic Engineering and Biotechnology "Georgi D. Efremov" , Macedonian Academy of Sciences and Arts | Aleksandar J. Dimovski, Dijana Plasheska-Karanfilska, Predrag Noveski, Gjorgji Bozinovski, Milena Jakimovska |
| EPI_ISL_949632, EPI_ISL_949634 | NHLS Universitas Academic | UFS Virology | PA Bester, MM Nyaga, P Nthiga, MT Mogotsi, Emmanuel Ogunbayo, D Goedhals, T de Oliveira |
| EPI_ISL_949747, EPI_ISL_950168 | University College London, Great Ormond Street Hospital for Children NHS Foundation Trust, Imperial College Healthcare NHS Trust | COVID-19 Genomics UK (COG-UK) Consortium | Sergi Castellano, Rachel Williams, Mark Kristiansen, Paola Resende Silva, Sunando Roy, Tony Brooks, Helena Tutill, Paola Niola, Patricia Dyal, Charlotte Williams, Leysa Forrest, Yasmin Panchbhaya, Jacqueline Findlay, Samuel Weeks, Julianne Brown, Kathryn Harris, Paul Randell, James Price, Alison Holmes, Judith Breuer |
| EPI_ISL_950545, EPI_ISL_950546, EPI_ISL_950547, EPI_ISL_950548, EPI_ISL_950549, EPI_ISL_950550, EPI_ISL_950552, EPI_ISL_950553, EPI_ISL_950556, EPI_ISL_950557, EPI_ISL_950558, EPI_ISL_950574, EPI_ISL_950575, EPI_ISL_950576, EPI_ISL_950580, EPI_ISL_950586, EPI_ISL_950588, EPI_ISL_950590, EPI_ISL_950591, EPI_ISL_950593, EPI_ISL_950594, EPI_ISL_950597, EPI_ISL_950598, EPI_ISL_950599, EPI_ISL_950600 | | | |
| see above | Quadram Institute Bioscience | COVID-19 Genomics UK (COG-UK) Consortium | Dave J. Baker, Gemma L. Kay, Alp Aydin, Thanh Le-Viet, Steven Rudder, Ana P. Tedim, Anastasia Kolyva, Maria Diaz, Leonardo de Oliveira Martins, Nabil-Fareed Alikhan, Lizzie Meadows, Rachael Stanley, Ngozi Elumogo, Muhammed Yasir, Nicholas M. Thomson, Alexander J Trotter, Rachel Gilroy, Samuel Bloomfield, Claire Stuart, Andrew Bell, Reenesh Prakash, Samir Dervisevic, Alison E. Mather, John Wain, Mark Webber, Andrew J. Page, Justin O'Grady |
| EPI_ISL_950706, EPI_ISL_950707, EPI_ISL_950708 | Lincolnshire Hospitals and DeepSeq Nottingham | COVID-19 Genomics UK (COG-UK) Consortium | Nichola Duckworth, Tim Sloan, Sarah Walsh, Jonathan Ball, Patrick McClure, Joeseph Chappell, Nadine Holmes, Matthew Carlisle, Christopher Moore, Fei Sang, Johnny Debebe, Victoria Wright, Matthew Loose |
| EPI_ISL_950763, EPI_ISL_950764, EPI_ISL_950765, EPI_ISL_950766, EPI_ISL_950767, EPI_ISL_950769, EPI_ISL_950772, EPI_ISL_950773, EPI_ISL_950774, EPI_ISL_950775, EPI_ISL_950776, EPI_ISL_950777, EPI_ISL_950778, EPI_ISL_950779, EPI_ISL_950785, EPI_ISL_950805, EPI_ISL_950806, EPI_ISL_950807, EPI_ISL_950808, EPI_ISL_950809, EPI_ISL_950810, EPI_ISL_950811, EPI_ISL_950814, EPI_ISL_950815, EPI_ISL_950816, EPI_ISL_950817, EPI_ISL_950820, EPI_ISL_950821, EPI_ISL_950822, EPI_ISL_950823, EPI_ISL_950825, EPI_ISL_950827, EPI_ISL_950828, EPI_ISL_950831, EPI_ISL_950833, EPI_ISL_950834, EPI_ISL_950835, EPI_ISL_950836, EPI_ISL_950837, EPI_ISL_950838, EPI_ISL_950840, EPI_ISL_950841, EPI_ISL_950842, EPI_ISL_950843, EPI_ISL_950845, EPI_ISL_950846, EPI_ISL_950848, EPI_ISL_950870, EPI_ISL_950873, EPI_ISL_950874, EPI_ISL_950875, EPI_ISL_950876, EPI_ISL_950877, EPI_ISL_950924, EPI_ISL_950925, EPI_ISL_950927, EPI_ISL_950929, EPI_ISL_950931, EPI_ISL_950933, EPI_ISL_950934, EPI_ISL_950935, EPI_ISL_950938, EPI_ISL_950939, EPI_ISL_950941, EPI_ISL_950943, EPI_ISL_950944, EPI_ISL_950946, EPI_ISL_950948, EPI_ISL_950949, EPI_ISL_950950 | | | |

| | | | |
|---|---|---|---|
| see above | Oxford Viromics, NDM, University of Oxford; Oxford University Hospitals; Basingstoke and North Hampshire Hospital | COVID-19 Genomics UK (COG-UK) Consortium | Tanya Golubchik, David Bonsall, George Macintyre, Amy Trebes, Mariateresa de Cesare, Catrin Moore, Alex Mobbs, Anita Justice, Robert Shaw, Monique Andersson, Timothy Peto, Emma Wise, Nathan Moore, Jessica Lynch, Nick Cortes, Matilde Mori, Stephen Kidd, David Buck, John Todd, Christophe Fraser |
| EPI_ISL_951750, EPI_ISL_951790, EPI_ISL_951791, EPI_ISL_951792, EPI_ISL_951793, EPI_ISL_951794, EPI_ISL_951795, EPI_ISL_951796, EPI_ISL_951797, EPI_ISL_951798, EPI_ISL_951799, EPI_ISL_951800, EPI_ISL_951801, EPI_ISL_951802, EPI_ISL_951803, EPI_ISL_951804, EPI_ISL_951805, EPI_ISL_951806, EPI_ISL_951807, EPI_ISL_951808, EPI_ISL_951809, EPI_ISL_951810, EPI_ISL_951811, EPI_ISL_951812, EPI_ISL_951813, EPI_ISL_951814 | | | |
| see above | Originating lab: Wales Specialist Virology Centre Sequencing lab: Pathogen Genomics Unit | Public Health Wales Microbiology Cardiff Wales Specialist Virology Centre | Catherine Moore, Johnathan Evans, Laura Gifford, Malorie Perry, Simon Cottrell, Angela Marchbank, Alec Birchley, Alexander Adams, Amy Gaskin, Bree Gatica-Wilcox, Jason Coombes, Joel Southgate, Lauren Gilbert, Lee Graham, Nicole Pacchiarini, Sara Kumziene-Summerhayes, Sarah Taylor, Sophie Jones, Sara Rey, Matthew Bull, Joanne Watkins, Sally Corden, Tom Connor |
| EPI_ISL_952817, EPI_ISL_952818 | Centre for Enzyme Innovation, University of Portsmouth / Translational Research Laboratory, Portsmouth Hospitals NHS Trust | COVID-19 Genomics UK (COG-UK) Consortium | Angela Beckett,Salman Goudarzi,Christopher Fearn,Kate Cook,Katie Loveson,Sharon Glaysher,Scott Elliott,Samuel Robson |
| EPI_ISL_953119, EPI_ISL_953120, EPI_ISL_953122, EPI_ISL_953125, EPI_ISL_953127, EPI_ISL_953128, EPI_ISL_953130, EPI_ISL_953137, EPI_ISL_953139, EPI_ISL_953140, EPI_ISL_953142, EPI_ISL_953144, EPI_ISL_953146, EPI_ISL_953154, EPI_ISL_953160, EPI_ISL_953164, EPI_ISL_953167, EPI_ISL_953171 | | | |
| see above | Bioinformatics and Biostatistics Lab, Advanced Sequencing Facility | COVID-19 Genomics UK (COG-UK) Consortium | Aengus Stewart,Jerome Nicod,Chelsea Sawyer,Laura Cubitt,Harshil Patel,Margaret Crawford |
| EPI_ISL_953939 | General Hospital - Veles | Research Center for Genetic Engineering and Biotechnology "Georgi D. Efremov" , Macedonian Academy of Sciences and Arts | Aleksandar J. Dimovski, Dijana Plasheska-Karanfilska, Predrag Noveski, Gjorgji Bozinovski, Milena Jakimovska |
| EPI_ISL_954139, EPI_ISL_954141 | Gerontology Institute - Skopje | Research Center for Genetic Engineering and Biotechnology "Georgi D. Efremov" , Macedonian Academy of Sciences and Arts | Aleksandar J. Dimovski, Dijana Plasheska-Karanfilska, Predrag Noveski, Gjorgji Bozinovski, Milena Jakimovska |
| EPI_ISL_954202, EPI_ISL_954203, EPI_ISL_954204, EPI_ISL_954205, EPI_ISL_954206, EPI_ISL_954207, EPI_ISL_954208, EPI_ISL_954209 | 1.AO Universitaria 'S. Giovanni di Dio e Ruggi D'Aragona, Scuola Medica Salernitana' Hospital / 2.UOC di Virologia e Microbiologia, Università della Campania 'L. Vanvitelli' / 3.AO Universitaria 'Federico II' Napoli Hospital / 4.AORN 'San Giuseppe Moscati' Avellino Hospital / 5.AO 'San Pio - presidio G. Rummo' Benevento Hospital / 6.AO 'Sant'Anna e San Sebastiano' Caserta Hospital / 7.PO 'Maria Santissima Addolorata' Eboli Hospital / 8.Biogem Istituto di Ricerche Genetiche | 1. Genome Research Center for Health (CRGS) / 2. Laboratory of Molecular Medicine and Genomics(LMMGe) / 3. Center for Research in Pure and Applied Mathematics (CRMPA) | Giorgio Giurato, Francesca Rizzo, Alessandro Weisz, Gianluigi Franci, Giovanni Nassa, Pasquale Pagliano, Roberta Tarallo, Elena Alexandrova, Ylenia D'Agostino, Carlo Ferravante, Jessica Lamberti, Viola Melone, Domenico Memoli, Valeria Mirici Cappa, Domenico Palumbo, Giovanni Pecoraro, Assunta Sellitto, Oriana Strianese, Ilaria Terenzi, Giuseppe Fenza, Aniello Gentile, Antonello Saccomanno, Sonia Amabile, Teresa Rocco, Annamaria Salvati, Emilia Vaccaro, Massimiliano Galdiero, Michele Cennamo, Giuseppe Portella, Maria Grazia Foti, Mariarosaria Ingino, Maria Landi, Maurizio Fumi, Vincenzo Rocco, Rita Greco, Vittoria Letizia, Arnolfo Petruzziello, Maddalena Schioppa, Gregorio Goffredi, Francesca Marciano, Michele Caraglia, Alessia Cossu, Marianna Scrima, Edmondo Adorisio, Morena D'Avenia, Michela Iacobellis, Rosanna Piluscio, Giorgio Dirani, Vittorio Sambri, Simona Semprini, Silvia Zanoli, Francesco Curcio, Stefania Marzinotto, Andreina Baj, Fausto Sessa. |
| EPI_ISL_954229, EPI_ISL_954255 | MRC/UVRI & LSHTM Uganda Research Unit | Where sequence data have been generated and submitted to GISAID | Matthew Cotten, Dan Lule Bugembe, My V.T. Phan, Isaac Sseeewanyana, Patrick Semanda, Susan Nabadda, Pontiano Kaleebu |
| EPI_ISL_954265 | General Hospital - Prilep | Research Center for Genetic Engineering and Biotechnology "Georgi D. Efremov" , Macedonian Academy of Sciences and Arts | Aleksandar J. Dimovski, Dijana Plasheska-Karanfilska, Predrag Noveski, Gjorgji Bozinovski, Milena Jakimovska |
| EPI_ISL_954301 | Institute for Lung Diseases in Children - Skopje | Research Center for Genetic Engineering and Biotechnology "Georgi D. Efremov" , Macedonian Academy of Sciences and Arts | Aleksandar J. Dimovski, Dijana Plasheska-Karanfilska, Predrag Noveski, Gjorgji Bozinovski, Milena Jakimovska |
| EPI_ISL_954611 | General Hospital - Struga | Research Center for Genetic Engineering and Biotechnology "Georgi D. Efremov" , Macedonian Academy of Sciences and Arts | Aleksandar J. Dimovski, Dijana Plasheska-Karanfilska, Predrag Noveski, Gjorgji Bozinovski, Milena Jakimovska |
| EPI_ISL_954745, EPI_ISL_954746 | General Hospital - Ohrid | Research Center for Genetic Engineering and Biotechnology "Georgi D. Efremov" , Macedonian Academy of Sciences and Arts | Aleksandar J. Dimovski, Dijana Plasheska-Karanfilska, Predrag Noveski, Gjorgji Bozinovski, Milena Jakimovska |
| EPI_ISL_954747 | Hospital Universitari Vall d'Hebron - Vall d'Hebron Institut de Rercerca | Hospital Universitari Vall d'Hebron - Vall d'Hebron Institut de Recerca | Cristina Andrés, Maria Piñana, Josep F Abril, Damir Garcia-Cehic, Ariadna Rando, Juliana Esperalba, Maria Gema Codina, Carla Castillo, Maria Carmen Martin, Tomàs Pumarola, Josep Quer, Andrés Antón |
| EPI_ISL_954894 | Colorado Department of Public Health and Environment | Colorado Department of Puplic Health and Environment | Laura Bankers, Molly C. Hetherington-Rauth, Diana Ir, Shannon Ely, Shannon R. Matzinger, Sarah Elizabeth Totten, Emily A. Travanty |
| EPI_ISL_955173 | University of Sarajevo, Veterinary Faculty, Laboratory for Molecular Diagnostic and Research Laboratory | University of Sarajevo, Veterinary Faculty, Laboratory for Molecular Diagnostic and Research Laboratory | Goleti Š., Goleti T., Softi A., Ali-Šeho A., Jaži A., Šabi E., Hodži A., Terzi I., Nicevi M. |
| EPI_ISL_955213, EPI_ISL_955214 | Indiana Animal Disease Diagnostic Laboratory | Carpi Laboratory - Purdue University | Jack Dorman, Ilinca I Ciubotariu, Lev Gorenstein, Abebe A Fola, G Kenitra Hendrix, Rebecca P Wilkes, Giovanna Carpi |
| EPI_ISL_955693, EPI_ISL_955694, EPI_ISL_955695, EPI_ISL_955696, EPI_ISL_955699, EPI_ISL_955701, EPI_ISL_955702, EPI_ISL_955703, EPI_ISL_955705 | Humboldt County Public Health Laboratory | Chan-Zuckerberg Biohub | CZB Cliahub Consortium |
| EPI_ISL_956325 | Laboratory Medicine | Department of Laboratory Medicine, Lin-Kou Chang Gung Memorial Hospital, Taoyuan, Taiwan | Kuo-Chien Tsao, Yu-Nong Gong, Shu-Li Yang, Yi-Chun Liu, Chung-Guei Huang, Mei-Jen Hsiao, Po-Wei Huang, Cheng-Ta Yang, Cheng-Hsun Chiu, Peng-Nien Huang, Kuo-Ming Lee, Guang-Wu Chen, Shin-Ru Shih |
| EPI_ISL_956340, EPI_ISL_956341, EPI_ISL_956342, EPI_ISL_956343, EPI_ISL_956344, EPI_ISL_956345, EPI_ISL_956346, EPI_ISL_956347, EPI_ISL_956348, EPI_ISL_956349, EPI_ISL_956350, EPI_ISL_956351 | | | |
| see above | Houston Health Department, Disease Prevention and Control | Houston Health Department, Disease Prevention and Control | Penn,R., Brown,P., Lara,A. |
| EPI_ISL_956367, EPI_ISL_956382, EPI_ISL_956383 | General Hospital - Prilep | Research Center for Genetic Engineering and Biotechnology "Georgi D. Efremov" , Macedonian Academy of Sciences and Arts | Aleksandar J. Dimovski, Dijana Plasheska-Karanfilska, Predrag Noveski, Gjorgji Bozinovski, Milena Jakimovska |
| EPI_ISL_960225 | QEII Health Sciences Centre | National Microbiology Laboratory (NML) | Anna Majer, Shari Tyson, Grace Seo, Philip Mabon, Elsie Grudeski, Rhiannon Huzarewich, Russell Mandes, Anneliese Landgraff, Jennifer Tanner, Natalie Knox, Morag Graham, Gary Van Domselaar, Todd Hatchette, Jason LeBlanc, Janice Pettipas, Dan Gaston, Nathalie Bastien, Yan Li, Timothy Booth, Darian Hole, Madison Chapel, Kirsten Biggar, CanCOGeN's metadata curation team, Public Health Agency of Canada CanCOGeN team |
| EPI_ISL_960435 | The National Institute of Public Health | State Veterinary Institute Prague | Nagy,A;Vecerova,J;Cernikova,L;Stara,M;Jirincova,H;Trnka,D |
| EPI_ISL_960480, EPI_ISL_960481, EPI_ISL_960492, EPI_ISL_960589, EPI_ISL_960658 | Istituto Zooprofilattico Sperimentale del Mezzogiorno | TIGEM | Patrizia Annunziata, Andrea Ballabio, Valentina Bouche, Davide Cacchiarelli, Pellegrino Cerino, Chiara Colantuono, Maria Concetta Cuomo, Denise Di Concilio, Lucio Di Filippo, Antonio Grimaldi, Antonio Limone, Anna Manfredi, Francesco Panariello, Biancamaria Pierri, Marcello Salvi |
| EPI_ISL_960664 | Germano de sousa | Instituto Gulbenkian de Ciencia | Maria Costa, Susana Ladeiro, Cathy Paulino, João Costa, João Sobral, Ricardo Leite |
| EPI_ISL_960665 | Germano de sousa | Instituto Gulbenkian de Ciencia | Cathy Paulino, João Costa, João Sobral, Maria Costa, Susana Ladeiro, Ricardo Leite |
| EPI_ISL_960666, EPI_ISL_960667 | Germano de sousa | Instituto Gulbenkian de Ciencia | João Costa, João Sobral, Maria Costa, Susana Ladeiro, Cathy Paulino, Ricardo Leite |
| EPI_ISL_960668 | Germano de sousa | Instituto Gulbenkian de Ciencia | Susana Ladeiro, Cathy Paulino, João Costa, João Sobral, Maria Costa, Ricardo Leite |
| EPI_ISL_960670 | Germano de sousa | Instituto Gulbenkian de Ciencia | João Costa, João Sobral, Maria Costa, Susana Ladeiro, Cathy Paulino, Ricardo Leite |

| | | | |
|---|---|---|---|
| EPI_ISL_960671 | Germano de sousa | Instituto Gulbenkian de Ciencia | Susana Ladeiro, Cathy Paulino, João Costa, João Sobral, Maria Costa, Ricardo Leite |
| EPI_ISL_960673 | Germano de sousa | Instituto Gulbenkian de Ciencia | Maria Costa, Susana Ladeiro, Cathy Paulino, João Costa, João Sobral, Ricardo Leite |
| EPI_ISL_960674 | Germano de sousa | Instituto Gulbenkian de Ciencia | João Costa, João Sobral, Maria Costa, Susana Ladeiro, Cathy Paulino, Ricardo Leite |
| EPI_ISL_960676 | Germano de sousa | Instituto Gulbenkian de Ciencia | Maria Costa, Susana Ladeiro, Cathy Paulino, João Costa, João Sobral, Ricardo Leite |
| EPI_ISL_960677 | Germano de sousa | Instituto Gulbenkian de Ciencia | Cathy Paulino, João Costa, João Sobral, Maria Costa, Susana Ladeiro, Ricardo Leite |
| EPI_ISL_960678, EPI_ISL_960679 | Germano de sousa | Instituto Gulbenkian de Ciencia | João Sobral, Maria Costa, Susana Ladeiro, Cathy Paulino, João Costa, Ricardo Leite |
| EPI_ISL_960680 | Germano de sousa | Instituto Gulbenkian de Ciencia | Maria Costa, Susana Ladeiro, Cathy Paulino, João Costa, João Sobral, Ricardo Leite |
| EPI_ISL_960683, EPI_ISL_960684 | Germano de sousa | Instituto Gulbenkian de Ciencia | Cathy Paulino, João Costa, João Sobral, Maria Costa, Susana Ladeiro, Ricardo Leite |
| EPI_ISL_960685 | Germano de sousa | Instituto Gulbenkian de Ciencia | João Costa, João Sobral, Maria Costa, Susana Ladeiro, Cathy Paulino, Ricardo Leite |
| EPI_ISL_960686 | Germano de sousa | Instituto Gulbenkian de Ciencia | Maria Costa, Susana Ladeiro, Cathy Paulino, João Costa, João Sobral, Ricardo Leite |
| EPI_ISL_960687, EPI_ISL_960688 | Germano de sousa | Instituto Gulbenkian de Ciencia | Cathy Paulino, João Costa, João Sobral, Maria Costa, Susana Ladeiro, Ricardo Leite |
| EPI_ISL_960689, EPI_ISL_960690, EPI_ISL_960691 | Germano de sousa | Instituto Gulbenkian de Ciencia | Maria Costa, Susana Ladeiro, Cathy Paulino, João Costa, João Sobral, Ricardo Leite |
| EPI_ISL_960694 | Germano de sousa | Instituto Gulbenkian de Ciencia | João Sobral, Maria Costa, Susana Ladeiro, Cathy Paulino, João Costa, Ricardo Leite |
| EPI_ISL_960695, EPI_ISL_960696 | Germano de sousa | Instituto Gulbenkian de Ciencia | Maria Costa, Susana Ladeiro, Cathy Paulino, João Costa, João Sobral, Ricardo Leite |
| EPI_ISL_960697, EPI_ISL_960698 | Germano de sousa | Instituto Gulbenkian de Ciencia | João Sobral, Maria Costa, Susana Ladeiro, Cathy Paulino, João Costa, Ricardo Leite |
| EPI_ISL_960699, EPI_ISL_960700, EPI_ISL_960701, EPI_ISL_960702 | Germano de sousa | Instituto Gulbenkian de Ciencia | João Costa, João Sobral, Maria Costa, Susana Ladeiro, Cathy Paulino, Ricardo Leite |
| EPI_ISL_960703, EPI_ISL_960704, EPI_ISL_960705 | Germano de sousa | Instituto Gulbenkian de Ciencia | Cathy Paulino, João Costa, João Sobral, Maria Costa, Susana Ladeiro, Ricardo Leite |
| EPI_ISL_960706 | Germano de sousa | Instituto Gulbenkian de Ciencia | João Costa, João Sobral, Maria Costa, Susana Ladeiro, Cathy Paulino, Ricardo Leite |
| EPI_ISL_960707, EPI_ISL_960709 | Germano de sousa | Instituto Gulbenkian de Ciencia | João Sobral, Maria Costa, Susana Ladeiro, Cathy Paulino, João Costa, Ricardo Leite |
| EPI_ISL_960710, EPI_ISL_960711, EPI_ISL_960712 | Germano de sousa | Instituto Gulbenkian de Ciencia | Cathy Paulino, João Costa, João Sobral, Maria Costa, Susana Ladeiro, Ricardo Leite |
| EPI_ISL_960713 | Germano de sousa | Instituto Gulbenkian de Ciencia | João Costa, João Sobral, Maria Costa, Susana Ladeiro, Cathy Paulino, Ricardo Leite |
| EPI_ISL_960714, EPI_ISL_960715 | Germano de sousa | Instituto Gulbenkian de Ciencia | João Sobral, Maria Costa, Susana Ladeiro, Cathy Paulino, João Costa, Ricardo Leite |
| EPI_ISL_960716, EPI_ISL_960717 | Germano de sousa | Instituto Gulbenkian de Ciencia | Susana Ladeiro, Cathy Paulino, João Costa, João Sobral, Maria Costa, Ricardo Leite |
| EPI_ISL_960720 | Germano de sousa | Instituto Gulbenkian de Ciencia | Cathy Paulino, João Costa, João Sobral, Maria Costa, Susana Ladeiro, Ricardo Leite |
| EPI_ISL_960721 | Germano de sousa | Instituto Gulbenkian de Ciencia | João Sobral, Maria Costa, Susana Ladeiro, Cathy Paulino, João Costa, Ricardo Leite |
| EPI_ISL_960722 | Germano de sousa | Instituto Gulbenkian de Ciencia | Maria Costa, Susana Ladeiro, Cathy Paulino, João Costa, João Sobral, Ricardo Leite |
| EPI_ISL_960724 | Germano de sousa | Instituto Gulbenkian de Ciencia | João Costa, João Sobral, Maria Costa, Susana Ladeiro, Cathy Paulino, Ricardo Leite |
| EPI_ISL_960725, EPI_ISL_960726, EPI_ISL_960727, EPI_ISL_960728, EPI_ISL_960729 | Germano de sousa | Instituto Gulbenkian de Ciencia | Cathy Paulino, João Costa, João Sobral, Maria Costa, Susana Ladeiro, Ricardo Leite |
| EPI_ISL_960730, EPI_ISL_960731, EPI_ISL_960732 | Germano de sousa | Instituto Gulbenkian de Ciencia | João Costa, João Sobral, Maria Costa, Susana Ladeiro, Cathy Paulino, Ricardo Leite |
| EPI_ISL_960733 | Germano de sousa | Instituto Gulbenkian de Ciencia | João Sobral, Maria Costa, Susana Ladeiro, Cathy Paulino, João Costa, Ricardo Leite |
| EPI_ISL_960734, EPI_ISL_960735, EPI_ISL_960736, EPI_ISL_960737, EPI_ISL_960738, EPI_ISL_960739, EPI_ISL_960740, EPI_ISL_960741, EPI_ISL_960742, EPI_ISL_960743, EPI_ISL_960744, EPI_ISL_960745, EPI_ISL_960746, EPI_ISL_960747 | | | |
| see above | Germano de sousa | Instituto Gulbenkian de Ciencia | Maria Costa, Susana Ladeiro, Cathy Paulino, João Costa, João Sobral, Ricardo Leite |
| EPI_ISL_960751, EPI_ISL_960752 | Germano de sousa | Instituto Gulbenkian de Ciencia | Cathy Paulino, João Costa, João Sobral, Maria Costa, Susana Ladeiro, Ricardo Leite |
| EPI_ISL_960753, EPI_ISL_960754, EPI_ISL_960755 | Germano de sousa | Instituto Gulbenkian de Ciencia | João Costa, João Sobral, Maria Costa, Susana Ladeiro, Cathy Paulino, Ricardo Leite |
| EPI_ISL_960756, EPI_ISL_960757 | Germano de sousa | Instituto Gulbenkian de Ciencia | João Sobral, Maria Costa, Susana Ladeiro, Cathy Paulino, João Costa, Ricardo Leite |
| EPI_ISL_960758, EPI_ISL_960759, EPI_ISL_960760, EPI_ISL_960761 | Germano de sousa | Instituto Gulbenkian de Ciencia | Susana Ladeiro, Cathy Paulino, João Costa, João Sobral, Maria Costa, Ricardo Leite |
| EPI_ISL_960763 | Germano de sousa | Instituto Gulbenkian de Ciencia | João Sobral, Maria Costa, Susana Ladeiro, Cathy Paulino, João Costa, Ricardo Leite |
| EPI_ISL_960764 | Germano de sousa | Instituto Gulbenkian de Ciencia | Cathy Paulino, João Costa, João Sobral, Maria Costa, Susana Ladeiro, Ricardo Leite |
| EPI_ISL_960765, EPI_ISL_960766, EPI_ISL_960767, EPI_ISL_960768 | Germano de sousa | Instituto Gulbenkian de Ciencia | João Sobral, Maria Costa, Susana Ladeiro, Cathy Paulino, João Costa, Ricardo Leite |
| EPI_ISL_960769 | Germano de sousa | Instituto Gulbenkian de Ciencia | Cathy Paulino, João Costa, João Sobral, Maria Costa, Susana Ladeiro, Ricardo Leite |
| EPI_ISL_960770, EPI_ISL_960771, EPI_ISL_960772 | Germano de sousa | Instituto Gulbenkian de Ciencia | João Sobral, Maria Costa, Susana Ladeiro, Cathy Paulino, João Costa, Ricardo Leite |
| EPI_ISL_960773 | Germano de sousa | Instituto Gulbenkian de Ciencia | Maria Costa, Susana Ladeiro, Cathy Paulino, João Costa, João Sobral, Ricardo Leite |
| EPI_ISL_960774 | Germano de sousa | Instituto Gulbenkian de Ciencia | João Costa, João Sobral, Maria Costa, Susana Ladeiro, Cathy Paulino, Ricardo Leite |
| EPI_ISL_960775 | Germano de sousa | Instituto Gulbenkian de Ciencia | Cathy Paulino, João Costa, João Sobral, Maria Costa, Susana Ladeiro, Ricardo Leite |
| EPI_ISL_960776 | Germano de sousa | Instituto Gulbenkian de Ciencia | João Costa, João Sobral, Maria Costa, Susana Ladeiro, Cathy Paulino, Ricardo Leite |
| EPI_ISL_960777 | Germano de sousa | Instituto Gulbenkian de Ciencia | João Sobral, Maria Costa, Susana Ladeiro, Cathy Paulino, João Costa, Ricardo Leite |
| EPI_ISL_960778 | Germano de sousa | Instituto Gulbenkian de Ciencia | Maria Costa, Susana Ladeiro, Cathy Paulino, João Costa, João Sobral, Ricardo Leite |
| EPI_ISL_960779 | Germano de sousa | Instituto Gulbenkian de Ciencia | João Sobral, Maria Costa, Susana Ladeiro, Cathy Paulino, João Costa, Ricardo Leite |
| EPI_ISL_960780 | Germano de sousa | Instituto Gulbenkian de Ciencia | Cathy Paulino, João Costa, João Sobral, Maria Costa, Susana Ladeiro, Ricardo Leite |
| EPI_ISL_960781, EPI_ISL_960782 | Germano de sousa | Instituto Gulbenkian de Ciencia | João Costa, João Sobral, Maria Costa, Susana Ladeiro, Cathy Paulino, Ricardo Leite |
| EPI_ISL_960784 | Germano de sousa | Instituto Gulbenkian de Ciencia | João Sobral, Maria Costa, Susana Ladeiro, Cathy Paulino, João Costa, Ricardo Leite |

| | | | |
|---|---|---|---|
| EPI_ISL_960785, EPI_ISL_960786, EPI_ISL_960787 | Germano de sousa | Instituto Gulbenkian de Ciencia | Susana Ladeiro, Cathy Paulino, João Costa, João Sobral, Maria Costa, Ricardo Leite |
| EPI_ISL_960788 | Germano de sousa | Instituto Gulbenkian de Ciencia | Maria Costa, Susana Ladeiro, Cathy Paulino, João Costa, João Sobral, Ricardo Leite |
| EPI_ISL_960789, EPI_ISL_960790, EPI_ISL_960791, EPI_ISL_960792 | Germano de sousa | Instituto Gulbenkian de Ciencia | Cathy Paulino, João Costa, João Sobral, Maria Costa, Susana Ladeiro, Ricardo Leite |
| EPI_ISL_960793, EPI_ISL_960794, EPI_ISL_960795 | Germano de sousa | Instituto Gulbenkian de Ciencia | João Costa, João Sobral, Maria Costa, Susana Ladeiro, Cathy Paulino, Ricardo Leite |
| EPI_ISL_960796, EPI_ISL_960797 | Germano de sousa | Instituto Gulbenkian de Ciencia | João Sobral, Maria Costa, Susana Ladeiro, Cathy Paulino, João Costa, Ricardo Leite |
| EPI_ISL_960798 | Germano de sousa | Instituto Gulbenkian de Ciencia | Maria Costa, Susana Ladeiro, Cathy Paulino, João Costa, João Sobral, Ricardo Leite |
| EPI_ISL_960799 | Germano de sousa | Instituto Gulbenkian de Ciencia | Cathy Paulino, João Costa, João Sobral, Maria Costa, Susana Ladeiro, Ricardo Leite |
| EPI_ISL_960800 | Germano de sousa | Instituto Gulbenkian de Ciencia | João Sobral, Maria Costa, Susana Ladeiro, Cathy Paulino, João Costa, Ricardo Leite |
| EPI_ISL_960801 | Germano de sousa | Instituto Gulbenkian de Ciencia | João Costa, João Sobral, Maria Costa, Susana Ladeiro, Cathy Paulino, Ricardo Leite |
| EPI_ISL_960802 | Germano de sousa | Instituto Gulbenkian de Ciencia | João Sobral, Maria Costa, Susana Ladeiro, Cathy Paulino, João Costa, Ricardo Leite |
| EPI_ISL_960803, EPI_ISL_960804 | Germano de sousa | Instituto Gulbenkian de Ciencia | João Costa, João Sobral, Maria Costa, Susana Ladeiro, Cathy Paulino, Ricardo Leite |
| EPI_ISL_960805, EPI_ISL_960806 | Germano de sousa | Instituto Gulbenkian de Ciencia | Susana Ladeiro, Cathy Paulino, João Costa, João Sobral, Maria Costa, Ricardo Leite |
| EPI_ISL_960807, EPI_ISL_960808 | Germano de sousa | Instituto Gulbenkian de Ciencia | Cathy Paulino, João Costa, João Sobral, Maria Costa, Susana Ladeiro, Ricardo Leite |
| EPI_ISL_960809, EPI_ISL_960810 | Germano de sousa | Instituto Gulbenkian de Ciencia | Maria Costa, Susana Ladeiro, Cathy Paulino, João Costa, João Sobral, Ricardo Leite |
| EPI_ISL_960811 | Germano de sousa | Instituto Gulbenkian de Ciencia | Cathy Paulino, João Costa, João Sobral, Maria Costa, Susana Ladeiro, Ricardo Leite |
| EPI_ISL_960812 | Germano de sousa | Instituto Gulbenkian de Ciencia | João Costa, João Sobral, Maria Costa, Susana Ladeiro, Cathy Paulino, Ricardo Leite |
| EPI_ISL_960813, EPI_ISL_960814, EPI_ISL_960815, EPI_ISL_960816 | Germano de sousa | Instituto Gulbenkian de Ciencia | João Sobral, Maria Costa, Susana Ladeiro, Cathy Paulino, João Costa, Ricardo Leite |
| EPI_ISL_960817, EPI_ISL_960818 | Germano de sousa | Instituto Gulbenkian de Ciencia | Maria Costa, Susana Ladeiro, Cathy Paulino, João Costa, João Sobral, Ricardo Leite |
| EPI_ISL_960819, EPI_ISL_960820, EPI_ISL_960821, EPI_ISL_960822 | Germano de sousa | Instituto Gulbenkian de Ciencia | Susana Ladeiro, Cathy Paulino, João Costa, João Sobral, Maria Costa, Ricardo Leite |
| EPI_ISL_960825 | Germano de sousa | Instituto Gulbenkian de Ciencia | João Sobral, Maria Costa, Susana Ladeiro, Cathy Paulino, João Costa, Ricardo Leite |
| EPI_ISL_960826 | Germano de sousa | Instituto Gulbenkian de Ciencia | Maria Costa, Susana Ladeiro, Cathy Paulino, João Costa, João Sobral, Ricardo Leite |
| EPI_ISL_960827, EPI_ISL_960828, EPI_ISL_960829 | Germano de sousa | Instituto Gulbenkian de Ciencia | João Costa, João Sobral, Maria Costa, Susana Ladeiro, Cathy Paulino, Ricardo Leite |
| EPI_ISL_960830 | Germano de sousa | Instituto Gulbenkian de Ciencia | Maria Costa, Susana Ladeiro, Cathy Paulino, João Costa, João Sobral, Ricardo Leite |
| EPI_ISL_960867 | Institute of Medical Microbiology and Hospital Hygiene | Institute of Medical Microbiology and Hospital Hygiene | Prof. Dr. Achim Kaasch, Aljoscha Tersteegen |
| EPI_ISL_961231 | Hospital General Universitario de Alicante - Instituto de Investigación Sanitaria y Biomédica de Alicante | SeqCOVID-SPAIN consortium/IBV(CSIC) | Maripaz Ventero Martín, Carmen Molina Pardines and SeqCOVID-SPAIN consortium |
| EPI_ISL_961661, EPI_ISL_961662, EPI_ISL_961663 | Hôpital Georges L. Dumont | National Microbiology Laboratory (NML) | Anna Majer, Shari Tyson, Grace Seo, Philip Mabon, Elsie Grudeski, Rhiannon Huzarewich, Russell Mandes, Anneliese Landgraff, Jennifer Tanner, Natalie Knox, Morag Graham, Gary Van Domselaar, Richard Garceau, Guillaume Desnoyers, Nathalie Bastien, Yan Li, Timothy Booth, Darian Hole, Madison Chapel, Kirsten Biggar, CanCOGeN's metadata curation team, Public Health Agency of Canada CanCOGeN team |
| EPI_ISL_962124 | Illinois Department of Public Health | Gagnon Lab, Southern Illinois University | Keith Gagnon |
| EPI_ISL_962507, EPI_ISL_962508, EPI_ISL_962514 | UCLA Clinical Micro Lab | Los Angeles County PHL | P. Hemarajata et al. |
| EPI_ISL_962811 | San Diego County Public Health Laboratory | Andersen lab at Scripps Research | SEARCH Alliance San Diego with Tracy Basler, Jovan Shephard, Brett Austin |
| EPI_ISL_962929, EPI_ISL_962943, EPI_ISL_962962, EPI_ISL_962964, EPI_ISL_962965, EPI_ISL_962966, EPI_ISL_962967, EPI_ISL_962968, EPI_ISL_962969, EPI_ISL_962971, EPI_ISL_962972 | | | |
| see above | Hospital Universitario de Gran Canaria Dr. Negrín | SeqCOVID-SPAIN consortium/IBV(CSIC) | M. Carmen Pérez González, Francisco J. Chamizo López, Ana Bordes Benítez and SeqCOVID-SPAIN consortium |
| EPI_ISL_964976 | Akershus University Hospital, Department for Microbiology and Infectious Disease Control | Norwegian Institute of Public Health, Department of Virology | Kathrine Stene-Johansen, Kamilla Heddeland Instefjord, Hilde Elshaug, Ignacio Garcia Llorente, Serina B Engebretsen, Atiya R Ali,Marie Paulsen Madsen, Rasmus Riis Kopperud, Hilde Vollan, Karoline Bragstad, Olav Hungnes |
| EPI_ISL_965218 | Virginia Division of Consolidated Laboratory Services | Virginia Division of Consolidated Laboratory Services | Virginia DCLS |
| EPI_ISL_965530, EPI_ISL_965602, EPI_ISL_965662, EPI_ISL_965724, EPI_ISL_965755, EPI_ISL_965758, EPI_ISL_965786 | Dutch COVID-19 response team | Medical Microbiology, Maastricht University Medical Centre | Jozef Dingemans*, Brian van der Veer*, Erik Beuken, Carmen Reumkens, Lieke van Alphen, Christian Hoebe, Paul Savelkoul |
| EPI_ISL_967586, EPI_ISL_967587, EPI_ISL_967613, EPI_ISL_967657, EPI_ISL_967661, EPI_ISL_967663, EPI_ISL_967685, EPI_ISL_967686, EPI_ISL_967747, EPI_ISL_967749 | State Laboratories Division, Hawaii State Department of Health | State Laboratories Division, Hawaii State Department of Health | Pamela O'Brien, Drew Kuwazaki, Ayana Garnet, Razvan Sultana, Edward Desmond |
| EPI_ISL_968082, EPI_ISL_968083, EPI_ISL_968084, EPI_ISL_968085, EPI_ISL_968086, EPI_ISL_968087, EPI_ISL_968088, EPI_ISL_968090, EPI_ISL_968091, EPI_ISL_968092, EPI_ISL_968093, EPI_ISL_968094, EPI_ISL_968095, EPI_ISL_968096, EPI_ISL_968097, EPI_ISL_968098, EPI_ISL_968099, EPI_ISL_968100, EPI_ISL_968101, EPI_ISL_968102, EPI_ISL_968103, EPI_ISL_968104, EPI_ISL_968105, EPI_ISL_968106, EPI_ISL_968107, EPI_ISL_968108, EPI_ISL_968109, EPI_ISL_968112, EPI_ISL_968113, EPI_ISL_968114, EPI_ISL_968115, EPI_ISL_968116, EPI_ISL_968117, EPI_ISL_968118, EPI_ISL_968119, EPI_ISL_968120, EPI_ISL_968121, EPI_ISL_968122, EPI_ISL_968123, EPI_ISL_968124, EPI_ISL_968125, EPI_ISL_968126, EPI_ISL_968127, EPI_ISL_968128, EPI_ISL_968129, EPI_ISL_968130, EPI_ISL_968131, EPI_ISL_968132, EPI_ISL_968133, EPI_ISL_968134, EPI_ISL_968135, EPI_ISL_968136 | | | |
| see above | Houston Health Department, Disease Prevention and Control | Houston Health Department, Disease Prevention and Control | Penn,R., Brown,P., Lara,A. |
| EPI_ISL_976691, EPI_ISL_976692, EPI_ISL_976693, EPI_ISL_976694, EPI_ISL_976695, EPI_ISL_976696, EPI_ISL_976697, EPI_ISL_976698, EPI_ISL_976699, EPI_ISL_976700, EPI_ISL_976701, EPI_ISL_976702, EPI_ISL_976703, EPI_ISL_976704, EPI_ISL_976705, EPI_ISL_976706, EPI_ISL_976707, EPI_ISL_976708, EPI_ISL_976709, EPI_ISL_976710, EPI_ISL_976711, EPI_ISL_976712, EPI_ISL_976713, EPI_ISL_976714, EPI_ISL_976715, EPI_ISL_976716, EPI_ISL_976717, EPI_ISL_976718, EPI_ISL_976719, EPI_ISL_976720, EPI_ISL_976721, EPI_ISL_976722, EPI_ISL_976723, EPI_ISL_976724, EPI_ISL_976725, EPI_ISL_976726, EPI_ISL_976727, EPI_ISL_976728, EPI_ISL_976729, EPI_ISL_976730, EPI_ISL_976731, EPI_ISL_976732, EPI_ISL_976733, EPI_ISL_976734, EPI_ISL_976735, EPI_ISL_976736, EPI_ISL_976737, EPI_ISL_976738, EPI_ISL_976739, EPI_ISL_976740, EPI_ISL_976741, EPI_ISL_976742, EPI_ISL_976743, EPI_ISL_976744, EPI_ISL_976745, EPI_ISL_976746, EPI_ISL_976747, EPI_ISL_976748, EPI_ISL_976749, EPI_ISL_976750, EPI_ISL_976751, EPI_ISL_976752, EPI_ISL_976753, EPI_ISL_976754, EPI_ISL_976755, EPI_ISL_976756, EPI_ISL_976757, EPI_ISL_976758, EPI_ISL_976759, EPI_ISL_976760, EPI_ISL_976761, EPI_ISL_976762, EPI_ISL_976763, EPI_ISL_976764, EPI_ISL_976765, EPI_ISL_976766, EPI_ISL_976767, EPI_ISL_976768, EPI_ISL_976769, EPI_ISL_976770, EPI_ISL_976771, EPI_ISL_976772, EPI_ISL_976773, EPI_ISL_976774, EPI_ISL_976775, EPI_ISL_976776, EPI_ISL_976777, EPI_ISL_976778, EPI_ISL_976779, EPI_ISL_976780, EPI_ISL_976781, EPI_ISL_976782, EPI_ISL_976783, EPI_ISL_976784, EPI_ISL_976785, EPI_ISL_976786, EPI_ISL_976787, EPI_ISL_976788, EPI_ISL_976789 | | | |
| see above | BCCDC Public Health Laboratory | BCCDC Public Health Laboratory | Prystajecky Natalie, Linda Hoang, Dan Fornika, John Tyson, Shannon Russell, Kim Macdonald, Kimia Kamelian, Ana Pacagnella, Corrinne Ng, Loretta Janz, Robert Azana Terry Snutch, Mel Krajden |
| EPI_ISL_977034 | Rhode Island Department of Health | Infectious Disease Program, Broad Institute of Harvard and MIT | Lemieux,J.E., Siddle,K.J., Huard,R., King,E., Azevedo,K., Miller,A., Adams,G., Gladden-Young,A., Lagerborg,K., Rudy,M., DeRuff,K., Carter,A., Normandin,E., Bauer,M., Reilly,S., Tomkins-Tinch,C., Loreth,C., Chaluvadi,S., Birren,B.W., Gallagher,G., Smole,S., Park,D.J., MacInnis,B.L., and Sabeti,P.C. |

| | | | |
|---|---|---|---|
| EPI_ISL_977079, EPI_ISL_977080, EPI_ISL_977081 | Massachusetts General Hospital | Infectious Disease Program, Broad Institute of Harvard and MIT | Lemieux,J.E., Siddle,K.J., Shaw,B., Adams,G., Pierce,V., Turbett,S., Anahtar,M., Branda,J., Slater,D., Harris,J., Lin,A.E., Gladden-Young,A., Lagerborg,K., Rudy,M., DeRuff,K., Carter,A., Normandin,E., Bauer,M., Reilly,S., Tomkins-Tinch,C., Loreth,C., Chaluvadi,S., Neumann,A., Cusick,C., Chapman,S.B., Gnirke,A., Flowers,K., Cerrato,F., Birren,B.W., Gallagher,G., Smole,S., Park,D.J., MacInnis,B.L., Ryan,E., LaRocque,R., Rosenberg,E. and Sabeti,P.C. |
| EPI_ISL_977194 | ULSS 8 Berica | Istituto Zooprofilattico Sperimentale delle Venezie | Adelaide Milani, Alessia Schivo, Annalisa Salviato, Erika Giorgia Quaranta, Ambra Pastori, Bianca Zecchin, Alice Fusaro, Isabella Monne, Calogero Terregino, Antonia Ricci |
| EPI_ISL_977207, EPI_ISL_977208, EPI_ISL_977209, EPI_ISL_977212 | Microbiologia e Virologia | Istituto Zooprofilattico Sperimentale delle Venezie | Adelaide Milani, Alessia Schivo, Annalisa Salviato, Erika Giorgia Quaranta, Ambra Pastori, Bianca Zecchin, Alice Fusaro, Isabella Monne, Calogero Terregino, Antonia Ricci |
| EPI_ISL_977287, EPI_ISL_977289, EPI_ISL_977344, EPI_ISL_977345 | University of Zambia, School of Veterinary Medicine | UNZAVET and PATH | Mulenga Mwenda-Chimfwembe, Ngonda Saasa, Daniel Bridges |
| EPI_ISL_977655 | Caribbean Public Health Agency | Carrington Lab, Department of PreClinical Sciences, Building 36, First Floor Biochemistry Unit, Faculty of Medical Sciences, The University of the West Indies | Nikita S. D. Sahadeo, Arianne Brown-Jordan, Vernie Ramkissoon, Sarah Hill, Naresh Nandram, Avery Hinds, Kenneth George, Jerome Foster, Stanley Giddings, Karla Georges, Marsha Ivey, Rahul Naidu, Risha Singh, SueMin Nathaniel, Rajini Haraksingh, Jaya Jayaraman, Chinna Chinnadurai, Adesh Ramsubhag, Nuno Faria, Oliver Pybus, Christopher Oura, Gabriel Escobar, Christine V. F. Carrington |
| EPI_ISL_977667, EPI_ISL_977668, EPI_ISL_977671, EPI_ISL_977672, EPI_ISL_977676, EPI_ISL_977679, EPI_ISL_977681, EPI_ISL_977682, EPI_ISL_977684, EPI_ISL_977685, EPI_ISL_977691, EPI_ISL_977698, EPI_ISL_977699, EPI_ISL_977700, EPI_ISL_977701, EPI_ISL_977703, EPI_ISL_977709, EPI_ISL_977712, EPI_ISL_977713, EPI_ISL_977716, EPI_ISL_977718, EPI_ISL_977731, EPI_ISL_977735, EPI_ISL_977739, EPI_ISL_977742 | | | |
| see above | California Department of Public Health | Chiu Laboratory, University of California, San Francisco | Charles Chiu, Xianding (Wayne) Deng, Candace Wang, Venice Servellita, Jill Hacker, Debra Wadford |
| EPI_ISL_978014, EPI_ISL_978015, EPI_ISL_978017, EPI_ISL_978018, EPI_ISL_978019, EPI_ISL_978020, EPI_ISL_978021, EPI_ISL_978022, EPI_ISL_978023, EPI_ISL_978024, EPI_ISL_978025, EPI_ISL_978026, EPI_ISL_978027, EPI_ISL_978028, EPI_ISL_978029, EPI_ISL_978030, EPI_ISL_978031, EPI_ISL_978032, EPI_ISL_978034, EPI_ISL_978035, EPI_ISL_978036, EPI_ISL_978037, EPI_ISL_978038, EPI_ISL_978039, EPI_ISL_978040, EPI_ISL_978041, EPI_ISL_978042, EPI_ISL_978043, EPI_ISL_978045, EPI_ISL_978049, EPI_ISL_978050, EPI_ISL_978057, EPI_ISL_978058, EPI_ISL_978059, EPI_ISL_978060, EPI_ISL_978061, EPI_ISL_978062, EPI_ISL_978131 | | | |
| see above | Chiu Laboratory, University of California, San Francisco | Chiu Laboratory, University of California, San Francisco | Charles Chiu, Xianding (Wayne) Deng, Candace Wang, Venice Servellita, Jill Hacker, Debra Wadford |
| EPI_ISL_978181, EPI_ISL_978182, EPI_ISL_978183, EPI_ISL_978184, EPI_ISL_978185, EPI_ISL_978186, EPI_ISL_978187, EPI_ISL_978188, EPI_ISL_978189, EPI_ISL_978190, EPI_ISL_978191, EPI_ISL_978192 | | | |
| see above | Virginia Division of Consolidated Laboratory Services | Virginia Division of Consolidated Laboratory Services | Virginia DCLS |
| EPI_ISL_978956, EPI_ISL_978957, EPI_ISL_978958, EPI_ISL_978959 | Chiu Laboratory, University of California, San Francisco | Chiu Laboratory, University of California, San Francisco | Charles Chiu, Xianding (Wayne) Deng, Candace Wang, Venice Servellita, Jill Hacker, Debra Wadford |
| EPI_ISL_978975, EPI_ISL_978976 | California Department of Public Health | Chiu Laboratory, University of California, San Francisco | Charles Chiu, Xianding (Wayne) Deng, Candace Wang, Venice Servellita, Jill Hacker, Debra Wadford |
| EPI_ISL_979228, EPI_ISL_979229, EPI_ISL_979230, EPI_ISL_979231, EPI_ISL_979232, EPI_ISL_979233, EPI_ISL_979235, EPI_ISL_979236, EPI_ISL_979237, EPI_ISL_979238, EPI_ISL_979239, EPI_ISL_979240, EPI_ISL_979241, EPI_ISL_979242, EPI_ISL_979243, EPI_ISL_979244, EPI_ISL_979245 | | | |
| see above | Humboldt County Public Health Laboratory | Chan-Zuckerberg Biohub | CZB Cliahub Consortium |
| EPI_ISL_979306, EPI_ISL_979307 | Cadham Provincial laboratory | National Microbiology Laboratory (NML) | Anna Majer, Shari Tyson, Grace Seo, Philip Mabon, Elsie Grudeski, Rhiannon Huzarewich, Russell Mandes, Anneliese Landgraff, Jennifer Tanner, Natalie Knox, Morag Graham, Gary Van Domselaar, Paul Van Caeseele, Jared Bullard, David Alexander, Kerry Dust, Nathalie Bastien, Yan Li, Timothy Booth, Darian Hole, Madison Chapel, Kirsten Biggar, CanCOGeN's metadata curation team, Public Health Agency of Canada CanCOGeN team |
| EPI_ISL_979330, EPI_ISL_979331, EPI_ISL_979332, EPI_ISL_979333, EPI_ISL_979334 | Laboratorio Estatal de Salud Pública de Nuevo León | Laboratorio de Infectología Molecular, Departamento de Bioquímica y Medicina Molecular,Facultad de Medicina - Universidad Autónoma de Nuevo León | Kame A. Galán-Huerta, María F. Herrera-Saldivar, Natalia Martínez-Acuña, Sonia A. Lozano-Sepúlveda, Daniel Arellanos-Soto, Ana M. Rivas-Estilla, Samuel Buentello-Wong, Else del Carmen García-García, Gloria A. Jasso-de-la-Peña, Roberto Montes-de-Oca, Consuelo Treviño-Garza, Manuel E. de-la-O-Cavazos |
| EPI_ISL_979594, EPI_ISL_979595, EPI_ISL_979607, EPI_ISL_979608, EPI_ISL_979609, EPI_ISL_979610, EPI_ISL_979611, EPI_ISL_979614, EPI_ISL_979615, EPI_ISL_979616, EPI_ISL_979617, EPI_ISL_979618, EPI_ISL_979619, EPI_ISL_979620, EPI_ISL_979621 | | | |
| see above | Santa Clara County Public Health Laboratory | Chan-Zuckerberg Biohub | CZB Cliahub Consortium |
| EPI_ISL_981968 | Microbiology Service, Hospital Universitario Clinico San Cecilio, Granada | Microbiology Service, Hospital Universitario Clinico San Cecilio, Granada | Adolfo de Salazar, Natalia Chueca, Laura Viñuela, Ana Fuentes, Federico García |
| EPI_ISL_983856, EPI_ISL_983857, EPI_ISL_983858, EPI_ISL_983859, EPI_ISL_983860 | Colorado Department of Public Health and Environment | Colorado Department of Puplic Health and Environment | Laura Bankers, Molly C. Hetherington-Rauth, Diana Ir, Shannon Ely, Shannon R. Matzinger, Sarah Elizabeth Totten, Emily A. Travanty |
