## Supplementary material for "Detection and characterization of the SARS-CoV-2 lineage B.1.526 in New York": Supp. Table 3 GISAID Acknowledgment part 2: gisaid_hcov-19_acknowledgement_table_2021_02_13_010-4.pdf

We gratefully acknowledge the following Authors from the Originating laboratories responsible for obtaining the specimens, as well as the Submitting laboratories where the genome data were generated and shared via GISAID, on which this research is based.

All Submitters of data may be contacted directly via www.gisaid.org

Authors are sorted alphabetically.

| Accession ID | Originating Laboratory | Submitting Laboratory | Authors |
| --- | --- | --- | --- |
| EPI_ISL_745029, EPI_ISL_745030, EPI_ISL_745031 | Area of Virology, Serology and Virology Division (SAViD), New South Wales Health Pathology Randwick | Area of Virology, Serology and Virology Division (SAViD), New South Wales Health Pathology Randwick | Rawlinson, W. |
| EPI_ISL_752599 | SA Pathology | SA Pathology | Lex Leong, Julien Soubrier, Chuan Kok Lim, Song Gao, Mark Turra, Karin Kassahn, Ivan Bastian, Geoff Higgins |
| EPI_ISL_754302 | Respiratory Virus Unit, National Infection Service, Public Health England | COVID-19 Genomics UK (COG-UK) Consortium | PHE Covid Sequencing Team |
| EPI_ISL_755574 | Center of Advanced Studies and Technology, CAST | Center of Advanced Studies and Technology, CAST | Ferrante,R., Mandatori,D., De Fabritiis,S. |
| EPI_ISL_755632, EPI_ISL_755633 | LabPLUS | Institute of Environmental Science and Research (ESR) | Xiaoyun Ren, Matt Storey, Nikki Freed, Muhammad Faisal, Jing Wang, Hermes Perez, Anja Werno, Antje van der Linden, Arlo Upton, Chris Mansell, David Hammer, Dragana Drinkovic, Gary McAuliffe, Hana Sofia Andersson, James Ussher, Jill Sherwood, Josh Freeman, Julia Howard, Juliet Elvy, Mary DeAlmeida, Matt Blakiston, Matthew Rogers, Max Bloomfield, Michael Addidle, Michelle Balm, Sally Roberts, Sarah Jefferies, Sharmini Muttaiyah, Susan Morpeth, Susan Taylor, Timothy Blackmore, Vani Sathyendran, Veronica Playle, Virginia Hope, Erasmus Smit, Lauren Jelly, Olin Silander, Joep de Ligt |
| EPI_ISL_755635, EPI_ISL_755636 | Middlemore Hospital | Institute of Environmental Science and Research (ESR) | Xiaoyun Ren, Matt Storey, Nikki Freed, Muhammad Faisal, Jing Wang, Hermes Perez, Anja Werno, Antje van der Linden, Arlo Upton, Chris Mansell, David Hammer, Dragana Drinkovic, Gary McAuliffe, Hana Sofia Andersson, James Ussher, Jill Sherwood, Josh Freeman, Julia Howard, Juliet Elvy, Mary DeAlmeida, Matt Blakiston, Matthew Rogers, Max Bloomfield, Michael Addidle, Michelle Balm, Sally Roberts, Sarah Jefferies, Sharmini Muttaiyah, Susan Morpeth, Susan Taylor, Timothy Blackmore, Vani Sathyendran, Veronica Playle, Virginia Hope, Erasmus Smit, Lauren Jelly, Olin Silander, Joep de Ligt |
| EPI_ISL_756274 | Hospital General Universitario Gregorio Marañón | Hospital General Universitario Gregorio Marañón | Sergio Buenestado Serrano, Pedro Sola Campoy, Pilar Catalán, Patricia Muñoz, Darío García de Viedma. |
| EPI_ISL_756375 | Siriraj Hospital, Mahidol University | National Institute of Health, Department of Medical Sciences, Ministry of Public Health, Thailand | Pilailuk Okada; Siripaporn Phuygun; Sittiporn Parnmen; Navin Horthongkham; Ratana Tacharoenmuang; Pakorn Piromtong; Natchaya Khiadsang; Thanutsapa Thanadachakul; Warawan Wongboot; Sirikanda wimol; Sunthareeya Waicharoen; Niracha Athipanyasilp; Archiraya Pattama; Chutikarn Chaimayo; Ruengpung Sutthent; Wannee Kantakamalakul. |
| EPI_ISL_757286 | Department of Virology, Public Health Laboratories Division | Department of Virology, Public Health Laboratories Division | Massab Umair, Aamer Ikram, Muhammad Salman |
| EPI_ISL_759723 | Instituto Nacional de Saude (INSA) | Instituto Nacional de Saude (INSA) | Borges et al |
| EPI_ISL_759724, EPI_ISL_759741, EPI_ISL_759743, EPI_ISL_759745, EPI_ISL_759746, EPI_ISL_759747, EPI_ISL_759748, EPI_ISL_759749, EPI_ISL_759750, EPI_ISL_759751 | University of Wisconsin-Madison AIDS Vaccine Research Laboratories | University of Wisconsin-Madison AIDS Vaccine Research Laboratories | Gage Moreno, Katarina Braun, et al. AIDS Vaccine Research Laboratories |
| EPI_ISL_763310 | Dutch COVID-19 response team | Erasmus Medical Center | Bas Oude Munnink, Reina Sikkema, David Nieuwenhuijse, Irina Chestakova, Anne van der Linden, Marjan Boter, Emmanuelle Munger, Corine GeurtsvanKessel, Annemiek van der Eijk, Richard Molenkamp, Marion Koopmans, on behalf of the Dutch national COVID-19 response team. |
| EPI_ISL_763888, EPI_ISL_764141 | Department of Pathology, University of Cambridge | COVID-19 Genomics UK (COG-UK) Consortium | Aminu S. Jahun, Yasmin Chaudhry, Grant Hall, Iliana Georgana, Myra Hosmillo, Martin D. Curran, Malte Pinckert, Surendra Parmar, Ian Goodfellow |
| EPI_ISL_764231 | Virology Department, Sheffield Teaching Hospitals NHS Foundation Trust/Department of Infection, Immunity and Cardiovascular Disease, The Medical School, University of Sheffield | COVID-19 Genomics UK (COG-UK) Consortium | Thushan de Silva, Matthew Parker, Nikki Smith, Adri Angyal, Rebecca Brown, Luke Green, Rachel Tucker, Paul Parsons, Danielle Groves, Katie Johnson, Laura Carrilero, Alex Keeley, Dave Partridge, Matthew Wyles, Benjamin Lindsey, Mehmet Yavuz, Mohammad Raza, Cariad Evans |
| EPI_ISL_764354 | Department of Pathology, University of Cambridge | COVID-19 Genomics UK (COG-UK) Consortium | Aminu S. Jahun, Yasmin Chaudhry, Grant Hall, Iliana Georgana, Myra Hosmillo, Martin D. Curran, Malte Pinckert, Surendra Parmar, Ian Goodfellow |
| EPI_ISL_765571 | National Institute for Infectious Diseases, INMI, "L. Spallanzani" IRCCS | National Institute for Infectious Diseases, INMI, "L. Spallanzani" IRCCS | E Giombini, B Bartolini, O Butera, C.E.M Gruber, M Rueca, F Messina, MR Capobianchi, A Di Caro |
| EPI_ISL_766022, EPI_ISL_766023, EPI_ISL_766027 | Department of Virology and Immunology, University of Helsinki and Helsinki University Hospital, Huslab Finland | Department of Virology, Faculty of Medicine, University of Helsinki, Helsinki, Finland | Teemu Smura, Olli Vapalahti, Maija Lappalainen, Satu Kurkela |
| EPI_ISL_766587 | Analytica Medizinische Laboratorien AG | Institute of Medical Virology, University of Zurich | Stefan Schmutz, Maryam Zaheri, Verena Kufner, Annette Audigé, Maria Grünberg, Kevin Steiner, Jon Huder, Cyril Shah, Riccarda Capaul, Jürg Böni, Michael Huber, Alexandra Trkola |
| EPI_ISL_766697 | Klinisk mikrobiologi, bakteriologi | The Public Health Agency of Sweden | Department of Microbiology, The Public Health Agency of Sweden |
| EPI_ISL_766699 | A05 Biomedicum | The Public Health Agency of Sweden | Department of Microbiology, The Public Health Agency of Sweden |
| EPI_ISL_766714 | Klinisk mikrobiologi | The Public Health Agency of Sweden | Department of Microbiology, The Public Health Agency of Sweden |
| EPI_ISL_766718, EPI_ISL_766719 | A05 Biomedicum | The Public Health Agency of Sweden | Department of Microbiology, The Public Health Agency of Sweden |
| EPI_ISL_766938 | Nevada State Health Lab | Nevada State Health Lab | Andrew Gorzalski |
| EPI_ISL_767013 | University of Bari Biomedical Sciences and Human Oncology | University of Bari Biomedical Sciences and Human Oncology | Maria Chironna, Anna Sallustio, Daniela Loconsole, Marisa Accogli |
| EPI_ISL_767044, EPI_ISL_767047, EPI_ISL_767048, EPI_ISL_767050, EPI_ISL_767064, EPI_ISL_767065, EPI_ISL_767066, EPI_ISL_767068, EPI_ISL_767069, EPI_ISL_767073, EPI_ISL_767074, EPI_ISL_767075, EPI_ISL_767076, EPI_ISL_767077, EPI_ISL_767078, EPI_ISL_767079, EPI_ISL_767080, EPI_ISL_767081, EPI_ISL_767082, EPI_ISL_767083, EPI_ISL_767084, EPI_ISL_767085, EPI_ISL_767086, EPI_ISL_767087, EPI_ISL_767088 | | | |
| see above | New Mexico Department of Health Scientific Laboratory | New Mexico Department of Health Scientific Laboratory | D'eldra Malone, Ellie Johnson, Anastacia Griego-Fisher |
| EPI_ISL_767613 | SARATOGA HOSPITAL LABORATORY | Wadsworth Center, New York State Department.of Health | Kirsten St. George, Daryl M. Lamson, Alexis Russel, Matthew Shudt, Melissa A Leisner, Jonathan Plitnick, Navjot Singh, John Kelly, Sara Griesemer, Erasmus Schneider, Erica Lasek-Nesselquist |
| EPI_ISL_767902 | Australian Clinical Labs | NSW Health Pathology - Institute of Clinical Pathology and Medical Research; Westmead Hospital; University of Sydney | CIDM-PH et al. |
| EPI_ISL_767903 | Laverty Pathology | NSW Health Pathology - Institute of Clinical Pathology and Medical Research; Westmead Hospital; University of Sydney | CIDM-PH et al. |
| EPI_ISL_767905 | Sydney South West Pathology Service (SSWPS) - Liverpool Hospital - NSW Health Pathology | NSW Health Pathology - Institute of Clinical Pathology and Medical Research; Westmead Hospital; University of Sydney | CIDM-PH et al. |
| EPI_ISL_767907 | Sydney South West Pathology Service (SSWPS) - Royal Prince Alfred Hospital - NSW Health Pathology | NSW Health Pathology - Institute of Clinical Pathology and Medical Research; Westmead Hospital; University of Sydney | CIDM-PH et al. |
| EPI_ISL_767916, EPI_ISL_767917, EPI_ISL_767918 | Australian Clinical Labs | NSW Health Pathology - Institute of Clinical Pathology and Medical Research; Westmead Hospital; University of Sydney | CIDM-PH et al. |

| | | | |
|---|---|---|---|
| EPI_ISL_767919 | Laverty Pathology | NSW Health Pathology - Institute of Clinical Pathology and Medical Research; Westmead Hospital; University of Sydney | CIDM-PH et al. |
| EPI_ISL_767920 | Pathology North - Hunter - NSW Health Pathology | NSW Health Pathology - Institute of Clinical Pathology and Medical Research; Westmead Hospital; University of Sydney | CIDM-PH et al. |
| EPI_ISL_767923, EPI_ISL_767924 | South Eastern Area Laboratory Services (SEALS) | NSW Health Pathology - Institute of Clinical Pathology and Medical Research; Westmead Hospital; University of Sydney | CIDM-PH et al. |
| EPI_ISL_767925, EPI_ISL_767926 | Histopath | NSW Health Pathology - Institute of Clinical Pathology and Medical Research; Westmead Hospital; University of Sydney | CIDM-PH et al. |
| EPI_ISL_767929 | Southern. IML Pathology | NSW Health Pathology - Institute of Clinical Pathology and Medical Research; Westmead Hospital; University of Sydney | CIDM-PH et al. |
| EPI_ISL_767930 | Sydney South West Pathology Service (SSWPS) - Royal Prince Alfred Hospital - NSW Health Pathology | NSW Health Pathology - Institute of Clinical Pathology and Medical Research; Westmead Hospital; University of Sydney | CIDM-PH et al. |
| EPI_ISL_768602 | South Eastern Area Laboratory Services (SEALS) | NSW Health Pathology - Institute of Clinical Pathology and Medical Research; Westmead Hospital; University of Sydney | CIDM-PH et al. |
| EPI_ISL_768619, EPI_ISL_768622, EPI_ISL_768625, EPI_ISL_768626, EPI_ISL_768627 | National Public Health Laboratory, National Centre for Infectious Diseases | National Public Health Laboratory, National Centre for Infectious Diseases | Tze Minn Mak, Sophie Octavia, Zhenyang Zhou, Lin Cui, Raymond Tzer Pin Lin |
| EPI_ISL_768824 | Laboratoire BIOLAB33 | CNR Virus des Infections Respiratoires - France SUD | Antonin Bal, Gregory Destras, Gwendolyne Burfin, Hadrien Règue, Quentin Semanas, Martine Valette, Bruno Lina, Laurence Josset |
| EPI_ISL_768829 | Hôpital d'Instruction des Armées Sainte-Anne | CNR Virus des Infections Respiratoires - France SUD | Antonin Bal, Gregory Destras, Gwendolyne Burfin, Hadrien Règue, Quentin Semanas, Martine Valette, Bruno Lina, Laurence Josset |
| EPI_ISL_768830 | Laboratoire BIOMED 05 | CNR Virus des Infections Respiratoires - France SUD | Antonin Bal, Gregory Destras, Gwendolyne Burfin, Hadrien Règue, Quentin Semanas, Martine Valette, Bruno Lina, Laurence Josset |
| EPI_ISL_769917, EPI_ISL_769918, EPI_ISL_769941, EPI_ISL_769946, EPI_ISL_769950, EPI_ISL_769951, EPI_ISL_769952, EPI_ISL_769953, EPI_ISL_769954, EPI_ISL_769956, EPI_ISL_769957, EPI_ISL_769958, EPI_ISL_769971, EPI_ISL_769972, EPI_ISL_769973, EPI_ISL_769974, EPI_ISL_769975, EPI_ISL_769976, EPI_ISL_769977, EPI_ISL_769978, EPI_ISL_769979, EPI_ISL_769980, EPI_ISL_769981, EPI_ISL_769982, EPI_ISL_769983, EPI_ISL_769984, EPI_ISL_769985 | | | |
| see above | Albany Medical Center Hospital Clinical Laboratories | Wadsworth Center, New York State Department.of Health | Kirsten St. George, Daryl M. Lamson, Alexis Russel, Matthew Shudt, Melissa A Leisner, Jonathan Plitnick, Navjot Singh, John Kelly, Sara Griesemer, Erasmus Schneider, Erica Lasek-Nesselquist |
| EPI_ISL_770442, EPI_ISL_770443, EPI_ISL_770444 | Wyoming Public Health Laboratory | Wyoming Public Health Laboratory | Noah Hull, Taylor Fearing, Lynette Gumbleton, Channing Weber, Ashley Norberg, Bailey Bowcutt, and Wanda Manley |
| EPI_ISL_770462, EPI_ISL_770463, EPI_ISL_770464, EPI_ISL_770465, EPI_ISL_770466, EPI_ISL_770467, EPI_ISL_770468, EPI_ISL_770469 | Respiratory Virus Unit, National Infection Service, Public Health England | COVID-19 Genomics UK (COG-UK) Consortium | PHE Covid Sequencing Team |
| EPI_ISL_770850, EPI_ISL_770851, EPI_ISL_770852, EPI_ISL_770855, EPI_ISL_770856, EPI_ISL_770858, EPI_ISL_770860, EPI_ISL_770868, EPI_ISL_770872, EPI_ISL_770881, EPI_ISL_770883, EPI_ISL_770889, EPI_ISL_770895, EPI_ISL_771051, EPI_ISL_771052, EPI_ISL_771060, EPI_ISL_771090, EPI_ISL_771092, EPI_ISL_771093, EPI_ISL_771095, EPI_ISL_771096, EPI_ISL_771112, EPI_ISL_771113, EPI_ISL_771114, EPI_ISL_771115, EPI_ISL_771116, EPI_ISL_771117, EPI_ISL_771118, EPI_ISL_771119, EPI_ISL_771120, EPI_ISL_771121, EPI_ISL_771122, EPI_ISL_771123, EPI_ISL_771124, EPI_ISL_771125, EPI_ISL_771126, EPI_ISL_771127, EPI_ISL_771133, EPI_ISL_771134, EPI_ISL_771141, EPI_ISL_771142, EPI_ISL_771143, EPI_ISL_771144, EPI_ISL_771145, EPI_ISL_771146 | | | |
| see above | Laboratoire national de santé, Microbiology, Virology | Laboratoire national de santé, Microbiology, Microbial Genomics Platform | Anke Wienecke-Baldacchino, Catherine Ragimbeau,Jessica Tapp, Fatu Djabi, Lise Pignon, Raoul Salmon, Tamir Abdelrahman |
| EPI_ISL_775428, EPI_ISL_775429, EPI_ISL_775432, EPI_ISL_775433, EPI_ISL_775436 | Department of Medical Microbiology - section Molde, Molde Hospital | Norwegian Institute of Public Health, Department of Virology | Kathrine Stene-Johansen, Kamilla Heddeland Instefjord, Hilde Elshaug, Atiya R Ali,Marie Paulsen Madsen, Rasmus Riis Kopperud, Hilde Vollan, Karoline Bragstad, Olav Hungnes |
| EPI_ISL_776569, EPI_ISL_776575, EPI_ISL_776590, EPI_ISL_776604, EPI_ISL_776609, EPI_ISL_776653, EPI_ISL_776654, EPI_ISL_776655, EPI_ISL_776656, EPI_ISL_776775, EPI_ISL_776776, EPI_ISL_776777, EPI_ISL_776778, EPI_ISL_776779, EPI_ISL_776780, EPI_ISL_776781, EPI_ISL_776782, EPI_ISL_776783, EPI_ISL_776784, EPI_ISL_776785, EPI_ISL_776786, EPI_ISL_776787, EPI_ISL_776788, EPI_ISL_776789, EPI_ISL_776790, EPI_ISL_776791, EPI_ISL_776792, EPI_ISL_776793, EPI_ISL_776794, EPI_ISL_776795, EPI_ISL_776796, EPI_ISL_776797, EPI_ISL_776798, EPI_ISL_776799, EPI_ISL_776800, EPI_ISL_776801, EPI_ISL_776802, EPI_ISL_776803, EPI_ISL_776804, EPI_ISL_776805, EPI_ISL_776806, EPI_ISL_776807, EPI_ISL_776808, EPI_ISL_776809, EPI_ISL_776810, EPI_ISL_776811, EPI_ISL_776812, EPI_ISL_776813, EPI_ISL_776814, EPI_ISL_776815, EPI_ISL_776816, EPI_ISL_776817, EPI_ISL_776818, EPI_ISL_776819, EPI_ISL_776820, EPI_ISL_776821, EPI_ISL_776822, EPI_ISL_776823, EPI_ISL_776824, EPI_ISL_776825, EPI_ISL_776826, EPI_ISL_776827, EPI_ISL_776828, EPI_ISL_776829, EPI_ISL_776830, EPI_ISL_776831, EPI_ISL_776832, EPI_ISL_776833, EPI_ISL_776834, EPI_ISL_776835, EPI_ISL_776836, EPI_ISL_776837, EPI_ISL_776838, EPI_ISL_776839, EPI_ISL_776840, EPI_ISL_776841, EPI_ISL_776842, EPI_ISL_776843, EPI_ISL_776844, EPI_ISL_776845, EPI_ISL_776846, EPI_ISL_776847, EPI_ISL_776848, EPI_ISL_776849, EPI_ISL_776850, EPI_ISL_776851, EPI_ISL_776852, EPI_ISL_776853, EPI_ISL_776854, EPI_ISL_776855, EPI_ISL_776856, EPI_ISL_776857, EPI_ISL_776858, EPI_ISL_776859, EPI_ISL_776860, EPI_ISL_776861, EPI_ISL_776862, EPI_ISL_776863, EPI_ISL_776864, EPI_ISL_776865, EPI_ISL_776866, EPI_ISL_776867, EPI_ISL_776868 | | | |
| see above | Israel Central Virology laboratory | Israel Central Virology laboratory | Neta Zuckerman, Efrat Dahan Bucris, Oran Erster, Michal Mandelboim, Orna Mor, Ella Mendelson |
| EPI_ISL_778821 | AIID | Irish Coronavirus Sequencing Consortium-Teagasc Grange | Matthew McCabe, Aljandro Abner Garcia Leon, Fiona Crispie, Calum Walsh, Michael Carr, John Kenny, Paul Cotter, Patrick Mallon, Gabriel Gonzalez |
| EPI_ISL_778842 | Clinical Pathology Labs | Centers for Disease Control and Prevention Division of Viral Diseases, Pathogen Discovery | Ying Tao Yan Li Jing Zhang Krista Queen Anna Uehara Peter Cook Clinton R. Paden Haibin Wang Suxiang Tong |
| EPI_ISL_779137, EPI_ISL_779141, EPI_ISL_779142, EPI_ISL_779143, EPI_ISL_779144, EPI_ISL_779145, EPI_ISL_779146 | Yale Clinical Virology Laboratory | Grubaugh Lab - Yale School of Public Health | Tara Alpert, Joseph Fauver, Anderson Brito, Mallery Breban, Anne Wyllie, Chantal Vogels, Mary Petrone, Chaney Kalinich, Isabel Ott, Arnau Casanovas, Catherine Muenker, Adam Moore, Alice Lu, Maria Tokuyama, Patrick Wong, Peiwen Lu, Saad Omer, Richard Martinello, Allison Nelson, Shelli Farhadian, Akiko Iwasaki, Charlese Dela Cruz, Albert Ko, Nathan Grubaugh |
| EPI_ISL_779356, EPI_ISL_779364, EPI_ISL_779369, EPI_ISL_779371 | University of Wisconsin-Madison AIDS Vaccine Research Laboratories | University of Wisconsin-Madison AIDS Vaccine Research Laboratories | Gage Moreno, Katarina Braun, et al. AIDS Vaccine Research Laboratories |
| EPI_ISL_779621 | Victorian Infectious Diseases Reference Laboratory (VIDRL) | VIDRL and MDU-PHL | Caly L., Seemann T., Sait, M.L., Druce J., Sherry, N.L. |
| EPI_ISL_779651 | Institute of Virology, Biomedical Research Center of the Slovak Academy of Sciences, Bratislava | Faculty of Natural Sciences, Comenius University, Bratislava | Viktória abanová, Kristína Boršová, Broa Brejová, Viktória Hodorová, Sabina Fumaová Havlíková, Juraj Kopáek, Martina Liková, ubomíra Lukáiková, Martina Neboháová, Monika Sláviková, Andrej Belák, Tomáš Vina, Jozef Nosek, Boris Klempa |
| EPI_ISL_779652 | Institute of Virology, Biomedical Research Center of the Slovak Academy of Sciences, Bratislava | Faculty of Natural Sciences, Comenius University, Bratislava | Kristína Boršová, Viktória abanová, Broa Brejová, Viktória Hodorová, Sabina Fumaová Havlíková, Juraj Kopáek, Martina Liková, ubomíra Lukáiková, Martina Neboháová, Monika Sláviková, Andrej Belák, Tomáš Vina, Boris Klempa, Jozef Nosek |
| EPI_ISL_779653 | Institute of Virology, Biomedical Research Center of the Slovak Academy of Sciences, Bratislava | Faculty of Natural Sciences, Comenius University, Bratislava | Viktória abanová, Kristína Boršová, Broa Brejová, Viktória Hodorová, Sabina Fumaová Havlíková, Juraj Kopáek, Martina Liková, ubomíra Lukáiková, Martina Neboháová, Monika Sláviková, Andrej Belák, Tomáš Vina, Jozef Nosek, Boris Klempa |
| EPI_ISL_779654 | Institute of Virology, Biomedical Research Center of the Slovak Academy of Sciences, Bratislava | Faculty of Natural Sciences, Comenius University, Bratislava | Kristína Boršová, Viktória abanová, Broa Brejová, Viktória Hodorová, Sabina Fumaová Havlíková, Juraj Kopáek, Martina Liková, ubomíra Lukáiková, Martina Neboháová, Monika Sláviková, Andrej Belák, Tomáš Vina, Boris Klempa, Jozef Nosek |
| EPI_ISL_779655 | Institute of Virology, Biomedical Research Center of the Slovak Academy of Sciences, Bratislava | Faculty of Natural Sciences, Comenius University, Bratislava | Viktória abanová, Kristína Boršová, Broa Brejová, Viktória Hodorová, Sabina Fumaová Havlíková, Juraj Kopáek, Martina Liková, ubomíra Lukáiková, Martina Neboháová, Monika Sláviková, Andrej Belák, Tomáš Vina, Jozef Nosek, Boris Klempa |
| EPI_ISL_779656 | Institute of Virology, Biomedical Research Center of the Slovak Academy of Sciences, Bratislava | Faculty of Natural Sciences, Comenius University, Bratislava | Kristína Boršová, Viktória abanová, Broa Brejová, Viktória Hodorová, Sabina Fumaová Havlíková, Juraj Kopáek, Martina Liková, ubomíra Lukáiková, Martina Neboháová, Monika Sláviková, Andrej Belák, Tomáš Vina, Boris Klempa, Jozef Nosek |
| EPI_ISL_779657, EPI_ISL_779658, EPI_ISL_779659 | Institute of Virology, Biomedical Research Center of the Slovak Academy of Sciences, Bratislava | Faculty of Natural Sciences, Comenius University, Bratislava | Broa Brejová, Viktória abanová, Kristína Boršová, Viktória Hodorová, Sabina Fumaová Havlíková, Juraj Kopáek, Martina Liková, ubomíra Lukáiková, Martina Neboháová, Monika Sláviková, Andrej Belák, Tomáš Vina, Jozef Nosek, Boris Klempa |
| EPI_ISL_779660 | Institute of Virology, Biomedical Research Center of the Slovak Academy of Sciences, Bratislava | Faculty of Natural Sciences, Comenius University, Bratislava | Viktória abanová, Kristína Boršová, Broa Brejová, Viktória Hodorová, Sabina Fumaová Havlíková, Juraj Kopáek, Martina Liková, ubomíra Lukáiková, Martina Neboháová, Monika Sláviková, Andrej Belák, Tomáš Vina, Jozef Nosek, Boris Klempa |

| | | | |
|---|---|---|---|
| EPI_ISL_779810, EPI_ISL_779811, EPI_ISL_779812 | CNR Virus des Infections Respiratoires - France SUD | CNR Virus des Infections Respiratoires - France SUD | Antonin Bal, Gregory Destras, Gwendolyne Burfin, Hadrien Règue, Quentin Semanas, Martine Valette, Bruno Lina, Laurence Josset |
| EPI_ISL_779836 | Laboratoire Mirialis | CNR Virus des Infections Respiratoires - France SUD | Antonin Bal, Gregory Destras, Gwendolyne Burfin, Hadrien Règue, Quentin Semanas, Martine Valette, Bruno Lina, Laurence Josset |
| EPI_ISL_788920 | University of Wisconsin-Madison AIDS Vaccine Research Laboratories | University of Wisconsin-Madison AIDS Vaccine Research Laboratories | Gage Moreno, Katarina Braun, et al. AIDS Vaccine Research Laboratories |
| EPI_ISL_789044 | Klinisk mikrobiologi | The Public Health Agency of Sweden | Department of Microbiology, The Public Health Agency of Sweden |
| EPI_ISL_789054 | Unilabs AB | The Public Health Agency of Sweden | Department of Microbiology, The Public Health Agency of Sweden |
| EPI_ISL_790588, EPI_ISL_790984, EPI_ISL_790985, EPI_ISL_791032 | Dutch COVID-19 response team | National Institute for Public Health and the Environment (RIVM) | Adam Meijer, Harry Vennema, Jeroen Cremer, Sharon van den Brink, Bas van der Veer, AnneMarie van den Brandt, Florian Zwagemaker, Dennis Schmitz, Chantal Reusken, on behalf of the national COVID-19 response team |
| EPI_ISL_791087, EPI_ISL_791088 | Instituto Nacional de Salud - Unidad de Secuenciación y Análisis Genómico | Instituto Nacional de Salud - Dirección de Investigación en Salud Pública | Katherine Laiton-Donato, Diego A. Álvarez-Díaz, Carlos Franco-Muñoz, Mauricio Pacheco-Montealegre, Jonathan Reales, Sheryl Corchuelo, Maria T. Herrera, Julian Naizaque, Gerardo Santamaría, Paola Muñoz-Laiton, Diego Andrés Prada, Magdalena Wiesner, Martha Lucia Ospina Martinez, Marcela Mercado-Reyes |
| EPI_ISL_791139 | University of Wisconsin-Madison AIDS Vaccine Research Laboratories | University of Wisconsin-Madison AIDS Vaccine Research Laboratories | Gage Moreno, Katarina Braun, et al. AIDS Vaccine Research Laboratories |
| EPI_ISL_791335, EPI_ISL_791337, EPI_ISL_791339, EPI_ISL_791424, EPI_ISL_791435, EPI_ISL_791436, EPI_ISL_791443, EPI_ISL_791444, EPI_ISL_791451, EPI_ISL_791463, EPI_ISL_791474, EPI_ISL_791477, EPI_ISL_791485, EPI_ISL_791486 | | | |
| see above | Johns Hopkins Hospital Department of Pathology | Johns Hopkins Hospital Department of Pathology | C. Paul Morris, Chun Huai Luo, Adannaya Amadi, Nicholas Gallagher, Heba H. Mostafa |
| EPI_ISL_792039 | hospital | National Reference Center for Viruses of Respiratory Infections, Institut Pasteur, Paris | Marion Barbet, Sylvie Behillil, Méline Bizard, Angela Brisebarre, Camille Capel, Etienne Simon-Lorière, Vincent Enouf, Maud Vanpeene, Sylvie van der Werf,Laurent Roudière |
| EPI_ISL_792054 | hospital | National Reference Center for Viruses of Respiratory Infections, Institut Pasteur, Paris | Marion Barbet, Sylvie Behillil, Méline Bizard, Angela Brisebarre, Camille Capel, Etienne Simon-Lorière, Vincent Enouf, Maud Vanpeene, Sylvie van der Werf,Esther Gyde |
| EPI_ISL_792055 | hospital | National Reference Center for Viruses of Respiratory Infections, Institut Pasteur, Paris | Marion Barbet, Sylvie Behillil, Méline Bizard, Angela Brisebarre, Camille Capel, Etienne Simon-Lorière, Vincent Enouf, Maud Vanpeene, Sylvie van der Werf,Patricia Stoessel |
| EPI_ISL_792084, EPI_ISL_792085 | Toronto Invasive Bacterial Diseases Network | McMaster University | Allison McGeer, Patryk Aftanas, Hooman Derakhshani, Angel Li, Kuganya Nirmalarajah, Emily Panousis, Ahmed Draia, Jalees Nasir, Michael Surette, Samira Mubareka, Andrew G. McArthur |
| EPI_ISL_792674, EPI_ISL_792675 | Los Angeles County PHL | Los Angeles County PHL | P. Hemarajata et al. |
| EPI_ISL_793030, EPI_ISL_793031, EPI_ISL_793032, EPI_ISL_793033, EPI_ISL_793034, EPI_ISL_793035, EPI_ISL_793036, EPI_ISL_793037, EPI_ISL_793038, EPI_ISL_793039, EPI_ISL_793040, EPI_ISL_793041, EPI_ISL_793042, EPI_ISL_793043, EPI_ISL_793044, EPI_ISL_793045, EPI_ISL_793046, EPI_ISL_793047, EPI_ISL_793048, EPI_ISL_793049, EPI_ISL_793050, EPI_ISL_793116, EPI_ISL_793117, EPI_ISL_793120, EPI_ISL_793130, EPI_ISL_793134, EPI_ISL_793142, EPI_ISL_793145, EPI_ISL_793150, EPI_ISL_793151, EPI_ISL_793159, EPI_ISL_793163, EPI_ISL_793164, EPI_ISL_793171, EPI_ISL_793179, EPI_ISL_793180, EPI_ISL_793181, EPI_ISL_793182, EPI_ISL_793183, EPI_ISL_793184, EPI_ISL_793185, EPI_ISL_793186, EPI_ISL_793187, EPI_ISL_793188, EPI_ISL_793189, EPI_ISL_793190, EPI_ISL_793191, EPI_ISL_793192, EPI_ISL_793193, EPI_ISL_793205, EPI_ISL_793206, EPI_ISL_793207, EPI_ISL_793208, EPI_ISL_793209, EPI_ISL_793210, EPI_ISL_793211, EPI_ISL_793212, EPI_ISL_793213, EPI_ISL_793214, EPI_ISL_793215, EPI_ISL_793216, EPI_ISL_793217, EPI_ISL_793296, EPI_ISL_793299, EPI_ISL_793301, EPI_ISL_793316, EPI_ISL_793317, EPI_ISL_793318, EPI_ISL_793319, EPI_ISL_793320, EPI_ISL_793321, EPI_ISL_793322, EPI_ISL_793323, EPI_ISL_793324, EPI_ISL_793325, EPI_ISL_793326, EPI_ISL_793327, EPI_ISL_793328, EPI_ISL_793329, EPI_ISL_793330, EPI_ISL_793331, EPI_ISL_793371, EPI_ISL_793374, EPI_ISL_793394, EPI_ISL_793396, EPI_ISL_793399, EPI_ISL_793400, EPI_ISL_793401, EPI_ISL_793402, EPI_ISL_793403, EPI_ISL_793406, EPI_ISL_793417, EPI_ISL_793421, EPI_ISL_793422, EPI_ISL_793423, EPI_ISL_793424, EPI_ISL_793425, EPI_ISL_793426, EPI_ISL_793427, EPI_ISL_793428, EPI_ISL_793429, EPI_ISL_793430, EPI_ISL_793431, EPI_ISL_793432, EPI_ISL_793433, EPI_ISL_793434, EPI_ISL_793435, EPI_ISL_793436, EPI_ISL_793437, EPI_ISL_793438, EPI_ISL_793439, EPI_ISL_793440, EPI_ISL_793441, EPI_ISL_793442, EPI_ISL_793443, EPI_ISL_793444, EPI_ISL_793445, EPI_ISL_793446, EPI_ISL_793447, EPI_ISL_793448, EPI_ISL_793449, EPI_ISL_793450, EPI_ISL_793451, EPI_ISL_793452, EPI_ISL_793453, EPI_ISL_793454, EPI_ISL_793455, EPI_ISL_793456, EPI_ISL_793457, EPI_ISL_793458, EPI_ISL_793459, EPI_ISL_793460, EPI_ISL_793461, EPI_ISL_793462, EPI_ISL_793463, EPI_ISL_793464, EPI_ISL_793465, EPI_ISL_793466, EPI_ISL_793467, EPI_ISL_793468, EPI_ISL_793469, EPI_ISL_793470, EPI_ISL_793471, EPI_ISL_793472, EPI_ISL_793473, EPI_ISL_793474, EPI_ISL_793475, EPI_ISL_793476, EPI_ISL_793477, EPI_ISL_793478, EPI_ISL_793479, EPI_ISL_793480, EPI_ISL_793481, EPI_ISL_793482, EPI_ISL_793483, EPI_ISL_793484, EPI_ISL_793485, EPI_ISL_793486, EPI_ISL_793487, EPI_ISL_793488, EPI_ISL_793489, EPI_ISL_793490, EPI_ISL_793491, EPI_ISL_793492, EPI_ISL_793493, EPI_ISL_793494, EPI_ISL_793495, EPI_ISL_793496, EPI_ISL_793497, EPI_ISL_793498, EPI_ISL_793499, EPI_ISL_793500, EPI_ISL_793501, EPI_ISL_793502, EPI_ISL_793503, EPI_ISL_793504, EPI_ISL_793505, EPI_ISL_793506, EPI_ISL_793507, EPI_ISL_793508, EPI_ISL_793509, EPI_ISL_793510, EPI_ISL_793511, EPI_ISL_793512, EPI_ISL_793513, EPI_ISL_793514, EPI_ISL_793515, EPI_ISL_793516, EPI_ISL_793517, EPI_ISL_793518, EPI_ISL_793519, EPI_ISL_793520, EPI_ISL_793521, EPI_ISL_793522, EPI_ISL_793523, EPI_ISL_793524, EPI_ISL_793525, EPI_ISL_793526, EPI_ISL_793527, EPI_ISL_793528, EPI_ISL_793529, EPI_ISL_793530, EPI_ISL_793531, EPI_ISL_793532, EPI_ISL_793533, EPI_ISL_793534, EPI_ISL_793535, EPI_ISL_793536, EPI_ISL_793537, EPI_ISL_793538, EPI_ISL_793539, EPI_ISL_793540, EPI_ISL_793541, EPI_ISL_793542, EPI_ISL_793543, EPI_ISL_793544, EPI_ISL_793545, EPI_ISL_793546, EPI_ISL_793547, EPI_ISL_793548, EPI_ISL_793549, EPI_ISL_793550, EPI_ISL_793551, EPI_ISL_793552, EPI_ISL_793553, EPI_ISL_793554, EPI_ISL_793555, EPI_ISL_793556, EPI_ISL_793557, EPI_ISL_793558, EPI_ISL_793559, EPI_ISL_793560, EPI_ISL_793561, EPI_ISL_793562, EPI_ISL_793563, EPI_ISL_793564, EPI_ISL_793565, EPI_ISL_793566, EPI_ISL_793567, EPI_ISL_793568, EPI_ISL_793569, EPI_ISL_793570, EPI_ISL_793571, EPI_ISL_793572, EPI_ISL_793573, EPI_ISL_793574, EPI_ISL_793575, EPI_ISL_793576, EPI_ISL_793577, EPI_ISL_793578, EPI_ISL_793579, EPI_ISL_793580, EPI_ISL_793581, EPI_ISL_793582, EPI_ISL_793583, EPI_ISL_793584, EPI_ISL_793585, EPI_ISL_793586, EPI_ISL_793587, EPI_ISL_793588, EPI_ISL_793589, EPI_ISL_793590, EPI_ISL_793591, EPI_ISL_793592, EPI_ISL_793593, EPI_ISL_793594, EPI_ISL_793595, EPI_ISL_793596, EPI_ISL_793597, EPI_ISL_793598, EPI_ISL_793599, EPI_ISL_793600, EPI_ISL_793601, EPI_ISL_793602, EPI_ISL_793603, EPI_ISL_793604, EPI_ISL_793605, EPI_ISL_793606, EPI_ISL_793607, EPI_ISL_793608, EPI_ISL_793609, EPI_ISL_793610, EPI_ISL_793611, EPI_ISL_793612, EPI_ISL_793613, EPI_ISL_793614, EPI_ISL_793615, EPI_ISL_793616, EPI_ISL_793617, EPI_ISL_793618, EPI_ISL_793619, EPI_ISL_793620, EPI_ISL_793621, EPI_ISL_793622, EPI_ISL_793623, EPI_ISL_793624, EPI_ISL_793625, EPI_ISL_793626, EPI_ISL_793627, EPI_ISL_793628, EPI_ISL_793629, EPI_ISL_793630, EPI_ISL_793631, EPI_ISL_793632, EPI_ISL_793633, EPI_ISL_793634, EPI_ISL_793635, EPI_ISL_793636, EPI_ISL_793637, EPI_ISL_793638, EPI_ISL_793639, EPI_ISL_793640, EPI_ISL_793641, EPI_ISL_793642, EPI_ISL_793643, EPI_ISL_793644, EPI_ISL_793645, EPI_ISL_793646, EPI_ISL_793647, EPI_ISL_793648, EPI_ISL_793649 | | | |
| see above | Department of Virus and Microbiological Special Diagnostics, Statens Serum Institut, Copenhagen, Denmark | Albertsen Lab, Department of Chemistry and Bioscience, Aalborg University, Denmark | Danish Covid-19 Genome Consortium |
| EPI_ISL_794083, EPI_ISL_794084, EPI_ISL_794085, EPI_ISL_794086, EPI_ISL_794087, EPI_ISL_794088, EPI_ISL_794089, EPI_ISL_794090, EPI_ISL_794091, EPI_ISL_794093, EPI_ISL_794094, EPI_ISL_794095, EPI_ISL_794096, EPI_ISL_794097, EPI_ISL_794098, EPI_ISL_794099, EPI_ISL_794100, EPI_ISL_794101, EPI_ISL_794102, EPI_ISL_794103, EPI_ISL_794105, EPI_ISL_794106 | | | |
| see above | URMC LABS | Wadsworth Center, New York State Department.of Health | Kirsten St. George, Daryl M. Lamson, Alexis Russel, Matthew Shudt, Melissa A Leisner, Jonathan Plitnick, Navjot Singh, John Kelly, Sara Griesemer, Erasmus Schneider, Erica Lasek-Nesselquist |
| EPI_ISL_794112, EPI_ISL_794113, EPI_ISL_794115, EPI_ISL_794118 | GLENS FALLS HOSPITAL LABORATORY | Wadsworth Center, New York State Department.of Health | Kirsten St. George, Daryl M. Lamson, Alexis Russel, Matthew Shudt, Melissa A Leisner, Jonathan Plitnick, Navjot Singh, John Kelly, Sara Griesemer, Erasmus Schneider, Erica Lasek-Nesselquist |
| EPI_ISL_794138, EPI_ISL_794139, EPI_ISL_794141, EPI_ISL_794143, EPI_ISL_794144, EPI_ISL_794145, EPI_ISL_794148, EPI_ISL_794149, EPI_ISL_794150, EPI_ISL_794151, EPI_ISL_794152, EPI_ISL_794153, EPI_ISL_794154, EPI_ISL_794155, EPI_ISL_794156, EPI_ISL_794157, EPI_ISL_794158 | | | |
| see above | NORTHWELL HEALTH LABORATORIES | Wadsworth Center, New York State Department.of Health | Kirsten St. George, Daryl M. Lamson, Alexis Russel, Matthew Shudt, Melissa A Leisner, Jonathan Plitnick, Navjot Singh, John Kelly, Sara Griesemer, Erasmus Schneider, Erica Lasek-Nesselquist |
| EPI_ISL_794166, EPI_ISL_794167 | Wadsworth Center, New York State Department.of Health | Wadsworth Center, New York State Department.of Health | Kirsten St. George, Daryl M. Lamson, Alexis Russel, Matthew Shudt, Melissa A Leisner, Jonathan Plitnick, Navjot Singh, John Kelly, Sara Griesemer, Erasmus Schneider, Erica Lasek-Nesselquist |
| EPI_ISL_794182, EPI_ISL_794183, EPI_ISL_794184, EPI_ISL_794185, EPI_ISL_794186, EPI_ISL_794187, EPI_ISL_794188, EPI_ISL_794189, EPI_ISL_794190, EPI_ISL_794191, EPI_ISL_794192, EPI_ISL_794193, EPI_ISL_794194, EPI_ISL_794195, EPI_ISL_794196, EPI_ISL_794197, EPI_ISL_794198, EPI_ISL_794199, EPI_ISL_794200, EPI_ISL_794201, EPI_ISL_794202, EPI_ISL_794203, EPI_ISL_794204, EPI_ISL_794205 | | | |
| see above | URMC LABS | Wadsworth Center, New York State Department.of Health | Kirsten St. George, Daryl M. Lamson, Alexis Russel, Matthew Shudt, Melissa A Leisner, Jonathan Plitnick, Navjot Singh, John Kelly, Sara Griesemer, Erasmus Schneider, Erica Lasek-Nesselquist |
| EPI_ISL_794219, EPI_ISL_794220, EPI_ISL_794221, EPI_ISL_794222, EPI_ISL_794223, EPI_ISL_794224, EPI_ISL_794225, EPI_ISL_794226, EPI_ISL_794227, EPI_ISL_794228, EPI_ISL_794229, EPI_ISL_794230, EPI_ISL_794231, EPI_ISL_794232, EPI_ISL_794233, EPI_ISL_794235, EPI_ISL_794238, EPI_ISL_794241, EPI_ISL_794242, EPI_ISL_794243 | | | |
| see above | WESTCHESTER MEDICAL CENTER | Wadsworth Center, New York State Department.of Health | Kirsten St. George, Daryl M. Lamson, Alexis Russel, Matthew Shudt, Melissa A Leisner, Jonathan Plitnick, Navjot Singh, John Kelly, Sara Griesemer, Erasmus Schneider, Erica Lasek-Nesselquist |
| EPI_ISL_794661 | DRECCION DE SANIDAD POLICIA NACIONAL | Instituto Nacional de Salud - Dirección de Investigación en Salud Pública | Katherine Laiton-Donato, Diego A. Álvarez-Díaz, Carlos Franco-Muñoz, Mauricio Pacheco-Montealegre, Jonathan Reales, Sheryl Corchuelo, Maria T. Herrera, Julian Naizaque, Gerardo Santamaría, Paola Muñoz-Laiton, Diego Andrés Prada, Magdalena Wiesner, Martha Lucia Ospina Martinez, Marcela Mercado-Reyes |
| EPI_ISL_795011, EPI_ISL_795012, EPI_ISL_795013, EPI_ISL_795014, EPI_ISL_795015, EPI_ISL_795016, EPI_ISL_795017, EPI_ISL_795018, EPI_ISL_795019, EPI_ISL_795021, EPI_ISL_795022, EPI_ISL_795023, EPI_ISL_795024, EPI_ISL_795025, EPI_ISL_795026, EPI_ISL_795027, EPI_ISL_795028, EPI_ISL_795029, EPI_ISL_795030, EPI_ISL_795031, EPI_ISL_795032, EPI_ISL_795033, EPI_ISL_795034, EPI_ISL_795035, EPI_ISL_795036, EPI_ISL_795037, EPI_ISL_795038, EPI_ISL_795040, EPI_ISL_795041, EPI_ISL_795042, EPI_ISL_795043, EPI_ISL_795044, EPI_ISL_795045, EPI_ISL_795046, EPI_ISL_795047, EPI_ISL_795048, EPI_ISL_795049, EPI_ISL_795050, EPI_ISL_795051, EPI_ISL_795052, EPI_ISL_795053, EPI_ISL_795054, EPI_ISL_795055, EPI_ISL_795056, EPI_ISL_795057, EPI_ISL_795058, EPI_ISL_795059, EPI_ISL_795060, EPI_ISL_795061, EPI_ISL_795062, EPI_ISL_795063, EPI_ISL_795064, EPI_ISL_795065, EPI_ISL_795066, EPI_ISL_795067, EPI_ISL_795068, EPI_ISL_795069, EPI_ISL_795070, EPI_ISL_795071, EPI_ISL_795072, EPI_ISL_795073, EPI_ISL_795074, EPI_ISL_795075, EPI_ISL_795076, EPI_ISL_795077, EPI_ISL_795078, EPI_ISL_795079, EPI_ISL_795080, EPI_ISL_795081, EPI_ISL_795082, EPI_ISL_795083, EPI_ISL_795084, EPI_ISL_795085, EPI_ISL_795086, EPI_ISL_795087, EPI_ISL_795088, EPI_ISL_795089, EPI_ISL_795090, EPI_ISL_795091, EPI_ISL_795092, EPI_ISL_795093, EPI_ISL_795094, EPI_ISL_795095, EPI_ISL_795096, EPI_ISL_795097, EPI_ISL_795098, EPI_ISL_795099, EPI_ISL_795100, EPI_ISL_795101, EPI_ISL_795102, EPI_ISL_795103, EPI_ISL_795104, EPI_ISL_795105, EPI_ISL_795106, EPI_ISL_795107, EPI_ISL_795108, EPI_ISL_795109, EPI_ISL_795110, EPI_ISL_795111, EPI_ISL_795112, EPI_ISL_795113, EPI_ISL_795114, EPI_ISL_795115, EPI_ISL_795116, EPI_ISL_795117, EPI_ISL_795118, EPI_ISL_795119, EPI_ISL_795120, | | | |

| | | | |
|---|---|---|---|
| EPI_ISL_795121, EPI_ISL_795122, EPI_ISL_795123, EPI_ISL_795124, EPI_ISL_795125, EPI_ISL_795126, EPI_ISL_795127, EPI_ISL_795128, EPI_ISL_795129, EPI_ISL_795130, EPI_ISL_795131, EPI_ISL_795132, EPI_ISL_795133, EPI_ISL_795134, EPI_ISL_795135, EPI_ISL_795136, EPI_ISL_795137, EPI_ISL_795138, EPI_ISL_795139, EPI_ISL_795140, EPI_ISL_795141, EPI_ISL_795142, EPI_ISL_795143, EPI_ISL_795144, EPI_ISL_795145, EPI_ISL_795146, EPI_ISL_795147, EPI_ISL_795148, EPI_ISL_795154, EPI_ISL_795155, EPI_ISL_795156, EPI_ISL_795157, EPI_ISL_795158, EPI_ISL_795159, EPI_ISL_795160, EPI_ISL_795161, EPI_ISL_795162, EPI_ISL_795163, EPI_ISL_795164, EPI_ISL_795165, EPI_ISL_795166, EPI_ISL_795167, EPI_ISL_795168, EPI_ISL_795169, EPI_ISL_795171, EPI_ISL_795172, EPI_ISL_795173, EPI_ISL_795175, EPI_ISL_795176, EPI_ISL_795177, EPI_ISL_795178, EPI_ISL_795179, EPI_ISL_795180, EPI_ISL_795181, EPI_ISL_795182, EPI_ISL_795183, EPI_ISL_795184, EPI_ISL_795185, EPI_ISL_795186, EPI_ISL_795187, EPI_ISL_795188, EPI_ISL_795189, EPI_ISL_795190, EPI_ISL_795191, EPI_ISL_795192, EPI_ISL_795193, EPI_ISL_795194, EPI_ISL_795195, EPI_ISL_795196, EPI_ISL_795197, EPI_ISL_795198, EPI_ISL_795199, EPI_ISL_795200, EPI_ISL_795201, EPI_ISL_795202, EPI_ISL_795203, EPI_ISL_795204, EPI_ISL_795205, EPI_ISL_795206, EPI_ISL_795207, EPI_ISL_795208, EPI_ISL_795209, EPI_ISL_795210, EPI_ISL_795211, EPI_ISL_795212, EPI_ISL_795213, EPI_ISL_795214, EPI_ISL_795215, EPI_ISL_795216, EPI_ISL_795217, EPI_ISL_795218, EPI_ISL_795219, EPI_ISL_795220, EPI_ISL_795221, EPI_ISL_795222, EPI_ISL_795223, EPI_ISL_795224, EPI_ISL_795225, EPI_ISL_795226, EPI_ISL_795227, EPI_ISL_795228, EPI_ISL_795229, EPI_ISL_795230, EPI_ISL_795231, EPI_ISL_795232, EPI_ISL_795233, EPI_ISL_795234, EPI_ISL_795235, EPI_ISL_795236, EPI_ISL_795237, EPI_ISL_795238, EPI_ISL_795242, EPI_ISL_795244, EPI_ISL_795245, EPI_ISL_795246, EPI_ISL_795247, EPI_ISL_795248, EPI_ISL_795249, EPI_ISL_795250, EPI_ISL_795252, EPI_ISL_795253, EPI_ISL_795254, EPI_ISL_795255, EPI_ISL_795256, EPI_ISL_795257, EPI_ISL_795258, EPI_ISL_795259, EPI_ISL_795260, EPI_ISL_795261, EPI_ISL_795262, EPI_ISL_795263, EPI_ISL_795264, EPI_ISL_795265, EPI_ISL_795266, EPI_ISL_795267, EPI_ISL_795268, EPI_ISL_795269, EPI_ISL_795270, EPI_ISL_795271, EPI_ISL_795272, EPI_ISL_795273, EPI_ISL_795274, EPI_ISL_795275, EPI_ISL_795276, EPI_ISL_795277, EPI_ISL_795281, EPI_ISL_795282, EPI_ISL_795283, EPI_ISL_795284, EPI_ISL_795285, EPI_ISL_795286, EPI_ISL_795287, EPI_ISL_795288, EPI_ISL_795289, EPI_ISL_795290, EPI_ISL_795291, EPI_ISL_795292, EPI_ISL_795293, EPI_ISL_795294, EPI_ISL_795295, EPI_ISL_795296, EPI_ISL_795297, EPI_ISL_795298, EPI_ISL_795299, EPI_ISL_795300, EPI_ISL_795301, EPI_ISL_795302, EPI_ISL_795303, EPI_ISL_795304, EPI_ISL_795305, EPI_ISL_795306, EPI_ISL_795307, EPI_ISL_795308, EPI_ISL_795309, EPI_ISL_795310, EPI_ISL_795311, EPI_ISL_795312, EPI_ISL_795313, EPI_ISL_795314, EPI_ISL_795315, EPI_ISL_795316, EPI_ISL_795317, EPI_ISL_795318, EPI_ISL_795319, EPI_ISL_795320, EPI_ISL_795321, EPI_ISL_795322, EPI_ISL_795323, EPI_ISL_795324, EPI_ISL_795325, EPI_ISL_795326, EPI_ISL_795327, EPI_ISL_795328, EPI_ISL_795329, EPI_ISL_795330, EPI_ISL_795331, EPI_ISL_795332, EPI_ISL_795333, EPI_ISL_795334, EPI_ISL_795335, EPI_ISL_795336, EPI_ISL_795337, EPI_ISL_795338, EPI_ISL_795339, EPI_ISL_795340, EPI_ISL_795341, EPI_ISL_795342, EPI_ISL_795343, EPI_ISL_795344, EPI_ISL_795345, EPI_ISL_795346, EPI_ISL_795347, EPI_ISL_795348, EPI_ISL_795349, EPI_ISL_795350, EPI_ISL_795351, EPI_ISL_795352, EPI_ISL_795353, EPI_ISL_795354, EPI_ISL_795355, EPI_ISL_795356, EPI_ISL_795357, EPI_ISL_795358, EPI_ISL_795359, EPI_ISL_795360, EPI_ISL_795361, EPI_ISL_795368, EPI_ISL_795369, EPI_ISL_795370, EPI_ISL_795371, EPI_ISL_795372, EPI_ISL_795373, EPI_ISL_795374, EPI_ISL_795375, EPI_ISL_795376, EPI_ISL_795377, EPI_ISL_795378, EPI_ISL_795379, EPI_ISL_795380, EPI_ISL_795381, EPI_ISL_795382, EPI_ISL_795384, EPI_ISL_795385, EPI_ISL_795386, EPI_ISL_795387, EPI_ISL_795388, EPI_ISL_795389, EPI_ISL_795390, EPI_ISL_795391, EPI_ISL_795392, EPI_ISL_795393, EPI_ISL_795394, EPI_ISL_795395, EPI_ISL_795396, EPI_ISL_795397, EPI_ISL_795398, EPI_ISL_795399, EPI_ISL_795400, EPI_ISL_795401, EPI_ISL_795402, EPI_ISL_795403, EPI_ISL_795404, EPI_ISL_795405, EPI_ISL_795406, EPI_ISL_795407, EPI_ISL_795408, EPI_ISL_795409, EPI_ISL_795410, EPI_ISL_795411, EPI_ISL_795412, EPI_ISL_795414, EPI_ISL_795415, EPI_ISL_795416, EPI_ISL_795417, EPI_ISL_795418, EPI_ISL_795419, EPI_ISL_795420, EPI_ISL_795421, EPI_ISL_795422, EPI_ISL_795423, EPI_ISL_795424, EPI_ISL_795426, EPI_ISL_795427, EPI_ISL_795428, EPI_ISL_795429, EPI_ISL_795430, EPI_ISL_795431, EPI_ISL_795432, EPI_ISL_795433, EPI_ISL_795434, EPI_ISL_795435, EPI_ISL_795436, EPI_ISL_795437, EPI_ISL_795438, EPI_ISL_795439, EPI_ISL_795440, EPI_ISL_795441, EPI_ISL_795442, EPI_ISL_795443, EPI_ISL_795444, EPI_ISL_795445, EPI_ISL_795446, EPI_ISL_795447, EPI_ISL_795448, EPI_ISL_795449, EPI_ISL_795450, EPI_ISL_795451, EPI_ISL_795452, EPI_ISL_795453, EPI_ISL_795454, EPI_ISL_795455, EPI_ISL_795456, EPI_ISL_795457, EPI_ISL_795458, EPI_ISL_795459, EPI_ISL_795460, EPI_ISL_795461, EPI_ISL_795462, EPI_ISL_795463, EPI_ISL_795464, EPI_ISL_795465, EPI_ISL_795466, EPI_ISL_795467, EPI_ISL_795468, EPI_ISL_795469, EPI_ISL_795470, EPI_ISL_795471, EPI_ISL_795472, EPI_ISL_795473, EPI_ISL_795474, EPI_ISL_795475, EPI_ISL_795476, EPI_ISL_795477, EPI_ISL_795478, EPI_ISL_795479, EPI_ISL_795480, EPI_ISL_795481, EPI_ISL_795482, EPI_ISL_795483, EPI_ISL_795484, EPI_ISL_795485, EPI_ISL_795486, EPI_ISL_795487, EPI_ISL_795488, EPI_ISL_795489, EPI_ISL_795490, EPI_ISL_795491, EPI_ISL_795492, EPI_ISL_795493, EPI_ISL_795494, EPI_ISL_795495, EPI_ISL_795496, EPI_ISL_795497, EPI_ISL_795498, EPI_ISL_795499, EPI_ISL_795500, EPI_ISL_795501, EPI_ISL_795502, EPI_ISL_795503, EPI_ISL_795504, EPI_ISL_795505, EPI_ISL_795506, EPI_ISL_795507, EPI_ISL_795508, EPI_ISL_795509, EPI_ISL_795510, EPI_ISL_795511, EPI_ISL_795512, EPI_ISL_795513, EPI_ISL_795514, EPI_ISL_795515, EPI_ISL_795516, EPI_ISL_795517, EPI_ISL_795518, EPI_ISL_795519, EPI_ISL_795520, EPI_ISL_795521, EPI_ISL_795522, EPI_ISL_795523, EPI_ISL_795524, EPI_ISL_795525, EPI_ISL_795526, EPI_ISL_795527, EPI_ISL_795528, EPI_ISL_795529, EPI_ISL_795530, EPI_ISL_795531, EPI_ISL_795532, EPI_ISL_795533, EPI_ISL_795534, EPI_ISL_795535, EPI_ISL_795536, EPI_ISL_795537, EPI_ISL_795538, EPI_ISL_795539, EPI_ISL_795540, EPI_ISL_795541, EPI_ISL_795542, EPI_ISL_795543, EPI_ISL_795544, EPI_ISL_795545, EPI_ISL_795546, EPI_ISL_795547, EPI_ISL_795548, EPI_ISL_795549, EPI_ISL_795550, EPI_ISL_795551, EPI_ISL_795552, EPI_ISL_795553, EPI_ISL_795554, EPI_ISL_795555, EPI_ISL_795556, EPI_ISL_795557, EPI_ISL_795558, EPI_ISL_795559, EPI_ISL_795560, EPI_ISL_795561, EPI_ISL_795562, EPI_ISL_795563, EPI_ISL_795564, EPI_ISL_795565, EPI_ISL_795566, EPI_ISL_795567, EPI_ISL_795568, EPI_ISL_795569, EPI_ISL_795570, EPI_ISL_795571, EPI_ISL_795572, EPI_ISL_795573, EPI_ISL_795574, EPI_ISL_795575, EPI_ISL_795576, EPI_ISL_795577, EPI_ISL_795578, EPI_ISL_795579, EPI_ISL_795580, EPI_ISL_795581, EPI_ISL_795582, EPI_ISL_795583, EPI_ISL_795584, EPI_ISL_795585, EPI_ISL_795586, EPI_ISL_795587, EPI_ISL_795588, EPI_ISL_795589, EPI_ISL_795590, EPI_ISL_795591, EPI_ISL_795592, EPI_ISL_795593, EPI_ISL_795594, EPI_ISL_795595, EPI_ISL_795596, EPI_ISL_795597, EPI_ISL_795598, EPI_ISL_795599, EPI_ISL_795600, EPI_ISL_795601, EPI_ISL_795602, EPI_ISL_795603, EPI_ISL_795604, EPI_ISL_795605, EPI_ISL_795606, EPI_ISL_795607, EPI_ISL_795608, EPI_ISL_795609, EPI_ISL_795610, EPI_ISL_795611, EPI_ISL_795612, EPI_ISL_795613, EPI_ISL_795614, EPI_ISL_795615, EPI_ISL_795616, EPI_ISL_795617, EPI_ISL_795618, EPI_ISL_795619, EPI_ISL_795620, EPI_ISL_795621, EPI_ISL_795622, EPI_ISL_795623, EPI_ISL_795624, EPI_ISL_795625, EPI_ISL_795626, EPI_ISL_795627, EPI_ISL_795628, EPI_ISL_795629, EPI_ISL_795630, EPI_ISL_795631, EPI_ISL_795632, EPI_ISL_795633, EPI_ISL_795634, EPI_ISL_795635, EPI_ISL_795636, EPI_ISL_795637, EPI_ISL_795638, EPI_ISL_795639, EPI_ISL_795640, EPI_ISL_795641, EPI_ISL_795642, EPI_ISL_795643, EPI_ISL_795644, EPI_ISL_795645, EPI_ISL_795646, EPI_ISL_795647, EPI_ISL_795648, EPI_ISL_795649, EPI_ISL_795650, EPI_ISL_795651, EPI_ISL_795652, EPI_ISL_795653, EPI_ISL_795654, EPI_ISL_795655, EPI_ISL_795656, EPI_ISL_795657, EPI_ISL_795658, EPI_ISL_795659, EPI_ISL_795660, EPI_ISL_795661, EPI_ISL_795662, EPI_ISL_795663, EPI_ISL_795664, EPI_ISL_795665, EPI_ISL_795666, EPI_ISL_795667, EPI_ISL_795668, EPI_ISL_795669, EPI_ISL_795670, EPI_ISL_795671, EPI_ISL_795672, EPI_ISL_795673, EPI_ISL_795674, EPI_ISL_795675, EPI_ISL_795676, EPI_ISL_795677, EPI_ISL_795678, EPI_ISL_795679, EPI_ISL_795680, EPI_ISL_795681, EPI_ISL_795682, EPI_ISL_795683, EPI_ISL_795684, EPI_ISL_795685, EPI_ISL_795686, EPI_ISL_795687, EPI_ISL_795688, EPI_ISL_795689, EPI_ISL_795690, EPI_ISL_795691, EPI_ISL_795692, EPI_ISL_795693, EPI_ISL_795694, EPI_ISL_795695, EPI_ISL_795696, EPI_ISL_795697, EPI_ISL_795698, EPI_ISL_795699, EPI_ISL_795700, EPI_ISL_795701, EPI_ISL_795702, EPI_ISL_795703, EPI_ISL_795704, EPI_ISL_795705, EPI_ISL_795706, EPI_ISL_795707, EPI_ISL_795708, EPI_ISL_795709, EPI_ISL_795710, EPI_ISL_795711, EPI_ISL_795712, EPI_ISL_795713, EPI_ISL_795714, EPI_ISL_795715, EPI_ISL_795716, EPI_ISL_795717, EPI_ISL_795718, EPI_ISL_795719, EPI_ISL_795720, EPI_ISL_795721, EPI_ISL_795722, EPI_ISL_795723, EPI_ISL_795724, EPI_ISL_795725, EPI_ISL_795726, EPI_ISL_795727, EPI_ISL_795728, EPI_ISL_795729, EPI_ISL_795730, EPI_ISL_795731, EPI_ISL_795732, EPI_ISL_795733, EPI_ISL_795734, EPI_ISL_795735, EPI_ISL_795736, EPI_ISL_795737, EPI_ISL_795738, EPI_ISL_795739, EPI_ISL_795740, EPI_ISL_795741, EPI_ISL_795742, EPI_ISL_795743, EPI_ISL_795744, EPI_ISL_795745, EPI_ISL_795746, EPI_ISL_795747, EPI_ISL_795748, EPI_ISL_795749, EPI_ISL_795750, EPI_ISL_795751, EPI_ISL_795752, EPI_ISL_795753, EPI_ISL_795754, EPI_ISL_795755, EPI_ISL_795756, EPI_ISL_795757, EPI_ISL_795758, EPI_ISL_795759, EPI_ISL_795760, EPI_ISL_795761, EPI_ISL_795762, EPI_ISL_795763, EPI_ISL_795764, EPI_ISL_795765, EPI_ISL_795766, EPI_ISL_795767, EPI_ISL_795768, EPI_ISL_795769, EPI_ISL_795770, EPI_ISL_795771, EPI_ISL_795772, EPI_ISL_795773, EPI_ISL_795774, EPI_ISL_795775, EPI_ISL_795776, EPI_ISL_795777, EPI_ISL_795778, EPI_ISL_795779, EPI_ISL_795780, EPI_ISL_795781, EPI_ISL_795782, EPI_ISL_795783, EPI_ISL_795784, EPI_ISL_795785, EPI_ISL_795786, EPI_ISL_795787, EPI_ISL_795788, EPI_ISL_795789, EPI_ISL_795790, EPI_ISL_795791, EPI_ISL_795792, EPI_ISL_795793, EPI_ISL_795794, EPI_ISL_795795, EPI_ISL_795796, EPI_ISL_795797, EPI_ISL_795798, EPI_ISL_795799, EPI_ISL_795800, EPI_ISL_795801, EPI_ISL_795802, EPI_ISL_795803, EPI_ISL_795804, EPI_ISL_795805, EPI_ISL_795806, EPI_ISL_795807, EPI_ISL_795808, EPI_ISL_795809, EPI_ISL_795810, EPI_ISL_795811, EPI_ISL_795812, EPI_ISL_795813, EPI_ISL_795814, EPI_ISL_795815, EPI_ISL_795816, EPI_ISL_795817, EPI_ISL_795818, EPI_ISL_795819, EPI_ISL_795820, EPI_ISL_795821, EPI_ISL_795822, EPI_ISL_795823, EPI_ISL_795824, EPI_ISL_795825, EPI_ISL_795826, EPI_ISL_795827, EPI_ISL_795828, EPI_ISL_795829, EPI_ISL_795830, EPI_ISL_795831, EPI_ISL_795832, EPI_ISL_795833, EPI_ISL_795834, EPI_ISL_795835, EPI_ISL_795836, EPI_ISL_795837, EPI_ISL_795838, EPI_ISL_795839, EPI_ISL_795840, EPI_ISL_795841, EPI_ISL_795842, EPI_ISL_795843, EPI_ISL_795844, EPI_ISL_795845, EPI_ISL_795846, EPI_ISL_795847, EPI_ISL_795848, EPI_ISL_795849, EPI_ISL_795850, EPI_ISL_795851, EPI_ISL_795852, EPI_ISL_795853, EPI_ISL_795854, EPI_ISL_795855, EPI_ISL_795856, EPI_ISL_795857, EPI_ISL_795858, EPI_ISL_795859, EPI_ISL_795860, EPI_ISL_795861, EPI_ISL_795862, EPI_ISL_795863, EPI_ISL_795864, EPI_ISL_795865, EPI_ISL_795866, EPI_ISL_795867, EPI_ISL_795868, EPI_ISL_795869, EPI_ISL_795870, EPI_ISL_795871, EPI_ISL_795872, EPI_ISL_795873, EPI_ISL_795874, EPI_ISL_795875, EPI_ISL_795876, EPI_ISL_795877, EPI_ISL_795878, EPI_ISL_795879, EPI_ISL_795880, EPI_ISL_795881, EPI_ISL_795882, EPI_ISL_795883, EPI_ISL_795884, EPI_ISL_795885, EPI_ISL_795886, EPI_ISL_795887, EPI_ISL_795888, EPI_ISL_795889, EPI_ISL_795890, EPI_ISL_795891, EPI_ISL_795892, EPI_ISL_795893, EPI_ISL_795894, EPI_ISL_795895, EPI_ISL_795896, EPI_ISL_795897, EPI_ISL_795898, EPI_ISL_795899, EPI_ISL_795900, EPI_ISL_795901, EPI_ISL_795902, EPI_ISL_795903, EPI_ISL_795904, EPI_ISL_795905, EPI_ISL_795906, EPI_ISL_795907, EPI_ISL_795908, EPI_ISL_795909, EPI_ISL_795910, EPI_ISL_795911, EPI_ISL_795912, EPI_ISL_795913, EPI_ISL_795914, EPI_ISL_795915, EPI_ISL_795916, EPI_ISL_795917, EPI_ISL_795918, EPI_ISL_795919, EPI_ISL_795920, EPI_ISL_795921, EPI_ISL_795922, EPI_ISL_795923, EPI_ISL_795924, EPI_ISL_795925, EPI_ISL_795926, EPI_ISL_795927, EPI_ISL_795928, EPI_ISL_795929, EPI_ISL_795930, EPI_ISL_795931, EPI_ISL_795932, EPI_ISL_795933, EPI_ISL_795934, EPI_ISL_795935, EPI_ISL_795936, EPI_ISL_795937, EPI_ISL_795938, EPI_ISL_795939, EPI_ISL_795940, EPI_ISL_795941, EPI_ISL_795942, EPI_ISL_795943, EPI_ISL_795944, EPI_ISL_795945, EPI_ISL_795946, EPI_ISL_795947, EPI_ISL_795948, EPI_ISL_795949, EPI_ISL_795950, EPI_ISL_795951, EPI_ISL_795952, EPI_ISL_795953, EPI_ISL_795954, EPI_ISL_795955, EPI_ISL_795956, EPI_ISL_795957, EPI_ISL_795958, EPI_ISL_795959, EPI_ISL_795960, EPI_ISL_795961, EPI_ISL_795962, EPI_ISL_795963, EPI_ISL_795964, EPI_ISL_795965, EPI_ISL_795966, EPI_ISL_795967, EPI_ISL_795968, EPI_ISL_795969, EPI_ISL_795970, EPI_ISL_795971, EPI_ISL_795972, EPI_ISL_795973, EPI_ISL_795974, EPI_ISL_795975, EPI_ISL_795976, EPI_ISL_795977, EPI_ISL_795978, EPI_ISL_795979, EPI_ISL_795980, EPI_ISL_795981, EPI_ISL_795982, EPI_ISL_795983 | | | |
| see above | Department of Virus and Microbiological Special Diagnostics, Statens Serum Institut, Copenhagen, Denmark | Albertsen Lab, Department of Chemistry and Bioscience, Aalborg University, Denmark | Danish Covid-19 Genome Consortium |
| EPI_ISL_796159, EPI_ISL_796160, EPI_ISL_796167, EPI_ISL_796171, EPI_ISL_796172, EPI_ISL_796173, EPI_ISL_796174, EPI_ISL_796177, EPI_ISL_796179, EPI_ISL_796180, EPI_ISL_796184, EPI_ISL_796188, EPI_ISL_796189, EPI_ISL_796190, EPI_ISL_796192, EPI_ISL_796194, EPI_ISL_796198, EPI_ISL_796199, EPI_ISL_796202, EPI_ISL_796204, EPI_ISL_796208, EPI_ISL_796218, EPI_ISL_796222, EPI_ISL_796223, EPI_ISL_796225, EPI_ISL_796226, EPI_ISL_796227, EPI_ISL_796229, EPI_ISL_796233, EPI_ISL_796242, EPI_ISL_796245, EPI_ISL_796246, EPI_ISL_796247, EPI_ISL_796249, EPI_ISL_796250, EPI_ISL_796252, EPI_ISL_796253, EPI_ISL_796256, EPI_ISL_796266, EPI_ISL_796268, EPI_ISL_796270, EPI_ISL_796271, EPI_ISL_796273, EPI_ISL_796275, EPI_ISL_796276, EPI_ISL_796279, EPI_ISL_796280, EPI_ISL_796282, EPI_ISL_796283, EPI_ISL_796284, EPI_ISL_796290, EPI_ISL_796293, EPI_ISL_796297, EPI_ISL_796298, EPI_ISL_796302, EPI_ISL_796303, EPI_ISL_796312, EPI_ISL_796316, EPI_ISL_796318, EPI_ISL_796319, EPI_ISL_796326, EPI_ISL_796333, EPI_ISL_796337, EPI_ISL_796340, EPI_ISL_796342, EPI_ISL_796348, EPI_ISL_796355, EPI_ISL_796359, EPI_ISL_796361, EPI_ISL_796362, EPI_ISL_796363, EPI_ISL_796364, EPI_ISL_796368, EPI_ISL_796371, EPI_ISL_796372, EPI_ISL_796374, EPI_ISL_796382, EPI_ISL_796396, EPI_ISL_796416, EPI_ISL_796418, EPI_ISL_796420, EPI_ISL_796421, EPI_ISL_796422, EPI_ISL_796423, EPI_ISL_796427, EPI_ISL_796428, EPI_ISL_796431, EPI_ISL_796436, EPI_ISL_796438, EPI_ISL_796443, EPI_ISL_796444, EPI_ISL_796446, EPI_ISL_796447, EPI_ISL_796449, EPI_ISL_796450, EPI_ISL_796452, EPI_ISL_796453, EPI_ISL_796454, EPI_ISL_796455, EPI_ISL_796458, EPI_ISL_796459, EPI_ISL_796463, EPI_ISL_796466, EPI_ISL_796467, EPI_ISL_796468, EPI_ISL_796469, EPI_ISL_796472, EPI_ISL_796473, EPI_ISL_796474, EPI_ISL_796475, EPI_ISL_796478, EPI_ISL_796484, EPI_ISL_796488, EPI_ISL_796491, EPI_ISL_796497, EPI_ISL_796501, EPI_ISL_796506, EPI_ISL_796515, EPI_ISL_796516, EPI_ISL_796528, EPI_ISL_796531, EPI_ISL_796536, EPI_ISL_796537, EPI_ISL_796549, EPI_ISL_796551, EPI_ISL_796553, EPI_ISL_796558, EPI_ISL_796564, EPI_ISL_796565, EPI_ISL_796575, EPI_ISL_796576, EPI_ISL_796581, EPI_ISL_796584, EPI_ISL_796595, EPI_ISL_796598, EPI_ISL_796599, EPI_ISL_796602, EPI_ISL_796604, EPI_ISL_796606, EPI_ISL_796608, EPI_ISL_796612, EPI_ISL_796622, EPI_ISL_796627, EPI_ISL_796628, EPI_ISL_796630, EPI_ISL_796633, EPI_ISL_796635 | | | |
| see above | Viollier AG | Department of Biosystems Science and Engineering, ETH Zürich | Chaoran Chen, Sarah Nadeau, Catharine Aquino, Ivan Topolsky, Philipp Jablonski, Lara Fuhrmann, David Dreifuss, Katharina Jahn, Andreia Cabral de Gouvea, Maria Domenica Moccia, Simon Grüter, Timothy Sykes, Lennart Opitz, Griffin White, Laura Neff, Doris Popovic, Andrea Patrignani, Jay Tracy, Ralph Schlapbach, Christiane Beckmann, Maurice Redondo, Olivier Kobel, Christoph Noppen, Sophie Seidel, Noemie Santamaria de Souza, Niko Beerenwinkel, Tanja Stadler |
| EPI_ISL_796733, EPI_ISL_796734, EPI_ISL_796752, EPI_ISL_796753, EPI_ISL_796764, EPI_ISL_796774, EPI_ISL_796775 | Instituto Nacional de Saude (INSA) | Instituto Nacional de Saude (INSA) | Borges et al |
| EPI_ISL_796795, EPI_ISL_796997 | Lighthouse Lab in Alderley Park | Wellcome Sanger Institute for the COVID-19 Genomics UK (COG-UK) Consortium | Jacquelyn Wynn, Mairead Hyland, The Lighthouse Lab in Alderley Park and Alex Alderton, Roberto Amato, Sonia Goncalves, Ewan Harrison, David K. Jackson, Ian Johnston, Dominic Kwiatkowski, Cordelia Langford, John Sillitoe on behalf of the Wellcome Sanger Institute COVID-19 Surveillance Team |
| EPI_ISL_797119, EPI_ISL_797122, EPI_ISL_797123, EPI_ISL_797124, EPI_ISL_797127, EPI_ISL_797130, EPI_ISL_797131, EPI_ISL_797132, EPI_ISL_797135, EPI_ISL_797136, EPI_ISL_797137, EPI_ISL_797140, EPI_ISL_797145, EPI_ISL_797147, EPI_ISL_797150, EPI_ISL_797152, EPI_ISL_797157, EPI_ISL_797158, EPI_ISL_797159, EPI_ISL_797162, EPI_ISL_797166, EPI_ISL_797167, EPI_ISL_797169, EPI_ISL_797170, EPI_ISL_797172, EPI_ISL_797174, EPI_ISL_797180, EPI_ISL_797181, EPI_ISL_797182, EPI_ISL_797183, EPI_ISL_797184, EPI_ISL_797186, EPI_ISL_797188, EPI_ISL_797189, EPI_ISL_797191, EPI_ISL_797192, EPI_ISL_797193, EPI_ISL_797195, EPI_ISL_797196, EPI_ISL_797197, EPI_ISL_797199, EPI_ISL_797200, EPI_ISL_797201, EPI_ISL_797206, EPI_ISL_797208, EPI_ISL_797209, EPI_ISL_797211, EPI_ISL_797213, EPI_ISL_797214, EPI_ISL_797215, EPI_ISL_797217, EPI_ISL_797219, EPI_ISL_797221, EPI_ISL_797222, EPI_ISL_797223, EPI_ISL_797224, EPI_ISL_797227, EPI_ISL_797230, EPI_ISL_797233, EPI_ISL_797235, EPI_ISL_797236, EPI_ISL_797237, EPI_ISL_797238, EPI_ISL_797241, EPI_ISL_797244, EPI_ISL_797245, EPI_ISL_797249, EPI_ISL_797250, EPI_ISL_797251, EPI_ISL_797255, EPI_ISL_797257, EPI_ISL_797260, EPI_ISL_797262, EPI_ISL_797263, EPI_ISL_797265, EPI_ISL_797267, EPI_ISL_797268, EPI_ISL_797270, EPI_ISL_797271, EPI_ISL_797273, EPI_ISL_797274, EPI_ISL_797275, EPI_ISL_797276, EPI_ISL_797278, EPI_ISL_797279, EPI_ISL_797280, EPI_ISL_797282, EPI_ISL_797283, EPI_ISL_797284, EPI_ISL_797285, EPI_ISL_797287, EPI_ISL_797288, EPI_ISL_797291, EPI_ISL_797292, EPI_ISL_797293, EPI_ISL_797294, EPI_ISL_797297, EPI_ISL_797298, EPI_ISL_797299, EPI_ISL_797300, EPI_ISL_797302, EPI_ISL_797303, EPI_ISL_797304, EPI_ISL_797306, EPI_ISL_797307, EPI_ISL_797308, EPI_ISL_797309, EPI_ISL_797310, EPI_ISL_797311, EPI_ISL_797312, EPI_ISL_797314, EPI_ISL_797316, EPI_ISL_797320, EPI_ISL_797321, EPI_ISL_797322, EPI_ISL_797324, EPI_ISL_797328, EPI_ISL_797329, EPI_ISL_797330, EPI_ISL_797331, EPI_ISL_797333, EPI_ISL_797334, EPI_ISL_797336, EPI_ISL_797340, EPI_ISL_797341, EPI_ISL_797343, EPI_ISL_797344, EPI_ISL_797347, EPI_ISL_797348, EPI_ISL_797350, EPI_ISL_797351, EPI_ISL_797352, EPI_ISL_797353, EPI_ISL_797355, EPI_ISL_797356, EPI_ISL_797358, EPI_ISL_797359, EPI_ISL_797360, EPI_ISL_797361, EPI_ISL_797364, EPI_ISL_797368, EPI_ISL_797369, EPI_ISL_797370, EPI_ISL_797371, EPI_ISL_797372, EPI_ISL_797373, EPI_ISL_797374, EPI_ISL_797375, EPI_ISL_797377, EPI_ISL_797378, EPI_ISL_797381, EPI_ISL_797382, EPI_ISL_797383, EPI_ISL_797384, EPI_ISL_797386, EPI_ISL_797387, EPI_ISL_797388, EPI_ISL_797392, EPI_ISL_797393, EPI_ISL_797394, EPI_ISL_797396 | | | |
| see above | Lighthouse Lab in Milton Keynes | Wellcome Sanger Institute for the COVID-19 Genomics UK (COG-UK) Consortium | The Lighthouse Lab in Milton Keynes and Alex Alderton, Roberto Amato, Sonia Goncalves, Ewan Harrison, David K. Jackson, Ian Johnston, Dominic Kwiatkowski, Cordelia Langford, John Sillitoe on behalf of the Wellcome Sanger Institute COVID-19 Surveillance Team |
| EPI_ISL_797400, EPI_ISL_797403, EPI_ISL_797434, EPI_ISL_797446, EPI_ISL_797465, EPI_ISL_797481, EPI_ISL_797483, EPI_ISL_797492, EPI_ISL_797497, EPI_ISL_797498, EPI_ISL_797502, EPI_ISL_797505, EPI_ISL_797510, EPI_ISL_797511, EPI_ISL_797532, EPI_ISL_797537, EPI_ISL_797548, EPI_ISL_797566, EPI_ISL_797578, EPI_ISL_797579, EPI_ISL_797580, EPI_ISL_797590, EPI_ISL_797593, EPI_ISL_797595, EPI_ISL_797597, EPI_ISL_797600, EPI_ISL_797625, EPI_ISL_797631, EPI_ISL_797643, EPI_ISL_797645, EPI_ISL_797646, EPI_ISL_797647, EPI_ISL_797648, EPI_ISL_797652, EPI_ISL_797657, EPI_ISL_797659, EPI_ISL_797664, EPI_ISL_797668, EPI_ISL_797676, EPI_ISL_797688, EPI_ISL_797690, EPI_ISL_797692, EPI_ISL_797694, EPI_ISL_797705 | | | |
| see above | Lighthouse Lab in Cambridge | Wellcome Sanger Institute for the COVID-19 Genomics UK | Rob Howes, The Lighthouse Lab in Cambridge and Alex Alderton, Roberto Amato, Sonia Goncalves, Ewan Harrison, David K. Jackson, Ian Johnston, |

| | | | |
|---|---|---|---|
| | | (COG-UK) Consortium | Dominic Kwiatkowski, Cordelia Langford, John Sillitoe on behalf of the Wellcome Sanger Institute COVID-19 Surveillance Team |
| EPI_ISL_797739, EPI_ISL_797775, EPI_ISL_797789, EPI_ISL_797797, EPI_ISL_797817, EPI_ISL_797831, EPI_ISL_797838, EPI_ISL_797851, EPI_ISL_797868, EPI_ISL_797884, EPI_ISL_797903, EPI_ISL_797916, EPI_ISL_797921, EPI_ISL_797937, EPI_ISL_797952, EPI_ISL_797965, EPI_ISL_797985 | | | |
| see above | Lighthouse Lab in Glasgow | Wellcome Sanger Institute for the COVID-19 Genomics UK (COG-UK) Consortium | Harper VanSteenhouse, Yumi Kasai, David Gray, Carol Clugston, Anna Dominiczak and Alex Alderton, Roberto Amato, Sonia Goncalves, Ewan Harrison, David K. Jackson, Ian Johnston, Dominic Kwiatkowski, Cordelia Langford, John Sillitoe on behalf of the Wellcome Sanger Institute COVID-19 Surveillance Team |
| EPI_ISL_797999, EPI_ISL_798002, EPI_ISL_798007, EPI_ISL_798009, EPI_ISL_798011, EPI_ISL_798013, EPI_ISL_798015 | Lighthouse Lab in Alderley Park | Wellcome Sanger Institute for the COVID-19 Genomics UK (COG-UK) Consortium | Jacquelyn Wynn, Mairead Hyland, The Lighthouse Lab in Alderley Park and Alex Alderton, Roberto Amato, Sonia Goncalves, Ewan Harrison, David K. Jackson, Ian Johnston, Dominic Kwiatkowski, Cordelia Langford, John Sillitoe on behalf of the Wellcome Sanger Institute COVID-19 Surveillance Team |
| EPI_ISL_798017, EPI_ISL_798022 | Lighthouse Lab in Glasgow | Wellcome Sanger Institute for the COVID-19 Genomics UK (COG-UK) Consortium | Harper VanSteenhouse, Yumi Kasai, David Gray, Carol Clugston, Anna Dominiczak and Alex Alderton, Roberto Amato, Sonia Goncalves, Ewan Harrison, David K. Jackson, Ian Johnston, Dominic Kwiatkowski, Cordelia Langford, John Sillitoe on behalf of the Wellcome Sanger Institute COVID-19 Surveillance Team |
| EPI_ISL_798025, EPI_ISL_798027, EPI_ISL_798029, EPI_ISL_798035 | Lighthouse Lab in Alderley Park | Wellcome Sanger Institute for the COVID-19 Genomics UK (COG-UK) Consortium | Jacquelyn Wynn, Mairead Hyland, The Lighthouse Lab in Alderley Park and Alex Alderton, Roberto Amato, Sonia Goncalves, Ewan Harrison, David K. Jackson, Ian Johnston, Dominic Kwiatkowski, Cordelia Langford, John Sillitoe on behalf of the Wellcome Sanger Institute COVID-19 Surveillance Team |
| EPI_ISL_798043, EPI_ISL_798051 | Lighthouse Lab in Glasgow | Wellcome Sanger Institute for the COVID-19 Genomics UK (COG-UK) Consortium | Harper VanSteenhouse, Yumi Kasai, David Gray, Carol Clugston, Anna Dominiczak and Alex Alderton, Roberto Amato, Sonia Goncalves, Ewan Harrison, David K. Jackson, Ian Johnston, Dominic Kwiatkowski, Cordelia Langford, John Sillitoe on behalf of the Wellcome Sanger Institute COVID-19 Surveillance Team |
| EPI_ISL_798054, EPI_ISL_798061, EPI_ISL_798062, EPI_ISL_798064, EPI_ISL_798066, EPI_ISL_798068, EPI_ISL_798073 | Lighthouse Lab in Alderley Park | Wellcome Sanger Institute for the COVID-19 Genomics UK (COG-UK) Consortium | Jacquelyn Wynn, Mairead Hyland, The Lighthouse Lab in Alderley Park and Alex Alderton, Roberto Amato, Sonia Goncalves, Ewan Harrison, David K. Jackson, Ian Johnston, Dominic Kwiatkowski, Cordelia Langford, John Sillitoe on behalf of the Wellcome Sanger Institute COVID-19 Surveillance Team |
| EPI_ISL_798080 | Lighthouse Lab in Glasgow | Wellcome Sanger Institute for the COVID-19 Genomics UK (COG-UK) Consortium | Harper VanSteenhouse, Yumi Kasai, David Gray, Carol Clugston, Anna Dominiczak and Alex Alderton, Roberto Amato, Sonia Goncalves, Ewan Harrison, David K. Jackson, Ian Johnston, Dominic Kwiatkowski, Cordelia Langford, John Sillitoe on behalf of the Wellcome Sanger Institute COVID-19 Surveillance Team |
| EPI_ISL_798083, EPI_ISL_798084, EPI_ISL_798096, EPI_ISL_798101, EPI_ISL_798103, EPI_ISL_798106 | Lighthouse Lab in Alderley Park | Wellcome Sanger Institute for the COVID-19 Genomics UK (COG-UK) Consortium | Jacquelyn Wynn, Mairead Hyland, The Lighthouse Lab in Alderley Park and Alex Alderton, Roberto Amato, Sonia Goncalves, Ewan Harrison, David K. Jackson, Ian Johnston, Dominic Kwiatkowski, Cordelia Langford, John Sillitoe on behalf of the Wellcome Sanger Institute COVID-19 Surveillance Team |
| EPI_ISL_798107 | Lighthouse Lab in Glasgow | Wellcome Sanger Institute for the COVID-19 Genomics UK (COG-UK) Consortium | Harper VanSteenhouse, Yumi Kasai, David Gray, Carol Clugston, Anna Dominiczak and Alex Alderton, Roberto Amato, Sonia Goncalves, Ewan Harrison, David K. Jackson, Ian Johnston, Dominic Kwiatkowski, Cordelia Langford, John Sillitoe on behalf of the Wellcome Sanger Institute COVID-19 Surveillance Team |
| EPI_ISL_798108, EPI_ISL_798119, EPI_ISL_798122, EPI_ISL_798123, EPI_ISL_798124, EPI_ISL_798125, EPI_ISL_798127, EPI_ISL_798131, EPI_ISL_798133, EPI_ISL_798134, EPI_ISL_798139 | | | |
| see above | Lighthouse Lab in Alderley Park | Wellcome Sanger Institute for the COVID-19 Genomics UK (COG-UK) Consortium | Jacquelyn Wynn, Mairead Hyland, The Lighthouse Lab in Alderley Park and Alex Alderton, Roberto Amato, Sonia Goncalves, Ewan Harrison, David K. Jackson, Ian Johnston, Dominic Kwiatkowski, Cordelia Langford, John Sillitoe on behalf of the Wellcome Sanger Institute COVID-19 Surveillance Team |
| EPI_ISL_798141 | Lighthouse Lab in Glasgow | Wellcome Sanger Institute for the COVID-19 Genomics UK (COG-UK) Consortium | Harper VanSteenhouse, Yumi Kasai, David Gray, Carol Clugston, Anna Dominiczak and Alex Alderton, Roberto Amato, Sonia Goncalves, Ewan Harrison, David K. Jackson, Ian Johnston, Dominic Kwiatkowski, Cordelia Langford, John Sillitoe on behalf of the Wellcome Sanger Institute COVID-19 Surveillance Team |
| EPI_ISL_798152, EPI_ISL_798154 | Lighthouse Lab in Alderley Park | Wellcome Sanger Institute for the COVID-19 Genomics UK (COG-UK) Consortium | Jacquelyn Wynn, Mairead Hyland, The Lighthouse Lab in Alderley Park and Alex Alderton, Roberto Amato, Sonia Goncalves, Ewan Harrison, David K. Jackson, Ian Johnston, Dominic Kwiatkowski, Cordelia Langford, John Sillitoe on behalf of the Wellcome Sanger Institute COVID-19 Surveillance Team |
| EPI_ISL_798159 | Lighthouse Lab in Glasgow | Wellcome Sanger Institute for the COVID-19 Genomics UK (COG-UK) Consortium | Harper VanSteenhouse, Yumi Kasai, David Gray, Carol Clugston, Anna Dominiczak and Alex Alderton, Roberto Amato, Sonia Goncalves, Ewan Harrison, David K. Jackson, Ian Johnston, Dominic Kwiatkowski, Cordelia Langford, John Sillitoe on behalf of the Wellcome Sanger Institute COVID-19 Surveillance Team |
| EPI_ISL_798169, EPI_ISL_798170 | Lighthouse Lab in Alderley Park | Wellcome Sanger Institute for the COVID-19 Genomics UK (COG-UK) Consortium | Jacquelyn Wynn, Mairead Hyland, The Lighthouse Lab in Alderley Park and Alex Alderton, Roberto Amato, Sonia Goncalves, Ewan Harrison, David K. Jackson, Ian Johnston, Dominic Kwiatkowski, Cordelia Langford, John Sillitoe on behalf of the Wellcome Sanger Institute COVID-19 Surveillance Team |
| EPI_ISL_798171 | Lighthouse Lab in Glasgow | Wellcome Sanger Institute for the COVID-19 Genomics UK (COG-UK) Consortium | Harper VanSteenhouse, Yumi Kasai, David Gray, Carol Clugston, Anna Dominiczak and Alex Alderton, Roberto Amato, Sonia Goncalves, Ewan Harrison, David K. Jackson, Ian Johnston, Dominic Kwiatkowski, Cordelia Langford, John Sillitoe on behalf of the Wellcome Sanger Institute COVID-19 Surveillance Team |
| EPI_ISL_798173, EPI_ISL_798174, EPI_ISL_798175, EPI_ISL_798176 | Lighthouse Lab in Alderley Park | Wellcome Sanger Institute for the COVID-19 Genomics UK (COG-UK) Consortium | Jacquelyn Wynn, Mairead Hyland, The Lighthouse Lab in Alderley Park and Alex Alderton, Roberto Amato, Sonia Goncalves, Ewan Harrison, David K. Jackson, Ian Johnston, Dominic Kwiatkowski, Cordelia Langford, John Sillitoe on behalf of the Wellcome Sanger Institute COVID-19 Surveillance Team |
| EPI_ISL_798178 | Lighthouse Lab in Glasgow | Wellcome Sanger Institute for the COVID-19 Genomics UK (COG-UK) Consortium | Harper VanSteenhouse, Yumi Kasai, David Gray, Carol Clugston, Anna Dominiczak and Alex Alderton, Roberto Amato, Sonia Goncalves, Ewan Harrison, David K. Jackson, Ian Johnston, Dominic Kwiatkowski, Cordelia Langford, John Sillitoe on behalf of the Wellcome Sanger Institute COVID-19 Surveillance Team |
| EPI_ISL_798181, EPI_ISL_798182 | Lighthouse Lab in Alderley Park | Wellcome Sanger Institute for the COVID-19 Genomics UK (COG-UK) Consortium | Jacquelyn Wynn, Mairead Hyland, The Lighthouse Lab in Alderley Park and Alex Alderton, Roberto Amato, Sonia Goncalves, Ewan Harrison, David K. Jackson, Ian Johnston, Dominic Kwiatkowski, Cordelia Langford, John Sillitoe on behalf of the Wellcome Sanger Institute COVID-19 Surveillance Team |
| EPI_ISL_798184, EPI_ISL_798192 | Lighthouse Lab in Glasgow | Wellcome Sanger Institute for the COVID-19 Genomics UK (COG-UK) Consortium | Harper VanSteenhouse, Yumi Kasai, David Gray, Carol Clugston, Anna Dominiczak and Alex Alderton, Roberto Amato, Sonia Goncalves, Ewan Harrison, David K. Jackson, Ian Johnston, Dominic Kwiatkowski, Cordelia Langford, John Sillitoe on behalf of the Wellcome Sanger Institute COVID-19 Surveillance Team |
| EPI_ISL_798202, EPI_ISL_798204, EPI_ISL_798208, EPI_ISL_798220, EPI_ISL_798221, EPI_ISL_798222 | Lighthouse Lab in Alderley Park | Wellcome Sanger Institute for the COVID-19 Genomics UK (COG-UK) Consortium | Jacquelyn Wynn, Mairead Hyland, The Lighthouse Lab in Alderley Park and Alex Alderton, Roberto Amato, Sonia Goncalves, Ewan Harrison, David K. Jackson, Ian Johnston, Dominic Kwiatkowski, Cordelia Langford, John Sillitoe on behalf of the Wellcome Sanger Institute COVID-19 Surveillance Team |
| EPI_ISL_798223 | Lighthouse Lab in Glasgow | Wellcome Sanger Institute for the COVID-19 Genomics UK (COG-UK) Consortium | Harper VanSteenhouse, Yumi Kasai, David Gray, Carol Clugston, Anna Dominiczak and Alex Alderton, Roberto Amato, Sonia Goncalves, Ewan Harrison, David K. Jackson, Ian Johnston, Dominic Kwiatkowski, Cordelia Langford, John Sillitoe on behalf of the Wellcome Sanger Institute COVID-19 Surveillance Team |
| EPI_ISL_798225, EPI_ISL_798229 | Lighthouse Lab in Alderley Park | Wellcome Sanger Institute for the COVID-19 Genomics UK (COG-UK) Consortium | Jacquelyn Wynn, Mairead Hyland, The Lighthouse Lab in Alderley Park and Alex Alderton, Roberto Amato, Sonia Goncalves, Ewan Harrison, David K. Jackson, Ian Johnston, Dominic Kwiatkowski, Cordelia Langford, John Sillitoe on behalf of the Wellcome Sanger Institute COVID-19 Surveillance Team |
| EPI_ISL_798231 | Lighthouse Lab in Glasgow | Wellcome Sanger Institute for the COVID-19 Genomics UK (COG-UK) Consortium | Harper VanSteenhouse, Yumi Kasai, David Gray, Carol Clugston, Anna Dominiczak and Alex Alderton, Roberto Amato, Sonia Goncalves, Ewan Harrison, David K. Jackson, Ian Johnston, Dominic Kwiatkowski, Cordelia Langford, John Sillitoe on behalf of the Wellcome Sanger Institute COVID-19 Surveillance Team |
| EPI_ISL_798234, EPI_ISL_798235, EPI_ISL_798236, EPI_ISL_798239, EPI_ISL_798241, EPI_ISL_798242, EPI_ISL_798245, EPI_ISL_798246, EPI_ISL_798248 | Lighthouse Lab in Alderley Park | Wellcome Sanger Institute for the COVID-19 Genomics UK (COG-UK) Consortium | Jacquelyn Wynn, Mairead Hyland, The Lighthouse Lab in Alderley Park and Alex Alderton, Roberto Amato, Sonia Goncalves, Ewan Harrison, David K. Jackson, Ian Johnston, Dominic Kwiatkowski, Cordelia Langford, John Sillitoe on behalf of the Wellcome Sanger Institute COVID-19 Surveillance Team |
| EPI_ISL_798249 | Lighthouse Lab in Glasgow | Wellcome Sanger Institute for the COVID-19 Genomics UK (COG-UK) Consortium | Harper VanSteenhouse, Yumi Kasai, David Gray, Carol Clugston, Anna Dominiczak and Alex Alderton, Roberto Amato, Sonia Goncalves, Ewan Harrison, David K. Jackson, Ian Johnston, Dominic Kwiatkowski, Cordelia Langford, John Sillitoe on behalf of the Wellcome Sanger Institute COVID-19 Surveillance |

| | | | |
|---|---|---|---|
| | | | Team |
| EPI_ISL_798250, EPI_ISL_798255 | Lighthouse Lab in Alderley Park | Wellcome Sanger Institute for the COVID-19 Genomics UK (COG-UK) Consortium | Jacquelyn Wynn, Mairead Hyland, The Lighthouse Lab in Alderley Park and Alex Alderton, Roberto Amato, Sonia Goncalves, Ewan Harrison, David K. Jackson, Ian Johnston, Dominic Kwiatkowski, Cordelia Langford, John Sillitoe on behalf of the Wellcome Sanger Institute COVID-19 Surveillance Team |
| EPI_ISL_798261 | Lighthouse Lab in Glasgow | Wellcome Sanger Institute for the COVID-19 Genomics UK (COG-UK) Consortium | Harper VanSteenhouse, Yumi Kasai, David Gray, Carol Clugston, Anna Dominiczak and Alex Alderton, Roberto Amato, Sonia Goncalves, Ewan Harrison, David K. Jackson, Ian Johnston, Dominic Kwiatkowski, Cordelia Langford, John Sillitoe on behalf of the Wellcome Sanger Institute COVID-19 Surveillance Team |
| EPI_ISL_798268 | Lighthouse Lab in Alderley Park | Wellcome Sanger Institute for the COVID-19 Genomics UK (COG-UK) Consortium | Jacquelyn Wynn, Mairead Hyland, The Lighthouse Lab in Alderley Park and Alex Alderton, Roberto Amato, Sonia Goncalves, Ewan Harrison, David K. Jackson, Ian Johnston, Dominic Kwiatkowski, Cordelia Langford, John Sillitoe on behalf of the Wellcome Sanger Institute COVID-19 Surveillance Team |
| EPI_ISL_798271 | Lighthouse Lab in Glasgow | Wellcome Sanger Institute for the COVID-19 Genomics UK (COG-UK) Consortium | Harper VanSteenhouse, Yumi Kasai, David Gray, Carol Clugston, Anna Dominiczak and Alex Alderton, Roberto Amato, Sonia Goncalves, Ewan Harrison, David K. Jackson, Ian Johnston, Dominic Kwiatkowski, Cordelia Langford, John Sillitoe on behalf of the Wellcome Sanger Institute COVID-19 Surveillance Team |
| EPI_ISL_798274, EPI_ISL_798275, EPI_ISL_798277 | Lighthouse Lab in Alderley Park | Wellcome Sanger Institute for the COVID-19 Genomics UK (COG-UK) Consortium | Jacquelyn Wynn, Mairead Hyland, The Lighthouse Lab in Alderley Park and Alex Alderton, Roberto Amato, Sonia Goncalves, Ewan Harrison, David K. Jackson, Ian Johnston, Dominic Kwiatkowski, Cordelia Langford, John Sillitoe on behalf of the Wellcome Sanger Institute COVID-19 Surveillance Team |
| EPI_ISL_798278 | Lighthouse Lab in Glasgow | Wellcome Sanger Institute for the COVID-19 Genomics UK (COG-UK) Consortium | Harper VanSteenhouse, Yumi Kasai, David Gray, Carol Clugston, Anna Dominiczak and Alex Alderton, Roberto Amato, Sonia Goncalves, Ewan Harrison, David K. Jackson, Ian Johnston, Dominic Kwiatkowski, Cordelia Langford, John Sillitoe on behalf of the Wellcome Sanger Institute COVID-19 Surveillance Team |
| EPI_ISL_798280, EPI_ISL_798282, EPI_ISL_798283, EPI_ISL_798284, EPI_ISL_798285 | Lighthouse Lab in Alderley Park | Wellcome Sanger Institute for the COVID-19 Genomics UK (COG-UK) Consortium | Jacquelyn Wynn, Mairead Hyland, The Lighthouse Lab in Alderley Park and Alex Alderton, Roberto Amato, Sonia Goncalves, Ewan Harrison, David K. Jackson, Ian Johnston, Dominic Kwiatkowski, Cordelia Langford, John Sillitoe on behalf of the Wellcome Sanger Institute COVID-19 Surveillance Team |
| EPI_ISL_798287 | Lighthouse Lab in Glasgow | Wellcome Sanger Institute for the COVID-19 Genomics UK (COG-UK) Consortium | Harper VanSteenhouse, Yumi Kasai, David Gray, Carol Clugston, Anna Dominiczak and Alex Alderton, Roberto Amato, Sonia Goncalves, Ewan Harrison, David K. Jackson, Ian Johnston, Dominic Kwiatkowski, Cordelia Langford, John Sillitoe on behalf of the Wellcome Sanger Institute COVID-19 Surveillance Team |
| EPI_ISL_798292, EPI_ISL_798295, EPI_ISL_798296, EPI_ISL_798299, EPI_ISL_798305 | Lighthouse Lab in Alderley Park | Wellcome Sanger Institute for the COVID-19 Genomics UK (COG-UK) Consortium | Jacquelyn Wynn, Mairead Hyland, The Lighthouse Lab in Alderley Park and Alex Alderton, Roberto Amato, Sonia Goncalves, Ewan Harrison, David K. Jackson, Ian Johnston, Dominic Kwiatkowski, Cordelia Langford, John Sillitoe on behalf of the Wellcome Sanger Institute COVID-19 Surveillance Team |
| EPI_ISL_798308 | Lighthouse Lab in Glasgow | Wellcome Sanger Institute for the COVID-19 Genomics UK (COG-UK) Consortium | Harper VanSteenhouse, Yumi Kasai, David Gray, Carol Clugston, Anna Dominiczak and Alex Alderton, Roberto Amato, Sonia Goncalves, Ewan Harrison, David K. Jackson, Ian Johnston, Dominic Kwiatkowski, Cordelia Langford, John Sillitoe on behalf of the Wellcome Sanger Institute COVID-19 Surveillance Team |
| EPI_ISL_798310, EPI_ISL_798314, EPI_ISL_798321, EPI_ISL_798323, EPI_ISL_798324 | Lighthouse Lab in Alderley Park | Wellcome Sanger Institute for the COVID-19 Genomics UK (COG-UK) Consortium | Jacquelyn Wynn, Mairead Hyland, The Lighthouse Lab in Alderley Park and Alex Alderton, Roberto Amato, Sonia Goncalves, Ewan Harrison, David K. Jackson, Ian Johnston, Dominic Kwiatkowski, Cordelia Langford, John Sillitoe on behalf of the Wellcome Sanger Institute COVID-19 Surveillance Team |
| EPI_ISL_798325 | Lighthouse Lab in Glasgow | Wellcome Sanger Institute for the COVID-19 Genomics UK (COG-UK) Consortium | Harper VanSteenhouse, Yumi Kasai, David Gray, Carol Clugston, Anna Dominiczak and Alex Alderton, Roberto Amato, Sonia Goncalves, Ewan Harrison, David K. Jackson, Ian Johnston, Dominic Kwiatkowski, Cordelia Langford, John Sillitoe on behalf of the Wellcome Sanger Institute COVID-19 Surveillance Team |
| EPI_ISL_798328, EPI_ISL_798329 | Lighthouse Lab in Alderley Park | Wellcome Sanger Institute for the COVID-19 Genomics UK (COG-UK) Consortium | Jacquelyn Wynn, Mairead Hyland, The Lighthouse Lab in Alderley Park and Alex Alderton, Roberto Amato, Sonia Goncalves, Ewan Harrison, David K. Jackson, Ian Johnston, Dominic Kwiatkowski, Cordelia Langford, John Sillitoe on behalf of the Wellcome Sanger Institute COVID-19 Surveillance Team |
| EPI_ISL_798331, EPI_ISL_798332 | Lighthouse Lab in Glasgow | Wellcome Sanger Institute for the COVID-19 Genomics UK (COG-UK) Consortium | Harper VanSteenhouse, Yumi Kasai, David Gray, Carol Clugston, Anna Dominiczak and Alex Alderton, Roberto Amato, Sonia Goncalves, Ewan Harrison, David K. Jackson, Ian Johnston, Dominic Kwiatkowski, Cordelia Langford, John Sillitoe on behalf of the Wellcome Sanger Institute COVID-19 Surveillance Team |
| EPI_ISL_798334, EPI_ISL_798336, EPI_ISL_798341, EPI_ISL_798345, EPI_ISL_798346, EPI_ISL_798349, EPI_ISL_798356, EPI_ISL_798362, EPI_ISL_798368, EPI_ISL_798369, EPI_ISL_798370, EPI_ISL_798378, EPI_ISL_798379, EPI_ISL_798384, EPI_ISL_798386, EPI_ISL_798388, EPI_ISL_798389, EPI_ISL_798396, EPI_ISL_798399, EPI_ISL_798404, EPI_ISL_798405, EPI_ISL_798413, EPI_ISL_798416, EPI_ISL_798417, EPI_ISL_798431, EPI_ISL_798436, EPI_ISL_798444, EPI_ISL_798455, EPI_ISL_798465, EPI_ISL_798468, EPI_ISL_798475, EPI_ISL_798479, EPI_ISL_798483, EPI_ISL_798487, EPI_ISL_798492, EPI_ISL_798493, EPI_ISL_798500, EPI_ISL_798502, EPI_ISL_798508, EPI_ISL_798510, EPI_ISL_798513, EPI_ISL_798516, EPI_ISL_798517, EPI_ISL_798519, EPI_ISL_798524, EPI_ISL_798531, EPI_ISL_798534, EPI_ISL_798539, EPI_ISL_798541, EPI_ISL_798542, EPI_ISL_798548, EPI_ISL_798550, EPI_ISL_798555, EPI_ISL_798561, EPI_ISL_798564, EPI_ISL_798579, EPI_ISL_798591, EPI_ISL_798593, EPI_ISL_798598, EPI_ISL_798601, EPI_ISL_798607, EPI_ISL_798609, EPI_ISL_798612, EPI_ISL_798622, EPI_ISL_798631, EPI_ISL_798632, EPI_ISL_798633, EPI_ISL_798635, EPI_ISL_798637, EPI_ISL_798639, EPI_ISL_798640, EPI_ISL_798641, EPI_ISL_798643, EPI_ISL_798652, EPI_ISL_798659, EPI_ISL_798661, EPI_ISL_798665, EPI_ISL_798667, EPI_ISL_798668, EPI_ISL_798669, EPI_ISL_798671 | | | |
| see above | Lighthouse Lab in Alderley Park | Wellcome Sanger Institute for the COVID-19 Genomics UK (COG-UK) Consortium | Jacquelyn Wynn, Mairead Hyland, The Lighthouse Lab in Alderley Park and Alex Alderton, Roberto Amato, Sonia Goncalves, Ewan Harrison, David K. Jackson, Ian Johnston, Dominic Kwiatkowski, Cordelia Langford, John Sillitoe on behalf of the Wellcome Sanger Institute COVID-19 Surveillance Team |
| EPI_ISL_798674 | Lighthouse Lab in Cambridge | Wellcome Sanger Institute for the COVID-19 Genomics UK (COG-UK) Consortium | Rob Howes, The Lighthouse Lab in Cambridge and Alex Alderton, Roberto Amato, Sonia Goncalves, Ewan Harrison, David K. Jackson, Ian Johnston, Dominic Kwiatkowski, Cordelia Langford, John Sillitoe on behalf of the Wellcome Sanger Institute COVID-19 Surveillance Team |
| EPI_ISL_798676 | Lighthouse Lab in Alderley Park | Wellcome Sanger Institute for the COVID-19 Genomics UK (COG-UK) Consortium | Jacquelyn Wynn, Mairead Hyland, The Lighthouse Lab in Alderley Park and Alex Alderton, Roberto Amato, Sonia Goncalves, Ewan Harrison, David K. Jackson, Ian Johnston, Dominic Kwiatkowski, Cordelia Langford, John Sillitoe on behalf of the Wellcome Sanger Institute COVID-19 Surveillance Team |
| EPI_ISL_798678, EPI_ISL_798687 | Lighthouse Lab in Cambridge | Wellcome Sanger Institute for the COVID-19 Genomics UK (COG-UK) Consortium | Rob Howes, The Lighthouse Lab in Cambridge and Alex Alderton, Roberto Amato, Sonia Goncalves, Ewan Harrison, David K. Jackson, Ian Johnston, Dominic Kwiatkowski, Cordelia Langford, John Sillitoe on behalf of the Wellcome Sanger Institute COVID-19 Surveillance Team |
| EPI_ISL_798688, EPI_ISL_798689, EPI_ISL_798690, EPI_ISL_798692, EPI_ISL_798695 | Lighthouse Lab in Alderley Park | Wellcome Sanger Institute for the COVID-19 Genomics UK (COG-UK) Consortium | Jacquelyn Wynn, Mairead Hyland, The Lighthouse Lab in Alderley Park and Alex Alderton, Roberto Amato, Sonia Goncalves, Ewan Harrison, David K. Jackson, Ian Johnston, Dominic Kwiatkowski, Cordelia Langford, John Sillitoe on behalf of the Wellcome Sanger Institute COVID-19 Surveillance Team |
| EPI_ISL_798700 | Lighthouse Lab in Cambridge | Wellcome Sanger Institute for the COVID-19 Genomics UK (COG-UK) Consortium | Rob Howes, The Lighthouse Lab in Cambridge and Alex Alderton, Roberto Amato, Sonia Goncalves, Ewan Harrison, David K. Jackson, Ian Johnston, Dominic Kwiatkowski, Cordelia Langford, John Sillitoe on behalf of the Wellcome Sanger Institute COVID-19 Surveillance Team |
| EPI_ISL_798704, EPI_ISL_798705, EPI_ISL_798722, EPI_ISL_798727, EPI_ISL_798735, EPI_ISL_798744, EPI_ISL_798749, EPI_ISL_798768, EPI_ISL_798771 | Lighthouse Lab in Alderley Park | Wellcome Sanger Institute for the COVID-19 Genomics UK (COG-UK) Consortium | Jacquelyn Wynn, Mairead Hyland, The Lighthouse Lab in Alderley Park and Alex Alderton, Roberto Amato, Sonia Goncalves, Ewan Harrison, David K. Jackson, Ian Johnston, Dominic Kwiatkowski, Cordelia Langford, John Sillitoe on behalf of the Wellcome Sanger Institute COVID-19 Surveillance Team |
| EPI_ISL_798778 | Lighthouse Lab in Cambridge | Wellcome Sanger Institute for the COVID-19 Genomics UK (COG-UK) Consortium | Rob Howes, The Lighthouse Lab in Cambridge and Alex Alderton, Roberto Amato, Sonia Goncalves, Ewan Harrison, David K. Jackson, Ian Johnston, Dominic Kwiatkowski, Cordelia Langford, John Sillitoe on behalf of the Wellcome Sanger Institute COVID-19 Surveillance Team |
| EPI_ISL_798784 | Lighthouse Lab in Alderley Park | Wellcome Sanger Institute for the COVID-19 Genomics UK (COG-UK) Consortium | Jacquelyn Wynn, Mairead Hyland, The Lighthouse Lab in Alderley Park and Alex Alderton, Roberto Amato, Sonia Goncalves, Ewan Harrison, David K. Jackson, Ian Johnston, Dominic Kwiatkowski, Cordelia Langford, John Sillitoe on behalf of the Wellcome Sanger Institute COVID-19 Surveillance Team |
| EPI_ISL_798786, EPI_ISL_798790 | Lighthouse Lab in Cambridge | Wellcome Sanger Institute for the COVID-19 Genomics UK (COG-UK) Consortium | Rob Howes, The Lighthouse Lab in Cambridge and Alex Alderton, Roberto Amato, Sonia Goncalves, Ewan Harrison, David K. Jackson, Ian Johnston, Dominic Kwiatkowski, Cordelia Langford, John Sillitoe on behalf of the Wellcome Sanger Institute COVID-19 Surveillance Team |
| EPI_ISL_798791, EPI_ISL_798793, EPI_ISL_798798, EPI_ISL_798804, EPI_ISL_798811, EPI_ISL_798814, EPI_ISL_798819, EPI_ISL_798824 | Lighthouse Lab in Alderley Park | Wellcome Sanger Institute for the COVID-19 Genomics UK (COG-UK) Consortium | Jacquelyn Wynn, Mairead Hyland, The Lighthouse Lab in Alderley Park and Alex Alderton, Roberto Amato, Sonia Goncalves, Ewan Harrison, David K. Jackson, Ian Johnston, Dominic Kwiatkowski, Cordelia Langford, John Sillitoe on behalf of the Wellcome Sanger Institute COVID-19 Surveillance Team |
| EPI_ISL_798825 | Lighthouse Lab in Cambridge | Wellcome Sanger Institute for the COVID-19 Genomics UK (COG-UK) Consortium | Rob Howes, The Lighthouse Lab in Cambridge and Alex Alderton, Roberto Amato, Sonia Goncalves, Ewan Harrison, David K. Jackson, Ian Johnston, Dominic Kwiatkowski, Cordelia Langford, John Sillitoe on behalf of the Wellcome Sanger Institute COVID-19 Surveillance Team |

| | | | |
|---|---|---|---|
| EPI_ISL_798826, EPI_ISL_798833 | Lighthouse Lab in Alderley Park | Wellcome Sanger Institute for the COVID-19 Genomics UK (COG-UK) Consortium | Jacquelyn Wynn, Mairead Hyland, The Lighthouse Lab in Alderley Park and Alex Alderton, Roberto Amato, Sonia Goncalves, Ewan Harrison, David K. Jackson, Ian Johnston, Dominic Kwiatkowski, Cordelia Langford, John Sillitoe on behalf of the Wellcome Sanger Institute COVID-19 Surveillance Team |
| EPI_ISL_798834, EPI_ISL_798841 | Lighthouse Lab in Cambridge | Wellcome Sanger Institute for the COVID-19 Genomics UK (COG-UK) Consortium | Rob Howes, The Lighthouse Lab in Cambridge and Alex Alderton, Roberto Amato, Sonia Goncalves, Ewan Harrison, David K. Jackson, Ian Johnston, Dominic Kwiatkowski, Cordelia Langford, John Sillitoe on behalf of the Wellcome Sanger Institute COVID-19 Surveillance Team |
| EPI_ISL_798845 | Lighthouse Lab in Alderley Park | Wellcome Sanger Institute for the COVID-19 Genomics UK (COG-UK) Consortium | Jacquelyn Wynn, Mairead Hyland, The Lighthouse Lab in Alderley Park and Alex Alderton, Roberto Amato, Sonia Goncalves, Ewan Harrison, David K. Jackson, Ian Johnston, Dominic Kwiatkowski, Cordelia Langford, John Sillitoe on behalf of the Wellcome Sanger Institute COVID-19 Surveillance Team |
| EPI_ISL_798848 | Lighthouse Lab in Cambridge | Wellcome Sanger Institute for the COVID-19 Genomics UK (COG-UK) Consortium | Rob Howes, The Lighthouse Lab in Cambridge and Alex Alderton, Roberto Amato, Sonia Goncalves, Ewan Harrison, David K. Jackson, Ian Johnston, Dominic Kwiatkowski, Cordelia Langford, John Sillitoe on behalf of the Wellcome Sanger Institute COVID-19 Surveillance Team |
| EPI_ISL_798850, EPI_ISL_798852, EPI_ISL_798869, EPI_ISL_798872, EPI_ISL_798878, EPI_ISL_798884, EPI_ISL_798888, EPI_ISL_798891, EPI_ISL_798898, EPI_ISL_798902, EPI_ISL_798906 | | | |
| see above | Lighthouse Lab in Alderley Park | Wellcome Sanger Institute for the COVID-19 Genomics UK (COG-UK) Consortium | Jacquelyn Wynn, Mairead Hyland, The Lighthouse Lab in Alderley Park and Alex Alderton, Roberto Amato, Sonia Goncalves, Ewan Harrison, David K. Jackson, Ian Johnston, Dominic Kwiatkowski, Cordelia Langford, John Sillitoe on behalf of the Wellcome Sanger Institute COVID-19 Surveillance Team |
| EPI_ISL_798907 | Lighthouse Lab in Cambridge | Wellcome Sanger Institute for the COVID-19 Genomics UK (COG-UK) Consortium | Rob Howes, The Lighthouse Lab in Cambridge and Alex Alderton, Roberto Amato, Sonia Goncalves, Ewan Harrison, David K. Jackson, Ian Johnston, Dominic Kwiatkowski, Cordelia Langford, John Sillitoe on behalf of the Wellcome Sanger Institute COVID-19 Surveillance Team |
| EPI_ISL_798914, EPI_ISL_798921, EPI_ISL_798924, EPI_ISL_798926, EPI_ISL_798928 | Lighthouse Lab in Alderley Park | Wellcome Sanger Institute for the COVID-19 Genomics UK (COG-UK) Consortium | Jacquelyn Wynn, Mairead Hyland, The Lighthouse Lab in Alderley Park and Alex Alderton, Roberto Amato, Sonia Goncalves, Ewan Harrison, David K. Jackson, Ian Johnston, Dominic Kwiatkowski, Cordelia Langford, John Sillitoe on behalf of the Wellcome Sanger Institute COVID-19 Surveillance Team |
| EPI_ISL_798931 | Lighthouse Lab in Cambridge | Wellcome Sanger Institute for the COVID-19 Genomics UK (COG-UK) Consortium | Rob Howes, The Lighthouse Lab in Cambridge and Alex Alderton, Roberto Amato, Sonia Goncalves, Ewan Harrison, David K. Jackson, Ian Johnston, Dominic Kwiatkowski, Cordelia Langford, John Sillitoe on behalf of the Wellcome Sanger Institute COVID-19 Surveillance Team |
| EPI_ISL_798934, EPI_ISL_798939 | Lighthouse Lab in Alderley Park | Wellcome Sanger Institute for the COVID-19 Genomics UK (COG-UK) Consortium | Jacquelyn Wynn, Mairead Hyland, The Lighthouse Lab in Alderley Park and Alex Alderton, Roberto Amato, Sonia Goncalves, Ewan Harrison, David K. Jackson, Ian Johnston, Dominic Kwiatkowski, Cordelia Langford, John Sillitoe on behalf of the Wellcome Sanger Institute COVID-19 Surveillance Team |
| EPI_ISL_798944 | Lighthouse Lab in Cambridge | Wellcome Sanger Institute for the COVID-19 Genomics UK (COG-UK) Consortium | Rob Howes, The Lighthouse Lab in Cambridge and Alex Alderton, Roberto Amato, Sonia Goncalves, Ewan Harrison, David K. Jackson, Ian Johnston, Dominic Kwiatkowski, Cordelia Langford, John Sillitoe on behalf of the Wellcome Sanger Institute COVID-19 Surveillance Team |
| EPI_ISL_798951, EPI_ISL_798952, EPI_ISL_798962, EPI_ISL_798963, EPI_ISL_798964, EPI_ISL_798968, EPI_ISL_798969, EPI_ISL_798984 | Lighthouse Lab in Alderley Park | Wellcome Sanger Institute for the COVID-19 Genomics UK (COG-UK) Consortium | Jacquelyn Wynn, Mairead Hyland, The Lighthouse Lab in Alderley Park and Alex Alderton, Roberto Amato, Sonia Goncalves, Ewan Harrison, David K. Jackson, Ian Johnston, Dominic Kwiatkowski, Cordelia Langford, John Sillitoe on behalf of the Wellcome Sanger Institute COVID-19 Surveillance Team |
| EPI_ISL_798985 | Lighthouse Lab in Cambridge | Wellcome Sanger Institute for the COVID-19 Genomics UK (COG-UK) Consortium | Rob Howes, The Lighthouse Lab in Cambridge and Alex Alderton, Roberto Amato, Sonia Goncalves, Ewan Harrison, David K. Jackson, Ian Johnston, Dominic Kwiatkowski, Cordelia Langford, John Sillitoe on behalf of the Wellcome Sanger Institute COVID-19 Surveillance Team |
| EPI_ISL_798986, EPI_ISL_798988, EPI_ISL_798991, EPI_ISL_799000, EPI_ISL_799003, EPI_ISL_799005, EPI_ISL_799010 | Lighthouse Lab in Alderley Park | Wellcome Sanger Institute for the COVID-19 Genomics UK (COG-UK) Consortium | Jacquelyn Wynn, Mairead Hyland, The Lighthouse Lab in Alderley Park and Alex Alderton, Roberto Amato, Sonia Goncalves, Ewan Harrison, David K. Jackson, Ian Johnston, Dominic Kwiatkowski, Cordelia Langford, John Sillitoe on behalf of the Wellcome Sanger Institute COVID-19 Surveillance Team |
| EPI_ISL_799020 | Lighthouse Lab in Cambridge | Wellcome Sanger Institute for the COVID-19 Genomics UK (COG-UK) Consortium | Rob Howes, The Lighthouse Lab in Cambridge and Alex Alderton, Roberto Amato, Sonia Goncalves, Ewan Harrison, David K. Jackson, Ian Johnston, Dominic Kwiatkowski, Cordelia Langford, John Sillitoe on behalf of the Wellcome Sanger Institute COVID-19 Surveillance Team |
| EPI_ISL_799021, EPI_ISL_799022, EPI_ISL_799023, EPI_ISL_799025, EPI_ISL_799026, EPI_ISL_799029 | Lighthouse Lab in Alderley Park | Wellcome Sanger Institute for the COVID-19 Genomics UK (COG-UK) Consortium | Jacquelyn Wynn, Mairead Hyland, The Lighthouse Lab in Alderley Park and Alex Alderton, Roberto Amato, Sonia Goncalves, Ewan Harrison, David K. Jackson, Ian Johnston, Dominic Kwiatkowski, Cordelia Langford, John Sillitoe on behalf of the Wellcome Sanger Institute COVID-19 Surveillance Team |
| EPI_ISL_799031 | Lighthouse Lab in Cambridge | Wellcome Sanger Institute for the COVID-19 Genomics UK (COG-UK) Consortium | Rob Howes, The Lighthouse Lab in Cambridge and Alex Alderton, Roberto Amato, Sonia Goncalves, Ewan Harrison, David K. Jackson, Ian Johnston, Dominic Kwiatkowski, Cordelia Langford, John Sillitoe on behalf of the Wellcome Sanger Institute COVID-19 Surveillance Team |
| EPI_ISL_799040, EPI_ISL_799045, EPI_ISL_799047, EPI_ISL_799053, EPI_ISL_799055, EPI_ISL_799057, EPI_ISL_799059, EPI_ISL_799069, EPI_ISL_799070 | Lighthouse Lab in Alderley Park | Wellcome Sanger Institute for the COVID-19 Genomics UK (COG-UK) Consortium | Jacquelyn Wynn, Mairead Hyland, The Lighthouse Lab in Alderley Park and Alex Alderton, Roberto Amato, Sonia Goncalves, Ewan Harrison, David K. Jackson, Ian Johnston, Dominic Kwiatkowski, Cordelia Langford, John Sillitoe on behalf of the Wellcome Sanger Institute COVID-19 Surveillance Team |
| EPI_ISL_799074 | Lighthouse Lab in Cambridge | Wellcome Sanger Institute for the COVID-19 Genomics UK (COG-UK) Consortium | Rob Howes, The Lighthouse Lab in Cambridge and Alex Alderton, Roberto Amato, Sonia Goncalves, Ewan Harrison, David K. Jackson, Ian Johnston, Dominic Kwiatkowski, Cordelia Langford, John Sillitoe on behalf of the Wellcome Sanger Institute COVID-19 Surveillance Team |
| EPI_ISL_799079 | Lighthouse Lab in Alderley Park | Wellcome Sanger Institute for the COVID-19 Genomics UK (COG-UK) Consortium | Jacquelyn Wynn, Mairead Hyland, The Lighthouse Lab in Alderley Park and Alex Alderton, Roberto Amato, Sonia Goncalves, Ewan Harrison, David K. Jackson, Ian Johnston, Dominic Kwiatkowski, Cordelia Langford, John Sillitoe on behalf of the Wellcome Sanger Institute COVID-19 Surveillance Team |
| EPI_ISL_799083 | Lighthouse Lab in Cambridge | Wellcome Sanger Institute for the COVID-19 Genomics UK (COG-UK) Consortium | Rob Howes, The Lighthouse Lab in Cambridge and Alex Alderton, Roberto Amato, Sonia Goncalves, Ewan Harrison, David K. Jackson, Ian Johnston, Dominic Kwiatkowski, Cordelia Langford, John Sillitoe on behalf of the Wellcome Sanger Institute COVID-19 Surveillance Team |
| EPI_ISL_799085, EPI_ISL_799086 | Lighthouse Lab in Alderley Park | Wellcome Sanger Institute for the COVID-19 Genomics UK (COG-UK) Consortium | Jacquelyn Wynn, Mairead Hyland, The Lighthouse Lab in Alderley Park and Alex Alderton, Roberto Amato, Sonia Goncalves, Ewan Harrison, David K. Jackson, Ian Johnston, Dominic Kwiatkowski, Cordelia Langford, John Sillitoe on behalf of the Wellcome Sanger Institute COVID-19 Surveillance Team |
| EPI_ISL_799087 | Lighthouse Lab in Cambridge | Wellcome Sanger Institute for the COVID-19 Genomics UK (COG-UK) Consortium | Rob Howes, The Lighthouse Lab in Cambridge and Alex Alderton, Roberto Amato, Sonia Goncalves, Ewan Harrison, David K. Jackson, Ian Johnston, Dominic Kwiatkowski, Cordelia Langford, John Sillitoe on behalf of the Wellcome Sanger Institute COVID-19 Surveillance Team |
| EPI_ISL_799099, EPI_ISL_799103, EPI_ISL_799104, EPI_ISL_799105, EPI_ISL_799106, EPI_ISL_799111, EPI_ISL_799113, EPI_ISL_799114, EPI_ISL_799122, EPI_ISL_799124, EPI_ISL_799137, EPI_ISL_799138, EPI_ISL_799139, EPI_ISL_799143, EPI_ISL_799145, EPI_ISL_799149 | | | |
| see above | Lighthouse Lab in Alderley Park | Wellcome Sanger Institute for the COVID-19 Genomics UK (COG-UK) Consortium | Jacquelyn Wynn, Mairead Hyland, The Lighthouse Lab in Alderley Park and Alex Alderton, Roberto Amato, Sonia Goncalves, Ewan Harrison, David K. Jackson, Ian Johnston, Dominic Kwiatkowski, Cordelia Langford, John Sillitoe on behalf of the Wellcome Sanger Institute COVID-19 Surveillance Team |
| EPI_ISL_799152, EPI_ISL_799153 | Lighthouse Lab in Cambridge | Wellcome Sanger Institute for the COVID-19 Genomics UK (COG-UK) Consortium | Rob Howes, The Lighthouse Lab in Cambridge and Alex Alderton, Roberto Amato, Sonia Goncalves, Ewan Harrison, David K. Jackson, Ian Johnston, Dominic Kwiatkowski, Cordelia Langford, John Sillitoe on behalf of the Wellcome Sanger Institute COVID-19 Surveillance Team |
| EPI_ISL_799154, EPI_ISL_799160, EPI_ISL_799161, EPI_ISL_799165, EPI_ISL_799166, EPI_ISL_799170, EPI_ISL_799171, EPI_ISL_799177, EPI_ISL_799178, EPI_ISL_799179, EPI_ISL_799182 | | | |
| see above | Lighthouse Lab in Alderley Park | Wellcome Sanger Institute for the COVID-19 Genomics UK (COG-UK) Consortium | Jacquelyn Wynn, Mairead Hyland, The Lighthouse Lab in Alderley Park and Alex Alderton, Roberto Amato, Sonia Goncalves, Ewan Harrison, David K. Jackson, Ian Johnston, Dominic Kwiatkowski, Cordelia Langford, John Sillitoe on behalf of the Wellcome Sanger Institute COVID-19 Surveillance Team |
| EPI_ISL_799186 | Lighthouse Lab in Cambridge | Wellcome Sanger Institute for the COVID-19 Genomics UK (COG-UK) Consortium | Rob Howes, The Lighthouse Lab in Cambridge and Alex Alderton, Roberto Amato, Sonia Goncalves, Ewan Harrison, David K. Jackson, Ian Johnston, Dominic Kwiatkowski, Cordelia Langford, John Sillitoe on behalf of the Wellcome Sanger Institute COVID-19 Surveillance Team |
| EPI_ISL_799188, EPI_ISL_799191, EPI_ISL_799198, EPI_ISL_799200, EPI_ISL_799202 | Lighthouse Lab in Alderley Park | Wellcome Sanger Institute for the COVID-19 Genomics UK (COG-UK) Consortium | Jacquelyn Wynn, Mairead Hyland, The Lighthouse Lab in Alderley Park and Alex Alderton, Roberto Amato, Sonia Goncalves, Ewan Harrison, David K. Jackson, Ian Johnston, Dominic Kwiatkowski, Cordelia Langford, John Sillitoe on behalf of the Wellcome Sanger Institute COVID-19 Surveillance Team |
| EPI_ISL_799208, EPI_ISL_799210 | Lighthouse Lab in Cambridge | Wellcome Sanger Institute for the COVID-19 Genomics UK (COG-UK) Consortium | Rob Howes, The Lighthouse Lab in Cambridge and Alex Alderton, Roberto Amato, Sonia Goncalves, Ewan Harrison, David K. Jackson, Ian Johnston, Dominic Kwiatkowski, Cordelia Langford, John Sillitoe on behalf of the Wellcome Sanger Institute COVID-19 Surveillance Team |
| EPI_ISL_799212, EPI_ISL_799213, EPI_ISL_799214, EPI_ISL_799219 | Lighthouse Lab in Alderley Park | Wellcome Sanger Institute for the COVID-19 Genomics UK (COG-UK) Consortium | Jacquelyn Wynn, Mairead Hyland, The Lighthouse Lab in Alderley Park and Alex Alderton, Roberto Amato, Sonia Goncalves, Ewan Harrison, David K. Jackson, Ian Johnston, Dominic Kwiatkowski, Cordelia Langford, John Sillitoe on behalf of the Wellcome Sanger Institute COVID-19 Surveillance Team |
| EPI_ISL_799226 | Lighthouse Lab in Cambridge | Wellcome Sanger Institute for the COVID-19 Genomics UK | Rob Howes, The Lighthouse Lab in Cambridge and Alex Alderton, Roberto Amato, Sonia Goncalves, Ewan Harrison, David K. Jackson, Ian Johnston, |

| | | | |
|---|---|---|---|
| | | (COG-UK) Consortium | Dominic Kwiatkowski, Cordelia Langford, John Sillitoe on behalf of the Wellcome Sanger Institute COVID-19 Surveillance Team |
| EPI_ISL_799227, EPI_ISL_799228, EPI_ISL_799229, EPI_ISL_799231, EPI_ISL_799233, EPI_ISL_799234, EPI_ISL_799235, EPI_ISL_799237, EPI_ISL_799244, EPI_ISL_799248, EPI_ISL_799252, EPI_ISL_799257, EPI_ISL_799261, EPI_ISL_799265, EPI_ISL_799269, EPI_ISL_799271, EPI_ISL_799276, EPI_ISL_799278, EPI_ISL_799281 | | | |
| see above | Lighthouse Lab in Alderley Park | Wellcome Sanger Institute for the COVID-19 Genomics UK (COG-UK) Consortium | Jacquelyn Wynn, Mairead Hyland, The Lighthouse Lab in Alderley Park and Alex Alderton, Roberto Amato, Sonia Goncalves, Ewan Harrison, David K. Jackson, Ian Johnston, Dominic Kwiatkowski, Cordelia Langford, John Sillitoe on behalf of the Wellcome Sanger Institute COVID-19 Surveillance Team |
| EPI_ISL_799291, EPI_ISL_799292 | Lighthouse Lab in Cambridge | Wellcome Sanger Institute for the COVID-19 Genomics UK (COG-UK) Consortium | Rob Howes, The Lighthouse Lab in Cambridge and Alex Alderton, Roberto Amato, Sonia Goncalves, Ewan Harrison, David K. Jackson, Ian Johnston, Dominic Kwiatkowski, Cordelia Langford, John Sillitoe on behalf of the Wellcome Sanger Institute COVID-19 Surveillance Team |
| EPI_ISL_799297, EPI_ISL_799300, EPI_ISL_799301, EPI_ISL_799304, EPI_ISL_799307, EPI_ISL_799309, EPI_ISL_799312, EPI_ISL_799313, EPI_ISL_799316, EPI_ISL_799321, EPI_ISL_799324, EPI_ISL_799328, EPI_ISL_799329, EPI_ISL_799330, EPI_ISL_799331, EPI_ISL_799332, EPI_ISL_799335, EPI_ISL_799336, EPI_ISL_799337, EPI_ISL_799338, EPI_ISL_799339, EPI_ISL_799340, EPI_ISL_799341, EPI_ISL_799342, EPI_ISL_799343, EPI_ISL_799344, EPI_ISL_799349, EPI_ISL_799358, EPI_ISL_799359, EPI_ISL_799361, EPI_ISL_799365, EPI_ISL_799367, EPI_ISL_799371, EPI_ISL_799374, EPI_ISL_799379, EPI_ISL_799381, EPI_ISL_799382, EPI_ISL_799383, EPI_ISL_799385, EPI_ISL_799389, EPI_ISL_799391, EPI_ISL_799393, EPI_ISL_799396, EPI_ISL_799402, EPI_ISL_799407, EPI_ISL_799408, EPI_ISL_799409, EPI_ISL_799414, EPI_ISL_799421, EPI_ISL_799423, EPI_ISL_799426, EPI_ISL_799428, EPI_ISL_799429, EPI_ISL_799431, EPI_ISL_799435, EPI_ISL_799442, EPI_ISL_799444, EPI_ISL_799449, EPI_ISL_799450, EPI_ISL_799451, EPI_ISL_799453, EPI_ISL_799455, EPI_ISL_799457, EPI_ISL_799459, EPI_ISL_799460, EPI_ISL_799462, EPI_ISL_799463, EPI_ISL_799464, EPI_ISL_799471, EPI_ISL_799472, EPI_ISL_799475, EPI_ISL_799478, EPI_ISL_799480, EPI_ISL_799481, EPI_ISL_799483, EPI_ISL_799484, EPI_ISL_799486, EPI_ISL_799489, EPI_ISL_799491, EPI_ISL_799493, EPI_ISL_799495, EPI_ISL_799499, EPI_ISL_799503, EPI_ISL_799504, EPI_ISL_799507, EPI_ISL_799511, EPI_ISL_799513, EPI_ISL_799516, EPI_ISL_799517, EPI_ISL_799518, EPI_ISL_799521, EPI_ISL_799524, EPI_ISL_799527, EPI_ISL_799532, EPI_ISL_799539, EPI_ISL_799544, EPI_ISL_799548, EPI_ISL_799552, EPI_ISL_799553, EPI_ISL_799554, EPI_ISL_799558, EPI_ISL_799561, EPI_ISL_799564, EPI_ISL_799568, EPI_ISL_799569, EPI_ISL_799571, EPI_ISL_799574, EPI_ISL_799576, EPI_ISL_799581, EPI_ISL_799584, EPI_ISL_799585, EPI_ISL_799589, EPI_ISL_799591, EPI_ISL_799592, EPI_ISL_799593, EPI_ISL_799594, EPI_ISL_799596, EPI_ISL_799599, EPI_ISL_799601, EPI_ISL_799605, EPI_ISL_799606, EPI_ISL_799609, EPI_ISL_799612, EPI_ISL_799613, EPI_ISL_799615, EPI_ISL_799634, EPI_ISL_799659, EPI_ISL_799669, EPI_ISL_799681, EPI_ISL_799682, EPI_ISL_799686 | | | |
| see above | Lighthouse Lab in Glasgow | Wellcome Sanger Institute for the COVID-19 Genomics UK (COG-UK) Consortium | Harper VanSteenhouse, Yumi Kasai, David Gray, Carol Clugston, Anna Dominiczak and Alex Alderton, Roberto Amato, Sonia Goncalves, Ewan Harrison, David K. Jackson, Ian Johnston, Dominic Kwiatkowski, Cordelia Langford, John Sillitoe on behalf of the Wellcome Sanger Institute COVID-19 Surveillance Team |
| EPI_ISL_799687, EPI_ISL_799693 | Lighthouse Lab in Alderley Park | Wellcome Sanger Institute for the COVID-19 Genomics UK (COG-UK) Consortium | Jacquelyn Wynn, Mairead Hyland, The Lighthouse Lab in Alderley Park and Alex Alderton, Roberto Amato, Sonia Goncalves, Ewan Harrison, David K. Jackson, Ian Johnston, Dominic Kwiatkowski, Cordelia Langford, John Sillitoe on behalf of the Wellcome Sanger Institute COVID-19 Surveillance Team |
| EPI_ISL_799699, EPI_ISL_799707, EPI_ISL_799726 | Lighthouse Lab in Glasgow | Wellcome Sanger Institute for the COVID-19 Genomics UK (COG-UK) Consortium | Harper VanSteenhouse, Yumi Kasai, David Gray, Carol Clugston, Anna Dominiczak and Alex Alderton, Roberto Amato, Sonia Goncalves, Ewan Harrison, David K. Jackson, Ian Johnston, Dominic Kwiatkowski, Cordelia Langford, John Sillitoe on behalf of the Wellcome Sanger Institute COVID-19 Surveillance Team |
| EPI_ISL_799734, EPI_ISL_799744, EPI_ISL_799747 | Lighthouse Lab in Alderley Park | Wellcome Sanger Institute for the COVID-19 Genomics UK (COG-UK) Consortium | Jacquelyn Wynn, Mairead Hyland, The Lighthouse Lab in Alderley Park and Alex Alderton, Roberto Amato, Sonia Goncalves, Ewan Harrison, David K. Jackson, Ian Johnston, Dominic Kwiatkowski, Cordelia Langford, John Sillitoe on behalf of the Wellcome Sanger Institute COVID-19 Surveillance Team |
| EPI_ISL_799748, EPI_ISL_799758, EPI_ISL_799763, EPI_ISL_799765 | Lighthouse Lab in Glasgow | Wellcome Sanger Institute for the COVID-19 Genomics UK (COG-UK) Consortium | Harper VanSteenhouse, Yumi Kasai, David Gray, Carol Clugston, Anna Dominiczak and Alex Alderton, Roberto Amato, Sonia Goncalves, Ewan Harrison, David K. Jackson, Ian Johnston, Dominic Kwiatkowski, Cordelia Langford, John Sillitoe on behalf of the Wellcome Sanger Institute COVID-19 Surveillance Team |
| EPI_ISL_799767 | Lighthouse Lab in Alderley Park | Wellcome Sanger Institute for the COVID-19 Genomics UK (COG-UK) Consortium | Jacquelyn Wynn, Mairead Hyland, The Lighthouse Lab in Alderley Park and Alex Alderton, Roberto Amato, Sonia Goncalves, Ewan Harrison, David K. Jackson, Ian Johnston, Dominic Kwiatkowski, Cordelia Langford, John Sillitoe on behalf of the Wellcome Sanger Institute COVID-19 Surveillance Team |
| EPI_ISL_799768 | Lighthouse Lab in Glasgow | Wellcome Sanger Institute for the COVID-19 Genomics UK (COG-UK) Consortium | Harper VanSteenhouse, Yumi Kasai, David Gray, Carol Clugston, Anna Dominiczak and Alex Alderton, Roberto Amato, Sonia Goncalves, Ewan Harrison, David K. Jackson, Ian Johnston, Dominic Kwiatkowski, Cordelia Langford, John Sillitoe on behalf of the Wellcome Sanger Institute COVID-19 Surveillance Team |
| EPI_ISL_799784, EPI_ISL_799787, EPI_ISL_799799 | Lighthouse Lab in Alderley Park | Wellcome Sanger Institute for the COVID-19 Genomics UK (COG-UK) Consortium | Jacquelyn Wynn, Mairead Hyland, The Lighthouse Lab in Alderley Park and Alex Alderton, Roberto Amato, Sonia Goncalves, Ewan Harrison, David K. Jackson, Ian Johnston, Dominic Kwiatkowski, Cordelia Langford, John Sillitoe on behalf of the Wellcome Sanger Institute COVID-19 Surveillance Team |
| EPI_ISL_799801, EPI_ISL_799805 | Lighthouse Lab in Glasgow | Wellcome Sanger Institute for the COVID-19 Genomics UK (COG-UK) Consortium | Harper VanSteenhouse, Yumi Kasai, David Gray, Carol Clugston, Anna Dominiczak and Alex Alderton, Roberto Amato, Sonia Goncalves, Ewan Harrison, David K. Jackson, Ian Johnston, Dominic Kwiatkowski, Cordelia Langford, John Sillitoe on behalf of the Wellcome Sanger Institute COVID-19 Surveillance Team |
| EPI_ISL_799807, EPI_ISL_799812 | Lighthouse Lab in Alderley Park | Wellcome Sanger Institute for the COVID-19 Genomics UK (COG-UK) Consortium | Jacquelyn Wynn, Mairead Hyland, The Lighthouse Lab in Alderley Park and Alex Alderton, Roberto Amato, Sonia Goncalves, Ewan Harrison, David K. Jackson, Ian Johnston, Dominic Kwiatkowski, Cordelia Langford, John Sillitoe on behalf of the Wellcome Sanger Institute COVID-19 Surveillance Team |
| EPI_ISL_799819 | Lighthouse Lab in Glasgow | Wellcome Sanger Institute for the COVID-19 Genomics UK (COG-UK) Consortium | Harper VanSteenhouse, Yumi Kasai, David Gray, Carol Clugston, Anna Dominiczak and Alex Alderton, Roberto Amato, Sonia Goncalves, Ewan Harrison, David K. Jackson, Ian Johnston, Dominic Kwiatkowski, Cordelia Langford, John Sillitoe on behalf of the Wellcome Sanger Institute COVID-19 Surveillance Team |
| EPI_ISL_799821, EPI_ISL_799822, EPI_ISL_799823 | Lighthouse Lab in Alderley Park | Wellcome Sanger Institute for the COVID-19 Genomics UK (COG-UK) Consortium | Jacquelyn Wynn, Mairead Hyland, The Lighthouse Lab in Alderley Park and Alex Alderton, Roberto Amato, Sonia Goncalves, Ewan Harrison, David K. Jackson, Ian Johnston, Dominic Kwiatkowski, Cordelia Langford, John Sillitoe on behalf of the Wellcome Sanger Institute COVID-19 Surveillance Team |
| EPI_ISL_799827 | Lighthouse Lab in Glasgow | Wellcome Sanger Institute for the COVID-19 Genomics UK (COG-UK) Consortium | Harper VanSteenhouse, Yumi Kasai, David Gray, Carol Clugston, Anna Dominiczak and Alex Alderton, Roberto Amato, Sonia Goncalves, Ewan Harrison, David K. Jackson, Ian Johnston, Dominic Kwiatkowski, Cordelia Langford, John Sillitoe on behalf of the Wellcome Sanger Institute COVID-19 Surveillance Team |
| EPI_ISL_799831, EPI_ISL_799832 | Lighthouse Lab in Alderley Park | Wellcome Sanger Institute for the COVID-19 Genomics UK (COG-UK) Consortium | Jacquelyn Wynn, Mairead Hyland, The Lighthouse Lab in Alderley Park and Alex Alderton, Roberto Amato, Sonia Goncalves, Ewan Harrison, David K. Jackson, Ian Johnston, Dominic Kwiatkowski, Cordelia Langford, John Sillitoe on behalf of the Wellcome Sanger Institute COVID-19 Surveillance Team |
| EPI_ISL_799838, EPI_ISL_799839 | Lighthouse Lab in Glasgow | Wellcome Sanger Institute for the COVID-19 Genomics UK (COG-UK) Consortium | Harper VanSteenhouse, Yumi Kasai, David Gray, Carol Clugston, Anna Dominiczak and Alex Alderton, Roberto Amato, Sonia Goncalves, Ewan Harrison, David K. Jackson, Ian Johnston, Dominic Kwiatkowski, Cordelia Langford, John Sillitoe on behalf of the Wellcome Sanger Institute COVID-19 Surveillance Team |
| EPI_ISL_799841, EPI_ISL_799844, EPI_ISL_799846 | Lighthouse Lab in Alderley Park | Wellcome Sanger Institute for the COVID-19 Genomics UK (COG-UK) Consortium | Jacquelyn Wynn, Mairead Hyland, The Lighthouse Lab in Alderley Park and Alex Alderton, Roberto Amato, Sonia Goncalves, Ewan Harrison, David K. Jackson, Ian Johnston, Dominic Kwiatkowski, Cordelia Langford, John Sillitoe on behalf of the Wellcome Sanger Institute COVID-19 Surveillance Team |
| EPI_ISL_799855 | Lighthouse Lab in Glasgow | Wellcome Sanger Institute for the COVID-19 Genomics UK (COG-UK) Consortium | Harper VanSteenhouse, Yumi Kasai, David Gray, Carol Clugston, Anna Dominiczak and Alex Alderton, Roberto Amato, Sonia Goncalves, Ewan Harrison, David K. Jackson, Ian Johnston, Dominic Kwiatkowski, Cordelia Langford, John Sillitoe on behalf of the Wellcome Sanger Institute COVID-19 Surveillance Team |
| EPI_ISL_799859 | Lighthouse Lab in Alderley Park | Wellcome Sanger Institute for the COVID-19 Genomics UK (COG-UK) Consortium | Jacquelyn Wynn, Mairead Hyland, The Lighthouse Lab in Alderley Park and Alex Alderton, Roberto Amato, Sonia Goncalves, Ewan Harrison, David K. Jackson, Ian Johnston, Dominic Kwiatkowski, Cordelia Langford, John Sillitoe on behalf of the Wellcome Sanger Institute COVID-19 Surveillance Team |
| EPI_ISL_799865, EPI_ISL_799888, EPI_ISL_799892, EPI_ISL_799894, EPI_ISL_799903, EPI_ISL_799906 | Lighthouse Lab in Glasgow | Wellcome Sanger Institute for the COVID-19 Genomics UK (COG-UK) Consortium | Harper VanSteenhouse, Yumi Kasai, David Gray, Carol Clugston, Anna Dominiczak and Alex Alderton, Roberto Amato, Sonia Goncalves, Ewan Harrison, David K. Jackson, Ian Johnston, Dominic Kwiatkowski, Cordelia Langford, John Sillitoe on behalf of the Wellcome Sanger Institute COVID-19 Surveillance Team |
| EPI_ISL_799920, EPI_ISL_799924, EPI_ISL_799930 | Lighthouse Lab in Alderley Park | Wellcome Sanger Institute for the COVID-19 Genomics UK (COG-UK) Consortium | Jacquelyn Wynn, Mairead Hyland, The Lighthouse Lab in Alderley Park and Alex Alderton, Roberto Amato, Sonia Goncalves, Ewan Harrison, David K. Jackson, Ian Johnston, Dominic Kwiatkowski, Cordelia Langford, John Sillitoe on behalf of the Wellcome Sanger Institute COVID-19 Surveillance Team |
| EPI_ISL_799932 | Lighthouse Lab in Glasgow | Wellcome Sanger Institute for the COVID-19 Genomics UK (COG-UK) Consortium | Harper VanSteenhouse, Yumi Kasai, David Gray, Carol Clugston, Anna Dominiczak and Alex Alderton, Roberto Amato, Sonia Goncalves, Ewan Harrison, David K. Jackson, Ian Johnston, Dominic Kwiatkowski, Cordelia Langford, John Sillitoe on behalf of the Wellcome Sanger Institute COVID-19 Surveillance Team |
| EPI_ISL_799933 | Lighthouse Lab in Alderley Park | Wellcome Sanger Institute for the COVID-19 Genomics UK (COG-UK) Consortium | Jacquelyn Wynn, Mairead Hyland, The Lighthouse Lab in Alderley Park and Alex Alderton, Roberto Amato, Sonia Goncalves, Ewan Harrison, David K. Jackson, Ian Johnston, Dominic Kwiatkowski, Cordelia Langford, John Sillitoe on behalf of the Wellcome Sanger Institute COVID-19 Surveillance Team |
| EPI_ISL_799937 | Lighthouse Lab in Glasgow | Wellcome Sanger Institute for the COVID-19 Genomics UK (COG-UK) Consortium | Harper VanSteenhouse, Yumi Kasai, David Gray, Carol Clugston, Anna Dominiczak and Alex Alderton, Roberto Amato, Sonia Goncalves, Ewan Harrison, David K. Jackson, Ian Johnston, Dominic Kwiatkowski, Cordelia Langford, John Sillitoe on behalf of the Wellcome Sanger Institute COVID-19 Surveillance Team |

| | | | |
|---|---|---|---|
| EPI_ISL_799940, EPI_ISL_799941, EPI_ISL_799943 | Lighthouse Lab in Alderley Park | Wellcome Sanger Institute for the COVID-19 Genomics UK (COG-UK) Consortium | Jacquelyn Wynn, Mairead Hyland, The Lighthouse Lab in Alderley Park and Alex Alderton, Roberto Amato, Sonia Goncalves, Ewan Harrison, David K. Jackson, Ian Johnston, Dominic Kwiatkowski, Cordelia Langford, John Sillitoe on behalf of the Wellcome Sanger Institute COVID-19 Surveillance Team |
| EPI_ISL_799946 | Lighthouse Lab in Glasgow | Wellcome Sanger Institute for the COVID-19 Genomics UK (COG-UK) Consortium | Harper VanSteenhouse, Yumi Kasai, David Gray, Carol Clugston, Anna Dominiczak and Alex Alderton, Roberto Amato, Sonia Goncalves, Ewan Harrison, David K. Jackson, Ian Johnston, Dominic Kwiatkowski, Cordelia Langford, John Sillitoe on behalf of the Wellcome Sanger Institute COVID-19 Surveillance Team |
| EPI_ISL_799948 | Lighthouse Lab in Alderley Park | Wellcome Sanger Institute for the COVID-19 Genomics UK (COG-UK) Consortium | Jacquelyn Wynn, Mairead Hyland, The Lighthouse Lab in Alderley Park and Alex Alderton, Roberto Amato, Sonia Goncalves, Ewan Harrison, David K. Jackson, Ian Johnston, Dominic Kwiatkowski, Cordelia Langford, John Sillitoe on behalf of the Wellcome Sanger Institute COVID-19 Surveillance Team |
| EPI_ISL_799949, EPI_ISL_799954 | Lighthouse Lab in Glasgow | Wellcome Sanger Institute for the COVID-19 Genomics UK (COG-UK) Consortium | Harper VanSteenhouse, Yumi Kasai, David Gray, Carol Clugston, Anna Dominiczak and Alex Alderton, Roberto Amato, Sonia Goncalves, Ewan Harrison, David K. Jackson, Ian Johnston, Dominic Kwiatkowski, Cordelia Langford, John Sillitoe on behalf of the Wellcome Sanger Institute COVID-19 Surveillance Team |
| EPI_ISL_799956, EPI_ISL_799961, EPI_ISL_799964 | Lighthouse Lab in Alderley Park | Wellcome Sanger Institute for the COVID-19 Genomics UK (COG-UK) Consortium | Jacquelyn Wynn, Mairead Hyland, The Lighthouse Lab in Alderley Park and Alex Alderton, Roberto Amato, Sonia Goncalves, Ewan Harrison, David K. Jackson, Ian Johnston, Dominic Kwiatkowski, Cordelia Langford, John Sillitoe on behalf of the Wellcome Sanger Institute COVID-19 Surveillance Team |
| EPI_ISL_799965 | Lighthouse Lab in Glasgow | Wellcome Sanger Institute for the COVID-19 Genomics UK (COG-UK) Consortium | Harper VanSteenhouse, Yumi Kasai, David Gray, Carol Clugston, Anna Dominiczak and Alex Alderton, Roberto Amato, Sonia Goncalves, Ewan Harrison, David K. Jackson, Ian Johnston, Dominic Kwiatkowski, Cordelia Langford, John Sillitoe on behalf of the Wellcome Sanger Institute COVID-19 Surveillance Team |
| EPI_ISL_799967 | Lighthouse Lab in Alderley Park | Wellcome Sanger Institute for the COVID-19 Genomics UK (COG-UK) Consortium | Jacquelyn Wynn, Mairead Hyland, The Lighthouse Lab in Alderley Park and Alex Alderton, Roberto Amato, Sonia Goncalves, Ewan Harrison, David K. Jackson, Ian Johnston, Dominic Kwiatkowski, Cordelia Langford, John Sillitoe on behalf of the Wellcome Sanger Institute COVID-19 Surveillance Team |
| EPI_ISL_799968, EPI_ISL_799969 | Lighthouse Lab in Glasgow | Wellcome Sanger Institute for the COVID-19 Genomics UK (COG-UK) Consortium | Harper VanSteenhouse, Yumi Kasai, David Gray, Carol Clugston, Anna Dominiczak and Alex Alderton, Roberto Amato, Sonia Goncalves, Ewan Harrison, David K. Jackson, Ian Johnston, Dominic Kwiatkowski, Cordelia Langford, John Sillitoe on behalf of the Wellcome Sanger Institute COVID-19 Surveillance Team |
| EPI_ISL_799970, EPI_ISL_799975, EPI_ISL_799977, EPI_ISL_799979 | Lighthouse Lab in Alderley Park | Wellcome Sanger Institute for the COVID-19 Genomics UK (COG-UK) Consortium | Jacquelyn Wynn, Mairead Hyland, The Lighthouse Lab in Alderley Park and Alex Alderton, Roberto Amato, Sonia Goncalves, Ewan Harrison, David K. Jackson, Ian Johnston, Dominic Kwiatkowski, Cordelia Langford, John Sillitoe on behalf of the Wellcome Sanger Institute COVID-19 Surveillance Team |
| EPI_ISL_799980 | Lighthouse Lab in Glasgow | Wellcome Sanger Institute for the COVID-19 Genomics UK (COG-UK) Consortium | Harper VanSteenhouse, Yumi Kasai, David Gray, Carol Clugston, Anna Dominiczak and Alex Alderton, Roberto Amato, Sonia Goncalves, Ewan Harrison, David K. Jackson, Ian Johnston, Dominic Kwiatkowski, Cordelia Langford, John Sillitoe on behalf of the Wellcome Sanger Institute COVID-19 Surveillance Team |
| EPI_ISL_799984 | Lighthouse Lab in Alderley Park | Wellcome Sanger Institute for the COVID-19 Genomics UK (COG-UK) Consortium | Jacquelyn Wynn, Mairead Hyland, The Lighthouse Lab in Alderley Park and Alex Alderton, Roberto Amato, Sonia Goncalves, Ewan Harrison, David K. Jackson, Ian Johnston, Dominic Kwiatkowski, Cordelia Langford, John Sillitoe on behalf of the Wellcome Sanger Institute COVID-19 Surveillance Team |
| EPI_ISL_799986, EPI_ISL_799987, EPI_ISL_799992 | Lighthouse Lab in Glasgow | Wellcome Sanger Institute for the COVID-19 Genomics UK (COG-UK) Consortium | Harper VanSteenhouse, Yumi Kasai, David Gray, Carol Clugston, Anna Dominiczak and Alex Alderton, Roberto Amato, Sonia Goncalves, Ewan Harrison, David K. Jackson, Ian Johnston, Dominic Kwiatkowski, Cordelia Langford, John Sillitoe on behalf of the Wellcome Sanger Institute COVID-19 Surveillance Team |
| EPI_ISL_799995, EPI_ISL_799999 | Lighthouse Lab in Alderley Park | Wellcome Sanger Institute for the COVID-19 Genomics UK (COG-UK) Consortium | Jacquelyn Wynn, Mairead Hyland, The Lighthouse Lab in Alderley Park and Alex Alderton, Roberto Amato, Sonia Goncalves, Ewan Harrison, David K. Jackson, Ian Johnston, Dominic Kwiatkowski, Cordelia Langford, John Sillitoe on behalf of the Wellcome Sanger Institute COVID-19 Surveillance Team |
| EPI_ISL_800000 | Lighthouse Lab in Glasgow | Wellcome Sanger Institute for the COVID-19 Genomics UK (COG-UK) Consortium | Harper VanSteenhouse, Yumi Kasai, David Gray, Carol Clugston, Anna Dominiczak and Alex Alderton, Roberto Amato, Sonia Goncalves, Ewan Harrison, David K. Jackson, Ian Johnston, Dominic Kwiatkowski, Cordelia Langford, John Sillitoe on behalf of the Wellcome Sanger Institute COVID-19 Surveillance Team |
| EPI_ISL_800002, EPI_ISL_800009, EPI_ISL_800011, EPI_ISL_800013, EPI_ISL_800014 | Lighthouse Lab in Alderley Park | Wellcome Sanger Institute for the COVID-19 Genomics UK (COG-UK) Consortium | Jacquelyn Wynn, Mairead Hyland, The Lighthouse Lab in Alderley Park and Alex Alderton, Roberto Amato, Sonia Goncalves, Ewan Harrison, David K. Jackson, Ian Johnston, Dominic Kwiatkowski, Cordelia Langford, John Sillitoe on behalf of the Wellcome Sanger Institute COVID-19 Surveillance Team |
| EPI_ISL_800015 | Lighthouse Lab in Glasgow | Wellcome Sanger Institute for the COVID-19 Genomics UK (COG-UK) Consortium | Harper VanSteenhouse, Yumi Kasai, David Gray, Carol Clugston, Anna Dominiczak and Alex Alderton, Roberto Amato, Sonia Goncalves, Ewan Harrison, David K. Jackson, Ian Johnston, Dominic Kwiatkowski, Cordelia Langford, John Sillitoe on behalf of the Wellcome Sanger Institute COVID-19 Surveillance Team |
| EPI_ISL_800016, EPI_ISL_800019, EPI_ISL_800020, EPI_ISL_800021, EPI_ISL_800023, EPI_ISL_800031 | Lighthouse Lab in Alderley Park | Wellcome Sanger Institute for the COVID-19 Genomics UK (COG-UK) Consortium | Jacquelyn Wynn, Mairead Hyland, The Lighthouse Lab in Alderley Park and Alex Alderton, Roberto Amato, Sonia Goncalves, Ewan Harrison, David K. Jackson, Ian Johnston, Dominic Kwiatkowski, Cordelia Langford, John Sillitoe on behalf of the Wellcome Sanger Institute COVID-19 Surveillance Team |
| EPI_ISL_800032 | Lighthouse Lab in Glasgow | Wellcome Sanger Institute for the COVID-19 Genomics UK (COG-UK) Consortium | Harper VanSteenhouse, Yumi Kasai, David Gray, Carol Clugston, Anna Dominiczak and Alex Alderton, Roberto Amato, Sonia Goncalves, Ewan Harrison, David K. Jackson, Ian Johnston, Dominic Kwiatkowski, Cordelia Langford, John Sillitoe on behalf of the Wellcome Sanger Institute COVID-19 Surveillance Team |
| EPI_ISL_800033, EPI_ISL_800035, EPI_ISL_800041, EPI_ISL_800042, EPI_ISL_800046 | Lighthouse Lab in Alderley Park | Wellcome Sanger Institute for the COVID-19 Genomics UK (COG-UK) Consortium | Jacquelyn Wynn, Mairead Hyland, The Lighthouse Lab in Alderley Park and Alex Alderton, Roberto Amato, Sonia Goncalves, Ewan Harrison, David K. Jackson, Ian Johnston, Dominic Kwiatkowski, Cordelia Langford, John Sillitoe on behalf of the Wellcome Sanger Institute COVID-19 Surveillance Team |
| EPI_ISL_800048 | Lighthouse Lab in Glasgow | Wellcome Sanger Institute for the COVID-19 Genomics UK (COG-UK) Consortium | Harper VanSteenhouse, Yumi Kasai, David Gray, Carol Clugston, Anna Dominiczak and Alex Alderton, Roberto Amato, Sonia Goncalves, Ewan Harrison, David K. Jackson, Ian Johnston, Dominic Kwiatkowski, Cordelia Langford, John Sillitoe on behalf of the Wellcome Sanger Institute COVID-19 Surveillance Team |
| EPI_ISL_800049 | Lighthouse Lab in Alderley Park | Wellcome Sanger Institute for the COVID-19 Genomics UK (COG-UK) Consortium | Jacquelyn Wynn, Mairead Hyland, The Lighthouse Lab in Alderley Park and Alex Alderton, Roberto Amato, Sonia Goncalves, Ewan Harrison, David K. Jackson, Ian Johnston, Dominic Kwiatkowski, Cordelia Langford, John Sillitoe on behalf of the Wellcome Sanger Institute COVID-19 Surveillance Team |
| EPI_ISL_800052 | Lighthouse Lab in Glasgow | Wellcome Sanger Institute for the COVID-19 Genomics UK (COG-UK) Consortium | Harper VanSteenhouse, Yumi Kasai, David Gray, Carol Clugston, Anna Dominiczak and Alex Alderton, Roberto Amato, Sonia Goncalves, Ewan Harrison, David K. Jackson, Ian Johnston, Dominic Kwiatkowski, Cordelia Langford, John Sillitoe on behalf of the Wellcome Sanger Institute COVID-19 Surveillance Team |
| EPI_ISL_800053 | Lighthouse Lab in Alderley Park | Wellcome Sanger Institute for the COVID-19 Genomics UK (COG-UK) Consortium | Jacquelyn Wynn, Mairead Hyland, The Lighthouse Lab in Alderley Park and Alex Alderton, Roberto Amato, Sonia Goncalves, Ewan Harrison, David K. Jackson, Ian Johnston, Dominic Kwiatkowski, Cordelia Langford, John Sillitoe on behalf of the Wellcome Sanger Institute COVID-19 Surveillance Team |
| EPI_ISL_800057 | Lighthouse Lab in Glasgow | Wellcome Sanger Institute for the COVID-19 Genomics UK (COG-UK) Consortium | Harper VanSteenhouse, Yumi Kasai, David Gray, Carol Clugston, Anna Dominiczak and Alex Alderton, Roberto Amato, Sonia Goncalves, Ewan Harrison, David K. Jackson, Ian Johnston, Dominic Kwiatkowski, Cordelia Langford, John Sillitoe on behalf of the Wellcome Sanger Institute COVID-19 Surveillance Team |
| EPI_ISL_800058, EPI_ISL_800060, EPI_ISL_800062, EPI_ISL_800063 | Lighthouse Lab in Alderley Park | Wellcome Sanger Institute for the COVID-19 Genomics UK (COG-UK) Consortium | Jacquelyn Wynn, Mairead Hyland, The Lighthouse Lab in Alderley Park and Alex Alderton, Roberto Amato, Sonia Goncalves, Ewan Harrison, David K. Jackson, Ian Johnston, Dominic Kwiatkowski, Cordelia Langford, John Sillitoe on behalf of the Wellcome Sanger Institute COVID-19 Surveillance Team |
| EPI_ISL_800064 | Lighthouse Lab in Glasgow | Wellcome Sanger Institute for the COVID-19 Genomics UK (COG-UK) Consortium | Harper VanSteenhouse, Yumi Kasai, David Gray, Carol Clugston, Anna Dominiczak and Alex Alderton, Roberto Amato, Sonia Goncalves, Ewan Harrison, David K. Jackson, Ian Johnston, Dominic Kwiatkowski, Cordelia Langford, John Sillitoe on behalf of the Wellcome Sanger Institute COVID-19 Surveillance Team |
| EPI_ISL_800066, EPI_ISL_800069 | Lighthouse Lab in Alderley Park | Wellcome Sanger Institute for the COVID-19 Genomics UK (COG-UK) Consortium | Jacquelyn Wynn, Mairead Hyland, The Lighthouse Lab in Alderley Park and Alex Alderton, Roberto Amato, Sonia Goncalves, Ewan Harrison, David K. Jackson, Ian Johnston, Dominic Kwiatkowski, Cordelia Langford, John Sillitoe on behalf of the Wellcome Sanger Institute COVID-19 Surveillance Team |
| EPI_ISL_800071 | Lighthouse Lab in Glasgow | Wellcome Sanger Institute for the COVID-19 Genomics UK (COG-UK) Consortium | Harper VanSteenhouse, Yumi Kasai, David Gray, Carol Clugston, Anna Dominiczak and Alex Alderton, Roberto Amato, Sonia Goncalves, Ewan Harrison, David K. Jackson, Ian Johnston, Dominic Kwiatkowski, Cordelia Langford, John Sillitoe on behalf of the Wellcome Sanger Institute COVID-19 Surveillance Team |

EPI_ISL_800073, EPI_ISL_800075, EPI_ISL_800076, EPI_ISL_800080, EPI_ISL_800081, EPI_ISL_800087, EPI_ISL_800090, EPI_ISL_800091, EPI_ISL_800097, EPI_ISL_800099, EPI_ISL_800104

| | | | |
|---|---|---|---|
| see above | Lighthouse Lab in Alderley Park | Wellcome Sanger Institute for the COVID-19 Genomics UK (COG-UK) Consortium | Jacquelyn Wynn, Mairead Hyland, The Lighthouse Lab in Alderley Park and Alex Alderton, Roberto Amato, Sonia Goncalves, Ewan Harrison, David K. Jackson, Ian Johnston, Dominic Kwiatkowski, Cordelia Langford, John Sillitoe on behalf of the Wellcome Sanger Institute COVID-19 Surveillance Team |
| EPI_ISL_800106 | Lighthouse Lab in Glasgow | Wellcome Sanger Institute for the COVID-19 Genomics UK (COG-UK) Consortium | Harper VanSteenhouse, Yumi Kasai, David Gray, Carol Clugston, Anna Dominiczak and Alex Alderton, Roberto Amato, Sonia Goncalves, Ewan Harrison, David K. Jackson, Ian Johnston, Dominic Kwiatkowski, Cordelia Langford, John Sillitoe on behalf of the Wellcome Sanger Institute COVID-19 Surveillance Team |
| EPI_ISL_800107, EPI_ISL_800108, EPI_ISL_800109 | Lighthouse Lab in Alderley Park | Wellcome Sanger Institute for the COVID-19 Genomics UK (COG-UK) Consortium | Jacquelyn Wynn, Mairead Hyland, The Lighthouse Lab in Alderley Park and Alex Alderton, Roberto Amato, Sonia Goncalves, Ewan Harrison, David K. Jackson, Ian Johnston, Dominic Kwiatkowski, Cordelia Langford, John Sillitoe on behalf of the Wellcome Sanger Institute COVID-19 Surveillance Team |
| EPI_ISL_800111, EPI_ISL_800113, EPI_ISL_800117, EPI_ISL_800120, EPI_ISL_800122 | Lighthouse Lab in Glasgow | Wellcome Sanger Institute for the COVID-19 Genomics UK (COG-UK) Consortium | Harper VanSteenhouse, Yumi Kasai, David Gray, Carol Clugston, Anna Dominiczak and Alex Alderton, Roberto Amato, Sonia Goncalves, Ewan Harrison, David K. Jackson, Ian Johnston, Dominic Kwiatkowski, Cordelia Langford, John Sillitoe on behalf of the Wellcome Sanger Institute COVID-19 Surveillance Team |
| EPI_ISL_800126, EPI_ISL_800129 | Lighthouse Lab in Alderley Park | Wellcome Sanger Institute for the COVID-19 Genomics UK (COG-UK) Consortium | Jacquelyn Wynn, Mairead Hyland, The Lighthouse Lab in Alderley Park and Alex Alderton, Roberto Amato, Sonia Goncalves, Ewan Harrison, David K. Jackson, Ian Johnston, Dominic Kwiatkowski, Cordelia Langford, John Sillitoe on behalf of the Wellcome Sanger Institute COVID-19 Surveillance Team |
| EPI_ISL_800131 | Lighthouse Lab in Glasgow | Wellcome Sanger Institute for the COVID-19 Genomics UK (COG-UK) Consortium | Harper VanSteenhouse, Yumi Kasai, David Gray, Carol Clugston, Anna Dominiczak and Alex Alderton, Roberto Amato, Sonia Goncalves, Ewan Harrison, David K. Jackson, Ian Johnston, Dominic Kwiatkowski, Cordelia Langford, John Sillitoe on behalf of the Wellcome Sanger Institute COVID-19 Surveillance Team |
| EPI_ISL_800135, EPI_ISL_800136 | Lighthouse Lab in Alderley Park | Wellcome Sanger Institute for the COVID-19 Genomics UK (COG-UK) Consortium | Jacquelyn Wynn, Mairead Hyland, The Lighthouse Lab in Alderley Park and Alex Alderton, Roberto Amato, Sonia Goncalves, Ewan Harrison, David K. Jackson, Ian Johnston, Dominic Kwiatkowski, Cordelia Langford, John Sillitoe on behalf of the Wellcome Sanger Institute COVID-19 Surveillance Team |
| EPI_ISL_800138 | Lighthouse Lab in Glasgow | Wellcome Sanger Institute for the COVID-19 Genomics UK (COG-UK) Consortium | Harper VanSteenhouse, Yumi Kasai, David Gray, Carol Clugston, Anna Dominiczak and Alex Alderton, Roberto Amato, Sonia Goncalves, Ewan Harrison, David K. Jackson, Ian Johnston, Dominic Kwiatkowski, Cordelia Langford, John Sillitoe on behalf of the Wellcome Sanger Institute COVID-19 Surveillance Team |
| EPI_ISL_800140 | Lighthouse Lab in Alderley Park | Wellcome Sanger Institute for the COVID-19 Genomics UK (COG-UK) Consortium | Jacquelyn Wynn, Mairead Hyland, The Lighthouse Lab in Alderley Park and Alex Alderton, Roberto Amato, Sonia Goncalves, Ewan Harrison, David K. Jackson, Ian Johnston, Dominic Kwiatkowski, Cordelia Langford, John Sillitoe on behalf of the Wellcome Sanger Institute COVID-19 Surveillance Team |
| EPI_ISL_800142 | Lighthouse Lab in Glasgow | Wellcome Sanger Institute for the COVID-19 Genomics UK (COG-UK) Consortium | Harper VanSteenhouse, Yumi Kasai, David Gray, Carol Clugston, Anna Dominiczak and Alex Alderton, Roberto Amato, Sonia Goncalves, Ewan Harrison, David K. Jackson, Ian Johnston, Dominic Kwiatkowski, Cordelia Langford, John Sillitoe on behalf of the Wellcome Sanger Institute COVID-19 Surveillance Team |
| EPI_ISL_800144, EPI_ISL_800145, EPI_ISL_800146 | Lighthouse Lab in Alderley Park | Wellcome Sanger Institute for the COVID-19 Genomics UK (COG-UK) Consortium | Jacquelyn Wynn, Mairead Hyland, The Lighthouse Lab in Alderley Park and Alex Alderton, Roberto Amato, Sonia Goncalves, Ewan Harrison, David K. Jackson, Ian Johnston, Dominic Kwiatkowski, Cordelia Langford, John Sillitoe on behalf of the Wellcome Sanger Institute COVID-19 Surveillance Team |
| EPI_ISL_800147, EPI_ISL_800158 | Lighthouse Lab in Glasgow | Wellcome Sanger Institute for the COVID-19 Genomics UK (COG-UK) Consortium | Harper VanSteenhouse, Yumi Kasai, David Gray, Carol Clugston, Anna Dominiczak and Alex Alderton, Roberto Amato, Sonia Goncalves, Ewan Harrison, David K. Jackson, Ian Johnston, Dominic Kwiatkowski, Cordelia Langford, John Sillitoe on behalf of the Wellcome Sanger Institute COVID-19 Surveillance Team |
| EPI_ISL_800159, EPI_ISL_800160, EPI_ISL_800162, EPI_ISL_800165, EPI_ISL_800171 | Lighthouse Lab in Alderley Park | Wellcome Sanger Institute for the COVID-19 Genomics UK (COG-UK) Consortium | Jacquelyn Wynn, Mairead Hyland, The Lighthouse Lab in Alderley Park and Alex Alderton, Roberto Amato, Sonia Goncalves, Ewan Harrison, David K. Jackson, Ian Johnston, Dominic Kwiatkowski, Cordelia Langford, John Sillitoe on behalf of the Wellcome Sanger Institute COVID-19 Surveillance Team |
| EPI_ISL_800174, EPI_ISL_800183 | Lighthouse Lab in Glasgow | Wellcome Sanger Institute for the COVID-19 Genomics UK (COG-UK) Consortium | Harper VanSteenhouse, Yumi Kasai, David Gray, Carol Clugston, Anna Dominiczak and Alex Alderton, Roberto Amato, Sonia Goncalves, Ewan Harrison, David K. Jackson, Ian Johnston, Dominic Kwiatkowski, Cordelia Langford, John Sillitoe on behalf of the Wellcome Sanger Institute COVID-19 Surveillance Team |
| EPI_ISL_800185 | Lighthouse Lab in Alderley Park | Wellcome Sanger Institute for the COVID-19 Genomics UK (COG-UK) Consortium | Jacquelyn Wynn, Mairead Hyland, The Lighthouse Lab in Alderley Park and Alex Alderton, Roberto Amato, Sonia Goncalves, Ewan Harrison, David K. Jackson, Ian Johnston, Dominic Kwiatkowski, Cordelia Langford, John Sillitoe on behalf of the Wellcome Sanger Institute COVID-19 Surveillance Team |
| EPI_ISL_800186, EPI_ISL_800189 | Lighthouse Lab in Glasgow | Wellcome Sanger Institute for the COVID-19 Genomics UK (COG-UK) Consortium | Harper VanSteenhouse, Yumi Kasai, David Gray, Carol Clugston, Anna Dominiczak and Alex Alderton, Roberto Amato, Sonia Goncalves, Ewan Harrison, David K. Jackson, Ian Johnston, Dominic Kwiatkowski, Cordelia Langford, John Sillitoe on behalf of the Wellcome Sanger Institute COVID-19 Surveillance Team |
| EPI_ISL_800193, EPI_ISL_800194, EPI_ISL_800197, EPI_ISL_800198, EPI_ISL_800199, EPI_ISL_800202 | Lighthouse Lab in Alderley Park | Wellcome Sanger Institute for the COVID-19 Genomics UK (COG-UK) Consortium | Jacquelyn Wynn, Mairead Hyland, The Lighthouse Lab in Alderley Park and Alex Alderton, Roberto Amato, Sonia Goncalves, Ewan Harrison, David K. Jackson, Ian Johnston, Dominic Kwiatkowski, Cordelia Langford, John Sillitoe on behalf of the Wellcome Sanger Institute COVID-19 Surveillance Team |
| EPI_ISL_800204 | Lighthouse Lab in Glasgow | Wellcome Sanger Institute for the COVID-19 Genomics UK (COG-UK) Consortium | Harper VanSteenhouse, Yumi Kasai, David Gray, Carol Clugston, Anna Dominiczak and Alex Alderton, Roberto Amato, Sonia Goncalves, Ewan Harrison, David K. Jackson, Ian Johnston, Dominic Kwiatkowski, Cordelia Langford, John Sillitoe on behalf of the Wellcome Sanger Institute COVID-19 Surveillance Team |
| EPI_ISL_800205, EPI_ISL_800206, EPI_ISL_800207, EPI_ISL_800208, EPI_ISL_800209 | Lighthouse Lab in Alderley Park | Wellcome Sanger Institute for the COVID-19 Genomics UK (COG-UK) Consortium | Jacquelyn Wynn, Mairead Hyland, The Lighthouse Lab in Alderley Park and Alex Alderton, Roberto Amato, Sonia Goncalves, Ewan Harrison, David K. Jackson, Ian Johnston, Dominic Kwiatkowski, Cordelia Langford, John Sillitoe on behalf of the Wellcome Sanger Institute COVID-19 Surveillance Team |
| EPI_ISL_800210 | Lighthouse Lab in Glasgow | Wellcome Sanger Institute for the COVID-19 Genomics UK (COG-UK) Consortium | Harper VanSteenhouse, Yumi Kasai, David Gray, Carol Clugston, Anna Dominiczak and Alex Alderton, Roberto Amato, Sonia Goncalves, Ewan Harrison, David K. Jackson, Ian Johnston, Dominic Kwiatkowski, Cordelia Langford, John Sillitoe on behalf of the Wellcome Sanger Institute COVID-19 Surveillance Team |
| EPI_ISL_800212, EPI_ISL_800213, EPI_ISL_800215, EPI_ISL_800216 | Lighthouse Lab in Alderley Park | Wellcome Sanger Institute for the COVID-19 Genomics UK (COG-UK) Consortium | Jacquelyn Wynn, Mairead Hyland, The Lighthouse Lab in Alderley Park and Alex Alderton, Roberto Amato, Sonia Goncalves, Ewan Harrison, David K. Jackson, Ian Johnston, Dominic Kwiatkowski, Cordelia Langford, John Sillitoe on behalf of the Wellcome Sanger Institute COVID-19 Surveillance Team |
| EPI_ISL_800217 | Lighthouse Lab in Glasgow | Wellcome Sanger Institute for the COVID-19 Genomics UK (COG-UK) Consortium | Harper VanSteenhouse, Yumi Kasai, David Gray, Carol Clugston, Anna Dominiczak and Alex Alderton, Roberto Amato, Sonia Goncalves, Ewan Harrison, David K. Jackson, Ian Johnston, Dominic Kwiatkowski, Cordelia Langford, John Sillitoe on behalf of the Wellcome Sanger Institute COVID-19 Surveillance Team |
| EPI_ISL_800219 | Lighthouse Lab in Alderley Park | Wellcome Sanger Institute for the COVID-19 Genomics UK (COG-UK) Consortium | Jacquelyn Wynn, Mairead Hyland, The Lighthouse Lab in Alderley Park and Alex Alderton, Roberto Amato, Sonia Goncalves, Ewan Harrison, David K. Jackson, Ian Johnston, Dominic Kwiatkowski, Cordelia Langford, John Sillitoe on behalf of the Wellcome Sanger Institute COVID-19 Surveillance Team |
| EPI_ISL_800221 | Lighthouse Lab in Glasgow | Wellcome Sanger Institute for the COVID-19 Genomics UK (COG-UK) Consortium | Harper VanSteenhouse, Yumi Kasai, David Gray, Carol Clugston, Anna Dominiczak and Alex Alderton, Roberto Amato, Sonia Goncalves, Ewan Harrison, David K. Jackson, Ian Johnston, Dominic Kwiatkowski, Cordelia Langford, John Sillitoe on behalf of the Wellcome Sanger Institute COVID-19 Surveillance Team |
| EPI_ISL_800223, EPI_ISL_800226, EPI_ISL_800229, EPI_ISL_800230, EPI_ISL_800231, EPI_ISL_800232 | Lighthouse Lab in Alderley Park | Wellcome Sanger Institute for the COVID-19 Genomics UK (COG-UK) Consortium | Jacquelyn Wynn, Mairead Hyland, The Lighthouse Lab in Alderley Park and Alex Alderton, Roberto Amato, Sonia Goncalves, Ewan Harrison, David K. Jackson, Ian Johnston, Dominic Kwiatkowski, Cordelia Langford, John Sillitoe on behalf of the Wellcome Sanger Institute COVID-19 Surveillance Team |
| EPI_ISL_800237 | Lighthouse Lab in Glasgow | Wellcome Sanger Institute for the COVID-19 Genomics UK (COG-UK) Consortium | Harper VanSteenhouse, Yumi Kasai, David Gray, Carol Clugston, Anna Dominiczak and Alex Alderton, Roberto Amato, Sonia Goncalves, Ewan Harrison, David K. Jackson, Ian Johnston, Dominic Kwiatkowski, Cordelia Langford, John Sillitoe on behalf of the Wellcome Sanger Institute COVID-19 Surveillance Team |
| EPI_ISL_800239, EPI_ISL_800242 | Lighthouse Lab in Alderley Park | Wellcome Sanger Institute for the COVID-19 Genomics UK (COG-UK) Consortium | Jacquelyn Wynn, Mairead Hyland, The Lighthouse Lab in Alderley Park and Alex Alderton, Roberto Amato, Sonia Goncalves, Ewan Harrison, David K. Jackson, Ian Johnston, Dominic Kwiatkowski, Cordelia Langford, John Sillitoe on behalf of the Wellcome Sanger Institute COVID-19 Surveillance Team |
| EPI_ISL_800245 | Lighthouse Lab in Glasgow | Wellcome Sanger Institute for the COVID-19 Genomics UK | Harper VanSteenhouse, Yumi Kasai, David Gray, Carol Clugston, Anna Dominiczak and Alex Alderton, Roberto Amato, Sonia Goncalves, Ewan Harrison, |

| | | | |
|---|---|---|---|
| | | (COG-UK) Consortium | David K. Jackson, Ian Johnston, Dominic Kwiatkowski, Cordelia Langford, John Sillitoe on behalf of the Wellcome Sanger Institute COVID-19 Surveillance Team |
| EPI_ISL_800247, EPI_ISL_800249 | Lighthouse Lab in Alderley Park | Wellcome Sanger Institute for the COVID-19 Genomics UK (COG-UK) Consortium | Jacquelyn Wynn, Mairead Hyland, The Lighthouse Lab in Alderley Park and Alex Alderton, Roberto Amato, Sonia Goncalves, Ewan Harrison, David K. Jackson, Ian Johnston, Dominic Kwiatkowski, Cordelia Langford, John Sillitoe on behalf of the Wellcome Sanger Institute COVID-19 Surveillance Team |
| EPI_ISL_800251, EPI_ISL_800253 | Lighthouse Lab in Glasgow | Wellcome Sanger Institute for the COVID-19 Genomics UK (COG-UK) Consortium | Harper VanSteenhouse, Yumi Kasai, David Gray, Carol Clugston, Anna Dominiczak and Alex Alderton, Roberto Amato, Sonia Goncalves, Ewan Harrison, David K. Jackson, Ian Johnston, Dominic Kwiatkowski, Cordelia Langford, John Sillitoe on behalf of the Wellcome Sanger Institute COVID-19 Surveillance Team |
| EPI_ISL_800267, EPI_ISL_800271, EPI_ISL_800274, EPI_ISL_800280, EPI_ISL_800281, EPI_ISL_800282, EPI_ISL_800286 | Lighthouse Lab in Alderley Park | Wellcome Sanger Institute for the COVID-19 Genomics UK (COG-UK) Consortium | Jacquelyn Wynn, Mairead Hyland, The Lighthouse Lab in Alderley Park and Alex Alderton, Roberto Amato, Sonia Goncalves, Ewan Harrison, David K. Jackson, Ian Johnston, Dominic Kwiatkowski, Cordelia Langford, John Sillitoe on behalf of the Wellcome Sanger Institute COVID-19 Surveillance Team |
| EPI_ISL_800297 | Lighthouse Lab in Cambridge | Wellcome Sanger Institute for the COVID-19 Genomics UK (COG-UK) Consortium | Rob Howes, The Lighthouse Lab in Cambridge and Alex Alderton, Roberto Amato, Sonia Goncalves, Ewan Harrison, David K. Jackson, Ian Johnston, Dominic Kwiatkowski, Cordelia Langford, John Sillitoe on behalf of the Wellcome Sanger Institute COVID-19 Surveillance Team |
| EPI_ISL_800299, EPI_ISL_800301, EPI_ISL_800303, EPI_ISL_800308, EPI_ISL_800317, EPI_ISL_800338, EPI_ISL_800340, EPI_ISL_800342, EPI_ISL_800347, EPI_ISL_800348, EPI_ISL_800350, EPI_ISL_800351, EPI_ISL_800354, EPI_ISL_800359, EPI_ISL_800360, EPI_ISL_800365, EPI_ISL_800367, EPI_ISL_800370 | | | |
| see above | Lighthouse Lab in Alderley Park | Wellcome Sanger Institute for the COVID-19 Genomics UK (COG-UK) Consortium | Jacquelyn Wynn, Mairead Hyland, The Lighthouse Lab in Alderley Park and Alex Alderton, Roberto Amato, Sonia Goncalves, Ewan Harrison, David K. Jackson, Ian Johnston, Dominic Kwiatkowski, Cordelia Langford, John Sillitoe on behalf of the Wellcome Sanger Institute COVID-19 Surveillance Team |
| EPI_ISL_800372 | Lighthouse Lab in Cambridge | Wellcome Sanger Institute for the COVID-19 Genomics UK (COG-UK) Consortium | Rob Howes, The Lighthouse Lab in Cambridge and Alex Alderton, Roberto Amato, Sonia Goncalves, Ewan Harrison, David K. Jackson, Ian Johnston, Dominic Kwiatkowski, Cordelia Langford, John Sillitoe on behalf of the Wellcome Sanger Institute COVID-19 Surveillance Team |
| EPI_ISL_800376, EPI_ISL_800384 | Lighthouse Lab in Alderley Park | Wellcome Sanger Institute for the COVID-19 Genomics UK (COG-UK) Consortium | Jacquelyn Wynn, Mairead Hyland, The Lighthouse Lab in Alderley Park and Alex Alderton, Roberto Amato, Sonia Goncalves, Ewan Harrison, David K. Jackson, Ian Johnston, Dominic Kwiatkowski, Cordelia Langford, John Sillitoe on behalf of the Wellcome Sanger Institute COVID-19 Surveillance Team |
| EPI_ISL_800389 | Lighthouse Lab in Cambridge | Wellcome Sanger Institute for the COVID-19 Genomics UK (COG-UK) Consortium | Rob Howes, The Lighthouse Lab in Cambridge and Alex Alderton, Roberto Amato, Sonia Goncalves, Ewan Harrison, David K. Jackson, Ian Johnston, Dominic Kwiatkowski, Cordelia Langford, John Sillitoe on behalf of the Wellcome Sanger Institute COVID-19 Surveillance Team |
| EPI_ISL_800392, EPI_ISL_800407, EPI_ISL_800420, EPI_ISL_800428, EPI_ISL_800441 | Lighthouse Lab in Alderley Park | Wellcome Sanger Institute for the COVID-19 Genomics UK (COG-UK) Consortium | Jacquelyn Wynn, Mairead Hyland, The Lighthouse Lab in Alderley Park and Alex Alderton, Roberto Amato, Sonia Goncalves, Ewan Harrison, David K. Jackson, Ian Johnston, Dominic Kwiatkowski, Cordelia Langford, John Sillitoe on behalf of the Wellcome Sanger Institute COVID-19 Surveillance Team |
| EPI_ISL_800442 | Lighthouse Lab in Cambridge | Wellcome Sanger Institute for the COVID-19 Genomics UK (COG-UK) Consortium | Rob Howes, The Lighthouse Lab in Cambridge and Alex Alderton, Roberto Amato, Sonia Goncalves, Ewan Harrison, David K. Jackson, Ian Johnston, Dominic Kwiatkowski, Cordelia Langford, John Sillitoe on behalf of the Wellcome Sanger Institute COVID-19 Surveillance Team |
| EPI_ISL_800444, EPI_ISL_800446, EPI_ISL_800452, EPI_ISL_800453, EPI_ISL_800454, EPI_ISL_800456, EPI_ISL_800457, EPI_ISL_800468, EPI_ISL_800474, EPI_ISL_800481, EPI_ISL_800485, EPI_ISL_800494, EPI_ISL_800495, EPI_ISL_800501, EPI_ISL_800502, EPI_ISL_800509, EPI_ISL_800516, EPI_ISL_800518, EPI_ISL_800524, EPI_ISL_800525, EPI_ISL_800528, EPI_ISL_800531, EPI_ISL_800558, EPI_ISL_800566 | | | |
| see above | Lighthouse Lab in Alderley Park | Wellcome Sanger Institute for the COVID-19 Genomics UK (COG-UK) Consortium | Jacquelyn Wynn, Mairead Hyland, The Lighthouse Lab in Alderley Park and Alex Alderton, Roberto Amato, Sonia Goncalves, Ewan Harrison, David K. Jackson, Ian Johnston, Dominic Kwiatkowski, Cordelia Langford, John Sillitoe on behalf of the Wellcome Sanger Institute COVID-19 Surveillance Team |
| EPI_ISL_801432, EPI_ISL_801433, EPI_ISL_801434, EPI_ISL_801435, EPI_ISL_801436, EPI_ISL_801439, EPI_ISL_801440, EPI_ISL_801442, EPI_ISL_801458, EPI_ISL_801496, EPI_ISL_801498, EPI_ISL_801499, EPI_ISL_801500, EPI_ISL_801501, EPI_ISL_801502, EPI_ISL_801503, EPI_ISL_801504, EPI_ISL_801505, EPI_ISL_801506, EPI_ISL_801507, EPI_ISL_801508, EPI_ISL_801509, EPI_ISL_801515 | | | |
| see above | Dutch COVID-19 response team | Erasmus Medical Center | Bas Oude Munnink, Reina Sikkema, David Nieuwenhuijse, Irina Chestakova, Anne van der Linden, Marjan Boter, Emmanuelle Munger, Corine GeurtsvanKessel, Annemiek van der Eijk, Richard Molenkamp, Marion Koopmans, on behalf of the Dutch national COVID-19 response team. |
| EPI_ISL_801521 | Instituto Nacional de Saude (INSA) | Instituto Nacional de Saude (INSA) | Borges et al |
| EPI_ISL_802430, EPI_ISL_802445, EPI_ISL_802446 | BIO-REFERENCE LABORATORIES | Wadsworth Center, New York State Department.of Health | Kirsten St. George, Daryl M. Lamson, Alexis Russel, Matthew Shudt, Melissa A Leisner, Jonathan Plitnick, Navjot Singh, John Kelly, Sara Griesemer, Erasmus Schneider, Erica Lasek-Nesselquist |
| EPI_ISL_802449, EPI_ISL_802450, EPI_ISL_802451, EPI_ISL_802452, EPI_ISL_802453, EPI_ISL_802454, EPI_ISL_802455, EPI_ISL_802456, EPI_ISL_802457, EPI_ISL_802458, EPI_ISL_802459, EPI_ISL_802460, EPI_ISL_802461, EPI_ISL_802462 | | | |
| see above | Wadsworth Center, New York State Department.of Health | Wadsworth Center, New York State Department.of Health | Kirsten St. George, Daryl M. Lamson, Alexis Russel, Matthew Shudt, Melissa A Leisner, Jonathan Plitnick, Navjot Singh, John Kelly, Sara Griesemer, Erasmus Schneider, Erica Lasek-Nesselquist |
| EPI_ISL_802575, EPI_ISL_802577, EPI_ISL_802579, EPI_ISL_802580, EPI_ISL_802585, EPI_ISL_802586, EPI_ISL_802587, EPI_ISL_802589, EPI_ISL_802590, EPI_ISL_802591, EPI_ISL_802592, EPI_ISL_802593, EPI_ISL_802594, EPI_ISL_802597 | | | |
| see above | Vault Health | Minnesota Department of Health, Public Health Laboratory | Alexandra Lorentz, Jacob Garfin, Matt Plumb, and Xiong Wang |
| EPI_ISL_802600, EPI_ISL_802605, EPI_ISL_802608, EPI_ISL_802609, EPI_ISL_802610, EPI_ISL_802611, EPI_ISL_802629, EPI_ISL_802630, EPI_ISL_802631, EPI_ISL_802637, EPI_ISL_802638, EPI_ISL_802639, EPI_ISL_802640, EPI_ISL_802648, EPI_ISL_802650, EPI_ISL_802654, EPI_ISL_802656, EPI_ISL_802657, EPI_ISL_802658, EPI_ISL_802659, EPI_ISL_802660, EPI_ISL_802669, EPI_ISL_802670, EPI_ISL_802671, EPI_ISL_802672, EPI_ISL_802673, EPI_ISL_802674, EPI_ISL_802679, EPI_ISL_802680, EPI_ISL_802681, EPI_ISL_802684, EPI_ISL_802685, EPI_ISL_802686, EPI_ISL_802688, EPI_ISL_802690, EPI_ISL_802699, EPI_ISL_802700, EPI_ISL_802706, EPI_ISL_802707, EPI_ISL_802708, EPI_ISL_802709, EPI_ISL_802710, EPI_ISL_802719, EPI_ISL_802722 | | | |
| see above | Helix/Illumina | Genomics and Discovery, Respiratory Viruses Branch, Division of Viral Diseases, Centers for Disease Control and Prevention | Peter W. Cook, Dhwani Batra, Ben L. Rambo-Martin Eileen de Feo, Jan Antico, Christine Tran, Matthew Tolentino, Shannon Wickline, Kim Gietzen, Brad Sickler, Jingtao Liu, Eric Allen, Phil Febbo, Summer Galloway, Nicole L. Washington, Simon White, Geraint Levan, Kelly Schiabor Barrett, Elizabeth Cirulli, Alexandre Bolze, Ary Ascencio, Charlotte Rivera-Garcia, Ryan Cho, Jason Nguyen, Sherry Wang, Jimmy Ramirez, Tyler Cassens, Efren Sandoval, Magnus Isaksson, William Lee, David Becker, Marc Laurent, James Lu, Clinton R. Paden, Suxiang Tong, Duncan MacCannell |
| EPI_ISL_802788, EPI_ISL_802789 | Columbia University Irving Medical Center | Wadsworth Center, New York State Department.of Health | Kirsten St. George, Daryl M. Lamson, Alexis Russel, Matthew Shudt, Melissa A Leisner, Jonathan Plitnick, Navjot Singh, John Kelly, Sara Griesemer, Erasmus Schneider, Erica Lasek-Nesselquist |
| EPI_ISL_803079 | Quest Diagnostics | Quest Diagnostics | Rosenthal,S.H., Gerasimova,A., Kagan,R.M., Anderson, B., Livingston, K.E., Hua, M., Liu Y., Shalhout, D.F., Owen, R., Lacbawan, F. |
| EPI_ISL_803894 | Robert Koch Institut | Robert Koch Institut | Annika Brinkmann, Janine Michel, Livia Schrick, Steven Uddin, Dominique Seifert, Alexander Dalpke, Leo Büttner, Kristina Hochauf-Stange, Dirk Lindemann, Lars Schaade, Andreas Nitsche |
| EPI_ISL_803900 | National Institute for Infectious Diseases, INMI, "L. Spallanzani" IRCCS | National Institute for Infectious Diseases, INMI, "L. Spallanzani" IRCCS | B Bartolini, O Butera, C.E.M Gruber, M Rueca, F Messina, E Giombini, MR Capobianchi, A Di Caro |
| EPI_ISL_803955, EPI_ISL_803956, EPI_ISL_803957 | Labor Dr. Wisplinghoff - Köln | Robert Koch Institute, Influenza and respiratory viruses FG17 & Bioinformatics MF1, Berlin, Germany | Dr. R. Grosser, Stephan Fuchs, Stefan Kroeger, Marianne Wedde, Oliver Drechsel, Aleksandar Radonic, Rene Kmiecinski, Ralf Duerrwald, Thorsten Wolff |
| EPI_ISL_803984, EPI_ISL_803985, EPI_ISL_803986, EPI_ISL_803987, EPI_ISL_803988 | National Public Health Laboratory, National Centre for Infectious Diseases | National Public Health Laboratory, National Centre for Infectious Diseases | Tze Minn Mak, Sophie Octavia, Zhenyang Zhou, Lin Cui, Raymond Tzer Pin Lin |
| EPI_ISL_804020, EPI_ISL_804021, EPI_ISL_804022, EPI_ISL_804023, EPI_ISL_804024, EPI_ISL_804025, EPI_ISL_804027, EPI_ISL_804028 | Maryland Public Health Laboratory | Maryland Public Health Laboratory | Maryland Department of Health Laboratories Administration |
| EPI_ISL_804056, EPI_ISL_804071, EPI_ISL_804072, EPI_ISL_804073, EPI_ISL_804074, EPI_ISL_804075, EPI_ISL_804076, EPI_ISL_804077, EPI_ISL_804078, EPI_ISL_804079, EPI_ISL_804080, EPI_ISL_804081, EPI_ISL_804082, EPI_ISL_804083, EPI_ISL_804092, EPI_ISL_804093, EPI_ISL_804094, EPI_ISL_804096, EPI_ISL_804098, EPI_ISL_804102, EPI_ISL_804103, EPI_ISL_804104, EPI_ISL_804106, EPI_ISL_804142, EPI_ISL_804143, EPI_ISL_804144, EPI_ISL_804145, EPI_ISL_804146, EPI_ISL_804147, EPI_ISL_804148, EPI_ISL_804149, EPI_ISL_804150, EPI_ISL_804151, EPI_ISL_804152, EPI_ISL_804153, EPI_ISL_804154, EPI_ISL_804155, EPI_ISL_804156, EPI_ISL_804157, EPI_ISL_804158, EPI_ISL_804159, EPI_ISL_804160, EPI_ISL_804161, EPI_ISL_804162, EPI_ISL_804163, EPI_ISL_804164, EPI_ISL_804165, EPI_ISL_804166, EPI_ISL_804167, EPI_ISL_804168, EPI_ISL_804169, EPI_ISL_804170, EPI_ISL_804171, EPI_ISL_804172, EPI_ISL_804173, EPI_ISL_804174, EPI_ISL_804175, EPI_ISL_804176, EPI_ISL_804177, EPI_ISL_804178, EPI_ISL_804179, EPI_ISL_804180, EPI_ISL_804181, EPI_ISL_804182, EPI_ISL_804183, EPI_ISL_804184, EPI_ISL_804185, EPI_ISL_804186, EPI_ISL_804187, EPI_ISL_804188, EPI_ISL_804191, EPI_ISL_804209, EPI_ISL_804210, EPI_ISL_804211, EPI_ISL_804212 | | | |
| see above | Israel Central Virology laboratory | Israel National Consortium for SARS-CoV-2 sequencing | Neta Zuckerman, Efrat Dahan Bucris, Michal Mandelboim, Dana Bar-Ilan, Oran Erster, Tzvia Mann, Omer Murik, David A. Zeevi, Assaf Rokney, Joseph |

| | | | |
|---|---|---|---|
| | | | Jaffe, Eva Nachum, Maya Davidovich Cohen, Ephraim Fass, Gal Zizelski Valenci, Mor Rubinstein, Efrat Rorman, Israel Nissan, Efrat Glick-Saar, Omri Nayshool, Gideon Rechavi, Ella Mendelson, Orna Mor |
| EPI_ISL_804223, EPI_ISL_804253, EPI_ISL_804274, EPI_ISL_804275, EPI_ISL_804277, EPI_ISL_804278, EPI_ISL_804279 | Respiratory Virus Unit, National Infection Service, Public Health England | COVID-19 Genomics UK (COG-UK) Consortium | PHE Covid Sequencing Team |
| EPI_ISL_804946 | BIO-REFERENCE LABORATORIES | Wadsworth Center, New York State Department.of Health | Kirsten St. George, Daryl M. Lamson, Alexis Russel, Matthew Shudt, Melissa A Leisner, Jonathan Plitnick, Navjot Singh, John Kelly, Sara Griesemer, Erasmus Schneider, Erica Lasek-Nesselquist |
| EPI_ISL_806721 | Histopath | NSW Health Pathology - Institute of Clinical Pathology and Medical Research; Westmead Hospital; University of Sydney | CIDM-PH et al. |
| EPI_ISL_806722 | South Eastern Area Laboratory Services (SEALS) | NSW Health Pathology - Institute of Clinical Pathology and Medical Research; Westmead Hospital; University of Sydney | CIDM-PH et al. |
| EPI_ISL_807156 | Deva County Emergency Hospital | National Institute of Infectious Diseases-Prof. Dr. Matei Bals Molecular Diagnostics Laboratory | Leontina Banica, Marius Surleac, Corina Casangiu, Petre Milu, Andreea Tudor, Simona Paraschiv, Dan Otelea |
| EPI_ISL_810804 | PathWest Laboratory Medicine WA | PathWest Laboratory Medicine WA Microbial Surveillance Unit | PathWest Laboratory Medicine WA Microbial Surveillance Unit |
| EPI_ISL_811122 | Respiratory Virus Unit, National Infection Service, Public Health England | COVID-19 Genomics UK (COG-UK) Consortium | PHE Covid Sequencing Team |
| EPI_ISL_811809, EPI_ISL_811810, EPI_ISL_811813, EPI_ISL_811815, EPI_ISL_811816, EPI_ISL_811820, EPI_ISL_811822, EPI_ISL_811829, EPI_ISL_811838, EPI_ISL_811842, EPI_ISL_811844, EPI_ISL_811863, EPI_ISL_811864, EPI_ISL_811866, EPI_ISL_811872, EPI_ISL_811876, EPI_ISL_811882, EPI_ISL_811895, EPI_ISL_811901, EPI_ISL_811903, EPI_ISL_811904, EPI_ISL_811906, EPI_ISL_811908, EPI_ISL_811911, EPI_ISL_811920, EPI_ISL_811926, EPI_ISL_811933, EPI_ISL_811934, EPI_ISL_811940, EPI_ISL_811947, EPI_ISL_811961, EPI_ISL_811963, EPI_ISL_811964, EPI_ISL_811966, EPI_ISL_811980, EPI_ISL_811982, EPI_ISL_811992, EPI_ISL_811995, EPI_ISL_812001, EPI_ISL_812004, EPI_ISL_812005, EPI_ISL_812006, EPI_ISL_812008, EPI_ISL_812011, EPI_ISL_812013, EPI_ISL_812016, EPI_ISL_812023, EPI_ISL_812029, EPI_ISL_812031, EPI_ISL_812032, EPI_ISL_812034, EPI_ISL_812039, EPI_ISL_812043, EPI_ISL_812044, EPI_ISL_812053, EPI_ISL_812056, EPI_ISL_812057, EPI_ISL_812060, EPI_ISL_812065, EPI_ISL_812071, EPI_ISL_812072, EPI_ISL_812081, EPI_ISL_812082, EPI_ISL_812087, EPI_ISL_812089, EPI_ISL_812095, EPI_ISL_812107, EPI_ISL_812108, EPI_ISL_812110, EPI_ISL_812111, EPI_ISL_812112 | | | |
| see above | Lighthouse Lab in Milton Keynes | Wellcome Sanger Institute for the COVID-19 Genomics UK (COG-UK) Consortium | The Lighthouse Lab in Milton Keynes and Alex Alderton, Roberto Amato, Sonia Goncalves, Ewan Harrison, David K. Jackson, Ian Johnston, Dominic Kwiatkowski, Cordelia Langford, John Sillitoe on behalf of the Wellcome Sanger Institute COVID-19 Surveillance Team |
| EPI_ISL_812114 | Arizona State University | Arizona State University | LaRinda A. Holland, Peter T. Skidmore, Emily A. Kaelin, Nicholas J. Mellor, Kristina Buss, Joy M. Blain, Valerie Harris, Joshua LaBaer, Vel Murugan, Efrem S. Lim |
| EPI_ISL_812286, EPI_ISL_812287, EPI_ISL_812288, EPI_ISL_812289, EPI_ISL_812297, EPI_ISL_812298, EPI_ISL_812299, EPI_ISL_812300, EPI_ISL_812301, EPI_ISL_812302, EPI_ISL_812303, EPI_ISL_812304, EPI_ISL_812305, EPI_ISL_812306, EPI_ISL_812307 | | | |
| see above | Delaware Public Health Lab | Delaware Public Health Lab | Gregory Hovan |
| EPI_ISL_812763, EPI_ISL_812764, EPI_ISL_812765, EPI_ISL_812766, EPI_ISL_812767 | Ministry of Health Turkey | Ministry of Health Turkey | Fatma Bayrakdar, Yasemin Cogun, Süleyman Yalcin, Aye Baak Alta, Gülay Korukluolu |
| EPI_ISL_813094, EPI_ISL_813095, EPI_ISL_813096, EPI_ISL_813097, EPI_ISL_813098, EPI_ISL_813099, EPI_ISL_813100, EPI_ISL_813101 | University of Birmingham | COVID-19 Genomics UK (COG-UK) Consortium | Institute of Microbiology, University of Birmingham: Claire McMurray, Joanne Stockton, Samuel Nicholls, Radoslaw Poplawski, Will Rowe, Josh Quick, Nicholas Loman. University of Birmingham Testing Laboratory: Celina M Whalley, Andrew Bosworth, Charlotte Poxon, Kasun Wanigasooriya, Oliver Pickles, Mike Kidd, Alex Richter, Andrew D Beggs PHE Heartlands Lab: Husam Osman, Andrew Bosworth. Queen Elizabeth Hospital: Anna Casey |
| EPI_ISL_813167, EPI_ISL_813171, EPI_ISL_813173, EPI_ISL_813175, EPI_ISL_813183, EPI_ISL_813227, EPI_ISL_813230, EPI_ISL_813250, EPI_ISL_813256, EPI_ISL_813265, EPI_ISL_813266, EPI_ISL_813267, EPI_ISL_813268, EPI_ISL_813269, EPI_ISL_813278, EPI_ISL_813281, EPI_ISL_813283, EPI_ISL_813285, EPI_ISL_813294 | | | |
| see above | Department of Pathology, University of Cambridge | COVID-19 Genomics UK (COG-UK) Consortium | Aminu S. Jahun, Yasmin Chaudhry, Grant Hall, Iliana Georgana, Myra Hosmillo, Martin D. Curran, Malte Pinckert, Surendra Parmar, Ian Goodfellow |
| EPI_ISL_813581, EPI_ISL_813582, EPI_ISL_813583, EPI_ISL_813587, EPI_ISL_813588, EPI_ISL_813589, EPI_ISL_813590, EPI_ISL_813591, EPI_ISL_813592, EPI_ISL_813593, EPI_ISL_813594, EPI_ISL_813595, EPI_ISL_813596, EPI_ISL_813597, EPI_ISL_813598, EPI_ISL_813600, EPI_ISL_813601, EPI_ISL_813602, EPI_ISL_813603, EPI_ISL_813604, EPI_ISL_813605, EPI_ISL_813606, EPI_ISL_813607, EPI_ISL_813609, EPI_ISL_813610, EPI_ISL_813611, EPI_ISL_813612, EPI_ISL_813613, EPI_ISL_813614, EPI_ISL_813615, EPI_ISL_813616, EPI_ISL_813617, EPI_ISL_813618, EPI_ISL_813619, EPI_ISL_813620, EPI_ISL_813621, EPI_ISL_813622, EPI_ISL_813624, EPI_ISL_813637 | | | |
| see above | University of Exeter | COVID-19 Genomics UK (COG-UK) Consortium | Ben Temperton,Aaron Jeffries,Michelle Michelsen,Joanna Warwick-Dugdale,Audrey Farbos,Robyn Manley,Stephen Michell,Jane Masoli |
| EPI_ISL_813773, EPI_ISL_813774, EPI_ISL_813775, EPI_ISL_813776, EPI_ISL_813777, EPI_ISL_813778, EPI_ISL_813779, EPI_ISL_813780, EPI_ISL_813781, EPI_ISL_813782, EPI_ISL_813784 | | | |
| see above | Liverpool Clinical Laboratories | COVID-19 Genomics UK (COG-UK) Consortium | Sam Haldenby, Anita Lucaci, Steve Paterson, Julian Hiscox, Alistair Darby, M Almsaud, A Alrezaihi, Muhannad Alruwaili, Stuart D Armstrong, Jones Benjamin, Eleanor G Bentley, Anu Chawla, Jordan J Clark, Angela Cowell, Richard Eccles, Isabel García-Dorival, Matthew Gemmell, Alessandro Gerada, PKF Gilmore, Richard Gregory, Ximeng Han, Catherine Hartley, Margaret Hughes, Miren Iturriza-Gomara, James Johnson, L Luu, Jenifer Manson, Charlotte Nelson, Elaine O'Toole, Cassie Olateju, Rebekah Penrice-Randal , Lucille Rainbow, N.P Randle, Trevor Ian Robinson, Parul Sharma, Ghada T Shawli, James P Stewart, Neil Swainston, Ecaterina Vamos, Joanne Watts, Mark Whitehead |
| EPI_ISL_813900, EPI_ISL_813901, EPI_ISL_813902, EPI_ISL_813903, EPI_ISL_813904, EPI_ISL_813905, EPI_ISL_813906, EPI_ISL_813956, EPI_ISL_813959, EPI_ISL_813962, EPI_ISL_813963, EPI_ISL_813964, EPI_ISL_813966 | | | |
| see above | University College London, Great Ormond Street Hospital for Children NHS Foundation Trust, Imperial College Healthcare NHS Trust | COVID-19 Genomics UK (COG-UK) Consortium | Sergi Castellano, Rachel Williams, Mark Kristiansen, Paola Resende Silva, Sunando Roy, Tony Brooks, Helena Tutill, Paola Niola, Patricia Dyal, Charlotte Williams, Leysa Forrest, Yasmin Panchbhaya, Jacqueline Findlay, Samuel Weeks, Julianne Brown, Kathryn Harris, Paul Randell, James Price, Alison Holmes, Judith Breuer |
| EPI_ISL_813975 | Akershus University Hospital, Department for Microbiology and Infectious Disease Control | Norwegian Institute of Public Health, Department of Virology | Kathrine Stene-Johansen, Kamilla Heddeland Instefjord, Hilde Elshaug, Atiya R Ali,Marie Paulsen Madsen, Rasmus Riis Kopperud, Hilde Vollan, Karoline Bragstad, Olav Hungnes |
| EPI_ISL_814068 | Ministry of Health Turkey | Ministry of Health Turkey | Fatma Bayrakdar, Yasemin Cogun, Süleyman Yalcin, Aye Baak Alta, Gülay Korukluolu |
| EPI_ISL_814092, EPI_ISL_814093, EPI_ISL_814094, EPI_ISL_814095, EPI_ISL_814096, EPI_ISL_814098, EPI_ISL_814099, EPI_ISL_814100, EPI_ISL_814101, EPI_ISL_814180, EPI_ISL_814181, EPI_ISL_814182, EPI_ISL_814183, EPI_ISL_814184, EPI_ISL_814185, EPI_ISL_814186, EPI_ISL_814187, EPI_ISL_814188, EPI_ISL_814189, EPI_ISL_814190, EPI_ISL_814191, EPI_ISL_814192, EPI_ISL_814193, EPI_ISL_814194, EPI_ISL_814195, EPI_ISL_814196, EPI_ISL_814197, EPI_ISL_814198, EPI_ISL_814199, EPI_ISL_814200, EPI_ISL_814201, EPI_ISL_814202, EPI_ISL_814203, EPI_ISL_814204, EPI_ISL_814205, EPI_ISL_814206, EPI_ISL_814207, EPI_ISL_814208, EPI_ISL_814209, EPI_ISL_814210, EPI_ISL_814211, EPI_ISL_814212, EPI_ISL_814213, EPI_ISL_814214, EPI_ISL_814215, EPI_ISL_814216, EPI_ISL_814217, EPI_ISL_814218, EPI_ISL_814219, EPI_ISL_814220, EPI_ISL_814221, EPI_ISL_814222, EPI_ISL_814223, EPI_ISL_814224, EPI_ISL_814225, EPI_ISL_814226, EPI_ISL_814227, EPI_ISL_814228, EPI_ISL_814229, EPI_ISL_814230, EPI_ISL_814231, EPI_ISL_814232, EPI_ISL_814233, EPI_ISL_814234, EPI_ISL_814235, EPI_ISL_814236, EPI_ISL_814237, EPI_ISL_814238, EPI_ISL_814239, EPI_ISL_814240, EPI_ISL_814241, EPI_ISL_814242, EPI_ISL_814243, EPI_ISL_814244, EPI_ISL_814245, EPI_ISL_814246, EPI_ISL_814247, EPI_ISL_814248, EPI_ISL_814249, EPI_ISL_814250, EPI_ISL_814251, EPI_ISL_814252, EPI_ISL_814253, EPI_ISL_814254, EPI_ISL_814255, EPI_ISL_814256, EPI_ISL_814257, EPI_ISL_814258, EPI_ISL_814259, EPI_ISL_814260, EPI_ISL_814261, EPI_ISL_814262, EPI_ISL_814263, EPI_ISL_814264, EPI_ISL_814265, EPI_ISL_814266, EPI_ISL_814267, EPI_ISL_814268, EPI_ISL_814269, EPI_ISL_814270, EPI_ISL_814271, EPI_ISL_814272, EPI_ISL_814273 | | | |
| see above | Israel Central Virology laboratory | Israel National Consortium for SARS-CoV-2 sequencing | Neta Zuckerman, Efrat Dahan Bucris, Michal Mandelboim, Dana Bar-Ilan, Oran Erster, Tzvia Mann, Omer Murik, David A. Zeevi, Assaf Rokney, Joseph Jaffe, Eva Nachum, Maya Davidovich Cohen, Ephraim Fass, Gal Zizelski Valenci, Mor Rubinstein, Efrat Rorman, Israel Nissan, Efrat Glick-Saar, Omri Nayshool, Gideon Rechavi, Ella Mendelson, Orna Mor |
| EPI_ISL_814350, EPI_ISL_814487, EPI_ISL_814504, EPI_ISL_814505, EPI_ISL_814506, EPI_ISL_814508, EPI_ISL_814509, EPI_ISL_814510, EPI_ISL_814541, EPI_ISL_814602, EPI_ISL_815045, EPI_ISL_815046, EPI_ISL_815068, EPI_ISL_815069, EPI_ISL_815070, EPI_ISL_815071, EPI_ISL_815233, EPI_ISL_815234, EPI_ISL_815235, EPI_ISL_815236, EPI_ISL_815237, EPI_ISL_815238, EPI_ISL_815239, EPI_ISL_815240, EPI_ISL_815241, EPI_ISL_815242, EPI_ISL_815243 | | | |
| see above | Wales Specialist Virology Centre Sequencing lab: Pathogen Genomics Unit | COVID-19 Genomics UK (COG-UK) Consortium | Catherine Moore, Johnathan Evans, Laura Gifford, Malorie Perry, Simon Cottrell, Angela Marchbank, Alec Birchley, Alexander Adams, Amy Gaskin, Bree Gatica-Wilcox, Jason Coombes, Joel Southgate, Lauren Gilbert, Lee Graham, Nicole Pacchiarini, Sara Kumziene-Summerhayes, Sarah Taylor, Sophie Jones, Sara Rey, Matthew Bull, Joanne Watkins, Sally Corden, Tom Connor |
| EPI_ISL_815402, EPI_ISL_815404, EPI_ISL_815405, EPI_ISL_815406, EPI_ISL_815408, EPI_ISL_815409, EPI_ISL_815412, EPI_ISL_815413, EPI_ISL_815414, EPI_ISL_815415, EPI_ISL_815470, EPI_ISL_815472, EPI_ISL_815476, EPI_ISL_815485, EPI_ISL_815488, EPI_ISL_815492, EPI_ISL_815502, EPI_ISL_815503, EPI_ISL_815504, EPI_ISL_815505, EPI_ISL_815506, EPI_ISL_815507, EPI_ISL_815537, EPI_ISL_815538, EPI_ISL_815547, EPI_ISL_815548, EPI_ISL_815549, EPI_ISL_815550, EPI_ISL_815559, EPI_ISL_815560, EPI_ISL_815561, EPI_ISL_815562, EPI_ISL_815605, EPI_ISL_815608, EPI_ISL_815609, EPI_ISL_815622, EPI_ISL_815627, EPI_ISL_815631, EPI_ISL_815632, EPI_ISL_815633, EPI_ISL_815637, EPI_ISL_815646, EPI_ISL_815661, EPI_ISL_815662, EPI_ISL_815663, EPI_ISL_815664, EPI_ISL_815665, EPI_ISL_815666, EPI_ISL_815667, EPI_ISL_815668, EPI_ISL_815669, EPI_ISL_815670, EPI_ISL_815671, EPI_ISL_815672, | | | |

| | | | |
|---|---|---|---|
| EPI_ISL_815673, EPI_ISL_815674, EPI_ISL_815675, EPI_ISL_815676, EPI_ISL_815677, EPI_ISL_815678, EPI_ISL_815679, EPI_ISL_815680, EPI_ISL_815681, EPI_ISL_815682, EPI_ISL_815683, EPI_ISL_815684, EPI_ISL_815685, EPI_ISL_815686, EPI_ISL_815687, EPI_ISL_815688, EPI_ISL_815689, EPI_ISL_815690, EPI_ISL_815691, EPI_ISL_815692, EPI_ISL_815693, EPI_ISL_815694, EPI_ISL_815695, EPI_ISL_815696, EPI_ISL_815697, EPI_ISL_815698, EPI_ISL_815699, EPI_ISL_815700, EPI_ISL_815701, EPI_ISL_815702, EPI_ISL_815703, EPI_ISL_815704, EPI_ISL_815705, EPI_ISL_815706, EPI_ISL_815707, EPI_ISL_815708, EPI_ISL_815709, EPI_ISL_815710, EPI_ISL_815711, EPI_ISL_815712, EPI_ISL_815713, EPI_ISL_815714, EPI_ISL_815715, EPI_ISL_815716, EPI_ISL_815717, EPI_ISL_815718, EPI_ISL_815719, EPI_ISL_815720, EPI_ISL_815721 | | | |
| see above | Department of Virus and Microbiological Special Diagnostics, Statens Serum Institut, Copenhagen, Denmark | Albertsen Lab, Department of Chemistry and Bioscience, Aalborg University, Denmark | Danish Covid-19 Genome Consortium |
| EPI_ISL_816194, EPI_ISL_816195, EPI_ISL_816196, EPI_ISL_816203, EPI_ISL_816224, EPI_ISL_816225 | Centre for Enzyme Innovation, University of Portsmouth / Translational Research Laboratory, Portsmouth Hospitals NHS Trust | COVID-19 Genomics UK (COG-UK) Consortium | Angela Beckett,Yann Bourgeois,Garry Scarlett,Sharon Glaysher,Scott Elliott,Kelly Bicknell,Robert Impey,Allyson Lloyd,Sarah Wyllie,Ethan Butcher,Anoop Chauhan,Samuel Robson |
| EPI_ISL_816230, EPI_ISL_816236, EPI_ISL_816239, EPI_ISL_816255, EPI_ISL_816262, EPI_ISL_816268, EPI_ISL_816290, EPI_ISL_816292, EPI_ISL_816302, EPI_ISL_816310, EPI_ISL_816313, EPI_ISL_816316, EPI_ISL_816320, EPI_ISL_816350, EPI_ISL_816357, EPI_ISL_816361, EPI_ISL_816365, EPI_ISL_816380, EPI_ISL_816382, EPI_ISL_816392, EPI_ISL_816421, EPI_ISL_816425, EPI_ISL_816430, EPI_ISL_816435, EPI_ISL_816447, EPI_ISL_816474, EPI_ISL_816480, EPI_ISL_816503, EPI_ISL_816505, EPI_ISL_816507, EPI_ISL_816513, EPI_ISL_816531, EPI_ISL_816533, EPI_ISL_816537, EPI_ISL_816542, EPI_ISL_816548, EPI_ISL_816560, EPI_ISL_816562, EPI_ISL_816577, EPI_ISL_816585, EPI_ISL_816589, EPI_ISL_816597, EPI_ISL_816599, EPI_ISL_816600, EPI_ISL_816605, EPI_ISL_816606, EPI_ISL_816607, EPI_ISL_816616, EPI_ISL_816619, EPI_ISL_816620, EPI_ISL_816621, EPI_ISL_816625, EPI_ISL_816635 | | | |
| see above | Virology Department, Sheffield Teaching Hospitals NHS Foundation Trust/Department of Infection, Immunity and Cardiovascular Disease, The Medical School, University of Sheffield | COVID-19 Genomics UK (COG-UK) Consortium | Thushan de Silva, Matthew Parker, Nikki Smith, Adri Angyal, Rebecca Brown, Luke Green, Rachel Tucker, Paul Parsons, Danielle Groves, Katie Johnson, Laura Carrilero, Alex Keeley, Dave Partridge, Matthew Wyles, Benjamin Lindsey, Mehmet Yavuz, Mohammad Raza, Cariad Evans |
| EPI_ISL_817123, EPI_ISL_817124, EPI_ISL_817125, EPI_ISL_817126, EPI_ISL_817127, EPI_ISL_817128, EPI_ISL_817129, EPI_ISL_817130, EPI_ISL_817131, EPI_ISL_817132, EPI_ISL_817133, EPI_ISL_817134, EPI_ISL_817136, EPI_ISL_817139, EPI_ISL_817140, EPI_ISL_817141, EPI_ISL_817142, EPI_ISL_817143, EPI_ISL_817144, EPI_ISL_817145, EPI_ISL_817148, EPI_ISL_817150, EPI_ISL_817151, EPI_ISL_817152, EPI_ISL_817153, EPI_ISL_817154, EPI_ISL_817155, EPI_ISL_817156, EPI_ISL_817157, EPI_ISL_817158, EPI_ISL_817177, EPI_ISL_817178, EPI_ISL_817179, EPI_ISL_817180, EPI_ISL_817181, EPI_ISL_817182, EPI_ISL_817183, EPI_ISL_817184, EPI_ISL_817185, EPI_ISL_817186, EPI_ISL_817187, EPI_ISL_817188, EPI_ISL_817189, EPI_ISL_817190, EPI_ISL_817191, EPI_ISL_817192, EPI_ISL_817193, EPI_ISL_817194, EPI_ISL_817195, EPI_ISL_817196, EPI_ISL_817197, EPI_ISL_817198, EPI_ISL_817199, EPI_ISL_817200, EPI_ISL_817201, EPI_ISL_817202, EPI_ISL_817203, EPI_ISL_817204, EPI_ISL_817205, EPI_ISL_817206, EPI_ISL_817207, EPI_ISL_817208, EPI_ISL_817209, EPI_ISL_817210, EPI_ISL_817211, EPI_ISL_817212, EPI_ISL_817213, EPI_ISL_817214, EPI_ISL_817215, EPI_ISL_817216, EPI_ISL_817217, EPI_ISL_817218, EPI_ISL_817219, EPI_ISL_817220, EPI_ISL_817221, EPI_ISL_817222, EPI_ISL_817223, EPI_ISL_817224, EPI_ISL_817225, EPI_ISL_817226, EPI_ISL_817227, EPI_ISL_817228, EPI_ISL_817229, EPI_ISL_817230, EPI_ISL_817231, EPI_ISL_817232, EPI_ISL_817233, EPI_ISL_817234, EPI_ISL_817235, EPI_ISL_817236, EPI_ISL_817237, EPI_ISL_817238, EPI_ISL_817239, EPI_ISL_817240, EPI_ISL_817241, EPI_ISL_817242, EPI_ISL_817243, EPI_ISL_817244, EPI_ISL_817245, EPI_ISL_817246, EPI_ISL_817247, EPI_ISL_817248, EPI_ISL_817249, EPI_ISL_817250, EPI_ISL_817251, EPI_ISL_817252, EPI_ISL_817253, EPI_ISL_817254, EPI_ISL_817255, EPI_ISL_817256, EPI_ISL_817257, EPI_ISL_817258, EPI_ISL_817259, EPI_ISL_817260, EPI_ISL_817261, EPI_ISL_817262, EPI_ISL_817263, EPI_ISL_817265, EPI_ISL_817266, EPI_ISL_817267, EPI_ISL_817268, EPI_ISL_817270, EPI_ISL_817271, EPI_ISL_817274, EPI_ISL_817275, EPI_ISL_817276, EPI_ISL_817277, EPI_ISL_817280, EPI_ISL_817283, EPI_ISL_817284, EPI_ISL_817285, EPI_ISL_817286, EPI_ISL_817287, EPI_ISL_817289, EPI_ISL_817293, EPI_ISL_817294, EPI_ISL_817295, EPI_ISL_817298, EPI_ISL_817300, EPI_ISL_817302, EPI_ISL_817304, EPI_ISL_817305, EPI_ISL_817307, EPI_ISL_817309, EPI_ISL_817310, EPI_ISL_817311, EPI_ISL_817313, EPI_ISL_817314, EPI_ISL_817318, EPI_ISL_817319, EPI_ISL_817320, EPI_ISL_817321, EPI_ISL_817322, EPI_ISL_817323, EPI_ISL_817324, EPI_ISL_817325, EPI_ISL_817326, EPI_ISL_817327, EPI_ISL_817328, EPI_ISL_817329, EPI_ISL_817330, EPI_ISL_817331, EPI_ISL_817332, EPI_ISL_817336, EPI_ISL_817337, EPI_ISL_817338, EPI_ISL_817339, EPI_ISL_817340, EPI_ISL_817341, EPI_ISL_817342, EPI_ISL_817343, EPI_ISL_817344, EPI_ISL_817345, EPI_ISL_817346, EPI_ISL_817347, EPI_ISL_817348, EPI_ISL_817349, EPI_ISL_817350, EPI_ISL_817351, EPI_ISL_817352, EPI_ISL_817353, EPI_ISL_817354, EPI_ISL_817355, EPI_ISL_817356, EPI_ISL_817357, EPI_ISL_817358, EPI_ISL_817359, EPI_ISL_817360, EPI_ISL_817361, EPI_ISL_817362, EPI_ISL_817363, EPI_ISL_817364, EPI_ISL_817365, EPI_ISL_817366, EPI_ISL_817367, EPI_ISL_817368, EPI_ISL_817369, EPI_ISL_817370, EPI_ISL_817371, EPI_ISL_817372, EPI_ISL_817373, EPI_ISL_817374, EPI_ISL_817377, EPI_ISL_817378, EPI_ISL_817379, EPI_ISL_817380, EPI_ISL_817381, EPI_ISL_817382, EPI_ISL_817387, EPI_ISL_817388, EPI_ISL_817389, EPI_ISL_817390, EPI_ISL_817391, EPI_ISL_817393, EPI_ISL_817394, EPI_ISL_817395, EPI_ISL_817396, EPI_ISL_817397, EPI_ISL_817400, EPI_ISL_817402, EPI_ISL_817403, EPI_ISL_817406, EPI_ISL_817407, EPI_ISL_817408, EPI_ISL_817409, EPI_ISL_817410, EPI_ISL_817411, EPI_ISL_817412, EPI_ISL_817420, EPI_ISL_817421, EPI_ISL_817422, EPI_ISL_817423, EPI_ISL_817424, EPI_ISL_817425, EPI_ISL_817426, EPI_ISL_817427, EPI_ISL_817428, EPI_ISL_817429, EPI_ISL_817430, EPI_ISL_817431, EPI_ISL_817432, EPI_ISL_817433, EPI_ISL_817434, EPI_ISL_817435, EPI_ISL_817436, EPI_ISL_817437, EPI_ISL_817438, EPI_ISL_817439, EPI_ISL_817440, EPI_ISL_817441, EPI_ISL_817442, EPI_ISL_817443, EPI_ISL_817444, EPI_ISL_817445, EPI_ISL_817446, EPI_ISL_817447, EPI_ISL_817448, EPI_ISL_817449, EPI_ISL_817450, EPI_ISL_817451, EPI_ISL_817452, EPI_ISL_817453, EPI_ISL_817454, EPI_ISL_817455, EPI_ISL_817456, EPI_ISL_817457, EPI_ISL_817458, EPI_ISL_817459, EPI_ISL_817460, EPI_ISL_817461, EPI_ISL_817462, EPI_ISL_817463, EPI_ISL_817464, EPI_ISL_817465, EPI_ISL_817466, EPI_ISL_817467, EPI_ISL_817468, EPI_ISL_817469, EPI_ISL_817470, EPI_ISL_817471, EPI_ISL_817472, EPI_ISL_817473, EPI_ISL_817474, EPI_ISL_817475, EPI_ISL_817476, EPI_ISL_817477, EPI_ISL_817478, EPI_ISL_817479, EPI_ISL_817480, EPI_ISL_817481, EPI_ISL_817482, EPI_ISL_817483, EPI_ISL_817484, EPI_ISL_817485, EPI_ISL_817486, EPI_ISL_817488, EPI_ISL_817489, EPI_ISL_817490, EPI_ISL_817491, EPI_ISL_817492, EPI_ISL_817493, EPI_ISL_817494, EPI_ISL_817495, EPI_ISL_817498, EPI_ISL_817500, EPI_ISL_817502, EPI_ISL_817503, EPI_ISL_817504, EPI_ISL_817505, EPI_ISL_817506, EPI_ISL_817507, EPI_ISL_817512, EPI_ISL_817513, EPI_ISL_817514, EPI_ISL_817515, EPI_ISL_817516, EPI_ISL_817517, EPI_ISL_817518, EPI_ISL_817519, EPI_ISL_817520, EPI_ISL_817521, EPI_ISL_817522, EPI_ISL_817523, EPI_ISL_817524, EPI_ISL_817525, EPI_ISL_817526, EPI_ISL_817527, EPI_ISL_817528, EPI_ISL_817529, EPI_ISL_817531, EPI_ISL_817533, EPI_ISL_817534, EPI_ISL_817535, EPI_ISL_817536, EPI_ISL_817539, EPI_ISL_817540, EPI_ISL_817541, EPI_ISL_817542, EPI_ISL_817543, EPI_ISL_817544, EPI_ISL_817545, EPI_ISL_817546, EPI_ISL_817547, EPI_ISL_817548, EPI_ISL_817549, EPI_ISL_817550, EPI_ISL_817551, EPI_ISL_817552, EPI_ISL_817553, EPI_ISL_817554, EPI_ISL_817653, EPI_ISL_817654, EPI_ISL_817655, EPI_ISL_817656, EPI_ISL_817657, EPI_ISL_817658, EPI_ISL_817659, EPI_ISL_817660, EPI_ISL_817661, EPI_ISL_817662, EPI_ISL_817663, EPI_ISL_817664, EPI_ISL_817665, EPI_ISL_817666, EPI_ISL_817667, EPI_ISL_817668, EPI_ISL_817669, EPI_ISL_817670, EPI_ISL_817671, EPI_ISL_817672, EPI_ISL_817673, EPI_ISL_817674, EPI_ISL_817675, EPI_ISL_817676, EPI_ISL_817677, EPI_ISL_817678, EPI_ISL_817679, EPI_ISL_817680, EPI_ISL_817681, EPI_ISL_817682, EPI_ISL_817683, EPI_ISL_817684, EPI_ISL_817685, EPI_ISL_817686, EPI_ISL_817687, EPI_ISL_817688, EPI_ISL_817689, EPI_ISL_817690, EPI_ISL_817691, EPI_ISL_817692, EPI_ISL_817693, EPI_ISL_817694, EPI_ISL_817695, EPI_ISL_817696, EPI_ISL_817697, EPI_ISL_817698, EPI_ISL_817699, EPI_ISL_817700, EPI_ISL_817701, EPI_ISL_817702, EPI_ISL_817703, EPI_ISL_817704, EPI_ISL_817705, EPI_ISL_817706, EPI_ISL_817707, EPI_ISL_817708, EPI_ISL_817709, EPI_ISL_817710, EPI_ISL_817711, EPI_ISL_817712, EPI_ISL_817713, EPI_ISL_817714, EPI_ISL_817715, EPI_ISL_817716, EPI_ISL_817717, EPI_ISL_817718, EPI_ISL_817719, EPI_ISL_817720, EPI_ISL_817721, EPI_ISL_817722, EPI_ISL_817723, EPI_ISL_817724, EPI_ISL_817725, EPI_ISL_817726, EPI_ISL_817727, EPI_ISL_817728, EPI_ISL_817729, EPI_ISL_817730, EPI_ISL_817731, EPI_ISL_817732, EPI_ISL_817733, EPI_ISL_817734, EPI_ISL_817735, EPI_ISL_817736, EPI_ISL_817737, EPI_ISL_817738, EPI_ISL_817739, EPI_ISL_817740, EPI_ISL_817741, EPI_ISL_817742, EPI_ISL_817743, EPI_ISL_817744, EPI_ISL_817745, EPI_ISL_817746, EPI_ISL_817747, EPI_ISL_817748, EPI_ISL_817749, EPI_ISL_817750, EPI_ISL_817751, EPI_ISL_817752, EPI_ISL_817753, EPI_ISL_817754, EPI_ISL_817755, EPI_ISL_817756, EPI_ISL_817757, EPI_ISL_817758, EPI_ISL_817759, EPI_ISL_817760, EPI_ISL_817761, EPI_ISL_817762, EPI_ISL_817763, EPI_ISL_817764, EPI_ISL_817765, EPI_ISL_817766, EPI_ISL_817767, EPI_ISL_817768, EPI_ISL_817769, EPI_ISL_817770, EPI_ISL_817771, EPI_ISL_817772, EPI_ISL_817773, EPI_ISL_817774, EPI_ISL_817775, EPI_ISL_817776, EPI_ISL_817777, EPI_ISL_817778, EPI_ISL_817779, EPI_ISL_817780, EPI_ISL_817781, EPI_ISL_817782, EPI_ISL_817783, EPI_ISL_817784, EPI_ISL_817785, EPI_ISL_817786, EPI_ISL_817787, EPI_ISL_817788, EPI_ISL_817789, EPI_ISL_817790, EPI_ISL_817791, EPI_ISL_817792, EPI_ISL_817793, EPI_ISL_817794, EPI_ISL_817795, EPI_ISL_817796, EPI_ISL_817797, EPI_ISL_817798, EPI_ISL_817799, EPI_ISL_817800, EPI_ISL_817801, EPI_ISL_817802, EPI_ISL_817803, EPI_ISL_817804, EPI_ISL_817805, EPI_ISL_817806, EPI_ISL_817807, EPI_ISL_817808, EPI_ISL_817809, EPI_ISL_817810, EPI_ISL_817811, EPI_ISL_817812, EPI_ISL_817813, EPI_ISL_817814, EPI_ISL_817815, EPI_ISL_817816, EPI_ISL_817817, EPI_ISL_817818, EPI_ISL_817819, EPI_ISL_817820, EPI_ISL_817821, EPI_ISL_817822, EPI_ISL_817823, EPI_ISL_817824, EPI_ISL_817825, EPI_ISL_817826, EPI_ISL_817827, EPI_ISL_817828, EPI_ISL_817829, EPI_ISL_817830, EPI_ISL_817831, EPI_ISL_817832, EPI_ISL_817833, EPI_ISL_817834, EPI_ISL_817835, EPI_ISL_817836, EPI_ISL_817837, EPI_ISL_817838, EPI_ISL_817839, EPI_ISL_817840, EPI_ISL_817841, EPI_ISL_817842, EPI_ISL_817843, EPI_ISL_817844, EPI_ISL_817845, EPI_ISL_817846, EPI_ISL_817847, EPI_ISL_817848, EPI_ISL_817849, EPI_ISL_817850, EPI_ISL_817851, EPI_ISL_817852, EPI_ISL_817853, EPI_ISL_817854, EPI_ISL_817855, EPI_ISL_817856, EPI_ISL_817857, EPI_ISL_817858, EPI_ISL_817859, EPI_ISL_817860, EPI_ISL_817861, EPI_ISL_817862, EPI_ISL_817863, EPI_ISL_817864, EPI_ISL_817865, EPI_ISL_817866, EPI_ISL_817867, EPI_ISL_817868, EPI_ISL_817869, EPI_ISL_817870, EPI_ISL_817871, EPI_ISL_817872, EPI_ISL_817873, EPI_ISL_817874, EPI_ISL_817875, EPI_ISL_817876, EPI_ISL_817877, EPI_ISL_817878, EPI_ISL_817879, EPI_ISL_817880, EPI_ISL_817881, EPI_ISL_817882, EPI_ISL_817883, EPI_ISL_817884, EPI_ISL_817885, EPI_ISL_817886, EPI_ISL_817887, EPI_ISL_817888, EPI_ISL_817889, EPI_ISL_817890, EPI_ISL_817891, EPI_ISL_817892, EPI_ISL_817893, EPI_ISL_817894, EPI_ISL_817895, EPI_ISL_817896, EPI_ISL_817897, EPI_ISL_817898, EPI_ISL_817899, EPI_ISL_817900, EPI_ISL_817901, EPI_ISL_817902, EPI_ISL_817903, EPI_ISL_817904, EPI_ISL_817905, EPI_ISL_817906, EPI_ISL_817907, EPI_ISL_817908, EPI_ISL_817909, EPI_ISL_817910, EPI_ISL_817911, EPI_ISL_817912, EPI_ISL_817913, EPI_ISL_817914, EPI_ISL_817915, EPI_ISL_817916, EPI_ISL_817917, EPI_ISL_817918, EPI_ISL_817919, EPI_ISL_817920, EPI_ISL_817921, EPI_ISL_817922, EPI_ISL_817923, EPI_ISL_817924, EPI_ISL_817925, EPI_ISL_817926, EPI_ISL_817927, EPI_ISL_817928, EPI_ISL_817929, EPI_ISL_817930, EPI_ISL_817931, EPI_ISL_817932, EPI_ISL_817933, EPI_ISL_817934, EPI_ISL_817935, EPI_ISL_817936, EPI_ISL_817937, EPI_ISL_817938, EPI_ISL_817939, EPI_ISL_817940, EPI_ISL_817941, EPI_ISL_817942, EPI_ISL_817943, EPI_ISL_817944, EPI_ISL_817945, EPI_ISL_817946, EPI_ISL_817947, EPI_ISL_817948, EPI_ISL_817949, EPI_ISL_817950, EPI_ISL_817951, EPI_ISL_817952, EPI_ISL_817953, EPI_ISL_817954, EPI_ISL_817955, EPI_ISL_817956, EPI_ISL_817957, EPI_ISL_817958, EPI_ISL_817959, EPI_ISL_817960, EPI_ISL_817961, EPI_ISL_817962, EPI_ISL_817963, EPI_ISL_817964, EPI_ISL_817965, EPI_ISL_817966, EPI_ISL_817967, EPI_ISL_817968, EPI_ISL_817969, EPI_ISL_817970, EPI_ISL_817971, EPI_ISL_817972, EPI_ISL_817973, EPI_ISL_817974, EPI_ISL_817975, EPI_ISL_817976, EPI_ISL_817977, EPI_ISL_817978, EPI_ISL_817979, EPI_ISL_817980, EPI_ISL_817981, EPI_ISL_817982, EPI_ISL_817983, EPI_ISL_817984, EPI_ISL_817985, EPI_ISL_817986, EPI_ISL_817987, EPI_ISL_817988, EPI_ISL_817989, EPI_ISL_817990, EPI_ISL_817991, EPI_ISL_817992, EPI_ISL_817993, EPI_ISL_817994, EPI_ISL_817995, EPI_ISL_817996, EPI_ISL_817997, EPI_ISL_817998, EPI_ISL_817999, EPI_ISL_818000, EPI_ISL_818001, EPI_ISL_818002, EPI_ISL_818003, EPI_ISL_818004, EPI_ISL_818005, EPI_ISL_818006, EPI_ISL_818007, EPI_ISL_818008, EPI_ISL_818009, EPI_ISL_818010, EPI_ISL_818011, EPI_ISL_818012, EPI_ISL_818013, EPI_ISL_818014, EPI_ISL_818015, EPI_ISL_818016, EPI_ISL_818017, EPI_ISL_818018, EPI_ISL_818019, EPI_ISL_818020, EPI_ISL_818021, EPI_ISL_818022, EPI_ISL_818023, EPI_ISL_818024, EPI_ISL_818025, EPI_ISL_818026, EPI_ISL_818027, EPI_ISL_818028, EPI_ISL_818029, EPI_ISL_818030, EPI_ISL_818031, EPI_ISL_818032, EPI_ISL_818033, EPI_ISL_818034, EPI_ISL_818035, EPI_ISL_818036, EPI_ISL_818037, EPI_ISL_818038, EPI_ISL_818039, EPI_ISL_818040, EPI_ISL_818041, EPI_ISL_818042, EPI_ISL_818043, EPI_ISL_818044, EPI_ISL_818045, EPI_ISL_818046, EPI_ISL_818047, EPI_ISL_818048, EPI_ISL_818049, EPI_ISL_818050, EPI_ISL_818051, EPI_ISL_818052, EPI_ISL_818053, EPI_ISL_818054, EPI_ISL_818055, EPI_ISL_818056, EPI_ISL_818057, EPI_ISL_818058, EPI_ISL_818059, EPI_ISL_818060, EPI_ISL_818061, EPI_ISL_818062, EPI_ISL_818063, EPI_ISL_818064, EPI_ISL_818065, EPI_ISL_818066, EPI_ISL_818067, EPI_ISL_818068, EPI_ISL_818069, EPI_ISL_818070, EPI_ISL_818071, EPI_ISL_818072, EPI_ISL_818073, EPI_ISL_818074, EPI_ISL_818075, EPI_ISL_818076, EPI_ISL_818077, EPI_ISL_818078, EPI_ISL_818079, EPI_ISL_818080, EPI_ISL_818081, EPI_ISL_818082, EPI_ISL_818083, EPI_ISL_818084, EPI_ISL_818085, EPI_ISL_818086, EPI_ISL_818087, EPI_ISL_818088, EPI_ISL_818089, EPI_ISL_818090, EPI_ISL_818091, EPI_ISL_818092, EPI_ISL_818093, EPI_ISL_818094, EPI_ISL_818095, EPI_ISL_818096, EPI_ISL_818097, EPI_ISL_818098, EPI_ISL_818099, EPI_ISL_818100, EPI_ISL_818101, EPI_ISL_818102, EPI_ISL_818103, EPI_ISL_818104, EPI_ISL_818105, EPI_ISL_818106, EPI_ISL_818107, EPI_ISL_818108, EPI_ISL_818109, EPI_ISL_818110, EPI_ISL_818111, EPI_ISL_818112, EPI_ISL_818113, EPI_ISL_818114, EPI_ISL_818115, EPI_ISL_818116, EPI_ISL_818117, EPI_ISL_818118, EPI_ISL_818119, EPI_ISL_818120, EPI_ISL_818121, EPI_ISL_818122 | | | |
| see above | Department of Virus and Microbiological Special Diagnostics, Statens Serum Institut, Copenhagen, Denmark | Albertsen Lab, Department of Chemistry and Bioscience, Aalborg University, Denmark | Danish Covid-19 Genome Consortium |
| EPI_ISL_819140, EPI_ISL_819144, EPI_ISL_819149, EPI_ISL_819180, EPI_ISL_819201 | Servicio de Microbiología, Hospital Universitario Son Espases | SeqCOVID-SPAIN consortium/IBV(CSIC) | Carla López-Causapé, Jordi Reina, Antonio Oliver and SeqCOVID-SPAIN consortium |
| EPI_ISL_819370, EPI_ISL_819437, EPI_ISL_819439 | Quadram Institute Bioscience | COVID-19 Genomics UK (COG-UK) Consortium | Dave J. Baker, Gemma L. Kay, Alp Aydin, Thanh Le-Viet, Steven Rudder, Ana P. Tedim, Anastasia Kolyva, Maria Diaz, Leonardo de Oliveira Martins, Nabil-Fareed Alikhan, Lizzie Meadows, Rachael Stanley, Ngozi Elumogo, Muhammed Yasir, Nicholas M. Thomson, Alexander J Trotter, Rachel Gilroy, Samuel Bloomfield, Claire Stuart, Andrew Bell, Reenesh Prakash, Samir Dervisevic, Alison E. Mather, John Wain, Mark Webber, Andrew J. Page, Justin O'Grady |
| EPI_ISL_820109 | University College London, Great Ormond Street Hospital for Children NHS Foundation Trust, Imperial College Healthcare NHS Trust | COVID-19 Genomics UK (COG-UK) Consortium | Sergi Castellano, Rachel Williams, Mark Kristiansen, Paola Resende Silva, Sunando Roy, Tony Brooks, Helena Tutill, Paola Niola, Patricia Dyal, Charlotte Williams, Leysa Forrest, Yasmin Panchbhaya, Jacqueline Findlay, Samuel Weeks, Julianne Brown, Kathryn Harris, Paul Randell, James Price, Alison Holmes, Judith Breuer |
| EPI_ISL_820110 | Quadram Institute Bioscience | COVID-19 Genomics UK (COG-UK) Consortium | Dave J. Baker, Gemma L. Kay, Alp Aydin, Thanh Le-Viet, Steven Rudder, Ana P. Tedim, Anastasia Kolyva, Maria Diaz, Leonardo de Oliveira Martins, Nabil-Fareed Alikhan, Lizzie Meadows, Rachael Stanley, Ngozi Elumogo, Muhammed Yasir, Nicholas M. Thomson, Alexander J Trotter, Rachel Gilroy, Samuel Bloomfield, Claire Stuart, Andrew Bell, Reenesh Prakash, Samir Dervisevic, Alison E. Mather, John Wain, Mark Webber, Andrew J. Page, Justin O'Grady |
| EPI_ISL_820111 | University College London, Great Ormond Street Hospital for Children NHS Foundation Trust, Imperial College Healthcare NHS Trust | COVID-19 Genomics UK (COG-UK) Consortium | Sergi Castellano, Rachel Williams, Mark Kristiansen, Paola Resende Silva, Sunando Roy, Tony Brooks, Helena Tutill, Paola Niola, Patricia Dyal, Charlotte Williams, Leysa Forrest, Yasmin Panchbhaya, Jacqueline Findlay, Samuel Weeks, Julianne Brown, Kathryn Harris, Paul Randell, James Price, Alison Holmes, Judith Breuer |
| EPI_ISL_820112 | Quadram Institute Bioscience | COVID-19 Genomics UK (COG-UK) Consortium | Dave J. Baker, Gemma L. Kay, Alp Aydin, Thanh Le-Viet, Steven Rudder, Ana P. Tedim, Anastasia Kolyva, Maria Diaz, Leonardo de Oliveira Martins, |

| | | | |
|---|---|---|---|
| | | | Nabil-Fareed Alikhan, Lizzie Meadows, Rachael Stanley, Ngozi Elumogo, Muhammed Yasir, Nicholas M. Thomson, Alexander J Trotter, Rachel Gilroy, Samuel Bloomfield, Claire Stuart, Andrew Bell, Reenesh Prakash, Samir Dervisevic, Alison E. Mather, John Wain, Mark Webber, Andrew J. Page, Justin O'Grady |
| EPI_ISL_820113, EPI_ISL_820114, EPI_ISL_820115, EPI_ISL_820116, EPI_ISL_820117, EPI_ISL_820118 | University College London, Great Ormond Street Hospital for Children NHS Foundation Trust, Imperial College Healthcare NHS Trust | COVID-19 Genomics UK (COG-UK) Consortium | Sergi Castellano, Rachel Williams, Mark Kristiansen, Paola Resende Silva, Sunando Roy, Tony Brooks, Helena Tutill, Paola Niola, Patricia Dyal, Charlotte Williams, Leysa Forrest, Yasmin Panchbhaya, Jacqueline Findlay, Samuel Weeks, Julianne Brown, Kathryn Harris, Paul Randell, James Price, Alison Holmes, Judith Breuer |
| EPI_ISL_820119 | Quadram Institute Bioscience | COVID-19 Genomics UK (COG-UK) Consortium | Dave J. Baker, Gemma L. Kay, Alp Aydin, Thanh Le-Viet, Steven Rudder, Ana P. Tedim, Anastasia Kolyva, Maria Diaz, Leonardo de Oliveira Martins, Nabil-Fareed Alikhan, Lizzie Meadows, Rachael Stanley, Ngozi Elumogo, Muhammed Yasir, Nicholas M. Thomson, Alexander J Trotter, Rachel Gilroy, Samuel Bloomfield, Claire Stuart, Andrew Bell, Reenesh Prakash, Samir Dervisevic, Alison E. Mather, John Wain, Mark Webber, Andrew J. Page, Justin O'Grady |
| EPI_ISL_820120, EPI_ISL_820121, EPI_ISL_820122, EPI_ISL_820123 | University College London, Great Ormond Street Hospital for Children NHS Foundation Trust, Imperial College Healthcare NHS Trust | COVID-19 Genomics UK (COG-UK) Consortium | Sergi Castellano, Rachel Williams, Mark Kristiansen, Paola Resende Silva, Sunando Roy, Tony Brooks, Helena Tutill, Paola Niola, Patricia Dyal, Charlotte Williams, Leysa Forrest, Yasmin Panchbhaya, Jacqueline Findlay, Samuel Weeks, Julianne Brown, Kathryn Harris, Paul Randell, James Price, Alison Holmes, Judith Breuer |
| EPI_ISL_820124 | Quadram Institute Bioscience | COVID-19 Genomics UK (COG-UK) Consortium | Dave J. Baker, Gemma L. Kay, Alp Aydin, Thanh Le-Viet, Steven Rudder, Ana P. Tedim, Anastasia Kolyva, Maria Diaz, Leonardo de Oliveira Martins, Nabil-Fareed Alikhan, Lizzie Meadows, Rachael Stanley, Ngozi Elumogo, Muhammed Yasir, Nicholas M. Thomson, Alexander J Trotter, Rachel Gilroy, Samuel Bloomfield, Claire Stuart, Andrew Bell, Reenesh Prakash, Samir Dervisevic, Alison E. Mather, John Wain, Mark Webber, Andrew J. Page, Justin O'Grady |
| EPI_ISL_820125 | University College London, Great Ormond Street Hospital for Children NHS Foundation Trust, Imperial College Healthcare NHS Trust | COVID-19 Genomics UK (COG-UK) Consortium | Sergi Castellano, Rachel Williams, Mark Kristiansen, Paola Resende Silva, Sunando Roy, Tony Brooks, Helena Tutill, Paola Niola, Patricia Dyal, Charlotte Williams, Leysa Forrest, Yasmin Panchbhaya, Jacqueline Findlay, Samuel Weeks, Julianne Brown, Kathryn Harris, Paul Randell, James Price, Alison Holmes, Judith Breuer |
| EPI_ISL_820126 | Quadram Institute Bioscience | COVID-19 Genomics UK (COG-UK) Consortium | Dave J. Baker, Gemma L. Kay, Alp Aydin, Thanh Le-Viet, Steven Rudder, Ana P. Tedim, Anastasia Kolyva, Maria Diaz, Leonardo de Oliveira Martins, Nabil-Fareed Alikhan, Lizzie Meadows, Rachael Stanley, Ngozi Elumogo, Muhammed Yasir, Nicholas M. Thomson, Alexander J Trotter, Rachel Gilroy, Samuel Bloomfield, Claire Stuart, Andrew Bell, Reenesh Prakash, Samir Dervisevic, Alison E. Mather, John Wain, Mark Webber, Andrew J. Page, Justin O'Grady |
| EPI_ISL_820127, EPI_ISL_820128 | University College London, Great Ormond Street Hospital for Children NHS Foundation Trust, Imperial College Healthcare NHS Trust | COVID-19 Genomics UK (COG-UK) Consortium | Sergi Castellano, Rachel Williams, Mark Kristiansen, Paola Resende Silva, Sunando Roy, Tony Brooks, Helena Tutill, Paola Niola, Patricia Dyal, Charlotte Williams, Leysa Forrest, Yasmin Panchbhaya, Jacqueline Findlay, Samuel Weeks, Julianne Brown, Kathryn Harris, Paul Randell, James Price, Alison Holmes, Judith Breuer |
| EPI_ISL_820131, EPI_ISL_820134, EPI_ISL_820136 | Quadram Institute Bioscience | COVID-19 Genomics UK (COG-UK) Consortium | Dave J. Baker, Gemma L. Kay, Alp Aydin, Thanh Le-Viet, Steven Rudder, Ana P. Tedim, Anastasia Kolyva, Maria Diaz, Leonardo de Oliveira Martins, Nabil-Fareed Alikhan, Lizzie Meadows, Rachael Stanley, Ngozi Elumogo, Muhammed Yasir, Nicholas M. Thomson, Alexander J Trotter, Rachel Gilroy, Samuel Bloomfield, Claire Stuart, Andrew Bell, Reenesh Prakash, Samir Dervisevic, Alison E. Mather, John Wain, Mark Webber, Andrew J. Page, Justin O'Grady |
| EPI_ISL_820138 | University College London, Great Ormond Street Hospital for Children NHS Foundation Trust, Imperial College Healthcare NHS Trust | COVID-19 Genomics UK (COG-UK) Consortium | Sergi Castellano, Rachel Williams, Mark Kristiansen, Paola Resende Silva, Sunando Roy, Tony Brooks, Helena Tutill, Paola Niola, Patricia Dyal, Charlotte Williams, Leysa Forrest, Yasmin Panchbhaya, Jacqueline Findlay, Samuel Weeks, Julianne Brown, Kathryn Harris, Paul Randell, James Price, Alison Holmes, Judith Breuer |
| EPI_ISL_820141, EPI_ISL_820144, EPI_ISL_820146, EPI_ISL_820149, EPI_ISL_820151, EPI_ISL_820154, EPI_ISL_820156, EPI_ISL_820159, EPI_ISL_820161, EPI_ISL_820164, EPI_ISL_820167, EPI_ISL_820169, EPI_ISL_820172, EPI_ISL_820175, EPI_ISL_820178, EPI_ISL_820180, EPI_ISL_820183, EPI_ISL_820186, EPI_ISL_820188, EPI_ISL_820191, EPI_ISL_820193, EPI_ISL_820196, EPI_ISL_820199, EPI_ISL_820202 | | | |
| see above | Quadram Institute Bioscience | COVID-19 Genomics UK (COG-UK) Consortium | Dave J. Baker, Gemma L. Kay, Alp Aydin, Thanh Le-Viet, Steven Rudder, Ana P. Tedim, Anastasia Kolyva, Maria Diaz, Leonardo de Oliveira Martins, Nabil-Fareed Alikhan, Lizzie Meadows, Rachael Stanley, Ngozi Elumogo, Muhammed Yasir, Nicholas M. Thomson, Alexander J Trotter, Rachel Gilroy, Samuel Bloomfield, Claire Stuart, Andrew Bell, Reenesh Prakash, Samir Dervisevic, Alison E. Mather, John Wain, Mark Webber, Andrew J. Page, Justin O'Grady |
| EPI_ISL_820204, EPI_ISL_820206, EPI_ISL_820209, EPI_ISL_820211, EPI_ISL_820213, EPI_ISL_820216, EPI_ISL_820218 | University College London, Great Ormond Street Hospital for Children NHS Foundation Trust, Imperial College Healthcare NHS Trust | COVID-19 Genomics UK (COG-UK) Consortium | Sergi Castellano, Rachel Williams, Mark Kristiansen, Paola Resende Silva, Sunando Roy, Tony Brooks, Helena Tutill, Paola Niola, Patricia Dyal, Charlotte Williams, Leysa Forrest, Yasmin Panchbhaya, Jacqueline Findlay, Samuel Weeks, Julianne Brown, Kathryn Harris, Paul Randell, James Price, Alison Holmes, Judith Breuer |
| EPI_ISL_820221, EPI_ISL_820223 | Quadram Institute Bioscience | COVID-19 Genomics UK (COG-UK) Consortium | Dave J. Baker, Gemma L. Kay, Alp Aydin, Thanh Le-Viet, Steven Rudder, Ana P. Tedim, Anastasia Kolyva, Maria Diaz, Leonardo de Oliveira Martins, Nabil-Fareed Alikhan, Lizzie Meadows, Rachael Stanley, Ngozi Elumogo, Muhammed Yasir, Nicholas M. Thomson, Alexander J Trotter, Rachel Gilroy, Samuel Bloomfield, Claire Stuart, Andrew Bell, Reenesh Prakash, Samir Dervisevic, Alison E. Mather, John Wain, Mark Webber, Andrew J. Page, Justin O'Grady |
| EPI_ISL_820226, EPI_ISL_820229 | University College London, Great Ormond Street Hospital for Children NHS Foundation Trust, Imperial College Healthcare NHS Trust | COVID-19 Genomics UK (COG-UK) Consortium | Sergi Castellano, Rachel Williams, Mark Kristiansen, Paola Resende Silva, Sunando Roy, Tony Brooks, Helena Tutill, Paola Niola, Patricia Dyal, Charlotte Williams, Leysa Forrest, Yasmin Panchbhaya, Jacqueline Findlay, Samuel Weeks, Julianne Brown, Kathryn Harris, Paul Randell, James Price, Alison Holmes, Judith Breuer |
| EPI_ISL_820231, EPI_ISL_820234 | Quadram Institute Bioscience | COVID-19 Genomics UK (COG-UK) Consortium | Dave J. Baker, Gemma L. Kay, Alp Aydin, Thanh Le-Viet, Steven Rudder, Ana P. Tedim, Anastasia Kolyva, Maria Diaz, Leonardo de Oliveira Martins, Nabil-Fareed Alikhan, Lizzie Meadows, Rachael Stanley, Ngozi Elumogo, Muhammed Yasir, Nicholas M. Thomson, Alexander J Trotter, Rachel Gilroy, Samuel Bloomfield, Claire Stuart, Andrew Bell, Reenesh Prakash, Samir Dervisevic, Alison E. Mather, John Wain, Mark Webber, Andrew J. Page, Justin O'Grady |
| EPI_ISL_820236 | University College London, Great Ormond Street Hospital for Children NHS Foundation Trust, Imperial College Healthcare NHS Trust | COVID-19 Genomics UK (COG-UK) Consortium | Sergi Castellano, Rachel Williams, Mark Kristiansen, Paola Resende Silva, Sunando Roy, Tony Brooks, Helena Tutill, Paola Niola, Patricia Dyal, Charlotte Williams, Leysa Forrest, Yasmin Panchbhaya, Jacqueline Findlay, Samuel Weeks, Julianne Brown, Kathryn Harris, Paul Randell, James Price, Alison Holmes, Judith Breuer |
| EPI_ISL_820238, EPI_ISL_820241, EPI_ISL_820244 | Quadram Institute Bioscience | COVID-19 Genomics UK (COG-UK) Consortium | Dave J. Baker, Gemma L. Kay, Alp Aydin, Thanh Le-Viet, Steven Rudder, Ana P. Tedim, Anastasia Kolyva, Maria Diaz, Leonardo de Oliveira Martins, Nabil-Fareed Alikhan, Lizzie Meadows, Rachael Stanley, Ngozi Elumogo, Muhammed Yasir, Nicholas M. Thomson, Alexander J Trotter, Rachel Gilroy, Samuel Bloomfield, Claire Stuart, Andrew Bell, Reenesh Prakash, Samir Dervisevic, Alison E. Mather, John Wain, Mark Webber, Andrew J. Page, Justin O'Grady |
| EPI_ISL_820246, EPI_ISL_820249, EPI_ISL_820251 | University College London, Great Ormond Street Hospital for Children NHS Foundation Trust, Imperial College Healthcare NHS Trust | COVID-19 Genomics UK (COG-UK) Consortium | Sergi Castellano, Rachel Williams, Mark Kristiansen, Paola Resende Silva, Sunando Roy, Tony Brooks, Helena Tutill, Paola Niola, Patricia Dyal, Charlotte Williams, Leysa Forrest, Yasmin Panchbhaya, Jacqueline Findlay, Samuel Weeks, Julianne Brown, Kathryn Harris, Paul Randell, James Price, Alison Holmes, Judith Breuer |
| EPI_ISL_820253 | Quadram Institute Bioscience | COVID-19 Genomics UK (COG-UK) Consortium | Dave J. Baker, Gemma L. Kay, Alp Aydin, Thanh Le-Viet, Steven Rudder, Ana P. Tedim, Anastasia Kolyva, Maria Diaz, Leonardo de Oliveira Martins, Nabil-Fareed Alikhan, Lizzie Meadows, Rachael Stanley, Ngozi Elumogo, Muhammed Yasir, Nicholas M. Thomson, Alexander J Trotter, Rachel Gilroy, Samuel Bloomfield, Claire Stuart, Andrew Bell, Reenesh Prakash, Samir Dervisevic, Alison E. Mather, John Wain, Mark Webber, Andrew J. Page, Justin O'Grady |
| EPI_ISL_820256, EPI_ISL_820259 | University College London, Great Ormond Street Hospital for Children NHS Foundation Trust, Imperial College Healthcare NHS Trust | COVID-19 Genomics UK (COG-UK) Consortium | Sergi Castellano, Rachel Williams, Mark Kristiansen, Paola Resende Silva, Sunando Roy, Tony Brooks, Helena Tutill, Paola Niola, Patricia Dyal, Charlotte Williams, Leysa Forrest, Yasmin Panchbhaya, Jacqueline Findlay, Samuel Weeks, Julianne Brown, Kathryn Harris, Paul Randell, James Price, Alison Holmes, Judith Breuer |
| EPI_ISL_820261 | Quadram Institute Bioscience | COVID-19 Genomics UK (COG-UK) Consortium | Dave J. Baker, Gemma L. Kay, Alp Aydin, Thanh Le-Viet, Steven Rudder, Ana P. Tedim, Anastasia Kolyva, Maria Diaz, Leonardo de Oliveira Martins, Nabil-Fareed Alikhan, Lizzie Meadows, Rachael Stanley, Ngozi Elumogo, Muhammed Yasir, Nicholas M. Thomson, Alexander J Trotter, Rachel Gilroy, Samuel Bloomfield, Claire Stuart, Andrew Bell, Reenesh Prakash, Samir Dervisevic, Alison E. Mather, John Wain, Mark Webber, Andrew J. Page, Justin O'Grady |

| | | | |
|---|---|---|---|
| EPI_ISL_820264, EPI_ISL_820266, EPI_ISL_820268, EPI_ISL_820271, EPI_ISL_820273, EPI_ISL_820276 | University College London, Great Ormond Street Hospital for Children NHS Foundation Trust, Imperial College Healthcare NHS Trust | COVID-19 Genomics UK (COG-UK) Consortium | Sergi Castellano, Rachel Williams, Mark Kristiansen, Paola Resende Silva, Sunando Roy, Tony Brooks, Helena Tutill, Paola Niola, Patricia Dyal, Charlotte Williams, Leysa Forrest, Yasmin Panchbhaya, Jacqueline Findlay, Samuel Weeks, Julianne Brown, Kathryn Harris, Paul Randell, James Price, Alison Holmes, Judith Breuer |
| EPI_ISL_820279 | Quadram Institute Bioscience | COVID-19 Genomics UK (COG-UK) Consortium | Dave J. Baker, Gemma L. Kay, Alp Aydin, Thanh Le-Viet, Steven Rudder, Ana P. Tedim, Anastasia Kolyva, Maria Diaz, Leonardo de Oliveira Martins, Nabil-Fareed Alikhan, Lizzie Meadows, Rachael Stanley, Ngozi Elumogo, Muhammed Yasir, Nicholas M. Thomson, Alexander J Trotter, Rachel Gilroy, Samuel Bloomfield, Claire Stuart, Andrew Bell, Reenesh Prakash, Samir Dervisevic, Alison E. Mather, John Wain, Mark Webber, Andrew J. Page, Justin O'Grady |
| EPI_ISL_820282 | University College London, Great Ormond Street Hospital for Children NHS Foundation Trust, Imperial College Healthcare NHS Trust | COVID-19 Genomics UK (COG-UK) Consortium | Sergi Castellano, Rachel Williams, Mark Kristiansen, Paola Resende Silva, Sunando Roy, Tony Brooks, Helena Tutill, Paola Niola, Patricia Dyal, Charlotte Williams, Leysa Forrest, Yasmin Panchbhaya, Jacqueline Findlay, Samuel Weeks, Julianne Brown, Kathryn Harris, Paul Randell, James Price, Alison Holmes, Judith Breuer |
| EPI_ISL_820284, EPI_ISL_820287, EPI_ISL_820289, EPI_ISL_820292, EPI_ISL_820294 | Quadram Institute Bioscience | COVID-19 Genomics UK (COG-UK) Consortium | Dave J. Baker, Gemma L. Kay, Alp Aydin, Thanh Le-Viet, Steven Rudder, Ana P. Tedim, Anastasia Kolyva, Maria Diaz, Leonardo de Oliveira Martins, Nabil-Fareed Alikhan, Lizzie Meadows, Rachael Stanley, Ngozi Elumogo, Muhammed Yasir, Nicholas M. Thomson, Alexander J Trotter, Rachel Gilroy, Samuel Bloomfield, Claire Stuart, Andrew Bell, Reenesh Prakash, Samir Dervisevic, Alison E. Mather, John Wain, Mark Webber, Andrew J. Page, Justin O'Grady |
| EPI_ISL_820297 | Queens Medical Centre, Clinical Microbiology Department / DeepSeq Nottingham | COVID-19 Genomics UK (COG-UK) Consortium | Gemma Clark, Wendy Smith, Manjinder Khakh, Vicki M Fleming, Michelle M Lister, Hannah Howson-Wells, Jonathan Ball, Patrick McClure, Joseph Chappell, Theocharis Tsoleridis, Nadine Holmes, Matthew Carlisle, Christopher Moore, Fei Sang, Johnny Debebe, Victoria Wright, Matthew Loose |
| EPI_ISL_820300, EPI_ISL_820302, EPI_ISL_820304, EPI_ISL_820306 | Quadram Institute Bioscience | COVID-19 Genomics UK (COG-UK) Consortium | Dave J. Baker, Gemma L. Kay, Alp Aydin, Thanh Le-Viet, Steven Rudder, Ana P. Tedim, Anastasia Kolyva, Maria Diaz, Leonardo de Oliveira Martins, Nabil-Fareed Alikhan, Lizzie Meadows, Rachael Stanley, Ngozi Elumogo, Muhammed Yasir, Nicholas M. Thomson, Alexander J Trotter, Rachel Gilroy, Samuel Bloomfield, Claire Stuart, Andrew Bell, Reenesh Prakash, Samir Dervisevic, Alison E. Mather, John Wain, Mark Webber, Andrew J. Page, Justin O'Grady |
| EPI_ISL_820309 | University College London, Great Ormond Street Hospital for Children NHS Foundation Trust, Imperial College Healthcare NHS Trust | COVID-19 Genomics UK (COG-UK) Consortium | Sergi Castellano, Rachel Williams, Mark Kristiansen, Paola Resende Silva, Sunando Roy, Tony Brooks, Helena Tutill, Paola Niola, Patricia Dyal, Charlotte Williams, Leysa Forrest, Yasmin Panchbhaya, Jacqueline Findlay, Samuel Weeks, Julianne Brown, Kathryn Harris, Paul Randell, James Price, Alison Holmes, Judith Breuer |
| EPI_ISL_820311 | Quadram Institute Bioscience | COVID-19 Genomics UK (COG-UK) Consortium | Dave J. Baker, Gemma L. Kay, Alp Aydin, Thanh Le-Viet, Steven Rudder, Ana P. Tedim, Anastasia Kolyva, Maria Diaz, Leonardo de Oliveira Martins, Nabil-Fareed Alikhan, Lizzie Meadows, Rachael Stanley, Ngozi Elumogo, Muhammed Yasir, Nicholas M. Thomson, Alexander J Trotter, Rachel Gilroy, Samuel Bloomfield, Claire Stuart, Andrew Bell, Reenesh Prakash, Samir Dervisevic, Alison E. Mather, John Wain, Mark Webber, Andrew J. Page, Justin O'Grady |
| EPI_ISL_820314, EPI_ISL_820316, EPI_ISL_820318, EPI_ISL_820321 | University College London, Great Ormond Street Hospital for Children NHS Foundation Trust, Imperial College Healthcare NHS Trust | COVID-19 Genomics UK (COG-UK) Consortium | Sergi Castellano, Rachel Williams, Mark Kristiansen, Paola Resende Silva, Sunando Roy, Tony Brooks, Helena Tutill, Paola Niola, Patricia Dyal, Charlotte Williams, Leysa Forrest, Yasmin Panchbhaya, Jacqueline Findlay, Samuel Weeks, Julianne Brown, Kathryn Harris, Paul Randell, James Price, Alison Holmes, Judith Breuer |
| EPI_ISL_820323 | Quadram Institute Bioscience | COVID-19 Genomics UK (COG-UK) Consortium | Dave J. Baker, Gemma L. Kay, Alp Aydin, Thanh Le-Viet, Steven Rudder, Ana P. Tedim, Anastasia Kolyva, Maria Diaz, Leonardo de Oliveira Martins, Nabil-Fareed Alikhan, Lizzie Meadows, Rachael Stanley, Ngozi Elumogo, Muhammed Yasir, Nicholas M. Thomson, Alexander J Trotter, Rachel Gilroy, Samuel Bloomfield, Claire Stuart, Andrew Bell, Reenesh Prakash, Samir Dervisevic, Alison E. Mather, John Wain, Mark Webber, Andrew J. Page, Justin O'Grady |
| EPI_ISL_821629, EPI_ISL_821630, EPI_ISL_821631, EPI_ISL_821633, EPI_ISL_821634, EPI_ISL_821635, EPI_ISL_821636, EPI_ISL_821637, EPI_ISL_821641, EPI_ISL_821648, EPI_ISL_821653 | | | |
| see above | Lighthouse Lab in Milton Keynes | Wellcome Sanger Institute for the COVID-19 Genomics UK (COG-UK) Consortium | The Lighthouse Lab in Milton Keynes and Alex Alderton, Roberto Amato, Sonia Goncalves, Ewan Harrison, David K. Jackson, Ian Johnston, Dominic Kwiatkowski, Cordelia Langford, John Sillitoe on behalf of the Wellcome Sanger Institute COVID-19 Surveillance Team |
| EPI_ISL_821667, EPI_ISL_821672, EPI_ISL_821677, EPI_ISL_821733, EPI_ISL_821787, EPI_ISL_821790, EPI_ISL_821827, EPI_ISL_821877, EPI_ISL_821889, EPI_ISL_821934, EPI_ISL_821943, EPI_ISL_821959 | | | |
| see above | Lighthouse Lab in Cambridge | Wellcome Sanger Institute for the COVID-19 Genomics UK (COG-UK) Consortium | Rob Howes, The Lighthouse Lab in Cambridge and Alex Alderton, Roberto Amato, Sonia Goncalves, Ewan Harrison, David K. Jackson, Ian Johnston, Dominic Kwiatkowski, Cordelia Langford, John Sillitoe on behalf of the Wellcome Sanger Institute COVID-19 Surveillance Team |
| EPI_ISL_822291, EPI_ISL_822294, EPI_ISL_822297, EPI_ISL_822298, EPI_ISL_822300, EPI_ISL_822303 | Lighthouse Lab in Alderley Park | Wellcome Sanger Institute for the COVID-19 Genomics UK (COG-UK) Consortium | Jacquelyn Wynn, Mairead Hyland, The Lighthouse Lab in Alderley Park and Alex Alderton, Roberto Amato, Sonia Goncalves, Ewan Harrison, David K. Jackson, Ian Johnston, Dominic Kwiatkowski, Cordelia Langford, John Sillitoe on behalf of the Wellcome Sanger Institute COVID-19 Surveillance Team |
| EPI_ISL_822307 | Lighthouse Lab in Glasgow | Wellcome Sanger Institute for the COVID-19 Genomics UK (COG-UK) Consortium | Harper VanSteenhouse, Yumi Kasai, David Gray, Carol Clugston, Anna Dominiczak and Alex Alderton, Roberto Amato, Sonia Goncalves, Ewan Harrison, David K. Jackson, Ian Johnston, Dominic Kwiatkowski, Cordelia Langford, John Sillitoe on behalf of the Wellcome Sanger Institute COVID-19 Surveillance Team |
| EPI_ISL_822529, EPI_ISL_822530, EPI_ISL_822531, EPI_ISL_822532, EPI_ISL_822533, EPI_ISL_822534, EPI_ISL_822535, EPI_ISL_822536, EPI_ISL_822537, EPI_ISL_822538, EPI_ISL_822539, EPI_ISL_822540, EPI_ISL_822541, EPI_ISL_822542, EPI_ISL_822543, EPI_ISL_822544, EPI_ISL_822545, EPI_ISL_822546, EPI_ISL_822547, EPI_ISL_822548, EPI_ISL_822549, EPI_ISL_822550, EPI_ISL_822551, EPI_ISL_822552, EPI_ISL_822553, EPI_ISL_822554, EPI_ISL_822555, EPI_ISL_822556, EPI_ISL_822557, EPI_ISL_822558, EPI_ISL_822559, EPI_ISL_822560, EPI_ISL_822561, EPI_ISL_822562, EPI_ISL_822563, EPI_ISL_822564, EPI_ISL_822565, EPI_ISL_822566, EPI_ISL_822567, EPI_ISL_822568, EPI_ISL_822569, EPI_ISL_822570, EPI_ISL_822571, EPI_ISL_822572, EPI_ISL_822573, EPI_ISL_822574, EPI_ISL_822575, EPI_ISL_822576, EPI_ISL_822577, EPI_ISL_822578, EPI_ISL_822579, EPI_ISL_822580, EPI_ISL_822581, EPI_ISL_822582, EPI_ISL_822583, EPI_ISL_822584, EPI_ISL_822585, EPI_ISL_822586, EPI_ISL_822587, EPI_ISL_822588, EPI_ISL_822589, EPI_ISL_822590, EPI_ISL_822591, EPI_ISL_822592, EPI_ISL_822593, EPI_ISL_822594, EPI_ISL_822595, EPI_ISL_822596, EPI_ISL_822597, EPI_ISL_822598, EPI_ISL_822599, EPI_ISL_822600, EPI_ISL_822601, EPI_ISL_822602, EPI_ISL_822603, EPI_ISL_822604, EPI_ISL_822605, EPI_ISL_822606, EPI_ISL_822607, EPI_ISL_822608, EPI_ISL_822609, EPI_ISL_822610, EPI_ISL_822611, EPI_ISL_822612, EPI_ISL_822613, EPI_ISL_822614, EPI_ISL_822615, EPI_ISL_822616, EPI_ISL_822617, EPI_ISL_822618, EPI_ISL_822619, EPI_ISL_822620, EPI_ISL_822621, EPI_ISL_822622, EPI_ISL_822623, EPI_ISL_822624, EPI_ISL_822625, EPI_ISL_822626, EPI_ISL_822627, EPI_ISL_822628, EPI_ISL_822629, EPI_ISL_822630, EPI_ISL_822631, EPI_ISL_822632, EPI_ISL_822633, EPI_ISL_822634, EPI_ISL_822635, EPI_ISL_822636, EPI_ISL_822637, EPI_ISL_822638, EPI_ISL_822639, EPI_ISL_822640, EPI_ISL_822641, EPI_ISL_822642, EPI_ISL_822643, EPI_ISL_822644, EPI_ISL_822645, EPI_ISL_822646, EPI_ISL_822647, EPI_ISL_822648, EPI_ISL_822649, EPI_ISL_822650, EPI_ISL_822651, EPI_ISL_822652, EPI_ISL_822653, EPI_ISL_822654, EPI_ISL_822655, EPI_ISL_822656, EPI_ISL_822657, EPI_ISL_822658, EPI_ISL_822839, EPI_ISL_822865, EPI_ISL_822885, EPI_ISL_822887, EPI_ISL_822921, EPI_ISL_822932, EPI_ISL_822949, EPI_ISL_822954, EPI_ISL_822963, EPI_ISL_822978, EPI_ISL_823019, EPI_ISL_823065, EPI_ISL_823093, EPI_ISL_823094, EPI_ISL_823095, EPI_ISL_823096, EPI_ISL_823097, EPI_ISL_823104, EPI_ISL_823106, EPI_ISL_823107, EPI_ISL_823108, EPI_ISL_823115, EPI_ISL_823141, EPI_ISL_823153, EPI_ISL_823643, EPI_ISL_823644, EPI_ISL_823645, EPI_ISL_823769, EPI_ISL_823770, EPI_ISL_823771, EPI_ISL_823772, EPI_ISL_823773, EPI_ISL_823774, EPI_ISL_823775, EPI_ISL_823776, EPI_ISL_823777, EPI_ISL_823778, EPI_ISL_823779, EPI_ISL_823800, EPI_ISL_823801, EPI_ISL_823803, EPI_ISL_823804, EPI_ISL_823808, EPI_ISL_823809, EPI_ISL_823810 | | | |
| see above | Wales Specialist Virology Centre Sequencing lab: Pathogen Genomics Unit | COVID-19 Genomics UK (COG-UK) Consortium | Catherine Moore, Johnathan Evans, Laura Gifford, Malorie Perry, Simon Cottrell, Angela Marchbank, Alec Birchley, Alexander Adams, Amy Gaskin, Bree Gatica-Wilcox, Jason Coombes, Joel Southgate, Lauren Gilbert, Lee Graham, Nicole Pacchiarini, Sara Kumziene-Summerhayes, Sarah Taylor, Sophie Jones, Sara Rey, Matthew Bull, Joanne Watkins, Sally Corden, Tom Connor |
| EPI_ISL_823979, EPI_ISL_823980, EPI_ISL_823992, EPI_ISL_824002, EPI_ISL_824041, EPI_ISL_824042, EPI_ISL_824043, EPI_ISL_824044, EPI_ISL_824045, EPI_ISL_824046, EPI_ISL_824056, EPI_ISL_824057, EPI_ISL_824058, EPI_ISL_824059, EPI_ISL_824069, EPI_ISL_824070, EPI_ISL_824072, EPI_ISL_824094, EPI_ISL_824110, EPI_ISL_824115, EPI_ISL_824117, EPI_ISL_824118, EPI_ISL_824119, EPI_ISL_824121, EPI_ISL_824122, EPI_ISL_824140, EPI_ISL_824163, EPI_ISL_824170, EPI_ISL_824171, EPI_ISL_824172, EPI_ISL_824173, EPI_ISL_824174, EPI_ISL_824175, EPI_ISL_824176, EPI_ISL_824177, EPI_ISL_824178, EPI_ISL_824179, EPI_ISL_824180, EPI_ISL_824195, EPI_ISL_824196, EPI_ISL_824210, EPI_ISL_824211, EPI_ISL_824212, EPI_ISL_824213, EPI_ISL_824230, EPI_ISL_824231, EPI_ISL_824248, EPI_ISL_824265, EPI_ISL_824269, EPI_ISL_824271, EPI_ISL_824274, EPI_ISL_824275, EPI_ISL_824277, EPI_ISL_824281, EPI_ISL_824282, EPI_ISL_824283 | | | |
| see above | Dutch COVID-19 response team | National Institute for Public Health and the Environment (RIVM) | Adam Meijer, Harry Vennema, Jeroen Cremer, Sharon van den Brink, Bas van der Veer, AnneMarie van den Brandt, Florian Zwagemaker, Dennis Schmitz, Chantal Reusken, on behalf of the national COVID-19 response team |
| EPI_ISL_824286, EPI_ISL_824287 | Institute of Microbiology, Universidad San Francisco de Quito | Institute of Microbiology, Universidad San Francisco de Quito | Belén Prado-Vivar, Sully Márquez, Juan José Guadalupe, Monica Becerra-Wong, Bernardo Gutiérrez, Andrea Cunguan, Nabih Dahik, Verónica Barragán, Patricio Rojas-Silva, Gabriel Trueba, Michelle Grunauer, Paúl Cárdenas |
| EPI_ISL_824288, EPI_ISL_824289 | Institute of Microbiology, Universidad San Francisco de Quito | Institute of Microbiology, Universidad San Francisco de Quito | Belén Prado-Vivar, Sully Márquez, Juan José Guadalupe, Monica Becerra-Wong, Bernardo Gutiérrez, Tania Guayasamin, Patricio Reyes, Verónica Barragán, Patricio Rojas-Silva, Gabriel Trueba, Michelle Grunauer, Paúl Cárdenas |
| EPI_ISL_824519, EPI_ISL_824520, EPI_ISL_824521, EPI_ISL_824522, EPI_ISL_824523, EPI_ISL_824536, EPI_ISL_824537 | New Mexico Department of Health Scientific Laboratory | New Mexico Department of Health Scientific Laboratory | Ellie Johnson, Anastacia Griego-Fisher, D'eldra Malone |
| EPI_ISL_824556, EPI_ISL_824557, EPI_ISL_824562, EPI_ISL_824563, EPI_ISL_824564, EPI_ISL_824566, EPI_ISL_824567, EPI_ISL_824568, EPI_ISL_824569, EPI_ISL_824574, EPI_ISL_824575, EPI_ISL_824576, EPI_ISL_824577, EPI_ISL_824578, EPI_ISL_824579, EPI_ISL_824584, EPI_ISL_824585, EPI_ISL_824586, EPI_ISL_824587, EPI_ISL_824588, EPI_ISL_824589, EPI_ISL_824596, EPI_ISL_824597, EPI_ISL_824598, EPI_ISL_824599, EPI_ISL_824600, EPI_ISL_824601, EPI_ISL_824607, EPI_ISL_824608, EPI_ISL_824609, EPI_ISL_824610, EPI_ISL_824611, EPI_ISL_824612, EPI_ISL_824618, EPI_ISL_824619, EPI_ISL_824620, EPI_ISL_824621, EPI_ISL_824622, EPI_ISL_824623, EPI_ISL_824625, EPI_ISL_824629, EPI_ISL_824630, EPI_ISL_824631, EPI_ISL_824632, EPI_ISL_824633, EPI_ISL_824634, EPI_ISL_824635, EPI_ISL_824637, EPI_ISL_824641, EPI_ISL_824642, EPI_ISL_824643, EPI_ISL_824644, EPI_ISL_824645, EPI_ISL_824646, | | | |

| | | | |
|---|---|---|---|
| EPI_ISL_824647, EPI_ISL_824663, EPI_ISL_824675, EPI_ISL_824695, EPI_ISL_824719 | | | |
| see above | Cedars-Sinai Medical Center, Department of Pathology & Laboratory Medicine, Molecular Pathology Laboratory | Cedars-Sinai Medical Center, Molecular Pathology Laboratory of Department of Pathology & Laboratory Medicine and Genomic Core | Wenjuan Zhang, Brian Davis, Stephanie Chen, Jorge Mario Sincuir Martinez, Jasmine T Plummer, Eric Vail |
| EPI_ISL_824800, EPI_ISL_824801, EPI_ISL_824802, EPI_ISL_824803, EPI_ISL_824804, EPI_ISL_824805, EPI_ISL_824806, EPI_ISL_824807, EPI_ISL_824808 | Department of Clinical Microbiology | GIGA Medical Genomics | Keith Durkin, Maria Artesi, Sébastien Bontems, Raphaël Boreux, Bouchra Boujemla, Cécile Meex, Pierrette Melin, Marie-Pierre Hayette, Vincent Bours |
| EPI_ISL_825011, EPI_ISL_825012 | Arizona State Public Health Laboratory | Arizona State Public Health Laboratory | Trung Huynh, Jessica Escobar, Katherine Fullerton, Nobuko Fukushima, Stacy White, Linda Getsinger, Victor Waddell |
| EPI_ISL_825152 | National Institute of Mental Health and Neurosciences (NIMHANS) | Department of Neurovirology, National Institute of Mental Health and Neurosciences (NIMHANS) | Chitra Pattabiraman, Pramada Prasad, Anita S Desai, V Ravi |
| EPI_ISL_825504, EPI_ISL_825532, EPI_ISL_825533, EPI_ISL_825556 | Respiratory Virus Unit, National Infection Service, Public Health England | COVID-19 Genomics UK (COG-UK) Consortium | PHE Covid Sequencing Team |
| EPI_ISL_826536 | Dutch COVID-19 response team | National Institute for Public Health and the Environment (RIVM) | Adam Meijer, Harry Vennema, Jeroen Cremer, Sharon van den Brink, Bas van der Veer, AnneMarie van den Brandt, Florian Zwagemaker, Dennis Schmitz, Chantal Reusken, on behalf of the national COVID-19 response team |
| EPI_ISL_827820, EPI_ISL_828111, EPI_ISL_828140, EPI_ISL_828809, EPI_ISL_828998, EPI_ISL_829136, EPI_ISL_829139, EPI_ISL_829310, EPI_ISL_829733, EPI_ISL_829805, EPI_ISL_830121, EPI_ISL_830122, EPI_ISL_830123, EPI_ISL_830124, EPI_ISL_830126 | | | |
| see above | deCODE genetics | deCODE genetics | Daniel F Gudbjartsson; Agnar Helgason; Hakon Jonsson; Olafur T Magnusson; Pall Melsted; Gudmundur L Norddahl; Jona Saemundsdottir; Asgeir Sigurdsson; Patrick Sulem; Arna B Agustsdottir; Hannes Eggertsson; Berglind Eiriksdottir; Run Fridriksdottir; Elisabet E Gardarsdottir; Gudmundur Georgsson; Olafia S Gretarsdottir; Kjartan R Gudmundsson; Thora R Gunnarsdottir; Arnaldur Gylfason; Hilma Holm; Brynjar O Jensson; Aslaug Jonasdottir; Kamilla S Josefsdottir; Thordur Kristjansson; Droplaug N Magnusdottir; Solvi Rognvaldsson; Louise le Roux; Gudrun Sigmundsdottir; Gardar Sveinbjornsson; Kristin E Sveinsdottir; Maney Sveinsdottir; Emil A Thorarensen; Bjarni Thorbjornsson; Gisli Masson; Ingileif Jonsdottir; Alma Moller; Thorolfur Gudnason; Karl G Kristinsson; Unnur Thorsteinsdottir; Kari Stefansson |
| EPI_ISL_830212, EPI_ISL_830243, EPI_ISL_830571, EPI_ISL_830573, EPI_ISL_830583, EPI_ISL_830584, EPI_ISL_830585, EPI_ISL_830586, EPI_ISL_830587, EPI_ISL_830588, EPI_ISL_830589, EPI_ISL_830590, EPI_ISL_830591, EPI_ISL_830592, EPI_ISL_830593, EPI_ISL_830594, EPI_ISL_830595, EPI_ISL_830596, EPI_ISL_830597, EPI_ISL_830598, EPI_ISL_830599, EPI_ISL_830600, EPI_ISL_830601, EPI_ISL_830602, EPI_ISL_830603, EPI_ISL_830604, EPI_ISL_830605, EPI_ISL_830606, EPI_ISL_830607 | | | |
| see above | KALEIDA CENTER FOR LABORATORY MEDICINE | Wadsworth Center, New York State Department of Health | Kirsten St. George, Daryl M. Lamson, Alexis Russel, Matthew Shudt, Melissa A Leisner, Jonathan Plitnick, Navjot Singh, John Kelly, Erasmus Schneider, Erica Lasek-Nesselquist |
| EPI_ISL_830720, EPI_ISL_830721 | BIO-REFERENCE LABORATORIES | Wadsworth Center, New York State Department of Health | Kirsten St. George, Daryl M. Lamson, Alexis Russel, Matthew Shudt, Melissa A Leisner, Jonathan Plitnick, Navjot Singh, John Kelly, Erasmus Schneider, Erica Lasek-Nesselquist |
| EPI_ISL_831021, EPI_ISL_831022, EPI_ISL_831023, EPI_ISL_831024, EPI_ISL_831025, EPI_ISL_831026 | Hospital Universitario La Paz (Madrid) | SeqCOVID-SPAIN consortium/IBV(CSIC) | Fernando Lázaro-Perona, María Rodríguez-Tejedor, Elias Dahdouh, Jesús Mingorance and SeqCOVID-SPAIN consortium |
| EPI_ISL_832019 | Southern Nevada Public Health Laboratory | Southern Nevada Public Health Laboratory | Michael Picker, Holly Hansen, Erin Buttery, Joseph Benson |
| EPI_ISL_832047, EPI_ISL_832048, EPI_ISL_832049, EPI_ISL_832050, EPI_ISL_832051, EPI_ISL_832052, EPI_ISL_832053, EPI_ISL_832054, EPI_ISL_832055, EPI_ISL_832056, EPI_ISL_832057, EPI_ISL_832058, EPI_ISL_832059, EPI_ISL_832060, EPI_ISL_832061, EPI_ISL_832062, EPI_ISL_832063, EPI_ISL_832064 | | | |
| see above | Wyoming Public Health Laboratory | Wyoming Public Health Laboratory | Noah Hull, Taylor Fearing, Lynette Gumbleton, Channing Weber, Ashley Norberg, Bailey Bowcutt, and Wanda Manley |
| EPI_ISL_832078 | Santa Clara County Public Health Laboratory | Santa Clara County Public Health Laboratory | Santa Clara County Public Health Department |
| EPI_ISL_832116 | MD Laboratories | Los Angeles County PHL | P. Hemarajata et al. |
| EPI_ISL_832197 | Hospital | National Reference Center for Viruses of Respiratory Infections, Institut Pasteur, Paris | Marion Barbet, Sylvie Behillil, Méline Bizard, Angela Brisebarre, Camille Capel, Etienne Simon-Lorière, Vincent Enouf, Maud Vanpeene, Sylvie van der Werf, Esther Gyde |
| EPI_ISL_832199 | Hospital | National Reference Center for Viruses of Respiratory Infections, Institut Pasteur, Paris | Marion Barbet, Sylvie Behillil, Méline Bizard, Angela Brisebarre, Camille Capel, Etienne Simon-Lorière, Vincent Enouf, Maud Vanpeene, Sylvie van der Werf, Patricia Stoessel-Thouvenin |
| EPI_ISL_832265 | DOHMH Jamaica | New York City Public Health Laboratory | Jade Wang, et al. |
| EPI_ISL_832355, EPI_ISL_832356 | DOHMH Corona | New York City Public Health Laboratory | Jade Wang, et al. |
| EPI_ISL_832357, EPI_ISL_832358 | DOHMH Jamaica | New York City Public Health Laboratory | Jade Wang, et al. |
| EPI_ISL_832359, EPI_ISL_832360 | DOHMH Corona | New York City Public Health Laboratory | Jade Wang, et al. |
| EPI_ISL_832361 | DOHMH Jamaica | New York City Public Health Laboratory | Jade Wang, et al. |
| EPI_ISL_832362, EPI_ISL_832363 | DOHMH Chelsea | New York City Public Health Laboratory | Jade Wang, et al. |
| EPI_ISL_832835, EPI_ISL_832836, EPI_ISL_832837, EPI_ISL_832838, EPI_ISL_832839, EPI_ISL_832840, EPI_ISL_832841, EPI_ISL_832842, EPI_ISL_832843, EPI_ISL_832844, EPI_ISL_832845, EPI_ISL_832846, EPI_ISL_832847, EPI_ISL_832848, EPI_ISL_832849, EPI_ISL_832850, EPI_ISL_832875, EPI_ISL_832911, EPI_ISL_832912, EPI_ISL_832914, EPI_ISL_832915, EPI_ISL_832916, EPI_ISL_832917, EPI_ISL_832918, EPI_ISL_832927, EPI_ISL_832928, EPI_ISL_832929, EPI_ISL_832933, EPI_ISL_832934 | | | |
| see above | Maine HETL | Tewhey Lab, The Jackson Laboratory | Matluk,N., Dewey,H., Iosue,F., Barter,M., Lynch,R., Munger,H. and Tewhey,R. |
| EPI_ISL_833149 | Platform BIS UZA/UAntwerpen, University Hospital Antwerp, Edegem, Belgium | UAntwerp, Laboratory of Medical Microbiology, Campus Drie Eiken S6.26, Universiteitsplein 1, 2610, Wilrijk, Belgium | Basil Britto Xavier, Jasmine Coppens, Sahaya Glingston Rajakani, Christine Lammens, Veerle Matheeussen, Herman Goossens |
| EPI_ISL_833183 | Microbiology Department. Complexo Hospitalario Universitario de Vigo | Microbiology Department. Complexo Hospitalario Universitario de Vigo | Microbiology Department, Complexo Hospitalario Universitario de Vigo. EPICOVIGAL. |
| EPI_ISL_833186 | Department of Clinical Microbiology | GIGA Medical Genomics | Keith Durkin, Maria Artesi, Sébastien Bontems, Raphaël Boreux, Bouchra Boujemla, Cécile Meex, Pierrette Melin, Marie-Pierre Hayette, Vincent Bours |
| EPI_ISL_833210, EPI_ISL_833227 | Department of Virology and Immunology, University of Helsinki and Helsinki University Hospital, Huslab Finland | Department of Virology, Faculty of Medicine, University of Helsinki, Helsinki, Finland | Teemu Smura, Ravi Kant, Phuoc Truong, Hussein Alburkat, Hannimari Kallio-Kokko, Jenni Virtanen, Maija Suvanto, Fathiah Zakham, Essi Korhonen, Sari Hannula, Harri Kangas, Pekka Ellonen, Olli Vapalahti |
| EPI_ISL_833251 | Ospedale Civile Atri-Medicina Interna | Istituto Zooprofilattico Sperimentale dell'Abruzzo e Molise "G. Caporale" | Lorusso A, Marcacci M, Di Domenico M, Ancora M, Curini V, Mangone I, Rinaldi A, Di Pasquale A, Cammà C, Puglia I, Calistri P, Savini G |
| EPI_ISL_833252 | Ospedale Civile Giulianova-Pronto Soccorso | Istituto Zooprofilattico Sperimentale dell'Abruzzo e Molise "G. Caporale" | Lorusso A, Marcacci M, Di Domenico M, Ancora M, Curini V, Mangone I, Rinaldi A, Di Pasquale A, Cammà C, Puglia I, Calistri P, Savini G |
| EPI_ISL_833253 | USCA-Avezzano | Istituto Zooprofilattico Sperimentale dell'Abruzzo e Molise "G. Caporale" | Lorusso A, Marcacci M, Di Domenico M, Ancora M, Curini V, Mangone I, Rinaldi A, Di Pasquale A, Cammà C, Puglia I, Calistri P, Savini G |
| EPI_ISL_833254 | USCA-Pescina | Istituto Zooprofilattico Sperimentale dell'Abruzzo e Molise "G. Caporale" | Lorusso A, Marcacci M, Di Domenico M, Ancora M, Curini V, Mangone I, Rinaldi A, Di Pasquale A, Cammà C, Puglia I, Calistri P, Savini G |
| EPI_ISL_833255, EPI_ISL_833256, EPI_ISL_833257 | Ospedale Civile Atri | Istituto Zooprofilattico Sperimentale dell'Abruzzo e Molise "G. Caporale" | Lorusso A, Marcacci M, Di Domenico M, Ancora M, Curini V, Mangone I, Rinaldi A, Di Pasquale A, Cammà C, Puglia I, Calistri P, Savini G |
| EPI_ISL_833258, EPI_ISL_833259 | SIESP DIPARTIMENTO DI PREVENZIONE TERAMO | Istituto Zooprofilattico Sperimentale dell'Abruzzo e Molise "G. Caporale" | Lorusso A, Marcacci M, Di Domenico M, Ancora M, Curini V, Mangone I, Rinaldi A, Di Pasquale A, Cammà C, Puglia I, Calistri P, Savini G |

| | | | |
|---|---|---|---|
| EPI_ISL_833260, EPI_ISL_833261 | Giulianova | Istituto Zooprofilattico Sperimentale dell'Abruzzo e Molise "G. Caporale" | Lorusso A, Marcacci M, Di Domenico M, Ancora M, Curini V, Mangone I, Rinaldi A, Di Pasquale A, Cammà C, Puglia I, Calistri P, Savini G |
| EPI_ISL_833262, EPI_ISL_833263, EPI_ISL_833264 | SIESP DIPARTIMENTO DI PREVENZIONE CHIETI | Istituto Zooprofilattico Sperimentale dell'Abruzzo e Molise "G. Caporale" | Lorusso A, Marcacci M, Di Domenico M, Ancora M, Curini V, Mangone I, Rinaldi A, Di Pasquale A, Cammà C, Puglia I, Calistri P, Savini G |
| EPI_ISL_833265, EPI_ISL_833266, EPI_ISL_833267, EPI_ISL_833268, EPI_ISL_833269 | Ospedale SS Annunziata Chieti | Istituto Zooprofilattico Sperimentale dell'Abruzzo e Molise "G. Caporale" | Lorusso A, Marcacci M, Di Domenico M, Ancora M, Curini V, Mangone I, Rinaldi A, Di Pasquale A, Cammà C, Puglia I, Calistri P, Savini G |
| EPI_ISL_833270, EPI_ISL_833272, EPI_ISL_833273, EPI_ISL_833274, EPI_ISL_833275 | SIESP CHIETI - DRIVE IN ORTONA | Istituto Zooprofilattico Sperimentale dell'Abruzzo e Molise "G. Caporale" | Lorusso A, Marcacci M, Di Domenico M, Ancora M, Curini V, Mangone I, Rinaldi A, Di Pasquale A, Cammà C, Puglia I, Calistri P, Savini G |
| EPI_ISL_833276 | Ospedale Civile G.Mazzini-Teramo | Istituto Zooprofilattico Sperimentale dell'Abruzzo e Molise "G. Caporale" | Lorusso A, Marcacci M, Di Domenico M, Ancora M, Curini V, Mangone I, Rinaldi A, Di Pasquale A, Cammà C, Puglia I, Calistri P, Savini G |
| EPI_ISL_833278, EPI_ISL_833279, EPI_ISL_833280 | SIESP CHIETI- DRIVE IN CHIETI | Istituto Zooprofilattico Sperimentale dell'Abruzzo e Molise "G. Caporale" | Lorusso A, Marcacci M, Di Domenico M, Ancora M, Curini V, Mangone I, Rinaldi A, Di Pasquale A, Cammà C, Puglia I, Calistri P, Savini G |
| EPI_ISL_833281, EPI_ISL_833282 | SIESP DIPARTIMENTO DI PREVENZIONE TERAMO | Istituto Zooprofilattico Sperimentale dell'Abruzzo e Molise "G. Caporale" | Lorusso A, Marcacci M, Di Domenico M, Ancora M, Curini V, Mangone I, Rinaldi A, Di Pasquale A, Cammà C, Puglia I, Calistri P, Savini G |
| EPI_ISL_833283 | DIPARTIMENTO PREVENZIONE AVEZZANO-SERVIZIO DI IGIENE EPIDEMIOLOGIA E SANITA' PUBBLICA | Istituto Zooprofilattico Sperimentale dell'Abruzzo e Molise "G. Caporale" | Lorusso A, Marcacci M, Di Domenico M, Ancora M, Curini V, Mangone I, Rinaldi A, Di Pasquale A, Cammà C, Puglia I, Calistri P, Savini G |
| EPI_ISL_833284, EPI_ISL_833285, EPI_ISL_833286, EPI_ISL_833287 | SIESP CHIETI - DRIVE IN ORTONA | Istituto Zooprofilattico Sperimentale dell'Abruzzo e Molise "G. Caporale" | Lorusso A, Marcacci M, Di Domenico M, Ancora M, Curini V, Mangone I, Rinaldi A, Di Pasquale A, Cammà C, Puglia I, Calistri P, Savini G |
| EPI_ISL_833288, EPI_ISL_833290 | PRESIDIO TERRITORIALE CASOLI CH | Istituto Zooprofilattico Sperimentale dell'Abruzzo e Molise "G. Caporale" | Lorusso A, Marcacci M, Di Domenico M, Ancora M, Curini V, Mangone I, Rinaldi A, Di Pasquale A, Cammà C, Puglia I, Calistri P, Savini G |
| EPI_ISL_833291 | SIESP DIPARTIMENTO DI PREVENZIONE DELL'AQUILA | Istituto Zooprofilattico Sperimentale dell'Abruzzo e Molise "G. Caporale" | Lorusso A, Marcacci M, Di Domenico M, Ancora M, Curini V, Mangone I, Rinaldi A, Di Pasquale A, Cammà C, Puglia I, Calistri P, Savini G |
| EPI_ISL_833292, EPI_ISL_833293, EPI_ISL_833294 | SIESP CHIETI - DRIVE IN LANCIANO | Istituto Zooprofilattico Sperimentale dell'Abruzzo e Molise "G. Caporale" | Lorusso A, Marcacci M, Di Domenico M, Ancora M, Curini V, Mangone I, Rinaldi A, Di Pasquale A, Cammà C, Puglia I, Calistri P, Savini G |
| EPI_ISL_833295, EPI_ISL_833296 | SIESP DIPARTIMENTO DI PREVENZIONE TERAMO | Istituto Zooprofilattico Sperimentale dell'Abruzzo e Molise "G. Caporale" | Lorusso A, Marcacci M, Di Domenico M, Ancora M, Curini V, Mangone I, Rinaldi A, Di Pasquale A, Cammà C, Puglia I, Calistri P, Savini G |
| EPI_ISL_833298, EPI_ISL_833300 | SIESP CHIETI - DRIVE IN LANCIANO | Istituto Zooprofilattico Sperimentale dell'Abruzzo e Molise "G. Caporale" | Lorusso A, Marcacci M, Di Domenico M, Ancora M, Curini V, Mangone I, Rinaldi A, Di Pasquale A, Cammà C, Puglia I, Calistri P, Savini G |
| EPI_ISL_833301 | DIPARTIMENTO PREVENZIONE AVEZZANO-SERVIZIO DI IGIENE EPIDEMIOLOGIA E SANITA' PUBBLICA | Istituto Zooprofilattico Sperimentale dell'Abruzzo e Molise "G. Caporale" | Lorusso A, Marcacci M, Di Domenico M, Ancora M, Curini V, Mangone I, Rinaldi A, Di Pasquale A, Cammà C, Puglia I, Calistri P, Savini G |
| EPI_ISL_833302, EPI_ISL_833303, EPI_ISL_833304, EPI_ISL_833305 | SIESP DIPARTIMENTO DI PREVENZIONE TERAMO | Istituto Zooprofilattico Sperimentale dell'Abruzzo e Molise "G. Caporale" | Lorusso A, Marcacci M, Di Domenico M, Ancora M, Curini V, Mangone I, Rinaldi A, Di Pasquale A, Cammà C, Puglia I, Calistri P, Savini G |
| EPI_ISL_833306, EPI_ISL_833307, EPI_ISL_833308, EPI_ISL_833309 | SIESP CHIETI - DRIVE IN ORTONA | Istituto Zooprofilattico Sperimentale dell'Abruzzo e Molise "G. Caporale" | Lorusso A, Marcacci M, Di Domenico M, Ancora M, Curini V, Mangone I, Rinaldi A, Di Pasquale A, Cammà C, Puglia I, Calistri P, Savini G |
| EPI_ISL_833311, EPI_ISL_833312 | SIESP CHIETI- DRIVE IN CHIETI | Istituto Zooprofilattico Sperimentale dell'Abruzzo e Molise "G. Caporale" | Lorusso A, Marcacci M, Di Domenico M, Ancora M, Curini V, Mangone I, Rinaldi A, Di Pasquale A, Cammà C, Puglia I, Calistri P, Savini G |
| EPI_ISL_833313, EPI_ISL_833314, EPI_ISL_833315 | SIESP CHIETI - DRIVE IN LANCIANO | Istituto Zooprofilattico Sperimentale dell'Abruzzo e Molise "G. Caporale" | Lorusso A, Marcacci M, Di Domenico M, Ancora M, Curini V, Mangone I, Rinaldi A, Di Pasquale A, Cammà C, Puglia I, Calistri P, Savini G |
| EPI_ISL_833316 | SIESP CHIETI - DRIVE IN GISSI | Istituto Zooprofilattico Sperimentale dell'Abruzzo e Molise "G. Caporale" | Lorusso A, Marcacci M, Di Domenico M, Ancora M, Curini V, Mangone I, Rinaldi A, Di Pasquale A, Cammà C, Puglia I, Calistri P, Savini G |
| EPI_ISL_833317, EPI_ISL_833318 | SIESP CHIETI - DRIVE IN CHIETI | Istituto Zooprofilattico Sperimentale dell'Abruzzo e Molise "G. Caporale" | Lorusso A, Marcacci M, Di Domenico M, Ancora M, Curini V, Mangone I, Rinaldi A, Di Pasquale A, Cammà C, Puglia I, Calistri P, Savini G |
| EPI_ISL_833319, EPI_ISL_833320, EPI_ISL_833322 | SIESP DIPARTIMENTO DI PREVENZIONE SULMONA | Istituto Zooprofilattico Sperimentale dell'Abruzzo e Molise "G. Caporale" | Lorusso A, Marcacci M, Di Domenico M, Ancora M, Curini V, Mangone I, Rinaldi A, Di Pasquale A, Cammà C, Puglia I, Calistri P, Savini G |
| EPI_ISL_833323 | SIESP CHIETI - DRIVE IN CHIETI | Istituto Zooprofilattico Sperimentale dell'Abruzzo e Molise "G. Caporale" | Lorusso A, Marcacci M, Di Domenico M, Ancora M, Curini V, Mangone I, Rinaldi A, Di Pasquale A, Cammà C, Puglia I, Calistri P, Savini G |
| EPI_ISL_833324 | SIESP DIPARTIMENTO DI PREVENZIONE CHIETI | Istituto Zooprofilattico Sperimentale dell'Abruzzo e Molise "G. Caporale" | Lorusso A, Marcacci M, Di Domenico M, Ancora M, Curini V, Mangone I, Rinaldi A, Di Pasquale A, Cammà C, Puglia I, Calistri P, Savini G |
| EPI_ISL_833325 | Ospedale SS Annunziata Chieti | Istituto Zooprofilattico Sperimentale dell'Abruzzo e Molise "G. Caporale" | Lorusso A, Marcacci M, Di Domenico M, Ancora M, Curini V, Mangone I, Rinaldi A, Di Pasquale A, Cammà C, Puglia I, Calistri P, Savini G |
| EPI_ISL_833326 | SIESP DIPARTIMENTO DI PREVENZIONE CHIETI | Istituto Zooprofilattico Sperimentale dell'Abruzzo e Molise "G. Caporale" | Lorusso A, Marcacci M, Di Domenico M, Ancora M, Curini V, Mangone I, Rinaldi A, Di Pasquale A, Cammà C, Puglia I, Calistri P, Savini G |
| EPI_ISL_833327 | Ospedale SS Annunziata Chieti | Istituto Zooprofilattico Sperimentale dell'Abruzzo e Molise "G. Caporale" | Lorusso A, Marcacci M, Di Domenico M, Ancora M, Curini V, Mangone I, Rinaldi A, Di Pasquale A, Cammà C, Puglia I, Calistri P, Savini G |
| EPI_ISL_833328, EPI_ISL_833329, EPI_ISL_833330 | SIESP CHIETI - DRIVE IN ORTONA | Istituto Zooprofilattico Sperimentale dell'Abruzzo e Molise "G. Caporale" | Lorusso A, Marcacci M, Di Domenico M, Ancora M, Curini V, Mangone I, Rinaldi A, Di Pasquale A, Cammà C, Puglia I, Calistri P, Savini G |
| EPI_ISL_833331 | Ospedale Civile Atri | Istituto Zooprofilattico Sperimentale dell'Abruzzo e Molise "G. Caporale" | Lorusso A, Marcacci M, Di Domenico M, Ancora M, Curini V, Mangone I, Rinaldi A, Di Pasquale A, Cammà C, Puglia I, Calistri P, Savini G |
| EPI_ISL_834578 | Lighthouse Lab in Glasgow | Wellcome Sanger Institute for the COVID-19 Genomics UK (COG-UK) Consortium | Harper VanSteenhouse, Yumi Kasai, David Gray, Carol Clugston, Anna Dominiczak and Alex Alderton, Roberto Amato, Sonia Goncalves, Ewan Harrison, David K. Jackson, Ian Johnston, Dominic Kwiatkowski, Cordelia Langford, John Sillitoe on behalf of the Wellcome Sanger Institute COVID-19 Surveillance Team |
| EPI_ISL_837225 | Respiratory Virus Unit, National Infection Service, Public Health England | COVID-19 Genomics UK (COG-UK) Consortium | PHE Covid Sequencing Team |
| EPI_ISL_837254, EPI_ISL_837326, EPI_ISL_837327, EPI_ISL_837328, EPI_ISL_837329, EPI_ISL_837330 | Istituto Zooprofilattico Sperimentale del Mezzogiorno | TIGEM | Patrizia Annunziata, Andrea Ballabio, Valentina Bouche, Davide Cacchiarelli (CorrespAuthor), Pellegrino Cerino, Chiara Colantuono, Lucio Di Filippo, Antonio Grimaldi, Antonio Limone, Gabriella Loconte, Anna Manfredi, Francesco Panariello, Biancamaria Pierri, Marcello Salvi, Lucia Vassallo |

EPI_ISL_837341, EPI_ISL_837343, EPI_ISL_837344, EPI_ISL_837346, EPI_ISL_837347, EPI_ISL_837349, EPI_ISL_837351, EPI_ISL_837352, EPI_ISL_837354, EPI_ISL_837355, EPI_ISL_837356, EPI_ISL_837357, EPI_ISL_837358, EPI_ISL_837359, EPI_ISL_837360, EPI_ISL_837361, EPI_ISL_837362, EPI_ISL_837363, EPI_ISL_837364, EPI_ISL_837365, EPI_ISL_837366, EPI_ISL_837367, EPI_ISL_837368, EPI_ISL_837369, EPI_ISL_837370, EPI_ISL_837371, EPI_ISL_837372, EPI_ISL_837373, EPI_ISL_837374, EPI_ISL_837375, EPI_ISL_837376, EPI_ISL_837377, EPI_ISL_837378, EPI_ISL_837379, EPI_ISL_837380, EPI_ISL_837381,

| | | | |
|---|---|---|---|
| EPI_ISL_837384, EPI_ISL_837385, EPI_ISL_837386, EPI_ISL_837388, EPI_ISL_837389, EPI_ISL_837390, EPI_ISL_837391, EPI_ISL_837392, EPI_ISL_837393, EPI_ISL_837396, EPI_ISL_837397, EPI_ISL_837398, EPI_ISL_837399, EPI_ISL_837400, EPI_ISL_837401, EPI_ISL_837402, EPI_ISL_837403, EPI_ISL_837404, EPI_ISL_837405, EPI_ISL_837406, EPI_ISL_837407, EPI_ISL_837408, EPI_ISL_837409, EPI_ISL_837410, EPI_ISL_837411, EPI_ISL_837412, EPI_ISL_837413, EPI_ISL_837414, EPI_ISL_837415, EPI_ISL_837417, EPI_ISL_837418, EPI_ISL_837419, EPI_ISL_837420, EPI_ISL_837421, EPI_ISL_837422, EPI_ISL_837423, EPI_ISL_837427, EPI_ISL_837428, EPI_ISL_837429, EPI_ISL_837431 | | | |
| see above | National Virus Reference Laboratory | National Virus Reference Laboratory | Michael Carr, Gabriel Gonzalez, Jonathan Dean, Cillian F De Gascun |
| EPI_ISL_837448, EPI_ISL_837483, EPI_ISL_837484, EPI_ISL_837485 | Istituto Zooprofilattico Sperimentale del Mezzogiorno | TIGEM | Patrizia Annunziata, Andrea Ballabio, Valentina Bouche, Davide Cacchiarelli (CorrespAuthor), Pellegrino Cerino, Chiara Colantuono, Lucio Di Filippo, Antonio Grimaldi, Antonio Limone, Gabriella Loconte, Anna Manfredi, Francesco Panariello, Biancamaria Pierri, Marcello Salvi, Lucia Vassallo |
| EPI_ISL_837518, EPI_ISL_837521, EPI_ISL_837522, EPI_ISL_837546 | UW Virology Lab | UW Virology Lab | Pavitra Roychoudhury, Hong Xie, Lasata Shrestha, Meei-Li Huang, Keith R Jerome, Alexander Greninger |
| EPI_ISL_837829, EPI_ISL_837830, EPI_ISL_837832 | Wyoming Public Health Laboratory | Wyoming Public Health Laboratory | Noah Hull, Taylor Fearing, Lynette Gumbleton, Channing Weber, Ashley Norberg, Bailey Bowcutt, and Wanda Manley |
| EPI_ISL_837879, EPI_ISL_837889, EPI_ISL_837891, EPI_ISL_837892, EPI_ISL_837893, EPI_ISL_837895, EPI_ISL_837898, EPI_ISL_837901 | Department of Pathology, University of Cambridge | COVID-19 Genomics UK (COG-UK) Consortium | Aminu S. Jahun, Yasmin Chaudhry, Grant Hall, Iliana Georgana, Myra Hosmillo, Martin D. Curran, Malte Pinckert, Surendra Parmar, Ian Goodfellow |
| EPI_ISL_838185, EPI_ISL_838188, EPI_ISL_838189, EPI_ISL_838190, EPI_ISL_838191, EPI_ISL_838192, EPI_ISL_838193, EPI_ISL_838194, EPI_ISL_838195, EPI_ISL_838196, EPI_ISL_838197 | | | |
| see above | West of Scotland Specialist Virology Centre, NHSGGC / MRC-University of Glasgow Centre for Virus Research | COVID-19 Genomics UK (COG-UK) Consortium | Ana da Silva Filipe, Natasha Johnson, Kathy Smollett, Daniel Mair, Stephen Carmichael, Alice Broos, Lily Tong, Jenna Nichols, Kyriaki Nomikou; Sarah McDonald; Richard Orton, Joseph Hughes, Sreenu Vattipally, David L Robertson; Alasdair MacLean, Rory Gunson; Sharif Shaaban, Matthew Holden; Rachel Blacow, Guy Mollett, Kathy Li, James Shepherd, Antonia Ho, Emma Thomson |
| EPI_ISL_838765, EPI_ISL_838766, EPI_ISL_838772, EPI_ISL_838774, EPI_ISL_838779, EPI_ISL_838780, EPI_ISL_838781, EPI_ISL_838782, EPI_ISL_838784, EPI_ISL_838785, EPI_ISL_838786, EPI_ISL_838787, EPI_ISL_838788, EPI_ISL_838789, EPI_ISL_838790, EPI_ISL_838791, EPI_ISL_838792, EPI_ISL_838793, EPI_ISL_838794, EPI_ISL_838795, EPI_ISL_838796, EPI_ISL_838797, EPI_ISL_839311, EPI_ISL_839313, EPI_ISL_839314, EPI_ISL_839324, EPI_ISL_839325, EPI_ISL_839326, EPI_ISL_839327, EPI_ISL_839328 | | | |
| see above | University College London, Great Ormond Street Hospital for Children NHS Foundation Trust, Imperial College Healthcare NHS Trust | COVID-19 Genomics UK (COG-UK) Consortium | Sergi Castellano, Rachel Williams, Mark Kristiansen, Paola Resende Silva, Sunando Roy, Tony Brooks, Helena Tutill, Paola Niola, Patricia Dyal, Charlotte Williams, Leysa Forrest, Yasmin Panchbhaya, Jacqueline Findlay, Samuel Weeks, Julianne Brown, Kathryn Harris, Paul Randell, James Price, Alison Holmes, Judith Breuer |
| EPI_ISL_839668, EPI_ISL_839669, EPI_ISL_839670, EPI_ISL_839671, EPI_ISL_839673, EPI_ISL_839674, EPI_ISL_839675, EPI_ISL_839677, EPI_ISL_839678, EPI_ISL_839679, EPI_ISL_839680, EPI_ISL_839681, EPI_ISL_839683, EPI_ISL_839684, EPI_ISL_839686, EPI_ISL_839687, EPI_ISL_839688, EPI_ISL_839689, EPI_ISL_839690, EPI_ISL_839691, EPI_ISL_839692, EPI_ISL_839693, EPI_ISL_839694, EPI_ISL_839695, EPI_ISL_839696, EPI_ISL_839697, EPI_ISL_839698, EPI_ISL_839701, EPI_ISL_839702, EPI_ISL_839705, EPI_ISL_839706, EPI_ISL_839707, EPI_ISL_839708 | | | |
| see above | Northumbria University / South Tees Hospitals NHS Foundation Trust / North Cumbria Integrated Care NHS Foundation Trust / North Tees and Hartlepool NHS Foundation Trust / Newcastle Hospitals NHS Foundation Trust | COVID-19 Genomics UK (COG-UK) Consortium | Darren L Smith,Andrew Nelson,Matthew Bashton,Greg R Young,Joshua Loh,John Allan,Mohammad A Tariq,Giles S Holt,Gary Black,Wen C Yew,Lynn Dover,Paul Baker,Steve Liggett,Sarah Essex,Jane Greenaway,Debra Padgett,Clive Graham,Garren Scott,Edward Barton,Emma Swindells,Brendan Payne,Jennifer Collins,Yusri Taha,Gary Eltringham |
| EPI_ISL_839992, EPI_ISL_839994 | Queens Medical Centre, Clinical Microbiology Department / DeepSeq Nottingham | COVID-19 Genomics UK (COG-UK) Consortium | Gemma Clark, Wendy Smith, Manjinder Khakh, Vicki M Fleming, Michelle M Lister, Hannah Howson-Wells, Jonathan Ball, Patrick McClure, Joseph Chappell, Theocharis Tsoleridis, Nadine Holmes, Matthew Carlisle, Christopher Moore, Fei Sang, Johnny Debebe, Victoria Wright, Matthew Loose |
| EPI_ISL_840104, EPI_ISL_840105, EPI_ISL_840106, EPI_ISL_840107, EPI_ISL_840111 | Lincolnshire Hospitals and DeepSeq Nottingham | COVID-19 Genomics UK (COG-UK) Consortium | Nichola Duckworth, Tim Sloan, Sarah Walsh, Jonathan Ball, Patrick McClure, Joeseph Chappell, Nadine Holmes, Matthew Carlisle, Christopher Moore, Fei Sang, Johnny Debebe, Victoria Wright, Matthew Loose |
| EPI_ISL_840332, EPI_ISL_840333, EPI_ISL_840334, EPI_ISL_840335, EPI_ISL_840336, EPI_ISL_840338, EPI_ISL_840343, EPI_ISL_840344, EPI_ISL_840345, EPI_ISL_840348 | Oxford Viromics, NDM, University of Oxford; Oxford University Hospitals; Basingstoke and North Hampshire Hospital | COVID-19 Genomics UK (COG-UK) Consortium | Tanya Golubchik, David Bonsall, George Macintyre, Amy Trebes, Mariateresa de Cesare, Catrin Moore, Alex Mobbs, Anita Justice, Robert Shaw, Monique Andersson, Timothy Peto, Emma Wise, Nathan Moore, Jessica Lynch, Nick Cortes, Matilde Mori, Stephen Kidd, David Buck, John Todd, Christophe Fraser |
| EPI_ISL_840370, EPI_ISL_840387, EPI_ISL_840388, EPI_ISL_840389, EPI_ISL_840513, EPI_ISL_840514, EPI_ISL_840519, EPI_ISL_840520, EPI_ISL_840521, EPI_ISL_840522, EPI_ISL_840523, EPI_ISL_840524, EPI_ISL_840525, EPI_ISL_840526, EPI_ISL_840527, EPI_ISL_840528, EPI_ISL_840529, EPI_ISL_840530, EPI_ISL_840531, EPI_ISL_840532, EPI_ISL_840533, EPI_ISL_840534, EPI_ISL_840535, EPI_ISL_840536, EPI_ISL_840537, EPI_ISL_840538, EPI_ISL_840539, EPI_ISL_840540, EPI_ISL_840541, EPI_ISL_840542, EPI_ISL_840543, EPI_ISL_840544, EPI_ISL_840545, EPI_ISL_840546, EPI_ISL_840547, EPI_ISL_840548, EPI_ISL_840549, EPI_ISL_840550, EPI_ISL_840551, EPI_ISL_840553, EPI_ISL_840555, EPI_ISL_840556, EPI_ISL_840557, EPI_ISL_840558, EPI_ISL_840559, EPI_ISL_840560, EPI_ISL_840561, EPI_ISL_840562, EPI_ISL_840563, EPI_ISL_840564, EPI_ISL_840565, EPI_ISL_840566, EPI_ISL_840567, EPI_ISL_840568, EPI_ISL_840569, EPI_ISL_840571, EPI_ISL_840572, EPI_ISL_840573, EPI_ISL_840574, EPI_ISL_840575, EPI_ISL_840576, EPI_ISL_840577, EPI_ISL_840578, EPI_ISL_840581, EPI_ISL_840582, EPI_ISL_840583, EPI_ISL_840585, EPI_ISL_840587, EPI_ISL_840588, EPI_ISL_840589, EPI_ISL_840591, EPI_ISL_840592, EPI_ISL_840593, EPI_ISL_840594, EPI_ISL_840595, EPI_ISL_840599 | | | |
| see above | Originating lab: Wales Specialist Virology Centre Sequencing lab: Pathogen Genomics Unit | Public Health Wales Microbiology Cardiff Wales Specialist Virology Centre | Catherine Moore, Johnathan Evans, Laura Gifford, Malorie Perry, Simon Cottrell, Angela Marchbank, Alec Birchley, Alexander Adams, Amy Gaskin, Bree Gatica-Wilcox, Jason Coombes, Joel Southgate, Lauren Gilbert, Lee Graham, Nicole Pacchiarini, Sara Kumziene-Summerhayes, Sarah Taylor, Sophie Jones, Sara Rey, Matthew Bull, Joanne Watkins, Sally Corden, Tom Connor |
| EPI_ISL_842010, EPI_ISL_842013 | Centre for Enzyme Innovation, University of Portsmouth / Translational Research Laboratory, Portsmouth Hospitals NHS Trust | COVID-19 Genomics UK (COG-UK) Consortium | Angela Beckett,Yann Bourgeois,Garry Scarlett,Sharon Glaysher,Scott Elliott,Kelly Bicknell,Robert Impey,Allyson Lloyd,Sarah Wyllie,Ethan Butcher,Anoop Chauhan,Samuel Robson |
| EPI_ISL_842230, EPI_ISL_842270, EPI_ISL_842297, EPI_ISL_842325 | Virology Department, Sheffield Teaching Hospitals NHS Foundation Trust/Department of Infection, Immunity and Cardiovascular Disease, The Medical School, University of Sheffield | COVID-19 Genomics UK (COG-UK) Consortium | Thushan de Silva, Matthew Parker, Nikki Smith, Adri Angyal, Rebecca Brown, Luke Green, Rachel Tucker, Paul Parsons, Danielle Groves, Katie Johnson, Laura Carrilero, Alex Keeley, Dave Partridge, Matthew Wyles, Benjamin Lindsey, Mehmet Yavuz, Mohammad Raza, Cariad Evans |
| EPI_ISL_842609, EPI_ISL_842613, EPI_ISL_842616, EPI_ISL_842620, EPI_ISL_842621, EPI_ISL_842622, EPI_ISL_842629, EPI_ISL_842630 | Wyoming Public Health Laboratory | Wyoming Public Health Laboratory | Noah Hull, Taylor Fearing, Lynette Gumbleton, Channing Weber, Ashley Norberg, Bailey Bowcutt, and Wanda Manley |
| EPI_ISL_842643 | Department of Medical Microbiology, St. Olavs hospital | Norwegian Institute of Public Health, Department of Virology | Kathrine Stene-Johansen, Kamilla Heddeland Instefjord, Hilde Elshaug, Atiya R Ali,Marie Paulsen Madsen, Rasmus Riis Kopperud, Hilde Vollan, Karoline Bragstad, Olav Hungnes |
| EPI_ISL_843144, EPI_ISL_843145 | Barts Health NHS Trust | COVID-19 Genomics UK (COG-UK) Consortium | CUTINO-MOGUEL, Maria-Teresa; HARRINGTON, David; OWOYEMI, Dola; SHYLINI, Raghavendran; BROAD, Claire; KELE, Beatrix |
| EPI_ISL_843151, EPI_ISL_843152, EPI_ISL_843153, EPI_ISL_843154, EPI_ISL_843163 | Regional Virus Laboratory, Belfast Health and Social Care Trust | COVID-19 Genomics UK (COG-UK) Consortium | Conall McCaughey, James McKenna, Tanya Curran, Susan Feeney, Alison Watt, Ciara Cox, Mairead Connor, Zoltan Molnar, David Simpson, Derek Fairley |
| EPI_ISL_843213, EPI_ISL_843216, EPI_ISL_843270, EPI_ISL_843302, EPI_ISL_843316, EPI_ISL_843341, EPI_ISL_843360, EPI_ISL_843370, EPI_ISL_843384, EPI_ISL_843547, EPI_ISL_843574, EPI_ISL_843591, EPI_ISL_843595, EPI_ISL_843598, EPI_ISL_843608, EPI_ISL_843615, EPI_ISL_843616, EPI_ISL_843617, EPI_ISL_843618, EPI_ISL_843619, EPI_ISL_843620, EPI_ISL_843621, EPI_ISL_843622, EPI_ISL_843623, EPI_ISL_843624, EPI_ISL_843625, EPI_ISL_843626, EPI_ISL_843627, EPI_ISL_843628, EPI_ISL_843629, EPI_ISL_843630, EPI_ISL_843631, EPI_ISL_843632, EPI_ISL_843633, EPI_ISL_843634, EPI_ISL_843635, EPI_ISL_843636, EPI_ISL_843637, EPI_ISL_843638, EPI_ISL_843639, EPI_ISL_843640, EPI_ISL_843641, EPI_ISL_843642, EPI_ISL_843643, EPI_ISL_843644, EPI_ISL_843645, EPI_ISL_843646, EPI_ISL_843647, EPI_ISL_843648, EPI_ISL_843649, EPI_ISL_843650, EPI_ISL_843651, EPI_ISL_843652, EPI_ISL_843653, EPI_ISL_843654, EPI_ISL_843655, EPI_ISL_843656, EPI_ISL_843657, EPI_ISL_843658, EPI_ISL_843659, EPI_ISL_843660, EPI_ISL_843661, EPI_ISL_843662, EPI_ISL_843663, EPI_ISL_843664, EPI_ISL_843665, EPI_ISL_843666, EPI_ISL_843667, EPI_ISL_843668, EPI_ISL_843669, EPI_ISL_843670, EPI_ISL_843671, EPI_ISL_843674, EPI_ISL_843675, EPI_ISL_843676, EPI_ISL_843677, EPI_ISL_843678, EPI_ISL_843679, EPI_ISL_843680, EPI_ISL_843681, EPI_ISL_843682, EPI_ISL_843683, EPI_ISL_843684, EPI_ISL_843685, EPI_ISL_843686, EPI_ISL_843687, EPI_ISL_843688, EPI_ISL_843689, EPI_ISL_843690, EPI_ISL_843691, EPI_ISL_843692, EPI_ISL_843693, EPI_ISL_843694, EPI_ISL_843695, EPI_ISL_843696, EPI_ISL_843697, EPI_ISL_843698, EPI_ISL_843699, EPI_ISL_843701, EPI_ISL_843702, EPI_ISL_843703, EPI_ISL_843704, EPI_ISL_843705, EPI_ISL_843706, EPI_ISL_843707, EPI_ISL_843708, EPI_ISL_843709, EPI_ISL_843710, EPI_ISL_843711, EPI_ISL_843712, EPI_ISL_843713, EPI_ISL_843714, EPI_ISL_843715, EPI_ISL_843716, EPI_ISL_843717, EPI_ISL_843718, EPI_ISL_843719, EPI_ISL_843720, EPI_ISL_843721, EPI_ISL_843722, EPI_ISL_843723, EPI_ISL_843724, EPI_ISL_843725, EPI_ISL_843726, EPI_ISL_843727, EPI_ISL_843728, EPI_ISL_843729, EPI_ISL_843730, EPI_ISL_843731, EPI_ISL_843732, EPI_ISL_843733, EPI_ISL_843734, EPI_ISL_843735, EPI_ISL_843736, EPI_ISL_843737, EPI_ISL_843738, EPI_ISL_843739, EPI_ISL_843740, EPI_ISL_843741, EPI_ISL_843742, EPI_ISL_843743, EPI_ISL_843744, EPI_ISL_843745, EPI_ISL_843746, EPI_ISL_843747, EPI_ISL_843748, EPI_ISL_843749, EPI_ISL_843750, EPI_ISL_843751, EPI_ISL_843752, EPI_ISL_843753, EPI_ISL_843754, EPI_ISL_843755, EPI_ISL_843756, EPI_ISL_843757, EPI_ISL_843758, EPI_ISL_843759, EPI_ISL_843760, EPI_ISL_843761, EPI_ISL_843762, EPI_ISL_843763, EPI_ISL_843764, EPI_ISL_843765, EPI_ISL_843766, EPI_ISL_843767, EPI_ISL_843768, EPI_ISL_843769, EPI_ISL_843770, EPI_ISL_843771, EPI_ISL_843772, EPI_ISL_843773, EPI_ISL_843774, EPI_ISL_843775, EPI_ISL_843776, EPI_ISL_843777, EPI_ISL_843778, EPI_ISL_843779, EPI_ISL_843780, EPI_ISL_843781, EPI_ISL_843782, EPI_ISL_843783, EPI_ISL_843784, EPI_ISL_843785, EPI_ISL_843786, EPI_ISL_843787, EPI_ISL_843788, EPI_ISL_843790, EPI_ISL_843791, EPI_ISL_843792, EPI_ISL_843793, EPI_ISL_843794, EPI_ISL_843795, EPI_ISL_843796, EPI_ISL_843797, EPI_ISL_843798, EPI_ISL_843799, EPI_ISL_843800, EPI_ISL_843801, EPI_ISL_843802, EPI_ISL_843804, EPI_ISL_843805, EPI_ISL_843806, EPI_ISL_843807, EPI_ISL_843808, EPI_ISL_843809, EPI_ISL_843810, EPI_ISL_843811, EPI_ISL_843812, EPI_ISL_843813, EPI_ISL_843814, EPI_ISL_843815, EPI_ISL_843816, EPI_ISL_843817, EPI_ISL_843818, EPI_ISL_843819, EPI_ISL_843820, | | | |

| | | | |
|---|---|---|---|
| EPI_ISL_843821, EPI_ISL_843822, EPI_ISL_843823, EPI_ISL_843824, EPI_ISL_843825, EPI_ISL_843826, EPI_ISL_843827, EPI_ISL_843828, EPI_ISL_843829, EPI_ISL_843830, EPI_ISL_843831, EPI_ISL_843832, EPI_ISL_843833, EPI_ISL_843834, EPI_ISL_843835, EPI_ISL_843836, EPI_ISL_843837, EPI_ISL_843838, EPI_ISL_843839, EPI_ISL_843840, EPI_ISL_843841, EPI_ISL_843842, EPI_ISL_843843, EPI_ISL_843844, EPI_ISL_843845, EPI_ISL_843846, EPI_ISL_843847, EPI_ISL_843848, EPI_ISL_843849, EPI_ISL_843850, EPI_ISL_843851, EPI_ISL_843852, EPI_ISL_843853, EPI_ISL_843854, EPI_ISL_843855, EPI_ISL_843856, EPI_ISL_843857, EPI_ISL_843858, EPI_ISL_843859, EPI_ISL_843860, EPI_ISL_843861, EPI_ISL_843862, EPI_ISL_843863, EPI_ISL_843864, EPI_ISL_843865, EPI_ISL_843866, EPI_ISL_843867, EPI_ISL_843868, EPI_ISL_843869, EPI_ISL_843870, EPI_ISL_843871, EPI_ISL_843872, EPI_ISL_843873, EPI_ISL_843874, EPI_ISL_843875, EPI_ISL_843876, EPI_ISL_843877, EPI_ISL_843878, EPI_ISL_843879, EPI_ISL_843880, EPI_ISL_843881, EPI_ISL_843882, EPI_ISL_843883, EPI_ISL_843884, EPI_ISL_843885, EPI_ISL_843886, EPI_ISL_843887, EPI_ISL_843888, EPI_ISL_843889, EPI_ISL_843890, EPI_ISL_843891, EPI_ISL_843892, EPI_ISL_843893, EPI_ISL_843894, EPI_ISL_843895, EPI_ISL_843896, EPI_ISL_843897, EPI_ISL_843898, EPI_ISL_843899, EPI_ISL_843900, EPI_ISL_843901, EPI_ISL_843902, EPI_ISL_843903, EPI_ISL_843904, EPI_ISL_843905, EPI_ISL_843906, EPI_ISL_843907, EPI_ISL_843908, EPI_ISL_843909, EPI_ISL_843910, EPI_ISL_843911, EPI_ISL_843913, EPI_ISL_843914, EPI_ISL_843915, EPI_ISL_843916, EPI_ISL_843917, EPI_ISL_843918, EPI_ISL_843919, EPI_ISL_843920, EPI_ISL_843921, EPI_ISL_843922, EPI_ISL_843923, EPI_ISL_843924, EPI_ISL_843926, EPI_ISL_843927, EPI_ISL_843928, EPI_ISL_843929, EPI_ISL_843930, EPI_ISL_843931, EPI_ISL_843932, EPI_ISL_843933, EPI_ISL_843934, EPI_ISL_843935, EPI_ISL_843936, EPI_ISL_843937, EPI_ISL_843938, EPI_ISL_843939, EPI_ISL_843940, EPI_ISL_843941, EPI_ISL_843942, EPI_ISL_843943, EPI_ISL_843944, EPI_ISL_843945, EPI_ISL_843946, EPI_ISL_843947, EPI_ISL_843948, EPI_ISL_843949, EPI_ISL_843950, EPI_ISL_843951, EPI_ISL_843952, EPI_ISL_843953, EPI_ISL_843954, EPI_ISL_843955, EPI_ISL_843956, EPI_ISL_843957, EPI_ISL_843958, EPI_ISL_843959, EPI_ISL_843960, EPI_ISL_843961, EPI_ISL_843962, EPI_ISL_843963, EPI_ISL_843964, EPI_ISL_843965, EPI_ISL_843966, EPI_ISL_843967, EPI_ISL_843968, EPI_ISL_843969, EPI_ISL_843970, EPI_ISL_843971, EPI_ISL_843972, EPI_ISL_843974, EPI_ISL_843975, EPI_ISL_843976, EPI_ISL_843977, EPI_ISL_843978, EPI_ISL_843979, EPI_ISL_843980, EPI_ISL_843981, EPI_ISL_843982, EPI_ISL_843983, EPI_ISL_843984, EPI_ISL_843985, EPI_ISL_843986, EPI_ISL_843987, EPI_ISL_843988, EPI_ISL_843989, EPI_ISL_843990, EPI_ISL_843991, EPI_ISL_843992, EPI_ISL_843993, EPI_ISL_843994, EPI_ISL_843995, EPI_ISL_843996, EPI_ISL_843997, EPI_ISL_843998, EPI_ISL_843999, EPI_ISL_844001, EPI_ISL_844002, EPI_ISL_844003, EPI_ISL_844004, EPI_ISL_844005, EPI_ISL_844006, EPI_ISL_844007, EPI_ISL_844009, EPI_ISL_844010, EPI_ISL_844011, EPI_ISL_844012, EPI_ISL_844013, EPI_ISL_844014, EPI_ISL_844016, EPI_ISL_844017, EPI_ISL_844018, EPI_ISL_844019, EPI_ISL_844020, EPI_ISL_844021, EPI_ISL_844022, EPI_ISL_844023, EPI_ISL_844024, EPI_ISL_844025, EPI_ISL_844027, EPI_ISL_844028, EPI_ISL_844029, EPI_ISL_844030, EPI_ISL_844031, EPI_ISL_844032, EPI_ISL_844033, EPI_ISL_844034, EPI_ISL_844035, EPI_ISL_844036, EPI_ISL_844037, EPI_ISL_844038, EPI_ISL_844039, EPI_ISL_844040, EPI_ISL_844041, EPI_ISL_844042, EPI_ISL_844043, EPI_ISL_844044, EPI_ISL_844045, EPI_ISL_844046, EPI_ISL_844047, EPI_ISL_844048, EPI_ISL_844049, EPI_ISL_844050, EPI_ISL_844051, EPI_ISL_844052, EPI_ISL_844053, EPI_ISL_844054, EPI_ISL_844055, EPI_ISL_844056, EPI_ISL_844057, EPI_ISL_844058, EPI_ISL_844059, EPI_ISL_844060, EPI_ISL_844061, EPI_ISL_844062, EPI_ISL_844063, EPI_ISL_844064, EPI_ISL_844065, EPI_ISL_844066, EPI_ISL_844068, EPI_ISL_844069, EPI_ISL_844070, EPI_ISL_844071, EPI_ISL_844072, EPI_ISL_844073, EPI_ISL_844074, EPI_ISL_844075, EPI_ISL_844076, EPI_ISL_844077, EPI_ISL_844078, EPI_ISL_844079, EPI_ISL_844080, EPI_ISL_844081, EPI_ISL_844082, EPI_ISL_844083, EPI_ISL_844084, EPI_ISL_844085, EPI_ISL_844086, EPI_ISL_844087, EPI_ISL_844088, EPI_ISL_844089, EPI_ISL_844090, EPI_ISL_844091, EPI_ISL_844092, EPI_ISL_844093, EPI_ISL_844094, EPI_ISL_844095, EPI_ISL_844096, EPI_ISL_844097, EPI_ISL_844098, EPI_ISL_844099, EPI_ISL_844100, EPI_ISL_844101, EPI_ISL_844102, EPI_ISL_844103, EPI_ISL_844104, EPI_ISL_844105, EPI_ISL_844106, EPI_ISL_844107, EPI_ISL_844109, EPI_ISL_844110, EPI_ISL_844111, EPI_ISL_844112, EPI_ISL_844114, EPI_ISL_844115, EPI_ISL_844116, EPI_ISL_844117, EPI_ISL_844118, EPI_ISL_844119, EPI_ISL_844121, EPI_ISL_844122, EPI_ISL_844123, EPI_ISL_844124, EPI_ISL_844125, EPI_ISL_844126, EPI_ISL_844127, EPI_ISL_844128, EPI_ISL_844130, EPI_ISL_844131, EPI_ISL_844132, EPI_ISL_844133, EPI_ISL_844134, EPI_ISL_844135, EPI_ISL_844136, EPI_ISL_844137, EPI_ISL_844138, EPI_ISL_844139, EPI_ISL_844140, EPI_ISL_844141, EPI_ISL_844142, EPI_ISL_844143, EPI_ISL_844144, EPI_ISL_844190, EPI_ISL_844196 | | | |
| see above | Department of Virus and Microbiological Special Diagnostics, Statens Serum Institut, Copenhagen, Denmark | Albertsen Lab, Department of Chemistry and Bioscience, Aalborg University, Denmark | Danish Covid-19 Genome Consortium |
| EPI_ISL_845592, EPI_ISL_845594, EPI_ISL_845596, EPI_ISL_845604 | KU Leuven, Rega Institute, Clinical and Epidemiological Virology | KU Leuven, Rega Institute, Clinical and Epidemiological Virology | Tony Wawina-Bokalanga, Bert Vanmechelen, Joan Marti-Carerras, Piet Maes |
| EPI_ISL_845620 | compensar calle 63 | Instituto Nacional de Salud - Dirección de Investigación en Salud Pública | Katherine Laiton-Donato, Diego A. Álvarez-Díaz, Carlos Franco-Muñoz, Mauricio Pacheco-Montealegre, Maria T. Herrera-Sepúlveda, Jonathan Reales, Sheryll Corchuelo, Julian Naizaque, Gerardo Santamaría, Paola Muñoz-Laiton, Diego Andrés Prada, Magdalena Wiesner, Martha Lucia Ospina Martinez, Marcela Mercado-Reyes |
| EPI_ISL_845622 | Dirección de Sanidad Ejército | Instituto Nacional de Salud - Dirección de Investigación en Salud Pública | Katherine Laiton-Donato, Diego A. Álvarez-Díaz, Carlos Franco-Muñoz, Mauricio Pacheco-Montealegre, Maria T. Herrera-Sepúlveda, Jonathan Reales, Sheryll Corchuelo, Julian Naizaque, Gerardo Santamaría, Paola Muñoz-Laiton, Diego Andrés Prada, Magdalena Wiesner, Martha Lucia Ospina Martinez, Marcela Mercado-Reyes |
| EPI_ISL_845630, EPI_ISL_845631, EPI_ISL_845632, EPI_ISL_845633 | IDIME S.A | Instituto Nacional de Salud - Dirección de Investigación en Salud Pública | Katherine Laiton-Donato, Diego A. Álvarez-Díaz, Carlos Franco-Muñoz, Mauricio Pacheco-Montealegre, Maria T. Herrera-Sepúlveda, Jonathan Reales, Sheryll Corchuelo, Julian Naizaque, Gerardo Santamaría, Paola Muñoz-Laiton, Diego Andrés Prada, Magdalena Wiesner, Martha Lucia Ospina Martinez, Marcela Mercado-Reyes |
| EPI_ISL_845650 | Laboratorio de Salud Pública - Secretaría Distrital de Salud | Instituto Nacional de Salud - Dirección de Investigación en Salud Pública | Katherine Laiton-Donato, Diego A. Álvarez-Díaz, Carlos Franco-Muñoz, Mauricio Pacheco-Montealegre, Maria T. Herrera-Sepúlveda, Jonathan Reales, Sheryll Corchuelo, Julian Naizaque, Gerardo Santamaría, Paola Muñoz-Laiton, Diego Andrés Prada, Magdalena Wiesner, Martha Lucia Ospina Martinez, Marcela Mercado-Reyes |
| EPI_ISL_845850, EPI_ISL_845851, EPI_ISL_845852, EPI_ISL_845854, EPI_ISL_845855, EPI_ISL_845856, EPI_ISL_845858, EPI_ISL_845859, EPI_ISL_845860 | TGen North | TGen North | Jolene Bowers, Megan Folkerts, Chris French, Hayley Yaglom, Ashlyn Pfeiffer, Darrin Lemmer, Dave Engelthaler, The Arizona COVID Genomics Union (ACGU) |
| EPI_ISL_845874, EPI_ISL_845875, EPI_ISL_845876, EPI_ISL_845883, EPI_ISL_845884, EPI_ISL_845889 | BBMP Urban PHC | Department of Neurovirology, National Institute of Mental Health and Neurosciences (NIMHANS) | Chitra Pattabiraman, Pramada Prasad, Risha Rasheed, Darshan Sreenivas, Nakka Vijay Kiran Reddy, Anita S Desai, V Ravi |
| EPI_ISL_846818, EPI_ISL_846819, EPI_ISL_846851, EPI_ISL_846852, EPI_ISL_846861, EPI_ISL_846866, EPI_ISL_846894, EPI_ISL_846895, EPI_ISL_846913, EPI_ISL_847168, EPI_ISL_847169, EPI_ISL_847170, EPI_ISL_847171, EPI_ISL_847172, EPI_ISL_847173, EPI_ISL_847174, EPI_ISL_847175, EPI_ISL_847176, EPI_ISL_847320, EPI_ISL_847348, EPI_ISL_847349, EPI_ISL_847368, EPI_ISL_847369, EPI_ISL_847370, EPI_ISL_847510, EPI_ISL_847511, EPI_ISL_847512, EPI_ISL_847513 | | | |
| see above | Department of Virus and Microbiological Special Diagnostics, Statens Serum Institut, Copenhagen, Denmark | Albertsen Lab, Department of Chemistry and Bioscience, Aalborg University, Denmark | Danish Covid-19 Genome Consortium |
| EPI_ISL_847532 | California Department of Public Health | Chiu Laboratory, University of California, San Francisco | Charles Chiu, Xianding (Wayne) Deng, Candace Wang, Brian Bushnell, Scot Federman, Jill Hacker, Debra Wadford |
| EPI_ISL_847536 | Chiu Laboratory, University of California, San Francisco | Chiu Laboratory, University of California, San Francisco | Charles Chiu, Xianding (Wayne) Deng, Candace Wang, Brian Bushnell, Scot Federman, Jill Hacker, Debra Wadford |
| EPI_ISL_847572, EPI_ISL_847574, EPI_ISL_847601, EPI_ISL_847602, EPI_ISL_847603, EPI_ISL_847604, EPI_ISL_847605, EPI_ISL_847606, EPI_ISL_847607 | California Department of Public Health | Chiu Laboratory, University of California, San Francisco | Charles Chiu, Xianding (Wayne) Deng, Candace Wang, Brian Bushnell, Scot Federman, Jill Hacker, Debra Wadford |
| EPI_ISL_847736, EPI_ISL_847741 | Chiu Laboratory, University of California, San Francisco | Chiu Laboratory, University of California, San Francisco | Charles Chiu, Xianding (Wayne) Deng, Candace Wang, Brian Bushnell, Scot Federman, Jill Hacker, Debra Wadford |
| EPI_ISL_847766, EPI_ISL_847767, EPI_ISL_847768, EPI_ISL_847769, EPI_ISL_847770, EPI_ISL_847771 | California Department of Public Health | Chiu Laboratory, University of California, San Francisco | Charles Chiu, Xianding (Wayne) Deng, Candace Wang, Brian Bushnell, Scot Federman, Jill Hacker, Debra Wadford |
| EPI_ISL_847830 | Tempus | Grubaugh Lab - Yale School of Public Health | Tara Alpert, Joseph Fauver, Anderson Brito, Mallery Breban, Anne Wyllie, Chantal Vogels, Mary Petrone, Chaney Kalinich, Isabel Ott, Nathan Grubaugh |
| EPI_ISL_847845, EPI_ISL_847848, EPI_ISL_847849, EPI_ISL_847890, EPI_ISL_847891, EPI_ISL_847892, EPI_ISL_847893, EPI_ISL_847894, EPI_ISL_847895, EPI_ISL_847896, EPI_ISL_847897, EPI_ISL_847898, EPI_ISL_847899, EPI_ISL_847900, EPI_ISL_847901, EPI_ISL_847925, EPI_ISL_847960, EPI_ISL_847961, EPI_ISL_847962, EPI_ISL_847963, EPI_ISL_847964, EPI_ISL_847965, EPI_ISL_847966, EPI_ISL_847967 | | | |
| see above | University Hospitals of Geneva, Laboratory of Virology | HUG, Laboratory of Virology and the Health2030 Genome Center | Samuel Cordey, Ana Rita Goncalves, Laurent Kaiser, Lorenzo Cerutti, Henri Pegeot, Melyssa Elies, Keith Harshman, Ioannis Xenarios, Emmanouil Dermitzakis |
| EPI_ISL_847989, EPI_ISL_847993, EPI_ISL_848000, EPI_ISL_848002, EPI_ISL_848012 | Michigan Department of Health and Human Services, Bureau of Laboratories | Michigan Department of Health and Human Services, Bureau of Laboratories | Blankenship HM, Riner D, Soehnlen MK |
| EPI_ISL_848063, EPI_ISL_848064 | CHU Purpan - Laboratoire de Virologie - Institut Fédératif de Biologie | CHU Purpan - Laboratoire de Virologie - Institut Fédératif de Biologie | Latour J., Ranger N., Dubois M., Carcenac R., Harter A., Boyer P., Tremeaux P., Izopet J. |
| EPI_ISL_849783, EPI_ISL_849787, EPI_ISL_849793, EPI_ISL_849881, EPI_ISL_849889, EPI_ISL_849890, EPI_ISL_849891, EPI_ISL_849892, EPI_ISL_849893, EPI_ISL_849894, EPI_ISL_849896, EPI_ISL_849897, EPI_ISL_849898, EPI_ISL_849899 | | | |
| see above | Utah Public Health Laboratory | Utah Public Health Laboratory | Erin L. Young, Kelly F. Oakeson, Tara Gallagher |
| EPI_ISL_849926 | UCSF Clinical Microbiology Laboratory | Chan-Zuckerberg Biohub | CZB Cliahub Consortium |
| EPI_ISL_850657, EPI_ISL_850662 | Division of Emerging Infectious Diseases, Bureau of Infectious Diseases Diagnosis Control, Korea Disease Control and Prevention Agency | Division of Emerging Infectious Diseases, Bureau of Infectious Diseases Diagnosis Control, Korea Disease Control and Prevention Agency | Ae Kyung Park, Il-Hwan Kim, Heui Man Kim, Jeong-Min Kim, Namjoo Lee, Chaeyoung Lee, Sang Hee Woo, Eun-Jin Kim |
| EPI_ISL_852961 | Hospital General Universitario Gregorio Marañón | SeqCOVID-SPAIN consortium/IBV(CSIC) | Darío García de Viedma, Laura Pérez-Lago, Pedro J Sola-Campoy, Sergio Buenestado-Serrano, Marta Herranz, Victor Manuel de la Cueva, Julia Suárez, Pilar Catalán, Patricia Muñoz and SeqCOVID-SPAIN consortium |

| | | | |
|---|---|---|---|
| EPI_ISL_853373, EPI_ISL_853374 | UPMC Clinical Microbiology Laboratory | Microbial Genome Sequencing Center; Microbial Genomic Epidemiology Laboratory | Mustapha M. Mustapha, Jane W. Marsh, Dan Snyder, Marissa P. Griffith, Stephanie L. Mitchell, Vatsala R. Srinivasa, Kady D. Waggle, Chinelo Ezeonwuku, Vaughn S. Cooper, Lee H. Harrison |
| EPI_ISL_853722, EPI_ISL_853743, EPI_ISL_853745, EPI_ISL_853956, EPI_ISL_854226 | Department of Microbiology, University Innsbruck | Bergthaler laboratory, CeMM Research Center for Molecular Medicine of the Austrian Academy of Sciences | Lukas Endler, Alexandra Popa, Benedikt Agerer, Jakob-Wendelin Genger, Alexander Lercher, Anna Schedl, Thomas Penz, Michael Schuster, Jan Laine, Martin Senekowitsch, Christoph Bock, Andreas Bergthaler |
| EPI_ISL_854416, EPI_ISL_854417, EPI_ISL_854418, EPI_ISL_854419, EPI_ISL_854420 | MONTEFIORE MEDICAL CENTER LABORATORIES | Wadsworth Center, New York State Department of Health | Kirsten St. George, Daryl M. Lamson, Alexis Russel, Matthew Shudt, Melissa A Leisner, Jonathan Plitnick, Navjot Singh, John Kelly, Erasmus Schneider, Erica Lasek-Nesselquist |
| EPI_ISL_854421 | SARATOGA HOSPITAL LABORATORY | Wadsworth Center, New York State Department of Health | Kirsten St. George, Daryl M. Lamson, Alexis Russel, Matthew Shudt, Melissa A Leisner, Jonathan Plitnick, Navjot Singh, John Kelly, Erasmus Schneider, Erica Lasek-Nesselquist |
| EPI_ISL_854422, EPI_ISL_854423, EPI_ISL_854424, EPI_ISL_854425, EPI_ISL_854426, EPI_ISL_854453, EPI_ISL_854456, EPI_ISL_854459 | MONTEFIORE MEDICAL CENTER LABORATORIES | Wadsworth Center, New York State Department of Health | Kirsten St. George, Daryl M. Lamson, Alexis Russel, Matthew Shudt, Melissa A Leisner, Jonathan Plitnick, Navjot Singh, John Kelly, Erasmus Schneider, Erica Lasek-Nesselquist |
| EPI_ISL_855016, EPI_ISL_855019, EPI_ISL_855023, EPI_ISL_855027, EPI_ISL_855030, EPI_ISL_855031, EPI_ISL_855033, EPI_ISL_855034, EPI_ISL_855037, EPI_ISL_855038, EPI_ISL_855039, EPI_ISL_855041, EPI_ISL_855046, EPI_ISL_855047, EPI_ISL_855049, EPI_ISL_855050, EPI_ISL_855052, EPI_ISL_855053, EPI_ISL_855054, EPI_ISL_855057, EPI_ISL_855059, EPI_ISL_855060, EPI_ISL_855061, EPI_ISL_855062, EPI_ISL_855063 | | | |
| see above | Quest Diagnostics | Quest Diagnostics | Rosenthal,S.H., Gerasimova,A., Kagan,R.M., Anderson, B., Hua, M., Liu Y., Bernstein, L.E., Livingston, K.E., Perez, A., Shalhout, D.F., Shlyakhter, I.A., Owen, R., Tanpaiboon, P., Lacbawan, F. |
| EPI_ISL_855401 | Chiu Laboratory, University of California, San Francisco | Chiu Laboratory, University of California, San Francisco | Charles Chiu, Xianding (Wayne) Deng, Candace Wang, Brian Bushnell, Scot Federman, Jill Hacker, Debra Wadford |
| EPI_ISL_855910, EPI_ISL_855912 | Lab voor klinische biologie | Onderzoeksgroep Virologie | Laurens Lambrechts, Nick Vereecke, Marthe Pauwels, Bruno Verhasselt, Linos Vandekerckhove, Hans Nauwynck, Sebastiaan Theuns |
| EPI_ISL_856612, EPI_ISL_856613, EPI_ISL_856614, EPI_ISL_856615, EPI_ISL_856616, EPI_ISL_856617, EPI_ISL_856618, EPI_ISL_856619, EPI_ISL_856620, EPI_ISL_856621, EPI_ISL_856622, EPI_ISL_856623, EPI_ISL_856624, EPI_ISL_856625, EPI_ISL_856626, EPI_ISL_856627, EPI_ISL_856628, EPI_ISL_856629, EPI_ISL_856630, EPI_ISL_856631, EPI_ISL_856632, EPI_ISL_856633, EPI_ISL_856634, EPI_ISL_856635, EPI_ISL_856636, EPI_ISL_856637, EPI_ISL_856638, EPI_ISL_856639, EPI_ISL_856640, EPI_ISL_856641, EPI_ISL_856642, EPI_ISL_856643, EPI_ISL_856644, EPI_ISL_856645, EPI_ISL_856646 | | | |
| see above | Department of Virus and Microbiological Special Diagnostics, Statens Serum Institut, Copenhagen, Denmark | Aalborg University | Danish Covid-19 Genome Consortium |
| EPI_ISL_856973, EPI_ISL_857040 | Platform BIS UZA/UAntwerpen, University Hospital Antwerp, Edegem, Belgium | UAntwerp, Laboratory of Medical Microbiology, Campus Drie Eiken S6.26, Universiteitsplein 1, 2610, Wilrijk, Antwerp, Belgium | Basil Britto Xavier, Jasmine Coppens, Christine Lammens, Veerle Matheeussen, Herman Goossens |
| EPI_ISL_857047 | Colorado Department of Public Health and Environment | Colorado Department of Puplic Health and Environment | Laura Bankers, Molly C. Hetherington-Rauth, Diana Ir, Shannon Ely, Shannon R. Matzinger, Sarah Elizabeth Totten, Emily A. Travanty |
| EPI_ISL_857187, EPI_ISL_857192 | DOHMH PHL | New York City Public Health Laboratory | Jade Wang, et al. |
| EPI_ISL_857198 | Department of Homeless Services | New York City Public Health Laboratory | Jade Wang, et al. |
| EPI_ISL_857279, EPI_ISL_857280 | DOHMH Fort Greene | New York City Public Health Laboratory | Jade Wang, et al. |
| EPI_ISL_857281 | DOHMH Riverside | New York City Public Health Laboratory | Jade Wang, et al. |
| EPI_ISL_857282, EPI_ISL_857283 | DOHMH Morrisania | New York City Public Health Laboratory | Jade Wang, et al. |
| EPI_ISL_857284 | DOHMH PHL | New York City Public Health Laboratory | Jade Wang, et al. |
| EPI_ISL_857285, EPI_ISL_857286 | DOHMH Central Harlem | New York City Public Health Laboratory | Jade Wang, et al. |
| EPI_ISL_857287, EPI_ISL_857288 | Department of Homeless Services | New York City Public Health Laboratory | Jade Wang, et al. |
| EPI_ISL_857289 | OCME Office Of Chief Medical Examiner | New York City Public Health Laboratory | Jade Wang, et al. |
| EPI_ISL_857290 | Department of Homeless Services | New York City Public Health Laboratory | Jade Wang, et al. |
| EPI_ISL_857515 | Swiss National Reference Centre for Influenza | Swiss National Reference Centre for Influenza | Ana Rita Goncalves,Samuel Cordey, Laurent Kaiser, Lorenzo Cerutti, Henri Pegeot, Melyssa Elies, Keith Harshman, Ioannis Xenarios, Emmanouil Dermitzakis |
| EPI_ISL_858590, EPI_ISL_858597, EPI_ISL_858619 | Lighthouse Lab in Glasgow | Wellcome Sanger Institute for the COVID-19 Genomics UK (COG-UK) Consortium | Harper VanSteenhouse, Yumi Kasai, David Gray, Carol Clugston, Anna Dominiczak and Alex Alderton, Roberto Amato, Sonia Goncalves, Ewan Harrison, David K. Jackson, Ian Johnston, Dominic Kwiatkowski, Cordelia Langford, John Sillitoe on behalf of the Wellcome Sanger Institute COVID-19 Surveillance Team |
| EPI_ISL_858701, EPI_ISL_858702, EPI_ISL_858704, EPI_ISL_858706, EPI_ISL_858707, EPI_ISL_858708, EPI_ISL_858709, EPI_ISL_858713, EPI_ISL_858714, EPI_ISL_858719, EPI_ISL_858720, EPI_ISL_858725, EPI_ISL_858732, EPI_ISL_858737, EPI_ISL_858739, EPI_ISL_858748, EPI_ISL_858752, EPI_ISL_858753 | | | |
| see above | Lighthouse Lab in Alderley Park | Wellcome Sanger Institute for the COVID-19 Genomics UK (COG-UK) Consortium | Jacquelyn Wynn, Mairead Hyland, The Lighthouse Lab in Alderley Park and Alex Alderton, Roberto Amato, Sonia Goncalves, Ewan Harrison, David K. Jackson, Ian Johnston, Dominic Kwiatkowski, Cordelia Langford, John Sillitoe on behalf of the Wellcome Sanger Institute COVID-19 Surveillance Team |
| EPI_ISL_858778 | Lighthouse Lab in Glasgow | Wellcome Sanger Institute for the COVID-19 Genomics UK (COG-UK) Consortium | Harper VanSteenhouse, Yumi Kasai, David Gray, Carol Clugston, Anna Dominiczak and Alex Alderton, Roberto Amato, Sonia Goncalves, Ewan Harrison, David K. Jackson, Ian Johnston, Dominic Kwiatkowski, Cordelia Langford, John Sillitoe on behalf of the Wellcome Sanger Institute COVID-19 Surveillance Team |
| EPI_ISL_858779, EPI_ISL_858780 | Lighthouse Lab in Alderley Park | Wellcome Sanger Institute for the COVID-19 Genomics UK (COG-UK) Consortium | Jacquelyn Wynn, Mairead Hyland, The Lighthouse Lab in Alderley Park and Alex Alderton, Roberto Amato, Sonia Goncalves, Ewan Harrison, David K. Jackson, Ian Johnston, Dominic Kwiatkowski, Cordelia Langford, John Sillitoe on behalf of the Wellcome Sanger Institute COVID-19 Surveillance Team |
| EPI_ISL_858782 | Lighthouse Lab in Glasgow | Wellcome Sanger Institute for the COVID-19 Genomics UK (COG-UK) Consortium | Harper VanSteenhouse, Yumi Kasai, David Gray, Carol Clugston, Anna Dominiczak and Alex Alderton, Roberto Amato, Sonia Goncalves, Ewan Harrison, David K. Jackson, Ian Johnston, Dominic Kwiatkowski, Cordelia Langford, John Sillitoe on behalf of the Wellcome Sanger Institute COVID-19 Surveillance Team |
| EPI_ISL_858784, EPI_ISL_858785, EPI_ISL_858786, EPI_ISL_858788, EPI_ISL_858789, EPI_ISL_858790, EPI_ISL_858791 | Lighthouse Lab in Alderley Park | Wellcome Sanger Institute for the COVID-19 Genomics UK (COG-UK) Consortium | Jacquelyn Wynn, Mairead Hyland, The Lighthouse Lab in Alderley Park and Alex Alderton, Roberto Amato, Sonia Goncalves, Ewan Harrison, David K. Jackson, Ian Johnston, Dominic Kwiatkowski, Cordelia Langford, John Sillitoe on behalf of the Wellcome Sanger Institute COVID-19 Surveillance Team |
| EPI_ISL_858793, EPI_ISL_858794, EPI_ISL_858796, EPI_ISL_858797, EPI_ISL_858801, EPI_ISL_858802, EPI_ISL_858806 | Lighthouse Lab in Glasgow | Wellcome Sanger Institute for the COVID-19 Genomics UK (COG-UK) Consortium | Harper VanSteenhouse, Yumi Kasai, David Gray, Carol Clugston, Anna Dominiczak and Alex Alderton, Roberto Amato, Sonia Goncalves, Ewan Harrison, David K. Jackson, Ian Johnston, Dominic Kwiatkowski, Cordelia Langford, John Sillitoe on behalf of the Wellcome Sanger Institute COVID-19 Surveillance Team |
| EPI_ISL_858809 | Lighthouse Lab in Alderley Park | Wellcome Sanger Institute for the COVID-19 Genomics UK (COG-UK) Consortium | Jacquelyn Wynn, Mairead Hyland, The Lighthouse Lab in Alderley Park and Alex Alderton, Roberto Amato, Sonia Goncalves, Ewan Harrison, David K. Jackson, Ian Johnston, Dominic Kwiatkowski, Cordelia Langford, John Sillitoe on behalf of the Wellcome Sanger Institute COVID-19 Surveillance Team |
| EPI_ISL_858810 | Lighthouse Lab in Glasgow | Wellcome Sanger Institute for the COVID-19 Genomics UK (COG-UK) Consortium | Harper VanSteenhouse, Yumi Kasai, David Gray, Carol Clugston, Anna Dominiczak and Alex Alderton, Roberto Amato, Sonia Goncalves, Ewan Harrison, David K. Jackson, Ian Johnston, Dominic Kwiatkowski, Cordelia Langford, John Sillitoe on behalf of the Wellcome Sanger Institute COVID-19 Surveillance Team |
| EPI_ISL_858811, EPI_ISL_858813, EPI_ISL_858814 | Lighthouse Lab in Alderley Park | Wellcome Sanger Institute for the COVID-19 Genomics UK (COG-UK) Consortium | Jacquelyn Wynn, Mairead Hyland, The Lighthouse Lab in Alderley Park and Alex Alderton, Roberto Amato, Sonia Goncalves, Ewan Harrison, David K. Jackson, Ian Johnston, Dominic Kwiatkowski, Cordelia Langford, John Sillitoe on behalf of the Wellcome Sanger Institute COVID-19 Surveillance Team |
| EPI_ISL_858815 | Lighthouse Lab in Glasgow | Wellcome Sanger Institute for the COVID-19 Genomics UK | Harper VanSteenhouse, Yumi Kasai, David Gray, Carol Clugston, Anna Dominiczak and Alex Alderton, Roberto Amato, Sonia Goncalves, Ewan Harrison, |

| | | | |
|---|---|---|---|
| | | (COG-UK) Consortium | David K. Jackson, Ian Johnston, Dominic Kwiatkowski, Cordelia Langford, John Sillitoe on behalf of the Wellcome Sanger Institute COVID-19 Surveillance Team |
| EPI_ISL_858816, EPI_ISL_858817, EPI_ISL_858821, EPI_ISL_858822, EPI_ISL_858824, EPI_ISL_858825, EPI_ISL_858827, EPI_ISL_858828 | Lighthouse Lab in Alderley Park | Wellcome Sanger Institute for the COVID-19 Genomics UK (COG-UK) Consortium | Jacquelyn Wynn, Mairead Hyland, The Lighthouse Lab in Alderley Park and Alex Alderton, Roberto Amato, Sonia Goncalves, Ewan Harrison, David K. Jackson, Ian Johnston, Dominic Kwiatkowski, Cordelia Langford, John Sillitoe on behalf of the Wellcome Sanger Institute COVID-19 Surveillance Team |
| EPI_ISL_858830 | Lighthouse Lab in Glasgow | Wellcome Sanger Institute for the COVID-19 Genomics UK (COG-UK) Consortium | Harper VanSteenhouse, Yumi Kasai, David Gray, Carol Clugston, Anna Dominiczak and Alex Alderton, Roberto Amato, Sonia Goncalves, Ewan Harrison, David K. Jackson, Ian Johnston, Dominic Kwiatkowski, Cordelia Langford, John Sillitoe on behalf of the Wellcome Sanger Institute COVID-19 Surveillance Team |
| EPI_ISL_858831, EPI_ISL_858832, EPI_ISL_858834, EPI_ISL_858835 | Lighthouse Lab in Alderley Park | Wellcome Sanger Institute for the COVID-19 Genomics UK (COG-UK) Consortium | Jacquelyn Wynn, Mairead Hyland, The Lighthouse Lab in Alderley Park and Alex Alderton, Roberto Amato, Sonia Goncalves, Ewan Harrison, David K. Jackson, Ian Johnston, Dominic Kwiatkowski, Cordelia Langford, John Sillitoe on behalf of the Wellcome Sanger Institute COVID-19 Surveillance Team |
| EPI_ISL_858842, EPI_ISL_858843, EPI_ISL_858844 | Lighthouse Lab in Glasgow | Wellcome Sanger Institute for the COVID-19 Genomics UK (COG-UK) Consortium | Harper VanSteenhouse, Yumi Kasai, David Gray, Carol Clugston, Anna Dominiczak and Alex Alderton, Roberto Amato, Sonia Goncalves, Ewan Harrison, David K. Jackson, Ian Johnston, Dominic Kwiatkowski, Cordelia Langford, John Sillitoe on behalf of the Wellcome Sanger Institute COVID-19 Surveillance Team |
| EPI_ISL_858847, EPI_ISL_858851, EPI_ISL_858854, EPI_ISL_858856 | Lighthouse Lab in Alderley Park | Wellcome Sanger Institute for the COVID-19 Genomics UK (COG-UK) Consortium | Jacquelyn Wynn, Mairead Hyland, The Lighthouse Lab in Alderley Park and Alex Alderton, Roberto Amato, Sonia Goncalves, Ewan Harrison, David K. Jackson, Ian Johnston, Dominic Kwiatkowski, Cordelia Langford, John Sillitoe on behalf of the Wellcome Sanger Institute COVID-19 Surveillance Team |
| EPI_ISL_858859, EPI_ISL_858860, EPI_ISL_858863, EPI_ISL_858865, EPI_ISL_858867 | Lighthouse Lab in Glasgow | Wellcome Sanger Institute for the COVID-19 Genomics UK (COG-UK) Consortium | Harper VanSteenhouse, Yumi Kasai, David Gray, Carol Clugston, Anna Dominiczak and Alex Alderton, Roberto Amato, Sonia Goncalves, Ewan Harrison, David K. Jackson, Ian Johnston, Dominic Kwiatkowski, Cordelia Langford, John Sillitoe on behalf of the Wellcome Sanger Institute COVID-19 Surveillance Team |
| EPI_ISL_858873 | Lighthouse Lab in Alderley Park | Wellcome Sanger Institute for the COVID-19 Genomics UK (COG-UK) Consortium | Jacquelyn Wynn, Mairead Hyland, The Lighthouse Lab in Alderley Park and Alex Alderton, Roberto Amato, Sonia Goncalves, Ewan Harrison, David K. Jackson, Ian Johnston, Dominic Kwiatkowski, Cordelia Langford, John Sillitoe on behalf of the Wellcome Sanger Institute COVID-19 Surveillance Team |
| EPI_ISL_858874 | Lighthouse Lab in Glasgow | Wellcome Sanger Institute for the COVID-19 Genomics UK (COG-UK) Consortium | Harper VanSteenhouse, Yumi Kasai, David Gray, Carol Clugston, Anna Dominiczak and Alex Alderton, Roberto Amato, Sonia Goncalves, Ewan Harrison, David K. Jackson, Ian Johnston, Dominic Kwiatkowski, Cordelia Langford, John Sillitoe on behalf of the Wellcome Sanger Institute COVID-19 Surveillance Team |
| EPI_ISL_858879, EPI_ISL_858887, EPI_ISL_858891, EPI_ISL_858895 | Lighthouse Lab in Alderley Park | Wellcome Sanger Institute for the COVID-19 Genomics UK (COG-UK) Consortium | Jacquelyn Wynn, Mairead Hyland, The Lighthouse Lab in Alderley Park and Alex Alderton, Roberto Amato, Sonia Goncalves, Ewan Harrison, David K. Jackson, Ian Johnston, Dominic Kwiatkowski, Cordelia Langford, John Sillitoe on behalf of the Wellcome Sanger Institute COVID-19 Surveillance Team |
| EPI_ISL_860027, EPI_ISL_860042, EPI_ISL_860043, EPI_ISL_860048, EPI_ISL_860049, EPI_ISL_860051, EPI_ISL_860052, EPI_ISL_860053, EPI_ISL_860054, EPI_ISL_860055, EPI_ISL_860056, EPI_ISL_860057, EPI_ISL_860063, EPI_ISL_860064, EPI_ISL_860066, EPI_ISL_860067, EPI_ISL_860068, EPI_ISL_860069, EPI_ISL_860071, EPI_ISL_860073, EPI_ISL_860075, EPI_ISL_860076, EPI_ISL_860079, EPI_ISL_860080, EPI_ISL_860089 | | | |
| see above | BTC, Khalifa University | BTC, Khalifa University | Al Safar et al |
| EPI_ISL_860203, EPI_ISL_860204, EPI_ISL_860205 | Lab. Microbiologia e Virologia, Cotugno, A.O. dei Colli | Lab. Microbiologia e Virologia, Cotugno, A.O. dei Colli | Luigi Atripaldi, Claudia Tiberio, Anna Perfetti |
| EPI_ISL_860229 | Ostfold Hospital Trust - Kalnes, Centre for Laboratory Medicine, Section for gene technology and infection serology | Norwegian Institute of Public Health, Department of Virology | Kathrine Stene-Johansen, Kamilla Heddeland Instefjord, Hilde Elshaug, Atiya R Ali,Marie Paulsen Madsen, Rasmus Riis Kopperud, Hilde Vollan, Karoline Bragstad, Olav Hungnes |
| EPI_ISL_860231 | Vestfold Hospital, Toensberg Department of Microbiology | Norwegian Institute of Public Health, Department of Virology | Kathrine Stene-Johansen, Kamilla Heddeland Instefjord, Hilde Elshaug, Atiya R Ali,Marie Paulsen Madsen, Rasmus Riis Kopperud, Hilde Vollan, Karoline Bragstad, Olav Hungnes |
| EPI_ISL_860247 | University Hospitals of Geneva, Laboratory of Virology | HUG, Laboratory of Virology and Universitätsspital Basel | Samuel Cordey, Ana Rita Goncalves, Laurent Kaiser, Tim Roloff, Madlen Stange, Helena MB Seth-Smith, Alfredo Mari, Karoline Leuzinger, Julia Bielicki, Manuel Battegay, Hans Hirsch, Adrian Egli |
| EPI_ISL_860257 | Lab. Microbiologia e Virologia, Cotugno, A.O. dei Colli | Lab. Microbiologia e Virologia, Cotugno, A.O. dei Colli | Luigi Atripaldi, Claudia Tiberio, Anna Perfetti |
| EPI_ISL_860298, EPI_ISL_860301 | Unit 17: Influenza & Other Respiratory Viruses, German National Influenza Center | Project group Epidemiology of Highly Pathogenic Microorganisms, Robert Koch-Institute | Andreas Sachse, Grit Schubert, Essia Belarbi, Sébastien Calvignac-Spencer, Thorsten Wolff, Ralf Dürrwald, Djin-Ye Oh, Marianne Wedde |
| EPI_ISL_860619 | NHLS-IALCH | KRISP, KZn Research Innovation and Sequencing Platform | Giandhari J, Pillay S, Lessells R, Mdlalose K, York D, Khan S, Tegally H, Wilkinson E, de Oliveira T |
| EPI_ISL_860641 | Respiratory Virus Unit, National Infection Service, Public Health England | COVID-19 Genomics UK (COG-UK) Consortium | PHE Covid Sequencing Team |
| EPI_ISL_860942, EPI_ISL_860945, EPI_ISL_861101, EPI_ISL_861103 | Johns Hopkins Hospital Department of Pathology | Johns Hopkins Hospital Department of Pathology | C. Paul Morris, Chun Huai Luo, Adannaya Amadi, Nicholas Gallagher, Heba H. Mostafa |
| EPI_ISL_861119 | New York Presbyterian Hospital | Wadsworth Center, New York State Department of Health | Kirsten St. George, Daryl M. Lamson, Alexis Russel, Matthew Shudt, Melissa A Leisner, Jonathan Plitnick, Navjot Singh, John Kelly, Erasmus Schneider, Erica Lasek-Nesselquist |
| EPI_ISL_861175, EPI_ISL_861176, EPI_ISL_861179, EPI_ISL_861185, EPI_ISL_861187, EPI_ISL_861210, EPI_ISL_861211, EPI_ISL_861212, EPI_ISL_861213, EPI_ISL_861214, EPI_ISL_861215, EPI_ISL_861216, EPI_ISL_861217, EPI_ISL_861218, EPI_ISL_861219, EPI_ISL_861220, EPI_ISL_861221, EPI_ISL_861222, EPI_ISL_861223, EPI_ISL_861224, EPI_ISL_861225, EPI_ISL_861226, EPI_ISL_861227, EPI_ISL_861228, EPI_ISL_861229, EPI_ISL_861230, EPI_ISL_861231, EPI_ISL_861232 | | | |
| see above | BIO-REFERENCE LABORATORIES | Wadsworth Center, New York State Department of Health | Kirsten St. George, Daryl M. Lamson, Alexis Russel, Matthew Shudt, Melissa A Leisner, Jonathan Plitnick, Navjot Singh, John Kelly, Erasmus Schneider, Erica Lasek-Nesselquist |
| EPI_ISL_861243 | Ohio Department of Health Laboratory | Ohio Department of Health Laboratory | Holmes, Jennifer; Eric Brandt, Keoni Omura, Glen McGillivary, Caitlin McDonnell, Kirtana Ramadugu, Erica Leasure, Kelsey Florek, Heather Blankenship, Quanta Brown, and Tammy Bannerman |
| EPI_ISL_861245, EPI_ISL_861246, EPI_ISL_861247, EPI_ISL_861249, EPI_ISL_861253, EPI_ISL_861256, EPI_ISL_861261, EPI_ISL_861267, EPI_ISL_861289, EPI_ISL_861291, EPI_ISL_861293, EPI_ISL_861294, EPI_ISL_861299, EPI_ISL_861301, EPI_ISL_861302, EPI_ISL_861308, EPI_ISL_861309, EPI_ISL_861311, EPI_ISL_861313, EPI_ISL_861315, EPI_ISL_861318, EPI_ISL_861319, EPI_ISL_861320, EPI_ISL_861321, EPI_ISL_861324, EPI_ISL_861326 | | | |
| see above | MONTEFIORE MEDICAL CENTER LABORATORIES | Wadsworth Center, New York State Department of Health | Kirsten St. George, Daryl M. Lamson, Alexis Russel, Matthew Shudt, Melissa A Leisner, Jonathan Plitnick, Navjot Singh, John Kelly, Erasmus Schneider, Erica Lasek-Nesselquist |
| EPI_ISL_861534, EPI_ISL_861537, EPI_ISL_861573, EPI_ISL_861574, EPI_ISL_861588 | Instituto Nacional de Saude (INSA) | Instituto Nacional de Saude (INSA) | Borges et al |
| EPI_ISL_861679 | Instituto Adolfo Lutz - Regional de Taubate | Instituto Adolfo Lutz, Interdiciplinary Procedures Center, Strategic Laboratory | Claudio Tavares Sacchi, Claudia Regina Gonçalves, Erica Valessa Ramos Gomes, Karoline Rodrigues Campos |
| EPI_ISL_861862 | Labormedizinisches Zentrum Dr Risch | University Hospital Basel, Clinical Bacteriology | Tim Roloff, Madlen Stange, Helena MB Seth-Smith, Alfredo Mari, Karoline Leuzinger, Julia Bielicki, Nadia Wohlwend,Martin Risch, Lorenz Risch, Manuel Battegay, Hans Hirsch, Adrian Egli |
| EPI_ISL_861865 | Bioanalytika AG | University Hospital Basel, Clinical Bacteriology | Tim Roloff, Madlen Stange, Helena MB Seth-Smith, Alfredo Mari, Karoline Leuzinger, Julia Bielicki, Adrian Härri, Manuel Battegay, Hans Hirsch, Adrian Egli |
| EPI_ISL_861997, EPI_ISL_861998, EPI_ISL_861999 | OHSU Lab Services Molecular Microbiology Lab | Oregon SARS-CoV-2 Genome Sequencing Center | Brendan L. O'Connell, Sally Grindstaff, Kayla Carter, Ruth V. Nichols, Alec J. Hirsch, Donna Hansel, Guang Fan, Xuan Qin, Daniel N. Streblow, William B. Messer, Andrew C. Adey, Benjamin N. Bimber, Brian J. O'Roak |
| EPI_ISL_862039 | RSUP Dr. Sardjito | Genetics Working Group (Pokja Genetik) Faculty of Medicine, Public Health and Nursing Universitas Gadjah Mada (FK-KMK UGM); Disease Investigation Center Wates Ministry of Agriculture Indonesia; Department of Microbiology FK-KMK | Gunadi, Hendra Wibawa, . Marcellus, Mohamad S. Hakim, Edwin W. Daniwijaya, Ludhang P. Rizki, Endah Supriyati, Eggi Arguni, Titik Nuryastuti, Tri Wibawa, Dwi AA Nugrahaningsih, Afiahayati, Siswanto, Kristy Iskandar, Nungki Anggorowati, Ika Trisnawati, Sumardi, Eko Budiono, Bambang Sigit Riyanto, Heni Retnowulan, Munawar Gani, Satria Maulana, Nur Rahmi Ananda, Riat El Khair, Yunika Puspadewi, Osman Sianipar, Umi Solekhah Intansari, Elizabeth Henny Herningtiyas, Ira Puspitawati, Nur Imma Fatimah Harahap,William Widitjiarso, Maria Patricia Inggriani |

| | | | |
|---|---|---|---|
| | | UGM; Laboratorium Diagnostik Yayasan Tahija World Mosquito Program (WMP) Yogyakarta Center for Tropical Medicine FK-KMK UGM; Integrated Research Center FK-KMK UGM; Department of Computer Science and Electronics FMIPA UGM; RSUP Dr. Sardjito | |
| EPI_ISL_862040 | RSUP Dr. Sardjito | Genetics Working Group (Pokja Genetik) Faculty of Medicine, Public Health and Nursing Universitas Gadjah Mada (FK-KMK UGM); Disease Investigation Center Wates Ministry of Agriculture Indonesia; Department of Microbiology FK-KMK UGM; Laboratorium Diagnostik Yayasan Tahija World Mosquito Program (WMP) Yogyakarta Center for Tropical Medicine FK-KMK UGM; Integrated Research Center FK-KMK UGM; Department of Computer Science and Electronics FMIPA UGM; RSUP Dr. Sardjito | Gunadi, Hendra Wibawa, . Marcellus, Mohamad S. Hakim, Edwin W. Daniwijaya, Ludhang P. Rizki, Endah Supriyati, Eggi Arguni, Titik Nuryastuti, Tri Wibawa, Dwi AA Nugrahaningsih, Afiahayati, Siswanto, Kristy Iskandar, Nungki Anggorowati, Ika Trisnawati, Sumardi, Eko Budiono, Bambang Sigit Riyanto, Heni Retnowulan, Munawar Gani, Satria Maulana, Nur Rahmi Ananda, Riat El Khair, Yunika Puspadewi, Osman Sianipar, Umi Solekhah Intansari, Elizabeth Henny Herningtiyas, Ira Puspitawati, Nur Imma Fatimah Harahap,Alvin Santoso Kalim, Untung Riawan |
| EPI_ISL_862041 | RSUP Dr. Sardjito | Genetics Working Group (Pokja Genetik) Faculty of Medicine, Public Health and Nursing Universitas Gadjah Mada (FK-KMK UGM); Disease Investigation Center Wates Ministry of Agriculture Indonesia; Department of Microbiology FK-KMK UGM; Laboratorium Diagnostik Yayasan Tahija World Mosquito Program (WMP) Yogyakarta Center for Tropical Medicine FK-KMK UGM; Integrated Research Center FK-KMK UGM; Department of Computer Science and Electronics FMIPA UGM; RSUP Dr. Sardjito | Gunadi, Hendra Wibawa, . Marcellus, Mohamad S. Hakim, Edwin W. Daniwijaya, Ludhang P. Rizki, Endah Supriyati, Eggi Arguni, Titik Nuryastuti, Tri Wibawa, Dwi AA Nugrahaningsih, Afiahayati, Siswanto, Kristy Iskandar, Nungki Anggorowati, Ika Trisnawati, Sumardi, Eko Budiono, Bambang Sigit Riyanto, Heni Retnowulan, Munawar Gani, Satria Maulana, Nur Rahmi Ananda, Riat El Khair, Yunika Puspadewi, Osman Sianipar, Umi Solekhah Intansari, Elizabeth Henny Herningtiyas, Ira Puspitawati, Nur Imma Fatimah Harahap, Dyah Ayu Puspitarani, Audric Kenny Tedja |
| EPI_ISL_862548 | Hospital Clínico San Carlos | Instituto de Salud Carlos III | Iglesias-Caballero, M. Camarero, S. Molinero Calamita, M. González-Esguevillas, M. Pozo, F. Casas, I. Jiménez, P. Jiménez, M. Zaballos, A. Monzón, S. Varona, S. Juliá, M. Cuesta, I. Rodríguez, I. |
| EPI_ISL_862551 | Hospital J.M. Morales Meseguer | Instituto de Salud Carlos III | Iglesias-Caballero, M. Camarero, S. Molinero Calamita, M. González-Esguevillas, M. . Pozo, F. Casas, I. Jiménez, P. Jiménez, M. Zaballos, A. Monzón, S. Varona, S. Juliá, M. Cuesta, I. Guerrero, C. |
| EPI_ISL_862552 | Complejo Hospitalario de Navarra | Instituto de Salud Carlos III | Iglesias-Caballero, M.Camarero, S. Molinero Calamita, M. González-Esguevillas, M. Pozo, F. Casas, I. Jiménez, P. Jiménez, M. Zaballos, A. Monzón, S. Varona, S. Juliá, M. Cuesta, I. Ezpeleta, C. |
| EPI_ISL_862557, EPI_ISL_862559, EPI_ISL_862560, EPI_ISL_862561, EPI_ISL_862566 | Consejería de Sanidad y Asuntos Sociales | Instituto de Salud Carlos III | Iglesias-Caballero, M. Camarero, S. Molinero Calamita, M. González-Esguevillas, M. Pozo, F. Casas, I. Jiménez, P. Jiménez, M. Zaballos, A. Monzón, S. Varona, S. Juliá, M. Cuesta, I. Gutiérrez, G. |
| EPI_ISL_862569 | Hospital Clínico San Carlos | Instituto de Salud Carlos III | Iglesias-Caballero, M. Camarero, S. Molinero Calamita, M. González-Esguevillas, M. Pozo, F. Casas, I. Jiménez, P. Jiménez, M. Zaballos, A. Monzón, S. Varona, S. Juliá, M. Cuesta, I. Rodríguez, I. |
| EPI_ISL_862573, EPI_ISL_862576, EPI_ISL_862577, EPI_ISL_862580, EPI_ISL_862585, EPI_ISL_862587 | Hospital Clínic | Instituto de Salud Carlos III | Iglesias-Caballero, M.Camarero, S. Molinero Calamita, M. González-Esguevillas, M. Pozo, F. Casas, I. Jiménez, P. Jiménez, M. Zaballos, A. Monzón, S. Varona, S. Juliá, M. Cuesta, I. Marcos, M.A. |
| EPI_ISL_862728, EPI_ISL_862748, EPI_ISL_862749, EPI_ISL_862754, EPI_ISL_862756, EPI_ISL_862758, EPI_ISL_862765, EPI_ISL_862775 | Utah Public Health Laboratory, Utah Public Health Laboratory Infectious Disease submission group | Utah Public Health Laboratory, Utah Public Health Laboratory Infectious Disease submission group | Young,E.L., Oakeson,K.F., Gallagher,T. |
| EPI_ISL_865079, EPI_ISL_865083, EPI_ISL_865084, EPI_ISL_865085, EPI_ISL_865086, EPI_ISL_865088, EPI_ISL_865142 | Virology Department, Royal Infirmary of Edinburgh, NHS Lothian / School of Biological Sciences, University of Edinburgh / Institute of Genetics and Molecular Medicine, University of Edinburgh | COVID-19 Genomics UK (COG-UK) Consortium | McHugh M, Dewar R, Rooke S, Gallagher M, Balcaza C, O'Toole Á, Scher E, Hill V, McCrone JT, Colquhoun R, Yu X, Jackson B, Rambaut A, Williams TC, Templeton K |
| EPI_ISL_865373, EPI_ISL_865374, EPI_ISL_865376, EPI_ISL_865398, EPI_ISL_865400, EPI_ISL_865405 | Liverpool Clinical Laboratories | COVID-19 Genomics UK (COG-UK) Consortium | Sam Haldenby, Anita Lucaci, Steve Paterson, Julian Hiscox, Alistair Darby, M Almsaud, A Alrezaihi, Muhannad Alruwaili, Stuart D Armstrong, Jones Benjamin, Eleanor G Bentley, Anu Chawla, Jordan J Clark, Angela Cowell, Richard Eccles, Isabel García-Dorival, Matthew Gemmell, Alessandro Gerada, PKF Gilmore, Richard Gregory, Ximeng Han, Catherine Hartley, Margaret Hughes, Miren Iturriza-Gomara, James Johnson, L Luu, Jenifer Manson, Charlotte Nelson, Elaine O'Toole, Cassie Olateju, Rebekah Penrice-Randal , Lucille Rainbow, N.P Randle, Trevor Ian Robinson, Parul Sharma, Ghada T Shawli, James P Stewart, Neil Swainston, Ecaterina Vamos, Joanne Watts, Mark Whitehead |
| EPI_ISL_865488, EPI_ISL_865489, EPI_ISL_865494, EPI_ISL_865495, EPI_ISL_865496, EPI_ISL_865497, EPI_ISL_865501, EPI_ISL_865514, EPI_ISL_865515, EPI_ISL_865516, EPI_ISL_865517, EPI_ISL_865518, EPI_ISL_865519, EPI_ISL_865520, EPI_ISL_865521, EPI_ISL_865522, EPI_ISL_865523, EPI_ISL_865524, EPI_ISL_865525, EPI_ISL_865526, EPI_ISL_865563, EPI_ISL_865564, EPI_ISL_865565, EPI_ISL_865566, EPI_ISL_865576, EPI_ISL_865577, EPI_ISL_865578 | | | |
| see above | Barts Health NHS Trust | COVID-19 Genomics UK (COG-UK) Consortium | CUTINO-MOGUEL, Maria-Teresa; HARRINGTON, David; OWOYEMI, Dola; KULASEGARAN-SHYLINI, Raghavendran; BROAD, Claire; KELE, Beatrix |
| EPI_ISL_865689, EPI_ISL_865797, EPI_ISL_865798, EPI_ISL_865799, EPI_ISL_865800, EPI_ISL_865801, EPI_ISL_865802, EPI_ISL_865803, EPI_ISL_865804, EPI_ISL_865805, EPI_ISL_865806, EPI_ISL_865807, EPI_ISL_865808, EPI_ISL_865809, EPI_ISL_865810, EPI_ISL_865811, EPI_ISL_865814, EPI_ISL_865815, EPI_ISL_865818, EPI_ISL_865837 | | | |
| see above | University College London, Great Ormond Street Hospital for Children NHS Foundation Trust, Imperial College Healthcare NHS Trust | COVID-19 Genomics UK (COG-UK) Consortium | Sergi Castellano, Rachel Williams, Mark Kristiansen, Paola Resende Silva, Sunando Roy, Tony Brooks, Helena Tutill, Paola Niola, Patricia Dyal, Charlotte Williams, Leysa Forrest, Yasmin Panchbhaya, Jacqueline Findlay, Samuel Weeks, Julianne Brown, Kathryn Harris, Paul Randell, James Price, Alison Holmes, Judith Breuer |
| EPI_ISL_866131, EPI_ISL_866183 | University College London Hospital | COVID-19 Genomics UK (COG-UK) Consortium | Judith Heaney, Matthew Byott, Catherine Houlihan, Dan Frampton, Stuart Kirk, Moira Spyer and Eleni Nastouli |
| EPI_ISL_867037, EPI_ISL_867038, EPI_ISL_867047, EPI_ISL_867051, EPI_ISL_867055 | Oxford Viromics, NDM, University of Oxford; Oxford University Hospitals; Basingstoke and North Hampshire Hospital | COVID-19 Genomics UK (COG-UK) Consortium | Tanya Golubchik, David Bonsall, George Macintyre, Amy Trebes, Mariateresa de Cesare, Catrin Moore, Alex Mobbs, Anita Justice, Robert Shaw, Monique Andersson, Timothy Peto, Emma Wise, Nathan Moore, Jessica Lynch, Nick Cortes, Matilde Mori, Stephen Kidd, David Buck, John Todd, Christophe Fraser |
| EPI_ISL_867218 | Originating lab: Wales Specialist Virology Centre Sequencing lab: Pathogen Genomics Unit | Public Health Wales Microbiology Cardiff Wales Specialist Virology Centre | Catherine Moore, Johnathan Evans, Laura Gifford, Malorie Perry, Simon Cottrell, Angela Marchbank, Alec Birchley, Alexander Adams, Amy Gaskin, Bree Gatica-Wilcox, Jason Coombes, Joel Southgate, Lauren Gilbert, Lee Graham, Nicole Pacchiarini, Sara Kumziene-Summerhayes, Sarah Taylor, Sophie Jones, Sara Rey, Matthew Bull, Joanne Watkins, Sally Corden, Tom Connor |
| EPI_ISL_868372, EPI_ISL_868438, EPI_ISL_868559, EPI_ISL_868570, EPI_ISL_868576 | Virology Department, Sheffield Teaching Hospitals NHS Foundation Trust/Department of Infection, Immunity and Cardiovascular Disease, The Medical School, University of Sheffield | COVID-19 Genomics UK (COG-UK) Consortium | Thushan de Silva, Matthew Parker, Nikki Smith, Adri Angyal, Rebecca Brown, Luke Green, Rachel Tucker, Paul Parsons, Danielle Groves, Katie Johnson, Laura Carrilero, Alex Keeley, Dave Partridge, Matthew Wyles, Benjamin Lindsey, Mehmet Yavuz, Mohammad Raza, Cariad Evans |
| EPI_ISL_869079, EPI_ISL_869080, EPI_ISL_869081, EPI_ISL_871325 | Ohio Department of Health Laboratory | Ohio Department of Health Laboratory | Holmes, Jennifer; Eric Brandt, Keoni Omura, Glen McGillivary, Caitlin McDonnell, Kirtana Ramadugu, Erica Leasure, Kelsey Florek, Heather Blankenship, Quanta Brown, and Tammy Bannerman |
| EPI_ISL_871458, EPI_ISL_871459, EPI_ISL_871460, EPI_ISL_871461, EPI_ISL_871462, EPI_ISL_871463, EPI_ISL_871464, EPI_ISL_871465, EPI_ISL_871466, EPI_ISL_871467, EPI_ISL_871468, EPI_ISL_871469, EPI_ISL_871470, EPI_ISL_871471, EPI_ISL_871472, EPI_ISL_871473, EPI_ISL_871474, EPI_ISL_871475, EPI_ISL_871476, EPI_ISL_871477, EPI_ISL_871478, EPI_ISL_871479, EPI_ISL_871480, EPI_ISL_871481, EPI_ISL_871482, EPI_ISL_871483, EPI_ISL_871484, EPI_ISL_871485, EPI_ISL_871486, EPI_ISL_871487, EPI_ISL_871488, EPI_ISL_871489, EPI_ISL_871490, EPI_ISL_871491, EPI_ISL_871492, EPI_ISL_871493, EPI_ISL_871494, EPI_ISL_871495, EPI_ISL_871496, EPI_ISL_871497, EPI_ISL_871498, EPI_ISL_871499, EPI_ISL_871500, EPI_ISL_871501, EPI_ISL_871502, EPI_ISL_871503, EPI_ISL_871504, EPI_ISL_871505, EPI_ISL_871506, EPI_ISL_871507, EPI_ISL_871508, EPI_ISL_871509, EPI_ISL_871510, EPI_ISL_871511, EPI_ISL_871512, EPI_ISL_871513, EPI_ISL_871514, EPI_ISL_871515, EPI_ISL_871516, EPI_ISL_871517, EPI_ISL_871518, EPI_ISL_871519, EPI_ISL_871520, EPI_ISL_871521, EPI_ISL_871522, EPI_ISL_871523, EPI_ISL_871524, EPI_ISL_871525, EPI_ISL_871526, EPI_ISL_871527, EPI_ISL_871528, EPI_ISL_871529, | | | |

| | | | |
|---|---|---|---|
| EPI_ISL_871530, EPI_ISL_871531, EPI_ISL_871532, EPI_ISL_871533, EPI_ISL_871534, EPI_ISL_871535, EPI_ISL_871536, EPI_ISL_871537, EPI_ISL_871538, EPI_ISL_871539, EPI_ISL_871540, EPI_ISL_871541, EPI_ISL_871542, EPI_ISL_871543, EPI_ISL_871544, EPI_ISL_871545, EPI_ISL_871546, EPI_ISL_871547, EPI_ISL_871548, EPI_ISL_871549, EPI_ISL_871550, EPI_ISL_871551, EPI_ISL_871552, EPI_ISL_871553, EPI_ISL_871554, EPI_ISL_871555, EPI_ISL_871556, EPI_ISL_871557, EPI_ISL_871558, EPI_ISL_871559, EPI_ISL_871560, EPI_ISL_871561, EPI_ISL_871562, EPI_ISL_871563, EPI_ISL_871564, EPI_ISL_871565, EPI_ISL_871566, EPI_ISL_871567, EPI_ISL_871568, EPI_ISL_871569, EPI_ISL_871570, EPI_ISL_871571, EPI_ISL_871572, EPI_ISL_871573, EPI_ISL_871574, EPI_ISL_871575, EPI_ISL_871576, EPI_ISL_871577 | | | |
| see above | Department of Virus and Microbiological Special Diagnostics, Statens Serum Institut, Copenhagen, Denmark | Aalborg University | Danish Covid-19 Genome Consortium |
| EPI_ISL_871788, EPI_ISL_871791 | Ohio Department of Health Laboratory | Ohio Department of Health Laboratory | Holmes, Jennifer; Eric Brandt, Keoni Omura, Glen McGillivary, Caitlin McDonnell, Kirtana Ramadugu, Erica Leasure, Kelsey Florek, Heather Blankenship, Quanta Brown, and Tammy Bannerman |
| EPI_ISL_871806, EPI_ISL_871810 | AIID | Irish Coronavirus Sequencing Consortium-Teagasc Grange | Matthew McCabe, Aljandro Abner Garcia Leon, Fiona Crispie, Calum Walsh, Michael Carr, John Kenny, Paul Cotter, Patrick Mallon, Gabriel Gonzalez |
| EPI_ISL_871893 | Hospital Universitario Marqués de Valdecilla - IDIVAL (Santander, Cantabria) | SeqCOVID-SPAIN consortium/IBV(CSIC) | Mónica Gozalo Margüello, María Eliecer Cano García, Jose Manuel Méndez Legaza, Daniel Pablo Marcos, Jesús Rodríguez Rodríguez, María Siller Ruiz and SeqCOVID-SPAIN consortium |
| EPI_ISL_871910 | Servicio de Microbiología Clínica (Complejo Hospitalario de Navarra, Pamplona), Instituto de Investigación Sanitaria de Navarra (IdiSNA) | SeqCOVID-SPAIN consortium/IBV(CSIC) | Carmen Ezpeleta Baquedano, Ana Navascués, Ana Miqueleiz and SeqCOVID-SPAIN consortium |
| EPI_ISL_872040, EPI_ISL_872058, EPI_ISL_872059, EPI_ISL_872061, EPI_ISL_872062 | Department of Clinical Microbiology | GIGA Medical Genomics | Keith Durkin, Maria Artesi, Sébastien Bontems, Raphaël Boreux, Bouchra Boujemla, Cécile Meex, Pierrette Melin, Marie-Pierre Hayette, Vincent Bours |
| EPI_ISL_872188 | RSUP Dr. Sardjito | Genetics Working Group (Pokja Genetik) Faculty of Medicine, Public Health and Nursing Universitas Gadjah Mada (FK-KMK UGM); Disease Investigation Center Wates Ministry of Agriculture Indonesia; Department of Microbiology FK-KMK UGM; Laboratorium Diagnostik Yayasan Tahija World Mosquito Program (WMP) Yogyakarta Center for Tropical Medicine FK-KMK UGM; Integrated Research Center FK-KMK UGM; Department of Computer Science and Electronics FMIPA UGM; RSUP Dr. Sardjito | Gunadi, Hendra Wibawa, . Marcellus, Mohamad S. Hakim, Edwin W. Daniwijaya, Ludhang P. Rizki, Endah Supriyati, Eggi Arguni, Titik Nuryastuti, Tri Wibawa, Dwi AA Nugrahaningsih, Afiahayati, Siswanto, Kristy Iskandar, Nungki Anggorowati, Ika Trisnawati, Sumardi, Eko Budiono, Bambang Sigit Riyanto, Heni Retnowulan, Munawar Gani, Satria Maulana, Nur Rahmi Ananda, Riat El Khair, Yunika Puspadewi, Osman Sianipar, Umi Solekhah Intansari, Elizabeth Henny Herningtiyas, Ira Puspitawati, Nur Imma Fatimah Harahap, Aditya Rifqi Fauzi, Fadil Fahri |
| EPI_ISL_872189 | RSUP Dr. Sardjito | Genetics Working Group (Pokja Genetik) Faculty of Medicine, Public Health and Nursing Universitas Gadjah Mada (FK-KMK UGM); Disease Investigation Center Wates Ministry of Agriculture Indonesia; Department of Microbiology FK-KMK UGM; Laboratorium Diagnostik Yayasan Tahija World Mosquito Program (WMP) Yogyakarta Center for Tropical Medicine FK-KMK UGM; Integrated Research Center FK-KMK UGM; Department of Computer Science and Electronics FMIPA UGM; RSUP Dr. Sardjito | Gunadi, Hendra Wibawa, . Marcellus, Mohamad S. Hakim, Edwin W. Daniwijaya, Ludhang P. Rizki, Endah Supriyati, Eggi Arguni, Titik Nuryastuti, Tri Wibawa, Dwi AA Nugrahaningsih, Afiahayati, Siswanto, Kristy Iskandar, Nungki Anggorowati, Ika Trisnawati, Sumardi, Eko Budiono, Bambang Sigit Riyanto, Heni Retnowulan, Munawar Gani, Satria Maulana, Nur Rahmi Ananda, Riat El Khair, Yunika Puspadewi, Osman Sianipar, Umi Solekhah Intansari, Elizabeth Henny Herningtiyas, Ira Puspitawati, Nur Imma Fatimah Harahap, Alvin S. Kalim, Dwiki afandy |
| EPI_ISL_872190 | RSUP Dr. Sardjito | Genetics Working Group (Pokja Genetik) Faculty of Medicine, Public Health and Nursing Universitas Gadjah Mada (FK-KMK UGM); Disease Investigation Center Wates Ministry of Agriculture Indonesia; Department of Microbiology FK-KMK UGM; Laboratorium Diagnostik Yayasan Tahija World Mosquito Program (WMP) Yogyakarta Center for Tropical Medicine FK-KMK UGM; Integrated Research Center FK-KMK UGM; Department of Computer Science and Electronics FMIPA UGM; RSUP Dr. Sardjito | Gunadi, Hendra Wibawa, . Marcellus, Mohamad S. Hakim, Edwin W. Daniwijaya, Ludhang P. Rizki, Endah Supriyati, Eggi Arguni, Titik Nuryastuti, Tri Wibawa, Dwi AA Nugrahaningsih, Afiahayati, Siswanto, Kristy Iskandar, Nungki Anggorowati, Ika Trisnawati, Sumardi, Eko Budiono, Bambang Sigit Riyanto, Heni Retnowulan, Munawar Gani, Satria Maulana, Nur Rahmi Ananda, Riat El Khair, Yunika Puspadewi, Osman Sianipar, Umi Solekhah Intansari, Elizabeth Henny Herningtiyas, Ira Puspitawati, Nur Imma Fatimah Harahap, Dyah Ayu Puspitarani, Kemala Athollah |
| EPI_ISL_872380, EPI_ISL_872381, EPI_ISL_872386, EPI_ISL_872387, EPI_ISL_872388, EPI_ISL_872389, EPI_ISL_872390, EPI_ISL_872391, EPI_ISL_872392, EPI_ISL_872393 | Texas Department of State Health Services (TXDSHS) | Texas Department of State Health Services (TXDSHS) | Bonnie Oh, Anita Pokharel, James Daniel Bonser, Myong Koag, Chung Wang, Rachel Lee, Grace Kubin, Rashmi Tuladhar, Mayela Pedrueza, Maliha Rahman, Jenny Zhang |
| EPI_ISL_872623, EPI_ISL_872624 | Nigeria Centre for Disease Control (NCDC) | African Centre of Excellence for Genomics of Infectious Diseases (ACEGID), Redeemer's University | Oluniyi P.E. et al |
| EPI_ISL_872741 | Rhode Island Department of Health | Infectious Disease Program, Broad Institute of Harvard and MIT | Lemieux,J.E., Siddle,K.J., Huard,R., King,E., Azevedo,K., Miller,A., Adams,G., Gladden-Young,A., Lagerborg,K., Rudy,M., DeRuff,K., Carter,A., Normandin,E., Bauer,M., Reilly,S., Tomkins-Tinch,C., Loreth,C., Chaluvadi,S., Birren,B.W., Gallagher,G., Smole,S., Park,D.J., MacInnis,B.L., and Sabeti,P.C. |
| EPI_ISL_873021, EPI_ISL_873022, EPI_ISL_873023, EPI_ISL_873024, EPI_ISL_873025, EPI_ISL_873026, EPI_ISL_873034 | HELIX LLC | WHO National Influenza Centre Russian Federation | Andrey Komissarov, Artem Fadeev, Anna Ivanova, Kseniya Komissarova, Dmitry Bazhenov, Mikhail Bakaev, Daria Danilenko, Ksenia Safina, Elena Nabieva, Georgii Bazykin, Dmitry Lioznov |
| EPI_ISL_873234 | M Health Fairview | Minnesota Department of Health, Public Health Laboratory | Alexandra Lorentz, Jacob Garfin, Matt Plumb, and Xiong Wang |
| EPI_ISL_875513 | National Virus Reference Laboratory | National Virus Reference Laboratory | Michael Carr, Gabriel Gonzalez, Jonathan Dean, Cillian F De Gascun |
| EPI_ISL_876058, EPI_ISL_876072, EPI_ISL_876088, EPI_ISL_876097, EPI_ISL_876132, EPI_ISL_876150, EPI_ISL_876151, EPI_ISL_876169, EPI_ISL_876182, EPI_ISL_876261, EPI_ISL_876262, EPI_ISL_876263, EPI_ISL_876264, EPI_ISL_876265, EPI_ISL_876266, EPI_ISL_876267, EPI_ISL_876268, EPI_ISL_876269, EPI_ISL_876270, EPI_ISL_876271, EPI_ISL_876272 | | | |
| see above | Massachusetts State Public Health Laboratory | Massachusetts State Public Health Laboratory | Andrew Lang, Timelia Fink, Glen Gallagher, Sandra Smole |
| EPI_ISL_876744, EPI_ISL_876745, EPI_ISL_876746, EPI_ISL_876747 | Istituto Zooprofilattico Sperimentale della Puglia e della Basilicata | Istituto Zooprofilattico Sperimentale della Puglia e della Basilicata | Parisi A., Bianco A., Capozzi L., Del Sambro L., Manzulli V, Rondinone V., Pace L., Cipolletta D., Galante D. |
| EPI_ISL_877186, EPI_ISL_877188 | Quest Diagnostics | Quest Diagnostics | Rosenthal,S.H., Gerasimova,A., Kagan,R.M., Anderson, B., Hua, M., Liu Y., Bernstein, L.E., Livingston, K.E., Perez, A., Shalhout, D.F., Shlyakhter, I.A., Owen, R., Tanpaiboon, P., Lacbawan, F. |
| EPI_ISL_877228 | Institute for Medical Research, Infectious Disease Research Centre, National Institutes of Health, Ministry of Health Malaysia | Institute for Medical Research, Infectious Disease Research Centre, National Institutes of Health, Ministry of Health Malaysia | Suppiah J, Kamel K, Azizan MA, Thayan R |
| EPI_ISL_877451, EPI_ISL_877452 | Institute of Microbiology and Immunology, Faculty of Medicine, University of Ljubljana | Institute of Microbiology and Immunology, Faculty of Medicine, University of Ljubljana | Samo Zakotnik, Tomaž Mark Zorec, Matic Brvar, Miša Korva, Mario Poljak, Tatjana Avši - Županc |
| EPI_ISL_877538 | Siti Khodijah Hospital | Institute of Tropical Disease, Universitas Airlangga | Krisnoadi Rahardjo, Aldise M Nastri, Jezzy R Dewantari, Rima R Prasetya, Muhammad Hamdan, Gatot Soegiarto, Laksmi Wulandari, Resti Yudhawati, Soetjipto, Yasuko Mori, Maria I Lusida, Kazufumi Shimizu |
| EPI_ISL_878779, EPI_ISL_880173, EPI_ISL_880195 | Rady's Childrens Hospital | Andersen lab at Scripps Research | SEARCH Alliance San Diego with Nanda Radamchar, David Dimmock, Linda Luo, Christina Clarke, Kathryn Bouic, Teresa Mueller, Denise Malicki |

| | | | |
|---|---|---|---|
| EPI_ISL_882314 | Lighthouse Lab in Alderley Park | Wellcome Sanger Institute for the COVID-19 Genomics UK (COG-UK) Consortium | Jacquelyn Wynn, Mairead Hyland, The Lighthouse Lab in Alderley Park and Alex Alderton, Roberto Amato, Sonia Goncalves, Ewan Harrison, David K. Jackson, Ian Johnston, Dominic Kwiatkowski, Cordelia Langford, John Sillitoe on behalf of the Wellcome Sanger Institute COVID-19 Surveillance Team |
| EPI_ISL_882771 | Siti Khodijah Hospital | Institute of Tropical Disease, Universitas Airlangga | Kazufumi Shimizu, Krisnoadi Rahardjo, Aldise M Nastri, Jezzy R Dewantari, Rima R Prasetya, Muhammad Hamdan, Gatot Soegiarto, Laksmi Wulandari, Resti Yudhawati, Yasuko Mori, Soetjipto, Maria I Lusida |
| EPI_ISL_882921 | Genomic Medicine Laboratory, IRCCS Santa Lucia Foundation | INMI Lazzaro Spallanzani IRCCS | M Rueca, E Giombini, C.E.M Gruber, B Bartolini, O Butera, F Messina, E Giardina, A Di Caro, MR Capobianchi |
| EPI_ISL_882924 | INMI Lazzaro Spallanzani IRCCS | INMI Lazzaro Spallanzani IRCCS | F Messina, O Butera, E Giombini, M Rueca, B Bartolini, C.E.M Gruber, MR Capobianchi, A Di Caro |
| EPI_ISL_882925 | INMI Lazzaro Spallanzani IRCCS | INMI Lazzaro Spallanzani IRCCS | C.E.M Gruber, B Bartolini, E Giombini, M Rueca, O Butera, F Messina, A Di Caro, MR Capobianchi |
| EPI_ISL_884260, EPI_ISL_884264, EPI_ISL_884272, EPI_ISL_884273, EPI_ISL_884275, EPI_ISL_884276, EPI_ISL_884288, EPI_ISL_884289, EPI_ISL_884290 | Institute of Medical Microbiology and Hospital Hygiene | Institute of Medical Microbiology and Hospital Hygiene | Prof. Dr. Achim Kaasch, Aljoscha Tersteegen |
| EPI_ISL_884983, EPI_ISL_884987, EPI_ISL_884988, EPI_ISL_884989, EPI_ISL_884990, EPI_ISL_884991, EPI_ISL_884992, EPI_ISL_884993, EPI_ISL_884994, EPI_ISL_885009, EPI_ISL_885010, EPI_ISL_885011 | | | |
| see above | Santa Clara County Public Health Laboratory | Chan-Zuckerberg Biohub | CZB Cliahub Consortium |
| EPI_ISL_885108, EPI_ISL_885109, EPI_ISL_885110, EPI_ISL_885111, EPI_ISL_885112, EPI_ISL_885113, EPI_ISL_885114, EPI_ISL_885115, EPI_ISL_885116, EPI_ISL_885117, EPI_ISL_885118, EPI_ISL_885119, EPI_ISL_885120, EPI_ISL_885121, EPI_ISL_885122, EPI_ISL_885123, EPI_ISL_885124, EPI_ISL_885125, EPI_ISL_885126, EPI_ISL_885127 | | | |
| see above | Orange County Public Health Lab | Chan-Zuckerberg Biohub | CZB Cliahub Consortium |
| EPI_ISL_885142 | RSUP Dr. Sardjito | Genetics Working Group (Pokja Genetik) Faculty of Medicine, Public Health and Nursing Universitas Gadjah Mada (FK-KMK UGM); Disease Investigation Center Wates Ministry of Agriculture Indonesia; Department of Microbiology FK-KMK UGM; Laboratorium Diagnostik Yayasan Tahija World Mosquito Program (WMP) Yogyakarta Center for Tropical Medicine FK-KMK UGM; Integrated Research Center FK-KMK UGM; Department of Computer Science and Electronics FMIPA UGM; RSUP Dr. Sardjito | Gunadi, Hendra Wibawa, . Marcellus, Mohamad S. Hakim, Edwin W. Daniwijaya, Ludhang P. Rizki, Endah Supriyati, Eggi Arguni, Titik Nuryastuti, Tri Wibawa, Dwi AA Nugrahaningsih, Afiahayati, Siswanto, Kristy Iskandar, Nungki Anggorowati, Ika Trisnawati, Sumardi, Eko Budiono, Bambang Sigit Riyanto, Heni Retnowulan, Munawar Gani, Satria Maulana, Nur Rahmi Ananda, Riat El Khair, Yunika Puspadewi, Osman Sianipar, Umi Solekhah Intansari, Elizabeth Henny Herningtiyas, Ira Puspitawati, Nur Imma Fatimah Harahap, Alvin Santoso Kalim, Susan Simanjaya |
| EPI_ISL_886825 | Labcorp | Genomics and Discovery, Respiratory Viruses Branch, Division of Viral Diseases, Centers for Disease Control and Prevention | Peter W. Cook,Dhwani Batra,Ben L. Rambo-Martin,Summer Galloway,Brian Krueger,Minoo Agarwal,Eyad Almasri,Debbie Boles,Ayla Burns,Nuthawin Charoensri,Oren Cohen,Susan Countryman,Mary Ann Cristobal,Bobbi Croy,Suzanne Dale,Hrushikesh Deshmukh,Amanda Douglas,Vincent Drouillon,Marcia Eisenberg,Howard Engler,Rama Ghatti,Prashant Gupta,Susan Hicks,Jake Humphrey,Lax Iyer,Manoj Jain,Mohan Kolli,Tim Kuphal,Stanley Letovsky,Michael Levandoski,Craig Lukasik,Jonathan Meltzer,Brian Norvell,Mindy Nye,Scott Parker,Christos Petropoulos,John Pruitt,Steven Ragan,Scott Ryan,Mike Sapeta,Jana Schroth,Suresh Babu Selvaraju,Goran Stevovic,Amanda Suchanek,Andrea Throop,Lyndon Tilson,Thomas Urban,Joe Voshell,Kimberly Wagner,Jonathan Williams,Mary Williamson,Qian Zeng,Tricia Zwiefelhofer,Clinton R. Paden,Suxiang Tong,Duncan MacCannell, |
| EPI_ISL_887440, EPI_ISL_887486, EPI_ISL_887487, EPI_ISL_887488, EPI_ISL_887489, EPI_ISL_887490, EPI_ISL_887491, EPI_ISL_887492 | Instituto Nacional de Saude (INS), Mozambique | KRISP, KZN Research Innovation and Sequencing Platform | Nalia Ismael, Nadia Sitoe, Paulo Arnaldo, Nedio Mabunda, Giandhari J, Pillay S, Tegally H, Wilkinson E, de Oliveira T |
| EPI_ISL_887593, EPI_ISL_887594, EPI_ISL_887596, EPI_ISL_887597, EPI_ISL_887601, EPI_ISL_887602, EPI_ISL_887603, EPI_ISL_887605, EPI_ISL_887607, EPI_ISL_887608, EPI_ISL_887609, EPI_ISL_887613, EPI_ISL_887615, EPI_ISL_887622, EPI_ISL_887627, EPI_ISL_887630, EPI_ISL_887631, EPI_ISL_887634, EPI_ISL_887640, EPI_ISL_887646, EPI_ISL_887649, EPI_ISL_887652, EPI_ISL_887661, EPI_ISL_887664, EPI_ISL_887666, EPI_ISL_887667, EPI_ISL_887675, EPI_ISL_887689, EPI_ISL_887690, EPI_ISL_887691, EPI_ISL_887693, EPI_ISL_887695, EPI_ISL_887704, EPI_ISL_887705, EPI_ISL_887707, EPI_ISL_887710, EPI_ISL_887712, EPI_ISL_887713, EPI_ISL_887717, EPI_ISL_887718, EPI_ISL_887720, EPI_ISL_887727, EPI_ISL_887734, EPI_ISL_887735, EPI_ISL_887736, EPI_ISL_887737, EPI_ISL_887741, EPI_ISL_887749, EPI_ISL_887750, EPI_ISL_887764, EPI_ISL_887769, EPI_ISL_887772, EPI_ISL_887774, EPI_ISL_887775, EPI_ISL_887783, EPI_ISL_887794, EPI_ISL_887795, EPI_ISL_887796, EPI_ISL_887799, EPI_ISL_887800, EPI_ISL_887801, EPI_ISL_887802, EPI_ISL_887803, EPI_ISL_887804, EPI_ISL_887805, EPI_ISL_887806, EPI_ISL_887809, EPI_ISL_887815, EPI_ISL_887818, EPI_ISL_887819, EPI_ISL_887820, EPI_ISL_887825, EPI_ISL_887827, EPI_ISL_887833, EPI_ISL_887835, EPI_ISL_887839, EPI_ISL_887843, EPI_ISL_887844, EPI_ISL_887846, EPI_ISL_887847, EPI_ISL_887848, EPI_ISL_887850, EPI_ISL_887854, EPI_ISL_887860, EPI_ISL_887863, EPI_ISL_887875, EPI_ISL_887876, EPI_ISL_887878, EPI_ISL_887881, EPI_ISL_887882, EPI_ISL_887888, EPI_ISL_887895, EPI_ISL_887896, EPI_ISL_887897, EPI_ISL_887899, EPI_ISL_887900, EPI_ISL_887904, EPI_ISL_887911, EPI_ISL_887914, EPI_ISL_887926, EPI_ISL_887929, EPI_ISL_887933, EPI_ISL_887941, EPI_ISL_887942, EPI_ISL_887943, EPI_ISL_887948, EPI_ISL_887952, EPI_ISL_887955, EPI_ISL_887958, EPI_ISL_887967, EPI_ISL_887969, EPI_ISL_887982, EPI_ISL_887983, EPI_ISL_887990, EPI_ISL_887996, EPI_ISL_887999, EPI_ISL_888005, EPI_ISL_888006, EPI_ISL_888011, EPI_ISL_888016, EPI_ISL_888017, EPI_ISL_888019, EPI_ISL_888020, EPI_ISL_888021, EPI_ISL_888025, EPI_ISL_888031, EPI_ISL_888034, EPI_ISL_888040, EPI_ISL_888046, EPI_ISL_888048, EPI_ISL_888052, EPI_ISL_888055, EPI_ISL_888056, EPI_ISL_888061, EPI_ISL_888067, EPI_ISL_888070, EPI_ISL_888072, EPI_ISL_888073, EPI_ISL_888077, EPI_ISL_888078, EPI_ISL_888079, EPI_ISL_888082, EPI_ISL_888086, EPI_ISL_888087, EPI_ISL_888089, EPI_ISL_888093, EPI_ISL_888098, EPI_ISL_888099, EPI_ISL_888106, EPI_ISL_888108, EPI_ISL_888115, EPI_ISL_888118, EPI_ISL_888119, EPI_ISL_888121, EPI_ISL_888124, EPI_ISL_888126, EPI_ISL_888129, EPI_ISL_888130, EPI_ISL_888131, EPI_ISL_888135, EPI_ISL_888136, EPI_ISL_888137, EPI_ISL_888138, EPI_ISL_888141, EPI_ISL_888142, EPI_ISL_888145, EPI_ISL_888146, EPI_ISL_888148, EPI_ISL_888163, EPI_ISL_888167, EPI_ISL_888168, EPI_ISL_888169, EPI_ISL_888170, EPI_ISL_888171, EPI_ISL_888172, EPI_ISL_888177, EPI_ISL_888178, EPI_ISL_888180, EPI_ISL_888181, EPI_ISL_888182, EPI_ISL_888190, EPI_ISL_888193, EPI_ISL_888194, EPI_ISL_888197, EPI_ISL_888198, EPI_ISL_888200, EPI_ISL_888203, EPI_ISL_888206, EPI_ISL_888212, EPI_ISL_888213, EPI_ISL_888214, EPI_ISL_888216, EPI_ISL_888224, EPI_ISL_888227, EPI_ISL_888230, EPI_ISL_888231, EPI_ISL_888233, EPI_ISL_888236, EPI_ISL_888239, EPI_ISL_888240, EPI_ISL_888244, EPI_ISL_888246, EPI_ISL_888264, EPI_ISL_888267, EPI_ISL_888269, EPI_ISL_888270, EPI_ISL_888279, EPI_ISL_888288, EPI_ISL_888289, EPI_ISL_888294, EPI_ISL_888295, EPI_ISL_888298, EPI_ISL_888303, EPI_ISL_888313, EPI_ISL_888317, EPI_ISL_888319, EPI_ISL_888321, EPI_ISL_888322, EPI_ISL_888331, EPI_ISL_888336, EPI_ISL_888338, EPI_ISL_888340, EPI_ISL_888343, EPI_ISL_888345, EPI_ISL_888346, EPI_ISL_888349, EPI_ISL_888355, EPI_ISL_888357, EPI_ISL_888358, EPI_ISL_888359, EPI_ISL_888361, EPI_ISL_888362, EPI_ISL_888364, EPI_ISL_888371, EPI_ISL_888374, EPI_ISL_888376, EPI_ISL_888383, EPI_ISL_888390, EPI_ISL_888398, EPI_ISL_888403, EPI_ISL_888412, EPI_ISL_888416, EPI_ISL_888418, EPI_ISL_888423, EPI_ISL_888425, EPI_ISL_888428, EPI_ISL_888430, EPI_ISL_888431, EPI_ISL_888433, EPI_ISL_888434, EPI_ISL_888438, EPI_ISL_888455, EPI_ISL_888456, EPI_ISL_888457, EPI_ISL_888459, EPI_ISL_888461, EPI_ISL_888465, EPI_ISL_888469, EPI_ISL_888470, EPI_ISL_888472, EPI_ISL_888473, EPI_ISL_888474, EPI_ISL_888477, EPI_ISL_888482, EPI_ISL_888485, EPI_ISL_888487, EPI_ISL_888489, EPI_ISL_888494, EPI_ISL_888495, EPI_ISL_888497, EPI_ISL_888499, EPI_ISL_888502, EPI_ISL_888503, EPI_ISL_888505, EPI_ISL_888506, EPI_ISL_888507, EPI_ISL_888508, EPI_ISL_888517, EPI_ISL_888518, EPI_ISL_888519, EPI_ISL_888520, EPI_ISL_888530, EPI_ISL_888531, EPI_ISL_888533, EPI_ISL_888542, EPI_ISL_888549, EPI_ISL_888550, EPI_ISL_888551, EPI_ISL_888554, EPI_ISL_888565, EPI_ISL_888569, EPI_ISL_888570, EPI_ISL_888575, EPI_ISL_888576, EPI_ISL_888577, EPI_ISL_888578, EPI_ISL_888579, EPI_ISL_888580, EPI_ISL_888584, EPI_ISL_888585, EPI_ISL_888586 | | | |
| see above | Labcorp | Genomics and Discovery, Respiratory Viruses Branch, Division of Viral Diseases, Centers for Disease Control and Prevention | Peter W. Cook,Dhwani Batra,Ben L. Rambo-Martin,Summer Galloway,Brian Krueger,Minoo Agarwal,Eyad Almasri,Debbie Boles,Ayla Burns,Nuthawin Charoensri,Oren Cohen,Susan Countryman,Mary Ann Cristobal,Bobbi Croy,Suzanne Dale,Hrushikesh Deshmukh,Amanda Douglas,Vincent Drouillon,Marcia Eisenberg,Howard Engler,Rama Ghatti,Prashant Gupta,Susan Hicks,Jake Humphrey,Lax Iyer,Manoj Jain,Mohan Kolli,Tim Kuphal,Stanley Letovsky,Michael Levandoski,Craig Lukasik,Jonathan Meltzer,Brian Norvell,Mindy Nye,Scott Parker,Christos Petropoulos,John Pruitt,Steven Ragan,Scott Ryan,Mike Sapeta,Jana Schroth,Suresh Babu Selvaraju,Goran Stevovic,Amanda Suchanek,Andrea Throop,Lyndon Tilson,Thomas Urban,Joe Voshell,Kimberly Wagner,Jonathan Williams,Mary Williamson,Qian Zeng,Tricia Zwiefelhofer,Clinton R. Paden,Suxiang Tong,Duncan MacCannell, |
| EPI_ISL_888676 | Center of Advanced Studies and Technology, Molecular Genetics Laboratory | Center of Advanced Studies and Technology, Molecular Genetics Laboratory | De Fabritiis Simone, Mandatori Domitilla, Ferrante Rossella |
| EPI_ISL_888786 | KU Leuven, Rega Institute, Clinical and Epidemiological Virology | KU Leuven, Rega Institute, Clinical and Epidemiological Virology | Tony Wawina-Bokalanga, Bert Vanmechelen, Joan Marti-Carerras, Piet Maes |
| EPI_ISL_888869 | Michigan Department of Health and Human Services, Bureau of Laboratories | Michigan Department of Health and Human Services, Bureau of Laboratories | Blankenship HM, Riner D, Soehnlen MK |
| EPI_ISL_889028, EPI_ISL_889029, EPI_ISL_889030, EPI_ISL_889031, EPI_ISL_889032, EPI_ISL_889033, EPI_ISL_889034, EPI_ISL_889035, EPI_ISL_889036, EPI_ISL_889037, EPI_ISL_889038, EPI_ISL_889039, EPI_ISL_889040, EPI_ISL_889041, EPI_ISL_889042, EPI_ISL_889043, EPI_ISL_889044, EPI_ISL_889045, EPI_ISL_889046, EPI_ISL_889047, EPI_ISL_889048, EPI_ISL_889049, EPI_ISL_889050, EPI_ISL_889051, EPI_ISL_889052, EPI_ISL_889053, EPI_ISL_889054, EPI_ISL_889055, EPI_ISL_889056, EPI_ISL_889057, EPI_ISL_889058, EPI_ISL_889059, EPI_ISL_889060, EPI_ISL_889061, EPI_ISL_889062, EPI_ISL_889063, EPI_ISL_889064, EPI_ISL_889065, EPI_ISL_889066, EPI_ISL_889067, EPI_ISL_889068, EPI_ISL_889069, EPI_ISL_889070, EPI_ISL_889154, EPI_ISL_889155, EPI_ISL_889156, EPI_ISL_889157, EPI_ISL_889162, EPI_ISL_889163, EPI_ISL_889164, EPI_ISL_889165, EPI_ISL_889166, EPI_ISL_889167, EPI_ISL_889168, EPI_ISL_889169, EPI_ISL_889170, EPI_ISL_889171, EPI_ISL_889172, EPI_ISL_889173, EPI_ISL_889174, EPI_ISL_889175 | | | |
| see above | Israel Central Virology laboratory | Israel National Consortium for SARS-CoV-2 sequencing | Neta Zuckerman, Efrat Dahan Bucris, Michal Mandelboim, Dana Bar-Ilan, Oran Erster, Tzvia Mann, Omer Murik, David A. Zeevi, Assaf Rokney, Joseph Jaffe, Eva Nachum, Maya Davidovich Cohen, Ephraim Fass, Gal Zizelski Valenci, Mor Rubinstein, Efrat Rorman, Israel Nissan, Efrat Glick-Saar, Omri Nayshool, Gideon Rechavi, Ella Mendelson, Orna Mor |
| EPI_ISL_889461, EPI_ISL_889462, EPI_ISL_889463, EPI_ISL_889464, EPI_ISL_889465, EPI_ISL_889466, EPI_ISL_889467, EPI_ISL_889468, EPI_ISL_889469, EPI_ISL_889470, EPI_ISL_889471, EPI_ISL_889472, EPI_ISL_889473, EPI_ISL_889474, EPI_ISL_889475, EPI_ISL_889476, EPI_ISL_889477, EPI_ISL_889478, EPI_ISL_889479, EPI_ISL_889480, EPI_ISL_889481, EPI_ISL_889483, EPI_ISL_889484, EPI_ISL_889485, EPI_ISL_889497 | | | |
| see above | LSUHS Emerging Viral Threat Laboratory | Microbial Genome Sequencing Center | Jeremy P. Kamil, Jennifer L. Carroll, Camille F. Abshire, Maarten Van Diest, Mohammed N.A. Siddiquey, Andrew D. Yurochko, Martin J. Sapp, Rona S. Scott, Christopher G. Kevil, Daniel J. Snyder, Vaughn S. Cooper, John A. Vanchiere |

| | | | |
|---|---|---|---|
| EPI_ISL_890236 | Center for Genome Sciences, US Army Medical Research Institute of Infectious Disease (USAMRIID) | Center for Genome Sciences, US Army Medical Research Institute of Infectious Disease (USAMRIID) | Kugelman,J., Di Paola,N., Richardson,J., Gibson,W., Bateman,S. and Jerke,K. |
| EPI_ISL_890248, EPI_ISL_890252, EPI_ISL_890257, EPI_ISL_890277, EPI_ISL_890328, EPI_ISL_890332 | KU Leuven, Rega Institute, Clinical and Epidemiological Virology | KU Leuven, Rega Institute, Clinical and Epidemiological Virology | Tony Wawina-Bokalanga, Bert Vanmechelen, Joan Marti-Carerras, Piet Maes |
| EPI_ISL_890894 | Hospital | National Reference Center for Viruses of Respiratory Infections, Institut Pasteur, Paris | Marion Barbet, Sylvie Behillil, Méline Bizard, Angela Brisebarre, Camille Capel, Etienne Simon-Lorière, Vincent Enouf, Maud Vanpeene, Sylvie van der Werf, Patricia Stoessel-Thouvenin |
| EPI_ISL_890998, EPI_ISL_890999, EPI_ISL_891000, EPI_ISL_891001, EPI_ISL_891002, EPI_ISL_891003, EPI_ISL_891004, EPI_ISL_891005, EPI_ISL_891006, EPI_ISL_891007, EPI_ISL_891008, EPI_ISL_891009, EPI_ISL_891010, EPI_ISL_891011, EPI_ISL_891020, EPI_ISL_891021, EPI_ISL_891022, EPI_ISL_891023, EPI_ISL_891024, EPI_ISL_891025, EPI_ISL_891026, EPI_ISL_891027, EPI_ISL_891028, EPI_ISL_891029, EPI_ISL_891030, EPI_ISL_891031, EPI_ISL_891032, EPI_ISL_891033, EPI_ISL_891034, EPI_ISL_891036, EPI_ISL_891037, EPI_ISL_891038, EPI_ISL_891039, EPI_ISL_891040, EPI_ISL_891041, EPI_ISL_891042, EPI_ISL_891043, EPI_ISL_891045, EPI_ISL_891048 | | | |
| see above | Washington State Department of Health | Seattle Flu Study | Deborah A. Nickerson, Chris D. Frazar, Jover Lee, Benjamin Pelle, Erica Ryke, Matthew Richardson, Amanda Adler, Elisabeth Brandstetter, Peter D. Han, Kairsten Fay, Misja Ilcisin, Kirsten Lacombe, Thomas R. Sibley, Melissa Truong, Caitlin R. Wolf, Romesh Gautom, Geoff Melly, Brian Hiatt, Philip Dykema, Scott Lindquist, Michael Boeckh, Janet A. Englund, Michael Famulare, Barry R. Lutz, Mark J. Rieder, Lea M. Starita, Matthew Thompson, Helen Y. Chu, Jay Shendure, Trevor Bedford |
| EPI_ISL_891211 | SMS Medical College Jaipur | SMS Medical College jaipur | Bharti Malhotra, Swati Gautam, Himanshu Sharma, Pratibha Sharma, Neha Bhomia, Nivedita Gupta, Pragya Yadav, Varsha Potdar |
| EPI_ISL_891213 | SMS Medical College Jaipur | SMS Medical College Jaipur | Bharti Malhotra, Swati Gautam, Himanshu Sharma, Pratibha Sharma, Neha Bhomia, Nivedita Gupta, Pragya Yadav, Varsha Potdar |
| EPI_ISL_891214 | SMS Medical College Jaipur | SMS Medical College Jaipur | Bharti Malhotra, Swati Gautam, Himanshu Sharma, Pratibha Sharma, Neha Bhomia, Nivedita Gupta, Pragya Yadav,Varsha Potdar |
| EPI_ISL_891219 | SMS Medical College Jaipur | SMS Medical College Jaipur | Bharti Malhotra, Swati Gautam, Himanshu Sharma, Pratibha Sharma, Neha Bhomia, Nivedita Gupta, Pragya Yadav, Varsha Potdar |
| EPI_ISL_892212 | Lighthouse Lab in Alderley Park | Wellcome Sanger Institute for the COVID-19 Genomics UK (COG-UK) Consortium | Jacquelyn Wynn, Mairead Hyland, The Lighthouse Lab in Alderley Park and Alex Alderton, Roberto Amato, Sonia Goncalves, Ewan Harrison, David K. Jackson, Ian Johnston, Dominic Kwiatkowski, Cordelia Langford, John Sillitoe on behalf of the Wellcome Sanger Institute COVID-19 Surveillance Team |
| EPI_ISL_893771, EPI_ISL_893773 | Institute of Virology, Medical Center, University of Freiburg, Freiburg, Germany | Institute of Virology, Clinial Virus Genomics, Medical Center, University of Freiburg, Freiburg, Germany | Jonas Fuchs, Lisa Kern, Sandra Reuter, Hajo Grundmann, Marcus Panning |
| EPI_ISL_894169 | Institute of Medical Microbiology and Hospital Hygiene | Institute of Medical Microbiology and Hospital Hygiene | Prof. Dr. Achim Kaasch, Aljoscha Tersteegen |
| EPI_ISL_896114 | Viollier AG | University Hospital Basel, Clinical Bacteriology | Tim Roloff, Madlen Stange, Helena MB Seth-Smith, Alfredo Mari, Karoline Leuzinger, Julia Bielicki, Christiane Beckmann, Manuel Battegay, Hans Hirsch, Adrian Egli |
| EPI_ISL_896204 | MEPHI, Aix Marseille University | MEPHI, Aix Marseille University | Anthony LEVASSEUR |
| EPI_ISL_899015, EPI_ISL_899019, EPI_ISL_899020, EPI_ISL_899033, EPI_ISL_899109, EPI_ISL_899118, EPI_ISL_899119, EPI_ISL_899120, EPI_ISL_899134, EPI_ISL_899161, EPI_ISL_899179, EPI_ISL_899185, EPI_ISL_899186, EPI_ISL_899187, EPI_ISL_899285, EPI_ISL_899286, EPI_ISL_899287, EPI_ISL_899288, EPI_ISL_899289, EPI_ISL_899290, EPI_ISL_899291, EPI_ISL_899292, EPI_ISL_899293, EPI_ISL_899294, EPI_ISL_899295, EPI_ISL_899296, EPI_ISL_899297, EPI_ISL_899405, EPI_ISL_899406, EPI_ISL_899407, EPI_ISL_899408, EPI_ISL_899409, EPI_ISL_899410, EPI_ISL_899411, EPI_ISL_899412, EPI_ISL_899413, EPI_ISL_899414, EPI_ISL_899415, EPI_ISL_899416, EPI_ISL_899417, EPI_ISL_899418, EPI_ISL_899419, EPI_ISL_899420, EPI_ISL_899517, EPI_ISL_899518, EPI_ISL_899519, EPI_ISL_899520, EPI_ISL_899521, EPI_ISL_899559, EPI_ISL_899597, EPI_ISL_899621, EPI_ISL_899623, EPI_ISL_899624, EPI_ISL_899625, EPI_ISL_899626, EPI_ISL_899664, EPI_ISL_899768, EPI_ISL_899769, EPI_ISL_899770, EPI_ISL_899771, EPI_ISL_899772, EPI_ISL_899774, EPI_ISL_899775, EPI_ISL_899776, EPI_ISL_899777, EPI_ISL_899778, EPI_ISL_899779, EPI_ISL_899780, EPI_ISL_899811, EPI_ISL_899812, EPI_ISL_899813, EPI_ISL_899814, EPI_ISL_899815, EPI_ISL_899816, EPI_ISL_899817, EPI_ISL_899818, EPI_ISL_899819, EPI_ISL_899820, EPI_ISL_899821, EPI_ISL_899822, EPI_ISL_899823, EPI_ISL_899824, EPI_ISL_899825, EPI_ISL_899826, EPI_ISL_899827, EPI_ISL_899863, EPI_ISL_899875, EPI_ISL_899876, EPI_ISL_899877, EPI_ISL_899878, EPI_ISL_899879, EPI_ISL_899880, EPI_ISL_899881, EPI_ISL_899933, EPI_ISL_899937, EPI_ISL_899938, EPI_ISL_899939, EPI_ISL_899940, EPI_ISL_899941, EPI_ISL_899944, EPI_ISL_899945, EPI_ISL_899946, EPI_ISL_899947, EPI_ISL_899948, EPI_ISL_899949, EPI_ISL_899950, EPI_ISL_899951 | | | |
| see above | Viollier AG | Department of Biosystems Science and Engineering, ETH Zürich | Christian Beisel, Sarah Nadeau, Chaoran Chen, Ivan Topolsky, Philipp Jablonski, Lara Fuhrmann, David Dreifuss, Katharina Jahn, Tobias Schär, Ina Nissen, Natascha Santacroce, Elodie Burcklen, Christiane Beckmann, Maurice Redondo, Olivier Kobel, Christoph Noppen, Sophie Seidel, Noemie Santamaria de Souza, Niko Beerenwinkel, Tanja Stadler |
| EPI_ISL_900512 | Centre hospitalier Emile Roux | CNR Virus des Infections Respiratoires - France SUD | Antonin Bal, Gregory Destras, Gwendolyne Burfin, Hadrien Règue, Quentin Semanas, Martine Valette, Bruno Lina, Laurence Josset |
| EPI_ISL_900520 | Bioesterel | CNR Virus des Infections Respiratoires - France SUD | Antonin Bal, Gregory Destras, Gwendolyne Burfin, Hadrien Règue, Quentin Semanas, Martine Valette, Bruno Lina, Laurence Josset |
| EPI_ISL_900533, EPI_ISL_900544 | CNR Virus des Infections Respiratoires - France SUD | CNR Virus des Infections Respiratoires - France SUD | Antonin Bal, Gregory Destras, Gwendolyne Burfin, Hadrien Règue, Quentin Semanas, Martine Valette, Bruno Lina, Laurence Josset |
| EPI_ISL_900583, EPI_ISL_900633, EPI_ISL_900634, EPI_ISL_900635, EPI_ISL_900636, EPI_ISL_900637, EPI_ISL_900638, EPI_ISL_900639, EPI_ISL_900640, EPI_ISL_900641, EPI_ISL_900642, EPI_ISL_900643, EPI_ISL_900644, EPI_ISL_900645, EPI_ISL_900646, EPI_ISL_900648, EPI_ISL_900649, EPI_ISL_900650, EPI_ISL_900685, EPI_ISL_900686, EPI_ISL_900687, EPI_ISL_900688, EPI_ISL_900689, EPI_ISL_900690 | | | |
| see above | IZSM | TIGEM | Patrizia Annunziata, Andrea Ballabio, Valentina Bouche, Davide Cacchiarelli (CorrespAuthor), Pellegrino Cerino, Chiara Colantuono, Maria Concetta Cuomo, Denise Di Concilio, Lucio Di Filippo, Antonio Grimaldi, Antonio Limone, Anna Manfredi, Francesco Panariello, Biancamaria Pierri, Marcello Salvi |
| EPI_ISL_903037, EPI_ISL_903038, EPI_ISL_903039, EPI_ISL_903040, EPI_ISL_903041, EPI_ISL_903042, EPI_ISL_903045, EPI_ISL_903046, EPI_ISL_903047, EPI_ISL_903082, EPI_ISL_903083 | | | |
| see above | Seattle Flu Study | Seattle Flu Study | Deborah A. Nickerson, Chris D. Frazar, Jover Lee, Benjamin Pelle, Erica Ryke, Matthew Richardson, Amanda Adler, Elisabeth Brandstetter, Peter D. Han, Kairsten Fay, Misja Ilcisin, Kirsten Lacombe, Thomas R. Sibley, Melissa Truong, Caitlin R. Wolf, Karen Cowgill, Stephanie Schrag, Jeff Duchin, Michael Boeckh, Janet A. Englund, Michael Famulare, Barry R. Lutz, Mark J. Rieder, Lea M. Starita, Matthew Thompson, Helen Y. Chu, Trevor Bedford, Jay Shendure |
| EPI_ISL_903106 | Washington State Department of Health | Seattle Flu Study | Deborah A. Nickerson, Chris D. Frazar, Jover Lee, Benjamin Pelle, Erica Ryke, Matthew Richardson, Amanda Adler, Elisabeth Brandstetter, Peter D. Han, Kairsten Fay, Misja Ilcisin, Kirsten Lacombe, Thomas R. Sibley, Melissa Truong, Caitlin R. Wolf, Romesh Gautom, Geoff Melly, Brian Hiatt, Philip Dykema, Scott Lindquist, Michael Boeckh, Janet A. Englund, Michael Famulare, Barry R. Lutz, Mark J. Rieder, Lea M. Starita, Matthew Thompson, Helen Y. Chu, Jay Shendure, Trevor Bedford |
| EPI_ISL_903264, EPI_ISL_903273 | M Health Fairview | Minnesota Department of Health, Public Health Laboratory | Alexandra Lorentz, Jacob Garfin, Matt Plumb, and Xiong Wang |
| EPI_ISL_903375, EPI_ISL_903377, EPI_ISL_903379, EPI_ISL_903380, EPI_ISL_903381, EPI_ISL_903388, EPI_ISL_903389 | MOH - Jaber Al-Ahmad Hospital (Innovation Research Laboratory) | MOH - Jaber Al-Ahmad Hospital (Innovation Research Laboratory) | Salman Al-Sabah , Mohammad Alghounaim |
| EPI_ISL_903574 | HI Dept. of Health, State Laboratories Division | Genomics and Discovery, Respiratory Viruses Branch, Division of Viral Diseases, Centers for Disease Control and Prevention | Krista Queen, Yan Li, Ying Tao, Jing Zhang, Anna Uehara, Anna Montmayeur, Clinton R. Paden, Peter W. Cook, Rachel Marine, Mili Sheth, Jasmine Padilla, Sarah Nobles, Mark Burroughs, Lori Rowe, Haibin Wang, Ben L. Rambo-Martin, Dhwani Batra, Justin Lee, Suxiang Tong |
| EPI_ISL_903580 | AZ SPHL, Arizona Department of Health Services | Genomics and Discovery, Respiratory Viruses Branch, Division of Viral Diseases, Centers for Disease Control and Prevention | Krista Queen, Yan Li, Ying Tao, Jing Zhang, Anna Uehara, Anna Montmayeur, Clinton R. Paden, Peter W. Cook, Rachel Marine, Mili Sheth, Jasmine Padilla, Sarah Nobles, Mark Burroughs, Lori Rowe, Haibin Wang, Ben L. Rambo-Martin, Dhwani Batra, Justin Lee, Suxiang Tong |
| EPI_ISL_903584 | OH Department of Health Laboratory | Genomics and Discovery, Respiratory Viruses Branch, Division of Viral Diseases, Centers for Disease Control and Prevention | Krista Queen, Yan Li, Ying Tao, Jing Zhang, Anna Uehara, Anna Montmayeur, Clinton R. Paden, Peter W. Cook, Rachel Marine, Mili Sheth, Jasmine Padilla, Sarah Nobles, Mark Burroughs, Lori Rowe, Haibin Wang, Ben L. Rambo-Martin, Dhwani Batra, Justin Lee, Suxiang Tong |
| EPI_ISL_903592 | PA Department of Health, Bureau of Laboratories | Genomics and Discovery, Respiratory Viruses Branch, Division of Viral Diseases, Centers for Disease Control and Prevention | Krista Queen, Yan Li, Ying Tao, Jing Zhang, Anna Uehara, Anna Montmayeur, Clinton R. Paden, Peter W. Cook, Rachel Marine, Mili Sheth, Jasmine Padilla, Sarah Nobles, Mark Burroughs, Lori Rowe, Haibin Wang, Ben L. Rambo-Martin, Dhwani Batra, Justin Lee, Suxiang Tong |
| EPI_ISL_903622 | NM Dept. Health, Scientific Laboratory Division | Genomics and Discovery, Respiratory Viruses Branch, Division of Viral Diseases, Centers for Disease Control and Prevention | Krista Queen, Yan Li, Ying Tao, Jing Zhang, Anna Uehara, Anna Montmayeur, Clinton R. Paden, Peter W. Cook, Rachel Marine, Mili Sheth, Jasmine Padilla, Sarah Nobles, Mark Burroughs, Lori Rowe, Haibin Wang, Ben L. Rambo-Martin, Dhwani Batra, Justin Lee, Suxiang Tong |
| EPI_ISL_903634 | AZ SPHL, Arizona Department of Health Services | Genomics and Discovery, Respiratory Viruses Branch, | Krista Queen, Yan Li, Ying Tao, Jing Zhang, Anna Uehara, Anna Montmayeur, Clinton R. Paden, Peter W. Cook, Rachel Marine, Mili Sheth, Jasmine |

| | | | |
|---|---|---|---|
| | | Division of Viral Diseases, Centers for Disease Control and Prevention | Padilla, Sarah Nobles, Mark Burroughs, Lori Rowe, Haibin Wang, Ben L. Rambo-Martin, Dhwani Batra, Justin Lee, Suxiang Tong |
| EPI_ISL_903677 | MN PHL Division, Minnesota Department of Health | Genomics and Discovery, Respiratory Viruses Branch, Division of Viral Diseases, Centers for Disease Control and Prevention | Krista Queen, Yan Li, Ying Tao, Jing Zhang, Anna Uehara, Anna Montmayeur, Clinton R. Paden, Peter W. Cook, Rachel Marine, Mili Sheth, Jasmine Padilla, Sarah Nobles, Mark Burroughs, Lori Rowe, Haibin Wang, Ben L. Rambo-Martin, Dhwani Batra, Justin Lee, Suxiang Tong |
| EPI_ISL_903715 | VT Dept. of Health Laboratory | Genomics and Discovery, Respiratory Viruses Branch, Division of Viral Diseases, Centers for Disease Control and Prevention | Krista Queen, Yan Li, Ying Tao, Jing Zhang, Anna Uehara, Anna Montmayeur, Clinton R. Paden, Peter W. Cook, Rachel Marine, Mili Sheth, Jasmine Padilla, Sarah Nobles, Mark Burroughs, Lori Rowe, Haibin Wang, Ben L. Rambo-Martin, Dhwani Batra, Justin Lee, Suxiang Tong |
| EPI_ISL_903740, EPI_ISL_903755 | NM Dept. Health, Scientific Laboratory Division | Genomics and Discovery, Respiratory Viruses Branch, Division of Viral Diseases, Centers for Disease Control and Prevention | Krista Queen, Yan Li, Ying Tao, Jing Zhang, Anna Uehara, Anna Montmayeur, Clinton R. Paden, Peter W. Cook, Rachel Marine, Mili Sheth, Jasmine Padilla, Sarah Nobles, Mark Burroughs, Lori Rowe, Haibin Wang, Ben L. Rambo-Martin, Dhwani Batra, Justin Lee, Suxiang Tong |
| EPI_ISL_903778 | PA Department of Health, Bureau of Laboratories | Genomics and Discovery, Respiratory Viruses Branch, Division of Viral Diseases, Centers for Disease Control and Prevention | Krista Queen, Yan Li, Ying Tao, Jing Zhang, Anna Uehara, Anna Montmayeur, Clinton R. Paden, Peter W. Cook, Rachel Marine, Mili Sheth, Jasmine Padilla, Sarah Nobles, Mark Burroughs, Lori Rowe, Haibin Wang, Ben L. Rambo-Martin, Dhwani Batra, Justin Lee, Suxiang Tong |
| EPI_ISL_903796, EPI_ISL_903800 | NM Dept. Health, Scientific Laboratory Division | Genomics and Discovery, Respiratory Viruses Branch, Division of Viral Diseases, Centers for Disease Control and Prevention | Krista Queen, Yan Li, Ying Tao, Jing Zhang, Anna Uehara, Anna Montmayeur, Clinton R. Paden, Peter W. Cook, Rachel Marine, Mili Sheth, Jasmine Padilla, Sarah Nobles, Mark Burroughs, Lori Rowe, Haibin Wang, Ben L. Rambo-Martin, Dhwani Batra, Justin Lee, Suxiang Tong |
| EPI_ISL_903808 | AZ SPHL, Arizona Department of Health Services | Genomics and Discovery, Respiratory Viruses Branch, Division of Viral Diseases, Centers for Disease Control and Prevention | Krista Queen, Yan Li, Ying Tao, Jing Zhang, Anna Uehara, Anna Montmayeur, Clinton R. Paden, Peter W. Cook, Rachel Marine, Mili Sheth, Jasmine Padilla, Sarah Nobles, Mark Burroughs, Lori Rowe, Haibin Wang, Ben L. Rambo-Martin, Dhwani Batra, Justin Lee, Suxiang Tong |
| EPI_ISL_903811 | NM Dept. Health, Scientific Laboratory Division | Genomics and Discovery, Respiratory Viruses Branch, Division of Viral Diseases, Centers for Disease Control and Prevention | Krista Queen, Yan Li, Ying Tao, Jing Zhang, Anna Uehara, Anna Montmayeur, Clinton R. Paden, Peter W. Cook, Rachel Marine, Mili Sheth, Jasmine Padilla, Sarah Nobles, Mark Burroughs, Lori Rowe, Haibin Wang, Ben L. Rambo-Martin, Dhwani Batra, Justin Lee, Suxiang Tong |
| EPI_ISL_903834 | PA Department of Health, Bureau of Laboratories | Genomics and Discovery, Respiratory Viruses Branch, Division of Viral Diseases, Centers for Disease Control and Prevention | Krista Queen, Yan Li, Ying Tao, Jing Zhang, Anna Uehara, Anna Montmayeur, Clinton R. Paden, Peter W. Cook, Rachel Marine, Mili Sheth, Jasmine Padilla, Sarah Nobles, Mark Burroughs, Lori Rowe, Haibin Wang, Ben L. Rambo-Martin, Dhwani Batra, Justin Lee, Suxiang Tong |
| EPI_ISL_903838, EPI_ISL_903855 | AZ SPHL, Arizona Department of Health Services | Genomics and Discovery, Respiratory Viruses Branch, Division of Viral Diseases, Centers for Disease Control and Prevention | Krista Queen, Yan Li, Ying Tao, Jing Zhang, Anna Uehara, Anna Montmayeur, Clinton R. Paden, Peter W. Cook, Rachel Marine, Mili Sheth, Jasmine Padilla, Sarah Nobles, Mark Burroughs, Lori Rowe, Haibin Wang, Ben L. Rambo-Martin, Dhwani Batra, Justin Lee, Suxiang Tong |
| EPI_ISL_903864 | DC Public Health Lab/ Dept. of Forensic Sciences | Genomics and Discovery, Respiratory Viruses Branch, Division of Viral Diseases, Centers for Disease Control and Prevention | Krista Queen, Yan Li, Ying Tao, Jing Zhang, Anna Uehara, Anna Montmayeur, Clinton R. Paden, Peter W. Cook, Rachel Marine, Mili Sheth, Jasmine Padilla, Sarah Nobles, Mark Burroughs, Lori Rowe, Haibin Wang, Ben L. Rambo-Martin, Dhwani Batra, Justin Lee, Suxiang Tong |
| EPI_ISL_903866 | MD DOH Laboratories Administration | Genomics and Discovery, Respiratory Viruses Branch, Division of Viral Diseases, Centers for Disease Control and Prevention | Krista Queen, Yan Li, Ying Tao, Jing Zhang, Anna Uehara, Anna Montmayeur, Clinton R. Paden, Peter W. Cook, Rachel Marine, Mili Sheth, Jasmine Padilla, Sarah Nobles, Mark Burroughs, Lori Rowe, Haibin Wang, Ben L. Rambo-Martin, Dhwani Batra, Justin Lee, Suxiang Tong |
| EPI_ISL_903870 | NM Dept. Health, Scientific Laboratory Division | Genomics and Discovery, Respiratory Viruses Branch, Division of Viral Diseases, Centers for Disease Control and Prevention | Krista Queen, Yan Li, Ying Tao, Jing Zhang, Anna Uehara, Anna Montmayeur, Clinton R. Paden, Peter W. Cook, Rachel Marine, Mili Sheth, Jasmine Padilla, Sarah Nobles, Mark Burroughs, Lori Rowe, Haibin Wang, Ben L. Rambo-Martin, Dhwani Batra, Justin Lee, Suxiang Tong |
| EPI_ISL_903876 | PA Department of Health, Bureau of Laboratories | Genomics and Discovery, Respiratory Viruses Branch, Division of Viral Diseases, Centers for Disease Control and Prevention | Krista Queen, Yan Li, Ying Tao, Jing Zhang, Anna Uehara, Anna Montmayeur, Clinton R. Paden, Peter W. Cook, Rachel Marine, Mili Sheth, Jasmine Padilla, Sarah Nobles, Mark Burroughs, Lori Rowe, Haibin Wang, Ben L. Rambo-Martin, Dhwani Batra, Justin Lee, Suxiang Tong |
| EPI_ISL_903885 | NE Public Health Laboratory | Genomics and Discovery, Respiratory Viruses Branch, Division of Viral Diseases, Centers for Disease Control and Prevention | Krista Queen, Yan Li, Ying Tao, Jing Zhang, Anna Uehara, Anna Montmayeur, Clinton R. Paden, Peter W. Cook, Rachel Marine, Mili Sheth, Jasmine Padilla, Sarah Nobles, Mark Burroughs, Lori Rowe, Haibin Wang, Ben L. Rambo-Martin, Dhwani Batra, Justin Lee, Suxiang Tong |
| EPI_ISL_903886 | IN State Department of Health Laboratory Services | Genomics and Discovery, Respiratory Viruses Branch, Division of Viral Diseases, Centers for Disease Control and Prevention | Krista Queen, Yan Li, Ying Tao, Jing Zhang, Anna Uehara, Anna Montmayeur, Clinton R. Paden, Peter W. Cook, Rachel Marine, Mili Sheth, Jasmine Padilla, Sarah Nobles, Mark Burroughs, Lori Rowe, Haibin Wang, Ben L. Rambo-Martin, Dhwani Batra, Justin Lee, Suxiang Tong |
| EPI_ISL_903902, EPI_ISL_903919, EPI_ISL_903935 | AZ SPHL, Arizona Department of Health Services | Genomics and Discovery, Respiratory Viruses Branch, Division of Viral Diseases, Centers for Disease Control and Prevention | Krista Queen, Yan Li, Ying Tao, Jing Zhang, Anna Uehara, Anna Montmayeur, Clinton R. Paden, Peter W. Cook, Rachel Marine, Mili Sheth, Jasmine Padilla, Sarah Nobles, Mark Burroughs, Lori Rowe, Haibin Wang, Ben L. Rambo-Martin, Dhwani Batra, Justin Lee, Suxiang Tong |
| EPI_ISL_903958 | MD DOH Laboratories Administration | Genomics and Discovery, Respiratory Viruses Branch, Division of Viral Diseases, Centers for Disease Control and Prevention | Krista Queen, Yan Li, Ying Tao, Jing Zhang, Anna Uehara, Anna Montmayeur, Clinton R. Paden, Peter W. Cook, Rachel Marine, Mili Sheth, Jasmine Padilla, Sarah Nobles, Mark Burroughs, Lori Rowe, Haibin Wang, Ben L. Rambo-Martin, Dhwani Batra, Justin Lee, Suxiang Tong |
| EPI_ISL_903959, EPI_ISL_903964 | DC Public Health Lab/ Dept. of Forensic Sciences | Genomics and Discovery, Respiratory Viruses Branch, Division of Viral Diseases, Centers for Disease Control and Prevention | Krista Queen, Yan Li, Ying Tao, Jing Zhang, Anna Uehara, Anna Montmayeur, Clinton R. Paden, Peter W. Cook, Rachel Marine, Mili Sheth, Jasmine Padilla, Sarah Nobles, Mark Burroughs, Lori Rowe, Haibin Wang, Ben L. Rambo-Martin, Dhwani Batra, Justin Lee, Suxiang Tong |
| EPI_ISL_903967 | MD DOH Laboratories Administration | Genomics and Discovery, Respiratory Viruses Branch, Division of Viral Diseases, Centers for Disease Control and Prevention | Krista Queen, Yan Li, Ying Tao, Jing Zhang, Anna Uehara, Anna Montmayeur, Clinton R. Paden, Peter W. Cook, Rachel Marine, Mili Sheth, Jasmine Padilla, Sarah Nobles, Mark Burroughs, Lori Rowe, Haibin Wang, Ben L. Rambo-Martin, Dhwani Batra, Justin Lee, Suxiang Tong |
| EPI_ISL_903976 | DC Public Health Lab/ Dept. of Forensic Sciences | Genomics and Discovery, Respiratory Viruses Branch, Division of Viral Diseases, Centers for Disease Control and Prevention | Krista Queen, Yan Li, Ying Tao, Jing Zhang, Anna Uehara, Anna Montmayeur, Clinton R. Paden, Peter W. Cook, Rachel Marine, Mili Sheth, Jasmine Padilla, Sarah Nobles, Mark Burroughs, Lori Rowe, Haibin Wang, Ben L. Rambo-Martin, Dhwani Batra, Justin Lee, Suxiang Tong |
| EPI_ISL_904170, EPI_ISL_904196, EPI_ISL_904197, EPI_ISL_904198, EPI_ISL_904300, EPI_ISL_904301, EPI_ISL_904302, EPI_ISL_904363, EPI_ISL_904506, EPI_ISL_904507, EPI_ISL_904508, EPI_ISL_904509, EPI_ISL_904510, EPI_ISL_904511, EPI_ISL_904512, EPI_ISL_904513, EPI_ISL_904514, EPI_ISL_904515, EPI_ISL_904516, EPI_ISL_904517, EPI_ISL_904518, EPI_ISL_904519, EPI_ISL_904618, EPI_ISL_904619, EPI_ISL_904620 | | | |
| see above | Dutch COVID-19 response team | Erasmus Medical Center | Bas Oude Munnink, Reina Sikkema, David Nieuwenhuijse, Irina Chestakova, Anne van der Linden, Marjan Boter, Emmanuelle Munger, Corine GeurtsvanKessel, Annemiek van der Eijk, Richard Molenkamp, Marion Koopmans, on behalf of the Dutch national COVID-19 response team. |
| EPI_ISL_904658, EPI_ISL_904746, EPI_ISL_904785, EPI_ISL_904887, EPI_ISL_904899, EPI_ISL_904903, EPI_ISL_905004, EPI_ISL_905099, EPI_ISL_905100, EPI_ISL_905128, EPI_ISL_905151, EPI_ISL_905156, EPI_ISL_905167, EPI_ISL_905174, EPI_ISL_905215, EPI_ISL_905224, EPI_ISL_905229, EPI_ISL_905232, EPI_ISL_905233, EPI_ISL_905237, EPI_ISL_905298, EPI_ISL_905370, EPI_ISL_905371, EPI_ISL_905377, EPI_ISL_905394, EPI_ISL_905395, EPI_ISL_905396, EPI_ISL_905397, EPI_ISL_905398, EPI_ISL_905399, EPI_ISL_905400, EPI_ISL_905401, EPI_ISL_905402, EPI_ISL_905403, EPI_ISL_905428, EPI_ISL_905433, EPI_ISL_905462, EPI_ISL_905463, EPI_ISL_905464, EPI_ISL_905465, EPI_ISL_905466, EPI_ISL_905467, EPI_ISL_905468, EPI_ISL_905469, EPI_ISL_905470, EPI_ISL_905471, EPI_ISL_905483, EPI_ISL_905491, EPI_ISL_905492, EPI_ISL_905493, EPI_ISL_905495, EPI_ISL_905496, EPI_ISL_905497, EPI_ISL_905498, EPI_ISL_905499, EPI_ISL_905500, EPI_ISL_905504, EPI_ISL_905505, EPI_ISL_905506, EPI_ISL_905508, EPI_ISL_905509, EPI_ISL_905510, EPI_ISL_905514, EPI_ISL_905560, EPI_ISL_905671, EPI_ISL_905672, EPI_ISL_905709, EPI_ISL_905710 | | | |
| see above | Dutch COVID-19 response team | National Institute for Public Health and the Environment (RIVM) | Adam Meijer, Harry Vennema, Dirk Eggink, Jeroen Cremer, Sharon van den Brink, Bas van der Veer, AnneMarie van den Brandt, Florian Zwagemaker, Dennis Schmitz, Chantal Reusken, on behalf of the national COVID-19 response team |
| EPI_ISL_906128, EPI_ISL_906129 | Institute of Microbiology and Immunology, Faculty of Medicine, University of Ljubljana | Institute of Microbiology and Immunology, Faculty of Medicine, University of Ljubljana | Samo Zakotnik, Tomaž Mark Zorec, Matic Brvar, Miša Korva, Mario Poljak, Tatjana Avši - Županc |

| | | | |
|---|---|---|---|
| EPI_ISL_906543 | FUNDACION HOSPITAL SAN PEDRO | Instituto Nacional de Salud- Dirección de Investigación en Salud Pública, Universidad de los Andes- Applied genomics research group, Vicerrectoria de Investigación y Creación, Universidad de los Andes- Systems and Computing Engineering Department | Katherine Laiton-Donato, Diego A. Álvarez-Díaz, Carlos Franco-Muñoz, Mauricio Pacheco-Montealegre, Héctor Alejandro Ruiz-Moreno, Maria T. Herrera-Sepúlveda, Diego Andrés Prada, Jhonnatan Reales-González, Sheryll Corchuelo, Julian Naizaque, Gerardo Santamaría Jorge Duitama, Laura Natalia Gonzalez, Jorge Ivan Diaz, Silvia Restrepo-Restrepo, Magdalena Wiesner, Martha Lucia Ospina Martinez, Marcela Mercado-Reyes |
| EPI_ISL_906553, EPI_ISL_906555 | Laboratorio Bienestar | Instituto Nacional de Salud- Dirección de Investigación en Salud Pública, Universidad de los Andes- Applied genomics research group, Vicerrectoria de Investigación y Creación, Universidad de los Andes- Systems and Computing Engineering Department | Katherine Laiton-Donato, Diego A. Álvarez-Díaz, Carlos Franco-Muñoz, Mauricio Pacheco-Montealegre, Héctor Alejandro Ruiz-Moreno, Maria T. Herrera-Sepúlveda, Diego Andrés Prada, Jhonnatan Reales-González, Sheryll Corchuelo, Julian Naizaque, Gerardo Santamaría Jorge Duitama, Laura Natalia Gonzalez, Jorge Ivan Diaz, Silvia Restrepo-Restrepo, Magdalena Wiesner, Martha Lucia Ospina Martinez, Marcela Mercado-Reyes |
| EPI_ISL_906571 | Maine Health and Environmental Testing Laboratory (Maine HETL) | Tewhey Lab, The Jackson Laboratory | Matluk,N., Dewey,H., Iosue,F., Barter,M., Lynch,R., Munger,H. and Tewhey,R. |
| EPI_ISL_906848, EPI_ISL_906851 | Respiratory Viruses Branch, Centers for Disease Control and Prevention | Respiratory Viruses Branch, Centers for Disease Control and Prevention | Tao,Y., Li,Y., Zhang,J., Queen,K., Uehara,A., Cook,P., Paden,C.R., Wang,H., Tong,S. |
| EPI_ISL_906918 | Bureau of Public Health Laboratories, Florida Department of Health (BPHL, FLDOH) | Bureau of Public Health Laboratories, Florida Department of Health (BPHL, FLDOH) | Schmedes,S., Blanton,J. |
| EPI_ISL_910547, EPI_ISL_910548, EPI_ISL_910550, EPI_ISL_910551, EPI_ISL_910552, EPI_ISL_910633, EPI_ISL_911241 | Laboratoire national de sante, Microbiology, Virology | Laboratoire national de sante, Microbiology, Microbial Genomics Platform | Anke Wienecke-Baldacchino, Catherine Ragimbeau,Jessica Tapp, Fatu Djabi, Lise Pignon, Raoul Salmon, Tamir Abdelrahman |
| EPI_ISL_912279, EPI_ISL_912297, EPI_ISL_912299, EPI_ISL_912315, EPI_ISL_912324, EPI_ISL_912336, EPI_ISL_912352 | Hospital General Universitario Gregorio Marañón | SeqCOVID-SPAIN consortium / IBV (CSIC) | Darío García de Viedma, Laura Pérez-Lago, Pedro J Sola-Campoy, Sergio Buenestado-Serrano, Marta Herranz, Victor Manuel de la Cueva, Julia Suárez, Pilar Catalán, Patricia Muñoz and SeqCOVID-SPAIN consortium |
| EPI_ISL_912407, EPI_ISL_912409, EPI_ISL_912417, EPI_ISL_912426 | KU Leuven, Rega Institute, Clinical and Epidemiological Virology | KU Leuven, Rega Institute, Clinical and Epidemiological Virology | Tony Wawina-Bokalanga, Bert Vanmechelen, Joan Marti-Carerras, Piet Maes |
| EPI_ISL_912508 | NHLS Universitas Academic | UFS Virology | PA Bester, MM Nyaga, P Nthiga, MT Mogotsi, D Goedhals, T de Oliveira |
| EPI_ISL_912892, EPI_ISL_912896, EPI_ISL_912906, EPI_ISL_912921, EPI_ISL_912929, EPI_ISL_912932, EPI_ISL_913012, EPI_ISL_913013, EPI_ISL_913014, EPI_ISL_913015, EPI_ISL_913016, EPI_ISL_913017, EPI_ISL_913018, EPI_ISL_913019 | | | |
| see above | Hôpital Henri Mondor | Department of Virology, Henri Mondor University Hospital, Assistance Publique Hôpitaux de Paris, Université Paris-Est Créteil, INSERM U955 | Christophe Rodriguez, Slim Fourati, Vanessa Demontant, Guillaume Gricourt, Melissa N'Debi, Alexandre Soulier, Elisabeth Trawinski, Jean-Michel Pawlotsky |
| EPI_ISL_913051 | Hospital Comarcal Sierrallana | Instituto de Salud Carlos III | Iglesias-Caballero, M. Camarero, S. Sandonís,V. Vázquez, S. Pozo, F. Casas, I. Jiménez, P. Zaballos, A. Monzón, S. Varona, S. Cuesta, I. De Benito, I. |
| EPI_ISL_913052, EPI_ISL_913053 | Complejo Asistencial Universitario de Salamanca | Instituto de Salud Carlos III | Iglesias-Caballero, M. Camarero, S. Sandonís,V. Vázquez, S. Pozo, F. Casas, I. Jiménez, P. Zaballos, A. Monzón, S. Varona, S. Cuesta, I. Ávila, A. |
| EPI_ISL_913276 | Synlab Medilab, Mikrobiologi | The Public Health Agency of Sweden | Anna-Malin Linde, Maria Lind Karlberg, Carlo Berg, Oskar Karlsson Lindsjo, Sofia Stamouli, Reza Advani, Mattias Haukland, Petra Holmstrom, Noura Walai, Petra Edquist, Mia Brytting, Anna Risberg, Karin Tegmark-Wisell |
| EPI_ISL_913280 | Klinisk mikrobiologi | The Public Health Agency of Sweden | Anna-Malin Linde, Maria Lind Karlberg, Carlo Berg, Oskar Karlsson Lindsjo, Sofia Stamouli, Reza Advani, Mattias Haukland, Petra Holmstrom, Noura Walai, Petra Edquist, Mia Brytting, Anna Risberg, Karin Tegmark-Wisell |
| EPI_ISL_913284 | Synlab Medilab, Mikrobiologi | The Public Health Agency of Sweden | Anna-Malin Linde, Maria Lind Karlberg, Carlo Berg, Oskar Karlsson Lindsjo, Sofia Stamouli, Reza Advani, Mattias Haukland, Petra Holmstrom, Noura Walai, Petra Edquist, Mia Brytting, Anna Risberg, Karin Tegmark-Wisell |
| EPI_ISL_913286, EPI_ISL_913314 | The Public Health Agency of Sweden | The Public Health Agency of Sweden | Anna-Malin Linde, Maria Lind Karlberg, Carlo Berg, Oskar Karlsson Lindsjo, Sofia Stamouli, Reza Advani, Mattias Haukland, Petra Holmstrom, Noura Walai, Petra Edquist, Mia Brytting, Anna Risberg, Karin Tegmark-Wisell |
| EPI_ISL_913327, EPI_ISL_913329 | Klinisk mikrobiologi | The Public Health Agency of Sweden | Anna-Malin Linde, Maria Lind Karlberg, Carlo Berg, Oskar Karlsson Lindsjo, Sofia Stamouli, Reza Advani, Mattias Haukland, Petra Holmstrom, Noura Walai, Petra Edquist, Mia Brytting, Anna Risberg, Karin Tegmark-Wisell |
| EPI_ISL_913340 | Unilabs, Mikrobiologiska laboratoriet | The Public Health Agency of Sweden | Anna-Malin Linde, Maria Lind Karlberg, Carlo Berg, Oskar Karlsson Lindsjo, Sofia Stamouli, Reza Advani, Mattias Haukland, Petra Holmstrom, Noura Walai, Petra Edquist, Mia Brytting, Anna Risberg, Karin Tegmark-Wisell |
| EPI_ISL_913378, EPI_ISL_913379 | Dynamic Code AB | The Public Health Agency of Sweden | Anna-Malin Linde, Maria Lind Karlberg, Carlo Berg, Oskar Karlsson Lindsjo, Sofia Stamouli, Reza Advani, Mattias Haukland, Petra Holmstrom, Noura Walai, Petra Edquist, Mia Brytting, Anna Risberg, Karin Tegmark-Wisell |
| EPI_ISL_913463 | Klinisk mikrobiologi | The Public Health Agency of Sweden | Anna-Malin Linde, Maria Lind Karlberg, Carlo Berg, Oskar Karlsson Lindsjo, Sofia Stamouli, Reza Advani, Mattias Haukland, Petra Holmstrom, Noura Walai, Petra Edquist, Mia Brytting, Anna Risberg, Karin Tegmark-Wisell |
| EPI_ISL_913464 | The Public Health Agency of Sweden | The Public Health Agency of Sweden | Anna-Malin Linde, Maria Lind Karlberg, Carlo Berg, Oskar Karlsson Lindsjo, Sofia Stamouli, Reza Advani, Mattias Haukland, Petra Holmstrom, Noura Walai, Petra Edquist, Mia Brytting, Anna Risberg, Karin Tegmark-Wisell |
| EPI_ISL_913485 | Klinisk mikrobiologi | The Public Health Agency of Sweden | Anna-Malin Linde, Maria Lind Karlberg, Carlo Berg, Oskar Karlsson Lindsjo, Sofia Stamouli, Reza Advani, Mattias Haukland, Petra Holmstrom, Noura Walai, Petra Edquist, Mia Brytting, Anna Risberg, Karin Tegmark-Wisell |
| EPI_ISL_913519, EPI_ISL_913520, EPI_ISL_913521, EPI_ISL_913522, EPI_ISL_913523, EPI_ISL_913524, EPI_ISL_913526, EPI_ISL_913527, EPI_ISL_913536 | M Health Fairview | Minnesota Department of Health, Public Health Laboratory | Alexandra Lorentz, Jacob Garfin, Matt Plumb, and Xiong Wang |
| EPI_ISL_913611, EPI_ISL_913612, EPI_ISL_913634, EPI_ISL_913635, EPI_ISL_913637, EPI_ISL_913638, EPI_ISL_913639 | Michigan Department of Health and Human Services, Bureau of Laboratories | Michigan Department of Health and Human Services, Bureau of Laboratories | Blankenship HM, Riner D, Soehnlen MK |
| EPI_ISL_913676, EPI_ISL_913677, EPI_ISL_913678, EPI_ISL_913679, EPI_ISL_913680, EPI_ISL_913681, EPI_ISL_913682, EPI_ISL_913683, EPI_ISL_913684, EPI_ISL_913685, EPI_ISL_913692 | | | |
| see above | Minnesota Department of Health, Public Health Laboratory | Minnesota Department of Health, Public Health Laboratory | Alexandra Lorentz, Jacob Garfin, Matt Plumb, and Xiong Wang |
| EPI_ISL_913811, EPI_ISL_913897 | TGen North | TGen North | "Jolene Bowers, Megan Folkerts, Chris French, Hayley Yaglom, Ashlyn Pfeiffer, Darrin Lemmer, Dave Engelthaler, The Arizona COVID Genomics Union (ACGU)" |
| EPI_ISL_914648, EPI_ISL_914649, EPI_ISL_914650, EPI_ISL_914651, EPI_ISL_914652, EPI_ISL_914653, EPI_ISL_914654, EPI_ISL_914655, EPI_ISL_914656, EPI_ISL_914657, EPI_ISL_914658, EPI_ISL_914659, EPI_ISL_914660, EPI_ISL_914661, EPI_ISL_914663, EPI_ISL_914664, EPI_ISL_914665, EPI_ISL_914666, EPI_ISL_914696, EPI_ISL_914697, EPI_ISL_914698, EPI_ISL_914699, EPI_ISL_914700, EPI_ISL_914701 | | | |
| see above | Utah Public Health Laboratory | Utah Public Health Laboratory | Erin L. Young, Kelly F. Oakeson, Tara Gallagher |
| EPI_ISL_914814 | HOSPITAL METROPOLITANO | Inciensa, Instituto Costarricense de Investigación y Enseñanza en Nutrición y Salud | Francisco Duarte, Hebleen Porras, Claudio Soto-Garita, Estela Cordero, Adriana Godínez, Melany Calderón & Margarita Lee-Lui |
| EPI_ISL_918172, EPI_ISL_918173 | Department of Infectious Diseases and Immunology, National | Clinical Research Center, National Hospital Organization | Yoshihiro Nakata, Hirotaka Ode, Mai Kubota, Masakazu Matsuda, Kazuhiro Matsuoka, Miho Nakasuji, Mikiko Mori, Mayumi Imahashi, Yoshiyuki |

| | | | |
|---|---|---|---|
| | Hospital Organization Nagoya Medical Center | Nagoya Medical Center | Yokomaku, Yasumasa Iwatani |
| EPI_ISL_918193, EPI_ISL_918241, EPI_ISL_918248, EPI_ISL_918250 | Innovative Genomics Institute, UC Berkeley | Innovative Genomics Institute, UC Berkeley | Stacia Wyman, Haridha Shivram, Phil Frankino, Liana Lareau, Shana McDevitt, Justin Choi |
| EPI_ISL_918349, EPI_ISL_918350 | Institute of Virology, Medical Center, University of Freiburg, Freiburg, Germany | Institute of Virology, Clinial Virus Genomics, Medical Center, University of Freiburg, Freiburg, Germany | Jonas Fuchs, Lisa Kern, Sandra Reuter, Hajo Grundmann, Marcus Panning |
| EPI_ISL_918408 | Ospedale Santa Caterina Novella | Istituto Zooprofilattico Sperimentale della Puglia e della Basilicata | Parisi A., Bianco A., Capozzi L., Del Sambro L., Simone D., Manzulli V, Rondinone V., Pace L., Cipolletta D., Galante D. |
| EPI_ISL_918500, EPI_ISL_918501, EPI_ISL_918503 | LACEN - Laboratório Central de Saúde Pública do Amazonas | Evandro Chagas Institute | Santos, M.C.; Silva, A.M.; Junior, W.D.C.; Barbagelata, L.S.; Ferreira, J.A.; Sousa, E.M.A.; da Silva, P.S.; Pinheiro, K.C.; L.C.; Sousa Junior, E.C. |
| EPI_ISL_918955 | University of Birmingham | COVID-19 Genomics UK (COG-UK) Consortium | Institute of Microbiology, University of Birmingham: Claire McMurray, Joanne Stockton, Samuel Nicholls, Radoslaw Poplawski, Will Rowe, Josh Quick, Nicholas Loman. University of Birmingham Testing Laboratory: Celina M Whalley, Andrew Bosworth, Charlotte Poxon, Kasun Wanigasooriya, Oliver Pickles, Mike Kidd, Alex Richter, Andrew D Beggs PHE Heartlands Lab: Husam Osman, Andrew Bosworth. Queen Elizabeth Hospital: Anna Casey |
| EPI_ISL_919406 | Virology Department, Royal Infirmary of Edinburgh, NHS Lothian / School of Biological Sciences, University of Edinburgh / Institute of Genetics and Molecular Medicine, University of Edinburgh | COVID-19 Genomics UK (COG-UK) Consortium | McHugh M, Dewar R, Rooke S, Gallagher M, Balcaza C, O'Toole Á, Scher E, Hill V, McCrone JT, Colquhoun R, Yu X, Jackson B, Rambaut A, Williams TC, Templeton K |
| EPI_ISL_919456 | Liverpool Clinical Laboratories | COVID-19 Genomics UK (COG-UK) Consortium | Sam Haldenby, Anita Lucaci, Steve Paterson, Julian Hiscox, Alistair Darby, M Almsaud, A Alrezaihi, Muhannad Alruwaili, Stuart D Armstrong, Jones Benjamin, Eleanor G Bentley, Anu Chawla, Jordan J Clark, Angela Cowell, Richard Eccles, Isabel García-Dorival, Matthew Gemmell, Alessandro Gerada, PKF Gilmore, Richard Gregory, Ximeng Han, Catherine Hartley, Margaret Hughes, Miren Iturriza-Gomara, James Johnson, L Luu, Jenifer Manson, Charlotte Nelson, Elaine O'Toole, Cassie Olateju, Rebekah Penrice-Randal , Lucille Rainbow, N.P Randle, Trevor Ian Robinson, Parul Sharma, Ghada T Shawli, James P Stewart, Neil Swainston, Ecaterina Vamos, Joanne Watts, Mark Whitehead |
| EPI_ISL_919760, EPI_ISL_919761, EPI_ISL_919762, EPI_ISL_919763, EPI_ISL_919764, EPI_ISL_919765, EPI_ISL_919766, EPI_ISL_919767 | University College London, Great Ormond Street Hospital for Children NHS Foundation Trust, Imperial College Healthcare NHS Trust | COVID-19 Genomics UK (COG-UK) Consortium | Sergi Castellano, Rachel Williams, Mark Kristiansen, Paola Resende Silva, Sunando Roy, Tony Brooks, Helena Tutill, Paola Niola, Patricia Dyal, Charlotte Williams, Leysa Forrest, Yasmin Panchbhaya, Jacqueline Findlay, Samuel Weeks, Julianne Brown, Kathryn Harris, Paul Randell, James Price, Alison Holmes, Judith Breuer |
| EPI_ISL_920189, EPI_ISL_920319, EPI_ISL_920320, EPI_ISL_920354, EPI_ISL_920370, EPI_ISL_920569, EPI_ISL_920629, EPI_ISL_920653, EPI_ISL_920697 | University College London Hospital | COVID-19 Genomics UK (COG-UK) Consortium | Judith Heaney, Matthew Byott, Catherine Houlihan, Dan Frampton, Stuart Kirk, Moira Spyer and Eleni Nastouli |
| EPI_ISL_920841, EPI_ISL_920854, EPI_ISL_920855 | University College London, Great Ormond Street Hospital for Children NHS Foundation Trust, Imperial College Healthcare NHS Trust | COVID-19 Genomics UK (COG-UK) Consortium | Sergi Castellano, Rachel Williams, Mark Kristiansen, Paola Resende Silva, Sunando Roy, Tony Brooks, Helena Tutill, Paola Niola, Patricia Dyal, Charlotte Williams, Leysa Forrest, Yasmin Panchbhaya, Jacqueline Findlay, Samuel Weeks, Julianne Brown, Kathryn Harris, Paul Randell, James Price, Alison Holmes, Judith Breuer |
| EPI_ISL_921090 | Regional Virus Laboratory, Belfast Health and Social Care Trust | COVID-19 Genomics UK (COG-UK) Consortium | Conall McCaughey, James McKenna, Tanya Curran, Susan Feeney, Alison Watt, Ciara Cox, Mairead Connor, Zoltan Molnar, David Simpson, Derek Fairley |
| EPI_ISL_921227, EPI_ISL_921230, EPI_ISL_921231, EPI_ISL_921232, EPI_ISL_921233, EPI_ISL_921234, EPI_ISL_921235, EPI_ISL_921237, EPI_ISL_921240, EPI_ISL_921364, EPI_ISL_921365, EPI_ISL_921366, EPI_ISL_921367, EPI_ISL_921368, EPI_ISL_921369, EPI_ISL_921370, EPI_ISL_921372, EPI_ISL_921374, EPI_ISL_921375, EPI_ISL_921376, EPI_ISL_921377, EPI_ISL_921670, EPI_ISL_921671, EPI_ISL_921771, EPI_ISL_921774, EPI_ISL_921778, EPI_ISL_921779 | | | |
| see above | Northumbria University / South Tees Hospitals NHS Foundation Trust / North Cumbria Integrated Care NHS Foundation Trust / North Tees and Hartlepool NHS Foundation Trust / Newcastle Hospitals NHS Foundation Trust | COVID-19 Genomics UK (COG-UK) Consortium | Darren L Smith,Andrew Nelson,Matthew Bashton,Greg R Young,Joshua Loh,John Allan,Mohammad A Tariq,Giles S Holt,Gary Black,Wen C Yew,Lynn Dover,Paul Baker,Steve Liggett,Sarah Essex,Jane Greenaway,Debra Padgett,Clive Graham,Garren Scott,Edward Barton,Emma Swindells,Brendan Payne,Jennifer Collins,Yusri Taha,Gary Eltringham |
| EPI_ISL_921789, EPI_ISL_921790, EPI_ISL_921791, EPI_ISL_921792, EPI_ISL_921796, EPI_ISL_921798, EPI_ISL_921799, EPI_ISL_921801, EPI_ISL_921802, EPI_ISL_921811, EPI_ISL_921815, EPI_ISL_921820, EPI_ISL_921821, EPI_ISL_921824, EPI_ISL_921826, EPI_ISL_921828, EPI_ISL_921839, EPI_ISL_921841, EPI_ISL_921842, EPI_ISL_921846, EPI_ISL_921847, EPI_ISL_921848, EPI_ISL_921849 | | | |
| see above | Quadram Institute Bioscience | COVID-19 Genomics UK (COG-UK) Consortium | Dave J. Baker, Gemma L. Kay, Alp Aydin, Thanh Le-Viet, Steven Rudder, Ana P. Tedim, Anastasia Kolyva, Maria Diaz, Leonardo de Oliveira Martins, Nabil-Fareed Alikhan, Lizzie Meadows, Rachael Stanley, Ngozi Elumogo, Muhammed Yasir, Nicholas M. Thomson, Alexander J Trotter, Rachel Gilroy, Samuel Bloomfield, Claire Stuart, Andrew Bell, Reenesh Prakash, Samir Dervisevic, Alison E. Mather, John Wain, Mark Webber, Andrew J. Page, Justin O'Grady |
| EPI_ISL_922268, EPI_ISL_922270, EPI_ISL_922276, EPI_ISL_922281, EPI_ISL_922282, EPI_ISL_922307, EPI_ISL_922313, EPI_ISL_922314 | Oxford Viromics, NDM, University of Oxford; Oxford University Hospitals; Basingstoke and North Hampshire Hospital | COVID-19 Genomics UK (COG-UK) Consortium | Tanya Golubchik, David Bonsall, George Macintyre, Amy Trebes, Mariateresa de Cesare, Catrin Moore, Alex Mobbs, Anita Justice, Robert Shaw, Monique Andersson, Timothy Peto, Emma Wise, Nathan Moore, Jessica Lynch, Nick Cortes, Matilde Mori, Stephen Kidd, David Buck, John Todd, Christophe Fraser |
| EPI_ISL_923258, EPI_ISL_923267, EPI_ISL_923268 | Centre for Enzyme Innovation, University of Portsmouth / Translational Research Laboratory, Portsmouth Hospitals NHS Trust | COVID-19 Genomics UK (COG-UK) Consortium | Angela Beckett,Salman Goudarzi,Christopher Fearn,Kate Cook,Katie Loveson,Sharon Glaysher,Scott Elliott,Samuel Robson |
| EPI_ISL_925923, EPI_ISL_925937, EPI_ISL_925953, EPI_ISL_926020, EPI_ISL_926024, EPI_ISL_926046, EPI_ISL_926059, EPI_ISL_926177, EPI_ISL_926192, EPI_ISL_926218, EPI_ISL_926224, EPI_ISL_926286, EPI_ISL_926289, EPI_ISL_926361, EPI_ISL_926438, EPI_ISL_926487, EPI_ISL_926539, EPI_ISL_926551, EPI_ISL_926575, EPI_ISL_926631, EPI_ISL_926661, EPI_ISL_926708, EPI_ISL_926770, EPI_ISL_926793, EPI_ISL_926809, EPI_ISL_926914, EPI_ISL_926937, EPI_ISL_926940, EPI_ISL_926942, EPI_ISL_926959, EPI_ISL_926960, EPI_ISL_926987, EPI_ISL_927099, EPI_ISL_927192, EPI_ISL_927194, EPI_ISL_927216, EPI_ISL_927314, EPI_ISL_927493, EPI_ISL_927505, EPI_ISL_927544, EPI_ISL_927565, EPI_ISL_927602, EPI_ISL_927626, EPI_ISL_927652, EPI_ISL_927657, EPI_ISL_927671, EPI_ISL_927721, EPI_ISL_927845, EPI_ISL_927981, EPI_ISL_928047, EPI_ISL_928048, EPI_ISL_928092, EPI_ISL_928149, EPI_ISL_928274, EPI_ISL_928395, EPI_ISL_928404, EPI_ISL_928446, EPI_ISL_928468, EPI_ISL_928532, EPI_ISL_928546, EPI_ISL_928560, EPI_ISL_928609, EPI_ISL_928636, EPI_ISL_928675, EPI_ISL_928750, EPI_ISL_928772, EPI_ISL_928796, EPI_ISL_928812, EPI_ISL_928814, EPI_ISL_928816, EPI_ISL_928843, EPI_ISL_928849, EPI_ISL_928935, EPI_ISL_928966, EPI_ISL_928984, EPI_ISL_929093, EPI_ISL_929141, EPI_ISL_929147, EPI_ISL_929169, EPI_ISL_929173, EPI_ISL_929175, EPI_ISL_929227, EPI_ISL_929319, EPI_ISL_929334, EPI_ISL_929387, EPI_ISL_929452, EPI_ISL_929458, EPI_ISL_929552, EPI_ISL_929570, EPI_ISL_929588, EPI_ISL_929605, EPI_ISL_929627, EPI_ISL_929706, EPI_ISL_929712, EPI_ISL_929745, EPI_ISL_929771, EPI_ISL_929802, EPI_ISL_929972, EPI_ISL_930001, EPI_ISL_930019, EPI_ISL_930044, EPI_ISL_930053, EPI_ISL_930062, EPI_ISL_930064, EPI_ISL_930208, EPI_ISL_930238, EPI_ISL_930257, EPI_ISL_930272, EPI_ISL_930300, EPI_ISL_930327, EPI_ISL_930343, EPI_ISL_930385, EPI_ISL_930465, EPI_ISL_930496 | | | |
| see above | Department of Virus and Microbiological Special Diagnostics, Statens Serum Institut, Copenhagen, Denmark | Aalborg University | Danish Covid-19 Genome Consortium |
| EPI_ISL_930665, EPI_ISL_930666, EPI_ISL_930667, EPI_ISL_930671, EPI_ISL_930672, EPI_ISL_930673, EPI_ISL_930674, EPI_ISL_930720, EPI_ISL_930722, EPI_ISL_930723, EPI_ISL_930724, EPI_ISL_930725, EPI_ISL_930726, EPI_ISL_930727, EPI_ISL_930728, EPI_ISL_930730, EPI_ISL_930731, EPI_ISL_930732, EPI_ISL_930733, EPI_ISL_930735, EPI_ISL_930736, EPI_ISL_930737, EPI_ISL_930738, EPI_ISL_930739, EPI_ISL_930740, EPI_ISL_930741, EPI_ISL_930742, EPI_ISL_930743, EPI_ISL_930744, EPI_ISL_930745, EPI_ISL_930746, EPI_ISL_930747, EPI_ISL_930748, EPI_ISL_930749, EPI_ISL_930750, EPI_ISL_930751, EPI_ISL_930752, EPI_ISL_930753, EPI_ISL_930754, EPI_ISL_930755, EPI_ISL_930756, EPI_ISL_930757, EPI_ISL_930758, EPI_ISL_930759, EPI_ISL_930761, EPI_ISL_930762, EPI_ISL_930763, EPI_ISL_930765, EPI_ISL_930767, EPI_ISL_930768, EPI_ISL_930777, EPI_ISL_930779, EPI_ISL_930781 | | | |
| see above | Utah Public Health Laboratory | Utah Public Health Laboratory | Erin L. Young, Kelly F. Oakeson, Tara Gallagher |
| EPI_ISL_931471 | Maryland Public Health Laboratory (MD PHL) | Maryland Public Health Laboratory (MD PHL) | Maryland Department of Health Laboratories Administration |
| EPI_ISL_931507, EPI_ISL_931516, EPI_ISL_931537, EPI_ISL_931539, EPI_ISL_931540 | Utah Public Health Laboratory | Utah Public Health Laboratory | Erin L. Young, Kelly F. Oakeson, Tara Gallagher |
| EPI_ISL_933426, EPI_ISL_933427, EPI_ISL_933428, EPI_ISL_933429, EPI_ISL_933431, EPI_ISL_933432, EPI_ISL_933433, EPI_ISL_933434, EPI_ISL_933436, EPI_ISL_933437, EPI_ISL_933438, EPI_ISL_933439, EPI_ISL_933440, EPI_ISL_933447, EPI_ISL_933453, EPI_ISL_933456, EPI_ISL_933463, EPI_ISL_933467, EPI_ISL_933469, EPI_ISL_933472, EPI_ISL_933474, EPI_ISL_933475 | | | |
| see above | Lighthouse Lab in Milton Keynes | Wellcome Sanger Institute for the COVID-19 Genomics UK (COG-UK) Consortium | The Lighthouse Lab in Milton Keynes and Alex Alderton, Roberto Amato, Sonia Goncalves, Ewan Harrison, David K. Jackson, Ian Johnston, Dominic Kwiatkowski, Cordelia Langford, John Sillitoe on behalf of the Wellcome Sanger Institute COVID-19 Surveillance Team |

| | | | |
|---|---|---|---|
| EPI_ISL_934349 | Klinisk mikrobiologi | The Public Health Agency of Sweden | Anna-Malin Linde, Maria Lind Karlberg, Carlo Berg, Oskar Karlsson Lindsjo, Sofia Stamouli, Reza Advani, Mattias Haukland, Petra Holmstrom, Noura Walai, Petra Edquist, Mia Brytting, Anna Risberg, Karin Tegmark-Wisell |
| EPI_ISL_934388 | Klinisk Mikrobiologi | The Public Health Agency of Sweden | Anna-Malin Linde, Maria Lind Karlberg, Carlo Berg, Oskar Karlsson Lindsjo, Sofia Stamouli, Reza Advani, Mattias Haukland, Petra Holmstrom, Noura Walai, Petra Edquist, Mia Brytting, Anna Risberg, Karin Tegmark-Wisell |
| EPI_ISL_934639, EPI_ISL_934642, EPI_ISL_934646 | Department of Microbiology, University Innsbruck | Bergthaler laboratory, CeMM Research Center for Molecular Medicine of the Austrian Academy of Sciences | Lukas Endler, Anna Schedl, Thomas Penz, Benedikt Agerer, Maelle Le Moing, Michael Schuster, Bekir Erguner, Jan Laine, Martin Senekowitsch, Christoph Bock, Andreas Bergthaler |
| EPI_ISL_935209 | KU Leuven, Rega Institute, Clinical and Epidemiological Virology | KU Leuven, Rega Institute, Clinical and Epidemiological Virology | Tony Wawina-Bokalanga, Bert Vanmechelen, Joan Marti-Carerras, Piet Maes |
| EPI_ISL_935708, EPI_ISL_935709, EPI_ISL_935710, EPI_ISL_935750, EPI_ISL_935751, EPI_ISL_935752 | Houston Health Dept. | Houston Health Dept. | Ryker Penn, Pamela Brown, Adolpho Lara |
| EPI_ISL_936036, EPI_ISL_936037, EPI_ISL_936038, EPI_ISL_936039, EPI_ISL_936040, EPI_ISL_936041, EPI_ISL_936042, EPI_ISL_936043 | Columbia University Irving Medical Center | Wadsworth Center, New York State Department of Health | Kirsten St. George, Daryl M. Lamson, Alexis Russel, Matthew Shudt, Melissa A Leisner, Jonathan Plitnick, Navjot Singh, John Kelly, Erasmus Schneider, Erica Lasek-Nesselquist |
| EPI_ISL_936180 | ALBANY MEDICAL CENTER | Wadsworth Center, New York State Department of Health | Kirsten St. George, Daryl M. Lamson, Alexis Russel, Matthew Shudt, Melissa A Leisner, Jonathan Plitnick, Navjot Singh, John Kelly, Erasmus Schneider, Erica Lasek-Nesselquist |
| EPI_ISL_936424 | TGen North | TGen North | Jolene Bowers, Megan Folkerts, Chris French, Hayley Yaglom, Ashlyn Pfeiffer, Darrin Lemmer, Dave Engelthaler, The Arizona COVID Genomics Union (ACGU) |
| EPI_ISL_936813, EPI_ISL_936814, EPI_ISL_936815, EPI_ISL_936816 | Northwestern Memorial Hospital | Ozer Lab | Ramon Lorenzo-Redondo, Lacy M. Simons, Chad J. Achenbach, Lawrence J. Jennings, Michael G. Ison, Judd F. Hultquist, Egon A. Ozer |
| EPI_ISL_940149 | NHLS Universitas Academic | UFS Virology | PA Bester, MM Nyaga, P Nthiga, MT Mogotsi, D Goedhals, T de Oliveira |
| EPI_ISL_940802, EPI_ISL_940819, EPI_ISL_940820 | Houston Health Dept. | Houston Health Dept. | Ryker Penn, Pamela Brown, Adolpho Lara |
| EPI_ISL_940861, EPI_ISL_940883 | Vaccines and Infectious Diseases Analytics Research Unit (VIDA) | KRISP, KZN Research Innovation and Sequencing Platform | Baillie Vicky, du Plessis Jeanine, Giandhari Jennifer, Pillay Sureshnee, Naidoo Yeshnee, Tegally Houriiyah, de Oliveira Tulio, Madhi Shabir |
| EPI_ISL_941233, EPI_ISL_941234, EPI_ISL_941235, EPI_ISL_941236, EPI_ISL_941237, EPI_ISL_941238, EPI_ISL_941239 | Hospital Universitario de La Ribera (Alzira, València) | SeqCOVID-SPAIN consortium/IBV(CSIC) | Olalla Martínez Macias, Julia González Cantó and SeqCOVID-SPAIN consortium |
| EPI_ISL_941269 | Virginia DCLS | Virginia DCLS | Virginia DCLS |
| EPI_ISL_941624 | Instituto Nacional de Saude (INSA) | Instituto Nacional de Saude (INSA) | Borges et al |
| EPI_ISL_941932 | Florida Bureau of Public Health Laboratories | Florida Bureau of Public Health Laboratories | Sarah Schmedes, Jason Blanton |
| EPI_ISL_942096, EPI_ISL_942142 | Wisconsin State Laboratory of Hygiene Communicable Disease Division | Wisconsin State Laboratory of Hygiene Communicable Disease Division | Kelsey R. Florek, Abigail C. Shockey |
| EPI_ISL_942753, EPI_ISL_942754, EPI_ISL_942755, EPI_ISL_942756, EPI_ISL_942757, EPI_ISL_942758, EPI_ISL_942759, EPI_ISL_942760, EPI_ISL_942761, EPI_ISL_942762 | Gundersen Molecular Diagnostics Laboratory | Kabara Cancer Research Institute | Craig S. Richmond, Paraic A. Kenny |
| EPI_ISL_943568 | Servizo de Microbioloxía. Complexo Hospitalario Universitario de Santiago de Compostela | Servizo de Microbioloxía. Complexo Hospitalario Universitario de Santiago de Compostela | Antonio Aguilera, Gema Barbeito, Amparo Coira, José Costa, Rocio Trastoy, María Luisa Pérez del Molino. |
| EPI_ISL_943818 | Utah Public Health Laboratory | Utah Public Health Laboratory | Erin L. Young, Kelly F. Oakeson, Tara Gallagher |
| EPI_ISL_945126, EPI_ISL_945136 | Lighthouse Lab in Glasgow | Wellcome Sanger Institute for the COVID-19 Genomics UK (COG-UK) Consortium | Harper VanSteenhouse, Yumi Kasai, David Gray, Carol Clugston, Anna Dominiczak and Alex Alderton, Roberto Amato, Sonia Goncalves, Ewan Harrison, David K. Jackson, Ian Johnston, Dominic Kwiatkowski, Cordelia Langford, John Sillitoe on behalf of the Wellcome Sanger Institute COVID-19 Surveillance Team |
| EPI_ISL_945231 | Lighthouse Lab in Milton Keynes | Wellcome Sanger Institute for the COVID-19 Genomics UK (COG-UK) Consortium | The Lighthouse Lab in Milton Keynes and Alex Alderton, Roberto Amato, Sonia Goncalves, Ewan Harrison, David K. Jackson, Ian Johnston, Dominic Kwiatkowski, Cordelia Langford, John Sillitoe on behalf of the Wellcome Sanger Institute COVID-19 Surveillance Team |
| EPI_ISL_945335 | Lighthouse Lab in Alderley Park | Wellcome Sanger Institute for the COVID-19 Genomics UK (COG-UK) Consortium | Jacquelyn Wynn, Mairead Hyland, The Lighthouse Lab in Alderley Park and Alex Alderton, Roberto Amato, Sonia Goncalves, Ewan Harrison, David K. Jackson, Ian Johnston, Dominic Kwiatkowski, Cordelia Langford, John Sillitoe on behalf of the Wellcome Sanger Institute COVID-19 Surveillance Team |
| EPI_ISL_949761 | Barts Health NHS Trust | COVID-19 Genomics UK (COG-UK) Consortium | CUTINO-MOGUEL, Maria-Teresa; HARRINGTON, David; OWOYEMI, Dola; KULASEGARAN-SHYLINI, Raghavendran; BROAD, Claire; KELE, Beatrix |
| EPI_ISL_950105, EPI_ISL_950138, EPI_ISL_950145 | University College London, Great Ormond Street Hospital for Children NHS Foundation Trust, Imperial College Healthcare NHS Trust | COVID-19 Genomics UK (COG-UK) Consortium | Sergi Castellano, Rachel Williams, Mark Kristiansen, Paola Resende Silva, Sunando Roy, Tony Brooks, Helena Tutill, Paola Niola, Patricia Dyal, Charlotte Williams, Leysa Forrest, Yasmin Panchbhaya, Jacqueline Findlay, Samuel Weeks, Julianne Brown, Kathryn Harris, Paul Randell, James Price, Alison Holmes, Judith Breuer |
| EPI_ISL_950240, EPI_ISL_950436, EPI_ISL_950438, EPI_ISL_950439, EPI_ISL_950440, EPI_ISL_950441, EPI_ISL_950442, EPI_ISL_950443 | Northumbria University / South Tees Hospitals NHS Foundation Trust / North Cumbria Integrated Care NHS Foundation Trust / North Tees and Hartlepool NHS Foundation Trust / Newcastle Hospitals NHS Foundation Trust | COVID-19 Genomics UK (COG-UK) Consortium | Darren L Smith,Andrew Nelson,Matthew Bashton,Greg R Young,Joshua Loh,John Allan,Mohammad A Tariq,Giles S Holt,Gary Black,Wen C Yew,Lynn Dover,Paul Baker,Steve Liggett,Sarah Essex,Jane Greenaway,Debra Padgett,Clive Graham,Garren Scott,Edward Barton,Emma Swindells,Brendan Payne,Jennifer Collins,Yusri Taha,Gary Eltringham |
| EPI_ISL_950544, EPI_ISL_950560, EPI_ISL_950561, EPI_ISL_950562, EPI_ISL_950572, EPI_ISL_950578, EPI_ISL_950579, EPI_ISL_950583, EPI_ISL_950584, EPI_ISL_950589, EPI_ISL_950595 | | | |
| see above | Quadram Institute Bioscience | COVID-19 Genomics UK (COG-UK) Consortium | Dave J. Baker, Gemma L. Kay, Alp Aydin, Thanh Le-Viet, Steven Rudder, Ana P. Tedim, Anastasia Kolyva, Maria Diaz, Leonardo de Oliveira Martins, Nabil-Fareed Alikhan, Lizzie Meadows, Rachael Stanley, Ngozi Elumogo, Muhammed Yasir, Nicholas M. Thomson, Alexander J Trotter, Rachel Gilroy, Samuel Bloomfield, Claire Stuart, Andrew Bell, Reenesh Prakash, Samir Dervisevic, Alison E. Mather, John Wain, Mark Webber, Andrew J. Page, Justin O'Grady |
| EPI_ISL_950849, EPI_ISL_950850, EPI_ISL_950851, EPI_ISL_950852, EPI_ISL_950853, EPI_ISL_950854, EPI_ISL_950855, EPI_ISL_950856, EPI_ISL_950857, EPI_ISL_950858, EPI_ISL_950859, EPI_ISL_950861, EPI_ISL_950862, EPI_ISL_950863, EPI_ISL_950864, EPI_ISL_950865, EPI_ISL_950868, EPI_ISL_950871, EPI_ISL_950879, EPI_ISL_950880, EPI_ISL_950881, EPI_ISL_950882, EPI_ISL_950884, EPI_ISL_950885, EPI_ISL_950886, EPI_ISL_950887, EPI_ISL_950889, EPI_ISL_950890, EPI_ISL_950891, EPI_ISL_950892, EPI_ISL_950893, EPI_ISL_950894, EPI_ISL_950895, EPI_ISL_950896, EPI_ISL_950897, EPI_ISL_950898, EPI_ISL_950899, EPI_ISL_950955, EPI_ISL_950961, EPI_ISL_950963, EPI_ISL_950964, EPI_ISL_950966, EPI_ISL_950968, EPI_ISL_950970, EPI_ISL_950971, EPI_ISL_950972, EPI_ISL_950973, EPI_ISL_950974, EPI_ISL_950975, EPI_ISL_950976, EPI_ISL_950977, EPI_ISL_950984, EPI_ISL_950997, EPI_ISL_951005, EPI_ISL_951006, EPI_ISL_951007, EPI_ISL_951012, EPI_ISL_951015, EPI_ISL_951343 | | | |
| see above | Oxford Viromics, NDM, University of Oxford; Oxford University Hospitals; Basingstoke and North Hampshire Hospital | COVID-19 Genomics UK (COG-UK) Consortium | Tanya Golubchik, David Bonsall, George Macintyre, Amy Trebes, Mariateresa de Cesare, Catrin Moore, Alex Mobbs, Anita Justice, Robert Shaw, Monique Andersson, Timothy Peto, Emma Wise, Nathan Moore, Jessica Lynch, Nick Cortes, Matilde Mori, Stephen Kidd, David Buck, John Todd, Christophe Fraser |
| EPI_ISL_954210 | 1.AO Universitaria 'S. Giovanni di Dio e Ruggi D'Aragona, Scuola Medica Salernitana' Hospital / 2.UOC di Virologia e Microbiologia, Università della Campania 'L. Vanvitelli' / 3.AO | 1. Genome Research Center for Health (CRGS) / 2. Laboratory of Molecular Medicine and Genomics(LMMGe) / 3. Center for Research in Pure and Applied Mathematics | Giorgio Giurato, Francesca Rizzo, Alessandro Weisz, Gianluigi Franci, Giovanni Nassa, Pasquale Pagliano, Roberta Tarallo, Elena Alexandrova, Ylenia D'Agostino, Carlo Ferravante, Jessica Lamberti, Viola Melone, Domenico Memoli, Valeria Mirici Cappa, Domenico Palumbo, Giovanni Pecoraro, Assunta Sellitto, Oriana Strianese, Ilaria Terenzi, Giuseppe Fenza, Aniello Gentile, Antonello Saccomanno, Sonia Amabile, Teresa Rocco, Annamaria Salvati, |

| | | | |
|---|---|---|---|
| | Universitaria 'Federico II' Napoli Hospital / 4.AORN 'San Giuseppe Moscati' Avellino Hospital / 5.AO 'San Pio - presidio G. Rummo' Benevento Hospital / 6.AO 'Sant'Anna e San Sebastiano' Caserta Hospital / 7.PO 'Maria Santissima Addolorata' Eboli Hospital / 8.Biogem Istituto di Ricerche Genetiche | (CRMPA) | Emilia Vaccaro, Massimiliano Galdiero, Michele Cennamo, Giuseppe Portella, Maria Grazia Foti, Mariarosaria Ingino, Maria Landi, Maurizio Fumi, Vincenzo Rocco, Rita Greco, Vittoria Letizia, Arnolfo Petruzziello, Maddalena Schioppa, Gregorio Goffredi, Francesca Marciano, Michele Caraglia, Alessia Cossu, Marianna Scrima, Edmondo Adorisio, Morena D'Avenia, Michela Iacobellis, Rosanna Piluscio, Giorgio Dirani, Vittorio Sambri, Simona Semprini, Silvia Zanoli, Francesco Curcio, Stefania Marzinotto, Andreina Baj, Fausto Sessa. |
| EPI_ISL_954227, EPI_ISL_954230, EPI_ISL_954256, EPI_ISL_954257, EPI_ISL_954258, EPI_ISL_954259, EPI_ISL_954260, EPI_ISL_954261, EPI_ISL_954262, EPI_ISL_954263, EPI_ISL_954264, EPI_ISL_954266, EPI_ISL_954267, EPI_ISL_954268, EPI_ISL_954269, EPI_ISL_954270, EPI_ISL_954271, EPI_ISL_954272, EPI_ISL_954273, EPI_ISL_954274, EPI_ISL_954275, EPI_ISL_954276, EPI_ISL_954277, EPI_ISL_954278, EPI_ISL_954279, EPI_ISL_954280 | | | |
| see above | MRC/UVRI & LSHTM Uganda Research Unit | Where sequence data have been generated and submitted to GISAID | Matthew Cotten, Dan Lule Bugembe, My V.T. Phan, Isaac Sseeewanyana, Patrick Semanda, Susan Nabadda, Pontiano Kaleebu |
| EPI_ISL_954869 | Colorado Department of Public Health and Environment | Colorado Department of Puplic Health and Environment | Laura Bankers, Molly C. Hetherington-Rauth, Diana Ir, Shannon Ely, Shannon R. Matzinger, Sarah Elizabeth Totten, Emily A. Travanty |
| EPI_ISL_955128 | Innovative Genomics Institute, UC Berkeley | Innovative Genomics Institute, UC Berkeley | Stacia Wyman, Haridha Shivram, Phil Frankino, Liana Lareau, Shana McDevitt, Justin Choi |
| EPI_ISL_955261, EPI_ISL_955263 | Fulgent Genetics | Fulgent Genetics | Harry Gao, Mickey Li, John Gao, Joseph Fierro, Benafsh Sapra, Becky Tsai, Yan Meng, Doreen Ng, James Xie |
| EPI_ISL_955389 | Alameda County Public Health Lab | Chan-Zuckerberg Biohub | CZB Cliahub Consortium |
| EPI_ISL_955697, EPI_ISL_955698, EPI_ISL_955700, EPI_ISL_955704, EPI_ISL_955706, EPI_ISL_955707, EPI_ISL_955708, EPI_ISL_955709, EPI_ISL_955710, EPI_ISL_955711, EPI_ISL_955712, EPI_ISL_955713, EPI_ISL_955714, EPI_ISL_955715, EPI_ISL_955716, EPI_ISL_955717, EPI_ISL_955718, EPI_ISL_955719, EPI_ISL_955720, EPI_ISL_955721, EPI_ISL_955722, EPI_ISL_955723, EPI_ISL_955724, EPI_ISL_955725, EPI_ISL_955726, EPI_ISL_955732, EPI_ISL_955733 | | | |
| see above | Humboldt County Public Health Laboratory | Chan-Zuckerberg Biohub | CZB Cliahub Consortium |
| EPI_ISL_956352, EPI_ISL_956353, EPI_ISL_956354 | Houston Health Department, Disease Prevention and Control | Houston Health Department, Disease Prevention and Control | Penn,R., Brown,P., Lara,A. |
| EPI_ISL_956402 | General Hospital - Ohrid | Research Center for Genetic Engineering and Biotechnology "Georgi D. Efremov" , Macedonian Academy of Sciences and Arts | Aleksandar J. Dimovski, Dijana Plasheska-Karanfilska, Predrag Noveski, Gjorgji Bozinovski, Milena Jakimovska |
| EPI_ISL_956403 | Institute for prevention, treatment and rehabilitation of cardiovascular diseases - Ohrid | Research Center for Genetic Engineering and Biotechnology "Georgi D. Efremov" , Macedonian Academy of Sciences and Arts | Aleksandar J. Dimovski, Dijana Plasheska-Karanfilska, Predrag Noveski, Gjorgji Bozinovski, Milena Jakimovska |
| EPI_ISL_960463, EPI_ISL_960464, EPI_ISL_960471, EPI_ISL_960560, EPI_ISL_960561, EPI_ISL_960562, EPI_ISL_960563, EPI_ISL_960564, EPI_ISL_960565, EPI_ISL_960566, EPI_ISL_960567, EPI_ISL_960649, EPI_ISL_960650, EPI_ISL_960651, EPI_ISL_960652, EPI_ISL_960653, EPI_ISL_960654, EPI_ISL_960655, EPI_ISL_960656, EPI_ISL_960657, EPI_ISL_960659 | | | |
| see above | Istituto Zooprofilattico Sperimentale del Mezzogiorno | TIGEM | Patrizia Annunziata, Andrea Ballabio, Valentina Bouche, Davide Cacchiarelli, Pellegrino Cerino, Chiara Colantuono, Maria Concetta Cuomo, Denise Di Concilio, Lucio Di Filippo, Antonio Grimaldi, Antonio Limone, Anna Manfredi, Francesco Panariello, Biancamaria Pierri, Marcello Salvi |
| EPI_ISL_960669, EPI_ISL_960672, EPI_ISL_960675, EPI_ISL_960681, EPI_ISL_960682, EPI_ISL_960692, EPI_ISL_960693, EPI_ISL_960708, EPI_ISL_960718, EPI_ISL_960719, EPI_ISL_960723, EPI_ISL_960748, EPI_ISL_960749, EPI_ISL_960750, EPI_ISL_960762, EPI_ISL_960783, EPI_ISL_960823, EPI_ISL_960824 | | | |
| see above | Germano de sousa | Instituto Gulbenkian de Ciencia | Susana Ladeiro, Cathy Paulino, João Costa, João Sobral, Maria Costa, Ricardo Leite |
| EPI_ISL_962013 | Illinois Department of Public Health | Gagnon Lab, Southern Illinois University | Keith Gagnon |
| EPI_ISL_962876 | UCLA Clinical Micro Lab | Los Angeles County PHL | P. Hemarajata et al. |
| EPI_ISL_962877, EPI_ISL_962878 | National Virology Reference Laboratory | National Public Health Laboratory, National Centre for Infectious Diseases | Tze Minn Mak, Zhenyang Zhou, Zaini Zainun, Taib Surita, Lin Cui, Raymond Tzer Pin Lin |
| EPI_ISL_962930, EPI_ISL_962931, EPI_ISL_962932, EPI_ISL_962942 | Hospital Universitario de Gran Canaria Dr. Negrín | SeqCOVID-SPAIN consortium/IBV(CSIC) | M. Carmen Pérez González, Francisco J. Chamizo López, Ana Bordes Benítez and SeqCOVID-SPAIN consortium |
| EPI_ISL_965217 | Virginia Division of Consolidated Laboratory Services | Virginia Division of Consolidated Laboratory Services | Virginia DCLS |
| EPI_ISL_965531, EPI_ISL_965534, EPI_ISL_965555, EPI_ISL_965588, EPI_ISL_965591, EPI_ISL_965642, EPI_ISL_965669, EPI_ISL_965701, EPI_ISL_965708, EPI_ISL_965726, EPI_ISL_965739, EPI_ISL_965742, EPI_ISL_965743, EPI_ISL_965744, EPI_ISL_965753, EPI_ISL_965757, EPI_ISL_965760, EPI_ISL_965768, EPI_ISL_965777 | | | |
| see above | Dutch COVID-19 response team | Medical Microbiology, Maastricht University Medical Centre | Jozef Dingemans*, Brian van der Veer*, Erik Beuken, Carmen Reumkens, Lieke van Alphen, Christian Hoebe, Paul Savelkoul |
| EPI_ISL_967591, EPI_ISL_967615, EPI_ISL_967616, EPI_ISL_967619, EPI_ISL_967622, EPI_ISL_967623, EPI_ISL_967626, EPI_ISL_967627, EPI_ISL_967633, EPI_ISL_967636, EPI_ISL_967640, EPI_ISL_967645, EPI_ISL_967646, EPI_ISL_967679, EPI_ISL_967687, EPI_ISL_967688, EPI_ISL_967725, EPI_ISL_967728, EPI_ISL_967730, EPI_ISL_967752 | | | |
| see above | State Laboratories Division, Hawaii State Department of Health | State Laboratories Division, Hawaii State Department of Health | Pamela O'Brien, Drew Kuwazaki, Ayana Garnet, Razvan Sultana, Edward Desmond |
| EPI_ISL_968110, EPI_ISL_968111, EPI_ISL_968137, EPI_ISL_968138, EPI_ISL_968139 | Houston Health Department, Disease Prevention and Control | Houston Health Department, Disease Prevention and Control | Penn,R., Brown,P., Lara,A. |
| EPI_ISL_970811, EPI_ISL_970821, EPI_ISL_970942, EPI_ISL_971140, EPI_ISL_971279, EPI_ISL_971532, EPI_ISL_971617, EPI_ISL_971844, EPI_ISL_971862, EPI_ISL_971872, EPI_ISL_971921, EPI_ISL_971945, EPI_ISL_972051, EPI_ISL_972263, EPI_ISL_972342, EPI_ISL_972479, EPI_ISL_972653, EPI_ISL_972714, EPI_ISL_972752, EPI_ISL_972805, EPI_ISL_972811, EPI_ISL_972871, EPI_ISL_972882, EPI_ISL_972929, EPI_ISL_972972, EPI_ISL_972996, EPI_ISL_973116, EPI_ISL_973213, EPI_ISL_973234, EPI_ISL_973342, EPI_ISL_973369, EPI_ISL_973415, EPI_ISL_973417, EPI_ISL_973463, EPI_ISL_973582 | | | |
| see above | Department of Virus and Microbiological Special Diagnostics, Statens Serum Institut, Copenhagen, Denmark | Aalborg University | Danish Covid-19 Genome Consortium |
| EPI_ISL_976790, EPI_ISL_976791, EPI_ISL_976792, EPI_ISL_976793, EPI_ISL_976794, EPI_ISL_976795, EPI_ISL_976796, EPI_ISL_976797, EPI_ISL_976798, EPI_ISL_976799, EPI_ISL_976800, EPI_ISL_976801, EPI_ISL_976802, EPI_ISL_976803, EPI_ISL_976804, EPI_ISL_976805, EPI_ISL_976806, EPI_ISL_976807, EPI_ISL_976808, EPI_ISL_976809, EPI_ISL_976810, EPI_ISL_976811, EPI_ISL_976812, EPI_ISL_976813, EPI_ISL_976814, EPI_ISL_976815, EPI_ISL_976817, EPI_ISL_976818, EPI_ISL_976819, EPI_ISL_976820, EPI_ISL_976821, EPI_ISL_976822, EPI_ISL_976823, EPI_ISL_976824, EPI_ISL_976825, EPI_ISL_976826, EPI_ISL_976827, EPI_ISL_976828, EPI_ISL_976829, EPI_ISL_976830, EPI_ISL_976831, EPI_ISL_976832, EPI_ISL_976833, EPI_ISL_976834, EPI_ISL_976835, EPI_ISL_976836, EPI_ISL_976837, EPI_ISL_976838, EPI_ISL_976839, EPI_ISL_976840, EPI_ISL_976841, EPI_ISL_976842, EPI_ISL_976843, EPI_ISL_976844, EPI_ISL_976845, EPI_ISL_976846, EPI_ISL_976847, EPI_ISL_976848, EPI_ISL_976849, EPI_ISL_976850 | | | |
| see above | BCCDC Public Health Laboratory | BCCDC Public Health Laboratory | Prystajecky Natalie, Linda Hoang, Dan Fornika, John Tyson, Shannon Russell, Kim Macdonald, Kimia Kamelian, Ana Pacagnella, Corrinne Ng, Loretta Janz, Robert Azana Terry Snutch, Mel Krajden |
| EPI_ISL_977082 | Massachusetts General Hospital | Infectious Disease Program, Broad Institute of Harvard and MIT | Lemieux,J.E., Siddle,K.J., Shaw,B., Adams,G., Pierce,V., Turbett,S., Anahtar,M., Branda,J., Slater,D., Harris,J., Lin,A.E., Gladden-Young,A., Lagerborg,K., Rudy,M., DeRuff,K., Carter,A., Normandin,E., Bauer,M., Reilly,S., Tomkins-Tinch,C., Loreth,C., Chaluvadi,S., Neumann,A., Cusick,C., Chapman,S.B., Gnirke,A., Flowers,K., Cerrato,F., Birren,B.W., Gallagher,G., Smole,S., Park,D.J., MacInnis,B.L., Ryan,E., LaRocque,R., Rosenberg,E. and Sabeti,P.C. |
| EPI_ISL_977156, EPI_ISL_977249, EPI_ISL_977250 | ULSS 03 Venezia | Istituto Zooprofilattico Sperimentale delle Venezie | Adelaide Milani, Alessia Schivo, Annalisa Salviato, Erika Giorgia Quaranta, Ambra Pastori, Bianca Zecchin, Alice Fusaro, Isabella Monne, Calogero Terregino, Antonia Ricci |
| EPI_ISL_977721, EPI_ISL_977722, EPI_ISL_977723, EPI_ISL_977726, EPI_ISL_977727, EPI_ISL_977728, EPI_ISL_977729, EPI_ISL_977730, EPI_ISL_977733, EPI_ISL_977734, EPI_ISL_977740 | | | |
| see above | California Department of Public Health | Chiu Laboratory, University of California, San Francisco | Charles Chiu, Xianding (Wayne) Deng, Candace Wang, Venice Servellita, Jill Hacker, Debra Wadford |
| EPI_ISL_978044, EPI_ISL_978048, EPI_ISL_978051, EPI_ISL_978052, EPI_ISL_978053, EPI_ISL_978054, EPI_ISL_978055, EPI_ISL_978056, EPI_ISL_978070, EPI_ISL_978071, EPI_ISL_978072, EPI_ISL_978073, EPI_ISL_978074 | | | |
| see above | Chiu Laboratory, University of California, San Francisco | Chiu Laboratory, University of California, San Francisco | Charles Chiu, Xianding (Wayne) Deng, Candace Wang, Venice Servellita, Jill Hacker, Debra Wadford |
| EPI_ISL_978417, EPI_ISL_978418, EPI_ISL_978419, EPI_ISL_978420, EPI_ISL_978421, EPI_ISL_978422, | Arizona State Public Health Laboratory | Arizona State Public Health Laboratory | Trung Huynh, Jessica Escobar, Katherine Fullerton, Nobuko Fukushima, Stacy White, Linda Getsinger, Victor Waddell |

| | | | |
|---|---|---|---|
| EPI_ISL_978423 | | | |
| EPI_ISL_978960 | Chiu Laboratory, University of California, San Francisco | Chiu Laboratory, University of California, San Francisco | Charles Chiu, Xianding (Wayne) Deng, Candace Wang, Venice Servellita, Jill Hacker, Debra Wadford |
| EPI_ISL_979060 | Santa Clara County Public Health Laboratory | Chan-Zuckerberg Biohub | CZB Cliahub Consortium |
| EPI_ISL_979246 | Institute of Microbiology and Immunology, Faculty of Medicine, University of Ljubljana | Institute of Microbiology and Immunology, Faculty of Medicine, University of Ljubljana | Samo Zakotnik, Tomaž Mark Zorec, Matic Brvar, Doroteja Vlaj, Patricija Pozvek,Špela Pleh, Miša Korva, Mario Poljak, Tatjana Avši - Županc |
| EPI_ISL_979335, EPI_ISL_979336, EPI_ISL_979337 | Laboratorio Estatal de Salud Pública de Nuevo León | Laboratorio de Infectología Molecular, Departamento de Bioquímica y Medicina Molecular,Facultad de Medicina - Universidad Autónoma de Nuevo León | Kame A. Galán-Huerta, María F. Herrera-Saldivar, Natalia Martínez-Acuña, Sonia A. Lozano-Sepúlveda, Daniel Arellanos-Soto, Ana M. Rivas-Estilla, Samuel Buentello-Wong, Else del Carmen García-García, Gloria A. Jasso-de-la-Peña, Roberto Montes-de-Oca, Consuelo Treviño-Garza, Manuel E. de-la-O-Cavazos |
| EPI_ISL_979596, EPI_ISL_979597, EPI_ISL_979598, EPI_ISL_979599, EPI_ISL_979600, EPI_ISL_979601, EPI_ISL_979602, EPI_ISL_979603, EPI_ISL_979604, EPI_ISL_979605, EPI_ISL_979606, EPI_ISL_979612, EPI_ISL_979613, EPI_ISL_979629, EPI_ISL_979630 | | | |
| see above | Santa Clara County Public Health Laboratory | Chan-Zuckerberg Biohub | CZB Cliahub Consortium |
| EPI_ISL_979656, EPI_ISL_979657 | Orange County Public Health Lab | Chan-Zuckerberg Biohub | CZB Cliahub Consortium |
| EPI_ISL_981062, EPI_ISL_981068 | Johns Hopkins Hospital Department of Pathology | Johns Hopkins Hospital Department of Pathology | C. Paul Morris, Chun Huai Luo, Adannaya Amadi, Matthew Schwartz, Nicholas Gallagher, Heba H. Mostafa |
| EPI_ISL_983323 | INMI Lazzaro Spallanzani IRCCS | INMI Lazzaro Spallanzani IRCCS | CEM Gruber, B Bartolini, E Giombini, M Rueca, O Butera, F Messina, A Di Caro, MR Capobianchi |
| EPI_ISL_983853, EPI_ISL_983854, EPI_ISL_983855 | Colorado Department of Public Health and Environment | Colorado Department of Puplic Health and Environment | Laura Bankers, Molly C. Hetherington-Rauth, Diana Ir, Shannon Ely, Shannon R. Matzinger, Sarah Elizabeth Totten, Emily A. Travanty |
