## Supplementary material for "Detection and characterization of the SARS-CoV-2 lineage B.1.526 in New York": Supp. Table 3 GISAID Acknowledgment part 2: gisaid_hcov-19_acknowledgement_table_2021_02_13_010-5.pdf

We gratefully acknowledge the following Authors from the Originating laboratories responsible for obtaining the specimens, as well as the Submitting laboratories where the genome data were generated and shared via GISAID, on which this research is based.

All Submitters of data may be contacted directly via www.gisaid.org

Authors are sorted alphabetically.

| Accession ID | Originating Laboratory | Submitting Laboratory | Authors |
|---|---|---|---|
| EPI_ISL_751801 | UCSD EXCITE | Andersen lab at Scripps Research | SEARCH Alliance San Diego |
| EPI_ISL_754238 | Laboratory for Respiratory Viruses, Cantacuzino National Military-Medical Institute for Research and Development | Cantacuzino Institute Virology | Luiza Ustea, Mühlemann Barbara, Mihaela Lazar |
| EPI_ISL_755568, EPI_ISL_755569, EPI_ISL_755570, EPI_ISL_755571 | SA Pathology | SA Pathology | Lex Leong, Julien Soubrier, Chuan Kok Lim, Song Gao, Mark Turra, Karin Kassahn, Ivan Bastian, Geoff Higgins |
| EPI_ISL_763769 | Quadram Institute Bioscience | COVID-19 Genomics UK (COG-UK) Consortium | Dave J. Baker, Gemma L. Kay, Alp Aydin, Thanh Le-Viet, Steven Rudder, Ana P. Tedim, Anastasia Kolyva, Maria Diaz, Leonardo de Oliveira Martins, Nabil-Fareed Alikhan, Lizzie Meadows, Rachael Stanley, Ngozi Elumogo, Muhammed Yasir, Nicholas M. Thomson, Alexander J Trotter, Rachel Gilroy, Samuel Bloomfield, Claire Stuart, Andrew Bell, Reenesh Prakash, Samir Dervisevic, Alison E. Mather, John Wain, Mark Webber, Andrew J. Page, Justin O'Grady |
| EPI_ISL_766221 | Respiratory Virus Unit, National Infection Service, Public Health England | COVID-19 Genomics UK (COG-UK) Consortium | PHE Covid Sequencing Team |
| EPI_ISL_766585 | Triemli Hospital | Institute of Medical Virology, University of Zurich | Stefan Schmutz, Maryam Zaheri, Verena Kufner, Annette Audigé, Maria Grünberg, Kevin Steiner, Jon Huder, Cyril Shah, Riccarda Capaul, Jürg Böni, Michael Huber, Alexandra Trkola |
| EPI_ISL_766694 | Klinisk mikrobiologi | The Public Health Agency of Sweden | Department of Microbiology, The Public Health Agency of Sweden |
| EPI_ISL_767906, EPI_ISL_767908, EPI_ISL_767909, EPI_ISL_767910 | Sydney South West Pathology Service (SSWPS) - Royal Prince Alfred Hospital - NSW Health Pathology | NSW Health Pathology - Institute of Clinical Pathology and Medical Research; Westmead Hospital; University of Sydney | CIDM-PH et al. |
| EPI_ISL_767911, EPI_ISL_767912, EPI_ISL_767913 | Sydney South West Pathology Service (SSWPS) - Liverpool Hospital - NSW Health Pathology | NSW Health Pathology - Institute of Clinical Pathology and Medical Research; Westmead Hospital; University of Sydney | CIDM-PH et al. |
| EPI_ISL_767914 | Sydney South West Pathology Service (SSWPS) - Royal Prince Alfred Hospital - NSW Health Pathology | NSW Health Pathology - Institute of Clinical Pathology and Medical Research; Westmead Hospital; University of Sydney | CIDM-PH et al. |
| EPI_ISL_767921 | South Eastern Area Laboratory Services (SEALS) | NSW Health Pathology - Institute of Clinical Pathology and Medical Research; Westmead Hospital; University of Sydney | CIDM-PH et al. |
| EPI_ISL_767927 | St Vincent's Pathology (SydPath) | NSW Health Pathology - Institute of Clinical Pathology and Medical Research; Westmead Hospital; University of Sydney | CIDM-PH et al. |
| EPI_ISL_767928 | South Eastern Area Laboratory Services (SEALS) | NSW Health Pathology - Institute of Clinical Pathology and Medical Research; Westmead Hospital; University of Sydney | CIDM-PH et al. |
| EPI_ISL_767933, EPI_ISL_767934, EPI_ISL_767935, EPI_ISL_767936 | Pathology West - NSW Health Pathology | NSW Health Pathology - Institute of Clinical Pathology and Medical Research; Westmead Hospital; University of Sydney | CIDM-PH et al. |
| EPI_ISL_768324 | Singapore General Hospital | Department of Microbiology | Nurdyana Abdul Rahman, Kun Lee Lim, Chenhao Li, Sui Sin Goh, Kenneth Xin Long Chan, Kian Sing Chan, Lynette Oon, Kern Rei Chng, Niranjan Nagarajan, Karrie Ko |
| EPI_ISL_769955 | Albany Medical Center Hospital Clinical Laboratories | Wadsworth Center, New York State Department.of Health | Kirsten St. George, Daryl M. Lamson, Alexis Russel, Matthew Shudt, Melissa A Leisner, Jonathan Plitnick, Navjot Singh, John Kelly, Sara Griesemer, Erasmus Schneider, Erica Lasek-Nesselquist |
| EPI_ISL_770445, EPI_ISL_770446, EPI_ISL_770447, EPI_ISL_770448, EPI_ISL_770449, EPI_ISL_770450, EPI_ISL_770451, EPI_ISL_770452, EPI_ISL_770453, EPI_ISL_770454, EPI_ISL_770455, EPI_ISL_770456, EPI_ISL_770457, EPI_ISL_770458, EPI_ISL_770459, EPI_ISL_770460, EPI_ISL_770461, EPI_ISL_770476, EPI_ISL_770478, EPI_ISL_770479, EPI_ISL_770480, EPI_ISL_770481, EPI_ISL_770482, EPI_ISL_770483, EPI_ISL_770484, EPI_ISL_770485, EPI_ISL_770486, EPI_ISL_770487, EPI_ISL_770488, EPI_ISL_770489, EPI_ISL_770490, EPI_ISL_770491, EPI_ISL_770492, EPI_ISL_770493, EPI_ISL_770494, EPI_ISL_770495 | | | |
| see above | Wyoming Public Health Laboratory | Wyoming Public Health Laboratory | Noah Hull, Taylor Fearing, Lynette Gumbleton, Channing Weber, Ashley Norberg, Bailey Bowcutt, and Wanda Manley |
| EPI_ISL_770820, EPI_ISL_770821 | South East Regional Hospital | NSW Health Pathology - Institute of Clinical Pathology and Medical Research; Westmead Hospital; University of Sydney | CIDM-PH et al. |
| EPI_ISL_770871, EPI_ISL_770880, EPI_ISL_770886, EPI_ISL_770887, EPI_ISL_770905, EPI_ISL_771097, EPI_ISL_771098, EPI_ISL_771099, EPI_ISL_771128, EPI_ISL_771129, EPI_ISL_771130, EPI_ISL_771131, EPI_ISL_771132, EPI_ISL_771135, EPI_ISL_771136, EPI_ISL_771137, EPI_ISL_771138, EPI_ISL_771139, EPI_ISL_771140, EPI_ISL_771147, EPI_ISL_771148, EPI_ISL_771149, EPI_ISL_771150 | | | |
| see above | Laboratoire national de santé, Microbiology, Virology | Laboratoire national de santé, Microbiology, Microbial Genomics Platform | Anke Wienecke-Baldacchino, Catherine Ragimbeau,Jessica Tapp, Fatu Djabi, Lise Pignon, Raoul Salmon, Tamir Abdelrahman |
| EPI_ISL_771367 | SA Pathology | SA Pathology | Lex Leong, Julien Soubrier, Chuan Kok Lim, Song Gao, Mark Turra, Karin Kassahn, Ivan Bastian, Geoff Higgins |
| EPI_ISL_775416 | Oslo University Hospital, Department of Medical Microbiology | Norwegian Institute of Public Health, Department of Virology | Kathrine Stene-Johansen, Kamilla Heddeland Instefjord, Hilde Elshaug, Atiya R Ali,Marie Paulsen Madsen, Rasmus Riis Kopperud, Hilde Vollan, Karoline Bragstad, Olav Hungnes |
| EPI_ISL_775550, EPI_ISL_775551, EPI_ISL_775552, EPI_ISL_775553, EPI_ISL_775554, EPI_ISL_775555, EPI_ISL_775556, EPI_ISL_775557, EPI_ISL_775558, EPI_ISL_775559, EPI_ISL_775560, EPI_ISL_775561, EPI_ISL_775562, EPI_ISL_775563, EPI_ISL_775564, EPI_ISL_775565, EPI_ISL_775566, EPI_ISL_775567, EPI_ISL_775568, EPI_ISL_775569, EPI_ISL_775570, EPI_ISL_775571 | | | |
| see above | ABC | The Public Health Agency of Sweden | Department of Microbiology, The Public Health Agency of Sweden |
| EPI_ISL_775572, EPI_ISL_775573, EPI_ISL_775574 | Klinisk mikrobiologi | The Public Health Agency of Sweden | Department of Microbiology, The Public Health Agency of Sweden |
| EPI_ISL_775575 | ABC | The Public Health Agency of Sweden | Department of Microbiology, The Public Health Agency of Sweden |
| EPI_ISL_779138, EPI_ISL_779139, EPI_ISL_779140, EPI_ISL_779147, EPI_ISL_779148, EPI_ISL_779149, EPI_ISL_779150, EPI_ISL_779152 | Yale Clinical Virology Laboratory | Grubaugh Lab - Yale School of Public Health | Tara Alpert, Joseph Fauver, Anderson Brito, Mallery Breban, Anne Wyllie, Chantal Vogels, Mary Petrone, Chaney Kalinich, Isabel Ott, Arnau Casanovas, Catherine Muenker, Adam Moore, Alice Lu, Maria Tokuyama, Patrick Wong, Peiwen Lu, Saad Omer, Richard Martinello, Allison Nelson, Shelli Farhadian, Akiko Iwasaki, Charlese Dela Cruz, Albert Ko, Nathan Grubaugh |
| EPI_ISL_779319, EPI_ISL_779320, EPI_ISL_779321, EPI_ISL_779322, EPI_ISL_779323, EPI_ISL_779324, EPI_ISL_779325, EPI_ISL_779326, EPI_ISL_779327, EPI_ISL_779328, EPI_ISL_779329, EPI_ISL_779330, EPI_ISL_779331, EPI_ISL_779332, EPI_ISL_779333, EPI_ISL_779334, EPI_ISL_779335, EPI_ISL_779336, EPI_ISL_779337, EPI_ISL_779338, EPI_ISL_779339, EPI_ISL_779340, EPI_ISL_779341, EPI_ISL_779342, EPI_ISL_779343, EPI_ISL_779344, EPI_ISL_779345, EPI_ISL_779346, EPI_ISL_779347, EPI_ISL_779348, EPI_ISL_779349, EPI_ISL_779350, EPI_ISL_779351, EPI_ISL_779352, EPI_ISL_779353, EPI_ISL_779354, EPI_ISL_779355, EPI_ISL_779357, EPI_ISL_779358, EPI_ISL_779359, EPI_ISL_779360, EPI_ISL_779361, EPI_ISL_779362, EPI_ISL_779363, EPI_ISL_779365, EPI_ISL_779366, EPI_ISL_779367, EPI_ISL_779368, EPI_ISL_779370, EPI_ISL_779372, EPI_ISL_779373, EPI_ISL_779374, EPI_ISL_779375, EPI_ISL_779376, EPI_ISL_779377, EPI_ISL_779378, EPI_ISL_779379, EPI_ISL_779380, EPI_ISL_779381, EPI_ISL_779382 | | | |
| see above | University of Wisconsin-Madison AIDS Vaccine Research Laboratories | University of Wisconsin-Madison AIDS Vaccine Research Laboratories | Gage Moreno, Katarina Braun, et al. AIDS Vaccine Research Laboratories |

| | | | |
|---|---|---|---|
| EPI_ISL_779399, EPI_ISL_779400 | South Eastern Area Laboratory Services (SEALS) | NSW Health Pathology - Institute of Clinical Pathology and Medical Research; Westmead Hospital; University of Sydney | CIDM-PH et al. |
| EPI_ISL_779413 | Royal Darwin Hospital Pathology | MDU-PHL | Meumann, E., Caly L., Seemann T., Sait, M.L., Druce J., Sherry, N.L. |
| EPI_ISL_779624 | Microbiological Diagnostic Unit - Public Health Laboratory (MDU-PHL) | MDU-PHL | Seemann T., Sait, M.L., Sherry, N.L. |
| EPI_ISL_779626 | Victorian Infectious Diseases Reference Laboratory (VIDRL) | VIDRL and MDU-PHL | Caly L., Seemann T., Sait, M.L., Druce J., Sherry, N.L. |
| EPI_ISL_779628, EPI_ISL_779630 | Microbiological Diagnostic Unit - Public Health Laboratory (MDU-PHL) | MDU-PHL | Seemann T., Sait, M.L., Sherry, N.L. |
| EPI_ISL_779631, EPI_ISL_779632, EPI_ISL_779633 | Victorian Infectious Diseases Reference Laboratory (VIDRL) | VIDRL and MDU-PHL | Caly L., Seemann T., Sait, M.L., Druce J., Sherry, N.L. |
| EPI_ISL_779641, EPI_ISL_779642, EPI_ISL_779643, EPI_ISL_779644, EPI_ISL_779645, EPI_ISL_779646 | Microbiological Diagnostic Unit - Public Health Laboratory (MDU-PHL) | MDU-PHL | Seemann T., Sait, M.L., Sherry, N.L. |
| EPI_ISL_779784 | CNR Virus des Infections Respiratoires - France SUD | CNR Virus des Infections Respiratoires - France SUD | Antonin Bal, Gregory Destras, Gwendolyne Burfin, Hadrien Règue, Quentin Semanas, Martine Valette, Bruno Lina, Laurence Josset |
| EPI_ISL_779848, EPI_ISL_779849 | Laboratoire BIOMED 05 | CNR Virus des Infections Respiratoires - France SUD | Antonin Bal, Gregory Destras, Gwendolyne Burfin, Hadrien Règue, Quentin Semanas, Martine Valette, Bruno Lina, Laurence Josset |
| EPI_ISL_788898, EPI_ISL_788899, EPI_ISL_788900, EPI_ISL_788901, EPI_ISL_788902, EPI_ISL_788903, EPI_ISL_788904, EPI_ISL_788905, EPI_ISL_788906, EPI_ISL_788907, EPI_ISL_788908, EPI_ISL_788919, EPI_ISL_788939, EPI_ISL_788940, EPI_ISL_788941 | | | |
| see above | University of Wisconsin-Madison AIDS Vaccine Research Laboratories | University of Wisconsin-Madison AIDS Vaccine Research Laboratories | Gage Moreno, Katarina Braun, et al. AIDS Vaccine Research Laboratories |
| EPI_ISL_788993, EPI_ISL_789000, EPI_ISL_789001, EPI_ISL_789002, EPI_ISL_789003, EPI_ISL_789004, EPI_ISL_789005, EPI_ISL_789006, EPI_ISL_789007, EPI_ISL_789008, EPI_ISL_789009, EPI_ISL_789010, EPI_ISL_789011, EPI_ISL_789018, EPI_ISL_789019, EPI_ISL_789020, EPI_ISL_789021, EPI_ISL_789022, EPI_ISL_789023, EPI_ISL_789024 | | | |
| see above | ABC | The Public Health Agency of Sweden | Department of Microbiology, The Public Health Agency of Sweden |
| EPI_ISL_789062, EPI_ISL_789063, EPI_ISL_789064, EPI_ISL_789065, EPI_ISL_789066, EPI_ISL_789073, EPI_ISL_789076, EPI_ISL_789079, EPI_ISL_789083, EPI_ISL_789086, EPI_ISL_789092, EPI_ISL_789095, EPI_ISL_789098 | | | |
| see above | National Virus Reference Laboratory | National Virus Reference Laboratory | Michael Carr, Gabriel Gonzalez, Dana Alalwan, Jonathan Dean, Daniel Hare, Cillian F De Gascun |
| EPI_ISL_790573, EPI_ISL_790574, EPI_ISL_790617, EPI_ISL_790618, EPI_ISL_790986, EPI_ISL_790987, EPI_ISL_790988, EPI_ISL_791002, EPI_ISL_791003, EPI_ISL_791004, EPI_ISL_791033, EPI_ISL_791038, EPI_ISL_791039, EPI_ISL_791040, EPI_ISL_791041, EPI_ISL_791054, EPI_ISL_791055, EPI_ISL_791056, EPI_ISL_791057, EPI_ISL_791058, EPI_ISL_791059, EPI_ISL_791060, EPI_ISL_791061, EPI_ISL_791062, EPI_ISL_791063 | | | |
| see above | Dutch COVID-19 response team | National Institute for Public Health and the Environment (RIVM) | Adam Meijer, Harry Vennema, Jeroen Cremer, Sharon van den Brink, Bas van der Veer, AnneMarie van den Brandt, Florian Zwagemaker, Dennis Schmitz, Chantal Reusken, on behalf of the national COVID-19 response team |
| EPI_ISL_791144, EPI_ISL_791150, EPI_ISL_791152, EPI_ISL_791153, EPI_ISL_791154, EPI_ISL_791155, EPI_ISL_791157, EPI_ISL_791158, EPI_ISL_791159, EPI_ISL_791160, EPI_ISL_791161, EPI_ISL_791163, EPI_ISL_791164, EPI_ISL_791165, EPI_ISL_791166, EPI_ISL_791167, EPI_ISL_791168, EPI_ISL_791169, EPI_ISL_791170, EPI_ISL_791171, EPI_ISL_791172, EPI_ISL_791173, EPI_ISL_791174, EPI_ISL_791175, EPI_ISL_791176, EPI_ISL_791177, EPI_ISL_791178, EPI_ISL_791179, EPI_ISL_791180, EPI_ISL_791181, EPI_ISL_791182 | | | |
| see above | University of Wisconsin-Madison AIDS Vaccine Research Laboratories | University of Wisconsin-Madison AIDS Vaccine Research Laboratories | Gage Moreno, Katarina Braun, et al. AIDS Vaccine Research Laboratories |
| EPI_ISL_791193 | Respiratory Virus Unit, National Infection Service, Public Health England | COVID-19 Genomics UK (COG-UK) Consortium | PHE Covid Sequencing Team |
| EPI_ISL_791330, EPI_ISL_791331 | University of Wisconsin-Madison AIDS Vaccine Research Laboratories | University of Wisconsin-Madison AIDS Vaccine Research Laboratories | Gage Moreno, Katarina Braun, et al. AIDS Vaccine Research Laboratories |
| EPI_ISL_791340, EPI_ISL_791346, EPI_ISL_791415, EPI_ISL_791421, EPI_ISL_791426, EPI_ISL_791437, EPI_ISL_791441, EPI_ISL_791442, EPI_ISL_791445, EPI_ISL_791446, EPI_ISL_791452, EPI_ISL_791453, EPI_ISL_791455, EPI_ISL_791456, EPI_ISL_791457, EPI_ISL_791461, EPI_ISL_791465, EPI_ISL_791466, EPI_ISL_791468, EPI_ISL_791469, EPI_ISL_791471, EPI_ISL_791472, EPI_ISL_791480, EPI_ISL_791487 | | | |
| see above | Johns Hopkins Hospital Department of Pathology | Johns Hopkins Hospital Department of Pathology | C. Paul Morris, Chun Huai Luo, Adannaya Amadi, Nicholas Gallagher, Heba H. Mostafa |
| EPI_ISL_792058 | Hopital | National Reference Center for Viruses of Respiratory Infections, Institut Pasteur, Paris | Marion Barbet, Sylvie Behillil, Méline Bizard, Angela Brisebarre, Camille Capel, Etienne Simon-Lorière, Vincent Enouf, Maud Vanpeene, Sylvie van der Werf,Patricia Stoessel |
| EPI_ISL_792088 | Toronto Invasive Bacterial Diseases Network | McMaster University | Allison McGeer, Patryk Aftanas, Hooman Derakhshani, Angel Li, Kuganya Nirmalarajah, Emily Panousis, Ahmed Draia, Jalees Nasir, Michael Surette, Samira Mubareka, Andrew G. McArthur |
| EPI_ISL_792550 | Centre for Dengue Research and AICBU, Department of Immunology and Molecular Medicine | Centre for Dengue Research and AICBU, Department of Immunology and Molecular Medicine | Chandima Jeewandara, Deshni Jayathilaka, Dinuka Ariyaratne, Diyanath Ranasinghe, Laksiri Gomes, Gathsaurie Neelika Malavige |
| EPI_ISL_792668, EPI_ISL_792669, EPI_ISL_792670, EPI_ISL_792671, EPI_ISL_792672, EPI_ISL_792673 | Los Angeles County PHL | Los Angeles County PHL | P. Hemarajata et al. |
| EPI_ISL_794092 | URMC LABS | Wadsworth Center, New York State Department.of Health | Kirsten St. George, Daryl M. Lamson, Alexis Russel, Matthew Shudt, Melissa A Leisner, Jonathan Plitnick, Navjot Singh, John Kelly, Sara Griesemer, Erasmus Schneider, Erica Lasek-Nesselquist |
| EPI_ISL_794116, EPI_ISL_794117 | GLENS FALLS HOSPITAL LABORATORY | Wadsworth Center, New York State Department.of Health | Kirsten St. George, Daryl M. Lamson, Alexis Russel, Matthew Shudt, Melissa A Leisner, Jonathan Plitnick, Navjot Singh, John Kelly, Sara Griesemer, Erasmus Schneider, Erica Lasek-Nesselquist |
| EPI_ISL_794159, EPI_ISL_794160, EPI_ISL_794161, EPI_ISL_794162, EPI_ISL_794163, EPI_ISL_794164, EPI_ISL_794165, EPI_ISL_794168, EPI_ISL_794169, EPI_ISL_794170, EPI_ISL_794171, EPI_ISL_794172, EPI_ISL_794173, EPI_ISL_794174, EPI_ISL_794175, EPI_ISL_794176, EPI_ISL_794177, EPI_ISL_794178, EPI_ISL_794179, EPI_ISL_794180, EPI_ISL_794181 | | | |
| see above | Wadsworth Center, New York State Department.of Health | Wadsworth Center, New York State Department.of Health | Kirsten St. George, Daryl M. Lamson, Alexis Russel, Matthew Shudt, Melissa A Leisner, Jonathan Plitnick, Navjot Singh, John Kelly, Sara Griesemer, Erasmus Schneider, Erica Lasek-Nesselquist |
| EPI_ISL_794234, EPI_ISL_794236, EPI_ISL_794237, EPI_ISL_794244, EPI_ISL_794245, EPI_ISL_794246, EPI_ISL_794247, EPI_ISL_794248, EPI_ISL_794249, EPI_ISL_794250, EPI_ISL_794251, EPI_ISL_794252, EPI_ISL_794253, EPI_ISL_794254, EPI_ISL_794255, EPI_ISL_794256, EPI_ISL_794257, EPI_ISL_794258, EPI_ISL_794259, EPI_ISL_794260, EPI_ISL_794281, EPI_ISL_794282 | | | |
| see above | WESTCHESTER MEDICAL CENTER | Wadsworth Center, New York State Department.of Health | Kirsten St. George, Daryl M. Lamson, Alexis Russel, Matthew Shudt, Melissa A Leisner, Jonathan Plitnick, Navjot Singh, John Kelly, Sara Griesemer, Erasmus Schneider, Erica Lasek-Nesselquist |
| EPI_ISL_794592 | Laboratorio Estatal de Salud Pública de Tamaulipas | Instituto de diagnóstico y Referencia Epidemiologicos (INDRE) | Abril Rodriguez-Maldonado, Claudia Wong-Arambula, Fabiola Garces-Ayala, Gisela Barrera-Badillo, Ana Maria Cortez-Calderon, Bernardita Reyes-Berrones, Hilda del Carmen-Selvera, Gloria Molina-Gamboa, Lucia Hernandez-Rivas, Irma Lopez-Martinez, Celia Alpuche-Aranda, Jose Luis Alomia-Zegarra, Hugo Lopez Gatell-Ramirez, Ernesto Ramirez-Gonzalez. |
| EPI_ISL_794610, EPI_ISL_794620, EPI_ISL_794621, EPI_ISL_794622 | LabPLUS | Institute of Environmental Science and Research (ESR) | Xiaoyun Ren, Matt Storey, Nikki Freed, Muhammad Faisal, Jing Wang, Hermes Perez, Anja Werno, Antje van der Linden, Arlo Upton, Chris Mansell, David Hammer, Dragana Drinkovic, Gary McAuliffe, Hana Sofia Andersson, James Ussher, Jill Sherwood, Josh Freeman, Julia Howard, Juliet Elvy, Mary DeAlmeida, Matt Blakiston, Matthew Rogers, Max Bloomfield, Michael Addidle, Michelle Balm, Sally Roberts, Sarah Jefferies, Sharmini Muttaiyah, Susan Morpeth, Susan Taylor, Timothy Blackmore, Vani Sathyendran, Veronica Playle, Virginia Hope, Erasmus Smit, Lauren Jelly, Olin Silander, Joep de Ligt |
| EPI_ISL_794624 | Middlemore Hospital | Institute of Environmental Science and Research (ESR) | Xiaoyun Ren, Matt Storey, Nikki Freed, Muhammad Faisal, Jing Wang, Hermes Perez, Anja Werno, Antje van der Linden, Arlo Upton, Chris Mansell, David Hammer, Dragana Drinkovic, Gary McAuliffe, Hana Sofia Andersson, James Ussher, Jill Sherwood, Josh Freeman, Julia Howard, Juliet Elvy, Mary DeAlmeida, Matt Blakiston, Matthew Rogers, Max Bloomfield, Michael Addidle, Michelle Balm, Sally Roberts, Sarah Jefferies, Sharmini Muttaiyah, Susan Morpeth, Susan Taylor, Timothy Blackmore, Vani Sathyendran, Veronica Playle, Virginia Hope, Erasmus Smit, Lauren Jelly, Olin Silander, Joep de Ligt |

| | | | |
|---|---|---|---|
| EPI_ISL_794627 | Canterbury Health Laboratories | Institute of Environmental Science and Research (ESR) | Xiaoyun Ren, Matt Storey, Nikki Freed, Muhammad Faisal, Jing Wang, Hermes Perez, Anja Werno, Antje van der Linden, Arlo Upton, Chris Mansell, David Hammer, Dragana Drinkovic, Gary McAuliffe, Hana Sofia Andersson, James Ussher, Jill Sherwood, Josh Freeman, Julia Howard, Juliet Elvy, Mary DeAlmeida, Matt Blakiston, Matthew Rogers, Max Bloomfield, Michael Addidle, Michelle Balm, Sally Roberts, Sarah Jefferies, Sharmini Muttaiyah, Susan Morpeth, Susan Taylor, Timothy Blackmore, Vani Sathyendran, Veronica Playle, Virginia Hope, Erasmus Smit, Lauren Jelly, Olin Silander, Joep de Ligt |
| EPI_ISL_794649, EPI_ISL_794650, EPI_ISL_794651 | Fundación Cardio Infantil | Instituto Nacional de Salud - Dirección de Investigación en Salud Pública | Katherine Laiton-Donato, Diego A. Álvarez-Díaz, Carlos Franco-Muñoz, Mauricio Pacheco-Montealegre, Jonathan Reales, Sheryl Corchuelo, Maria T. Herrera, Julian Naizaque, Gerardo Santamaría, Paola Muñoz-Laiton, Diego Andrés Prada, Magdalena Wiesner, Martha Lucia Ospina Martinez, Marcela Mercado-Reyes |
| EPI_ISL_794652, EPI_ISL_794653 | LSP DEL TOLIMA | Instituto Nacional de Salud - Dirección de Investigación en Salud Pública | Katherine Laiton-Donato, Diego A. Álvarez-Díaz, Carlos Franco-Muñoz, Mauricio Pacheco-Montealegre, Jonathan Reales, Sheryl Corchuelo, Maria T. Herrera, Julian Naizaque, Gerardo Santamaría, Paola Muñoz-Laiton, Diego Andrés Prada, Magdalena Wiesner, Martha Lucia Ospina Martinez, Marcela Mercado-Reyes |
| EPI_ISL_794654, EPI_ISL_794655 | Fundación Valle del Lili | Instituto Nacional de Salud - Dirección de Investigación en Salud Pública | Katherine Laiton-Donato, Diego A. Álvarez-Díaz, Carlos Franco-Muñoz, Mauricio Pacheco-Montealegre, Jonathan Reales, Sheryl Corchuelo, Maria T. Herrera, Julian Naizaque, Gerardo Santamaría, Paola Muñoz-Laiton, Diego Andrés Prada, Magdalena Wiesner, Martha Lucia Ospina Martinez, Marcela Mercado-Reyes |
| EPI_ISL_794656, EPI_ISL_794657 | UNIDAD HEMATOLOGICA ESPECIALIZADA | Instituto Nacional de Salud - Dirección de Investigación en Salud Pública | Katherine Laiton-Donato, Diego A. Álvarez-Díaz, Carlos Franco-Muñoz, Mauricio Pacheco-Montealegre, Jonathan Reales, Sheryl Corchuelo, Maria T. Herrera, Julian Naizaque, Gerardo Santamaría, Paola Muñoz-Laiton, Diego Andrés Prada, Magdalena Wiesner, Martha Lucia Ospina Martinez, Marcela Mercado-Reyes |
| EPI_ISL_794658 | DIRECCION DE SANIDAD POLICIA NACIONAL | Instituto Nacional de Salud - Dirección de Investigación en Salud Pública | Katherine Laiton-Donato, Diego A. Álvarez-Díaz, Carlos Franco-Muñoz, Mauricio Pacheco-Montealegre, Jonathan Reales, Sheryl Corchuelo, Maria T. Herrera, Julian Naizaque, Gerardo Santamaría, Paola Muñoz-Laiton, Diego Andrés Prada, Magdalena Wiesner, Martha Lucia Ospina Martinez, Marcela Mercado-Reyes |
| EPI_ISL_794659, EPI_ISL_794660 | HOSPITAL UNIVERSITARIO SAN IGNACIO | Instituto Nacional de Salud - Dirección de Investigación en Salud Pública | Katherine Laiton-Donato, Diego A. Álvarez-Díaz, Carlos Franco-Muñoz, Mauricio Pacheco-Montealegre, Jonathan Reales, Sheryl Corchuelo, Maria T. Herrera, Julian Naizaque, Gerardo Santamaría, Paola Muñoz-Laiton, Diego Andrés Prada, Magdalena Wiesner, Martha Lucia Ospina Martinez, Marcela Mercado-Reyes |
| EPI_ISL_794662 | Fundación Cardio Infantil | Instituto Nacional de Salud - Dirección de Investigación en Salud Pública | Katherine Laiton-Donato, Diego A. Álvarez-Díaz, Carlos Franco-Muñoz, Mauricio Pacheco-Montealegre, Jonathan Reales, Sheryl Corchuelo, Maria T. Herrera, Julian Naizaque, Gerardo Santamaría, Paola Muñoz-Laiton, Diego Andrés Prada, Magdalena Wiesner, Martha Lucia Ospina Martinez, Marcela Mercado-Reyes |
| EPI_ISL_794663 | Carvajal Laboratorios | Instituto Nacional de Salud - Dirección de Investigación en Salud Pública | Katherine Laiton-Donato, Diego A. Álvarez-Díaz, Carlos Franco-Muñoz, Mauricio Pacheco-Montealegre, Jonathan Reales, Sheryl Corchuelo, Maria T. Herrera, Julian Naizaque, Gerardo Santamaría, Paola Muñoz-Laiton, Diego Andrés Prada, Magdalena Wiesner, Martha Lucia Ospina Martinez, Marcela Mercado-Reyes |
| EPI_ISL_794664, EPI_ISL_794665 | Fundación Valle del Lili | Instituto Nacional de Salud - Dirección de Investigación en Salud Pública | Katherine Laiton-Donato, Diego A. Álvarez-Díaz, Carlos Franco-Muñoz, Mauricio Pacheco-Montealegre, Jonathan Reales, Sheryl Corchuelo, Maria T. Herrera, Julian Naizaque, Gerardo Santamaría, Paola Muñoz-Laiton, Diego Andrés Prada, Magdalena Wiesner, Martha Lucia Ospina Martinez, Marcela Mercado-Reyes |
| EPI_ISL_794672, EPI_ISL_794673, EPI_ISL_794674 | PathWest Laboratory Medicine WA | PathWest Laboratory Medicine WA Microbial Surveillance Unit | PathWest Laboratory Medicine WA Microbial Surveillance Unit |
| EPI_ISL_794818, EPI_ISL_794819, EPI_ISL_794820, EPI_ISL_794822, EPI_ISL_794823, EPI_ISL_794824 | Greek Genome Center, Biomedical Research Foundation of the Academy of Athens (BRFAA) | Greek Genome Center, Biomedical Research Foundation of the Academy of Athens (BRFAA) | Emmanouil Athanasiadis, Ioannis Vatsellas, Thodoris Loupis, Christina Maria Kravvari, Katerina Zoi, Dimitrios Thanos |
| EPI_ISL_796161, EPI_ISL_796162, EPI_ISL_796166, EPI_ISL_796168, EPI_ISL_796169, EPI_ISL_796175, EPI_ISL_796176, EPI_ISL_796182, EPI_ISL_796183, EPI_ISL_796185, EPI_ISL_796191, EPI_ISL_796195, EPI_ISL_796196, EPI_ISL_796197, EPI_ISL_796201, EPI_ISL_796203, EPI_ISL_796206, EPI_ISL_796209, EPI_ISL_796210, EPI_ISL_796211, EPI_ISL_796212, EPI_ISL_796214, EPI_ISL_796215, EPI_ISL_796216, EPI_ISL_796217, EPI_ISL_796219, EPI_ISL_796220, EPI_ISL_796224, EPI_ISL_796228, EPI_ISL_796234, EPI_ISL_796235, EPI_ISL_796236, EPI_ISL_796237, EPI_ISL_796240, EPI_ISL_796243, EPI_ISL_796251, EPI_ISL_796255, EPI_ISL_796259, EPI_ISL_796264, EPI_ISL_796265, EPI_ISL_796267, EPI_ISL_796269, EPI_ISL_796285, EPI_ISL_796286, EPI_ISL_796287, EPI_ISL_796288, EPI_ISL_796289, EPI_ISL_796291, EPI_ISL_796294, EPI_ISL_796295, EPI_ISL_796300, EPI_ISL_796301, EPI_ISL_796304, EPI_ISL_796305, EPI_ISL_796306, EPI_ISL_796307, EPI_ISL_796309, EPI_ISL_796310, EPI_ISL_796317, EPI_ISL_796320, EPI_ISL_796321, EPI_ISL_796322, EPI_ISL_796324, EPI_ISL_796325, EPI_ISL_796327, EPI_ISL_796334, EPI_ISL_796351, EPI_ISL_796353, EPI_ISL_796356, EPI_ISL_796369, EPI_ISL_796373, EPI_ISL_796376, EPI_ISL_796377, EPI_ISL_796378, EPI_ISL_796384, EPI_ISL_796385, EPI_ISL_796386, EPI_ISL_796390, EPI_ISL_796391, EPI_ISL_796393, EPI_ISL_796395, EPI_ISL_796398, EPI_ISL_796399, EPI_ISL_796400, EPI_ISL_796401, EPI_ISL_796403, EPI_ISL_796407, EPI_ISL_796411, EPI_ISL_796413, EPI_ISL_796417, EPI_ISL_796426, EPI_ISL_796437, EPI_ISL_796439, EPI_ISL_796440, EPI_ISL_796441, EPI_ISL_796448, EPI_ISL_796451, EPI_ISL_796456, EPI_ISL_796457, EPI_ISL_796461, EPI_ISL_796462, EPI_ISL_796464, EPI_ISL_796465, EPI_ISL_796470, EPI_ISL_796471, EPI_ISL_796480, EPI_ISL_796481, EPI_ISL_796482, EPI_ISL_796486, EPI_ISL_796489, EPI_ISL_796493, EPI_ISL_796498, EPI_ISL_796504, EPI_ISL_796505, EPI_ISL_796511, EPI_ISL_796514, EPI_ISL_796517, EPI_ISL_796518, EPI_ISL_796523, EPI_ISL_796535, EPI_ISL_796538, EPI_ISL_796539, EPI_ISL_796540, EPI_ISL_796541, EPI_ISL_796542, EPI_ISL_796543, EPI_ISL_796544, EPI_ISL_796545, EPI_ISL_796546, EPI_ISL_796547, EPI_ISL_796548, EPI_ISL_796550, EPI_ISL_796561, EPI_ISL_796567, EPI_ISL_796571, EPI_ISL_796572, EPI_ISL_796573, EPI_ISL_796574, EPI_ISL_796577, EPI_ISL_796578, EPI_ISL_796579, EPI_ISL_796582, EPI_ISL_796585, EPI_ISL_796586, EPI_ISL_796587, EPI_ISL_796589, EPI_ISL_796591, EPI_ISL_796592, EPI_ISL_796600, EPI_ISL_796601, EPI_ISL_796607, EPI_ISL_796609, EPI_ISL_796610, EPI_ISL_796611, EPI_ISL_796613, EPI_ISL_796614, EPI_ISL_796615, EPI_ISL_796616, EPI_ISL_796617, EPI_ISL_796619, EPI_ISL_796620, EPI_ISL_796621, EPI_ISL_796623, EPI_ISL_796624, EPI_ISL_796625, EPI_ISL_796626, EPI_ISL_796629, EPI_ISL_796631, EPI_ISL_796632, EPI_ISL_796634, EPI_ISL_796636, EPI_ISL_796637, EPI_ISL_796638, EPI_ISL_796639, EPI_ISL_796640, EPI_ISL_796643 | | | |
| see above | Viollier AG | Department of Biosystems Science and Engineering, ETH Zürich | Chaoran Chen, Sarah Nadeau, Catharine Aquino, Ivan Topolsky, Philipp Jablonski, Lara Fuhrmann, David Dreifuss, Katharina Jahn, Andreia Cabral de Gouvea, Maria Domenica Moccia, Simon Grüter, Timothy Sykes, Lennart Opitz, Griffin White, Laura Neff, Doris Popovic, Andrea Patrignani, Jay Tracy, Ralph Schlapbach, Christiane Beckmann, Maurice Redondo, Olivier Kobel, Christoph Noppen, Sophie Seidel, Noemie Santamaria de Souza, Niko Beerenwinkel, Tanja Stadler |
| EPI_ISL_796661, EPI_ISL_796675 | Norwegian Institute of Public Health, Department of Virology | Norwegian Institute of Public Health, Department of Virology | Kathrine Stene-Johansen, Kamilla Heddeland Instefjord, Hilde Elshaug, Atiya R Ali,Marie Paulsen Madsen, Rasmus Riis Kopperud, Hilde Vollan, Karoline Bragstad, Olav Hungnes |
| EPI_ISL_796737, EPI_ISL_796738, EPI_ISL_796754, EPI_ISL_796761, EPI_ISL_796768, EPI_ISL_796771, EPI_ISL_796773 | Instituto Nacional de Saude (INSA) | Instituto Nacional de Saude (INSA) | Borges et al |
| EPI_ISL_796784, EPI_ISL_796785, EPI_ISL_796786, EPI_ISL_796787, EPI_ISL_796788, EPI_ISL_796789, EPI_ISL_796790, EPI_ISL_796791, EPI_ISL_796792, EPI_ISL_796793, EPI_ISL_796794, EPI_ISL_796796, EPI_ISL_796797, EPI_ISL_796798, EPI_ISL_796799, EPI_ISL_796800, EPI_ISL_796801, EPI_ISL_796802, EPI_ISL_796803, EPI_ISL_796804, EPI_ISL_796805, EPI_ISL_796806, EPI_ISL_796807, EPI_ISL_796808, EPI_ISL_796809, EPI_ISL_796810, EPI_ISL_796811, EPI_ISL_796812, EPI_ISL_796813, EPI_ISL_796814, EPI_ISL_796815, EPI_ISL_796816, EPI_ISL_796817, EPI_ISL_796818, EPI_ISL_796819, EPI_ISL_796820, EPI_ISL_796821, EPI_ISL_796822, EPI_ISL_796823, EPI_ISL_796824, EPI_ISL_796825, EPI_ISL_796826, EPI_ISL_796827, EPI_ISL_796828, EPI_ISL_796829, EPI_ISL_796830, EPI_ISL_796831, EPI_ISL_796832, EPI_ISL_796833, EPI_ISL_796834, EPI_ISL_796835, EPI_ISL_796836, EPI_ISL_796837, EPI_ISL_796838, EPI_ISL_796839, EPI_ISL_796840, EPI_ISL_796841, EPI_ISL_796842, EPI_ISL_796843, EPI_ISL_796844, EPI_ISL_796845, EPI_ISL_796846, EPI_ISL_796847, EPI_ISL_796848, EPI_ISL_796849, EPI_ISL_796850, EPI_ISL_796851, EPI_ISL_796852, EPI_ISL_796853, EPI_ISL_796854, EPI_ISL_796855, EPI_ISL_796856, EPI_ISL_796857, EPI_ISL_796858, EPI_ISL_796859, EPI_ISL_796860, EPI_ISL_796861, EPI_ISL_796862, EPI_ISL_796863, EPI_ISL_796864, EPI_ISL_796865, EPI_ISL_796866, EPI_ISL_796867, EPI_ISL_796868, EPI_ISL_796869, EPI_ISL_796870, EPI_ISL_796871, EPI_ISL_796872, EPI_ISL_796873, EPI_ISL_796874, EPI_ISL_796875, EPI_ISL_796876, EPI_ISL_796877, EPI_ISL_796878, EPI_ISL_796879, EPI_ISL_796880, EPI_ISL_796881, EPI_ISL_796882, EPI_ISL_796883, EPI_ISL_796884, EPI_ISL_796885, EPI_ISL_796886, EPI_ISL_796887, EPI_ISL_796888, EPI_ISL_796889, EPI_ISL_796890, EPI_ISL_796891, EPI_ISL_796892, EPI_ISL_796893, EPI_ISL_796894, EPI_ISL_796895, EPI_ISL_796896, EPI_ISL_796897, EPI_ISL_796898, EPI_ISL_796899, EPI_ISL_796900, EPI_ISL_796901, EPI_ISL_796902, EPI_ISL_796903, EPI_ISL_796904, EPI_ISL_796905, EPI_ISL_796906, EPI_ISL_796907, EPI_ISL_796908, EPI_ISL_796909, EPI_ISL_796910, EPI_ISL_796911, EPI_ISL_796912, EPI_ISL_796913, EPI_ISL_796914, EPI_ISL_796915, EPI_ISL_796916, EPI_ISL_796917, EPI_ISL_796918, EPI_ISL_796919, EPI_ISL_796920, EPI_ISL_796921, EPI_ISL_796922, EPI_ISL_796923, EPI_ISL_796924, EPI_ISL_796925, EPI_ISL_796926, EPI_ISL_796927, EPI_ISL_796928, EPI_ISL_796929, EPI_ISL_796930, EPI_ISL_796931, EPI_ISL_796932, EPI_ISL_796933, EPI_ISL_796934, EPI_ISL_796935, EPI_ISL_796936, EPI_ISL_796937, EPI_ISL_796938, EPI_ISL_796939, EPI_ISL_796940, EPI_ISL_796941, EPI_ISL_796942, EPI_ISL_796943, EPI_ISL_796944, EPI_ISL_796945, EPI_ISL_796946, EPI_ISL_796947, EPI_ISL_796948, EPI_ISL_796949, EPI_ISL_796950, EPI_ISL_796951, EPI_ISL_796952, EPI_ISL_796953, EPI_ISL_796954, EPI_ISL_796955, EPI_ISL_796956, EPI_ISL_796957, EPI_ISL_796958, EPI_ISL_796959, EPI_ISL_796960, EPI_ISL_796961, EPI_ISL_796962, EPI_ISL_796963, EPI_ISL_796964, EPI_ISL_796965, EPI_ISL_796966, EPI_ISL_796967, EPI_ISL_796968, EPI_ISL_796969, EPI_ISL_796970, EPI_ISL_796971, EPI_ISL_796972, EPI_ISL_796973, EPI_ISL_796974, EPI_ISL_796975, EPI_ISL_796976, EPI_ISL_796977, EPI_ISL_796978, EPI_ISL_796979, EPI_ISL_796980, EPI_ISL_796981, EPI_ISL_796982, EPI_ISL_796983, EPI_ISL_796984, EPI_ISL_796985, EPI_ISL_796986, EPI_ISL_796987, EPI_ISL_796988, EPI_ISL_796989, EPI_ISL_796990, EPI_ISL_796991, EPI_ISL_796992, EPI_ISL_796993, EPI_ISL_796994, EPI_ISL_796995, EPI_ISL_796996, EPI_ISL_796998, EPI_ISL_796999, EPI_ISL_797000, EPI_ISL_797001, EPI_ISL_797002, EPI_ISL_797003, EPI_ISL_797004, EPI_ISL_797005, EPI_ISL_797006, EPI_ISL_797007, EPI_ISL_797008, EPI_ISL_797009, EPI_ISL_797010, EPI_ISL_797011, EPI_ISL_797012, EPI_ISL_797013, EPI_ISL_797014, EPI_ISL_797015, EPI_ISL_797016, EPI_ISL_797017, EPI_ISL_797018, EPI_ISL_797019, EPI_ISL_797020, EPI_ISL_797021, EPI_ISL_797022, EPI_ISL_797023, EPI_ISL_797024, EPI_ISL_797025, EPI_ISL_797026, EPI_ISL_797027, EPI_ISL_797028, EPI_ISL_797029, EPI_ISL_797030, EPI_ISL_797031, EPI_ISL_797032, EPI_ISL_797033, EPI_ISL_797034, EPI_ISL_797035, EPI_ISL_797036, EPI_ISL_797037, EPI_ISL_797038, EPI_ISL_797039, EPI_ISL_797040, EPI_ISL_797041, EPI_ISL_797042, EPI_ISL_797043, EPI_ISL_797044, EPI_ISL_797045, EPI_ISL_797046, EPI_ISL_797047, EPI_ISL_797048, EPI_ISL_797049, EPI_ISL_797050, EPI_ISL_797051, EPI_ISL_797052, EPI_ISL_797053, EPI_ISL_797054, EPI_ISL_797055, EPI_ISL_797056, EPI_ISL_797057, EPI_ISL_797058, EPI_ISL_797059, EPI_ISL_797060, EPI_ISL_797061, EPI_ISL_797062, EPI_ISL_797063, EPI_ISL_797064, EPI_ISL_797065, EPI_ISL_797066, EPI_ISL_797067, EPI_ISL_797068, EPI_ISL_797069, EPI_ISL_797070, EPI_ISL_797071, EPI_ISL_797072, EPI_ISL_797073, EPI_ISL_797074, EPI_ISL_797075, EPI_ISL_797076, EPI_ISL_797077, EPI_ISL_797078, EPI_ISL_797079, EPI_ISL_797080, EPI_ISL_797081, EPI_ISL_797082, EPI_ISL_797083, EPI_ISL_797084, EPI_ISL_797085, EPI_ISL_797086, EPI_ISL_797088, EPI_ISL_797089, EPI_ISL_797090, EPI_ISL_797091, EPI_ISL_797092, EPI_ISL_797093, EPI_ISL_797094, EPI_ISL_797095, EPI_ISL_797096, EPI_ISL_797097, EPI_ISL_797098, EPI_ISL_797099, EPI_ISL_797100, EPI_ISL_797101, EPI_ISL_797102, EPI_ISL_797103, EPI_ISL_797104, EPI_ISL_797105, EPI_ISL_797106, EPI_ISL_797107, EPI_ISL_797108, EPI_ISL_797109, EPI_ISL_797110, EPI_ISL_797111, EPI_ISL_797112, EPI_ISL_797113, EPI_ISL_797114, EPI_ISL_797115, EPI_ISL_797116, EPI_ISL_797117 | | | |
| see above | Lighthouse Lab in Alderley Park | Wellcome Sanger Institute for the COVID-19 Genomics UK (COG-UK) Consortium | Jacquelyn Wynn, Mairead Hyland, The Lighthouse Lab in Alderley Park and Alex Alderton, Roberto Amato, Sonia Goncalves, Ewan Harrison, David K. Jackson, Ian Johnston, Dominic Kwiatkowski, Cordelia Langford, John Sillitoe on behalf of the Wellcome Sanger Institute COVID-19 Surveillance Team |
| EPI_ISL_797118, EPI_ISL_797120, EPI_ISL_797121, EPI_ISL_797125, EPI_ISL_797126, EPI_ISL_797128, EPI_ISL_797129, EPI_ISL_797133, EPI_ISL_797134, EPI_ISL_797138, EPI_ISL_797139, EPI_ISL_797141, EPI_ISL_797142, EPI_ISL_797143, EPI_ISL_797144, EPI_ISL_797146, EPI_ISL_797148, EPI_ISL_797149, | | | |

| | | | |
|---|---|---|---|
| EPI_ISL_797151, EPI_ISL_797153, EPI_ISL_797154, EPI_ISL_797155, EPI_ISL_797156, EPI_ISL_797160, EPI_ISL_797161, EPI_ISL_797163, EPI_ISL_797164, EPI_ISL_797165, EPI_ISL_797168, EPI_ISL_797171, EPI_ISL_797173, EPI_ISL_797175, EPI_ISL_797176, EPI_ISL_797177, EPI_ISL_797178, EPI_ISL_797179, EPI_ISL_797185, EPI_ISL_797187, EPI_ISL_797190, EPI_ISL_797194, EPI_ISL_797198, EPI_ISL_797202, EPI_ISL_797203, EPI_ISL_797204, EPI_ISL_797205, EPI_ISL_797207, EPI_ISL_797210, EPI_ISL_797212, EPI_ISL_797216, EPI_ISL_797218, EPI_ISL_797220, EPI_ISL_797225, EPI_ISL_797226, EPI_ISL_797228, EPI_ISL_797229, EPI_ISL_797231, EPI_ISL_797232, EPI_ISL_797234, EPI_ISL_797239, EPI_ISL_797240, EPI_ISL_797242, EPI_ISL_797243, EPI_ISL_797246, EPI_ISL_797247, EPI_ISL_797248, EPI_ISL_797252, EPI_ISL_797253, EPI_ISL_797254, EPI_ISL_797256, EPI_ISL_797258, EPI_ISL_797259, EPI_ISL_797261, EPI_ISL_797264, EPI_ISL_797266, EPI_ISL_797269, EPI_ISL_797272, EPI_ISL_797277, EPI_ISL_797281, EPI_ISL_797286, EPI_ISL_797289, EPI_ISL_797290, EPI_ISL_797295, EPI_ISL_797296, EPI_ISL_797301, EPI_ISL_797305, EPI_ISL_797313, EPI_ISL_797315, EPI_ISL_797317, EPI_ISL_797318, EPI_ISL_797319, EPI_ISL_797323, EPI_ISL_797325, EPI_ISL_797326, EPI_ISL_797327, EPI_ISL_797332, EPI_ISL_797335, EPI_ISL_797337, EPI_ISL_797338, EPI_ISL_797339, EPI_ISL_797342, EPI_ISL_797345, EPI_ISL_797346, EPI_ISL_797349, EPI_ISL_797354, EPI_ISL_797357, EPI_ISL_797362, EPI_ISL_797363, EPI_ISL_797365, EPI_ISL_797366, EPI_ISL_797367, EPI_ISL_797376, EPI_ISL_797379, EPI_ISL_797380, EPI_ISL_797385, EPI_ISL_797389, EPI_ISL_797390, EPI_ISL_797391, EPI_ISL_797395 | | | |
| see above | Lighthouse Lab in Milton Keynes | Wellcome Sanger Institute for the COVID-19 Genomics UK (COG-UK) Consortium | The Lighthouse Lab in Milton Keynes and Alex Alderton, Roberto Amato, Sonia Goncalves, Ewan Harrison, David K. Jackson, Ian Johnston, Dominic Kwiatkowski, Cordelia Langford, John Sillitoe on behalf of the Wellcome Sanger Institute COVID-19 Surveillance Team |
| EPI_ISL_797397, EPI_ISL_797398 | Lighthouse Lab in Cambridge | Wellcome Sanger Institute for the COVID-19 Genomics UK (COG-UK) Consortium | Rob Howes, The Lighthouse Lab in Cambridge and Alex Alderton, Roberto Amato, Sonia Goncalves, Ewan Harrison, David K. Jackson, Ian Johnston, Dominic Kwiatkowski, Cordelia Langford, John Sillitoe on behalf of the Wellcome Sanger Institute COVID-19 Surveillance Team |
| EPI_ISL_797399 | Lighthouse Lab in Alderley Park | Wellcome Sanger Institute for the COVID-19 Genomics UK (COG-UK) Consortium | Jacquelyn Wynn, Mairead Hyland, The Lighthouse Lab in Alderley Park and Alex Alderton, Roberto Amato, Sonia Goncalves, Ewan Harrison, David K. Jackson, Ian Johnston, Dominic Kwiatkowski, Cordelia Langford, John Sillitoe on behalf of the Wellcome Sanger Institute COVID-19 Surveillance Team |
| EPI_ISL_797401 | Lighthouse Lab in Cambridge | Wellcome Sanger Institute for the COVID-19 Genomics UK (COG-UK) Consortium | Rob Howes, The Lighthouse Lab in Cambridge and Alex Alderton, Roberto Amato, Sonia Goncalves, Ewan Harrison, David K. Jackson, Ian Johnston, Dominic Kwiatkowski, Cordelia Langford, John Sillitoe on behalf of the Wellcome Sanger Institute COVID-19 Surveillance Team |
| EPI_ISL_797402 | Lighthouse Lab in Alderley Park | Wellcome Sanger Institute for the COVID-19 Genomics UK (COG-UK) Consortium | Jacquelyn Wynn, Mairead Hyland, The Lighthouse Lab in Alderley Park and Alex Alderton, Roberto Amato, Sonia Goncalves, Ewan Harrison, David K. Jackson, Ian Johnston, Dominic Kwiatkowski, Cordelia Langford, John Sillitoe on behalf of the Wellcome Sanger Institute COVID-19 Surveillance Team |
| EPI_ISL_797404, EPI_ISL_797406, EPI_ISL_797407 | Lighthouse Lab in Cambridge | Wellcome Sanger Institute for the COVID-19 Genomics UK (COG-UK) Consortium | Rob Howes, The Lighthouse Lab in Cambridge and Alex Alderton, Roberto Amato, Sonia Goncalves, Ewan Harrison, David K. Jackson, Ian Johnston, Dominic Kwiatkowski, Cordelia Langford, John Sillitoe on behalf of the Wellcome Sanger Institute COVID-19 Surveillance Team |
| EPI_ISL_797409 | Lighthouse Lab in Alderley Park | Wellcome Sanger Institute for the COVID-19 Genomics UK (COG-UK) Consortium | Jacquelyn Wynn, Mairead Hyland, The Lighthouse Lab in Alderley Park and Alex Alderton, Roberto Amato, Sonia Goncalves, Ewan Harrison, David K. Jackson, Ian Johnston, Dominic Kwiatkowski, Cordelia Langford, John Sillitoe on behalf of the Wellcome Sanger Institute COVID-19 Surveillance Team |
| EPI_ISL_797410, EPI_ISL_797412 | Lighthouse Lab in Cambridge | Wellcome Sanger Institute for the COVID-19 Genomics UK (COG-UK) Consortium | Rob Howes, The Lighthouse Lab in Cambridge and Alex Alderton, Roberto Amato, Sonia Goncalves, Ewan Harrison, David K. Jackson, Ian Johnston, Dominic Kwiatkowski, Cordelia Langford, John Sillitoe on behalf of the Wellcome Sanger Institute COVID-19 Surveillance Team |
| EPI_ISL_797413 | Lighthouse Lab in Alderley Park | Wellcome Sanger Institute for the COVID-19 Genomics UK (COG-UK) Consortium | Jacquelyn Wynn, Mairead Hyland, The Lighthouse Lab in Alderley Park and Alex Alderton, Roberto Amato, Sonia Goncalves, Ewan Harrison, David K. Jackson, Ian Johnston, Dominic Kwiatkowski, Cordelia Langford, John Sillitoe on behalf of the Wellcome Sanger Institute COVID-19 Surveillance Team |
| EPI_ISL_797414, EPI_ISL_797415 | Lighthouse Lab in Cambridge | Wellcome Sanger Institute for the COVID-19 Genomics UK (COG-UK) Consortium | Rob Howes, The Lighthouse Lab in Cambridge and Alex Alderton, Roberto Amato, Sonia Goncalves, Ewan Harrison, David K. Jackson, Ian Johnston, Dominic Kwiatkowski, Cordelia Langford, John Sillitoe on behalf of the Wellcome Sanger Institute COVID-19 Surveillance Team |
| EPI_ISL_797416 | Lighthouse Lab in Alderley Park | Wellcome Sanger Institute for the COVID-19 Genomics UK (COG-UK) Consortium | Jacquelyn Wynn, Mairead Hyland, The Lighthouse Lab in Alderley Park and Alex Alderton, Roberto Amato, Sonia Goncalves, Ewan Harrison, David K. Jackson, Ian Johnston, Dominic Kwiatkowski, Cordelia Langford, John Sillitoe on behalf of the Wellcome Sanger Institute COVID-19 Surveillance Team |
| EPI_ISL_797417, EPI_ISL_797418, EPI_ISL_797419 | Lighthouse Lab in Cambridge | Wellcome Sanger Institute for the COVID-19 Genomics UK (COG-UK) Consortium | Rob Howes, The Lighthouse Lab in Cambridge and Alex Alderton, Roberto Amato, Sonia Goncalves, Ewan Harrison, David K. Jackson, Ian Johnston, Dominic Kwiatkowski, Cordelia Langford, John Sillitoe on behalf of the Wellcome Sanger Institute COVID-19 Surveillance Team |
| EPI_ISL_797420, EPI_ISL_797421 | Lighthouse Lab in Alderley Park | Wellcome Sanger Institute for the COVID-19 Genomics UK (COG-UK) Consortium | Jacquelyn Wynn, Mairead Hyland, The Lighthouse Lab in Alderley Park and Alex Alderton, Roberto Amato, Sonia Goncalves, Ewan Harrison, David K. Jackson, Ian Johnston, Dominic Kwiatkowski, Cordelia Langford, John Sillitoe on behalf of the Wellcome Sanger Institute COVID-19 Surveillance Team |
| EPI_ISL_797422, EPI_ISL_797423 | Lighthouse Lab in Cambridge | Wellcome Sanger Institute for the COVID-19 Genomics UK (COG-UK) Consortium | Rob Howes, The Lighthouse Lab in Cambridge and Alex Alderton, Roberto Amato, Sonia Goncalves, Ewan Harrison, David K. Jackson, Ian Johnston, Dominic Kwiatkowski, Cordelia Langford, John Sillitoe on behalf of the Wellcome Sanger Institute COVID-19 Surveillance Team |
| EPI_ISL_797424 | Lighthouse Lab in Alderley Park | Wellcome Sanger Institute for the COVID-19 Genomics UK (COG-UK) Consortium | Jacquelyn Wynn, Mairead Hyland, The Lighthouse Lab in Alderley Park and Alex Alderton, Roberto Amato, Sonia Goncalves, Ewan Harrison, David K. Jackson, Ian Johnston, Dominic Kwiatkowski, Cordelia Langford, John Sillitoe on behalf of the Wellcome Sanger Institute COVID-19 Surveillance Team |
| EPI_ISL_797425, EPI_ISL_797426, EPI_ISL_797427 | Lighthouse Lab in Cambridge | Wellcome Sanger Institute for the COVID-19 Genomics UK (COG-UK) Consortium | Rob Howes, The Lighthouse Lab in Cambridge and Alex Alderton, Roberto Amato, Sonia Goncalves, Ewan Harrison, David K. Jackson, Ian Johnston, Dominic Kwiatkowski, Cordelia Langford, John Sillitoe on behalf of the Wellcome Sanger Institute COVID-19 Surveillance Team |
| EPI_ISL_797428 | Lighthouse Lab in Alderley Park | Wellcome Sanger Institute for the COVID-19 Genomics UK (COG-UK) Consortium | Jacquelyn Wynn, Mairead Hyland, The Lighthouse Lab in Alderley Park and Alex Alderton, Roberto Amato, Sonia Goncalves, Ewan Harrison, David K. Jackson, Ian Johnston, Dominic Kwiatkowski, Cordelia Langford, John Sillitoe on behalf of the Wellcome Sanger Institute COVID-19 Surveillance Team |
| EPI_ISL_797429 | Lighthouse Lab in Cambridge | Wellcome Sanger Institute for the COVID-19 Genomics UK (COG-UK) Consortium | Rob Howes, The Lighthouse Lab in Cambridge and Alex Alderton, Roberto Amato, Sonia Goncalves, Ewan Harrison, David K. Jackson, Ian Johnston, Dominic Kwiatkowski, Cordelia Langford, John Sillitoe on behalf of the Wellcome Sanger Institute COVID-19 Surveillance Team |
| EPI_ISL_797430 | Lighthouse Lab in Alderley Park | Wellcome Sanger Institute for the COVID-19 Genomics UK (COG-UK) Consortium | Jacquelyn Wynn, Mairead Hyland, The Lighthouse Lab in Alderley Park and Alex Alderton, Roberto Amato, Sonia Goncalves, Ewan Harrison, David K. Jackson, Ian Johnston, Dominic Kwiatkowski, Cordelia Langford, John Sillitoe on behalf of the Wellcome Sanger Institute COVID-19 Surveillance Team |
| EPI_ISL_797431, EPI_ISL_797433 | Lighthouse Lab in Cambridge | Wellcome Sanger Institute for the COVID-19 Genomics UK (COG-UK) Consortium | Rob Howes, The Lighthouse Lab in Cambridge and Alex Alderton, Roberto Amato, Sonia Goncalves, Ewan Harrison, David K. Jackson, Ian Johnston, Dominic Kwiatkowski, Cordelia Langford, John Sillitoe on behalf of the Wellcome Sanger Institute COVID-19 Surveillance Team |
| EPI_ISL_797435 | Lighthouse Lab in Alderley Park | Wellcome Sanger Institute for the COVID-19 Genomics UK (COG-UK) Consortium | Jacquelyn Wynn, Mairead Hyland, The Lighthouse Lab in Alderley Park and Alex Alderton, Roberto Amato, Sonia Goncalves, Ewan Harrison, David K. Jackson, Ian Johnston, Dominic Kwiatkowski, Cordelia Langford, John Sillitoe on behalf of the Wellcome Sanger Institute COVID-19 Surveillance Team |
| EPI_ISL_797436, EPI_ISL_797437, EPI_ISL_797439, EPI_ISL_797440, EPI_ISL_797442, EPI_ISL_797443, EPI_ISL_797444, EPI_ISL_797447, EPI_ISL_797448, EPI_ISL_797449, EPI_ISL_797450, EPI_ISL_797452, EPI_ISL_797453, EPI_ISL_797454, EPI_ISL_797455 | | | |
| see above | Lighthouse Lab in Cambridge | Wellcome Sanger Institute for the COVID-19 Genomics UK (COG-UK) Consortium | Rob Howes, The Lighthouse Lab in Cambridge and Alex Alderton, Roberto Amato, Sonia Goncalves, Ewan Harrison, David K. Jackson, Ian Johnston, Dominic Kwiatkowski, Cordelia Langford, John Sillitoe on behalf of the Wellcome Sanger Institute COVID-19 Surveillance Team |
| EPI_ISL_797456 | Lighthouse Lab in Alderley Park | Wellcome Sanger Institute for the COVID-19 Genomics UK (COG-UK) Consortium | Jacquelyn Wynn, Mairead Hyland, The Lighthouse Lab in Alderley Park and Alex Alderton, Roberto Amato, Sonia Goncalves, Ewan Harrison, David K. Jackson, Ian Johnston, Dominic Kwiatkowski, Cordelia Langford, John Sillitoe on behalf of the Wellcome Sanger Institute COVID-19 Surveillance Team |
| EPI_ISL_797457, EPI_ISL_797458, EPI_ISL_797460, EPI_ISL_797463, EPI_ISL_797464, EPI_ISL_797466, EPI_ISL_797467 | Lighthouse Lab in Cambridge | Wellcome Sanger Institute for the COVID-19 Genomics UK (COG-UK) Consortium | Rob Howes, The Lighthouse Lab in Cambridge and Alex Alderton, Roberto Amato, Sonia Goncalves, Ewan Harrison, David K. Jackson, Ian Johnston, Dominic Kwiatkowski, Cordelia Langford, John Sillitoe on behalf of the Wellcome Sanger Institute COVID-19 Surveillance Team |
| EPI_ISL_797468 | Lighthouse Lab in Alderley Park | Wellcome Sanger Institute for the COVID-19 Genomics UK (COG-UK) Consortium | Jacquelyn Wynn, Mairead Hyland, The Lighthouse Lab in Alderley Park and Alex Alderton, Roberto Amato, Sonia Goncalves, Ewan Harrison, David K. Jackson, Ian Johnston, Dominic Kwiatkowski, Cordelia Langford, John Sillitoe on behalf of the Wellcome Sanger Institute COVID-19 Surveillance Team |
| EPI_ISL_797469, EPI_ISL_797470, EPI_ISL_797471, EPI_ISL_797472, EPI_ISL_797473, EPI_ISL_797474, EPI_ISL_797476, EPI_ISL_797477, EPI_ISL_797478, EPI_ISL_797479 | Lighthouse Lab in Cambridge | Wellcome Sanger Institute for the COVID-19 Genomics UK (COG-UK) Consortium | Rob Howes, The Lighthouse Lab in Cambridge and Alex Alderton, Roberto Amato, Sonia Goncalves, Ewan Harrison, David K. Jackson, Ian Johnston, Dominic Kwiatkowski, Cordelia Langford, John Sillitoe on behalf of the Wellcome Sanger Institute COVID-19 Surveillance Team |
| EPI_ISL_797480 | Lighthouse Lab in Alderley Park | Wellcome Sanger Institute for the COVID-19 Genomics UK (COG-UK) Consortium | Jacquelyn Wynn, Mairead Hyland, The Lighthouse Lab in Alderley Park and Alex Alderton, Roberto Amato, Sonia Goncalves, Ewan Harrison, David K. Jackson, Ian Johnston, Dominic Kwiatkowski, Cordelia Langford, John Sillitoe on behalf of the Wellcome Sanger Institute COVID-19 Surveillance Team |
| EPI_ISL_797482, EPI_ISL_797485, EPI_ISL_797486, EPI_ISL_797487, EPI_ISL_797488 | Lighthouse Lab in Cambridge | Wellcome Sanger Institute for the COVID-19 Genomics UK (COG-UK) Consortium | Rob Howes, The Lighthouse Lab in Cambridge and Alex Alderton, Roberto Amato, Sonia Goncalves, Ewan Harrison, David K. Jackson, Ian Johnston, Dominic Kwiatkowski, Cordelia Langford, John Sillitoe on behalf of the Wellcome Sanger Institute COVID-19 Surveillance Team |
| EPI_ISL_797489 | Lighthouse Lab in Alderley Park | Wellcome Sanger Institute for the COVID-19 Genomics UK (COG-UK) Consortium | Jacquelyn Wynn, Mairead Hyland, The Lighthouse Lab in Alderley Park and Alex Alderton, Roberto Amato, Sonia Goncalves, Ewan Harrison, David K. Jackson, Ian Johnston, Dominic Kwiatkowski, Cordelia Langford, John Sillitoe on behalf of the Wellcome Sanger Institute COVID-19 Surveillance Team |
| EPI_ISL_797490 | Lighthouse Lab in Cambridge | Wellcome Sanger Institute for the COVID-19 Genomics UK | Rob Howes, The Lighthouse Lab in Cambridge and Alex Alderton, Roberto Amato, Sonia Goncalves, Ewan Harrison, David K. Jackson, Ian Johnston, |

| | | | |
|---|---|---|---|
| | | (COG-UK) Consortium | Dominic Kwiatkowski, Cordelia Langford, John Sillitoe on behalf of the Wellcome Sanger Institute COVID-19 Surveillance Team |
| EPI_ISL_797491 | Lighthouse Lab in Alderley Park | Wellcome Sanger Institute for the COVID-19 Genomics UK (COG-UK) Consortium | Jacquelyn Wynn, Mairead Hyland, The Lighthouse Lab in Alderley Park and Alex Alderton, Roberto Amato, Sonia Goncalves, Ewan Harrison, David K. Jackson, Ian Johnston, Dominic Kwiatkowski, Cordelia Langford, John Sillitoe on behalf of the Wellcome Sanger Institute COVID-19 Surveillance Team |
| EPI_ISL_797493, EPI_ISL_797494, EPI_ISL_797495, EPI_ISL_797496 | Lighthouse Lab in Cambridge | Wellcome Sanger Institute for the COVID-19 Genomics UK (COG-UK) Consortium | Rob Howes, The Lighthouse Lab in Cambridge and Alex Alderton, Roberto Amato, Sonia Goncalves, Ewan Harrison, David K. Jackson, Ian Johnston, Dominic Kwiatkowski, Cordelia Langford, John Sillitoe on behalf of the Wellcome Sanger Institute COVID-19 Surveillance Team |
| EPI_ISL_797499 | Lighthouse Lab in Alderley Park | Wellcome Sanger Institute for the COVID-19 Genomics UK (COG-UK) Consortium | Jacquelyn Wynn, Mairead Hyland, The Lighthouse Lab in Alderley Park and Alex Alderton, Roberto Amato, Sonia Goncalves, Ewan Harrison, David K. Jackson, Ian Johnston, Dominic Kwiatkowski, Cordelia Langford, John Sillitoe on behalf of the Wellcome Sanger Institute COVID-19 Surveillance Team |
| EPI_ISL_797501, EPI_ISL_797503, EPI_ISL_797506, EPI_ISL_797507, EPI_ISL_797508 | Lighthouse Lab in Cambridge | Wellcome Sanger Institute for the COVID-19 Genomics UK (COG-UK) Consortium | Rob Howes, The Lighthouse Lab in Cambridge and Alex Alderton, Roberto Amato, Sonia Goncalves, Ewan Harrison, David K. Jackson, Ian Johnston, Dominic Kwiatkowski, Cordelia Langford, John Sillitoe on behalf of the Wellcome Sanger Institute COVID-19 Surveillance Team |
| EPI_ISL_797512 | Lighthouse Lab in Alderley Park | Wellcome Sanger Institute for the COVID-19 Genomics UK (COG-UK) Consortium | Jacquelyn Wynn, Mairead Hyland, The Lighthouse Lab in Alderley Park and Alex Alderton, Roberto Amato, Sonia Goncalves, Ewan Harrison, David K. Jackson, Ian Johnston, Dominic Kwiatkowski, Cordelia Langford, John Sillitoe on behalf of the Wellcome Sanger Institute COVID-19 Surveillance Team |
| EPI_ISL_797513, EPI_ISL_797515, EPI_ISL_797516, EPI_ISL_797517, EPI_ISL_797518, EPI_ISL_797519, EPI_ISL_797520, EPI_ISL_797521, EPI_ISL_797522, EPI_ISL_797523, EPI_ISL_797524, EPI_ISL_797525, EPI_ISL_797526, EPI_ISL_797527, EPI_ISL_797528, EPI_ISL_797529, EPI_ISL_797530 | | | |
| see above | Lighthouse Lab in Cambridge | Wellcome Sanger Institute for the COVID-19 Genomics UK (COG-UK) Consortium | Rob Howes, The Lighthouse Lab in Cambridge and Alex Alderton, Roberto Amato, Sonia Goncalves, Ewan Harrison, David K. Jackson, Ian Johnston, Dominic Kwiatkowski, Cordelia Langford, John Sillitoe on behalf of the Wellcome Sanger Institute COVID-19 Surveillance Team |
| EPI_ISL_797533 | Lighthouse Lab in Alderley Park | Wellcome Sanger Institute for the COVID-19 Genomics UK (COG-UK) Consortium | Jacquelyn Wynn, Mairead Hyland, The Lighthouse Lab in Alderley Park and Alex Alderton, Roberto Amato, Sonia Goncalves, Ewan Harrison, David K. Jackson, Ian Johnston, Dominic Kwiatkowski, Cordelia Langford, John Sillitoe on behalf of the Wellcome Sanger Institute COVID-19 Surveillance Team |
| EPI_ISL_797534 | Lighthouse Lab in Cambridge | Wellcome Sanger Institute for the COVID-19 Genomics UK (COG-UK) Consortium | Rob Howes, The Lighthouse Lab in Cambridge and Alex Alderton, Roberto Amato, Sonia Goncalves, Ewan Harrison, David K. Jackson, Ian Johnston, Dominic Kwiatkowski, Cordelia Langford, John Sillitoe on behalf of the Wellcome Sanger Institute COVID-19 Surveillance Team |
| EPI_ISL_797535 | Lighthouse Lab in Alderley Park | Wellcome Sanger Institute for the COVID-19 Genomics UK (COG-UK) Consortium | Jacquelyn Wynn, Mairead Hyland, The Lighthouse Lab in Alderley Park and Alex Alderton, Roberto Amato, Sonia Goncalves, Ewan Harrison, David K. Jackson, Ian Johnston, Dominic Kwiatkowski, Cordelia Langford, John Sillitoe on behalf of the Wellcome Sanger Institute COVID-19 Surveillance Team |
| EPI_ISL_797536, EPI_ISL_797538, EPI_ISL_797539 | Lighthouse Lab in Cambridge | Wellcome Sanger Institute for the COVID-19 Genomics UK (COG-UK) Consortium | Rob Howes, The Lighthouse Lab in Cambridge and Alex Alderton, Roberto Amato, Sonia Goncalves, Ewan Harrison, David K. Jackson, Ian Johnston, Dominic Kwiatkowski, Cordelia Langford, John Sillitoe on behalf of the Wellcome Sanger Institute COVID-19 Surveillance Team |
| EPI_ISL_797540, EPI_ISL_797541 | Lighthouse Lab in Alderley Park | Wellcome Sanger Institute for the COVID-19 Genomics UK (COG-UK) Consortium | Jacquelyn Wynn, Mairead Hyland, The Lighthouse Lab in Alderley Park and Alex Alderton, Roberto Amato, Sonia Goncalves, Ewan Harrison, David K. Jackson, Ian Johnston, Dominic Kwiatkowski, Cordelia Langford, John Sillitoe on behalf of the Wellcome Sanger Institute COVID-19 Surveillance Team |
| EPI_ISL_797542 | Lighthouse Lab in Cambridge | Wellcome Sanger Institute for the COVID-19 Genomics UK (COG-UK) Consortium | Rob Howes, The Lighthouse Lab in Cambridge and Alex Alderton, Roberto Amato, Sonia Goncalves, Ewan Harrison, David K. Jackson, Ian Johnston, Dominic Kwiatkowski, Cordelia Langford, John Sillitoe on behalf of the Wellcome Sanger Institute COVID-19 Surveillance Team |
| EPI_ISL_797543 | Lighthouse Lab in Alderley Park | Wellcome Sanger Institute for the COVID-19 Genomics UK (COG-UK) Consortium | Jacquelyn Wynn, Mairead Hyland, The Lighthouse Lab in Alderley Park and Alex Alderton, Roberto Amato, Sonia Goncalves, Ewan Harrison, David K. Jackson, Ian Johnston, Dominic Kwiatkowski, Cordelia Langford, John Sillitoe on behalf of the Wellcome Sanger Institute COVID-19 Surveillance Team |
| EPI_ISL_797544, EPI_ISL_797545, EPI_ISL_797546 | Lighthouse Lab in Cambridge | Wellcome Sanger Institute for the COVID-19 Genomics UK (COG-UK) Consortium | Rob Howes, The Lighthouse Lab in Cambridge and Alex Alderton, Roberto Amato, Sonia Goncalves, Ewan Harrison, David K. Jackson, Ian Johnston, Dominic Kwiatkowski, Cordelia Langford, John Sillitoe on behalf of the Wellcome Sanger Institute COVID-19 Surveillance Team |
| EPI_ISL_797547, EPI_ISL_797549 | Lighthouse Lab in Alderley Park | Wellcome Sanger Institute for the COVID-19 Genomics UK (COG-UK) Consortium | Jacquelyn Wynn, Mairead Hyland, The Lighthouse Lab in Alderley Park and Alex Alderton, Roberto Amato, Sonia Goncalves, Ewan Harrison, David K. Jackson, Ian Johnston, Dominic Kwiatkowski, Cordelia Langford, John Sillitoe on behalf of the Wellcome Sanger Institute COVID-19 Surveillance Team |
| EPI_ISL_797550, EPI_ISL_797551, EPI_ISL_797552, EPI_ISL_797555, EPI_ISL_797556, EPI_ISL_797557, EPI_ISL_797558, EPI_ISL_797561, EPI_ISL_797562, EPI_ISL_797565, EPI_ISL_797567 | | | |
| see above | Lighthouse Lab in Cambridge | Wellcome Sanger Institute for the COVID-19 Genomics UK (COG-UK) Consortium | Rob Howes, The Lighthouse Lab in Cambridge and Alex Alderton, Roberto Amato, Sonia Goncalves, Ewan Harrison, David K. Jackson, Ian Johnston, Dominic Kwiatkowski, Cordelia Langford, John Sillitoe on behalf of the Wellcome Sanger Institute COVID-19 Surveillance Team |
| EPI_ISL_797568 | Lighthouse Lab in Alderley Park | Wellcome Sanger Institute for the COVID-19 Genomics UK (COG-UK) Consortium | Jacquelyn Wynn, Mairead Hyland, The Lighthouse Lab in Alderley Park and Alex Alderton, Roberto Amato, Sonia Goncalves, Ewan Harrison, David K. Jackson, Ian Johnston, Dominic Kwiatkowski, Cordelia Langford, John Sillitoe on behalf of the Wellcome Sanger Institute COVID-19 Surveillance Team |
| EPI_ISL_797569, EPI_ISL_797570, EPI_ISL_797571, EPI_ISL_797572, EPI_ISL_797573, EPI_ISL_797574, EPI_ISL_797577, EPI_ISL_797581, EPI_ISL_797582, EPI_ISL_797583, EPI_ISL_797584, EPI_ISL_797585 | | | |
| see above | Lighthouse Lab in Cambridge | Wellcome Sanger Institute for the COVID-19 Genomics UK (COG-UK) Consortium | Rob Howes, The Lighthouse Lab in Cambridge and Alex Alderton, Roberto Amato, Sonia Goncalves, Ewan Harrison, David K. Jackson, Ian Johnston, Dominic Kwiatkowski, Cordelia Langford, John Sillitoe on behalf of the Wellcome Sanger Institute COVID-19 Surveillance Team |
| EPI_ISL_797587 | Lighthouse Lab in Alderley Park | Wellcome Sanger Institute for the COVID-19 Genomics UK (COG-UK) Consortium | Jacquelyn Wynn, Mairead Hyland, The Lighthouse Lab in Alderley Park and Alex Alderton, Roberto Amato, Sonia Goncalves, Ewan Harrison, David K. Jackson, Ian Johnston, Dominic Kwiatkowski, Cordelia Langford, John Sillitoe on behalf of the Wellcome Sanger Institute COVID-19 Surveillance Team |
| EPI_ISL_797588, EPI_ISL_797589, EPI_ISL_797591, EPI_ISL_797592, EPI_ISL_797594, EPI_ISL_797596, EPI_ISL_797598 | Lighthouse Lab in Cambridge | Wellcome Sanger Institute for the COVID-19 Genomics UK (COG-UK) Consortium | Rob Howes, The Lighthouse Lab in Cambridge and Alex Alderton, Roberto Amato, Sonia Goncalves, Ewan Harrison, David K. Jackson, Ian Johnston, Dominic Kwiatkowski, Cordelia Langford, John Sillitoe on behalf of the Wellcome Sanger Institute COVID-19 Surveillance Team |
| EPI_ISL_797599 | Lighthouse Lab in Alderley Park | Wellcome Sanger Institute for the COVID-19 Genomics UK (COG-UK) Consortium | Jacquelyn Wynn, Mairead Hyland, The Lighthouse Lab in Alderley Park and Alex Alderton, Roberto Amato, Sonia Goncalves, Ewan Harrison, David K. Jackson, Ian Johnston, Dominic Kwiatkowski, Cordelia Langford, John Sillitoe on behalf of the Wellcome Sanger Institute COVID-19 Surveillance Team |
| EPI_ISL_797601, EPI_ISL_797602, EPI_ISL_797603, EPI_ISL_797604 | Lighthouse Lab in Cambridge | Wellcome Sanger Institute for the COVID-19 Genomics UK (COG-UK) Consortium | Rob Howes, The Lighthouse Lab in Cambridge and Alex Alderton, Roberto Amato, Sonia Goncalves, Ewan Harrison, David K. Jackson, Ian Johnston, Dominic Kwiatkowski, Cordelia Langford, John Sillitoe on behalf of the Wellcome Sanger Institute COVID-19 Surveillance Team |
| EPI_ISL_797606 | Lighthouse Lab in Alderley Park | Wellcome Sanger Institute for the COVID-19 Genomics UK (COG-UK) Consortium | Jacquelyn Wynn, Mairead Hyland, The Lighthouse Lab in Alderley Park and Alex Alderton, Roberto Amato, Sonia Goncalves, Ewan Harrison, David K. Jackson, Ian Johnston, Dominic Kwiatkowski, Cordelia Langford, John Sillitoe on behalf of the Wellcome Sanger Institute COVID-19 Surveillance Team |
| EPI_ISL_797607, EPI_ISL_797608, EPI_ISL_797609 | Lighthouse Lab in Cambridge | Wellcome Sanger Institute for the COVID-19 Genomics UK (COG-UK) Consortium | Rob Howes, The Lighthouse Lab in Cambridge and Alex Alderton, Roberto Amato, Sonia Goncalves, Ewan Harrison, David K. Jackson, Ian Johnston, Dominic Kwiatkowski, Cordelia Langford, John Sillitoe on behalf of the Wellcome Sanger Institute COVID-19 Surveillance Team |
| EPI_ISL_797610 | Lighthouse Lab in Alderley Park | Wellcome Sanger Institute for the COVID-19 Genomics UK (COG-UK) Consortium | Jacquelyn Wynn, Mairead Hyland, The Lighthouse Lab in Alderley Park and Alex Alderton, Roberto Amato, Sonia Goncalves, Ewan Harrison, David K. Jackson, Ian Johnston, Dominic Kwiatkowski, Cordelia Langford, John Sillitoe on behalf of the Wellcome Sanger Institute COVID-19 Surveillance Team |
| EPI_ISL_797611, EPI_ISL_797612 | Lighthouse Lab in Cambridge | Wellcome Sanger Institute for the COVID-19 Genomics UK (COG-UK) Consortium | Rob Howes, The Lighthouse Lab in Cambridge and Alex Alderton, Roberto Amato, Sonia Goncalves, Ewan Harrison, David K. Jackson, Ian Johnston, Dominic Kwiatkowski, Cordelia Langford, John Sillitoe on behalf of the Wellcome Sanger Institute COVID-19 Surveillance Team |
| EPI_ISL_797613 | Lighthouse Lab in Alderley Park | Wellcome Sanger Institute for the COVID-19 Genomics UK (COG-UK) Consortium | Jacquelyn Wynn, Mairead Hyland, The Lighthouse Lab in Alderley Park and Alex Alderton, Roberto Amato, Sonia Goncalves, Ewan Harrison, David K. Jackson, Ian Johnston, Dominic Kwiatkowski, Cordelia Langford, John Sillitoe on behalf of the Wellcome Sanger Institute COVID-19 Surveillance Team |
| EPI_ISL_797614 | Lighthouse Lab in Cambridge | Wellcome Sanger Institute for the COVID-19 Genomics UK (COG-UK) Consortium | Rob Howes, The Lighthouse Lab in Cambridge and Alex Alderton, Roberto Amato, Sonia Goncalves, Ewan Harrison, David K. Jackson, Ian Johnston, Dominic Kwiatkowski, Cordelia Langford, John Sillitoe on behalf of the Wellcome Sanger Institute COVID-19 Surveillance Team |
| EPI_ISL_797616 | Lighthouse Lab in Alderley Park | Wellcome Sanger Institute for the COVID-19 Genomics UK (COG-UK) Consortium | Jacquelyn Wynn, Mairead Hyland, The Lighthouse Lab in Alderley Park and Alex Alderton, Roberto Amato, Sonia Goncalves, Ewan Harrison, David K. Jackson, Ian Johnston, Dominic Kwiatkowski, Cordelia Langford, John Sillitoe on behalf of the Wellcome Sanger Institute COVID-19 Surveillance Team |
| EPI_ISL_797617, EPI_ISL_797618 | Lighthouse Lab in Cambridge | Wellcome Sanger Institute for the COVID-19 Genomics UK (COG-UK) Consortium | Rob Howes, The Lighthouse Lab in Cambridge and Alex Alderton, Roberto Amato, Sonia Goncalves, Ewan Harrison, David K. Jackson, Ian Johnston, Dominic Kwiatkowski, Cordelia Langford, John Sillitoe on behalf of the Wellcome Sanger Institute COVID-19 Surveillance Team |
| EPI_ISL_797619 | Lighthouse Lab in Alderley Park | Wellcome Sanger Institute for the COVID-19 Genomics UK (COG-UK) Consortium | Jacquelyn Wynn, Mairead Hyland, The Lighthouse Lab in Alderley Park and Alex Alderton, Roberto Amato, Sonia Goncalves, Ewan Harrison, David K. Jackson, Ian Johnston, Dominic Kwiatkowski, Cordelia Langford, John Sillitoe on behalf of the Wellcome Sanger Institute COVID-19 Surveillance Team |
| EPI_ISL_797621, EPI_ISL_797622, EPI_ISL_797623, EPI_ISL_797624, EPI_ISL_797626, EPI_ISL_797628, EPI_ISL_797629, EPI_ISL_797630, EPI_ISL_797633, EPI_ISL_797634, EPI_ISL_797636 | | | |

| | | | |
|---|---|---|---|
| see above | Lighthouse Lab in Cambridge | Wellcome Sanger Institute for the COVID-19 Genomics UK (COG-UK) Consortium | Rob Howes, The Lighthouse Lab in Cambridge and Alex Alderton, Roberto Amato, Sonia Goncalves, Ewan Harrison, David K. Jackson, Ian Johnston, Dominic Kwiatkowski, Cordelia Langford, John Sillitoe on behalf of the Wellcome Sanger Institute COVID-19 Surveillance Team |
| EPI_ISL_797637 | Lighthouse Lab in Alderley Park | Wellcome Sanger Institute for the COVID-19 Genomics UK (COG-UK) Consortium | Jacquelyn Wynn, Mairead Hyland, The Lighthouse Lab in Alderley Park and Alex Alderton, Roberto Amato, Sonia Goncalves, Ewan Harrison, David K. Jackson, Ian Johnston, Dominic Kwiatkowski, Cordelia Langford, John Sillitoe on behalf of the Wellcome Sanger Institute COVID-19 Surveillance Team |
| EPI_ISL_797638, EPI_ISL_797639, EPI_ISL_797640 | Lighthouse Lab in Cambridge | Wellcome Sanger Institute for the COVID-19 Genomics UK (COG-UK) Consortium | Rob Howes, The Lighthouse Lab in Cambridge and Alex Alderton, Roberto Amato, Sonia Goncalves, Ewan Harrison, David K. Jackson, Ian Johnston, Dominic Kwiatkowski, Cordelia Langford, John Sillitoe on behalf of the Wellcome Sanger Institute COVID-19 Surveillance Team |
| EPI_ISL_797641, EPI_ISL_797642, EPI_ISL_797644 | Lighthouse Lab in Alderley Park | Wellcome Sanger Institute for the COVID-19 Genomics UK (COG-UK) Consortium | Jacquelyn Wynn, Mairead Hyland, The Lighthouse Lab in Alderley Park and Alex Alderton, Roberto Amato, Sonia Goncalves, Ewan Harrison, David K. Jackson, Ian Johnston, Dominic Kwiatkowski, Cordelia Langford, John Sillitoe on behalf of the Wellcome Sanger Institute COVID-19 Surveillance Team |
| EPI_ISL_797649, EPI_ISL_797650, EPI_ISL_797651, EPI_ISL_797653 | Lighthouse Lab in Cambridge | Wellcome Sanger Institute for the COVID-19 Genomics UK (COG-UK) Consortium | Rob Howes, The Lighthouse Lab in Cambridge and Alex Alderton, Roberto Amato, Sonia Goncalves, Ewan Harrison, David K. Jackson, Ian Johnston, Dominic Kwiatkowski, Cordelia Langford, John Sillitoe on behalf of the Wellcome Sanger Institute COVID-19 Surveillance Team |
| EPI_ISL_797654 | Lighthouse Lab in Alderley Park | Wellcome Sanger Institute for the COVID-19 Genomics UK (COG-UK) Consortium | Jacquelyn Wynn, Mairead Hyland, The Lighthouse Lab in Alderley Park and Alex Alderton, Roberto Amato, Sonia Goncalves, Ewan Harrison, David K. Jackson, Ian Johnston, Dominic Kwiatkowski, Cordelia Langford, John Sillitoe on behalf of the Wellcome Sanger Institute COVID-19 Surveillance Team |
| EPI_ISL_797655, EPI_ISL_797656, EPI_ISL_797658 | Lighthouse Lab in Cambridge | Wellcome Sanger Institute for the COVID-19 Genomics UK (COG-UK) Consortium | Rob Howes, The Lighthouse Lab in Cambridge and Alex Alderton, Roberto Amato, Sonia Goncalves, Ewan Harrison, David K. Jackson, Ian Johnston, Dominic Kwiatkowski, Cordelia Langford, John Sillitoe on behalf of the Wellcome Sanger Institute COVID-19 Surveillance Team |
| EPI_ISL_797660 | Lighthouse Lab in Alderley Park | Wellcome Sanger Institute for the COVID-19 Genomics UK (COG-UK) Consortium | Jacquelyn Wynn, Mairead Hyland, The Lighthouse Lab in Alderley Park and Alex Alderton, Roberto Amato, Sonia Goncalves, Ewan Harrison, David K. Jackson, Ian Johnston, Dominic Kwiatkowski, Cordelia Langford, John Sillitoe on behalf of the Wellcome Sanger Institute COVID-19 Surveillance Team |
| EPI_ISL_797661, EPI_ISL_797662 | Lighthouse Lab in Cambridge | Wellcome Sanger Institute for the COVID-19 Genomics UK (COG-UK) Consortium | Rob Howes, The Lighthouse Lab in Cambridge and Alex Alderton, Roberto Amato, Sonia Goncalves, Ewan Harrison, David K. Jackson, Ian Johnston, Dominic Kwiatkowski, Cordelia Langford, John Sillitoe on behalf of the Wellcome Sanger Institute COVID-19 Surveillance Team |
| EPI_ISL_797663 | Lighthouse Lab in Alderley Park | Wellcome Sanger Institute for the COVID-19 Genomics UK (COG-UK) Consortium | Jacquelyn Wynn, Mairead Hyland, The Lighthouse Lab in Alderley Park and Alex Alderton, Roberto Amato, Sonia Goncalves, Ewan Harrison, David K. Jackson, Ian Johnston, Dominic Kwiatkowski, Cordelia Langford, John Sillitoe on behalf of the Wellcome Sanger Institute COVID-19 Surveillance Team |
| EPI_ISL_797665 | Lighthouse Lab in Cambridge | Wellcome Sanger Institute for the COVID-19 Genomics UK (COG-UK) Consortium | Rob Howes, The Lighthouse Lab in Cambridge and Alex Alderton, Roberto Amato, Sonia Goncalves, Ewan Harrison, David K. Jackson, Ian Johnston, Dominic Kwiatkowski, Cordelia Langford, John Sillitoe on behalf of the Wellcome Sanger Institute COVID-19 Surveillance Team |
| EPI_ISL_797666 | Lighthouse Lab in Alderley Park | Wellcome Sanger Institute for the COVID-19 Genomics UK (COG-UK) Consortium | Jacquelyn Wynn, Mairead Hyland, The Lighthouse Lab in Alderley Park and Alex Alderton, Roberto Amato, Sonia Goncalves, Ewan Harrison, David K. Jackson, Ian Johnston, Dominic Kwiatkowski, Cordelia Langford, John Sillitoe on behalf of the Wellcome Sanger Institute COVID-19 Surveillance Team |
| EPI_ISL_797667, EPI_ISL_797669, EPI_ISL_797670, EPI_ISL_797671, EPI_ISL_797672, EPI_ISL_797674, EPI_ISL_797675, EPI_ISL_797679, EPI_ISL_797680, EPI_ISL_797681, EPI_ISL_797682, EPI_ISL_797683, EPI_ISL_797684, EPI_ISL_797687, EPI_ISL_797689, EPI_ISL_797691, EPI_ISL_797693, EPI_ISL_797695, EPI_ISL_797696, EPI_ISL_797697, EPI_ISL_797698 | | | |
| see above | Lighthouse Lab in Cambridge | Wellcome Sanger Institute for the COVID-19 Genomics UK (COG-UK) Consortium | Rob Howes, The Lighthouse Lab in Cambridge and Alex Alderton, Roberto Amato, Sonia Goncalves, Ewan Harrison, David K. Jackson, Ian Johnston, Dominic Kwiatkowski, Cordelia Langford, John Sillitoe on behalf of the Wellcome Sanger Institute COVID-19 Surveillance Team |
| EPI_ISL_797699 | Lighthouse Lab in Alderley Park | Wellcome Sanger Institute for the COVID-19 Genomics UK (COG-UK) Consortium | Jacquelyn Wynn, Mairead Hyland, The Lighthouse Lab in Alderley Park and Alex Alderton, Roberto Amato, Sonia Goncalves, Ewan Harrison, David K. Jackson, Ian Johnston, Dominic Kwiatkowski, Cordelia Langford, John Sillitoe on behalf of the Wellcome Sanger Institute COVID-19 Surveillance Team |
| EPI_ISL_797700 | Lighthouse Lab in Cambridge | Wellcome Sanger Institute for the COVID-19 Genomics UK (COG-UK) Consortium | Rob Howes, The Lighthouse Lab in Cambridge and Alex Alderton, Roberto Amato, Sonia Goncalves, Ewan Harrison, David K. Jackson, Ian Johnston, Dominic Kwiatkowski, Cordelia Langford, John Sillitoe on behalf of the Wellcome Sanger Institute COVID-19 Surveillance Team |
| EPI_ISL_797701 | Lighthouse Lab in Alderley Park | Wellcome Sanger Institute for the COVID-19 Genomics UK (COG-UK) Consortium | Jacquelyn Wynn, Mairead Hyland, The Lighthouse Lab in Alderley Park and Alex Alderton, Roberto Amato, Sonia Goncalves, Ewan Harrison, David K. Jackson, Ian Johnston, Dominic Kwiatkowski, Cordelia Langford, John Sillitoe on behalf of the Wellcome Sanger Institute COVID-19 Surveillance Team |
| EPI_ISL_797702, EPI_ISL_797703, EPI_ISL_797706, EPI_ISL_797707, EPI_ISL_797708 | Lighthouse Lab in Cambridge | Wellcome Sanger Institute for the COVID-19 Genomics UK (COG-UK) Consortium | Rob Howes, The Lighthouse Lab in Cambridge and Alex Alderton, Roberto Amato, Sonia Goncalves, Ewan Harrison, David K. Jackson, Ian Johnston, Dominic Kwiatkowski, Cordelia Langford, John Sillitoe on behalf of the Wellcome Sanger Institute COVID-19 Surveillance Team |
| EPI_ISL_797709 | Lighthouse Lab in Alderley Park | Wellcome Sanger Institute for the COVID-19 Genomics UK (COG-UK) Consortium | Jacquelyn Wynn, Mairead Hyland, The Lighthouse Lab in Alderley Park and Alex Alderton, Roberto Amato, Sonia Goncalves, Ewan Harrison, David K. Jackson, Ian Johnston, Dominic Kwiatkowski, Cordelia Langford, John Sillitoe on behalf of the Wellcome Sanger Institute COVID-19 Surveillance Team |
| EPI_ISL_797710, EPI_ISL_797711 | Lighthouse Lab in Cambridge | Wellcome Sanger Institute for the COVID-19 Genomics UK (COG-UK) Consortium | Rob Howes, The Lighthouse Lab in Cambridge and Alex Alderton, Roberto Amato, Sonia Goncalves, Ewan Harrison, David K. Jackson, Ian Johnston, Dominic Kwiatkowski, Cordelia Langford, John Sillitoe on behalf of the Wellcome Sanger Institute COVID-19 Surveillance Team |
| EPI_ISL_797712, EPI_ISL_797714, EPI_ISL_797715, EPI_ISL_797717 | Lighthouse Lab in Alderley Park | Wellcome Sanger Institute for the COVID-19 Genomics UK (COG-UK) Consortium | Jacquelyn Wynn, Mairead Hyland, The Lighthouse Lab in Alderley Park and Alex Alderton, Roberto Amato, Sonia Goncalves, Ewan Harrison, David K. Jackson, Ian Johnston, Dominic Kwiatkowski, Cordelia Langford, John Sillitoe on behalf of the Wellcome Sanger Institute COVID-19 Surveillance Team |
| EPI_ISL_797718 | Lighthouse Lab in Cambridge | Wellcome Sanger Institute for the COVID-19 Genomics UK (COG-UK) Consortium | Rob Howes, The Lighthouse Lab in Cambridge and Alex Alderton, Roberto Amato, Sonia Goncalves, Ewan Harrison, David K. Jackson, Ian Johnston, Dominic Kwiatkowski, Cordelia Langford, John Sillitoe on behalf of the Wellcome Sanger Institute COVID-19 Surveillance Team |
| EPI_ISL_797719, EPI_ISL_797720, EPI_ISL_797721, EPI_ISL_797722, EPI_ISL_797723, EPI_ISL_797724, EPI_ISL_797725, EPI_ISL_797726, EPI_ISL_797727, EPI_ISL_797728, EPI_ISL_797729, EPI_ISL_797730, EPI_ISL_797731, EPI_ISL_797732, EPI_ISL_797733, EPI_ISL_797734, EPI_ISL_797735, EPI_ISL_797736, EPI_ISL_797737, EPI_ISL_797738, EPI_ISL_797740, EPI_ISL_797741, EPI_ISL_797742, EPI_ISL_797743, EPI_ISL_797744, EPI_ISL_797745, EPI_ISL_797746, EPI_ISL_797747, EPI_ISL_797748, EPI_ISL_797749, EPI_ISL_797750, EPI_ISL_797751, EPI_ISL_797752, EPI_ISL_797753, EPI_ISL_797754, EPI_ISL_797755, EPI_ISL_797756, EPI_ISL_797757, EPI_ISL_797758, EPI_ISL_797759, EPI_ISL_797760, EPI_ISL_797761, EPI_ISL_797762, EPI_ISL_797763, EPI_ISL_797764, EPI_ISL_797765, EPI_ISL_797767, EPI_ISL_797768, EPI_ISL_797769, EPI_ISL_797770, EPI_ISL_797771, EPI_ISL_797772, EPI_ISL_797773, EPI_ISL_797774, EPI_ISL_797776, EPI_ISL_797777, EPI_ISL_797778, EPI_ISL_797779, EPI_ISL_797780, EPI_ISL_797781, EPI_ISL_797782, EPI_ISL_797783, EPI_ISL_797784, EPI_ISL_797785, EPI_ISL_797786, EPI_ISL_797787, EPI_ISL_797788, EPI_ISL_797790, EPI_ISL_797791, EPI_ISL_797792, EPI_ISL_797793, EPI_ISL_797794, EPI_ISL_797795, EPI_ISL_797796, EPI_ISL_797798, EPI_ISL_797799, EPI_ISL_797800, EPI_ISL_797801, EPI_ISL_797802, EPI_ISL_797803, EPI_ISL_797804, EPI_ISL_797805, EPI_ISL_797806, EPI_ISL_797807, EPI_ISL_797808, EPI_ISL_797809, EPI_ISL_797810, EPI_ISL_797811, EPI_ISL_797812, EPI_ISL_797813, EPI_ISL_797814, EPI_ISL_797815, EPI_ISL_797816, EPI_ISL_797818, EPI_ISL_797819, EPI_ISL_797820, EPI_ISL_797821, EPI_ISL_797822, EPI_ISL_797823, EPI_ISL_797824, EPI_ISL_797826, EPI_ISL_797827, EPI_ISL_797828, EPI_ISL_797832, EPI_ISL_797833, EPI_ISL_797834, EPI_ISL_797835, EPI_ISL_797836, EPI_ISL_797837, EPI_ISL_797839, EPI_ISL_797840, EPI_ISL_797841, EPI_ISL_797842, EPI_ISL_797843, EPI_ISL_797844, EPI_ISL_797845, EPI_ISL_797846, EPI_ISL_797848, EPI_ISL_797849, EPI_ISL_797852, EPI_ISL_797854, EPI_ISL_797855, EPI_ISL_797856, EPI_ISL_797857, EPI_ISL_797858, EPI_ISL_797859, EPI_ISL_797860, EPI_ISL_797861, EPI_ISL_797862, EPI_ISL_797863, EPI_ISL_797864, EPI_ISL_797865, EPI_ISL_797866, EPI_ISL_797869, EPI_ISL_797871, EPI_ISL_797872, EPI_ISL_797873, EPI_ISL_797874, EPI_ISL_797875, EPI_ISL_797876, EPI_ISL_797877, EPI_ISL_797878, EPI_ISL_797879, EPI_ISL_797880, EPI_ISL_797881, EPI_ISL_797882, EPI_ISL_797883, EPI_ISL_797886, EPI_ISL_797887, EPI_ISL_797888, EPI_ISL_797889, EPI_ISL_797890, EPI_ISL_797891, EPI_ISL_797892, EPI_ISL_797893, EPI_ISL_797894, EPI_ISL_797895, EPI_ISL_797896, EPI_ISL_797897, EPI_ISL_797898, EPI_ISL_797899, EPI_ISL_797900, EPI_ISL_797901, EPI_ISL_797902, EPI_ISL_797904, EPI_ISL_797905, EPI_ISL_797906, EPI_ISL_797907, EPI_ISL_797908, EPI_ISL_797909, EPI_ISL_797910, EPI_ISL_797911, EPI_ISL_797912, EPI_ISL_797913, EPI_ISL_797915, EPI_ISL_797917, EPI_ISL_797918, EPI_ISL_797919, EPI_ISL_797920, EPI_ISL_797922, EPI_ISL_797923, EPI_ISL_797924, EPI_ISL_797925, EPI_ISL_797926, EPI_ISL_797927, EPI_ISL_797928, EPI_ISL_797930, EPI_ISL_797931, EPI_ISL_797932, EPI_ISL_797933, EPI_ISL_797934, EPI_ISL_797935, EPI_ISL_797936, EPI_ISL_797938, EPI_ISL_797939, EPI_ISL_797940, EPI_ISL_797941, EPI_ISL_797942, EPI_ISL_797943, EPI_ISL_797944, EPI_ISL_797946, EPI_ISL_797947, EPI_ISL_797948, EPI_ISL_797949, EPI_ISL_797950, EPI_ISL_797951, EPI_ISL_797953, EPI_ISL_797954, EPI_ISL_797955, EPI_ISL_797956, EPI_ISL_797957, EPI_ISL_797958, EPI_ISL_797959, EPI_ISL_797960, EPI_ISL_797962, EPI_ISL_797963, EPI_ISL_797964, EPI_ISL_797966, EPI_ISL_797967, EPI_ISL_797968, EPI_ISL_797969, EPI_ISL_797971, EPI_ISL_797972, EPI_ISL_797973, EPI_ISL_797974, EPI_ISL_797975, EPI_ISL_797976, EPI_ISL_797977, EPI_ISL_797978, EPI_ISL_797979, EPI_ISL_797981, EPI_ISL_797982, EPI_ISL_797983, EPI_ISL_797984, EPI_ISL_797986, EPI_ISL_797987, EPI_ISL_797988, EPI_ISL_797989, EPI_ISL_797990, EPI_ISL_797991, EPI_ISL_797992, EPI_ISL_797993, EPI_ISL_797995, EPI_ISL_797997, EPI_ISL_797998, EPI_ISL_798001, EPI_ISL_798004, EPI_ISL_798005, EPI_ISL_798006, EPI_ISL_798008, EPI_ISL_798010, EPI_ISL_798012, EPI_ISL_798014, EPI_ISL_798016, EPI_ISL_798019, EPI_ISL_798020, EPI_ISL_798021, EPI_ISL_798023, EPI_ISL_798024, EPI_ISL_798026, EPI_ISL_798028, EPI_ISL_798030, EPI_ISL_798031, EPI_ISL_798033, EPI_ISL_798034, EPI_ISL_798036, EPI_ISL_798037, EPI_ISL_798039, EPI_ISL_798041, EPI_ISL_798042, EPI_ISL_798044, EPI_ISL_798045, EPI_ISL_798047, EPI_ISL_798049, EPI_ISL_798050, EPI_ISL_798052, EPI_ISL_798056, EPI_ISL_798057, EPI_ISL_798059, EPI_ISL_798063, EPI_ISL_798065, EPI_ISL_798067, EPI_ISL_798070, EPI_ISL_798071, EPI_ISL_798074, EPI_ISL_798075, EPI_ISL_798081, EPI_ISL_798082, EPI_ISL_798085, EPI_ISL_798087, EPI_ISL_798089, EPI_ISL_798090, EPI_ISL_798091, EPI_ISL_798092, EPI_ISL_798098, EPI_ISL_798099, EPI_ISL_798100, EPI_ISL_798102, EPI_ISL_798104, EPI_ISL_798109, EPI_ISL_798114, EPI_ISL_798116, EPI_ISL_798118, EPI_ISL_798120, EPI_ISL_798129, EPI_ISL_798135, EPI_ISL_798136, EPI_ISL_798138, EPI_ISL_798142, EPI_ISL_798144, EPI_ISL_798146, EPI_ISL_798147, EPI_ISL_798150, EPI_ISL_798155, EPI_ISL_798156, EPI_ISL_798157, EPI_ISL_798158, EPI_ISL_798162, EPI_ISL_798163, EPI_ISL_798164, EPI_ISL_798165, EPI_ISL_798172, EPI_ISL_798177, EPI_ISL_798179, EPI_ISL_798180, EPI_ISL_798187, EPI_ISL_798188, EPI_ISL_798189, EPI_ISL_798190, EPI_ISL_798191, EPI_ISL_798194, EPI_ISL_798195, EPI_ISL_798196, EPI_ISL_798198, EPI_ISL_798200, EPI_ISL_798205, EPI_ISL_798206, EPI_ISL_798207, EPI_ISL_798209, EPI_ISL_798210, EPI_ISL_798212, EPI_ISL_798215, EPI_ISL_798219, EPI_ISL_798226, EPI_ISL_798227, EPI_ISL_798228, EPI_ISL_798232, EPI_ISL_798233, EPI_ISL_798237, EPI_ISL_798238, EPI_ISL_798240, EPI_ISL_798247, EPI_ISL_798257, EPI_ISL_798259, EPI_ISL_798260, EPI_ISL_798262, EPI_ISL_798263, EPI_ISL_798264, EPI_ISL_798265, EPI_ISL_798267, EPI_ISL_798269, EPI_ISL_798276, EPI_ISL_798279, EPI_ISL_798286, EPI_ISL_798290, EPI_ISL_798291, EPI_ISL_798293, EPI_ISL_798297, EPI_ISL_798301, EPI_ISL_798302, EPI_ISL_798303, EPI_ISL_798304, EPI_ISL_798306, EPI_ISL_798309, EPI_ISL_798313, EPI_ISL_798317, EPI_ISL_798318, EPI_ISL_798320, EPI_ISL_798330, EPI_ISL_799006, EPI_ISL_799009, EPI_ISL_799011, EPI_ISL_799012, EPI_ISL_799014, EPI_ISL_799015, EPI_ISL_799016, EPI_ISL_799019, EPI_ISL_799024, EPI_ISL_799027, EPI_ISL_799028, EPI_ISL_799030, EPI_ISL_799033, EPI_ISL_799034, EPI_ISL_799038, EPI_ISL_799041, EPI_ISL_799042, EPI_ISL_799043, EPI_ISL_799044, EPI_ISL_799046, EPI_ISL_799049, EPI_ISL_799050, EPI_ISL_799051, EPI_ISL_799054, EPI_ISL_799056, EPI_ISL_799060, EPI_ISL_799062, EPI_ISL_799063, EPI_ISL_799064, EPI_ISL_799065, EPI_ISL_799066, EPI_ISL_799067, EPI_ISL_799068, EPI_ISL_799071, EPI_ISL_799072, EPI_ISL_799073, EPI_ISL_799075, EPI_ISL_799077, EPI_ISL_799080, EPI_ISL_799081, EPI_ISL_799082, EPI_ISL_799084, EPI_ISL_799089, EPI_ISL_799090, EPI_ISL_799096, EPI_ISL_799098, EPI_ISL_799100, EPI_ISL_799101, EPI_ISL_799102, EPI_ISL_799107, EPI_ISL_799108, EPI_ISL_799112, EPI_ISL_799115, EPI_ISL_799118, EPI_ISL_799120, EPI_ISL_799123, EPI_ISL_799125, EPI_ISL_799126, EPI_ISL_799127, EPI_ISL_799129, EPI_ISL_799130, EPI_ISL_799131, EPI_ISL_799132, EPI_ISL_799134, EPI_ISL_799140, EPI_ISL_799144, EPI_ISL_799147, EPI_ISL_799150, EPI_ISL_799151, EPI_ISL_799155, EPI_ISL_799156, EPI_ISL_799157, EPI_ISL_799158, EPI_ISL_799162, EPI_ISL_799163, EPI_ISL_799164, EPI_ISL_799167, EPI_ISL_799169, EPI_ISL_799172, EPI_ISL_799175, EPI_ISL_799176, EPI_ISL_799180, EPI_ISL_799183, EPI_ISL_799189, EPI_ISL_799190, EPI_ISL_799192, EPI_ISL_799193, EPI_ISL_799194, EPI_ISL_799196, EPI_ISL_799197, EPI_ISL_799201, EPI_ISL_799203, EPI_ISL_799204, EPI_ISL_799205, EPI_ISL_799207, EPI_ISL_799209, EPI_ISL_799215, EPI_ISL_799216, EPI_ISL_799218, EPI_ISL_799221, EPI_ISL_799222, EPI_ISL_799224, EPI_ISL_799230, EPI_ISL_799232, EPI_ISL_799236, EPI_ISL_799240, EPI_ISL_799243, EPI_ISL_799245, EPI_ISL_799246, EPI_ISL_799249, EPI_ISL_799250, EPI_ISL_799255, EPI_ISL_799256, EPI_ISL_799258, EPI_ISL_799260, EPI_ISL_799262, EPI_ISL_799263, EPI_ISL_799264, EPI_ISL_799267, EPI_ISL_799268, EPI_ISL_799270, EPI_ISL_799272, EPI_ISL_799273, EPI_ISL_799274, EPI_ISL_799275, EPI_ISL_799277, EPI_ISL_799279, EPI_ISL_799280, EPI_ISL_799282, EPI_ISL_799283, EPI_ISL_799285, EPI_ISL_799286, EPI_ISL_799287, EPI_ISL_799288, EPI_ISL_799289, EPI_ISL_799290, EPI_ISL_799294 | | | |
| see above | Lighthouse Lab in Glasgow | Wellcome Sanger Institute for the COVID-19 Genomics UK (COG-UK) Consortium | Harper VanSteenhouse, Yumi Kasai, David Gray, Carol Clugston, Anna Dominiczak and Alex Alderton, Roberto Amato, Sonia Goncalves, Ewan Harrison, David K. Jackson, Ian Johnston, Dominic Kwiatkowski, Cordelia Langford, John Sillitoe on behalf of the Wellcome Sanger Institute COVID-19 Surveillance |

| | | | |
|---|---|---|---|
| | | | Team |
| EPI_ISL_801412, EPI_ISL_801437, EPI_ISL_801511 | Dutch COVID-19 response team | Erasmus Medical Center | Bas Oude Munnink, Reina Sikkema, David Nieuwenhuijse, Irina Chestakova, Anne van der Linden, Marjan Boter, Emmanuelle Munger, Corine GeurtsvanKessel, Annemiek van der Eijk, Richard Molenkamp, Marion Koopmans, on behalf of the Dutch national COVID-19 response team. |
| EPI_ISL_801517, EPI_ISL_801518 | Instituto Nacional de Saude (INSA) | Instituto Nacional de Saude (INSA) | Borges et al |
| EPI_ISL_802429, EPI_ISL_802431 | Wadsworth Center, New York State Department.of Health | Wadsworth Center, New York State Department.of Health | Kirsten St. George, Daryl M. Lamson, Alexis Russel, Matthew Shudt, Melissa A Leisner, Jonathan Plitnick, Navjot Singh, John Kelly, Sara Griesemer, Erasmus Schneider, Erica Lasek-Nesselquist |
| EPI_ISL_802434, EPI_ISL_802435 | WESTCHESTER MEDICAL CENTER | Wadsworth Center, New York State Department.of Health | Kirsten St. George, Daryl M. Lamson, Alexis Russel, Matthew Shudt, Melissa A Leisner, Jonathan Plitnick, Navjot Singh, John Kelly, Sara Griesemer, Erasmus Schneider, Erica Lasek-Nesselquist |
| EPI_ISL_802439 | SARATOGA HOSPITAL LABORATORY | Wadsworth Center, New York State Department.of Health | Kirsten St. George, Daryl M. Lamson, Alexis Russel, Matthew Shudt, Melissa A Leisner, Jonathan Plitnick, Navjot Singh, John Kelly, Sara Griesemer, Erasmus Schneider, Erica Lasek-Nesselquist |
| EPI_ISL_802440, EPI_ISL_802441 | WESTCHESTER MEDICAL CENTER | Wadsworth Center, New York State Department.of Health | Kirsten St. George, Daryl M. Lamson, Alexis Russel, Matthew Shudt, Melissa A Leisner, Jonathan Plitnick, Navjot Singh, John Kelly, Sara Griesemer, Erasmus Schneider, Erica Lasek-Nesselquist |
| EPI_ISL_802442 | Wadsworth Center, New York State Department.of Health | Wadsworth Center, New York State Department.of Health | Kirsten St. George, Daryl M. Lamson, Alexis Russel, Matthew Shudt, Melissa A Leisner, Jonathan Plitnick, Navjot Singh, John Kelly, Sara Griesemer, Erasmus Schneider, Erica Lasek-Nesselquist |
| EPI_ISL_802443 | WESTCHESTER MEDICAL CENTER | Wadsworth Center, New York State Department.of Health | Kirsten St. George, Daryl M. Lamson, Alexis Russel, Matthew Shudt, Melissa A Leisner, Jonathan Plitnick, Navjot Singh, John Kelly, Sara Griesemer, Erasmus Schneider, Erica Lasek-Nesselquist |
| EPI_ISL_802447 | BIO-REFERENCE LABORATORIES | Wadsworth Center, New York State Department.of Health | Kirsten St. George, Daryl M. Lamson, Alexis Russel, Matthew Shudt, Melissa A Leisner, Jonathan Plitnick, Navjot Singh, John Kelly, Sara Griesemer, Erasmus Schneider, Erica Lasek-Nesselquist |
| EPI_ISL_802448 | WESTCHESTER MEDICAL CENTER | Wadsworth Center, New York State Department.of Health | Kirsten St. George, Daryl M. Lamson, Alexis Russel, Matthew Shudt, Melissa A Leisner, Jonathan Plitnick, Navjot Singh, John Kelly, Sara Griesemer, Erasmus Schneider, Erica Lasek-Nesselquist |
| EPI_ISL_802463, EPI_ISL_802464, EPI_ISL_802465, EPI_ISL_802466 | Wadsworth Center, New York State Department.of Health | Wadsworth Center, New York State Department.of Health | Kirsten St. George, Daryl M. Lamson, Alexis Russel, Matthew Shudt, Melissa A Leisner, Jonathan Plitnick, Navjot Singh, John Kelly, Sara Griesemer, Erasmus Schneider, Erica Lasek-Nesselquist |
| EPI_ISL_802467, EPI_ISL_802468 | SARATOGA HOSPITAL LABORATORY | Wadsworth Center, New York State Department.of Health | Kirsten St. George, Daryl M. Lamson, Alexis Russel, Matthew Shudt, Melissa A Leisner, Jonathan Plitnick, Navjot Singh, John Kelly, Sara Griesemer, Erasmus Schneider, Erica Lasek-Nesselquist |
| EPI_ISL_802469, EPI_ISL_802470, EPI_ISL_802471, EPI_ISL_802472, EPI_ISL_802473, EPI_ISL_802474, EPI_ISL_802475, EPI_ISL_802476, EPI_ISL_802477, EPI_ISL_802478, EPI_ISL_802479, EPI_ISL_802480, EPI_ISL_802481, EPI_ISL_802482, EPI_ISL_802483, EPI_ISL_802484, EPI_ISL_802485, EPI_ISL_802486, EPI_ISL_802487, EPI_ISL_802488, EPI_ISL_802489, EPI_ISL_802490 | | | |
| see above | WESTCHESTER MEDICAL CENTER | Wadsworth Center, New York State Department.of Health | Kirsten St. George, Daryl M. Lamson, Alexis Russel, Matthew Shudt, Melissa A Leisner, Jonathan Plitnick, Navjot Singh, John Kelly, Sara Griesemer, Erasmus Schneider, Erica Lasek-Nesselquist |
| EPI_ISL_802506, EPI_ISL_802509, EPI_ISL_802510, EPI_ISL_802511, EPI_ISL_802517, EPI_ISL_802518, EPI_ISL_802519, EPI_ISL_802526, EPI_ISL_802527, EPI_ISL_802528, EPI_ISL_802529, EPI_ISL_802530, EPI_ISL_802536, EPI_ISL_802542 | | | |
| see above | Dutch COVID-19 response team | Erasmus Medical Center | Bas Oude Munnink, Reina Sikkema, David Nieuwenhuijse, Irina Chestakova, Anne van der Linden, Marjan Boter, Emmanuelle Munger, Corine GeurtsvanKessel, Annemiek van der Eijk, Richard Molenkamp, Marion Koopmans, on behalf of the Dutch national COVID-19 response team. |
| EPI_ISL_802599, EPI_ISL_802601, EPI_ISL_802604, EPI_ISL_802606, EPI_ISL_802607, EPI_ISL_802618, EPI_ISL_802619, EPI_ISL_802620, EPI_ISL_802621, EPI_ISL_802622, EPI_ISL_802623, EPI_ISL_802624, EPI_ISL_802625, EPI_ISL_802626, EPI_ISL_802627, EPI_ISL_802628, EPI_ISL_802641, EPI_ISL_802642, EPI_ISL_802643, EPI_ISL_802644, EPI_ISL_802645, EPI_ISL_802646, EPI_ISL_802649, EPI_ISL_802652, EPI_ISL_802662, EPI_ISL_802663, EPI_ISL_802664, EPI_ISL_802665, EPI_ISL_802666, EPI_ISL_802667, EPI_ISL_802668, EPI_ISL_802682, EPI_ISL_802683, EPI_ISL_802687, EPI_ISL_802689, EPI_ISL_802691, EPI_ISL_802692, EPI_ISL_802693, EPI_ISL_802694, EPI_ISL_802695, EPI_ISL_802696, EPI_ISL_802711, EPI_ISL_802712, EPI_ISL_802713, EPI_ISL_802715, EPI_ISL_802716, EPI_ISL_802717 | | | |
| see above | Helix/Illumina | Genomics and Discovery, Respiratory Viruses Branch, Division of Viral Diseases, Centers for Disease Control and Prevention | Peter W. Cook, Dhwani Batra, Ben L. Rambo-Martin Eileen de Feo, Jan Antico, Christine Tran, Matthew Tolentino, Shannon Wickline, Kim Gietzen, Brad Sickler, Jingtao Liu, Eric Allen, Phil Febbo, Summer Galloway, Nicole L. Washington, Simon White, Geraint Levan, Kelly Schiabor Barrett, Elizabeth Cirulli, Alexandre Bolze, Ary Ascencio, Charlotte Rivera-Garcia, Ryan Cho, Jason Nguyen, Sherry Wang, Jimmy Ramirez, Tyler Cassens, Efren Sandoval, Magnus Isaksson, William Lee, David Becker, Marc Laurent, James Lu, Clinton R. Paden, Suxiang Tong, Duncan MacCannell |
| EPI_ISL_802725 | Florida Bureau of Public Health Laboratories | Florida Bureau of Public Health Laboratories | Sarah Schmedes, Jason Blanton |
| EPI_ISL_802790, EPI_ISL_802791, EPI_ISL_802792, EPI_ISL_802793 | BIO-REFERENCE LABORATORIES | Wadsworth Center, New York State Department.of Health | Kirsten St. George, Daryl M. Lamson, Alexis Russel, Matthew Shudt, Melissa A Leisner, Jonathan Plitnick, Navjot Singh, John Kelly, Sara Griesemer, Erasmus Schneider, Erica Lasek-Nesselquist |
| EPI_ISL_803855 | Hospital General Universitario Gregorio Marañón | Hospital General Universitario Gregorio Marañón | Sergio Buenestado Serrano, Pedro Sola Campoy, Laura Pérez-Lago, Pilar Catalán, Patricia Muñoz, Darío García de Viedma |
| EPI_ISL_803856, EPI_ISL_803857, EPI_ISL_803858, EPI_ISL_803859, EPI_ISL_803860, EPI_ISL_803861, EPI_ISL_803862, EPI_ISL_803863, EPI_ISL_803864 | Hospital General Universitario Gregorio Marañón | Hospital General Universitario Gregorio Marañón | Sergio Buenestado Serrano, Pedro Sola Campoy, Laura Pérez-Lago, Pilar Catalán, Patricia Muñoz, Darío García de Viedma. |
| EPI_ISL_803959, EPI_ISL_803960, EPI_ISL_803961, EPI_ISL_803966, EPI_ISL_803969, EPI_ISL_803970, EPI_ISL_803971, EPI_ISL_803972, EPI_ISL_803973, EPI_ISL_803974, EPI_ISL_803975, EPI_ISL_803976, EPI_ISL_803977, EPI_ISL_803978, EPI_ISL_803979, EPI_ISL_803980, EPI_ISL_803981, EPI_ISL_803982, EPI_ISL_803983 | | | |
| see above | National Public Health Laboratory, National Centre for Infectious Diseases | National Public Health Laboratory, National Centre for Infectious Diseases | Tze Minn Mak, Sophie Octavia, Zhenyang Zhou, Lin Cui, Raymond Tzer Pin Lin |
| EPI_ISL_804026, EPI_ISL_804029, EPI_ISL_804030, EPI_ISL_804031, EPI_ISL_804032, EPI_ISL_804033, EPI_ISL_804034, EPI_ISL_804035, EPI_ISL_804036 | Maryland Public Health Laboratory | Maryland Public Health Laboratory | Maryland Department of Health Laboratories Administration |
| EPI_ISL_804040 | SC (UCO) Igiene e Sanità Pubblica (funzione integrata con SC Microbiologia e Virologia) e Laboratory of Molecular Virology of the International Centre for Genetic Engineering and Biotechnology (ICGEB) | ARGO Laboratorio Genomica ed Epigenomica | Licastro D, Dal Monego S, Degasperi M, Marcello A, D'Agaro P |
| EPI_ISL_804057, EPI_ISL_804058, EPI_ISL_804059, EPI_ISL_804060, EPI_ISL_804061, EPI_ISL_804062, EPI_ISL_804063, EPI_ISL_804064, EPI_ISL_804065, EPI_ISL_804066, EPI_ISL_804067, EPI_ISL_804068, EPI_ISL_804069, EPI_ISL_804070, EPI_ISL_804085, EPI_ISL_804086, EPI_ISL_804087, EPI_ISL_804088, EPI_ISL_804089, EPI_ISL_804090, EPI_ISL_804091, EPI_ISL_804095, EPI_ISL_804097, EPI_ISL_804099, EPI_ISL_804100, EPI_ISL_804101, EPI_ISL_804105, EPI_ISL_804107, EPI_ISL_804108, EPI_ISL_804109, EPI_ISL_804110, EPI_ISL_804111, EPI_ISL_804112, EPI_ISL_804113, EPI_ISL_804114, EPI_ISL_804115, EPI_ISL_804116, EPI_ISL_804117, EPI_ISL_804118, EPI_ISL_804119, EPI_ISL_804120, EPI_ISL_804121, EPI_ISL_804122, EPI_ISL_804123, EPI_ISL_804124, EPI_ISL_804125, EPI_ISL_804126, EPI_ISL_804127, EPI_ISL_804128, EPI_ISL_804129, EPI_ISL_804130, EPI_ISL_804131, EPI_ISL_804132, EPI_ISL_804133, EPI_ISL_804134, EPI_ISL_804135, EPI_ISL_804136, EPI_ISL_804137, EPI_ISL_804138, EPI_ISL_804139, EPI_ISL_804140, EPI_ISL_804141, EPI_ISL_804189, EPI_ISL_804190, EPI_ISL_804192, EPI_ISL_804193, EPI_ISL_804194, EPI_ISL_804195, EPI_ISL_804196, EPI_ISL_804197, EPI_ISL_804198, EPI_ISL_804199, EPI_ISL_804200, EPI_ISL_804201, EPI_ISL_804202, EPI_ISL_804203, EPI_ISL_804204, EPI_ISL_804205, EPI_ISL_804206, EPI_ISL_804207, EPI_ISL_804208, EPI_ISL_804213 | | | |
| see above | Israel Central Virology laboratory | Israel National Consortium for SARS-CoV-2 sequencing | Neta Zuckerman, Efrat Dahan Bucris, Michal Mandelboim, Dana Bar-Ilan, Oran Erster, Tzvia Mann, Omer Murik, David A. Zeevi, Assaf Rokney, Joseph Jaffe, Eva Nachum, Maya Davidovich Cohen, Ephraim Fass, Gal Zizelski Valenci, Mor Rubinstein, Efrat Rorman, Israel Nissan, Efrat Glick-Saar, Omri Nayshool, Gideon Rechavi, Ella Mendelson, Orna Mor |
| EPI_ISL_804222, EPI_ISL_804224, EPI_ISL_804242, EPI_ISL_804243, EPI_ISL_804244, EPI_ISL_804245, EPI_ISL_804246, EPI_ISL_804247, EPI_ISL_804248, EPI_ISL_804251, EPI_ISL_804252, EPI_ISL_804260, EPI_ISL_804280, EPI_ISL_804281, EPI_ISL_804282, EPI_ISL_804283, EPI_ISL_804284, EPI_ISL_804285, EPI_ISL_804286, EPI_ISL_804287, EPI_ISL_804288, EPI_ISL_804289, EPI_ISL_804290, EPI_ISL_804291, EPI_ISL_804292, EPI_ISL_804293 | | | |
| see above | Respiratory Virus Unit, National Infection Service, Public Health England | COVID-19 Genomics UK (COG-UK) Consortium | PHE Covid Sequencing Team |

| | | | |
|---|---|---|---|
| EPI_ISL_804463, EPI_ISL_804464, EPI_ISL_804465 | Maryland Public Health Laboratory | Maryland Public Health Laboratory | Maryland Department of Health Laboratories Administration |
| EPI_ISL_804982, EPI_ISL_804983, EPI_ISL_804984 | BIO-REFERENCE LABORATORIES | Wadsworth Center, New York State Department of Health | Kirsten St. George, Daryl M. Lamson, Alexis Russel, Matthew Shudt, Melissa A Leisner, Jonathan Plitnick, Navjot Singh, John Kelly, Erasmus Schneider, Erica Lasek-Nesselquist |
| EPI_ISL_806723 | 4Cyte Pathology | NSW Health Pathology - Institute of Clinical Pathology and Medical Research; Westmead Hospital; University of Sydney | CIDM-PH et al. |
| EPI_ISL_806724 | Sydney South West Pathology Service (SSWPS) - Royal Prince Alfred Hospital - NSW Health Pathology | NSW Health Pathology - Institute of Clinical Pathology and Medical Research; Westmead Hospital; University of Sydney | CIDM-PH et al. |
| EPI_ISL_806727 | 4Cyte Pathology | NSW Health Pathology - Institute of Clinical Pathology and Medical Research; Westmead Hospital; University of Sydney | CIDM-PH et al. |
| EPI_ISL_811040, EPI_ISL_811041, EPI_ISL_811042, EPI_ISL_811043, EPI_ISL_811044, EPI_ISL_811045, EPI_ISL_811046, EPI_ISL_811047, EPI_ISL_811048, EPI_ISL_811049, EPI_ISL_811050, EPI_ISL_811051, EPI_ISL_811052, EPI_ISL_811053, EPI_ISL_811054, EPI_ISL_811055, EPI_ISL_811056, EPI_ISL_811057, EPI_ISL_811058, EPI_ISL_811059, EPI_ISL_811060, EPI_ISL_811061, EPI_ISL_811062, EPI_ISL_811063, EPI_ISL_811064, EPI_ISL_811065, EPI_ISL_811066, EPI_ISL_811067, EPI_ISL_811068, EPI_ISL_811069, EPI_ISL_811070, EPI_ISL_811071, EPI_ISL_811073, EPI_ISL_811074, EPI_ISL_811075, EPI_ISL_811076, EPI_ISL_811077, EPI_ISL_811078, EPI_ISL_811079, EPI_ISL_811080, EPI_ISL_811081, EPI_ISL_811082, EPI_ISL_811083, EPI_ISL_811084, EPI_ISL_811085, EPI_ISL_811086, EPI_ISL_811087, EPI_ISL_811088, EPI_ISL_811089, EPI_ISL_811090, EPI_ISL_811091, EPI_ISL_811092, EPI_ISL_811093, EPI_ISL_811094, EPI_ISL_811095, EPI_ISL_811096, EPI_ISL_811097, EPI_ISL_811098, EPI_ISL_811099, EPI_ISL_811100, EPI_ISL_811101, EPI_ISL_811102, EPI_ISL_811103, EPI_ISL_811104, EPI_ISL_811105, EPI_ISL_811115, EPI_ISL_811117, EPI_ISL_811118, EPI_ISL_811119 | | | |
| see above | Respiratory Virus Unit, National Infection Service, Public Health England | COVID-19 Genomics UK (COG-UK) Consortium | PHE Covid Sequencing Team |
| EPI_ISL_811137, EPI_ISL_811138, EPI_ISL_811139, EPI_ISL_811143 | Ministry of Health Turkey | Ministry of Health Turkey | Fatma Bayrakdar, Yasemin Cogun, Süleyman Yalcin, Aye Baak Alta, Gülay Korukluolu |
| EPI_ISL_811149 | Laboratorio de Ecologia de Doencas Transmissiveis na Amazonia, Instituto Leonidas e Maria Deane - Fiocruz Amazonia | Laboratorio de Ecologia de Doencas Transmissiveis na Amazonia, Instituto Leonidas e Maria Deane - Fiocruz Amazonia | Valdinete Nascimento, Victor Souza, André Corado, Fernanda Nascimento, George Silva, Ágatha Costa, Debora Duarte, Luciana Gonçalves, Matilde Mejía, Karina Pessoa, Maria Júlia Brandão, Michele Jesus, Felipe Naveca |
| EPI_ISL_811214, EPI_ISL_811234, EPI_ISL_811236, EPI_ISL_811241, EPI_ISL_811247, EPI_ISL_811275, EPI_ISL_811293, EPI_ISL_811310, EPI_ISL_811312, EPI_ISL_811321, EPI_ISL_811345, EPI_ISL_811350, EPI_ISL_811353, EPI_ISL_811366, EPI_ISL_811377, EPI_ISL_811414, EPI_ISL_811422, EPI_ISL_811472, EPI_ISL_811490, EPI_ISL_811494, EPI_ISL_811500, EPI_ISL_811516, EPI_ISL_811519, EPI_ISL_811524, EPI_ISL_811530, EPI_ISL_811548, EPI_ISL_811563, EPI_ISL_811573, EPI_ISL_811580, EPI_ISL_811584, EPI_ISL_811594, EPI_ISL_811617, EPI_ISL_811618, EPI_ISL_811620, EPI_ISL_811621, EPI_ISL_811627, EPI_ISL_811645, EPI_ISL_811650, EPI_ISL_811651, EPI_ISL_811661, EPI_ISL_811669, EPI_ISL_811670, EPI_ISL_811671, EPI_ISL_811677, EPI_ISL_811678, EPI_ISL_811682, EPI_ISL_811688 | | | |
| see above | Lighthouse Lab in Milton Keynes | Wellcome Sanger Institute for the COVID-19 Genomics UK (COG-UK) Consortium | The Lighthouse Lab in Milton Keynes and Alex Alderton, Roberto Amato, Sonia Goncalves, Ewan Harrison, David K. Jackson, Ian Johnston, Dominic Kwiatkowski, Cordelia Langford, John Sillitoe on behalf of the Wellcome Sanger Institute COVID-19 Surveillance Team |
| EPI_ISL_811713 | Lighthouse Lab in Cambridge | Wellcome Sanger Institute for the COVID-19 Genomics UK (COG-UK) Consortium | Rob Howes, The Lighthouse Lab in Cambridge and Alex Alderton, Roberto Amato, Sonia Goncalves, Ewan Harrison, David K. Jackson, Ian Johnston, Dominic Kwiatkowski, Cordelia Langford, John Sillitoe on behalf of the Wellcome Sanger Institute COVID-19 Surveillance Team |
| EPI_ISL_811720, EPI_ISL_811727, EPI_ISL_811750, EPI_ISL_811753, EPI_ISL_811756, EPI_ISL_811759, EPI_ISL_811767, EPI_ISL_811785, EPI_ISL_811808, EPI_ISL_811811, EPI_ISL_811812, EPI_ISL_811814, EPI_ISL_811817, EPI_ISL_811818, EPI_ISL_811819, EPI_ISL_811821, EPI_ISL_811823, EPI_ISL_811824, EPI_ISL_811825, EPI_ISL_811826, EPI_ISL_811827, EPI_ISL_811828, EPI_ISL_811830, EPI_ISL_811831, EPI_ISL_811832, EPI_ISL_811833, EPI_ISL_811834, EPI_ISL_811835, EPI_ISL_811836, EPI_ISL_811837, EPI_ISL_811839, EPI_ISL_811840, EPI_ISL_811841, EPI_ISL_811843, EPI_ISL_811845, EPI_ISL_811846, EPI_ISL_811847, EPI_ISL_811848, EPI_ISL_811849, EPI_ISL_811850, EPI_ISL_811851, EPI_ISL_811852, EPI_ISL_811853, EPI_ISL_811854, EPI_ISL_811855, EPI_ISL_811856, EPI_ISL_811857, EPI_ISL_811858, EPI_ISL_811859, EPI_ISL_811860, EPI_ISL_811861, EPI_ISL_811862, EPI_ISL_811865, EPI_ISL_811867, EPI_ISL_811868, EPI_ISL_811869, EPI_ISL_811870, EPI_ISL_811871, EPI_ISL_811873, EPI_ISL_811874, EPI_ISL_811875, EPI_ISL_811877, EPI_ISL_811878, EPI_ISL_811879, EPI_ISL_811880, EPI_ISL_811881, EPI_ISL_811883, EPI_ISL_811884, EPI_ISL_811885, EPI_ISL_811886, EPI_ISL_811887, EPI_ISL_811888, EPI_ISL_811889, EPI_ISL_811890, EPI_ISL_811891, EPI_ISL_811892, EPI_ISL_811893, EPI_ISL_811894, EPI_ISL_811896, EPI_ISL_811897, EPI_ISL_811898, EPI_ISL_811899, EPI_ISL_811900, EPI_ISL_811902, EPI_ISL_811905, EPI_ISL_811907, EPI_ISL_811909, EPI_ISL_811910, EPI_ISL_811912, EPI_ISL_811913, EPI_ISL_811914, EPI_ISL_811915, EPI_ISL_811916, EPI_ISL_811917, EPI_ISL_811918, EPI_ISL_811919, EPI_ISL_811921, EPI_ISL_811922, EPI_ISL_811923, EPI_ISL_811924, EPI_ISL_811925, EPI_ISL_811927, EPI_ISL_811928, EPI_ISL_811929, EPI_ISL_811930, EPI_ISL_811931, EPI_ISL_811932, EPI_ISL_811935, EPI_ISL_811936, EPI_ISL_811937, EPI_ISL_811938, EPI_ISL_811939, EPI_ISL_811941, EPI_ISL_811942, EPI_ISL_811943, EPI_ISL_811944, EPI_ISL_811945, EPI_ISL_811946, EPI_ISL_811948, EPI_ISL_811949, EPI_ISL_811950, EPI_ISL_811951, EPI_ISL_811952, EPI_ISL_811953, EPI_ISL_811954, EPI_ISL_811955, EPI_ISL_811956, EPI_ISL_811957, EPI_ISL_811958, EPI_ISL_811959, EPI_ISL_811960, EPI_ISL_811962, EPI_ISL_811965, EPI_ISL_811967, EPI_ISL_811968, EPI_ISL_811969, EPI_ISL_811970, EPI_ISL_811971, EPI_ISL_811972, EPI_ISL_811973, EPI_ISL_811974, EPI_ISL_811975, EPI_ISL_811976, EPI_ISL_811977, EPI_ISL_811978, EPI_ISL_811979, EPI_ISL_811981, EPI_ISL_811983, EPI_ISL_811984, EPI_ISL_811985, EPI_ISL_811986, EPI_ISL_811987, EPI_ISL_811988, EPI_ISL_811989, EPI_ISL_811990, EPI_ISL_811991, EPI_ISL_811993, EPI_ISL_811994, EPI_ISL_811996, EPI_ISL_811997, EPI_ISL_811998, EPI_ISL_811999, EPI_ISL_812000, EPI_ISL_812002, EPI_ISL_812003, EPI_ISL_812007, EPI_ISL_812009, EPI_ISL_812010, EPI_ISL_812012, EPI_ISL_812014, EPI_ISL_812015, EPI_ISL_812017, EPI_ISL_812018, EPI_ISL_812019, EPI_ISL_812020, EPI_ISL_812021, EPI_ISL_812022, EPI_ISL_812024, EPI_ISL_812025, EPI_ISL_812026, EPI_ISL_812027, EPI_ISL_812028, EPI_ISL_812030, EPI_ISL_812033, EPI_ISL_812035, EPI_ISL_812036, EPI_ISL_812037, EPI_ISL_812038, EPI_ISL_812040, EPI_ISL_812041, EPI_ISL_812042, EPI_ISL_812045, EPI_ISL_812046, EPI_ISL_812047, EPI_ISL_812048, EPI_ISL_812049, EPI_ISL_812050, EPI_ISL_812051, EPI_ISL_812052, EPI_ISL_812054, EPI_ISL_812055, EPI_ISL_812058, EPI_ISL_812059, EPI_ISL_812061, EPI_ISL_812062, EPI_ISL_812063, EPI_ISL_812064, EPI_ISL_812066, EPI_ISL_812067, EPI_ISL_812068, EPI_ISL_812069, EPI_ISL_812070, EPI_ISL_812073, EPI_ISL_812074, EPI_ISL_812075, EPI_ISL_812076, EPI_ISL_812077, EPI_ISL_812078, EPI_ISL_812079, EPI_ISL_812080, EPI_ISL_812083, EPI_ISL_812084, EPI_ISL_812085, EPI_ISL_812086, EPI_ISL_812088, EPI_ISL_812090, EPI_ISL_812091, EPI_ISL_812092, EPI_ISL_812093, EPI_ISL_812094, EPI_ISL_812096, EPI_ISL_812097, EPI_ISL_812098, EPI_ISL_812099, EPI_ISL_812100, EPI_ISL_812101, EPI_ISL_812102, EPI_ISL_812103, EPI_ISL_812104, EPI_ISL_812105, EPI_ISL_812106, EPI_ISL_812109, EPI_ISL_812113 | | | |
| see above | Lighthouse Lab in Milton Keynes | Wellcome Sanger Institute for the COVID-19 Genomics UK (COG-UK) Consortium | The Lighthouse Lab in Milton Keynes and Alex Alderton, Roberto Amato, Sonia Goncalves, Ewan Harrison, David K. Jackson, Ian Johnston, Dominic Kwiatkowski, Cordelia Langford, John Sillitoe on behalf of the Wellcome Sanger Institute COVID-19 Surveillance Team |
| EPI_ISL_812250 | Invenimus AG | Institute of Medical Virology, University of Zurich | Stefan Schmutz, Maryam Zaheri, Verena Kufner, Annette Audigé, Maria Grünberg, Kevin Steiner, Jon Huder, Cyril Shah, Riccarda Capaul, Guido Bloemberg, Jürg Böni, Michael Huber, Alexandra Trkola |
| EPI_ISL_812348 | University of Wisconsin-Madison AIDS Vaccine Research Laboratories | University of Wisconsin-Madison AIDS Vaccine Research Laboratories | Gage Moreno, Katarina Braun, et al. AIDS Vaccine Research Laboratories |
| EPI_ISL_812424, EPI_ISL_812425, EPI_ISL_812426, EPI_ISL_812427, EPI_ISL_812428, EPI_ISL_812429, EPI_ISL_812430, EPI_ISL_812431, EPI_ISL_812432, EPI_ISL_812433 | Microbiological Diagnostic Unit - Public Health Laboratory (MDU-PHL) | MDU-PHL | Seemann T., Sait, M.L., Sherry, N.L. |
| EPI_ISL_812437 | Victorian Infectious Diseases Reference Laboratory (VIDRL) | VIDRL and MDU-PHL | Caly L., Seemann T., Sait, M.L., Druce J., Sherry, N.L. |
| EPI_ISL_812768, EPI_ISL_812769, EPI_ISL_812770, EPI_ISL_812771, EPI_ISL_812777, EPI_ISL_812779, EPI_ISL_812780, EPI_ISL_812781, EPI_ISL_812874, EPI_ISL_812875, EPI_ISL_812876 | | | |
| see above | Ministry of Health Turkey | Ministry of Health Turkey | Fatma Bayrakdar, Yasemin Cogun, Süleyman Yalcin, Aye Baak Alta, Gülay Korukluolu |
| EPI_ISL_813091, EPI_ISL_813092, EPI_ISL_813093, EPI_ISL_813116 | University of Birmingham | COVID-19 Genomics UK (COG-UK) Consortium | Institute of Microbiology, University of Birmingham: Claire McMurray, Joanne Stockton, Samuel Nicholls, Radoslaw Poplawski, Will Rowe, Josh Quick, Nicholas Loman. University of Birmingham Testing Laboratory: Celina M Whalley, Andrew Bosworth, Charlotte Poxon, Kasun Wanigasooriya, Oliver Pickles, Mike Kidd, Alex Richter, Andrew D Beggs PHE Heartlands Lab: Husam Osman, Andrew Bosworth. Queen Elizabeth Hospital: Anna Casey |
| EPI_ISL_813198, EPI_ISL_813201, EPI_ISL_813202, EPI_ISL_813203, EPI_ISL_813204, EPI_ISL_813205, EPI_ISL_813206, EPI_ISL_813207, EPI_ISL_813209, EPI_ISL_813210, EPI_ISL_813211, EPI_ISL_813212, EPI_ISL_813213, EPI_ISL_813214, EPI_ISL_813215, EPI_ISL_813216, EPI_ISL_813217, EPI_ISL_813219, EPI_ISL_813220, EPI_ISL_813221, EPI_ISL_813222, EPI_ISL_813223, EPI_ISL_813224 | | | |
| see above | Department of Pathology, University of Cambridge | COVID-19 Genomics UK (COG-UK) Consortium | Aminu S. Jahun, Yasmin Chaudhry, Grant Hall, Iliana Georgana, Myra Hosmillo, Martin D. Curran, Malte Pinckert, Surendra Parmar, Ian Goodfellow |
| EPI_ISL_813608, EPI_ISL_813623, EPI_ISL_813625, EPI_ISL_813626, EPI_ISL_813627, EPI_ISL_813628, EPI_ISL_813629, EPI_ISL_813630, EPI_ISL_813631, EPI_ISL_813632, EPI_ISL_813633, EPI_ISL_813634, EPI_ISL_813635, EPI_ISL_813636, EPI_ISL_813638, EPI_ISL_813639, EPI_ISL_813640, EPI_ISL_813641, EPI_ISL_813642, EPI_ISL_813643, EPI_ISL_813644, EPI_ISL_813645, EPI_ISL_813646, EPI_ISL_813647, EPI_ISL_813648, EPI_ISL_813649, EPI_ISL_813650, EPI_ISL_813651, EPI_ISL_813652, EPI_ISL_813653, EPI_ISL_813654 | | | |
| see above | University of Exeter | COVID-19 Genomics UK (COG-UK) Consortium | Ben Temperton,Aaron Jeffries,Michelle Michelsen,Joanna Warwick-Dugdale,Audrey Farbos,Robyn Manley,Stephen Michell,Jane Masoli |
| EPI_ISL_813810, EPI_ISL_813811, EPI_ISL_813812, EPI_ISL_813813, EPI_ISL_813814, EPI_ISL_813815 | Liverpool Clinical Laboratories | COVID-19 Genomics UK (COG-UK) Consortium | Sam Haldenby, Anita Lucaci, Steve Paterson, Julian Hiscox, Alistair Darby, M Almsaud, A Alrezaihi, Muhannad Alruwaili, Stuart D Armstrong, Jones Benjamin, Eleanor G Bentley, Anu Chawla, Jordan J Clark, Angela Cowell, Richard Eccles, Isabel García-Dorival, Matthew Gemmell, Alessandro Gerada, PKF Gilmore, Richard Gregory, Ximeng Han, Catherine Hartley, Margaret Hughes, Miren Iturriza-Gomara, James Johnson, L Luu, Jenifer Manson, Charlotte Nelson, Elaine O'Toole, Cassie Olateju, Rebekah Penrice-Randal , Lucille Rainbow, N.P Randle, Trevor Ian Robinson, Parul Sharma, Ghada T Shawli, James P Stewart, Neil Swainston, Ecaterina Vamos, Joanne Watts, Mark Whitehead |
| EPI_ISL_813958 | University College London, Great Ormond Street Hospital for Children NHS Foundation Trust, Imperial College Healthcare NHS Trust | COVID-19 Genomics UK (COG-UK) Consortium | Sergi Castellano, Rachel Williams, Mark Kristiansen, Paola Resende Silva, Sunando Roy, Tony Brooks, Helena Tutill, Paola Niola, Patricia Dyal, Charlotte Williams, Leysa Forrest, Yasmin Panchbhaya, Jacqueline Findlay, Samuel Weeks, Julianne Brown, Kathryn Harris, Paul Randell, James Price, Alison Holmes, Judith Breuer |
| EPI_ISL_813980, EPI_ISL_813981, EPI_ISL_813982, EPI_ISL_813984, EPI_ISL_813985, EPI_ISL_813986, EPI_ISL_813989, EPI_ISL_813990, EPI_ISL_813991, EPI_ISL_813992, EPI_ISL_813994, EPI_ISL_813997, EPI_ISL_813998, EPI_ISL_814001, EPI_ISL_814002, EPI_ISL_814003, EPI_ISL_814004, EPI_ISL_814005, | | | |

| | | | |
|---|---|---|---|
| EPI_ISL_814006, EPI_ISL_814008, EPI_ISL_814009, EPI_ISL_814015, EPI_ISL_814018, EPI_ISL_814019, EPI_ISL_814020, EPI_ISL_814027, EPI_ISL_814028, EPI_ISL_814029, EPI_ISL_814030, EPI_ISL_814031, EPI_ISL_814032, EPI_ISL_814033, EPI_ISL_814037, EPI_ISL_814038, EPI_ISL_814039, EPI_ISL_814040, EPI_ISL_814051, EPI_ISL_814052, EPI_ISL_814053, EPI_ISL_814054, EPI_ISL_814055, EPI_ISL_814056, EPI_ISL_814057, EPI_ISL_814058, EPI_ISL_814059, EPI_ISL_814060, EPI_ISL_814061 | | | |
| see above | Hospital General Universitario Gregorio Marañón | SeqCOVID-SPAIN consortium/IBV(CSIC) | Darío García de Viedma, Laura Pérez-Lago, Marta Herranz, Jon Sicilia, Julia Suárez, Pilar Catalán, Patricia Muñoz and SeqCOVID-SPAIN consortium |
| EPI_ISL_814358, EPI_ISL_814429, EPI_ISL_814433, EPI_ISL_814434 | Bioinformatics and Biostatistics Lab, Advanced Sequencing Facility | COVID-19 Genomics UK (COG-UK) Consortium | Aengus Stewart,Jerome Nicod,Chelsea Sawyer,Laura Cubitt,Harshil Patel,Margaret Crawford |
| EPI_ISL_816213, EPI_ISL_816214, EPI_ISL_816215, EPI_ISL_816220, EPI_ISL_816221, EPI_ISL_816222, EPI_ISL_816223 | Centre for Enzyme Innovation, University of Portsmouth / Translational Research Laboratory, Portsmouth Hospitals NHS Trust | COVID-19 Genomics UK (COG-UK) Consortium | Angela Beckett,Yann Bourgeois,Garry Scarlett,Sharon Glaysher,Scott Elliott,Kelly Bicknell,Robert Impey,Allyson Lloyd,Sarah Wyllie,Ethan Butcher,Anoop Chauhan,Samuel Robson |
| EPI_ISL_816226, EPI_ISL_816227, EPI_ISL_816228, EPI_ISL_816229, EPI_ISL_816232, EPI_ISL_816233, EPI_ISL_816235, EPI_ISL_816246, EPI_ISL_816248, EPI_ISL_816256, EPI_ISL_816261, EPI_ISL_816275, EPI_ISL_816276, EPI_ISL_816277, EPI_ISL_816285, EPI_ISL_816291, EPI_ISL_816294, EPI_ISL_816296, EPI_ISL_816297, EPI_ISL_816305, EPI_ISL_816311, EPI_ISL_816314, EPI_ISL_816318, EPI_ISL_816319, EPI_ISL_816323, EPI_ISL_816327, EPI_ISL_816331, EPI_ISL_816348, EPI_ISL_816353, EPI_ISL_816364, EPI_ISL_816368, EPI_ISL_816383, EPI_ISL_816390, EPI_ISL_816396, EPI_ISL_816405, EPI_ISL_816406, EPI_ISL_816407, EPI_ISL_816409, EPI_ISL_816415, EPI_ISL_816423, EPI_ISL_816426, EPI_ISL_816427, EPI_ISL_816431, EPI_ISL_816434, EPI_ISL_816437, EPI_ISL_816438, EPI_ISL_816441, EPI_ISL_816442, EPI_ISL_816444, EPI_ISL_816448, EPI_ISL_816449, EPI_ISL_816462, EPI_ISL_816469, EPI_ISL_816472, EPI_ISL_816475, EPI_ISL_816477, EPI_ISL_816492, EPI_ISL_816494, EPI_ISL_816495, EPI_ISL_816496, EPI_ISL_816501, EPI_ISL_816508, EPI_ISL_816525, EPI_ISL_816536, EPI_ISL_816539, EPI_ISL_816555, EPI_ISL_816561, EPI_ISL_816566, EPI_ISL_816571, EPI_ISL_816576, EPI_ISL_816581, EPI_ISL_816582, EPI_ISL_816584, EPI_ISL_816601, EPI_ISL_816608, EPI_ISL_816611, EPI_ISL_816613, EPI_ISL_816629, EPI_ISL_816633, EPI_ISL_816634, EPI_ISL_816641, EPI_ISL_816642, EPI_ISL_816646 | | | |
| see above | Virology Department, Sheffield Teaching Hospitals NHS Foundation Trust/Department of Infection, Immunity and Cardiovascular Disease, The Medical School, University of Sheffield | COVID-19 Genomics UK (COG-UK) Consortium | Thushan de Silva, Matthew Parker, Nikki Smith, Adri Angyal, Rebecca Brown, Luke Green, Rachel Tucker, Paul Parsons, Danielle Groves, Katie Johnson, Laura Carrilero, Alex Keeley, Dave Partridge, Matthew Wyles, Benjamin Lindsey, Mehmet Yavuz, Mohammad Raza, Cariad Evans |
| EPI_ISL_816825, EPI_ISL_816826, EPI_ISL_816827, EPI_ISL_816828, EPI_ISL_816829, EPI_ISL_816830, EPI_ISL_816831, EPI_ISL_816832, EPI_ISL_816833, EPI_ISL_816834, EPI_ISL_816835, EPI_ISL_816836, EPI_ISL_816837, EPI_ISL_816838, EPI_ISL_816839, EPI_ISL_816840, EPI_ISL_816841, EPI_ISL_816842, EPI_ISL_816843, EPI_ISL_816844, EPI_ISL_816845, EPI_ISL_816846, EPI_ISL_816847, EPI_ISL_816848, EPI_ISL_816849, EPI_ISL_816850, EPI_ISL_816851, EPI_ISL_816852, EPI_ISL_816853, EPI_ISL_816854, EPI_ISL_816855, EPI_ISL_816856, EPI_ISL_816857, EPI_ISL_816858, EPI_ISL_816859, EPI_ISL_816860, EPI_ISL_816892, EPI_ISL_816897, EPI_ISL_816898, EPI_ISL_816903, EPI_ISL_816959, EPI_ISL_816964, EPI_ISL_816965, EPI_ISL_816970, EPI_ISL_817039 | | | |
| see above | Bioinformatics and Biostatistics Lab, Advanced Sequencing Facility | COVID-19 Genomics UK (COG-UK) Consortium | Aengus Stewart,Jerome Nicod,Chelsea Sawyer,Laura Cubitt,Harshil Patel,Margaret Crawford |
| EPI_ISL_818342 | Charité Universitätsmedizin Berlin, Institute of Virology | Charité Universitätsmedizin Berlin, Institute of Virology | Victor M Corman, Julia Schneider, Jörn Beheim-Schwarzbach, Barbara Mühlemann, Talitha Veith, Terry Jones, Christian Drosten |
| EPI_ISL_819133, EPI_ISL_819135, EPI_ISL_819142, EPI_ISL_819156, EPI_ISL_819157, EPI_ISL_819163, EPI_ISL_819169, EPI_ISL_819171, EPI_ISL_819175, EPI_ISL_819186, EPI_ISL_819187, EPI_ISL_819191, EPI_ISL_819195 | | | |
| see above | Servicio de Microbiología, Hospital Universitario Son Espases | SeqCOVID-SPAIN consortium/IBV(CSIC) | Carla López-Causapé, Jordi Reina, Antonio Oliver and SeqCOVID-SPAIN consortium |
| EPI_ISL_819378, EPI_ISL_819387, EPI_ISL_819388, EPI_ISL_819389, EPI_ISL_819390, EPI_ISL_819398, EPI_ISL_819399, EPI_ISL_819400, EPI_ISL_819401, EPI_ISL_819402, EPI_ISL_819403, EPI_ISL_819404, EPI_ISL_819428, EPI_ISL_819431, EPI_ISL_819433, EPI_ISL_819438 | | | |
| see above | Quadram Institute Bioscience | COVID-19 Genomics UK (COG-UK) Consortium | Dave J. Baker, Gemma L. Kay, Alp Aydin, Thanh Le-Viet, Steven Rudder, Ana P. Tedim, Anastasia Kolyva, Maria Diaz, Leonardo de Oliveira Martins, Nabil-Fareed Alikhan, Lizzie Meadows, Rachael Stanley, Ngozi Elumogo, Muhammed Yasir, Nicholas M. Thomson, Alexander J Trotter, Rachel Gilroy, Samuel Bloomfield, Claire Stuart, Andrew Bell, Reenesh Prakash, Samir Dervisevic, Alison E. Mather, John Wain, Mark Webber, Andrew J. Page, Justin O'Grady |
| EPI_ISL_820326, EPI_ISL_820328, EPI_ISL_820331, EPI_ISL_820333, EPI_ISL_820335, EPI_ISL_820337, EPI_ISL_820340, EPI_ISL_820343, EPI_ISL_820345, EPI_ISL_820347, EPI_ISL_820350 | | | |
| see above | University College London, Great Ormond Street Hospital for Children NHS Foundation Trust, Imperial College Healthcare NHS Trust | COVID-19 Genomics UK (COG-UK) Consortium | Sergi Castellano, Rachel Williams, Mark Kristiansen, Paola Resende Silva, Sunando Roy, Tony Brooks, Helena Tutill, Paola Niola, Patricia Dyal, Charlotte Williams, Leysa Forrest, Yasmin Panchbhaya, Jacqueline Findlay, Samuel Weeks, Julianne Brown, Kathryn Harris, Paul Randell, James Price, Alison Holmes, Judith Breuer |
| EPI_ISL_820353, EPI_ISL_820355, EPI_ISL_820358, EPI_ISL_820360, EPI_ISL_820363, EPI_ISL_820365, EPI_ISL_820368, EPI_ISL_820371 | Quadram Institute Bioscience | COVID-19 Genomics UK (COG-UK) Consortium | Dave J. Baker, Gemma L. Kay, Alp Aydin, Thanh Le-Viet, Steven Rudder, Ana P. Tedim, Anastasia Kolyva, Maria Diaz, Leonardo de Oliveira Martins, Nabil-Fareed Alikhan, Lizzie Meadows, Rachael Stanley, Ngozi Elumogo, Muhammed Yasir, Nicholas M. Thomson, Alexander J Trotter, Rachel Gilroy, Samuel Bloomfield, Claire Stuart, Andrew Bell, Reenesh Prakash, Samir Dervisevic, Alison E. Mather, John Wain, Mark Webber, Andrew J. Page, Justin O'Grady |
| EPI_ISL_820373, EPI_ISL_820376 | University College London, Great Ormond Street Hospital for Children NHS Foundation Trust, Imperial College Healthcare NHS Trust | COVID-19 Genomics UK (COG-UK) Consortium | Sergi Castellano, Rachel Williams, Mark Kristiansen, Paola Resende Silva, Sunando Roy, Tony Brooks, Helena Tutill, Paola Niola, Patricia Dyal, Charlotte Williams, Leysa Forrest, Yasmin Panchbhaya, Jacqueline Findlay, Samuel Weeks, Julianne Brown, Kathryn Harris, Paul Randell, James Price, Alison Holmes, Judith Breuer |
| EPI_ISL_820378 | Quadram Institute Bioscience | COVID-19 Genomics UK (COG-UK) Consortium | Dave J. Baker, Gemma L. Kay, Alp Aydin, Thanh Le-Viet, Steven Rudder, Ana P. Tedim, Anastasia Kolyva, Maria Diaz, Leonardo de Oliveira Martins, Nabil-Fareed Alikhan, Lizzie Meadows, Rachael Stanley, Ngozi Elumogo, Muhammed Yasir, Nicholas M. Thomson, Alexander J Trotter, Rachel Gilroy, Samuel Bloomfield, Claire Stuart, Andrew Bell, Reenesh Prakash, Samir Dervisevic, Alison E. Mather, John Wain, Mark Webber, Andrew J. Page, Justin O'Grady |
| EPI_ISL_820381, EPI_ISL_820383, EPI_ISL_820385 | University College London, Great Ormond Street Hospital for Children NHS Foundation Trust, Imperial College Healthcare NHS Trust | COVID-19 Genomics UK (COG-UK) Consortium | Sergi Castellano, Rachel Williams, Mark Kristiansen, Paola Resende Silva, Sunando Roy, Tony Brooks, Helena Tutill, Paola Niola, Patricia Dyal, Charlotte Williams, Leysa Forrest, Yasmin Panchbhaya, Jacqueline Findlay, Samuel Weeks, Julianne Brown, Kathryn Harris, Paul Randell, James Price, Alison Holmes, Judith Breuer |
| EPI_ISL_820388, EPI_ISL_820391 | Quadram Institute Bioscience | COVID-19 Genomics UK (COG-UK) Consortium | Dave J. Baker, Gemma L. Kay, Alp Aydin, Thanh Le-Viet, Steven Rudder, Ana P. Tedim, Anastasia Kolyva, Maria Diaz, Leonardo de Oliveira Martins, Nabil-Fareed Alikhan, Lizzie Meadows, Rachael Stanley, Ngozi Elumogo, Muhammed Yasir, Nicholas M. Thomson, Alexander J Trotter, Rachel Gilroy, Samuel Bloomfield, Claire Stuart, Andrew Bell, Reenesh Prakash, Samir Dervisevic, Alison E. Mather, John Wain, Mark Webber, Andrew J. Page, Justin O'Grady |
| EPI_ISL_820394 | Queens Medical Centre, Clinical Microbiology Department / DeepSeq Nottingham | COVID-19 Genomics UK (COG-UK) Consortium | Gemma Clark, Wendy Smith, Manjinder Khakh, Vicki M Fleming, Michelle M Lister, Hannah Howson-Wells, Jonathan Ball, Patrick McClure, Joseph Chappell, Theocharis Tsoleridis, Nadine Holmes, Matthew Carlisle, Christopher Moore, Fei Sang, Johnny Debebe, Victoria Wright, Matthew Loose |
| EPI_ISL_820396 | University College London, Great Ormond Street Hospital for Children NHS Foundation Trust, Imperial College Healthcare NHS Trust | COVID-19 Genomics UK (COG-UK) Consortium | Sergi Castellano, Rachel Williams, Mark Kristiansen, Paola Resende Silva, Sunando Roy, Tony Brooks, Helena Tutill, Paola Niola, Patricia Dyal, Charlotte Williams, Leysa Forrest, Yasmin Panchbhaya, Jacqueline Findlay, Samuel Weeks, Julianne Brown, Kathryn Harris, Paul Randell, James Price, Alison Holmes, Judith Breuer |
| EPI_ISL_820399 | Quadram Institute Bioscience | COVID-19 Genomics UK (COG-UK) Consortium | Dave J. Baker, Gemma L. Kay, Alp Aydin, Thanh Le-Viet, Steven Rudder, Ana P. Tedim, Anastasia Kolyva, Maria Diaz, Leonardo de Oliveira Martins, Nabil-Fareed Alikhan, Lizzie Meadows, Rachael Stanley, Ngozi Elumogo, Muhammed Yasir, Nicholas M. Thomson, Alexander J Trotter, Rachel Gilroy, Samuel Bloomfield, Claire Stuart, Andrew Bell, Reenesh Prakash, Samir Dervisevic, Alison E. Mather, John Wain, Mark Webber, Andrew J. Page, Justin O'Grady |
| EPI_ISL_820402 | Queens Medical Centre, Clinical Microbiology Department / DeepSeq Nottingham | COVID-19 Genomics UK (COG-UK) Consortium | Gemma Clark, Wendy Smith, Manjinder Khakh, Vicki M Fleming, Michelle M Lister, Hannah Howson-Wells, Jonathan Ball, Patrick McClure, Joseph Chappell, Theocharis Tsoleridis, Nadine Holmes, Matthew Carlisle, Christopher Moore, Fei Sang, Johnny Debebe, Victoria Wright, Matthew Loose |
| EPI_ISL_820404, EPI_ISL_820407, EPI_ISL_820410, EPI_ISL_820412, EPI_ISL_820415 | Quadram Institute Bioscience | COVID-19 Genomics UK (COG-UK) Consortium | Dave J. Baker, Gemma L. Kay, Alp Aydin, Thanh Le-Viet, Steven Rudder, Ana P. Tedim, Anastasia Kolyva, Maria Diaz, Leonardo de Oliveira Martins, Nabil-Fareed Alikhan, Lizzie Meadows, Rachael Stanley, Ngozi Elumogo, Muhammed Yasir, Nicholas M. Thomson, Alexander J Trotter, Rachel Gilroy, Samuel Bloomfield, Claire Stuart, Andrew Bell, Reenesh Prakash, Samir Dervisevic, Alison E. Mather, John Wain, Mark Webber, Andrew J. Page, Justin O'Grady |
| EPI_ISL_820418 | Queens Medical Centre, Clinical Microbiology Department / DeepSeq Nottingham | COVID-19 Genomics UK (COG-UK) Consortium | Gemma Clark, Wendy Smith, Manjinder Khakh, Vicki M Fleming, Michelle M Lister, Hannah Howson-Wells, Jonathan Ball, Patrick McClure, Joseph Chappell, Theocharis Tsoleridis, Nadine Holmes, Matthew Carlisle, Christopher Moore, Fei Sang, Johnny Debebe, Victoria Wright, Matthew Loose |
| EPI_ISL_820420 | University College London, Great Ormond Street Hospital for Children NHS Foundation Trust, Imperial College Healthcare NHS Trust | COVID-19 Genomics UK (COG-UK) Consortium | Sergi Castellano, Rachel Williams, Mark Kristiansen, Paola Resende Silva, Sunando Roy, Tony Brooks, Helena Tutill, Paola Niola, Patricia Dyal, Charlotte Williams, Leysa Forrest, Yasmin Panchbhaya, Jacqueline Findlay, Samuel Weeks, Julianne Brown, Kathryn Harris, Paul Randell, James Price, Alison Holmes, Judith Breuer |
| EPI_ISL_820423 | Queens Medical Centre, Clinical Microbiology Department / | COVID-19 Genomics UK (COG-UK) Consortium | Gemma Clark, Wendy Smith, Manjinder Khakh, Vicki M Fleming, Michelle M Lister, Hannah Howson-Wells, Jonathan Ball, Patrick McClure, Joseph |

| | | | |
|---|---|---|---|
| | DeepSeq Nottingham | | Chappell, Theocharis Tsoleridis, Nadine Holmes, Matthew Carlisle, Christopher Moore, Fei Sang, Johnny Debebe, Victoria Wright, Matthew Loose |
| EPI_ISL_820426 | Quadram Institute Bioscience | COVID-19 Genomics UK (COG-UK) Consortium | Dave J. Baker, Gemma L. Kay, Alp Aydin, Thanh Le-Viet, Steven Rudder, Ana P. Tedim, Anastasia Kolyva, Maria Diaz, Leonardo de Oliveira Martins, Nabil-Fareed Alikhan, Lizzie Meadows, Rachael Stanley, Ngozi Elumogo, Muhammed Yasir, Nicholas M. Thomson, Alexander J Trotter, Rachel Gilroy, Samuel Bloomfield, Claire Stuart, Andrew Bell, Reenesh Prakash, Samir Dervisevic, Alison E. Mather, John Wain, Mark Webber, Andrew J. Page, Justin O'Grady |
| EPI_ISL_820428, EPI_ISL_820431, EPI_ISL_820433 | Queens Medical Centre, Clinical Microbiology Department / DeepSeq Nottingham | COVID-19 Genomics UK (COG-UK) Consortium | Gemma Clark, Wendy Smith, Manjinder Khakh, Vicki M Fleming, Michelle M Lister, Rachael Howson-Wells, Jonathan Ball, Patrick McClure, Joseph Chappell, Theocharis Tsoleridis, Nadine Holmes, Matthew Carlisle, Christopher Moore, Fei Sang, Johnny Debebe, Victoria Wright, Matthew Loose |
| EPI_ISL_820436, EPI_ISL_820439, EPI_ISL_820441, EPI_ISL_820444, EPI_ISL_820446, EPI_ISL_820449, EPI_ISL_820451, EPI_ISL_820454, EPI_ISL_820456, EPI_ISL_820459, EPI_ISL_820462, EPI_ISL_820464, EPI_ISL_820467, EPI_ISL_820470, EPI_ISL_820472, EPI_ISL_820475, EPI_ISL_820478, EPI_ISL_820480, EPI_ISL_820483, EPI_ISL_820485, EPI_ISL_820488, EPI_ISL_820490, EPI_ISL_820493, EPI_ISL_820496, EPI_ISL_820498, EPI_ISL_820501, EPI_ISL_820504, EPI_ISL_820506, EPI_ISL_820509, EPI_ISL_820511, EPI_ISL_820514, EPI_ISL_820516, EPI_ISL_820519, EPI_ISL_820521, EPI_ISL_820524, EPI_ISL_820526, EPI_ISL_820528, EPI_ISL_820531, EPI_ISL_820533, EPI_ISL_820536, EPI_ISL_820539, EPI_ISL_820541, EPI_ISL_820543, EPI_ISL_820546, EPI_ISL_820549, EPI_ISL_820552, EPI_ISL_820554, EPI_ISL_820557 | | | |
| see above | Quadram Institute Bioscience | COVID-19 Genomics UK (COG-UK) Consortium | Dave J. Baker, Gemma L. Kay, Alp Aydin, Thanh Le-Viet, Steven Rudder, Ana P. Tedim, Anastasia Kolyva, Maria Diaz, Leonardo de Oliveira Martins, Nabil-Fareed Alikhan, Lizzie Meadows, Rachael Stanley, Ngozi Elumogo, Muhammed Yasir, Nicholas M. Thomson, Alexander J Trotter, Rachel Gilroy, Samuel Bloomfield, Claire Stuart, Andrew Bell, Reenesh Prakash, Samir Dervisevic, Alison E. Mather, John Wain, Mark Webber, Andrew J. Page, Justin O'Grady |
| EPI_ISL_821039, EPI_ISL_821128, EPI_ISL_821140, EPI_ISL_821163, EPI_ISL_821165, EPI_ISL_821172, EPI_ISL_821197, EPI_ISL_821220, EPI_ISL_821235, EPI_ISL_821263, EPI_ISL_821270, EPI_ISL_821271 | | | |
| see above | Lighthouse Lab in Alderley Park | Wellcome Sanger Institute for the COVID-19 Genomics UK (COG-UK) Consortium | Jacquelyn Wynn, Mairead Hyland, The Lighthouse Lab in Alderley Park and Alex Alderton, Roberto Amato, Sonia Goncalves, Ewan Harrison, David K. Jackson, Ian Johnston, Dominic Kwiatkowski, Cordelia Langford, John Sillitoe on behalf of the Wellcome Sanger Institute COVID-19 Surveillance Team |
| EPI_ISL_821275, EPI_ISL_821277 | Lighthouse Lab in Milton Keynes | Wellcome Sanger Institute for the COVID-19 Genomics UK (COG-UK) Consortium | The Lighthouse Lab in Milton Keynes and Alex Alderton, Roberto Amato, Sonia Goncalves, Ewan Harrison, David K. Jackson, Ian Johnston, Dominic Kwiatkowski, Cordelia Langford, John Sillitoe on behalf of the Wellcome Sanger Institute COVID-19 Surveillance Team |
| EPI_ISL_821278 | Lighthouse Lab in Cambridge | Wellcome Sanger Institute for the COVID-19 Genomics UK (COG-UK) Consortium | Rob Howes, The Lighthouse Lab in Cambridge and Alex Alderton, Roberto Amato, Sonia Goncalves, Ewan Harrison, David K. Jackson, Ian Johnston, Dominic Kwiatkowski, Cordelia Langford, John Sillitoe on behalf of the Wellcome Sanger Institute COVID-19 Surveillance Team |
| EPI_ISL_821291, EPI_ISL_821299, EPI_ISL_821302, EPI_ISL_821305 | Lighthouse Lab in Milton Keynes | Wellcome Sanger Institute for the COVID-19 Genomics UK (COG-UK) Consortium | The Lighthouse Lab in Milton Keynes and Alex Alderton, Roberto Amato, Sonia Goncalves, Ewan Harrison, David K. Jackson, Ian Johnston, Dominic Kwiatkowski, Cordelia Langford, John Sillitoe on behalf of the Wellcome Sanger Institute COVID-19 Surveillance Team |
| EPI_ISL_821311 | Lighthouse Lab in Cambridge | Wellcome Sanger Institute for the COVID-19 Genomics UK (COG-UK) Consortium | Rob Howes, The Lighthouse Lab in Cambridge and Alex Alderton, Roberto Amato, Sonia Goncalves, Ewan Harrison, David K. Jackson, Ian Johnston, Dominic Kwiatkowski, Cordelia Langford, John Sillitoe on behalf of the Wellcome Sanger Institute COVID-19 Surveillance Team |
| EPI_ISL_821312, EPI_ISL_821313, EPI_ISL_821315 | Lighthouse Lab in Milton Keynes | Wellcome Sanger Institute for the COVID-19 Genomics UK (COG-UK) Consortium | The Lighthouse Lab in Milton Keynes and Alex Alderton, Roberto Amato, Sonia Goncalves, Ewan Harrison, David K. Jackson, Ian Johnston, Dominic Kwiatkowski, Cordelia Langford, John Sillitoe on behalf of the Wellcome Sanger Institute COVID-19 Surveillance Team |
| EPI_ISL_821316 | Lighthouse Lab in Cambridge | Wellcome Sanger Institute for the COVID-19 Genomics UK (COG-UK) Consortium | Rob Howes, The Lighthouse Lab in Cambridge and Alex Alderton, Roberto Amato, Sonia Goncalves, Ewan Harrison, David K. Jackson, Ian Johnston, Dominic Kwiatkowski, Cordelia Langford, John Sillitoe on behalf of the Wellcome Sanger Institute COVID-19 Surveillance Team |
| EPI_ISL_821318 | Lighthouse Lab in Alderley Park | Wellcome Sanger Institute for the COVID-19 Genomics UK (COG-UK) Consortium | Jacquelyn Wynn, Mairead Hyland, The Lighthouse Lab in Alderley Park and Alex Alderton, Roberto Amato, Sonia Goncalves, Ewan Harrison, David K. Jackson, Ian Johnston, Dominic Kwiatkowski, Cordelia Langford, John Sillitoe on behalf of the Wellcome Sanger Institute COVID-19 Surveillance Team |
| EPI_ISL_821320 | Lighthouse Lab in Cambridge | Wellcome Sanger Institute for the COVID-19 Genomics UK (COG-UK) Consortium | Rob Howes, The Lighthouse Lab in Cambridge and Alex Alderton, Roberto Amato, Sonia Goncalves, Ewan Harrison, David K. Jackson, Ian Johnston, Dominic Kwiatkowski, Cordelia Langford, John Sillitoe on behalf of the Wellcome Sanger Institute COVID-19 Surveillance Team |
| EPI_ISL_821321, EPI_ISL_821322, EPI_ISL_821325, EPI_ISL_821326, EPI_ISL_821330, EPI_ISL_821332, EPI_ISL_821337, EPI_ISL_821345, EPI_ISL_821349, EPI_ISL_821355, EPI_ISL_821357, EPI_ISL_821361, EPI_ISL_821364 | | | |
| see above | Lighthouse Lab in Milton Keynes | Wellcome Sanger Institute for the COVID-19 Genomics UK (COG-UK) Consortium | The Lighthouse Lab in Milton Keynes and Alex Alderton, Roberto Amato, Sonia Goncalves, Ewan Harrison, David K. Jackson, Ian Johnston, Dominic Kwiatkowski, Cordelia Langford, John Sillitoe on behalf of the Wellcome Sanger Institute COVID-19 Surveillance Team |
| EPI_ISL_821368 | Lighthouse Lab in Cambridge | Wellcome Sanger Institute for the COVID-19 Genomics UK (COG-UK) Consortium | Rob Howes, The Lighthouse Lab in Cambridge and Alex Alderton, Roberto Amato, Sonia Goncalves, Ewan Harrison, David K. Jackson, Ian Johnston, Dominic Kwiatkowski, Cordelia Langford, John Sillitoe on behalf of the Wellcome Sanger Institute COVID-19 Surveillance Team |
| EPI_ISL_821369 | Lighthouse Lab in Milton Keynes | Wellcome Sanger Institute for the COVID-19 Genomics UK (COG-UK) Consortium | The Lighthouse Lab in Milton Keynes and Alex Alderton, Roberto Amato, Sonia Goncalves, Ewan Harrison, David K. Jackson, Ian Johnston, Dominic Kwiatkowski, Cordelia Langford, John Sillitoe on behalf of the Wellcome Sanger Institute COVID-19 Surveillance Team |
| EPI_ISL_821376, EPI_ISL_821393 | Lighthouse Lab in Cambridge | Wellcome Sanger Institute for the COVID-19 Genomics UK (COG-UK) Consortium | Rob Howes, The Lighthouse Lab in Cambridge and Alex Alderton, Roberto Amato, Sonia Goncalves, Ewan Harrison, David K. Jackson, Ian Johnston, Dominic Kwiatkowski, Cordelia Langford, John Sillitoe on behalf of the Wellcome Sanger Institute COVID-19 Surveillance Team |
| EPI_ISL_821396 | Lighthouse Lab in Milton Keynes | Wellcome Sanger Institute for the COVID-19 Genomics UK (COG-UK) Consortium | The Lighthouse Lab in Milton Keynes and Alex Alderton, Roberto Amato, Sonia Goncalves, Ewan Harrison, David K. Jackson, Ian Johnston, Dominic Kwiatkowski, Cordelia Langford, John Sillitoe on behalf of the Wellcome Sanger Institute COVID-19 Surveillance Team |
| EPI_ISL_821399 | Lighthouse Lab in Cambridge | Wellcome Sanger Institute for the COVID-19 Genomics UK (COG-UK) Consortium | Rob Howes, The Lighthouse Lab in Cambridge and Alex Alderton, Roberto Amato, Sonia Goncalves, Ewan Harrison, David K. Jackson, Ian Johnston, Dominic Kwiatkowski, Cordelia Langford, John Sillitoe on behalf of the Wellcome Sanger Institute COVID-19 Surveillance Team |
| EPI_ISL_821413, EPI_ISL_821420 | Lighthouse Lab in Milton Keynes | Wellcome Sanger Institute for the COVID-19 Genomics UK (COG-UK) Consortium | The Lighthouse Lab in Milton Keynes and Alex Alderton, Roberto Amato, Sonia Goncalves, Ewan Harrison, David K. Jackson, Ian Johnston, Dominic Kwiatkowski, Cordelia Langford, John Sillitoe on behalf of the Wellcome Sanger Institute COVID-19 Surveillance Team |
| EPI_ISL_821421, EPI_ISL_821427 | Lighthouse Lab in Cambridge | Wellcome Sanger Institute for the COVID-19 Genomics UK (COG-UK) Consortium | Rob Howes, The Lighthouse Lab in Cambridge and Alex Alderton, Roberto Amato, Sonia Goncalves, Ewan Harrison, David K. Jackson, Ian Johnston, Dominic Kwiatkowski, Cordelia Langford, John Sillitoe on behalf of the Wellcome Sanger Institute COVID-19 Surveillance Team |
| EPI_ISL_821429, EPI_ISL_821430 | Lighthouse Lab in Milton Keynes | Wellcome Sanger Institute for the COVID-19 Genomics UK (COG-UK) Consortium | The Lighthouse Lab in Milton Keynes and Alex Alderton, Roberto Amato, Sonia Goncalves, Ewan Harrison, David K. Jackson, Ian Johnston, Dominic Kwiatkowski, Cordelia Langford, John Sillitoe on behalf of the Wellcome Sanger Institute COVID-19 Surveillance Team |
| EPI_ISL_821442, EPI_ISL_821444 | Lighthouse Lab in Cambridge | Wellcome Sanger Institute for the COVID-19 Genomics UK (COG-UK) Consortium | Rob Howes, The Lighthouse Lab in Cambridge and Alex Alderton, Roberto Amato, Sonia Goncalves, Ewan Harrison, David K. Jackson, Ian Johnston, Dominic Kwiatkowski, Cordelia Langford, John Sillitoe on behalf of the Wellcome Sanger Institute COVID-19 Surveillance Team |
| EPI_ISL_821450 | Lighthouse Lab in Milton Keynes | Wellcome Sanger Institute for the COVID-19 Genomics UK (COG-UK) Consortium | The Lighthouse Lab in Milton Keynes and Alex Alderton, Roberto Amato, Sonia Goncalves, Ewan Harrison, David K. Jackson, Ian Johnston, Dominic Kwiatkowski, Cordelia Langford, John Sillitoe on behalf of the Wellcome Sanger Institute COVID-19 Surveillance Team |
| EPI_ISL_821454 | Lighthouse Lab in Cambridge | Wellcome Sanger Institute for the COVID-19 Genomics UK (COG-UK) Consortium | Rob Howes, The Lighthouse Lab in Cambridge and Alex Alderton, Roberto Amato, Sonia Goncalves, Ewan Harrison, David K. Jackson, Ian Johnston, Dominic Kwiatkowski, Cordelia Langford, John Sillitoe on behalf of the Wellcome Sanger Institute COVID-19 Surveillance Team |
| EPI_ISL_821455 | Lighthouse Lab in Milton Keynes | Wellcome Sanger Institute for the COVID-19 Genomics UK (COG-UK) Consortium | The Lighthouse Lab in Milton Keynes and Alex Alderton, Roberto Amato, Sonia Goncalves, Ewan Harrison, David K. Jackson, Ian Johnston, Dominic Kwiatkowski, Cordelia Langford, John Sillitoe on behalf of the Wellcome Sanger Institute COVID-19 Surveillance Team |
| EPI_ISL_821456 | Lighthouse Lab in Alderley Park | Wellcome Sanger Institute for the COVID-19 Genomics UK (COG-UK) Consortium | Jacquelyn Wynn, Mairead Hyland, The Lighthouse Lab in Alderley Park and Alex Alderton, Roberto Amato, Sonia Goncalves, Ewan Harrison, David K. Jackson, Ian Johnston, Dominic Kwiatkowski, Cordelia Langford, John Sillitoe on behalf of the Wellcome Sanger Institute COVID-19 Surveillance Team |
| EPI_ISL_821463 | Lighthouse Lab in Cambridge | Wellcome Sanger Institute for the COVID-19 Genomics UK (COG-UK) Consortium | Rob Howes, The Lighthouse Lab in Cambridge and Alex Alderton, Roberto Amato, Sonia Goncalves, Ewan Harrison, David K. Jackson, Ian Johnston, Dominic Kwiatkowski, Cordelia Langford, John Sillitoe on behalf of the Wellcome Sanger Institute COVID-19 Surveillance Team |
| EPI_ISL_821464, EPI_ISL_821471 | Lighthouse Lab in Milton Keynes | Wellcome Sanger Institute for the COVID-19 Genomics UK (COG-UK) Consortium | The Lighthouse Lab in Milton Keynes and Alex Alderton, Roberto Amato, Sonia Goncalves, Ewan Harrison, David K. Jackson, Ian Johnston, Dominic Kwiatkowski, Cordelia Langford, John Sillitoe on behalf of the Wellcome Sanger Institute COVID-19 Surveillance Team |
| EPI_ISL_821473 | Lighthouse Lab in Cambridge | Wellcome Sanger Institute for the COVID-19 Genomics UK (COG-UK) Consortium | Rob Howes, The Lighthouse Lab in Cambridge and Alex Alderton, Roberto Amato, Sonia Goncalves, Ewan Harrison, David K. Jackson, Ian Johnston, Dominic Kwiatkowski, Cordelia Langford, John Sillitoe on behalf of the Wellcome Sanger Institute COVID-19 Surveillance Team |
| EPI_ISL_821481 | Lighthouse Lab in Milton Keynes | Wellcome Sanger Institute for the COVID-19 Genomics UK (COG-UK) Consortium | The Lighthouse Lab in Milton Keynes and Alex Alderton, Roberto Amato, Sonia Goncalves, Ewan Harrison, David K. Jackson, Ian Johnston, Dominic Kwiatkowski, Cordelia Langford, John Sillitoe on behalf of the Wellcome Sanger Institute COVID-19 Surveillance Team |
| EPI_ISL_821485 | Lighthouse Lab in Cambridge | Wellcome Sanger Institute for the COVID-19 Genomics UK | Rob Howes, The Lighthouse Lab in Cambridge and Alex Alderton, Roberto Amato, Sonia Goncalves, Ewan Harrison, David K. Jackson, Ian Johnston, |

| | | | |
|---|---|---|---|
| | | (COG-UK) Consortium | Dominic Kwiatkowski, Cordelia Langford, John Sillitoe on behalf of the Wellcome Sanger Institute COVID-19 Surveillance Team |
| EPI_ISL_821489, EPI_ISL_821495, EPI_ISL_821499, EPI_ISL_821505, EPI_ISL_821508, EPI_ISL_821512 | Lighthouse Lab in Milton Keynes | Wellcome Sanger Institute for the COVID-19 Genomics UK (COG-UK) Consortium | The Lighthouse Lab in Milton Keynes and Alex Alderton, Roberto Amato, Sonia Goncalves, Ewan Harrison, David K. Jackson, Ian Johnston, Dominic Kwiatkowski, Cordelia Langford, John Sillitoe on behalf of the Wellcome Sanger Institute COVID-19 Surveillance Team |
| EPI_ISL_821513 | Lighthouse Lab in Cambridge | Wellcome Sanger Institute for the COVID-19 Genomics UK (COG-UK) Consortium | Rob Howes, The Lighthouse Lab in Cambridge and Alex Alderton, Roberto Amato, Sonia Goncalves, Ewan Harrison, David K. Jackson, Ian Johnston, Dominic Kwiatkowski, Cordelia Langford, John Sillitoe on behalf of the Wellcome Sanger Institute COVID-19 Surveillance Team |
| EPI_ISL_821525 | Lighthouse Lab in Alderley Park | Wellcome Sanger Institute for the COVID-19 Genomics UK (COG-UK) Consortium | Jacquelyn Wynn, Mairead Hyland, The Lighthouse Lab in Alderley Park and Alex Alderton, Roberto Amato, Sonia Goncalves, Ewan Harrison, David K. Jackson, Ian Johnston, Dominic Kwiatkowski, Cordelia Langford, John Sillitoe on behalf of the Wellcome Sanger Institute COVID-19 Surveillance Team |
| EPI_ISL_821531 | Lighthouse Lab in Milton Keynes | Wellcome Sanger Institute for the COVID-19 Genomics UK (COG-UK) Consortium | The Lighthouse Lab in Milton Keynes and Alex Alderton, Roberto Amato, Sonia Goncalves, Ewan Harrison, David K. Jackson, Ian Johnston, Dominic Kwiatkowski, Cordelia Langford, John Sillitoe on behalf of the Wellcome Sanger Institute COVID-19 Surveillance Team |
| EPI_ISL_821546 | Lighthouse Lab in Cambridge | Wellcome Sanger Institute for the COVID-19 Genomics UK (COG-UK) Consortium | Rob Howes, The Lighthouse Lab in Cambridge and Alex Alderton, Roberto Amato, Sonia Goncalves, Ewan Harrison, David K. Jackson, Ian Johnston, Dominic Kwiatkowski, Cordelia Langford, John Sillitoe on behalf of the Wellcome Sanger Institute COVID-19 Surveillance Team |
| EPI_ISL_821557, EPI_ISL_821578, EPI_ISL_821581, EPI_ISL_821591 | Lighthouse Lab in Milton Keynes | Wellcome Sanger Institute for the COVID-19 Genomics UK (COG-UK) Consortium | The Lighthouse Lab in Milton Keynes and Alex Alderton, Roberto Amato, Sonia Goncalves, Ewan Harrison, David K. Jackson, Ian Johnston, Dominic Kwiatkowski, Cordelia Langford, John Sillitoe on behalf of the Wellcome Sanger Institute COVID-19 Surveillance Team |
| EPI_ISL_821602 | Lighthouse Lab in Cambridge | Wellcome Sanger Institute for the COVID-19 Genomics UK (COG-UK) Consortium | Rob Howes, The Lighthouse Lab in Cambridge and Alex Alderton, Roberto Amato, Sonia Goncalves, Ewan Harrison, David K. Jackson, Ian Johnston, Dominic Kwiatkowski, Cordelia Langford, John Sillitoe on behalf of the Wellcome Sanger Institute COVID-19 Surveillance Team |
| EPI_ISL_821611 | Lighthouse Lab in Milton Keynes | Wellcome Sanger Institute for the COVID-19 Genomics UK (COG-UK) Consortium | The Lighthouse Lab in Milton Keynes and Alex Alderton, Roberto Amato, Sonia Goncalves, Ewan Harrison, David K. Jackson, Ian Johnston, Dominic Kwiatkowski, Cordelia Langford, John Sillitoe on behalf of the Wellcome Sanger Institute COVID-19 Surveillance Team |
| EPI_ISL_821613 | Lighthouse Lab in Cambridge | Wellcome Sanger Institute for the COVID-19 Genomics UK (COG-UK) Consortium | Rob Howes, The Lighthouse Lab in Cambridge and Alex Alderton, Roberto Amato, Sonia Goncalves, Ewan Harrison, David K. Jackson, Ian Johnston, Dominic Kwiatkowski, Cordelia Langford, John Sillitoe on behalf of the Wellcome Sanger Institute COVID-19 Surveillance Team |
| EPI_ISL_821617 | Lighthouse Lab in Alderley Park | Wellcome Sanger Institute for the COVID-19 Genomics UK (COG-UK) Consortium | Jacquelyn Wynn, Mairead Hyland, The Lighthouse Lab in Alderley Park and Alex Alderton, Roberto Amato, Sonia Goncalves, Ewan Harrison, David K. Jackson, Ian Johnston, Dominic Kwiatkowski, Cordelia Langford, John Sillitoe on behalf of the Wellcome Sanger Institute COVID-19 Surveillance Team |
| EPI_ISL_821619 | Lighthouse Lab in Milton Keynes | Wellcome Sanger Institute for the COVID-19 Genomics UK (COG-UK) Consortium | The Lighthouse Lab in Milton Keynes and Alex Alderton, Roberto Amato, Sonia Goncalves, Ewan Harrison, David K. Jackson, Ian Johnston, Dominic Kwiatkowski, Cordelia Langford, John Sillitoe on behalf of the Wellcome Sanger Institute COVID-19 Surveillance Team |
| EPI_ISL_821626, EPI_ISL_821627, EPI_ISL_821628 | Lighthouse Lab in Alderley Park | Wellcome Sanger Institute for the COVID-19 Genomics UK (COG-UK) Consortium | Jacquelyn Wynn, Mairead Hyland, The Lighthouse Lab in Alderley Park and Alex Alderton, Roberto Amato, Sonia Goncalves, Ewan Harrison, David K. Jackson, Ian Johnston, Dominic Kwiatkowski, Cordelia Langford, John Sillitoe on behalf of the Wellcome Sanger Institute COVID-19 Surveillance Team |
| EPI_ISL_821632, EPI_ISL_821638, EPI_ISL_821639, EPI_ISL_821640, EPI_ISL_821642, EPI_ISL_821643, EPI_ISL_821644, EPI_ISL_821645, EPI_ISL_821646, EPI_ISL_821647, EPI_ISL_821649, EPI_ISL_821650, EPI_ISL_821651, EPI_ISL_821652, EPI_ISL_821654, EPI_ISL_821655, EPI_ISL_821656, EPI_ISL_821657, EPI_ISL_821658 | | | |
| see above | Lighthouse Lab in Milton Keynes | Wellcome Sanger Institute for the COVID-19 Genomics UK (COG-UK) Consortium | The Lighthouse Lab in Milton Keynes and Alex Alderton, Roberto Amato, Sonia Goncalves, Ewan Harrison, David K. Jackson, Ian Johnston, Dominic Kwiatkowski, Cordelia Langford, John Sillitoe on behalf of the Wellcome Sanger Institute COVID-19 Surveillance Team |
| EPI_ISL_821659, EPI_ISL_821660 | Lighthouse Lab in Glasgow | Wellcome Sanger Institute for the COVID-19 Genomics UK (COG-UK) Consortium | Harper VanSteenhouse, Yumi Kasai, David Gray, Carol Clugston, Anna Dominiczak and Alex Alderton, Roberto Amato, Sonia Goncalves, Ewan Harrison, David K. Jackson, Ian Johnston, Dominic Kwiatkowski, Cordelia Langford, John Sillitoe on behalf of the Wellcome Sanger Institute COVID-19 Surveillance Team |
| EPI_ISL_821661, EPI_ISL_821662, EPI_ISL_821663, EPI_ISL_821664, EPI_ISL_821665, EPI_ISL_821666, EPI_ISL_821668, EPI_ISL_821670, EPI_ISL_821671, EPI_ISL_821673, EPI_ISL_821674, EPI_ISL_821675, EPI_ISL_821676, EPI_ISL_821678, EPI_ISL_821679, EPI_ISL_821680, EPI_ISL_821681, EPI_ISL_821682, EPI_ISL_821683, EPI_ISL_821684, EPI_ISL_821685, EPI_ISL_821686, EPI_ISL_821687, EPI_ISL_821688, EPI_ISL_821689, EPI_ISL_821690, EPI_ISL_821691, EPI_ISL_821692, EPI_ISL_821693, EPI_ISL_821694, EPI_ISL_821695, EPI_ISL_821696, EPI_ISL_821697, EPI_ISL_821698, EPI_ISL_821699, EPI_ISL_821700, EPI_ISL_821701, EPI_ISL_821702, EPI_ISL_821703, EPI_ISL_821704, EPI_ISL_821705, EPI_ISL_821706, EPI_ISL_821707, EPI_ISL_821708, EPI_ISL_821709, EPI_ISL_821710, EPI_ISL_821711, EPI_ISL_821712, EPI_ISL_821713, EPI_ISL_821714, EPI_ISL_821715, EPI_ISL_821716, EPI_ISL_821717, EPI_ISL_821718, EPI_ISL_821719, EPI_ISL_821720, EPI_ISL_821721, EPI_ISL_821722, EPI_ISL_821723, EPI_ISL_821724, EPI_ISL_821725, EPI_ISL_821726, EPI_ISL_821727, EPI_ISL_821728, EPI_ISL_821729, EPI_ISL_821730, EPI_ISL_821731, EPI_ISL_821732, EPI_ISL_821734, EPI_ISL_821735, EPI_ISL_821736, EPI_ISL_821737, EPI_ISL_821738, EPI_ISL_821739, EPI_ISL_821740, EPI_ISL_821741, EPI_ISL_821742, EPI_ISL_821743, EPI_ISL_821744, EPI_ISL_821745, EPI_ISL_821746, EPI_ISL_821747, EPI_ISL_821748, EPI_ISL_821749, EPI_ISL_821750, EPI_ISL_821751, EPI_ISL_821752, EPI_ISL_821753, EPI_ISL_821754, EPI_ISL_821755, EPI_ISL_821756, EPI_ISL_821757, EPI_ISL_821758, EPI_ISL_821759, EPI_ISL_821760, EPI_ISL_821761, EPI_ISL_821762, EPI_ISL_821763, EPI_ISL_821764, EPI_ISL_821765, EPI_ISL_821766, EPI_ISL_821767, EPI_ISL_821768, EPI_ISL_821769, EPI_ISL_821770, EPI_ISL_821771, EPI_ISL_821772, EPI_ISL_821773, EPI_ISL_821774, EPI_ISL_821775, EPI_ISL_821776, EPI_ISL_821777, EPI_ISL_821778, EPI_ISL_821779, EPI_ISL_821780, EPI_ISL_821781, EPI_ISL_821782, EPI_ISL_821783, EPI_ISL_821784, EPI_ISL_821785, EPI_ISL_821786, EPI_ISL_821788, EPI_ISL_821789, EPI_ISL_821791, EPI_ISL_821792, EPI_ISL_821793, EPI_ISL_821794, EPI_ISL_821795, EPI_ISL_821796, EPI_ISL_821797, EPI_ISL_821798, EPI_ISL_821799, EPI_ISL_821800, EPI_ISL_821801, EPI_ISL_821802, EPI_ISL_821803, EPI_ISL_821804, EPI_ISL_821805, EPI_ISL_821806, EPI_ISL_821807, EPI_ISL_821808, EPI_ISL_821809, EPI_ISL_821810, EPI_ISL_821811, EPI_ISL_821812, EPI_ISL_821813, EPI_ISL_821814, EPI_ISL_821815, EPI_ISL_821816, EPI_ISL_821817, EPI_ISL_821818, EPI_ISL_821819, EPI_ISL_821820, EPI_ISL_821821, EPI_ISL_821822, EPI_ISL_821823, EPI_ISL_821824, EPI_ISL_821825, EPI_ISL_821826, EPI_ISL_821828, EPI_ISL_821829, EPI_ISL_821830, EPI_ISL_821831, EPI_ISL_821832, EPI_ISL_821833, EPI_ISL_821835, EPI_ISL_821836, EPI_ISL_821837, EPI_ISL_821838, EPI_ISL_821839, EPI_ISL_821840, EPI_ISL_821841, EPI_ISL_821843, EPI_ISL_821844, EPI_ISL_821845, EPI_ISL_821846, EPI_ISL_821847, EPI_ISL_821848, EPI_ISL_821849, EPI_ISL_821850, EPI_ISL_821851, EPI_ISL_821852, EPI_ISL_821853, EPI_ISL_821854, EPI_ISL_821855, EPI_ISL_821856, EPI_ISL_821857, EPI_ISL_821859, EPI_ISL_821860, EPI_ISL_821862, EPI_ISL_821863, EPI_ISL_821864, EPI_ISL_821865, EPI_ISL_821866, EPI_ISL_821867, EPI_ISL_821868, EPI_ISL_821869, EPI_ISL_821870, EPI_ISL_821871, EPI_ISL_821872, EPI_ISL_821873, EPI_ISL_821874, EPI_ISL_821875, EPI_ISL_821876, EPI_ISL_821878, EPI_ISL_821879, EPI_ISL_821880, EPI_ISL_821881, EPI_ISL_821882, EPI_ISL_821883, EPI_ISL_821884, EPI_ISL_821885, EPI_ISL_821886, EPI_ISL_821887, EPI_ISL_821888, EPI_ISL_821890, EPI_ISL_821891, EPI_ISL_821892, EPI_ISL_821893, EPI_ISL_821894, EPI_ISL_821896, EPI_ISL_821897, EPI_ISL_821898, EPI_ISL_821899, EPI_ISL_821900, EPI_ISL_821901, EPI_ISL_821902, EPI_ISL_821904, EPI_ISL_821905, EPI_ISL_821906, EPI_ISL_821907, EPI_ISL_821908, EPI_ISL_821909, EPI_ISL_821910, EPI_ISL_821911, EPI_ISL_821912, EPI_ISL_821913, EPI_ISL_821914, EPI_ISL_821915, EPI_ISL_821916, EPI_ISL_821917, EPI_ISL_821918, EPI_ISL_821919, EPI_ISL_821920, EPI_ISL_821921, EPI_ISL_821922, EPI_ISL_821923, EPI_ISL_821924, EPI_ISL_821925, EPI_ISL_821926, EPI_ISL_821927, EPI_ISL_821928, EPI_ISL_821929, EPI_ISL_821930, EPI_ISL_821931, EPI_ISL_821932, EPI_ISL_821933, EPI_ISL_821935, EPI_ISL_821936, EPI_ISL_821937, EPI_ISL_821938, EPI_ISL_821939, EPI_ISL_821940, EPI_ISL_821941, EPI_ISL_821942, EPI_ISL_821944, EPI_ISL_821945, EPI_ISL_821946, EPI_ISL_821947, EPI_ISL_821948, EPI_ISL_821949, EPI_ISL_821951, EPI_ISL_821952, EPI_ISL_821953, EPI_ISL_821954, EPI_ISL_821955, EPI_ISL_821956, EPI_ISL_821957, EPI_ISL_821958, EPI_ISL_821960, EPI_ISL_821961, EPI_ISL_821962, EPI_ISL_821963, EPI_ISL_821964, EPI_ISL_821965, EPI_ISL_821966, EPI_ISL_821968, EPI_ISL_821969, EPI_ISL_821970 | | | |
| see above | Lighthouse Lab in Cambridge | Wellcome Sanger Institute for the COVID-19 Genomics UK (COG-UK) Consortium | Rob Howes, The Lighthouse Lab in Cambridge and Alex Alderton, Roberto Amato, Sonia Goncalves, Ewan Harrison, David K. Jackson, Ian Johnston, Dominic Kwiatkowski, Cordelia Langford, John Sillitoe on behalf of the Wellcome Sanger Institute COVID-19 Surveillance Team |
| EPI_ISL_821971, EPI_ISL_821974, EPI_ISL_821976, EPI_ISL_821977, EPI_ISL_821978, EPI_ISL_821979, EPI_ISL_821980, EPI_ISL_821981, EPI_ISL_821982, EPI_ISL_821983, EPI_ISL_821984, EPI_ISL_821985, EPI_ISL_821986, EPI_ISL_821989, EPI_ISL_821990, EPI_ISL_821994, EPI_ISL_821998, EPI_ISL_821999, EPI_ISL_822000, EPI_ISL_822002, EPI_ISL_822003, EPI_ISL_822007, EPI_ISL_822008, EPI_ISL_822009, EPI_ISL_822010, EPI_ISL_822012, EPI_ISL_822014, EPI_ISL_822015, EPI_ISL_822016, EPI_ISL_822017, EPI_ISL_822018, EPI_ISL_822020, EPI_ISL_822021, EPI_ISL_822024, EPI_ISL_822025, EPI_ISL_822026, EPI_ISL_822027, EPI_ISL_822028, EPI_ISL_822031, EPI_ISL_822032, EPI_ISL_822034, EPI_ISL_822036, EPI_ISL_822037, EPI_ISL_822038, EPI_ISL_822039, EPI_ISL_822042, EPI_ISL_822045, EPI_ISL_822046, EPI_ISL_822049, EPI_ISL_822050, EPI_ISL_822052, EPI_ISL_822054, EPI_ISL_822055, EPI_ISL_822056, EPI_ISL_822057, EPI_ISL_822058, EPI_ISL_822061, EPI_ISL_822062, EPI_ISL_822064, EPI_ISL_822066, EPI_ISL_822068, EPI_ISL_822070, EPI_ISL_822071, EPI_ISL_822073, EPI_ISL_822074, EPI_ISL_822076, EPI_ISL_822077, EPI_ISL_822079, EPI_ISL_822082, EPI_ISL_822084, EPI_ISL_822085, EPI_ISL_822087, EPI_ISL_822088, EPI_ISL_822089, EPI_ISL_822090, EPI_ISL_822091, EPI_ISL_822092, EPI_ISL_822095, EPI_ISL_822096, EPI_ISL_822100, EPI_ISL_822101, EPI_ISL_822102, EPI_ISL_822103, EPI_ISL_822106, EPI_ISL_822107, EPI_ISL_822108, EPI_ISL_822109, EPI_ISL_822110, EPI_ISL_822111, EPI_ISL_822112, EPI_ISL_822114, EPI_ISL_822115, EPI_ISL_822116, EPI_ISL_822118, EPI_ISL_822120, EPI_ISL_822121, EPI_ISL_822123, EPI_ISL_822124, EPI_ISL_822125, EPI_ISL_822126, EPI_ISL_822128, EPI_ISL_822129, EPI_ISL_822131, EPI_ISL_822132, EPI_ISL_822133, EPI_ISL_822134, EPI_ISL_822136, EPI_ISL_822137, EPI_ISL_822138, EPI_ISL_822143, EPI_ISL_822144, EPI_ISL_822145, EPI_ISL_822146, EPI_ISL_822147, EPI_ISL_822149, EPI_ISL_822151, EPI_ISL_822152, EPI_ISL_822154, EPI_ISL_822155, EPI_ISL_822157, EPI_ISL_822160, EPI_ISL_822161, EPI_ISL_822162, EPI_ISL_822163, EPI_ISL_822164, EPI_ISL_822166, EPI_ISL_822167, EPI_ISL_822168, EPI_ISL_822171, EPI_ISL_822172, EPI_ISL_822174, EPI_ISL_822175, EPI_ISL_822176, EPI_ISL_822177, EPI_ISL_822179, EPI_ISL_822180, EPI_ISL_822181, EPI_ISL_822185, EPI_ISL_822186, EPI_ISL_822187, EPI_ISL_822188, EPI_ISL_822189, EPI_ISL_822190, EPI_ISL_822192, EPI_ISL_822194, EPI_ISL_822195, EPI_ISL_822196, EPI_ISL_822198, EPI_ISL_822200, EPI_ISL_822203, EPI_ISL_822204, EPI_ISL_822205, EPI_ISL_822206, EPI_ISL_822207, EPI_ISL_822208, EPI_ISL_822210, EPI_ISL_822211, EPI_ISL_822212, EPI_ISL_822213, EPI_ISL_822215, EPI_ISL_822216, EPI_ISL_822217, EPI_ISL_822218, EPI_ISL_822219, EPI_ISL_822220, EPI_ISL_822221, EPI_ISL_822222, EPI_ISL_822225, EPI_ISL_822226, EPI_ISL_822227, EPI_ISL_822228, EPI_ISL_822229, EPI_ISL_822231, EPI_ISL_822232, EPI_ISL_822233, EPI_ISL_822234, EPI_ISL_822235, EPI_ISL_822236, EPI_ISL_822238, EPI_ISL_822239, EPI_ISL_822241, EPI_ISL_822246, EPI_ISL_822251, EPI_ISL_822252, EPI_ISL_822254, EPI_ISL_822255, EPI_ISL_822257, EPI_ISL_822258, EPI_ISL_822259, EPI_ISL_822261, EPI_ISL_822262, EPI_ISL_822263, EPI_ISL_822266, EPI_ISL_822269, EPI_ISL_822270, EPI_ISL_822271, EPI_ISL_822273, EPI_ISL_822274, EPI_ISL_822276, EPI_ISL_822277, EPI_ISL_822278, EPI_ISL_822279, EPI_ISL_822280, EPI_ISL_822284, EPI_ISL_822285, EPI_ISL_822286, EPI_ISL_822287, EPI_ISL_822289, EPI_ISL_822290 | | | |
| see above | Lighthouse Lab in Alderley Park | Wellcome Sanger Institute for the COVID-19 Genomics UK (COG-UK) Consortium | Jacquelyn Wynn, Mairead Hyland, The Lighthouse Lab in Alderley Park and Alex Alderton, Roberto Amato, Sonia Goncalves, Ewan Harrison, David K. Jackson, Ian Johnston, Dominic Kwiatkowski, Cordelia Langford, John Sillitoe on behalf of the Wellcome Sanger Institute COVID-19 Surveillance Team |
| EPI_ISL_822292, EPI_ISL_822293, EPI_ISL_822295, EPI_ISL_822296, EPI_ISL_822302, EPI_ISL_822304 | Lighthouse Lab in Glasgow | Wellcome Sanger Institute for the COVID-19 Genomics UK (COG-UK) Consortium | Harper VanSteenhouse, Yumi Kasai, David Gray, Carol Clugston, Anna Dominiczak and Alex Alderton, Roberto Amato, Sonia Goncalves, Ewan Harrison, David K. Jackson, Ian Johnston, Dominic Kwiatkowski, Cordelia Langford, John Sillitoe on behalf of the Wellcome Sanger Institute COVID-19 Surveillance Team |
| EPI_ISL_822659, EPI_ISL_822660, EPI_ISL_822661, EPI_ISL_822662, EPI_ISL_822663, EPI_ISL_822664, EPI_ISL_822665, EPI_ISL_822666, EPI_ISL_822667, EPI_ISL_822668, EPI_ISL_822669, EPI_ISL_822670, EPI_ISL_822671, EPI_ISL_822672, EPI_ISL_822673, EPI_ISL_822674, EPI_ISL_822675, EPI_ISL_822676, EPI_ISL_822677, EPI_ISL_822678, EPI_ISL_822679, EPI_ISL_822680, EPI_ISL_822681, EPI_ISL_822682, EPI_ISL_822683, EPI_ISL_822684, EPI_ISL_822685, EPI_ISL_822686, EPI_ISL_822687, EPI_ISL_822688, EPI_ISL_822689, EPI_ISL_822690, EPI_ISL_822691, EPI_ISL_822692, EPI_ISL_822693, EPI_ISL_822694, EPI_ISL_822695, EPI_ISL_822696, EPI_ISL_822697, EPI_ISL_822698, EPI_ISL_822699, EPI_ISL_822700, EPI_ISL_822701, EPI_ISL_822702, EPI_ISL_822703, EPI_ISL_822704, EPI_ISL_822705, EPI_ISL_822706, EPI_ISL_822707, EPI_ISL_822708, EPI_ISL_822709, EPI_ISL_822710, EPI_ISL_822711, EPI_ISL_822712, EPI_ISL_822713, EPI_ISL_822714, EPI_ISL_822715, EPI_ISL_822716, EPI_ISL_822717, EPI_ISL_822718, EPI_ISL_822719, EPI_ISL_822720, EPI_ISL_822721, EPI_ISL_822722, EPI_ISL_822723, EPI_ISL_822724, EPI_ISL_822725, EPI_ISL_822726, EPI_ISL_822727, EPI_ISL_822728, EPI_ISL_822729, EPI_ISL_822730, EPI_ISL_822731, EPI_ISL_822732, EPI_ISL_822733, EPI_ISL_822734, EPI_ISL_822735, EPI_ISL_822736, EPI_ISL_822737, EPI_ISL_822738, EPI_ISL_822739, EPI_ISL_822740, EPI_ISL_822741, EPI_ISL_822742, EPI_ISL_822743, EPI_ISL_822744, EPI_ISL_822745, EPI_ISL_822746, EPI_ISL_822747, EPI_ISL_822748, EPI_ISL_822749, EPI_ISL_822750, EPI_ISL_822751, EPI_ISL_822752, EPI_ISL_822753, EPI_ISL_822754, EPI_ISL_822755, EPI_ISL_822756, EPI_ISL_822757, EPI_ISL_822758, EPI_ISL_822759, EPI_ISL_822760, EPI_ISL_822761, EPI_ISL_822762, EPI_ISL_822763, EPI_ISL_822764, EPI_ISL_822765, EPI_ISL_822766, EPI_ISL_822767, EPI_ISL_822768, EPI_ISL_822769, EPI_ISL_822770, EPI_ISL_822771, EPI_ISL_822772, EPI_ISL_822773, EPI_ISL_822774, EPI_ISL_822775, EPI_ISL_822776, EPI_ISL_822777, EPI_ISL_822778, EPI_ISL_822779, EPI_ISL_822780, EPI_ISL_822781, EPI_ISL_822782, EPI_ISL_822783, EPI_ISL_822784, EPI_ISL_822785, EPI_ISL_822786, EPI_ISL_822787, EPI_ISL_822788, EPI_ISL_822789, EPI_ISL_822790, EPI_ISL_822791, EPI_ISL_822792, EPI_ISL_822793, EPI_ISL_822794, EPI_ISL_822795, EPI_ISL_822796, EPI_ISL_822797, EPI_ISL_822798, EPI_ISL_822799, EPI_ISL_822800, EPI_ISL_822801, EPI_ISL_822802, | | | |

| | | | |
|---|---|---|---|
| EPI_ISL_822803, EPI_ISL_822804, EPI_ISL_822805, EPI_ISL_822844, EPI_ISL_822880, EPI_ISL_822925, EPI_ISL_822940, EPI_ISL_822947, EPI_ISL_822951, EPI_ISL_822952, EPI_ISL_822955, EPI_ISL_822957, EPI_ISL_822971, EPI_ISL_823003, EPI_ISL_823066, EPI_ISL_823098, EPI_ISL_823099, EPI_ISL_823100, EPI_ISL_823101, EPI_ISL_823102, EPI_ISL_823103, EPI_ISL_823112, EPI_ISL_823149, EPI_ISL_823152, EPI_ISL_823154, EPI_ISL_823155, EPI_ISL_823646, EPI_ISL_823647, EPI_ISL_823648, EPI_ISL_823649, EPI_ISL_823780, EPI_ISL_823781, EPI_ISL_823782, EPI_ISL_823783, EPI_ISL_823784, EPI_ISL_823785, EPI_ISL_823786, EPI_ISL_823787, EPI_ISL_823788, EPI_ISL_823789, EPI_ISL_823790, EPI_ISL_823791, EPI_ISL_823792, EPI_ISL_823793, EPI_ISL_823794, EPI_ISL_823795 | | | |
| see above | Wales Specialist Virology Centre Sequencing lab: Pathogen Genomics Unit | COVID-19 Genomics UK (COG-UK) Consortium | Catherine Moore, Johnathan Evans, Laura Gifford, Malorie Perry, Simon Cottrell, Angela Marchbank, Alec Birchley, Alexander Adams, Amy Gaskin, Bree Gatica-Wilcox, Jason Coombes, Joel Southgate, Lauren Gilbert, Lee Graham, Nicole Pacchiarini, Sara Kumziene-Summerhayes, Sarah Taylor, Sophie Jones, Sara Rey, Matthew Bull, Joanne Watkins, Sally Corden, Tom Connor |
| EPI_ISL_823976, EPI_ISL_823981, EPI_ISL_823983, EPI_ISL_823984, EPI_ISL_823985, EPI_ISL_823986, EPI_ISL_823987, EPI_ISL_823988, EPI_ISL_823990, EPI_ISL_823991, EPI_ISL_823994, EPI_ISL_823995, EPI_ISL_823996, EPI_ISL_824020, EPI_ISL_824025, EPI_ISL_824026, EPI_ISL_824038, EPI_ISL_824039, EPI_ISL_824040, EPI_ISL_824047, EPI_ISL_824048, EPI_ISL_824051, EPI_ISL_824052, EPI_ISL_824068, EPI_ISL_824071, EPI_ISL_824088, EPI_ISL_824109, EPI_ISL_824111, EPI_ISL_824112, EPI_ISL_824113, EPI_ISL_824114, EPI_ISL_824116, EPI_ISL_824120, EPI_ISL_824123, EPI_ISL_824164, EPI_ISL_824165, EPI_ISL_824166, EPI_ISL_824167, EPI_ISL_824168, EPI_ISL_824181, EPI_ISL_824182, EPI_ISL_824199, EPI_ISL_824200, EPI_ISL_824201, EPI_ISL_824202, EPI_ISL_824203, EPI_ISL_824204, EPI_ISL_824207, EPI_ISL_824208, EPI_ISL_824214, EPI_ISL_824215, EPI_ISL_824216, EPI_ISL_824217, EPI_ISL_824242, EPI_ISL_824243, EPI_ISL_824244, EPI_ISL_824245, EPI_ISL_824246, EPI_ISL_824247, EPI_ISL_824249, EPI_ISL_824250, EPI_ISL_824251, EPI_ISL_824252, EPI_ISL_824259, EPI_ISL_824260, EPI_ISL_824266, EPI_ISL_824272, EPI_ISL_824273, EPI_ISL_824276, EPI_ISL_824278 | | | |
| see above | Dutch COVID-19 response team | National Institute for Public Health and the Environment (RIVM) | Adam Meijer, Harry Vennema, Jeroen Cremer, Sharon van den Brink, Bas van der Veer, AnneMarie van den Brandt, Florian Zwagemaker, Dennis Schmitz, Chantal Reusken, on behalf of the national COVID-19 response team |
| EPI_ISL_824290, EPI_ISL_824291 | Institute of Microbiology, Universidad San Francisco de Quito | Institute of Microbiology, Universidad San Francisco de Quito | Belén Prado-Vivar, Sully Márquez, Juan José Guadalupe, Monica Becerra-Wong, Bernardo Gutiérrez, Andrea Cunguan, Nabih Dahik, Verónica Barragán, Patricio Rojas-Silva, Gabriel Trueba, Michelle Grunauer, Paúl Cárdenas |
| EPI_ISL_824342 | Michigan Department of Health and Human Services, Bureau of Laboratories | Michigan Department of Health and Human Services, Bureau of Laboratories | Blankenship HM, Riner D, Soehnlen MK |
| EPI_ISL_824524, EPI_ISL_824525, EPI_ISL_824526, EPI_ISL_824527, EPI_ISL_824528, EPI_ISL_824529, EPI_ISL_824530, EPI_ISL_824531, EPI_ISL_824532, EPI_ISL_824533, EPI_ISL_824534, EPI_ISL_824535, EPI_ISL_824538, EPI_ISL_824539, EPI_ISL_824540, EPI_ISL_824541, EPI_ISL_824542, EPI_ISL_824543, EPI_ISL_824544, EPI_ISL_824545, EPI_ISL_824546, EPI_ISL_824547, EPI_ISL_824548, EPI_ISL_824549, EPI_ISL_824550, EPI_ISL_824551, EPI_ISL_824552, EPI_ISL_824553 | | | |
| see above | New Mexico Department of Health Scientific Laboratory | New Mexico Department of Health Scientific Laboratory | Ellie Johnson, Anastacia Griego-Fisher, D'eldra Malone |
| EPI_ISL_824790, EPI_ISL_824791, EPI_ISL_824792, EPI_ISL_824793, EPI_ISL_824794, EPI_ISL_824795, EPI_ISL_824796, EPI_ISL_824797, EPI_ISL_824798, EPI_ISL_824809, EPI_ISL_824810, EPI_ISL_824811, EPI_ISL_824812, EPI_ISL_824813, EPI_ISL_824814, EPI_ISL_824815, EPI_ISL_824816, EPI_ISL_824817, EPI_ISL_824818, EPI_ISL_824819, EPI_ISL_824820, EPI_ISL_824821, EPI_ISL_824822, EPI_ISL_824823, EPI_ISL_824824, EPI_ISL_824825, EPI_ISL_824826, EPI_ISL_824827, EPI_ISL_824828, EPI_ISL_824829, EPI_ISL_824830, EPI_ISL_824831, EPI_ISL_824834 | | | |
| see above | Department of Clinical Microbiology | GIGA Medical Genomics | Keith Durkin, Maria Artesi, Sébastien Bontems, Raphaël Boreux, Bouchra Boujemla, Cécile Meex, Pierrette Melin, Marie-Pierre Hayette, Vincent Bours |
| EPI_ISL_824921, EPI_ISL_824944 | Charité Universitätsmedizin Berlin, Institute of Virology, Charitéplatz 1, 10117 Berlin, Germany | Charité Universitätsmedizin Berlin, Institute of Virology, Charitéplatz 1, 10117 Berlin, Germany | Victor M Corman, Jörn Beheim-Schwarzbach, Tobias Bleicker, Julia Tesch, Barbara Mühlemann, Talitha Veith, Julia Schneider, Terry Jones, Christian Drosten |
| EPI_ISL_824945, EPI_ISL_824946, EPI_ISL_824947, EPI_ISL_824948, EPI_ISL_824949, EPI_ISL_824950, EPI_ISL_824951, EPI_ISL_824952 | Maryland Public Health Laboratory | Maryland Public Health Laboratory | Maryland Department of Health Laboratories Administration |
| EPI_ISL_824987 | Arizona State Public Health Laboratory | Arizona State Public Health Laboratory | Trung Huynh, Jessica Escobar, Katherine Fullerton, Nobuko Fukushima, Stacy White, Linda Getsinger, Victor Waddell |
| EPI_ISL_825014 | Northwestern Memorial Hospital | Ozer Lab | Ramon Lorenzo-Redondo, Lacy M. Simons, Lawrence J. Jennings, Michael G. Ison, Judd F. Hultquist, Egon A. Ozer |
| EPI_ISL_825061 | GMERS Medical College and Hospital, Gotri, Vadodara | Gujarat Biotechnology Research Centre | Apurvasinh Puvar, Ramesh Pandit, Janvi Raval, Zarna Patel, Nitin Savaliya, Dinesh Kumar, Zuber Saiyed, Afzal Ansari, Nikha Trivedi, Mitesh Kamothi, Bithika Duttaroy, Kalpesh Mistry, Chaitanya Joshi, Madhvi Joshi |
| EPI_ISL_825062 | GMERS Medical College and Hospital, Gotri, Vadodara | Gujarat Biotechnology Research Centre | Ramesh Pandit, Janvi Raval, Zarna Patel, Nitin Savaliya, Dinesh Kumar, Zuber Saiyed, Afzal Ansari, Nikha Trivedi, Apurvasinh Puvar, Mitesh Kamothi, Bithika Duttaroy, Kalpesh Mistry, Chaitanya Joshi, Madhvi Joshi |
| EPI_ISL_825063 | NHL Municipal Medical College, Ahmedbad | Gujarat Biotechnology Research Centre | Janvi Raval, Zarna Patel, Nitin Savaliya, Dinesh Kumar, Zuber Saiyed, Afzal Ansari, Nikha Trivedi, Apurvasinh Puvar, Ramesh Pandit, Jayshri Pethani, Monila Patel, Atit Shah, NM Shaikh, Bimal Chauhan, Tanmay Mehta, Bhavin Prajapati, Chaitanya Joshi, Madhvi Joshi |
| EPI_ISL_825156 | Microbiology and Virology Unit, Florence Careggi University Hospital | Microbiology and Virology Unit, Florence Careggi University Hospital | Vincenzo Di Pilato, Marco Coppi, Fabio Morecchiato, Alberto Antonelli, Emanuele Gori, Gian Maria Rossolini |
| EPI_ISL_825165, EPI_ISL_825169, EPI_ISL_825170, EPI_ISL_825171, EPI_ISL_825172 | Robert Koch Institute, ZBS1 Highly Pathogenic Viruses, Berlin, Germany | Robert Koch Institute, ZBS1 Highly Pathogenic Viruses & Bioinformatics MF1, Berlin, Germany | Annika Brinkmann, Janine Michel, Livia Schrick, Steven Uddin, Dominique Seifert, Alexander Dalpke, Leo Büttner, Kristina Hochauf-Stange, Dirk Lindemann, Lars Schaade, Andreas Nitsche |
| EPI_ISL_825499, EPI_ISL_825500, EPI_ISL_825534, EPI_ISL_825535, EPI_ISL_825536, EPI_ISL_825537, EPI_ISL_825538, EPI_ISL_825539, EPI_ISL_825540, EPI_ISL_825541, EPI_ISL_825542, EPI_ISL_825543, EPI_ISL_825544, EPI_ISL_825545, EPI_ISL_825568, EPI_ISL_825569, EPI_ISL_825570, EPI_ISL_825610, EPI_ISL_825611, EPI_ISL_825612, EPI_ISL_825613, EPI_ISL_825614, EPI_ISL_825615 | | | |
| see above | Respiratory Virus Unit, National Infection Service, Public Health England | COVID-19 Genomics UK (COG-UK) Consortium | PHE Covid Sequencing Team |
| EPI_ISL_825626, EPI_ISL_825627, EPI_ISL_825628, EPI_ISL_825629, EPI_ISL_825630, EPI_ISL_825631 | Hospital Universitari Vall d'Hebron - Vall d'Hebron Institut de Rercerca | Hospital Universitari Vall d'Hebron | Cristina Andrés, Maria Piñana, Josep F Abril, Damir Garcia-Cehic, Ariadna Rando, Juliana Esperalba, Maria Gema Codina, Carla Castillo, Maria Carmen Martin, Tomàs Pumarola, Josep Quer, Andrés Antón |
| EPI_ISL_826458 | University of Bari Biomedical Sciences and Human Oncology | University of Bari Biomedical Sciences and Human Oncology | Chironna Maria, Sallustio Anna, Loconsole Daniela, Accogli Marisa |
| EPI_ISL_826463 | The Ohio State University | OSU Polaris Molecular Laboratory | Huolin Tu, Matthew R Avenarius, Laura Kubatko, Matthew Hunt, Xiaokang Pan, Peng Ru, Jason Garee, Keelie Thomas, Peter Mohler, Preeti Pancholi, Dan Jones |
| EPI_ISL_826472 | Lighthouse Lab in Alderley Park | Wellcome Sanger Institute for the COVID-19 Genomics UK (COG-UK) Consortium | Jacquelyn Wynn, Mairead Hyland, The Lighthouse Lab in Alderley Park and Alex Alderton, Roberto Amato, Sonia Goncalves, Ewan Harrison, David K. Jackson, Ian Johnston, Dominic Kwiatkowski, Cordelia Langford, John Sillitoe on behalf of the Wellcome Sanger Institute COVID-19 Surveillance Team |
| EPI_ISL_826475 | Lighthouse Lab in Cambridge | Wellcome Sanger Institute for the COVID-19 Genomics UK (COG-UK) Consortium | Rob Howes, The Lighthouse Lab in Cambridge and Alex Alderton, Roberto Amato, Sonia Goncalves, Ewan Harrison, David K. Jackson, Ian Johnston, Dominic Kwiatkowski, Cordelia Langford, John Sillitoe on behalf of the Wellcome Sanger Institute COVID-19 Surveillance Team |
| EPI_ISL_826476 | Lighthouse Lab in Alderley Park | Wellcome Sanger Institute for the COVID-19 Genomics UK (COG-UK) Consortium | Jacquelyn Wynn, Mairead Hyland, The Lighthouse Lab in Alderley Park and Alex Alderton, Roberto Amato, Sonia Goncalves, Ewan Harrison, David K. Jackson, Ian Johnston, Dominic Kwiatkowski, Cordelia Langford, John Sillitoe on behalf of the Wellcome Sanger Institute COVID-19 Surveillance Team |
| EPI_ISL_826521 | The Ohio State University | James Molecular Laboratory | Huolin Tu, Matthew R Avenarius, Laura Kubatko, Matthew Hunt, Xiaokang Pan, Peng Ru, Jason Garee, Keelie Thomas, Peter Mohler, Preeti Pancholi, Dan Jones |
| EPI_ISL_826523, EPI_ISL_826524, EPI_ISL_826525, EPI_ISL_826526, EPI_ISL_826527, EPI_ISL_826528, EPI_ISL_826529, EPI_ISL_826530, EPI_ISL_826531, EPI_ISL_826532, EPI_ISL_826533, EPI_ISL_826534, EPI_ISL_826535 | | | |
| see above | Dutch COVID-19 response team | National Institute for Public Health and the Environment (RIVM) | Adam Meijer, Harry Vennema, Jeroen Cremer, Sharon van den Brink, Bas van der Veer, AnneMarie van den Brandt, Florian Zwagemaker, Dennis Schmitz, Chantal Reusken, on behalf of the national COVID-19 response team |
| EPI_ISL_827042, EPI_ISL_827043 | Institute of Virology, University of Cologne | Institute of Virology, University of Cologne | Saleta Sierra, Gibran Rubio, Zevanya Tessalonica, Dominik Aschenmeier, Eva Heger, Elena Knops, Rolf Kaiser, Martin Däumer, Alex Thielen |
| EPI_ISL_827146, EPI_ISL_827927, EPI_ISL_827929, EPI_ISL_827932, EPI_ISL_828221, EPI_ISL_828335 | deCODE genetics | deCODE genetics | Daniel F Gudbjartsson; Agnar Helgason; Hakon Jonsson; Olafur T Magnusson; Pall Melsted; Gudmundur L Norddahl; Jona Saemundsdottir; Asgeir Sigurdsson; Patrick Sulem; Arna B Agustsdottir; Hannes Eggertsson; Berglind Eiriksdottir; Run Fridriksdottir; Elisabet E Gardarsdottir; Gudmundur Georgsson; Olafia S Gretarsdottir; Kjartan R Gudmundsson; Thora R Gunnarsdottir; Arnaldur Gylfason; Hilma Holm; Brynjar O Jensson; Aslaug Jonasdottir; Kamilla S Josefsdottir; Thordur Kristjansson; Droplaug N Magnusdottir; Solvi Rognvaldsson; Louise le Roux; Gudrun Sigmundsdottir; Gardar Sveinbjornsson; Kristin E Sveinsdottir; Maney Sveinsdottir; Emil A Thorarensen; Bjarni Thorbjornsson; Gisli Masson; Ingileif Jonsdottir; Alma Moller; Thorolfur Gudnason; Karl G Kristinsson; Unnur Thorsteinsdottir; Kari Stefansson |
| EPI_ISL_828336 | The National University Hospital of Iceland | deCODE genetics | Daniel F Gudbjartsson; Agnar Helgason; Hakon Jonsson; Olafur T Magnusson; Pall Melsted; Gudmundur L Norddahl; Jona Saemundsdottir; Asgeir Sigurdsson; Patrick Sulem; Arna B Agustsdottir; Hannes Eggertsson; Berglind Eiriksdottir; Run Fridriksdottir; Elisabet E Gardarsdottir; Gudmundur Georgsson; Olafia S Gretarsdottir; Kjartan R Gudmundsson; Thora R Gunnarsdottir; Arnaldur Gylfason; Hilma Holm; Brynjar O Jensson; Aslaug |

| | | | |
|---|---|---|---|
| | | | Jonasdottir; Kamilla S Josefsdottir; Thordur Kristjansson; Droplaug N Magnusdottir; Solvi Rognvaldsson; Louise le Roux; Gudrun Sigmundsdottir; Gardar Sveinbjornsson; Kristin E Sveinsdottir; Maney Sveinsdottir; Emil A Thorarensen; Bjarni Thorbjornsson; Gisli Masson; Ingileif Jonsdottir; Alma Moller; Thorolfur Gudnason; Karl G Kristinsson; Unnur Thorsteinsdottir; Kari Stefansson |
| EPI_ISL_828414, EPI_ISL_828786, EPI_ISL_829305, EPI_ISL_829306, EPI_ISL_829307, EPI_ISL_829311 | deCODE genetics | deCODE genetics | Daniel F Gudbjartsson; Agnar Helgason; Hakon Jonsson; Olafur T Magnusson; Pall Melsted; Gudmundur L Norddahl; Jona Saemundsdottir; Asgeir Sigurdsson; Patrick Sulem; Arna B Agustsdottir; Hannes Eggertsson; Berglind Eiriksdottir; Run Fridriksdottir; Elisabet E Gardarsdottir; Gudmundur Georgsson; Olafia S Gretarsdottir; Kjartan R Gudmundsson; Thora R Gunnarsdottir; Arnaldur Gylfason; Hilma Holm; Brynjar O Jensson; Aslaug Jonasdottir; Kamilla S Josefsdottir; Thordur Kristjansson; Droplaug N Magnusdottir; Solvi Rognvaldsson; Louise le Roux; Gudrun Sigmundsdottir; Gardar Sveinbjornsson; Kristin E Sveinsdottir; Maney Sveinsdottir; Emil A Thorarensen; Bjarni Thorbjornsson; Gisli Masson; Ingileif Jonsdottir; Alma Moller; Thorolfur Gudnason; Karl G Kristinsson; Unnur Thorsteinsdottir; Kari Stefansson |
| EPI_ISL_829543 | The National University Hospital of Iceland | deCODE genetics | Daniel F Gudbjartsson; Agnar Helgason; Hakon Jonsson; Olafur T Magnusson; Pall Melsted; Gudmundur L Norddahl; Jona Saemundsdottir; Asgeir Sigurdsson; Patrick Sulem; Arna B Agustsdottir; Hannes Eggertsson; Berglind Eiriksdottir; Run Fridriksdottir; Elisabet E Gardarsdottir; Gudmundur Georgsson; Olafia S Gretarsdottir; Kjartan R Gudmundsson; Thora R Gunnarsdottir; Arnaldur Gylfason; Hilma Holm; Brynjar O Jensson; Aslaug Jonasdottir; Kamilla S Josefsdottir; Thordur Kristjansson; Droplaug N Magnusdottir; Solvi Rognvaldsson; Louise le Roux; Gudrun Sigmundsdottir; Gardar Sveinbjornsson; Kristin E Sveinsdottir; Maney Sveinsdottir; Emil A Thorarensen; Bjarni Thorbjornsson; Gisli Masson; Ingileif Jonsdottir; Alma Moller; Thorolfur Gudnason; Karl G Kristinsson; Unnur Thorsteinsdottir; Kari Stefansson |
| EPI_ISL_829545, EPI_ISL_829546, EPI_ISL_829547 | deCODE genetics | deCODE genetics | Daniel F Gudbjartsson; Agnar Helgason; Hakon Jonsson; Olafur T Magnusson; Pall Melsted; Gudmundur L Norddahl; Jona Saemundsdottir; Asgeir Sigurdsson; Patrick Sulem; Arna B Agustsdottir; Hannes Eggertsson; Berglind Eiriksdottir; Run Fridriksdottir; Elisabet E Gardarsdottir; Gudmundur Georgsson; Olafia S Gretarsdottir; Kjartan R Gudmundsson; Thora R Gunnarsdottir; Arnaldur Gylfason; Hilma Holm; Brynjar O Jensson; Aslaug Jonasdottir; Kamilla S Josefsdottir; Thordur Kristjansson; Droplaug N Magnusdottir; Solvi Rognvaldsson; Louise le Roux; Gudrun Sigmundsdottir; Gardar Sveinbjornsson; Kristin E Sveinsdottir; Maney Sveinsdottir; Emil A Thorarensen; Bjarni Thorbjornsson; Gisli Masson; Ingileif Jonsdottir; Alma Moller; Thorolfur Gudnason; Karl G Kristinsson; Unnur Thorsteinsdottir; Kari Stefansson |
| EPI_ISL_830214 | BOSTON HEART DIAGNOSTICS CORP | Wadsworth Center, New York State Department of Health | Kirsten St. George, Daryl M. Lamson, Alexis Russel, Matthew Shudt, Melissa A Leisner, Jonathan Plitnick, Navjot Singh, John Kelly, Erasmus Schneider, Erica Lasek-Nesselquist |
| EPI_ISL_830608, EPI_ISL_830609, EPI_ISL_830610, EPI_ISL_830611, EPI_ISL_830612, EPI_ISL_830613, EPI_ISL_830614, EPI_ISL_830615, EPI_ISL_830616, EPI_ISL_830617, EPI_ISL_830618, EPI_ISL_830619, EPI_ISL_830620, EPI_ISL_830621, EPI_ISL_830622, EPI_ISL_830623, EPI_ISL_830624, EPI_ISL_830625, EPI_ISL_830626, EPI_ISL_830627 | | | |
| see above | KALEIDA CENTER FOR LABORATORY MEDICINE | Wadsworth Center, New York State Department of Health | Kirsten St. George, Daryl M. Lamson, Alexis Russel, Matthew Shudt, Melissa A Leisner, Jonathan Plitnick, Navjot Singh, John Kelly, Erasmus Schneider, Erica Lasek-Nesselquist |
| EPI_ISL_830649 | Gundersen Molecular Diagnostics Laboratory | Kabara Cancer Research Institute | Craig S. Richmond, Paraic A. Kenny |
| EPI_ISL_830650, EPI_ISL_830651, EPI_ISL_830652, EPI_ISL_830653, EPI_ISL_830654, EPI_ISL_830655, EPI_ISL_830656, EPI_ISL_830657, EPI_ISL_830658, EPI_ISL_830659, EPI_ISL_830660, EPI_ISL_830661, EPI_ISL_830662, EPI_ISL_830663, EPI_ISL_830665, EPI_ISL_830666, EPI_ISL_830667, EPI_ISL_830668, EPI_ISL_830670, EPI_ISL_830671, EPI_ISL_830672, EPI_ISL_830673, EPI_ISL_830674, EPI_ISL_830675, EPI_ISL_830676, EPI_ISL_830677, EPI_ISL_830678, EPI_ISL_830679, EPI_ISL_830680, EPI_ISL_830681 | | | |
| see above | SUNY UPSTATE MEDICAL UNIVERSITY | Wadsworth Center, New York State Department of Health | Kirsten St. George, Daryl M. Lamson, Alexis Russel, Matthew Shudt, Melissa A Leisner, Jonathan Plitnick, Navjot Singh, John Kelly, Erasmus Schneider, Erica Lasek-Nesselquist |
| EPI_ISL_830722, EPI_ISL_830723 | BIO-REFERENCE LABORATORIES | Wadsworth Center, New York State Department of Health | Kirsten St. George, Daryl M. Lamson, Alexis Russel, Matthew Shudt, Melissa A Leisner, Jonathan Plitnick, Navjot Singh, John Kelly, Erasmus Schneider, Erica Lasek-Nesselquist |
| EPI_ISL_831490, EPI_ISL_831498, EPI_ISL_831499, EPI_ISL_831572, EPI_ISL_831573, EPI_ISL_831575, EPI_ISL_831577, EPI_ISL_831578, EPI_ISL_831579, EPI_ISL_831580, EPI_ISL_831581, EPI_ISL_831582, EPI_ISL_831583, EPI_ISL_831584, EPI_ISL_831585, EPI_ISL_831586 | | | |
| see above | University of Wisconsin-Madison AIDS Vaccine Research Laboratories | University of Wisconsin-Madison AIDS Vaccine Research Laboratories | Gage Moreno, Katarina Braun, et al. AIDS Vaccine Research Laboratories |
| EPI_ISL_831649 | Institute for Infectious Diseases, University of Bern, Switzerland | Institute for Infectious Diseases, University of Bern, Switzerland | Michel C Koch, Christian Baumann, Miguel A Terrazos Miani, Cora Sägesser, Pascal Bittel, Stephen L Leib, Peter Keller, Franziska Suter-Riniker, Alban Ramette |
| EPI_ISL_831944, EPI_ISL_831947, EPI_ISL_831948, EPI_ISL_831949, EPI_ISL_831950, EPI_ISL_831951, EPI_ISL_831952 | ABC | The Public Health Agency of Sweden | Department of Microbiology, The Public Health Agency of Sweden |
| EPI_ISL_831980, EPI_ISL_831981 | Klinisk mikrobiologi | The Public Health Agency of Sweden | Department of Microbiology, The Public Health Agency of Sweden |
| EPI_ISL_832008 | Unilabs, Mikrobiologiska laboratoriet | The Public Health Agency of Sweden | Department of Microbiology, The Public Health Agency of Sweden |
| EPI_ISL_832014 | Utah Public Health Laboratory | Utah Public Health Laboratory | Erin L. Young, Kelly F. Oakeson, Tara Gallagher |
| EPI_ISL_832017 | University of Wisconsin-Madison AIDS Vaccine Research Laboratories | University of Wisconsin-Madison AIDS Vaccine Research Laboratories | Gage Moreno, Katarina Braun, et al. AIDS Vaccine Research Laboratories |
| EPI_ISL_832079, EPI_ISL_832080, EPI_ISL_832081 | Santa Clara County Public Health Laboratory | Santa Clara County Public Health Laboratory | Santa Clara County Public Health Department |
| EPI_ISL_832207 | Department of Clinical Microbiology | GIGA Medical Genomics | Keith Durkin, Maria Artesi, Sébastien Bontems, Raphaël Boreux, Bouchra Boujemla, Cécile Meex, Pierrette Melin, Marie-Pierre Hayette, Vincent Bours |
| EPI_ISL_832823, EPI_ISL_832824, EPI_ISL_832825, EPI_ISL_832826, EPI_ISL_832827, EPI_ISL_832828, EPI_ISL_832829, EPI_ISL_832830, EPI_ISL_832831, EPI_ISL_832832, EPI_ISL_832833, EPI_ISL_832834, EPI_ISL_832852, EPI_ISL_832853, EPI_ISL_832854, EPI_ISL_832855, EPI_ISL_832856, EPI_ISL_832857, EPI_ISL_832858, EPI_ISL_832859, EPI_ISL_832860, EPI_ISL_832861, EPI_ISL_832862, EPI_ISL_832863, EPI_ISL_832864, EPI_ISL_832865, EPI_ISL_832866, EPI_ISL_832867, EPI_ISL_832868, EPI_ISL_832869, EPI_ISL_832870, EPI_ISL_832873, EPI_ISL_832874, EPI_ISL_832885, EPI_ISL_832886, EPI_ISL_832887, EPI_ISL_832888, EPI_ISL_832889, EPI_ISL_832890, EPI_ISL_832891, EPI_ISL_832892, EPI_ISL_832893, EPI_ISL_832894, EPI_ISL_832895, EPI_ISL_832896, EPI_ISL_832897, EPI_ISL_832926, EPI_ISL_832935, EPI_ISL_832936, EPI_ISL_832937, EPI_ISL_832938, EPI_ISL_832939, EPI_ISL_832940, EPI_ISL_832941, EPI_ISL_832942, EPI_ISL_832943, EPI_ISL_832944, EPI_ISL_832947, EPI_ISL_832948, EPI_ISL_832949, EPI_ISL_832950, EPI_ISL_832951, EPI_ISL_832952, EPI_ISL_832953, EPI_ISL_832954, EPI_ISL_832955, EPI_ISL_832956, EPI_ISL_832957, EPI_ISL_832958, EPI_ISL_832959, EPI_ISL_832960, EPI_ISL_832961, EPI_ISL_832962, EPI_ISL_832963, EPI_ISL_832964, EPI_ISL_832965, EPI_ISL_832966, EPI_ISL_832967 | | | |
| see above | Maine HETL | Tewhey Lab, The Jackson Laboratory | Matluk,N., Dewey,H., Iosue,F., Barter,M., Lynch,R., Munger,H. and Tewhey,R. |
| EPI_ISL_833038 | Laboratorio de Referencia Nacional de Virus Respiratorio. Instituto Nacional de Salud Perú | Laboratorio de Referencia Nacional de Biotecnología y Biología Molecular. Instituto Nacional de Salud Perú | Carlos Padilla Rojas, Luis Barcena, Karolyn Vega Chozo, Priscila Lope Pari, Omar Caceres Rey, Marco Galarza Perez, Maribel Huaringa Nuñez, Johanna Balbuena Torrez, Henri Bailon Calderon, Nancy Rojas Serrano |
| EPI_ISL_833136 | Laboratorio de Ecologia de Doencas Transmissiveis na Amazonia, Instituto Leonidas e Maria Deane - Fiocruz Amazonia | Laboratorio de Ecologia de Doencas Transmissiveis na Amazonia, Instituto Leonidas e Maria Deane - Fiocruz Amazonia | Valdinete Nascimento, Victor Souza, André Corado, Fernanda Nascimento, George Silva, Ágatha Costa, Debora Duarte, Karina Pessoa, Matilde Mejía, Luciana Gonçalves, Maria Júlia Brandão, Michele Jesus, Felipe Naveca |
| EPI_ISL_833146 | Genomic Laboratory (GLAB) (Conjoint lab of Health Directorate of Istanbul and Istanbul Technical University) | Genomic Laboratory (GLAB), Istanbul Technical University | Ilker Karacan, Tugba Kizilboga Akgun, Payam Zolfagharian, Nisan Denizce Can, Pari Sharifli, Levent Doganay, Gizem Dinler Doganay |
| EPI_ISL_833187 | Department of Clinical Microbiology | GIGA Medical Genomics | Keith Durkin, Maria Artesi, Sébastien Bontems, Raphaël Boreux, Bouchra Boujemla, Cécile Meex, Pierrette Melin, Marie-Pierre Hayette, Vincent Bours |
| EPI_ISL_833202, EPI_ISL_833204, EPI_ISL_833207, EPI_ISL_833211, EPI_ISL_833212, EPI_ISL_833216, EPI_ISL_833217, EPI_ISL_833218, EPI_ISL_833219 | Department of Virology and Immunology, University of Helsinki and Helsinki University Hospital, Huslab Finland | Department of Virology, Faculty of Medicine, University of Helsinki, Helsinki, Finland | Teemu Smura, Ravi Kant, Phuoc Truong, Hussein Alburkat, Hannimari Kallio-Kokko, Jenni Virtanen, Maija Suvanto, Fathiah Zakham, Essi Korhonen, Sari Hannula, Harri Kangas, Pekka Ellonen, Olli Vapalahti |
| EPI_ISL_833228, EPI_ISL_833229 | R.P. GUARDIAGRELE Ospedale di Comunità | Istituto Zooprofilattico Sperimentale dell'Abruzzo e Molise "G.Caporale" | Lorusso A, Marcacci M, Di Domenico M, Curini V, Ancora M, Cammà C, Rinaldi A, Mangone I, Di Pasquale A, Puglia I, Calistri P, Savini G. |
| EPI_ISL_833230 | OSPEDALE S.S. ANNUNZIATA CHIETI - CLINICA MEDICA | Istituto Zooprofilattico Sperimentale dell'Abruzzo e Molise | Lorusso A, Marcacci M, Di Domenico M, Curini V, Ancora M, Cammà C, Rinaldi A, Mangone I, Di Pasquale A, Puglia I, Calistri P, Savini G. |

| | | | |
|---|---|---|---|
| | (MEDICINA GENERALE 1) | "G.Caporale" | |
| EPI_ISL_833231 | SIESP CHIETI - DRIVE IN LANCIANO | Istituto Zooprofilattico Sperimentale dell'Abruzzo e Molise "G.Caporale" | Lorusso A, Marcacci M, Di Domenico M, Curini V, Ancora M, Cammà C, Rinaldi A, Mangone I, Di Pasquale A, Puglia I, Calistri P, Savini G. |
| EPI_ISL_833232 | SIESP CHIETI - DRIVE IN CHIETI | Istituto Zooprofilattico Sperimentale dell'Abruzzo e Molise "G.Caporale" | Lorusso A, Marcacci M, Di Domenico M, Curini V, Ancora M, Cammà C, Rinaldi A, Mangone I, Di Pasquale A, Puglia I, Calistri P, Savini G. |
| EPI_ISL_833233, EPI_ISL_833234, EPI_ISL_833235, EPI_ISL_833236, EPI_ISL_833237, EPI_ISL_833238 | SIESP DIPARTIMENTO DI PREVENZIONE CHIETI | Istituto Zooprofilattico Sperimentale dell'Abruzzo e Molise "G.Caporale" | Lorusso A, Marcacci M, Di Domenico M, Curini V, Ancora M, Cammà C, Rinaldi A, Mangone I, Di Pasquale A, Puglia I, Calistri P, Savini G. |
| EPI_ISL_833239, EPI_ISL_833240 | SIESP DIPARTIMENTO DI PREVENZIONE TERAMO | Istituto Zooprofilattico Sperimentale dell'Abruzzo e Molise "G.Caporale" | Lorusso A, Marcacci M, Di Domenico M, Curini V, Ancora M, Cammà C, Rinaldi A, Mangone I, Di Pasquale A, Puglia I, Calistri P, Savini G. |
| EPI_ISL_833241 | SIESP DIPARTIMENTO DI PREVENZIONE CHIETI | Istituto Zooprofilattico Sperimentale dell'Abruzzo e Molise "G.Caporale" | Lorusso A, Marcacci M, Di Domenico M, Curini V, Ancora M, Cammà C, Rinaldi A, Mangone I, Di Pasquale A, Puglia I, Calistri P, Savini G. |
| EPI_ISL_833242 | SIESP CHIETI - DRIVE IN LANCIANO | Istituto Zooprofilattico Sperimentale dell'Abruzzo e Molise "G.Caporale" | Lorusso A, Marcacci M, Di Domenico M, Curini V, Ancora M, Cammà C, Rinaldi A, Mangone I, Di Pasquale A, Puglia I, Calistri P, Savini G. |
| EPI_ISL_833243, EPI_ISL_833244, EPI_ISL_833245, EPI_ISL_833246, EPI_ISL_833247 | SIESP DIPARTIMENTO DI PREVENZIONE CHIETI | Istituto Zooprofilattico Sperimentale dell'Abruzzo e Molise "G.Caporale" | Lorusso A, Marcacci M, Di Domenico M, Curini V, Ancora M, Cammà C, Rinaldi A, Mangone I, Di Pasquale A, Puglia I, Calistri P, Savini G. |
| EPI_ISL_833297 | Ospedale Civile Giulianova | Istituto Zooprofilattico Sperimentale dell'Abruzzo e Molise "G. Caporale" | Lorusso A, Marcacci M, Di Domenico M, Ancora M, Curini V, Mangone I, Rinaldi A, Di Pasquale A, Cammà C, Puglia I, Calistri P, Savini G |
| EPI_ISL_833435, EPI_ISL_833436, EPI_ISL_833437, EPI_ISL_833438, EPI_ISL_833439, EPI_ISL_833440, EPI_ISL_833441, EPI_ISL_833442, EPI_ISL_833443, EPI_ISL_833444, EPI_ISL_833445, EPI_ISL_833446 | | | |
| see above | SC (UCO) Igiene e Sanità Pubblica (funzione integrata con SC Microbiologia e Virologia) e Laboratory of Molecular Virology of the International Centre for Genetic Engineering and Biotechnology (ICGEB) | ARGO Laboratorio Genomica ed Epigenomica | Licastro D, Dal Monego S, Degasperi M, Marcello A, D'Agaro P |
| EPI_ISL_834146, EPI_ISL_834211, EPI_ISL_834212, EPI_ISL_834223, EPI_ISL_834226, EPI_ISL_834228, EPI_ISL_834231, EPI_ISL_834233, EPI_ISL_834236, EPI_ISL_834237, EPI_ISL_834248, EPI_ISL_834250, EPI_ISL_834256, EPI_ISL_834264, EPI_ISL_834271, EPI_ISL_834280, EPI_ISL_834283, EPI_ISL_834288, EPI_ISL_834290, EPI_ISL_834293, EPI_ISL_834298, EPI_ISL_834299, EPI_ISL_834311, EPI_ISL_834312, EPI_ISL_834313, EPI_ISL_834344, EPI_ISL_834352, EPI_ISL_834353, EPI_ISL_834360, EPI_ISL_834373, EPI_ISL_834376, EPI_ISL_834377, EPI_ISL_834378, EPI_ISL_834380, EPI_ISL_834381, EPI_ISL_834382, EPI_ISL_834388, EPI_ISL_834404, EPI_ISL_834407, EPI_ISL_834409, EPI_ISL_834417, EPI_ISL_834423, EPI_ISL_834427, EPI_ISL_834440, EPI_ISL_834443, EPI_ISL_834445, EPI_ISL_834448, EPI_ISL_834449, EPI_ISL_834459, EPI_ISL_834466, EPI_ISL_834467, EPI_ISL_834468, EPI_ISL_834485, EPI_ISL_834486, EPI_ISL_834488, EPI_ISL_834494, EPI_ISL_834497, EPI_ISL_834499, EPI_ISL_834500, EPI_ISL_834503, EPI_ISL_834511, EPI_ISL_834517, EPI_ISL_834529, EPI_ISL_834541, EPI_ISL_834545, EPI_ISL_834804, EPI_ISL_834814, EPI_ISL_834823, EPI_ISL_834825, EPI_ISL_834826, EPI_ISL_834833, EPI_ISL_834834, EPI_ISL_834837, EPI_ISL_834848, EPI_ISL_834865, EPI_ISL_834870, EPI_ISL_834874, EPI_ISL_834903, EPI_ISL_834909, EPI_ISL_834910, EPI_ISL_834920, EPI_ISL_834925, EPI_ISL_834931, EPI_ISL_834933, EPI_ISL_834949, EPI_ISL_834959, EPI_ISL_834972, EPI_ISL_834979, EPI_ISL_834981, EPI_ISL_834983, EPI_ISL_834989, EPI_ISL_834991, EPI_ISL_835001, EPI_ISL_835004, EPI_ISL_835008, EPI_ISL_835017, EPI_ISL_835038, EPI_ISL_835047, EPI_ISL_835055, EPI_ISL_835058, EPI_ISL_835060, EPI_ISL_835067, EPI_ISL_835112, EPI_ISL_835114, EPI_ISL_835115, EPI_ISL_835127, EPI_ISL_835130, EPI_ISL_835132, EPI_ISL_835140, EPI_ISL_835483, EPI_ISL_835492, EPI_ISL_835495, EPI_ISL_835532, EPI_ISL_835546, EPI_ISL_835550, EPI_ISL_835562, EPI_ISL_835615, EPI_ISL_835639, EPI_ISL_835655, EPI_ISL_835680, EPI_ISL_835698 | | | |
| see above | Lighthouse Lab in Alderley Park | Wellcome Sanger Institute for the COVID-19 Genomics UK (COG-UK) Consortium | Jacquelyn Wynn, Mairead Hyland, The Lighthouse Lab in Alderley Park and Alex Alderton, Roberto Amato, Sonia Goncalves, Ewan Harrison, David K. Jackson, Ian Johnston, Dominic Kwiatkowski, Cordelia Langford, John Sillitoe on behalf of the Wellcome Sanger Institute COVID-19 Surveillance Team |
| EPI_ISL_837041, EPI_ISL_837042, EPI_ISL_837043, EPI_ISL_837044, EPI_ISL_837045, EPI_ISL_837046, EPI_ISL_837047, EPI_ISL_837048, EPI_ISL_837049, EPI_ISL_837050, EPI_ISL_837051, EPI_ISL_837064, EPI_ISL_837070, EPI_ISL_837080, EPI_ISL_837086, EPI_ISL_837092, EPI_ISL_837093, EPI_ISL_837094, EPI_ISL_837096, EPI_ISL_837135, EPI_ISL_837136, EPI_ISL_837137, EPI_ISL_837138, EPI_ISL_837139, EPI_ISL_837140, EPI_ISL_837141, EPI_ISL_837142, EPI_ISL_837143, EPI_ISL_837144, EPI_ISL_837145, EPI_ISL_837146, EPI_ISL_837147, EPI_ISL_837148, EPI_ISL_837149, EPI_ISL_837150, EPI_ISL_837151, EPI_ISL_837152, EPI_ISL_837153, EPI_ISL_837154, EPI_ISL_837155, EPI_ISL_837156, EPI_ISL_837157, EPI_ISL_837158, EPI_ISL_837159, EPI_ISL_837160, EPI_ISL_837161, EPI_ISL_837162, EPI_ISL_837163, EPI_ISL_837164, EPI_ISL_837165, EPI_ISL_837166, EPI_ISL_837167, EPI_ISL_837168, EPI_ISL_837169, EPI_ISL_837170, EPI_ISL_837171, EPI_ISL_837184, EPI_ISL_837188, EPI_ISL_837190, EPI_ISL_837191, EPI_ISL_837192, EPI_ISL_837193, EPI_ISL_837194, EPI_ISL_837226, EPI_ISL_837227, EPI_ISL_837228, EPI_ISL_837229, EPI_ISL_837230, EPI_ISL_837231, EPI_ISL_837232, EPI_ISL_837233, EPI_ISL_837234, EPI_ISL_837235, EPI_ISL_837236, EPI_ISL_837237, EPI_ISL_837238 | | | |
| see above | Respiratory Virus Unit, National Infection Service, Public Health England | COVID-19 Genomics UK (COG-UK) Consortium | PHE Covid Sequencing Team |
| EPI_ISL_837345, EPI_ISL_837348, EPI_ISL_837353, EPI_ISL_837382, EPI_ISL_837383, EPI_ISL_837394, EPI_ISL_837416, EPI_ISL_837424, EPI_ISL_837425, EPI_ISL_837430, EPI_ISL_837432 | | | |
| see above | National Virus Reference Laboratory | National Virus Reference Laboratory | Michael Carr, Gabriel Gonzalez, Jonathan Dean, Cillian F De Gascun |
| EPI_ISL_837525, EPI_ISL_837527 | UW Virology Lab | UW Virology Lab | Pavitra Roychoudhury, Hong Xie, Lasata Shrestha, Meei-Li Huang, Keith R Jerome, Alexander Greninger |
| EPI_ISL_837817, EPI_ISL_837820, EPI_ISL_837821, EPI_ISL_837826, EPI_ISL_837828, EPI_ISL_837831 | Wyoming Public Health Laboratory | Wyoming Public Health Laboratory | Noah Hull, Taylor Fearing, Lynette Gumbleton, Channing Weber, Ashley Norberg, Bailey Bowcutt, and Wanda Manley |
| EPI_ISL_837851, EPI_ISL_837853, EPI_ISL_837858, EPI_ISL_837866, EPI_ISL_837867, EPI_ISL_837868, EPI_ISL_837872, EPI_ISL_837873, EPI_ISL_837874, EPI_ISL_837875, EPI_ISL_837878, EPI_ISL_837880, EPI_ISL_837881, EPI_ISL_837883, EPI_ISL_837884, EPI_ISL_837885, EPI_ISL_837888, EPI_ISL_837890, EPI_ISL_837894, EPI_ISL_837897, EPI_ISL_837899, EPI_ISL_837900 | | | |
| see above | Department of Pathology, University of Cambridge | COVID-19 Genomics UK (COG-UK) Consortium | Aminu S. Jahun, Yasmin Chaudhry, Grant Hall, Iliana Georgana, Myra Hosmillo, Martin D. Curran, Malte Pinckert, Surendra Parmar, Ian Goodfellow |
| EPI_ISL_838268, EPI_ISL_838269, EPI_ISL_838270, EPI_ISL_838271, EPI_ISL_838273, EPI_ISL_838277, EPI_ISL_838278, EPI_ISL_838279 | Virology Department, Royal Infirmary of Edinburgh, NHS Lothian / School of Biological Sciences, University of Edinburgh / Institute of Genetics and Molecular Medicine, University of Edinburgh | COVID-19 Genomics UK (COG-UK) Consortium | McHugh M, Dewar R, Rooke S, Gallagher M, Balcaza C, O'Toole Á, Scher E, Hill V, McCrone JT, Colquhoun R, Yu X, Jackson B, Rambaut A, Williams TC, Templeton K |
| EPI_ISL_838330, EPI_ISL_838332, EPI_ISL_838333 | University of Exeter | COVID-19 Genomics UK (COG-UK) Consortium | Ben Temperton,Aaron Jeffries,Michelle Michelsen,Joanna Warwick-Dugdale,Audrey Farbos,Robyn Manley,Stephen Michell,Jane Masoli |
| EPI_ISL_838580, EPI_ISL_838581, EPI_ISL_838582, EPI_ISL_838583, EPI_ISL_838584, EPI_ISL_838589, EPI_ISL_838590, EPI_ISL_838591, EPI_ISL_838594, EPI_ISL_838595, EPI_ISL_838596, EPI_ISL_838601, EPI_ISL_838604, EPI_ISL_838606, EPI_ISL_838607, EPI_ISL_838608, EPI_ISL_838609, EPI_ISL_838611, EPI_ISL_838614, EPI_ISL_838615, EPI_ISL_838616, EPI_ISL_838621, EPI_ISL_838623, EPI_ISL_838624, EPI_ISL_838629, EPI_ISL_838630, EPI_ISL_838631, EPI_ISL_838632, EPI_ISL_838634, EPI_ISL_838635, EPI_ISL_838636, EPI_ISL_838637, EPI_ISL_838639, EPI_ISL_838642, EPI_ISL_838643, EPI_ISL_838644, EPI_ISL_838647, EPI_ISL_838648, EPI_ISL_838649, EPI_ISL_838650, EPI_ISL_838651, EPI_ISL_838654, EPI_ISL_838656, EPI_ISL_838657, EPI_ISL_838658, EPI_ISL_838659, EPI_ISL_838660, EPI_ISL_838661, EPI_ISL_838663 | | | |
| see above | Liverpool Clinical Laboratories | COVID-19 Genomics UK (COG-UK) Consortium | Sam Haldenby, Anita Lucaci, Steve Paterson, Julian Hiscox, Alistair Darby, M Almsaud, A Alrezaihi, Muhannad Alruwaili, Stuart D Armstrong, Jones Benjamin, Eleanor G Bentley, Anu Chawla, Jordan J Clark, Angela Cowell, Richard Eccles, Isabel García-Dorival, Matthew Gemmell, Alessandro Gerada, PKF Gilmore, Richard Gregory, Ximeng Han, Catherine Hartley, Margaret Hughes, Miren Iturriza-Gomara, James Johnson, L Luu, Jenifer Manson, Charlotte Nelson, Elaine O'Toole, Cassie Olateju, Rebekah Penrice-Randal , Lucille Rainbow, N.P Randle, Trevor Ian Robinson, Parul Sharma, Ghada T Shawli, James P Stewart, Neil Swainston, Ecaterina Vamos, Joanne Watts, Mark Whitehead |
| EPI_ISL_838767, EPI_ISL_838768, EPI_ISL_838769, EPI_ISL_838770, EPI_ISL_838771, EPI_ISL_838773, EPI_ISL_838775, EPI_ISL_838776, EPI_ISL_838777, EPI_ISL_838799, EPI_ISL_838800, EPI_ISL_838801, EPI_ISL_838802, EPI_ISL_838803, EPI_ISL_838804, EPI_ISL_838805, EPI_ISL_838806, EPI_ISL_838808, EPI_ISL_838809, EPI_ISL_838810, EPI_ISL_838811, EPI_ISL_838812, EPI_ISL_838813, EPI_ISL_838814, EPI_ISL_838815, EPI_ISL_838816, EPI_ISL_838817, EPI_ISL_838818, EPI_ISL_838819, EPI_ISL_838820, EPI_ISL_838821, EPI_ISL_838822, EPI_ISL_838823, EPI_ISL_838824, EPI_ISL_838825, EPI_ISL_838826, EPI_ISL_838827, EPI_ISL_839041, EPI_ISL_839042, EPI_ISL_839043, EPI_ISL_839063, EPI_ISL_839064, EPI_ISL_839065, EPI_ISL_839066, EPI_ISL_839067, EPI_ISL_839068, EPI_ISL_839069, EPI_ISL_839080, EPI_ISL_839081, EPI_ISL_839082, EPI_ISL_839083, EPI_ISL_839084, EPI_ISL_839101, EPI_ISL_839102, EPI_ISL_839103, EPI_ISL_839104, EPI_ISL_839105, EPI_ISL_839106, EPI_ISL_839107, EPI_ISL_839108, EPI_ISL_839109, EPI_ISL_839110, EPI_ISL_839111, EPI_ISL_839112, EPI_ISL_839113, EPI_ISL_839114, EPI_ISL_839116, EPI_ISL_839215, EPI_ISL_839218, EPI_ISL_839219, EPI_ISL_839220, EPI_ISL_839222, EPI_ISL_839223, EPI_ISL_839226, EPI_ISL_839227, EPI_ISL_839305, EPI_ISL_839306, EPI_ISL_839307, EPI_ISL_839308, EPI_ISL_839310, EPI_ISL_839316, EPI_ISL_839317, EPI_ISL_839330 | | | |
| see above | University College London, Great Ormond Street Hospital for Children NHS Foundation Trust, Imperial College Healthcare NHS Trust | COVID-19 Genomics UK (COG-UK) Consortium | Sergi Castellano, Rachel Williams, Mark Kristiansen, Paola Resende Silva, Sunando Roy, Tony Brooks, Helena Tutill, Paola Niola, Patricia Dyal, Charlotte Williams, Leysa Forrest, Yasmin Panchbhaya, Jacqueline Findlay, Samuel Weeks, Julianne Brown, Kathryn Harris, Paul Randell, James Price, Alison Holmes, Judith Breuer |
| EPI_ISL_839710, EPI_ISL_839711, EPI_ISL_839712, EPI_ISL_839714, EPI_ISL_839715, EPI_ISL_839716, EPI_ISL_839717, EPI_ISL_839718, EPI_ISL_839719, EPI_ISL_839720, EPI_ISL_839721, EPI_ISL_839722, EPI_ISL_839723, EPI_ISL_839724, EPI_ISL_839725, EPI_ISL_839726, EPI_ISL_839727, EPI_ISL_839728, EPI_ISL_839729, EPI_ISL_839730 | | | |

| | | | |
|---|---|---|---|
| see above | Northumbria University / South Tees Hospitals NHS Foundation Trust / North Cumbria Integrated Care NHS Foundation Trust / North Tees and Hartlepool NHS Foundation Trust / Newcastle Hospitals NHS Foundation Trust | COVID-19 Genomics UK (COG-UK) Consortium | Darren L Smith,Andrew Nelson,Matthew Bashton,Greg R Young,Joshua Loh,John Allan,Mohammad A Tariq,Giles S Holt,Gary Black,Wen C Yew,Lynn Dover,Paul Baker,Steve Liggett,Sarah Essex,Jane Greenaway,Debra Padgett,Clive Graham,Garren Scott,Edward Barton,Emma Swindells,Brendan Payne,Jennifer Collins,Yusri Taha,Gary Eltringham |
| EPI_ISL_840108, EPI_ISL_840109, EPI_ISL_840110, EPI_ISL_840112, EPI_ISL_840113, EPI_ISL_840114, EPI_ISL_840115, EPI_ISL_840116, EPI_ISL_840117, EPI_ISL_840118, EPI_ISL_840119, EPI_ISL_840120 | | | |
| see above | Lincolnshire Hospitals and DeepSeq Nottingham | COVID-19 Genomics UK (COG-UK) Consortium | Nichola Duckworth, Tim Sloan, Sarah Walsh, Jonathan Ball, Patrick McClure, Joeseph Chappell, Nadine Holmes, Matthew Carlisle, Christopher Moore, Fei Sang, Johnny Debebe, Victoria Wright, Matthew Loose |
| EPI_ISL_840371, EPI_ISL_840372, EPI_ISL_840390, EPI_ISL_840391, EPI_ISL_840392, EPI_ISL_840393, EPI_ISL_840394, EPI_ISL_840395, EPI_ISL_840396, EPI_ISL_840397, EPI_ISL_840399, EPI_ISL_840600, EPI_ISL_840603, EPI_ISL_840604, EPI_ISL_840606, EPI_ISL_840609, EPI_ISL_840610, EPI_ISL_840611, EPI_ISL_840612, EPI_ISL_840615, EPI_ISL_840617, EPI_ISL_840618, EPI_ISL_840619, EPI_ISL_840620, EPI_ISL_840621, EPI_ISL_840622, EPI_ISL_840623, EPI_ISL_840624, EPI_ISL_840625, EPI_ISL_840626, EPI_ISL_840627, EPI_ISL_840628, EPI_ISL_840629, EPI_ISL_840630, EPI_ISL_840631, EPI_ISL_840632, EPI_ISL_840633, EPI_ISL_840634, EPI_ISL_840635, EPI_ISL_840636 | | | |
| see above | Originating lab: Wales Specialist Virology Centre Sequencing lab: Pathogen Genomics Unit | Public Health Wales Microbiology Cardiff Wales Specialist Virology Centre | Catherine Moore, Johnathan Evans, Laura Gifford, Malorie Perry, Simon Cottrell, Angela Marchbank, Alec Birchley, Alexander Adams, Amy Gaskin, Bree Gatica-Wilcox, Jason Coombes, Joel Southgate, Lauren Gilbert, Lee Graham, Nicole Pacchiarini, Sara Kumziene-Summerhayes, Sarah Taylor, Sophie Jones, Sara Rey, Matthew Bull, Joanne Watkins, Sally Corden, Tom Connor |
| EPI_ISL_840637 | Wyoming Public Health Laboratory | Wyoming Public Health Laboratory | Noah Hull, Taylor Fearing, Lynette Gumbleton, Channing Weber, Ashley Norberg, Bailey Bowcutt, and Wanda Manley |
| EPI_ISL_840638, EPI_ISL_840639, EPI_ISL_840640, EPI_ISL_840641, EPI_ISL_840643, EPI_ISL_840644, EPI_ISL_840645, EPI_ISL_840646, EPI_ISL_840647, EPI_ISL_840648, EPI_ISL_840649, EPI_ISL_840651, EPI_ISL_840653, EPI_ISL_840655, EPI_ISL_840656, EPI_ISL_840657, EPI_ISL_840658, EPI_ISL_840671, EPI_ISL_840672, EPI_ISL_840695, EPI_ISL_840696, EPI_ISL_840697, EPI_ISL_840698, EPI_ISL_840699, EPI_ISL_840700, EPI_ISL_840701, EPI_ISL_840702, EPI_ISL_840703, EPI_ISL_840774, EPI_ISL_840775, EPI_ISL_840776, EPI_ISL_840777, EPI_ISL_840778, EPI_ISL_840779, EPI_ISL_840781, EPI_ISL_840782, EPI_ISL_840783, EPI_ISL_840785, EPI_ISL_840787, EPI_ISL_840790, EPI_ISL_840794, EPI_ISL_840795, EPI_ISL_840796, EPI_ISL_840798, EPI_ISL_840799, EPI_ISL_840801, EPI_ISL_840802, EPI_ISL_840803, EPI_ISL_840804, EPI_ISL_840805, EPI_ISL_840806, EPI_ISL_840808, EPI_ISL_840810 | | | |
| see above | Originating lab: Wales Specialist Virology Centre Sequencing lab: Pathogen Genomics Unit | Public Health Wales Microbiology Cardiff Wales Specialist Virology Centre | Catherine Moore, Johnathan Evans, Laura Gifford, Malorie Perry, Simon Cottrell, Angela Marchbank, Alec Birchley, Alexander Adams, Amy Gaskin, Bree Gatica-Wilcox, Jason Coombes, Joel Southgate, Lauren Gilbert, Lee Graham, Nicole Pacchiarini, Sara Kumziene-Summerhayes, Sarah Taylor, Sophie Jones, Sara Rey, Matthew Bull, Joanne Watkins, Sally Corden, Tom Connor |
| EPI_ISL_840837 | Wales Specialist Virology Centre Sequencing lab: Pathogen Genomics Unit | Public Health Wales Microbiology Cardiff Wales Specialist Virology Centre | Catherine Moore, Johnathan Evans, Laura Gifford, Malorie Perry, Simon Cottrell, Angela Marchbank, Alec Birchley, Alexander Adams, Amy Gaskin, Bree Gatica-Wilcox, Jason Coombes, Joel Southgate, Lauren Gilbert, Lee Graham, Nicole Pacchiarini, Sara Kumziene-Summerhayes, Sarah Taylor, Sophie Jones, Sara Rey, Matthew Bull, Joanne Watkins, Sally Corden, Tom Connor |
| EPI_ISL_840850 | Wyoming Public Health Laboratory | Wyoming Public Health Laboratory | Noah Hull, Taylor Fearing, Lynette Gumbleton, Channing Weber, Ashley Norberg, Bailey Bowcutt, and Wanda Manley |
| EPI_ISL_840895, EPI_ISL_840897, EPI_ISL_840898, EPI_ISL_840899, EPI_ISL_840900, EPI_ISL_841057, EPI_ISL_841058, EPI_ISL_841083, EPI_ISL_841087, EPI_ISL_841088, EPI_ISL_841089, EPI_ISL_841090, EPI_ISL_841091, EPI_ISL_841092, EPI_ISL_841093, EPI_ISL_841094, EPI_ISL_841095, EPI_ISL_841096, EPI_ISL_841097, EPI_ISL_841099, EPI_ISL_841100, EPI_ISL_841101, EPI_ISL_841102, EPI_ISL_841103, EPI_ISL_841104, EPI_ISL_841105, EPI_ISL_841106, EPI_ISL_841107, EPI_ISL_841109, EPI_ISL_841110, EPI_ISL_841111, EPI_ISL_841112, EPI_ISL_841113, EPI_ISL_841302, EPI_ISL_841303, EPI_ISL_841304, EPI_ISL_841305 | | | |
| see above | Wales Specialist Virology Centre Sequencing lab: Pathogen Genomics Unit | Public Health Wales Microbiology Cardiff Wales Specialist Virology Centre | Catherine Moore, Johnathan Evans, Laura Gifford, Malorie Perry, Simon Cottrell, Angela Marchbank, Alec Birchley, Alexander Adams, Amy Gaskin, Bree Gatica-Wilcox, Jason Coombes, Joel Southgate, Lauren Gilbert, Lee Graham, Nicole Pacchiarini, Sara Kumziene-Summerhayes, Sarah Taylor, Sophie Jones, Sara Rey, Matthew Bull, Joanne Watkins, Sally Corden, Tom Connor |
| EPI_ISL_841321, EPI_ISL_841355, EPI_ISL_841359, EPI_ISL_841360, EPI_ISL_841361, EPI_ISL_841521, EPI_ISL_841522, EPI_ISL_841580, EPI_ISL_841581, EPI_ISL_841584, EPI_ISL_841587 | | | |
| see above | Originating lab: Wales Specialist Virology Centre Sequencing lab: Pathogen Genomics Unit | Public Health Wales Microbiology Cardiff Wales Specialist Virology Centre | Catherine Moore, Johnathan Evans, Laura Gifford, Malorie Perry, Simon Cottrell, Angela Marchbank, Alec Birchley, Alexander Adams, Amy Gaskin, Bree Gatica-Wilcox, Jason Coombes, Joel Southgate, Lauren Gilbert, Lee Graham, Nicole Pacchiarini, Sara Kumziene-Summerhayes, Sarah Taylor, Sophie Jones, Sara Rey, Matthew Bull, Joanne Watkins, Sally Corden, Tom Connor |
| EPI_ISL_841978, EPI_ISL_842004, EPI_ISL_842014 | Centre for Enzyme Innovation, University of Portsmouth / Translational Research Laboratory, Portsmouth Hospitals NHS Trust | COVID-19 Genomics UK (COG-UK) Consortium | Angela Beckett,Yann Bourgeois,Garry Scarlett,Sharon Glaysher,Scott Elliott,Kelly Bicknell,Robert Impey,Allyson Lloyd,Sarah Wyllie,Ethan Butcher,Anoop Chauhan,Samuel Robson |
| EPI_ISL_842209, EPI_ISL_842265, EPI_ISL_842273, EPI_ISL_842293, EPI_ISL_842312, EPI_ISL_842338, EPI_ISL_842341 | Virology Department, Sheffield Teaching Hospitals NHS Foundation Trust/Department of Infection, Immunity and Cardiovascular Disease, The Medical School, University of Sheffield | COVID-19 Genomics UK (COG-UK) Consortium | Thushan de Silva, Matthew Parker, Nikki Smith, Adri Angyal, Rebecca Brown, Luke Green, Rachel Tucker, Paul Parsons, Danielle Groves, Katie Johnson, Laura Carrilero, Alex Keeley, Dave Partridge, Matthew Wyles, Benjamin Lindsey, Mehmet Yavuz, Mohammad Raza, Cariad Evans |
| EPI_ISL_842356, EPI_ISL_842361, EPI_ISL_842362, EPI_ISL_842367, EPI_ISL_842422, EPI_ISL_842423, EPI_ISL_842424, EPI_ISL_842425, EPI_ISL_842426, EPI_ISL_842427, EPI_ISL_842428, EPI_ISL_842429, EPI_ISL_842430, EPI_ISL_842431, EPI_ISL_842432, EPI_ISL_842433, EPI_ISL_842434, EPI_ISL_842435, EPI_ISL_842436, EPI_ISL_842437, EPI_ISL_842438, EPI_ISL_842439, EPI_ISL_842440, EPI_ISL_842441, EPI_ISL_842442, EPI_ISL_842443, EPI_ISL_842444, EPI_ISL_842445, EPI_ISL_842447, EPI_ISL_842448, EPI_ISL_842449, EPI_ISL_842450, EPI_ISL_842451, EPI_ISL_842452, EPI_ISL_842453, EPI_ISL_842454, EPI_ISL_842455, EPI_ISL_842456, EPI_ISL_842457, EPI_ISL_842458, EPI_ISL_842459, EPI_ISL_842460, EPI_ISL_842461, EPI_ISL_842462, EPI_ISL_842463, EPI_ISL_842464, EPI_ISL_842465, EPI_ISL_842466, EPI_ISL_842467, EPI_ISL_842468, EPI_ISL_842469, EPI_ISL_842470, EPI_ISL_842472, EPI_ISL_842473, EPI_ISL_842474, EPI_ISL_842475, EPI_ISL_842476, EPI_ISL_842477, EPI_ISL_842478, EPI_ISL_842479, EPI_ISL_842480, EPI_ISL_842481, EPI_ISL_842482, EPI_ISL_842483, EPI_ISL_842484, EPI_ISL_842485, EPI_ISL_842486, EPI_ISL_842487, EPI_ISL_842488, EPI_ISL_842489, EPI_ISL_842490, EPI_ISL_842491, EPI_ISL_842492, EPI_ISL_842493, EPI_ISL_842494, EPI_ISL_842496, EPI_ISL_842497, EPI_ISL_842498, EPI_ISL_842499, EPI_ISL_842500, EPI_ISL_842501, EPI_ISL_842502, EPI_ISL_842503, EPI_ISL_842504, EPI_ISL_842505, EPI_ISL_842506, EPI_ISL_842507, EPI_ISL_842508, EPI_ISL_842509, EPI_ISL_842510, EPI_ISL_842511, EPI_ISL_842513, EPI_ISL_842583, EPI_ISL_842585, EPI_ISL_842586, EPI_ISL_842587, EPI_ISL_842588, EPI_ISL_842589, EPI_ISL_842590, EPI_ISL_842591, EPI_ISL_842592, EPI_ISL_842593, EPI_ISL_842594, EPI_ISL_842596, EPI_ISL_842598, EPI_ISL_842599, EPI_ISL_842600, EPI_ISL_842601, EPI_ISL_842602, EPI_ISL_842603, EPI_ISL_842604, EPI_ISL_842605, EPI_ISL_842606, EPI_ISL_842607, EPI_ISL_842608 | | | |
| see above | Bioinformatics and Biostatistics Lab, Advanced Sequencing Facility | COVID-19 Genomics UK (COG-UK) Consortium | Aengus Stewart,Jerome Nicod,Chelsea Sawyer,Laura Cubitt,Harshil Patel,Margaret Crawford |
| EPI_ISL_842610, EPI_ISL_842611, EPI_ISL_842612, EPI_ISL_842614, EPI_ISL_842615, EPI_ISL_842617, EPI_ISL_842618, EPI_ISL_842619, EPI_ISL_842623, EPI_ISL_842624, EPI_ISL_842625, EPI_ISL_842626, EPI_ISL_842627, EPI_ISL_842628, EPI_ISL_842631, EPI_ISL_842632 | | | |
| see above | Wyoming Public Health Laboratory | Wyoming Public Health Laboratory | Noah Hull, Taylor Fearing, Lynette Gumbleton, Channing Weber, Ashley Norberg, Bailey Bowcutt, and Wanda Manley |
| EPI_ISL_843147, EPI_ISL_843148, EPI_ISL_843149, EPI_ISL_843150 | Barts Health NHS Trust | COVID-19 Genomics UK (COG-UK) Consortium | CUTINO-MOGUEL, Maria-Teresa; HARRINGTON, David; OWOYEMI, Dola; SHYLINI, Raghavendran; BROAD, Claire; KELE, Beatrix |
| EPI_ISL_843155, EPI_ISL_843156, EPI_ISL_843157, EPI_ISL_843158, EPI_ISL_843159, EPI_ISL_843160, EPI_ISL_843161, EPI_ISL_843164, EPI_ISL_843165, EPI_ISL_843166 | Regional Virus Laboratory, Belfast Health and Social Care Trust | COVID-19 Genomics UK (COG-UK) Consortium | Conall McCaughey, James McKenna, Tanya Curran, Susan Feeney, Alison Watt, Ciara Cox, Mairead Connor, Zoltan Molnar, David Simpson, Derek Fairley |
| EPI_ISL_843196 | LabPLUS | Institute of Environmental Science and Research (ESR) | Xiaoyun Ren, Matt Storey, Nikki Freed, Muhammad Faisal, Jing Wang, Hermes Perez, Anja Werno, Antje van der Linden, Arlo Upton, Chris Mansell, David Hammer, Dragana Drinkovic, Gary McAuliffe, Hana Sofia Andersson, James Ussher, Jill Sherwood, Josh Freeman, Julia Howard, Juliet Elvy, Mary DeAlmeida, Matt Blakiston, Matthew Rogers, Max Bloomfield, Michael Addidle, Michelle Balm, Sally Roberts, Sarah Jefferies, Sharmini Muttaiyah, Susan Morpeth, Susan Taylor, Timothy Blackmore, Vani Sathyendran, Veronica Playle, Virginia Hope, Erasmus Smit, Lauren Jelly, Olin Silander, Joep de Ligt |
| EPI_ISL_845543, EPI_ISL_845544 | Santa Clara County Public Health Laboratory | Santa Clara County Public Health Laboratory | Santa Clara County Public Health Department |
| EPI_ISL_845588, EPI_ISL_845590, EPI_ISL_845593, EPI_ISL_845606, EPI_ISL_845608, EPI_ISL_845610, EPI_ISL_845611, EPI_ISL_845614, EPI_ISL_845615 | KU Leuven, Rega Institute, Clinical and Epidemiological Virology | KU Leuven, Rega Institute, Clinical and Epidemiological Virology | Tony Wawina-Bokalanga, Bert Vanmechelen, Joan Marti-Carerras, Piet Maes |
| EPI_ISL_845626 | FUNDACION CARDIOINFANTIL | Instituto Nacional de Salud - Dirección de Investigación en Salud Pública | Katherine Laiton-Donato, Diego A. Álvarez-Díaz, Carlos Franco-Muñoz, Mauricio Pacheco-Montealegre, Maria T. Herrera-Sepúlveda, Jonathan Reales, Sheryll Corchuelo, Julian Naizaque, Gerardo Santamaría, Paola Muñoz-Laiton, Diego Andrés Prada, Magdalena Wiesner, Martha Lucia Ospina Martinez, Marcela Mercado-Reyes |
| EPI_ISL_845634 | Instituto Nacional de Cancerologia | Instituto Nacional de Salud - Dirección de Investigación en Salud Pública | Katherine Laiton-Donato, Diego A. Álvarez-Díaz, Carlos Franco-Muñoz, Mauricio Pacheco-Montealegre, Maria T. Herrera-Sepúlveda, Jonathan Reales, Sheryll Corchuelo, Julian Naizaque, Gerardo Santamaría, Paola Muñoz-Laiton, Diego Andrés Prada, Magdalena Wiesner, Martha Lucia Ospina Martinez, |

| | | | |
|---|---|---|---|
| | | | Marcela Mercado-Reyes |
| EPI_ISL_845636 | Laboratorio de Salud Pública - Secretaría Distrital de Salud | Instituto Nacional de Salud - Dirección de Investigación en Salud Pública | Katherine Laiton-Donato, Diego A. Álvarez-Díaz, Carlos Franco-Muñoz, Mauricio Pacheco-Montealegre, Maria T. Herrera-Sepúlveda, Jonathan Reales, Sheryll Corchuelo, Julian Naizaque, Gerardo Santamaría, Paola Muñoz-Laiton, Diego Andrés Prada, Magdalena Wiesner, Martha Lucia Ospina Martinez, Marcela Mercado-Reyes |
| EPI_ISL_845770 | Emory Molecular Diagnostics Laboratory, Emory Healthcare | Piantadosi Lab, Emory Department of Pathology | Ahmed Babiker, Anne Piantadosi |
| EPI_ISL_845825, EPI_ISL_845826, EPI_ISL_845827, EPI_ISL_845828, EPI_ISL_845829, EPI_ISL_845830, EPI_ISL_845831, EPI_ISL_845832, EPI_ISL_845833, EPI_ISL_845834, EPI_ISL_845835, EPI_ISL_845836, EPI_ISL_845837, EPI_ISL_845838, EPI_ISL_845839, EPI_ISL_845840, EPI_ISL_845841, EPI_ISL_845842, EPI_ISL_845843, EPI_ISL_845844, EPI_ISL_845845, EPI_ISL_845846, EPI_ISL_845847, EPI_ISL_845848, EPI_ISL_845849, EPI_ISL_845853, EPI_ISL_845857, EPI_ISL_845872, EPI_ISL_845873 | | | |
| see above | TGen North | TGen North | Jolene Bowers, Megan Folkerts, Chris French, Hayley Yaglom, Ashlyn Pfeiffer, Darrin Lemmer, Dave Engelthaler, The Arizona COVID Genomics Union (ACGU) |
| EPI_ISL_845877 | National Institute of Mental Health and Neurosciences (NIMHANS) | Department of Neurovirology, National Institute of Mental Health and Neurosciences (NIMHANS) | Chitra Pattabiraman, Pramada Prasad, Risha Rasheed, Darshan Sreenivas, Nakka Vijay Kiran Reddy, Anita S Desai, V Ravi |
| EPI_ISL_845878, EPI_ISL_845879, EPI_ISL_845880 | BBMP Urban PHC | Department of Neurovirology, National Institute of Mental Health and Neurosciences (NIMHANS) | Chitra Pattabiraman, Pramada Prasad, Risha Rasheed, Darshan Sreenivas, Nakka Vijay Kiran Reddy, Anita S Desai, V Ravi |
| EPI_ISL_845881 | Kidwai Memorial Institute of Oncology | Department of Neurovirology, National Institute of Mental Health and Neurosciences (NIMHANS) | Chitra Pattabiraman, Pramada Prasad, Risha Rasheed, Darshan Sreenivas, Nakka Vijay Kiran Reddy, Anita S Desai, V Ravi |
| EPI_ISL_845882 | VRDL, Mysore Medical College | Department of Neurovirology, National Institute of Mental Health and Neurosciences (NIMHANS) | Chitra Pattabiraman, Pramada Prasad, Risha Rasheed, Darshan Sreenivas, Nakka Vijay Kiran Reddy, Anita S Desai, V Ravi |
| EPI_ISL_845885, EPI_ISL_845886 | Railway Hospital | Department of Neurovirology, National Institute of Mental Health and Neurosciences (NIMHANS) | Chitra Pattabiraman, Pramada Prasad, Risha Rasheed, Darshan Sreenivas, Nakka Vijay Kiran Reddy, Anita S Desai, V Ravi |
| EPI_ISL_845887, EPI_ISL_845888, EPI_ISL_845890 | BBMP Urban PHC | Department of Neurovirology, National Institute of Mental Health and Neurosciences (NIMHANS) | Chitra Pattabiraman, Pramada Prasad, Risha Rasheed, Darshan Sreenivas, Nakka Vijay Kiran Reddy, Anita S Desai, V Ravi |
| EPI_ISL_846526, EPI_ISL_846536 | Lighthouse Lab in Alderley Park | Wellcome Sanger Institute for the COVID-19 Genomics UK (COG-UK) Consortium | Jacquelyn Wynn, Mairead Hyland, The Lighthouse Lab in Alderley Park and Alex Alderton, Roberto Amato, Sonia Goncalves, Ewan Harrison, David K. Jackson, Ian Johnston, Dominic Kwiatkowski, Cordelia Langford, John Sillitoe on behalf of the Wellcome Sanger Institute COVID-19 Surveillance Team |
| EPI_ISL_847594, EPI_ISL_847595, EPI_ISL_847596, EPI_ISL_847597, EPI_ISL_847598, EPI_ISL_847599, EPI_ISL_847600 | California Department of Public Health | Chiu Laboratory, University of California, San Francisco | Charles Chiu, Xianding (Wayne) Deng, Candace Wang, Brian Bushnell, Scot Federman, Jill Hacker, Debra Wadford |
| EPI_ISL_847730, EPI_ISL_847743 | Chiu Laboratory, University of California, San Francisco | Chiu Laboratory, University of California, San Francisco | Charles Chiu, Xianding (Wayne) Deng, Candace Wang, Brian Bushnell, Scot Federman, Jill Hacker, Debra Wadford |
| EPI_ISL_847756, EPI_ISL_847757, EPI_ISL_847760, EPI_ISL_847761, EPI_ISL_847762, EPI_ISL_847763, EPI_ISL_847764, EPI_ISL_847765 | California Department of Public Health | Chiu Laboratory, University of California, San Francisco | Charles Chiu, Xianding (Wayne) Deng, Candace Wang, Brian Bushnell, Scot Federman, Jill Hacker, Debra Wadford |
| EPI_ISL_847836, EPI_ISL_847839 | Tempus | Grubaugh Lab - Yale School of Public Health | Tara Alpert, Joseph Fauver, Anderson Brito, Mallery Breban, Anne Wyllie, Chantal Vogels, Mary Petrone, Chaney Kalinich, Isabel Ott, Nathan Grubaugh |
| EPI_ISL_847841 | Yale New Haven Hospital | Grubaugh Lab - Yale School of Public Health | Tara Alpert, Joseph Fauver, Anderson Brito, Mallery Breban, Anne Wyllie, Chantal Vogels, Mary Petrone, Chaney Kalinich, Isabel Ott, Nathan Grubaugh |
| EPI_ISL_847873, EPI_ISL_847874, EPI_ISL_847875, EPI_ISL_847876, EPI_ISL_847877, EPI_ISL_847878, EPI_ISL_847879, EPI_ISL_847880, EPI_ISL_847881, EPI_ISL_847882, EPI_ISL_847883, EPI_ISL_847884, EPI_ISL_847885, EPI_ISL_847886, EPI_ISL_847887, EPI_ISL_847888, EPI_ISL_847889, EPI_ISL_847921, EPI_ISL_847924, EPI_ISL_847927, EPI_ISL_847928, EPI_ISL_847937, EPI_ISL_847938, EPI_ISL_847953, EPI_ISL_847954, EPI_ISL_847955, EPI_ISL_847956, EPI_ISL_847957, EPI_ISL_847958, EPI_ISL_847959 | | | |
| see above | University Hospitals of Geneva, Laboratory of Virology | HUG, Laboratory of Virology and the Health2030 Genome Center | Samuel Cordey, Ana Rita Goncalves, Laurent Kaiser, Lorenzo Cerutti, Henri Pegeot, Melyssa Elies, Keith Harshman, Ioannis Xenarios, Emmanouil Dermitzakis |
| EPI_ISL_847996, EPI_ISL_848013, EPI_ISL_848044 | Michigan Department of Health and Human Services, Bureau of Laboratories | Michigan Department of Health and Human Services, Bureau of Laboratories | Blankenship HM, Riner D, Soehnlen MK |
| EPI_ISL_848633, EPI_ISL_848642 | Montana Public Health Laboratory | Wyoming Public Health Laboratory | Noah Hull, Joy Ritter, Taylor Fearing, Lynette Gumbleton, Channing Weber, Ashley Norberg, Bailey Bowcutt, Wanda Manley, Deborah Gibson |
| EPI_ISL_848643 | Florida Bureau of Public Health Laboratories | Florida Bureau of Public Health Laboratories | Sarah Schmedes, Jason Blanton |
| EPI_ISL_849322, EPI_ISL_849323, EPI_ISL_849324, EPI_ISL_849325 | Delaware Public Health Lab | Delaware Public Health Lab | Gregory Hovan |
| EPI_ISL_849634 | SIESP DIPARTIMENTO DI PREVENZIONE CHIETI | Istituto Zooprofilattico Sperimentale dell'Abruzzo e Molise "G.Caporale" | Lorusso A, Marcacci M, Di Domenico M, Curini V, Ancora M, Cammà C, Rinaldi A, Mangone I, Di Pasquale A, Puglia I, Savini G. |
| EPI_ISL_849635, EPI_ISL_849636 | DIPARTIMENTO PREVENZIONE AVEZZANO-SERVIZIO DI IGIENE EPIDEMIOLOGIA E SANITA' PUBBLICA | Istituto Zooprofilattico Sperimentale dell'Abruzzo e Molise "G.Caporale" | Lorusso A, Marcacci M, Di Domenico M, Curini V, Ancora M, Cammà C, Rinaldi A, Mangone I, Di Pasquale A, Puglia I, Savini G. |
| EPI_ISL_849637, EPI_ISL_849638, EPI_ISL_849639, EPI_ISL_849640, EPI_ISL_849641, EPI_ISL_849642, EPI_ISL_849643, EPI_ISL_849644, EPI_ISL_849645, EPI_ISL_849646 | SIESP DIPARTIMENTO DI PREVENZIONE CHIETI | Istituto Zooprofilattico Sperimentale dell'Abruzzo e Molise "G.Caporale" | Lorusso A, Marcacci M, Di Domenico M, Curini V, Ancora M, Cammà C, Rinaldi A, Mangone I, Di Pasquale A, Puglia I, Savini G. |
| EPI_ISL_849647, EPI_ISL_849648, EPI_ISL_849649 | SIESP- DRIVE IN CHIETI | Istituto Zooprofilattico Sperimentale dell'Abruzzo e Molise "G.Caporale" | Lorusso A, Marcacci M, Di Domenico M, Curini V, Ancora M, Cammà C, Rinaldi A, Mangone I, Di Pasquale A, Puglia I, Savini G. |
| EPI_ISL_849865, EPI_ISL_849866 | A. Krumbholz, Labor Dr. Krause und Kollegen MVZ GmbH, Kiel | Charité Universitätsmedizin Berlin, Institut für Virologie | Victor M Corman, Jörn Beheim-Schwarzbach, Talitha Veith, Julia Schneider, Tobias Bleicker, Julia Tesch, Barbara Mühlemann, Terry Jones, Christian Drosten |
| EPI_ISL_849895 | Utah Public Health Laboratory | Utah Public Health Laboratory | Erin L. Young, Kelly F. Oakeson, Tara Gallagher |
| EPI_ISL_849925, EPI_ISL_849927 | UCSF Clinical Microbiology Laboratory | Chan-Zuckerberg Biohub | CZB Cliahub Consortium |
| EPI_ISL_850508 | Hospital de Tortosa Verge de la Cinta | Hospital Universitari Vall d'Hebron - Vall d'Hebron Institut de Rercerca | Cristina Andrés, Maria Piñana, Josep F Abril, Damir Garcia-Cehic, Ariadna Rando, Juliana Esperalba, Maria Gema Codina, Carla Castillo, Maria Carmen Martin, Tomàs Pumarola, Josep Quer, Andrés Antón |
| EPI_ISL_850601, EPI_ISL_850602, EPI_ISL_850603, EPI_ISL_850604, EPI_ISL_850605, EPI_ISL_850606, EPI_ISL_850607, EPI_ISL_850608, EPI_ISL_850609, EPI_ISL_850610, EPI_ISL_850611, EPI_ISL_850612, EPI_ISL_850613, EPI_ISL_850614, EPI_ISL_850615, EPI_ISL_850616, EPI_ISL_850617, EPI_ISL_850618, EPI_ISL_850619, EPI_ISL_850620, EPI_ISL_850621, EPI_ISL_850622, EPI_ISL_850623, EPI_ISL_850624, EPI_ISL_850625, EPI_ISL_850626, EPI_ISL_850627, EPI_ISL_850628, EPI_ISL_850629, EPI_ISL_850630, EPI_ISL_850631, EPI_ISL_850632, EPI_ISL_850633, EPI_ISL_850634, EPI_ISL_850635, EPI_ISL_850636, EPI_ISL_850637, EPI_ISL_850638, EPI_ISL_850639, EPI_ISL_850640, EPI_ISL_850641, EPI_ISL_850642, EPI_ISL_850643, EPI_ISL_850644, EPI_ISL_850645, EPI_ISL_850646, EPI_ISL_850647, EPI_ISL_850648, EPI_ISL_850649, EPI_ISL_850650, EPI_ISL_850651 | | | |
| see above | Helix/Illumina | Genomics and Discovery, Respiratory Viruses Branch, Division of Viral Diseases, Centers for Disease Control and Prevention | Peter W. Cook, Dhwani Batra, Ben L. Rambo-Martin Eileen de Feo, Jan Antico, Christine Tran, Matthew Tolentino, Shannon Wickline, Kim Gietzen, Brad Sickler, Jingtao Liu, Eric Allen, Phil Febbo, Summer Galloway, Nicole L. Washington, Simon White, Geraint Levan, Kelly Schiabor Barrett, Elizabeth Cirulli, Alexandre Bolze, Ary Ascencio, Charlotte Rivera-Garcia, Ryan Cho, Jason Nguyen, Sherry Wang, Jimmy Ramirez, Tyler Cassens, Efren Sandoval, Magnus Isaksson, William Lee, David Becker, Marc Laurent, James Lu, Clinton R. Paden, Suxiang Tong, Duncan MacCannell |
| EPI_ISL_850660, EPI_ISL_850661 | Division of Emerging Infectious Diseases, Bureau of Infectious Diseases Diagnosis Control, Korea Disease Control and Prevention Agency | Division of Emerging Infectious Diseases, Bureau of Infectious Diseases Diagnosis Control, Korea Disease Control and Prevention Agency | Ae Kyung Park, Il-Hwan Kim, Heui Man Kim, Jeong-Min Kim, Namjoo Lee, Chaeyoung Lee, Sang Hee Woo, Eun-Jin Kim |

| | | | |
|---|---|---|---|
| EPI_ISL_850942, EPI_ISL_850943, EPI_ISL_850944 | Helix / Illumina | Genomics and Discovery, Respiratory Viruses Branch, Division of Viral Diseases, Centers for Disease Control and Prevention | Peter W. Cook, Dhwani Batra, Ben L. Rambo-Martin Eileen de Feo, Jan Antico, Christine Tran, Matthew Tolentino, Shannon Wickline, Kim Gietzen, Brad Sickler, Jingtao Liu, Eric Allen, Phil Febbo, Summer Galloway, Nicole L. Washington, Simon White, Geraint Levan, Kelly Schiabor Barrett, Elizabeth Cirulli, Alexandre Bolze, Ary Ascencio, Charlotte Rivera-Garcia, Ryan Cho, Jason Nguyen, Sherry Wang, Jimmy Ramirez, Tyler Cassens, Efren Sandoval, Magnus Isaksson, William Lee, David Becker, Marc Laurent, James Lu, Clinton R. Paden, Suxiang Tong, Duncan MacCannell |
| EPI_ISL_851007, EPI_ISL_851008, EPI_ISL_851009, EPI_ISL_851010, EPI_ISL_851011, EPI_ISL_851012, EPI_ISL_851013, EPI_ISL_851014, EPI_ISL_851015, EPI_ISL_851016, EPI_ISL_851017, EPI_ISL_851018, EPI_ISL_851019, EPI_ISL_851020, EPI_ISL_851026, EPI_ISL_851027, EPI_ISL_851028, EPI_ISL_851033, EPI_ISL_851034, EPI_ISL_851035, EPI_ISL_851036, EPI_ISL_851037, EPI_ISL_851038, EPI_ISL_851039, EPI_ISL_851040 | | | |
| see above | Helix/Illumina | Genomics and Discovery, Respiratory Viruses Branch, Division of Viral Diseases, Centers for Disease Control and Prevention | Peter W. Cook, Dhwani Batra, Ben L. Rambo-Martin Eileen de Feo, Jan Antico, Christine Tran, Matthew Tolentino, Shannon Wickline, Kim Gietzen, Brad Sickler, Jingtao Liu, Eric Allen, Phil Febbo, Summer Galloway, Nicole L. Washington, Simon White, Geraint Levan, Kelly Schiabor Barrett, Elizabeth Cirulli, Alexandre Bolze, Ary Ascencio, Charlotte Rivera-Garcia, Ryan Cho, Jason Nguyen, Sherry Wang, Jimmy Ramirez, Tyler Cassens, Efren Sandoval, Magnus Isaksson, William Lee, David Becker, Marc Laurent, James Lu, Clinton R. Paden, Suxiang Tong, Duncan MacCannell |
| EPI_ISL_852017 | Lighthouse Lab in Milton Keynes | Wellcome Sanger Institute for the COVID-19 Genomics UK (COG-UK) Consortium | The Lighthouse Lab in Milton Keynes and Alex Alderton, Roberto Amato, Sonia Goncalves, Ewan Harrison, David K. Jackson, Ian Johnston, Dominic Kwiatkowski, Cordelia Langford, John Sillitoe on behalf of the Wellcome Sanger Institute COVID-19 Surveillance Team |
| EPI_ISL_852191 | Lighthouse Lab in Alderley Park | Wellcome Sanger Institute for the COVID-19 Genomics UK (COG-UK) Consortium | Jacquelyn Wynn, Mairead Hyland, The Lighthouse Lab in Alderley Park and Alex Alderton, Roberto Amato, Sonia Goncalves, Ewan Harrison, David K. Jackson, Ian Johnston, Dominic Kwiatkowski, Cordelia Langford, John Sillitoe on behalf of the Wellcome Sanger Institute COVID-19 Surveillance Team |
| EPI_ISL_852565, EPI_ISL_852579 | Max von Pettenkofer Institute, Virology, National Reference Center for Retroviruses, LMU München | Laboratory for Functional Genome Analysis, Dept. Genomics, Gene Center of the LMU Munich | Max Muenchhoff, Stefan Krebs, Alexander Graf, Oliver Keppler, Helmut Blum |
| EPI_ISL_852619, EPI_ISL_852622, EPI_ISL_852628 | Lab. Microbiologia e Virologia, Cotugno, A.O. dei Colli | Lab. Microbiologia e Virologia, Cotugno, A.O. dei Colli | Luigi Atripaldi, Claudia Tiberio, Anna Perfetti |
| EPI_ISL_852818 | CHU Purpan - Laboratoire de Virologie - Institut Fédératif de Biologie | CHU Purpan - Laboratoire de Virologie - Institut Fédératif de Biologie | Latour J., Ranger N., Dubois M., Carcenac R., Harter A., Boyer P., Tremeaux P., Izopet J. |
| EPI_ISL_852974, EPI_ISL_853002 | Hospital General Universitario Gregorio Marañón | SeqCOVID-SPAIN consortium/IBV(CSIC) | Darío García de Viedma, Laura Pérez-Lago, Pedro J Sola-Campoy, Sergio Buenestado-Serrano, Marta Herranz, Victor Manuel de la Cueva, Julia Suárez, Pilar Catalán, Patricia Muñoz and SeqCOVID-SPAIN consortium |
| EPI_ISL_853401, EPI_ISL_853402, EPI_ISL_853403 | Charité Universitätsmedizin Berlin, Institut für Virologie/Labor Berlin | Charité Universitätsmedizin Berlin, Institut für Virologie | Victor M Corman, Julia Schneider, Barbara Mühlemann, Jörn Beheim-Schwarzbach, Talitha Veith, Julia Tesch, Tobias Bleicker, Terry Jones, Christian Drosten |
| EPI_ISL_853688 | THE MARY IMOGENE BASSETT HOSPITAL | Wadsworth Center, New York State Department of Health | Kirsten St. George, Daryl M. Lamson, Alexis Russel, Matthew Shudt, Melissa A Leisner, Jonathan Plitnick, Navjot Singh, John Kelly, Erasmus Schneider, Erica Lasek-Nesselquist |
| EPI_ISL_853689 | SUNY UPSTATE MEDICAL UNIVERSITY | Wadsworth Center, New York State Department of Health | Kirsten St. George, Daryl M. Lamson, Alexis Russel, Matthew Shudt, Melissa A Leisner, Jonathan Plitnick, Navjot Singh, John Kelly, Erasmus Schneider, Erica Lasek-Nesselquist |
| EPI_ISL_853690, EPI_ISL_853691, EPI_ISL_853692, EPI_ISL_853693, EPI_ISL_853694, EPI_ISL_853695, EPI_ISL_853696, EPI_ISL_853697, EPI_ISL_853698, EPI_ISL_853699, EPI_ISL_853700, EPI_ISL_853701, EPI_ISL_853702, EPI_ISL_853703, EPI_ISL_853704 | | | |
| see above | ALBANY MEDICAL CENTER HOSPITAL CLINICAL LABORATORIES | Wadsworth Center, New York State Department of Health | Kirsten St. George, Daryl M. Lamson, Alexis Russel, Matthew Shudt, Melissa A Leisner, Jonathan Plitnick, Navjot Singh, John Kelly, Erasmus Schneider, Erica Lasek-Nesselquist |
| EPI_ISL_853705 | NORTHWELL HEALTH LABORATORIES | Wadsworth Center, New York State Department of Health | Kirsten St. George, Daryl M. Lamson, Alexis Russel, Matthew Shudt, Melissa A Leisner, Jonathan Plitnick, Navjot Singh, John Kelly, Erasmus Schneider, Erica Lasek-Nesselquist |
| EPI_ISL_853706, EPI_ISL_853707, EPI_ISL_853708 | SUNY UPSTATE MEDICAL UNIVERSITY | Wadsworth Center, New York State Department of Health | Kirsten St. George, Daryl M. Lamson, Alexis Russel, Matthew Shudt, Melissa A Leisner, Jonathan Plitnick, Navjot Singh, John Kelly, Erasmus Schneider, Erica Lasek-Nesselquist |
| EPI_ISL_853709, EPI_ISL_853710, EPI_ISL_853711 | ALBANY MEDICAL CENTER HOSPITAL CLINICAL LABORATORIES | Wadsworth Center, New York State Department of Health | Kirsten St. George, Daryl M. Lamson, Alexis Russel, Matthew Shudt, Melissa A Leisner, Jonathan Plitnick, Navjot Singh, John Kelly, Erasmus Schneider, Erica Lasek-Nesselquist |
| EPI_ISL_853715 | SUNY UPSTATE MEDICAL UNIVERSITY | Wadsworth Center, New York State Department of Health | Kirsten St. George, Daryl M. Lamson, Alexis Russel, Matthew Shudt, Melissa A Leisner, Jonathan Plitnick, Navjot Singh, John Kelly, Erasmus Schneider, Erica Lasek-Nesselquist |
| EPI_ISL_853742 | Department of Microbiology, University Innsbruck | Bergthaler laboratory, CeMM Research Center for Molecular Medicine of the Austrian Academy of Sciences | Lukas Endler, Alexandra Popa, Benedikt Agerer, Jakob-Wendelin Genger, Alexander Lercher, Anna Schedl, Thomas Penz, Michael Schuster, Jan Laine, Martin Senekowitsch, Christoph Bock, Andreas Bergthaler |
| EPI_ISL_854350, EPI_ISL_854351, EPI_ISL_854352, EPI_ISL_854353, EPI_ISL_854354, EPI_ISL_854355, EPI_ISL_854356, EPI_ISL_854357, EPI_ISL_854358, EPI_ISL_854359, EPI_ISL_854360, EPI_ISL_854361, EPI_ISL_854362, EPI_ISL_854363, EPI_ISL_854364, EPI_ISL_854365, EPI_ISL_854366, EPI_ISL_854367, EPI_ISL_854369, EPI_ISL_854372 | | | |
| see above | BIO-REFERENCE LABORATORIES | Wadsworth Center, New York State Department of Health | Kirsten St. George, Daryl M. Lamson, Alexis Russel, Matthew Shudt, Melissa A Leisner, Jonathan Plitnick, Navjot Singh, John Kelly, Erasmus Schneider, Erica Lasek-Nesselquist |
| EPI_ISL_854405, EPI_ISL_854406, EPI_ISL_854407, EPI_ISL_854408, EPI_ISL_854409, EPI_ISL_854410, EPI_ISL_854411, EPI_ISL_854412, EPI_ISL_854413, EPI_ISL_854414, EPI_ISL_854415, EPI_ISL_854450, EPI_ISL_854452, EPI_ISL_854454, EPI_ISL_854455, EPI_ISL_854457, EPI_ISL_854458 | | | |
| see above | MONTEFIORE MEDICAL CENTER LABORATORIES | Wadsworth Center, New York State Department of Health | Kirsten St. George, Daryl M. Lamson, Alexis Russel, Matthew Shudt, Melissa A Leisner, Jonathan Plitnick, Navjot Singh, John Kelly, Erasmus Schneider, Erica Lasek-Nesselquist |
| EPI_ISL_854748 | Victorian Infectious Diseases Reference Laboratory (VIDRL) | VIDRL and MDU-PHL | Caly L., Seemann T., Sait, M.L., Druce J., Sherry, N.L. |
| EPI_ISL_855388 | Hospital | National Reference Center for Viruses of Respiratory Infections, Institut Pasteur, Paris | Marion Barbet, Sylvie Behillil, Méline Bizard, Angela Brisebarre, Camille Capel, Etienne Simon-Lorière, Vincent Enouf, Maud Vanpeene, Sylvie van der Werf,Ducancelle Alexandra |
| EPI_ISL_855393, EPI_ISL_855394 | California Department of Public Health | Chiu Laboratory, University of California, San Francisco | Charles Chiu, Xianding (Wayne) Deng, Candace Wang, Brian Bushnell, Scot Federman, Jill Hacker, Debra Wadford |
| EPI_ISL_855556 | Lab. Microbiologia e Virologia, Cotugno, A.O. dei Colli | Lab. Microbiologia e Virologia, Cotugno, A.O. dei Colli | Luigi Atripaldi, Claudia Tiberio, Anna Perfetti |
| EPI_ISL_855908 | Lab voor klinische biologie | Onderzoeksgroep Virologie | Nick Vereecke, Laurens Lambrechts, Marthe Pauwels, Bruno Verhasselt, Linos Vandekerckhove, Hans Nauwynck, Sebastiaan Theuns |
| EPI_ISL_855923, EPI_ISL_855925, EPI_ISL_855927, EPI_ISL_855928, EPI_ISL_855934 | Lab voor klinische biologie | Onderzoeksgroep Virologie | Laurens Lambrechts, Nick Vereecke, Marthe Pauwels, Bruno Verhasselt, Linos Vandekerckhove, Hans Nauwynck, Sebastiaan Theuns |
| EPI_ISL_855940, EPI_ISL_855942 | Lab voor klinische biologie | Onderzoeksgroep Virologie | Nick Vereecke, Laurens Lambrechts, Marthe Pauwels, Bruno Verhasselt, Linos Vandekerckhove, Hans Nauwynck, Sebastiaan Theuns |
| EPI_ISL_856680 | Charité Universitätsmedizin Berlin, Institute of Virology, Charitéplatz 1, 10117 Berlin, Germany | Charité Universitätsmedizin Berlin, Institute of Virology, Charitéplatz 1, 10117 Berlin, Germany | Victor M Corman, Julia Schneider, Jörn Beheim-Schwarzbach, Tobias Bleicker, Julia Tesch, Barbara Mühlemann, Talitha Veith, Terry Jones, Christian Drosten |
| EPI_ISL_857052, EPI_ISL_857055 | Platform BIS UZA/UAntwerpen, University Hospital Antwerp, Edegem, Belgium | UAntwerp, Laboratory of Medical Microbiology, Campus Drie Eiken S6.26, Universiteitsplein 1, 2610, Wilrijk, Antwerp, Belgium | Basil Britto Xavier, Jasmine Coppens, Christine Lammens, Veerle Matheeussen, Herman Goossens |
| EPI_ISL_857056, EPI_ISL_857057, EPI_ISL_857058 | OCME Office Of Chief Medical Examiner | New York City Public Health Laboratory | Jade Wang, et al. |
| EPI_ISL_857189 | DOHMH Corona | New York City Public Health Laboratory | Jade Wang, et al. |
| EPI_ISL_857190 | DOHMH PHL | New York City Public Health Laboratory | Jade Wang, et al. |
| EPI_ISL_857191 | DOHMH Morrisania | New York City Public Health Laboratory | Jade Wang, et al. |
| EPI_ISL_857195 | DOHMH Riverside | New York City Public Health Laboratory | Jade Wang, et al. |

| | | | |
|---|---|---|---|
| EPI_ISL_857196 | DOHMH Crown Heights | New York City Public Health Laboratory | Jade Wang, et al. |
| EPI_ISL_857197 | DOHMH Jamaica | New York City Public Health Laboratory | Jade Wang, et al. |
| EPI_ISL_857202 | DOHMH PHL | New York City Public Health Laboratory | Jade Wang, et al. |
| EPI_ISL_857204, EPI_ISL_857205 | DOHMH Central Harlem | New York City Public Health Laboratory | Jade Wang, et al. |
| EPI_ISL_857227, EPI_ISL_857228 | DOHMH Riverside | New York City Public Health Laboratory | Jade Wang, et al. |
| EPI_ISL_857229 | DOHMH Morrisania | New York City Public Health Laboratory | Jade Wang, et al. |
| EPI_ISL_857230 | DOHMH Chelsea | New York City Public Health Laboratory | Jade Wang, et al. |
| EPI_ISL_857231, EPI_ISL_857232, EPI_ISL_857233 | DOHMH Central Harlem | New York City Public Health Laboratory | Jade Wang, et al. |
| EPI_ISL_857234, EPI_ISL_857235, EPI_ISL_857236, EPI_ISL_857237 | DOHMH Jamaica | New York City Public Health Laboratory | Jade Wang, et al. |
| EPI_ISL_857238, EPI_ISL_857239, EPI_ISL_857240 | DOHMH Corona | New York City Public Health Laboratory | Jade Wang, et al. |
| EPI_ISL_857241, EPI_ISL_857242, EPI_ISL_857243, EPI_ISL_857244, EPI_ISL_857245, EPI_ISL_857246, EPI_ISL_857247, EPI_ISL_857248, EPI_ISL_857249, EPI_ISL_857250 | DOHMH Jamaica | New York City Public Health Laboratory | Jade Wang, et al. |
| EPI_ISL_857251, EPI_ISL_857252 | DOHMH Morrisania | New York City Public Health Laboratory | Jade Wang, et al. |
| EPI_ISL_857253, EPI_ISL_857254 | DOHMH Chelsea | New York City Public Health Laboratory | Jade Wang, et al. |
| EPI_ISL_857255 | DOHMH Central Harlem | New York City Public Health Laboratory | Jade Wang, et al. |
| EPI_ISL_857256 | DOHMH Fort Greene | New York City Public Health Laboratory | Jade Wang, et al. |
| EPI_ISL_857257 | DOHMH PHL | New York City Public Health Laboratory | Jade Wang, et al. |
| EPI_ISL_857258, EPI_ISL_857259 | DOHMH Central Harlem | New York City Public Health Laboratory | Jade Wang, et al. |
| EPI_ISL_857260, EPI_ISL_857261, EPI_ISL_857262 | DOHMH Chelsea | New York City Public Health Laboratory | Jade Wang, et al. |
| EPI_ISL_857263, EPI_ISL_857264 | DOHMH Corona | New York City Public Health Laboratory | Jade Wang, et al. |
| EPI_ISL_857265 | DOHMH Fort Greene | New York City Public Health Laboratory | Jade Wang, et al. |
| EPI_ISL_857266 | DOHMH PHL | New York City Public Health Laboratory | Jade Wang, et al. |
| EPI_ISL_857267, EPI_ISL_857268, EPI_ISL_857269, EPI_ISL_857270 | DOHMH Jamaica | New York City Public Health Laboratory | Jade Wang, et al. |
| EPI_ISL_857271, EPI_ISL_857272, EPI_ISL_857273 | DOHMH Morrisania | New York City Public Health Laboratory | Jade Wang, et al. |
| EPI_ISL_857274, EPI_ISL_857275, EPI_ISL_857276 | DOHMH Jamaica | New York City Public Health Laboratory | Jade Wang, et al. |
| EPI_ISL_857277 | DOHMH Corona | New York City Public Health Laboratory | Jade Wang, et al. |
| EPI_ISL_857278 | DOHMH PHL | New York City Public Health Laboratory | Jade Wang, et al. |
| EPI_ISL_858274, EPI_ISL_858313 | Lighthouse Lab in Glasgow | Wellcome Sanger Institute for the COVID-19 Genomics UK (COG-UK) Consortium | Harper VanSteenhouse, Yumi Kasai, David Gray, Carol Clugston, Anna Dominiczak and Alex Alderton, Roberto Amato, Sonia Goncalves, Ewan Harrison, David K. Jackson, Ian Johnston, Dominic Kwiatkowski, Cordelia Langford, John Sillitoe on behalf of the Wellcome Sanger Institute COVID-19 Surveillance Team |
| EPI_ISL_858573, EPI_ISL_858575, EPI_ISL_858577, EPI_ISL_858578, EPI_ISL_858579, EPI_ISL_858582 | Lighthouse Lab in Alderley Park | Wellcome Sanger Institute for the COVID-19 Genomics UK (COG-UK) Consortium | Jacquelyn Wynn, Mairead Hyland, The Lighthouse Lab in Alderley Park and Alex Alderton, Roberto Amato, Sonia Goncalves, Ewan Harrison, David K. Jackson, Ian Johnston, Dominic Kwiatkowski, Cordelia Langford, John Sillitoe on behalf of the Wellcome Sanger Institute COVID-19 Surveillance Team |
| EPI_ISL_858589, EPI_ISL_858591, EPI_ISL_858593, EPI_ISL_858595, EPI_ISL_858596, EPI_ISL_858599, EPI_ISL_858602, EPI_ISL_858603, EPI_ISL_858605, EPI_ISL_858606, EPI_ISL_858611, EPI_ISL_858617, EPI_ISL_858620, EPI_ISL_858622, EPI_ISL_858625, EPI_ISL_858629, EPI_ISL_858631, EPI_ISL_858632, EPI_ISL_858639 | | | |
| see above | Lighthouse Lab in Glasgow | Wellcome Sanger Institute for the COVID-19 Genomics UK (COG-UK) Consortium | Harper VanSteenhouse, Yumi Kasai, David Gray, Carol Clugston, Anna Dominiczak and Alex Alderton, Roberto Amato, Sonia Goncalves, Ewan Harrison, David K. Jackson, Ian Johnston, Dominic Kwiatkowski, Cordelia Langford, John Sillitoe on behalf of the Wellcome Sanger Institute COVID-19 Surveillance Team |
| EPI_ISL_858644, EPI_ISL_858647, EPI_ISL_858648, EPI_ISL_858649, EPI_ISL_858653, EPI_ISL_858654, EPI_ISL_858655, EPI_ISL_858662, EPI_ISL_858663, EPI_ISL_858666, EPI_ISL_858667, EPI_ISL_858669, EPI_ISL_858670, EPI_ISL_858671, EPI_ISL_858672, EPI_ISL_858673, EPI_ISL_858674, EPI_ISL_858675, EPI_ISL_858676, EPI_ISL_858677, EPI_ISL_858678, EPI_ISL_858679, EPI_ISL_858680, EPI_ISL_858681, EPI_ISL_858682, EPI_ISL_858683, EPI_ISL_858684, EPI_ISL_858685, EPI_ISL_858686, EPI_ISL_858687, EPI_ISL_858688, EPI_ISL_858689, EPI_ISL_858690, EPI_ISL_858691, EPI_ISL_858692, EPI_ISL_858693, EPI_ISL_858694, EPI_ISL_858695, EPI_ISL_858696, EPI_ISL_858697, EPI_ISL_858698, EPI_ISL_858699, EPI_ISL_858700 | | | |
| see above | Lighthouse Lab in Alderley Park | Wellcome Sanger Institute for the COVID-19 Genomics UK (COG-UK) Consortium | Jacquelyn Wynn, Mairead Hyland, The Lighthouse Lab in Alderley Park and Alex Alderton, Roberto Amato, Sonia Goncalves, Ewan Harrison, David K. Jackson, Ian Johnston, Dominic Kwiatkowski, Cordelia Langford, John Sillitoe on behalf of the Wellcome Sanger Institute COVID-19 Surveillance Team |
| EPI_ISL_858711, EPI_ISL_858712, EPI_ISL_858715, EPI_ISL_858716, EPI_ISL_858717, EPI_ISL_858718, EPI_ISL_858721, EPI_ISL_858722, EPI_ISL_858723, EPI_ISL_858724, EPI_ISL_858726, EPI_ISL_858727, EPI_ISL_858728, EPI_ISL_858729, EPI_ISL_858730, EPI_ISL_858731, EPI_ISL_858733, EPI_ISL_858734, EPI_ISL_858735, EPI_ISL_858736, EPI_ISL_858738, EPI_ISL_858740, EPI_ISL_858741, EPI_ISL_858742, EPI_ISL_858743, EPI_ISL_858744, EPI_ISL_858745, EPI_ISL_858746, EPI_ISL_858747, EPI_ISL_858749, EPI_ISL_858750, EPI_ISL_858751, EPI_ISL_858754, EPI_ISL_858755, EPI_ISL_858756, EPI_ISL_858757, EPI_ISL_858758, EPI_ISL_858759, EPI_ISL_858760, EPI_ISL_858761, EPI_ISL_858762, EPI_ISL_858763, EPI_ISL_858764, EPI_ISL_858765, EPI_ISL_858766, EPI_ISL_858767, EPI_ISL_858768, EPI_ISL_858769, EPI_ISL_858770, EPI_ISL_858771, EPI_ISL_858772, EPI_ISL_858773, EPI_ISL_858774, EPI_ISL_858775, EPI_ISL_858776, EPI_ISL_858777, EPI_ISL_858781, EPI_ISL_858783 | | | |
| see above | Lighthouse Lab in Glasgow | Wellcome Sanger Institute for the COVID-19 Genomics UK (COG-UK) Consortium | Harper VanSteenhouse, Yumi Kasai, David Gray, Carol Clugston, Anna Dominiczak and Alex Alderton, Roberto Amato, Sonia Goncalves, Ewan Harrison, David K. Jackson, Ian Johnston, Dominic Kwiatkowski, Cordelia Langford, John Sillitoe on behalf of the Wellcome Sanger Institute COVID-19 Surveillance Team |
| EPI_ISL_860026, EPI_ISL_860028, EPI_ISL_860029, EPI_ISL_860030, EPI_ISL_860031, EPI_ISL_860032, EPI_ISL_860033, EPI_ISL_860034, EPI_ISL_860035, EPI_ISL_860036, EPI_ISL_860037, EPI_ISL_860038, EPI_ISL_860039, EPI_ISL_860040, EPI_ISL_860041, EPI_ISL_860044, EPI_ISL_860045, EPI_ISL_860046, EPI_ISL_860047, EPI_ISL_860070, EPI_ISL_860072, EPI_ISL_860074, EPI_ISL_860078 | | | |
| see above | BTC, Khalifa University | BTC, Khalifa University | Al Safar et al |
| EPI_ISL_860182, EPI_ISL_860183 | Bangalore Medical College and Research Institute | Department of Neurovirology, National Institute of Mental Health and Neurosciences (NIMHANS) | Chitra Pattabiraman, Pramada Prasad, Anson Kunjumon George, Risha Rasheed, Darshan Sreenivas, Nakka Vijay Kiran Reddy, Anita S Desai, V Ravi |
| EPI_ISL_860208, EPI_ISL_860211 | Haukeland University Hospital, Dept. of Microbiology | Norwegian Institute of Public Health, Department of Virology | Kathrine Stene-Johansen, Kamilla Heddeland Instefjord, Hilde Elshaug, Atiya R Ali,Marie Paulsen Madsen, Rasmus Riis Kopperud, Hilde Vollan, Karoline Bragstad, Olav Hungnes |
| EPI_ISL_860213 | Furst Medical Laboratory | Norwegian Institute of Public Health, Department of Virology | Kathrine Stene-Johansen, Kamilla Heddeland Instefjord, Hilde Elshaug, Atiya R Ali,Marie Paulsen Madsen, Rasmus Riis Kopperud, Hilde Vollan, Karoline Bragstad, Olav Hungnes |

| | | | |
|---|---|---|---|
| EPI_ISL_860230 | Ostfold Hospital Trust - Kalnes, Centre for Laboratory Medicine, Section for gene technology and infection serology | Norwegian Institute of Public Health, Department of Virology | Kathrine Stene-Johansen, Kamilla Heddeland Instefjord, Hilde Elshaug, Atiya R Ali,Marie Paulsen Madsen, Rasmus Riis Kopperud, Hilde Vollan, Karoline Bragstad, Olav Hungnes |
| EPI_ISL_860236 | Hospital of Southern Norway - Kristiansand, Department of Medical Microbiology | Norwegian Institute of Public Health, Department of Virology | Kathrine Stene-Johansen, Kamilla Heddeland Instefjord, Hilde Elshaug, Atiya R Ali,Marie Paulsen Madsen, Rasmus Riis Kopperud, Hilde Vollan, Karoline Bragstad, Olav Hungnes |
| EPI_ISL_860289 | Dept. of Medical Microbiology, Stavanger University Hospital, Helse Stavanger HF | Norwegian Institute of Public Health, Department of Virology | Kathrine Stene-Johansen, Kamilla Heddeland Instefjord, Hilde Elshaug, Atiya R Ali,Marie Paulsen Madsen, Rasmus Riis Kopperud, Hilde Vollan, Karoline Bragstad, Olav Hungnes |
| EPI_ISL_860302, EPI_ISL_860303 | Unit 17: Influenza & Other Respiratory Viruses, German National Influenza Center | Project group Epidemiology of Highly Pathogenic Microorganisms, Robert Koch-Institute | Andreas Sachse, Grit Schubert, Essia Belarbi, Sébastien Calvignac-Spencer, Thorsten Wolff, Ralf Dürrwald, Djin-Ye Oh, Marianne Wedde |
| EPI_ISL_860559, EPI_ISL_860576, EPI_ISL_860579, EPI_ISL_860580, EPI_ISL_860581, EPI_ISL_860583, EPI_ISL_860584, EPI_ISL_860586 | NHLS-IALCH | KRISP, KZn Research Innovation and Sequencing Platform | Giandhari J, Pillay S, Lessells R, Mdlalose K, York D, Khan S, Tegally H, Wilkinson E, de Oliveira T |
| EPI_ISL_860642, EPI_ISL_860643, EPI_ISL_860644, EPI_ISL_860645, EPI_ISL_860646, EPI_ISL_860647, EPI_ISL_860648, EPI_ISL_860649, EPI_ISL_860650, EPI_ISL_860651, EPI_ISL_860653, EPI_ISL_860654, EPI_ISL_860655, EPI_ISL_860656, EPI_ISL_860657, EPI_ISL_860658, EPI_ISL_860659, EPI_ISL_860660, EPI_ISL_860661, EPI_ISL_860662, EPI_ISL_860663, EPI_ISL_860664, EPI_ISL_860665, EPI_ISL_860782 | | | |
| see above | Respiratory Virus Unit, National Infection Service, Public Health England | COVID-19 Genomics UK (COG-UK) Consortium | PHE Covid Sequencing Team |
| EPI_ISL_860864, EPI_ISL_860869, EPI_ISL_860870, EPI_ISL_860871, EPI_ISL_860882, EPI_ISL_860883, EPI_ISL_860884 | Labo Analyses Med | National Reference Center for Viruses of Respiratory Infections, Institut Pasteur, Paris | Marion Barbet, Sylvie Behillil, Méline Bizard, Angela Brisebarre, Camille Capel, Etienne Simon-Lorière, Vincent Enouf, Maud Vanpeene, Sylvie van der Werf,Lefaure Brieuc |
| EPI_ISL_860898 | Centre Hospitalier Compiègne-Noyon - Laboratoire | National Reference Center for Viruses of Respiratory Infections, Institut Pasteur, Paris | Marion Barbet, Sylvie Behillil, Méline Bizard, Angela Brisebarre, Camille Capel, Etienne Simon-Lorière, Vincent Enouf, Maud Vanpeene, Sylvie van der Werf,Emond Jean-Philippe |
| EPI_ISL_860938, EPI_ISL_860946, EPI_ISL_860950, EPI_ISL_860957, EPI_ISL_860960, EPI_ISL_860969, EPI_ISL_861097, EPI_ISL_861102 | Johns Hopkins Hospital Department of Pathology | Johns Hopkins Hospital Department of Pathology | C. Paul Morris, Chun Huai Luo, Adannaya Amadi, Nicholas Gallagher, Heba H. Mostafa |
| EPI_ISL_861109 | New York Presbyterian Hospital | Wadsworth Center, New York State Department of Health | Kirsten St. George, Daryl M. Lamson, Alexis Russel, Matthew Shudt, Melissa A Leisner, Jonathan Plitnick, Navjot Singh, John Kelly, Erasmus Schneider, Erica Lasek-Nesselquist |
| EPI_ISL_861135, EPI_ISL_861136, EPI_ISL_861137, EPI_ISL_861138, EPI_ISL_861139, EPI_ISL_861140, EPI_ISL_861141, EPI_ISL_861142, EPI_ISL_861143 | ADIRONDACK MEDICAL CENTER | Wadsworth Center, New York State Department of Health | Kirsten St. George, Daryl M. Lamson, Alexis Russel, Matthew Shudt, Melissa A Leisner, Jonathan Plitnick, Navjot Singh, John Kelly, Erasmus Schneider, Erica Lasek-Nesselquist |
| EPI_ISL_861191 | NORTH SHORE UNIVERSITY HOSPITAL | Wadsworth Center, New York State Department of Health | Kirsten St. George, Daryl M. Lamson, Alexis Russel, Matthew Shudt, Melissa A Leisner, Jonathan Plitnick, Navjot Singh, John Kelly, Erasmus Schneider, Erica Lasek-Nesselquist |
| EPI_ISL_861290, EPI_ISL_861292, EPI_ISL_861295, EPI_ISL_861296, EPI_ISL_861297, EPI_ISL_861298, EPI_ISL_861300, EPI_ISL_861305, EPI_ISL_861306, EPI_ISL_861310, EPI_ISL_861312, EPI_ISL_861314, EPI_ISL_861317, EPI_ISL_861323, EPI_ISL_861325, EPI_ISL_861327, EPI_ISL_861329 | | | |
| see above | MONTEFIORE MEDICAL CENTER LABORATORIES | Wadsworth Center, New York State Department of Health | Kirsten St. George, Daryl M. Lamson, Alexis Russel, Matthew Shudt, Melissa A Leisner, Jonathan Plitnick, Navjot Singh, John Kelly, Erasmus Schneider, Erica Lasek-Nesselquist |
| EPI_ISL_861413 | NORTH SHORE UNIVERSITY HOSPITAL | Wadsworth Center, New York State Department of Health | Kirsten St. George, Daryl M. Lamson, Alexis Russel, Matthew Shudt, Melissa A Leisner, Jonathan Plitnick, Navjot Singh, John Kelly, Erasmus Schneider, Erica Lasek-Nesselquist |
| EPI_ISL_861434, EPI_ISL_861435 | Ohio State University | James Molecular Laboratory | Huolin Tu, Matthew R Avenarius, Laura Kubatko, Matthew Hunt, Xiaokang Pan, Peng Ru, Jason Garee, Keelie Thomas, Peter Mohler, Preeti Pancholi, Dan Jones |
| EPI_ISL_861439, EPI_ISL_861440, EPI_ISL_861441, EPI_ISL_861442 | The Ohio State University | James Molecular Laboratory | Huolin Tu, Matthew R Avenarius, Laura Kubatko, Matthew Hunt, Xiaokang Pan, Peng Ru, Jason Garee, Keelie Thomas, Peter Mohler, Preeti Pancholi, Dan Jones |
| EPI_ISL_861538, EPI_ISL_861552, EPI_ISL_861568, EPI_ISL_861570, EPI_ISL_861589 | Instituto Nacional de Saude (INSA) | Instituto Nacional de Saude (INSA) | Borges et al |
| EPI_ISL_861699 | Los Angeles County PHL | Los Angeles County PHL | P. Hemarajata et al. |
| EPI_ISL_861866 | Bioanalytika AG | University Hospital Basel, Clinical Bacteriology | Tim Roloff, Madlen Stange, Helena MB Seth-Smith, Alfredo Mari, Karoline Leuzinger, Julia Bielicki, Adrian Härri, Manuel Battegay, Hans Hirsch, Adrian Egli |
| EPI_ISL_862000, EPI_ISL_862001, EPI_ISL_862002, EPI_ISL_862003, EPI_ISL_862004, EPI_ISL_862005, EPI_ISL_862006, EPI_ISL_862007, EPI_ISL_862008, EPI_ISL_862009, EPI_ISL_862010, EPI_ISL_862011, EPI_ISL_862012, EPI_ISL_862013, EPI_ISL_862014, EPI_ISL_862015, EPI_ISL_862016, EPI_ISL_862017, EPI_ISL_862018, EPI_ISL_862019, EPI_ISL_862020, EPI_ISL_862021, EPI_ISL_862022, EPI_ISL_862023, EPI_ISL_862024, EPI_ISL_862025, EPI_ISL_862026, EPI_ISL_862027, EPI_ISL_862028, EPI_ISL_862029, EPI_ISL_862030, EPI_ISL_862031, EPI_ISL_862032, EPI_ISL_862033, EPI_ISL_862034, EPI_ISL_862035 | | | |
| see above | OHSU Lab Services Molecular Microbiology Lab | Oregon SARS-CoV-2 Genome Sequencing Center | Brendan L. O'Connell, Sally Grindstaff, Kayla Carter, Ruth V. Nichols, Alec J. Hirsch, Donna Hansel, Guang Fan, Xuan Qin, Daniel N. Streblow, William B. Messer, Andrew C. Adey, Benjamin N. Bimber, Brian J. O'Roak |
| EPI_ISL_862125, EPI_ISL_862132 | Charité Universitätsmedizin Berlin, Institut fur Virologie/Labor Berlin | Charité Universitätsmedizin Berlin, Institut für Virologie | Victor M Corman, Barbara Mühlemann, Jörn Beheim-Schwarzbach, Tobias Bleicker, Julia Tesch, Talitha Veith, Julia Schneider, Terry Jones, Christian Drosten |
| EPI_ISL_862556 | Gerencia de Asistencia Sanitaria de Soria | Instituto de Salud Carlos III | Iglesias-Caballero, M.Camarero, S. Molinero Calamita, M. González-Esguevillas, M. Pozo, F. Casas, I. Jiménez, P. Jiménez, M. Zaballos, A. Monzón, S. Varona, S. Juliá, M. Cuesta, I. Aldea, C. |
| EPI_ISL_862575 | Hospital Clínic | Instituto de Salud Carlos III | Iglesias-Caballero, M.Camarero, S. Molinero Calamita, M. González-Esguevillas, M. Pozo, F. Casas, I. Jiménez, P. Jiménez, M. Zaballos, A. Monzón, S. Varona, S. Juliá, M. Cuesta, I. Marcos, M.A. |
| EPI_ISL_862759 | Utah Public Health Laboratory, Utah Public Health Laboratory Infectious Disease submission group | Utah Public Health Laboratory, Utah Public Health Laboratory Infectious Disease submission group | Young,E.L., Oakeson,K.F., Gallagher,T. |
| EPI_ISL_862832 | National Institute of Infectious Diseases-Prof. Dr. Matei Bals Molecular Diagnostics Laboratory | National Institute of Infectious Diseases-Prof. Dr. Matei Bals Molecular Diagnostics Laboratory | Leontina Banica, Marius Surleac, Corina Casangiu, Petre Milu, Andreea Tudor, Simona Paraschiv, Dan Otelea |
| EPI_ISL_863532, EPI_ISL_863590, EPI_ISL_863600, EPI_ISL_863631, EPI_ISL_863635, EPI_ISL_863639, EPI_ISL_863645, EPI_ISL_863651, EPI_ISL_863707, EPI_ISL_863721, EPI_ISL_863729, EPI_ISL_863746, EPI_ISL_863803, EPI_ISL_863811, EPI_ISL_863830, EPI_ISL_863841, EPI_ISL_863853 | | | |
| see above | Lighthouse Lab in Glasgow | Wellcome Sanger Institute for the COVID-19 Genomics UK (COG-UK) Consortium | Harper VanSteenhouse, Yumi Kasai, David Gray, Carol Clugston, Anna Dominiczak and Alex Alderton, Roberto Amato, Sonia Goncalves, Ewan Harrison, David K. Jackson, Ian Johnston, Dominic Kwiatkowski, Cordelia Langford, John Sillitoe on behalf of the Wellcome Sanger Institute COVID-19 Surveillance Team |
| EPI_ISL_864738, EPI_ISL_864739, EPI_ISL_864740, EPI_ISL_864741, EPI_ISL_864742, EPI_ISL_864743, EPI_ISL_864744, EPI_ISL_864745, EPI_ISL_864752, EPI_ISL_864753, EPI_ISL_864787, EPI_ISL_864869, EPI_ISL_864871, EPI_ISL_864873 | | | |
| see above | Department of Pathology, University of Cambridge | COVID-19 Genomics UK (COG-UK) Consortium | Aminu S. Jahun, Yasmin Chaudhry, Grant Hall, Iliana Georgana, Myra Hosmillo, Martin D. Curran, Malte Pinckert, Surendra Parmar, Ian Goodfellow |
| EPI_ISL_864961, EPI_ISL_864962, EPI_ISL_864963, EPI_ISL_864964, EPI_ISL_864965, EPI_ISL_864966, EPI_ISL_864967, EPI_ISL_864968, EPI_ISL_864969, EPI_ISL_864970, EPI_ISL_864971, EPI_ISL_864972, EPI_ISL_864973, EPI_ISL_864974, EPI_ISL_864975, EPI_ISL_864976, EPI_ISL_864977, EPI_ISL_864978, EPI_ISL_864979, EPI_ISL_864980, EPI_ISL_864982, EPI_ISL_864983, EPI_ISL_864992, EPI_ISL_864993 | | | |

| | | | |
|---|---|---|---|
| see above | West of Scotland Specialist Virology Centre, NHSGGC / MRC-University of Glasgow Centre for Virus Research | COVID-19 Genomics UK (COG-UK) Consortium | Ana da Silva Filipe, Natasha Johnson, Kathy Smollett, Daniel Mair, Stephen Carmichael, Alice Broos, Lily Tong, Jenna Nichols, Kyriaki Nomikou; Sarah McDonald; Richard Orton, Joseph Hughes, Sreenu Vattipally, David L Robertson; Alasdair MacLean, Rory Gunson; Sharif Shaaban, Matthew Holden; Rachel Blacow, Guy Mollett, Kathy Li, James Shepherd, Antonia Ho, Emma Thomson |
| EPI_ISL_865000, EPI_ISL_865001, EPI_ISL_865002, EPI_ISL_865003, EPI_ISL_865008 | Lighthouse Lab in Glasgow / MRC-University of Glasgow Centre for Virus Research | COVID-19 Genomics UK (COG-UK) Consortium | Ana da Silva Filipe, Natasha Johnson, Kathy Smollett, Daniel Mair, Stephen Carmichael, Alice Broos, Lily Tong, Jenna Nichols, Kyriaki Nomikou; Sarah McDonald; Harper VanSteenhouse, Yumi Kasai, David Gray, Carol Clugston, Anna Dominiczak; Alasdair MacLean, Rory Gunson; Richard Orton, Joseph Hughes, Sreenu Vattipally, David L Robertson; Sharif Shaaban, Matthew Holden; Kathy Li, James Shepherd, Antonia Ho, Emma Thomson |
| EPI_ISL_865080, EPI_ISL_865081, EPI_ISL_865082, EPI_ISL_865087 | Virology Department, Royal Infirmary of Edinburgh, NHS Lothian / School of Biological Sciences, University of Edinburgh / Institute of Genetics and Molecular Medicine, University of Edinburgh | COVID-19 Genomics UK (COG-UK) Consortium | McHugh M, Dewar R, Rooke S, Gallagher M, Balcaza C, O'Toole Á, Scher E, Hill V, McCrone JT, Colquhoun R, Yu X, Jackson B, Rambaut A, Williams TC, Templeton K |
| EPI_ISL_865177, EPI_ISL_865354, EPI_ISL_865355, EPI_ISL_865356, EPI_ISL_865357, EPI_ISL_865358, EPI_ISL_865359, EPI_ISL_865360, EPI_ISL_865361, EPI_ISL_865362, EPI_ISL_865363, EPI_ISL_865371, EPI_ISL_865372, EPI_ISL_865375, EPI_ISL_865377, EPI_ISL_865378, EPI_ISL_865379, EPI_ISL_865380, EPI_ISL_865381, EPI_ISL_865382, EPI_ISL_865386 | | | |
| see above | Liverpool Clinical Laboratories | COVID-19 Genomics UK (COG-UK) Consortium | Sam Haldenby, Anita Lucaci, Steve Paterson, Julian Hiscox, Alistair Darby, M Almsaud, A Alrezaihi, Muhannad Alruwaili, Stuart D Armstrong, Jones Benjamin, Eleanor G Bentley, Anu Chawla, Jordan J Clark, Angela Cowell, Richard Eccles, Isabel García-Dorival, Matthew Gemmell, Alessandro Gerada, PKF Gilmore, Richard Gregory, Ximeng Han, Catherine Hartley, Margaret Hughes, Miren Iturriza-Gomara, James Johnson, L Luu, Jenifer Manson, Charlotte Nelson, Elaine O'Toole, Cassie Olateju, Rebekah Penrice-Randal , Lucille Rainbow, N.P Randle, Trevor Ian Robinson, Parul Sharma, Ghada T Shawli, James P Stewart, Neil Swainston, Ecaterina Vamos, Joanne Watts, Mark Whitehead |
| EPI_ISL_865498, EPI_ISL_865502, EPI_ISL_865527, EPI_ISL_865529, EPI_ISL_865530, EPI_ISL_865531, EPI_ISL_865532, EPI_ISL_865533, EPI_ISL_865534, EPI_ISL_865535, EPI_ISL_865536, EPI_ISL_865537, EPI_ISL_865538, EPI_ISL_865539, EPI_ISL_865540, EPI_ISL_865541, EPI_ISL_865542, EPI_ISL_865546, EPI_ISL_865547, EPI_ISL_865548, EPI_ISL_865549, EPI_ISL_865550, EPI_ISL_865551, EPI_ISL_865552, EPI_ISL_865553, EPI_ISL_865554, EPI_ISL_865555, EPI_ISL_865556, EPI_ISL_865557, EPI_ISL_865558, EPI_ISL_865561, EPI_ISL_865562, EPI_ISL_865567, EPI_ISL_865568, EPI_ISL_865569, EPI_ISL_865570, EPI_ISL_865579, EPI_ISL_865580, EPI_ISL_865582, EPI_ISL_865583 | | | |
| see above | Barts Health NHS Trust | COVID-19 Genomics UK (COG-UK) Consortium | CUTINO-MOGUEL, Maria-Teresa; HARRINGTON, David; OWOYEMI, Dola; KULASEGARAN-SHYLINI, Raghavendran; BROAD, Claire; KELE, Beatrix |
| EPI_ISL_865682, EPI_ISL_865691, EPI_ISL_865692, EPI_ISL_865693, EPI_ISL_865694, EPI_ISL_865812, EPI_ISL_865816, EPI_ISL_865817, EPI_ISL_865819, EPI_ISL_865820, EPI_ISL_865821, EPI_ISL_865822, EPI_ISL_865823, EPI_ISL_865824, EPI_ISL_865825, EPI_ISL_865826, EPI_ISL_865827, EPI_ISL_865828, EPI_ISL_865829, EPI_ISL_865830, EPI_ISL_865831, EPI_ISL_865832, EPI_ISL_865833, EPI_ISL_865834, EPI_ISL_865835, EPI_ISL_865836, EPI_ISL_865838, EPI_ISL_865839, EPI_ISL_865840, EPI_ISL_865841, EPI_ISL_865843, EPI_ISL_865845, EPI_ISL_865846 | | | |
| see above | University College London, Great Ormond Street Hospital for Children NHS Foundation Trust, Imperial College Healthcare NHS Trust | COVID-19 Genomics UK (COG-UK) Consortium | Sergi Castellano, Rachel Williams, Mark Kristiansen, Paola Resende Silva, Sunando Roy, Tony Brooks, Helena Tutill, Paola Niola, Patricia Dyal, Charlotte Williams, Leysa Forrest, Yasmin Panchbhaya, Jacqueline Findlay, Samuel Weeks, Julianne Brown, Kathryn Harris, Paul Randell, James Price, Alison Holmes, Judith Breuer |
| EPI_ISL_866071, EPI_ISL_866072, EPI_ISL_866073, EPI_ISL_866074, EPI_ISL_866115, EPI_ISL_866122, EPI_ISL_866127, EPI_ISL_866135, EPI_ISL_866140, EPI_ISL_866162, EPI_ISL_866168 | | | |
| see above | University College London Hospital | COVID-19 Genomics UK (COG-UK) Consortium | Judith Heaney, Matthew Byott, Catherine Houlihan, Dan Frampton, Stuart Kirk, Moira Spyer and Eleni Nastouli |
| EPI_ISL_866349, EPI_ISL_866350, EPI_ISL_866351, EPI_ISL_866352, EPI_ISL_866364, EPI_ISL_866365, EPI_ISL_866372 | Regional Virus Laboratory, Belfast Health and Social Care Trust | COVID-19 Genomics UK (COG-UK) Consortium | Conall McCaughey, James McKenna, Tanya Curran, Susan Feeney, Alison Watt, Ciara Cox, Mairead Connor, Zoltan Molnar, David Simpson, Derek Fairley |
| EPI_ISL_866486, EPI_ISL_866487, EPI_ISL_866489, EPI_ISL_866490, EPI_ISL_866491, EPI_ISL_866492, EPI_ISL_866493, EPI_ISL_866494, EPI_ISL_866495, EPI_ISL_866496, EPI_ISL_866497, EPI_ISL_866498, EPI_ISL_866499, EPI_ISL_866501, EPI_ISL_866502, EPI_ISL_866503, EPI_ISL_866504, EPI_ISL_866506, EPI_ISL_866507, EPI_ISL_866508, EPI_ISL_866510, EPI_ISL_866511, EPI_ISL_866512, EPI_ISL_866515, EPI_ISL_866516, EPI_ISL_866517, EPI_ISL_866518, EPI_ISL_866519, EPI_ISL_866522, EPI_ISL_866524, EPI_ISL_866525 | | | |
| see above | Northumbria University / South Tees Hospitals NHS Foundation Trust / North Cumbria Integrated Care NHS Foundation Trust / North Tees and Hartlepool NHS Foundation Trust / Newcastle Hospitals NHS Foundation Trust | COVID-19 Genomics UK (COG-UK) Consortium | Darren L Smith,Andrew Nelson,Matthew Bashton,Greg R Young,Joshua Loh,John Allan,Mohammad A Tariq,Giles S Holt,Gary Black,Wen C Yew,Lynn Dover,Paul Baker,Steve Liggett,Sarah Essex,Jane Greenaway,Debra Padgett,Clive Graham,Garren Scott,Edward Barton,Emma Swindells,Brendan Payne,Jennifer Collins,Yusri Taha,Gary Eltringham |
| EPI_ISL_866905 | Queens Medical Centre, Clinical Microbiology Department / DeepSeq Nottingham | COVID-19 Genomics UK (COG-UK) Consortium | Gemma Clark, Wendy Smith, Manjinder Khakh, Vicki M Fleming, Michelle M Lister, Hannah Howson-Wells, Jonathan Ball, Patrick McClure, Joseph Chappell, Theocharis Tsoleridis, Nadine Holmes, Matthew Carlisle, Christopher Moore, Fei Sang, Johnny Debebe, Victoria Wright, Matthew Loose |
| EPI_ISL_866992, EPI_ISL_866993, EPI_ISL_866994, EPI_ISL_866995, EPI_ISL_866996, EPI_ISL_866997, EPI_ISL_866998, EPI_ISL_866999, EPI_ISL_867000, EPI_ISL_867001, EPI_ISL_867002, EPI_ISL_867003, EPI_ISL_867004, EPI_ISL_867005, EPI_ISL_867006, EPI_ISL_867007, EPI_ISL_867010 | | | |
| see above | Lincolnshire Hospitals and DeepSeq Nottingham | COVID-19 Genomics UK (COG-UK) Consortium | Nichola Duckworth, Tim Sloan, Sarah Walsh, Jonathan Ball, Patrick McClure, Joeseph Chappell, Nadine Holmes, Matthew Carlisle, Christopher Moore, Fei Sang, Johnny Debebe, Victoria Wright, Matthew Loose |
| EPI_ISL_867039, EPI_ISL_867040, EPI_ISL_867041, EPI_ISL_867044, EPI_ISL_867045, EPI_ISL_867046, EPI_ISL_867048, EPI_ISL_867049, EPI_ISL_867050, EPI_ISL_867052, EPI_ISL_867053, EPI_ISL_867054, EPI_ISL_867056, EPI_ISL_867057, EPI_ISL_867058, EPI_ISL_867059, EPI_ISL_867060, EPI_ISL_867061, EPI_ISL_867062, EPI_ISL_867063, EPI_ISL_867064, EPI_ISL_867065, EPI_ISL_867066, EPI_ISL_867067, EPI_ISL_867068, EPI_ISL_867069, EPI_ISL_867071, EPI_ISL_867072, EPI_ISL_867073, EPI_ISL_867074, EPI_ISL_867075, EPI_ISL_867077, EPI_ISL_867078, EPI_ISL_867079, EPI_ISL_867080, EPI_ISL_867081, EPI_ISL_867082, EPI_ISL_867084, EPI_ISL_867087, EPI_ISL_867088, EPI_ISL_867089, EPI_ISL_867090, EPI_ISL_867093, EPI_ISL_867094, EPI_ISL_867095, EPI_ISL_867096, EPI_ISL_867097, EPI_ISL_867098, EPI_ISL_867130, EPI_ISL_867155, EPI_ISL_867159, EPI_ISL_867161, EPI_ISL_867162, EPI_ISL_867168, EPI_ISL_867169, EPI_ISL_867170, EPI_ISL_867171, EPI_ISL_867181 | | | |
| see above | Oxford Viromics, NDM, University of Oxford; Oxford University Hospitals; Basingstoke and North Hampshire Hospital | COVID-19 Genomics UK (COG-UK) Consortium | Tanya Golubchik, David Bonsall, George Macintyre, Amy Trebes, Mariateresa de Cesare, Catrin Moore, Alex Mobbs, Anita Justice, Robert Shaw, Monique Andersson, Timothy Peto, Emma Wise, Nathan Moore, Jessica Lynch, Nick Cortes, Matilde Mori, Stephen Kidd, David Buck, John Todd, Christophe Fraser |
| EPI_ISL_867216, EPI_ISL_867217, EPI_ISL_867437, EPI_ISL_867438, EPI_ISL_867439, EPI_ISL_867440, EPI_ISL_867441, EPI_ISL_867442, EPI_ISL_867443 | Originating lab: Wales Specialist Virology Centre Sequencing lab: Pathogen Genomics Unit | Public Health Wales Microbiology Cardiff Wales Specialist Virology Centre | Catherine Moore, Johnathan Evans, Laura Gifford, Malorie Perry, Simon Cottrell, Angela Marchbank, Alec Birchley, Alexander Adams, Amy Gaskin, Bree Gatica-Wilcox, Jason Coombes, Joel Southgate, Lauren Gilbert, Lee Graham, Nicole Pacchiarini, Sara Kumziene-Summerhayes, Sarah Taylor, Sophie Jones, Sara Rey, Matthew Bull, Joanne Watkins, Sally Corden, Tom Connor |
| EPI_ISL_868420, EPI_ISL_868423, EPI_ISL_868485, EPI_ISL_868629 | Virology Department, Sheffield Teaching Hospitals NHS Foundation Trust/Department of Infection, Immunity and Cardiovascular Disease, The Medical School, University of Sheffield | COVID-19 Genomics UK (COG-UK) Consortium | Thushan de Silva, Matthew Parker, Nikki Smith, Adri Angyal, Rebecca Brown, Luke Green, Rachel Tucker, Paul Parsons, Danielle Groves, Katie Johnson, Laura Carrilero, Alex Keeley, Dave Partridge, Matthew Wyles, Benjamin Lindsey, Mehmet Yavuz, Mohammad Raza, Cariad Evans |
| EPI_ISL_868716, EPI_ISL_868717, EPI_ISL_868718, EPI_ISL_868719, EPI_ISL_868720, EPI_ISL_868721, EPI_ISL_868722, EPI_ISL_868723, EPI_ISL_868724, EPI_ISL_868725, EPI_ISL_868726, EPI_ISL_868727, EPI_ISL_868728, EPI_ISL_868729, EPI_ISL_868730, EPI_ISL_868731, EPI_ISL_868732, EPI_ISL_868733, EPI_ISL_868734, EPI_ISL_868735, EPI_ISL_868736, EPI_ISL_868737, EPI_ISL_868738, EPI_ISL_868739, EPI_ISL_868740, EPI_ISL_868741, EPI_ISL_868742, EPI_ISL_868743, EPI_ISL_868744, EPI_ISL_868745, EPI_ISL_868747, EPI_ISL_868748, EPI_ISL_868749, EPI_ISL_868750, EPI_ISL_868751, EPI_ISL_868752, EPI_ISL_868753, EPI_ISL_868754, EPI_ISL_868755, EPI_ISL_868756, EPI_ISL_868757, EPI_ISL_868842, EPI_ISL_868846, EPI_ISL_868847, EPI_ISL_868849, EPI_ISL_868852, EPI_ISL_868857, EPI_ISL_868858, EPI_ISL_868862, EPI_ISL_868870, EPI_ISL_868871, EPI_ISL_868874, EPI_ISL_868876, EPI_ISL_868877, EPI_ISL_868901, EPI_ISL_868902, EPI_ISL_868903, EPI_ISL_868904, EPI_ISL_868905, EPI_ISL_868906, EPI_ISL_868907, EPI_ISL_868908, EPI_ISL_868909, EPI_ISL_868910, EPI_ISL_868911, EPI_ISL_868912, EPI_ISL_868913, EPI_ISL_868914, EPI_ISL_868915, EPI_ISL_868916, EPI_ISL_868917, EPI_ISL_868918, EPI_ISL_868919, EPI_ISL_868920, EPI_ISL_868921, EPI_ISL_868922, EPI_ISL_868923, EPI_ISL_868924, EPI_ISL_868925, EPI_ISL_868926, EPI_ISL_868927, EPI_ISL_868928, EPI_ISL_868929, EPI_ISL_868930, EPI_ISL_868931, EPI_ISL_868932, EPI_ISL_868933, EPI_ISL_868934, EPI_ISL_868935, EPI_ISL_868936, EPI_ISL_868937 | | | |
| see above | Bioinformatics and Biostatistics Lab, Advanced Sequencing Facility | COVID-19 Genomics UK (COG-UK) Consortium | Aengus Stewart,Jerome Nicod,Chelsea Sawyer,Laura Cubitt,Harshil Patel,Margaret Crawford |
| EPI_ISL_868949 | Charité Universitätsmedizin Berlin, Institute of Virology, Charitéplatz 1, 10117 Berlin, Germany | Charité Universitätsmedizin Berlin, Institute of Virology, Charitéplatz 1, 10117 Berlin, Germany | Victor M Corman, Julia Schneider, Jörn Beheim-Schwarzbach, Tobias Bleicker, Julia Tesch, Barbara Mühlemann, Talitha Veith, Terry Jones, Christian Drosten |
| EPI_ISL_869014, EPI_ISL_869015, EPI_ISL_869016, EPI_ISL_869017, EPI_ISL_869018, EPI_ISL_869019, EPI_ISL_869020, EPI_ISL_869021, EPI_ISL_869022, EPI_ISL_869023, EPI_ISL_869024, EPI_ISL_869025, EPI_ISL_869027, EPI_ISL_869028, EPI_ISL_869029, EPI_ISL_869030, EPI_ISL_869031, EPI_ISL_869032, EPI_ISL_869033, EPI_ISL_869034, EPI_ISL_869035, EPI_ISL_869036, EPI_ISL_869037, EPI_ISL_869038, EPI_ISL_869039, EPI_ISL_869040, EPI_ISL_869041, EPI_ISL_869042, EPI_ISL_869043, EPI_ISL_869044, EPI_ISL_869051, EPI_ISL_869052, EPI_ISL_869053, EPI_ISL_869054, EPI_ISL_869056, EPI_ISL_869058, EPI_ISL_869061, EPI_ISL_869062, EPI_ISL_869067, EPI_ISL_869069, EPI_ISL_869071, EPI_ISL_869072, EPI_ISL_869073, EPI_ISL_869074, EPI_ISL_869075, EPI_ISL_869077, EPI_ISL_869078 | | | |
| see above | Bioinformatics and Biostatistics Lab, Advanced Sequencing Facility | COVID-19 Genomics UK (COG-UK) Consortium | Aengus Stewart,Jerome Nicod,Chelsea Sawyer,Laura Cubitt,Harshil Patel,Margaret Crawford |
| EPI_ISL_870065 | Ohio Department of Health Laboratory | Ohio Department of Health Laboratory | Holmes, Jennifer; Eric Brandt, Keoni Omura, Glen McGillivary, Caitlin McDonnell, Kirtana Ramadugu, Erica Leasure, Kelsey Florek, Heather Blankenship, |

| | | | |
|---|---|---|---|
| | | | Quanta Brown, and Tammy Bannerman |
| EPI_ISL_871794, EPI_ISL_871795, EPI_ISL_871797, EPI_ISL_871800, EPI_ISL_871801, EPI_ISL_871802, EPI_ISL_871803, EPI_ISL_871804, EPI_ISL_871807, EPI_ISL_871808, EPI_ISL_871811, EPI_ISL_871812 | | | |
| see above | AIID | Irish Coronavirus Sequencing Consortium-Teagasc Grange | Matthew McCabe, Aljandro Abner Garcia Leon, Fiona Crispie, Calum Walsh, Michael Carr, John Kenny, Paul Cotter, Patrick Mallon, Gabriel Gonzalez |
| EPI_ISL_871816, EPI_ISL_871818, EPI_ISL_871819 | Ohio Department of Health Laboratory | Ohio Department of Health Laboratory | Holmes, Jennifer; Eric Brandt, Keoni Omura, Glen McGillivary, Caitlin McDonnell, Kirtana Ramadugu, Erica Leasure, Kelsey Florek, Heather Blankenship, Quanta Brown, and Tammy Bannerman |
| EPI_ISL_871908, EPI_ISL_871909 | Servicio de Microbiología Clínica (Complejo Hospitalario de Navarra, Pamplona), Instituto de Investigación Sanitaria de Navarra (IdiSNA) | SeqCOVID-SPAIN consortium/IBV(CSIC) | Carmen Ezpeleta Baquedano, Ana Navascués, Ana Miqueleiz and SeqCOVID-SPAIN consortium |
| EPI_ISL_871911 | Hospital Universitario Marqués de Valdecilla - IDIVAL (Santander, Cantabria) | SeqCOVID-SPAIN consortium/IBV(CSIC) | Mónica Gozalo Margüello, María Eliecer Cano García, Jose Manuel Méndez Legaza, Daniel Pablo Marcos, Jesús Rodríguez Rodríguez, María Siller Ruiz and SeqCOVID-SPAIN consortium |
| EPI_ISL_871956, EPI_ISL_871961 | Servicio de Microbiología Clínica (Complejo Hospitalario de Navarra, Pamplona), Instituto de Investigación Sanitaria de Navarra (IdiSNA) | SeqCOVID-SPAIN consortium/IBV(CSIC) | Carmen Ezpeleta Baquedano, Ana Navascués, Ana Miqueleiz and SeqCOVID-SPAIN consortium |
| EPI_ISL_871965, EPI_ISL_871966, EPI_ISL_871967, EPI_ISL_871968, EPI_ISL_871969, EPI_ISL_872002 | Ohio Department of Health Laboratory | Ohio Department of Health Laboratory | Holmes, Jennifer; Eric Brandt, Keoni Omura, Glen McGillivary, Caitlin McDonnell, Kirtana Ramadugu, Erica Leasure, Kelsey Florek, Heather Blankenship, Quanta Brown, and Tammy Bannerman |
| EPI_ISL_872041, EPI_ISL_872063, EPI_ISL_872064, EPI_ISL_872153, EPI_ISL_872154, EPI_ISL_872155 | Department of Clinical Microbiology | GIGA Medical Genomics | Keith Durkin, Maria Artesi, Sébastien Bontems, Raphaël Boreux, Bouchra Boujemla, Cécile Meex, Pierrette Melin, Marie-Pierre Hayette, Vincent Bours |
| EPI_ISL_872209, EPI_ISL_872259, EPI_ISL_872290 | Labo Analyses Med | National Reference Center for Viruses of Respiratory Infections, Institut Pasteur, Paris | Marion Barbet, Sylvie Behillil, Méline Bizard, Angela Brisebarre, Camille Capel, Etienne Simon-Lorière, Vincent Enouf, Maud Vanpeene, Sylvie van der Werf |
| EPI_ISL_872313 | Labo analyses med | National Reference Center for Viruses of Respiratory Infections, Institut Pasteur, Paris | Marion Barbet, Sylvie Behillil, Méline Bizard, Angela Brisebarre, Camille Capel, Etienne Simon-Lorière, Vincent Enouf, Maud Vanpeene, Sylvie van der Werf,Amzalag Jonas |
| EPI_ISL_872323 | Labo Analyses Med | National Reference Center for Viruses of Respiratory Infections, Institut Pasteur, Paris | Marion Barbet, Sylvie Behillil, Méline Bizard, Angela Brisebarre, Camille Capel, Etienne Simon-Lorière, Vincent Enouf, Maud Vanpeene, Sylvie van der Werf,Rouah Raquel |
| EPI_ISL_872382, EPI_ISL_872383, EPI_ISL_872384, EPI_ISL_872385 | Texas Department of State Health Services (TXDSHS) | Texas Department of State Health Services (TXDSHS) | Bonnie Oh, Anita Pokharel, James Daniel Bonser, Myong Koag, Chung Wang, Rachel Lee, Grace Kubin, Rashmi Tuladhar, Mayela Pedrueza, Maliha Rahman, Jenny Zhang |
| EPI_ISL_872572 | Sydney South West Pathology Service (SSWPS) - Liverpool Hospital - NSW Health Pathology | NSW Health Pathology - Institute of Clinical Pathology and Medical Research; Westmead Hospital; University of Sydney | CIDM-PH et al. |
| EPI_ISL_872625, EPI_ISL_872626, EPI_ISL_872627 | Nigeria Centre for Disease Control (NCDC) | African Centre of Excellence for Genomics of Infectious Diseases (ACEGID), Redeemer's University | Oluniyi P.E. et al |
| EPI_ISL_872742, EPI_ISL_872743 | Rhode Island Department of Health | Infectious Disease Program, Broad Institute of Harvard and MIT | Lemieux,J.E., Siddle,K.J., Huard,R., King,E., Azevedo,K., Miller,A., Adams,G., Gladden-Young,A., Lagerborg,K., Rudy,M., DeRuff,K., Carter,A., Normandin,E., Bauer,M., Reilly,S., Tomkins-Tinch,C., Loreth,C., Chaluvadi,S., Birren,B.W., Gallagher,G., Smole,S., Park,D.J., MacInnis,B.L., and Sabeti,P.C. |
| EPI_ISL_875347, EPI_ISL_875348 | Ohio Department of Health Laboratory | Ohio Department of Health Laboratory | Holmes, Jennifer; Eric Brandt, Keoni Omura, Glen McGillivary, Caitlin McDonnell, Kirtana Ramadugu, Erica Leasure, Kelsey Florek, Heather Blankenship, Quanta Brown, and Tammy Bannerman |
| EPI_ISL_875516 | Institute of Virology, Biomedical Research Center of the Slovak Academy of Sciences, Bratislava | Faculty of Natural Sciences, Comenius University, Bratislava | Kristína Boršová, Viktória abanová, Broa Brejová, Viktória Hodorová, Sabina Fumaová Havlíková, Juraj Kopáek, Martina Liková, ubomíra Lukáiková, Martina Neboháová, Monika Sláviková, Tomáš Vina, Boris Klempa, Jozef Nosek |
| EPI_ISL_876054, EPI_ISL_876076, EPI_ISL_876078, EPI_ISL_876084, EPI_ISL_876106, EPI_ISL_876122, EPI_ISL_876141, EPI_ISL_876145, EPI_ISL_876171, EPI_ISL_876204, EPI_ISL_876205, EPI_ISL_876252, EPI_ISL_876253, EPI_ISL_876254, EPI_ISL_876255, EPI_ISL_876256, EPI_ISL_876257, EPI_ISL_876258, EPI_ISL_876259, EPI_ISL_876260 | | | |
| see above | Massachusetts State Public Health Laboratory | Massachusetts State Public Health Laboratory | Andrew Lang, Timelia Fink, Glen Gallagher, Sandra Smole |
| EPI_ISL_876738 | Istituto Zooprofilattico Sperimentale della Puglia e della Basilicata | Istituto Zooprofilattico Sperimentale della Puglia e della Basilicata | Parisi A., Bianco A., Capozzi L., Del Sambro L., Manzulli V, Rondinone V., Pace L., Cipolletta D., Galante D. |
| EPI_ISL_876742 | Ospedale Santa Caterina Novella | Istituto Zooprofilattico Sperimentale della Puglia e della Basilicata | Parisi A., Bianco A., Capozzi L., Del Sambro L., Manzulli V, Rondinone V., Pace L., Cipolletta D., Galante D. |
| EPI_ISL_876743 | Presidio di Brindisi Di Summa - Perrino | Istituto Zooprofilattico Sperimentale della Puglia e della Basilicata | Parisi A., Bianco A., Capozzi L., Del Sambro L., Manzulli V, Rondinone V., Pace L., Cipolletta D., Galante D. |
| EPI_ISL_878734, EPI_ISL_878750, EPI_ISL_878752, EPI_ISL_878771, EPI_ISL_878773, EPI_ISL_878789, EPI_ISL_880171 | Rady's Childrens Hospital | Andersen lab at Scripps Research | SEARCH Alliance San Diego with Nanda Radamchar, David Dimmock, Linda Luo, Christina Clarke, Kathryn Bouic, Teresa Mueller, Denise Malicki |
| EPI_ISL_882303, EPI_ISL_882305, EPI_ISL_882307, EPI_ISL_882308, EPI_ISL_882309, EPI_ISL_882316, EPI_ISL_882320, EPI_ISL_882323, EPI_ISL_882327, EPI_ISL_882332, EPI_ISL_882343, EPI_ISL_882352, EPI_ISL_882361, EPI_ISL_882363, EPI_ISL_882364, EPI_ISL_882371, EPI_ISL_882372, EPI_ISL_882375, EPI_ISL_882376, EPI_ISL_882377, EPI_ISL_882385, EPI_ISL_882388, EPI_ISL_882390, EPI_ISL_882393, EPI_ISL_882397, EPI_ISL_882398, EPI_ISL_882402, EPI_ISL_882404, EPI_ISL_882409, EPI_ISL_882413, EPI_ISL_882415, EPI_ISL_882418, EPI_ISL_882420, EPI_ISL_882421, EPI_ISL_882427, EPI_ISL_882437, EPI_ISL_882438, EPI_ISL_882440, EPI_ISL_882441, EPI_ISL_882442, EPI_ISL_882452, EPI_ISL_882471, EPI_ISL_882473, EPI_ISL_882475, EPI_ISL_882476, EPI_ISL_882480, EPI_ISL_882481, EPI_ISL_882484, EPI_ISL_882491, EPI_ISL_882506, EPI_ISL_882512, EPI_ISL_882518, EPI_ISL_882520, EPI_ISL_882525, EPI_ISL_882534, EPI_ISL_882538, EPI_ISL_882539, EPI_ISL_882542, EPI_ISL_882548, EPI_ISL_882550, EPI_ISL_882559, EPI_ISL_882562, EPI_ISL_882563, EPI_ISL_882571, EPI_ISL_882579, EPI_ISL_882580, EPI_ISL_882585, EPI_ISL_882586, EPI_ISL_882589, EPI_ISL_882595 | | | |
| see above | Lighthouse Lab in Alderley Park | Wellcome Sanger Institute for the COVID-19 Genomics UK (COG-UK) Consortium | Jacquelyn Wynn, Mairead Hyland, The Lighthouse Lab in Alderley Park and Alex Alderton, Roberto Amato, Sonia Goncalves, Ewan Harrison, David K. Jackson, Ian Johnston, Dominic Kwiatkowski, Cordelia Langford, John Sillitoe on behalf of the Wellcome Sanger Institute COVID-19 Surveillance Team |
| EPI_ISL_882617, EPI_ISL_882621, EPI_ISL_882623 | Hospital Ramón y Cajal | Hospital Ramón y Cajal | José M Gonzalez-Alba, Concepción Rodríguez, Melanie Abreu, Laura Martínez, Val F Lanza, Luz Leticia Olavarrieta, Rafael Cantón, JC Galán |
| EPI_ISL_882635 | SC (UCO) Igiene e Sanità Pubblica (funzione integrata con SC Microbiologia e Virologia) e Laboratory of Molecular Virology of the International Centre for Genetic Engineering and Biotechnology (ICGEB) | ARGO Laboratorio Genomica ed EpigenomicA | Licastro D, Dal Monego S, Degasperi M, Marcello A, D'Agaro P |
| EPI_ISL_882636 | SC (UCO) Igiene e Sanità Pubblica (funzione integrata con SC Microbiologia e Virologia) e Laboratory of Molecular Virology of the International Centre for Genetic Engineering and Biotechnology (ICGEB) | ARGO Laboratorio Genomica ed Epigenomica | Licastro D, Dal Monego S, Degasperi M, Marcello A, D'Agaro P |
| EPI_ISL_882663, EPI_ISL_882664 | LACEN do Estado do Piaui, Dr. Costa Alvarenga | Instituto Adolfo Lutz, Interdiciplinary Procedures Center, Strategic Laboratory | Claudio Tavares Sacchi, Claudia Regina Gonçalves, Erica Valessa Ramos Gomes, Karoline Rodrigues Campos |
| EPI_ISL_882779, EPI_ISL_882781, EPI_ISL_882800, EPI_ISL_882801, EPI_ISL_882802, EPI_ISL_882803, EPI_ISL_882804 | Institute for Urban Disease Control and Prevention | COVID-19 Network Investigations (CONI) Alliance | Kamolthip Atsawawaranunt, Elizabeth Batty, Wasun Chantratita, Thanat Chookajorn, Stefan Fernandez, Angkana Huang, Anthony R. Jones, Khajohn Joonsalak, Chonticha Klungtong, Theerarat Kochakarn, Prayuth Kaewmalang, Amornmas Kongklieng, Namfon Kotanan, Krittikorn Kumpornsin, Duangkamon Loesbanluechai, Wudtichai Manasatienkij, Anek Mungaomklang, Bhakbhoom Panthan, Pukkaporn Parnwijitkul, Ekawat Pasomsub, Vichan Pawun, Kingkan Rakmanee, Insee Sensorn, Janjira Thaipadungpanit, Arporn Wangwiwatsin, Treewat Watthanachockchai |

| | | | |
|---|---|---|---|
| EPI_ISL_882955 | State Veterinary Institute Prague | State Veterinary Institute Prague | Nagy,A;Cernikova,L;Stara,M |
| EPI_ISL_883033, EPI_ISL_883034 | Maryland Public Health Laboratory | Maryland Public Health Laboratory | Maryland Department of Health Laboratories Administration |
| EPI_ISL_883350 | OCME Office Of Chief Medical Examiner | New York City Public Health Laboratory | Jade Wang, et al. |
| EPI_ISL_884265, EPI_ISL_884268, EPI_ISL_884269, EPI_ISL_884270, EPI_ISL_884274, EPI_ISL_884277, EPI_ISL_884279, EPI_ISL_884286, EPI_ISL_884287 | Institute of Medical Microbiology and Hospital Hygiene | Institute of Medical Microbiology and Hospital Hygiene | Prof. Dr. Achim Kaasch, Aljoscha Tersteegen |
| EPI_ISL_884984, EPI_ISL_884985, EPI_ISL_884986, EPI_ISL_885013, EPI_ISL_885014, EPI_ISL_885015, EPI_ISL_885016, EPI_ISL_885017, EPI_ISL_885018, EPI_ISL_885019, EPI_ISL_885020, EPI_ISL_885021, EPI_ISL_885023, EPI_ISL_885024, EPI_ISL_885025, EPI_ISL_885033, EPI_ISL_885034, EPI_ISL_885036, EPI_ISL_885039, EPI_ISL_885040, EPI_ISL_885043 | | | |
| see above | Santa Clara County Public Health Laboratory | Chan-Zuckerberg Biohub | CZB Cliahub Consortium |
| EPI_ISL_885139 | PathWest Laboratory Medicine WA | PathWest Laboratory Medicine WA Microbial Surveillance Unit | PathWest Laboratory Medicine WA Microbial Surveillance Unit |
| EPI_ISL_886232, EPI_ISL_886279, EPI_ISL_886361, EPI_ISL_886455, EPI_ISL_886472, EPI_ISL_886576, EPI_ISL_886600, EPI_ISL_886610, EPI_ISL_886702, EPI_ISL_886863, EPI_ISL_887054, EPI_ISL_887103, EPI_ISL_887635, EPI_ISL_887648, EPI_ISL_887672, EPI_ISL_887676, EPI_ISL_887682, EPI_ISL_887685, EPI_ISL_887688, EPI_ISL_887714, EPI_ISL_887719, EPI_ISL_887722, EPI_ISL_887730, EPI_ISL_887760, EPI_ISL_887761, EPI_ISL_887781, EPI_ISL_887790, EPI_ISL_887814, EPI_ISL_887836, EPI_ISL_887840, EPI_ISL_887842, EPI_ISL_887853, EPI_ISL_887855, EPI_ISL_887858, EPI_ISL_887865, EPI_ISL_887879, EPI_ISL_887883, EPI_ISL_887884, EPI_ISL_887889, EPI_ISL_887894, EPI_ISL_887906, EPI_ISL_887907, EPI_ISL_887919, EPI_ISL_887954, EPI_ISL_887956, EPI_ISL_887962, EPI_ISL_887963, EPI_ISL_887977, EPI_ISL_888033, EPI_ISL_888035, EPI_ISL_888038, EPI_ISL_888039, EPI_ISL_888045, EPI_ISL_888047, EPI_ISL_888049, EPI_ISL_888090, EPI_ISL_888092, EPI_ISL_888096, EPI_ISL_888134, EPI_ISL_888154, EPI_ISL_888161, EPI_ISL_888282, EPI_ISL_888285, EPI_ISL_888335, EPI_ISL_888504, EPI_ISL_888513, EPI_ISL_888521, EPI_ISL_888529, EPI_ISL_888532, EPI_ISL_888534, EPI_ISL_888535, EPI_ISL_888536, EPI_ISL_888537, EPI_ISL_888539, EPI_ISL_888540, EPI_ISL_888552, EPI_ISL_888555, EPI_ISL_888558, EPI_ISL_888561, EPI_ISL_888573, EPI_ISL_888574, EPI_ISL_888583 | | | |
| see above | Labcorp | Genomics and Discovery, Respiratory Viruses Branch, Division of Viral Diseases, Centers for Disease Control and Prevention | Peter W. Cook,Dhwani Batra,Ben L. Rambo-Martin,Summer Galloway,Brian Krueger,Minoo Agarwal,Eyad Almasri,Debbie Boles,Ayla Burns,Nuthawin Charoensri,Oren Cohen,Susan Countryman,Mary Ann Cristobal,Bobbi Croy,Suzanne Dale,Hrushikesh Deshmukh,Amanda Douglas,Vincent Drouillon,Marcia Eisenberg,Howard Engler,Rama Ghatti,Prashant Gupta,Susan Hicks,Jake Humphrey,Lax Iyer,Manoj Jain,Mohan Kolli,Tim Kuphal,Stanley Letovsky,Michael Levandoski,Craig Lukasik,Jonathan Meltzer,Brian Norvell,Mindy Nye,Scott Parker,Christos Petropoulos,John Pruitt,Steven Ragan,Scott Ryan,Mike Sapeta,Jana Schroth,Suresh Babu Selvaraju,Goran Stevovic,Amanda Suchanek,Andrea Throop,Lyndon Tilson,Thomas Urban,Joe Voshell,Kimberly Wagner,Jonathan Williams,Mary Williamson,Qian Zeng,Tricia Zwiefelhofer,Clinton R. Paden,Suxiang Tong,Duncan MacCannell, |
| EPI_ISL_888785, EPI_ISL_888789, EPI_ISL_888790 | KU Leuven, Rega Institute, Clinical and Epidemiological Virology | KU Leuven, Rega Institute, Clinical and Epidemiological Virology | Tony Wawina-Bokalanga, Bert Vanmechelen, Joan Marti-Carerras, Piet Maes |
| EPI_ISL_888832, EPI_ISL_888834 | National Virus Reference Laboratory | National Virus Reference Laboratory | Michael Carr, Gabriel Gonzalez, Jonathan Dean, Cillian F De Gascun |
| EPI_ISL_888845, EPI_ISL_888846, EPI_ISL_888847, EPI_ISL_888852 | Michigan Department of Health and Human Services, Bureau of Laboratories | Michigan Department of Health and Human Services, Bureau of Laboratories | Blankenship HM, Riner D, Soehnlen MK |
| EPI_ISL_888994 | RSU MMC | Eijkman Institute for Molecular Biology, Ministry of Research and Technology/National Agency for Research and Innovation | Lydia V. Panggalo, Sukma Oktavianthi, Willy Agustine, Edison Johar, Hidayat Trimarsanto, Iskandar Adnan, Frilasita A Yudhaputri, Safarina G Malik, Khin Saw Myint, Amin Soebandrio |
| EPI_ISL_888995 | RSU Sumber Waras | Eijkman Institute for Molecular Biology, Ministry of Research and Technology/National Agency for Research and Innovation | Lydia V. Panggalo, Sukma Oktavianthi, Willy Agustine, Edison Johar, Hidayat Trimarsanto, Iskandar Adnan, Frilasita A Yudhaputri, Safarina G Malik, Khin Saw Myint, Amin Soebandrio |
| EPI_ISL_888996 | RSU Graha Juanda | Eijkman Institute for Molecular Biology, Ministry of Research and Technology/National Agency for Research and Innovation | Lydia V. Panggalo, Sukma Oktavianthi, Willy Agustine, Edison Johar, Hidayat Trimarsanto, Iskandar Adnan, Frilasita A Yudhaputri, Safarina G Malik, Khin Saw Myint, Amin Soebandrio |
| EPI_ISL_889003 | RS Mitra Keluarga Gading Serpong | Eijkman Institute for Molecular Biology, Ministry of Research and Technology/National Agency for Research and Innovation | Willy Agustine, Edison Johar, Hidayat Trimarsanto, Iskandar Adnan, Lydia V. Panggalo, Sukma Oktavianthi, Frilasita A Yudhaputri, Safarina G Malik, Khin Saw Myint, Amin Soebandrio |
| EPI_ISL_889017 | RS Hermina Tangerang | Eijkman Institute for Molecular Biology, Ministry of Research and Technology/National Agency for Research and Innovation | Lydia V. Panggalo, Sukma Oktavianthi, Willy Agustine, Edison Johar, Hidayat Trimarsanto, Iskandar Adnan, Frilasita A Yudhaputri, Safarina G Malik, Khin Saw Myint, Amin Soebandrio |
| EPI_ISL_889023, EPI_ISL_889024, EPI_ISL_889025, EPI_ISL_889026, EPI_ISL_889027, EPI_ISL_889071, EPI_ISL_889072, EPI_ISL_889073, EPI_ISL_889074, EPI_ISL_889075, EPI_ISL_889076, EPI_ISL_889077, EPI_ISL_889078, EPI_ISL_889079, EPI_ISL_889080, EPI_ISL_889081, EPI_ISL_889082, EPI_ISL_889083, EPI_ISL_889084, EPI_ISL_889085, EPI_ISL_889086, EPI_ISL_889087, EPI_ISL_889088, EPI_ISL_889089, EPI_ISL_889090, EPI_ISL_889091, EPI_ISL_889092, EPI_ISL_889093, EPI_ISL_889094, EPI_ISL_889095, EPI_ISL_889096, EPI_ISL_889097, EPI_ISL_889098, EPI_ISL_889099, EPI_ISL_889100, EPI_ISL_889101, EPI_ISL_889102, EPI_ISL_889103, EPI_ISL_889104, EPI_ISL_889105, EPI_ISL_889106, EPI_ISL_889107, EPI_ISL_889108, EPI_ISL_889109, EPI_ISL_889110, EPI_ISL_889111, EPI_ISL_889112, EPI_ISL_889113, EPI_ISL_889114, EPI_ISL_889115, EPI_ISL_889116, EPI_ISL_889117, EPI_ISL_889118, EPI_ISL_889119, EPI_ISL_889120, EPI_ISL_889121, EPI_ISL_889122, EPI_ISL_889123, EPI_ISL_889124, EPI_ISL_889125, EPI_ISL_889126, EPI_ISL_889127, EPI_ISL_889128, EPI_ISL_889129, EPI_ISL_889130, EPI_ISL_889131, EPI_ISL_889132, EPI_ISL_889133, EPI_ISL_889134, EPI_ISL_889135, EPI_ISL_889136, EPI_ISL_889137, EPI_ISL_889138, EPI_ISL_889139, EPI_ISL_889140, EPI_ISL_889141, EPI_ISL_889142, EPI_ISL_889143, EPI_ISL_889144, EPI_ISL_889145, EPI_ISL_889146, EPI_ISL_889147, EPI_ISL_889148, EPI_ISL_889149, EPI_ISL_889150, EPI_ISL_889151, EPI_ISL_889152, EPI_ISL_889153, EPI_ISL_889158, EPI_ISL_889159, EPI_ISL_889160, EPI_ISL_889161, EPI_ISL_889176, EPI_ISL_889177, EPI_ISL_889178, EPI_ISL_889179, EPI_ISL_889180, EPI_ISL_889181, EPI_ISL_889182, EPI_ISL_889183, EPI_ISL_889184, EPI_ISL_889185, EPI_ISL_889186, EPI_ISL_889187, EPI_ISL_889188, EPI_ISL_889189, EPI_ISL_889190, EPI_ISL_889191, EPI_ISL_889192, EPI_ISL_889193, EPI_ISL_889194, EPI_ISL_889195, EPI_ISL_889196, EPI_ISL_889197, EPI_ISL_889198, EPI_ISL_889199, EPI_ISL_889200, EPI_ISL_889201, EPI_ISL_889202, EPI_ISL_889203, EPI_ISL_889204, EPI_ISL_889205, EPI_ISL_889206, EPI_ISL_889207, EPI_ISL_889208, EPI_ISL_889209, EPI_ISL_889210, EPI_ISL_889211, EPI_ISL_889212, EPI_ISL_889213, EPI_ISL_889214, EPI_ISL_889215, EPI_ISL_889216, EPI_ISL_889217, EPI_ISL_889218, EPI_ISL_889219, EPI_ISL_889220, EPI_ISL_889221, EPI_ISL_889222, EPI_ISL_889223, EPI_ISL_889224, EPI_ISL_889225, EPI_ISL_889226, EPI_ISL_889227, EPI_ISL_889228, EPI_ISL_889229, EPI_ISL_889230, EPI_ISL_889231, EPI_ISL_889232, EPI_ISL_889233, EPI_ISL_889234, EPI_ISL_889235, EPI_ISL_889236, EPI_ISL_889237, EPI_ISL_889238, EPI_ISL_889239, EPI_ISL_889240, EPI_ISL_889241, EPI_ISL_889242, EPI_ISL_889243, EPI_ISL_889244, EPI_ISL_889245, EPI_ISL_889246, EPI_ISL_889247, EPI_ISL_889248, EPI_ISL_889249, EPI_ISL_889250, EPI_ISL_889251, EPI_ISL_889252, EPI_ISL_889253, EPI_ISL_889254, EPI_ISL_889255, EPI_ISL_889256, EPI_ISL_889257, EPI_ISL_889258, EPI_ISL_889259, EPI_ISL_889260, EPI_ISL_889261, EPI_ISL_889262, EPI_ISL_889263, EPI_ISL_889264, EPI_ISL_889265, EPI_ISL_889266, EPI_ISL_889267, EPI_ISL_889268, EPI_ISL_889269, EPI_ISL_889270, EPI_ISL_889271, EPI_ISL_889272, EPI_ISL_889273, EPI_ISL_889274, EPI_ISL_889275, EPI_ISL_889276, EPI_ISL_889277, EPI_ISL_889278, EPI_ISL_889279, EPI_ISL_889280, EPI_ISL_889281, EPI_ISL_889282, EPI_ISL_889283, EPI_ISL_889284, EPI_ISL_889285, EPI_ISL_889286, EPI_ISL_889287, EPI_ISL_889288, EPI_ISL_889289, EPI_ISL_889290, EPI_ISL_889291, EPI_ISL_889292, EPI_ISL_889293, EPI_ISL_889294, EPI_ISL_889295, EPI_ISL_889296, EPI_ISL_889297, EPI_ISL_889298, EPI_ISL_889299, EPI_ISL_889300, EPI_ISL_889301, EPI_ISL_889302, EPI_ISL_889303, EPI_ISL_889304, EPI_ISL_889305, EPI_ISL_889306 | | | |
| see above | Israel Central Virology laboratory | Israel National Consortium for SARS-CoV-2 sequencing | Neta Zuckerman, Efrat Dahan Bucris, Michal Mandelboim, Dana Bar-Ilan, Oran Erster, Tzvia Mann, Omer Murik, David A. Zeevi, Assaf Rokney, Joseph Jaffe, Eva Nachum, Maya Davidovich Cohen, Ephraim Fass, Gal Zizelski Valenci, Mor Rubinstein, Efrat Rorman, Israel Nissan, Efrat Glick-Saar, Omri Nayshool, Gideon Rechavi, Ella Mendelson, Orna Mor |
| EPI_ISL_889482, EPI_ISL_889486, EPI_ISL_889487, EPI_ISL_889488, EPI_ISL_889489, EPI_ISL_889490, EPI_ISL_889491, EPI_ISL_889492, EPI_ISL_889493, EPI_ISL_889494, EPI_ISL_889495, EPI_ISL_889496, EPI_ISL_889498, EPI_ISL_889499, EPI_ISL_889500, EPI_ISL_889501, EPI_ISL_889502, EPI_ISL_889503, EPI_ISL_889504, EPI_ISL_889505, EPI_ISL_889506, EPI_ISL_889507, EPI_ISL_889508, EPI_ISL_889509, EPI_ISL_889510, EPI_ISL_889511, EPI_ISL_889512, EPI_ISL_889513, EPI_ISL_889514, EPI_ISL_889515, EPI_ISL_889516, EPI_ISL_889517, EPI_ISL_889518, EPI_ISL_889519, EPI_ISL_889520, EPI_ISL_889521, EPI_ISL_889522, EPI_ISL_889523, EPI_ISL_889524, EPI_ISL_889525, EPI_ISL_889526, EPI_ISL_889527, EPI_ISL_889528, EPI_ISL_889529, EPI_ISL_889530, EPI_ISL_889531, EPI_ISL_889532, EPI_ISL_889533, EPI_ISL_889534, EPI_ISL_889535, EPI_ISL_889536, EPI_ISL_889537, EPI_ISL_889538, EPI_ISL_889539, EPI_ISL_889540, EPI_ISL_889541, EPI_ISL_889543, EPI_ISL_889544, EPI_ISL_889545, EPI_ISL_889546, EPI_ISL_889547, EPI_ISL_889548, EPI_ISL_889549, EPI_ISL_889550, EPI_ISL_889551 | | | |
| see above | LSUHS Emerging Viral Threat Laboratory | Microbial Genome Sequencing Center | Jeremy P. Kamil, Jennifer L. Carroll, Camille F. Abshire, Maarten Van Diest, Mohammed N.A. Siddiquey, Andrew D. Yurochko, Martin J. Sapp, Rona S. Scott, Christopher G. Kevil, Daniel J. Snyder, Vaughn S. Cooper, John A. Vanchiere |
| EPI_ISL_890238, EPI_ISL_890243, EPI_ISL_890295, EPI_ISL_890336, EPI_ISL_890340, EPI_ISL_890343, EPI_ISL_890346, EPI_ISL_890350 | KU Leuven, Rega Institute, Clinical and Epidemiological Virology | KU Leuven, Rega Institute, Clinical and Epidemiological Virology | Tony Wawina-Bokalanga, Bert Vanmechelen, Joan Marti-Carerras, Piet Maes |
| EPI_ISL_890361, EPI_ISL_890362 | LSUHS Emerging Viral Threat Laboratory | Microbial Genome Sequencing Center | Jeremy P. Kamil, Jennifer L. Carroll, Camille F. Abshire, Maarten Van Diest, Mohammed N.A. Siddiquey, Andrew D. Yurochko, Martin J. Sapp, Rona S. Scott, Christopher G. Kevil, Daniel J. Snyder, Vaughn S. Cooper, John A. Vanchiere |
| EPI_ISL_891035, EPI_ISL_891044, EPI_ISL_891046, EPI_ISL_891047, EPI_ISL_891049, EPI_ISL_891050, EPI_ISL_891051, EPI_ISL_891052, EPI_ISL_891053, EPI_ISL_891054, EPI_ISL_891055, EPI_ISL_891056, EPI_ISL_891057 | | | |
| see above | Washington State Department of Health | Seattle Flu Study | Deborah A. Nickerson, Chris D. Frazar, Jover Lee, Benjamin Pelle, Erica Ryke, Matthew Richardson, Amanda Adler, Elisabeth Brandstetter, Peter D. Han, Kairsten Fay, Misja Ilcisin, Kirsten Lacombe, Thomas R. Sibley, Melissa Truong, Caitlin R. Wolf, Romesh Gautom, Geoff Melly, Brian Hiatt, Philip Dykema, Scott Lindquist, Michael Boeckh, Janet A. Englund, Michael Famulare, Barry R. Lutz, Mark J. Rieder, Lea M. Starita, Matthew Thompson, Helen Y. Chu, Jay Shendure, Trevor Bedford |
| EPI_ISL_891136 | The Jackson Laboratory | The Jackson Laboratory | Lloyd M, Sanderson B, Srivastava A, Maurya R, Renzette N, Omerza G, Kelly K, Li L, Wei C L, Adams M |
| EPI_ISL_891198, EPI_ISL_891201 | DPH, Massachusetts State Public Health Lab | DPH, Massachusetts State Public Health Lab | Lang,A.S., Fink,T., Gallagher,G.R., Smole,S.C. |

| | | | |
|---|---|---|---|
| EPI_ISL_892244 | National Laboratory for Health, Environment and Food | National Laboratory for Health, Environment and Food | Aleksander Mahnic, Sandra Janezic, Maja Rupnik |
| EPI_ISL_893744, EPI_ISL_893774 | Institute of Virology, Medical Center, University of Freiburg, Freiburg, Germany | Institute of Virology, Clinial Virus Genomics, Medical Center, University of Freiburg, Freiburg, Germany | Jonas Fuchs, Lisa Kern, Sandra Reuter, Hajo Grundmann, Marcus Panning |
| EPI_ISL_894168 | Institute of Medical Microbiology and Hospital Hygiene | Institute of Medical Microbiology and Hospital Hygiene | Prof. Dr. Achim Kaasch, Aljoscha Tersteegen |
| EPI_ISL_896072 | Viollier AG | University Hospital Basel, Clinical Bacteriology | Tim Roloff, Madlen Stange, Helena MB Seth-Smith, Alfredo Mari, Karoline Leuzinger, Julia Bielicki, Christiane Beckmann, Manuel Battegay, Hans Hirsch, Adrian Egli |
| EPI_ISL_896087, EPI_ISL_896088 | Labormedizinisches Zentrum Dr Risch | University Hospital Basel, Clinical Bacteriology | Tim Roloff, Madlen Stange, Helena MB Seth-Smith, Alfredo Mari, Karoline Leuzinger, Julia Bielicki, Nadia Wohlwend,Martin Risch, Lorenz Risch, Manuel Battegay, Hans Hirsch, Adrian Egli |
| EPI_ISL_896095 | Bioanalytika AG | University Hospital Basel, Clinical Bacteriology | Tim Roloff, Madlen Stange, Helena MB Seth-Smith, Alfredo Mari, Karoline Leuzinger, Julia Bielicki, Adrian Härri, Manuel Battegay, Hans Hirsch, Adrian Egli |
| EPI_ISL_896101, EPI_ISL_896102 | Kantonsspital Aarau, Institut für Labormedizin | University Hospital Basel, Clinical Bacteriology | Tim Roloff, Madlen Stange, Helena MB Seth-Smith, Alfredo Mari, Karoline Leuzinger, Julia Bielicki, Michael Oberle, Manuel Battegay, Hans Hirsch, Adrian Egli |
| EPI_ISL_896103, EPI_ISL_896104 | Labormedizinisches Zentrum Dr Risch | University Hospital Basel, Clinical Bacteriology | Tim Roloff, Madlen Stange, Helena MB Seth-Smith, Alfredo Mari, Karoline Leuzinger, Julia Bielicki, Nadia Wohlwend,Martin Risch, Lorenz Risch, Manuel Battegay, Hans Hirsch, Adrian Egli |
| EPI_ISL_898010, EPI_ISL_898011, EPI_ISL_898025, EPI_ISL_898026, EPI_ISL_898027, EPI_ISL_898028 | KU Leuven, Rega Institute, Clinical and Epidemiological Virology | KU Leuven, Rega Institute, Clinical and Epidemiological Virology | Tony Wawina-Bokalanga, Bert Vanmechelen, Joan Marti-Carerras, Piet Maes |
| EPI_ISL_899008, EPI_ISL_899021, EPI_ISL_899045, EPI_ISL_899046, EPI_ISL_899049, EPI_ISL_899066, EPI_ISL_899067, EPI_ISL_899068, EPI_ISL_899091, EPI_ISL_899111, EPI_ISL_899121, EPI_ISL_899122, EPI_ISL_899135, EPI_ISL_899143, EPI_ISL_899144, EPI_ISL_899155, EPI_ISL_899168, EPI_ISL_899170, EPI_ISL_899173, EPI_ISL_899175, EPI_ISL_899188, EPI_ISL_899189, EPI_ISL_899190, EPI_ISL_899191, EPI_ISL_899192, EPI_ISL_899193, EPI_ISL_899194, EPI_ISL_899195, EPI_ISL_899196, EPI_ISL_899298, EPI_ISL_899299, EPI_ISL_899300, EPI_ISL_899301, EPI_ISL_899421, EPI_ISL_899422, EPI_ISL_899423, EPI_ISL_899424, EPI_ISL_899425, EPI_ISL_899426, EPI_ISL_899427, EPI_ISL_899428, EPI_ISL_899522, EPI_ISL_899523, EPI_ISL_899537, EPI_ISL_899547, EPI_ISL_899548, EPI_ISL_899578, EPI_ISL_899579, EPI_ISL_899580, EPI_ISL_899581, EPI_ISL_899582, EPI_ISL_899598, EPI_ISL_899622, EPI_ISL_899627, EPI_ISL_899628, EPI_ISL_899629, EPI_ISL_899630, EPI_ISL_899665, EPI_ISL_899666, EPI_ISL_899667, EPI_ISL_899773, EPI_ISL_899828, EPI_ISL_899829, EPI_ISL_899830, EPI_ISL_899831, EPI_ISL_899832, EPI_ISL_899833, EPI_ISL_899834, EPI_ISL_899835, EPI_ISL_899836, EPI_ISL_899837, EPI_ISL_899921, EPI_ISL_899942, EPI_ISL_899943, EPI_ISL_899952, EPI_ISL_899953, EPI_ISL_899954, EPI_ISL_899955, EPI_ISL_899956, EPI_ISL_899957, EPI_ISL_899958, EPI_ISL_899959, EPI_ISL_899960, EPI_ISL_899961, EPI_ISL_899962, EPI_ISL_899963, EPI_ISL_899964, EPI_ISL_899965, EPI_ISL_899966, EPI_ISL_899967, EPI_ISL_899968, EPI_ISL_899969, EPI_ISL_899970, EPI_ISL_899971, EPI_ISL_899972, EPI_ISL_899973 | | | |
| see above | Viollier AG | Department of Biosystems Science and Engineering, ETH Zürich | Christian Beisel, Sarah Nadeau, Chaoran Chen, Ivan Topolsky, Philipp Jablonski, Lara Fuhrmann, David Dreifuss, Katharina Jahn, Tobias Schär, Ina Nissen, Natascha Santacroce, Elodie Burcklen, Christiane Beckmann, Maurice Redondo, Olivier Kobel, Christoph Noppen, Sophie Seidel, Noemie Santamaria de Souza, Niko Beerenwinkel, Tanja Stadler |
| EPI_ISL_900502 | Gen-Bio | CNR Virus des Infections Respiratoires - France SUD | Antonin Bal, Gregory Destras, Gwendolyne Burfin, Hadrien Règue, Quentin Semanas, Martine Valette, Bruno Lina, Sylvie Larrat, Laurence Josset |
| EPI_ISL_900504, EPI_ISL_900507, EPI_ISL_900508, EPI_ISL_900509 | Mirialis | CNR Virus des Infections Respiratoires - France SUD | Antonin Bal, Gregory Destras, Gwendolyne Burfin, Hadrien Règue, Quentin Semanas, Martine Valette, Bruno Lina, Laurence Josset |
| EPI_ISL_900510 | Laboratoire de virologie, CHU de Grenoble - CS 10217 - 38043 Grenoble cedex 9 | CNR Virus des Infections Respiratoires - France SUD | Antonin Bal, Gregory Destras, Gwendolyne Burfin, Hadrien Règue, Quentin Semanas, Martine Valette, Bruno Lina, Sylvie Larrat, Laurence Josset |
| EPI_ISL_900513 | Labosud | CNR Virus des Infections Respiratoires - France SUD | Antonin Bal, Gregory Destras, Gwendolyne Burfin, Hadrien Règue, Quentin Semanas, Martine Valette, Bruno Lina, Laurence Josset |
| EPI_ISL_900515, EPI_ISL_900516, EPI_ISL_900517 | Bioesterel | CNR Virus des Infections Respiratoires - France SUD | Antonin Bal, Gregory Destras, Gwendolyne Burfin, Hadrien Règue, Quentin Semanas, Martine Valette, Bruno Lina, Laurence Josset |
| EPI_ISL_900521 | Novabio | CNR Virus des Infections Respiratoires - France SUD | Antonin Bal, Gregory Destras, Gwendolyne Burfin, Hadrien Règue, Quentin Semanas, Martine Valette, Bruno Lina, Laurence Josset |
| EPI_ISL_900523 | Novabio | CNR Virus des Infections Respiratoires - France SUD | Antonin Bal, Gregory Destras, Gwendolyne Burfin, Hadrien Règue, Quentin Semanas, Martine Valette, Bruno Lina, Sylvie Larrat, Laurence Josset |
| EPI_ISL_900525 | Astralab | CNR Virus des Infections Respiratoires - France SUD | Antonin Bal, Gregory Destras, Gwendolyne Burfin, Hadrien Règue, Quentin Semanas, Martine Valette, Bruno Lina, Laurence Josset |
| EPI_ISL_900526 | CNR Virus des Infections Respiratoires - France SUD | CNR Virus des Infections Respiratoires - France SUD | Antonin Bal, Gregory Destras, Gwendolyne Burfin, Hadrien Règue, Quentin Semanas, Martine Valette, Bruno Lina, Laurence Josset |
| EPI_ISL_900527 | Cerballiance | CNR Virus des Infections Respiratoires - France SUD | Antonin Bal, Gregory Destras, Gwendolyne Burfin, Hadrien Règue, Quentin Semanas, Martine Valette, Bruno Lina, Laurence Josset |
| EPI_ISL_900529 | Biosterel | CNR Virus des Infections Respiratoires - France SUD | Antonin Bal, Gregory Destras, Gwendolyne Burfin, Hadrien Règue, Quentin Semanas, Martine Valette, Bruno Lina, Laurence Josset |
| EPI_ISL_900530 | Centre Hospitalier de Guéret | CNR Virus des Infections Respiratoires - France SUD | Antonin Bal, Gregory Destras, Gwendolyne Burfin, Hadrien Règue, Quentin Semanas, Martine Valette, Bruno Lina, Sylvie Larrat, Laurence Josset |
| EPI_ISL_900531 | CHU Poitiers | CNR Virus des Infections Respiratoires - France SUD | Antonin Bal, Gregory Destras, Gwendolyne Burfin, Hadrien Règue, Quentin Semanas, Martine Valette, Bruno Lina, Agnès Beby-Defaux, Magali Garcia, Clément Jousselin, Nicolas Lévêque, Laurence Josset |
| EPI_ISL_900532 | CNR Virus des Infections Respiratoires - France SUD | CNR Virus des Infections Respiratoires - France SUD | Antonin Bal, Gregory Destras, Gwendolyne Burfin, Hadrien Règue, Quentin Semanas, Martine Valette, Bruno Lina, Laurence Josset |
| EPI_ISL_900536, EPI_ISL_900537, EPI_ISL_900538, EPI_ISL_900539 | Gen-Bio | CNR Virus des Infections Respiratoires - France SUD | Antonin Bal, Gregory Destras, Gwendolyne Burfin, Hadrien Règue, Quentin Semanas, Martine Valette, Bruno Lina, Laurence Josset |
| EPI_ISL_900540, EPI_ISL_900541, EPI_ISL_900542 | Gen-Bio | CNR Virus des Infections Respiratoires - France SUD | Antonin Bal, Gregory Destras, Gwendolyne Burfin, Hadrien Règue, Quentin Semanas, Martine Valette, Bruno Lina, Sylvie Larrat, Laurence Josset |
| EPI_ISL_900545, EPI_ISL_900546, EPI_ISL_900547, EPI_ISL_900548, EPI_ISL_900549, EPI_ISL_900550, EPI_ISL_900551, EPI_ISL_900552, EPI_ISL_900553, EPI_ISL_900554, EPI_ISL_900555, EPI_ISL_900556, EPI_ISL_900557, EPI_ISL_900558, EPI_ISL_900559, EPI_ISL_900560, EPI_ISL_900561, EPI_ISL_900562, EPI_ISL_900563, EPI_ISL_900564, EPI_ISL_900566, EPI_ISL_900567 | | | |
| see above | CNR Virus des Infections Respiratoires - France SUD | CNR Virus des Infections Respiratoires - France SUD | Antonin Bal, Gregory Destras, Gwendolyne Burfin, Hadrien Règue, Quentin Semanas, Martine Valette, Bruno Lina, Laurence Josset |
| EPI_ISL_902856, EPI_ISL_902857, EPI_ISL_902858, EPI_ISL_902859, EPI_ISL_902860, EPI_ISL_902861, EPI_ISL_902862, EPI_ISL_902863, EPI_ISL_902864, EPI_ISL_902865, EPI_ISL_902866, EPI_ISL_902867, EPI_ISL_902868, EPI_ISL_902869, EPI_ISL_902870, EPI_ISL_902871, EPI_ISL_902872, EPI_ISL_902873, EPI_ISL_902874, EPI_ISL_902875, EPI_ISL_902876, EPI_ISL_902877, EPI_ISL_902878, EPI_ISL_902879, EPI_ISL_902880, EPI_ISL_902881, EPI_ISL_902882, EPI_ISL_902883, EPI_ISL_902884, EPI_ISL_902885, EPI_ISL_902886, EPI_ISL_902887, EPI_ISL_902888 | | | |
| see above | Department of Virology and Immunology, University of Helsinki and Helsinki University Hospital, Huslab Finland | Department of Virology, Faculty of Medicine, University of Helsinki, Helsinki, Finland | Teemu Smura, Ravi Kant, Phuoc Truong, Hussein Alburkat, Hannimari Kallio-Kokko, Jenni Virtanen, Maija Suvanto, Essi Korhonen, Sari Hannula, Harri Kangas, Hanna Liimatainen, Satu Kurkela, Hanna Jarva, Maija Lappalainen, Pekka Ellonen, Olli Vapalahti |
| EPI_ISL_903043, EPI_ISL_903044, EPI_ISL_903048 | Seattle Flu Study | Seattle Flu Study | Deborah A. Nickerson, Chris D. Frazar, Jover Lee, Benjamin Pelle, Erica Ryke, Matthew Richardson, Amanda Adler, Elisabeth Brandstetter, Peter D. Han, Kairsten Fay, Misja Ilcisin, Kirsten Lacombe, Thomas R. Sibley, Melissa Truong, Caitlin R. Wolf, Michael Boeckh, Janet A. Englund, Michael Famulare, Barry R. Lutz, Mark J. Rieder, Lea M. Starita, Matthew Thompson, Jay Shendure, Trevor Bedford, Helen Y. Chu |
| EPI_ISL_903049, EPI_ISL_903050, EPI_ISL_903051, EPI_ISL_903052, EPI_ISL_903053 | Seattle Flu Study | Seattle Flu Study | Deborah A. Nickerson, Chris D. Frazar, Jover Lee, Benjamin Pelle, Erica Ryke, Matthew Richardson, Amanda Adler, Elisabeth Brandstetter, Peter D. Han, Kairsten Fay, Misja Ilcisin, Kirsten Lacombe, Thomas R. Sibley, Melissa Truong, Caitlin R. Wolf, Karen Cowgill, Stephanie Schrag, Jeff Duchin, Michael Boeckh, Janet A. Englund, Michael Famulare, Barry R. Lutz, Mark J. Rieder, Lea M. Starita, Matthew Thompson, Helen Y. Chu, Trevor Bedford, Jay Shendure |
| EPI_ISL_903054 | Seattle Flu Study | Seattle Flu Study | Deborah A. Nickerson, Chris D. Frazar, Jover Lee, Benjamin Pelle, Erica Ryke, Matthew Richardson, Amanda Adler, Elisabeth Brandstetter, Peter D. Han, Kairsten Fay, Misja Ilcisin, Kirsten Lacombe, Thomas R. Sibley, Melissa Truong, Caitlin R. Wolf, Michael Boeckh, Janet A. Englund, Michael Famulare, Barry R. Lutz, Mark J. Rieder, Lea M. Starita, Matthew Thompson, Jay Shendure, Trevor Bedford, Helen Y. Chu |
| EPI_ISL_903055 | Seattle Flu Study | Seattle Flu Study | Deborah A. Nickerson, Chris D. Frazar, Jover Lee, Benjamin Pelle, Erica Ryke, Matthew Richardson, Amanda Adler, Elisabeth Brandstetter, Peter D. Han, Kairsten Fay, Misja Ilcisin, Kirsten Lacombe, Thomas R. Sibley, Melissa Truong, Caitlin R. Wolf, Karen Cowgill, Stephanie Schrag, Jeff Duchin, Michael Boeckh, Janet A. Englund, Michael Famulare, Barry R. Lutz, Mark J. Rieder, Lea M. Starita, Matthew Thompson, Helen Y. Chu, Trevor Bedford, Jay Shendure |
| EPI_ISL_903056 | Seattle Flu Study | Seattle Flu Study | Deborah A. Nickerson, Chris D. Frazar, Jover Lee, Benjamin Pelle, Erica Ryke, Matthew Richardson, Amanda Adler, Elisabeth Brandstetter, Peter D. Han, Kairsten Fay, Misja Ilcisin, Kirsten Lacombe, Thomas R. Sibley, Melissa Truong, Caitlin R. Wolf, Michael Boeckh, Janet A. Englund, Michael Famulare, |

| | | | |
|---|---|---|---|
| | | | Barry R. Lutz, Mark J. Rieder, Lea M. Starita, Matthew Thompson, Jay Shendure, Trevor Bedford, Helen Y. Chu |
| EPI_ISL_903057, EPI_ISL_903058, EPI_ISL_903059, EPI_ISL_903060, EPI_ISL_903087, EPI_ISL_903088, EPI_ISL_903089, EPI_ISL_903090, EPI_ISL_903091 | Seattle Flu Study | Seattle Flu Study | Deborah A. Nickerson, Chris D. Frazar, Jover Lee, Benjamin Pelle, Erica Ryke, Matthew Richardson, Amanda Adler, Elisabeth Brandstetter, Peter D. Han, Kairsten Fay, Misja Ilcisin, Kirsten Lacombe, Thomas R. Sibley, Melissa Truong, Caitlin R. Wolf, Karen Cowgill, Stephanie Schrag, Jeff Duchin, Michael Boeckh, Janet A. Englund, Michael Famulare, Barry R. Lutz, Mark J. Rieder, Lea M. Starita, Matthew Thompson, Helen Y. Chu, Trevor Bedford, Jay Shendure |
| EPI_ISL_903105, EPI_ISL_903107, EPI_ISL_903109, EPI_ISL_903112, EPI_ISL_903114, EPI_ISL_903115, EPI_ISL_903116, EPI_ISL_903117, EPI_ISL_903118, EPI_ISL_903119, EPI_ISL_903124, EPI_ISL_903125, EPI_ISL_903126, EPI_ISL_903127, EPI_ISL_903128 | | | |
| see above | Washington State Department of Health | Seattle Flu Study | Deborah A. Nickerson, Chris D. Frazar, Jover Lee, Benjamin Pelle, Erica Ryke, Matthew Richardson, Amanda Adler, Elisabeth Brandstetter, Peter D. Han, Kairsten Fay, Misja Ilcisin, Kirsten Lacombe, Thomas R. Sibley, Melissa Truong, Caitlin R. Wolf, Romesh Gautom, Geoff Melly, Brian Hiatt, Philip Dykema, Scott Lindquist, Michael Boeckh, Janet A. Englund, Michael Famulare, Barry R. Lutz, Mark J. Rieder, Lea M. Starita, Matthew Thompson, Helen Y. Chu, Jay Shendure, Trevor Bedford |
| EPI_ISL_903241, EPI_ISL_903274, EPI_ISL_903275, EPI_ISL_903276, EPI_ISL_903277, EPI_ISL_903278, EPI_ISL_903279, EPI_ISL_903280, EPI_ISL_903282, EPI_ISL_903283, EPI_ISL_903286, EPI_ISL_903288 | | | |
| see above | M Health Fairview | Minnesota Department of Health, Public Health Laboratory | Alexandra Lorentz, Jacob Garfin, Matt Plumb, and Xiong Wang |
| EPI_ISL_903378, EPI_ISL_903383 | MOH - Jaber Al-Ahmad Hospital (Innovation Research Laboratory) | MOH - Jaber Al-Ahmad Hospital (Innovation Research Laboratory) | Salman Al-Sabah , Mohammad Alghounaim |
| EPI_ISL_903577 | NJ Public Health and Environmental Laboratories | Genomics and Discovery, Respiratory Viruses Branch, Division of Viral Diseases, Centers for Disease Control and Prevention | Krista Queen, Yan Li, Ying Tao, Jing Zhang, Anna Uehara, Anna Montmayeur, Clinton R. Paden, Peter W. Cook, Rachel Marine, Mili Sheth, Jasmine Padilla, Sarah Nobles, Mark Burroughs, Lori Rowe, Haibin Wang, Ben L. Rambo-Martin, Dhwani Batra, Justin Lee, Suxiang Tong |
| EPI_ISL_903585 | MD DOH Laboratories Administration | Genomics and Discovery, Respiratory Viruses Branch, Division of Viral Diseases, Centers for Disease Control and Prevention | Krista Queen, Yan Li, Ying Tao, Jing Zhang, Anna Uehara, Anna Montmayeur, Clinton R. Paden, Peter W. Cook, Rachel Marine, Mili Sheth, Jasmine Padilla, Sarah Nobles, Mark Burroughs, Lori Rowe, Haibin Wang, Ben L. Rambo-Martin, Dhwani Batra, Justin Lee, Suxiang Tong |
| EPI_ISL_903591 | CDPH, Viral and Rickettsial Disease Laboratory | Genomics and Discovery, Respiratory Viruses Branch, Division of Viral Diseases, Centers for Disease Control and Prevention | Krista Queen, Yan Li, Ying Tao, Jing Zhang, Anna Uehara, Anna Montmayeur, Clinton R. Paden, Peter W. Cook, Rachel Marine, Mili Sheth, Jasmine Padilla, Sarah Nobles, Mark Burroughs, Lori Rowe, Haibin Wang, Ben L. Rambo-Martin, Dhwani Batra, Justin Lee, Suxiang Tong |
| EPI_ISL_903594, EPI_ISL_903595 | NJ Public Health and Environmental Laboratories | Genomics and Discovery, Respiratory Viruses Branch, Division of Viral Diseases, Centers for Disease Control and Prevention | Krista Queen, Yan Li, Ying Tao, Jing Zhang, Anna Uehara, Anna Montmayeur, Clinton R. Paden, Peter W. Cook, Rachel Marine, Mili Sheth, Jasmine Padilla, Sarah Nobles, Mark Burroughs, Lori Rowe, Haibin Wang, Ben L. Rambo-Martin, Dhwani Batra, Justin Lee, Suxiang Tong |
| EPI_ISL_903596 | OK Public Health Laboratory, Oklahoma State DOH | Genomics and Discovery, Respiratory Viruses Branch, Division of Viral Diseases, Centers for Disease Control and Prevention | Krista Queen, Yan Li, Ying Tao, Jing Zhang, Anna Uehara, Anna Montmayeur, Clinton R. Paden, Peter W. Cook, Rachel Marine, Mili Sheth, Jasmine Padilla, Sarah Nobles, Mark Burroughs, Lori Rowe, Haibin Wang, Ben L. Rambo-Martin, Dhwani Batra, Justin Lee, Suxiang Tong |
| EPI_ISL_903607 | IL Dept. of Public Health Springfield Laboratory | Genomics and Discovery, Respiratory Viruses Branch, Division of Viral Diseases, Centers for Disease Control and Prevention | Krista Queen, Yan Li, Ying Tao, Jing Zhang, Anna Uehara, Anna Montmayeur, Clinton R. Paden, Peter W. Cook, Rachel Marine, Mili Sheth, Jasmine Padilla, Sarah Nobles, Mark Burroughs, Lori Rowe, Haibin Wang, Ben L. Rambo-Martin, Dhwani Batra, Justin Lee, Suxiang Tong |
| EPI_ISL_903609 | NJ Public Health and Environmental Laboratories | Genomics and Discovery, Respiratory Viruses Branch, Division of Viral Diseases, Centers for Disease Control and Prevention | Krista Queen, Yan Li, Ying Tao, Jing Zhang, Anna Uehara, Anna Montmayeur, Clinton R. Paden, Peter W. Cook, Rachel Marine, Mili Sheth, Jasmine Padilla, Sarah Nobles, Mark Burroughs, Lori Rowe, Haibin Wang, Ben L. Rambo-Martin, Dhwani Batra, Justin Lee, Suxiang Tong |
| EPI_ISL_903610 | NV State Public Health Laboratory | Genomics and Discovery, Respiratory Viruses Branch, Division of Viral Diseases, Centers for Disease Control and Prevention | Krista Queen, Yan Li, Ying Tao, Jing Zhang, Anna Uehara, Anna Montmayeur, Clinton R. Paden, Peter W. Cook, Rachel Marine, Mili Sheth, Jasmine Padilla, Sarah Nobles, Mark Burroughs, Lori Rowe, Haibin Wang, Ben L. Rambo-Martin, Dhwani Batra, Justin Lee, Suxiang Tong |
| EPI_ISL_903616 | IN State Department of Health Laboratory Services | Genomics and Discovery, Respiratory Viruses Branch, Division of Viral Diseases, Centers for Disease Control and Prevention | Krista Queen, Yan Li, Ying Tao, Jing Zhang, Anna Uehara, Anna Montmayeur, Clinton R. Paden, Peter W. Cook, Rachel Marine, Mili Sheth, Jasmine Padilla, Sarah Nobles, Mark Burroughs, Lori Rowe, Haibin Wang, Ben L. Rambo-Martin, Dhwani Batra, Justin Lee, Suxiang Tong |
| EPI_ISL_903637, EPI_ISL_903638 | NJ Public Health and Environmental Laboratories | Genomics and Discovery, Respiratory Viruses Branch, Division of Viral Diseases, Centers for Disease Control and Prevention | Krista Queen, Yan Li, Ying Tao, Jing Zhang, Anna Uehara, Anna Montmayeur, Clinton R. Paden, Peter W. Cook, Rachel Marine, Mili Sheth, Jasmine Padilla, Sarah Nobles, Mark Burroughs, Lori Rowe, Haibin Wang, Ben L. Rambo-Martin, Dhwani Batra, Justin Lee, Suxiang Tong |
| EPI_ISL_903642 | PA Department of Health, Bureau of Laboratories | Genomics and Discovery, Respiratory Viruses Branch, Division of Viral Diseases, Centers for Disease Control and Prevention | Krista Queen, Yan Li, Ying Tao, Jing Zhang, Anna Uehara, Anna Montmayeur, Clinton R. Paden, Peter W. Cook, Rachel Marine, Mili Sheth, Jasmine Padilla, Sarah Nobles, Mark Burroughs, Lori Rowe, Haibin Wang, Ben L. Rambo-Martin, Dhwani Batra, Justin Lee, Suxiang Tong |
| EPI_ISL_903651 | LA Office of Public Health Laboratories | Genomics and Discovery, Respiratory Viruses Branch, Division of Viral Diseases, Centers for Disease Control and Prevention | Krista Queen, Yan Li, Ying Tao, Jing Zhang, Anna Uehara, Anna Montmayeur, Clinton R. Paden, Peter W. Cook, Rachel Marine, Mili Sheth, Jasmine Padilla, Sarah Nobles, Mark Burroughs, Lori Rowe, Haibin Wang, Ben L. Rambo-Martin, Dhwani Batra, Justin Lee, Suxiang Tong |
| EPI_ISL_903654, EPI_ISL_903655, EPI_ISL_903658 | NJ Public Health and Environmental Laboratories | Genomics and Discovery, Respiratory Viruses Branch, Division of Viral Diseases, Centers for Disease Control and Prevention | Krista Queen, Yan Li, Ying Tao, Jing Zhang, Anna Uehara, Anna Montmayeur, Clinton R. Paden, Peter W. Cook, Rachel Marine, Mili Sheth, Jasmine Padilla, Sarah Nobles, Mark Burroughs, Lori Rowe, Haibin Wang, Ben L. Rambo-Martin, Dhwani Batra, Justin Lee, Suxiang Tong |
| EPI_ISL_903663 | IL Dept. of Public Health Springfield Laboratory | Genomics and Discovery, Respiratory Viruses Branch, Division of Viral Diseases, Centers for Disease Control and Prevention | Krista Queen, Yan Li, Ying Tao, Jing Zhang, Anna Uehara, Anna Montmayeur, Clinton R. Paden, Peter W. Cook, Rachel Marine, Mili Sheth, Jasmine Padilla, Sarah Nobles, Mark Burroughs, Lori Rowe, Haibin Wang, Ben L. Rambo-Martin, Dhwani Batra, Justin Lee, Suxiang Tong |
| EPI_ISL_903664 | NV State Public Health Laboratory | Genomics and Discovery, Respiratory Viruses Branch, Division of Viral Diseases, Centers for Disease Control and Prevention | Krista Queen, Yan Li, Ying Tao, Jing Zhang, Anna Uehara, Anna Montmayeur, Clinton R. Paden, Peter W. Cook, Rachel Marine, Mili Sheth, Jasmine Padilla, Sarah Nobles, Mark Burroughs, Lori Rowe, Haibin Wang, Ben L. Rambo-Martin, Dhwani Batra, Justin Lee, Suxiang Tong |
| EPI_ISL_903673, EPI_ISL_903674, EPI_ISL_903678, EPI_ISL_903680 | NJ Public Health and Environmental Laboratories | Genomics and Discovery, Respiratory Viruses Branch, Division of Viral Diseases, Centers for Disease Control and Prevention | Krista Queen, Yan Li, Ying Tao, Jing Zhang, Anna Uehara, Anna Montmayeur, Clinton R. Paden, Peter W. Cook, Rachel Marine, Mili Sheth, Jasmine Padilla, Sarah Nobles, Mark Burroughs, Lori Rowe, Haibin Wang, Ben L. Rambo-Martin, Dhwani Batra, Justin Lee, Suxiang Tong |
| EPI_ISL_903682 | IL Dept. of Public Health Springfield Laboratory | Genomics and Discovery, Respiratory Viruses Branch, Division of Viral Diseases, Centers for Disease Control and Prevention | Krista Queen, Yan Li, Ying Tao, Jing Zhang, Anna Uehara, Anna Montmayeur, Clinton R. Paden, Peter W. Cook, Rachel Marine, Mili Sheth, Jasmine Padilla, Sarah Nobles, Mark Burroughs, Lori Rowe, Haibin Wang, Ben L. Rambo-Martin, Dhwani Batra, Justin Lee, Suxiang Tong |
| EPI_ISL_903683, EPI_ISL_903686 | NJ Public Health and Environmental Laboratories | Genomics and Discovery, Respiratory Viruses Branch, Division of Viral Diseases, Centers for Disease Control and Prevention | Krista Queen, Yan Li, Ying Tao, Jing Zhang, Anna Uehara, Anna Montmayeur, Clinton R. Paden, Peter W. Cook, Rachel Marine, Mili Sheth, Jasmine Padilla, Sarah Nobles, Mark Burroughs, Lori Rowe, Haibin Wang, Ben L. Rambo-Martin, Dhwani Batra, Justin Lee, Suxiang Tong |
| EPI_ISL_903689 | LA Office of Public Health Laboratories | Genomics and Discovery, Respiratory Viruses Branch, Division of Viral Diseases, Centers for Disease Control and Prevention | Krista Queen, Yan Li, Ying Tao, Jing Zhang, Anna Uehara, Anna Montmayeur, Clinton R. Paden, Peter W. Cook, Rachel Marine, Mili Sheth, Jasmine Padilla, Sarah Nobles, Mark Burroughs, Lori Rowe, Haibin Wang, Ben L. Rambo-Martin, Dhwani Batra, Justin Lee, Suxiang Tong |
| EPI_ISL_903690 | NV State Public Health Laboratory | Genomics and Discovery, Respiratory Viruses Branch, Division of Viral Diseases, Centers for Disease Control and | Krista Queen, Yan Li, Ying Tao, Jing Zhang, Anna Uehara, Anna Montmayeur, Clinton R. Paden, Peter W. Cook, Rachel Marine, Mili Sheth, Jasmine Padilla, Sarah Nobles, Mark Burroughs, Lori Rowe, Haibin Wang, Ben L. Rambo-Martin, Dhwani Batra, Justin Lee, Suxiang Tong |

| | | | |
|---|---|---|---|
| | | Prevention | |
| EPI_ISL_903692 | NJ Public Health and Environmental Laboratories | Genomics and Discovery, Respiratory Viruses Branch, Division of Viral Diseases, Centers for Disease Control and Prevention | Krista Queen, Yan Li, Ying Tao, Jing Zhang, Anna Uehara, Anna Montmayeur, Clinton R. Paden, Peter W. Cook, Rachel Marine, Mili Sheth, Jasmine Padilla, Sarah Nobles, Mark Burroughs, Lori Rowe, Haibin Wang, Ben L. Rambo-Martin, Dhwani Batra, Justin Lee, Suxiang Tong |
| EPI_ISL_903695 | OK Public Health Laboratory, Oklahoma State DOH | Genomics and Discovery, Respiratory Viruses Branch, Division of Viral Diseases, Centers for Disease Control and Prevention | Krista Queen, Yan Li, Ying Tao, Jing Zhang, Anna Uehara, Anna Montmayeur, Clinton R. Paden, Peter W. Cook, Rachel Marine, Mili Sheth, Jasmine Padilla, Sarah Nobles, Mark Burroughs, Lori Rowe, Haibin Wang, Ben L. Rambo-Martin, Dhwani Batra, Justin Lee, Suxiang Tong |
| EPI_ISL_903699 | NJ Public Health and Environmental Laboratories | Genomics and Discovery, Respiratory Viruses Branch, Division of Viral Diseases, Centers for Disease Control and Prevention | Krista Queen, Yan Li, Ying Tao, Jing Zhang, Anna Uehara, Anna Montmayeur, Clinton R. Paden, Peter W. Cook, Rachel Marine, Mili Sheth, Jasmine Padilla, Sarah Nobles, Mark Burroughs, Lori Rowe, Haibin Wang, Ben L. Rambo-Martin, Dhwani Batra, Justin Lee, Suxiang Tong |
| EPI_ISL_903700, EPI_ISL_903702 | NV State Public Health Laboratory | Genomics and Discovery, Respiratory Viruses Branch, Division of Viral Diseases, Centers for Disease Control and Prevention | Krista Queen, Yan Li, Ying Tao, Jing Zhang, Anna Uehara, Anna Montmayeur, Clinton R. Paden, Peter W. Cook, Rachel Marine, Mili Sheth, Jasmine Padilla, Sarah Nobles, Mark Burroughs, Lori Rowe, Haibin Wang, Ben L. Rambo-Martin, Dhwani Batra, Justin Lee, Suxiang Tong |
| EPI_ISL_903704 | NJ Public Health and Environmental Laboratories | Genomics and Discovery, Respiratory Viruses Branch, Division of Viral Diseases, Centers for Disease Control and Prevention | Krista Queen, Yan Li, Ying Tao, Jing Zhang, Anna Uehara, Anna Montmayeur, Clinton R. Paden, Peter W. Cook, Rachel Marine, Mili Sheth, Jasmine Padilla, Sarah Nobles, Mark Burroughs, Lori Rowe, Haibin Wang, Ben L. Rambo-Martin, Dhwani Batra, Justin Lee, Suxiang Tong |
| EPI_ISL_903709 | LA Office of Public Health Laboratories | Genomics and Discovery, Respiratory Viruses Branch, Division of Viral Diseases, Centers for Disease Control and Prevention | Krista Queen, Yan Li, Ying Tao, Jing Zhang, Anna Uehara, Anna Montmayeur, Clinton R. Paden, Peter W. Cook, Rachel Marine, Mili Sheth, Jasmine Padilla, Sarah Nobles, Mark Burroughs, Lori Rowe, Haibin Wang, Ben L. Rambo-Martin, Dhwani Batra, Justin Lee, Suxiang Tong |
| EPI_ISL_903711, EPI_ISL_903714 | NJ Public Health and Environmental Laboratories | Genomics and Discovery, Respiratory Viruses Branch, Division of Viral Diseases, Centers for Disease Control and Prevention | Krista Queen, Yan Li, Ying Tao, Jing Zhang, Anna Uehara, Anna Montmayeur, Clinton R. Paden, Peter W. Cook, Rachel Marine, Mili Sheth, Jasmine Padilla, Sarah Nobles, Mark Burroughs, Lori Rowe, Haibin Wang, Ben L. Rambo-Martin, Dhwani Batra, Justin Lee, Suxiang Tong |
| EPI_ISL_903716 | OK Public Health Laboratory, Oklahoma State DOH | Genomics and Discovery, Respiratory Viruses Branch, Division of Viral Diseases, Centers for Disease Control and Prevention | Krista Queen, Yan Li, Ying Tao, Jing Zhang, Anna Uehara, Anna Montmayeur, Clinton R. Paden, Peter W. Cook, Rachel Marine, Mili Sheth, Jasmine Padilla, Sarah Nobles, Mark Burroughs, Lori Rowe, Haibin Wang, Ben L. Rambo-Martin, Dhwani Batra, Justin Lee, Suxiang Tong |
| EPI_ISL_903721, EPI_ISL_903723 | NV State Public Health Laboratory | Genomics and Discovery, Respiratory Viruses Branch, Division of Viral Diseases, Centers for Disease Control and Prevention | Krista Queen, Yan Li, Ying Tao, Jing Zhang, Anna Uehara, Anna Montmayeur, Clinton R. Paden, Peter W. Cook, Rachel Marine, Mili Sheth, Jasmine Padilla, Sarah Nobles, Mark Burroughs, Lori Rowe, Haibin Wang, Ben L. Rambo-Martin, Dhwani Batra, Justin Lee, Suxiang Tong |
| EPI_ISL_903726 | NJ Public Health and Environmental Laboratories | Genomics and Discovery, Respiratory Viruses Branch, Division of Viral Diseases, Centers for Disease Control and Prevention | Krista Queen, Yan Li, Ying Tao, Jing Zhang, Anna Uehara, Anna Montmayeur, Clinton R. Paden, Peter W. Cook, Rachel Marine, Mili Sheth, Jasmine Padilla, Sarah Nobles, Mark Burroughs, Lori Rowe, Haibin Wang, Ben L. Rambo-Martin, Dhwani Batra, Justin Lee, Suxiang Tong |
| EPI_ISL_903727 | NV State Public Health Laboratory | Genomics and Discovery, Respiratory Viruses Branch, Division of Viral Diseases, Centers for Disease Control and Prevention | Krista Queen, Yan Li, Ying Tao, Jing Zhang, Anna Uehara, Anna Montmayeur, Clinton R. Paden, Peter W. Cook, Rachel Marine, Mili Sheth, Jasmine Padilla, Sarah Nobles, Mark Burroughs, Lori Rowe, Haibin Wang, Ben L. Rambo-Martin, Dhwani Batra, Justin Lee, Suxiang Tong |
| EPI_ISL_903728 | LA Office of Public Health Laboratories | Genomics and Discovery, Respiratory Viruses Branch, Division of Viral Diseases, Centers for Disease Control and Prevention | Krista Queen, Yan Li, Ying Tao, Jing Zhang, Anna Uehara, Anna Montmayeur, Clinton R. Paden, Peter W. Cook, Rachel Marine, Mili Sheth, Jasmine Padilla, Sarah Nobles, Mark Burroughs, Lori Rowe, Haibin Wang, Ben L. Rambo-Martin, Dhwani Batra, Justin Lee, Suxiang Tong |
| EPI_ISL_903733 | NV State Public Health Laboratory | Genomics and Discovery, Respiratory Viruses Branch, Division of Viral Diseases, Centers for Disease Control and Prevention | Krista Queen, Yan Li, Ying Tao, Jing Zhang, Anna Uehara, Anna Montmayeur, Clinton R. Paden, Peter W. Cook, Rachel Marine, Mili Sheth, Jasmine Padilla, Sarah Nobles, Mark Burroughs, Lori Rowe, Haibin Wang, Ben L. Rambo-Martin, Dhwani Batra, Justin Lee, Suxiang Tong |
| EPI_ISL_903734 | NJ Public Health and Environmental Laboratories | Genomics and Discovery, Respiratory Viruses Branch, Division of Viral Diseases, Centers for Disease Control and Prevention | Krista Queen, Yan Li, Ying Tao, Jing Zhang, Anna Uehara, Anna Montmayeur, Clinton R. Paden, Peter W. Cook, Rachel Marine, Mili Sheth, Jasmine Padilla, Sarah Nobles, Mark Burroughs, Lori Rowe, Haibin Wang, Ben L. Rambo-Martin, Dhwani Batra, Justin Lee, Suxiang Tong |
| EPI_ISL_903751 | PA Department of Health, Bureau of Laboratories | Genomics and Discovery, Respiratory Viruses Branch, Division of Viral Diseases, Centers for Disease Control and Prevention | Krista Queen, Yan Li, Ying Tao, Jing Zhang, Anna Uehara, Anna Montmayeur, Clinton R. Paden, Peter W. Cook, Rachel Marine, Mili Sheth, Jasmine Padilla, Sarah Nobles, Mark Burroughs, Lori Rowe, Haibin Wang, Ben L. Rambo-Martin, Dhwani Batra, Justin Lee, Suxiang Tong |
| EPI_ISL_903754 | IN State Department of Health Laboratory Services | Genomics and Discovery, Respiratory Viruses Branch, Division of Viral Diseases, Centers for Disease Control and Prevention | Krista Queen, Yan Li, Ying Tao, Jing Zhang, Anna Uehara, Anna Montmayeur, Clinton R. Paden, Peter W. Cook, Rachel Marine, Mili Sheth, Jasmine Padilla, Sarah Nobles, Mark Burroughs, Lori Rowe, Haibin Wang, Ben L. Rambo-Martin, Dhwani Batra, Justin Lee, Suxiang Tong |
| EPI_ISL_903759 | NM Dept. Health, Scientific Laboratory Division | Genomics and Discovery, Respiratory Viruses Branch, Division of Viral Diseases, Centers for Disease Control and Prevention | Krista Queen, Yan Li, Ying Tao, Jing Zhang, Anna Uehara, Anna Montmayeur, Clinton R. Paden, Peter W. Cook, Rachel Marine, Mili Sheth, Jasmine Padilla, Sarah Nobles, Mark Burroughs, Lori Rowe, Haibin Wang, Ben L. Rambo-Martin, Dhwani Batra, Justin Lee, Suxiang Tong |
| EPI_ISL_903761 | CDPH, Viral and Rickettsial Disease Laboratory | Genomics and Discovery, Respiratory Viruses Branch, Division of Viral Diseases, Centers for Disease Control and Prevention | Krista Queen, Yan Li, Ying Tao, Jing Zhang, Anna Uehara, Anna Montmayeur, Clinton R. Paden, Peter W. Cook, Rachel Marine, Mili Sheth, Jasmine Padilla, Sarah Nobles, Mark Burroughs, Lori Rowe, Haibin Wang, Ben L. Rambo-Martin, Dhwani Batra, Justin Lee, Suxiang Tong |
| EPI_ISL_903762 | MD DOH Laboratories Administration | Genomics and Discovery, Respiratory Viruses Branch, Division of Viral Diseases, Centers for Disease Control and Prevention | Krista Queen, Yan Li, Ying Tao, Jing Zhang, Anna Uehara, Anna Montmayeur, Clinton R. Paden, Peter W. Cook, Rachel Marine, Mili Sheth, Jasmine Padilla, Sarah Nobles, Mark Burroughs, Lori Rowe, Haibin Wang, Ben L. Rambo-Martin, Dhwani Batra, Justin Lee, Suxiang Tong |
| EPI_ISL_903773 | IN State Department of Health Laboratory Services | Genomics and Discovery, Respiratory Viruses Branch, Division of Viral Diseases, Centers for Disease Control and Prevention | Krista Queen, Yan Li, Ying Tao, Jing Zhang, Anna Uehara, Anna Montmayeur, Clinton R. Paden, Peter W. Cook, Rachel Marine, Mili Sheth, Jasmine Padilla, Sarah Nobles, Mark Burroughs, Lori Rowe, Haibin Wang, Ben L. Rambo-Martin, Dhwani Batra, Justin Lee, Suxiang Tong |
| EPI_ISL_903779, EPI_ISL_903789 | IL Dept. of Public Health Springfield Laboratory | Genomics and Discovery, Respiratory Viruses Branch, Division of Viral Diseases, Centers for Disease Control and Prevention | Krista Queen, Yan Li, Ying Tao, Jing Zhang, Anna Uehara, Anna Montmayeur, Clinton R. Paden, Peter W. Cook, Rachel Marine, Mili Sheth, Jasmine Padilla, Sarah Nobles, Mark Burroughs, Lori Rowe, Haibin Wang, Ben L. Rambo-Martin, Dhwani Batra, Justin Lee, Suxiang Tong |
| EPI_ISL_903790 | PA Department of Health, Bureau of Laboratories | Genomics and Discovery, Respiratory Viruses Branch, Division of Viral Diseases, Centers for Disease Control and Prevention | Krista Queen, Yan Li, Ying Tao, Jing Zhang, Anna Uehara, Anna Montmayeur, Clinton R. Paden, Peter W. Cook, Rachel Marine, Mili Sheth, Jasmine Padilla, Sarah Nobles, Mark Burroughs, Lori Rowe, Haibin Wang, Ben L. Rambo-Martin, Dhwani Batra, Justin Lee, Suxiang Tong |
| EPI_ISL_903810 | CDPH, Viral and Rickettsial Disease Laboratory | Genomics and Discovery, Respiratory Viruses Branch, Division of Viral Diseases, Centers for Disease Control and Prevention | Krista Queen, Yan Li, Ying Tao, Jing Zhang, Anna Uehara, Anna Montmayeur, Clinton R. Paden, Peter W. Cook, Rachel Marine, Mili Sheth, Jasmine Padilla, Sarah Nobles, Mark Burroughs, Lori Rowe, Haibin Wang, Ben L. Rambo-Martin, Dhwani Batra, Justin Lee, Suxiang Tong |
| EPI_ISL_903823 | OK Public Health Laboratory, Oklahoma State DOH | Genomics and Discovery, Respiratory Viruses Branch, Division of Viral Diseases, Centers for Disease Control and Prevention | Krista Queen, Yan Li, Ying Tao, Jing Zhang, Anna Uehara, Anna Montmayeur, Clinton R. Paden, Peter W. Cook, Rachel Marine, Mili Sheth, Jasmine Padilla, Sarah Nobles, Mark Burroughs, Lori Rowe, Haibin Wang, Ben L. Rambo-Martin, Dhwani Batra, Justin Lee, Suxiang Tong |
| EPI_ISL_903828 | IL Dept. of Public Health Springfield Laboratory | Genomics and Discovery, Respiratory Viruses Branch, Division of Viral Diseases, Centers for Disease Control and | Krista Queen, Yan Li, Ying Tao, Jing Zhang, Anna Uehara, Anna Montmayeur, Clinton R. Paden, Peter W. Cook, Rachel Marine, Mili Sheth, Jasmine Padilla, Sarah Nobles, Mark Burroughs, Lori Rowe, Haibin Wang, Ben L. Rambo-Martin, Dhwani Batra, Justin Lee, Suxiang Tong |

| | | | |
|---|---|---|---|
| | | Prevention | |
| EPI_ISL_903832 | LA Office of Public Health Laboratories | Genomics and Discovery, Respiratory Viruses Branch, Division of Viral Diseases, Centers for Disease Control and Prevention | Krista Queen, Yan Li, Ying Tao, Jing Zhang, Anna Uehara, Anna Montmayeur, Clinton R. Paden, Peter W. Cook, Rachel Marine, Mili Sheth, Jasmine Padilla, Sarah Nobles, Mark Burroughs, Lori Rowe, Haibin Wang, Ben L. Rambo-Martin, Dhwani Batra, Justin Lee, Suxiang Tong |
| EPI_ISL_903837 | PA Department of Health, Bureau of Laboratories | Genomics and Discovery, Respiratory Viruses Branch, Division of Viral Diseases, Centers for Disease Control and Prevention | Krista Queen, Yan Li, Ying Tao, Jing Zhang, Anna Uehara, Anna Montmayeur, Clinton R. Paden, Peter W. Cook, Rachel Marine, Mili Sheth, Jasmine Padilla, Sarah Nobles, Mark Burroughs, Lori Rowe, Haibin Wang, Ben L. Rambo-Martin, Dhwani Batra, Justin Lee, Suxiang Tong |
| EPI_ISL_903848 | MD DOH Laboratories Administration | Genomics and Discovery, Respiratory Viruses Branch, Division of Viral Diseases, Centers for Disease Control and Prevention | Krista Queen, Yan Li, Ying Tao, Jing Zhang, Anna Uehara, Anna Montmayeur, Clinton R. Paden, Peter W. Cook, Rachel Marine, Mili Sheth, Jasmine Padilla, Sarah Nobles, Mark Burroughs, Lori Rowe, Haibin Wang, Ben L. Rambo-Martin, Dhwani Batra, Justin Lee, Suxiang Tong |
| EPI_ISL_903852, EPI_ISL_903859 | PA Department of Health, Bureau of Laboratories | Genomics and Discovery, Respiratory Viruses Branch, Division of Viral Diseases, Centers for Disease Control and Prevention | Krista Queen, Yan Li, Ying Tao, Jing Zhang, Anna Uehara, Anna Montmayeur, Clinton R. Paden, Peter W. Cook, Rachel Marine, Mili Sheth, Jasmine Padilla, Sarah Nobles, Mark Burroughs, Lori Rowe, Haibin Wang, Ben L. Rambo-Martin, Dhwani Batra, Justin Lee, Suxiang Tong |
| EPI_ISL_903862 | NM Dept. Health, Scientific Laboratory Division | Genomics and Discovery, Respiratory Viruses Branch, Division of Viral Diseases, Centers for Disease Control and Prevention | Krista Queen, Yan Li, Ying Tao, Jing Zhang, Anna Uehara, Anna Montmayeur, Clinton R. Paden, Peter W. Cook, Rachel Marine, Mili Sheth, Jasmine Padilla, Sarah Nobles, Mark Burroughs, Lori Rowe, Haibin Wang, Ben L. Rambo-Martin, Dhwani Batra, Justin Lee, Suxiang Tong |
| EPI_ISL_903875 | IN State Department of Health Laboratory Services | Genomics and Discovery, Respiratory Viruses Branch, Division of Viral Diseases, Centers for Disease Control and Prevention | Krista Queen, Yan Li, Ying Tao, Jing Zhang, Anna Uehara, Anna Montmayeur, Clinton R. Paden, Peter W. Cook, Rachel Marine, Mili Sheth, Jasmine Padilla, Sarah Nobles, Mark Burroughs, Lori Rowe, Haibin Wang, Ben L. Rambo-Martin, Dhwani Batra, Justin Lee, Suxiang Tong |
| EPI_ISL_903877 | LA Office of Public Health Laboratories | Genomics and Discovery, Respiratory Viruses Branch, Division of Viral Diseases, Centers for Disease Control and Prevention | Krista Queen, Yan Li, Ying Tao, Jing Zhang, Anna Uehara, Anna Montmayeur, Clinton R. Paden, Peter W. Cook, Rachel Marine, Mili Sheth, Jasmine Padilla, Sarah Nobles, Mark Burroughs, Lori Rowe, Haibin Wang, Ben L. Rambo-Martin, Dhwani Batra, Justin Lee, Suxiang Tong |
| EPI_ISL_903883 | IN State Department of Health Laboratory Services | Genomics and Discovery, Respiratory Viruses Branch, Division of Viral Diseases, Centers for Disease Control and Prevention | Krista Queen, Yan Li, Ying Tao, Jing Zhang, Anna Uehara, Anna Montmayeur, Clinton R. Paden, Peter W. Cook, Rachel Marine, Mili Sheth, Jasmine Padilla, Sarah Nobles, Mark Burroughs, Lori Rowe, Haibin Wang, Ben L. Rambo-Martin, Dhwani Batra, Justin Lee, Suxiang Tong |
| EPI_ISL_903892 | CDPH, Viral and Rickettsial Disease Laboratory | Genomics and Discovery, Respiratory Viruses Branch, Division of Viral Diseases, Centers for Disease Control and Prevention | Krista Queen, Yan Li, Ying Tao, Jing Zhang, Anna Uehara, Anna Montmayeur, Clinton R. Paden, Peter W. Cook, Rachel Marine, Mili Sheth, Jasmine Padilla, Sarah Nobles, Mark Burroughs, Lori Rowe, Haibin Wang, Ben L. Rambo-Martin, Dhwani Batra, Justin Lee, Suxiang Tong |
| EPI_ISL_903896 | IL Dept. of Public Health Springfield Laboratory | Genomics and Discovery, Respiratory Viruses Branch, Division of Viral Diseases, Centers for Disease Control and Prevention | Krista Queen, Yan Li, Ying Tao, Jing Zhang, Anna Uehara, Anna Montmayeur, Clinton R. Paden, Peter W. Cook, Rachel Marine, Mili Sheth, Jasmine Padilla, Sarah Nobles, Mark Burroughs, Lori Rowe, Haibin Wang, Ben L. Rambo-Martin, Dhwani Batra, Justin Lee, Suxiang Tong |
| EPI_ISL_903904 | IN State Department of Health Laboratory Services | Genomics and Discovery, Respiratory Viruses Branch, Division of Viral Diseases, Centers for Disease Control and Prevention | Krista Queen, Yan Li, Ying Tao, Jing Zhang, Anna Uehara, Anna Montmayeur, Clinton R. Paden, Peter W. Cook, Rachel Marine, Mili Sheth, Jasmine Padilla, Sarah Nobles, Mark Burroughs, Lori Rowe, Haibin Wang, Ben L. Rambo-Martin, Dhwani Batra, Justin Lee, Suxiang Tong |
| EPI_ISL_903909 | PA Department of Health, Bureau of Laboratories | Genomics and Discovery, Respiratory Viruses Branch, Division of Viral Diseases, Centers for Disease Control and Prevention | Krista Queen, Yan Li, Ying Tao, Jing Zhang, Anna Uehara, Anna Montmayeur, Clinton R. Paden, Peter W. Cook, Rachel Marine, Mili Sheth, Jasmine Padilla, Sarah Nobles, Mark Burroughs, Lori Rowe, Haibin Wang, Ben L. Rambo-Martin, Dhwani Batra, Justin Lee, Suxiang Tong |
| EPI_ISL_903915 | CDPH, Viral and Rickettsial Disease Laboratory | Genomics and Discovery, Respiratory Viruses Branch, Division of Viral Diseases, Centers for Disease Control and Prevention | Krista Queen, Yan Li, Ying Tao, Jing Zhang, Anna Uehara, Anna Montmayeur, Clinton R. Paden, Peter W. Cook, Rachel Marine, Mili Sheth, Jasmine Padilla, Sarah Nobles, Mark Burroughs, Lori Rowe, Haibin Wang, Ben L. Rambo-Martin, Dhwani Batra, Justin Lee, Suxiang Tong |
| EPI_ISL_903933 | PA Department of Health, Bureau of Laboratories | Genomics and Discovery, Respiratory Viruses Branch, Division of Viral Diseases, Centers for Disease Control and Prevention | Krista Queen, Yan Li, Ying Tao, Jing Zhang, Anna Uehara, Anna Montmayeur, Clinton R. Paden, Peter W. Cook, Rachel Marine, Mili Sheth, Jasmine Padilla, Sarah Nobles, Mark Burroughs, Lori Rowe, Haibin Wang, Ben L. Rambo-Martin, Dhwani Batra, Justin Lee, Suxiang Tong |
| EPI_ISL_903963 | MD DOH Laboratories Administration | Genomics and Discovery, Respiratory Viruses Branch, Division of Viral Diseases, Centers for Disease Control and Prevention | Krista Queen, Yan Li, Ying Tao, Jing Zhang, Anna Uehara, Anna Montmayeur, Clinton R. Paden, Peter W. Cook, Rachel Marine, Mili Sheth, Jasmine Padilla, Sarah Nobles, Mark Burroughs, Lori Rowe, Haibin Wang, Ben L. Rambo-Martin, Dhwani Batra, Justin Lee, Suxiang Tong |
| EPI_ISL_904036, EPI_ISL_904037, EPI_ISL_904038, EPI_ISL_904039, EPI_ISL_904040, EPI_ISL_904041, EPI_ISL_904042 | New Mexico Department of Health Scientific Laboratory | New Mexico Department of Health Scientific Laboratory | Ellie Johnson, Anastacia Griego-Fisher, D'eldra Malone |
| EPI_ISL_904148, EPI_ISL_904153, EPI_ISL_904164, EPI_ISL_904186, EPI_ISL_904187, EPI_ISL_904190, EPI_ISL_904199, EPI_ISL_904265, EPI_ISL_904266, EPI_ISL_904267, EPI_ISL_904303, EPI_ISL_904304, EPI_ISL_904364, EPI_ISL_904365, EPI_ISL_904520, EPI_ISL_904521, EPI_ISL_904522, EPI_ISL_904523, EPI_ISL_904524, EPI_ISL_904525, EPI_ISL_904526, EPI_ISL_904527, EPI_ISL_904528, EPI_ISL_904621, EPI_ISL_904622, EPI_ISL_904623, EPI_ISL_904624 | | | |
| see above | Dutch COVID-19 response team | Erasmus Medical Center | Bas Oude Munnink, Reina Sikkema, David Nieuwenhuijse, Irina Chestakova, Anne van der Linden, Marjan Boter, Emmanuelle Munger, Corine GeurtsvanKessel, Annemiek van der Eijk, Richard Molenkamp, Marion Koopmans, on behalf of the Dutch national COVID-19 response team. |
| EPI_ISL_904767, EPI_ISL_904784, EPI_ISL_904891, EPI_ISL_904904, EPI_ISL_904972, EPI_ISL_904992, EPI_ISL_904994, EPI_ISL_905011, EPI_ISL_905033, EPI_ISL_905069, EPI_ISL_905101, EPI_ISL_905102, EPI_ISL_905103, EPI_ISL_905104, EPI_ISL_905149, EPI_ISL_905153, EPI_ISL_905157, EPI_ISL_905164, EPI_ISL_905169, EPI_ISL_905187, EPI_ISL_905201, EPI_ISL_905213, EPI_ISL_905214, EPI_ISL_905218, EPI_ISL_905219, EPI_ISL_905225, EPI_ISL_905226, EPI_ISL_905227, EPI_ISL_905352, EPI_ISL_905353, EPI_ISL_905354, EPI_ISL_905372, EPI_ISL_905373, EPI_ISL_905374, EPI_ISL_905375, EPI_ISL_905378, EPI_ISL_905379, EPI_ISL_905380, EPI_ISL_905381, EPI_ISL_905382, EPI_ISL_905404, EPI_ISL_905418, EPI_ISL_905429, EPI_ISL_905430, EPI_ISL_905431, EPI_ISL_905434, EPI_ISL_905435, EPI_ISL_905472, EPI_ISL_905473, EPI_ISL_905486, EPI_ISL_905502, EPI_ISL_905503, EPI_ISL_905507, EPI_ISL_905515, EPI_ISL_905536, EPI_ISL_905538, EPI_ISL_905539, EPI_ISL_905540, EPI_ISL_905551, EPI_ISL_905553, EPI_ISL_905554, EPI_ISL_905555, EPI_ISL_905556, EPI_ISL_905557, EPI_ISL_905558, EPI_ISL_905559, EPI_ISL_905561, EPI_ISL_905562, EPI_ISL_905563, EPI_ISL_905564, EPI_ISL_905565, EPI_ISL_905662, EPI_ISL_905673, EPI_ISL_905674, EPI_ISL_905675, EPI_ISL_905688, EPI_ISL_905711, EPI_ISL_905712, EPI_ISL_905713, EPI_ISL_905725, EPI_ISL_905729 | | | |
| see above | Dutch COVID-19 response team | National Institute for Public Health and the Environment (RIVM) | Adam Meijer, Harry Vennema, Dirk Eggink, Jeroen Cremer, Sharon van den Brink, Bas van der Veer, AnneMarie van den Brandt, Florian Zwagemaker, Dennis Schmitz, Chantal Reusken, on behalf of the national COVID-19 response team |
| EPI_ISL_905849, EPI_ISL_905850, EPI_ISL_905851, EPI_ISL_905852, EPI_ISL_905853, EPI_ISL_905854, EPI_ISL_905855, EPI_ISL_905856, EPI_ISL_905857, EPI_ISL_905858, EPI_ISL_905859, EPI_ISL_905860, EPI_ISL_905861, EPI_ISL_905862, EPI_ISL_905863, EPI_ISL_905864, EPI_ISL_905865, EPI_ISL_905900, EPI_ISL_905901 | | | |
| see above | OHSU Lab Services Molecular Microbiology Lab | Oregon SARS-CoV-2 Genome Sequencing Center | Brendan L. O'Connell, Sally Grindstaff, Kayla Carter, Ruth V. Nichols, Alec J. Hirsch, Donna Hansel, Guang Fan, Xuan, Qin, Daniel N. Streblow, William B. Messer, Andrew C. Adey, Benjamin N. Bimber, Brian J. O'Roak |
| EPI_ISL_906058, EPI_ISL_906062 | Tilia Laboratories s.r.o. | Tilia Laboratories s.r.o. | Sona Pekova, MD, PhD. |
| EPI_ISL_906091 | Shimantik Pathology and Diagnostic Center | Child Health Research Foundation | Senjuti Saha, Syed Muktadir Al Sium, Arif Mohammad Tanmoy, Afroza Akter Tanni, Sharmistha Goswami, Roly Malaker Md Hafizur Rahman, Md. Parvej Alam, Md. Mobarok Karim, Samir K Saha |
| EPI_ISL_906116 | Child Health Research Foundation | Child Health Research Foundation | Senjuti Saha, Afroza Akter Tanni, Sharmistha Goswami, Syed Muktadir Al Sium, Arif Mohammad Tanmoy, Roly Malaker, Md Hafizur Rahman, Samir K Saha |
| EPI_ISL_906536 | CLINICA OCCIDENTE | Instituto Nacional de Salud- Dirección de Investigación en Salud Pública, Universidad de los Andes- Applied genomics | Katherine Laiton-Donato, Diego A. Álvarez-Díaz, Carlos Franco-Muñoz, Mauricio Pacheco-Montealegre, Héctor Alejandro Ruiz-Moreno, Maria T. Herrera-Sepúlveda, Diego Andrés Prada, Jhonnatan Reales-González, Sheryll Corchuelo, Julian Naizaque, Gerardo Santamaría Jorge Duitama, Laura Natalia Gonzalez, Jorge Ivan Diaz, Silvia Restrepo-Restrepo, Magdalena Wiesner, Martha Lucia Ospina Martinez, Marcela Mercado-Reyes |

| | | | |
|---|---|---|---|
| | | research group, Vicerrectoria de Investigación y Creación, Universidad de los Andes- Systems and Computing Engineering Department | |
| EPI_ISL_906540, EPI_ISL_906544 | CLINICA DE OCCIDENTE | Instituto Nacional de Salud- Dirección de Investigación en Salud Pública, Universidad de los Andes- Applied genomics research group, Vicerrectoria de Investigación y Creación, Universidad de los Andes- Systems and Computing Engineering Department | Katherine Laiton-Donato, Diego A. Álvarez-Díaz, Carlos Franco-Muñoz, Mauricio Pacheco-Montealegre, Héctor Alejandro Ruiz-Moreno, Maria T. Herrera-Sepúlveda, Diego Andrés Prada, Jhonnatan Reales-González, Sheryll Corchuelo, Julian Naizaque, Gerardo Santamaría Jorge Duitama, Laura Natalia Gonzalez, Jorge Ivan Diaz, Silvia Restrepo-Restrepo, Magdalena Wiesner, Martha Lucia Ospina Martinez, Marcela Mercado-Reyes |
| EPI_ISL_906572 | Maine Health and Environmental Testing Laboratory (Maine HETL) | Tewhey Lab, The Jackson Laboratory | Matluk,N., Dewey,H., Iosue,F., Barter,M., Lynch,R., Munger,H. and Tewhey,R. |
| EPI_ISL_906777, EPI_ISL_906778 | New Mexico Department of Health Scientific Laboratory | New Mexico Department of Health Scientific Laboratory | Ellie Johnson, Anastacia Griego-Fisher, D'eldra Malone |
| EPI_ISL_909686 | GH Cochin-Broca Hôtel Dieu/Lab Virologie | National Reference Center for Viruses of Respiratory Infections, Institut Pasteur, Paris | Marion Barbet, Sylvie Behillil, Méline Bizard, Angela Brisebarre, Camille Capel, Etienne Simon-Lorière, Vincent Enouf, Maud Vanpeene, Sylvie van der Werf,L'Honneur Anne-Sophie |
| EPI_ISL_910030, EPI_ISL_910222, EPI_ISL_910324 | Viollier AG | University Hospital Basel, Clinical Bacteriology | Tim Roloff, Madlen Stange, Helena MB Seth-Smith, Alfredo Mari, Karoline Leuzinger, Julia Bielicki, Christiane Beckmann, Manuel Battegay, Hans Hirsch, Adrian Egli |
| EPI_ISL_910336 | Laboratory for Respiratory Viruses, Cantacuzino National Military-Medical Institute for Research and Development | Cantacuzino Institute Virology | Luiza Ustea, Nicoleta Paraschiv, Mihaela Lazar |
| EPI_ISL_910553, EPI_ISL_910554, EPI_ISL_910555, EPI_ISL_910556, EPI_ISL_910557, EPI_ISL_910558, EPI_ISL_910559, EPI_ISL_910560, EPI_ISL_910561, EPI_ISL_910562, EPI_ISL_910563, EPI_ISL_910564, EPI_ISL_910565, EPI_ISL_910566, EPI_ISL_910567, EPI_ISL_910568, EPI_ISL_910569, EPI_ISL_910571, EPI_ISL_910572, EPI_ISL_910573, EPI_ISL_910574, EPI_ISL_910575, EPI_ISL_910576, EPI_ISL_910577, EPI_ISL_910580, EPI_ISL_910581, EPI_ISL_910582, EPI_ISL_910583, EPI_ISL_910584, EPI_ISL_911233 | | | |
| see above | Laboratoire national de sante, Microbiology, Virology | Laboratoire national de sante, Microbiology, Microbial Genomics Platform | Anke Wienecke-Baldacchino, Catherine Ragimbeau,Jessica Tapp, Fatu Djabi, Lise Pignon, Raoul Salmon, Tamir Abdelrahman |
| EPI_ISL_911331 | Servicio de Microbiología, Hospital Universitario Son Espases | SeqCOVID-SPAIN consortium/IBV(CSIC) | Carla López-Causapé, Jordi Reina, Antonio Oliver and SeqCOVID-SPAIN consortium |
| EPI_ISL_911524 | Microbiology and Virology Unit, Florence Careggi University Hospital | Microbiology and Virology Unit, Florence Careggi University Hospital | Vincenzo Di Pilato, Marco Coppi, Fabio Morecchiato, Noemi Aiezza, Ilaria Baccani, Alberto Antonelli, Emanuele Gori, Gian Maria Rossolini |
| EPI_ISL_911967 | LabCorp | Wyoming Public Health Laboratory | Noah Hull, Taylor Fearing, Lynette Gumbleton, Channing Weber, Ashley Norberg, Bailey Bowcutt, and Wanda Manley |
| EPI_ISL_912165 | Connecticut Department of Health | Grubaugh Lab - Yale School of Public Health | Tara Alpert, Joseph Fauver, Anderson Brito, Mallery Breban, Anne Wyllie, Chantal Vogels, Mary Petrone, Annie Watkins, Chaney Kalinich, Isabel Ott, Nathan Grubaugh |
| EPI_ISL_912269, EPI_ISL_912306 | Hospital General Universitario Gregorio Marañón | SeqCOVID-SPAIN consortium / IBV (CSIC) | Darío García de Viedma, Laura Pérez-Lago, Pedro J Sola-Campoy, Sergio Buenestado-Serrano, Marta Herranz, Victor Manuel de la Cueva, Julia Suárez, Pilar Catalán, Patricia Muñoz and SeqCOVID-SPAIN consortium |
| EPI_ISL_912374, EPI_ISL_912381, EPI_ISL_912382, EPI_ISL_912383, EPI_ISL_912384, EPI_ISL_912385 | Fondation Congolaise pour la recherche medicale (FCRM), Francine Ntoumi | NGS Competence Center Tuebingen, Institut für Medizinische Mikrobiologie und Hygiene, Universitaetsklinikum Tübingen | Angel Angelov |
| EPI_ISL_912405, EPI_ISL_912411, EPI_ISL_912412, EPI_ISL_912419 | KU Leuven, Rega Institute, Clinical and Epidemiological Virology | KU Leuven, Rega Institute, Clinical and Epidemiological Virology | Tony Wawina-Bokalanga, Bert Vanmechelen, Joan Marti-Carerras, Piet Maes |
| EPI_ISL_912458, EPI_ISL_912461, EPI_ISL_912491, EPI_ISL_912519 | NHLS Universitas Academic | UFS Virology | PA Bester, MM Nyaga, P Nthiga, MT Mogotsi, D Goedhals, T de Oliveira |
| EPI_ISL_912900, EPI_ISL_912922, EPI_ISL_912937, EPI_ISL_912962, EPI_ISL_912963, EPI_ISL_912964, EPI_ISL_912965, EPI_ISL_912966, EPI_ISL_912967, EPI_ISL_912968, EPI_ISL_912969, EPI_ISL_912970, EPI_ISL_912971, EPI_ISL_912972, EPI_ISL_912973, EPI_ISL_912974, EPI_ISL_912975, EPI_ISL_912976, EPI_ISL_912977 | | | |
| see above | Hôpital Henri Mondor | Department of Virology, Henri Mondor University Hospital, Assistance Publique Hôpitaux de Paris, Université Paris-Est Créteil, INSERM U955 | Christophe Rodriguez, Slim Fourati, Vanessa Demontant, Guillaume Gricourt, Melissa N'Debi, Alexandre Soulier, Elisabeth Trawinski, Jean-Michel Pawlotsky |
| EPI_ISL_913021, EPI_ISL_913022, EPI_ISL_913023 | Hospital San Pedro de Alcántara | Instituto de Salud Carlos III | Iglesias-Caballero, M. Camarero, S. Sandonís,V. Vázquez, S. Pozo, F. Casas, I. Jiménez, P. Zaballos, A. Monzón, S. Varona, S. Cuesta, I. Rodríguez, G. |
| EPI_ISL_913024 | Complejo Hospitalario Universitario de Santiago | Instituto de Salud Carlos III | Iglesias-Caballero, M. Camarero, S. Sandonís,V. Vázquez, S. Pozo, F. Casas, I. Jiménez, P. Zaballos, A. Monzón, S. Varona, S. Cuesta, I. Peña,F. |
| EPI_ISL_913033 | Hospital San Pedro de Alcántara | Instituto de Salud Carlos III | Iglesias-Caballero, M. Camarero, S. Sandonís,V. Vázquez, S. Pozo, F. Casas, I. Jiménez, P. Zaballos, A. Monzón, S. Varona, S. Cuesta, I. Rodríguez, G. |
| EPI_ISL_913040 | Hospital Universitario de Guadalajara | Instituto de Salud Carlos III | Iglesias-Caballero, M. Camarero, S. Sandonís,V. Vázquez, S. Pozo, F. Casas, I. Jiménez, P. Zaballos, A. Monzón, S. Varona, S. Cuesta, I. González, A. |
| EPI_ISL_913041 | Consejería de Sanidad y Asuntos Sociales | Instituto de Salud Carlos III | Iglesias-Caballero, M. Camarero, S. Sandonís,V. Vázquez, S. Pozo, F. Casas, I. Jiménez, P. Zaballos, A. Monzón, S. Varona, S. Cuesta, I. Gutiérrez, G. |
| EPI_ISL_913048, EPI_ISL_913049, EPI_ISL_913050 | Hospital Comarcal Sierrallana | Instituto de Salud Carlos III | Iglesias-Caballero, M. Camarero, S. Sandonís,V. Vázquez, S. Pozo, F. Casas, I. Jiménez, P. Zaballos, A. Monzón, S. Varona, S. Cuesta, I. De Benito, I. |
| EPI_ISL_913076, EPI_ISL_913085 | Center for Virology | Center for Virology | Jeremy V. Camp, Irene Goerzer, Monika Redlberger-Fritz, Stephan W. Aberle |
| EPI_ISL_913102 | CHU Purpan - Laboratoire de Virologie - Institut Fédératif de Biologie | CHU Purpan - Laboratoire de Virologie - Institut Fédératif de Biologie | Latour J., Ranger N., Dubois M., Carcenac R., Harter A., Boyer P., Tremeaux P., Izopet J. |
| EPI_ISL_913269, EPI_ISL_913274 | Klinisk mikrobiologi | The Public Health Agency of Sweden | Anna-Malin Linde, Maria Lind Karlberg, Carlo Berg, Oskar Karlsson Lindsjo, Sofia Stamouli, Reza Advani, Mattias Haukland, Petra Holmstrom, Noura Walai, Petra Edquist, Mia Brytting, Anna Risberg, Karin Tegmark-Wisell |
| EPI_ISL_913291, EPI_ISL_913292, EPI_ISL_913293 | ABC Labs | The Public Health Agency of Sweden | Anna-Malin Linde, Maria Lind Karlberg, Carlo Berg, Oskar Karlsson Lindsjo, Sofia Stamouli, Reza Advani, Mattias Haukland, Petra Holmstrom, Noura Walai, Petra Edquist, Mia Brytting, Anna Risberg, Karin Tegmark-Wisell |
| EPI_ISL_913296 | Klinisk mikrobiologi | The Public Health Agency of Sweden | Anna-Malin Linde, Maria Lind Karlberg, Carlo Berg, Oskar Karlsson Lindsjo, Sofia Stamouli, Reza Advani, Mattias Haukland, Petra Holmstrom, Noura Walai, Petra Edquist, Mia Brytting, Anna Risberg, Karin Tegmark-Wisell |
| EPI_ISL_913305 | Synlab Medilab, Mikrobiologi | The Public Health Agency of Sweden | Anna-Malin Linde, Maria Lind Karlberg, Carlo Berg, Oskar Karlsson Lindsjo, Sofia Stamouli, Reza Advani, Mattias Haukland, Petra Holmstrom, Noura Walai, Petra Edquist, Mia Brytting, Anna Risberg, Karin Tegmark-Wisell |
| EPI_ISL_913311 | Klinisk mikrobiologi | The Public Health Agency of Sweden | Anna-Malin Linde, Maria Lind Karlberg, Carlo Berg, Oskar Karlsson Lindsjo, Sofia Stamouli, Reza Advani, Mattias Haukland, Petra Holmstrom, Noura Walai, Petra Edquist, Mia Brytting, Anna Risberg, Karin Tegmark-Wisell |
| EPI_ISL_913339, EPI_ISL_913341, EPI_ISL_913342 | Unilabs, Mikrobiologiska laboratoriet | The Public Health Agency of Sweden | Anna-Malin Linde, Maria Lind Karlberg, Carlo Berg, Oskar Karlsson Lindsjo, Sofia Stamouli, Reza Advani, Mattias Haukland, Petra Holmstrom, Noura Walai, Petra Edquist, Mia Brytting, Anna Risberg, Karin Tegmark-Wisell |
| EPI_ISL_913367, EPI_ISL_913369 | Klinisk mikrobiologi | The Public Health Agency of Sweden | Anna-Malin Linde, Maria Lind Karlberg, Carlo Berg, Oskar Karlsson Lindsjo, Sofia Stamouli, Reza Advani, Mattias Haukland, Petra Holmstrom, Noura Walai, Petra Edquist, Mia Brytting, Anna Risberg, Karin Tegmark-Wisell |
| EPI_ISL_913380, EPI_ISL_913381, EPI_ISL_913382, EPI_ISL_913383, EPI_ISL_913384 | Dynamic Code AB | The Public Health Agency of Sweden | Anna-Malin Linde, Maria Lind Karlberg, Carlo Berg, Oskar Karlsson Lindsjo, Sofia Stamouli, Reza Advani, Mattias Haukland, Petra Holmstrom, Noura Walai, Petra Edquist, Mia Brytting, Anna Risberg, Karin Tegmark-Wisell |
| EPI_ISL_913450 | Unilabs AB Skovde | The Public Health Agency of Sweden | Anna-Malin Linde, Maria Lind Karlberg, Carlo Berg, Oskar Karlsson Lindsjo, Sofia Stamouli, Reza Advani, Mattias Haukland, Petra Holmstrom, Noura |

| | | | |
|---|---|---|---|
| | | | Walai, Petra Edquist, Mia Brytting, Anna Risberg, Karin Tegmark-Wisell |
| EPI_ISL_913486 | ABC Labs | The Public Health Agency of Sweden | Anna-Malin Linde, Maria Lind Karlberg, Carlo Berg, Oskar Karlsson Lindsjo, Sofia Stamouli, Reza Advani, Mattias Haukland, Petra Holmstrom, Noura Walai, Petra Edquist, Mia Brytting, Anna Risberg, Karin Tegmark-Wisell |
| EPI_ISL_913487, EPI_ISL_913488, EPI_ISL_913489 | Klinisk mikrobiologi | The Public Health Agency of Sweden | Anna-Malin Linde, Maria Lind Karlberg, Carlo Berg, Oskar Karlsson Lindsjo, Sofia Stamouli, Reza Advani, Mattias Haukland, Petra Holmstrom, Noura Walai, Petra Edquist, Mia Brytting, Anna Risberg, Karin Tegmark-Wisell |
| EPI_ISL_913495 | Complejo Hospitalario Universitario de Santiago | Instituto de Salud Carlos III | Iglesias-Caballero, M. Camarero, S. Sandonís,V. Vázquez, S. Pozo, F. Casas, I. Jiménez, P. Zaballos, A. Monzón, S. Varona, S. Cuesta, I. Peña,F. |
| EPI_ISL_913525, EPI_ISL_913528, EPI_ISL_913529, EPI_ISL_913530, EPI_ISL_913531, EPI_ISL_913532, EPI_ISL_913533, EPI_ISL_913534, EPI_ISL_913535, EPI_ISL_913537, EPI_ISL_913538, EPI_ISL_913539, EPI_ISL_913540, EPI_ISL_913541, EPI_ISL_913542, EPI_ISL_913543, EPI_ISL_913544, EPI_ISL_913545, EPI_ISL_913546, EPI_ISL_913547, EPI_ISL_913548, EPI_ISL_913549, EPI_ISL_913550, EPI_ISL_913551, EPI_ISL_913552, EPI_ISL_913553, EPI_ISL_913554, EPI_ISL_913555, EPI_ISL_913556, EPI_ISL_913557, EPI_ISL_913558, EPI_ISL_913559, EPI_ISL_913560, EPI_ISL_913561, EPI_ISL_913562 | | | |
| see above | M Health Fairview | Minnesota Department of Health, Public Health Laboratory | Alexandra Lorentz, Jacob Garfin, Matt Plumb, and Xiong Wang |
| EPI_ISL_913613, EPI_ISL_913614, EPI_ISL_913640, EPI_ISL_913641, EPI_ISL_913642, EPI_ISL_913643, EPI_ISL_913644, EPI_ISL_913645, EPI_ISL_913646 | Michigan Department of Health and Human Services, Bureau of Laboratories | Michigan Department of Health and Human Services, Bureau of Laboratories | Blankenship HM, Riner D, Soehnlen MK |
| EPI_ISL_913686, EPI_ISL_913687, EPI_ISL_913688, EPI_ISL_913689, EPI_ISL_913690, EPI_ISL_913691, EPI_ISL_913693, EPI_ISL_913694, EPI_ISL_913695, EPI_ISL_913696, EPI_ISL_913697, EPI_ISL_913698, EPI_ISL_913699, EPI_ISL_913700, EPI_ISL_913701, EPI_ISL_913702, EPI_ISL_913703, EPI_ISL_913704, EPI_ISL_913705, EPI_ISL_913706, EPI_ISL_913707, EPI_ISL_913708, EPI_ISL_913709, EPI_ISL_913710 | | | |
| see above | Minnesota Department of Health, Public Health Laboratory | Minnesota Department of Health, Public Health Laboratory | Alexandra Lorentz, Jacob Garfin, Matt Plumb, and Xiong Wang |
| EPI_ISL_913726 | Missouri State Public Health Laboratory | Minnesota Department of Health, Public Health Laboratory | Alexandra Lorentz, Jacob Garfin, Matt Plumb, and Xiong Wang |
| EPI_ISL_913791, EPI_ISL_913808, EPI_ISL_913822, EPI_ISL_913832, EPI_ISL_913839, EPI_ISL_913846, EPI_ISL_913862, EPI_ISL_913877, EPI_ISL_913883, EPI_ISL_913884, EPI_ISL_913891, EPI_ISL_913892, EPI_ISL_913893, EPI_ISL_913898, EPI_ISL_913902, EPI_ISL_913903, EPI_ISL_913904, EPI_ISL_913906, EPI_ISL_913909 | | | |
| see above | TGen North | TGen North | "Jolene Bowers, Megan Folkerts, Chris French, Hayley Yaglom, Ashlyn Pfeiffer, Darrin Lemmer, Dave Engelthaler, The Arizona COVID Genomics Union (ACGU)" |
| EPI_ISL_913999 | TX DSHS, Lab Services Section MC 1947 | Pathogen Discovery, Respiratory Viruses Branch, Division of Viral Diseases, Centers for Disease Control and Prevention | Ying Tao, Yan Li, Jing Zhang, Krista Queen, Anna Uehara, Peter Cook, Clinton R. Paden, Haibin Wang, Suxiang Tong |
| EPI_ISL_914038 | CT-Dr. Katherine A. Kelley State Public Health Lab | Pathogen Discovery, Respiratory Viruses Branch, Division of Viral Diseases, Centers for Disease Control and Prevention | Ying Tao, Yan Li, Jing Zhang, Krista Queen, Anna Uehara, Peter Cook, Clinton R. Paden, Haibin Wang, Suxiang Tong |
| EPI_ISL_914644 | Santa Clara County Public Health Laboratory | Santa Clara County Public Health Laboratory | Santa Clara County Public Health Department |
| EPI_ISL_914817, EPI_ISL_914819 | HOSPITAL DE NIÑOS DR. CARLOS SAENZ HERRERA | Inciensa, Instituto Costarricense de Investigación y Enseñanza en Nutrición y Salud | Francisco Duarte, Hebleen Porras, Claudio Soto-Garita, Estela Cordero, Adriana Godínez, Melany Calderón & Cristian Pérez-Corrales |
| EPI_ISL_918189, EPI_ISL_918194, EPI_ISL_918224, EPI_ISL_918225, EPI_ISL_918227, EPI_ISL_918228, EPI_ISL_918233, EPI_ISL_918239, EPI_ISL_918240, EPI_ISL_918242, EPI_ISL_918244, EPI_ISL_918261 | | | |
| see above | Innovative Genomics Institute, UC Berkeley | Innovative Genomics Institute, UC Berkeley | Stacia Wyman, Haridha Shivram, Phil Frankino, Liana Lareau, Shana McDevitt, Justin Choi |
| EPI_ISL_918351 | Institute of Virology, Medical Center, University of Freiburg, Freiburg, Germany | Institute of Virology, Clinial Virus Genomics, Medical Center, University of Freiburg, Freiburg, Germany | Jonas Fuchs, Lisa Kern, Sandra Reuter, Hajo Grundmann, Marcus Panning |
| EPI_ISL_918409 | Ospedale Santa Caterina Novella | Istituto Zooprofilattico Sperimentale della Puglia e della Basilicata | Parisi A., Bianco A., Capozzi L., Del Sambro L., Simone D., Manzulli V, Rondinone V., Pace L., Cipolletta D., Galante D. |
| EPI_ISL_918444, EPI_ISL_918445, EPI_ISL_918446, EPI_ISL_918447, EPI_ISL_918448, EPI_ISL_918449, EPI_ISL_918450, EPI_ISL_918451, EPI_ISL_918452, EPI_ISL_918453, EPI_ISL_918454, EPI_ISL_918455, EPI_ISL_918456 | | | |
| see above | AIID | Irish Coronavirus Sequencing Consortium-Teagasc Grange | Matthew McCabe, Aljandro Abner Garcia Leon, Fiona Crispie, Calum Walsh, Michael Carr, John Kenny, Paul Cotter, Patrick Mallon, Gabriel Gonzalez |
| EPI_ISL_918502, EPI_ISL_918505, EPI_ISL_918506, EPI_ISL_918511 | LACEN - Laboratório Central de Saúde Pública do Amazonas | Evandro Chagas Institute | Santos, M.C.; Silva, A.M.; Junior, W.D.C.; Barbagelata, L.S.; Ferreira, J.A.; Sousa, E.M.A.; da Silva, P.S.; Pinheiro, K.C.; L.C.; Sousa Junior, E.C. |
| EPI_ISL_918748, EPI_ISL_918749, EPI_ISL_918750, EPI_ISL_918751, EPI_ISL_918752, EPI_ISL_918753, EPI_ISL_918754, EPI_ISL_918755, EPI_ISL_918756, EPI_ISL_918757, EPI_ISL_918760, EPI_ISL_918761, EPI_ISL_918762, EPI_ISL_918763, EPI_ISL_918940, EPI_ISL_918941, EPI_ISL_918942, EPI_ISL_918943, EPI_ISL_918944, EPI_ISL_918945, EPI_ISL_918946, EPI_ISL_918947, EPI_ISL_918948, EPI_ISL_918949, EPI_ISL_918950, EPI_ISL_918951, EPI_ISL_918952, EPI_ISL_918953, EPI_ISL_918954, EPI_ISL_918959, EPI_ISL_918961, EPI_ISL_918962, EPI_ISL_918963, EPI_ISL_918965 | | | |
| see above | University of Birmingham | COVID-19 Genomics UK (COG-UK) Consortium | Institute of Microbiology, University of Birmingham: Claire McMurray, Joanne Stockton, Samuel Nicholls, Radoslaw Poplawski, Will Rowe, Josh Quick, Nicholas Loman. University of Birmingham Testing Laboratory: Celina M Whalley, Andrew Bosworth, Charlotte Poxon, Kasun Wanigasooriya, Oliver Pickles, Mike Kidd, Alex Richter, Andrew D Beggs PHE Heartlands Lab: Husam Osman, Andrew Bosworth. Queen Elizabeth Hospital: Anna Casey |
| EPI_ISL_919257 | West of Scotland Specialist Virology Centre, NHSGGC / MRC-University of Glasgow Centre for Virus Research | COVID-19 Genomics UK (COG-UK) Consortium | Ana da Silva Filipe, Natasha Johnson, Kathy Smollett, Daniel Mair, Stephen Carmichael, Alice Broos, Lily Tong, Jenna Nichols, Kyriaki Nomikou; Sarah McDonald; Richard Orton, Joseph Hughes, Sreenu Vattipally, David L Robertson; Alasdair MacLean, Rory Gunson; Sharif Shaaban, Matthew Holden; Rachel Blacow, Guy Mollett, Kathy Li, James Shepherd, Antonia Ho, Emma Thomson |
| EPI_ISL_919321, EPI_ISL_919322, EPI_ISL_919323, EPI_ISL_919324, EPI_ISL_919325, EPI_ISL_919326, EPI_ISL_919327, EPI_ISL_919328, EPI_ISL_919329, EPI_ISL_919330, EPI_ISL_919331, EPI_ISL_919332, EPI_ISL_919333, EPI_ISL_919334, EPI_ISL_919335 | | | |
| see above | Virology Department, Royal Infirmary of Edinburgh, NHS Lothian / School of Biological Sciences, University of Edinburgh / Institute of Genetics and Molecular Medicine, University of Edinburgh | COVID-19 Genomics UK (COG-UK) Consortium | McHugh M, Dewar R, Rooke S, Gallagher M, Balcaza C, O'Toole Á, Scher E, Hill V, McCrone JT, Colquhoun R, Yu X, Jackson B, Rambaut A, Williams TC, Templeton K |
| EPI_ISL_919452, EPI_ISL_919453, EPI_ISL_919476 | Liverpool Clinical Laboratories | COVID-19 Genomics UK (COG-UK) Consortium | Sam Haldenby, Anita Lucaci, Steve Paterson, Julian Hiscox, Alistair Darby, M Almsaud, A Alrezaihi, Muhannad Alruwaili, Stuart D Armstrong, Jones Benjamin, Eleanor G Bentley, Anu Chawla, Jordan J Clark, Angela Cowell, Richard Eccles, Isabel García-Dorival, Matthew Gemmell, Alessandro Gerada, PKF Gilmore, Richard Gregory, Ximeng Han, Catherine Hartley, Margaret Hughes, Miren Iturriza-Gomara, James Johnson, L Luu, Jenifer Manson, Charlotte Nelson, Elaine O'Toole, Cassie Olateju, Rebekah Penrice-Randal , Lucille Rainbow, N.P Randle, Trevor Ian Robinson, Parul Sharma, Ghada T Shawli, James P Stewart, Neil Swainston, Ecaterina Vamos, Joanne Watts, Mark Whitehead |
| EPI_ISL_919781, EPI_ISL_919782, EPI_ISL_919783, EPI_ISL_919784, EPI_ISL_919785, EPI_ISL_919786, EPI_ISL_919787, EPI_ISL_919788, EPI_ISL_919789, EPI_ISL_919790, EPI_ISL_919791, EPI_ISL_919792, EPI_ISL_919793, EPI_ISL_919794, EPI_ISL_919795, EPI_ISL_919796 | | | |
| see above | Barts Health NHS Trust | COVID-19 Genomics UK (COG-UK) Consortium | CUTINO-MOGUEL, Maria-Teresa; HARRINGTON, David; OWOYEMI, Dola; KULASEGARAN-SHYLINI, Raghavendran; BROAD, Claire; KELE, Beatrix |
| EPI_ISL_920499 | University College London Hospital | COVID-19 Genomics UK (COG-UK) Consortium | Judith Heaney, Matthew Byott, Catherine Houlihan, Dan Frampton, Stuart Kirk, Moira Spyer and Eleni Nastouli |
| EPI_ISL_920843, EPI_ISL_920844, EPI_ISL_920845, EPI_ISL_920846, EPI_ISL_920847 | University College London, Great Ormond Street Hospital for Children NHS Foundation Trust, Imperial College Healthcare NHS Trust | COVID-19 Genomics UK (COG-UK) Consortium | Sergi Castellano, Rachel Williams, Mark Kristiansen, Paola Resende Silva, Sunando Roy, Tony Brooks, Helena Tutill, Paola Niola, Patricia Dyal, Charlotte Williams, Leysa Forrest, Yasmin Panchbhaya, Jacqueline Findlay, Samuel Weeks, Julianne Brown, Kathryn Harris, Paul Randell, James Price, Alison Holmes, Judith Breuer |
| EPI_ISL_921022, EPI_ISL_921027, EPI_ISL_921028, EPI_ISL_921029, EPI_ISL_921030, EPI_ISL_921031, EPI_ISL_921032, EPI_ISL_921034, EPI_ISL_921035, EPI_ISL_921036, EPI_ISL_921042, EPI_ISL_921043, EPI_ISL_921044, EPI_ISL_921045, EPI_ISL_921085, EPI_ISL_921087, EPI_ISL_921088, EPI_ISL_921089, EPI_ISL_921118, EPI_ISL_921119, EPI_ISL_921120, EPI_ISL_921133, EPI_ISL_921135, EPI_ISL_921136 | | | |
| see above | Regional Virus Laboratory, Belfast Health and Social Care Trust | COVID-19 Genomics UK (COG-UK) Consortium | Conall McCaughey, James McKenna, Tanya Curran, Susan Feeney, Alison Watt, Ciara Cox, Mairead Connor, Zoltan Molnar, David Simpson, Derek Fairley |
| EPI_ISL_921253, EPI_ISL_921255, EPI_ISL_921258, EPI_ISL_921259, EPI_ISL_921261, EPI_ISL_921262, EPI_ISL_921264, EPI_ISL_921265, EPI_ISL_921271, EPI_ISL_921272, EPI_ISL_921275, EPI_ISL_921276, EPI_ISL_921371, EPI_ISL_921379, EPI_ISL_921380, EPI_ISL_921381, EPI_ISL_921382, EPI_ISL_921384, EPI_ISL_921385, EPI_ISL_921386, EPI_ISL_921387, EPI_ISL_921388, EPI_ISL_921389, EPI_ISL_921390, EPI_ISL_921392, EPI_ISL_921393, EPI_ISL_921394, EPI_ISL_921395, EPI_ISL_921396, EPI_ISL_921397, EPI_ISL_921398, EPI_ISL_921399, EPI_ISL_921400, EPI_ISL_921401, EPI_ISL_921402, EPI_ISL_921403, EPI_ISL_921404, EPI_ISL_921405, EPI_ISL_921406, EPI_ISL_921414, EPI_ISL_921415, EPI_ISL_921416, EPI_ISL_921417, EPI_ISL_921418, EPI_ISL_921420, EPI_ISL_921422 | | | |

| | | | |
|---|---|---|---|
| see above | Northumbria University / South Tees Hospitals NHS Foundation Trust / North Cumbria Integrated Care NHS Foundation Trust / North Tees and Hartlepool NHS Foundation Trust / Newcastle Hospitals NHS Foundation Trust | COVID-19 Genomics UK (COG-UK) Consortium | Darren L Smith,Andrew Nelson,Matthew Bashton,Greg R Young,Joshua Loh,John Allan,Mohammad A Tariq,Giles S Holt,Gary Black,Wen C Yew,Lynn Dover,Paul Baker,Steve Liggett,Sarah Essex,Jane Greenaway,Debra Padgett,Clive Graham,Garren Scott,Edward Barton,Emma Swindells,Brendan Payne,Jennifer Collins,Yusri Taha,Gary Eltringham |
| EPI_ISL_921786, EPI_ISL_921787, EPI_ISL_921788, EPI_ISL_921793, EPI_ISL_921800, EPI_ISL_921805, EPI_ISL_921808, EPI_ISL_921809, EPI_ISL_921810, EPI_ISL_921813, EPI_ISL_921819, EPI_ISL_921825 | | | |
| see above | Quadram Institute Bioscience | COVID-19 Genomics UK (COG-UK) Consortium | Dave J. Baker, Gemma L. Kay, Alp Aydin, Thanh Le-Viet, Steven Rudder, Ana P. Tedim, Anastasia Kolyva, Maria Diaz, Leonardo de Oliveira Martins, Nabil-Fareed Alikhan, Lizzie Meadows, Rachael Stanley, Ngozi Elumogo, Muhammed Yasir, Nicholas M. Thomson, Alexander J Trotter, Rachel Gilroy, Samuel Bloomfield, Claire Stuart, Andrew Bell, Reenesh Prakash, Samir Dervisevic, Alison E. Mather, John Wain, Mark Webber, Andrew J. Page, Justin O'Grady |
| EPI_ISL_922241, EPI_ISL_922260, EPI_ISL_922261, EPI_ISL_922262, EPI_ISL_922263, EPI_ISL_922264, EPI_ISL_922267, EPI_ISL_922271, EPI_ISL_922272, EPI_ISL_922274, EPI_ISL_922275, EPI_ISL_922277, EPI_ISL_922279, EPI_ISL_922287, EPI_ISL_922288, EPI_ISL_922290, EPI_ISL_922291, EPI_ISL_922292, EPI_ISL_922293, EPI_ISL_922296, EPI_ISL_922297, EPI_ISL_922299, EPI_ISL_922300, EPI_ISL_922302, EPI_ISL_922305 | | | |
| see above | Oxford Viromics, NDM, University of Oxford; Oxford University Hospitals; Basingstoke and North Hampshire Hospital | COVID-19 Genomics UK (COG-UK) Consortium | Tanya Golubchik, David Bonsall, George Macintyre, Amy Trebes, Mariateresa de Cesare, Catrin Moore, Alex Mobbs, Anita Justice, Robert Shaw, Monique Andersson, Timothy Peto, Emma Wise, Nathan Moore, Jessica Lynch, Nick Cortes, Matilde Mori, Stephen Kidd, David Buck, John Todd, Christophe Fraser |
| EPI_ISL_923244, EPI_ISL_923265 | Centre for Enzyme Innovation, University of Portsmouth / Translational Research Laboratory, Portsmouth Hospitals NHS Trust | COVID-19 Genomics UK (COG-UK) Consortium | Angela Beckett,Salman Goudarzi,Christopher Fearn,Kate Cook,Katie Loveson,Sharon Glaysher,Scott Elliott,Samuel Robson |
| EPI_ISL_924094, EPI_ISL_924168, EPI_ISL_924213, EPI_ISL_924237, EPI_ISL_924250, EPI_ISL_924280, EPI_ISL_924283 | Virology Department, Sheffield Teaching Hospitals NHS Foundation Trust/Department of Infection, Immunity and Cardiovascular Disease, The Medical School, University of Sheffield | COVID-19 Genomics UK (COG-UK) Consortium | Thushan de Silva, Matthew Parker, Nikki Smith, Adri Angyal, Rebecca Brown, Luke Green, Rachel Tucker, Paul Parsons, Danielle Groves, Katie Johnson, Laura Carrilero, Alex Keeley, Dave Partridge, Matthew Wyles, Benjamin Lindsey, Mehmet Yavuz, Mohammad Raza, Cariad Evans |
| EPI_ISL_924430, EPI_ISL_924431, EPI_ISL_924432, EPI_ISL_924433, EPI_ISL_924434, EPI_ISL_924435, EPI_ISL_924436, EPI_ISL_924437, EPI_ISL_924438, EPI_ISL_924439, EPI_ISL_924440, EPI_ISL_924441, EPI_ISL_924443, EPI_ISL_924444, EPI_ISL_924445, EPI_ISL_924446, EPI_ISL_924447, EPI_ISL_924448, EPI_ISL_924449, EPI_ISL_924450, EPI_ISL_924451, EPI_ISL_924452, EPI_ISL_924453, EPI_ISL_924454, EPI_ISL_924455, EPI_ISL_924456, EPI_ISL_924457, EPI_ISL_924458, EPI_ISL_924459, EPI_ISL_924460, EPI_ISL_924467, EPI_ISL_924468, EPI_ISL_924469, EPI_ISL_924470, EPI_ISL_924472, EPI_ISL_924474, EPI_ISL_924476, EPI_ISL_924477, EPI_ISL_924478, EPI_ISL_924479, EPI_ISL_924483, EPI_ISL_924485, EPI_ISL_924487, EPI_ISL_924488, EPI_ISL_924489, EPI_ISL_924490, EPI_ISL_924491, EPI_ISL_924493, EPI_ISL_924494, EPI_ISL_924497, EPI_ISL_924498, EPI_ISL_924500, EPI_ISL_924501, EPI_ISL_924502, EPI_ISL_924503, EPI_ISL_924504, EPI_ISL_924505, EPI_ISL_924506, EPI_ISL_924507, EPI_ISL_924508, EPI_ISL_924509, EPI_ISL_924510, EPI_ISL_924511, EPI_ISL_924512, EPI_ISL_924513, EPI_ISL_924514, EPI_ISL_924515, EPI_ISL_924516, EPI_ISL_924519, EPI_ISL_924520, EPI_ISL_924522, EPI_ISL_924523, EPI_ISL_924524, EPI_ISL_924525, EPI_ISL_924526, EPI_ISL_924527, EPI_ISL_924532, EPI_ISL_924534, EPI_ISL_924535, EPI_ISL_924537, EPI_ISL_924538, EPI_ISL_924549, EPI_ISL_924551, EPI_ISL_924554, EPI_ISL_924561, EPI_ISL_924569 | | | |
| see above | Bioinformatics and Biostatistics Lab, Advanced Sequencing Facility | COVID-19 Genomics UK (COG-UK) Consortium | Aengus Stewart,Jerome Nicod,Chelsea Sawyer,Laura Cubitt,Harshil Patel,Margaret Crawford |
| EPI_ISL_925395, EPI_ISL_925396 | Department of Clinical Microbiology | GIGA Medical Genomics | Keith Durkin, Maria Artesi, Sébastien Bontems, Raphaël Boreux, Bouchra Boujemla, Cécile Meex, Pierrette Melin, Marie-Pierre Hayette, Vincent Bours |
| EPI_ISL_925528 | Arizona State Public Health Laboratory | Arizona State Public Health Laboratory | Trung Huynh, Jessica Escobar, Katherine Fullerton, Nobuko Fukushima, Stacy White, Linda Getsinger, Victor Waddell |
| EPI_ISL_925913 | Nucleic Acid Testing, National Reference Laboratory | GIGA Medical Genomics | Yvan Butera, Keith Durkin, Maria Artesi, Bouchra Boujemla, Robert Rutayisire, Patrick Tuyisenge, Esperence Umumararungu, Sébastien Bontems, Marie-Pierre Hayette, Nathalie Renotte, Swaibu Gatare, Jacob Souopgui, Sabin Nsanzimana, Vincent Bours, Léon Mutesa |
| EPI_ISL_930668, EPI_ISL_930669, EPI_ISL_930670, EPI_ISL_930675, EPI_ISL_930676, EPI_ISL_930677, EPI_ISL_930678, EPI_ISL_930679, EPI_ISL_930680, EPI_ISL_930681, EPI_ISL_930682, EPI_ISL_930683, EPI_ISL_930684, EPI_ISL_930685, EPI_ISL_930686, EPI_ISL_930687, EPI_ISL_930688, EPI_ISL_930689, EPI_ISL_930690, EPI_ISL_930691, EPI_ISL_930692, EPI_ISL_930693, EPI_ISL_930694, EPI_ISL_930695, EPI_ISL_930696, EPI_ISL_930697, EPI_ISL_930698, EPI_ISL_930699, EPI_ISL_930700, EPI_ISL_930701, EPI_ISL_930702, EPI_ISL_930703, EPI_ISL_930704, EPI_ISL_930705, EPI_ISL_930706, EPI_ISL_930707, EPI_ISL_930708, EPI_ISL_930709, EPI_ISL_930710, EPI_ISL_930711, EPI_ISL_930712, EPI_ISL_930713, EPI_ISL_930714, EPI_ISL_930715, EPI_ISL_930716, EPI_ISL_930717, EPI_ISL_930718, EPI_ISL_930719, EPI_ISL_930721, EPI_ISL_930729, EPI_ISL_930760, EPI_ISL_930764, EPI_ISL_930766, EPI_ISL_930769, EPI_ISL_930770, EPI_ISL_930771, EPI_ISL_930772, EPI_ISL_930773, EPI_ISL_930774, EPI_ISL_930775, EPI_ISL_930776, EPI_ISL_930778, EPI_ISL_930780, EPI_ISL_930782, EPI_ISL_931508, EPI_ISL_931509, EPI_ISL_931510, EPI_ISL_931511, EPI_ISL_931512, EPI_ISL_931513, EPI_ISL_931514, EPI_ISL_931515, EPI_ISL_931517, EPI_ISL_931518, EPI_ISL_931519, EPI_ISL_931520, EPI_ISL_931521, EPI_ISL_931522, EPI_ISL_931523, EPI_ISL_931524, EPI_ISL_931525, EPI_ISL_931526, EPI_ISL_931527, EPI_ISL_931528, EPI_ISL_931529, EPI_ISL_931530, EPI_ISL_931531, EPI_ISL_931532, EPI_ISL_931533, EPI_ISL_931534, EPI_ISL_931535, EPI_ISL_931536, EPI_ISL_931538, EPI_ISL_931541, EPI_ISL_931542, EPI_ISL_931543, EPI_ISL_931544, EPI_ISL_931545, EPI_ISL_931546, EPI_ISL_931547, EPI_ISL_931548, EPI_ISL_931549, EPI_ISL_931550, EPI_ISL_931551, EPI_ISL_931552, EPI_ISL_931553, EPI_ISL_931554, EPI_ISL_931555, EPI_ISL_931556, EPI_ISL_931557, EPI_ISL_931558, EPI_ISL_931559, EPI_ISL_931560, EPI_ISL_931561, EPI_ISL_931562, EPI_ISL_931563, EPI_ISL_931564, EPI_ISL_931565, EPI_ISL_931566, EPI_ISL_931567, EPI_ISL_931568, EPI_ISL_931569, EPI_ISL_931571, EPI_ISL_931572, EPI_ISL_931573 | | | |
| see above | Utah Public Health Laboratory | Utah Public Health Laboratory | Erin L. Young, Kelly F. Oakeson, Tara Gallagher |
| EPI_ISL_933441, EPI_ISL_933442, EPI_ISL_933443, EPI_ISL_933444, EPI_ISL_933445, EPI_ISL_933446, EPI_ISL_933448, EPI_ISL_933449, EPI_ISL_933450, EPI_ISL_933451, EPI_ISL_933452, EPI_ISL_933454, EPI_ISL_933455, EPI_ISL_933457, EPI_ISL_933458, EPI_ISL_933459, EPI_ISL_933460, EPI_ISL_933461, EPI_ISL_933462, EPI_ISL_933464, EPI_ISL_933465, EPI_ISL_933466, EPI_ISL_933468, EPI_ISL_933470, EPI_ISL_933471, EPI_ISL_933473, EPI_ISL_933476, EPI_ISL_933477, EPI_ISL_933478 | | | |
| see above | Lighthouse Lab in Milton Keynes | Wellcome Sanger Institute for the COVID-19 Genomics UK (COG-UK) Consortium | The Lighthouse Lab in Milton Keynes and Alex Alderton, Roberto Amato, Sonia Goncalves, Ewan Harrison, David K. Jackson, Ian Johnston, Dominic Kwiatkowski, Cordelia Langford, John Sillitoe on behalf of the Wellcome Sanger Institute COVID-19 Surveillance Team |
| EPI_ISL_933679 | Instituto de Diagnostico y Referencia Epidemiologicos INDRE_RNLSP | Instituto de Diagnostico y Referencia Epidemiologicos (INDRE) | Claudia Wong-Arambula, Abril Rodriguez-Maldonado, Fabiola Garces-Ayala, Adnan Araiza-Rodriguez, David Fragoso-Fonseca, Sergio Rangel-Guerrero, Mayra Jimenez-Morales, Nancy Munoz-Hernandez, Natividad Cruz-Ortiz, Tatiana Nunez-Garcia, Gisela Barrera-Badillo, Lucia Hernandez-Rivas, Irma Lopez-Martinez, Ernesto Ramirez-Gonzalez. |
| EPI_ISL_933780, EPI_ISL_933781, EPI_ISL_933782, EPI_ISL_933783, EPI_ISL_933784 | Virology Unit, Institut Pasteur du Cambodge | Virology Unit, Institut Pasteur du Cambodge | Sokhoun Yann, Ly Sovann, Kraing Sidonn, Yi Sengdoeurn, Chin Savuth, Chau Darapheak, Etienne Simon-Loriere, Veasna Duong, Erik A Karlsson |
| EPI_ISL_934348, EPI_ISL_934358, EPI_ISL_934359, EPI_ISL_934361, EPI_ISL_934362, EPI_ISL_934363 | Klinisk mikrobiologi | The Public Health Agency of Sweden | Anna-Malin Linde, Maria Lind Karlberg, Carlo Berg, Oskar Karlsson Lindsjo, Sofia Stamouli, Reza Advani, Mattias Haukland, Petra Holmstrom, Noura Walai, Petra Edquist, Mia Brytting, Anna Risberg, Karin Tegmark-Wisell |
| EPI_ISL_934390, EPI_ISL_934407 | Klinisk Mikrobiologi | The Public Health Agency of Sweden | Anna-Malin Linde, Maria Lind Karlberg, Carlo Berg, Oskar Karlsson Lindsjo, Sofia Stamouli, Reza Advani, Mattias Haukland, Petra Holmstrom, Noura Walai, Petra Edquist, Mia Brytting, Anna Risberg, Karin Tegmark-Wisell |
| EPI_ISL_934569 | Department of Laboratory Medicine, Division of Clinical Virology, University of Medicine, Vienna | Bergthaler laboratory, CeMM Research Center for Molecular Medicine of the Austrian Academy of Sciences | Lukas Endler, Anna Schedl, Thomas Penz, Benedikt Agerer, Maelle Le Moing, Michael Schuster, Bekir Erguner, Jan Laine, Martin Senekowitsch, Christoph Bock, Andreas Bergthaler |
| EPI_ISL_934640 | Department of Microbiology, University Innsbruck | Bergthaler laboratory, CeMM Research Center for Molecular Medicine of the Austrian Academy of Sciences | Lukas Endler, Anna Schedl, Thomas Penz, Benedikt Agerer, Maelle Le Moing, Michael Schuster, Bekir Erguner, Jan Laine, Martin Senekowitsch, Christoph Bock, Andreas Bergthaler |
| EPI_ISL_935214, EPI_ISL_935217, EPI_ISL_935218, EPI_ISL_935219 | KU Leuven, Rega Institute, Clinical and Epidemiological Virology | KU Leuven, Rega Institute, Clinical and Epidemiological Virology | Tony Wawina-Bokalanga, Bert Vanmechelen, Joan Marti-Carerras, Piet Maes |
| EPI_ISL_935982, EPI_ISL_935983, EPI_ISL_935984, EPI_ISL_935985 | GLENS FALLS HOSPITAL LABORATORY | Wadsworth Center, New York State Department of Health | Kirsten St. George, Daryl M. Lamson, Alexis Russel, Matthew Shudt, Melissa A Leisner, Jonathan Plitnick, Navjot Singh, John Kelly, Erasmus Schneider, Erica Lasek-Nesselquist |
| EPI_ISL_936044, EPI_ISL_936045 | Columbia University Irving Medical Center | Wadsworth Center, New York State Department of Health | Kirsten St. George, Daryl M. Lamson, Alexis Russel, Matthew Shudt, Melissa A Leisner, Jonathan Plitnick, Navjot Singh, John Kelly, Erasmus Schneider, Erica Lasek-Nesselquist |
| EPI_ISL_936060 | KALEIDA CENTER FOR LABORATORY MEDICINE | Wadsworth Center, New York State Department of Health | Kirsten St. George, Daryl M. Lamson, Alexis Russel, Matthew Shudt, Melissa A Leisner, Jonathan Plitnick, Navjot Singh, John Kelly, Erasmus Schneider, Erica Lasek-Nesselquist |
| EPI_ISL_936063, EPI_ISL_936064, EPI_ISL_936065, EPI_ISL_936066, EPI_ISL_936069, EPI_ISL_936082, EPI_ISL_936086, EPI_ISL_936089, EPI_ISL_936092, EPI_ISL_936093 | Columbia University Irving Medical Center | Wadsworth Center, New York State Department of Health | Kirsten St. George, Daryl M. Lamson, Alexis Russel, Matthew Shudt, Melissa A Leisner, Jonathan Plitnick, Navjot Singh, John Kelly, Erasmus Schneider, Erica Lasek-Nesselquist |
| EPI_ISL_936147 | New York Presbyterian Hospital | Wadsworth Center, New York State Department of Health | Kirsten St. George, Daryl M. Lamson, Alexis Russel, Matthew Shudt, Melissa A Leisner, Jonathan Plitnick, Navjot Singh, John Kelly, Erasmus Schneider, Erica Lasek-Nesselquist |

| | | | |
|---|---|---|---|
| EPI_ISL_936390, EPI_ISL_936396, EPI_ISL_936397, EPI_ISL_936400, EPI_ISL_936401, EPI_ISL_936402, EPI_ISL_936405, EPI_ISL_936406, EPI_ISL_936407, EPI_ISL_936408, EPI_ISL_936410, EPI_ISL_936414, EPI_ISL_936415, EPI_ISL_936419, EPI_ISL_936420, EPI_ISL_936423, EPI_ISL_936437, EPI_ISL_936439, EPI_ISL_936450, EPI_ISL_936451, EPI_ISL_936453, EPI_ISL_936456, EPI_ISL_936458, EPI_ISL_936459, EPI_ISL_936460, EPI_ISL_936463 | | | |
| see above | TGen North | TGen North | Jolene Bowers, Megan Folkerts, Chris French, Hayley Yaglom, Ashlyn Pfeiffer, Darrin Lemmer, Dave Engelthaler, The Arizona COVID Genomics Union (ACGU) |
| EPI_ISL_936817, EPI_ISL_936818, EPI_ISL_936819, EPI_ISL_936820, EPI_ISL_936821, EPI_ISL_936822, EPI_ISL_936823, EPI_ISL_936824, EPI_ISL_936825, EPI_ISL_936826, EPI_ISL_936827 | | | |
| see above | Northwestern Memorial Hospital | Ozer Lab | Ramon Lorenzo-Redondo, Lacy M. Simons, Chad J. Achenbach, Lawrence J. Jennings, Michael G. Ison, Judd F. Hultquist, Egon A. Ozer |
| EPI_ISL_940148 | NHLS Universitas Academic | UFS Virology | PA Bester, MM Nyaga, P Nthiga, MT Mogotsi, D Goedhals, T de Oliveira |
| EPI_ISL_940769, EPI_ISL_940775 | INSPI-CRN de Influenza y otros virus respiratorios | INSPI-Centro de Investigación Multidisciplinaria de la DTIDI | Leandro Patiño, Doménica de Mora, Maritza Olmedo, Andrés Carrazco-Montalvo, Orson Mestanza, Mary Regato-Arrata, Melissa Zambrano, Manuel González, Alfredo Bruno, Alberto Orlando. |
| EPI_ISL_940863, EPI_ISL_940873, EPI_ISL_940884, EPI_ISL_940885 | Vaccines and Infectious Diseases Analytics Research Unit (VIDA) | KRISP, KZN Research Innovation and Sequencing Platform | Baillie Vicky, du Plessis Jeanine, Giandhari Jennifer, Pillay Sureshnee, Naidoo Yeshnee, Tegally Houriiyah, de Oliveira Tulio, Madhi Shabir |
| EPI_ISL_941223, EPI_ISL_941225 | Laboratorio de Microbiología. Hospital General Universitario de Elda, Alicante | SeqCOVID-SPAIN consortium/IBV(CSIC) | Mª Isabel Gascón Ros, Cristina Torregrosa Hetland, Eva Pastor Boix, Paloma Cascales Ramos and SeqCOVID-SPAIN consortium |
| EPI_ISL_941285, EPI_ISL_941286, EPI_ISL_941287, EPI_ISL_941289, EPI_ISL_941290, EPI_ISL_941291 | Nigeria Centre for Disease Control (NCDC) | African Centre of Excellence for Genomics of Infectious Diseases (ACEGID), Redeemer's University | Oluniyi P.E. et al |
| EPI_ISL_941370, EPI_ISL_941623 | Instituto Nacional de Saude (INSA) | Instituto Nacional de Saude (INSA) | Borges et al |
| EPI_ISL_942763, EPI_ISL_942764, EPI_ISL_942765, EPI_ISL_942766, EPI_ISL_942767, EPI_ISL_942768, EPI_ISL_942769, EPI_ISL_942771 | Gundersen Molecular Diagnostics Laboratory | Kabara Cancer Research Institute | Craig S. Richmond, Paraic A. Kenny |
| EPI_ISL_943564, EPI_ISL_943565, EPI_ISL_943567 | Servizo de Microbioloxía. Complexo Hospitalario Universitario de Santiago de Compostela | Servizo de Microbioloxía. Complexo Hospitalario Universitario de Santiago de Compostela | Antonio Aguilera, Gema Barbeito, Amparo Coira, José Costa, Rocio Trastoy, María Luisa Pérez del Molino. |
| EPI_ISL_943819, EPI_ISL_943820, EPI_ISL_943821, EPI_ISL_943931 | Utah Public Health Laboratory | Utah Public Health Laboratory | Erin L. Young, Kelly F. Oakeson, Tara Gallagher |
| EPI_ISL_944126, EPI_ISL_944129, EPI_ISL_944140, EPI_ISL_944141, EPI_ISL_944142, EPI_ISL_944143, EPI_ISL_944149 | National Health Laboratory Service, South Africa | KRISP, KZN Research Innovation and Sequencing Platform | Laguda-Akingba O, Giandhari J, Pillay S, Lessells R, Mdlalose K, York D, Khan S, Emmanuel SJ, Tegally H, Wilkinson E, de Oliveira T |
| EPI_ISL_944202, EPI_ISL_944225, EPI_ISL_944458 | Israel Central Virology laboratory | Israel National Consortium for SARS-CoV-2 sequencing | Neta Zuckerman, Efrat Dahan Bucris, Michal Mandelboim, Dana Bar-Ilan, Oran Erster, Tzvia Mann, Omer Murik, David A. Zeevi, Assaf Rokney, Joseph Jaffe, Eva Nachum, Maya Davidovich Cohen, Ephraim Fass, Gal Zizelski Valenci, Mor Rubinstein, Efrat Rorman, Israel Nissan, Efrat Glick-Saar, Omri Nayshool, Gideon Rechavi, Ella Mendelson, Orna Mor |
| EPI_ISL_944737 | unknown | Public Health Virology-Forensic and Scientific Services (PHV-FSS) | Son Nguyen et al. |
| EPI_ISL_945065 | Lighthouse Lab in Glasgow | Wellcome Sanger Institute for the COVID-19 Genomics UK (COG-UK) Consortium | Harper VanSteenhouse, Yumi Kasai, David Gray, Carol Clugston, Anna Dominiczak and Alex Alderton, Roberto Amato, Sonia Goncalves, Ewan Harrison, David K. Jackson, Ian Johnston, Dominic Kwiatkowski, Cordelia Langford, John Sillitoe on behalf of the Wellcome Sanger Institute COVID-19 Surveillance Team |
| EPI_ISL_945273 | Lighthouse Lab in Milton Keynes | Wellcome Sanger Institute for the COVID-19 Genomics UK (COG-UK) Consortium | The Lighthouse Lab in Milton Keynes and Alex Alderton, Roberto Amato, Sonia Goncalves, Ewan Harrison, David K. Jackson, Ian Johnston, Dominic Kwiatkowski, Cordelia Langford, John Sillitoe on behalf of the Wellcome Sanger Institute COVID-19 Surveillance Team |
| EPI_ISL_945301 | Lighthouse Lab in Glasgow | Wellcome Sanger Institute for the COVID-19 Genomics UK (COG-UK) Consortium | Harper VanSteenhouse, Yumi Kasai, David Gray, Carol Clugston, Anna Dominiczak and Alex Alderton, Roberto Amato, Sonia Goncalves, Ewan Harrison, David K. Jackson, Ian Johnston, Dominic Kwiatkowski, Cordelia Langford, John Sillitoe on behalf of the Wellcome Sanger Institute COVID-19 Surveillance Team |
| EPI_ISL_945339 | Lighthouse Lab in Milton Keynes | Wellcome Sanger Institute for the COVID-19 Genomics UK (COG-UK) Consortium | The Lighthouse Lab in Milton Keynes and Alex Alderton, Roberto Amato, Sonia Goncalves, Ewan Harrison, David K. Jackson, Ian Johnston, Dominic Kwiatkowski, Cordelia Langford, John Sillitoe on behalf of the Wellcome Sanger Institute COVID-19 Surveillance Team |
| EPI_ISL_949959, EPI_ISL_950131, EPI_ISL_950132, EPI_ISL_950137, EPI_ISL_950139, EPI_ISL_950140, EPI_ISL_950141, EPI_ISL_950143, EPI_ISL_950144, EPI_ISL_950149, EPI_ISL_950167 | | | |
| see above | University College London, Great Ormond Street Hospital for Children NHS Foundation Trust, Imperial College Healthcare NHS Trust | COVID-19 Genomics UK (COG-UK) Consortium | Sergi Castellano, Rachel Williams, Mark Kristiansen, Paola Resende Silva, Sunando Roy, Tony Brooks, Helena Tutill, Paola Niola, Patricia Dyal, Charlotte Williams, Leysa Forrest, Yasmin Panchbhaya, Jacqueline Findlay, Samuel Weeks, Julianne Brown, Kathryn Harris, Paul Randell, James Price, Alison Holmes, Judith Breuer |
| EPI_ISL_950234, EPI_ISL_950237, EPI_ISL_950238, EPI_ISL_950249, EPI_ISL_950444, EPI_ISL_950445, EPI_ISL_950446, EPI_ISL_950447, EPI_ISL_950448, EPI_ISL_950449, EPI_ISL_950450, EPI_ISL_950451, EPI_ISL_950452, EPI_ISL_950453, EPI_ISL_950454, EPI_ISL_950455, EPI_ISL_950456, EPI_ISL_950457, EPI_ISL_950458 | | | |
| see above | Northumbria University / South Tees Hospitals NHS Foundation Trust / North Cumbria Integrated Care NHS Foundation Trust / North Tees and Hartlepool NHS Foundation Trust / Newcastle Hospitals NHS Foundation Trust | COVID-19 Genomics UK (COG-UK) Consortium | Darren L Smith,Andrew Nelson,Matthew Bashton,Greg R Young,Joshua Loh,John Allan,Mohammad A Tariq,Giles S Holt,Gary Black,Wen C Yew,Lynn Dover,Paul Baker,Steve Liggett,Sarah Essex,Jane Greenaway,Debra Padgett,Clive Graham,Garren Scott,Edward Barton,Emma Swindells,Brendan Payne,Jennifer Collins,Yusri Taha,Gary Eltringham |
| EPI_ISL_950555 | Quadram Institute Bioscience | COVID-19 Genomics UK (COG-UK) Consortium | Dave J. Baker, Gemma L. Kay, Alp Aydin, Thanh Le-Viet, Steven Rudder, Ana P. Tedim, Anastasia Kolyva, Maria Diaz, Leonardo de Oliveira Martins, Nabil-Fareed Alikhan, Lizzie Meadows, Rachael Stanley, Ngozi Elumogo, Muhammed Yasir, Nicholas M. Thomson, Alexander J Trotter, Rachel Gilroy, Samuel Bloomfield, Claire Stuart, Andrew Bell, Reenesh Prakash, Samir Dervisevic, Alison E. Mather, John Wain, Mark Webber, Andrew J. Page, Justin O'Grady |
| EPI_ISL_950710 | Lincolnshire Hospitals and DeepSeq Nottingham | COVID-19 Genomics UK (COG-UK) Consortium | Nichola Duckworth, Tim Sloan, Sarah Walsh, Jonathan Ball, Patrick McClure, Joeseph Chappell, Nadine Holmes, Matthew Carlisle, Christopher Moore, Fei Sang, Johnny Debebe, Victoria Wright, Matthew Loose |
| EPI_ISL_950953, EPI_ISL_950954, EPI_ISL_950956, EPI_ISL_950957, EPI_ISL_950958, EPI_ISL_950959, EPI_ISL_950965, EPI_ISL_950979, EPI_ISL_950980, EPI_ISL_950981, EPI_ISL_950982, EPI_ISL_950983, EPI_ISL_950985, EPI_ISL_950987, EPI_ISL_950988, EPI_ISL_950989, EPI_ISL_950990, EPI_ISL_950991, EPI_ISL_950992, EPI_ISL_950993, EPI_ISL_950994, EPI_ISL_950995, EPI_ISL_950996, EPI_ISL_950999, EPI_ISL_951000, EPI_ISL_951002, EPI_ISL_951003, EPI_ISL_951004, EPI_ISL_951008, EPI_ISL_951009, EPI_ISL_951010, EPI_ISL_951011, EPI_ISL_951013, EPI_ISL_951014, EPI_ISL_951016, EPI_ISL_951018, EPI_ISL_951020, EPI_ISL_951021, EPI_ISL_951023, EPI_ISL_951024, EPI_ISL_951025, EPI_ISL_951026, EPI_ISL_951027, EPI_ISL_951028, EPI_ISL_951029, EPI_ISL_951030, EPI_ISL_951031, EPI_ISL_951032, EPI_ISL_951033, EPI_ISL_951034, EPI_ISL_951036, EPI_ISL_951037, EPI_ISL_951038, EPI_ISL_951042, EPI_ISL_951043, EPI_ISL_951044, EPI_ISL_951045, EPI_ISL_951046, EPI_ISL_951049, EPI_ISL_951050, EPI_ISL_951052, EPI_ISL_951053, EPI_ISL_951054, EPI_ISL_951055, EPI_ISL_951056, EPI_ISL_951057, EPI_ISL_951058, EPI_ISL_951059, EPI_ISL_951061, EPI_ISL_951062, EPI_ISL_951064, EPI_ISL_951066, EPI_ISL_951079, EPI_ISL_951080, EPI_ISL_951081, EPI_ISL_951083, EPI_ISL_951094, EPI_ISL_951097, EPI_ISL_951098, EPI_ISL_951163, EPI_ISL_951164, EPI_ISL_951165, EPI_ISL_951166, EPI_ISL_951169, EPI_ISL_951170, EPI_ISL_951171, EPI_ISL_951249 | | | |
| see above | Oxford Viromics, NDM, University of Oxford; Oxford University Hospitals; Basingstoke and North Hampshire Hospital | COVID-19 Genomics UK (COG-UK) Consortium | Tanya Golubchik, David Bonsall, George Macintyre, Amy Trebes, Mariateresa de Cesare, Catrin Moore, Alex Mobbs, Anita Justice, Robert Shaw, Monique Andersson, Timothy Peto, Emma Wise, Nathan Moore, Jessica Lynch, Nick Cortes, Matilde Mori, Stephen Kidd, David Buck, John Todd, Christophe Fraser |
| EPI_ISL_953665 | University Hospitals of Geneva, Laboratory of Virology | HUG, Laboratory of Virology and the Health2030 Genome Center | Samuel Cordey, Ana Rita Goncalves, Laurent Kaiser, Lorenzo Cerutti, Henri Pegeot, Melyssa Elies, Deborah Penet, Keith Harshman, Ioannis Xenarios, Emmanouil Dermitzakis |
| EPI_ISL_954211, EPI_ISL_954212, EPI_ISL_954213, EPI_ISL_954214, EPI_ISL_954215 | 1.AO Universitaria 'S. Giovanni di Dio e Ruggi D'Aragona, Scuola Medica Salernitana' Hospital / 2.UOC di Virologia e Microbiologia, Università della Campania 'L. Vanvitelli' / 3.AO Universitaria 'Federico II' Napoli Hospital / 4.AORN 'San | 1. Genome Research Center for Health (CRGS) / 2. Laboratory of Molecular Medicine and Genomics(LMMGe) / 3. Center for Research in Pure and Applied Mathematics (CRMPA) | Giorgio Giurato, Francesca Rizzo, Alessandro Weisz, Gianluigi Franci, Giovanni Nassa, Pasquale Pagliano, Roberta Tarallo, Elena Alexandrova, Ylenia D'Agostino, Carlo Ferravante, Jessica Lamberti, Viola Melone, Domenico Memoli, Valeria Mirici Cappa, Domenico Palumbo, Giovanni Pecoraro, Assunta Sellitto, Oriana Strianese, Ilaria Terenzi, Giuseppe Fenza, Aniello Gentile, Antonello Saccomanno, Sonia Amabile, Teresa Rocco, Annamaria Salvati, Emilia Vaccaro, Massimiliano Galdiero, Michele Cennamo, Giuseppe Portella, Maria Grazia Foti, Mariarosaria Ingino, Maria Landi, Maurizio Fumi, Vincenzo |

| | | | |
|---|---|---|---|
| | Giuseppe Moscati' Avellino Hospital / 5.AO 'San Pio - presidio G. Rummo' Benevento Hospital / 6.AO 'Sant'Anna e San Sebastiano' Caserta Hospital / 7.PO 'Maria Santissima Addolorata' Eboli Hospital / 8.Biogem Istituto di Ricerche Genetiche | | Rocco, Rita Greco, Vittoria Letizia, Arnolfo Petruzziello, Maddalena Schioppa, Gregorio Goffredi, Francesca Marciano, Michele Caraglia, Alessia Cossu, Marianna Scrima, Edmondo Adorisio, Morena D'Avenia, Michela Iacobellis, Rosanna Piluscio, Giorgio Dirani, Vittorio Sambri, Simona Semprini, Silvia Zanoli, Francesco Curcio, Stefania Marzinotto, Andreina Baj, Fausto Sessa. |
| EPI_ISL_954281, EPI_ISL_954282, EPI_ISL_954283, EPI_ISL_954284, EPI_ISL_954285, EPI_ISL_954286, EPI_ISL_954287, EPI_ISL_954288, EPI_ISL_954289, EPI_ISL_954290, EPI_ISL_954291, EPI_ISL_954292, EPI_ISL_954293, EPI_ISL_954294, EPI_ISL_954295, EPI_ISL_954296 | | | |
| see above | MRC/UVRI & LSHTM Uganda Research Unit | Where sequence data have been generated and submitted to GISAID | Matthew Cotten, Dan Lule Bugembe, My V.T. Phan, Isaac Sseeewanyana, Patrick Semanda, Susan Nabadda, Pontiano Kaleebu |
| EPI_ISL_955126 | Innovative Genomics Institute, UC Berkeley | Innovative Genomics Institute, UC Berkeley | Stacia Wyman, Haridha Shivram, Phil Frankino, Liana Lareau, Shana McDevitt, Justin Choi |
| EPI_ISL_955136 | MRC/UVRI & LSHTM Uganda Research Unit | Where sequence data have been generated and submitted to GISAID | Matthew Cotten, Dan Lule Bugembe, My V.T. Phan, Isaac Sseeewanyana, Patrick Semanda, Susan Nabadda, Pontiano Kaleebu |
| EPI_ISL_955387, EPI_ISL_955388, EPI_ISL_955390 | Alameda County Public Health Lab | Chan-Zuckerberg Biohub | CZB Cliahub Consortium |
| EPI_ISL_955727, EPI_ISL_955728, EPI_ISL_955729, EPI_ISL_955730, EPI_ISL_955731, EPI_ISL_955734, EPI_ISL_955735, EPI_ISL_955736, EPI_ISL_955737, EPI_ISL_955738, EPI_ISL_955739, EPI_ISL_955740, EPI_ISL_955741, EPI_ISL_955742, EPI_ISL_955743, EPI_ISL_955744, EPI_ISL_955745, EPI_ISL_955746, EPI_ISL_955747, EPI_ISL_955748, EPI_ISL_955749, EPI_ISL_955750, EPI_ISL_955751, EPI_ISL_955752, EPI_ISL_955753, EPI_ISL_955754, EPI_ISL_955755, EPI_ISL_955756, EPI_ISL_955757, EPI_ISL_955758, EPI_ISL_955759, EPI_ISL_955760, EPI_ISL_955761, EPI_ISL_955762, EPI_ISL_955763, EPI_ISL_955765, EPI_ISL_955766 | | | |
| see above | Humboldt County Public Health Laboratory | Chan-Zuckerberg Biohub | CZB Cliahub Consortium |
| EPI_ISL_956279 | Siti Khodijah Hospital | Institute of Tropical Disease, Universitas Airlangga | Jezzy R Dewantari, Rima R Prasetya, Krisnoadi Rahardjo, Aldise M Nastri, Muhammad Hamdan, Gatot Soegiarto, Laksmi Wulandari, Resti Yudhawati, Soetjipto, Yasuko Mori, Maria I Lusida, Kazufumi Shimizu |
| EPI_ISL_956326 | Laboratory Medicine | Department of Laboratory Medicine, Lin-Kou Chang Gung Memorial Hospital, Taoyuan, Taiwan | Kuo-Chien Tsao, Yu-Nong Gong, Shu-Li Yang, Yi-Chun Liu, Chung-Guei Huang, Mei-Jen Hsiao, Po-Wei Huang, Cheng-Ta Yang, Cheng-Hsun Chiu, Peng-Nien Huang, Kuo-Ming Lee, Guang-Wu Chen, Shin-Ru Shih |
| EPI_ISL_960224 | QEII Health Sciences Centre | National Microbiology Laboratory (NML) | Anna Majer, Shari Tyson, Grace Seo, Philip Mabon, Elsie Grudeski, Rhiannon Huzarewich, Russell Mandes, Anneliese Landgraff, Jennifer Tanner, Natalie Knox, Morag Graham, Gary Van Domselaar, Todd Hatchette, Jason LeBlanc, Janice Pettipas, Dan Gaston, Nathalie Bastien, Yan Li, Timothy Booth, Darian Hole, Madison Chapel, Kirsten Biggar, CanCOGeN's metadata curation team, Public Health Agency of Canada CanCOGeN team |
| EPI_ISL_960425, EPI_ISL_960438 | The National Institute of Public Health | State Veterinary Institute Prague | Nagy,A;Vecerova,J;Cernikova,L;Stara,M;Jirincova,H;Trnka,D |
| EPI_ISL_960460, EPI_ISL_960466, EPI_ISL_960467, EPI_ISL_960479, EPI_ISL_960571 | Istituto Zooprofilattico Sperimentale del Mezzogiorno | TIGEM | Patrizia Annunziata, Andrea Ballabio, Valentina Bouche, Davide Cacchiarelli, Pellegrino Cerino, Chiara Colantuono, Maria Concetta Cuomo, Denise Di Concilio, Lucio Di Filippo, Antonio Grimaldi, Antonio Limone, Anna Manfredi, Francesco Panariello, Biancamaria Pierri, Marcello Salvi |
| EPI_ISL_961270, EPI_ISL_961271 | Hospital General Universitario de Alicante - Instituto de Investigación Sanitaria y Biomédica de Alicante | SeqCOVID-SPAIN consortium/IBV(CSIC) | Maripaz Ventero Martín, Carmen Molina Pardines and SeqCOVID-SPAIN consortium |
| EPI_ISL_961570, EPI_ISL_961571, EPI_ISL_961572, EPI_ISL_961573, EPI_ISL_961574, EPI_ISL_961575, EPI_ISL_961576, EPI_ISL_961577, EPI_ISL_961578, EPI_ISL_961579, EPI_ISL_961607, EPI_ISL_961608 | | | |
| see above | Hôpital Georges L. Dumont | National Microbiology Laboratory (NML) | Anna Majer, Shari Tyson, Grace Seo, Philip Mabon, Elsie Grudeski, Rhiannon Huzarewich, Russell Mandes, Anneliese Landgraff, Jennifer Tanner, Natalie Knox, Morag Graham, Gary Van Domselaar, Richard Garceau, Guillaume Desnoyers, Nathalie Bastien, Yan Li, Timothy Booth, Darian Hole, Madison Chapel, Kirsten Biggar, CanCOGeN's metadata curation team, Public Health Agency of Canada CanCOGeN team |
| EPI_ISL_961891, EPI_ISL_961936, EPI_ISL_962004, EPI_ISL_962108, EPI_ISL_962109, EPI_ISL_962110, EPI_ISL_962111, EPI_ISL_962112, EPI_ISL_962113, EPI_ISL_962114, EPI_ISL_962115, EPI_ISL_962116, EPI_ISL_962117, EPI_ISL_962118, EPI_ISL_962119, EPI_ISL_962120, EPI_ISL_962128 | | | |
| see above | Illinois Department of Public Health | Gagnon Lab, Southern Illinois University | Keith Gagnon |
| EPI_ISL_962611, EPI_ISL_962613 | San Diego County Public Health Laboratory | Andersen lab at Scripps Research | SEARCH Alliance San Diego with Tracy Basler, Jovan Shephard, Brett Austin |
| EPI_ISL_962815 | Robert Garry lab | Andersen lab at Scripps Research | Allison Smither, Gilberto Sabino-Santos, Patricia Snarski, Lilia Melnik, Antoinette Bell, Kaylynn Genemaras, Arnaud Drouin, Dahlene Fusco, Robert Garry with SEARCH Alliance San Diego |
| EPI_ISL_962907 | Ostfold Hospital Trust - Kalnes, Centre for Laboratory Medicine, Section for gene technology and infection serology | Norwegian Institute of Public Health, Department of Virology | Kathrine Stene-Johansen, Kamilla Heddeland Instefjord, Hilde Elshaug, Ignacio Garcia Llorente, Serina B Engebretsen, Atiya R Ali,Marie Paulsen Madsen, Rasmus Riis Kopperud, Hilde Vollan, Karoline Bragstad, Olav Hungnes |
| EPI_ISL_962936, EPI_ISL_962940, EPI_ISL_962941 | Hospital Universitario de Gran Canaria Dr. Negrín | SeqCOVID-SPAIN consortium/IBV(CSIC) | M. Carmen Pérez González, Francisco J. Chamizo López, Ana Bordes Benítez and SeqCOVID-SPAIN consortium |
| EPI_ISL_965053 | Wyoming Public Health Laboratory | Wyoming Public Health Laboratory | Noah Hull, Taylor Fearing, Lynette Gumbleton, Channing Weber, Ashley Norberg, Bailey Bowcutt, and Wanda Manley |
| EPI_ISL_965189, EPI_ISL_965214, EPI_ISL_965215, EPI_ISL_965216 | Virginia Division of Consolidated Laboratory Services | Virginia Division of Consolidated Laboratory Services | Virginia DCLS |
| EPI_ISL_965528 | NYU Langone Health | Departments of Pathology and Medicine, New York University School of Medicine | Adriana Heguy, Dacia Dimartino, Emily Guzman, Christian Marier, Peter Meyn, Sitharam Ramaswami, Gael Westby, Paul Zappile, Yutong Zhang, Paolo Cotzia, Guiqing Wang |
| EPI_ISL_965533, EPI_ISL_965536, EPI_ISL_965547, EPI_ISL_965549, EPI_ISL_965556, EPI_ISL_965560, EPI_ISL_965561, EPI_ISL_965563, EPI_ISL_965589, EPI_ISL_965594, EPI_ISL_965601, EPI_ISL_965606, EPI_ISL_965607, EPI_ISL_965620, EPI_ISL_965667, EPI_ISL_965720, EPI_ISL_965729, EPI_ISL_965731, EPI_ISL_965741, EPI_ISL_965747, EPI_ISL_965748, EPI_ISL_965749, EPI_ISL_965756, EPI_ISL_965759, EPI_ISL_965761, EPI_ISL_965762 | | | |
| see above | Dutch COVID-19 response team | Medical Microbiology, Maastricht University Medical Centre | Jozef Dingemans*, Brian van der Veer*, Erik Beuken, Carmen Reumkens, Lieke van Alphen, Christian Hoebe, Paul Savelkoul |
| EPI_ISL_965770 | NYU Langone Health | Departments of Pathology and Medicine, New York University School of Medicine | Adriana Heguy, Dacia Dimartino, Emily Guzman, Christian Marier, Peter Meyn, Sitharam Ramaswami, Gael Westby, Paul Zappile, Yutong Zhang, Paolo Cotzia, Guiqing Wang |
| EPI_ISL_965773, EPI_ISL_965774, EPI_ISL_965776, EPI_ISL_965778, EPI_ISL_965779, EPI_ISL_965785, EPI_ISL_965805 | Dutch COVID-19 response team | Medical Microbiology, Maastricht University Medical Centre | Jozef Dingemans*, Brian van der Veer*, Erik Beuken, Carmen Reumkens, Lieke van Alphen, Christian Hoebe, Paul Savelkoul |
| EPI_ISL_965812, EPI_ISL_965814, EPI_ISL_965826, EPI_ISL_965837, EPI_ISL_965838, EPI_ISL_965839, EPI_ISL_965910, EPI_ISL_965948, EPI_ISL_965962, EPI_ISL_965963, EPI_ISL_965964, EPI_ISL_965965, EPI_ISL_965968, EPI_ISL_965969, EPI_ISL_965970, EPI_ISL_965971, EPI_ISL_965972, EPI_ISL_965973, EPI_ISL_965977, EPI_ISL_965980 | | | |
| see above | NYU Langone Health | Departments of Pathology and Medicine, New York University School of Medicine | Adriana Heguy, Dacia Dimartino, Emily Guzman, Christian Marier, Peter Meyn, Sitharam Ramaswami, Gael Westby, Paul Zappile, Yutong Zhang, Paolo Cotzia, Guiqing Wang |
| EPI_ISL_967618, EPI_ISL_967620, EPI_ISL_967621, EPI_ISL_967624, EPI_ISL_967625, EPI_ISL_967628, EPI_ISL_967630, EPI_ISL_967631, EPI_ISL_967632, EPI_ISL_967634, EPI_ISL_967635, EPI_ISL_967637, EPI_ISL_967639, EPI_ISL_967643, EPI_ISL_967689, EPI_ISL_967690, EPI_ISL_967691, EPI_ISL_967692, EPI_ISL_967706, EPI_ISL_967707, EPI_ISL_967708, EPI_ISL_967722, EPI_ISL_967723, EPI_ISL_967724, EPI_ISL_967727, EPI_ISL_967729, EPI_ISL_967750, EPI_ISL_967751, EPI_ISL_967758, EPI_ISL_967759, EPI_ISL_967760, EPI_ISL_967762, EPI_ISL_967766 | | | |
| see above | State Laboratories Division, Hawaii State Department of Health | State Laboratories Division, Hawaii State Department of Health | Pamela O'Brien, Drew Kuwazaki, Ayana Garnet, Razvan Sultana, Edward Desmond |
| EPI_ISL_968081 | Public Health Virology-Forensic and Scientific Services (PHV-FSS) | Public Health Virology-Forensic and Scientific Services (PHV-FSS) | Alyssa T. Pyke et al. |
| EPI_ISL_968854, EPI_ISL_968878 | KEMRI-Wellcome Trust Research Programme/KEMRI-CGMR-C Kilifi | KEMRI-Wellcome Trust Research Programme/KEMRI-CGMR-C Kilifi | Githinji et al |
| EPI_ISL_976851, EPI_ISL_976852, EPI_ISL_976853, EPI_ISL_976854, EPI_ISL_976855, EPI_ISL_976856, EPI_ISL_976857, EPI_ISL_976858, EPI_ISL_976859, EPI_ISL_976860, EPI_ISL_976861, EPI_ISL_976862, EPI_ISL_976863, EPI_ISL_976864, EPI_ISL_976865, EPI_ISL_976866, EPI_ISL_976867, EPI_ISL_976868, | | | |

| | | | |
|---|---|---|---|
| EPI_ISL_976869, EPI_ISL_976870, EPI_ISL_976871, EPI_ISL_976872, EPI_ISL_976873, EPI_ISL_976874, EPI_ISL_976875, EPI_ISL_976876, EPI_ISL_976877, EPI_ISL_976878, EPI_ISL_976879, EPI_ISL_976880, EPI_ISL_976881, EPI_ISL_976882 | | | |
| see above | BCCDC Public Health Laboratory | BCCDC Public Health Laboratory | Prystajecky Natalie, Linda Hoang, Dan Fornika, John Tyson, Shannon Russell, Kim Macdonald, Kimia Kamelian, Ana Pacagnella, Corrinne Ng, Loretta Janz, Robert Azana Terry Snutch, Mel Krajden |
| EPI_ISL_977078 | Massachusetts General Hospital | Infectious Disease Program, Broad Institute of Harvard and MIT | Lemieux,J.E., Siddle,K.J., Shaw,B., Adams,G., Pierce,V., Turbett,S., Anahtar,M., Branda,J., Slater,D., Harris,J., Lin,A.E., Gladden-Young,A., Lagerborg,K., Rudy,M., DeRuff,K., Carter,A., Normandin,E., Bauer,M., Reilly,S., Tomkins-Tinch,C., Loreth,C., Chaluvadi,S., Neumann,A., Cusick,C., Chapman,S.B., Gnirke,A., Flowers,K., Cerrato,F., Birren,B.W., Gallagher,G., Smole,S., Park,D.J., MacInnis,B.L., Ryan,E., LaRocque,R., Rosenberg,E. and Sabeti,P.C. |
| EPI_ISL_977210 | Microbiologia e Virologia | Istituto Zooprofilattico Sperimentale delle Venezie | Adelaide Milani, Alessia Schivo, Annalisa Salviato, Erika Giorgia Quaranta, Ambra Pastori, Bianca Zecchin, Alice Fusaro, Isabella Monne, Calogero Terregino, Antonia Ricci |
| EPI_ISL_977656, EPI_ISL_977657, EPI_ISL_977658 | Caribbean Public Health Agency | Carrington Lab, Department of PreClinical Sciences, Building 36, First Floor Biochemistry Unit, Faculty of Medical Sciences, The University of the West Indies | Nikita S. D. Sahadeo, Arianne Brown-Jordan, Vernie Ramkissoon, Sarah Hill, Naresh Nandram, Avery Hinds, Kenneth George, Jerome Foster, Stanley Giddings, Karla Georges, Marsha Ivey, Rahul Naidu, Risha Singh, SueMin Nathaniel, Rajini Haraksingh, Jaya Jayaraman, Chinna Chinnadurai, Adesh Ramsubhag, Nuno Faria, Oliver Pybus, Christopher Oura, Gabriel Escobar, Christine V. F. Carrington |
| EPI_ISL_977686, EPI_ISL_977693, EPI_ISL_977694, EPI_ISL_977697, EPI_ISL_977705, EPI_ISL_977708, EPI_ISL_977711, EPI_ISL_977719, EPI_ISL_977720 | California Department of Public Health | Chiu Laboratory, University of California, San Francisco | Charles Chiu, Xianding (Wayne) Deng, Candace Wang, Venice Servellita, Jill Hacker, Debra Wadford |
| EPI_ISL_978063, EPI_ISL_978064, EPI_ISL_978065, EPI_ISL_978066, EPI_ISL_978067, EPI_ISL_978068, EPI_ISL_978069, EPI_ISL_978075, EPI_ISL_978076, EPI_ISL_978077, EPI_ISL_978078, EPI_ISL_978079, EPI_ISL_978080, EPI_ISL_978081, EPI_ISL_978082, EPI_ISL_978083, EPI_ISL_978084, EPI_ISL_978085, EPI_ISL_978086, EPI_ISL_978087, EPI_ISL_978088, EPI_ISL_978119, EPI_ISL_978120, EPI_ISL_978121, EPI_ISL_978122, EPI_ISL_978123, EPI_ISL_978124, EPI_ISL_978125, EPI_ISL_978126, EPI_ISL_978127, EPI_ISL_978128, EPI_ISL_978129, EPI_ISL_978130 | | | |
| see above | Chiu Laboratory, University of California, San Francisco | Chiu Laboratory, University of California, San Francisco | Charles Chiu, Xianding (Wayne) Deng, Candace Wang, Venice Servellita, Jill Hacker, Debra Wadford |
| EPI_ISL_978411, EPI_ISL_978412, EPI_ISL_978413, EPI_ISL_978414, EPI_ISL_978415, EPI_ISL_978416 | Arizona State Public Health Laboratory | Arizona State Public Health Laboratory | Trung Huynh, Jessica Escobar, Katherine Fullerton, Nobuko Fukushima, Stacy White, Linda Getsinger, Victor Waddell |
| EPI_ISL_978920, EPI_ISL_978921, EPI_ISL_978922, EPI_ISL_978923 | Centre for Dengue Research and AICBU, Department of Immunology and Molecular Medicine | Centre for Dengue Research and AICBU, Department of Immunology and Molecular Medicine | Chandima Jeewandara, Deshni Jayathilaka, Dinuka Ariyaratne, Tibutius Thanesh Pramanayagam, Diyanath Ranasinghe, Laksiri Gomes, Gathsaurie Neelika Malavige |
| EPI_ISL_978961 | Chiu Laboratory, University of California, San Francisco | Chiu Laboratory, University of California, San Francisco | Charles Chiu, Xianding (Wayne) Deng, Candace Wang, Venice Servellita, Jill Hacker, Debra Wadford |
| EPI_ISL_979141 | Santa Clara County Public Health Laboratory | Chan-Zuckerberg Biohub | CZB Cliahub Consortium |
| EPI_ISL_979250 | Institute of Microbiology and Immunology, Faculty of Medicine, University of Ljubljana | Institute of Microbiology and Immunology, Faculty of Medicine, University of Ljubljana | Samo Zakotnik, Tomaž Mark Zorec, Matic Brvar, Doroteja Vlaj, Patricija Pozvek,Špela Pleh, Miša Korva, Mario Poljak, Tatjana Avši - Županc |
| EPI_ISL_979338 | Laboratorio Estatal de Salud Pública de Nuevo León | Laboratorio de Infectología Molecular, Departamento de Bioquímica y Medicina Molecular,Facultad de Medicina - Universidad Autónoma de Nuevo León | Kame A. Galán-Huerta, María F. Herrera-Saldivar, Natalia Martínez-Acuña, Sonia A. Lozano-Sepúlveda, Daniel Arellanos-Soto, Ana M. Rivas-Estilla, Samuel Buentello-Wong, Else del Carmen García-García, Gloria A. Jasso-de-la-Peña, Roberto Montes-de-Oca, Consuelo Treviño-Garza, Manuel E. de-la-O-Cavazos |
| EPI_ISL_979623, EPI_ISL_979624, EPI_ISL_979625, EPI_ISL_979626, EPI_ISL_979627, EPI_ISL_979628, EPI_ISL_979631, EPI_ISL_979632, EPI_ISL_979633 | Santa Clara County Public Health Laboratory | Chan-Zuckerberg Biohub | CZB Cliahub Consortium |
| EPI_ISL_979643, EPI_ISL_979644, EPI_ISL_979645, EPI_ISL_979646, EPI_ISL_979647, EPI_ISL_979648, EPI_ISL_979649, EPI_ISL_979650, EPI_ISL_979651, EPI_ISL_979652, EPI_ISL_979653, EPI_ISL_979654, EPI_ISL_979655, EPI_ISL_979658, EPI_ISL_979659, EPI_ISL_979660, EPI_ISL_979661, EPI_ISL_979662, EPI_ISL_979663, EPI_ISL_979664, EPI_ISL_979665, EPI_ISL_979666, EPI_ISL_979667, EPI_ISL_979668, EPI_ISL_979669, EPI_ISL_979670, EPI_ISL_979671, EPI_ISL_979672, EPI_ISL_979673, EPI_ISL_979674, EPI_ISL_979675, EPI_ISL_979676, EPI_ISL_979677, EPI_ISL_979678, EPI_ISL_979679, EPI_ISL_979680, EPI_ISL_979681, EPI_ISL_979682, EPI_ISL_979683, EPI_ISL_979684, EPI_ISL_979685, EPI_ISL_979686, EPI_ISL_979687, EPI_ISL_979688, EPI_ISL_979689, EPI_ISL_979690, EPI_ISL_979691 | | | |
| see above | Orange County Public Health Lab | Chan-Zuckerberg Biohub | CZB Cliahub Consortium |
| EPI_ISL_980974 | Innovative Genomics Institute, UC Berkeley | Innovative Genomics Institute, UC Berkeley | Stacia Wyman, Haridha Shivram, Phil Frankino, Liana Lareau |
| EPI_ISL_981069 | Johns Hopkins Hospital Department of Pathology | Johns Hopkins Hospital Department of Pathology | C. Paul Morris, Chun Huai Luo, Adannaya Amadi, Matthew Schwartz, Nicholas Gallagher, Heba H. Mostafa |
| EPI_ISL_981954, EPI_ISL_981955, EPI_ISL_981957 | Microbiology Service, Hospital Universitario Clinico San Cecilio, Granada | Microbiology Service, Hospital Universitario Clinico San Cecilio, Granada | Adolfo de Salazar, Natalia Chueca, Laura Viñuela, Ana Fuentes, Federico García |
| EPI_ISL_982239 | Lab voor klinische biologie | Lab voor klinische biologie | Hannelore Hamerlinck, Marija Janevska, Bruno Verhasselt |
| EPI_ISL_982355 | M Health Fairview | Minnesota Department of Health, Public Health Laboratory | Alexandra Lorentz, Jacob Garfin, Matt Plumb, and Xiong Wang |
| EPI_ISL_982442 | MONTEFIORE MEDICAL CENTER LABORATORIES | Wadsworth Center, New York State Department of Health | Kirsten St. George, Daryl M. Lamson, Alexis Russel, Matthew Shudt, Melissa A Leisner, Jonathan Plitnick, Navjot Singh, John Kelly, Erasmus Schneider, Erica Lasek-Nesselquist |
| EPI_ISL_982613 | Landstuhl Regional Medical Center | US Air Force School of Aerospace Medicine | Anthony Fries, Jennifer Meyer, William Gruner, William Buggele, Amanda Javorina, Sarah Purves, Fritz Castillo, Cole Anderson, Clarise Starr, Elizabeth Macias |
| EPI_ISL_982853 | Kentucky State Public Health Lab | Kentucky State Public Health Lab | Stephanie Lunn, Karim George, Joshua Tobias, William Grooms, Vaneet Arora, Matthew Johnson, Rachel Zinner, Rhonda Lucas |
| EPI_ISL_983408 | Utah Public Health Laboratory | Utah Public Health Laboratory | Erin L. Young, Kelly F. Oakeson, Tara Gallagher |
| EPI_ISL_983836, EPI_ISL_983837, EPI_ISL_983838, EPI_ISL_983839, EPI_ISL_983840, EPI_ISL_983841, EPI_ISL_983842, EPI_ISL_983843, EPI_ISL_983844, EPI_ISL_983845, EPI_ISL_983846, EPI_ISL_983847, EPI_ISL_983848, EPI_ISL_983849, EPI_ISL_983850, EPI_ISL_983851, EPI_ISL_983852 | | | |
| see above | Colorado Department of Public Health and Environment | Colorado Department of Puplic Health and Environment | Laura Bankers, Molly C. Hetherington-Rauth, Diana Ir, Shannon Ely, Shannon R. Matzinger, Sarah Elizabeth Totten, Emily A. Travanty |
