## Supplementary material for "Detection and characterization of the SARS-CoV-2 lineage B.1.526 in New York": Supp. Table 3 GISAID Acknowledgment part 2: gisaid_hcov-19_acknowledgement_table_2021_02_13_010-6.pdf

We gratefully acknowledge the following Authors from the Originating laboratories responsible for obtaining the specimens, as well as the Submitting laboratories where the genome data were generated and shared via GISAID, on which this research is based.

All Submitters of data may be contacted directly via www.gisaid.org

Authors are sorted alphabetically.

| Accession ID | Originating Laboratory | Submitting Laboratory | Authors |
|---|---|---|---|
| EPI_ISL_767937 | Pathology West - NSW Health Pathology | NSW Health Pathology - Institute of Clinical Pathology and Medical Research; Westmead Hospital; University of Sydney | CIDM-PH et al. |
| EPI_ISL_767938 | Laverty Pathology | NSW Health Pathology - Institute of Clinical Pathology and Medical Research; Westmead Hospital; University of Sydney | CIDM-PH et al. |
| EPI_ISL_767939, EPI_ISL_767940 | Sydney South West Pathology Service (SSWPS) - Liverpool Hospital - NSW Health Pathology | NSW Health Pathology - Institute of Clinical Pathology and Medical Research; Westmead Hospital; University of Sydney | CIDM-PH et al. |
| EPI_ISL_768815, EPI_ISL_768816 | Respiratory Virus Unit, National Infection Service, Public Health England | COVID-19 Genomics UK (COG-UK) Consortium | PHE Covid Sequencing Team |
| EPI_ISL_770477, EPI_ISL_770496, EPI_ISL_770497 | Wyoming Public Health Laboratory | Wyoming Public Health Laboratory | Noah Hull, Taylor Fearing, Lynette Gumbleton, Channing Weber, Ashley Norberg, Bailey Bowcutt, and Wanda Manley |
| EPI_ISL_771368, EPI_ISL_771369 | SA Pathology | SA Pathology | Lex Leong, Julien Soubrier, Chuan Kok Lim, Song Gao, Mark Turra, Karin Kassahn, Ivan Bastian, Geoff Higgins |
| EPI_ISL_779151 | Yale Clinical Virology Laboratory | Grubaugh Lab - Yale School of Public Health | Tara Alpert, Joseph Fauver, Anderson Brito, Mallery Breban, Anne Wyllie, Chantal Vogels, Mary Petrone, Chaney Kalinich, Isabel Ott, Arnau Casanovas, Catherine Muenker, Adam Moore, Alice Lu, Maria Tokuyama, Patrick Wong, Peiwen Lu, Saad Omer, Richard Martinello, Allison Nelson, Shelli Farhadian, Akiko Iwasaki, Charlese Dela Cruz, Albert Ko, Nathan Grubaugh |
| EPI_ISL_779153 | Yale Pathology Lab | Grubaugh Lab - Yale School of Public Health | Tara Alpert, Joseph Fauver, Chen Liu, Pei Hui, Jianhui Wang, Susan Bell and Han Zhou, Anderson Brito, Mallery Breban, Anne Wyllie, Chantal Vogels, Mary Petrone, Chaney Kalinich, Isabel Ott, Arnau Casanovas, Catherine Muenker, Adam Moore, Alice Lu, Maria Tokuyama, Patrick Wong, Peiwen Lu, Saad Omer, Richard Martinello, Allison Nelson, Shelli Farhadian, Akiko Iwasaki, Charlese Dela Cruz, Albert Ko, Nathan Grubaugh |
| EPI_ISL_779401, EPI_ISL_779402 | South Eastern Area Laboratory Services (SEALS) | NSW Health Pathology - Institute of Clinical Pathology and Medical Research; Westmead Hospital; University of Sydney | CIDM-PH et al. |
| EPI_ISL_779409, EPI_ISL_779410, EPI_ISL_779411 | Royal Darwin Hospital Pathology | MDU-PHL | Meumann, E., Caly L., Seemann T., Sait, M.L., Druce J., Sherry, N.L. |
| EPI_ISL_779647 | Microbiological Diagnostic Unit - Public Health Laboratory (MDU-PHL) | MDU-PHL | Seemann T., Sait, M.L., Sherry, N.L. |
| EPI_ISL_779648, EPI_ISL_779649 | Victorian Infectious Diseases Reference Laboratory (VIDRL) | VIDRL and MDU-PHL | Caly L., Seemann T., Sait, M.L., Druce J., Sherry, N.L. |
| EPI_ISL_789061, EPI_ISL_789074, EPI_ISL_789075, EPI_ISL_789077, EPI_ISL_789078, EPI_ISL_789080, EPI_ISL_789081, EPI_ISL_789082, EPI_ISL_789094, EPI_ISL_789096, EPI_ISL_789097, EPI_ISL_789099, EPI_ISL_789100, EPI_ISL_789101 | | | |
| see above | National Virus Reference Laboratory | National Virus Reference Laboratory | Michael Carr, Gabriel Gonzalez, Dana Alalwan, Jonathan Dean, Daniel Hare, Cillian F De Gascun |
| EPI_ISL_791107, EPI_ISL_791108, EPI_ISL_791109, EPI_ISL_791110, EPI_ISL_791111, EPI_ISL_791112, EPI_ISL_791113, EPI_ISL_791114, EPI_ISL_791115, EPI_ISL_791116, EPI_ISL_791117, EPI_ISL_791118, EPI_ISL_791119, EPI_ISL_791120, EPI_ISL_791121, EPI_ISL_791122, EPI_ISL_791123, EPI_ISL_791124, EPI_ISL_791125, EPI_ISL_791126, EPI_ISL_791127, EPI_ISL_791128, EPI_ISL_791129, EPI_ISL_791130, EPI_ISL_791131, EPI_ISL_791132, EPI_ISL_791133, EPI_ISL_791134, EPI_ISL_791135, EPI_ISL_791136, EPI_ISL_791137, EPI_ISL_791138, EPI_ISL_791140, EPI_ISL_791141, EPI_ISL_791142, EPI_ISL_791143, EPI_ISL_791145, EPI_ISL_791146, EPI_ISL_791147, EPI_ISL_791148, EPI_ISL_791149, EPI_ISL_791151, EPI_ISL_791156, EPI_ISL_791162 | | | |
| see above | University of Wisconsin-Madison AIDS Vaccine Research Laboratories | University of Wisconsin-Madison AIDS Vaccine Research Laboratories | Gage Moreno, Katarina Braun, et al. AIDS Vaccine Research Laboratories |
| EPI_ISL_791279 | South Eastern Area Laboratory Services (SEALS) | NSW Health Pathology - Institute of Clinical Pathology and Medical Research; Westmead Hospital; University of Sydney | CIDM-PH et al. |
| EPI_ISL_791327, EPI_ISL_791328, EPI_ISL_791329 | University of Wisconsin-Madison AIDS Vaccine Research Laboratories | University of Wisconsin-Madison AIDS Vaccine Research Laboratories | Gage Moreno, Katarina Braun, et al. AIDS Vaccine Research Laboratories |
| EPI_ISL_791427 | Johns Hopkins Hospital Department of Pathology | Johns Hopkins Hospital Department of Pathology | C. Paul Morris, Chun Huai Luo, Adannaya Amadi, Nicholas Gallagher, Heba H. Mostafa |
| EPI_ISL_792037 | South Eastern Area Laboratory Services (SEALS) | NSW Health Pathology - Institute of Clinical Pathology and Medical Research; Westmead Hospital; University of Sydney | CIDM-PH et al. |
| EPI_ISL_792555 | Centre for Dengue Research and AICBU, Department of Immunology and Molecular Medicine | Centre for Dengue Research and AICBU, Department of Immunology and Molecular Medicine | Chandima Jeewandara, Deshni Jayathilaka, Dinuka Ariyaratne, Diyanath Ranasinghe, Laksiri Gomes, Gathsaurie Neelika Malavige |
| EPI_ISL_792680, EPI_ISL_792681, EPI_ISL_792682, EPI_ISL_792683 | Pathogen Genomics Center, National Institute of Infectious Diseases | Pathogen Genomics Center, National Institute of Infectious Diseases | Tsuyoshi Sekizuka, Kentaro Itokawa, Rina Tanaka, Masanori Hashino, Makoto Kuroda |
| EPI_ISL_794206, EPI_ISL_794207 | Wadsworth Center, New York State Department.of Health | Wadsworth Center, New York State Department.of Health | Kirsten St. George, Daryl M. Lamson, Alexis Russel, Matthew Shudt, Melissa A Leisner, Jonathan Plitnick, Navjot Singh, John Kelly, Sara Griesemer, Erasmus Schneider, Erica Lasek-Nesselquist |
| EPI_ISL_794261, EPI_ISL_794262, EPI_ISL_794263, EPI_ISL_794264, EPI_ISL_794265, EPI_ISL_794266, EPI_ISL_794267, EPI_ISL_794268, EPI_ISL_794269, EPI_ISL_794270, EPI_ISL_794271, EPI_ISL_794272, EPI_ISL_794273, EPI_ISL_794274, EPI_ISL_794275, EPI_ISL_794276, EPI_ISL_794277, EPI_ISL_794278, EPI_ISL_794279, EPI_ISL_794280 | | | |
| see above | WESTCHESTER MEDICAL CENTER | Wadsworth Center, New York State Department.of Health | Kirsten St. George, Daryl M. Lamson, Alexis Russel, Matthew Shudt, Melissa A Leisner, Jonathan Plitnick, Navjot Singh, John Kelly, Sara Griesemer, Erasmus Schneider, Erica Lasek-Nesselquist |
| EPI_ISL_794611, EPI_ISL_794612, EPI_ISL_794613, EPI_ISL_794614, EPI_ISL_794615, EPI_ISL_794616, EPI_ISL_794617, EPI_ISL_794618, EPI_ISL_794619, EPI_ISL_794623 | LabPLUS | Institute of Environmental Science and Research (ESR) | Xiaoyun Ren, Matt Storey, Nikki Freed, Muhammad Faisal, Jing Wang, Hermes Perez, Anja Werno, Antje van der Linden, Arlo Upton, Chris Mansell, David Hammer, Dragana Drinkovic, Gary McAuliffe, Hana Sofia Andersson, James Ussher, Jill Sherwood, Josh Freeman, Julia Howard, Juliet Elvy, Mary DeAlmeida, Matt Blakiston, Matthew Rogers, Max Bloomfield, Michael Addidle, Michelle Balm, Sally Roberts, Sarah Jefferies, Sharmini Muttaiyah, Susan Morpeth, Susan Taylor, Timothy Blackmore, Vani Sathyendran, Veronica Playle, Virginia Hope, Erasmus Smit, Lauren Jelly, Olin Silander, Joep de Ligt |
| EPI_ISL_794625 | Middlemore Hospital | Institute of Environmental Science and Research (ESR) | Xiaoyun Ren, Matt Storey, Nikki Freed, Muhammad Faisal, Jing Wang, Hermes Perez, Anja Werno, Antje van der Linden, Arlo Upton, Chris Mansell, David Hammer, Dragana Drinkovic, Gary McAuliffe, Hana Sofia Andersson, James Ussher, Jill Sherwood, Josh Freeman, Julia Howard, Juliet Elvy, Mary DeAlmeida, Matt Blakiston, Matthew Rogers, Max Bloomfield, Michael Addidle, Michelle Balm, Sally Roberts, Sarah Jefferies, Sharmini Muttaiyah, Susan Morpeth, Susan Taylor, Timothy Blackmore, Vani Sathyendran, Veronica Playle, Virginia Hope, Erasmus Smit, Lauren Jelly, Olin Silander, Joep de Ligt |
| EPI_ISL_794635, EPI_ISL_794636, EPI_ISL_794637, EPI_ISL_794638 | USC Clinical Lab | Los Angeles County PHL | P. Hemarajata et al. |
| EPI_ISL_794675, EPI_ISL_794676 | PathWest Laboratory Medicine WA | PathWest Laboratory Medicine WA Microbial Surveillance Unit | PathWest Laboratory Medicine WA Microbial Surveillance Unit |
| EPI_ISL_796012, EPI_ISL_796013, EPI_ISL_796014, EPI_ISL_796015, EPI_ISL_796016, EPI_ISL_796017, EPI_ISL_796018, EPI_ISL_796019, EPI_ISL_796020, EPI_ISL_796021, EPI_ISL_796022 | | | |

| | | | |
|---|---|---|---|
| see above | Hebei Provincial Center for Disease Control and Prevention, Shijiazhuang, Hebei Province; National Institute for Viral Disease Control and Prevention, China CDC | Hebei Provincial Center for Disease Control and Prevention, Shijiazhuang, Hebei Province; National Institute for Viral Disease Control and Prevention, China CDC | Shunxiang Qi, Xiang Zhao, Nankun Liu, George F. Gao, Yang Song, Wenbo Xu, Qi Li |
| EPI_ISL_796755, EPI_ISL_796756, EPI_ISL_796757, EPI_ISL_796758, EPI_ISL_796762, EPI_ISL_796767 | Instituto Nacional de Saude (INSA) | Instituto Nacional de Saude (INSA) | Borges et al |
| EPI_ISL_797405, EPI_ISL_797408 | Lighthouse Lab in Glasgow | Wellcome Sanger Institute for the COVID-19 Genomics UK (COG-UK) Consortium | Harper VanSteenhouse, Yumi Kasai, David Gray, Carol Clugston, Anna Dominiczak and Alex Alderton, Roberto Amato, Sonia Goncalves, Ewan Harrison, David K. Jackson, Ian Johnston, Dominic Kwiatkowski, Cordelia Langford, John Sillitoe on behalf of the Wellcome Sanger Institute COVID-19 Surveillance Team |
| EPI_ISL_797411 | Lighthouse Lab in Alderley Park | Wellcome Sanger Institute for the COVID-19 Genomics UK (COG-UK) Consortium | Jacquelyn Wynn, Mairead Hyland, The Lighthouse Lab in Alderley Park and Alex Alderton, Roberto Amato, Sonia Goncalves, Ewan Harrison, David K. Jackson, Ian Johnston, Dominic Kwiatkowski, Cordelia Langford, John Sillitoe on behalf of the Wellcome Sanger Institute COVID-19 Surveillance Team |
| EPI_ISL_797432 | Lighthouse Lab in Glasgow | Wellcome Sanger Institute for the COVID-19 Genomics UK (COG-UK) Consortium | Harper VanSteenhouse, Yumi Kasai, David Gray, Carol Clugston, Anna Dominiczak and Alex Alderton, Roberto Amato, Sonia Goncalves, Ewan Harrison, David K. Jackson, Ian Johnston, Dominic Kwiatkowski, Cordelia Langford, John Sillitoe on behalf of the Wellcome Sanger Institute COVID-19 Surveillance Team |
| EPI_ISL_797438, EPI_ISL_797441 | Lighthouse Lab in Alderley Park | Wellcome Sanger Institute for the COVID-19 Genomics UK (COG-UK) Consortium | Jacquelyn Wynn, Mairead Hyland, The Lighthouse Lab in Alderley Park and Alex Alderton, Roberto Amato, Sonia Goncalves, Ewan Harrison, David K. Jackson, Ian Johnston, Dominic Kwiatkowski, Cordelia Langford, John Sillitoe on behalf of the Wellcome Sanger Institute COVID-19 Surveillance Team |
| EPI_ISL_797445, EPI_ISL_797451, EPI_ISL_797459 | Lighthouse Lab in Glasgow | Wellcome Sanger Institute for the COVID-19 Genomics UK (COG-UK) Consortium | Harper VanSteenhouse, Yumi Kasai, David Gray, Carol Clugston, Anna Dominiczak and Alex Alderton, Roberto Amato, Sonia Goncalves, Ewan Harrison, David K. Jackson, Ian Johnston, Dominic Kwiatkowski, Cordelia Langford, John Sillitoe on behalf of the Wellcome Sanger Institute COVID-19 Surveillance Team |
| EPI_ISL_797461, EPI_ISL_797462, EPI_ISL_797475, EPI_ISL_797484 | Lighthouse Lab in Alderley Park | Wellcome Sanger Institute for the COVID-19 Genomics UK (COG-UK) Consortium | Jacquelyn Wynn, Mairead Hyland, The Lighthouse Lab in Alderley Park and Alex Alderton, Roberto Amato, Sonia Goncalves, Ewan Harrison, David K. Jackson, Ian Johnston, Dominic Kwiatkowski, Cordelia Langford, John Sillitoe on behalf of the Wellcome Sanger Institute COVID-19 Surveillance Team |
| EPI_ISL_797500 | Lighthouse Lab in Glasgow | Wellcome Sanger Institute for the COVID-19 Genomics UK (COG-UK) Consortium | Harper VanSteenhouse, Yumi Kasai, David Gray, Carol Clugston, Anna Dominiczak and Alex Alderton, Roberto Amato, Sonia Goncalves, Ewan Harrison, David K. Jackson, Ian Johnston, Dominic Kwiatkowski, Cordelia Langford, John Sillitoe on behalf of the Wellcome Sanger Institute COVID-19 Surveillance Team |
| EPI_ISL_797504, EPI_ISL_797509 | Lighthouse Lab in Alderley Park | Wellcome Sanger Institute for the COVID-19 Genomics UK (COG-UK) Consortium | Jacquelyn Wynn, Mairead Hyland, The Lighthouse Lab in Alderley Park and Alex Alderton, Roberto Amato, Sonia Goncalves, Ewan Harrison, David K. Jackson, Ian Johnston, Dominic Kwiatkowski, Cordelia Langford, John Sillitoe on behalf of the Wellcome Sanger Institute COVID-19 Surveillance Team |
| EPI_ISL_797514, EPI_ISL_797531 | Lighthouse Lab in Glasgow | Wellcome Sanger Institute for the COVID-19 Genomics UK (COG-UK) Consortium | Harper VanSteenhouse, Yumi Kasai, David Gray, Carol Clugston, Anna Dominiczak and Alex Alderton, Roberto Amato, Sonia Goncalves, Ewan Harrison, David K. Jackson, Ian Johnston, Dominic Kwiatkowski, Cordelia Langford, John Sillitoe on behalf of the Wellcome Sanger Institute COVID-19 Surveillance Team |
| EPI_ISL_797553 | Lighthouse Lab in Alderley Park | Wellcome Sanger Institute for the COVID-19 Genomics UK (COG-UK) Consortium | Jacquelyn Wynn, Mairead Hyland, The Lighthouse Lab in Alderley Park and Alex Alderton, Roberto Amato, Sonia Goncalves, Ewan Harrison, David K. Jackson, Ian Johnston, Dominic Kwiatkowski, Cordelia Langford, John Sillitoe on behalf of the Wellcome Sanger Institute COVID-19 Surveillance Team |
| EPI_ISL_797554 | Lighthouse Lab in Glasgow | Wellcome Sanger Institute for the COVID-19 Genomics UK (COG-UK) Consortium | Harper VanSteenhouse, Yumi Kasai, David Gray, Carol Clugston, Anna Dominiczak and Alex Alderton, Roberto Amato, Sonia Goncalves, Ewan Harrison, David K. Jackson, Ian Johnston, Dominic Kwiatkowski, Cordelia Langford, John Sillitoe on behalf of the Wellcome Sanger Institute COVID-19 Surveillance Team |
| EPI_ISL_797559, EPI_ISL_797560 | Lighthouse Lab in Alderley Park | Wellcome Sanger Institute for the COVID-19 Genomics UK (COG-UK) Consortium | Jacquelyn Wynn, Mairead Hyland, The Lighthouse Lab in Alderley Park and Alex Alderton, Roberto Amato, Sonia Goncalves, Ewan Harrison, David K. Jackson, Ian Johnston, Dominic Kwiatkowski, Cordelia Langford, John Sillitoe on behalf of the Wellcome Sanger Institute COVID-19 Surveillance Team |
| EPI_ISL_797563, EPI_ISL_797564 | Lighthouse Lab in Glasgow | Wellcome Sanger Institute for the COVID-19 Genomics UK (COG-UK) Consortium | Harper VanSteenhouse, Yumi Kasai, David Gray, Carol Clugston, Anna Dominiczak and Alex Alderton, Roberto Amato, Sonia Goncalves, Ewan Harrison, David K. Jackson, Ian Johnston, Dominic Kwiatkowski, Cordelia Langford, John Sillitoe on behalf of the Wellcome Sanger Institute COVID-19 Surveillance Team |
| EPI_ISL_797575, EPI_ISL_797576, EPI_ISL_797586, EPI_ISL_797605 | Lighthouse Lab in Alderley Park | Wellcome Sanger Institute for the COVID-19 Genomics UK (COG-UK) Consortium | Jacquelyn Wynn, Mairead Hyland, The Lighthouse Lab in Alderley Park and Alex Alderton, Roberto Amato, Sonia Goncalves, Ewan Harrison, David K. Jackson, Ian Johnston, Dominic Kwiatkowski, Cordelia Langford, John Sillitoe on behalf of the Wellcome Sanger Institute COVID-19 Surveillance Team |
| EPI_ISL_797615, EPI_ISL_797620, EPI_ISL_797627, EPI_ISL_797632 | Lighthouse Lab in Glasgow | Wellcome Sanger Institute for the COVID-19 Genomics UK (COG-UK) Consortium | Harper VanSteenhouse, Yumi Kasai, David Gray, Carol Clugston, Anna Dominiczak and Alex Alderton, Roberto Amato, Sonia Goncalves, Ewan Harrison, David K. Jackson, Ian Johnston, Dominic Kwiatkowski, Cordelia Langford, John Sillitoe on behalf of the Wellcome Sanger Institute COVID-19 Surveillance Team |
| EPI_ISL_797635, EPI_ISL_797673 | Lighthouse Lab in Alderley Park | Wellcome Sanger Institute for the COVID-19 Genomics UK (COG-UK) Consortium | Jacquelyn Wynn, Mairead Hyland, The Lighthouse Lab in Alderley Park and Alex Alderton, Roberto Amato, Sonia Goncalves, Ewan Harrison, David K. Jackson, Ian Johnston, Dominic Kwiatkowski, Cordelia Langford, John Sillitoe on behalf of the Wellcome Sanger Institute COVID-19 Surveillance Team |
| EPI_ISL_797677, EPI_ISL_797678, EPI_ISL_797685 | Lighthouse Lab in Glasgow | Wellcome Sanger Institute for the COVID-19 Genomics UK (COG-UK) Consortium | Harper VanSteenhouse, Yumi Kasai, David Gray, Carol Clugston, Anna Dominiczak and Alex Alderton, Roberto Amato, Sonia Goncalves, Ewan Harrison, David K. Jackson, Ian Johnston, Dominic Kwiatkowski, Cordelia Langford, John Sillitoe on behalf of the Wellcome Sanger Institute COVID-19 Surveillance Team |
| EPI_ISL_797686 | Lighthouse Lab in Alderley Park | Wellcome Sanger Institute for the COVID-19 Genomics UK (COG-UK) Consortium | Jacquelyn Wynn, Mairead Hyland, The Lighthouse Lab in Alderley Park and Alex Alderton, Roberto Amato, Sonia Goncalves, Ewan Harrison, David K. Jackson, Ian Johnston, Dominic Kwiatkowski, Cordelia Langford, John Sillitoe on behalf of the Wellcome Sanger Institute COVID-19 Surveillance Team |
| EPI_ISL_797704, EPI_ISL_797713, EPI_ISL_797716, EPI_ISL_797847, EPI_ISL_797867, EPI_ISL_797870, EPI_ISL_797914, EPI_ISL_797929, EPI_ISL_797945, EPI_ISL_797961, EPI_ISL_797980, EPI_ISL_797996 | | | |
| see above | Lighthouse Lab in Glasgow | Wellcome Sanger Institute for the COVID-19 Genomics UK (COG-UK) Consortium | Harper VanSteenhouse, Yumi Kasai, David Gray, Carol Clugston, Anna Dominiczak and Alex Alderton, Roberto Amato, Sonia Goncalves, Ewan Harrison, David K. Jackson, Ian Johnston, Dominic Kwiatkowski, Cordelia Langford, John Sillitoe on behalf of the Wellcome Sanger Institute COVID-19 Surveillance Team |
| EPI_ISL_801519, EPI_ISL_801520 | Instituto Nacional de Saude (INSA) | Instituto Nacional de Saude (INSA) | Borges et al |
| EPI_ISL_802428, EPI_ISL_802432, EPI_ISL_802437, EPI_ISL_802491, EPI_ISL_802492, EPI_ISL_802493, EPI_ISL_802494, EPI_ISL_802495, EPI_ISL_802496, EPI_ISL_802497 | SARATOGA HOSPITAL LABORATORY | Wadsworth Center, New York State Department.of Health | Kirsten St. George, Daryl M. Lamson, Alexis Russel, Matthew Shudt, Melissa A Leisner, Jonathan Plitnick, Navjot Singh, John Kelly, Sara Griesemer, Erasmus Schneider, Erica Lasek-Nesselquist |
| EPI_ISL_803958, EPI_ISL_803967, EPI_ISL_803968, EPI_ISL_803989, EPI_ISL_803990, EPI_ISL_803991, EPI_ISL_803992, EPI_ISL_803993, EPI_ISL_803998 | National Public Health Laboratory, National Centre for Infectious Diseases | National Public Health Laboratory, National Centre for Infectious Diseases | Tze Minn Mak, Sophie Octavia, Zhenyang Zhou, Lin Cui, Raymond Tzer Pin Lin |
| EPI_ISL_804249 | Respiratory Virus Unit, National Infection Service, Public Health England | COVID-19 Genomics UK (COG-UK) Consortium | PHE Covid Sequencing Team |
| EPI_ISL_804981 | BIO-REFERENCE LABORATORIES | Wadsworth Center, New York State Department of Health | Kirsten St. George, Daryl M. Lamson, Alexis Russel, Matthew Shudt, Melissa A Leisner, Jonathan Plitnick, Navjot Singh, John Kelly, Erasmus Schneider, Erica Lasek-Nesselquist |
| EPI_ISL_806725 | Medlab Pathology | NSW Health Pathology - Institute of Clinical Pathology and Medical Research; Westmead Hospital; University of Sydney | CIDM-PH et al. |
| EPI_ISL_806726 | 4Cyte Pathology | NSW Health Pathology - Institute of Clinical Pathology and | CIDM-PH et al. |

| | | | |
|---|---|---|---|
| | | Medical Research; Westmead Hospital; University of Sydney | |
| EPI_ISL_806728 | Sydney South West Pathology Service (SSWPS) - Liverpool Hospital - NSW Health Pathology | NSW Health Pathology - Institute of Clinical Pathology and Medical Research; Westmead Hospital; University of Sydney | CIDM-PH et al. |
| EPI_ISL_806807 | Gundersen Molecular Diagnostics Laboratory | Kabara Cancer Research Institute | Craig S. Richmond, Paraic A. Kenny |
| EPI_ISL_810964 | PathWest Laboratory Medicine WA | PathWest Laboratory Medicine WA Microbial Surveillance Unit | PathWest Laboratory Medicine WA Microbial Surveillance Unit |
| EPI_ISL_811072, EPI_ISL_811106, EPI_ISL_811107, EPI_ISL_811108, EPI_ISL_811109, EPI_ISL_811110, EPI_ISL_811111, EPI_ISL_811112, EPI_ISL_811113, EPI_ISL_811114, EPI_ISL_811116 | | | |
| see above | Respiratory Virus Unit, National Infection Service, Public Health England | COVID-19 Genomics UK (COG-UK) Consortium | PHE Covid Sequencing Team |
| EPI_ISL_811194, EPI_ISL_811195, EPI_ISL_811196 | Lighthouse Lab in Glasgow | Wellcome Sanger Institute for the COVID-19 Genomics UK (COG-UK) Consortium | Harper VanSteenhouse, Yumi Kasai, David Gray, Carol Clugston, Anna Dominiczak and Alex Alderton, Roberto Amato, Sonia Goncalves, Ewan Harrison, David K. Jackson, Ian Johnston, Dominic Kwiatkowski, Cordelia Langford, John Sillitoe on behalf of the Wellcome Sanger Institute COVID-19 Surveillance Team |
| EPI_ISL_811197 | Lighthouse Lab in Milton Keynes | Wellcome Sanger Institute for the COVID-19 Genomics UK (COG-UK) Consortium | The Lighthouse Lab in Milton Keynes and Alex Alderton, Roberto Amato, Sonia Goncalves, Ewan Harrison, David K. Jackson, Ian Johnston, Dominic Kwiatkowski, Cordelia Langford, John Sillitoe on behalf of the Wellcome Sanger Institute COVID-19 Surveillance Team |
| EPI_ISL_811198, EPI_ISL_811199, EPI_ISL_811201, EPI_ISL_811202, EPI_ISL_811203, EPI_ISL_811204 | Lighthouse Lab in Glasgow | Wellcome Sanger Institute for the COVID-19 Genomics UK (COG-UK) Consortium | Harper VanSteenhouse, Yumi Kasai, David Gray, Carol Clugston, Anna Dominiczak and Alex Alderton, Roberto Amato, Sonia Goncalves, Ewan Harrison, David K. Jackson, Ian Johnston, Dominic Kwiatkowski, Cordelia Langford, John Sillitoe on behalf of the Wellcome Sanger Institute COVID-19 Surveillance Team |
| EPI_ISL_811205 | Lighthouse Lab in Milton Keynes | Wellcome Sanger Institute for the COVID-19 Genomics UK (COG-UK) Consortium | The Lighthouse Lab in Milton Keynes and Alex Alderton, Roberto Amato, Sonia Goncalves, Ewan Harrison, David K. Jackson, Ian Johnston, Dominic Kwiatkowski, Cordelia Langford, John Sillitoe on behalf of the Wellcome Sanger Institute COVID-19 Surveillance Team |
| EPI_ISL_811206 | Lighthouse Lab in Glasgow | Wellcome Sanger Institute for the COVID-19 Genomics UK (COG-UK) Consortium | Harper VanSteenhouse, Yumi Kasai, David Gray, Carol Clugston, Anna Dominiczak and Alex Alderton, Roberto Amato, Sonia Goncalves, Ewan Harrison, David K. Jackson, Ian Johnston, Dominic Kwiatkowski, Cordelia Langford, John Sillitoe on behalf of the Wellcome Sanger Institute COVID-19 Surveillance Team |
| EPI_ISL_811207 | Lighthouse Lab in Milton Keynes | Wellcome Sanger Institute for the COVID-19 Genomics UK (COG-UK) Consortium | The Lighthouse Lab in Milton Keynes and Alex Alderton, Roberto Amato, Sonia Goncalves, Ewan Harrison, David K. Jackson, Ian Johnston, Dominic Kwiatkowski, Cordelia Langford, John Sillitoe on behalf of the Wellcome Sanger Institute COVID-19 Surveillance Team |
| EPI_ISL_811208, EPI_ISL_811209, EPI_ISL_811210 | Lighthouse Lab in Glasgow | Wellcome Sanger Institute for the COVID-19 Genomics UK (COG-UK) Consortium | Harper VanSteenhouse, Yumi Kasai, David Gray, Carol Clugston, Anna Dominiczak and Alex Alderton, Roberto Amato, Sonia Goncalves, Ewan Harrison, David K. Jackson, Ian Johnston, Dominic Kwiatkowski, Cordelia Langford, John Sillitoe on behalf of the Wellcome Sanger Institute COVID-19 Surveillance Team |
| EPI_ISL_811211, EPI_ISL_811212, EPI_ISL_811215, EPI_ISL_811217 | Lighthouse Lab in Milton Keynes | Wellcome Sanger Institute for the COVID-19 Genomics UK (COG-UK) Consortium | The Lighthouse Lab in Milton Keynes and Alex Alderton, Roberto Amato, Sonia Goncalves, Ewan Harrison, David K. Jackson, Ian Johnston, Dominic Kwiatkowski, Cordelia Langford, John Sillitoe on behalf of the Wellcome Sanger Institute COVID-19 Surveillance Team |
| EPI_ISL_811218, EPI_ISL_811219, EPI_ISL_811220, EPI_ISL_811223, EPI_ISL_811224, EPI_ISL_811225, EPI_ISL_811227, EPI_ISL_811228 | Lighthouse Lab in Glasgow | Wellcome Sanger Institute for the COVID-19 Genomics UK (COG-UK) Consortium | Harper VanSteenhouse, Yumi Kasai, David Gray, Carol Clugston, Anna Dominiczak and Alex Alderton, Roberto Amato, Sonia Goncalves, Ewan Harrison, David K. Jackson, Ian Johnston, Dominic Kwiatkowski, Cordelia Langford, John Sillitoe on behalf of the Wellcome Sanger Institute COVID-19 Surveillance Team |
| EPI_ISL_811229 | Lighthouse Lab in Milton Keynes | Wellcome Sanger Institute for the COVID-19 Genomics UK (COG-UK) Consortium | The Lighthouse Lab in Milton Keynes and Alex Alderton, Roberto Amato, Sonia Goncalves, Ewan Harrison, David K. Jackson, Ian Johnston, Dominic Kwiatkowski, Cordelia Langford, John Sillitoe on behalf of the Wellcome Sanger Institute COVID-19 Surveillance Team |
| EPI_ISL_811230, EPI_ISL_811231, EPI_ISL_811232, EPI_ISL_811233, EPI_ISL_811237, EPI_ISL_811238, EPI_ISL_811239, EPI_ISL_811240, EPI_ISL_811242, EPI_ISL_811243, EPI_ISL_811244 | | | |
| see above | Lighthouse Lab in Glasgow | Wellcome Sanger Institute for the COVID-19 Genomics UK (COG-UK) Consortium | Harper VanSteenhouse, Yumi Kasai, David Gray, Carol Clugston, Anna Dominiczak and Alex Alderton, Roberto Amato, Sonia Goncalves, Ewan Harrison, David K. Jackson, Ian Johnston, Dominic Kwiatkowski, Cordelia Langford, John Sillitoe on behalf of the Wellcome Sanger Institute COVID-19 Surveillance Team |
| EPI_ISL_811245 | Lighthouse Lab in Milton Keynes | Wellcome Sanger Institute for the COVID-19 Genomics UK (COG-UK) Consortium | The Lighthouse Lab in Milton Keynes and Alex Alderton, Roberto Amato, Sonia Goncalves, Ewan Harrison, David K. Jackson, Ian Johnston, Dominic Kwiatkowski, Cordelia Langford, John Sillitoe on behalf of the Wellcome Sanger Institute COVID-19 Surveillance Team |
| EPI_ISL_811246 | Lighthouse Lab in Glasgow | Wellcome Sanger Institute for the COVID-19 Genomics UK (COG-UK) Consortium | Harper VanSteenhouse, Yumi Kasai, David Gray, Carol Clugston, Anna Dominiczak and Alex Alderton, Roberto Amato, Sonia Goncalves, Ewan Harrison, David K. Jackson, Ian Johnston, Dominic Kwiatkowski, Cordelia Langford, John Sillitoe on behalf of the Wellcome Sanger Institute COVID-19 Surveillance Team |
| EPI_ISL_811248 | Lighthouse Lab in Milton Keynes | Wellcome Sanger Institute for the COVID-19 Genomics UK (COG-UK) Consortium | The Lighthouse Lab in Milton Keynes and Alex Alderton, Roberto Amato, Sonia Goncalves, Ewan Harrison, David K. Jackson, Ian Johnston, Dominic Kwiatkowski, Cordelia Langford, John Sillitoe on behalf of the Wellcome Sanger Institute COVID-19 Surveillance Team |
| EPI_ISL_811249, EPI_ISL_811250 | Lighthouse Lab in Glasgow | Wellcome Sanger Institute for the COVID-19 Genomics UK (COG-UK) Consortium | Harper VanSteenhouse, Yumi Kasai, David Gray, Carol Clugston, Anna Dominiczak and Alex Alderton, Roberto Amato, Sonia Goncalves, Ewan Harrison, David K. Jackson, Ian Johnston, Dominic Kwiatkowski, Cordelia Langford, John Sillitoe on behalf of the Wellcome Sanger Institute COVID-19 Surveillance Team |
| EPI_ISL_811251 | Lighthouse Lab in Milton Keynes | Wellcome Sanger Institute for the COVID-19 Genomics UK (COG-UK) Consortium | The Lighthouse Lab in Milton Keynes and Alex Alderton, Roberto Amato, Sonia Goncalves, Ewan Harrison, David K. Jackson, Ian Johnston, Dominic Kwiatkowski, Cordelia Langford, John Sillitoe on behalf of the Wellcome Sanger Institute COVID-19 Surveillance Team |
| EPI_ISL_811252, EPI_ISL_811253 | Lighthouse Lab in Glasgow | Wellcome Sanger Institute for the COVID-19 Genomics UK (COG-UK) Consortium | Harper VanSteenhouse, Yumi Kasai, David Gray, Carol Clugston, Anna Dominiczak and Alex Alderton, Roberto Amato, Sonia Goncalves, Ewan Harrison, David K. Jackson, Ian Johnston, Dominic Kwiatkowski, Cordelia Langford, John Sillitoe on behalf of the Wellcome Sanger Institute COVID-19 Surveillance Team |
| EPI_ISL_811255 | Lighthouse Lab in Milton Keynes | Wellcome Sanger Institute for the COVID-19 Genomics UK (COG-UK) Consortium | The Lighthouse Lab in Milton Keynes and Alex Alderton, Roberto Amato, Sonia Goncalves, Ewan Harrison, David K. Jackson, Ian Johnston, Dominic Kwiatkowski, Cordelia Langford, John Sillitoe on behalf of the Wellcome Sanger Institute COVID-19 Surveillance Team |
| EPI_ISL_811259 | Lighthouse Lab in Glasgow | Wellcome Sanger Institute for the COVID-19 Genomics UK (COG-UK) Consortium | Harper VanSteenhouse, Yumi Kasai, David Gray, Carol Clugston, Anna Dominiczak and Alex Alderton, Roberto Amato, Sonia Goncalves, Ewan Harrison, David K. Jackson, Ian Johnston, Dominic Kwiatkowski, Cordelia Langford, John Sillitoe on behalf of the Wellcome Sanger Institute COVID-19 Surveillance Team |
| EPI_ISL_811262 | Lighthouse Lab in Milton Keynes | Wellcome Sanger Institute for the COVID-19 Genomics UK (COG-UK) Consortium | The Lighthouse Lab in Milton Keynes and Alex Alderton, Roberto Amato, Sonia Goncalves, Ewan Harrison, David K. Jackson, Ian Johnston, Dominic Kwiatkowski, Cordelia Langford, John Sillitoe on behalf of the Wellcome Sanger Institute COVID-19 Surveillance Team |
| EPI_ISL_811264, EPI_ISL_811265, EPI_ISL_811266, EPI_ISL_811267 | Lighthouse Lab in Glasgow | Wellcome Sanger Institute for the COVID-19 Genomics UK (COG-UK) Consortium | Harper VanSteenhouse, Yumi Kasai, David Gray, Carol Clugston, Anna Dominiczak and Alex Alderton, Roberto Amato, Sonia Goncalves, Ewan Harrison, David K. Jackson, Ian Johnston, Dominic Kwiatkowski, Cordelia Langford, John Sillitoe on behalf of the Wellcome Sanger Institute COVID-19 Surveillance Team |
| EPI_ISL_811269 | Lighthouse Lab in Milton Keynes | Wellcome Sanger Institute for the COVID-19 Genomics UK (COG-UK) Consortium | The Lighthouse Lab in Milton Keynes and Alex Alderton, Roberto Amato, Sonia Goncalves, Ewan Harrison, David K. Jackson, Ian Johnston, Dominic Kwiatkowski, Cordelia Langford, John Sillitoe on behalf of the Wellcome Sanger Institute COVID-19 Surveillance Team |
| EPI_ISL_811270, EPI_ISL_811271, EPI_ISL_811272, EPI_ISL_811274, EPI_ISL_811276, EPI_ISL_811277, EPI_ISL_811278, EPI_ISL_811279 | Lighthouse Lab in Glasgow | Wellcome Sanger Institute for the COVID-19 Genomics UK (COG-UK) Consortium | Harper VanSteenhouse, Yumi Kasai, David Gray, Carol Clugston, Anna Dominiczak and Alex Alderton, Roberto Amato, Sonia Goncalves, Ewan Harrison, David K. Jackson, Ian Johnston, Dominic Kwiatkowski, Cordelia Langford, John Sillitoe on behalf of the Wellcome Sanger Institute COVID-19 Surveillance Team |
| EPI_ISL_811280 | Lighthouse Lab in Milton Keynes | Wellcome Sanger Institute for the COVID-19 Genomics UK (COG-UK) Consortium | The Lighthouse Lab in Milton Keynes and Alex Alderton, Roberto Amato, Sonia Goncalves, Ewan Harrison, David K. Jackson, Ian Johnston, Dominic Kwiatkowski, Cordelia Langford, John Sillitoe on behalf of the Wellcome Sanger Institute COVID-19 Surveillance Team |
| EPI_ISL_811281 | Lighthouse Lab in Glasgow | Wellcome Sanger Institute for the COVID-19 Genomics UK | Harper VanSteenhouse, Yumi Kasai, David Gray, Carol Clugston, Anna Dominiczak and Alex Alderton, Roberto Amato, Sonia Goncalves, Ewan Harrison, |

| | | | |
|---|---|---|---|
| | | (COG-UK) Consortium | David K. Jackson, Ian Johnston, Dominic Kwiatkowski, Cordelia Langford, John Sillitoe on behalf of the Wellcome Sanger Institute COVID-19 Surveillance Team |
| EPI_ISL_811282 | Lighthouse Lab in Milton Keynes | Wellcome Sanger Institute for the COVID-19 Genomics UK (COG-UK) Consortium | The Lighthouse Lab in Milton Keynes and Alex Alderton, Roberto Amato, Sonia Goncalves, Ewan Harrison, David K. Jackson, Ian Johnston, Dominic Kwiatkowski, Cordelia Langford, John Sillitoe on behalf of the Wellcome Sanger Institute COVID-19 Surveillance Team |
| EPI_ISL_811283 | Lighthouse Lab in Glasgow | Wellcome Sanger Institute for the COVID-19 Genomics UK (COG-UK) Consortium | Harper VanSteenhouse, Yumi Kasai, David Gray, Carol Clugston, Anna Dominiczak and Alex Alderton, Roberto Amato, Sonia Goncalves, Ewan Harrison, David K. Jackson, Ian Johnston, Dominic Kwiatkowski, Cordelia Langford, John Sillitoe on behalf of the Wellcome Sanger Institute COVID-19 Surveillance Team |
| EPI_ISL_811284, EPI_ISL_811285 | Lighthouse Lab in Milton Keynes | Wellcome Sanger Institute for the COVID-19 Genomics UK (COG-UK) Consortium | The Lighthouse Lab in Milton Keynes and Alex Alderton, Roberto Amato, Sonia Goncalves, Ewan Harrison, David K. Jackson, Ian Johnston, Dominic Kwiatkowski, Cordelia Langford, John Sillitoe on behalf of the Wellcome Sanger Institute COVID-19 Surveillance Team |
| EPI_ISL_811286, EPI_ISL_811287, EPI_ISL_811288, EPI_ISL_811290, EPI_ISL_811291, EPI_ISL_811292, EPI_ISL_811294, EPI_ISL_811296, EPI_ISL_811298, EPI_ISL_811299 | Lighthouse Lab in Glasgow | Wellcome Sanger Institute for the COVID-19 Genomics UK (COG-UK) Consortium | Harper VanSteenhouse, Yumi Kasai, David Gray, Carol Clugston, Anna Dominiczak and Alex Alderton, Roberto Amato, Sonia Goncalves, Ewan Harrison, David K. Jackson, Ian Johnston, Dominic Kwiatkowski, Cordelia Langford, John Sillitoe on behalf of the Wellcome Sanger Institute COVID-19 Surveillance Team |
| EPI_ISL_811300 | Lighthouse Lab in Milton Keynes | Wellcome Sanger Institute for the COVID-19 Genomics UK (COG-UK) Consortium | The Lighthouse Lab in Milton Keynes and Alex Alderton, Roberto Amato, Sonia Goncalves, Ewan Harrison, David K. Jackson, Ian Johnston, Dominic Kwiatkowski, Cordelia Langford, John Sillitoe on behalf of the Wellcome Sanger Institute COVID-19 Surveillance Team |
| EPI_ISL_811301 | Lighthouse Lab in Glasgow | Wellcome Sanger Institute for the COVID-19 Genomics UK (COG-UK) Consortium | Harper VanSteenhouse, Yumi Kasai, David Gray, Carol Clugston, Anna Dominiczak and Alex Alderton, Roberto Amato, Sonia Goncalves, Ewan Harrison, David K. Jackson, Ian Johnston, Dominic Kwiatkowski, Cordelia Langford, John Sillitoe on behalf of the Wellcome Sanger Institute COVID-19 Surveillance Team |
| EPI_ISL_811302 | Lighthouse Lab in Milton Keynes | Wellcome Sanger Institute for the COVID-19 Genomics UK (COG-UK) Consortium | The Lighthouse Lab in Milton Keynes and Alex Alderton, Roberto Amato, Sonia Goncalves, Ewan Harrison, David K. Jackson, Ian Johnston, Dominic Kwiatkowski, Cordelia Langford, John Sillitoe on behalf of the Wellcome Sanger Institute COVID-19 Surveillance Team |
| EPI_ISL_811303 | Lighthouse Lab in Glasgow | Wellcome Sanger Institute for the COVID-19 Genomics UK (COG-UK) Consortium | Harper VanSteenhouse, Yumi Kasai, David Gray, Carol Clugston, Anna Dominiczak and Alex Alderton, Roberto Amato, Sonia Goncalves, Ewan Harrison, David K. Jackson, Ian Johnston, Dominic Kwiatkowski, Cordelia Langford, John Sillitoe on behalf of the Wellcome Sanger Institute COVID-19 Surveillance Team |
| EPI_ISL_811304 | Lighthouse Lab in Milton Keynes | Wellcome Sanger Institute for the COVID-19 Genomics UK (COG-UK) Consortium | The Lighthouse Lab in Milton Keynes and Alex Alderton, Roberto Amato, Sonia Goncalves, Ewan Harrison, David K. Jackson, Ian Johnston, Dominic Kwiatkowski, Cordelia Langford, John Sillitoe on behalf of the Wellcome Sanger Institute COVID-19 Surveillance Team |
| EPI_ISL_811305 | Lighthouse Lab in Glasgow | Wellcome Sanger Institute for the COVID-19 Genomics UK (COG-UK) Consortium | Harper VanSteenhouse, Yumi Kasai, David Gray, Carol Clugston, Anna Dominiczak and Alex Alderton, Roberto Amato, Sonia Goncalves, Ewan Harrison, David K. Jackson, Ian Johnston, Dominic Kwiatkowski, Cordelia Langford, John Sillitoe on behalf of the Wellcome Sanger Institute COVID-19 Surveillance Team |
| EPI_ISL_811306, EPI_ISL_811307 | Lighthouse Lab in Milton Keynes | Wellcome Sanger Institute for the COVID-19 Genomics UK (COG-UK) Consortium | The Lighthouse Lab in Milton Keynes and Alex Alderton, Roberto Amato, Sonia Goncalves, Ewan Harrison, David K. Jackson, Ian Johnston, Dominic Kwiatkowski, Cordelia Langford, John Sillitoe on behalf of the Wellcome Sanger Institute COVID-19 Surveillance Team |
| EPI_ISL_811308 | Lighthouse Lab in Glasgow | Wellcome Sanger Institute for the COVID-19 Genomics UK (COG-UK) Consortium | Harper VanSteenhouse, Yumi Kasai, David Gray, Carol Clugston, Anna Dominiczak and Alex Alderton, Roberto Amato, Sonia Goncalves, Ewan Harrison, David K. Jackson, Ian Johnston, Dominic Kwiatkowski, Cordelia Langford, John Sillitoe on behalf of the Wellcome Sanger Institute COVID-19 Surveillance Team |
| EPI_ISL_811309, EPI_ISL_811311 | Lighthouse Lab in Milton Keynes | Wellcome Sanger Institute for the COVID-19 Genomics UK (COG-UK) Consortium | The Lighthouse Lab in Milton Keynes and Alex Alderton, Roberto Amato, Sonia Goncalves, Ewan Harrison, David K. Jackson, Ian Johnston, Dominic Kwiatkowski, Cordelia Langford, John Sillitoe on behalf of the Wellcome Sanger Institute COVID-19 Surveillance Team |
| EPI_ISL_811313, EPI_ISL_811314 | Lighthouse Lab in Glasgow | Wellcome Sanger Institute for the COVID-19 Genomics UK (COG-UK) Consortium | Harper VanSteenhouse, Yumi Kasai, David Gray, Carol Clugston, Anna Dominiczak and Alex Alderton, Roberto Amato, Sonia Goncalves, Ewan Harrison, David K. Jackson, Ian Johnston, Dominic Kwiatkowski, Cordelia Langford, John Sillitoe on behalf of the Wellcome Sanger Institute COVID-19 Surveillance Team |
| EPI_ISL_811315 | Lighthouse Lab in Milton Keynes | Wellcome Sanger Institute for the COVID-19 Genomics UK (COG-UK) Consortium | The Lighthouse Lab in Milton Keynes and Alex Alderton, Roberto Amato, Sonia Goncalves, Ewan Harrison, David K. Jackson, Ian Johnston, Dominic Kwiatkowski, Cordelia Langford, John Sillitoe on behalf of the Wellcome Sanger Institute COVID-19 Surveillance Team |
| EPI_ISL_811318, EPI_ISL_811319, EPI_ISL_811320 | Lighthouse Lab in Glasgow | Wellcome Sanger Institute for the COVID-19 Genomics UK (COG-UK) Consortium | Harper VanSteenhouse, Yumi Kasai, David Gray, Carol Clugston, Anna Dominiczak and Alex Alderton, Roberto Amato, Sonia Goncalves, Ewan Harrison, David K. Jackson, Ian Johnston, Dominic Kwiatkowski, Cordelia Langford, John Sillitoe on behalf of the Wellcome Sanger Institute COVID-19 Surveillance Team |
| EPI_ISL_811322 | Lighthouse Lab in Milton Keynes | Wellcome Sanger Institute for the COVID-19 Genomics UK (COG-UK) Consortium | The Lighthouse Lab in Milton Keynes and Alex Alderton, Roberto Amato, Sonia Goncalves, Ewan Harrison, David K. Jackson, Ian Johnston, Dominic Kwiatkowski, Cordelia Langford, John Sillitoe on behalf of the Wellcome Sanger Institute COVID-19 Surveillance Team |
| EPI_ISL_811323, EPI_ISL_811324 | Lighthouse Lab in Glasgow | Wellcome Sanger Institute for the COVID-19 Genomics UK (COG-UK) Consortium | Harper VanSteenhouse, Yumi Kasai, David Gray, Carol Clugston, Anna Dominiczak and Alex Alderton, Roberto Amato, Sonia Goncalves, Ewan Harrison, David K. Jackson, Ian Johnston, Dominic Kwiatkowski, Cordelia Langford, John Sillitoe on behalf of the Wellcome Sanger Institute COVID-19 Surveillance Team |
| EPI_ISL_811325 | Lighthouse Lab in Milton Keynes | Wellcome Sanger Institute for the COVID-19 Genomics UK (COG-UK) Consortium | The Lighthouse Lab in Milton Keynes and Alex Alderton, Roberto Amato, Sonia Goncalves, Ewan Harrison, David K. Jackson, Ian Johnston, Dominic Kwiatkowski, Cordelia Langford, John Sillitoe on behalf of the Wellcome Sanger Institute COVID-19 Surveillance Team |
| EPI_ISL_811326, EPI_ISL_811327, EPI_ISL_811328 | Lighthouse Lab in Glasgow | Wellcome Sanger Institute for the COVID-19 Genomics UK (COG-UK) Consortium | Harper VanSteenhouse, Yumi Kasai, David Gray, Carol Clugston, Anna Dominiczak and Alex Alderton, Roberto Amato, Sonia Goncalves, Ewan Harrison, David K. Jackson, Ian Johnston, Dominic Kwiatkowski, Cordelia Langford, John Sillitoe on behalf of the Wellcome Sanger Institute COVID-19 Surveillance Team |
| EPI_ISL_811329 | Lighthouse Lab in Milton Keynes | Wellcome Sanger Institute for the COVID-19 Genomics UK (COG-UK) Consortium | The Lighthouse Lab in Milton Keynes and Alex Alderton, Roberto Amato, Sonia Goncalves, Ewan Harrison, David K. Jackson, Ian Johnston, Dominic Kwiatkowski, Cordelia Langford, John Sillitoe on behalf of the Wellcome Sanger Institute COVID-19 Surveillance Team |
| EPI_ISL_811330, EPI_ISL_811331 | Lighthouse Lab in Glasgow | Wellcome Sanger Institute for the COVID-19 Genomics UK (COG-UK) Consortium | Harper VanSteenhouse, Yumi Kasai, David Gray, Carol Clugston, Anna Dominiczak and Alex Alderton, Roberto Amato, Sonia Goncalves, Ewan Harrison, David K. Jackson, Ian Johnston, Dominic Kwiatkowski, Cordelia Langford, John Sillitoe on behalf of the Wellcome Sanger Institute COVID-19 Surveillance Team |
| EPI_ISL_811332 | Lighthouse Lab in Milton Keynes | Wellcome Sanger Institute for the COVID-19 Genomics UK (COG-UK) Consortium | The Lighthouse Lab in Milton Keynes and Alex Alderton, Roberto Amato, Sonia Goncalves, Ewan Harrison, David K. Jackson, Ian Johnston, Dominic Kwiatkowski, Cordelia Langford, John Sillitoe on behalf of the Wellcome Sanger Institute COVID-19 Surveillance Team |
| EPI_ISL_811333 | Lighthouse Lab in Glasgow | Wellcome Sanger Institute for the COVID-19 Genomics UK (COG-UK) Consortium | Harper VanSteenhouse, Yumi Kasai, David Gray, Carol Clugston, Anna Dominiczak and Alex Alderton, Roberto Amato, Sonia Goncalves, Ewan Harrison, David K. Jackson, Ian Johnston, Dominic Kwiatkowski, Cordelia Langford, John Sillitoe on behalf of the Wellcome Sanger Institute COVID-19 Surveillance Team |
| EPI_ISL_811334, EPI_ISL_811335, EPI_ISL_811336 | Lighthouse Lab in Milton Keynes | Wellcome Sanger Institute for the COVID-19 Genomics UK (COG-UK) Consortium | The Lighthouse Lab in Milton Keynes and Alex Alderton, Roberto Amato, Sonia Goncalves, Ewan Harrison, David K. Jackson, Ian Johnston, Dominic Kwiatkowski, Cordelia Langford, John Sillitoe on behalf of the Wellcome Sanger Institute COVID-19 Surveillance Team |
| EPI_ISL_811337 | Lighthouse Lab in Glasgow | Wellcome Sanger Institute for the COVID-19 Genomics UK (COG-UK) Consortium | Harper VanSteenhouse, Yumi Kasai, David Gray, Carol Clugston, Anna Dominiczak and Alex Alderton, Roberto Amato, Sonia Goncalves, Ewan Harrison, David K. Jackson, Ian Johnston, Dominic Kwiatkowski, Cordelia Langford, John Sillitoe on behalf of the Wellcome Sanger Institute COVID-19 Surveillance Team |
| EPI_ISL_811339 | Lighthouse Lab in Milton Keynes | Wellcome Sanger Institute for the COVID-19 Genomics UK (COG-UK) Consortium | The Lighthouse Lab in Milton Keynes and Alex Alderton, Roberto Amato, Sonia Goncalves, Ewan Harrison, David K. Jackson, Ian Johnston, Dominic Kwiatkowski, Cordelia Langford, John Sillitoe on behalf of the Wellcome Sanger Institute COVID-19 Surveillance Team |
| EPI_ISL_811340, EPI_ISL_811341 | Lighthouse Lab in Glasgow | Wellcome Sanger Institute for the COVID-19 Genomics UK (COG-UK) Consortium | Harper VanSteenhouse, Yumi Kasai, David Gray, Carol Clugston, Anna Dominiczak and Alex Alderton, Roberto Amato, Sonia Goncalves, Ewan Harrison, David K. Jackson, Ian Johnston, Dominic Kwiatkowski, Cordelia Langford, John Sillitoe on behalf of the Wellcome Sanger Institute COVID-19 Surveillance |

| | | | |
|---|---|---|---|
| | | | Team |
| EPI_ISL_811342 | Lighthouse Lab in Milton Keynes | Wellcome Sanger Institute for the COVID-19 Genomics UK (COG-UK) Consortium | The Lighthouse Lab in Milton Keynes and Alex Alderton, Roberto Amato, Sonia Goncalves, Ewan Harrison, David K. Jackson, Ian Johnston, Dominic Kwiatkowski, Cordelia Langford, John Sillitoe on behalf of the Wellcome Sanger Institute COVID-19 Surveillance Team |
| EPI_ISL_811343 | Lighthouse Lab in Glasgow | Wellcome Sanger Institute for the COVID-19 Genomics UK (COG-UK) Consortium | Harper VanSteenhouse, Yumi Kasai, David Gray, Carol Clugston, Anna Dominiczak and Alex Alderton, Roberto Amato, Sonia Goncalves, Ewan Harrison, David K. Jackson, Ian Johnston, Dominic Kwiatkowski, Cordelia Langford, John Sillitoe on behalf of the Wellcome Sanger Institute COVID-19 Surveillance Team |
| EPI_ISL_811344 | Lighthouse Lab in Milton Keynes | Wellcome Sanger Institute for the COVID-19 Genomics UK (COG-UK) Consortium | The Lighthouse Lab in Milton Keynes and Alex Alderton, Roberto Amato, Sonia Goncalves, Ewan Harrison, David K. Jackson, Ian Johnston, Dominic Kwiatkowski, Cordelia Langford, John Sillitoe on behalf of the Wellcome Sanger Institute COVID-19 Surveillance Team |
| EPI_ISL_811346, EPI_ISL_811349, EPI_ISL_811351, EPI_ISL_811352, EPI_ISL_811354, EPI_ISL_811357 | Lighthouse Lab in Glasgow | Wellcome Sanger Institute for the COVID-19 Genomics UK (COG-UK) Consortium | Harper VanSteenhouse, Yumi Kasai, David Gray, Carol Clugston, Anna Dominiczak and Alex Alderton, Roberto Amato, Sonia Goncalves, Ewan Harrison, David K. Jackson, Ian Johnston, Dominic Kwiatkowski, Cordelia Langford, John Sillitoe on behalf of the Wellcome Sanger Institute COVID-19 Surveillance Team |
| EPI_ISL_811358, EPI_ISL_811359 | Lighthouse Lab in Milton Keynes | Wellcome Sanger Institute for the COVID-19 Genomics UK (COG-UK) Consortium | The Lighthouse Lab in Milton Keynes and Alex Alderton, Roberto Amato, Sonia Goncalves, Ewan Harrison, David K. Jackson, Ian Johnston, Dominic Kwiatkowski, Cordelia Langford, John Sillitoe on behalf of the Wellcome Sanger Institute COVID-19 Surveillance Team |
| EPI_ISL_811360, EPI_ISL_811361 | Lighthouse Lab in Glasgow | Wellcome Sanger Institute for the COVID-19 Genomics UK (COG-UK) Consortium | Harper VanSteenhouse, Yumi Kasai, David Gray, Carol Clugston, Anna Dominiczak and Alex Alderton, Roberto Amato, Sonia Goncalves, Ewan Harrison, David K. Jackson, Ian Johnston, Dominic Kwiatkowski, Cordelia Langford, John Sillitoe on behalf of the Wellcome Sanger Institute COVID-19 Surveillance Team |
| EPI_ISL_811362 | Lighthouse Lab in Milton Keynes | Wellcome Sanger Institute for the COVID-19 Genomics UK (COG-UK) Consortium | The Lighthouse Lab in Milton Keynes and Alex Alderton, Roberto Amato, Sonia Goncalves, Ewan Harrison, David K. Jackson, Ian Johnston, Dominic Kwiatkowski, Cordelia Langford, John Sillitoe on behalf of the Wellcome Sanger Institute COVID-19 Surveillance Team |
| EPI_ISL_811363 | Lighthouse Lab in Glasgow | Wellcome Sanger Institute for the COVID-19 Genomics UK (COG-UK) Consortium | Harper VanSteenhouse, Yumi Kasai, David Gray, Carol Clugston, Anna Dominiczak and Alex Alderton, Roberto Amato, Sonia Goncalves, Ewan Harrison, David K. Jackson, Ian Johnston, Dominic Kwiatkowski, Cordelia Langford, John Sillitoe on behalf of the Wellcome Sanger Institute COVID-19 Surveillance Team |
| EPI_ISL_811367, EPI_ISL_811368, EPI_ISL_811369 | Lighthouse Lab in Milton Keynes | Wellcome Sanger Institute for the COVID-19 Genomics UK (COG-UK) Consortium | The Lighthouse Lab in Milton Keynes and Alex Alderton, Roberto Amato, Sonia Goncalves, Ewan Harrison, David K. Jackson, Ian Johnston, Dominic Kwiatkowski, Cordelia Langford, John Sillitoe on behalf of the Wellcome Sanger Institute COVID-19 Surveillance Team |
| EPI_ISL_811370, EPI_ISL_811371, EPI_ISL_811372 | Lighthouse Lab in Glasgow | Wellcome Sanger Institute for the COVID-19 Genomics UK (COG-UK) Consortium | Harper VanSteenhouse, Yumi Kasai, David Gray, Carol Clugston, Anna Dominiczak and Alex Alderton, Roberto Amato, Sonia Goncalves, Ewan Harrison, David K. Jackson, Ian Johnston, Dominic Kwiatkowski, Cordelia Langford, John Sillitoe on behalf of the Wellcome Sanger Institute COVID-19 Surveillance Team |
| EPI_ISL_811373 | Lighthouse Lab in Milton Keynes | Wellcome Sanger Institute for the COVID-19 Genomics UK (COG-UK) Consortium | The Lighthouse Lab in Milton Keynes and Alex Alderton, Roberto Amato, Sonia Goncalves, Ewan Harrison, David K. Jackson, Ian Johnston, Dominic Kwiatkowski, Cordelia Langford, John Sillitoe on behalf of the Wellcome Sanger Institute COVID-19 Surveillance Team |
| EPI_ISL_811375, EPI_ISL_811378, EPI_ISL_811379 | Lighthouse Lab in Glasgow | Wellcome Sanger Institute for the COVID-19 Genomics UK (COG-UK) Consortium | Harper VanSteenhouse, Yumi Kasai, David Gray, Carol Clugston, Anna Dominiczak and Alex Alderton, Roberto Amato, Sonia Goncalves, Ewan Harrison, David K. Jackson, Ian Johnston, Dominic Kwiatkowski, Cordelia Langford, John Sillitoe on behalf of the Wellcome Sanger Institute COVID-19 Surveillance Team |
| EPI_ISL_811380, EPI_ISL_811381, EPI_ISL_811382, EPI_ISL_811383, EPI_ISL_811385 | Lighthouse Lab in Milton Keynes | Wellcome Sanger Institute for the COVID-19 Genomics UK (COG-UK) Consortium | The Lighthouse Lab in Milton Keynes and Alex Alderton, Roberto Amato, Sonia Goncalves, Ewan Harrison, David K. Jackson, Ian Johnston, Dominic Kwiatkowski, Cordelia Langford, John Sillitoe on behalf of the Wellcome Sanger Institute COVID-19 Surveillance Team |
| EPI_ISL_811386 | Lighthouse Lab in Glasgow | Wellcome Sanger Institute for the COVID-19 Genomics UK (COG-UK) Consortium | Harper VanSteenhouse, Yumi Kasai, David Gray, Carol Clugston, Anna Dominiczak and Alex Alderton, Roberto Amato, Sonia Goncalves, Ewan Harrison, David K. Jackson, Ian Johnston, Dominic Kwiatkowski, Cordelia Langford, John Sillitoe on behalf of the Wellcome Sanger Institute COVID-19 Surveillance Team |
| EPI_ISL_811389 | Lighthouse Lab in Milton Keynes | Wellcome Sanger Institute for the COVID-19 Genomics UK (COG-UK) Consortium | The Lighthouse Lab in Milton Keynes and Alex Alderton, Roberto Amato, Sonia Goncalves, Ewan Harrison, David K. Jackson, Ian Johnston, Dominic Kwiatkowski, Cordelia Langford, John Sillitoe on behalf of the Wellcome Sanger Institute COVID-19 Surveillance Team |
| EPI_ISL_811390 | Lighthouse Lab in Glasgow | Wellcome Sanger Institute for the COVID-19 Genomics UK (COG-UK) Consortium | Harper VanSteenhouse, Yumi Kasai, David Gray, Carol Clugston, Anna Dominiczak and Alex Alderton, Roberto Amato, Sonia Goncalves, Ewan Harrison, David K. Jackson, Ian Johnston, Dominic Kwiatkowski, Cordelia Langford, John Sillitoe on behalf of the Wellcome Sanger Institute COVID-19 Surveillance Team |
| EPI_ISL_811392 | Lighthouse Lab in Milton Keynes | Wellcome Sanger Institute for the COVID-19 Genomics UK (COG-UK) Consortium | The Lighthouse Lab in Milton Keynes and Alex Alderton, Roberto Amato, Sonia Goncalves, Ewan Harrison, David K. Jackson, Ian Johnston, Dominic Kwiatkowski, Cordelia Langford, John Sillitoe on behalf of the Wellcome Sanger Institute COVID-19 Surveillance Team |
| EPI_ISL_811393 | Lighthouse Lab in Glasgow | Wellcome Sanger Institute for the COVID-19 Genomics UK (COG-UK) Consortium | Harper VanSteenhouse, Yumi Kasai, David Gray, Carol Clugston, Anna Dominiczak and Alex Alderton, Roberto Amato, Sonia Goncalves, Ewan Harrison, David K. Jackson, Ian Johnston, Dominic Kwiatkowski, Cordelia Langford, John Sillitoe on behalf of the Wellcome Sanger Institute COVID-19 Surveillance Team |
| EPI_ISL_811394, EPI_ISL_811395 | Lighthouse Lab in Milton Keynes | Wellcome Sanger Institute for the COVID-19 Genomics UK (COG-UK) Consortium | The Lighthouse Lab in Milton Keynes and Alex Alderton, Roberto Amato, Sonia Goncalves, Ewan Harrison, David K. Jackson, Ian Johnston, Dominic Kwiatkowski, Cordelia Langford, John Sillitoe on behalf of the Wellcome Sanger Institute COVID-19 Surveillance Team |
| EPI_ISL_811396 | Lighthouse Lab in Glasgow | Wellcome Sanger Institute for the COVID-19 Genomics UK (COG-UK) Consortium | Harper VanSteenhouse, Yumi Kasai, David Gray, Carol Clugston, Anna Dominiczak and Alex Alderton, Roberto Amato, Sonia Goncalves, Ewan Harrison, David K. Jackson, Ian Johnston, Dominic Kwiatkowski, Cordelia Langford, John Sillitoe on behalf of the Wellcome Sanger Institute COVID-19 Surveillance Team |
| EPI_ISL_811397 | Lighthouse Lab in Milton Keynes | Wellcome Sanger Institute for the COVID-19 Genomics UK (COG-UK) Consortium | The Lighthouse Lab in Milton Keynes and Alex Alderton, Roberto Amato, Sonia Goncalves, Ewan Harrison, David K. Jackson, Ian Johnston, Dominic Kwiatkowski, Cordelia Langford, John Sillitoe on behalf of the Wellcome Sanger Institute COVID-19 Surveillance Team |
| EPI_ISL_811398, EPI_ISL_811399 | Lighthouse Lab in Glasgow | Wellcome Sanger Institute for the COVID-19 Genomics UK (COG-UK) Consortium | Harper VanSteenhouse, Yumi Kasai, David Gray, Carol Clugston, Anna Dominiczak and Alex Alderton, Roberto Amato, Sonia Goncalves, Ewan Harrison, David K. Jackson, Ian Johnston, Dominic Kwiatkowski, Cordelia Langford, John Sillitoe on behalf of the Wellcome Sanger Institute COVID-19 Surveillance Team |
| EPI_ISL_811400 | Lighthouse Lab in Milton Keynes | Wellcome Sanger Institute for the COVID-19 Genomics UK (COG-UK) Consortium | The Lighthouse Lab in Milton Keynes and Alex Alderton, Roberto Amato, Sonia Goncalves, Ewan Harrison, David K. Jackson, Ian Johnston, Dominic Kwiatkowski, Cordelia Langford, John Sillitoe on behalf of the Wellcome Sanger Institute COVID-19 Surveillance Team |
| EPI_ISL_811401, EPI_ISL_811402, EPI_ISL_811403 | Lighthouse Lab in Glasgow | Wellcome Sanger Institute for the COVID-19 Genomics UK (COG-UK) Consortium | Harper VanSteenhouse, Yumi Kasai, David Gray, Carol Clugston, Anna Dominiczak and Alex Alderton, Roberto Amato, Sonia Goncalves, Ewan Harrison, David K. Jackson, Ian Johnston, Dominic Kwiatkowski, Cordelia Langford, John Sillitoe on behalf of the Wellcome Sanger Institute COVID-19 Surveillance Team |
| EPI_ISL_811404, EPI_ISL_811406 | Lighthouse Lab in Milton Keynes | Wellcome Sanger Institute for the COVID-19 Genomics UK (COG-UK) Consortium | The Lighthouse Lab in Milton Keynes and Alex Alderton, Roberto Amato, Sonia Goncalves, Ewan Harrison, David K. Jackson, Ian Johnston, Dominic Kwiatkowski, Cordelia Langford, John Sillitoe on behalf of the Wellcome Sanger Institute COVID-19 Surveillance Team |
| EPI_ISL_811407, EPI_ISL_811408 | Lighthouse Lab in Glasgow | Wellcome Sanger Institute for the COVID-19 Genomics UK (COG-UK) Consortium | Harper VanSteenhouse, Yumi Kasai, David Gray, Carol Clugston, Anna Dominiczak and Alex Alderton, Roberto Amato, Sonia Goncalves, Ewan Harrison, David K. Jackson, Ian Johnston, Dominic Kwiatkowski, Cordelia Langford, John Sillitoe on behalf of the Wellcome Sanger Institute COVID-19 Surveillance Team |
| EPI_ISL_811409, EPI_ISL_811410 | Lighthouse Lab in Milton Keynes | Wellcome Sanger Institute for the COVID-19 Genomics UK (COG-UK) Consortium | The Lighthouse Lab in Milton Keynes and Alex Alderton, Roberto Amato, Sonia Goncalves, Ewan Harrison, David K. Jackson, Ian Johnston, Dominic Kwiatkowski, Cordelia Langford, John Sillitoe on behalf of the Wellcome Sanger Institute COVID-19 Surveillance Team |
| EPI_ISL_811411, EPI_ISL_811412, EPI_ISL_811413, EPI_ISL_811416, EPI_ISL_811418, EPI_ISL_811419 | Lighthouse Lab in Glasgow | Wellcome Sanger Institute for the COVID-19 Genomics UK (COG-UK) Consortium | Harper VanSteenhouse, Yumi Kasai, David Gray, Carol Clugston, Anna Dominiczak and Alex Alderton, Roberto Amato, Sonia Goncalves, Ewan Harrison, David K. Jackson, Ian Johnston, Dominic Kwiatkowski, Cordelia Langford, John Sillitoe on behalf of the Wellcome Sanger Institute COVID-19 Surveillance Team |
| EPI_ISL_811420 | Lighthouse Lab in Milton Keynes | Wellcome Sanger Institute for the COVID-19 Genomics UK | The Lighthouse Lab in Milton Keynes and Alex Alderton, Roberto Amato, Sonia Goncalves, Ewan Harrison, David K. Jackson, Ian Johnston, Dominic |

| | | | |
|---|---|---|---|
| | | (COG-UK) Consortium | Kwiatkowski, Cordelia Langford, John Sillitoe on behalf of the Wellcome Sanger Institute COVID-19 Surveillance Team |
| EPI_ISL_811421, EPI_ISL_811425, EPI_ISL_811428 | Lighthouse Lab in Glasgow | Wellcome Sanger Institute for the COVID-19 Genomics UK (COG-UK) Consortium | Harper VanSteenhouse, Yumi Kasai, David Gray, Carol Clugston, Anna Dominiczak and Alex Alderton, Roberto Amato, Sonia Goncalves, Ewan Harrison, David K. Jackson, Ian Johnston, Dominic Kwiatkowski, Cordelia Langford, John Sillitoe on behalf of the Wellcome Sanger Institute COVID-19 Surveillance Team |
| EPI_ISL_811429, EPI_ISL_811430, EPI_ISL_811431, EPI_ISL_811433 | Lighthouse Lab in Milton Keynes | Wellcome Sanger Institute for the COVID-19 Genomics UK (COG-UK) Consortium | The Lighthouse Lab in Milton Keynes and Alex Alderton, Roberto Amato, Sonia Goncalves, Ewan Harrison, David K. Jackson, Ian Johnston, Dominic Kwiatkowski, Cordelia Langford, John Sillitoe on behalf of the Wellcome Sanger Institute COVID-19 Surveillance Team |
| EPI_ISL_811434, EPI_ISL_811435 | Lighthouse Lab in Glasgow | Wellcome Sanger Institute for the COVID-19 Genomics UK (COG-UK) Consortium | Harper VanSteenhouse, Yumi Kasai, David Gray, Carol Clugston, Anna Dominiczak and Alex Alderton, Roberto Amato, Sonia Goncalves, Ewan Harrison, David K. Jackson, Ian Johnston, Dominic Kwiatkowski, Cordelia Langford, John Sillitoe on behalf of the Wellcome Sanger Institute COVID-19 Surveillance Team |
| EPI_ISL_811436 | Lighthouse Lab in Milton Keynes | Wellcome Sanger Institute for the COVID-19 Genomics UK (COG-UK) Consortium | The Lighthouse Lab in Milton Keynes and Alex Alderton, Roberto Amato, Sonia Goncalves, Ewan Harrison, David K. Jackson, Ian Johnston, Dominic Kwiatkowski, Cordelia Langford, John Sillitoe on behalf of the Wellcome Sanger Institute COVID-19 Surveillance Team |
| EPI_ISL_811437, EPI_ISL_811438, EPI_ISL_811439, EPI_ISL_811440, EPI_ISL_811441, EPI_ISL_811442, EPI_ISL_811443 | Lighthouse Lab in Glasgow | Wellcome Sanger Institute for the COVID-19 Genomics UK (COG-UK) Consortium | Harper VanSteenhouse, Yumi Kasai, David Gray, Carol Clugston, Anna Dominiczak and Alex Alderton, Roberto Amato, Sonia Goncalves, Ewan Harrison, David K. Jackson, Ian Johnston, Dominic Kwiatkowski, Cordelia Langford, John Sillitoe on behalf of the Wellcome Sanger Institute COVID-19 Surveillance Team |
| EPI_ISL_811444, EPI_ISL_811445 | Lighthouse Lab in Milton Keynes | Wellcome Sanger Institute for the COVID-19 Genomics UK (COG-UK) Consortium | The Lighthouse Lab in Milton Keynes and Alex Alderton, Roberto Amato, Sonia Goncalves, Ewan Harrison, David K. Jackson, Ian Johnston, Dominic Kwiatkowski, Cordelia Langford, John Sillitoe on behalf of the Wellcome Sanger Institute COVID-19 Surveillance Team |
| EPI_ISL_811446, EPI_ISL_811447, EPI_ISL_811448, EPI_ISL_811449, EPI_ISL_811450, EPI_ISL_811451 | Lighthouse Lab in Glasgow | Wellcome Sanger Institute for the COVID-19 Genomics UK (COG-UK) Consortium | Harper VanSteenhouse, Yumi Kasai, David Gray, Carol Clugston, Anna Dominiczak and Alex Alderton, Roberto Amato, Sonia Goncalves, Ewan Harrison, David K. Jackson, Ian Johnston, Dominic Kwiatkowski, Cordelia Langford, John Sillitoe on behalf of the Wellcome Sanger Institute COVID-19 Surveillance Team |
| EPI_ISL_811452, EPI_ISL_811454, EPI_ISL_811455, EPI_ISL_811456 | Lighthouse Lab in Milton Keynes | Wellcome Sanger Institute for the COVID-19 Genomics UK (COG-UK) Consortium | The Lighthouse Lab in Milton Keynes and Alex Alderton, Roberto Amato, Sonia Goncalves, Ewan Harrison, David K. Jackson, Ian Johnston, Dominic Kwiatkowski, Cordelia Langford, John Sillitoe on behalf of the Wellcome Sanger Institute COVID-19 Surveillance Team |
| EPI_ISL_811457, EPI_ISL_811458, EPI_ISL_811459, EPI_ISL_811460, EPI_ISL_811461, EPI_ISL_811462, EPI_ISL_811463, EPI_ISL_811464, EPI_ISL_811465, EPI_ISL_811466, EPI_ISL_811467, EPI_ISL_811468 | | | |
| see above | Lighthouse Lab in Cambridge | Wellcome Sanger Institute for the COVID-19 Genomics UK (COG-UK) Consortium | Rob Howes, The Lighthouse Lab in Cambridge and Alex Alderton, Roberto Amato, Sonia Goncalves, Ewan Harrison, David K. Jackson, Ian Johnston, Dominic Kwiatkowski, Cordelia Langford, John Sillitoe on behalf of the Wellcome Sanger Institute COVID-19 Surveillance Team |
| EPI_ISL_811469 | Lighthouse Lab in Milton Keynes | Wellcome Sanger Institute for the COVID-19 Genomics UK (COG-UK) Consortium | The Lighthouse Lab in Milton Keynes and Alex Alderton, Roberto Amato, Sonia Goncalves, Ewan Harrison, David K. Jackson, Ian Johnston, Dominic Kwiatkowski, Cordelia Langford, John Sillitoe on behalf of the Wellcome Sanger Institute COVID-19 Surveillance Team |
| EPI_ISL_811470, EPI_ISL_811471, EPI_ISL_811473, EPI_ISL_811474, EPI_ISL_811475, EPI_ISL_811476, EPI_ISL_811477 | Lighthouse Lab in Cambridge | Wellcome Sanger Institute for the COVID-19 Genomics UK (COG-UK) Consortium | Rob Howes, The Lighthouse Lab in Cambridge and Alex Alderton, Roberto Amato, Sonia Goncalves, Ewan Harrison, David K. Jackson, Ian Johnston, Dominic Kwiatkowski, Cordelia Langford, John Sillitoe on behalf of the Wellcome Sanger Institute COVID-19 Surveillance Team |
| EPI_ISL_811478 | Lighthouse Lab in Milton Keynes | Wellcome Sanger Institute for the COVID-19 Genomics UK (COG-UK) Consortium | The Lighthouse Lab in Milton Keynes and Alex Alderton, Roberto Amato, Sonia Goncalves, Ewan Harrison, David K. Jackson, Ian Johnston, Dominic Kwiatkowski, Cordelia Langford, John Sillitoe on behalf of the Wellcome Sanger Institute COVID-19 Surveillance Team |
| EPI_ISL_811479, EPI_ISL_811480, EPI_ISL_811481 | Lighthouse Lab in Cambridge | Wellcome Sanger Institute for the COVID-19 Genomics UK (COG-UK) Consortium | Rob Howes, The Lighthouse Lab in Cambridge and Alex Alderton, Roberto Amato, Sonia Goncalves, Ewan Harrison, David K. Jackson, Ian Johnston, Dominic Kwiatkowski, Cordelia Langford, John Sillitoe on behalf of the Wellcome Sanger Institute COVID-19 Surveillance Team |
| EPI_ISL_811482 | Lighthouse Lab in Milton Keynes | Wellcome Sanger Institute for the COVID-19 Genomics UK (COG-UK) Consortium | The Lighthouse Lab in Milton Keynes and Alex Alderton, Roberto Amato, Sonia Goncalves, Ewan Harrison, David K. Jackson, Ian Johnston, Dominic Kwiatkowski, Cordelia Langford, John Sillitoe on behalf of the Wellcome Sanger Institute COVID-19 Surveillance Team |
| EPI_ISL_811483 | Lighthouse Lab in Cambridge | Wellcome Sanger Institute for the COVID-19 Genomics UK (COG-UK) Consortium | Rob Howes, The Lighthouse Lab in Cambridge and Alex Alderton, Roberto Amato, Sonia Goncalves, Ewan Harrison, David K. Jackson, Ian Johnston, Dominic Kwiatkowski, Cordelia Langford, John Sillitoe on behalf of the Wellcome Sanger Institute COVID-19 Surveillance Team |
| EPI_ISL_811484 | Lighthouse Lab in Milton Keynes | Wellcome Sanger Institute for the COVID-19 Genomics UK (COG-UK) Consortium | The Lighthouse Lab in Milton Keynes and Alex Alderton, Roberto Amato, Sonia Goncalves, Ewan Harrison, David K. Jackson, Ian Johnston, Dominic Kwiatkowski, Cordelia Langford, John Sillitoe on behalf of the Wellcome Sanger Institute COVID-19 Surveillance Team |
| EPI_ISL_811485, EPI_ISL_811486 | Lighthouse Lab in Cambridge | Wellcome Sanger Institute for the COVID-19 Genomics UK (COG-UK) Consortium | Rob Howes, The Lighthouse Lab in Cambridge and Alex Alderton, Roberto Amato, Sonia Goncalves, Ewan Harrison, David K. Jackson, Ian Johnston, Dominic Kwiatkowski, Cordelia Langford, John Sillitoe on behalf of the Wellcome Sanger Institute COVID-19 Surveillance Team |
| EPI_ISL_811487 | Lighthouse Lab in Milton Keynes | Wellcome Sanger Institute for the COVID-19 Genomics UK (COG-UK) Consortium | The Lighthouse Lab in Milton Keynes and Alex Alderton, Roberto Amato, Sonia Goncalves, Ewan Harrison, David K. Jackson, Ian Johnston, Dominic Kwiatkowski, Cordelia Langford, John Sillitoe on behalf of the Wellcome Sanger Institute COVID-19 Surveillance Team |
| EPI_ISL_811488, EPI_ISL_811489, EPI_ISL_811491, EPI_ISL_811492, EPI_ISL_811493, EPI_ISL_811495, EPI_ISL_811496, EPI_ISL_811497 | Lighthouse Lab in Cambridge | Wellcome Sanger Institute for the COVID-19 Genomics UK (COG-UK) Consortium | Rob Howes, The Lighthouse Lab in Cambridge and Alex Alderton, Roberto Amato, Sonia Goncalves, Ewan Harrison, David K. Jackson, Ian Johnston, Dominic Kwiatkowski, Cordelia Langford, John Sillitoe on behalf of the Wellcome Sanger Institute COVID-19 Surveillance Team |
| EPI_ISL_811498 | Lighthouse Lab in Milton Keynes | Wellcome Sanger Institute for the COVID-19 Genomics UK (COG-UK) Consortium | The Lighthouse Lab in Milton Keynes and Alex Alderton, Roberto Amato, Sonia Goncalves, Ewan Harrison, David K. Jackson, Ian Johnston, Dominic Kwiatkowski, Cordelia Langford, John Sillitoe on behalf of the Wellcome Sanger Institute COVID-19 Surveillance Team |
| EPI_ISL_811499, EPI_ISL_811501 | Lighthouse Lab in Cambridge | Wellcome Sanger Institute for the COVID-19 Genomics UK (COG-UK) Consortium | Rob Howes, The Lighthouse Lab in Cambridge and Alex Alderton, Roberto Amato, Sonia Goncalves, Ewan Harrison, David K. Jackson, Ian Johnston, Dominic Kwiatkowski, Cordelia Langford, John Sillitoe on behalf of the Wellcome Sanger Institute COVID-19 Surveillance Team |
| EPI_ISL_811502 | Lighthouse Lab in Milton Keynes | Wellcome Sanger Institute for the COVID-19 Genomics UK (COG-UK) Consortium | The Lighthouse Lab in Milton Keynes and Alex Alderton, Roberto Amato, Sonia Goncalves, Ewan Harrison, David K. Jackson, Ian Johnston, Dominic Kwiatkowski, Cordelia Langford, John Sillitoe on behalf of the Wellcome Sanger Institute COVID-19 Surveillance Team |
| EPI_ISL_811503, EPI_ISL_811504, EPI_ISL_811505, EPI_ISL_811506, EPI_ISL_811507, EPI_ISL_811508, EPI_ISL_811509, EPI_ISL_811510 | Lighthouse Lab in Cambridge | Wellcome Sanger Institute for the COVID-19 Genomics UK (COG-UK) Consortium | Rob Howes, The Lighthouse Lab in Cambridge and Alex Alderton, Roberto Amato, Sonia Goncalves, Ewan Harrison, David K. Jackson, Ian Johnston, Dominic Kwiatkowski, Cordelia Langford, John Sillitoe on behalf of the Wellcome Sanger Institute COVID-19 Surveillance Team |
| EPI_ISL_811511 | Lighthouse Lab in Milton Keynes | Wellcome Sanger Institute for the COVID-19 Genomics UK (COG-UK) Consortium | The Lighthouse Lab in Milton Keynes and Alex Alderton, Roberto Amato, Sonia Goncalves, Ewan Harrison, David K. Jackson, Ian Johnston, Dominic Kwiatkowski, Cordelia Langford, John Sillitoe on behalf of the Wellcome Sanger Institute COVID-19 Surveillance Team |
| EPI_ISL_811512, EPI_ISL_811513, EPI_ISL_811514, EPI_ISL_811515, EPI_ISL_811517, EPI_ISL_811518, EPI_ISL_811520, EPI_ISL_811521, EPI_ISL_811522, EPI_ISL_811523 | Lighthouse Lab in Cambridge | Wellcome Sanger Institute for the COVID-19 Genomics UK (COG-UK) Consortium | Rob Howes, The Lighthouse Lab in Cambridge and Alex Alderton, Roberto Amato, Sonia Goncalves, Ewan Harrison, David K. Jackson, Ian Johnston, Dominic Kwiatkowski, Cordelia Langford, John Sillitoe on behalf of the Wellcome Sanger Institute COVID-19 Surveillance Team |
| EPI_ISL_811525 | Lighthouse Lab in Milton Keynes | Wellcome Sanger Institute for the COVID-19 Genomics UK (COG-UK) Consortium | The Lighthouse Lab in Milton Keynes and Alex Alderton, Roberto Amato, Sonia Goncalves, Ewan Harrison, David K. Jackson, Ian Johnston, Dominic Kwiatkowski, Cordelia Langford, John Sillitoe on behalf of the Wellcome Sanger Institute COVID-19 Surveillance Team |
| EPI_ISL_811526, EPI_ISL_811527, EPI_ISL_811528, EPI_ISL_811529, EPI_ISL_811531, EPI_ISL_811532, EPI_ISL_811533, EPI_ISL_811534, EPI_ISL_811535, EPI_ISL_811536, EPI_ISL_811537 | | | |
| see above | Lighthouse Lab in Cambridge | Wellcome Sanger Institute for the COVID-19 Genomics UK (COG-UK) Consortium | Rob Howes, The Lighthouse Lab in Cambridge and Alex Alderton, Roberto Amato, Sonia Goncalves, Ewan Harrison, David K. Jackson, Ian Johnston, Dominic Kwiatkowski, Cordelia Langford, John Sillitoe on behalf of the Wellcome Sanger Institute COVID-19 Surveillance Team |
| EPI_ISL_811538 | Lighthouse Lab in Milton Keynes | Wellcome Sanger Institute for the COVID-19 Genomics UK (COG-UK) Consortium | The Lighthouse Lab in Milton Keynes and Alex Alderton, Roberto Amato, Sonia Goncalves, Ewan Harrison, David K. Jackson, Ian Johnston, Dominic Kwiatkowski, Cordelia Langford, John Sillitoe on behalf of the Wellcome Sanger Institute COVID-19 Surveillance Team |

| | | | |
|---|---|---|---|
| EPI_ISL_811539, EPI_ISL_811540 | Lighthouse Lab in Cambridge | Wellcome Sanger Institute for the COVID-19 Genomics UK (COG-UK) Consortium | Rob Howes, The Lighthouse Lab in Cambridge and Alex Alderton, Roberto Amato, Sonia Goncalves, Ewan Harrison, David K. Jackson, Ian Johnston, Dominic Kwiatkowski, Cordelia Langford, John Sillitoe on behalf of the Wellcome Sanger Institute COVID-19 Surveillance Team |
| EPI_ISL_811541 | Lighthouse Lab in Milton Keynes | Wellcome Sanger Institute for the COVID-19 Genomics UK (COG-UK) Consortium | The Lighthouse Lab in Milton Keynes and Alex Alderton, Roberto Amato, Sonia Goncalves, Ewan Harrison, David K. Jackson, Ian Johnston, Dominic Kwiatkowski, Cordelia Langford, John Sillitoe on behalf of the Wellcome Sanger Institute COVID-19 Surveillance Team |
| EPI_ISL_811542, EPI_ISL_811543, EPI_ISL_811544 | Lighthouse Lab in Cambridge | Wellcome Sanger Institute for the COVID-19 Genomics UK (COG-UK) Consortium | Rob Howes, The Lighthouse Lab in Cambridge and Alex Alderton, Roberto Amato, Sonia Goncalves, Ewan Harrison, David K. Jackson, Ian Johnston, Dominic Kwiatkowski, Cordelia Langford, John Sillitoe on behalf of the Wellcome Sanger Institute COVID-19 Surveillance Team |
| EPI_ISL_811545 | Lighthouse Lab in Milton Keynes | Wellcome Sanger Institute for the COVID-19 Genomics UK (COG-UK) Consortium | The Lighthouse Lab in Milton Keynes and Alex Alderton, Roberto Amato, Sonia Goncalves, Ewan Harrison, David K. Jackson, Ian Johnston, Dominic Kwiatkowski, Cordelia Langford, John Sillitoe on behalf of the Wellcome Sanger Institute COVID-19 Surveillance Team |
| EPI_ISL_811546, EPI_ISL_811547 | Lighthouse Lab in Cambridge | Wellcome Sanger Institute for the COVID-19 Genomics UK (COG-UK) Consortium | Rob Howes, The Lighthouse Lab in Cambridge and Alex Alderton, Roberto Amato, Sonia Goncalves, Ewan Harrison, David K. Jackson, Ian Johnston, Dominic Kwiatkowski, Cordelia Langford, John Sillitoe on behalf of the Wellcome Sanger Institute COVID-19 Surveillance Team |
| EPI_ISL_811549 | Lighthouse Lab in Milton Keynes | Wellcome Sanger Institute for the COVID-19 Genomics UK (COG-UK) Consortium | The Lighthouse Lab in Milton Keynes and Alex Alderton, Roberto Amato, Sonia Goncalves, Ewan Harrison, David K. Jackson, Ian Johnston, Dominic Kwiatkowski, Cordelia Langford, John Sillitoe on behalf of the Wellcome Sanger Institute COVID-19 Surveillance Team |
| EPI_ISL_811550, EPI_ISL_811551, EPI_ISL_811552, EPI_ISL_811553, EPI_ISL_811554, EPI_ISL_811555 | Lighthouse Lab in Cambridge | Wellcome Sanger Institute for the COVID-19 Genomics UK (COG-UK) Consortium | Rob Howes, The Lighthouse Lab in Cambridge and Alex Alderton, Roberto Amato, Sonia Goncalves, Ewan Harrison, David K. Jackson, Ian Johnston, Dominic Kwiatkowski, Cordelia Langford, John Sillitoe on behalf of the Wellcome Sanger Institute COVID-19 Surveillance Team |
| EPI_ISL_811556 | Lighthouse Lab in Milton Keynes | Wellcome Sanger Institute for the COVID-19 Genomics UK (COG-UK) Consortium | The Lighthouse Lab in Milton Keynes and Alex Alderton, Roberto Amato, Sonia Goncalves, Ewan Harrison, David K. Jackson, Ian Johnston, Dominic Kwiatkowski, Cordelia Langford, John Sillitoe on behalf of the Wellcome Sanger Institute COVID-19 Surveillance Team |
| EPI_ISL_811557, EPI_ISL_811558 | Lighthouse Lab in Cambridge | Wellcome Sanger Institute for the COVID-19 Genomics UK (COG-UK) Consortium | Rob Howes, The Lighthouse Lab in Cambridge and Alex Alderton, Roberto Amato, Sonia Goncalves, Ewan Harrison, David K. Jackson, Ian Johnston, Dominic Kwiatkowski, Cordelia Langford, John Sillitoe on behalf of the Wellcome Sanger Institute COVID-19 Surveillance Team |
| EPI_ISL_811559 | Lighthouse Lab in Milton Keynes | Wellcome Sanger Institute for the COVID-19 Genomics UK (COG-UK) Consortium | The Lighthouse Lab in Milton Keynes and Alex Alderton, Roberto Amato, Sonia Goncalves, Ewan Harrison, David K. Jackson, Ian Johnston, Dominic Kwiatkowski, Cordelia Langford, John Sillitoe on behalf of the Wellcome Sanger Institute COVID-19 Surveillance Team |
| EPI_ISL_811560, EPI_ISL_811561, EPI_ISL_811562, EPI_ISL_811564 | Lighthouse Lab in Cambridge | Wellcome Sanger Institute for the COVID-19 Genomics UK (COG-UK) Consortium | Rob Howes, The Lighthouse Lab in Cambridge and Alex Alderton, Roberto Amato, Sonia Goncalves, Ewan Harrison, David K. Jackson, Ian Johnston, Dominic Kwiatkowski, Cordelia Langford, John Sillitoe on behalf of the Wellcome Sanger Institute COVID-19 Surveillance Team |
| EPI_ISL_811565 | Lighthouse Lab in Milton Keynes | Wellcome Sanger Institute for the COVID-19 Genomics UK (COG-UK) Consortium | The Lighthouse Lab in Milton Keynes and Alex Alderton, Roberto Amato, Sonia Goncalves, Ewan Harrison, David K. Jackson, Ian Johnston, Dominic Kwiatkowski, Cordelia Langford, John Sillitoe on behalf of the Wellcome Sanger Institute COVID-19 Surveillance Team |
| EPI_ISL_811566, EPI_ISL_811567, EPI_ISL_811568 | Lighthouse Lab in Cambridge | Wellcome Sanger Institute for the COVID-19 Genomics UK (COG-UK) Consortium | Rob Howes, The Lighthouse Lab in Cambridge and Alex Alderton, Roberto Amato, Sonia Goncalves, Ewan Harrison, David K. Jackson, Ian Johnston, Dominic Kwiatkowski, Cordelia Langford, John Sillitoe on behalf of the Wellcome Sanger Institute COVID-19 Surveillance Team |
| EPI_ISL_811569 | Lighthouse Lab in Milton Keynes | Wellcome Sanger Institute for the COVID-19 Genomics UK (COG-UK) Consortium | The Lighthouse Lab in Milton Keynes and Alex Alderton, Roberto Amato, Sonia Goncalves, Ewan Harrison, David K. Jackson, Ian Johnston, Dominic Kwiatkowski, Cordelia Langford, John Sillitoe on behalf of the Wellcome Sanger Institute COVID-19 Surveillance Team |
| EPI_ISL_811570, EPI_ISL_811571, EPI_ISL_811572, EPI_ISL_811574, EPI_ISL_811575, EPI_ISL_811576, EPI_ISL_811577, EPI_ISL_811578, EPI_ISL_811579, EPI_ISL_811581, EPI_ISL_811582, EPI_ISL_811583, EPI_ISL_811585, EPI_ISL_811586, EPI_ISL_811587, EPI_ISL_811588, EPI_ISL_811589, EPI_ISL_811590 | | | |
| see above | Lighthouse Lab in Cambridge | Wellcome Sanger Institute for the COVID-19 Genomics UK (COG-UK) Consortium | Rob Howes, The Lighthouse Lab in Cambridge and Alex Alderton, Roberto Amato, Sonia Goncalves, Ewan Harrison, David K. Jackson, Ian Johnston, Dominic Kwiatkowski, Cordelia Langford, John Sillitoe on behalf of the Wellcome Sanger Institute COVID-19 Surveillance Team |
| EPI_ISL_811591 | Lighthouse Lab in Milton Keynes | Wellcome Sanger Institute for the COVID-19 Genomics UK (COG-UK) Consortium | The Lighthouse Lab in Milton Keynes and Alex Alderton, Roberto Amato, Sonia Goncalves, Ewan Harrison, David K. Jackson, Ian Johnston, Dominic Kwiatkowski, Cordelia Langford, John Sillitoe on behalf of the Wellcome Sanger Institute COVID-19 Surveillance Team |
| EPI_ISL_811592, EPI_ISL_811593, EPI_ISL_811595, EPI_ISL_811596, EPI_ISL_811597, EPI_ISL_811598, EPI_ISL_811599, EPI_ISL_811600, EPI_ISL_811601, EPI_ISL_811602 | Lighthouse Lab in Cambridge | Wellcome Sanger Institute for the COVID-19 Genomics UK (COG-UK) Consortium | Rob Howes, The Lighthouse Lab in Cambridge and Alex Alderton, Roberto Amato, Sonia Goncalves, Ewan Harrison, David K. Jackson, Ian Johnston, Dominic Kwiatkowski, Cordelia Langford, John Sillitoe on behalf of the Wellcome Sanger Institute COVID-19 Surveillance Team |
| EPI_ISL_811603 | Lighthouse Lab in Milton Keynes | Wellcome Sanger Institute for the COVID-19 Genomics UK (COG-UK) Consortium | The Lighthouse Lab in Milton Keynes and Alex Alderton, Roberto Amato, Sonia Goncalves, Ewan Harrison, David K. Jackson, Ian Johnston, Dominic Kwiatkowski, Cordelia Langford, John Sillitoe on behalf of the Wellcome Sanger Institute COVID-19 Surveillance Team |
| EPI_ISL_811604, EPI_ISL_811605 | Lighthouse Lab in Cambridge | Wellcome Sanger Institute for the COVID-19 Genomics UK (COG-UK) Consortium | Rob Howes, The Lighthouse Lab in Cambridge and Alex Alderton, Roberto Amato, Sonia Goncalves, Ewan Harrison, David K. Jackson, Ian Johnston, Dominic Kwiatkowski, Cordelia Langford, John Sillitoe on behalf of the Wellcome Sanger Institute COVID-19 Surveillance Team |
| EPI_ISL_811606 | Lighthouse Lab in Milton Keynes | Wellcome Sanger Institute for the COVID-19 Genomics UK (COG-UK) Consortium | The Lighthouse Lab in Milton Keynes and Alex Alderton, Roberto Amato, Sonia Goncalves, Ewan Harrison, David K. Jackson, Ian Johnston, Dominic Kwiatkowski, Cordelia Langford, John Sillitoe on behalf of the Wellcome Sanger Institute COVID-19 Surveillance Team |
| EPI_ISL_811607, EPI_ISL_811608, EPI_ISL_811609 | Lighthouse Lab in Cambridge | Wellcome Sanger Institute for the COVID-19 Genomics UK (COG-UK) Consortium | Rob Howes, The Lighthouse Lab in Cambridge and Alex Alderton, Roberto Amato, Sonia Goncalves, Ewan Harrison, David K. Jackson, Ian Johnston, Dominic Kwiatkowski, Cordelia Langford, John Sillitoe on behalf of the Wellcome Sanger Institute COVID-19 Surveillance Team |
| EPI_ISL_811610, EPI_ISL_811611 | Lighthouse Lab in Milton Keynes | Wellcome Sanger Institute for the COVID-19 Genomics UK (COG-UK) Consortium | The Lighthouse Lab in Milton Keynes and Alex Alderton, Roberto Amato, Sonia Goncalves, Ewan Harrison, David K. Jackson, Ian Johnston, Dominic Kwiatkowski, Cordelia Langford, John Sillitoe on behalf of the Wellcome Sanger Institute COVID-19 Surveillance Team |
| EPI_ISL_811612, EPI_ISL_811613, EPI_ISL_811614, EPI_ISL_811615, EPI_ISL_811616, EPI_ISL_811619 | Lighthouse Lab in Cambridge | Wellcome Sanger Institute for the COVID-19 Genomics UK (COG-UK) Consortium | Rob Howes, The Lighthouse Lab in Cambridge and Alex Alderton, Roberto Amato, Sonia Goncalves, Ewan Harrison, David K. Jackson, Ian Johnston, Dominic Kwiatkowski, Cordelia Langford, John Sillitoe on behalf of the Wellcome Sanger Institute COVID-19 Surveillance Team |
| EPI_ISL_811622 | Lighthouse Lab in Milton Keynes | Wellcome Sanger Institute for the COVID-19 Genomics UK (COG-UK) Consortium | The Lighthouse Lab in Milton Keynes and Alex Alderton, Roberto Amato, Sonia Goncalves, Ewan Harrison, David K. Jackson, Ian Johnston, Dominic Kwiatkowski, Cordelia Langford, John Sillitoe on behalf of the Wellcome Sanger Institute COVID-19 Surveillance Team |
| EPI_ISL_811623, EPI_ISL_811624 | Lighthouse Lab in Cambridge | Wellcome Sanger Institute for the COVID-19 Genomics UK (COG-UK) Consortium | Rob Howes, The Lighthouse Lab in Cambridge and Alex Alderton, Roberto Amato, Sonia Goncalves, Ewan Harrison, David K. Jackson, Ian Johnston, Dominic Kwiatkowski, Cordelia Langford, John Sillitoe on behalf of the Wellcome Sanger Institute COVID-19 Surveillance Team |
| EPI_ISL_811625, EPI_ISL_811626 | Lighthouse Lab in Milton Keynes | Wellcome Sanger Institute for the COVID-19 Genomics UK (COG-UK) Consortium | The Lighthouse Lab in Milton Keynes and Alex Alderton, Roberto Amato, Sonia Goncalves, Ewan Harrison, David K. Jackson, Ian Johnston, Dominic Kwiatkowski, Cordelia Langford, John Sillitoe on behalf of the Wellcome Sanger Institute COVID-19 Surveillance Team |
| EPI_ISL_811628, EPI_ISL_811629, EPI_ISL_811630, EPI_ISL_811631, EPI_ISL_811632, EPI_ISL_811633 | Lighthouse Lab in Cambridge | Wellcome Sanger Institute for the COVID-19 Genomics UK (COG-UK) Consortium | Rob Howes, The Lighthouse Lab in Cambridge and Alex Alderton, Roberto Amato, Sonia Goncalves, Ewan Harrison, David K. Jackson, Ian Johnston, Dominic Kwiatkowski, Cordelia Langford, John Sillitoe on behalf of the Wellcome Sanger Institute COVID-19 Surveillance Team |
| EPI_ISL_811634 | Lighthouse Lab in Milton Keynes | Wellcome Sanger Institute for the COVID-19 Genomics UK (COG-UK) Consortium | The Lighthouse Lab in Milton Keynes and Alex Alderton, Roberto Amato, Sonia Goncalves, Ewan Harrison, David K. Jackson, Ian Johnston, Dominic Kwiatkowski, Cordelia Langford, John Sillitoe on behalf of the Wellcome Sanger Institute COVID-19 Surveillance Team |
| EPI_ISL_811635, EPI_ISL_811636, EPI_ISL_811637, EPI_ISL_811638, EPI_ISL_811639, EPI_ISL_811640, EPI_ISL_811641, EPI_ISL_811642, EPI_ISL_811643 | Lighthouse Lab in Cambridge | Wellcome Sanger Institute for the COVID-19 Genomics UK (COG-UK) Consortium | Rob Howes, The Lighthouse Lab in Cambridge and Alex Alderton, Roberto Amato, Sonia Goncalves, Ewan Harrison, David K. Jackson, Ian Johnston, Dominic Kwiatkowski, Cordelia Langford, John Sillitoe on behalf of the Wellcome Sanger Institute COVID-19 Surveillance Team |
| EPI_ISL_811644 | Lighthouse Lab in Milton Keynes | Wellcome Sanger Institute for the COVID-19 Genomics UK (COG-UK) Consortium | The Lighthouse Lab in Milton Keynes and Alex Alderton, Roberto Amato, Sonia Goncalves, Ewan Harrison, David K. Jackson, Ian Johnston, Dominic Kwiatkowski, Cordelia Langford, John Sillitoe on behalf of the Wellcome Sanger Institute COVID-19 Surveillance Team |
| EPI_ISL_811646, EPI_ISL_811647, EPI_ISL_811648, EPI_ISL_811649, | Lighthouse Lab in Cambridge | Wellcome Sanger Institute for the COVID-19 Genomics UK (COG-UK) Consortium | Rob Howes, The Lighthouse Lab in Cambridge and Alex Alderton, Roberto Amato, Sonia Goncalves, Ewan Harrison, David K. Jackson, Ian Johnston, Dominic Kwiatkowski, Cordelia Langford, John Sillitoe on behalf of the Wellcome Sanger Institute COVID-19 Surveillance Team |

| | | | |
|---|---|---|---|
| EPI_ISL_811652, EPI_ISL_811653, EPI_ISL_811654, EPI_ISL_811655, EPI_ISL_811656 | | | |
| EPI_ISL_811657 | Lighthouse Lab in Milton Keynes | Wellcome Sanger Institute for the COVID-19 Genomics UK (COG-UK) Consortium | The Lighthouse Lab in Milton Keynes and Alex Alderton, Roberto Amato, Sonia Goncalves, Ewan Harrison, David K. Jackson, Ian Johnston, Dominic Kwiatkowski, Cordelia Langford, John Sillitoe on behalf of the Wellcome Sanger Institute COVID-19 Surveillance Team |
| EPI_ISL_811658, EPI_ISL_811659 | Lighthouse Lab in Cambridge | Wellcome Sanger Institute for the COVID-19 Genomics UK (COG-UK) Consortium | Rob Howes, The Lighthouse Lab in Cambridge and Alex Alderton, Roberto Amato, Sonia Goncalves, Ewan Harrison, David K. Jackson, Ian Johnston, Dominic Kwiatkowski, Cordelia Langford, John Sillitoe on behalf of the Wellcome Sanger Institute COVID-19 Surveillance Team |
| EPI_ISL_811660 | Lighthouse Lab in Milton Keynes | Wellcome Sanger Institute for the COVID-19 Genomics UK (COG-UK) Consortium | The Lighthouse Lab in Milton Keynes and Alex Alderton, Roberto Amato, Sonia Goncalves, Ewan Harrison, David K. Jackson, Ian Johnston, Dominic Kwiatkowski, Cordelia Langford, John Sillitoe on behalf of the Wellcome Sanger Institute COVID-19 Surveillance Team |
| EPI_ISL_811662, EPI_ISL_811663 | Lighthouse Lab in Cambridge | Wellcome Sanger Institute for the COVID-19 Genomics UK (COG-UK) Consortium | Rob Howes, The Lighthouse Lab in Cambridge and Alex Alderton, Roberto Amato, Sonia Goncalves, Ewan Harrison, David K. Jackson, Ian Johnston, Dominic Kwiatkowski, Cordelia Langford, John Sillitoe on behalf of the Wellcome Sanger Institute COVID-19 Surveillance Team |
| EPI_ISL_811664 | Lighthouse Lab in Milton Keynes | Wellcome Sanger Institute for the COVID-19 Genomics UK (COG-UK) Consortium | The Lighthouse Lab in Milton Keynes and Alex Alderton, Roberto Amato, Sonia Goncalves, Ewan Harrison, David K. Jackson, Ian Johnston, Dominic Kwiatkowski, Cordelia Langford, John Sillitoe on behalf of the Wellcome Sanger Institute COVID-19 Surveillance Team |
| EPI_ISL_811665 | Lighthouse Lab in Cambridge | Wellcome Sanger Institute for the COVID-19 Genomics UK (COG-UK) Consortium | Rob Howes, The Lighthouse Lab in Cambridge and Alex Alderton, Roberto Amato, Sonia Goncalves, Ewan Harrison, David K. Jackson, Ian Johnston, Dominic Kwiatkowski, Cordelia Langford, John Sillitoe on behalf of the Wellcome Sanger Institute COVID-19 Surveillance Team |
| EPI_ISL_811666 | Lighthouse Lab in Milton Keynes | Wellcome Sanger Institute for the COVID-19 Genomics UK (COG-UK) Consortium | The Lighthouse Lab in Milton Keynes and Alex Alderton, Roberto Amato, Sonia Goncalves, Ewan Harrison, David K. Jackson, Ian Johnston, Dominic Kwiatkowski, Cordelia Langford, John Sillitoe on behalf of the Wellcome Sanger Institute COVID-19 Surveillance Team |
| EPI_ISL_811667, EPI_ISL_811668 | Lighthouse Lab in Cambridge | Wellcome Sanger Institute for the COVID-19 Genomics UK (COG-UK) Consortium | Rob Howes, The Lighthouse Lab in Cambridge and Alex Alderton, Roberto Amato, Sonia Goncalves, Ewan Harrison, David K. Jackson, Ian Johnston, Dominic Kwiatkowski, Cordelia Langford, John Sillitoe on behalf of the Wellcome Sanger Institute COVID-19 Surveillance Team |
| EPI_ISL_811672, EPI_ISL_811673 | Lighthouse Lab in Milton Keynes | Wellcome Sanger Institute for the COVID-19 Genomics UK (COG-UK) Consortium | The Lighthouse Lab in Milton Keynes and Alex Alderton, Roberto Amato, Sonia Goncalves, Ewan Harrison, David K. Jackson, Ian Johnston, Dominic Kwiatkowski, Cordelia Langford, John Sillitoe on behalf of the Wellcome Sanger Institute COVID-19 Surveillance Team |
| EPI_ISL_811674, EPI_ISL_811675, EPI_ISL_811676, EPI_ISL_811679, EPI_ISL_811680, EPI_ISL_811681, EPI_ISL_811683, EPI_ISL_811684, EPI_ISL_811685, EPI_ISL_811686, EPI_ISL_811687, EPI_ISL_811689, EPI_ISL_811690, EPI_ISL_811691, EPI_ISL_811692, EPI_ISL_811693, EPI_ISL_811694, EPI_ISL_811695 | | | |
| see above | Lighthouse Lab in Cambridge | Wellcome Sanger Institute for the COVID-19 Genomics UK (COG-UK) Consortium | Rob Howes, The Lighthouse Lab in Cambridge and Alex Alderton, Roberto Amato, Sonia Goncalves, Ewan Harrison, David K. Jackson, Ian Johnston, Dominic Kwiatkowski, Cordelia Langford, John Sillitoe on behalf of the Wellcome Sanger Institute COVID-19 Surveillance Team |
| EPI_ISL_811696 | Lighthouse Lab in Milton Keynes | Wellcome Sanger Institute for the COVID-19 Genomics UK (COG-UK) Consortium | The Lighthouse Lab in Milton Keynes and Alex Alderton, Roberto Amato, Sonia Goncalves, Ewan Harrison, David K. Jackson, Ian Johnston, Dominic Kwiatkowski, Cordelia Langford, John Sillitoe on behalf of the Wellcome Sanger Institute COVID-19 Surveillance Team |
| EPI_ISL_811697 | Lighthouse Lab in Cambridge | Wellcome Sanger Institute for the COVID-19 Genomics UK (COG-UK) Consortium | Rob Howes, The Lighthouse Lab in Cambridge and Alex Alderton, Roberto Amato, Sonia Goncalves, Ewan Harrison, David K. Jackson, Ian Johnston, Dominic Kwiatkowski, Cordelia Langford, John Sillitoe on behalf of the Wellcome Sanger Institute COVID-19 Surveillance Team |
| EPI_ISL_811698 | Lighthouse Lab in Milton Keynes | Wellcome Sanger Institute for the COVID-19 Genomics UK (COG-UK) Consortium | The Lighthouse Lab in Milton Keynes and Alex Alderton, Roberto Amato, Sonia Goncalves, Ewan Harrison, David K. Jackson, Ian Johnston, Dominic Kwiatkowski, Cordelia Langford, John Sillitoe on behalf of the Wellcome Sanger Institute COVID-19 Surveillance Team |
| EPI_ISL_811699, EPI_ISL_811700 | Lighthouse Lab in Cambridge | Wellcome Sanger Institute for the COVID-19 Genomics UK (COG-UK) Consortium | Rob Howes, The Lighthouse Lab in Cambridge and Alex Alderton, Roberto Amato, Sonia Goncalves, Ewan Harrison, David K. Jackson, Ian Johnston, Dominic Kwiatkowski, Cordelia Langford, John Sillitoe on behalf of the Wellcome Sanger Institute COVID-19 Surveillance Team |
| EPI_ISL_811701, EPI_ISL_811702, EPI_ISL_811703 | Lighthouse Lab in Milton Keynes | Wellcome Sanger Institute for the COVID-19 Genomics UK (COG-UK) Consortium | The Lighthouse Lab in Milton Keynes and Alex Alderton, Roberto Amato, Sonia Goncalves, Ewan Harrison, David K. Jackson, Ian Johnston, Dominic Kwiatkowski, Cordelia Langford, John Sillitoe on behalf of the Wellcome Sanger Institute COVID-19 Surveillance Team |
| EPI_ISL_811704, EPI_ISL_811705, EPI_ISL_811706, EPI_ISL_811707 | Lighthouse Lab in Cambridge | Wellcome Sanger Institute for the COVID-19 Genomics UK (COG-UK) Consortium | Rob Howes, The Lighthouse Lab in Cambridge and Alex Alderton, Roberto Amato, Sonia Goncalves, Ewan Harrison, David K. Jackson, Ian Johnston, Dominic Kwiatkowski, Cordelia Langford, John Sillitoe on behalf of the Wellcome Sanger Institute COVID-19 Surveillance Team |
| EPI_ISL_811708 | Lighthouse Lab in Milton Keynes | Wellcome Sanger Institute for the COVID-19 Genomics UK (COG-UK) Consortium | The Lighthouse Lab in Milton Keynes and Alex Alderton, Roberto Amato, Sonia Goncalves, Ewan Harrison, David K. Jackson, Ian Johnston, Dominic Kwiatkowski, Cordelia Langford, John Sillitoe on behalf of the Wellcome Sanger Institute COVID-19 Surveillance Team |
| EPI_ISL_811709, EPI_ISL_811710, EPI_ISL_811711, EPI_ISL_811712, EPI_ISL_811714, EPI_ISL_811715, EPI_ISL_811716, EPI_ISL_811717, EPI_ISL_811718, EPI_ISL_811719, EPI_ISL_811721, EPI_ISL_811722, EPI_ISL_811723, EPI_ISL_811724, EPI_ISL_811725 | | | |
| see above | Lighthouse Lab in Cambridge | Wellcome Sanger Institute for the COVID-19 Genomics UK (COG-UK) Consortium | Rob Howes, The Lighthouse Lab in Cambridge and Alex Alderton, Roberto Amato, Sonia Goncalves, Ewan Harrison, David K. Jackson, Ian Johnston, Dominic Kwiatkowski, Cordelia Langford, John Sillitoe on behalf of the Wellcome Sanger Institute COVID-19 Surveillance Team |
| EPI_ISL_811726 | Lighthouse Lab in Milton Keynes | Wellcome Sanger Institute for the COVID-19 Genomics UK (COG-UK) Consortium | The Lighthouse Lab in Milton Keynes and Alex Alderton, Roberto Amato, Sonia Goncalves, Ewan Harrison, David K. Jackson, Ian Johnston, Dominic Kwiatkowski, Cordelia Langford, John Sillitoe on behalf of the Wellcome Sanger Institute COVID-19 Surveillance Team |
| EPI_ISL_811728, EPI_ISL_811729, EPI_ISL_811730, EPI_ISL_811731, EPI_ISL_811732, EPI_ISL_811733, EPI_ISL_811734, EPI_ISL_811735 | Lighthouse Lab in Cambridge | Wellcome Sanger Institute for the COVID-19 Genomics UK (COG-UK) Consortium | Rob Howes, The Lighthouse Lab in Cambridge and Alex Alderton, Roberto Amato, Sonia Goncalves, Ewan Harrison, David K. Jackson, Ian Johnston, Dominic Kwiatkowski, Cordelia Langford, John Sillitoe on behalf of the Wellcome Sanger Institute COVID-19 Surveillance Team |
| EPI_ISL_811736 | Lighthouse Lab in Milton Keynes | Wellcome Sanger Institute for the COVID-19 Genomics UK (COG-UK) Consortium | The Lighthouse Lab in Milton Keynes and Alex Alderton, Roberto Amato, Sonia Goncalves, Ewan Harrison, David K. Jackson, Ian Johnston, Dominic Kwiatkowski, Cordelia Langford, John Sillitoe on behalf of the Wellcome Sanger Institute COVID-19 Surveillance Team |
| EPI_ISL_811737, EPI_ISL_811738, EPI_ISL_811739 | Lighthouse Lab in Cambridge | Wellcome Sanger Institute for the COVID-19 Genomics UK (COG-UK) Consortium | Rob Howes, The Lighthouse Lab in Cambridge and Alex Alderton, Roberto Amato, Sonia Goncalves, Ewan Harrison, David K. Jackson, Ian Johnston, Dominic Kwiatkowski, Cordelia Langford, John Sillitoe on behalf of the Wellcome Sanger Institute COVID-19 Surveillance Team |
| EPI_ISL_811740, EPI_ISL_811741 | Lighthouse Lab in Milton Keynes | Wellcome Sanger Institute for the COVID-19 Genomics UK (COG-UK) Consortium | The Lighthouse Lab in Milton Keynes and Alex Alderton, Roberto Amato, Sonia Goncalves, Ewan Harrison, David K. Jackson, Ian Johnston, Dominic Kwiatkowski, Cordelia Langford, John Sillitoe on behalf of the Wellcome Sanger Institute COVID-19 Surveillance Team |
| EPI_ISL_811742, EPI_ISL_811743, EPI_ISL_811744, EPI_ISL_811745, EPI_ISL_811746, EPI_ISL_811747, EPI_ISL_811748, EPI_ISL_811749, EPI_ISL_811751, EPI_ISL_811752 | Lighthouse Lab in Cambridge | Wellcome Sanger Institute for the COVID-19 Genomics UK (COG-UK) Consortium | Rob Howes, The Lighthouse Lab in Cambridge and Alex Alderton, Roberto Amato, Sonia Goncalves, Ewan Harrison, David K. Jackson, Ian Johnston, Dominic Kwiatkowski, Cordelia Langford, John Sillitoe on behalf of the Wellcome Sanger Institute COVID-19 Surveillance Team |
| EPI_ISL_811754 | Lighthouse Lab in Milton Keynes | Wellcome Sanger Institute for the COVID-19 Genomics UK (COG-UK) Consortium | The Lighthouse Lab in Milton Keynes and Alex Alderton, Roberto Amato, Sonia Goncalves, Ewan Harrison, David K. Jackson, Ian Johnston, Dominic Kwiatkowski, Cordelia Langford, John Sillitoe on behalf of the Wellcome Sanger Institute COVID-19 Surveillance Team |
| EPI_ISL_811755, EPI_ISL_811757, EPI_ISL_811758, EPI_ISL_811760, EPI_ISL_811761 | Lighthouse Lab in Cambridge | Wellcome Sanger Institute for the COVID-19 Genomics UK (COG-UK) Consortium | Rob Howes, The Lighthouse Lab in Cambridge and Alex Alderton, Roberto Amato, Sonia Goncalves, Ewan Harrison, David K. Jackson, Ian Johnston, Dominic Kwiatkowski, Cordelia Langford, John Sillitoe on behalf of the Wellcome Sanger Institute COVID-19 Surveillance Team |
| EPI_ISL_811762 | Lighthouse Lab in Milton Keynes | Wellcome Sanger Institute for the COVID-19 Genomics UK (COG-UK) Consortium | The Lighthouse Lab in Milton Keynes and Alex Alderton, Roberto Amato, Sonia Goncalves, Ewan Harrison, David K. Jackson, Ian Johnston, Dominic Kwiatkowski, Cordelia Langford, John Sillitoe on behalf of the Wellcome Sanger Institute COVID-19 Surveillance Team |
| EPI_ISL_811763, EPI_ISL_811764, EPI_ISL_811765, EPI_ISL_811766, EPI_ISL_811768, EPI_ISL_811769, EPI_ISL_811770, EPI_ISL_811771, EPI_ISL_811772, EPI_ISL_811773, EPI_ISL_811774, EPI_ISL_811775 | | | |
| see above | Lighthouse Lab in Cambridge | Wellcome Sanger Institute for the COVID-19 Genomics UK (COG-UK) Consortium | Rob Howes, The Lighthouse Lab in Cambridge and Alex Alderton, Roberto Amato, Sonia Goncalves, Ewan Harrison, David K. Jackson, Ian Johnston, Dominic Kwiatkowski, Cordelia Langford, John Sillitoe on behalf of the Wellcome Sanger Institute COVID-19 Surveillance Team |
| EPI_ISL_811776 | Lighthouse Lab in Milton Keynes | Wellcome Sanger Institute for the COVID-19 Genomics UK (COG-UK) Consortium | The Lighthouse Lab in Milton Keynes and Alex Alderton, Roberto Amato, Sonia Goncalves, Ewan Harrison, David K. Jackson, Ian Johnston, Dominic Kwiatkowski, Cordelia Langford, John Sillitoe on behalf of the Wellcome Sanger Institute COVID-19 Surveillance Team |
| EPI_ISL_811777, EPI_ISL_811778, | Lighthouse Lab in Cambridge | Wellcome Sanger Institute for the COVID-19 Genomics UK | Rob Howes, The Lighthouse Lab in Cambridge and Alex Alderton, Roberto Amato, Sonia Goncalves, Ewan Harrison, David K. Jackson, Ian Johnston, |

| | | | |
|---|---|---|---|
| EPI_ISL_811779, EPI_ISL_811780, EPI_ISL_811781, EPI_ISL_811782, EPI_ISL_811783, EPI_ISL_811784, EPI_ISL_811786 | | (COG-UK) Consortium | Dominic Kwiatkowski, Cordelia Langford, John Sillitoe on behalf of the Wellcome Sanger Institute COVID-19 Surveillance Team |
| EPI_ISL_811787 | Lighthouse Lab in Milton Keynes | Wellcome Sanger Institute for the COVID-19 Genomics UK (COG-UK) Consortium | The Lighthouse Lab in Milton Keynes and Alex Alderton, Roberto Amato, Sonia Goncalves, Ewan Harrison, David K. Jackson, Ian Johnston, Dominic Kwiatkowski, Cordelia Langford, John Sillitoe on behalf of the Wellcome Sanger Institute COVID-19 Surveillance Team |
| EPI_ISL_811788, EPI_ISL_811789, EPI_ISL_811790, EPI_ISL_811791, EPI_ISL_811792, EPI_ISL_811793, EPI_ISL_811794, EPI_ISL_811795, EPI_ISL_811796, EPI_ISL_811797, EPI_ISL_811798, EPI_ISL_811799, EPI_ISL_811800, EPI_ISL_811801 | | | |
| see above | Lighthouse Lab in Cambridge | Wellcome Sanger Institute for the COVID-19 Genomics UK (COG-UK) Consortium | Rob Howes, The Lighthouse Lab in Cambridge and Alex Alderton, Roberto Amato, Sonia Goncalves, Ewan Harrison, David K. Jackson, Ian Johnston, Dominic Kwiatkowski, Cordelia Langford, John Sillitoe on behalf of the Wellcome Sanger Institute COVID-19 Surveillance Team |
| EPI_ISL_811802 | Lighthouse Lab in Milton Keynes | Wellcome Sanger Institute for the COVID-19 Genomics UK (COG-UK) Consortium | The Lighthouse Lab in Milton Keynes and Alex Alderton, Roberto Amato, Sonia Goncalves, Ewan Harrison, David K. Jackson, Ian Johnston, Dominic Kwiatkowski, Cordelia Langford, John Sillitoe on behalf of the Wellcome Sanger Institute COVID-19 Surveillance Team |
| EPI_ISL_811803, EPI_ISL_811804, EPI_ISL_811805, EPI_ISL_811806 | Lighthouse Lab in Cambridge | Wellcome Sanger Institute for the COVID-19 Genomics UK (COG-UK) Consortium | Rob Howes, The Lighthouse Lab in Cambridge and Alex Alderton, Roberto Amato, Sonia Goncalves, Ewan Harrison, David K. Jackson, Ian Johnston, Dominic Kwiatkowski, Cordelia Langford, John Sillitoe on behalf of the Wellcome Sanger Institute COVID-19 Surveillance Team |
| EPI_ISL_811807 | Lighthouse Lab in Milton Keynes | Wellcome Sanger Institute for the COVID-19 Genomics UK (COG-UK) Consortium | The Lighthouse Lab in Milton Keynes and Alex Alderton, Roberto Amato, Sonia Goncalves, Ewan Harrison, David K. Jackson, Ian Johnston, Dominic Kwiatkowski, Cordelia Langford, John Sillitoe on behalf of the Wellcome Sanger Institute COVID-19 Surveillance Team |
| EPI_ISL_812249 | Cantonal Hospital Baden | Institute of Medical Virology, University of Zurich | Stefan Schmutz, Maryam Zaheri, Verena Kufner, Annette Audigé, Maria Grünberg, Kevin Steiner, Jon Huder, Cyril Shah, Riccarda Capaul, Guido Bloemberg, Jürg Böni, Michael Huber, Alexandra Trkola |
| EPI_ISL_812251, EPI_ISL_812252, EPI_ISL_812253, EPI_ISL_812254 | Kantonsarztamt Bern | Institute of Medical Virology, University of Zurich | Stefan Schmutz, Maryam Zaheri, Verena Kufner, Annette Audigé, Maria Grünberg, Kevin Steiner, Jon Huder, Cyril Shah, Riccarda Capaul, Guido Bloemberg, Jürg Böni, Michael Huber, Alexandra Trkola |
| EPI_ISL_812255 | Kantonsärztlicher Dienst Aargau | Institute of Medical Virology, University of Zurich | Stefan Schmutz, Maryam Zaheri, Verena Kufner, Annette Audigé, Maria Grünberg, Kevin Steiner, Jon Huder, Cyril Shah, Riccarda Capaul, Guido Bloemberg, Jürg Böni, Michael Huber, Alexandra Trkola |
| EPI_ISL_812434 | Microbiological Diagnostic Unit - Public Health Laboratory (MDU-PHL) | MDU-PHL | Seemann T., Sait, M.L., Sherry, N.L. |
| EPI_ISL_812435, EPI_ISL_812436, EPI_ISL_812438, EPI_ISL_812439 | Victorian Infectious Diseases Reference Laboratory (VIDRL) | VIDRL and MDU-PHL | Caly L., Seemann T., Sait, M.L., Druce J., Sherry, N.L. |
| EPI_ISL_813199 | Department of Pathology, University of Cambridge | COVID-19 Genomics UK (COG-UK) Consortium | Aminu S. Jahun, Yasmin Chaudhry, Grant Hall, Iliana Georgana, Myra Hosmillo, Martin D. Curran, Malte Pinckert, Surendra Parmar, Ian Goodfellow |
| EPI_ISL_813816, EPI_ISL_813817, EPI_ISL_813818, EPI_ISL_813819, EPI_ISL_813820, EPI_ISL_813821, EPI_ISL_813822, EPI_ISL_813823, EPI_ISL_813824, EPI_ISL_813825, EPI_ISL_813827, EPI_ISL_813828 | | | |
| see above | Liverpool Clinical Laboratories | COVID-19 Genomics UK (COG-UK) Consortium | Sam Haldenby, Anita Lucaci, Steve Paterson, Julian Hiscox, Alistair Darby, M Almsaud, A Alrezaihi, Muhannad Alruwaili, Stuart D Armstrong, Jones Benjamin, Eleanor G Bentley, Anu Chawla, Jordan J Clark, Angela Cowell, Richard Eccles, Isabel García-Dorival, Matthew Gemmell, Alessandro Gerada, PKF Gilmore, Richard Gregory, Ximeng Han, Catherine Hartley, Margaret Hughes, Miren Iturriza-Gomara, James Johnson, L Luu, Jenifer Manson, Charlotte Nelson, Elaine O'Toole, Cassie Olateju, Rebekah Penrice-Randal , Lucille Rainbow, N.P Randle, Trevor Ian Robinson, Parul Sharma, Ghada T Shawli, James P Stewart, Neil Swainston, Ecaterina Vamos, Joanne Watts, Mark Whitehead |
| EPI_ISL_813987, EPI_ISL_813988, EPI_ISL_813996, EPI_ISL_814000, EPI_ISL_814007, EPI_ISL_814011, EPI_ISL_814012, EPI_ISL_814013, EPI_ISL_814017, EPI_ISL_814023, EPI_ISL_814024, EPI_ISL_814025, EPI_ISL_814036, EPI_ISL_814041, EPI_ISL_814042, EPI_ISL_814043, EPI_ISL_814044, EPI_ISL_814045, EPI_ISL_814046, EPI_ISL_814047, EPI_ISL_814048, EPI_ISL_814049, EPI_ISL_814050 | | | |
| see above | Hospital General Universitario Gregorio Marañón | SeqCOVID-SPAIN consortium/IBV(CSIC) | Darío García de Viedma, Laura Pérez-Lago, Marta Herranz, Jon Sicilia, Julia Suárez, Pilar Catalán, Patricia Muñoz and SeqCOVID-SPAIN consortium |
| EPI_ISL_816210, EPI_ISL_816211, EPI_ISL_816212, EPI_ISL_816216, EPI_ISL_816217, EPI_ISL_816218, EPI_ISL_816219 | Centre for Enzyme Innovation, University of Portsmouth / Translational Research Laboratory, Portsmouth Hospitals NHS Trust | COVID-19 Genomics UK (COG-UK) Consortium | Angela Beckett,Yann Bourgeois,Garry Scarlett,Sharon Glaysher,Scott Elliott,Kelly Bicknell,Robert Impey,Allyson Lloyd,Sarah Wyllie,Ethan Butcher,Anoop Chauhan,Samuel Robson |
| EPI_ISL_816237, EPI_ISL_816243, EPI_ISL_816251, EPI_ISL_816260, EPI_ISL_816264, EPI_ISL_816267, EPI_ISL_816281, EPI_ISL_816286, EPI_ISL_816287, EPI_ISL_816293, EPI_ISL_816303, EPI_ISL_816309, EPI_ISL_816312, EPI_ISL_816315, EPI_ISL_816322, EPI_ISL_816326, EPI_ISL_816333, EPI_ISL_816334, EPI_ISL_816336, EPI_ISL_816338, EPI_ISL_816341, EPI_ISL_816355, EPI_ISL_816367, EPI_ISL_816394, EPI_ISL_816395, EPI_ISL_816399, EPI_ISL_816402, EPI_ISL_816408, EPI_ISL_816416, EPI_ISL_816436, EPI_ISL_816439, EPI_ISL_816450, EPI_ISL_816452, EPI_ISL_816458, EPI_ISL_816463, EPI_ISL_816470, EPI_ISL_816471, EPI_ISL_816498, EPI_ISL_816502, EPI_ISL_816509, EPI_ISL_816517, EPI_ISL_816520, EPI_ISL_816523, EPI_ISL_816526, EPI_ISL_816527, EPI_ISL_816543, EPI_ISL_816546, EPI_ISL_816552, EPI_ISL_816564, EPI_ISL_816565, EPI_ISL_816572, EPI_ISL_816573, EPI_ISL_816583, EPI_ISL_816588, EPI_ISL_816590, EPI_ISL_816591, EPI_ISL_816596, EPI_ISL_816603, EPI_ISL_816618, EPI_ISL_816622, EPI_ISL_816624, EPI_ISL_816632, EPI_ISL_816636, EPI_ISL_816649, EPI_ISL_816655 | | | |
| see above | Virology Department, Sheffield Teaching Hospitals NHS Foundation Trust/Department of Infection, Immunity and Cardiovascular Disease, The Medical School, University of Sheffield | COVID-19 Genomics UK (COG-UK) Consortium | Thushan de Silva, Matthew Parker, Nikki Smith, Adri Angyal, Rebecca Brown, Luke Green, Rachel Tucker, Paul Parsons, Danielle Groves, Katie Johnson, Laura Carrilero, Alex Keeley, Dave Partridge, Matthew Wyles, Benjamin Lindsey, Mehmet Yavuz, Mohammad Raza, Cariad Evans |
| EPI_ISL_819267, EPI_ISL_819269, EPI_ISL_819270, EPI_ISL_819271 | Wyoming Public Health Laboratory | Wyoming Public Health Laboratory | Noah Hull, Taylor Fearing, Lynette Gumbleton, Channing Weber, Ashley Norberg, Bailey Bowcutt, and Wanda Manley |
| EPI_ISL_819366, EPI_ISL_819375, EPI_ISL_819377, EPI_ISL_819391, EPI_ISL_819392, EPI_ISL_819393, EPI_ISL_819394, EPI_ISL_819395, EPI_ISL_819405, EPI_ISL_819406, EPI_ISL_819407, EPI_ISL_819408, EPI_ISL_819409, EPI_ISL_819410, EPI_ISL_819411, EPI_ISL_819412, EPI_ISL_819413, EPI_ISL_819415, EPI_ISL_819416, EPI_ISL_819418, EPI_ISL_819419, EPI_ISL_819421, EPI_ISL_819424, EPI_ISL_819425, EPI_ISL_819427, EPI_ISL_819440, EPI_ISL_819441, EPI_ISL_819442, EPI_ISL_819443, EPI_ISL_819444 | | | |
| see above | Quadram Institute Bioscience | COVID-19 Genomics UK (COG-UK) Consortium | Dave J. Baker, Gemma L. Kay, Alp Aydin, Thanh Le-Viet, Steven Rudder, Ana P. Tedim, Anastasia Kolyva, Maria Diaz, Leonardo de Oliveira Martins, Nabil-Fareed Alikhan, Lizzie Meadows, Rachael Stanley, Ngozi Elumogo, Muhammed Yasir, Nicholas M. Thomson, Alexander J Trotter, Rachel Gilroy, Samuel Bloomfield, Claire Stuart, Andrew Bell, Reenesh Prakash, Samir Dervisevic, Alison E. Mather, John Wain, Mark Webber, Andrew J. Page, Justin O'Grady |
| EPI_ISL_820560 | Queens Medical Centre, Clinical Microbiology Department / DeepSeq Nottingham | COVID-19 Genomics UK (COG-UK) Consortium | Gemma Clark, Wendy Smith, Manjinder Khakh, Vicki M Fleming, Michelle M Lister, Hannah Howson-Wells, Jonathan Ball, Patrick McClure, Joseph Chappell, Theocharis Tsoleridis, Nadine Holmes, Matthew Carlisle, Christopher Moore, Fei Sang, Johnny Debebe, Victoria Wright, Matthew Loose |
| EPI_ISL_820563, EPI_ISL_820565, EPI_ISL_820568, EPI_ISL_820570 | Quadram Institute Bioscience | COVID-19 Genomics UK (COG-UK) Consortium | Dave J. Baker, Gemma L. Kay, Alp Aydin, Thanh Le-Viet, Steven Rudder, Ana P. Tedim, Anastasia Kolyva, Maria Diaz, Leonardo de Oliveira Martins, Nabil-Fareed Alikhan, Lizzie Meadows, Rachael Stanley, Ngozi Elumogo, Muhammed Yasir, Nicholas M. Thomson, Alexander J Trotter, Rachel Gilroy, Samuel Bloomfield, Claire Stuart, Andrew Bell, Reenesh Prakash, Samir Dervisevic, Alison E. Mather, John Wain, Mark Webber, Andrew J. Page, Justin O'Grady |
| EPI_ISL_820573, EPI_ISL_820575, EPI_ISL_820578, EPI_ISL_820580, EPI_ISL_820583, EPI_ISL_820586 | Queens Medical Centre, Clinical Microbiology Department / DeepSeq Nottingham | COVID-19 Genomics UK (COG-UK) Consortium | Gemma Clark, Wendy Smith, Manjinder Khakh, Vicki M Fleming, Michelle M Lister, Hannah Howson-Wells, Jonathan Ball, Patrick McClure, Joseph Chappell, Theocharis Tsoleridis, Nadine Holmes, Matthew Carlisle, Christopher Moore, Fei Sang, Johnny Debebe, Victoria Wright, Matthew Loose |
| EPI_ISL_820648, EPI_ISL_820649, EPI_ISL_820650, EPI_ISL_820651, EPI_ISL_820652, EPI_ISL_820653, EPI_ISL_820654, EPI_ISL_820655, EPI_ISL_820656, EPI_ISL_820657, EPI_ISL_820658, EPI_ISL_820659, EPI_ISL_820660, EPI_ISL_820661, EPI_ISL_820662, EPI_ISL_820663, EPI_ISL_820664, EPI_ISL_820665, EPI_ISL_820666, EPI_ISL_820667, EPI_ISL_820668, EPI_ISL_820669, EPI_ISL_820670, EPI_ISL_820671, EPI_ISL_820672, EPI_ISL_820673, EPI_ISL_820674, EPI_ISL_820675, EPI_ISL_820676, EPI_ISL_820677, EPI_ISL_820678, EPI_ISL_820679, EPI_ISL_820680, EPI_ISL_820681, EPI_ISL_820682, EPI_ISL_820683, EPI_ISL_820684, EPI_ISL_820685, EPI_ISL_820686, EPI_ISL_820687, EPI_ISL_820688, EPI_ISL_820689, EPI_ISL_820690, EPI_ISL_820691, EPI_ISL_820692, EPI_ISL_820693, EPI_ISL_820694, EPI_ISL_820695, EPI_ISL_820696, EPI_ISL_820697, EPI_ISL_820698, EPI_ISL_820699, EPI_ISL_820700, EPI_ISL_820701, EPI_ISL_820702, EPI_ISL_820703, EPI_ISL_820704, EPI_ISL_820705, EPI_ISL_820706, EPI_ISL_820707, EPI_ISL_820708, EPI_ISL_820709, EPI_ISL_820710, EPI_ISL_820711, EPI_ISL_820712, EPI_ISL_820713, EPI_ISL_820714, EPI_ISL_820715, EPI_ISL_820716, EPI_ISL_820717, EPI_ISL_820718, EPI_ISL_820719, EPI_ISL_820720, EPI_ISL_820721, EPI_ISL_820722, EPI_ISL_820723, EPI_ISL_820724, EPI_ISL_820725, EPI_ISL_820726, EPI_ISL_820727, EPI_ISL_820728, EPI_ISL_820729, EPI_ISL_820730, EPI_ISL_820731, EPI_ISL_820732, EPI_ISL_820733, EPI_ISL_820734, EPI_ISL_820735, EPI_ISL_820736, EPI_ISL_820737, EPI_ISL_820738, EPI_ISL_820739, EPI_ISL_820740, EPI_ISL_820741, EPI_ISL_820742, EPI_ISL_820743, EPI_ISL_820744, EPI_ISL_820745, EPI_ISL_820746, EPI_ISL_820747, EPI_ISL_820748, EPI_ISL_820749, EPI_ISL_820750, EPI_ISL_820751, EPI_ISL_820752, EPI_ISL_820753, EPI_ISL_820754, EPI_ISL_820755, EPI_ISL_820756, EPI_ISL_820757, EPI_ISL_820758, EPI_ISL_820759, EPI_ISL_820760, EPI_ISL_820761, EPI_ISL_820762, EPI_ISL_820763, EPI_ISL_820764, EPI_ISL_820765, EPI_ISL_820766, EPI_ISL_820767, EPI_ISL_820768, EPI_ISL_820769, EPI_ISL_820770, EPI_ISL_820771, EPI_ISL_820772, EPI_ISL_820773, EPI_ISL_820774, EPI_ISL_820775, EPI_ISL_820776, EPI_ISL_820777, EPI_ISL_820778, EPI_ISL_820779, EPI_ISL_820780, EPI_ISL_820781, EPI_ISL_820782, EPI_ISL_820783, EPI_ISL_820784, EPI_ISL_820785, EPI_ISL_820786, EPI_ISL_820787, EPI_ISL_820788, EPI_ISL_820789, EPI_ISL_820790, EPI_ISL_820791, EPI_ISL_820792, EPI_ISL_820793, EPI_ISL_820794, EPI_ISL_820795, EPI_ISL_820796, EPI_ISL_820797, EPI_ISL_820798, EPI_ISL_820799, EPI_ISL_820800, EPI_ISL_820801, EPI_ISL_820802, EPI_ISL_820803, EPI_ISL_820804, EPI_ISL_820805, EPI_ISL_820806, EPI_ISL_820807, EPI_ISL_820808, EPI_ISL_820809, EPI_ISL_820810, EPI_ISL_820811, EPI_ISL_820812, EPI_ISL_820813, EPI_ISL_820814, EPI_ISL_820815, EPI_ISL_820816, EPI_ISL_820817, EPI_ISL_820818, EPI_ISL_820819, EPI_ISL_820820, EPI_ISL_820821, EPI_ISL_820822, EPI_ISL_820823, EPI_ISL_820824, EPI_ISL_820825, EPI_ISL_820826, EPI_ISL_820827, | | | |

EPI_ISL_820828, EPI_ISL_820829, EPI_ISL_820830, EPI_ISL_820831, EPI_ISL_820832, EPI_ISL_820833, EPI_ISL_820834, EPI_ISL_820835, EPI_ISL_820836, EPI_ISL_820837, EPI_ISL_820838, EPI_ISL_820839, EPI_ISL_820840, EPI_ISL_820841, EPI_ISL_820842, EPI_ISL_820843, EPI_ISL_820844, EPI_ISL_820845, EPI_ISL_820846, EPI_ISL_820847, EPI_ISL_820848, EPI_ISL_820849, EPI_ISL_820850, EPI_ISL_820851, EPI_ISL_820852, EPI_ISL_820853, EPI_ISL_820854, EPI_ISL_820855, EPI_ISL_820856, EPI_ISL_820857, EPI_ISL_820858, EPI_ISL_820859, EPI_ISL_820860, EPI_ISL_820861, EPI_ISL_820862, EPI_ISL_820863, EPI_ISL_820864, EPI_ISL_820865, EPI_ISL_820866, EPI_ISL_820867, EPI_ISL_820868, EPI_ISL_820869, EPI_ISL_820870, EPI_ISL_820871, EPI_ISL_820872, EPI_ISL_820873, EPI_ISL_820874, EPI_ISL_820875, EPI_ISL_820876, EPI_ISL_820877, EPI_ISL_820878, EPI_ISL_820879, EPI_ISL_820880, EPI_ISL_820881, EPI_ISL_820882, EPI_ISL_820883, EPI_ISL_820884, EPI_ISL_820885, EPI_ISL_820886, EPI_ISL_820887, EPI_ISL_820888, EPI_ISL_820889, EPI_ISL_820890, EPI_ISL_820891, EPI_ISL_820892, EPI_ISL_820893, EPI_ISL_820894, EPI_ISL_820895, EPI_ISL_820896, EPI_ISL_820897, EPI_ISL_820898, EPI_ISL_820899, EPI_ISL_820900, EPI_ISL_820901, EPI_ISL_820902, EPI_ISL_820903, EPI_ISL_820904, EPI_ISL_820905, EPI_ISL_820906, EPI_ISL_820907, EPI_ISL_820908, EPI_ISL_820909, EPI_ISL_820910, EPI_ISL_820911, EPI_ISL_820912, EPI_ISL_820913, EPI_ISL_820914, EPI_ISL_820915, EPI_ISL_820916, EPI_ISL_820917, EPI_ISL_820918, EPI_ISL_820919, EPI_ISL_820920, EPI_ISL_820921, EPI_ISL_820922, EPI_ISL_820923, EPI_ISL_820924, EPI_ISL_820925, EPI_ISL_820926, EPI_ISL_820927, EPI_ISL_820928, EPI_ISL_820929, EPI_ISL_820930, EPI_ISL_820931, EPI_ISL_820932, EPI_ISL_820933, EPI_ISL_820934, EPI_ISL_820935, EPI_ISL_820936, EPI_ISL_820937, EPI_ISL_820938, EPI_ISL_820939, EPI_ISL_820940, EPI_ISL_820941, EPI_ISL_820942, EPI_ISL_820943, EPI_ISL_820944, EPI_ISL_820945, EPI_ISL_820946, EPI_ISL_820947, EPI_ISL_820948, EPI_ISL_820949, EPI_ISL_820950, EPI_ISL_820951, EPI_ISL_820952, EPI_ISL_820953, EPI_ISL_820954, EPI_ISL_820955, EPI_ISL_820956, EPI_ISL_820957, EPI_ISL_820958, EPI_ISL_820959, EPI_ISL_820960, EPI_ISL_820961, EPI_ISL_820962, EPI_ISL_820963, EPI_ISL_820964, EPI_ISL_820965, EPI_ISL_820966, EPI_ISL_820967, EPI_ISL_820968, EPI_ISL_820969, EPI_ISL_820970, EPI_ISL_820971, EPI_ISL_820972, EPI_ISL_820973, EPI_ISL_820974, EPI_ISL_820975, EPI_ISL_820976, EPI_ISL_820977, EPI_ISL_820978, EPI_ISL_820979, EPI_ISL_820980, EPI_ISL_820981, EPI_ISL_820982, EPI_ISL_820983, EPI_ISL_820984, EPI_ISL_820985, EPI_ISL_820986, EPI_ISL_820987, EPI_ISL_820988, EPI_ISL_820989, EPI_ISL_820990, EPI_ISL_820991, EPI_ISL_820992, EPI_ISL_820993, EPI_ISL_820994, EPI_ISL_820995, EPI_ISL_820996, EPI_ISL_820997, EPI_ISL_820998, EPI_ISL_820999, EPI_ISL_821000, EPI_ISL_821001

| | | | |
|---|---|---|---|
| see above | Lighthouse Lab in Milton Keynes | Wellcome Sanger Institute for the COVID-19 Genomics UK (COG-UK) Consortium | The Lighthouse Lab in Milton Keynes and Alex Alderton, Roberto Amato, Sonia Goncalves, Ewan Harrison, David K. Jackson, Ian Johnston, Dominic Kwiatkowski, Cordelia Langford, John Sillitoe on behalf of the Wellcome Sanger Institute COVID-19 Surveillance Team |

EPI_ISL_821002, EPI_ISL_821003, EPI_ISL_821004, EPI_ISL_821005, EPI_ISL_821006, EPI_ISL_821007, EPI_ISL_821008, EPI_ISL_821009, EPI_ISL_821010, EPI_ISL_821011, EPI_ISL_821012, EPI_ISL_821013, EPI_ISL_821014, EPI_ISL_821015, EPI_ISL_821016, EPI_ISL_821017, EPI_ISL_821018, EPI_ISL_821019, EPI_ISL_821020, EPI_ISL_821021, EPI_ISL_821022, EPI_ISL_821023, EPI_ISL_821024, EPI_ISL_821025, EPI_ISL_821026, EPI_ISL_821027, EPI_ISL_821028, EPI_ISL_821029, EPI_ISL_821030, EPI_ISL_821031, EPI_ISL_821033, EPI_ISL_821034, EPI_ISL_821035, EPI_ISL_821036, EPI_ISL_821038, EPI_ISL_821040, EPI_ISL_821041, EPI_ISL_821042, EPI_ISL_821043, EPI_ISL_821044, EPI_ISL_821045, EPI_ISL_821046, EPI_ISL_821047, EPI_ISL_821048, EPI_ISL_821049, EPI_ISL_821050, EPI_ISL_821051, EPI_ISL_821052, EPI_ISL_821053, EPI_ISL_821054, EPI_ISL_821055, EPI_ISL_821057, EPI_ISL_821058, EPI_ISL_821059, EPI_ISL_821060, EPI_ISL_821063, EPI_ISL_821064, EPI_ISL_821065, EPI_ISL_821066, EPI_ISL_821067, EPI_ISL_821068, EPI_ISL_821070, EPI_ISL_821071, EPI_ISL_821072, EPI_ISL_821073, EPI_ISL_821074, EPI_ISL_821075, EPI_ISL_821076, EPI_ISL_821078, EPI_ISL_821079, EPI_ISL_821080, EPI_ISL_821081, EPI_ISL_821082, EPI_ISL_821083, EPI_ISL_821084, EPI_ISL_821085, EPI_ISL_821086, EPI_ISL_821087, EPI_ISL_821089, EPI_ISL_821090, EPI_ISL_821091, EPI_ISL_821092, EPI_ISL_821093, EPI_ISL_821094, EPI_ISL_821095, EPI_ISL_821096, EPI_ISL_821097, EPI_ISL_821099, EPI_ISL_821100, EPI_ISL_821101, EPI_ISL_821102, EPI_ISL_821103, EPI_ISL_821104, EPI_ISL_821106, EPI_ISL_821107, EPI_ISL_821108, EPI_ISL_821109, EPI_ISL_821110, EPI_ISL_821111, EPI_ISL_821112, EPI_ISL_821113, EPI_ISL_821114, EPI_ISL_821115, EPI_ISL_821116, EPI_ISL_821117, EPI_ISL_821118, EPI_ISL_821119, EPI_ISL_821120, EPI_ISL_821121, EPI_ISL_821122, EPI_ISL_821123, EPI_ISL_821124, EPI_ISL_821125, EPI_ISL_821126, EPI_ISL_821127, EPI_ISL_821129, EPI_ISL_821130, EPI_ISL_821131, EPI_ISL_821132, EPI_ISL_821133, EPI_ISL_821134, EPI_ISL_821135, EPI_ISL_821136, EPI_ISL_821137, EPI_ISL_821138, EPI_ISL_821141, EPI_ISL_821143, EPI_ISL_821144, EPI_ISL_821145, EPI_ISL_821146, EPI_ISL_821147, EPI_ISL_821148, EPI_ISL_821149, EPI_ISL_821151, EPI_ISL_821152, EPI_ISL_821153, EPI_ISL_821154, EPI_ISL_821155, EPI_ISL_821156, EPI_ISL_821157, EPI_ISL_821158, EPI_ISL_821159, EPI_ISL_821161, EPI_ISL_821162, EPI_ISL_821164, EPI_ISL_821166, EPI_ISL_821167, EPI_ISL_821168, EPI_ISL_821169, EPI_ISL_821170, EPI_ISL_821171, EPI_ISL_821173, EPI_ISL_821174, EPI_ISL_821175, EPI_ISL_821176, EPI_ISL_821177, EPI_ISL_821178, EPI_ISL_821179, EPI_ISL_821180, EPI_ISL_821181, EPI_ISL_821182, EPI_ISL_821183, EPI_ISL_821184, EPI_ISL_821185, EPI_ISL_821186, EPI_ISL_821187, EPI_ISL_821188, EPI_ISL_821189, EPI_ISL_821190, EPI_ISL_821191, EPI_ISL_821192, EPI_ISL_821193, EPI_ISL_821194, EPI_ISL_821196, EPI_ISL_821199, EPI_ISL_821200, EPI_ISL_821201, EPI_ISL_821202, EPI_ISL_821203, EPI_ISL_821204, EPI_ISL_821205, EPI_ISL_821206, EPI_ISL_821208, EPI_ISL_821209, EPI_ISL_821210, EPI_ISL_821211, EPI_ISL_821212, EPI_ISL_821213, EPI_ISL_821214, EPI_ISL_821215, EPI_ISL_821216, EPI_ISL_821217, EPI_ISL_821218, EPI_ISL_821219, EPI_ISL_821221, EPI_ISL_821222, EPI_ISL_821223, EPI_ISL_821224, EPI_ISL_821226, EPI_ISL_821228, EPI_ISL_821229, EPI_ISL_821230, EPI_ISL_821231, EPI_ISL_821232, EPI_ISL_821233, EPI_ISL_821234, EPI_ISL_821236, EPI_ISL_821237, EPI_ISL_821238, EPI_ISL_821239, EPI_ISL_821240, EPI_ISL_821242, EPI_ISL_821243, EPI_ISL_821244, EPI_ISL_821246, EPI_ISL_821247, EPI_ISL_821248, EPI_ISL_821250, EPI_ISL_821251, EPI_ISL_821252, EPI_ISL_821253, EPI_ISL_821254, EPI_ISL_821255, EPI_ISL_821256, EPI_ISL_821259, EPI_ISL_821260, EPI_ISL_821261, EPI_ISL_821262, EPI_ISL_821264, EPI_ISL_821265, EPI_ISL_821267, EPI_ISL_821268, EPI_ISL_821272, EPI_ISL_821273

| | | | |
|---|---|---|---|
| see above | Lighthouse Lab in Alderley Park | Wellcome Sanger Institute for the COVID-19 Genomics UK (COG-UK) Consortium | Jacquelyn Wynn, Mairead Hyland, The Lighthouse Lab in Alderley Park and Alex Alderton, Roberto Amato, Sonia Goncalves, Ewan Harrison, David K. Jackson, Ian Johnston, Dominic Kwiatkowski, Cordelia Langford, John Sillitoe on behalf of the Wellcome Sanger Institute COVID-19 Surveillance Team |
| EPI_ISL_821274 | Lighthouse Lab in Milton Keynes | Wellcome Sanger Institute for the COVID-19 Genomics UK (COG-UK) Consortium | The Lighthouse Lab in Milton Keynes and Alex Alderton, Roberto Amato, Sonia Goncalves, Ewan Harrison, David K. Jackson, Ian Johnston, Dominic Kwiatkowski, Cordelia Langford, John Sillitoe on behalf of the Wellcome Sanger Institute COVID-19 Surveillance Team |
| EPI_ISL_821276, EPI_ISL_821279 | Lighthouse Lab in Cambridge | Wellcome Sanger Institute for the COVID-19 Genomics UK (COG-UK) Consortium | Rob Howes, The Lighthouse Lab in Cambridge and Alex Alderton, Roberto Amato, Sonia Goncalves, Ewan Harrison, David K. Jackson, Ian Johnston, Dominic Kwiatkowski, Cordelia Langford, John Sillitoe on behalf of the Wellcome Sanger Institute COVID-19 Surveillance Team |
| EPI_ISL_821280, EPI_ISL_821281, EPI_ISL_821282 | Lighthouse Lab in Milton Keynes | Wellcome Sanger Institute for the COVID-19 Genomics UK (COG-UK) Consortium | The Lighthouse Lab in Milton Keynes and Alex Alderton, Roberto Amato, Sonia Goncalves, Ewan Harrison, David K. Jackson, Ian Johnston, Dominic Kwiatkowski, Cordelia Langford, John Sillitoe on behalf of the Wellcome Sanger Institute COVID-19 Surveillance Team |
| EPI_ISL_821283 | Lighthouse Lab in Alderley Park | Wellcome Sanger Institute for the COVID-19 Genomics UK (COG-UK) Consortium | Jacquelyn Wynn, Mairead Hyland, The Lighthouse Lab in Alderley Park and Alex Alderton, Roberto Amato, Sonia Goncalves, Ewan Harrison, David K. Jackson, Ian Johnston, Dominic Kwiatkowski, Cordelia Langford, John Sillitoe on behalf of the Wellcome Sanger Institute COVID-19 Surveillance Team |
| EPI_ISL_821284 | Lighthouse Lab in Cambridge | Wellcome Sanger Institute for the COVID-19 Genomics UK (COG-UK) Consortium | Rob Howes, The Lighthouse Lab in Cambridge and Alex Alderton, Roberto Amato, Sonia Goncalves, Ewan Harrison, David K. Jackson, Ian Johnston, Dominic Kwiatkowski, Cordelia Langford, John Sillitoe on behalf of the Wellcome Sanger Institute COVID-19 Surveillance Team |
| EPI_ISL_821285 | Lighthouse Lab in Alderley Park | Wellcome Sanger Institute for the COVID-19 Genomics UK (COG-UK) Consortium | Jacquelyn Wynn, Mairead Hyland, The Lighthouse Lab in Alderley Park and Alex Alderton, Roberto Amato, Sonia Goncalves, Ewan Harrison, David K. Jackson, Ian Johnston, Dominic Kwiatkowski, Cordelia Langford, John Sillitoe on behalf of the Wellcome Sanger Institute COVID-19 Surveillance Team |
| EPI_ISL_821286 | Lighthouse Lab in Cambridge | Wellcome Sanger Institute for the COVID-19 Genomics UK (COG-UK) Consortium | Rob Howes, The Lighthouse Lab in Cambridge and Alex Alderton, Roberto Amato, Sonia Goncalves, Ewan Harrison, David K. Jackson, Ian Johnston, Dominic Kwiatkowski, Cordelia Langford, John Sillitoe on behalf of the Wellcome Sanger Institute COVID-19 Surveillance Team |
| EPI_ISL_821287 | Lighthouse Lab in Milton Keynes | Wellcome Sanger Institute for the COVID-19 Genomics UK (COG-UK) Consortium | The Lighthouse Lab in Milton Keynes and Alex Alderton, Roberto Amato, Sonia Goncalves, Ewan Harrison, David K. Jackson, Ian Johnston, Dominic Kwiatkowski, Cordelia Langford, John Sillitoe on behalf of the Wellcome Sanger Institute COVID-19 Surveillance Team |
| EPI_ISL_821288 | Lighthouse Lab in Alderley Park | Wellcome Sanger Institute for the COVID-19 Genomics UK (COG-UK) Consortium | Jacquelyn Wynn, Mairead Hyland, The Lighthouse Lab in Alderley Park and Alex Alderton, Roberto Amato, Sonia Goncalves, Ewan Harrison, David K. Jackson, Ian Johnston, Dominic Kwiatkowski, Cordelia Langford, John Sillitoe on behalf of the Wellcome Sanger Institute COVID-19 Surveillance Team |
| EPI_ISL_821289, EPI_ISL_821290 | Lighthouse Lab in Cambridge | Wellcome Sanger Institute for the COVID-19 Genomics UK (COG-UK) Consortium | Rob Howes, The Lighthouse Lab in Cambridge and Alex Alderton, Roberto Amato, Sonia Goncalves, Ewan Harrison, David K. Jackson, Ian Johnston, Dominic Kwiatkowski, Cordelia Langford, John Sillitoe on behalf of the Wellcome Sanger Institute COVID-19 Surveillance Team |
| EPI_ISL_821292 | Lighthouse Lab in Alderley Park | Wellcome Sanger Institute for the COVID-19 Genomics UK (COG-UK) Consortium | Jacquelyn Wynn, Mairead Hyland, The Lighthouse Lab in Alderley Park and Alex Alderton, Roberto Amato, Sonia Goncalves, Ewan Harrison, David K. Jackson, Ian Johnston, Dominic Kwiatkowski, Cordelia Langford, John Sillitoe on behalf of the Wellcome Sanger Institute COVID-19 Surveillance Team |
| EPI_ISL_821293 | Lighthouse Lab in Milton Keynes | Wellcome Sanger Institute for the COVID-19 Genomics UK (COG-UK) Consortium | The Lighthouse Lab in Milton Keynes and Alex Alderton, Roberto Amato, Sonia Goncalves, Ewan Harrison, David K. Jackson, Ian Johnston, Dominic Kwiatkowski, Cordelia Langford, John Sillitoe on behalf of the Wellcome Sanger Institute COVID-19 Surveillance Team |
| EPI_ISL_821294 | Lighthouse Lab in Alderley Park | Wellcome Sanger Institute for the COVID-19 Genomics UK (COG-UK) Consortium | Jacquelyn Wynn, Mairead Hyland, The Lighthouse Lab in Alderley Park and Alex Alderton, Roberto Amato, Sonia Goncalves, Ewan Harrison, David K. Jackson, Ian Johnston, Dominic Kwiatkowski, Cordelia Langford, John Sillitoe on behalf of the Wellcome Sanger Institute COVID-19 Surveillance Team |
| EPI_ISL_821295 | Lighthouse Lab in Cambridge | Wellcome Sanger Institute for the COVID-19 Genomics UK (COG-UK) Consortium | Rob Howes, The Lighthouse Lab in Cambridge and Alex Alderton, Roberto Amato, Sonia Goncalves, Ewan Harrison, David K. Jackson, Ian Johnston, Dominic Kwiatkowski, Cordelia Langford, John Sillitoe on behalf of the Wellcome Sanger Institute COVID-19 Surveillance Team |
| EPI_ISL_821296, EPI_ISL_821297 | Lighthouse Lab in Milton Keynes | Wellcome Sanger Institute for the COVID-19 Genomics UK (COG-UK) Consortium | The Lighthouse Lab in Milton Keynes and Alex Alderton, Roberto Amato, Sonia Goncalves, Ewan Harrison, David K. Jackson, Ian Johnston, Dominic Kwiatkowski, Cordelia Langford, John Sillitoe on behalf of the Wellcome Sanger Institute COVID-19 Surveillance Team |
| EPI_ISL_821298 | Lighthouse Lab in Alderley Park | Wellcome Sanger Institute for the COVID-19 Genomics UK (COG-UK) Consortium | Jacquelyn Wynn, Mairead Hyland, The Lighthouse Lab in Alderley Park and Alex Alderton, Roberto Amato, Sonia Goncalves, Ewan Harrison, David K. Jackson, Ian Johnston, Dominic Kwiatkowski, Cordelia Langford, John Sillitoe on behalf of the Wellcome Sanger Institute COVID-19 Surveillance Team |
| EPI_ISL_821300, EPI_ISL_821301 | Lighthouse Lab in Milton Keynes | Wellcome Sanger Institute for the COVID-19 Genomics UK (COG-UK) Consortium | The Lighthouse Lab in Milton Keynes and Alex Alderton, Roberto Amato, Sonia Goncalves, Ewan Harrison, David K. Jackson, Ian Johnston, Dominic Kwiatkowski, Cordelia Langford, John Sillitoe on behalf of the Wellcome Sanger Institute COVID-19 Surveillance Team |
| EPI_ISL_821303 | Lighthouse Lab in Alderley Park | Wellcome Sanger Institute for the COVID-19 Genomics UK (COG-UK) Consortium | Jacquelyn Wynn, Mairead Hyland, The Lighthouse Lab in Alderley Park and Alex Alderton, Roberto Amato, Sonia Goncalves, Ewan Harrison, David K. Jackson, Ian Johnston, Dominic Kwiatkowski, Cordelia Langford, John Sillitoe on behalf of the Wellcome Sanger Institute COVID-19 Surveillance Team |
| EPI_ISL_821304, EPI_ISL_821306, EPI_ISL_821307 | Lighthouse Lab in Milton Keynes | Wellcome Sanger Institute for the COVID-19 Genomics UK (COG-UK) Consortium | The Lighthouse Lab in Milton Keynes and Alex Alderton, Roberto Amato, Sonia Goncalves, Ewan Harrison, David K. Jackson, Ian Johnston, Dominic Kwiatkowski, Cordelia Langford, John Sillitoe on behalf of the Wellcome Sanger Institute COVID-19 Surveillance Team |
| EPI_ISL_821308 | Lighthouse Lab in Alderley Park | Wellcome Sanger Institute for the COVID-19 Genomics UK (COG-UK) Consortium | Jacquelyn Wynn, Mairead Hyland, The Lighthouse Lab in Alderley Park and Alex Alderton, Roberto Amato, Sonia Goncalves, Ewan Harrison, David K. Jackson, Ian Johnston, Dominic Kwiatkowski, Cordelia Langford, John Sillitoe on behalf of the Wellcome Sanger Institute COVID-19 Surveillance Team |
| EPI_ISL_821309, EPI_ISL_821310 | Lighthouse Lab in Cambridge | Wellcome Sanger Institute for the COVID-19 Genomics UK (COG-UK) Consortium | Rob Howes, The Lighthouse Lab in Cambridge and Alex Alderton, Roberto Amato, Sonia Goncalves, Ewan Harrison, David K. Jackson, Ian Johnston, Dominic Kwiatkowski, Cordelia Langford, John Sillitoe on behalf of the Wellcome Sanger Institute COVID-19 Surveillance Team |
| EPI_ISL_821314 | Lighthouse Lab in Alderley Park | Wellcome Sanger Institute for the COVID-19 Genomics UK (COG-UK) Consortium | Jacquelyn Wynn, Mairead Hyland, The Lighthouse Lab in Alderley Park and Alex Alderton, Roberto Amato, Sonia Goncalves, Ewan Harrison, David K. Jackson, Ian Johnston, Dominic Kwiatkowski, Cordelia Langford, John Sillitoe on behalf of the Wellcome Sanger Institute COVID-19 Surveillance Team |
| EPI_ISL_821317, EPI_ISL_821319, EPI_ISL_821323, EPI_ISL_821324 | Lighthouse Lab in Milton Keynes | Wellcome Sanger Institute for the COVID-19 Genomics UK (COG-UK) Consortium | The Lighthouse Lab in Milton Keynes and Alex Alderton, Roberto Amato, Sonia Goncalves, Ewan Harrison, David K. Jackson, Ian Johnston, Dominic Kwiatkowski, Cordelia Langford, John Sillitoe on behalf of the Wellcome Sanger Institute COVID-19 Surveillance Team |
| EPI_ISL_821327 | Lighthouse Lab in Alderley Park | Wellcome Sanger Institute for the COVID-19 Genomics UK | Jacquelyn Wynn, Mairead Hyland, The Lighthouse Lab in Alderley Park and Alex Alderton, Roberto Amato, Sonia Goncalves, Ewan Harrison, David K. |

| | | | |
|---|---|---|---|
| | | (COG-UK) Consortium | Jackson, Ian Johnston, Dominic Kwiatkowski, Cordelia Langford, John Sillitoe on behalf of the Wellcome Sanger Institute COVID-19 Surveillance Team |
| EPI_ISL_821328 | Lighthouse Lab in Milton Keynes | Wellcome Sanger Institute for the COVID-19 Genomics UK (COG-UK) Consortium | The Lighthouse Lab in Milton Keynes and Alex Alderton, Roberto Amato, Sonia Goncalves, Ewan Harrison, David K. Jackson, Ian Johnston, Dominic Kwiatkowski, Cordelia Langford, John Sillitoe on behalf of the Wellcome Sanger Institute COVID-19 Surveillance Team |
| EPI_ISL_821329 | Lighthouse Lab in Alderley Park | Wellcome Sanger Institute for the COVID-19 Genomics UK (COG-UK) Consortium | Jacquelyn Wynn, Mairead Hyland, The Lighthouse Lab in Alderley Park and Alex Alderton, Roberto Amato, Sonia Goncalves, Ewan Harrison, David K. Jackson, Ian Johnston, Dominic Kwiatkowski, Cordelia Langford, John Sillitoe on behalf of the Wellcome Sanger Institute COVID-19 Surveillance Team |
| EPI_ISL_821331 | Lighthouse Lab in Milton Keynes | Wellcome Sanger Institute for the COVID-19 Genomics UK (COG-UK) Consortium | The Lighthouse Lab in Milton Keynes and Alex Alderton, Roberto Amato, Sonia Goncalves, Ewan Harrison, David K. Jackson, Ian Johnston, Dominic Kwiatkowski, Cordelia Langford, John Sillitoe on behalf of the Wellcome Sanger Institute COVID-19 Surveillance Team |
| EPI_ISL_821333 | Lighthouse Lab in Cambridge | Wellcome Sanger Institute for the COVID-19 Genomics UK (COG-UK) Consortium | Rob Howes, The Lighthouse Lab in Cambridge and Alex Alderton, Roberto Amato, Sonia Goncalves, Ewan Harrison, David K. Jackson, Ian Johnston, Dominic Kwiatkowski, Cordelia Langford, John Sillitoe on behalf of the Wellcome Sanger Institute COVID-19 Surveillance Team |
| EPI_ISL_821334 | Lighthouse Lab in Milton Keynes | Wellcome Sanger Institute for the COVID-19 Genomics UK (COG-UK) Consortium | The Lighthouse Lab in Milton Keynes and Alex Alderton, Roberto Amato, Sonia Goncalves, Ewan Harrison, David K. Jackson, Ian Johnston, Dominic Kwiatkowski, Cordelia Langford, John Sillitoe on behalf of the Wellcome Sanger Institute COVID-19 Surveillance Team |
| EPI_ISL_821335 | Lighthouse Lab in Alderley Park | Wellcome Sanger Institute for the COVID-19 Genomics UK (COG-UK) Consortium | Jacquelyn Wynn, Mairead Hyland, The Lighthouse Lab in Alderley Park and Alex Alderton, Roberto Amato, Sonia Goncalves, Ewan Harrison, David K. Jackson, Ian Johnston, Dominic Kwiatkowski, Cordelia Langford, John Sillitoe on behalf of the Wellcome Sanger Institute COVID-19 Surveillance Team |
| EPI_ISL_821336 | Lighthouse Lab in Milton Keynes | Wellcome Sanger Institute for the COVID-19 Genomics UK (COG-UK) Consortium | The Lighthouse Lab in Milton Keynes and Alex Alderton, Roberto Amato, Sonia Goncalves, Ewan Harrison, David K. Jackson, Ian Johnston, Dominic Kwiatkowski, Cordelia Langford, John Sillitoe on behalf of the Wellcome Sanger Institute COVID-19 Surveillance Team |
| EPI_ISL_821338, EPI_ISL_821339 | Lighthouse Lab in Cambridge | Wellcome Sanger Institute for the COVID-19 Genomics UK (COG-UK) Consortium | Rob Howes, The Lighthouse Lab in Cambridge and Alex Alderton, Roberto Amato, Sonia Goncalves, Ewan Harrison, David K. Jackson, Ian Johnston, Dominic Kwiatkowski, Cordelia Langford, John Sillitoe on behalf of the Wellcome Sanger Institute COVID-19 Surveillance Team |
| EPI_ISL_821340, EPI_ISL_821341 | Lighthouse Lab in Milton Keynes | Wellcome Sanger Institute for the COVID-19 Genomics UK (COG-UK) Consortium | The Lighthouse Lab in Milton Keynes and Alex Alderton, Roberto Amato, Sonia Goncalves, Ewan Harrison, David K. Jackson, Ian Johnston, Dominic Kwiatkowski, Cordelia Langford, John Sillitoe on behalf of the Wellcome Sanger Institute COVID-19 Surveillance Team |
| EPI_ISL_821342, EPI_ISL_821343 | Lighthouse Lab in Cambridge | Wellcome Sanger Institute for the COVID-19 Genomics UK (COG-UK) Consortium | Rob Howes, The Lighthouse Lab in Cambridge and Alex Alderton, Roberto Amato, Sonia Goncalves, Ewan Harrison, David K. Jackson, Ian Johnston, Dominic Kwiatkowski, Cordelia Langford, John Sillitoe on behalf of the Wellcome Sanger Institute COVID-19 Surveillance Team |
| EPI_ISL_821344, EPI_ISL_821346 | Lighthouse Lab in Milton Keynes | Wellcome Sanger Institute for the COVID-19 Genomics UK (COG-UK) Consortium | The Lighthouse Lab in Milton Keynes and Alex Alderton, Roberto Amato, Sonia Goncalves, Ewan Harrison, David K. Jackson, Ian Johnston, Dominic Kwiatkowski, Cordelia Langford, John Sillitoe on behalf of the Wellcome Sanger Institute COVID-19 Surveillance Team |
| EPI_ISL_821347 | Lighthouse Lab in Alderley Park | Wellcome Sanger Institute for the COVID-19 Genomics UK (COG-UK) Consortium | Jacquelyn Wynn, Mairead Hyland, The Lighthouse Lab in Alderley Park and Alex Alderton, Roberto Amato, Sonia Goncalves, Ewan Harrison, David K. Jackson, Ian Johnston, Dominic Kwiatkowski, Cordelia Langford, John Sillitoe on behalf of the Wellcome Sanger Institute COVID-19 Surveillance Team |
| EPI_ISL_821348 | Lighthouse Lab in Cambridge | Wellcome Sanger Institute for the COVID-19 Genomics UK (COG-UK) Consortium | Rob Howes, The Lighthouse Lab in Cambridge and Alex Alderton, Roberto Amato, Sonia Goncalves, Ewan Harrison, David K. Jackson, Ian Johnston, Dominic Kwiatkowski, Cordelia Langford, John Sillitoe on behalf of the Wellcome Sanger Institute COVID-19 Surveillance Team |
| EPI_ISL_821350 | Lighthouse Lab in Milton Keynes | Wellcome Sanger Institute for the COVID-19 Genomics UK (COG-UK) Consortium | The Lighthouse Lab in Milton Keynes and Alex Alderton, Roberto Amato, Sonia Goncalves, Ewan Harrison, David K. Jackson, Ian Johnston, Dominic Kwiatkowski, Cordelia Langford, John Sillitoe on behalf of the Wellcome Sanger Institute COVID-19 Surveillance Team |
| EPI_ISL_821351 | Lighthouse Lab in Alderley Park | Wellcome Sanger Institute for the COVID-19 Genomics UK (COG-UK) Consortium | Jacquelyn Wynn, Mairead Hyland, The Lighthouse Lab in Alderley Park and Alex Alderton, Roberto Amato, Sonia Goncalves, Ewan Harrison, David K. Jackson, Ian Johnston, Dominic Kwiatkowski, Cordelia Langford, John Sillitoe on behalf of the Wellcome Sanger Institute COVID-19 Surveillance Team |
| EPI_ISL_821352, EPI_ISL_821353 | Lighthouse Lab in Milton Keynes | Wellcome Sanger Institute for the COVID-19 Genomics UK (COG-UK) Consortium | The Lighthouse Lab in Milton Keynes and Alex Alderton, Roberto Amato, Sonia Goncalves, Ewan Harrison, David K. Jackson, Ian Johnston, Dominic Kwiatkowski, Cordelia Langford, John Sillitoe on behalf of the Wellcome Sanger Institute COVID-19 Surveillance Team |
| EPI_ISL_821354 | Lighthouse Lab in Cambridge | Wellcome Sanger Institute for the COVID-19 Genomics UK (COG-UK) Consortium | Rob Howes, The Lighthouse Lab in Cambridge and Alex Alderton, Roberto Amato, Sonia Goncalves, Ewan Harrison, David K. Jackson, Ian Johnston, Dominic Kwiatkowski, Cordelia Langford, John Sillitoe on behalf of the Wellcome Sanger Institute COVID-19 Surveillance Team |
| EPI_ISL_821356, EPI_ISL_821358, EPI_ISL_821359 | Lighthouse Lab in Milton Keynes | Wellcome Sanger Institute for the COVID-19 Genomics UK (COG-UK) Consortium | The Lighthouse Lab in Milton Keynes and Alex Alderton, Roberto Amato, Sonia Goncalves, Ewan Harrison, David K. Jackson, Ian Johnston, Dominic Kwiatkowski, Cordelia Langford, John Sillitoe on behalf of the Wellcome Sanger Institute COVID-19 Surveillance Team |
| EPI_ISL_821360 | Lighthouse Lab in Cambridge | Wellcome Sanger Institute for the COVID-19 Genomics UK (COG-UK) Consortium | Rob Howes, The Lighthouse Lab in Cambridge and Alex Alderton, Roberto Amato, Sonia Goncalves, Ewan Harrison, David K. Jackson, Ian Johnston, Dominic Kwiatkowski, Cordelia Langford, John Sillitoe on behalf of the Wellcome Sanger Institute COVID-19 Surveillance Team |
| EPI_ISL_821362 | Lighthouse Lab in Milton Keynes | Wellcome Sanger Institute for the COVID-19 Genomics UK (COG-UK) Consortium | The Lighthouse Lab in Milton Keynes and Alex Alderton, Roberto Amato, Sonia Goncalves, Ewan Harrison, David K. Jackson, Ian Johnston, Dominic Kwiatkowski, Cordelia Langford, John Sillitoe on behalf of the Wellcome Sanger Institute COVID-19 Surveillance Team |
| EPI_ISL_821363, EPI_ISL_821365 | Lighthouse Lab in Cambridge | Wellcome Sanger Institute for the COVID-19 Genomics UK (COG-UK) Consortium | Rob Howes, The Lighthouse Lab in Cambridge and Alex Alderton, Roberto Amato, Sonia Goncalves, Ewan Harrison, David K. Jackson, Ian Johnston, Dominic Kwiatkowski, Cordelia Langford, John Sillitoe on behalf of the Wellcome Sanger Institute COVID-19 Surveillance Team |
| EPI_ISL_821366 | Lighthouse Lab in Alderley Park | Wellcome Sanger Institute for the COVID-19 Genomics UK (COG-UK) Consortium | Jacquelyn Wynn, Mairead Hyland, The Lighthouse Lab in Alderley Park and Alex Alderton, Roberto Amato, Sonia Goncalves, Ewan Harrison, David K. Jackson, Ian Johnston, Dominic Kwiatkowski, Cordelia Langford, John Sillitoe on behalf of the Wellcome Sanger Institute COVID-19 Surveillance Team |
| EPI_ISL_821367 | Lighthouse Lab in Cambridge | Wellcome Sanger Institute for the COVID-19 Genomics UK (COG-UK) Consortium | Rob Howes, The Lighthouse Lab in Cambridge and Alex Alderton, Roberto Amato, Sonia Goncalves, Ewan Harrison, David K. Jackson, Ian Johnston, Dominic Kwiatkowski, Cordelia Langford, John Sillitoe on behalf of the Wellcome Sanger Institute COVID-19 Surveillance Team |
| EPI_ISL_821370, EPI_ISL_821371, EPI_ISL_821372 | Lighthouse Lab in Alderley Park | Wellcome Sanger Institute for the COVID-19 Genomics UK (COG-UK) Consortium | Jacquelyn Wynn, Mairead Hyland, The Lighthouse Lab in Alderley Park and Alex Alderton, Roberto Amato, Sonia Goncalves, Ewan Harrison, David K. Jackson, Ian Johnston, Dominic Kwiatkowski, Cordelia Langford, John Sillitoe on behalf of the Wellcome Sanger Institute COVID-19 Surveillance Team |
| EPI_ISL_821373 | Lighthouse Lab in Milton Keynes | Wellcome Sanger Institute for the COVID-19 Genomics UK (COG-UK) Consortium | The Lighthouse Lab in Milton Keynes and Alex Alderton, Roberto Amato, Sonia Goncalves, Ewan Harrison, David K. Jackson, Ian Johnston, Dominic Kwiatkowski, Cordelia Langford, John Sillitoe on behalf of the Wellcome Sanger Institute COVID-19 Surveillance Team |
| EPI_ISL_821374, EPI_ISL_821375 | Lighthouse Lab in Cambridge | Wellcome Sanger Institute for the COVID-19 Genomics UK (COG-UK) Consortium | Rob Howes, The Lighthouse Lab in Cambridge and Alex Alderton, Roberto Amato, Sonia Goncalves, Ewan Harrison, David K. Jackson, Ian Johnston, Dominic Kwiatkowski, Cordelia Langford, John Sillitoe on behalf of the Wellcome Sanger Institute COVID-19 Surveillance Team |
| EPI_ISL_821377, EPI_ISL_821378, EPI_ISL_821379, EPI_ISL_821380, EPI_ISL_821381 | Lighthouse Lab in Milton Keynes | Wellcome Sanger Institute for the COVID-19 Genomics UK (COG-UK) Consortium | The Lighthouse Lab in Milton Keynes and Alex Alderton, Roberto Amato, Sonia Goncalves, Ewan Harrison, David K. Jackson, Ian Johnston, Dominic Kwiatkowski, Cordelia Langford, John Sillitoe on behalf of the Wellcome Sanger Institute COVID-19 Surveillance Team |
| EPI_ISL_821382 | Lighthouse Lab in Alderley Park | Wellcome Sanger Institute for the COVID-19 Genomics UK (COG-UK) Consortium | Jacquelyn Wynn, Mairead Hyland, The Lighthouse Lab in Alderley Park and Alex Alderton, Roberto Amato, Sonia Goncalves, Ewan Harrison, David K. Jackson, Ian Johnston, Dominic Kwiatkowski, Cordelia Langford, John Sillitoe on behalf of the Wellcome Sanger Institute COVID-19 Surveillance Team |
| EPI_ISL_821383 | Lighthouse Lab in Milton Keynes | Wellcome Sanger Institute for the COVID-19 Genomics UK (COG-UK) Consortium | The Lighthouse Lab in Milton Keynes and Alex Alderton, Roberto Amato, Sonia Goncalves, Ewan Harrison, David K. Jackson, Ian Johnston, Dominic Kwiatkowski, Cordelia Langford, John Sillitoe on behalf of the Wellcome Sanger Institute COVID-19 Surveillance Team |
| EPI_ISL_821384 | Lighthouse Lab in Alderley Park | Wellcome Sanger Institute for the COVID-19 Genomics UK (COG-UK) Consortium | Jacquelyn Wynn, Mairead Hyland, The Lighthouse Lab in Alderley Park and Alex Alderton, Roberto Amato, Sonia Goncalves, Ewan Harrison, David K. Jackson, Ian Johnston, Dominic Kwiatkowski, Cordelia Langford, John Sillitoe on behalf of the Wellcome Sanger Institute COVID-19 Surveillance Team |
| EPI_ISL_821385 | Lighthouse Lab in Milton Keynes | Wellcome Sanger Institute for the COVID-19 Genomics UK (COG-UK) Consortium | The Lighthouse Lab in Milton Keynes and Alex Alderton, Roberto Amato, Sonia Goncalves, Ewan Harrison, David K. Jackson, Ian Johnston, Dominic Kwiatkowski, Cordelia Langford, John Sillitoe on behalf of the Wellcome Sanger Institute COVID-19 Surveillance Team |
| EPI_ISL_821386 | Lighthouse Lab in Cambridge | Wellcome Sanger Institute for the COVID-19 Genomics UK (COG-UK) Consortium | Rob Howes, The Lighthouse Lab in Cambridge and Alex Alderton, Roberto Amato, Sonia Goncalves, Ewan Harrison, David K. Jackson, Ian Johnston, Dominic Kwiatkowski, Cordelia Langford, John Sillitoe on behalf of the Wellcome Sanger Institute COVID-19 Surveillance Team |
| EPI_ISL_821387, EPI_ISL_821388 | Lighthouse Lab in Milton Keynes | Wellcome Sanger Institute for the COVID-19 Genomics UK (COG-UK) Consortium | The Lighthouse Lab in Milton Keynes and Alex Alderton, Roberto Amato, Sonia Goncalves, Ewan Harrison, David K. Jackson, Ian Johnston, Dominic Kwiatkowski, Cordelia Langford, John Sillitoe on behalf of the Wellcome Sanger Institute COVID-19 Surveillance Team |
| EPI_ISL_821389, EPI_ISL_821390 | Lighthouse Lab in Alderley Park | Wellcome Sanger Institute for the COVID-19 Genomics UK (COG-UK) Consortium | Jacquelyn Wynn, Mairead Hyland, The Lighthouse Lab in Alderley Park and Alex Alderton, Roberto Amato, Sonia Goncalves, Ewan Harrison, David K. Jackson, Ian Johnston, Dominic Kwiatkowski, Cordelia Langford, John Sillitoe on behalf of the Wellcome Sanger Institute COVID-19 Surveillance Team |

| | | | |
|---|---|---|---|
| EPI_ISL_821391 | Lighthouse Lab in Milton Keynes | Wellcome Sanger Institute for the COVID-19 Genomics UK (COG-UK) Consortium | The Lighthouse Lab in Milton Keynes and Alex Alderton, Roberto Amato, Sonia Goncalves, Ewan Harrison, David K. Jackson, Ian Johnston, Dominic Kwiatkowski, Cordelia Langford, John Sillitoe on behalf of the Wellcome Sanger Institute COVID-19 Surveillance Team |
| EPI_ISL_821392, EPI_ISL_821394 | Lighthouse Lab in Cambridge | Wellcome Sanger Institute for the COVID-19 Genomics UK (COG-UK) Consortium | Rob Howes, The Lighthouse Lab in Cambridge and Alex Alderton, Roberto Amato, Sonia Goncalves, Ewan Harrison, David K. Jackson, Ian Johnston, Dominic Kwiatkowski, Cordelia Langford, John Sillitoe on behalf of the Wellcome Sanger Institute COVID-19 Surveillance Team |
| EPI_ISL_821395, EPI_ISL_821397 | Lighthouse Lab in Milton Keynes | Wellcome Sanger Institute for the COVID-19 Genomics UK (COG-UK) Consortium | The Lighthouse Lab in Milton Keynes and Alex Alderton, Roberto Amato, Sonia Goncalves, Ewan Harrison, David K. Jackson, Ian Johnston, Dominic Kwiatkowski, Cordelia Langford, John Sillitoe on behalf of the Wellcome Sanger Institute COVID-19 Surveillance Team |
| EPI_ISL_821398 | Lighthouse Lab in Alderley Park | Wellcome Sanger Institute for the COVID-19 Genomics UK (COG-UK) Consortium | Jacquelyn Wynn, Mairead Hyland, The Lighthouse Lab in Alderley Park and Alex Alderton, Roberto Amato, Sonia Goncalves, Ewan Harrison, David K. Jackson, Ian Johnston, Dominic Kwiatkowski, Cordelia Langford, John Sillitoe on behalf of the Wellcome Sanger Institute COVID-19 Surveillance Team |
| EPI_ISL_821400 | Lighthouse Lab in Milton Keynes | Wellcome Sanger Institute for the COVID-19 Genomics UK (COG-UK) Consortium | The Lighthouse Lab in Milton Keynes and Alex Alderton, Roberto Amato, Sonia Goncalves, Ewan Harrison, David K. Jackson, Ian Johnston, Dominic Kwiatkowski, Cordelia Langford, John Sillitoe on behalf of the Wellcome Sanger Institute COVID-19 Surveillance Team |
| EPI_ISL_821401 | Lighthouse Lab in Alderley Park | Wellcome Sanger Institute for the COVID-19 Genomics UK (COG-UK) Consortium | Jacquelyn Wynn, Mairead Hyland, The Lighthouse Lab in Alderley Park and Alex Alderton, Roberto Amato, Sonia Goncalves, Ewan Harrison, David K. Jackson, Ian Johnston, Dominic Kwiatkowski, Cordelia Langford, John Sillitoe on behalf of the Wellcome Sanger Institute COVID-19 Surveillance Team |
| EPI_ISL_821402 | Lighthouse Lab in Milton Keynes | Wellcome Sanger Institute for the COVID-19 Genomics UK (COG-UK) Consortium | The Lighthouse Lab in Milton Keynes and Alex Alderton, Roberto Amato, Sonia Goncalves, Ewan Harrison, David K. Jackson, Ian Johnston, Dominic Kwiatkowski, Cordelia Langford, John Sillitoe on behalf of the Wellcome Sanger Institute COVID-19 Surveillance Team |
| EPI_ISL_821403 | Lighthouse Lab in Alderley Park | Wellcome Sanger Institute for the COVID-19 Genomics UK (COG-UK) Consortium | Jacquelyn Wynn, Mairead Hyland, The Lighthouse Lab in Alderley Park and Alex Alderton, Roberto Amato, Sonia Goncalves, Ewan Harrison, David K. Jackson, Ian Johnston, Dominic Kwiatkowski, Cordelia Langford, John Sillitoe on behalf of the Wellcome Sanger Institute COVID-19 Surveillance Team |
| EPI_ISL_821404 | Lighthouse Lab in Cambridge | Wellcome Sanger Institute for the COVID-19 Genomics UK (COG-UK) Consortium | Rob Howes, The Lighthouse Lab in Cambridge and Alex Alderton, Roberto Amato, Sonia Goncalves, Ewan Harrison, David K. Jackson, Ian Johnston, Dominic Kwiatkowski, Cordelia Langford, John Sillitoe on behalf of the Wellcome Sanger Institute COVID-19 Surveillance Team |
| EPI_ISL_821405, EPI_ISL_821406, EPI_ISL_821407 | Lighthouse Lab in Milton Keynes | Wellcome Sanger Institute for the COVID-19 Genomics UK (COG-UK) Consortium | The Lighthouse Lab in Milton Keynes and Alex Alderton, Roberto Amato, Sonia Goncalves, Ewan Harrison, David K. Jackson, Ian Johnston, Dominic Kwiatkowski, Cordelia Langford, John Sillitoe on behalf of the Wellcome Sanger Institute COVID-19 Surveillance Team |
| EPI_ISL_821408 | Lighthouse Lab in Cambridge | Wellcome Sanger Institute for the COVID-19 Genomics UK (COG-UK) Consortium | Rob Howes, The Lighthouse Lab in Cambridge and Alex Alderton, Roberto Amato, Sonia Goncalves, Ewan Harrison, David K. Jackson, Ian Johnston, Dominic Kwiatkowski, Cordelia Langford, John Sillitoe on behalf of the Wellcome Sanger Institute COVID-19 Surveillance Team |
| EPI_ISL_821409, EPI_ISL_821410 | Lighthouse Lab in Milton Keynes | Wellcome Sanger Institute for the COVID-19 Genomics UK (COG-UK) Consortium | The Lighthouse Lab in Milton Keynes and Alex Alderton, Roberto Amato, Sonia Goncalves, Ewan Harrison, David K. Jackson, Ian Johnston, Dominic Kwiatkowski, Cordelia Langford, John Sillitoe on behalf of the Wellcome Sanger Institute COVID-19 Surveillance Team |
| EPI_ISL_821411 | Lighthouse Lab in Cambridge | Wellcome Sanger Institute for the COVID-19 Genomics UK (COG-UK) Consortium | Rob Howes, The Lighthouse Lab in Cambridge and Alex Alderton, Roberto Amato, Sonia Goncalves, Ewan Harrison, David K. Jackson, Ian Johnston, Dominic Kwiatkowski, Cordelia Langford, John Sillitoe on behalf of the Wellcome Sanger Institute COVID-19 Surveillance Team |
| EPI_ISL_821412, EPI_ISL_821414, EPI_ISL_821415 | Lighthouse Lab in Alderley Park | Wellcome Sanger Institute for the COVID-19 Genomics UK (COG-UK) Consortium | Jacquelyn Wynn, Mairead Hyland, The Lighthouse Lab in Alderley Park and Alex Alderton, Roberto Amato, Sonia Goncalves, Ewan Harrison, David K. Jackson, Ian Johnston, Dominic Kwiatkowski, Cordelia Langford, John Sillitoe on behalf of the Wellcome Sanger Institute COVID-19 Surveillance Team |
| EPI_ISL_821416 | Lighthouse Lab in Cambridge | Wellcome Sanger Institute for the COVID-19 Genomics UK (COG-UK) Consortium | Rob Howes, The Lighthouse Lab in Cambridge and Alex Alderton, Roberto Amato, Sonia Goncalves, Ewan Harrison, David K. Jackson, Ian Johnston, Dominic Kwiatkowski, Cordelia Langford, John Sillitoe on behalf of the Wellcome Sanger Institute COVID-19 Surveillance Team |
| EPI_ISL_821417 | Lighthouse Lab in Milton Keynes | Wellcome Sanger Institute for the COVID-19 Genomics UK (COG-UK) Consortium | The Lighthouse Lab in Milton Keynes and Alex Alderton, Roberto Amato, Sonia Goncalves, Ewan Harrison, David K. Jackson, Ian Johnston, Dominic Kwiatkowski, Cordelia Langford, John Sillitoe on behalf of the Wellcome Sanger Institute COVID-19 Surveillance Team |
| EPI_ISL_821418, EPI_ISL_821419 | Lighthouse Lab in Alderley Park | Wellcome Sanger Institute for the COVID-19 Genomics UK (COG-UK) Consortium | Jacquelyn Wynn, Mairead Hyland, The Lighthouse Lab in Alderley Park and Alex Alderton, Roberto Amato, Sonia Goncalves, Ewan Harrison, David K. Jackson, Ian Johnston, Dominic Kwiatkowski, Cordelia Langford, John Sillitoe on behalf of the Wellcome Sanger Institute COVID-19 Surveillance Team |
| EPI_ISL_821422, EPI_ISL_821423, EPI_ISL_821424 | Lighthouse Lab in Milton Keynes | Wellcome Sanger Institute for the COVID-19 Genomics UK (COG-UK) Consortium | The Lighthouse Lab in Milton Keynes and Alex Alderton, Roberto Amato, Sonia Goncalves, Ewan Harrison, David K. Jackson, Ian Johnston, Dominic Kwiatkowski, Cordelia Langford, John Sillitoe on behalf of the Wellcome Sanger Institute COVID-19 Surveillance Team |
| EPI_ISL_821425, EPI_ISL_821426 | Lighthouse Lab in Alderley Park | Wellcome Sanger Institute for the COVID-19 Genomics UK (COG-UK) Consortium | Jacquelyn Wynn, Mairead Hyland, The Lighthouse Lab in Alderley Park and Alex Alderton, Roberto Amato, Sonia Goncalves, Ewan Harrison, David K. Jackson, Ian Johnston, Dominic Kwiatkowski, Cordelia Langford, John Sillitoe on behalf of the Wellcome Sanger Institute COVID-19 Surveillance Team |
| EPI_ISL_821428, EPI_ISL_821431, EPI_ISL_821432 | Lighthouse Lab in Milton Keynes | Wellcome Sanger Institute for the COVID-19 Genomics UK (COG-UK) Consortium | The Lighthouse Lab in Milton Keynes and Alex Alderton, Roberto Amato, Sonia Goncalves, Ewan Harrison, David K. Jackson, Ian Johnston, Dominic Kwiatkowski, Cordelia Langford, John Sillitoe on behalf of the Wellcome Sanger Institute COVID-19 Surveillance Team |
| EPI_ISL_821433 | Lighthouse Lab in Alderley Park | Wellcome Sanger Institute for the COVID-19 Genomics UK (COG-UK) Consortium | Jacquelyn Wynn, Mairead Hyland, The Lighthouse Lab in Alderley Park and Alex Alderton, Roberto Amato, Sonia Goncalves, Ewan Harrison, David K. Jackson, Ian Johnston, Dominic Kwiatkowski, Cordelia Langford, John Sillitoe on behalf of the Wellcome Sanger Institute COVID-19 Surveillance Team |
| EPI_ISL_821434 | Lighthouse Lab in Milton Keynes | Wellcome Sanger Institute for the COVID-19 Genomics UK (COG-UK) Consortium | The Lighthouse Lab in Milton Keynes and Alex Alderton, Roberto Amato, Sonia Goncalves, Ewan Harrison, David K. Jackson, Ian Johnston, Dominic Kwiatkowski, Cordelia Langford, John Sillitoe on behalf of the Wellcome Sanger Institute COVID-19 Surveillance Team |
| EPI_ISL_821435 | Lighthouse Lab in Cambridge | Wellcome Sanger Institute for the COVID-19 Genomics UK (COG-UK) Consortium | Rob Howes, The Lighthouse Lab in Cambridge and Alex Alderton, Roberto Amato, Sonia Goncalves, Ewan Harrison, David K. Jackson, Ian Johnston, Dominic Kwiatkowski, Cordelia Langford, John Sillitoe on behalf of the Wellcome Sanger Institute COVID-19 Surveillance Team |
| EPI_ISL_821436, EPI_ISL_821437 | Lighthouse Lab in Alderley Park | Wellcome Sanger Institute for the COVID-19 Genomics UK (COG-UK) Consortium | Jacquelyn Wynn, Mairead Hyland, The Lighthouse Lab in Alderley Park and Alex Alderton, Roberto Amato, Sonia Goncalves, Ewan Harrison, David K. Jackson, Ian Johnston, Dominic Kwiatkowski, Cordelia Langford, John Sillitoe on behalf of the Wellcome Sanger Institute COVID-19 Surveillance Team |
| EPI_ISL_821438 | Lighthouse Lab in Milton Keynes | Wellcome Sanger Institute for the COVID-19 Genomics UK (COG-UK) Consortium | The Lighthouse Lab in Milton Keynes and Alex Alderton, Roberto Amato, Sonia Goncalves, Ewan Harrison, David K. Jackson, Ian Johnston, Dominic Kwiatkowski, Cordelia Langford, John Sillitoe on behalf of the Wellcome Sanger Institute COVID-19 Surveillance Team |
| EPI_ISL_821439 | Lighthouse Lab in Cambridge | Wellcome Sanger Institute for the COVID-19 Genomics UK (COG-UK) Consortium | Rob Howes, The Lighthouse Lab in Cambridge and Alex Alderton, Roberto Amato, Sonia Goncalves, Ewan Harrison, David K. Jackson, Ian Johnston, Dominic Kwiatkowski, Cordelia Langford, John Sillitoe on behalf of the Wellcome Sanger Institute COVID-19 Surveillance Team |
| EPI_ISL_821440 | Lighthouse Lab in Alderley Park | Wellcome Sanger Institute for the COVID-19 Genomics UK (COG-UK) Consortium | Jacquelyn Wynn, Mairead Hyland, The Lighthouse Lab in Alderley Park and Alex Alderton, Roberto Amato, Sonia Goncalves, Ewan Harrison, David K. Jackson, Ian Johnston, Dominic Kwiatkowski, Cordelia Langford, John Sillitoe on behalf of the Wellcome Sanger Institute COVID-19 Surveillance Team |
| EPI_ISL_821441 | Lighthouse Lab in Cambridge | Wellcome Sanger Institute for the COVID-19 Genomics UK (COG-UK) Consortium | Rob Howes, The Lighthouse Lab in Cambridge and Alex Alderton, Roberto Amato, Sonia Goncalves, Ewan Harrison, David K. Jackson, Ian Johnston, Dominic Kwiatkowski, Cordelia Langford, John Sillitoe on behalf of the Wellcome Sanger Institute COVID-19 Surveillance Team |
| EPI_ISL_821443, EPI_ISL_821445 | Lighthouse Lab in Milton Keynes | Wellcome Sanger Institute for the COVID-19 Genomics UK (COG-UK) Consortium | The Lighthouse Lab in Milton Keynes and Alex Alderton, Roberto Amato, Sonia Goncalves, Ewan Harrison, David K. Jackson, Ian Johnston, Dominic Kwiatkowski, Cordelia Langford, John Sillitoe on behalf of the Wellcome Sanger Institute COVID-19 Surveillance Team |
| EPI_ISL_821446 | Lighthouse Lab in Alderley Park | Wellcome Sanger Institute for the COVID-19 Genomics UK (COG-UK) Consortium | Jacquelyn Wynn, Mairead Hyland, The Lighthouse Lab in Alderley Park and Alex Alderton, Roberto Amato, Sonia Goncalves, Ewan Harrison, David K. Jackson, Ian Johnston, Dominic Kwiatkowski, Cordelia Langford, John Sillitoe on behalf of the Wellcome Sanger Institute COVID-19 Surveillance Team |
| EPI_ISL_821447 | Lighthouse Lab in Cambridge | Wellcome Sanger Institute for the COVID-19 Genomics UK (COG-UK) Consortium | Rob Howes, The Lighthouse Lab in Cambridge and Alex Alderton, Roberto Amato, Sonia Goncalves, Ewan Harrison, David K. Jackson, Ian Johnston, Dominic Kwiatkowski, Cordelia Langford, John Sillitoe on behalf of the Wellcome Sanger Institute COVID-19 Surveillance Team |
| EPI_ISL_821448, EPI_ISL_821449 | Lighthouse Lab in Milton Keynes | Wellcome Sanger Institute for the COVID-19 Genomics UK (COG-UK) Consortium | The Lighthouse Lab in Milton Keynes and Alex Alderton, Roberto Amato, Sonia Goncalves, Ewan Harrison, David K. Jackson, Ian Johnston, Dominic Kwiatkowski, Cordelia Langford, John Sillitoe on behalf of the Wellcome Sanger Institute COVID-19 Surveillance Team |
| EPI_ISL_821451 | Lighthouse Lab in Alderley Park | Wellcome Sanger Institute for the COVID-19 Genomics UK (COG-UK) Consortium | Jacquelyn Wynn, Mairead Hyland, The Lighthouse Lab in Alderley Park and Alex Alderton, Roberto Amato, Sonia Goncalves, Ewan Harrison, David K. Jackson, Ian Johnston, Dominic Kwiatkowski, Cordelia Langford, John Sillitoe on behalf of the Wellcome Sanger Institute COVID-19 Surveillance Team |
| EPI_ISL_821452 | Lighthouse Lab in Milton Keynes | Wellcome Sanger Institute for the COVID-19 Genomics UK (COG-UK) Consortium | The Lighthouse Lab in Milton Keynes and Alex Alderton, Roberto Amato, Sonia Goncalves, Ewan Harrison, David K. Jackson, Ian Johnston, Dominic Kwiatkowski, Cordelia Langford, John Sillitoe on behalf of the Wellcome Sanger Institute COVID-19 Surveillance Team |
| EPI_ISL_821453 | Lighthouse Lab in Alderley Park | Wellcome Sanger Institute for the COVID-19 Genomics UK (COG-UK) Consortium | Jacquelyn Wynn, Mairead Hyland, The Lighthouse Lab in Alderley Park and Alex Alderton, Roberto Amato, Sonia Goncalves, Ewan Harrison, David K. Jackson, Ian Johnston, Dominic Kwiatkowski, Cordelia Langford, John Sillitoe on behalf of the Wellcome Sanger Institute COVID-19 Surveillance Team |

| | | | |
|---|---|---|---|
| EPI_ISL_821457, EPI_ISL_821458 | Lighthouse Lab in Milton Keynes | Wellcome Sanger Institute for the COVID-19 Genomics UK (COG-UK) Consortium | The Lighthouse Lab in Milton Keynes and Alex Alderton, Roberto Amato, Sonia Goncalves, Ewan Harrison, David K. Jackson, Ian Johnston, Dominic Kwiatkowski, Cordelia Langford, John Sillitoe on behalf of the Wellcome Sanger Institute COVID-19 Surveillance Team |
| EPI_ISL_821459 | Lighthouse Lab in Alderley Park | Wellcome Sanger Institute for the COVID-19 Genomics UK (COG-UK) Consortium | Jacquelyn Wynn, Mairead Hyland, The Lighthouse Lab in Alderley Park and Alex Alderton, Roberto Amato, Sonia Goncalves, Ewan Harrison, David K. Jackson, Ian Johnston, Dominic Kwiatkowski, Cordelia Langford, John Sillitoe on behalf of the Wellcome Sanger Institute COVID-19 Surveillance Team |
| EPI_ISL_821460, EPI_ISL_821461, EPI_ISL_821462 | Lighthouse Lab in Milton Keynes | Wellcome Sanger Institute for the COVID-19 Genomics UK (COG-UK) Consortium | The Lighthouse Lab in Milton Keynes and Alex Alderton, Roberto Amato, Sonia Goncalves, Ewan Harrison, David K. Jackson, Ian Johnston, Dominic Kwiatkowski, Cordelia Langford, John Sillitoe on behalf of the Wellcome Sanger Institute COVID-19 Surveillance Team |
| EPI_ISL_821465, EPI_ISL_821466 | Lighthouse Lab in Alderley Park | Wellcome Sanger Institute for the COVID-19 Genomics UK (COG-UK) Consortium | Jacquelyn Wynn, Mairead Hyland, The Lighthouse Lab in Alderley Park and Alex Alderton, Roberto Amato, Sonia Goncalves, Ewan Harrison, David K. Jackson, Ian Johnston, Dominic Kwiatkowski, Cordelia Langford, John Sillitoe on behalf of the Wellcome Sanger Institute COVID-19 Surveillance Team |
| EPI_ISL_821467, EPI_ISL_821468 | Lighthouse Lab in Milton Keynes | Wellcome Sanger Institute for the COVID-19 Genomics UK (COG-UK) Consortium | The Lighthouse Lab in Milton Keynes and Alex Alderton, Roberto Amato, Sonia Goncalves, Ewan Harrison, David K. Jackson, Ian Johnston, Dominic Kwiatkowski, Cordelia Langford, John Sillitoe on behalf of the Wellcome Sanger Institute COVID-19 Surveillance Team |
| EPI_ISL_821469, EPI_ISL_821470 | Lighthouse Lab in Alderley Park | Wellcome Sanger Institute for the COVID-19 Genomics UK (COG-UK) Consortium | Jacquelyn Wynn, Mairead Hyland, The Lighthouse Lab in Alderley Park and Alex Alderton, Roberto Amato, Sonia Goncalves, Ewan Harrison, David K. Jackson, Ian Johnston, Dominic Kwiatkowski, Cordelia Langford, John Sillitoe on behalf of the Wellcome Sanger Institute COVID-19 Surveillance Team |
| EPI_ISL_821472, EPI_ISL_821474, EPI_ISL_821475 | Lighthouse Lab in Milton Keynes | Wellcome Sanger Institute for the COVID-19 Genomics UK (COG-UK) Consortium | The Lighthouse Lab in Milton Keynes and Alex Alderton, Roberto Amato, Sonia Goncalves, Ewan Harrison, David K. Jackson, Ian Johnston, Dominic Kwiatkowski, Cordelia Langford, John Sillitoe on behalf of the Wellcome Sanger Institute COVID-19 Surveillance Team |
| EPI_ISL_821476 | Lighthouse Lab in Alderley Park | Wellcome Sanger Institute for the COVID-19 Genomics UK (COG-UK) Consortium | Jacquelyn Wynn, Mairead Hyland, The Lighthouse Lab in Alderley Park and Alex Alderton, Roberto Amato, Sonia Goncalves, Ewan Harrison, David K. Jackson, Ian Johnston, Dominic Kwiatkowski, Cordelia Langford, John Sillitoe on behalf of the Wellcome Sanger Institute COVID-19 Surveillance Team |
| EPI_ISL_821477, EPI_ISL_821478 | Lighthouse Lab in Milton Keynes | Wellcome Sanger Institute for the COVID-19 Genomics UK (COG-UK) Consortium | The Lighthouse Lab in Milton Keynes and Alex Alderton, Roberto Amato, Sonia Goncalves, Ewan Harrison, David K. Jackson, Ian Johnston, Dominic Kwiatkowski, Cordelia Langford, John Sillitoe on behalf of the Wellcome Sanger Institute COVID-19 Surveillance Team |
| EPI_ISL_821479 | Lighthouse Lab in Cambridge | Wellcome Sanger Institute for the COVID-19 Genomics UK (COG-UK) Consortium | Rob Howes, The Lighthouse Lab in Cambridge and Alex Alderton, Roberto Amato, Sonia Goncalves, Ewan Harrison, David K. Jackson, Ian Johnston, Dominic Kwiatkowski, Cordelia Langford, John Sillitoe on behalf of the Wellcome Sanger Institute COVID-19 Surveillance Team |
| EPI_ISL_821480 | Lighthouse Lab in Milton Keynes | Wellcome Sanger Institute for the COVID-19 Genomics UK (COG-UK) Consortium | The Lighthouse Lab in Milton Keynes and Alex Alderton, Roberto Amato, Sonia Goncalves, Ewan Harrison, David K. Jackson, Ian Johnston, Dominic Kwiatkowski, Cordelia Langford, John Sillitoe on behalf of the Wellcome Sanger Institute COVID-19 Surveillance Team |
| EPI_ISL_821482, EPI_ISL_821483 | Lighthouse Lab in Alderley Park | Wellcome Sanger Institute for the COVID-19 Genomics UK (COG-UK) Consortium | Jacquelyn Wynn, Mairead Hyland, The Lighthouse Lab in Alderley Park and Alex Alderton, Roberto Amato, Sonia Goncalves, Ewan Harrison, David K. Jackson, Ian Johnston, Dominic Kwiatkowski, Cordelia Langford, John Sillitoe on behalf of the Wellcome Sanger Institute COVID-19 Surveillance Team |
| EPI_ISL_821484 | Lighthouse Lab in Cambridge | Wellcome Sanger Institute for the COVID-19 Genomics UK (COG-UK) Consortium | Rob Howes, The Lighthouse Lab in Cambridge and Alex Alderton, Roberto Amato, Sonia Goncalves, Ewan Harrison, David K. Jackson, Ian Johnston, Dominic Kwiatkowski, Cordelia Langford, John Sillitoe on behalf of the Wellcome Sanger Institute COVID-19 Surveillance Team |
| EPI_ISL_821486, EPI_ISL_821487 | Lighthouse Lab in Milton Keynes | Wellcome Sanger Institute for the COVID-19 Genomics UK (COG-UK) Consortium | The Lighthouse Lab in Milton Keynes and Alex Alderton, Roberto Amato, Sonia Goncalves, Ewan Harrison, David K. Jackson, Ian Johnston, Dominic Kwiatkowski, Cordelia Langford, John Sillitoe on behalf of the Wellcome Sanger Institute COVID-19 Surveillance Team |
| EPI_ISL_821488 | Lighthouse Lab in Cambridge | Wellcome Sanger Institute for the COVID-19 Genomics UK (COG-UK) Consortium | Rob Howes, The Lighthouse Lab in Cambridge and Alex Alderton, Roberto Amato, Sonia Goncalves, Ewan Harrison, David K. Jackson, Ian Johnston, Dominic Kwiatkowski, Cordelia Langford, John Sillitoe on behalf of the Wellcome Sanger Institute COVID-19 Surveillance Team |
| EPI_ISL_821490 | Lighthouse Lab in Alderley Park | Wellcome Sanger Institute for the COVID-19 Genomics UK (COG-UK) Consortium | Jacquelyn Wynn, Mairead Hyland, The Lighthouse Lab in Alderley Park and Alex Alderton, Roberto Amato, Sonia Goncalves, Ewan Harrison, David K. Jackson, Ian Johnston, Dominic Kwiatkowski, Cordelia Langford, John Sillitoe on behalf of the Wellcome Sanger Institute COVID-19 Surveillance Team |
| EPI_ISL_821491, EPI_ISL_821492 | Lighthouse Lab in Cambridge | Wellcome Sanger Institute for the COVID-19 Genomics UK (COG-UK) Consortium | Rob Howes, The Lighthouse Lab in Cambridge and Alex Alderton, Roberto Amato, Sonia Goncalves, Ewan Harrison, David K. Jackson, Ian Johnston, Dominic Kwiatkowski, Cordelia Langford, John Sillitoe on behalf of the Wellcome Sanger Institute COVID-19 Surveillance Team |
| EPI_ISL_821493 | Lighthouse Lab in Alderley Park | Wellcome Sanger Institute for the COVID-19 Genomics UK (COG-UK) Consortium | Jacquelyn Wynn, Mairead Hyland, The Lighthouse Lab in Alderley Park and Alex Alderton, Roberto Amato, Sonia Goncalves, Ewan Harrison, David K. Jackson, Ian Johnston, Dominic Kwiatkowski, Cordelia Langford, John Sillitoe on behalf of the Wellcome Sanger Institute COVID-19 Surveillance Team |
| EPI_ISL_821494 | Lighthouse Lab in Milton Keynes | Wellcome Sanger Institute for the COVID-19 Genomics UK (COG-UK) Consortium | The Lighthouse Lab in Milton Keynes and Alex Alderton, Roberto Amato, Sonia Goncalves, Ewan Harrison, David K. Jackson, Ian Johnston, Dominic Kwiatkowski, Cordelia Langford, John Sillitoe on behalf of the Wellcome Sanger Institute COVID-19 Surveillance Team |
| EPI_ISL_821496, EPI_ISL_821497 | Lighthouse Lab in Alderley Park | Wellcome Sanger Institute for the COVID-19 Genomics UK (COG-UK) Consortium | Jacquelyn Wynn, Mairead Hyland, The Lighthouse Lab in Alderley Park and Alex Alderton, Roberto Amato, Sonia Goncalves, Ewan Harrison, David K. Jackson, Ian Johnston, Dominic Kwiatkowski, Cordelia Langford, John Sillitoe on behalf of the Wellcome Sanger Institute COVID-19 Surveillance Team |
| EPI_ISL_821498 | Lighthouse Lab in Milton Keynes | Wellcome Sanger Institute for the COVID-19 Genomics UK (COG-UK) Consortium | The Lighthouse Lab in Milton Keynes and Alex Alderton, Roberto Amato, Sonia Goncalves, Ewan Harrison, David K. Jackson, Ian Johnston, Dominic Kwiatkowski, Cordelia Langford, John Sillitoe on behalf of the Wellcome Sanger Institute COVID-19 Surveillance Team |
| EPI_ISL_821500 | Lighthouse Lab in Alderley Park | Wellcome Sanger Institute for the COVID-19 Genomics UK (COG-UK) Consortium | Jacquelyn Wynn, Mairead Hyland, The Lighthouse Lab in Alderley Park and Alex Alderton, Roberto Amato, Sonia Goncalves, Ewan Harrison, David K. Jackson, Ian Johnston, Dominic Kwiatkowski, Cordelia Langford, John Sillitoe on behalf of the Wellcome Sanger Institute COVID-19 Surveillance Team |
| EPI_ISL_821501 | Lighthouse Lab in Milton Keynes | Wellcome Sanger Institute for the COVID-19 Genomics UK (COG-UK) Consortium | The Lighthouse Lab in Milton Keynes and Alex Alderton, Roberto Amato, Sonia Goncalves, Ewan Harrison, David K. Jackson, Ian Johnston, Dominic Kwiatkowski, Cordelia Langford, John Sillitoe on behalf of the Wellcome Sanger Institute COVID-19 Surveillance Team |
| EPI_ISL_821502, EPI_ISL_821503 | Lighthouse Lab in Alderley Park | Wellcome Sanger Institute for the COVID-19 Genomics UK (COG-UK) Consortium | Jacquelyn Wynn, Mairead Hyland, The Lighthouse Lab in Alderley Park and Alex Alderton, Roberto Amato, Sonia Goncalves, Ewan Harrison, David K. Jackson, Ian Johnston, Dominic Kwiatkowski, Cordelia Langford, John Sillitoe on behalf of the Wellcome Sanger Institute COVID-19 Surveillance Team |
| EPI_ISL_821504 | Lighthouse Lab in Milton Keynes | Wellcome Sanger Institute for the COVID-19 Genomics UK (COG-UK) Consortium | The Lighthouse Lab in Milton Keynes and Alex Alderton, Roberto Amato, Sonia Goncalves, Ewan Harrison, David K. Jackson, Ian Johnston, Dominic Kwiatkowski, Cordelia Langford, John Sillitoe on behalf of the Wellcome Sanger Institute COVID-19 Surveillance Team |
| EPI_ISL_821506 | Lighthouse Lab in Cambridge | Wellcome Sanger Institute for the COVID-19 Genomics UK (COG-UK) Consortium | Rob Howes, The Lighthouse Lab in Cambridge and Alex Alderton, Roberto Amato, Sonia Goncalves, Ewan Harrison, David K. Jackson, Ian Johnston, Dominic Kwiatkowski, Cordelia Langford, John Sillitoe on behalf of the Wellcome Sanger Institute COVID-19 Surveillance Team |
| EPI_ISL_821507, EPI_ISL_821509 | Lighthouse Lab in Milton Keynes | Wellcome Sanger Institute for the COVID-19 Genomics UK (COG-UK) Consortium | The Lighthouse Lab in Milton Keynes and Alex Alderton, Roberto Amato, Sonia Goncalves, Ewan Harrison, David K. Jackson, Ian Johnston, Dominic Kwiatkowski, Cordelia Langford, John Sillitoe on behalf of the Wellcome Sanger Institute COVID-19 Surveillance Team |
| EPI_ISL_821510 | Lighthouse Lab in Cambridge | Wellcome Sanger Institute for the COVID-19 Genomics UK (COG-UK) Consortium | Rob Howes, The Lighthouse Lab in Cambridge and Alex Alderton, Roberto Amato, Sonia Goncalves, Ewan Harrison, David K. Jackson, Ian Johnston, Dominic Kwiatkowski, Cordelia Langford, John Sillitoe on behalf of the Wellcome Sanger Institute COVID-19 Surveillance Team |
| EPI_ISL_821511 | Lighthouse Lab in Alderley Park | Wellcome Sanger Institute for the COVID-19 Genomics UK (COG-UK) Consortium | Jacquelyn Wynn, Mairead Hyland, The Lighthouse Lab in Alderley Park and Alex Alderton, Roberto Amato, Sonia Goncalves, Ewan Harrison, David K. Jackson, Ian Johnston, Dominic Kwiatkowski, Cordelia Langford, John Sillitoe on behalf of the Wellcome Sanger Institute COVID-19 Surveillance Team |
| EPI_ISL_821514, EPI_ISL_821515 | Lighthouse Lab in Milton Keynes | Wellcome Sanger Institute for the COVID-19 Genomics UK (COG-UK) Consortium | The Lighthouse Lab in Milton Keynes and Alex Alderton, Roberto Amato, Sonia Goncalves, Ewan Harrison, David K. Jackson, Ian Johnston, Dominic Kwiatkowski, Cordelia Langford, John Sillitoe on behalf of the Wellcome Sanger Institute COVID-19 Surveillance Team |
| EPI_ISL_821516 | Lighthouse Lab in Cambridge | Wellcome Sanger Institute for the COVID-19 Genomics UK (COG-UK) Consortium | Rob Howes, The Lighthouse Lab in Cambridge and Alex Alderton, Roberto Amato, Sonia Goncalves, Ewan Harrison, David K. Jackson, Ian Johnston, Dominic Kwiatkowski, Cordelia Langford, John Sillitoe on behalf of the Wellcome Sanger Institute COVID-19 Surveillance Team |
| EPI_ISL_821517 | Lighthouse Lab in Milton Keynes | Wellcome Sanger Institute for the COVID-19 Genomics UK (COG-UK) Consortium | The Lighthouse Lab in Milton Keynes and Alex Alderton, Roberto Amato, Sonia Goncalves, Ewan Harrison, David K. Jackson, Ian Johnston, Dominic Kwiatkowski, Cordelia Langford, John Sillitoe on behalf of the Wellcome Sanger Institute COVID-19 Surveillance Team |
| EPI_ISL_821518 | Lighthouse Lab in Alderley Park | Wellcome Sanger Institute for the COVID-19 Genomics UK (COG-UK) Consortium | Jacquelyn Wynn, Mairead Hyland, The Lighthouse Lab in Alderley Park and Alex Alderton, Roberto Amato, Sonia Goncalves, Ewan Harrison, David K. Jackson, Ian Johnston, Dominic Kwiatkowski, Cordelia Langford, John Sillitoe on behalf of the Wellcome Sanger Institute COVID-19 Surveillance Team |
| EPI_ISL_821519 | Lighthouse Lab in Cambridge | Wellcome Sanger Institute for the COVID-19 Genomics UK (COG-UK) Consortium | Rob Howes, The Lighthouse Lab in Cambridge and Alex Alderton, Roberto Amato, Sonia Goncalves, Ewan Harrison, David K. Jackson, Ian Johnston, Dominic Kwiatkowski, Cordelia Langford, John Sillitoe on behalf of the Wellcome Sanger Institute COVID-19 Surveillance Team |
| EPI_ISL_821520, EPI_ISL_821521, EPI_ISL_821522 | Lighthouse Lab in Milton Keynes | Wellcome Sanger Institute for the COVID-19 Genomics UK (COG-UK) Consortium | The Lighthouse Lab in Milton Keynes and Alex Alderton, Roberto Amato, Sonia Goncalves, Ewan Harrison, David K. Jackson, Ian Johnston, Dominic Kwiatkowski, Cordelia Langford, John Sillitoe on behalf of the Wellcome Sanger Institute COVID-19 Surveillance Team |

| | | | |
|---|---|---|---|
| EPI_ISL_821523 | Lighthouse Lab in Cambridge | Wellcome Sanger Institute for the COVID-19 Genomics UK (COG-UK) Consortium | Rob Howes, The Lighthouse Lab in Cambridge and Alex Alderton, Roberto Amato, Sonia Goncalves, Ewan Harrison, David K. Jackson, Ian Johnston, Dominic Kwiatkowski, Cordelia Langford, John Sillitoe on behalf of the Wellcome Sanger Institute COVID-19 Surveillance Team |
| EPI_ISL_821524 | Lighthouse Lab in Milton Keynes | Wellcome Sanger Institute for the COVID-19 Genomics UK (COG-UK) Consortium | The Lighthouse Lab in Milton Keynes and Alex Alderton, Roberto Amato, Sonia Goncalves, Ewan Harrison, David K. Jackson, Ian Johnston, Dominic Kwiatkowski, Cordelia Langford, John Sillitoe on behalf of the Wellcome Sanger Institute COVID-19 Surveillance Team |
| EPI_ISL_821526 | Lighthouse Lab in Alderley Park | Wellcome Sanger Institute for the COVID-19 Genomics UK (COG-UK) Consortium | Jacquelyn Wynn, Mairead Hyland, The Lighthouse Lab in Alderley Park and Alex Alderton, Roberto Amato, Sonia Goncalves, Ewan Harrison, David K. Jackson, Ian Johnston, Dominic Kwiatkowski, Cordelia Langford, John Sillitoe on behalf of the Wellcome Sanger Institute COVID-19 Surveillance Team |
| EPI_ISL_821527 | Lighthouse Lab in Milton Keynes | Wellcome Sanger Institute for the COVID-19 Genomics UK (COG-UK) Consortium | The Lighthouse Lab in Milton Keynes and Alex Alderton, Roberto Amato, Sonia Goncalves, Ewan Harrison, David K. Jackson, Ian Johnston, Dominic Kwiatkowski, Cordelia Langford, John Sillitoe on behalf of the Wellcome Sanger Institute COVID-19 Surveillance Team |
| EPI_ISL_821528 | Lighthouse Lab in Alderley Park | Wellcome Sanger Institute for the COVID-19 Genomics UK (COG-UK) Consortium | Jacquelyn Wynn, Mairead Hyland, The Lighthouse Lab in Alderley Park and Alex Alderton, Roberto Amato, Sonia Goncalves, Ewan Harrison, David K. Jackson, Ian Johnston, Dominic Kwiatkowski, Cordelia Langford, John Sillitoe on behalf of the Wellcome Sanger Institute COVID-19 Surveillance Team |
| EPI_ISL_821529 | Lighthouse Lab in Cambridge | Wellcome Sanger Institute for the COVID-19 Genomics UK (COG-UK) Consortium | Rob Howes, The Lighthouse Lab in Cambridge and Alex Alderton, Roberto Amato, Sonia Goncalves, Ewan Harrison, David K. Jackson, Ian Johnston, Dominic Kwiatkowski, Cordelia Langford, John Sillitoe on behalf of the Wellcome Sanger Institute COVID-19 Surveillance Team |
| EPI_ISL_821530, EPI_ISL_821532 | Lighthouse Lab in Milton Keynes | Wellcome Sanger Institute for the COVID-19 Genomics UK (COG-UK) Consortium | The Lighthouse Lab in Milton Keynes and Alex Alderton, Roberto Amato, Sonia Goncalves, Ewan Harrison, David K. Jackson, Ian Johnston, Dominic Kwiatkowski, Cordelia Langford, John Sillitoe on behalf of the Wellcome Sanger Institute COVID-19 Surveillance Team |
| EPI_ISL_821533, EPI_ISL_821534 | Lighthouse Lab in Alderley Park | Wellcome Sanger Institute for the COVID-19 Genomics UK (COG-UK) Consortium | Jacquelyn Wynn, Mairead Hyland, The Lighthouse Lab in Alderley Park and Alex Alderton, Roberto Amato, Sonia Goncalves, Ewan Harrison, David K. Jackson, Ian Johnston, Dominic Kwiatkowski, Cordelia Langford, John Sillitoe on behalf of the Wellcome Sanger Institute COVID-19 Surveillance Team |
| EPI_ISL_821535, EPI_ISL_821536, EPI_ISL_821537 | Lighthouse Lab in Milton Keynes | Wellcome Sanger Institute for the COVID-19 Genomics UK (COG-UK) Consortium | The Lighthouse Lab in Milton Keynes and Alex Alderton, Roberto Amato, Sonia Goncalves, Ewan Harrison, David K. Jackson, Ian Johnston, Dominic Kwiatkowski, Cordelia Langford, John Sillitoe on behalf of the Wellcome Sanger Institute COVID-19 Surveillance Team |
| EPI_ISL_821538 | Lighthouse Lab in Alderley Park | Wellcome Sanger Institute for the COVID-19 Genomics UK (COG-UK) Consortium | Jacquelyn Wynn, Mairead Hyland, The Lighthouse Lab in Alderley Park and Alex Alderton, Roberto Amato, Sonia Goncalves, Ewan Harrison, David K. Jackson, Ian Johnston, Dominic Kwiatkowski, Cordelia Langford, John Sillitoe on behalf of the Wellcome Sanger Institute COVID-19 Surveillance Team |
| EPI_ISL_821539, EPI_ISL_821540, EPI_ISL_821541, EPI_ISL_821542, EPI_ISL_821543 | Lighthouse Lab in Milton Keynes | Wellcome Sanger Institute for the COVID-19 Genomics UK (COG-UK) Consortium | The Lighthouse Lab in Milton Keynes and Alex Alderton, Roberto Amato, Sonia Goncalves, Ewan Harrison, David K. Jackson, Ian Johnston, Dominic Kwiatkowski, Cordelia Langford, John Sillitoe on behalf of the Wellcome Sanger Institute COVID-19 Surveillance Team |
| EPI_ISL_821544 | Lighthouse Lab in Cambridge | Wellcome Sanger Institute for the COVID-19 Genomics UK (COG-UK) Consortium | Rob Howes, The Lighthouse Lab in Cambridge and Alex Alderton, Roberto Amato, Sonia Goncalves, Ewan Harrison, David K. Jackson, Ian Johnston, Dominic Kwiatkowski, Cordelia Langford, John Sillitoe on behalf of the Wellcome Sanger Institute COVID-19 Surveillance Team |
| EPI_ISL_821545 | Lighthouse Lab in Milton Keynes | Wellcome Sanger Institute for the COVID-19 Genomics UK (COG-UK) Consortium | The Lighthouse Lab in Milton Keynes and Alex Alderton, Roberto Amato, Sonia Goncalves, Ewan Harrison, David K. Jackson, Ian Johnston, Dominic Kwiatkowski, Cordelia Langford, John Sillitoe on behalf of the Wellcome Sanger Institute COVID-19 Surveillance Team |
| EPI_ISL_821547 | Lighthouse Lab in Cambridge | Wellcome Sanger Institute for the COVID-19 Genomics UK (COG-UK) Consortium | Rob Howes, The Lighthouse Lab in Cambridge and Alex Alderton, Roberto Amato, Sonia Goncalves, Ewan Harrison, David K. Jackson, Ian Johnston, Dominic Kwiatkowski, Cordelia Langford, John Sillitoe on behalf of the Wellcome Sanger Institute COVID-19 Surveillance Team |
| EPI_ISL_821548 | Lighthouse Lab in Alderley Park | Wellcome Sanger Institute for the COVID-19 Genomics UK (COG-UK) Consortium | Jacquelyn Wynn, Mairead Hyland, The Lighthouse Lab in Alderley Park and Alex Alderton, Roberto Amato, Sonia Goncalves, Ewan Harrison, David K. Jackson, Ian Johnston, Dominic Kwiatkowski, Cordelia Langford, John Sillitoe on behalf of the Wellcome Sanger Institute COVID-19 Surveillance Team |
| EPI_ISL_821549 | Lighthouse Lab in Cambridge | Wellcome Sanger Institute for the COVID-19 Genomics UK (COG-UK) Consortium | Rob Howes, The Lighthouse Lab in Cambridge and Alex Alderton, Roberto Amato, Sonia Goncalves, Ewan Harrison, David K. Jackson, Ian Johnston, Dominic Kwiatkowski, Cordelia Langford, John Sillitoe on behalf of the Wellcome Sanger Institute COVID-19 Surveillance Team |
| EPI_ISL_821550, EPI_ISL_821551, EPI_ISL_821552, EPI_ISL_821553 | Lighthouse Lab in Alderley Park | Wellcome Sanger Institute for the COVID-19 Genomics UK (COG-UK) Consortium | Jacquelyn Wynn, Mairead Hyland, The Lighthouse Lab in Alderley Park and Alex Alderton, Roberto Amato, Sonia Goncalves, Ewan Harrison, David K. Jackson, Ian Johnston, Dominic Kwiatkowski, Cordelia Langford, John Sillitoe on behalf of the Wellcome Sanger Institute COVID-19 Surveillance Team |
| EPI_ISL_821554 | Lighthouse Lab in Milton Keynes | Wellcome Sanger Institute for the COVID-19 Genomics UK (COG-UK) Consortium | The Lighthouse Lab in Milton Keynes and Alex Alderton, Roberto Amato, Sonia Goncalves, Ewan Harrison, David K. Jackson, Ian Johnston, Dominic Kwiatkowski, Cordelia Langford, John Sillitoe on behalf of the Wellcome Sanger Institute COVID-19 Surveillance Team |
| EPI_ISL_821555 | Lighthouse Lab in Cambridge | Wellcome Sanger Institute for the COVID-19 Genomics UK (COG-UK) Consortium | Rob Howes, The Lighthouse Lab in Cambridge and Alex Alderton, Roberto Amato, Sonia Goncalves, Ewan Harrison, David K. Jackson, Ian Johnston, Dominic Kwiatkowski, Cordelia Langford, John Sillitoe on behalf of the Wellcome Sanger Institute COVID-19 Surveillance Team |
| EPI_ISL_821556, EPI_ISL_821558 | Lighthouse Lab in Alderley Park | Wellcome Sanger Institute for the COVID-19 Genomics UK (COG-UK) Consortium | Jacquelyn Wynn, Mairead Hyland, The Lighthouse Lab in Alderley Park and Alex Alderton, Roberto Amato, Sonia Goncalves, Ewan Harrison, David K. Jackson, Ian Johnston, Dominic Kwiatkowski, Cordelia Langford, John Sillitoe on behalf of the Wellcome Sanger Institute COVID-19 Surveillance Team |
| EPI_ISL_821559, EPI_ISL_821560, EPI_ISL_821561, EPI_ISL_821562 | Lighthouse Lab in Milton Keynes | Wellcome Sanger Institute for the COVID-19 Genomics UK (COG-UK) Consortium | The Lighthouse Lab in Milton Keynes and Alex Alderton, Roberto Amato, Sonia Goncalves, Ewan Harrison, David K. Jackson, Ian Johnston, Dominic Kwiatkowski, Cordelia Langford, John Sillitoe on behalf of the Wellcome Sanger Institute COVID-19 Surveillance Team |
| EPI_ISL_821563, EPI_ISL_821564, EPI_ISL_821565 | Lighthouse Lab in Cambridge | Wellcome Sanger Institute for the COVID-19 Genomics UK (COG-UK) Consortium | Rob Howes, The Lighthouse Lab in Cambridge and Alex Alderton, Roberto Amato, Sonia Goncalves, Ewan Harrison, David K. Jackson, Ian Johnston, Dominic Kwiatkowski, Cordelia Langford, John Sillitoe on behalf of the Wellcome Sanger Institute COVID-19 Surveillance Team |
| EPI_ISL_821566 | Lighthouse Lab in Alderley Park | Wellcome Sanger Institute for the COVID-19 Genomics UK (COG-UK) Consortium | Jacquelyn Wynn, Mairead Hyland, The Lighthouse Lab in Alderley Park and Alex Alderton, Roberto Amato, Sonia Goncalves, Ewan Harrison, David K. Jackson, Ian Johnston, Dominic Kwiatkowski, Cordelia Langford, John Sillitoe on behalf of the Wellcome Sanger Institute COVID-19 Surveillance Team |
| EPI_ISL_821567 | Lighthouse Lab in Milton Keynes | Wellcome Sanger Institute for the COVID-19 Genomics UK (COG-UK) Consortium | The Lighthouse Lab in Milton Keynes and Alex Alderton, Roberto Amato, Sonia Goncalves, Ewan Harrison, David K. Jackson, Ian Johnston, Dominic Kwiatkowski, Cordelia Langford, John Sillitoe on behalf of the Wellcome Sanger Institute COVID-19 Surveillance Team |
| EPI_ISL_821568 | Lighthouse Lab in Cambridge | Wellcome Sanger Institute for the COVID-19 Genomics UK (COG-UK) Consortium | Rob Howes, The Lighthouse Lab in Cambridge and Alex Alderton, Roberto Amato, Sonia Goncalves, Ewan Harrison, David K. Jackson, Ian Johnston, Dominic Kwiatkowski, Cordelia Langford, John Sillitoe on behalf of the Wellcome Sanger Institute COVID-19 Surveillance Team |
| EPI_ISL_821569 | Lighthouse Lab in Milton Keynes | Wellcome Sanger Institute for the COVID-19 Genomics UK (COG-UK) Consortium | The Lighthouse Lab in Milton Keynes and Alex Alderton, Roberto Amato, Sonia Goncalves, Ewan Harrison, David K. Jackson, Ian Johnston, Dominic Kwiatkowski, Cordelia Langford, John Sillitoe on behalf of the Wellcome Sanger Institute COVID-19 Surveillance Team |
| EPI_ISL_821570 | Lighthouse Lab in Cambridge | Wellcome Sanger Institute for the COVID-19 Genomics UK (COG-UK) Consortium | Rob Howes, The Lighthouse Lab in Cambridge and Alex Alderton, Roberto Amato, Sonia Goncalves, Ewan Harrison, David K. Jackson, Ian Johnston, Dominic Kwiatkowski, Cordelia Langford, John Sillitoe on behalf of the Wellcome Sanger Institute COVID-19 Surveillance Team |
| EPI_ISL_821571, EPI_ISL_821572, EPI_ISL_821573, EPI_ISL_821574, EPI_ISL_821575 | Lighthouse Lab in Milton Keynes | Wellcome Sanger Institute for the COVID-19 Genomics UK (COG-UK) Consortium | The Lighthouse Lab in Milton Keynes and Alex Alderton, Roberto Amato, Sonia Goncalves, Ewan Harrison, David K. Jackson, Ian Johnston, Dominic Kwiatkowski, Cordelia Langford, John Sillitoe on behalf of the Wellcome Sanger Institute COVID-19 Surveillance Team |
| EPI_ISL_821576, EPI_ISL_821577, EPI_ISL_821579 | Lighthouse Lab in Cambridge | Wellcome Sanger Institute for the COVID-19 Genomics UK (COG-UK) Consortium | Rob Howes, The Lighthouse Lab in Cambridge and Alex Alderton, Roberto Amato, Sonia Goncalves, Ewan Harrison, David K. Jackson, Ian Johnston, Dominic Kwiatkowski, Cordelia Langford, John Sillitoe on behalf of the Wellcome Sanger Institute COVID-19 Surveillance Team |
| EPI_ISL_821580, EPI_ISL_821582 | Lighthouse Lab in Milton Keynes | Wellcome Sanger Institute for the COVID-19 Genomics UK (COG-UK) Consortium | The Lighthouse Lab in Milton Keynes and Alex Alderton, Roberto Amato, Sonia Goncalves, Ewan Harrison, David K. Jackson, Ian Johnston, Dominic Kwiatkowski, Cordelia Langford, John Sillitoe on behalf of the Wellcome Sanger Institute COVID-19 Surveillance Team |
| EPI_ISL_821583 | Lighthouse Lab in Cambridge | Wellcome Sanger Institute for the COVID-19 Genomics UK (COG-UK) Consortium | Rob Howes, The Lighthouse Lab in Cambridge and Alex Alderton, Roberto Amato, Sonia Goncalves, Ewan Harrison, David K. Jackson, Ian Johnston, Dominic Kwiatkowski, Cordelia Langford, John Sillitoe on behalf of the Wellcome Sanger Institute COVID-19 Surveillance Team |
| EPI_ISL_821584, EPI_ISL_821585, EPI_ISL_821586 | Lighthouse Lab in Alderley Park | Wellcome Sanger Institute for the COVID-19 Genomics UK (COG-UK) Consortium | Jacquelyn Wynn, Mairead Hyland, The Lighthouse Lab in Alderley Park and Alex Alderton, Roberto Amato, Sonia Goncalves, Ewan Harrison, David K. Jackson, Ian Johnston, Dominic Kwiatkowski, Cordelia Langford, John Sillitoe on behalf of the Wellcome Sanger Institute COVID-19 Surveillance Team |
| EPI_ISL_821587 | Lighthouse Lab in Milton Keynes | Wellcome Sanger Institute for the COVID-19 Genomics UK (COG-UK) Consortium | The Lighthouse Lab in Milton Keynes and Alex Alderton, Roberto Amato, Sonia Goncalves, Ewan Harrison, David K. Jackson, Ian Johnston, Dominic Kwiatkowski, Cordelia Langford, John Sillitoe on behalf of the Wellcome Sanger Institute COVID-19 Surveillance Team |
| EPI_ISL_821588 | Lighthouse Lab in Cambridge | Wellcome Sanger Institute for the COVID-19 Genomics UK (COG-UK) Consortium | Rob Howes, The Lighthouse Lab in Cambridge and Alex Alderton, Roberto Amato, Sonia Goncalves, Ewan Harrison, David K. Jackson, Ian Johnston, Dominic Kwiatkowski, Cordelia Langford, John Sillitoe on behalf of the Wellcome Sanger Institute COVID-19 Surveillance Team |
| EPI_ISL_821589 | Lighthouse Lab in Milton Keynes | Wellcome Sanger Institute for the COVID-19 Genomics UK | The Lighthouse Lab in Milton Keynes and Alex Alderton, Roberto Amato, Sonia Goncalves, Ewan Harrison, David K. Jackson, Ian Johnston, Dominic |

| | | | |
|---|---|---|---|
| | | (COG-UK) Consortium | Kwiatkowski, Cordelia Langford, John Sillitoe on behalf of the Wellcome Sanger Institute COVID-19 Surveillance Team |
| EPI_ISL_821590, EPI_ISL_821592, EPI_ISL_821593 | Lighthouse Lab in Cambridge | Wellcome Sanger Institute for the COVID-19 Genomics UK (COG-UK) Consortium | Rob Howes, The Lighthouse Lab in Cambridge and Alex Alderton, Roberto Amato, Sonia Goncalves, Ewan Harrison, David K. Jackson, Ian Johnston, Dominic Kwiatkowski, Cordelia Langford, John Sillitoe on behalf of the Wellcome Sanger Institute COVID-19 Surveillance Team |
| EPI_ISL_821594, EPI_ISL_821595 | Lighthouse Lab in Milton Keynes | Wellcome Sanger Institute for the COVID-19 Genomics UK (COG-UK) Consortium | The Lighthouse Lab in Milton Keynes and Alex Alderton, Roberto Amato, Sonia Goncalves, Ewan Harrison, David K. Jackson, Ian Johnston, Dominic Kwiatkowski, Cordelia Langford, John Sillitoe on behalf of the Wellcome Sanger Institute COVID-19 Surveillance Team |
| EPI_ISL_821596 | Lighthouse Lab in Cambridge | Wellcome Sanger Institute for the COVID-19 Genomics UK (COG-UK) Consortium | Rob Howes, The Lighthouse Lab in Cambridge and Alex Alderton, Roberto Amato, Sonia Goncalves, Ewan Harrison, David K. Jackson, Ian Johnston, Dominic Kwiatkowski, Cordelia Langford, John Sillitoe on behalf of the Wellcome Sanger Institute COVID-19 Surveillance Team |
| EPI_ISL_821597, EPI_ISL_821598 | Lighthouse Lab in Milton Keynes | Wellcome Sanger Institute for the COVID-19 Genomics UK (COG-UK) Consortium | The Lighthouse Lab in Milton Keynes and Alex Alderton, Roberto Amato, Sonia Goncalves, Ewan Harrison, David K. Jackson, Ian Johnston, Dominic Kwiatkowski, Cordelia Langford, John Sillitoe on behalf of the Wellcome Sanger Institute COVID-19 Surveillance Team |
| EPI_ISL_821599, EPI_ISL_821600 | Lighthouse Lab in Alderley Park | Wellcome Sanger Institute for the COVID-19 Genomics UK (COG-UK) Consortium | Jacquelyn Wynn, Mairead Hyland, The Lighthouse Lab in Alderley Park and Alex Alderton, Roberto Amato, Sonia Goncalves, Ewan Harrison, David K. Jackson, Ian Johnston, Dominic Kwiatkowski, Cordelia Langford, John Sillitoe on behalf of the Wellcome Sanger Institute COVID-19 Surveillance Team |
| EPI_ISL_821601, EPI_ISL_821603 | Lighthouse Lab in Milton Keynes | Wellcome Sanger Institute for the COVID-19 Genomics UK (COG-UK) Consortium | The Lighthouse Lab in Milton Keynes and Alex Alderton, Roberto Amato, Sonia Goncalves, Ewan Harrison, David K. Jackson, Ian Johnston, Dominic Kwiatkowski, Cordelia Langford, John Sillitoe on behalf of the Wellcome Sanger Institute COVID-19 Surveillance Team |
| EPI_ISL_821604, EPI_ISL_821605 | Lighthouse Lab in Cambridge | Wellcome Sanger Institute for the COVID-19 Genomics UK (COG-UK) Consortium | Rob Howes, The Lighthouse Lab in Cambridge and Alex Alderton, Roberto Amato, Sonia Goncalves, Ewan Harrison, David K. Jackson, Ian Johnston, Dominic Kwiatkowski, Cordelia Langford, John Sillitoe on behalf of the Wellcome Sanger Institute COVID-19 Surveillance Team |
| EPI_ISL_821606, EPI_ISL_821607 | Lighthouse Lab in Alderley Park | Wellcome Sanger Institute for the COVID-19 Genomics UK (COG-UK) Consortium | Jacquelyn Wynn, Mairead Hyland, The Lighthouse Lab in Alderley Park and Alex Alderton, Roberto Amato, Sonia Goncalves, Ewan Harrison, David K. Jackson, Ian Johnston, Dominic Kwiatkowski, Cordelia Langford, John Sillitoe on behalf of the Wellcome Sanger Institute COVID-19 Surveillance Team |
| EPI_ISL_821608 | Lighthouse Lab in Cambridge | Wellcome Sanger Institute for the COVID-19 Genomics UK (COG-UK) Consortium | Rob Howes, The Lighthouse Lab in Cambridge and Alex Alderton, Roberto Amato, Sonia Goncalves, Ewan Harrison, David K. Jackson, Ian Johnston, Dominic Kwiatkowski, Cordelia Langford, John Sillitoe on behalf of the Wellcome Sanger Institute COVID-19 Surveillance Team |
| EPI_ISL_821609, EPI_ISL_821610, EPI_ISL_821612 | Lighthouse Lab in Milton Keynes | Wellcome Sanger Institute for the COVID-19 Genomics UK (COG-UK) Consortium | The Lighthouse Lab in Milton Keynes and Alex Alderton, Roberto Amato, Sonia Goncalves, Ewan Harrison, David K. Jackson, Ian Johnston, Dominic Kwiatkowski, Cordelia Langford, John Sillitoe on behalf of the Wellcome Sanger Institute COVID-19 Surveillance Team |
| EPI_ISL_821614 | Lighthouse Lab in Alderley Park | Wellcome Sanger Institute for the COVID-19 Genomics UK (COG-UK) Consortium | Jacquelyn Wynn, Mairead Hyland, The Lighthouse Lab in Alderley Park and Alex Alderton, Roberto Amato, Sonia Goncalves, Ewan Harrison, David K. Jackson, Ian Johnston, Dominic Kwiatkowski, Cordelia Langford, John Sillitoe on behalf of the Wellcome Sanger Institute COVID-19 Surveillance Team |
| EPI_ISL_821615 | Lighthouse Lab in Milton Keynes | Wellcome Sanger Institute for the COVID-19 Genomics UK (COG-UK) Consortium | The Lighthouse Lab in Milton Keynes and Alex Alderton, Roberto Amato, Sonia Goncalves, Ewan Harrison, David K. Jackson, Ian Johnston, Dominic Kwiatkowski, Cordelia Langford, John Sillitoe on behalf of the Wellcome Sanger Institute COVID-19 Surveillance Team |
| EPI_ISL_821616, EPI_ISL_821618, EPI_ISL_821620, EPI_ISL_821622, EPI_ISL_821623, EPI_ISL_821624 | Lighthouse Lab in Cambridge | Wellcome Sanger Institute for the COVID-19 Genomics UK (COG-UK) Consortium | Rob Howes, The Lighthouse Lab in Cambridge and Alex Alderton, Roberto Amato, Sonia Goncalves, Ewan Harrison, David K. Jackson, Ian Johnston, Dominic Kwiatkowski, Cordelia Langford, John Sillitoe on behalf of the Wellcome Sanger Institute COVID-19 Surveillance Team |
| EPI_ISL_821625 | Lighthouse Lab in Milton Keynes | Wellcome Sanger Institute for the COVID-19 Genomics UK (COG-UK) Consortium | The Lighthouse Lab in Milton Keynes and Alex Alderton, Roberto Amato, Sonia Goncalves, Ewan Harrison, David K. Jackson, Ian Johnston, Dominic Kwiatkowski, Cordelia Langford, John Sillitoe on behalf of the Wellcome Sanger Institute COVID-19 Surveillance Team |
| EPI_ISL_821669, EPI_ISL_821834, EPI_ISL_821842, EPI_ISL_821858, EPI_ISL_821861, EPI_ISL_821895, EPI_ISL_821903, EPI_ISL_821950, EPI_ISL_821967 | Lighthouse Lab in Cambridge | Wellcome Sanger Institute for the COVID-19 Genomics UK (COG-UK) Consortium | Rob Howes, The Lighthouse Lab in Cambridge and Alex Alderton, Roberto Amato, Sonia Goncalves, Ewan Harrison, David K. Jackson, Ian Johnston, Dominic Kwiatkowski, Cordelia Langford, John Sillitoe on behalf of the Wellcome Sanger Institute COVID-19 Surveillance Team |
| EPI_ISL_821972, EPI_ISL_821973 | Lighthouse Lab in Alderley Park | Wellcome Sanger Institute for the COVID-19 Genomics UK (COG-UK) Consortium | Jacquelyn Wynn, Mairead Hyland, The Lighthouse Lab in Alderley Park and Alex Alderton, Roberto Amato, Sonia Goncalves, Ewan Harrison, David K. Jackson, Ian Johnston, Dominic Kwiatkowski, Cordelia Langford, John Sillitoe on behalf of the Wellcome Sanger Institute COVID-19 Surveillance Team |
| EPI_ISL_821975 | Lighthouse Lab in Glasgow | Wellcome Sanger Institute for the COVID-19 Genomics UK (COG-UK) Consortium | Harper VanSteenhouse, Yumi Kasai, David Gray, Carol Clugston, Anna Dominiczak and Alex Alderton, Roberto Amato, Sonia Goncalves, Ewan Harrison, David K. Jackson, Ian Johnston, Dominic Kwiatkowski, Cordelia Langford, John Sillitoe on behalf of the Wellcome Sanger Institute COVID-19 Surveillance Team |
| EPI_ISL_821987, EPI_ISL_821988, EPI_ISL_821991, EPI_ISL_821992 | Lighthouse Lab in Alderley Park | Wellcome Sanger Institute for the COVID-19 Genomics UK (COG-UK) Consortium | Jacquelyn Wynn, Mairead Hyland, The Lighthouse Lab in Alderley Park and Alex Alderton, Roberto Amato, Sonia Goncalves, Ewan Harrison, David K. Jackson, Ian Johnston, Dominic Kwiatkowski, Cordelia Langford, John Sillitoe on behalf of the Wellcome Sanger Institute COVID-19 Surveillance Team |
| EPI_ISL_821993, EPI_ISL_821995 | Lighthouse Lab in Glasgow | Wellcome Sanger Institute for the COVID-19 Genomics UK (COG-UK) Consortium | Harper VanSteenhouse, Yumi Kasai, David Gray, Carol Clugston, Anna Dominiczak and Alex Alderton, Roberto Amato, Sonia Goncalves, Ewan Harrison, David K. Jackson, Ian Johnston, Dominic Kwiatkowski, Cordelia Langford, John Sillitoe on behalf of the Wellcome Sanger Institute COVID-19 Surveillance Team |
| EPI_ISL_821996, EPI_ISL_821997 | Lighthouse Lab in Alderley Park | Wellcome Sanger Institute for the COVID-19 Genomics UK (COG-UK) Consortium | Jacquelyn Wynn, Mairead Hyland, The Lighthouse Lab in Alderley Park and Alex Alderton, Roberto Amato, Sonia Goncalves, Ewan Harrison, David K. Jackson, Ian Johnston, Dominic Kwiatkowski, Cordelia Langford, John Sillitoe on behalf of the Wellcome Sanger Institute COVID-19 Surveillance Team |
| EPI_ISL_822001, EPI_ISL_822004, EPI_ISL_822005, EPI_ISL_822006, EPI_ISL_822011 | Lighthouse Lab in Glasgow | Wellcome Sanger Institute for the COVID-19 Genomics UK (COG-UK) Consortium | Harper VanSteenhouse, Yumi Kasai, David Gray, Carol Clugston, Anna Dominiczak and Alex Alderton, Roberto Amato, Sonia Goncalves, Ewan Harrison, David K. Jackson, Ian Johnston, Dominic Kwiatkowski, Cordelia Langford, John Sillitoe on behalf of the Wellcome Sanger Institute COVID-19 Surveillance Team |
| EPI_ISL_822013, EPI_ISL_822019 | Lighthouse Lab in Alderley Park | Wellcome Sanger Institute for the COVID-19 Genomics UK (COG-UK) Consortium | Jacquelyn Wynn, Mairead Hyland, The Lighthouse Lab in Alderley Park and Alex Alderton, Roberto Amato, Sonia Goncalves, Ewan Harrison, David K. Jackson, Ian Johnston, Dominic Kwiatkowski, Cordelia Langford, John Sillitoe on behalf of the Wellcome Sanger Institute COVID-19 Surveillance Team |
| EPI_ISL_822022, EPI_ISL_822023, EPI_ISL_822029 | Lighthouse Lab in Glasgow | Wellcome Sanger Institute for the COVID-19 Genomics UK (COG-UK) Consortium | Harper VanSteenhouse, Yumi Kasai, David Gray, Carol Clugston, Anna Dominiczak and Alex Alderton, Roberto Amato, Sonia Goncalves, Ewan Harrison, David K. Jackson, Ian Johnston, Dominic Kwiatkowski, Cordelia Langford, John Sillitoe on behalf of the Wellcome Sanger Institute COVID-19 Surveillance Team |
| EPI_ISL_822030 | Lighthouse Lab in Alderley Park | Wellcome Sanger Institute for the COVID-19 Genomics UK (COG-UK) Consortium | Jacquelyn Wynn, Mairead Hyland, The Lighthouse Lab in Alderley Park and Alex Alderton, Roberto Amato, Sonia Goncalves, Ewan Harrison, David K. Jackson, Ian Johnston, Dominic Kwiatkowski, Cordelia Langford, John Sillitoe on behalf of the Wellcome Sanger Institute COVID-19 Surveillance Team |
| EPI_ISL_822033, EPI_ISL_822035, EPI_ISL_822040, EPI_ISL_822041 | Lighthouse Lab in Glasgow | Wellcome Sanger Institute for the COVID-19 Genomics UK (COG-UK) Consortium | Harper VanSteenhouse, Yumi Kasai, David Gray, Carol Clugston, Anna Dominiczak and Alex Alderton, Roberto Amato, Sonia Goncalves, Ewan Harrison, David K. Jackson, Ian Johnston, Dominic Kwiatkowski, Cordelia Langford, John Sillitoe on behalf of the Wellcome Sanger Institute COVID-19 Surveillance Team |
| EPI_ISL_822043 | Lighthouse Lab in Alderley Park | Wellcome Sanger Institute for the COVID-19 Genomics UK (COG-UK) Consortium | Jacquelyn Wynn, Mairead Hyland, The Lighthouse Lab in Alderley Park and Alex Alderton, Roberto Amato, Sonia Goncalves, Ewan Harrison, David K. Jackson, Ian Johnston, Dominic Kwiatkowski, Cordelia Langford, John Sillitoe on behalf of the Wellcome Sanger Institute COVID-19 Surveillance Team |
| EPI_ISL_822044 | Lighthouse Lab in Glasgow | Wellcome Sanger Institute for the COVID-19 Genomics UK (COG-UK) Consortium | Harper VanSteenhouse, Yumi Kasai, David Gray, Carol Clugston, Anna Dominiczak and Alex Alderton, Roberto Amato, Sonia Goncalves, Ewan Harrison, David K. Jackson, Ian Johnston, Dominic Kwiatkowski, Cordelia Langford, John Sillitoe on behalf of the Wellcome Sanger Institute COVID-19 Surveillance Team |
| EPI_ISL_822047 | Lighthouse Lab in Alderley Park | Wellcome Sanger Institute for the COVID-19 Genomics UK (COG-UK) Consortium | Jacquelyn Wynn, Mairead Hyland, The Lighthouse Lab in Alderley Park and Alex Alderton, Roberto Amato, Sonia Goncalves, Ewan Harrison, David K. Jackson, Ian Johnston, Dominic Kwiatkowski, Cordelia Langford, John Sillitoe on behalf of the Wellcome Sanger Institute COVID-19 Surveillance Team |
| EPI_ISL_822048, EPI_ISL_822051, EPI_ISL_822053, EPI_ISL_822059 | Lighthouse Lab in Glasgow | Wellcome Sanger Institute for the COVID-19 Genomics UK (COG-UK) Consortium | Harper VanSteenhouse, Yumi Kasai, David Gray, Carol Clugston, Anna Dominiczak and Alex Alderton, Roberto Amato, Sonia Goncalves, Ewan Harrison, David K. Jackson, Ian Johnston, Dominic Kwiatkowski, Cordelia Langford, John Sillitoe on behalf of the Wellcome Sanger Institute COVID-19 Surveillance Team |
| EPI_ISL_822060 | Lighthouse Lab in Alderley Park | Wellcome Sanger Institute for the COVID-19 Genomics UK (COG-UK) Consortium | Jacquelyn Wynn, Mairead Hyland, The Lighthouse Lab in Alderley Park and Alex Alderton, Roberto Amato, Sonia Goncalves, Ewan Harrison, David K. Jackson, Ian Johnston, Dominic Kwiatkowski, Cordelia Langford, John Sillitoe on behalf of the Wellcome Sanger Institute COVID-19 Surveillance Team |

| | | | |
|---|---|---|---|
| EPI_ISL_822063 | Lighthouse Lab in Glasgow | Wellcome Sanger Institute for the COVID-19 Genomics UK (COG-UK) Consortium | Harper VanSteenhouse, Yumi Kasai, David Gray, Carol Clugston, Anna Dominiczak and Alex Alderton, Roberto Amato, Sonia Goncalves, Ewan Harrison, David K. Jackson, Ian Johnston, Dominic Kwiatkowski, Cordelia Langford, John Sillitoe on behalf of the Wellcome Sanger Institute COVID-19 Surveillance Team |
| EPI_ISL_822065 | Lighthouse Lab in Alderley Park | Wellcome Sanger Institute for the COVID-19 Genomics UK (COG-UK) Consortium | Jacquelyn Wynn, Mairead Hyland, The Lighthouse Lab in Alderley Park and Alex Alderton, Roberto Amato, Sonia Goncalves, Ewan Harrison, David K. Jackson, Ian Johnston, Dominic Kwiatkowski, Cordelia Langford, John Sillitoe on behalf of the Wellcome Sanger Institute COVID-19 Surveillance Team |
| EPI_ISL_822067, EPI_ISL_822069, EPI_ISL_822072, EPI_ISL_822075, EPI_ISL_822078 | Lighthouse Lab in Glasgow | Wellcome Sanger Institute for the COVID-19 Genomics UK (COG-UK) Consortium | Harper VanSteenhouse, Yumi Kasai, David Gray, Carol Clugston, Anna Dominiczak and Alex Alderton, Roberto Amato, Sonia Goncalves, Ewan Harrison, David K. Jackson, Ian Johnston, Dominic Kwiatkowski, Cordelia Langford, John Sillitoe on behalf of the Wellcome Sanger Institute COVID-19 Surveillance Team |
| EPI_ISL_822080, EPI_ISL_822081 | Lighthouse Lab in Alderley Park | Wellcome Sanger Institute for the COVID-19 Genomics UK (COG-UK) Consortium | Jacquelyn Wynn, Mairead Hyland, The Lighthouse Lab in Alderley Park and Alex Alderton, Roberto Amato, Sonia Goncalves, Ewan Harrison, David K. Jackson, Ian Johnston, Dominic Kwiatkowski, Cordelia Langford, John Sillitoe on behalf of the Wellcome Sanger Institute COVID-19 Surveillance Team |
| EPI_ISL_822083 | Lighthouse Lab in Glasgow | Wellcome Sanger Institute for the COVID-19 Genomics UK (COG-UK) Consortium | Harper VanSteenhouse, Yumi Kasai, David Gray, Carol Clugston, Anna Dominiczak and Alex Alderton, Roberto Amato, Sonia Goncalves, Ewan Harrison, David K. Jackson, Ian Johnston, Dominic Kwiatkowski, Cordelia Langford, John Sillitoe on behalf of the Wellcome Sanger Institute COVID-19 Surveillance Team |
| EPI_ISL_822086, EPI_ISL_822094, EPI_ISL_822097 | Lighthouse Lab in Alderley Park | Wellcome Sanger Institute for the COVID-19 Genomics UK (COG-UK) Consortium | Jacquelyn Wynn, Mairead Hyland, The Lighthouse Lab in Alderley Park and Alex Alderton, Roberto Amato, Sonia Goncalves, Ewan Harrison, David K. Jackson, Ian Johnston, Dominic Kwiatkowski, Cordelia Langford, John Sillitoe on behalf of the Wellcome Sanger Institute COVID-19 Surveillance Team |
| EPI_ISL_822098, EPI_ISL_822099 | Lighthouse Lab in Glasgow | Wellcome Sanger Institute for the COVID-19 Genomics UK (COG-UK) Consortium | Harper VanSteenhouse, Yumi Kasai, David Gray, Carol Clugston, Anna Dominiczak and Alex Alderton, Roberto Amato, Sonia Goncalves, Ewan Harrison, David K. Jackson, Ian Johnston, Dominic Kwiatkowski, Cordelia Langford, John Sillitoe on behalf of the Wellcome Sanger Institute COVID-19 Surveillance Team |
| EPI_ISL_822104, EPI_ISL_822105 | Lighthouse Lab in Alderley Park | Wellcome Sanger Institute for the COVID-19 Genomics UK (COG-UK) Consortium | Jacquelyn Wynn, Mairead Hyland, The Lighthouse Lab in Alderley Park and Alex Alderton, Roberto Amato, Sonia Goncalves, Ewan Harrison, David K. Jackson, Ian Johnston, Dominic Kwiatkowski, Cordelia Langford, John Sillitoe on behalf of the Wellcome Sanger Institute COVID-19 Surveillance Team |
| EPI_ISL_822113, EPI_ISL_822117, EPI_ISL_822119, EPI_ISL_822122, EPI_ISL_822127, EPI_ISL_822130 | Lighthouse Lab in Glasgow | Wellcome Sanger Institute for the COVID-19 Genomics UK (COG-UK) Consortium | Harper VanSteenhouse, Yumi Kasai, David Gray, Carol Clugston, Anna Dominiczak and Alex Alderton, Roberto Amato, Sonia Goncalves, Ewan Harrison, David K. Jackson, Ian Johnston, Dominic Kwiatkowski, Cordelia Langford, John Sillitoe on behalf of the Wellcome Sanger Institute COVID-19 Surveillance Team |
| EPI_ISL_822135, EPI_ISL_822139 | Lighthouse Lab in Alderley Park | Wellcome Sanger Institute for the COVID-19 Genomics UK (COG-UK) Consortium | Jacquelyn Wynn, Mairead Hyland, The Lighthouse Lab in Alderley Park and Alex Alderton, Roberto Amato, Sonia Goncalves, Ewan Harrison, David K. Jackson, Ian Johnston, Dominic Kwiatkowski, Cordelia Langford, John Sillitoe on behalf of the Wellcome Sanger Institute COVID-19 Surveillance Team |
| EPI_ISL_822140 | Lighthouse Lab in Glasgow | Wellcome Sanger Institute for the COVID-19 Genomics UK (COG-UK) Consortium | Harper VanSteenhouse, Yumi Kasai, David Gray, Carol Clugston, Anna Dominiczak and Alex Alderton, Roberto Amato, Sonia Goncalves, Ewan Harrison, David K. Jackson, Ian Johnston, Dominic Kwiatkowski, Cordelia Langford, John Sillitoe on behalf of the Wellcome Sanger Institute COVID-19 Surveillance Team |
| EPI_ISL_822141, EPI_ISL_822142 | Lighthouse Lab in Alderley Park | Wellcome Sanger Institute for the COVID-19 Genomics UK (COG-UK) Consortium | Jacquelyn Wynn, Mairead Hyland, The Lighthouse Lab in Alderley Park and Alex Alderton, Roberto Amato, Sonia Goncalves, Ewan Harrison, David K. Jackson, Ian Johnston, Dominic Kwiatkowski, Cordelia Langford, John Sillitoe on behalf of the Wellcome Sanger Institute COVID-19 Surveillance Team |
| EPI_ISL_822148 | Lighthouse Lab in Glasgow | Wellcome Sanger Institute for the COVID-19 Genomics UK (COG-UK) Consortium | Harper VanSteenhouse, Yumi Kasai, David Gray, Carol Clugston, Anna Dominiczak and Alex Alderton, Roberto Amato, Sonia Goncalves, Ewan Harrison, David K. Jackson, Ian Johnston, Dominic Kwiatkowski, Cordelia Langford, John Sillitoe on behalf of the Wellcome Sanger Institute COVID-19 Surveillance Team |
| EPI_ISL_822150 | Lighthouse Lab in Alderley Park | Wellcome Sanger Institute for the COVID-19 Genomics UK (COG-UK) Consortium | Jacquelyn Wynn, Mairead Hyland, The Lighthouse Lab in Alderley Park and Alex Alderton, Roberto Amato, Sonia Goncalves, Ewan Harrison, David K. Jackson, Ian Johnston, Dominic Kwiatkowski, Cordelia Langford, John Sillitoe on behalf of the Wellcome Sanger Institute COVID-19 Surveillance Team |
| EPI_ISL_822153, EPI_ISL_822156, EPI_ISL_822158, EPI_ISL_822159, EPI_ISL_822165, EPI_ISL_822169, EPI_ISL_822170 | Lighthouse Lab in Glasgow | Wellcome Sanger Institute for the COVID-19 Genomics UK (COG-UK) Consortium | Harper VanSteenhouse, Yumi Kasai, David Gray, Carol Clugston, Anna Dominiczak and Alex Alderton, Roberto Amato, Sonia Goncalves, Ewan Harrison, David K. Jackson, Ian Johnston, Dominic Kwiatkowski, Cordelia Langford, John Sillitoe on behalf of the Wellcome Sanger Institute COVID-19 Surveillance Team |
| EPI_ISL_822173 | Lighthouse Lab in Alderley Park | Wellcome Sanger Institute for the COVID-19 Genomics UK (COG-UK) Consortium | Jacquelyn Wynn, Mairead Hyland, The Lighthouse Lab in Alderley Park and Alex Alderton, Roberto Amato, Sonia Goncalves, Ewan Harrison, David K. Jackson, Ian Johnston, Dominic Kwiatkowski, Cordelia Langford, John Sillitoe on behalf of the Wellcome Sanger Institute COVID-19 Surveillance Team |
| EPI_ISL_822178, EPI_ISL_822182 | Lighthouse Lab in Glasgow | Wellcome Sanger Institute for the COVID-19 Genomics UK (COG-UK) Consortium | Harper VanSteenhouse, Yumi Kasai, David Gray, Carol Clugston, Anna Dominiczak and Alex Alderton, Roberto Amato, Sonia Goncalves, Ewan Harrison, David K. Jackson, Ian Johnston, Dominic Kwiatkowski, Cordelia Langford, John Sillitoe on behalf of the Wellcome Sanger Institute COVID-19 Surveillance Team |
| EPI_ISL_822183, EPI_ISL_822184, EPI_ISL_822191, EPI_ISL_822193, EPI_ISL_822197, EPI_ISL_822199 | Lighthouse Lab in Alderley Park | Wellcome Sanger Institute for the COVID-19 Genomics UK (COG-UK) Consortium | Jacquelyn Wynn, Mairead Hyland, The Lighthouse Lab in Alderley Park and Alex Alderton, Roberto Amato, Sonia Goncalves, Ewan Harrison, David K. Jackson, Ian Johnston, Dominic Kwiatkowski, Cordelia Langford, John Sillitoe on behalf of the Wellcome Sanger Institute COVID-19 Surveillance Team |
| EPI_ISL_822201 | Lighthouse Lab in Glasgow | Wellcome Sanger Institute for the COVID-19 Genomics UK (COG-UK) Consortium | Harper VanSteenhouse, Yumi Kasai, David Gray, Carol Clugston, Anna Dominiczak and Alex Alderton, Roberto Amato, Sonia Goncalves, Ewan Harrison, David K. Jackson, Ian Johnston, Dominic Kwiatkowski, Cordelia Langford, John Sillitoe on behalf of the Wellcome Sanger Institute COVID-19 Surveillance Team |
| EPI_ISL_822202, EPI_ISL_822209 | Lighthouse Lab in Alderley Park | Wellcome Sanger Institute for the COVID-19 Genomics UK (COG-UK) Consortium | Jacquelyn Wynn, Mairead Hyland, The Lighthouse Lab in Alderley Park and Alex Alderton, Roberto Amato, Sonia Goncalves, Ewan Harrison, David K. Jackson, Ian Johnston, Dominic Kwiatkowski, Cordelia Langford, John Sillitoe on behalf of the Wellcome Sanger Institute COVID-19 Surveillance Team |
| EPI_ISL_822214, EPI_ISL_822223 | Lighthouse Lab in Glasgow | Wellcome Sanger Institute for the COVID-19 Genomics UK (COG-UK) Consortium | Harper VanSteenhouse, Yumi Kasai, David Gray, Carol Clugston, Anna Dominiczak and Alex Alderton, Roberto Amato, Sonia Goncalves, Ewan Harrison, David K. Jackson, Ian Johnston, Dominic Kwiatkowski, Cordelia Langford, John Sillitoe on behalf of the Wellcome Sanger Institute COVID-19 Surveillance Team |
| EPI_ISL_822224 | Lighthouse Lab in Alderley Park | Wellcome Sanger Institute for the COVID-19 Genomics UK (COG-UK) Consortium | Jacquelyn Wynn, Mairead Hyland, The Lighthouse Lab in Alderley Park and Alex Alderton, Roberto Amato, Sonia Goncalves, Ewan Harrison, David K. Jackson, Ian Johnston, Dominic Kwiatkowski, Cordelia Langford, John Sillitoe on behalf of the Wellcome Sanger Institute COVID-19 Surveillance Team |
| EPI_ISL_822230, EPI_ISL_822237, EPI_ISL_822240, EPI_ISL_822242, EPI_ISL_822243, EPI_ISL_822244, EPI_ISL_822245, EPI_ISL_822247 | Lighthouse Lab in Glasgow | Wellcome Sanger Institute for the COVID-19 Genomics UK (COG-UK) Consortium | Harper VanSteenhouse, Yumi Kasai, David Gray, Carol Clugston, Anna Dominiczak and Alex Alderton, Roberto Amato, Sonia Goncalves, Ewan Harrison, David K. Jackson, Ian Johnston, Dominic Kwiatkowski, Cordelia Langford, John Sillitoe on behalf of the Wellcome Sanger Institute COVID-19 Surveillance Team |
| EPI_ISL_822248 | Lighthouse Lab in Alderley Park | Wellcome Sanger Institute for the COVID-19 Genomics UK (COG-UK) Consortium | Jacquelyn Wynn, Mairead Hyland, The Lighthouse Lab in Alderley Park and Alex Alderton, Roberto Amato, Sonia Goncalves, Ewan Harrison, David K. Jackson, Ian Johnston, Dominic Kwiatkowski, Cordelia Langford, John Sillitoe on behalf of the Wellcome Sanger Institute COVID-19 Surveillance Team |
| EPI_ISL_822249 | Lighthouse Lab in Glasgow | Wellcome Sanger Institute for the COVID-19 Genomics UK (COG-UK) Consortium | Harper VanSteenhouse, Yumi Kasai, David Gray, Carol Clugston, Anna Dominiczak and Alex Alderton, Roberto Amato, Sonia Goncalves, Ewan Harrison, David K. Jackson, Ian Johnston, Dominic Kwiatkowski, Cordelia Langford, John Sillitoe on behalf of the Wellcome Sanger Institute COVID-19 Surveillance Team |
| EPI_ISL_822250 | Lighthouse Lab in Alderley Park | Wellcome Sanger Institute for the COVID-19 Genomics UK (COG-UK) Consortium | Jacquelyn Wynn, Mairead Hyland, The Lighthouse Lab in Alderley Park and Alex Alderton, Roberto Amato, Sonia Goncalves, Ewan Harrison, David K. Jackson, Ian Johnston, Dominic Kwiatkowski, Cordelia Langford, John Sillitoe on behalf of the Wellcome Sanger Institute COVID-19 Surveillance Team |
| EPI_ISL_822253, EPI_ISL_822256, EPI_ISL_822260, EPI_ISL_822264, EPI_ISL_822265, EPI_ISL_822267, EPI_ISL_822268, EPI_ISL_822272, EPI_ISL_822275, EPI_ISL_822281, EPI_ISL_822282 | | | |
| see above | Lighthouse Lab in Glasgow | Wellcome Sanger Institute for the COVID-19 Genomics UK (COG-UK) Consortium | Harper VanSteenhouse, Yumi Kasai, David Gray, Carol Clugston, Anna Dominiczak and Alex Alderton, Roberto Amato, Sonia Goncalves, Ewan Harrison, David K. Jackson, Ian Johnston, Dominic Kwiatkowski, Cordelia Langford, John Sillitoe on behalf of the Wellcome Sanger Institute COVID-19 Surveillance Team |
| EPI_ISL_822283 | Lighthouse Lab in Alderley Park | Wellcome Sanger Institute for the COVID-19 Genomics UK | Jacquelyn Wynn, Mairead Hyland, The Lighthouse Lab in Alderley Park and Alex Alderton, Roberto Amato, Sonia Goncalves, Ewan Harrison, David K. |

| | | | |
|---|---|---|---|
| | | (COG-UK) Consortium | Jackson, Ian Johnston, Dominic Kwiatkowski, Cordelia Langford, John Sillitoe on behalf of the Wellcome Sanger Institute COVID-19 Surveillance Team |
| EPI_ISL_822288 | Lighthouse Lab in Glasgow | Wellcome Sanger Institute for the COVID-19 Genomics UK (COG-UK) Consortium | Harper VanSteenhouse, Yumi Kasai, David Gray, Carol Clugston, Anna Dominiczak and Alex Alderton, Roberto Amato, Sonia Goncalves, Ewan Harrison, David K. Jackson, Ian Johnston, Dominic Kwiatkowski, Cordelia Langford, John Sillitoe on behalf of the Wellcome Sanger Institute COVID-19 Surveillance Team |
| EPI_ISL_822806, EPI_ISL_822807, EPI_ISL_822808, EPI_ISL_822809, EPI_ISL_822810, EPI_ISL_822811, EPI_ISL_822812, EPI_ISL_822813, EPI_ISL_822814, EPI_ISL_822815, EPI_ISL_822816, EPI_ISL_822817, EPI_ISL_822818, EPI_ISL_822819, EPI_ISL_822820, EPI_ISL_822821, EPI_ISL_822822, EPI_ISL_822823, EPI_ISL_822824, EPI_ISL_822825, EPI_ISL_822826, EPI_ISL_822827, EPI_ISL_822828, EPI_ISL_822829, EPI_ISL_822830, EPI_ISL_822831, EPI_ISL_822832, EPI_ISL_822833, EPI_ISL_822834, EPI_ISL_822835, EPI_ISL_822836 | | | |
| see above | Wales Specialist Virology Centre Sequencing lab: Pathogen Genomics Unit | COVID-19 Genomics UK (COG-UK) Consortium | Catherine Moore, Johnathan Evans, Laura Gifford, Malorie Perry, Simon Cottrell, Angela Marchbank, Alec Birchley, Alexander Adams, Amy Gaskin, Bree Gatica-Wilcox, Jason Coombes, Joel Southgate, Lauren Gilbert, Lee Graham, Nicole Pacchiarini, Sara Kumziene-Summerhayes, Sarah Taylor, Sophie Jones, Sara Rey, Matthew Bull, Joanne Watkins, Sally Corden, Tom Connor |
| EPI_ISL_824169, EPI_ISL_824209, EPI_ISL_824264, EPI_ISL_824267, EPI_ISL_824268, EPI_ISL_824270 | Dutch COVID-19 response team | National Institute for Public Health and the Environment (RIVM) | Adam Meijer, Harry Vennema, Jeroen Cremer, Sharon van den Brink, Bas van der Veer, AnneMarie van den Brandt, Florian Zwagemaker, Dennis Schmitz, Chantal Reusken, on behalf of the national COVID-19 response team |
| EPI_ISL_824554 | New Mexico Department of Health Scientific Laboratory | New Mexico Department of Health Scientific Laboratory | Ellie Johnson, Anastacia Griego-Fisher, D'eldra Malone |
| EPI_ISL_824832, EPI_ISL_824835, EPI_ISL_824836, EPI_ISL_824837, EPI_ISL_824838, EPI_ISL_824839, EPI_ISL_824840, EPI_ISL_824841 | Department of Clinical Microbiology | GIGA Medical Genomics | Keith Durkin, Maria Artesi, Sébastien Bontems, Raphaël Boreux, Bouchra Boujemla, Cécile Meex, Pierrette Melin, Marie-Pierre Hayette, Vincent Bours |
| EPI_ISL_824943 | Charité Universitätsmedizin Berlin, Institute of Virology, Charitéplatz 1, 10117 Berlin, Germany | Charité Universitätsmedizin Berlin, Institute of Virology, Charitéplatz 1, 10117 Berlin, Germany | Victor M Corman, Jörn Beheim-Schwarzbach, Tobias Bleicker, Julia Tesch, Barbara Mühlemann, Talitha Veith, Julia Schneider, Terry Jones, Christian Drosten |
| EPI_ISL_826466 | Lighthouse Lab in Milton Keynes | Wellcome Sanger Institute for the COVID-19 Genomics UK (COG-UK) Consortium | The Lighthouse Lab in Milton Keynes and Alex Alderton, Roberto Amato, Sonia Goncalves, Ewan Harrison, David K. Jackson, Ian Johnston, Dominic Kwiatkowski, Cordelia Langford, John Sillitoe on behalf of the Wellcome Sanger Institute COVID-19 Surveillance Team |
| EPI_ISL_826467, EPI_ISL_826468, EPI_ISL_826469, EPI_ISL_826470, EPI_ISL_826471 | Lighthouse Lab in Alderley Park | Wellcome Sanger Institute for the COVID-19 Genomics UK (COG-UK) Consortium | Jacquelyn Wynn, Mairead Hyland, The Lighthouse Lab in Alderley Park and Alex Alderton, Roberto Amato, Sonia Goncalves, Ewan Harrison, David K. Jackson, Ian Johnston, Dominic Kwiatkowski, Cordelia Langford, John Sillitoe on behalf of the Wellcome Sanger Institute COVID-19 Surveillance Team |
| EPI_ISL_826473 | Lighthouse Lab in Cambridge | Wellcome Sanger Institute for the COVID-19 Genomics UK (COG-UK) Consortium | Rob Howes, The Lighthouse Lab in Cambridge and Alex Alderton, Roberto Amato, Sonia Goncalves, Ewan Harrison, David K. Jackson, Ian Johnston, Dominic Kwiatkowski, Cordelia Langford, John Sillitoe on behalf of the Wellcome Sanger Institute COVID-19 Surveillance Team |
| EPI_ISL_826474 | Lighthouse Lab in Alderley Park | Wellcome Sanger Institute for the COVID-19 Genomics UK (COG-UK) Consortium | Jacquelyn Wynn, Mairead Hyland, The Lighthouse Lab in Alderley Park and Alex Alderton, Roberto Amato, Sonia Goncalves, Ewan Harrison, David K. Jackson, Ian Johnston, Dominic Kwiatkowski, Cordelia Langford, John Sillitoe on behalf of the Wellcome Sanger Institute COVID-19 Surveillance Team |
| EPI_ISL_826477 | Lighthouse Lab in Glasgow | Wellcome Sanger Institute for the COVID-19 Genomics UK (COG-UK) Consortium | Harper VanSteenhouse, Yumi Kasai, David Gray, Carol Clugston, Anna Dominiczak and Alex Alderton, Roberto Amato, Sonia Goncalves, Ewan Harrison, David K. Jackson, Ian Johnston, Dominic Kwiatkowski, Cordelia Langford, John Sillitoe on behalf of the Wellcome Sanger Institute COVID-19 Surveillance Team |
| EPI_ISL_826478 | Lighthouse Lab in Alderley Park | Wellcome Sanger Institute for the COVID-19 Genomics UK (COG-UK) Consortium | Jacquelyn Wynn, Mairead Hyland, The Lighthouse Lab in Alderley Park and Alex Alderton, Roberto Amato, Sonia Goncalves, Ewan Harrison, David K. Jackson, Ian Johnston, Dominic Kwiatkowski, Cordelia Langford, John Sillitoe on behalf of the Wellcome Sanger Institute COVID-19 Surveillance Team |
| EPI_ISL_826479 | Lighthouse Lab in Glasgow | Wellcome Sanger Institute for the COVID-19 Genomics UK (COG-UK) Consortium | Harper VanSteenhouse, Yumi Kasai, David Gray, Carol Clugston, Anna Dominiczak and Alex Alderton, Roberto Amato, Sonia Goncalves, Ewan Harrison, David K. Jackson, Ian Johnston, Dominic Kwiatkowski, Cordelia Langford, John Sillitoe on behalf of the Wellcome Sanger Institute COVID-19 Surveillance Team |
| EPI_ISL_826726, EPI_ISL_826727, EPI_ISL_826728, EPI_ISL_826729, EPI_ISL_826730, EPI_ISL_826731, EPI_ISL_826740, EPI_ISL_827038, EPI_ISL_827041, EPI_ISL_827044, EPI_ISL_827896, EPI_ISL_827897, EPI_ISL_827898 | | | |
| see above | deCODE genetics | deCODE genetics | Daniel F Gudbjartsson; Agnar Helgason; Hakon Jonsson; Olafur T Magnusson; Pall Melsted; Gudmundur L Norddahl; Jona Saemundsdottir; Asgeir Sigurdsson; Patrick Sulem; Arna B Agustsdottir; Hannes Eggertsson; Berglind Eiriksdottir; Run Fridriksdottir; Elisabet E Gardarsdottir; Gudmundur Georgsson; Olafia S Gretarsdottir; Kjartan R Gudmundsson; Thora R Gunnarsdottir; Arnaldur Gylfason; Hilma Holm; Brynjar O Jensson; Aslaug Jonasdottir; Kamilla S Josefsdottir; Thordur Kristjansson; Droplaug N Magnusdottir; Solvi Rognvaldsson; Louise le Roux; Gudrun Sigmundsdottir; Gardar Sveinbjornsson; Kristin E Sveinsdottir; Maney Sveinsdottir; Emil A Thorarensen; Bjarni Thorbjornsson; Gisli Masson; Ingileif Jonsdottir; Alma Moller; Thorolfur Gudnason; Karl G Kristinsson; Unnur Thorsteinsdottir; Kari Stefansson |
| EPI_ISL_827899 | The National University Hospital of Iceland | deCODE genetics | Daniel F Gudbjartsson; Agnar Helgason; Hakon Jonsson; Olafur T Magnusson; Pall Melsted; Gudmundur L Norddahl; Jona Saemundsdottir; Asgeir Sigurdsson; Patrick Sulem; Arna B Agustsdottir; Hannes Eggertsson; Berglind Eiriksdottir; Run Fridriksdottir; Elisabet E Gardarsdottir; Gudmundur Georgsson; Olafia S Gretarsdottir; Kjartan R Gudmundsson; Thora R Gunnarsdottir; Arnaldur Gylfason; Hilma Holm; Brynjar O Jensson; Aslaug Jonasdottir; Kamilla S Josefsdottir; Thordur Kristjansson; Droplaug N Magnusdottir; Solvi Rognvaldsson; Louise le Roux; Gudrun Sigmundsdottir; Gardar Sveinbjornsson; Kristin E Sveinsdottir; Maney Sveinsdottir; Emil A Thorarensen; Bjarni Thorbjornsson; Gisli Masson; Ingileif Jonsdottir; Alma Moller; Thorolfur Gudnason; Karl G Kristinsson; Unnur Thorsteinsdottir; Kari Stefansson |
| EPI_ISL_827930, EPI_ISL_827931, EPI_ISL_827933, EPI_ISL_828523, EPI_ISL_828538, EPI_ISL_828785, EPI_ISL_828873, EPI_ISL_829062, EPI_ISL_829080, EPI_ISL_829316, EPI_ISL_829323, EPI_ISL_829325, EPI_ISL_829326, EPI_ISL_829743, EPI_ISL_830186, EPI_ISL_830318, EPI_ISL_830396 | | | |
| see above | deCODE genetics | deCODE genetics | Daniel F Gudbjartsson; Agnar Helgason; Hakon Jonsson; Olafur T Magnusson; Pall Melsted; Gudmundur L Norddahl; Jona Saemundsdottir; Asgeir Sigurdsson; Patrick Sulem; Arna B Agustsdottir; Hannes Eggertsson; Berglind Eiriksdottir; Run Fridriksdottir; Elisabet E Gardarsdottir; Gudmundur Georgsson; Olafia S Gretarsdottir; Kjartan R Gudmundsson; Thora R Gunnarsdottir; Arnaldur Gylfason; Hilma Holm; Brynjar O Jensson; Aslaug Jonasdottir; Kamilla S Josefsdottir; Thordur Kristjansson; Droplaug N Magnusdottir; Solvi Rognvaldsson; Louise le Roux; Gudrun Sigmundsdottir; Gardar Sveinbjornsson; Kristin E Sveinsdottir; Maney Sveinsdottir; Emil A Thorarensen; Bjarni Thorbjornsson; Gisli Masson; Ingileif Jonsdottir; Alma Moller; Thorolfur Gudnason; Karl G Kristinsson; Unnur Thorsteinsdottir; Kari Stefansson |
| EPI_ISL_830628 | NORTHWELL HEALTH LABORATORIES | Wadsworth Center, New York State Department of Health | Kirsten St. George, Daryl M. Lamson, Alexis Russel, Matthew Shudt, Melissa A Leisner, Jonathan Plitnick, Navjot Singh, John Kelly, Erasmus Schneider, Erica Lasek-Nesselquist |
| EPI_ISL_830648 | Institute for Infectious Diseases, University of Bern, Switzerland | Institute for Infectious Diseases, University of Bern, Switzerland | Michel C Koch, Christian Baumann, Miguel A Terrazos Miani, Cora Sägesser, Pascal Bittel, Stephen L Leib, Peter Keller, Franziska Suter-Riniker, Alban Ramette |
| EPI_ISL_830669 | SUNY UPSTATE MEDICAL UNIVERSITY | Wadsworth Center, New York State Department of Health | Kirsten St. George, Daryl M. Lamson, Alexis Russel, Matthew Shudt, Melissa A Leisner, Jonathan Plitnick, Navjot Singh, John Kelly, Erasmus Schneider, Erica Lasek-Nesselquist |
| EPI_ISL_830683, EPI_ISL_830684, EPI_ISL_830685, EPI_ISL_830686, EPI_ISL_830687, EPI_ISL_830688, EPI_ISL_830689, EPI_ISL_830690, EPI_ISL_830691, EPI_ISL_830692, EPI_ISL_830693, EPI_ISL_830694, EPI_ISL_830695, EPI_ISL_830696, EPI_ISL_830697, EPI_ISL_830698, EPI_ISL_830709, EPI_ISL_830710, EPI_ISL_830711, EPI_ISL_830712, EPI_ISL_830713, EPI_ISL_830714, EPI_ISL_830715, EPI_ISL_830716 | | | |
| see above | ALBANY MEDICAL CENTER HOSPITAL CLINICAL LABORATORIES | Wadsworth Center, New York State Department of Health | Kirsten St. George, Daryl M. Lamson, Alexis Russel, Matthew Shudt, Melissa A Leisner, Jonathan Plitnick, Navjot Singh, John Kelly, Erasmus Schneider, Erica Lasek-Nesselquist |
| EPI_ISL_831552, EPI_ISL_831553, EPI_ISL_831554 | University of Wisconsin-Madison AIDS Vaccine Research Laboratories | University of Wisconsin-Madison AIDS Vaccine Research Laboratories | Gage Moreno, Katarina Braun, et al. AIDS Vaccine Research Laboratories |
| EPI_ISL_832006, EPI_ISL_832007 | Unilabs AB Skovde | The Public Health Agency of Sweden | Department of Microbiology, The Public Health Agency of Sweden |
| EPI_ISL_832090, EPI_ISL_832095 | Santa Clara County Public Health Laboratory | Santa Clara County Public Health Laboratory | Santa Clara County Public Health Department |
| EPI_ISL_832851, EPI_ISL_832871, EPI_ISL_832872, EPI_ISL_832876, EPI_ISL_832877, EPI_ISL_832878, EPI_ISL_832879, EPI_ISL_832880, EPI_ISL_832881, EPI_ISL_832882, EPI_ISL_832883, EPI_ISL_832884, EPI_ISL_832913, EPI_ISL_832945, EPI_ISL_832946 | | | |
| see above | Maine HETL | Tewhey Lab, The Jackson Laboratory | Matluk,N., Dewey,H., Iosue,F., Barter,M., Lynch,R., Munger,H. and Tewhey,R. |
| EPI_ISL_833208, EPI_ISL_833209, | Department of Virology and Immunology, University of Helsinki | Department of Virology, Faculty of Medicine, University of | Teemu Smura, Ravi Kant, Phuoc Truong, Hussein Alburkat, Hannimari Kallio-Kokko, Jenni Virtanen, Maija Suvanto, Fathiah Zakham, Essi Korhonen, Sari |

| | | | |
|---|---|---|---|
| EPI_ISL_833220, EPI_ISL_833221, EPI_ISL_833222, EPI_ISL_833223, EPI_ISL_833224, EPI_ISL_833225, EPI_ISL_833226 | and Helsinki University Hospital, Huslab Finland | Helsinki, Helsinki, Finland | Hannula, Harri Kangas, Pekka Ellonen, Olli Vapalahti |
| EPI_ISL_833344, EPI_ISL_833345, EPI_ISL_833346, EPI_ISL_833347, EPI_ISL_833348, EPI_ISL_833349, EPI_ISL_833350, EPI_ISL_833351, EPI_ISL_833352, EPI_ISL_833353, EPI_ISL_833354, EPI_ISL_833355 | | | |
| see above | Microbiology Division, South Carolina Department of Health and Environmental Control (SC DHEC) | Microbiology Division, South Carolina Department of Health and Environmental Control (SC DHEC) | Flores,H., Freeman,J. |
| EPI_ISL_833394, EPI_ISL_833395, EPI_ISL_833396, EPI_ISL_833397, EPI_ISL_833398, EPI_ISL_833399, EPI_ISL_833400 | St. Francis Medical Center | Los Angeles County PHL | P. Hemarajata et al. |
| EPI_ISL_833414 | USC Clinical Lab | Los Angeles County PHL | P. Hemarajata et al. |
| EPI_ISL_834213, EPI_ISL_834215, EPI_ISL_834217, EPI_ISL_834218, EPI_ISL_834220, EPI_ISL_834222, EPI_ISL_834224, EPI_ISL_834227, EPI_ISL_834230, EPI_ISL_834234, EPI_ISL_834235, EPI_ISL_834238, EPI_ISL_834239, EPI_ISL_834240, EPI_ISL_834241, EPI_ISL_834242, EPI_ISL_834243, EPI_ISL_834244, EPI_ISL_834245, EPI_ISL_834246, EPI_ISL_834247, EPI_ISL_834249, EPI_ISL_834251, EPI_ISL_834252, EPI_ISL_834253, EPI_ISL_834254, EPI_ISL_834257, EPI_ISL_834258, EPI_ISL_834259, EPI_ISL_834261, EPI_ISL_834262, EPI_ISL_834263, EPI_ISL_834265, EPI_ISL_834266, EPI_ISL_834267, EPI_ISL_834268, EPI_ISL_834269, EPI_ISL_834270, EPI_ISL_834272, EPI_ISL_834274, EPI_ISL_834277, EPI_ISL_834278, EPI_ISL_834279, EPI_ISL_834281, EPI_ISL_834284, EPI_ISL_834285, EPI_ISL_834286, EPI_ISL_834287, EPI_ISL_834289, EPI_ISL_834291, EPI_ISL_834292, EPI_ISL_834294, EPI_ISL_834295, EPI_ISL_834296, EPI_ISL_834297, EPI_ISL_834300, EPI_ISL_834301, EPI_ISL_834302, EPI_ISL_834303, EPI_ISL_834305, EPI_ISL_834306, EPI_ISL_834307, EPI_ISL_834310, EPI_ISL_834315, EPI_ISL_834316, EPI_ISL_834318, EPI_ISL_834319, EPI_ISL_834320, EPI_ISL_834322, EPI_ISL_834323, EPI_ISL_834324, EPI_ISL_834325, EPI_ISL_834326, EPI_ISL_834328, EPI_ISL_834329, EPI_ISL_834331, EPI_ISL_834332, EPI_ISL_834334, EPI_ISL_834337, EPI_ISL_834339, EPI_ISL_834340, EPI_ISL_834341, EPI_ISL_834342, EPI_ISL_834346, EPI_ISL_834347, EPI_ISL_834350, EPI_ISL_834351, EPI_ISL_834354, EPI_ISL_834356, EPI_ISL_834357, EPI_ISL_834358, EPI_ISL_834361, EPI_ISL_834362, EPI_ISL_834363, EPI_ISL_834364, EPI_ISL_834365, EPI_ISL_834366, EPI_ISL_834367, EPI_ISL_834368, EPI_ISL_834369, EPI_ISL_834370, EPI_ISL_834371, EPI_ISL_834372, EPI_ISL_834374, EPI_ISL_834379, EPI_ISL_834383, EPI_ISL_834384, EPI_ISL_834385, EPI_ISL_834386, EPI_ISL_834389, EPI_ISL_834390, EPI_ISL_834391, EPI_ISL_834393, EPI_ISL_834394, EPI_ISL_834396, EPI_ISL_834397, EPI_ISL_834400, EPI_ISL_834401, EPI_ISL_834402, EPI_ISL_834405, EPI_ISL_834406, EPI_ISL_834408, EPI_ISL_834410, EPI_ISL_834412, EPI_ISL_834413, EPI_ISL_834416, EPI_ISL_834419, EPI_ISL_834420, EPI_ISL_834422, EPI_ISL_834424, EPI_ISL_834425, EPI_ISL_834428, EPI_ISL_834429, EPI_ISL_834430, EPI_ISL_834431, EPI_ISL_834432, EPI_ISL_834433, EPI_ISL_834435, EPI_ISL_834436, EPI_ISL_834437, EPI_ISL_834439, EPI_ISL_834441, EPI_ISL_834442, EPI_ISL_834444, EPI_ISL_834446, EPI_ISL_834447, EPI_ISL_834451, EPI_ISL_834452, EPI_ISL_834453, EPI_ISL_834454, EPI_ISL_834455, EPI_ISL_834456, EPI_ISL_834457, EPI_ISL_834458, EPI_ISL_834460, EPI_ISL_834462, EPI_ISL_834464, EPI_ISL_834465, EPI_ISL_834469, EPI_ISL_834470, EPI_ISL_834473, EPI_ISL_834474, EPI_ISL_834475, EPI_ISL_834477, EPI_ISL_834478, EPI_ISL_834479, EPI_ISL_834483, EPI_ISL_834484, EPI_ISL_834487, EPI_ISL_834490, EPI_ISL_834491, EPI_ISL_834492, EPI_ISL_834495, EPI_ISL_834496, EPI_ISL_834501, EPI_ISL_834502, EPI_ISL_834504, EPI_ISL_834507, EPI_ISL_834508, EPI_ISL_834512, EPI_ISL_834513, EPI_ISL_834514, EPI_ISL_834515, EPI_ISL_834516, EPI_ISL_834518, EPI_ISL_834519, EPI_ISL_834520, EPI_ISL_834521, EPI_ISL_834522, EPI_ISL_834523, EPI_ISL_834524, EPI_ISL_834525, EPI_ISL_834526, EPI_ISL_834528, EPI_ISL_834530, EPI_ISL_834531, EPI_ISL_834532, EPI_ISL_834533, EPI_ISL_834534, EPI_ISL_834535, EPI_ISL_834537, EPI_ISL_834539, EPI_ISL_834540, EPI_ISL_834543, EPI_ISL_834546, EPI_ISL_834547, EPI_ISL_834548, EPI_ISL_834549, EPI_ISL_834551 | | | |
| see above | Lighthouse Lab in Alderley Park | Wellcome Sanger Institute for the COVID-19 Genomics UK (COG-UK) Consortium | Jacquelyn Wynn, Mairead Hyland, The Lighthouse Lab in Alderley Park and Alex Alderton, Roberto Amato, Sonia Goncalves, Ewan Harrison, David K. Jackson, Ian Johnston, Dominic Kwiatkowski, Cordelia Langford, John Sillitoe on behalf of the Wellcome Sanger Institute COVID-19 Surveillance Team |
| EPI_ISL_834562, EPI_ISL_834572, EPI_ISL_834583, EPI_ISL_834584, EPI_ISL_834640, EPI_ISL_834644, EPI_ISL_834646, EPI_ISL_834650, EPI_ISL_834729, EPI_ISL_834746, EPI_ISL_834761, EPI_ISL_834783, EPI_ISL_834791 | | | |
| see above | Lighthouse Lab in Glasgow | Wellcome Sanger Institute for the COVID-19 Genomics UK (COG-UK) Consortium | Harper VanSteenhouse, Yumi Kasai, David Gray, Carol Clugston, Anna Dominiczak and Alex Alderton, Roberto Amato, Sonia Goncalves, Ewan Harrison, David K. Jackson, Ian Johnston, Dominic Kwiatkowski, Cordelia Langford, John Sillitoe on behalf of the Wellcome Sanger Institute COVID-19 Surveillance Team |
| EPI_ISL_834798, EPI_ISL_834800, EPI_ISL_834802, EPI_ISL_834803, EPI_ISL_834805, EPI_ISL_834806, EPI_ISL_834807, EPI_ISL_834808, EPI_ISL_834809, EPI_ISL_834810, EPI_ISL_834812, EPI_ISL_834815, EPI_ISL_834817, EPI_ISL_834818, EPI_ISL_834819, EPI_ISL_834820, EPI_ISL_834821, EPI_ISL_834822, EPI_ISL_834824, EPI_ISL_834829, EPI_ISL_834831, EPI_ISL_834832, EPI_ISL_834835, EPI_ISL_834839, EPI_ISL_834840, EPI_ISL_834841, EPI_ISL_834842, EPI_ISL_834843, EPI_ISL_834845, EPI_ISL_834846, EPI_ISL_834847, EPI_ISL_834849, EPI_ISL_834850, EPI_ISL_834851, EPI_ISL_834852, EPI_ISL_834853, EPI_ISL_834854, EPI_ISL_834855, EPI_ISL_834857, EPI_ISL_834859, EPI_ISL_834860, EPI_ISL_834861, EPI_ISL_834862, EPI_ISL_834866, EPI_ISL_834867, EPI_ISL_834869, EPI_ISL_834872, EPI_ISL_834873, EPI_ISL_834875, EPI_ISL_834876, EPI_ISL_834877, EPI_ISL_834878, EPI_ISL_834879, EPI_ISL_834880, EPI_ISL_834881, EPI_ISL_834883, EPI_ISL_834885, EPI_ISL_834886, EPI_ISL_834888, EPI_ISL_834889, EPI_ISL_834891, EPI_ISL_834892, EPI_ISL_834893, EPI_ISL_834894, EPI_ISL_834896, EPI_ISL_834900, EPI_ISL_834901, EPI_ISL_834904, EPI_ISL_834906, EPI_ISL_834907, EPI_ISL_834908, EPI_ISL_834913, EPI_ISL_834914, EPI_ISL_834917, EPI_ISL_834919, EPI_ISL_834921, EPI_ISL_834922, EPI_ISL_834923, EPI_ISL_834924, EPI_ISL_834926, EPI_ISL_834927, EPI_ISL_834929, EPI_ISL_834930, EPI_ISL_834932, EPI_ISL_834934, EPI_ISL_834935, EPI_ISL_834936, EPI_ISL_834937, EPI_ISL_834938, EPI_ISL_834939, EPI_ISL_834940, EPI_ISL_834941, EPI_ISL_834944, EPI_ISL_834945, EPI_ISL_834947, EPI_ISL_834948, EPI_ISL_834950, EPI_ISL_834954, EPI_ISL_834955, EPI_ISL_834956, EPI_ISL_834957, EPI_ISL_834958, EPI_ISL_834960, EPI_ISL_834961, EPI_ISL_834962, EPI_ISL_834963, EPI_ISL_834964, EPI_ISL_834966, EPI_ISL_834967, EPI_ISL_834969, EPI_ISL_834970, EPI_ISL_834971, EPI_ISL_834973, EPI_ISL_834975, EPI_ISL_834977, EPI_ISL_834980, EPI_ISL_834982, EPI_ISL_834986, EPI_ISL_834987, EPI_ISL_834990, EPI_ISL_834992, EPI_ISL_834993, EPI_ISL_834995, EPI_ISL_834997, EPI_ISL_834998, EPI_ISL_834999, EPI_ISL_835000, EPI_ISL_835002, EPI_ISL_835006, EPI_ISL_835007, EPI_ISL_835011, EPI_ISL_835012, EPI_ISL_835014, EPI_ISL_835015, EPI_ISL_835019, EPI_ISL_835020, EPI_ISL_835021, EPI_ISL_835022, EPI_ISL_835023, EPI_ISL_835025, EPI_ISL_835027, EPI_ISL_835029, EPI_ISL_835030, EPI_ISL_835031, EPI_ISL_835032, EPI_ISL_835034, EPI_ISL_835035, EPI_ISL_835037, EPI_ISL_835039, EPI_ISL_835040, EPI_ISL_835042, EPI_ISL_835043, EPI_ISL_835044, EPI_ISL_835045, EPI_ISL_835046, EPI_ISL_835048, EPI_ISL_835049, EPI_ISL_835050, EPI_ISL_835051, EPI_ISL_835052, EPI_ISL_835054, EPI_ISL_835056, EPI_ISL_835057, EPI_ISL_835059, EPI_ISL_835061, EPI_ISL_835062, EPI_ISL_835063, EPI_ISL_835064, EPI_ISL_835070, EPI_ISL_835071, EPI_ISL_835072, EPI_ISL_835073, EPI_ISL_835074, EPI_ISL_835075, EPI_ISL_835076, EPI_ISL_835078, EPI_ISL_835079, EPI_ISL_835081, EPI_ISL_835082, EPI_ISL_835084, EPI_ISL_835086, EPI_ISL_835087, EPI_ISL_835089, EPI_ISL_835090, EPI_ISL_835091, EPI_ISL_835092, EPI_ISL_835093, EPI_ISL_835094, EPI_ISL_835095, EPI_ISL_835096, EPI_ISL_835097, EPI_ISL_835098, EPI_ISL_835099, EPI_ISL_835100, EPI_ISL_835101, EPI_ISL_835104, EPI_ISL_835105, EPI_ISL_835106, EPI_ISL_835111, EPI_ISL_835113, EPI_ISL_835116, EPI_ISL_835118, EPI_ISL_835119, EPI_ISL_835123, EPI_ISL_835124, EPI_ISL_835125, EPI_ISL_835126, EPI_ISL_835128, EPI_ISL_835129, EPI_ISL_835131, EPI_ISL_835134, EPI_ISL_835135, EPI_ISL_835136, EPI_ISL_835137, EPI_ISL_835139, EPI_ISL_835141, EPI_ISL_835142 | | | |
| see above | Lighthouse Lab in Alderley Park | Wellcome Sanger Institute for the COVID-19 Genomics UK (COG-UK) Consortium | Jacquelyn Wynn, Mairead Hyland, The Lighthouse Lab in Alderley Park and Alex Alderton, Roberto Amato, Sonia Goncalves, Ewan Harrison, David K. Jackson, Ian Johnston, Dominic Kwiatkowski, Cordelia Langford, John Sillitoe on behalf of the Wellcome Sanger Institute COVID-19 Surveillance Team |
| EPI_ISL_835145, EPI_ISL_835155, EPI_ISL_835177, EPI_ISL_835212, EPI_ISL_835235, EPI_ISL_835241, EPI_ISL_835247, EPI_ISL_835261, EPI_ISL_835282, EPI_ISL_835307, EPI_ISL_835319, EPI_ISL_835330, EPI_ISL_835333, EPI_ISL_835368, EPI_ISL_835378, EPI_ISL_835390 | | | |
| see above | Lighthouse Lab in Glasgow | Wellcome Sanger Institute for the COVID-19 Genomics UK (COG-UK) Consortium | Harper VanSteenhouse, Yumi Kasai, David Gray, Carol Clugston, Anna Dominiczak and Alex Alderton, Roberto Amato, Sonia Goncalves, Ewan Harrison, David K. Jackson, Ian Johnston, Dominic Kwiatkowski, Cordelia Langford, John Sillitoe on behalf of the Wellcome Sanger Institute COVID-19 Surveillance Team |
| EPI_ISL_835426, EPI_ISL_835427, EPI_ISL_835428, EPI_ISL_835429, EPI_ISL_835430, EPI_ISL_835431, EPI_ISL_835432, EPI_ISL_835434, EPI_ISL_835435, EPI_ISL_835436, EPI_ISL_835437, EPI_ISL_835438, EPI_ISL_835440, EPI_ISL_835441, EPI_ISL_835442, EPI_ISL_835443, EPI_ISL_835444, EPI_ISL_835445, EPI_ISL_835446, EPI_ISL_835449, EPI_ISL_835450, EPI_ISL_835451, EPI_ISL_835452, EPI_ISL_835453, EPI_ISL_835455, EPI_ISL_835456, EPI_ISL_835458, EPI_ISL_835459, EPI_ISL_835460, EPI_ISL_835461, EPI_ISL_835462, EPI_ISL_835463, EPI_ISL_835466, EPI_ISL_835467, EPI_ISL_835468, EPI_ISL_835469, EPI_ISL_835471, EPI_ISL_835472, EPI_ISL_835473, EPI_ISL_835476, EPI_ISL_835477, EPI_ISL_835478, EPI_ISL_835479, EPI_ISL_835480, EPI_ISL_835482, EPI_ISL_835484, EPI_ISL_835485, EPI_ISL_835489, EPI_ISL_835490, EPI_ISL_835491, EPI_ISL_835494, EPI_ISL_835496, EPI_ISL_835498, EPI_ISL_835501, EPI_ISL_835504, EPI_ISL_835508, EPI_ISL_835509, EPI_ISL_835510, EPI_ISL_835511, EPI_ISL_835512, EPI_ISL_835513, EPI_ISL_835514, EPI_ISL_835516, EPI_ISL_835517, EPI_ISL_835519, EPI_ISL_835520, EPI_ISL_835521, EPI_ISL_835522, EPI_ISL_835523, EPI_ISL_835524, EPI_ISL_835525, EPI_ISL_835526, EPI_ISL_835528, EPI_ISL_835529, EPI_ISL_835534, EPI_ISL_835536, EPI_ISL_835537, EPI_ISL_835538, EPI_ISL_835542, EPI_ISL_835543, EPI_ISL_835544, EPI_ISL_835545, EPI_ISL_835549, EPI_ISL_835551, EPI_ISL_835552, EPI_ISL_835554, EPI_ISL_835556, EPI_ISL_835557, EPI_ISL_835558, EPI_ISL_835559, EPI_ISL_835560, EPI_ISL_835561, EPI_ISL_835563, EPI_ISL_835565, EPI_ISL_835566, EPI_ISL_835567, EPI_ISL_835568, EPI_ISL_835569, EPI_ISL_835571, EPI_ISL_835572, EPI_ISL_835575, EPI_ISL_835578, EPI_ISL_835580, EPI_ISL_835581, EPI_ISL_835583, EPI_ISL_835584, EPI_ISL_835585, EPI_ISL_835586, EPI_ISL_835587, EPI_ISL_835588, EPI_ISL_835591, EPI_ISL_835593, EPI_ISL_835594, EPI_ISL_835595, EPI_ISL_835596, EPI_ISL_835598, EPI_ISL_835599, EPI_ISL_835601, EPI_ISL_835602, EPI_ISL_835604, EPI_ISL_835605, EPI_ISL_835606, EPI_ISL_835607, EPI_ISL_835608, EPI_ISL_835609, EPI_ISL_835611, EPI_ISL_835612, EPI_ISL_835613, EPI_ISL_835617, EPI_ISL_835619, EPI_ISL_835620, EPI_ISL_835621, EPI_ISL_835624, EPI_ISL_835625, EPI_ISL_835626, EPI_ISL_835627, EPI_ISL_835628, EPI_ISL_835629, EPI_ISL_835630, EPI_ISL_835632, EPI_ISL_835633, EPI_ISL_835634, EPI_ISL_835635, EPI_ISL_835636, EPI_ISL_835637, EPI_ISL_835638, EPI_ISL_835640, EPI_ISL_835641, EPI_ISL_835642, EPI_ISL_835644, EPI_ISL_835645, EPI_ISL_835648, EPI_ISL_835649, EPI_ISL_835650, EPI_ISL_835652, EPI_ISL_835653, EPI_ISL_835654, EPI_ISL_835656, EPI_ISL_835657, EPI_ISL_835658, EPI_ISL_835660, EPI_ISL_835661, EPI_ISL_835662, EPI_ISL_835663, EPI_ISL_835665, EPI_ISL_835666, EPI_ISL_835667, EPI_ISL_835668, EPI_ISL_835669, EPI_ISL_835670, EPI_ISL_835672, EPI_ISL_835674, EPI_ISL_835675, EPI_ISL_835676, EPI_ISL_835679, EPI_ISL_835681, EPI_ISL_835684, EPI_ISL_835685, EPI_ISL_835689, EPI_ISL_835690, EPI_ISL_835691, EPI_ISL_835693, EPI_ISL_835694, EPI_ISL_835696, EPI_ISL_835697, EPI_ISL_835699, EPI_ISL_835700, EPI_ISL_835702, EPI_ISL_835704, EPI_ISL_835705, EPI_ISL_835706, EPI_ISL_835708, EPI_ISL_835709 | | | |
| see above | Lighthouse Lab in Alderley Park | Wellcome Sanger Institute for the COVID-19 Genomics UK (COG-UK) Consortium | Jacquelyn Wynn, Mairead Hyland, The Lighthouse Lab in Alderley Park and Alex Alderton, Roberto Amato, Sonia Goncalves, Ewan Harrison, David K. Jackson, Ian Johnston, Dominic Kwiatkowski, Cordelia Langford, John Sillitoe on behalf of the Wellcome Sanger Institute COVID-19 Surveillance Team |
| EPI_ISL_835710, EPI_ISL_835711, EPI_ISL_835712, EPI_ISL_835713, EPI_ISL_835714, EPI_ISL_835715, EPI_ISL_835716, EPI_ISL_835717, EPI_ISL_835718, EPI_ISL_835719, EPI_ISL_835720, EPI_ISL_835721, EPI_ISL_835722, EPI_ISL_835723, EPI_ISL_835724, EPI_ISL_835725, EPI_ISL_835726, EPI_ISL_835727, EPI_ISL_835728, EPI_ISL_835729, EPI_ISL_835730, EPI_ISL_835731, EPI_ISL_835732, EPI_ISL_835733, EPI_ISL_835734, EPI_ISL_835735, EPI_ISL_835736, EPI_ISL_835737, EPI_ISL_835738, EPI_ISL_835739, EPI_ISL_835740, EPI_ISL_835741, EPI_ISL_835742, EPI_ISL_835743, EPI_ISL_835744, EPI_ISL_835745, EPI_ISL_835746, EPI_ISL_835747, EPI_ISL_835748, EPI_ISL_835749, EPI_ISL_835750, EPI_ISL_835751, EPI_ISL_835752, EPI_ISL_835753, EPI_ISL_835754, EPI_ISL_835755, EPI_ISL_835756, EPI_ISL_835757, EPI_ISL_835758, EPI_ISL_835759, EPI_ISL_835760, EPI_ISL_835761, EPI_ISL_835762, EPI_ISL_835763, EPI_ISL_835764, EPI_ISL_835765, EPI_ISL_835766, EPI_ISL_835767, EPI_ISL_835768, EPI_ISL_835769, EPI_ISL_835770, EPI_ISL_835771, EPI_ISL_835772, EPI_ISL_835773, EPI_ISL_835774, EPI_ISL_835775, EPI_ISL_835776, EPI_ISL_835777, EPI_ISL_835778, EPI_ISL_835779, EPI_ISL_835780, EPI_ISL_835781, EPI_ISL_835782, EPI_ISL_835783, EPI_ISL_835784, EPI_ISL_835785, EPI_ISL_835786, EPI_ISL_835787, EPI_ISL_835788, EPI_ISL_835789, EPI_ISL_835790, EPI_ISL_835791, EPI_ISL_835792, EPI_ISL_835793, EPI_ISL_835794, EPI_ISL_835795, EPI_ISL_835796, EPI_ISL_835797, EPI_ISL_835798, EPI_ISL_835799, EPI_ISL_835800, EPI_ISL_835801, EPI_ISL_835802, EPI_ISL_835803, EPI_ISL_835804, EPI_ISL_835805, EPI_ISL_835806, EPI_ISL_835807, EPI_ISL_835808, EPI_ISL_835809, EPI_ISL_835810, EPI_ISL_835811, EPI_ISL_835812, EPI_ISL_835813, EPI_ISL_835814, EPI_ISL_835815, EPI_ISL_835816, EPI_ISL_835817, EPI_ISL_835818, EPI_ISL_835819, EPI_ISL_835820, EPI_ISL_835821, EPI_ISL_835822, EPI_ISL_835823, EPI_ISL_835824, EPI_ISL_835825, EPI_ISL_835826, EPI_ISL_835827, EPI_ISL_835828, EPI_ISL_835829, EPI_ISL_835830, EPI_ISL_835831, EPI_ISL_835832, EPI_ISL_835833, EPI_ISL_835834, EPI_ISL_835835, EPI_ISL_835836, EPI_ISL_835837, EPI_ISL_835838, EPI_ISL_835839, EPI_ISL_835840, EPI_ISL_835841, EPI_ISL_835842, EPI_ISL_835843, EPI_ISL_835844, EPI_ISL_835845, EPI_ISL_835846, EPI_ISL_835847, EPI_ISL_835848, EPI_ISL_835849, EPI_ISL_835850, EPI_ISL_835851, EPI_ISL_835852, EPI_ISL_835853, EPI_ISL_835854, EPI_ISL_835855, EPI_ISL_835856, EPI_ISL_835857, EPI_ISL_835858, EPI_ISL_835859, EPI_ISL_835860, EPI_ISL_835861, EPI_ISL_835862, EPI_ISL_835863, EPI_ISL_835864, EPI_ISL_835865, EPI_ISL_835866, EPI_ISL_835867, EPI_ISL_835868, EPI_ISL_835869, EPI_ISL_835870, EPI_ISL_835871, EPI_ISL_835872, EPI_ISL_835873, EPI_ISL_835874, EPI_ISL_835875, EPI_ISL_835876, EPI_ISL_835877, EPI_ISL_835878, EPI_ISL_835879, EPI_ISL_835880, EPI_ISL_835881, EPI_ISL_835882, EPI_ISL_835883, EPI_ISL_835884, EPI_ISL_835885, EPI_ISL_835886, EPI_ISL_835887, EPI_ISL_835888, EPI_ISL_835889, EPI_ISL_835890, EPI_ISL_835891, EPI_ISL_835892, EPI_ISL_835893, EPI_ISL_835894, EPI_ISL_835895, EPI_ISL_835896, EPI_ISL_835897, EPI_ISL_835898, EPI_ISL_835899, EPI_ISL_835900, EPI_ISL_835901, EPI_ISL_835902, EPI_ISL_835903, EPI_ISL_835904, EPI_ISL_835905, EPI_ISL_835906, EPI_ISL_835907, EPI_ISL_835908, EPI_ISL_835909, EPI_ISL_835910, EPI_ISL_835911, EPI_ISL_835912, EPI_ISL_835913, EPI_ISL_835914, EPI_ISL_835915, EPI_ISL_835916, EPI_ISL_835917, EPI_ISL_835918, EPI_ISL_835919, EPI_ISL_835920, EPI_ISL_835921, EPI_ISL_835922, EPI_ISL_835923, EPI_ISL_835924, EPI_ISL_835925, EPI_ISL_835926, EPI_ISL_835927, EPI_ISL_835928, EPI_ISL_835929, EPI_ISL_835930, EPI_ISL_835931, EPI_ISL_835932, EPI_ISL_835933, EPI_ISL_835934, EPI_ISL_835935, EPI_ISL_835936, EPI_ISL_835937, EPI_ISL_835938, EPI_ISL_835939, EPI_ISL_835940, EPI_ISL_835941, EPI_ISL_835942, EPI_ISL_835943, EPI_ISL_835944, EPI_ISL_835945, EPI_ISL_835946, EPI_ISL_835947, EPI_ISL_835948, EPI_ISL_835949, EPI_ISL_835950, EPI_ISL_835951, EPI_ISL_835952, EPI_ISL_835953, EPI_ISL_835954, EPI_ISL_835955, EPI_ISL_835956, EPI_ISL_835957, EPI_ISL_835958, EPI_ISL_835959, EPI_ISL_835960, EPI_ISL_835961, EPI_ISL_835962, EPI_ISL_835963, EPI_ISL_835964, EPI_ISL_835965, EPI_ISL_835966, EPI_ISL_835967, EPI_ISL_835968, EPI_ISL_835969, EPI_ISL_835970, EPI_ISL_835971, EPI_ISL_835972, EPI_ISL_835973, EPI_ISL_835974, EPI_ISL_835975, EPI_ISL_835976, EPI_ISL_835977, EPI_ISL_835978, EPI_ISL_835979, EPI_ISL_835980, EPI_ISL_835981, EPI_ISL_835982, EPI_ISL_835983, EPI_ISL_835984, EPI_ISL_835985, EPI_ISL_835986, EPI_ISL_835987, EPI_ISL_835988, EPI_ISL_835989, EPI_ISL_835990, EPI_ISL_835991, EPI_ISL_835992, EPI_ISL_835993, EPI_ISL_835994, EPI_ISL_835995, EPI_ISL_835996, EPI_ISL_835997, EPI_ISL_835998, EPI_ISL_835999, EPI_ISL_836000, EPI_ISL_836001, EPI_ISL_836002, EPI_ISL_836003, EPI_ISL_836004, EPI_ISL_836005, EPI_ISL_836006, EPI_ISL_836007, EPI_ISL_836008, EPI_ISL_836009, EPI_ISL_836010, EPI_ISL_836011, EPI_ISL_836012, EPI_ISL_836013, EPI_ISL_836014, EPI_ISL_836015, EPI_ISL_836016, EPI_ISL_836017, EPI_ISL_836018, EPI_ISL_836019, EPI_ISL_836020, EPI_ISL_836021, EPI_ISL_836022, EPI_ISL_836023, EPI_ISL_836024, EPI_ISL_836025, EPI_ISL_836026, EPI_ISL_836027, EPI_ISL_836028, EPI_ISL_836029, EPI_ISL_836030, EPI_ISL_836031, EPI_ISL_836032, EPI_ISL_836033, | | | |

| | | | |
|---|---|---|---|
| EPI_ISL_836034, EPI_ISL_836035, EPI_ISL_836036, EPI_ISL_836037, EPI_ISL_836038, EPI_ISL_836039, EPI_ISL_836040, EPI_ISL_836041, EPI_ISL_836042, EPI_ISL_836043, EPI_ISL_836044, EPI_ISL_836045, EPI_ISL_836046, EPI_ISL_836047, EPI_ISL_836048, EPI_ISL_836049, EPI_ISL_836050, EPI_ISL_836051, EPI_ISL_836052, EPI_ISL_836053, EPI_ISL_836054, EPI_ISL_836055, EPI_ISL_836056, EPI_ISL_836057, EPI_ISL_836058, EPI_ISL_836059, EPI_ISL_836060, EPI_ISL_836061, EPI_ISL_836062, EPI_ISL_836063, EPI_ISL_836064, EPI_ISL_836065, EPI_ISL_836066, EPI_ISL_836067, EPI_ISL_836068, EPI_ISL_836069, EPI_ISL_836070, EPI_ISL_836071, EPI_ISL_836072, EPI_ISL_836073, EPI_ISL_836074, EPI_ISL_836076, EPI_ISL_836080, EPI_ISL_836081, EPI_ISL_836082, EPI_ISL_836085, EPI_ISL_836086, EPI_ISL_836087, EPI_ISL_836090, EPI_ISL_836091, EPI_ISL_836092, EPI_ISL_836093, EPI_ISL_836094, EPI_ISL_836096, EPI_ISL_836097, EPI_ISL_836098, EPI_ISL_836099, EPI_ISL_836100, EPI_ISL_836102, EPI_ISL_836103, EPI_ISL_836104, EPI_ISL_836109, EPI_ISL_836110, EPI_ISL_836113, EPI_ISL_836114, EPI_ISL_836116, EPI_ISL_836117, EPI_ISL_836118, EPI_ISL_836119, EPI_ISL_836121, EPI_ISL_836122, EPI_ISL_836123, EPI_ISL_836124, EPI_ISL_836125, EPI_ISL_836126, EPI_ISL_836127, EPI_ISL_836130, EPI_ISL_836133, EPI_ISL_836134, EPI_ISL_836135, EPI_ISL_836136, EPI_ISL_836139, EPI_ISL_836140, EPI_ISL_836141, EPI_ISL_836145, EPI_ISL_836146, EPI_ISL_836148, EPI_ISL_836150, EPI_ISL_836151, EPI_ISL_836152, EPI_ISL_836153, EPI_ISL_836157, EPI_ISL_836158, EPI_ISL_836159, EPI_ISL_836161, EPI_ISL_836163, EPI_ISL_836164, EPI_ISL_836166, EPI_ISL_836167, EPI_ISL_836170, EPI_ISL_836171, EPI_ISL_836172, EPI_ISL_836173, EPI_ISL_836175, EPI_ISL_836176, EPI_ISL_836177, EPI_ISL_836179, EPI_ISL_836181, EPI_ISL_836182, EPI_ISL_836183, EPI_ISL_836185, EPI_ISL_836186, EPI_ISL_836187, EPI_ISL_836188, EPI_ISL_836189, EPI_ISL_836191, EPI_ISL_836193, EPI_ISL_836194, EPI_ISL_836195, EPI_ISL_836196, EPI_ISL_836199, EPI_ISL_836200, EPI_ISL_836202, EPI_ISL_836204, EPI_ISL_836205, EPI_ISL_836206, EPI_ISL_836207, EPI_ISL_836208, EPI_ISL_836209, EPI_ISL_836210, EPI_ISL_836213, EPI_ISL_836214, EPI_ISL_836215, EPI_ISL_836216, EPI_ISL_836217, EPI_ISL_836219, EPI_ISL_836220, EPI_ISL_836221, EPI_ISL_836222, EPI_ISL_836224, EPI_ISL_836226, EPI_ISL_836227, EPI_ISL_836229, EPI_ISL_836232, EPI_ISL_836235, EPI_ISL_836236, EPI_ISL_836237, EPI_ISL_836239, EPI_ISL_836240, EPI_ISL_836242, EPI_ISL_836243, EPI_ISL_836244, EPI_ISL_836245, EPI_ISL_836246, EPI_ISL_836249, EPI_ISL_836253, EPI_ISL_836254, EPI_ISL_836255, EPI_ISL_836256, EPI_ISL_836257, EPI_ISL_836258, EPI_ISL_836259, EPI_ISL_836260, EPI_ISL_836263, EPI_ISL_836264, EPI_ISL_836265, EPI_ISL_836267, EPI_ISL_836269, EPI_ISL_836270, EPI_ISL_836271, EPI_ISL_836272, EPI_ISL_836273, EPI_ISL_836276, EPI_ISL_836277, EPI_ISL_836278, EPI_ISL_836279, EPI_ISL_836280, EPI_ISL_836281, EPI_ISL_836283, EPI_ISL_836286, EPI_ISL_836287, EPI_ISL_836288, EPI_ISL_836289, EPI_ISL_836290, EPI_ISL_836294, EPI_ISL_836295, EPI_ISL_836296, EPI_ISL_836297, EPI_ISL_836299, EPI_ISL_836300, EPI_ISL_836301, EPI_ISL_836302, EPI_ISL_836303, EPI_ISL_836304, EPI_ISL_836309, EPI_ISL_836312, EPI_ISL_836316, EPI_ISL_836317, EPI_ISL_836322, EPI_ISL_836324, EPI_ISL_836325, EPI_ISL_836326, EPI_ISL_836327, EPI_ISL_836330, EPI_ISL_836331, EPI_ISL_836332, EPI_ISL_836334, EPI_ISL_836335, EPI_ISL_836337, EPI_ISL_836339, EPI_ISL_836340, EPI_ISL_836341, EPI_ISL_836344, EPI_ISL_836345, EPI_ISL_836346, EPI_ISL_836347, EPI_ISL_836348, EPI_ISL_836349, EPI_ISL_836350, EPI_ISL_836353, EPI_ISL_836354, EPI_ISL_836355, EPI_ISL_836359, EPI_ISL_836360, EPI_ISL_836361, EPI_ISL_836364, EPI_ISL_836365, EPI_ISL_836368, EPI_ISL_836369, EPI_ISL_836370, EPI_ISL_836372, EPI_ISL_836374, EPI_ISL_836375, EPI_ISL_836376, EPI_ISL_836379, EPI_ISL_836380, EPI_ISL_836381, EPI_ISL_836382, EPI_ISL_836383, EPI_ISL_836384, EPI_ISL_836385, EPI_ISL_836386, EPI_ISL_836388, EPI_ISL_836389, EPI_ISL_836391, EPI_ISL_836392, EPI_ISL_836393, EPI_ISL_836394, EPI_ISL_836397, EPI_ISL_836398, EPI_ISL_836399, EPI_ISL_836400, EPI_ISL_836401, EPI_ISL_836402, EPI_ISL_836404, EPI_ISL_836405, EPI_ISL_836408, EPI_ISL_836411, EPI_ISL_836412, EPI_ISL_836413, EPI_ISL_836414, EPI_ISL_836417, EPI_ISL_836419, EPI_ISL_836420 | | | |
| see above | Lighthouse Lab in Milton Keynes | Wellcome Sanger Institute for the COVID-19 Genomics UK (COG-UK) Consortium | The Lighthouse Lab in Milton Keynes and Alex Alderton, Roberto Amato, Sonia Goncalves, Ewan Harrison, David K. Jackson, Ian Johnston, Dominic Kwiatkowski, Cordelia Langford, John Sillitoe on behalf of the Wellcome Sanger Institute COVID-19 Surveillance Team |
| EPI_ISL_836754, EPI_ISL_836765, EPI_ISL_836772, EPI_ISL_836777, EPI_ISL_836781, EPI_ISL_836788, EPI_ISL_836792, EPI_ISL_836795, EPI_ISL_836797, EPI_ISL_836803, EPI_ISL_836806, EPI_ISL_836807, EPI_ISL_836811, EPI_ISL_836813, EPI_ISL_836814, EPI_ISL_836822, EPI_ISL_836836, EPI_ISL_836837, EPI_ISL_836842, EPI_ISL_836843, EPI_ISL_836848, EPI_ISL_836851, EPI_ISL_836858, EPI_ISL_836859, EPI_ISL_836862, EPI_ISL_836863, EPI_ISL_836866, EPI_ISL_836867, EPI_ISL_836877, EPI_ISL_836883, EPI_ISL_836887, EPI_ISL_836890, EPI_ISL_836891, EPI_ISL_836897, EPI_ISL_836903 | | | |
| see above | Lighthouse Lab in Alderley Park | Wellcome Sanger Institute for the COVID-19 Genomics UK (COG-UK) Consortium | Jacquelyn Wynn, Mairead Hyland, The Lighthouse Lab in Alderley Park and Alex Alderton, Roberto Amato, Sonia Goncalves, Ewan Harrison, David K. Jackson, Ian Johnston, Dominic Kwiatkowski, Cordelia Langford, John Sillitoe on behalf of the Wellcome Sanger Institute COVID-19 Surveillance Team |
| EPI_ISL_836907 | Lighthouse Lab in Glasgow | Wellcome Sanger Institute for the COVID-19 Genomics UK (COG-UK) Consortium | Harper VanSteenhouse, Yumi Kasai, David Gray, Carol Clugston, Anna Dominiczak and Alex Alderton, Roberto Amato, Sonia Goncalves, Ewan Harrison, David K. Jackson, Ian Johnston, Dominic Kwiatkowski, Cordelia Langford, John Sillitoe on behalf of the Wellcome Sanger Institute COVID-19 Surveillance Team |
| EPI_ISL_836909, EPI_ISL_836910, EPI_ISL_836913, EPI_ISL_836920, EPI_ISL_836921, EPI_ISL_836922, EPI_ISL_836926, EPI_ISL_836928, EPI_ISL_836929, EPI_ISL_836936, EPI_ISL_836940, EPI_ISL_836946, EPI_ISL_836947, EPI_ISL_836948, EPI_ISL_836955, EPI_ISL_836975, EPI_ISL_836976 | | | |
| see above | Lighthouse Lab in Alderley Park | Wellcome Sanger Institute for the COVID-19 Genomics UK (COG-UK) Consortium | Jacquelyn Wynn, Mairead Hyland, The Lighthouse Lab in Alderley Park and Alex Alderton, Roberto Amato, Sonia Goncalves, Ewan Harrison, David K. Jackson, Ian Johnston, Dominic Kwiatkowski, Cordelia Langford, John Sillitoe on behalf of the Wellcome Sanger Institute COVID-19 Surveillance Team |
| EPI_ISL_837052, EPI_ISL_837067, EPI_ISL_837077, EPI_ISL_837173, EPI_ISL_837174, EPI_ISL_837175, EPI_ISL_837239, EPI_ISL_837240 | Respiratory Virus Unit, National Infection Service, Public Health England | COVID-19 Genomics UK (COG-UK) Consortium | PHE Covid Sequencing Team |
| EPI_ISL_837510, EPI_ISL_837526 | UW Virology Lab | UW Virology Lab | Pavitra Roychoudhury, Hong Xie, Lasata Shrestha, Meei-Li Huang, Keith R Jerome, Alexander Greninger |
| EPI_ISL_837843, EPI_ISL_837846, EPI_ISL_837849, EPI_ISL_837852, EPI_ISL_837854, EPI_ISL_837855, EPI_ISL_837856, EPI_ISL_837857, EPI_ISL_837859, EPI_ISL_837860, EPI_ISL_837861, EPI_ISL_837862, EPI_ISL_837863, EPI_ISL_837864, EPI_ISL_837865, EPI_ISL_837869, EPI_ISL_837870, EPI_ISL_837871, EPI_ISL_837905, EPI_ISL_837908, EPI_ISL_837909, EPI_ISL_837911, EPI_ISL_837913, EPI_ISL_837914, EPI_ISL_837915, EPI_ISL_837916, EPI_ISL_837917, EPI_ISL_837918, EPI_ISL_837919, EPI_ISL_837923, EPI_ISL_837924, EPI_ISL_837925, EPI_ISL_837926, EPI_ISL_837927, EPI_ISL_837928, EPI_ISL_837930, EPI_ISL_837932, EPI_ISL_837933 | | | |
| see above | Department of Pathology, University of Cambridge | COVID-19 Genomics UK (COG-UK) Consortium | Aminu S. Jahun, Yasmin Chaudhry, Grant Hall, Iliana Georgana, Myra Hosmillo, Martin D. Curran, Malte Pinckert, Surendra Parmar, Ian Goodfellow |
| EPI_ISL_838331, EPI_ISL_838334, EPI_ISL_838335, EPI_ISL_838336, EPI_ISL_838337, EPI_ISL_838338, EPI_ISL_838339 | University of Exeter | COVID-19 Genomics UK (COG-UK) Consortium | Ben Temperton,Aaron Jeffries,Michelle Michelsen,Joanna Warwick-Dugdale,Audrey Farbos,Robyn Manley,Stephen Michell,Jane Masoli |
| EPI_ISL_838586, EPI_ISL_838587, EPI_ISL_838592, EPI_ISL_838593, EPI_ISL_838597, EPI_ISL_838598, EPI_ISL_838600, EPI_ISL_838602, EPI_ISL_838603, EPI_ISL_838605, EPI_ISL_838610, EPI_ISL_838612, EPI_ISL_838613, EPI_ISL_838618, EPI_ISL_838620, EPI_ISL_838622, EPI_ISL_838628, EPI_ISL_838633, EPI_ISL_838640, EPI_ISL_838641, EPI_ISL_838652, EPI_ISL_838653, EPI_ISL_838655, EPI_ISL_838667, EPI_ISL_838668, EPI_ISL_838669, EPI_ISL_838670, EPI_ISL_838671 | | | |
| see above | Liverpool Clinical Laboratories | COVID-19 Genomics UK (COG-UK) Consortium | Sam Haldenby, Anita Lucaci, Steve Paterson, Julian Hiscox, Alistair Darby, M Almsaud, A Alrezaihi, Muhannad Alruwaili, Stuart D Armstrong, Jones Benjamin, Eleanor G Bentley, Anu Chawla, Jordan J Clark, Angela Cowell, Richard Eccles, Isabel García-Dorival, Matthew Gemmell, Alessandro Gerada, PKF Gilmore, Richard Gregory, Ximeng Han, Catherine Hartley, Margaret Hughes, Miren Iturriza-Gomara, James Johnson, L Luu, Jenifer Manson, Charlotte Nelson, Elaine O'Toole, Cassie Olateju, Rebekah Penrice-Randal , Lucille Rainbow, N.P Randle, Trevor Ian Robinson, Parul Sharma, Ghada T Shawli, James P Stewart, Neil Swainston, Ecaterina Vamos, Joanne Watts, Mark Whitehead |
| EPI_ISL_838807, EPI_ISL_838828, EPI_ISL_838829, EPI_ISL_838830, EPI_ISL_838831, EPI_ISL_838832, EPI_ISL_838833, EPI_ISL_838834, EPI_ISL_838835, EPI_ISL_838836, EPI_ISL_838837, EPI_ISL_838838, EPI_ISL_838839, EPI_ISL_838840, EPI_ISL_838841, EPI_ISL_838842, EPI_ISL_838843, EPI_ISL_838844, EPI_ISL_838846, EPI_ISL_838847, EPI_ISL_838848, EPI_ISL_838849, EPI_ISL_838850, EPI_ISL_838851, EPI_ISL_838852, EPI_ISL_838853, EPI_ISL_838855, EPI_ISL_838856, EPI_ISL_838857, EPI_ISL_838858, EPI_ISL_838859, EPI_ISL_838860, EPI_ISL_838861, EPI_ISL_838862, EPI_ISL_838863, EPI_ISL_838922, EPI_ISL_838923, EPI_ISL_839040, EPI_ISL_839045, EPI_ISL_839047, EPI_ISL_839048, EPI_ISL_839049, EPI_ISL_839050, EPI_ISL_839051, EPI_ISL_839052, EPI_ISL_839053, EPI_ISL_839054, EPI_ISL_839055, EPI_ISL_839056, EPI_ISL_839057, EPI_ISL_839058, EPI_ISL_839059, EPI_ISL_839060, EPI_ISL_839061, EPI_ISL_839062, EPI_ISL_839074, EPI_ISL_839075, EPI_ISL_839076, EPI_ISL_839077, EPI_ISL_839078, EPI_ISL_839079, EPI_ISL_839085, EPI_ISL_839086, EPI_ISL_839087, EPI_ISL_839088, EPI_ISL_839089, EPI_ISL_839090, EPI_ISL_839091, EPI_ISL_839092, EPI_ISL_839093, EPI_ISL_839094, EPI_ISL_839095, EPI_ISL_839096, EPI_ISL_839097, EPI_ISL_839098, EPI_ISL_839099, EPI_ISL_839100, EPI_ISL_839115, EPI_ISL_839118, EPI_ISL_839119, EPI_ISL_839120, EPI_ISL_839121, EPI_ISL_839122, EPI_ISL_839123, EPI_ISL_839124, EPI_ISL_839125, EPI_ISL_839184, EPI_ISL_839185, EPI_ISL_839198, EPI_ISL_839199, EPI_ISL_839201, EPI_ISL_839203, EPI_ISL_839204, EPI_ISL_839205, EPI_ISL_839206, EPI_ISL_839207, EPI_ISL_839208, EPI_ISL_839209, EPI_ISL_839211, EPI_ISL_839213, EPI_ISL_839216, EPI_ISL_839217, EPI_ISL_839221, EPI_ISL_839224, EPI_ISL_839225, EPI_ISL_839228, EPI_ISL_839229, EPI_ISL_839232 | | | |
| see above | University College London, Great Ormond Street Hospital for Children NHS Foundation Trust, Imperial College Healthcare NHS Trust | COVID-19 Genomics UK (COG-UK) Consortium | Sergi Castellano, Rachel Williams, Mark Kristiansen, Paola Resende Silva, Sunando Roy, Tony Brooks, Helena Tutill, Paola Niola, Patricia Dyal, Charlotte Williams, Leysa Forrest, Yasmin Panchbhaya, Jacqueline Findlay, Samuel Weeks, Julianne Brown, Kathryn Harris, Paul Randell, James Price, Alison Holmes, Judith Breuer |
| EPI_ISL_839733, EPI_ISL_839734 | Northumbria University / South Tees Hospitals NHS Foundation Trust / North Cumbria Integrated Care NHS Foundation Trust / North Tees and Hartlepool NHS Foundation Trust / Newcastle Hospitals NHS Foundation Trust | COVID-19 Genomics UK (COG-UK) Consortium | Darren L Smith,Andrew Nelson,Matthew Bashton,Greg R Young,Joshua Loh,John Allan,Mohammad A Tariq,Giles S Holt,Gary Black,Wen C Yew,Lynn Dover,Paul Baker,Steve Liggett,Sarah Essex,Jane Greenaway,Debra Padgett,Clive Graham,Garren Scott,Edward Barton,Emma Swindells,Brendan Payne,Jennifer Collins,Yusri Taha,Gary Eltringham |
| EPI_ISL_839825, EPI_ISL_839826, EPI_ISL_839827, EPI_ISL_839828, EPI_ISL_839829, EPI_ISL_839830, EPI_ISL_839831, EPI_ISL_839832, EPI_ISL_839833, EPI_ISL_839834, EPI_ISL_839835, EPI_ISL_839836, EPI_ISL_839837, EPI_ISL_839838, EPI_ISL_839839, EPI_ISL_839901, EPI_ISL_839902, EPI_ISL_839903, EPI_ISL_839904, EPI_ISL_839905, EPI_ISL_839906, EPI_ISL_839909, EPI_ISL_839910, EPI_ISL_839912, EPI_ISL_839915, EPI_ISL_839917, EPI_ISL_839919, EPI_ISL_839920, EPI_ISL_839922, EPI_ISL_839923, EPI_ISL_839924, EPI_ISL_839925, EPI_ISL_839926, EPI_ISL_839927, EPI_ISL_839928, EPI_ISL_839929, EPI_ISL_839930, EPI_ISL_839931 | | | |
| see above | Quadram Institute Bioscience | COVID-19 Genomics UK (COG-UK) Consortium | Dave J. Baker, Gemma L. Kay, Alp Aydin, Thanh Le-Viet, Steven Rudder, Ana P. Tedim, Anastasia Kolyva, Maria Diaz, Leonardo de Oliveira Martins, Nabil-Fareed Alikhan, Lizzie Meadows, Rachael Stanley, Ngozi Elumogo, Muhammed Yasir, Nicholas M. Thomson, Alexander J Trotter, Rachel Gilroy, Samuel Bloomfield, Claire Stuart, Andrew Bell, Reenesh Prakash, Samir Dervisevic, Alison E. Mather, John Wain, Mark Webber, Andrew J. Page, Justin O'Grady |
| EPI_ISL_839995, EPI_ISL_839996 | Queens Medical Centre, Clinical Microbiology Department / DeepSeq Nottingham | COVID-19 Genomics UK (COG-UK) Consortium | Gemma Clark, Wendy Smith, Manjinder Khakh, Vicki M Fleming, Michelle M Lister, Hannah Howson-Wells, Jonathan Ball, Patrick McClure, Joseph Chappell, Theocharis Tsoleridis, Nadine Holmes, Matthew Carlisle, Christopher Moore, Fei Sang, Johnny Debebe, Victoria Wright, Matthew Loose |
| EPI_ISL_840121, EPI_ISL_840122, EPI_ISL_840123, EPI_ISL_840124, EPI_ISL_840125, EPI_ISL_840126, EPI_ISL_840127, EPI_ISL_840128, EPI_ISL_840129, EPI_ISL_840130, EPI_ISL_840131, EPI_ISL_840132, EPI_ISL_840133, EPI_ISL_840134 | | | |
| see above | Lincolnshire Hospitals and DeepSeq Nottingham | COVID-19 Genomics UK (COG-UK) Consortium | Nichola Duckworth, Tim Sloan, Sarah Walsh, Jonathan Ball, Patrick McClure, Joeseph Chappell, Nadine Holmes, Matthew Carlisle, Christopher Moore, Fei Sang, Johnny Debebe, Victoria Wright, Matthew Loose |
| EPI_ISL_840650, EPI_ISL_840652, EPI_ISL_840660, EPI_ISL_840661, EPI_ISL_840663, EPI_ISL_840664, EPI_ISL_840666, EPI_ISL_840673, EPI_ISL_840674, EPI_ISL_840675, EPI_ISL_840676, EPI_ISL_840677, EPI_ISL_840678, EPI_ISL_840679, EPI_ISL_840680, EPI_ISL_840681, EPI_ISL_840682, EPI_ISL_840683, EPI_ISL_840684, EPI_ISL_840685, EPI_ISL_840686, EPI_ISL_840687, EPI_ISL_840688, EPI_ISL_840689, EPI_ISL_840690, EPI_ISL_840691, EPI_ISL_840692, EPI_ISL_840693, EPI_ISL_840811, EPI_ISL_840812, EPI_ISL_840813, EPI_ISL_840814, EPI_ISL_840815 | | | |
| see above | Originating lab: Wales Specialist Virology Centre Sequencing lab: Pathogen Genomics Unit | Public Health Wales Microbiology Cardiff Wales Specialist Virology Centre | Catherine Moore, Johnathan Evans, Laura Gifford, Malorie Perry, Simon Cottrell, Angela Marchbank, Alec Birchley, Alexander Adams, Amy Gaskin, Bree Gatica-Wilcox, Jason Coombes, Joel Southgate, Lauren Gilbert, Lee Graham, Nicole Pacchiarini, Sara Kumziene-Summerhayes, Sarah Taylor, Sophie |

| | | | |
|---|---|---|---|
| | | | Jones, Sara Rey, Matthew Bull, Joanne Watkins, Sally Corden, Tom Connor |
| EPI_ISL_840816, EPI_ISL_840819, EPI_ISL_840820, EPI_ISL_840821, EPI_ISL_840822, EPI_ISL_840824, EPI_ISL_840826, EPI_ISL_840827, EPI_ISL_840839, EPI_ISL_840840, EPI_ISL_840841, EPI_ISL_840842, EPI_ISL_840845, EPI_ISL_840846, EPI_ISL_840847, EPI_ISL_840848, EPI_ISL_840853, EPI_ISL_840916, EPI_ISL_840917, EPI_ISL_840918, EPI_ISL_840919, EPI_ISL_840922, EPI_ISL_840925, EPI_ISL_840926, EPI_ISL_840928, EPI_ISL_840929, EPI_ISL_840930, EPI_ISL_840932, EPI_ISL_840933, EPI_ISL_840934, EPI_ISL_840935, EPI_ISL_840936, EPI_ISL_840942, EPI_ISL_840944, EPI_ISL_840945, EPI_ISL_840947, EPI_ISL_840952, EPI_ISL_840954, EPI_ISL_840955, EPI_ISL_840956, EPI_ISL_840957, EPI_ISL_840958, EPI_ISL_840959, EPI_ISL_840960, EPI_ISL_840962, EPI_ISL_840966, EPI_ISL_840967, EPI_ISL_841041, EPI_ISL_841042, EPI_ISL_841059, EPI_ISL_841062, EPI_ISL_841063, EPI_ISL_841064, EPI_ISL_841081, EPI_ISL_841114, EPI_ISL_841115, EPI_ISL_841116, EPI_ISL_841117, EPI_ISL_841118, EPI_ISL_841119, EPI_ISL_841120, EPI_ISL_841121, EPI_ISL_841122, EPI_ISL_841123, EPI_ISL_841124, EPI_ISL_841125, EPI_ISL_841126, EPI_ISL_841127, EPI_ISL_841128, EPI_ISL_841129, EPI_ISL_841130, EPI_ISL_841131, EPI_ISL_841133, EPI_ISL_841134, EPI_ISL_841137, EPI_ISL_841138, EPI_ISL_841139, EPI_ISL_841140, EPI_ISL_841141, EPI_ISL_841142, EPI_ISL_841143, EPI_ISL_841144, EPI_ISL_841145, EPI_ISL_841146, EPI_ISL_841156, EPI_ISL_841157, EPI_ISL_841158, EPI_ISL_841159, EPI_ISL_841161, EPI_ISL_841162, EPI_ISL_841163, EPI_ISL_841164, EPI_ISL_841165, EPI_ISL_841167, EPI_ISL_841306 | | | |
| see above | Wales Specialist Virology Centre Sequencing lab: Pathogen Genomics Unit | Public Health Wales Microbiology Cardiff Wales Specialist Virology Centre | Catherine Moore, Johnathan Evans, Laura Gifford, Malorie Perry, Simon Cottrell, Angela Marchbank, Alec Birchley, Alexander Adams, Amy Gaskin, Bree Gatica-Wilcox, Jason Coombes, Joel Southgate, Lauren Gilbert, Lee Graham, Nicole Pacchiarini, Sara Kumziene-Summerhayes, Sarah Taylor, Sophie Jones, Sara Rey, Matthew Bull, Joanne Watkins, Sally Corden, Tom Connor |
| EPI_ISL_841333, EPI_ISL_841362, EPI_ISL_841363, EPI_ISL_841364, EPI_ISL_841365, EPI_ISL_841366, EPI_ISL_841367, EPI_ISL_841368, EPI_ISL_841369, EPI_ISL_841370, EPI_ISL_841371, EPI_ISL_841372, EPI_ISL_841374, EPI_ISL_841508, EPI_ISL_841509, EPI_ISL_841510, EPI_ISL_841511, EPI_ISL_841591, EPI_ISL_841608, EPI_ISL_841610 | | | |
| see above | Originating lab: Wales Specialist Virology Centre Sequencing lab: Pathogen Genomics Unit | Public Health Wales Microbiology Cardiff Wales Specialist Virology Centre | Catherine Moore, Johnathan Evans, Laura Gifford, Malorie Perry, Simon Cottrell, Angela Marchbank, Alec Birchley, Alexander Adams, Amy Gaskin, Bree Gatica-Wilcox, Jason Coombes, Joel Southgate, Lauren Gilbert, Lee Graham, Nicole Pacchiarini, Sara Kumziene-Summerhayes, Sarah Taylor, Sophie Jones, Sara Rey, Matthew Bull, Joanne Watkins, Sally Corden, Tom Connor |
| EPI_ISL_841922, EPI_ISL_841927, EPI_ISL_841945, EPI_ISL_842008, EPI_ISL_842015 | Centre for Enzyme Innovation, University of Portsmouth / Translational Research Laboratory, Portsmouth Hospitals NHS Trust | COVID-19 Genomics UK (COG-UK) Consortium | Angela Beckett,Yann Bourgeois,Garry Scarlett,Sharon Glaysher,Scott Elliott,Kelly Bicknell,Robert Impey,Allyson Lloyd,Sarah Wyllie,Ethan Butcher,Anoop Chauhan,Samuel Robson |
| EPI_ISL_842285, EPI_ISL_842315, EPI_ISL_842322 | Virology Department, Sheffield Teaching Hospitals NHS Foundation Trust/Department of Infection, Immunity and Cardiovascular Disease, The Medical School, University of Sheffield | COVID-19 Genomics UK (COG-UK) Consortium | Thushan de Silva, Matthew Parker, Nikki Smith, Adri Angyal, Rebecca Brown, Luke Green, Rachel Tucker, Paul Parsons, Danielle Groves, Katie Johnson, Laura Carrilero, Alex Keeley, Dave Partridge, Matthew Wyles, Benjamin Lindsey, Mehmet Yavuz, Mohammad Raza, Cariad Evans |
| EPI_ISL_843193 | LabPLUS | Institute of Environmental Science and Research (ESR) | Xiaoyun Ren, Matt Storey, Nikki Freed, Muhammad Faisal, Jing Wang, Hermes Perez, Anja Werno, Antje van der Linden, Arlo Upton, Chris Mansell, David Hammer, Dragana Drinkovic, Gary McAuliffe, Hana Sofia Andersson, James Ussher, Jill Sherwood, Josh Freeman, Julia Howard, Juliet Elvy, Mary DeAlmeida, Matt Blakiston, Matthew Rogers, Max Bloomfield, Michael Addidle, Michelle Balm, Sally Roberts, Sarah Jefferies, Sharmini Muttaiyah, Susan Morpeth, Susan Taylor, Timothy Blackmore, Vani Sathyendran, Veronica Playle, Virginia Hope, Erasmus Smit, Lauren Jelly, Olin Silander, Joep de Ligt |
| EPI_ISL_845566, EPI_ISL_845567, EPI_ISL_845571, EPI_ISL_845574, EPI_ISL_845577, EPI_ISL_845581, EPI_ISL_845584, EPI_ISL_845591, EPI_ISL_845598, EPI_ISL_845600, EPI_ISL_845605 | | | |
| see above | KU Leuven, Rega Institute, Clinical and Epidemiological Virology | KU Leuven, Rega Institute, Clinical and Epidemiological Virology | Tony Wawina-Bokalanga, Bert Vanmechelen, Joan Marti-Carerras, Piet Maes |
| EPI_ISL_845641, EPI_ISL_845642, EPI_ISL_845643, EPI_ISL_845644, EPI_ISL_845645, EPI_ISL_845648, EPI_ISL_845649, EPI_ISL_845651 | Laboratorio de Salud Pública - Secretaría Distrital de Salud | Instituto Nacional de Salud - Dirección de Investigación en Salud Pública | Katherine Laiton-Donato, Diego A. Álvarez-Díaz, Carlos Franco-Muñoz, Mauricio Pacheco-Montealegre, Maria T. Herrera-Sepúlveda, Jonathan Reales, Sheryll Corchuelo, Julian Naizaque, Gerardo Santamaría, Paola Muñoz-Laiton, Diego Andrés Prada, Magdalena Wiesner, Martha Lucia Ospina Martinez, Marcela Mercado-Reyes |
| EPI_ISL_845751 | Toronto Invasive Bacterial Diseases Network | McMaster University | Allison McGeer, Patryk Aftanas, Hooman Derakhshani, Angel Li, Kuganya Nirmalarajah, Emily Panousis, Ahmed Draia, Jalees Nasir, Michael Surette, Samira Mubareka, Andrew G. McArthur |
| EPI_ISL_845771 | Emory Molecular Diagnostics Laboratory, Emory Healthcare | Piantadosi Lab, Emory Department of Pathology | Ahmed Babiker, Anne Piantadosi |
| EPI_ISL_846523, EPI_ISL_846524, EPI_ISL_846525 | Lighthouse Lab in Alderley Park | Wellcome Sanger Institute for the COVID-19 Genomics UK (COG-UK) Consortium | Jacquelyn Wynn, Mairead Hyland, The Lighthouse Lab in Alderley Park and Alex Alderton, Roberto Amato, Sonia Goncalves, Ewan Harrison, David K. Jackson, Ian Johnston, Dominic Kwiatkowski, Cordelia Langford, John Sillitoe on behalf of the Wellcome Sanger Institute COVID-19 Surveillance Team |
| EPI_ISL_846527 | Lighthouse Lab in Glasgow | Wellcome Sanger Institute for the COVID-19 Genomics UK (COG-UK) Consortium | Harper VanSteenhouse, Yumi Kasai, David Gray, Carol Clugston, Anna Dominiczak and Alex Alderton, Roberto Amato, Sonia Goncalves, Ewan Harrison, David K. Jackson, Ian Johnston, Dominic Kwiatkowski, Cordelia Langford, John Sillitoe on behalf of the Wellcome Sanger Institute COVID-19 Surveillance Team |
| EPI_ISL_846543 | Lab voor klinische biologie | Lab voor klinische biologie | Hannelore Hamerlinck, Bruno Verhasselt |
| EPI_ISL_846595 | Respiratory Virus Unit, National Infection Service, Public Health England | COVID-19 Genomics UK (COG-UK) Consortium | PHE Covid Sequencing Team |
| EPI_ISL_846631, EPI_ISL_846632, EPI_ISL_846633, EPI_ISL_846634, EPI_ISL_846635, EPI_ISL_846649 | Hopital | National Reference Center for Viruses of Respiratory Infections, Institut Pasteur, Paris | Marion Barbet, Sylvie Behillil, Méline Bizard, Angela Brisebarre, Camille Capel, Etienne Simon-Lorière, Vincent Enouf, Maud Vanpeene, Sylvie van der Werf,Farfour Eric |
| EPI_ISL_847568, EPI_ISL_847590, EPI_ISL_847591, EPI_ISL_847592, EPI_ISL_847593, EPI_ISL_847758, EPI_ISL_847759 | California Department of Public Health | Chiu Laboratory, University of California, San Francisco | Charles Chiu, Xianding (Wayne) Deng, Candace Wang, Brian Bushnell, Scot Federman, Jill Hacker, Debra Wadford |
| EPI_ISL_847825 | Lab. Microbiologia e Virologia, Cotugno, A.O. dei Colli | Lab. Microbiologia e Virologia, Cotugno, A.O. dei Colli | Luigi Atripaldi, Claudia Tiberio, Anna Perfetti |
| EPI_ISL_847829 | Tempus | Grubaugh Lab - Yale School of Public Health | Tara Alpert, Joseph Fauver, Anderson Brito, Mallery Breban, Anne Wyllie, Chantal Vogels, Mary Petrone, Chaney Kalinich, Isabel Ott, Nathan Grubaugh |
| EPI_ISL_847832, EPI_ISL_847833, EPI_ISL_847837 | Yale New Haven Hospital | Grubaugh Lab - Yale School of Public Health | Tara Alpert, Joseph Fauver, Anderson Brito, Mallery Breban, Anne Wyllie, Chantal Vogels, Mary Petrone, Chaney Kalinich, Isabel Ott, Nathan Grubaugh |
| EPI_ISL_847844, EPI_ISL_847847, EPI_ISL_847852, EPI_ISL_847853, EPI_ISL_847855, EPI_ISL_847856, EPI_ISL_847857, EPI_ISL_847858, EPI_ISL_847859, EPI_ISL_847860, EPI_ISL_847861, EPI_ISL_847862, EPI_ISL_847863, EPI_ISL_847864, EPI_ISL_847865, EPI_ISL_847866, EPI_ISL_847867, EPI_ISL_847868, EPI_ISL_847869, EPI_ISL_847870, EPI_ISL_847871, EPI_ISL_847872, EPI_ISL_847920, EPI_ISL_847929, EPI_ISL_847930, EPI_ISL_847931, EPI_ISL_847932, EPI_ISL_847933, EPI_ISL_847934, EPI_ISL_847935, EPI_ISL_847936, EPI_ISL_847940, EPI_ISL_847941, EPI_ISL_847942, EPI_ISL_847943, EPI_ISL_847944, EPI_ISL_847945, EPI_ISL_847946, EPI_ISL_847947, EPI_ISL_847948, EPI_ISL_847949, EPI_ISL_847950, EPI_ISL_847951, EPI_ISL_847952 | | | |
| see above | University Hospitals of Geneva, Laboratory of Virology | HUG, Laboratory of Virology and the Health2030 Genome Center | Samuel Cordey, Ana Rita Goncalves, Laurent Kaiser, Lorenzo Cerutti, Henri Pegeot, Melyssa Elies, Keith Harshman, Ioannis Xenarios, Emmanouil Dermitzakis |
| EPI_ISL_847998, EPI_ISL_848056 | Michigan Department of Health and Human Services, Bureau of Laboratories | Michigan Department of Health and Human Services, Bureau of Laboratories | Blankenship HM, Riner D, Soehnlen MK |
| EPI_ISL_849326, EPI_ISL_849327, EPI_ISL_849328, EPI_ISL_849329, EPI_ISL_849330, EPI_ISL_849331, EPI_ISL_849332, EPI_ISL_849333, EPI_ISL_849334, EPI_ISL_849335 | Delaware Public Health Lab | Delaware Public Health Lab | Gregory Hovan |
| EPI_ISL_849750, EPI_ISL_849751, EPI_ISL_849752 | unknown | PHV-FSS | Son Nguyen et al. |
| EPI_ISL_849780 | Utah Public Health Laboratory | Utah Public Health Laboratory | Erin L. Young, Kelly F. Oakeson, Tara Gallagher |
| EPI_ISL_849924 | UCSF Clinical Microbiology Laboratory | Chan-Zuckerberg Biohub | CZB Cliahub Consortium |
| EPI_ISL_850519, EPI_ISL_850520, EPI_ISL_850521, EPI_ISL_850522, EPI_ISL_850523, EPI_ISL_850524, EPI_ISL_850525, EPI_ISL_850526, EPI_ISL_850527, EPI_ISL_850528, EPI_ISL_850529, EPI_ISL_850530, EPI_ISL_850531, EPI_ISL_850532, EPI_ISL_850533, EPI_ISL_850534, EPI_ISL_850535, EPI_ISL_850536, EPI_ISL_850537, EPI_ISL_850538, EPI_ISL_850539, EPI_ISL_850540, EPI_ISL_850541, EPI_ISL_850542, EPI_ISL_850543, EPI_ISL_850544, EPI_ISL_850545, EPI_ISL_850546, EPI_ISL_850547, EPI_ISL_850548, EPI_ISL_850549, EPI_ISL_850550, EPI_ISL_850551, EPI_ISL_850552, EPI_ISL_850553, EPI_ISL_850554, | | | |

| | | | |
|---|---|---|---|
| EPI_ISL_850555, EPI_ISL_850556, EPI_ISL_850557, EPI_ISL_850558, EPI_ISL_850559, EPI_ISL_850560, EPI_ISL_850561, EPI_ISL_850562, EPI_ISL_850563, EPI_ISL_850564, EPI_ISL_850565, EPI_ISL_850566, EPI_ISL_850567, EPI_ISL_850568, EPI_ISL_850569, EPI_ISL_850570, EPI_ISL_850571, EPI_ISL_850572, EPI_ISL_850573, EPI_ISL_850574, EPI_ISL_850575, EPI_ISL_850576, EPI_ISL_850577, EPI_ISL_850578, EPI_ISL_850579, EPI_ISL_850580, EPI_ISL_850581, EPI_ISL_850582, EPI_ISL_850583, EPI_ISL_850584, EPI_ISL_850585, EPI_ISL_850586, EPI_ISL_850587, EPI_ISL_850588, EPI_ISL_850589, EPI_ISL_850590, EPI_ISL_850591, EPI_ISL_850592, EPI_ISL_850593, EPI_ISL_850594, EPI_ISL_850595, EPI_ISL_850596, EPI_ISL_850597, EPI_ISL_850598, EPI_ISL_850599, EPI_ISL_850600 | | | |
| see above | Helix/Illumina | Genomics and Discovery, Respiratory Viruses Branch, Division of Viral Diseases, Centers for Disease Control and Prevention | Peter W. Cook, Dhwani Batra, Ben L. Rambo-Martin Eileen de Feo, Jan Antico, Christine Tran, Matthew Tolentino, Shannon Wickline, Kim Gietzen, Brad Sickler, Jingtao Liu, Eric Allen, Phil Febbo, Summer Galloway, Nicole L. Washington, Simon White, Geraint Levan, Kelly Schiabor Barrett, Elizabeth Cirulli, Alexandre Bolze, Ary Ascencio, Charlotte Rivera-Garcia, Ryan Cho, Jason Nguyen, Sherry Wang, Jimmy Ramirez, Tyler Cassens, Efren Sandoval, Magnus Isaksson, William Lee, David Becker, Marc Laurent, James Lu, Clinton R. Paden, Suxiang Tong, Duncan MacCannell |
| EPI_ISL_850671, EPI_ISL_850672 | The National Institute of Public Health | State Veterinary Institute Prague | Nagy,A;Jirincova,H;Trnka,D;Vecerova,J;Trinklova,M |
| EPI_ISL_850894, EPI_ISL_850895, EPI_ISL_850896, EPI_ISL_850897, EPI_ISL_850898, EPI_ISL_850899, EPI_ISL_850900, EPI_ISL_850901, EPI_ISL_850902, EPI_ISL_850903, EPI_ISL_850904, EPI_ISL_850905, EPI_ISL_850906, EPI_ISL_850907, EPI_ISL_850908, EPI_ISL_850909, EPI_ISL_850910, EPI_ISL_850911, EPI_ISL_850912, EPI_ISL_850913, EPI_ISL_850914, EPI_ISL_850915, EPI_ISL_850916, EPI_ISL_850917, EPI_ISL_850918, EPI_ISL_850919, EPI_ISL_850920, EPI_ISL_850921, EPI_ISL_850922, EPI_ISL_850923, EPI_ISL_850924, EPI_ISL_850925, EPI_ISL_850926, EPI_ISL_850927, EPI_ISL_850928, EPI_ISL_850929, EPI_ISL_850930, EPI_ISL_850931, EPI_ISL_850932, EPI_ISL_850933, EPI_ISL_850934, EPI_ISL_850935, EPI_ISL_850936, EPI_ISL_850937, EPI_ISL_850938, EPI_ISL_850939, EPI_ISL_850940, EPI_ISL_850941 | | | |
| see above | Helix / Illumina | Genomics and Discovery, Respiratory Viruses Branch, Division of Viral Diseases, Centers for Disease Control and Prevention | Peter W. Cook, Dhwani Batra, Ben L. Rambo-Martin Eileen de Feo, Jan Antico, Christine Tran, Matthew Tolentino, Shannon Wickline, Kim Gietzen, Brad Sickler, Jingtao Liu, Eric Allen, Phil Febbo, Summer Galloway, Nicole L. Washington, Simon White, Geraint Levan, Kelly Schiabor Barrett, Elizabeth Cirulli, Alexandre Bolze, Ary Ascencio, Charlotte Rivera-Garcia, Ryan Cho, Jason Nguyen, Sherry Wang, Jimmy Ramirez, Tyler Cassens, Efren Sandoval, Magnus Isaksson, William Lee, David Becker, Marc Laurent, James Lu, Clinton R. Paden, Suxiang Tong, Duncan MacCannell |
| EPI_ISL_850961, EPI_ISL_850962, EPI_ISL_850963, EPI_ISL_850964, EPI_ISL_850965, EPI_ISL_850966, EPI_ISL_850967, EPI_ISL_850968, EPI_ISL_850969, EPI_ISL_850970, EPI_ISL_850971, EPI_ISL_850972, EPI_ISL_850973, EPI_ISL_850974, EPI_ISL_850975, EPI_ISL_850976, EPI_ISL_850977, EPI_ISL_850978, EPI_ISL_850979, EPI_ISL_850980, EPI_ISL_850981, EPI_ISL_850982, EPI_ISL_850983, EPI_ISL_850984, EPI_ISL_850985, EPI_ISL_850986, EPI_ISL_850987, EPI_ISL_850988, EPI_ISL_850989, EPI_ISL_850990, EPI_ISL_850991, EPI_ISL_850992, EPI_ISL_850993, EPI_ISL_850994, EPI_ISL_850995, EPI_ISL_850996, EPI_ISL_850997, EPI_ISL_850998, EPI_ISL_850999, EPI_ISL_851000, EPI_ISL_851001, EPI_ISL_851002, EPI_ISL_851003, EPI_ISL_851004, EPI_ISL_851005, EPI_ISL_851006, EPI_ISL_851021, EPI_ISL_851022, EPI_ISL_851023, EPI_ISL_851024, EPI_ISL_851025, EPI_ISL_851029, EPI_ISL_851030, EPI_ISL_851031, EPI_ISL_851032 | | | |
| see above | Helix/Illumina | Genomics and Discovery, Respiratory Viruses Branch, Division of Viral Diseases, Centers for Disease Control and Prevention | Peter W. Cook, Dhwani Batra, Ben L. Rambo-Martin Eileen de Feo, Jan Antico, Christine Tran, Matthew Tolentino, Shannon Wickline, Kim Gietzen, Brad Sickler, Jingtao Liu, Eric Allen, Phil Febbo, Summer Galloway, Nicole L. Washington, Simon White, Geraint Levan, Kelly Schiabor Barrett, Elizabeth Cirulli, Alexandre Bolze, Ary Ascencio, Charlotte Rivera-Garcia, Ryan Cho, Jason Nguyen, Sherry Wang, Jimmy Ramirez, Tyler Cassens, Efren Sandoval, Magnus Isaksson, William Lee, David Becker, Marc Laurent, James Lu, Clinton R. Paden, Suxiang Tong, Duncan MacCannell |
| EPI_ISL_852047, EPI_ISL_852225 | Lighthouse Lab in Alderley Park | Wellcome Sanger Institute for the COVID-19 Genomics UK (COG-UK) Consortium | Jacquelyn Wynn, Mairead Hyland, The Lighthouse Lab in Alderley Park and Alex Alderton, Roberto Amato, Sonia Goncalves, Ewan Harrison, David K. Jackson, Ian Johnston, Dominic Kwiatkowski, Cordelia Langford, John Sillitoe on behalf of the Wellcome Sanger Institute COVID-19 Surveillance Team |
| EPI_ISL_852567, EPI_ISL_852570 | Max von Pettenkofer Institute, Virology, National Reference Center for Retroviruses, LMU München | Laboratory for Functional Genome Analysis, Dept. Genomics, Gene Center of the LMU Munich | Max Muenchhoff, Stefan Krebs, Alexander Graf, Oliver Keppler, Helmut Blum |
| EPI_ISL_852629, EPI_ISL_852630, EPI_ISL_852631, EPI_ISL_852632, EPI_ISL_852813, EPI_ISL_852819, EPI_ISL_852821, EPI_ISL_852823, EPI_ISL_852825, EPI_ISL_852828 | Lab. Microbiologia e Virologia, Cotugno, A.O. dei Colli | Lab. Microbiologia e Virologia, Cotugno, A.O. dei Colli | Luigi Atripaldi, Claudia Tiberio, Anna Perfetti |
| EPI_ISL_853400 | Charité Universitätsmedizin Berlin, Institut für Virologie/Labor Berlin | Charité Universitätsmedizin Berlin, Institut für Virologie | Victor M Corman, Julia Schneider, Barbara Mühlemann, Jörn Beheim-Schwarzbach, Talitha Veith, Julia Tesch, Tobias Bleicker, Terry Jones, Christian Drosten |
| EPI_ISL_853683 | NORTHWELL HEALTH LABORATORIES | Wadsworth Center, New York State Department of Health | Kirsten St. George, Daryl M. Lamson, Alexis Russel, Matthew Shudt, Melissa A Leisner, Jonathan Plitnick, Navjot Singh, John Kelly, Erasmus Schneider, Erica Lasek-Nesselquist |
| EPI_ISL_853684, EPI_ISL_853685, EPI_ISL_853686, EPI_ISL_853687 | ALBANY MEDICAL CENTER HOSPITAL CLINICAL LABORATORIES | Wadsworth Center, New York State Department of Health | Kirsten St. George, Daryl M. Lamson, Alexis Russel, Matthew Shudt, Melissa A Leisner, Jonathan Plitnick, Navjot Singh, John Kelly, Erasmus Schneider, Erica Lasek-Nesselquist |
| EPI_ISL_853727, EPI_ISL_853748, EPI_ISL_853760 | Department of Microbiology, University Innsbruck | Bergthaler laboratory, CeMM Research Center for Molecular Medicine of the Austrian Academy of Sciences | Lukas Endler, Alexandra Popa, Benedikt Agerer, Jakob-Wendelin Genger, Alexander Lercher, Anna Schedl, Thomas Penz, Michael Schuster, Jan Laine, Martin Senekowitsch, Christoph Bock, Andreas Bergthaler |
| EPI_ISL_855353 | Hospital | National Reference Center for Viruses of Respiratory Infections, Institut Pasteur, Paris | Marion Barbet, Sylvie Behillil, Méline Bizard, Angela Brisebarre, Camille Capel, Etienne Simon-Lorière, Vincent Enouf, Maud Vanpeene, Sylvie van der Werf,Pilorge Léa |
| EPI_ISL_855383, EPI_ISL_855384 | Hospital | National Reference Center for Viruses of Respiratory Infections, Institut Pasteur, Paris | Marion Barbet, Sylvie Behillil, Méline Bizard, Angela Brisebarre, Camille Capel, Etienne Simon-Lorière, Vincent Enouf, Maud Vanpeene, Sylvie van der Werf |
| EPI_ISL_855387 | Labo Analyses Med | National Reference Center for Viruses of Respiratory Infections, Institut Pasteur, Paris | Marion Barbet, Sylvie Behillil, Méline Bizard, Angela Brisebarre, Camille Capel, Etienne Simon-Lorière, Vincent Enouf, Maud Vanpeene, Sylvie van der Werf |
| EPI_ISL_856930, EPI_ISL_856931, EPI_ISL_856941, EPI_ISL_856942, EPI_ISL_856968, EPI_ISL_856969, EPI_ISL_856970 | Wyoming Public Health Laboratory | Wyoming Public Health Laboratory | Noah Hull, Taylor Fearing, Lynette Gumbleton, Channing Weber, Ashley Norberg, Bailey Bowcutt, and Wanda Manley |
| EPI_ISL_857059, EPI_ISL_857060, EPI_ISL_857061 | OCME Office Of Chief Medical Examiner | New York City Public Health Laboratory | Jade Wang, et al. |
| EPI_ISL_857314 | Centers for Disease Control, R.O.C. (Taiwan) | Centers for Disease Control, R.O.C. (Taiwan) | Ji-Rong Yang, Yu-Chi Lin, Jung-Jung-Mu, Ming-Tsan Liu |
| EPI_ISL_858077, EPI_ISL_858085, EPI_ISL_858101, EPI_ISL_858142, EPI_ISL_858145, EPI_ISL_858148, EPI_ISL_858151, EPI_ISL_858207, EPI_ISL_858208, EPI_ISL_858209, EPI_ISL_858210, EPI_ISL_858211, EPI_ISL_858212, EPI_ISL_858213, EPI_ISL_858214, EPI_ISL_858215, EPI_ISL_858217, EPI_ISL_858218, EPI_ISL_858219, EPI_ISL_858220, EPI_ISL_858221, EPI_ISL_858222 | | | |
| see above | Lighthouse Lab in Alderley Park | Wellcome Sanger Institute for the COVID-19 Genomics UK (COG-UK) Consortium | Jacquelyn Wynn, Mairead Hyland, The Lighthouse Lab in Alderley Park and Alex Alderton, Roberto Amato, Sonia Goncalves, Ewan Harrison, David K. Jackson, Ian Johnston, Dominic Kwiatkowski, Cordelia Langford, John Sillitoe on behalf of the Wellcome Sanger Institute COVID-19 Surveillance Team |
| EPI_ISL_858267 | Lighthouse Lab in Glasgow | Wellcome Sanger Institute for the COVID-19 Genomics UK (COG-UK) Consortium | Harper VanSteenhouse, Yumi Kasai, David Gray, Carol Clugston, Anna Dominiczak and Alex Alderton, Roberto Amato, Sonia Goncalves, Ewan Harrison, David K. Jackson, Ian Johnston, Dominic Kwiatkowski, Cordelia Langford, John Sillitoe on behalf of the Wellcome Sanger Institute COVID-19 Surveillance Team |
| EPI_ISL_858268 | Lighthouse Lab in Milton Keynes | Wellcome Sanger Institute for the COVID-19 Genomics UK (COG-UK) Consortium | The Lighthouse Lab in Milton Keynes and Alex Alderton, Roberto Amato, Sonia Goncalves, Ewan Harrison, David K. Jackson, Ian Johnston, Dominic Kwiatkowski, Cordelia Langford, John Sillitoe on behalf of the Wellcome Sanger Institute COVID-19 Surveillance Team |
| EPI_ISL_858269, EPI_ISL_858270, EPI_ISL_858296, EPI_ISL_858306, EPI_ISL_858311, EPI_ISL_858326, EPI_ISL_858328, EPI_ISL_858357, EPI_ISL_858381, EPI_ISL_858382 | Lighthouse Lab in Alderley Park | Wellcome Sanger Institute for the COVID-19 Genomics UK (COG-UK) Consortium | Jacquelyn Wynn, Mairead Hyland, The Lighthouse Lab in Alderley Park and Alex Alderton, Roberto Amato, Sonia Goncalves, Ewan Harrison, David K. Jackson, Ian Johnston, Dominic Kwiatkowski, Cordelia Langford, John Sillitoe on behalf of the Wellcome Sanger Institute COVID-19 Surveillance Team |
| EPI_ISL_858383 | Lighthouse Lab in Glasgow | Wellcome Sanger Institute for the COVID-19 Genomics UK (COG-UK) Consortium | Harper VanSteenhouse, Yumi Kasai, David Gray, Carol Clugston, Anna Dominiczak and Alex Alderton, Roberto Amato, Sonia Goncalves, Ewan Harrison, David K. Jackson, Ian Johnston, Dominic Kwiatkowski, Cordelia Langford, John Sillitoe on behalf of the Wellcome Sanger Institute COVID-19 Surveillance Team |
| EPI_ISL_858387, EPI_ISL_858388, EPI_ISL_858389, EPI_ISL_858393 | Lighthouse Lab in Alderley Park | Wellcome Sanger Institute for the COVID-19 Genomics UK (COG-UK) Consortium | Jacquelyn Wynn, Mairead Hyland, The Lighthouse Lab in Alderley Park and Alex Alderton, Roberto Amato, Sonia Goncalves, Ewan Harrison, David K. Jackson, Ian Johnston, Dominic Kwiatkowski, Cordelia Langford, John Sillitoe on behalf of the Wellcome Sanger Institute COVID-19 Surveillance Team |
| EPI_ISL_858394 | Lighthouse Lab in Glasgow | Wellcome Sanger Institute for the COVID-19 Genomics UK (COG-UK) Consortium | Harper VanSteenhouse, Yumi Kasai, David Gray, Carol Clugston, Anna Dominiczak and Alex Alderton, Roberto Amato, Sonia Goncalves, Ewan Harrison, David K. Jackson, Ian Johnston, Dominic Kwiatkowski, Cordelia Langford, John Sillitoe on behalf of the Wellcome Sanger Institute COVID-19 Surveillance |

| | | | |
|---|---|---|---|
| | | | Team |
| EPI_ISL_858397 | Lighthouse Lab in Alderley Park | Wellcome Sanger Institute for the COVID-19 Genomics UK (COG-UK) Consortium | Jacquelyn Wynn, Mairead Hyland, The Lighthouse Lab in Alderley Park and Alex Alderton, Roberto Amato, Sonia Goncalves, Ewan Harrison, David K. Jackson, Ian Johnston, Dominic Kwiatkowski, Cordelia Langford, John Sillitoe on behalf of the Wellcome Sanger Institute COVID-19 Surveillance Team |
| EPI_ISL_858426, EPI_ISL_858428, EPI_ISL_858438, EPI_ISL_858439, EPI_ISL_858442, EPI_ISL_858443, EPI_ISL_858445, EPI_ISL_858449, EPI_ISL_858450, EPI_ISL_858458, EPI_ISL_858461, EPI_ISL_858464, EPI_ISL_858469, EPI_ISL_858471, EPI_ISL_858473, EPI_ISL_858477, EPI_ISL_858478, EPI_ISL_858479, EPI_ISL_858481, EPI_ISL_858485, EPI_ISL_858486, EPI_ISL_858491, EPI_ISL_858501 | | | |
| see above | Lighthouse Lab in Glasgow | Wellcome Sanger Institute for the COVID-19 Genomics UK (COG-UK) Consortium | Harper VanSteenhouse, Yumi Kasai, David Gray, Carol Clugston, Anna Dominiczak and Alex Alderton, Roberto Amato, Sonia Goncalves, Ewan Harrison, David K. Jackson, Ian Johnston, Dominic Kwiatkowski, Cordelia Langford, John Sillitoe on behalf of the Wellcome Sanger Institute COVID-19 Surveillance Team |
| EPI_ISL_858516, EPI_ISL_858521, EPI_ISL_858523, EPI_ISL_858531, EPI_ISL_858557, EPI_ISL_858563, EPI_ISL_858572, EPI_ISL_858574, EPI_ISL_858576, EPI_ISL_858580, EPI_ISL_858581 | | | |
| see above | Lighthouse Lab in Alderley Park | Wellcome Sanger Institute for the COVID-19 Genomics UK (COG-UK) Consortium | Jacquelyn Wynn, Mairead Hyland, The Lighthouse Lab in Alderley Park and Alex Alderton, Roberto Amato, Sonia Goncalves, Ewan Harrison, David K. Jackson, Ian Johnston, Dominic Kwiatkowski, Cordelia Langford, John Sillitoe on behalf of the Wellcome Sanger Institute COVID-19 Surveillance Team |
| EPI_ISL_858583, EPI_ISL_858584, EPI_ISL_858585, EPI_ISL_858586, EPI_ISL_858587, EPI_ISL_858588, EPI_ISL_858592, EPI_ISL_858594, EPI_ISL_858598, EPI_ISL_858600, EPI_ISL_858601, EPI_ISL_858604, EPI_ISL_858607, EPI_ISL_858608, EPI_ISL_858609, EPI_ISL_858610, EPI_ISL_858612, EPI_ISL_858613, EPI_ISL_858614, EPI_ISL_858615, EPI_ISL_858616, EPI_ISL_858618, EPI_ISL_858621, EPI_ISL_858623, EPI_ISL_858624, EPI_ISL_858626, EPI_ISL_858627, EPI_ISL_858628, EPI_ISL_858630, EPI_ISL_858633, EPI_ISL_858634, EPI_ISL_858635, EPI_ISL_858636, EPI_ISL_858637, EPI_ISL_858638, EPI_ISL_858640, EPI_ISL_858641, EPI_ISL_858642, EPI_ISL_858643 | | | |
| see above | Lighthouse Lab in Glasgow | Wellcome Sanger Institute for the COVID-19 Genomics UK (COG-UK) Consortium | Harper VanSteenhouse, Yumi Kasai, David Gray, Carol Clugston, Anna Dominiczak and Alex Alderton, Roberto Amato, Sonia Goncalves, Ewan Harrison, David K. Jackson, Ian Johnston, Dominic Kwiatkowski, Cordelia Langford, John Sillitoe on behalf of the Wellcome Sanger Institute COVID-19 Surveillance Team |
| EPI_ISL_858645 | Lighthouse Lab in Alderley Park | Wellcome Sanger Institute for the COVID-19 Genomics UK (COG-UK) Consortium | Jacquelyn Wynn, Mairead Hyland, The Lighthouse Lab in Alderley Park and Alex Alderton, Roberto Amato, Sonia Goncalves, Ewan Harrison, David K. Jackson, Ian Johnston, Dominic Kwiatkowski, Cordelia Langford, John Sillitoe on behalf of the Wellcome Sanger Institute COVID-19 Surveillance Team |
| EPI_ISL_858646, EPI_ISL_858650, EPI_ISL_858651, EPI_ISL_858652, EPI_ISL_858656, EPI_ISL_858657, EPI_ISL_858658, EPI_ISL_858659, EPI_ISL_858660, EPI_ISL_858661, EPI_ISL_858664, EPI_ISL_858665, EPI_ISL_858668 | | | |
| see above | Lighthouse Lab in Glasgow | Wellcome Sanger Institute for the COVID-19 Genomics UK (COG-UK) Consortium | Harper VanSteenhouse, Yumi Kasai, David Gray, Carol Clugston, Anna Dominiczak and Alex Alderton, Roberto Amato, Sonia Goncalves, Ewan Harrison, David K. Jackson, Ian Johnston, Dominic Kwiatkowski, Cordelia Langford, John Sillitoe on behalf of the Wellcome Sanger Institute COVID-19 Surveillance Team |
| EPI_ISL_860184, EPI_ISL_860185 | Bangalore Medical College and Research Institute | Department of Neurovirology, National Institute of Mental Health and Neurosciences (NIMHANS) | Chitra Pattabiraman, Pramada Prasad, Anson Kunjumon George, Risha Rasheed, Darshan Sreenivas, Nakka Vijay Kiran Reddy, Anita S Desai, V Ravi |
| EPI_ISL_860187, EPI_ISL_860188 | Shimoga Institute of Medical Sciences | Department of Neurovirology, National Institute of Mental Health and Neurosciences (NIMHANS) | Chitra Pattabiraman, Pramada Prasad, Anson Kunjumon George, Risha Rasheed, Darshan Sreenivas, Nakka Vijay Kiran Reddy, Anita S Desai, V Ravi |
| EPI_ISL_860189 | BBMP Urban PHC | Department of Neurovirology, National Institute of Mental Health and Neurosciences (NIMHANS) | Chitra Pattabiraman, Pramada Prasad, Anson Kunjumon George, Risha Rasheed, Darshan Sreenivas, Nakka Vijay Kiran Reddy, Anita S Desai, V Ravi |
| EPI_ISL_860190 | Railway Hospital | Department of Neurovirology, National Institute of Mental Health and Neurosciences (NIMHANS) | Chitra Pattabiraman, Pramada Prasad, Anson Kunjumon George, Risha Rasheed, Darshan Sreenivas, Nakka Vijay Kiran Reddy, Anita S Desai, V Ravi |
| EPI_ISL_860191 | National Institute of Mental Health and Neurosciences (NIMHANS) | Department of Neurovirology, National Institute of Mental Health and Neurosciences (NIMHANS) | Chitra Pattabiraman, Pramada Prasad, Anson Kunjumon George, Risha Rasheed, Darshan Sreenivas, Nakka Vijay Kiran Reddy, Anita S Desai, V Ravi |
| EPI_ISL_860192 | Railway Hospital | Department of Neurovirology, National Institute of Mental Health and Neurosciences (NIMHANS) | Chitra Pattabiraman, Pramada Prasad, Anson Kunjumon George, Risha Rasheed, Darshan Sreenivas, Nakka Vijay Kiran Reddy, Anita S Desai, V Ravi |
| EPI_ISL_860193 | Jayanagar General Hospital | Department of Neurovirology, National Institute of Mental Health and Neurosciences (NIMHANS) | Chitra Pattabiraman, Pramada Prasad, Anson Kunjumon George, Risha Rasheed, Darshan Sreenivas, Nakka Vijay Kiran Reddy, Anita S Desai, V Ravi |
| EPI_ISL_860194, EPI_ISL_860195 | BBMP Urban PHC | Department of Neurovirology, National Institute of Mental Health and Neurosciences (NIMHANS) | Chitra Pattabiraman, Pramada Prasad, Anson Kunjumon George, Risha Rasheed, Darshan Sreenivas, Nakka Vijay Kiran Reddy, Anita S Desai, V Ravi |
| EPI_ISL_860210, EPI_ISL_860212 | Akershus University Hospital, Department for Microbiology and Infectious Disease Control | Norwegian Institute of Public Health, Department of Virology | Kathrine Stene-Johansen, Kamilla Heddeland Instefjord, Hilde Elshaug, Atiya R Ali,Marie Paulsen Madsen, Rasmus Riis Kopperud, Hilde Vollan, Karoline Bragstad, Olav Hungnes |
| EPI_ISL_860241, EPI_ISL_860242, EPI_ISL_860248, EPI_ISL_860254 | University Hospitals of Geneva, Laboratory of Virology | HUG, Laboratory of Virology and Universitätsspital Basel | Samuel Cordey, Ana Rita Goncalves, Laurent Kaiser, Tim Roloff, Madlen Stange, Helena MB Seth-Smith, Alfredo Mari, Karoline Leuzinger, Julia Bielicki, Manuel Battegay, Hans Hirsch, Adrian Egli |
| EPI_ISL_860259 | Akershus University Hospital, Department for Microbiology and Infectious Disease Control | Norwegian Institute of Public Health, Department of Virology | Kathrine Stene-Johansen, Kamilla Heddeland Instefjord, Hilde Elshaug, Atiya R Ali,Marie Paulsen Madsen, Rasmus Riis Kopperud, Hilde Vollan, Karoline Bragstad, Olav Hungnes |
| EPI_ISL_860276 | Foerde Hospital, Department of Microbiology | Norwegian Institute of Public Health, Department of Virology | Kathrine Stene-Johansen, Kamilla Heddeland Instefjord, Hilde Elshaug, Atiya R Ali,Marie Paulsen Madsen, Rasmus Riis Kopperud, Hilde Vollan, Karoline Bragstad, Olav Hungnes |
| EPI_ISL_860277 | Akershus University Hospital, Department for Microbiology and Infectious Disease Control | Norwegian Institute of Public Health, Department of Virology | Kathrine Stene-Johansen, Kamilla Heddeland Instefjord, Hilde Elshaug, Atiya R Ali,Marie Paulsen Madsen, Rasmus Riis Kopperud, Hilde Vollan, Karoline Bragstad, Olav Hungnes |
| EPI_ISL_860285, EPI_ISL_860288 | Department of Medical Microbiology, St. Olavs hospital | Norwegian Institute of Public Health, Department of Virology | Kathrine Stene-Johansen, Kamilla Heddeland Instefjord, Hilde Elshaug, Atiya R Ali,Marie Paulsen Madsen, Rasmus Riis Kopperud, Hilde Vollan, Karoline Bragstad, Olav Hungnes |
| EPI_ISL_860560, EPI_ISL_860561, EPI_ISL_860562, EPI_ISL_860563, EPI_ISL_860564, EPI_ISL_860565, EPI_ISL_860567, EPI_ISL_860568, EPI_ISL_860569, EPI_ISL_860570, EPI_ISL_860571, EPI_ISL_860572, EPI_ISL_860573, EPI_ISL_860574, EPI_ISL_860575, EPI_ISL_860577, EPI_ISL_860578, EPI_ISL_860582, EPI_ISL_860585, EPI_ISL_860587, EPI_ISL_860588, EPI_ISL_860589, EPI_ISL_860590, EPI_ISL_860591, EPI_ISL_860592, EPI_ISL_860593, EPI_ISL_860594 | | | |
| see above | NHLS-IALCH | KRISP, KZn Research Innovation and Sequencing Platform | Giandhari J, Pillay S, Lessells R, Mdlalose K, York D, Khan S, Tegally H, Wilkinson E, de Oliveira T |
| EPI_ISL_860666, EPI_ISL_860667 | Respiratory Virus Unit, National Infection Service, Public Health England | COVID-19 Genomics UK (COG-UK) Consortium | PHE Covid Sequencing Team |
| EPI_ISL_860881, EPI_ISL_860888 | Labo Analyses Med | National Reference Center for Viruses of Respiratory Infections, Institut Pasteur, Paris | Marion Barbet, Sylvie Behillil, Méline Bizard, Angela Brisebarre, Camille Capel, Etienne Simon-Lorière, Vincent Enouf, Maud Vanpeene, Sylvie van der Werf,Lefaure Brieuc |
| EPI_ISL_860940, EPI_ISL_860943, EPI_ISL_860948, EPI_ISL_860951, EPI_ISL_860955, EPI_ISL_860962, EPI_ISL_860971, EPI_ISL_860976, EPI_ISL_860979, EPI_ISL_860981, EPI_ISL_860989, EPI_ISL_860992, EPI_ISL_860995, EPI_ISL_861018, EPI_ISL_861021, EPI_ISL_861023, EPI_ISL_861025, EPI_ISL_861056, EPI_ISL_861059, EPI_ISL_861060, EPI_ISL_861061, EPI_ISL_861062, EPI_ISL_861077, EPI_ISL_861079 | | | |
| see above | Johns Hopkins Hospital Department of Pathology | Johns Hopkins Hospital Department of Pathology | C. Paul Morris, Chun Huai Luo, Adannaya Amadi, Nicholas Gallagher, Heba H. Mostafa |
| EPI_ISL_861110, EPI_ISL_861111, EPI_ISL_861112, EPI_ISL_861113, EPI_ISL_861114, EPI_ISL_861118, EPI_ISL_861120, EPI_ISL_861121, EPI_ISL_861122, EPI_ISL_861123, EPI_ISL_861124, EPI_ISL_861125, EPI_ISL_861126, EPI_ISL_861127, EPI_ISL_861128, EPI_ISL_861129, EPI_ISL_861130, EPI_ISL_861131, EPI_ISL_861132, EPI_ISL_861133 | | | |
| see above | New York Presbyterian Hospital | Wadsworth Center, New York State Department of Health | Kirsten St. George, Daryl M. Lamson, Alexis Russel, Matthew Shudt, Melissa A Leisner, Jonathan Plitnick, Navjot Singh, John Kelly, Erasmus Schneider, Erica Lasek-Nesselquist |
| EPI_ISL_861177, EPI_ISL_861178, EPI_ISL_861181, EPI_ISL_861182 | BIO-REFERENCE LABORATORIES | Wadsworth Center, New York State Department of Health | Kirsten St. George, Daryl M. Lamson, Alexis Russel, Matthew Shudt, Melissa A Leisner, Jonathan Plitnick, Navjot Singh, John Kelly, Erasmus Schneider, Erica Lasek-Nesselquist |
| EPI_ISL_861186, EPI_ISL_861188, EPI_ISL_861189, EPI_ISL_861190, EPI_ISL_861192, EPI_ISL_861193, EPI_ISL_861194, EPI_ISL_861195, EPI_ISL_861196, EPI_ISL_861197, EPI_ISL_861198 | | | |
| see above | NORTH SHORE UNIVERSITY HOSPITAL | Wadsworth Center, New York State Department of Health | Kirsten St. George, Daryl M. Lamson, Alexis Russel, Matthew Shudt, Melissa A Leisner, Jonathan Plitnick, Navjot Singh, John Kelly, Erasmus Schneider, Erica Lasek-Nesselquist |

| | | | |
|---|---|---|---|
| EPI_ISL_861369 | WESTCHESTER MEDICAL CENTER | Wadsworth Center, New York State Department of Health | Kirsten St. George, Daryl M. Lamson, Alexis Russel, Matthew Shudt, Melissa A Leisner, Jonathan Plitnick, Navjot Singh, John Kelly, Erasmus Schneider, Erica Lasek-Nesselquist |
| EPI_ISL_861414, EPI_ISL_861415, EPI_ISL_861416, EPI_ISL_861418 | NORTH SHORE UNIVERSITY HOSPITAL | Wadsworth Center, New York State Department of Health | Kirsten St. George, Daryl M. Lamson, Alexis Russel, Matthew Shudt, Melissa A Leisner, Jonathan Plitnick, Navjot Singh, John Kelly, Erasmus Schneider, Erica Lasek-Nesselquist |
| EPI_ISL_861536, EPI_ISL_861539, EPI_ISL_861540, EPI_ISL_861557, EPI_ISL_861577 | Instituto Nacional de Saude (INSA) | Instituto Nacional de Saude (INSA) | Borges et al |
| EPI_ISL_861681 | Hospital Municipal Dr. Waldemar Tebaldi | Instituto Adolfo Lutz, Interdiciplinary Procedures Center, Strategic Laboratory | Claudio Tavares Sacchi, Claudia Regina Gonçalves, Erica Valessa Ramos Gomes, Karoline Rodrigues Campos |
| EPI_ISL_861684, EPI_ISL_861685 | Hospital Pronto Socorro Itaquera | Instituto Adolfo Lutz, Interdiciplinary Procedures Center, Strategic Laboratory | Claudio Tavares Sacchi, Claudia Regina Gonçalves, Erica Valessa Ramos Gomes, Karoline Rodrigues Campos |
| EPI_ISL_861689, EPI_ISL_861692, EPI_ISL_861694, EPI_ISL_861695, EPI_ISL_861696, EPI_ISL_861697 | Los Angeles County PHL | Los Angeles County PHL | P. Hemarajata et al. |
| EPI_ISL_861750, EPI_ISL_861771 | Yale Clinical Virology Lab | Grubaugh Lab - Yale School of Public Health | Tara Alpert, Joseph Fauver, Anderson Brito, Mallery Breban, Anne Wyllie, Chantal Vogels, Mary Petrone, Chaney Kalinich, Isabel Ott, Nathan Grubaugh |
| EPI_ISL_862660 | Charité Universitätsmedizin Berlin, Institut fur Virologie/Labor Berlin | Charité Universitätsmedizin Berlin, Institut für Virologie | Victor M Corman, Barbara Mühlemann, Jörn Beheim-Schwarzbach, Tobias Bleicker, Julia Tesch, Talitha Veith, Julia Schneider, Terry Jones, Christian Drosten |
| EPI_ISL_862665, EPI_ISL_862666, EPI_ISL_862667, EPI_ISL_862668, EPI_ISL_862714, EPI_ISL_862715 | University of Michigan Clinical Microbiology Laboratory | Lauring Lab, University of Michigan, Department of Microbiology and Immunology | Valesano |
| EPI_ISL_862742, EPI_ISL_862743, EPI_ISL_862744, EPI_ISL_862751, EPI_ISL_862760, EPI_ISL_862763, EPI_ISL_862771, EPI_ISL_862773, EPI_ISL_862777 | Utah Public Health Laboratory, Utah Public Health Laboratory Infectious Disease submission group | Utah Public Health Laboratory, Utah Public Health Laboratory Infectious Disease submission group | Young,E.L., Oakeson,K.F., Gallagher,T. |
| EPI_ISL_862786, EPI_ISL_862787, EPI_ISL_862788, EPI_ISL_862789, EPI_ISL_862790, EPI_ISL_862791, EPI_ISL_862792, EPI_ISL_862793, EPI_ISL_862794, EPI_ISL_862795, EPI_ISL_862796, EPI_ISL_862797, EPI_ISL_862798, EPI_ISL_862799, EPI_ISL_862800, EPI_ISL_862801, EPI_ISL_862802, EPI_ISL_862803, EPI_ISL_862804, EPI_ISL_862805, EPI_ISL_862806, EPI_ISL_862807, EPI_ISL_862808, EPI_ISL_862809, EPI_ISL_862810, EPI_ISL_862811, EPI_ISL_862812, EPI_ISL_862813 | | | |
| see above | Cancer Biology Department, National Cancer Institute | Cancer Biology Department, National Cancer Institute | Zekri,A.N., Sedawy,M.G., Ahmed,O.S., Hafez,M.M., Soliman,H.K., Bahnassy,A.A., Elhosieny,F.W., Gad,A.E., Hamdy,M.S., Soliman,M.S., Soliman,L., Abouelhoda,M. |
| EPI_ISL_863534, EPI_ISL_863535, EPI_ISL_863536, EPI_ISL_863538, EPI_ISL_863541, EPI_ISL_863545, EPI_ISL_863548, EPI_ISL_863550, EPI_ISL_863554, EPI_ISL_863559, EPI_ISL_863563, EPI_ISL_863564, EPI_ISL_863568, EPI_ISL_863569, EPI_ISL_863571, EPI_ISL_863573, EPI_ISL_863575, EPI_ISL_863585, EPI_ISL_863596, EPI_ISL_863597, EPI_ISL_863598, EPI_ISL_863602, EPI_ISL_863607, EPI_ISL_863611, EPI_ISL_863613, EPI_ISL_863615, EPI_ISL_863617, EPI_ISL_863620, EPI_ISL_863624, EPI_ISL_863632, EPI_ISL_863634, EPI_ISL_863637, EPI_ISL_863638, EPI_ISL_863642, EPI_ISL_863646, EPI_ISL_863653, EPI_ISL_863657, EPI_ISL_863660, EPI_ISL_863669, EPI_ISL_863677, EPI_ISL_863678, EPI_ISL_863679, EPI_ISL_863681, EPI_ISL_863696, EPI_ISL_863699, EPI_ISL_863700, EPI_ISL_863705, EPI_ISL_863709, EPI_ISL_863711, EPI_ISL_863723, EPI_ISL_863725, EPI_ISL_863726, EPI_ISL_863733, EPI_ISL_863734, EPI_ISL_863741, EPI_ISL_863742, EPI_ISL_863747, EPI_ISL_863759, EPI_ISL_863761, EPI_ISL_863766, EPI_ISL_863773, EPI_ISL_863776, EPI_ISL_863781, EPI_ISL_863784, EPI_ISL_863785, EPI_ISL_863788, EPI_ISL_863789, EPI_ISL_863792, EPI_ISL_863793, EPI_ISL_863795, EPI_ISL_863796, EPI_ISL_863797, EPI_ISL_863802, EPI_ISL_863804, EPI_ISL_863806, EPI_ISL_863810, EPI_ISL_863812, EPI_ISL_863838, EPI_ISL_863842, EPI_ISL_863845, EPI_ISL_863847, EPI_ISL_863848, EPI_ISL_863850, EPI_ISL_863851, EPI_ISL_863856, EPI_ISL_863858, EPI_ISL_863860, EPI_ISL_863863, EPI_ISL_863865 | | | |
| see above | Lighthouse Lab in Milton Keynes | Wellcome Sanger Institute for the COVID-19 Genomics UK (COG-UK) Consortium | The Lighthouse Lab in Milton Keynes and Alex Alderton, Roberto Amato, Sonia Goncalves, Ewan Harrison, David K. Jackson, Ian Johnston, Dominic Kwiatkowski, Cordelia Langford, John Sillitoe on behalf of the Wellcome Sanger Institute COVID-19 Surveillance Team |
| EPI_ISL_864543 | Lighthouse Lab in Glasgow | Wellcome Sanger Institute for the COVID-19 Genomics UK (COG-UK) Consortium | Harper VanSteenhouse, Yumi Kasai, David Gray, Carol Clugston, Anna Dominiczak and Alex Alderton, Roberto Amato, Sonia Goncalves, Ewan Harrison, David K. Jackson, Ian Johnston, Dominic Kwiatkowski, Cordelia Langford, John Sillitoe on behalf of the Wellcome Sanger Institute COVID-19 Surveillance Team |
| EPI_ISL_864746, EPI_ISL_864747, EPI_ISL_864748, EPI_ISL_864749, EPI_ISL_864750, EPI_ISL_864751, EPI_ISL_864766, EPI_ISL_864768, EPI_ISL_864771, EPI_ISL_864773, EPI_ISL_864775, EPI_ISL_864777, EPI_ISL_864779 | | | |
| see above | Department of Pathology, University of Cambridge | COVID-19 Genomics UK (COG-UK) Consortium | Aminu S. Jahun, Yasmin Chaudhry, Grant Hall, Iliana Georgana, Myra Hosmillo, Martin D. Curran, Malte Pinckert, Surendra Parmar, Ian Goodfellow |
| EPI_ISL_864981, EPI_ISL_864984, EPI_ISL_864985, EPI_ISL_864986, EPI_ISL_864987, EPI_ISL_864988, EPI_ISL_864989, EPI_ISL_864990, EPI_ISL_864991, EPI_ISL_864994, EPI_ISL_864995 | | | |
| see above | West of Scotland Specialist Virology Centre, NHSGGC / MRC-University of Glasgow Centre for Virus Research | COVID-19 Genomics UK (COG-UK) Consortium | Ana da Silva Filipe, Natasha Johnson, Kathy Smollett, Daniel Mair, Stephen Carmichael, Alice Broos, Lily Tong, Jenna Nichols, Kyriaki Nomikou; Sarah McDonald; Richard Orton, Joseph Hughes, Sreenu Vattipally, David L Robertson; Alasdair MacLean, Rory Gunson; Sharif Shaaban, Matthew Holden; Rachel Blacow, Guy Mollett, Kathy Li, James Shepherd, Antonia Ho, Emma Thomson |
| EPI_ISL_865004, EPI_ISL_865005, EPI_ISL_865006, EPI_ISL_865009 | Lighthouse Lab in Glasgow / MRC-University of Glasgow Centre for Virus Research | COVID-19 Genomics UK (COG-UK) Consortium | Ana da Silva Filipe, Natasha Johnson, Kathy Smollett, Daniel Mair, Stephen Carmichael, Alice Broos, Lily Tong, Jenna Nichols, Kyriaki Nomikou; Sarah McDonald; Harper VanSteenhouse, Yumi Kasai, David Gray, Carol Clugston, Anna Dominiczak; Alasdair MacLean, Rory Gunson; Richard Orton, Joseph Hughes, Sreenu Vattipally, David L Robertson; Sharif Shaaban, Matthew Holden; Kathy Li, James Shepherd, Antonia Ho, Emma Thomson |
| EPI_ISL_865013 | West of Scotland Specialist Virology Centre, NHSGGC / MRC-University of Glasgow Centre for Virus Research | COVID-19 Genomics UK (COG-UK) Consortium | Ana da Silva Filipe, Natasha Johnson, Kathy Smollett, Daniel Mair, Stephen Carmichael, Alice Broos, Lily Tong, Jenna Nichols, Kyriaki Nomikou; Sarah McDonald; Richard Orton, Joseph Hughes, Sreenu Vattipally, David L Robertson; Alasdair MacLean, Rory Gunson; Sharif Shaaban, Matthew Holden; Rachel Blacow, Guy Mollett, Kathy Li, James Shepherd, Antonia Ho, Emma Thomson |
| EPI_ISL_865073, EPI_ISL_865094, EPI_ISL_865096, EPI_ISL_865097, EPI_ISL_865098, EPI_ISL_865099, EPI_ISL_865100, EPI_ISL_865101, EPI_ISL_865103, EPI_ISL_865104 | Virology Department, Royal Infirmary of Edinburgh, NHS Lothian / School of Biological Sciences, University of Edinburgh / Institute of Genetics and Molecular Medicine, University of Edinburgh | COVID-19 Genomics UK (COG-UK) Consortium | McHugh M, Dewar R, Rooke S, Gallagher M, Balcaza C, O'Toole Á, Scher E, Hill V, McCrone JT, Colquhoun R, Yu X, Jackson B, Rambaut A, Williams TC, Templeton K |
| EPI_ISL_865178, EPI_ISL_865179, EPI_ISL_865183, EPI_ISL_865184, EPI_ISL_865190, EPI_ISL_865191, EPI_ISL_865198, EPI_ISL_865364, EPI_ISL_865365, EPI_ISL_865366, EPI_ISL_865367, EPI_ISL_865368, EPI_ISL_865369, EPI_ISL_865370, EPI_ISL_865383, EPI_ISL_865384, EPI_ISL_865385, EPI_ISL_865387, EPI_ISL_865388, EPI_ISL_865389, EPI_ISL_865390, EPI_ISL_865481 | | | |
| see above | Liverpool Clinical Laboratories | COVID-19 Genomics UK (COG-UK) Consortium | Sam Haldenby, Anita Lucaci, Steve Paterson, Julian Hiscox, Alistair Darby, M Almsaud, A Alrezaihi, Muhannad Alruwaili, Stuart D Armstrong, Jones Benjamin, Eleanor G Bentley, Anu Chawla, Jordan J Clark, Angela Cowell, Richard Eccles, Isabel García-Dorival, Matthew Gemmell, Alessandro Gerada, PKF Gilmore, Richard Gregory, Ximeng Han, Catherine Hartley, Margaret Hughes, Miren Iturriza-Gomara, James Johnson, L Luu, Jenifer Manson, Charlotte Nelson, Elaine O'Toole, Cassie Olateju, Rebekah Penrice-Randal , Lucille Rainbow, N.P Randle, Trevor Ian Robinson, Parul Sharma, Ghada T Shawli, James P Stewart, Neil Swainston, Ecaterina Vamos, Joanne Watts, Mark Whitehead |
| EPI_ISL_865499, EPI_ISL_865571, EPI_ISL_865572, EPI_ISL_865574, EPI_ISL_865575 | Barts Health NHS Trust | COVID-19 Genomics UK (COG-UK) Consortium | CUTINO-MOGUEL, Maria-Teresa; HARRINGTON, David; OWOYEMI, Dola; KULASEGARAN-SHYLINI, Raghavendran; BROAD, Claire; KELE, Beatrix |
| EPI_ISL_866147 | University College London Hospital | COVID-19 Genomics UK (COG-UK) Consortium | Judith Heaney, Matthew Byott, Catherine Houlihan, Dan Frampton, Stuart Kirk, Moira Spyer and Eleni Nastouli |
| EPI_ISL_866188, EPI_ISL_866189, EPI_ISL_866190 | University College London, Great Ormond Street Hospital for Children NHS Foundation Trust, Imperial College Healthcare NHS Trust | COVID-19 Genomics UK (COG-UK) Consortium | Sergi Castellano, Rachel Williams, Mark Kristiansen, Paola Resende Silva, Sunando Roy, Tony Brooks, Helena Tutill, Paola Niola, Patricia Dyal, Charlotte Williams, Leysa Forrest, Yasmin Panchbhaya, Jacqueline Findlay, Samuel Weeks, Julianne Brown, Kathryn Harris, Paul Randell, James Price, Alison Holmes, Judith Breuer |
| EPI_ISL_866346, EPI_ISL_866347, EPI_ISL_866348 | Regional Virus Laboratory, Belfast Health and Social Care Trust | COVID-19 Genomics UK (COG-UK) Consortium | Conall McCaughey, James McKenna, Tanya Curran, Susan Feeney, Alison Watt, Ciara Cox, Mairead Connor, Zoltan Molnar, David Simpson, Derek Fairley |
| EPI_ISL_866520, EPI_ISL_866523, EPI_ISL_866528, EPI_ISL_866529, EPI_ISL_866531, EPI_ISL_866533, EPI_ISL_866534, EPI_ISL_866535, EPI_ISL_866536, EPI_ISL_866537, EPI_ISL_866538, EPI_ISL_866539, EPI_ISL_866540 | | | |

| | | | |
|---|---|---|---|
| see above | Northumbria University / South Tees Hospitals NHS Foundation Trust / North Cumbria Integrated Care NHS Foundation Trust / North Tees and Hartlepool NHS Foundation Trust / Newcastle Hospitals NHS Foundation Trust | COVID-19 Genomics UK (COG-UK) Consortium | Darren L Smith,Andrew Nelson,Matthew Bashton,Greg R Young,Joshua Loh,John Allan,Mohammad A Tariq,Giles S Holt,Gary Black,Wen C Yew,Lynn Dover,Paul Baker,Steve Liggett,Sarah Essex,Jane Greenaway,Debra Padgett,Clive Graham,Garren Scott,Edward Barton,Emma Swindells,Brendan Payne,Jennifer Collins,Yusri Taha,Gary Eltringham |
| EPI_ISL_866588, EPI_ISL_866589, EPI_ISL_866590, EPI_ISL_866591, EPI_ISL_866592, EPI_ISL_866593, EPI_ISL_866594, EPI_ISL_866595, EPI_ISL_866596, EPI_ISL_866597, EPI_ISL_866598, EPI_ISL_866599, EPI_ISL_866600, EPI_ISL_866601, EPI_ISL_866602 | | | |
| see above | Quadram Institute Bioscience | COVID-19 Genomics UK (COG-UK) Consortium | Dave J. Baker, Gemma L. Kay, Alp Aydin, Thanh Le-Viet, Steven Rudder, Ana P. Tedim, Anastasia Kolyva, Maria Diaz, Leonardo de Oliveira Martins, Nabil-Fareed Alikhan, Lizzie Meadows, Rachael Stanley, Ngozi Elumogo, Muhammad Yasir, Nicholas M. Thomson, Alexander J Trotter, Rachel Gilroy, Samuel Bloomfield, Claire Stuart, Andrew Bell, Reenesh Prakash, Samir Dervisevic, Alison E. Mather, John Wain, Mark Webber, Andrew J. Page, Justin O'Grady |
| EPI_ISL_866906 | Queens Medical Centre, Clinical Microbiology Department / DeepSeq Nottingham | COVID-19 Genomics UK (COG-UK) Consortium | Gemma Clark, Wendy Smith, Manjinder Khakh, Vicki M Fleming, Michelle M Lister, Hannah Howson-Wells, Jonathan Ball, Patrick McClure, Joseph Chappell, Theocharis Tsoleridis, Nadine Holmes, Matthew Carlisle, Christopher Moore, Fei Sang, Johnny Debebe, Victoria Wright, Matthew Loose |
| EPI_ISL_867008, EPI_ISL_867009, EPI_ISL_867011, EPI_ISL_867012, EPI_ISL_867013, EPI_ISL_867014, EPI_ISL_867015, EPI_ISL_867016, EPI_ISL_867017, EPI_ISL_867018, EPI_ISL_867019, EPI_ISL_867020, EPI_ISL_867021, EPI_ISL_867022, EPI_ISL_867023, EPI_ISL_867024, EPI_ISL_867025, EPI_ISL_867026, EPI_ISL_867027, EPI_ISL_867028, EPI_ISL_867029, EPI_ISL_867031 | | | |
| see above | Lincolnshire Hospitals and DeepSeq Nottingham | COVID-19 Genomics UK (COG-UK) Consortium | Nichola Duckworth, Tim Sloan, Sarah Walsh, Jonathan Ball, Patrick McClure, Joeseph Chappell, Nadine Holmes, Matthew Carlisle, Christopher Moore, Fei Sang, Johnny Debebe, Victoria Wright, Matthew Loose |
| EPI_ISL_867091, EPI_ISL_867092, EPI_ISL_867099, EPI_ISL_867103, EPI_ISL_867105, EPI_ISL_867106, EPI_ISL_867107, EPI_ISL_867108, EPI_ISL_867110, EPI_ISL_867111, EPI_ISL_867112, EPI_ISL_867113, EPI_ISL_867114, EPI_ISL_867118, EPI_ISL_867119, EPI_ISL_867120, EPI_ISL_867121, EPI_ISL_867123, EPI_ISL_867125, EPI_ISL_867127, EPI_ISL_867128, EPI_ISL_867129, EPI_ISL_867131, EPI_ISL_867132, EPI_ISL_867139, EPI_ISL_867151, EPI_ISL_867152, EPI_ISL_867163 | | | |
| see above | Oxford Viromics, NDM, University of Oxford; Oxford University Hospitals; Basingstoke and North Hampshire Hospital | COVID-19 Genomics UK (COG-UK) Consortium | Tanya Golubchik, David Bonsall, George Macintyre, Amy Trebes, Mariateresa de Cesare, Catrin Moore, Alex Mobbs, Anita Justice, Robert Shaw, Monique Andersson, Timothy Peto, Emma Wise, Nathan Moore, Jessica Lynch, Nick Cortes, Matilde Mori, Stephen Kidd, David Buck, John Todd, Christophe Fraser |
| EPI_ISL_867203, EPI_ISL_867204, EPI_ISL_867205, EPI_ISL_867431, EPI_ISL_867432, EPI_ISL_867433, EPI_ISL_867434 | Originating lab: Wales Specialist Virology Centre Sequencing lab: Pathogen Genomics Unit | Public Health Wales Microbiology Cardiff Wales Specialist Virology Centre | Catherine Moore, Johnathan Evans, Laura Gifford, Malorie Perry, Simon Cottrell, Angela Marchbank, Alec Birchley, Alexander Adams, Amy Gaskin, Bree Gatica-Wilcox, Jason Coombes, Joel Southgate, Lauren Gilbert, Lee Graham, Nicole Pacchiarini, Sara Kumziene-Summerhayes, Sarah Taylor, Sophie Jones, Sara Rey, Matthew Bull, Joanne Watkins, Sally Corden, Tom Connor |
| EPI_ISL_868311, EPI_ISL_868314, EPI_ISL_868317, EPI_ISL_868318, EPI_ISL_868319, EPI_ISL_868320, EPI_ISL_868321, EPI_ISL_868328, EPI_ISL_868351, EPI_ISL_868352 | Centre for Enzyme Innovation, University of Portsmouth / Translational Research Laboratory, Portsmouth Hospitals NHS Trust | COVID-19 Genomics UK (COG-UK) Consortium | Angela Beckett,Yann Bourgeois,Garry Scarlett,Sharon Glaysher,Scott Elliott,Kelly Bicknell,Robert Impey,Allyson Lloyd,Sarah Wyllie,Ethan Butcher,Anoop Chauhan,Samuel Robson |
| EPI_ISL_868479 | Virology Department, Sheffield Teaching Hospitals NHS Foundation Trust/Department of Infection, Immunity and Cardiovascular Disease, The Medical School, University of Sheffield | COVID-19 Genomics UK (COG-UK) Consortium | Thushan de Silva, Matthew Parker, Nikki Smith, Adri Angyal, Rebecca Brown, Luke Green, Rachel Tucker, Paul Parsons, Danielle Groves, Katie Johnson, Laura Carrilero, Alex Keeley, Dave Partridge, Matthew Wyles, Benjamin Lindsey, Mehmet Yavuz, Mohammad Raza, Cariad Evans |
| EPI_ISL_869045, EPI_ISL_869046, EPI_ISL_869048, EPI_ISL_869049, EPI_ISL_869050, EPI_ISL_869055, EPI_ISL_869057, EPI_ISL_869059, EPI_ISL_869060, EPI_ISL_869063, EPI_ISL_869064, EPI_ISL_869065, EPI_ISL_869066, EPI_ISL_869068, EPI_ISL_869070, EPI_ISL_869076 | | | |
| see above | Bioinformatics and Biostatistics Lab, Advanced Sequencing Facility | COVID-19 Genomics UK (COG-UK) Consortium | Aengus Stewart,Jerome Nicod,Chelsea Sawyer,Laura Cubitt,Harshil Patel,Margaret Crawford |
| EPI_ISL_869133 | Charité Universitätsmedizin Berlin, Institut für Virologie/Labor Berlin | Charité Universitätsmedizin Berlin, Institut für Virologie | Victor M Corman, Barbara Mühlemann, Jörn Beheim-Schwarzbach, Tobias Bleicker, Julia Tesch, Talitha Veith, Julia Schneider, Terry Jones, Christian Drosten |
| EPI_ISL_871832, EPI_ISL_871833, EPI_ISL_871834, EPI_ISL_871835, EPI_ISL_871836, EPI_ISL_871837, EPI_ISL_871838, EPI_ISL_871839 | Jessa | Jessa | Brigitte Maes, Bert Cruys |
| EPI_ISL_872204, EPI_ISL_872282 | Labo Analyses Med | National Reference Center for Viruses of Respiratory Infections, Institut Pasteur, Paris | Marion Barbet, Sylvie Behillil, Méline Bizard, Angela Brisebarre, Camille Capel, Etienne Simon-Lorière, Vincent Enouf, Maud Vanpeene, Sylvie van der Werf,Girard Sophie |
| EPI_ISL_872575 | South Eastern Area Laboratory Services (SEALS) | NSW Health Pathology - Institute of Clinical Pathology and Medical Research; Westmead Hospital; University of Sydney | CIDM-PH et al. |
| EPI_ISL_872744 | Rhode Island Department of Health | Infectious Disease Program, Broad Institute of Harvard and MIT | Lemieux,J.E., Siddle,K.J., Huard,R., King,E., Azevedo,K., Miller,A., Adams,G., Gladden-Young,A., Lagerborg,K., Rudy,M., DeRuff,K., Carter,A., Normandin,E., Bauer,M., Reilly,S., Tomkins-Tinch,C., Loreth,C., Chaluvadi,S., Birren,B.W., Gallagher,G., Smole,S., Park,D.J., MacInnis,B.L., and Sabeti,P.C. |
| EPI_ISL_872972, EPI_ISL_872973, EPI_ISL_872974, EPI_ISL_872975, EPI_ISL_872976, EPI_ISL_872977, EPI_ISL_872978, EPI_ISL_872979, EPI_ISL_872980, EPI_ISL_872981, EPI_ISL_872982, EPI_ISL_872983, EPI_ISL_872984, EPI_ISL_873027, EPI_ISL_873028, EPI_ISL_873029, EPI_ISL_873030, EPI_ISL_873031, EPI_ISL_873032, EPI_ISL_873033, EPI_ISL_873035, EPI_ISL_873036, EPI_ISL_873037, EPI_ISL_873038, EPI_ISL_873039 | | | |
| see above | HELIX LLC | WHO National Influenza Centre Russian Federation | Andrey Komissarov, Artem Fadeev, Anna Ivanova, Kseniya Komissarova, Dmitry Bazhenov, Mikhail Bakaev, Daria Danilenko, Ksenia Safina, Elena Nabieva, Georgii Bazykin, Dmitry Lioznov |
| EPI_ISL_873041, EPI_ISL_873042, EPI_ISL_873043, EPI_ISL_873044, EPI_ISL_873045, EPI_ISL_873046, EPI_ISL_873047, EPI_ISL_873048 | University of Michigan Clinical Microbiology Laboratory | Lauring Lab, University of Michigan, Department of Microbiology and Immunology | Valesano |
| EPI_ISL_873184, EPI_ISL_873186, EPI_ISL_873187, EPI_ISL_873188, EPI_ISL_873189, EPI_ISL_873190, EPI_ISL_873191, EPI_ISL_873192, EPI_ISL_873193 | Microbiology Division, South Carolina Department of Health and Environmental Control (SC DHEC) | Microbiology Division, South Carolina Department of Health and Environmental Control (SC DHEC) | Flores,H., Freeman,J. |
| EPI_ISL_873204, EPI_ISL_873205 | North Dakota Department of Health, Public Health Laboratory | North Dakota Department of Health, Public Health Laboratory | Lisa Wingerter |
| EPI_ISL_875512 | National Virus Reference Laboratory | National Virus Reference Laboratory | Michael Carr, Gabriel Gonzalez, Jonathan Dean, Cillian F De Gascun |
| EPI_ISL_876101 | Massachusetts State Public Health Laboratory | Massachusetts State Public Health Laboratory | Andrew Lang, Timelia Fink, Glen Gallagher, Sandra Smole |
| EPI_ISL_876631, EPI_ISL_876667, EPI_ISL_876679, EPI_ISL_876697, EPI_ISL_876707, EPI_ISL_876710, EPI_ISL_876717 | Helix/Illumina | Genomics and Discovery, Respiratory Viruses Branch, Division of Viral Diseases, Centers for Disease Control and Prevention | Peter W. Cook,Dhwani Batra,Ben L. Rambo-Martin,Eileen de Feo,Jan Antico,Christine Tran,Matthew Tolentino,Shannon Wickline,Kim Gietzen,Brad Sickler,Jingtao Liu,Eric Allen,Phil Febbo,Summer Galloway,Nicole L. Washington,Simon White,Geraint Levan,Kelly Schiabor Barrett,Elizabeth Cirulli,Alexandre Bolze,Ary Ascencio,Charlotte Rivera-Garcia,Ryan Cho,Jason Nguyen,Sherry Wang,Jimmy Ramirez,Tyler Cassens,Efren Sandoval,Magnus Isaksson,William Lee,David Becker,Marc Laurent,James Lu,Clinton R. Paden,Suxiang Tong,Duncan MacCannell, |
| EPI_ISL_878495, EPI_ISL_878498, EPI_ISL_878500, EPI_ISL_878576, EPI_ISL_878579, EPI_ISL_878581 | Biolab Diagnostic Laboratories | Andersen lab at Scripps Research | Issa Abu-Dayyeh, Ahmad Tibi, Lama Hussein, Lina Mohammad, Zein Naber, Amid Abdelnour with SEARCH Alliance San Diego |
| EPI_ISL_878622, EPI_ISL_878627 | San Diego County Public Health Laboratory | Andersen lab at Scripps Research | SEARCH Alliance San Diego with Tracy Basler, Jovan Shephard, Brett Austin |
| EPI_ISL_878695, EPI_ISL_878744, | Rady's Childrens Hospital | Andersen lab at Scripps Research | SEARCH Alliance San Diego with Nanda Radamchar, David Dimmock, Linda Luo, Christina Clarke, Kathryn Bouic, Teresa Mueller, Denise Malicki |

| | | | |
|---|---|---|---|
| EPI_ISL_880176, EPI_ISL_880177, EPI_ISL_880178 | | | |
| EPI_ISL_882301, EPI_ISL_882302, EPI_ISL_882304, EPI_ISL_882306, EPI_ISL_882310, EPI_ISL_882311, EPI_ISL_882312, EPI_ISL_882313, EPI_ISL_882317, EPI_ISL_882318, EPI_ISL_882319, EPI_ISL_882321, EPI_ISL_882322, EPI_ISL_882325, EPI_ISL_882326, EPI_ISL_882328, EPI_ISL_882330, EPI_ISL_882331, EPI_ISL_882334, EPI_ISL_882335, EPI_ISL_882336, EPI_ISL_882337, EPI_ISL_882339, EPI_ISL_882341, EPI_ISL_882342, EPI_ISL_882344, EPI_ISL_882345, EPI_ISL_882346, EPI_ISL_882348, EPI_ISL_882349, EPI_ISL_882350, EPI_ISL_882351, EPI_ISL_882353, EPI_ISL_882354, EPI_ISL_882355, EPI_ISL_882356, EPI_ISL_882358, EPI_ISL_882359, EPI_ISL_882360, EPI_ISL_882362, EPI_ISL_882365, EPI_ISL_882366, EPI_ISL_882367, EPI_ISL_882368, EPI_ISL_882369, EPI_ISL_882373, EPI_ISL_882374, EPI_ISL_882379, EPI_ISL_882380, EPI_ISL_882381, EPI_ISL_882382, EPI_ISL_882383, EPI_ISL_882384, EPI_ISL_882386, EPI_ISL_882387, EPI_ISL_882389, EPI_ISL_882391, EPI_ISL_882392, EPI_ISL_882394, EPI_ISL_882395, EPI_ISL_882396, EPI_ISL_882399, EPI_ISL_882400, EPI_ISL_882401, EPI_ISL_882403, EPI_ISL_882406, EPI_ISL_882407, EPI_ISL_882408, EPI_ISL_882410, EPI_ISL_882411, EPI_ISL_882412, EPI_ISL_882414, EPI_ISL_882416, EPI_ISL_882417, EPI_ISL_882419, EPI_ISL_882422, EPI_ISL_882423, EPI_ISL_882424, EPI_ISL_882425, EPI_ISL_882426, EPI_ISL_882428, EPI_ISL_882429, EPI_ISL_882431, EPI_ISL_882432, EPI_ISL_882433, EPI_ISL_882434, EPI_ISL_882435, EPI_ISL_882436, EPI_ISL_882439, EPI_ISL_882443, EPI_ISL_882444, EPI_ISL_882445, EPI_ISL_882446, EPI_ISL_882447, EPI_ISL_882448, EPI_ISL_882449, EPI_ISL_882450, EPI_ISL_882451, EPI_ISL_882453, EPI_ISL_882454, EPI_ISL_882455, EPI_ISL_882456, EPI_ISL_882457, EPI_ISL_882458, EPI_ISL_882459, EPI_ISL_882460, EPI_ISL_882461, EPI_ISL_882462, EPI_ISL_882463, EPI_ISL_882464, EPI_ISL_882465, EPI_ISL_882466, EPI_ISL_882467, EPI_ISL_882468, EPI_ISL_882469, EPI_ISL_882470, EPI_ISL_882472, EPI_ISL_882474, EPI_ISL_882477, EPI_ISL_882478, EPI_ISL_882479, EPI_ISL_882482, EPI_ISL_882483, EPI_ISL_882485, EPI_ISL_882486, EPI_ISL_882488, EPI_ISL_882489, EPI_ISL_882490, EPI_ISL_882492, EPI_ISL_882493, EPI_ISL_882494, EPI_ISL_882495, EPI_ISL_882496, EPI_ISL_882497, EPI_ISL_882498, EPI_ISL_882499, EPI_ISL_882500, EPI_ISL_882501, EPI_ISL_882502, EPI_ISL_882503, EPI_ISL_882504, EPI_ISL_882505, EPI_ISL_882507, EPI_ISL_882508, EPI_ISL_882509, EPI_ISL_882510, EPI_ISL_882511, EPI_ISL_882513, EPI_ISL_882514, EPI_ISL_882515, EPI_ISL_882516, EPI_ISL_882517, EPI_ISL_882519, EPI_ISL_882521, EPI_ISL_882522, EPI_ISL_882523, EPI_ISL_882524, EPI_ISL_882526, EPI_ISL_882527, EPI_ISL_882528, EPI_ISL_882529, EPI_ISL_882531, EPI_ISL_882532, EPI_ISL_882533, EPI_ISL_882535, EPI_ISL_882536, EPI_ISL_882537, EPI_ISL_882541, EPI_ISL_882543, EPI_ISL_882544, EPI_ISL_882545, EPI_ISL_882546, EPI_ISL_882547, EPI_ISL_882549, EPI_ISL_882551, EPI_ISL_882552, EPI_ISL_882553, EPI_ISL_882554, EPI_ISL_882555, EPI_ISL_882557, EPI_ISL_882558, EPI_ISL_882560, EPI_ISL_882561, EPI_ISL_882564, EPI_ISL_882565, EPI_ISL_882566, EPI_ISL_882568, EPI_ISL_882569, EPI_ISL_882570, EPI_ISL_882572, EPI_ISL_882573, EPI_ISL_882574, EPI_ISL_882576, EPI_ISL_882577, EPI_ISL_882578, EPI_ISL_882581, EPI_ISL_882582, EPI_ISL_882583, EPI_ISL_882584, EPI_ISL_882587, EPI_ISL_882588, EPI_ISL_882592, EPI_ISL_882593, EPI_ISL_882594, EPI_ISL_882596, EPI_ISL_882598, EPI_ISL_882599, EPI_ISL_882600, EPI_ISL_882601, EPI_ISL_882602, EPI_ISL_882603, EPI_ISL_882604 | | | |
| see above | Lighthouse Lab in Alderley Park | Wellcome Sanger Institute for the COVID-19 Genomics UK (COG-UK) Consortium | Jacquelyn Wynn, Mairead Hyland, The Lighthouse Lab in Alderley Park and Alex Alderton, Roberto Amato, Sonia Goncalves, Ewan Harrison, David K. Jackson, Ian Johnston, Dominic Kwiatkowski, Cordelia Langford, John Sillitoe on behalf of the Wellcome Sanger Institute COVID-19 Surveillance Team |
| EPI_ISL_882612, EPI_ISL_882613, EPI_ISL_882615 | Hospital Ramón y Cajal | Hospital Ramón y Cajal | José M Gonzalez-Alba, Concepción Rodríguez, Melanie Abreu, Laura Martínez, Val F Lanza, Luz Leticia Olavarrieta, Rafael Cantón, JC Galán |
| EPI_ISL_882637, EPI_ISL_882638, EPI_ISL_882639 | Azerbaijan National Hematology Center Division of Medical Genetics | Azerbaijan National Hematology Center Division of Medical Genetics | Aghayev Agha Rza, Bayraml Ramin |
| EPI_ISL_882918 | INMI Lazzaro Spallanzani IRCCS | INMI Lazzaro Spallanzani IRCCS | E Giombini, M. Rueca, B Bartolini, O Butera, C.E.M Gruber, F Messina, MR Capobianchi, A Di Caro |
| EPI_ISL_882920 | INMI Lazzaro Spallanzani IRCCS | INMI Lazzaro Spallanzani IRCCS | M Rueca, E Giombini, C.E.M Gruber, B Bartolini, O Butera, F Messina, A Di Caro, MR Capobianchi |
| EPI_ISL_882926 | INMI Lazzaro Spallanzani IRCCS | INMI Lazzaro Spallanzani IRCCS | B Bartolini, O Butera, C.E.M Gruber, M Rueca, F Messina, E Giombini, MR Capobianchi, A Di Caro |
| EPI_ISL_882928 | INMI Lazzaro Spallanzani IRCCS | INMI Lazzaro Spallanzani IRCCS | F Messina, O Butera, E Giombini, M Rueca, B Bartolini, C.E.M Gruber, MR Capobianchi, A Di Caro |
| EPI_ISL_882929 | INMI Lazzaro Spallanzani IRCCS | INMI Lazzaro Spallanzani IRCCS | C.E.M Gruber, B Bartolini, E Giombini, M Rueca, O Butera, F Messina, A Di Caro, MR Capobianchi |
| EPI_ISL_882930 | INMI Lazzaro Spallanzani IRCCS | INMI Lazzaro Spallanzani IRCCS | E Giombini, M. Rueca, B Bartolini, O Butera, C.E.M Gruber, F Messina, MR Capobianchi, A Di Caro |
| EPI_ISL_882931 | INMI Lazzaro Spallanzani IRCCS | INMI Lazzaro Spallanzani IRCCS | B Bartolini, O Butera, C.E.M Gruber, M Rueca, F Messina, E Giombini, MR Capobianchi, A Di Caro |
| EPI_ISL_882945, EPI_ISL_882946, EPI_ISL_882947, EPI_ISL_882948, EPI_ISL_882949 | The National Institute of Public Health | State Veterinary Institute Prague | Nagy,A;Jirincova,H;Vecerova,JTrnka,D |
| EPI_ISL_883286, EPI_ISL_883287 | SIESP DIPARTIMENTO DI PREVENZIONE CHIE | Istituto Zooprofilattico Sperimentale dell'Abruzzo e Molise "G. Caporale" | Lorusso A, Marcacci M, Di Domenico M, Ancora M, Curini V, Mangone I, Rinaldi A, Scialabba S, Di Pasquale A, Cammà C, Puglia I, Calistri P, Savini G |
| EPI_ISL_883288, EPI_ISL_883289, EPI_ISL_883296 | SIESP CHIETI-DRIVE IN ORTONA | Istituto Zooprofilattico Sperimentale dell'Abruzzo e Molise "G. Caporale" | Lorusso A, Marcacci M, Di Domenico M, Ancora M, Curini V, Mangone I, Rinaldi A, Scialabba S, Di Pasquale A, Cammà C, Puglia I, Calistri P, Savini G |
| EPI_ISL_883332 | Department of Homeless Services | New York City Public Health Laboratory | Jade Wang, et al. |
| EPI_ISL_883351, EPI_ISL_883352, EPI_ISL_883353 | OCME Office Of Chief Medical Examiner | New York City Public Health Laboratory | Jade Wang, et al. |
| EPI_ISL_885022, EPI_ISL_885026, EPI_ISL_885027, EPI_ISL_885028, EPI_ISL_885029, EPI_ISL_885030, EPI_ISL_885031, EPI_ISL_885032, EPI_ISL_885035, EPI_ISL_885037, EPI_ISL_885038, EPI_ISL_885041, EPI_ISL_885042, EPI_ISL_885044, EPI_ISL_885045, EPI_ISL_885046, EPI_ISL_885047, EPI_ISL_885048, EPI_ISL_885050 | | | |
| see above | Santa Clara County Public Health Laboratory | Chan-Zuckerberg Biohub | CZB Cliahub Consortium |
| EPI_ISL_886162, EPI_ISL_886163, EPI_ISL_886164, EPI_ISL_886166, EPI_ISL_886167, EPI_ISL_886173, EPI_ISL_886175, EPI_ISL_886183, EPI_ISL_886190, EPI_ISL_886196, EPI_ISL_886197, EPI_ISL_886198, EPI_ISL_886203, EPI_ISL_886222, EPI_ISL_886230, EPI_ISL_886234, EPI_ISL_886236, EPI_ISL_886239, EPI_ISL_886256, EPI_ISL_886278, EPI_ISL_886282, EPI_ISL_886296, EPI_ISL_886304, EPI_ISL_886307, EPI_ISL_886313, EPI_ISL_886315, EPI_ISL_886316, EPI_ISL_886323, EPI_ISL_886325, EPI_ISL_886328, EPI_ISL_886329, EPI_ISL_886334, EPI_ISL_886335, EPI_ISL_886368, EPI_ISL_886382, EPI_ISL_886389, EPI_ISL_886394, EPI_ISL_886408, EPI_ISL_886410, EPI_ISL_886415, EPI_ISL_886419, EPI_ISL_886425, EPI_ISL_886428, EPI_ISL_886441, EPI_ISL_886449, EPI_ISL_886453, EPI_ISL_886456, EPI_ISL_886458, EPI_ISL_886459, EPI_ISL_886462, EPI_ISL_886468, EPI_ISL_886480, EPI_ISL_886481, EPI_ISL_886484, EPI_ISL_886486, EPI_ISL_886490, EPI_ISL_886493, EPI_ISL_886500, EPI_ISL_886501, EPI_ISL_886507, EPI_ISL_886512, EPI_ISL_886517, EPI_ISL_886524, EPI_ISL_886536, EPI_ISL_886548, EPI_ISL_886549, EPI_ISL_886556, EPI_ISL_886573, EPI_ISL_886578, EPI_ISL_886585, EPI_ISL_886598, EPI_ISL_886601, EPI_ISL_886604, EPI_ISL_886605, EPI_ISL_886607, EPI_ISL_886611, EPI_ISL_886615, EPI_ISL_886621, EPI_ISL_886623, EPI_ISL_886629, EPI_ISL_886639, EPI_ISL_886641, EPI_ISL_886642, EPI_ISL_886643, EPI_ISL_886644, EPI_ISL_886648, EPI_ISL_886660, EPI_ISL_886664, EPI_ISL_886667, EPI_ISL_886670, EPI_ISL_886673, EPI_ISL_886679, EPI_ISL_886680, EPI_ISL_886681, EPI_ISL_886686, EPI_ISL_886698, EPI_ISL_886704, EPI_ISL_886708, EPI_ISL_886724, EPI_ISL_886733, EPI_ISL_886738, EPI_ISL_886739, EPI_ISL_886755, EPI_ISL_886760, EPI_ISL_886762, EPI_ISL_886769, EPI_ISL_886770, EPI_ISL_886780, EPI_ISL_886785, EPI_ISL_886787, EPI_ISL_886791, EPI_ISL_886792, EPI_ISL_886793, EPI_ISL_886795, EPI_ISL_886809, EPI_ISL_886811, EPI_ISL_886812, EPI_ISL_886831, EPI_ISL_886832, EPI_ISL_886845, EPI_ISL_886868, EPI_ISL_886877, EPI_ISL_886913, EPI_ISL_886922, EPI_ISL_886929, EPI_ISL_886930, EPI_ISL_886939, EPI_ISL_886943, EPI_ISL_886947, EPI_ISL_886962, EPI_ISL_886982, EPI_ISL_886985, EPI_ISL_886989, EPI_ISL_886995, EPI_ISL_887009, EPI_ISL_887014, EPI_ISL_887015, EPI_ISL_887021, EPI_ISL_887030, EPI_ISL_887040, EPI_ISL_887051, EPI_ISL_887056, EPI_ISL_887062, EPI_ISL_887069, EPI_ISL_887076, EPI_ISL_887090, EPI_ISL_887096, EPI_ISL_887100, EPI_ISL_887101, EPI_ISL_887104 | | | |
| see above | Labcorp | Genomics and Discovery, Respiratory Viruses Branch, Division of Viral Diseases, Centers for Disease Control and Prevention | Peter W. Cook,Dhwani Batra,Ben L. Rambo-Martin,Summer Galloway,Brian Krueger,Minoo Agarwal,Eyad Almasri,Debbie Boles,Ayla Burns,Nuthawin Charoensri,Oren Cohen,Susan Countryman,Mary Ann Cristobal,Bobbi Croy,Suzanne Dale,Hrushikesh Deshmukh,Amanda Douglas,Vincent Drouillon,Marcia Eisenberg,Howard Engler,Rama Ghatti,Prashant Gupta,Susan Hicks,Jake Humphrey,Lax Iyer,Manoj Jain,Mohan Kolli,Tim Kuphal,Stanley Letovsky,Michael Levandoski,Craig Lukasik,Jonathan Meltzer,Brian Norvell,Mindy Nye,Scott Parker,Christos Petropoulos,John Pruitt,Steven Ragan,Scott Ryan,Mike Sapeta,Jana Schroth,Suresh Babu Selvaraju,Goran Stevovic,Amanda Suchanek,Andrea Throop,Lyndon Tilson,Thomas Urban,Joe Voshell,Kimberly Wagner,Jonathan Williams,Mary Williamson,Qian Zeng,Tricia Zwiefelhofer,Clinton R. Paden,Suxiang Tong,Duncan MacCannell, |
| EPI_ISL_887115, EPI_ISL_887118, EPI_ISL_887126, EPI_ISL_887127, EPI_ISL_887135, EPI_ISL_887142, EPI_ISL_887143 | Institute of Medical Microbiology and Hospital Hygiene | Institute of Medical Microbiology and Hospital Hygiene | Prof. Dr. Achim Kaasch, Aljoscha Tersteegen |
| EPI_ISL_887531, EPI_ISL_887535, EPI_ISL_887536 | Johns Hopkins Hospital Department of Pathology | Johns Hopkins Hospital Department of Pathology | C. Paul Morris, Chun Huai Luo, Adannaya Amadi, Matthew Schwartz, Nicholas Gallagher, Heba H. Mostafa |
| EPI_ISL_887595, EPI_ISL_887617, EPI_ISL_887619, EPI_ISL_887628, EPI_ISL_887641, EPI_ISL_887647, EPI_ISL_887651, EPI_ISL_887671, EPI_ISL_887674, EPI_ISL_887677, EPI_ISL_887679, EPI_ISL_887702, EPI_ISL_887716, EPI_ISL_887725, EPI_ISL_887729, EPI_ISL_887733, EPI_ISL_887739, EPI_ISL_887748, EPI_ISL_887753, EPI_ISL_887754, EPI_ISL_887756, EPI_ISL_887757, EPI_ISL_887758, EPI_ISL_887765, EPI_ISL_887784, EPI_ISL_887798, EPI_ISL_887807, EPI_ISL_887810, EPI_ISL_887817, EPI_ISL_887829, EPI_ISL_887841, EPI_ISL_887845, EPI_ISL_887852, EPI_ISL_887857, EPI_ISL_887866, EPI_ISL_887910, EPI_ISL_887917, EPI_ISL_887921, EPI_ISL_887923, EPI_ISL_887928, EPI_ISL_887945, EPI_ISL_887947, EPI_ISL_887949, EPI_ISL_887950, EPI_ISL_887966, EPI_ISL_887974, EPI_ISL_887975, EPI_ISL_887981, EPI_ISL_887984, EPI_ISL_887985, EPI_ISL_887993, EPI_ISL_887995, EPI_ISL_888007, EPI_ISL_888009, EPI_ISL_888012, EPI_ISL_888028, EPI_ISL_888032, EPI_ISL_888041, EPI_ISL_888054, EPI_ISL_888059, EPI_ISL_888060, EPI_ISL_888065, EPI_ISL_888071, EPI_ISL_888094, EPI_ISL_888100, EPI_ISL_888104, EPI_ISL_888112, EPI_ISL_888123, EPI_ISL_888125, EPI_ISL_888127, EPI_ISL_888157, EPI_ISL_888159, EPI_ISL_888173, EPI_ISL_888186, EPI_ISL_888196, EPI_ISL_888211, EPI_ISL_888219, EPI_ISL_888223, EPI_ISL_888234, EPI_ISL_888238, EPI_ISL_888304, EPI_ISL_888318, EPI_ISL_888324, EPI_ISL_888401, EPI_ISL_888450, EPI_ISL_888511, EPI_ISL_888515, EPI_ISL_888516, EPI_ISL_888522, EPI_ISL_888523, EPI_ISL_888524, EPI_ISL_888525, EPI_ISL_888526, EPI_ISL_888543, EPI_ISL_888545, EPI_ISL_888546, EPI_ISL_888553, EPI_ISL_888556, EPI_ISL_888559, EPI_ISL_888560, EPI_ISL_888563, EPI_ISL_888572, EPI_ISL_888587 | | | |
| see above | Labcorp | Genomics and Discovery, Respiratory Viruses Branch, Division of Viral Diseases, Centers for Disease Control and Prevention | Peter W. Cook,Dhwani Batra,Ben L. Rambo-Martin,Summer Galloway,Brian Krueger,Minoo Agarwal,Eyad Almasri,Debbie Boles,Ayla Burns,Nuthawin Charoensri,Oren Cohen,Susan Countryman,Mary Ann Cristobal,Bobbi Croy,Suzanne Dale,Hrushikesh Deshmukh,Amanda Douglas,Vincent Drouillon,Marcia Eisenberg,Howard Engler,Rama Ghatti,Prashant Gupta,Susan Hicks,Jake Humphrey,Lax Iyer,Manoj Jain,Mohan Kolli,Tim Kuphal,Stanley Letovsky,Michael Levandoski,Craig Lukasik,Jonathan Meltzer,Brian Norvell,Mindy Nye,Scott Parker,Christos Petropoulos,John Pruitt,Steven Ragan,Scott Ryan,Mike Sapeta,Jana Schroth,Suresh Babu Selvaraju,Goran Stevovic,Amanda Suchanek,Andrea Throop,Lyndon Tilson,Thomas Urban,Joe Voshell,Kimberly Wagner,Jonathan Williams,Mary Williamson,Qian Zeng,Tricia Zwiefelhofer,Clinton R. Paden,Suxiang Tong,Duncan MacCannell, |
| EPI_ISL_889004 | RS Mitra Keluarga Gading Serpong | Eijkman Institute for Molecular Biology, Ministry of Research and Technology/National Agency for Research and Innovation | Willy Agustine, Edison Johar, Hidayat Trimarsanto, Iskandar Adnan, Lydia V. Panggalo, Sukma Oktavianthi, Frilasita A Yudhaputri, Safarina G Malik, Khin Saw Myint, Amin Soebandrio |

| | | | |
|---|---|---|---|
| EPI_ISL_889552, EPI_ISL_889553, EPI_ISL_889554, EPI_ISL_889555, EPI_ISL_889556, EPI_ISL_889557, EPI_ISL_889558, EPI_ISL_889559, EPI_ISL_889560, EPI_ISL_889561, EPI_ISL_889562, EPI_ISL_889563, EPI_ISL_889564, EPI_ISL_889565, EPI_ISL_889566, EPI_ISL_889567 | | | |
| see above | LSUHS Emerging Viral Threat Laboratory | Microbial Genome Sequencing Center | Jeremy P. Kamil, Jennifer L. Carroll, Camille F. Abshire, Maarten Van Diest, Mohammed N.A. Siddiquey, Andrew D. Yurochko, Martin J. Sapp, Rona S. Scott, Christopher G. Kevil, Daniel J. Snyder, Vaughn S. Cooper, John A. Vanchiere |
| EPI_ISL_890280, EPI_ISL_890283, EPI_ISL_890284, EPI_ISL_890287, EPI_ISL_890288, EPI_ISL_890291, EPI_ISL_890294, EPI_ISL_890298, EPI_ISL_890299, EPI_ISL_890304, EPI_ISL_890305, EPI_ISL_890306, EPI_ISL_890320, EPI_ISL_890322, EPI_ISL_890329, EPI_ISL_890333 | | | |
| see above | KU Leuven, Rega Institute, Clinical and Epidemiological Virology | KU Leuven, Rega Institute, Clinical and Epidemiological Virology | Tony Wawina-Bokalanga, Bert Vanmechelen, Joan Marti-Carerras, Piet Maes |
| EPI_ISL_890352 | Labo Analyses Med | National Reference Center for Viruses of Respiratory Infections, Institut Pasteur, Paris | Marion Barbet, Sylvie Behillil, Méline Bizard, Angela Brisebarre, Camille Capel, Etienne Simon-Lorière, Vincent Enouf, Maud Vanpeene, Sylvie van der Werf,Rousset Dominique |
| EPI_ISL_891137 | The Jackson Laboratory | The Jackson Laboratory | Lloyd M, Sanderson B, Srivastava A, Maurya R, Renzette N, Omerza G, Kelly K, Li L, Wei C L, Adams M |
| EPI_ISL_892059, EPI_ISL_892061, EPI_ISL_892062, EPI_ISL_892063, EPI_ISL_892064, EPI_ISL_892065, EPI_ISL_892066 | Lighthouse Lab in Alderley Park | Wellcome Sanger Institute for the COVID-19 Genomics UK (COG-UK) Consortium | Jacquelyn Wynn, Mairead Hyland, The Lighthouse Lab in Alderley Park and Alex Alderton, Roberto Amato, Sonia Goncalves, Ewan Harrison, David K. Jackson, Ian Johnston, Dominic Kwiatkowski, Cordelia Langford, John Sillitoe on behalf of the Wellcome Sanger Institute COVID-19 Surveillance Team |
| EPI_ISL_893746, EPI_ISL_893775 | Institute of Virology, Medical Center, University of Freiburg, Freiburg, Germany | Institute of Virology, Clinial Virus Genomics, Medical Center, University of Freiburg, Freiburg, Germany | Jonas Fuchs, Lisa Kern, Sandra Reuter, Hajo Grundmann, Marcus Panning |
| EPI_ISL_894171 | KU Leuven, Rega Institute, Clinical and Epidemiological Virology | KU Leuven, Rega Institute, Clinical and Epidemiological Virology | Tony Wawina-Bokalanga, Bert Vanmechelen, Joan Marti-Carerras, Piet Maes |
| EPI_ISL_896073 | Labormedizinisches Zentrum Dr Risch | University Hospital Basel, Clinical Bacteriology | Tim Roloff, Madlen Stange, Helena MB Seth-Smith, Alfredo Mari, Karoline Leuzinger, Julia Bielicki, Nadia Wohlwend,Martin Risch, Lorenz Risch, Manuel Battegay, Hans Hirsch, Adrian Egli |
| EPI_ISL_896074, EPI_ISL_896075, EPI_ISL_896076 | Viollier AG | University Hospital Basel, Clinical Bacteriology | Tim Roloff, Madlen Stange, Helena MB Seth-Smith, Alfredo Mari, Karoline Leuzinger, Julia Bielicki, Christiane Beckmann, Manuel Battegay, Hans Hirsch, Adrian Egli |
| EPI_ISL_896096, EPI_ISL_896108 | Labormedizinisches Zentrum Dr Risch | University Hospital Basel, Clinical Bacteriology | Tim Roloff, Madlen Stange, Helena MB Seth-Smith, Alfredo Mari, Karoline Leuzinger, Julia Bielicki, Nadia Wohlwend,Martin Risch, Lorenz Risch, Manuel Battegay, Hans Hirsch, Adrian Egli |
| EPI_ISL_896110 | Viollier AG | University Hospital Basel, Clinical Bacteriology | Tim Roloff, Madlen Stange, Helena MB Seth-Smith, Alfredo Mari, Karoline Leuzinger, Julia Bielicki, Christiane Beckmann, Manuel Battegay, Hans Hirsch, Adrian Egli |
| EPI_ISL_896167, EPI_ISL_896209 | MEPHI, Aix Marseille University | MEPHI, Aix Marseille University | Anthony LEVASSEUR |
| EPI_ISL_896422, EPI_ISL_896423, EPI_ISL_896424, EPI_ISL_896425, EPI_ISL_896426, EPI_ISL_896427, EPI_ISL_896428, EPI_ISL_896429 | Columbia University Irving Medical Center | Wadsworth Center, New York State Department of Health | Kirsten St. George, Daryl M. Lamson, Alexis Russel, Matthew Shudt, Melissa A Leisner, Jonathan Plitnick, Navjot Singh, John Kelly, Erasmus Schneider, Erica Lasek-Nesselquist |
| EPI_ISL_899006, EPI_ISL_899017, EPI_ISL_899050, EPI_ISL_899051, EPI_ISL_899085, EPI_ISL_899092, EPI_ISL_899101, EPI_ISL_899107, EPI_ISL_899108, EPI_ISL_899112, EPI_ISL_899113, EPI_ISL_899114, EPI_ISL_899127, EPI_ISL_899128, EPI_ISL_899131, EPI_ISL_899145, EPI_ISL_899146, EPI_ISL_899156, EPI_ISL_899162, EPI_ISL_899172, EPI_ISL_899197, EPI_ISL_899198, EPI_ISL_899199, EPI_ISL_899200, EPI_ISL_899201, EPI_ISL_899202, EPI_ISL_899203, EPI_ISL_899204, EPI_ISL_899205, EPI_ISL_899206, EPI_ISL_899207, EPI_ISL_899208, EPI_ISL_899209, EPI_ISL_899210, EPI_ISL_899211, EPI_ISL_899212, EPI_ISL_899213, EPI_ISL_899214, EPI_ISL_899215, EPI_ISL_899216, EPI_ISL_899217, EPI_ISL_899302, EPI_ISL_899303, EPI_ISL_899304, EPI_ISL_899305, EPI_ISL_899306, EPI_ISL_899307, EPI_ISL_899308, EPI_ISL_899309, EPI_ISL_899310, EPI_ISL_899311, EPI_ISL_899312, EPI_ISL_899313, EPI_ISL_899314, EPI_ISL_899315, EPI_ISL_899316, EPI_ISL_899317, EPI_ISL_899429, EPI_ISL_899430, EPI_ISL_899431, EPI_ISL_899432, EPI_ISL_899433, EPI_ISL_899434, EPI_ISL_899435, EPI_ISL_899436, EPI_ISL_899437, EPI_ISL_899438, EPI_ISL_899524, EPI_ISL_899525, EPI_ISL_899526, EPI_ISL_899527, EPI_ISL_899528, EPI_ISL_899549, EPI_ISL_899550, EPI_ISL_899551, EPI_ISL_899563, EPI_ISL_899631, EPI_ISL_899632, EPI_ISL_899633, EPI_ISL_899634, EPI_ISL_899635, EPI_ISL_899636, EPI_ISL_899637, EPI_ISL_899638, EPI_ISL_899639, EPI_ISL_899668, EPI_ISL_899669, EPI_ISL_899670, EPI_ISL_899671, EPI_ISL_899672, EPI_ISL_899673, EPI_ISL_899674, EPI_ISL_899675, EPI_ISL_899676, EPI_ISL_899677, EPI_ISL_899678, EPI_ISL_899679, EPI_ISL_899680, EPI_ISL_899681, EPI_ISL_899682, EPI_ISL_899683, EPI_ISL_899684, EPI_ISL_899685, EPI_ISL_899781, EPI_ISL_899782, EPI_ISL_899838, EPI_ISL_899839, EPI_ISL_899882, EPI_ISL_899883, EPI_ISL_899975, EPI_ISL_899976, EPI_ISL_899977, EPI_ISL_899978, EPI_ISL_899979, EPI_ISL_899980, EPI_ISL_899981, EPI_ISL_899982, EPI_ISL_899983, EPI_ISL_899984 | | | |
| see above | Viollier AG | Department of Biosystems Science and Engineering, ETH Zürich | Christian Beisel, Sarah Nadeau, Chaoran Chen, Ivan Topolsky, Philipp Jablonski, Lara Fuhrmann, David Dreifuss, Katharina Jahn, Tobias Schär, Ina Nissen, Natascha Santacroce, Elodie Burcklen, Christiane Beckmann, Maurice Redondo, Olivier Kobel, Christoph Noppen, Sophie Seidel, Noemie Santamaria de Souza, Niko Beerenwinkel, Tanja Stadler |
| EPI_ISL_900511 | Oriade | CNR Virus des Infections Respiratoires - France SUD | Antonin Bal, Gregory Destras, Gwendolyne Burfin, Hadrien Règue, Quentin Semanas, Martine Valette, Bruno Lina, Sylvie Larrat, Laurence Josset |
| EPI_ISL_900518 | Bioesterel | CNR Virus des Infections Respiratoires - France SUD | Antonin Bal, Gregory Destras, Gwendolyne Burfin, Hadrien Règue, Quentin Semanas, Martine Valette, Bruno Lina, Sylvie Larrat, Laurence Josset |
| EPI_ISL_900519 | Bioesterel | CNR Virus des Infections Respiratoires - France SUD | Antonin Bal, Géraldine Gonfrier, Gregory Destras, Gwendolyne Burfin, Hadrien Règue, Quentin Semanas, Martine Valette, Bruno Lina, Valérie Giordanengo, Laurence Josset |
| EPI_ISL_900524 | Labosud | CNR Virus des Infections Respiratoires - France SUD | Antonin Bal, Gregory Destras, Gwendolyne Burfin, Hadrien Règue, Quentin Semanas, Martine Valette, Bruno Lina, Laurence Josset |
| EPI_ISL_900565 | CNR Virus des Infections Respiratoires - France SUD | CNR Virus des Infections Respiratoires - France SUD | Antonin Bal, Gregory Destras, Gwendolyne Burfin, Hadrien Règue, Quentin Semanas, Martine Valette, Bruno Lina, Laurence Josset |
| EPI_ISL_902759, EPI_ISL_902760, EPI_ISL_902761, EPI_ISL_902762, EPI_ISL_902763, EPI_ISL_902764, EPI_ISL_902765, EPI_ISL_902766, EPI_ISL_902767 | Department of Virology and Immunology, University of Helsinki and Helsinki University Hospital, Huslab Finland | Department of Virology, Faculty of Medicine, University of Helsinki, Helsinki, Finland | Teemu Smura, Ravi Kant, Phuoc Truong, Hussein Alburkat, Hannimari Kallio-Kokko, Jenni Virtanen, Maija Suvanto, Essi Korhonen, Sari Hannula, Harri Kangas, Hanna Liimatainen, Satu Kurkela, Hanna Jarva, Maija Lappalainen, Pekka Ellonen, Olli Vapalahti |
| EPI_ISL_903061, EPI_ISL_903062, EPI_ISL_903063, EPI_ISL_903092, EPI_ISL_903093, EPI_ISL_903094, EPI_ISL_903095, EPI_ISL_903096, EPI_ISL_903097, EPI_ISL_903098, EPI_ISL_903099, EPI_ISL_903100, EPI_ISL_903101, EPI_ISL_903102, EPI_ISL_903103 | | | |
| see above | Seattle Flu Study | Seattle Flu Study | Deborah A. Nickerson, Chris D. Frazar, Jover Lee, Benjamin Pelle, Erica Ryke, Matthew Richardson, Amanda Adler, Elisabeth Brandstetter, Peter D. Han, Kairsten Fay, Misja Ilcisin, Kirsten Lacombe, Thomas R. Sibley, Melissa Truong, Caitlin R. Wolf, Karen Cowgill, Stephanie Schrag, Jeff Duchin, Michael Boeckh, Janet A. Englund, Michael Famulare, Barry R. Lutz, Mark J. Rieder, Lea M. Starita, Matthew Thompson, Helen Y. Chu, Trevor Bedford, Jay Shendure |
| EPI_ISL_903108, EPI_ISL_903110, EPI_ISL_903111, EPI_ISL_903113, EPI_ISL_903120, EPI_ISL_903121, EPI_ISL_903122, EPI_ISL_903123, EPI_ISL_903129, EPI_ISL_903130, EPI_ISL_903131, EPI_ISL_903132, EPI_ISL_903133, EPI_ISL_903134, EPI_ISL_903135, EPI_ISL_903149 | | | |
| see above | Washington State Department of Health | Seattle Flu Study | Deborah A. Nickerson, Chris D. Frazar, Jover Lee, Benjamin Pelle, Erica Ryke, Matthew Richardson, Amanda Adler, Elisabeth Brandstetter, Peter D. Han, Kairsten Fay, Misja Ilcisin, Kirsten Lacombe, Thomas R. Sibley, Melissa Truong, Caitlin R. Wolf, Romesh Gautom, Geoff Melly, Brian Hiatt, Philip Dykema, Scott Lindquist, Michael Boeckh, Janet A. Englund, Michael Famulare, Barry R. Lutz, Mark J. Rieder, Lea M. Starita, Matthew Thompson, Helen Y. Chu, Jay Shendure, Trevor Bedford |
| EPI_ISL_903231, EPI_ISL_903238, EPI_ISL_903239, EPI_ISL_903281, EPI_ISL_903284, EPI_ISL_903285, EPI_ISL_903287, EPI_ISL_903289, EPI_ISL_903290, EPI_ISL_903292, EPI_ISL_903294, EPI_ISL_903299, EPI_ISL_903303, EPI_ISL_903304, EPI_ISL_903305, EPI_ISL_903306, EPI_ISL_903309, EPI_ISL_903311 | | | |
| see above | M Health Fairview | Minnesota Department of Health, Public Health Laboratory | Alexandra Lorentz, Jacob Garfin, Matt Plumb, and Xiong Wang |
| EPI_ISL_903567, EPI_ISL_903614, EPI_ISL_903626 | NM Dept. Health, Scientific Laboratory Division | Genomics and Discovery, Respiratory Viruses Branch, Division of Viral Diseases, Centers for Disease Control and Prevention | Krista Queen, Yan Li, Ying Tao, Jing Zhang, Anna Uehara, Anna Montmayeur, Clinton R. Paden, Peter W. Cook, Rachel Marine, Mili Sheth, Jasmine Padilla, Sarah Nobles, Mark Burroughs, Lori Rowe, Haibin Wang, Ben L. Rambo-Martin, Dhwani Batra, Justin Lee, Suxiang Tong |
| EPI_ISL_903668, EPI_ISL_903671, EPI_ISL_903672, EPI_ISL_903688, EPI_ISL_903717 | SC Dept of Health and Env. Control-Bureau of Laboratories | Genomics and Discovery, Respiratory Viruses Branch, Division of Viral Diseases, Centers for Disease Control and Prevention | Krista Queen, Yan Li, Ying Tao, Jing Zhang, Anna Uehara, Anna Montmayeur, Clinton R. Paden, Peter W. Cook, Rachel Marine, Mili Sheth, Jasmine Padilla, Sarah Nobles, Mark Burroughs, Lori Rowe, Haibin Wang, Ben L. Rambo-Martin, Dhwani Batra, Justin Lee, Suxiang Tong |
| EPI_ISL_903752 | LA Office of Public Health Laboratories | Genomics and Discovery, Respiratory Viruses Branch, Division of Viral Diseases, Centers for Disease Control and Prevention | Krista Queen, Yan Li, Ying Tao, Jing Zhang, Anna Uehara, Anna Montmayeur, Clinton R. Paden, Peter W. Cook, Rachel Marine, Mili Sheth, Jasmine Padilla, Sarah Nobles, Mark Burroughs, Lori Rowe, Haibin Wang, Ben L. Rambo-Martin, Dhwani Batra, Justin Lee, Suxiang Tong |

| | | | |
|---|---|---|---|
| EPI_ISL_903769, EPI_ISL_903804 | NM Dept. Health, Scientific Laboratory Division | Genomics and Discovery, Respiratory Viruses Branch, Division of Viral Diseases, Centers for Disease Control and Prevention | Krista Queen, Yan Li, Ying Tao, Jing Zhang, Anna Uehara, Anna Montmayeur, Clinton R. Paden, Peter W. Cook, Rachel Marine, Mili Sheth, Jasmine Padilla, Sarah Nobles, Mark Burroughs, Lori Rowe, Haibin Wang, Ben L. Rambo-Martin, Dhwani Batra, Justin Lee, Suxiang Tong |
| EPI_ISL_903819 | SC Dept of Health and Env. Control-Bureau of Laboratories | Genomics and Discovery, Respiratory Viruses Branch, Division of Viral Diseases, Centers for Disease Control and Prevention | Krista Queen, Yan Li, Ying Tao, Jing Zhang, Anna Uehara, Anna Montmayeur, Clinton R. Paden, Peter W. Cook, Rachel Marine, Mili Sheth, Jasmine Padilla, Sarah Nobles, Mark Burroughs, Lori Rowe, Haibin Wang, Ben L. Rambo-Martin, Dhwani Batra, Justin Lee, Suxiang Tong |
| EPI_ISL_903833 | CDPH, Viral and Rickettsial Disease Laboratory | Genomics and Discovery, Respiratory Viruses Branch, Division of Viral Diseases, Centers for Disease Control and Prevention | Krista Queen, Yan Li, Ying Tao, Jing Zhang, Anna Uehara, Anna Montmayeur, Clinton R. Paden, Peter W. Cook, Rachel Marine, Mili Sheth, Jasmine Padilla, Sarah Nobles, Mark Burroughs, Lori Rowe, Haibin Wang, Ben L. Rambo-Martin, Dhwani Batra, Justin Lee, Suxiang Tong |
| EPI_ISL_903839 | NM Dept. Health, Scientific Laboratory Division | Genomics and Discovery, Respiratory Viruses Branch, Division of Viral Diseases, Centers for Disease Control and Prevention | Krista Queen, Yan Li, Ying Tao, Jing Zhang, Anna Uehara, Anna Montmayeur, Clinton R. Paden, Peter W. Cook, Rachel Marine, Mili Sheth, Jasmine Padilla, Sarah Nobles, Mark Burroughs, Lori Rowe, Haibin Wang, Ben L. Rambo-Martin, Dhwani Batra, Justin Lee, Suxiang Tong |
| EPI_ISL_903844, EPI_ISL_903856 | CDPH, Viral and Rickettsial Disease Laboratory | Genomics and Discovery, Respiratory Viruses Branch, Division of Viral Diseases, Centers for Disease Control and Prevention | Krista Queen, Yan Li, Ying Tao, Jing Zhang, Anna Uehara, Anna Montmayeur, Clinton R. Paden, Peter W. Cook, Rachel Marine, Mili Sheth, Jasmine Padilla, Sarah Nobles, Mark Burroughs, Lori Rowe, Haibin Wang, Ben L. Rambo-Martin, Dhwani Batra, Justin Lee, Suxiang Tong |
| EPI_ISL_903872 | SC Dept of Health and Env. Control-Bureau of Laboratories | Genomics and Discovery, Respiratory Viruses Branch, Division of Viral Diseases, Centers for Disease Control and Prevention | Krista Queen, Yan Li, Ying Tao, Jing Zhang, Anna Uehara, Anna Montmayeur, Clinton R. Paden, Peter W. Cook, Rachel Marine, Mili Sheth, Jasmine Padilla, Sarah Nobles, Mark Burroughs, Lori Rowe, Haibin Wang, Ben L. Rambo-Martin, Dhwani Batra, Justin Lee, Suxiang Tong |
| EPI_ISL_903907 | NM Dept. Health, Scientific Laboratory Division | Genomics and Discovery, Respiratory Viruses Branch, Division of Viral Diseases, Centers for Disease Control and Prevention | Krista Queen, Yan Li, Ying Tao, Jing Zhang, Anna Uehara, Anna Montmayeur, Clinton R. Paden, Peter W. Cook, Rachel Marine, Mili Sheth, Jasmine Padilla, Sarah Nobles, Mark Burroughs, Lori Rowe, Haibin Wang, Ben L. Rambo-Martin, Dhwani Batra, Justin Lee, Suxiang Tong |
| EPI_ISL_903913, EPI_ISL_903941 | CDPH, Viral and Rickettsial Disease Laboratory | Genomics and Discovery, Respiratory Viruses Branch, Division of Viral Diseases, Centers for Disease Control and Prevention | Krista Queen, Yan Li, Ying Tao, Jing Zhang, Anna Uehara, Anna Montmayeur, Clinton R. Paden, Peter W. Cook, Rachel Marine, Mili Sheth, Jasmine Padilla, Sarah Nobles, Mark Burroughs, Lori Rowe, Haibin Wang, Ben L. Rambo-Martin, Dhwani Batra, Justin Lee, Suxiang Tong |
| EPI_ISL_904043, EPI_ISL_904044, EPI_ISL_904045, EPI_ISL_904046, EPI_ISL_904047, EPI_ISL_904048, EPI_ISL_904049 | New Mexico Department of Health Scientific Laboratory | New Mexico Department of Health Scientific Laboratory | Ellie Johnson, Anastacia Griego-Fisher, D'eldra Malone |
| EPI_ISL_904143, EPI_ISL_904145, EPI_ISL_904149, EPI_ISL_904160, EPI_ISL_904165, EPI_ISL_904180, EPI_ISL_904191, EPI_ISL_904192, EPI_ISL_904200, EPI_ISL_904223, EPI_ISL_904224, EPI_ISL_904268, EPI_ISL_904269, EPI_ISL_904280, EPI_ISL_904376, EPI_ISL_904529, EPI_ISL_904530, EPI_ISL_904531, EPI_ISL_904532, EPI_ISL_904533, EPI_ISL_904534, EPI_ISL_904535, EPI_ISL_904536, EPI_ISL_904537, EPI_ISL_904538, EPI_ISL_904539 | | | |
| see above | Dutch COVID-19 response team | Erasmus Medical Center | Bas Oude Munnink, Reina Sikkema, David Nieuwenhuijse, Irina Chestakova, Anne van der Linden, Marjan Boter, Emmanuelle Munger, Corine GeurtsvanKessel, Annemiek van der Eijk, Richard Molenkamp, Marion Koopmans, on behalf of the Dutch national COVID-19 response team. |
| EPI_ISL_904626, EPI_ISL_904630, EPI_ISL_904644, EPI_ISL_904649, EPI_ISL_904653 | Servicio de Microbiología, Laboratori Clínic Metropolitana Nord. Hospital Universitari Germans Trias i Pujol. Institut d'Investigació en Ciències de la Salut Germans Trias i Pujol (IGTP) | IrsiCaixa - Can Ruti CovidSeq | Marc Noguera-Julian, Mariona Parera, Maria Casadellà, Pilar Armengol, Francesc Catala-Moll, Roger Paredes, Bonaventura Clotet Elisa Martró, Verónica Saludes, Anna Not, Ana Pérez, Montserrat Giménez, Ignacio Blanco, Cristina Casañ, Antoni E. Bordoy, Adrián Antuori |
| EPI_ISL_904743, EPI_ISL_904776, EPI_ISL_904973, EPI_ISL_905005, EPI_ISL_905006, EPI_ISL_905008, EPI_ISL_905009, EPI_ISL_905034, EPI_ISL_905049, EPI_ISL_905070, EPI_ISL_905105, EPI_ISL_905106, EPI_ISL_905235, EPI_ISL_905238, EPI_ISL_905339, EPI_ISL_905340, EPI_ISL_905351, EPI_ISL_905360, EPI_ISL_905376, EPI_ISL_905419, EPI_ISL_905420, EPI_ISL_905421, EPI_ISL_905422, EPI_ISL_905423, EPI_ISL_905432, EPI_ISL_905436, EPI_ISL_905437, EPI_ISL_905438, EPI_ISL_905488, EPI_ISL_905494, EPI_ISL_905524, EPI_ISL_905534, EPI_ISL_905541, EPI_ISL_905542, EPI_ISL_905543, EPI_ISL_905544, EPI_ISL_905566, EPI_ISL_905567, EPI_ISL_905661 | | | |
| see above | Dutch COVID-19 response team | National Institute for Public Health and the Environment (RIVM) | Adam Meijer, Harry Vennema, Dirk Eggink, Jeroen Cremer, Sharon van den Brink, Bas van der Veer, AnneMarie van den Brandt, Florian Zwagemaker, Dennis Schmitz, Chantal Reusken, on behalf of the national COVID-19 response team |
| EPI_ISL_905866, EPI_ISL_905867, EPI_ISL_905868, EPI_ISL_905869, EPI_ISL_905870, EPI_ISL_905871, EPI_ISL_905872, EPI_ISL_905873, EPI_ISL_905874, EPI_ISL_905875, EPI_ISL_905876, EPI_ISL_905877, EPI_ISL_905878, EPI_ISL_905879, EPI_ISL_905880, EPI_ISL_905881, EPI_ISL_905882, EPI_ISL_905883, EPI_ISL_905884, EPI_ISL_905885, EPI_ISL_905886, EPI_ISL_905887, EPI_ISL_905888, EPI_ISL_905889, EPI_ISL_905890, EPI_ISL_905891, EPI_ISL_905892, EPI_ISL_905893, EPI_ISL_905894, EPI_ISL_905895, EPI_ISL_905896, EPI_ISL_905897, EPI_ISL_905898, EPI_ISL_905899, EPI_ISL_905907, EPI_ISL_905908, EPI_ISL_905909, EPI_ISL_905910, EPI_ISL_905911, EPI_ISL_905912, EPI_ISL_905913 | | | |
| see above | OHSU Lab Services Molecular Microbiology Lab | Oregon SARS-CoV-2 Genome Sequencing Center | Brendan L. O'Connell, Sally Grindstaff, Kayla Carter, Ruth V. Nichols, Alec J. Hirsch, Donna Hansel, Guang Fan, Xuan, Qin, Daniel N. Streblow, William B. Messer, Andrew C. Adey, Benjamin N. Bimber, Brian J. O'Roak |
| EPI_ISL_906539 | HOSPITAL DEPARTAMENTAL DE VILLAVICENCIO | Instituto Nacional de Salud- Dirección de Investigación en Salud Pública, Universidad de los Andes- Applied genomics research group, Vicerrectoria de Investigación y Creación, Universidad de los Andes- Systems and Computing Engineering Department | Katherine Laiton-Donato, Diego A. Álvarez-Díaz, Carlos Franco-Muñoz, Mauricio Pacheco-Montealegre, Héctor Alejandro Ruiz-Moreno, Maria T. Herrera-Sepúlveda, Diego Andrés Prada, Jhonnatan Reales-González, Sheryll Corchuelo, Julian Naizaque, Gerardo Santamaría Jorge Duitama, Laura Natalia Gonzalez, Jorge Ivan Diaz, Silvia Restrepo-Restrepo, Magdalena Wiesner, Martha Lucia Ospina Martinez, Marcela Mercado-Reyes |
| EPI_ISL_906712, EPI_ISL_906714, EPI_ISL_906725, EPI_ISL_906744, EPI_ISL_906746 | Hematology Laboratory, Section of Molecular Diagnostics, University Clinical Centre, Medical University of Gdansk | Laboratory of Recombinant Vaccines | Lukasz Rabalski, Maciej Kosinski, Maciej Grzybek, Adam Sodol, Aneta Szulc, Krzysztof Lewandowski, Ewa Milosz, Marlena Robakowska, Boguslaw Szewczyk, Krystyna Bienkowska-Szewczyk |
| EPI_ISL_907093, EPI_ISL_907101, EPI_ISL_907102, EPI_ISL_907103, EPI_ISL_907104, EPI_ISL_907105, EPI_ISL_907106, EPI_ISL_907107, EPI_ISL_907108, EPI_ISL_907109, EPI_ISL_907110, EPI_ISL_907111, EPI_ISL_907112, EPI_ISL_907113, EPI_ISL_907114, EPI_ISL_907115, EPI_ISL_907116, EPI_ISL_907117, EPI_ISL_907118 | | | |
| see above | Cancer Biology Department, National Cancer Institute | Cancer Biology Department, National Cancer Institute | Zekri,A.N., Sedawy,M.G., Ahmed,O.S., Hafez,M.M., Soliman,H.K., Bahnassy,A.A., Elhosieny,F.W., Gad,A.E., Hamdy,M.S., Soliman,M.S., Soliman,L., Abouelhoda,M. |
| EPI_ISL_910337 | Laboratory for Respiratory Viruses, Cantacuzino National Military-Medical Institute for Research and Development | Cantacuzino Institute Virology | Luiza Ustea, Nicoleta Paraschiv, Mihaela Lazar |
| EPI_ISL_910585, EPI_ISL_910586, EPI_ISL_910587, EPI_ISL_910588, EPI_ISL_910589, EPI_ISL_910590, EPI_ISL_910591, EPI_ISL_910592, EPI_ISL_910593, EPI_ISL_910594, EPI_ISL_910595, EPI_ISL_910596, EPI_ISL_910597, EPI_ISL_910598, EPI_ISL_910599, EPI_ISL_910600, EPI_ISL_910601, EPI_ISL_910602, EPI_ISL_910603, EPI_ISL_910604, EPI_ISL_910605, EPI_ISL_910606, EPI_ISL_910607, EPI_ISL_910608, EPI_ISL_910609, EPI_ISL_910611, EPI_ISL_910612, EPI_ISL_910613, EPI_ISL_910614, EPI_ISL_910615, EPI_ISL_910616, EPI_ISL_910617, EPI_ISL_910618, EPI_ISL_910619, EPI_ISL_910620, EPI_ISL_910621, EPI_ISL_910622, EPI_ISL_910623, EPI_ISL_910624, EPI_ISL_910625, EPI_ISL_910626, EPI_ISL_910627, EPI_ISL_910628, EPI_ISL_910629, EPI_ISL_910630, EPI_ISL_910631, EPI_ISL_910632, EPI_ISL_910881, EPI_ISL_910882, EPI_ISL_910883, EPI_ISL_910884, EPI_ISL_910885, EPI_ISL_910886, EPI_ISL_910887 | | | |
| see above | Laboratoire national de sante, Microbiology, Virology | Laboratoire national de sante, Microbiology, Microbial Genomics Platform | Anke Wienecke-Baldacchino, Catherine Ragimbeau,Jessica Tapp, Fatu Djabi, Lise Pignon, Raoul Salmon, Tamir Abdelrahman |
| EPI_ISL_911530, EPI_ISL_911531 | M Health Fairview | Minnesota Department of Health, Public Health Laboratory | Alexandra Lorentz, Jacob Garfin, Matt Plumb, and Xiong Wang |
| EPI_ISL_911658, EPI_ISL_911659, EPI_ISL_911660, EPI_ISL_911661 | Texas Department of State Health Services (TXDSHS) | Texas Department of State Health Services (TXDSHS) | Bonnie Oh, Anita Pokharel, James Daniel Bonser, Myong Koag, Chung Wang, Rachel Lee, Grace Kubin, Rashmi Tuladhar, Mayela Pedrueza, Maliha Rahman, Jenny Zhang |
| EPI_ISL_911826, EPI_ISL_911866, EPI_ISL_911867 | Johns Hopkins Hospital Department of Pathology | Johns Hopkins Hospital Department of Pathology | C. Paul Morris, Chun Huai Luo, Adannaya Amadi, Matthew Schwartz, Nicholas Gallagher, Heba H. Mostafa |
| EPI_ISL_912170 | Tempus | Grubaugh Lab - Yale School of Public Health | Tara Alpert, Joseph Fauver, Anderson Brito, Mallery Breban, Anne Wyllie, Chantal Vogels, Mary Petrone, Annie Watkins, Chaney Kalinich, Isabel Ott, Nathan Grubaugh |
| EPI_ISL_912375 | Fondation Congolaise pour la recherche medicale (FCRM), Francine Ntoumi | NGS Competence Center Tuebingen, Institut für Medizinische Mikrobiologie und Hygiene, Universitaetsklinikum Tübingen | Angel Angelov |

| | | | |
|---|---|---|---|
| EPI_ISL_912454 | KU Leuven, Rega Institute, Clinical and Epidemiological Virology | KU Leuven, Rega Institute, Clinical and Epidemiological Virology | Tony Wawina-Bokalanga, Bert Vanmechelen, Joan Marti-Carerras, Piet Maes |
| EPI_ISL_912484, EPI_ISL_912526 | NHLS Universitas Academic | UFS Virology | PA Bester, MM Nyaga, P Nthiga, MT Mogotsi, D Goedhals, T de Oliveira |
| EPI_ISL_912691, EPI_ISL_912721, EPI_ISL_912901 | Hôpital Henri Mondor | Department of Virology, Henri Mondor University Hospital, Assistance Publique Hôpitaux de Paris, Université Paris-Est Créteil, INSERM U955 | Christophe Rodriguez, Slim Fourati, Vanessa Demontant, Guillaume Gricourt, Melissa N'Debi, Alexandre Soulier, Elisabeth Trawinski, Jean-Michel Pawlotsky |
| EPI_ISL_912924 | Hôpital Pitié-Salpêtrière | Department of Virology, Henri Mondor University Hospital, Assistance Publique Hôpitaux de Paris, Université Paris-Est Créteil, INSERM U955 | Christophe Rodriguez, Slim Fourati, Vanessa Demontant, Guillaume Gricourt, Melissa N'Debi, Alexandre Soulier, Elisabeth Trawinski, Jean-Michel Pawlotsky |
| EPI_ISL_912943, EPI_ISL_912978, EPI_ISL_912979, EPI_ISL_912980, EPI_ISL_912981 | Hôpital Henri Mondor | Department of Virology, Henri Mondor University Hospital, Assistance Publique Hôpitaux de Paris, Université Paris-Est Créteil, INSERM U955 | Christophe Rodriguez, Slim Fourati, Vanessa Demontant, Guillaume Gricourt, Melissa N'Debi, Alexandre Soulier, Elisabeth Trawinski, Jean-Michel Pawlotsky |
| EPI_ISL_913034, EPI_ISL_913035 | Hospital San Pedro de Alcántara | Instituto de Salud Carlos III | Iglesias-Caballero, M. Camarero, S. Sandonís,V. Vázquez, S. Pozo, F. Casas, I. Jiménez, P. Zaballos, A. Monzón, S. Varona, S. Cuesta, I. Rodríguez, G. |
| EPI_ISL_913044, EPI_ISL_913045, EPI_ISL_913047 | Hospital Comarcal Sierrallana | Instituto de Salud Carlos III | Iglesias-Caballero, M. Camarero, S. Sandonís,V. Vázquez, S. Pozo, F. Casas, I. Jiménez, P. Zaballos, A. Monzón, S. Varona, S. Cuesta, I. De Benito, I. |
| EPI_ISL_913065 | Center for Virology | Center for Virology | Jeremy V. Camp, Irene Goerzer, Monika Redlberger-Fritz, Stephan W. Aberle |
| EPI_ISL_913256, EPI_ISL_913257 | Synlab Medilab, Mikrobiologi | The Public Health Agency of Sweden | Anna-Malin Linde, Maria Lind Karlberg, Carlo Berg, Oskar Karlsson Lindsjo, Sofia Stamouli, Reza Advani, Mattias Haukland, Petra Holmstrom, Noura Walai, Petra Edquist, Mia Brytting, Anna Risberg, Karin Tegmark-Wisell |
| EPI_ISL_913275, EPI_ISL_913277, EPI_ISL_913281 | Dynamic Code AB | The Public Health Agency of Sweden | Anna-Malin Linde, Maria Lind Karlberg, Carlo Berg, Oskar Karlsson Lindsjo, Sofia Stamouli, Reza Advani, Mattias Haukland, Petra Holmstrom, Noura Walai, Petra Edquist, Mia Brytting, Anna Risberg, Karin Tegmark-Wisell |
| EPI_ISL_913297 | Klinisk mikrobiologi | The Public Health Agency of Sweden | Anna-Malin Linde, Maria Lind Karlberg, Carlo Berg, Oskar Karlsson Lindsjo, Sofia Stamouli, Reza Advani, Mattias Haukland, Petra Holmstrom, Noura Walai, Petra Edquist, Mia Brytting, Anna Risberg, Karin Tegmark-Wisell |
| EPI_ISL_913301, EPI_ISL_913303, EPI_ISL_913304 | Synlab Medilab, Mikrobiologi | The Public Health Agency of Sweden | Anna-Malin Linde, Maria Lind Karlberg, Carlo Berg, Oskar Karlsson Lindsjo, Sofia Stamouli, Reza Advani, Mattias Haukland, Petra Holmstrom, Noura Walai, Petra Edquist, Mia Brytting, Anna Risberg, Karin Tegmark-Wisell |
| EPI_ISL_913309 | Dynamic Code AB | The Public Health Agency of Sweden | Anna-Malin Linde, Maria Lind Karlberg, Carlo Berg, Oskar Karlsson Lindsjo, Sofia Stamouli, Reza Advani, Mattias Haukland, Petra Holmstrom, Noura Walai, Petra Edquist, Mia Brytting, Anna Risberg, Karin Tegmark-Wisell |
| EPI_ISL_913328 | Klinisk mikrobiologi | The Public Health Agency of Sweden | Anna-Malin Linde, Maria Lind Karlberg, Carlo Berg, Oskar Karlsson Lindsjo, Sofia Stamouli, Reza Advani, Mattias Haukland, Petra Holmstrom, Noura Walai, Petra Edquist, Mia Brytting, Anna Risberg, Karin Tegmark-Wisell |
| EPI_ISL_913331, EPI_ISL_913332, EPI_ISL_913333 | Mikrobiologen | The Public Health Agency of Sweden | Anna-Malin Linde, Maria Lind Karlberg, Carlo Berg, Oskar Karlsson Lindsjo, Sofia Stamouli, Reza Advani, Mattias Haukland, Petra Holmstrom, Noura Walai, Petra Edquist, Mia Brytting, Anna Risberg, Karin Tegmark-Wisell |
| EPI_ISL_913337, EPI_ISL_913338 | Unilabs AB Skovde | The Public Health Agency of Sweden | Anna-Malin Linde, Maria Lind Karlberg, Carlo Berg, Oskar Karlsson Lindsjo, Sofia Stamouli, Reza Advani, Mattias Haukland, Petra Holmstrom, Noura Walai, Petra Edquist, Mia Brytting, Anna Risberg, Karin Tegmark-Wisell |
| EPI_ISL_913385, EPI_ISL_913386, EPI_ISL_913387, EPI_ISL_913388, EPI_ISL_913389, EPI_ISL_913390, EPI_ISL_913391, EPI_ISL_913392, EPI_ISL_913459 | Dynamic Code AB | The Public Health Agency of Sweden | Anna-Malin Linde, Maria Lind Karlberg, Carlo Berg, Oskar Karlsson Lindsjo, Sofia Stamouli, Reza Advani, Mattias Haukland, Petra Holmstrom, Noura Walai, Petra Edquist, Mia Brytting, Anna Risberg, Karin Tegmark-Wisell |
| EPI_ISL_913465, EPI_ISL_913490, EPI_ISL_913491, EPI_ISL_913492 | Klinisk mikrobiologi | The Public Health Agency of Sweden | Anna-Malin Linde, Maria Lind Karlberg, Carlo Berg, Oskar Karlsson Lindsjo, Sofia Stamouli, Reza Advani, Mattias Haukland, Petra Holmstrom, Noura Walai, Petra Edquist, Mia Brytting, Anna Risberg, Karin Tegmark-Wisell |
| EPI_ISL_913563, EPI_ISL_913564, EPI_ISL_913565, EPI_ISL_913566 | M Health Fairview | Minnesota Department of Health, Public Health Laboratory | Alexandra Lorentz, Jacob Garfin, Matt Plumb, and Xiong Wang |
| EPI_ISL_913648 | Michigan Department of Health and Human Services, Bureau of Laboratories | Michigan Department of Health and Human Services, Bureau of Laboratories | Blankenship HM, Riner D, Soehnlen MK |
| EPI_ISL_913711, EPI_ISL_913712 | Minnesota Department of Health, Public Health Laboratory | Minnesota Department of Health, Public Health Laboratory | Alexandra Lorentz, Jacob Garfin, Matt Plumb, and Xiong Wang |
| EPI_ISL_914815, EPI_ISL_914816, EPI_ISL_914818 | HOSPITAL DE NIÑOS DR. CARLOS SAENZ HERRERA | Inciensa, Instituto Costarricense de Investigación y Enseñanza en Nutrición y Salud | Francisco Duarte, Hebleen Porras, Claudio Soto-Garita, Estela Cordero, Adriana Godínez, Melany Calderón & Cristian Pérez-Corrales |
| EPI_ISL_914823 | HOSPITAL DR. ENRIQUE BALTODANO BRICEÑO | Inciensa, Instituto Costarricense de Investigación y Enseñanza en Nutrición y Salud | Francisco Duarte, Hebleen Porras, Claudio Soto-Garita, Estela Cordero, Adriana Godínez, Melany Calderón & Mariel López |
| EPI_ISL_917936, EPI_ISL_917938 | Los Angeles County PHL | Los Angeles County PHL | P. Hemarajata et al. |
| EPI_ISL_918435, EPI_ISL_918438, EPI_ISL_918439, EPI_ISL_918440 | AIID | Irish Coronavirus Sequencing Consortium-Teagasc Grange | Matthew McCabe, Aljandro Abner Garcia Leon, Fiona Crispie, Calum Walsh, Michael Carr, John Kenny, Paul Cotter, Patrick Mallon, Gabriel Gonzalez |
| EPI_ISL_918557, EPI_ISL_918558, EPI_ISL_918559 | LACEN - Laboratório Central de Saúde Pública do Amapa | Evandro Chagas Institute | Santos, M.C.; Silva, A.M.; Junior, W.D.C.; Barbagelata, L.S.; Ferreira, J.A.; Sousa, E.M.A.; da Silva, P.S.; Pinheiro, K.C.; L.C.; Sousa Junior, E.C. |
| EPI_ISL_918721, EPI_ISL_918745, EPI_ISL_918901, EPI_ISL_918902, EPI_ISL_918903, EPI_ISL_918906, EPI_ISL_918909, EPI_ISL_918910, EPI_ISL_918911, EPI_ISL_918958, EPI_ISL_918960, EPI_ISL_918964, EPI_ISL_918966, EPI_ISL_918967, EPI_ISL_918968 | | | |
| see above | University of Birmingham | COVID-19 Genomics UK (COG-UK) Consortium | Institute of Microbiology, University of Birmingham: Claire McMurray, Joanne Stockton, Samuel Nicholls, Radoslaw Poplawski, Will Rowe, Josh Quick, Nicholas Loman. University of Birmingham Testing Laboratory: Celina M Whalley, Andrew Bosworth, Charlotte Poxon, Kasun Wanigasooriya, Oliver Pickles, Mike Kidd, Alex Richter, Andrew D Beggs PHE Heartlands Lab: Husam Osman, Andrew Bosworth. Queen Elizabeth Hospital: Anna Casey |
| EPI_ISL_919336 | Virology Department, Royal Infirmary of Edinburgh, NHS Lothian / School of Biological Sciences, University of Edinburgh / Institute of Genetics and Molecular Medicine, University of Edinburgh | COVID-19 Genomics UK (COG-UK) Consortium | McHugh M, Dewar R, Rooke S, Gallagher M, Balcaza C, O'Toole Á, Scher E, Hill V, McCrone JT, Colquhoun R, Yu X, Jackson B, Rambaut A, Williams TC, Templeton K |
| EPI_ISL_919449, EPI_ISL_919450, EPI_ISL_919451, EPI_ISL_919454 | Liverpool Clinical Laboratories | COVID-19 Genomics UK (COG-UK) Consortium | Sam Haldenby, Anita Lucaci, Steve Paterson, Julian Hiscox, Alistair Darby, M Almsaud, A Alrezaihi, Muhannad Alruwaili, Stuart D Armstrong, Jones Benjamin, Eleanor G Bentley, Anu Chawla, Jordan J Clark, Angela Cowell, Richard Eccles, Isabel García-Dorival, Matthew Gemmell, Alessandro Gerada, PKF Gilmore, Richard Gregory, Ximeng Han, Catherine Hartley, Margaret Hughes, Miren Iturriza-Gomara, James Johnson, L Luu, Jenifer Manson, Charlotte Nelson, Elaine O'Toole, Cassie Olateju, Rebekah Penrice-Randal , Lucille Rainbow, N.P Randle, Trevor Ian Robinson, Parul Sharma, Ghada T Shawli, James P Stewart, Neil Swainston, Ecaterina Vamos, Joanne Watts, Mark Whitehead |
| EPI_ISL_919806, EPI_ISL_919807, EPI_ISL_919808, EPI_ISL_919809, EPI_ISL_919810, EPI_ISL_919811, EPI_ISL_919812, EPI_ISL_919813, EPI_ISL_919842 | Barts Health NHS Trust | COVID-19 Genomics UK (COG-UK) Consortium | CUTINO-MOGUEL, Maria-Teresa; HARRINGTON, David; OWOYEMI, Dola; KULASEGARAN-SHYLINI, Raghavendran; BROAD, Claire; KELE, Beatrix |
| EPI_ISL_919961, EPI_ISL_919962, EPI_ISL_919963, EPI_ISL_919964, EPI_ISL_919965, EPI_ISL_919966, EPI_ISL_919967, EPI_ISL_919968, EPI_ISL_919969, EPI_ISL_919971, EPI_ISL_919972, EPI_ISL_919974, EPI_ISL_919975, EPI_ISL_919976, EPI_ISL_919977 | | | |

| | | | |
|---|---|---|---|
| see above | University College London, Great Ormond Street Hospital for Children NHS Foundation Trust, Imperial College Healthcare NHS Trust | COVID-19 Genomics UK (COG-UK) Consortium | Sergi Castellano, Rachel Williams, Mark Kristiansen, Paola Resende Silva, Sunando Roy, Tony Brooks, Helena Tutill, Paola Niola, Patricia Dyal, Charlotte Williams, Leysa Forrest, Yasmin Panchbhaya, Jacqueline Findlay, Samuel Weeks, Julianne Brown, Kathryn Harris, Paul Randell, James Price, Alison Holmes, Judith Breuer |
| EPI_ISL_920344, EPI_ISL_920457 | University College London Hospital | COVID-19 Genomics UK (COG-UK) Consortium | Judith Heaney, Matthew Byott, Catherine Houlihan, Dan Frampton, Stuart Kirk, Moira Spyer and Eleni Nastouli |
| EPI_ISL_920840, EPI_ISL_920848 | University College London, Great Ormond Street Hospital for Children NHS Foundation Trust, Imperial College Healthcare NHS Trust | COVID-19 Genomics UK (COG-UK) Consortium | Sergi Castellano, Rachel Williams, Mark Kristiansen, Paola Resende Silva, Sunando Roy, Tony Brooks, Helena Tutill, Paola Niola, Patricia Dyal, Charlotte Williams, Leysa Forrest, Yasmin Panchbhaya, Jacqueline Findlay, Samuel Weeks, Julianne Brown, Kathryn Harris, Paul Randell, James Price, Alison Holmes, Judith Breuer |
| EPI_ISL_921024, EPI_ISL_921033, EPI_ISL_921038, EPI_ISL_921039, EPI_ISL_921040, EPI_ISL_921041, EPI_ISL_921046, EPI_ISL_921047, EPI_ISL_921048, EPI_ISL_921049, EPI_ISL_921050, EPI_ISL_921051, EPI_ISL_921052, EPI_ISL_921053, EPI_ISL_921054, EPI_ISL_921055, EPI_ISL_921056, EPI_ISL_921057, EPI_ISL_921058, EPI_ISL_921059, EPI_ISL_921060, EPI_ISL_921061, EPI_ISL_921062, EPI_ISL_921063, EPI_ISL_921064, EPI_ISL_921065, EPI_ISL_921066, EPI_ISL_921067, EPI_ISL_921068, EPI_ISL_921069, EPI_ISL_921070, EPI_ISL_921071, EPI_ISL_921072, EPI_ISL_921073, EPI_ISL_921074, EPI_ISL_921075, EPI_ISL_921076, EPI_ISL_921077, EPI_ISL_921091, EPI_ISL_921092, EPI_ISL_921093, EPI_ISL_921094, EPI_ISL_921095, EPI_ISL_921098, EPI_ISL_921100, EPI_ISL_921102, EPI_ISL_921106, EPI_ISL_921107, EPI_ISL_921109, EPI_ISL_921110, EPI_ISL_921111, EPI_ISL_921113, EPI_ISL_921114, EPI_ISL_921115, EPI_ISL_921116, EPI_ISL_921117, EPI_ISL_921125, EPI_ISL_921131, EPI_ISL_921134 | | | |
| see above | Regional Virus Laboratory, Belfast Health and Social Care Trust | COVID-19 Genomics UK (COG-UK) Consortium | Conall McCaughey, James McKenna, Tanya Curran, Susan Feeney, Alison Watt, Ciara Cox, Mairead Connor, Zoltan Molnar, David Simpson, Derek Fairley |
| EPI_ISL_921246, EPI_ISL_921250, EPI_ISL_921251, EPI_ISL_921283, EPI_ISL_921284, EPI_ISL_921285, EPI_ISL_921286, EPI_ISL_921419, EPI_ISL_921424, EPI_ISL_921425, EPI_ISL_921426, EPI_ISL_921427, EPI_ISL_921429, EPI_ISL_921430, EPI_ISL_921431, EPI_ISL_921432, EPI_ISL_921434 | | | |
| see above | Northumbria University / South Tees Hospitals NHS Foundation Trust / North Cumbria Integrated Care NHS Foundation Trust / North Tees and Hartlepool NHS Foundation Trust / Newcastle Hospitals NHS Foundation Trust | COVID-19 Genomics UK (COG-UK) Consortium | Darren L Smith,Andrew Nelson,Matthew Bashton,Greg R Young,Joshua Loh,John Allan,Mohammad A Tariq,Giles S Holt,Gary Black,Wen C Yew,Lynn Dover,Paul Baker,Steve Liggett,Sarah Essex,Jane Greenaway,Debra Padgett,Clive Graham,Garren Scott,Edward Barton,Emma Swindells,Brendan Payne,Jennifer Collins,Yusri Taha,Gary Eltringham |
| EPI_ISL_922180, EPI_ISL_922242, EPI_ISL_922255, EPI_ISL_922286, EPI_ISL_922289, EPI_ISL_922308, EPI_ISL_922311, EPI_ISL_922327, EPI_ISL_922336, EPI_ISL_922337 | Oxford Viromics, NDM, University of Oxford; Oxford University Hospitals; Basingstoke and North Hampshire Hospital | COVID-19 Genomics UK (COG-UK) Consortium | Tanya Golubchik, David Bonsall, George Macintyre, Amy Trebes, Mariateresa de Cesare, Catrin Moore, Alex Mobbs, Anita Justice, Robert Shaw, Monique Andersson, Timothy Peto, Emma Wise, Nathan Moore, Jessica Lynch, Nick Cortes, Matilde Mori, Stephen Kidd, David Buck, John Todd, Christophe Fraser |
| EPI_ISL_923242 | Centre for Enzyme Innovation, University of Portsmouth / Translational Research Laboratory, Portsmouth Hospitals NHS Trust | COVID-19 Genomics UK (COG-UK) Consortium | Angela Beckett,Salman Goudarzi,Christopher Fearn,Kate Cook,Katie Loveson,Sharon Glaysher,Scott Elliott,Samuel Robson |
| EPI_ISL_924157, EPI_ISL_924333, EPI_ISL_924387, EPI_ISL_924402, EPI_ISL_924412 | Virology Department, Sheffield Teaching Hospitals NHS Foundation Trust/Department of Infection, Immunity and Cardiovascular Disease, The Medical School, University of Sheffield | COVID-19 Genomics UK (COG-UK) Consortium | Thushan de Silva, Matthew Parker, Nikki Smith, Adri Angyal, Rebecca Brown, Luke Green, Rachel Tucker, Paul Parsons, Danielle Groves, Katie Johnson, Laura Carrilero, Alex Keeley, Dave Partridge, Matthew Wyles, Benjamin Lindsey, Mehmet Yavuz, Mohammad Raza, Cariad Evans |
| EPI_ISL_924461, EPI_ISL_924462, EPI_ISL_924464, EPI_ISL_924465, EPI_ISL_924466, EPI_ISL_924471, EPI_ISL_924473, EPI_ISL_924475, EPI_ISL_924480, EPI_ISL_924481, EPI_ISL_924482, EPI_ISL_924484, EPI_ISL_924486, EPI_ISL_924492, EPI_ISL_924495, EPI_ISL_924517, EPI_ISL_924518, EPI_ISL_924521, EPI_ISL_924528, EPI_ISL_924529, EPI_ISL_924530, EPI_ISL_924531, EPI_ISL_924533, EPI_ISL_924539, EPI_ISL_924540, EPI_ISL_924542, EPI_ISL_924544, EPI_ISL_924545, EPI_ISL_924547, EPI_ISL_924548, EPI_ISL_924550, EPI_ISL_924552, EPI_ISL_924553, EPI_ISL_924555, EPI_ISL_924556, EPI_ISL_924557, EPI_ISL_924558, EPI_ISL_924559, EPI_ISL_924560, EPI_ISL_924562, EPI_ISL_924563, EPI_ISL_924564, EPI_ISL_924565, EPI_ISL_924566, EPI_ISL_924567, EPI_ISL_924568, EPI_ISL_924570, EPI_ISL_924571, EPI_ISL_924572, EPI_ISL_924573, EPI_ISL_924574, EPI_ISL_924575, EPI_ISL_924576, EPI_ISL_924577, EPI_ISL_924578, EPI_ISL_924579, EPI_ISL_924580, EPI_ISL_924581, EPI_ISL_924582, EPI_ISL_924583, EPI_ISL_924584, EPI_ISL_924585, EPI_ISL_924586, EPI_ISL_924587, EPI_ISL_924588, EPI_ISL_924589, EPI_ISL_924590, EPI_ISL_924591, EPI_ISL_924592, EPI_ISL_924593, EPI_ISL_924594, EPI_ISL_924595, EPI_ISL_924596, EPI_ISL_924597, EPI_ISL_924598, EPI_ISL_924599, EPI_ISL_924600, EPI_ISL_924601, EPI_ISL_924602, EPI_ISL_924603, EPI_ISL_924604, EPI_ISL_924738 | | | |
| see above | Bioinformatics and Biostatistics Lab, Advanced Sequencing Facility | COVID-19 Genomics UK (COG-UK) Consortium | Aengus Stewart,Jerome Nicod,Chelsea Sawyer,Laura Cubitt,Harshil Patel,Margaret Crawford |
| EPI_ISL_925397 | Department of Clinical Microbiology | GIGA Medical Genomics | Keith Durkin, Maria Artesi, Sébastien Bontems, Raphaël Boreux, Bouchra Boujemla, Cécile Meex, Pierrette Melin, Marie-Pierre Hayette, Vincent Bours |
| EPI_ISL_925514, EPI_ISL_925520, EPI_ISL_925521 | Arizona State Public Health Laboratory | Arizona State Public Health Laboratory | Trung Huynh, Jessica Escobar, Katherine Fullerton, Nobuko Fukushima, Stacy White, Linda Getsinger, Victor Waddell |
| EPI_ISL_925915 | Nucleic Acid Testing, National Reference Laboratory | GIGA Medical Genomics | Yvan Butera, Keith Durkin, Maria Artesi, Bouchra Boujemla, Robert Rutayisire, Patrick Tuyisenge, Esperence Umumararungu, Sébastien Bontems, Marie-Pierre Hayette, Nathalie Renotte, Swaibu Gatare, Jacob Souopgui, Sabin Nsanzimana, Vincent Bours, Léon Mutesa |
| EPI_ISL_931570 | Utah Public Health Laboratory | Utah Public Health Laboratory | Erin L. Young, Kelly F. Oakeson, Tara Gallagher |
| EPI_ISL_934174, EPI_ISL_934175, EPI_ISL_934176, EPI_ISL_934177, EPI_ISL_934178, EPI_ISL_934179, EPI_ISL_934187, EPI_ISL_934203, EPI_ISL_934222 | Vilnius university hospital Santaros Klinikos, Center of Laboratory Medicine | Vilnius university hospital Santaros Klinikos, Center of Laboratory Medicine | Ingrida Olendraite, Daniel Naumovas, Rimvydas Norvilas, Dovile Ezerskyte, Justinas Slikas, Gytis Dudas |
| EPI_ISL_934345, EPI_ISL_934346, EPI_ISL_934347 | Klinisk mikrobiologi | The Public Health Agency of Sweden | Anna-Malin Linde, Maria Lind Karlberg, Carlo Berg, Oskar Karlsson Lindsjo, Sofia Stamouli, Reza Advani, Mattias Haukland, Petra Holmstrom, Noura Walai, Petra Edquist, Mia Brytting, Anna Risberg, Karin Tegmark-Wisell |
| EPI_ISL_934377 | Dynamic Code AB | The Public Health Agency of Sweden | Anna-Malin Linde, Maria Lind Karlberg, Carlo Berg, Oskar Karlsson Lindsjo, Sofia Stamouli, Reza Advani, Mattias Haukland, Petra Holmstrom, Noura Walai, Petra Edquist, Mia Brytting, Anna Risberg, Karin Tegmark-Wisell |
| EPI_ISL_934384 | Klinisk mikrobiologi | The Public Health Agency of Sweden | Anna-Malin Linde, Maria Lind Karlberg, Carlo Berg, Oskar Karlsson Lindsjo, Sofia Stamouli, Reza Advani, Mattias Haukland, Petra Holmstrom, Noura Walai, Petra Edquist, Mia Brytting, Anna Risberg, Karin Tegmark-Wisell |
| EPI_ISL_934593, EPI_ISL_934594, EPI_ISL_934595, EPI_ISL_934596, EPI_ISL_934597, EPI_ISL_934598, EPI_ISL_934599, EPI_ISL_934600, EPI_ISL_934601, EPI_ISL_934602, EPI_ISL_934603, EPI_ISL_934604, EPI_ISL_934605, EPI_ISL_934606, EPI_ISL_934607, EPI_ISL_934608, EPI_ISL_934609, EPI_ISL_934610, EPI_ISL_934611, EPI_ISL_934612, EPI_ISL_934613, EPI_ISL_934614, EPI_ISL_934615, EPI_ISL_934616, EPI_ISL_934617, EPI_ISL_934618, EPI_ISL_934619, EPI_ISL_934620, EPI_ISL_934621, EPI_ISL_934622, EPI_ISL_934623, EPI_ISL_934624, EPI_ISL_934625, EPI_ISL_934626, EPI_ISL_934627, EPI_ISL_934628, EPI_ISL_934629, EPI_ISL_934630, EPI_ISL_934631, EPI_ISL_934632 | | | |
| see above | Austrian Agency for Health and Food Safety (AGES) | Bergthaler laboratory, CeMM Research Center for Molecular Medicine of the Austrian Academy of Sciences | Lukas Endler, Anna Schedl, Thomas Penz, Benedikt Agerer, Maelle Le Moing, Michael Schuster, Bekir Erguner, Jan Laine, Martin Senekowitsch, Christoph Bock, Andreas Bergthaler |
| EPI_ISL_934637, EPI_ISL_934643 | Department of Microbiology, University Innsbruck | Bergthaler laboratory, CeMM Research Center for Molecular Medicine of the Austrian Academy of Sciences | Lukas Endler, Anna Schedl, Thomas Penz, Benedikt Agerer, Maelle Le Moing, Michael Schuster, Bekir Erguner, Jan Laine, Martin Senekowitsch, Christoph Bock, Andreas Bergthaler |
| EPI_ISL_934650 | Institute for Water Quality and Resource Management, Technical University Vienna | Bergthaler laboratory, CeMM Research Center for Molecular Medicine of the Austrian Academy of Sciences | Lukas Endler, Anna Schedl, Thomas Penz, Benedikt Agerer, Maelle Le Moing, Michael Schuster, Bekir Erguner, Jan Laine, Martin Senekowitsch, Christoph Bock, Andreas Bergthaler |
| EPI_ISL_935220, EPI_ISL_935223, EPI_ISL_935224 | KU Leuven, Rega Institute, Clinical and Epidemiological Virology | KU Leuven, Rega Institute, Clinical and Epidemiological Virology | Tony Wawina-Bokalanga, Bert Vanmechelen, Joan Marti-Carerras, Piet Maes |
| EPI_ISL_935361, EPI_ISL_935363, EPI_ISL_935364 | Florida Bureau of Public Health Laboratories | Florida Bureau of Public Health Laboratories | Sarah Schmedes, Jason Blanton |
| EPI_ISL_935711, EPI_ISL_935712, EPI_ISL_935713, EPI_ISL_935714, EPI_ISL_935715, EPI_ISL_935716, EPI_ISL_935717, EPI_ISL_935718, EPI_ISL_935719, EPI_ISL_935720, EPI_ISL_935721, EPI_ISL_935722 | | | |
| see above | Houston Health Dept. | Houston Health Dept. | Ryker Penn, Pamela Brown, Adolpho Lara |
| EPI_ISL_935986, EPI_ISL_935987, EPI_ISL_935992 | GLENS FALLS HOSPITAL LABORATORY | Wadsworth Center, New York State Department of Health | Kirsten St. George, Daryl M. Lamson, Alexis Russel, Matthew Shudt, Melissa A Leisner, Jonathan Plitnick, Navjot Singh, John Kelly, Erasmus Schneider, Erica Lasek-Nesselquist |
| EPI_ISL_936061, EPI_ISL_936067, EPI_ISL_936068, EPI_ISL_936070, EPI_ISL_936071, EPI_ISL_936072, EPI_ISL_936073, EPI_ISL_936074, EPI_ISL_936075, EPI_ISL_936076, EPI_ISL_936077, EPI_ISL_936084, EPI_ISL_936085, EPI_ISL_936087, EPI_ISL_936088, EPI_ISL_936090, EPI_ISL_936091, EPI_ISL_936094 | | | |
| see above | KALEIDA CENTER FOR LABORATORY MEDICINE | Wadsworth Center, New York State Department of Health | Kirsten St. George, Daryl M. Lamson, Alexis Russel, Matthew Shudt, Melissa A Leisner, Jonathan Plitnick, Navjot Singh, John Kelly, Erasmus Schneider, |

| | | | |
|---|---|---|---|
| | | | Erica Lasek-Nesselquist |
| EPI_ISL_936861, EPI_ISL_936862, EPI_ISL_936863, EPI_ISL_936864, EPI_ISL_936865, EPI_ISL_936866, EPI_ISL_936868, EPI_ISL_936869, EPI_ISL_936870, EPI_ISL_936871, EPI_ISL_936872, EPI_ISL_936873, EPI_ISL_936874, EPI_ISL_936882, EPI_ISL_936883, EPI_ISL_936884, EPI_ISL_936896, EPI_ISL_936919, EPI_ISL_936920, EPI_ISL_936921, EPI_ISL_936922 | | | |
| see above | Northwestern Memorial Hospital | Ozer Lab | Ramon Lorenzo-Redondo, Lacy M. Simons, Chad J. Achenbach, Lawrence J. Jennings, Michael G. Ison, Judd F. Hultquist, Egon A. Ozer |
| EPI_ISL_940715 | City of Milwaukee Health Department Laboratory | City of Milwaukee Health Department Laboratory | Sanjib Bhattacharyya |
| EPI_ISL_940773 | INSPI-CRN de Influenza y otros virus respiratorios | INSPI-Centro de Investigación Multidisciplinaria de la DTIDI | Leandro Patiño, Doménica de Mora, Maritza Olmedo, Andrés Carrazco-Montalvo, Orson Mestanza, Mary Regato-Arrata, Melissa Zambrano, Manuel González, Alfredo Bruno, Alberto Orlando. |
| EPI_ISL_941371, EPI_ISL_941372, EPI_ISL_941442, EPI_ISL_941443, EPI_ISL_941444, EPI_ISL_941445, EPI_ISL_941625, EPI_ISL_941626, EPI_ISL_941627 | Instituto Nacional de Saude (INSA) | Instituto Nacional de Saude (INSA) | Borges et al |
| EPI_ISL_942770, EPI_ISL_942772, EPI_ISL_942773, EPI_ISL_942774, EPI_ISL_942775, EPI_ISL_942776, EPI_ISL_942777, EPI_ISL_942778, EPI_ISL_942779, EPI_ISL_942780, EPI_ISL_942781, EPI_ISL_942782, EPI_ISL_942783, EPI_ISL_942784, EPI_ISL_942785, EPI_ISL_942786, EPI_ISL_942787 | | | |
| see above | Gundersen Molecular Diagnostics Laboratory | Kabara Cancer Research Institute | Craig S. Richmond, Paraic A. Kenny |
| EPI_ISL_942900, EPI_ISL_942901, EPI_ISL_942902, EPI_ISL_942903, EPI_ISL_942904, EPI_ISL_942905, EPI_ISL_942906, EPI_ISL_942907, EPI_ISL_942908, EPI_ISL_942909, EPI_ISL_942910, EPI_ISL_942911, EPI_ISL_942912, EPI_ISL_942913, EPI_ISL_942914, EPI_ISL_942915 | | | |
| see above | Houston Health Dept. | Houston Health Dept. | Ryker Penn, Pamela Brown, Adolpho Lara |
| EPI_ISL_943153, EPI_ISL_943154, EPI_ISL_943200, EPI_ISL_943201, EPI_ISL_943202, EPI_ISL_943203, EPI_ISL_943204, EPI_ISL_943205, EPI_ISL_943245, EPI_ISL_943328, EPI_ISL_943408, EPI_ISL_943451, EPI_ISL_943452, EPI_ISL_943465, EPI_ISL_943489, EPI_ISL_943502, EPI_ISL_943514, EPI_ISL_943515 | | | |
| see above | Dutch COVID-19 response team | National Institute for Public Health and the Environment (RIVM) | Adam Meijer, Harry Vennema, Dirk Eggink, Jeroen Cremer, Sharon van den Brink, Bas van der Veer, AnneMarie van den Brandt, Florian Zwagemaker, Dennis Schmitz, Chantal Reusken, on behalf of the national COVID-19 response team |
| EPI_ISL_943573 | Servizo de Microbioloxía. Complexo Hospitalario Universitario de Santiago de Compostela | Servizo de Microbioloxía. Complexo Hospitalario Universitario de Santiago de Compostela | Antonio Aguilera, Gema Barbeito, Amparo Coira, José Costa, Rocio Trastoy, María Luisa Pérez del Molino. |
| EPI_ISL_943822 | Utah Public Health Laboratory | Utah Public Health Laboratory | Erin L. Young, Kelly F. Oakeson, Tara Gallagher |
| EPI_ISL_944125, EPI_ISL_944133, EPI_ISL_944151, EPI_ISL_944156, EPI_ISL_944164, EPI_ISL_944174 | National Health Laboratory Service, South Africa | KRISP, KZN Research Innovation and Sequencing Platform | Laguda-Akingba O, Giandhari J, Pillay S, Lessells R, Mdlalose K, York D, Khan S, Emmanuel SJ, Tegally H, Wilkinson E, de Oliveira T |
| EPI_ISL_944197, EPI_ISL_944198, EPI_ISL_944205, EPI_ISL_944213, EPI_ISL_944222, EPI_ISL_944241, EPI_ISL_944276, EPI_ISL_944277, EPI_ISL_944278, EPI_ISL_944378, EPI_ISL_944379, EPI_ISL_944382, EPI_ISL_944421, EPI_ISL_944422, EPI_ISL_944459, EPI_ISL_944460, EPI_ISL_944461, EPI_ISL_944462, EPI_ISL_944463 | | | |
| see above | Israel Central Virology laboratory | Israel National Consortium for SARS-CoV-2 sequencing | Neta Zuckerman, Efrat Dahan Bucris, Michal Mandelboim, Dana Bar-Ilan, Oran Erster, Tzvia Mann, Omer Murik, David A. Zeevi, Assaf Rokney, Joseph Jaffe, Eva Nachum, Maya Davidovich Cohen, Ephraim Fass, Gal Zizelski Valenci, Mor Rubinstein, Efrat Rorman, Israel Nissan, Efrat Glick-Saar, Omri Nayshool, Gideon Rechavi, Ella Mendelson, Orna Mor |
| EPI_ISL_944652, EPI_ISL_944673, EPI_ISL_944683, EPI_ISL_944719, EPI_ISL_944726, EPI_ISL_944727 | Department of Biochemistry, Cell and Molecular Biology, West African Centre for Cell Biology of Infectious Pathogens (WACCBIP), University of Ghana | Department of Biochemistry, Cell and Molecular Biology, West African Centre for Cell Biology of Infectious Pathogens (WACCBIP), University of Ghana | Morang'a,C.M., Ngoi,J.M., Quansah,E.B., Saiid,S., Amuzu,D.S., Asante,I., Bonney,J.H., Bonney,E., Odoom,J.K., Ndam,N.T., Tei-Maya,F., Adusei-Poku,M., Ofori-Boadu,L., Ampofo,W.K., Amenga-Etego,L.N., Quashie,P., Bediako,Y., Awandare,G.A. |
| EPI_ISL_945064 | Lighthouse Lab in Milton Keynes | Wellcome Sanger Institute for the COVID-19 Genomics UK (COG-UK) Consortium | The Lighthouse Lab in Milton Keynes and Alex Alderton, Roberto Amato, Sonia Goncalves, Ewan Harrison, David K. Jackson, Ian Johnston, Dominic Kwiatkowski, Cordelia Langford, John Sillitoe on behalf of the Wellcome Sanger Institute COVID-19 Surveillance Team |
| EPI_ISL_945068, EPI_ISL_945070 | Lighthouse Lab in Cambridge | Wellcome Sanger Institute for the COVID-19 Genomics UK (COG-UK) Consortium | Rob Howes, The Lighthouse Lab in Cambridge and Alex Alderton, Roberto Amato, Sonia Goncalves, Ewan Harrison, David K. Jackson, Ian Johnston, Dominic Kwiatkowski, Cordelia Langford, John Sillitoe on behalf of the Wellcome Sanger Institute COVID-19 Surveillance Team |
| EPI_ISL_945105, EPI_ISL_945159, EPI_ISL_945174 | Lighthouse Lab in Milton Keynes | Wellcome Sanger Institute for the COVID-19 Genomics UK (COG-UK) Consortium | The Lighthouse Lab in Milton Keynes and Alex Alderton, Roberto Amato, Sonia Goncalves, Ewan Harrison, David K. Jackson, Ian Johnston, Dominic Kwiatkowski, Cordelia Langford, John Sillitoe on behalf of the Wellcome Sanger Institute COVID-19 Surveillance Team |
| EPI_ISL_945180 | Lighthouse Lab in Cambridge | Wellcome Sanger Institute for the COVID-19 Genomics UK (COG-UK) Consortium | Rob Howes, The Lighthouse Lab in Cambridge and Alex Alderton, Roberto Amato, Sonia Goncalves, Ewan Harrison, David K. Jackson, Ian Johnston, Dominic Kwiatkowski, Cordelia Langford, John Sillitoe on behalf of the Wellcome Sanger Institute COVID-19 Surveillance Team |
| EPI_ISL_945295 | Lighthouse Lab in Milton Keynes | Wellcome Sanger Institute for the COVID-19 Genomics UK (COG-UK) Consortium | The Lighthouse Lab in Milton Keynes and Alex Alderton, Roberto Amato, Sonia Goncalves, Ewan Harrison, David K. Jackson, Ian Johnston, Dominic Kwiatkowski, Cordelia Langford, John Sillitoe on behalf of the Wellcome Sanger Institute COVID-19 Surveillance Team |
| EPI_ISL_947326 | RS Mitra Keluarga Gading Serpong | Eijkman Institute for Molecular Biology, Ministry of Research and Technology/National Agency for Research and Innovation | Willy Agustine, Edison Johar, Hidayat Trimarsanto, Iskandar Adnan, Lydia V. Panggalo, Sukma Oktavianthi, Frilasita A Yudhaputri, Safarina G Malik, Khin Saw Myint, Amin Soebandrio |
| EPI_ISL_947649 | Lighthouse Lab in Alderley Park | Wellcome Sanger Institute for the COVID-19 Genomics UK (COG-UK) Consortium | Jacquelyn Wynn, Mairead Hyland, The Lighthouse Lab in Alderley Park and Alex Alderton, Roberto Amato, Sonia Goncalves, Ewan Harrison, David K. Jackson, Ian Johnston, Dominic Kwiatkowski, Cordelia Langford, John Sillitoe on behalf of the Wellcome Sanger Institute COVID-19 Surveillance Team |
| EPI_ISL_949748, EPI_ISL_949784 | Barts Health NHS Trust | COVID-19 Genomics UK (COG-UK) Consortium | CUTINO-MOGUEL, Maria-Teresa; HARRINGTON, David; OWOYEMI, Dola; KULASEGARAN-SHYLINI, Raghavendran; BROAD, Claire; KELE, Beatrix |
| EPI_ISL_950048, EPI_ISL_950078, EPI_ISL_950082, EPI_ISL_950086, EPI_ISL_950121, EPI_ISL_950127, EPI_ISL_950134, EPI_ISL_950135, EPI_ISL_950153, EPI_ISL_950165 | University College London, Great Ormond Street Hospital for Children NHS Foundation Trust, Imperial College Healthcare NHS Trust | COVID-19 Genomics UK (COG-UK) Consortium | Sergi Castellano, Rachel Williams, Mark Kristiansen, Paola Resende Silva, Sunando Roy, Tony Brooks, Helena Tutill, Paola Niola, Patricia Dyal, Charlotte Williams, Leysa Forrest, Yasmin Panchbhaya, Jacqueline Findlay, Samuel Weeks, Julianne Brown, Kathryn Harris, Paul Randell, James Price, Alison Holmes, Judith Breuer |
| EPI_ISL_950241, EPI_ISL_950243, EPI_ISL_950244, EPI_ISL_950245, EPI_ISL_950247, EPI_ISL_950248, EPI_ISL_950250, EPI_ISL_950253, EPI_ISL_950255, EPI_ISL_950256, EPI_ISL_950258, EPI_ISL_950259, EPI_ISL_950260, EPI_ISL_950262, EPI_ISL_950263, EPI_ISL_950265, EPI_ISL_950267, EPI_ISL_950460, EPI_ISL_950461, EPI_ISL_950462, EPI_ISL_950463, EPI_ISL_950465, EPI_ISL_950466, EPI_ISL_950467, EPI_ISL_950468, EPI_ISL_950469, EPI_ISL_950470, EPI_ISL_950471, EPI_ISL_950472, EPI_ISL_950473, EPI_ISL_950474, EPI_ISL_950475, EPI_ISL_950476, EPI_ISL_950477, EPI_ISL_950478, EPI_ISL_950480, EPI_ISL_950481 | | | |
| see above | Northumbria University / South Tees Hospitals NHS Foundation Trust / North Cumbria Integrated Care NHS Foundation Trust / North Tees and Hartlepool NHS Foundation Trust / Newcastle Hospitals NHS Foundation Trust | COVID-19 Genomics UK (COG-UK) Consortium | Darren L Smith,Andrew Nelson,Matthew Bashton,Greg R Young,Joshua Loh,John Allan,Mohammad A Tariq,Giles S Holt,Gary Black,Wen C Yew,Lynn Dover,Paul Baker,Steve Liggett,Sarah Essex,Jane Greenaway,Debra Padgett,Clive Graham,Garren Scott,Edward Barton,Emma Swindells,Brendan Payne,Jennifer Collins,Yusri Taha,Gary Eltringham |
| EPI_ISL_950551, EPI_ISL_950582 | Quadram Institute Bioscience | COVID-19 Genomics UK (COG-UK) Consortium | Dave J. Baker, Gemma L. Kay, Alp Aydin, Thanh Le-Viet, Steven Rudder, Ana P. Tedim, Anastasia Kolyva, Maria Diaz, Leonardo de Oliveira Martins, Nabil-Fareed Alikhan, Lizzie Meadows, Rachael Stanley, Ngozi Elumogo, Muhammed Yasir, Nicholas M. Thomson, Alexander J Trotter, Rachel Gilroy, Samuel Bloomfield, Claire Stuart, Andrew Bell, Reenesh Prakash, Samir Dervisevic, Alison E. Mather, John Wain, Mark Webber, Andrew J. Page, Justin O'Grady |
| EPI_ISL_950711 | Lincolnshire Hospitals and DeepSeq Nottingham | COVID-19 Genomics UK (COG-UK) Consortium | Nichola Duckworth, Tim Sloan, Sarah Walsh, Jonathan Ball, Patrick McClure, Joeseph Chappell, Nadine Holmes, Matthew Carlisle, Christopher Moore, Fei Sang, Johnny Debebe, Victoria Wright, Matthew Loose |
| EPI_ISL_951068, EPI_ISL_951069, EPI_ISL_951070, EPI_ISL_951071, EPI_ISL_951072, EPI_ISL_951073, EPI_ISL_951074, EPI_ISL_951075, EPI_ISL_951077, EPI_ISL_951078, EPI_ISL_951082, EPI_ISL_951091, EPI_ISL_951101, EPI_ISL_951102, EPI_ISL_951103, EPI_ISL_951104, EPI_ISL_951105, EPI_ISL_951106, EPI_ISL_951107, EPI_ISL_951108, EPI_ISL_951109, EPI_ISL_951110, EPI_ISL_951111, EPI_ISL_951112, EPI_ISL_951114, EPI_ISL_951115, EPI_ISL_951116, EPI_ISL_951117, EPI_ISL_951118, EPI_ISL_951119, EPI_ISL_951120, EPI_ISL_951121, EPI_ISL_951122, EPI_ISL_951135, EPI_ISL_951137, EPI_ISL_951143, EPI_ISL_951161, EPI_ISL_951162, EPI_ISL_951188, EPI_ISL_951338, EPI_ISL_951381 | | | |
| see above | Oxford Viromics, NDM, University of Oxford; Oxford University Hospitals; Basingstoke and North Hampshire Hospital | COVID-19 Genomics UK (COG-UK) Consortium | Tanya Golubchik, David Bonsall, George Macintyre, Amy Trebes, Mariateresa de Cesare, Catrin Moore, Alex Mobbs, Anita Justice, Robert Shaw, Monique Andersson, Timothy Peto, Emma Wise, Nathan Moore, Jessica Lynch, Nick Cortes, Matilde Mori, Stephen Kidd, David Buck, John Todd, Christophe Fraser |

| | | | |
|---|---|---|---|
| EPI_ISL_952430, EPI_ISL_952435 | Centre for Enzyme Innovation, University of Portsmouth / Translational Research Laboratory, Portsmouth Hospitals NHS Trust | COVID-19 Genomics UK (COG-UK) Consortium | Angela Beckett,Salman Goudarzi,Christopher Fearn,Kate Cook,Katie Loveson,Sharon Glaysher,Scott Elliott,Samuel Robson |
| EPI_ISL_953174, EPI_ISL_953175, EPI_ISL_953179, EPI_ISL_953183, EPI_ISL_953186, EPI_ISL_953191, EPI_ISL_953194, EPI_ISL_953195, EPI_ISL_953196, EPI_ISL_953197, EPI_ISL_953198, EPI_ISL_953203, EPI_ISL_953204, EPI_ISL_953206, EPI_ISL_953208, EPI_ISL_953209, EPI_ISL_953211, EPI_ISL_953213, EPI_ISL_953215, EPI_ISL_953218, EPI_ISL_953220, EPI_ISL_953221, EPI_ISL_953224, EPI_ISL_953227, EPI_ISL_953228, EPI_ISL_953234, EPI_ISL_953235, EPI_ISL_953237 | | | |
| see above | Bioinformatics and Biostatistics Lab, Advanced Sequencing Facility | COVID-19 Genomics UK (COG-UK) Consortium | Aengus Stewart,Jerome Nicod,Chelsea Sawyer,Laura Cubitt,Harshil Patel,Margaret Crawford |
| EPI_ISL_953427 | National Institute of Health Research and Development | National Institute of Health Research and Development | Subangkit, Hana Apsari Pawestri, Kartika Dewi Puspa, Arie Ardiansyah Nugraha, Hartanti Dian Ikawati, Krisna Nur Andriana Pangesti, Yuni Rukminiati, Ririn Ramadhany, Agustiningsih, Kindi Adam, Holy Arif Wibowo, Triyani Soekarso, Ni Ketut Susilarini, Nurika Hariastuti, Ully Alfi Nikmah, Reni Herman, Nike Susanti, Herna, Tati Febriyanti, Natalie Laurencia Kipuw, Fauzul Muna, Irene Lorinda Indalao, Nelly Puspandari, Vivi Setiawaty |
| EPI_ISL_954218, EPI_ISL_954219 | 1.AO Universitaria 'S. Giovanni di Dio e Ruggi D'Aragona, Scuola Medica Salernitana' Hospital / 2.UOC di Virologia e Microbiologia, Università della Campania 'L. Vanvitelli' / 3.AO Universitaria 'Federico II' Napoli Hospital / 4.AORN 'San Giuseppe Moscati' Avellino Hospital / 5.AO 'San Pio - presidio G. Rummo' Benevento Hospital / 6.AO 'Sant'Anna e San Sebastiano' Caserta Hospital / 7.PO 'Maria Santissima Addolorata' Eboli Hospital / 8.Biogem Istituto di Ricerche Genetiche | 1. Genome Research Center for Health (CRGS) / 2. Laboratory of Molecular Medicine and Genomics(LMMGe) / 3. Center for Research in Pure and Applied Mathematics (CRMPA) | Giorgio Giurato, Francesca Rizzo, Alessandro Weisz, Gianluigi Franci, Giovanni Nassa, Pasquale Pagliano, Roberta Tarallo, Elena Alexandrova, Ylenia D'Agostino, Carlo Ferravante, Jessica Lamberti, Viola Melone, Domenico Memoli, Valeria Mirici Cappa, Domenico Palumbo, Giovanni Pecoraro, Assunta Sellitto, Oriana Strianese, Ilaria Terenzi, Giuseppe Fenza, Aniello Gentile, Antonello Saccomanno, Sonia Amabile, Teresa Rocco, Annamaria Salvati, Emilia Vaccaro, Massimiliano Galdiero, Michele Cennamo, Giuseppe Portella, Maria Grazia Foti, Mariarosaria Ingino, Maria Landi, Maurizio Fumi, Vincenzo Rocco, Rita Greco, Vittoria Letizia, Arnolfo Petruzziello, Maddalena Schioppa, Gregorio Goffredi, Francesca Marciano, Michele Caraglia, Alessia Cossu, Marianna Scrima, Edmondo Adorisio, Morena D'Avenia, Michela Iacobellis, Rosanna Piluscio, Giorgio Dirani, Vittorio Sambri, Simona Semprini, Silvia Zanoli, Francesco Curcio, Stefania Marzinotto, Andreina Baj, Fausto Sessa. |
| EPI_ISL_954226, EPI_ISL_954297, EPI_ISL_954298, EPI_ISL_954299, EPI_ISL_954300 | MRC/UVRI & LSHTM Uganda Research Unit | Where sequence data have been generated and submitted to GISAID | Matthew Cotten, Dan Lule Bugembe, My V.T. Phan, Isaac Sseeewanyana, Patrick Semanda, Susan Nabadda, Pontiano Kaleebu |
| EPI_ISL_954825, EPI_ISL_954829, EPI_ISL_954832, EPI_ISL_954835, EPI_ISL_954865, EPI_ISL_954867, EPI_ISL_954878, EPI_ISL_954881, EPI_ISL_954884, EPI_ISL_954895, EPI_ISL_954896, EPI_ISL_954897, EPI_ISL_954898, EPI_ISL_954899, EPI_ISL_954900, EPI_ISL_954901 | | | |
| see above | Colorado Department of Public Health and Environment | Colorado Department of Puplic Health and Environment | Laura Bankers, Molly C. Hetherington-Rauth, Diana Ir, Shannon Ely, Shannon R. Matzinger, Sarah Elizabeth Totten, Emily A. Travanty |
| EPI_ISL_955231, EPI_ISL_955232, EPI_ISL_955258, EPI_ISL_955259, EPI_ISL_955260 | BioReference Lab | NJ Public Health and Environmental Laboratories | Lindsey Bodnar, Shiv Verma, Dana Woell, Byeong Jeong |
| EPI_ISL_955392 | Alameda County Public Health Lab | Chan-Zuckerberg Biohub | CZB Cliahub Consortium |
| EPI_ISL_955427, EPI_ISL_955428, EPI_ISL_955429, EPI_ISL_955430, EPI_ISL_955431, EPI_ISL_955432, EPI_ISL_955433 | Orange County Public Health Lab | Chan-Zuckerberg Biohub | CZB Cliahub Consortium |
| EPI_ISL_955768 | Humboldt County Public Health Laboratory | Chan-Zuckerberg Biohub | CZB Cliahub Consortium |
| EPI_ISL_955905 | Pamela Youde Nethersole Eastern Hospital | Hong Kong Department of Health | Alan K.L. Tsang, Peter C.W. Yip, Edman T.K. Lam, Rickjason C.W. Chan, Dominic N.C. Tsang |
| EPI_ISL_956312 | Institute of Tropical Disease | Institute of Tropical Disease, Universitas Airlangga | Krisnoadi Rahardjo, Aldise M Nastri, Jezzy R Dewantari, Rima R Prasetya, Gatot Soegiarto, Laksmi Wulandari, Resti Yudhawati, Yasuko Mori, Soetjipto, Kazufumi Shimizu, Maria I Lusida |
| EPI_ISL_956327, EPI_ISL_956328 | Laboratory Medicine | Department of Laboratory Medicine, Lin-Kou Chang Gung Memorial Hospital, Taoyuan, Taiwan | Kuo-Chien Tsao, Yu-Nong Gong, Shu-Li Yang, Yi-Chun Liu, Chung-Guei Huang, Mei-Jen Hsiao, Po-Wei Huang, Cheng-Ta Yang, Cheng-Hsun Chiu, Peng-Nien Huang, Kuo-Ming Lee, Guang-Wu Chen, Shin-Ru Shih |
| EPI_ISL_956355, EPI_ISL_956356, EPI_ISL_956357, EPI_ISL_956358, EPI_ISL_956359, EPI_ISL_956360, EPI_ISL_956361, EPI_ISL_956362, EPI_ISL_956363, EPI_ISL_956364, EPI_ISL_956365, EPI_ISL_956366 | | | |
| see above | Houston Health Department, Disease Prevention and Control | Houston Health Department, Disease Prevention and Control | Penn,R., Brown,P., Lara,A. |
| EPI_ISL_956405 | General Hospital - Veles | Research Center for Genetic Engineering and Biotechnology "Georgi D. Efremov" , Macedonian Academy of Sciences and Arts | Aleksandar J. Dimovski, Dijana Plasheska-Karanfilska, Predrag Noveski, Gjorgji Bozinovski, Milena Jakimovska |
| EPI_ISL_961486, EPI_ISL_961509, EPI_ISL_961513, EPI_ISL_961518, EPI_ISL_961527, EPI_ISL_961554 | Michigan Department of Health and Human Services, Bureau of Laboratories | Michigan Department of Health and Human Services, Bureau of Laboratories | Blankenship HM, Riner D, Soehnlen MK |
| EPI_ISL_961561, EPI_ISL_961562, EPI_ISL_961563, EPI_ISL_961564, EPI_ISL_961565, EPI_ISL_961566, EPI_ISL_961567, EPI_ISL_961568, EPI_ISL_961569, EPI_ISL_961604, EPI_ISL_961605, EPI_ISL_961606, EPI_ISL_961653, EPI_ISL_961654, EPI_ISL_961655, EPI_ISL_961656, EPI_ISL_961657, EPI_ISL_961658, EPI_ISL_961659, EPI_ISL_961660 | | | |
| see above | Hôpital Georges L. Dumont | National Microbiology Laboratory (NML) | Anna Majer, Shari Tyson, Grace Seo, Philip Mabon, Elsie Grudeski, Rhiannon Huzarewich, Russell Mandes, Anneliese Landgraff, Jennifer Tanner, Natalie Knox, Morag Graham, Gary Van Domselaar, Richard Garceau, Guillaume Desnoyers, Nathalie Bastien, Yan Li, Timothy Booth, Darian Hole, Madison Chapel, Kirsten Biggar, CanCOGeN's metadata curation team, Public Health Agency of Canada CanCOGeN team |
| EPI_ISL_961977, EPI_ISL_961978, EPI_ISL_961992, EPI_ISL_962121, EPI_ISL_962129 | Illinois Department of Public Health | Gagnon Lab, Southern Illinois University | Keith Gagnon |
| EPI_ISL_962509 | UCLA Clinical Micro Lab | Los Angeles County PHL | P. Hemarajata et al. |
| EPI_ISL_962818, EPI_ISL_962819, EPI_ISL_962820, EPI_ISL_962821, EPI_ISL_962822 | Microbiological Diagnostic Unit - Public Health Laboratory (MDU-PHL) | MDU-PHL | Seemann T., Sait, M.L., Sherry, N.L. |
| EPI_ISL_962823, EPI_ISL_962824 | Victorian Infectious Diseases Reference Laboratory (VIDRL) | VIDRL and MDU-PHL | Caly L., Seemann T., Sait, M.L., Druce J., Sherry, N.L. |
| EPI_ISL_962895 | Akershus University Hospital, Department for Microbiology and Infectious Disease Control | Norwegian Institute of Public Health, Department of Virology | Kathrine Stene-Johansen, Kamilla Heddeland Instefjord, Hilde Elshaug, Ignacio Garcia Llorente, Serina B Engebretsen, Atiya R Ali,Marie Paulsen Madsen, Rasmus Riis Kopperud, Hilde Vollan, Karoline Bragstad, Olav Hungnes |
| EPI_ISL_964916 | Hopital | National Reference Center for Viruses of Respiratory Infections, Institut Pasteur, Paris | Marion Barbet, Sylvie Behillil, Méline Bizard, Angela Brisebarre, Camille Capel, Etienne Simon-Lorière, Vincent Enouf, Maud Vanpeene, Sylvie van der Werf,Smati-Lafarge Mounira |
| EPI_ISL_964949 | Laboratorio de Virologia del HUCA | Laboratorio de Virologia del HUCA | Castelló C, Gómez de Oña J, Boga JA, Rojo S, Alvarez-Arguelles ME, Abreu F, Costales I, Sandoval M, Perez-Martinez Z, Martín-Rodriguez G, Coto E, Melon S |
| EPI_ISL_965211, EPI_ISL_965212, EPI_ISL_965213 | Virginia Division of Consolidated Laboratory Services | Virginia Division of Consolidated Laboratory Services | Virginia DCLS |
| EPI_ISL_965548, EPI_ISL_965568, EPI_ISL_965608, EPI_ISL_965627, EPI_ISL_965633, EPI_ISL_965653, EPI_ISL_965661, EPI_ISL_965674, EPI_ISL_965802 | Dutch COVID-19 response team | Medical Microbiology, Maastricht University Medical Centre | Jozef Dingemans*, Brian van der Veer*, Erik Beuken, Carmen Reumkens, Lieke van Alphen, Christian Hoebe, Paul Savelkoul |
| EPI_ISL_965967, EPI_ISL_965989, EPI_ISL_966020, EPI_ISL_966044, EPI_ISL_966062, EPI_ISL_966084, EPI_ISL_966108, EPI_ISL_966160, EPI_ISL_966184, EPI_ISL_966204, EPI_ISL_966227, EPI_ISL_966264, EPI_ISL_966289, EPI_ISL_966308, EPI_ISL_966311, EPI_ISL_966314, EPI_ISL_966333, EPI_ISL_966338, | | | |

| | | | |
|---|---|---|---|
| EPI_ISL_966357, EPI_ISL_966375, EPI_ISL_966376, EPI_ISL_966377, EPI_ISL_966378, EPI_ISL_966380, EPI_ISL_966406, EPI_ISL_966428, EPI_ISL_966452, EPI_ISL_966472, EPI_ISL_966489, EPI_ISL_966496, EPI_ISL_966549, EPI_ISL_966551, EPI_ISL_966552, EPI_ISL_966565, EPI_ISL_966591, EPI_ISL_966605, EPI_ISL_966623, EPI_ISL_966646, EPI_ISL_966663, EPI_ISL_966682, EPI_ISL_966707 | | | |
| see above | NYU Langone Health | Departments of Pathology and Medicine, New York University School of Medicine | Adriana Heguy, Dacia Dimartino, Emily Guzman, Christian Marier, Peter Meyn, Sitharam Ramaswami, Gael Westby, Paul Zappile, Yutong Zhang, Paolo Cotzia, Guiqing Wang |
| EPI_ISL_966766, EPI_ISL_966767, EPI_ISL_966768 | Maine HETL | Tewhey Lab, The Jackson Laboratory | Matluk,N., Dewey,H., Iosue,F., Barter,M., Lynch,R., Munger,H. and Tewhey,R. |
| EPI_ISL_967592, EPI_ISL_967594, EPI_ISL_967595, EPI_ISL_967596, EPI_ISL_967597, EPI_ISL_967598, EPI_ISL_967641, EPI_ISL_967644, EPI_ISL_967647, EPI_ISL_967648, EPI_ISL_967650, EPI_ISL_967651, EPI_ISL_967652, EPI_ISL_967665, EPI_ISL_967676, EPI_ISL_967698, EPI_ISL_967699 | | | |
| see above | State Laboratories Division, Hawaii State Department of Health | State Laboratories Division, Hawaii State Department of Health | Pamela O'Brien, Drew Kuwazaki, Ayana Garnet, Razvan Sultana, Edward Desmond |
| EPI_ISL_967892, EPI_ISL_967903, EPI_ISL_967915, EPI_ISL_967920, EPI_ISL_967940, EPI_ISL_967958, EPI_ISL_967972, EPI_ISL_967981, EPI_ISL_967984, EPI_ISL_968006, EPI_ISL_968010, EPI_ISL_968018, EPI_ISL_968021, EPI_ISL_968022, EPI_ISL_968026, EPI_ISL_968029, EPI_ISL_968040, EPI_ISL_968042, EPI_ISL_968047, EPI_ISL_968048, EPI_ISL_968054 | | | |
| see above | TGen North | Sonora Quest Laboratories | "Jolene Bowers, Megan Folkerts, Chris French, Hayley Yaglom, Ashlyn Pfeiffer, Darrin Lemmer, Dave Engelthaler, The Arizona COVID Genomics Union (ACGU)" |
| EPI_ISL_968855 | KEMRI-Wellcome Trust Research Programme/KEMRI-CGMR-C Kilifi | KEMRI-Wellcome Trust Research Programme/KEMRI-CGMR-C Kilifi | Githinji et al |
| EPI_ISL_976883, EPI_ISL_976884, EPI_ISL_976885, EPI_ISL_976886, EPI_ISL_976887, EPI_ISL_976888, EPI_ISL_976889, EPI_ISL_976890, EPI_ISL_976891, EPI_ISL_976892 | BCCDC Public Health Laboratory | BCCDC Public Health Laboratory | Prystajecky Natalie, Linda Hoang, Dan Fornika, John Tyson, Shannon Russell, Kim Macdonald, Kimia Kamelian, Ana Pacagnella, Corrinne Ng, Loretta Janz, Robert Azana Terry Snutch, Mel Krajden |
| EPI_ISL_977211 | Microbiologia e Virologia | Istituto Zooprofilattico Sperimentale delle Venezie | Adelaide Milani, Alessia Schivo, Annalisa Salviato, Erika Giorgia Quaranta, Ambra Pastori, Bianca Zecchin, Alice Fusaro, Isabella Monne, Calogero Terregino, Antonia Ricci |
| EPI_ISL_977240, EPI_ISL_977241 | ULSS 5 Polesana | Istituto Zooprofilattico Sperimentale delle Venezie | Adelaide Milani, Alessia Schivo, Annalisa Salviato, Erika Giorgia Quaranta, Ambra Pastori, Bianca Zecchin, Alice Fusaro, Isabella Monne, Calogero Terregino, Antonia Ricci |
| EPI_ISL_977659 | Caribbean Public Health Agency | Carrington Lab, Department of PreClinical Sciences, Building 36, First Floor Biochemistry Unit, Faculty of Medical Sciences, The University of the West Indies | Nikita S. D. Sahadeo, Arianne Brown-Jordan, Vernie Ramkissoon, Sarah Hill, Naresh Nandram, Avery Hinds, Kenneth George, Jerome Foster, Stanley Giddings, Karla Georges, Marsha Ivey, Rahul Naidu, Risha Singh, SueMin Nathaniel, Rajini Haraksingh, Jaya Jayaraman, Chinna Chinnadurai, Adesh Ramsubhag, Nuno Faria, Oliver Pybus, Christopher Oura, Gabriel Escobar, Christine V. F. Carrington |
| EPI_ISL_978089, EPI_ISL_978090, EPI_ISL_978091, EPI_ISL_978092, EPI_ISL_978093, EPI_ISL_978094, EPI_ISL_978095, EPI_ISL_978096, EPI_ISL_978097, EPI_ISL_978098, EPI_ISL_978099, EPI_ISL_978100, EPI_ISL_978101, EPI_ISL_978102, EPI_ISL_978103, EPI_ISL_978104, EPI_ISL_978105, EPI_ISL_978106, EPI_ISL_978107, EPI_ISL_978108, EPI_ISL_978109, EPI_ISL_978110, EPI_ISL_978111, EPI_ISL_978112, EPI_ISL_978113, EPI_ISL_978114, EPI_ISL_978115, EPI_ISL_978116, EPI_ISL_978117, EPI_ISL_978118 | | | |
| see above | Chiu Laboratory, University of California, San Francisco | Chiu Laboratory, University of California, San Francisco | Charles Chiu, Xianding (Wayne) Deng, Candace Wang, Venice Servellita, Jill Hacker, Debra Wadford |
| EPI_ISL_979136, EPI_ISL_979138, EPI_ISL_979139, EPI_ISL_979140, EPI_ISL_979142 | Santa Clara County Public Health Laboratory | Chan-Zuckerberg Biohub | CZB Cliahub Consortium |
| EPI_ISL_979339, EPI_ISL_979340, EPI_ISL_979341, EPI_ISL_979342 | Laboratorio Estatal de Salud Pública de Nuevo León | Laboratorio de Infectología Molecular, Departamento de Bioquímica y Medicina Molecular,Facultad de Medicina - Universidad Autónoma de Nuevo León | Kame A. Galán-Huerta, María F. Herrera-Saldivar, Natalia Martínez-Acuña, Sonia A. Lozano-Sepúlveda, Daniel Arellanos-Soto, Ana M. Rivas-Estilla, Samuel Buentello-Wong, Else del Carmen García-García, Gloria A. Jasso-de-la-Peña, Roberto Montes-de-Oca, Consuelo Treviño-Garza, Manuel E. de-la-O-Cavazos |
| EPI_ISL_979634, EPI_ISL_979635, EPI_ISL_979636, EPI_ISL_979637 | Santa Clara County Public Health Laboratory | Chan-Zuckerberg Biohub | CZB Cliahub Consortium |
| EPI_ISL_979692, EPI_ISL_979693, EPI_ISL_979694, EPI_ISL_979696, EPI_ISL_979697, EPI_ISL_979705, EPI_ISL_979706, EPI_ISL_979708, EPI_ISL_979709 | Orange County Public Health Lab | Chan-Zuckerberg Biohub | CZB Cliahub Consortium |
| EPI_ISL_981058 | Johns Hopkins Hospital Department of Pathology | Johns Hopkins Hospital Department of Pathology | C. Paul Morris, Chun Huai Luo, Adannaya Amadi, Matthew Schwartz, Nicholas Gallagher, Heba H. Mostafa |
| EPI_ISL_981956, EPI_ISL_981958, EPI_ISL_981959, EPI_ISL_981960 | Microbiology Service, Hospital Universitario Clinico San Cecilio, Granada | Microbiology Service, Hospital Universitario Clinico San Cecilio, Granada | Adolfo de Salazar, Natalia Chueca, Laura Viñuela, Ana Fuentes, Federico García |
| EPI_ISL_981997, EPI_ISL_982034, EPI_ISL_982075, EPI_ISL_982080, EPI_ISL_982103, EPI_ISL_982110 | TGen North | Sonora Quest Laboratories | "Jolene Bowers, Megan Folkerts, Chris French, Hayley Yaglom, Ashlyn Pfeiffer, Darrin Lemmer, Dave Engelthaler, The Arizona COVID Genomics Union (ACGU)" |
| EPI_ISL_982243 | Lab voor klinische biologie | Lab voor klinische biologie | Hannelore Hamerlinck, Marija Janevska, Bruno Verhasselt |
| EPI_ISL_982348, EPI_ISL_982350, EPI_ISL_982351, EPI_ISL_982356, EPI_ISL_982357, EPI_ISL_982358, EPI_ISL_982359, EPI_ISL_982360, EPI_ISL_982361, EPI_ISL_982362, EPI_ISL_982363, EPI_ISL_982364, EPI_ISL_982365, EPI_ISL_982366, EPI_ISL_982367, EPI_ISL_982368, EPI_ISL_982370, EPI_ISL_982374, EPI_ISL_982384 | | | |
| see above | M Health Fairview | Minnesota Department of Health, Public Health Laboratory | Alexandra Lorentz, Jacob Garfin, Matt Plumb, and Xiong Wang |
| EPI_ISL_982426, EPI_ISL_982437, EPI_ISL_982438, EPI_ISL_982439, EPI_ISL_982443, EPI_ISL_982444, EPI_ISL_982447, EPI_ISL_982451, EPI_ISL_982452, EPI_ISL_982454, EPI_ISL_982455, EPI_ISL_982458, EPI_ISL_982461, EPI_ISL_982469, EPI_ISL_982474, EPI_ISL_982475, EPI_ISL_982482, EPI_ISL_982483, EPI_ISL_982484, EPI_ISL_982485, EPI_ISL_982488, EPI_ISL_982491, EPI_ISL_982496, EPI_ISL_982497, EPI_ISL_982498 | | | |
| see above | MONTEFIORE MEDICAL CENTER LABORATORIES | Wadsworth Center, New York State Department of Health | Kirsten St. George, Daryl M. Lamson, Alexis Russel, Matthew Shudt, Melissa A Leisner, Jonathan Plitnick, Navjot Singh, John Kelly, Erasmus Schneider, Erica Lasek-Nesselquist |
| EPI_ISL_982530, EPI_ISL_982608, EPI_ISL_982653 | US Air Force School of Aerospace Medicine | US Air Force School of Aerospace Medicine | Anthony Fries, Jennifer Meyer, William Gruner, William Buggele, Amanda Javorina, Sarah Purves, Clarise Starr, Elizabeth Macias |
| EPI_ISL_983101, EPI_ISL_983103, EPI_ISL_983104, EPI_ISL_983105, EPI_ISL_983106, EPI_ISL_983108, EPI_ISL_983113, EPI_ISL_983115, EPI_ISL_983134, EPI_ISL_983138, EPI_ISL_983142, EPI_ISL_983144, EPI_ISL_983146, EPI_ISL_983150, EPI_ISL_983152, EPI_ISL_983154, EPI_ISL_983157, EPI_ISL_983166, EPI_ISL_983168, EPI_ISL_983169 | | | |
| see above | MONTEFIORE MEDICAL CENTER LABORATORIES | Wadsworth Center, New York State Department of Health | Kirsten St. George, Daryl M. Lamson, Alexis Russel, Matthew Shudt, Melissa A Leisner, Jonathan Plitnick, Navjot Singh, John Kelly, Erasmus Schneider, Erica Lasek-Nesselquist |
