## Supplementary material for "Detection and characterization of the SARS-CoV-2 lineage B.1.526 in New York": Supp. Table 3 GISAID Acknowledgment part 2: gisaid_hcov-19_acknowledgement_table_2021_02_13_010-7.pdf

We gratefully acknowledge the following Authors from the Originating laboratories responsible for obtaining the specimens, as well as the Submitting laboratories where the genome data were generated and shared via GISAID, on which this research is based.

All Submitters of data may be contacted directly via www.gisaid.org

Authors are sorted alphabetically.

| Accession ID | Originating Laboratory | Submitting Laboratory | Authors |
|---|---|---|---|
| EPI_ISL_771370 | SA Pathology | SA Pathology | Lex Leong, Julien Soubrier, Chuan Kok Lim, Song Gao, Mark Turra, Karin Kassahn, Ivan Bastian, Geoff Higgins |
| EPI_ISL_775242, EPI_ISL_775245 | Area of Virology, Serology and Virology Division (SAViD), New South Wales Health Pathology Randwick | Area of Virology, Serology and Virology Division (SAViD), New South Wales Health Pathology Randwick | Rawlinson, W., Bull, R., Deveson, I. |
| EPI_ISL_779154 | Yale Pathology Lab | Grubaugh Lab - Yale School of Public Health | Tara Alpert, Joseph Fauver, Chen Liu, Pei Hui, Jianhui Wang, Susan Bell and Han Zhou, Anderson Brito, Mallery Breban, Anne Wyllie, Chantal Vogels, Mary Petrone, Chaney Kalinich, Isabel Ott, Arnau Casanovas, Catherine Muenker, Adam Moore, Alice Lu, Maria Tokuyama, Patrick Wong, Peiwen Lu, Saad Omer, Richard Martinello, Allison Nelson, Shelli Farhadian, Akiko Iwasaki, Charlese Dela Cruz, Albert Ko, Nathan Grubaugh |
| EPI_ISL_779398 | Douglass Hanly Moir Pathology | NSW Health Pathology - Institute of Clinical Pathology and Medical Research; Westmead Hospital; University of Sydney | CIDM-PH et al. |
| EPI_ISL_788909, EPI_ISL_788910, EPI_ISL_788911, EPI_ISL_788912, EPI_ISL_788913, EPI_ISL_788914, EPI_ISL_788915, EPI_ISL_788916, EPI_ISL_788917, EPI_ISL_788918, EPI_ISL_788942, EPI_ISL_791097, EPI_ISL_791098, EPI_ISL_791099, EPI_ISL_791100, EPI_ISL_791101, EPI_ISL_791102, EPI_ISL_791103, EPI_ISL_791104, EPI_ISL_791105, EPI_ISL_791106 | | | |
| see above | University of Wisconsin-Madison AIDS Vaccine Research Laboratories | University of Wisconsin-Madison AIDS Vaccine Research Laboratories | Gage Moreno, Katarina Braun, et al. AIDS Vaccine Research Laboratories |
| EPI_ISL_791194, EPI_ISL_791197, EPI_ISL_791198, EPI_ISL_791223, EPI_ISL_791224, EPI_ISL_791225, EPI_ISL_791226, EPI_ISL_791227, EPI_ISL_791228, EPI_ISL_791229, EPI_ISL_791230, EPI_ISL_791231, EPI_ISL_791232, EPI_ISL_791233, EPI_ISL_791234, EPI_ISL_791235, EPI_ISL_791236, EPI_ISL_791237, EPI_ISL_791238, EPI_ISL_791239, EPI_ISL_791240, EPI_ISL_791241, EPI_ISL_791242, EPI_ISL_791243, EPI_ISL_791244, EPI_ISL_791245, EPI_ISL_791246, EPI_ISL_791247, EPI_ISL_791248, EPI_ISL_791254, EPI_ISL_791255, EPI_ISL_791256, EPI_ISL_791257, EPI_ISL_791258, EPI_ISL_791259, EPI_ISL_791260, EPI_ISL_791261, EPI_ISL_791262, EPI_ISL_791263, EPI_ISL_791264, EPI_ISL_791265, EPI_ISL_791266, EPI_ISL_791267, EPI_ISL_791268 | | | |
| see above | Respiratory Virus Unit, National Infection Service, Public Health England | COVID-19 Genomics UK (COG-UK) Consortium | PHE Covid Sequencing Team |
| EPI_ISL_791430 | Johns Hopkins Hospital Department of Pathology | Johns Hopkins Hospital Department of Pathology | C. Paul Morris, Chun Huai Luo, Adannaya Amadi, Nicholas Gallagher, Heba H. Mostafa |
| EPI_ISL_792089 | Toronto Invasive Bacterial Diseases Network | McMaster University | Allison McGeer, Patryk Aftanas, Hooman Derakhshani, Angel Li, Kuganya Nirmalarajah, Emily Panousis, Ahmed Draia, Jalees Nasir, Michael Surette, Samira Mubareka, Andrew G. McArthur |
| EPI_ISL_792551 | Centre for Dengue Research and AICBU, Department of Immunology and Molecular Medicine | Centre for Dengue Research and AICBU, Department of Immunology and Molecular Medicine | Chandima Jeewandara, Deshni Jayathilaka, Dinuka Ariyaratne, Diyanath Ranasinghe, Laksiri Gomes, Gathsaurie Neelika Malavige |
| EPI_ISL_794308 | Wadsworth Center, New York State Department.of Health | Wadsworth Center, New York State Department.of Health | Kirsten St. George, Daryl M. Lamson, Alexis Russel, Matthew Shudt, Melissa A Leisner, Jonathan Plitnick, Navjot Singh, John Kelly, Sara Griesemer, Erasmus Schneider, Erica Lasek-Nesselquist |
| EPI_ISL_794626 | Middlemore Hospital | Institute of Environmental Science and Research (ESR) | Xiaoyun Ren, Matt Storey, Nikki Freed, Muhammad Faisal, Jing Wang, Hermes Perez, Anja Werno, Antje van der Linden, Arlo Upton, Chris Mansell, David Hammer, Dragana Drinkovic, Gary McAuliffe, Hana Sofia Andersson, James Ussher, Jill Sherwood, Josh Freeman, Julia Howard, Juliet Elvy, Mary DeAlmeida, Matt Blakiston, Matthew Rogers, Max Bloomfield, Michael Addidle, Michelle Balm, Sally Roberts, Sarah Jefferies, Sharmini Muttaiyah, Susan Morpeth, Susan Taylor, Timothy Blackmore, Vani Sathyendran, Veronica Playle, Virginia Hope, Erasmus Smit, Lauren Jelly, Olin Silander, Joep de Ligt |
| EPI_ISL_794639, EPI_ISL_794669 | USC Clinical Lab | Los Angeles County PHL | P. Hemarajata et al. |
| EPI_ISL_794744 | National Institute of Infectious Diseases-Prof. Dr. Matei Bals Molecular Diagnostics Laboratory | National Institute of Infectious Diseases-Prof. Dr. Matei Bals Molecular Diagnostics Laboratory | Leontina Banica, Marius Surleac, Corina Casangiu, Petre Milu, Andreea Tudor, Simona Paraschiv, Dan Otelea |
| EPI_ISL_796023, EPI_ISL_796024, EPI_ISL_796025, EPI_ISL_796026 | Hebei Provincial Center for Disease Control and Prevention, Shijiazhuang, Hebei Province; National Institute for Viral Disease Control and Prevention, China CDC | Hebei Provincial Center for Disease Control and Prevention, Shijiazhuang, Hebei Province; National Institute for Viral Disease Control and Prevention, China CDC | Shunxiang Qi, Xiang Zhao, Nankun Liu, George F. Gao, Yang Song, Wenbo Xu, Qi Li |
| EPI_ISL_796736, EPI_ISL_796765 | Instituto Nacional de Saude (INSA) | Instituto Nacional de Saude (INSA) | Borges et al |
| EPI_ISL_797970 | Lighthouse Lab in Glasgow | Wellcome Sanger Institute for the COVID-19 Genomics UK (COG-UK) Consortium | Harper VanSteenhouse, Yumi Kasai, David Gray, Carol Clugston, Anna Dominiczak and Alex Alderton, Roberto Amato, Sonia Goncalves, Ewan Harrison, David K. Jackson, Ian Johnston, Dominic Kwiatkowski, Cordelia Langford, John Sillitoe on behalf of the Wellcome Sanger Institute COVID-19 Surveillance Team |
| EPI_ISL_802498, EPI_ISL_802499, EPI_ISL_802500 | Wadsworth Center, New York State Department.of Health | Wadsworth Center, New York State Department.of Health | Kirsten St. George, Daryl M. Lamson, Alexis Russel, Matthew Shudt, Melissa A Leisner, Jonathan Plitnick, Navjot Singh, John Kelly, Sara Griesemer, Erasmus Schneider, Erica Lasek-Nesselquist |
| EPI_ISL_803994, EPI_ISL_803995, EPI_ISL_803996, EPI_ISL_803997, EPI_ISL_804000, EPI_ISL_804001, EPI_ISL_804002, EPI_ISL_804003, EPI_ISL_804004, EPI_ISL_804005 | National Public Health Laboratory, National Centre for Infectious Diseases | National Public Health Laboratory, National Centre for Infectious Diseases | Tze Minn Mak, Sophie Octavia, Zhenyang Zhou, Lin Cui, Raymond Tzer Pin Lin |
| EPI_ISL_804217, EPI_ISL_804225, EPI_ISL_804276 | Respiratory Virus Unit, National Infection Service, Public Health England | COVID-19 Genomics UK (COG-UK) Consortium | PHE Covid Sequencing Team |
| EPI_ISL_804945, EPI_ISL_804948 | Wadsworth Center, New York State Department.of Health | Wadsworth Center, New York State Department.of Health | Kirsten St. George, Daryl M. Lamson, Alexis Russel, Matthew Shudt, Melissa A Leisner, Jonathan Plitnick, Navjot Singh, John Kelly, Sara Griesemer, Erasmus Schneider, Erica Lasek-Nesselquist |
| EPI_ISL_810965, EPI_ISL_810966 | PathWest Laboratory Medicine WA | PathWest Laboratory Medicine WA Microbial Surveillance Unit | PathWest Laboratory Medicine WA Microbial Surveillance Unit |
| EPI_ISL_811121 | Respiratory Virus Unit, National Infection Service, Public Health England | COVID-19 Genomics UK (COG-UK) Consortium | PHE Covid Sequencing Team |
| EPI_ISL_811140, EPI_ISL_811141, EPI_ISL_811142 | Ministry of Health Turkey | Ministry of Health Turkey | Fatma Bayrakdar, Yasemin Cogun, Süleyman Yalcin, Aye Baak Alta, Gülay Korukluolu |
| EPI_ISL_811200, EPI_ISL_811213, EPI_ISL_811216, EPI_ISL_811221, EPI_ISL_811222, EPI_ISL_811226, EPI_ISL_811235, EPI_ISL_811254, EPI_ISL_811256, EPI_ISL_811257, EPI_ISL_811258, EPI_ISL_811260, EPI_ISL_811261, EPI_ISL_811263, EPI_ISL_811268, EPI_ISL_811273, EPI_ISL_811289, EPI_ISL_811295, EPI_ISL_811297, EPI_ISL_811316, EPI_ISL_811317, EPI_ISL_811338, EPI_ISL_811347, EPI_ISL_811348, EPI_ISL_811355, EPI_ISL_811356, EPI_ISL_811364, EPI_ISL_811365, EPI_ISL_811374, EPI_ISL_811376, EPI_ISL_811384, EPI_ISL_811387, EPI_ISL_811388, EPI_ISL_811391, EPI_ISL_811405, EPI_ISL_811415, EPI_ISL_811417, EPI_ISL_811423, EPI_ISL_811424, EPI_ISL_811426, EPI_ISL_811427, EPI_ISL_811432, EPI_ISL_811453 | | | |
| see above | Lighthouse Lab in Glasgow | Wellcome Sanger Institute for the COVID-19 Genomics UK (COG-UK) Consortium | Harper VanSteenhouse, Yumi Kasai, David Gray, Carol Clugston, Anna Dominiczak and Alex Alderton, Roberto Amato, Sonia Goncalves, Ewan Harrison, David K. Jackson, Ian Johnston, Dominic Kwiatkowski, Cordelia Langford, John Sillitoe on behalf of the Wellcome Sanger Institute COVID-19 Surveillance Team |
| EPI_ISL_812257 | Analytica Medizinische Laboratorien AG | Institute of Medical Virology, University of Zurich | Stefan Schmutz, Maryam Zaheri, Verena Kufner, Annette Audigé, Maria Grünberg, Kevin Steiner, Jon Huder, Cyril Shah, Riccarda Capaul, Guido Bloemberg, Jürg Böni, Michael Huber, Alexandra Trkola |

| | | | |
|---|---|---|---|
| EPI_ISL_812423 | Royal Darwin Hospital Pathology | MDU-PHL | Meumann, E., Caly L., Seemann T., Sait, M.L., Druce J., Sherry, N.L. |
| EPI_ISL_812772, EPI_ISL_812773 | Ministry of Health Turkey | Ministry of Health Turkey | Fatma Bayrakdar, Yasemin Cogun, Süleyman Yalcin, Aye Baak Alta, Gülay Korukluolu |
| EPI_ISL_813826, EPI_ISL_813829, EPI_ISL_813830, EPI_ISL_813831 | Liverpool Clinical Laboratories | COVID-19 Genomics UK (COG-UK) Consortium | Sam Haldenby, Anita Lucaci, Steve Paterson, Julian Hiscox, Alistair Darby, M Almsaud, A Alrezaihi, Muhannad Alruwaili, Stuart D Armstrong, Jones Benjamin, Eleanor G Bentley, Anu Chawla, Jordan J Clark, Angela Cowell, Richard Eccles, Isabel García-Dorival, Matthew Gemmell, Alessandro Gerada, PKF Gilmore, Richard Gregory, Ximeng Han, Catherine Hartley, Margaret Hughes, Miren Iturriza-Gomara, James Johnson, L Luu, Jenifer Manson, Charlotte Nelson, Elaine O'Toole, Cassie Olateju, Rebekah Penrice-Randal , Lucille Rainbow, N.P Randle, Trevor Ian Robinson, Parul Sharma, Ghada T Shawli, James P Stewart, Neil Swainston, Ecaterina Vamos, Joanne Watts, Mark Whitehead |
| EPI_ISL_813995, EPI_ISL_814010, EPI_ISL_814014, EPI_ISL_814021, EPI_ISL_814034, EPI_ISL_814035 | Hospital General Universitario Gregorio Marañón | SeqCOVID-SPAIN consortium/IBV(CSIC) | Darío García de Viedma, Laura Pérez-Lago, Marta Herranz, Jon Sicilia, Julia Suárez, Pilar Catalán, Patricia Muñoz and SeqCOVID-SPAIN consortium |
| EPI_ISL_816244, EPI_ISL_816253, EPI_ISL_816284, EPI_ISL_816299, EPI_ISL_816307, EPI_ISL_816358, EPI_ISL_816369, EPI_ISL_816376, EPI_ISL_816378, EPI_ISL_816387, EPI_ISL_816389, EPI_ISL_816391, EPI_ISL_816433, EPI_ISL_816455, EPI_ISL_816493, EPI_ISL_816518, EPI_ISL_816524, EPI_ISL_816559, EPI_ISL_816623, EPI_ISL_816639, EPI_ISL_816648, EPI_ISL_816652 | | | |
| see above | Virology Department, Sheffield Teaching Hospitals NHS Foundation Trust/Department of Infection, Immunity and Cardiovascular Disease, The Medical School, University of Sheffield | COVID-19 Genomics UK (COG-UK) Consortium | Thushan de Silva, Matthew Parker, Nikki Smith, Adri Angyal, Rebecca Brown, Luke Green, Rachel Tucker, Paul Parsons, Danielle Groves, Katie Johnson, Laura Carrilero, Alex Keeley, Dave Partridge, Matthew Wyles, Benjamin Lindsey, Mehmet Yavuz, Mohammad Raza, Cariad Evans |
| EPI_ISL_817135, EPI_ISL_817137, EPI_ISL_817138, EPI_ISL_817146, EPI_ISL_817147, EPI_ISL_817149, EPI_ISL_817159, EPI_ISL_817160, EPI_ISL_817161, EPI_ISL_817162, EPI_ISL_817163, EPI_ISL_817164, EPI_ISL_817165, EPI_ISL_817166, EPI_ISL_817167, EPI_ISL_817168, EPI_ISL_817169, EPI_ISL_817170, EPI_ISL_817171, EPI_ISL_817172, EPI_ISL_817173, EPI_ISL_817174, EPI_ISL_817175, EPI_ISL_817176, EPI_ISL_817264, EPI_ISL_817269, EPI_ISL_817272, EPI_ISL_817273, EPI_ISL_817278, EPI_ISL_817279, EPI_ISL_817281, EPI_ISL_817282, EPI_ISL_817288, EPI_ISL_817290, EPI_ISL_817291, EPI_ISL_817292, EPI_ISL_817296, EPI_ISL_817297, EPI_ISL_817299, EPI_ISL_817301, EPI_ISL_817303, EPI_ISL_817306, EPI_ISL_817308, EPI_ISL_817312, EPI_ISL_817315, EPI_ISL_817316, EPI_ISL_817317, EPI_ISL_817333, EPI_ISL_817334, EPI_ISL_817335, EPI_ISL_817375, EPI_ISL_817376, EPI_ISL_817383, EPI_ISL_817384, EPI_ISL_817385, EPI_ISL_817386, EPI_ISL_817392, EPI_ISL_817398, EPI_ISL_817399, EPI_ISL_817401, EPI_ISL_817404, EPI_ISL_817405, EPI_ISL_817413, EPI_ISL_817414, EPI_ISL_817415, EPI_ISL_817416, EPI_ISL_817417, EPI_ISL_817418, EPI_ISL_817419, EPI_ISL_817487, EPI_ISL_817496, EPI_ISL_817497, EPI_ISL_817499, EPI_ISL_817501, EPI_ISL_817508, EPI_ISL_817509, EPI_ISL_817510, EPI_ISL_817511, EPI_ISL_817530, EPI_ISL_817532, EPI_ISL_817537, EPI_ISL_817538, EPI_ISL_817555, EPI_ISL_817556, EPI_ISL_817557, EPI_ISL_817558, EPI_ISL_817559, EPI_ISL_817560, EPI_ISL_817561, EPI_ISL_817562, EPI_ISL_817563, EPI_ISL_817564, EPI_ISL_817565, EPI_ISL_817566, EPI_ISL_817567, EPI_ISL_817568, EPI_ISL_817569, EPI_ISL_817570, EPI_ISL_817571, EPI_ISL_817572, EPI_ISL_817573, EPI_ISL_817574, EPI_ISL_817575, EPI_ISL_817576, EPI_ISL_817577, EPI_ISL_817578, EPI_ISL_817579, EPI_ISL_817580, EPI_ISL_817581, EPI_ISL_817582, EPI_ISL_817583, EPI_ISL_817584, EPI_ISL_817585, EPI_ISL_817586, EPI_ISL_817587, EPI_ISL_817588, EPI_ISL_817589, EPI_ISL_817590, EPI_ISL_817591, EPI_ISL_817592, EPI_ISL_817593, EPI_ISL_817594, EPI_ISL_817595, EPI_ISL_817596, EPI_ISL_817597, EPI_ISL_817598, EPI_ISL_817599, EPI_ISL_817600, EPI_ISL_817601, EPI_ISL_817602, EPI_ISL_817603, EPI_ISL_817604, EPI_ISL_817605, EPI_ISL_817606, EPI_ISL_817607, EPI_ISL_817608, EPI_ISL_817609, EPI_ISL_817610, EPI_ISL_817611, EPI_ISL_817612, EPI_ISL_817613, EPI_ISL_817614, EPI_ISL_817615, EPI_ISL_817616, EPI_ISL_817617, EPI_ISL_817618, EPI_ISL_817619, EPI_ISL_817620, EPI_ISL_817621, EPI_ISL_817622, EPI_ISL_817623, EPI_ISL_817624, EPI_ISL_817625, EPI_ISL_817626, EPI_ISL_817627, EPI_ISL_817628, EPI_ISL_817629, EPI_ISL_817630, EPI_ISL_817631, EPI_ISL_817632, EPI_ISL_817633, EPI_ISL_817634, EPI_ISL_817635, EPI_ISL_817636, EPI_ISL_817637, EPI_ISL_817638, EPI_ISL_817639, EPI_ISL_817640, EPI_ISL_817641, EPI_ISL_817642, EPI_ISL_817643, EPI_ISL_817644, EPI_ISL_817645, EPI_ISL_817646, EPI_ISL_817647, EPI_ISL_817648, EPI_ISL_817649, EPI_ISL_817650, EPI_ISL_817651, EPI_ISL_817652, EPI_ISL_818123, EPI_ISL_818124, EPI_ISL_818125, EPI_ISL_818126, EPI_ISL_818127, EPI_ISL_818128, EPI_ISL_818129, EPI_ISL_818130, EPI_ISL_818131, EPI_ISL_818132, EPI_ISL_818133, EPI_ISL_818134, EPI_ISL_818135, EPI_ISL_818136, EPI_ISL_818137, EPI_ISL_818138, EPI_ISL_818139, EPI_ISL_818140, EPI_ISL_818141, EPI_ISL_818142, EPI_ISL_818143, EPI_ISL_818144, EPI_ISL_818145, EPI_ISL_818146, EPI_ISL_818147, EPI_ISL_818148, EPI_ISL_818149, EPI_ISL_818150, EPI_ISL_818151, EPI_ISL_818152, EPI_ISL_818153, EPI_ISL_818154, EPI_ISL_818155, EPI_ISL_818156, EPI_ISL_818157, EPI_ISL_818158, EPI_ISL_818159, EPI_ISL_818160, EPI_ISL_818161, EPI_ISL_818162, EPI_ISL_818163, EPI_ISL_818164, EPI_ISL_818165, EPI_ISL_818166, EPI_ISL_818167, EPI_ISL_818168, EPI_ISL_818169, EPI_ISL_818170, EPI_ISL_818171, EPI_ISL_818172, EPI_ISL_818173, EPI_ISL_818174, EPI_ISL_818175, EPI_ISL_818176, EPI_ISL_818177, EPI_ISL_818178, EPI_ISL_818179, EPI_ISL_818180, EPI_ISL_818181, EPI_ISL_818182, EPI_ISL_818183, EPI_ISL_818184, EPI_ISL_818185, EPI_ISL_818186, EPI_ISL_818187, EPI_ISL_818188, EPI_ISL_818189, EPI_ISL_818190, EPI_ISL_818191, EPI_ISL_818192, EPI_ISL_818193, EPI_ISL_818194, EPI_ISL_818195, EPI_ISL_818196, EPI_ISL_818197, EPI_ISL_818198, EPI_ISL_818199, EPI_ISL_818200, EPI_ISL_818201, EPI_ISL_818202, EPI_ISL_818203, EPI_ISL_818204, EPI_ISL_818205, EPI_ISL_818206, EPI_ISL_818207, EPI_ISL_818208, EPI_ISL_818209, EPI_ISL_818210, EPI_ISL_818211, EPI_ISL_818212, EPI_ISL_818213, EPI_ISL_818214, EPI_ISL_818215, EPI_ISL_818216, EPI_ISL_818217, EPI_ISL_818218, EPI_ISL_818219, EPI_ISL_818220, EPI_ISL_818221, EPI_ISL_818222, EPI_ISL_818223, EPI_ISL_818224, EPI_ISL_818225, EPI_ISL_818226, EPI_ISL_818227, EPI_ISL_818228, EPI_ISL_818229, EPI_ISL_818230, EPI_ISL_818231, EPI_ISL_818232, EPI_ISL_818233, EPI_ISL_818234, EPI_ISL_818235, EPI_ISL_818236, EPI_ISL_818237, EPI_ISL_818238, EPI_ISL_818239, EPI_ISL_818240, EPI_ISL_818241, EPI_ISL_818242, EPI_ISL_818243, EPI_ISL_818244, EPI_ISL_818245, EPI_ISL_818246, EPI_ISL_818247, EPI_ISL_818248, EPI_ISL_818249, EPI_ISL_818250, EPI_ISL_818251, EPI_ISL_818252, EPI_ISL_818253, EPI_ISL_818254, EPI_ISL_818255, EPI_ISL_818256, EPI_ISL_818257, EPI_ISL_818258, EPI_ISL_818259, EPI_ISL_818260, EPI_ISL_818261, EPI_ISL_818262, EPI_ISL_818263, EPI_ISL_818264, EPI_ISL_818265, EPI_ISL_818266, EPI_ISL_818267, EPI_ISL_818268, EPI_ISL_818269, EPI_ISL_818270, EPI_ISL_818271, EPI_ISL_818272, EPI_ISL_818273, EPI_ISL_818274, EPI_ISL_818275, EPI_ISL_818276, EPI_ISL_818277, EPI_ISL_818278, EPI_ISL_818279, EPI_ISL_818280, EPI_ISL_818281, EPI_ISL_818282, EPI_ISL_818283, EPI_ISL_818284, EPI_ISL_818285, EPI_ISL_818286, EPI_ISL_818287, EPI_ISL_818288, EPI_ISL_818289, EPI_ISL_818290, EPI_ISL_818291, EPI_ISL_818292, EPI_ISL_818293, EPI_ISL_818294, EPI_ISL_818295, EPI_ISL_818296, EPI_ISL_818297, EPI_ISL_818298, EPI_ISL_818299, EPI_ISL_818300, EPI_ISL_818301, EPI_ISL_818302, EPI_ISL_818303, EPI_ISL_818304, EPI_ISL_818305, EPI_ISL_818306, EPI_ISL_818307, EPI_ISL_818308, EPI_ISL_818309, EPI_ISL_818310, EPI_ISL_818311, EPI_ISL_818312, EPI_ISL_818313, EPI_ISL_818314, EPI_ISL_818315, EPI_ISL_818316, EPI_ISL_818317, EPI_ISL_818318, EPI_ISL_818319, EPI_ISL_818320, EPI_ISL_818321, EPI_ISL_818322, EPI_ISL_818323, EPI_ISL_818324, EPI_ISL_818325, EPI_ISL_818326, EPI_ISL_818327, EPI_ISL_818328, EPI_ISL_818329, EPI_ISL_818330, EPI_ISL_818331, EPI_ISL_818332, EPI_ISL_818333, EPI_ISL_818334, EPI_ISL_818335, EPI_ISL_818336, EPI_ISL_818337, EPI_ISL_818338, EPI_ISL_818339, EPI_ISL_818340, EPI_ISL_818341, EPI_ISL_818343, EPI_ISL_818344, EPI_ISL_818345, EPI_ISL_818346, EPI_ISL_818347, EPI_ISL_818348, EPI_ISL_818349, EPI_ISL_818350, EPI_ISL_818351, EPI_ISL_818352, EPI_ISL_818353, EPI_ISL_818354, EPI_ISL_818355, EPI_ISL_818356, EPI_ISL_818357, EPI_ISL_818358, EPI_ISL_818359, EPI_ISL_818360, EPI_ISL_818361, EPI_ISL_818362, EPI_ISL_818363, EPI_ISL_818364, EPI_ISL_818365, EPI_ISL_818366, EPI_ISL_818367, EPI_ISL_818368, EPI_ISL_818369, EPI_ISL_818370, EPI_ISL_818371, EPI_ISL_818372, EPI_ISL_818373, EPI_ISL_818374, EPI_ISL_818375, EPI_ISL_818376, EPI_ISL_818377, EPI_ISL_818378, EPI_ISL_818379, EPI_ISL_818380, EPI_ISL_818381, EPI_ISL_818382, EPI_ISL_818383, EPI_ISL_818384, EPI_ISL_818385, EPI_ISL_818386, EPI_ISL_818387, EPI_ISL_818388, EPI_ISL_818389, EPI_ISL_818390, EPI_ISL_818391, EPI_ISL_818392, EPI_ISL_818393, EPI_ISL_818394, EPI_ISL_818395, EPI_ISL_818396, EPI_ISL_818397, EPI_ISL_818398, EPI_ISL_818399, EPI_ISL_818400, EPI_ISL_818401, EPI_ISL_818402, EPI_ISL_818403, EPI_ISL_818404, EPI_ISL_818405, EPI_ISL_818406, EPI_ISL_818407, EPI_ISL_818408, EPI_ISL_818409, EPI_ISL_818410, EPI_ISL_818411, EPI_ISL_818412, EPI_ISL_818413, EPI_ISL_818414, EPI_ISL_818415, EPI_ISL_818416, EPI_ISL_818417, EPI_ISL_818418, EPI_ISL_818419, EPI_ISL_818420, EPI_ISL_818421, EPI_ISL_818422, EPI_ISL_818423, EPI_ISL_818424, EPI_ISL_818425, EPI_ISL_818426, EPI_ISL_818427, EPI_ISL_818428, EPI_ISL_818429, EPI_ISL_818430, EPI_ISL_818431, EPI_ISL_818432, EPI_ISL_818433, EPI_ISL_818434, EPI_ISL_818435, EPI_ISL_818436, EPI_ISL_818437, EPI_ISL_818438, EPI_ISL_818439, EPI_ISL_818440, EPI_ISL_818441, EPI_ISL_818442, EPI_ISL_818443, EPI_ISL_818444, EPI_ISL_818445, EPI_ISL_818446, EPI_ISL_818447, EPI_ISL_818448, EPI_ISL_818449, EPI_ISL_818450, EPI_ISL_818451, EPI_ISL_818452, EPI_ISL_818453, EPI_ISL_818454, EPI_ISL_818455, EPI_ISL_818456, EPI_ISL_818457, EPI_ISL_818458, EPI_ISL_818459, EPI_ISL_818460, EPI_ISL_818461, EPI_ISL_818462, EPI_ISL_818463, EPI_ISL_818464, EPI_ISL_818465, EPI_ISL_818466, EPI_ISL_818467, EPI_ISL_818468, EPI_ISL_818469, EPI_ISL_818470, EPI_ISL_818471, EPI_ISL_818472, EPI_ISL_818473, EPI_ISL_818474, EPI_ISL_818475, EPI_ISL_818476, EPI_ISL_818477, EPI_ISL_818478, EPI_ISL_818479, EPI_ISL_818480, EPI_ISL_818481, EPI_ISL_818482, EPI_ISL_818483, EPI_ISL_818484, EPI_ISL_818485, EPI_ISL_818486, EPI_ISL_818487, EPI_ISL_818488, EPI_ISL_818489, EPI_ISL_818490, EPI_ISL_818491, EPI_ISL_818492, EPI_ISL_818493, EPI_ISL_818494, EPI_ISL_818495, EPI_ISL_818496, EPI_ISL_818497, EPI_ISL_818498, EPI_ISL_818499, EPI_ISL_818500, EPI_ISL_818501, EPI_ISL_818502, EPI_ISL_818503, EPI_ISL_818504, EPI_ISL_818505, EPI_ISL_818506, EPI_ISL_818507, EPI_ISL_818508, EPI_ISL_818509, EPI_ISL_818510, EPI_ISL_818511, EPI_ISL_818512, EPI_ISL_818513, EPI_ISL_818514, EPI_ISL_818515, EPI_ISL_818516, EPI_ISL_818517, EPI_ISL_818518, EPI_ISL_818519, EPI_ISL_818520, EPI_ISL_818521, EPI_ISL_818522, EPI_ISL_818523, EPI_ISL_818524, EPI_ISL_818525, EPI_ISL_818526, EPI_ISL_818527, EPI_ISL_818528, EPI_ISL_818529, EPI_ISL_818530, EPI_ISL_818531, EPI_ISL_818532, EPI_ISL_818533, EPI_ISL_818534, EPI_ISL_818535, EPI_ISL_818536, EPI_ISL_818537, EPI_ISL_818538, EPI_ISL_818539, EPI_ISL_818540, EPI_ISL_818541, EPI_ISL_818542, EPI_ISL_818543, EPI_ISL_818544, EPI_ISL_818545, EPI_ISL_818546, EPI_ISL_818547, EPI_ISL_818548, EPI_ISL_818549, EPI_ISL_818550, EPI_ISL_818551, EPI_ISL_818552, EPI_ISL_818553, EPI_ISL_818554, EPI_ISL_818555, EPI_ISL_818556, EPI_ISL_818557, EPI_ISL_818558, EPI_ISL_818559, EPI_ISL_818560, EPI_ISL_818561, EPI_ISL_818562, EPI_ISL_818563, EPI_ISL_818564, EPI_ISL_818565, EPI_ISL_818566, EPI_ISL_818567, EPI_ISL_818568, EPI_ISL_818569, EPI_ISL_818570, EPI_ISL_818571, EPI_ISL_818572, EPI_ISL_818573, EPI_ISL_818574, EPI_ISL_818575, EPI_ISL_818576, EPI_ISL_818577, EPI_ISL_818578, EPI_ISL_818579, EPI_ISL_818580, EPI_ISL_818581, EPI_ISL_818582, EPI_ISL_818583, EPI_ISL_818584, EPI_ISL_818585, EPI_ISL_818586, EPI_ISL_818587, EPI_ISL_818588, EPI_ISL_818589, EPI_ISL_818590, EPI_ISL_818591, EPI_ISL_818592, EPI_ISL_818593, EPI_ISL_818594, EPI_ISL_818595, EPI_ISL_818596, EPI_ISL_818597, EPI_ISL_818598, EPI_ISL_818599, EPI_ISL_818600, EPI_ISL_818601, EPI_ISL_818602, EPI_ISL_818603, EPI_ISL_818604, EPI_ISL_818605, EPI_ISL_818606, EPI_ISL_818607, EPI_ISL_818608, EPI_ISL_818609, EPI_ISL_818610, EPI_ISL_818611, EPI_ISL_818612, EPI_ISL_818613, EPI_ISL_818614, EPI_ISL_818615, EPI_ISL_818616, EPI_ISL_818617, EPI_ISL_818618, EPI_ISL_818619, EPI_ISL_818620, EPI_ISL_818621, EPI_ISL_818622, EPI_ISL_818623, EPI_ISL_818624, EPI_ISL_818625, EPI_ISL_818626, EPI_ISL_818627, EPI_ISL_818628, EPI_ISL_818629, EPI_ISL_818630, EPI_ISL_818631, EPI_ISL_818632, EPI_ISL_818633, EPI_ISL_818634, EPI_ISL_818635, EPI_ISL_818636, EPI_ISL_818637, EPI_ISL_818638, EPI_ISL_818639, EPI_ISL_818640, EPI_ISL_818641, EPI_ISL_818642, EPI_ISL_818643, EPI_ISL_818644, EPI_ISL_818645, EPI_ISL_818646, EPI_ISL_818647, EPI_ISL_818648, EPI_ISL_818649, EPI_ISL_818650, EPI_ISL_818651, EPI_ISL_818652, EPI_ISL_818653, EPI_ISL_818654, EPI_ISL_818655, EPI_ISL_818656, EPI_ISL_818657, EPI_ISL_818658, EPI_ISL_818659, EPI_ISL_818660, EPI_ISL_818661, EPI_ISL_818662, EPI_ISL_818663, EPI_ISL_818664, EPI_ISL_818665, EPI_ISL_818666, EPI_ISL_818667, EPI_ISL_818668, EPI_ISL_818669, EPI_ISL_818670, EPI_ISL_818671, EPI_ISL_818672, EPI_ISL_818673, EPI_ISL_818674, EPI_ISL_818675, EPI_ISL_818676, EPI_ISL_818677, EPI_ISL_818678, EPI_ISL_818679, EPI_ISL_818680, EPI_ISL_818681, EPI_ISL_818682, EPI_ISL_818683, EPI_ISL_818684, EPI_ISL_818685, EPI_ISL_818686, EPI_ISL_818687, EPI_ISL_818688, EPI_ISL_818689, EPI_ISL_818690, EPI_ISL_818691, EPI_ISL_818692, EPI_ISL_818693, EPI_ISL_818694, EPI_ISL_818695, EPI_ISL_818696, EPI_ISL_818697, EPI_ISL_818698, EPI_ISL_818699, EPI_ISL_818700, EPI_ISL_818701, EPI_ISL_818702, EPI_ISL_818703, EPI_ISL_818704, EPI_ISL_818705, EPI_ISL_818706, EPI_ISL_818707, EPI_ISL_818708, EPI_ISL_818709, EPI_ISL_818710, EPI_ISL_818711, EPI_ISL_818712, EPI_ISL_818713, EPI_ISL_818714, EPI_ISL_818715, EPI_ISL_818716, EPI_ISL_818717, EPI_ISL_818718, EPI_ISL_818719, EPI_ISL_818720, EPI_ISL_818721, EPI_ISL_818722, EPI_ISL_818723, EPI_ISL_818724, EPI_ISL_818725, EPI_ISL_818726, EPI_ISL_818727, EPI_ISL_818728, EPI_ISL_818729, EPI_ISL_818730, EPI_ISL_818731, EPI_ISL_818732, EPI_ISL_818733, EPI_ISL_818734, EPI_ISL_818735, EPI_ISL_818736, EPI_ISL_818737, EPI_ISL_818738, EPI_ISL_818739, EPI_ISL_818740, EPI_ISL_818741, EPI_ISL_818742, EPI_ISL_818743, EPI_ISL_818744, EPI_ISL_818745, EPI_ISL_818746, EPI_ISL_818747, EPI_ISL_818748, EPI_ISL_818749, EPI_ISL_818750, EPI_ISL_818751, EPI_ISL_818752, EPI_ISL_818753, EPI_ISL_818754, EPI_ISL_818755, EPI_ISL_818756, EPI_ISL_818757, EPI_ISL_818758, EPI_ISL_818759, EPI_ISL_818760, EPI_ISL_818761, EPI_ISL_818762, EPI_ISL_818763, EPI_ISL_818764, EPI_ISL_818765, EPI_ISL_818766, EPI_ISL_818767, EPI_ISL_818768, EPI_ISL_818769, EPI_ISL_818770, EPI_ISL_818771, EPI_ISL_818772, EPI_ISL_818773, EPI_ISL_818774, EPI_ISL_818775, EPI_ISL_818776, EPI_ISL_818777, EPI_ISL_818778, EPI_ISL_818779, EPI_ISL_818780, EPI_ISL_818781, EPI_ISL_818782, EPI_ISL_818783, EPI_ISL_818784, EPI_ISL_818785, EPI_ISL_818786, EPI_ISL_818787, EPI_ISL_818788, EPI_ISL_818789, EPI_ISL_818790, EPI_ISL_818791, EPI_ISL_818792, EPI_ISL_818793, EPI_ISL_818794, EPI_ISL_818795, EPI_ISL_818796, EPI_ISL_818797, EPI_ISL_818798, EPI_ISL_818799, EPI_ISL_818800, EPI_ISL_818801, EPI_ISL_818802, EPI_ISL_818803, EPI_ISL_818804, EPI_ISL_818805, EPI_ISL_818806, EPI_ISL_818807, EPI_ISL_818808, EPI_ISL_818809, EPI_ISL_818810, EPI_ISL_818811, EPI_ISL_818812, EPI_ISL_818813, EPI_ISL_818814, EPI_ISL_818815, EPI_ISL_818816, EPI_ISL_818817, EPI_ISL_818818, EPI_ISL_818819, EPI_ISL_818820, EPI_ISL_818821, EPI_ISL_818822, EPI_ISL_818823, EPI_ISL_818824, EPI_ISL_818825, EPI_ISL_818826, EPI_ISL_818827, EPI_ISL_818828, EPI_ISL_818829, EPI_ISL_818830, EPI_ISL_818831, EPI_ISL_818832, EPI_ISL_818833, EPI_ISL_818834, EPI_ISL_818835, EPI_ISL_818836, EPI_ISL_818837, EPI_ISL_818838, EPI_ISL_818839, EPI_ISL_818840, EPI_ISL_818841, EPI_ISL_818842, EPI_ISL_818843, EPI_ISL_818844, EPI_ISL_818845, EPI_ISL_818846, EPI_ISL_818847, EPI_ISL_818848, EPI_ISL_818849, EPI_ISL_818850, EPI_ISL_818851, EPI_ISL_818852, EPI_ISL_818853, EPI_ISL_818854, EPI_ISL_818855, EPI_ISL_818856, EPI_ISL_818857, EPI_ISL_818858, EPI_ISL_818859, EPI_ISL_818860, EPI_ISL_818861, EPI_ISL_818862, EPI_ISL_818863, EPI_ISL_818864, EPI_ISL_818865, EPI_ISL_818866, EPI_ISL_818867, EPI_ISL_818868, EPI_ISL_818869, EPI_ISL_818870, EPI_ISL_818871, EPI_ISL_818872, EPI_ISL_818873, EPI_ISL_818874, EPI_ISL_818875, EPI_ISL_818876, EPI_ISL_818877, EPI_ISL_818878, EPI_ISL_818879, EPI_ISL_818880, EPI_ISL_818881, EPI_ISL_818882, EPI_ISL_818883, EPI_ISL_818884, EPI_ISL_818885, EPI_ISL_818886, EPI_ISL_818887, EPI_ISL_818888, EPI_ISL_818889, EPI_ISL_818890, EPI_ISL_818891, EPI_ISL_818892, EPI_ISL_818893, EPI_ISL_818894, EPI_ISL_818895, EPI_ISL_818896, EPI_ISL_818897, EPI_ISL_818898, EPI_ISL_818899, EPI_ISL_818900, EPI_ISL_818901, EPI_ISL_818902, EPI_ISL_818903, EPI_ISL_818904, EPI_ISL_818905, EPI_ISL_818906, EPI_ISL_818907, EPI_ISL_818908, EPI_ISL_818909, EPI_ISL_818910, EPI_ISL_818911, EPI_ISL_818912, EPI_ISL_818913, EPI_ISL_818914, EPI_ISL_818915, EPI_ISL_818916, EPI_ISL_818917, EPI_ISL_818918, EPI_ISL_818919, EPI_ISL_818920, EPI_ISL_818921, EPI_ISL_818922, EPI_ISL_818923, EPI_ISL_818924, EPI_ISL_818925, EPI_ISL_818926, EPI_ISL_818927, EPI_ISL_818928, EPI_ISL_818929, EPI_ISL_818930, EPI_ISL_818931, EPI_ISL_818932, EPI_ISL_818933, EPI_ISL_818934, EPI_ISL_818935, EPI_ISL_818936, EPI_ISL_818937, EPI_ISL_818938, EPI_ISL_818939, EPI_ISL_818940, EPI_ISL_818941, EPI_ISL_818942, EPI_ISL_818943, EPI_ISL_818944, EPI_ISL_818945, EPI_ISL_818946, EPI_ISL_818947, EPI_ISL_818948, EPI_ISL_818949, EPI_ISL_818950, EPI_ISL_818951, EPI_ISL_818952, EPI_ISL_818953, EPI_ISL_818954, EPI_ISL_818955, EPI_ISL_818956, EPI_ISL_818957, EPI_ISL_818958, EPI_ISL_818959, EPI_ISL_818960, EPI_ISL_818961, EPI_ISL_818962, EPI_ISL_818963, EPI_ISL_818964, EPI_ISL_818965, EPI_ISL_818966, EPI_ISL_818967, EPI_ISL_818968, EPI_ISL_818969, EPI_ISL_818970, EPI_ISL_818971, EPI_ISL_818972, EPI_ISL_818973, EPI_ISL_818974, EPI_ISL_818975, EPI_ISL_818976, EPI_ISL_818977, EPI_ISL_818978, EPI_ISL_818979, EPI_ISL_818980, EPI_ISL_818981, EPI_ISL_818982, EPI_ISL_818983, EPI_ISL_818984, EPI_ISL_818985, EPI_ISL_818986, EPI_ISL_818987, EPI_ISL_818988, EPI_ISL_818989, EPI_ISL_818990, EPI_ISL_818991, EPI_ISL_818992, EPI_ISL_818993, EPI_ISL_818994, EPI_ISL_818995, EPI_ISL_818996, EPI_ISL_818997, EPI_ISL_818998, EPI_ISL_818999, EPI_ISL_819000, EPI_ISL_819001, EPI_ISL_819002, EPI_ISL_819003, EPI_ISL_819004, EPI_ISL_819005, EPI_ISL_819006, EPI_ISL_819007, EPI_ISL_819008, EPI_ISL_819009, EPI_ISL_819010, EPI_ISL_819011, EPI_ISL_819012, EPI_ISL_819013, EPI_ISL_819014, EPI_ISL_819015, EPI_ISL_819016, EPI_ISL_819017, EPI_ISL_819018, EPI_ISL_819019, EPI_ISL_819020, EPI_ISL_819021, EPI_ISL_819022, EPI_ISL_819023, EPI_ISL_819024, EPI_ISL_819025, EPI_ISL_819026, EPI_ISL_819027, EPI_ISL_819028, EPI_ISL_819029, EPI_ISL_819030, EPI_ISL_819031, EPI_ISL_819032, EPI_ISL_819033, EPI_ISL_819034, EPI_ISL_819035, EPI_ISL_819036, EPI_ISL_819037, EPI_ISL_819038, EPI_ISL_819039, EPI_ISL_819040, EPI_ISL_819041, EPI_ISL_819042, EPI_ISL_819043, EPI_ISL_819044, EPI_ISL_819045, EPI_ISL_819046, EPI_ISL_819047, EPI_ISL_819048, EPI_ISL_819049, EPI_ISL_819050, EPI_ISL_819051, EPI_ISL_819052, EPI_ISL_819053, EPI_ISL_819054, EPI_ISL_819055, EPI_ISL_819056, EPI_ISL_819057, EPI_ISL_819058, EPI_ISL_819059, EPI_ISL_819060, EPI_ISL_819061, EPI_ISL_819062, EPI_ISL_819063, EPI_ISL_819064, EPI_ISL_819065, EPI_ISL_819066, EPI_ISL_819067, EPI_ISL_819068, EPI_ISL_819069, EPI_ISL_819070, EPI_ISL_819071, EPI_ISL_819072, EPI_ISL_819073, EPI_ISL_819074, EPI_ISL_819075, EPI_ISL_819076, EPI_ISL_819077, EPI_ISL_819078, EPI_ISL_819079, EPI_ISL_819080, EPI_ISL_819081, EPI_ISL_819082, EPI_ISL_819083, EPI_ISL_819084, EPI_ISL_819085, EPI_ISL_819086, EPI_ISL_819087, EPI_ISL_819088, EPI_ISL_819089, EPI_ISL_819090, EPI_ISL_819091, EPI_ISL_819092, EPI_ISL_819093, EPI_ISL_819094, EPI_ISL_819095, EPI_ISL_819096, EPI_ISL_819097, EPI_ISL_819098, EPI_ISL_819099, EPI_ISL_819100, EPI_ISL_819101, EPI_ISL_819102, EPI_ISL_819103, EPI_ISL_819104, EPI_ISL_819105, EPI_ISL_819106, EPI_ISL_819107, EPI_ISL_819108, EPI_ISL_819109, EPI_ISL_819110, EPI_ISL_819111, EPI_ISL_819112, EPI_ISL_819113, EPI_ISL_819114, EPI_ISL_819115, EPI_ISL_819116, EPI_ISL_819117, EPI_ISL_819118, EPI_ISL_819119, EPI_ISL_819120, EPI_ISL_819121, EPI_ISL_819122, EPI_ISL_819123 | | | |
| see above | Department of Virus and Microbiological Special Diagnostics, Statens Serum Institut, Copenhagen, Denmark | Albertsen Lab, Department of Chemistry and Bioscience, Aalborg University, Denmark | Danish Covid-19 Genome Consortium |

| | | | |
|---|---|---|---|
| EPI_ISL_819166 | Servicio de Microbiología, Hospital Universitario Son Espases | SeqCOVID-SPAIN consortium/IBV(CSIC) | Carla López-Causapé, Jordi Reina, Antonio Oliver and SeqCOVID-SPAIN consortium |
| EPI_ISL_819205, EPI_ISL_819207, EPI_ISL_819209, EPI_ISL_819210, EPI_ISL_819213, EPI_ISL_819215, EPI_ISL_819216, EPI_ISL_819217, EPI_ISL_819218, EPI_ISL_819219, EPI_ISL_819220, EPI_ISL_819221, EPI_ISL_819222, EPI_ISL_819223, EPI_ISL_819236, EPI_ISL_819237, EPI_ISL_819238, EPI_ISL_819239, EPI_ISL_819241, EPI_ISL_819242, EPI_ISL_819243, EPI_ISL_819244, EPI_ISL_819245, EPI_ISL_819268, EPI_ISL_819272, EPI_ISL_819273, EPI_ISL_819274, EPI_ISL_819275, EPI_ISL_819276, EPI_ISL_819277, EPI_ISL_819278, EPI_ISL_819279, EPI_ISL_819280, EPI_ISL_819281, EPI_ISL_819282, EPI_ISL_819283, EPI_ISL_819284, EPI_ISL_819285, EPI_ISL_819286, EPI_ISL_819287, EPI_ISL_819288, EPI_ISL_819289, EPI_ISL_819290, EPI_ISL_819291, EPI_ISL_819292 | | | |
| see above | Wyoming Public Health Laboratory | Wyoming Public Health Laboratory | Noah Hull, Taylor Fearing, Lynette Gumbleton, Channing Weber, Ashley Norberg, Bailey Bowcutt, and Wanda Manley |
| EPI_ISL_819384, EPI_ISL_819414, EPI_ISL_819417, EPI_ISL_819420, EPI_ISL_819429, EPI_ISL_819445, EPI_ISL_819446 | Quadram Institute Bioscience | COVID-19 Genomics UK (COG-UK) Consortium | Dave J. Baker, Gemma L. Kay, Alp Aydin, Thanh Le-Viet, Steven Rudder, Ana P. Tedim, Anastasia Kolyva, Maria Diaz, Leonardo de Oliveira Martins, Nabil-Fareed Alikhan, Lizzie Meadows, Rachael Stanley, Ngozi Elumogo, Muhammed Yasir, Nicholas M. Thomson, Alexander J Trotter, Rachel Gilroy, Samuel Bloomfield, Claire Stuart, Andrew Bell, Reenesh Prakash, Samir Dervisevic, Alison E. Mather, John Wain, Mark Webber, Andrew J. Page, Justin O'Grady |
| EPI_ISL_820588, EPI_ISL_820591, EPI_ISL_820593 | Queens Medical Centre, Clinical Microbiology Department / DeepSeq Nottingham | COVID-19 Genomics UK (COG-UK) Consortium | Gemma Clark, Wendy Smith, Manjinder Khakh, Vicki M Fleming, Michelle M Lister, Hannah Howson-Wells, Jonathan Ball, Patrick McClure, Joseph Chappell, Theocharis Tsoleridis, Nadine Holmes, Matthew Carlisle, Christopher Moore, Fei Sang, Johnny Debebe, Victoria Wright, Matthew Loose |
| EPI_ISL_821032, EPI_ISL_821037, EPI_ISL_821056, EPI_ISL_821061, EPI_ISL_821062, EPI_ISL_821069, EPI_ISL_821077, EPI_ISL_821098, EPI_ISL_821105, EPI_ISL_821142, EPI_ISL_821150, EPI_ISL_821160, EPI_ISL_821195, EPI_ISL_821198, EPI_ISL_821207, EPI_ISL_821225, EPI_ISL_821227, EPI_ISL_821241, EPI_ISL_821245, EPI_ISL_821249, EPI_ISL_821257, EPI_ISL_821258, EPI_ISL_821266, EPI_ISL_821269, EPI_ISL_821621 | | | |
| see above | Lighthouse Lab in Glasgow | Wellcome Sanger Institute for the COVID-19 Genomics UK (COG-UK) Consortium | Harper VanSteenhouse, Yumi Kasai, David Gray, Carol Clugston, Anna Dominiczak and Alex Alderton, Roberto Amato, Sonia Goncalves, Ewan Harrison, David K. Jackson, Ian Johnston, Dominic Kwiatkowski, Cordelia Langford, John Sillitoe on behalf of the Wellcome Sanger Institute COVID-19 Surveillance Team |
| EPI_ISL_824096, EPI_ISL_824097, EPI_ISL_824098, EPI_ISL_824099, EPI_ISL_824100, EPI_ISL_824141, EPI_ISL_824145, EPI_ISL_824146, EPI_ISL_824147, EPI_ISL_824148, EPI_ISL_824149, EPI_ISL_824150, EPI_ISL_824153, EPI_ISL_824154, EPI_ISL_824253, EPI_ISL_824254 | | | |
| see above | Dutch COVID-19 response team | National Institute for Public Health and the Environment (RIVM) | Adam Meijer, Harry Vennema, Jeroen Cremer, Sharon van den Brink, Bas van der Veer, AnneMarie van den Brandt, Florian Zwagemaker, Dennis Schmitz, Chantal Reusken, on behalf of the national COVID-19 response team |
| EPI_ISL_824842, EPI_ISL_824843, EPI_ISL_824844, EPI_ISL_824845, EPI_ISL_824846, EPI_ISL_824847, EPI_ISL_824848, EPI_ISL_824849, EPI_ISL_824850, EPI_ISL_824851 | Department of Clinical Microbiology | GIGA Medical Genomics | Keith Durkin, Maria Artesi, Sébastien Bontems, Raphaël Boreux, Bouchra Boujemla, Cécile Meex, Pierrette Melin, Marie-Pierre Hayette, Vincent Bours |
| EPI_ISL_824939, EPI_ISL_824940 | Arizona State Public Health Laboratory | Arizona State Public Health Laboratory | Trung Huynh, Jessica Escobar, Katherine Fullerton, Nobuko Fukushima, Stacy White, Linda Getsinger, Victor Waddell |
| EPI_ISL_824953, EPI_ISL_824954, EPI_ISL_824955, EPI_ISL_824956, EPI_ISL_824957, EPI_ISL_824958, EPI_ISL_824959, EPI_ISL_824960, EPI_ISL_824961, EPI_ISL_824962, EPI_ISL_824963, EPI_ISL_824964, EPI_ISL_824965, EPI_ISL_824974, EPI_ISL_824975, EPI_ISL_824976, EPI_ISL_824977 | | | |
| see above | Maryland Public Health Laboratory | Maryland Public Health Laboratory | Maryland Department of Health Laboratories Administration |
| EPI_ISL_827396 | The National University Hospital of Iceland | deCODE genetics | Daniel F Gudbjartsson; Agnar Helgason; Hakon Jonsson; Olafur T Magnusson; Pall Melsted; Gudmundur L Norddahl; Jona Saemundsdottir; Asgeir Sigurdsson; Patrick Sulem; Arna B Agustsdottir; Hannes Eggertsson; Berglind Eiriksdottir; Run Fridriksdottir; Elisabet E Gardarsdottir; Gudmundur Georgsson; Olafia S Gretarsdottir; Kjartan R Gudmundsson; Thora R Gunnarsdottir; Arnaldur Gylfason; Hilma Holm; Brynjar O Jensson; Aslaug Jonasdottir; Kamilla S Josefsdottir; Thordur Kristjansson; Droplaug N Magnusdottir; Solvi Rognvaldsson; Louise le Roux; Gudrun Sigmundsdottir; Gardar Sveinbjornsson; Kristin E Sveinsdottir; Maney Sveinsdottir; Emil A Thorarensen; Bjarni Thorbjornsson; Gisli Masson; Ingileif Jonsdottir; Alma Moller; Thorolfur Gudnason; Karl G Kristinsson; Unnur Thorsteinsdottir; Kari Stefansson |
| EPI_ISL_827545, EPI_ISL_827546, EPI_ISL_827554, EPI_ISL_827862, EPI_ISL_827921, EPI_ISL_827922, EPI_ISL_827923, EPI_ISL_827925, EPI_ISL_828255, EPI_ISL_828256, EPI_ISL_829703, EPI_ISL_829972, EPI_ISL_830194 | | | |
| see above | deCODE genetics | deCODE genetics | Daniel F Gudbjartsson; Agnar Helgason; Hakon Jonsson; Olafur T Magnusson; Pall Melsted; Gudmundur L Norddahl; Jona Saemundsdottir; Asgeir Sigurdsson; Patrick Sulem; Arna B Agustsdottir; Hannes Eggertsson; Berglind Eiriksdottir; Run Fridriksdottir; Elisabet E Gardarsdottir; Gudmundur Georgsson; Olafia S Gretarsdottir; Kjartan R Gudmundsson; Thora R Gunnarsdottir; Arnaldur Gylfason; Hilma Holm; Brynjar O Jensson; Aslaug Jonasdottir; Kamilla S Josefsdottir; Thordur Kristjansson; Droplaug N Magnusdottir; Solvi Rognvaldsson; Louise le Roux; Gudrun Sigmundsdottir; Gardar Sveinbjornsson; Kristin E Sveinsdottir; Maney Sveinsdottir; Emil A Thorarensen; Bjarni Thorbjornsson; Gisli Masson; Ingileif Jonsdottir; Alma Moller; Thorolfur Gudnason; Karl G Kristinsson; Unnur Thorsteinsdottir; Kari Stefansson |
| EPI_ISL_830244, EPI_ISL_830246, EPI_ISL_830248, EPI_ISL_830250, EPI_ISL_830252, EPI_ISL_830253, EPI_ISL_830255, EPI_ISL_830257, EPI_ISL_830259, EPI_ISL_830261 | Wadsworth Center, New York State Department of Health | Wadsworth Center, New York State Department of Health | Kirsten St. George, Daryl M. Lamson, Alexis Russel, Matthew Shudt, Melissa A Leisner, Jonathan Plitnick, Navjot Singh, John Kelly, Erasmus Schneider, Erica Lasek-Nesselquist |
| EPI_ISL_830567 | deCODE genetics | deCODE genetics | Daniel F Gudbjartsson; Agnar Helgason; Hakon Jonsson; Olafur T Magnusson; Pall Melsted; Gudmundur L Norddahl; Jona Saemundsdottir; Asgeir Sigurdsson; Patrick Sulem; Arna B Agustsdottir; Hannes Eggertsson; Berglind Eiriksdottir; Run Fridriksdottir; Elisabet E Gardarsdottir; Gudmundur Georgsson; Olafia S Gretarsdottir; Kjartan R Gudmundsson; Thora R Gunnarsdottir; Arnaldur Gylfason; Hilma Holm; Brynjar O Jensson; Aslaug Jonasdottir; Kamilla S Josefsdottir; Thordur Kristjansson; Droplaug N Magnusdottir; Solvi Rognvaldsson; Louise le Roux; Gudrun Sigmundsdottir; Gardar Sveinbjornsson; Kristin E Sveinsdottir; Maney Sveinsdottir; Emil A Thorarensen; Bjarni Thorbjornsson; Gisli Masson; Ingileif Jonsdottir; Alma Moller; Thorolfur Gudnason; Karl G Kristinsson; Unnur Thorsteinsdottir; Kari Stefansson |
| EPI_ISL_830572, EPI_ISL_830574, EPI_ISL_830576, EPI_ISL_830577, EPI_ISL_830578, EPI_ISL_830629, EPI_ISL_830630, EPI_ISL_830631, EPI_ISL_830632, EPI_ISL_830633 | NORTHWELL HEALTH LABORATORIES | Wadsworth Center, New York State Department of Health | Kirsten St. George, Daryl M. Lamson, Alexis Russel, Matthew Shudt, Melissa A Leisner, Jonathan Plitnick, Navjot Singh, John Kelly, Erasmus Schneider, Erica Lasek-Nesselquist |
| EPI_ISL_830634 | MARY IMOGENE BASSETT HOSPITAL | Wadsworth Center, New York State Department of Health | Kirsten St. George, Daryl M. Lamson, Alexis Russel, Matthew Shudt, Melissa A Leisner, Jonathan Plitnick, Navjot Singh, John Kelly, Erasmus Schneider, Erica Lasek-Nesselquist |
| EPI_ISL_830635, EPI_ISL_830636, EPI_ISL_830637, EPI_ISL_830638, EPI_ISL_830639, EPI_ISL_830640, EPI_ISL_830641, EPI_ISL_830642 | NORTHWELL HEALTH LABORATORIES | Wadsworth Center, New York State Department of Health | Kirsten St. George, Daryl M. Lamson, Alexis Russel, Matthew Shudt, Melissa A Leisner, Jonathan Plitnick, Navjot Singh, John Kelly, Erasmus Schneider, Erica Lasek-Nesselquist |
| EPI_ISL_830664 | SUNY UPSTATE MEDICAL UNIVERSITY | Wadsworth Center, New York State Department of Health | Kirsten St. George, Daryl M. Lamson, Alexis Russel, Matthew Shudt, Melissa A Leisner, Jonathan Plitnick, Navjot Singh, John Kelly, Erasmus Schneider, Erica Lasek-Nesselquist |
| EPI_ISL_830699, EPI_ISL_830700, EPI_ISL_830701, EPI_ISL_830702, EPI_ISL_830703, EPI_ISL_830707, EPI_ISL_830708, EPI_ISL_830717, EPI_ISL_830718, EPI_ISL_830719 | ALBANY MEDICAL CENTER HOSPITAL CLINICAL LABORATORIES | Wadsworth Center, New York State Department of Health | Kirsten St. George, Daryl M. Lamson, Alexis Russel, Matthew Shudt, Melissa A Leisner, Jonathan Plitnick, Navjot Singh, John Kelly, Erasmus Schneider, Erica Lasek-Nesselquist |
| EPI_ISL_830724, EPI_ISL_830725 | ACUTIS DIAGNOSTICS | Wadsworth Center, New York State Department of Health | Kirsten St. George, Daryl M. Lamson, Alexis Russel, Matthew Shudt, Melissa A Leisner, Jonathan Plitnick, Navjot Singh, John Kelly, Erasmus Schneider, Erica Lasek-Nesselquist |
| EPI_ISL_831289, EPI_ISL_831295, EPI_ISL_831296, EPI_ISL_831310, EPI_ISL_831324, EPI_ISL_831327, EPI_ISL_831333 | Texas Department of State Health Services | Texas Department of State Health Services | Anita Pokharel, Bonnie Oh, James Daniel Bonser, Rashmi Tuladhar, Mayela Pedrueza, Jenny Zhang, Maliha Rahman, Myong Koag, Chung Wang, Rachel Lee, Grace Kubin |

| | | | |
|---|---|---|---|
| EPI_ISL_831475, EPI_ISL_831477, EPI_ISL_831482, EPI_ISL_831485, EPI_ISL_831488, EPI_ISL_831491, EPI_ISL_831492, EPI_ISL_831493, EPI_ISL_831494, EPI_ISL_831497, EPI_ISL_831551, EPI_ISL_831555, EPI_ISL_831556, EPI_ISL_831557, EPI_ISL_831558, EPI_ISL_831559, EPI_ISL_831560, EPI_ISL_831561, EPI_ISL_831562, EPI_ISL_831563, EPI_ISL_831564, EPI_ISL_831565, EPI_ISL_831566, EPI_ISL_831567, EPI_ISL_831568, EPI_ISL_831569, EPI_ISL_831570, EPI_ISL_831571, EPI_ISL_831574, EPI_ISL_831576, EPI_ISL_831587, EPI_ISL_831588, EPI_ISL_831589, EPI_ISL_831590, EPI_ISL_831591, EPI_ISL_831592, EPI_ISL_831593, EPI_ISL_831594, EPI_ISL_831595, EPI_ISL_831596, EPI_ISL_831597, EPI_ISL_831598, EPI_ISL_831599, EPI_ISL_831600, EPI_ISL_831601, EPI_ISL_831602, EPI_ISL_831603, EPI_ISL_831604, EPI_ISL_831605, EPI_ISL_831606, EPI_ISL_831607, EPI_ISL_831608, EPI_ISL_831609, EPI_ISL_831610, EPI_ISL_831611, EPI_ISL_831612, EPI_ISL_831613, EPI_ISL_831614, EPI_ISL_831615, EPI_ISL_831616, EPI_ISL_831617, EPI_ISL_831618, EPI_ISL_831619, EPI_ISL_831620, EPI_ISL_831622 | | | |
| see above | University of Wisconsin-Madison AIDS Vaccine Research Laboratories | University of Wisconsin-Madison AIDS Vaccine Research Laboratories | Gage Moreno, Katarina Braun, et al. AIDS Vaccine Research Laboratories |
| EPI_ISL_831909 | New Mexico Department of Health Scientific Laboratory | New Mexico Department of Health Scientific Laboratory | Ellie Johnson, Anastacia Griego-Fisher, D'eldra Malone |
| EPI_ISL_831943 | Unilabs, Mikrobiologiska laboratoriet | The Public Health Agency of Sweden | Department of Microbiology, The Public Health Agency of Sweden |
| EPI_ISL_832003, EPI_ISL_832004 | Mikrobiologen | The Public Health Agency of Sweden | Department of Microbiology, The Public Health Agency of Sweden |
| EPI_ISL_832005 | Klinisk Mikrobiologi | The Public Health Agency of Sweden | Department of Microbiology, The Public Health Agency of Sweden |
| EPI_ISL_832018 | University of Wisconsin-Madison AIDS Vaccine Research Laboratories | University of Wisconsin-Madison AIDS Vaccine Research Laboratories | Gage Moreno, Katarina Braun, et al. AIDS Vaccine Research Laboratories |
| EPI_ISL_832121, EPI_ISL_832124 | Clinical Molecular Microbiology Laboratory, UNC Hospitals | Jeremy Wang | Jeremy Wang, Alexander Rubinsteyn, Colleen Rice, Jason Smedberg, Melissa Miller, Corbin Jones, Robert Hagan |
| EPI_ISL_832208, EPI_ISL_832209 | Department of Clinical Microbiology | GIGA Medical Genomics | Keith Durkin, Maria Artesi, Sébastien Bontems, Raphaël Boreux, Bouchra Boujemla, Cécile Meex, Pierrette Melin, Marie-Pierre Hayette, Vincent Bours |
| EPI_ISL_832978, EPI_ISL_832979, EPI_ISL_832980, EPI_ISL_832981, EPI_ISL_832982, EPI_ISL_832983, EPI_ISL_832984, EPI_ISL_832985 | Maine HETL | Tewhey Lab, The Jackson Laboratory | Matluk,N., Dewey,H., Iosue,F., Barter,M., Lynch,R., Munger,H. and Tewhey,R. |
| EPI_ISL_833041 | Defence Services Medical Research Center, Biological Research Laboratory | Defence Services Medical Research Center, Biological Research Laboratory | Khine Zaw Oo, Nay Myo Aung, Ko Ko Win, Phyo Kyaw Aung, Zaw Win Htun, Sat Paing Htoo, Kyaw Wanna, Thein Zaw, Kyee Myint, Ko Ko Lwin |
| EPI_ISL_833188 | Department of Clinical Microbiology | GIGA Medical Genomics | Keith Durkin, Maria Artesi, Sébastien Bontems, Raphaël Boreux, Bouchra Boujemla, Cécile Meex, Pierrette Melin, Marie-Pierre Hayette, Vincent Bours |
| EPI_ISL_833206, EPI_ISL_833213 | Department of Virology and Immunology, University of Helsinki and Helsinki University Hospital, Huslab Finland | Department of Virology, Faculty of Medicine, University of Helsinki, Helsinki, Finland | Teemu Smura, Ravi Kant, Phuoc Truong, Hussein Alburkat, Hannimari Kallio-Kokko, Jenni Virtanen, Maija Suvanto, Fathiah Zakham, Essi Korhonen, Sari Hannula, Harri Kangas, Pekka Ellonen, Olli Vapalahti |
| EPI_ISL_833415, EPI_ISL_833416 | USC Clinical Lab | Los Angeles County PHL | P. Hemarajata et al. |
| EPI_ISL_833944, EPI_ISL_833956, EPI_ISL_833966, EPI_ISL_833968, EPI_ISL_833995, EPI_ISL_833997, EPI_ISL_834009, EPI_ISL_834022, EPI_ISL_834029, EPI_ISL_834042, EPI_ISL_834069, EPI_ISL_834082, EPI_ISL_834104, EPI_ISL_834134, EPI_ISL_834144, EPI_ISL_834197, EPI_ISL_834210, EPI_ISL_834214, EPI_ISL_834216, EPI_ISL_834219, EPI_ISL_834221, EPI_ISL_834225, EPI_ISL_834229, EPI_ISL_834232, EPI_ISL_834255, EPI_ISL_834260, EPI_ISL_834273, EPI_ISL_834275, EPI_ISL_834276, EPI_ISL_834282, EPI_ISL_834304, EPI_ISL_834308, EPI_ISL_834309, EPI_ISL_834314, EPI_ISL_834317, EPI_ISL_834321, EPI_ISL_834327, EPI_ISL_834330, EPI_ISL_834333, EPI_ISL_834335, EPI_ISL_834336, EPI_ISL_834338, EPI_ISL_834345, EPI_ISL_834348, EPI_ISL_834349, EPI_ISL_834355, EPI_ISL_834359, EPI_ISL_834375, EPI_ISL_834387, EPI_ISL_834392, EPI_ISL_834395, EPI_ISL_834398, EPI_ISL_834399, EPI_ISL_834403, EPI_ISL_834411, EPI_ISL_834414, EPI_ISL_834415, EPI_ISL_834418, EPI_ISL_834421, EPI_ISL_834426, EPI_ISL_834434, EPI_ISL_834438, EPI_ISL_834450, EPI_ISL_834461, EPI_ISL_834463, EPI_ISL_834471, EPI_ISL_834472, EPI_ISL_834476, EPI_ISL_834480, EPI_ISL_834481, EPI_ISL_834482, EPI_ISL_834489, EPI_ISL_834493, EPI_ISL_834498, EPI_ISL_834505, EPI_ISL_834506, EPI_ISL_834509, EPI_ISL_834510, EPI_ISL_834527, EPI_ISL_834536, EPI_ISL_834538, EPI_ISL_834542, EPI_ISL_834544, EPI_ISL_834550, EPI_ISL_834552 | | | |
| see above | Lighthouse Lab in Alderley Park | Wellcome Sanger Institute for the COVID-19 Genomics UK (COG-UK) Consortium | Jacquelyn Wynn, Mairead Hyland, The Lighthouse Lab in Alderley Park and Alex Alderton, Roberto Amato, Sonia Goncalves, Ewan Harrison, David K. Jackson, Ian Johnston, Dominic Kwiatkowski, Cordelia Langford, John Sillitoe on behalf of the Wellcome Sanger Institute COVID-19 Surveillance Team |
| EPI_ISL_834558, EPI_ISL_834561, EPI_ISL_834568, EPI_ISL_834579, EPI_ISL_834581, EPI_ISL_834588, EPI_ISL_834592, EPI_ISL_834593, EPI_ISL_834599, EPI_ISL_834600, EPI_ISL_834603, EPI_ISL_834604, EPI_ISL_834608, EPI_ISL_834612, EPI_ISL_834619, EPI_ISL_834623, EPI_ISL_834629, EPI_ISL_834632, EPI_ISL_834639, EPI_ISL_834642, EPI_ISL_834645, EPI_ISL_834648, EPI_ISL_834658, EPI_ISL_834670, EPI_ISL_834672, EPI_ISL_834673, EPI_ISL_834683, EPI_ISL_834686, EPI_ISL_834687, EPI_ISL_834689, EPI_ISL_834690, EPI_ISL_834691, EPI_ISL_834692, EPI_ISL_834695, EPI_ISL_834699, EPI_ISL_834703, EPI_ISL_834708, EPI_ISL_834710, EPI_ISL_834714, EPI_ISL_834717, EPI_ISL_834718, EPI_ISL_834726, EPI_ISL_834733, EPI_ISL_834734, EPI_ISL_834738, EPI_ISL_834741, EPI_ISL_834751, EPI_ISL_834754, EPI_ISL_834756, EPI_ISL_834757, EPI_ISL_834760, EPI_ISL_834764, EPI_ISL_834765, EPI_ISL_834767, EPI_ISL_834782, EPI_ISL_834793 | | | |
| see above | Lighthouse Lab in Glasgow | Wellcome Sanger Institute for the COVID-19 Genomics UK (COG-UK) Consortium | Harper VanSteenhouse, Yumi Kasai, David Gray, Carol Clugston, Anna Dominiczak and Alex Alderton, Roberto Amato, Sonia Goncalves, Ewan Harrison, David K. Jackson, Ian Johnston, Dominic Kwiatkowski, Cordelia Langford, John Sillitoe on behalf of the Wellcome Sanger Institute COVID-19 Surveillance Team |
| EPI_ISL_834799, EPI_ISL_834811, EPI_ISL_834813, EPI_ISL_834816, EPI_ISL_834827, EPI_ISL_834828, EPI_ISL_834830, EPI_ISL_834836, EPI_ISL_834838, EPI_ISL_834844, EPI_ISL_834856, EPI_ISL_834858, EPI_ISL_834863, EPI_ISL_834864, EPI_ISL_834871, EPI_ISL_834882, EPI_ISL_834884, EPI_ISL_834887, EPI_ISL_834890, EPI_ISL_834895, EPI_ISL_834897, EPI_ISL_834898, EPI_ISL_834899, EPI_ISL_834902, EPI_ISL_834905, EPI_ISL_834911, EPI_ISL_834912, EPI_ISL_834915, EPI_ISL_834916, EPI_ISL_834918, EPI_ISL_834928, EPI_ISL_834942, EPI_ISL_834943, EPI_ISL_834946, EPI_ISL_834951, EPI_ISL_834952, EPI_ISL_834953, EPI_ISL_834965, EPI_ISL_834974, EPI_ISL_834978, EPI_ISL_834984, EPI_ISL_834988, EPI_ISL_834994, EPI_ISL_834996, EPI_ISL_835003, EPI_ISL_835005, EPI_ISL_835009, EPI_ISL_835013, EPI_ISL_835018, EPI_ISL_835024, EPI_ISL_835026, EPI_ISL_835028, EPI_ISL_835033, EPI_ISL_835036, EPI_ISL_835053, EPI_ISL_835066, EPI_ISL_835069, EPI_ISL_835077, EPI_ISL_835080, EPI_ISL_835083, EPI_ISL_835085, EPI_ISL_835088, EPI_ISL_835102, EPI_ISL_835103, EPI_ISL_835107, EPI_ISL_835109, EPI_ISL_835117, EPI_ISL_835120, EPI_ISL_835122, EPI_ISL_835133, EPI_ISL_835138 | | | |
| see above | Lighthouse Lab in Alderley Park | Wellcome Sanger Institute for the COVID-19 Genomics UK (COG-UK) Consortium | Jacquelyn Wynn, Mairead Hyland, The Lighthouse Lab in Alderley Park and Alex Alderton, Roberto Amato, Sonia Goncalves, Ewan Harrison, David K. Jackson, Ian Johnston, Dominic Kwiatkowski, Cordelia Langford, John Sillitoe on behalf of the Wellcome Sanger Institute COVID-19 Surveillance Team |
| EPI_ISL_835149, EPI_ISL_835153, EPI_ISL_835154, EPI_ISL_835174, EPI_ISL_835179, EPI_ISL_835181, EPI_ISL_835189, EPI_ISL_835192, EPI_ISL_835198, EPI_ISL_835200, EPI_ISL_835202, EPI_ISL_835205, EPI_ISL_835214, EPI_ISL_835219, EPI_ISL_835222, EPI_ISL_835223, EPI_ISL_835231, EPI_ISL_835243, EPI_ISL_835246, EPI_ISL_835253, EPI_ISL_835257, EPI_ISL_835259, EPI_ISL_835260, EPI_ISL_835268, EPI_ISL_835289, EPI_ISL_835298, EPI_ISL_835300, EPI_ISL_835303, EPI_ISL_835310, EPI_ISL_835313, EPI_ISL_835316, EPI_ISL_835320, EPI_ISL_835321, EPI_ISL_835334, EPI_ISL_835338, EPI_ISL_835339, EPI_ISL_835354, EPI_ISL_835357, EPI_ISL_835359, EPI_ISL_835361, EPI_ISL_835362, EPI_ISL_835365, EPI_ISL_835366, EPI_ISL_835369, EPI_ISL_835375, EPI_ISL_835384, EPI_ISL_835385, EPI_ISL_835389, EPI_ISL_835400, EPI_ISL_835403, EPI_ISL_835409, EPI_ISL_835410, EPI_ISL_835415, EPI_ISL_835416, EPI_ISL_835420, EPI_ISL_835421, EPI_ISL_835433, EPI_ISL_835439, EPI_ISL_835447, EPI_ISL_835448, EPI_ISL_835454, EPI_ISL_835457, EPI_ISL_835464, EPI_ISL_835465, EPI_ISL_835470, EPI_ISL_835475, EPI_ISL_835481, EPI_ISL_835486, EPI_ISL_835487, EPI_ISL_835488, EPI_ISL_835493, EPI_ISL_835497, EPI_ISL_835499, EPI_ISL_835500, EPI_ISL_835502, EPI_ISL_835503, EPI_ISL_835505, EPI_ISL_835506, EPI_ISL_835507, EPI_ISL_835515, EPI_ISL_835518, EPI_ISL_835527, EPI_ISL_835530, EPI_ISL_835531, EPI_ISL_835533, EPI_ISL_835535, EPI_ISL_835539, EPI_ISL_835540, EPI_ISL_835541, EPI_ISL_835547, EPI_ISL_835548, EPI_ISL_835553, EPI_ISL_835555, EPI_ISL_835564, EPI_ISL_835570, EPI_ISL_835573, EPI_ISL_835574, EPI_ISL_835576, EPI_ISL_835577, EPI_ISL_835579, EPI_ISL_835582, EPI_ISL_835589, EPI_ISL_835590, EPI_ISL_835592, EPI_ISL_835597, EPI_ISL_835600, EPI_ISL_835603, EPI_ISL_835610, EPI_ISL_835614, EPI_ISL_835616, EPI_ISL_835618, EPI_ISL_835622, EPI_ISL_835623, EPI_ISL_835631, EPI_ISL_835643, EPI_ISL_835646, EPI_ISL_835647, EPI_ISL_835651, EPI_ISL_835659, EPI_ISL_835664, EPI_ISL_835671, EPI_ISL_835673, EPI_ISL_835677, EPI_ISL_835678, EPI_ISL_835682, EPI_ISL_835683, EPI_ISL_835686, EPI_ISL_835687, EPI_ISL_835688, EPI_ISL_835692, EPI_ISL_835695, EPI_ISL_835701, EPI_ISL_835703, EPI_ISL_835707 | | | |
| see above | Lighthouse Lab in Glasgow | Wellcome Sanger Institute for the COVID-19 Genomics UK (COG-UK) Consortium | Harper VanSteenhouse, Yumi Kasai, David Gray, Carol Clugston, Anna Dominiczak and Alex Alderton, Roberto Amato, Sonia Goncalves, Ewan Harrison, David K. Jackson, Ian Johnston, Dominic Kwiatkowski, Cordelia Langford, John Sillitoe on behalf of the Wellcome Sanger Institute COVID-19 Surveillance Team |
| EPI_ISL_836075, EPI_ISL_836077, EPI_ISL_836078, EPI_ISL_836079, EPI_ISL_836083, EPI_ISL_836084, EPI_ISL_836088, EPI_ISL_836089, EPI_ISL_836095, EPI_ISL_836101, EPI_ISL_836105, EPI_ISL_836106, EPI_ISL_836107, EPI_ISL_836108, EPI_ISL_836111, EPI_ISL_836112, EPI_ISL_836115, EPI_ISL_836120, EPI_ISL_836128, EPI_ISL_836129, EPI_ISL_836131, EPI_ISL_836132, EPI_ISL_836137, EPI_ISL_836138, EPI_ISL_836142, EPI_ISL_836144, EPI_ISL_836147, EPI_ISL_836149, EPI_ISL_836154, EPI_ISL_836155, EPI_ISL_836156, EPI_ISL_836160, EPI_ISL_836162, EPI_ISL_836165, EPI_ISL_836168, EPI_ISL_836169, EPI_ISL_836174, EPI_ISL_836178, EPI_ISL_836180, EPI_ISL_836184, EPI_ISL_836190, EPI_ISL_836192, EPI_ISL_836197, EPI_ISL_836198, EPI_ISL_836201, EPI_ISL_836203, EPI_ISL_836211, EPI_ISL_836212, EPI_ISL_836218, EPI_ISL_836223, EPI_ISL_836225, EPI_ISL_836228, EPI_ISL_836230, EPI_ISL_836231, EPI_ISL_836233, EPI_ISL_836234, EPI_ISL_836238, EPI_ISL_836241, EPI_ISL_836247, EPI_ISL_836248, EPI_ISL_836250, EPI_ISL_836251, EPI_ISL_836252, EPI_ISL_836261, EPI_ISL_836262, EPI_ISL_836266, EPI_ISL_836268, EPI_ISL_836274, EPI_ISL_836275, EPI_ISL_836282, EPI_ISL_836284, EPI_ISL_836285, EPI_ISL_836291, EPI_ISL_836292, EPI_ISL_836293, EPI_ISL_836298, EPI_ISL_836305, EPI_ISL_836306, EPI_ISL_836307, EPI_ISL_836308, EPI_ISL_836310, EPI_ISL_836311, EPI_ISL_836313, EPI_ISL_836314, EPI_ISL_836315, EPI_ISL_836318, EPI_ISL_836319, EPI_ISL_836320, EPI_ISL_836321, EPI_ISL_836323, EPI_ISL_836328, EPI_ISL_836329, EPI_ISL_836333, EPI_ISL_836336, EPI_ISL_836338, EPI_ISL_836342, EPI_ISL_836343, EPI_ISL_836351, EPI_ISL_836352, EPI_ISL_836356, EPI_ISL_836357, EPI_ISL_836358, EPI_ISL_836362, EPI_ISL_836363, EPI_ISL_836366, EPI_ISL_836367, EPI_ISL_836371, EPI_ISL_836373, EPI_ISL_836377, EPI_ISL_836378, EPI_ISL_836387, EPI_ISL_836390, EPI_ISL_836395, EPI_ISL_836396, EPI_ISL_836403, EPI_ISL_836406, EPI_ISL_836407, EPI_ISL_836409, EPI_ISL_836410, EPI_ISL_836415, EPI_ISL_836416, EPI_ISL_836418 | | | |
| see above | Lighthouse Lab in Milton Keynes | Wellcome Sanger Institute for the COVID-19 Genomics UK (COG-UK) Consortium | The Lighthouse Lab in Milton Keynes and Alex Alderton, Roberto Amato, Sonia Goncalves, Ewan Harrison, David K. Jackson, Ian Johnston, Dominic Kwiatkowski, Cordelia Langford, John Sillitoe on behalf of the Wellcome Sanger Institute COVID-19 Surveillance Team |
| EPI_ISL_836421, EPI_ISL_836422, EPI_ISL_836425, EPI_ISL_836426, EPI_ISL_836427, EPI_ISL_836428, EPI_ISL_836429, EPI_ISL_836430, EPI_ISL_836433, EPI_ISL_836434, EPI_ISL_836435, EPI_ISL_836436, EPI_ISL_836437, EPI_ISL_836438, EPI_ISL_836440, EPI_ISL_836441, EPI_ISL_836442, EPI_ISL_836443, EPI_ISL_836446, EPI_ISL_836447, EPI_ISL_836448, EPI_ISL_836449, EPI_ISL_836450, EPI_ISL_836452, EPI_ISL_836453, EPI_ISL_836454, EPI_ISL_836455, EPI_ISL_836456, EPI_ISL_836457, EPI_ISL_836458, EPI_ISL_836459, EPI_ISL_836460, EPI_ISL_836461, EPI_ISL_836462, EPI_ISL_836463, EPI_ISL_836464, EPI_ISL_836465, EPI_ISL_836467, EPI_ISL_836468, EPI_ISL_836469, EPI_ISL_836470, EPI_ISL_836472, EPI_ISL_836473, EPI_ISL_836474, EPI_ISL_836475, EPI_ISL_836476, EPI_ISL_836477, EPI_ISL_836478, EPI_ISL_836479, EPI_ISL_836480, EPI_ISL_836481, EPI_ISL_836484, EPI_ISL_836485, EPI_ISL_836486, EPI_ISL_836487, EPI_ISL_836488, EPI_ISL_836489, EPI_ISL_836490, EPI_ISL_836492, EPI_ISL_836493, EPI_ISL_836495, EPI_ISL_836497, EPI_ISL_836498, EPI_ISL_836499, EPI_ISL_836502, EPI_ISL_836503, EPI_ISL_836505, EPI_ISL_836507, EPI_ISL_836508, EPI_ISL_836509, EPI_ISL_836510, EPI_ISL_836511, EPI_ISL_836512, EPI_ISL_836513, EPI_ISL_836514, EPI_ISL_836515, EPI_ISL_836516, EPI_ISL_836518, EPI_ISL_836519, EPI_ISL_836520, EPI_ISL_836521, EPI_ISL_836522, EPI_ISL_836523, EPI_ISL_836524, EPI_ISL_836525, EPI_ISL_836526, EPI_ISL_836529, EPI_ISL_836531, EPI_ISL_836532, EPI_ISL_836533, EPI_ISL_836534, EPI_ISL_836535, EPI_ISL_836537, EPI_ISL_836538, EPI_ISL_836539, EPI_ISL_836540, EPI_ISL_836541, EPI_ISL_836542, EPI_ISL_836543, EPI_ISL_836544, EPI_ISL_836545, EPI_ISL_836546, EPI_ISL_836547, EPI_ISL_836549, EPI_ISL_836550, EPI_ISL_836551, EPI_ISL_836552, EPI_ISL_836553, EPI_ISL_836554, EPI_ISL_836555, EPI_ISL_836557, EPI_ISL_836558, EPI_ISL_836559, EPI_ISL_836560, EPI_ISL_836561, EPI_ISL_836563, EPI_ISL_836564, EPI_ISL_836565, EPI_ISL_836566, EPI_ISL_836568, EPI_ISL_836569, EPI_ISL_836570, EPI_ISL_836571, EPI_ISL_836573, EPI_ISL_836574, EPI_ISL_836575, EPI_ISL_836577, EPI_ISL_836578, EPI_ISL_836579, EPI_ISL_836580, EPI_ISL_836581, EPI_ISL_836582, EPI_ISL_836583, EPI_ISL_836584, EPI_ISL_836586, EPI_ISL_836587, EPI_ISL_836588, EPI_ISL_836589, EPI_ISL_836590, EPI_ISL_836591, EPI_ISL_836593, EPI_ISL_836594, EPI_ISL_836598, EPI_ISL_836599, EPI_ISL_836600, EPI_ISL_836601, EPI_ISL_836602, EPI_ISL_836604, EPI_ISL_836605, EPI_ISL_836606, EPI_ISL_836607, EPI_ISL_836608, EPI_ISL_836609, EPI_ISL_836610, EPI_ISL_836612, EPI_ISL_836613, EPI_ISL_836614, EPI_ISL_836615, EPI_ISL_836616, EPI_ISL_836618, EPI_ISL_836619, EPI_ISL_836621, EPI_ISL_836622, EPI_ISL_836624, EPI_ISL_836625, EPI_ISL_836628, EPI_ISL_836629, EPI_ISL_836630, EPI_ISL_836631, EPI_ISL_836632, EPI_ISL_836633, EPI_ISL_836634, EPI_ISL_836635, EPI_ISL_836637, EPI_ISL_836638, EPI_ISL_836639, EPI_ISL_836640, EPI_ISL_836641, EPI_ISL_836642, EPI_ISL_836643, EPI_ISL_836646, EPI_ISL_836647, EPI_ISL_836648, EPI_ISL_836649, EPI_ISL_836650, EPI_ISL_836653, EPI_ISL_836654, EPI_ISL_836658, EPI_ISL_836659, EPI_ISL_836660, EPI_ISL_836661, EPI_ISL_836662, EPI_ISL_836663, EPI_ISL_836664, EPI_ISL_836666, EPI_ISL_836667, EPI_ISL_836668, EPI_ISL_836669, | | | |

| | | | |
|---|---|---|---|
| EPI_ISL_836670, EPI_ISL_836671, EPI_ISL_836672, EPI_ISL_836673, EPI_ISL_836674, EPI_ISL_836675, EPI_ISL_836676, EPI_ISL_836677, EPI_ISL_836678, EPI_ISL_836680, EPI_ISL_836684, EPI_ISL_836685, EPI_ISL_836687, EPI_ISL_836690, EPI_ISL_836691, EPI_ISL_836692, EPI_ISL_836693, EPI_ISL_836694, EPI_ISL_836696, EPI_ISL_836698, EPI_ISL_836699, EPI_ISL_836700, EPI_ISL_836701, EPI_ISL_836702, EPI_ISL_836703, EPI_ISL_836704, EPI_ISL_836705, EPI_ISL_836706, EPI_ISL_836707, EPI_ISL_836708, EPI_ISL_836709, EPI_ISL_836710, EPI_ISL_836711, EPI_ISL_836714, EPI_ISL_836715, EPI_ISL_836716, EPI_ISL_836717, EPI_ISL_836718, EPI_ISL_836719, EPI_ISL_836720, EPI_ISL_836721, EPI_ISL_836723, EPI_ISL_836724, EPI_ISL_836726, EPI_ISL_836727, EPI_ISL_836728, EPI_ISL_836729, EPI_ISL_836730, EPI_ISL_836731, EPI_ISL_836732, EPI_ISL_836733, EPI_ISL_836734, EPI_ISL_836735, EPI_ISL_836736, EPI_ISL_836737, EPI_ISL_836738, EPI_ISL_836739, EPI_ISL_836740, EPI_ISL_836741, EPI_ISL_836742, EPI_ISL_836743, EPI_ISL_836744, EPI_ISL_836745, EPI_ISL_836746, EPI_ISL_836747 | | | |
| see above | Lighthouse Lab in Cambridge | Wellcome Sanger Institute for the COVID-19 Genomics UK (COG-UK) Consortium | Rob Howes, The Lighthouse Lab in Cambridge and Alex Alderton, Roberto Amato, Sonia Goncalves, Ewan Harrison, David K. Jackson, Ian Johnston, Dominic Kwiatkowski, Cordelia Langford, John Sillitoe on behalf of the Wellcome Sanger Institute COVID-19 Surveillance Team |
| EPI_ISL_836748, EPI_ISL_836749, EPI_ISL_836750, EPI_ISL_836752, EPI_ISL_836753, EPI_ISL_836755, EPI_ISL_836756, EPI_ISL_836757, EPI_ISL_836758, EPI_ISL_836760, EPI_ISL_836761, EPI_ISL_836762, EPI_ISL_836763, EPI_ISL_836764 | | | |
| see above | Lighthouse Lab in Glasgow | Wellcome Sanger Institute for the COVID-19 Genomics UK (COG-UK) Consortium | Harper VanSteenhouse, Yumi Kasai, David Gray, Carol Clugston, Anna Dominiczak and Alex Alderton, Roberto Amato, Sonia Goncalves, Ewan Harrison, David K. Jackson, Ian Johnston, Dominic Kwiatkowski, Cordelia Langford, John Sillitoe on behalf of the Wellcome Sanger Institute COVID-19 Surveillance Team |
| EPI_ISL_836766 | Lighthouse Lab in Alderley Park | Wellcome Sanger Institute for the COVID-19 Genomics UK (COG-UK) Consortium | Jacquelyn Wynn, Mairead Hyland, The Lighthouse Lab in Alderley Park and Alex Alderton, Roberto Amato, Sonia Goncalves, Ewan Harrison, David K. Jackson, Ian Johnston, Dominic Kwiatkowski, Cordelia Langford, John Sillitoe on behalf of the Wellcome Sanger Institute COVID-19 Surveillance Team |
| EPI_ISL_836767, EPI_ISL_836768 | Lighthouse Lab in Glasgow | Wellcome Sanger Institute for the COVID-19 Genomics UK (COG-UK) Consortium | Harper VanSteenhouse, Yumi Kasai, David Gray, Carol Clugston, Anna Dominiczak and Alex Alderton, Roberto Amato, Sonia Goncalves, Ewan Harrison, David K. Jackson, Ian Johnston, Dominic Kwiatkowski, Cordelia Langford, John Sillitoe on behalf of the Wellcome Sanger Institute COVID-19 Surveillance Team |
| EPI_ISL_836769 | Lighthouse Lab in Alderley Park | Wellcome Sanger Institute for the COVID-19 Genomics UK (COG-UK) Consortium | Jacquelyn Wynn, Mairead Hyland, The Lighthouse Lab in Alderley Park and Alex Alderton, Roberto Amato, Sonia Goncalves, Ewan Harrison, David K. Jackson, Ian Johnston, Dominic Kwiatkowski, Cordelia Langford, John Sillitoe on behalf of the Wellcome Sanger Institute COVID-19 Surveillance Team |
| EPI_ISL_836771, EPI_ISL_836773, EPI_ISL_836774, EPI_ISL_836775, EPI_ISL_836776, EPI_ISL_836779, EPI_ISL_836780, EPI_ISL_836782, EPI_ISL_836783, EPI_ISL_836785, EPI_ISL_836786, EPI_ISL_836787, EPI_ISL_836789, EPI_ISL_836790, EPI_ISL_836791, EPI_ISL_836793, EPI_ISL_836794, EPI_ISL_836796, EPI_ISL_836798, EPI_ISL_836799, EPI_ISL_836800, EPI_ISL_836801 | | | |
| see above | Lighthouse Lab in Glasgow | Wellcome Sanger Institute for the COVID-19 Genomics UK (COG-UK) Consortium | Harper VanSteenhouse, Yumi Kasai, David Gray, Carol Clugston, Anna Dominiczak and Alex Alderton, Roberto Amato, Sonia Goncalves, Ewan Harrison, David K. Jackson, Ian Johnston, Dominic Kwiatkowski, Cordelia Langford, John Sillitoe on behalf of the Wellcome Sanger Institute COVID-19 Surveillance Team |
| EPI_ISL_836802 | Lighthouse Lab in Alderley Park | Wellcome Sanger Institute for the COVID-19 Genomics UK (COG-UK) Consortium | Jacquelyn Wynn, Mairead Hyland, The Lighthouse Lab in Alderley Park and Alex Alderton, Roberto Amato, Sonia Goncalves, Ewan Harrison, David K. Jackson, Ian Johnston, Dominic Kwiatkowski, Cordelia Langford, John Sillitoe on behalf of the Wellcome Sanger Institute COVID-19 Surveillance Team |
| EPI_ISL_836804, EPI_ISL_836805, EPI_ISL_836808, EPI_ISL_836809, EPI_ISL_836810, EPI_ISL_836812, EPI_ISL_836815, EPI_ISL_836816 | Lighthouse Lab in Glasgow | Wellcome Sanger Institute for the COVID-19 Genomics UK (COG-UK) Consortium | Harper VanSteenhouse, Yumi Kasai, David Gray, Carol Clugston, Anna Dominiczak and Alex Alderton, Roberto Amato, Sonia Goncalves, Ewan Harrison, David K. Jackson, Ian Johnston, Dominic Kwiatkowski, Cordelia Langford, John Sillitoe on behalf of the Wellcome Sanger Institute COVID-19 Surveillance Team |
| EPI_ISL_836817 | Lighthouse Lab in Alderley Park | Wellcome Sanger Institute for the COVID-19 Genomics UK (COG-UK) Consortium | Jacquelyn Wynn, Mairead Hyland, The Lighthouse Lab in Alderley Park and Alex Alderton, Roberto Amato, Sonia Goncalves, Ewan Harrison, David K. Jackson, Ian Johnston, Dominic Kwiatkowski, Cordelia Langford, John Sillitoe on behalf of the Wellcome Sanger Institute COVID-19 Surveillance Team |
| EPI_ISL_836818 | Lighthouse Lab in Glasgow | Wellcome Sanger Institute for the COVID-19 Genomics UK (COG-UK) Consortium | Harper VanSteenhouse, Yumi Kasai, David Gray, Carol Clugston, Anna Dominiczak and Alex Alderton, Roberto Amato, Sonia Goncalves, Ewan Harrison, David K. Jackson, Ian Johnston, Dominic Kwiatkowski, Cordelia Langford, John Sillitoe on behalf of the Wellcome Sanger Institute COVID-19 Surveillance Team |
| EPI_ISL_836819 | Lighthouse Lab in Alderley Park | Wellcome Sanger Institute for the COVID-19 Genomics UK (COG-UK) Consortium | Jacquelyn Wynn, Mairead Hyland, The Lighthouse Lab in Alderley Park and Alex Alderton, Roberto Amato, Sonia Goncalves, Ewan Harrison, David K. Jackson, Ian Johnston, Dominic Kwiatkowski, Cordelia Langford, John Sillitoe on behalf of the Wellcome Sanger Institute COVID-19 Surveillance Team |
| EPI_ISL_836821, EPI_ISL_836823, EPI_ISL_836824, EPI_ISL_836826, EPI_ISL_836828, EPI_ISL_836830, EPI_ISL_836832, EPI_ISL_836833, EPI_ISL_836835, EPI_ISL_836838, EPI_ISL_836839 | | | |
| see above | Lighthouse Lab in Glasgow | Wellcome Sanger Institute for the COVID-19 Genomics UK (COG-UK) Consortium | Harper VanSteenhouse, Yumi Kasai, David Gray, Carol Clugston, Anna Dominiczak and Alex Alderton, Roberto Amato, Sonia Goncalves, Ewan Harrison, David K. Jackson, Ian Johnston, Dominic Kwiatkowski, Cordelia Langford, John Sillitoe on behalf of the Wellcome Sanger Institute COVID-19 Surveillance Team |
| EPI_ISL_836840 | Lighthouse Lab in Alderley Park | Wellcome Sanger Institute for the COVID-19 Genomics UK (COG-UK) Consortium | Jacquelyn Wynn, Mairead Hyland, The Lighthouse Lab in Alderley Park and Alex Alderton, Roberto Amato, Sonia Goncalves, Ewan Harrison, David K. Jackson, Ian Johnston, Dominic Kwiatkowski, Cordelia Langford, John Sillitoe on behalf of the Wellcome Sanger Institute COVID-19 Surveillance Team |
| EPI_ISL_836841, EPI_ISL_836844, EPI_ISL_836845, EPI_ISL_836846, EPI_ISL_836852, EPI_ISL_836853, EPI_ISL_836854, EPI_ISL_836855, EPI_ISL_836856, EPI_ISL_836860, EPI_ISL_836861, EPI_ISL_836865, EPI_ISL_836868, EPI_ISL_836869, EPI_ISL_836871, EPI_ISL_836872, EPI_ISL_836873, EPI_ISL_836874, EPI_ISL_836875 | | | |
| see above | Lighthouse Lab in Glasgow | Wellcome Sanger Institute for the COVID-19 Genomics UK (COG-UK) Consortium | Harper VanSteenhouse, Yumi Kasai, David Gray, Carol Clugston, Anna Dominiczak and Alex Alderton, Roberto Amato, Sonia Goncalves, Ewan Harrison, David K. Jackson, Ian Johnston, Dominic Kwiatkowski, Cordelia Langford, John Sillitoe on behalf of the Wellcome Sanger Institute COVID-19 Surveillance Team |
| EPI_ISL_836876, EPI_ISL_836878 | Lighthouse Lab in Alderley Park | Wellcome Sanger Institute for the COVID-19 Genomics UK (COG-UK) Consortium | Jacquelyn Wynn, Mairead Hyland, The Lighthouse Lab in Alderley Park and Alex Alderton, Roberto Amato, Sonia Goncalves, Ewan Harrison, David K. Jackson, Ian Johnston, Dominic Kwiatkowski, Cordelia Langford, John Sillitoe on behalf of the Wellcome Sanger Institute COVID-19 Surveillance Team |
| EPI_ISL_836879, EPI_ISL_836880, EPI_ISL_836881, EPI_ISL_836882, EPI_ISL_836884, EPI_ISL_836885, EPI_ISL_836886, EPI_ISL_836888, EPI_ISL_836889, EPI_ISL_836892, EPI_ISL_836894, EPI_ISL_836896, EPI_ISL_836899, EPI_ISL_836901, EPI_ISL_836904, EPI_ISL_836905 | | | |
| see above | Lighthouse Lab in Glasgow | Wellcome Sanger Institute for the COVID-19 Genomics UK (COG-UK) Consortium | Harper VanSteenhouse, Yumi Kasai, David Gray, Carol Clugston, Anna Dominiczak and Alex Alderton, Roberto Amato, Sonia Goncalves, Ewan Harrison, David K. Jackson, Ian Johnston, Dominic Kwiatkowski, Cordelia Langford, John Sillitoe on behalf of the Wellcome Sanger Institute COVID-19 Surveillance Team |
| EPI_ISL_836906 | Lighthouse Lab in Alderley Park | Wellcome Sanger Institute for the COVID-19 Genomics UK (COG-UK) Consortium | Jacquelyn Wynn, Mairead Hyland, The Lighthouse Lab in Alderley Park and Alex Alderton, Roberto Amato, Sonia Goncalves, Ewan Harrison, David K. Jackson, Ian Johnston, Dominic Kwiatkowski, Cordelia Langford, John Sillitoe on behalf of the Wellcome Sanger Institute COVID-19 Surveillance Team |
| EPI_ISL_836911, EPI_ISL_836912 | Lighthouse Lab in Glasgow | Wellcome Sanger Institute for the COVID-19 Genomics UK (COG-UK) Consortium | Harper VanSteenhouse, Yumi Kasai, David Gray, Carol Clugston, Anna Dominiczak and Alex Alderton, Roberto Amato, Sonia Goncalves, Ewan Harrison, David K. Jackson, Ian Johnston, Dominic Kwiatkowski, Cordelia Langford, John Sillitoe on behalf of the Wellcome Sanger Institute COVID-19 Surveillance Team |
| EPI_ISL_836914 | Lighthouse Lab in Alderley Park | Wellcome Sanger Institute for the COVID-19 Genomics UK (COG-UK) Consortium | Jacquelyn Wynn, Mairead Hyland, The Lighthouse Lab in Alderley Park and Alex Alderton, Roberto Amato, Sonia Goncalves, Ewan Harrison, David K. Jackson, Ian Johnston, Dominic Kwiatkowski, Cordelia Langford, John Sillitoe on behalf of the Wellcome Sanger Institute COVID-19 Surveillance Team |
| EPI_ISL_836915, EPI_ISL_836916, EPI_ISL_836917, EPI_ISL_836918, EPI_ISL_836919, EPI_ISL_836923, EPI_ISL_836925, EPI_ISL_836927, EPI_ISL_836932, EPI_ISL_836933, EPI_ISL_836934 | | | |
| see above | Lighthouse Lab in Glasgow | Wellcome Sanger Institute for the COVID-19 Genomics UK (COG-UK) Consortium | Harper VanSteenhouse, Yumi Kasai, David Gray, Carol Clugston, Anna Dominiczak and Alex Alderton, Roberto Amato, Sonia Goncalves, Ewan Harrison, David K. Jackson, Ian Johnston, Dominic Kwiatkowski, Cordelia Langford, John Sillitoe on behalf of the Wellcome Sanger Institute COVID-19 Surveillance Team |
| EPI_ISL_836935 | Lighthouse Lab in Alderley Park | Wellcome Sanger Institute for the COVID-19 Genomics UK (COG-UK) Consortium | Jacquelyn Wynn, Mairead Hyland, The Lighthouse Lab in Alderley Park and Alex Alderton, Roberto Amato, Sonia Goncalves, Ewan Harrison, David K. Jackson, Ian Johnston, Dominic Kwiatkowski, Cordelia Langford, John Sillitoe on behalf of the Wellcome Sanger Institute COVID-19 Surveillance Team |
| EPI_ISL_836937, EPI_ISL_836938, EPI_ISL_836939, EPI_ISL_836941, EPI_ISL_836943, EPI_ISL_836945, EPI_ISL_836950, EPI_ISL_836951, EPI_ISL_836952, EPI_ISL_836953, EPI_ISL_836958, EPI_ISL_836959, EPI_ISL_836960, EPI_ISL_836961 | | | |
| see above | Lighthouse Lab in Glasgow | Wellcome Sanger Institute for the COVID-19 Genomics UK (COG-UK) Consortium | Harper VanSteenhouse, Yumi Kasai, David Gray, Carol Clugston, Anna Dominiczak and Alex Alderton, Roberto Amato, Sonia Goncalves, Ewan Harrison, David K. Jackson, Ian Johnston, Dominic Kwiatkowski, Cordelia Langford, John Sillitoe on behalf of the Wellcome Sanger Institute COVID-19 Surveillance Team |
| EPI_ISL_836962 | Lighthouse Lab in Alderley Park | Wellcome Sanger Institute for the COVID-19 Genomics UK (COG-UK) Consortium | Jacquelyn Wynn, Mairead Hyland, The Lighthouse Lab in Alderley Park and Alex Alderton, Roberto Amato, Sonia Goncalves, Ewan Harrison, David K. Jackson, Ian Johnston, Dominic Kwiatkowski, Cordelia Langford, John Sillitoe on behalf of the Wellcome Sanger Institute COVID-19 Surveillance Team |
| EPI_ISL_836963, EPI_ISL_836964, | Lighthouse Lab in Glasgow | Wellcome Sanger Institute for the COVID-19 Genomics UK | Harper VanSteenhouse, Yumi Kasai, David Gray, Carol Clugston, Anna Dominiczak and Alex Alderton, Roberto Amato, Sonia Goncalves, Ewan Harrison, |

| | | | |
|---|---|---|---|
| EPI_ISL_836965, EPI_ISL_836966, EPI_ISL_836967, EPI_ISL_836968, EPI_ISL_836970, EPI_ISL_836972 | | (COG-UK) Consortium | David K. Jackson, Ian Johnston, Dominic Kwiatkowski, Cordelia Langford, John Sillitoe on behalf of the Wellcome Sanger Institute COVID-19 Surveillance Team |
| EPI_ISL_837241, EPI_ISL_837242 | Respiratory Virus Unit, National Infection Service, Public Health England | COVID-19 Genomics UK (COG-UK) Consortium | PHE Covid Sequencing Team |
| EPI_ISL_837844, EPI_ISL_837845, EPI_ISL_837850, EPI_ISL_837903, EPI_ISL_837906, EPI_ISL_837907, EPI_ISL_837910, EPI_ISL_837912, EPI_ISL_837920, EPI_ISL_837921, EPI_ISL_837922, EPI_ISL_837929, EPI_ISL_837931, EPI_ISL_837934, EPI_ISL_837935, EPI_ISL_837936, EPI_ISL_837940, EPI_ISL_837942, EPI_ISL_837943, EPI_ISL_837955, EPI_ISL_837971 | | | |
| see above | Department of Pathology, University of Cambridge | COVID-19 Genomics UK (COG-UK) Consortium | Aminu S. Jahun, Yasmin Chaudhry, Grant Hall, Iliana Georgana, Myra Hosmillo, Martin D. Curran, Malte Pinckert, Surendra Parmar, Ian Goodfellow |
| EPI_ISL_838340 | University of Exeter | COVID-19 Genomics UK (COG-UK) Consortium | Ben Temperton,Aaron Jeffries,Michelle Michelsen,Joanna Warwick-Dugdale,Audrey Farbos,Robyn Manley,Stephen Michell,Jane Masoli |
| EPI_ISL_838864, EPI_ISL_838865, EPI_ISL_838866, EPI_ISL_838867, EPI_ISL_838868, EPI_ISL_838869, EPI_ISL_838870, EPI_ISL_838871, EPI_ISL_838872, EPI_ISL_838873, EPI_ISL_838874, EPI_ISL_838875, EPI_ISL_838876, EPI_ISL_838877, EPI_ISL_838878, EPI_ISL_838879, EPI_ISL_838880, EPI_ISL_838881, EPI_ISL_838882, EPI_ISL_838883, EPI_ISL_838884, EPI_ISL_838885, EPI_ISL_838886, EPI_ISL_838887, EPI_ISL_838888, EPI_ISL_838889, EPI_ISL_838890, EPI_ISL_838891, EPI_ISL_838892, EPI_ISL_838893, EPI_ISL_838894, EPI_ISL_838906, EPI_ISL_838907, EPI_ISL_838908, EPI_ISL_838924, EPI_ISL_838925, EPI_ISL_839044, EPI_ISL_839070, EPI_ISL_839071, EPI_ISL_839072, EPI_ISL_839073, EPI_ISL_839117, EPI_ISL_839156, EPI_ISL_839172, EPI_ISL_839176, EPI_ISL_839177, EPI_ISL_839178, EPI_ISL_839179, EPI_ISL_839180, EPI_ISL_839181, EPI_ISL_839182, EPI_ISL_839183, EPI_ISL_839186, EPI_ISL_839187, EPI_ISL_839189, EPI_ISL_839192, EPI_ISL_839195, EPI_ISL_839210, EPI_ISL_839214, EPI_ISL_839231 | | | |
| see above | University College London, Great Ormond Street Hospital for Children NHS Foundation Trust, Imperial College Healthcare NHS Trust | COVID-19 Genomics UK (COG-UK) Consortium | Sergi Castellano, Rachel Williams, Mark Kristiansen, Paola Resende Silva, Sunando Roy, Tony Brooks, Helena Tutill, Paola Niola, Patricia Dyal, Charlotte Williams, Leysa Forrest, Yasmin Panchbhaya, Jacqueline Findlay, Samuel Weeks, Julianne Brown, Kathryn Harris, Paul Randell, James Price, Alison Holmes, Judith Breuer |
| EPI_ISL_839823, EPI_ISL_839824, EPI_ISL_839840, EPI_ISL_839841, EPI_ISL_839842, EPI_ISL_839843, EPI_ISL_839844, EPI_ISL_839845, EPI_ISL_839846, EPI_ISL_839847, EPI_ISL_839848, EPI_ISL_839849, EPI_ISL_839850, EPI_ISL_839851, EPI_ISL_839852, EPI_ISL_839853, EPI_ISL_839854, EPI_ISL_839866, EPI_ISL_839899, EPI_ISL_839907, EPI_ISL_839908, EPI_ISL_839913, EPI_ISL_839914, EPI_ISL_839916, EPI_ISL_839918, EPI_ISL_839921, EPI_ISL_839932, EPI_ISL_839933, EPI_ISL_839934, EPI_ISL_839935, EPI_ISL_839936, EPI_ISL_839937, EPI_ISL_839938, EPI_ISL_839974, EPI_ISL_839975, EPI_ISL_839976, EPI_ISL_839977, EPI_ISL_839978, EPI_ISL_839979, EPI_ISL_839980 | | | |
| see above | Quadram Institute Bioscience | COVID-19 Genomics UK (COG-UK) Consortium | Dave J. Baker, Gemma L. Kay, Alp Aydin, Thanh Le-Viet, Steven Rudder, Ana P. Tedim, Anastasia Kolyva, Maria Diaz, Leonardo de Oliveira Martins, Nabil-Fareed Alikhan, Lizzie Meadows, Rachael Stanley, Ngozi Elumogo, Muhammed Yasir, Nicholas M. Thomson, Alexander J Trotter, Rachel Gilroy, Samuel Bloomfield, Claire Stuart, Andrew Bell, Reenesh Prakash, Samir Dervisevic, Alison E. Mather, John Wain, Mark Webber, Andrew J. Page, Justin O'Grady |
| EPI_ISL_839997 | Queens Medical Centre, Clinical Microbiology Department / DeepSeq Nottingham | COVID-19 Genomics UK (COG-UK) Consortium | Gemma Clark, Wendy Smith, Manjinder Khakh, Vicki M Fleming, Michelle M Lister, Hannah Howson-Wells, Jonathan Ball, Patrick McClure, Joseph Chappell, Theocharis Tsoleridis, Nadine Holmes, Matthew Carlisle, Christopher Moore, Fei Sang, Johnny Debebe, Victoria Wright, Matthew Loose |
| EPI_ISL_840135, EPI_ISL_840136, EPI_ISL_840137, EPI_ISL_840138, EPI_ISL_840140, EPI_ISL_840141, EPI_ISL_840142, EPI_ISL_840143, EPI_ISL_840144 | Lincolnshire Hospitals and DeepSeq Nottingham | COVID-19 Genomics UK (COG-UK) Consortium | Nichola Duckworth, Tim Sloan, Sarah Walsh, Jonathan Ball, Patrick McClure, Joeseph Chappell, Nadine Holmes, Matthew Carlisle, Christopher Moore, Fei Sang, Johnny Debebe, Victoria Wright, Matthew Loose |
| EPI_ISL_840694 | Originating lab: Wales Specialist Virology Centre Sequencing lab: Pathogen Genomics Unit | Public Health Wales Microbiology Cardiff Wales Specialist Virology Centre | Catherine Moore, Johnathan Evans, Laura Gifford, Malorie Perry, Simon Cottrell, Angela Marchbank, Alec Birchley, Alexander Adams, Amy Gaskin, Bree Gatica-Wilcox, Jason Coombes, Joel Southgate, Lauren Gilbert, Lee Graham, Nicole Pacchiarini, Sara Kumziene-Summerhayes, Sarah Taylor, Sophie Jones, Sara Rey, Matthew Bull, Joanne Watkins, Sally Corden, Tom Connor |
| EPI_ISL_840843, EPI_ISL_840849, EPI_ISL_840852, EPI_ISL_840855, EPI_ISL_840856, EPI_ISL_840857, EPI_ISL_840858, EPI_ISL_840859, EPI_ISL_840881, EPI_ISL_840907, EPI_ISL_840908, EPI_ISL_840951, EPI_ISL_840961, EPI_ISL_840963, EPI_ISL_840964, EPI_ISL_840965, EPI_ISL_840968, EPI_ISL_840969, EPI_ISL_840970, EPI_ISL_840971, EPI_ISL_840974, EPI_ISL_840975, EPI_ISL_840976, EPI_ISL_840977, EPI_ISL_840978, EPI_ISL_840979, EPI_ISL_840981, EPI_ISL_840982, EPI_ISL_840983, EPI_ISL_840984, EPI_ISL_840985, EPI_ISL_840989, EPI_ISL_840990, EPI_ISL_840991, EPI_ISL_840992, EPI_ISL_840993, EPI_ISL_840994, EPI_ISL_840995, EPI_ISL_840996, EPI_ISL_840997, EPI_ISL_840998, EPI_ISL_841000, EPI_ISL_841002, EPI_ISL_841004, EPI_ISL_841032, EPI_ISL_841040, EPI_ISL_841043, EPI_ISL_841044, EPI_ISL_841051, EPI_ISL_841052, EPI_ISL_841060, EPI_ISL_841086, EPI_ISL_841098, EPI_ISL_841166, EPI_ISL_841168, EPI_ISL_841169, EPI_ISL_841170, EPI_ISL_841172, EPI_ISL_841174, EPI_ISL_841175, EPI_ISL_841176, EPI_ISL_841178, EPI_ISL_841179, EPI_ISL_841183, EPI_ISL_841184, EPI_ISL_841185, EPI_ISL_841186, EPI_ISL_841193, EPI_ISL_841195, EPI_ISL_841204, EPI_ISL_841208, EPI_ISL_841214, EPI_ISL_841307, EPI_ISL_841308, EPI_ISL_841309, EPI_ISL_841310, EPI_ISL_841311, EPI_ISL_841312, EPI_ISL_841313, EPI_ISL_841316 | | | |
| see above | Wales Specialist Virology Centre Sequencing lab: Pathogen Genomics Unit | Public Health Wales Microbiology Cardiff Wales Specialist Virology Centre | Catherine Moore, Johnathan Evans, Laura Gifford, Malorie Perry, Simon Cottrell, Angela Marchbank, Alec Birchley, Alexander Adams, Amy Gaskin, Bree Gatica-Wilcox, Jason Coombes, Joel Southgate, Lauren Gilbert, Lee Graham, Nicole Pacchiarini, Sara Kumziene-Summerhayes, Sarah Taylor, Sophie Jones, Sara Rey, Matthew Bull, Joanne Watkins, Sally Corden, Tom Connor |
| EPI_ISL_841332, EPI_ISL_841334, EPI_ISL_841375, EPI_ISL_841376, EPI_ISL_841377, EPI_ISL_841387, EPI_ISL_841388, EPI_ISL_841389, EPI_ISL_841390, EPI_ISL_841498, EPI_ISL_841506, EPI_ISL_841512, EPI_ISL_841586, EPI_ISL_841588, EPI_ISL_841590, EPI_ISL_841596, EPI_ISL_841611 | | | |
| see above | Originating lab: Wales Specialist Virology Centre Sequencing lab: Pathogen Genomics Unit | Public Health Wales Microbiology Cardiff Wales Specialist Virology Centre | Catherine Moore, Johnathan Evans, Laura Gifford, Malorie Perry, Simon Cottrell, Angela Marchbank, Alec Birchley, Alexander Adams, Amy Gaskin, Bree Gatica-Wilcox, Jason Coombes, Joel Southgate, Lauren Gilbert, Lee Graham, Nicole Pacchiarini, Sara Kumziene-Summerhayes, Sarah Taylor, Sophie Jones, Sara Rey, Matthew Bull, Joanne Watkins, Sally Corden, Tom Connor |
| EPI_ISL_842208, EPI_ISL_842215, EPI_ISL_842216, EPI_ISL_842221, EPI_ISL_842224, EPI_ISL_842226, EPI_ISL_842239, EPI_ISL_842241, EPI_ISL_842246, EPI_ISL_842251, EPI_ISL_842258, EPI_ISL_842261, EPI_ISL_842280, EPI_ISL_842287, EPI_ISL_842288, EPI_ISL_842289, EPI_ISL_842294, EPI_ISL_842300, EPI_ISL_842301, EPI_ISL_842303, EPI_ISL_842305, EPI_ISL_842324, EPI_ISL_842329, EPI_ISL_842330, EPI_ISL_842334, EPI_ISL_842340 | | | |
| see above | Virology Department, Sheffield Teaching Hospitals NHS Foundation Trust/Department of Infection, Immunity and Cardiovascular Disease, The Medical School, University of Sheffield | COVID-19 Genomics UK (COG-UK) Consortium | Thushan de Silva, Matthew Parker, Nikki Smith, Adri Angyal, Rebecca Brown, Luke Green, Rachel Tucker, Paul Parsons, Danielle Groves, Katie Johnson, Laura Carrilero, Alex Keeley, Dave Partridge, Matthew Wyles, Benjamin Lindsey, Mehmet Yavuz, Mohammad Raza, Cariad Evans |
| EPI_ISL_842638, EPI_ISL_842640 | University Hospital of Northern Norway, Department for Microbiology and Infectious Disease Control | Norwegian Institute of Public Health, Department of Virology | Kathrine Stene-Johansen, Kamilla Heddeland Instefjord, Hilde Elshaug, Atiya R Ali,Marie Paulsen Madsen, Rasmus Riis Kopperud, Hilde Vollan, Karoline Bragstad, Olav Hungnes |
| EPI_ISL_842641 | Unilabs Laboratory Medicine | Norwegian Institute of Public Health, Department of Virology | Kathrine Stene-Johansen, Kamilla Heddeland Instefjord, Hilde Elshaug, Atiya R Ali,Marie Paulsen Madsen, Rasmus Riis Kopperud, Hilde Vollan, Karoline Bragstad, Olav Hungnes |
| EPI_ISL_842645 | Hospital of Southern Norway - Kristiansand, Department of Medical Microbiology | Norwegian Institute of Public Health, Department of Virology | Kathrine Stene-Johansen, Kamilla Heddeland Instefjord, Hilde Elshaug, Atiya R Ali,Marie Paulsen Madsen, Rasmus Riis Kopperud, Hilde Vollan, Karoline Bragstad, Olav Hungnes |
| EPI_ISL_843201, EPI_ISL_843202, EPI_ISL_843203, EPI_ISL_843204, EPI_ISL_843205, EPI_ISL_843206, EPI_ISL_843207, EPI_ISL_843208, EPI_ISL_843209, EPI_ISL_843210, EPI_ISL_843211, EPI_ISL_843212, EPI_ISL_843214, EPI_ISL_843215, EPI_ISL_843217, EPI_ISL_843218, EPI_ISL_843219, EPI_ISL_843220, EPI_ISL_843221, EPI_ISL_843222, EPI_ISL_843223, EPI_ISL_843224, EPI_ISL_843225, EPI_ISL_843226, EPI_ISL_843227, EPI_ISL_843228, EPI_ISL_843229, EPI_ISL_843230, EPI_ISL_843231, EPI_ISL_843232, EPI_ISL_843233, EPI_ISL_843234, EPI_ISL_843235, EPI_ISL_843236, EPI_ISL_843237, EPI_ISL_843238, EPI_ISL_843239, EPI_ISL_843240, EPI_ISL_843241, EPI_ISL_843242, EPI_ISL_843243, EPI_ISL_843244, EPI_ISL_843245, EPI_ISL_843246, EPI_ISL_843247, EPI_ISL_843248, EPI_ISL_843249, EPI_ISL_843250, EPI_ISL_843251, EPI_ISL_843252, EPI_ISL_843253, EPI_ISL_843254, EPI_ISL_843255, EPI_ISL_843256, EPI_ISL_843257, EPI_ISL_843258, EPI_ISL_843259, EPI_ISL_843260, EPI_ISL_843261, EPI_ISL_843262, EPI_ISL_843263, EPI_ISL_843264, EPI_ISL_843265, EPI_ISL_843266, EPI_ISL_843267, EPI_ISL_843268, EPI_ISL_843269, EPI_ISL_843271, EPI_ISL_843272, EPI_ISL_843273, EPI_ISL_843274, EPI_ISL_843275, EPI_ISL_843276, EPI_ISL_843277, EPI_ISL_843278, EPI_ISL_843279, EPI_ISL_843280, EPI_ISL_843281, EPI_ISL_843282, EPI_ISL_843283, EPI_ISL_843284, EPI_ISL_843285, EPI_ISL_843286, EPI_ISL_843287, EPI_ISL_843288, EPI_ISL_843289, EPI_ISL_843290, EPI_ISL_843291, EPI_ISL_843292, EPI_ISL_843293, EPI_ISL_843294, EPI_ISL_843295, EPI_ISL_843296, EPI_ISL_843297, EPI_ISL_843298, EPI_ISL_843299, EPI_ISL_843300, EPI_ISL_843301, EPI_ISL_843303, EPI_ISL_843304, EPI_ISL_843305, EPI_ISL_843306, EPI_ISL_843307, EPI_ISL_843308, EPI_ISL_843309, EPI_ISL_843310, EPI_ISL_843311, EPI_ISL_843312, EPI_ISL_843313, EPI_ISL_843314, EPI_ISL_843315, EPI_ISL_843317, EPI_ISL_843318, EPI_ISL_843319, EPI_ISL_843320, EPI_ISL_843321, EPI_ISL_843322, EPI_ISL_843323, EPI_ISL_843324, EPI_ISL_843325, EPI_ISL_843326, EPI_ISL_843327, EPI_ISL_843328, EPI_ISL_843329, EPI_ISL_843330, EPI_ISL_843331, EPI_ISL_843332, EPI_ISL_843333, EPI_ISL_843334, EPI_ISL_843335, EPI_ISL_843336, EPI_ISL_843337, EPI_ISL_843338, EPI_ISL_843339, EPI_ISL_843340, EPI_ISL_843342, EPI_ISL_843343, EPI_ISL_843344, EPI_ISL_843345, EPI_ISL_843346, EPI_ISL_843347, EPI_ISL_843348, EPI_ISL_843349, EPI_ISL_843350, EPI_ISL_843351, EPI_ISL_843352, EPI_ISL_843353, EPI_ISL_843354, EPI_ISL_843355, EPI_ISL_843356, EPI_ISL_843357, EPI_ISL_843358, EPI_ISL_843359, EPI_ISL_843361, EPI_ISL_843362, EPI_ISL_843363, EPI_ISL_843364, EPI_ISL_843365, EPI_ISL_843366, EPI_ISL_843367, EPI_ISL_843368, EPI_ISL_843369, EPI_ISL_843371, EPI_ISL_843372, EPI_ISL_843373, EPI_ISL_843374, EPI_ISL_843375, EPI_ISL_843376, EPI_ISL_843377, EPI_ISL_843378, EPI_ISL_843379, EPI_ISL_843380, EPI_ISL_843381, EPI_ISL_843382, EPI_ISL_843383, EPI_ISL_843385, EPI_ISL_843386, EPI_ISL_843387, EPI_ISL_843388, EPI_ISL_843389, EPI_ISL_843390, EPI_ISL_843391, EPI_ISL_843392, EPI_ISL_843393, EPI_ISL_843394, EPI_ISL_843395, EPI_ISL_843396, EPI_ISL_843397, EPI_ISL_843398, EPI_ISL_843399, EPI_ISL_843400, EPI_ISL_843401, EPI_ISL_843402, EPI_ISL_843403, EPI_ISL_843404, EPI_ISL_843405, EPI_ISL_843406, EPI_ISL_843407, EPI_ISL_843408, EPI_ISL_843409, EPI_ISL_843410, EPI_ISL_843411, EPI_ISL_843412, EPI_ISL_843413, EPI_ISL_843414, EPI_ISL_843415, EPI_ISL_843416, EPI_ISL_843417, EPI_ISL_843418, EPI_ISL_843419, EPI_ISL_843420, EPI_ISL_843421, EPI_ISL_843422, EPI_ISL_843423, EPI_ISL_843424, EPI_ISL_843425, EPI_ISL_843426, EPI_ISL_843427, EPI_ISL_843428, EPI_ISL_843429, EPI_ISL_843430, EPI_ISL_843431, EPI_ISL_843432, EPI_ISL_843433, EPI_ISL_843434, EPI_ISL_843435, EPI_ISL_843436, EPI_ISL_843437, EPI_ISL_843438, EPI_ISL_843439, EPI_ISL_843440, EPI_ISL_843441, EPI_ISL_843442, EPI_ISL_843443, EPI_ISL_843444, EPI_ISL_843445, EPI_ISL_843446, EPI_ISL_843447, EPI_ISL_843448, EPI_ISL_843449, EPI_ISL_843450, EPI_ISL_843451, EPI_ISL_843452, EPI_ISL_843453, EPI_ISL_843454, EPI_ISL_843455, EPI_ISL_843456, EPI_ISL_843457, EPI_ISL_843458, EPI_ISL_843459, EPI_ISL_843460, EPI_ISL_843461, EPI_ISL_843462, EPI_ISL_843463, EPI_ISL_843464, EPI_ISL_843465, EPI_ISL_843466, EPI_ISL_843467, EPI_ISL_843468, EPI_ISL_843469, EPI_ISL_843470, EPI_ISL_843471, EPI_ISL_843472, EPI_ISL_843473, EPI_ISL_843474, EPI_ISL_843475, EPI_ISL_843476, EPI_ISL_843477, EPI_ISL_843478, EPI_ISL_843479, EPI_ISL_843480, EPI_ISL_843481, EPI_ISL_843482, EPI_ISL_843483, EPI_ISL_843484, EPI_ISL_843485, EPI_ISL_843486, EPI_ISL_843487, EPI_ISL_843488, EPI_ISL_843489, EPI_ISL_843490, EPI_ISL_843491, EPI_ISL_843492, EPI_ISL_843493, EPI_ISL_843494, EPI_ISL_843495, EPI_ISL_843496, EPI_ISL_843497, EPI_ISL_843498, EPI_ISL_843499, EPI_ISL_843500, EPI_ISL_843501, EPI_ISL_843502, EPI_ISL_843503, EPI_ISL_843504, EPI_ISL_843505, EPI_ISL_843506, EPI_ISL_843507, EPI_ISL_843508, EPI_ISL_843509, EPI_ISL_843510, EPI_ISL_843511, EPI_ISL_843512, EPI_ISL_843513, EPI_ISL_843514, EPI_ISL_843515, EPI_ISL_843516, EPI_ISL_843517, EPI_ISL_843518, EPI_ISL_843519, EPI_ISL_843520, EPI_ISL_843521, EPI_ISL_843522, EPI_ISL_843523, EPI_ISL_843524, EPI_ISL_843525, EPI_ISL_843526, EPI_ISL_843527, EPI_ISL_843528, EPI_ISL_843529, EPI_ISL_843530, EPI_ISL_843531, EPI_ISL_843532, EPI_ISL_843533, EPI_ISL_843534, EPI_ISL_843535, EPI_ISL_843536, EPI_ISL_843537, EPI_ISL_843538, EPI_ISL_843539, EPI_ISL_843540, EPI_ISL_843541, EPI_ISL_843542, EPI_ISL_843543, EPI_ISL_843544, EPI_ISL_843545, EPI_ISL_843546, EPI_ISL_843548, EPI_ISL_843549, EPI_ISL_843550, EPI_ISL_843551, EPI_ISL_843552, EPI_ISL_843553, EPI_ISL_843554, EPI_ISL_843555, EPI_ISL_843556, EPI_ISL_843557, EPI_ISL_843558, EPI_ISL_843559, EPI_ISL_843560, EPI_ISL_843561, EPI_ISL_843562, EPI_ISL_843563, EPI_ISL_843564, EPI_ISL_843565, EPI_ISL_843566, EPI_ISL_843567, EPI_ISL_843568, EPI_ISL_843569, EPI_ISL_843570, | | | |

EPI_ISL_843571, EPI_ISL_843572, EPI_ISL_843573, EPI_ISL_843575, EPI_ISL_843576, EPI_ISL_843577, EPI_ISL_843578, EPI_ISL_843579, EPI_ISL_843580, EPI_ISL_843581, EPI_ISL_843582, EPI_ISL_843583, EPI_ISL_843584, EPI_ISL_843585, EPI_ISL_843586, EPI_ISL_843587, EPI_ISL_843588, EPI_ISL_843589, EPI_ISL_843590, EPI_ISL_843592, EPI_ISL_843593, EPI_ISL_843594, EPI_ISL_843596, EPI_ISL_843597, EPI_ISL_843599, EPI_ISL_843600, EPI_ISL_843601, EPI_ISL_843602, EPI_ISL_843603, EPI_ISL_843604, EPI_ISL_843605, EPI_ISL_843606, EPI_ISL_843607, EPI_ISL_843609, EPI_ISL_843610, EPI_ISL_843611, EPI_ISL_843612, EPI_ISL_843613, EPI_ISL_843614, EPI_ISL_843672, EPI_ISL_843700, EPI_ISL_843789, EPI_ISL_843803, EPI_ISL_843912, EPI_ISL_843925, EPI_ISL_843973, EPI_ISL_844000, EPI_ISL_844008, EPI_ISL_844026, EPI_ISL_844067, EPI_ISL_844108, EPI_ISL_844113, EPI_ISL_844120, EPI_ISL_844129, EPI_ISL_844544, EPI_ISL_844546, EPI_ISL_844547, EPI_ISL_844548, EPI_ISL_844549, EPI_ISL_844550, EPI_ISL_844551, EPI_ISL_844554, EPI_ISL_844555, EPI_ISL_844578, EPI_ISL_844579, EPI_ISL_844580, EPI_ISL_844581, EPI_ISL_844582, EPI_ISL_844583, EPI_ISL_844584, EPI_ISL_844585, EPI_ISL_844586, EPI_ISL_844587, EPI_ISL_844588, EPI_ISL_844589, EPI_ISL_844590, EPI_ISL_844591, EPI_ISL_844592, EPI_ISL_844593, EPI_ISL_844594, EPI_ISL_844595, EPI_ISL_844596, EPI_ISL_844597, EPI_ISL_844598, EPI_ISL_844599, EPI_ISL_844600, EPI_ISL_844601, EPI_ISL_844602, EPI_ISL_844603, EPI_ISL_844604, EPI_ISL_844605, EPI_ISL_844606, EPI_ISL_844607, EPI_ISL_844608, EPI_ISL_844609, EPI_ISL_844610, EPI_ISL_844611, EPI_ISL_844612, EPI_ISL_844613, EPI_ISL_844614, EPI_ISL_844615, EPI_ISL_844616, EPI_ISL_844617, EPI_ISL_844618, EPI_ISL_844619, EPI_ISL_844620, EPI_ISL_844621, EPI_ISL_844622, EPI_ISL_844623, EPI_ISL_844624, EPI_ISL_844625, EPI_ISL_844626, EPI_ISL_844627, EPI_ISL_844628, EPI_ISL_844629, EPI_ISL_844630, EPI_ISL_844631, EPI_ISL_844632, EPI_ISL_844633, EPI_ISL_844634, EPI_ISL_844640, EPI_ISL_844641, EPI_ISL_844642, EPI_ISL_844643, EPI_ISL_844644, EPI_ISL_844645, EPI_ISL_844646, EPI_ISL_844647, EPI_ISL_844648, EPI_ISL_844649, EPI_ISL_844650, EPI_ISL_844651, EPI_ISL_844652, EPI_ISL_844656, EPI_ISL_844657, EPI_ISL_844658, EPI_ISL_844659, EPI_ISL_844660, EPI_ISL_844661, EPI_ISL_844662, EPI_ISL_844663, EPI_ISL_844664, EPI_ISL_844665, EPI_ISL_844668, EPI_ISL_844671, EPI_ISL_844672, EPI_ISL_844673, EPI_ISL_844674, EPI_ISL_844675, EPI_ISL_844677, EPI_ISL_844678, EPI_ISL_844679, EPI_ISL_844682, EPI_ISL_844683, EPI_ISL_844684, EPI_ISL_844685, EPI_ISL_844686, EPI_ISL_844687, EPI_ISL_844688, EPI_ISL_844689, EPI_ISL_844690, EPI_ISL_844691, EPI_ISL_844692, EPI_ISL_844695, EPI_ISL_844696, EPI_ISL_844697, EPI_ISL_844698, EPI_ISL_844699, EPI_ISL_844700, EPI_ISL_844701, EPI_ISL_844703, EPI_ISL_844704, EPI_ISL_844705, EPI_ISL_844709, EPI_ISL_844710, EPI_ISL_844711, EPI_ISL_844712, EPI_ISL_844713, EPI_ISL_844714, EPI_ISL_844715, EPI_ISL_844719, EPI_ISL_844720, EPI_ISL_844721, EPI_ISL_844722, EPI_ISL_844723, EPI_ISL_844724, EPI_ISL_844725, EPI_ISL_844726, EPI_ISL_844727, EPI_ISL_844728, EPI_ISL_844729, EPI_ISL_844739, EPI_ISL_844740, EPI_ISL_844741, EPI_ISL_844742, EPI_ISL_844743, EPI_ISL_844744, EPI_ISL_844745, EPI_ISL_844746, EPI_ISL_844748, EPI_ISL_844749, EPI_ISL_844750, EPI_ISL_844751, EPI_ISL_844752, EPI_ISL_844753, EPI_ISL_844754, EPI_ISL_844755, EPI_ISL_844756, EPI_ISL_844757, EPI_ISL_844760, EPI_ISL_844761, EPI_ISL_844762, EPI_ISL_844764, EPI_ISL_844765, EPI_ISL_844769, EPI_ISL_844770, EPI_ISL_844771, EPI_ISL_844772, EPI_ISL_844773, EPI_ISL_844774, EPI_ISL_844775, EPI_ISL_844776, EPI_ISL_844777, EPI_ISL_844778, EPI_ISL_844779, EPI_ISL_844780, EPI_ISL_844781, EPI_ISL_844782, EPI_ISL_844783, EPI_ISL_844784, EPI_ISL_844785, EPI_ISL_844786, EPI_ISL_844787, EPI_ISL_844789, EPI_ISL_844790, EPI_ISL_844791, EPI_ISL_844792, EPI_ISL_844793, EPI_ISL_844794, EPI_ISL_844795, EPI_ISL_844796, EPI_ISL_844797, EPI_ISL_844798, EPI_ISL_844799, EPI_ISL_844800, EPI_ISL_844801, EPI_ISL_844802, EPI_ISL_844803, EPI_ISL_844804, EPI_ISL_844806, EPI_ISL_844809, EPI_ISL_844810, EPI_ISL_844811, EPI_ISL_844812, EPI_ISL_844813, EPI_ISL_844814, EPI_ISL_844815, EPI_ISL_844816, EPI_ISL_844817, EPI_ISL_844819, EPI_ISL_844820, EPI_ISL_844822, EPI_ISL_844823, EPI_ISL_844825, EPI_ISL_844826, EPI_ISL_844827, EPI_ISL_844828, EPI_ISL_844829, EPI_ISL_844830, EPI_ISL_844831, EPI_ISL_844832, EPI_ISL_844833, EPI_ISL_844835, EPI_ISL_844838, EPI_ISL_844839, EPI_ISL_844840, EPI_ISL_844841, EPI_ISL_844850, EPI_ISL_844851, EPI_ISL_844852, EPI_ISL_844853, EPI_ISL_844854, EPI_ISL_844855, EPI_ISL_844856, EPI_ISL_844857, EPI_ISL_844858, EPI_ISL_844859, EPI_ISL_844866, EPI_ISL_844867, EPI_ISL_844868, EPI_ISL_844869, EPI_ISL_844870, EPI_ISL_844871, EPI_ISL_844872, EPI_ISL_844873, EPI_ISL_844874, EPI_ISL_844875, EPI_ISL_844876, EPI_ISL_844877, EPI_ISL_844878, EPI_ISL_844879, EPI_ISL_844880, EPI_ISL_844881, EPI_ISL_844882, EPI_ISL_844883, EPI_ISL_844884, EPI_ISL_844885, EPI_ISL_844886, EPI_ISL_844887, EPI_ISL_844888, EPI_ISL_844889, EPI_ISL_844890, EPI_ISL_844891, EPI_ISL_844892, EPI_ISL_844893, EPI_ISL_844894, EPI_ISL_844895, EPI_ISL_844896, EPI_ISL_844897, EPI_ISL_844898, EPI_ISL_844899, EPI_ISL_844900, EPI_ISL_844901, EPI_ISL_844902, EPI_ISL_844903, EPI_ISL_844904, EPI_ISL_844905, EPI_ISL_844906, EPI_ISL_844907, EPI_ISL_844908, EPI_ISL_844909, EPI_ISL_844910, EPI_ISL_844913, EPI_ISL_844914, EPI_ISL_844915, EPI_ISL_844916, EPI_ISL_844917, EPI_ISL_844918, EPI_ISL_844919, EPI_ISL_844920, EPI_ISL_844921, EPI_ISL_844922, EPI_ISL_844923, EPI_ISL_844924, EPI_ISL_844925, EPI_ISL_844926, EPI_ISL_844927, EPI_ISL_844928, EPI_ISL_844929, EPI_ISL_844930, EPI_ISL_844933, EPI_ISL_844934, EPI_ISL_844935, EPI_ISL_844936, EPI_ISL_844937, EPI_ISL_844938, EPI_ISL_844939, EPI_ISL_844940, EPI_ISL_844941, EPI_ISL_844942, EPI_ISL_844943, EPI_ISL_844944, EPI_ISL_844945, EPI_ISL_844946, EPI_ISL_844947, EPI_ISL_844949, EPI_ISL_844950, EPI_ISL_844952, EPI_ISL_844959, EPI_ISL_844960, EPI_ISL_844961, EPI_ISL_844962, EPI_ISL_844963, EPI_ISL_844964, EPI_ISL_844965, EPI_ISL_844966, EPI_ISL_844967, EPI_ISL_844968, EPI_ISL_844969, EPI_ISL_844970, EPI_ISL_844971, EPI_ISL_844972, EPI_ISL_844973, EPI_ISL_844974, EPI_ISL_844975, EPI_ISL_844976, EPI_ISL_844977, EPI_ISL_844979, EPI_ISL_844980, EPI_ISL_844982, EPI_ISL_844983, EPI_ISL_844984, EPI_ISL_844985, EPI_ISL_844986, EPI_ISL_844987, EPI_ISL_844989, EPI_ISL_844990, EPI_ISL_844994, EPI_ISL_844995, EPI_ISL_844996, EPI_ISL_844997, EPI_ISL_845081, EPI_ISL_845082, EPI_ISL_845083, EPI_ISL_845084, EPI_ISL_845085, EPI_ISL_845086, EPI_ISL_845087, EPI_ISL_845088, EPI_ISL_845089, EPI_ISL_845090, EPI_ISL_845091, EPI_ISL_845092, EPI_ISL_845093, EPI_ISL_845094, EPI_ISL_845095, EPI_ISL_845096, EPI_ISL_845097, EPI_ISL_845098, EPI_ISL_845099, EPI_ISL_845100, EPI_ISL_845101, EPI_ISL_845102, EPI_ISL_845103, EPI_ISL_845104, EPI_ISL_845105, EPI_ISL_845106, EPI_ISL_845107, EPI_ISL_845108, EPI_ISL_845109, EPI_ISL_845110, EPI_ISL_845111, EPI_ISL_845112, EPI_ISL_845113, EPI_ISL_845114, EPI_ISL_845115, EPI_ISL_845116, EPI_ISL_845117, EPI_ISL_845118, EPI_ISL_845119, EPI_ISL_845120, EPI_ISL_845121, EPI_ISL_845122, EPI_ISL_845123, EPI_ISL_845124, EPI_ISL_845125, EPI_ISL_845126, EPI_ISL_845127, EPI_ISL_845128, EPI_ISL_845129, EPI_ISL_845130, EPI_ISL_845131, EPI_ISL_845132, EPI_ISL_845133, EPI_ISL_845134, EPI_ISL_845135, EPI_ISL_845136, EPI_ISL_845137, EPI_ISL_845138, EPI_ISL_845139, EPI_ISL_845140, EPI_ISL_845141, EPI_ISL_845142, EPI_ISL_845143, EPI_ISL_845144, EPI_ISL_845145, EPI_ISL_845146, EPI_ISL_845147, EPI_ISL_845148, EPI_ISL_845149, EPI_ISL_845150, EPI_ISL_845151, EPI_ISL_845152, EPI_ISL_845153, EPI_ISL_845154, EPI_ISL_845155, EPI_ISL_845156, EPI_ISL_845157, EPI_ISL_845158, EPI_ISL_845159, EPI_ISL_845160, EPI_ISL_845161, EPI_ISL_845162, EPI_ISL_845163, EPI_ISL_845164, EPI_ISL_845165, EPI_ISL_845166, EPI_ISL_845167, EPI_ISL_845168, EPI_ISL_845169, EPI_ISL_845170, EPI_ISL_845171, EPI_ISL_845172, EPI_ISL_845173, EPI_ISL_845174, EPI_ISL_845175, EPI_ISL_845176, EPI_ISL_845177, EPI_ISL_845178, EPI_ISL_845179, EPI_ISL_845180, EPI_ISL_845181, EPI_ISL_845182, EPI_ISL_845183, EPI_ISL_845184, EPI_ISL_845185, EPI_ISL_845186, EPI_ISL_845187, EPI_ISL_845188, EPI_ISL_845189, EPI_ISL_845190, EPI_ISL_845191, EPI_ISL_845192, EPI_ISL_845193, EPI_ISL_845194, EPI_ISL_845195, EPI_ISL_845196, EPI_ISL_845197, EPI_ISL_845198, EPI_ISL_845199, EPI_ISL_845200, EPI_ISL_845201, EPI_ISL_845202, EPI_ISL_845203, EPI_ISL_845204, EPI_ISL_845205, EPI_ISL_845206, EPI_ISL_845207, EPI_ISL_845208, EPI_ISL_845209, EPI_ISL_845210, EPI_ISL_845211, EPI_ISL_845212, EPI_ISL_845213, EPI_ISL_845214, EPI_ISL_845215, EPI_ISL_845216, EPI_ISL_845217, EPI_ISL_845218, EPI_ISL_845219, EPI_ISL_845220, EPI_ISL_845221, EPI_ISL_845222, EPI_ISL_845223, EPI_ISL_845224, EPI_ISL_845225, EPI_ISL_845226, EPI_ISL_845227, EPI_ISL_845228, EPI_ISL_845229, EPI_ISL_845230, EPI_ISL_845231, EPI_ISL_845232, EPI_ISL_845233, EPI_ISL_845234, EPI_ISL_845235, EPI_ISL_845236, EPI_ISL_845237, EPI_ISL_845238, EPI_ISL_845239, EPI_ISL_845240, EPI_ISL_845241, EPI_ISL_845242, EPI_ISL_845243, EPI_ISL_845244, EPI_ISL_845245, EPI_ISL_845246, EPI_ISL_845247, EPI_ISL_845248, EPI_ISL_845249, EPI_ISL_845250, EPI_ISL_845251, EPI_ISL_845252, EPI_ISL_845253, EPI_ISL_845254, EPI_ISL_845255, EPI_ISL_845256, EPI_ISL_845257, EPI_ISL_845258, EPI_ISL_845259, EPI_ISL_845260, EPI_ISL_845261, EPI_ISL_845262, EPI_ISL_845263, EPI_ISL_845264, EPI_ISL_845265, EPI_ISL_845266, EPI_ISL_845267, EPI_ISL_845268, EPI_ISL_845269, EPI_ISL_845270, EPI_ISL_845271, EPI_ISL_845272, EPI_ISL_845273, EPI_ISL_845274, EPI_ISL_845281, EPI_ISL_845282, EPI_ISL_845283, EPI_ISL_845284, EPI_ISL_845285, EPI_ISL_845286, EPI_ISL_845287, EPI_ISL_845288, EPI_ISL_845289, EPI_ISL_845290, EPI_ISL_845291, EPI_ISL_845292, EPI_ISL_845293, EPI_ISL_845294, EPI_ISL_845295, EPI_ISL_845296, EPI_ISL_845297, EPI_ISL_845298, EPI_ISL_845299, EPI_ISL_845300, EPI_ISL_845301, EPI_ISL_845302, EPI_ISL_845303, EPI_ISL_845304, EPI_ISL_845305, EPI_ISL_845306, EPI_ISL_845307, EPI_ISL_845308, EPI_ISL_845309, EPI_ISL_845310, EPI_ISL_845311, EPI_ISL_845312, EPI_ISL_845313, EPI_ISL_845314, EPI_ISL_845315, EPI_ISL_845316, EPI_ISL_845317, EPI_ISL_845318, EPI_ISL_845319, EPI_ISL_845320, EPI_ISL_845321, EPI_ISL_845322, EPI_ISL_845323, EPI_ISL_845324, EPI_ISL_845325, EPI_ISL_845326, EPI_ISL_845327, EPI_ISL_845328, EPI_ISL_845329, EPI_ISL_845330, EPI_ISL_845331, EPI_ISL_845332, EPI_ISL_845333, EPI_ISL_845334, EPI_ISL_845335, EPI_ISL_845336, EPI_ISL_845337, EPI_ISL_845338, EPI_ISL_845339, EPI_ISL_845340, EPI_ISL_845341, EPI_ISL_845342, EPI_ISL_845343, EPI_ISL_845344, EPI_ISL_845345, EPI_ISL_845346, EPI_ISL_845347, EPI_ISL_845348, EPI_ISL_845349, EPI_ISL_845350, EPI_ISL_845351, EPI_ISL_845352, EPI_ISL_845353, EPI_ISL_845354, EPI_ISL_845355, EPI_ISL_845356, EPI_ISL_845357, EPI_ISL_845358, EPI_ISL_845359, EPI_ISL_845360, EPI_ISL_845361, EPI_ISL_845362, EPI_ISL_845363, EPI_ISL_845364, EPI_ISL_845365, EPI_ISL_845366, EPI_ISL_845367, EPI_ISL_845368, EPI_ISL_845369, EPI_ISL_845370, EPI_ISL_845371, EPI_ISL_845372, EPI_ISL_845373, EPI_ISL_845374, EPI_ISL_845375, EPI_ISL_845376, EPI_ISL_845400, EPI_ISL_845401, EPI_ISL_845402, EPI_ISL_845403, EPI_ISL_845404, EPI_ISL_845405, EPI_ISL_845406, EPI_ISL_845407, EPI_ISL_845408, EPI_ISL_845409, EPI_ISL_845410, EPI_ISL_845411, EPI_ISL_845412, EPI_ISL_845413, EPI_ISL_845414, EPI_ISL_845415, EPI_ISL_845416, EPI_ISL_845417, EPI_ISL_845418, EPI_ISL_845419, EPI_ISL_845420, EPI_ISL_845421, EPI_ISL_845422, EPI_ISL_845423, EPI_ISL_845424, EPI_ISL_845425, EPI_ISL_845426, EPI_ISL_845427, EPI_ISL_845428, EPI_ISL_845429, EPI_ISL_845430, EPI_ISL_845431, EPI_ISL_845432, EPI_ISL_845433, EPI_ISL_845434, EPI_ISL_845435, EPI_ISL_845436, EPI_ISL_845437, EPI_ISL_845438, EPI_ISL_845439, EPI_ISL_845440, EPI_ISL_845441, EPI_ISL_845442, EPI_ISL_845443, EPI_ISL_845444, EPI_ISL_845445, EPI_ISL_845446, EPI_ISL_845447, EPI_ISL_845448, EPI_ISL_845449, EPI_ISL_845450, EPI_ISL_845451, EPI_ISL_845452, EPI_ISL_845453, EPI_ISL_845454, EPI_ISL_845455, EPI_ISL_845456, EPI_ISL_845457, EPI_ISL_845458, EPI_ISL_845459, EPI_ISL_845460, EPI_ISL_845461, EPI_ISL_845462, EPI_ISL_845463, EPI_ISL_845464, EPI_ISL_845465, EPI_ISL_845466, EPI_ISL_845467, EPI_ISL_845468, EPI_ISL_845469, EPI_ISL_845470, EPI_ISL_845471, EPI_ISL_845472, EPI_ISL_845473, EPI_ISL_845474, EPI_ISL_845475, EPI_ISL_845476, EPI_ISL_845477, EPI_ISL_845478, EPI_ISL_845479, EPI_ISL_845480, EPI_ISL_845481, EPI_ISL_845482, EPI_ISL_845483, EPI_ISL_845484, EPI_ISL_845485, EPI_ISL_845486, EPI_ISL_845487, EPI_ISL_845488, EPI_ISL_845489, EPI_ISL_845490, EPI_ISL_845491, EPI_ISL_845492, EPI_ISL_845493, EPI_ISL_845494, EPI_ISL_845495, EPI_ISL_845496, EPI_ISL_845497, EPI_ISL_845501, EPI_ISL_845502, EPI_ISL_845503, EPI_ISL_845504, EPI_ISL_845505, EPI_ISL_845506, EPI_ISL_845507, EPI_ISL_845508, EPI_ISL_845509, EPI_ISL_845510, EPI_ISL_845511, EPI_ISL_845512, EPI_ISL_845513, EPI_ISL_845514, EPI_ISL_845515, EPI_ISL_845516, EPI_ISL_845517, EPI_ISL_845518, EPI_ISL_845519, EPI_ISL_845520, EPI_ISL_845521, EPI_ISL_845522, EPI_ISL_845523, EPI_ISL_845524, EPI_ISL_845525, EPI_ISL_845526, EPI_ISL_845527, EPI_ISL_845528, EPI_ISL_845529, EPI_ISL_845530, EPI_ISL_845531, EPI_ISL_845532, EPI_ISL_845533, EPI_ISL_845534, EPI_ISL_845535, EPI_ISL_845536, EPI_ISL_845537, EPI_ISL_845538, EPI_ISL_845539, EPI_ISL_845540, EPI_ISL_845541, EPI_ISL_845542

| | | | |
|---|---|---|---|
| see above | Department of Virus and Microbiological Special Diagnostics, Statens Serum Institut, Copenhagen, Denmark | Albertsen Lab, Department of Chemistry and Bioscience, Aalborg University, Denmark | Danish Covid-19 Genome Consortium |

EPI_ISL_845573, EPI_ISL_845576, EPI_ISL_845579, EPI_ISL_845582, EPI_ISL_845595, EPI_ISL_845597, EPI_ISL_845599, EPI_ISL_845601, EPI_ISL_845603, EPI_ISL_845607, EPI_ISL_845609

| | | | |
|---|---|---|---|
| see above | KU Leuven, Rega Institute, Clinical and Epidemiological Virology | KU Leuven, Rega Institute, Clinical and Epidemiological Virology | Tony Wawina-Bokalanga, Bert Vanmechelen, Joan Marti-Carerras, Piet Maes |
| EPI_ISL_845652, EPI_ISL_845653 | Laboratorio de Salud Pública - Secretaría Distrital de Salud | Instituto Nacional de Salud - Dirección de Investigación en Salud Pública | Katherine Laiton-Donato, Diego A. Álvarez-Díaz, Carlos Franco-Muñoz, Mauricio Pacheco-Montealegre, Maria T. Herrera-Sepúlveda, Jonathan Reales, Sheryll Corchuelo, Julian Naizaque, Gerardo Santamaría, Paola Muñoz-Laiton, Diego Andrés Prada, Magdalena Wiesner, Martha Lucia Ospina Martinez, Marcela Mercado-Reyes |
| EPI_ISL_845760 | Emory Molecular Diagnostics Laboratory, Emory Healthcare | Piantadosi Lab, Emory Department of Pathology | Ahmed Babiker, Anne Piantadosi |

EPI_ISL_845861, EPI_ISL_845862, EPI_ISL_845863, EPI_ISL_845864, EPI_ISL_845865, EPI_ISL_845866, EPI_ISL_845867, EPI_ISL_845868, EPI_ISL_845869, EPI_ISL_845870, EPI_ISL_845871

| | | | |
|---|---|---|---|
| see above | TGen North | TGen North | Jolene Bowers, Megan Folkerts, Chris French, Hayley Yaglom, Ashlyn Pfeiffer, Darrin Lemmer, Dave Engelthaler, The Arizona COVID Genomics Union (ACGU) |
| EPI_ISL_846528, EPI_ISL_846529, EPI_ISL_846530, EPI_ISL_846531, EPI_ISL_846532, EPI_ISL_846533, EPI_ISL_846535 | Lighthouse Lab in Glasgow | Wellcome Sanger Institute for the COVID-19 Genomics UK (COG-UK) Consortium | Harper VanSteenhouse, Yumi Kasai, David Gray, Carol Clugston, Anna Dominiczak and Alex Alderton, Roberto Amato, Sonia Goncalves, Ewan Harrison, David K. Jackson, Ian Johnston, Dominic Kwiatkowski, Cordelia Langford, John Sillitoe on behalf of the Wellcome Sanger Institute COVID-19 Surveillance Team |
| EPI_ISL_846537 | Lighthouse Lab in Alderley Park | Wellcome Sanger Institute for the COVID-19 Genomics UK (COG-UK) Consortium | Jacquelyn Wynn, Mairead Hyland, The Lighthouse Lab in Alderley Park and Alex Alderton, Roberto Amato, Sonia Goncalves, Ewan Harrison, David K. Jackson, Ian Johnston, Dominic Kwiatkowski, Cordelia Langford, John Sillitoe on behalf of the Wellcome Sanger Institute COVID-19 Surveillance Team |
| EPI_ISL_846541, EPI_ISL_846552 | Lab voor klinische biologie | Lab voor klinische biologie | Hannelore Hamerlinck, Bruno Verhasselt |
| EPI_ISL_846596 | Respiratory Virus Unit, National Infection Service, Public Health England | COVID-19 Genomics UK (COG-UK) Consortium | PHE Covid Sequencing Team |
| EPI_ISL_846636 | Hopital | National Reference Center for Viruses of Respiratory Infections, Institut Pasteur, Paris | Marion Barbet, Sylvie Behillil, Méline Bizard, Angela Brisebarre, Camille Capel, Etienne Simon-Lorière, Vincent Enouf, Maud Vanpeene, Sylvie van der Werf,Farfour Eric |
| EPI_ISL_846642, EPI_ISL_846643, EPI_ISL_846645 | Labo Analyses Med | National Reference Center for Viruses of Respiratory Infections, Institut Pasteur, Paris | Marion Barbet, Sylvie Behillil, Méline Bizard, Angela Brisebarre, Camille Capel, Etienne Simon-Lorière, Vincent Enouf, Maud Vanpeene, Sylvie van der Werf |
| EPI_ISL_846646, EPI_ISL_846647 | Labo Analyses Med | National Reference Center for Viruses of Respiratory Infections, Institut Pasteur, Paris | Marion Barbet, Sylvie Behillil, Méline Bizard, Angela Brisebarre, Camille Capel, Etienne Simon-Lorière, Vincent Enouf, Maud Vanpeene, Sylvie van der Werf,Coignard Catherine |
| EPI_ISL_846650 | Hopital | National Reference Center for Viruses of Respiratory Infections, Institut Pasteur, Paris | Marion Barbet, Sylvie Behillil, Méline Bizard, Angela Brisebarre, Camille Capel, Etienne Simon-Lorière, Vincent Enouf, Maud Vanpeene, Sylvie van der Werf,Farfour Eric |

EPI_ISL_846779, EPI_ISL_846781, EPI_ISL_846812, EPI_ISL_846813, EPI_ISL_846814, EPI_ISL_846815, EPI_ISL_846816, EPI_ISL_846817, EPI_ISL_846860, EPI_ISL_846877, EPI_ISL_846884, EPI_ISL_846886, EPI_ISL_846888, EPI_ISL_846889, EPI_ISL_846893, EPI_ISL_846898, EPI_ISL_846904, EPI_ISL_846910, EPI_ISL_847125, EPI_ISL_847126, EPI_ISL_847127, EPI_ISL_847128, EPI_ISL_847129, EPI_ISL_847130, EPI_ISL_847131, EPI_ISL_847132, EPI_ISL_847133, EPI_ISL_847134, EPI_ISL_847135, EPI_ISL_847136, EPI_ISL_847137, EPI_ISL_847138, EPI_ISL_847139, EPI_ISL_847140, EPI_ISL_847141, EPI_ISL_847142, EPI_ISL_847143, EPI_ISL_847144, EPI_ISL_847145, EPI_ISL_847146, EPI_ISL_847147, EPI_ISL_847148, EPI_ISL_847149, EPI_ISL_847150, EPI_ISL_847151, EPI_ISL_847152, EPI_ISL_847153, EPI_ISL_847154, EPI_ISL_847155, EPI_ISL_847156, EPI_ISL_847157, EPI_ISL_847158, EPI_ISL_847159, EPI_ISL_847160,

| | | | |
|---|---|---|---|
| EPI_ISL_847161, EPI_ISL_847162, EPI_ISL_847163, EPI_ISL_847164, EPI_ISL_847165, EPI_ISL_847166, EPI_ISL_847167, EPI_ISL_847199, EPI_ISL_847200, EPI_ISL_847201, EPI_ISL_847202, EPI_ISL_847203, EPI_ISL_847204, EPI_ISL_847233, EPI_ISL_847234, EPI_ISL_847235, EPI_ISL_847236, EPI_ISL_847237, EPI_ISL_847244, EPI_ISL_847245, EPI_ISL_847246, EPI_ISL_847247, EPI_ISL_847248, EPI_ISL_847249, EPI_ISL_847252, EPI_ISL_847253, EPI_ISL_847254, EPI_ISL_847255, EPI_ISL_847262, EPI_ISL_847263, EPI_ISL_847264, EPI_ISL_847266, EPI_ISL_847270, EPI_ISL_847280, EPI_ISL_847332, EPI_ISL_847335, EPI_ISL_847337, EPI_ISL_847351, EPI_ISL_847352, EPI_ISL_847353, EPI_ISL_847358, EPI_ISL_847366, EPI_ISL_847377, EPI_ISL_847486, EPI_ISL_847487, EPI_ISL_847488, EPI_ISL_847489, EPI_ISL_847490, EPI_ISL_847491, EPI_ISL_847492, EPI_ISL_847493, EPI_ISL_847494, EPI_ISL_847495, EPI_ISL_847496, EPI_ISL_847497, EPI_ISL_847498, EPI_ISL_847499, EPI_ISL_847500, EPI_ISL_847501, EPI_ISL_847502, EPI_ISL_847503, EPI_ISL_847504, EPI_ISL_847505, EPI_ISL_847506, EPI_ISL_847507, EPI_ISL_847508, EPI_ISL_847509 | | | |
| see above | Department of Virus and Microbiological Special Diagnostics, Statens Serum Institut, Copenhagen, Denmark | Albertsen Lab, Department of Chemistry and Bioscience, Aalborg University, Denmark | Danish Covid-19 Genome Consortium |
| EPI_ISL_848003 | Michigan Department of Health and Human Services, Bureau of Laboratories | Michigan Department of Health and Human Services, Bureau of Laboratories | Blankenship HM, Riner D, Soehnlen MK |
| EPI_ISL_849336, EPI_ISL_849337, EPI_ISL_849338, EPI_ISL_849339, EPI_ISL_849340, EPI_ISL_849341, EPI_ISL_849342, EPI_ISL_849343, EPI_ISL_849344, EPI_ISL_849345, EPI_ISL_849346, EPI_ISL_849347, EPI_ISL_849348, EPI_ISL_849349 | | | |
| see above | Delaware Public Health Lab | Delaware Public Health Lab | Gregory Hovan |
| EPI_ISL_849778, EPI_ISL_849779, EPI_ISL_849781, EPI_ISL_849782, EPI_ISL_849784, EPI_ISL_849785, EPI_ISL_849786, EPI_ISL_849789, EPI_ISL_849791 | Utah Public Health Laboratory | Utah Public Health Laboratory | Erin L. Young, Kelly F. Oakeson, Tara Gallagher |
| EPI_ISL_849922 | Orange County Public Health Lab | Chan-Zuckerberg Biohub | CZB Cliahub Consortium |
| EPI_ISL_850517, EPI_ISL_850518 | Helix/Illumina | Genomics and Discovery, Respiratory Viruses Branch, Division of Viral Diseases, Centers for Disease Control and Prevention | Peter W. Cook, Dhwani Batra, Ben L. Rambo-Martin Eileen de Feo, Jan Antico, Christine Tran, Matthew Tolentino, Shannon Wickline, Kim Gietzen, Brad Sickler, Jingtao Liu, Eric Allen, Phil Febbo, Summer Galloway, Nicole L. Washington, Simon White, Geraint Levan, Kelly Schiabor Barrett, Elizabeth Cirulli, Alexandre Bolze, Ary Ascencio, Charlotte Rivera-Garcia, Ryan Cho, Jason Nguyen, Sherry Wang, Jimmy Ramirez, Tyler Cassens, Efren Sandoval, Magnus Isaksson, William Lee, David Becker, Marc Laurent, James Lu, Clinton R. Paden, Suxiang Tong, Duncan MacCannell |
| EPI_ISL_850673 | The National Institute of Public Health | State Veterinary Institute Prague | Nagy,A;Jirincova,H;Trnka,D;Vecerova,J;Trinklova,M |
| EPI_ISL_850678, EPI_ISL_850680, EPI_ISL_850681, EPI_ISL_850682, EPI_ISL_850683 | The National Institute of Public Health | State Veterinary Institute Prague | Nagy,A;Jirincova,H;Trnka,D;Vecerova,J |
| EPI_ISL_850817, EPI_ISL_850818, EPI_ISL_850819, EPI_ISL_850820, EPI_ISL_850821, EPI_ISL_850822, EPI_ISL_850823, EPI_ISL_850824, EPI_ISL_850825, EPI_ISL_850826, EPI_ISL_850827, EPI_ISL_850828, EPI_ISL_850829, EPI_ISL_850830, EPI_ISL_850831, EPI_ISL_850832, EPI_ISL_850833, EPI_ISL_850834, EPI_ISL_850835, EPI_ISL_850836, EPI_ISL_850837, EPI_ISL_850838, EPI_ISL_850839, EPI_ISL_850840, EPI_ISL_850841, EPI_ISL_850842, EPI_ISL_850843, EPI_ISL_850844, EPI_ISL_850845, EPI_ISL_850846, EPI_ISL_850847, EPI_ISL_850848, EPI_ISL_850849, EPI_ISL_850850, EPI_ISL_850851, EPI_ISL_850852, EPI_ISL_850853, EPI_ISL_850854, EPI_ISL_850855, EPI_ISL_850856, EPI_ISL_850857, EPI_ISL_850858, EPI_ISL_850859, EPI_ISL_850860, EPI_ISL_850861, EPI_ISL_850862, EPI_ISL_850863, EPI_ISL_850864, EPI_ISL_850865, EPI_ISL_850866, EPI_ISL_850867, EPI_ISL_850868, EPI_ISL_850869, EPI_ISL_850870, EPI_ISL_850871, EPI_ISL_850872, EPI_ISL_850873, EPI_ISL_850874, EPI_ISL_850875, EPI_ISL_850876, EPI_ISL_850877, EPI_ISL_850878, EPI_ISL_850879, EPI_ISL_850880, EPI_ISL_850881, EPI_ISL_850882, EPI_ISL_850883, EPI_ISL_850884, EPI_ISL_850885, EPI_ISL_850886, EPI_ISL_850887, EPI_ISL_850888, EPI_ISL_850889, EPI_ISL_850890, EPI_ISL_850891, EPI_ISL_850892, EPI_ISL_850893 | | | |
| see above | Helix / Illumina | Genomics and Discovery, Respiratory Viruses Branch, Division of Viral Diseases, Centers for Disease Control and Prevention | Peter W. Cook, Dhwani Batra, Ben L. Rambo-Martin Eileen de Feo, Jan Antico, Christine Tran, Matthew Tolentino, Shannon Wickline, Kim Gietzen, Brad Sickler, Jingtao Liu, Eric Allen, Phil Febbo, Summer Galloway, Nicole L. Washington, Simon White, Geraint Levan, Kelly Schiabor Barrett, Elizabeth Cirulli, Alexandre Bolze, Ary Ascencio, Charlotte Rivera-Garcia, Ryan Cho, Jason Nguyen, Sherry Wang, Jimmy Ramirez, Tyler Cassens, Efren Sandoval, Magnus Isaksson, William Lee, David Becker, Marc Laurent, James Lu, Clinton R. Paden, Suxiang Tong, Duncan MacCannell |
| EPI_ISL_850952, EPI_ISL_850953, EPI_ISL_850954, EPI_ISL_850955, EPI_ISL_850956, EPI_ISL_850957, EPI_ISL_850958, EPI_ISL_850959, EPI_ISL_850960 | Helix/Illumina | Genomics and Discovery, Respiratory Viruses Branch, Division of Viral Diseases, Centers for Disease Control and Prevention | Peter W. Cook, Dhwani Batra, Ben L. Rambo-Martin Eileen de Feo, Jan Antico, Christine Tran, Matthew Tolentino, Shannon Wickline, Kim Gietzen, Brad Sickler, Jingtao Liu, Eric Allen, Phil Febbo, Summer Galloway, Nicole L. Washington, Simon White, Geraint Levan, Kelly Schiabor Barrett, Elizabeth Cirulli, Alexandre Bolze, Ary Ascencio, Charlotte Rivera-Garcia, Ryan Cho, Jason Nguyen, Sherry Wang, Jimmy Ramirez, Tyler Cassens, Efren Sandoval, Magnus Isaksson, William Lee, David Becker, Marc Laurent, James Lu, Clinton R. Paden, Suxiang Tong, Duncan MacCannell |
| EPI_ISL_852383 | Lighthouse Lab in Glasgow | Wellcome Sanger Institute for the COVID-19 Genomics UK (COG-UK) Consortium | Harper VanSteenhouse, Yumi Kasai, David Gray, Carol Clugston, Anna Dominiczak and Alex Alderton, Roberto Amato, Sonia Goncalves, Ewan Harrison, David K. Jackson, Ian Johnston, Dominic Kwiatkowski, Cordelia Langford, John Sillitoe on behalf of the Wellcome Sanger Institute COVID-19 Surveillance Team |
| EPI_ISL_852858, EPI_ISL_852859, EPI_ISL_852860 | Florida Bureau of Public Health Laboratories | Florida Bureau of Public Health Laboratories | Sarah Schmedes, Jason Blanton |
| EPI_ISL_853004, EPI_ISL_853005 | Hospital General Universitario Gregorio Marañón | SeqCOVID-SPAIN consortium/IBV(CSIC) | Darío García de Viedma, Laura Pérez-Lago, Pedro J Sola-Campoy, Sergio Buenestado-Serrano, Marta Herranz, Victor Manuel de la Cueva, Julia Suárez, Pilar Catalán, Patricia Muñoz and SeqCOVID-SPAIN consortium |
| EPI_ISL_853648, EPI_ISL_853668, EPI_ISL_853669 | THE MARY IMOGENE BASSETT HOSPITAL | Wadsworth Center, New York State Department of Health | Kirsten St. George, Daryl M. Lamson, Alexis Russel, Matthew Shudt, Melissa A Leisner, Jonathan Plitnick, Navjot Singh, John Kelly, Erasmus Schneider, Erica Lasek-Nesselquist |
| EPI_ISL_853670, EPI_ISL_853671, EPI_ISL_853672, EPI_ISL_853673, EPI_ISL_853674, EPI_ISL_853675 | NORTHWELL HEALTH LABORATORIES | Wadsworth Center, New York State Department of Health | Kirsten St. George, Daryl M. Lamson, Alexis Russel, Matthew Shudt, Melissa A Leisner, Jonathan Plitnick, Navjot Singh, John Kelly, Erasmus Schneider, Erica Lasek-Nesselquist |
| EPI_ISL_853676, EPI_ISL_853677, EPI_ISL_853678, EPI_ISL_853679, EPI_ISL_853680, EPI_ISL_853681, EPI_ISL_853682 | ACUTIS DIAGNOSTICS | Wadsworth Center, New York State Department of Health | Kirsten St. George, Daryl M. Lamson, Alexis Russel, Matthew Shudt, Melissa A Leisner, Jonathan Plitnick, Navjot Singh, John Kelly, Erasmus Schneider, Erica Lasek-Nesselquist |
| EPI_ISL_853712 | NORTHWELL HEALTH LABORATORIES | Wadsworth Center, New York State Department of Health | Kirsten St. George, Daryl M. Lamson, Alexis Russel, Matthew Shudt, Melissa A Leisner, Jonathan Plitnick, Navjot Singh, John Kelly, Erasmus Schneider, Erica Lasek-Nesselquist |
| EPI_ISL_854306, EPI_ISL_854321, EPI_ISL_854322, EPI_ISL_854323, EPI_ISL_854324, EPI_ISL_854325, EPI_ISL_854326, EPI_ISL_854327, EPI_ISL_854328, EPI_ISL_854329, EPI_ISL_854330, EPI_ISL_854331, EPI_ISL_854332, EPI_ISL_854333, EPI_ISL_854334, EPI_ISL_854335, EPI_ISL_854336, EPI_ISL_854337, EPI_ISL_854338, EPI_ISL_854339, EPI_ISL_854340, EPI_ISL_854341, EPI_ISL_854342, EPI_ISL_854343, EPI_ISL_854344, EPI_ISL_854345, EPI_ISL_854346, EPI_ISL_854347, EPI_ISL_854348, EPI_ISL_854349, EPI_ISL_854370, EPI_ISL_854371, EPI_ISL_854404 | | | |
| see above | URMC LABS | Wadsworth Center, New York State Department of Health | Kirsten St. George, Daryl M. Lamson, Alexis Russel, Matthew Shudt, Melissa A Leisner, Jonathan Plitnick, Navjot Singh, John Kelly, Erasmus Schneider, Erica Lasek-Nesselquist |
| EPI_ISL_854843, EPI_ISL_854850, EPI_ISL_854862, EPI_ISL_854867, EPI_ISL_854885, EPI_ISL_854890, EPI_ISL_854892, EPI_ISL_854898, EPI_ISL_854943, EPI_ISL_854963, EPI_ISL_854964, EPI_ISL_854965, EPI_ISL_854967, EPI_ISL_855115, EPI_ISL_855129, EPI_ISL_855164 | | | |
| see above | Quest Diagnostics | Quest Diagnostics | Rosenthal,S.H., Gerasimova,A., Kagan,R.M., Anderson, B., Hua, M., Liu Y., Bernstein, L.E., Livingston, K.E., Perez, A., Shalhout, D.F., Shlyakhter, I.A., Owen, R., Tanpaiboon, P., Lacbawan, F. |
| EPI_ISL_855386 | Hospital | National Reference Center for Viruses of Respiratory Infections, Institut Pasteur, Paris | Marion Barbet, Sylvie Behillil, Méline Bizard, Angela Brisebarre, Camille Capel, Etienne Simon-Lorière, Vincent Enouf, Maud Vanpeene, Sylvie van der Werf |
| EPI_ISL_855588 | Respiratory Virus Unit, National Infection Service, Public Health England | COVID-19 Genomics UK (COG-UK) Consortium | PHE Covid Sequencing Team |
| EPI_ISL_855949, EPI_ISL_855951, EPI_ISL_855953, EPI_ISL_855955, EPI_ISL_855957, EPI_ISL_855959, EPI_ISL_855960, EPI_ISL_855962, EPI_ISL_855964, EPI_ISL_855966, EPI_ISL_855968 | | | |
| see above | Lab voor klinische biologie | Onderzoeksgroep Virologie | Nick Vereecke, Laurens Lambrechts, Marthe Pauwels, Bruno Verhasselt, Linos Vandekerckhove, Hans Nauwynck, Sebastiaan Theuns |
| EPI_ISL_856292, EPI_ISL_856293, EPI_ISL_856294, EPI_ISL_856295, EPI_ISL_856296, EPI_ISL_856297, EPI_ISL_856298, EPI_ISL_856299, EPI_ISL_856300, EPI_ISL_856301, EPI_ISL_856302, EPI_ISL_856303, EPI_ISL_856304, EPI_ISL_856305, EPI_ISL_856306, EPI_ISL_856307, EPI_ISL_856308, EPI_ISL_856309, EPI_ISL_856310, EPI_ISL_856311, EPI_ISL_856312, EPI_ISL_856313, EPI_ISL_856314, EPI_ISL_856315, EPI_ISL_856316, EPI_ISL_856317, EPI_ISL_856318, EPI_ISL_856319, EPI_ISL_856320, EPI_ISL_856321, EPI_ISL_856322, EPI_ISL_856323, EPI_ISL_856324, EPI_ISL_856325, EPI_ISL_856326, EPI_ISL_856327, EPI_ISL_856328, EPI_ISL_856329, EPI_ISL_856330, EPI_ISL_856331, EPI_ISL_856332, EPI_ISL_856333, EPI_ISL_856334, EPI_ISL_856335, EPI_ISL_856336, EPI_ISL_856337, EPI_ISL_856338, EPI_ISL_856339, EPI_ISL_856340, EPI_ISL_856341, EPI_ISL_856342, EPI_ISL_856343, EPI_ISL_856344, EPI_ISL_856345, EPI_ISL_856346, EPI_ISL_856347, EPI_ISL_856348, EPI_ISL_856349, EPI_ISL_856350, EPI_ISL_856351, EPI_ISL_856352, EPI_ISL_856353, EPI_ISL_856354, EPI_ISL_856355, EPI_ISL_856356, EPI_ISL_856357, EPI_ISL_856358, EPI_ISL_856359, EPI_ISL_856360, EPI_ISL_856361, EPI_ISL_856362, EPI_ISL_856363, | | | |

| | | | |
|---|---|---|---|
| EPI_ISL_856364, EPI_ISL_856365, EPI_ISL_856366, EPI_ISL_856367, EPI_ISL_856368, EPI_ISL_856369, EPI_ISL_856370, EPI_ISL_856371, EPI_ISL_856372, EPI_ISL_856373, EPI_ISL_856374, EPI_ISL_856375, EPI_ISL_856376, EPI_ISL_856377, EPI_ISL_856378, EPI_ISL_856379, EPI_ISL_856380, EPI_ISL_856381, EPI_ISL_856382, EPI_ISL_856383, EPI_ISL_856384, EPI_ISL_856385, EPI_ISL_856386, EPI_ISL_856387, EPI_ISL_856388, EPI_ISL_856389, EPI_ISL_856390, EPI_ISL_856391, EPI_ISL_856392, EPI_ISL_856393, EPI_ISL_856394, EPI_ISL_856395, EPI_ISL_856396, EPI_ISL_856397, EPI_ISL_856398, EPI_ISL_856399, EPI_ISL_856400, EPI_ISL_856401, EPI_ISL_856402, EPI_ISL_856403, EPI_ISL_856404, EPI_ISL_856405, EPI_ISL_856406, EPI_ISL_856407, EPI_ISL_856408, EPI_ISL_856409, EPI_ISL_856410, EPI_ISL_856411, EPI_ISL_856412, EPI_ISL_856413, EPI_ISL_856414, EPI_ISL_856415, EPI_ISL_856416, EPI_ISL_856417, EPI_ISL_856418, EPI_ISL_856419, EPI_ISL_856420, EPI_ISL_856421, EPI_ISL_856422, EPI_ISL_856423, EPI_ISL_856424, EPI_ISL_856425, EPI_ISL_856426, EPI_ISL_856427, EPI_ISL_856428, EPI_ISL_856429, EPI_ISL_856430, EPI_ISL_856431, EPI_ISL_856432, EPI_ISL_856433, EPI_ISL_856434, EPI_ISL_856435, EPI_ISL_856436, EPI_ISL_856437, EPI_ISL_856438, EPI_ISL_856439, EPI_ISL_856440, EPI_ISL_856441, EPI_ISL_856442, EPI_ISL_856443, EPI_ISL_856444, EPI_ISL_856445, EPI_ISL_856446, EPI_ISL_856447, EPI_ISL_856448, EPI_ISL_856449, EPI_ISL_856450, EPI_ISL_856451, EPI_ISL_856452, EPI_ISL_856453, EPI_ISL_856454, EPI_ISL_856455, EPI_ISL_856456, EPI_ISL_856457, EPI_ISL_856458, EPI_ISL_856459, EPI_ISL_856460, EPI_ISL_856461, EPI_ISL_856462, EPI_ISL_856463, EPI_ISL_856464, EPI_ISL_856465, EPI_ISL_856466, EPI_ISL_856467, EPI_ISL_856468, EPI_ISL_856469, EPI_ISL_856470, EPI_ISL_856471, EPI_ISL_856472, EPI_ISL_856473, EPI_ISL_856474, EPI_ISL_856475, EPI_ISL_856476, EPI_ISL_856477, EPI_ISL_856478, EPI_ISL_856479, EPI_ISL_856480, EPI_ISL_856481, EPI_ISL_856482, EPI_ISL_856483, EPI_ISL_856484, EPI_ISL_856485, EPI_ISL_856486, EPI_ISL_856487, EPI_ISL_856488, EPI_ISL_856489, EPI_ISL_856490, EPI_ISL_856491, EPI_ISL_856492, EPI_ISL_856493, EPI_ISL_856494, EPI_ISL_856495, EPI_ISL_856496, EPI_ISL_856497, EPI_ISL_856498, EPI_ISL_856499, EPI_ISL_856500, EPI_ISL_856501, EPI_ISL_856502, EPI_ISL_856503, EPI_ISL_856504, EPI_ISL_856505, EPI_ISL_856506, EPI_ISL_856507, EPI_ISL_856508, EPI_ISL_856509, EPI_ISL_856510, EPI_ISL_856511, EPI_ISL_856512, EPI_ISL_856513, EPI_ISL_856514, EPI_ISL_856515, EPI_ISL_856516, EPI_ISL_856517, EPI_ISL_856518, EPI_ISL_856519, EPI_ISL_856520, EPI_ISL_856521, EPI_ISL_856522, EPI_ISL_856523, EPI_ISL_856524, EPI_ISL_856525, EPI_ISL_856526, EPI_ISL_856527, EPI_ISL_856528, EPI_ISL_856529, EPI_ISL_856530, EPI_ISL_856531, EPI_ISL_856532, EPI_ISL_856533, EPI_ISL_856534, EPI_ISL_856535, EPI_ISL_856536, EPI_ISL_856537, EPI_ISL_856538, EPI_ISL_856539, EPI_ISL_856540, EPI_ISL_856541, EPI_ISL_856542, EPI_ISL_856543, EPI_ISL_856544, EPI_ISL_856545, EPI_ISL_856546, EPI_ISL_856547, EPI_ISL_856548, EPI_ISL_856549, EPI_ISL_856550, EPI_ISL_856551, EPI_ISL_856552, EPI_ISL_856553, EPI_ISL_856554, EPI_ISL_856555, EPI_ISL_856556, EPI_ISL_856557, EPI_ISL_856558, EPI_ISL_856559, EPI_ISL_856560, EPI_ISL_856561, EPI_ISL_856562, EPI_ISL_856563, EPI_ISL_856564, EPI_ISL_856565, EPI_ISL_856566, EPI_ISL_856567, EPI_ISL_856568, EPI_ISL_856569, EPI_ISL_856570, EPI_ISL_856571, EPI_ISL_856572, EPI_ISL_856573, EPI_ISL_856574, EPI_ISL_856575, EPI_ISL_856576, EPI_ISL_856577, EPI_ISL_856578, EPI_ISL_856579, EPI_ISL_856580, EPI_ISL_856581, EPI_ISL_856582, EPI_ISL_856583, EPI_ISL_856584, EPI_ISL_856585, EPI_ISL_856586, EPI_ISL_856587, EPI_ISL_856588, EPI_ISL_856589, EPI_ISL_856590, EPI_ISL_856591, EPI_ISL_856592, EPI_ISL_856593, EPI_ISL_856594, EPI_ISL_856595, EPI_ISL_856596, EPI_ISL_856597, EPI_ISL_856598, EPI_ISL_856599, EPI_ISL_856600, EPI_ISL_856601, EPI_ISL_856602, EPI_ISL_856603, EPI_ISL_856604, EPI_ISL_856605, EPI_ISL_856606, EPI_ISL_856607, EPI_ISL_856608, EPI_ISL_856609, EPI_ISL_856610, EPI_ISL_856611 | | | |
| see above | Department of Virus and Microbiological Special Diagnostics, Statens Serum Institut, Copenhagen, Denmark | Aalborg University | Danish Covid-19 Genome Consortium |
| EPI_ISL_856934, EPI_ISL_856938, EPI_ISL_856939, EPI_ISL_856940, EPI_ISL_856943, EPI_ISL_856944, EPI_ISL_856961, EPI_ISL_856962, EPI_ISL_856963, EPI_ISL_856964, EPI_ISL_856965, EPI_ISL_856966, EPI_ISL_856967 | | | |
| see above | Wyoming Public Health Laboratory | Wyoming Public Health Laboratory | Noah Hull, Taylor Fearing, Lynette Gumbleton, Channing Weber, Ashley Norberg, Bailey Bowcutt, and Wanda Manley |
| EPI_ISL_857046 | Colorado Department of Public Health and Environment | Colorado Department of Puplic Health and Environment | Laura Bankers, Molly C. Hetherington-Rauth, Diana Ir, Shannon Ely, Shannon R. Matzinger, Sarah Elizabeth Totten, Emily A. Travanty |
| EPI_ISL_857062 | OCME Office Of Chief Medical Examiner | New York City Public Health Laboratory | Jade Wang, et al. |
| EPI_ISL_857063, EPI_ISL_857064, EPI_ISL_857065 | DOHMH PHL | New York City Public Health Laboratory | Jade Wang, et al. |
| EPI_ISL_857066 | OCME Office Of Chief Medical Examiner | New York City Public Health Laboratory | Jade Wang, et al. |
| EPI_ISL_857194, EPI_ISL_857200 | DOHMH Central Harlem | New York City Public Health Laboratory | Jade Wang, et al. |
| EPI_ISL_857201 | DOHMH Jamaica | New York City Public Health Laboratory | Jade Wang, et al. |
| EPI_ISL_857203 | DOHMH Riverside | New York City Public Health Laboratory | Jade Wang, et al. |
| EPI_ISL_857206 | DOHMH PHL | New York City Public Health Laboratory | Jade Wang, et al. |
| EPI_ISL_857207, EPI_ISL_857208 | DOHMH Morrisania | New York City Public Health Laboratory | Jade Wang, et al. |
| EPI_ISL_857209 | DOHMH Central Harlem | New York City Public Health Laboratory | Jade Wang, et al. |
| EPI_ISL_857210, EPI_ISL_857211, EPI_ISL_857212, EPI_ISL_857213, EPI_ISL_857214, EPI_ISL_857215, EPI_ISL_857216, EPI_ISL_857217 | DOHMH Jamaica | New York City Public Health Laboratory | Jade Wang, et al. |
| EPI_ISL_857218, EPI_ISL_857219, EPI_ISL_857220 | DOHMH Corona | New York City Public Health Laboratory | Jade Wang, et al. |
| EPI_ISL_857221 | DOHMH Fort Greene | New York City Public Health Laboratory | Jade Wang, et al. |
| EPI_ISL_857222, EPI_ISL_857223, EPI_ISL_857224 | DOHMH Jamaica | New York City Public Health Laboratory | Jade Wang, et al. |
| EPI_ISL_857225 | DOHMH Corona | New York City Public Health Laboratory | Jade Wang, et al. |
| EPI_ISL_857226 | DOHMH Jamaica | New York City Public Health Laboratory | Jade Wang, et al. |
| EPI_ISL_858086, EPI_ISL_858122, EPI_ISL_858136, EPI_ISL_858143, EPI_ISL_858144, EPI_ISL_858146, EPI_ISL_858147, EPI_ISL_858149, EPI_ISL_858150, EPI_ISL_858152, EPI_ISL_858153, EPI_ISL_858154, EPI_ISL_858155, EPI_ISL_858156, EPI_ISL_858157 | | | |
| see above | Lighthouse Lab in Alderley Park | Wellcome Sanger Institute for the COVID-19 Genomics UK (COG-UK) Consortium | Jacquelyn Wynn, Mairead Hyland, The Lighthouse Lab in Alderley Park and Alex Alderton, Roberto Amato, Sonia Goncalves, Ewan Harrison, David K. Jackson, Ian Johnston, Dominic Kwiatkowski, Cordelia Langford, John Sillitoe on behalf of the Wellcome Sanger Institute COVID-19 Surveillance Team |
| EPI_ISL_858160 | Lighthouse Lab in Glasgow | Wellcome Sanger Institute for the COVID-19 Genomics UK (COG-UK) Consortium | Harper VanSteenhouse, Yumi Kasai, David Gray, Carol Clugston, Anna Dominiczak and Alex Alderton, Roberto Amato, Sonia Goncalves, Ewan Harrison, David K. Jackson, Ian Johnston, Dominic Kwiatkowski, Cordelia Langford, John Sillitoe on behalf of the Wellcome Sanger Institute COVID-19 Surveillance Team |
| EPI_ISL_858216 | Lighthouse Lab in Alderley Park | Wellcome Sanger Institute for the COVID-19 Genomics UK (COG-UK) Consortium | Jacquelyn Wynn, Mairead Hyland, The Lighthouse Lab in Alderley Park and Alex Alderton, Roberto Amato, Sonia Goncalves, Ewan Harrison, David K. Jackson, Ian Johnston, Dominic Kwiatkowski, Cordelia Langford, John Sillitoe on behalf of the Wellcome Sanger Institute COVID-19 Surveillance Team |
| EPI_ISL_858241, EPI_ISL_858246, EPI_ISL_858263, EPI_ISL_858272, EPI_ISL_858278, EPI_ISL_858285 | Lighthouse Lab in Glasgow | Wellcome Sanger Institute for the COVID-19 Genomics UK (COG-UK) Consortium | Harper VanSteenhouse, Yumi Kasai, David Gray, Carol Clugston, Anna Dominiczak and Alex Alderton, Roberto Amato, Sonia Goncalves, Ewan Harrison, David K. Jackson, Ian Johnston, Dominic Kwiatkowski, Cordelia Langford, John Sillitoe on behalf of the Wellcome Sanger Institute COVID-19 Surveillance Team |
| EPI_ISL_858287 | Lighthouse Lab in Alderley Park | Wellcome Sanger Institute for the COVID-19 Genomics UK (COG-UK) Consortium | Jacquelyn Wynn, Mairead Hyland, The Lighthouse Lab in Alderley Park and Alex Alderton, Roberto Amato, Sonia Goncalves, Ewan Harrison, David K. Jackson, Ian Johnston, Dominic Kwiatkowski, Cordelia Langford, John Sillitoe on behalf of the Wellcome Sanger Institute COVID-19 Surveillance Team |
| EPI_ISL_858290, EPI_ISL_858303 | Lighthouse Lab in Glasgow | Wellcome Sanger Institute for the COVID-19 Genomics UK (COG-UK) Consortium | Harper VanSteenhouse, Yumi Kasai, David Gray, Carol Clugston, Anna Dominiczak and Alex Alderton, Roberto Amato, Sonia Goncalves, Ewan Harrison, David K. Jackson, Ian Johnston, Dominic Kwiatkowski, Cordelia Langford, John Sillitoe on behalf of the Wellcome Sanger Institute COVID-19 Surveillance Team |
| EPI_ISL_858307, EPI_ISL_858308, EPI_ISL_858309 | Lighthouse Lab in Alderley Park | Wellcome Sanger Institute for the COVID-19 Genomics UK (COG-UK) Consortium | Jacquelyn Wynn, Mairead Hyland, The Lighthouse Lab in Alderley Park and Alex Alderton, Roberto Amato, Sonia Goncalves, Ewan Harrison, David K. Jackson, Ian Johnston, Dominic Kwiatkowski, Cordelia Langford, John Sillitoe on behalf of the Wellcome Sanger Institute COVID-19 Surveillance Team |
| EPI_ISL_858310, EPI_ISL_858312, EPI_ISL_858318 | Lighthouse Lab in Glasgow | Wellcome Sanger Institute for the COVID-19 Genomics UK (COG-UK) Consortium | Harper VanSteenhouse, Yumi Kasai, David Gray, Carol Clugston, Anna Dominiczak and Alex Alderton, Roberto Amato, Sonia Goncalves, Ewan Harrison, David K. Jackson, Ian Johnston, Dominic Kwiatkowski, Cordelia Langford, John Sillitoe on behalf of the Wellcome Sanger Institute COVID-19 Surveillance Team |
| EPI_ISL_858324, EPI_ISL_858325, EPI_ISL_858329 | Lighthouse Lab in Alderley Park | Wellcome Sanger Institute for the COVID-19 Genomics UK (COG-UK) Consortium | Jacquelyn Wynn, Mairead Hyland, The Lighthouse Lab in Alderley Park and Alex Alderton, Roberto Amato, Sonia Goncalves, Ewan Harrison, David K. Jackson, Ian Johnston, Dominic Kwiatkowski, Cordelia Langford, John Sillitoe on behalf of the Wellcome Sanger Institute COVID-19 Surveillance Team |
| EPI_ISL_858330 | Lighthouse Lab in Glasgow | Wellcome Sanger Institute for the COVID-19 Genomics UK (COG-UK) Consortium | Harper VanSteenhouse, Yumi Kasai, David Gray, Carol Clugston, Anna Dominiczak and Alex Alderton, Roberto Amato, Sonia Goncalves, Ewan Harrison, David K. Jackson, Ian Johnston, Dominic Kwiatkowski, Cordelia Langford, John Sillitoe on behalf of the Wellcome Sanger Institute COVID-19 Surveillance Team |
| EPI_ISL_858332, EPI_ISL_858334 | Lighthouse Lab in Alderley Park | Wellcome Sanger Institute for the COVID-19 Genomics UK (COG-UK) Consortium | Jacquelyn Wynn, Mairead Hyland, The Lighthouse Lab in Alderley Park and Alex Alderton, Roberto Amato, Sonia Goncalves, Ewan Harrison, David K. Jackson, Ian Johnston, Dominic Kwiatkowski, Cordelia Langford, John Sillitoe on behalf of the Wellcome Sanger Institute COVID-19 Surveillance Team |
| EPI_ISL_858339, EPI_ISL_858341 | Lighthouse Lab in Glasgow | Wellcome Sanger Institute for the COVID-19 Genomics UK (COG-UK) Consortium | Harper VanSteenhouse, Yumi Kasai, David Gray, Carol Clugston, Anna Dominiczak and Alex Alderton, Roberto Amato, Sonia Goncalves, Ewan Harrison, David K. Jackson, Ian Johnston, Dominic Kwiatkowski, Cordelia Langford, John Sillitoe on behalf of the Wellcome Sanger Institute COVID-19 Surveillance Team |

| | | | |
|---|---|---|---|
| EPI_ISL_858343 | Lighthouse Lab in Alderley Park | Wellcome Sanger Institute for the COVID-19 Genomics UK (COG-UK) Consortium | Jacquelyn Wynn, Mairead Hyland, The Lighthouse Lab in Alderley Park and Alex Alderton, Roberto Amato, Sonia Goncalves, Ewan Harrison, David K. Jackson, Ian Johnston, Dominic Kwiatkowski, Cordelia Langford, John Sillitoe on behalf of the Wellcome Sanger Institute COVID-19 Surveillance Team |
| EPI_ISL_858350, EPI_ISL_858351 | Lighthouse Lab in Glasgow | Wellcome Sanger Institute for the COVID-19 Genomics UK (COG-UK) Consortium | Harper VanSteenhouse, Yumi Kasai, David Gray, Carol Clugston, Anna Dominiczak and Alex Alderton, Roberto Amato, Sonia Goncalves, Ewan Harrison, David K. Jackson, Ian Johnston, Dominic Kwiatkowski, Cordelia Langford, John Sillitoe on behalf of the Wellcome Sanger Institute COVID-19 Surveillance Team |
| EPI_ISL_858354 | Lighthouse Lab in Alderley Park | Wellcome Sanger Institute for the COVID-19 Genomics UK (COG-UK) Consortium | Jacquelyn Wynn, Mairead Hyland, The Lighthouse Lab in Alderley Park and Alex Alderton, Roberto Amato, Sonia Goncalves, Ewan Harrison, David K. Jackson, Ian Johnston, Dominic Kwiatkowski, Cordelia Langford, John Sillitoe on behalf of the Wellcome Sanger Institute COVID-19 Surveillance Team |
| EPI_ISL_858363 | Lighthouse Lab in Glasgow | Wellcome Sanger Institute for the COVID-19 Genomics UK (COG-UK) Consortium | Harper VanSteenhouse, Yumi Kasai, David Gray, Carol Clugston, Anna Dominiczak and Alex Alderton, Roberto Amato, Sonia Goncalves, Ewan Harrison, David K. Jackson, Ian Johnston, Dominic Kwiatkowski, Cordelia Langford, John Sillitoe on behalf of the Wellcome Sanger Institute COVID-19 Surveillance Team |
| EPI_ISL_858368, EPI_ISL_858370 | Lighthouse Lab in Alderley Park | Wellcome Sanger Institute for the COVID-19 Genomics UK (COG-UK) Consortium | Jacquelyn Wynn, Mairead Hyland, The Lighthouse Lab in Alderley Park and Alex Alderton, Roberto Amato, Sonia Goncalves, Ewan Harrison, David K. Jackson, Ian Johnston, Dominic Kwiatkowski, Cordelia Langford, John Sillitoe on behalf of the Wellcome Sanger Institute COVID-19 Surveillance Team |
| EPI_ISL_858371 | Lighthouse Lab in Glasgow | Wellcome Sanger Institute for the COVID-19 Genomics UK (COG-UK) Consortium | Harper VanSteenhouse, Yumi Kasai, David Gray, Carol Clugston, Anna Dominiczak and Alex Alderton, Roberto Amato, Sonia Goncalves, Ewan Harrison, David K. Jackson, Ian Johnston, Dominic Kwiatkowski, Cordelia Langford, John Sillitoe on behalf of the Wellcome Sanger Institute COVID-19 Surveillance Team |
| EPI_ISL_858375 | Lighthouse Lab in Alderley Park | Wellcome Sanger Institute for the COVID-19 Genomics UK (COG-UK) Consortium | Jacquelyn Wynn, Mairead Hyland, The Lighthouse Lab in Alderley Park and Alex Alderton, Roberto Amato, Sonia Goncalves, Ewan Harrison, David K. Jackson, Ian Johnston, Dominic Kwiatkowski, Cordelia Langford, John Sillitoe on behalf of the Wellcome Sanger Institute COVID-19 Surveillance Team |
| EPI_ISL_858378, EPI_ISL_858385, EPI_ISL_858386, EPI_ISL_858391, EPI_ISL_858392, EPI_ISL_858395, EPI_ISL_858398, EPI_ISL_858399 | Lighthouse Lab in Glasgow | Wellcome Sanger Institute for the COVID-19 Genomics UK (COG-UK) Consortium | Harper VanSteenhouse, Yumi Kasai, David Gray, Carol Clugston, Anna Dominiczak and Alex Alderton, Roberto Amato, Sonia Goncalves, Ewan Harrison, David K. Jackson, Ian Johnston, Dominic Kwiatkowski, Cordelia Langford, John Sillitoe on behalf of the Wellcome Sanger Institute COVID-19 Surveillance Team |
| EPI_ISL_858400, EPI_ISL_858403 | Lighthouse Lab in Alderley Park | Wellcome Sanger Institute for the COVID-19 Genomics UK (COG-UK) Consortium | Jacquelyn Wynn, Mairead Hyland, The Lighthouse Lab in Alderley Park and Alex Alderton, Roberto Amato, Sonia Goncalves, Ewan Harrison, David K. Jackson, Ian Johnston, Dominic Kwiatkowski, Cordelia Langford, John Sillitoe on behalf of the Wellcome Sanger Institute COVID-19 Surveillance Team |
| EPI_ISL_858425, EPI_ISL_858427, EPI_ISL_858429, EPI_ISL_858430, EPI_ISL_858431, EPI_ISL_858432, EPI_ISL_858433, EPI_ISL_858434, EPI_ISL_858435, EPI_ISL_858436, EPI_ISL_858437, EPI_ISL_858440, EPI_ISL_858441, EPI_ISL_858444, EPI_ISL_858446, EPI_ISL_858447, EPI_ISL_858448, EPI_ISL_858451, EPI_ISL_858452, EPI_ISL_858453, EPI_ISL_858454, EPI_ISL_858455, EPI_ISL_858456, EPI_ISL_858457, EPI_ISL_858459, EPI_ISL_858460, EPI_ISL_858462, EPI_ISL_858463, EPI_ISL_858465, EPI_ISL_858466, EPI_ISL_858467, EPI_ISL_858468, EPI_ISL_858470, EPI_ISL_858472, EPI_ISL_858474, EPI_ISL_858475, EPI_ISL_858476, EPI_ISL_858480, EPI_ISL_858482, EPI_ISL_858483, EPI_ISL_858484, EPI_ISL_858487, EPI_ISL_858488, EPI_ISL_858489, EPI_ISL_858490, EPI_ISL_858492, EPI_ISL_858493, EPI_ISL_858494, EPI_ISL_858495, EPI_ISL_858496, EPI_ISL_858497, EPI_ISL_858498, EPI_ISL_858499, EPI_ISL_858500, EPI_ISL_858502, EPI_ISL_858503, EPI_ISL_858504, EPI_ISL_858505, EPI_ISL_858506, EPI_ISL_858507, EPI_ISL_858508, EPI_ISL_858509, EPI_ISL_858510, EPI_ISL_858511, EPI_ISL_858512, EPI_ISL_858513, EPI_ISL_858514, EPI_ISL_858515, EPI_ISL_858517, EPI_ISL_858518, EPI_ISL_858519, EPI_ISL_858520, EPI_ISL_858522, EPI_ISL_858524, EPI_ISL_858525, EPI_ISL_858526, EPI_ISL_858527, EPI_ISL_858528, EPI_ISL_858529, EPI_ISL_858530, EPI_ISL_858532, EPI_ISL_858533, EPI_ISL_858534, EPI_ISL_858535, EPI_ISL_858536, EPI_ISL_858537, EPI_ISL_858538, EPI_ISL_858539, EPI_ISL_858540, EPI_ISL_858541, EPI_ISL_858542, EPI_ISL_858543, EPI_ISL_858544, EPI_ISL_858545, EPI_ISL_858546, EPI_ISL_858547, EPI_ISL_858548, EPI_ISL_858549, EPI_ISL_858550, EPI_ISL_858551, EPI_ISL_858552, EPI_ISL_858553, EPI_ISL_858554, EPI_ISL_858555, EPI_ISL_858556, EPI_ISL_858558, EPI_ISL_858559, EPI_ISL_858560, EPI_ISL_858561, EPI_ISL_858562, EPI_ISL_858564, EPI_ISL_858565, EPI_ISL_858566, EPI_ISL_858567, EPI_ISL_858568, EPI_ISL_858569, EPI_ISL_858570, EPI_ISL_858571 | | | |
| see above | Lighthouse Lab in Glasgow | Wellcome Sanger Institute for the COVID-19 Genomics UK (COG-UK) Consortium | Harper VanSteenhouse, Yumi Kasai, David Gray, Carol Clugston, Anna Dominiczak and Alex Alderton, Roberto Amato, Sonia Goncalves, Ewan Harrison, David K. Jackson, Ian Johnston, Dominic Kwiatkowski, Cordelia Langford, John Sillitoe on behalf of the Wellcome Sanger Institute COVID-19 Surveillance Team |
| EPI_ISL_860186 | Bangalore Medical College and Research Institute | Department of Neurovirology, National Institute of Mental Health and Neurosciences (NIMHANS) | Chitra Pattabiraman, Pramada Prasad, Anson Kunjumon George, Risha Rasheed, Darshan Sreenivas, Nakka Vijay Kiran Reddy, Anita S Desai, V Ravi |
| EPI_ISL_860196 | National Institute of Mental Health and Neurosciences (NIMHANS) | Department of Neurovirology, National Institute of Mental Health and Neurosciences (NIMHANS) | Chitra Pattabiraman, Pramada Prasad, Anson Kunjumon George, Risha Rasheed, Darshan Sreenivas, Nakka Vijay Kiran Reddy, Anita S Desai, V Ravi |
| EPI_ISL_860197 | BBMP Urban PHC | Department of Neurovirology, National Institute of Mental Health and Neurosciences (NIMHANS) | Chitra Pattabiraman, Pramada Prasad, Anson Kunjumon George, Risha Rasheed, Darshan Sreenivas, Nakka Vijay Kiran Reddy, Anita S Desai, V Ravi |
| EPI_ISL_860215 | Norwegian Institute of Public Health, Department of Virology | Norwegian Institute of Public Health, Department of Virology | Kathrine Stene-Johansen, Kamilla Heddeland Instefjord, Hilde Elshaug, Atiya R Ali,Marie Paulsen Madsen, Rasmus Riis Kopperud, Hilde Vollan, Karoline Bragstad, Olav Hungnes |
| EPI_ISL_860219, EPI_ISL_860232 | Akershus University Hospital, Department for Microbiology and Infectious Disease Control | Norwegian Institute of Public Health, Department of Virology | Kathrine Stene-Johansen, Kamilla Heddeland Instefjord, Hilde Elshaug, Atiya R Ali,Marie Paulsen Madsen, Rasmus Riis Kopperud, Hilde Vollan, Karoline Bragstad, Olav Hungnes |
| EPI_ISL_860243, EPI_ISL_860249, EPI_ISL_860250, EPI_ISL_860252 | University Hospitals of Geneva, Laboratory of Virology | HUG, Laboratory of Virology and Universitätsspital Basel | Samuel Cordey, Ana Rita Goncalves, Laurent Kaiser, Tim Roloff, Madlen Stange, Helena MB Seth-Smith, Alfredo Mari, Karoline Leuzinger, Julia Bielicki, Manuel Battegay, Hans Hirsch, Adrian Egli |
| EPI_ISL_860261 | Nordland Hospital - Bodo, Laboratory Department, Molecular Biology Unit | Norwegian Institute of Public Health, Department of Virology | Kathrine Stene-Johansen, Kamilla Heddeland Instefjord, Hilde Elshaug, Atiya R Ali,Marie Paulsen Madsen, Rasmus Riis Kopperud, Hilde Vollan, Karoline Bragstad, Olav Hungnes |
| EPI_ISL_860264, EPI_ISL_860265, EPI_ISL_860266, EPI_ISL_860267 | Vestfold Hospital, Toensberg Department of Microbiology | Norwegian Institute of Public Health, Department of Virology | Kathrine Stene-Johansen, Kamilla Heddeland Instefjord, Hilde Elshaug, Atiya R Ali,Marie Paulsen Madsen, Rasmus Riis Kopperud, Hilde Vollan, Karoline Bragstad, Olav Hungnes |
| EPI_ISL_860272 | Ostfold Hospital Trust - Kalnes, Centre for Laboratory Medicine, Section for gene technology and infection serology | Norwegian Institute of Public Health, Department of Virology | Kathrine Stene-Johansen, Kamilla Heddeland Instefjord, Hilde Elshaug, Atiya R Ali,Marie Paulsen Madsen, Rasmus Riis Kopperud, Hilde Vollan, Karoline Bragstad, Olav Hungnes |
| EPI_ISL_860275 | Foerde Hospital, Department of Microbiology | Norwegian Institute of Public Health, Department of Virology | Kathrine Stene-Johansen, Kamilla Heddeland Instefjord, Hilde Elshaug, Atiya R Ali,Marie Paulsen Madsen, Rasmus Riis Kopperud, Hilde Vollan, Karoline Bragstad, Olav Hungnes |
| EPI_ISL_860284, EPI_ISL_860286, EPI_ISL_860287 | Department of Medical Microbiology, St. Olavs hospital | Norwegian Institute of Public Health, Department of Virology | Kathrine Stene-Johansen, Kamilla Heddeland Instefjord, Hilde Elshaug, Atiya R Ali,Marie Paulsen Madsen, Rasmus Riis Kopperud, Hilde Vollan, Karoline Bragstad, Olav Hungnes |
| EPI_ISL_860295 | Hospital of Southern Norway - Kristiansand, Department of Medical Microbiology | Norwegian Institute of Public Health, Department of Virology | Kathrine Stene-Johansen, Kamilla Heddeland Instefjord, Hilde Elshaug, Atiya R Ali,Marie Paulsen Madsen, Rasmus Riis Kopperud, Hilde Vollan, Karoline Bragstad, Olav Hungnes |
| EPI_ISL_860304, EPI_ISL_860305, EPI_ISL_860306, EPI_ISL_860307, EPI_ISL_860309, EPI_ISL_860310, EPI_ISL_860311, EPI_ISL_860312 | Unit 17: Influenza & Other Respiratory Viruses, German National Influenza Center | Project group Epidemiology of Highly Pathogenic Microorganisms, Robert Koch-Institute | Andreas Sachse, Grit Schubert, Essia Belarbi, Sébastien Calvignac-Spencer, Thorsten Wolff, Ralf Dürrwald, Djin-Ye Oh, Marianne Wedde |
| EPI_ISL_860668, EPI_ISL_860669, EPI_ISL_860670, EPI_ISL_860672, EPI_ISL_860673, EPI_ISL_860678, EPI_ISL_860679 | Respiratory Virus Unit, National Infection Service, Public Health England | COVID-19 Genomics UK (COG-UK) Consortium | PHE Covid Sequencing Team |
| EPI_ISL_860879, EPI_ISL_860880, EPI_ISL_860885 | Labo Analyses Med | National Reference Center for Viruses of Respiratory Infections, Institut Pasteur, Paris | Marion Barbet, Sylvie Behillil, Méline Bizard, Angela Brisebarre, Camille Capel, Etienne Simon-Lorière, Vincent Enouf, Maud Vanpeene, Sylvie van der Werf,Lefaure Brieuc |
| EPI_ISL_860937, EPI_ISL_860947, EPI_ISL_860949, EPI_ISL_860964, EPI_ISL_860987, EPI_ISL_860996, EPI_ISL_860998, EPI_ISL_861028, EPI_ISL_861031, EPI_ISL_861037, EPI_ISL_861068, EPI_ISL_861087, EPI_ISL_861098, EPI_ISL_861099 | | | |
| see above | Johns Hopkins Hospital Department of Pathology | Johns Hopkins Hospital Department of Pathology | C. Paul Morris, Chun Huai Luo, Adannaya Amadi, Nicholas Gallagher, Heba H. Mostafa |
| EPI_ISL_861115, EPI_ISL_861116, EPI_ISL_861117 | Wadsworth Center, New York State Department of Health | Wadsworth Center, New York State Department of Health | Kirsten St. George, Daryl M. Lamson, Alexis Russel, Matthew Shudt, Melissa A Leisner, Jonathan Plitnick, Navjot Singh, John Kelly, Erasmus Schneider, Erica Lasek-Nesselquist |
| EPI_ISL_861180 | BIO-REFERENCE LABORATORIES | Wadsworth Center, New York State Department of Health | Kirsten St. George, Daryl M. Lamson, Alexis Russel, Matthew Shudt, Melissa A Leisner, Jonathan Plitnick, Navjot Singh, John Kelly, Erasmus Schneider, Erica Lasek-Nesselquist |

| | | | |
|---|---|---|---|
| EPI_ISL_861360, EPI_ISL_861361, EPI_ISL_861362, EPI_ISL_861363, EPI_ISL_861364, EPI_ISL_861365, EPI_ISL_861366, EPI_ISL_861367, EPI_ISL_861368, EPI_ISL_861370, EPI_ISL_861371, EPI_ISL_861372, EPI_ISL_861381, EPI_ISL_861388, EPI_ISL_861390, EPI_ISL_861391, EPI_ISL_861398 | | | |
| see above | WESTCHESTER MEDICAL CENTER | Wadsworth Center, New York State Department of Health | Kirsten St. George, Daryl M. Lamson, Alexis Russel, Matthew Shudt, Melissa A Leisner, Jonathan Plitnick, Navjot Singh, John Kelly, Erasmus Schneider, Erica Lasek-Nesselquist |
| EPI_ISL_861427, EPI_ISL_861429, EPI_ISL_861431 | Wyoming Public Health Laboratory | Wyoming Public Health Laboratory | Noah Hull, Taylor Fearing, Lynette Gumbleton, Channing Weber, Ashley Norberg, Bailey Bowcutt, and Wanda Manley |
| EPI_ISL_861472, EPI_ISL_861473 | Gundersen Molecular Diagnostics Laboratory | Kabara Cancer Research Institute | Craig S. Richmond, Paraic A. Kenny |
| EPI_ISL_861550, EPI_ISL_861572 | Instituto Nacional de Saude (INSA) | Instituto Nacional de Saude (INSA) | Borges et al |
| EPI_ISL_861683 | Complexo Hospitalar Padre Bento de Guarulhos | Instituto Adolfo Lutz, Interdiciplinary Procedures Center, Strategic Laboratory | Claudio Tavares Sacchi, Claudia Regina Gonçalves, Erica Valessa Ramos Gomes, Karoline Rodrigues Campos |
| EPI_ISL_861686, EPI_ISL_861690, EPI_ISL_861693 | Los Angeles County PHL | Los Angeles County PHL | P. Hemarajata et al. |
| EPI_ISL_861742, EPI_ISL_861744 | Tempus | Grubaugh Lab - Yale School of Public Health | Tara Alpert, Joseph Fauver, Anderson Brito, Mallery Breban, Anne Wyllie, Chantal Vogels, Mary Petrone, Annie Watkins, Chaney Kalinich, Isabel Ott, Nathan Grubaugh |
| EPI_ISL_862037 | Johns Hopkins Hospital Department of Pathology | Johns Hopkins Hospital Department of Pathology | C. Paul Morris, Chun Huai Luo, Adannaya Amadi, Matthew Schwartz, Nicholas Gallagher, Heba H. Mostafa |
| EPI_ISL_862051 | Multidisciplinary Research Unit, DHR-ICMR, Institute of Medical Sciences, Banaras Hindu University | Multidisciplinary Research Unit, DHR-ICMR, Institute of Medical Sciences, Banaras Hindu University | Priyoneel Basu, Ashish, Nitish Kumar Singh, Abhay Kumar Yadav, Manpreet Kaur, Arup Acharjee, Deepa Devadas, Chetan Sahni, Sanjay Kumar, Tribhuwan Mohan Mahapatra, Richa Arya, Prashant Singh, Jay Prakash Maurya, Surendra Pratap Mishra, Royana Singh |
| EPI_ISL_862153, EPI_ISL_862155, EPI_ISL_862156 | Charité Universitätsmedizin Berlin, Institut fur Virologie/Labor Berlin | Charité Universitätsmedizin Berlin, Institut für Virologie | Victor M Corman, Barbara Mühlemann, Jörn Beheim-Schwarzbach, Tobias Bleicker, Julia Tesch, Talitha Veith, Julia Schneider, Terry Jones, Christian Drosten |
| EPI_ISL_862746, EPI_ISL_862747, EPI_ISL_862750, EPI_ISL_862753, EPI_ISL_862755, EPI_ISL_862762, EPI_ISL_862764, EPI_ISL_862767, EPI_ISL_862768, EPI_ISL_862770, EPI_ISL_862778 | | | |
| see above | Utah Public Health Laboratory, Utah Public Health Laboratory Infectious Disease submission group | Utah Public Health Laboratory, Utah Public Health Laboratory Infectious Disease submission group | Young,E.L., Oakeson,K.F., Gallagher,T. |
| EPI_ISL_862831 | National Institute of Infectious Diseases-Prof. Dr. Matei Bals Molecular Diagnostics Laboratory | National Institute of Infectious Diseases-Prof. Dr. Matei Bals Molecular Diagnostics Laboratory | Leontina Banica, Marius Surleac, Corina Casangiu, Petre Milu, Andreea Tudor, Simona Paraschiv, Dan Otelea |
| EPI_ISL_864368 | Lighthouse Lab in Milton Keynes | Wellcome Sanger Institute for the COVID-19 Genomics UK (COG-UK) Consortium | The Lighthouse Lab in Milton Keynes and Alex Alderton, Roberto Amato, Sonia Goncalves, Ewan Harrison, David K. Jackson, Ian Johnston, Dominic Kwiatkowski, Cordelia Langford, John Sillitoe on behalf of the Wellcome Sanger Institute COVID-19 Surveillance Team |
| EPI_ISL_864755, EPI_ISL_864756, EPI_ISL_864758, EPI_ISL_864762, EPI_ISL_864764, EPI_ISL_864774, EPI_ISL_864776, EPI_ISL_864778, EPI_ISL_864780, EPI_ISL_864805, EPI_ISL_864819, EPI_ISL_864821, EPI_ISL_864828, EPI_ISL_864832, EPI_ISL_864834, EPI_ISL_864838, EPI_ISL_864850, EPI_ISL_864856 | | | |
| see above | Department of Pathology, University of Cambridge | COVID-19 Genomics UK (COG-UK) Consortium | Aminu S. Jahun, Yasmin Chaudhry, Grant Hall, Iliana Georgana, Myra Hosmillo, Martin D. Curran, Malte Pinckert, Surendra Parmar, Ian Goodfellow |
| EPI_ISL_864996, EPI_ISL_864997, EPI_ISL_864998, EPI_ISL_864999 | West of Scotland Specialist Virology Centre, NHSGGC / MRC-University of Glasgow Centre for Virus Research | COVID-19 Genomics UK (COG-UK) Consortium | Ana da Silva Filipe, Natasha Johnson, Kathy Smollett, Daniel Mair, Stephen Carmichael, Alice Broos, Lily Tong, Jenna Nichols, Kyriaki Nomikou; Sarah McDonald; Richard Orton, Joseph Hughes, Sreenu Vattipally, David L Robertson; Alasdair MacLean, Rory Gunson; Sharif Shaaban, Matthew Holden; Rachel Blacow, Guy Mollett, Kathy Li, James Shepherd, Antonia Ho, Emma Thomson |
| EPI_ISL_865007, EPI_ISL_865010, EPI_ISL_865011 | Lighthouse Lab in Glasgow / MRC-University of Glasgow Centre for Virus Research | COVID-19 Genomics UK (COG-UK) Consortium | Ana da Silva Filipe, Natasha Johnson, Kathy Smollett, Daniel Mair, Stephen Carmichael, Alice Broos, Lily Tong, Jenna Nichols, Kyriaki Nomikou; Sarah McDonald; Harper VanSteenhouse, Yumi Kasai, David Gray, Carol Clugston, Anna Dominiczak; Alasdair MacLean, Rory Gunson; Richard Orton, Joseph Hughes, Sreenu Vattipally, David L Robertson; Sharif Shaaban, Matthew Holden; Kathy Li, James Shepherd, Antonia Ho, Emma Thomson |
| EPI_ISL_865095, EPI_ISL_865102 | Virology Department, Royal Infirmary of Edinburgh, NHS Lothian / School of Biological Sciences, University of Edinburgh / Institute of Genetics and Molecular Medicine, University of Edinburgh | COVID-19 Genomics UK (COG-UK) Consortium | McHugh M, Dewar R, Rooke S, Gallagher M, Balcaza C, O'Toole Á, Scher E, Hill V, McCrone JT, Colquhoun R, Yu X, Jackson B, Rambaut A, Williams TC, Templeton K |
| EPI_ISL_865181, EPI_ISL_865182, EPI_ISL_865185, EPI_ISL_865186, EPI_ISL_865187, EPI_ISL_865188, EPI_ISL_865189, EPI_ISL_865192, EPI_ISL_865193, EPI_ISL_865194, EPI_ISL_865196, EPI_ISL_865197, EPI_ISL_865199, EPI_ISL_865200, EPI_ISL_865201, EPI_ISL_865202, EPI_ISL_865334, EPI_ISL_865335, EPI_ISL_865337, EPI_ISL_865450, EPI_ISL_865478, EPI_ISL_865482, EPI_ISL_865483 | | | |
| see above | Liverpool Clinical Laboratories | COVID-19 Genomics UK (COG-UK) Consortium | Sam Haldenby, Anita Lucaci, Steve Paterson, Julian Hiscox, Alistair Darby, M Almsaud, A Alrezaihi, Muhannad Alruwaili, Stuart D Armstrong, Jones Benjamin, Eleanor G Bentley, Anu Chawla, Jordan J Clark, Angela Cowell, Richard Eccles, Isabel García-Dorival, Matthew Gemmell, Alessandro Gerada, PKF Gilmore, Richard Gregory, Ximeng Han, Catherine Hartley, Margaret Hughes, Miren Iturriza-Gomara, James Johnson, L Luu, Jenifer Manson, Charlotte Nelson, Elaine O'Toole, Cassie Olateju, Rebekah Penrice-Randal , Lucille Rainbow, N.P Randle, Trevor Ian Robinson, Parul Sharma, Ghada T Shawli, James P Stewart, Neil Swainston, Ecaterina Vamos, Joanne Watts, Mark Whitehead |
| EPI_ISL_865500 | Barts Health NHS Trust | COVID-19 Genomics UK (COG-UK) Consortium | CUTINO-MOGUEL, Maria-Teresa; HARRINGTON, David; OWOYEMI, Dola; KULASEGARAN-SHYLINI, Raghavendran; BROAD, Claire; KELE, Beatrix |
| EPI_ISL_866076 | University College London Hospital | COVID-19 Genomics UK (COG-UK) Consortium | Judith Heaney, Matthew Byott, Catherine Houlihan, Dan Frampton, Stuart Kirk, Moira Spyer and Eleni Nastouli |
| EPI_ISL_866187 | University College London, Great Ormond Street Hospital for Children NHS Foundation Trust, Imperial College Healthcare NHS Trust | COVID-19 Genomics UK (COG-UK) Consortium | Sergi Castellano, Rachel Williams, Mark Kristiansen, Paola Resende Silva, Sunando Roy, Tony Brooks, Helena Tutill, Paola Niola, Patricia Dyal, Charlotte Williams, Leysa Forrest, Yasmin Panchbhaya, Jacqueline Findlay, Samuel Weeks, Julianne Brown, Kathryn Harris, Paul Randell, James Price, Alison Holmes, Judith Breuer |
| EPI_ISL_866353, EPI_ISL_866360, EPI_ISL_866361, EPI_ISL_866370, EPI_ISL_866389, EPI_ISL_866390, EPI_ISL_866391, EPI_ISL_866392 | Regional Virus Laboratory, Belfast Health and Social Care Trust | COVID-19 Genomics UK (COG-UK) Consortium | Conall McCaughey, James McKenna, Tanya Curran, Susan Feeney, Alison Watt, Ciara Cox, Mairead Connor, Zoltan Molnar, David Simpson, Derek Fairley |
| EPI_ISL_867030, EPI_ISL_867032, EPI_ISL_867033 | Lincolnshire Hospitals and DeepSeq Nottingham | COVID-19 Genomics UK (COG-UK) Consortium | Nichola Duckworth, Tim Sloan, Sarah Walsh, Jonathan Ball, Patrick McClure, Joeseph Chappell, Nadine Holmes, Matthew Carlisle, Christopher Moore, Fei Sang, Johnny Debebe, Victoria Wright, Matthew Loose |
| EPI_ISL_867100, EPI_ISL_867101, EPI_ISL_867133, EPI_ISL_867134, EPI_ISL_867137, EPI_ISL_867138, EPI_ISL_867140, EPI_ISL_867141, EPI_ISL_867142, EPI_ISL_867144, EPI_ISL_867145, EPI_ISL_867147, EPI_ISL_867148, EPI_ISL_867150, EPI_ISL_867166 | | | |
| see above | Oxford Viromics, NDM, University of Oxford; Oxford University Hospitals; Basingstoke and North Hampshire Hospital | COVID-19 Genomics UK (COG-UK) Consortium | Tanya Golubchik, David Bonsall, George Macintyre, Amy Trebes, Mariateresa de Cesare, Catrin Moore, Alex Mobbs, Anita Justice, Robert Shaw, Monique Andersson, Timothy Peto, Emma Wise, Nathan Moore, Jessica Lynch, Nick Cortes, Matilde Mori, Stephen Kidd, David Buck, John Todd, Christophe Fraser |
| EPI_ISL_867186, EPI_ISL_867206, EPI_ISL_867207 | Originating lab: Wales Specialist Virology Centre Sequencing lab: Pathogen Genomics Unit | Public Health Wales Microbiology Cardiff Wales Specialist Virology Centre | Catherine Moore, Johnathan Evans, Laura Gifford, Malorie Perry, Simon Cottrell, Angela Marchbank, Alec Birchley, Alexander Adams, Amy Gaskin, Bree Gatica-Wilcox, Jason Coombes, Joel Southgate, Lauren Gilbert, Lee Graham, Nicole Pacchiarini, Sara Kumziene-Summerhayes, Sarah Taylor, Sophie Jones, Sara Rey, Matthew Bull, Joanne Watkins, Sally Corden, Tom Connor |
| EPI_ISL_867990, EPI_ISL_868312, EPI_ISL_868313, EPI_ISL_868315, EPI_ISL_868316, EPI_ISL_868326, EPI_ISL_868327, EPI_ISL_868349, EPI_ISL_868350 | Centre for Enzyme Innovation, University of Portsmouth / Translational Research Laboratory, Portsmouth Hospitals NHS Trust | COVID-19 Genomics UK (COG-UK) Consortium | Angela Beckett,Yann Bourgeois,Garry Scarlett,Sharon Glaysher,Scott Elliott,Kelly Bicknell,Robert Impey,Allyson Lloyd,Sarah Wyllie,Ethan Butcher,Anoop Chauhan,Samuel Robson |
| EPI_ISL_868357, EPI_ISL_868607, EPI_ISL_868704 | Virology Department, Sheffield Teaching Hospitals NHS Foundation Trust/Department of Infection, Immunity and Cardiovascular Disease, The Medical School, University of Sheffield | COVID-19 Genomics UK (COG-UK) Consortium | Thushan de Silva, Matthew Parker, Nikki Smith, Adri Angyal, Rebecca Brown, Luke Green, Rachel Tucker, Paul Parsons, Danielle Groves, Katie Johnson, Laura Carrilero, Alex Keeley, Dave Partridge, Matthew Wyles, Benjamin Lindsey, Mehmet Yavuz, Mohammad Raza, Cariad Evans |
| EPI_ISL_869083, EPI_ISL_869124, EPI_ISL_869125, EPI_ISL_869126, | Charité Universitätsmedizin Berlin, Institut für Virologie/Labor Berlin | Charité Universitätsmedizin Berlin, Institut für Virologie | Victor M Corman, Barbara Mühlemann, Jörn Beheim-Schwarzbach, Tobias Bleicker, Julia Tesch, Talitha Veith, Julia Schneider, Terry Jones, Christian Drosten |

| | | | |
|---|---|---|---|
| EPI_ISL_869127, EPI_ISL_869128, EPI_ISL_869129, EPI_ISL_869130 | | | |
| EPI_ISL_869182, EPI_ISL_869183 | New Mexico Department of Health Scientific Laboratory | Center for Global Health, University of New Mexico Health Sciences Center | Daryl Domman, Kurt Schwalm, Twila Kunde, Joseph Hicks, Anastacia Griego, Michael Edwards, Darrell Dinwiddie |
| EPI_ISL_871133, EPI_ISL_871134, EPI_ISL_871135, EPI_ISL_871136, EPI_ISL_871137, EPI_ISL_871138, EPI_ISL_871139, EPI_ISL_871140, EPI_ISL_871141, EPI_ISL_871142, EPI_ISL_871143, EPI_ISL_871144, EPI_ISL_871145, EPI_ISL_871146, EPI_ISL_871147, EPI_ISL_871148, EPI_ISL_871149, EPI_ISL_871150, EPI_ISL_871151, EPI_ISL_871152, EPI_ISL_871153, EPI_ISL_871154, EPI_ISL_871155, EPI_ISL_871156, EPI_ISL_871157, EPI_ISL_871158, EPI_ISL_871159, EPI_ISL_871160, EPI_ISL_871161, EPI_ISL_871162, EPI_ISL_871163, EPI_ISL_871164, EPI_ISL_871165, EPI_ISL_871166, EPI_ISL_871167, EPI_ISL_871168, EPI_ISL_871169, EPI_ISL_871170, EPI_ISL_871171, EPI_ISL_871172, EPI_ISL_871173, EPI_ISL_871174, EPI_ISL_871175, EPI_ISL_871176, EPI_ISL_871177, EPI_ISL_871178, EPI_ISL_871179, EPI_ISL_871180, EPI_ISL_871181, EPI_ISL_871182, EPI_ISL_871183, EPI_ISL_871184, EPI_ISL_871185, EPI_ISL_871186, EPI_ISL_871187, EPI_ISL_871188, EPI_ISL_871189, EPI_ISL_871190, EPI_ISL_871191, EPI_ISL_871192, EPI_ISL_871193, EPI_ISL_871194, EPI_ISL_871195, EPI_ISL_871196, EPI_ISL_871197, EPI_ISL_871198, EPI_ISL_871199, EPI_ISL_871200, EPI_ISL_871201, EPI_ISL_871202, EPI_ISL_871203, EPI_ISL_871204, EPI_ISL_871205, EPI_ISL_871206, EPI_ISL_871207, EPI_ISL_871208, EPI_ISL_871209, EPI_ISL_871210, EPI_ISL_871211, EPI_ISL_871212, EPI_ISL_871213, EPI_ISL_871214, EPI_ISL_871215, EPI_ISL_871216, EPI_ISL_871217, EPI_ISL_871218, EPI_ISL_871219, EPI_ISL_871220, EPI_ISL_871221, EPI_ISL_871222, EPI_ISL_871223, EPI_ISL_871224, EPI_ISL_871225, EPI_ISL_871226, EPI_ISL_871227, EPI_ISL_871228, EPI_ISL_871229, EPI_ISL_871230, EPI_ISL_871231, EPI_ISL_871232, EPI_ISL_871233, EPI_ISL_871234, EPI_ISL_871235, EPI_ISL_871236, EPI_ISL_871237, EPI_ISL_871238, EPI_ISL_871239, EPI_ISL_871240, EPI_ISL_871241, EPI_ISL_871242, EPI_ISL_871243, EPI_ISL_871244, EPI_ISL_871245, EPI_ISL_871246, EPI_ISL_871247, EPI_ISL_871248, EPI_ISL_871249, EPI_ISL_871250, EPI_ISL_871251, EPI_ISL_871252, EPI_ISL_871253, EPI_ISL_871254, EPI_ISL_871255, EPI_ISL_871256, EPI_ISL_871257, EPI_ISL_871258, EPI_ISL_871259, EPI_ISL_871260, EPI_ISL_871261, EPI_ISL_871262, EPI_ISL_871263, EPI_ISL_871264, EPI_ISL_871265, EPI_ISL_871266, EPI_ISL_871267, EPI_ISL_871268, EPI_ISL_871269, EPI_ISL_871270, EPI_ISL_871271, EPI_ISL_871272, EPI_ISL_871273, EPI_ISL_871274, EPI_ISL_871275, EPI_ISL_871276, EPI_ISL_871277, EPI_ISL_871278, EPI_ISL_871279, EPI_ISL_871280, EPI_ISL_871281, EPI_ISL_871282, EPI_ISL_871283, EPI_ISL_871284, EPI_ISL_871285, EPI_ISL_871286, EPI_ISL_871287, EPI_ISL_871288, EPI_ISL_871289, EPI_ISL_871290, EPI_ISL_871291, EPI_ISL_871292, EPI_ISL_871293, EPI_ISL_871294, EPI_ISL_871295, EPI_ISL_871296, EPI_ISL_871297, EPI_ISL_871298, EPI_ISL_871299, EPI_ISL_871300, EPI_ISL_871301, EPI_ISL_871302, EPI_ISL_871303, EPI_ISL_871304, EPI_ISL_871305, EPI_ISL_871306, EPI_ISL_871307, EPI_ISL_871308, EPI_ISL_871309, EPI_ISL_871310, EPI_ISL_871311, EPI_ISL_871312, EPI_ISL_871313, EPI_ISL_871314, EPI_ISL_871315, EPI_ISL_871316, EPI_ISL_871317, EPI_ISL_871318, EPI_ISL_871319, EPI_ISL_871320, EPI_ISL_871321, EPI_ISL_871322, EPI_ISL_871323, EPI_ISL_871324, EPI_ISL_871326, EPI_ISL_871327, EPI_ISL_871328, EPI_ISL_871329, EPI_ISL_871330, EPI_ISL_871331, EPI_ISL_871332, EPI_ISL_871333, EPI_ISL_871334, EPI_ISL_871335, EPI_ISL_871336, EPI_ISL_871337, EPI_ISL_871338, EPI_ISL_871339, EPI_ISL_871340, EPI_ISL_871341, EPI_ISL_871342, EPI_ISL_871343, EPI_ISL_871344, EPI_ISL_871345, EPI_ISL_871346, EPI_ISL_871347, EPI_ISL_871348, EPI_ISL_871349, EPI_ISL_871350, EPI_ISL_871351, EPI_ISL_871352, EPI_ISL_871353, EPI_ISL_871354, EPI_ISL_871355, EPI_ISL_871356, EPI_ISL_871357, EPI_ISL_871358, EPI_ISL_871359, EPI_ISL_871360, EPI_ISL_871361, EPI_ISL_871362, EPI_ISL_871363, EPI_ISL_871364, EPI_ISL_871365, EPI_ISL_871366, EPI_ISL_871367, EPI_ISL_871368, EPI_ISL_871369, EPI_ISL_871370, EPI_ISL_871371, EPI_ISL_871372, EPI_ISL_871373, EPI_ISL_871374, EPI_ISL_871375, EPI_ISL_871376, EPI_ISL_871377, EPI_ISL_871378, EPI_ISL_871379, EPI_ISL_871380, EPI_ISL_871381, EPI_ISL_871382, EPI_ISL_871383, EPI_ISL_871384, EPI_ISL_871385, EPI_ISL_871386, EPI_ISL_871387, EPI_ISL_871388, EPI_ISL_871389, EPI_ISL_871390, EPI_ISL_871391, EPI_ISL_871392, EPI_ISL_871393, EPI_ISL_871394, EPI_ISL_871395, EPI_ISL_871396, EPI_ISL_871397, EPI_ISL_871398, EPI_ISL_871399, EPI_ISL_871400, EPI_ISL_871401, EPI_ISL_871402, EPI_ISL_871403, EPI_ISL_871404, EPI_ISL_871405, EPI_ISL_871406, EPI_ISL_871407, EPI_ISL_871408, EPI_ISL_871409, EPI_ISL_871410, EPI_ISL_871411, EPI_ISL_871412, EPI_ISL_871413, EPI_ISL_871414, EPI_ISL_871415, EPI_ISL_871416, EPI_ISL_871417, EPI_ISL_871418, EPI_ISL_871419, EPI_ISL_871420, EPI_ISL_871421, EPI_ISL_871422, EPI_ISL_871423, EPI_ISL_871424, EPI_ISL_871425, EPI_ISL_871426, EPI_ISL_871427, EPI_ISL_871428, EPI_ISL_871429, EPI_ISL_871430, EPI_ISL_871431, EPI_ISL_871432, EPI_ISL_871433, EPI_ISL_871434, EPI_ISL_871435, EPI_ISL_871436, EPI_ISL_871437, EPI_ISL_871438, EPI_ISL_871439, EPI_ISL_871440, EPI_ISL_871441, EPI_ISL_871442, EPI_ISL_871443, EPI_ISL_871444, EPI_ISL_871445, EPI_ISL_871446, EPI_ISL_871447, EPI_ISL_871448, EPI_ISL_871449, EPI_ISL_871450, EPI_ISL_871451, EPI_ISL_871452, EPI_ISL_871453, EPI_ISL_871454, EPI_ISL_871455, EPI_ISL_871456, EPI_ISL_871457 | | | |
| see above | Department of Virus and Microbiological Special Diagnostics, Statens Serum Institut, Copenhagen, Denmark | Aalborg University | Danish Covid-19 Genome Consortium |
| EPI_ISL_871841, EPI_ISL_871871 | Wyoming Public Health Laboratory | Wyoming Public Health Laboratory | Noah Hull, Taylor Fearing, Lynette Gumbleton, Channing Weber, Ashley Norberg, Bailey Bowcutt, and Wanda Manley |
| EPI_ISL_871890 | Hospital Universitario Marqués de Valdecilla - IDIVAL (Santander, Cantabria) | SeqCOVID-SPAIN consortium/IBV(CSIC) | Mónica Gozalo Margüello, María Eliecer Cano García, Jose Manuel Méndez Legaza, Daniel Pablo Marcos, Jesús Rodríguez Rodríguez, María Siller Ruiz and SeqCOVID-SPAIN consortium |
| EPI_ISL_871959 | Servicio de Microbiología Clínica (Complejo Hospitalario de Navarra, Pamplona), Instituto de Investigación Sanitaria de Navarra (IdiSNA) | SeqCOVID-SPAIN consortium/IBV(CSIC) | Carmen Ezpeleta Baquedano, Ana Navascués, Ana Miqueleiz and SeqCOVID-SPAIN consortium |
| EPI_ISL_872095, EPI_ISL_872096 | Wyoming Public Health Laboratory | Wyoming Public Health Laboratory | Noah Hull, Taylor Fearing, Lynette Gumbleton, Channing Weber, Ashley Norberg, Bailey Bowcutt, and Wanda Manley |
| EPI_ISL_872239 | Labo Analyses Med | National Reference Center for Viruses of Respiratory Infections, Institut Pasteur, Paris | Marion Barbet, Sylvie Behillil, Méline Bizard, Angela Brisebarre, Camille Capel, Etienne Simon-Lorière, Vincent Enouf, Maud Vanpeene, Sylvie van der Werf,Rouah Raquel |
| EPI_ISL_872298 | Hopital | National Reference Center for Viruses of Respiratory Infections, Institut Pasteur, Paris | Marion Barbet, Sylvie Behillil, Méline Bizard, Angela Brisebarre, Camille Capel, Etienne Simon-Lorière, Vincent Enouf, Maud Vanpeene, Sylvie van der Werf,Florin CéCile |
| EPI_ISL_872573 | Laverty Pathology | NSW Health Pathology - Institute of Clinical Pathology and Medical Research; Westmead Hospital; University of Sydney | CIDM-PH et al. |
| EPI_ISL_872577 | South Eastern Area Laboratory Services (SEALS) | NSW Health Pathology - Institute of Clinical Pathology and Medical Research; Westmead Hospital; University of Sydney | CIDM-PH et al. |
| EPI_ISL_873049, EPI_ISL_873050, EPI_ISL_873051, EPI_ISL_873052, EPI_ISL_873053, EPI_ISL_873054, EPI_ISL_873056, EPI_ISL_873058, EPI_ISL_873059, EPI_ISL_873060, EPI_ISL_873061, EPI_ISL_873064 | | | |
| see above | University of Michigan Clinical Microbiology Laboratory | Lauring Lab, University of Michigan, Department of Microbiology and Immunology | Valesano |
| EPI_ISL_873164 | Medical Laboratory Sciences, Arab American University | Medical Laboratory Sciences, Arab American University | Dumaidi,k., Al-Jawabreh,A., Ereqat,S., Al-Jawabreh,H., Nasereddin,A. |
| EPI_ISL_873185 | Microbiology Division, South Carolina Department of Health and Environmental Control (SC DHEC) | Microbiology Division, South Carolina Department of Health and Environmental Control (SC DHEC) | Flores,H., Freeman,J. |
| EPI_ISL_873197, EPI_ISL_873201, EPI_ISL_873207, EPI_ISL_873208 | North Dakota Department of Health, Public Health Laboratory | North Dakota Department of Health, Public Health Laboratory | Lisa Wingerter |
| EPI_ISL_875511 | National Virus Reference Laboratory | National Virus Reference Laboratory | Michael Carr, Gabriel Gonzalez, Jonathan Dean, Cillian F De Gascun |
| EPI_ISL_875517 | Institute of Virology, Biomedical Research Center of the Slovak Academy of Sciences, Bratislava | Faculty of Natural Sciences, Comenius University, Bratislava | Broa Brejová, Viktória abanová, Kristína Boršová, Viktória Hodorová, Sabina Fumaová Havlíková, Juraj Kopáek, Martina Liková, ubomíra Lukáiková, Martina Neboháová, Monika Sláviková, Tomáš Vina, Jozef Nosek, Boris Klempa |
| EPI_ISL_875533 | Institute of Virology, Biomedical Research Center of the Slovak Academy of Sciences, Bratislava | Faculty of Natural Sciences, Comenius University, Bratislava | Viktória abanová, Kristína Boršová, Broa Brejová, Viktória Hodorová, Sabina Fumaová Havlíková, Juraj Kopáek, Martina Liková, ubomíra Lukáiková, Martina Neboháová, Monika Sláviková, Tomáš Vina, Jozef Nosek, Boris Klempa |
| EPI_ISL_875534 | Institute of Virology, Biomedical Research Center of the Slovak Academy of Sciences, Bratislava | Faculty of Natural Sciences, Comenius University, Bratislava | Kristína Boršová, Viktória abanová, Broa Brejová, Viktória Hodorová, Sabina Fumaová Havlíková, Juraj Kopáek, Martina Liková, ubomíra Lukáiková, Martina Neboháová, Monika Sláviková, Tomáš Vina, Boris Klempa, Jozef Nosek |
| EPI_ISL_875561, EPI_ISL_875562, EPI_ISL_875563, EPI_ISL_875565 | Ohio Department of Health Laboratory | Ohio Department of Health Laboratory | Holmes, Jennifer; Eric Brandt, Keoni Omura, Glen McGillivary, Caitlin McDonnell, Kirtana Ramadugu, Erica Leasure, Kelsey Florek, Heather Blankenship, Quanta Brown, and Tammy Bannerman |
| EPI_ISL_875688 | National Influenza Center - Instituto Adolfo Lutz | Instituto Adolfo Lutz, Interdiciplinary Procedures Center, Strategic Laboratory | Claudio Tavares Sacchi, Claudia Regina Gonçalves, Erica Valessa Ramos Gomes, Karoline Rodrigues Campos, Katia Correa de Oliveira Santos, Ana Lucia de Carvalho Avelino, Clovis Roberto Abe Constantino |
| EPI_ISL_875871, EPI_ISL_875872 | Ohio Department of Health Laboratory | Ohio Department of Health Laboratory | Holmes, Jennifer; Eric Brandt, Keoni Omura, Glen McGillivary, Caitlin McDonnell, Kirtana Ramadugu, Erica Leasure, Kelsey Florek, Heather Blankenship, Quanta Brown, and Tammy Bannerman |
| EPI_ISL_876046, EPI_ISL_876114, EPI_ISL_876176, EPI_ISL_876193, EPI_ISL_876197, EPI_ISL_876199 | Massachusetts State Public Health Laboratory | Massachusetts State Public Health Laboratory | Andrew Lang, Timelia Fink, Glen Gallagher, Sandra Smole |
| EPI_ISL_876519 | DOHMH Jamaica | New York City Public Health Laboratory | Jade Wang, et al. |
| EPI_ISL_876627, EPI_ISL_876642, EPI_ISL_876659, EPI_ISL_876669, EPI_ISL_876690, EPI_ISL_876694 | Helix/Illumina | Genomics and Discovery, Respiratory Viruses Branch, Division of Viral Diseases, Centers for Disease Control and Prevention | Peter W. Cook,Dhwani Batra,Ben L. Rambo-Martin,Eileen de Feo,Jan Antico,Christine Tran,Matthew Tolentino,Shannon Wickline,Kim Gietzen,Brad Sickler,Jingtao Liu,Eric Allen,Phil Febbo,Summer Galloway,Nicole L. Washington,Simon White,Geraint Levan,Kelly Schiabor Barrett,Elizabeth Cirulli,Alexandre Bolze,Ary Ascencio,Charlotte Rivera-Garcia,Ryan Cho,Jason Nguyen,Sherry Wang,Jimmy Ramirez,Tyler Cassens,Efren Sandoval,Magnus Isaksson,William Lee,David Becker,Marc Laurent,James Lu,Clinton R. Paden,Suxiang Tong,Duncan MacCannell, |
| EPI_ISL_878440, EPI_ISL_878443, EPI_ISL_878445, EPI_ISL_878448, EPI_ISL_878450, EPI_ISL_878452, EPI_ISL_878455, EPI_ISL_878457, EPI_ISL_878460, EPI_ISL_878463, EPI_ISL_878465, EPI_ISL_878468, EPI_ISL_878470, EPI_ISL_878472, EPI_ISL_878474 | | | |
| see above | Biolab Diagnostic Laboratories | Andersen lab at Scripps Research | Issa Abu-Dayyeh, Ahmad Tibi, Lama Hussein, Lina Mohammad, Zein Naber, Amid Abdelnour with SEARCH Alliance San Diego |
| EPI_ISL_878784, EPI_ISL_878794 | Rady's Childrens Hospital | Andersen lab at Scripps Research | SEARCH Alliance San Diego with Nanda Radamchar, David Dimmock, Linda Luo, Christina Clarke, Kathryn Bouic, Teresa Mueller, Denise Malicki |
| EPI_ISL_882315, EPI_ISL_882324, EPI_ISL_882329, EPI_ISL_882338, EPI_ISL_882340, EPI_ISL_882347, EPI_ISL_882357, EPI_ISL_882370, EPI_ISL_882378, EPI_ISL_882405, EPI_ISL_882430, EPI_ISL_882487, EPI_ISL_882530, EPI_ISL_882540, EPI_ISL_882556, EPI_ISL_882567, EPI_ISL_882575, EPI_ISL_882591, EPI_ISL_882597 | | | |

| | | | |
|---|---|---|---|
| see above | Lighthouse Lab in Alderley Park | Wellcome Sanger Institute for the COVID-19 Genomics UK (COG-UK) Consortium | Jacquelyn Wynn, Mairead Hyland, The Lighthouse Lab in Alderley Park and Alex Alderton, Roberto Amato, Sonia Goncalves, Ewan Harrison, David K. Jackson, Ian Johnston, Dominic Kwiatkowski, Cordelia Langford, John Sillitoe on behalf of the Wellcome Sanger Institute COVID-19 Surveillance Team |
| EPI_ISL_882640 | Azerbaijan National Hematology Center Division of Medical Genetics | Azerbaijan National Hematology Center Division of Medical Genetics | Aghayev Agha Rza |
| EPI_ISL_882641 | Azerbaijan National Hematology Center Division of Medical Genetics | Azerbaijan National Hematology Center Division of Medical Genetics | Aghayev Agha Rza, Bayraml Ramin |
| EPI_ISL_882645 | Medical Research Center, Faculty of Medicine, Syarif Hidayatullah State Islamic University Jakarta | Medical Research Center, Faculty of Medicine, Syarif Hidayatullah State Islamic University Jakarta | Chris Adhiyanto, Erike A Suwarsono, Laifa Hendarmin, Zeti Harriyati, Endah Wulandari, Flori Ratna Sari, Fika Ekayanti, Hari Hendarto |
| EPI_ISL_882772 | Office of Diseases Prevention and Control Region 4 Saraburi | COVID-19 Network Investigations (CONI) Alliance | Kamolthip Atsawawaranunt, Elizabeth Batty, Wasun Chantratita, Thanat Chookajorn, Stefan Fernandez, Angkana Huang, Anthony R. Jones, Khajohn Joonsalak, Chonticha Klungtong, Theerarat Kochakarn, Prayuth Kaewmalang, Amornmas Kongklieng, Namfon Kotanan, Krittikorn Kumpornsin, Duangkamon Loesbanluechai, Wudtichai Manasatienkij, Anek Mungaomklang, Bhakbhoom Panthan, Pukkaporn Parnwijitkul, Ekawat Pasomsub, Vichan Pawun, Kingkan Rakmanee, Insee Sensorn, Janjira Thaipadungpanit, Arporn Wangwiwatsin, Treewat Watthanachockchai |
| EPI_ISL_882792, EPI_ISL_882793, EPI_ISL_882794 | Institute for Urban Disease Control and Prevention | COVID-19 Network Investigations (CONI) Alliance | Kamolthip Atsawawaranunt, Elizabeth Batty, Wasun Chantratita, Thanat Chookajorn, Stefan Fernandez, Angkana Huang, Anthony R. Jones, Khajohn Joonsalak, Chonticha Klungtong, Theerarat Kochakarn, Prayuth Kaewmalang, Amornmas Kongklieng, Namfon Kotanan, Krittikorn Kumpornsin, Duangkamon Loesbanluechai, Wudtichai Manasatienkij, Anek Mungaomklang, Bhakbhoom Panthan, Pukkaporn Parnwijitkul, Ekawat Pasomsub, Vichan Pawun, Kingkan Rakmanee, Insee Sensorn, Janjira Thaipadungpanit, Arporn Wangwiwatsin, Treewat Watthanachockchai |
| EPI_ISL_882795, EPI_ISL_882796, EPI_ISL_882797, EPI_ISL_882798, EPI_ISL_882799 | Office of Diseases Prevention and Control Region 4 Saraburi | COVID-19 Network Investigations (CONI) Alliance | Kamolthip Atsawawaranunt, Elizabeth Batty, Wasun Chantratita, Thanat Chookajorn, Stefan Fernandez, Angkana Huang, Anthony R. Jones, Khajohn Joonsalak, Chonticha Klungtong, Theerarat Kochakarn, Prayuth Kaewmalang, Amornmas Kongklieng, Namfon Kotanan, Krittikorn Kumpornsin, Duangkamon Loesbanluechai, Wudtichai Manasatienkij, Anek Mungaomklang, Bhakbhoom Panthan, Pukkaporn Parnwijitkul, Ekawat Pasomsub, Vichan Pawun, Kingkan Rakmanee, Insee Sensorn, Janjira Thaipadungpanit, Arporn Wangwiwatsin, Treewat Watthanachockchai |
| EPI_ISL_882927 | INMI Lazzaro Spallanzani IRCCS | INMI Lazzaro Spallanzani IRCCS | M Rueca, E Giombini, C.E.M Gruber, B Bartolini, O Butera, F Messina, A Di Caro, MR Capobianchi |
| EPI_ISL_882932 | INMI Lazzaro Spallanzani IRCCS | INMI Lazzaro Spallanzani IRCCS | F Messina, O Butera, E Giombini, M Rueca, B Bartolini, C.E.M Gruber, MR Capobianchi, A Di Caro |
| EPI_ISL_882943 | The National Institute of Public Health | State Veterinary Institute Prague | Nagy,A;Jirincova,H;Vecerova,JTrnka,D |
| EPI_ISL_883003 | Maryland Public Health Laboratory | Maryland Public Health Laboratory | Maryland Department of Health Laboratories Administration |
| EPI_ISL_883290 | SIESP CHIETI-DRIVE IN CHIETI | Istituto Zooprofilattico Sperimentale dell'Abruzzo e Molise "G. Caporale" | Lorusso A, Marcacci M, Di Domenico M, Ancora M, Curini V, Mangone I, Rinaldi A, Scialabba S, Di Pasquale A, Cammà C, Puglia I, Calistri P, Savini G |
| EPI_ISL_883291 | RP Guardiagrele-Ospedale di Comunità | Istituto Zooprofilattico Sperimentale dell'Abruzzo e Molise "G. Caporale" | Lorusso A, Marcacci M, Di Domenico M, Ancora M, Curini V, Mangone I, Rinaldi A, Scialabba S, Di Pasquale A, Cammà C, Puglia I, Calistri P, Savini G |
| EPI_ISL_883292 | SIESP CHIETI-DRIVE IN ORTONA | Istituto Zooprofilattico Sperimentale dell'Abruzzo e Molise "G. Caporale" | Lorusso A, Marcacci M, Di Domenico M, Ancora M, Curini V, Mangone I, Rinaldi A, Scialabba S, Di Pasquale A, Cammà C, Puglia I, Calistri P, Savini G |
| EPI_ISL_883297 | RP Guardiagrele-Ospedale di Comunità | Istituto Zooprofilattico Sperimentale dell'Abruzzo e Molise "G. Caporale" | Lorusso A, Marcacci M, Di Domenico M, Ancora M, Curini V, Mangone I, Rinaldi A, Scialabba S, Di Pasquale A, Cammà C, Puglia I, Calistri P, Savini G |
| EPI_ISL_883298, EPI_ISL_883299 | SIESP DIPARTIMENTO DI PREVENZIONE CHIE | Istituto Zooprofilattico Sperimentale dell'Abruzzo e Molise "G. Caporale" | Lorusso A, Marcacci M, Di Domenico M, Ancora M, Curini V, Mangone I, Rinaldi A, Scialabba S, Di Pasquale A, Cammà C, Puglia I, Calistri P, Savini G |
| EPI_ISL_883300 | SIESP CHIETI-DRIVE IN CHIETI | Istituto Zooprofilattico Sperimentale dell'Abruzzo e Molise "G. Caporale" | Lorusso A, Marcacci M, Di Domenico M, Ancora M, Curini V, Mangone I, Rinaldi A, Scialabba S, Di Pasquale A, Cammà C, Puglia I, Calistri P, Savini G |
| EPI_ISL_883448, EPI_ISL_883449, EPI_ISL_883450, EPI_ISL_883451, EPI_ISL_883454, EPI_ISL_883455, EPI_ISL_883457 | NORTHWELL HEALTH LABORATORIES | Wadsworth Center, New York State Department of Health | Kirsten St. George, Daryl M. Lamson, Alexis Russel, Matthew Shudt, Melissa A Leisner, Jonathan Plitnick, Navjot Singh, John Kelly, Erasmus Schneider, Erica Lasek-Nesselquist |
| EPI_ISL_884000 | Labo Analyses Med | National Reference Center for Viruses of Respiratory Infections, Institut Pasteur, Paris | Marion Barbet, Sylvie Behillil, Méline Bizard, Angela Brisebarre, Camille Capel, Etienne Simon-Lorière, Vincent Enouf, Maud Vanpeene, Sylvie van der Werf,Rousset Dominique |
| EPI_ISL_884254 | Maryland Public Health Laboratory | Maryland Public Health Laboratory | Maryland Department of Health Laboratories Administration |
| EPI_ISL_884860 | Bacteriology, Georgia Public Health Laboratory (GPHL) | Bacteriology, Georgia Public Health Laboratory (GPHL) | Reeves,S., Edwards,J., Dixey,C., Parrott,T. |
| EPI_ISL_885049 | Santa Clara County Public Health Laboratory | Chan-Zuckerberg Biohub | CZB Cliahub Consortium |
| EPI_ISL_886161, EPI_ISL_886168, EPI_ISL_886187, EPI_ISL_886193, EPI_ISL_886206, EPI_ISL_886208, EPI_ISL_886209, EPI_ISL_886211, EPI_ISL_886217, EPI_ISL_886229, EPI_ISL_886231, EPI_ISL_886250, EPI_ISL_886251, EPI_ISL_886257, EPI_ISL_886258, EPI_ISL_886268, EPI_ISL_886272, EPI_ISL_886277, EPI_ISL_886290, EPI_ISL_886291, EPI_ISL_886299, EPI_ISL_886300, EPI_ISL_886301, EPI_ISL_886309, EPI_ISL_886311, EPI_ISL_886317, EPI_ISL_886320, EPI_ISL_886326, EPI_ISL_886332, EPI_ISL_886340, EPI_ISL_886346, EPI_ISL_886349, EPI_ISL_886373, EPI_ISL_886377, EPI_ISL_886379, EPI_ISL_886381, EPI_ISL_886383, EPI_ISL_886390, EPI_ISL_886395, EPI_ISL_886397, EPI_ISL_886398, EPI_ISL_886399, EPI_ISL_886400, EPI_ISL_886413, EPI_ISL_886417, EPI_ISL_886421, EPI_ISL_886422, EPI_ISL_886429, EPI_ISL_886444, EPI_ISL_886446, EPI_ISL_886450, EPI_ISL_886454, EPI_ISL_886477, EPI_ISL_886485, EPI_ISL_886491, EPI_ISL_886492, EPI_ISL_886497, EPI_ISL_886503, EPI_ISL_886508, EPI_ISL_886513, EPI_ISL_886526, EPI_ISL_886527, EPI_ISL_886543, EPI_ISL_886544, EPI_ISL_886550, EPI_ISL_886551, EPI_ISL_886559, EPI_ISL_886560, EPI_ISL_886561, EPI_ISL_886563, EPI_ISL_886581, EPI_ISL_886586, EPI_ISL_886589, EPI_ISL_886593, EPI_ISL_886596, EPI_ISL_886612, EPI_ISL_886619, EPI_ISL_886627, EPI_ISL_886631, EPI_ISL_886651, EPI_ISL_886654, EPI_ISL_886658, EPI_ISL_886665, EPI_ISL_886675, EPI_ISL_886691, EPI_ISL_886692, EPI_ISL_886693, EPI_ISL_886694, EPI_ISL_886712, EPI_ISL_886714, EPI_ISL_886715, EPI_ISL_886717, EPI_ISL_886722, EPI_ISL_886727, EPI_ISL_886731, EPI_ISL_886732, EPI_ISL_886742, EPI_ISL_886744, EPI_ISL_886752, EPI_ISL_886754, EPI_ISL_886756, EPI_ISL_886763, EPI_ISL_886766, EPI_ISL_886772, EPI_ISL_886782, EPI_ISL_886786, EPI_ISL_886797, EPI_ISL_886798, EPI_ISL_886801, EPI_ISL_886803, EPI_ISL_886804, EPI_ISL_886805, EPI_ISL_886806, EPI_ISL_886810, EPI_ISL_886815, EPI_ISL_886826, EPI_ISL_886843, EPI_ISL_886844, EPI_ISL_886846, EPI_ISL_886862, EPI_ISL_886872, EPI_ISL_886878, EPI_ISL_886889, EPI_ISL_886891, EPI_ISL_886893, EPI_ISL_886899, EPI_ISL_886909, EPI_ISL_886916, EPI_ISL_886919, EPI_ISL_886920, EPI_ISL_886921, EPI_ISL_886924, EPI_ISL_886925, EPI_ISL_886928, EPI_ISL_886931, EPI_ISL_886937, EPI_ISL_886938, EPI_ISL_886942, EPI_ISL_886945, EPI_ISL_886951, EPI_ISL_886960, EPI_ISL_886977, EPI_ISL_886979, EPI_ISL_886984, EPI_ISL_886987, EPI_ISL_886998, EPI_ISL_886999, EPI_ISL_887000, EPI_ISL_887002, EPI_ISL_887003, EPI_ISL_887004, EPI_ISL_887005, EPI_ISL_887024, EPI_ISL_887027, EPI_ISL_887031, EPI_ISL_887035, EPI_ISL_887036, EPI_ISL_887039, EPI_ISL_887044, EPI_ISL_887047, EPI_ISL_887075, EPI_ISL_887078, EPI_ISL_887079, EPI_ISL_887081, EPI_ISL_887102 | | | |
| see above | Labcorp | Genomics and Discovery, Respiratory Viruses Branch, Division of Viral Diseases, Centers for Disease Control and Prevention | Peter W. Cook,Dhwani Batra,Ben L. Rambo-Martin,Summer Galloway,Brian Krueger,Minoo Agarwal,Eyad Almasri,Debbie Boles,Ayla Burns,Nuthawin Charoensri,Oren Cohen,Susan Countryman,Mary Ann Cristobal,Bobbi Croy,Suzanne Dale,Hrushikesh Deshmukh,Amanda Douglas,Vincent Drouillon,Marcia Eisenberg,Howard Engler,Rama Ghatti,Prashant Gupta,Susan Hicks,Jake Humphrey,Lax Iyer,Manoj Jain,Mohan Kolli,Tim Kuphal,Stanley Letovsky,Michael Levandoski,Craig Lukasik,Jonathan Meltzer,Brian Norvell,Mindy Nye,Scott Parker,Christos Petropoulos,John Pruitt,Steven Ragan,Scott Ryan,Mike Sapeta,Jana Schroth,Suresh Babu Selvaraju,Goran Stevovic,Amanda Suchanek,Andrea Throop,Lyndon Tilson,Thomas Urban,Joe Voshell,Kimberly Wagner,Jonathan Williams,Mary Williamson,Qian Zeng,Tricia Zwiefelhofer,Clinton R. Paden,Suxiang Tong,Duncan MacCannell, |
| EPI_ISL_887534 | Johns Hopkins Hospital Department of Pathology | Johns Hopkins Hospital Department of Pathology | C. Paul Morris, Chun Huai Luo, Adannaya Amadi, Matthew Schwartz, Nicholas Gallagher, Heba H. Mostafa |
| EPI_ISL_887591, EPI_ISL_887618, EPI_ISL_887620, EPI_ISL_887724, EPI_ISL_887746, EPI_ISL_887773, EPI_ISL_887792, EPI_ISL_887824, EPI_ISL_887909, EPI_ISL_887913, EPI_ISL_887936, EPI_ISL_887939, EPI_ISL_887970, EPI_ISL_887978, EPI_ISL_887980, EPI_ISL_888004, EPI_ISL_888023, EPI_ISL_888053, EPI_ISL_888091, EPI_ISL_888097, EPI_ISL_888160, EPI_ISL_888176, EPI_ISL_888188, EPI_ISL_888207, EPI_ISL_888254, EPI_ISL_888263, EPI_ISL_888286, EPI_ISL_888509, EPI_ISL_888548, EPI_ISL_888557, EPI_ISL_888564, EPI_ISL_888581, EPI_ISL_888588 | | | |
| see above | Labcorp | Genomics and Discovery, Respiratory Viruses Branch, Division of Viral Diseases, Centers for Disease Control and Prevention | Peter W. Cook,Dhwani Batra,Ben L. Rambo-Martin,Summer Galloway,Brian Krueger,Minoo Agarwal,Eyad Almasri,Debbie Boles,Ayla Burns,Nuthawin Charoensri,Oren Cohen,Susan Countryman,Mary Ann Cristobal,Bobbi Croy,Suzanne Dale,Hrushikesh Deshmukh,Amanda Douglas,Vincent Drouillon,Marcia Eisenberg,Howard Engler,Rama Ghatti,Prashant Gupta,Susan Hicks,Jake Humphrey,Lax Iyer,Manoj Jain,Mohan Kolli,Tim Kuphal,Stanley Letovsky,Michael Levandoski,Craig Lukasik,Jonathan Meltzer,Brian Norvell,Mindy Nye,Scott Parker,Christos Petropoulos,John Pruitt,Steven Ragan,Scott Ryan,Mike Sapeta,Jana Schroth,Suresh Babu Selvaraju,Goran Stevovic,Amanda Suchanek,Andrea Throop,Lyndon Tilson,Thomas Urban,Joe Voshell,Kimberly Wagner,Jonathan Williams,Mary Williamson,Qian Zeng,Tricia Zwiefelhofer,Clinton R. Paden,Suxiang Tong,Duncan MacCannell, |
| EPI_ISL_888853 | Michigan Department of Health and Human Services, Bureau of Laboratories | Michigan Department of Health and Human Services, Bureau of Laboratories | Blankenship HM, Riner D, Soehnlen MK |
| EPI_ISL_888997 | RSU Asysyifaa | Eijkman Institute for Molecular Biology, Ministry of Research | Sukma Oktavianthi, Willy Agustine, Edison Johar, Hidayat Trimarsanto, Iskandar Adnan, Lydia V. Panggalo, Frilasita A Yudhaputri, Safarina G Malik, Khin |

| | | | |
|---|---|---|---|
| | | and Technology/National Agency for Research and Innovation | Saw Myint, Amin Soebandrio |
| EPI_ISL_888998 | RSUD Cileungsi | Eijkman Institute for Molecular Biology, Ministry of Research and Technology/National Agency for Research and Innovation | Sukma Oktavianthi, Willy Agustine, Edison Johar, Hidayat Trimarsanto, Iskandar Adnan, Lydia V. Panggalo, Frilasita A Yudhaputri, Safarina G Malik, Khin Saw Myint, Amin Soebandrio |
| EPI_ISL_888999 | RS Mitra Keluarga Cibubur | Eijkman Institute for Molecular Biology, Ministry of Research and Technology/National Agency for Research and Innovation | Sukma Oktavianthi, Willy Agustine, Edison Johar, Hidayat Trimarsanto, Iskandar Adnan, Lydia V. Panggalo, Frilasita A Yudhaputri, Safarina G Malik, Khin Saw Myint, Amin Soebandrio |
| EPI_ISL_889018 | RSU Medika Dramaga | Eijkman Institute for Molecular Biology, Ministry of Research and Technology/National Agency for Research and Innovation | Lydia V. Panggalo, Sukma Oktavianthi, Willy Agustine, Edison Johar, Hidayat Trimarsanto, Iskandar Adnan, Frilasita A Yudhaputri, Safarina G Malik, Khin Saw Myint, Amin Soebandrio |
| EPI_ISL_889019 | RSIA Sayyidah | Eijkman Institute for Molecular Biology, Ministry of Research and Technology/National Agency for Research and Innovation | Lydia V. Panggalo, Sukma Oktavianthi, Willy Agustine, Edison Johar, Hidayat Trimarsanto, Iskandar Adnan, Frilasita A Yudhaputri, Safarina G Malik, Khin Saw Myint, Amin Soebandrio |
| EPI_ISL_889568, EPI_ISL_889569, EPI_ISL_889570, EPI_ISL_889571, EPI_ISL_889572, EPI_ISL_889573, EPI_ISL_889574, EPI_ISL_889575, EPI_ISL_889576, EPI_ISL_889577, EPI_ISL_889578, EPI_ISL_889579, EPI_ISL_889580, EPI_ISL_889581, EPI_ISL_889582, EPI_ISL_889583, EPI_ISL_889584, EPI_ISL_889585, EPI_ISL_889586, EPI_ISL_889587, EPI_ISL_889588, EPI_ISL_889589, EPI_ISL_889590, EPI_ISL_889591, EPI_ISL_889592, EPI_ISL_889593, EPI_ISL_889594, EPI_ISL_889595, EPI_ISL_889596, EPI_ISL_889597, EPI_ISL_889598, EPI_ISL_889599, EPI_ISL_889600, EPI_ISL_889601, EPI_ISL_889602, EPI_ISL_889603, EPI_ISL_889604, EPI_ISL_889605, EPI_ISL_889606 | | | |
| see above | LSUHS Emerging Viral Threat Laboratory | Microbial Genome Sequencing Center | Jeremy P. Kamil, Jennifer L. Carroll, Camille F. Abshire, Maarten Van Diest, Mohammed N.A. Siddiquey, Andrew D. Yurochko, Martin J. Sapp, Rona S. Scott, Christopher G. Kevil, Daniel J. Snyder, Vaughn S. Cooper, John A. Vanchiere |
| EPI_ISL_890307, EPI_ISL_890310, EPI_ISL_890337, EPI_ISL_890341 | KU Leuven, Rega Institute, Clinical and Epidemiological Virology | KU Leuven, Rega Institute, Clinical and Epidemiological Virology | Tony Wawina-Bokalanga, Bert Vanmechelen, Joan Marti-Carerras, Piet Maes |
| EPI_ISL_891138 | The Jackson Laboratory | The Jackson Laboratory | Lloyd M, Sanderson B, Srivastava A, Maurya R, Renzette N, Omerza G, Kelly K, Li L, Wei C L, Adams M |
| EPI_ISL_891254 | IHU Méditerranée Infection | IHU Méditerranée Infection | Philippe Colson et al. |
| EPI_ISL_891255 | IHU Méditerranée Infection | 19-21, boulevard Jean Moulin, 13005 Marseille | Philippe Colson et al. |
| EPI_ISL_892060 | Lighthouse Lab in Alderley Park | Wellcome Sanger Institute for the COVID-19 Genomics UK (COG-UK) Consortium | Jacquelyn Wynn, Mairead Hyland, The Lighthouse Lab in Alderley Park and Alex Alderton, Roberto Amato, Sonia Goncalves, Ewan Harrison, David K. Jackson, Ian Johnston, Dominic Kwiatkowski, Cordelia Langford, John Sillitoe on behalf of the Wellcome Sanger Institute COVID-19 Surveillance Team |
| EPI_ISL_892211 | Lighthouse Lab in Glasgow | Wellcome Sanger Institute for the COVID-19 Genomics UK (COG-UK) Consortium | Harper VanSteenhouse, Yumi Kasai, David Gray, Carol Clugston, Anna Dominiczak and Alex Alderton, Roberto Amato, Sonia Goncalves, Ewan Harrison, David K. Jackson, Ian Johnston, Dominic Kwiatkowski, Cordelia Langford, John Sillitoe on behalf of the Wellcome Sanger Institute COVID-19 Surveillance Team |
| EPI_ISL_893748, EPI_ISL_893777 | Institute of Virology, Medical Center, University of Freiburg, Freiburg, Germany | Institute of Virology, Clinial Virus Genomics, Medical Center, University of Freiburg, Freiburg, Germany | Jonas Fuchs, Lisa Kern, Sandra Reuter, Hajo Grundmann, Marcus Panning |
| EPI_ISL_894172, EPI_ISL_894173, EPI_ISL_894174, EPI_ISL_894183, EPI_ISL_894187 | KU Leuven, Rega Institute, Clinical and Epidemiological Virology | KU Leuven, Rega Institute, Clinical and Epidemiological Virology | Tony Wawina-Bokalanga, Bert Vanmechelen, Joan Marti-Carerras, Piet Maes |
| EPI_ISL_896077 | Labormedizinisches Zentrum Dr Risch | University Hospital Basel, Clinical Bacteriology | Tim Roloff, Madlen Stange, Helena MB Seth-Smith, Alfredo Mari, Karoline Leuzinger, Julia Bielicki, Nadia Wohlwend,Martin Risch, Lorenz Risch, Manuel Battegay, Hans Hirsch, Adrian Egli |
| EPI_ISL_896078 | Viollier AG | University Hospital Basel, Clinical Bacteriology | Tim Roloff, Madlen Stange, Helena MB Seth-Smith, Alfredo Mari, Karoline Leuzinger, Julia Bielicki, Christiane Beckmann, Manuel Battegay, Hans Hirsch, Adrian Egli |
| EPI_ISL_896079, EPI_ISL_896080, EPI_ISL_896081, EPI_ISL_896082, EPI_ISL_896083, EPI_ISL_896092, EPI_ISL_896097 | Labormedizinisches Zentrum Dr Risch | University Hospital Basel, Clinical Bacteriology | Tim Roloff, Madlen Stange, Helena MB Seth-Smith, Alfredo Mari, Karoline Leuzinger, Julia Bielicki, Nadia Wohlwend,Martin Risch, Lorenz Risch, Manuel Battegay, Hans Hirsch, Adrian Egli |
| EPI_ISL_896098 | Rothen Medizinische Laboratorien AG | University Hospital Basel, Clinical Bacteriology | Tim Roloff, Madlen Stange, Helena MB Seth-Smith, Alfredo Mari, Karoline Leuzinger, Julia Bielicki, Ingrid Steffen, Manuel Battegay, Hans Hirsch, Adrian Egli |
| EPI_ISL_896106, EPI_ISL_896115 | Labormedizinisches Zentrum Dr Risch | University Hospital Basel, Clinical Bacteriology | Tim Roloff, Madlen Stange, Helena MB Seth-Smith, Alfredo Mari, Karoline Leuzinger, Julia Bielicki, Nadia Wohlwend,Martin Risch, Lorenz Risch, Manuel Battegay, Hans Hirsch, Adrian Egli |
| EPI_ISL_896155, EPI_ISL_896168, EPI_ISL_896205 | MEPHI, Aix Marseille University | MEPHI, Aix Marseille University | Anthony LEVASSEUR |
| EPI_ISL_896294, EPI_ISL_896298 | New York Presbyterian Hospital | Wadsworth Center, New York State Department of Health | Kirsten St. George, Daryl M. Lamson, Alexis Russel, Matthew Shudt, Melissa A Leisner, Jonathan Plitnick, Navjot Singh, John Kelly, Erasmus Schneider, Erica Lasek-Nesselquist |
| EPI_ISL_896301 | STONY BROOK UNIVERSITY HOSPITAL | Wadsworth Center, New York State Department of Health | Kirsten St. George, Daryl M. Lamson, Alexis Russel, Matthew Shudt, Melissa A Leisner, Jonathan Plitnick, Navjot Singh, John Kelly, Erasmus Schneider, Erica Lasek-Nesselquist |
| EPI_ISL_896304, EPI_ISL_896338, EPI_ISL_896339, EPI_ISL_896340, EPI_ISL_896341, EPI_ISL_896342, EPI_ISL_896347, EPI_ISL_896348, EPI_ISL_896350, EPI_ISL_896351, EPI_ISL_896352, EPI_ISL_896353, EPI_ISL_896354, EPI_ISL_896355, EPI_ISL_896356, EPI_ISL_896357, EPI_ISL_896358, EPI_ISL_896359, EPI_ISL_896360, EPI_ISL_896361, EPI_ISL_896362, EPI_ISL_896363, EPI_ISL_896364, EPI_ISL_896365 | | | |
| see above | New York Presbyterian Hospital | Wadsworth Center, New York State Department of Health | Kirsten St. George, Daryl M. Lamson, Alexis Russel, Matthew Shudt, Melissa A Leisner, Jonathan Plitnick, Navjot Singh, John Kelly, Erasmus Schneider, Erica Lasek-Nesselquist |
| EPI_ISL_896382, EPI_ISL_896383, EPI_ISL_896384, EPI_ISL_896386, EPI_ISL_896389, EPI_ISL_896391, EPI_ISL_896392, EPI_ISL_896393, EPI_ISL_896394, EPI_ISL_896395, EPI_ISL_896430, EPI_ISL_896431, EPI_ISL_896432, EPI_ISL_896433, EPI_ISL_896434, EPI_ISL_896435, EPI_ISL_896436, EPI_ISL_896437, EPI_ISL_896438, EPI_ISL_896439, EPI_ISL_896440, EPI_ISL_896441, EPI_ISL_896442, EPI_ISL_896443, EPI_ISL_896444, EPI_ISL_896445, EPI_ISL_896446 | | | |
| see above | Columbia University Irving Medical Center | Wadsworth Center, New York State Department of Health | Kirsten St. George, Daryl M. Lamson, Alexis Russel, Matthew Shudt, Melissa A Leisner, Jonathan Plitnick, Navjot Singh, John Kelly, Erasmus Schneider, Erica Lasek-Nesselquist |
| EPI_ISL_899027, EPI_ISL_899047, EPI_ISL_899048, EPI_ISL_899062, EPI_ISL_899095, EPI_ISL_899115, EPI_ISL_899129, EPI_ISL_899218, EPI_ISL_899219, EPI_ISL_899220, EPI_ISL_899221, EPI_ISL_899222, EPI_ISL_899223, EPI_ISL_899224, EPI_ISL_899225, EPI_ISL_899226, EPI_ISL_899227, EPI_ISL_899228, EPI_ISL_899229, EPI_ISL_899230, EPI_ISL_899231, EPI_ISL_899232, EPI_ISL_899233, EPI_ISL_899234, EPI_ISL_899235, EPI_ISL_899237, EPI_ISL_899318, EPI_ISL_899319, EPI_ISL_899320, EPI_ISL_899321, EPI_ISL_899322, EPI_ISL_899323, EPI_ISL_899324, EPI_ISL_899439, EPI_ISL_899440, EPI_ISL_899441, EPI_ISL_899564, EPI_ISL_899565, EPI_ISL_899566, EPI_ISL_899567, EPI_ISL_899568, EPI_ISL_899686, EPI_ISL_899687, EPI_ISL_899688, EPI_ISL_899689, EPI_ISL_899690, EPI_ISL_899691, EPI_ISL_899692, EPI_ISL_899693, EPI_ISL_899694, EPI_ISL_899695, EPI_ISL_899696, EPI_ISL_899697, EPI_ISL_899698, EPI_ISL_899699, EPI_ISL_899783, EPI_ISL_899784, EPI_ISL_899785, EPI_ISL_899786, EPI_ISL_899974, EPI_ISL_899985, EPI_ISL_899986, EPI_ISL_899987 | | | |
| see above | Viollier AG | Department of Biosystems Science and Engineering, ETH Zürich | Christian Beisel, Sarah Nadeau, Chaoran Chen, Ivan Topolsky, Philipp Jablonski, Lara Fuhrmann, David Dreifuss, Katharina Jahn, Tobias Schär, Ina Nissen, Natascha Santacroce, Elodie Burcklen, Christiane Beckmann, Maurice Redondo, Olivier Kobel, Christoph Noppen, Sophie Seidel, Noemie Santamaria de Souza, Niko Beerenwinkel, Tanja Stadler |
| EPI_ISL_900110, EPI_ISL_900317, EPI_ISL_900468 | MEPHI, Aix Marseille University | MEPHI, Aix Marseille University | Anthony LEVASSEUR |
| EPI_ISL_900522 | Novabio | CNR Virus des Infections Respiratoires - France SUD | Antonin Bal, Gregory Destras, Gwendolyne Burfin, Hadrien Règue, Quentin Semanas, Martine Valette, Bruno Lina, Sylvie Larrat, Laurence Josset |
| EPI_ISL_903064, EPI_ISL_903065 | Seattle Flu Study | Seattle Flu Study | Deborah A. Nickerson, Chris D. Frazar, Jover Lee, Benjamin Pelle, Erica Ryke, Matthew Richardson, Amanda Adler, Elisabeth Brandstetter, Peter D. Han, Kairsten Fay, Misja Ilcisin, Kirsten Lacombe, Thomas R. Sibley, Melissa Truong, Caitlin R. Wolf, Michael Boeckh, Janet A. Englund, Michael Famulare, Barry R. Lutz, Mark J. Rieder, Lea M. Starita, Matthew Thompson, Jay Shendure, Trevor Bedford, Helen Y. Chu |
| EPI_ISL_903136, EPI_ISL_903137, EPI_ISL_903138, EPI_ISL_903139, EPI_ISL_903140, EPI_ISL_903141, EPI_ISL_903142, EPI_ISL_903143, EPI_ISL_903144, EPI_ISL_903145, EPI_ISL_903146, EPI_ISL_903147, EPI_ISL_903148, EPI_ISL_903150, EPI_ISL_903152, EPI_ISL_903163, EPI_ISL_903165, EPI_ISL_903166, EPI_ISL_903167, EPI_ISL_903168, EPI_ISL_903169 | | | |
| see above | Washington State Department of Health | Seattle Flu Study | Deborah A. Nickerson, Chris D. Frazar, Jover Lee, Benjamin Pelle, Erica Ryke, Matthew Richardson, Amanda Adler, Elisabeth Brandstetter, Peter D. Han, |

| | | | |
|---|---|---|---|
| | | | Kairsten Fay, Misja Ilcisin, Kirsten Lacombe, Thomas R. Sibley, Melissa Truong, Caitlin R. Wolf, Romesh Gautom, Geoff Melly, Brian Hiatt, Philip Dykema, Scott Lindquist, Michael Boeckh, Janet A. Englund, Michael Famulare, Barry R. Lutz, Mark J. Rieder, Lea M. Starita, Matthew Thompson, Helen Y. Chu, Jay Shendure, Trevor Bedford |
| EPI_ISL_903180, EPI_ISL_903181, EPI_ISL_903182, EPI_ISL_903183, EPI_ISL_903184 | Seattle Flu Study | Seattle Flu Study | Deborah A. Nickerson, Chris D. Frazar, Jover Lee, Benjamin Pelle, Erica Ryke, Matthew Richardson, Amanda Adler, Elisabeth Brandstetter, Peter D. Han, Kairsten Fay, Misja Ilcisin, Kirsten Lacombe, Thomas R. Sibley, Melissa Truong, Caitlin R. Wolf, Karen Cowgill, Stephanie Schrag, Jeff Duchin, Michael Boeckh, Janet A. Englund, Michael Famulare, Barry R. Lutz, Mark J. Rieder, Lea M. Starita, Matthew Thompson, Helen Y. Chu, Trevor Bedford, Jay Shendure |
| EPI_ISL_903237, EPI_ISL_903291, EPI_ISL_903293, EPI_ISL_903295, EPI_ISL_903296, EPI_ISL_903297, EPI_ISL_903298, EPI_ISL_903300, EPI_ISL_903301, EPI_ISL_903302, EPI_ISL_903307, EPI_ISL_903308, EPI_ISL_903310 | | | |
| see above | M Health Fairview | Minnesota Department of Health, Public Health Laboratory | Alexandra Lorentz, Jacob Garfin, Matt Plumb, and Xiong Wang |
| EPI_ISL_903670, EPI_ISL_903685 | GA Department of Public Health Laboratory | Genomics and Discovery, Respiratory Viruses Branch, Division of Viral Diseases, Centers for Disease Control and Prevention | Krista Queen, Yan Li, Ying Tao, Jing Zhang, Anna Uehara, Anna Montmayeur, Clinton R. Paden, Peter W. Cook, Rachel Marine, Mili Sheth, Jasmine Padilla, Sarah Nobles, Mark Burroughs, Lori Rowe, Haibin Wang, Ben L. Rambo-Martin, Dhwani Batra, Justin Lee, Suxiang Tong |
| EPI_ISL_903822, EPI_ISL_903865 | WI State Laboratory of Hygiene | Genomics and Discovery, Respiratory Viruses Branch, Division of Viral Diseases, Centers for Disease Control and Prevention | Krista Queen, Yan Li, Ying Tao, Jing Zhang, Anna Uehara, Anna Montmayeur, Clinton R. Paden, Peter W. Cook, Rachel Marine, Mili Sheth, Jasmine Padilla, Sarah Nobles, Mark Burroughs, Lori Rowe, Haibin Wang, Ben L. Rambo-Martin, Dhwani Batra, Justin Lee, Suxiang Tong |
| EPI_ISL_904050, EPI_ISL_904051, EPI_ISL_904052, EPI_ISL_904053, EPI_ISL_904054, EPI_ISL_904055, EPI_ISL_904056, EPI_ISL_904057 | New Mexico Department of Health Scientific Laboratory | New Mexico Department of Health Scientific Laboratory | Ellie Johnson, Anastacia Griego-Fisher, D'eldra Malone |
| EPI_ISL_904270, EPI_ISL_904373, EPI_ISL_904381, EPI_ISL_904540, EPI_ISL_904541, EPI_ISL_904542, EPI_ISL_904543, EPI_ISL_904544, EPI_ISL_904545, EPI_ISL_904546, EPI_ISL_904547 | | | |
| see above | Dutch COVID-19 response team | Erasmus Medical Center | Bas Oude Munnink, Reina Sikkema, David Nieuwenhuijse, Irina Chestakova, Anne van der Linden, Marjan Boter, Emmanuelle Munger, Corine GeurtsvanKessel, Annemiek van der Eijk, Richard Molenkamp, Marion Koopmans, on behalf of the Dutch national COVID-19 response team. |
| EPI_ISL_904629, EPI_ISL_904632, EPI_ISL_904633, EPI_ISL_904648, EPI_ISL_904655 | Servicio de Microbiología, Laboratori Clínic Metropolitana Nord. Hospital Universitari Germans Trias i Pujol. Institut d'Investigació en Ciències de la Salut Germans Trias i Pujol (IGTP) | IrsiCaixa - Can Ruti CovidSeq | Marc Noguera-Julian, Mariona Parera, Maria Casadellà, Pilar Armengol, Francesc Catala-Moll, Roger Paredes, Bonaventura Clotet Elisa Martró, Verónica Saludes, Anna Not, Ana Pérez, Montserrat Giménez, Ignacio Blanco, Cristina Casañ, Antoni E. Bordoy, Adrián Antuori |
| EPI_ISL_904982, EPI_ISL_904984, EPI_ISL_904985, EPI_ISL_904991, EPI_ISL_904993, EPI_ISL_904996, EPI_ISL_905010, EPI_ISL_905012, EPI_ISL_905013, EPI_ISL_905014, EPI_ISL_905015, EPI_ISL_905016, EPI_ISL_905018, EPI_ISL_905019, EPI_ISL_905020, EPI_ISL_905021, EPI_ISL_905022, EPI_ISL_905023, EPI_ISL_905024, EPI_ISL_905025, EPI_ISL_905026, EPI_ISL_905027, EPI_ISL_905028, EPI_ISL_905030, EPI_ISL_905268, EPI_ISL_905275, EPI_ISL_905276, EPI_ISL_905282, EPI_ISL_905284, EPI_ISL_905285, EPI_ISL_905286, EPI_ISL_905287, EPI_ISL_905288, EPI_ISL_905361, EPI_ISL_905362, EPI_ISL_905384, EPI_ISL_905385, EPI_ISL_905386, EPI_ISL_905388, EPI_ISL_905391, EPI_ISL_905393, EPI_ISL_905424, EPI_ISL_905439, EPI_ISL_905440, EPI_ISL_905441, EPI_ISL_905442, EPI_ISL_905443, EPI_ISL_905444, EPI_ISL_905445, EPI_ISL_905446, EPI_ISL_905447, EPI_ISL_905448, EPI_ISL_905449, EPI_ISL_905459, EPI_ISL_905460, EPI_ISL_905461, EPI_ISL_905481, EPI_ISL_905501, EPI_ISL_905513, EPI_ISL_905516, EPI_ISL_905519, EPI_ISL_905520, EPI_ISL_905521, EPI_ISL_905522, EPI_ISL_905533, EPI_ISL_905568, EPI_ISL_905592, EPI_ISL_905593, EPI_ISL_905594, EPI_ISL_905595, EPI_ISL_905596, EPI_ISL_905610, EPI_ISL_905641, EPI_ISL_905642, EPI_ISL_905714, EPI_ISL_905715, EPI_ISL_905718, EPI_ISL_905728 | | | |
| see above | Dutch COVID-19 response team | National Institute for Public Health and the Environment (RIVM) | Adam Meijer, Harry Vennema, Dirk Eggink, Jeroen Cremer, Sharon van den Brink, Bas van der Veer, AnneMarie van den Brandt, Florian Zwagemaker, Dennis Schmitz, Chantal Reusken, on behalf of the national COVID-19 response team |
| EPI_ISL_905740 | NZOZ LM Diagnostyka Czstochowa | National Institute of Public Health - National Institute of Hygiene | Wokowicz Tomasz, Zacharczuk Katarzyna |
| EPI_ISL_905914, EPI_ISL_905915, EPI_ISL_905916, EPI_ISL_905917, EPI_ISL_905918, EPI_ISL_905919, EPI_ISL_905920, EPI_ISL_905921, EPI_ISL_905922, EPI_ISL_905923, EPI_ISL_905924, EPI_ISL_905925, EPI_ISL_905926 | | | |
| see above | OHSU Lab Services Molecular Microbiology Lab | Oregon SARS-CoV-2 Genome Sequencing Center | Brendan L. O'Connell, Sally Grindstaff, Kayla Carter, Ruth V. Nichols, Alec J. Hirsch, Donna Hansel, Guang Fan, Xuan, Qin, Daniel N. Streblow, William B. Messer, Andrew C. Adey, Benjamin N. Bimber, Brian J. O'Roak |
| EPI_ISL_906098 | Shimantik Pathology and Diagnostic Center | Child Health Research Foundation | Senjuti Saha, Arif Mohammad Tanmoy, Syed Muktadir Al Sium, Afroza Akter Tanni, Sharmistha Goswami, Roly Malaker, Md Hafizur Rahman, Md. Parvej Alam, Md. Mobarok Karim, Samir K Saha |
| EPI_ISL_906547 | HOSPITAL DEPARTAMENTAL DE VILLAVICENCIO E.S.E. | Instituto Nacional de Salud- Dirección de Investigación en Salud Pública, Universidad de los Andes- Applied genomics research group, Vicerrectoria de Investigación y Creación, Universidad de los Andes- Systems and Computing Engineering Department | Katherine Laiton-Donato, Diego A. Álvarez-Díaz, Carlos Franco-Muñoz, Mauricio Pacheco-Montealegre, Héctor Alejandro Ruiz-Moreno, Maria T. Herrera-Sepúlveda, Diego Andrés Prada, Jhonnatan Reales-González, Sheryll Corchuelo, Julian Naizaque, Gerardo Santamaría Jorge Duitama, Laura Natalia Gonzalez, Jorge Ivan Diaz, Silvia Restrepo-Restrepo, Magdalena Wiesner, Martha Lucia Ospina Martinez, Marcela Mercado-Reyes |
| EPI_ISL_906573, EPI_ISL_906574 | Maine Health and Environmental Testing Laboratory (Maine HETL) | Tewhey Lab, The Jackson Laboratory | Matluk,N., Dewey,H., Iosue,F., Barter,M., Lynch,R., Munger,H. and Tewhey,R. |
| EPI_ISL_906728 | Hematology Laboratory, Section of Molecular Diagnostics, University Clinical Centre, Medical University of Gdansk | Laboratory of Recombinant Vaccines | Lukasz Rabalski, Maciej Kosinski, Maciej Grzybek, Adam Sodol, Aneta Szulc, Krzysztof Lewandowski, Ewa Milosz, Marlena Robakowska, Boguslaw Szewczyk, Krystyna Bienkowska-Szewczyk |
| EPI_ISL_906780, EPI_ISL_906781, EPI_ISL_906790, EPI_ISL_906792 | Center for Genome Research and Biocomputing | Center for Genome Research and Biocomputing | Oregon State University TRACE Project |
| EPI_ISL_906800, EPI_ISL_906802 | Nordland Hospital - Bodo, Laboratory Department, Molecular Biology Unit | Norwegian Institute of Public Health, Department of Virology | Kathrine Stene-Johansen, Kamilla Heddeland Instefjord, Hilde Elshaug, Atiya R Ali,Marie Paulsen Madsen, Rasmus Riis Kopperud, Hilde Vollan, Karoline Bragstad, Olav Hungnes |
| EPI_ISL_906824 | Department of Medical Microbiology - section Molde, Molde Hospital | Norwegian Institute of Public Health, Department of Virology | Kathrine Stene-Johansen, Kamilla Heddeland Instefjord, Hilde Elshaug, Atiya R Ali,Marie Paulsen Madsen, Rasmus Riis Kopperud, Hilde Vollan, Karoline Bragstad, Olav Hungnes |
| EPI_ISL_906830 | Furst Medical Laboratory | Norwegian Institute of Public Health, Department of Virology | Kathrine Stene-Johansen, Kamilla Heddeland Instefjord, Hilde Elshaug, Atiya R Ali,Marie Paulsen Madsen, Rasmus Riis Kopperud, Hilde Vollan, Karoline Bragstad, Olav Hungnes |
| EPI_ISL_906831 | Hospital of Southern Norway - Kristiansand, Department of Medical Microbiology | Norwegian Institute of Public Health, Department of Virology | Kathrine Stene-Johansen, Kamilla Heddeland Instefjord, Hilde Elshaug, Atiya R Ali,Marie Paulsen Madsen, Rasmus Riis Kopperud, Hilde Vollan, Karoline Bragstad, Olav Hungnes |
| EPI_ISL_906832 | Oslo University Hospital, Department of Medical Microbiology | Norwegian Institute of Public Health, Department of Virology | Kathrine Stene-Johansen, Kamilla Heddeland Instefjord, Hilde Elshaug, Atiya R Ali,Marie Paulsen Madsen, Rasmus Riis Kopperud, Hilde Vollan, Karoline Bragstad, Olav Hungnes |
| EPI_ISL_910016, EPI_ISL_910338, EPI_ISL_910339 | Laboratory for Respiratory Viruses, Cantacuzino National Military-Medical Institute for Research and Development | Cantacuzino Institute Virology | Luiza Ustea, Nicoleta Paraschiv, Mihaela Lazar |
| EPI_ISL_910888, EPI_ISL_910889, EPI_ISL_910890, EPI_ISL_910891, EPI_ISL_910892, EPI_ISL_910893, EPI_ISL_910894, EPI_ISL_910895, EPI_ISL_910896, EPI_ISL_910897, EPI_ISL_910898, EPI_ISL_910899, EPI_ISL_910900, EPI_ISL_910901 | | | |
| see above | Laboratoire national de sante, Microbiology, Virology | Laboratoire national de sante, Microbiology, Microbial Genomics Platform | Anke Wienecke-Baldacchino, Catherine Ragimbeau,Jessica Tapp, Fatu Djabi, Lise Pignon, Raoul Salmon, Tamir Abdelrahman |
| EPI_ISL_911650, EPI_ISL_911651, EPI_ISL_911652, EPI_ISL_911653, EPI_ISL_911654, EPI_ISL_911655, EPI_ISL_911656, EPI_ISL_911657, EPI_ISL_911665, EPI_ISL_911666, EPI_ISL_911667, EPI_ISL_911668, EPI_ISL_911669 | | | |
| see above | Texas Department of State Health Services (TXDSHS) | Texas Department of State Health Services (TXDSHS) | Bonnie Oh, Anita Pokharel, James Daniel Bonser, Myong Koag, Chung Wang, Rachel Lee, Grace Kubin, Rashmi Tuladhar, Mayela Pedrueza, Maliha Rahman, Jenny Zhang |
| EPI_ISL_911923 | IL Department of Public Health Chicago Laboratory/Tempus Labs | Pathogen Discovery, Respiratory Viruses Branch, Division of Viral Diseases, Centers for Disease Control and Prevention | Ying Tao, Yan Li, Jing Zhang, Krista Queen, Anna Uehara, Peter Cook, Clinton R. Paden, Haibin Wang, Suxiang Tong |
| EPI_ISL_912163, EPI_ISL_912164 | Connecticut Department of Health | Grubaugh Lab - Yale School of Public Health | Tara Alpert, Joseph Fauver, Anderson Brito, Mallery Breban, Anne Wyllie, Chantal Vogels, Mary Petrone, Annie Watkins, Chaney Kalinich, Isabel Ott, |

| | | | |
|---|---|---|---|
| | | | Nathan Grubaugh |
| EPI_ISL_912272, EPI_ISL_912293, EPI_ISL_912318, EPI_ISL_912325, EPI_ISL_912330, EPI_ISL_912339, EPI_ISL_912349, EPI_ISL_912351 | Hospital General Universitario Gregorio Marañón | SeqCOVID-SPAIN consortium / IBV (CSIC) | Darío García de Viedma, Laura Pérez-Lago, Pedro J Sola-Campoy, Sergio Buenestado-Serrano, Marta Herranz, Victor Manuel de la Cueva, Julia Suárez, Pilar Catalán, Patricia Muñoz and SeqCOVID-SPAIN consortium |
| EPI_ISL_912376, EPI_ISL_912377, EPI_ISL_912378 | Fondation Congolaise pour la recherche medicale (FCRM), Francine Ntoumi | NGS Competence Center Tuebingen, Institut für Medizinische Mikrobiologie und Hygiene, Universitaetsklinikum Tübingen | Angel Angelov |
| EPI_ISL_912406, EPI_ISL_912410, EPI_ISL_912418, EPI_ISL_912421, EPI_ISL_912422, EPI_ISL_912427, EPI_ISL_912429, EPI_ISL_912453 | KU Leuven, Rega Institute, Clinical and Epidemiological Virology | KU Leuven, Rega Institute, Clinical and Epidemiological Virology | Tony Wawina-Bokalanga, Bert Vanmechelen, Joan Marti-Carerras, Piet Maes |
| EPI_ISL_912645, EPI_ISL_912650, EPI_ISL_912658, EPI_ISL_912669, EPI_ISL_912706, EPI_ISL_912718, EPI_ISL_912722, EPI_ISL_912723, EPI_ISL_912724, EPI_ISL_912727, EPI_ISL_912728, EPI_ISL_912729, EPI_ISL_912730, EPI_ISL_912731, EPI_ISL_912732, EPI_ISL_912733 | | | |
| see above | Hôpital Henri Mondor | Department of Virology, Henri Mondor University Hospital, Assistance Publique Hôpitaux de Paris, Université Paris-Est Créteil, INSERM U955 | Christophe Rodriguez, Slim Fourati, Vanessa Demontant, Guillaume Gricourt, Melissa N'Debi, Alexandre Soulier, Elisabeth Trawinski, Jean-Michel Pawlotsky |
| EPI_ISL_913059 | Consejería de Sanidad y Asuntos Sociales | Instituto de Salud Carlos III | Iglesias-Caballero, M. Camarero, S. Sandonís,V. Vázquez, S. Pozo, F. Casas, I. Jiménez, P. Zaballos, A. Monzón, S. Varona, S. Cuesta, I. Gutiérrez, G. |
| EPI_ISL_913298, EPI_ISL_913299 | Klinisk Mikrobiologi | The Public Health Agency of Sweden | Anna-Malin Linde, Maria Lind Karlberg, Carlo Berg, Oskar Karlsson Lindsjo, Sofia Stamouli, Reza Advani, Mattias Haukland, Petra Holmstrom, Noura Walai, Petra Edquist, Mia Brytting, Anna Risberg, Karin Tegmark-Wisell |
| EPI_ISL_913312, EPI_ISL_913330 | Mikrobiologen | The Public Health Agency of Sweden | Anna-Malin Linde, Maria Lind Karlberg, Carlo Berg, Oskar Karlsson Lindsjo, Sofia Stamouli, Reza Advani, Mattias Haukland, Petra Holmstrom, Noura Walai, Petra Edquist, Mia Brytting, Anna Risberg, Karin Tegmark-Wisell |
| EPI_ISL_913334, EPI_ISL_913335, EPI_ISL_913336 | Klinisk Mikrobiologi | The Public Health Agency of Sweden | Anna-Malin Linde, Maria Lind Karlberg, Carlo Berg, Oskar Karlsson Lindsjo, Sofia Stamouli, Reza Advani, Mattias Haukland, Petra Holmstrom, Noura Walai, Petra Edquist, Mia Brytting, Anna Risberg, Karin Tegmark-Wisell |
| EPI_ISL_913410 | Massachusetts State Public Health Laboratory | Massachusetts State Public Health Laboratory | Andrew Lang, Timelia Fink, Glen Gallagher, Sandra Smole |
| EPI_ISL_913568 | M Health Fairview | Minnesota Department of Health, Public Health Laboratory | Alexandra Lorentz, Jacob Garfin, Matt Plumb, and Xiong Wang |
| EPI_ISL_913647, EPI_ISL_913649, EPI_ISL_913650, EPI_ISL_913655, EPI_ISL_913656, EPI_ISL_913657, EPI_ISL_913658, EPI_ISL_913659, EPI_ISL_913660, EPI_ISL_913661, EPI_ISL_913662, EPI_ISL_913663, EPI_ISL_913664, EPI_ISL_913665 | | | |
| see above | Michigan Department of Health and Human Services, Bureau of Laboratories | Michigan Department of Health and Human Services, Bureau of Laboratories | Blankenship HM, Riner D, Soehnlen MK |
| EPI_ISL_913713, EPI_ISL_913714, EPI_ISL_913715, EPI_ISL_913716, EPI_ISL_913717, EPI_ISL_913718, EPI_ISL_913719 | Minnesota Department of Health, Public Health Laboratory | Minnesota Department of Health, Public Health Laboratory | Alexandra Lorentz, Jacob Garfin, Matt Plumb, and Xiong Wang |
| EPI_ISL_913732, EPI_ISL_913733, EPI_ISL_913734, EPI_ISL_913735, EPI_ISL_913736 | M Health Fairview | Minnesota Department of Health, Public Health Laboratory | Alexandra Lorentz, Jacob Garfin, Matt Plumb, and Xiong Wang |
| EPI_ISL_913778, EPI_ISL_913818, EPI_ISL_913842, EPI_ISL_913859, EPI_ISL_913866, EPI_ISL_913870, EPI_ISL_913882, EPI_ISL_913888, EPI_ISL_913889, EPI_ISL_913905, EPI_ISL_913910 | | | |
| see above | TGen North | TGen North | "Jolene Bowers, Megan Folkerts, Chris French, Hayley Yaglom, Ashlyn Pfeiffer, Darrin Lemmer, Dave Engelthaler, The Arizona COVID Genomics Union (ACGU)" |
| EPI_ISL_913990 | IL Department of Public Health Chicago Laboratory | Pathogen Discovery, Respiratory Viruses Branch, Division of Viral Diseases, Centers for Disease Control and Prevention | Ying Tao, Yan Li, Jing Zhang, Krista Queen, Anna Uehara, Peter Cook, Clinton R. Paden, Haibin Wang, Suxiang Tong |
| EPI_ISL_914036, EPI_ISL_914037 | CT-Dr. Katherine A. Kelley State Public Health Lab | Pathogen Discovery, Respiratory Viruses Branch, Division of Viral Diseases, Centers for Disease Control and Prevention | Ying Tao, Yan Li, Jing Zhang, Krista Queen, Anna Uehara, Peter Cook, Clinton R. Paden, Haibin Wang, Suxiang Tong |
| EPI_ISL_914820 | AREA DE SALUD SAN JUAN-SAN DIEGO-CONCEPCION 2 | Inciensa, Instituto Costarricense de Investigación y Enseñanza en Nutrición y Salud | Francisco Duarte, Hebleen Porras, Claudio Soto-Garita, Estela Cordero, Adriana Godínez, Melany Calderón & Mariel López |
| EPI_ISL_914824 | TAMIZAJE COMUNITARIO - PASO CANOAS | Inciensa, Instituto Costarricense de Investigación y Enseñanza en Nutrición y Salud | Francisco Duarte, Hebleen Porras, Claudio Soto-Garita, Estela Cordero, Adriana Godínez, Melany Calderón & Mariel López |
| EPI_ISL_918174 | Department of Infectious Diseases and Immunology, National Hospital Organization Nagoya Medical Center | Clinical Research Center, National Hospital Organization Nagoya Medical Center | Yoshihiro Nakata, Hirotaka Ode, Mai Kubota, Masakazu Matsuda, Kazuhiro Matsuoka, Miho Nakasuji, Mikiko Mori, Mayumi Imahashi, Yoshiyuki Yokomaku, Yasumasa Iwatani |
| EPI_ISL_918273, EPI_ISL_918274, EPI_ISL_918275, EPI_ISL_918276 | Hospital Universitari Vall d'Hebron - Vall d'Hebron Institut de Rercerca | Hospital Universitari Vall d'Hebron | Cristina Andrés, Maria Piñana, Josep F Abril, Damir Garcia-Cehic, Ariadna Rando, Juliana Esperalba, Maria Gema Codina, Carla Castillo, Maria Carmen Martin, Tomàs Pumarola, Josep Quer, Andrés Antón |
| EPI_ISL_918348 | Institute of Virology, Medical Center, University of Freiburg, Freiburg, Germany | Institute of Virology, Clinial Virus Genomics, Medical Center, University of Freiburg, Freiburg, Germany | Jonas Fuchs, Lisa Kern, Sandra Reuter, Hajo Grundmann, Marcus Panning |
| EPI_ISL_918434, EPI_ISL_918436, EPI_ISL_918437, EPI_ISL_918441, EPI_ISL_918442 | AIID | Irish Coronavirus Sequencing Consortium-Teagasc Grange | Matthew McCabe, Aljandro Abner Garcia Leon, Fiona Crispie, Calum Walsh, Michael Carr, John Kenny, Paul Cotter, Patrick Mallon, Gabriel Gonzalez |
| EPI_ISL_918541, EPI_ISL_918542 | LACEN - Laboratório Central de Saúde Pública do Ceara | Evandro Chagas Institute | Santos, M.C.; Silva, A.M.; Junior, W.D.C.; Barbagelata, L.S.; Ferreira, J.A.; Sousa, E.M.A.; da Silva, P.S.; Pinheiro, K.C.; L.C.; Sousa Junior, E.C. |
| EPI_ISL_918584, EPI_ISL_918738, EPI_ISL_918739, EPI_ISL_918741, EPI_ISL_918742, EPI_ISL_918743, EPI_ISL_918744, EPI_ISL_918746, EPI_ISL_918747, EPI_ISL_918907, EPI_ISL_918908 | | | |
| see above | University of Birmingham | COVID-19 Genomics UK (COG-UK) Consortium | Institute of Microbiology, University of Birmingham: Claire McMurray, Joanne Stockton, Sam Nicholls, Radoslaw Poplawski, Will Rowe, Josh Quick, Nicholas Loman. University of Birmingham Testing Laboratory: Celina M Whalley, Andrew Bosworth, Charlotte Poxon, Kasun Wanigasooriya, Oliver Pickles, Mike Kidd, Alex Richter, Andrew D Beggs PHE Heartlands Lab: Husam Osman, Andrew Bosworth. Queen Elizabeth Hospital: Anna Casey |
| EPI_ISL_919455, EPI_ISL_919459, EPI_ISL_919461 | Liverpool Clinical Laboratories | COVID-19 Genomics UK (COG-UK) Consortium | Sam Haldenby, Anita Lucaci, Steve Paterson, Julian Hiscox, Alistair Darby, M Almsaud, A Alrezaihi, Muhannad Alruwaili, Stuart D Armstrong, Jones Benjamin, Eleanor G Bentley, Anu Chawla, Jordan J Clark, Angela Cowell, Richard Eccles, Isabel García-Dorival, Matthew Gemmell, Alessandro Gerada, PKF Gilmore, Richard Gregory, Ximeng Han, Catherine Hartley, Margaret Hughes, Miren Iturriza-Gomara, James Johnson, L Luu, Jenifer Manson, Charlotte Nelson, Elaine O'Toole, Cassie Olateju, Rebekah Penrice-Randal , Lucille Rainbow, N.P Randle, Trevor Ian Robinson, Parul Sharma, Ghada T Shawli, James P Stewart, Neil Swainston, Ecaterina Vamos, Joanne Watts, Mark Whitehead |
| EPI_ISL_919798, EPI_ISL_919799, EPI_ISL_919800, EPI_ISL_919801, EPI_ISL_919802, EPI_ISL_919805, EPI_ISL_919814, EPI_ISL_919815, EPI_ISL_919816, EPI_ISL_919817, EPI_ISL_919818, EPI_ISL_919819, EPI_ISL_919821, EPI_ISL_919822, EPI_ISL_919823, EPI_ISL_919824, EPI_ISL_919825, EPI_ISL_919843, EPI_ISL_919844 | | | |
| see above | Barts Health NHS Trust | COVID-19 Genomics UK (COG-UK) Consortium | CUTINO-MOGUEL, Maria-Teresa; HARRINGTON, David; OWOYEMI, Dola; KULASEGARAN-SHYLINI, Raghavendran; BROAD, Claire; KELE, Beatrix |
| EPI_ISL_919970, EPI_ISL_919973, EPI_ISL_919978, EPI_ISL_919979, EPI_ISL_919980, EPI_ISL_919981, EPI_ISL_919985, EPI_ISL_919987, EPI_ISL_920040, EPI_ISL_920047 | University College London, Great Ormond Street Hospital for Children NHS Foundation Trust, Imperial College Healthcare NHS Trust | COVID-19 Genomics UK (COG-UK) Consortium | Sergi Castellano, Rachel Williams, Mark Kristiansen, Paola Resende Silva, Sunando Roy, Tony Brooks, Helena Tutill, Paola Niola, Patricia Dyal, Charlotte Williams, Leysa Forrest, Yasmin Panchbhaya, Jacqueline Findlay, Samuel Weeks, Julianne Brown, Kathryn Harris, Paul Randell, James Price, Alison Holmes, Judith Breuer |

| | | | |
|---|---|---|---|
| EPI_ISL_920711 | University College London Hospital | COVID-19 Genomics UK (COG-UK) Consortium | Judith Heaney, Matthew Byott, Catherine Houlihan, Dan Frampton, Stuart Kirk, Moira Spyer and Eleni Nastouli |
| EPI_ISL_920849 | University College London, Great Ormond Street Hospital for Children NHS Foundation Trust, Imperial College Healthcare NHS Trust | COVID-19 Genomics UK (COG-UK) Consortium | Sergi Castellano, Rachel Williams, Mark Kristiansen, Paola Resende Silva, Sunando Roy, Tony Brooks, Helena Tutill, Paola Niola, Patricia Dyal, Charlotte Williams, Leysa Forrest, Yasmin Panchbhaya, Jacqueline Findlay, Samuel Weeks, Julianne Brown, Kathryn Harris, Paul Randell, James Price, Alison Holmes, Judith Breuer |
| EPI_ISL_921020, EPI_ISL_921021, EPI_ISL_921023, EPI_ISL_921080, EPI_ISL_921081, EPI_ISL_921082, EPI_ISL_921083, EPI_ISL_921084, EPI_ISL_921099, EPI_ISL_921103, EPI_ISL_921104, EPI_ISL_921108, EPI_ISL_921112, EPI_ISL_921123, EPI_ISL_921124 | | | |
| see above | Regional Virus Laboratory, Belfast Health and Social Care Trust | COVID-19 Genomics UK (COG-UK) Consortium | Conall McCaughey, James McKenna, Tanya Curran, Susan Feeney, Alison Watt, Ciara Cox, Mairead Connor, Zoltan Molnar, David Simpson, Derek Fairley |
| EPI_ISL_921277, EPI_ISL_921287, EPI_ISL_921289, EPI_ISL_921290, EPI_ISL_921291, EPI_ISL_921292, EPI_ISL_921294, EPI_ISL_921296, EPI_ISL_921299, EPI_ISL_921407, EPI_ISL_921408, EPI_ISL_921409, EPI_ISL_921410, EPI_ISL_921411, EPI_ISL_921412, EPI_ISL_921413, EPI_ISL_921438, EPI_ISL_921440, EPI_ISL_921441, EPI_ISL_921443, EPI_ISL_921444, EPI_ISL_921445, EPI_ISL_921447, EPI_ISL_921448, EPI_ISL_921449 | | | |
| see above | Northumbria University / South Tees Hospitals NHS Foundation Trust / North Cumbria Integrated Care NHS Foundation Trust / North Tees and Hartlepool NHS Foundation Trust / Newcastle Hospitals NHS Foundation Trust | COVID-19 Genomics UK (COG-UK) Consortium | Darren L Smith,Andrew Nelson,Matthew Bashton,Greg R Young,Joshua Loh,John Allan,Mohammad A Tariq,Giles S Holt,Gary Black,Wen C Yew,Lynn Dover,Paul Baker,Steve Liggett,Sarah Essex,Jane Greenaway,Debra Padgett,Clive Graham,Garren Scott,Edward Barton,Emma Swindells,Brendan Payne,Jennifer Collins,Yusri Taha,Gary Eltringham |
| EPI_ISL_922175, EPI_ISL_922178, EPI_ISL_922183, EPI_ISL_922188, EPI_ISL_922192, EPI_ISL_922249, EPI_ISL_922283, EPI_ISL_922285, EPI_ISL_922304, EPI_ISL_922309, EPI_ISL_922310, EPI_ISL_922317, EPI_ISL_922321, EPI_ISL_922339, EPI_ISL_922340, EPI_ISL_922341 | | | |
| see above | Oxford Viromics, NDM, University of Oxford; Oxford University Hospitals; Basingstoke and North Hampshire Hospital | COVID-19 Genomics UK (COG-UK) Consortium | Tanya Golubchik, David Bonsall, George Macintyre, Amy Trebes, Mariateresa de Cesare, Catrin Moore, Alex Mobbs, Anita Justice, Robert Shaw, Monique Andersson, Timothy Peto, Emma Wise, Nathan Moore, Jessica Lynch, Nick Cortes, Matilde Mori, Stephen Kidd, David Buck, John Todd, Christophe Fraser |
| EPI_ISL_923577, EPI_ISL_923580 | Centre for Enzyme Innovation, University of Portsmouth / Translational Research Laboratory, Portsmouth Hospitals NHS Trust | COVID-19 Genomics UK (COG-UK) Consortium | Angela Beckett,Salman Goudarzi,Christopher Fearn,Kate Cook,Katie Loveson,Sharon Glaysher,Scott Elliott,Samuel Robson |
| EPI_ISL_924079, EPI_ISL_924107, EPI_ISL_924132, EPI_ISL_924292, EPI_ISL_924342, EPI_ISL_924400 | Virology Department, Sheffield Teaching Hospitals NHS Foundation Trust/Department of Infection, Immunity and Cardiovascular Disease, The Medical School, University of Sheffield | COVID-19 Genomics UK (COG-UK) Consortium | Thushan de Silva, Matthew Parker, Nikki Smith, Adri Angyal, Rebecca Brown, Luke Green, Rachel Tucker, Paul Parsons, Danielle Groves, Katie Johnson, Laura Carrilero, Alex Keeley, Dave Partridge, Matthew Wyles, Benjamin Lindsey, Mehmet Yavuz, Mohammad Raza, Cariad Evans |
| EPI_ISL_924622, EPI_ISL_924630, EPI_ISL_924634, EPI_ISL_924893, EPI_ISL_924894, EPI_ISL_924896, EPI_ISL_924900, EPI_ISL_924906 | Bioinformatics and Biostatistics Lab, Advanced Sequencing Facility | COVID-19 Genomics UK (COG-UK) Consortium | Aengus Stewart,Jerome Nicod,Chelsea Sawyer,Laura Cubitt,Harshil Patel,Margaret Crawford |
| EPI_ISL_925064 | TXDSHS | TXDSHS | Bonnie Oh, Anita Pokharel, James Daniel Bonser, Myong Koag, Chung Wang, Rachel Lee, Grace Kubin, Rashmi Tuladhar, Mayela Pedrueza, Maliha Rahman, Jenny Zhang |
| EPI_ISL_925398 | Department of Clinical Microbiology | GIGA Medical Genomics | Keith Durkin, Maria Artesi, Sébastien Bontems, Raphaël Boreux, Bouchra Boujemla, Cécile Meex, Pierrette Melin, Marie-Pierre Hayette, Vincent Bours |
| EPI_ISL_925513, EPI_ISL_925515, EPI_ISL_925516, EPI_ISL_925517, EPI_ISL_925518, EPI_ISL_925519, EPI_ISL_925527, EPI_ISL_925529, EPI_ISL_925530, EPI_ISL_925531, EPI_ISL_925532, EPI_ISL_925533, EPI_ISL_925832 | | | |
| see above | Arizona State Public Health Laboratory | Arizona State Public Health Laboratory | Trung Huynh, Jessica Escobar, Katherine Fullerton, Nobuko Fukushima, Stacy White, Linda Getsinger, Victor Waddell |
| EPI_ISL_925924, EPI_ISL_925933, EPI_ISL_925949, EPI_ISL_926010, EPI_ISL_926025, EPI_ISL_926041, EPI_ISL_926082, EPI_ISL_926104, EPI_ISL_926151, EPI_ISL_926237, EPI_ISL_926238, EPI_ISL_926265, EPI_ISL_926321, EPI_ISL_926410, EPI_ISL_926442, EPI_ISL_926505, EPI_ISL_926546, EPI_ISL_926593, EPI_ISL_926596, EPI_ISL_926609, EPI_ISL_926620, EPI_ISL_926632, EPI_ISL_926667, EPI_ISL_926716, EPI_ISL_926791, EPI_ISL_926795, EPI_ISL_926818, EPI_ISL_926844, EPI_ISL_926885, EPI_ISL_926936, EPI_ISL_927056, EPI_ISL_927075, EPI_ISL_927103, EPI_ISL_927125, EPI_ISL_927338, EPI_ISL_927356, EPI_ISL_927366, EPI_ISL_927432, EPI_ISL_927480, EPI_ISL_927543, EPI_ISL_927546, EPI_ISL_927601, EPI_ISL_927614, EPI_ISL_927639, EPI_ISL_927668, EPI_ISL_927698, EPI_ISL_927709, EPI_ISL_927740, EPI_ISL_927774, EPI_ISL_927792, EPI_ISL_927830, EPI_ISL_927832, EPI_ISL_927858, EPI_ISL_927878, EPI_ISL_927933, EPI_ISL_927978, EPI_ISL_927979, EPI_ISL_927987, EPI_ISL_928003, EPI_ISL_928026, EPI_ISL_928038, EPI_ISL_928062, EPI_ISL_928064, EPI_ISL_928085, EPI_ISL_928100, EPI_ISL_928137, EPI_ISL_928213, EPI_ISL_928218, EPI_ISL_928237, EPI_ISL_928298, EPI_ISL_928365, EPI_ISL_928434, EPI_ISL_928480, EPI_ISL_928481, EPI_ISL_928483, EPI_ISL_928529, EPI_ISL_928537, EPI_ISL_928720, EPI_ISL_928727, EPI_ISL_928768, EPI_ISL_928834, EPI_ISL_928867, EPI_ISL_928894, EPI_ISL_928932, EPI_ISL_928983, EPI_ISL_929020, EPI_ISL_929064, EPI_ISL_929066, EPI_ISL_929119, EPI_ISL_929202, EPI_ISL_929262, EPI_ISL_929279, EPI_ISL_929309, EPI_ISL_929370, EPI_ISL_929371, EPI_ISL_929390, EPI_ISL_929449, EPI_ISL_929469, EPI_ISL_929533, EPI_ISL_929677, EPI_ISL_929722, EPI_ISL_929726, EPI_ISL_929734, EPI_ISL_929744, EPI_ISL_929748, EPI_ISL_929827, EPI_ISL_929861, EPI_ISL_929881, EPI_ISL_929887, EPI_ISL_929917, EPI_ISL_929923, EPI_ISL_929940, EPI_ISL_929966, EPI_ISL_930004, EPI_ISL_930021, EPI_ISL_930040, EPI_ISL_930090, EPI_ISL_930096, EPI_ISL_930173, EPI_ISL_930259, EPI_ISL_930270, EPI_ISL_930280, EPI_ISL_930303, EPI_ISL_930356, EPI_ISL_930390, EPI_ISL_930455, EPI_ISL_930461, EPI_ISL_930494, EPI_ISL_930509, EPI_ISL_930510 | | | |
| see above | Department of Virus and Microbiological Special Diagnostics, Statens Serum Institut, Copenhagen, Denmark | Aalborg University | Danish Covid-19 Genome Consortium |
| EPI_ISL_931574, EPI_ISL_931575, EPI_ISL_931576, EPI_ISL_931577, EPI_ISL_931578, EPI_ISL_931579, EPI_ISL_931580, EPI_ISL_931581, EPI_ISL_931582, EPI_ISL_931583 | Utah Public Health Laboratory | Utah Public Health Laboratory | Erin L. Young, Kelly F. Oakeson, Tara Gallagher |
| EPI_ISL_932832, EPI_ISL_933425 | Lighthouse Lab in Glasgow | Wellcome Sanger Institute for the COVID-19 Genomics UK (COG-UK) Consortium | Harper VanSteenhouse, Yumi Kasai, David Gray, Carol Clugston, Anna Dominiczak and Alex Alderton, Roberto Amato, Sonia Goncalves, Ewan Harrison, David K. Jackson, Ian Johnston, Dominic Kwiatkowski, Cordelia Langford, John Sillitoe on behalf of the Wellcome Sanger Institute COVID-19 Surveillance Team |
| EPI_ISL_934180, EPI_ISL_934181, EPI_ISL_934188, EPI_ISL_934189, EPI_ISL_934190, EPI_ISL_934191, EPI_ISL_934192, EPI_ISL_934193, EPI_ISL_934194, EPI_ISL_934202, EPI_ISL_934204, EPI_ISL_934205, EPI_ISL_934206, EPI_ISL_934207, EPI_ISL_934208, EPI_ISL_934209, EPI_ISL_934210, EPI_ISL_934211, EPI_ISL_934212, EPI_ISL_934213, EPI_ISL_934214, EPI_ISL_934215, EPI_ISL_934216, EPI_ISL_934217, EPI_ISL_934218, EPI_ISL_934219, EPI_ISL_934220, EPI_ISL_934221, EPI_ISL_934223, EPI_ISL_934224, EPI_ISL_934225, EPI_ISL_934226, EPI_ISL_934227, EPI_ISL_934228 | | | |
| see above | Vilnius university hospital Santaros Klinikos, Center of Laboratory Medicine | Vilnius university hospital Santaros Klinikos, Center of Laboratory Medicine | Ingrida Olendraite, Daniel Naumovas, Rimvydas Norvilas, Dovile Ezerskyte, Justinas Slikas, Gytis Dudas |
| EPI_ISL_934364, EPI_ISL_934365 | Synlab Medilab, Mikrobiologi | The Public Health Agency of Sweden | Anna-Malin Linde, Maria Lind Karlberg, Carlo Berg, Oskar Karlsson Lindsjo, Sofia Stamouli, Reza Advani, Mattias Haukland, Petra Holmstrom, Noura Walai, Petra Edquist, Mia Brytting, Anna Risberg, Karin Tegmark-Wisell |
| EPI_ISL_935211 | KU Leuven, Rega Institute, Clinical and Epidemiological Virology | KU Leuven, Rega Institute, Clinical and Epidemiological Virology | Tony Wawina-Bokalanga, Bert Vanmechelen, Joan Marti-Carerras, Piet Maes |
| EPI_ISL_935359, EPI_ISL_935360 | Florida Bureau of Public Health Laboratories | Florida Bureau of Public Health Laboratories | Sarah Schmedes, Jason Blanton |
| EPI_ISL_935988, EPI_ISL_935989, EPI_ISL_935990 | GLENS FALLS HOSPITAL LABORATORY | Wadsworth Center, New York State Department of Health | Kirsten St. George, Daryl M. Lamson, Alexis Russel, Matthew Shudt, Melissa A Leisner, Jonathan Plitnick, Navjot Singh, John Kelly, Erasmus Schneider, Erica Lasek-Nesselquist |
| EPI_ISL_936048 | ADIRONDACK MEDICAL CENTER | Wadsworth Center, New York State Department of Health | Kirsten St. George, Daryl M. Lamson, Alexis Russel, Matthew Shudt, Melissa A Leisner, Jonathan Plitnick, Navjot Singh, John Kelly, Erasmus Schneider, Erica Lasek-Nesselquist |
| EPI_ISL_936078, EPI_ISL_936079, EPI_ISL_936080, EPI_ISL_936081, EPI_ISL_936083, EPI_ISL_936095, EPI_ISL_936096, EPI_ISL_936097, EPI_ISL_936101 | KALEIDA CENTER FOR LABORATORY MEDICINE | Wadsworth Center, New York State Department of Health | Kirsten St. George, Daryl M. Lamson, Alexis Russel, Matthew Shudt, Melissa A Leisner, Jonathan Plitnick, Navjot Singh, John Kelly, Erasmus Schneider, Erica Lasek-Nesselquist |
| EPI_ISL_936142 | SUNY UPSTATE MEDICAL UNIVERSITY | Wadsworth Center, New York State Department of Health | Kirsten St. George, Daryl M. Lamson, Alexis Russel, Matthew Shudt, Melissa A Leisner, Jonathan Plitnick, Navjot Singh, John Kelly, Erasmus Schneider, Erica Lasek-Nesselquist |
| EPI_ISL_936144 | KALEIDA CENTER FOR LABORATORY MEDICINE | Wadsworth Center, New York State Department of Health | Kirsten St. George, Daryl M. Lamson, Alexis Russel, Matthew Shudt, Melissa A Leisner, Jonathan Plitnick, Navjot Singh, John Kelly, Erasmus Schneider, Erica Lasek-Nesselquist |
| EPI_ISL_936391, EPI_ISL_936395, EPI_ISL_936426, EPI_ISL_936431, EPI_ISL_936433, EPI_ISL_936435, EPI_ISL_936436, EPI_ISL_936442, EPI_ISL_936445, EPI_ISL_936447, EPI_ISL_936449 | | | |

| | | | |
|---|---|---|---|
| see above | TGen North | TGen North | Jolene Bowers, Megan Folkerts, Chris French, Hayley Yaglom, Ashlyn Pfeiffer, Darrin Lemmer, Dave Engelthaler, The Arizona COVID Genomics Union (ACGU) |
| EPI_ISL_936493 | Santa Clara County Public Health Laboratory | Santa Clara County Public Health Laboratory | Santa Clara County Public Health Department |
| EPI_ISL_936867, EPI_ISL_936875, EPI_ISL_936876, EPI_ISL_936877, EPI_ISL_936878, EPI_ISL_936879, EPI_ISL_936880, EPI_ISL_936881, EPI_ISL_936885, EPI_ISL_936886, EPI_ISL_936887, EPI_ISL_936889, EPI_ISL_936893, EPI_ISL_936895, EPI_ISL_936897, EPI_ISL_936898, EPI_ISL_936899, EPI_ISL_936900, EPI_ISL_936901, EPI_ISL_936902, EPI_ISL_936903, EPI_ISL_936904 | | | |
| see above | Northwestern Memorial Hospital | Ozer Lab | Ramon Lorenzo-Redondo, Lacy M. Simons, Chad J. Achenbach, Lawrence J. Jennings, Michael G. Ison, Judd F. Hultquist, Egon A. Ozer |
| EPI_ISL_940716, EPI_ISL_940717, EPI_ISL_940718, EPI_ISL_940719, EPI_ISL_940720, EPI_ISL_940721, EPI_ISL_940722, EPI_ISL_940733 | City of Milwaukee Health Department Laboratory | City of Milwaukee Health Department Laboratory | Sanjib Bhattacharyya |
| EPI_ISL_940777 | NSPI-CRN de Influenza y otros virus respiratorios | INSPI-Centro de Investigación Multidisciplinaria de la DTIDI | Leandro Patiño, Doménica de Mora, Maritza Olmedo, Andrés Carrazco-Montalvo, Orson Mestanza, Mary Regato-Arrata, Melissa Zambrano, Manuel González, Alfredo Bruno, Alberto Orlando. |
| EPI_ISL_940781 | INSPI-CRN de Influenza y otros virus respiratorios | INSPI-Centro de Investigación Multidisciplinaria de la DTIDI | Leandro Patiño, Doménica de Mora, Maritza Olmedo, Andrés Carrazco-Montalvo, Orson Mestanza, Mary Regato-Arrata, Melissa Zambrano, Manuel González, Alfredo Bruno, Alberto Orlando |
| EPI_ISL_940821 | INSPI-CRN de Influenza y otros virus respiratorios | INSPI-Centro de Investigación Multidisciplinaria de la DTIDI | Leandro Patiño, Doménica de Mora, Maritza Olmedo, Andrés Carrazco-Montalvo, Orson Mestanza, Mary Regato-Arrata, Melissa Zambrano, Manuel González, Alfredo Bruno, Alberto Orlando. |
| EPI_ISL_941226, EPI_ISL_941227 | Laboratorio de Microbiología. Hospital General Universitario de Elda, Alicante | SeqCOVID-SPAIN consortium/IBV(CSIC) | Mª Isabel Gascón Ros, Cristina Torregrosa Hetland, Eva Pastor Boix, Paloma Cascales Ramos and SeqCOVID-SPAIN consortium |
| EPI_ISL_941288, EPI_ISL_941292 | Nigeria Centre for Disease Control (NCDC) | African Centre of Excellence for Genomics of Infectious Diseases (ACEGID), Redeemer's University | Oluniyi P.E. et al |
| EPI_ISL_941373, EPI_ISL_941374, EPI_ISL_941446, EPI_ISL_941447, EPI_ISL_941448, EPI_ISL_941449, EPI_ISL_941450, EPI_ISL_941628 | Instituto Nacional de Saude (INSA) | Instituto Nacional de Saude (INSA) | Borges et al |
| EPI_ISL_941926 | Nigeria Centre for Disease Control (NCDC) | African Centre of Excellence for Genomics of Infectious Diseases (ACEGID), Redeemer's University | Oluniyi P.E. et al |
| EPI_ISL_942013 | Wyoming Public Health Laboratory | Wyoming Public Health Laboratory | Noah Hull, Taylor Fearing, Lynette Gumbleton, Channing Weber, Ashley Norberg, Bailey Bowcutt, and Wanda Manley |
| EPI_ISL_942788, EPI_ISL_942789, EPI_ISL_942790, EPI_ISL_942791, EPI_ISL_942792, EPI_ISL_942793, EPI_ISL_942794 | Gundersen Molecular Diagnostics Laboratory | Kabara Cancer Research Institute | Craig S. Richmond, Paraic A. Kenny |
| EPI_ISL_942960, EPI_ISL_942971 | National Institute of Health Research and Development | National Institute of Health Research and Development | Subangkit, Hana Apsari Pawestri, Kartika Dewi Puspa, Arie Ardiansyah Nugraha, Hartanti Dian Ikawati, Krisna Nur Andriana Pangesti, Yuni Rukminiati, Ririn Ramadhany, Agustiningsih, Kindi Adam, Holy Arif Wibowo, Triyani Soekarso, Ni Ketut Susilarini, Nurika Hariastuti, Ully Alfi Nikmah, Reni Herman, Nike Susanti, Herna, Tati Febriyanti, Natalie Laurencia Kipuw, Fauzul Muna, Irene Lorinda Indalao, Nelly Puspandari, Vivi Setiawaty. |
| EPI_ISL_942999, EPI_ISL_943183, EPI_ISL_943184, EPI_ISL_943242, EPI_ISL_943395, EPI_ISL_943396, EPI_ISL_943397, EPI_ISL_943478 | Dutch COVID-19 response team | National Institute for Public Health and the Environment (RIVM) | Adam Meijer, Harry Vennema, Dirk Eggink, Jeroen Cremer, Sharon van den Brink, Bas van der Veer, AnneMarie van den Brandt, Florian Zwagemaker, Dennis Schmitz, Chantal Reusken, on behalf of the national COVID-19 response team |
| EPI_ISL_943823 | Utah Public Health Laboratory | Utah Public Health Laboratory | Erin L. Young, Kelly F. Oakeson, Tara Gallagher |
| EPI_ISL_944124, EPI_ISL_944127, EPI_ISL_944128, EPI_ISL_944131, EPI_ISL_944135, EPI_ISL_944137, EPI_ISL_944145, EPI_ISL_944148, EPI_ISL_944152, EPI_ISL_944153, EPI_ISL_944154, EPI_ISL_944155, EPI_ISL_944157, EPI_ISL_944158, EPI_ISL_944159, EPI_ISL_944162, EPI_ISL_944163, EPI_ISL_944166, EPI_ISL_944167, EPI_ISL_944168, EPI_ISL_944169, EPI_ISL_944170, EPI_ISL_944171, EPI_ISL_944173, EPI_ISL_944175, EPI_ISL_944176 | | | |
| see above | National Health Laboratory Service, South Africa | KRISP, KZN Research Innovation and Sequencing Platform | Laguda-Akingba O, Giandhari J, Pillay S, Lessells R, Mdlalose K, York D, Khan S, Emmanuel SJ, Tegally H, Wilkinson E, de Oliveira T |
| EPI_ISL_944208 | Israel Central Virology laboratory | Israel National Consortium for SARS-CoV-2 sequencing | Neta Zuckerman, Efrat Dahan Bucris, Michal Mandelboim, Dana Bar-Ilan, Oran Erster, Tzvia Mann, Omer Murik, David A. Zeevi, Assaf Rokney, Joseph Jaffe, Eva Nachum, Maya Davidovich Cohen, Ephraim Fass, Gal Zizelski Valenci, Mor Rubinstein, Efrat Rorman, Israel Nissan, Efrat Glick-Saar, Omri Nayshool, Gideon Rechavi, Ella Mendelson, Orna Mor |
| EPI_ISL_944614 | Instituto Nacional de Medicina Genomica | Instituto Nacional de Medicina Genomica | Hidalgo-Miranda A, Mendoza-Vargas A, Reyes-Grajeda JP, Cisneros-Villanueva M, Cedro-Tanda A,Peñaloza-Figueroa F, Herrera-Montalvo LA |
| EPI_ISL_944691, EPI_ISL_944713 | Department of Biochemistry, Cell and Molecular Biology, West African Centre for Cell Biology of Infectious Pathogens (WACCBIP), University of Ghana | Department of Biochemistry, Cell and Molecular Biology, West African Centre for Cell Biology of Infectious Pathogens (WACCBIP), University of Ghana | Morang'a,C.M., Ngoi,J.M., Quansah,E.B., Saiid,S., Amuzu,D.S., Asante,I., Bonney,J.H., Bonney,E., Odoom,J.K., Ndam,N.T., Tei-Maya,F., Adusei-Poku,M., Ofori-Boadu,L., Ampofo,W.K., Amenga-Etego,L.N., Quashie,P., Bediako,Y., Awandare,G.A. |
| EPI_ISL_945057, EPI_ISL_945071, EPI_ISL_945074, EPI_ISL_945075, EPI_ISL_945077, EPI_ISL_945078, EPI_ISL_945079, EPI_ISL_945081, EPI_ISL_945086, EPI_ISL_945097, EPI_ISL_945104 | | | |
| see above | Lighthouse Lab in Milton Keynes | Wellcome Sanger Institute for the COVID-19 Genomics UK (COG-UK) Consortium | The Lighthouse Lab in Milton Keynes and Alex Alderton, Roberto Amato, Sonia Goncalves, Ewan Harrison, David K. Jackson, Ian Johnston, Dominic Kwiatkowski, Cordelia Langford, John Sillitoe on behalf of the Wellcome Sanger Institute COVID-19 Surveillance Team |
| EPI_ISL_945107 | Lighthouse Lab in Cambridge | Wellcome Sanger Institute for the COVID-19 Genomics UK (COG-UK) Consortium | Rob Howes, The Lighthouse Lab in Cambridge and Alex Alderton, Roberto Amato, Sonia Goncalves, Ewan Harrison, David K. Jackson, Ian Johnston, Dominic Kwiatkowski, Cordelia Langford, John Sillitoe on behalf of the Wellcome Sanger Institute COVID-19 Surveillance Team |
| EPI_ISL_945110, EPI_ISL_945127, EPI_ISL_945132, EPI_ISL_945135, EPI_ISL_945137 | Lighthouse Lab in Milton Keynes | Wellcome Sanger Institute for the COVID-19 Genomics UK (COG-UK) Consortium | The Lighthouse Lab in Milton Keynes and Alex Alderton, Roberto Amato, Sonia Goncalves, Ewan Harrison, David K. Jackson, Ian Johnston, Dominic Kwiatkowski, Cordelia Langford, John Sillitoe on behalf of the Wellcome Sanger Institute COVID-19 Surveillance Team |
| EPI_ISL_945144 | Lighthouse Lab in Glasgow | Wellcome Sanger Institute for the COVID-19 Genomics UK (COG-UK) Consortium | Harper VanSteenhouse, Yumi Kasai, David Gray, Carol Clugston, Anna Dominiczak and Alex Alderton, Roberto Amato, Sonia Goncalves, Ewan Harrison, David K. Jackson, Ian Johnston, Dominic Kwiatkowski, Cordelia Langford, John Sillitoe on behalf of the Wellcome Sanger Institute COVID-19 Surveillance Team |
| EPI_ISL_945151, EPI_ISL_945154, EPI_ISL_945158 | Lighthouse Lab in Milton Keynes | Wellcome Sanger Institute for the COVID-19 Genomics UK (COG-UK) Consortium | The Lighthouse Lab in Milton Keynes and Alex Alderton, Roberto Amato, Sonia Goncalves, Ewan Harrison, David K. Jackson, Ian Johnston, Dominic Kwiatkowski, Cordelia Langford, John Sillitoe on behalf of the Wellcome Sanger Institute COVID-19 Surveillance Team |
| EPI_ISL_945162 | Lighthouse Lab in Glasgow | Wellcome Sanger Institute for the COVID-19 Genomics UK (COG-UK) Consortium | Harper VanSteenhouse, Yumi Kasai, David Gray, Carol Clugston, Anna Dominiczak and Alex Alderton, Roberto Amato, Sonia Goncalves, Ewan Harrison, David K. Jackson, Ian Johnston, Dominic Kwiatkowski, Cordelia Langford, John Sillitoe on behalf of the Wellcome Sanger Institute COVID-19 Surveillance Team |
| EPI_ISL_945165, EPI_ISL_945167, EPI_ISL_945168, EPI_ISL_945173, EPI_ISL_945182, EPI_ISL_945187, EPI_ISL_945194, EPI_ISL_945196, EPI_ISL_945200, EPI_ISL_945206, EPI_ISL_945211, EPI_ISL_945213, EPI_ISL_945214, EPI_ISL_945215, EPI_ISL_945216, EPI_ISL_945219, EPI_ISL_945225, EPI_ISL_945227, EPI_ISL_945229, EPI_ISL_945233, EPI_ISL_945242, EPI_ISL_945245, EPI_ISL_945259, EPI_ISL_945260, EPI_ISL_945262, EPI_ISL_945263, EPI_ISL_945264, EPI_ISL_945265, EPI_ISL_945267, EPI_ISL_945272, EPI_ISL_945274, EPI_ISL_945280, EPI_ISL_945281, EPI_ISL_945283, EPI_ISL_945284, EPI_ISL_945286, EPI_ISL_945290, EPI_ISL_945291, EPI_ISL_945294, EPI_ISL_945296, EPI_ISL_945298, EPI_ISL_945300, EPI_ISL_945302, EPI_ISL_945303, EPI_ISL_945308, EPI_ISL_945312, EPI_ISL_945316, EPI_ISL_945317, EPI_ISL_945318, EPI_ISL_945326, EPI_ISL_945327, EPI_ISL_945328, EPI_ISL_945330, EPI_ISL_945348, EPI_ISL_945349, EPI_ISL_945353, EPI_ISL_945354, EPI_ISL_945355, EPI_ISL_945359, EPI_ISL_945369, EPI_ISL_945370, EPI_ISL_945371 | | | |
| see above | Lighthouse Lab in Milton Keynes | Wellcome Sanger Institute for the COVID-19 Genomics UK (COG-UK) Consortium | The Lighthouse Lab in Milton Keynes and Alex Alderton, Roberto Amato, Sonia Goncalves, Ewan Harrison, David K. Jackson, Ian Johnston, Dominic Kwiatkowski, Cordelia Langford, John Sillitoe on behalf of the Wellcome Sanger Institute COVID-19 Surveillance Team |
| EPI_ISL_947323 | RS Siloam Kebon Jeruk | Eijkman Institute for Molecular Biology, Ministry of Research | Sukma Oktavianthi, Willy Agustine, Edison Johar, Hidayat Trimarsanto, Iskandar Adnan, Lydia V. Panggalo, Frilasita A Yudhaputri, Safarina G Malik, Khin |

| | | | |
|---|---|---|---|
| | | and Technology/National Agency for Research and Innovation | Saw Myint, Amin Soebandrio |
| EPI_ISL_949406 | University of Birmingham | COVID-19 Genomics UK (COG-UK) Consortium | Institute of Microbiology, University of Birmingham: Claire McMurray, Joanne Stockton, Samuel Nicholls, Radoslaw Poplawski, Will Rowe, Josh Quick, Nicholas Loman. University of Birmingham Testing Laboratory: Celina M Whalley, Andrew Bosworth, Charlotte Poxon, Kasun Wanigasooriya, Oliver Pickles, Mike Kidd, Alex Richter, Andrew D Beggs PHE Heartlands Lab: Husam Osman, Andrew Bosworth. Queen Elizabeth Hospital: Anna Casey |
| EPI_ISL_949755, EPI_ISL_949767 | Barts Health NHS Trust | COVID-19 Genomics UK (COG-UK) Consortium | CUTINO-MOGUEL, Maria-Teresa; HARRINGTON, David; OWOYEMI, Dola; KULASEGARAN-SHYLINI, Raghavendran; BROAD, Claire; KELE, Beatrix |
| EPI_ISL_950094, EPI_ISL_950111, EPI_ISL_950146, EPI_ISL_950151 | University College London, Great Ormond Street Hospital for Children NHS Foundation Trust, Imperial College Healthcare NHS Trust | COVID-19 Genomics UK (COG-UK) Consortium | Sergi Castellano, Rachel Williams, Mark Kristiansen, Paola Resende Silva, Sunando Roy, Tony Brooks, Helena Tutill, Paola Niola, Patricia Dyal, Charlotte Williams, Leysa Forrest, Yasmin Panchbhaya, Jacqueline Findlay, Samuel Weeks, Julianne Brown, Kathryn Harris, Paul Randell, James Price, Alison Holmes, Judith Breuer |
| EPI_ISL_950288, EPI_ISL_950482, EPI_ISL_950483, EPI_ISL_950484, EPI_ISL_950485, EPI_ISL_950486, EPI_ISL_950487, EPI_ISL_950489, EPI_ISL_950490, EPI_ISL_950491, EPI_ISL_950492, EPI_ISL_950493 | | | |
| see above | Northumbria University / South Tees Hospitals NHS Foundation Trust / North Cumbria Integrated Care NHS Foundation Trust / North Tees and Hartlepool NHS Foundation Trust / Newcastle Hospitals NHS Foundation Trust | COVID-19 Genomics UK (COG-UK) Consortium | Darren L Smith,Andrew Nelson,Matthew Bashton,Greg R Young,Joshua Loh,John Allan,Mohammad A Tariq,Giles S Holt,Gary Black,Wen C Yew,Lynn Dover,Paul Baker,Steve Liggett,Sarah Essex,Jane Greenaway,Debra Padgett,Clive Graham,Garren Scott,Edward Barton,Emma Swindells,Brendan Payne,Jennifer Collins,Yusri Taha,Gary Eltringham |
| EPI_ISL_951084, EPI_ISL_951085, EPI_ISL_951087, EPI_ISL_951088, EPI_ISL_951089, EPI_ISL_951090, EPI_ISL_951093, EPI_ISL_951126, EPI_ISL_951131, EPI_ISL_951132, EPI_ISL_951134, EPI_ISL_951136, EPI_ISL_951138, EPI_ISL_951139, EPI_ISL_951144, EPI_ISL_951145, EPI_ISL_951147, EPI_ISL_951148, EPI_ISL_951149, EPI_ISL_951153, EPI_ISL_951155, EPI_ISL_951159, EPI_ISL_951173, EPI_ISL_951178, EPI_ISL_951183, EPI_ISL_951185, EPI_ISL_951187, EPI_ISL_951191, EPI_ISL_951192, EPI_ISL_951198, EPI_ISL_951200, EPI_ISL_951214, EPI_ISL_951229, EPI_ISL_951230, EPI_ISL_951319, EPI_ISL_951384, EPI_ISL_951386 | | | |
| see above | Oxford Viromics, NDM, University of Oxford; Oxford University Hospitals; Basingstoke and North Hampshire Hospital | COVID-19 Genomics UK (COG-UK) Consortium | Tanya Golubchik, David Bonsall, George Macintyre, Amy Trebes, Mariateresa de Cesare, Catrin Moore, Alex Mobbs, Anita Justice, Robert Shaw, Monique Andersson, Timothy Peto, Emma Wise, Nathan Moore, Jessica Lynch, Nick Cortes, Matilde Mori, Stephen Kidd, David Buck, John Todd, Christophe Fraser |
| EPI_ISL_952420, EPI_ISL_952423 | Centre for Enzyme Innovation, University of Portsmouth / Translational Research Laboratory, Portsmouth Hospitals NHS Trust | COVID-19 Genomics UK (COG-UK) Consortium | Angela Beckett,Salman Goudarzi,Christopher Fearn,Kate Cook,Katie Loveson,Sharon Glaysher,Scott Elliott,Samuel Robson |
| EPI_ISL_953176, EPI_ISL_953184, EPI_ISL_953188, EPI_ISL_953189, EPI_ISL_953190, EPI_ISL_953192, EPI_ISL_953200, EPI_ISL_953201, EPI_ISL_953202, EPI_ISL_953212, EPI_ISL_953223, EPI_ISL_953225, EPI_ISL_953229 | | | |
| see above | Bioinformatics and Biostatistics Lab, Advanced Sequencing Facility | COVID-19 Genomics UK (COG-UK) Consortium | Aengus Stewart,Jerome Nicod,Chelsea Sawyer,Laura Cubitt,Harshil Patel,Margaret Crawford |
| EPI_ISL_954805 | Hospital General Universitario de Ciudad Real | Instituto de Salud Carlos III | Iglesias-Caballero, M. Camarero, S. Sandonís,V. Vázquez, S. Pozo, F. Casas, I. Jiménez, P. Zaballos, A. Monzón, S. Varona, S. Cuesta, I. Illescas, S. |
| EPI_ISL_954820, EPI_ISL_954877, EPI_ISL_954885 | Colorado Department of Public Health and Environment | Colorado Department of Puplic Health and Environment | Laura Bankers, Molly C. Hetherington-Rauth, Diana Ir, Shannon Ely, Shannon R. Matzinger, Sarah Elizabeth Totten, Emily A. Travanty |
| EPI_ISL_955271 | American Esoteric Laboratory | Pathogen Discovery, Respiratory Viruses Branch, Division of Viral Diseases, Centers for Disease Control and Prevention | Ying Tao, Jing Zhang, Yan Li, Krista Queen, Anna Uehara, Peter Cook, Clinton R. Paden, Haibin Wang, Suxiang Tong |
| EPI_ISL_955393, EPI_ISL_955394, EPI_ISL_955395, EPI_ISL_955396, EPI_ISL_955397, EPI_ISL_955398, EPI_ISL_955399, EPI_ISL_955400, EPI_ISL_955401, EPI_ISL_955402, EPI_ISL_955403, EPI_ISL_955404, EPI_ISL_955405, EPI_ISL_955406, EPI_ISL_955407, EPI_ISL_955408, EPI_ISL_955409, EPI_ISL_955410, EPI_ISL_955412, EPI_ISL_955413, EPI_ISL_955414, EPI_ISL_955415, EPI_ISL_955416 | | | |
| see above | Alameda County Public Health Lab | Chan-Zuckerberg Biohub | CZB Cliahub Consortium |
| EPI_ISL_955434, EPI_ISL_955435, EPI_ISL_955436, EPI_ISL_955437, EPI_ISL_955438, EPI_ISL_955439, EPI_ISL_955440, EPI_ISL_955441, EPI_ISL_955442, EPI_ISL_955443, EPI_ISL_955444, EPI_ISL_955445, EPI_ISL_955446, EPI_ISL_955448, EPI_ISL_955449, EPI_ISL_955450, EPI_ISL_955451, EPI_ISL_955454, EPI_ISL_955455, EPI_ISL_955456, EPI_ISL_955457, EPI_ISL_955458, EPI_ISL_955459, EPI_ISL_955460, EPI_ISL_955461, EPI_ISL_955462, EPI_ISL_955463, EPI_ISL_955468, EPI_ISL_955469 | | | |
| see above | Orange County Public Health Lab | Chan-Zuckerberg Biohub | CZB Cliahub Consortium |
| EPI_ISL_955764, EPI_ISL_955767, EPI_ISL_955769, EPI_ISL_955770, EPI_ISL_955771, EPI_ISL_955772, EPI_ISL_955773, EPI_ISL_955774, EPI_ISL_955775, EPI_ISL_955776 | Humboldt County Public Health Laboratory | Chan-Zuckerberg Biohub | CZB Cliahub Consortium |
| EPI_ISL_956280 | Institute of Tropical Disease | Institute of Tropical Disease, Universitas Airlangga | Rima R Prasetya, Krisnoadi Rahardjo, Aldise M Nastri, Jezzy R Dewantari, Gatot Soegiarto, Laksmi Wulandari, Resti Yudhawati, Soetjipto, Yasuko Mori, Maria I Lusida, Kazufumi Shimizu |
| EPI_ISL_956281 | Siti Khodijah Hospital | Institute of Tropical Disease, Universitas Airlangga | Maria I Lusida, Krisnoadi Rahardjo, Aldise M Nastri, Jezzy R Dewantari, Rima R Prasetya, Muhammad Hamdan, Gatot Soegiarto, Laksmi Wulandari, Resti Yudhawati, Soetjipto, Yasuko Mori, Kazufumi Shimizu |
| EPI_ISL_956297 | Instituto Nacional de Salud- Dirección de Redes de Laboratorios de Salud Pública | Instituto Nacional de Salud- Dirección de Investigación en Salud Pública | Katherine Laiton-Donato, Diego A. Álvarez-Díaz, Carlos Franco-Muñoz, Mauricio Pacheco-Montealegre, Hector Alejandro Ruiz-Moreno, Maria T. Herrera-Sepúlveda, Diego Andrés Prada, Jhonnatan Reales-González, Sheryll Corchuelo, Julian Naizaque, Gerardo Santamaría, Magdalena Wiesner, Martha Lucia Ospina Martinez, Marcela Mercado-Reyes |
| EPI_ISL_956315 | Siti Khodijah Hospital | Institute of Tropical Disease, Universitas Airlangga | Kazufumi Shimizu, Krisnoadi Rahardjo, Aldise M Nastri, Jezzy R Dewantari, Rima R Prasetya, Muhammad Hamdan, Gatot Soegiarto, Laksmi Wulandari, Resti Yudhawati, Yasuko Mori, Soetjipto, Maria I Lusida |
| EPI_ISL_959369 | Viollier AG | University Hospital Basel, Clinical Bacteriology | Tim Roloff, Madlen Stange, Helena MB Seth-Smith, Alfredo Mari, Karoline Leuzinger, Julia Bielicki, Christiane Beckmann, Manuel Battegay, Hans Hirsch, Adrian Egli |
| EPI_ISL_960408, EPI_ISL_960410, EPI_ISL_960412, EPI_ISL_960437, EPI_ISL_960439 | The National Institute of Public Health | State Veterinary Institute Prague | Nagy,A;Vecerova,J;Cernikova,L;Stara,M;Jirincova,H;Trnka,D |
| EPI_ISL_960459, EPI_ISL_960462 | Istituto Zooprofilattico Sperimentale del Mezzogiorno | TIGEM | Patrizia Annunziata, Andrea Ballabio, Valentina Bouche, Davide Cacchiarelli, Pellegrino Cerino, Chiara Colantuono, Maria Concetta Cuomo, Denise Di Concilio, Lucio Di Filippo, Antonio Grimaldi, Antonio Limone, Anna Manfredi, Francesco Panariello, Biancamaria Pierri, Marcello Salvi |
| EPI_ISL_961207, EPI_ISL_961253 | Hospital General Universitario de Alicante - Instituto de Investigación Sanitaria y Biomédica de Alicante | SeqCOVID-SPAIN consortium/IBV(CSIC) | Maripaz Ventero Martín, Carmen Molina Pardines and SeqCOVID-SPAIN consortium |
| EPI_ISL_961482, EPI_ISL_961492, EPI_ISL_961552, EPI_ISL_961553 | Michigan Department of Health and Human Services, Bureau of Laboratories | Michigan Department of Health and Human Services, Bureau of Laboratories | Blankenship HM, Riner D, Soehnlen MK |
| EPI_ISL_961583, EPI_ISL_961652 | Hôpital Georges L. Dumont | National Microbiology Laboratory (NML) | Anna Majer, Shari Tyson, Grace Seo, Philip Mabon, Elsie Grudeski, Rhiannon Huzarewich, Russell Mandes, Anneliese Landgraff, Jennifer Tanner, Natalie Knox, Morag Graham, Gary Van Domselaar, Richard Garceau, Guillaume Desnoyers, Nathalie Bastien, Yan Li, Timothy Booth, Darian Hole, Madison Chapel, Kirsten Biggar, CanCOGeN's metadata curation team, Public Health Agency of Canada CanCOGeN team |
| EPI_ISL_961898, EPI_ISL_961927, EPI_ISL_961937, EPI_ISL_961967, EPI_ISL_962130, EPI_ISL_962131, EPI_ISL_962132, EPI_ISL_962133, EPI_ISL_962134, EPI_ISL_962135, EPI_ISL_962136, EPI_ISL_962137, EPI_ISL_962140 | | | |
| see above | Illinois Department of Public Health | Gagnon Lab, Southern Illinois University | Keith Gagnon |
| EPI_ISL_962918 | Servicio de Microbiología, Laboratori Clínic Metropolitana Nord. Hospital Universitari Germans Trias i Pujol. Institut d'Investigació en Ciències de la Salut Germans Trias i Pujol (IGTP) | SeqCOVID-SPAIN consortium/IBV(CSIC) | Elisa Martró, Antoni E. Bordoy, Anna Not, Adrián Antuori, Anabel Fernández, Nona Romaní, Verónica Saludes, Cristina Casañ and SeqCOVID-SPAIN consortium |
| EPI_ISL_965029 | Laboratorio de Virologia HUCA | Laboratorio de Virologia HUCA | Castelló C, Gómez de Oña J, Boga JA, Rojo S, Alvarez-Arguelles ME, Abreu F, Costales I, Sandoval M, Perez-Martínez Z, Martín-Rodríguez G, Coto E, Melón S |

| | | | |
|---|---|---|---|
| EPI_ISL_965033 | Laboratorio de Virología HUCA | Laboratorio de Virología HUCA | Castelló C, Gómez de Oña J, Boga JA, Rojo S, Alvarez-Arguelles ME, Abreu F, Costales I, Sandoval M, Perez-Martínez Z, Martín-Rodríguez G, Coto E, Melón S |
| EPI_ISL_965186, EPI_ISL_965188, EPI_ISL_965198, EPI_ISL_965199, EPI_ISL_965200, EPI_ISL_965201, EPI_ISL_965202, EPI_ISL_965203, EPI_ISL_965204, EPI_ISL_965205, EPI_ISL_965206, EPI_ISL_965207, EPI_ISL_965208, EPI_ISL_965209, EPI_ISL_965210 | | | |
| see above | Virginia Division of Consolidated Laboratory Services | Virginia Division of Consolidated Laboratory Services | Virginia DCLS |
| EPI_ISL_965572, EPI_ISL_965573, EPI_ISL_965605, EPI_ISL_965618, EPI_ISL_965641, EPI_ISL_965652, EPI_ISL_965659, EPI_ISL_965671, EPI_ISL_965692, EPI_ISL_965699, EPI_ISL_965721, EPI_ISL_965798, EPI_ISL_965809 | | | |
| see above | Dutch COVID-19 response team | Medical Microbiology, Maastricht University Medical Centre | Jozef Dingemans*, Brian van der Veer*, Erik Beuken, Carmen Reumkens, Lieke van Alphen, Christian Hoebe, Paul Savelkoul |
| EPI_ISL_966769, EPI_ISL_966770, EPI_ISL_966771, EPI_ISL_966772, EPI_ISL_966773, EPI_ISL_966774, EPI_ISL_966775, EPI_ISL_966776 | Maine HETL | Tewhey Lab, The Jackson Laboratory | Matluk,N., Dewey,H., Iosue,F., Barter,M., Lynch,R., Munger,H. and Tewhey,R. |
| EPI_ISL_967588, EPI_ISL_967593, EPI_ISL_967649, EPI_ISL_967680, EPI_ISL_967681, EPI_ISL_967701, EPI_ISL_967731, EPI_ISL_967732, EPI_ISL_967733 | State Laboratories Division, Hawaii State Department of Health | State Laboratories Division, Hawaii State Department of Health | Pamela O'Brien, Drew Kuwazaki, Ayana Garnet, Razvan Sultana, Edward Desmond |
| EPI_ISL_967929, EPI_ISL_967946, EPI_ISL_967954, EPI_ISL_967968, EPI_ISL_967996, EPI_ISL_968019 | TGen North | Sonora Quest Laboratories | "Jolene Bowers, Megan Folkerts, Chris French, Hayley Yaglom, Ashlyn Pfeiffer, Darrin Lemmer, Dave Engelthaler, The Arizona COVID Genomics Union (ACGU)" |
| EPI_ISL_968853, EPI_ISL_968856, EPI_ISL_968858 | KEMRI-Wellcome Trust Research Programme/KEMRI-CGMR-C Kilifi | KEMRI-Wellcome Trust Research Programme/KEMRI-CGMR-C Kilifi | Githinji et al |
| EPI_ISL_970868, EPI_ISL_970901, EPI_ISL_970984, EPI_ISL_971051, EPI_ISL_971210, EPI_ISL_971217, EPI_ISL_971226, EPI_ISL_971280, EPI_ISL_971341, EPI_ISL_971366, EPI_ISL_971387, EPI_ISL_971477, EPI_ISL_971518, EPI_ISL_971527, EPI_ISL_971553, EPI_ISL_971579, EPI_ISL_971614, EPI_ISL_971642, EPI_ISL_971655, EPI_ISL_971706, EPI_ISL_971738, EPI_ISL_971773, EPI_ISL_971784, EPI_ISL_971790, EPI_ISL_971805, EPI_ISL_971819, EPI_ISL_971834, EPI_ISL_971885, EPI_ISL_971914, EPI_ISL_971920, EPI_ISL_971939, EPI_ISL_971951, EPI_ISL_971987, EPI_ISL_971991, EPI_ISL_971995, EPI_ISL_971997, EPI_ISL_972027, EPI_ISL_972058, EPI_ISL_972116, EPI_ISL_972117, EPI_ISL_972118, EPI_ISL_972153, EPI_ISL_972207, EPI_ISL_972212, EPI_ISL_972213, EPI_ISL_972236, EPI_ISL_972302, EPI_ISL_972307, EPI_ISL_972341, EPI_ISL_972452, EPI_ISL_972459, EPI_ISL_972501, EPI_ISL_972525, EPI_ISL_972544, EPI_ISL_972559, EPI_ISL_972587, EPI_ISL_972602, EPI_ISL_972662, EPI_ISL_972668, EPI_ISL_972697, EPI_ISL_972702, EPI_ISL_972766, EPI_ISL_972779, EPI_ISL_972810, EPI_ISL_972846, EPI_ISL_972861, EPI_ISL_972891, EPI_ISL_972905, EPI_ISL_972924, EPI_ISL_972954, EPI_ISL_973009, EPI_ISL_973019, EPI_ISL_973080, EPI_ISL_973105, EPI_ISL_973124, EPI_ISL_973145, EPI_ISL_973165, EPI_ISL_973252, EPI_ISL_973277, EPI_ISL_973278, EPI_ISL_973332, EPI_ISL_973346, EPI_ISL_973443, EPI_ISL_973544, EPI_ISL_973578, EPI_ISL_973601, EPI_ISL_973607, EPI_ISL_973635, EPI_ISL_973656, EPI_ISL_973704, EPI_ISL_973780, EPI_ISL_973799, EPI_ISL_973827 | | | |
| see above | Department of Virus and Microbiological Special Diagnostics, Statens Serum Institut, Copenhagen, Denmark | Aalborg University | Danish Covid-19 Genome Consortium |
| EPI_ISL_977653 | Michigan Department of Health and Human Services, Bureau of Laboratories | Michigan Department of Health and Human Services, Bureau of Laboratories | Blankenship HM, Riner D, Soehnlen MK |
| EPI_ISL_978152, EPI_ISL_978153, EPI_ISL_978154, EPI_ISL_978155, EPI_ISL_978156, EPI_ISL_978157 | Chiu Laboratory, University of California, San Francisco | Chiu Laboratory, University of California, San Francisco | Charles Chiu, Xianding (Wayne) Deng, Candace Wang, Venice Servellita, Jill Hacker, Debra Wadford |
| EPI_ISL_978409, EPI_ISL_978410 | Arizona State Public Health Laboratory | Arizona State Public Health Laboratory | Trung Huynh, Jessica Escobar, Katherine Fullerton, Nobuko Fukushima, Stacy White, Linda Getsinger, Victor Waddell |
| EPI_ISL_979695, EPI_ISL_979698, EPI_ISL_979699, EPI_ISL_979700, EPI_ISL_979701, EPI_ISL_979702, EPI_ISL_979703, EPI_ISL_979704, EPI_ISL_979707 | Orange County Public Health Lab | Chan-Zuckerberg Biohub | CZB Cliahub Consortium |
| EPI_ISL_981953 | Microbiology Service, Hospital Universitario Clinico San Cecilio, Granada | Microbiology Service, Hospital Universitario Clinico San Cecilio, Granada | Adolfo de Salazar, Natalia Chueca, Laura Viñuela, Ana Fuentes, Federico García |
| EPI_ISL_982005, EPI_ISL_982072, EPI_ISL_982093, EPI_ISL_982102, EPI_ISL_982107 | TGen North | Sonora Quest Laboratories | "Jolene Bowers, Megan Folkerts, Chris French, Hayley Yaglom, Ashlyn Pfeiffer, Darrin Lemmer, Dave Engelthaler, The Arizona COVID Genomics Union (ACGU)" |
| EPI_ISL_982369, EPI_ISL_982371, EPI_ISL_982372, EPI_ISL_982373, EPI_ISL_982375, EPI_ISL_982376, EPI_ISL_982377, EPI_ISL_982378, EPI_ISL_982379, EPI_ISL_982380, EPI_ISL_982381, EPI_ISL_982382, EPI_ISL_982383 | | | |
| see above | M Health Fairview | Minnesota Department of Health, Public Health Laboratory | Alexandra Lorentz, Jacob Garfin, Matt Plumb, and Xiong Wang |
| EPI_ISL_982427, EPI_ISL_982428, EPI_ISL_982429, EPI_ISL_982430, EPI_ISL_982431, EPI_ISL_982432, EPI_ISL_982433, EPI_ISL_982434, EPI_ISL_982435, EPI_ISL_982440, EPI_ISL_982445, EPI_ISL_982446, EPI_ISL_982448, EPI_ISL_982456, EPI_ISL_982459, EPI_ISL_982462, EPI_ISL_982465, EPI_ISL_982467, EPI_ISL_982468, EPI_ISL_982470, EPI_ISL_982472, EPI_ISL_982478, EPI_ISL_982480, EPI_ISL_982481, EPI_ISL_982486, EPI_ISL_982487, EPI_ISL_982489, EPI_ISL_982492, EPI_ISL_982495 | | | |
| see above | MONTEFIORE MEDICAL CENTER LABORATORIES | Wadsworth Center, New York State Department of Health | Kirsten St. George, Daryl M. Lamson, Alexis Russel, Matthew Shudt, Melissa A Leisner, Jonathan Plitnick, Navjot Singh, John Kelly, Erasmus Schneider, Erica Lasek-Nesselquist |
| EPI_ISL_982534, EPI_ISL_982542, EPI_ISL_982566, EPI_ISL_982571, EPI_ISL_982574, EPI_ISL_982575, EPI_ISL_982586, EPI_ISL_982592, EPI_ISL_982593, EPI_ISL_982594, EPI_ISL_982601, EPI_ISL_982602, EPI_ISL_982605, EPI_ISL_982606, EPI_ISL_982616, EPI_ISL_982650, EPI_ISL_982669, EPI_ISL_982678, EPI_ISL_982696, EPI_ISL_982697, EPI_ISL_982698, EPI_ISL_982699, EPI_ISL_982700, EPI_ISL_982701, EPI_ISL_982728, EPI_ISL_982736, EPI_ISL_982739, EPI_ISL_982741, EPI_ISL_982748, EPI_ISL_982749, EPI_ISL_982750, EPI_ISL_982759 | | | |
| see above | US Air Force School of Aerospace Medicine | US Air Force School of Aerospace Medicine | Anthony Fries, Jennifer Meyer, William Gruner, William Buggele, Amanda Javorina, Sarah Purves, Clarise Starr, Elizabeth Macias |
| EPI_ISL_982854, EPI_ISL_982855, EPI_ISL_982856 | Kentucky State Public Health Lab | Kentucky State Public Health Lab | Stephanie Lunn, Karim George, Joshua Tobias, William Grooms, Vaneet Arora, Matthew Johnson, Rachel Zinner, Rhonda Lucas |
| EPI_ISL_982857, EPI_ISL_982858, EPI_ISL_982859, EPI_ISL_983081 | Gravity Diagnostics | Kentucky State Public Health Lab | Stephanie Lunn, Karim George, Joshua Tobias, William Grooms, Vaneet Arora, Matthew Johnson, Rachel Zinner, Rhonda Lucas |
| EPI_ISL_983099, EPI_ISL_983102, EPI_ISL_983111, EPI_ISL_983112, EPI_ISL_983132, EPI_ISL_983133, EPI_ISL_983135, EPI_ISL_983137, EPI_ISL_983140, EPI_ISL_983141, EPI_ISL_983147, EPI_ISL_983149, EPI_ISL_983156, EPI_ISL_983162, EPI_ISL_983163, EPI_ISL_983164, EPI_ISL_983167, EPI_ISL_983170 | | | |
| see above | MONTEFIORE MEDICAL CENTER LABORATORIES | Wadsworth Center, New York State Department of Health | Kirsten St. George, Daryl M. Lamson, Alexis Russel, Matthew Shudt, Melissa A Leisner, Jonathan Plitnick, Navjot Singh, John Kelly, Erasmus Schneider, Erica Lasek-Nesselquist |
| EPI_ISL_983245 | SUNY UPSTATE MEDICAL UNIVERSITY | Wadsworth Center, New York State Department of Health | Kirsten St. George, Daryl M. Lamson, Alexis Russel, Matthew Shudt, Melissa A Leisner, Jonathan Plitnick, Navjot Singh, John Kelly, Erasmus Schneider, Erica Lasek-Nesselquist |
| EPI_ISL_983247 | KALEIDA CENTER FOR LABORATORY MEDICINE | Wadsworth Center, New York State Department of Health | Kirsten St. George, Daryl M. Lamson, Alexis Russel, Matthew Shudt, Melissa A Leisner, Jonathan Plitnick, Navjot Singh, John Kelly, Erasmus Schneider, Erica Lasek-Nesselquist |
| EPI_ISL_983248 | SUNY UPSTATE MEDICAL UNIVERSITY | Wadsworth Center, New York State Department of Health | Kirsten St. George, Daryl M. Lamson, Alexis Russel, Matthew Shudt, Melissa A Leisner, Jonathan Plitnick, Navjot Singh, John Kelly, Erasmus Schneider, Erica Lasek-Nesselquist |
| EPI_ISL_983296, EPI_ISL_983297, EPI_ISL_983298, EPI_ISL_983299 | KALEIDA CENTER FOR LABORATORY MEDICINE | Wadsworth Center, New York State Department of Health | Kirsten St. George, Daryl M. Lamson, Alexis Russel, Matthew Shudt, Melissa A Leisner, Jonathan Plitnick, Navjot Singh, John Kelly, Erasmus Schneider, Erica Lasek-Nesselquist |
| EPI_ISL_983300, EPI_ISL_983301, EPI_ISL_983302, EPI_ISL_983303, EPI_ISL_983304, EPI_ISL_983305, EPI_ISL_983306, EPI_ISL_983307, EPI_ISL_983308, EPI_ISL_983309, EPI_ISL_983310, EPI_ISL_983311, EPI_ISL_983312, EPI_ISL_983313, EPI_ISL_983314, EPI_ISL_983315, EPI_ISL_983316, EPI_ISL_983317, EPI_ISL_983318, EPI_ISL_983319 | | | |
| see above | SUNY UPSTATE MEDICAL UNIVERSITY | Wadsworth Center, New York State Department of Health | Kirsten St. George, Daryl M. Lamson, Alexis Russel, Matthew Shudt, Melissa A Leisner, Jonathan Plitnick, Navjot Singh, John Kelly, Erasmus Schneider, Erica Lasek-Nesselquist |

| | | | |
|---|---|---|---|
| EPI_ISL_983375, EPI_ISL_983376, EPI_ISL_983377, EPI_ISL_983378, EPI_ISL_983379, EPI_ISL_983380, EPI_ISL_983381 | Utah Public Health Laboratory | Utah Public Health Laboratory | Erin L. Young, Kelly F. Oakeson, Tara Gallagher |
| EPI_ISL_983826, EPI_ISL_983827, EPI_ISL_983828, EPI_ISL_983829, EPI_ISL_983830, EPI_ISL_983831, EPI_ISL_983832, EPI_ISL_983833, EPI_ISL_983834, EPI_ISL_983835 | Colorado Department of Public Health and Environment | Colorado Department of Puplic Health and Environment | Laura Bankers, Molly C. Hetherington-Rauth, Diana Ir, Shannon Ely, Shannon R. Matzinger, Sarah Elizabeth Totten, Emily A. Travanty |
