## Supplementary material for "Detection and characterization of the SARS-CoV-2 lineage B.1.526 in New York": Supp. Table 3 GISAID Acknowledgment part 2: gisaid_hcov-19_acknowledgement_table_2021_02_13_010-8.pdf

We gratefully acknowledge the following Authors from the Originating laboratories responsible for obtaining the specimens, as well as the Submitting laboratories where the genome data were generated and shared via GISAID, on which this research is based.

All Submitters of data may be contacted directly via www.gisaid.org

Authors are sorted alphabetically.

| Accession ID | Originating Laboratory | Submitting Laboratory | Authors |
|---|---|---|---|
| EPI_ISL_791422, EPI_ISL_791423, EPI_ISL_791425, EPI_ISL_791429 | Johns Hopkins Hospital Department of Pathology | Johns Hopkins Hospital Department of Pathology | C. Paul Morris, Chun Huai Luo, Adannaya Amadi, Nicholas Gallagher, Heba H. Mostafa |
| EPI_ISL_792090 | Toronto Invasive Bacterial Diseases Network | McMaster University | Allison McGeer, Patryk Aftanas, Hooman Derakhshani, Angel Li, Kuganya Nirmalarajah, Emily Panousis, Ahmed Draia, Jalees Nasir, Michael Surette, Samira Mubareka, Andrew G. McArthur |
| EPI_ISL_794309, EPI_ISL_794310, EPI_ISL_794311, EPI_ISL_794312, EPI_ISL_794313, EPI_ISL_794314, EPI_ISL_794315, EPI_ISL_794316, EPI_ISL_794317, EPI_ISL_794318, EPI_ISL_794319, EPI_ISL_794320 | | | |
| see above | Wadsworth Center, New York State Department.of Health | Wadsworth Center, New York State Department.of Health | Kirsten St. George, Daryl M. Lamson, Alexis Russel, Matthew Shudt, Melissa A Leisner, Jonathan Plitnick, Navjot Singh, John Kelly, Sara Griesemer, Erasmus Schneider, Erica Lasek-Nesselquist |
| EPI_ISL_794640 | USC Clinical Lab | Los Angeles County PHL | P. Hemarajata et al. |
| EPI_ISL_794641, EPI_ISL_794642, EPI_ISL_794643, EPI_ISL_794644, EPI_ISL_794645, EPI_ISL_794646, EPI_ISL_794647, EPI_ISL_794648 | UCLA Clinical Micro Lab | Los Angeles County PHL | P. Hemarajata et al. |
| EPI_ISL_794666, EPI_ISL_794667 | SYNLAB REGIONAL NOROCCIDENTE | Instituto Nacional de Salud - Dirección de Investigación en Salud Pública | Katherine Laiton-Donato, Diego A. Álvarez-Díaz, Carlos Franco-Muñoz, Mauricio Pacheco-Montealegre, Jonathan Reales, Sheryl Corchuelo, Maria T. Herrera, Julian Naizaque, Gerardo Santamaría, Paola Muñoz-Laiton, Diego Andrés Prada, Magdalena Wiesner, Martha Lucia Ospina Martinez, Marcela Mercado-Reyes |
| EPI_ISL_794670 | UCLA Clinical Micro Lab | Los Angeles County PHL | P. Hemarajata et al. |
| EPI_ISL_794671 | Area of Virology, Serology and Virology Division (SAViD), New South Wales Health Pathology Randwick | Area of Virology, Serology and Virology Division (SAViD), New South Wales Health Pathology Randwick | Rawlinson, W., Deveson, I., Van Haal, S., Bull, R. |
| EPI_ISL_794735, EPI_ISL_794736, EPI_ISL_794737, EPI_ISL_794738 | 1-Laboratory of Microbiology, National Reference Lab, Charles Nicolle Hospital; 2-University of Tunis ElManar, Faculty of Medicine of Tunis, LR99ES09, Tunis, Tunisia | 1-Clinical and Experimental Pharmacology Lab, LR16SP02, National Center of Pharmacovigilance, University of Tunis El Manar, Tunis, Tunisia. 2-Neurodegenerative diseases and psychiatric troubles, LR18SP03, Razi Hospital, University of Tunis El Manar, Tunis, Tunisia. 3- Ministry of Health, National Observatory of New and Emerging Diseases, 1006, Tunis, Tunisia | Ilhem Boutiba-Ben Boubaker, Sameh Trabelsi, Nissaf Ben Alaya, Maher Kharrat, Alia BenKahla, Jalila Ben Khelil, Salma Abid, Sana Ferjani, Mouna Ben Sassi, Mouna Safer, Zaineb Hamzaoui, Guedi Ali Barreh, Habiba Ben Romdhane, Souissi Amira, Sarra Chamman, Hanen El Jebari, Asma Ferjani, Gaies Emna, Riadh Daghfous, Riadh Gouider. |
| EPI_ISL_803098 | Institute of Microbiology, Universidad San Francisco de Quito | Institute of Microbiology, Universidad San Francisco de Quito | Sully Márquez, Belén Prado-Vivar, Juan José Guadalupe, Monica Becerra-Wong, Bernardo Gutiérrez, Eulalia Pazmiño, Katalina Pacheco, Verónica Barragán, Patricio Rojas-Silva, Gabriel Trueba, Michelle Grunauer, Paúl Cárdenas |
| EPI_ISL_803120, EPI_ISL_803430 | Laboratory of Microbiology, National Reference Lab, Charles Nicolle Hospital; 2-University of Tunis ElManar, Faculty of Medicine of Tunis, LR99ES09, Tunis, Tunisia | Clinical and Experimental Pharmacology Lab, LR16SP02, National Center of Pharmacovigilance, University of Tunis El Manar, Tunis, Tunisia. 2-Neurodegenerative diseases and psychiatric troubles, LR18SP03, Razi Hospital, University of Tunis El Manar, Tunis, Tunisia. 3- Ministry of Health, National Observatory of New and Emerging Diseases, 1006, Tunis, Tunisia | Ilhem Boutiba-Ben Boubaker, Sameh Trabelsi, Nissaf Ben Alaya, Maher Kharrat, Alia BenKahla, Jalila Ben Khelil, Salma Abid, Sana Ferjani, Mouna Ben Sassi, Mouna Safer, Zaineb Hamzaoui, Habiba Ben Romdhane, Souissi Amira, Sarra Chamman, Hanen El Jebari, Ahmed Fakhfakh, Gaies Emna, Riadh Daghfous, Riadh Gouider. |
| EPI_ISL_803962, EPI_ISL_803963, EPI_ISL_803964, EPI_ISL_803965, EPI_ISL_803999, EPI_ISL_804006 | National Public Health Laboratory, National Centre for Infectious Diseases | National Public Health Laboratory, National Centre for Infectious Diseases | Tze Minn Mak, Sophie Octavia, Zhenyang Zhou, Lin Cui, Raymond Tzer Pin Lin |
| EPI_ISL_804259, EPI_ISL_804369 | Respiratory Virus Unit, National Infection Service, Public Health England | COVID-19 Genomics UK (COG-UK) Consortium | PHE Covid Sequencing Team |
| EPI_ISL_804947 | Wadsworth Center, New York State Department.of Health | Wadsworth Center, New York State Department.of Health | Kirsten St. George, Daryl M. Lamson, Alexis Russel, Matthew Shudt, Melissa A Leisner, Jonathan Plitnick, Navjot Singh, John Kelly, Sara Griesemer, Erasmus Schneider, Erica Lasek-Nesselquist |
| EPI_ISL_806544 | INSPI Instituto Nacional de Investigación en Salud Pública | Av. Julián Coronel 905 entre Esmeraldas y José Mascote Av. Juan Tanca Marengo No. 100 y Av. de las Américas | Leandro Patiño, Doménica de Mora, Maritza Olmedo, Andrés Carrazco, Orson Mestanza, Mary Regato, Melissa Zambrano, Manuel González, Alfredo Bruno, Alberto Orlando. |
| EPI_ISL_806729 | Sydney South West Pathology Service (SSWPS) - Royal Prince Alfred Hospital - NSW Health Pathology | NSW Health Pathology - Institute of Clinical Pathology and Medical Research; Westmead Hospital; University of Sydney | CIDM-PH et al. |
| EPI_ISL_806808 | Gundersen Molecular Diagnostics Laboratory | Kabara Cancer Research Institute | Craig S. Richmond, Paraic A. Kenny |
| EPI_ISL_807148, EPI_ISL_807149 | Department of Virology | Department of Virology | Massab Umair, Aamer Ikram, Muhammad Salman, Sana Tamim, Nazish Badar, Adnan Khurshid, Salmaan Sharif, Zaira Rehman, Abdul Ahad, Hamza Ahmad, Samee Ullah |
| EPI_ISL_810967 | PathWest Laboratory Medicine WA | PathWest Laboratory Medicine WA Microbial Surveillance Unit | PathWest Laboratory Medicine WA Microbial Surveillance Unit |
| EPI_ISL_811127 | Cantonal Hospital Winterthur | Institute of Medical Virology, University of Zurich | Stefan Schmutz, Maryam Zaheri, Verena Kufner, Annette Audigé, Maria Grünberg, Kevin Steiner, Jon Huder, Cyril Shah, Riccarda Capaul, Guido Bloemberg, Jürg Böni, Michael Huber, Alexandra Trkola, Alexander Wepf, Urs Karrer |
| EPI_ISL_811147 | WHO National Influenza Centre Russian Federation | WHO National Influenza Centre Russian Federation | Andrey Komissarov, Artem Fadeev, Maxim Burlakov, Anna Ivanova, Kseniya Komissarova, Dmitry Bazhenov, Daria Danilenko, Ksenia Safina, Elena Nabieva, Georgii Bazykin, Dmitry Lioznov |
| EPI_ISL_812122 | Department of Clinical Microbiology | GIGA Medical Genomics | Keith Durkin, Maria Artesi, Sébastien Bontems, Raphaël Boreux, Bouchra Boujemla, Cécile Meex, Pierrette Melin, Marie-Pierre Hayette, Vincent Bours |
| EPI_ISL_812356, EPI_ISL_812357 | SA Pathology | SA Pathology | Lex Leong, Julien Soubrier, Chuan Kok Lim, Song Gao, Mark Turra, Karin Kassahn, Ivan Bastian, Geoff Higgins |
| EPI_ISL_812440 | Victorian Infectious Diseases Reference Laboratory (VIDRL) | VIDRL and MDU-PHL | Caly L., Seemann T., Sait, M.L., Druce J., Sherry, N.L. |
| EPI_ISL_812778 | Ministry of Health Turkey | Ministry of Health Turkey | Fatma Bayrakdar, Yasemin Cogun, Süleyman Yalcin, Aye Baak Alta, Gülay Korukluolu |
| EPI_ISL_816258, EPI_ISL_816328, EPI_ISL_816363, EPI_ISL_816371, EPI_ISL_816373, EPI_ISL_816377, EPI_ISL_816413, EPI_ISL_816424, EPI_ISL_816451, EPI_ISL_816479, EPI_ISL_816506, EPI_ISL_816519, EPI_ISL_816614, EPI_ISL_816653 | | | |
| see above | Virology Department, Sheffield Teaching Hospitals NHS Foundation Trust/Department of Infection, Immunity and Cardiovascular Disease, The Medical School, University of Sheffield | COVID-19 Genomics UK (COG-UK) Consortium | Thushan de Silva, Matthew Parker, Nikki Smith, Adri Angyal, Rebecca Brown, Luke Green, Rachel Tucker, Paul Parsons, Danielle Groves, Katie Johnson, Laura Carrilero, Alex Keeley, Dave Partridge, Matthew Wyles, Benjamin Lindsey, Mehmet Yavuz, Mohammad Raza, Cariad Evans |

| | | | |
|---|---|---|---|
| EPI_ISL_819125 | Wyoming Public Health Laboratory | Wyoming Public Health Laboratory | Noah Hull, Taylor Fearing, Lynette Gumbleton, Channing Weber, Ashley Norberg, Bailey Bowcutt, and Wanda Manley |
| EPI_ISL_819134, EPI_ISL_819139, EPI_ISL_819141, EPI_ISL_819145, EPI_ISL_819147, EPI_ISL_819151, EPI_ISL_819152, EPI_ISL_819153, EPI_ISL_819154, EPI_ISL_819155, EPI_ISL_819158, EPI_ISL_819159, EPI_ISL_819160, EPI_ISL_819161, EPI_ISL_819162, EPI_ISL_819164, EPI_ISL_819165, EPI_ISL_819177 | | | |
| see above | Servicio de Microbiología, Hospital Universitario Son Espases | SeqCOVID-SPAIN consortium/IBV(CSIC) | Carla López-Causapé, Jordi Reina, Antonio Oliver and SeqCOVID-SPAIN consortium |
| EPI_ISL_819204, EPI_ISL_819206, EPI_ISL_819208, EPI_ISL_819211, EPI_ISL_819212, EPI_ISL_819214, EPI_ISL_819224, EPI_ISL_819225, EPI_ISL_819226, EPI_ISL_819227, EPI_ISL_819228, EPI_ISL_819229, EPI_ISL_819230, EPI_ISL_819231, EPI_ISL_819232, EPI_ISL_819233, EPI_ISL_819234, EPI_ISL_819235, EPI_ISL_819240, EPI_ISL_819246, EPI_ISL_819247, EPI_ISL_819248, EPI_ISL_819249, EPI_ISL_819250, EPI_ISL_819251, EPI_ISL_819252, EPI_ISL_819253, EPI_ISL_819254, EPI_ISL_819255, EPI_ISL_819256, EPI_ISL_819257, EPI_ISL_819258, EPI_ISL_819259, EPI_ISL_819260, EPI_ISL_819261, EPI_ISL_819262, EPI_ISL_819263, EPI_ISL_819264, EPI_ISL_819265, EPI_ISL_819266 | | | |
| see above | Wyoming Public Health Laboratory | Wyoming Public Health Laboratory | Noah Hull, Taylor Fearing, Lynette Gumbleton, Channing Weber, Ashley Norberg, Bailey Bowcutt, and Wanda Manley |
| EPI_ISL_820596, EPI_ISL_820598, EPI_ISL_820601, EPI_ISL_820603, EPI_ISL_820605, EPI_ISL_820607, EPI_ISL_820610 | Queens Medical Centre, Clinical Microbiology Department / DeepSeq Nottingham | COVID-19 Genomics UK (COG-UK) Consortium | Gemma Clark, Wendy Smith, Manjinder Khakh, Vicki M Fleming, Michelle M Lister, Hannah Howson-Wells, Jonathan Ball, Patrick McClure, Joseph Chappell, Theocharis Tsoleridis, Nadine Holmes, Matthew Carlisle, Christopher Moore, Fei Sang, Johnny Debebe, Victoria Wright, Matthew Loose |
| EPI_ISL_821088, EPI_ISL_821139 | Lighthouse Lab in Alderley Park | Wellcome Sanger Institute for the COVID-19 Genomics UK (COG-UK) Consortium | Jacquelyn Wynn, Mairead Hyland, The Lighthouse Lab in Alderley Park and Alex Alderton, Roberto Amato, Sonia Goncalves, Ewan Harrison, David K. Jackson, Ian Johnston, Dominic Kwiatkowski, Cordelia Langford, John Sillitoe on behalf of the Wellcome Sanger Institute COVID-19 Surveillance Team |
| EPI_ISL_823989, EPI_ISL_824261, EPI_ISL_824262, EPI_ISL_824263, EPI_ISL_824279, EPI_ISL_824280 | Dutch COVID-19 response team | National Institute for Public Health and the Environment (RIVM) | Adam Meijer, Harry Vennema, Jeroen Cremer, Sharon van den Brink, Bas van der Veer, AnneMarie van den Brandt, Florian Zwagemaker, Dennis Schmitz, Chantal Reusken, on behalf of the national COVID-19 response team |
| EPI_ISL_824292 | Institute of Microbiology, Universidad San Francisco de Quito | Institute of Microbiology, Universidad San Francisco de Quito | Belén Prado-Vivar, Sully Márquez, Juan José Guadalupe, Monica Becerra-Wong, Bernardo Gutiérrez, Tania Guayasamin, Patricio Reyes, Verónica Barragán, Patricio Rojas-Silva, Gabriel Trueba, Michelle Grunauer, Paúl Cárdenas |
| EPI_ISL_824293, EPI_ISL_824294 | Institute of Microbiology, Universidad San Francisco de Quito | Institute of Microbiology, Universidad San Francisco de Quito | Belén Prado-Vivar, Sully Márquez, Juan José Guadalupe, Monica Becerra-Wong, Bernardo Gutiérrez, Eulalia Pazmiño, Katalina Pacheco, Verónica Barragán, Patricio Rojas-Silva, Gabriel Trueba, Michelle Grunauer, Paúl Cárdenas |
| EPI_ISL_824852, EPI_ISL_824853, EPI_ISL_824854, EPI_ISL_824855, EPI_ISL_824856, EPI_ISL_824857, EPI_ISL_824858, EPI_ISL_824859, EPI_ISL_824860, EPI_ISL_824861, EPI_ISL_824862, EPI_ISL_824863 | | | |
| see above | Department of Clinical Microbiology | GIGA Medical Genomics | Keith Durkin, Maria Artesi, Sébastien Bontems, Raphaël Boreux, Bouchra Boujemla, Cécile Meex, Pierrette Melin, Marie-Pierre Hayette, Vincent Bours |
| EPI_ISL_824966, EPI_ISL_824967, EPI_ISL_824968, EPI_ISL_824969, EPI_ISL_824970, EPI_ISL_824971, EPI_ISL_824972, EPI_ISL_824973, EPI_ISL_824978, EPI_ISL_824979 | Maryland Public Health Laboratory | Maryland Public Health Laboratory | Maryland Department of Health Laboratories Administration |
| EPI_ISL_824985, EPI_ISL_824986 | Charité Universitätsmedizin Berlin, Institute of Virology, Charitéplatz 1, 10117 Berlin, Germany | Charité Universitätsmedizin Berlin, Institute of Virology, Charitéplatz 1, 10117 Berlin, Germany | Victor M Corman, Jörn Beheim-Schwarzbach, Tobias Bleicker, Julia Tesch, Barbara Mühlemann, Talitha Veith, Julia Schneider, Terry Jones, Christian Drosten |
| EPI_ISL_824995 | Arizona State Public Health Laboratory | Arizona State Public Health Laboratory | Trung Huynh, Jessica Escobar, Katherine Fullerton, Nobuko Fukushima, Stacy White, Linda Getsinger, Victor Waddell |
| EPI_ISL_825013 | Johns Hopkins Pathology | Johns Hopkins Hospital Department of Pathology | C. Paul Morris, Chun Huai Luo, Adannaya Amadi, Nicholas Gallagher, Heba H. Mostafa |
| EPI_ISL_825069, EPI_ISL_825074, EPI_ISL_825076, EPI_ISL_825080 | National Public Health Laboratory, National Centre for Infectious Diseases | National Public Health Laboratory, National Centre for Infectious Diseases | Tze Minn Mak, Sophie Octavia, Zhenyang Zhou, Lin Cui, Raymond Tzer Pin Lin |
| EPI_ISL_826269 | University of Debrecen, Department of Medical Microbiology | National Laboratory of Virology, Szentágothai Research Centre | Endre Gábor Tóth, Balázs Somogyi, Brigitta Zana, Eszter Csoma, Ferenc Jakab, Gábor Kemenesi |
| EPI_ISL_826304, EPI_ISL_826306 | Wyoming Public Health Laboratory | Wyoming Public Health Laboratory | Noah Hull, Taylor Fearing, Lynette Gumbleton, Channing Weber, Ashley Norberg, Bailey Bowcutt, and Wanda Manley |
| EPI_ISL_826461 | Montana Public Health Laboratory | Wyoming Public Health Laboratory | Noah Hull, Joy Ritter, Taylor Fearing, Lynette Gumbleton, Channing Weber, Ashley Norberg, Bailey Bowcutt, Wanda Manley, Deborah Gibson |
| EPI_ISL_826896, EPI_ISL_826918, EPI_ISL_827031, EPI_ISL_827120, EPI_ISL_827307, EPI_ISL_827345, EPI_ISL_827358, EPI_ISL_827416, EPI_ISL_827689, EPI_ISL_827690, EPI_ISL_827757, EPI_ISL_827780, EPI_ISL_827871, EPI_ISL_827951, EPI_ISL_827992, EPI_ISL_828262, EPI_ISL_828497, EPI_ISL_829977, EPI_ISL_829982 | | | |
| see above | deCODE genetics | deCODE genetics | Daniel F Gudbjartsson; Agnar Helgason; Hakon Jonsson; Olafur T Magnusson; Pall Melsted; Gudmundur L Norddahl; Jona Saemundsdottir; Asgeir Sigurdsson; Patrick Sulem; Arna B Agustsdottir; Hannes Eggertsson; Berglind Eiriksdottir; Run Fridriksdottir; Elisabet E Gardarsdottir; Gudmundur Georgsson; Olafia S Gretarsdottir; Kjartan R Gudmundsson; Thora R Gunnarsdottir; Arnaldur Gylfason; Hilma Holm; Brynjar O Jensson; Aslaug Jonasdottir; Kamilla S Josefsdottir; Thordur Kristjansson; Droplaug N Magnusdottir; Solvi Rognvaldsson; Louise le Roux; Gudrun Sigmundsdottir; Gardar Sveinbjornsson; Kristin E Sveinsdottir; Maney Sveinsdottir; Emil A Thorarensen; Bjarni Thorbjornsson; Gisli Masson; Ingileif Jonsdottir; Alma Moller; Thorolfur Gudnason; Karl G Kristinsson; Unnur Thorsteinsdottir; Kari Stefansson |
| EPI_ISL_830216, EPI_ISL_830218, EPI_ISL_830220, EPI_ISL_830221, EPI_ISL_830263, EPI_ISL_830265, EPI_ISL_830267, EPI_ISL_830269, EPI_ISL_830270, EPI_ISL_830272, EPI_ISL_830274, EPI_ISL_830276, EPI_ISL_830278, EPI_ISL_830280, EPI_ISL_830282, EPI_ISL_830284, EPI_ISL_830286, EPI_ISL_830288, EPI_ISL_830289, EPI_ISL_830291, EPI_ISL_830293, EPI_ISL_830294, EPI_ISL_830296, EPI_ISL_830298, EPI_ISL_830300, EPI_ISL_830302, EPI_ISL_830304, EPI_ISL_830306, EPI_ISL_830308, EPI_ISL_830310, EPI_ISL_830312, EPI_ISL_830313, EPI_ISL_830315, EPI_ISL_830317, EPI_ISL_830319, EPI_ISL_830321, EPI_ISL_830323, EPI_ISL_830325, EPI_ISL_830326, EPI_ISL_830329, EPI_ISL_830331, EPI_ISL_830333, EPI_ISL_830334, EPI_ISL_830335, EPI_ISL_830337, EPI_ISL_830338, EPI_ISL_830340, EPI_ISL_830342, EPI_ISL_830344, EPI_ISL_830346, EPI_ISL_830348, EPI_ISL_830349 | | | |
| see above | Wadsworth Center, New York State Department of Health | Wadsworth Center, New York State Department of Health | Kirsten St. George, Daryl M. Lamson, Alexis Russel, Matthew Shudt, Melissa A Leisner, Jonathan Plitnick, Navjot Singh, John Kelly, Erasmus Schneider, Erica Lasek-Nesselquist |
| EPI_ISL_830682 | SUNY UPSTATE MEDICAL UNIVERSITY | Wadsworth Center, New York State Department of Health | Kirsten St. George, Daryl M. Lamson, Alexis Russel, Matthew Shudt, Melissa A Leisner, Jonathan Plitnick, Navjot Singh, John Kelly, Erasmus Schneider, Erica Lasek-Nesselquist |
| EPI_ISL_830704, EPI_ISL_830705, EPI_ISL_830706 | ALBANY MEDICAL CENTER HOSPITAL CLINICAL LABORATORIES | Wadsworth Center, New York State Department of Health | Kirsten St. George, Daryl M. Lamson, Alexis Russel, Matthew Shudt, Melissa A Leisner, Jonathan Plitnick, Navjot Singh, John Kelly, Erasmus Schneider, Erica Lasek-Nesselquist |
| EPI_ISL_831366, EPI_ISL_831367 | Clinical Molecular Microbiology Laboratory, UNC Hospitals | Jeremy Wang | Jeremy Wang, Alexander Rubinsteyn, Colleen Rice, Jason Smedberg, Melissa Miller, Corbin Jones, Robert Hagan |
| EPI_ISL_831479, EPI_ISL_831480, EPI_ISL_831481, EPI_ISL_831483, EPI_ISL_831486, EPI_ISL_831487, EPI_ISL_831496, EPI_ISL_831511, EPI_ISL_831521, EPI_ISL_831527, EPI_ISL_831529, EPI_ISL_831530, EPI_ISL_831533, EPI_ISL_831534, EPI_ISL_831536, EPI_ISL_831538, EPI_ISL_831539, EPI_ISL_831540, EPI_ISL_831542, EPI_ISL_831543, EPI_ISL_831544, EPI_ISL_831545, EPI_ISL_831546, EPI_ISL_831548, EPI_ISL_831550, EPI_ISL_831623, EPI_ISL_831624, EPI_ISL_831625, EPI_ISL_831626, EPI_ISL_831627, EPI_ISL_831628, EPI_ISL_831629, EPI_ISL_831630, EPI_ISL_831631, EPI_ISL_831632, EPI_ISL_831633, EPI_ISL_831634, EPI_ISL_831635, EPI_ISL_831636, EPI_ISL_831637, EPI_ISL_831638, EPI_ISL_831639, EPI_ISL_831640, EPI_ISL_831641, EPI_ISL_831642, EPI_ISL_831643 | | | |
| see above | University of Wisconsin-Madison AIDS Vaccine Research Laboratories | University of Wisconsin-Madison AIDS Vaccine Research Laboratories | Gage Moreno, Katarina Braun, et al. AIDS Vaccine Research Laboratories |
| EPI_ISL_831667 | Institute of Virology, Biomedical Research Center of the Slovak Academy of Sciences, Bratislava | Faculty of Natural Sciences, Comenius University, Bratislava | Viktória abanová, Kristína Boršová, Broa Brejová, Viktória Hodorová, Sabina Fumaová Havlíková, Juraj Kopáek, Martina Liková, ubomíra Lukáiková, Martina Neboháová, Monika Sláviková, Tomáš Vina, Jozef Nosek, Boris Klempa |
| EPI_ISL_831668 | Institute of Virology, Biomedical Research Center of the Slovak Academy of Sciences, Bratislava | Faculty of Natural Sciences, Comenius University, Bratislava | Kristína Boršová, Viktória abanová, Broa Brejová, Viktória Hodorová, Sabina Fumaová Havlíková, Juraj Kopáek, Martina Liková, ubomíra Lukáiková, Martina Neboháová, Monika Sláviková, Tomáš Vina, Boris Klempa, Jozef Nosek |
| EPI_ISL_831672 | Institute of Virology, Biomedical Research Center of the Slovak Academy of Sciences, Bratislava | Faculty of Natural Sciences, Comenius University, Bratislava | Viktória abanová, Kristína Boršová, Broa Brejová, Viktória Hodorová, Sabina Fumaová Havlíková, Juraj Kopáek, Martina Liková, ubomíra Lukáiková, Martina Neboháová, Monika Sláviková, Tomáš Vina, Jozef Nosek, Boris Klempa |
| EPI_ISL_831906, EPI_ISL_831907, EPI_ISL_831911, EPI_ISL_831928, EPI_ISL_831929, EPI_ISL_831930 | New Mexico Department of Health Scientific Laboratory | New Mexico Department of Health Scientific Laboratory | Ellie Johnson, Anastacia Griego-Fisher, D'eldra Malone |
| EPI_ISL_831941, EPI_ISL_831942 | Klinisk mikrobiologi | The Public Health Agency of Sweden | Department of Microbiology, The Public Health Agency of Sweden |

| | | | |
|---|---|---|---|
| EPI_ISL_832022, EPI_ISL_832065, EPI_ISL_832096, EPI_ISL_832097, EPI_ISL_832098, EPI_ISL_832100 | Santa Clara County Public Health Laboratory | Santa Clara County Public Health Laboratory | Santa Clara County Public Health Department |
| EPI_ISL_832122, EPI_ISL_832123 | Clinical Molecular Microbiology Laboratory, UNC Hospitals | Jeremy Wang | Jeremy Wang, Alexander Rubinsteyn, Colleen Rice, Jason Smedberg, Melissa Miller, Corbin Jones, Robert Hagan |
| EPI_ISL_832210, EPI_ISL_832211, EPI_ISL_832212, EPI_ISL_832213, EPI_ISL_832214, EPI_ISL_832215, EPI_ISL_832216, EPI_ISL_832217, EPI_ISL_832218, EPI_ISL_832219, EPI_ISL_832220, EPI_ISL_832221, EPI_ISL_832222, EPI_ISL_832223, EPI_ISL_832224, EPI_ISL_832225, EPI_ISL_832226, EPI_ISL_832227, EPI_ISL_832228, EPI_ISL_832229, EPI_ISL_832230, EPI_ISL_832231 | | | |
| see above | Department of Clinical Microbiology | GIGA Medical Genomics | Keith Durkin, Maria Artesi, Sébastien Bontems, Raphaël Boreux, Bouchra Boujemla, Cécile Meex, Pierrette Melin, Marie-Pierre Hayette, Vincent Bours |
| EPI_ISL_832378 | Ohio State | James Molecular Lab | Huolin Tu, Matthew R Avenarius, Laura Kubatko, Matthew Hunt, Xiaokang Pan, Peng Ru, Jason Garee, Keelie Thomas, Peter Mohler, Preeti Pancholi, Dan Jones |
| EPI_ISL_832968, EPI_ISL_832969, EPI_ISL_832970, EPI_ISL_832971, EPI_ISL_832972, EPI_ISL_832973, EPI_ISL_832974, EPI_ISL_832975, EPI_ISL_832976, EPI_ISL_832977, EPI_ISL_832986, EPI_ISL_832987, EPI_ISL_832988, EPI_ISL_832989, EPI_ISL_832990, EPI_ISL_832991, EPI_ISL_832992, EPI_ISL_832993, EPI_ISL_832994, EPI_ISL_832995, EPI_ISL_832996, EPI_ISL_832997, EPI_ISL_832998, EPI_ISL_832999, EPI_ISL_833000, EPI_ISL_833001, EPI_ISL_833002, EPI_ISL_833003, EPI_ISL_833004, EPI_ISL_833005, EPI_ISL_833006, EPI_ISL_833007, EPI_ISL_833008, EPI_ISL_833009, EPI_ISL_833010, EPI_ISL_833011, EPI_ISL_833012, EPI_ISL_833013, EPI_ISL_833014, EPI_ISL_833015, EPI_ISL_833016, EPI_ISL_833017, EPI_ISL_833018, EPI_ISL_833019, EPI_ISL_833020, EPI_ISL_833021, EPI_ISL_833022, EPI_ISL_833023, EPI_ISL_833024, EPI_ISL_833025, EPI_ISL_833026, EPI_ISL_833027, EPI_ISL_833029, EPI_ISL_833030, EPI_ISL_833031, EPI_ISL_833032, EPI_ISL_833033, EPI_ISL_833034, EPI_ISL_833035, EPI_ISL_833036, EPI_ISL_833037 | | | |
| see above | Maine HETL | Tewhey Lab, The Jackson Laboratory | Matluk,N., Dewey,H., Iosue,F., Barter,M., Lynch,R., Munger,H. and Tewhey,R. |
| EPI_ISL_833147 | Genomic Laboratory (GLAB) (Conjoint lab of Health Directorate of Istanbul and Istanbul Technical University) | Genomic Laboratory (GLAB), Istanbul Technical University | Ilker Karacan, Tugba Kizilboga Akgun, Payam Zolfagharian, Nisan Denizce Can, Pari Sharifli, Levent Doganay, Gizem Dinler Doganay |
| EPI_ISL_833165, EPI_ISL_833166 | Hospital Samaritano | Instituto Adolfo Lutz, Interdiciplinary Procedures Center, Strategic Laboratory | Claudio Tavares Sacchi, Claudia Regina Gonçalves, Erica Valessa Ramos Gomes, Karoline Rodrigues Campos |
| EPI_ISL_833185, EPI_ISL_833189, EPI_ISL_833190 | Department of Clinical Microbiology | GIGA Medical Genomics | Keith Durkin, Maria Artesi, Sébastien Bontems, Raphaël Boreux, Bouchra Boujemla, Cécile Meex, Pierrette Melin, Marie-Pierre Hayette, Vincent Bours |
| EPI_ISL_833401, EPI_ISL_833402, EPI_ISL_833403 | St. Francis Medical Center | Los Angeles County PHL | P. Hemarajata et al. |
| EPI_ISL_833405, EPI_ISL_833406, EPI_ISL_833407, EPI_ISL_833408, EPI_ISL_833409, EPI_ISL_833410, EPI_ISL_833411, EPI_ISL_833412 | Beverly Hospital | Los Angeles County PHL | P. Hemarajata et al. |
| EPI_ISL_833413 | City of Hope Comprehensive Cancer Center | Los Angeles County PHL | P. Hemarajata et al. |
| EPI_ISL_833417, EPI_ISL_833419, EPI_ISL_833420, EPI_ISL_833421, EPI_ISL_833422, EPI_ISL_833423, EPI_ISL_833424, EPI_ISL_833425, EPI_ISL_833426, EPI_ISL_833427 | MD Laboratories | Los Angeles County PHL | P. Hemarajata et al. |
| EPI_ISL_833428, EPI_ISL_833429, EPI_ISL_833430, EPI_ISL_833431 | UCLA Clinical Micro Lab | Los Angeles County PHL | P. Hemarajata et al. |
| EPI_ISL_833489 | St. Francis Medical Center | Los Angeles County PHL | P. Hemarajata et al. |
| EPI_ISL_833586, EPI_ISL_833590, EPI_ISL_833593, EPI_ISL_833596, EPI_ISL_833597, EPI_ISL_833599, EPI_ISL_833602, EPI_ISL_833604, EPI_ISL_833606, EPI_ISL_833607 | Lighthouse Lab in Milton Keynes | Wellcome Sanger Institute for the COVID-19 Genomics UK (COG-UK) Consortium | The Lighthouse Lab in Milton Keynes and Alex Alderton, Roberto Amato, Sonia Goncalves, Ewan Harrison, David K. Jackson, Ian Johnston, Dominic Kwiatkowski, Cordelia Langford, John Sillitoe on behalf of the Wellcome Sanger Institute COVID-19 Surveillance Team |
| EPI_ISL_833609, EPI_ISL_833610 | Lighthouse Lab in Cambridge | Wellcome Sanger Institute for the COVID-19 Genomics UK (COG-UK) Consortium | Rob Howes, The Lighthouse Lab in Cambridge and Alex Alderton, Roberto Amato, Sonia Goncalves, Ewan Harrison, David K. Jackson, Ian Johnston, Dominic Kwiatkowski, Cordelia Langford, John Sillitoe on behalf of the Wellcome Sanger Institute COVID-19 Surveillance Team |
| EPI_ISL_833611, EPI_ISL_833612, EPI_ISL_833614, EPI_ISL_833615 | Lighthouse Lab in Milton Keynes | Wellcome Sanger Institute for the COVID-19 Genomics UK (COG-UK) Consortium | The Lighthouse Lab in Milton Keynes and Alex Alderton, Roberto Amato, Sonia Goncalves, Ewan Harrison, David K. Jackson, Ian Johnston, Dominic Kwiatkowski, Cordelia Langford, John Sillitoe on behalf of the Wellcome Sanger Institute COVID-19 Surveillance Team |
| EPI_ISL_833616 | Lighthouse Lab in Cambridge | Wellcome Sanger Institute for the COVID-19 Genomics UK (COG-UK) Consortium | Rob Howes, The Lighthouse Lab in Cambridge and Alex Alderton, Roberto Amato, Sonia Goncalves, Ewan Harrison, David K. Jackson, Ian Johnston, Dominic Kwiatkowski, Cordelia Langford, John Sillitoe on behalf of the Wellcome Sanger Institute COVID-19 Surveillance Team |
| EPI_ISL_833617, EPI_ISL_833618, EPI_ISL_833619, EPI_ISL_833620, EPI_ISL_833621 | Lighthouse Lab in Milton Keynes | Wellcome Sanger Institute for the COVID-19 Genomics UK (COG-UK) Consortium | The Lighthouse Lab in Milton Keynes and Alex Alderton, Roberto Amato, Sonia Goncalves, Ewan Harrison, David K. Jackson, Ian Johnston, Dominic Kwiatkowski, Cordelia Langford, John Sillitoe on behalf of the Wellcome Sanger Institute COVID-19 Surveillance Team |
| EPI_ISL_833626 | Lighthouse Lab in Cambridge | Wellcome Sanger Institute for the COVID-19 Genomics UK (COG-UK) Consortium | Rob Howes, The Lighthouse Lab in Cambridge and Alex Alderton, Roberto Amato, Sonia Goncalves, Ewan Harrison, David K. Jackson, Ian Johnston, Dominic Kwiatkowski, Cordelia Langford, John Sillitoe on behalf of the Wellcome Sanger Institute COVID-19 Surveillance Team |
| EPI_ISL_833629 | Lighthouse Lab in Milton Keynes | Wellcome Sanger Institute for the COVID-19 Genomics UK (COG-UK) Consortium | The Lighthouse Lab in Milton Keynes and Alex Alderton, Roberto Amato, Sonia Goncalves, Ewan Harrison, David K. Jackson, Ian Johnston, Dominic Kwiatkowski, Cordelia Langford, John Sillitoe on behalf of the Wellcome Sanger Institute COVID-19 Surveillance Team |
| EPI_ISL_833630, EPI_ISL_833631 | Lighthouse Lab in Cambridge | Wellcome Sanger Institute for the COVID-19 Genomics UK (COG-UK) Consortium | Rob Howes, The Lighthouse Lab in Cambridge and Alex Alderton, Roberto Amato, Sonia Goncalves, Ewan Harrison, David K. Jackson, Ian Johnston, Dominic Kwiatkowski, Cordelia Langford, John Sillitoe on behalf of the Wellcome Sanger Institute COVID-19 Surveillance Team |
| EPI_ISL_833632, EPI_ISL_833634 | Lighthouse Lab in Milton Keynes | Wellcome Sanger Institute for the COVID-19 Genomics UK (COG-UK) Consortium | The Lighthouse Lab in Milton Keynes and Alex Alderton, Roberto Amato, Sonia Goncalves, Ewan Harrison, David K. Jackson, Ian Johnston, Dominic Kwiatkowski, Cordelia Langford, John Sillitoe on behalf of the Wellcome Sanger Institute COVID-19 Surveillance Team |
| EPI_ISL_833635, EPI_ISL_833636, EPI_ISL_833640 | Lighthouse Lab in Cambridge | Wellcome Sanger Institute for the COVID-19 Genomics UK (COG-UK) Consortium | Rob Howes, The Lighthouse Lab in Cambridge and Alex Alderton, Roberto Amato, Sonia Goncalves, Ewan Harrison, David K. Jackson, Ian Johnston, Dominic Kwiatkowski, Cordelia Langford, John Sillitoe on behalf of the Wellcome Sanger Institute COVID-19 Surveillance Team |
| EPI_ISL_833643 | Lighthouse Lab in Milton Keynes | Wellcome Sanger Institute for the COVID-19 Genomics UK (COG-UK) Consortium | The Lighthouse Lab in Milton Keynes and Alex Alderton, Roberto Amato, Sonia Goncalves, Ewan Harrison, David K. Jackson, Ian Johnston, Dominic Kwiatkowski, Cordelia Langford, John Sillitoe on behalf of the Wellcome Sanger Institute COVID-19 Surveillance Team |
| EPI_ISL_833646 | Lighthouse Lab in Cambridge | Wellcome Sanger Institute for the COVID-19 Genomics UK (COG-UK) Consortium | Rob Howes, The Lighthouse Lab in Cambridge and Alex Alderton, Roberto Amato, Sonia Goncalves, Ewan Harrison, David K. Jackson, Ian Johnston, Dominic Kwiatkowski, Cordelia Langford, John Sillitoe on behalf of the Wellcome Sanger Institute COVID-19 Surveillance Team |
| EPI_ISL_833647, EPI_ISL_833648 | Lighthouse Lab in Milton Keynes | Wellcome Sanger Institute for the COVID-19 Genomics UK (COG-UK) Consortium | The Lighthouse Lab in Milton Keynes and Alex Alderton, Roberto Amato, Sonia Goncalves, Ewan Harrison, David K. Jackson, Ian Johnston, Dominic Kwiatkowski, Cordelia Langford, John Sillitoe on behalf of the Wellcome Sanger Institute COVID-19 Surveillance Team |
| EPI_ISL_833650, EPI_ISL_833651 | Lighthouse Lab in Cambridge | Wellcome Sanger Institute for the COVID-19 Genomics UK (COG-UK) Consortium | Rob Howes, The Lighthouse Lab in Cambridge and Alex Alderton, Roberto Amato, Sonia Goncalves, Ewan Harrison, David K. Jackson, Ian Johnston, Dominic Kwiatkowski, Cordelia Langford, John Sillitoe on behalf of the Wellcome Sanger Institute COVID-19 Surveillance Team |
| EPI_ISL_833652 | Lighthouse Lab in Milton Keynes | Wellcome Sanger Institute for the COVID-19 Genomics UK (COG-UK) Consortium | The Lighthouse Lab in Milton Keynes and Alex Alderton, Roberto Amato, Sonia Goncalves, Ewan Harrison, David K. Jackson, Ian Johnston, Dominic Kwiatkowski, Cordelia Langford, John Sillitoe on behalf of the Wellcome Sanger Institute COVID-19 Surveillance Team |
| EPI_ISL_833654 | Lighthouse Lab in Cambridge | Wellcome Sanger Institute for the COVID-19 Genomics UK (COG-UK) Consortium | Rob Howes, The Lighthouse Lab in Cambridge and Alex Alderton, Roberto Amato, Sonia Goncalves, Ewan Harrison, David K. Jackson, Ian Johnston, Dominic Kwiatkowski, Cordelia Langford, John Sillitoe on behalf of the Wellcome Sanger Institute COVID-19 Surveillance Team |
| EPI_ISL_833656, EPI_ISL_833658 | Lighthouse Lab in Milton Keynes | Wellcome Sanger Institute for the COVID-19 Genomics UK | The Lighthouse Lab in Milton Keynes and Alex Alderton, Roberto Amato, Sonia Goncalves, Ewan Harrison, David K. Jackson, Ian Johnston, Dominic |

| | | | |
|---|---|---|---|
| | | (COG-UK) Consortium | Kwiatkowski, Cordelia Langford, John Sillitoe on behalf of the Wellcome Sanger Institute COVID-19 Surveillance Team |
| EPI_ISL_833662, EPI_ISL_833665 | Lighthouse Lab in Cambridge | Wellcome Sanger Institute for the COVID-19 Genomics UK (COG-UK) Consortium | Rob Howes, The Lighthouse Lab in Cambridge and Alex Alderton, Roberto Amato, Sonia Goncalves, Ewan Harrison, David K. Jackson, Ian Johnston, Dominic Kwiatkowski, Cordelia Langford, John Sillitoe on behalf of the Wellcome Sanger Institute COVID-19 Surveillance Team |
| EPI_ISL_833667 | Lighthouse Lab in Milton Keynes | Wellcome Sanger Institute for the COVID-19 Genomics UK (COG-UK) Consortium | The Lighthouse Lab in Milton Keynes and Alex Alderton, Roberto Amato, Sonia Goncalves, Ewan Harrison, David K. Jackson, Ian Johnston, Dominic Kwiatkowski, Cordelia Langford, John Sillitoe on behalf of the Wellcome Sanger Institute COVID-19 Surveillance Team |
| EPI_ISL_833668, EPI_ISL_833670, EPI_ISL_833671, EPI_ISL_833673 | Lighthouse Lab in Cambridge | Wellcome Sanger Institute for the COVID-19 Genomics UK (COG-UK) Consortium | Rob Howes, The Lighthouse Lab in Cambridge and Alex Alderton, Roberto Amato, Sonia Goncalves, Ewan Harrison, David K. Jackson, Ian Johnston, Dominic Kwiatkowski, Cordelia Langford, John Sillitoe on behalf of the Wellcome Sanger Institute COVID-19 Surveillance Team |
| EPI_ISL_833676, EPI_ISL_833678 | Lighthouse Lab in Milton Keynes | Wellcome Sanger Institute for the COVID-19 Genomics UK (COG-UK) Consortium | The Lighthouse Lab in Milton Keynes and Alex Alderton, Roberto Amato, Sonia Goncalves, Ewan Harrison, David K. Jackson, Ian Johnston, Dominic Kwiatkowski, Cordelia Langford, John Sillitoe on behalf of the Wellcome Sanger Institute COVID-19 Surveillance Team |
| EPI_ISL_833679, EPI_ISL_833680, EPI_ISL_833683, EPI_ISL_833684 | Lighthouse Lab in Cambridge | Wellcome Sanger Institute for the COVID-19 Genomics UK (COG-UK) Consortium | Rob Howes, The Lighthouse Lab in Cambridge and Alex Alderton, Roberto Amato, Sonia Goncalves, Ewan Harrison, David K. Jackson, Ian Johnston, Dominic Kwiatkowski, Cordelia Langford, John Sillitoe on behalf of the Wellcome Sanger Institute COVID-19 Surveillance Team |
| EPI_ISL_833685 | Lighthouse Lab in Milton Keynes | Wellcome Sanger Institute for the COVID-19 Genomics UK (COG-UK) Consortium | The Lighthouse Lab in Milton Keynes and Alex Alderton, Roberto Amato, Sonia Goncalves, Ewan Harrison, David K. Jackson, Ian Johnston, Dominic Kwiatkowski, Cordelia Langford, John Sillitoe on behalf of the Wellcome Sanger Institute COVID-19 Surveillance Team |
| EPI_ISL_833686, EPI_ISL_833687, EPI_ISL_833689, EPI_ISL_833691 | Lighthouse Lab in Cambridge | Wellcome Sanger Institute for the COVID-19 Genomics UK (COG-UK) Consortium | Rob Howes, The Lighthouse Lab in Cambridge and Alex Alderton, Roberto Amato, Sonia Goncalves, Ewan Harrison, David K. Jackson, Ian Johnston, Dominic Kwiatkowski, Cordelia Langford, John Sillitoe on behalf of the Wellcome Sanger Institute COVID-19 Surveillance Team |
| EPI_ISL_833693 | Lighthouse Lab in Milton Keynes | Wellcome Sanger Institute for the COVID-19 Genomics UK (COG-UK) Consortium | The Lighthouse Lab in Milton Keynes and Alex Alderton, Roberto Amato, Sonia Goncalves, Ewan Harrison, David K. Jackson, Ian Johnston, Dominic Kwiatkowski, Cordelia Langford, John Sillitoe on behalf of the Wellcome Sanger Institute COVID-19 Surveillance Team |
| EPI_ISL_833695, EPI_ISL_833696 | Lighthouse Lab in Cambridge | Wellcome Sanger Institute for the COVID-19 Genomics UK (COG-UK) Consortium | Rob Howes, The Lighthouse Lab in Cambridge and Alex Alderton, Roberto Amato, Sonia Goncalves, Ewan Harrison, David K. Jackson, Ian Johnston, Dominic Kwiatkowski, Cordelia Langford, John Sillitoe on behalf of the Wellcome Sanger Institute COVID-19 Surveillance Team |
| EPI_ISL_833700 | Lighthouse Lab in Milton Keynes | Wellcome Sanger Institute for the COVID-19 Genomics UK (COG-UK) Consortium | The Lighthouse Lab in Milton Keynes and Alex Alderton, Roberto Amato, Sonia Goncalves, Ewan Harrison, David K. Jackson, Ian Johnston, Dominic Kwiatkowski, Cordelia Langford, John Sillitoe on behalf of the Wellcome Sanger Institute COVID-19 Surveillance Team |
| EPI_ISL_833702, EPI_ISL_833703, EPI_ISL_833704, EPI_ISL_833705, EPI_ISL_833706, EPI_ISL_833707, EPI_ISL_833708, EPI_ISL_833709, EPI_ISL_833710 | Lighthouse Lab in Cambridge | Wellcome Sanger Institute for the COVID-19 Genomics UK (COG-UK) Consortium | Rob Howes, The Lighthouse Lab in Cambridge and Alex Alderton, Roberto Amato, Sonia Goncalves, Ewan Harrison, David K. Jackson, Ian Johnston, Dominic Kwiatkowski, Cordelia Langford, John Sillitoe on behalf of the Wellcome Sanger Institute COVID-19 Surveillance Team |
| EPI_ISL_833712 | Lighthouse Lab in Milton Keynes | Wellcome Sanger Institute for the COVID-19 Genomics UK (COG-UK) Consortium | The Lighthouse Lab in Milton Keynes and Alex Alderton, Roberto Amato, Sonia Goncalves, Ewan Harrison, David K. Jackson, Ian Johnston, Dominic Kwiatkowski, Cordelia Langford, John Sillitoe on behalf of the Wellcome Sanger Institute COVID-19 Surveillance Team |
| EPI_ISL_833714 | Lighthouse Lab in Cambridge | Wellcome Sanger Institute for the COVID-19 Genomics UK (COG-UK) Consortium | Rob Howes, The Lighthouse Lab in Cambridge and Alex Alderton, Roberto Amato, Sonia Goncalves, Ewan Harrison, David K. Jackson, Ian Johnston, Dominic Kwiatkowski, Cordelia Langford, John Sillitoe on behalf of the Wellcome Sanger Institute COVID-19 Surveillance Team |
| EPI_ISL_833716, EPI_ISL_833719, EPI_ISL_833721 | Lighthouse Lab in Milton Keynes | Wellcome Sanger Institute for the COVID-19 Genomics UK (COG-UK) Consortium | The Lighthouse Lab in Milton Keynes and Alex Alderton, Roberto Amato, Sonia Goncalves, Ewan Harrison, David K. Jackson, Ian Johnston, Dominic Kwiatkowski, Cordelia Langford, John Sillitoe on behalf of the Wellcome Sanger Institute COVID-19 Surveillance Team |
| EPI_ISL_833724 | Lighthouse Lab in Cambridge | Wellcome Sanger Institute for the COVID-19 Genomics UK (COG-UK) Consortium | Rob Howes, The Lighthouse Lab in Cambridge and Alex Alderton, Roberto Amato, Sonia Goncalves, Ewan Harrison, David K. Jackson, Ian Johnston, Dominic Kwiatkowski, Cordelia Langford, John Sillitoe on behalf of the Wellcome Sanger Institute COVID-19 Surveillance Team |
| EPI_ISL_833725, EPI_ISL_833726 | Lighthouse Lab in Milton Keynes | Wellcome Sanger Institute for the COVID-19 Genomics UK (COG-UK) Consortium | The Lighthouse Lab in Milton Keynes and Alex Alderton, Roberto Amato, Sonia Goncalves, Ewan Harrison, David K. Jackson, Ian Johnston, Dominic Kwiatkowski, Cordelia Langford, John Sillitoe on behalf of the Wellcome Sanger Institute COVID-19 Surveillance Team |
| EPI_ISL_833727, EPI_ISL_833728, EPI_ISL_833733 | Lighthouse Lab in Cambridge | Wellcome Sanger Institute for the COVID-19 Genomics UK (COG-UK) Consortium | Rob Howes, The Lighthouse Lab in Cambridge and Alex Alderton, Roberto Amato, Sonia Goncalves, Ewan Harrison, David K. Jackson, Ian Johnston, Dominic Kwiatkowski, Cordelia Langford, John Sillitoe on behalf of the Wellcome Sanger Institute COVID-19 Surveillance Team |
| EPI_ISL_833734, EPI_ISL_833735, EPI_ISL_833736, EPI_ISL_833738 | Lighthouse Lab in Milton Keynes | Wellcome Sanger Institute for the COVID-19 Genomics UK (COG-UK) Consortium | The Lighthouse Lab in Milton Keynes and Alex Alderton, Roberto Amato, Sonia Goncalves, Ewan Harrison, David K. Jackson, Ian Johnston, Dominic Kwiatkowski, Cordelia Langford, John Sillitoe on behalf of the Wellcome Sanger Institute COVID-19 Surveillance Team |
| EPI_ISL_833741, EPI_ISL_833742 | Lighthouse Lab in Cambridge | Wellcome Sanger Institute for the COVID-19 Genomics UK (COG-UK) Consortium | Rob Howes, The Lighthouse Lab in Cambridge and Alex Alderton, Roberto Amato, Sonia Goncalves, Ewan Harrison, David K. Jackson, Ian Johnston, Dominic Kwiatkowski, Cordelia Langford, John Sillitoe on behalf of the Wellcome Sanger Institute COVID-19 Surveillance Team |
| EPI_ISL_833743 | Lighthouse Lab in Milton Keynes | Wellcome Sanger Institute for the COVID-19 Genomics UK (COG-UK) Consortium | The Lighthouse Lab in Milton Keynes and Alex Alderton, Roberto Amato, Sonia Goncalves, Ewan Harrison, David K. Jackson, Ian Johnston, Dominic Kwiatkowski, Cordelia Langford, John Sillitoe on behalf of the Wellcome Sanger Institute COVID-19 Surveillance Team |
| EPI_ISL_833746, EPI_ISL_833748, EPI_ISL_833749, EPI_ISL_833750, EPI_ISL_833753 | Lighthouse Lab in Cambridge | Wellcome Sanger Institute for the COVID-19 Genomics UK (COG-UK) Consortium | Rob Howes, The Lighthouse Lab in Cambridge and Alex Alderton, Roberto Amato, Sonia Goncalves, Ewan Harrison, David K. Jackson, Ian Johnston, Dominic Kwiatkowski, Cordelia Langford, John Sillitoe on behalf of the Wellcome Sanger Institute COVID-19 Surveillance Team |
| EPI_ISL_833755, EPI_ISL_833756 | Lighthouse Lab in Milton Keynes | Wellcome Sanger Institute for the COVID-19 Genomics UK (COG-UK) Consortium | The Lighthouse Lab in Milton Keynes and Alex Alderton, Roberto Amato, Sonia Goncalves, Ewan Harrison, David K. Jackson, Ian Johnston, Dominic Kwiatkowski, Cordelia Langford, John Sillitoe on behalf of the Wellcome Sanger Institute COVID-19 Surveillance Team |
| EPI_ISL_833758 | Lighthouse Lab in Cambridge | Wellcome Sanger Institute for the COVID-19 Genomics UK (COG-UK) Consortium | Rob Howes, The Lighthouse Lab in Cambridge and Alex Alderton, Roberto Amato, Sonia Goncalves, Ewan Harrison, David K. Jackson, Ian Johnston, Dominic Kwiatkowski, Cordelia Langford, John Sillitoe on behalf of the Wellcome Sanger Institute COVID-19 Surveillance Team |
| EPI_ISL_833759 | Lighthouse Lab in Milton Keynes | Wellcome Sanger Institute for the COVID-19 Genomics UK (COG-UK) Consortium | The Lighthouse Lab in Milton Keynes and Alex Alderton, Roberto Amato, Sonia Goncalves, Ewan Harrison, David K. Jackson, Ian Johnston, Dominic Kwiatkowski, Cordelia Langford, John Sillitoe on behalf of the Wellcome Sanger Institute COVID-19 Surveillance Team |
| EPI_ISL_833762 | Lighthouse Lab in Cambridge | Wellcome Sanger Institute for the COVID-19 Genomics UK (COG-UK) Consortium | Rob Howes, The Lighthouse Lab in Cambridge and Alex Alderton, Roberto Amato, Sonia Goncalves, Ewan Harrison, David K. Jackson, Ian Johnston, Dominic Kwiatkowski, Cordelia Langford, John Sillitoe on behalf of the Wellcome Sanger Institute COVID-19 Surveillance Team |
| EPI_ISL_833766, EPI_ISL_833767, EPI_ISL_833768 | Lighthouse Lab in Milton Keynes | Wellcome Sanger Institute for the COVID-19 Genomics UK (COG-UK) Consortium | The Lighthouse Lab in Milton Keynes and Alex Alderton, Roberto Amato, Sonia Goncalves, Ewan Harrison, David K. Jackson, Ian Johnston, Dominic Kwiatkowski, Cordelia Langford, John Sillitoe on behalf of the Wellcome Sanger Institute COVID-19 Surveillance Team |
| EPI_ISL_833770, EPI_ISL_833771, EPI_ISL_833773, EPI_ISL_833774, EPI_ISL_833778 | Lighthouse Lab in Cambridge | Wellcome Sanger Institute for the COVID-19 Genomics UK (COG-UK) Consortium | Rob Howes, The Lighthouse Lab in Cambridge and Alex Alderton, Roberto Amato, Sonia Goncalves, Ewan Harrison, David K. Jackson, Ian Johnston, Dominic Kwiatkowski, Cordelia Langford, John Sillitoe on behalf of the Wellcome Sanger Institute COVID-19 Surveillance Team |
| EPI_ISL_833779, EPI_ISL_833780 | Lighthouse Lab in Milton Keynes | Wellcome Sanger Institute for the COVID-19 Genomics UK (COG-UK) Consortium | The Lighthouse Lab in Milton Keynes and Alex Alderton, Roberto Amato, Sonia Goncalves, Ewan Harrison, David K. Jackson, Ian Johnston, Dominic Kwiatkowski, Cordelia Langford, John Sillitoe on behalf of the Wellcome Sanger Institute COVID-19 Surveillance Team |
| EPI_ISL_833784 | Lighthouse Lab in Cambridge | Wellcome Sanger Institute for the COVID-19 Genomics UK (COG-UK) Consortium | Rob Howes, The Lighthouse Lab in Cambridge and Alex Alderton, Roberto Amato, Sonia Goncalves, Ewan Harrison, David K. Jackson, Ian Johnston, Dominic Kwiatkowski, Cordelia Langford, John Sillitoe on behalf of the Wellcome Sanger Institute COVID-19 Surveillance Team |
| EPI_ISL_833789, EPI_ISL_833791 | Lighthouse Lab in Milton Keynes | Wellcome Sanger Institute for the COVID-19 Genomics UK (COG-UK) Consortium | The Lighthouse Lab in Milton Keynes and Alex Alderton, Roberto Amato, Sonia Goncalves, Ewan Harrison, David K. Jackson, Ian Johnston, Dominic Kwiatkowski, Cordelia Langford, John Sillitoe on behalf of the Wellcome Sanger Institute COVID-19 Surveillance Team |
| EPI_ISL_833792, EPI_ISL_833794 | Lighthouse Lab in Cambridge | Wellcome Sanger Institute for the COVID-19 Genomics UK (COG-UK) Consortium | Rob Howes, The Lighthouse Lab in Cambridge and Alex Alderton, Roberto Amato, Sonia Goncalves, Ewan Harrison, David K. Jackson, Ian Johnston, Dominic Kwiatkowski, Cordelia Langford, John Sillitoe on behalf of the Wellcome Sanger Institute COVID-19 Surveillance Team |
| EPI_ISL_833795, EPI_ISL_833796, EPI_ISL_833797 | Lighthouse Lab in Milton Keynes | Wellcome Sanger Institute for the COVID-19 Genomics UK (COG-UK) Consortium | The Lighthouse Lab in Milton Keynes and Alex Alderton, Roberto Amato, Sonia Goncalves, Ewan Harrison, David K. Jackson, Ian Johnston, Dominic Kwiatkowski, Cordelia Langford, John Sillitoe on behalf of the Wellcome Sanger Institute COVID-19 Surveillance Team |
| EPI_ISL_833800 | Lighthouse Lab in Cambridge | Wellcome Sanger Institute for the COVID-19 Genomics UK | Rob Howes, The Lighthouse Lab in Cambridge and Alex Alderton, Roberto Amato, Sonia Goncalves, Ewan Harrison, David K. Jackson, Ian Johnston, |

| | | | |
|---|---|---|---|
| | | (COG-UK) Consortium | Dominic Kwiatkowski, Cordelia Langford, John Sillitoe on behalf of the Wellcome Sanger Institute COVID-19 Surveillance Team |
| EPI_ISL_833801, EPI_ISL_833803, EPI_ISL_833804 | Lighthouse Lab in Milton Keynes | Wellcome Sanger Institute for the COVID-19 Genomics UK (COG-UK) Consortium | The Lighthouse Lab in Milton Keynes and Alex Alderton, Roberto Amato, Sonia Goncalves, Ewan Harrison, David K. Jackson, Ian Johnston, Dominic Kwiatkowski, Cordelia Langford, John Sillitoe on behalf of the Wellcome Sanger Institute COVID-19 Surveillance Team |
| EPI_ISL_833806, EPI_ISL_833807 | Lighthouse Lab in Cambridge | Wellcome Sanger Institute for the COVID-19 Genomics UK (COG-UK) Consortium | Rob Howes, The Lighthouse Lab in Cambridge and Alex Alderton, Roberto Amato, Sonia Goncalves, Ewan Harrison, David K. Jackson, Ian Johnston, Dominic Kwiatkowski, Cordelia Langford, John Sillitoe on behalf of the Wellcome Sanger Institute COVID-19 Surveillance Team |
| EPI_ISL_833809 | Lighthouse Lab in Milton Keynes | Wellcome Sanger Institute for the COVID-19 Genomics UK (COG-UK) Consortium | The Lighthouse Lab in Milton Keynes and Alex Alderton, Roberto Amato, Sonia Goncalves, Ewan Harrison, David K. Jackson, Ian Johnston, Dominic Kwiatkowski, Cordelia Langford, John Sillitoe on behalf of the Wellcome Sanger Institute COVID-19 Surveillance Team |
| EPI_ISL_833810, EPI_ISL_833812, EPI_ISL_833813 | Lighthouse Lab in Cambridge | Wellcome Sanger Institute for the COVID-19 Genomics UK (COG-UK) Consortium | Rob Howes, The Lighthouse Lab in Cambridge and Alex Alderton, Roberto Amato, Sonia Goncalves, Ewan Harrison, David K. Jackson, Ian Johnston, Dominic Kwiatkowski, Cordelia Langford, John Sillitoe on behalf of the Wellcome Sanger Institute COVID-19 Surveillance Team |
| EPI_ISL_833814 | Lighthouse Lab in Milton Keynes | Wellcome Sanger Institute for the COVID-19 Genomics UK (COG-UK) Consortium | The Lighthouse Lab in Milton Keynes and Alex Alderton, Roberto Amato, Sonia Goncalves, Ewan Harrison, David K. Jackson, Ian Johnston, Dominic Kwiatkowski, Cordelia Langford, John Sillitoe on behalf of the Wellcome Sanger Institute COVID-19 Surveillance Team |
| EPI_ISL_833815, EPI_ISL_833816 | Lighthouse Lab in Cambridge | Wellcome Sanger Institute for the COVID-19 Genomics UK (COG-UK) Consortium | Rob Howes, The Lighthouse Lab in Cambridge and Alex Alderton, Roberto Amato, Sonia Goncalves, Ewan Harrison, David K. Jackson, Ian Johnston, Dominic Kwiatkowski, Cordelia Langford, John Sillitoe on behalf of the Wellcome Sanger Institute COVID-19 Surveillance Team |
| EPI_ISL_833817, EPI_ISL_833818, EPI_ISL_833821, EPI_ISL_833823 | Lighthouse Lab in Milton Keynes | Wellcome Sanger Institute for the COVID-19 Genomics UK (COG-UK) Consortium | The Lighthouse Lab in Milton Keynes and Alex Alderton, Roberto Amato, Sonia Goncalves, Ewan Harrison, David K. Jackson, Ian Johnston, Dominic Kwiatkowski, Cordelia Langford, John Sillitoe on behalf of the Wellcome Sanger Institute COVID-19 Surveillance Team |
| EPI_ISL_833829 | Lighthouse Lab in Cambridge | Wellcome Sanger Institute for the COVID-19 Genomics UK (COG-UK) Consortium | Rob Howes, The Lighthouse Lab in Cambridge and Alex Alderton, Roberto Amato, Sonia Goncalves, Ewan Harrison, David K. Jackson, Ian Johnston, Dominic Kwiatkowski, Cordelia Langford, John Sillitoe on behalf of the Wellcome Sanger Institute COVID-19 Surveillance Team |
| EPI_ISL_833830 | Lighthouse Lab in Milton Keynes | Wellcome Sanger Institute for the COVID-19 Genomics UK (COG-UK) Consortium | The Lighthouse Lab in Milton Keynes and Alex Alderton, Roberto Amato, Sonia Goncalves, Ewan Harrison, David K. Jackson, Ian Johnston, Dominic Kwiatkowski, Cordelia Langford, John Sillitoe on behalf of the Wellcome Sanger Institute COVID-19 Surveillance Team |
| EPI_ISL_833831, EPI_ISL_833832, EPI_ISL_833833 | Lighthouse Lab in Cambridge | Wellcome Sanger Institute for the COVID-19 Genomics UK (COG-UK) Consortium | Rob Howes, The Lighthouse Lab in Cambridge and Alex Alderton, Roberto Amato, Sonia Goncalves, Ewan Harrison, David K. Jackson, Ian Johnston, Dominic Kwiatkowski, Cordelia Langford, John Sillitoe on behalf of the Wellcome Sanger Institute COVID-19 Surveillance Team |
| EPI_ISL_833834, EPI_ISL_833835 | Lighthouse Lab in Milton Keynes | Wellcome Sanger Institute for the COVID-19 Genomics UK (COG-UK) Consortium | The Lighthouse Lab in Milton Keynes and Alex Alderton, Roberto Amato, Sonia Goncalves, Ewan Harrison, David K. Jackson, Ian Johnston, Dominic Kwiatkowski, Cordelia Langford, John Sillitoe on behalf of the Wellcome Sanger Institute COVID-19 Surveillance Team |
| EPI_ISL_833840 | Lighthouse Lab in Cambridge | Wellcome Sanger Institute for the COVID-19 Genomics UK (COG-UK) Consortium | Rob Howes, The Lighthouse Lab in Cambridge and Alex Alderton, Roberto Amato, Sonia Goncalves, Ewan Harrison, David K. Jackson, Ian Johnston, Dominic Kwiatkowski, Cordelia Langford, John Sillitoe on behalf of the Wellcome Sanger Institute COVID-19 Surveillance Team |
| EPI_ISL_833841 | Lighthouse Lab in Milton Keynes | Wellcome Sanger Institute for the COVID-19 Genomics UK (COG-UK) Consortium | The Lighthouse Lab in Milton Keynes and Alex Alderton, Roberto Amato, Sonia Goncalves, Ewan Harrison, David K. Jackson, Ian Johnston, Dominic Kwiatkowski, Cordelia Langford, John Sillitoe on behalf of the Wellcome Sanger Institute COVID-19 Surveillance Team |
| EPI_ISL_833842, EPI_ISL_833844 | Lighthouse Lab in Cambridge | Wellcome Sanger Institute for the COVID-19 Genomics UK (COG-UK) Consortium | Rob Howes, The Lighthouse Lab in Cambridge and Alex Alderton, Roberto Amato, Sonia Goncalves, Ewan Harrison, David K. Jackson, Ian Johnston, Dominic Kwiatkowski, Cordelia Langford, John Sillitoe on behalf of the Wellcome Sanger Institute COVID-19 Surveillance Team |
| EPI_ISL_833845, EPI_ISL_833846, EPI_ISL_833848 | Lighthouse Lab in Milton Keynes | Wellcome Sanger Institute for the COVID-19 Genomics UK (COG-UK) Consortium | The Lighthouse Lab in Milton Keynes and Alex Alderton, Roberto Amato, Sonia Goncalves, Ewan Harrison, David K. Jackson, Ian Johnston, Dominic Kwiatkowski, Cordelia Langford, John Sillitoe on behalf of the Wellcome Sanger Institute COVID-19 Surveillance Team |
| EPI_ISL_833849, EPI_ISL_833852, EPI_ISL_833853, EPI_ISL_833854, EPI_ISL_833855 | Lighthouse Lab in Cambridge | Wellcome Sanger Institute for the COVID-19 Genomics UK (COG-UK) Consortium | Rob Howes, The Lighthouse Lab in Cambridge and Alex Alderton, Roberto Amato, Sonia Goncalves, Ewan Harrison, David K. Jackson, Ian Johnston, Dominic Kwiatkowski, Cordelia Langford, John Sillitoe on behalf of the Wellcome Sanger Institute COVID-19 Surveillance Team |
| EPI_ISL_833856 | Lighthouse Lab in Milton Keynes | Wellcome Sanger Institute for the COVID-19 Genomics UK (COG-UK) Consortium | The Lighthouse Lab in Milton Keynes and Alex Alderton, Roberto Amato, Sonia Goncalves, Ewan Harrison, David K. Jackson, Ian Johnston, Dominic Kwiatkowski, Cordelia Langford, John Sillitoe on behalf of the Wellcome Sanger Institute COVID-19 Surveillance Team |
| EPI_ISL_833859, EPI_ISL_833860 | Lighthouse Lab in Cambridge | Wellcome Sanger Institute for the COVID-19 Genomics UK (COG-UK) Consortium | Rob Howes, The Lighthouse Lab in Cambridge and Alex Alderton, Roberto Amato, Sonia Goncalves, Ewan Harrison, David K. Jackson, Ian Johnston, Dominic Kwiatkowski, Cordelia Langford, John Sillitoe on behalf of the Wellcome Sanger Institute COVID-19 Surveillance Team |
| EPI_ISL_833861 | Lighthouse Lab in Milton Keynes | Wellcome Sanger Institute for the COVID-19 Genomics UK (COG-UK) Consortium | The Lighthouse Lab in Milton Keynes and Alex Alderton, Roberto Amato, Sonia Goncalves, Ewan Harrison, David K. Jackson, Ian Johnston, Dominic Kwiatkowski, Cordelia Langford, John Sillitoe on behalf of the Wellcome Sanger Institute COVID-19 Surveillance Team |
| EPI_ISL_833862, EPI_ISL_833863, EPI_ISL_833864 | Lighthouse Lab in Cambridge | Wellcome Sanger Institute for the COVID-19 Genomics UK (COG-UK) Consortium | Rob Howes, The Lighthouse Lab in Cambridge and Alex Alderton, Roberto Amato, Sonia Goncalves, Ewan Harrison, David K. Jackson, Ian Johnston, Dominic Kwiatkowski, Cordelia Langford, John Sillitoe on behalf of the Wellcome Sanger Institute COVID-19 Surveillance Team |
| EPI_ISL_833865 | Lighthouse Lab in Milton Keynes | Wellcome Sanger Institute for the COVID-19 Genomics UK (COG-UK) Consortium | The Lighthouse Lab in Milton Keynes and Alex Alderton, Roberto Amato, Sonia Goncalves, Ewan Harrison, David K. Jackson, Ian Johnston, Dominic Kwiatkowski, Cordelia Langford, John Sillitoe on behalf of the Wellcome Sanger Institute COVID-19 Surveillance Team |
| EPI_ISL_833866 | Lighthouse Lab in Cambridge | Wellcome Sanger Institute for the COVID-19 Genomics UK (COG-UK) Consortium | Rob Howes, The Lighthouse Lab in Cambridge and Alex Alderton, Roberto Amato, Sonia Goncalves, Ewan Harrison, David K. Jackson, Ian Johnston, Dominic Kwiatkowski, Cordelia Langford, John Sillitoe on behalf of the Wellcome Sanger Institute COVID-19 Surveillance Team |
| EPI_ISL_833868, EPI_ISL_833872 | Lighthouse Lab in Milton Keynes | Wellcome Sanger Institute for the COVID-19 Genomics UK (COG-UK) Consortium | The Lighthouse Lab in Milton Keynes and Alex Alderton, Roberto Amato, Sonia Goncalves, Ewan Harrison, David K. Jackson, Ian Johnston, Dominic Kwiatkowski, Cordelia Langford, John Sillitoe on behalf of the Wellcome Sanger Institute COVID-19 Surveillance Team |
| EPI_ISL_833876, EPI_ISL_833877 | Lighthouse Lab in Cambridge | Wellcome Sanger Institute for the COVID-19 Genomics UK (COG-UK) Consortium | Rob Howes, The Lighthouse Lab in Cambridge and Alex Alderton, Roberto Amato, Sonia Goncalves, Ewan Harrison, David K. Jackson, Ian Johnston, Dominic Kwiatkowski, Cordelia Langford, John Sillitoe on behalf of the Wellcome Sanger Institute COVID-19 Surveillance Team |
| EPI_ISL_833879, EPI_ISL_833881 | Lighthouse Lab in Milton Keynes | Wellcome Sanger Institute for the COVID-19 Genomics UK (COG-UK) Consortium | The Lighthouse Lab in Milton Keynes and Alex Alderton, Roberto Amato, Sonia Goncalves, Ewan Harrison, David K. Jackson, Ian Johnston, Dominic Kwiatkowski, Cordelia Langford, John Sillitoe on behalf of the Wellcome Sanger Institute COVID-19 Surveillance Team |
| EPI_ISL_833882 | Lighthouse Lab in Cambridge | Wellcome Sanger Institute for the COVID-19 Genomics UK (COG-UK) Consortium | Rob Howes, The Lighthouse Lab in Cambridge and Alex Alderton, Roberto Amato, Sonia Goncalves, Ewan Harrison, David K. Jackson, Ian Johnston, Dominic Kwiatkowski, Cordelia Langford, John Sillitoe on behalf of the Wellcome Sanger Institute COVID-19 Surveillance Team |
| EPI_ISL_833884 | Lighthouse Lab in Milton Keynes | Wellcome Sanger Institute for the COVID-19 Genomics UK (COG-UK) Consortium | The Lighthouse Lab in Milton Keynes and Alex Alderton, Roberto Amato, Sonia Goncalves, Ewan Harrison, David K. Jackson, Ian Johnston, Dominic Kwiatkowski, Cordelia Langford, John Sillitoe on behalf of the Wellcome Sanger Institute COVID-19 Surveillance Team |
| EPI_ISL_833886, EPI_ISL_833888 | Lighthouse Lab in Cambridge | Wellcome Sanger Institute for the COVID-19 Genomics UK (COG-UK) Consortium | Rob Howes, The Lighthouse Lab in Cambridge and Alex Alderton, Roberto Amato, Sonia Goncalves, Ewan Harrison, David K. Jackson, Ian Johnston, Dominic Kwiatkowski, Cordelia Langford, John Sillitoe on behalf of the Wellcome Sanger Institute COVID-19 Surveillance Team |
| EPI_ISL_833889, EPI_ISL_833890 | Lighthouse Lab in Milton Keynes | Wellcome Sanger Institute for the COVID-19 Genomics UK (COG-UK) Consortium | The Lighthouse Lab in Milton Keynes and Alex Alderton, Roberto Amato, Sonia Goncalves, Ewan Harrison, David K. Jackson, Ian Johnston, Dominic Kwiatkowski, Cordelia Langford, John Sillitoe on behalf of the Wellcome Sanger Institute COVID-19 Surveillance Team |
| EPI_ISL_833892 | Lighthouse Lab in Cambridge | Wellcome Sanger Institute for the COVID-19 Genomics UK (COG-UK) Consortium | Rob Howes, The Lighthouse Lab in Cambridge and Alex Alderton, Roberto Amato, Sonia Goncalves, Ewan Harrison, David K. Jackson, Ian Johnston, Dominic Kwiatkowski, Cordelia Langford, John Sillitoe on behalf of the Wellcome Sanger Institute COVID-19 Surveillance Team |
| EPI_ISL_833894, EPI_ISL_833895 | Lighthouse Lab in Milton Keynes | Wellcome Sanger Institute for the COVID-19 Genomics UK (COG-UK) Consortium | The Lighthouse Lab in Milton Keynes and Alex Alderton, Roberto Amato, Sonia Goncalves, Ewan Harrison, David K. Jackson, Ian Johnston, Dominic Kwiatkowski, Cordelia Langford, John Sillitoe on behalf of the Wellcome Sanger Institute COVID-19 Surveillance Team |
| EPI_ISL_833896, EPI_ISL_833898 | Lighthouse Lab in Cambridge | Wellcome Sanger Institute for the COVID-19 Genomics UK (COG-UK) Consortium | Rob Howes, The Lighthouse Lab in Cambridge and Alex Alderton, Roberto Amato, Sonia Goncalves, Ewan Harrison, David K. Jackson, Ian Johnston, Dominic Kwiatkowski, Cordelia Langford, John Sillitoe on behalf of the Wellcome Sanger Institute COVID-19 Surveillance Team |
| EPI_ISL_833901 | Lighthouse Lab in Milton Keynes | Wellcome Sanger Institute for the COVID-19 Genomics UK (COG-UK) Consortium | The Lighthouse Lab in Milton Keynes and Alex Alderton, Roberto Amato, Sonia Goncalves, Ewan Harrison, David K. Jackson, Ian Johnston, Dominic Kwiatkowski, Cordelia Langford, John Sillitoe on behalf of the Wellcome Sanger Institute COVID-19 Surveillance Team |
| EPI_ISL_833903, EPI_ISL_833907 | Lighthouse Lab in Cambridge | Wellcome Sanger Institute for the COVID-19 Genomics UK (COG-UK) Consortium | Rob Howes, The Lighthouse Lab in Cambridge and Alex Alderton, Roberto Amato, Sonia Goncalves, Ewan Harrison, David K. Jackson, Ian Johnston, Dominic Kwiatkowski, Cordelia Langford, John Sillitoe on behalf of the Wellcome Sanger Institute COVID-19 Surveillance Team |

| | | | |
|---|---|---|---|
| EPI_ISL_833908 | Lighthouse Lab in Milton Keynes | Wellcome Sanger Institute for the COVID-19 Genomics UK (COG-UK) Consortium | The Lighthouse Lab in Milton Keynes and Alex Alderton, Roberto Amato, Sonia Goncalves, Ewan Harrison, David K. Jackson, Ian Johnston, Dominic Kwiatkowski, Cordelia Langford, John Sillitoe on behalf of the Wellcome Sanger Institute COVID-19 Surveillance Team |
| EPI_ISL_833909, EPI_ISL_833910, EPI_ISL_833912 | Lighthouse Lab in Cambridge | Wellcome Sanger Institute for the COVID-19 Genomics UK (COG-UK) Consortium | Rob Howes, The Lighthouse Lab in Cambridge and Alex Alderton, Roberto Amato, Sonia Goncalves, Ewan Harrison, David K. Jackson, Ian Johnston, Dominic Kwiatkowski, Cordelia Langford, John Sillitoe on behalf of the Wellcome Sanger Institute COVID-19 Surveillance Team |
| EPI_ISL_833916 | Lighthouse Lab in Milton Keynes | Wellcome Sanger Institute for the COVID-19 Genomics UK (COG-UK) Consortium | The Lighthouse Lab in Milton Keynes and Alex Alderton, Roberto Amato, Sonia Goncalves, Ewan Harrison, David K. Jackson, Ian Johnston, Dominic Kwiatkowski, Cordelia Langford, John Sillitoe on behalf of the Wellcome Sanger Institute COVID-19 Surveillance Team |
| EPI_ISL_833917, EPI_ISL_833918, EPI_ISL_833919 | Lighthouse Lab in Cambridge | Wellcome Sanger Institute for the COVID-19 Genomics UK (COG-UK) Consortium | Rob Howes, The Lighthouse Lab in Cambridge and Alex Alderton, Roberto Amato, Sonia Goncalves, Ewan Harrison, David K. Jackson, Ian Johnston, Dominic Kwiatkowski, Cordelia Langford, John Sillitoe on behalf of the Wellcome Sanger Institute COVID-19 Surveillance Team |
| EPI_ISL_833922, EPI_ISL_833924, EPI_ISL_833930, EPI_ISL_833931 | Lighthouse Lab in Milton Keynes | Wellcome Sanger Institute for the COVID-19 Genomics UK (COG-UK) Consortium | The Lighthouse Lab in Milton Keynes and Alex Alderton, Roberto Amato, Sonia Goncalves, Ewan Harrison, David K. Jackson, Ian Johnston, Dominic Kwiatkowski, Cordelia Langford, John Sillitoe on behalf of the Wellcome Sanger Institute COVID-19 Surveillance Team |
| EPI_ISL_833932 | Lighthouse Lab in Cambridge | Wellcome Sanger Institute for the COVID-19 Genomics UK (COG-UK) Consortium | Rob Howes, The Lighthouse Lab in Cambridge and Alex Alderton, Roberto Amato, Sonia Goncalves, Ewan Harrison, David K. Jackson, Ian Johnston, Dominic Kwiatkowski, Cordelia Langford, John Sillitoe on behalf of the Wellcome Sanger Institute COVID-19 Surveillance Team |
| EPI_ISL_833933, EPI_ISL_833934, EPI_ISL_833935, EPI_ISL_833936, EPI_ISL_833937, EPI_ISL_833938, EPI_ISL_833939, EPI_ISL_833940, EPI_ISL_833941, EPI_ISL_833942, EPI_ISL_833943, EPI_ISL_833945, EPI_ISL_833946, EPI_ISL_833947, EPI_ISL_833948, EPI_ISL_833949, EPI_ISL_833950, EPI_ISL_833951, EPI_ISL_833952, EPI_ISL_833953, EPI_ISL_833954, EPI_ISL_833955, EPI_ISL_833957, EPI_ISL_833958, EPI_ISL_833959, EPI_ISL_833960, EPI_ISL_833961, EPI_ISL_833962, EPI_ISL_833963, EPI_ISL_833964, EPI_ISL_833965, EPI_ISL_833967, EPI_ISL_833969, EPI_ISL_833970, EPI_ISL_833971, EPI_ISL_833972, EPI_ISL_833973, EPI_ISL_833974, EPI_ISL_833975, EPI_ISL_833976, EPI_ISL_833977, EPI_ISL_833978, EPI_ISL_833979, EPI_ISL_833980, EPI_ISL_833981, EPI_ISL_833982, EPI_ISL_833983, EPI_ISL_833984, EPI_ISL_833985, EPI_ISL_833986, EPI_ISL_833987, EPI_ISL_833988, EPI_ISL_833989, EPI_ISL_833990, EPI_ISL_833991, EPI_ISL_833992, EPI_ISL_833993, EPI_ISL_833994, EPI_ISL_833996, EPI_ISL_833998, EPI_ISL_833999, EPI_ISL_834000, EPI_ISL_834001, EPI_ISL_834002, EPI_ISL_834003, EPI_ISL_834004, EPI_ISL_834005, EPI_ISL_834006, EPI_ISL_834007, EPI_ISL_834008, EPI_ISL_834010, EPI_ISL_834011, EPI_ISL_834012, EPI_ISL_834013, EPI_ISL_834014, EPI_ISL_834015, EPI_ISL_834016, EPI_ISL_834017, EPI_ISL_834018, EPI_ISL_834019, EPI_ISL_834020, EPI_ISL_834021, EPI_ISL_834023, EPI_ISL_834024, EPI_ISL_834025, EPI_ISL_834026, EPI_ISL_834027, EPI_ISL_834028, EPI_ISL_834030, EPI_ISL_834031, EPI_ISL_834032, EPI_ISL_834033, EPI_ISL_834034, EPI_ISL_834035, EPI_ISL_834036, EPI_ISL_834037, EPI_ISL_834038, EPI_ISL_834039, EPI_ISL_834040, EPI_ISL_834041, EPI_ISL_834043, EPI_ISL_834044, EPI_ISL_834045, EPI_ISL_834046, EPI_ISL_834047, EPI_ISL_834048, EPI_ISL_834049, EPI_ISL_834050, EPI_ISL_834051, EPI_ISL_834052, EPI_ISL_834053, EPI_ISL_834054, EPI_ISL_834055, EPI_ISL_834056, EPI_ISL_834057, EPI_ISL_834058, EPI_ISL_834059, EPI_ISL_834060, EPI_ISL_834061, EPI_ISL_834062, EPI_ISL_834063, EPI_ISL_834064, EPI_ISL_834065, EPI_ISL_834066, EPI_ISL_834067, EPI_ISL_834068, EPI_ISL_834070, EPI_ISL_834071, EPI_ISL_834072, EPI_ISL_834073, EPI_ISL_834074, EPI_ISL_834075, EPI_ISL_834076, EPI_ISL_834077, EPI_ISL_834078, EPI_ISL_834079, EPI_ISL_834080, EPI_ISL_834081, EPI_ISL_834083, EPI_ISL_834084, EPI_ISL_834085, EPI_ISL_834086, EPI_ISL_834087, EPI_ISL_834088, EPI_ISL_834089, EPI_ISL_834090, EPI_ISL_834091, EPI_ISL_834092, EPI_ISL_834093, EPI_ISL_834094, EPI_ISL_834095, EPI_ISL_834096, EPI_ISL_834097, EPI_ISL_834098, EPI_ISL_834099, EPI_ISL_834100, EPI_ISL_834101, EPI_ISL_834102, EPI_ISL_834103, EPI_ISL_834105, EPI_ISL_834106, EPI_ISL_834107, EPI_ISL_834108, EPI_ISL_834109, EPI_ISL_834110, EPI_ISL_834111, EPI_ISL_834112, EPI_ISL_834113, EPI_ISL_834114, EPI_ISL_834115, EPI_ISL_834116, EPI_ISL_834117, EPI_ISL_834118, EPI_ISL_834119, EPI_ISL_834120, EPI_ISL_834121, EPI_ISL_834122, EPI_ISL_834123, EPI_ISL_834124, EPI_ISL_834125, EPI_ISL_834126, EPI_ISL_834127, EPI_ISL_834128, EPI_ISL_834129, EPI_ISL_834130, EPI_ISL_834131, EPI_ISL_834132, EPI_ISL_834133, EPI_ISL_834135, EPI_ISL_834136, EPI_ISL_834137, EPI_ISL_834138, EPI_ISL_834139, EPI_ISL_834140, EPI_ISL_834141, EPI_ISL_834142, EPI_ISL_834143, EPI_ISL_834145, EPI_ISL_834147, EPI_ISL_834148, EPI_ISL_834149, EPI_ISL_834150, EPI_ISL_834151, EPI_ISL_834152, EPI_ISL_834153, EPI_ISL_834154, EPI_ISL_834155, EPI_ISL_834156, EPI_ISL_834157, EPI_ISL_834158, EPI_ISL_834159, EPI_ISL_834160, EPI_ISL_834161, EPI_ISL_834162, EPI_ISL_834163, EPI_ISL_834164, EPI_ISL_834165, EPI_ISL_834166, EPI_ISL_834167, EPI_ISL_834168, EPI_ISL_834169, EPI_ISL_834170, EPI_ISL_834171, EPI_ISL_834172, EPI_ISL_834173, EPI_ISL_834174, EPI_ISL_834175, EPI_ISL_834176, EPI_ISL_834177, EPI_ISL_834178, EPI_ISL_834179, EPI_ISL_834180, EPI_ISL_834181, EPI_ISL_834182, EPI_ISL_834183, EPI_ISL_834184, EPI_ISL_834185, EPI_ISL_834186, EPI_ISL_834187, EPI_ISL_834188, EPI_ISL_834189, EPI_ISL_834190, EPI_ISL_834191, EPI_ISL_834192, EPI_ISL_834193, EPI_ISL_834194, EPI_ISL_834195, EPI_ISL_834196, EPI_ISL_834198, EPI_ISL_834199, EPI_ISL_834200, EPI_ISL_834201, EPI_ISL_834202, EPI_ISL_834203, EPI_ISL_834204, EPI_ISL_834205, EPI_ISL_834206, EPI_ISL_834207, EPI_ISL_834208, EPI_ISL_834209 | | | |
| see above | Lighthouse Lab in Alderley Park | Wellcome Sanger Institute for the COVID-19 Genomics UK (COG-UK) Consortium | Jacquelyn Wynn, Mairead Hyland, The Lighthouse Lab in Alderley Park and Alex Alderton, Roberto Amato, Sonia Goncalves, Ewan Harrison, David K. Jackson, Ian Johnston, Dominic Kwiatkowski, Cordelia Langford, John Sillitoe on behalf of the Wellcome Sanger Institute COVID-19 Surveillance Team |
| EPI_ISL_834553, EPI_ISL_834554, EPI_ISL_834555, EPI_ISL_834556, EPI_ISL_834557, EPI_ISL_834559, EPI_ISL_834560, EPI_ISL_834563, EPI_ISL_834564, EPI_ISL_834565, EPI_ISL_834566, EPI_ISL_834567, EPI_ISL_834569, EPI_ISL_834570, EPI_ISL_834571, EPI_ISL_834573, EPI_ISL_834574, EPI_ISL_834575, EPI_ISL_834576, EPI_ISL_834577, EPI_ISL_834580, EPI_ISL_834582, EPI_ISL_834585, EPI_ISL_834586, EPI_ISL_834587, EPI_ISL_834589, EPI_ISL_834590, EPI_ISL_834591, EPI_ISL_834594, EPI_ISL_834595, EPI_ISL_834596, EPI_ISL_834597, EPI_ISL_834598, EPI_ISL_834601, EPI_ISL_834602, EPI_ISL_834605, EPI_ISL_834606, EPI_ISL_834607, EPI_ISL_834609, EPI_ISL_834610, EPI_ISL_834611, EPI_ISL_834613, EPI_ISL_834614, EPI_ISL_834615, EPI_ISL_834616, EPI_ISL_834617, EPI_ISL_834618, EPI_ISL_834620, EPI_ISL_834621, EPI_ISL_834622, EPI_ISL_834624, EPI_ISL_834625, EPI_ISL_834626, EPI_ISL_834627, EPI_ISL_834628, EPI_ISL_834630, EPI_ISL_834631, EPI_ISL_834633, EPI_ISL_834634, EPI_ISL_834635, EPI_ISL_834636, EPI_ISL_834637, EPI_ISL_834638, EPI_ISL_834641, EPI_ISL_834643, EPI_ISL_834647, EPI_ISL_834649, EPI_ISL_834651, EPI_ISL_834652, EPI_ISL_834653, EPI_ISL_834654, EPI_ISL_834655, EPI_ISL_834656, EPI_ISL_834657, EPI_ISL_834659, EPI_ISL_834660, EPI_ISL_834661, EPI_ISL_834662, EPI_ISL_834663, EPI_ISL_834664, EPI_ISL_834665, EPI_ISL_834666, EPI_ISL_834667, EPI_ISL_834668, EPI_ISL_834669, EPI_ISL_834671, EPI_ISL_834674, EPI_ISL_834675, EPI_ISL_834676, EPI_ISL_834677, EPI_ISL_834678, EPI_ISL_834679, EPI_ISL_834680, EPI_ISL_834681, EPI_ISL_834682, EPI_ISL_834684, EPI_ISL_834685, EPI_ISL_834688, EPI_ISL_834693, EPI_ISL_834694, EPI_ISL_834696, EPI_ISL_834697, EPI_ISL_834698, EPI_ISL_834700, EPI_ISL_834701, EPI_ISL_834702, EPI_ISL_834704, EPI_ISL_834705, EPI_ISL_834706, EPI_ISL_834707, EPI_ISL_834709, EPI_ISL_834711, EPI_ISL_834712, EPI_ISL_834713, EPI_ISL_834715, EPI_ISL_834716, EPI_ISL_834719, EPI_ISL_834720, EPI_ISL_834721, EPI_ISL_834722, EPI_ISL_834723, EPI_ISL_834724, EPI_ISL_834725, EPI_ISL_834727, EPI_ISL_834728, EPI_ISL_834730, EPI_ISL_834731, EPI_ISL_834732, EPI_ISL_834735, EPI_ISL_834736, EPI_ISL_834737, EPI_ISL_834739, EPI_ISL_834740, EPI_ISL_834742, EPI_ISL_834743, EPI_ISL_834744, EPI_ISL_834745, EPI_ISL_834747, EPI_ISL_834748, EPI_ISL_834749, EPI_ISL_834750, EPI_ISL_834752, EPI_ISL_834753, EPI_ISL_834755, EPI_ISL_834758, EPI_ISL_834759, EPI_ISL_834762, EPI_ISL_834763, EPI_ISL_834766, EPI_ISL_834768, EPI_ISL_834769, EPI_ISL_834770, EPI_ISL_834771, EPI_ISL_834772, EPI_ISL_834773, EPI_ISL_834774, EPI_ISL_834775, EPI_ISL_834776, EPI_ISL_834777, EPI_ISL_834778, EPI_ISL_834779, EPI_ISL_834780, EPI_ISL_834781, EPI_ISL_834784, EPI_ISL_834785, EPI_ISL_834786, EPI_ISL_834787, EPI_ISL_834788, EPI_ISL_834789, EPI_ISL_834790, EPI_ISL_834792, EPI_ISL_834794, EPI_ISL_834795, EPI_ISL_834796, EPI_ISL_834797 | | | |
| see above | Lighthouse Lab in Glasgow | Wellcome Sanger Institute for the COVID-19 Genomics UK (COG-UK) Consortium | Harper VanSteenhouse, Yumi Kasai, David Gray, Carol Clugston, Anna Dominiczak and Alex Alderton, Roberto Amato, Sonia Goncalves, Ewan Harrison, David K. Jackson, Ian Johnston, Dominic Kwiatkowski, Cordelia Langford, John Sillitoe on behalf of the Wellcome Sanger Institute COVID-19 Surveillance Team |
| EPI_ISL_834801, EPI_ISL_834868, EPI_ISL_834968, EPI_ISL_834976, EPI_ISL_834985, EPI_ISL_835010, EPI_ISL_835016, EPI_ISL_835065, EPI_ISL_835068, EPI_ISL_835108, EPI_ISL_835110, EPI_ISL_835121 | | | |
| see above | Lighthouse Lab in Alderley Park | Wellcome Sanger Institute for the COVID-19 Genomics UK (COG-UK) Consortium | Jacquelyn Wynn, Mairead Hyland, The Lighthouse Lab in Alderley Park and Alex Alderton, Roberto Amato, Sonia Goncalves, Ewan Harrison, David K. Jackson, Ian Johnston, Dominic Kwiatkowski, Cordelia Langford, John Sillitoe on behalf of the Wellcome Sanger Institute COVID-19 Surveillance Team |
| EPI_ISL_835143, EPI_ISL_835144, EPI_ISL_835146, EPI_ISL_835147, EPI_ISL_835148, EPI_ISL_835150, EPI_ISL_835151, EPI_ISL_835152, EPI_ISL_835156, EPI_ISL_835157, EPI_ISL_835158, EPI_ISL_835159, EPI_ISL_835160, EPI_ISL_835161, EPI_ISL_835162, EPI_ISL_835163, EPI_ISL_835164, EPI_ISL_835165, EPI_ISL_835166, EPI_ISL_835167, EPI_ISL_835168, EPI_ISL_835169, EPI_ISL_835170, EPI_ISL_835171, EPI_ISL_835172, EPI_ISL_835173, EPI_ISL_835175, EPI_ISL_835176, EPI_ISL_835178, EPI_ISL_835180, EPI_ISL_835182, EPI_ISL_835183, EPI_ISL_835184, EPI_ISL_835185, EPI_ISL_835186, EPI_ISL_835187, EPI_ISL_835188, EPI_ISL_835190, EPI_ISL_835191, EPI_ISL_835193, EPI_ISL_835194, EPI_ISL_835195, EPI_ISL_835196, EPI_ISL_835197, EPI_ISL_835199, EPI_ISL_835201, EPI_ISL_835203, EPI_ISL_835204, EPI_ISL_835206, EPI_ISL_835207, EPI_ISL_835208, EPI_ISL_835209, EPI_ISL_835210, EPI_ISL_835211, EPI_ISL_835213, EPI_ISL_835215, EPI_ISL_835216, EPI_ISL_835217, EPI_ISL_835218, EPI_ISL_835220, EPI_ISL_835221, EPI_ISL_835224, EPI_ISL_835225, EPI_ISL_835226, EPI_ISL_835227, EPI_ISL_835228, EPI_ISL_835229, EPI_ISL_835230, EPI_ISL_835232, EPI_ISL_835233, EPI_ISL_835234, EPI_ISL_835236, EPI_ISL_835237, EPI_ISL_835238, EPI_ISL_835239, EPI_ISL_835240, EPI_ISL_835242, EPI_ISL_835244, EPI_ISL_835245, EPI_ISL_835248, EPI_ISL_835249, EPI_ISL_835250, EPI_ISL_835251, EPI_ISL_835252, EPI_ISL_835254, EPI_ISL_835255, EPI_ISL_835256, EPI_ISL_835258, EPI_ISL_835262, EPI_ISL_835263, EPI_ISL_835264, EPI_ISL_835265, EPI_ISL_835266, EPI_ISL_835267, EPI_ISL_835269, EPI_ISL_835270, EPI_ISL_835271, EPI_ISL_835272, EPI_ISL_835273, EPI_ISL_835274, EPI_ISL_835275, EPI_ISL_835276, EPI_ISL_835277, EPI_ISL_835278, EPI_ISL_835279, EPI_ISL_835280, EPI_ISL_835281, EPI_ISL_835283, EPI_ISL_835284, EPI_ISL_835285, EPI_ISL_835286, EPI_ISL_835287, EPI_ISL_835288, EPI_ISL_835290, EPI_ISL_835291, EPI_ISL_835292, EPI_ISL_835293, EPI_ISL_835294, EPI_ISL_835295, EPI_ISL_835296, EPI_ISL_835297, EPI_ISL_835299, EPI_ISL_835301, EPI_ISL_835302, EPI_ISL_835304, EPI_ISL_835305, EPI_ISL_835306, EPI_ISL_835308, EPI_ISL_835309, EPI_ISL_835311, EPI_ISL_835312, EPI_ISL_835314, EPI_ISL_835315, EPI_ISL_835317, EPI_ISL_835318, EPI_ISL_835322, EPI_ISL_835323, EPI_ISL_835324, EPI_ISL_835325, EPI_ISL_835326, EPI_ISL_835327, EPI_ISL_835328, EPI_ISL_835329, EPI_ISL_835331, EPI_ISL_835332, EPI_ISL_835335, EPI_ISL_835336, EPI_ISL_835337, EPI_ISL_835340, EPI_ISL_835341, EPI_ISL_835342, EPI_ISL_835343, EPI_ISL_835344, EPI_ISL_835345, EPI_ISL_835346, EPI_ISL_835347, EPI_ISL_835348, EPI_ISL_835349, EPI_ISL_835350, EPI_ISL_835351, EPI_ISL_835352, EPI_ISL_835353, EPI_ISL_835355, EPI_ISL_835356, EPI_ISL_835358, EPI_ISL_835360, EPI_ISL_835363, EPI_ISL_835364, EPI_ISL_835367, EPI_ISL_835370, EPI_ISL_835371, EPI_ISL_835372, EPI_ISL_835373, EPI_ISL_835374, EPI_ISL_835376, EPI_ISL_835377, EPI_ISL_835379, EPI_ISL_835380, EPI_ISL_835381, EPI_ISL_835382, EPI_ISL_835383, EPI_ISL_835386, EPI_ISL_835387, EPI_ISL_835388, EPI_ISL_835391, EPI_ISL_835392, EPI_ISL_835393, EPI_ISL_835394, EPI_ISL_835395, EPI_ISL_835396, EPI_ISL_835397, EPI_ISL_835398, EPI_ISL_835399, EPI_ISL_835401, EPI_ISL_835402, EPI_ISL_835404, EPI_ISL_835405, EPI_ISL_835406, EPI_ISL_835407, EPI_ISL_835408, EPI_ISL_835411, EPI_ISL_835412, EPI_ISL_835413, EPI_ISL_835414, EPI_ISL_835417, EPI_ISL_835418, EPI_ISL_835419, EPI_ISL_835422, EPI_ISL_835423, EPI_ISL_835424, EPI_ISL_835425 | | | |
| see above | Lighthouse Lab in Glasgow | Wellcome Sanger Institute for the COVID-19 Genomics UK (COG-UK) Consortium | Harper VanSteenhouse, Yumi Kasai, David Gray, Carol Clugston, Anna Dominiczak and Alex Alderton, Roberto Amato, Sonia Goncalves, Ewan Harrison, David K. Jackson, Ian Johnston, Dominic Kwiatkowski, Cordelia Langford, John Sillitoe on behalf of the Wellcome Sanger Institute COVID-19 Surveillance Team |
| EPI_ISL_836423, EPI_ISL_836424, EPI_ISL_836431, EPI_ISL_836432, EPI_ISL_836439, EPI_ISL_836444, EPI_ISL_836445, EPI_ISL_836451, EPI_ISL_836466, EPI_ISL_836471, EPI_ISL_836482, EPI_ISL_836483, EPI_ISL_836491, EPI_ISL_836494, EPI_ISL_836496, EPI_ISL_836500, EPI_ISL_836501, EPI_ISL_836504, EPI_ISL_836506, EPI_ISL_836517, EPI_ISL_836527, EPI_ISL_836528, EPI_ISL_836530, EPI_ISL_836536, EPI_ISL_836548, EPI_ISL_836556, EPI_ISL_836562, EPI_ISL_836567, EPI_ISL_836572, EPI_ISL_836576, EPI_ISL_836585, EPI_ISL_836592, EPI_ISL_836595, EPI_ISL_836596, EPI_ISL_836597, EPI_ISL_836603, EPI_ISL_836611, EPI_ISL_836617, EPI_ISL_836620, EPI_ISL_836623, EPI_ISL_836626, EPI_ISL_836627, EPI_ISL_836636, EPI_ISL_836644, EPI_ISL_836645, EPI_ISL_836651, EPI_ISL_836652, EPI_ISL_836655, EPI_ISL_836656, EPI_ISL_836657, EPI_ISL_836665, EPI_ISL_836679, EPI_ISL_836681, EPI_ISL_836682, EPI_ISL_836683, EPI_ISL_836686, EPI_ISL_836688, EPI_ISL_836689, EPI_ISL_836695, EPI_ISL_836697, EPI_ISL_836712, EPI_ISL_836713, EPI_ISL_836722, EPI_ISL_836725 | | | |
| see above | Lighthouse Lab in Cambridge | Wellcome Sanger Institute for the COVID-19 Genomics UK (COG-UK) Consortium | Rob Howes, The Lighthouse Lab in Cambridge and Alex Alderton, Roberto Amato, Sonia Goncalves, Ewan Harrison, David K. Jackson, Ian Johnston, Dominic Kwiatkowski, Cordelia Langford, John Sillitoe on behalf of the Wellcome Sanger Institute COVID-19 Surveillance Team |
| EPI_ISL_836751, EPI_ISL_836759, EPI_ISL_836770, EPI_ISL_836778, EPI_ISL_836784, EPI_ISL_836820, EPI_ISL_836825, EPI_ISL_836827, EPI_ISL_836829, EPI_ISL_836831, EPI_ISL_836834, EPI_ISL_836847, EPI_ISL_836849, EPI_ISL_836850, EPI_ISL_836857, EPI_ISL_836864, EPI_ISL_836870, EPI_ISL_836893, EPI_ISL_836895, EPI_ISL_836898, EPI_ISL_836900, EPI_ISL_836902, EPI_ISL_836908, EPI_ISL_836924, EPI_ISL_836930, EPI_ISL_836931, EPI_ISL_836942, EPI_ISL_836944, EPI_ISL_836949, EPI_ISL_836954, EPI_ISL_836956, EPI_ISL_836957, EPI_ISL_836969, EPI_ISL_836971, EPI_ISL_836973, EPI_ISL_836974 | | | |
| see above | Lighthouse Lab in Glasgow | Wellcome Sanger Institute for the COVID-19 Genomics UK (COG-UK) Consortium | Harper VanSteenhouse, Yumi Kasai, David Gray, Carol Clugston, Anna Dominiczak and Alex Alderton, Roberto Amato, Sonia Goncalves, Ewan Harrison, David K. Jackson, Ian Johnston, Dominic Kwiatkowski, Cordelia Langford, John Sillitoe on behalf of the Wellcome Sanger Institute COVID-19 Surveillance Team |
| EPI_ISL_837055, EPI_ISL_837176, EPI_ISL_837177, EPI_ISL_837178, EPI_ISL_837179, EPI_ISL_837180 | Respiratory Virus Unit, National Infection Service, Public Health England | COVID-19 Genomics UK (COG-UK) Consortium | PHE Covid Sequencing Team |
| EPI_ISL_837847, EPI_ISL_837848, EPI_ISL_837937, EPI_ISL_837938, EPI_ISL_837939, EPI_ISL_837941, EPI_ISL_837944, EPI_ISL_837945, EPI_ISL_837946, EPI_ISL_837947, EPI_ISL_837948, EPI_ISL_837949, EPI_ISL_837950, EPI_ISL_837951, EPI_ISL_837952, EPI_ISL_837953, EPI_ISL_837954, EPI_ISL_837956, EPI_ISL_837958, EPI_ISL_837960, EPI_ISL_837961, EPI_ISL_837962, EPI_ISL_837963, EPI_ISL_837964, EPI_ISL_837965, EPI_ISL_837966, EPI_ISL_837967, EPI_ISL_837968, EPI_ISL_837969, EPI_ISL_837970 | | | |
| see above | Department of Pathology, University of Cambridge | COVID-19 Genomics UK (COG-UK) Consortium | Aminu S. Jahun, Yasmin Chaudhry, Grant Hall, Iliana Georgana, Myra Hosmillo, Martin D. Curran, Malte Pinckert, Surendra Parmar, Ian Goodfellow |

| | | | |
|---|---|---|---|
| EPI_ISL_838290, EPI_ISL_838291, EPI_ISL_838293, EPI_ISL_838294, EPI_ISL_838295, EPI_ISL_838296, EPI_ISL_838297, EPI_ISL_838298, EPI_ISL_838299, EPI_ISL_838300, EPI_ISL_838301, EPI_ISL_838302, EPI_ISL_838303, EPI_ISL_838304, EPI_ISL_838305, EPI_ISL_838306, EPI_ISL_838307, EPI_ISL_838308, EPI_ISL_838309, EPI_ISL_838310, EPI_ISL_838311, EPI_ISL_838312 | | | |
| see above | Virology Department, Royal Infirmary of Edinburgh, NHS Lothian / School of Biological Sciences, University of Edinburgh / Institute of Genetics and Molecular Medicine, University of Edinburgh | COVID-19 Genomics UK (COG-UK) Consortium | McHugh M, Dewar R, Rooke S, Gallagher M, Balcaza C, O'Toole Á, Scher E, Hill V, McCrone JT, Colquhoun R, Yu X, Jackson B, Rambaut A, Williams TC, Templeton K |
| EPI_ISL_838341 | University of Exeter | COVID-19 Genomics UK (COG-UK) Consortium | Ben Temperton,Aaron Jeffries,Michelle Michelsen,Joanna Warwick-Dugdale,Audrey Farbos,Robyn Manley,Stephen Michell,Jane Masoli |
| EPI_ISL_838895, EPI_ISL_838896, EPI_ISL_838897, EPI_ISL_838899, EPI_ISL_838900, EPI_ISL_838901, EPI_ISL_838902, EPI_ISL_838903, EPI_ISL_838904, EPI_ISL_838905, EPI_ISL_838909, EPI_ISL_838910, EPI_ISL_838911, EPI_ISL_838912, EPI_ISL_838913, EPI_ISL_838914, EPI_ISL_838915, EPI_ISL_838917, EPI_ISL_838919, EPI_ISL_838920, EPI_ISL_838926, EPI_ISL_838927, EPI_ISL_838928, EPI_ISL_838929, EPI_ISL_838930, EPI_ISL_838931, EPI_ISL_838932, EPI_ISL_838933, EPI_ISL_838934, EPI_ISL_838935, EPI_ISL_838936, EPI_ISL_838937, EPI_ISL_838938, EPI_ISL_838939, EPI_ISL_838940, EPI_ISL_838941, EPI_ISL_838942, EPI_ISL_838943, EPI_ISL_838944, EPI_ISL_838945, EPI_ISL_838946, EPI_ISL_838947, EPI_ISL_838949, EPI_ISL_838950, EPI_ISL_838951, EPI_ISL_838952, EPI_ISL_838953, EPI_ISL_838954, EPI_ISL_838955, EPI_ISL_838956, EPI_ISL_838957, EPI_ISL_838958, EPI_ISL_838959, EPI_ISL_838969, EPI_ISL_838988, EPI_ISL_838989, EPI_ISL_838990, EPI_ISL_839013, EPI_ISL_839015, EPI_ISL_839026, EPI_ISL_839027, EPI_ISL_839028, EPI_ISL_839029, EPI_ISL_839030, EPI_ISL_839031, EPI_ISL_839032, EPI_ISL_839033, EPI_ISL_839034, EPI_ISL_839035, EPI_ISL_839036, EPI_ISL_839037, EPI_ISL_839038, EPI_ISL_839039, EPI_ISL_839128, EPI_ISL_839133, EPI_ISL_839134, EPI_ISL_839135, EPI_ISL_839136, EPI_ISL_839137, EPI_ISL_839138, EPI_ISL_839139, EPI_ISL_839140, EPI_ISL_839141, EPI_ISL_839142, EPI_ISL_839143, EPI_ISL_839144, EPI_ISL_839145, EPI_ISL_839146, EPI_ISL_839147, EPI_ISL_839148, EPI_ISL_839149, EPI_ISL_839150, EPI_ISL_839151, EPI_ISL_839152, EPI_ISL_839153, EPI_ISL_839154, EPI_ISL_839155, EPI_ISL_839157, EPI_ISL_839158, EPI_ISL_839159, EPI_ISL_839160, EPI_ISL_839161, EPI_ISL_839162, EPI_ISL_839163, EPI_ISL_839165, EPI_ISL_839166, EPI_ISL_839167, EPI_ISL_839168, EPI_ISL_839169, EPI_ISL_839170, EPI_ISL_839171, EPI_ISL_839173, EPI_ISL_839174, EPI_ISL_839175, EPI_ISL_839188, EPI_ISL_839190, EPI_ISL_839191, EPI_ISL_839193, EPI_ISL_839194, EPI_ISL_839196, EPI_ISL_839197, EPI_ISL_839200, EPI_ISL_839212, EPI_ISL_839250, EPI_ISL_839251, EPI_ISL_839252, EPI_ISL_839253, EPI_ISL_839254, EPI_ISL_839255, EPI_ISL_839256, EPI_ISL_839257, EPI_ISL_839258, EPI_ISL_839260, EPI_ISL_839261, EPI_ISL_839262, EPI_ISL_839264, EPI_ISL_839266, EPI_ISL_839267, EPI_ISL_839268, EPI_ISL_839269, EPI_ISL_839270, EPI_ISL_839271, EPI_ISL_839272, EPI_ISL_839273, EPI_ISL_839274, EPI_ISL_839275, EPI_ISL_839276 | | | |
| see above | University College London, Great Ormond Street Hospital for Children NHS Foundation Trust, Imperial College Healthcare NHS Trust | COVID-19 Genomics UK (COG-UK) Consortium | Sergi Castellano, Rachel Williams, Mark Kristiansen, Paola Resende Silva, Sunando Roy, Tony Brooks, Helena Tutill, Paola Niola, Patricia Dyal, Charlotte Williams, Leysa Forrest, Yasmin Panchbhaya, Jacqueline Findlay, Samuel Weeks, Julianne Brown, Kathryn Harris, Paul Randell, James Price, Alison Holmes, Judith Breuer |
| EPI_ISL_839855, EPI_ISL_839856, EPI_ISL_839857, EPI_ISL_839858, EPI_ISL_839859, EPI_ISL_839860, EPI_ISL_839861, EPI_ISL_839862, EPI_ISL_839863, EPI_ISL_839864, EPI_ISL_839865, EPI_ISL_839867, EPI_ISL_839868, EPI_ISL_839869, EPI_ISL_839870, EPI_ISL_839871, EPI_ISL_839872, EPI_ISL_839873, EPI_ISL_839874, EPI_ISL_839875, EPI_ISL_839876, EPI_ISL_839877, EPI_ISL_839878, EPI_ISL_839879, EPI_ISL_839880, EPI_ISL_839881, EPI_ISL_839882, EPI_ISL_839883, EPI_ISL_839884, EPI_ISL_839885, EPI_ISL_839886, EPI_ISL_839887, EPI_ISL_839888, EPI_ISL_839889, EPI_ISL_839890, EPI_ISL_839891, EPI_ISL_839892, EPI_ISL_839893, EPI_ISL_839894, EPI_ISL_839895, EPI_ISL_839896, EPI_ISL_839897, EPI_ISL_839898, EPI_ISL_839900, EPI_ISL_839939, EPI_ISL_839940, EPI_ISL_839941, EPI_ISL_839942, EPI_ISL_839943, EPI_ISL_839944, EPI_ISL_839945, EPI_ISL_839946, EPI_ISL_839947, EPI_ISL_839948, EPI_ISL_839949, EPI_ISL_839950, EPI_ISL_839951, EPI_ISL_839952, EPI_ISL_839953, EPI_ISL_839954, EPI_ISL_839955, EPI_ISL_839956, EPI_ISL_839957, EPI_ISL_839958, EPI_ISL_839959, EPI_ISL_839960, EPI_ISL_839961, EPI_ISL_839962, EPI_ISL_839963, EPI_ISL_839964, EPI_ISL_839965, EPI_ISL_839966, EPI_ISL_839967, EPI_ISL_839968, EPI_ISL_839969, EPI_ISL_839970, EPI_ISL_839971, EPI_ISL_839972, EPI_ISL_839973 | | | |
| see above | Quadram Institute Bioscience | COVID-19 Genomics UK (COG-UK) Consortium | Dave J. Baker, Gemma L. Kay, Alp Aydin, Thanh Le-Viet, Steven Rudder, Ana P. Tedim, Anastasia Kolyva, Maria Diaz, Leonardo de Oliveira Martins, Nabil-Fareed Alikhan, Lizzie Meadows, Rachael Stanley, Ngozi Elumogo, Muhammed Yasir, Nicholas M. Thomson, Alexander J Trotter, Rachel Gilroy, Samuel Bloomfield, Claire Stuart, Andrew Bell, Reenesh Prakash, Samir Dervisevic, Alison E. Mather, John Wain, Mark Webber, Andrew J. Page, Justin O'Grady |
| EPI_ISL_839998, EPI_ISL_839999, EPI_ISL_840000, EPI_ISL_840001, EPI_ISL_840002, EPI_ISL_840003, EPI_ISL_840004, EPI_ISL_840005, EPI_ISL_840006, EPI_ISL_840007, EPI_ISL_840008, EPI_ISL_840009, EPI_ISL_840010, EPI_ISL_840011, EPI_ISL_840012, EPI_ISL_840013, EPI_ISL_840014, EPI_ISL_840015, EPI_ISL_840016, EPI_ISL_840051, EPI_ISL_840052 | | | |
| see above | Queens Medical Centre, Clinical Microbiology Department / DeepSeq Nottingham | COVID-19 Genomics UK (COG-UK) Consortium | Gemma Clark, Wendy Smith, Manjinder Khakh, Vicki M Fleming, Michelle M Lister, Hannah Howson-Wells, Jonathan Ball, Patrick McClure, Joseph Chappell, Theocharis Tsoleridis, Nadine Holmes, Matthew Carlisle, Christopher Moore, Fei Sang, Johnny Debebe, Victoria Wright, Matthew Loose |
| EPI_ISL_840139, EPI_ISL_840145 | Lincolnshire Hospitals and DeepSeq Nottingham | COVID-19 Genomics UK (COG-UK) Consortium | Nichola Duckworth, Tim Sloan, Sarah Walsh, Jonathan Ball, Patrick McClure, Joeseph Chappell, Nadine Holmes, Matthew Carlisle, Christopher Moore, Fei Sang, Johnny Debebe, Victoria Wright, Matthew Loose |
| EPI_ISL_840861, EPI_ISL_840862, EPI_ISL_840863, EPI_ISL_840864, EPI_ISL_840865, EPI_ISL_840866, EPI_ISL_840867, EPI_ISL_840869, EPI_ISL_840870, EPI_ISL_840871, EPI_ISL_840872, EPI_ISL_840874, EPI_ISL_840875, EPI_ISL_840876, EPI_ISL_840877, EPI_ISL_840878, EPI_ISL_840880, EPI_ISL_840882, EPI_ISL_840883, EPI_ISL_840884, EPI_ISL_841008, EPI_ISL_841010, EPI_ISL_841012, EPI_ISL_841013, EPI_ISL_841014, EPI_ISL_841015, EPI_ISL_841016, EPI_ISL_841017, EPI_ISL_841018, EPI_ISL_841019, EPI_ISL_841020, EPI_ISL_841021, EPI_ISL_841027, EPI_ISL_841029, EPI_ISL_841030, EPI_ISL_841031, EPI_ISL_841036, EPI_ISL_841037, EPI_ISL_841038, EPI_ISL_841039, EPI_ISL_841045, EPI_ISL_841046, EPI_ISL_841047, EPI_ISL_841049, EPI_ISL_841050, EPI_ISL_841053, EPI_ISL_841054, EPI_ISL_841055, EPI_ISL_841056, EPI_ISL_841067, EPI_ISL_841068, EPI_ISL_841069, EPI_ISL_841070, EPI_ISL_841071, EPI_ISL_841072, EPI_ISL_841073, EPI_ISL_841074, EPI_ISL_841075, EPI_ISL_841076, EPI_ISL_841077, EPI_ISL_841078, EPI_ISL_841079, EPI_ISL_841080, EPI_ISL_841108, EPI_ISL_841147, EPI_ISL_841148, EPI_ISL_841149, EPI_ISL_841150, EPI_ISL_841151, EPI_ISL_841152, EPI_ISL_841153, EPI_ISL_841154, EPI_ISL_841155, EPI_ISL_841182, EPI_ISL_841191, EPI_ISL_841192, EPI_ISL_841194, EPI_ISL_841196, EPI_ISL_841197, EPI_ISL_841200, EPI_ISL_841201, EPI_ISL_841202, EPI_ISL_841203, EPI_ISL_841205, EPI_ISL_841206, EPI_ISL_841207, EPI_ISL_841209, EPI_ISL_841210, EPI_ISL_841213, EPI_ISL_841216, EPI_ISL_841217, EPI_ISL_841220, EPI_ISL_841221, EPI_ISL_841222, EPI_ISL_841223, EPI_ISL_841224, EPI_ISL_841226, EPI_ISL_841227, EPI_ISL_841229, EPI_ISL_841230, EPI_ISL_841231, EPI_ISL_841232, EPI_ISL_841233, EPI_ISL_841234, EPI_ISL_841235, EPI_ISL_841236, EPI_ISL_841237, EPI_ISL_841238, EPI_ISL_841240, EPI_ISL_841242, EPI_ISL_841244, EPI_ISL_841245, EPI_ISL_841246, EPI_ISL_841247, EPI_ISL_841248, EPI_ISL_841250, EPI_ISL_841251, EPI_ISL_841252, EPI_ISL_841253, EPI_ISL_841255, EPI_ISL_841256, EPI_ISL_841257, EPI_ISL_841258, EPI_ISL_841259, EPI_ISL_841261, EPI_ISL_841262, EPI_ISL_841263, EPI_ISL_841264, EPI_ISL_841265, EPI_ISL_841266, EPI_ISL_841267, EPI_ISL_841268, EPI_ISL_841269, EPI_ISL_841270, EPI_ISL_841271, EPI_ISL_841273, EPI_ISL_841275, EPI_ISL_841276, EPI_ISL_841277, EPI_ISL_841278, EPI_ISL_841280, EPI_ISL_841281, EPI_ISL_841283, EPI_ISL_841284, EPI_ISL_841287, EPI_ISL_841288, EPI_ISL_841290, EPI_ISL_841291, EPI_ISL_841292, EPI_ISL_841293, EPI_ISL_841294, EPI_ISL_841295, EPI_ISL_841296, EPI_ISL_841297, EPI_ISL_841298, EPI_ISL_841299, EPI_ISL_841300, EPI_ISL_841314, EPI_ISL_841315 | | | |
| see above | Wales Specialist Virology Centre Sequencing lab: Pathogen Genomics Unit | Public Health Wales Microbiology Cardiff Wales Specialist Virology Centre | Catherine Moore, Johnathan Evans, Laura Gifford, Malorie Perry, Simon Cottrell, Angela Marchbank, Alec Birchley, Alexander Adams, Amy Gaskin, Bree Gatica-Wilcox, Jason Coombes, Joel Southgate, Lauren Gilbert, Lee Graham, Nicole Pacchiarini, Sara Kumziene-Summerhayes, Sarah Taylor, Sophie Jones, Sara Rey, Matthew Bull, Joanne Watkins, Sally Corden, Tom Connor |
| EPI_ISL_841317, EPI_ISL_841318, EPI_ISL_841319, EPI_ISL_841320, EPI_ISL_841322, EPI_ISL_841323, EPI_ISL_841324, EPI_ISL_841327, EPI_ISL_841335, EPI_ISL_841336, EPI_ISL_841337, EPI_ISL_841338, EPI_ISL_841339, EPI_ISL_841341, EPI_ISL_841342, EPI_ISL_841343, EPI_ISL_841344, EPI_ISL_841345, EPI_ISL_841346, EPI_ISL_841347, EPI_ISL_841348, EPI_ISL_841353, EPI_ISL_841354, EPI_ISL_841358, EPI_ISL_841378, EPI_ISL_841379, EPI_ISL_841380, EPI_ISL_841381, EPI_ISL_841382, EPI_ISL_841383, EPI_ISL_841384, EPI_ISL_841385, EPI_ISL_841386, EPI_ISL_841391, EPI_ISL_841392, EPI_ISL_841393, EPI_ISL_841394, EPI_ISL_841395, EPI_ISL_841396, EPI_ISL_841397, EPI_ISL_841399, EPI_ISL_841400, EPI_ISL_841401, EPI_ISL_841402, EPI_ISL_841403, EPI_ISL_841404, EPI_ISL_841405, EPI_ISL_841406, EPI_ISL_841413, EPI_ISL_841414, EPI_ISL_841416, EPI_ISL_841417, EPI_ISL_841418, EPI_ISL_841419, EPI_ISL_841420, EPI_ISL_841421, EPI_ISL_841422, EPI_ISL_841423, EPI_ISL_841424, EPI_ISL_841425, EPI_ISL_841428, EPI_ISL_841429, EPI_ISL_841431, EPI_ISL_841432, EPI_ISL_841434, EPI_ISL_841435, EPI_ISL_841436, EPI_ISL_841437, EPI_ISL_841440, EPI_ISL_841443, EPI_ISL_841444, EPI_ISL_841445, EPI_ISL_841446, EPI_ISL_841448, EPI_ISL_841451, EPI_ISL_841453, EPI_ISL_841454, EPI_ISL_841455, EPI_ISL_841456, EPI_ISL_841458, EPI_ISL_841460, EPI_ISL_841461, EPI_ISL_841462, EPI_ISL_841464, EPI_ISL_841465, EPI_ISL_841467, EPI_ISL_841468, EPI_ISL_841469, EPI_ISL_841470, EPI_ISL_841472, EPI_ISL_841473, EPI_ISL_841474, EPI_ISL_841475, EPI_ISL_841476, EPI_ISL_841477, EPI_ISL_841478, EPI_ISL_841480, EPI_ISL_841481, EPI_ISL_841502, EPI_ISL_841513, EPI_ISL_841514, EPI_ISL_841515, EPI_ISL_841516, EPI_ISL_841517, EPI_ISL_841518, EPI_ISL_841519, EPI_ISL_841520, EPI_ISL_841548, EPI_ISL_841594, EPI_ISL_841595, EPI_ISL_841597, EPI_ISL_841598, EPI_ISL_841599, EPI_ISL_841600, EPI_ISL_841601, EPI_ISL_841602, EPI_ISL_841603, EPI_ISL_841604, EPI_ISL_841605, EPI_ISL_841606, EPI_ISL_841607, EPI_ISL_841613, EPI_ISL_841615, EPI_ISL_841616, EPI_ISL_841621, EPI_ISL_841624, EPI_ISL_841628, EPI_ISL_841629, EPI_ISL_841631, EPI_ISL_841632, EPI_ISL_841642, EPI_ISL_841644, EPI_ISL_841645 | | | |
| see above | Originating lab: Wales Specialist Virology Centre Sequencing lab: Pathogen Genomics Unit | Public Health Wales Microbiology Cardiff Wales Specialist Virology Centre | Catherine Moore, Johnathan Evans, Laura Gifford, Malorie Perry, Simon Cottrell, Angela Marchbank, Alec Birchley, Alexander Adams, Amy Gaskin, Bree Gatica-Wilcox, Jason Coombes, Joel Southgate, Lauren Gilbert, Lee Graham, Nicole Pacchiarini, Sara Kumziene-Summerhayes, Sarah Taylor, Sophie Jones, Sara Rey, Matthew Bull, Joanne Watkins, Sally Corden, Tom Connor |
| EPI_ISL_842202, EPI_ISL_842203, EPI_ISL_842205, EPI_ISL_842206, EPI_ISL_842219, EPI_ISL_842220, EPI_ISL_842227, EPI_ISL_842231, EPI_ISL_842237, EPI_ISL_842240, EPI_ISL_842244, EPI_ISL_842245, EPI_ISL_842253, EPI_ISL_842257, EPI_ISL_842260, EPI_ISL_842262, EPI_ISL_842268, EPI_ISL_842269, EPI_ISL_842279, EPI_ISL_842284, EPI_ISL_842286, EPI_ISL_842290, EPI_ISL_842298, EPI_ISL_842299, EPI_ISL_842307, EPI_ISL_842308, EPI_ISL_842311, EPI_ISL_842326, EPI_ISL_842332, EPI_ISL_842344, EPI_ISL_842345, EPI_ISL_842346 | | | |
| see above | Virology Department, Sheffield Teaching Hospitals NHS Foundation Trust/Department of Infection, Immunity and Cardiovascular Disease, The Medical School, University of Sheffield | COVID-19 Genomics UK (COG-UK) Consortium | Thushan de Silva, Matthew Parker, Nikki Smith, Adri Angyal, Rebecca Brown, Luke Green, Rachel Tucker, Paul Parsons, Danielle Groves, Katie Johnson, Laura Carrilero, Alex Keeley, Dave Partridge, Matthew Wyles, Benjamin Lindsey, Mehmet Yavuz, Mohammad Raza, Cariad Evans |
| EPI_ISL_842634, EPI_ISL_842635, EPI_ISL_842636 | Medical Microbiology Unit, Department for Laboratory Medicine, Drammen Hospital, Vestre Viken Health Trust, | Norwegian Institute of Public Health, Department of Virology | Kathrine Stene-Johansen, Kamilla Heddeland Instefjord, Hilde Elshaug, Atiya R Ali,Marie Paulsen Madsen, Rasmus Riis Kopperud, Hilde Vollan, Karoline Bragstad, Olav Hungnes |
| EPI_ISL_842639 | Oslo University Hospital, Department of Medical Microbiology | Norwegian Institute of Public Health, Department of Virology | Kathrine Stene-Johansen, Kamilla Heddeland Instefjord, Hilde Elshaug, Atiya R Ali,Marie Paulsen Madsen, Rasmus Riis Kopperud, Hilde Vollan, Karoline Bragstad, Olav Hungnes |
| EPI_ISL_842644 | Furst Medical Laboratory | Norwegian Institute of Public Health, Department of Virology | Kathrine Stene-Johansen, Kamilla Heddeland Instefjord, Hilde Elshaug, Atiya R Ali,Marie Paulsen Madsen, Rasmus Riis Kopperud, Hilde Vollan, Karoline Bragstad, Olav Hungnes |
| EPI_ISL_843170, EPI_ISL_843171, EPI_ISL_843172, EPI_ISL_843173, EPI_ISL_843174, EPI_ISL_843175, EPI_ISL_843176, EPI_ISL_843177, EPI_ISL_843178, EPI_ISL_843179, EPI_ISL_843180, EPI_ISL_843181, EPI_ISL_843182, EPI_ISL_843183 | | | |
| see above | Maryland Public Health Laboratory | Maryland Public Health Laboratory | Maryland Department of Health Laboratories Administration |
| EPI_ISL_843195, EPI_ISL_843197, EPI_ISL_843198 | Canterbury Health Laboratories | Institute of Environmental Science and Research (ESR) | Xiaoyun Ren, Matt Storey, Nikki Freed, Muhammad Faisal, Jing Wang, Hermes Perez, Anja Werno, Antje van der Linden, Arlo Upton, Chris Mansell, David Hammer, Dragana Drinkovic, Gary McAuliffe, Hana Sofia Andersson, James Ussher, Jill Sherwood, Josh Freeman, Julia Howard, Juliet Elvy, Mary DeAlmeida, Matt Blakiston, Matthew Rogers, Max Bloomfield, Michael Addidle, Michelle Balm, Sally Roberts, Sarah Jefferies, Sharmini Muttaiyah, Susan Morpeth, Susan Taylor, Timothy Blackmore, Vani Sathyendran, Veronica Playle, Virginia Hope, Erasmus Smit, Lauren Jelly, Olin Silander, Joep de Ligt |
| EPI_ISL_845575, EPI_ISL_845586, | KU Leuven, Rega Institute, Clinical and Epidemiological | KU Leuven, Rega Institute, Clinical and Epidemiological | Tony Wawina-Bokalanga, Bert Vanmechelen, Joan Marti-Carerras, Piet Maes |

| | | | |
|---|---|---|---|
| EPI_ISL_845612, EPI_ISL_845613, EPI_ISL_845616 | Virology | Virology | |
| EPI_ISL_845752 | Toronto Invasive Bacterial Diseases Network | McMaster University | Allison McGeer, Patryk Aftanas, Hooman Derakhshani, Angel Li, Kuganya Nirmalarajah, Emily Panousis, Ahmed Draia, Jalees Nasir, Michael Surette, Samira Mubareka, Andrew G. McArthur |
| EPI_ISL_845761, EPI_ISL_845762, EPI_ISL_845763, EPI_ISL_845764, EPI_ISL_845765, EPI_ISL_845766, EPI_ISL_845767, EPI_ISL_845768, EPI_ISL_845769, EPI_ISL_845772, EPI_ISL_845773, EPI_ISL_845774, EPI_ISL_845775, EPI_ISL_845776, EPI_ISL_845777, EPI_ISL_845778, EPI_ISL_845779, EPI_ISL_845780, EPI_ISL_845781, EPI_ISL_845782 | | | |
| see above | Emory Molecular Diagnostics Laboratory, Emory Healthcare | Piantadosi Lab, Emory Department of Pathology | Ahmed Babiker, Anne Piantadosi |
| EPI_ISL_845893, EPI_ISL_845896, EPI_ISL_845973, EPI_ISL_846007, EPI_ISL_846022, EPI_ISL_846023, EPI_ISL_846052, EPI_ISL_846059, EPI_ISL_846070, EPI_ISL_846084, EPI_ISL_846139, EPI_ISL_846147, EPI_ISL_846152, EPI_ISL_846168 | | | |
| see above | Lighthouse Lab in Cambridge | Wellcome Sanger Institute for the COVID-19 Genomics UK (COG-UK) Consortium | Rob Howes, The Lighthouse Lab in Cambridge and Alex Alderton, Roberto Amato, Sonia Goncalves, Ewan Harrison, David K. Jackson, Ian Johnston, Dominic Kwiatkowski, Cordelia Langford, John Sillitoe on behalf of the Wellcome Sanger Institute COVID-19 Surveillance Team |
| EPI_ISL_846283, EPI_ISL_846287, EPI_ISL_846299, EPI_ISL_846315, EPI_ISL_846319, EPI_ISL_846321, EPI_ISL_846354, EPI_ISL_846399, EPI_ISL_846400, EPI_ISL_846421, EPI_ISL_846429, EPI_ISL_846431, EPI_ISL_846443, EPI_ISL_846491 | | | |
| see above | Lighthouse Lab in Alderley Park | Wellcome Sanger Institute for the COVID-19 Genomics UK (COG-UK) Consortium | Jacquelyn Wynn, Mairead Hyland, The Lighthouse Lab in Alderley Park and Alex Alderton, Roberto Amato, Sonia Goncalves, Ewan Harrison, David K. Jackson, Ian Johnston, Dominic Kwiatkowski, Cordelia Langford, John Sillitoe on behalf of the Wellcome Sanger Institute COVID-19 Surveillance Team |
| EPI_ISL_846509 | Lighthouse Lab in Cambridge | Wellcome Sanger Institute for the COVID-19 Genomics UK (COG-UK) Consortium | Rob Howes, The Lighthouse Lab in Cambridge and Alex Alderton, Roberto Amato, Sonia Goncalves, Ewan Harrison, David K. Jackson, Ian Johnston, Dominic Kwiatkowski, Cordelia Langford, John Sillitoe on behalf of the Wellcome Sanger Institute COVID-19 Surveillance Team |
| EPI_ISL_846510, EPI_ISL_846511, EPI_ISL_846513, EPI_ISL_846514, EPI_ISL_846515, EPI_ISL_846516, EPI_ISL_846517, EPI_ISL_846518, EPI_ISL_846519, EPI_ISL_846520, EPI_ISL_846521, EPI_ISL_846522 | | | |
| see above | Lighthouse Lab in Alderley Park | Wellcome Sanger Institute for the COVID-19 Genomics UK (COG-UK) Consortium | Jacquelyn Wynn, Mairead Hyland, The Lighthouse Lab in Alderley Park and Alex Alderton, Roberto Amato, Sonia Goncalves, Ewan Harrison, David K. Jackson, Ian Johnston, Dominic Kwiatkowski, Cordelia Langford, John Sillitoe on behalf of the Wellcome Sanger Institute COVID-19 Surveillance Team |
| EPI_ISL_846534, EPI_ISL_846538, EPI_ISL_846539 | Lighthouse Lab in Glasgow | Wellcome Sanger Institute for the COVID-19 Genomics UK (COG-UK) Consortium | Harper VanSteenhouse, Yumi Kasai, David Gray, Carol Clugston, Anna Dominiczak and Alex Alderton, Roberto Amato, Sonia Goncalves, Ewan Harrison, David K. Jackson, Ian Johnston, Dominic Kwiatkowski, Cordelia Langford, John Sillitoe on behalf of the Wellcome Sanger Institute COVID-19 Surveillance Team |
| EPI_ISL_846540 | National Laboratory for Health, Environment and Food | National Laboratory for Health, Environment and Food | Aleksander Mahnic, Sandra Janezic, Maja Rupnik |
| EPI_ISL_846545, EPI_ISL_846546 | Lab voor klinische biologie | Lab voor klinische biologie | Hannelore Hamerlinck, Bruno Verhasselt |
| EPI_ISL_846598 | Respiratory Virus Unit, National Infection Service, Public Health England | COVID-19 Genomics UK (COG-UK) Consortium | PHE Covid Sequencing Team |
| EPI_ISL_846644 | Hopital | National Reference Center for Viruses of Respiratory Infections, Institut Pasteur, Paris | Marion Barbet, Sylvie Behillil, Méline Bizard, Angela Brisebarre, Camille Capel, Etienne Simon-Lorière, Vincent Enouf, Maud Vanpeene, Sylvie van der Werf |
| EPI_ISL_846648 | Labo Analyses Med | National Reference Center for Viruses of Respiratory Infections, Institut Pasteur, Paris | Marion Barbet, Sylvie Behillil, Méline Bizard, Angela Brisebarre, Camille Capel, Etienne Simon-Lorière, Vincent Enouf, Maud Vanpeene, Sylvie van der Werf |
| EPI_ISL_847835 | Oak Ridge National Laboratory | Grubaugh Lab - Yale School of Public Health | Tara Alpert, Joseph Fauver, Anderson Brito, Mallery Breban, Anne Wyllie, Chantal Vogels, Mary Petrone, Chaney Kalinich, Isabel Ott, Nathan Grubaugh |
| EPI_ISL_847987, EPI_ISL_848050, EPI_ISL_848055 | Michigan Department of Health and Human Services, Bureau of Laboratories | Michigan Department of Health and Human Services, Bureau of Laboratories | Blankenship HM, Riner D, Soehnlen MK |
| EPI_ISL_848065, EPI_ISL_848066, EPI_ISL_848067, EPI_ISL_848068 | CHU Purpan - Laboratoire de Virologie - Institut Fédératif de Biologie | CHU Purpan - Laboratoire de Virologie - Institut Fédératif de Biologie | Latour J., Ranger N., Dubois M., Carcenac R., Harter A., Boyer P., Tremeaux P., Izopet J. |
| EPI_ISL_848069 | Tulane Medical Center | Centers for Disease Control and Prevention Division of Viral Diseases, Pathogen Discovery | Ying Tao, Yan Li, Jing Zhang, Krista Queen, Anna Uehara, Peter Cook, Clinton R. Paden, Haibin Wang, Suxiang Tong |
| EPI_ISL_848630, EPI_ISL_848631, EPI_ISL_848632 | Montana Public Health Laboratory | Wyoming Public Health Laboratory | Noah Hull, Joy Ritter, Taylor Fearing, Lynette Gumbleton, Channing Weber, Ashley Norberg, Bailey Bowcutt, Wanda Manley, Deborah Gibson |
| EPI_ISL_849753, EPI_ISL_849754, EPI_ISL_849755 | unknown | PHV-FSS | Son Nguyen et al. |
| EPI_ISL_849906, EPI_ISL_849907, EPI_ISL_849908, EPI_ISL_849909, EPI_ISL_849910, EPI_ISL_849911, EPI_ISL_849912, EPI_ISL_849921 | Utah Public Health Laboratory | Utah Public Health Laboratory | Erin L. Young, Kelly F. Oakeson, Tara Gallagher |
| EPI_ISL_849923 | UCSF Clinical Microbiology Laboratory | Chan-Zuckerberg Biohub | CZB Cliahub Consortium |
| EPI_ISL_850677, EPI_ISL_850679, EPI_ISL_850684, EPI_ISL_850685, EPI_ISL_850686, EPI_ISL_850688 | The National Institute of Public Health | State Veterinary Institute Prague | Nagy,A;Jirincova,H;Trnka,D;Vecerova,J |
| EPI_ISL_850689, EPI_ISL_850690, EPI_ISL_850691, EPI_ISL_850692, EPI_ISL_850693, EPI_ISL_850694, EPI_ISL_850695, EPI_ISL_850696, EPI_ISL_850697, EPI_ISL_850698, EPI_ISL_850699, EPI_ISL_850700, EPI_ISL_850701, EPI_ISL_850702, EPI_ISL_850703, EPI_ISL_850704, EPI_ISL_850705, EPI_ISL_850706, EPI_ISL_850707, EPI_ISL_850708, EPI_ISL_850709, EPI_ISL_850710, EPI_ISL_850711, EPI_ISL_850712, EPI_ISL_850713, EPI_ISL_850714, EPI_ISL_850715, EPI_ISL_850716, EPI_ISL_850717, EPI_ISL_850718, EPI_ISL_850719, EPI_ISL_850720, EPI_ISL_850721, EPI_ISL_850722, EPI_ISL_850723, EPI_ISL_850724, EPI_ISL_850725, EPI_ISL_850726, EPI_ISL_850727, EPI_ISL_850728, EPI_ISL_850729, EPI_ISL_850730, EPI_ISL_850731, EPI_ISL_850732, EPI_ISL_850733, EPI_ISL_850734, EPI_ISL_850735, EPI_ISL_850736, EPI_ISL_850737, EPI_ISL_850738, EPI_ISL_850739, EPI_ISL_850740, EPI_ISL_850741, EPI_ISL_850742, EPI_ISL_850743, EPI_ISL_850744, EPI_ISL_850745, EPI_ISL_850746, EPI_ISL_850747, EPI_ISL_850748, EPI_ISL_850749, EPI_ISL_850750, EPI_ISL_850751, EPI_ISL_850752, EPI_ISL_850753, EPI_ISL_850754, EPI_ISL_850755, EPI_ISL_850756, EPI_ISL_850757, EPI_ISL_850758, EPI_ISL_850759, EPI_ISL_850760, EPI_ISL_850761, EPI_ISL_850762, EPI_ISL_850763, EPI_ISL_850764, EPI_ISL_850765, EPI_ISL_850766, EPI_ISL_850767, EPI_ISL_850768, EPI_ISL_850769, EPI_ISL_850770, EPI_ISL_850771, EPI_ISL_850772, EPI_ISL_850773, EPI_ISL_850774, EPI_ISL_850775, EPI_ISL_850776, EPI_ISL_850777, EPI_ISL_850778, EPI_ISL_850779, EPI_ISL_850780, EPI_ISL_850781, EPI_ISL_850782, EPI_ISL_850783, EPI_ISL_850784, EPI_ISL_850785, EPI_ISL_850786, EPI_ISL_850787, EPI_ISL_850788, EPI_ISL_850789, EPI_ISL_850790, EPI_ISL_850791, EPI_ISL_850792, EPI_ISL_850793, EPI_ISL_850794, EPI_ISL_850795, EPI_ISL_850796, EPI_ISL_850797, EPI_ISL_850798, EPI_ISL_850799, EPI_ISL_850800, EPI_ISL_850801, EPI_ISL_850802, EPI_ISL_850803, EPI_ISL_850804, EPI_ISL_850805, EPI_ISL_850806, EPI_ISL_850807, EPI_ISL_850808, EPI_ISL_850809, EPI_ISL_850810, EPI_ISL_850811, EPI_ISL_850812, EPI_ISL_850813, EPI_ISL_850814, EPI_ISL_850815, EPI_ISL_850816 | | | |
| see above | Helix / Illumina | Genomics and Discovery, Respiratory Viruses Branch, Division of Viral Diseases, Centers for Disease Control and Prevention | Peter W. Cook, Dhwani Batra, Ben L. Rambo-Martin Eileen de Feo, Jan Antico, Christine Tran, Matthew Tolentino, Shannon Wickline, Kim Gietzen, Brad Sickler, Jingtao Liu, Eric Allen, Phil Febbo, Summer Galloway, Nicole L. Washington, Simon White, Geraint Levan, Kelly Schiabor Barrett, Elizabeth Cirulli, Alexandre Bolze, Ary Ascencio, Charlotte Rivera-Garcia, Ryan Cho, Jason Nguyen, Sherry Wang, Jimmy Ramirez, Tyler Cassens, Efren Sandoval, Magnus Isaksson, William Lee, David Becker, Marc Laurent, James Lu, Clinton R. Paden, Suxiang Tong, Duncan MacCannell |
| EPI_ISL_851062, EPI_ISL_851063, EPI_ISL_851079, EPI_ISL_851083, EPI_ISL_851110, EPI_ISL_851115, EPI_ISL_851133, EPI_ISL_851143, EPI_ISL_851150, EPI_ISL_851166, EPI_ISL_851170, EPI_ISL_851175, EPI_ISL_851189, EPI_ISL_851193, EPI_ISL_851194, EPI_ISL_851202, EPI_ISL_851208, EPI_ISL_851213, EPI_ISL_851214, EPI_ISL_851251, EPI_ISL_851254, EPI_ISL_851269, EPI_ISL_851273, EPI_ISL_851287, EPI_ISL_851298, EPI_ISL_851332, EPI_ISL_851338, EPI_ISL_851342, EPI_ISL_851343, EPI_ISL_851353, EPI_ISL_851354, EPI_ISL_851370, EPI_ISL_851377 | | | |
| see above | Lighthouse Lab in Cambridge | Wellcome Sanger Institute for the COVID-19 Genomics UK (COG-UK) Consortium | Rob Howes, The Lighthouse Lab in Cambridge and Alex Alderton, Roberto Amato, Sonia Goncalves, Ewan Harrison, David K. Jackson, Ian Johnston, Dominic Kwiatkowski, Cordelia Langford, John Sillitoe on behalf of the Wellcome Sanger Institute COVID-19 Surveillance Team |
| EPI_ISL_851390, EPI_ISL_851395, EPI_ISL_851413, EPI_ISL_851423, EPI_ISL_851436, EPI_ISL_851448, EPI_ISL_851454, EPI_ISL_851455, EPI_ISL_851461, EPI_ISL_851467, EPI_ISL_851484, EPI_ISL_851514, EPI_ISL_851559, EPI_ISL_851567, EPI_ISL_851572, EPI_ISL_851582, EPI_ISL_851590, EPI_ISL_851594, EPI_ISL_851598, EPI_ISL_851601, EPI_ISL_851614, EPI_ISL_851617, EPI_ISL_851629, EPI_ISL_851633, EPI_ISL_851635, EPI_ISL_851646, EPI_ISL_851648, EPI_ISL_851657, EPI_ISL_851664 | | | |
| see above | Lighthouse Lab in Glasgow | Wellcome Sanger Institute for the COVID-19 Genomics UK (COG-UK) Consortium | Harper VanSteenhouse, Yumi Kasai, David Gray, Carol Clugston, Anna Dominiczak and Alex Alderton, Roberto Amato, Sonia Goncalves, Ewan Harrison, David K. Jackson, Ian Johnston, Dominic Kwiatkowski, Cordelia Langford, John Sillitoe on behalf of the Wellcome Sanger Institute COVID-19 Surveillance Team |
| EPI_ISL_851750, EPI_ISL_851875 | Lighthouse Lab in Alderley Park | Wellcome Sanger Institute for the COVID-19 Genomics UK (COG-UK) Consortium | Jacquelyn Wynn, Mairead Hyland, The Lighthouse Lab in Alderley Park and Alex Alderton, Roberto Amato, Sonia Goncalves, Ewan Harrison, David K. Jackson, Ian Johnston, Dominic Kwiatkowski, Cordelia Langford, John Sillitoe on behalf of the Wellcome Sanger Institute COVID-19 Surveillance Team |

| | | | |
|---|---|---|---|
| EPI_ISL_851954 | Lighthouse Lab in Glasgow | Wellcome Sanger Institute for the COVID-19 Genomics UK (COG-UK) Consortium | Harper VanSteenhouse, Yumi Kasai, David Gray, Carol Clugston, Anna Dominiczak and Alex Alderton, Roberto Amato, Sonia Goncalves, Ewan Harrison, David K. Jackson, Ian Johnston, Dominic Kwiatkowski, Cordelia Langford, John Sillitoe on behalf of the Wellcome Sanger Institute COVID-19 Surveillance Team |
| EPI_ISL_851963 | Lighthouse Lab in Milton Keynes | Wellcome Sanger Institute for the COVID-19 Genomics UK (COG-UK) Consortium | The Lighthouse Lab in Milton Keynes and Alex Alderton, Roberto Amato, Sonia Goncalves, Ewan Harrison, David K. Jackson, Ian Johnston, Dominic Kwiatkowski, Cordelia Langford, John Sillitoe on behalf of the Wellcome Sanger Institute COVID-19 Surveillance Team |
| EPI_ISL_851964 | Lighthouse Lab in Alderley Park | Wellcome Sanger Institute for the COVID-19 Genomics UK (COG-UK) Consortium | Jacquelyn Wynn, Mairead Hyland, The Lighthouse Lab in Alderley Park and Alex Alderton, Roberto Amato, Sonia Goncalves, Ewan Harrison, David K. Jackson, Ian Johnston, Dominic Kwiatkowski, Cordelia Langford, John Sillitoe on behalf of the Wellcome Sanger Institute COVID-19 Surveillance Team |
| EPI_ISL_851968, EPI_ISL_851972, EPI_ISL_851974, EPI_ISL_851976, EPI_ISL_851979, EPI_ISL_851985, EPI_ISL_851993, EPI_ISL_852002, EPI_ISL_852009 | Lighthouse Lab in Milton Keynes | Wellcome Sanger Institute for the COVID-19 Genomics UK (COG-UK) Consortium | The Lighthouse Lab in Milton Keynes and Alex Alderton, Roberto Amato, Sonia Goncalves, Ewan Harrison, David K. Jackson, Ian Johnston, Dominic Kwiatkowski, Cordelia Langford, John Sillitoe on behalf of the Wellcome Sanger Institute COVID-19 Surveillance Team |
| EPI_ISL_852025 | Lighthouse Lab in Alderley Park | Wellcome Sanger Institute for the COVID-19 Genomics UK (COG-UK) Consortium | Jacquelyn Wynn, Mairead Hyland, The Lighthouse Lab in Alderley Park and Alex Alderton, Roberto Amato, Sonia Goncalves, Ewan Harrison, David K. Jackson, Ian Johnston, Dominic Kwiatkowski, Cordelia Langford, John Sillitoe on behalf of the Wellcome Sanger Institute COVID-19 Surveillance Team |
| EPI_ISL_852029, EPI_ISL_852042, EPI_ISL_852062, EPI_ISL_852077, EPI_ISL_852086, EPI_ISL_852093, EPI_ISL_852100, EPI_ISL_852114, EPI_ISL_852118, EPI_ISL_852124, EPI_ISL_852127, EPI_ISL_852132, EPI_ISL_852135, EPI_ISL_852165, EPI_ISL_852174 | | | |
| see above | Lighthouse Lab in Milton Keynes | Wellcome Sanger Institute for the COVID-19 Genomics UK (COG-UK) Consortium | The Lighthouse Lab in Milton Keynes and Alex Alderton, Roberto Amato, Sonia Goncalves, Ewan Harrison, David K. Jackson, Ian Johnston, Dominic Kwiatkowski, Cordelia Langford, John Sillitoe on behalf of the Wellcome Sanger Institute COVID-19 Surveillance Team |
| EPI_ISL_852183 | Lighthouse Lab in Alderley Park | Wellcome Sanger Institute for the COVID-19 Genomics UK (COG-UK) Consortium | Jacquelyn Wynn, Mairead Hyland, The Lighthouse Lab in Alderley Park and Alex Alderton, Roberto Amato, Sonia Goncalves, Ewan Harrison, David K. Jackson, Ian Johnston, Dominic Kwiatkowski, Cordelia Langford, John Sillitoe on behalf of the Wellcome Sanger Institute COVID-19 Surveillance Team |
| EPI_ISL_852184, EPI_ISL_852190, EPI_ISL_852203, EPI_ISL_852206, EPI_ISL_852213 | Lighthouse Lab in Milton Keynes | Wellcome Sanger Institute for the COVID-19 Genomics UK (COG-UK) Consortium | The Lighthouse Lab in Milton Keynes and Alex Alderton, Roberto Amato, Sonia Goncalves, Ewan Harrison, David K. Jackson, Ian Johnston, Dominic Kwiatkowski, Cordelia Langford, John Sillitoe on behalf of the Wellcome Sanger Institute COVID-19 Surveillance Team |
| EPI_ISL_852214 | Lighthouse Lab in Alderley Park | Wellcome Sanger Institute for the COVID-19 Genomics UK (COG-UK) Consortium | Jacquelyn Wynn, Mairead Hyland, The Lighthouse Lab in Alderley Park and Alex Alderton, Roberto Amato, Sonia Goncalves, Ewan Harrison, David K. Jackson, Ian Johnston, Dominic Kwiatkowski, Cordelia Langford, John Sillitoe on behalf of the Wellcome Sanger Institute COVID-19 Surveillance Team |
| EPI_ISL_852218, EPI_ISL_852234, EPI_ISL_852236 | Lighthouse Lab in Milton Keynes | Wellcome Sanger Institute for the COVID-19 Genomics UK (COG-UK) Consortium | The Lighthouse Lab in Milton Keynes and Alex Alderton, Roberto Amato, Sonia Goncalves, Ewan Harrison, David K. Jackson, Ian Johnston, Dominic Kwiatkowski, Cordelia Langford, John Sillitoe on behalf of the Wellcome Sanger Institute COVID-19 Surveillance Team |
| EPI_ISL_852256, EPI_ISL_852257, EPI_ISL_852258, EPI_ISL_852259, EPI_ISL_852260, EPI_ISL_852261, EPI_ISL_852262, EPI_ISL_852263, EPI_ISL_852264, EPI_ISL_852265, EPI_ISL_852266, EPI_ISL_852267, EPI_ISL_852269, EPI_ISL_852270, EPI_ISL_852271, EPI_ISL_852272, EPI_ISL_852273, EPI_ISL_852274, EPI_ISL_852275, EPI_ISL_852276, EPI_ISL_852277, EPI_ISL_852278, EPI_ISL_852279, EPI_ISL_852280, EPI_ISL_852281, EPI_ISL_852282, EPI_ISL_852283, EPI_ISL_852284, EPI_ISL_852285, EPI_ISL_852287, EPI_ISL_852288, EPI_ISL_852289, EPI_ISL_852290, EPI_ISL_852291, EPI_ISL_852292, EPI_ISL_852293, EPI_ISL_852294, EPI_ISL_852295, EPI_ISL_852296, EPI_ISL_852297, EPI_ISL_852298, EPI_ISL_852299, EPI_ISL_852300, EPI_ISL_852301, EPI_ISL_852302, EPI_ISL_852304, EPI_ISL_852305, EPI_ISL_852306, EPI_ISL_852307, EPI_ISL_852308, EPI_ISL_852309, EPI_ISL_852310, EPI_ISL_852311, EPI_ISL_852312, EPI_ISL_852313, EPI_ISL_852314, EPI_ISL_852315, EPI_ISL_852316, EPI_ISL_852317, EPI_ISL_852318, EPI_ISL_852320, EPI_ISL_852321, EPI_ISL_852322, EPI_ISL_852323, EPI_ISL_852324, EPI_ISL_852325, EPI_ISL_852326, EPI_ISL_852327, EPI_ISL_852328, EPI_ISL_852329, EPI_ISL_852330, EPI_ISL_852331, EPI_ISL_852332, EPI_ISL_852333, EPI_ISL_852334, EPI_ISL_852335, EPI_ISL_852336, EPI_ISL_852337, EPI_ISL_852339, EPI_ISL_852340, EPI_ISL_852341, EPI_ISL_852342, EPI_ISL_852343, EPI_ISL_852344, EPI_ISL_852345, EPI_ISL_852346, EPI_ISL_852347, EPI_ISL_852348, EPI_ISL_852349, EPI_ISL_852350, EPI_ISL_852351, EPI_ISL_852352, EPI_ISL_852353, EPI_ISL_852354, EPI_ISL_852355, EPI_ISL_852356, EPI_ISL_852357, EPI_ISL_852358, EPI_ISL_852359, EPI_ISL_852360, EPI_ISL_852362, EPI_ISL_852363, EPI_ISL_852364, EPI_ISL_852365, EPI_ISL_852366, EPI_ISL_852367, EPI_ISL_852368, EPI_ISL_852369, EPI_ISL_852370, EPI_ISL_852371, EPI_ISL_852372, EPI_ISL_852373, EPI_ISL_852374, EPI_ISL_852376, EPI_ISL_852377, EPI_ISL_852378, EPI_ISL_852379, EPI_ISL_852380, EPI_ISL_852381, EPI_ISL_852382, EPI_ISL_852384, EPI_ISL_852385, EPI_ISL_852386, EPI_ISL_852387, EPI_ISL_852388, EPI_ISL_852390, EPI_ISL_852391, EPI_ISL_852392, EPI_ISL_852393, EPI_ISL_852394, EPI_ISL_852395, EPI_ISL_852396, EPI_ISL_852397, EPI_ISL_852398, EPI_ISL_852399, EPI_ISL_852400, EPI_ISL_852401, EPI_ISL_852402, EPI_ISL_852403, EPI_ISL_852404, EPI_ISL_852405, EPI_ISL_852408, EPI_ISL_852409, EPI_ISL_852410, EPI_ISL_852411, EPI_ISL_852412, EPI_ISL_852415, EPI_ISL_852417, EPI_ISL_852418, EPI_ISL_852419, EPI_ISL_852420, EPI_ISL_852421, EPI_ISL_852422, EPI_ISL_852423, EPI_ISL_852424, EPI_ISL_852425, EPI_ISL_852426, EPI_ISL_852427, EPI_ISL_852428, EPI_ISL_852429, EPI_ISL_852430, EPI_ISL_852431, EPI_ISL_852432, EPI_ISL_852433, EPI_ISL_852434, EPI_ISL_852435, EPI_ISL_852436, EPI_ISL_852437, EPI_ISL_852439, EPI_ISL_852440, EPI_ISL_852441, EPI_ISL_852442, EPI_ISL_852443, EPI_ISL_852444, EPI_ISL_852445, EPI_ISL_852446, EPI_ISL_852447, EPI_ISL_852449, EPI_ISL_852450, EPI_ISL_852451, EPI_ISL_852452, EPI_ISL_852453, EPI_ISL_852454, EPI_ISL_852455, EPI_ISL_852456, EPI_ISL_852457, EPI_ISL_852458, EPI_ISL_852460, EPI_ISL_852461, EPI_ISL_852462, EPI_ISL_852463, EPI_ISL_852464, EPI_ISL_852465, EPI_ISL_852466, EPI_ISL_852467, EPI_ISL_852468, EPI_ISL_852469, EPI_ISL_852470, EPI_ISL_852471, EPI_ISL_852472, EPI_ISL_852473, EPI_ISL_852474, EPI_ISL_852475, EPI_ISL_852476, EPI_ISL_852477, EPI_ISL_852478, EPI_ISL_852479, EPI_ISL_852480, EPI_ISL_852481, EPI_ISL_852482, EPI_ISL_852483, EPI_ISL_852484, EPI_ISL_852485, EPI_ISL_852486, EPI_ISL_852487, EPI_ISL_852488, EPI_ISL_852489, EPI_ISL_852490, EPI_ISL_852491, EPI_ISL_852492, EPI_ISL_852493, EPI_ISL_852494, EPI_ISL_852495, EPI_ISL_852496, EPI_ISL_852497, EPI_ISL_852498, EPI_ISL_852499, EPI_ISL_852501, EPI_ISL_852502, EPI_ISL_852503, EPI_ISL_852504, EPI_ISL_852505, EPI_ISL_852506, EPI_ISL_852507, EPI_ISL_852508, EPI_ISL_852509, EPI_ISL_852510, EPI_ISL_852511, EPI_ISL_852512, EPI_ISL_852513, EPI_ISL_852514, EPI_ISL_852515, EPI_ISL_852516, EPI_ISL_852518, EPI_ISL_852519, EPI_ISL_852520, EPI_ISL_852521, EPI_ISL_852522, EPI_ISL_852523, EPI_ISL_852525, EPI_ISL_852526, EPI_ISL_852528, EPI_ISL_852529, EPI_ISL_852530, EPI_ISL_852531, EPI_ISL_852532, EPI_ISL_852533, EPI_ISL_852534, EPI_ISL_852537, EPI_ISL_852538, EPI_ISL_852539, EPI_ISL_852540, EPI_ISL_852541, EPI_ISL_852542, EPI_ISL_852543, EPI_ISL_852545, EPI_ISL_852546, EPI_ISL_852547, EPI_ISL_852548, EPI_ISL_852550, EPI_ISL_852551, EPI_ISL_852552, EPI_ISL_852553, EPI_ISL_852554, EPI_ISL_852555, EPI_ISL_852556, EPI_ISL_852557, EPI_ISL_852558, EPI_ISL_852559, EPI_ISL_852560, EPI_ISL_852561, EPI_ISL_852562, EPI_ISL_852563 | | | |
| see above | Lighthouse Lab in Glasgow | Wellcome Sanger Institute for the COVID-19 Genomics UK (COG-UK) Consortium | Harper VanSteenhouse, Yumi Kasai, David Gray, Carol Clugston, Anna Dominiczak and Alex Alderton, Roberto Amato, Sonia Goncalves, Ewan Harrison, David K. Jackson, Ian Johnston, Dominic Kwiatkowski, Cordelia Langford, John Sillitoe on behalf of the Wellcome Sanger Institute COVID-19 Surveillance Team |
| EPI_ISL_852861 | Florida Bureau of Public Health Laboratories | Florida Bureau of Public Health Laboratories | Sarah Schmedes, Jason Blanton |
| EPI_ISL_852947, EPI_ISL_852948, EPI_ISL_852950, EPI_ISL_852954, EPI_ISL_852964, EPI_ISL_853003, EPI_ISL_853009, EPI_ISL_853010, EPI_ISL_853011, EPI_ISL_853012, EPI_ISL_853024, EPI_ISL_853025, EPI_ISL_853026, EPI_ISL_853027, EPI_ISL_853028, EPI_ISL_853029, EPI_ISL_853032 | | | |
| see above | Hospital General Universitario Gregorio Marañón | SeqCOVID-SPAIN consortium/IBV(CSIC) | Darío García de Viedma, Laura Pérez-Lago, Pedro J Sola-Campoy, Sergio Buenestado-Serrano, Marta Herranz, Victor Manuel de la Cueva, Julia Suárez, Pilar Catalán, Patricia Muñoz and SeqCOVID-SPAIN consortium |
| EPI_ISL_853397, EPI_ISL_853398, EPI_ISL_853399 | Charité Universitätsmedizin Berlin, Institut für Virologie/Labor Berlin | Charité Universitätsmedizin Berlin, Institut für Virologie | Victor M Corman, Julia Schneider, Barbara Mühlemann, Jörn Beheim-Schwarzbach, Talitha Veith, Julia Tesch, Tobias Bleicker, Terry Jones, Christian Drosten |
| EPI_ISL_853649 | THE MARY IMOGENE BASSETT HOSPITAL | Wadsworth Center, New York State Department of Health | Kirsten St. George, Daryl M. Lamson, Alexis Russel, Matthew Shudt, Melissa A Leisner, Jonathan Plitnick, Navjot Singh, John Kelly, Erasmus Schneider, Erica Lasek-Nesselquist |
| EPI_ISL_853650, EPI_ISL_853651 | NORTHWELL HEALTH LABORATORIES | Wadsworth Center, New York State Department of Health | Kirsten St. George, Daryl M. Lamson, Alexis Russel, Matthew Shudt, Melissa A Leisner, Jonathan Plitnick, Navjot Singh, John Kelly, Erasmus Schneider, Erica Lasek-Nesselquist |
| EPI_ISL_853652 | THE MARY IMOGENE BASSETT HOSPITAL | Wadsworth Center, New York State Department of Health | Kirsten St. George, Daryl M. Lamson, Alexis Russel, Matthew Shudt, Melissa A Leisner, Jonathan Plitnick, Navjot Singh, John Kelly, Erasmus Schneider, Erica Lasek-Nesselquist |
| EPI_ISL_853653, EPI_ISL_853654, EPI_ISL_853655, EPI_ISL_853656, EPI_ISL_853657, EPI_ISL_853658, EPI_ISL_853659, EPI_ISL_853660, EPI_ISL_853661, EPI_ISL_853662, EPI_ISL_853663, EPI_ISL_853664, EPI_ISL_853665, EPI_ISL_853666 | | | |
| see above | NORTHWELL HEALTH LABORATORIES | Wadsworth Center, New York State Department of Health | Kirsten St. George, Daryl M. Lamson, Alexis Russel, Matthew Shudt, Melissa A Leisner, Jonathan Plitnick, Navjot Singh, John Kelly, Erasmus Schneider, Erica Lasek-Nesselquist |
| EPI_ISL_853667 | THE MARY IMOGENE BASSETT HOSPITAL | Wadsworth Center, New York State Department of Health | Kirsten St. George, Daryl M. Lamson, Alexis Russel, Matthew Shudt, Melissa A Leisner, Jonathan Plitnick, Navjot Singh, John Kelly, Erasmus Schneider, Erica Lasek-Nesselquist |
| EPI_ISL_853713, EPI_ISL_853714, EPI_ISL_853716, EPI_ISL_853717 | NORTHWELL HEALTH LABORATORIES | Wadsworth Center, New York State Department of Health | Kirsten St. George, Daryl M. Lamson, Alexis Russel, Matthew Shudt, Melissa A Leisner, Jonathan Plitnick, Navjot Singh, John Kelly, Erasmus Schneider, Erica Lasek-Nesselquist |
| EPI_ISL_854305 | ESOTERIX GENETIC LABORATORIES LLC | Wadsworth Center, New York State Department of Health | Kirsten St. George, Daryl M. Lamson, Alexis Russel, Matthew Shudt, Melissa A Leisner, Jonathan Plitnick, Navjot Singh, John Kelly, Erasmus Schneider, Erica Lasek-Nesselquist |
| EPI_ISL_854307, EPI_ISL_854308, EPI_ISL_854309, EPI_ISL_854310, EPI_ISL_854311, EPI_ISL_854312, EPI_ISL_854313, EPI_ISL_854314, EPI_ISL_854315, EPI_ISL_854316, EPI_ISL_854317, EPI_ISL_854318, EPI_ISL_854319, EPI_ISL_854320 | | | |
| see above | URMC LABS | Wadsworth Center, New York State Department of Health | Kirsten St. George, Daryl M. Lamson, Alexis Russel, Matthew Shudt, Melissa A Leisner, Jonathan Plitnick, Navjot Singh, John Kelly, Erasmus Schneider, Erica Lasek-Nesselquist |
| EPI_ISL_854368 | Wadsworth Center, New York State Department of Health | Wadsworth Center, New York State Department of Health | Kirsten St. George, Daryl M. Lamson, Alexis Russel, Matthew Shudt, Melissa A Leisner, Jonathan Plitnick, Navjot Singh, John Kelly, Erasmus Schneider, Erica Lasek-Nesselquist |
| EPI_ISL_854381, EPI_ISL_854382, EPI_ISL_854383, EPI_ISL_854384, EPI_ISL_854385, EPI_ISL_854386, EPI_ISL_854387, EPI_ISL_854388, EPI_ISL_854389, EPI_ISL_854390, EPI_ISL_854391, EPI_ISL_854392, EPI_ISL_854393, EPI_ISL_854394, EPI_ISL_854395, EPI_ISL_854396, EPI_ISL_854397, EPI_ISL_854398 | | | |
| see above | WESTCHESTER MEDICAL CENTER | Wadsworth Center, New York State Department of Health | Kirsten St. George, Daryl M. Lamson, Alexis Russel, Matthew Shudt, Melissa A Leisner, Jonathan Plitnick, Navjot Singh, John Kelly, Erasmus Schneider, |

| | | | |
|---|---|---|---|
| | | | Erica Lasek-Nesselquist |
| EPI_ISL_854399 | URMC LABS | Wadsworth Center, New York State Department of Health | Kirsten St. George, Daryl M. Lamson, Alexis Russel, Matthew Shudt, Melissa A Leisner, Jonathan Plitnick, Navjot Singh, John Kelly, Erasmus Schneider, Erica Lasek-Nesselquist |
| EPI_ISL_854400, EPI_ISL_854401, EPI_ISL_854402, EPI_ISL_854403 | WESTCHESTER MEDICAL CENTER | Wadsworth Center, New York State Department of Health | Kirsten St. George, Daryl M. Lamson, Alexis Russel, Matthew Shudt, Melissa A Leisner, Jonathan Plitnick, Navjot Singh, John Kelly, Erasmus Schneider, Erica Lasek-Nesselquist |
| EPI_ISL_854446 | BOSTON HEART DIAGNOSTICS CORP | Wadsworth Center, New York State Department of Health | Kirsten St. George, Daryl M. Lamson, Alexis Russel, Matthew Shudt, Melissa A Leisner, Jonathan Plitnick, Navjot Singh, John Kelly, Erasmus Schneider, Erica Lasek-Nesselquist |
| EPI_ISL_854447, EPI_ISL_854448 | TEMPUS LABS INC | Wadsworth Center, New York State Department of Health | Kirsten St. George, Daryl M. Lamson, Alexis Russel, Matthew Shudt, Melissa A Leisner, Jonathan Plitnick, Navjot Singh, John Kelly, Erasmus Schneider, Erica Lasek-Nesselquist |
| EPI_ISL_854449 | WESTCHESTER MEDICAL CENTER | Wadsworth Center, New York State Department of Health | Kirsten St. George, Daryl M. Lamson, Alexis Russel, Matthew Shudt, Melissa A Leisner, Jonathan Plitnick, Navjot Singh, John Kelly, Erasmus Schneider, Erica Lasek-Nesselquist |
| EPI_ISL_854460 | Northwestern Memorial Hospital | Ozer Lab | Ramon Lorenzo-Redondo, Lacy M. Simons, Lawrence J. Jennings, Michael G. Ison, Judd F. Hultquist, Egon A. Ozer |
| EPI_ISL_854745 | Royal Darwin Hospital Pathology | MDU-PHL | Meumann, E., Caly L., Seemann T., Sait, M.L., Druce J., Sherry, N.L. |
| EPI_ISL_854810, EPI_ISL_854813, EPI_ISL_854816, EPI_ISL_854817, EPI_ISL_854818, EPI_ISL_854820, EPI_ISL_854824, EPI_ISL_854828, EPI_ISL_854829, EPI_ISL_854830, EPI_ISL_854831, EPI_ISL_854832, EPI_ISL_854833, EPI_ISL_854834, EPI_ISL_854842, EPI_ISL_854844, EPI_ISL_854845, EPI_ISL_854846, EPI_ISL_854847, EPI_ISL_854848, EPI_ISL_854849, EPI_ISL_854851, EPI_ISL_854852, EPI_ISL_854853, EPI_ISL_854854, EPI_ISL_854855, EPI_ISL_854856, EPI_ISL_854857, EPI_ISL_854858, EPI_ISL_854859, EPI_ISL_854860, EPI_ISL_854861, EPI_ISL_854863, EPI_ISL_854864, EPI_ISL_854865, EPI_ISL_854866, EPI_ISL_854868, EPI_ISL_854869, EPI_ISL_854870, EPI_ISL_854871, EPI_ISL_854872, EPI_ISL_854873, EPI_ISL_854874, EPI_ISL_854875, EPI_ISL_854876, EPI_ISL_854877, EPI_ISL_854878, EPI_ISL_854879, EPI_ISL_854880, EPI_ISL_854881, EPI_ISL_854882, EPI_ISL_854883, EPI_ISL_854884, EPI_ISL_854886, EPI_ISL_854887, EPI_ISL_854888, EPI_ISL_854889, EPI_ISL_854891, EPI_ISL_854893, EPI_ISL_854894, EPI_ISL_854895, EPI_ISL_854896, EPI_ISL_854897, EPI_ISL_854899, EPI_ISL_854900, EPI_ISL_854901, EPI_ISL_854902, EPI_ISL_854903, EPI_ISL_854904, EPI_ISL_854905, EPI_ISL_854906, EPI_ISL_854907, EPI_ISL_854908, EPI_ISL_854909, EPI_ISL_854910, EPI_ISL_854911, EPI_ISL_854912, EPI_ISL_854913, EPI_ISL_854914, EPI_ISL_854915, EPI_ISL_854916, EPI_ISL_854917, EPI_ISL_854918, EPI_ISL_854919, EPI_ISL_854920, EPI_ISL_854921, EPI_ISL_854922, EPI_ISL_854923, EPI_ISL_854924, EPI_ISL_854925, EPI_ISL_854926, EPI_ISL_854927, EPI_ISL_854928, EPI_ISL_854929, EPI_ISL_854930, EPI_ISL_854931, EPI_ISL_854932, EPI_ISL_854933, EPI_ISL_854934, EPI_ISL_854935, EPI_ISL_854936, EPI_ISL_854937, EPI_ISL_854938, EPI_ISL_854939, EPI_ISL_854940, EPI_ISL_854941, EPI_ISL_854942, EPI_ISL_854944, EPI_ISL_854945, EPI_ISL_854946, EPI_ISL_854947, EPI_ISL_854948, EPI_ISL_854949, EPI_ISL_854950, EPI_ISL_854951, EPI_ISL_854952, EPI_ISL_854953, EPI_ISL_854954, EPI_ISL_854955, EPI_ISL_854956, EPI_ISL_854957, EPI_ISL_854958, EPI_ISL_854959, EPI_ISL_854960, EPI_ISL_854961, EPI_ISL_854962, EPI_ISL_854966, EPI_ISL_854968, EPI_ISL_854969, EPI_ISL_854970, EPI_ISL_854971, EPI_ISL_854972, EPI_ISL_854973, EPI_ISL_854974, EPI_ISL_854975, EPI_ISL_854976, EPI_ISL_855015, EPI_ISL_855018, EPI_ISL_855020, EPI_ISL_855021, EPI_ISL_855024, EPI_ISL_855025, EPI_ISL_855026, EPI_ISL_855028, EPI_ISL_855029, EPI_ISL_855032, EPI_ISL_855036, EPI_ISL_855040, EPI_ISL_855042, EPI_ISL_855043, EPI_ISL_855044, EPI_ISL_855045, EPI_ISL_855048, EPI_ISL_855051, EPI_ISL_855056, EPI_ISL_855064, EPI_ISL_855065, EPI_ISL_855066, EPI_ISL_855067, EPI_ISL_855068, EPI_ISL_855069, EPI_ISL_855071, EPI_ISL_855072, EPI_ISL_855073, EPI_ISL_855074, EPI_ISL_855075, EPI_ISL_855081, EPI_ISL_855082, EPI_ISL_855083, EPI_ISL_855084, EPI_ISL_855085, EPI_ISL_855086, EPI_ISL_855087, EPI_ISL_855088, EPI_ISL_855089, EPI_ISL_855090, EPI_ISL_855091, EPI_ISL_855092, EPI_ISL_855093, EPI_ISL_855094, EPI_ISL_855095, EPI_ISL_855096, EPI_ISL_855097, EPI_ISL_855098, EPI_ISL_855099, EPI_ISL_855100, EPI_ISL_855101, EPI_ISL_855102, EPI_ISL_855103, EPI_ISL_855104, EPI_ISL_855105, EPI_ISL_855106, EPI_ISL_855107, EPI_ISL_855108, EPI_ISL_855109, EPI_ISL_855110, EPI_ISL_855111, EPI_ISL_855112, EPI_ISL_855113, EPI_ISL_855114, EPI_ISL_855116, EPI_ISL_855117, EPI_ISL_855118, EPI_ISL_855119, EPI_ISL_855120, EPI_ISL_855121, EPI_ISL_855122, EPI_ISL_855123, EPI_ISL_855124, EPI_ISL_855125, EPI_ISL_855126, EPI_ISL_855127, EPI_ISL_855128, EPI_ISL_855130, EPI_ISL_855131, EPI_ISL_855132, EPI_ISL_855133, EPI_ISL_855134, EPI_ISL_855135, EPI_ISL_855136, EPI_ISL_855137, EPI_ISL_855138, EPI_ISL_855139, EPI_ISL_855140, EPI_ISL_855141, EPI_ISL_855142, EPI_ISL_855143, EPI_ISL_855144, EPI_ISL_855145, EPI_ISL_855146, EPI_ISL_855147, EPI_ISL_855148, EPI_ISL_855149, EPI_ISL_855150, EPI_ISL_855151, EPI_ISL_855152, EPI_ISL_855153, EPI_ISL_855155, EPI_ISL_855156, EPI_ISL_855157, EPI_ISL_855158, EPI_ISL_855160, EPI_ISL_855161, EPI_ISL_855162, EPI_ISL_855163, EPI_ISL_855165, EPI_ISL_855166, EPI_ISL_855167, EPI_ISL_855168, EPI_ISL_855169, EPI_ISL_855170, EPI_ISL_855171, EPI_ISL_855172, EPI_ISL_855173, EPI_ISL_855174, EPI_ISL_855175, EPI_ISL_855176, EPI_ISL_855177, EPI_ISL_855178, EPI_ISL_855179, EPI_ISL_855180, EPI_ISL_855181, EPI_ISL_855182, EPI_ISL_855183, EPI_ISL_855185 | | | |
| see above | Quest Diagnostics | Quest Diagnostics | Rosenthal,S.H., Gerasimova,A., Kagan,R.M., Anderson, B., Hua, M., Liu Y., Bernstein, L.E., Livingston, K.E., Perez, A., Shalhout, D.F., Shlyakhter, I.A., Owen, R., Tanpaiboon, P., Lacbawan, F. |
| EPI_ISL_855358 | Hospital | National Reference Center for Viruses of Respiratory Infections, Institut Pasteur, Paris | Marion Barbet, Sylvie Behillil, Méline Bizard, Angela Brisebarre, Camille Capel, Etienne Simon-Lorière, Vincent Enouf, Maud Vanpeene, Sylvie van der Werf,Griscelli Franck |
| EPI_ISL_855359 | Hospital | National Reference Center for Viruses of Respiratory Infections, Institut Pasteur, Paris | Marion Barbet, Sylvie Behillil, Méline Bizard, Angela Brisebarre, Camille Capel, Etienne Simon-Lorière, Vincent Enouf, Maud Vanpeene, Sylvie van der Werf,Fourgeaud Jacques |
| EPI_ISL_855360, EPI_ISL_855361, EPI_ISL_855362, EPI_ISL_855363, EPI_ISL_855364, EPI_ISL_855365, EPI_ISL_855366, EPI_ISL_855367, EPI_ISL_855368, EPI_ISL_855369, EPI_ISL_855370, EPI_ISL_855371, EPI_ISL_855372, EPI_ISL_855373, EPI_ISL_855374, EPI_ISL_855375, EPI_ISL_855376, EPI_ISL_855377, EPI_ISL_855378, EPI_ISL_855379, EPI_ISL_855380 | | | |
| see above | Hospital | National Reference Center for Viruses of Respiratory Infections, Institut Pasteur, Paris | Marion Barbet, Sylvie Behillil, Méline Bizard, Angela Brisebarre, Camille Capel, Etienne Simon-Lorière, Vincent Enouf, Maud Vanpeene, Sylvie van der Werf,Combe Patrice |
| EPI_ISL_855381 | Labo Analyses Med | National Reference Center for Viruses of Respiratory Infections, Institut Pasteur, Paris | Marion Barbet, Sylvie Behillil, Méline Bizard, Angela Brisebarre, Camille Capel, Etienne Simon-Lorière, Vincent Enouf, Maud Vanpeene, Sylvie van der Werf |
| EPI_ISL_855385 | Hospital | National Reference Center for Viruses of Respiratory Infections, Institut Pasteur, Paris | Marion Barbet, Sylvie Behillil, Méline Bizard, Angela Brisebarre, Camille Capel, Etienne Simon-Lorière, Vincent Enouf, Maud Vanpeene, Sylvie van der Werf |
| EPI_ISL_855551 | Cerballiance Wilson | CERBA LAB | Roquebert B; Merah K; Olivi M; Herhira S; Lecorche E; Trombert S; Verdurme L; Malek R; Zimmer S; Costa JM; Haïm-Boukobza S |
| EPI_ISL_855574, EPI_ISL_855575, EPI_ISL_855577, EPI_ISL_855578, EPI_ISL_855579, EPI_ISL_855581, EPI_ISL_855582, EPI_ISL_855584, EPI_ISL_855589, EPI_ISL_855590, EPI_ISL_855591, EPI_ISL_855592, EPI_ISL_855593, EPI_ISL_855594, EPI_ISL_855595, EPI_ISL_855596, EPI_ISL_855597, EPI_ISL_855598, EPI_ISL_855599, EPI_ISL_855600, EPI_ISL_855601, EPI_ISL_855602, EPI_ISL_855606, EPI_ISL_855607, EPI_ISL_855608, EPI_ISL_855609 | | | |
| see above | Respiratory Virus Unit, National Infection Service, Public Health England | COVID-19 Genomics UK (COG-UK) Consortium | PHE Covid Sequencing Team |
| EPI_ISL_855904, EPI_ISL_855930 | Lab voor klinische biologie | Onderzoeksgroep Virologie | Laurens Lambrechts, Nick Vereecke, Marthe Pauwels, Bruno Verhasselt, Linos Vandekerckhove, Hans Nauwynck, Sebastiaan Theuns |
| EPI_ISL_855936, EPI_ISL_855938, EPI_ISL_855944, EPI_ISL_855945, EPI_ISL_855947 | Lab voor klinische biologie | Onderzoeksgroep Virologie | Nick Vereecke, Laurens Lambrechts, Marthe Pauwels, Bruno Verhasselt, Linos Vandekerckhove, Hans Nauwynck, Sebastiaan Theuns |
| EPI_ISL_856703, EPI_ISL_856704, EPI_ISL_856705, EPI_ISL_856706 | Respiratory Virus Unit, National Infection Service, Public Health England | COVID-19 Genomics UK (COG-UK) Consortium | PHE Covid Sequencing Team |
| EPI_ISL_856711 | Wyoming Public Health Laboratory | Wyoming Public Health Laboratory | Noah Hull, Kevin Libuit, Joel Sevinsky, Taylor Fearing, Lynette Gumbleton, Channing Weber, Ashley Norberg, Bailey Bowcutt, and Wanda Manley |
| EPI_ISL_856913, EPI_ISL_856914, EPI_ISL_856915, EPI_ISL_856916, EPI_ISL_856917, EPI_ISL_856918, EPI_ISL_856919, EPI_ISL_856932, EPI_ISL_856933, EPI_ISL_856935, EPI_ISL_856936, EPI_ISL_856937, EPI_ISL_856945, EPI_ISL_856946, EPI_ISL_856947, EPI_ISL_856948, EPI_ISL_856949, EPI_ISL_856950, EPI_ISL_856951, EPI_ISL_856952, EPI_ISL_856953, EPI_ISL_856954, EPI_ISL_856955, EPI_ISL_856956, EPI_ISL_856957, EPI_ISL_856958, EPI_ISL_856959, EPI_ISL_856960 | | | |
| see above | Wyoming Public Health Laboratory | Wyoming Public Health Laboratory | Noah Hull, Taylor Fearing, Lynette Gumbleton, Channing Weber, Ashley Norberg, Bailey Bowcutt, and Wanda Manley |
| EPI_ISL_857042 | Platform BIS UZA/UAntwerpen, University Hospital Antwerp, Edegem, Belgium | UAntwerp, Laboratory of Medical Microbiology, Campus Drie Eiken S6.26, Universiteitsplein 1, 2610, Wilrijk, Antwerp, Belgium | Basil Britto Xavier, Jasmine Coppens, Christine Lammens, Veerle Matheeussen, Herman Goossens |
| EPI_ISL_857043, EPI_ISL_857044, EPI_ISL_857045 | Colorado Department of Public Health and Environment | Colorado Department of Puplic Health and Environment | Laura Bankers, Molly C. Hetherington-Rauth, Diana Ir, Shannon Ely, Shannon R. Matzinger, Sarah Elizabeth Totten, Emily A. Travanty |
| EPI_ISL_857053, EPI_ISL_857054 | Platform BIS UZA/UAntwerpen, University Hospital Antwerp, Edegem, Belgium | UAntwerp, Laboratory of Medical Microbiology, Campus Drie Eiken S6.26, Universiteitsplein 1, 2610, Wilrijk, Antwerp, Belgium | Basil Britto Xavier, Jasmine Coppens, Christine Lammens, Veerle Matheeussen, Herman Goossens |
| EPI_ISL_857067 | DOHMH Corona | New York City Public Health Laboratory | Jade Wang, et al. |
| EPI_ISL_857068 | DOHMH PHL | New York City Public Health Laboratory | Jade Wang, et al. |
| EPI_ISL_857069 | DOHMH Jamaica | New York City Public Health Laboratory | Jade Wang, et al. |
| EPI_ISL_857070 | DOHMH PHL | New York City Public Health Laboratory | Jade Wang, et al. |
| EPI_ISL_857071 | DOHMH Morrisania | New York City Public Health Laboratory | Jade Wang, et al. |
| EPI_ISL_857072 | DOHMH PHL | New York City Public Health Laboratory | Jade Wang, et al. |

| | | | |
|---|---|---|---|
| EPI_ISL_857073 | DOHMH Corona | New York City Public Health Laboratory | Jade Wang, et al. |
| EPI_ISL_857074 | DOHMH Morrisania | New York City Public Health Laboratory | Jade Wang, et al. |
| EPI_ISL_857075 | DOHMH Jamaica | New York City Public Health Laboratory | Jade Wang, et al. |
| EPI_ISL_857076 | DOHMH Chelsea | New York City Public Health Laboratory | Jade Wang, et al. |
| EPI_ISL_857077, EPI_ISL_857078, EPI_ISL_857079, EPI_ISL_857080 | DOHMH Central Harlem | New York City Public Health Laboratory | Jade Wang, et al. |
| EPI_ISL_857081 | DOHMH Morrisania | New York City Public Health Laboratory | Jade Wang, et al. |
| EPI_ISL_857082, EPI_ISL_857083, EPI_ISL_857084 | DOHMH Corona | New York City Public Health Laboratory | Jade Wang, et al. |
| EPI_ISL_857085, EPI_ISL_857086 | DOHMH Central Harlem | New York City Public Health Laboratory | Jade Wang, et al. |
| EPI_ISL_857087, EPI_ISL_857088 | DOHMH PHL | New York City Public Health Laboratory | Jade Wang, et al. |
| EPI_ISL_857089 | DOHMH Crown Heights | New York City Public Health Laboratory | Jade Wang, et al. |
| EPI_ISL_857090 | DOHMH Riverside | New York City Public Health Laboratory | Jade Wang, et al. |
| EPI_ISL_857091 | DOHMH Jamaica | New York City Public Health Laboratory | Jade Wang, et al. |
| EPI_ISL_857092 | DOHMH Riverside | New York City Public Health Laboratory | Jade Wang, et al. |
| EPI_ISL_857093 | DOHMH Morrisania | New York City Public Health Laboratory | Jade Wang, et al. |
| EPI_ISL_857094 | DOHMH Central Harlem | New York City Public Health Laboratory | Jade Wang, et al. |
| EPI_ISL_857095 | DOHMH Fort Greene | New York City Public Health Laboratory | Jade Wang, et al. |
| EPI_ISL_857096 | DOHMH Jamaica | New York City Public Health Laboratory | Jade Wang, et al. |
| EPI_ISL_857097 | DOHMH Crown Heights | New York City Public Health Laboratory | Jade Wang, et al. |
| EPI_ISL_857098, EPI_ISL_857099 | DOHMH Morrisania | New York City Public Health Laboratory | Jade Wang, et al. |
| EPI_ISL_857100 | DOHMH Central Harlem | New York City Public Health Laboratory | Jade Wang, et al. |
| EPI_ISL_857101, EPI_ISL_857102, EPI_ISL_857103, EPI_ISL_857104, EPI_ISL_857105 | DOHMH Corona | New York City Public Health Laboratory | Jade Wang, et al. |
| EPI_ISL_857106, EPI_ISL_857107, EPI_ISL_857108, EPI_ISL_857109, EPI_ISL_857110 | DOHMH Chelsea | New York City Public Health Laboratory | Jade Wang, et al. |
| EPI_ISL_857111 | DOHMH Riverside | New York City Public Health Laboratory | Jade Wang, et al. |
| EPI_ISL_857112, EPI_ISL_857113, EPI_ISL_857114, EPI_ISL_857115 | DOHMH Fort Greene | New York City Public Health Laboratory | Jade Wang, et al. |
| EPI_ISL_857116 | DOHMH Central Harlem | New York City Public Health Laboratory | Jade Wang, et al. |
| EPI_ISL_857117 | DOHMH Chelsea | New York City Public Health Laboratory | Jade Wang, et al. |
| EPI_ISL_857118 | DOHMH Central Harlem | New York City Public Health Laboratory | Jade Wang, et al. |
| EPI_ISL_857119 | DOHMH Morrisania | New York City Public Health Laboratory | Jade Wang, et al. |
| EPI_ISL_857120 | DOHMH Central Harlem | New York City Public Health Laboratory | Jade Wang, et al. |
| EPI_ISL_857121 | DOHMH PHL | New York City Public Health Laboratory | Jade Wang, et al. |
| EPI_ISL_857122, EPI_ISL_857123, EPI_ISL_857124, EPI_ISL_857125 | DOHMH Morrisania | New York City Public Health Laboratory | Jade Wang, et al. |
| EPI_ISL_857126 | DOHMH Central Harlem | New York City Public Health Laboratory | Jade Wang, et al. |
| EPI_ISL_857127, EPI_ISL_857128, EPI_ISL_857129 | DOHMH PHL | New York City Public Health Laboratory | Jade Wang, et al. |
| EPI_ISL_857130, EPI_ISL_857131 | DOHMH Chelsea | New York City Public Health Laboratory | Jade Wang, et al. |
| EPI_ISL_857132, EPI_ISL_857133 | DOHMH Central Harlem | New York City Public Health Laboratory | Jade Wang, et al. |
| EPI_ISL_857349, EPI_ISL_857350, EPI_ISL_857351, EPI_ISL_857352, EPI_ISL_857353, EPI_ISL_857354, EPI_ISL_857355, EPI_ISL_857356, EPI_ISL_857357, EPI_ISL_857358, EPI_ISL_857359, EPI_ISL_857360, EPI_ISL_857361, EPI_ISL_857362, EPI_ISL_857363, EPI_ISL_857364, EPI_ISL_857365 | | | |
| see above | Texas Department of State Health Services (TXDSHS) | Texas Department of State Health Services (TXDSHS) | Anita Pokharel, Bonnie Oh, James Daniel Bonser, Rashmi Tuladhar, Mayela Pedrueza, Jenny Zhang, Maliha Rahman, Myong Koag, Chung Wang, Rachel Lee, Grace Kubin |
| EPI_ISL_857402, EPI_ISL_857454, EPI_ISL_857456, EPI_ISL_857458, EPI_ISL_857459 | Maine HETL | Tewhey Lab, The Jackson Laboratory | Matluk,N., Dewey,H., Iosue,F., Barter,M., Lynch,R., Munger,H. and Tewhey,R. |
| EPI_ISL_857478 | National Public Health Laboratory, National Centre for Infectious Diseases | National Public Health Laboratory, National Centre for Infectious Diseases | Tze Minn Mak, Sophie Octavia, Zhenyang Zhou, Lin Cui, Raymond Tzer Pin Lin |
| EPI_ISL_857546, EPI_ISL_857548, EPI_ISL_857550, EPI_ISL_857553, EPI_ISL_857562, EPI_ISL_857571, EPI_ISL_857584, EPI_ISL_857599, EPI_ISL_857603, EPI_ISL_857604, EPI_ISL_857606, EPI_ISL_857618, EPI_ISL_857642, EPI_ISL_857645, EPI_ISL_857654, EPI_ISL_857656, EPI_ISL_857669, EPI_ISL_857680, EPI_ISL_857682, EPI_ISL_857685, EPI_ISL_857699, EPI_ISL_857701, EPI_ISL_857715, EPI_ISL_857733, EPI_ISL_857752, EPI_ISL_857759, EPI_ISL_857781, EPI_ISL_857790, EPI_ISL_857802, EPI_ISL_857818, EPI_ISL_857819, EPI_ISL_857821, EPI_ISL_857823, EPI_ISL_857852 | | | |
| see above | Lighthouse Lab in Cambridge | Wellcome Sanger Institute for the COVID-19 Genomics UK (COG-UK) Consortium | Rob Howes, The Lighthouse Lab in Cambridge and Alex Alderton, Roberto Amato, Sonia Goncalves, Ewan Harrison, David K. Jackson, Ian Johnston, Dominic Kwiatkowski, Cordelia Langford, John Sillitoe on behalf of the Wellcome Sanger Institute COVID-19 Surveillance Team |
| EPI_ISL_857875, EPI_ISL_857879, EPI_ISL_857880, EPI_ISL_857882, EPI_ISL_857890, EPI_ISL_857960, EPI_ISL_857965, EPI_ISL_857966, EPI_ISL_857984, EPI_ISL_858000 | Lighthouse Lab in Alderley Park | Wellcome Sanger Institute for the COVID-19 Genomics UK (COG-UK) Consortium | Jacquelyn Wynn, Mairead Hyland, The Lighthouse Lab in Alderley Park and Alex Alderton, Roberto Amato, Sonia Goncalves, Ewan Harrison, David K. Jackson, Ian Johnston, Dominic Kwiatkowski, Cordelia Langford, John Sillitoe on behalf of the Wellcome Sanger Institute COVID-19 Surveillance Team |
| EPI_ISL_858012, EPI_ISL_858013, EPI_ISL_858014, EPI_ISL_858015, EPI_ISL_858016, EPI_ISL_858017, EPI_ISL_858018, EPI_ISL_858019, EPI_ISL_858020, EPI_ISL_858021, EPI_ISL_858023, EPI_ISL_858024, EPI_ISL_858026, EPI_ISL_858027, EPI_ISL_858028, EPI_ISL_858029, EPI_ISL_858030, EPI_ISL_858031, EPI_ISL_858032, EPI_ISL_858033, EPI_ISL_858034, EPI_ISL_858036, EPI_ISL_858037, EPI_ISL_858038, EPI_ISL_858041, EPI_ISL_858042, EPI_ISL_858044, EPI_ISL_858045, EPI_ISL_858046, EPI_ISL_858047, EPI_ISL_858048, EPI_ISL_858049, EPI_ISL_858050, EPI_ISL_858051, EPI_ISL_858052, EPI_ISL_858053, EPI_ISL_858055, EPI_ISL_858056, EPI_ISL_858057, EPI_ISL_858058, EPI_ISL_858059, EPI_ISL_858060, EPI_ISL_858061, EPI_ISL_858062, EPI_ISL_858063 | | | |
| see above | Lighthouse Lab in Glasgow | Wellcome Sanger Institute for the COVID-19 Genomics UK (COG-UK) Consortium | Harper VanSteenhouse, Yumi Kasai, David Gray, Carol Clugston, Anna Dominiczak and Alex Alderton, Roberto Amato, Sonia Goncalves, Ewan Harrison, David K. Jackson, Ian Johnston, Dominic Kwiatkowski, Cordelia Langford, John Sillitoe on behalf of the Wellcome Sanger Institute COVID-19 Surveillance |

| | | | |
|---|---|---|---|
| | | | Team |
| EPI_ISL_858064, EPI_ISL_858065, EPI_ISL_858066, EPI_ISL_858067, EPI_ISL_858068, EPI_ISL_858069, EPI_ISL_858070, EPI_ISL_858071, EPI_ISL_858072, EPI_ISL_858073, EPI_ISL_858074, EPI_ISL_858075, EPI_ISL_858076, EPI_ISL_858078, EPI_ISL_858079, EPI_ISL_858080, EPI_ISL_858081, EPI_ISL_858082, EPI_ISL_858083, EPI_ISL_858084, EPI_ISL_858087, EPI_ISL_858088, EPI_ISL_858089, EPI_ISL_858090, EPI_ISL_858091, EPI_ISL_858092, EPI_ISL_858093, EPI_ISL_858094, EPI_ISL_858095, EPI_ISL_858096, EPI_ISL_858097, EPI_ISL_858098, EPI_ISL_858099, EPI_ISL_858100, EPI_ISL_858102, EPI_ISL_858103, EPI_ISL_858104, EPI_ISL_858105, EPI_ISL_858106, EPI_ISL_858107, EPI_ISL_858108, EPI_ISL_858109, EPI_ISL_858110, EPI_ISL_858111, EPI_ISL_858112, EPI_ISL_858113, EPI_ISL_858114, EPI_ISL_858115, EPI_ISL_858116, EPI_ISL_858117, EPI_ISL_858118, EPI_ISL_858119, EPI_ISL_858120, EPI_ISL_858121, EPI_ISL_858123, EPI_ISL_858124, EPI_ISL_858125, EPI_ISL_858126, EPI_ISL_858127, EPI_ISL_858128, EPI_ISL_858129, EPI_ISL_858130, EPI_ISL_858131, EPI_ISL_858132, EPI_ISL_858133, EPI_ISL_858134, EPI_ISL_858135, EPI_ISL_858137, EPI_ISL_858138, EPI_ISL_858139, EPI_ISL_858140, EPI_ISL_858141 | | | |
| see above | Lighthouse Lab in Alderley Park | Wellcome Sanger Institute for the COVID-19 Genomics UK (COG-UK) Consortium | Jacquelyn Wynn, Mairead Hyland, The Lighthouse Lab in Alderley Park and Alex Alderton, Roberto Amato, Sonia Goncalves, Ewan Harrison, David K. Jackson, Ian Johnston, Dominic Kwiatkowski, Cordelia Langford, John Sillitoe on behalf of the Wellcome Sanger Institute COVID-19 Surveillance Team |
| EPI_ISL_858158, EPI_ISL_858159, EPI_ISL_858161, EPI_ISL_858162, EPI_ISL_858163, EPI_ISL_858164, EPI_ISL_858165, EPI_ISL_858166, EPI_ISL_858167, EPI_ISL_858168, EPI_ISL_858169, EPI_ISL_858170, EPI_ISL_858171, EPI_ISL_858172, EPI_ISL_858173, EPI_ISL_858174, EPI_ISL_858175, EPI_ISL_858176, EPI_ISL_858177, EPI_ISL_858178, EPI_ISL_858179, EPI_ISL_858180, EPI_ISL_858181, EPI_ISL_858182, EPI_ISL_858183, EPI_ISL_858184, EPI_ISL_858185, EPI_ISL_858186, EPI_ISL_858187, EPI_ISL_858188, EPI_ISL_858189, EPI_ISL_858190, EPI_ISL_858191, EPI_ISL_858192, EPI_ISL_858193, EPI_ISL_858194, EPI_ISL_858195, EPI_ISL_858196, EPI_ISL_858197, EPI_ISL_858198, EPI_ISL_858199, EPI_ISL_858200, EPI_ISL_858201, EPI_ISL_858202, EPI_ISL_858203, EPI_ISL_858204, EPI_ISL_858205, EPI_ISL_858206, EPI_ISL_858223, EPI_ISL_858224, EPI_ISL_858225, EPI_ISL_858226, EPI_ISL_858227, EPI_ISL_858228, EPI_ISL_858229, EPI_ISL_858230, EPI_ISL_858231, EPI_ISL_858232, EPI_ISL_858233, EPI_ISL_858234, EPI_ISL_858235, EPI_ISL_858236, EPI_ISL_858237, EPI_ISL_858238, EPI_ISL_858239, EPI_ISL_858240, EPI_ISL_858242, EPI_ISL_858243, EPI_ISL_858244, EPI_ISL_858245, EPI_ISL_858247, EPI_ISL_858248, EPI_ISL_858249, EPI_ISL_858250, EPI_ISL_858251, EPI_ISL_858252, EPI_ISL_858253, EPI_ISL_858254, EPI_ISL_858255, EPI_ISL_858256, EPI_ISL_858257, EPI_ISL_858258, EPI_ISL_858259, EPI_ISL_858260, EPI_ISL_858261, EPI_ISL_858262, EPI_ISL_858264, EPI_ISL_858265, EPI_ISL_858266, EPI_ISL_858271, EPI_ISL_858273, EPI_ISL_858275, EPI_ISL_858276, EPI_ISL_858277, EPI_ISL_858279, EPI_ISL_858280, EPI_ISL_858281, EPI_ISL_858282, EPI_ISL_858283, EPI_ISL_858284, EPI_ISL_858286, EPI_ISL_858288, EPI_ISL_858289, EPI_ISL_858291, EPI_ISL_858292, EPI_ISL_858293, EPI_ISL_858294, EPI_ISL_858295, EPI_ISL_858297, EPI_ISL_858298, EPI_ISL_858299, EPI_ISL_858300, EPI_ISL_858301, EPI_ISL_858302, EPI_ISL_858304, EPI_ISL_858305, EPI_ISL_858314, EPI_ISL_858315, EPI_ISL_858316, EPI_ISL_858317, EPI_ISL_858319, EPI_ISL_858320, EPI_ISL_858321, EPI_ISL_858322, EPI_ISL_858323, EPI_ISL_858327, EPI_ISL_858331, EPI_ISL_858333, EPI_ISL_858335, EPI_ISL_858336, EPI_ISL_858337, EPI_ISL_858338, EPI_ISL_858340, EPI_ISL_858342, EPI_ISL_858344, EPI_ISL_858345, EPI_ISL_858346, EPI_ISL_858347, EPI_ISL_858348, EPI_ISL_858349, EPI_ISL_858352, EPI_ISL_858353, EPI_ISL_858355, EPI_ISL_858356, EPI_ISL_858358, EPI_ISL_858359, EPI_ISL_858360, EPI_ISL_858361, EPI_ISL_858362, EPI_ISL_858364, EPI_ISL_858365, EPI_ISL_858366, EPI_ISL_858367, EPI_ISL_858369, EPI_ISL_858372, EPI_ISL_858373, EPI_ISL_858374, EPI_ISL_858376, EPI_ISL_858377, EPI_ISL_858379, EPI_ISL_858380, EPI_ISL_858384, EPI_ISL_858390, EPI_ISL_858396, EPI_ISL_858401, EPI_ISL_858402, EPI_ISL_858404, EPI_ISL_858405, EPI_ISL_858406, EPI_ISL_858407 | | | |
| see above | Lighthouse Lab in Glasgow | Wellcome Sanger Institute for the COVID-19 Genomics UK (COG-UK) Consortium | Harper VanSteenhouse, Yumi Kasai, David Gray, Carol Clugston, Anna Dominiczak and Alex Alderton, Roberto Amato, Sonia Goncalves, Ewan Harrison, David K. Jackson, Ian Johnston, Dominic Kwiatkowski, Cordelia Langford, John Sillitoe on behalf of the Wellcome Sanger Institute COVID-19 Surveillance Team |
| EPI_ISL_860198 | Railway Hospital | Department of Neurovirology, National Institute of Mental Health and Neurosciences (NIMHANS) | Chitra Pattabiraman, Pramada Prasad, Anson Kunjumon George, Risha Rasheed, Darshan Sreenivas, Nakka Vijay Kiran Reddy, Anita S Desai, V Ravi |
| EPI_ISL_860240, EPI_ISL_860244, EPI_ISL_860245, EPI_ISL_860246, EPI_ISL_860251, EPI_ISL_860253, EPI_ISL_860255 | University Hospitals of Geneva, Laboratory of Virology | HUG, Laboratory of Virology and Universitätsspital Basel | Samuel Cordey, Ana Rita Goncalves, Laurent Kaiser, Tim Roloff, Madlen Stange, Helena MB Seth-Smith, Alfredo Mari, Karoline Leuzinger, Julia Bielicki, Manuel Battegay, Hans Hirsch, Adrian Egli |
| EPI_ISL_860271, EPI_ISL_860273 | Ostfold Hospital Trust - Kalnes, Centre for Laboratory Medicine, Section for gene technology and infection serology | Norwegian Institute of Public Health, Department of Virology | Kathrine Stene-Johansen, Kamilla Heddeland Instefjord, Hilde Elshaug, Atiya R Ali,Marie Paulsen Madsen, Rasmus Riis Kopperud, Hilde Vollan, Karoline Bragstad, Olav Hungnes |
| EPI_ISL_860274 | Foerde Hospital, Department of Microbiology | Norwegian Institute of Public Health, Department of Virology | Kathrine Stene-Johansen, Kamilla Heddeland Instefjord, Hilde Elshaug, Atiya R Ali,Marie Paulsen Madsen, Rasmus Riis Kopperud, Hilde Vollan, Karoline Bragstad, Olav Hungnes |
| EPI_ISL_860279 | Ostfold Hospital Trust - Kalnes, Centre for Laboratory Medicine, Section for gene technology and infection serology | Norwegian Institute of Public Health, Department of Virology | Kathrine Stene-Johansen, Kamilla Heddeland Instefjord, Hilde Elshaug, Atiya R Ali,Marie Paulsen Madsen, Rasmus Riis Kopperud, Hilde Vollan, Karoline Bragstad, Olav Hungnes |
| EPI_ISL_860294 | Norwegian Institute of Public Health, Department of Virology | Norwegian Institute of Public Health, Department of Virology | Kathrine Stene-Johansen, Kamilla Heddeland Instefjord, Hilde Elshaug, Atiya R Ali,Marie Paulsen Madsen, Rasmus Riis Kopperud, Hilde Vollan, Karoline Bragstad, Olav Hungnes |
| EPI_ISL_860313, EPI_ISL_860314, EPI_ISL_860315 | Unit 17: Influenza & Other Respiratory Viruses, German National Influenza Center | Project group Epidemiology of Highly Pathogenic Microorganisms, Robert Koch-Institute | Andreas Sachse, Grit Schubert, Essia Belarbi, Sébastien Calvignac-Spencer, Thorsten Wolff, Ralf Dürrwald, Djin-Ye Oh, Marianne Wedde |
| EPI_ISL_860317 | Instituto Adolfo Lutz - Regional de Sorocaba | Instituto Adolfo Lutz, Interdiciplinary Procedures Center, Strategic Laboratory | Claudio Tavares Sacchi, Claudia Regina Gonçalves, Erica Valessa Ramos Gomes, Karoline Rodrigues Campos |
| EPI_ISL_860553 | Ampath-Netcare | KRISP, KZn Research Innovation and Sequencing Platform | Giandhari J, Pillay S, Lessells R, Mdlalose K, York D, Khan S, Tegally H, Wilkinson E, de Oliveira T |
| EPI_ISL_860554, EPI_ISL_860555, EPI_ISL_860556, EPI_ISL_860557, EPI_ISL_860558 | NHLS-IALCH | KRISP, KZn Research Innovation and Sequencing Platform | Giandhari J, Pillay S, Lessells R, Mdlalose K, York D, Khan S, Tegally H, Wilkinson E, de Oliveira T |
| EPI_ISL_860633 | Instituto Adolfo Lutz - Regional de Sorocaba | Instituto Adolfo Lutz, Interdiciplinary Procedures Center, Strategic Laboratory | Claudio Tavares Sacchi, Claudia Regina Gonçalves, Erica Valessa Ramos Gomes, Karoline Rodrigues Campos |
| EPI_ISL_860671, EPI_ISL_860674, EPI_ISL_860675, EPI_ISL_860676, EPI_ISL_860680, EPI_ISL_860681, EPI_ISL_860682, EPI_ISL_860683, EPI_ISL_860684 | Respiratory Virus Unit, National Infection Service, Public Health England | COVID-19 Genomics UK (COG-UK) Consortium | PHE Covid Sequencing Team |
| EPI_ISL_860719 | Institute for Infectious Diseases, University of Bern, Switzerland | Institute for Infectious Diseases, University of Bern, Switzerland | Michel C Koch, Christian Baumann, Miguel A Terrazos Miani, Cora Sägesser, Pascal Bittel, Stephen L Leib, Peter Keller, Franziska Suter-Riniker, Alban Ramette |
| EPI_ISL_860784, EPI_ISL_860786, EPI_ISL_860788 | Charité Universitätsmedizin Berlin, Institute of Virology, Charitéplatz 1, 10117 Berlin, Germany | Charité Universitätsmedizin Berlin, Institute of Virology, Charitéplatz 1, 10117 Berlin, Germany | Victor M Corman, Julia Schneider, Jörn Beheim-Schwarzbach, Tobias Bleicker, Julia Tesch, Barbara Mühlemann, Talitha Veith, Terry Jones, Christian Drosten |
| EPI_ISL_860821 | CHU de Nantes - Hôtel Dieu - Labo. Virologie | National Reference Center for Viruses of Respiratory Infections, Institut Pasteur, Paris | Marion Barbet, Sylvie Behillil, Méline Bizard, Angela Brisebarre, Camille Capel, Etienne Simon-Lorière, Vincent Enouf, Maud Vanpeene, Sylvie van der Werf,Andre-Garnier Elisabeth |
| EPI_ISL_860822 | CHU Tours - Virologie | National Reference Center for Viruses of Respiratory Infections, Institut Pasteur, Paris | Marion Barbet, Sylvie Behillil, Méline Bizard, Angela Brisebarre, Camille Capel, Etienne Simon-Lorière, Vincent Enouf, Maud Vanpeene, Sylvie van der Werf,Gaudy Graffin Catherine |
| EPI_ISL_860824 | Labo Analyses Med | National Reference Center for Viruses of Respiratory Infections, Institut Pasteur, Paris | Marion Barbet, Sylvie Behillil, Méline Bizard, Angela Brisebarre, Camille Capel, Etienne Simon-Lorière, Vincent Enouf, Maud Vanpeene, Sylvie van der Werf,Amzalag Jonas |
| EPI_ISL_860826 | Labo Analyses med | National Reference Center for Viruses of Respiratory Infections, Institut Pasteur, Paris | Marion Barbet, Sylvie Behillil, Méline Bizard, Angela Brisebarre, Camille Capel, Etienne Simon-Lorière, Vincent Enouf, Maud Vanpeene, Sylvie van der Werf,Amzalag Jonas |
| EPI_ISL_860828 | CHU de Nantes - Hôtel Dieu - Labo. Virologie | National Reference Center for Viruses of Respiratory Infections, Institut Pasteur, Paris | Marion Barbet, Sylvie Behillil, Méline Bizard, Angela Brisebarre, Camille Capel, Etienne Simon-Lorière, Vincent Enouf, Maud Vanpeene, Sylvie van der Werf,Andre-Garnier Elisabeth |
| EPI_ISL_860831, EPI_ISL_860833, EPI_ISL_860836, EPI_ISL_860838, EPI_ISL_860840 | Labo Analyses Med | National Reference Center for Viruses of Respiratory Infections, Institut Pasteur, Paris | Marion Barbet, Sylvie Behillil, Méline Bizard, Angela Brisebarre, Camille Capel, Etienne Simon-Lorière, Vincent Enouf, Maud Vanpeene, Sylvie van der Werf,Amzalag Jonas |
| EPI_ISL_860841 | Labo Analyses Med | National Reference Center for Viruses of Respiratory Infections, Institut Pasteur, Paris | Marion Barbet, Sylvie Behillil, Méline Bizard, Angela Brisebarre, Camille Capel, Etienne Simon-Lorière, Vincent Enouf, Maud Vanpeene, Sylvie van der Werf, |
| EPI_ISL_860842, EPI_ISL_860843, EPI_ISL_860844 | Labo Analyses med | National Reference Center for Viruses of Respiratory Infections, Institut Pasteur, Paris | Marion Barbet, Sylvie Behillil, Méline Bizard, Angela Brisebarre, Camille Capel, Etienne Simon-Lorière, Vincent Enouf, Maud Vanpeene, Sylvie van der Werf,Amzalag Jonas |
| EPI_ISL_860849 | HIA BÉGIN Service de Biologie | National Reference Center for Viruses of Respiratory Infections, Institut Pasteur, Paris | Marion Barbet, Sylvie Behillil, Méline Bizard, Angela Brisebarre, Camille Capel, Etienne Simon-Lorière, Vincent Enouf, Maud Vanpeene, Sylvie van der Werf,Merens Audrey |
| EPI_ISL_860856 | Labo Analyses Med | National Reference Center for Viruses of Respiratory | Marion Barbet, Sylvie Behillil, Méline Bizard, Angela Brisebarre, Camille Capel, Etienne Simon-Lorière, Vincent Enouf, Maud Vanpeene, Sylvie van der |

| | | | |
|---|---|---|---|
| | | Infections, Institut Pasteur, Paris | Werf,Amzalag Jonas |
| EPI_ISL_860860, EPI_ISL_860861, EPI_ISL_860862 | Labo Analyses med | National Reference Center for Viruses of Respiratory Infections, Institut Pasteur, Paris | Marion Barbet, Sylvie Behillil, Méline Bizard, Angela Brisebarre, Camille Capel, Etienne Simon-Lorière, Vincent Enouf, Maud Vanpeene, Sylvie van der Werf,Amzalag Jonas |
| EPI_ISL_860863, EPI_ISL_860865 | CHU de Nantes - Hôtel Dieu - Labo. Virologie | National Reference Center for Viruses of Respiratory Infections, Institut Pasteur, Paris | Marion Barbet, Sylvie Behillil, Méline Bizard, Angela Brisebarre, Camille Capel, Etienne Simon-Lorière, Vincent Enouf, Maud Vanpeene, Sylvie van der Werf,Andre-Garnier Elisabeth |
| EPI_ISL_860866 | Labo Analyses Med | National Reference Center for Viruses of Respiratory Infections, Institut Pasteur, Paris | Marion Barbet, Sylvie Behillil, Méline Bizard, Angela Brisebarre, Camille Capel, Etienne Simon-Lorière, Vincent Enouf, Maud Vanpeene, Sylvie van der Werf,Lefaure Brieuc |
| EPI_ISL_860867 | CHU Tours - Virologie | National Reference Center for Viruses of Respiratory Infections, Institut Pasteur, Paris | Marion Barbet, Sylvie Behillil, Méline Bizard, Angela Brisebarre, Camille Capel, Etienne Simon-Lorière, Vincent Enouf, Maud Vanpeene, Sylvie van der Werf,Gaudy Graffin Catherine |
| EPI_ISL_860868 | CHU de Nantes - Hôtel Dieu - Labo. Virologie | National Reference Center for Viruses of Respiratory Infections, Institut Pasteur, Paris | Marion Barbet, Sylvie Behillil, Méline Bizard, Angela Brisebarre, Camille Capel, Etienne Simon-Lorière, Vincent Enouf, Maud Vanpeene, Sylvie van der Werf,Andre-Garnier Elisabeth |
| EPI_ISL_860872, EPI_ISL_860873, EPI_ISL_860874, EPI_ISL_860875, EPI_ISL_860876, EPI_ISL_860877, EPI_ISL_860878, EPI_ISL_860886, EPI_ISL_860887 | Labo Analyses Med | National Reference Center for Viruses of Respiratory Infections, Institut Pasteur, Paris | Marion Barbet, Sylvie Behillil, Méline Bizard, Angela Brisebarre, Camille Capel, Etienne Simon-Lorière, Vincent Enouf, Maud Vanpeene, Sylvie van der Werf,Lefaure Brieuc |
| EPI_ISL_860889 | Labo Analyses Med | National Reference Center for Viruses of Respiratory Infections, Institut Pasteur, Paris | Marion Barbet, Sylvie Behillil, Méline Bizard, Angela Brisebarre, Camille Capel, Etienne Simon-Lorière, Vincent Enouf, Maud Vanpeene, Sylvie van der Werf,Amzalag Jonas |
| EPI_ISL_860891, EPI_ISL_860893, EPI_ISL_860894, EPI_ISL_860896 | CHU Tours - Virologie | National Reference Center for Viruses of Respiratory Infections, Institut Pasteur, Paris | Marion Barbet, Sylvie Behillil, Méline Bizard, Angela Brisebarre, Camille Capel, Etienne Simon-Lorière, Vincent Enouf, Maud Vanpeene, Sylvie van der Werf,Gaudy Graffin Catherine |
| EPI_ISL_860905 | CHU Angers - Dpt des Agents Infectieux | National Reference Center for Viruses of Respiratory Infections, Institut Pasteur, Paris | Marion Barbet, Sylvie Behillil, Méline Bizard, Angela Brisebarre, Camille Capel, Etienne Simon-Lorière, Vincent Enouf, Maud Vanpeene, Sylvie van der Werf,Ducancelle Alexandra |
| EPI_ISL_860907 | CHU Tours - Virologie | National Reference Center for Viruses of Respiratory Infections, Institut Pasteur, Paris | Marion Barbet, Sylvie Behillil, Méline Bizard, Angela Brisebarre, Camille Capel, Etienne Simon-Lorière, Vincent Enouf, Maud Vanpeene, Sylvie van der Werf,Gaudy Graffin Catherine |
| EPI_ISL_860911, EPI_ISL_860912 | CHU Angers - Dpt des Agents Infectieux | National Reference Center for Viruses of Respiratory Infections, Institut Pasteur, Paris | Marion Barbet, Sylvie Behillil, Méline Bizard, Angela Brisebarre, Camille Capel, Etienne Simon-Lorière, Vincent Enouf, Maud Vanpeene, Sylvie van der Werf,Ducancelle Alexandra |
| EPI_ISL_860913, EPI_ISL_860918, EPI_ISL_860919, EPI_ISL_860920, EPI_ISL_860921, EPI_ISL_860926, EPI_ISL_860927, EPI_ISL_860928, EPI_ISL_860929, EPI_ISL_860930 | CHU de Nantes - Hôtel Dieu - Labo. Virologie | National Reference Center for Viruses of Respiratory Infections, Institut Pasteur, Paris | Marion Barbet, Sylvie Behillil, Méline Bizard, Angela Brisebarre, Camille Capel, Etienne Simon-Lorière, Vincent Enouf, Maud Vanpeene, Sylvie van der Werf,Andre-Garnier Elisabeth |
| EPI_ISL_860952, EPI_ISL_860953, EPI_ISL_860965, EPI_ISL_860977, EPI_ISL_860984, EPI_ISL_860988, EPI_ISL_860999, EPI_ISL_861002, EPI_ISL_861008, EPI_ISL_861016, EPI_ISL_861030, EPI_ISL_861041, EPI_ISL_861042, EPI_ISL_861044, EPI_ISL_861047, EPI_ISL_861050, EPI_ISL_861074, EPI_ISL_861076 | | | |
| see above | Johns Hopkins Hospital Department of Pathology | Johns Hopkins Hospital Department of Pathology | C. Paul Morris, Chun Huai Luo, Adannaya Amadi, Nicholas Gallagher, Heba H. Mostafa |
| EPI_ISL_861235 | CHU Tours - Virologie | National Reference Center for Viruses of Respiratory Infections, Institut Pasteur, Paris | Marion Barbet, Sylvie Behillil, Méline Bizard, Angela Brisebarre, Camille Capel, Etienne Simon-Lorière, Vincent Enouf, Maud Vanpeene, Sylvie van der Werf,Gaudy Graffin Catherine |
| EPI_ISL_861236 | CHU Angers - Dpt des Agents Infectieux | National Reference Center for Viruses of Respiratory Infections, Institut Pasteur, Paris | Marion Barbet, Sylvie Behillil, Méline Bizard, Angela Brisebarre, Camille Capel, Etienne Simon-Lorière, Vincent Enouf, Maud Vanpeene, Sylvie van der Werf,Ducancelle Alexandra |
| EPI_ISL_861239 | Labo Analyses med | National Reference Center for Viruses of Respiratory Infections, Institut Pasteur, Paris | Marion Barbet, Sylvie Behillil, Méline Bizard, Angela Brisebarre, Camille Capel, Etienne Simon-Lorière, Vincent Enouf, Maud Vanpeene, Sylvie van der Werf,Amzalag Jonas |
| EPI_ISL_861333, EPI_ISL_861337, EPI_ISL_861340, EPI_ISL_861348, EPI_ISL_861355, EPI_ISL_861356, EPI_ISL_861373, EPI_ISL_861374, EPI_ISL_861375, EPI_ISL_861376, EPI_ISL_861377, EPI_ISL_861378, EPI_ISL_861379, EPI_ISL_861380, EPI_ISL_861382, EPI_ISL_861383, EPI_ISL_861384, EPI_ISL_861385, EPI_ISL_861386, EPI_ISL_861387, EPI_ISL_861389, EPI_ISL_861392, EPI_ISL_861393, EPI_ISL_861394, EPI_ISL_861395, EPI_ISL_861396, EPI_ISL_861397, EPI_ISL_861399, EPI_ISL_861400, EPI_ISL_861401, EPI_ISL_861402, EPI_ISL_861403, EPI_ISL_861404, EPI_ISL_861405, EPI_ISL_861406, EPI_ISL_861407, EPI_ISL_861408, EPI_ISL_861409, EPI_ISL_861410, EPI_ISL_861411, EPI_ISL_861412 | | | |
| see above | WESTCHESTER MEDICAL CENTER | Wadsworth Center, New York State Department of Health | Kirsten St. George, Daryl M. Lamson, Alexis Russel, Matthew Shudt, Melissa A Leisner, Jonathan Plitnick, Navjot Singh, John Kelly, Erasmus Schneider, Erica Lasek-Nesselquist |
| EPI_ISL_861417 | NORTH SHORE UNIVERSITY HOSPITAL | Wadsworth Center, New York State Department of Health | Kirsten St. George, Daryl M. Lamson, Alexis Russel, Matthew Shudt, Melissa A Leisner, Jonathan Plitnick, Navjot Singh, John Kelly, Erasmus Schneider, Erica Lasek-Nesselquist |
| EPI_ISL_861428, EPI_ISL_861430, EPI_ISL_861432, EPI_ISL_861476, EPI_ISL_861477, EPI_ISL_861478, EPI_ISL_861479, EPI_ISL_861482, EPI_ISL_861483, EPI_ISL_861484, EPI_ISL_861485, EPI_ISL_861486 | | | |
| see above | Wyoming Public Health Laboratory | Wyoming Public Health Laboratory | Noah Hull, Taylor Fearing, Lynette Gumbleton, Channing Weber, Ashley Norberg, Bailey Bowcutt, and Wanda Manley |
| EPI_ISL_861496 | Cerballiance Côte d'Azur | CERBA LAB | Bausset O; Juvet O; Prots L. |
| EPI_ISL_861503 | Cerballiance Alpes Durance | CERBA LAB | Dyen P; Prots L. |
| EPI_ISL_861505 | Cerballiance Provence | CERBA LAB | Delaunay E; Roig JC; Prots L. |
| EPI_ISL_861506 | Cerballiance Provence | CERBA LAB | Roig JC; Delaunay E; Prots L. |
| EPI_ISL_861508 | Cerballiance Haut de France | CERBA LAB | Felloni C; Mainardi A; Lahmidi S. |
| EPI_ISL_861535, EPI_ISL_861541, EPI_ISL_861542, EPI_ISL_861543, EPI_ISL_861544, EPI_ISL_861545, EPI_ISL_861546, EPI_ISL_861547, EPI_ISL_861548, EPI_ISL_861549, EPI_ISL_861551, EPI_ISL_861558, EPI_ISL_861559, EPI_ISL_861565, EPI_ISL_861566, EPI_ISL_861578, EPI_ISL_861580, EPI_ISL_861592, EPI_ISL_861593, EPI_ISL_861594, EPI_ISL_861595, EPI_ISL_861596, EPI_ISL_861597, EPI_ISL_861598, EPI_ISL_861599, EPI_ISL_861600, EPI_ISL_861601, EPI_ISL_861602, EPI_ISL_861603, EPI_ISL_861604, EPI_ISL_861605, EPI_ISL_861606, EPI_ISL_861607, EPI_ISL_861608, EPI_ISL_861609, EPI_ISL_861610, EPI_ISL_861611, EPI_ISL_861612, EPI_ISL_861613, EPI_ISL_861614, EPI_ISL_861615, EPI_ISL_861616, EPI_ISL_861617, EPI_ISL_861618, EPI_ISL_861619, EPI_ISL_861620, EPI_ISL_861621, EPI_ISL_861622, EPI_ISL_861623 | | | |
| see above | Instituto Nacional de Saude (INSA) | Instituto Nacional de Saude (INSA) | Borges et al |
| EPI_ISL_861687, EPI_ISL_861688, EPI_ISL_861691 | Los Angeles County PHL | Los Angeles County PHL | P. Hemarajata et al. |
| EPI_ISL_861700, EPI_ISL_861701, EPI_ISL_861702, EPI_ISL_861703 | MD Laboratories | Los Angeles County PHL | P. Hemarajata et al. |
| EPI_ISL_861704, EPI_ISL_861705, EPI_ISL_861706 | UCLA Clinical Micro Lab | Los Angeles County PHL | P. Hemarajata et al. |
| EPI_ISL_861707, EPI_ISL_861708, EPI_ISL_861709 | Instituto Nacional de Saude (INSA) | Instituto Nacional de Saude (INSA) | Borges et al |
| EPI_ISL_861714 | Cerba | CERBA LAB | Roquebert B; Costa JM; Hedbaut E; Trombert S; Lecorche E; Verdurme L; Malek Ramdane, Olivi M; Haïm-Boukobza S. |
| EPI_ISL_861715 | CERBA | CERBA LAB | Roquebert B; Costa JM; Hedbaut E; Trombert S; Lecorche E; Verdurme L; Malek Ramdane, Olivi M; Haïm-Boukobza S. |
| EPI_ISL_861727 | CERBA | CERBA LAB | Roquebert B; Costa JM; Hedbaut E; Trombert S; Lecorche E; Verdurme L; Malek Ramdane; Olivi M; Haïm-Boukobza S. |
| EPI_ISL_861734 | Yale Pathology Lab | Grubaugh Lab - Yale School of Public Health | Tara Alpert, Joseph Fauver, Chen Liu, Pei Hui, Jianhui Wang, Susan Bell and Han Zhou, Anderson Brito, Mallery Breban, Anne Wyllie, Chantal Vogels, |

| | | | |
|---|---|---|---|
| | | | Mary Petrone, Chaney Kalinich, Isabel Ott, Arnau Casanovas, Catherine Muenker, Adam Moore, Alice Lu, Maria Tokuyama, Patrick Wong, Peiwen Lu, Saad Omer, Richard Martinello, Allison Nelson, Shelli Farhadian, Akiko Iwasaki, Charlese Dela Cruz, Albert Ko, Nathan Grubaugh |
| EPI_ISL_861739, EPI_ISL_861740, EPI_ISL_861743, EPI_ISL_861747 | Tempus | Grubaugh Lab - Yale School of Public Health | Tara Alpert, Joseph Fauver, Anderson Brito, Mallery Breban, Anne Wyllie, Chantal Vogels, Mary Petrone, Annie Watkins, Chaney Kalinich, Isabel Ott, Nathan Grubaugh |
| EPI_ISL_861752, EPI_ISL_861753 | Yale Clinical Virology Lab | Grubaugh Lab - Yale School of Public Health | Tara Alpert, Joseph Fauver, Anderson Brito, Mallery Breban, Anne Wyllie, Chantal Vogels, Mary Petrone, Chaney Kalinich, Isabel Ott, Nathan Grubaugh |
| EPI_ISL_861762, EPI_ISL_861763 | Tempus | Grubaugh Lab - Yale School of Public Health | Tara Alpert, Joseph Fauver, Anderson Brito, Mallery Breban, Anne Wyllie, Chantal Vogels, Mary Petrone, Annie Watkins, Chaney Kalinich, Isabel Ott, Nathan Grubaugh |
| EPI_ISL_861764 | Yale Pathology Lab | Grubaugh Lab - Yale School of Public Health | Tara Alpert, Joseph Fauver, Chen Liu, Pei Hui, Jianhui Wang, Susan Bell and Han Zhou, Anderson Brito, Mallery Breban, Anne Wyllie, Chantal Vogels, Mary Petrone, Chaney Kalinich, Isabel Ott, Arnau Casanovas, Catherine Muenker, Adam Moore, Alice Lu, Maria Tokuyama, Patrick Wong, Peiwen Lu, Saad Omer, Richard Martinello, Allison Nelson, Shelli Farhadian, Akiko Iwasaki, Charlese Dela Cruz, Albert Ko, Nathan Grubaugh |
| EPI_ISL_861765 | Tempus | Grubaugh Lab - Yale School of Public Health | Tara Alpert, Joseph Fauver, Anderson Brito, Mallery Breban, Anne Wyllie, Chantal Vogels, Mary Petrone, Annie Watkins, Chaney Kalinich, Isabel Ott, Nathan Grubaugh |
| EPI_ISL_862043, EPI_ISL_862044, EPI_ISL_862045 | CERBA | CERBA LAB | Roquebert B; Costa JM; Hedbaut E; Trombert S; Lecorche E; Verdurme L; Malek Ramdane, Olivi M; Haïm-Boukobza S. |
| EPI_ISL_862046, EPI_ISL_862047 | Cerballiance Val de Loire | CERBA LAB | Abs G. |
| EPI_ISL_862121 | MD Laboratories | Los Angeles County PHL | P. Hemarajata et al. |
| EPI_ISL_862122, EPI_ISL_862123 | UCLA Clinical Micro Lab | Los Angeles County PHL | P. Hemarajata et al. |
| EPI_ISL_862131, EPI_ISL_862148, EPI_ISL_862154, EPI_ISL_862157, EPI_ISL_862158, EPI_ISL_862172, EPI_ISL_862173, EPI_ISL_862174, EPI_ISL_862182, EPI_ISL_862184, EPI_ISL_862185 | | | |
| see above | Charité Universitätsmedizin Berlin, Institut fur Virologie/Labor Berlin | Charité Universitätsmedizin Berlin, Institut für Virologie | Victor M Corman, Barbara Mühlemann, Jörn Beheim-Schwarzbach, Tobias Bleicker, Julia Tesch, Talitha Veith, Julia Schneider, Terry Jones, Christian Drosten |
| EPI_ISL_862676 | New Mexico Department of Health Scientific Laboratory | New Mexico Department of Health Scientific Laboratory | D'eldra Malone, Ellie Johnson, Anastacia Griego-Fisher |
| EPI_ISL_862730, EPI_ISL_862731, EPI_ISL_862733, EPI_ISL_862737, EPI_ISL_862745, EPI_ISL_862752, EPI_ISL_862772, EPI_ISL_862774, EPI_ISL_862779 | Utah Public Health Laboratory, Utah Public Health Laboratory Infectious Disease submission group | Utah Public Health Laboratory, Utah Public Health Laboratory Infectious Disease submission group | Young,E.L., Oakeson,K.F., Gallagher,T. |
| EPI_ISL_863809 | Lighthouse Lab in Milton Keynes | Wellcome Sanger Institute for the COVID-19 Genomics UK (COG-UK) Consortium | The Lighthouse Lab in Milton Keynes and Alex Alderton, Roberto Amato, Sonia Goncalves, Ewan Harrison, David K. Jackson, Ian Johnston, Dominic Kwiatkowski, Cordelia Langford, John Sillitoe on behalf of the Wellcome Sanger Institute COVID-19 Surveillance Team |
| EPI_ISL_864039 | Lighthouse Lab in Alderley Park | Wellcome Sanger Institute for the COVID-19 Genomics UK (COG-UK) Consortium | Jacquelyn Wynn, Mairead Hyland, The Lighthouse Lab in Alderley Park and Alex Alderton, Roberto Amato, Sonia Goncalves, Ewan Harrison, David K. Jackson, Ian Johnston, Dominic Kwiatkowski, Cordelia Langford, John Sillitoe on behalf of the Wellcome Sanger Institute COVID-19 Surveillance Team |
| EPI_ISL_864204, EPI_ISL_864211, EPI_ISL_864213, EPI_ISL_864216, EPI_ISL_864217, EPI_ISL_864221, EPI_ISL_864227, EPI_ISL_864228, EPI_ISL_864230, EPI_ISL_864239, EPI_ISL_864244, EPI_ISL_864255, EPI_ISL_864262, EPI_ISL_864266, EPI_ISL_864274, EPI_ISL_864276, EPI_ISL_864281, EPI_ISL_864288, EPI_ISL_864291, EPI_ISL_864297, EPI_ISL_864301, EPI_ISL_864305, EPI_ISL_864309, EPI_ISL_864311, EPI_ISL_864313, EPI_ISL_864317, EPI_ISL_864321, EPI_ISL_864324, EPI_ISL_864328, EPI_ISL_864336, EPI_ISL_864352, EPI_ISL_864354, EPI_ISL_864356, EPI_ISL_864361, EPI_ISL_864365, EPI_ISL_864370, EPI_ISL_864383, EPI_ISL_864384, EPI_ISL_864390, EPI_ISL_864396, EPI_ISL_864399, EPI_ISL_864400, EPI_ISL_864402, EPI_ISL_864403, EPI_ISL_864409, EPI_ISL_864411, EPI_ISL_864414, EPI_ISL_864422, EPI_ISL_864436, EPI_ISL_864439, EPI_ISL_864440, EPI_ISL_864443, EPI_ISL_864445, EPI_ISL_864447, EPI_ISL_864452, EPI_ISL_864461, EPI_ISL_864465, EPI_ISL_864467, EPI_ISL_864469, EPI_ISL_864472, EPI_ISL_864475, EPI_ISL_864478, EPI_ISL_864483, EPI_ISL_864484, EPI_ISL_864485, EPI_ISL_864487, EPI_ISL_864493, EPI_ISL_864501, EPI_ISL_864513, EPI_ISL_864515, EPI_ISL_864516, EPI_ISL_864520, EPI_ISL_864527, EPI_ISL_864528, EPI_ISL_864529, EPI_ISL_864534, EPI_ISL_864539, EPI_ISL_864540, EPI_ISL_864542 | | | |
| see above | Lighthouse Lab in Milton Keynes | Wellcome Sanger Institute for the COVID-19 Genomics UK (COG-UK) Consortium | The Lighthouse Lab in Milton Keynes and Alex Alderton, Roberto Amato, Sonia Goncalves, Ewan Harrison, David K. Jackson, Ian Johnston, Dominic Kwiatkowski, Cordelia Langford, John Sillitoe on behalf of the Wellcome Sanger Institute COVID-19 Surveillance Team |
| EPI_ISL_864582 | CHU Purpan - Laboratoire de Virologie - Institut Fédératif de Biologie | CHU Purpan - Laboratoire de Virologie - Institut Fédératif de Biologie | Latour J., Ranger N., Dubois M., Carcenac R., Harter A., Boyer P., Tremeaux P., Izopet J. |
| EPI_ISL_864719, EPI_ISL_864720, EPI_ISL_864734, EPI_ISL_864737 | University Hospitals of Geneva, Laboratory of Virology | HUG, Laboratory of Virology and the Health2030 Genome Center | Samuel Cordey, Ana Rita Goncalves, Laurent Kaiser, Lorenzo Cerutti, Henri Pegeot, Melyssa Elies, Deborah Penet, Keith Harshman, Ioannis Xenarios, Emmanouil Dermitzakis |
| EPI_ISL_864754, EPI_ISL_864760, EPI_ISL_864797, EPI_ISL_864799, EPI_ISL_864803, EPI_ISL_864807, EPI_ISL_864809, EPI_ISL_864810, EPI_ISL_864811, EPI_ISL_864813, EPI_ISL_864814, EPI_ISL_864815, EPI_ISL_864818, EPI_ISL_864829, EPI_ISL_864846, EPI_ISL_864875, EPI_ISL_864883, EPI_ISL_864889, EPI_ISL_864893 | | | |
| see above | Department of Pathology, University of Cambridge | COVID-19 Genomics UK (COG-UK) Consortium | Aminu S. Jahun, Yasmin Chaudhry, Grant Hall, Iliana Georgana, Myra Hosmillo, Martin D. Curran, Malte Pinckert, Surendra Parmar, Ian Goodfellow |
| EPI_ISL_865012 | Lighthouse Lab in Glasgow / MRC-University of Glasgow Centre for Virus Research | COVID-19 Genomics UK (COG-UK) Consortium | Ana da Silva Filipe, Natasha Johnson, Kathy Smollett, Daniel Mair, Stephen Carmichael, Alice Broos, Lily Tong, Jenna Nichols, Kyriaki Nomikou; Sarah McDonald; Harper VanSteenhouse, Yumi Kasai, David Gray, Carol Clugston, Anna Dominiczak; Alasdair MacLean, Rory Gunson; Richard Orton, Joseph Hughes, Sreenu Vattipally, David L Robertson; Sharif Shaaban, Matthew Holden; Kathy Li, James Shepherd, Antonia Ho, Emma Thomson |
| EPI_ISL_865014, EPI_ISL_865015, EPI_ISL_865016, EPI_ISL_865017, EPI_ISL_865018, EPI_ISL_865020, EPI_ISL_865041, EPI_ISL_865042, EPI_ISL_865043, EPI_ISL_865044, EPI_ISL_865045, EPI_ISL_865046, EPI_ISL_865047, EPI_ISL_865048, EPI_ISL_865049, EPI_ISL_865050, EPI_ISL_865051, EPI_ISL_865052, EPI_ISL_865054, EPI_ISL_865055, EPI_ISL_865066, EPI_ISL_865067 | | | |
| see above | West of Scotland Specialist Virology Centre, NHSGGC / MRC-University of Glasgow Centre for Virus Research | COVID-19 Genomics UK (COG-UK) Consortium | Ana da Silva Filipe, Natasha Johnson, Kathy Smollett, Daniel Mair, Stephen Carmichael, Alice Broos, Lily Tong, Jenna Nichols, Kyriaki Nomikou; Sarah McDonald; Richard Orton, Joseph Hughes, Sreenu Vattipally, David L Robertson; Alasdair MacLean, Rory Gunson; Sharif Shaaban, Matthew Holden; Rachel Blacow, Guy Mollett, Kathy Li, James Shepherd, Antonia Ho, Emma Thomson |
| EPI_ISL_865074, EPI_ISL_865075, EPI_ISL_865076, EPI_ISL_865077, EPI_ISL_865089, EPI_ISL_865115, EPI_ISL_865117, EPI_ISL_865143, EPI_ISL_865149, EPI_ISL_865150, EPI_ISL_865151, EPI_ISL_865152 | | | |
| see above | Virology Department, Royal Infirmary of Edinburgh, NHS Lothian / School of Biological Sciences, University of Edinburgh / Institute of Genetics and Molecular Medicine, University of Edinburgh | COVID-19 Genomics UK (COG-UK) Consortium | McHugh M, Dewar R, Rooke S, Gallagher M, Balcaza C, O'Toole Á, Scher E, Hill V, McCrone JT, Colquhoun R, Yu X, Jackson B, Rambaut A, Williams TC, Templeton K |
| EPI_ISL_865203, EPI_ISL_865204, EPI_ISL_865205, EPI_ISL_865206, EPI_ISL_865270, EPI_ISL_865271, EPI_ISL_865322, EPI_ISL_865324, EPI_ISL_865331, EPI_ISL_865332, EPI_ISL_865333, EPI_ISL_865336, EPI_ISL_865338, EPI_ISL_865339, EPI_ISL_865340, EPI_ISL_865479, EPI_ISL_865480, EPI_ISL_865484 | | | |
| see above | Liverpool Clinical Laboratories | COVID-19 Genomics UK (COG-UK) Consortium | Sam Haldenby, Anita Lucaci, Steve Paterson, Julian Hiscox, Alistair Darby, M Almsaud, A Alrezaihi, Muhannad Alruwaili, Stuart D Armstrong, Jones Benjamin, Eleanor G Bentley, Anu Chawla, Jordan J Clark, Angela Cowell, Richard Eccles, Isabel García-Dorival, Matthew Gemmell, Alessandro Gerada, PKF Gilmore, Richard Gregory, Ximeng Han, Catherine Hartley, Margaret Hughes, Miren Iturriza-Gomara, James Johnson, L Luu, Jenifer Manson, Charlotte Nelson, Elaine O'Toole, Cassie Olateju, Rebekah Penrice-Randal , Lucille Rainbow, N.P Randle, Trevor Ian Robinson, Parul Sharma, Ghada T Shawli, James P Stewart, Neil Swainston, Ecaterina Vamos, Joanne Watts, Mark Whitehead |
| EPI_ISL_866077, EPI_ISL_866078, EPI_ISL_866079, EPI_ISL_866080, EPI_ISL_866109, EPI_ISL_866110, EPI_ISL_866111 | University College London Hospital | COVID-19 Genomics UK (COG-UK) Consortium | Judith Heaney, Matthew Byott, Catherine Houlihan, Dan Frampton, Stuart Kirk, Moira Spyer and Eleni Nastouli |
| EPI_ISL_866191, EPI_ISL_866192, EPI_ISL_866193 | University College London, Great Ormond Street Hospital for Children NHS Foundation Trust, Imperial College Healthcare NHS Trust | COVID-19 Genomics UK (COG-UK) Consortium | Sergi Castellano, Rachel Williams, Mark Kristiansen, Paola Resende Silva, Sunando Roy, Tony Brooks, Helena Tutill, Paola Niola, Patricia Dyal, Charlotte Williams, Leysa Forrest, Yasmin Panchbhaya, Jacqueline Findlay, Samuel Weeks, Julianne Brown, Kathryn Harris, Paul Randell, James Price, Alison Holmes, Judith Breuer |
| EPI_ISL_866354, EPI_ISL_866355, EPI_ISL_866357, EPI_ISL_866358, EPI_ISL_866359, EPI_ISL_866362, EPI_ISL_866366, EPI_ISL_866367, EPI_ISL_866373, EPI_ISL_866374, EPI_ISL_866375, EPI_ISL_866376, EPI_ISL_866377, EPI_ISL_866378, EPI_ISL_866379, EPI_ISL_866380, EPI_ISL_866381, EPI_ISL_866382, EPI_ISL_866384, EPI_ISL_866385, EPI_ISL_866386, EPI_ISL_866387, EPI_ISL_866388, EPI_ISL_866395, EPI_ISL_866396, EPI_ISL_866397, EPI_ISL_866398, EPI_ISL_866399 | | | |

| | | | |
|---|---|---|---|
| see above | Regional Virus Laboratory, Belfast Health and Social Care Trust | COVID-19 Genomics UK (COG-UK) Consortium | Conall McCaughey, James McKenna, Tanya Curran, Susan Feeney, Alison Watt, Ciara Cox, Mairead Connor, Zoltan Molnar, David Simpson, Derek Fairley |
| EPI_ISL_866662, EPI_ISL_866675, EPI_ISL_866676, EPI_ISL_866678, EPI_ISL_866680, EPI_ISL_866685, EPI_ISL_866687, EPI_ISL_866690, EPI_ISL_866693, EPI_ISL_866694 | Quadram Institute Bioscience | COVID-19 Genomics UK (COG-UK) Consortium | Dave J. Baker, Gemma L. Kay, Alp Aydin, Thanh Le-Viet, Steven Rudder, Ana P. Tedim, Anastasia Kolyva, Maria Diaz, Leonardo de Oliveira Martins, Nabil-Fareed Alikhan, Lizzie Meadows, Rachael Stanley, Ngozi Elumogo, Muhammed Yasir, Nicholas M. Thomson, Alexander J Trotter, Rachel Gilroy, Samuel Bloomfield, Claire Stuart, Andrew Bell, Reenesh Prakash, Samir Dervisevic, Alison E. Mather, John Wain, Mark Webber, Andrew J. Page, Justin O'Grady |
| EPI_ISL_866907, EPI_ISL_866908 | Queens Medical Centre, Clinical Microbiology Department / DeepSeq Nottingham | COVID-19 Genomics UK (COG-UK) Consortium | Gemma Clark, Wendy Smith, Manjinder Khakh, Vicki M Fleming, Michelle M Lister, Hannah Howson-Wells, Jonathan Ball, Patrick McClure, Joseph Chappell, Theocharis Tsoleridis, Nadine Holmes, Matthew Carlisle, Christopher Moore, Fei Sang, Johnny Debebe, Victoria Wright, Matthew Loose |
| EPI_ISL_867034 | Lincolnshire Hospitals and DeepSeq Nottingham | COVID-19 Genomics UK (COG-UK) Consortium | Nichola Duckworth, Tim Sloan, Sarah Walsh, Jonathan Ball, Patrick McClure, Joeseph Chappell, Nadine Holmes, Matthew Carlisle, Christopher Moore, Fei Sang, Johnny Debebe, Victoria Wright, Matthew Loose |
| EPI_ISL_867164, EPI_ISL_867165, EPI_ISL_867167 | Oxford Viromics, NDM, University of Oxford; Oxford University Hospitals; Basingstoke and North Hampshire Hospital | COVID-19 Genomics UK (COG-UK) Consortium | Tanya Golubchik, David Bonsall, George Macintyre, Amy Trebes, Mariateresa de Cesare, Catrin Moore, Alex Mobbs, Anita Justice, Robert Shaw, Monique Andersson, Timothy Peto, Emma Wise, Nathan Moore, Jessica Lynch, Nick Cortes, Matilde Mori, Stephen Kidd, David Buck, John Todd, Christophe Fraser |
| EPI_ISL_867208, EPI_ISL_867209, EPI_ISL_867210, EPI_ISL_867211, EPI_ISL_867219, EPI_ISL_867220, EPI_ISL_867313, EPI_ISL_867314, EPI_ISL_867315, EPI_ISL_867416, EPI_ISL_867456, EPI_ISL_867503, EPI_ISL_867511, EPI_ISL_867512, EPI_ISL_867515, EPI_ISL_867751, EPI_ISL_867758, EPI_ISL_867759, EPI_ISL_867762, EPI_ISL_867763, EPI_ISL_867764, EPI_ISL_867766, EPI_ISL_867768, EPI_ISL_867771, EPI_ISL_867773, EPI_ISL_867776, EPI_ISL_867779 | | | |
| see above | Originating lab: Wales Specialist Virology Centre Sequencing lab: Pathogen Genomics Unit | Public Health Wales Microbiology Cardiff Wales Specialist Virology Centre | Catherine Moore, Johnathan Evans, Laura Gifford, Malorie Perry, Simon Cottrell, Angela Marchbank, Alec Birchley, Alexander Adams, Amy Gaskin, Bree Gatica-Wilcox, Jason Coombes, Joel Southgate, Lauren Gilbert, Lee Graham, Nicole Pacchiarini, Sara Kumziene-Summerhayes, Sarah Taylor, Sophie Jones, Sara Rey, Matthew Bull, Joanne Watkins, Sally Corden, Tom Connor |
| EPI_ISL_868001, EPI_ISL_868008, EPI_ISL_868309, EPI_ISL_868310, EPI_ISL_868322, EPI_ISL_868323, EPI_ISL_868324, EPI_ISL_868325, EPI_ISL_868329, EPI_ISL_868330, EPI_ISL_868331, EPI_ISL_868339, EPI_ISL_868340, EPI_ISL_868341, EPI_ISL_868342, EPI_ISL_868343, EPI_ISL_868344, EPI_ISL_868345, EPI_ISL_868346, EPI_ISL_868347, EPI_ISL_868348, EPI_ISL_868353 | | | |
| see above | Centre for Enzyme Innovation, University of Portsmouth / Translational Research Laboratory, Portsmouth Hospitals NHS Trust | COVID-19 Genomics UK (COG-UK) Consortium | Angela Beckett,Yann Bourgeois,Garry Scarlett,Sharon Glaysher,Scott Elliott,Kelly Bicknell,Robert Impey,Allyson Lloyd,Sarah Wyllie,Ethan Butcher,Anoop Chauhan,Samuel Robson |
| EPI_ISL_868370, EPI_ISL_868390, EPI_ISL_868392, EPI_ISL_868400, EPI_ISL_868402, EPI_ISL_868406, EPI_ISL_868416, EPI_ISL_868429, EPI_ISL_868431, EPI_ISL_868444, EPI_ISL_868450, EPI_ISL_868453, EPI_ISL_868454, EPI_ISL_868455, EPI_ISL_868468, EPI_ISL_868470, EPI_ISL_868474, EPI_ISL_868478, EPI_ISL_868481, EPI_ISL_868491, EPI_ISL_868498, EPI_ISL_868499, EPI_ISL_868502, EPI_ISL_868507, EPI_ISL_868515, EPI_ISL_868516, EPI_ISL_868518, EPI_ISL_868525, EPI_ISL_868531, EPI_ISL_868537, EPI_ISL_868542, EPI_ISL_868543, EPI_ISL_868545, EPI_ISL_868561, EPI_ISL_868565, EPI_ISL_868572, EPI_ISL_868590, EPI_ISL_868598, EPI_ISL_868601, EPI_ISL_868603, EPI_ISL_868608, EPI_ISL_868609, EPI_ISL_868612, EPI_ISL_868613, EPI_ISL_868620, EPI_ISL_868622, EPI_ISL_868625, EPI_ISL_868627, EPI_ISL_868630, EPI_ISL_868644, EPI_ISL_868654, EPI_ISL_868659, EPI_ISL_868664, EPI_ISL_868665, EPI_ISL_868675, EPI_ISL_868678, EPI_ISL_868680, EPI_ISL_868683, EPI_ISL_868699 | | | |
| see above | Virology Department, Sheffield Teaching Hospitals NHS Foundation Trust/Department of Infection, Immunity and Cardiovascular Disease, The Medical School, University of Sheffield | COVID-19 Genomics UK (COG-UK) Consortium | Thushan de Silva, Matthew Parker, Nikki Smith, Adri Angyal, Rebecca Brown, Luke Green, Rachel Tucker, Paul Parsons, Danielle Groves, Katie Johnson, Laura Carrilero, Alex Keeley, Dave Partridge, Matthew Wyles, Benjamin Lindsey, Mehmet Yavuz, Mohammad Raza, Cariad Evans |
| EPI_ISL_869167, EPI_ISL_869168, EPI_ISL_869170, EPI_ISL_869171, EPI_ISL_869174, EPI_ISL_869175, EPI_ISL_869176, EPI_ISL_869180, EPI_ISL_869181, EPI_ISL_869184, EPI_ISL_869185, EPI_ISL_869186, EPI_ISL_869187, EPI_ISL_869188, EPI_ISL_869189, EPI_ISL_869190, EPI_ISL_869191, EPI_ISL_869192, EPI_ISL_869193, EPI_ISL_869194, EPI_ISL_869195, EPI_ISL_869196, EPI_ISL_869197, EPI_ISL_869198, EPI_ISL_869199, EPI_ISL_869200, EPI_ISL_869201, EPI_ISL_869202, EPI_ISL_869203, EPI_ISL_869204, EPI_ISL_869205, EPI_ISL_869206, EPI_ISL_869207, EPI_ISL_869208, EPI_ISL_869209, EPI_ISL_869210, EPI_ISL_869211, EPI_ISL_869212, EPI_ISL_869213, EPI_ISL_869214, EPI_ISL_869215, EPI_ISL_869216, EPI_ISL_869225, EPI_ISL_869227 | | | |
| see above | New Mexico Department of Health Scientific Laboratory | Center for Global Health, University of New Mexico Health Sciences Center | Daryl Domman, Kurt Schwalm, Twila Kunde, Joseph Hicks, Anastacia Griego, Michael Edwards, Darrell Dinwiddie |
| EPI_ISL_871789 | Cerballiance Normandie | CERBA LAB | LAFOREST D, BRIAND G, LEPIGEON, technicien V; COUDRAY J; MARTINS-AMARAL F; COULON C, THOMAS Johan, VITEL R, HAMEL A. |
| EPI_ISL_871827, EPI_ISL_871828, EPI_ISL_871829, EPI_ISL_871830, EPI_ISL_871831 | Jessa | Jessa | Brigitte Maes, Bert Cruys |
| EPI_ISL_871844, EPI_ISL_871845, EPI_ISL_871846, EPI_ISL_871848, EPI_ISL_871849, EPI_ISL_871870, EPI_ISL_871875 | Wyoming Public Health Laboratory | Wyoming Public Health Laboratory | Noah Hull, Taylor Fearing, Lynette Gumbleton, Channing Weber, Ashley Norberg, Bailey Bowcutt, and Wanda Manley |
| EPI_ISL_871876 | Cerballiance Côte d'Azur | CERBA LAB | Prots L; Barrieu-Moussat S; Roquebert B; Costa JM; Hedbaut E; Trombert S; Lecorche E; Verdurme L; Malek Ramdane, Olivi M; Haïm-Boukobza S. |
| EPI_ISL_871879, EPI_ISL_871880, EPI_ISL_871881, EPI_ISL_871882, EPI_ISL_871883, EPI_ISL_871884, EPI_ISL_871885, EPI_ISL_871886, EPI_ISL_871887, EPI_ISL_871888, EPI_ISL_871889, EPI_ISL_871892, EPI_ISL_871894, EPI_ISL_871895, EPI_ISL_871897, EPI_ISL_871898, EPI_ISL_871899, EPI_ISL_871900, EPI_ISL_871901, EPI_ISL_871902, EPI_ISL_871903, EPI_ISL_871904, EPI_ISL_871905, EPI_ISL_871906, EPI_ISL_871907 | | | |
| see above | Hospital Universitario Marqués de Valdecilla - IDIVAL (Santander, Cantabria) | SeqCOVID-SPAIN consortium/IBV(CSIC) | Mónica Gozalo Margüello, María Eliecer Cano García, Jose Manuel Méndez Legaza, Daniel Pablo Marcos, Jesús Rodríguez Rodríguez, María Siller Ruiz and SeqCOVID-SPAIN consortium |
| EPI_ISL_871913, EPI_ISL_871938, EPI_ISL_871942, EPI_ISL_871943, EPI_ISL_871944, EPI_ISL_871945, EPI_ISL_871946, EPI_ISL_871947, EPI_ISL_871948, EPI_ISL_871949, EPI_ISL_871950, EPI_ISL_871951, EPI_ISL_871952, EPI_ISL_871953, EPI_ISL_871955, EPI_ISL_871957, EPI_ISL_871958 | | | |
| see above | Servicio de Microbiología Clínica (Complejo Hospitalario de Navarra, Pamplona), Instituto de Investigación Sanitaria de Navarra (IdiSNA) | SeqCOVID-SPAIN consortium/IBV(CSIC) | Carmen Ezpeleta Baquedano, Ana Navascués, Ana Miqueleiz and SeqCOVID-SPAIN consortium |
| EPI_ISL_871999, EPI_ISL_872001, EPI_ISL_872094 | Wyoming Public Health Laboratory | Wyoming Public Health Laboratory | Noah Hull, Taylor Fearing, Lynette Gumbleton, Channing Weber, Ashley Norberg, Bailey Bowcutt, and Wanda Manley |
| EPI_ISL_872104 | Chu Tivoli | GIGA Medical Genomics | Keith Durkin, Maria Artesi, Sébastien Bontems, Raphaël Boreux, Bouchra Boujemla, Cécile Meex, Pierrette Melin, Marie-Pierre Hayette, Vincent Bours |
| EPI_ISL_872198 | Hopital | National Reference Center for Viruses of Respiratory Infections, Institut Pasteur, Paris | Marion Barbet, Sylvie Behillil, Méline Bizard, Angela Brisebarre, Camille Capel, Etienne Simon-Lorière, Vincent Enouf, Maud Vanpeene, Sylvie van der Werf |
| EPI_ISL_872201, EPI_ISL_872205 | Labo Analyses Med | National Reference Center for Viruses of Respiratory Infections, Institut Pasteur, Paris | Marion Barbet, Sylvie Behillil, Méline Bizard, Angela Brisebarre, Camille Capel, Etienne Simon-Lorière, Vincent Enouf, Maud Vanpeene, Sylvie van der Werf,Girard Sophie |
| EPI_ISL_872206 | Hopital | National Reference Center for Viruses of Respiratory Infections, Institut Pasteur, Paris | Marion Barbet, Sylvie Behillil, Méline Bizard, Angela Brisebarre, Camille Capel, Etienne Simon-Lorière, Vincent Enouf, Maud Vanpeene, Sylvie van der Werf,Morvan Odile |
| EPI_ISL_872208 | Labo Analyses Med | National Reference Center for Viruses of Respiratory Infections, Institut Pasteur, Paris | Marion Barbet, Sylvie Behillil, Méline Bizard, Angela Brisebarre, Camille Capel, Etienne Simon-Lorière, Vincent Enouf, Maud Vanpeene, Sylvie van der Werf |
| EPI_ISL_872212 | Labo analyses med | National Reference Center for Viruses of Respiratory Infections, Institut Pasteur, Paris | Marion Barbet, Sylvie Behillil, Méline Bizard, Angela Brisebarre, Camille Capel, Etienne Simon-Lorière, Vincent Enouf, Maud Vanpeene, Sylvie van der Werf,Amzalag Jonas |
| EPI_ISL_872213, EPI_ISL_872214, EPI_ISL_872215, EPI_ISL_872216, EPI_ISL_872217, EPI_ISL_872218, EPI_ISL_872219, EPI_ISL_872233 | Hopital | National Reference Center for Viruses of Respiratory Infections, Institut Pasteur, Paris | Marion Barbet, Sylvie Behillil, Méline Bizard, Angela Brisebarre, Camille Capel, Etienne Simon-Lorière, Vincent Enouf, Maud Vanpeene, Sylvie van der Werf,Fourgeaud Jacques |
| EPI_ISL_872234 | Labo Analyses Med | National Reference Center for Viruses of Respiratory Infections, Institut Pasteur, Paris | Marion Barbet, Sylvie Behillil, Méline Bizard, Angela Brisebarre, Camille Capel, Etienne Simon-Lorière, Vincent Enouf, Maud Vanpeene, Sylvie van der Werf,Girard Sophie |
| EPI_ISL_872236 | Hopital | National Reference Center for Viruses of Respiratory | Marion Barbet, Sylvie Behillil, Méline Bizard, Angela Brisebarre, Camille Capel, Etienne Simon-Lorière, Vincent Enouf, Maud Vanpeene, Sylvie van der |

| | | | |
|---|---|---|---|
| | | Infections, Institut Pasteur, Paris | Werf,Guigon AuréLie |
| EPI_ISL_872238 | Labo analyses med | National Reference Center for Viruses of Respiratory Infections, Institut Pasteur, Paris | Marion Barbet, Sylvie Behillil, Méline Bizard, Angela Brisebarre, Camille Capel, Etienne Simon-Lorière, Vincent Enouf, Maud Vanpeene, Sylvie van der Werf,Amzalag Jonas |
| EPI_ISL_872254 | hopital | National Reference Center for Viruses of Respiratory Infections, Institut Pasteur, Paris | Marion Barbet, Sylvie Behillil, Méline Bizard, Angela Brisebarre, Camille Capel, Etienne Simon-Lorière, Vincent Enouf, Maud Vanpeene, Sylvie van der Werf,Ducancelle Alexandra |
| EPI_ISL_872256 | Labo Analyses Med | National Reference Center for Viruses of Respiratory Infections, Institut Pasteur, Paris | Marion Barbet, Sylvie Behillil, Méline Bizard, Angela Brisebarre, Camille Capel, Etienne Simon-Lorière, Vincent Enouf, Maud Vanpeene, Sylvie van der Werf |
| EPI_ISL_872264, EPI_ISL_872265 | Labo analyses med | National Reference Center for Viruses of Respiratory Infections, Institut Pasteur, Paris | Marion Barbet, Sylvie Behillil, Méline Bizard, Angela Brisebarre, Camille Capel, Etienne Simon-Lorière, Vincent Enouf, Maud Vanpeene, Sylvie van der Werf,Amzalag Jonas |
| EPI_ISL_872266, EPI_ISL_872267, EPI_ISL_872268, EPI_ISL_872270 | Hopital | National Reference Center for Viruses of Respiratory Infections, Institut Pasteur, Paris | Marion Barbet, Sylvie Behillil, Méline Bizard, Angela Brisebarre, Camille Capel, Etienne Simon-Lorière, Vincent Enouf, Maud Vanpeene, Sylvie van der Werf,Fourgeaud Jacques |
| EPI_ISL_872274 | Hopital | National Reference Center for Viruses of Respiratory Infections, Institut Pasteur, Paris | Marion Barbet, Sylvie Behillil, Méline Bizard, Angela Brisebarre, Camille Capel, Etienne Simon-Lorière, Vincent Enouf, Maud Vanpeene, Sylvie van der Werf,Irimia Alix |
| EPI_ISL_872278 | Hopital | National Reference Center for Viruses of Respiratory Infections, Institut Pasteur, Paris | Marion Barbet, Sylvie Behillil, Méline Bizard, Angela Brisebarre, Camille Capel, Etienne Simon-Lorière, Vincent Enouf, Maud Vanpeene, Sylvie van der Werf,Ramanantsoa CéLine |
| EPI_ISL_872279 | Labo Analyses Med | National Reference Center for Viruses of Respiratory Infections, Institut Pasteur, Paris | Marion Barbet, Sylvie Behillil, Méline Bizard, Angela Brisebarre, Camille Capel, Etienne Simon-Lorière, Vincent Enouf, Maud Vanpeene, Sylvie van der Werf,Bour Jean Baptiste |
| EPI_ISL_872291, EPI_ISL_872292 | Hopital | National Reference Center for Viruses of Respiratory Infections, Institut Pasteur, Paris | Marion Barbet, Sylvie Behillil, Méline Bizard, Angela Brisebarre, Camille Capel, Etienne Simon-Lorière, Vincent Enouf, Maud Vanpeene, Sylvie van der Werf,Guigon AuréLie |
| EPI_ISL_872309, EPI_ISL_872310, EPI_ISL_872311, EPI_ISL_872312, EPI_ISL_872314 | Labo analyses med | National Reference Center for Viruses of Respiratory Infections, Institut Pasteur, Paris | Marion Barbet, Sylvie Behillil, Méline Bizard, Angela Brisebarre, Camille Capel, Etienne Simon-Lorière, Vincent Enouf, Maud Vanpeene, Sylvie van der Werf,Amzalag Jonas |
| EPI_ISL_872315, EPI_ISL_872316, EPI_ISL_872317, EPI_ISL_872318, EPI_ISL_872319 | Hopital | National Reference Center for Viruses of Respiratory Infections, Institut Pasteur, Paris | Marion Barbet, Sylvie Behillil, Méline Bizard, Angela Brisebarre, Camille Capel, Etienne Simon-Lorière, Vincent Enouf, Maud Vanpeene, Sylvie van der Werf,Fourgeaud Jacques |
| EPI_ISL_872339, EPI_ISL_872340 | Hopital | National Reference Center for Viruses of Respiratory Infections, Institut Pasteur, Paris | Marion Barbet, Sylvie Behillil, Méline Bizard, Angela Brisebarre, Camille Capel, Etienne Simon-Lorière, Vincent Enouf, Maud Vanpeene, Sylvie van der Werf,Ducancelle Alexandra |
| EPI_ISL_872347 | Hopital | National Reference Center for Viruses of Respiratory Infections, Institut Pasteur, Paris | Marion Barbet, Sylvie Behillil, Méline Bizard, Angela Brisebarre, Camille Capel, Etienne Simon-Lorière, Vincent Enouf, Maud Vanpeene, Sylvie van der Werf,Irimia Alix |
| EPI_ISL_872512 | New Mexico Department of Health Scientific Laboratory | Center for Global Health, University of New Mexico Health Sciences Center | Daryl Domman, Kurt Schwalm, Twila Kunde, Joseph Hicks, Anastacia Griego, Michael Edwards, Darrell Dinwiddie |
| EPI_ISL_872576 | Laverty Pathology | NSW Health Pathology - Institute of Clinical Pathology and Medical Research; Westmead Hospital; University of Sydney | CIDM-PH et al. |
| EPI_ISL_872578 | Sydney South West Pathology Service (SSWPS) - Liverpool Hospital - NSW Health Pathology | NSW Health Pathology - Institute of Clinical Pathology and Medical Research; Westmead Hospital; University of Sydney | CIDM-PH et al. |
| EPI_ISL_872825, EPI_ISL_872826, EPI_ISL_872827, EPI_ISL_872830, EPI_ISL_872831, EPI_ISL_872832, EPI_ISL_872834, EPI_ISL_872836, EPI_ISL_872838, EPI_ISL_872844, EPI_ISL_872846, EPI_ISL_872848, EPI_ISL_872855, EPI_ISL_872857, EPI_ISL_872859, EPI_ISL_872860, EPI_ISL_872861, EPI_ISL_872863, EPI_ISL_872865, EPI_ISL_872866, EPI_ISL_872868, EPI_ISL_872871, EPI_ISL_872873, EPI_ISL_872875, EPI_ISL_872876, EPI_ISL_872877, EPI_ISL_872878, EPI_ISL_872879, EPI_ISL_872881, EPI_ISL_872883, EPI_ISL_872884, EPI_ISL_872885, EPI_ISL_872887, EPI_ISL_872889, EPI_ISL_872892, EPI_ISL_872895, EPI_ISL_872896 | | | |
| see above | University of Wisconsin-Madison AIDS Vaccine Research Laboratories | University of Wisconsin-Madison AIDS Vaccine Research Laboratories | Gage Moreno, Katarina Braun, et al. AIDS Vaccine Research Laboratories |
| EPI_ISL_873055, EPI_ISL_873057, EPI_ISL_873062, EPI_ISL_873063, EPI_ISL_873065, EPI_ISL_873066, EPI_ISL_873067, EPI_ISL_873068, EPI_ISL_873069, EPI_ISL_873070, EPI_ISL_873071, EPI_ISL_873072, EPI_ISL_873073, EPI_ISL_873074, EPI_ISL_873075, EPI_ISL_873076, EPI_ISL_873077, EPI_ISL_873078, EPI_ISL_873079, EPI_ISL_873080, EPI_ISL_873081, EPI_ISL_873082, EPI_ISL_873083, EPI_ISL_873084, EPI_ISL_873085, EPI_ISL_873086, EPI_ISL_873087, EPI_ISL_873088, EPI_ISL_873089, EPI_ISL_873090, EPI_ISL_873091, EPI_ISL_873092, EPI_ISL_873093, EPI_ISL_873094, EPI_ISL_873095, EPI_ISL_873096, EPI_ISL_873097, EPI_ISL_873098, EPI_ISL_873099, EPI_ISL_873100, EPI_ISL_873101, EPI_ISL_873102, EPI_ISL_873103, EPI_ISL_873104, EPI_ISL_873105, EPI_ISL_873106, EPI_ISL_873107, EPI_ISL_873108, EPI_ISL_873109, EPI_ISL_873111, EPI_ISL_873113, EPI_ISL_873116, EPI_ISL_873117, EPI_ISL_873119, EPI_ISL_873120, EPI_ISL_873122, EPI_ISL_873156, EPI_ISL_873159 | | | |
| see above | University of Michigan Clinical Microbiology Laboratory | Lauring Lab, University of Michigan, Department of Microbiology and Immunology | Valesano |
| EPI_ISL_873165 | Medical Laboratory Sciences, Arab American University | Medical Laboratory Sciences, Arab American University | Dumaidi,k., Al-Jawabreh,A., Ereqat,S., Al-Jawabreh,H., Nasereddin,A. |
| EPI_ISL_873176, EPI_ISL_873183 | Microbiology Division, South Carolina Department of Health and Environmental Control (SC DHEC) | Microbiology Division, South Carolina Department of Health and Environmental Control (SC DHEC) | Flores,H., Freeman,J. |
| EPI_ISL_873195, EPI_ISL_873196, EPI_ISL_873198, EPI_ISL_873199, EPI_ISL_873200, EPI_ISL_873202, EPI_ISL_873203, EPI_ISL_873206 | North Dakota Department of Health, Public Health Laboratory | North Dakota Department of Health, Public Health Laboratory | Lisa Wingerter |
| EPI_ISL_873238, EPI_ISL_873241, EPI_ISL_873243, EPI_ISL_873245, EPI_ISL_873246, EPI_ISL_873249, EPI_ISL_873250, EPI_ISL_873252, EPI_ISL_873254 | M Health Fairview | Minnesota Department of Health, Public Health Laboratory | Alexandra Lorentz, Jacob Garfin, Matt Plumb, and Xiong Wang |
| EPI_ISL_873273, EPI_ISL_873274, EPI_ISL_873275 | Vault Health | Minnesota Department of Health, Public Health Laboratory | Alexandra Lorentz, Jacob Garfin, Matt Plumb, and Xiong Wang |
| EPI_ISL_873277 | Environmental and Global Health, University of Florida | Environmental and Global Health, University of Florida | Iovine,N.M., Waltzek,T.B., Subramaniam,K., Loeb,J.C.,Stephenson,C.J., Lessard,K., Reeves,M., Wilkerson,G., Morris,J.G. and Lednicky,J.A. |
| EPI_ISL_873624, EPI_ISL_873746 | Lighthouse Lab in Cambridge | Wellcome Sanger Institute for the COVID-19 Genomics UK (COG-UK) Consortium | Rob Howes, The Lighthouse Lab in Cambridge and Alex Alderton, Roberto Amato, Sonia Goncalves, Ewan Harrison, David K. Jackson, Ian Johnston, Dominic Kwiatkowski, Cordelia Langford, John Sillitoe on behalf of the Wellcome Sanger Institute COVID-19 Surveillance Team |
| EPI_ISL_874170, EPI_ISL_874281 | Lighthouse Lab in Alderley Park | Wellcome Sanger Institute for the COVID-19 Genomics UK (COG-UK) Consortium | Jacquelyn Wynn, Mairead Hyland, The Lighthouse Lab in Alderley Park and Alex Alderton, Roberto Amato, Sonia Goncalves, Ewan Harrison, David K. Jackson, Ian Johnston, Dominic Kwiatkowski, Cordelia Langford, John Sillitoe on behalf of the Wellcome Sanger Institute COVID-19 Surveillance Team |
| EPI_ISL_874990, EPI_ISL_874991, EPI_ISL_874992, EPI_ISL_874993, EPI_ISL_874994, EPI_ISL_874995, EPI_ISL_874996, EPI_ISL_874997, EPI_ISL_874998, EPI_ISL_874999, EPI_ISL_875000, EPI_ISL_875001, EPI_ISL_875002, EPI_ISL_875003, EPI_ISL_875004, EPI_ISL_875005, EPI_ISL_875006, EPI_ISL_875007, EPI_ISL_875008, EPI_ISL_875009, EPI_ISL_875010, EPI_ISL_875011, EPI_ISL_875012, EPI_ISL_875013, EPI_ISL_875014, EPI_ISL_875015, EPI_ISL_875016, EPI_ISL_875017, EPI_ISL_875018, EPI_ISL_875019, EPI_ISL_875020, EPI_ISL_875021, EPI_ISL_875022, EPI_ISL_875023, EPI_ISL_875024, EPI_ISL_875025, EPI_ISL_875026, EPI_ISL_875027, EPI_ISL_875028, EPI_ISL_875029, EPI_ISL_875030, EPI_ISL_875031, EPI_ISL_875032, EPI_ISL_875033, EPI_ISL_875034, EPI_ISL_875035, EPI_ISL_875036, EPI_ISL_875037, EPI_ISL_875038, EPI_ISL_875039, EPI_ISL_875040, EPI_ISL_875041, EPI_ISL_875042, EPI_ISL_875043, EPI_ISL_875044, EPI_ISL_875045, EPI_ISL_875046, EPI_ISL_875047, EPI_ISL_875048, EPI_ISL_875049, EPI_ISL_875050, EPI_ISL_875051, EPI_ISL_875052, EPI_ISL_875053, EPI_ISL_875054, EPI_ISL_875055, EPI_ISL_875056, EPI_ISL_875057, EPI_ISL_875058, EPI_ISL_875059, EPI_ISL_875060, EPI_ISL_875061, EPI_ISL_875062, EPI_ISL_875063, EPI_ISL_875064, EPI_ISL_875065, EPI_ISL_875066, EPI_ISL_875067, EPI_ISL_875068, EPI_ISL_875069, EPI_ISL_875070, EPI_ISL_875071, EPI_ISL_875072, EPI_ISL_875073, EPI_ISL_875074, EPI_ISL_875075, EPI_ISL_875076, EPI_ISL_875077, EPI_ISL_875078, EPI_ISL_875079, EPI_ISL_875080, EPI_ISL_875081, EPI_ISL_875082, EPI_ISL_875083, EPI_ISL_875084, EPI_ISL_875085, EPI_ISL_875086, EPI_ISL_875087, EPI_ISL_875088, EPI_ISL_875089, EPI_ISL_875090, EPI_ISL_875091, EPI_ISL_875092, EPI_ISL_875093, EPI_ISL_875094, EPI_ISL_875095, EPI_ISL_875096, EPI_ISL_875097, EPI_ISL_875098, EPI_ISL_875099, EPI_ISL_875100, EPI_ISL_875101, EPI_ISL_875102, EPI_ISL_875103, EPI_ISL_875104, EPI_ISL_875105, EPI_ISL_875106, EPI_ISL_875107, EPI_ISL_875108, EPI_ISL_875109, EPI_ISL_875110, EPI_ISL_875111, EPI_ISL_875112, EPI_ISL_875113, EPI_ISL_875114, EPI_ISL_875115, EPI_ISL_875116, EPI_ISL_875117, EPI_ISL_875118, EPI_ISL_875119, EPI_ISL_875120, EPI_ISL_875121, EPI_ISL_875122, EPI_ISL_875123, EPI_ISL_875124, EPI_ISL_875125, EPI_ISL_875126, EPI_ISL_875127, EPI_ISL_875128, EPI_ISL_875129, EPI_ISL_875130, EPI_ISL_875131, EPI_ISL_875132, EPI_ISL_875133, | | | |

| | | | |
|---|---|---|---|
| EPI_ISL_875134, EPI_ISL_875135, EPI_ISL_875136, EPI_ISL_875137, EPI_ISL_875138, EPI_ISL_875139, EPI_ISL_875140, EPI_ISL_875141, EPI_ISL_875142, EPI_ISL_875143, EPI_ISL_875144, EPI_ISL_875145, EPI_ISL_875146, EPI_ISL_875147, EPI_ISL_875148, EPI_ISL_875149, EPI_ISL_875150, EPI_ISL_875151, EPI_ISL_875152, EPI_ISL_875153, EPI_ISL_875154, EPI_ISL_875155, EPI_ISL_875156, EPI_ISL_875157, EPI_ISL_875158, EPI_ISL_875159, EPI_ISL_875160, EPI_ISL_875161, EPI_ISL_875162, EPI_ISL_875163, EPI_ISL_875164, EPI_ISL_875165, EPI_ISL_875166, EPI_ISL_875167, EPI_ISL_875168, EPI_ISL_875169, EPI_ISL_875170, EPI_ISL_875171, EPI_ISL_875172, EPI_ISL_875173, EPI_ISL_875174, EPI_ISL_875175, EPI_ISL_875176, EPI_ISL_875177, EPI_ISL_875178, EPI_ISL_875179, EPI_ISL_875180, EPI_ISL_875181, EPI_ISL_875182, EPI_ISL_875183, EPI_ISL_875184, EPI_ISL_875185, EPI_ISL_875186, EPI_ISL_875187, EPI_ISL_875188, EPI_ISL_875189, EPI_ISL_875190, EPI_ISL_875191, EPI_ISL_875192, EPI_ISL_875193, EPI_ISL_875194, EPI_ISL_875195, EPI_ISL_875196, EPI_ISL_875197, EPI_ISL_875198, EPI_ISL_875199, EPI_ISL_875200, EPI_ISL_875201, EPI_ISL_875202, EPI_ISL_875203, EPI_ISL_875204, EPI_ISL_875205, EPI_ISL_875206, EPI_ISL_875207, EPI_ISL_875208, EPI_ISL_875209, EPI_ISL_875210, EPI_ISL_875211, EPI_ISL_875212, EPI_ISL_875213, EPI_ISL_875214, EPI_ISL_875215, EPI_ISL_875216, EPI_ISL_875217, EPI_ISL_875218, EPI_ISL_875219, EPI_ISL_875220, EPI_ISL_875221, EPI_ISL_875222, EPI_ISL_875223, EPI_ISL_875224, EPI_ISL_875225, EPI_ISL_875226, EPI_ISL_875227, EPI_ISL_875228, EPI_ISL_875229, EPI_ISL_875230, EPI_ISL_875231, EPI_ISL_875232, EPI_ISL_875233, EPI_ISL_875234, EPI_ISL_875235, EPI_ISL_875236, EPI_ISL_875237, EPI_ISL_875238, EPI_ISL_875239, EPI_ISL_875240, EPI_ISL_875241, EPI_ISL_875242, EPI_ISL_875243, EPI_ISL_875244, EPI_ISL_875245, EPI_ISL_875246, EPI_ISL_875247, EPI_ISL_875248, EPI_ISL_875249, EPI_ISL_875250, EPI_ISL_875251, EPI_ISL_875252, EPI_ISL_875253, EPI_ISL_875254, EPI_ISL_875255, EPI_ISL_875256, EPI_ISL_875257, EPI_ISL_875258, EPI_ISL_875259, EPI_ISL_875260, EPI_ISL_875261, EPI_ISL_875262, EPI_ISL_875263, EPI_ISL_875264, EPI_ISL_875265, EPI_ISL_875266, EPI_ISL_875267, EPI_ISL_875268, EPI_ISL_875269, EPI_ISL_875270, EPI_ISL_875271, EPI_ISL_875272, EPI_ISL_875273, EPI_ISL_875274, EPI_ISL_875275, EPI_ISL_875276, EPI_ISL_875277, EPI_ISL_875278, EPI_ISL_875279, EPI_ISL_875280, EPI_ISL_875281, EPI_ISL_875282, EPI_ISL_875283, EPI_ISL_875284, EPI_ISL_875285, EPI_ISL_875286, EPI_ISL_875287, EPI_ISL_875288, EPI_ISL_875289, EPI_ISL_875290, EPI_ISL_875291, EPI_ISL_875292, EPI_ISL_875293, EPI_ISL_875294, EPI_ISL_875295, EPI_ISL_875296, EPI_ISL_875297, EPI_ISL_875298, EPI_ISL_875299, EPI_ISL_875300, EPI_ISL_875301, EPI_ISL_875302, EPI_ISL_875303, EPI_ISL_875304, EPI_ISL_875305, EPI_ISL_875306, EPI_ISL_875307, EPI_ISL_875308, EPI_ISL_875309, EPI_ISL_875310, EPI_ISL_875311, EPI_ISL_875312, EPI_ISL_875313, EPI_ISL_875314, EPI_ISL_875315, EPI_ISL_875316, EPI_ISL_875317, EPI_ISL_875318, EPI_ISL_875319, EPI_ISL_875320, EPI_ISL_875321, EPI_ISL_875322, EPI_ISL_875323, EPI_ISL_875324, EPI_ISL_875325, EPI_ISL_875326, EPI_ISL_875327, EPI_ISL_875328, EPI_ISL_875329, EPI_ISL_875330, EPI_ISL_875331, EPI_ISL_875332, EPI_ISL_875333, EPI_ISL_875334, EPI_ISL_875335, EPI_ISL_875336, EPI_ISL_875337, EPI_ISL_875338, EPI_ISL_875339, EPI_ISL_875340, EPI_ISL_875341 | | | |
| see above | Lighthouse Lab in Milton Keynes | Wellcome Sanger Institute for the COVID-19 Genomics UK (COG-UK) Consortium | The Lighthouse Lab in Milton Keynes and Alex Alderton, Roberto Amato, Sonia Goncalves, Ewan Harrison, David K. Jackson, Ian Johnston, Dominic Kwiatkowski, Cordelia Langford, John Sillitoe on behalf of the Wellcome Sanger Institute COVID-19 Surveillance Team |
| EPI_ISL_875346 | National Institute of Infectious Diseases-Prof. Dr. Matei Bals Molecular Diagnostics Laboratory | National Institute of Infectious Diseases-Prof. Dr. Matei Bals Molecular Diagnostics Laboratory | Leontina Banica, Marius Surleac, Corina Casangiu, Petre Milu, Andreea Tudor, Simona Paraschiv, Dan Otelea |
| EPI_ISL_875460, EPI_ISL_875461, EPI_ISL_875462, EPI_ISL_875463, EPI_ISL_875464, EPI_ISL_875465, EPI_ISL_875466, EPI_ISL_875467, EPI_ISL_875468, EPI_ISL_875469, EPI_ISL_875470, EPI_ISL_875471, EPI_ISL_875472, EPI_ISL_875473, EPI_ISL_875474, EPI_ISL_875475, EPI_ISL_875476, EPI_ISL_875477, EPI_ISL_875478, EPI_ISL_875479, EPI_ISL_875480, EPI_ISL_875481, EPI_ISL_875482, EPI_ISL_875483, EPI_ISL_875484, EPI_ISL_875485, EPI_ISL_875486, EPI_ISL_875487, EPI_ISL_875488, EPI_ISL_875489, EPI_ISL_875490, EPI_ISL_875491, EPI_ISL_875492, EPI_ISL_875493, EPI_ISL_875494, EPI_ISL_875495, EPI_ISL_875496, EPI_ISL_875497, EPI_ISL_875498, EPI_ISL_875499, EPI_ISL_875500, EPI_ISL_875501, EPI_ISL_875502, EPI_ISL_875503, EPI_ISL_875504, EPI_ISL_875505, EPI_ISL_875506, EPI_ISL_875507, EPI_ISL_875508, EPI_ISL_875509, EPI_ISL_875510 | | | |
| see above | National Virus Reference Laboratory | National Virus Reference Laboratory | Michael Carr, Gabriel Gonzalez, Jonathan Dean, Cillian F De Gascun |
| EPI_ISL_875518 | Institute of Virology, Biomedical Research Center of the Slovak Academy of Sciences, Bratislava | Faculty of Natural Sciences, Comenius University, Bratislava | Broa Brejová, Viktória abanová, Kristína Boršová, Viktória Hodorová, Sabina Fumaová Havlíková, Juraj Kopáek, Martina Liková, ubomíra Lukáiková, Martina Neboháová, Monika Sláviková, Tomáš Vina, Jozef Nosek, Boris Klempa |
| EPI_ISL_875520 | Institute of Virology, Biomedical Research Center of the Slovak Academy of Sciences, Bratislava | Faculty of Natural Sciences, Comenius University, Bratislava | Kristína Boršová, Viktória abanová, Broa Brejová, Viktória Hodorová, Sabina Fumaová Havlíková, Juraj Kopáek, Martina Liková, ubomíra Lukáiková, Martina Neboháová, Monika Sláviková, Tomáš Vina, Boris Klempa, Jozef Nosek |
| EPI_ISL_875521, EPI_ISL_875522 | Institute of Virology, Biomedical Research Center of the Slovak Academy of Sciences, Bratislava | Faculty of Natural Sciences, Comenius University, Bratislava | Viktória abanová, Kristína Boršová, Broa Brejová, Viktória Hodorová, Sabina Fumaová Havlíková, Juraj Kopáek, Martina Liková, ubomíra Lukáiková, Martina Neboháová, Monika Sláviková, Tomáš Vina, Jozef Nosek, Boris Klempa |
| EPI_ISL_875523 | Institute of Virology, Biomedical Research Center of the Slovak Academy of Sciences, Bratislava | Faculty of Natural Sciences, Comenius University, Bratislava | Kristína Boršová, Viktória abanová, Broa Brejová, Viktória Hodorová, Sabina Fumaová Havlíková, Juraj Kopáek, Martina Liková, ubomíra Lukáiková, Martina Neboháová, Monika Sláviková, Tomáš Vina, Boris Klempa, Jozef Nosek |
| EPI_ISL_875524 | Institute of Virology, Biomedical Research Center of the Slovak Academy of Sciences, Bratislava | Faculty of Natural Sciences, Comenius University, Bratislava | Viktória abanová, Kristína Boršová, Broa Brejová, Viktória Hodorová, Sabina Fumaová Havlíková, Juraj Kopáek, Martina Liková, ubomíra Lukáiková, Martina Neboháová, Monika Sláviková, Tomáš Vina, Jozef Nosek, Boris Klempa |
| EPI_ISL_875525 | Institute of Virology, Biomedical Research Center of the Slovak Academy of Sciences, Bratislava | Faculty of Natural Sciences, Comenius University, Bratislava | Kristína Boršová, Viktória abanová, Broa Brejová, Viktória Hodorová, Sabina Fumaová Havlíková, Juraj Kopáek, Martina Liková, ubomíra Lukáiková, Martina Neboháová, Monika Sláviková, Tomáš Vina, Boris Klempa, Jozef Nosek |
| EPI_ISL_875526, EPI_ISL_875527 | Institute of Virology, Biomedical Research Center of the Slovak Academy of Sciences, Bratislava | Faculty of Natural Sciences, Comenius University, Bratislava | Broa Brejová, Viktória abanová, Kristína Boršová, Viktória Hodorová, Sabina Fumaová Havlíková, Juraj Kopáek, Martina Liková, ubomíra Lukáiková, Martina Neboháová, Monika Sláviková, Tomáš Vina, Jozef Nosek, Boris Klempa |
| EPI_ISL_875528 | Institute of Virology, Biomedical Research Center of the Slovak Academy of Sciences, Bratislava | Faculty of Natural Sciences, Comenius University, Bratislava | Kristína Boršová, Viktória abanová, Broa Brejová, Viktória Hodorová, Sabina Fumaová Havlíková, Juraj Kopáek, Martina Liková, ubomíra Lukáiková, Martina Neboháová, Monika Sláviková, Tomáš Vina, Boris Klempa, Jozef Nosek |
| EPI_ISL_875530 | Institute of Virology, Biomedical Research Center of the Slovak Academy of Sciences, Bratislava | Faculty of Natural Sciences, Comenius University, Bratislava | Viktória abanová, Kristína Boršová, Broa Brejová, Viktória Hodorová, Sabina Fumaová Havlíková, Juraj Kopáek, Martina Liková, ubomíra Lukáiková, Martina Neboháová, Monika Sláviková, Tomáš Vina, Jozef Nosek, Boris Klempa |
| EPI_ISL_875660, EPI_ISL_875661 | University of Michigan Clinical Microbiology Laboratory | Lauring Lab, University of Michigan, Department of Microbiology and Immunology | Valesano |
| EPI_ISL_875686 | National Virus Reference Laboratory | National Virus Reference Laboratory | Michael Carr, Gabriel Gonzalez, Jonathan Dean, Cillian F De Gascun |
| EPI_ISL_876052, EPI_ISL_876053, EPI_ISL_876056, EPI_ISL_876059, EPI_ISL_876060, EPI_ISL_876062, EPI_ISL_876081, EPI_ISL_876127, EPI_ISL_876128, EPI_ISL_876131, EPI_ISL_876137, EPI_ISL_876146, EPI_ISL_876147, EPI_ISL_876149, EPI_ISL_876163, EPI_ISL_876172, EPI_ISL_876173, EPI_ISL_876178, EPI_ISL_876180, EPI_ISL_876184, EPI_ISL_876186, EPI_ISL_876196, EPI_ISL_876198, EPI_ISL_876200, EPI_ISL_876203, EPI_ISL_876208, EPI_ISL_876245, EPI_ISL_876246, EPI_ISL_876247, EPI_ISL_876248, EPI_ISL_876249, EPI_ISL_876250, EPI_ISL_876251 | | | |
| see above | Massachusetts State Public Health Laboratory | Massachusetts State Public Health Laboratory | Andrew Lang, Timelia Fink, Glen Gallagher, Sandra Smole |
| EPI_ISL_876588 | Blackhawk Genomics | Pathogen Discovery, Respiratory Viruses Branch, Division of Viral Diseases, Centers for Disease Control and Prevention | Ying Tao, Yan Li, Jing Zhang, Krista Queen, Anna Uehara, Peter Cook, Clinton R. Paden, Haibin Wang, Suxiang Tong |
| EPI_ISL_876594, EPI_ISL_876596 | TN Division of Laboratory Services | Pathogen Discovery, Respiratory Viruses Branch, Division of Viral Diseases, Centers for Disease Control and Prevention | Ying Tao, Yan Li, Jing Zhang, Krista Queen, Anna Uehara, Peter Cook, Clinton R. Paden, Haibin Wang, Suxiang Tong |
| EPI_ISL_876625, EPI_ISL_876626, EPI_ISL_876628, EPI_ISL_876629, EPI_ISL_876630, EPI_ISL_876632, EPI_ISL_876633, EPI_ISL_876634, EPI_ISL_876635, EPI_ISL_876636, EPI_ISL_876637, EPI_ISL_876638, EPI_ISL_876639, EPI_ISL_876640, EPI_ISL_876641, EPI_ISL_876643, EPI_ISL_876644, EPI_ISL_876645, EPI_ISL_876646, EPI_ISL_876647, EPI_ISL_876648, EPI_ISL_876649, EPI_ISL_876650, EPI_ISL_876651, EPI_ISL_876652, EPI_ISL_876653, EPI_ISL_876654, EPI_ISL_876655, EPI_ISL_876656, EPI_ISL_876657, EPI_ISL_876658, EPI_ISL_876660, EPI_ISL_876661, EPI_ISL_876662, EPI_ISL_876663, EPI_ISL_876664, EPI_ISL_876665, EPI_ISL_876666, EPI_ISL_876668, EPI_ISL_876670, EPI_ISL_876671, EPI_ISL_876672, EPI_ISL_876673, EPI_ISL_876674, EPI_ISL_876675, EPI_ISL_876676, EPI_ISL_876677, EPI_ISL_876678, EPI_ISL_876680, EPI_ISL_876681, EPI_ISL_876682, EPI_ISL_876683, EPI_ISL_876684, EPI_ISL_876685, EPI_ISL_876686, EPI_ISL_876687, EPI_ISL_876688, EPI_ISL_876689, EPI_ISL_876691, EPI_ISL_876692, EPI_ISL_876693, EPI_ISL_876695, EPI_ISL_876696, EPI_ISL_876698, EPI_ISL_876699, EPI_ISL_876700, EPI_ISL_876701, EPI_ISL_876702, EPI_ISL_876703, EPI_ISL_876704, EPI_ISL_876705, EPI_ISL_876706, EPI_ISL_876708, EPI_ISL_876709, EPI_ISL_876711, EPI_ISL_876712, EPI_ISL_876713, EPI_ISL_876714, EPI_ISL_876715, EPI_ISL_876716, EPI_ISL_876718, EPI_ISL_876719, EPI_ISL_876720, EPI_ISL_876721, EPI_ISL_876722, EPI_ISL_876723, EPI_ISL_876724, EPI_ISL_876725, EPI_ISL_876726, EPI_ISL_876727, EPI_ISL_876728, EPI_ISL_876729, EPI_ISL_876730, EPI_ISL_876731, EPI_ISL_876732, EPI_ISL_876733, EPI_ISL_876734, EPI_ISL_876735, EPI_ISL_876736 | | | |
| see above | Helix/Illumina | Genomics and Discovery, Respiratory Viruses Branch, Division of Viral Diseases, Centers for Disease Control and Prevention | Peter W. Cook,Dhwani Batra,Ben L. Rambo-Martin,Eileen de Feo,Jan Antico,Christine Tran,Matthew Tolentino,Shannon Wickline,Kim Gietzen,Brad Sickler,Jingtao Liu,Eric Allen,Phil Febbo,Summer Galloway,Nicole L. Washington,Simon White,Geraint Levan,Kelly Schiabor Barrett,Elizabeth Cirulli,Alexandre Bolze,Ary Ascencio,Charlotte Rivera-Garcia,Ryan Cho,Jason Nguyen,Sherry Wang,Jimmy Ramirez,Tyler Cassens,Efren Sandoval,Magnus Isaksson,William Lee,David Becker,Marc Laurent,James Lu,Clinton R. Paden,Suxiang Tong,Duncan MacCannell, |
| EPI_ISL_877187, EPI_ISL_877189, EPI_ISL_877190, EPI_ISL_877191, EPI_ISL_877192, EPI_ISL_877193, EPI_ISL_877194, EPI_ISL_877195, EPI_ISL_877196, EPI_ISL_877197, EPI_ISL_877198, EPI_ISL_877199, EPI_ISL_877200, EPI_ISL_877201, EPI_ISL_877202, EPI_ISL_877203, EPI_ISL_877204 | | | |
| see above | Quest Diagnostics | Quest Diagnostics | Rosenthal,S.H., Gerasimova,A., Kagan,R.M., Anderson, B., Hua, M., Liu Y., Bernstein, L.E., Livingston, K.E., Perez, A., Shalhout, D.F., Shlyakhter, I.A., Owen, R., Tanpaiboon, P., Lacbawan, F. |
| EPI_ISL_877205, EPI_ISL_877206, EPI_ISL_877207, EPI_ISL_877208, EPI_ISL_877209 | LabPLUS | Institute of Environmental Science and Research (ESR) | Xiaoyun Ren, Matt Storey, Nikki Freed, Muhammad Faisal, Jing Wang, Hermes Perez, Anja Werno, Antje van der Linden, Arlo Upton, Chris Mansell, David Hammer, Dragana Drinkovic, Gary McAuliffe, Hana Sofia Andersson, James Ussher, Jill Sherwood, Josh Freeman, Julia Howard, Juliet Elvy, Mary DeAlmeida, Matt Blakiston, Matthew Rogers, Max Bloomfield, Michael Addidle, Michelle Balm, Sally Roberts, Sarah Jefferies, Sharmini Muttaiyah, Susan Morpeth, Susan Taylor, Timothy Blackmore, Vani Sathyendran, Veronica Playle, Virginia Hope, Erasmus Smit, Lauren Jelly, Olin Silander, Joep de Ligt |
| EPI_ISL_877214 | Middlemore Hospital | Institute of Environmental Science and Research (ESR) | Xiaoyun Ren, Matt Storey, Nikki Freed, Muhammad Faisal, Jing Wang, Hermes Perez, Anja Werno, Antje van der Linden, Arlo Upton, Chris Mansell, David Hammer, Dragana Drinkovic, Gary McAuliffe, Hana Sofia Andersson, James Ussher, Jill Sherwood, Josh Freeman, Julia Howard, Juliet Elvy, Mary DeAlmeida, Matt Blakiston, Matthew Rogers, Max Bloomfield, Michael Addidle, Michelle Balm, Sally Roberts, Sarah Jefferies, Sharmini Muttaiyah, Susan Morpeth, Susan Taylor, Timothy Blackmore, Vani Sathyendran, Veronica Playle, Virginia Hope, Erasmus Smit, Lauren Jelly, Olin Silander, Joep de Ligt |
| EPI_ISL_877453 | Institute of Microbiology and Immunology, Faculty of Medicine, University of Ljubljana | Institute of Microbiology and Immunology, Faculty of Medicine, University of Ljubljana | Samo Zakotnik, Tomaž Mark Zorec, Matic Brvar, Miša Korva, Mario Poljak, Tatjana Avši - Županc |
| EPI_ISL_878477, EPI_ISL_878480, EPI_ISL_878483, EPI_ISL_878485, EPI_ISL_878488, EPI_ISL_878491, EPI_ISL_878492, EPI_ISL_878503, EPI_ISL_878506, EPI_ISL_878509, EPI_ISL_878511, EPI_ISL_878514, EPI_ISL_878516, EPI_ISL_878519, EPI_ISL_878522, EPI_ISL_878524, EPI_ISL_878526 | | | |
| see above | Biolab Diagnostic Laboratories | Andersen lab at Scripps Research | Issa Abu-Dayyeh, Ahmad Tibi, Lama Hussein, Lina Mohammad, Zein Naber, Amid Abdelnour with SEARCH Alliance San Diego |

| | | | |
|---|---|---|---|
| EPI_ISL_880037, EPI_ISL_880045, EPI_ISL_880048, EPI_ISL_880050, EPI_ISL_880053, EPI_ISL_880056, EPI_ISL_880061, EPI_ISL_880063, EPI_ISL_880071, EPI_ISL_880074, EPI_ISL_880079, EPI_ISL_880082, EPI_ISL_880090, EPI_ISL_880092, EPI_ISL_880095, EPI_ISL_880101, EPI_ISL_880103, EPI_ISL_880106, EPI_ISL_880117, EPI_ISL_880119, EPI_ISL_880122, EPI_ISL_880125, EPI_ISL_880128, EPI_ISL_880130, EPI_ISL_880133, EPI_ISL_880135, EPI_ISL_880138, EPI_ISL_880140, EPI_ISL_880141, EPI_ISL_880143, EPI_ISL_880144, EPI_ISL_880146, EPI_ISL_880147, EPI_ISL_880149, EPI_ISL_880150, EPI_ISL_880151, EPI_ISL_880152, EPI_ISL_880156, EPI_ISL_880157, EPI_ISL_880159, EPI_ISL_880161, EPI_ISL_880162, EPI_ISL_880164, EPI_ISL_880165, EPI_ISL_880168, EPI_ISL_880169 | | | |
| see above | Sharp HealthCare Laboratory | Andersen lab at Scripps Research | SEARCH Alliance San Diego with Aaron Harding, Jacquelyn Berumen, Cathy Woerle, Liam McGinnis, Art Mendoza, Omid Bakhtar |
| EPI_ISL_880182, EPI_ISL_880183, EPI_ISL_880192 | Rady's Childrens Hospital | Andersen lab at Scripps Research | SEARCH Alliance San Diego with Nanda Radamchar, David Dimmock, Linda Luo, Christina Clarke, Kathryn Bouic, Teresa Mueller, Denise Malicki |
| EPI_ISL_880201, EPI_ISL_880202, EPI_ISL_880203, EPI_ISL_880204, EPI_ISL_880205, EPI_ISL_880206, EPI_ISL_880207, EPI_ISL_880208, EPI_ISL_880209, EPI_ISL_880212, EPI_ISL_880215, EPI_ISL_880218, EPI_ISL_880220, EPI_ISL_880223, EPI_ISL_880225, EPI_ISL_880228, EPI_ISL_880231, EPI_ISL_880233, EPI_ISL_880236, EPI_ISL_880239, EPI_ISL_880241, EPI_ISL_880244, EPI_ISL_880246, EPI_ISL_880249, EPI_ISL_880251, EPI_ISL_880253, EPI_ISL_880256, EPI_ISL_880259, EPI_ISL_880261, EPI_ISL_880263, EPI_ISL_880266, EPI_ISL_880268, EPI_ISL_880271, EPI_ISL_880274, EPI_ISL_880277, EPI_ISL_880279, EPI_ISL_880282, EPI_ISL_880284, EPI_ISL_880287, EPI_ISL_880290, EPI_ISL_880292, EPI_ISL_880294, EPI_ISL_880297, EPI_ISL_880300, EPI_ISL_880303, EPI_ISL_880305, EPI_ISL_880308, EPI_ISL_880310, EPI_ISL_880313, EPI_ISL_880316, EPI_ISL_880319, EPI_ISL_880321, EPI_ISL_880324, EPI_ISL_880326, EPI_ISL_880329 | | | |
| see above | Scripps Medical Laboratory | Andersen lab at Scripps Research | SEARCH Alliance San Diego with Michael Quigley, Ellen Stefanski, Ian Mchardy |
| EPI_ISL_882610 | Lighthouse Lab in Milton Keynes | Wellcome Sanger Institute for the COVID-19 Genomics UK (COG-UK) Consortium | The Lighthouse Lab in Milton Keynes and Alex Alderton, Roberto Amato, Sonia Goncalves, Ewan Harrison, David K. Jackson, Ian Johnston, Dominic Kwiatkowski, Cordelia Langford, John Sillitoe on behalf of the Wellcome Sanger Institute COVID-19 Surveillance Team |
| EPI_ISL_882616, EPI_ISL_882618, EPI_ISL_882628, EPI_ISL_882629 | Hospital Ramón y Cajal | Hospital Ramón y Cajal | José M Gonzalez-Alba, Concepción Rodríguez, Melanie Abreu, Laura Martínez, Val F Lanza, Luz Leticia Olavarrieta, Rafael Cantón, JC Galán |
| EPI_ISL_882642, EPI_ISL_882643, EPI_ISL_882644 | Azerbaijan National Hematology Center Division of Medical Genetics | Azerbaijan National Hematology Center Division of Medical Genetics | Aghayev Agha Rza |
| EPI_ISL_882646 | Medical Research Center, Faculty of Medicine, Syarif Hidayatullah State Islamic University Jakarta | Medical Research Center, Faculty of Medicine, Syarif Hidayatullah State Islamic University Jakarta | Chris Adhiyanto, Laifa Hendarmin, Zeti Harriyati, Endah Wulandari, Flori Ratna Sari, Fika Ekayanti, Hari Hendarto |
| EPI_ISL_882656 | Medical Research Center, Faculty of Medicine, Syarif Hidayatullah State Islamic University Jakarta | Medical Research Center, Faculty of Medicine, Syarif Hidayatullah State Islamic University Jakarta | Erike A Suwarsono, Chris Adhiyanto, Laifa Hendarmin, Zeti Harriyati, Flori Ratna Sari, Endah Wulandari, Fika Ekayanti |
| EPI_ISL_882665 | Unidade de Pronto Atendimento Dra Zilda Arns | Instituto Adolfo Lutz, Interdiciplinary Procedures Center, Strategic Laboratory | Claudio Tavares Sacchi, Claudia Regina Gonçalves, Erica Valessa Ramos Gomes, Karoline Rodrigues Campos |
| EPI_ISL_882773, EPI_ISL_882774, EPI_ISL_882775, EPI_ISL_882777, EPI_ISL_882778 | Institute for Urban Disease Control and Prevention | COVID-19 Network Investigations (CONI) Alliance | Kamolthip Atsawawaranunt, Elizabeth Batty, Wasun Chantratita, Thanat Chookajorn, Stefan Fernandez, Angkana Huang, Anthony R. Jones, Khajohn Joonsalak, Chonticha Klungtong, Theerarat Kochakarn, Prayuth Kaewmalang, Amornmas Kongklieng, Namfon Kotanan, Krittikorn Kumpornsin, Duangkamon Loesbanluechai, Wudtichai Manasatienkij, Anek Mungaomklang, Bhakbhoom Panthan, Pukkaporn Parnwijitkul, Ekawat Pasomsub, Vichan Pawun, Kingkan Rakmanee, Insee Sensorn, Janjira Thaipadungpanit, Arporn Wangwiwatsin, Treewat Watthanachockchai |
| EPI_ISL_882780 | Ramathibodi Chakri Naruebodindra Hospital | COVID-19 Network Investigations (CONI) Alliance | Elizabeth Batty, Wasun Chantratita, Thanat Chookajorn, Stefan Fernandez, Angkana Huang, Anthony R. Jones, Khajohn Joonsalak, Chonticha Klungtong, Theerarat Kochakarn, Namfon Kotanan, Krittikorn Kumpornsin, Wudtichai Manasatienkij, Bhakbhoom Panthan, Ekawat Pasomsub, Kingkan Rakmanee, Insee Sensorn, Janjira Thaipadungpanit, Arporn Wangwiwatsin,Treewat Watthanachockchai |
| EPI_ISL_882783, EPI_ISL_882784, EPI_ISL_882785, EPI_ISL_882786, EPI_ISL_882787, EPI_ISL_882788, EPI_ISL_882789, EPI_ISL_882790, EPI_ISL_882791 | Institute for Urban Disease Control and Prevention | COVID-19 Network Investigations (CONI) Alliance | Kamolthip Atsawawaranunt, Elizabeth Batty, Wasun Chantratita, Thanat Chookajorn, Stefan Fernandez, Angkana Huang, Anthony R. Jones, Khajohn Joonsalak, Chonticha Klungtong, Theerarat Kochakarn, Prayuth Kaewmalang, Amornmas Kongklieng, Namfon Kotanan, Krittikorn Kumpornsin, Duangkamon Loesbanluechai, Wudtichai Manasatienkij, Anek Mungaomklang, Bhakbhoom Panthan, Pukkaporn Parnwijitkul, Ekawat Pasomsub, Vichan Pawun, Kingkan Rakmanee, Insee Sensorn, Janjira Thaipadungpanit, Arporn Wangwiwatsin, Treewat Watthanachockchai |
| EPI_ISL_882805 | MSHS Clinical Microbiology Laboratories | MSHS Pathogen Surveillance Program | Ana S. Gonzalez-Reiche, Hala Alshammary, Mitchell J. Sullivan, Brianne Ciferri, Ajay Obla, Angela Amoako, Mahmoud Awawda, Elena Hirsch, Ashley S. Salimbangon, Levy Sominsky, Katherine Beach, Kayla Russo, Charles Gleason, Shelcie Fabre, Giulio Kleiner, Zenab Khan, Bremy Alburquerque, Adriana van de Guchte, Komal Srivastava, Matthew M. Hernandez, Jayeeta Dutta, Denise Jurczyszak, Emily Ferreri, Rachel Chernet, Nancy Francoeur, Betsaida Salom Melo, Irina Oussenko, Gintaras Deikus, Juan Soto, Shwetha Hara Sridhar, Ying-Chih Wang, Kathryn Twyman, Andrew Kasarskis, Deena R. Altman, Robert Sebra, Adolfo García-Sastre, Marta Luksza, Gopi Patel, Sarah Schaefer, Melissa Gitman, Michael D. Nowak, Alberto Paniz-Mondolfi, Emilia Mia Sordillo, Viviana Simon, Harm van Bakel |
| EPI_ISL_882919 | INMI Lazzaro Spallanzani IRCCS | INMI Lazzaro Spallanzani IRCCS | C.E.M Gruber, B Bartolini, E Giombini, M Rueca, O Butera, F Messina, A Di Caro, MR Capobianchi |
| EPI_ISL_882944, EPI_ISL_882950, EPI_ISL_882951, EPI_ISL_882952, EPI_ISL_882953 | The National Institute of Public Health | State Veterinary Institute Prague | Nagy,A;Jirincova,H;Vecerova,JTrnka,D |
| EPI_ISL_882987 | MD Laboratories | Los Angeles County PHL | P. Hemarajata et al. |
| EPI_ISL_883035 | Maryland Public Health Laboratory | Maryland Public Health Laboratory | Maryland Department of Health Laboratories Administration |
| EPI_ISL_883293 | SIESP CHIETI-DRIVE IN LANCIANO | Istituto Zooprofilattico Sperimentale dell'Abruzzo e Molise "G. Caporale" | Lorusso A, Marcacci M, Di Domenico M, Ancora M, Curini V, Mangone I, Rinaldi A, Scialabba S, Di Pasquale A, Cammà C, Puglia I, Calistri P, Savini G |
| EPI_ISL_883294, EPI_ISL_883295, EPI_ISL_883301, EPI_ISL_883302 | RP Guardiagrele-Ospedale di Comunità | Istituto Zooprofilattico Sperimentale dell'Abruzzo e Molise "G. Caporale" | Lorusso A, Marcacci M, Di Domenico M, Ancora M, Curini V, Mangone I, Rinaldi A, Scialabba S, Di Pasquale A, Cammà C, Puglia I, Calistri P, Savini G |
| EPI_ISL_883303, EPI_ISL_883304, EPI_ISL_883305 | SIESP DIPARTIMENTO DI PREVENZIONE CHIE | Istituto Zooprofilattico Sperimentale dell'Abruzzo e Molise "G. Caporale" | Lorusso A, Marcacci M, Di Domenico M, Ancora M, Curini V, Mangone I, Rinaldi A, Scialabba S, Di Pasquale A, Cammà C, Puglia I, Calistri P, Savini G |
| EPI_ISL_883427, EPI_ISL_883452, EPI_ISL_883453, EPI_ISL_883456, EPI_ISL_883458, EPI_ISL_883459, EPI_ISL_883460, EPI_ISL_883461, EPI_ISL_883462, EPI_ISL_883463, EPI_ISL_883464, EPI_ISL_883465, EPI_ISL_883466, EPI_ISL_883467, EPI_ISL_883468, EPI_ISL_883469, EPI_ISL_883470, EPI_ISL_883481 | | | |
| see above | NORTHWELL HEALTH LABORATORIES | Wadsworth Center, New York State Department of Health | Kirsten St. George, Daryl M. Lamson, Alexis Russel, Matthew Shudt, Melissa A Leisner, Jonathan Plitnick, Navjot Singh, John Kelly, Erasmus Schneider, Erica Lasek-Nesselquist |
| EPI_ISL_883957 | Medical Research Center, Faculty of Medicine, Syarif Hidayatullah State Islamic University Jakarta | Medical Research Center, Faculty of Medicine, Syarif Hidayatullah State Islamic University Jakarta | Erike A Suwarsono, Chris Adhiyanto, Laifa Hendarmin, Zeti Harriyati, Endah Wulandari, Flori Ratna Sari, Fika Ekayanti, Hari Hendarto |
| EPI_ISL_883961 | Labo Analyses Med | National Reference Center for Viruses of Respiratory Infections, Institut Pasteur, Paris | Marion Barbet, Sylvie Behillil, Méline Bizard, Angela Brisebarre, Camille Capel, Etienne Simon-Lorière, Vincent Enouf, Maud Vanpeene, Sylvie van der Werf,Jacques Alexandra |
| EPI_ISL_883962, EPI_ISL_883963 | Labo Analyses Med | National Reference Center for Viruses of Respiratory Infections, Institut Pasteur, Paris | Marion Barbet, Sylvie Behillil, Méline Bizard, Angela Brisebarre, Camille Capel, Etienne Simon-Lorière, Vincent Enouf, Maud Vanpeene, Sylvie van der Werf,Amzalag Jonas |
| EPI_ISL_883964, EPI_ISL_883967, EPI_ISL_883968, EPI_ISL_883969, EPI_ISL_883970 | Labo Analyses med | National Reference Center for Viruses of Respiratory Infections, Institut Pasteur, Paris | Marion Barbet, Sylvie Behillil, Méline Bizard, Angela Brisebarre, Camille Capel, Etienne Simon-Lorière, Vincent Enouf, Maud Vanpeene, Sylvie van der Werf,Amzalag Jonas |
| EPI_ISL_883972, EPI_ISL_883973, EPI_ISL_883974, EPI_ISL_883975, EPI_ISL_883976, EPI_ISL_883979, EPI_ISL_883980, EPI_ISL_883981, EPI_ISL_883985 | Hopital | National Reference Center for Viruses of Respiratory Infections, Institut Pasteur, Paris | Marion Barbet, Sylvie Behillil, Méline Bizard, Angela Brisebarre, Camille Capel, Etienne Simon-Lorière, Vincent Enouf, Maud Vanpeene, Sylvie van der Werf,Fourgeaud Jacques |
| EPI_ISL_883986, EPI_ISL_883987, EPI_ISL_883988, EPI_ISL_883989 | Hopital | National Reference Center for Viruses of Respiratory Infections, Institut Pasteur, Paris | Marion Barbet, Sylvie Behillil, Méline Bizard, Angela Brisebarre, Camille Capel, Etienne Simon-Lorière, Vincent Enouf, Maud Vanpeene, Sylvie van der Werf |
| EPI_ISL_883990, EPI_ISL_883991, | hopital | National Reference Center for Viruses of Respiratory | Marion Barbet, Sylvie Behillil, Méline Bizard, Angela Brisebarre, Camille Capel, Etienne Simon-Lorière, Vincent Enouf, Maud Vanpeene, Sylvie van der |

| | | | |
|---|---|---|---|
| EPI_ISL_883992, EPI_ISL_883993, EPI_ISL_883994, EPI_ISL_883995, EPI_ISL_883996 | | Infections, Institut Pasteur, Paris | Werf,Andre-Garnier Elisabeth |
| EPI_ISL_884001, EPI_ISL_884003 | Labo Analyses Med | National Reference Center for Viruses of Respiratory Infections, Institut Pasteur, Paris | Marion Barbet, Sylvie Behillil, Méline Bizard, Angela Brisebarre, Camille Capel, Etienne Simon-Lorière, Vincent Enouf, Maud Vanpeene, Sylvie van der Werf,Rousset Dominique |
| EPI_ISL_884080 | Wadsworth Center, New York State Department of Health | Wadsworth Center, New York State Department of Health | Kirsten St. George, Daryl M. Lamson, Alexis Russel, Matthew Shudt, Melissa A Leisner, Jonathan Plitnick, Navjot Singh, John Kelly, Erasmus Schneider, Erica Lasek-Nesselquist |
| EPI_ISL_884081, EPI_ISL_884082, EPI_ISL_884083, EPI_ISL_884084, EPI_ISL_884085 | ALBANY MEDICAL CENTER HOSPITAL CLINICAL LABORATORIES | Wadsworth Center, New York State Department of Health | Kirsten St. George, Daryl M. Lamson, Alexis Russel, Matthew Shudt, Melissa A Leisner, Jonathan Plitnick, Navjot Singh, John Kelly, Erasmus Schneider, Erica Lasek-Nesselquist |
| EPI_ISL_884086 | Wadsworth Center, New York State Department of Health | Wadsworth Center, New York State Department of Health | Kirsten St. George, Daryl M. Lamson, Alexis Russel, Matthew Shudt, Melissa A Leisner, Jonathan Plitnick, Navjot Singh, John Kelly, Erasmus Schneider, Erica Lasek-Nesselquist |
| EPI_ISL_884089, EPI_ISL_884090, EPI_ISL_884091, EPI_ISL_884092, EPI_ISL_884093, EPI_ISL_884094, EPI_ISL_884095, EPI_ISL_884096, EPI_ISL_884097, EPI_ISL_884098, EPI_ISL_884099, EPI_ISL_884100, EPI_ISL_884101, EPI_ISL_884102, EPI_ISL_884103, EPI_ISL_884104, EPI_ISL_884105, EPI_ISL_884106, EPI_ISL_884107, EPI_ISL_884108, EPI_ISL_884109, EPI_ISL_884110, EPI_ISL_884111, EPI_ISL_884112, EPI_ISL_884113, EPI_ISL_884114, EPI_ISL_884115, EPI_ISL_884116, EPI_ISL_884117, EPI_ISL_884118, EPI_ISL_884119, EPI_ISL_884120, EPI_ISL_884121, EPI_ISL_884122, EPI_ISL_884123, EPI_ISL_884124, EPI_ISL_884125, EPI_ISL_884126, EPI_ISL_884127, EPI_ISL_884128, EPI_ISL_884129, EPI_ISL_884130, EPI_ISL_884131, EPI_ISL_884132, EPI_ISL_884133, EPI_ISL_884134, EPI_ISL_884135, EPI_ISL_884136, EPI_ISL_884137, EPI_ISL_884138, EPI_ISL_884139, EPI_ISL_884140, EPI_ISL_884141, EPI_ISL_884142, EPI_ISL_884143, EPI_ISL_884144, EPI_ISL_884145, EPI_ISL_884146, EPI_ISL_884147, EPI_ISL_884148, EPI_ISL_884149, EPI_ISL_884150, EPI_ISL_884151, EPI_ISL_884152, EPI_ISL_884153, EPI_ISL_884154, EPI_ISL_884155, EPI_ISL_884156, EPI_ISL_884157, EPI_ISL_884158, EPI_ISL_884159, EPI_ISL_884160, EPI_ISL_884161, EPI_ISL_884162, EPI_ISL_884163, EPI_ISL_884164, EPI_ISL_884165, EPI_ISL_884166, EPI_ISL_884167, EPI_ISL_884168, EPI_ISL_884169, EPI_ISL_884170, EPI_ISL_884171, EPI_ISL_884172, EPI_ISL_884173, EPI_ISL_884174, EPI_ISL_884175, EPI_ISL_884176, EPI_ISL_884177, EPI_ISL_884178, EPI_ISL_884179, EPI_ISL_884180, EPI_ISL_884181 | | | |
| see above | Eurofins Diatherix | Hudsonalpha Genome Sequencing Center | Jane Grimwood, Melissa Williams, Lori H. Handley, Joshua Stough, Leslie Malone, Stefan Brzezinski, Ada Stewart, Teresa Jones, Jenell Webber, John Lovell, Jennifer Cart, and Jeremy Schmutz |
| EPI_ISL_884248 | Kansas Health and Environmental Lab | Kansas Health and Environmental Lab | Mike Grose, Paige Drury, Carissa Robertson, Ben Olsen, and Phil Adam |
| EPI_ISL_884296, EPI_ISL_884297, EPI_ISL_884298, EPI_ISL_884299, EPI_ISL_884300, EPI_ISL_884301, EPI_ISL_884302 | Santa Clara County Public Health Laboratory | Santa Clara County Public Health Laboratory | Santa Clara County Public Health Department |
| EPI_ISL_886159, EPI_ISL_886165, EPI_ISL_886178, EPI_ISL_886179, EPI_ISL_886182, EPI_ISL_886192, EPI_ISL_886202, EPI_ISL_886210, EPI_ISL_886216, EPI_ISL_886225, EPI_ISL_886226, EPI_ISL_886260, EPI_ISL_886262, EPI_ISL_886273, EPI_ISL_886275, EPI_ISL_886284, EPI_ISL_886287, EPI_ISL_886314, EPI_ISL_886319, EPI_ISL_886327, EPI_ISL_886343, EPI_ISL_886356, EPI_ISL_886386, EPI_ISL_886387, EPI_ISL_886396, EPI_ISL_886427, EPI_ISL_886439, EPI_ISL_886440, EPI_ISL_886457, EPI_ISL_886461, EPI_ISL_886475, EPI_ISL_886478, EPI_ISL_886502, EPI_ISL_886509, EPI_ISL_886516, EPI_ISL_886521, EPI_ISL_886532, EPI_ISL_886533, EPI_ISL_886537, EPI_ISL_886538, EPI_ISL_886539, EPI_ISL_886540, EPI_ISL_886541, EPI_ISL_886542, EPI_ISL_886552, EPI_ISL_886555, EPI_ISL_886579, EPI_ISL_886582, EPI_ISL_886590, EPI_ISL_886603, EPI_ISL_886613, EPI_ISL_886614, EPI_ISL_886624, EPI_ISL_886628, EPI_ISL_886640, EPI_ISL_886652, EPI_ISL_886674, EPI_ISL_886676, EPI_ISL_886678, EPI_ISL_886682, EPI_ISL_886683, EPI_ISL_886687, EPI_ISL_886695, EPI_ISL_886709, EPI_ISL_886713, EPI_ISL_886718, EPI_ISL_886737, EPI_ISL_886740, EPI_ISL_886741, EPI_ISL_886768, EPI_ISL_886781, EPI_ISL_886800, EPI_ISL_886820, EPI_ISL_886823, EPI_ISL_886859, EPI_ISL_886865, EPI_ISL_886879, EPI_ISL_886881, EPI_ISL_886882, EPI_ISL_886884, EPI_ISL_886887, EPI_ISL_886892, EPI_ISL_886903, EPI_ISL_886908, EPI_ISL_886911, EPI_ISL_886912, EPI_ISL_886917, EPI_ISL_886923, EPI_ISL_886932, EPI_ISL_886949, EPI_ISL_886954, EPI_ISL_886955, EPI_ISL_886956, EPI_ISL_887001, EPI_ISL_887008, EPI_ISL_887010, EPI_ISL_887019, EPI_ISL_887029, EPI_ISL_887034, EPI_ISL_887038, EPI_ISL_887042, EPI_ISL_887045, EPI_ISL_887058, EPI_ISL_887071, EPI_ISL_887077, EPI_ISL_887080, EPI_ISL_887083, EPI_ISL_887084 | | | |
| see above | Labcorp | Genomics and Discovery, Respiratory Viruses Branch, Division of Viral Diseases, Centers for Disease Control and Prevention | Peter W. Cook,Dhwani Batra,Ben L. Rambo-Martin,Summer Galloway,Brian Krueger,Minoo Agarwal,Eyad Almasri,Debbie Boles,Ayla Burns,Nuthawin Charoensri,Oren Cohen,Susan Countryman,Mary Ann Cristobal,Bobbi Croy,Suzanne Dale,Hrushikesh Deshmukh,Amanda Douglas,Vincent Drouillon,Marcia Eisenberg,Howard Engler,Rama Ghatti,Prashant Gupta,Susan Hicks,Jake Humphrey,Lax Iyer,Manoj Jain,Mohan Kolli,Tim Kuphal,Stanley Letovsky,Michael Levandoski,Craig Lukasik,Jonathan Meltzer,Brian Norvell,Mindy Nye,Scott Parker,Christos Petropoulos,John Pruitt,Steven Ragan,Scott Ryan,Mike Sapeta,Jana Schroth,Suresh Babu Selvaraju,Goran Stevovic,Amanda Suchanek,Andrea Throop,Lyndon Tilson,Thomas Urban,Joe Voshell,Kimberly Wagner,Jonathan Williams,Mary Williamson,Qian Zeng,Tricia Zwiefelhofer,Clinton R. Paden,Suxiang Tong,Duncan MacCannell, |
| EPI_ISL_887105 | Institute for Urban Disease Control and Prevention | COVID-19 Network Investigations (CONI) Alliance | Kamolthip Atsawawaranunt, Elizabeth Batty, Wasun Chantratita, Thanat Chookajorn, Stefan Fernandez, Angkana Huang, Anthony R. Jones, Khajohn Joonsalak, Chonticha Klungtong, Theerarat Kochakarn, Prayuth Kaewmalang, Amornmas Kongklieng, Namfon Kotanan, Krittikorn Kumpornsin, Duangkamon Loesbanluechai, Wudtichai Manasatienkij, Anek Mungaomklang, Bhakbhoom Panthan, Pukkaporn Parnwijitkul, Ekawat Pasomsub, Vichan Pawun, Kingkan Rakmanee, Insee Sensorn, Janjira Thaipadungpanit, Arporn Wangwiwatsin, Treewat Watthanachockchai |
| EPI_ISL_887117, EPI_ISL_887123, EPI_ISL_887129, EPI_ISL_887134 | Institute of Medical Microbiology and Hospital Hygiene | Institute of Medical Microbiology and Hospital Hygiene | Prof. Dr. Achim Kaasch, Aljoscha Tersteegen |
| EPI_ISL_887419, EPI_ISL_887422, EPI_ISL_887423, EPI_ISL_887427, EPI_ISL_887430, EPI_ISL_887432, EPI_ISL_887433, EPI_ISL_887448, EPI_ISL_887449, EPI_ISL_887450, EPI_ISL_887451, EPI_ISL_887452, EPI_ISL_887453, EPI_ISL_887454, EPI_ISL_887504, EPI_ISL_887505 | | | |
| see above | Instituto Nacional de Saude (INS), Mozambique | KRISP, KZN Research Innovation and Sequencing Platform | Nalia Ismael, Nadia Sitoe, Paulo Arnaldo, Nedio Mabunda, Giandhari J, Pillay S, Tegally H, Wilkinson E, de Oliveira T |
| EPI_ISL_887514, EPI_ISL_887515, EPI_ISL_887519, EPI_ISL_887520, EPI_ISL_887521, EPI_ISL_887525, EPI_ISL_887526, EPI_ISL_887527, EPI_ISL_887537, EPI_ISL_887561, EPI_ISL_887562, EPI_ISL_887564, EPI_ISL_887565, EPI_ISL_887566, EPI_ISL_887584, EPI_ISL_887585, EPI_ISL_887586 | | | |
| see above | Johns Hopkins Hospital Department of Pathology | Johns Hopkins Hospital Department of Pathology | C. Paul Morris, Chun Huai Luo, Adannaya Amadi, Matthew Schwartz, Nicholas Gallagher, Heba H. Mostafa |
| EPI_ISL_887606, EPI_ISL_887723, EPI_ISL_887965, EPI_ISL_888062 | Labcorp | Genomics and Discovery, Respiratory Viruses Branch, Division of Viral Diseases, Centers for Disease Control and Prevention | Peter W. Cook,Dhwani Batra,Ben L. Rambo-Martin,Summer Galloway,Brian Krueger,Minoo Agarwal,Eyad Almasri,Debbie Boles,Ayla Burns,Nuthawin Charoensri,Oren Cohen,Susan Countryman,Mary Ann Cristobal,Bobbi Croy,Suzanne Dale,Hrushikesh Deshmukh,Amanda Douglas,Vincent Drouillon,Marcia Eisenberg,Howard Engler,Rama Ghatti,Prashant Gupta,Susan Hicks,Jake Humphrey,Lax Iyer,Manoj Jain,Mohan Kolli,Tim Kuphal,Stanley Letovsky,Michael Levandoski,Craig Lukasik,Jonathan Meltzer,Brian Norvell,Mindy Nye,Scott Parker,Christos Petropoulos,John Pruitt,Steven Ragan,Scott Ryan,Mike Sapeta,Jana Schroth,Suresh Babu Selvaraju,Goran Stevovic,Amanda Suchanek,Andrea Throop,Lyndon Tilson,Thomas Urban,Joe Voshell,Kimberly Wagner,Jonathan Williams,Mary Williamson,Qian Zeng,Tricia Zwiefelhofer,Clinton R. Paden,Suxiang Tong,Duncan MacCannell, |
| EPI_ISL_888628, EPI_ISL_888629, EPI_ISL_888630, EPI_ISL_888631, EPI_ISL_888632, EPI_ISL_888633, EPI_ISL_888634, EPI_ISL_888635, EPI_ISL_888636, EPI_ISL_888637, EPI_ISL_888638, EPI_ISL_888639, EPI_ISL_888640, EPI_ISL_888641, EPI_ISL_888642, EPI_ISL_888643, EPI_ISL_888649, EPI_ISL_888650 | | | |
| see above | Univeristy of New Mexico Hospital | Center for Global Health, University of New Mexico Health Sciences Center | Daryl Domman, Kurt Schwalm, Justin Bacca, Jon Femling, Darrell Dinwiddie |
| EPI_ISL_888854, EPI_ISL_888870, EPI_ISL_888871, EPI_ISL_888879, EPI_ISL_888880, EPI_ISL_888894, EPI_ISL_888895 | Michigan Department of Health and Human Services, Bureau of Laboratories | Michigan Department of Health and Human Services, Bureau of Laboratories | Blankenship HM, Riner D, Soehnlen MK |
| EPI_ISL_888979 | RS Bhakti Mulia | Eijkman Institute for Molecular Biology, Ministry of Research and Technology/National Agency for Research and Innovation | Willy Agustine, Edison Johar, Hidayat Trimarsanto, Iskandar Adnan, Lydia V. Panggalo, Sukma Oktavianthi, Frilasita A Yudhaputri, Safarina G Malik, Khin Saw Myint, Amin Soebandrio |
| EPI_ISL_889000 | RSUD Cileungsi | Eijkman Institute for Molecular Biology, Ministry of Research and Technology/National Agency for Research and Innovation | Sukma Oktavianthi, Willy Agustine, Edison Johar, Hidayat Trimarsanto, Iskandar Adnan, Lydia V. Panggalo, Frilasita A Yudhaputri, Safarina G Malik, Khin Saw Myint, Amin Soebandrio |
| EPI_ISL_889001 | RSU Azra | Eijkman Institute for Molecular Biology, Ministry of Research and Technology/National Agency for Research and Innovation | Sukma Oktavianthi, Willy Agustine, Edison Johar, Hidayat Trimarsanto, Iskandar Adnan, Lydia V. Panggalo, Frilasita A Yudhaputri, Safarina G Malik, Khin Saw Myint, Amin Soebandrio |
| EPI_ISL_889002 | RSU Sumber Waras | Eijkman Institute for Molecular Biology, Ministry of Research and Technology/National Agency for Research and Innovation | Sukma Oktavianthi, Willy Agustine, Edison Johar, Hidayat Trimarsanto, Iskandar Adnan, Lydia V. Panggalo, Frilasita A Yudhaputri, Safarina G Malik, Khin Saw Myint, Amin Soebandrio |
| EPI_ISL_889607, EPI_ISL_889608, EPI_ISL_889609, EPI_ISL_889610, EPI_ISL_889611, EPI_ISL_889612, EPI_ISL_889613, EPI_ISL_889614, EPI_ISL_889615, EPI_ISL_889616, EPI_ISL_889617, EPI_ISL_889618, EPI_ISL_889619, EPI_ISL_889620, EPI_ISL_889621, EPI_ISL_889622, EPI_ISL_889623, EPI_ISL_889624, EPI_ISL_889625, EPI_ISL_889626, EPI_ISL_889627, EPI_ISL_889628, EPI_ISL_889629, EPI_ISL_889630, EPI_ISL_889631, EPI_ISL_889632, EPI_ISL_889633, EPI_ISL_889634, EPI_ISL_889635, EPI_ISL_889636, EPI_ISL_889637, EPI_ISL_889638, EPI_ISL_889639, EPI_ISL_889640, EPI_ISL_889641, EPI_ISL_889642, EPI_ISL_889643, EPI_ISL_889644, EPI_ISL_889645, EPI_ISL_889646, EPI_ISL_889647, EPI_ISL_889648, EPI_ISL_889649, EPI_ISL_889650, EPI_ISL_889651, EPI_ISL_889652, EPI_ISL_889653, EPI_ISL_889654, EPI_ISL_889655, EPI_ISL_889656, EPI_ISL_889657, EPI_ISL_889658, EPI_ISL_889659, EPI_ISL_889660, EPI_ISL_889661, EPI_ISL_889662, EPI_ISL_889663, EPI_ISL_889664, EPI_ISL_889665, EPI_ISL_889666, EPI_ISL_889667, EPI_ISL_889668, EPI_ISL_889669, EPI_ISL_889670 | | | |
| see above | LSUHS Emerging Viral Threat Laboratory | Microbial Genome Sequencing Center | Jeremy P. Kamil, Jennifer L. Carroll, Camille F. Abshire, Maarten Van Diest, Mohammed N.A. Siddiquey, Andrew D. Yurochko, Martin J. Sapp, Rona S. Scott, Christopher G. Kevil, Daniel J. Snyder, Vaughn S. Cooper, John A. Vanchiere |

| | | | |
|---|---|---|---|
| EPI_ISL_890235 | Respiratory Viruses Branch, Centers for Disease Control and Prevention | Centers for Disease Control and Prevention | Tao,Y., Li,Y., Zhang,J., Queen,K., Uehara,A., Cook,P., Paden,C.R., Wang,H. and Tong,S. |
| EPI_ISL_890237 | Virology, International Centre for Diarrhoeal Disease Research, Bangladesh (ICDDR,B) | International Centre for Diarrhoeal Disease Research (ICDDR,B) | Hossain,M.E., Rahman,M.M., Sumiya,M.K., Alam,M.S., Karim,M.Y.,Hoque,A.F., Rahman,M.Z. and Rahman,M. |
| EPI_ISL_890282, EPI_ISL_890286, EPI_ISL_890290, EPI_ISL_890293, EPI_ISL_890296, EPI_ISL_890300, EPI_ISL_890308, EPI_ISL_890311, EPI_ISL_890313, EPI_ISL_890314, EPI_ISL_890316, EPI_ISL_890317, EPI_ISL_890319, EPI_ISL_890321, EPI_ISL_890323, EPI_ISL_890326, EPI_ISL_890330, EPI_ISL_890334, EPI_ISL_890338, EPI_ISL_890347, EPI_ISL_890351 | | | |
| see above | KU Leuven, Rega Institute, Clinical and Epidemiological Virology | KU Leuven, Rega Institute, Clinical and Epidemiological Virology | Tony Wawina-Bokalanga, Bert Vanmechelen, Joan Marti-Carerras, Piet Maes |
| EPI_ISL_890354 | Labo Analyses Med | National Reference Center for Viruses of Respiratory Infections, Institut Pasteur, Paris | Marion Barbet, Sylvie Behillil, Méline Bizard, Angela Brisebarre, Camille Capel, Etienne Simon-Lorière, Vincent Enouf, Maud Vanpeene, Sylvie van der Werf,Coignard Catherine |
| EPI_ISL_890359 | Hospital | National Reference Center for Viruses of Respiratory Infections, Institut Pasteur, Paris | Marion Barbet, Sylvie Behillil, Méline Bizard, Angela Brisebarre, Camille Capel, Etienne Simon-Lorière, Vincent Enouf, Maud Vanpeene, Sylvie van der Werf,Guigon AuréLie |
| EPI_ISL_890363, EPI_ISL_890364 | LSUHS Emerging Viral Threat Laboratory | Microbial Genome Sequencing Center | Jeremy P. Kamil, Jennifer L. Carroll, Camille F. Abshire, Maarten Van Diest, Mohammed N.A. Siddiquey, Andrew D. Yurochko, Martin J. Sapp, Rona S. Scott, Christopher G. Kevil, Daniel J. Snyder, Vaughn S. Cooper, John A. Vanchiere |
| EPI_ISL_891139, EPI_ISL_891140 | The Jackson Laboratory | The Jackson Laboratory | Lloyd M, Sanderson B, Srivastava A, Maurya R, Renzette N, Omerza G, Kelly K, Li L, Wei C L, Adams M |
| EPI_ISL_891200, EPI_ISL_891202, EPI_ISL_891203 | DPH, Massachusetts State Public Health Lab | DPH, Massachusetts State Public Health Lab | Lang,A.S., Fink,T., Gallagher,G.R., Smole,S.C. |
| EPI_ISL_891212 | Population Medicine and Diagnostic Sciences, Cornell University | Population Medicine and Diagnostic Sciences, Cornell University | Cardia Caserta,L., Mitchel,P.K., Plachorzyk,E. and Diel,D.G. |
| EPI_ISL_892207, EPI_ISL_892208, EPI_ISL_892209 | Lighthouse Lab in Alderley Park | Wellcome Sanger Institute for the COVID-19 Genomics UK (COG-UK) Consortium | Jacquelyn Wynn, Mairead Hyland, The Lighthouse Lab in Alderley Park and Alex Alderton, Roberto Amato, Sonia Goncalves, Ewan Harrison, David K. Jackson, Ian Johnston, Dominic Kwiatkowski, Cordelia Langford, John Sillitoe on behalf of the Wellcome Sanger Institute COVID-19 Surveillance Team |
| EPI_ISL_892210 | Lighthouse Lab in Glasgow | Wellcome Sanger Institute for the COVID-19 Genomics UK (COG-UK) Consortium | Harper VanSteenhouse, Yumi Kasai, David Gray, Carol Clugston, Anna Dominiczak and Alex Alderton, Roberto Amato, Sonia Goncalves, Ewan Harrison, David K. Jackson, Ian Johnston, Dominic Kwiatkowski, Cordelia Langford, John Sillitoe on behalf of the Wellcome Sanger Institute COVID-19 Surveillance Team |
| EPI_ISL_892266 | MD Laboratories | Los Angeles County PHL | P. Hemarajata et al. |
| EPI_ISL_892267, EPI_ISL_892268 | UCLA Clinical Micro Lab | Los Angeles County PHL | P. Hemarajata et al. |
| EPI_ISL_893750, EPI_ISL_893752, EPI_ISL_893755, EPI_ISL_893757, EPI_ISL_893759, EPI_ISL_893761, EPI_ISL_893778 | Institute of Virology, Medical Center, University of Freiburg, Freiburg, Germany | Institute of Virology, Clinial Virus Genomics, Medical Center, University of Freiburg, Freiburg, Germany | Jonas Fuchs, Lisa Kern, Sandra Reuter, Hajo Grundmann, Marcus Panning |
| EPI_ISL_894176, EPI_ISL_894178, EPI_ISL_894179, EPI_ISL_894180, EPI_ISL_894181, EPI_ISL_894185, EPI_ISL_894186, EPI_ISL_894192, EPI_ISL_894193, EPI_ISL_894196, EPI_ISL_894197, EPI_ISL_894209, EPI_ISL_894211, EPI_ISL_894212, EPI_ISL_894213, EPI_ISL_894216 | | | |
| see above | KU Leuven, Rega Institute, Clinical and Epidemiological Virology | KU Leuven, Rega Institute, Clinical and Epidemiological Virology | Tony Wawina-Bokalanga, Bert Vanmechelen, Joan Marti-Carerras, Piet Maes |
| EPI_ISL_894219 | Hopital | National Reference Center for Viruses of Respiratory Infections, Institut Pasteur, Paris | Marion Barbet, Sylvie Behillil, Méline Bizard, Angela Brisebarre, Camille Capel, Etienne Simon-Lorière, Vincent Enouf, Maud Vanpeene, Sylvie van der Werf,Guigon AuréLie |
| EPI_ISL_894223, EPI_ISL_894232, EPI_ISL_894244, EPI_ISL_894245 | Labo Analyses med | National Reference Center for Viruses of Respiratory Infections, Institut Pasteur, Paris | Marion Barbet, Sylvie Behillil, Méline Bizard, Angela Brisebarre, Camille Capel, Etienne Simon-Lorière, Vincent Enouf, Maud Vanpeene, Sylvie van der Werf,Amzalag Jonas |
| EPI_ISL_894246, EPI_ISL_894247 | Ministry of Health Turkey | Ministry of Health Turkey | Fatma Bayrakdar, Yasemin Cogun, Süleyman Yalcin, Aye Baak Alta, Gülay Korukluolu |
| EPI_ISL_896071, EPI_ISL_896084 | Labormedizinisches Zentrum Dr Risch | University Hospital Basel, Clinical Bacteriology | Tim Roloff, Madlen Stange, Helena MB Seth-Smith, Alfredo Mari, Karoline Leuzinger, Julia Bielicki, Nadia Wohlwend,Martin Risch, Lorenz Risch, Manuel Battegay, Hans Hirsch, Adrian Egli |
| EPI_ISL_896089 | Rothen Medizinische Laboratorien AG | University Hospital Basel, Clinical Bacteriology | Tim Roloff, Madlen Stange, Helena MB Seth-Smith, Alfredo Mari, Karoline Leuzinger, Julia Bielicki, Ingrid Steffen, Manuel Battegay, Hans Hirsch, Adrian Egli |
| EPI_ISL_896090 | Labormedizinisches Zentrum Dr Risch | University Hospital Basel, Clinical Bacteriology | Tim Roloff, Madlen Stange, Helena MB Seth-Smith, Alfredo Mari, Karoline Leuzinger, Julia Bielicki, Nadia Wohlwend,Martin Risch, Lorenz Risch, Manuel Battegay, Hans Hirsch, Adrian Egli |
| EPI_ISL_896091 | Kantonsspital Aarau, Institut für Labormedizin | University Hospital Basel, Clinical Bacteriology | Tim Roloff, Madlen Stange, Helena MB Seth-Smith, Alfredo Mari, Karoline Leuzinger, Julia Bielicki, Michael Oberle, Manuel Battegay, Hans Hirsch, Adrian Egli |
| EPI_ISL_896112 | Labormedizinisches Zentrum Dr Risch | University Hospital Basel, Clinical Bacteriology | Tim Roloff, Madlen Stange, Helena MB Seth-Smith, Alfredo Mari, Karoline Leuzinger, Julia Bielicki, Nadia Wohlwend,Martin Risch, Lorenz Risch, Manuel Battegay, Hans Hirsch, Adrian Egli |
| EPI_ISL_896113 | Viollier AG | University Hospital Basel, Clinical Bacteriology | Tim Roloff, Madlen Stange, Helena MB Seth-Smith, Alfredo Mari, Karoline Leuzinger, Julia Bielicki, Christiane Beckmann, Manuel Battegay, Hans Hirsch, Adrian Egli |
| EPI_ISL_896210 | MEPHI, Aix Marseille University | MEPHI, Aix Marseille University | Anthony LEVASSEUR |
| EPI_ISL_896296 | New York Presbyterian Hospital | Wadsworth Center, New York State Department of Health | Kirsten St. George, Daryl M. Lamson, Alexis Russel, Matthew Shudt, Melissa A Leisner, Jonathan Plitnick, Navjot Singh, John Kelly, Erasmus Schneider, Erica Lasek-Nesselquist |
| EPI_ISL_896297 | BIO-REFERENCE LABORATORIES | Wadsworth Center, New York State Department of Health | Kirsten St. George, Daryl M. Lamson, Alexis Russel, Matthew Shudt, Melissa A Leisner, Jonathan Plitnick, Navjot Singh, John Kelly, Erasmus Schneider, Erica Lasek-Nesselquist |
| EPI_ISL_896302, EPI_ISL_896303, EPI_ISL_896305, EPI_ISL_896306, EPI_ISL_896366, EPI_ISL_896367, EPI_ISL_896368, EPI_ISL_896369, EPI_ISL_896370, EPI_ISL_896371, EPI_ISL_896372 | | | |
| see above | New York Presbyterian Hospital | Wadsworth Center, New York State Department of Health | Kirsten St. George, Daryl M. Lamson, Alexis Russel, Matthew Shudt, Melissa A Leisner, Jonathan Plitnick, Navjot Singh, John Kelly, Erasmus Schneider, Erica Lasek-Nesselquist |
| EPI_ISL_896487 | KU Leuven, Rega Institute, Clinical and Epidemiological Virology | KU Leuven, Rega Institute, Clinical and Epidemiological Virology | Tony Wawina-Bokalanga, Bert Vanmechelen, Joan Marti-Carerras, Piet Maes |
| EPI_ISL_896504, EPI_ISL_896505, EPI_ISL_896506, EPI_ISL_896507, EPI_ISL_896525, EPI_ISL_896526, EPI_ISL_896527, EPI_ISL_896528, EPI_ISL_896529, EPI_ISL_896531, EPI_ISL_896532, EPI_ISL_896534, EPI_ISL_896537, EPI_ISL_896538, EPI_ISL_896539, EPI_ISL_896540, EPI_ISL_896549, EPI_ISL_896550, EPI_ISL_896551, EPI_ISL_896553, EPI_ISL_896554, EPI_ISL_896556, EPI_ISL_896557, EPI_ISL_896558, EPI_ISL_896559, EPI_ISL_896560, EPI_ISL_896561 | | | |
| see above | New York Presbyterian Hospital | Wadsworth Center, New York State Department of Health | Kirsten St. George, Daryl M. Lamson, Alexis Russel, Matthew Shudt, Melissa A Leisner, Jonathan Plitnick, Navjot Singh, John Kelly, Erasmus Schneider, Erica Lasek-Nesselquist |
| EPI_ISL_897987, EPI_ISL_898008, EPI_ISL_898029 | KU Leuven, Rega Institute, Clinical and Epidemiological Virology | KU Leuven, Rega Institute, Clinical and Epidemiological Virology | Tony Wawina-Bokalanga, Bert Vanmechelen, Joan Marti-Carerras, Piet Maes |
| EPI_ISL_899063, EPI_ISL_899065, EPI_ISL_899097, EPI_ISL_899102, EPI_ISL_899136, EPI_ISL_899139, EPI_ISL_899147, EPI_ISL_899148, EPI_ISL_899149, EPI_ISL_899150, EPI_ISL_899157, EPI_ISL_899163, EPI_ISL_899178, EPI_ISL_899180, EPI_ISL_899181, EPI_ISL_899236, EPI_ISL_899238, EPI_ISL_899239, EPI_ISL_899240, EPI_ISL_899325, EPI_ISL_899326, EPI_ISL_899327, EPI_ISL_899328, EPI_ISL_899329, EPI_ISL_899330, EPI_ISL_899331, EPI_ISL_899332, EPI_ISL_899333, EPI_ISL_899334, EPI_ISL_899335, EPI_ISL_899336, EPI_ISL_899337, EPI_ISL_899338, EPI_ISL_899339, EPI_ISL_899442, EPI_ISL_899443, EPI_ISL_899444, EPI_ISL_899445, EPI_ISL_899446, EPI_ISL_899447, EPI_ISL_899448, EPI_ISL_899449, EPI_ISL_899450, EPI_ISL_899451, EPI_ISL_899452, EPI_ISL_899453, EPI_ISL_899454, EPI_ISL_899455, EPI_ISL_899529, EPI_ISL_899539, EPI_ISL_899540, EPI_ISL_899541, EPI_ISL_899552, EPI_ISL_899553, EPI_ISL_899569, EPI_ISL_899570, EPI_ISL_899700, EPI_ISL_899701, EPI_ISL_899702, EPI_ISL_899703, EPI_ISL_899704, EPI_ISL_899705, EPI_ISL_899706, EPI_ISL_899707, EPI_ISL_899708, EPI_ISL_899709, EPI_ISL_899710, EPI_ISL_899712, EPI_ISL_899713, EPI_ISL_899714, EPI_ISL_899715, EPI_ISL_899716, | | | |

| | | | |
|---|---|---|---|
| EPI_ISL_899717, EPI_ISL_899718, EPI_ISL_899719, EPI_ISL_899720, EPI_ISL_899840, EPI_ISL_899841, EPI_ISL_899842, EPI_ISL_899884, EPI_ISL_899907, EPI_ISL_899908, EPI_ISL_899909, EPI_ISL_899910, EPI_ISL_899911, EPI_ISL_899988, EPI_ISL_899989, EPI_ISL_899990, EPI_ISL_899991, EPI_ISL_899992, EPI_ISL_899993, EPI_ISL_899994, EPI_ISL_899995, EPI_ISL_899996, EPI_ISL_899997, EPI_ISL_899998, EPI_ISL_899999, EPI_ISL_900000, EPI_ISL_900001, EPI_ISL_900002, EPI_ISL_900003, EPI_ISL_900004, EPI_ISL_900005, EPI_ISL_900006, EPI_ISL_900007 | | | |
| see above | Viollier AG | Department of Biosystems Science and Engineering, ETH Zürich | Christian Beisel, Sarah Nadeau, Chaoran Chen, Ivan Topolsky, Philipp Jablonski, Lara Fuhrmann, David Dreifuss, Katharina Jahn, Tobias Schär, Ina Nissen, Natascha Santacroce, Elodie Burcklen, Christiane Beckmann, Maurice Redondo, Olivier Kobel, Christoph Noppen, Sophie Seidel, Noemie Santamaria de Souza, Niko Beerenwinkel, Tanja Stadler |
| EPI_ISL_900023 | Viollier AG | Department of Biosystems Science and Engineering, ETH Zürich | Chaoran Chen, Sarah Nadeau, Ivan Topolsky, Emmanouil Dermitzakis, Keith Harshman, Ioannis Xenarios, Henri Pegeot, Lorenzo Cerutti, Deborah Penet, Philipp Jablonski, Lara Fuhrmann, David Dreifuss, Katharina Jahn, Christiane Beckmann, Maurice Redondo, Olivier Kobel, Christoph Noppen, Sophie Seidel, Noemie Santamaria de Souza, Niko Beerenwinkel, Tanja Stadler |
| EPI_ISL_900568 | Synlab | CNR Virus des Infections Respiratoires - France SUD | Antonin Bal, Gregory Destras, Gwendolyne Burfin, Hadrien Règue, Quentin Semanas, Martine Valette, Bruno Lina, Sylvie Larrat, Laurence Josset |
| EPI_ISL_902736 | Dirección Departamental de Salud de Pública Leticia | Instituto Nacional de Salud- Dirección de Investigación en Salud Pública, Universidad de los Andes- Applied genomics research group, Vicerrectoria de Investigación y Creación, Universidad de los Andes- Systems and Computing Engineering Department | Katherine Laiton-Donato, Diego A. Álvarez-Díaz, Carlos Franco-Muñoz, Mauricio Pacheco-Montealegre, Maria T. Herrera-Sepúlveda, Diego Andrés Prada, Jorge Duitama, Laura Natalia Gonzalez, Jorge Ivan Diaz, Silvia Restrepo-Restrepo, Magdalena Wiesner, Martha Lucia Ospina Martinez, Marcela Mercado-Reyes |
| EPI_ISL_902889, EPI_ISL_902890, EPI_ISL_902891, EPI_ISL_902892, EPI_ISL_902893 | Department of Virology and Immunology, University of Helsinki and Helsinki University Hospital, Huslab Finland | Department of Virology, Faculty of Medicine, University of Helsinki, Helsinki, Finland | Teemu Smura, Ravi Kant, Phuoc Truong, Hussein Alburkat, Hannimari Kallio-Kokko, Jenni Virtanen, Maija Suvanto, Essi Korhonen, Sari Hannula, Harri Kangas, Hanna Liimatainen, Satu Kurkela, Hanna Jarva, Maija Lappalainen, Pekka Ellonen, Olli Vapalahti |
| EPI_ISL_903066, EPI_ISL_903067, EPI_ISL_903068, EPI_ISL_903069 | Seattle Flu Study | Seattle Flu Study | Deborah A. Nickerson, Chris D. Frazar, Jover Lee, Benjamin Pelle, Erica Ryke, Matthew Richardson, Amanda Adler, Elisabeth Brandstetter, Peter D. Han, Kairsten Fay, Misja Ilcisin, Kirsten Lacombe, Thomas R. Sibley, Melissa Truong, Caitlin R. Wolf, Michael Boeckh, Janet A. Englund, Michael Famulare, Barry R. Lutz, Mark J. Rieder, Lea M. Starita, Matthew Thompson, Jay Shendure, Trevor Bedford, Helen Y. Chu |
| EPI_ISL_903073 | Seattle Flu Study | Seattle Flu Study | Deborah A. Nickerson, Chris D. Frazar, Jover Lee, Benjamin Pelle, Erica Ryke, Matthew Richardson, Amanda Adler, Elisabeth Brandstetter, Peter D. Han, Kairsten Fay, Misja Ilcisin, Kirsten Lacombe, Thomas R. Sibley, Melissa Truong, Caitlin R. Wolf, Karen Cowgill, Stephanie Schrag, Jeff Duchin, Michael Boeckh, Janet A. Englund, Michael Famulare, Barry R. Lutz, Mark J. Rieder, Lea M. Starita, Matthew Thompson, Helen Y. Chu, Trevor Bedford, Jay Shendure |
| EPI_ISL_903074, EPI_ISL_903075, EPI_ISL_903076, EPI_ISL_903077 | Seattle Flu Study | Seattle Flu Study | Deborah A. Nickerson, Chris D. Frazar, Jover Lee, Benjamin Pelle, Erica Ryke, Matthew Richardson, Amanda Adler, Elisabeth Brandstetter, Peter D. Han, Kairsten Fay, Misja Ilcisin, Kirsten Lacombe, Thomas R. Sibley, Melissa Truong, Caitlin R. Wolf, Michael Boeckh, Janet A. Englund, Michael Famulare, Barry R. Lutz, Mark J. Rieder, Lea M. Starita, Matthew Thompson, Jay Shendure, Trevor Bedford, Helen Y. Chu |
| EPI_ISL_903078, EPI_ISL_903079, EPI_ISL_903080, EPI_ISL_903104 | Seattle Flu Study | Seattle Flu Study | Deborah A. Nickerson, Chris D. Frazar, Jover Lee, Benjamin Pelle, Erica Ryke, Matthew Richardson, Amanda Adler, Elisabeth Brandstetter, Peter D. Han, Kairsten Fay, Misja Ilcisin, Kirsten Lacombe, Thomas R. Sibley, Melissa Truong, Caitlin R. Wolf, Karen Cowgill, Stephanie Schrag, Jeff Duchin, Michael Boeckh, Janet A. Englund, Michael Famulare, Barry R. Lutz, Mark J. Rieder, Lea M. Starita, Matthew Thompson, Helen Y. Chu, Trevor Bedford, Jay Shendure |
| EPI_ISL_903151, EPI_ISL_903153, EPI_ISL_903154, EPI_ISL_903155, EPI_ISL_903156, EPI_ISL_903157, EPI_ISL_903158, EPI_ISL_903159, EPI_ISL_903160, EPI_ISL_903161, EPI_ISL_903162, EPI_ISL_903164, EPI_ISL_903170, EPI_ISL_903171, EPI_ISL_903172, EPI_ISL_903173, EPI_ISL_903174, EPI_ISL_903175, EPI_ISL_903176, EPI_ISL_903177, EPI_ISL_903178, EPI_ISL_903179 | | | |
| see above | Washington State Department of Health | Seattle Flu Study | Deborah A. Nickerson, Chris D. Frazar, Jover Lee, Benjamin Pelle, Erica Ryke, Matthew Richardson, Amanda Adler, Elisabeth Brandstetter, Peter D. Han, Kairsten Fay, Misja Ilcisin, Kirsten Lacombe, Thomas R. Sibley, Melissa Truong, Caitlin R. Wolf, Romesh Gautom, Geoff Melly, Brian Hiatt, Philip Dykema, Scott Lindquist, Michael Boeckh, Janet A. Englund, Michael Famulare, Barry R. Lutz, Mark J. Rieder, Lea M. Starita, Matthew Thompson, Helen Y. Chu, Jay Shendure, Trevor Bedford |
| EPI_ISL_903185, EPI_ISL_903186, EPI_ISL_903187, EPI_ISL_903188, EPI_ISL_903189, EPI_ISL_903190, EPI_ISL_903191, EPI_ISL_903192, EPI_ISL_903193, EPI_ISL_903194, EPI_ISL_903195, EPI_ISL_903196, EPI_ISL_903197, EPI_ISL_903198, EPI_ISL_903199, EPI_ISL_903200, EPI_ISL_903201, EPI_ISL_903202, EPI_ISL_903203, EPI_ISL_903205 | | | |
| see above | Seattle Flu Study | Seattle Flu Study | Deborah A. Nickerson, Chris D. Frazar, Jover Lee, Benjamin Pelle, Erica Ryke, Matthew Richardson, Amanda Adler, Elisabeth Brandstetter, Peter D. Han, Kairsten Fay, Misja Ilcisin, Kirsten Lacombe, Thomas R. Sibley, Melissa Truong, Caitlin R. Wolf, Karen Cowgill, Stephanie Schrag, Jeff Duchin, Michael Boeckh, Janet A. Englund, Michael Famulare, Barry R. Lutz, Mark J. Rieder, Lea M. Starita, Matthew Thompson, Helen Y. Chu, Trevor Bedford, Jay Shendure |
| EPI_ISL_903361, EPI_ISL_903362, EPI_ISL_903363 | Arizona State University | Arizona State University | Emily A. Kaelin, LaRinda A. Holland, Peter T. Skidmore, Nicholas J. Mellor, Kristina Buss, Joy M. Blain, Valerie Harris, Joshua LaBaer, Vel Murugan, Efrem S. Lim |
| EPI_ISL_903394, EPI_ISL_903398, EPI_ISL_903400, EPI_ISL_903406, EPI_ISL_903417, EPI_ISL_903421, EPI_ISL_903423, EPI_ISL_903425, EPI_ISL_903437, EPI_ISL_903444, EPI_ISL_903459, EPI_ISL_903467, EPI_ISL_903473, EPI_ISL_903478, EPI_ISL_903483, EPI_ISL_903498, EPI_ISL_903504, EPI_ISL_903526, EPI_ISL_903527, EPI_ISL_903528, EPI_ISL_903529, EPI_ISL_903530, EPI_ISL_903531, EPI_ISL_903532, EPI_ISL_903533, EPI_ISL_903534, EPI_ISL_903535, EPI_ISL_903536, EPI_ISL_903537, EPI_ISL_903538, EPI_ISL_903539, EPI_ISL_903540, EPI_ISL_903541, EPI_ISL_903542, EPI_ISL_903543, EPI_ISL_903544, EPI_ISL_903545, EPI_ISL_903546, EPI_ISL_903547, EPI_ISL_903548, EPI_ISL_903549, EPI_ISL_903550, EPI_ISL_903551, EPI_ISL_903553, EPI_ISL_903558, EPI_ISL_903563, EPI_ISL_903564 | | | |
| see above | Quest Diagnostics | Quest Diagnostics | Rosenthal,S.H., Gerasimova,A., Kagan,R.M., Anderson, B., Hua, M., Liu Y., Bernstein, L.E., Livingston, K.E., Perez, A., Shalhout, D.F., Shlyakhter, I.A., Owen, R., Tanpaiboon, P., Lacbawan, F. |
| EPI_ISL_904058, EPI_ISL_904059, EPI_ISL_904060, EPI_ISL_904061, EPI_ISL_904062, EPI_ISL_904063 | New Mexico Department of Health Scientific Laboratory | New Mexico Department of Health Scientific Laboratory | Ellie Johnson, Anastacia Griego-Fisher, D'eldra Malone |
| EPI_ISL_904122 | National Institute of Public Health - National Institute of Hygiene | National Institute of Public Health - National Institute of Hygiene | Wokowicz Tomasz, Zacharczuk Katarzyna |
| EPI_ISL_904125 | Laboratorium Analityczne Szpital Specjalistyczny w Chorzowie | National Institute of Public Health - National Institute of Hygiene | Wokowicz Tomasz, Zacharczuk Katarzyna |
| EPI_ISL_904128, EPI_ISL_904130, EPI_ISL_904131, EPI_ISL_904132, EPI_ISL_904133, EPI_ISL_904134 | Quest Diagnostics | Quest Diagnostics | Rosenthal,S.H., Gerasimova,A., Kagan,R.M., Anderson, B., Hua, M., Liu Y., Bernstein, L.E., Livingston, K.E., Perez, A., Shalhout, D.F., Shlyakhter, I.A., Owen, R., Tanpaiboon, P., Lacbawan, F. |
| EPI_ISL_904144, EPI_ISL_904161, EPI_ISL_904201, EPI_ISL_904202, EPI_ISL_904203, EPI_ISL_904219, EPI_ISL_904220, EPI_ISL_904237, EPI_ISL_904246, EPI_ISL_904271, EPI_ISL_904272, EPI_ISL_904305, EPI_ISL_904306, EPI_ISL_904307, EPI_ISL_904308, EPI_ISL_904309, EPI_ISL_904310, EPI_ISL_904318, EPI_ISL_904366, EPI_ISL_904367, EPI_ISL_904548, EPI_ISL_904549, EPI_ISL_904550, EPI_ISL_904551, EPI_ISL_904552, EPI_ISL_904553, EPI_ISL_904554, EPI_ISL_904555, EPI_ISL_904556, EPI_ISL_904557, EPI_ISL_904558, EPI_ISL_904559, EPI_ISL_904560, EPI_ISL_904561, EPI_ISL_904562, EPI_ISL_904563, EPI_ISL_904564 | | | |
| see above | Dutch COVID-19 response team | Erasmus Medical Center | Bas Oude Munnink, Reina Sikkema, David Nieuwenhuijse, Irina Chestakova, Anne van der Linden, Marjan Boter, Emmanuelle Munger, Corine GeurtsvanKessel, Annemiek van der Eijk, Richard Molenkamp, Marion Koopmans, on behalf of the Dutch national COVID-19 response team. |
| EPI_ISL_904631, EPI_ISL_904634, EPI_ISL_904636, EPI_ISL_904637, EPI_ISL_904638, EPI_ISL_904640, EPI_ISL_904641, EPI_ISL_904642, EPI_ISL_904647, EPI_ISL_904650, EPI_ISL_904651, EPI_ISL_904652, EPI_ISL_904654, EPI_ISL_904656, EPI_ISL_904657 | | | |
| see above | Servicio de Microbiología, Laboratori Clínic Metropolitana Nord. Hospital Universitari Germans Trias i Pujol. Institut d'Investigació en Ciències de la Salut Germans Trias i Pujol (IGTP) | IrsiCaixa - Can Ruti CovidSeq | Marc Noguera-Julian, Mariona Parera, Maria Casadellà, Pilar Armengol, Francesc Catala-Moll, Roger Paredes, Bonaventura Clotet Elisa Martró, Verónica Saludes, Anna Not, Ana Pérez, Montserrat Giménez, Ignacio Blanco, Cristina Casañ, Antoni E. Bordoy, Adrián Antuori |
| EPI_ISL_904747, EPI_ISL_904781, EPI_ISL_904892, EPI_ISL_904975, EPI_ISL_904983, EPI_ISL_904987, EPI_ISL_904989, EPI_ISL_905000, EPI_ISL_905003, EPI_ISL_905007, EPI_ISL_905029, EPI_ISL_905035, EPI_ISL_905036, EPI_ISL_905037, EPI_ISL_905038, EPI_ISL_905039, EPI_ISL_905040, EPI_ISL_905041, EPI_ISL_905042, EPI_ISL_905043, EPI_ISL_905044, EPI_ISL_905045, EPI_ISL_905046, EPI_ISL_905047, EPI_ISL_905072, EPI_ISL_905073, EPI_ISL_905074, EPI_ISL_905075, EPI_ISL_905107, EPI_ISL_905108, EPI_ISL_905109, EPI_ISL_905111, EPI_ISL_905120, EPI_ISL_905121, EPI_ISL_905175, EPI_ISL_905190, EPI_ISL_905202, EPI_ISL_905203, EPI_ISL_905263, EPI_ISL_905269, EPI_ISL_905270, EPI_ISL_905271, EPI_ISL_905272, EPI_ISL_905273, EPI_ISL_905277, EPI_ISL_905278, EPI_ISL_905279, EPI_ISL_905289, EPI_ISL_905290, EPI_ISL_905349, EPI_ISL_905363, EPI_ISL_905364, EPI_ISL_905365, EPI_ISL_905366, EPI_ISL_905367, EPI_ISL_905368, EPI_ISL_905369, EPI_ISL_905387, EPI_ISL_905389, EPI_ISL_905390, EPI_ISL_905392, EPI_ISL_905405, EPI_ISL_905406, EPI_ISL_905407, EPI_ISL_905408, EPI_ISL_905409, EPI_ISL_905410, EPI_ISL_905411, EPI_ISL_905412, EPI_ISL_905413, EPI_ISL_905415, EPI_ISL_905416, EPI_ISL_905417, EPI_ISL_905425, EPI_ISL_905426, EPI_ISL_905427, EPI_ISL_905450, EPI_ISL_905451, EPI_ISL_905452, EPI_ISL_905453, EPI_ISL_905454, EPI_ISL_905455, EPI_ISL_905456, EPI_ISL_905457, EPI_ISL_905458, EPI_ISL_905474, EPI_ISL_905475, EPI_ISL_905476, EPI_ISL_905477, EPI_ISL_905478, | | | |

| | | | |
|---|---|---|---|
| EPI_ISL_905479, EPI_ISL_905480, EPI_ISL_905482, EPI_ISL_905485, EPI_ISL_905511, EPI_ISL_905512, EPI_ISL_905517, EPI_ISL_905518, EPI_ISL_905523, EPI_ISL_905525, EPI_ISL_905526, EPI_ISL_905527, EPI_ISL_905528, EPI_ISL_905529, EPI_ISL_905530, EPI_ISL_905531, EPI_ISL_905545, EPI_ISL_905546, EPI_ISL_905547, EPI_ISL_905569, EPI_ISL_905570, EPI_ISL_905571, EPI_ISL_905572, EPI_ISL_905573, EPI_ISL_905574, EPI_ISL_905576, EPI_ISL_905582, EPI_ISL_905585, EPI_ISL_905587, EPI_ISL_905588, EPI_ISL_905597, EPI_ISL_905598, EPI_ISL_905599, EPI_ISL_905600, EPI_ISL_905601, EPI_ISL_905602, EPI_ISL_905609, EPI_ISL_905615, EPI_ISL_905616, EPI_ISL_905622, EPI_ISL_905629, EPI_ISL_905630, EPI_ISL_905631, EPI_ISL_905632, EPI_ISL_905633, EPI_ISL_905634, EPI_ISL_905635, EPI_ISL_905636, EPI_ISL_905643, EPI_ISL_905644, EPI_ISL_905645, EPI_ISL_905646, EPI_ISL_905664, EPI_ISL_905665, EPI_ISL_905666, EPI_ISL_905676, EPI_ISL_905681, EPI_ISL_905682, EPI_ISL_905683, EPI_ISL_905684, EPI_ISL_905697, EPI_ISL_905698, EPI_ISL_905699, EPI_ISL_905719, EPI_ISL_905720 | | | |
| see above | Dutch COVID-19 response team | National Institute for Public Health and the Environment (RIVM) | Adam Meijer, Harry Vennema, Dirk Eggink, Jeroen Cremer, Sharon van den Brink, Bas van der Veer, AnneMarie van den Brandt, Florian Zwagemaker, Dennis Schmitz, Chantal Reusken, on behalf of the national COVID-19 response team |
| EPI_ISL_905732, EPI_ISL_905733, EPI_ISL_905734, EPI_ISL_905735, EPI_ISL_905736, EPI_ISL_905737, EPI_ISL_905738 | National Institute of Public Health - National Institute of Hygiene | National Institute of Public Health - National Institute of Hygiene | Wokowicz Tomasz, Zacharczuk Katarzyna |
| EPI_ISL_905927, EPI_ISL_905928, EPI_ISL_905929, EPI_ISL_905930, EPI_ISL_905931, EPI_ISL_905932, EPI_ISL_905933, EPI_ISL_905934, EPI_ISL_905935, EPI_ISL_905936, EPI_ISL_905937, EPI_ISL_905938, EPI_ISL_905939, EPI_ISL_905940, EPI_ISL_905941, EPI_ISL_905942, EPI_ISL_905943, EPI_ISL_905944, EPI_ISL_905945, EPI_ISL_905946, EPI_ISL_905947, EPI_ISL_905948, EPI_ISL_905949, EPI_ISL_905950, EPI_ISL_905951, EPI_ISL_905952, EPI_ISL_905953, EPI_ISL_905954, EPI_ISL_905955, EPI_ISL_905956, EPI_ISL_905957, EPI_ISL_905958, EPI_ISL_905959, EPI_ISL_905960, EPI_ISL_905961, EPI_ISL_905962, EPI_ISL_905963, EPI_ISL_905964, EPI_ISL_905965, EPI_ISL_905966, EPI_ISL_905967, EPI_ISL_905968, EPI_ISL_905969, EPI_ISL_905970, EPI_ISL_905971, EPI_ISL_905972, EPI_ISL_905973, EPI_ISL_905974, EPI_ISL_905975, EPI_ISL_905976, EPI_ISL_905977, EPI_ISL_905978, EPI_ISL_905979, EPI_ISL_905980, EPI_ISL_905981, EPI_ISL_905982, EPI_ISL_905983, EPI_ISL_905984, EPI_ISL_905985, EPI_ISL_905986, EPI_ISL_905987, EPI_ISL_905988, EPI_ISL_905989, EPI_ISL_905990, EPI_ISL_905991, EPI_ISL_905992, EPI_ISL_905993, EPI_ISL_905994, EPI_ISL_905995, EPI_ISL_905996, EPI_ISL_905997, EPI_ISL_905998, EPI_ISL_905999, EPI_ISL_906000, EPI_ISL_906001, EPI_ISL_906002, EPI_ISL_906003, EPI_ISL_906004, EPI_ISL_906005, EPI_ISL_906006, EPI_ISL_906007, EPI_ISL_906008, EPI_ISL_906009, EPI_ISL_906010, EPI_ISL_906011, EPI_ISL_906012, EPI_ISL_906013, EPI_ISL_906014, EPI_ISL_906015, EPI_ISL_906016, EPI_ISL_906017, EPI_ISL_906018, EPI_ISL_906019, EPI_ISL_906020, EPI_ISL_906021, EPI_ISL_906022, EPI_ISL_906023, EPI_ISL_906024, EPI_ISL_906025, EPI_ISL_906026, EPI_ISL_906027, EPI_ISL_906028, EPI_ISL_906029, EPI_ISL_906030, EPI_ISL_906031, EPI_ISL_906032, EPI_ISL_906033, EPI_ISL_906034, EPI_ISL_906035, EPI_ISL_906036, EPI_ISL_906037, EPI_ISL_906038, EPI_ISL_906039, EPI_ISL_906040, EPI_ISL_906041, EPI_ISL_906042, EPI_ISL_906043, EPI_ISL_906044, EPI_ISL_906046 | | | |
| see above | OHSU Lab Services Molecular Microbiology Lab | Oregon SARS-CoV-2 Genome Sequencing Center | Brendan L. O'Connell, Sally Grindstaff, Kayla Carter, Ruth V. Nichols, Alec J. Hirsch, Donna Hansel, Guang Fan, Xuan, Qin, Daniel N. Streblow, William B. Messer, Andrew C. Adey, Benjamin N. Bimber, Brian J. O'Roak |
| EPI_ISL_906086 | Child Health Research Foundation | Child Health Research Foundation | Senjuti Saha, Syed Muktadir Al Sium, Sharmistha Goswami, Afroza Akter Tanni, Arif Mohammad Tanmoy, Roly Malaker, Md Hafizur Rahman, Samir K Saha |
| EPI_ISL_906145 | Laboratorio de Salud Publica de Amazonas | Instituto Nacional de Salud- Dirección de Investigación en Salud Pública, Universidad de los Andes- Applied genomics research group, Vicerrectoria de Investigación y Creación, Universidad de los Andes- Systems and Computing Engineering Department | Katherine Laiton-Donato, Diego A. Álvarez-Díaz, Carlos Franco-Muñoz, Mauricio Pacheco-Montealegre, Héctor Alejandro Ruiz-Moreno, Maria T. Herrera-Sepúlveda, Diego Andrés Prada, Jhonnatan Reales-González, Sheryll Corchuelo, Julian Naizaque, Gerardo Santamaría Jorge Duitama, Laura Natalia Gonzalez, Jorge Ivan Diaz, Silvia Restrepo-Restrepo, Magdalena Wiesner, Martha Lucia Ospina Martinez, Marcela Mercado-Reyes |
| EPI_ISL_906147, EPI_ISL_906148, EPI_ISL_906150, EPI_ISL_906153, EPI_ISL_906158, EPI_ISL_906159, EPI_ISL_906161, EPI_ISL_906164, EPI_ISL_906167, EPI_ISL_906169, EPI_ISL_906172, EPI_ISL_906174, EPI_ISL_906176, EPI_ISL_906178, EPI_ISL_906181, EPI_ISL_906182, EPI_ISL_906185, EPI_ISL_906187 | | | |
| see above | University of Wisconsin-Madison AIDS Vaccine Research Laboratories | University of Wisconsin-Madison AIDS Vaccine Research Laboratories | Gage Moreno, Katarina Braun, et al. AIDS Vaccine Research Laboratories |
| EPI_ISL_906277, EPI_ISL_906278 | Nigeria Centre for Disease Control (NCDC) | African Centre of Excellence for Genomics of Infectious Diseases (ACEGID), Redeemer's University | Oluniyi P.E. et al |
| EPI_ISL_906315, EPI_ISL_906339, EPI_ISL_906342, EPI_ISL_906365, EPI_ISL_906369, EPI_ISL_906389, EPI_ISL_906394, EPI_ISL_906396, EPI_ISL_906403, EPI_ISL_906424, EPI_ISL_906425, EPI_ISL_906426, EPI_ISL_906427, EPI_ISL_906428, EPI_ISL_906429, EPI_ISL_906430, EPI_ISL_906431, EPI_ISL_906432, EPI_ISL_906434, EPI_ISL_906435, EPI_ISL_906437, EPI_ISL_906438, EPI_ISL_906439, EPI_ISL_906440, EPI_ISL_906441, EPI_ISL_906442, EPI_ISL_906443, EPI_ISL_906444, EPI_ISL_906445, EPI_ISL_906446, EPI_ISL_906447, EPI_ISL_906449, EPI_ISL_906450, EPI_ISL_906451, EPI_ISL_906452, EPI_ISL_906453, EPI_ISL_906459, EPI_ISL_906460, EPI_ISL_906464 | | | |
| see above | Quest Diagnostics | Quest Diagnostics | Rosenthal,S.H., Gerasimova,A., Kagan,R.M., Anderson, B., Hua, M., Liu Y., Bernstein, L.E., Livingston, K.E., Perez, A., Shalhout, D.F., Shlyakhter, I.A., Owen, R., Tanpaiboon, P., Lacbawan, F. |
| EPI_ISL_906529 | Direccion de partamental de salud del Amazonas | Instituto Nacional de Salud- Dirección de Investigación en Salud Pública, Universidad de los Andes- Applied genomics research group, Vicerrectoria de Investigación y Creación, Universidad de los Andes- Systems and Computing Engineering Department | Katherine Laiton-Donato, Diego A. Álvarez-Díaz, Carlos Franco-Muñoz, Mauricio Pacheco-Montealegre, Héctor Alejandro Ruiz-Moreno, Maria T. Herrera-Sepúlveda, Diego Andrés Prada, Jhonnatan Reales-González, Sheryll Corchuelo, Julian Naizaque, Gerardo Santamaría Jorge Duitama, Laura Natalia Gonzalez, Jorge Ivan Diaz, Silvia Restrepo-Restrepo, Magdalena Wiesner, Martha Lucia Ospina Martinez, Marcela Mercado-Reyes |
| EPI_ISL_906535, EPI_ISL_906537 | HOSPITAL DEPARTAMENTAL DE VILLAVICENCIO E.S.E. | Instituto Nacional de Salud- Dirección de Investigación en Salud Pública, Universidad de los Andes- Applied genomics research group, Vicerrectoria de Investigación y Creación, Universidad de los Andes- Systems and Computing Engineering Department | Katherine Laiton-Donato, Diego A. Álvarez-Díaz, Carlos Franco-Muñoz, Mauricio Pacheco-Montealegre, Héctor Alejandro Ruiz-Moreno, Maria T. Herrera-Sepúlveda, Diego Andrés Prada, Jhonnatan Reales-González, Sheryll Corchuelo, Julian Naizaque, Gerardo Santamaría Jorge Duitama, Laura Natalia Gonzalez, Jorge Ivan Diaz, Silvia Restrepo-Restrepo, Magdalena Wiesner, Martha Lucia Ospina Martinez, Marcela Mercado-Reyes |
| EPI_ISL_906550, EPI_ISL_906551 | DIRECCION DEPARTAMENTAL DE SALUD DE AMAZONAS | Instituto Nacional de Salud- Dirección de Investigación en Salud Pública, Universidad de los Andes- Applied genomics research group, Vicerrectoria de Investigación y Creación, Universidad de los Andes- Systems and Computing Engineering Department | Katherine Laiton-Donato, Diego A. Álvarez-Díaz, Carlos Franco-Muñoz, Mauricio Pacheco-Montealegre, Héctor Alejandro Ruiz-Moreno, Maria T. Herrera-Sepúlveda, Diego Andrés Prada, Jhonnatan Reales-González, Sheryll Corchuelo, Julian Naizaque, Gerardo Santamaría Jorge Duitama, Laura Natalia Gonzalez, Jorge Ivan Diaz, Silvia Restrepo-Restrepo, Magdalena Wiesner, Martha Lucia Ospina Martinez, Marcela Mercado-Reyes |
| EPI_ISL_906716, EPI_ISL_906718, EPI_ISL_906719, EPI_ISL_906722, EPI_ISL_906732, EPI_ISL_906734, EPI_ISL_906737, EPI_ISL_906739, EPI_ISL_906741, EPI_ISL_906742, EPI_ISL_906745, EPI_ISL_906747, EPI_ISL_906750, EPI_ISL_906751, EPI_ISL_906753, EPI_ISL_906754 | | | |
| see above | Hematology Laboratory, Section of Molecular Diagnostics, University Clinical Centre, Medical University of Gdansk | Laboratory of Recombinant Vaccines | Lukasz Rabalski, Maciej Kosinski, Maciej Grzybek, Adam Sodol, Aneta Szulc, Krzysztof Lewandowski, Ewa Milosz, Marlena Robakowska, Boguslaw Szewczyk, Krystyna Bienkowska-Szewczyk |
| EPI_ISL_906798, EPI_ISL_906799, EPI_ISL_906801, EPI_ISL_906803 | Nordland Hospital - Bodo, Laboratory Department, Molecular Biology Unit | Norwegian Institute of Public Health, Department of Virology | Kathrine Stene-Johansen, Kamilla Heddeland Instefjord, Hilde Elshaug, Atiya R Ali,Marie Paulsen Madsen, Rasmus Riis Kopperud, Hilde Vollan, Karoline Bragstad, Olav Hungnes |
| EPI_ISL_906822, EPI_ISL_906823 | Department of Medical Microbiology - section Molde, Molde Hospital | Norwegian Institute of Public Health, Department of Virology | Kathrine Stene-Johansen, Kamilla Heddeland Instefjord, Hilde Elshaug, Atiya R Ali,Marie Paulsen Madsen, Rasmus Riis Kopperud, Hilde Vollan, Karoline Bragstad, Olav Hungnes |
| EPI_ISL_906826, EPI_ISL_906827, EPI_ISL_906828, EPI_ISL_906829 | Akershus University Hospital, Department for Microbiology and Infectious Disease Control | Norwegian Institute of Public Health, Department of Virology | Kathrine Stene-Johansen, Kamilla Heddeland Instefjord, Hilde Elshaug, Atiya R Ali,Marie Paulsen Madsen, Rasmus Riis Kopperud, Hilde Vollan, Karoline Bragstad, Olav Hungnes |
| EPI_ISL_906884, EPI_ISL_906885, EPI_ISL_906886, EPI_ISL_906887, EPI_ISL_906888, EPI_ISL_906889, EPI_ISL_906890, EPI_ISL_906891, EPI_ISL_906892, EPI_ISL_906893, EPI_ISL_906894, EPI_ISL_906895, EPI_ISL_906896, EPI_ISL_906897, EPI_ISL_906898, EPI_ISL_906899, EPI_ISL_906900, EPI_ISL_906901, EPI_ISL_906902, EPI_ISL_906903, EPI_ISL_906904 | | | |
| see above | Bureau of Public Health Laboratories, Florida Department of Health (BPHL, FLDOH) | Bureau of Public Health Laboratories, Florida Department of Health (BPHL, FLDOH) | Schmedes,S., Blanton,J. |
| EPI_ISL_906924, EPI_ISL_906930, EPI_ISL_906954, EPI_ISL_906964, EPI_ISL_906978, EPI_ISL_907064, EPI_ISL_907066, EPI_ISL_907068, EPI_ISL_907070, EPI_ISL_907074 | Infectious Diseases, Quest Diagnostics | Infectious Diseases, Quest Diagnostics | Rosenthal,S.H., Gerasimova,A., Kagan,R.M., Anderson,B., Bernstein,L.E., Livingston,K.E., Hua,M., Liu,Y., Shalhout,D.F., Owen,R., Lacbawan,F. |
| EPI_ISL_909946, EPI_ISL_909947 | CUSL/UCLouvain COVID testing federal platform | UCLouvain/IREC/MBLG | Jean Ruelle, Lysa Pinsmaye, Benoit Kabamba Mukadi |
| EPI_ISL_910029, EPI_ISL_910335 | Laboratory for Respiratory Viruses, Cantacuzino National Military-Medical Institute for Research and Development | Cantacuzino Institute Virology | Luiza Ustea, Nicoleta Paraschiv, Mihaela Lazar |
| EPI_ISL_910902, EPI_ISL_910903, EPI_ISL_910904, EPI_ISL_910905, EPI_ISL_910906, EPI_ISL_910907, EPI_ISL_910908, EPI_ISL_910909, EPI_ISL_910910, EPI_ISL_910911, EPI_ISL_910912, EPI_ISL_910913, EPI_ISL_910914, EPI_ISL_910915, EPI_ISL_910916, EPI_ISL_910917, EPI_ISL_910918, EPI_ISL_910919, EPI_ISL_910920, EPI_ISL_910921, EPI_ISL_910922, EPI_ISL_910923, EPI_ISL_910924, EPI_ISL_910925, EPI_ISL_910926, EPI_ISL_910927, EPI_ISL_910928, EPI_ISL_910929, EPI_ISL_910930, EPI_ISL_910931, EPI_ISL_910932, EPI_ISL_910933, EPI_ISL_910934, EPI_ISL_910935, EPI_ISL_910936, EPI_ISL_910937, EPI_ISL_910938, EPI_ISL_910939, EPI_ISL_910940, EPI_ISL_910942 | | | |

| | | | |
|---|---|---|---|
| see above | Laboratoire national de sante, Microbiology, Virology | Laboratoire national de sante, Microbiology, Microbial Genomics Platform | Anke Wienecke-Baldacchino, Catherine Ragimbeau,Jessica Tapp, Fatu Djabi, Lise Pignon, Raoul Salmon, Tamir Abdelrahman |
| EPI_ISL_911292, EPI_ISL_911302, EPI_ISL_911311, EPI_ISL_911321, EPI_ISL_911329, EPI_ISL_911346 | Servicio de Microbiología, Hospital Universitario Son Espases | SeqCOVID-SPAIN consortium/IBV(CSIC) | Carla López-Causapé, Jordi Reina, Antonio Oliver and SeqCOVID-SPAIN consortium |
| EPI_ISL_911518, EPI_ISL_911519, EPI_ISL_911520, EPI_ISL_911521, EPI_ISL_911522, EPI_ISL_911523 | Florida Bureau of Public Health Laboratories | Florida Bureau of Public Health Laboratories | Sarah Schmedes, Jason Blanton |
| EPI_ISL_911526, EPI_ISL_911527 | Microbiology and Virology Unit, Florence Careggi University Hospital | Microbiology and Virology Unit, Florence Careggi University Hospital | Vincenzo Di Pilato, Marco Coppi, Fabio Morecchiato, Noemi Aiezza, Ilaria Baccani, Alberto Antonelli, Emanuele Gori, Gian Maria Rossolini |
| EPI_ISL_911635, EPI_ISL_911636, EPI_ISL_911637, EPI_ISL_911638, EPI_ISL_911641, EPI_ISL_911642, EPI_ISL_911643, EPI_ISL_911644, EPI_ISL_911645, EPI_ISL_911646, EPI_ISL_911647, EPI_ISL_911648, EPI_ISL_911649, EPI_ISL_911670, EPI_ISL_911671, EPI_ISL_911672, EPI_ISL_911673 | | | |
| see above | Texas Department of State Health Services (TXDSHS) | Texas Department of State Health Services (TXDSHS) | Bonnie Oh, Anita Pokharel, James Daniel Bonser, Myong Koag, Chung Wang, Rachel Lee, Grace Kubin, Rashmi Tuladhar, Mayela Pedrueza, Maliha Rahman, Jenny Zhang |
| EPI_ISL_911789, EPI_ISL_911824, EPI_ISL_911825 | Johns Hopkins Hospital Department of Pathology | Johns Hopkins Hospital Department of Pathology | C. Paul Morris, Chun Huai Luo, Adannaya Amadi, Matthew Schwartz, Nicholas Gallagher, Heba H. Mostafa |
| EPI_ISL_912035, EPI_ISL_912036, EPI_ISL_912037, EPI_ISL_912038, EPI_ISL_912039, EPI_ISL_912040, EPI_ISL_912041, EPI_ISL_912042, EPI_ISL_912043, EPI_ISL_912044, EPI_ISL_912045, EPI_ISL_912046 | | | |
| see above | Washington State Department of Health | Seattle Flu Study | Deborah A. Nickerson, Chris D. Frazar, Jover Lee, Benjamin Pelle, Erica Ryke, Matthew Richardson, Amanda Adler, Elisabeth Brandstetter, Peter D. Han, Kairsten Fay, Misja Ilcisin, Kirsten Lacombe, Thomas R. Sibley, Melissa Truong, Caitlin R. Wolf, Romesh Gautom, Geoff Melly, Brian Hiatt, Philip Dykema, Scott Lindquist, Michael Boeckh, Janet A. Englund, Michael Famulare, Barry R. Lutz, Mark J. Rieder, Lea M. Starita, Matthew Thompson, Helen Y. Chu, Jay Shendure, Trevor Bedford |
| EPI_ISL_912162 | Baptist Medical Center | Grubaugh Lab - Yale School of Public Health | Tara Alpert, Joseph Fauver, Anderson Brito, Mallery Breban, Anne Wyllie, Chantal Vogels, Mary Petrone, Annie Watkins, Chaney Kalinich, Isabel Ott, Nathan Grubaugh |
| EPI_ISL_912169 | East Gene | Grubaugh Lab - Yale School of Public Health | Tara Alpert, Joseph Fauver, Anderson Brito, Mallery Breban, Anne Wyllie, Chantal Vogels, Mary Petrone, Annie Watkins, Chaney Kalinich, Isabel Ott, Nathan Grubaugh |
| EPI_ISL_912185 | Connecticut Department of Health | Grubaugh Lab - Yale School of Public Health | Tara Alpert, Joseph Fauver, Anderson Brito, Mallery Breban, Anne Wyllie, Chantal Vogels, Mary Petrone, Annie Watkins, Chaney Kalinich, Isabel Ott, Nathan Grubaugh |
| EPI_ISL_912280 | Hospital General Universitario Gregorio Marañón | SeqCOVID-SPAIN consortium / IBV (CSIC) | Darío García de Viedma, Laura Pérez-Lago, Pedro J Sola-Campoy, Sergio Buenestado-Serrano, Marta Herranz, Victor Manuel de la Cueva, Julia Suárez, Pilar Catalán, Patricia Muñoz and SeqCOVID-SPAIN consortium |
| EPI_ISL_912305, EPI_ISL_912309, EPI_ISL_912312 | Hospital General Universitario Gregorio Marañón | SeqCOVID-SPAIN consortium / IBV (CSIC) | Sergio Buenestado Serrano, Pedro J. Sola Campoy, Laura Pérez-Lago, Pilar Catalán, Arturo fraile Torres, Andrés Von Wernitz, Carmen del Arco, Patricia Muñoz, Laura Cardeñoso, Darío García de Viedma and SeqCOVID-SPAIN consortium |
| EPI_ISL_912317, EPI_ISL_912319 | Hospital General Universitario Gregorio Marañón | SeqCOVID-SPAIN consortium / IBV (CSIC) | Darío García de Viedma, Laura Pérez-Lago, Pedro J Sola-Campoy, Sergio Buenestado-Serrano, Marta Herranz, Victor Manuel de la Cueva, Julia Suárez, Pilar Catalán, Patricia Muñoz and SeqCOVID-SPAIN consortium |
| EPI_ISL_912332 | Hospital General Universitario Gregorio Marañón | SeqCOVID-SPAIN consortium / IBV (CSIC) | Sergio Buenestado Serrano, Pedro J. Sola Campoy, Laura Pérez-Lago, Pilar Catalán, Arturo fraile Torres, Andrés Von Wernitz, Carmen del Arco, Patricia Muñoz, Laura Cardeñoso, Darío García de Viedma and SeqCOVID-SPAIN consortium |
| EPI_ISL_912333, EPI_ISL_912335 | Hospital General Universitario Gregorio Marañón | SeqCOVID-SPAIN consortium / IBV (CSIC) | Darío García de Viedma, Laura Pérez-Lago, Pedro J Sola-Campoy, Sergio Buenestado-Serrano, Marta Herranz, Victor Manuel de la Cueva, Julia Suárez, Pilar Catalán, Patricia Muñoz and SeqCOVID-SPAIN consortium |
| EPI_ISL_912379, EPI_ISL_912380, EPI_ISL_912386, EPI_ISL_912387, EPI_ISL_912388, EPI_ISL_912389, EPI_ISL_912390 | Fondation Congolaise pour la recherche medicale (FCRM), Francine Ntoumi | NGS Competence Center Tuebingen, Institut für Medizinische Mikrobiologie und Hygiene, Universitaetsklinikum Tübingen | Angel Angelov |
| EPI_ISL_912404, EPI_ISL_912420, EPI_ISL_912449, EPI_ISL_912450, EPI_ISL_912451, EPI_ISL_912452, EPI_ISL_912455 | KU Leuven, Rega Institute, Clinical and Epidemiological Virology | KU Leuven, Rega Institute, Clinical and Epidemiological Virology | Tony Wawina-Bokalanga, Bert Vanmechelen, Joan Marti-Carerras, Piet Maes |
| EPI_ISL_912521, EPI_ISL_912524 | NHLS Universitas Academic | UFS Virology | PA Bester, MM Nyaga, P Nthiga, MT Mogotsi, D Goedhals, T de Oliveira |
| EPI_ISL_912597 | Labo Analyses Med | National Reference Center for Viruses of Respiratory Infections, Institut Pasteur, Paris | Marion Barbet, Sylvie Behillil, Méline Bizard, Angela Brisebarre, Camille Capel, Etienne Simon-Lorière, Vincent Enouf, Maud Vanpeene, Sylvie van der Werf,Le Vicky |
| EPI_ISL_912647, EPI_ISL_912652, EPI_ISL_912680, EPI_ISL_912688, EPI_ISL_912689, EPI_ISL_912693, EPI_ISL_912701, EPI_ISL_912705, EPI_ISL_912708, EPI_ISL_912714, EPI_ISL_912715, EPI_ISL_912725, EPI_ISL_912726, EPI_ISL_912735, EPI_ISL_912736, EPI_ISL_912737, EPI_ISL_912738, EPI_ISL_912739, EPI_ISL_912740, EPI_ISL_912741, EPI_ISL_912850 | | | |
| see above | Hôpital Henri Mondor | Department of Virology, Henri Mondor University Hospital, Assistance Publique Hôpitaux de Paris, Université Paris-Est Créteil, INSERM U955 | Christophe Rodriguez, Slim Fourati, Vanessa Demontant, Guillaume Gricourt, Melissa N'Debi, Alexandre Soulier, Elisabeth Trawinski, Jean-Michel Pawlotsky |
| EPI_ISL_912919, EPI_ISL_912920 | Hôpital Pitié-Salpêtrière | Department of Virology, Henri Mondor University Hospital, Assistance Publique Hôpitaux de Paris, Université Paris-Est Créteil, INSERM U955 | Christophe Rodriguez, Slim Fourati, Vanessa Demontant, Guillaume Gricourt, Melissa N'Debi, Alexandre Soulier, Elisabeth Trawinski, Jean-Michel Pawlotsky |
| EPI_ISL_913020 | Hospital Universitario de Ceuta | Instituto de Salud Carlos III | Iglesias-Caballero, M. Camarero, S. Sandonís,V. Vázquez, S. Pozo, F. Casas, I. Jiménez, P. Zaballos, A. Monzón, S. Varona, S. Cuesta, I. López, J. |
| EPI_ISL_913038 | Complejo Asistencial Universitario de Burgos | Instituto de Salud Carlos III | Iglesias-Caballero, M. Camarero, S. Sandonís,V. Vázquez, S. Pozo, F. Casas, I. Jiménez, P. Zaballos, A. Monzón, S. Varona, S. Cuesta, I. Megías, G. |
| EPI_ISL_913042, EPI_ISL_913043, EPI_ISL_913046 | Hospital Comarcal Sierrallana | Instituto de Salud Carlos III | Iglesias-Caballero, M. Camarero, S. Sandonís,V. Vázquez, S. Pozo, F. Casas, I. Jiménez, P. Zaballos, A. Monzón, S. Varona, S. Cuesta, I. De Benito, I. |
| EPI_ISL_913054, EPI_ISL_913055, EPI_ISL_913056, EPI_ISL_913058 | Hospital Universitario de Ceuta | Instituto de Salud Carlos III | Iglesias-Caballero, M. Camarero, S. Sandonís,V. Vázquez, S. Pozo, F. Casas, I. Jiménez, P. Zaballos, A. Monzón, S. Varona, S. Cuesta, I. López, J. |
| EPI_ISL_913077, EPI_ISL_913078 | Center for Virology | Center for Virology | Jeremy V. Camp, Irene Goerzer, Monika Redlberger-Fritz, Stephan W. Aberle |
| EPI_ISL_913127, EPI_ISL_913134 | University of Michigan Clinical Microbiology Laboratory | Lauring Lab, University of Michigan, Department of Microbiology and Immunology | Valesano |
| EPI_ISL_913259 | Klinisk mikrobiologi | The Public Health Agency of Sweden | Anna-Malin Linde, Maria Lind Karlberg, Carlo Berg, Oskar Karlsson Lindsjo, Sofia Stamouli, Reza Advani, Mattias Haukland, Petra Holmstrom, Noura Walai, Petra Edquist, Mia Brytting, Anna Risberg, Karin Tegmark-Wisell |
| EPI_ISL_913263, EPI_ISL_913267 | The Public Health Agency of Sweden | The Public Health Agency of Sweden | Anna-Malin Linde, Maria Lind Karlberg, Carlo Berg, Oskar Karlsson Lindsjo, Sofia Stamouli, Reza Advani, Mattias Haukland, Petra Holmstrom, Noura Walai, Petra Edquist, Mia Brytting, Anna Risberg, Karin Tegmark-Wisell |
| EPI_ISL_913268, EPI_ISL_913272, EPI_ISL_913300 | Klinisk mikrobiologi | The Public Health Agency of Sweden | Anna-Malin Linde, Maria Lind Karlberg, Carlo Berg, Oskar Karlsson Lindsjo, Sofia Stamouli, Reza Advani, Mattias Haukland, Petra Holmstrom, Noura Walai, Petra Edquist, Mia Brytting, Anna Risberg, Karin Tegmark-Wisell |
| EPI_ISL_913370 | Klinisk Mikrobiologi | The Public Health Agency of Sweden | Anna-Malin Linde, Maria Lind Karlberg, Carlo Berg, Oskar Karlsson Lindsjo, Sofia Stamouli, Reza Advani, Mattias Haukland, Petra Holmstrom, Noura |

| | | | |
|---|---|---|---|
| | | | Walai, Petra Edquist, Mia Brytting, Anna Risberg, Karin Tegmark-Wisell |
| EPI_ISL_913371, EPI_ISL_913372, EPI_ISL_913374 | Klinisk mikrobiologi | The Public Health Agency of Sweden | Anna-Malin Linde, Maria Lind Karlberg, Carlo Berg, Oskar Karlsson Lindsjo, Sofia Stamouli, Reza Advani, Mattias Haukland, Petra Holmstrom, Noura Walai, Petra Edquist, Mia Brytting, Anna Risberg, Karin Tegmark-Wisell |
| EPI_ISL_913375, EPI_ISL_913376, EPI_ISL_913377 | The Public Health Agency of Sweden | The Public Health Agency of Sweden | Anna-Malin Linde, Maria Lind Karlberg, Carlo Berg, Oskar Karlsson Lindsjo, Sofia Stamouli, Reza Advani, Mattias Haukland, Petra Holmstrom, Noura Walai, Petra Edquist, Mia Brytting, Anna Risberg, Karin Tegmark-Wisell |
| EPI_ISL_913399, EPI_ISL_913417, EPI_ISL_913430 | Massachusetts State Public Health Laboratory | Massachusetts State Public Health Laboratory | Andrew Lang, Timelia Fink, Glen Gallagher, Sandra Smole |
| EPI_ISL_913451, EPI_ISL_913452 | The Public Health Agency of Sweden | The Public Health Agency of Sweden | Anna-Malin Linde, Maria Lind Karlberg, Carlo Berg, Oskar Karlsson Lindsjo, Sofia Stamouli, Reza Advani, Mattias Haukland, Petra Holmstrom, Noura Walai, Petra Edquist, Mia Brytting, Anna Risberg, Karin Tegmark-Wisell |
| EPI_ISL_913466 | Synlab Medilab, Mikrobiologi | The Public Health Agency of Sweden | Anna-Malin Linde, Maria Lind Karlberg, Carlo Berg, Oskar Karlsson Lindsjo, Sofia Stamouli, Reza Advani, Mattias Haukland, Petra Holmstrom, Noura Walai, Petra Edquist, Mia Brytting, Anna Risberg, Karin Tegmark-Wisell |
| EPI_ISL_913471, EPI_ISL_913493 | Klinisk mikrobiologi | The Public Health Agency of Sweden | Anna-Malin Linde, Maria Lind Karlberg, Carlo Berg, Oskar Karlsson Lindsjo, Sofia Stamouli, Reza Advani, Mattias Haukland, Petra Holmstrom, Noura Walai, Petra Edquist, Mia Brytting, Anna Risberg, Karin Tegmark-Wisell |
| EPI_ISL_913567, EPI_ISL_913569, EPI_ISL_913570, EPI_ISL_913571, EPI_ISL_913572, EPI_ISL_913573, EPI_ISL_913574, EPI_ISL_913575, EPI_ISL_913576, EPI_ISL_913577, EPI_ISL_913578, EPI_ISL_913579, EPI_ISL_913580, EPI_ISL_913581, EPI_ISL_913582, EPI_ISL_913583, EPI_ISL_913585 | | | |
| see above | M Health Fairview | Minnesota Department of Health, Public Health Laboratory | Alexandra Lorentz, Jacob Garfin, Matt Plumb, and Xiong Wang |
| EPI_ISL_913651, EPI_ISL_913652, EPI_ISL_913653, EPI_ISL_913654 | Michigan Department of Health and Human Services, Bureau of Laboratories | Michigan Department of Health and Human Services, Bureau of Laboratories | Blankenship HM, Riner D, Soehnlen MK |
| EPI_ISL_913720, EPI_ISL_913721, EPI_ISL_913722, EPI_ISL_913723, EPI_ISL_913724, EPI_ISL_913725 | Minnesota Department of Health, Public Health Laboratory | Minnesota Department of Health, Public Health Laboratory | Alexandra Lorentz, Jacob Garfin, Matt Plumb, and Xiong Wang |
| EPI_ISL_913727, EPI_ISL_913728, EPI_ISL_913729, EPI_ISL_913730, EPI_ISL_913731 | M Health Fairview | Minnesota Department of Health, Public Health Laboratory | Alexandra Lorentz, Jacob Garfin, Matt Plumb, and Xiong Wang |
| EPI_ISL_913737, EPI_ISL_913738, EPI_ISL_913739 | Minnesota Department of Health, Public Health Laboratory | Minnesota Department of Health, Public Health Laboratory | Alexandra Lorentz, Jacob Garfin, Matt Plumb, and Xiong Wang |
| EPI_ISL_913771, EPI_ISL_913774, EPI_ISL_913782, EPI_ISL_913788, EPI_ISL_913795, EPI_ISL_913798, EPI_ISL_913805, EPI_ISL_913815, EPI_ISL_913829, EPI_ISL_913836, EPI_ISL_913849, EPI_ISL_913852, EPI_ISL_913856, EPI_ISL_913873, EPI_ISL_913879, EPI_ISL_913881, EPI_ISL_913885, EPI_ISL_913886, EPI_ISL_913890, EPI_ISL_913894, EPI_ISL_913895, EPI_ISL_913896, EPI_ISL_913899, EPI_ISL_913900, EPI_ISL_913901, EPI_ISL_913907, EPI_ISL_913908 | | | |
| see above | TGen North | TGen North | "Jolene Bowers, Megan Folkerts, Chris French, Hayley Yaglom, Ashlyn Pfeiffer, Darrin Lemmer, Dave Engelthaler, The Arizona COVID Genomics Union (ACGU)" |
| EPI_ISL_914000, EPI_ISL_914001 | Quest Diagnostics Tucker GA | Pathogen Discovery, Respiratory Viruses Branch, Division of Viral Diseases, Centers for Disease Control and Prevention | Ying Tao, Yan Li, Jing Zhang, Krista Queen, Anna Uehara, Peter Cook, Clinton R. Paden, Haibin Wang, Suxiang Tong |
| EPI_ISL_914013, EPI_ISL_914014 | IL Department of Public Health Chicago Laboratory | Pathogen Discovery, Respiratory Viruses Branch, Division of Viral Diseases, Centers for Disease Control and Prevention | Ying Tao, Yan Li, Jing Zhang, Krista Queen, Anna Uehara, Peter Cook, Clinton R. Paden, Haibin Wang, Suxiang Tong |
| EPI_ISL_914641, EPI_ISL_914645 | Santa Clara County Public Health Laboratory | Santa Clara County Public Health Laboratory | Santa Clara County Public Health Department |
| EPI_ISL_914825 | AREA DE SALUD MORAVIA | Inciensa, Instituto Costarricense de Investigación y Enseñanza en Nutrición y Salud | Francisco Duarte, Hebleen Porras, Claudio Soto-Garita, Estela Cordero, Adriana Godínez, Melany Calderón & Mariel López |
| EPI_ISL_914826 | HLE - MANOS ABIERTAS | Inciensa, Instituto Costarricense de Investigación y Enseñanza en Nutrición y Salud | Francisco Duarte, Hebleen Porras, Claudio Soto-Garita, Estela Cordero, Adriana Godínez, Melany Calderón & Mariel López |
| EPI_ISL_914827 | HLE - ASOCIACION DE ALBERGUE PARA ANCIANOS DE GOLFITO | Inciensa, Instituto Costarricense de Investigación y Enseñanza en Nutrición y Salud | Francisco Duarte, Hebleen Porras, Claudio Soto-Garita, Estela Cordero, Adriana Godínez, Melany Calderón & Mariel López |
| EPI_ISL_914830 | AREA DE SALUD CARRILLO | Inciensa, Instituto Costarricense de Investigación y Enseñanza en Nutrición y Salud | Francisco Duarte, Hebleen Porras, Claudio Soto-Garita, Estela Cordero, Adriana Godínez, Melany Calderón & Adriana Bermúdez-Espinoza |
| EPI_ISL_914970, EPI_ISL_915192 | MRCG at LSHTM Genomics lab | MRCG at LSHTM Genomics lab | Abdul Karim sesay, Abdoulie Kanteh, Jarra Manneh, Mariama Kujabi, Bakary Sanyang |
| EPI_ISL_915326, EPI_ISL_915333, EPI_ISL_915334, EPI_ISL_915341 | Quest Diagnostics | Quest Diagnostics | Rosenthal,S.H., Gerasimova,A., Kagan,R.M., Anderson, B., Hua, M., Liu Y., Bernstein, L.E., Livingston, K.E., Perez, A., Shalhout, D.F., Shlyakhter, I.A., Owen, R., Tanpaiboon, P., Lacbawan, F. |
| EPI_ISL_915434 | MD Laboratories | Los Angeles County PHL | P. Hemarajata et al. |
| EPI_ISL_917941 | Los Angeles County PHL | Los Angeles County PHL | P. Hemarajata et al. |
| EPI_ISL_918176 | Department of Infectious Diseases and Immunology, National Hospital Organization Nagoya Medical Center | Clinical Research Center, National Hospital Organization Nagoya Medical Center | Yoshihiro Nakata, Hirotaka Ode, Mai Kubota, Masakazu Matsuda, Kazuhiro Matsuoka, Miho Nakasuji, Mikiko Mori, Mayumi Imahashi, Yoshiyuki Yokomaku, Yasumasa Iwatani |
| EPI_ISL_918177, EPI_ISL_918180, EPI_ISL_918183, EPI_ISL_918185, EPI_ISL_918187, EPI_ISL_918200, EPI_ISL_918237, EPI_ISL_918243, EPI_ISL_918249, EPI_ISL_918251, EPI_ISL_918254, EPI_ISL_918255 | | | |
| see above | Innovative Genomics Institute, UC Berkeley | Innovative Genomics Institute, UC Berkeley | Stacia Wyman, Haridha Shivram, Phil Frankino, Liana Lareau, Shana McDevitt, Justin Choi |
| EPI_ISL_918263 | SIESP DIPARTIMENTO DI PREVENZIONE CHIE | Istituto Zooprofilattico Sperimentale dell'Abruzzo e Molise "G. Caporale" | Lorusso A, Marcacci M, Di Domenico M, Ancora M, Curini V, Mangone I, Rinaldi A, Scialabba S, Di Pasquale A, Cammà C, Puglia I, Calistri P, Savini G |
| EPI_ISL_918266 | SIESP CHIETI-Drive in Lanciano | Istituto Zooprofilattico Sperimentale dell'Abruzzo e Molise "G. Caporale" | Lorusso A, Marcacci M, Di Domenico M, Ancora M, Curini V, Mangone I, Rinaldi A, Scialabba S, Di Pasquale A, Cammà C, Puglia I, Calistri P, Savini G |
| EPI_ISL_918267 | SIESP CHIETI - DRIVE IN CHIETI | Istituto Zooprofilattico Sperimentale dell'Abruzzo e Molise "G. Caporale" | Lorusso A, Marcacci M, Di Domenico M, Ancora M, Curini V, Mangone I, Rinaldi A, Scialabba S, Di Pasquale A, Cammà C, Puglia I, Calistri P, Savini G |
| EPI_ISL_918268 | SIESP CHIETI - DRIVE IN ORTONA | Istituto Zooprofilattico Sperimentale dell'Abruzzo e Molise "G. Caporale" | Lorusso A, Marcacci M, Di Domenico M, Ancora M, Curini V, Mangone I, Rinaldi A, Scialabba S, Di Pasquale A, Cammà C, Puglia I, Calistri P, Savini G |
| EPI_ISL_918271 | SIESP DIPARTIMENTO DI PREVENZIONE CHIETI | Istituto Zooprofilattico Sperimentale dell'Abruzzo e Molise "G. Caporale" | Lorusso A, Marcacci M, Di Domenico M, Ancora M, Curini V, Mangone I, Rinaldi A, Scialabba S, Di Pasquale A, Cammà C, Puglia I, Calistri P, Savini G |
| EPI_ISL_918277, EPI_ISL_918278, EPI_ISL_918279 | Hospital Universitari Vall d'Hebron - Vall d'Hebron Institut de Rercerca | Hospital Universitari Vall d'Hebron | Cristina Andrés, Maria Piñana, Josep F Abril, Damir Garcia-Cehic, Ariadna Rando, Juliana Esperalba, Maria Gema Codina, Carla Castillo, Maria Carmen Martin, Tomàs Pumarola, Josep Quer, Andrés Antón |
| EPI_ISL_918489 | CUSL/UCLouvain COVID testing federal platform | UCLouvain/IREC/MBLG | Jean Ruelle, Lysa Pinsmaye, Benoit Kabamba Mukadi |
| EPI_ISL_918507, EPI_ISL_918508, EPI_ISL_918509, EPI_ISL_918510, EPI_ISL_918535 | LACEN - Laboratório Central de Saúde Pública do Amazonas | Evandro Chagas Institute | Santos, M.C.; Silva, A.M.; Junior, W.D.C.; Barbagelata, L.S.; Ferreira, J.A.; Sousa, E.M.A.; da Silva, P.S.; Pinheiro, K.C.; L.C.; Sousa Junior, E.C. |
| EPI_ISL_918537, EPI_ISL_918538 | LACEN - Laboratório Central de Saúde Pública do Ceara | Evandro Chagas Institute | Santos, M.C.; Silva, A.M.; Junior, W.D.C.; Barbagelata, L.S.; Ferreira, J.A.; Sousa, E.M.A.; da Silva, P.S.; Pinheiro, K.C.; L.C.; Sousa Junior, E.C. |
| EPI_ISL_918545, EPI_ISL_918546, | LACEN - Laboratório Central de Saúde Pública do Para | Evandro Chagas Institute | Santos, M.C.; Silva, A.M.; Junior, W.D.C.; Barbagelata, L.S.; Ferreira, J.A.; Sousa, E.M.A.; da Silva, P.S.; Pinheiro, K.C.; L.C.; Sousa Junior, E.C. |

| | | | |
|---|---|---|---|
| EPI_ISL_918547, EPI_ISL_918548, EPI_ISL_918549 | | | |
| EPI_ISL_918697, EPI_ISL_918698, EPI_ISL_918699, EPI_ISL_918700, EPI_ISL_918702, EPI_ISL_918703, EPI_ISL_918704, EPI_ISL_918705, EPI_ISL_918706, EPI_ISL_918707, EPI_ISL_918708, EPI_ISL_918709, EPI_ISL_918710, EPI_ISL_918712, EPI_ISL_918713, EPI_ISL_918714, EPI_ISL_918715, EPI_ISL_918716, EPI_ISL_918717, EPI_ISL_918718, EPI_ISL_918719, EPI_ISL_918720, EPI_ISL_918722, EPI_ISL_918723, EPI_ISL_918724, EPI_ISL_918725, EPI_ISL_918726, EPI_ISL_918727, EPI_ISL_918728, EPI_ISL_918729, EPI_ISL_918730, EPI_ISL_918731, EPI_ISL_918732, EPI_ISL_918735, EPI_ISL_918736, EPI_ISL_918737, EPI_ISL_918740, EPI_ISL_918904, EPI_ISL_918905 | | | |
| see above | University of Birmingham | COVID-19 Genomics UK (COG-UK) Consortium | Institute of Microbiology, University of Birmingham: Claire McMurray, Joanne Stockton, Samuel Nicholls, Radoslaw Poplawski, Will Rowe, Josh Quick, Nicholas Loman. University of Birmingham Testing Laboratory: Celina M Whalley, Andrew Bosworth, Charlotte Poxon, Kasun Wanigasooriya, Oliver Pickles, Mike Kidd, Alex Richter, Andrew D Beggs PHE Heartlands Lab: Husam Osman, Andrew Bosworth. Queen Elizabeth Hospital: Anna Casey |
| EPI_ISL_919071, EPI_ISL_919073, EPI_ISL_919077, EPI_ISL_919081, EPI_ISL_919104 | Department of Pathology, University of Cambridge | COVID-19 Genomics UK (COG-UK) Consortium | Aminu S. Jahun, Yasmin Chaudhry, Iliana Georgana, Myra Hosmillo, Rhys Izu, Martin D. Curran, Surendra Parmar, Ian Goodfellow |
| EPI_ISL_919196 | West of Scotland Specialist Virology Centre, NHSGGC / MRC-University of Glasgow Centre for Virus Research | COVID-19 Genomics UK (COG-UK) Consortium | Ana da Silva Filipe, Natasha Johnson, Kathy Smollett, Daniel Mair, Stephen Carmichael, Alice Broos, Lily Tong, Jenna Nichols, Kyriaki Nomikou; Sarah McDonald; Richard Orton, Joseph Hughes, Sreenu Vattipally, David L Robertson; Alasdair MacLean, Rory Gunson; Sharif Shaaban, Matthew Holden; Rachel Blacow, Guy Mollett, Kathy Li, James Shepherd, Antonia Ho, Emma Thomson |
| EPI_ISL_919460, EPI_ISL_919462, EPI_ISL_919463, EPI_ISL_919465, EPI_ISL_919594, EPI_ISL_919595 | Liverpool Clinical Laboratories | COVID-19 Genomics UK (COG-UK) Consortium | Sam Haldenby, Anita Lucaci, Steve Paterson, Julian Hiscox, Alistair Darby, M Almsaud, A Alrezaihi, Muhannad Alruwaili, Stuart D Armstrong, Jones Benjamin, Eleanor G Bentley, Anu Chawla, Jordan J Clark, Angela Cowell, Richard Eccles, Isabel García-Dorival, Matthew Gemmell, Alessandro Gerada, PKF Gilmore, Richard Gregory, Ximeng Han, Catherine Hartley, Margaret Hughes, Miren Iturriza-Gomara, James Johnson, L Luu, Jenifer Manson, Charlotte Nelson, Elaine O'Toole, Cassie Olateju, Rebekah Penrice-Randal , Lucille Rainbow, N.P Randle, Trevor Ian Robinson, Parul Sharma, Ghada T Shawli, James P Stewart, Neil Swainston, Ecaterina Vamos, Joanne Watts, Mark Whitehead |
| EPI_ISL_919779, EPI_ISL_919780, EPI_ISL_919797, EPI_ISL_919803, EPI_ISL_919804, EPI_ISL_919826, EPI_ISL_919827, EPI_ISL_919828, EPI_ISL_919829, EPI_ISL_919830, EPI_ISL_919831, EPI_ISL_919832, EPI_ISL_919833, EPI_ISL_919834, EPI_ISL_919835, EPI_ISL_919836, EPI_ISL_919837, EPI_ISL_919838, EPI_ISL_919839, EPI_ISL_919840, EPI_ISL_919841, EPI_ISL_919845, EPI_ISL_919846, EPI_ISL_919847, EPI_ISL_919848, EPI_ISL_919849, EPI_ISL_919850, EPI_ISL_919851 | | | |
| see above | Barts Health NHS Trust | COVID-19 Genomics UK (COG-UK) Consortium | CUTINO-MOGUEL, Maria-Teresa; HARRINGTON, David; OWOYEMI, Dola; KULASEGARAN-SHYLINI, Raghavendran; BROAD, Claire; KELE, Beatrix |
| EPI_ISL_919982, EPI_ISL_919983, EPI_ISL_919984, EPI_ISL_919986, EPI_ISL_919988, EPI_ISL_919989, EPI_ISL_919990, EPI_ISL_919991, EPI_ISL_919992, EPI_ISL_919993, EPI_ISL_919994, EPI_ISL_919995, EPI_ISL_919996, EPI_ISL_919997, EPI_ISL_919998, EPI_ISL_919999, EPI_ISL_920041, EPI_ISL_920048 | | | |
| see above | University College London, Great Ormond Street Hospital for Children NHS Foundation Trust, Imperial College Healthcare NHS Trust | COVID-19 Genomics UK (COG-UK) Consortium | Sergi Castellano, Rachel Williams, Mark Kristiansen, Paola Resende Silva, Sunando Roy, Tony Brooks, Helena Tutill, Paola Niola, Patricia Dyal, Charlotte Williams, Leysa Forrest, Yasmin Panchbhaya, Jacqueline Findlay, Samuel Weeks, Julianne Brown, Kathryn Harris, Paul Randell, James Price, Alison Holmes, Judith Breuer |
| EPI_ISL_920179, EPI_ISL_920288, EPI_ISL_920297, EPI_ISL_920311, EPI_ISL_920689 | University College London Hospital | COVID-19 Genomics UK (COG-UK) Consortium | Judith Heaney, Matthew Byott, Catherine Houlihan, Dan Frampton, Stuart Kirk, Moira Spyer and Eleni Nastouli |
| EPI_ISL_920839, EPI_ISL_920850, EPI_ISL_920852, EPI_ISL_920853, EPI_ISL_920856, EPI_ISL_920857, EPI_ISL_920858, EPI_ISL_920859 | University College London, Great Ormond Street Hospital for Children NHS Foundation Trust, Imperial College Healthcare NHS Trust | COVID-19 Genomics UK (COG-UK) Consortium | Sergi Castellano, Rachel Williams, Mark Kristiansen, Paola Resende Silva, Sunando Roy, Tony Brooks, Helena Tutill, Paola Niola, Patricia Dyal, Charlotte Williams, Leysa Forrest, Yasmin Panchbhaya, Jacqueline Findlay, Samuel Weeks, Julianne Brown, Kathryn Harris, Paul Randell, James Price, Alison Holmes, Judith Breuer |
| EPI_ISL_920980, EPI_ISL_921019, EPI_ISL_921026, EPI_ISL_921126, EPI_ISL_921127, EPI_ISL_921129, EPI_ISL_921130 | Regional Virus Laboratory, Belfast Health and Social Care Trust | COVID-19 Genomics UK (COG-UK) Consortium | Conall McCaughey, James McKenna, Tanya Curran, Susan Feeney, Alison Watt, Ciara Cox, Mairead Connor, Zoltan Molnar, David Simpson, Derek Fairley |
| EPI_ISL_921300, EPI_ISL_921302, EPI_ISL_921307, EPI_ISL_921308, EPI_ISL_921309, EPI_ISL_921310, EPI_ISL_921311, EPI_ISL_921312, EPI_ISL_921315, EPI_ISL_921319, EPI_ISL_921320, EPI_ISL_921452, EPI_ISL_921453, EPI_ISL_921454, EPI_ISL_921455, EPI_ISL_921457, EPI_ISL_921458, EPI_ISL_921459, EPI_ISL_921462, EPI_ISL_921464, EPI_ISL_921465, EPI_ISL_921468, EPI_ISL_921469, EPI_ISL_921470, EPI_ISL_921471, EPI_ISL_921472, EPI_ISL_921473, EPI_ISL_921474, EPI_ISL_921475, EPI_ISL_921476, EPI_ISL_921477, EPI_ISL_921478, EPI_ISL_921479, EPI_ISL_921480, EPI_ISL_921481, EPI_ISL_921482, EPI_ISL_921483, EPI_ISL_921484, EPI_ISL_921494 | | | |
| see above | Northumbria University / South Tees Hospitals NHS Foundation Trust / North Cumbria Integrated Care NHS Foundation Trust / North Tees and Hartlepool NHS Foundation Trust / Newcastle Hospitals NHS Foundation Trust | COVID-19 Genomics UK (COG-UK) Consortium | Darren L Smith,Andrew Nelson,Matthew Bashton,Greg R Young,Joshua Loh,John Allan,Mohammad A Tariq,Giles S Holt,Gary Black,Wen C Yew,Lynn Dover,Paul Baker,Steve Liggett,Sarah Essex,Jane Greenaway,Debra Padgett,Clive Graham,Garren Scott,Edward Barton,Emma Swindells,Brendan Payne,Jennifer Collins,Yusri Taha,Gary Eltringham |
| EPI_ISL_922041, EPI_ISL_922042, EPI_ISL_922043, EPI_ISL_922044, EPI_ISL_922045, EPI_ISL_922046, EPI_ISL_922047, EPI_ISL_922048, EPI_ISL_922049, EPI_ISL_922050, EPI_ISL_922051, EPI_ISL_922052, EPI_ISL_922053, EPI_ISL_922054, EPI_ISL_922055, EPI_ISL_922056, EPI_ISL_922057, EPI_ISL_922058, EPI_ISL_922059, EPI_ISL_922060, EPI_ISL_922061, EPI_ISL_922062, EPI_ISL_922063, EPI_ISL_922064, EPI_ISL_922065, EPI_ISL_922066, EPI_ISL_922067, EPI_ISL_922068, EPI_ISL_922069, EPI_ISL_922070, EPI_ISL_922073 | | | |
| see above | Lincolnshire Hospitals and DeepSeq Nottingham | COVID-19 Genomics UK (COG-UK) Consortium | Nichola Duckworth, Tim Sloan, Sarah Walsh, Jonathan Ball, Patrick McClure, Joeseph Chappell, Nadine Holmes, Matthew Carlisle, Christopher Moore, Fei Sang, Johnny Debebe, Victoria Wright, Matthew Loose |
| EPI_ISL_922174, EPI_ISL_922177, EPI_ISL_922182, EPI_ISL_922184, EPI_ISL_922185, EPI_ISL_922189, EPI_ISL_922190, EPI_ISL_922191, EPI_ISL_922194, EPI_ISL_922197, EPI_ISL_922198, EPI_ISL_922199, EPI_ISL_922200, EPI_ISL_922201, EPI_ISL_922203, EPI_ISL_922208, EPI_ISL_922209, EPI_ISL_922244, EPI_ISL_922246, EPI_ISL_922252, EPI_ISL_922254, EPI_ISL_922258, EPI_ISL_922284, EPI_ISL_922295, EPI_ISL_922323, EPI_ISL_922324, EPI_ISL_922325, EPI_ISL_922326, EPI_ISL_922328, EPI_ISL_922329, EPI_ISL_922330, EPI_ISL_922331, EPI_ISL_922332, EPI_ISL_922333, EPI_ISL_922334, EPI_ISL_922335, EPI_ISL_922338, EPI_ISL_922342, EPI_ISL_922343, EPI_ISL_922345, EPI_ISL_922346, EPI_ISL_922347, EPI_ISL_922348 | | | |
| see above | Oxford Viromics, NDM, University of Oxford; Oxford University Hospitals; Basingstoke and North Hampshire Hospital | COVID-19 Genomics UK (COG-UK) Consortium | Tanya Golubchik, David Bonsall, George Macintyre, Amy Trebes, Mariateresa de Cesare, Catrin Moore, Alex Mobbs, Anita Justice, Robert Shaw, Monique Andersson, Timothy Peto, Emma Wise, Nathan Moore, Jessica Lynch, Nick Cortes, Matilde Mori, Stephen Kidd, David Buck, John Todd, Christophe Fraser |
| EPI_ISL_923256, EPI_ISL_923257, EPI_ISL_923260, EPI_ISL_923261, EPI_ISL_923263, EPI_ISL_923266, EPI_ISL_923275, EPI_ISL_923296, EPI_ISL_923300, EPI_ISL_923372, EPI_ISL_923374, EPI_ISL_923377, EPI_ISL_923379, EPI_ISL_923380, EPI_ISL_923381, EPI_ISL_923382, EPI_ISL_923383, EPI_ISL_923384, EPI_ISL_923385, EPI_ISL_923386, EPI_ISL_923387, EPI_ISL_923388, EPI_ISL_923430, EPI_ISL_923443, EPI_ISL_923663 | | | |
| see above | Centre for Enzyme Innovation, University of Portsmouth / Translational Research Laboratory, Portsmouth Hospitals NHS Trust | COVID-19 Genomics UK (COG-UK) Consortium | Angela Beckett,Salman Goudarzi,Christopher Fearn,Kate Cook,Katie Loveson,Sharon Glaysher,Scott Elliott,Samuel Robson |
| EPI_ISL_924285, EPI_ISL_924369, EPI_ISL_924383 | Virology Department, Sheffield Teaching Hospitals NHS Foundation Trust/Department of Infection, Immunity and Cardiovascular Disease, The Medical School, University of Sheffield | COVID-19 Genomics UK (COG-UK) Consortium | Thushan de Silva, Matthew Parker, Nikki Smith, Adri Angyal, Rebecca Brown, Luke Green, Rachel Tucker, Paul Parsons, Danielle Groves, Katie Johnson, Laura Carrilero, Alex Keeley, Dave Partridge, Matthew Wyles, Benjamin Lindsey, Mehmet Yavuz, Mohammad Raza, Cariad Evans |
| EPI_ISL_924623, EPI_ISL_924625, EPI_ISL_924626, EPI_ISL_924627, EPI_ISL_924628, EPI_ISL_924629, EPI_ISL_924631, EPI_ISL_924632, EPI_ISL_924633, EPI_ISL_924635, EPI_ISL_924636, EPI_ISL_924637, EPI_ISL_924638, EPI_ISL_924639, EPI_ISL_924640, EPI_ISL_924641, EPI_ISL_924642, EPI_ISL_924644, EPI_ISL_924645, EPI_ISL_924646, EPI_ISL_924647, EPI_ISL_924648, EPI_ISL_924649, EPI_ISL_924650, EPI_ISL_924651, EPI_ISL_924723, EPI_ISL_924724, EPI_ISL_924725, EPI_ISL_924726, EPI_ISL_924727, EPI_ISL_924728, EPI_ISL_924730, EPI_ISL_924732, EPI_ISL_924733, EPI_ISL_924734, EPI_ISL_924735, EPI_ISL_924736, EPI_ISL_924737, EPI_ISL_924739, EPI_ISL_924740, EPI_ISL_924741, EPI_ISL_924742, EPI_ISL_924743, EPI_ISL_924744, EPI_ISL_924745, EPI_ISL_924747, EPI_ISL_924748, EPI_ISL_924750, EPI_ISL_924752, EPI_ISL_924753, EPI_ISL_924754, EPI_ISL_924755, EPI_ISL_924756, EPI_ISL_924757, EPI_ISL_924758, EPI_ISL_924759, EPI_ISL_924760, EPI_ISL_924761, EPI_ISL_924762, EPI_ISL_924763, EPI_ISL_924765, EPI_ISL_924766, EPI_ISL_924768, EPI_ISL_924769, EPI_ISL_924771, EPI_ISL_924772, EPI_ISL_924773, EPI_ISL_924775, EPI_ISL_924776, EPI_ISL_924777, EPI_ISL_924778, EPI_ISL_924779, EPI_ISL_924780, EPI_ISL_924781, EPI_ISL_924782, EPI_ISL_924897 | | | |
| see above | Bioinformatics and Biostatistics Lab, Advanced Sequencing Facility | COVID-19 Genomics UK (COG-UK) Consortium | Aengus Stewart,Jerome Nicod,Chelsea Sawyer,Laura Cubitt,Harshil Patel,Margaret Crawford |
| EPI_ISL_925065, EPI_ISL_925066, EPI_ISL_925067, EPI_ISL_925068, EPI_ISL_925069, EPI_ISL_925070, EPI_ISL_925071, EPI_ISL_925072, EPI_ISL_925073, EPI_ISL_925074 | TXDSHS | TXDSHS | Bonnie Oh, Anita Pokharel, James Daniel Bonser, Myong Koag, Chung Wang, Rachel Lee, Grace Kubin, Rashmi Tuladhar, Mayela Pedrueza, Maliha Rahman, Jenny Zhang |
| EPI_ISL_925290, EPI_ISL_925304 | Wyoming Public Health Laboratory | Wyoming Public Health Laboratory | Noah Hull, Taylor Fearing, Lynette Gumbleton, Channing Weber, Ashley Norberg, Bailey Bowcutt, and Wanda Manley |

| | | | |
|---|---|---|---|
| EPI_ISL_925509, EPI_ISL_925510, EPI_ISL_925511, EPI_ISL_925512, EPI_ISL_925522, EPI_ISL_925523, EPI_ISL_925525, EPI_ISL_925526, EPI_ISL_925534, EPI_ISL_925824, EPI_ISL_925825, EPI_ISL_925826, EPI_ISL_925827, EPI_ISL_925828, EPI_ISL_925829, EPI_ISL_925830, EPI_ISL_925831, EPI_ISL_925833, EPI_ISL_925834, EPI_ISL_925835, EPI_ISL_925836, EPI_ISL_925837, EPI_ISL_925838, EPI_ISL_925839, EPI_ISL_925840, EPI_ISL_925841, EPI_ISL_925842, EPI_ISL_925843, EPI_ISL_925844, EPI_ISL_925845 | | | |
| see above | Arizona State Public Health Laboratory | Arizona State Public Health Laboratory | Trung Huynh, Jessica Escobar, Katherine Fullerton, Nobuko Fukushima, Stacy White, Linda Getsinger, Victor Waddell |
| EPI_ISL_925887, EPI_ISL_925908, EPI_ISL_925912 | Nucleic Acid Testing, National Reference Laboratory | GIGA Medical Genomics | Yvan Butera, Keith Durkin, Maria Artesi, Bouchra Boujemla, Robert Rutayisire, Patrick Tuyisenge, Esperence Umumararungu, Sébastien Bontems, Marie-Pierre Hayette, Nathalie Renotte, Swaibu Gatare, Jacob Souopgui, Sabin Nsanzimana, Vincent Bours, Léon Mutesa |
| EPI_ISL_930636, EPI_ISL_930649, EPI_ISL_930650, EPI_ISL_930661 | Arizona State Public Health Laboratory | Arizona State Public Health Laboratory | Trung Huynh, Jessica Escobar, Katherine Fullerton, Nobuko Fukushima, Stacy White, Linda Getsinger, Victor Waddell |
| EPI_ISL_931489, EPI_ISL_931496, EPI_ISL_931497, EPI_ISL_931498, EPI_ISL_931499, EPI_ISL_931500 | Maryland Public Health Laboratory (MD PHL) | Maryland Public Health Laboratory (MD PHL) | Maryland Department of Health Laboratories Administration |
| EPI_ISL_931818 | Lighthouse Lab in Alderley Park | Wellcome Sanger Institute for the COVID-19 Genomics UK (COG-UK) Consortium | Jacquelyn Wynn, Mairead Hyland, The Lighthouse Lab in Alderley Park and Alex Alderton, Roberto Amato, Sonia Goncalves, Ewan Harrison, David K. Jackson, Ian Johnston, Dominic Kwiatkowski, Cordelia Langford, John Sillitoe on behalf of the Wellcome Sanger Institute COVID-19 Surveillance Team |
| EPI_ISL_933661, EPI_ISL_933662, EPI_ISL_933680, EPI_ISL_933681, EPI_ISL_933682, EPI_ISL_933683, EPI_ISL_933684, EPI_ISL_933685, EPI_ISL_933686, EPI_ISL_933687, EPI_ISL_933688, EPI_ISL_933689, EPI_ISL_933704 | | | |
| see above | Instituto de Diagnostico y Referencia Epidemiologicos INDRE_RNLSP | Instituto de Diagnostico y Referencia Epidemiologicos (INDRE) | Claudia Wong-Arambula, Abril Rodriguez-Maldonado, Fabiola Garces-Ayala, Adnan Araiza-Rodriguez, David Fragoso-Fonseca, Sergio Rangel-Guerrero, Mayra Jimenez-Morales, Nancy Munoz-Hernandez, Natividad Cruz-Ortiz, Tatiana Nunez-Garcia, Gisela Barrera-Badillo, Lucia Hernandez-Rivas, Irma Lopez-Martinez, Ernesto Ramirez-Gonzalez. |
| EPI_ISL_934182, EPI_ISL_934183, EPI_ISL_934184, EPI_ISL_934185, EPI_ISL_934186, EPI_ISL_934195, EPI_ISL_934196, EPI_ISL_934197, EPI_ISL_934198, EPI_ISL_934199, EPI_ISL_934200, EPI_ISL_934201, EPI_ISL_934229, EPI_ISL_934230, EPI_ISL_934231, EPI_ISL_934232, EPI_ISL_934233 | | | |
| see above | Vilnius university hospital Santaros Klinikos, Center of Laboratory Medicine | Vilnius university hospital Santaros Klinikos, Center of Laboratory Medicine | Ingrida Olendraite, Daniel Naumovas, Rimvydas Norvilas, Dovile Ezerskyte, Justinas Slikas, Gytis Dudas |
| EPI_ISL_934343, EPI_ISL_934344, EPI_ISL_934355, EPI_ISL_934356, EPI_ISL_934357, EPI_ISL_934360 | Klinisk mikrobiologi | The Public Health Agency of Sweden | Anna-Malin Linde, Maria Lind Karlberg, Carlo Berg, Oskar Karlsson Lindsjo, Sofia Stamouli, Reza Advani, Mattias Haukland, Petra Holmstrom, Noura Walai, Petra Edquist, Mia Brytting, Anna Risberg, Karin Tegmark-Wisell |
| EPI_ISL_934366, EPI_ISL_934367, EPI_ISL_934368, EPI_ISL_934369 | Synlab Medilab, Mikrobiologi | The Public Health Agency of Sweden | Anna-Malin Linde, Maria Lind Karlberg, Carlo Berg, Oskar Karlsson Lindsjo, Sofia Stamouli, Reza Advani, Mattias Haukland, Petra Holmstrom, Noura Walai, Petra Edquist, Mia Brytting, Anna Risberg, Karin Tegmark-Wisell |
| EPI_ISL_934431, EPI_ISL_934432, EPI_ISL_934433 | ILV Kärnten | Bergthaler laboratory, CeMM Research Center for Molecular Medicine of the Austrian Academy of Sciences | Lukas Endler, Anna Schedl, Thomas Penz, Benedikt Agerer, Maelle Le Moing, Michael Schuster, Bekir Erguner, Jan Laine, Martin Senekowitsch, Christoph Bock, Andreas Bergthaler |
| EPI_ISL_934456, EPI_ISL_934460, EPI_ISL_934486, EPI_ISL_934510, EPI_ISL_934511, EPI_ISL_934512, EPI_ISL_934513, EPI_ISL_934514, EPI_ISL_934515, EPI_ISL_934516, EPI_ISL_934517, EPI_ISL_934518, EPI_ISL_934519, EPI_ISL_934520, EPI_ISL_934521, EPI_ISL_934522, EPI_ISL_934523, EPI_ISL_934524, EPI_ISL_934525, EPI_ISL_934526, EPI_ISL_934527, EPI_ISL_934528, EPI_ISL_934529, EPI_ISL_934530, EPI_ISL_934531, EPI_ISL_934532, EPI_ISL_934533, EPI_ISL_934534, EPI_ISL_934535, EPI_ISL_934536, EPI_ISL_934537, EPI_ISL_934538 | | | |
| see above | Austrian Agency for Health and Food Safety (AGES) | Bergthaler laboratory, CeMM Research Center for Molecular Medicine of the Austrian Academy of Sciences | Lukas Endler, Anna Schedl, Thomas Penz, Benedikt Agerer, Maelle Le Moing, Michael Schuster, Bekir Erguner, Jan Laine, Martin Senekowitsch, Christoph Bock, Andreas Bergthaler |
| EPI_ISL_934634, EPI_ISL_934641 | Department of Microbiology, University Innsbruck | Bergthaler laboratory, CeMM Research Center for Molecular Medicine of the Austrian Academy of Sciences | Lukas Endler, Anna Schedl, Thomas Penz, Benedikt Agerer, Maelle Le Moing, Michael Schuster, Bekir Erguner, Jan Laine, Martin Senekowitsch, Christoph Bock, Andreas Bergthaler |
| EPI_ISL_935026 | Biolab Diagnostic Laboratories | Princess Haya Biotechnology Center, Jordan University of Science and Technology | Saied Jaradat, Hazem Haddad, Areej Alquran, Maha Karam, Shereen Issa,Suha Hasan, Amid Abdelnour, Issa Abu-Dayyeh, Mustafa Ababneh, Moh"D Al-Zghoul, Mohammad Alboom |
| EPI_ISL_935038, EPI_ISL_935039, EPI_ISL_935041 | Biolab Diagnostic Laboratories | Princess Haya Biotechnology Center, Jordan University of Science and Technology | Saied Jaradat, Hazem Haddad, Areej Alquran, Maha Karam, Shereen Issa,Suha Hasan, Amid Abdelnour, Issa Abu-Dayyeh |
| EPI_ISL_935043 | Biolab Diagnostic Laboratories | Princess Haya Biotechnology Center/ Jordan University of Science & Technology | Saied Jaradat, Hazem Haddad, Areej Alquran, Maha Karam, Shereen Issa,Suha Hasan, Amid Abdelnour, Issa Abu-Dayyeh |
| EPI_ISL_935046 | Biolab Diagnostic Laboratories | Princess Haya Biotechnology Center/ Jordan University of Science & Technology | Saied Jaradat, Hazem Haddad, Areej Alquran, Maha Karam, Shereen Issa,Suha Hasan, Amid Abdelnour, Issa Abu-Dayyeh, Mustafa Ababneh, Moh"D Al-Zghoul, Mohammad Alboom |
| EPI_ISL_935157 | SIESP CHIETI-DRIVE IN ORTONA | Istituto Zooprofilattico Sperimentale dell'Abruzzo e Molise "G. Caporale" | Lorusso A, Marcacci M, Di Domenico M, Ancora M, Curini V, Mangone I, Rinaldi A, Scialabba S, Di Pasquale A, Cammà C, Puglia I, Calistri P, Savini G |
| EPI_ISL_935158 | SIESP CHIETI | Istituto Zooprofilattico Sperimentale dell'Abruzzo e Molise "G. Caporale" | Lorusso A, Marcacci M, Di Domenico M, Ancora M, Curini V, Mangone I, Rinaldi A, Scialabba S, Di Pasquale A, Cammà C, Puglia I, Calistri P, Savini G |
| EPI_ISL_935226 | KU Leuven, Rega Institute, Clinical and Epidemiological Virology | KU Leuven, Rega Institute, Clinical and Epidemiological Virology | Tony Wawina-Bokalanga, Bert Vanmechelen, Joan Marti-Carerras, Piet Maes |
| EPI_ISL_935362, EPI_ISL_935365, EPI_ISL_935366 | Florida Bureau of Public Health Laboratories | Florida Bureau of Public Health Laboratories | Sarah Schmedes, Jason Blanton |
| EPI_ISL_935974, EPI_ISL_935976, EPI_ISL_935977, EPI_ISL_935978, EPI_ISL_935979, EPI_ISL_935980, EPI_ISL_935981 | GLENS FALLS HOSPITAL LABORATORY | Wadsworth Center, New York State Department of Health | Kirsten St. George, Daryl M. Lamson, Alexis Russel, Matthew Shudt, Melissa A Leisner, Jonathan Plitnick, Navjot Singh, John Kelly, Erasmus Schneider, Erica Lasek-Nesselquist |
| EPI_ISL_935993, EPI_ISL_935994, EPI_ISL_935995, EPI_ISL_935996, EPI_ISL_935997, EPI_ISL_935998, EPI_ISL_935999, EPI_ISL_936000, EPI_ISL_936001 | SUNY UPSTATE MEDICAL UNIVERSITY | Wadsworth Center, New York State Department of Health | Kirsten St. George, Daryl M. Lamson, Alexis Russel, Matthew Shudt, Melissa A Leisner, Jonathan Plitnick, Navjot Singh, John Kelly, Erasmus Schneider, Erica Lasek-Nesselquist |
| EPI_ISL_936023, EPI_ISL_936024, EPI_ISL_936029, EPI_ISL_936030, EPI_ISL_936032, EPI_ISL_936033 | THE MARY IMOGENE BASSETT HOSPITAL | Wadsworth Center, New York State Department of Health | Kirsten St. George, Daryl M. Lamson, Alexis Russel, Matthew Shudt, Melissa A Leisner, Jonathan Plitnick, Navjot Singh, John Kelly, Erasmus Schneider, Erica Lasek-Nesselquist |
| EPI_ISL_936046, EPI_ISL_936047, EPI_ISL_936049, EPI_ISL_936052, EPI_ISL_936053, EPI_ISL_936054, EPI_ISL_936056 | ADIRONDACK MEDICAL CENTER | Wadsworth Center, New York State Department of Health | Kirsten St. George, Daryl M. Lamson, Alexis Russel, Matthew Shudt, Melissa A Leisner, Jonathan Plitnick, Navjot Singh, John Kelly, Erasmus Schneider, Erica Lasek-Nesselquist |
| EPI_ISL_936062, EPI_ISL_936098, EPI_ISL_936099, EPI_ISL_936100, EPI_ISL_936102 | KALEIDA CENTER FOR LABORATORY MEDICINE | Wadsworth Center, New York State Department of Health | Kirsten St. George, Daryl M. Lamson, Alexis Russel, Matthew Shudt, Melissa A Leisner, Jonathan Plitnick, Navjot Singh, John Kelly, Erasmus Schneider, Erica Lasek-Nesselquist |
| EPI_ISL_936139 | THE MARY IMOGENE BASSETT HOSPITAL | Wadsworth Center, New York State Department of Health | Kirsten St. George, Daryl M. Lamson, Alexis Russel, Matthew Shudt, Melissa A Leisner, Jonathan Plitnick, Navjot Singh, John Kelly, Erasmus Schneider, Erica Lasek-Nesselquist |
| EPI_ISL_936143 | SUNY UPSTATE MEDICAL UNIVERSITY | Wadsworth Center, New York State Department of Health | Kirsten St. George, Daryl M. Lamson, Alexis Russel, Matthew Shudt, Melissa A Leisner, Jonathan Plitnick, Navjot Singh, John Kelly, Erasmus Schneider, Erica Lasek-Nesselquist |
| EPI_ISL_936145 | New York Presbyterian Hospital | Wadsworth Center, New York State Department of Health | Kirsten St. George, Daryl M. Lamson, Alexis Russel, Matthew Shudt, Melissa A Leisner, Jonathan Plitnick, Navjot Singh, John Kelly, Erasmus Schneider, |

| | | | |
|---|---|---|---|
| | | | Erica Lasek-Nesselquist |
| EPI_ISL_936212, EPI_ISL_936213 | Wadsworth Center, New York State Department of Health | Wadsworth Center, New York State Department of Health | Kirsten St. George, Daryl M. Lamson, Alexis Russel, Matthew Shudt, Melissa A Leisner, Jonathan Plitnick, Navjot Singh, John Kelly, Erasmus Schneider, Erica Lasek-Nesselquist |
| EPI_ISL_936387, EPI_ISL_936389, EPI_ISL_936393, EPI_ISL_936394, EPI_ISL_936399, EPI_ISL_936403, EPI_ISL_936404, EPI_ISL_936409, EPI_ISL_936411, EPI_ISL_936412, EPI_ISL_936413, EPI_ISL_936416, EPI_ISL_936417, EPI_ISL_936418, EPI_ISL_936422, EPI_ISL_936425, EPI_ISL_936427, EPI_ISL_936428, EPI_ISL_936429, EPI_ISL_936430, EPI_ISL_936434, EPI_ISL_936438, EPI_ISL_936440, EPI_ISL_936441, EPI_ISL_936443, EPI_ISL_936444, EPI_ISL_936446, EPI_ISL_936448, EPI_ISL_936452, EPI_ISL_936454, EPI_ISL_936455, EPI_ISL_936457, EPI_ISL_936461, EPI_ISL_936462 | | | |
| see above | TGen North | TGen North | Jolene Bowers, Megan Folkerts, Chris French, Hayley Yaglom, Ashlyn Pfeiffer, Darrin Lemmer, Dave Engelthaler, The Arizona COVID Genomics Union (ACGU) |
| EPI_ISL_936469 | DPH, Massachusetts State Public Health Lab | DPH, Massachusetts State Public Health Lab | Lang,A.S., Fink,T., Gallagher,G.R., Smole,S.C. |
| EPI_ISL_936888, EPI_ISL_936890, EPI_ISL_936891, EPI_ISL_936892, EPI_ISL_936894, EPI_ISL_936905, EPI_ISL_936906, EPI_ISL_936907, EPI_ISL_936908, EPI_ISL_936909, EPI_ISL_936910, EPI_ISL_936911, EPI_ISL_936912, EPI_ISL_936913, EPI_ISL_936914, EPI_ISL_936915, EPI_ISL_936916, EPI_ISL_936917, EPI_ISL_936918, EPI_ISL_936923, EPI_ISL_936924, EPI_ISL_936925, EPI_ISL_936926, EPI_ISL_936927, EPI_ISL_936928, EPI_ISL_936929, EPI_ISL_936930, EPI_ISL_936931, EPI_ISL_936932, EPI_ISL_936933, EPI_ISL_936934, EPI_ISL_936935, EPI_ISL_936936, EPI_ISL_936937, EPI_ISL_936938, EPI_ISL_936939, EPI_ISL_936940, EPI_ISL_936941, EPI_ISL_936942, EPI_ISL_936943, EPI_ISL_936944, EPI_ISL_936945, EPI_ISL_936946, EPI_ISL_936947, EPI_ISL_936948, EPI_ISL_936949, EPI_ISL_936950, EPI_ISL_936951, EPI_ISL_936952, EPI_ISL_936953, EPI_ISL_936954, EPI_ISL_936955, EPI_ISL_936956, EPI_ISL_936957, EPI_ISL_936958, EPI_ISL_936959, EPI_ISL_936960, EPI_ISL_936961, EPI_ISL_936962, EPI_ISL_936963, EPI_ISL_936964, EPI_ISL_936965, EPI_ISL_936966 | | | |
| see above | Northwestern Memorial Hospital | Ozer Lab | Ramon Lorenzo-Redondo, Lacy M. Simons, Chad J. Achenbach, Lawrence J. Jennings, Michael G. Ison, Judd F. Hultquist, Egon A. Ozer |
| EPI_ISL_937259 | NYC HH Elmhurst Hospital Medical Center | New York City Public Health Laboratory | Jade Wang, et al. |
| EPI_ISL_940723, EPI_ISL_940724, EPI_ISL_940725, EPI_ISL_940726, EPI_ISL_940734 | City of Milwaukee Health Department Laboratory | City of Milwaukee Health Department Laboratory | Sanjib Bhattacharyya |
| EPI_ISL_940852, EPI_ISL_940856, EPI_ISL_940862, EPI_ISL_940865, EPI_ISL_940866, EPI_ISL_940867, EPI_ISL_940869, EPI_ISL_940870, EPI_ISL_940874, EPI_ISL_940876, EPI_ISL_940886, EPI_ISL_940887 | | | |
| see above | Vaccines and Infectious Diseases Analytics Research Unit (VIDA) | KRISP, KZN Research Innovation and Sequencing Platform | Baillie Vicky, du Plessis Jeanine, Giandhari Jennifer, Pillay Sureshnee, Naidoo Yeshnee, Tegally Houriiyah, de Oliveira Tulio, Madhi Shabir |
| EPI_ISL_941005, EPI_ISL_941006, EPI_ISL_941009, EPI_ISL_941010, EPI_ISL_941011, EPI_ISL_941012, EPI_ISL_941014 | Labo Analyses Med | National Reference Center for Viruses of Respiratory Infections, Institut Pasteur, Paris | Marion Barbet, Sylvie Behillil, Méline Bizard, Angela Brisebarre, Camille Capel, Etienne Simon-Lorière, Vincent Enouf, Maud Vanpeene, Sylvie van der Werf, Amzalag Jonas |
| EPI_ISL_941016, EPI_ISL_941019, EPI_ISL_941021, EPI_ISL_941022 | Hopital | National Reference Center for Viruses of Respiratory Infections, Institut Pasteur, Paris | Marion Barbet, Sylvie Behillil, Méline Bizard, Angela Brisebarre, Camille Capel, Etienne Simon-Lorière, Vincent Enouf, Maud Vanpeene, Sylvie van der Werf, Bour Jean Baptiste |
| EPI_ISL_941089, EPI_ISL_941092, EPI_ISL_941094 | Labo Analyses Med | National Reference Center for Viruses of Respiratory Infections, Institut Pasteur, Paris | Marion Barbet, Sylvie Behillil, Méline Bizard, Angela Brisebarre, Camille Capel, Etienne Simon-Lorière, Vincent Enouf, Maud Vanpeene, Sylvie van der Werf, Merah Kader |
| EPI_ISL_941164 | Servicio de Microbiología. Hospital Arnau de Vilanova | SeqCOVID-SPAIN consortium/IBV(CSIC) | Victoria Domínguez, Rocío Falcón, Amparo Farga and SeqCOVID-SPAIN consortium |
| EPI_ISL_941220, EPI_ISL_941221, EPI_ISL_941222, EPI_ISL_941224 | Laboratorio de Microbiología. Hospital General Universitario de Elda, Alicante | SeqCOVID-SPAIN consortium/IBV(CSIC) | Mª Isabel Gascón Ros, Cristina Torregrosa Hetland, Eva Pastor Boix, Paloma Cascales Ramos and SeqCOVID-SPAIN consortium |
| EPI_ISL_941375, EPI_ISL_941451, EPI_ISL_941452, EPI_ISL_941453, EPI_ISL_941454, EPI_ISL_941629, EPI_ISL_941630 | Instituto Nacional de Saude (INSA) | Instituto Nacional de Saude (INSA) | Borges et al |
| EPI_ISL_941901 | Virginia DCLS | Virginia DCLS | Virginia DCLS |
| EPI_ISL_942795, EPI_ISL_942796, EPI_ISL_942797, EPI_ISL_942798, EPI_ISL_942799, EPI_ISL_942800, EPI_ISL_942801, EPI_ISL_942802, EPI_ISL_942803, EPI_ISL_942804, EPI_ISL_942805, EPI_ISL_942806, EPI_ISL_942807, EPI_ISL_942808, EPI_ISL_942944 | | | |
| see above | Gundersen Molecular Diagnostics Laboratory | Kabara Cancer Research Institute | Craig S. Richmond, Paraic A. Kenny |
| EPI_ISL_942996, EPI_ISL_943010, EPI_ISL_943013, EPI_ISL_943014, EPI_ISL_943019, EPI_ISL_943067, EPI_ISL_943148, EPI_ISL_943149, EPI_ISL_943185, EPI_ISL_943186, EPI_ISL_943187, EPI_ISL_943188, EPI_ISL_943189, EPI_ISL_943190, EPI_ISL_943191, EPI_ISL_943192, EPI_ISL_943292, EPI_ISL_943337, EPI_ISL_943358, EPI_ISL_943380, EPI_ISL_943398, EPI_ISL_943399, EPI_ISL_943400, EPI_ISL_943401, EPI_ISL_943402, EPI_ISL_943403, EPI_ISL_943404, EPI_ISL_943437, EPI_ISL_943497, EPI_ISL_943508, EPI_ISL_943511, EPI_ISL_943512 | | | |
| see above | Dutch COVID-19 response team | National Institute for Public Health and the Environment (RIVM) | Adam Meijer, Harry Vennema, Dirk Eggink, Jeroen Cremer, Sharon van den Brink, Bas van der Veer, AnneMarie van den Brandt, Florian Zwagemaker, Dennis Schmitz, Chantal Reusken, on behalf of the national COVID-19 response team |
| EPI_ISL_943548 | National Institute of Health Research and Development | National Institute of Health Research and Development | Subangkit, Hana Apsari Pawestri, Kartika Dewi Puspa, Arie Ardiansyah Nugraha, Hartanti Dian Ikawati, Krisna Nur Andriana Pangesti, Yuni Rukminiati, Ririn Ramadhany, Agustiningsih, Kindi Adam, Holy Arif Wibowo, Triyani Soekarso, Ni Ketut Susilarini, Nurika Hariastuti, Ully Alfi Nikmah, Reni Herman, Nike Susanti, Herna, Tati Febriyanti, Natalie Laurencia Kipuw, Fauzul Muna, Irene Lorinda Indalao, Nelly Puspandari, Vivi Setiawaty |
| EPI_ISL_943824, EPI_ISL_943825, EPI_ISL_943826, EPI_ISL_943827, EPI_ISL_943828, EPI_ISL_943829, EPI_ISL_943830, EPI_ISL_943836, EPI_ISL_943950 | Utah Public Health Laboratory | Utah Public Health Laboratory | Erin L. Young, Kelly F. Oakeson, Tara Gallagher |
| EPI_ISL_944130, EPI_ISL_944132, EPI_ISL_944134, EPI_ISL_944136, EPI_ISL_944138, EPI_ISL_944139, EPI_ISL_944144, EPI_ISL_944146, EPI_ISL_944147, EPI_ISL_944150, EPI_ISL_944160, EPI_ISL_944161, EPI_ISL_944165, EPI_ISL_944172, EPI_ISL_944177 | | | |
| see above | National Health Laboratory Service, South Africa | KRISP, KZN Research Innovation and Sequencing Platform | Laguda-Akingba O, Giandhari J, Pillay S, Lessells R, Mdlalose K, York D, Khan S, Emmanuel SJ, Tegally H, Wilkinson E, de Oliveira T |
| EPI_ISL_944234, EPI_ISL_944235, EPI_ISL_944246, EPI_ISL_944249, EPI_ISL_944250, EPI_ISL_944279, EPI_ISL_944280, EPI_ISL_944281, EPI_ISL_944282, EPI_ISL_944283, EPI_ISL_944284, EPI_ISL_944285, EPI_ISL_944286, EPI_ISL_944287, EPI_ISL_944288, EPI_ISL_944289, EPI_ISL_944290, EPI_ISL_944291, EPI_ISL_944292, EPI_ISL_944293, EPI_ISL_944294, EPI_ISL_944295, EPI_ISL_944296, EPI_ISL_944297, EPI_ISL_944298, EPI_ISL_944299, EPI_ISL_944300, EPI_ISL_944301, EPI_ISL_944302, EPI_ISL_944303, EPI_ISL_944304, EPI_ISL_944305, EPI_ISL_944306, EPI_ISL_944307, EPI_ISL_944308, EPI_ISL_944309, EPI_ISL_944310, EPI_ISL_944311, EPI_ISL_944312, EPI_ISL_944313, EPI_ISL_944314, EPI_ISL_944315, EPI_ISL_944316, EPI_ISL_944317, EPI_ISL_944318, EPI_ISL_944319, EPI_ISL_944320, EPI_ISL_944321, EPI_ISL_944396, EPI_ISL_944399, EPI_ISL_944400, EPI_ISL_944416 | | | |
| see above | Israel Central Virology laboratory | Israel National Consortium for SARS-CoV-2 sequencing | Neta Zuckerman, Efrat Dahan Bucris, Michal Mandelboim, Dana Bar-Ilan, Oran Erster, Tzvia Mann, Omer Murik, David A. Zeevi, Assaf Rokney, Joseph Jaffe, Eva Nachum, Maya Davidovich Cohen, Ephraim Fass, Gal Zizelski Valenci, Mor Rubinstein, Efrat Rorman, Israel Nissan, Efrat Glick-Saar, Omri Nayshool, Gideon Rechavi, Ella Mendelson, Orna Mor |
| EPI_ISL_944644 | Public Health Virology-Forensic and Scientific Services (PHV-FSS) | Public Health Virology-Forensic and Scientific Services (PHV-FSS) | Alyssa T. Pyke et al. |
| EPI_ISL_944646, EPI_ISL_944647, EPI_ISL_944648, EPI_ISL_944649, EPI_ISL_944650, EPI_ISL_944651, EPI_ISL_944653, EPI_ISL_944654, EPI_ISL_944655, EPI_ISL_944658, EPI_ISL_944659, EPI_ISL_944661, EPI_ISL_944662, EPI_ISL_944663, EPI_ISL_944664, EPI_ISL_944665, EPI_ISL_944666, EPI_ISL_944667, EPI_ISL_944668, EPI_ISL_944670, EPI_ISL_944671, EPI_ISL_944675, EPI_ISL_944677, EPI_ISL_944678, EPI_ISL_944684, EPI_ISL_944689, EPI_ISL_944692, EPI_ISL_944694, EPI_ISL_944695, EPI_ISL_944696, EPI_ISL_944700, EPI_ISL_944701, EPI_ISL_944702, EPI_ISL_944703, EPI_ISL_944704, EPI_ISL_944705, EPI_ISL_944709, EPI_ISL_944710, EPI_ISL_944712, EPI_ISL_944715, EPI_ISL_944716, EPI_ISL_944717, EPI_ISL_944718, EPI_ISL_944720, EPI_ISL_944721, EPI_ISL_944723, EPI_ISL_944724, EPI_ISL_944725, EPI_ISL_944728, EPI_ISL_944729, EPI_ISL_944730, EPI_ISL_944731, EPI_ISL_944732, EPI_ISL_944733, EPI_ISL_944735 | | | |
| see above | Department of Biochemistry, Cell and Molecular Biology, West African Centre for Cell Biology of Infectious Pathogens (WACCBIP), University of Ghana | Department of Biochemistry, Cell and Molecular Biology, West African Centre for Cell Biology of Infectious Pathogens (WACCBIP), University of Ghana | Morang'a,C.M., Ngoi,J.M., Quansah,E.B., Saiid,S., Amuzu,D.S., Asante,I., Bonney,J.H., Bonney,E., Odoom,J.K., Ndam,N.T., Tei-Maya,F., Adusei-Poku,M., Ofori-Boadu,L., Ampofo,W.K., Amenga-Etego,L.N., Quashie,P., Bediako,Y., Awandare,G.A. |
| EPI_ISL_945050, EPI_ISL_945053, EPI_ISL_945055, EPI_ISL_945058, EPI_ISL_945059, EPI_ISL_945060, EPI_ISL_945062 | Lighthouse Lab in Milton Keynes | Wellcome Sanger Institute for the COVID-19 Genomics UK (COG-UK) Consortium | The Lighthouse Lab in Milton Keynes and Alex Alderton, Roberto Amato, Sonia Goncalves, Ewan Harrison, David K. Jackson, Ian Johnston, Dominic Kwiatkowski, Cordelia Langford, John Sillitoe on behalf of the Wellcome Sanger Institute COVID-19 Surveillance Team |
| EPI_ISL_945063 | Lighthouse Lab in Cambridge | Wellcome Sanger Institute for the COVID-19 Genomics UK | Rob Howes, The Lighthouse Lab in Cambridge and Alex Alderton, Roberto Amato, Sonia Goncalves, Ewan Harrison, David K. Jackson, Ian Johnston, |

| | | | |
|---|---|---|---|
| | | (COG-UK) Consortium | Dominic Kwiatkowski, Cordelia Langford, John Sillitoe on behalf of the Wellcome Sanger Institute COVID-19 Surveillance Team |
| EPI_ISL_945066 | Lighthouse Lab in Milton Keynes | Wellcome Sanger Institute for the COVID-19 Genomics UK (COG-UK) Consortium | The Lighthouse Lab in Milton Keynes and Alex Alderton, Roberto Amato, Sonia Goncalves, Ewan Harrison, David K. Jackson, Ian Johnston, Dominic Kwiatkowski, Cordelia Langford, John Sillitoe on behalf of the Wellcome Sanger Institute COVID-19 Surveillance Team |
| EPI_ISL_945067 | Lighthouse Lab in Cambridge | Wellcome Sanger Institute for the COVID-19 Genomics UK (COG-UK) Consortium | Rob Howes, The Lighthouse Lab in Cambridge and Alex Alderton, Roberto Amato, Sonia Goncalves, Ewan Harrison, David K. Jackson, Ian Johnston, Dominic Kwiatkowski, Cordelia Langford, John Sillitoe on behalf of the Wellcome Sanger Institute COVID-19 Surveillance Team |
| EPI_ISL_945069 | Lighthouse Lab in Milton Keynes | Wellcome Sanger Institute for the COVID-19 Genomics UK (COG-UK) Consortium | The Lighthouse Lab in Milton Keynes and Alex Alderton, Roberto Amato, Sonia Goncalves, Ewan Harrison, David K. Jackson, Ian Johnston, Dominic Kwiatkowski, Cordelia Langford, John Sillitoe on behalf of the Wellcome Sanger Institute COVID-19 Surveillance Team |
| EPI_ISL_945076 | Lighthouse Lab in Cambridge | Wellcome Sanger Institute for the COVID-19 Genomics UK (COG-UK) Consortium | Rob Howes, The Lighthouse Lab in Cambridge and Alex Alderton, Roberto Amato, Sonia Goncalves, Ewan Harrison, David K. Jackson, Ian Johnston, Dominic Kwiatkowski, Cordelia Langford, John Sillitoe on behalf of the Wellcome Sanger Institute COVID-19 Surveillance Team |
| EPI_ISL_945080, EPI_ISL_945082 | Lighthouse Lab in Milton Keynes | Wellcome Sanger Institute for the COVID-19 Genomics UK (COG-UK) Consortium | The Lighthouse Lab in Milton Keynes and Alex Alderton, Roberto Amato, Sonia Goncalves, Ewan Harrison, David K. Jackson, Ian Johnston, Dominic Kwiatkowski, Cordelia Langford, John Sillitoe on behalf of the Wellcome Sanger Institute COVID-19 Surveillance Team |
| EPI_ISL_945083 | Lighthouse Lab in Cambridge | Wellcome Sanger Institute for the COVID-19 Genomics UK (COG-UK) Consortium | Rob Howes, The Lighthouse Lab in Cambridge and Alex Alderton, Roberto Amato, Sonia Goncalves, Ewan Harrison, David K. Jackson, Ian Johnston, Dominic Kwiatkowski, Cordelia Langford, John Sillitoe on behalf of the Wellcome Sanger Institute COVID-19 Surveillance Team |
| EPI_ISL_945087, EPI_ISL_945090, EPI_ISL_945091, EPI_ISL_945092, EPI_ISL_945093, EPI_ISL_945094, EPI_ISL_945096 | Lighthouse Lab in Milton Keynes | Wellcome Sanger Institute for the COVID-19 Genomics UK (COG-UK) Consortium | The Lighthouse Lab in Milton Keynes and Alex Alderton, Roberto Amato, Sonia Goncalves, Ewan Harrison, David K. Jackson, Ian Johnston, Dominic Kwiatkowski, Cordelia Langford, John Sillitoe on behalf of the Wellcome Sanger Institute COVID-19 Surveillance Team |
| EPI_ISL_945100 | Lighthouse Lab in Cambridge | Wellcome Sanger Institute for the COVID-19 Genomics UK (COG-UK) Consortium | Rob Howes, The Lighthouse Lab in Cambridge and Alex Alderton, Roberto Amato, Sonia Goncalves, Ewan Harrison, David K. Jackson, Ian Johnston, Dominic Kwiatkowski, Cordelia Langford, John Sillitoe on behalf of the Wellcome Sanger Institute COVID-19 Surveillance Team |
| EPI_ISL_945101 | Lighthouse Lab in Milton Keynes | Wellcome Sanger Institute for the COVID-19 Genomics UK (COG-UK) Consortium | The Lighthouse Lab in Milton Keynes and Alex Alderton, Roberto Amato, Sonia Goncalves, Ewan Harrison, David K. Jackson, Ian Johnston, Dominic Kwiatkowski, Cordelia Langford, John Sillitoe on behalf of the Wellcome Sanger Institute COVID-19 Surveillance Team |
| EPI_ISL_945103, EPI_ISL_945106 | Lighthouse Lab in Cambridge | Wellcome Sanger Institute for the COVID-19 Genomics UK (COG-UK) Consortium | Rob Howes, The Lighthouse Lab in Cambridge and Alex Alderton, Roberto Amato, Sonia Goncalves, Ewan Harrison, David K. Jackson, Ian Johnston, Dominic Kwiatkowski, Cordelia Langford, John Sillitoe on behalf of the Wellcome Sanger Institute COVID-19 Surveillance Team |
| EPI_ISL_945108, EPI_ISL_945112 | Lighthouse Lab in Milton Keynes | Wellcome Sanger Institute for the COVID-19 Genomics UK (COG-UK) Consortium | The Lighthouse Lab in Milton Keynes and Alex Alderton, Roberto Amato, Sonia Goncalves, Ewan Harrison, David K. Jackson, Ian Johnston, Dominic Kwiatkowski, Cordelia Langford, John Sillitoe on behalf of the Wellcome Sanger Institute COVID-19 Surveillance Team |
| EPI_ISL_945116 | Lighthouse Lab in Cambridge | Wellcome Sanger Institute for the COVID-19 Genomics UK (COG-UK) Consortium | Rob Howes, The Lighthouse Lab in Cambridge and Alex Alderton, Roberto Amato, Sonia Goncalves, Ewan Harrison, David K. Jackson, Ian Johnston, Dominic Kwiatkowski, Cordelia Langford, John Sillitoe on behalf of the Wellcome Sanger Institute COVID-19 Surveillance Team |
| EPI_ISL_945118, EPI_ISL_945120, EPI_ISL_945121, EPI_ISL_945124, EPI_ISL_945128, EPI_ISL_945129 | Lighthouse Lab in Milton Keynes | Wellcome Sanger Institute for the COVID-19 Genomics UK (COG-UK) Consortium | The Lighthouse Lab in Milton Keynes and Alex Alderton, Roberto Amato, Sonia Goncalves, Ewan Harrison, David K. Jackson, Ian Johnston, Dominic Kwiatkowski, Cordelia Langford, John Sillitoe on behalf of the Wellcome Sanger Institute COVID-19 Surveillance Team |
| EPI_ISL_945131 | Lighthouse Lab in Cambridge | Wellcome Sanger Institute for the COVID-19 Genomics UK (COG-UK) Consortium | Rob Howes, The Lighthouse Lab in Cambridge and Alex Alderton, Roberto Amato, Sonia Goncalves, Ewan Harrison, David K. Jackson, Ian Johnston, Dominic Kwiatkowski, Cordelia Langford, John Sillitoe on behalf of the Wellcome Sanger Institute COVID-19 Surveillance Team |
| EPI_ISL_945133, EPI_ISL_945134 | Lighthouse Lab in Milton Keynes | Wellcome Sanger Institute for the COVID-19 Genomics UK (COG-UK) Consortium | The Lighthouse Lab in Milton Keynes and Alex Alderton, Roberto Amato, Sonia Goncalves, Ewan Harrison, David K. Jackson, Ian Johnston, Dominic Kwiatkowski, Cordelia Langford, John Sillitoe on behalf of the Wellcome Sanger Institute COVID-19 Surveillance Team |
| EPI_ISL_945139 | Lighthouse Lab in Cambridge | Wellcome Sanger Institute for the COVID-19 Genomics UK (COG-UK) Consortium | Rob Howes, The Lighthouse Lab in Cambridge and Alex Alderton, Roberto Amato, Sonia Goncalves, Ewan Harrison, David K. Jackson, Ian Johnston, Dominic Kwiatkowski, Cordelia Langford, John Sillitoe on behalf of the Wellcome Sanger Institute COVID-19 Surveillance Team |
| EPI_ISL_945141, EPI_ISL_945145 | Lighthouse Lab in Milton Keynes | Wellcome Sanger Institute for the COVID-19 Genomics UK (COG-UK) Consortium | The Lighthouse Lab in Milton Keynes and Alex Alderton, Roberto Amato, Sonia Goncalves, Ewan Harrison, David K. Jackson, Ian Johnston, Dominic Kwiatkowski, Cordelia Langford, John Sillitoe on behalf of the Wellcome Sanger Institute COVID-19 Surveillance Team |
| EPI_ISL_945148 | Lighthouse Lab in Cambridge | Wellcome Sanger Institute for the COVID-19 Genomics UK (COG-UK) Consortium | Rob Howes, The Lighthouse Lab in Cambridge and Alex Alderton, Roberto Amato, Sonia Goncalves, Ewan Harrison, David K. Jackson, Ian Johnston, Dominic Kwiatkowski, Cordelia Langford, John Sillitoe on behalf of the Wellcome Sanger Institute COVID-19 Surveillance Team |
| EPI_ISL_945149, EPI_ISL_945150, EPI_ISL_945153, EPI_ISL_945156 | Lighthouse Lab in Milton Keynes | Wellcome Sanger Institute for the COVID-19 Genomics UK (COG-UK) Consortium | The Lighthouse Lab in Milton Keynes and Alex Alderton, Roberto Amato, Sonia Goncalves, Ewan Harrison, David K. Jackson, Ian Johnston, Dominic Kwiatkowski, Cordelia Langford, John Sillitoe on behalf of the Wellcome Sanger Institute COVID-19 Surveillance Team |
| EPI_ISL_945157 | Lighthouse Lab in Cambridge | Wellcome Sanger Institute for the COVID-19 Genomics UK (COG-UK) Consortium | Rob Howes, The Lighthouse Lab in Cambridge and Alex Alderton, Roberto Amato, Sonia Goncalves, Ewan Harrison, David K. Jackson, Ian Johnston, Dominic Kwiatkowski, Cordelia Langford, John Sillitoe on behalf of the Wellcome Sanger Institute COVID-19 Surveillance Team |
| EPI_ISL_945161, EPI_ISL_945164 | Lighthouse Lab in Milton Keynes | Wellcome Sanger Institute for the COVID-19 Genomics UK (COG-UK) Consortium | The Lighthouse Lab in Milton Keynes and Alex Alderton, Roberto Amato, Sonia Goncalves, Ewan Harrison, David K. Jackson, Ian Johnston, Dominic Kwiatkowski, Cordelia Langford, John Sillitoe on behalf of the Wellcome Sanger Institute COVID-19 Surveillance Team |
| EPI_ISL_945166 | Lighthouse Lab in Cambridge | Wellcome Sanger Institute for the COVID-19 Genomics UK (COG-UK) Consortium | Rob Howes, The Lighthouse Lab in Cambridge and Alex Alderton, Roberto Amato, Sonia Goncalves, Ewan Harrison, David K. Jackson, Ian Johnston, Dominic Kwiatkowski, Cordelia Langford, John Sillitoe on behalf of the Wellcome Sanger Institute COVID-19 Surveillance Team |
| EPI_ISL_945172, EPI_ISL_945175 | Lighthouse Lab in Milton Keynes | Wellcome Sanger Institute for the COVID-19 Genomics UK (COG-UK) Consortium | The Lighthouse Lab in Milton Keynes and Alex Alderton, Roberto Amato, Sonia Goncalves, Ewan Harrison, David K. Jackson, Ian Johnston, Dominic Kwiatkowski, Cordelia Langford, John Sillitoe on behalf of the Wellcome Sanger Institute COVID-19 Surveillance Team |
| EPI_ISL_945176 | Lighthouse Lab in Cambridge | Wellcome Sanger Institute for the COVID-19 Genomics UK (COG-UK) Consortium | Rob Howes, The Lighthouse Lab in Cambridge and Alex Alderton, Roberto Amato, Sonia Goncalves, Ewan Harrison, David K. Jackson, Ian Johnston, Dominic Kwiatkowski, Cordelia Langford, John Sillitoe on behalf of the Wellcome Sanger Institute COVID-19 Surveillance Team |
| EPI_ISL_945177, EPI_ISL_945178, EPI_ISL_945179, EPI_ISL_945183, EPI_ISL_945185 | Lighthouse Lab in Milton Keynes | Wellcome Sanger Institute for the COVID-19 Genomics UK (COG-UK) Consortium | The Lighthouse Lab in Milton Keynes and Alex Alderton, Roberto Amato, Sonia Goncalves, Ewan Harrison, David K. Jackson, Ian Johnston, Dominic Kwiatkowski, Cordelia Langford, John Sillitoe on behalf of the Wellcome Sanger Institute COVID-19 Surveillance Team |
| EPI_ISL_945186 | Lighthouse Lab in Cambridge | Wellcome Sanger Institute for the COVID-19 Genomics UK (COG-UK) Consortium | Rob Howes, The Lighthouse Lab in Cambridge and Alex Alderton, Roberto Amato, Sonia Goncalves, Ewan Harrison, David K. Jackson, Ian Johnston, Dominic Kwiatkowski, Cordelia Langford, John Sillitoe on behalf of the Wellcome Sanger Institute COVID-19 Surveillance Team |
| EPI_ISL_945189 | Lighthouse Lab in Milton Keynes | Wellcome Sanger Institute for the COVID-19 Genomics UK (COG-UK) Consortium | The Lighthouse Lab in Milton Keynes and Alex Alderton, Roberto Amato, Sonia Goncalves, Ewan Harrison, David K. Jackson, Ian Johnston, Dominic Kwiatkowski, Cordelia Langford, John Sillitoe on behalf of the Wellcome Sanger Institute COVID-19 Surveillance Team |
| EPI_ISL_945191, EPI_ISL_945192 | Lighthouse Lab in Cambridge | Wellcome Sanger Institute for the COVID-19 Genomics UK (COG-UK) Consortium | Rob Howes, The Lighthouse Lab in Cambridge and Alex Alderton, Roberto Amato, Sonia Goncalves, Ewan Harrison, David K. Jackson, Ian Johnston, Dominic Kwiatkowski, Cordelia Langford, John Sillitoe on behalf of the Wellcome Sanger Institute COVID-19 Surveillance Team |
| EPI_ISL_945193, EPI_ISL_945198 | Lighthouse Lab in Milton Keynes | Wellcome Sanger Institute for the COVID-19 Genomics UK (COG-UK) Consortium | The Lighthouse Lab in Milton Keynes and Alex Alderton, Roberto Amato, Sonia Goncalves, Ewan Harrison, David K. Jackson, Ian Johnston, Dominic Kwiatkowski, Cordelia Langford, John Sillitoe on behalf of the Wellcome Sanger Institute COVID-19 Surveillance Team |
| EPI_ISL_945199, EPI_ISL_945203 | Lighthouse Lab in Cambridge | Wellcome Sanger Institute for the COVID-19 Genomics UK (COG-UK) Consortium | Rob Howes, The Lighthouse Lab in Cambridge and Alex Alderton, Roberto Amato, Sonia Goncalves, Ewan Harrison, David K. Jackson, Ian Johnston, Dominic Kwiatkowski, Cordelia Langford, John Sillitoe on behalf of the Wellcome Sanger Institute COVID-19 Surveillance Team |
| EPI_ISL_945205, EPI_ISL_945210, EPI_ISL_945218, EPI_ISL_945220 | Lighthouse Lab in Milton Keynes | Wellcome Sanger Institute for the COVID-19 Genomics UK (COG-UK) Consortium | The Lighthouse Lab in Milton Keynes and Alex Alderton, Roberto Amato, Sonia Goncalves, Ewan Harrison, David K. Jackson, Ian Johnston, Dominic Kwiatkowski, Cordelia Langford, John Sillitoe on behalf of the Wellcome Sanger Institute COVID-19 Surveillance Team |
| EPI_ISL_945222 | Lighthouse Lab in Cambridge | Wellcome Sanger Institute for the COVID-19 Genomics UK (COG-UK) Consortium | Rob Howes, The Lighthouse Lab in Cambridge and Alex Alderton, Roberto Amato, Sonia Goncalves, Ewan Harrison, David K. Jackson, Ian Johnston, Dominic Kwiatkowski, Cordelia Langford, John Sillitoe on behalf of the Wellcome Sanger Institute COVID-19 Surveillance Team |
| EPI_ISL_945228, EPI_ISL_945230 | Lighthouse Lab in Milton Keynes | Wellcome Sanger Institute for the COVID-19 Genomics UK (COG-UK) Consortium | The Lighthouse Lab in Milton Keynes and Alex Alderton, Roberto Amato, Sonia Goncalves, Ewan Harrison, David K. Jackson, Ian Johnston, Dominic Kwiatkowski, Cordelia Langford, John Sillitoe on behalf of the Wellcome Sanger Institute COVID-19 Surveillance Team |

| | | | |
|---|---|---|---|
| EPI_ISL_945232, EPI_ISL_945237 | Lighthouse Lab in Cambridge | Wellcome Sanger Institute for the COVID-19 Genomics UK (COG-UK) Consortium | Rob Howes, The Lighthouse Lab in Cambridge and Alex Alderton, Roberto Amato, Sonia Goncalves, Ewan Harrison, David K. Jackson, Ian Johnston, Dominic Kwiatkowski, Cordelia Langford, John Sillitoe on behalf of the Wellcome Sanger Institute COVID-19 Surveillance Team |
| EPI_ISL_945239, EPI_ISL_945246, EPI_ISL_945251, EPI_ISL_945253, EPI_ISL_945256 | Lighthouse Lab in Milton Keynes | Wellcome Sanger Institute for the COVID-19 Genomics UK (COG-UK) Consortium | The Lighthouse Lab in Milton Keynes and Alex Alderton, Roberto Amato, Sonia Goncalves, Ewan Harrison, David K. Jackson, Ian Johnston, Dominic Kwiatkowski, Cordelia Langford, John Sillitoe on behalf of the Wellcome Sanger Institute COVID-19 Surveillance Team |
| EPI_ISL_945257 | Lighthouse Lab in Cambridge | Wellcome Sanger Institute for the COVID-19 Genomics UK (COG-UK) Consortium | Rob Howes, The Lighthouse Lab in Cambridge and Alex Alderton, Roberto Amato, Sonia Goncalves, Ewan Harrison, David K. Jackson, Ian Johnston, Dominic Kwiatkowski, Cordelia Langford, John Sillitoe on behalf of the Wellcome Sanger Institute COVID-19 Surveillance Team |
| EPI_ISL_945258 | Lighthouse Lab in Milton Keynes | Wellcome Sanger Institute for the COVID-19 Genomics UK (COG-UK) Consortium | The Lighthouse Lab in Milton Keynes and Alex Alderton, Roberto Amato, Sonia Goncalves, Ewan Harrison, David K. Jackson, Ian Johnston, Dominic Kwiatkowski, Cordelia Langford, John Sillitoe on behalf of the Wellcome Sanger Institute COVID-19 Surveillance Team |
| EPI_ISL_945261 | Lighthouse Lab in Cambridge | Wellcome Sanger Institute for the COVID-19 Genomics UK (COG-UK) Consortium | Rob Howes, The Lighthouse Lab in Cambridge and Alex Alderton, Roberto Amato, Sonia Goncalves, Ewan Harrison, David K. Jackson, Ian Johnston, Dominic Kwiatkowski, Cordelia Langford, John Sillitoe on behalf of the Wellcome Sanger Institute COVID-19 Surveillance Team |
| EPI_ISL_945266 | Lighthouse Lab in Milton Keynes | Wellcome Sanger Institute for the COVID-19 Genomics UK (COG-UK) Consortium | The Lighthouse Lab in Milton Keynes and Alex Alderton, Roberto Amato, Sonia Goncalves, Ewan Harrison, David K. Jackson, Ian Johnston, Dominic Kwiatkowski, Cordelia Langford, John Sillitoe on behalf of the Wellcome Sanger Institute COVID-19 Surveillance Team |
| EPI_ISL_945268, EPI_ISL_945271 | Lighthouse Lab in Cambridge | Wellcome Sanger Institute for the COVID-19 Genomics UK (COG-UK) Consortium | Rob Howes, The Lighthouse Lab in Cambridge and Alex Alderton, Roberto Amato, Sonia Goncalves, Ewan Harrison, David K. Jackson, Ian Johnston, Dominic Kwiatkowski, Cordelia Langford, John Sillitoe on behalf of the Wellcome Sanger Institute COVID-19 Surveillance Team |
| EPI_ISL_945276, EPI_ISL_945277, EPI_ISL_945278, EPI_ISL_945287, EPI_ISL_945292 | Lighthouse Lab in Milton Keynes | Wellcome Sanger Institute for the COVID-19 Genomics UK (COG-UK) Consortium | The Lighthouse Lab in Milton Keynes and Alex Alderton, Roberto Amato, Sonia Goncalves, Ewan Harrison, David K. Jackson, Ian Johnston, Dominic Kwiatkowski, Cordelia Langford, John Sillitoe on behalf of the Wellcome Sanger Institute COVID-19 Surveillance Team |
| EPI_ISL_945309 | Lighthouse Lab in Cambridge | Wellcome Sanger Institute for the COVID-19 Genomics UK (COG-UK) Consortium | Rob Howes, The Lighthouse Lab in Cambridge and Alex Alderton, Roberto Amato, Sonia Goncalves, Ewan Harrison, David K. Jackson, Ian Johnston, Dominic Kwiatkowski, Cordelia Langford, John Sillitoe on behalf of the Wellcome Sanger Institute COVID-19 Surveillance Team |
| EPI_ISL_945313, EPI_ISL_945314, EPI_ISL_945315 | Lighthouse Lab in Milton Keynes | Wellcome Sanger Institute for the COVID-19 Genomics UK (COG-UK) Consortium | The Lighthouse Lab in Milton Keynes and Alex Alderton, Roberto Amato, Sonia Goncalves, Ewan Harrison, David K. Jackson, Ian Johnston, Dominic Kwiatkowski, Cordelia Langford, John Sillitoe on behalf of the Wellcome Sanger Institute COVID-19 Surveillance Team |
| EPI_ISL_945321 | Lighthouse Lab in Cambridge | Wellcome Sanger Institute for the COVID-19 Genomics UK (COG-UK) Consortium | Rob Howes, The Lighthouse Lab in Cambridge and Alex Alderton, Roberto Amato, Sonia Goncalves, Ewan Harrison, David K. Jackson, Ian Johnston, Dominic Kwiatkowski, Cordelia Langford, John Sillitoe on behalf of the Wellcome Sanger Institute COVID-19 Surveillance Team |
| EPI_ISL_945322, EPI_ISL_945325, EPI_ISL_945329, EPI_ISL_945331, EPI_ISL_945333, EPI_ISL_945336 | Lighthouse Lab in Milton Keynes | Wellcome Sanger Institute for the COVID-19 Genomics UK (COG-UK) Consortium | The Lighthouse Lab in Milton Keynes and Alex Alderton, Roberto Amato, Sonia Goncalves, Ewan Harrison, David K. Jackson, Ian Johnston, Dominic Kwiatkowski, Cordelia Langford, John Sillitoe on behalf of the Wellcome Sanger Institute COVID-19 Surveillance Team |
| EPI_ISL_945341 | Lighthouse Lab in Cambridge | Wellcome Sanger Institute for the COVID-19 Genomics UK (COG-UK) Consortium | Rob Howes, The Lighthouse Lab in Cambridge and Alex Alderton, Roberto Amato, Sonia Goncalves, Ewan Harrison, David K. Jackson, Ian Johnston, Dominic Kwiatkowski, Cordelia Langford, John Sillitoe on behalf of the Wellcome Sanger Institute COVID-19 Surveillance Team |
| EPI_ISL_945344 | Lighthouse Lab in Milton Keynes | Wellcome Sanger Institute for the COVID-19 Genomics UK (COG-UK) Consortium | The Lighthouse Lab in Milton Keynes and Alex Alderton, Roberto Amato, Sonia Goncalves, Ewan Harrison, David K. Jackson, Ian Johnston, Dominic Kwiatkowski, Cordelia Langford, John Sillitoe on behalf of the Wellcome Sanger Institute COVID-19 Surveillance Team |
| EPI_ISL_945346 | Lighthouse Lab in Cambridge | Wellcome Sanger Institute for the COVID-19 Genomics UK (COG-UK) Consortium | Rob Howes, The Lighthouse Lab in Cambridge and Alex Alderton, Roberto Amato, Sonia Goncalves, Ewan Harrison, David K. Jackson, Ian Johnston, Dominic Kwiatkowski, Cordelia Langford, John Sillitoe on behalf of the Wellcome Sanger Institute COVID-19 Surveillance Team |
| EPI_ISL_945352 | Lighthouse Lab in Milton Keynes | Wellcome Sanger Institute for the COVID-19 Genomics UK (COG-UK) Consortium | The Lighthouse Lab in Milton Keynes and Alex Alderton, Roberto Amato, Sonia Goncalves, Ewan Harrison, David K. Jackson, Ian Johnston, Dominic Kwiatkowski, Cordelia Langford, John Sillitoe on behalf of the Wellcome Sanger Institute COVID-19 Surveillance Team |
| EPI_ISL_945356 | Lighthouse Lab in Cambridge | Wellcome Sanger Institute for the COVID-19 Genomics UK (COG-UK) Consortium | Rob Howes, The Lighthouse Lab in Cambridge and Alex Alderton, Roberto Amato, Sonia Goncalves, Ewan Harrison, David K. Jackson, Ian Johnston, Dominic Kwiatkowski, Cordelia Langford, John Sillitoe on behalf of the Wellcome Sanger Institute COVID-19 Surveillance Team |
| EPI_ISL_945362 | Lighthouse Lab in Milton Keynes | Wellcome Sanger Institute for the COVID-19 Genomics UK (COG-UK) Consortium | The Lighthouse Lab in Milton Keynes and Alex Alderton, Roberto Amato, Sonia Goncalves, Ewan Harrison, David K. Jackson, Ian Johnston, Dominic Kwiatkowski, Cordelia Langford, John Sillitoe on behalf of the Wellcome Sanger Institute COVID-19 Surveillance Team |
| EPI_ISL_945365 | Lighthouse Lab in Cambridge | Wellcome Sanger Institute for the COVID-19 Genomics UK (COG-UK) Consortium | Rob Howes, The Lighthouse Lab in Cambridge and Alex Alderton, Roberto Amato, Sonia Goncalves, Ewan Harrison, David K. Jackson, Ian Johnston, Dominic Kwiatkowski, Cordelia Langford, John Sillitoe on behalf of the Wellcome Sanger Institute COVID-19 Surveillance Team |
| EPI_ISL_945368, EPI_ISL_945372, EPI_ISL_945373 | Lighthouse Lab in Milton Keynes | Wellcome Sanger Institute for the COVID-19 Genomics UK (COG-UK) Consortium | The Lighthouse Lab in Milton Keynes and Alex Alderton, Roberto Amato, Sonia Goncalves, Ewan Harrison, David K. Jackson, Ian Johnston, Dominic Kwiatkowski, Cordelia Langford, John Sillitoe on behalf of the Wellcome Sanger Institute COVID-19 Surveillance Team |
| EPI_ISL_946746 | Lighthouse Lab in Alderley Park | Wellcome Sanger Institute for the COVID-19 Genomics UK (COG-UK) Consortium | Jacquelyn Wynn, Mairead Hyland, The Lighthouse Lab in Alderley Park and Alex Alderton, Roberto Amato, Sonia Goncalves, Ewan Harrison, David K. Jackson, Ian Johnston, Dominic Kwiatkowski, Cordelia Langford, John Sillitoe on behalf of the Wellcome Sanger Institute COVID-19 Surveillance Team |
| EPI_ISL_947322 | RS Mitra Keluarga Kelapa Gading | Eijkman Institute for Molecular Biology, Ministry of Research and Technology/National Agency for Research and Innovation | Sukma Oktavianthi, Willy Agustine, Edison Johar, Hidayat Trimarsanto, Iskandar Adnan, Lydia V. Panggalo, Frilasita A Yudhaputri, Safarina G Malik, Khin Saw Myint, Amin Soebandrio |
| EPI_ISL_947324 | RSU Medirossa | Eijkman Institute for Molecular Biology, Ministry of Research and Technology/National Agency for Research and Innovation | Sukma Oktavianthi, Willy Agustine, Edison Johar, Hidayat Trimarsanto, Iskandar Adnan, Lydia V. Panggalo, Frilasita A Yudhaputri, Safarina G Malik, Khin Saw Myint, Amin Soebandrio |
| EPI_ISL_947325 | RSIA PKU Muhammadiyah | Eijkman Institute for Molecular Biology, Ministry of Research and Technology/National Agency for Research and Innovation | Sukma Oktavianthi, Willy Agustine, Edison Johar, Hidayat Trimarsanto, Iskandar Adnan, Lydia V. Panggalo, Frilasita A Yudhaputri, Safarina G Malik, Khin Saw Myint, Amin Soebandrio |
| EPI_ISL_947327 | RS Mitra Keluarga Gading Serpong | Eijkman Institute for Molecular Biology, Ministry of Research and Technology/National Agency for Research and Innovation | Willy Agustine, Edison Johar, Hidayat Trimarsanto, Iskandar Adnan, Lydia V. Panggalo, Sukma Oktavianthi, Frilasita A Yudhaputri, Safarina G Malik, Khin Saw Myint, Amin Soebandrio |
| EPI_ISL_949183 | National Institute of Health Research and Development | National Institute of Health Research and Development | Kindi Adam; Holy Arif Wibowo; Ririn Ramadhany; Yuni Rukminiati; Agustiningsih; Hana Apsari Pawestri; Subangkit; Kartika Dewi Puspa; Arie Ardiansyah Nugraha; Hartanti Dian Ikawati; Krisna Nur Andriana Pangesti; Triyani Soekarso; Ni Ketut Susilarini; Nur Ika Hariastuti; Uly Alfi Nikmah; Mursinah; Asri Febriyani; Reni Herman; Nike Susanti; Herna; Tati Febriyanti; Natalie Laurencia Kipuw; Fauzul Muna; Irene Lorinda Indalao; Aulia Rizki; Nelly Puspandari; Vivi Setiawaty |
| EPI_ISL_949411 | University of Birmingham | COVID-19 Genomics UK (COG-UK) Consortium | Institute of Microbiology, University of Birmingham: Claire McMurray, Joanne Stockton, Samuel Nicholls, Radoslaw Poplawski, Will Rowe, Josh Quick, Nicholas Loman. University of Birmingham Testing Laboratory: Celina M Whalley, Andrew Bosworth, Charlotte Poxon, Kasun Wanigasooriya, Oliver Pickles, Mike Kidd, Alex Richter, Andrew D Beggs PHE Heartlands Lab: Husam Osman, Andrew Bosworth. Queen Elizabeth Hospital: Anna Casey |
| EPI_ISL_949754, EPI_ISL_949756, EPI_ISL_949764, EPI_ISL_949769, EPI_ISL_949771, EPI_ISL_949773, EPI_ISL_949775, EPI_ISL_949781 | Barts Health NHS Trust | COVID-19 Genomics UK (COG-UK) Consortium | CUTINO-MOGUEL, Maria-Teresa; HARRINGTON, David; OWOYEMI, Dola; KULASEGARAN-SHYLINI, Raghavendran; BROAD, Claire; KELE, Beatrix |
| EPI_ISL_949788, EPI_ISL_949927, EPI_ISL_950068, EPI_ISL_950073, EPI_ISL_950076, EPI_ISL_950081, EPI_ISL_950087, EPI_ISL_950089, EPI_ISL_950090, EPI_ISL_950091, EPI_ISL_950093, EPI_ISL_950095, EPI_ISL_950096, EPI_ISL_950099, EPI_ISL_950103, EPI_ISL_950104, EPI_ISL_950106, EPI_ISL_950107, EPI_ISL_950108, EPI_ISL_950109, EPI_ISL_950113, EPI_ISL_950116, EPI_ISL_950117, EPI_ISL_950118, EPI_ISL_950119, EPI_ISL_950120, EPI_ISL_950122, EPI_ISL_950123, EPI_ISL_950124, EPI_ISL_950125, EPI_ISL_950128, EPI_ISL_950129, EPI_ISL_950130, EPI_ISL_950133, EPI_ISL_950136, EPI_ISL_950142, EPI_ISL_950147, EPI_ISL_950148, EPI_ISL_950152, EPI_ISL_950154 | | | |
| see above | University College London, Great Ormond Street Hospital for Children NHS Foundation Trust, Imperial College Healthcare NHS Trust | COVID-19 Genomics UK (COG-UK) Consortium | Sergi Castellano, Rachel Williams, Mark Kristiansen, Paola Resende Silva, Sunando Roy, Tony Brooks, Helena Tutill, Paola Niola, Patricia Dyal, Charlotte Williams, Leysa Forrest, Yasmin Panchbhaya, Jacqueline Findlay, Samuel Weeks, Julianne Brown, Kathryn Harris, Paul Randell, James Price, Alison Holmes, Judith Breuer |
| EPI_ISL_950268, EPI_ISL_950270 | Northumbria University / South Tees Hospitals NHS Foundation Trust / North Cumbria Integrated Care NHS Foundation Trust / North Tees and Hartlepool NHS | COVID-19 Genomics UK (COG-UK) Consortium | Darren L Smith,Andrew Nelson,Matthew Bashton,Greg R Young,Joshua Loh,John Allan,Mohammad A Tariq,Giles S Holt,Gary Black,Wen C Yew,Lynn Dover,Paul Baker,Steve Liggett,Sarah Essex,Jane Greenaway,Debra Padgett,Clive Graham,Garren Scott,Edward Barton,Emma Swindells,Brendan Payne,Jennifer Collins,Yusri Taha,Gary Eltringham |

| | | | |
|---|---|---|---|
| | Foundation Trust / Newcastle Hospitals NHS Foundation Trust | | |
| EPI_ISL_950712 | Lincolnshire Hospitals and DeepSeq Nottingham | COVID-19 Genomics UK (COG-UK) Consortium | Nichola Duckworth, Tim Sloan, Sarah Walsh, Jonathan Ball, Patrick McClure, Joeseph Chappell, Nadine Holmes, Matthew Carlisle, Christopher Moore, Fei Sang, Johnny Debebe, Victoria Wright, Matthew Loose |
| EPI_ISL_951125, EPI_ISL_951141, EPI_ISL_951152, EPI_ISL_951154, EPI_ISL_951158, EPI_ISL_951160, EPI_ISL_951172, EPI_ISL_951174, EPI_ISL_951175, EPI_ISL_951176, EPI_ISL_951177, EPI_ISL_951180, EPI_ISL_951181, EPI_ISL_951182, EPI_ISL_951184, EPI_ISL_951186, EPI_ISL_951189, EPI_ISL_951193, EPI_ISL_951195, EPI_ISL_951196, EPI_ISL_951197, EPI_ISL_951199, EPI_ISL_951201, EPI_ISL_951202, EPI_ISL_951204, EPI_ISL_951205, EPI_ISL_951206, EPI_ISL_951207, EPI_ISL_951209, EPI_ISL_951210, EPI_ISL_951211, EPI_ISL_951212, EPI_ISL_951213, EPI_ISL_951215, EPI_ISL_951216, EPI_ISL_951218, EPI_ISL_951220, EPI_ISL_951221, EPI_ISL_951223, EPI_ISL_951224, EPI_ISL_951225, EPI_ISL_951226, EPI_ISL_951227, EPI_ISL_951228, EPI_ISL_951233, EPI_ISL_951234, EPI_ISL_951235, EPI_ISL_951237, EPI_ISL_951238, EPI_ISL_951241, EPI_ISL_951242, EPI_ISL_951264, EPI_ISL_951267, EPI_ISL_951301, EPI_ISL_951320, EPI_ISL_951323, EPI_ISL_951324, EPI_ISL_951326, EPI_ISL_951327, EPI_ISL_951328, EPI_ISL_951329, EPI_ISL_951331, EPI_ISL_951333, EPI_ISL_951334, EPI_ISL_951335, EPI_ISL_951336, EPI_ISL_951337, EPI_ISL_951339, EPI_ISL_951340, EPI_ISL_951341, EPI_ISL_951342, EPI_ISL_951344, EPI_ISL_951345, EPI_ISL_951346, EPI_ISL_951347, EPI_ISL_951348, EPI_ISL_951349, EPI_ISL_951350, EPI_ISL_951351, EPI_ISL_951352, EPI_ISL_951353, EPI_ISL_951354, EPI_ISL_951382, EPI_ISL_951397, EPI_ISL_951398, EPI_ISL_951400, EPI_ISL_951401, EPI_ISL_951402, EPI_ISL_951404, EPI_ISL_951405, EPI_ISL_951410 | | | |
| see above | Oxford Viromics, NDM, University of Oxford; Oxford University Hospitals; Basingstoke and North Hampshire Hospital | COVID-19 Genomics UK (COG-UK) Consortium | Tanya Golubchik, David Bonsall, George Macintyre, Amy Trebes, Mariateresa de Cesare, Catrin Moore, Alex Mobbs, Anita Justice, Robert Shaw, Monique Andersson, Timothy Peto, Emma Wise, Nathan Moore, Jessica Lynch, Nick Cortes, Matilde Mori, Stephen Kidd, David Buck, John Todd, Christophe Fraser |
| EPI_ISL_952401, EPI_ISL_952407, EPI_ISL_952408, EPI_ISL_952413, EPI_ISL_952415, EPI_ISL_952785, EPI_ISL_952813, EPI_ISL_952814, EPI_ISL_952815 | Centre for Enzyme Innovation, University of Portsmouth / Translational Research Laboratory, Portsmouth Hospitals NHS Trust | COVID-19 Genomics UK (COG-UK) Consortium | Angela Beckett,Salman Goudarzi,Christopher Fearn,Kate Cook,Katie Loveson,Sharon Glaysher,Scott Elliott,Samuel Robson |
| EPI_ISL_953178, EPI_ISL_953210, EPI_ISL_953214, EPI_ISL_953242, EPI_ISL_953260, EPI_ISL_953273, EPI_ISL_953274, EPI_ISL_953278 | Bioinformatics and Biostatistics Lab, Advanced Sequencing Facility | COVID-19 Genomics UK (COG-UK) Consortium | Aengus Stewart,Jerome Nicod,Chelsea Sawyer,Laura Cubitt,Harshil Patel,Margaret Crawford |
| EPI_ISL_953997 | Hopital | National Reference Center for Viruses of Respiratory Infections, Institut Pasteur, Paris | Marion Barbet, Sylvie Behillil, Méline Bizard, Angela Brisebarre, Camille Capel, Etienne Simon-Lorière, Vincent Enouf, Maud Vanpeene, Sylvie van der Werf,Lagathu GisèLe |
| EPI_ISL_954220, EPI_ISL_954221, EPI_ISL_954222 | 1.AO Universitaria 'S. Giovanni di Dio e Ruggi D'Aragona, Scuola Medica Salernitana' Hospital / 2.UOC di Virologia e Microbiologia, Università della Campania 'L. Vanvitelli' / 3.AO Universitaria 'Federico II' Napoli Hospital / 4.AORN 'San Giuseppe Moscati' Avellino Hospital / 5.AO 'San Pio - presidio G. Rummo' Benevento Hospital / 6.AO 'Sant'Anna e San Sebastiano' Caserta Hospital / 7.PO 'Maria Santissima Addolorata' Eboli Hospital / 8.Biogem Istituto di Ricerche Genetiche | 1. Genome Research Center for Health (CRGS) / 2. Laboratory of Molecular Medicine and Genomics(LMMGe) / 3. Center for Research in Pure and Applied Mathematics (CRMPA) | Giorgio Giurato, Francesca Rizzo, Alessandro Weisz, Gianluigi Franci, Giovanni Nassa, Pasquale Pagliano, Roberta Tarallo, Elena Alexandrova, Ylenia D'Agostino, Carlo Ferravante, Jessica Lamberti, Viola Melone, Domenico Memoli, Valeria Mirici Cappa, Domenico Palumbo, Giovanni Pecoraro, Assunta Sellitto, Oriana Strianese, Ilaria Terenzi, Giuseppe Fenza, Aniello Gentile, Antonello Saccomanno, Sonia Amabile, Teresa Rocco, Annamaria Salvati, Emilia Vaccaro, Massimiliano Galdiero, Michele Cennamo, Giuseppe Portella, Maria Grazia Foti, Mariarosaria Ingino, Maria Landi, Maurizio Fumi, Vincenzo Rocco, Rita Greco, Vittoria Letizia, Arnolfo Petruzziello, Maddalena Schioppa, Gregorio Goffredi, Francesca Marciano, Michele Caraglia, Alessia Cossu, Marianna Scrima, Edmondo Adorisio, Morena D'Avenia, Michela Iacobellis, Rosanna Piluscio, Giorgio Dirani, Vittorio Sambri, Simona Semprini, Silvia Zanoli, Francesco Curcio, Stefania Marzinotto, Andreina Baj, Fausto Sessa. |
| EPI_ISL_954831, EPI_ISL_954844, EPI_ISL_954848, EPI_ISL_954852, EPI_ISL_954856, EPI_ISL_954857, EPI_ISL_954862, EPI_ISL_954864, EPI_ISL_954886, EPI_ISL_954902, EPI_ISL_954903, EPI_ISL_954904, EPI_ISL_954905, EPI_ISL_954906, EPI_ISL_954907, EPI_ISL_954908, EPI_ISL_954909, EPI_ISL_954910, EPI_ISL_954911, EPI_ISL_954912, EPI_ISL_954913, EPI_ISL_954914, EPI_ISL_954915, EPI_ISL_954916, EPI_ISL_954917, EPI_ISL_954918 | | | |
| see above | Colorado Department of Public Health and Environment | Colorado Department of Puplic Health and Environment | Laura Bankers, Molly C. Hetherington-Rauth, Diana Ir, Shannon Ely, Shannon R. Matzinger, Sarah Elizabeth Totten, Emily A. Travanty |
| EPI_ISL_955127 | Innovative Genomics Institute, UC Berkeley | Innovative Genomics Institute, UC Berkeley | Stacia Wyman, Haridha Shivram, Phil Frankino, Liana Lareau, Shana McDevitt, Justin Choi |
| EPI_ISL_955184 | University of Sarajevo, Veterinary Faculty, Laboratory for Molecular Diagnostic and Research Laboratory | University of Sarajevo, Veterinary Faculty, Laboratory for Molecular Diagnostic and Research Laboratory | Goleti Š., Goleti T., Softi A., Ali-Šeho A., Nicevi M., Hodži A., Terzi I., Šabi E., Jaži A. |
| EPI_ISL_955185 | University of Sarajevo, Veterinary Faculty, Laboratory for Molecular Diagnostic and Research Laboratory | University of Sarajevo, Veterinary Faculty, Laboratory for Molecular Diagnostic and Research Laboratory | Goleti Š., Goleti T., Ali-Šeho A., Softi A., Terzi I., Šabi E., Jaži A., Nicevi M., Hodži A. |
| EPI_ISL_955211 | University of Sarajevo, Veterinary Faculty, Laboratory for Molecular Diagnostic and Research Laboratory | University of Sarajevo, Veterinary Faculty, Laboratory for Molecular Diagnostic and Research Laboratory | Goleti Š., Goleti T., Ali-Šeho A., Softi A., Jaži A., Terzi I., Šabi E., Nicevi M., Hodži A. |
| EPI_ISL_955274, EPI_ISL_955285 | American Esoteric Laboratory | Pathogen Discovery, Respiratory Viruses Branch, Division of Viral Diseases, Centers for Disease Control and Prevention | Ying Tao, Jing Zhang, Yan Li, Krista Queen, Anna Uehara, Peter Cook, Clinton R. Paden, Haibin Wang, Suxiang Tong |
| EPI_ISL_955311 | TX DSHS, Lab Services Section MC 1947 | Pathogen Discovery, Respiratory Viruses Branch, Division of Viral Diseases, Centers for Disease Control and Prevention | Ying Tao, Jing Zhang, Yan Li, Krista Queen, Anna Uehara, Peter Cook, Clinton R. Paden, Haibin Wang, Suxiang Tong |
| EPI_ISL_955324, EPI_ISL_955325, EPI_ISL_955326, EPI_ISL_955327 | American Esoteric Laboratory | Pathogen Discovery, Respiratory Viruses Branch, Division of Viral Diseases, Centers for Disease Control and Prevention | Ying Tao, Jing Zhang, Yan Li, Krista Queen, Anna Uehara, Peter Cook, Clinton R. Paden, Haibin Wang, Suxiang Tong |
| EPI_ISL_955330, EPI_ISL_955335 | GA Department of Public Health Laboratory | Pathogen Discovery, Respiratory Viruses Branch, Division of Viral Diseases, Centers for Disease Control and Prevention | Ying Tao, Jing Zhang, Yan Li, Krista Queen, Anna Uehara, Peter Cook, Clinton R. Paden, Haibin Wang, Suxiang Tong |
| EPI_ISL_955411, EPI_ISL_955417, EPI_ISL_955418, EPI_ISL_955419, EPI_ISL_955420, EPI_ISL_955421, EPI_ISL_955422, EPI_ISL_955423, EPI_ISL_955424, EPI_ISL_955425 | Alameda County Public Health Lab | Chan-Zuckerberg Biohub | CZB Cliahub Consortium |
| EPI_ISL_955447, EPI_ISL_955452, EPI_ISL_955453, EPI_ISL_955464, EPI_ISL_955465, EPI_ISL_955466, EPI_ISL_955467, EPI_ISL_955470, EPI_ISL_955471, EPI_ISL_955472, EPI_ISL_955473, EPI_ISL_955474, EPI_ISL_955475, EPI_ISL_955476, EPI_ISL_955477, EPI_ISL_955478, EPI_ISL_955479, EPI_ISL_955480, EPI_ISL_955481, EPI_ISL_955482, EPI_ISL_955483, EPI_ISL_955484, EPI_ISL_955485, EPI_ISL_955486, EPI_ISL_955487, EPI_ISL_955488, EPI_ISL_955489, EPI_ISL_955490, EPI_ISL_955491, EPI_ISL_955492, EPI_ISL_955493, EPI_ISL_955494, EPI_ISL_955495 | | | |
| see above | Orange County Public Health Lab | Chan-Zuckerberg Biohub | CZB Cliahub Consortium |
| EPI_ISL_955777 | Humboldt County Public Health Laboratory | Chan-Zuckerberg Biohub | CZB Cliahub Consortium |
| EPI_ISL_956300 | Laboratorio de salud publica Arauca | Instituto Nacional de Salud- Dirección de Investigación en Salud Pública | Katherine Laiton-Donato, Diego A. Álvarez-Díaz, Carlos Franco-Muñoz, Mauricio Pacheco-Montealegre, Hector Alejandro Ruiz-Moreno, Maria T. Herrera-Sepúlveda, Diego Andrés Prada, Jhonnatan Reales-González, Sheryll Corchuelo, Julian Naizaque, Gerardo Santamaría, Magdalena Wiesner, Martha Lucia Ospina Martinez, Marcela Mercado-Reyes |
| EPI_ISL_956316 | Institute of Tropical Disease | Institute of Tropical Disease, Universitas Airlangga | Krisnoadi Rahardjo, Aldise M Nastri, Jezzy R Dewantari, Rima R Prasetya, Gatot Soegiarto, Laksmi Wulandari, Resti Yudhawati, Yasuko Mori, Soetjipto, Kazufumi Shimizu, Maria I Lusida |
| EPI_ISL_956421, EPI_ISL_956668 | General Hospital - Kumanovo | Research Center for Genetic Engineering and Biotechnology "Georgi D. Efremov" , Macedonian Academy of Sciences and Arts | Aleksandar J. Dimovski, Dijana Plasheska-Karanfilska, Predrag Noveski, Gjorgji Bozinovski, Milena Jakimovska |
| EPI_ISL_957423 | General Hospital - Bitola | Research Center for Genetic Engineering and Biotechnology "Georgi D. Efremov" , Macedonian Academy of Sciences and Arts | Aleksandar J. Dimovski, Dijana Plasheska-Karanfilska, Predrag Noveski, Gjorgji Bozinovski, Milena Jakimovska |
| EPI_ISL_957966 | General Hospital - Tetovo | Research Center for Genetic Engineering and Biotechnology "Georgi D. Efremov" , Macedonian Academy of Sciences and Arts | Aleksandar J. Dimovski, Dijana Plasheska-Karanfilska, Predrag Noveski, Gjorgji Bozinovski, Milena Jakimovska |
| EPI_ISL_958564, EPI_ISL_958565 | Lighthouse Lab in Milton Keynes | Wellcome Sanger Institute for the COVID-19 Genomics UK (COG-UK) Consortium | The Lighthouse Lab in Milton Keynes and Alex Alderton, Roberto Amato, Sonia Goncalves, Ewan Harrison, David K. Jackson, Ian Johnston, Dominic Kwiatkowski, Cordelia Langford, John Sillitoe on behalf of the Wellcome Sanger Institute COVID-19 Surveillance Team |

| | | | |
|---|---|---|---|
| EPI_ISL_958751 | General Hospital - Bitola | Research Center for Genetic Engineering and Biotechnology "Georgi D. Efremov" , Macedonian Academy of Sciences and Arts | Aleksandar J. Dimovski, Dijana Plasheska-Karanfilska, Predrag Noveski, Gjorgji Bozinovski, Milena Jakimovska |
| EPI_ISL_959851 | National Virus Reference Laboratory | National Virus Reference Laboratory | Michael Carr, Gabriel Gonzalez, Jonathan Dean, Cillian F De Gascun |
| EPI_ISL_960313 | University of Wisconsin-Madison AIDS Vaccine Research Laboratories | University of Wisconsin-Madison AIDS Vaccine Research Laboratories | Gage Moreno, Katarina Braun, et al. AIDS Vaccine Research Laboratories |
| EPI_ISL_960409, EPI_ISL_960411, EPI_ISL_960416, EPI_ISL_960426, EPI_ISL_960432 | The National Institute of Public Health | State Veterinary Institute Prague | Nagy,A;Vecerova,J;Cernikova,L;Stara,M;Jirincova,H;Trnka,D |
| EPI_ISL_960465 | Istituto Zooprofilattico Sperimentale del Mezzogiorno | TIGEM | Patrizia Annunziata, Andrea Ballabio, Valentina Bouche, Davide Cacchiarelli, Pellegrino Cerino, Chiara Colantuono, Maria Concetta Cuomo, Denise Di Concilio, Lucio Di Filippo, Antonio Grimaldi, Antonio Limone, Anna Manfredi, Francesco Panariello, Biancamaria Pierri, Marcello Salvi |
| EPI_ISL_960853, EPI_ISL_960868 | Institute of Medical Microbiology and Hospital Hygiene | Institute of Medical Microbiology and Hospital Hygiene | Prof. Dr. Achim Kaasch, Aljoscha Tersteegen |
| EPI_ISL_961127, EPI_ISL_961128 | Delaware Public Health Laboratory | Delaware Public Health Lab | Gregory Hovan |
| EPI_ISL_961223, EPI_ISL_961230 | Hospital General Universitario de Alicante - Instituto de Investigación Sanitaria y Biomédica de Alicante | SeqCOVID-SPAIN consortium/IBV(CSIC) | Maripaz Ventero Martín, Carmen Molina Pardines and SeqCOVID-SPAIN consortium |
| EPI_ISL_961470, EPI_ISL_961476, EPI_ISL_961493, EPI_ISL_961496, EPI_ISL_961498, EPI_ISL_961510, EPI_ISL_961512, EPI_ISL_961519, EPI_ISL_961525, EPI_ISL_961548, EPI_ISL_961549, EPI_ISL_961550, EPI_ISL_961551 | | | |
| see above | Michigan Department of Health and Human Services, Bureau of Laboratories | Michigan Department of Health and Human Services, Bureau of Laboratories | Blankenship HM, Riner D, Soehnlen MK |
| EPI_ISL_961560, EPI_ISL_961599, EPI_ISL_961600, EPI_ISL_961601, EPI_ISL_961602, EPI_ISL_961603, EPI_ISL_961649, EPI_ISL_961650, EPI_ISL_961651 | Hôpital Georges L. Dumont | National Microbiology Laboratory (NML) | Anna Majer, Shari Tyson, Grace Seo, Philip Mabon, Elsie Grudeski, Rhiannon Huzarewich, Russell Mandes, Anneliese Landgraff, Jennifer Tanner, Natalie Knox, Morag Graham, Gary Van Domselaar, Richard Garceau, Guillaume Desnoyers, Nathalie Bastien, Yan Li, Timothy Booth, Darian Hole, Madison Chapel, Kirsten Biggar, CanCOGeN's metadata curation team, Public Health Agency of Canada CanCOGeN team |
| EPI_ISL_961803 | E. Gulbja laboratorija | Latvian Biomedical Research and Study Centre | Janis Pjalkovskis, Nikita Zrelovs, Monta Ustinova, Ivars Silamikelis, Liga Birzniece, Kaspars Megnis, Vita Rovite, Lauma Freimane, Laila Silamikele, Laura Ansone, Davids Fridmanis, Mikus Gavars, Dmitrijs Perminovs, Jurijs Perevoscikovs, Uga Dumpis, Janis Klovins |
| EPI_ISL_961888, EPI_ISL_961889, EPI_ISL_961896, EPI_ISL_961900, EPI_ISL_961918, EPI_ISL_961968, EPI_ISL_961985, EPI_ISL_961990, EPI_ISL_961996, EPI_ISL_962138, EPI_ISL_962139, EPI_ISL_962141, EPI_ISL_962142, EPI_ISL_962143, EPI_ISL_962144, EPI_ISL_962145, EPI_ISL_962146, EPI_ISL_962147, EPI_ISL_962148, EPI_ISL_962149, EPI_ISL_962150, EPI_ISL_962151, EPI_ISL_962152, EPI_ISL_962153, EPI_ISL_962160 | | | |
| see above | Illinois Department of Public Health | Gagnon Lab, Southern Illinois University | Keith Gagnon |
| EPI_ISL_962510 | UCLA Clinical Micro Lab | Los Angeles County PHL | P. Hemarajata et al. |
| EPI_ISL_962585, EPI_ISL_962586, EPI_ISL_962587, EPI_ISL_962588, EPI_ISL_962589, EPI_ISL_962590, EPI_ISL_962591, EPI_ISL_962592, EPI_ISL_962593, EPI_ISL_962594, EPI_ISL_962595, EPI_ISL_962596, EPI_ISL_962597, EPI_ISL_962598, EPI_ISL_962599, EPI_ISL_962600, EPI_ISL_962601 | | | |
| see above | Scripps Medical Laboratory | Andersen lab at Scripps Research | SEARCH Alliance San Diego with Michael Quigley, Ellen Stefanski, Ian Mchardy |
| EPI_ISL_962904, EPI_ISL_962905 | Ostfold Hospital Trust - Kalnes, Centre for Laboratory Medicine, Section for gene technology and infection serology | Norwegian Institute of Public Health, Department of Virology | Kathrine Stene-Johansen, Kamilla Heddeland Instefjord, Hilde Elshaug, Ignacio Garcia Llorente, Serina B Engebretsen, Atiya R Ali,Marie Paulsen Madsen, Rasmus Riis Kopperud, Hilde Vollan, Karoline Bragstad, Olav Hungnes |
| EPI_ISL_962919, EPI_ISL_962920, EPI_ISL_962921 | Servicio de Microbiología, Laboratori Clínic Metropolitana Nord. Hospital Universitari Germans Trias i Pujol. Institut d'Investigació en Ciències de la Salut Germans Trias i Pujol (IGTP) | SeqCOVID-SPAIN consortium/IBV(CSIC) | Elisa Martró, Antoni E. Bordoy, Anna Not, Adrián Antuori, Anabel Fernández, Nona Romaní, Verónica Saludes, Cristina Casañ and SeqCOVID-SPAIN consortium |
| EPI_ISL_962935 | Hospital Universitario de Gran Canaria Dr. Negrín | SeqCOVID-SPAIN consortium/IBV(CSIC) | M. Carmen Pérez González, Francisco J. Chamizo López, Ana Bordes Benítez and SeqCOVID-SPAIN consortium |
| EPI_ISL_963106, EPI_ISL_963980, EPI_ISL_964107, EPI_ISL_964219, EPI_ISL_964234 | Lighthouse Lab in Alderley Park | Wellcome Sanger Institute for the COVID-19 Genomics UK (COG-UK) Consortium | Jacquelyn Wynn, Mairead Hyland, The Lighthouse Lab in Alderley Park and Alex Alderton, Roberto Amato, Sonia Goncalves, Ewan Harrison, David K. Jackson, Ian Johnston, Dominic Kwiatkowski, Cordelia Langford, John Sillitoe on behalf of the Wellcome Sanger Institute COVID-19 Surveillance Team |
| EPI_ISL_964281 | Oslo University Hospital, Department of Medical Microbiology | Norwegian Institute of Public Health, Department of Virology | Kathrine Stene-Johansen, Kamilla Heddeland Instefjord, Hilde Elshaug, Ignacio Garcia Llorente, Serina B Engebretsen, Atiya R Ali, Marie Paulsen Madsen, Rasmus Riis Kopperud, Hilde Vollan, Karoline Bragstad, Olav Hungnes |
| EPI_ISL_964303 | Akershus University Hospital, Department for Microbiology and Infectious Disease Control | Norwegian Institute of Public Health, Department of Virology | Kathrine Stene-Johansen, Kamilla Heddeland Instefjord, Hilde Elshaug, Ignacio Garcia Llorente, Serina B Engebretsen, Atiya R Ali, Marie Paulsen Madsen, Rasmus Riis Kopperud, Hilde Vollan, Karoline Bragstad, Olav Hungnes |
| EPI_ISL_965191, EPI_ISL_965192, EPI_ISL_965193, EPI_ISL_965194, EPI_ISL_965195, EPI_ISL_965196, EPI_ISL_965197 | Virginia Division of Consolidated Laboratory Services | Virginia Division of Consolidated Laboratory Services | Virginia DCLS |
| EPI_ISL_965579, EPI_ISL_965585, EPI_ISL_965614, EPI_ISL_965615, EPI_ISL_965625, EPI_ISL_965632, EPI_ISL_965640, EPI_ISL_965668, EPI_ISL_965676, EPI_ISL_965686, EPI_ISL_965752, EPI_ISL_965765, EPI_ISL_965795, EPI_ISL_965810, EPI_ISL_965811 | | | |
| see above | Dutch COVID-19 response team | Medical Microbiology, Maastricht University Medical Centre | Jozef Dingemans*, Brian van der Veer*, Erik Beuken, Carmen Reumkens, Lieke van Alphen, Christian Hoebe, Paul Savelkoul |
| EPI_ISL_966777, EPI_ISL_966778, EPI_ISL_966779, EPI_ISL_966780, EPI_ISL_966781 | Maine HETL | Tewhey Lab, The Jackson Laboratory | Matluk,N., Dewey,H., Iosue,F., Barter,M., Lynch,R., Munger,H. and Tewhey,R. |
| EPI_ISL_967589, EPI_ISL_967638, EPI_ISL_967653, EPI_ISL_967654, EPI_ISL_967666, EPI_ISL_967667, EPI_ISL_967668, EPI_ISL_967669, EPI_ISL_967670, EPI_ISL_967673, EPI_ISL_967674, EPI_ISL_967682, EPI_ISL_967694, EPI_ISL_967695, EPI_ISL_967709, EPI_ISL_967711, EPI_ISL_967712, EPI_ISL_967734, EPI_ISL_967735, EPI_ISL_967737, EPI_ISL_967738, EPI_ISL_967739, EPI_ISL_967740, EPI_ISL_967741, EPI_ISL_967742, EPI_ISL_967763, EPI_ISL_967765 | | | |
| see above | State Laboratories Division, Hawaii State Department of Health | State Laboratories Division, Hawaii State Department of Health | Pamela O'Brien, Drew Kuwazaki, Ayana Garnet, Razvan Sultana, Edward Desmond |
| EPI_ISL_968860, EPI_ISL_968861, EPI_ISL_968863, EPI_ISL_968864, EPI_ISL_968865, EPI_ISL_968868, EPI_ISL_968869, EPI_ISL_968871, EPI_ISL_968873, EPI_ISL_968877 | KEMRI-Wellcome Trust Research Programme/KEMRI-CGMR-C Kilifi | KEMRI-Wellcome Trust Research Programme/KEMRI-CGMR-C Kilifi | Githinji et al |
| EPI_ISL_976945, EPI_ISL_976953, EPI_ISL_976961, EPI_ISL_976969, EPI_ISL_976977, EPI_ISL_976982, EPI_ISL_976986, EPI_ISL_976994, EPI_ISL_977001, EPI_ISL_977008, EPI_ISL_977014 | | | |
| see above | Broad Institute Clinical Research Sequencing Platform | Infectious Disease Program, Broad Institute of Harvard and MIT | Lemieux,J.E., Siddle,K.J., Adams,G., Gladden-Young,A., Lagerborg,K., Rudy,M., DeRuff,K., Carter,A., Normandin,E., Bauer,M., Reilly,S., Tomkins-Tinch,C., Loreth,C., Chaluvadi,S., Birren,B.W., Gallagher,G., Smole,S., Park,D.J., MacInnis,B.L., and Sabeti,P.C. |
| EPI_ISL_977130 | Flow Health | Infectious Disease Program, Broad Institute of Harvard and MIT | Lemieux,J.E., Siddle,K.J., Adams,G., Gladden-Young,A., Lagerborg,K., Rudy,M., DeRuff,K., Carter,A., Normandin,E., Bauer,M., Reilly,S., Tomkins-Tinch,C., Loreth,C., Chaluvadi,S., Birren,B.W., Gallagher,G., Smole,S., Park,D.J., MacInnis,B.L., and Sabeti,P.C. |
| EPI_ISL_977158 | ULSS 03 Venezia | Istituto Zooprofilattico Sperimentale delle Venezie | Adelaide Milani, Alessia Schivo, Annalisa Salviato, Erika Giorgia Quaranta, Ambra Pastori, Bianca Zecchin, Alice Fusaro, Isabella Monne, Calogero Terregino, Antonia Ricci |
| EPI_ISL_977544, EPI_ISL_977549, | Nigeria Centre of Disease Control (NCDC) | African Centre of Excellence for Genomics of Infectious | Olawoye I. B. et al |

| | | | |
|---|---|---|---|
| EPI_ISL_977555, EPI_ISL_977556, EPI_ISL_977560, EPI_ISL_977561, EPI_ISL_977562, EPI_ISL_977564 | | Diseases (ACEGID), Redeemer's University | |
| EPI_ISL_977661, EPI_ISL_977662, EPI_ISL_977663 | Labo analyses med | National Reference Center for Viruses of Respiratory Infections, Institut Pasteur, Paris | Marion Barbet, Sylvie Behillil, Méline Bizard, Angela Brisebarre, Camille Capel, Etienne Simon-Lorière, Vincent Enouf, Maud Vanpeene, Sylvie van der Werf,Amzalag Jonas |
| EPI_ISL_978132, EPI_ISL_978133, EPI_ISL_978134, EPI_ISL_978135, EPI_ISL_978136, EPI_ISL_978137, EPI_ISL_978138, EPI_ISL_978139, EPI_ISL_978140, EPI_ISL_978141, EPI_ISL_978142, EPI_ISL_978143, EPI_ISL_978144, EPI_ISL_978145, EPI_ISL_978146, EPI_ISL_978147, EPI_ISL_978148, EPI_ISL_978149, EPI_ISL_978150, EPI_ISL_978151 | | | |
| see above | Chiu Laboratory, University of California, San Francisco | Chiu Laboratory, University of California, San Francisco | Charles Chiu, Xianding (Wayne) Deng, Candace Wang, Venice Servellita, Jill Hacker, Debra Wadford |
| EPI_ISL_978371, EPI_ISL_978372, EPI_ISL_978397, EPI_ISL_978398, EPI_ISL_978399, EPI_ISL_978400, EPI_ISL_978401, EPI_ISL_978402, EPI_ISL_978403, EPI_ISL_978404, EPI_ISL_978405, EPI_ISL_978406, EPI_ISL_978407, EPI_ISL_978408 | | | |
| see above | Arizona State Public Health Laboratory | Arizona State Public Health Laboratory | Trung Huynh, Jessica Escobar, Katherine Fullerton, Nobuko Fukushima, Stacy White, Linda Getsinger, Victor Waddell |
| EPI_ISL_978741, EPI_ISL_978742, EPI_ISL_978743, EPI_ISL_978744, EPI_ISL_978745, EPI_ISL_978746, EPI_ISL_978747, EPI_ISL_978748, EPI_ISL_978749, EPI_ISL_978750, EPI_ISL_978751, EPI_ISL_978752, EPI_ISL_978753, EPI_ISL_978754, EPI_ISL_978755, EPI_ISL_978756, EPI_ISL_978757, EPI_ISL_978758, EPI_ISL_978759, EPI_ISL_978760, EPI_ISL_978761, EPI_ISL_978762, EPI_ISL_978763, EPI_ISL_978764, EPI_ISL_978765, EPI_ISL_978766, EPI_ISL_978767, EPI_ISL_978768, EPI_ISL_978769, EPI_ISL_978770, EPI_ISL_978771, EPI_ISL_978772, EPI_ISL_978773, EPI_ISL_978774, EPI_ISL_978775, EPI_ISL_978776, EPI_ISL_978777, EPI_ISL_978778, EPI_ISL_978779, EPI_ISL_978780, EPI_ISL_978781, EPI_ISL_978782, EPI_ISL_978783, EPI_ISL_978784, EPI_ISL_978785, EPI_ISL_978786, EPI_ISL_978787 | | | |
| see above | Helix/Illumina | Respiratory Viruses Branch, Division of Viral Diseases, Centers for Disease Control and Prevention | Peter W. Cook,Dakota Howard,Dhwani Batra,Ben L. Rambo-Martin,Eileen de Feo,Jan Antico,Christine Tran,Matthew Tolentino,Shannon Wickline,Kim Gietzen,Brad Sickler,Jingtao Liu,Eric Allen,Phil Febbo,Summer Galloway,Nicole L. Washington,Simon White,Geraint Levan,Kelly Schiabor Barrett,Elizabeth Cirulli,Alexandre Bolze,Ary Ascencio,Charlotte Rivera-Garcia,Ryan Cho,Jason Nguyen,Sherry Wang,Jimmy Ramirez,Tyler Cassens,Efren Sandoval,Magnus Isaksson,William Lee,David Becker,Marc Laurent,James Lu,Clinton R. Paden,Suxiang Tong,Duncan MacCannell, |
| EPI_ISL_978924 | Centre for Dengue Research and AICBU, Department of Immunology and Molecular Medicine | Centre for Dengue Research and AICBU, Department of Immunology and Molecular Medicine | Chandima Jeewandara, Deshni Jayathilaka, Dinuka Ariyaratne, Tibutius Thanesh Pramanayagam, Diyanath Ranasinghe, Laksiri Gomes, Gathsaurie Neelika Malavige |
| EPI_ISL_978931, EPI_ISL_978933, EPI_ISL_978935, EPI_ISL_978936 | Caribbean Public Health Agency | Carrington Lab, Department of PreClinical Sciences, Building 36, First Floor Biochemistry Unit, Faculty of Medical Sciences, The University of the West Indies | Nikita S. D. Sahadeo, Arianne Brown-Jordan, Vernie Ramkissoon, Sarah Hill, Naresh Nandram, Avery Hinds, Irad Potter, Jerome Foster, Stanley Giddings, Karla Georges, Marsha Ivey, Rahul Naidu, Risha Singh, SueMin Nathaniel, Rajini Haraksingh, Jaya Jayaraman, Chinna Chinnadurai, Adesh Ramsubhag, Nuno Faria, Oliver Pybus, Christopher Oura, Gabriel Escobar, Christine V. F. Carrington |
| EPI_ISL_978962, EPI_ISL_978963, EPI_ISL_978964, EPI_ISL_978965, EPI_ISL_978966 | Chiu Laboratory, University of California, San Francisco | Chiu Laboratory, University of California, San Francisco | Charles Chiu, Xianding (Wayne) Deng, Candace Wang, Venice Servellita, Jill Hacker, Debra Wadford |
| EPI_ISL_979137 | Santa Clara County Public Health Laboratory | Chan-Zuckerberg Biohub | CZB Cliahub Consortium |
| EPI_ISL_979159 | County of San Luis Obispo Public Health Laboratory | Chan-Zuckerberg Biohub | CZB Cliahub Consortium |
| EPI_ISL_979454, EPI_ISL_979456 | University of Iowa Hospitals and Clinics | State Hygienic Laboratory at the University of Iowa | Valerie Reeb, Wes Hottel, Alankar Kampoowale |
| EPI_ISL_981136 | Johns Hopkins Hospital Department of Pathology | Johns Hopkins Hospital Department of Pathology | C. Paul Morris, Chun Huai Luo, Adannaya Amadi, Matthew Schwartz, Nicholas Gallagher, Heba H. Mostafa |
| EPI_ISL_981370 | AZ Klina | AZ Klina | Dr. C. Vael |
| EPI_ISL_981946 | Microbiology Service, Hospital Universitario Clinico San Cecilio, Granada | Microbiology Service, Hospital Universitario Clinico San Cecilio, Granada | Adolfo de Salazar, Natalia Chueca, Laura Viñuela, Ana Fuentes, Federico García |
| EPI_ISL_981976, EPI_ISL_981980, EPI_ISL_981984, EPI_ISL_981986, EPI_ISL_981991, EPI_ISL_981993, EPI_ISL_981994 | TGen North | Sonora Quest Laboratories | "Jolene Bowers, Megan Folkerts, Chris French, Hayley Yaglom, Ashlyn Pfeiffer, Darrin Lemmer, Dave Engelthaler, The Arizona COVID Genomics Union (ACGU)" |
| EPI_ISL_982008 | TGen North | TGen North | "Jolene Bowers, Megan Folkerts, Chris French, Hayley Yaglom, Ashlyn Pfeiffer, Darrin Lemmer, Dave Engelthaler, The Arizona COVID Genomics Union (ACGU)" |
| EPI_ISL_982009 | TGen North | Sonora Quest Laboratories | "Jolene Bowers, Megan Folkerts, Chris French, Hayley Yaglom, Ashlyn Pfeiffer, Darrin Lemmer, Dave Engelthaler, The Arizona COVID Genomics Union (ACGU)" |
| EPI_ISL_982010, EPI_ISL_982020 | TGen North | TGen North | "Jolene Bowers, Megan Folkerts, Chris French, Hayley Yaglom, Ashlyn Pfeiffer, Darrin Lemmer, Dave Engelthaler, The Arizona COVID Genomics Union (ACGU)" |
| EPI_ISL_982025, EPI_ISL_982028, EPI_ISL_982030, EPI_ISL_982031, EPI_ISL_982033, EPI_ISL_982036, EPI_ISL_982037, EPI_ISL_982039, EPI_ISL_982040, EPI_ISL_982042, EPI_ISL_982051, EPI_ISL_982054, EPI_ISL_982058, EPI_ISL_982062, EPI_ISL_982069, EPI_ISL_982076, EPI_ISL_982091, EPI_ISL_982095, EPI_ISL_982096, EPI_ISL_982098, EPI_ISL_982101 | | | |
| see above | TGen North | Sonora Quest Laboratories | "Jolene Bowers, Megan Folkerts, Chris French, Hayley Yaglom, Ashlyn Pfeiffer, Darrin Lemmer, Dave Engelthaler, The Arizona COVID Genomics Union (ACGU)" |
| EPI_ISL_982105 | TGen North | TGen North | "Jolene Bowers, Megan Folkerts, Chris French, Hayley Yaglom, Ashlyn Pfeiffer, Darrin Lemmer, Dave Engelthaler, The Arizona COVID Genomics Union (ACGU)" |
| EPI_ISL_982240, EPI_ISL_982242 | Lab voor klinische biologie | Lab voor klinische biologie | Hannelore Hamerlinck, Marija Janevska, Bruno Verhasselt |
| EPI_ISL_982425, EPI_ISL_982436, EPI_ISL_982441, EPI_ISL_982449, EPI_ISL_982450, EPI_ISL_982453, EPI_ISL_982457, EPI_ISL_982460, EPI_ISL_982463, EPI_ISL_982464, EPI_ISL_982466, EPI_ISL_982471, EPI_ISL_982473, EPI_ISL_982476, EPI_ISL_982477, EPI_ISL_982479, EPI_ISL_982490, EPI_ISL_982493, EPI_ISL_982494 | | | |
| see above | MONTEFIORE MEDICAL CENTER LABORATORIES | Wadsworth Center, New York State Department of Health | Kirsten St. George, Daryl M. Lamson, Alexis Russel, Matthew Shudt, Melissa A Leisner, Jonathan Plitnick, Navjot Singh, John Kelly, Erasmus Schneider, Erica Lasek-Nesselquist |
| EPI_ISL_982527, EPI_ISL_982528, EPI_ISL_982531, EPI_ISL_982532, EPI_ISL_982533, EPI_ISL_982536, EPI_ISL_982543, EPI_ISL_982549, EPI_ISL_982554, EPI_ISL_982556, EPI_ISL_982559, EPI_ISL_982562, EPI_ISL_982564, EPI_ISL_982565, EPI_ISL_982570, EPI_ISL_982572, EPI_ISL_982577, EPI_ISL_982580, EPI_ISL_982581, EPI_ISL_982582, EPI_ISL_982584, EPI_ISL_982587, EPI_ISL_982588, EPI_ISL_982590, EPI_ISL_982595, EPI_ISL_982598, EPI_ISL_982599, EPI_ISL_982604, EPI_ISL_982607, EPI_ISL_982610, EPI_ISL_982612, EPI_ISL_982614, EPI_ISL_982618, EPI_ISL_982619, EPI_ISL_982620, EPI_ISL_982621, EPI_ISL_982622, EPI_ISL_982631, EPI_ISL_982632, EPI_ISL_982633, EPI_ISL_982636, EPI_ISL_982638, EPI_ISL_982639, EPI_ISL_982640, EPI_ISL_982644, EPI_ISL_982645, EPI_ISL_982646, EPI_ISL_982648, EPI_ISL_982649, EPI_ISL_982651, EPI_ISL_982654, EPI_ISL_982655, EPI_ISL_982656, EPI_ISL_982657, EPI_ISL_982658, EPI_ISL_982659, EPI_ISL_982660, EPI_ISL_982671, EPI_ISL_982681, EPI_ISL_982682, EPI_ISL_982685, EPI_ISL_982686, EPI_ISL_982687, EPI_ISL_982688, EPI_ISL_982689, EPI_ISL_982690, EPI_ISL_982694, EPI_ISL_982702, EPI_ISL_982703, EPI_ISL_982704, EPI_ISL_982705, EPI_ISL_982710, EPI_ISL_982711, EPI_ISL_982712, EPI_ISL_982713, EPI_ISL_982714, EPI_ISL_982715, EPI_ISL_982716, EPI_ISL_982717, EPI_ISL_982718, EPI_ISL_982729, EPI_ISL_982737, EPI_ISL_982740, EPI_ISL_982743, EPI_ISL_982744, EPI_ISL_982745, EPI_ISL_982751, EPI_ISL_982752, EPI_ISL_982753, EPI_ISL_982754, EPI_ISL_982755, EPI_ISL_982756, EPI_ISL_982757 | | | |
| see above | US Air Force School of Aerospace Medicine | US Air Force School of Aerospace Medicine | Anthony Fries, Jennifer Meyer, William Gruner, William Buggele, Amanda Javorina, Sarah Purves, Clarise Starr, Elizabeth Macias |
| EPI_ISL_983066, EPI_ISL_983067, EPI_ISL_983068, EPI_ISL_983069, EPI_ISL_983070, EPI_ISL_983071, EPI_ISL_983072 | Gravity Diagnostics | Kentucky State Public Health Lab | Stephanie Lunn, Karim George, Joshua Tobias, William Grooms, Vaneet Arora, Matthew Johnson, Rachel Zinner, Rhonda Lucas |
| EPI_ISL_983073, EPI_ISL_983074 | Kentucky State Public Health Lab | Kentucky State Public Health Lab | Stephanie Lunn, Karim George, Joshua Tobias, William Grooms, Vaneet Arora, Matthew Johnson, Rachel Zinner, Rhonda Lucas |
| EPI_ISL_983075, EPI_ISL_983076, EPI_ISL_983077, EPI_ISL_983078, EPI_ISL_983079, EPI_ISL_983080 | Gravity Diagnostics | Kentucky State Public Health Lab | Stephanie Lunn, Karim George, Joshua Tobias, William Grooms, Vaneet Arora, Matthew Johnson, Rachel Zinner, Rhonda Lucas |
| EPI_ISL_983100, EPI_ISL_983107, EPI_ISL_983109, EPI_ISL_983110, EPI_ISL_983114, EPI_ISL_983136, EPI_ISL_983139, EPI_ISL_983143, EPI_ISL_983145, EPI_ISL_983148, EPI_ISL_983151, EPI_ISL_983153, EPI_ISL_983155, EPI_ISL_983158, EPI_ISL_983159, EPI_ISL_983160, EPI_ISL_983161, EPI_ISL_983165 | | | |
| see above | MONTEFIORE MEDICAL CENTER LABORATORIES | Wadsworth Center, New York State Department of Health | Kirsten St. George, Daryl M. Lamson, Alexis Russel, Matthew Shudt, Melissa A Leisner, Jonathan Plitnick, Navjot Singh, John Kelly, Erasmus Schneider, Erica Lasek-Nesselquist |

| | | | |
|---|---|---|---|
| EPI_ISL_983240, EPI_ISL_983241, EPI_ISL_983242 | KALEIDA CENTER FOR LABORATORY MEDICINE | Wadsworth Center, New York State Department of Health | Kirsten St. George, Daryl M. Lamson, Alexis Russel, Matthew Shudt, Melissa A Leisner, Jonathan Plitnick, Navjot Singh, John Kelly, Erasmus Schneider, Erica Lasek-Nesselquist |
| EPI_ISL_983243 | SUNY UPSTATE MEDICAL UNIVERSITY | Wadsworth Center, New York State Department of Health | Kirsten St. George, Daryl M. Lamson, Alexis Russel, Matthew Shudt, Melissa A Leisner, Jonathan Plitnick, Navjot Singh, John Kelly, Erasmus Schneider, Erica Lasek-Nesselquist |
| EPI_ISL_983244, EPI_ISL_983246 | KALEIDA CENTER FOR LABORATORY MEDICINE | Wadsworth Center, New York State Department of Health | Kirsten St. George, Daryl M. Lamson, Alexis Russel, Matthew Shudt, Melissa A Leisner, Jonathan Plitnick, Navjot Singh, John Kelly, Erasmus Schneider, Erica Lasek-Nesselquist |
| EPI_ISL_983249 | SUNY UPSTATE MEDICAL UNIVERSITY | Wadsworth Center, New York State Department of Health | Kirsten St. George, Daryl M. Lamson, Alexis Russel, Matthew Shudt, Melissa A Leisner, Jonathan Plitnick, Navjot Singh, John Kelly, Erasmus Schneider, Erica Lasek-Nesselquist |
| EPI_ISL_983250, EPI_ISL_983263, EPI_ISL_983264, EPI_ISL_983265, EPI_ISL_983266, EPI_ISL_983267, EPI_ISL_983268, EPI_ISL_983269, EPI_ISL_983270, EPI_ISL_983271, EPI_ISL_983272, EPI_ISL_983273, EPI_ISL_983274, EPI_ISL_983275, EPI_ISL_983276, EPI_ISL_983277, EPI_ISL_983278, EPI_ISL_983279, EPI_ISL_983280, EPI_ISL_983281, EPI_ISL_983282 | | | |
| see above | KALEIDA CENTER FOR LABORATORY MEDICINE | Wadsworth Center, New York State Department of Health | Kirsten St. George, Daryl M. Lamson, Alexis Russel, Matthew Shudt, Melissa A Leisner, Jonathan Plitnick, Navjot Singh, John Kelly, Erasmus Schneider, Erica Lasek-Nesselquist |
| EPI_ISL_983283 | SUNY UPSTATE MEDICAL UNIVERSITY | Wadsworth Center, New York State Department of Health | Kirsten St. George, Daryl M. Lamson, Alexis Russel, Matthew Shudt, Melissa A Leisner, Jonathan Plitnick, Navjot Singh, John Kelly, Erasmus Schneider, Erica Lasek-Nesselquist |
| EPI_ISL_983284, EPI_ISL_983285, EPI_ISL_983286, EPI_ISL_983287, EPI_ISL_983288 | KALEIDA CENTER FOR LABORATORY MEDICINE | Wadsworth Center, New York State Department of Health | Kirsten St. George, Daryl M. Lamson, Alexis Russel, Matthew Shudt, Melissa A Leisner, Jonathan Plitnick, Navjot Singh, John Kelly, Erasmus Schneider, Erica Lasek-Nesselquist |
| EPI_ISL_983289, EPI_ISL_983290, EPI_ISL_983291, EPI_ISL_983292, EPI_ISL_983293, EPI_ISL_983294, EPI_ISL_983295 | SUNY UPSTATE MEDICAL UNIVERSITY | Wadsworth Center, New York State Department of Health | Kirsten St. George, Daryl M. Lamson, Alexis Russel, Matthew Shudt, Melissa A Leisner, Jonathan Plitnick, Navjot Singh, John Kelly, Erasmus Schneider, Erica Lasek-Nesselquist |
| EPI_ISL_983324 | INMI Lazzaro Spallanzani IRCCS | INMI Lazzaro Spallanzani IRCCS | B Bartolini, O Butera, C.E.M Gruber, M Rueca, F Messina, E Giombini, MR Capobianchi, A Di Caro |
| EPI_ISL_983382, EPI_ISL_983383, EPI_ISL_983384, EPI_ISL_983385, EPI_ISL_983386, EPI_ISL_983387, EPI_ISL_983388, EPI_ISL_983389, EPI_ISL_983390, EPI_ISL_983391, EPI_ISL_983392, EPI_ISL_983393, EPI_ISL_983394, EPI_ISL_983395, EPI_ISL_983396, EPI_ISL_983397, EPI_ISL_983398, EPI_ISL_983399, EPI_ISL_983400, EPI_ISL_983401, EPI_ISL_983402, EPI_ISL_983403, EPI_ISL_983404, EPI_ISL_983405, EPI_ISL_983406, EPI_ISL_983407 | | | |
| see above | Utah Public Health Laboratory | Utah Public Health Laboratory | Erin L. Young, Kelly F. Oakeson, Tara Gallagher |
| EPI_ISL_983720, EPI_ISL_983735, EPI_ISL_983813, EPI_ISL_983814, EPI_ISL_983815, EPI_ISL_983816, EPI_ISL_983817, EPI_ISL_983818, EPI_ISL_983819, EPI_ISL_983820, EPI_ISL_983821, EPI_ISL_983822, EPI_ISL_983823, EPI_ISL_983824, EPI_ISL_983825 | | | |
| see above | Colorado Department of Public Health and Environment | Colorado Department of Puplic Health and Environment | Laura Bankers, Molly C. Hetherington-Rauth, Diana Ir, Shannon Ely, Shannon R. Matzinger, Sarah Elizabeth Totten, Emily A. Travanty |
