## Supplementary material for "Detection and characterization of the SARS-CoV-2 lineage B.1.526 in New York": Supp. Table 3 GISAID Acknowledgment part 2: gisaid_hcov-19_acknowledgement_table_2021_02_13_010-9.pdf

We gratefully acknowledge the following Authors from the Originating laboratories responsible for obtaining the specimens, as well as the Submitting laboratories where the genome data were generated and shared via GISAID, on which this research is based.

All Submitters of data may be contacted directly via www.gisaid.org

Authors are sorted alphabetically.

| Accession ID | Originating Laboratory | Submitting Laboratory | Authors |
|---|---|---|---|
| EPI_ISL_803108 | Pathology West - NSW Health Pathology | NSW Health Pathology - Institute of Clinical Pathology and Medical Research; Westmead Hospital; University of Sydney | CIDM-PH et al. |
| EPI_ISL_803110, EPI_ISL_803111, EPI_ISL_803112, EPI_ISL_803113, EPI_ISL_803114 | Sydney South West Pathology Service (SSWPS) - Royal Prince Alfred Hospital - NSW Health Pathology | NSW Health Pathology - Institute of Clinical Pathology and Medical Research; Westmead Hospital; University of Sydney | CIDM-PH et al. |
| EPI_ISL_803119, EPI_ISL_803851 | Laboratory of Microbiology, National Reference Lab, Charles Nicolle Hospital; 2-University of Tunis ElManar, Faculty of Medicine of Tunis, LR99ES09, Tunis, Tunisia | Clinical and Experimental Pharmacology Lab, LR16SP02, National Center of Pharmacovigilance, University of Tunis El Manar, Tunis, Tunisia. 2-Neurodegenerative diseases and psychiatric troubles, LR18SP03, Razi Hospital, University of Tunis El Manar, Tunis, Tunisia. 3- Ministry of Health, National Observatory of New and Emerging Diseases, 1006, Tunis, Tunisia | Ilhem Boutiba-Ben Boubaker, Sameh Trabelsi, Nissaf Ben Alaya, Maher Kharrat, Alia BenKahla, Jalila Ben Khelil, Salma Abid, Sana Ferjani, Mouna Ben Sassi, Mouna Safer, Zaineb Hamzaoui, Habiba Ben Romdhane, Souissi Amira, Sarra Chamman, Hanen El Jebari, Ahmed Fakhfakh, Gaies Emna, Riadh Daghfous, Riadh Gouider. |
| EPI_ISL_810968, EPI_ISL_810969 | PathWest Laboratory Medicine WA | PathWest Laboratory Medicine WA Microbial Surveillance Unit | PathWest Laboratory Medicine WA Microbial Surveillance Unit |
| EPI_ISL_811123, EPI_ISL_811124, EPI_ISL_811125, EPI_ISL_811126, EPI_ISL_811150 | Respiratory Virus Unit, National Infection Service, Public Health England | COVID-19 Genomics UK (COG-UK) Consortium | PHE Covid Sequencing Team |
| EPI_ISL_812256 | University Hospital Zürich | Institute of Medical Virology, University of Zurich | Stefan Schmutz, Maryam Zaheri, Verena Kufner, Annette Audigé, Maria Grünberg, Kevin Steiner, Jon Huder, Cyril Shah, Riccarda Capaul, Guido Bloemberg, Jürg Böni, Michael Huber, Alexandra Trkola |
| EPI_ISL_812346 | Oklahoma Animal Disease Diagnostic Laboratory | Oklahoma Animal Disease Diagnostic Laboratory | Sai Narayanan, Girish Patil, Sunil More, Jeremiah Saliki, Akhilesh Ramachandran |
| EPI_ISL_812347 | Area of Virology, Serology and Virology Division (SAViD), New South Wales Health Pathology Randwick | Virology Research Laboratory; Area of Virology, Serology and Virology Division (SAViD), New South Wales Health Pathology Randwick | Foster, C.; Au, J.; Ruiz Silva, M.; Deveson, I.; Bull, R.; Van Hal, S.; Rawlinson, W. |
| EPI_ISL_812358 | SA Pathology | SA Pathology | Lex Leong, Julien Soubrier, Chuan Kok Lim, Song Gao, Mark Turra, Karin Kassahn, Ivan Bastian, Geoff Higgins |
| EPI_ISL_812441, EPI_ISL_812442 | Victorian Infectious Diseases Reference Laboratory (VIDRL) | VIDRL and MDU-PHL | Caly L., Seemann T., Sait, M.L., Druce J., Sherry, N.L. |
| EPI_ISL_812521 | Pathology North - Royal North Shore Hospital - NSW Health Pathology | NSW Health Pathology - Institute of Clinical Pathology and Medical Research; Westmead Hospital; University of Sydney | CIDM-PH et al. |
| EPI_ISL_825016 | Sydney South West Pathology Service (SSWPS) - Royal Prince Alfred Hospital - NSW Health Pathology | NSW Health Pathology - Institute of Clinical Pathology and Medical Research; Westmead Hospital; University of Sydney | CIDM-PH et al. |
| EPI_ISL_825064, EPI_ISL_825065, EPI_ISL_825066, EPI_ISL_825067, EPI_ISL_825068, EPI_ISL_825070, EPI_ISL_825071, EPI_ISL_825072, EPI_ISL_825073, EPI_ISL_825075, EPI_ISL_825077, EPI_ISL_825078, EPI_ISL_825079, EPI_ISL_825081, EPI_ISL_825082, EPI_ISL_825083 | | | |
| see above | National Public Health Laboratory, National Centre for Infectious Diseases | National Public Health Laboratory, National Centre for Infectious Diseases | Tze Minn Mak, Sophie Octavia, Zhenyang Zhou, Lin Cui, Raymond Tzer Pin Lin |
| EPI_ISL_825491, EPI_ISL_825616 | Respiratory Virus Unit, National Infection Service, Public Health England | COVID-19 Genomics UK (COG-UK) Consortium | PHE Covid Sequencing Team |
| EPI_ISL_826270, EPI_ISL_826271, EPI_ISL_826272, EPI_ISL_826273 | University of Debrecen, Department of Medical Microbiology | National Laboratory of Virology, Szentágothai Research Centre | Endre Gábor Tóth, Balázs Somogyi, Brigitta Zana, Eszter Csoma, Ferenc Jakab, Gábor Kemenesi |
| EPI_ISL_826282 | Respiratory Virus Unit, National Infection Service, Public Health England | COVID-19 Genomics UK (COG-UK) Consortium | PHE Covid Sequencing Team |
| EPI_ISL_826297, EPI_ISL_826298, EPI_ISL_826299, EPI_ISL_826300, EPI_ISL_826301, EPI_ISL_826302, EPI_ISL_826303, EPI_ISL_826305, EPI_ISL_826307, EPI_ISL_826308, EPI_ISL_826309, EPI_ISL_826310, EPI_ISL_826311, EPI_ISL_826312, EPI_ISL_826313, EPI_ISL_826314, EPI_ISL_826315, EPI_ISL_826316, EPI_ISL_826317, EPI_ISL_826318 | | | |
| see above | Wyoming Public Health Laboratory | Wyoming Public Health Laboratory | Noah Hull, Taylor Fearing, Lynette Gumbleton, Channing Weber, Ashley Norberg, Bailey Bowcutt, and Wanda Manley |
| EPI_ISL_830643, EPI_ISL_830644, EPI_ISL_830645, EPI_ISL_830646, EPI_ISL_830647 | Wadsworth Center, New York State Department of Health | Wadsworth Center, New York State Department of Health | Kirsten St. George, Daryl M. Lamson, Alexis Russel, Matthew Shudt, Melissa A Leisner, Jonathan Plitnick, Navjot Singh, John Kelly, Erasmus Schneider, Erica Lasek-Nesselquist |
| EPI_ISL_831361, EPI_ISL_831362, EPI_ISL_831363, EPI_ISL_831364, EPI_ISL_831365, EPI_ISL_831368, EPI_ISL_831369 | Clinical Molecular Microbiology Laboratory, UNC Hospitals | Jeremy Wang | Jeremy Wang, Alexander Rubinsteyn, Colleen Rice, Jason Smedberg, Melissa Miller, Corbin Jones, Robert Hagan |
| EPI_ISL_831476, EPI_ISL_831478, EPI_ISL_831484, EPI_ISL_831489, EPI_ISL_831495, EPI_ISL_831504, EPI_ISL_831505, EPI_ISL_831506, EPI_ISL_831507, EPI_ISL_831508, EPI_ISL_831509, EPI_ISL_831510, EPI_ISL_831512, EPI_ISL_831513, EPI_ISL_831514, EPI_ISL_831515, EPI_ISL_831516, EPI_ISL_831517, EPI_ISL_831518, EPI_ISL_831519, EPI_ISL_831520, EPI_ISL_831522, EPI_ISL_831523, EPI_ISL_831524, EPI_ISL_831525, EPI_ISL_831526, EPI_ISL_831528, EPI_ISL_831531, EPI_ISL_831532, EPI_ISL_831535, EPI_ISL_831537, EPI_ISL_831541, EPI_ISL_831547, EPI_ISL_831549 | | | |
| see above | University of Wisconsin-Madison AIDS Vaccine Research Laboratories | University of Wisconsin-Madison AIDS Vaccine Research Laboratories | Gage Moreno, Katarina Braun, et al. AIDS Vaccine Research Laboratories |
| EPI_ISL_831674 | Institute of Virology, Biomedical Research Center of the Slovak Academy of Sciences, Bratislava | Faculty of Natural Sciences, Comenius University, Bratislava | Kristína Boršová, Viktória abanová, Broa Brejová, Viktória Hodorová, Sabina Fumaová Havlíková, Juraj Kopáek, Martina Liková, ubomíra Lukáiková, Martina Neboháová, Monika Sláviková, Tomáš Vina, Boris Klempa, Jozef Nosek |
| EPI_ISL_831675 | Institute of Virology, Biomedical Research Center of the Slovak Academy of Sciences, Bratislava | Faculty of Natural Sciences, Comenius University, Bratislava | Viktória abanová, Kristína Boršová, Broa Brejová, Viktória Hodorová, Sabina Fumaová Havlíková, Juraj Kopáek, Martina Liková, ubomíra Lukáiková, Martina Neboháová, Monika Sláviková, Tomáš Vina, Jozef Nosek, Boris Klempa |
| EPI_ISL_831676 | Institute of Virology, Biomedical Research Center of the Slovak Academy of Sciences, Bratislava | Faculty of Natural Sciences, Comenius University, Bratislava | Broa Brejová, Viktória abanová, Kristína Boršová, Viktória Hodorová, Sabina Fumaová Havlíková, Juraj Kopáek, Martina Liková, ubomíra Lukáiková, Martina Neboháová, Monika Sláviková, Tomáš Vina, Jozef Nosek, Boris Klempa |
| EPI_ISL_831900, EPI_ISL_831901, EPI_ISL_831903, EPI_ISL_831905, EPI_ISL_831908, EPI_ISL_831910, EPI_ISL_831920, EPI_ISL_831921, EPI_ISL_831922, EPI_ISL_831923, EPI_ISL_831924, EPI_ISL_831925, EPI_ISL_831926, EPI_ISL_831927, EPI_ISL_831931, EPI_ISL_831932, EPI_ISL_831933, EPI_ISL_831934, EPI_ISL_831935, EPI_ISL_831936, EPI_ISL_831937 | | | |
| see above | New Mexico Department of Health Scientific Laboratory | New Mexico Department of Health Scientific Laboratory | Ellie Johnson, Anastacia Griego-Fisher, D'eldra Malone |
| EPI_ISL_832015, EPI_ISL_832016 | University of Wisconsin-Madison AIDS Vaccine Research | University of Wisconsin-Madison AIDS Vaccine Research | Gage Moreno, Katarina Braun, et al. AIDS Vaccine Research Laboratories |

| | Laboratories | Laboratories | |
|---|---|---|---|
| EPI_ISL_832125, EPI_ISL_832126 | Clinical Molecular Microbiology Laboratory, UNC Hospitals | Jeremy Wang | Jeremy Wang, Alexander Rubinsteyn, Colleen Rice, Jason Smedberg, Melissa Miller, Corbin Jones, Robert Hagan |
| EPI_ISL_832127 | Michigan Department of Health and Human Services, Bureau of Laboratories | Michigan Department of Health and Human Services, Bureau of Laboratories | Blankenship HM, Riner D, Soehnlen MK |
| EPI_ISL_832203, EPI_ISL_832204 | Northwest Pathology | UW Virology Lab | Pavitra Roychoudhury, Hong Xie, Lasata Shrestha, Michelle Lin, Meei-Li Huang, Keith R Jerome, Alexander Greninger |
| EPI_ISL_832232, EPI_ISL_832233, EPI_ISL_832234, EPI_ISL_832235, EPI_ISL_832236, EPI_ISL_832237, EPI_ISL_832238, EPI_ISL_832239, EPI_ISL_832240, EPI_ISL_832241 | Department of Clinical Microbiology | GIGA Medical Genomics | Keith Durkin, Maria Artesi, Sébastien Bontems, Raphaël Boreux, Bouchra Boujemla, Cécile Meex, Pierrette Melin, Marie-Pierre Hayette, Vincent Bours |
| EPI_ISL_833028 | Maine HETL | Tewhey Lab, The Jackson Laboratory | Matluk,N., Dewey,H., Iosue,F., Barter,M., Lynch,R., Munger,H. and Tewhey,R. |
| EPI_ISL_833148 | Genomic Laboratory (GLAB) (Conjoint lab of Health Directorate of Istanbul and Istanbul Technical University) | Genomic Laboratory (GLAB), Istanbul Technical University | Ilker Karacan, Tugba Kizilboga Akgun, Payam Zolfagharian, Nisan Denizce Can, Pari Sharifli, Levent Doganay, Gizem Dinler Doganay |
| EPI_ISL_833249 | Division of Emerging Infectious Diseases, Bureau of Infectious Diseases Diagnosis Control, Korea Disease Control and Prevention Agency | Division of Emerging Infectious Diseases, Bureau of Infectious Diseases Diagnosis Control, Korea Disease Control and Prevention Agency | Ae Kyung Park, Il-Hwan Kim, Heui Man Kim, Jeong-Min Kim, Namjoo Lee, Chaeyoung Lee, Sang Hee Woo, Eun-Jin Kim |
| EPI_ISL_833404 | St. Francis Medical Center | Los Angeles County PHL | P. Hemarajata et al. |
| EPI_ISL_833418, EPI_ISL_833490, EPI_ISL_833491 | MD Laboratories | Los Angeles County PHL | P. Hemarajata et al. |
| EPI_ISL_833578 | National Reference Laboratory for COVID-19, Pasteur Institute of Iran | National Reference Laboratory for COVID-19, Pasteur Institute of Iran | Zahra Ahmadi, Marzieh Sadjadi, Tahmineh Jalali, Mohammad Hassan Pouriayevali, Mahsa Tavakoli, Zahra Fereydouni, Setareh Kashanian, Sanam Azad-Manjiri, Tahereh Mohammadi, Kayhan Azadmanesh, Zabihollah Shoja, Parastoo Yekta, Farideh Niknam, Hessam Nemati, Ahmad Ghasemi, Sahar Khakifirouz, Sepideh Gerdooei,Maryam Rostamtabar,Sana Eybpoosh,Mohammad Mehdi Mortazavipour, Mohamad Sadegh Shams Nosrati, Zeynab VeisiZadeh, Mostafa Salehi-Vaziri |
| EPI_ISL_833585, EPI_ISL_833587, EPI_ISL_833588, EPI_ISL_833589, EPI_ISL_833591, EPI_ISL_833592, EPI_ISL_833594, EPI_ISL_833595, EPI_ISL_833598, EPI_ISL_833600, EPI_ISL_833601, EPI_ISL_833603, EPI_ISL_833605, EPI_ISL_833608, EPI_ISL_833613, EPI_ISL_833622, EPI_ISL_833623, EPI_ISL_833624, EPI_ISL_833625, EPI_ISL_833627, EPI_ISL_833628, EPI_ISL_833633, EPI_ISL_833637, EPI_ISL_833638, EPI_ISL_833639, EPI_ISL_833641, EPI_ISL_833642, EPI_ISL_833644, EPI_ISL_833645, EPI_ISL_833649, EPI_ISL_833653, EPI_ISL_833655, EPI_ISL_833657, EPI_ISL_833659, EPI_ISL_833660, EPI_ISL_833661, EPI_ISL_833663, EPI_ISL_833664, EPI_ISL_833666, EPI_ISL_833669, EPI_ISL_833672, EPI_ISL_833674, EPI_ISL_833675, EPI_ISL_833677, EPI_ISL_833681, EPI_ISL_833682, EPI_ISL_833688, EPI_ISL_833690, EPI_ISL_833692, EPI_ISL_833694, EPI_ISL_833697, EPI_ISL_833698, EPI_ISL_833699, EPI_ISL_833701, EPI_ISL_833711, EPI_ISL_833713, EPI_ISL_833715, EPI_ISL_833717, EPI_ISL_833718, EPI_ISL_833720, EPI_ISL_833722, EPI_ISL_833723, EPI_ISL_833729, EPI_ISL_833730, EPI_ISL_833731, EPI_ISL_833732, EPI_ISL_833737, EPI_ISL_833739, EPI_ISL_833740, EPI_ISL_833744, EPI_ISL_833745, EPI_ISL_833747, EPI_ISL_833751, EPI_ISL_833752, EPI_ISL_833754, EPI_ISL_833757, EPI_ISL_833760, EPI_ISL_833761, EPI_ISL_833763, EPI_ISL_833764, EPI_ISL_833765, EPI_ISL_833769, EPI_ISL_833772, EPI_ISL_833775, EPI_ISL_833776, EPI_ISL_833777, EPI_ISL_833781, EPI_ISL_833782, EPI_ISL_833783, EPI_ISL_833785, EPI_ISL_833786, EPI_ISL_833787, EPI_ISL_833788, EPI_ISL_833790, EPI_ISL_833793, EPI_ISL_833798, EPI_ISL_833799, EPI_ISL_833802, EPI_ISL_833805, EPI_ISL_833808, EPI_ISL_833811, EPI_ISL_833819, EPI_ISL_833820, EPI_ISL_833822, EPI_ISL_833824, EPI_ISL_833825, EPI_ISL_833826, EPI_ISL_833827, EPI_ISL_833828, EPI_ISL_833836, EPI_ISL_833837, EPI_ISL_833838, EPI_ISL_833839, EPI_ISL_833843, EPI_ISL_833847, EPI_ISL_833850, EPI_ISL_833851, EPI_ISL_833857, EPI_ISL_833858, EPI_ISL_833867, EPI_ISL_833869, EPI_ISL_833870, EPI_ISL_833871, EPI_ISL_833873, EPI_ISL_833874, EPI_ISL_833875, EPI_ISL_833878, EPI_ISL_833880, EPI_ISL_833883, EPI_ISL_833885, EPI_ISL_833887, EPI_ISL_833891, EPI_ISL_833893, EPI_ISL_833897, EPI_ISL_833899, EPI_ISL_833900, EPI_ISL_833902, EPI_ISL_833904, EPI_ISL_833905, EPI_ISL_833906, EPI_ISL_833911, EPI_ISL_833913, EPI_ISL_833914, EPI_ISL_833915, EPI_ISL_833920, EPI_ISL_833921, EPI_ISL_833923, EPI_ISL_833925, EPI_ISL_833926, EPI_ISL_833927, EPI_ISL_833928, EPI_ISL_833929 | | | |
| see above | Lighthouse Lab in Cambridge | Wellcome Sanger Institute for the COVID-19 Genomics UK (COG-UK) Consortium | Rob Howes, The Lighthouse Lab in Cambridge and Alex Alderton, Roberto Amato, Sonia Goncalves, Ewan Harrison, David K. Jackson, Ian Johnston, Dominic Kwiatkowski, Cordelia Langford, John Sillitoe on behalf of the Wellcome Sanger Institute COVID-19 Surveillance Team |
| EPI_ISL_835041 | Lighthouse Lab in Alderley Park | Wellcome Sanger Institute for the COVID-19 Genomics UK (COG-UK) Consortium | Jacquelyn Wynn, Mairead Hyland, The Lighthouse Lab in Alderley Park and Alex Alderton, Roberto Amato, Sonia Goncalves, Ewan Harrison, David K. Jackson, Ian Johnston, Dominic Kwiatkowski, Cordelia Langford, John Sillitoe on behalf of the Wellcome Sanger Institute COVID-19 Surveillance Team |
| EPI_ISL_837056, EPI_ISL_837059, EPI_ISL_837060, EPI_ISL_837069, EPI_ISL_837181, EPI_ISL_837243, EPI_ISL_837244 | Respiratory Virus Unit, National Infection Service, Public Health England | COVID-19 Genomics UK (COG-UK) Consortium | PHE Covid Sequencing Team |
| EPI_ISL_837387 | National Virus Reference Laboratory | National Virus Reference Laboratory | Michael Carr, Gabriel Gonzalez, Jonathan Dean, Cillian F De Gascun |
| EPI_ISL_837957, EPI_ISL_837959, EPI_ISL_837974, EPI_ISL_837976, EPI_ISL_837977, EPI_ISL_837979, EPI_ISL_837980, EPI_ISL_837982, EPI_ISL_837983, EPI_ISL_837986, EPI_ISL_837987, EPI_ISL_837988, EPI_ISL_837991, EPI_ISL_837993, EPI_ISL_837997, EPI_ISL_838000, EPI_ISL_838001, EPI_ISL_838003, EPI_ISL_838004, EPI_ISL_838005, EPI_ISL_838006, EPI_ISL_838007, EPI_ISL_838008, EPI_ISL_838009, EPI_ISL_838011, EPI_ISL_838012, EPI_ISL_838013, EPI_ISL_838016, EPI_ISL_838026, EPI_ISL_838032, EPI_ISL_838049, EPI_ISL_838050, EPI_ISL_838051, EPI_ISL_838052, EPI_ISL_838053, EPI_ISL_838054 | | | |
| see above | Department of Pathology, University of Cambridge | COVID-19 Genomics UK (COG-UK) Consortium | Aminu S. Jahun, Yasmin Chaudhry, Grant Hall, Iliana Georgana, Myra Hosmillo, Martin D. Curran, Malte Pinckert, Surendra Parmar, Ian Goodfellow |
| EPI_ISL_838315 | University of Exeter | COVID-19 Genomics UK (COG-UK) Consortium | Ben Temperton,Aaron Jeffries,Michelle Michelsen,Joanna Warwick-Dugdale,Audrey Farbos,Robyn Manley,Stephen Michell,Jane Masoli |
| EPI_ISL_838960, EPI_ISL_838961, EPI_ISL_838962, EPI_ISL_838963, EPI_ISL_838964, EPI_ISL_838965, EPI_ISL_838966, EPI_ISL_838967, EPI_ISL_838968, EPI_ISL_838970, EPI_ISL_838971, EPI_ISL_838972, EPI_ISL_838973, EPI_ISL_838974, EPI_ISL_838975, EPI_ISL_838976, EPI_ISL_838977, EPI_ISL_838978, EPI_ISL_838979, EPI_ISL_838980, EPI_ISL_838982, EPI_ISL_838983, EPI_ISL_838984, EPI_ISL_838985, EPI_ISL_838986, EPI_ISL_838987, EPI_ISL_838991, EPI_ISL_838992, EPI_ISL_838993, EPI_ISL_838994, EPI_ISL_838995, EPI_ISL_838996, EPI_ISL_838999, EPI_ISL_839000, EPI_ISL_839001, EPI_ISL_839002, EPI_ISL_839003, EPI_ISL_839004, EPI_ISL_839005, EPI_ISL_839007, EPI_ISL_839008, EPI_ISL_839009, EPI_ISL_839010, EPI_ISL_839011, EPI_ISL_839012, EPI_ISL_839126, EPI_ISL_839127, EPI_ISL_839129, EPI_ISL_839130, EPI_ISL_839131, EPI_ISL_839132, EPI_ISL_839233, EPI_ISL_839234, EPI_ISL_839235, EPI_ISL_839236, EPI_ISL_839237, EPI_ISL_839238, EPI_ISL_839239, EPI_ISL_839240, EPI_ISL_839241, EPI_ISL_839242, EPI_ISL_839243, EPI_ISL_839244, EPI_ISL_839245, EPI_ISL_839246, EPI_ISL_839247, EPI_ISL_839248, EPI_ISL_839249, EPI_ISL_839259, EPI_ISL_839263, EPI_ISL_839277, EPI_ISL_839278, EPI_ISL_839279, EPI_ISL_839280, EPI_ISL_839281, EPI_ISL_839282, EPI_ISL_839283, EPI_ISL_839284, EPI_ISL_839285, EPI_ISL_839286, EPI_ISL_839287, EPI_ISL_839288, EPI_ISL_839289, EPI_ISL_839290, EPI_ISL_839291, EPI_ISL_839292, EPI_ISL_839293, EPI_ISL_839294, EPI_ISL_839295, EPI_ISL_839296, EPI_ISL_839297, EPI_ISL_839298, EPI_ISL_839299, EPI_ISL_839300, EPI_ISL_839301, EPI_ISL_839302, EPI_ISL_839303, EPI_ISL_839304 | | | |
| see above | University College London, Great Ormond Street Hospital for Children NHS Foundation Trust, Imperial College Healthcare NHS Trust | COVID-19 Genomics UK (COG-UK) Consortium | Sergi Castellano, Rachel Williams, Mark Kristiansen, Paola Resende Silva, Sunando Roy, Tony Brooks, Helena Tutill, Paola Niola, Patricia Dyal, Charlotte Williams, Leysa Forrest, Yasmin Panchbhaya, Jacqueline Findlay, Samuel Weeks, Julianne Brown, Kathryn Harris, Paul Randell, James Price, Alison Holmes, Judith Breuer |
| EPI_ISL_840053, EPI_ISL_840054, EPI_ISL_840055, EPI_ISL_840056, EPI_ISL_840057, EPI_ISL_840058, EPI_ISL_840059, EPI_ISL_840060 | Queens Medical Centre, Clinical Microbiology Department / DeepSeq Nottingham | COVID-19 Genomics UK (COG-UK) Consortium | Gemma Clark, Wendy Smith, Manjinder Khakh, Vicki M Fleming, Michelle M Lister, Hannah Howson-Wells, Jonathan Ball, Patrick McClure, Joseph Chappell, Theocharis Tsoleridis, Nadine Holmes, Matthew Carlisle, Christopher Moore, Fei Sang, Johnny Debebe, Victoria Wright, Matthew Loose |
| EPI_ISL_841326, EPI_ISL_841328, EPI_ISL_841330, EPI_ISL_841331, EPI_ISL_841340, EPI_ISL_841349, EPI_ISL_841350, EPI_ISL_841351, EPI_ISL_841352, EPI_ISL_841426, EPI_ISL_841427, EPI_ISL_841430, EPI_ISL_841457, EPI_ISL_841484, EPI_ISL_841487, EPI_ISL_841488, EPI_ISL_841489, EPI_ISL_841492, EPI_ISL_841493, EPI_ISL_841494, EPI_ISL_841495, EPI_ISL_841496, EPI_ISL_841497, EPI_ISL_841523, EPI_ISL_841524, EPI_ISL_841525, EPI_ISL_841527, EPI_ISL_841528, EPI_ISL_841530, EPI_ISL_841532, EPI_ISL_841533, EPI_ISL_841534, EPI_ISL_841535, EPI_ISL_841536, EPI_ISL_841537, EPI_ISL_841538, EPI_ISL_841539, EPI_ISL_841540, EPI_ISL_841541, EPI_ISL_841542, EPI_ISL_841547, EPI_ISL_841549, EPI_ISL_841550, EPI_ISL_841551, EPI_ISL_841553, EPI_ISL_841554, EPI_ISL_841557, EPI_ISL_841558, EPI_ISL_841614, EPI_ISL_841625, EPI_ISL_841659, EPI_ISL_841661, EPI_ISL_841665, EPI_ISL_841667, EPI_ISL_841669, EPI_ISL_841670, EPI_ISL_841673, EPI_ISL_841676, EPI_ISL_841677, EPI_ISL_841679, EPI_ISL_841684, EPI_ISL_841685, EPI_ISL_841686, EPI_ISL_841687, EPI_ISL_841688, EPI_ISL_841689, EPI_ISL_841690, EPI_ISL_841694, EPI_ISL_841695, EPI_ISL_841700, EPI_ISL_841702, EPI_ISL_841704, EPI_ISL_841705, EPI_ISL_841706, EPI_ISL_841707, EPI_ISL_841709, EPI_ISL_841710, EPI_ISL_841711, EPI_ISL_841712, EPI_ISL_841713, EPI_ISL_841714, EPI_ISL_841715, EPI_ISL_841716, EPI_ISL_841717, EPI_ISL_841718, EPI_ISL_841719, EPI_ISL_841720, EPI_ISL_841721, EPI_ISL_841722, EPI_ISL_841723, EPI_ISL_841724, EPI_ISL_841725, EPI_ISL_841726, EPI_ISL_841727, EPI_ISL_841728, EPI_ISL_841729, EPI_ISL_841730, EPI_ISL_841733, EPI_ISL_841734, EPI_ISL_841735, EPI_ISL_841736, EPI_ISL_841738, EPI_ISL_841739, EPI_ISL_841740 | | | |
| see above | Originating lab: Wales Specialist Virology Centre Sequencing lab: Pathogen Genomics Unit | Public Health Wales Microbiology Cardiff Wales Specialist Virology Centre | Catherine Moore, Johnathan Evans, Laura Gifford, Malorie Perry, Simon Cottrell, Angela Marchbank, Alec Birchley, Alexander Adams, Amy Gaskin, Bree Gatica-Wilcox, Jason Coombes, Joel Southgate, Lauren Gilbert, Lee Graham, Nicole Pacchiarini, Sara Kumziene-Summerhayes, Sarah Taylor, Sophie Jones, Sara Rey, Matthew Bull, Joanne Watkins, Sally Corden, Tom Connor |
| EPI_ISL_841921 | Centre for Enzyme Innovation, University of Portsmouth / Translational Research Laboratory, Portsmouth Hospitals NHS Trust | COVID-19 Genomics UK (COG-UK) Consortium | Angela Beckett,Yann Bourgeois,Garry Scarlett,Sharon Glaysher,Scott Elliott,Kelly Bicknell,Robert Impey,Allyson Lloyd,Sarah Wyllie,Ethan Butcher,Anoop Chauhan,Samuel Robson |
| EPI_ISL_842207, EPI_ISL_842213, EPI_ISL_842222, EPI_ISL_842223, EPI_ISL_842225, EPI_ISL_842229, EPI_ISL_842234, EPI_ISL_842242, EPI_ISL_842243, EPI_ISL_842247, EPI_ISL_842249, EPI_ISL_842252, EPI_ISL_842259, EPI_ISL_842267, EPI_ISL_842271, EPI_ISL_842275, EPI_ISL_842276, EPI_ISL_842277, EPI_ISL_842296, EPI_ISL_842302, EPI_ISL_842309, EPI_ISL_842310, EPI_ISL_842314, EPI_ISL_842320, EPI_ISL_842321, EPI_ISL_842323, EPI_ISL_842328, EPI_ISL_842331, EPI_ISL_842333, EPI_ISL_842336, EPI_ISL_842337, EPI_ISL_842343 | | | |
| see above | Virology Department, Sheffield Teaching Hospitals NHS Foundation Trust/Department of Infection, Immunity and | COVID-19 Genomics UK (COG-UK) Consortium | Thushan de Silva, Matthew Parker, Nikki Smith, Adri Angyal, Rebecca Brown, Luke Green, Rachel Tucker, Paul Parsons, Danielle Groves, Katie Johnson, Laura Carrilero, Alex Keeley, Dave Partridge, Matthew Wyles, Benjamin Lindsey, Mehmet Yavuz, Mohammad Raza, Cariad Evans |

| | | | |
|---|---|---|---|
| | Cardiovascular Disease, The Medical School, University of Sheffield | | |
| EPI_ISL_842637 | Medical Microbiology Unit, Department for Laboratory Medicine, Drammen Hospital, Vestre Viken Health Trust, | Norwegian Institute of Public Health, Department of Virology | Kathrine Stene-Johansen, Kamilla Heddeland Instefjord, Hilde Elshaug, Atiya R Ali,Marie Paulsen Madsen, Rasmus Riis Kopperud, Hilde Vollan, Karoline Bragstad, Olav Hungnes |
| EPI_ISL_842642 | Norwegian Institute of Public Health, Department of Virology | Norwegian Institute of Public Health, Department of Virology | Kathrine Stene-Johansen, Kamilla Heddeland Instefjord, Hilde Elshaug, Atiya R Ali,Marie Paulsen Madsen, Rasmus Riis Kopperud, Hilde Vollan, Karoline Bragstad, Olav Hungnes |
| EPI_ISL_843184, EPI_ISL_843185, EPI_ISL_843186, EPI_ISL_843187 | Maryland Public Health Laboratory | Maryland Public Health Laboratory | Maryland Department of Health Laboratories Administration |
| EPI_ISL_843194, EPI_ISL_843199, EPI_ISL_843200 | Canterbury Health Laboratories | Institute of Environmental Science and Research (ESR) | Xiaoyun Ren, Matt Storey, Nikki Freed, Muhammad Faisal, Jing Wang, Hermes Perez, Anja Werno, Antje van der Linden, Arlo Upton, Chris Mansell, David Hammer, Dragana Drinkovic, Gary McAuliffe, Hana Sofia Andersson, James Ussher, Jill Sherwood, Josh Freeman, Julia Howard, Juliet Elvy, Mary DeAlmeida, Matt Blakiston, Matthew Rogers, Max Bloomfield, Michael Addidle, Michelle Balm, Sally Roberts, Sarah Jefferies, Sharmini Muttaiyah, Susan Morpeth, Susan Taylor, Timothy Blackmore, Vani Sathyendran, Veronica Playle, Virginia Hope, Erasmus Smit, Lauren Jelly, Olin Silander, Joep de Ligt |
| EPI_ISL_845638, EPI_ISL_845639, EPI_ISL_845640, EPI_ISL_845646, EPI_ISL_845647, EPI_ISL_845654 | Laboratorio de Salud Pública - Secretaría Distrital de Salud | Instituto Nacional de Salud - Dirección de Investigación en Salud Pública | Katherine Laiton-Donato, Diego A. Álvarez-Díaz, Carlos Franco-Muñoz, Mauricio Pacheco-Montealegre, Maria T. Herrera-Sepúlveda, Jonathan Reales, Sheryll Corchuelo, Julian Naizaque, Gerardo Santamaría, Paola Muñoz-Laiton, Diego Andrés Prada, Magdalena Wiesner, Martha Lucia Ospina Martinez, Marcela Mercado-Reyes |
| EPI_ISL_845753 | Toronto Invasive Bacterial Diseases Network | McMaster University | Allison McGeer, Patryk Aftanas, Hooman Derakhshani, Angel Li, Kuganya Nirmalarajah, Emily Panousis, Ahmed Draia, Jalees Nasir, Michael Surette, Samira Mubareka, Andrew G. McArthur |
| EPI_ISL_845797 | South Eastern Area Laboratory Services (SEALS) | NSW Health Pathology - Institute of Clinical Pathology and Medical Research; Westmead Hospital; University of Sydney | CIDM-PH et al. |
| EPI_ISL_845798, EPI_ISL_845799 | Sydney South West Pathology Service (SSWPS) - Royal Prince Alfred Hospital - NSW Health Pathology | NSW Health Pathology - Institute of Clinical Pathology and Medical Research; Westmead Hospital; University of Sydney | CIDM-PH et al. |
| EPI_ISL_845894, EPI_ISL_845895, EPI_ISL_845897, EPI_ISL_845898, EPI_ISL_845899, EPI_ISL_845900, EPI_ISL_845901, EPI_ISL_845902, EPI_ISL_845903, EPI_ISL_845904, EPI_ISL_845905, EPI_ISL_845906, EPI_ISL_845907, EPI_ISL_845908, EPI_ISL_845909, EPI_ISL_845910, EPI_ISL_845911, EPI_ISL_845912, EPI_ISL_845913, EPI_ISL_845914, EPI_ISL_845915, EPI_ISL_845916, EPI_ISL_845917, EPI_ISL_845918, EPI_ISL_845919, EPI_ISL_845920, EPI_ISL_845921, EPI_ISL_845922, EPI_ISL_845923, EPI_ISL_845924, EPI_ISL_845925, EPI_ISL_845926, EPI_ISL_845927, EPI_ISL_845928, EPI_ISL_845929, EPI_ISL_845930, EPI_ISL_845931, EPI_ISL_845932, EPI_ISL_845933, EPI_ISL_845934, EPI_ISL_845935, EPI_ISL_845936, EPI_ISL_845937, EPI_ISL_845938, EPI_ISL_845939, EPI_ISL_845940, EPI_ISL_845941, EPI_ISL_845942, EPI_ISL_845943, EPI_ISL_845944, EPI_ISL_845945, EPI_ISL_845946, EPI_ISL_845947, EPI_ISL_845948, EPI_ISL_845949, EPI_ISL_845950, EPI_ISL_845951, EPI_ISL_845952, EPI_ISL_845953, EPI_ISL_845954, EPI_ISL_845955, EPI_ISL_845956, EPI_ISL_845957, EPI_ISL_845958, EPI_ISL_845959, EPI_ISL_845960, EPI_ISL_845961, EPI_ISL_845962, EPI_ISL_845963, EPI_ISL_845964, EPI_ISL_845965, EPI_ISL_845966, EPI_ISL_845967, EPI_ISL_845968, EPI_ISL_845969, EPI_ISL_845970, EPI_ISL_845971, EPI_ISL_845972, EPI_ISL_845974, EPI_ISL_845975, EPI_ISL_845976, EPI_ISL_845977, EPI_ISL_845978, EPI_ISL_845979, EPI_ISL_845980, EPI_ISL_845981, EPI_ISL_845982, EPI_ISL_845983, EPI_ISL_845984, EPI_ISL_845985, EPI_ISL_845986, EPI_ISL_845987, EPI_ISL_845988, EPI_ISL_845989, EPI_ISL_845990, EPI_ISL_845991, EPI_ISL_845992, EPI_ISL_845993, EPI_ISL_845994, EPI_ISL_845995, EPI_ISL_845996, EPI_ISL_845997, EPI_ISL_845998, EPI_ISL_845999, EPI_ISL_846000, EPI_ISL_846001, EPI_ISL_846002, EPI_ISL_846003, EPI_ISL_846004, EPI_ISL_846005, EPI_ISL_846006, EPI_ISL_846008, EPI_ISL_846009, EPI_ISL_846010, EPI_ISL_846011, EPI_ISL_846012, EPI_ISL_846013, EPI_ISL_846014, EPI_ISL_846015, EPI_ISL_846016, EPI_ISL_846017, EPI_ISL_846018, EPI_ISL_846019, EPI_ISL_846020, EPI_ISL_846021, EPI_ISL_846024, EPI_ISL_846025, EPI_ISL_846026, EPI_ISL_846027, EPI_ISL_846028, EPI_ISL_846029, EPI_ISL_846030, EPI_ISL_846031, EPI_ISL_846032, EPI_ISL_846033, EPI_ISL_846034, EPI_ISL_846035, EPI_ISL_846036, EPI_ISL_846037, EPI_ISL_846038, EPI_ISL_846039, EPI_ISL_846040, EPI_ISL_846041, EPI_ISL_846042, EPI_ISL_846043, EPI_ISL_846044, EPI_ISL_846045, EPI_ISL_846046, EPI_ISL_846047, EPI_ISL_846048, EPI_ISL_846049, EPI_ISL_846050, EPI_ISL_846051, EPI_ISL_846053, EPI_ISL_846054, EPI_ISL_846055, EPI_ISL_846056, EPI_ISL_846057, EPI_ISL_846058, EPI_ISL_846060, EPI_ISL_846061, EPI_ISL_846062, EPI_ISL_846063, EPI_ISL_846064, EPI_ISL_846065, EPI_ISL_846066, EPI_ISL_846067, EPI_ISL_846068, EPI_ISL_846069, EPI_ISL_846071, EPI_ISL_846072, EPI_ISL_846073, EPI_ISL_846074, EPI_ISL_846075, EPI_ISL_846076, EPI_ISL_846077, EPI_ISL_846078, EPI_ISL_846079, EPI_ISL_846080, EPI_ISL_846081, EPI_ISL_846082, EPI_ISL_846083, EPI_ISL_846085, EPI_ISL_846086, EPI_ISL_846087, EPI_ISL_846088, EPI_ISL_846089, EPI_ISL_846090, EPI_ISL_846091, EPI_ISL_846092, EPI_ISL_846093, EPI_ISL_846094, EPI_ISL_846095, EPI_ISL_846096, EPI_ISL_846097, EPI_ISL_846098, EPI_ISL_846099, EPI_ISL_846100, EPI_ISL_846101, EPI_ISL_846102, EPI_ISL_846103, EPI_ISL_846104, EPI_ISL_846105, EPI_ISL_846106, EPI_ISL_846107, EPI_ISL_846108, EPI_ISL_846109, EPI_ISL_846110, EPI_ISL_846111, EPI_ISL_846112, EPI_ISL_846113, EPI_ISL_846114, EPI_ISL_846115, EPI_ISL_846116, EPI_ISL_846117, EPI_ISL_846118, EPI_ISL_846119, EPI_ISL_846120, EPI_ISL_846121, EPI_ISL_846122, EPI_ISL_846123, EPI_ISL_846124, EPI_ISL_846125, EPI_ISL_846126, EPI_ISL_846127, EPI_ISL_846128, EPI_ISL_846129, EPI_ISL_846130, EPI_ISL_846131, EPI_ISL_846132, EPI_ISL_846133, EPI_ISL_846134, EPI_ISL_846135, EPI_ISL_846136, EPI_ISL_846137, EPI_ISL_846138, EPI_ISL_846140, EPI_ISL_846141, EPI_ISL_846142, EPI_ISL_846143, EPI_ISL_846144, EPI_ISL_846145, EPI_ISL_846146, EPI_ISL_846148, EPI_ISL_846149, EPI_ISL_846150, EPI_ISL_846151, EPI_ISL_846153, EPI_ISL_846154, EPI_ISL_846155, EPI_ISL_846156, EPI_ISL_846157, EPI_ISL_846158, EPI_ISL_846159, EPI_ISL_846160, EPI_ISL_846161, EPI_ISL_846162, EPI_ISL_846163, EPI_ISL_846164, EPI_ISL_846165, EPI_ISL_846166, EPI_ISL_846167, EPI_ISL_846169, EPI_ISL_846170, EPI_ISL_846171, EPI_ISL_846172, EPI_ISL_846173, EPI_ISL_846174, EPI_ISL_846175, EPI_ISL_846176, EPI_ISL_846177, EPI_ISL_846178, EPI_ISL_846179, EPI_ISL_846180, EPI_ISL_846181, EPI_ISL_846182, EPI_ISL_846183, EPI_ISL_846184, EPI_ISL_846185, EPI_ISL_846186, EPI_ISL_846187, EPI_ISL_846188, EPI_ISL_846189, EPI_ISL_846190, EPI_ISL_846191, EPI_ISL_846192, EPI_ISL_846193, EPI_ISL_846194, EPI_ISL_846195, EPI_ISL_846196, EPI_ISL_846197, EPI_ISL_846198, EPI_ISL_846199 | | | |
| see above | Lighthouse Lab in Cambridge | Wellcome Sanger Institute for the COVID-19 Genomics UK (COG-UK) Consortium | Rob Howes, The Lighthouse Lab in Cambridge and Alex Alderton, Roberto Amato, Sonia Goncalves, Ewan Harrison, David K. Jackson, Ian Johnston, Dominic Kwiatkowski, Cordelia Langford, John Sillitoe on behalf of the Wellcome Sanger Institute COVID-19 Surveillance Team |
| EPI_ISL_846200, EPI_ISL_846201, EPI_ISL_846202, EPI_ISL_846203, EPI_ISL_846204, EPI_ISL_846205, EPI_ISL_846206, EPI_ISL_846207, EPI_ISL_846208, EPI_ISL_846209, EPI_ISL_846210, EPI_ISL_846211, EPI_ISL_846212, EPI_ISL_846213, EPI_ISL_846214, EPI_ISL_846215, EPI_ISL_846216, EPI_ISL_846217, EPI_ISL_846218, EPI_ISL_846219, EPI_ISL_846220, EPI_ISL_846221, EPI_ISL_846222, EPI_ISL_846223, EPI_ISL_846224, EPI_ISL_846225, EPI_ISL_846226, EPI_ISL_846227, EPI_ISL_846228, EPI_ISL_846229, EPI_ISL_846230, EPI_ISL_846231, EPI_ISL_846232, EPI_ISL_846233, EPI_ISL_846234, EPI_ISL_846235, EPI_ISL_846236, EPI_ISL_846237, EPI_ISL_846238, EPI_ISL_846239, EPI_ISL_846240, EPI_ISL_846241, EPI_ISL_846242, EPI_ISL_846243, EPI_ISL_846244, EPI_ISL_846245, EPI_ISL_846246, EPI_ISL_846247, EPI_ISL_846248, EPI_ISL_846249, EPI_ISL_846250, EPI_ISL_846251, EPI_ISL_846252, EPI_ISL_846253, EPI_ISL_846254, EPI_ISL_846255, EPI_ISL_846256, EPI_ISL_846257, EPI_ISL_846258, EPI_ISL_846259, EPI_ISL_846260, EPI_ISL_846261, EPI_ISL_846262, EPI_ISL_846263, EPI_ISL_846264, EPI_ISL_846265, EPI_ISL_846266, EPI_ISL_846267, EPI_ISL_846268, EPI_ISL_846269, EPI_ISL_846270, EPI_ISL_846271, EPI_ISL_846272, EPI_ISL_846273, EPI_ISL_846274, EPI_ISL_846275, EPI_ISL_846276, EPI_ISL_846277, EPI_ISL_846278, EPI_ISL_846279, EPI_ISL_846280, EPI_ISL_846281, EPI_ISL_846282, EPI_ISL_846284, EPI_ISL_846285, EPI_ISL_846286, EPI_ISL_846288, EPI_ISL_846289, EPI_ISL_846290, EPI_ISL_846291, EPI_ISL_846292, EPI_ISL_846293, EPI_ISL_846294, EPI_ISL_846295, EPI_ISL_846296, EPI_ISL_846297, EPI_ISL_846298, EPI_ISL_846300, EPI_ISL_846301, EPI_ISL_846302, EPI_ISL_846303, EPI_ISL_846304, EPI_ISL_846305, EPI_ISL_846306, EPI_ISL_846307, EPI_ISL_846308, EPI_ISL_846309, EPI_ISL_846310, EPI_ISL_846311, EPI_ISL_846312, EPI_ISL_846313, EPI_ISL_846314, EPI_ISL_846316, EPI_ISL_846317, EPI_ISL_846318, EPI_ISL_846320, EPI_ISL_846322, EPI_ISL_846323, EPI_ISL_846324, EPI_ISL_846325, EPI_ISL_846326, EPI_ISL_846327, EPI_ISL_846328, EPI_ISL_846329, EPI_ISL_846330, EPI_ISL_846331, EPI_ISL_846332, EPI_ISL_846333, EPI_ISL_846334, EPI_ISL_846335, EPI_ISL_846336, EPI_ISL_846337, EPI_ISL_846338, EPI_ISL_846339, EPI_ISL_846340, EPI_ISL_846341, EPI_ISL_846342, EPI_ISL_846343, EPI_ISL_846344, EPI_ISL_846345, EPI_ISL_846346, EPI_ISL_846347, EPI_ISL_846348, EPI_ISL_846349, EPI_ISL_846350, EPI_ISL_846351, EPI_ISL_846352, EPI_ISL_846353, EPI_ISL_846355, EPI_ISL_846356, EPI_ISL_846357, EPI_ISL_846358, EPI_ISL_846359, EPI_ISL_846360, EPI_ISL_846361, EPI_ISL_846362, EPI_ISL_846363, EPI_ISL_846364, EPI_ISL_846365, EPI_ISL_846366, EPI_ISL_846367, EPI_ISL_846368, EPI_ISL_846369, EPI_ISL_846370, EPI_ISL_846371, EPI_ISL_846372, EPI_ISL_846373, EPI_ISL_846374, EPI_ISL_846375, EPI_ISL_846376, EPI_ISL_846377, EPI_ISL_846378, EPI_ISL_846379, EPI_ISL_846380, EPI_ISL_846381, EPI_ISL_846382, EPI_ISL_846383, EPI_ISL_846384, EPI_ISL_846385, EPI_ISL_846386, EPI_ISL_846387, EPI_ISL_846388, EPI_ISL_846389, EPI_ISL_846390, EPI_ISL_846391, EPI_ISL_846392, EPI_ISL_846393, EPI_ISL_846394, EPI_ISL_846395, EPI_ISL_846396, EPI_ISL_846397, EPI_ISL_846398, EPI_ISL_846401, EPI_ISL_846402, EPI_ISL_846403, EPI_ISL_846404, EPI_ISL_846405, EPI_ISL_846406, EPI_ISL_846407, EPI_ISL_846408, EPI_ISL_846409, EPI_ISL_846410, EPI_ISL_846411, EPI_ISL_846412, EPI_ISL_846413, EPI_ISL_846414, EPI_ISL_846415, EPI_ISL_846416, EPI_ISL_846417, EPI_ISL_846418, EPI_ISL_846419, EPI_ISL_846420, EPI_ISL_846422, EPI_ISL_846423, EPI_ISL_846424, EPI_ISL_846425, EPI_ISL_846426, EPI_ISL_846427, EPI_ISL_846428, EPI_ISL_846430, EPI_ISL_846432, EPI_ISL_846433, EPI_ISL_846434, EPI_ISL_846435, EPI_ISL_846436, EPI_ISL_846437, EPI_ISL_846438, EPI_ISL_846439, EPI_ISL_846440, EPI_ISL_846441, EPI_ISL_846442, EPI_ISL_846444, EPI_ISL_846445, EPI_ISL_846446, EPI_ISL_846447, EPI_ISL_846448, EPI_ISL_846449, EPI_ISL_846450, EPI_ISL_846451, EPI_ISL_846452, EPI_ISL_846453, EPI_ISL_846454, EPI_ISL_846455, EPI_ISL_846456, EPI_ISL_846457, EPI_ISL_846458, EPI_ISL_846459, EPI_ISL_846460, EPI_ISL_846461, EPI_ISL_846462, EPI_ISL_846463, EPI_ISL_846464, EPI_ISL_846465, EPI_ISL_846466, EPI_ISL_846467, EPI_ISL_846468, EPI_ISL_846469, EPI_ISL_846470, EPI_ISL_846471, EPI_ISL_846472, EPI_ISL_846473, EPI_ISL_846474, EPI_ISL_846475, EPI_ISL_846476, EPI_ISL_846477, EPI_ISL_846478, EPI_ISL_846479, EPI_ISL_846480, EPI_ISL_846481, EPI_ISL_846482, EPI_ISL_846483, EPI_ISL_846484, EPI_ISL_846485, EPI_ISL_846486, EPI_ISL_846487, EPI_ISL_846488, EPI_ISL_846489, EPI_ISL_846490, EPI_ISL_846492, EPI_ISL_846493, EPI_ISL_846494, EPI_ISL_846495, EPI_ISL_846496, EPI_ISL_846497, EPI_ISL_846498, EPI_ISL_846499, EPI_ISL_846500, EPI_ISL_846501, EPI_ISL_846502, EPI_ISL_846503, EPI_ISL_846504, EPI_ISL_846505, EPI_ISL_846506, EPI_ISL_846507, EPI_ISL_846508, EPI_ISL_846512 | | | |
| see above | Lighthouse Lab in Alderley Park | Wellcome Sanger Institute for the COVID-19 Genomics UK (COG-UK) Consortium | Jacquelyn Wynn, Mairead Hyland, The Lighthouse Lab in Alderley Park and Alex Alderton, Roberto Amato, Sonia Goncalves, Ewan Harrison, David K. Jackson, Ian Johnston, Dominic Kwiatkowski, Cordelia Langford, John Sillitoe on behalf of the Wellcome Sanger Institute COVID-19 Surveillance Team |
| EPI_ISL_846542, EPI_ISL_846547, EPI_ISL_846548, EPI_ISL_846549, EPI_ISL_846559, EPI_ISL_846560, EPI_ISL_846575, EPI_ISL_846576 | Lab voor klinische biologie | Lab voor klinische biologie | Hannelore Hamerlinck, Bruno Verhasselt |
| EPI_ISL_846597 | Respiratory Virus Unit, National Infection Service, Public Health England | COVID-19 Genomics UK (COG-UK) Consortium | PHE Covid Sequencing Team |
| EPI_ISL_847589 | California Department of Public Health | Chiu Laboratory, University of California, San Francisco | Charles Chiu, Xianding (Wayne) Deng, Candace Wang, Brian Bushnell, Scot Federman, Jill Hacker, Debra Wadford |
| EPI_ISL_848192 | National Laboratory for Health, Environment and Food | National Laboratory for Health, Environment and Food | Aleksander Mahnic, Sandra Janezic, Maja Rupnik |
| EPI_ISL_849736, EPI_ISL_849737, EPI_ISL_849738, EPI_ISL_849739, EPI_ISL_849740, EPI_ISL_849741, EPI_ISL_849742, EPI_ISL_849743 | Special Operations Medical Research Division, Defence Services Medical Research Centre | Special Operations Medical Research Division, Defence Services Medical Research Centre | Oo,K.Z., Aung,N.M., Win,K.K., Aung,P.K., Htun,Z.W., Zaw,T., Myint,K., Lwin,K.K. |
| EPI_ISL_849756, EPI_ISL_849757 | unknown | PHV-FSS | Son Nguyen et al. |
| EPI_ISL_849763 | Molecular Biology, IZS Sicilia | Molecular Biology, IZS Sicilia | Reale,S., Seidita,G., Di Gaudio,F., Scibetta,S. and Vitale,F. |
| EPI_ISL_849913, EPI_ISL_849914, EPI_ISL_849915, EPI_ISL_849916, EPI_ISL_849917, EPI_ISL_849918, EPI_ISL_849919, EPI_ISL_849920 | Utah Public Health Laboratory | Utah Public Health Laboratory | Erin L. Young, Kelly F. Oakeson, Tara Gallagher |

| | | | |
|---|---|---|---|
| EPI_ISL_850511, EPI_ISL_850512, EPI_ISL_850513, EPI_ISL_850515 | Hospital Universitari de Bellvitge | Hospital Universitari Vall d'Hebron - Vall d'Hebron Institut de Rercerca | Cristina Andrés, Maria Piñana, Josep F Abril, Damir Garcia-Cehic, Ariadna Rando, Juliana Esperalba, Maria Gema Codina, Carla Castillo, Maria Carmen Martin, Tomàs Pumarola, Josep Quer, Andrés Antón |
| EPI_ISL_850516 | Hospital Universitari Vall d'Hebron - Vall d'Hebron Institut de Rercerca | Hospital Universitari Vall d'Hebron - Vall d'Hebron Institut de Rercerca | Cristina Andrés, Maria Piñana, Josep F Abril, Damir Garcia-Cehic, Ariadna Rando, Juliana Esperalba, Maria Gema Codina, Carla Castillo, Maria Carmen Martin, Tomàs Pumarola, Josep Quer, Andrés Antón |
| EPI_ISL_851064, EPI_ISL_851065, EPI_ISL_851066, EPI_ISL_851067, EPI_ISL_851068, EPI_ISL_851069 | Lighthouse Lab in Cambridge | Wellcome Sanger Institute for the COVID-19 Genomics UK (COG-UK) Consortium | Rob Howes, The Lighthouse Lab in Cambridge and Alex Alderton, Roberto Amato, Sonia Goncalves, Ewan Harrison, David K. Jackson, Ian Johnston, Dominic Kwiatkowski, Cordelia Langford, John Sillitoe on behalf of the Wellcome Sanger Institute COVID-19 Surveillance Team |
| EPI_ISL_851070 | Lighthouse Lab in Glasgow | Wellcome Sanger Institute for the COVID-19 Genomics UK (COG-UK) Consortium | Harper VanSteenhouse, Yumi Kasai, David Gray, Carol Clugston, Anna Dominiczak and Alex Alderton, Roberto Amato, Sonia Goncalves, Ewan Harrison, David K. Jackson, Ian Johnston, Dominic Kwiatkowski, Cordelia Langford, John Sillitoe on behalf of the Wellcome Sanger Institute COVID-19 Surveillance Team |
| EPI_ISL_851071, EPI_ISL_851072, EPI_ISL_851073, EPI_ISL_851074, EPI_ISL_851075, EPI_ISL_851076 | Lighthouse Lab in Cambridge | Wellcome Sanger Institute for the COVID-19 Genomics UK (COG-UK) Consortium | Rob Howes, The Lighthouse Lab in Cambridge and Alex Alderton, Roberto Amato, Sonia Goncalves, Ewan Harrison, David K. Jackson, Ian Johnston, Dominic Kwiatkowski, Cordelia Langford, John Sillitoe on behalf of the Wellcome Sanger Institute COVID-19 Surveillance Team |
| EPI_ISL_851077 | Lighthouse Lab in Glasgow | Wellcome Sanger Institute for the COVID-19 Genomics UK (COG-UK) Consortium | Harper VanSteenhouse, Yumi Kasai, David Gray, Carol Clugston, Anna Dominiczak and Alex Alderton, Roberto Amato, Sonia Goncalves, Ewan Harrison, David K. Jackson, Ian Johnston, Dominic Kwiatkowski, Cordelia Langford, John Sillitoe on behalf of the Wellcome Sanger Institute COVID-19 Surveillance Team |
| EPI_ISL_851078, EPI_ISL_851080 | Lighthouse Lab in Cambridge | Wellcome Sanger Institute for the COVID-19 Genomics UK (COG-UK) Consortium | Rob Howes, The Lighthouse Lab in Cambridge and Alex Alderton, Roberto Amato, Sonia Goncalves, Ewan Harrison, David K. Jackson, Ian Johnston, Dominic Kwiatkowski, Cordelia Langford, John Sillitoe on behalf of the Wellcome Sanger Institute COVID-19 Surveillance Team |
| EPI_ISL_851081, EPI_ISL_851082 | Lighthouse Lab in Glasgow | Wellcome Sanger Institute for the COVID-19 Genomics UK (COG-UK) Consortium | Harper VanSteenhouse, Yumi Kasai, David Gray, Carol Clugston, Anna Dominiczak and Alex Alderton, Roberto Amato, Sonia Goncalves, Ewan Harrison, David K. Jackson, Ian Johnston, Dominic Kwiatkowski, Cordelia Langford, John Sillitoe on behalf of the Wellcome Sanger Institute COVID-19 Surveillance Team |
| EPI_ISL_851084 | Lighthouse Lab in Cambridge | Wellcome Sanger Institute for the COVID-19 Genomics UK (COG-UK) Consortium | Rob Howes, The Lighthouse Lab in Cambridge and Alex Alderton, Roberto Amato, Sonia Goncalves, Ewan Harrison, David K. Jackson, Ian Johnston, Dominic Kwiatkowski, Cordelia Langford, John Sillitoe on behalf of the Wellcome Sanger Institute COVID-19 Surveillance Team |
| EPI_ISL_851085 | Lighthouse Lab in Glasgow | Wellcome Sanger Institute for the COVID-19 Genomics UK (COG-UK) Consortium | Harper VanSteenhouse, Yumi Kasai, David Gray, Carol Clugston, Anna Dominiczak and Alex Alderton, Roberto Amato, Sonia Goncalves, Ewan Harrison, David K. Jackson, Ian Johnston, Dominic Kwiatkowski, Cordelia Langford, John Sillitoe on behalf of the Wellcome Sanger Institute COVID-19 Surveillance Team |
| EPI_ISL_851086, EPI_ISL_851087, EPI_ISL_851088, EPI_ISL_851089, EPI_ISL_851090, EPI_ISL_851091, EPI_ISL_851092, EPI_ISL_851093 | Lighthouse Lab in Cambridge | Wellcome Sanger Institute for the COVID-19 Genomics UK (COG-UK) Consortium | Rob Howes, The Lighthouse Lab in Cambridge and Alex Alderton, Roberto Amato, Sonia Goncalves, Ewan Harrison, David K. Jackson, Ian Johnston, Dominic Kwiatkowski, Cordelia Langford, John Sillitoe on behalf of the Wellcome Sanger Institute COVID-19 Surveillance Team |
| EPI_ISL_851094, EPI_ISL_851095 | Lighthouse Lab in Glasgow | Wellcome Sanger Institute for the COVID-19 Genomics UK (COG-UK) Consortium | Harper VanSteenhouse, Yumi Kasai, David Gray, Carol Clugston, Anna Dominiczak and Alex Alderton, Roberto Amato, Sonia Goncalves, Ewan Harrison, David K. Jackson, Ian Johnston, Dominic Kwiatkowski, Cordelia Langford, John Sillitoe on behalf of the Wellcome Sanger Institute COVID-19 Surveillance Team |
| EPI_ISL_851096, EPI_ISL_851097, EPI_ISL_851098, EPI_ISL_851099, EPI_ISL_851100 | Lighthouse Lab in Cambridge | Wellcome Sanger Institute for the COVID-19 Genomics UK (COG-UK) Consortium | Rob Howes, The Lighthouse Lab in Cambridge and Alex Alderton, Roberto Amato, Sonia Goncalves, Ewan Harrison, David K. Jackson, Ian Johnston, Dominic Kwiatkowski, Cordelia Langford, John Sillitoe on behalf of the Wellcome Sanger Institute COVID-19 Surveillance Team |
| EPI_ISL_851102 | Lighthouse Lab in Glasgow | Wellcome Sanger Institute for the COVID-19 Genomics UK (COG-UK) Consortium | Harper VanSteenhouse, Yumi Kasai, David Gray, Carol Clugston, Anna Dominiczak and Alex Alderton, Roberto Amato, Sonia Goncalves, Ewan Harrison, David K. Jackson, Ian Johnston, Dominic Kwiatkowski, Cordelia Langford, John Sillitoe on behalf of the Wellcome Sanger Institute COVID-19 Surveillance Team |
| EPI_ISL_851103 | Lighthouse Lab in Cambridge | Wellcome Sanger Institute for the COVID-19 Genomics UK (COG-UK) Consortium | Rob Howes, The Lighthouse Lab in Cambridge and Alex Alderton, Roberto Amato, Sonia Goncalves, Ewan Harrison, David K. Jackson, Ian Johnston, Dominic Kwiatkowski, Cordelia Langford, John Sillitoe on behalf of the Wellcome Sanger Institute COVID-19 Surveillance Team |
| EPI_ISL_851104 | Lighthouse Lab in Glasgow | Wellcome Sanger Institute for the COVID-19 Genomics UK (COG-UK) Consortium | Harper VanSteenhouse, Yumi Kasai, David Gray, Carol Clugston, Anna Dominiczak and Alex Alderton, Roberto Amato, Sonia Goncalves, Ewan Harrison, David K. Jackson, Ian Johnston, Dominic Kwiatkowski, Cordelia Langford, John Sillitoe on behalf of the Wellcome Sanger Institute COVID-19 Surveillance Team |
| EPI_ISL_851105, EPI_ISL_851106 | Lighthouse Lab in Cambridge | Wellcome Sanger Institute for the COVID-19 Genomics UK (COG-UK) Consortium | Rob Howes, The Lighthouse Lab in Cambridge and Alex Alderton, Roberto Amato, Sonia Goncalves, Ewan Harrison, David K. Jackson, Ian Johnston, Dominic Kwiatkowski, Cordelia Langford, John Sillitoe on behalf of the Wellcome Sanger Institute COVID-19 Surveillance Team |
| EPI_ISL_851107 | Lighthouse Lab in Glasgow | Wellcome Sanger Institute for the COVID-19 Genomics UK (COG-UK) Consortium | Harper VanSteenhouse, Yumi Kasai, David Gray, Carol Clugston, Anna Dominiczak and Alex Alderton, Roberto Amato, Sonia Goncalves, Ewan Harrison, David K. Jackson, Ian Johnston, Dominic Kwiatkowski, Cordelia Langford, John Sillitoe on behalf of the Wellcome Sanger Institute COVID-19 Surveillance Team |
| EPI_ISL_851108, EPI_ISL_851109, EPI_ISL_851111, EPI_ISL_851112, EPI_ISL_851113, EPI_ISL_851114 | Lighthouse Lab in Cambridge | Wellcome Sanger Institute for the COVID-19 Genomics UK (COG-UK) Consortium | Rob Howes, The Lighthouse Lab in Cambridge and Alex Alderton, Roberto Amato, Sonia Goncalves, Ewan Harrison, David K. Jackson, Ian Johnston, Dominic Kwiatkowski, Cordelia Langford, John Sillitoe on behalf of the Wellcome Sanger Institute COVID-19 Surveillance Team |
| EPI_ISL_851116 | Lighthouse Lab in Glasgow | Wellcome Sanger Institute for the COVID-19 Genomics UK (COG-UK) Consortium | Harper VanSteenhouse, Yumi Kasai, David Gray, Carol Clugston, Anna Dominiczak and Alex Alderton, Roberto Amato, Sonia Goncalves, Ewan Harrison, David K. Jackson, Ian Johnston, Dominic Kwiatkowski, Cordelia Langford, John Sillitoe on behalf of the Wellcome Sanger Institute COVID-19 Surveillance Team |
| EPI_ISL_851117, EPI_ISL_851118 | Lighthouse Lab in Cambridge | Wellcome Sanger Institute for the COVID-19 Genomics UK (COG-UK) Consortium | Rob Howes, The Lighthouse Lab in Cambridge and Alex Alderton, Roberto Amato, Sonia Goncalves, Ewan Harrison, David K. Jackson, Ian Johnston, Dominic Kwiatkowski, Cordelia Langford, John Sillitoe on behalf of the Wellcome Sanger Institute COVID-19 Surveillance Team |
| EPI_ISL_851119 | Lighthouse Lab in Glasgow | Wellcome Sanger Institute for the COVID-19 Genomics UK (COG-UK) Consortium | Harper VanSteenhouse, Yumi Kasai, David Gray, Carol Clugston, Anna Dominiczak and Alex Alderton, Roberto Amato, Sonia Goncalves, Ewan Harrison, David K. Jackson, Ian Johnston, Dominic Kwiatkowski, Cordelia Langford, John Sillitoe on behalf of the Wellcome Sanger Institute COVID-19 Surveillance Team |
| EPI_ISL_851120, EPI_ISL_851121, EPI_ISL_851122, EPI_ISL_851123 | Lighthouse Lab in Cambridge | Wellcome Sanger Institute for the COVID-19 Genomics UK (COG-UK) Consortium | Rob Howes, The Lighthouse Lab in Cambridge and Alex Alderton, Roberto Amato, Sonia Goncalves, Ewan Harrison, David K. Jackson, Ian Johnston, Dominic Kwiatkowski, Cordelia Langford, John Sillitoe on behalf of the Wellcome Sanger Institute COVID-19 Surveillance Team |
| EPI_ISL_851124, EPI_ISL_851125 | Lighthouse Lab in Glasgow | Wellcome Sanger Institute for the COVID-19 Genomics UK (COG-UK) Consortium | Harper VanSteenhouse, Yumi Kasai, David Gray, Carol Clugston, Anna Dominiczak and Alex Alderton, Roberto Amato, Sonia Goncalves, Ewan Harrison, David K. Jackson, Ian Johnston, Dominic Kwiatkowski, Cordelia Langford, John Sillitoe on behalf of the Wellcome Sanger Institute COVID-19 Surveillance Team |
| EPI_ISL_851127, EPI_ISL_851128, EPI_ISL_851129 | Lighthouse Lab in Cambridge | Wellcome Sanger Institute for the COVID-19 Genomics UK (COG-UK) Consortium | Rob Howes, The Lighthouse Lab in Cambridge and Alex Alderton, Roberto Amato, Sonia Goncalves, Ewan Harrison, David K. Jackson, Ian Johnston, Dominic Kwiatkowski, Cordelia Langford, John Sillitoe on behalf of the Wellcome Sanger Institute COVID-19 Surveillance Team |
| EPI_ISL_851130 | Lighthouse Lab in Glasgow | Wellcome Sanger Institute for the COVID-19 Genomics UK (COG-UK) Consortium | Harper VanSteenhouse, Yumi Kasai, David Gray, Carol Clugston, Anna Dominiczak and Alex Alderton, Roberto Amato, Sonia Goncalves, Ewan Harrison, David K. Jackson, Ian Johnston, Dominic Kwiatkowski, Cordelia Langford, John Sillitoe on behalf of the Wellcome Sanger Institute COVID-19 Surveillance Team |
| EPI_ISL_851131, EPI_ISL_851132 | Lighthouse Lab in Cambridge | Wellcome Sanger Institute for the COVID-19 Genomics UK (COG-UK) Consortium | Rob Howes, The Lighthouse Lab in Cambridge and Alex Alderton, Roberto Amato, Sonia Goncalves, Ewan Harrison, David K. Jackson, Ian Johnston, Dominic Kwiatkowski, Cordelia Langford, John Sillitoe on behalf of the Wellcome Sanger Institute COVID-19 Surveillance Team |
| EPI_ISL_851134, EPI_ISL_851135 | Lighthouse Lab in Glasgow | Wellcome Sanger Institute for the COVID-19 Genomics UK (COG-UK) Consortium | Harper VanSteenhouse, Yumi Kasai, David Gray, Carol Clugston, Anna Dominiczak and Alex Alderton, Roberto Amato, Sonia Goncalves, Ewan Harrison, David K. Jackson, Ian Johnston, Dominic Kwiatkowski, Cordelia Langford, John Sillitoe on behalf of the Wellcome Sanger Institute COVID-19 Surveillance |

| | | | |
|---|---|---|---|
| | | | Team |
| EPI_ISL_851136 | Lighthouse Lab in Cambridge | Wellcome Sanger Institute for the COVID-19 Genomics UK (COG-UK) Consortium | Rob Howes, The Lighthouse Lab in Cambridge and Alex Alderton, Roberto Amato, Sonia Goncalves, Ewan Harrison, David K. Jackson, Ian Johnston, Dominic Kwiatkowski, Cordelia Langford, John Sillitoe on behalf of the Wellcome Sanger Institute COVID-19 Surveillance Team |
| EPI_ISL_851137 | Lighthouse Lab in Glasgow | Wellcome Sanger Institute for the COVID-19 Genomics UK (COG-UK) Consortium | Harper VanSteenhouse, Yumi Kasai, David Gray, Carol Clugston, Anna Dominiczak and Alex Alderton, Roberto Amato, Sonia Goncalves, Ewan Harrison, David K. Jackson, Ian Johnston, Dominic Kwiatkowski, Cordelia Langford, John Sillitoe on behalf of the Wellcome Sanger Institute COVID-19 Surveillance Team |
| EPI_ISL_851138, EPI_ISL_851139, EPI_ISL_851140, EPI_ISL_851141, EPI_ISL_851144, EPI_ISL_851145 | Lighthouse Lab in Cambridge | Wellcome Sanger Institute for the COVID-19 Genomics UK (COG-UK) Consortium | Rob Howes, The Lighthouse Lab in Cambridge and Alex Alderton, Roberto Amato, Sonia Goncalves, Ewan Harrison, David K. Jackson, Ian Johnston, Dominic Kwiatkowski, Cordelia Langford, John Sillitoe on behalf of the Wellcome Sanger Institute COVID-19 Surveillance Team |
| EPI_ISL_851146, EPI_ISL_851147, EPI_ISL_851148 | Lighthouse Lab in Glasgow | Wellcome Sanger Institute for the COVID-19 Genomics UK (COG-UK) Consortium | Harper VanSteenhouse, Yumi Kasai, David Gray, Carol Clugston, Anna Dominiczak and Alex Alderton, Roberto Amato, Sonia Goncalves, Ewan Harrison, David K. Jackson, Ian Johnston, Dominic Kwiatkowski, Cordelia Langford, John Sillitoe on behalf of the Wellcome Sanger Institute COVID-19 Surveillance Team |
| EPI_ISL_851149, EPI_ISL_851151, EPI_ISL_851152, EPI_ISL_851153 | Lighthouse Lab in Cambridge | Wellcome Sanger Institute for the COVID-19 Genomics UK (COG-UK) Consortium | Rob Howes, The Lighthouse Lab in Cambridge and Alex Alderton, Roberto Amato, Sonia Goncalves, Ewan Harrison, David K. Jackson, Ian Johnston, Dominic Kwiatkowski, Cordelia Langford, John Sillitoe on behalf of the Wellcome Sanger Institute COVID-19 Surveillance Team |
| EPI_ISL_851154 | Lighthouse Lab in Glasgow | Wellcome Sanger Institute for the COVID-19 Genomics UK (COG-UK) Consortium | Harper VanSteenhouse, Yumi Kasai, David Gray, Carol Clugston, Anna Dominiczak and Alex Alderton, Roberto Amato, Sonia Goncalves, Ewan Harrison, David K. Jackson, Ian Johnston, Dominic Kwiatkowski, Cordelia Langford, John Sillitoe on behalf of the Wellcome Sanger Institute COVID-19 Surveillance Team |
| EPI_ISL_851155, EPI_ISL_851156, EPI_ISL_851157, EPI_ISL_851158, EPI_ISL_851159, EPI_ISL_851160, EPI_ISL_851161, EPI_ISL_851162, EPI_ISL_851163, EPI_ISL_851164, EPI_ISL_851165 | | | |
| see above | Lighthouse Lab in Cambridge | Wellcome Sanger Institute for the COVID-19 Genomics UK (COG-UK) Consortium | Rob Howes, The Lighthouse Lab in Cambridge and Alex Alderton, Roberto Amato, Sonia Goncalves, Ewan Harrison, David K. Jackson, Ian Johnston, Dominic Kwiatkowski, Cordelia Langford, John Sillitoe on behalf of the Wellcome Sanger Institute COVID-19 Surveillance Team |
| EPI_ISL_851167 | Lighthouse Lab in Glasgow | Wellcome Sanger Institute for the COVID-19 Genomics UK (COG-UK) Consortium | Harper VanSteenhouse, Yumi Kasai, David Gray, Carol Clugston, Anna Dominiczak and Alex Alderton, Roberto Amato, Sonia Goncalves, Ewan Harrison, David K. Jackson, Ian Johnston, Dominic Kwiatkowski, Cordelia Langford, John Sillitoe on behalf of the Wellcome Sanger Institute COVID-19 Surveillance Team |
| EPI_ISL_851168, EPI_ISL_851169, EPI_ISL_851171, EPI_ISL_851172, EPI_ISL_851173, EPI_ISL_851174, EPI_ISL_851176, EPI_ISL_851177 | Lighthouse Lab in Cambridge | Wellcome Sanger Institute for the COVID-19 Genomics UK (COG-UK) Consortium | Rob Howes, The Lighthouse Lab in Cambridge and Alex Alderton, Roberto Amato, Sonia Goncalves, Ewan Harrison, David K. Jackson, Ian Johnston, Dominic Kwiatkowski, Cordelia Langford, John Sillitoe on behalf of the Wellcome Sanger Institute COVID-19 Surveillance Team |
| EPI_ISL_851178 | Lighthouse Lab in Glasgow | Wellcome Sanger Institute for the COVID-19 Genomics UK (COG-UK) Consortium | Harper VanSteenhouse, Yumi Kasai, David Gray, Carol Clugston, Anna Dominiczak and Alex Alderton, Roberto Amato, Sonia Goncalves, Ewan Harrison, David K. Jackson, Ian Johnston, Dominic Kwiatkowski, Cordelia Langford, John Sillitoe on behalf of the Wellcome Sanger Institute COVID-19 Surveillance Team |
| EPI_ISL_851179, EPI_ISL_851181 | Lighthouse Lab in Cambridge | Wellcome Sanger Institute for the COVID-19 Genomics UK (COG-UK) Consortium | Rob Howes, The Lighthouse Lab in Cambridge and Alex Alderton, Roberto Amato, Sonia Goncalves, Ewan Harrison, David K. Jackson, Ian Johnston, Dominic Kwiatkowski, Cordelia Langford, John Sillitoe on behalf of the Wellcome Sanger Institute COVID-19 Surveillance Team |
| EPI_ISL_851182 | Lighthouse Lab in Glasgow | Wellcome Sanger Institute for the COVID-19 Genomics UK (COG-UK) Consortium | Harper VanSteenhouse, Yumi Kasai, David Gray, Carol Clugston, Anna Dominiczak and Alex Alderton, Roberto Amato, Sonia Goncalves, Ewan Harrison, David K. Jackson, Ian Johnston, Dominic Kwiatkowski, Cordelia Langford, John Sillitoe on behalf of the Wellcome Sanger Institute COVID-19 Surveillance Team |
| EPI_ISL_851183, EPI_ISL_851184, EPI_ISL_851185, EPI_ISL_851186, EPI_ISL_851188, EPI_ISL_851191, EPI_ISL_851192, EPI_ISL_851195, EPI_ISL_851197, EPI_ISL_851198, EPI_ISL_851199, EPI_ISL_851200, EPI_ISL_851201, EPI_ISL_851203, EPI_ISL_851204, EPI_ISL_851205 | | | |
| see above | Lighthouse Lab in Cambridge | Wellcome Sanger Institute for the COVID-19 Genomics UK (COG-UK) Consortium | Rob Howes, The Lighthouse Lab in Cambridge and Alex Alderton, Roberto Amato, Sonia Goncalves, Ewan Harrison, David K. Jackson, Ian Johnston, Dominic Kwiatkowski, Cordelia Langford, John Sillitoe on behalf of the Wellcome Sanger Institute COVID-19 Surveillance Team |
| EPI_ISL_851206, EPI_ISL_851207 | Lighthouse Lab in Glasgow | Wellcome Sanger Institute for the COVID-19 Genomics UK (COG-UK) Consortium | Harper VanSteenhouse, Yumi Kasai, David Gray, Carol Clugston, Anna Dominiczak and Alex Alderton, Roberto Amato, Sonia Goncalves, Ewan Harrison, David K. Jackson, Ian Johnston, Dominic Kwiatkowski, Cordelia Langford, John Sillitoe on behalf of the Wellcome Sanger Institute COVID-19 Surveillance Team |
| EPI_ISL_851209, EPI_ISL_851210, EPI_ISL_851211, EPI_ISL_851212 | Lighthouse Lab in Cambridge | Wellcome Sanger Institute for the COVID-19 Genomics UK (COG-UK) Consortium | Rob Howes, The Lighthouse Lab in Cambridge and Alex Alderton, Roberto Amato, Sonia Goncalves, Ewan Harrison, David K. Jackson, Ian Johnston, Dominic Kwiatkowski, Cordelia Langford, John Sillitoe on behalf of the Wellcome Sanger Institute COVID-19 Surveillance Team |
| EPI_ISL_851215 | Lighthouse Lab in Glasgow | Wellcome Sanger Institute for the COVID-19 Genomics UK (COG-UK) Consortium | Harper VanSteenhouse, Yumi Kasai, David Gray, Carol Clugston, Anna Dominiczak and Alex Alderton, Roberto Amato, Sonia Goncalves, Ewan Harrison, David K. Jackson, Ian Johnston, Dominic Kwiatkowski, Cordelia Langford, John Sillitoe on behalf of the Wellcome Sanger Institute COVID-19 Surveillance Team |
| EPI_ISL_851216 | Lighthouse Lab in Cambridge | Wellcome Sanger Institute for the COVID-19 Genomics UK (COG-UK) Consortium | Rob Howes, The Lighthouse Lab in Cambridge and Alex Alderton, Roberto Amato, Sonia Goncalves, Ewan Harrison, David K. Jackson, Ian Johnston, Dominic Kwiatkowski, Cordelia Langford, John Sillitoe on behalf of the Wellcome Sanger Institute COVID-19 Surveillance Team |
| EPI_ISL_851218 | Lighthouse Lab in Glasgow | Wellcome Sanger Institute for the COVID-19 Genomics UK (COG-UK) Consortium | Harper VanSteenhouse, Yumi Kasai, David Gray, Carol Clugston, Anna Dominiczak and Alex Alderton, Roberto Amato, Sonia Goncalves, Ewan Harrison, David K. Jackson, Ian Johnston, Dominic Kwiatkowski, Cordelia Langford, John Sillitoe on behalf of the Wellcome Sanger Institute COVID-19 Surveillance Team |
| EPI_ISL_851219, EPI_ISL_851220, EPI_ISL_851221, EPI_ISL_851222, EPI_ISL_851223, EPI_ISL_851224 | Lighthouse Lab in Cambridge | Wellcome Sanger Institute for the COVID-19 Genomics UK (COG-UK) Consortium | Rob Howes, The Lighthouse Lab in Cambridge and Alex Alderton, Roberto Amato, Sonia Goncalves, Ewan Harrison, David K. Jackson, Ian Johnston, Dominic Kwiatkowski, Cordelia Langford, John Sillitoe on behalf of the Wellcome Sanger Institute COVID-19 Surveillance Team |
| EPI_ISL_851225 | Lighthouse Lab in Glasgow | Wellcome Sanger Institute for the COVID-19 Genomics UK (COG-UK) Consortium | Harper VanSteenhouse, Yumi Kasai, David Gray, Carol Clugston, Anna Dominiczak and Alex Alderton, Roberto Amato, Sonia Goncalves, Ewan Harrison, David K. Jackson, Ian Johnston, Dominic Kwiatkowski, Cordelia Langford, John Sillitoe on behalf of the Wellcome Sanger Institute COVID-19 Surveillance Team |
| EPI_ISL_851226, EPI_ISL_851227, EPI_ISL_851228, EPI_ISL_851230, EPI_ISL_851231, EPI_ISL_851232, EPI_ISL_851233, EPI_ISL_851234, EPI_ISL_851235 | Lighthouse Lab in Cambridge | Wellcome Sanger Institute for the COVID-19 Genomics UK (COG-UK) Consortium | Rob Howes, The Lighthouse Lab in Cambridge and Alex Alderton, Roberto Amato, Sonia Goncalves, Ewan Harrison, David K. Jackson, Ian Johnston, Dominic Kwiatkowski, Cordelia Langford, John Sillitoe on behalf of the Wellcome Sanger Institute COVID-19 Surveillance Team |
| EPI_ISL_851237 | Lighthouse Lab in Glasgow | Wellcome Sanger Institute for the COVID-19 Genomics UK (COG-UK) Consortium | Harper VanSteenhouse, Yumi Kasai, David Gray, Carol Clugston, Anna Dominiczak and Alex Alderton, Roberto Amato, Sonia Goncalves, Ewan Harrison, David K. Jackson, Ian Johnston, Dominic Kwiatkowski, Cordelia Langford, John Sillitoe on behalf of the Wellcome Sanger Institute COVID-19 Surveillance Team |
| EPI_ISL_851238, EPI_ISL_851239, EPI_ISL_851240, EPI_ISL_851242, EPI_ISL_851243, EPI_ISL_851244, EPI_ISL_851245 | Lighthouse Lab in Cambridge | Wellcome Sanger Institute for the COVID-19 Genomics UK (COG-UK) Consortium | Rob Howes, The Lighthouse Lab in Cambridge and Alex Alderton, Roberto Amato, Sonia Goncalves, Ewan Harrison, David K. Jackson, Ian Johnston, Dominic Kwiatkowski, Cordelia Langford, John Sillitoe on behalf of the Wellcome Sanger Institute COVID-19 Surveillance Team |
| EPI_ISL_851246 | Lighthouse Lab in Glasgow | Wellcome Sanger Institute for the COVID-19 Genomics UK (COG-UK) Consortium | Harper VanSteenhouse, Yumi Kasai, David Gray, Carol Clugston, Anna Dominiczak and Alex Alderton, Roberto Amato, Sonia Goncalves, Ewan Harrison, David K. Jackson, Ian Johnston, Dominic Kwiatkowski, Cordelia Langford, John Sillitoe on behalf of the Wellcome Sanger Institute COVID-19 Surveillance Team |
| EPI_ISL_851247, EPI_ISL_851248, EPI_ISL_851249, EPI_ISL_851250 | Lighthouse Lab in Cambridge | Wellcome Sanger Institute for the COVID-19 Genomics UK (COG-UK) Consortium | Rob Howes, The Lighthouse Lab in Cambridge and Alex Alderton, Roberto Amato, Sonia Goncalves, Ewan Harrison, David K. Jackson, Ian Johnston, Dominic Kwiatkowski, Cordelia Langford, John Sillitoe on behalf of the Wellcome Sanger Institute COVID-19 Surveillance Team |

| | | | |
|---|---|---|---|
| EPI_ISL_851252, EPI_ISL_851253 | Lighthouse Lab in Glasgow | Wellcome Sanger Institute for the COVID-19 Genomics UK (COG-UK) Consortium | Harper VanSteenhouse, Yumi Kasai, David Gray, Carol Clugston, Anna Dominiczak and Alex Alderton, Roberto Amato, Sonia Goncalves, Ewan Harrison, David K. Jackson, Ian Johnston, Dominic Kwiatkowski, Cordelia Langford, John Sillitoe on behalf of the Wellcome Sanger Institute COVID-19 Surveillance Team |
| EPI_ISL_851255, EPI_ISL_851256 | Lighthouse Lab in Cambridge | Wellcome Sanger Institute for the COVID-19 Genomics UK (COG-UK) Consortium | Rob Howes, The Lighthouse Lab in Cambridge and Alex Alderton, Roberto Amato, Sonia Goncalves, Ewan Harrison, David K. Jackson, Ian Johnston, Dominic Kwiatkowski, Cordelia Langford, John Sillitoe on behalf of the Wellcome Sanger Institute COVID-19 Surveillance Team |
| EPI_ISL_851257, EPI_ISL_851259 | Lighthouse Lab in Glasgow | Wellcome Sanger Institute for the COVID-19 Genomics UK (COG-UK) Consortium | Harper VanSteenhouse, Yumi Kasai, David Gray, Carol Clugston, Anna Dominiczak and Alex Alderton, Roberto Amato, Sonia Goncalves, Ewan Harrison, David K. Jackson, Ian Johnston, Dominic Kwiatkowski, Cordelia Langford, John Sillitoe on behalf of the Wellcome Sanger Institute COVID-19 Surveillance Team |
| EPI_ISL_851260, EPI_ISL_851261, EPI_ISL_851262, EPI_ISL_851263 | Lighthouse Lab in Cambridge | Wellcome Sanger Institute for the COVID-19 Genomics UK (COG-UK) Consortium | Rob Howes, The Lighthouse Lab in Cambridge and Alex Alderton, Roberto Amato, Sonia Goncalves, Ewan Harrison, David K. Jackson, Ian Johnston, Dominic Kwiatkowski, Cordelia Langford, John Sillitoe on behalf of the Wellcome Sanger Institute COVID-19 Surveillance Team |
| EPI_ISL_851264 | Lighthouse Lab in Glasgow | Wellcome Sanger Institute for the COVID-19 Genomics UK (COG-UK) Consortium | Harper VanSteenhouse, Yumi Kasai, David Gray, Carol Clugston, Anna Dominiczak and Alex Alderton, Roberto Amato, Sonia Goncalves, Ewan Harrison, David K. Jackson, Ian Johnston, Dominic Kwiatkowski, Cordelia Langford, John Sillitoe on behalf of the Wellcome Sanger Institute COVID-19 Surveillance Team |
| EPI_ISL_851265, EPI_ISL_851266, EPI_ISL_851267, EPI_ISL_851268 | Lighthouse Lab in Cambridge | Wellcome Sanger Institute for the COVID-19 Genomics UK (COG-UK) Consortium | Rob Howes, The Lighthouse Lab in Cambridge and Alex Alderton, Roberto Amato, Sonia Goncalves, Ewan Harrison, David K. Jackson, Ian Johnston, Dominic Kwiatkowski, Cordelia Langford, John Sillitoe on behalf of the Wellcome Sanger Institute COVID-19 Surveillance Team |
| EPI_ISL_851270 | Lighthouse Lab in Glasgow | Wellcome Sanger Institute for the COVID-19 Genomics UK (COG-UK) Consortium | Harper VanSteenhouse, Yumi Kasai, David Gray, Carol Clugston, Anna Dominiczak and Alex Alderton, Roberto Amato, Sonia Goncalves, Ewan Harrison, David K. Jackson, Ian Johnston, Dominic Kwiatkowski, Cordelia Langford, John Sillitoe on behalf of the Wellcome Sanger Institute COVID-19 Surveillance Team |
| EPI_ISL_851271, EPI_ISL_851272, EPI_ISL_851274, EPI_ISL_851275 | Lighthouse Lab in Cambridge | Wellcome Sanger Institute for the COVID-19 Genomics UK (COG-UK) Consortium | Rob Howes, The Lighthouse Lab in Cambridge and Alex Alderton, Roberto Amato, Sonia Goncalves, Ewan Harrison, David K. Jackson, Ian Johnston, Dominic Kwiatkowski, Cordelia Langford, John Sillitoe on behalf of the Wellcome Sanger Institute COVID-19 Surveillance Team |
| EPI_ISL_851276 | Lighthouse Lab in Glasgow | Wellcome Sanger Institute for the COVID-19 Genomics UK (COG-UK) Consortium | Harper VanSteenhouse, Yumi Kasai, David Gray, Carol Clugston, Anna Dominiczak and Alex Alderton, Roberto Amato, Sonia Goncalves, Ewan Harrison, David K. Jackson, Ian Johnston, Dominic Kwiatkowski, Cordelia Langford, John Sillitoe on behalf of the Wellcome Sanger Institute COVID-19 Surveillance Team |
| EPI_ISL_851277, EPI_ISL_851278, EPI_ISL_851279, EPI_ISL_851280, EPI_ISL_851281, EPI_ISL_851282, EPI_ISL_851283, EPI_ISL_851284 | Lighthouse Lab in Cambridge | Wellcome Sanger Institute for the COVID-19 Genomics UK (COG-UK) Consortium | Rob Howes, The Lighthouse Lab in Cambridge and Alex Alderton, Roberto Amato, Sonia Goncalves, Ewan Harrison, David K. Jackson, Ian Johnston, Dominic Kwiatkowski, Cordelia Langford, John Sillitoe on behalf of the Wellcome Sanger Institute COVID-19 Surveillance Team |
| EPI_ISL_851285 | Lighthouse Lab in Glasgow | Wellcome Sanger Institute for the COVID-19 Genomics UK (COG-UK) Consortium | Harper VanSteenhouse, Yumi Kasai, David Gray, Carol Clugston, Anna Dominiczak and Alex Alderton, Roberto Amato, Sonia Goncalves, Ewan Harrison, David K. Jackson, Ian Johnston, Dominic Kwiatkowski, Cordelia Langford, John Sillitoe on behalf of the Wellcome Sanger Institute COVID-19 Surveillance Team |
| EPI_ISL_851286, EPI_ISL_851288, EPI_ISL_851289, EPI_ISL_851290, EPI_ISL_851291, EPI_ISL_851292, EPI_ISL_851293, EPI_ISL_851294, EPI_ISL_851295 | Lighthouse Lab in Cambridge | Wellcome Sanger Institute for the COVID-19 Genomics UK (COG-UK) Consortium | Rob Howes, The Lighthouse Lab in Cambridge and Alex Alderton, Roberto Amato, Sonia Goncalves, Ewan Harrison, David K. Jackson, Ian Johnston, Dominic Kwiatkowski, Cordelia Langford, John Sillitoe on behalf of the Wellcome Sanger Institute COVID-19 Surveillance Team |
| EPI_ISL_851296 | Lighthouse Lab in Glasgow | Wellcome Sanger Institute for the COVID-19 Genomics UK (COG-UK) Consortium | Harper VanSteenhouse, Yumi Kasai, David Gray, Carol Clugston, Anna Dominiczak and Alex Alderton, Roberto Amato, Sonia Goncalves, Ewan Harrison, David K. Jackson, Ian Johnston, Dominic Kwiatkowski, Cordelia Langford, John Sillitoe on behalf of the Wellcome Sanger Institute COVID-19 Surveillance Team |
| EPI_ISL_851297, EPI_ISL_851299, EPI_ISL_851300 | Lighthouse Lab in Cambridge | Wellcome Sanger Institute for the COVID-19 Genomics UK (COG-UK) Consortium | Rob Howes, The Lighthouse Lab in Cambridge and Alex Alderton, Roberto Amato, Sonia Goncalves, Ewan Harrison, David K. Jackson, Ian Johnston, Dominic Kwiatkowski, Cordelia Langford, John Sillitoe on behalf of the Wellcome Sanger Institute COVID-19 Surveillance Team |
| EPI_ISL_851301 | Lighthouse Lab in Glasgow | Wellcome Sanger Institute for the COVID-19 Genomics UK (COG-UK) Consortium | Harper VanSteenhouse, Yumi Kasai, David Gray, Carol Clugston, Anna Dominiczak and Alex Alderton, Roberto Amato, Sonia Goncalves, Ewan Harrison, David K. Jackson, Ian Johnston, Dominic Kwiatkowski, Cordelia Langford, John Sillitoe on behalf of the Wellcome Sanger Institute COVID-19 Surveillance Team |
| EPI_ISL_851302, EPI_ISL_851303, EPI_ISL_851304 | Lighthouse Lab in Cambridge | Wellcome Sanger Institute for the COVID-19 Genomics UK (COG-UK) Consortium | Rob Howes, The Lighthouse Lab in Cambridge and Alex Alderton, Roberto Amato, Sonia Goncalves, Ewan Harrison, David K. Jackson, Ian Johnston, Dominic Kwiatkowski, Cordelia Langford, John Sillitoe on behalf of the Wellcome Sanger Institute COVID-19 Surveillance Team |
| EPI_ISL_851305, EPI_ISL_851306, EPI_ISL_851307 | Lighthouse Lab in Glasgow | Wellcome Sanger Institute for the COVID-19 Genomics UK (COG-UK) Consortium | Harper VanSteenhouse, Yumi Kasai, David Gray, Carol Clugston, Anna Dominiczak and Alex Alderton, Roberto Amato, Sonia Goncalves, Ewan Harrison, David K. Jackson, Ian Johnston, Dominic Kwiatkowski, Cordelia Langford, John Sillitoe on behalf of the Wellcome Sanger Institute COVID-19 Surveillance Team |
| EPI_ISL_851308, EPI_ISL_851309 | Lighthouse Lab in Cambridge | Wellcome Sanger Institute for the COVID-19 Genomics UK (COG-UK) Consortium | Rob Howes, The Lighthouse Lab in Cambridge and Alex Alderton, Roberto Amato, Sonia Goncalves, Ewan Harrison, David K. Jackson, Ian Johnston, Dominic Kwiatkowski, Cordelia Langford, John Sillitoe on behalf of the Wellcome Sanger Institute COVID-19 Surveillance Team |
| EPI_ISL_851310 | Lighthouse Lab in Glasgow | Wellcome Sanger Institute for the COVID-19 Genomics UK (COG-UK) Consortium | Harper VanSteenhouse, Yumi Kasai, David Gray, Carol Clugston, Anna Dominiczak and Alex Alderton, Roberto Amato, Sonia Goncalves, Ewan Harrison, David K. Jackson, Ian Johnston, Dominic Kwiatkowski, Cordelia Langford, John Sillitoe on behalf of the Wellcome Sanger Institute COVID-19 Surveillance Team |
| EPI_ISL_851311, EPI_ISL_851312, EPI_ISL_851313, EPI_ISL_851314, EPI_ISL_851315, EPI_ISL_851316, EPI_ISL_851317, EPI_ISL_851318, EPI_ISL_851319, EPI_ISL_851320, EPI_ISL_851321, EPI_ISL_851322, EPI_ISL_851323, EPI_ISL_851324 | | | |
| see above | Lighthouse Lab in Cambridge | Wellcome Sanger Institute for the COVID-19 Genomics UK (COG-UK) Consortium | Rob Howes, The Lighthouse Lab in Cambridge and Alex Alderton, Roberto Amato, Sonia Goncalves, Ewan Harrison, David K. Jackson, Ian Johnston, Dominic Kwiatkowski, Cordelia Langford, John Sillitoe on behalf of the Wellcome Sanger Institute COVID-19 Surveillance Team |
| EPI_ISL_851325 | Lighthouse Lab in Glasgow | Wellcome Sanger Institute for the COVID-19 Genomics UK (COG-UK) Consortium | Harper VanSteenhouse, Yumi Kasai, David Gray, Carol Clugston, Anna Dominiczak and Alex Alderton, Roberto Amato, Sonia Goncalves, Ewan Harrison, David K. Jackson, Ian Johnston, Dominic Kwiatkowski, Cordelia Langford, John Sillitoe on behalf of the Wellcome Sanger Institute COVID-19 Surveillance Team |
| EPI_ISL_851326 | Lighthouse Lab in Cambridge | Wellcome Sanger Institute for the COVID-19 Genomics UK (COG-UK) Consortium | Rob Howes, The Lighthouse Lab in Cambridge and Alex Alderton, Roberto Amato, Sonia Goncalves, Ewan Harrison, David K. Jackson, Ian Johnston, Dominic Kwiatkowski, Cordelia Langford, John Sillitoe on behalf of the Wellcome Sanger Institute COVID-19 Surveillance Team |
| EPI_ISL_851327 | Lighthouse Lab in Glasgow | Wellcome Sanger Institute for the COVID-19 Genomics UK (COG-UK) Consortium | Harper VanSteenhouse, Yumi Kasai, David Gray, Carol Clugston, Anna Dominiczak and Alex Alderton, Roberto Amato, Sonia Goncalves, Ewan Harrison, David K. Jackson, Ian Johnston, Dominic Kwiatkowski, Cordelia Langford, John Sillitoe on behalf of the Wellcome Sanger Institute COVID-19 Surveillance Team |
| EPI_ISL_851328, EPI_ISL_851330 | Lighthouse Lab in Cambridge | Wellcome Sanger Institute for the COVID-19 Genomics UK (COG-UK) Consortium | Rob Howes, The Lighthouse Lab in Cambridge and Alex Alderton, Roberto Amato, Sonia Goncalves, Ewan Harrison, David K. Jackson, Ian Johnston, Dominic Kwiatkowski, Cordelia Langford, John Sillitoe on behalf of the Wellcome Sanger Institute COVID-19 Surveillance Team |
| EPI_ISL_851331 | Lighthouse Lab in Glasgow | Wellcome Sanger Institute for the COVID-19 Genomics UK (COG-UK) Consortium | Harper VanSteenhouse, Yumi Kasai, David Gray, Carol Clugston, Anna Dominiczak and Alex Alderton, Roberto Amato, Sonia Goncalves, Ewan Harrison, David K. Jackson, Ian Johnston, Dominic Kwiatkowski, Cordelia Langford, John Sillitoe on behalf of the Wellcome Sanger Institute COVID-19 Surveillance Team |
| EPI_ISL_851333, EPI_ISL_851334 | Lighthouse Lab in Cambridge | Wellcome Sanger Institute for the COVID-19 Genomics UK (COG-UK) Consortium | Rob Howes, The Lighthouse Lab in Cambridge and Alex Alderton, Roberto Amato, Sonia Goncalves, Ewan Harrison, David K. Jackson, Ian Johnston, Dominic Kwiatkowski, Cordelia Langford, John Sillitoe on behalf of the Wellcome Sanger Institute COVID-19 Surveillance Team |
| EPI_ISL_851335 | Lighthouse Lab in Glasgow | Wellcome Sanger Institute for the COVID-19 Genomics UK (COG-UK) Consortium | Harper VanSteenhouse, Yumi Kasai, David Gray, Carol Clugston, Anna Dominiczak and Alex Alderton, Roberto Amato, Sonia Goncalves, Ewan Harrison, David K. Jackson, Ian Johnston, Dominic Kwiatkowski, Cordelia Langford, John Sillitoe on behalf of the Wellcome Sanger Institute COVID-19 Surveillance Team |

| | | | |
|---|---|---|---|
| EPI_ISL_851336, EPI_ISL_851337, EPI_ISL_851339, EPI_ISL_851340 | Lighthouse Lab in Cambridge | Wellcome Sanger Institute for the COVID-19 Genomics UK (COG-UK) Consortium | Rob Howes, The Lighthouse Lab in Cambridge and Alex Alderton, Roberto Amato, Sonia Goncalves, Ewan Harrison, David K. Jackson, Ian Johnston, Dominic Kwiatkowski, Cordelia Langford, John Sillitoe on behalf of the Wellcome Sanger Institute COVID-19 Surveillance Team |
| EPI_ISL_851341 | Lighthouse Lab in Glasgow | Wellcome Sanger Institute for the COVID-19 Genomics UK (COG-UK) Consortium | Harper VanSteenhouse, Yumi Kasai, David Gray, Carol Clugston, Anna Dominiczak and Alex Alderton, Roberto Amato, Sonia Goncalves, Ewan Harrison, David K. Jackson, Ian Johnston, Dominic Kwiatkowski, Cordelia Langford, John Sillitoe on behalf of the Wellcome Sanger Institute COVID-19 Surveillance Team |
| EPI_ISL_851344, EPI_ISL_851345 | Lighthouse Lab in Cambridge | Wellcome Sanger Institute for the COVID-19 Genomics UK (COG-UK) Consortium | Rob Howes, The Lighthouse Lab in Cambridge and Alex Alderton, Roberto Amato, Sonia Goncalves, Ewan Harrison, David K. Jackson, Ian Johnston, Dominic Kwiatkowski, Cordelia Langford, John Sillitoe on behalf of the Wellcome Sanger Institute COVID-19 Surveillance Team |
| EPI_ISL_851346 | Lighthouse Lab in Glasgow | Wellcome Sanger Institute for the COVID-19 Genomics UK (COG-UK) Consortium | Harper VanSteenhouse, Yumi Kasai, David Gray, Carol Clugston, Anna Dominiczak and Alex Alderton, Roberto Amato, Sonia Goncalves, Ewan Harrison, David K. Jackson, Ian Johnston, Dominic Kwiatkowski, Cordelia Langford, John Sillitoe on behalf of the Wellcome Sanger Institute COVID-19 Surveillance Team |
| EPI_ISL_851347, EPI_ISL_851348, EPI_ISL_851349, EPI_ISL_851350 | Lighthouse Lab in Cambridge | Wellcome Sanger Institute for the COVID-19 Genomics UK (COG-UK) Consortium | Rob Howes, The Lighthouse Lab in Cambridge and Alex Alderton, Roberto Amato, Sonia Goncalves, Ewan Harrison, David K. Jackson, Ian Johnston, Dominic Kwiatkowski, Cordelia Langford, John Sillitoe on behalf of the Wellcome Sanger Institute COVID-19 Surveillance Team |
| EPI_ISL_851351 | Lighthouse Lab in Glasgow | Wellcome Sanger Institute for the COVID-19 Genomics UK (COG-UK) Consortium | Harper VanSteenhouse, Yumi Kasai, David Gray, Carol Clugston, Anna Dominiczak and Alex Alderton, Roberto Amato, Sonia Goncalves, Ewan Harrison, David K. Jackson, Ian Johnston, Dominic Kwiatkowski, Cordelia Langford, John Sillitoe on behalf of the Wellcome Sanger Institute COVID-19 Surveillance Team |
| EPI_ISL_851352, EPI_ISL_851355, EPI_ISL_851356, EPI_ISL_851357 | Lighthouse Lab in Cambridge | Wellcome Sanger Institute for the COVID-19 Genomics UK (COG-UK) Consortium | Rob Howes, The Lighthouse Lab in Cambridge and Alex Alderton, Roberto Amato, Sonia Goncalves, Ewan Harrison, David K. Jackson, Ian Johnston, Dominic Kwiatkowski, Cordelia Langford, John Sillitoe on behalf of the Wellcome Sanger Institute COVID-19 Surveillance Team |
| EPI_ISL_851358 | Lighthouse Lab in Glasgow | Wellcome Sanger Institute for the COVID-19 Genomics UK (COG-UK) Consortium | Harper VanSteenhouse, Yumi Kasai, David Gray, Carol Clugston, Anna Dominiczak and Alex Alderton, Roberto Amato, Sonia Goncalves, Ewan Harrison, David K. Jackson, Ian Johnston, Dominic Kwiatkowski, Cordelia Langford, John Sillitoe on behalf of the Wellcome Sanger Institute COVID-19 Surveillance Team |
| EPI_ISL_851359, EPI_ISL_851360, EPI_ISL_851361, EPI_ISL_851362, EPI_ISL_851363, EPI_ISL_851364, EPI_ISL_851366 | Lighthouse Lab in Cambridge | Wellcome Sanger Institute for the COVID-19 Genomics UK (COG-UK) Consortium | Rob Howes, The Lighthouse Lab in Cambridge and Alex Alderton, Roberto Amato, Sonia Goncalves, Ewan Harrison, David K. Jackson, Ian Johnston, Dominic Kwiatkowski, Cordelia Langford, John Sillitoe on behalf of the Wellcome Sanger Institute COVID-19 Surveillance Team |
| EPI_ISL_851367 | Lighthouse Lab in Glasgow | Wellcome Sanger Institute for the COVID-19 Genomics UK (COG-UK) Consortium | Harper VanSteenhouse, Yumi Kasai, David Gray, Carol Clugston, Anna Dominiczak and Alex Alderton, Roberto Amato, Sonia Goncalves, Ewan Harrison, David K. Jackson, Ian Johnston, Dominic Kwiatkowski, Cordelia Langford, John Sillitoe on behalf of the Wellcome Sanger Institute COVID-19 Surveillance Team |
| EPI_ISL_851368, EPI_ISL_851369 | Lighthouse Lab in Cambridge | Wellcome Sanger Institute for the COVID-19 Genomics UK (COG-UK) Consortium | Rob Howes, The Lighthouse Lab in Cambridge and Alex Alderton, Roberto Amato, Sonia Goncalves, Ewan Harrison, David K. Jackson, Ian Johnston, Dominic Kwiatkowski, Cordelia Langford, John Sillitoe on behalf of the Wellcome Sanger Institute COVID-19 Surveillance Team |
| EPI_ISL_851371 | Lighthouse Lab in Glasgow | Wellcome Sanger Institute for the COVID-19 Genomics UK (COG-UK) Consortium | Harper VanSteenhouse, Yumi Kasai, David Gray, Carol Clugston, Anna Dominiczak and Alex Alderton, Roberto Amato, Sonia Goncalves, Ewan Harrison, David K. Jackson, Ian Johnston, Dominic Kwiatkowski, Cordelia Langford, John Sillitoe on behalf of the Wellcome Sanger Institute COVID-19 Surveillance Team |
| EPI_ISL_851372, EPI_ISL_851373, EPI_ISL_851375, EPI_ISL_851376, EPI_ISL_851378, EPI_ISL_851379 | Lighthouse Lab in Cambridge | Wellcome Sanger Institute for the COVID-19 Genomics UK (COG-UK) Consortium | Rob Howes, The Lighthouse Lab in Cambridge and Alex Alderton, Roberto Amato, Sonia Goncalves, Ewan Harrison, David K. Jackson, Ian Johnston, Dominic Kwiatkowski, Cordelia Langford, John Sillitoe on behalf of the Wellcome Sanger Institute COVID-19 Surveillance Team |
| EPI_ISL_851380 | Lighthouse Lab in Glasgow | Wellcome Sanger Institute for the COVID-19 Genomics UK (COG-UK) Consortium | Harper VanSteenhouse, Yumi Kasai, David Gray, Carol Clugston, Anna Dominiczak and Alex Alderton, Roberto Amato, Sonia Goncalves, Ewan Harrison, David K. Jackson, Ian Johnston, Dominic Kwiatkowski, Cordelia Langford, John Sillitoe on behalf of the Wellcome Sanger Institute COVID-19 Surveillance Team |
| EPI_ISL_851381 | Lighthouse Lab in Cambridge | Wellcome Sanger Institute for the COVID-19 Genomics UK (COG-UK) Consortium | Rob Howes, The Lighthouse Lab in Cambridge and Alex Alderton, Roberto Amato, Sonia Goncalves, Ewan Harrison, David K. Jackson, Ian Johnston, Dominic Kwiatkowski, Cordelia Langford, John Sillitoe on behalf of the Wellcome Sanger Institute COVID-19 Surveillance Team |
| EPI_ISL_851382, EPI_ISL_851383, EPI_ISL_851384, EPI_ISL_851385, EPI_ISL_851386, EPI_ISL_851387, EPI_ISL_851388, EPI_ISL_851389, EPI_ISL_851391, EPI_ISL_851392, EPI_ISL_851393, EPI_ISL_851394, EPI_ISL_851396, EPI_ISL_851397, EPI_ISL_851398, EPI_ISL_851399, EPI_ISL_851400, EPI_ISL_851401, EPI_ISL_851402, EPI_ISL_851403, EPI_ISL_851404, EPI_ISL_851405, EPI_ISL_851406, EPI_ISL_851407, EPI_ISL_851408, EPI_ISL_851409, EPI_ISL_851410, EPI_ISL_851411, EPI_ISL_851412, EPI_ISL_851414, EPI_ISL_851415, EPI_ISL_851416, EPI_ISL_851417, EPI_ISL_851418, EPI_ISL_851419, EPI_ISL_851420, EPI_ISL_851421, EPI_ISL_851422, EPI_ISL_851424, EPI_ISL_851425, EPI_ISL_851426, EPI_ISL_851427, EPI_ISL_851428, EPI_ISL_851429, EPI_ISL_851430, EPI_ISL_851431, EPI_ISL_851432, EPI_ISL_851433, EPI_ISL_851434, EPI_ISL_851435, EPI_ISL_851437, EPI_ISL_851438, EPI_ISL_851439, EPI_ISL_851440, EPI_ISL_851441, EPI_ISL_851442, EPI_ISL_851443, EPI_ISL_851444, EPI_ISL_851445, EPI_ISL_851446, EPI_ISL_851447, EPI_ISL_851449, EPI_ISL_851450, EPI_ISL_851451, EPI_ISL_851452, EPI_ISL_851453, EPI_ISL_851456, EPI_ISL_851457, EPI_ISL_851458, EPI_ISL_851459, EPI_ISL_851460, EPI_ISL_851462, EPI_ISL_851463, EPI_ISL_851464, EPI_ISL_851465, EPI_ISL_851466, EPI_ISL_851468, EPI_ISL_851469, EPI_ISL_851470, EPI_ISL_851471, EPI_ISL_851472, EPI_ISL_851473, EPI_ISL_851474, EPI_ISL_851475, EPI_ISL_851476, EPI_ISL_851477, EPI_ISL_851478, EPI_ISL_851479, EPI_ISL_851480, EPI_ISL_851481, EPI_ISL_851482, EPI_ISL_851483, EPI_ISL_851485, EPI_ISL_851486, EPI_ISL_851487, EPI_ISL_851488, EPI_ISL_851489, EPI_ISL_851490, EPI_ISL_851491, EPI_ISL_851492, EPI_ISL_851493, EPI_ISL_851494, EPI_ISL_851495, EPI_ISL_851496, EPI_ISL_851497, EPI_ISL_851498, EPI_ISL_851499, EPI_ISL_851500, EPI_ISL_851501, EPI_ISL_851502, EPI_ISL_851503, EPI_ISL_851504, EPI_ISL_851505, EPI_ISL_851506, EPI_ISL_851507, EPI_ISL_851508, EPI_ISL_851509, EPI_ISL_851510, EPI_ISL_851511, EPI_ISL_851512, EPI_ISL_851513, EPI_ISL_851515, EPI_ISL_851516, EPI_ISL_851517, EPI_ISL_851518, EPI_ISL_851519, EPI_ISL_851520, EPI_ISL_851521, EPI_ISL_851522, EPI_ISL_851523, EPI_ISL_851524, EPI_ISL_851525, EPI_ISL_851526, EPI_ISL_851527, EPI_ISL_851528, EPI_ISL_851529, EPI_ISL_851530, EPI_ISL_851531, EPI_ISL_851532, EPI_ISL_851533, EPI_ISL_851534, EPI_ISL_851535, EPI_ISL_851536, EPI_ISL_851537, EPI_ISL_851538, EPI_ISL_851539, EPI_ISL_851540, EPI_ISL_851541, EPI_ISL_851542, EPI_ISL_851543, EPI_ISL_851544, EPI_ISL_851545, EPI_ISL_851546, EPI_ISL_851547, EPI_ISL_851548, EPI_ISL_851549, EPI_ISL_851550, EPI_ISL_851551, EPI_ISL_851552, EPI_ISL_851553, EPI_ISL_851554, EPI_ISL_851555, EPI_ISL_851556, EPI_ISL_851557, EPI_ISL_851558, EPI_ISL_851560, EPI_ISL_851561, EPI_ISL_851562, EPI_ISL_851563, EPI_ISL_851564, EPI_ISL_851565, EPI_ISL_851566, EPI_ISL_851568, EPI_ISL_851569, EPI_ISL_851570, EPI_ISL_851571, EPI_ISL_851573, EPI_ISL_851574, EPI_ISL_851575, EPI_ISL_851576, EPI_ISL_851577, EPI_ISL_851578, EPI_ISL_851579, EPI_ISL_851580, EPI_ISL_851581, EPI_ISL_851583, EPI_ISL_851584, EPI_ISL_851585, EPI_ISL_851586, EPI_ISL_851587, EPI_ISL_851588, EPI_ISL_851589, EPI_ISL_851591, EPI_ISL_851592, EPI_ISL_851593, EPI_ISL_851595, EPI_ISL_851596, EPI_ISL_851597, EPI_ISL_851599, EPI_ISL_851600, EPI_ISL_851602, EPI_ISL_851603, EPI_ISL_851604, EPI_ISL_851605, EPI_ISL_851606, EPI_ISL_851607, EPI_ISL_851608, EPI_ISL_851609, EPI_ISL_851610, EPI_ISL_851611, EPI_ISL_851612, EPI_ISL_851613, EPI_ISL_851615, EPI_ISL_851616, EPI_ISL_851618, EPI_ISL_851619, EPI_ISL_851620, EPI_ISL_851621, EPI_ISL_851622, EPI_ISL_851623, EPI_ISL_851624, EPI_ISL_851625, EPI_ISL_851626, EPI_ISL_851627, EPI_ISL_851628, EPI_ISL_851630, EPI_ISL_851631, EPI_ISL_851632, EPI_ISL_851634, EPI_ISL_851636, EPI_ISL_851637, EPI_ISL_851638, EPI_ISL_851639, EPI_ISL_851640, EPI_ISL_851641, EPI_ISL_851642, EPI_ISL_851643, EPI_ISL_851644, EPI_ISL_851645, EPI_ISL_851647, EPI_ISL_851649, EPI_ISL_851650, EPI_ISL_851651, EPI_ISL_851652, EPI_ISL_851653, EPI_ISL_851654, EPI_ISL_851655, EPI_ISL_851656, EPI_ISL_851658, EPI_ISL_851659, EPI_ISL_851660, EPI_ISL_851661, EPI_ISL_851662, EPI_ISL_851663, EPI_ISL_851665, EPI_ISL_851666, EPI_ISL_851667, EPI_ISL_851668, EPI_ISL_851669, EPI_ISL_851670, EPI_ISL_851671, EPI_ISL_851672 | | | |
| see above | Lighthouse Lab in Glasgow | Wellcome Sanger Institute for the COVID-19 Genomics UK (COG-UK) Consortium | Harper VanSteenhouse, Yumi Kasai, David Gray, Carol Clugston, Anna Dominiczak and Alex Alderton, Roberto Amato, Sonia Goncalves, Ewan Harrison, David K. Jackson, Ian Johnston, Dominic Kwiatkowski, Cordelia Langford, John Sillitoe on behalf of the Wellcome Sanger Institute COVID-19 Surveillance Team |
| EPI_ISL_851673, EPI_ISL_851675 | Lighthouse Lab in Alderley Park | Wellcome Sanger Institute for the COVID-19 Genomics UK (COG-UK) Consortium | Jacquelyn Wynn, Mairead Hyland, The Lighthouse Lab in Alderley Park and Alex Alderton, Roberto Amato, Sonia Goncalves, Ewan Harrison, David K. Jackson, Ian Johnston, Dominic Kwiatkowski, Cordelia Langford, John Sillitoe on behalf of the Wellcome Sanger Institute COVID-19 Surveillance Team |
| EPI_ISL_851676 | Lighthouse Lab in Glasgow | Wellcome Sanger Institute for the COVID-19 Genomics UK (COG-UK) Consortium | Harper VanSteenhouse, Yumi Kasai, David Gray, Carol Clugston, Anna Dominiczak and Alex Alderton, Roberto Amato, Sonia Goncalves, Ewan Harrison, David K. Jackson, Ian Johnston, Dominic Kwiatkowski, Cordelia Langford, John Sillitoe on behalf of the Wellcome Sanger Institute COVID-19 Surveillance Team |
| EPI_ISL_851677, EPI_ISL_851678, EPI_ISL_851679, EPI_ISL_851680 | Lighthouse Lab in Alderley Park | Wellcome Sanger Institute for the COVID-19 Genomics UK (COG-UK) Consortium | Jacquelyn Wynn, Mairead Hyland, The Lighthouse Lab in Alderley Park and Alex Alderton, Roberto Amato, Sonia Goncalves, Ewan Harrison, David K. Jackson, Ian Johnston, Dominic Kwiatkowski, Cordelia Langford, John Sillitoe on behalf of the Wellcome Sanger Institute COVID-19 Surveillance Team |
| EPI_ISL_851681 | Lighthouse Lab in Glasgow | Wellcome Sanger Institute for the COVID-19 Genomics UK (COG-UK) Consortium | Harper VanSteenhouse, Yumi Kasai, David Gray, Carol Clugston, Anna Dominiczak and Alex Alderton, Roberto Amato, Sonia Goncalves, Ewan Harrison, David K. Jackson, Ian Johnston, Dominic Kwiatkowski, Cordelia Langford, John Sillitoe on behalf of the Wellcome Sanger Institute COVID-19 Surveillance Team |
| EPI_ISL_851682, EPI_ISL_851683, EPI_ISL_851684 | Lighthouse Lab in Alderley Park | Wellcome Sanger Institute for the COVID-19 Genomics UK (COG-UK) Consortium | Jacquelyn Wynn, Mairead Hyland, The Lighthouse Lab in Alderley Park and Alex Alderton, Roberto Amato, Sonia Goncalves, Ewan Harrison, David K. Jackson, Ian Johnston, Dominic Kwiatkowski, Cordelia Langford, John Sillitoe on behalf of the Wellcome Sanger Institute COVID-19 Surveillance Team |
| EPI_ISL_851685, EPI_ISL_851686, EPI_ISL_851687 | Lighthouse Lab in Glasgow | Wellcome Sanger Institute for the COVID-19 Genomics UK (COG-UK) Consortium | Harper VanSteenhouse, Yumi Kasai, David Gray, Carol Clugston, Anna Dominiczak and Alex Alderton, Roberto Amato, Sonia Goncalves, Ewan Harrison, David K. Jackson, Ian Johnston, Dominic Kwiatkowski, Cordelia Langford, John Sillitoe on behalf of the Wellcome Sanger Institute COVID-19 Surveillance Team |
| EPI_ISL_851688 | Lighthouse Lab in Alderley Park | Wellcome Sanger Institute for the COVID-19 Genomics UK (COG-UK) Consortium | Jacquelyn Wynn, Mairead Hyland, The Lighthouse Lab in Alderley Park and Alex Alderton, Roberto Amato, Sonia Goncalves, Ewan Harrison, David K. Jackson, Ian Johnston, Dominic Kwiatkowski, Cordelia Langford, John Sillitoe on behalf of the Wellcome Sanger Institute COVID-19 Surveillance Team |

| | | | |
|---|---|---|---|
| EPI_ISL_851689, EPI_ISL_851690 | Lighthouse Lab in Glasgow | Wellcome Sanger Institute for the COVID-19 Genomics UK (COG-UK) Consortium | Harper VanSteenhouse, Yumi Kasai, David Gray, Carol Clugston, Anna Dominiczak and Alex Alderton, Roberto Amato, Sonia Goncalves, Ewan Harrison, David K. Jackson, Ian Johnston, Dominic Kwiatkowski, Cordelia Langford, John Sillitoe on behalf of the Wellcome Sanger Institute COVID-19 Surveillance Team |
| EPI_ISL_851691, EPI_ISL_851692 | Lighthouse Lab in Alderley Park | Wellcome Sanger Institute for the COVID-19 Genomics UK (COG-UK) Consortium | Jacquelyn Wynn, Mairead Hyland, The Lighthouse Lab in Alderley Park and Alex Alderton, Roberto Amato, Sonia Goncalves, Ewan Harrison, David K. Jackson, Ian Johnston, Dominic Kwiatkowski, Cordelia Langford, John Sillitoe on behalf of the Wellcome Sanger Institute COVID-19 Surveillance Team |
| EPI_ISL_851693 | Lighthouse Lab in Glasgow | Wellcome Sanger Institute for the COVID-19 Genomics UK (COG-UK) Consortium | Harper VanSteenhouse, Yumi Kasai, David Gray, Carol Clugston, Anna Dominiczak and Alex Alderton, Roberto Amato, Sonia Goncalves, Ewan Harrison, David K. Jackson, Ian Johnston, Dominic Kwiatkowski, Cordelia Langford, John Sillitoe on behalf of the Wellcome Sanger Institute COVID-19 Surveillance Team |
| EPI_ISL_851694, EPI_ISL_851695, EPI_ISL_851696, EPI_ISL_851697, EPI_ISL_851698, EPI_ISL_851699, EPI_ISL_851700 | Lighthouse Lab in Alderley Park | Wellcome Sanger Institute for the COVID-19 Genomics UK (COG-UK) Consortium | Jacquelyn Wynn, Mairead Hyland, The Lighthouse Lab in Alderley Park and Alex Alderton, Roberto Amato, Sonia Goncalves, Ewan Harrison, David K. Jackson, Ian Johnston, Dominic Kwiatkowski, Cordelia Langford, John Sillitoe on behalf of the Wellcome Sanger Institute COVID-19 Surveillance Team |
| EPI_ISL_851701 | Lighthouse Lab in Glasgow | Wellcome Sanger Institute for the COVID-19 Genomics UK (COG-UK) Consortium | Harper VanSteenhouse, Yumi Kasai, David Gray, Carol Clugston, Anna Dominiczak and Alex Alderton, Roberto Amato, Sonia Goncalves, Ewan Harrison, David K. Jackson, Ian Johnston, Dominic Kwiatkowski, Cordelia Langford, John Sillitoe on behalf of the Wellcome Sanger Institute COVID-19 Surveillance Team |
| EPI_ISL_851702, EPI_ISL_851703, EPI_ISL_851704, EPI_ISL_851705, EPI_ISL_851706, EPI_ISL_851707, EPI_ISL_851708, EPI_ISL_851709, EPI_ISL_851710, EPI_ISL_851711, EPI_ISL_851712, EPI_ISL_851713 | | | |
| see above | Lighthouse Lab in Alderley Park | Wellcome Sanger Institute for the COVID-19 Genomics UK (COG-UK) Consortium | Jacquelyn Wynn, Mairead Hyland, The Lighthouse Lab in Alderley Park and Alex Alderton, Roberto Amato, Sonia Goncalves, Ewan Harrison, David K. Jackson, Ian Johnston, Dominic Kwiatkowski, Cordelia Langford, John Sillitoe on behalf of the Wellcome Sanger Institute COVID-19 Surveillance Team |
| EPI_ISL_851714 | Lighthouse Lab in Glasgow | Wellcome Sanger Institute for the COVID-19 Genomics UK (COG-UK) Consortium | Harper VanSteenhouse, Yumi Kasai, David Gray, Carol Clugston, Anna Dominiczak and Alex Alderton, Roberto Amato, Sonia Goncalves, Ewan Harrison, David K. Jackson, Ian Johnston, Dominic Kwiatkowski, Cordelia Langford, John Sillitoe on behalf of the Wellcome Sanger Institute COVID-19 Surveillance Team |
| EPI_ISL_851715, EPI_ISL_851716, EPI_ISL_851717, EPI_ISL_851718, EPI_ISL_851719 | Lighthouse Lab in Alderley Park | Wellcome Sanger Institute for the COVID-19 Genomics UK (COG-UK) Consortium | Jacquelyn Wynn, Mairead Hyland, The Lighthouse Lab in Alderley Park and Alex Alderton, Roberto Amato, Sonia Goncalves, Ewan Harrison, David K. Jackson, Ian Johnston, Dominic Kwiatkowski, Cordelia Langford, John Sillitoe on behalf of the Wellcome Sanger Institute COVID-19 Surveillance Team |
| EPI_ISL_851720 | Lighthouse Lab in Glasgow | Wellcome Sanger Institute for the COVID-19 Genomics UK (COG-UK) Consortium | Harper VanSteenhouse, Yumi Kasai, David Gray, Carol Clugston, Anna Dominiczak and Alex Alderton, Roberto Amato, Sonia Goncalves, Ewan Harrison, David K. Jackson, Ian Johnston, Dominic Kwiatkowski, Cordelia Langford, John Sillitoe on behalf of the Wellcome Sanger Institute COVID-19 Surveillance Team |
| EPI_ISL_851721, EPI_ISL_851722, EPI_ISL_851723, EPI_ISL_851724 | Lighthouse Lab in Alderley Park | Wellcome Sanger Institute for the COVID-19 Genomics UK (COG-UK) Consortium | Jacquelyn Wynn, Mairead Hyland, The Lighthouse Lab in Alderley Park and Alex Alderton, Roberto Amato, Sonia Goncalves, Ewan Harrison, David K. Jackson, Ian Johnston, Dominic Kwiatkowski, Cordelia Langford, John Sillitoe on behalf of the Wellcome Sanger Institute COVID-19 Surveillance Team |
| EPI_ISL_851725 | Lighthouse Lab in Glasgow | Wellcome Sanger Institute for the COVID-19 Genomics UK (COG-UK) Consortium | Harper VanSteenhouse, Yumi Kasai, David Gray, Carol Clugston, Anna Dominiczak and Alex Alderton, Roberto Amato, Sonia Goncalves, Ewan Harrison, David K. Jackson, Ian Johnston, Dominic Kwiatkowski, Cordelia Langford, John Sillitoe on behalf of the Wellcome Sanger Institute COVID-19 Surveillance Team |
| EPI_ISL_851726 | Lighthouse Lab in Alderley Park | Wellcome Sanger Institute for the COVID-19 Genomics UK (COG-UK) Consortium | Jacquelyn Wynn, Mairead Hyland, The Lighthouse Lab in Alderley Park and Alex Alderton, Roberto Amato, Sonia Goncalves, Ewan Harrison, David K. Jackson, Ian Johnston, Dominic Kwiatkowski, Cordelia Langford, John Sillitoe on behalf of the Wellcome Sanger Institute COVID-19 Surveillance Team |
| EPI_ISL_851727 | Lighthouse Lab in Glasgow | Wellcome Sanger Institute for the COVID-19 Genomics UK (COG-UK) Consortium | Harper VanSteenhouse, Yumi Kasai, David Gray, Carol Clugston, Anna Dominiczak and Alex Alderton, Roberto Amato, Sonia Goncalves, Ewan Harrison, David K. Jackson, Ian Johnston, Dominic Kwiatkowski, Cordelia Langford, John Sillitoe on behalf of the Wellcome Sanger Institute COVID-19 Surveillance Team |
| EPI_ISL_851728 | Lighthouse Lab in Alderley Park | Wellcome Sanger Institute for the COVID-19 Genomics UK (COG-UK) Consortium | Jacquelyn Wynn, Mairead Hyland, The Lighthouse Lab in Alderley Park and Alex Alderton, Roberto Amato, Sonia Goncalves, Ewan Harrison, David K. Jackson, Ian Johnston, Dominic Kwiatkowski, Cordelia Langford, John Sillitoe on behalf of the Wellcome Sanger Institute COVID-19 Surveillance Team |
| EPI_ISL_851729 | Lighthouse Lab in Glasgow | Wellcome Sanger Institute for the COVID-19 Genomics UK (COG-UK) Consortium | Harper VanSteenhouse, Yumi Kasai, David Gray, Carol Clugston, Anna Dominiczak and Alex Alderton, Roberto Amato, Sonia Goncalves, Ewan Harrison, David K. Jackson, Ian Johnston, Dominic Kwiatkowski, Cordelia Langford, John Sillitoe on behalf of the Wellcome Sanger Institute COVID-19 Surveillance Team |
| EPI_ISL_851730, EPI_ISL_851731, EPI_ISL_851732, EPI_ISL_851733 | Lighthouse Lab in Alderley Park | Wellcome Sanger Institute for the COVID-19 Genomics UK (COG-UK) Consortium | Jacquelyn Wynn, Mairead Hyland, The Lighthouse Lab in Alderley Park and Alex Alderton, Roberto Amato, Sonia Goncalves, Ewan Harrison, David K. Jackson, Ian Johnston, Dominic Kwiatkowski, Cordelia Langford, John Sillitoe on behalf of the Wellcome Sanger Institute COVID-19 Surveillance Team |
| EPI_ISL_851734 | Lighthouse Lab in Glasgow | Wellcome Sanger Institute for the COVID-19 Genomics UK (COG-UK) Consortium | Harper VanSteenhouse, Yumi Kasai, David Gray, Carol Clugston, Anna Dominiczak and Alex Alderton, Roberto Amato, Sonia Goncalves, Ewan Harrison, David K. Jackson, Ian Johnston, Dominic Kwiatkowski, Cordelia Langford, John Sillitoe on behalf of the Wellcome Sanger Institute COVID-19 Surveillance Team |
| EPI_ISL_851735, EPI_ISL_851736, EPI_ISL_851737 | Lighthouse Lab in Alderley Park | Wellcome Sanger Institute for the COVID-19 Genomics UK (COG-UK) Consortium | Jacquelyn Wynn, Mairead Hyland, The Lighthouse Lab in Alderley Park and Alex Alderton, Roberto Amato, Sonia Goncalves, Ewan Harrison, David K. Jackson, Ian Johnston, Dominic Kwiatkowski, Cordelia Langford, John Sillitoe on behalf of the Wellcome Sanger Institute COVID-19 Surveillance Team |
| EPI_ISL_851738 | Lighthouse Lab in Glasgow | Wellcome Sanger Institute for the COVID-19 Genomics UK (COG-UK) Consortium | Harper VanSteenhouse, Yumi Kasai, David Gray, Carol Clugston, Anna Dominiczak and Alex Alderton, Roberto Amato, Sonia Goncalves, Ewan Harrison, David K. Jackson, Ian Johnston, Dominic Kwiatkowski, Cordelia Langford, John Sillitoe on behalf of the Wellcome Sanger Institute COVID-19 Surveillance Team |
| EPI_ISL_851739, EPI_ISL_851740, EPI_ISL_851741, EPI_ISL_851742 | Lighthouse Lab in Alderley Park | Wellcome Sanger Institute for the COVID-19 Genomics UK (COG-UK) Consortium | Jacquelyn Wynn, Mairead Hyland, The Lighthouse Lab in Alderley Park and Alex Alderton, Roberto Amato, Sonia Goncalves, Ewan Harrison, David K. Jackson, Ian Johnston, Dominic Kwiatkowski, Cordelia Langford, John Sillitoe on behalf of the Wellcome Sanger Institute COVID-19 Surveillance Team |
| EPI_ISL_851743 | Lighthouse Lab in Glasgow | Wellcome Sanger Institute for the COVID-19 Genomics UK (COG-UK) Consortium | Harper VanSteenhouse, Yumi Kasai, David Gray, Carol Clugston, Anna Dominiczak and Alex Alderton, Roberto Amato, Sonia Goncalves, Ewan Harrison, David K. Jackson, Ian Johnston, Dominic Kwiatkowski, Cordelia Langford, John Sillitoe on behalf of the Wellcome Sanger Institute COVID-19 Surveillance Team |
| EPI_ISL_851744, EPI_ISL_851745, EPI_ISL_851746 | Lighthouse Lab in Alderley Park | Wellcome Sanger Institute for the COVID-19 Genomics UK (COG-UK) Consortium | Jacquelyn Wynn, Mairead Hyland, The Lighthouse Lab in Alderley Park and Alex Alderton, Roberto Amato, Sonia Goncalves, Ewan Harrison, David K. Jackson, Ian Johnston, Dominic Kwiatkowski, Cordelia Langford, John Sillitoe on behalf of the Wellcome Sanger Institute COVID-19 Surveillance Team |
| EPI_ISL_851747 | Lighthouse Lab in Glasgow | Wellcome Sanger Institute for the COVID-19 Genomics UK (COG-UK) Consortium | Harper VanSteenhouse, Yumi Kasai, David Gray, Carol Clugston, Anna Dominiczak and Alex Alderton, Roberto Amato, Sonia Goncalves, Ewan Harrison, David K. Jackson, Ian Johnston, Dominic Kwiatkowski, Cordelia Langford, John Sillitoe on behalf of the Wellcome Sanger Institute COVID-19 Surveillance Team |
| EPI_ISL_851748 | Lighthouse Lab in Alderley Park | Wellcome Sanger Institute for the COVID-19 Genomics UK (COG-UK) Consortium | Jacquelyn Wynn, Mairead Hyland, The Lighthouse Lab in Alderley Park and Alex Alderton, Roberto Amato, Sonia Goncalves, Ewan Harrison, David K. Jackson, Ian Johnston, Dominic Kwiatkowski, Cordelia Langford, John Sillitoe on behalf of the Wellcome Sanger Institute COVID-19 Surveillance Team |
| EPI_ISL_851749 | Lighthouse Lab in Glasgow | Wellcome Sanger Institute for the COVID-19 Genomics UK (COG-UK) Consortium | Harper VanSteenhouse, Yumi Kasai, David Gray, Carol Clugston, Anna Dominiczak and Alex Alderton, Roberto Amato, Sonia Goncalves, Ewan Harrison, David K. Jackson, Ian Johnston, Dominic Kwiatkowski, Cordelia Langford, John Sillitoe on behalf of the Wellcome Sanger Institute COVID-19 Surveillance Team |
| EPI_ISL_851751, EPI_ISL_851752, EPI_ISL_851753 | Lighthouse Lab in Alderley Park | Wellcome Sanger Institute for the COVID-19 Genomics UK (COG-UK) Consortium | Jacquelyn Wynn, Mairead Hyland, The Lighthouse Lab in Alderley Park and Alex Alderton, Roberto Amato, Sonia Goncalves, Ewan Harrison, David K. Jackson, Ian Johnston, Dominic Kwiatkowski, Cordelia Langford, John Sillitoe on behalf of the Wellcome Sanger Institute COVID-19 Surveillance Team |
| EPI_ISL_851754 | Lighthouse Lab in Glasgow | Wellcome Sanger Institute for the COVID-19 Genomics UK (COG-UK) Consortium | Harper VanSteenhouse, Yumi Kasai, David Gray, Carol Clugston, Anna Dominiczak and Alex Alderton, Roberto Amato, Sonia Goncalves, Ewan Harrison, David K. Jackson, Ian Johnston, Dominic Kwiatkowski, Cordelia Langford, John Sillitoe on behalf of the Wellcome Sanger Institute COVID-19 Surveillance Team |
| EPI_ISL_851755, EPI_ISL_851756 | Lighthouse Lab in Alderley Park | Wellcome Sanger Institute for the COVID-19 Genomics UK | Jacquelyn Wynn, Mairead Hyland, The Lighthouse Lab in Alderley Park and Alex Alderton, Roberto Amato, Sonia Goncalves, Ewan Harrison, David K. |

| | | | |
|---|---|---|---|
| | | (COG-UK) Consortium | Jackson, Ian Johnston, Dominic Kwiatkowski, Cordelia Langford, John Sillitoe on behalf of the Wellcome Sanger Institute COVID-19 Surveillance Team |
| EPI_ISL_851757 | Lighthouse Lab in Glasgow | Wellcome Sanger Institute for the COVID-19 Genomics UK (COG-UK) Consortium | Harper VanSteenhouse, Yumi Kasai, David Gray, Carol Clugston, Anna Dominiczak and Alex Alderton, Roberto Amato, Sonia Goncalves, Ewan Harrison, David K. Jackson, Ian Johnston, Dominic Kwiatkowski, Cordelia Langford, John Sillitoe on behalf of the Wellcome Sanger Institute COVID-19 Surveillance Team |
| EPI_ISL_851758, EPI_ISL_851759 | Lighthouse Lab in Alderley Park | Wellcome Sanger Institute for the COVID-19 Genomics UK (COG-UK) Consortium | Jacquelyn Wynn, Mairead Hyland, The Lighthouse Lab in Alderley Park and Alex Alderton, Roberto Amato, Sonia Goncalves, Ewan Harrison, David K. Jackson, Ian Johnston, Dominic Kwiatkowski, Cordelia Langford, John Sillitoe on behalf of the Wellcome Sanger Institute COVID-19 Surveillance Team |
| EPI_ISL_851760 | Lighthouse Lab in Glasgow | Wellcome Sanger Institute for the COVID-19 Genomics UK (COG-UK) Consortium | Harper VanSteenhouse, Yumi Kasai, David Gray, Carol Clugston, Anna Dominiczak and Alex Alderton, Roberto Amato, Sonia Goncalves, Ewan Harrison, David K. Jackson, Ian Johnston, Dominic Kwiatkowski, Cordelia Langford, John Sillitoe on behalf of the Wellcome Sanger Institute COVID-19 Surveillance Team |
| EPI_ISL_851761, EPI_ISL_851762 | Lighthouse Lab in Alderley Park | Wellcome Sanger Institute for the COVID-19 Genomics UK (COG-UK) Consortium | Jacquelyn Wynn, Mairead Hyland, The Lighthouse Lab in Alderley Park and Alex Alderton, Roberto Amato, Sonia Goncalves, Ewan Harrison, David K. Jackson, Ian Johnston, Dominic Kwiatkowski, Cordelia Langford, John Sillitoe on behalf of the Wellcome Sanger Institute COVID-19 Surveillance Team |
| EPI_ISL_851763 | Lighthouse Lab in Glasgow | Wellcome Sanger Institute for the COVID-19 Genomics UK (COG-UK) Consortium | Harper VanSteenhouse, Yumi Kasai, David Gray, Carol Clugston, Anna Dominiczak and Alex Alderton, Roberto Amato, Sonia Goncalves, Ewan Harrison, David K. Jackson, Ian Johnston, Dominic Kwiatkowski, Cordelia Langford, John Sillitoe on behalf of the Wellcome Sanger Institute COVID-19 Surveillance Team |
| EPI_ISL_851764, EPI_ISL_851765, EPI_ISL_851766 | Lighthouse Lab in Alderley Park | Wellcome Sanger Institute for the COVID-19 Genomics UK (COG-UK) Consortium | Jacquelyn Wynn, Mairead Hyland, The Lighthouse Lab in Alderley Park and Alex Alderton, Roberto Amato, Sonia Goncalves, Ewan Harrison, David K. Jackson, Ian Johnston, Dominic Kwiatkowski, Cordelia Langford, John Sillitoe on behalf of the Wellcome Sanger Institute COVID-19 Surveillance Team |
| EPI_ISL_851767 | Lighthouse Lab in Glasgow | Wellcome Sanger Institute for the COVID-19 Genomics UK (COG-UK) Consortium | Harper VanSteenhouse, Yumi Kasai, David Gray, Carol Clugston, Anna Dominiczak and Alex Alderton, Roberto Amato, Sonia Goncalves, Ewan Harrison, David K. Jackson, Ian Johnston, Dominic Kwiatkowski, Cordelia Langford, John Sillitoe on behalf of the Wellcome Sanger Institute COVID-19 Surveillance Team |
| EPI_ISL_851768, EPI_ISL_851769 | Lighthouse Lab in Alderley Park | Wellcome Sanger Institute for the COVID-19 Genomics UK (COG-UK) Consortium | Jacquelyn Wynn, Mairead Hyland, The Lighthouse Lab in Alderley Park and Alex Alderton, Roberto Amato, Sonia Goncalves, Ewan Harrison, David K. Jackson, Ian Johnston, Dominic Kwiatkowski, Cordelia Langford, John Sillitoe on behalf of the Wellcome Sanger Institute COVID-19 Surveillance Team |
| EPI_ISL_851770 | Lighthouse Lab in Glasgow | Wellcome Sanger Institute for the COVID-19 Genomics UK (COG-UK) Consortium | Harper VanSteenhouse, Yumi Kasai, David Gray, Carol Clugston, Anna Dominiczak and Alex Alderton, Roberto Amato, Sonia Goncalves, Ewan Harrison, David K. Jackson, Ian Johnston, Dominic Kwiatkowski, Cordelia Langford, John Sillitoe on behalf of the Wellcome Sanger Institute COVID-19 Surveillance Team |
| EPI_ISL_851771 | Lighthouse Lab in Alderley Park | Wellcome Sanger Institute for the COVID-19 Genomics UK (COG-UK) Consortium | Jacquelyn Wynn, Mairead Hyland, The Lighthouse Lab in Alderley Park and Alex Alderton, Roberto Amato, Sonia Goncalves, Ewan Harrison, David K. Jackson, Ian Johnston, Dominic Kwiatkowski, Cordelia Langford, John Sillitoe on behalf of the Wellcome Sanger Institute COVID-19 Surveillance Team |
| EPI_ISL_851772 | Lighthouse Lab in Glasgow | Wellcome Sanger Institute for the COVID-19 Genomics UK (COG-UK) Consortium | Harper VanSteenhouse, Yumi Kasai, David Gray, Carol Clugston, Anna Dominiczak and Alex Alderton, Roberto Amato, Sonia Goncalves, Ewan Harrison, David K. Jackson, Ian Johnston, Dominic Kwiatkowski, Cordelia Langford, John Sillitoe on behalf of the Wellcome Sanger Institute COVID-19 Surveillance Team |
| EPI_ISL_851773 | Lighthouse Lab in Alderley Park | Wellcome Sanger Institute for the COVID-19 Genomics UK (COG-UK) Consortium | Jacquelyn Wynn, Mairead Hyland, The Lighthouse Lab in Alderley Park and Alex Alderton, Roberto Amato, Sonia Goncalves, Ewan Harrison, David K. Jackson, Ian Johnston, Dominic Kwiatkowski, Cordelia Langford, John Sillitoe on behalf of the Wellcome Sanger Institute COVID-19 Surveillance Team |
| EPI_ISL_851774 | Lighthouse Lab in Glasgow | Wellcome Sanger Institute for the COVID-19 Genomics UK (COG-UK) Consortium | Harper VanSteenhouse, Yumi Kasai, David Gray, Carol Clugston, Anna Dominiczak and Alex Alderton, Roberto Amato, Sonia Goncalves, Ewan Harrison, David K. Jackson, Ian Johnston, Dominic Kwiatkowski, Cordelia Langford, John Sillitoe on behalf of the Wellcome Sanger Institute COVID-19 Surveillance Team |
| EPI_ISL_851775, EPI_ISL_851776, EPI_ISL_851777, EPI_ISL_851778, EPI_ISL_851779 | Lighthouse Lab in Alderley Park | Wellcome Sanger Institute for the COVID-19 Genomics UK (COG-UK) Consortium | Jacquelyn Wynn, Mairead Hyland, The Lighthouse Lab in Alderley Park and Alex Alderton, Roberto Amato, Sonia Goncalves, Ewan Harrison, David K. Jackson, Ian Johnston, Dominic Kwiatkowski, Cordelia Langford, John Sillitoe on behalf of the Wellcome Sanger Institute COVID-19 Surveillance Team |
| EPI_ISL_851780, EPI_ISL_851781 | Lighthouse Lab in Glasgow | Wellcome Sanger Institute for the COVID-19 Genomics UK (COG-UK) Consortium | Harper VanSteenhouse, Yumi Kasai, David Gray, Carol Clugston, Anna Dominiczak and Alex Alderton, Roberto Amato, Sonia Goncalves, Ewan Harrison, David K. Jackson, Ian Johnston, Dominic Kwiatkowski, Cordelia Langford, John Sillitoe on behalf of the Wellcome Sanger Institute COVID-19 Surveillance Team |
| EPI_ISL_851782, EPI_ISL_851783, EPI_ISL_851784 | Lighthouse Lab in Alderley Park | Wellcome Sanger Institute for the COVID-19 Genomics UK (COG-UK) Consortium | Jacquelyn Wynn, Mairead Hyland, The Lighthouse Lab in Alderley Park and Alex Alderton, Roberto Amato, Sonia Goncalves, Ewan Harrison, David K. Jackson, Ian Johnston, Dominic Kwiatkowski, Cordelia Langford, John Sillitoe on behalf of the Wellcome Sanger Institute COVID-19 Surveillance Team |
| EPI_ISL_851785 | Lighthouse Lab in Glasgow | Wellcome Sanger Institute for the COVID-19 Genomics UK (COG-UK) Consortium | Harper VanSteenhouse, Yumi Kasai, David Gray, Carol Clugston, Anna Dominiczak and Alex Alderton, Roberto Amato, Sonia Goncalves, Ewan Harrison, David K. Jackson, Ian Johnston, Dominic Kwiatkowski, Cordelia Langford, John Sillitoe on behalf of the Wellcome Sanger Institute COVID-19 Surveillance Team |
| EPI_ISL_851786, EPI_ISL_851787, EPI_ISL_851788, EPI_ISL_851789, EPI_ISL_851790, EPI_ISL_851791 | Lighthouse Lab in Alderley Park | Wellcome Sanger Institute for the COVID-19 Genomics UK (COG-UK) Consortium | Jacquelyn Wynn, Mairead Hyland, The Lighthouse Lab in Alderley Park and Alex Alderton, Roberto Amato, Sonia Goncalves, Ewan Harrison, David K. Jackson, Ian Johnston, Dominic Kwiatkowski, Cordelia Langford, John Sillitoe on behalf of the Wellcome Sanger Institute COVID-19 Surveillance Team |
| EPI_ISL_851792 | Lighthouse Lab in Glasgow | Wellcome Sanger Institute for the COVID-19 Genomics UK (COG-UK) Consortium | Harper VanSteenhouse, Yumi Kasai, David Gray, Carol Clugston, Anna Dominiczak and Alex Alderton, Roberto Amato, Sonia Goncalves, Ewan Harrison, David K. Jackson, Ian Johnston, Dominic Kwiatkowski, Cordelia Langford, John Sillitoe on behalf of the Wellcome Sanger Institute COVID-19 Surveillance Team |
| EPI_ISL_851793 | Lighthouse Lab in Alderley Park | Wellcome Sanger Institute for the COVID-19 Genomics UK (COG-UK) Consortium | Jacquelyn Wynn, Mairead Hyland, The Lighthouse Lab in Alderley Park and Alex Alderton, Roberto Amato, Sonia Goncalves, Ewan Harrison, David K. Jackson, Ian Johnston, Dominic Kwiatkowski, Cordelia Langford, John Sillitoe on behalf of the Wellcome Sanger Institute COVID-19 Surveillance Team |
| EPI_ISL_851794 | Lighthouse Lab in Glasgow | Wellcome Sanger Institute for the COVID-19 Genomics UK (COG-UK) Consortium | Harper VanSteenhouse, Yumi Kasai, David Gray, Carol Clugston, Anna Dominiczak and Alex Alderton, Roberto Amato, Sonia Goncalves, Ewan Harrison, David K. Jackson, Ian Johnston, Dominic Kwiatkowski, Cordelia Langford, John Sillitoe on behalf of the Wellcome Sanger Institute COVID-19 Surveillance Team |
| EPI_ISL_851795 | Lighthouse Lab in Alderley Park | Wellcome Sanger Institute for the COVID-19 Genomics UK (COG-UK) Consortium | Jacquelyn Wynn, Mairead Hyland, The Lighthouse Lab in Alderley Park and Alex Alderton, Roberto Amato, Sonia Goncalves, Ewan Harrison, David K. Jackson, Ian Johnston, Dominic Kwiatkowski, Cordelia Langford, John Sillitoe on behalf of the Wellcome Sanger Institute COVID-19 Surveillance Team |
| EPI_ISL_851796, EPI_ISL_851797, EPI_ISL_851798, EPI_ISL_851799 | Lighthouse Lab in Glasgow | Wellcome Sanger Institute for the COVID-19 Genomics UK (COG-UK) Consortium | Harper VanSteenhouse, Yumi Kasai, David Gray, Carol Clugston, Anna Dominiczak and Alex Alderton, Roberto Amato, Sonia Goncalves, Ewan Harrison, David K. Jackson, Ian Johnston, Dominic Kwiatkowski, Cordelia Langford, John Sillitoe on behalf of the Wellcome Sanger Institute COVID-19 Surveillance Team |
| EPI_ISL_851800, EPI_ISL_851801, EPI_ISL_851802 | Lighthouse Lab in Alderley Park | Wellcome Sanger Institute for the COVID-19 Genomics UK (COG-UK) Consortium | Jacquelyn Wynn, Mairead Hyland, The Lighthouse Lab in Alderley Park and Alex Alderton, Roberto Amato, Sonia Goncalves, Ewan Harrison, David K. Jackson, Ian Johnston, Dominic Kwiatkowski, Cordelia Langford, John Sillitoe on behalf of the Wellcome Sanger Institute COVID-19 Surveillance Team |
| EPI_ISL_851803 | Lighthouse Lab in Glasgow | Wellcome Sanger Institute for the COVID-19 Genomics UK (COG-UK) Consortium | Harper VanSteenhouse, Yumi Kasai, David Gray, Carol Clugston, Anna Dominiczak and Alex Alderton, Roberto Amato, Sonia Goncalves, Ewan Harrison, David K. Jackson, Ian Johnston, Dominic Kwiatkowski, Cordelia Langford, John Sillitoe on behalf of the Wellcome Sanger Institute COVID-19 Surveillance Team |
| EPI_ISL_851804 | Lighthouse Lab in Alderley Park | Wellcome Sanger Institute for the COVID-19 Genomics UK (COG-UK) Consortium | Jacquelyn Wynn, Mairead Hyland, The Lighthouse Lab in Alderley Park and Alex Alderton, Roberto Amato, Sonia Goncalves, Ewan Harrison, David K. Jackson, Ian Johnston, Dominic Kwiatkowski, Cordelia Langford, John Sillitoe on behalf of the Wellcome Sanger Institute COVID-19 Surveillance Team |
| EPI_ISL_851805 | Lighthouse Lab in Glasgow | Wellcome Sanger Institute for the COVID-19 Genomics UK (COG-UK) Consortium | Harper VanSteenhouse, Yumi Kasai, David Gray, Carol Clugston, Anna Dominiczak and Alex Alderton, Roberto Amato, Sonia Goncalves, Ewan Harrison, David K. Jackson, Ian Johnston, Dominic Kwiatkowski, Cordelia Langford, John Sillitoe on behalf of the Wellcome Sanger Institute COVID-19 Surveillance Team |
| EPI_ISL_851806 | Lighthouse Lab in Alderley Park | Wellcome Sanger Institute for the COVID-19 Genomics UK (COG-UK) Consortium | Jacquelyn Wynn, Mairead Hyland, The Lighthouse Lab in Alderley Park and Alex Alderton, Roberto Amato, Sonia Goncalves, Ewan Harrison, David K. Jackson, Ian Johnston, Dominic Kwiatkowski, Cordelia Langford, John Sillitoe on behalf of the Wellcome Sanger Institute COVID-19 Surveillance Team |

| | | | |
|---|---|---|---|
| EPI_ISL_851807 | Lighthouse Lab in Glasgow | Wellcome Sanger Institute for the COVID-19 Genomics UK (COG-UK) Consortium | Harper VanSteenhouse, Yumi Kasai, David Gray, Carol Clugston, Anna Dominiczak and Alex Alderton, Roberto Amato, Sonia Goncalves, Ewan Harrison, David K. Jackson, Ian Johnston, Dominic Kwiatkowski, Cordelia Langford, John Sillitoe on behalf of the Wellcome Sanger Institute COVID-19 Surveillance Team |
| EPI_ISL_851808, EPI_ISL_851809, EPI_ISL_851810, EPI_ISL_851811 | Lighthouse Lab in Alderley Park | Wellcome Sanger Institute for the COVID-19 Genomics UK (COG-UK) Consortium | Jacquelyn Wynn, Mairead Hyland, The Lighthouse Lab in Alderley Park and Alex Alderton, Roberto Amato, Sonia Goncalves, Ewan Harrison, David K. Jackson, Ian Johnston, Dominic Kwiatkowski, Cordelia Langford, John Sillitoe on behalf of the Wellcome Sanger Institute COVID-19 Surveillance Team |
| EPI_ISL_851812 | Lighthouse Lab in Glasgow | Wellcome Sanger Institute for the COVID-19 Genomics UK (COG-UK) Consortium | Harper VanSteenhouse, Yumi Kasai, David Gray, Carol Clugston, Anna Dominiczak and Alex Alderton, Roberto Amato, Sonia Goncalves, Ewan Harrison, David K. Jackson, Ian Johnston, Dominic Kwiatkowski, Cordelia Langford, John Sillitoe on behalf of the Wellcome Sanger Institute COVID-19 Surveillance Team |
| EPI_ISL_851813 | Lighthouse Lab in Alderley Park | Wellcome Sanger Institute for the COVID-19 Genomics UK (COG-UK) Consortium | Jacquelyn Wynn, Mairead Hyland, The Lighthouse Lab in Alderley Park and Alex Alderton, Roberto Amato, Sonia Goncalves, Ewan Harrison, David K. Jackson, Ian Johnston, Dominic Kwiatkowski, Cordelia Langford, John Sillitoe on behalf of the Wellcome Sanger Institute COVID-19 Surveillance Team |
| EPI_ISL_851814 | Lighthouse Lab in Glasgow | Wellcome Sanger Institute for the COVID-19 Genomics UK (COG-UK) Consortium | Harper VanSteenhouse, Yumi Kasai, David Gray, Carol Clugston, Anna Dominiczak and Alex Alderton, Roberto Amato, Sonia Goncalves, Ewan Harrison, David K. Jackson, Ian Johnston, Dominic Kwiatkowski, Cordelia Langford, John Sillitoe on behalf of the Wellcome Sanger Institute COVID-19 Surveillance Team |
| EPI_ISL_851815, EPI_ISL_851816, EPI_ISL_851817, EPI_ISL_851818, EPI_ISL_851819, EPI_ISL_851820, EPI_ISL_851821, EPI_ISL_851822, EPI_ISL_851823, EPI_ISL_851824, EPI_ISL_851825, EPI_ISL_851826, EPI_ISL_851827, EPI_ISL_851828 | | | |
| see above | Lighthouse Lab in Alderley Park | Wellcome Sanger Institute for the COVID-19 Genomics UK (COG-UK) Consortium | Jacquelyn Wynn, Mairead Hyland, The Lighthouse Lab in Alderley Park and Alex Alderton, Roberto Amato, Sonia Goncalves, Ewan Harrison, David K. Jackson, Ian Johnston, Dominic Kwiatkowski, Cordelia Langford, John Sillitoe on behalf of the Wellcome Sanger Institute COVID-19 Surveillance Team |
| EPI_ISL_851829 | Lighthouse Lab in Glasgow | Wellcome Sanger Institute for the COVID-19 Genomics UK (COG-UK) Consortium | Harper VanSteenhouse, Yumi Kasai, David Gray, Carol Clugston, Anna Dominiczak and Alex Alderton, Roberto Amato, Sonia Goncalves, Ewan Harrison, David K. Jackson, Ian Johnston, Dominic Kwiatkowski, Cordelia Langford, John Sillitoe on behalf of the Wellcome Sanger Institute COVID-19 Surveillance Team |
| EPI_ISL_851830, EPI_ISL_851831, EPI_ISL_851832, EPI_ISL_851833, EPI_ISL_851834, EPI_ISL_851835, EPI_ISL_851836 | Lighthouse Lab in Alderley Park | Wellcome Sanger Institute for the COVID-19 Genomics UK (COG-UK) Consortium | Jacquelyn Wynn, Mairead Hyland, The Lighthouse Lab in Alderley Park and Alex Alderton, Roberto Amato, Sonia Goncalves, Ewan Harrison, David K. Jackson, Ian Johnston, Dominic Kwiatkowski, Cordelia Langford, John Sillitoe on behalf of the Wellcome Sanger Institute COVID-19 Surveillance Team |
| EPI_ISL_851837 | Lighthouse Lab in Glasgow | Wellcome Sanger Institute for the COVID-19 Genomics UK (COG-UK) Consortium | Harper VanSteenhouse, Yumi Kasai, David Gray, Carol Clugston, Anna Dominiczak and Alex Alderton, Roberto Amato, Sonia Goncalves, Ewan Harrison, David K. Jackson, Ian Johnston, Dominic Kwiatkowski, Cordelia Langford, John Sillitoe on behalf of the Wellcome Sanger Institute COVID-19 Surveillance Team |
| EPI_ISL_851838, EPI_ISL_851839 | Lighthouse Lab in Alderley Park | Wellcome Sanger Institute for the COVID-19 Genomics UK (COG-UK) Consortium | Jacquelyn Wynn, Mairead Hyland, The Lighthouse Lab in Alderley Park and Alex Alderton, Roberto Amato, Sonia Goncalves, Ewan Harrison, David K. Jackson, Ian Johnston, Dominic Kwiatkowski, Cordelia Langford, John Sillitoe on behalf of the Wellcome Sanger Institute COVID-19 Surveillance Team |
| EPI_ISL_851840 | Lighthouse Lab in Glasgow | Wellcome Sanger Institute for the COVID-19 Genomics UK (COG-UK) Consortium | Harper VanSteenhouse, Yumi Kasai, David Gray, Carol Clugston, Anna Dominiczak and Alex Alderton, Roberto Amato, Sonia Goncalves, Ewan Harrison, David K. Jackson, Ian Johnston, Dominic Kwiatkowski, Cordelia Langford, John Sillitoe on behalf of the Wellcome Sanger Institute COVID-19 Surveillance Team |
| EPI_ISL_851841, EPI_ISL_851842, EPI_ISL_851843, EPI_ISL_851844, EPI_ISL_851845 | Lighthouse Lab in Alderley Park | Wellcome Sanger Institute for the COVID-19 Genomics UK (COG-UK) Consortium | Jacquelyn Wynn, Mairead Hyland, The Lighthouse Lab in Alderley Park and Alex Alderton, Roberto Amato, Sonia Goncalves, Ewan Harrison, David K. Jackson, Ian Johnston, Dominic Kwiatkowski, Cordelia Langford, John Sillitoe on behalf of the Wellcome Sanger Institute COVID-19 Surveillance Team |
| EPI_ISL_851846, EPI_ISL_851847 | Lighthouse Lab in Glasgow | Wellcome Sanger Institute for the COVID-19 Genomics UK (COG-UK) Consortium | Harper VanSteenhouse, Yumi Kasai, David Gray, Carol Clugston, Anna Dominiczak and Alex Alderton, Roberto Amato, Sonia Goncalves, Ewan Harrison, David K. Jackson, Ian Johnston, Dominic Kwiatkowski, Cordelia Langford, John Sillitoe on behalf of the Wellcome Sanger Institute COVID-19 Surveillance Team |
| EPI_ISL_851848, EPI_ISL_851849, EPI_ISL_851850 | Lighthouse Lab in Alderley Park | Wellcome Sanger Institute for the COVID-19 Genomics UK (COG-UK) Consortium | Jacquelyn Wynn, Mairead Hyland, The Lighthouse Lab in Alderley Park and Alex Alderton, Roberto Amato, Sonia Goncalves, Ewan Harrison, David K. Jackson, Ian Johnston, Dominic Kwiatkowski, Cordelia Langford, John Sillitoe on behalf of the Wellcome Sanger Institute COVID-19 Surveillance Team |
| EPI_ISL_851851 | Lighthouse Lab in Glasgow | Wellcome Sanger Institute for the COVID-19 Genomics UK (COG-UK) Consortium | Harper VanSteenhouse, Yumi Kasai, David Gray, Carol Clugston, Anna Dominiczak and Alex Alderton, Roberto Amato, Sonia Goncalves, Ewan Harrison, David K. Jackson, Ian Johnston, Dominic Kwiatkowski, Cordelia Langford, John Sillitoe on behalf of the Wellcome Sanger Institute COVID-19 Surveillance Team |
| EPI_ISL_851852, EPI_ISL_851853, EPI_ISL_851854, EPI_ISL_851855 | Lighthouse Lab in Alderley Park | Wellcome Sanger Institute for the COVID-19 Genomics UK (COG-UK) Consortium | Jacquelyn Wynn, Mairead Hyland, The Lighthouse Lab in Alderley Park and Alex Alderton, Roberto Amato, Sonia Goncalves, Ewan Harrison, David K. Jackson, Ian Johnston, Dominic Kwiatkowski, Cordelia Langford, John Sillitoe on behalf of the Wellcome Sanger Institute COVID-19 Surveillance Team |
| EPI_ISL_851856, EPI_ISL_851857 | Lighthouse Lab in Glasgow | Wellcome Sanger Institute for the COVID-19 Genomics UK (COG-UK) Consortium | Harper VanSteenhouse, Yumi Kasai, David Gray, Carol Clugston, Anna Dominiczak and Alex Alderton, Roberto Amato, Sonia Goncalves, Ewan Harrison, David K. Jackson, Ian Johnston, Dominic Kwiatkowski, Cordelia Langford, John Sillitoe on behalf of the Wellcome Sanger Institute COVID-19 Surveillance Team |
| EPI_ISL_851858 | Lighthouse Lab in Alderley Park | Wellcome Sanger Institute for the COVID-19 Genomics UK (COG-UK) Consortium | Jacquelyn Wynn, Mairead Hyland, The Lighthouse Lab in Alderley Park and Alex Alderton, Roberto Amato, Sonia Goncalves, Ewan Harrison, David K. Jackson, Ian Johnston, Dominic Kwiatkowski, Cordelia Langford, John Sillitoe on behalf of the Wellcome Sanger Institute COVID-19 Surveillance Team |
| EPI_ISL_851859 | Lighthouse Lab in Glasgow | Wellcome Sanger Institute for the COVID-19 Genomics UK (COG-UK) Consortium | Harper VanSteenhouse, Yumi Kasai, David Gray, Carol Clugston, Anna Dominiczak and Alex Alderton, Roberto Amato, Sonia Goncalves, Ewan Harrison, David K. Jackson, Ian Johnston, Dominic Kwiatkowski, Cordelia Langford, John Sillitoe on behalf of the Wellcome Sanger Institute COVID-19 Surveillance Team |
| EPI_ISL_851860, EPI_ISL_851861, EPI_ISL_851862 | Lighthouse Lab in Alderley Park | Wellcome Sanger Institute for the COVID-19 Genomics UK (COG-UK) Consortium | Jacquelyn Wynn, Mairead Hyland, The Lighthouse Lab in Alderley Park and Alex Alderton, Roberto Amato, Sonia Goncalves, Ewan Harrison, David K. Jackson, Ian Johnston, Dominic Kwiatkowski, Cordelia Langford, John Sillitoe on behalf of the Wellcome Sanger Institute COVID-19 Surveillance Team |
| EPI_ISL_851863, EPI_ISL_851864, EPI_ISL_851865, EPI_ISL_851866 | Lighthouse Lab in Glasgow | Wellcome Sanger Institute for the COVID-19 Genomics UK (COG-UK) Consortium | Harper VanSteenhouse, Yumi Kasai, David Gray, Carol Clugston, Anna Dominiczak and Alex Alderton, Roberto Amato, Sonia Goncalves, Ewan Harrison, David K. Jackson, Ian Johnston, Dominic Kwiatkowski, Cordelia Langford, John Sillitoe on behalf of the Wellcome Sanger Institute COVID-19 Surveillance Team |
| EPI_ISL_851867, EPI_ISL_851868, EPI_ISL_851869 | Lighthouse Lab in Alderley Park | Wellcome Sanger Institute for the COVID-19 Genomics UK (COG-UK) Consortium | Jacquelyn Wynn, Mairead Hyland, The Lighthouse Lab in Alderley Park and Alex Alderton, Roberto Amato, Sonia Goncalves, Ewan Harrison, David K. Jackson, Ian Johnston, Dominic Kwiatkowski, Cordelia Langford, John Sillitoe on behalf of the Wellcome Sanger Institute COVID-19 Surveillance Team |
| EPI_ISL_851870, EPI_ISL_851871 | Lighthouse Lab in Glasgow | Wellcome Sanger Institute for the COVID-19 Genomics UK (COG-UK) Consortium | Harper VanSteenhouse, Yumi Kasai, David Gray, Carol Clugston, Anna Dominiczak and Alex Alderton, Roberto Amato, Sonia Goncalves, Ewan Harrison, David K. Jackson, Ian Johnston, Dominic Kwiatkowski, Cordelia Langford, John Sillitoe on behalf of the Wellcome Sanger Institute COVID-19 Surveillance Team |
| EPI_ISL_851872, EPI_ISL_851873, EPI_ISL_851874 | Lighthouse Lab in Alderley Park | Wellcome Sanger Institute for the COVID-19 Genomics UK (COG-UK) Consortium | Jacquelyn Wynn, Mairead Hyland, The Lighthouse Lab in Alderley Park and Alex Alderton, Roberto Amato, Sonia Goncalves, Ewan Harrison, David K. Jackson, Ian Johnston, Dominic Kwiatkowski, Cordelia Langford, John Sillitoe on behalf of the Wellcome Sanger Institute COVID-19 Surveillance Team |
| EPI_ISL_851876, EPI_ISL_851877 | Lighthouse Lab in Glasgow | Wellcome Sanger Institute for the COVID-19 Genomics UK (COG-UK) Consortium | Harper VanSteenhouse, Yumi Kasai, David Gray, Carol Clugston, Anna Dominiczak and Alex Alderton, Roberto Amato, Sonia Goncalves, Ewan Harrison, David K. Jackson, Ian Johnston, Dominic Kwiatkowski, Cordelia Langford, John Sillitoe on behalf of the Wellcome Sanger Institute COVID-19 Surveillance Team |
| EPI_ISL_851878, EPI_ISL_851879, EPI_ISL_851880, EPI_ISL_851881, EPI_ISL_851882, EPI_ISL_851883, EPI_ISL_851884 | Lighthouse Lab in Alderley Park | Wellcome Sanger Institute for the COVID-19 Genomics UK (COG-UK) Consortium | Jacquelyn Wynn, Mairead Hyland, The Lighthouse Lab in Alderley Park and Alex Alderton, Roberto Amato, Sonia Goncalves, Ewan Harrison, David K. Jackson, Ian Johnston, Dominic Kwiatkowski, Cordelia Langford, John Sillitoe on behalf of the Wellcome Sanger Institute COVID-19 Surveillance Team |
| EPI_ISL_851885 | Lighthouse Lab in Glasgow | Wellcome Sanger Institute for the COVID-19 Genomics UK (COG-UK) Consortium | Harper VanSteenhouse, Yumi Kasai, David Gray, Carol Clugston, Anna Dominiczak and Alex Alderton, Roberto Amato, Sonia Goncalves, Ewan Harrison, David K. Jackson, Ian Johnston, Dominic Kwiatkowski, Cordelia Langford, John Sillitoe on behalf of the Wellcome Sanger Institute COVID-19 Surveillance Team |

| | | | |
|---|---|---|---|
| EPI_ISL_851886, EPI_ISL_851887 | Lighthouse Lab in Alderley Park | Wellcome Sanger Institute for the COVID-19 Genomics UK (COG-UK) Consortium | Jacquelyn Wynn, Mairead Hyland, The Lighthouse Lab in Alderley Park and Alex Alderton, Roberto Amato, Sonia Goncalves, Ewan Harrison, David K. Jackson, Ian Johnston, Dominic Kwiatkowski, Cordelia Langford, John Sillitoe on behalf of the Wellcome Sanger Institute COVID-19 Surveillance Team |
| EPI_ISL_851888 | Lighthouse Lab in Glasgow | Wellcome Sanger Institute for the COVID-19 Genomics UK (COG-UK) Consortium | Harper VanSteenhouse, Yumi Kasai, David Gray, Carol Clugston, Anna Dominiczak and Alex Alderton, Roberto Amato, Sonia Goncalves, Ewan Harrison, David K. Jackson, Ian Johnston, Dominic Kwiatkowski, Cordelia Langford, John Sillitoe on behalf of the Wellcome Sanger Institute COVID-19 Surveillance Team |
| EPI_ISL_851889, EPI_ISL_851890 | Lighthouse Lab in Alderley Park | Wellcome Sanger Institute for the COVID-19 Genomics UK (COG-UK) Consortium | Jacquelyn Wynn, Mairead Hyland, The Lighthouse Lab in Alderley Park and Alex Alderton, Roberto Amato, Sonia Goncalves, Ewan Harrison, David K. Jackson, Ian Johnston, Dominic Kwiatkowski, Cordelia Langford, John Sillitoe on behalf of the Wellcome Sanger Institute COVID-19 Surveillance Team |
| EPI_ISL_851891, EPI_ISL_851892 | Lighthouse Lab in Glasgow | Wellcome Sanger Institute for the COVID-19 Genomics UK (COG-UK) Consortium | Harper VanSteenhouse, Yumi Kasai, David Gray, Carol Clugston, Anna Dominiczak and Alex Alderton, Roberto Amato, Sonia Goncalves, Ewan Harrison, David K. Jackson, Ian Johnston, Dominic Kwiatkowski, Cordelia Langford, John Sillitoe on behalf of the Wellcome Sanger Institute COVID-19 Surveillance Team |
| EPI_ISL_851893, EPI_ISL_851894 | Lighthouse Lab in Alderley Park | Wellcome Sanger Institute for the COVID-19 Genomics UK (COG-UK) Consortium | Jacquelyn Wynn, Mairead Hyland, The Lighthouse Lab in Alderley Park and Alex Alderton, Roberto Amato, Sonia Goncalves, Ewan Harrison, David K. Jackson, Ian Johnston, Dominic Kwiatkowski, Cordelia Langford, John Sillitoe on behalf of the Wellcome Sanger Institute COVID-19 Surveillance Team |
| EPI_ISL_851895 | Lighthouse Lab in Glasgow | Wellcome Sanger Institute for the COVID-19 Genomics UK (COG-UK) Consortium | Harper VanSteenhouse, Yumi Kasai, David Gray, Carol Clugston, Anna Dominiczak and Alex Alderton, Roberto Amato, Sonia Goncalves, Ewan Harrison, David K. Jackson, Ian Johnston, Dominic Kwiatkowski, Cordelia Langford, John Sillitoe on behalf of the Wellcome Sanger Institute COVID-19 Surveillance Team |
| EPI_ISL_851896, EPI_ISL_851897 | Lighthouse Lab in Alderley Park | Wellcome Sanger Institute for the COVID-19 Genomics UK (COG-UK) Consortium | Jacquelyn Wynn, Mairead Hyland, The Lighthouse Lab in Alderley Park and Alex Alderton, Roberto Amato, Sonia Goncalves, Ewan Harrison, David K. Jackson, Ian Johnston, Dominic Kwiatkowski, Cordelia Langford, John Sillitoe on behalf of the Wellcome Sanger Institute COVID-19 Surveillance Team |
| EPI_ISL_851898 | Lighthouse Lab in Glasgow | Wellcome Sanger Institute for the COVID-19 Genomics UK (COG-UK) Consortium | Harper VanSteenhouse, Yumi Kasai, David Gray, Carol Clugston, Anna Dominiczak and Alex Alderton, Roberto Amato, Sonia Goncalves, Ewan Harrison, David K. Jackson, Ian Johnston, Dominic Kwiatkowski, Cordelia Langford, John Sillitoe on behalf of the Wellcome Sanger Institute COVID-19 Surveillance Team |
| EPI_ISL_851899, EPI_ISL_851900 | Lighthouse Lab in Alderley Park | Wellcome Sanger Institute for the COVID-19 Genomics UK (COG-UK) Consortium | Jacquelyn Wynn, Mairead Hyland, The Lighthouse Lab in Alderley Park and Alex Alderton, Roberto Amato, Sonia Goncalves, Ewan Harrison, David K. Jackson, Ian Johnston, Dominic Kwiatkowski, Cordelia Langford, John Sillitoe on behalf of the Wellcome Sanger Institute COVID-19 Surveillance Team |
| EPI_ISL_851901 | Lighthouse Lab in Glasgow | Wellcome Sanger Institute for the COVID-19 Genomics UK (COG-UK) Consortium | Harper VanSteenhouse, Yumi Kasai, David Gray, Carol Clugston, Anna Dominiczak and Alex Alderton, Roberto Amato, Sonia Goncalves, Ewan Harrison, David K. Jackson, Ian Johnston, Dominic Kwiatkowski, Cordelia Langford, John Sillitoe on behalf of the Wellcome Sanger Institute COVID-19 Surveillance Team |
| EPI_ISL_851902, EPI_ISL_851903, EPI_ISL_851904 | Lighthouse Lab in Alderley Park | Wellcome Sanger Institute for the COVID-19 Genomics UK (COG-UK) Consortium | Jacquelyn Wynn, Mairead Hyland, The Lighthouse Lab in Alderley Park and Alex Alderton, Roberto Amato, Sonia Goncalves, Ewan Harrison, David K. Jackson, Ian Johnston, Dominic Kwiatkowski, Cordelia Langford, John Sillitoe on behalf of the Wellcome Sanger Institute COVID-19 Surveillance Team |
| EPI_ISL_851905, EPI_ISL_851906 | Lighthouse Lab in Glasgow | Wellcome Sanger Institute for the COVID-19 Genomics UK (COG-UK) Consortium | Harper VanSteenhouse, Yumi Kasai, David Gray, Carol Clugston, Anna Dominiczak and Alex Alderton, Roberto Amato, Sonia Goncalves, Ewan Harrison, David K. Jackson, Ian Johnston, Dominic Kwiatkowski, Cordelia Langford, John Sillitoe on behalf of the Wellcome Sanger Institute COVID-19 Surveillance Team |
| EPI_ISL_851907, EPI_ISL_851908, EPI_ISL_851909, EPI_ISL_851910, EPI_ISL_851911, EPI_ISL_851912, EPI_ISL_851913, EPI_ISL_851914, EPI_ISL_851915 | Lighthouse Lab in Alderley Park | Wellcome Sanger Institute for the COVID-19 Genomics UK (COG-UK) Consortium | Jacquelyn Wynn, Mairead Hyland, The Lighthouse Lab in Alderley Park and Alex Alderton, Roberto Amato, Sonia Goncalves, Ewan Harrison, David K. Jackson, Ian Johnston, Dominic Kwiatkowski, Cordelia Langford, John Sillitoe on behalf of the Wellcome Sanger Institute COVID-19 Surveillance Team |
| EPI_ISL_851916 | Lighthouse Lab in Glasgow | Wellcome Sanger Institute for the COVID-19 Genomics UK (COG-UK) Consortium | Harper VanSteenhouse, Yumi Kasai, David Gray, Carol Clugston, Anna Dominiczak and Alex Alderton, Roberto Amato, Sonia Goncalves, Ewan Harrison, David K. Jackson, Ian Johnston, Dominic Kwiatkowski, Cordelia Langford, John Sillitoe on behalf of the Wellcome Sanger Institute COVID-19 Surveillance Team |
| EPI_ISL_851917, EPI_ISL_851918, EPI_ISL_851919 | Lighthouse Lab in Alderley Park | Wellcome Sanger Institute for the COVID-19 Genomics UK (COG-UK) Consortium | Jacquelyn Wynn, Mairead Hyland, The Lighthouse Lab in Alderley Park and Alex Alderton, Roberto Amato, Sonia Goncalves, Ewan Harrison, David K. Jackson, Ian Johnston, Dominic Kwiatkowski, Cordelia Langford, John Sillitoe on behalf of the Wellcome Sanger Institute COVID-19 Surveillance Team |
| EPI_ISL_851920 | Lighthouse Lab in Glasgow | Wellcome Sanger Institute for the COVID-19 Genomics UK (COG-UK) Consortium | Harper VanSteenhouse, Yumi Kasai, David Gray, Carol Clugston, Anna Dominiczak and Alex Alderton, Roberto Amato, Sonia Goncalves, Ewan Harrison, David K. Jackson, Ian Johnston, Dominic Kwiatkowski, Cordelia Langford, John Sillitoe on behalf of the Wellcome Sanger Institute COVID-19 Surveillance Team |
| EPI_ISL_851921 | Lighthouse Lab in Alderley Park | Wellcome Sanger Institute for the COVID-19 Genomics UK (COG-UK) Consortium | Jacquelyn Wynn, Mairead Hyland, The Lighthouse Lab in Alderley Park and Alex Alderton, Roberto Amato, Sonia Goncalves, Ewan Harrison, David K. Jackson, Ian Johnston, Dominic Kwiatkowski, Cordelia Langford, John Sillitoe on behalf of the Wellcome Sanger Institute COVID-19 Surveillance Team |
| EPI_ISL_851922 | Lighthouse Lab in Glasgow | Wellcome Sanger Institute for the COVID-19 Genomics UK (COG-UK) Consortium | Harper VanSteenhouse, Yumi Kasai, David Gray, Carol Clugston, Anna Dominiczak and Alex Alderton, Roberto Amato, Sonia Goncalves, Ewan Harrison, David K. Jackson, Ian Johnston, Dominic Kwiatkowski, Cordelia Langford, John Sillitoe on behalf of the Wellcome Sanger Institute COVID-19 Surveillance Team |
| EPI_ISL_851923, EPI_ISL_851924, EPI_ISL_851925, EPI_ISL_851926, EPI_ISL_851927, EPI_ISL_851928 | Lighthouse Lab in Alderley Park | Wellcome Sanger Institute for the COVID-19 Genomics UK (COG-UK) Consortium | Jacquelyn Wynn, Mairead Hyland, The Lighthouse Lab in Alderley Park and Alex Alderton, Roberto Amato, Sonia Goncalves, Ewan Harrison, David K. Jackson, Ian Johnston, Dominic Kwiatkowski, Cordelia Langford, John Sillitoe on behalf of the Wellcome Sanger Institute COVID-19 Surveillance Team |
| EPI_ISL_851929 | Lighthouse Lab in Glasgow | Wellcome Sanger Institute for the COVID-19 Genomics UK (COG-UK) Consortium | Harper VanSteenhouse, Yumi Kasai, David Gray, Carol Clugston, Anna Dominiczak and Alex Alderton, Roberto Amato, Sonia Goncalves, Ewan Harrison, David K. Jackson, Ian Johnston, Dominic Kwiatkowski, Cordelia Langford, John Sillitoe on behalf of the Wellcome Sanger Institute COVID-19 Surveillance Team |
| EPI_ISL_851930, EPI_ISL_851931, EPI_ISL_851932, EPI_ISL_851933 | Lighthouse Lab in Alderley Park | Wellcome Sanger Institute for the COVID-19 Genomics UK (COG-UK) Consortium | Jacquelyn Wynn, Mairead Hyland, The Lighthouse Lab in Alderley Park and Alex Alderton, Roberto Amato, Sonia Goncalves, Ewan Harrison, David K. Jackson, Ian Johnston, Dominic Kwiatkowski, Cordelia Langford, John Sillitoe on behalf of the Wellcome Sanger Institute COVID-19 Surveillance Team |
| EPI_ISL_851934 | Lighthouse Lab in Glasgow | Wellcome Sanger Institute for the COVID-19 Genomics UK (COG-UK) Consortium | Harper VanSteenhouse, Yumi Kasai, David Gray, Carol Clugston, Anna Dominiczak and Alex Alderton, Roberto Amato, Sonia Goncalves, Ewan Harrison, David K. Jackson, Ian Johnston, Dominic Kwiatkowski, Cordelia Langford, John Sillitoe on behalf of the Wellcome Sanger Institute COVID-19 Surveillance Team |
| EPI_ISL_851935 | Lighthouse Lab in Alderley Park | Wellcome Sanger Institute for the COVID-19 Genomics UK (COG-UK) Consortium | Jacquelyn Wynn, Mairead Hyland, The Lighthouse Lab in Alderley Park and Alex Alderton, Roberto Amato, Sonia Goncalves, Ewan Harrison, David K. Jackson, Ian Johnston, Dominic Kwiatkowski, Cordelia Langford, John Sillitoe on behalf of the Wellcome Sanger Institute COVID-19 Surveillance Team |
| EPI_ISL_851936 | Lighthouse Lab in Glasgow | Wellcome Sanger Institute for the COVID-19 Genomics UK (COG-UK) Consortium | Harper VanSteenhouse, Yumi Kasai, David Gray, Carol Clugston, Anna Dominiczak and Alex Alderton, Roberto Amato, Sonia Goncalves, Ewan Harrison, David K. Jackson, Ian Johnston, Dominic Kwiatkowski, Cordelia Langford, John Sillitoe on behalf of the Wellcome Sanger Institute COVID-19 Surveillance Team |
| EPI_ISL_851937, EPI_ISL_851938, EPI_ISL_851939 | Lighthouse Lab in Alderley Park | Wellcome Sanger Institute for the COVID-19 Genomics UK (COG-UK) Consortium | Jacquelyn Wynn, Mairead Hyland, The Lighthouse Lab in Alderley Park and Alex Alderton, Roberto Amato, Sonia Goncalves, Ewan Harrison, David K. Jackson, Ian Johnston, Dominic Kwiatkowski, Cordelia Langford, John Sillitoe on behalf of the Wellcome Sanger Institute COVID-19 Surveillance Team |
| EPI_ISL_851940 | Lighthouse Lab in Glasgow | Wellcome Sanger Institute for the COVID-19 Genomics UK (COG-UK) Consortium | Harper VanSteenhouse, Yumi Kasai, David Gray, Carol Clugston, Anna Dominiczak and Alex Alderton, Roberto Amato, Sonia Goncalves, Ewan Harrison, David K. Jackson, Ian Johnston, Dominic Kwiatkowski, Cordelia Langford, John Sillitoe on behalf of the Wellcome Sanger Institute COVID-19 Surveillance Team |
| EPI_ISL_851941, EPI_ISL_851942, EPI_ISL_851943, EPI_ISL_851944, EPI_ISL_851945, EPI_ISL_851946, EPI_ISL_851947, EPI_ISL_851948, EPI_ISL_851949 | Lighthouse Lab in Alderley Park | Wellcome Sanger Institute for the COVID-19 Genomics UK (COG-UK) Consortium | Jacquelyn Wynn, Mairead Hyland, The Lighthouse Lab in Alderley Park and Alex Alderton, Roberto Amato, Sonia Goncalves, Ewan Harrison, David K. Jackson, Ian Johnston, Dominic Kwiatkowski, Cordelia Langford, John Sillitoe on behalf of the Wellcome Sanger Institute COVID-19 Surveillance Team |

| | | | |
|---|---|---|---|
| EPI_ISL_851950 | Lighthouse Lab in Glasgow | Wellcome Sanger Institute for the COVID-19 Genomics UK (COG-UK) Consortium | Harper VanSteenhouse, Yumi Kasai, David Gray, Carol Clugston, Anna Dominiczak and Alex Alderton, Roberto Amato, Sonia Goncalves, Ewan Harrison, David K. Jackson, Ian Johnston, Dominic Kwiatkowski, Cordelia Langford, John Sillitoe on behalf of the Wellcome Sanger Institute COVID-19 Surveillance Team |
| EPI_ISL_851951, EPI_ISL_851952, EPI_ISL_851953 | Lighthouse Lab in Alderley Park | Wellcome Sanger Institute for the COVID-19 Genomics UK (COG-UK) Consortium | Jacquelyn Wynn, Mairead Hyland, The Lighthouse Lab in Alderley Park and Alex Alderton, Roberto Amato, Sonia Goncalves, Ewan Harrison, David K. Jackson, Ian Johnston, Dominic Kwiatkowski, Cordelia Langford, John Sillitoe on behalf of the Wellcome Sanger Institute COVID-19 Surveillance Team |
| EPI_ISL_851955, EPI_ISL_851956 | Lighthouse Lab in Glasgow | Wellcome Sanger Institute for the COVID-19 Genomics UK (COG-UK) Consortium | Harper VanSteenhouse, Yumi Kasai, David Gray, Carol Clugston, Anna Dominiczak and Alex Alderton, Roberto Amato, Sonia Goncalves, Ewan Harrison, David K. Jackson, Ian Johnston, Dominic Kwiatkowski, Cordelia Langford, John Sillitoe on behalf of the Wellcome Sanger Institute COVID-19 Surveillance Team |
| EPI_ISL_851957, EPI_ISL_851958, EPI_ISL_851959, EPI_ISL_851960, EPI_ISL_851961, EPI_ISL_851962, EPI_ISL_851965, EPI_ISL_851966, EPI_ISL_851967, EPI_ISL_851969, EPI_ISL_851970, EPI_ISL_851971, EPI_ISL_851973, EPI_ISL_851975, EPI_ISL_851977, EPI_ISL_851978, EPI_ISL_851980, EPI_ISL_851981, EPI_ISL_851982, EPI_ISL_851983, EPI_ISL_851984, EPI_ISL_851986, EPI_ISL_851987, EPI_ISL_851988, EPI_ISL_851990, EPI_ISL_851991, EPI_ISL_851992, EPI_ISL_851994, EPI_ISL_851995, EPI_ISL_851996, EPI_ISL_851997, EPI_ISL_851998, EPI_ISL_851999, EPI_ISL_852000, EPI_ISL_852001, EPI_ISL_852003, EPI_ISL_852004, EPI_ISL_852005, EPI_ISL_852006, EPI_ISL_852007, EPI_ISL_852008, EPI_ISL_852010, EPI_ISL_852011, EPI_ISL_852012, EPI_ISL_852013, EPI_ISL_852015, EPI_ISL_852016, EPI_ISL_852018, EPI_ISL_852019, EPI_ISL_852020, EPI_ISL_852021, EPI_ISL_852022, EPI_ISL_852023, EPI_ISL_852024, EPI_ISL_852026, EPI_ISL_852027, EPI_ISL_852028, EPI_ISL_852030, EPI_ISL_852031, EPI_ISL_852032, EPI_ISL_852033, EPI_ISL_852034, EPI_ISL_852035, EPI_ISL_852036, EPI_ISL_852037, EPI_ISL_852038, EPI_ISL_852039, EPI_ISL_852040, EPI_ISL_852041, EPI_ISL_852043, EPI_ISL_852044, EPI_ISL_852045, EPI_ISL_852046, EPI_ISL_852048, EPI_ISL_852049, EPI_ISL_852050, EPI_ISL_852051, EPI_ISL_852052, EPI_ISL_852053, EPI_ISL_852054, EPI_ISL_852055, EPI_ISL_852056, EPI_ISL_852057, EPI_ISL_852058, EPI_ISL_852059, EPI_ISL_852060, EPI_ISL_852061, EPI_ISL_852063, EPI_ISL_852064, EPI_ISL_852065, EPI_ISL_852066, EPI_ISL_852067, EPI_ISL_852068, EPI_ISL_852069, EPI_ISL_852070, EPI_ISL_852071, EPI_ISL_852072, EPI_ISL_852073, EPI_ISL_852075, EPI_ISL_852076, EPI_ISL_852078, EPI_ISL_852079, EPI_ISL_852080, EPI_ISL_852081, EPI_ISL_852082, EPI_ISL_852084, EPI_ISL_852085, EPI_ISL_852087, EPI_ISL_852088, EPI_ISL_852089, EPI_ISL_852090 | | | |
| see above | Lighthouse Lab in Milton Keynes | Wellcome Sanger Institute for the COVID-19 Genomics UK (COG-UK) Consortium | The Lighthouse Lab in Milton Keynes and Alex Alderton, Roberto Amato, Sonia Goncalves, Ewan Harrison, David K. Jackson, Ian Johnston, Dominic Kwiatkowski, Cordelia Langford, John Sillitoe on behalf of the Wellcome Sanger Institute COVID-19 Surveillance Team |
| EPI_ISL_852091 | Lighthouse Lab in Alderley Park | Wellcome Sanger Institute for the COVID-19 Genomics UK (COG-UK) Consortium | Jacquelyn Wynn, Mairead Hyland, The Lighthouse Lab in Alderley Park and Alex Alderton, Roberto Amato, Sonia Goncalves, Ewan Harrison, David K. Jackson, Ian Johnston, Dominic Kwiatkowski, Cordelia Langford, John Sillitoe on behalf of the Wellcome Sanger Institute COVID-19 Surveillance Team |
| EPI_ISL_852092, EPI_ISL_852094, EPI_ISL_852095, EPI_ISL_852096, EPI_ISL_852097, EPI_ISL_852098, EPI_ISL_852099, EPI_ISL_852101, EPI_ISL_852102, EPI_ISL_852103, EPI_ISL_852104, EPI_ISL_852105, EPI_ISL_852106 | | | |
| see above | Lighthouse Lab in Milton Keynes | Wellcome Sanger Institute for the COVID-19 Genomics UK (COG-UK) Consortium | The Lighthouse Lab in Milton Keynes and Alex Alderton, Roberto Amato, Sonia Goncalves, Ewan Harrison, David K. Jackson, Ian Johnston, Dominic Kwiatkowski, Cordelia Langford, John Sillitoe on behalf of the Wellcome Sanger Institute COVID-19 Surveillance Team |
| EPI_ISL_852107 | Lighthouse Lab in Alderley Park | Wellcome Sanger Institute for the COVID-19 Genomics UK (COG-UK) Consortium | Jacquelyn Wynn, Mairead Hyland, The Lighthouse Lab in Alderley Park and Alex Alderton, Roberto Amato, Sonia Goncalves, Ewan Harrison, David K. Jackson, Ian Johnston, Dominic Kwiatkowski, Cordelia Langford, John Sillitoe on behalf of the Wellcome Sanger Institute COVID-19 Surveillance Team |
| EPI_ISL_852108, EPI_ISL_852109, EPI_ISL_852110, EPI_ISL_852111, EPI_ISL_852112, EPI_ISL_852113, EPI_ISL_852115, EPI_ISL_852116, EPI_ISL_852117, EPI_ISL_852119, EPI_ISL_852120, EPI_ISL_852121, EPI_ISL_852122, EPI_ISL_852123, EPI_ISL_852125, EPI_ISL_852126, EPI_ISL_852128, EPI_ISL_852129, EPI_ISL_852130, EPI_ISL_852131, EPI_ISL_852133, EPI_ISL_852134, EPI_ISL_852136, EPI_ISL_852137, EPI_ISL_852138, EPI_ISL_852139, EPI_ISL_852140, EPI_ISL_852141, EPI_ISL_852142, EPI_ISL_852143, EPI_ISL_852144, EPI_ISL_852145, EPI_ISL_852146, EPI_ISL_852147, EPI_ISL_852148, EPI_ISL_852149, EPI_ISL_852150, EPI_ISL_852151, EPI_ISL_852152, EPI_ISL_852153, EPI_ISL_852154, EPI_ISL_852155, EPI_ISL_852156, EPI_ISL_852157, EPI_ISL_852158, EPI_ISL_852159, EPI_ISL_852160, EPI_ISL_852161, EPI_ISL_852162, EPI_ISL_852163, EPI_ISL_852164, EPI_ISL_852166, EPI_ISL_852167, EPI_ISL_852168, EPI_ISL_852169, EPI_ISL_852170, EPI_ISL_852171, EPI_ISL_852172, EPI_ISL_852173, EPI_ISL_852175, EPI_ISL_852176, EPI_ISL_852177, EPI_ISL_852178, EPI_ISL_852179, EPI_ISL_852180, EPI_ISL_852181, EPI_ISL_852182, EPI_ISL_852185, EPI_ISL_852186, EPI_ISL_852187, EPI_ISL_852188, EPI_ISL_852189, EPI_ISL_852192, EPI_ISL_852193, EPI_ISL_852194, EPI_ISL_852195, EPI_ISL_852196, EPI_ISL_852197, EPI_ISL_852198, EPI_ISL_852199, EPI_ISL_852200, EPI_ISL_852201, EPI_ISL_852202, EPI_ISL_852204, EPI_ISL_852205, EPI_ISL_852207, EPI_ISL_852208, EPI_ISL_852209, EPI_ISL_852210, EPI_ISL_852211, EPI_ISL_852212, EPI_ISL_852215, EPI_ISL_852216, EPI_ISL_852217, EPI_ISL_852219, EPI_ISL_852220, EPI_ISL_852221, EPI_ISL_852222, EPI_ISL_852223, EPI_ISL_852224, EPI_ISL_852226, EPI_ISL_852227, EPI_ISL_852228, EPI_ISL_852229, EPI_ISL_852230, EPI_ISL_852231, EPI_ISL_852232, EPI_ISL_852233, EPI_ISL_852235, EPI_ISL_852237, EPI_ISL_852238, EPI_ISL_852239, EPI_ISL_852240, EPI_ISL_852241, EPI_ISL_852242, EPI_ISL_852243, EPI_ISL_852244, EPI_ISL_852245, EPI_ISL_852246, EPI_ISL_852247, EPI_ISL_852248, EPI_ISL_852249, EPI_ISL_852250, EPI_ISL_852251, EPI_ISL_852252, EPI_ISL_852253, EPI_ISL_852254, EPI_ISL_852255 | | | |
| see above | Lighthouse Lab in Milton Keynes | Wellcome Sanger Institute for the COVID-19 Genomics UK (COG-UK) Consortium | The Lighthouse Lab in Milton Keynes and Alex Alderton, Roberto Amato, Sonia Goncalves, Ewan Harrison, David K. Jackson, Ian Johnston, Dominic Kwiatkowski, Cordelia Langford, John Sillitoe on behalf of the Wellcome Sanger Institute COVID-19 Surveillance Team |
| EPI_ISL_852268, EPI_ISL_852286, EPI_ISL_852303, EPI_ISL_852319, EPI_ISL_852338, EPI_ISL_852361, EPI_ISL_852375, EPI_ISL_852389, EPI_ISL_852406, EPI_ISL_852407, EPI_ISL_852413, EPI_ISL_852414, EPI_ISL_852416, EPI_ISL_852438, EPI_ISL_852448, EPI_ISL_852459, EPI_ISL_852500, EPI_ISL_852517, EPI_ISL_852524, EPI_ISL_852527, EPI_ISL_852535, EPI_ISL_852536, EPI_ISL_852544, EPI_ISL_852549 | | | |
| see above | Lighthouse Lab in Glasgow | Wellcome Sanger Institute for the COVID-19 Genomics UK (COG-UK) Consortium | Harper VanSteenhouse, Yumi Kasai, David Gray, Carol Clugston, Anna Dominiczak and Alex Alderton, Roberto Amato, Sonia Goncalves, Ewan Harrison, David K. Jackson, Ian Johnston, Dominic Kwiatkowski, Cordelia Langford, John Sillitoe on behalf of the Wellcome Sanger Institute COVID-19 Surveillance Team |
| EPI_ISL_852952, EPI_ISL_852955, EPI_ISL_852962, EPI_ISL_852963, EPI_ISL_852992, EPI_ISL_852993, EPI_ISL_852994, EPI_ISL_852995, EPI_ISL_852996, EPI_ISL_852997, EPI_ISL_852999, EPI_ISL_853001, EPI_ISL_853007, EPI_ISL_853008, EPI_ISL_853018, EPI_ISL_853019, EPI_ISL_853020, EPI_ISL_853021, EPI_ISL_853022, EPI_ISL_853023 | | | |
| see above | Hospital General Universitario Gregorio Marañón | SeqCOVID-SPAIN consortium/IBV(CSIC) | Darío García de Viedma, Laura Pérez-Lago, Pedro J Sola-Campoy, Sergio Buenestado-Serrano, Marta Herranz, Victor Manuel de la Cueva, Julia Suárez, Pilar Catalán, Patricia Muñoz and SeqCOVID-SPAIN consortium |
| EPI_ISL_853394, EPI_ISL_853395, EPI_ISL_853396 | Charité Universitätsmedizin Berlin, Institut für Virologie/Labor Berlin | Charité Universitätsmedizin Berlin, Institut für Virologie | Victor M Corman, Julia Schneider, Barbara Mühlemann, Jörn Beheim-Schwarzbach, Talitha Veith, Julia Tesch, Tobias Bleicker, Terry Jones, Christian Drosten |
| EPI_ISL_853757, EPI_ISL_853808, EPI_ISL_853811, EPI_ISL_853812, EPI_ISL_853813, EPI_ISL_854187, EPI_ISL_854188, EPI_ISL_854189, EPI_ISL_854190, EPI_ISL_854191, EPI_ISL_854192, EPI_ISL_854193, EPI_ISL_854194, EPI_ISL_854195, EPI_ISL_854196, EPI_ISL_854197, EPI_ISL_854198, EPI_ISL_854199, EPI_ISL_854200, EPI_ISL_854201, EPI_ISL_854202, EPI_ISL_854203, EPI_ISL_854204, EPI_ISL_854205, EPI_ISL_854206, EPI_ISL_854207, EPI_ISL_854208 | | | |
| see above | Austrian Agency for Health and Food Safety (AGES) | Bergthaler laboratory, CeMM Research Center for Molecular Medicine of the Austrian Academy of Sciences | Lukas Endler, Alexandra Popa, Benedikt Agerer, Jakob-Wendelin Genger, Alexander Lercher, Anna Schedl, Thomas Penz, Michael Schuster, Jan Laine, Martin Senekowitsch, Christoph Bock, Andreas Bergthaler |
| EPI_ISL_854374, EPI_ISL_854375, EPI_ISL_854376, EPI_ISL_854377, EPI_ISL_854378, EPI_ISL_854379, EPI_ISL_854380, EPI_ISL_854451 | WESTCHESTER MEDICAL CENTER | Wadsworth Center, New York State Department of Health | Kirsten St. George, Daryl M. Lamson, Alexis Russel, Matthew Shudt, Melissa A Leisner, Jonathan Plitnick, Navjot Singh, John Kelly, Erasmus Schneider, Erica Lasek-Nesselquist |
| EPI_ISL_854750 | Microbiological Diagnostic Unit - Public Health Laboratory (MDU-PHL) | MDU-PHL | Seemann T., Sait, M.L., Sherry, N.L. |
| EPI_ISL_854751, EPI_ISL_854752, EPI_ISL_854753 | Victorian Infectious Diseases Reference Laboratory (VIDRL) | VIDRL and MDU-PHL | Caly L., Seemann T., Sait, M.L., Druce J., Sherry, N.L. |
| EPI_ISL_854755, EPI_ISL_854756 | Microbiological Diagnostic Unit - Public Health Laboratory (MDU-PHL) | MDU-PHL | Seemann T., Sait, M.L., Sherry, N.L. |
| EPI_ISL_854757 | Victorian Infectious Diseases Reference Laboratory (VIDRL) | VIDRL and MDU-PHL | Caly L., Seemann T., Sait, M.L., Druce J., Sherry, N.L. |
| EPI_ISL_854811, EPI_ISL_854812, EPI_ISL_854814, EPI_ISL_854815, EPI_ISL_854819, EPI_ISL_854821, EPI_ISL_854822, EPI_ISL_854823, EPI_ISL_854825, EPI_ISL_854826, EPI_ISL_854827, EPI_ISL_854835, EPI_ISL_854836, EPI_ISL_854837, EPI_ISL_854838, EPI_ISL_854839, EPI_ISL_854840, EPI_ISL_854841, EPI_ISL_855014, EPI_ISL_855017, EPI_ISL_855022, EPI_ISL_855035, EPI_ISL_855055, EPI_ISL_855058, EPI_ISL_855070, EPI_ISL_855076, EPI_ISL_855077, EPI_ISL_855078, EPI_ISL_855079, EPI_ISL_855080, EPI_ISL_855154, EPI_ISL_855159, EPI_ISL_855184 | | | |
| see above | Quest Diagnostics | Quest Diagnostics | Rosenthal,S.H., Gerasimova,A., Kagan,R.M., Anderson, B., Hua, M., Liu Y., Bernstein, L.E., Livingston, K.E., Perez, A., Shalhout, D.F., Shlyakhter, I.A., Owen, R., Tanpaiboon, P., Lacbawan, F. |
| EPI_ISL_855357 | Labo Analyses Med | National Reference Center for Viruses of Respiratory Infections, Institut Pasteur, Paris | Marion Barbet, Sylvie Behillil, Méline Bizard, Angela Brisebarre, Camille Capel, Etienne Simon-Lorière, Vincent Enouf, Maud Vanpeene, Sylvie van der Werf,Harich |
| EPI_ISL_856678 | Charité Universitätsmedizin Berlin, Institute of Virology, Charitéplatz 1, 10117 Berlin, Germany | Charité Universitätsmedizin Berlin, Institute of Virology, Charitéplatz 1, 10117 Berlin, Germany | Victor M Corman, Julia Schneider, Jörn Beheim-Schwarzbach, Tobias Bleicker, Julia Tesch, Barbara Mühlemann, Talitha Veith, Terry Jones, Christian Drosten |
| EPI_ISL_856722, EPI_ISL_856730, EPI_ISL_856731 | Department of Clinical Microbiology | GIGA Medical Genomics | Keith Durkin, Maria Artesi, Sébastien Bontems, Raphaël Boreux, Bouchra Boujemla, Cécile Meex, Pierrette Melin, Marie-Pierre Hayette, Vincent Bours |
| EPI_ISL_856912, EPI_ISL_856920, EPI_ISL_856921, EPI_ISL_856922, EPI_ISL_856923, EPI_ISL_856924, EPI_ISL_856925, EPI_ISL_856926, EPI_ISL_856927, EPI_ISL_856928, EPI_ISL_856929 | | | |
| see above | Wyoming Public Health Laboratory | Wyoming Public Health Laboratory | Noah Hull, Taylor Fearing, Lynette Gumbleton, Channing Weber, Ashley Norberg, Bailey Bowcutt, and Wanda Manley |
| EPI_ISL_857041 | Platform BIS UZA/UAntwerpen, University Hospital Antwerp, | UAntwerp, Laboratory of Medical Microbiology, Campus Drie | Basil Britto Xavier, Jasmine Coppens, Christine Lammens, Veerle Matheeussen, Herman Goossens |

| | | | |
|---|---|---|---|
| | Edegem, Belgium | Eiken S6.26, Universiteitsplein 1, 2610, Wilrijk, Antwerp, Belgium | |
| EPI_ISL_857134 | DOHMH PHL | New York City Public Health Laboratory | Jade Wang, et al. |
| EPI_ISL_857135 | DOHMH Corona | New York City Public Health Laboratory | Jade Wang, et al. |
| EPI_ISL_857136, EPI_ISL_857137, EPI_ISL_857138, EPI_ISL_857139 | DOHMH Chelsea | New York City Public Health Laboratory | Jade Wang, et al. |
| EPI_ISL_857140, EPI_ISL_857141 | DOHMH Morrisania | New York City Public Health Laboratory | Jade Wang, et al. |
| EPI_ISL_857142 | DOHMH Central Harlem | New York City Public Health Laboratory | Jade Wang, et al. |
| EPI_ISL_857143, EPI_ISL_857144 | DOHMH Riverside | New York City Public Health Laboratory | Jade Wang, et al. |
| EPI_ISL_857145, EPI_ISL_857146, EPI_ISL_857147 | DOHMH Corona | New York City Public Health Laboratory | Jade Wang, et al. |
| EPI_ISL_857148 | DOHMH Fort Greene | New York City Public Health Laboratory | Jade Wang, et al. |
| EPI_ISL_857149, EPI_ISL_857150, EPI_ISL_857151, EPI_ISL_857152 | DOHMH Jamaica | New York City Public Health Laboratory | Jade Wang, et al. |
| EPI_ISL_857153 | DOHMH Corona | New York City Public Health Laboratory | Jade Wang, et al. |
| EPI_ISL_857154, EPI_ISL_857155 | DOHMH Jamaica | New York City Public Health Laboratory | Jade Wang, et al. |
| EPI_ISL_857156 | DOHMH Corona | New York City Public Health Laboratory | Jade Wang, et al. |
| EPI_ISL_857157 | DOHMH PHL | New York City Public Health Laboratory | Jade Wang, et al. |
| EPI_ISL_857308 | South Eastern Area Laboratory Services (SEALS) | NSW Health Pathology - Institute of Clinical Pathology and Medical Research; Westmead Hospital; University of Sydney | CIDM-PH et al. |
| EPI_ISL_857390, EPI_ISL_857391, EPI_ISL_857392, EPI_ISL_857393, EPI_ISL_857394 | Texas Department of State Health Services (TXDSHS) | Texas Department of State Health Services (TXDSHS) | Anita Pokharel, Bonnie Oh, James Daniel Bonser, Rashmi Tuladhar, Mayela Pedrueza, Jenny Zhang, Maliha Rahman, Myong Koag, Chung Wang, Rachel Lee, Grace Kubin |
| EPI_ISL_857403, EPI_ISL_857404, EPI_ISL_857405, EPI_ISL_857406, EPI_ISL_857445, EPI_ISL_857448, EPI_ISL_857449, EPI_ISL_857450, EPI_ISL_857451, EPI_ISL_857452, EPI_ISL_857453, EPI_ISL_857455, EPI_ISL_857457, EPI_ISL_857460, EPI_ISL_857461, EPI_ISL_857462, EPI_ISL_857463, EPI_ISL_857464, EPI_ISL_857465 | | | |
| see above | Maine HETL | Tewhey Lab, The Jackson Laboratory | Matluk,N., Dewey,H., Iosue,F., Barter,M., Lynch,R., Munger,H. and Tewhey,R. |
| EPI_ISL_857542, EPI_ISL_857544, EPI_ISL_857545, EPI_ISL_857547, EPI_ISL_857549, EPI_ISL_857551, EPI_ISL_857552, EPI_ISL_857555, EPI_ISL_857556, EPI_ISL_857557, EPI_ISL_857558, EPI_ISL_857560, EPI_ISL_857561, EPI_ISL_857563, EPI_ISL_857565, EPI_ISL_857567, EPI_ISL_857568, EPI_ISL_857569, EPI_ISL_857570, EPI_ISL_857573, EPI_ISL_857574, EPI_ISL_857575, EPI_ISL_857576, EPI_ISL_857577, EPI_ISL_857578, EPI_ISL_857579, EPI_ISL_857580, EPI_ISL_857582, EPI_ISL_857583, EPI_ISL_857585, EPI_ISL_857586, EPI_ISL_857588, EPI_ISL_857589, EPI_ISL_857590, EPI_ISL_857591, EPI_ISL_857592, EPI_ISL_857594, EPI_ISL_857597, EPI_ISL_857598, EPI_ISL_857601, EPI_ISL_857608, EPI_ISL_857609, EPI_ISL_857610, EPI_ISL_857612, EPI_ISL_857615, EPI_ISL_857617, EPI_ISL_857620, EPI_ISL_857623, EPI_ISL_857624, EPI_ISL_857627, EPI_ISL_857628, EPI_ISL_857629, EPI_ISL_857630, EPI_ISL_857634, EPI_ISL_857636, EPI_ISL_857637, EPI_ISL_857639, EPI_ISL_857640, EPI_ISL_857643, EPI_ISL_857644, EPI_ISL_857647, EPI_ISL_857648, EPI_ISL_857650, EPI_ISL_857651, EPI_ISL_857652, EPI_ISL_857655, EPI_ISL_857657, EPI_ISL_857658, EPI_ISL_857661, EPI_ISL_857663, EPI_ISL_857664, EPI_ISL_857665, EPI_ISL_857666, EPI_ISL_857667, EPI_ISL_857668, EPI_ISL_857670, EPI_ISL_857671, EPI_ISL_857672, EPI_ISL_857673, EPI_ISL_857674, EPI_ISL_857676, EPI_ISL_857678, EPI_ISL_857681, EPI_ISL_857683, EPI_ISL_857684, EPI_ISL_857686, EPI_ISL_857688, EPI_ISL_857689, EPI_ISL_857690, EPI_ISL_857692, EPI_ISL_857693, EPI_ISL_857694, EPI_ISL_857695, EPI_ISL_857696, EPI_ISL_857697, EPI_ISL_857698, EPI_ISL_857702, EPI_ISL_857703, EPI_ISL_857704, EPI_ISL_857706, EPI_ISL_857707, EPI_ISL_857708, EPI_ISL_857710, EPI_ISL_857711, EPI_ISL_857712, EPI_ISL_857713, EPI_ISL_857714, EPI_ISL_857716, EPI_ISL_857718, EPI_ISL_857721, EPI_ISL_857722, EPI_ISL_857724, EPI_ISL_857725, EPI_ISL_857726, EPI_ISL_857727, EPI_ISL_857729, EPI_ISL_857730, EPI_ISL_857732, EPI_ISL_857735, EPI_ISL_857737, EPI_ISL_857741, EPI_ISL_857742, EPI_ISL_857745, EPI_ISL_857746, EPI_ISL_857747, EPI_ISL_857748, EPI_ISL_857750, EPI_ISL_857753, EPI_ISL_857755, EPI_ISL_857757, EPI_ISL_857758, EPI_ISL_857760, EPI_ISL_857761, EPI_ISL_857762, EPI_ISL_857764, EPI_ISL_857765, EPI_ISL_857766, EPI_ISL_857767, EPI_ISL_857768, EPI_ISL_857770, EPI_ISL_857771, EPI_ISL_857772, EPI_ISL_857774, EPI_ISL_857775, EPI_ISL_857777, EPI_ISL_857779, EPI_ISL_857784, EPI_ISL_857785, EPI_ISL_857788, EPI_ISL_857789, EPI_ISL_857791, EPI_ISL_857793, EPI_ISL_857796, EPI_ISL_857797, EPI_ISL_857800, EPI_ISL_857801, EPI_ISL_857804, EPI_ISL_857805, EPI_ISL_857806, EPI_ISL_857807, EPI_ISL_857808, EPI_ISL_857809, EPI_ISL_857810, EPI_ISL_857812, EPI_ISL_857813, EPI_ISL_857814, EPI_ISL_857815, EPI_ISL_857816, EPI_ISL_857817, EPI_ISL_857822, EPI_ISL_857824, EPI_ISL_857826, EPI_ISL_857827, EPI_ISL_857828, EPI_ISL_857830, EPI_ISL_857831, EPI_ISL_857832, EPI_ISL_857833, EPI_ISL_857834, EPI_ISL_857836, EPI_ISL_857837, EPI_ISL_857838, EPI_ISL_857841, EPI_ISL_857842, EPI_ISL_857843, EPI_ISL_857844, EPI_ISL_857848, EPI_ISL_857849, EPI_ISL_857850, EPI_ISL_857851, EPI_ISL_857854, EPI_ISL_857856, EPI_ISL_857858, EPI_ISL_857859, EPI_ISL_857860, EPI_ISL_857861, EPI_ISL_857862, EPI_ISL_857864, EPI_ISL_857865, EPI_ISL_857867, EPI_ISL_857868, EPI_ISL_857869 | | | |
| see above | Lighthouse Lab in Cambridge | Wellcome Sanger Institute for the COVID-19 Genomics UK (COG-UK) Consortium | Rob Howes, The Lighthouse Lab in Cambridge and Alex Alderton, Roberto Amato, Sonia Goncalves, Ewan Harrison, David K. Jackson, Ian Johnston, Dominic Kwiatkowski, Cordelia Langford, John Sillitoe on behalf of the Wellcome Sanger Institute COVID-19 Surveillance Team |
| EPI_ISL_857870, EPI_ISL_857871, EPI_ISL_857872, EPI_ISL_857873 | Lighthouse Lab in Glasgow | Wellcome Sanger Institute for the COVID-19 Genomics UK (COG-UK) Consortium | Harper VanSteenhouse, Yumi Kasai, David Gray, Carol Clugston, Anna Dominiczak and Alex Alderton, Roberto Amato, Sonia Goncalves, Ewan Harrison, David K. Jackson, Ian Johnston, Dominic Kwiatkowski, Cordelia Langford, John Sillitoe on behalf of the Wellcome Sanger Institute COVID-19 Surveillance Team |
| EPI_ISL_857874 | Lighthouse Lab in Alderley Park | Wellcome Sanger Institute for the COVID-19 Genomics UK (COG-UK) Consortium | Jacquelyn Wynn, Mairead Hyland, The Lighthouse Lab in Alderley Park and Alex Alderton, Roberto Amato, Sonia Goncalves, Ewan Harrison, David K. Jackson, Ian Johnston, Dominic Kwiatkowski, Cordelia Langford, John Sillitoe on behalf of the Wellcome Sanger Institute COVID-19 Surveillance Team |
| EPI_ISL_857876 | Lighthouse Lab in Glasgow | Wellcome Sanger Institute for the COVID-19 Genomics UK (COG-UK) Consortium | Harper VanSteenhouse, Yumi Kasai, David Gray, Carol Clugston, Anna Dominiczak and Alex Alderton, Roberto Amato, Sonia Goncalves, Ewan Harrison, David K. Jackson, Ian Johnston, Dominic Kwiatkowski, Cordelia Langford, John Sillitoe on behalf of the Wellcome Sanger Institute COVID-19 Surveillance Team |
| EPI_ISL_857877 | Lighthouse Lab in Alderley Park | Wellcome Sanger Institute for the COVID-19 Genomics UK (COG-UK) Consortium | Jacquelyn Wynn, Mairead Hyland, The Lighthouse Lab in Alderley Park and Alex Alderton, Roberto Amato, Sonia Goncalves, Ewan Harrison, David K. Jackson, Ian Johnston, Dominic Kwiatkowski, Cordelia Langford, John Sillitoe on behalf of the Wellcome Sanger Institute COVID-19 Surveillance Team |
| EPI_ISL_857878, EPI_ISL_857881 | Lighthouse Lab in Glasgow | Wellcome Sanger Institute for the COVID-19 Genomics UK (COG-UK) Consortium | Harper VanSteenhouse, Yumi Kasai, David Gray, Carol Clugston, Anna Dominiczak and Alex Alderton, Roberto Amato, Sonia Goncalves, Ewan Harrison, David K. Jackson, Ian Johnston, Dominic Kwiatkowski, Cordelia Langford, John Sillitoe on behalf of the Wellcome Sanger Institute COVID-19 Surveillance Team |
| EPI_ISL_857883, EPI_ISL_857884 | Lighthouse Lab in Alderley Park | Wellcome Sanger Institute for the COVID-19 Genomics UK (COG-UK) Consortium | Jacquelyn Wynn, Mairead Hyland, The Lighthouse Lab in Alderley Park and Alex Alderton, Roberto Amato, Sonia Goncalves, Ewan Harrison, David K. Jackson, Ian Johnston, Dominic Kwiatkowski, Cordelia Langford, John Sillitoe on behalf of the Wellcome Sanger Institute COVID-19 Surveillance Team |
| EPI_ISL_857885 | Lighthouse Lab in Glasgow | Wellcome Sanger Institute for the COVID-19 Genomics UK (COG-UK) Consortium | Harper VanSteenhouse, Yumi Kasai, David Gray, Carol Clugston, Anna Dominiczak and Alex Alderton, Roberto Amato, Sonia Goncalves, Ewan Harrison, David K. Jackson, Ian Johnston, Dominic Kwiatkowski, Cordelia Langford, John Sillitoe on behalf of the Wellcome Sanger Institute COVID-19 Surveillance Team |
| EPI_ISL_857886 | Lighthouse Lab in Alderley Park | Wellcome Sanger Institute for the COVID-19 Genomics UK (COG-UK) Consortium | Jacquelyn Wynn, Mairead Hyland, The Lighthouse Lab in Alderley Park and Alex Alderton, Roberto Amato, Sonia Goncalves, Ewan Harrison, David K. Jackson, Ian Johnston, Dominic Kwiatkowski, Cordelia Langford, John Sillitoe on behalf of the Wellcome Sanger Institute COVID-19 Surveillance Team |
| EPI_ISL_857887, EPI_ISL_857888, EPI_ISL_857889, EPI_ISL_857894, EPI_ISL_857911, EPI_ISL_857912, EPI_ISL_857913, EPI_ISL_857914, EPI_ISL_857915, EPI_ISL_857916, EPI_ISL_857917, EPI_ISL_857918, EPI_ISL_857919, EPI_ISL_857920, EPI_ISL_857921, EPI_ISL_857922, EPI_ISL_857923, EPI_ISL_857924, EPI_ISL_857925, EPI_ISL_857926, EPI_ISL_857927, EPI_ISL_857928, EPI_ISL_857929, EPI_ISL_857930, EPI_ISL_857931, EPI_ISL_857932, EPI_ISL_857933, EPI_ISL_857934, EPI_ISL_857935, EPI_ISL_857936, EPI_ISL_857937, EPI_ISL_857938, EPI_ISL_857939, EPI_ISL_857940, EPI_ISL_857941, EPI_ISL_857942, EPI_ISL_857943, EPI_ISL_857944, EPI_ISL_857945, EPI_ISL_857946, EPI_ISL_857947, EPI_ISL_857948, EPI_ISL_857949, EPI_ISL_857950, EPI_ISL_857951, EPI_ISL_857952, EPI_ISL_857953, EPI_ISL_857954, EPI_ISL_857955, EPI_ISL_857956, EPI_ISL_857957, EPI_ISL_857958, EPI_ISL_857959 | | | |
| see above | Lighthouse Lab in Glasgow | Wellcome Sanger Institute for the COVID-19 Genomics UK (COG-UK) Consortium | Harper VanSteenhouse, Yumi Kasai, David Gray, Carol Clugston, Anna Dominiczak and Alex Alderton, Roberto Amato, Sonia Goncalves, Ewan Harrison, David K. Jackson, Ian Johnston, Dominic Kwiatkowski, Cordelia Langford, John Sillitoe on behalf of the Wellcome Sanger Institute COVID-19 Surveillance Team |
| EPI_ISL_857961, EPI_ISL_857962, EPI_ISL_857963, EPI_ISL_857964, EPI_ISL_857967, EPI_ISL_857968, EPI_ISL_857969, EPI_ISL_857970, EPI_ISL_857971, EPI_ISL_857972, EPI_ISL_857973, EPI_ISL_857974, EPI_ISL_857975, EPI_ISL_857976, EPI_ISL_857977, EPI_ISL_857978, EPI_ISL_857979, EPI_ISL_857980, EPI_ISL_857981, EPI_ISL_857982, EPI_ISL_857983, EPI_ISL_857985, EPI_ISL_857986, EPI_ISL_857987, EPI_ISL_857988, EPI_ISL_857989, EPI_ISL_857990, EPI_ISL_857991, EPI_ISL_857992, EPI_ISL_857993, EPI_ISL_857994, EPI_ISL_857995, EPI_ISL_857996, EPI_ISL_857997, EPI_ISL_857998, EPI_ISL_857999, EPI_ISL_858001, EPI_ISL_858002, EPI_ISL_858003, EPI_ISL_858004, EPI_ISL_858005, EPI_ISL_858006, EPI_ISL_858007, EPI_ISL_858008, EPI_ISL_858009, EPI_ISL_858010 | | | |
| see above | Lighthouse Lab in Alderley Park | Wellcome Sanger Institute for the COVID-19 Genomics UK (COG-UK) Consortium | Jacquelyn Wynn, Mairead Hyland, The Lighthouse Lab in Alderley Park and Alex Alderton, Roberto Amato, Sonia Goncalves, Ewan Harrison, David K. Jackson, Ian Johnston, Dominic Kwiatkowski, Cordelia Langford, John Sillitoe on behalf of the Wellcome Sanger Institute COVID-19 Surveillance Team |
| EPI_ISL_858011, EPI_ISL_858022, | Lighthouse Lab in Glasgow | Wellcome Sanger Institute for the COVID-19 Genomics UK | Harper VanSteenhouse, Yumi Kasai, David Gray, Carol Clugston, Anna Dominiczak and Alex Alderton, Roberto Amato, Sonia Goncalves, Ewan Harrison, |

| | | | |
|---|---|---|---|
| EPI_ISL_858025, EPI_ISL_858035, EPI_ISL_858039, EPI_ISL_858040, EPI_ISL_858043, EPI_ISL_858054 | | (COG-UK) Consortium | David K. Jackson, Ian Johnston, Dominic Kwiatkowski, Cordelia Langford, John Sillitoe on behalf of the Wellcome Sanger Institute COVID-19 Surveillance Team |
| EPI_ISL_860260 | Akershus University Hospital, Department for Microbiology and Infectious Disease Control | Norwegian Institute of Public Health, Department of Virology | Kathrine Stene-Johansen, Kamilla Heddeland Instefjord, Hilde Elshaug, Atiya R Ali,Marie Paulsen Madsen, Rasmus Riis Kopperud, Hilde Vollan, Karoline Bragstad, Olav Hungnes |
| EPI_ISL_860263 | Oslo University Hospital, Department of Medical Microbiology | Norwegian Institute of Public Health, Department of Virology | Kathrine Stene-Johansen, Kamilla Heddeland Instefjord, Hilde Elshaug, Atiya R Ali,Marie Paulsen Madsen, Rasmus Riis Kopperud, Hilde Vollan, Karoline Bragstad, Olav Hungnes |
| EPI_ISL_860268 | Vestfold Hospital, Toensberg Department of Microbiology | Norwegian Institute of Public Health, Department of Virology | Kathrine Stene-Johansen, Kamilla Heddeland Instefjord, Hilde Elshaug, Atiya R Ali,Marie Paulsen Madsen, Rasmus Riis Kopperud, Hilde Vollan, Karoline Bragstad, Olav Hungnes |
| EPI_ISL_860269 | Akershus University Hospital, Department for Microbiology and Infectious Disease Control | Norwegian Institute of Public Health, Department of Virology | Kathrine Stene-Johansen, Kamilla Heddeland Instefjord, Hilde Elshaug, Atiya R Ali,Marie Paulsen Madsen, Rasmus Riis Kopperud, Hilde Vollan, Karoline Bragstad, Olav Hungnes |
| EPI_ISL_860281 | Furst Medical Laboratory | Norwegian Institute of Public Health, Department of Virology | Kathrine Stene-Johansen, Kamilla Heddeland Instefjord, Hilde Elshaug, Atiya R Ali,Marie Paulsen Madsen, Rasmus Riis Kopperud, Hilde Vollan, Karoline Bragstad, Olav Hungnes |
| EPI_ISL_860688 | Respiratory Virus Unit, National Infection Service, Public Health England | COVID-19 Genomics UK (COG-UK) Consortium | PHE Covid Sequencing Team |
| EPI_ISL_860716 | Cerballiance Côte d'Azur | CERBA LAB | Roquebert B; Prots.L; Barrieu-Moussat S;Haïm-Boukobza S. |
| EPI_ISL_860724, EPI_ISL_860725 | Institute for Infectious Diseases, University of Bern, Switzerland | Institute for Infectious Diseases, University of Bern, Switzerland | Michel C Koch, Christian Baumann, Miguel A Terrazos Miani, Cora Sägesser, Pascal Bittel, Stephen L Leib, Peter Keller, Franziska Suter-Riniker, Alban Ramette |
| EPI_ISL_860787 | Charité Universitätsmedizin Berlin, Institute of Virology, Chariteplatz 1, 10117 Berlin, Germany | Charité Universitätsmedizin Berlin, Institute of Virology, Charitéplatz 1, 10117 Berlin, Germany | Victor M Corman, Julia Schneider, Jörn Beheim-Schwarzbach, Tobias Bleicker, Julia Tesch, Barbara Mühlemann, Talitha Veith, Terry Jones, Christian Drosten |
| EPI_ISL_860820, EPI_ISL_860823, EPI_ISL_860825, EPI_ISL_860827 | CHU Tours - Virologie | National Reference Center for Viruses of Respiratory Infections, Institut Pasteur, Paris | Marion Barbet, Sylvie Behillil, Méline Bizard, Angela Brisebarre, Camille Capel, Etienne Simon-Lorière, Vincent Enouf, Maud Vanpeene, Sylvie van der Werf,Gaudy Graffin Catherine |
| EPI_ISL_860829, EPI_ISL_860830 | Labo Analyses Med | National Reference Center for Viruses of Respiratory Infections, Institut Pasteur, Paris | Marion Barbet, Sylvie Behillil, Méline Bizard, Angela Brisebarre, Camille Capel, Etienne Simon-Lorière, Vincent Enouf, Maud Vanpeene, Sylvie van der Werf,Amzalag Jonas |
| EPI_ISL_860832 | CHU Tours - Virologie | National Reference Center for Viruses of Respiratory Infections, Institut Pasteur, Paris | Marion Barbet, Sylvie Behillil, Méline Bizard, Angela Brisebarre, Camille Capel, Etienne Simon-Lorière, Vincent Enouf, Maud Vanpeene, Sylvie van der Werf,Gaudy Graffin Catherine |
| EPI_ISL_860834, EPI_ISL_860835, EPI_ISL_860837, EPI_ISL_860839 | Labo Analyses Med | National Reference Center for Viruses of Respiratory Infections, Institut Pasteur, Paris | Marion Barbet, Sylvie Behillil, Méline Bizard, Angela Brisebarre, Camille Capel, Etienne Simon-Lorière, Vincent Enouf, Maud Vanpeene, Sylvie van der Werf,Amzalag Jonas |
| EPI_ISL_860845 | Labo Analyses med | National Reference Center for Viruses of Respiratory Infections, Institut Pasteur, Paris | Marion Barbet, Sylvie Behillil, Méline Bizard, Angela Brisebarre, Camille Capel, Etienne Simon-Lorière, Vincent Enouf, Maud Vanpeene, Sylvie van der Werf,Amzalag Jonas |
| EPI_ISL_860847 | Centre Hospitalier Compiègne-Noyon - Laboratoire | National Reference Center for Viruses of Respiratory Infections, Institut Pasteur, Paris | Marion Barbet, Sylvie Behillil, Méline Bizard, Angela Brisebarre, Camille Capel, Etienne Simon-Lorière, Vincent Enouf, Maud Vanpeene, Sylvie van der Werf,Foissaud Vincent |
| EPI_ISL_860848 | CH Dunkerque Laboratoire de Biologie | National Reference Center for Viruses of Respiratory Infections, Institut Pasteur, Paris | Marion Barbet, Sylvie Behillil, Méline Bizard, Angela Brisebarre, Camille Capel, Etienne Simon-Lorière, Vincent Enouf, Maud Vanpeene, Sylvie van der Werf,Joly Isabelle |
| EPI_ISL_860852, EPI_ISL_860853, EPI_ISL_860854, EPI_ISL_860855, EPI_ISL_860857, EPI_ISL_860858, EPI_ISL_860859 | Labo Analyses Med | National Reference Center for Viruses of Respiratory Infections, Institut Pasteur, Paris | Marion Barbet, Sylvie Behillil, Méline Bizard, Angela Brisebarre, Camille Capel, Etienne Simon-Lorière, Vincent Enouf, Maud Vanpeene, Sylvie van der Werf,Amzalag Jonas |
| EPI_ISL_860890 | CHU de Nantes - Hôtel Dieu - Labo. Virologie | National Reference Center for Viruses of Respiratory Infections, Institut Pasteur, Paris | Marion Barbet, Sylvie Behillil, Méline Bizard, Angela Brisebarre, Camille Capel, Etienne Simon-Lorière, Vincent Enouf, Maud Vanpeene, Sylvie van der Werf,Andre-Garnier Elisabeth |
| EPI_ISL_860892, EPI_ISL_860895, EPI_ISL_860897 | CHU Tours - Virologie | National Reference Center for Viruses of Respiratory Infections, Institut Pasteur, Paris | Marion Barbet, Sylvie Behillil, Méline Bizard, Angela Brisebarre, Camille Capel, Etienne Simon-Lorière, Vincent Enouf, Maud Vanpeene, Sylvie van der Werf,Gaudy Graffin Catherine |
| EPI_ISL_860900, EPI_ISL_860901, EPI_ISL_860902, EPI_ISL_860903, EPI_ISL_860904 | CH du MANS - Lab. Bio. Moléculaire | National Reference Center for Viruses of Respiratory Infections, Institut Pasteur, Paris | Marion Barbet, Sylvie Behillil, Méline Bizard, Angela Brisebarre, Camille Capel, Etienne Simon-Lorière, Vincent Enouf, Maud Vanpeene, Sylvie van der Werf,Ramanantsoa CéLine |
| EPI_ISL_860906 | CHU Angers - Dpt des Agents Infectieux | National Reference Center for Viruses of Respiratory Infections, Institut Pasteur, Paris | Marion Barbet, Sylvie Behillil, Méline Bizard, Angela Brisebarre, Camille Capel, Etienne Simon-Lorière, Vincent Enouf, Maud Vanpeene, Sylvie van der Werf,Ducancelle Alexandra |
| EPI_ISL_860908, EPI_ISL_860909 | CHU Tours - Virologie | National Reference Center for Viruses of Respiratory Infections, Institut Pasteur, Paris | Marion Barbet, Sylvie Behillil, Méline Bizard, Angela Brisebarre, Camille Capel, Etienne Simon-Lorière, Vincent Enouf, Maud Vanpeene, Sylvie van der Werf,Gaudy Graffin Catherine |
| EPI_ISL_860910 | CH du MANS - Lab. Bio. Moléculaire | National Reference Center for Viruses of Respiratory Infections, Institut Pasteur, Paris | Marion Barbet, Sylvie Behillil, Méline Bizard, Angela Brisebarre, Camille Capel, Etienne Simon-Lorière, Vincent Enouf, Maud Vanpeene, Sylvie van der Werf,Ramanantsoa CéLine |
| EPI_ISL_860914, EPI_ISL_860915, EPI_ISL_860916, EPI_ISL_860917, EPI_ISL_860922, EPI_ISL_860923, EPI_ISL_860924, EPI_ISL_860925 | CHU de Nantes - Hôtel Dieu - Labo. Virologie | National Reference Center for Viruses of Respiratory Infections, Institut Pasteur, Paris | Marion Barbet, Sylvie Behillil, Méline Bizard, Angela Brisebarre, Camille Capel, Etienne Simon-Lorière, Vincent Enouf, Maud Vanpeene, Sylvie van der Werf,Andre-Garnier Elisabeth |
| EPI_ISL_860931 | Labo Analyses Med | National Reference Center for Viruses of Respiratory Infections, Institut Pasteur, Paris | Marion Barbet, Sylvie Behillil, Méline Bizard, Angela Brisebarre, Camille Capel, Etienne Simon-Lorière, Vincent Enouf, Maud Vanpeene, Sylvie van der Werf |
| EPI_ISL_860956, EPI_ISL_860970, EPI_ISL_860972, EPI_ISL_860973, EPI_ISL_860974, EPI_ISL_860978, EPI_ISL_860980, EPI_ISL_860982, EPI_ISL_860985, EPI_ISL_860994, EPI_ISL_860997, EPI_ISL_861000, EPI_ISL_861017, EPI_ISL_861019, EPI_ISL_861020, EPI_ISL_861022, EPI_ISL_861024, EPI_ISL_861029, EPI_ISL_861032, EPI_ISL_861033, EPI_ISL_861039, EPI_ISL_861040, EPI_ISL_861045, EPI_ISL_861051, EPI_ISL_861058, EPI_ISL_861064, EPI_ISL_861065, EPI_ISL_861078, EPI_ISL_861081, EPI_ISL_861083, EPI_ISL_861086, EPI_ISL_861088, EPI_ISL_861100 | | | |
| see above | Johns Hopkins Hospital Department of Pathology | Johns Hopkins Hospital Department of Pathology | C. Paul Morris, Chun Huai Luo, Adannaya Amadi, Nicholas Gallagher, Heba H. Mostafa |
| EPI_ISL_861205 | URMC LABS | Wadsworth Center, New York State Department of Health | Kirsten St. George, Daryl M. Lamson, Alexis Russel, Matthew Shudt, Melissa A Leisner, Jonathan Plitnick, Navjot Singh, John Kelly, Erasmus Schneider, Erica Lasek-Nesselquist |
| EPI_ISL_861234, EPI_ISL_861237 | Labo Analyses Med | National Reference Center for Viruses of Respiratory Infections, Institut Pasteur, Paris | Marion Barbet, Sylvie Behillil, Méline Bizard, Angela Brisebarre, Camille Capel, Etienne Simon-Lorière, Vincent Enouf, Maud Vanpeene, Sylvie van der Werf |
| EPI_ISL_861238 | CHU Tours - Virologie | National Reference Center for Viruses of Respiratory Infections, Institut Pasteur, Paris | Marion Barbet, Sylvie Behillil, Méline Bizard, Angela Brisebarre, Camille Capel, Etienne Simon-Lorière, Vincent Enouf, Maud Vanpeene, Sylvie van der Werf,Gaudy Graffin Catherine |
| EPI_ISL_861334, EPI_ISL_861335, EPI_ISL_861336, EPI_ISL_861338, EPI_ISL_861339, EPI_ISL_861341, EPI_ISL_861342, EPI_ISL_861343, EPI_ISL_861344, EPI_ISL_861345, EPI_ISL_861346, EPI_ISL_861347, EPI_ISL_861349, EPI_ISL_861350, EPI_ISL_861351, EPI_ISL_861352, EPI_ISL_861353, EPI_ISL_861354, EPI_ISL_861357, EPI_ISL_861358, EPI_ISL_861359 | | | |
| see above | WESTCHESTER MEDICAL CENTER | Wadsworth Center, New York State Department of Health | Kirsten St. George, Daryl M. Lamson, Alexis Russel, Matthew Shudt, Melissa A Leisner, Jonathan Plitnick, Navjot Singh, John Kelly, Erasmus Schneider, Erica Lasek-Nesselquist |
| EPI_ISL_861422, EPI_ISL_861423, | Oklahoma Animal Disease Diagnostic Laboratory | Oklahoma Animal Disease Diagnostic Laboratory | Sai Narayanan, Girish Patil, Sunil More, Jeremiah Saliki, Akhilesh Ramachandran |

| | | | |
|---|---|---|---|
| EPI_ISL_861424, EPI_ISL_861425, EPI_ISL_861426 | | | |
| EPI_ISL_861445 | Clinical Molecular Microbiology Laboratory, UNC Hospitals | Jeremy Wang | Jeremy Wang, Alexander Rubinsteyn, Colleen Rice, Jason Smedberg, Melissa Miller, Corbin Jones, Robert Hagan |
| EPI_ISL_861474, EPI_ISL_861475 | Gundersen Molecular Diagnostics Laboratory | Kabara Cancer Research Institute | Craig S. Richmond, Paraic A. Kenny |
| EPI_ISL_861480 | Wyoming Public Health Laboratory | Wyoming Public Health Laboratory | Noah Hull, Taylor Fearing, Lynette Gumbleton, Channing Weber, Ashley Norberg, Bailey Bowcutt, and Wanda Manley |
| EPI_ISL_861504 | Cerballiance Provence | CERBA LAB | Delaunay E; Roig JC; Prots L. |
| EPI_ISL_861529, EPI_ISL_861553, EPI_ISL_861554, EPI_ISL_861555, EPI_ISL_861560, EPI_ISL_861561, EPI_ISL_861562, EPI_ISL_861563, EPI_ISL_861564, EPI_ISL_861576, EPI_ISL_861579, EPI_ISL_861624 | | | |
| see above | Instituto Nacional de Saude (INSA) | Instituto Nacional de Saude (INSA) | Borges et al |
| EPI_ISL_861716, EPI_ISL_861728 | CERBA | CERBA LAB | Roquebert B; Costa JM; Hedbaut E; Trombert S; Lecorche E; Verdurme L; Malek Ramdane; Olivi M; Haïm-Boukobza S. |
| EPI_ISL_861732, EPI_ISL_861733, EPI_ISL_861748, EPI_ISL_861755 | Tempus | Grubaugh Lab - Yale School of Public Health | Tara Alpert, Joseph Fauver, Anderson Brito, Mallery Breban, Anne Wyllie, Chantal Vogels, Mary Petrone, Annie Watkins, Chaney Kalinich, Isabel Ott, Nathan Grubaugh |
| EPI_ISL_861758 | Murphy Medical Associates | Grubaugh Lab - Yale School of Public Health | Tara Alpert, Joseph Fauver, Anderson Brito, Mallery Breban, Anne Wyllie, Chantal Vogels, Mary Petrone, Annie Watkins, Chaney Kalinich, Isabel Ott, Nathan Grubaugh |
| EPI_ISL_861761, EPI_ISL_861766 | Tempus | Grubaugh Lab - Yale School of Public Health | Tara Alpert, Joseph Fauver, Anderson Brito, Mallery Breban, Anne Wyllie, Chantal Vogels, Mary Petrone, Annie Watkins, Chaney Kalinich, Isabel Ott, Nathan Grubaugh |
| EPI_ISL_862042 | CERBA | CERBA LAB | Roquebert B; Costa JM; Hedbaut E; Trombert S; Lecorche E; Verdurme L; Malek Ramdane, Olivi M; Haïm-Boukobza S. |
| EPI_ISL_862126, EPI_ISL_862143, EPI_ISL_862149, EPI_ISL_862175, EPI_ISL_862186 | Charité Universitätsmedizin Berlin, Institut fur Virologie/Labor Berlin | Charité Universitätsmedizin Berlin, Institut für Virologie | Victor M Corman, Barbara Mühlemann, Jörn Beheim-Schwarzbach, Tobias Bleicker, Julia Tesch, Talitha Veith, Julia Schneider, Terry Jones, Christian Drosten |
| EPI_ISL_862669, EPI_ISL_862710, EPI_ISL_862711, EPI_ISL_862712 | New Mexico Department of Health Scientific Laboratory | New Mexico Department of Health Scientific Laboratory | D'eldra Malone, Ellie Johnson, Anastacia Griego-Fisher |
| EPI_ISL_862722 | San Diego County Public Health Laboratory | Andersen lab at Scripps Research | SEARCH Alliance San Diego with Tracy Basler, Jovan Shephard, Brett Austin |
| EPI_ISL_862729, EPI_ISL_862732, EPI_ISL_862734 | Utah Public Health Laboratory, Utah Public Health Laboratory Infectious Disease submission group | Utah Public Health Laboratory, Utah Public Health Laboratory Infectious Disease submission group | Young,E.L., Oakeson,K.F., Gallagher,T. |
| EPI_ISL_862886, EPI_ISL_862887, EPI_ISL_862888, EPI_ISL_862900, EPI_ISL_862915, EPI_ISL_862938, EPI_ISL_862948, EPI_ISL_862983, EPI_ISL_862990, EPI_ISL_863013, EPI_ISL_863023, EPI_ISL_863046, EPI_ISL_863058, EPI_ISL_863061, EPI_ISL_863074, EPI_ISL_863084, EPI_ISL_863089, EPI_ISL_863091, EPI_ISL_863101, EPI_ISL_863103, EPI_ISL_863106, EPI_ISL_863112, EPI_ISL_863122, EPI_ISL_863126, EPI_ISL_863136, EPI_ISL_863137, EPI_ISL_863138, EPI_ISL_863140, EPI_ISL_863151, EPI_ISL_863156, EPI_ISL_863162, EPI_ISL_863179, EPI_ISL_863180, EPI_ISL_863188, EPI_ISL_863192, EPI_ISL_863193, EPI_ISL_863210, EPI_ISL_863218, EPI_ISL_863238 | | | |
| see above | Lighthouse Lab in Alderley Park | Wellcome Sanger Institute for the COVID-19 Genomics UK (COG-UK) Consortium | Jacquelyn Wynn, Mairead Hyland, The Lighthouse Lab in Alderley Park and Alex Alderton, Roberto Amato, Sonia Goncalves, Ewan Harrison, David K. Jackson, Ian Johnston, Dominic Kwiatkowski, Cordelia Langford, John Sillitoe on behalf of the Wellcome Sanger Institute COVID-19 Surveillance Team |
| EPI_ISL_863537, EPI_ISL_863543, EPI_ISL_863547, EPI_ISL_863553, EPI_ISL_863561, EPI_ISL_863562, EPI_ISL_863567, EPI_ISL_863577, EPI_ISL_863580, EPI_ISL_863587, EPI_ISL_863588, EPI_ISL_863589, EPI_ISL_863599, EPI_ISL_863601, EPI_ISL_863605, EPI_ISL_863606, EPI_ISL_863622, EPI_ISL_863623, EPI_ISL_863628, EPI_ISL_863633, EPI_ISL_863647, EPI_ISL_863649, EPI_ISL_863650, EPI_ISL_863655, EPI_ISL_863663, EPI_ISL_863666, EPI_ISL_863670, EPI_ISL_863683, EPI_ISL_863684, EPI_ISL_863686, EPI_ISL_863693, EPI_ISL_863703, EPI_ISL_863714, EPI_ISL_863716, EPI_ISL_863717, EPI_ISL_863718, EPI_ISL_863719, EPI_ISL_863724, EPI_ISL_863732, EPI_ISL_863735, EPI_ISL_863737, EPI_ISL_863739, EPI_ISL_863745, EPI_ISL_863753, EPI_ISL_863768, EPI_ISL_863775, EPI_ISL_863783, EPI_ISL_863805, EPI_ISL_863807, EPI_ISL_863808, EPI_ISL_863814, EPI_ISL_863815, EPI_ISL_863817, EPI_ISL_863825, EPI_ISL_863829, EPI_ISL_863833, EPI_ISL_863834, EPI_ISL_863835, EPI_ISL_863855, EPI_ISL_863857, EPI_ISL_863859, EPI_ISL_863867, EPI_ISL_863899, EPI_ISL_863901, EPI_ISL_863912, EPI_ISL_863913, EPI_ISL_863921, EPI_ISL_863922, EPI_ISL_863956, EPI_ISL_863969, EPI_ISL_863996, EPI_ISL_864005, EPI_ISL_864009, EPI_ISL_864026, EPI_ISL_864036, EPI_ISL_864043, EPI_ISL_864098 | | | |
| see above | Lighthouse Lab in Milton Keynes | Wellcome Sanger Institute for the COVID-19 Genomics UK (COG-UK) Consortium | The Lighthouse Lab in Milton Keynes and Alex Alderton, Roberto Amato, Sonia Goncalves, Ewan Harrison, David K. Jackson, Ian Johnston, Dominic Kwiatkowski, Cordelia Langford, John Sillitoe on behalf of the Wellcome Sanger Institute COVID-19 Surveillance Team |
| EPI_ISL_864110 | Lighthouse Lab in Alderley Park | Wellcome Sanger Institute for the COVID-19 Genomics UK (COG-UK) Consortium | Jacquelyn Wynn, Mairead Hyland, The Lighthouse Lab in Alderley Park and Alex Alderton, Roberto Amato, Sonia Goncalves, Ewan Harrison, David K. Jackson, Ian Johnston, Dominic Kwiatkowski, Cordelia Langford, John Sillitoe on behalf of the Wellcome Sanger Institute COVID-19 Surveillance Team |
| EPI_ISL_864124, EPI_ISL_864133, EPI_ISL_864165, EPI_ISL_864169, EPI_ISL_864173, EPI_ISL_864175, EPI_ISL_864176, EPI_ISL_864191, EPI_ISL_864193, EPI_ISL_864203, EPI_ISL_864205, EPI_ISL_864206, EPI_ISL_864207, EPI_ISL_864208, EPI_ISL_864209, EPI_ISL_864210, EPI_ISL_864212, EPI_ISL_864214, EPI_ISL_864215, EPI_ISL_864218, EPI_ISL_864219, EPI_ISL_864220, EPI_ISL_864222, EPI_ISL_864223, EPI_ISL_864224, EPI_ISL_864225 | | | |
| see above | Lighthouse Lab in Milton Keynes | Wellcome Sanger Institute for the COVID-19 Genomics UK (COG-UK) Consortium | The Lighthouse Lab in Milton Keynes and Alex Alderton, Roberto Amato, Sonia Goncalves, Ewan Harrison, David K. Jackson, Ian Johnston, Dominic Kwiatkowski, Cordelia Langford, John Sillitoe on behalf of the Wellcome Sanger Institute COVID-19 Surveillance Team |
| EPI_ISL_864226 | Lighthouse Lab in Alderley Park | Wellcome Sanger Institute for the COVID-19 Genomics UK (COG-UK) Consortium | Jacquelyn Wynn, Mairead Hyland, The Lighthouse Lab in Alderley Park and Alex Alderton, Roberto Amato, Sonia Goncalves, Ewan Harrison, David K. Jackson, Ian Johnston, Dominic Kwiatkowski, Cordelia Langford, John Sillitoe on behalf of the Wellcome Sanger Institute COVID-19 Surveillance Team |
| EPI_ISL_864229, EPI_ISL_864231, EPI_ISL_864232, EPI_ISL_864233, EPI_ISL_864234, EPI_ISL_864235, EPI_ISL_864236, EPI_ISL_864237, EPI_ISL_864238, EPI_ISL_864240, EPI_ISL_864241, EPI_ISL_864242, EPI_ISL_864243, EPI_ISL_864245, EPI_ISL_864246, EPI_ISL_864247, EPI_ISL_864248, EPI_ISL_864249, EPI_ISL_864250, EPI_ISL_864251, EPI_ISL_864252, EPI_ISL_864253, EPI_ISL_864254, EPI_ISL_864256, EPI_ISL_864257, EPI_ISL_864258, EPI_ISL_864259, EPI_ISL_864260, EPI_ISL_864261, EPI_ISL_864263, EPI_ISL_864264, EPI_ISL_864265 | | | |
| see above | Lighthouse Lab in Milton Keynes | Wellcome Sanger Institute for the COVID-19 Genomics UK (COG-UK) Consortium | The Lighthouse Lab in Milton Keynes and Alex Alderton, Roberto Amato, Sonia Goncalves, Ewan Harrison, David K. Jackson, Ian Johnston, Dominic Kwiatkowski, Cordelia Langford, John Sillitoe on behalf of the Wellcome Sanger Institute COVID-19 Surveillance Team |
| EPI_ISL_864267 | Lighthouse Lab in Alderley Park | Wellcome Sanger Institute for the COVID-19 Genomics UK (COG-UK) Consortium | Jacquelyn Wynn, Mairead Hyland, The Lighthouse Lab in Alderley Park and Alex Alderton, Roberto Amato, Sonia Goncalves, Ewan Harrison, David K. Jackson, Ian Johnston, Dominic Kwiatkowski, Cordelia Langford, John Sillitoe on behalf of the Wellcome Sanger Institute COVID-19 Surveillance Team |
| EPI_ISL_864268, EPI_ISL_864269, EPI_ISL_864270, EPI_ISL_864271, EPI_ISL_864272, EPI_ISL_864273, EPI_ISL_864275, EPI_ISL_864277, EPI_ISL_864278, EPI_ISL_864279, EPI_ISL_864280, EPI_ISL_864282, EPI_ISL_864283, EPI_ISL_864284, EPI_ISL_864285, EPI_ISL_864286, EPI_ISL_864287, EPI_ISL_864289, EPI_ISL_864290, EPI_ISL_864292, EPI_ISL_864293, EPI_ISL_864294, EPI_ISL_864295, EPI_ISL_864296, EPI_ISL_864298, EPI_ISL_864299, EPI_ISL_864300, EPI_ISL_864302, EPI_ISL_864303, EPI_ISL_864304, EPI_ISL_864306, EPI_ISL_864307 | | | |
| see above | Lighthouse Lab in Milton Keynes | Wellcome Sanger Institute for the COVID-19 Genomics UK (COG-UK) Consortium | The Lighthouse Lab in Milton Keynes and Alex Alderton, Roberto Amato, Sonia Goncalves, Ewan Harrison, David K. Jackson, Ian Johnston, Dominic Kwiatkowski, Cordelia Langford, John Sillitoe on behalf of the Wellcome Sanger Institute COVID-19 Surveillance Team |
| EPI_ISL_864308 | Lighthouse Lab in Alderley Park | Wellcome Sanger Institute for the COVID-19 Genomics UK (COG-UK) Consortium | Jacquelyn Wynn, Mairead Hyland, The Lighthouse Lab in Alderley Park and Alex Alderton, Roberto Amato, Sonia Goncalves, Ewan Harrison, David K. Jackson, Ian Johnston, Dominic Kwiatkowski, Cordelia Langford, John Sillitoe on behalf of the Wellcome Sanger Institute COVID-19 Surveillance Team |
| EPI_ISL_864310, EPI_ISL_864312, EPI_ISL_864314, EPI_ISL_864315, EPI_ISL_864316, EPI_ISL_864318, EPI_ISL_864319, EPI_ISL_864320, EPI_ISL_864322, EPI_ISL_864323, EPI_ISL_864325, EPI_ISL_864326, EPI_ISL_864327, EPI_ISL_864329, EPI_ISL_864330, EPI_ISL_864331, EPI_ISL_864332, EPI_ISL_864333, EPI_ISL_864334, EPI_ISL_864335, EPI_ISL_864337, EPI_ISL_864338, EPI_ISL_864339, EPI_ISL_864340, EPI_ISL_864341, EPI_ISL_864342, EPI_ISL_864343, EPI_ISL_864344, EPI_ISL_864345, EPI_ISL_864346, EPI_ISL_864347, EPI_ISL_864348, EPI_ISL_864349, EPI_ISL_864350, EPI_ISL_864351, EPI_ISL_864353, EPI_ISL_864355, EPI_ISL_864357, EPI_ISL_864358, EPI_ISL_864359, EPI_ISL_864360, EPI_ISL_864362, EPI_ISL_864363, EPI_ISL_864364, EPI_ISL_864366, EPI_ISL_864367, EPI_ISL_864369, EPI_ISL_864371, EPI_ISL_864372, EPI_ISL_864373, EPI_ISL_864374, EPI_ISL_864375, EPI_ISL_864376, EPI_ISL_864377, EPI_ISL_864378, EPI_ISL_864379, EPI_ISL_864380, EPI_ISL_864381, EPI_ISL_864382, EPI_ISL_864385, EPI_ISL_864386, EPI_ISL_864387, EPI_ISL_864388, EPI_ISL_864389, EPI_ISL_864391, EPI_ISL_864392, EPI_ISL_864393, EPI_ISL_864394, EPI_ISL_864395, EPI_ISL_864397, EPI_ISL_864398, EPI_ISL_864401, EPI_ISL_864404, EPI_ISL_864405, EPI_ISL_864406, EPI_ISL_864407, EPI_ISL_864408, EPI_ISL_864410, EPI_ISL_864412, EPI_ISL_864413, EPI_ISL_864415, EPI_ISL_864416, EPI_ISL_864417, EPI_ISL_864418, EPI_ISL_864419, EPI_ISL_864420, EPI_ISL_864421, EPI_ISL_864423, EPI_ISL_864424, EPI_ISL_864425, EPI_ISL_864426, EPI_ISL_864427, EPI_ISL_864428, EPI_ISL_864429, EPI_ISL_864430, EPI_ISL_864431, EPI_ISL_864432, EPI_ISL_864433, EPI_ISL_864434, EPI_ISL_864435, EPI_ISL_864437, EPI_ISL_864438, EPI_ISL_864441, EPI_ISL_864442, EPI_ISL_864444, EPI_ISL_864446, EPI_ISL_864448, EPI_ISL_864449, EPI_ISL_864450, EPI_ISL_864451, EPI_ISL_864453, EPI_ISL_864454, EPI_ISL_864455, EPI_ISL_864456, EPI_ISL_864457, EPI_ISL_864458, EPI_ISL_864459, EPI_ISL_864460, EPI_ISL_864462, EPI_ISL_864463, EPI_ISL_864464, EPI_ISL_864466, EPI_ISL_864468, EPI_ISL_864470, EPI_ISL_864471, EPI_ISL_864473, EPI_ISL_864474, EPI_ISL_864476, EPI_ISL_864477, EPI_ISL_864479, EPI_ISL_864480, EPI_ISL_864481, EPI_ISL_864482, EPI_ISL_864486, EPI_ISL_864488, EPI_ISL_864489, EPI_ISL_864490, EPI_ISL_864491, EPI_ISL_864492, EPI_ISL_864494, EPI_ISL_864495, EPI_ISL_864496, EPI_ISL_864497, EPI_ISL_864498, EPI_ISL_864499, EPI_ISL_864500, EPI_ISL_864502, EPI_ISL_864503, EPI_ISL_864504, EPI_ISL_864505, EPI_ISL_864506, EPI_ISL_864507, EPI_ISL_864508, EPI_ISL_864509, EPI_ISL_864510, EPI_ISL_864511, EPI_ISL_864512, EPI_ISL_864514, EPI_ISL_864517, EPI_ISL_864518, EPI_ISL_864519, EPI_ISL_864521, EPI_ISL_864522, EPI_ISL_864523, EPI_ISL_864524, EPI_ISL_864525, EPI_ISL_864526, EPI_ISL_864530, EPI_ISL_864531, EPI_ISL_864532, EPI_ISL_864533, EPI_ISL_864535, EPI_ISL_864536, EPI_ISL_864537, EPI_ISL_864538, EPI_ISL_864541 | | | |
| see above | Lighthouse Lab in Milton Keynes | Wellcome Sanger Institute for the COVID-19 Genomics UK (COG-UK) Consortium | The Lighthouse Lab in Milton Keynes and Alex Alderton, Roberto Amato, Sonia Goncalves, Ewan Harrison, David K. Jackson, Ian Johnston, Dominic Kwiatkowski, Cordelia Langford, John Sillitoe on behalf of the Wellcome Sanger Institute COVID-19 Surveillance Team |
| EPI_ISL_864594, EPI_ISL_864596, EPI_ISL_864597, EPI_ISL_864600, EPI_ISL_864601, EPI_ISL_864602, EPI_ISL_864603, EPI_ISL_864604, EPI_ISL_864605, EPI_ISL_864606, EPI_ISL_864613, EPI_ISL_864614, EPI_ISL_864615, EPI_ISL_864621, EPI_ISL_864622, EPI_ISL_864623, EPI_ISL_864626, EPI_ISL_864627, EPI_ISL_864631, EPI_ISL_864632, EPI_ISL_864633, EPI_ISL_864636, EPI_ISL_864637, EPI_ISL_864638, EPI_ISL_864639, EPI_ISL_864640, EPI_ISL_864641, EPI_ISL_864642, EPI_ISL_864643, EPI_ISL_864644, EPI_ISL_864645, EPI_ISL_864646, EPI_ISL_864647, EPI_ISL_864648, EPI_ISL_864649, EPI_ISL_864650, | | | |

| | | | |
|---|---|---|---|
| EPI_ISL_864651, EPI_ISL_864652, EPI_ISL_864653, EPI_ISL_864654, EPI_ISL_864655, EPI_ISL_864656, EPI_ISL_864657, EPI_ISL_864658, EPI_ISL_864659, EPI_ISL_864660, EPI_ISL_864661, EPI_ISL_864662, EPI_ISL_864663, EPI_ISL_864664, EPI_ISL_864665, EPI_ISL_864666, EPI_ISL_864667, EPI_ISL_864668, EPI_ISL_864669, EPI_ISL_864670, EPI_ISL_864671, EPI_ISL_864672, EPI_ISL_864673, EPI_ISL_864674, EPI_ISL_864675, EPI_ISL_864676, EPI_ISL_864677, EPI_ISL_864678, EPI_ISL_864679, EPI_ISL_864680, EPI_ISL_864681, EPI_ISL_864682, EPI_ISL_864683, EPI_ISL_864684, EPI_ISL_864685, EPI_ISL_864686, EPI_ISL_864687, EPI_ISL_864688, EPI_ISL_864717, EPI_ISL_864718, EPI_ISL_864721, EPI_ISL_864722, EPI_ISL_864723, EPI_ISL_864724, EPI_ISL_864735, EPI_ISL_864736 | | | |
| see above | University Hospitals of Geneva, Laboratory of Virology | HUG, Laboratory of Virology and the Health2030 Genome Center | Samuel Cordey, Ana Rita Goncalves, Laurent Kaiser, Lorenzo Cerutti, Henri Pegeot, Melyssa Elies, Deborah Penet, Keith Harshman, Ioannis Xenarios, Emmanouil Dermitzakis |
| EPI_ISL_865019, EPI_ISL_865021, EPI_ISL_865022, EPI_ISL_865023, EPI_ISL_865024, EPI_ISL_865025, EPI_ISL_865026, EPI_ISL_865027, EPI_ISL_865028, EPI_ISL_865029, EPI_ISL_865030, EPI_ISL_865031, EPI_ISL_865032, EPI_ISL_865033, EPI_ISL_865034, EPI_ISL_865037, EPI_ISL_865038, EPI_ISL_865039, EPI_ISL_865040, EPI_ISL_865053, EPI_ISL_865056, EPI_ISL_865057, EPI_ISL_865058, EPI_ISL_865059, EPI_ISL_865060, EPI_ISL_865061, EPI_ISL_865062, EPI_ISL_865063, EPI_ISL_865064, EPI_ISL_865065 | | | |
| see above | West of Scotland Specialist Virology Centre, NHSGGC / MRC-University of Glasgow Centre for Virus Research | COVID-19 Genomics UK (COG-UK) Consortium | Ana da Silva Filipe, Natasha Johnson, Kathy Smollett, Daniel Mair, Stephen Carmichael, Alice Broos, Lily Tong, Jenna Nichols, Kyriaki Nomikou; Sarah McDonald; Richard Orton, Joseph Hughes, Sreenu Vattipally, David L Robertson; Alasdair MacLean, Rory Gunson; Sharif Shaaban, Matthew Holden; Rachel Blacow, Guy Mollett, Kathy Li, James Shepherd, Antonia Ho, Emma Thomson |
| EPI_ISL_865111, EPI_ISL_865112, EPI_ISL_865113, EPI_ISL_865114, EPI_ISL_865116, EPI_ISL_865118, EPI_ISL_865119, EPI_ISL_865120, EPI_ISL_865121, EPI_ISL_865122, EPI_ISL_865123, EPI_ISL_865124, EPI_ISL_865125, EPI_ISL_865126, EPI_ISL_865127, EPI_ISL_865128, EPI_ISL_865129, EPI_ISL_865130, EPI_ISL_865132, EPI_ISL_865133, EPI_ISL_865134, EPI_ISL_865135, EPI_ISL_865145 | | | |
| see above | Virology Department, Royal Infirmary of Edinburgh, NHS Lothian / School of Biological Sciences, University of Edinburgh / Institute of Genetics and Molecular Medicine, University of Edinburgh | COVID-19 Genomics UK (COG-UK) Consortium | McHugh M, Dewar R, Rooke S, Gallagher M, Balcaza C, O'Toole Á, Scher E, Hill V, McCrone JT, Colquhoun R, Yu X, Jackson B, Rambaut A, Williams TC, Templeton K |
| EPI_ISL_865272, EPI_ISL_865273, EPI_ISL_865274, EPI_ISL_865275, EPI_ISL_865276, EPI_ISL_865277, EPI_ISL_865280, EPI_ISL_865281, EPI_ISL_865282, EPI_ISL_865284, EPI_ISL_865285, EPI_ISL_865286, EPI_ISL_865287, EPI_ISL_865288, EPI_ISL_865289, EPI_ISL_865290, EPI_ISL_865291, EPI_ISL_865292, EPI_ISL_865293, EPI_ISL_865294, EPI_ISL_865295, EPI_ISL_865296, EPI_ISL_865297, EPI_ISL_865298, EPI_ISL_865300, EPI_ISL_865301, EPI_ISL_865302, EPI_ISL_865303, EPI_ISL_865304, EPI_ISL_865305, EPI_ISL_865306, EPI_ISL_865307, EPI_ISL_865308, EPI_ISL_865309, EPI_ISL_865310, EPI_ISL_865311, EPI_ISL_865312, EPI_ISL_865313, EPI_ISL_865314, EPI_ISL_865315, EPI_ISL_865317, EPI_ISL_865318, EPI_ISL_865319, EPI_ISL_865320, EPI_ISL_865321, EPI_ISL_865326, EPI_ISL_865327, EPI_ISL_865328, EPI_ISL_865341, EPI_ISL_865342, EPI_ISL_865346, EPI_ISL_865347, EPI_ISL_865350 | | | |
| see above | Liverpool Clinical Laboratories | COVID-19 Genomics UK (COG-UK) Consortium | Sam Haldenby, Anita Lucaci, Steve Paterson, Julian Hiscox, Alistair Darby, M Almsaud, A Alrezaihi, Muhannad Alruwaili, Stuart D Armstrong, Jones Benjamin, Eleanor G Bentley, Anu Chawla, Jordan J Clark, Angela Cowell, Richard Eccles, Isabel García-Dorival, Matthew Gemmell, Alessandro Gerada, PKF Gilmore, Richard Gregory, Ximeng Han, Catherine Hartley, Margaret Hughes, Miren Iturriza-Gomara, James Johnson, L Luu, Jenifer Manson, Charlotte Nelson, Elaine O'Toole, Cassie Olateju, Rebekah Penrice-Randal , Lucille Rainbow, N.P Randle, Trevor Ian Robinson, Parul Sharma, Ghada T Shawli, James P Stewart, Neil Swainston, Ecaterina Vamos, Joanne Watts, Mark Whitehead |
| EPI_ISL_866082, EPI_ISL_866083, EPI_ISL_866085, EPI_ISL_866086, EPI_ISL_866087, EPI_ISL_866088, EPI_ISL_866089, EPI_ISL_866090, EPI_ISL_866091, EPI_ISL_866092, EPI_ISL_866093, EPI_ISL_866094, EPI_ISL_866095, EPI_ISL_866096, EPI_ISL_866097, EPI_ISL_866098, EPI_ISL_866099, EPI_ISL_866100, EPI_ISL_866101, EPI_ISL_866102, EPI_ISL_866112, EPI_ISL_866113, EPI_ISL_866114 | | | |
| see above | University College London Hospital | COVID-19 Genomics UK (COG-UK) Consortium | Judith Heaney, Matthew Byott, Catherine Houlihan, Dan Frampton, Stuart Kirk, Moira Spyer and Eleni Nastouli |
| EPI_ISL_866194, EPI_ISL_866196, EPI_ISL_866202, EPI_ISL_866206, EPI_ISL_866207, EPI_ISL_866208, EPI_ISL_866209, EPI_ISL_866210, EPI_ISL_866211, EPI_ISL_866212, EPI_ISL_866213, EPI_ISL_866222, EPI_ISL_866223, EPI_ISL_866225, EPI_ISL_866228, EPI_ISL_866229, EPI_ISL_866230, EPI_ISL_866232, EPI_ISL_866235, EPI_ISL_866236, EPI_ISL_866237, EPI_ISL_866238, EPI_ISL_866239, EPI_ISL_866240, EPI_ISL_866241, EPI_ISL_866242, EPI_ISL_866243, EPI_ISL_866244, EPI_ISL_866247, EPI_ISL_866250, EPI_ISL_866253, EPI_ISL_866255, EPI_ISL_866256, EPI_ISL_866257, EPI_ISL_866258, EPI_ISL_866259, EPI_ISL_866260, EPI_ISL_866261 | | | |
| see above | University College London, Great Ormond Street Hospital for Children NHS Foundation Trust, Imperial College Healthcare NHS Trust | COVID-19 Genomics UK (COG-UK) Consortium | Sergi Castellano, Rachel Williams, Mark Kristiansen, Paola Resende Silva, Sunando Roy, Tony Brooks, Helena Tutill, Paola Niola, Patricia Dyal, Charlotte Williams, Leysa Forrest, Yasmin Panchbhaya, Jacqueline Findlay, Samuel Weeks, Julianne Brown, Kathryn Harris, Paul Randell, James Price, Alison Holmes, Judith Breuer |
| EPI_ISL_866393, EPI_ISL_866394 | Regional Virus Laboratory, Belfast Health and Social Care Trust | COVID-19 Genomics UK (COG-UK) Consortium | Conall McCaughey, James McKenna, Tanya Curran, Susan Feeney, Alison Watt, Ciara Cox, Mairead Connor, Zoltan Molnar, David Simpson, Derek Fairley |
| EPI_ISL_866545, EPI_ISL_866573, EPI_ISL_866618, EPI_ISL_866621, EPI_ISL_866622, EPI_ISL_866627, EPI_ISL_866628, EPI_ISL_866629, EPI_ISL_866631, EPI_ISL_866635, EPI_ISL_866642, EPI_ISL_866643, EPI_ISL_866651, EPI_ISL_866652, EPI_ISL_866653, EPI_ISL_866654, EPI_ISL_866655, EPI_ISL_866656, EPI_ISL_866657, EPI_ISL_866658, EPI_ISL_866659, EPI_ISL_866660, EPI_ISL_866661, EPI_ISL_866663, EPI_ISL_866664, EPI_ISL_866665, EPI_ISL_866666, EPI_ISL_866667, EPI_ISL_866668, EPI_ISL_866669, EPI_ISL_866670, EPI_ISL_866671, EPI_ISL_866672, EPI_ISL_866673, EPI_ISL_866674, EPI_ISL_866677, EPI_ISL_866679, EPI_ISL_866681, EPI_ISL_866682, EPI_ISL_866683, EPI_ISL_866684, EPI_ISL_866686, EPI_ISL_866688, EPI_ISL_866689, EPI_ISL_866691, EPI_ISL_866692, EPI_ISL_866695, EPI_ISL_866696, EPI_ISL_866697, EPI_ISL_866698, EPI_ISL_866699, EPI_ISL_866700, EPI_ISL_866701, EPI_ISL_866702, EPI_ISL_866703, EPI_ISL_866704, EPI_ISL_866705, EPI_ISL_866706, EPI_ISL_866707, EPI_ISL_866708, EPI_ISL_866709, EPI_ISL_866710, EPI_ISL_866711, EPI_ISL_866712, EPI_ISL_866713, EPI_ISL_866714, EPI_ISL_866715, EPI_ISL_866716, EPI_ISL_866717, EPI_ISL_866718, EPI_ISL_866719, EPI_ISL_866720, EPI_ISL_866721, EPI_ISL_866722, EPI_ISL_866723, EPI_ISL_866724, EPI_ISL_866725, EPI_ISL_866726, EPI_ISL_866727, EPI_ISL_866728, EPI_ISL_866729, EPI_ISL_866730, EPI_ISL_866731, EPI_ISL_866732, EPI_ISL_866733, EPI_ISL_866734, EPI_ISL_866735, EPI_ISL_866736, EPI_ISL_866737, EPI_ISL_866738, EPI_ISL_866740, EPI_ISL_866741, EPI_ISL_866742, EPI_ISL_866744, EPI_ISL_866745, EPI_ISL_866746, EPI_ISL_866747, EPI_ISL_866831, EPI_ISL_866832, EPI_ISL_866833, EPI_ISL_866834 | | | |
| see above | Quadram Institute Bioscience | COVID-19 Genomics UK (COG-UK) Consortium | Dave J. Baker, Gemma L. Kay, Alp Aydin, Thanh Le-Viet, Steven Rudder, Ana P. Tedim, Anastasia Kolyva, Maria Diaz, Leonardo de Oliveira Martins, Nabil-Fareed Alikhan, Lizzie Meadows, Rachael Stanley, Ngozi Elumogo, Muhammed Yasir, Nicholas M. Thomson, Alexander J Trotter, Rachel Gilroy, Samuel Bloomfield, Claire Stuart, Andrew Bell, Reenesh Prakash, Samir Dervisevic, Alison E. Mather, John Wain, Mark Webber, Andrew J. Page, Justin O'Grady |
| EPI_ISL_866909, EPI_ISL_866910, EPI_ISL_866911, EPI_ISL_866912, EPI_ISL_866913, EPI_ISL_866914, EPI_ISL_866915, EPI_ISL_866916, EPI_ISL_866917, EPI_ISL_866918, EPI_ISL_866919, EPI_ISL_866920 | | | |
| see above | Queens Medical Centre, Clinical Microbiology Department / DeepSeq Nottingham | COVID-19 Genomics UK (COG-UK) Consortium | Gemma Clark, Wendy Smith, Manjinder Khakh, Vicki M Fleming, Michelle M Lister, Hannah Howson-Wells, Jonathan Ball, Patrick McClure, Joseph Chappell, Theocharis Tsoleridis, Nadine Holmes, Matthew Carlisle, Christopher Moore, Fei Sang, Johnny Debebe, Victoria Wright, Matthew Loose |
| EPI_ISL_867190, EPI_ISL_867191, EPI_ISL_867196, EPI_ISL_867197, EPI_ISL_867212, EPI_ISL_867213, EPI_ISL_867214, EPI_ISL_867221, EPI_ISL_867222, EPI_ISL_867223, EPI_ISL_867224, EPI_ISL_867225, EPI_ISL_867226, EPI_ISL_867237, EPI_ISL_867238, EPI_ISL_867239, EPI_ISL_867240, EPI_ISL_867241, EPI_ISL_867242, EPI_ISL_867243, EPI_ISL_867244, EPI_ISL_867245, EPI_ISL_867246, EPI_ISL_867247, EPI_ISL_867248, EPI_ISL_867249, EPI_ISL_867250, EPI_ISL_867251, EPI_ISL_867270, EPI_ISL_867305, EPI_ISL_867306, EPI_ISL_867307, EPI_ISL_867308, EPI_ISL_867309, EPI_ISL_867310, EPI_ISL_867311, EPI_ISL_867312, EPI_ISL_867317, EPI_ISL_867318, EPI_ISL_867319, EPI_ISL_867320, EPI_ISL_867321, EPI_ISL_867322, EPI_ISL_867323, EPI_ISL_867324, EPI_ISL_867325, EPI_ISL_867326, EPI_ISL_867327, EPI_ISL_867328, EPI_ISL_867329, EPI_ISL_867330, EPI_ISL_867331, EPI_ISL_867332, EPI_ISL_867333, EPI_ISL_867334, EPI_ISL_867335, EPI_ISL_867336, EPI_ISL_867337, EPI_ISL_867338, EPI_ISL_867339, EPI_ISL_867340, EPI_ISL_867341, EPI_ISL_867342, EPI_ISL_867343, EPI_ISL_867344, EPI_ISL_867345, EPI_ISL_867346, EPI_ISL_867348, EPI_ISL_867349, EPI_ISL_867350, EPI_ISL_867351, EPI_ISL_867352, EPI_ISL_867354, EPI_ISL_867355, EPI_ISL_867357, EPI_ISL_867359, EPI_ISL_867361, EPI_ISL_867368, EPI_ISL_867371, EPI_ISL_867417, EPI_ISL_867429, EPI_ISL_867457, EPI_ISL_867458, EPI_ISL_867497, EPI_ISL_867498, EPI_ISL_867506, EPI_ISL_867509, EPI_ISL_867510, EPI_ISL_867518, EPI_ISL_867519, EPI_ISL_867520, EPI_ISL_867521, EPI_ISL_867522, EPI_ISL_867523, EPI_ISL_867542, EPI_ISL_867543, EPI_ISL_867547, EPI_ISL_867550, EPI_ISL_867552, EPI_ISL_867553, EPI_ISL_867708, EPI_ISL_867717, EPI_ISL_867718, EPI_ISL_867719, EPI_ISL_867720, EPI_ISL_867721, EPI_ISL_867722, EPI_ISL_867723, EPI_ISL_867724, EPI_ISL_867725, EPI_ISL_867730, EPI_ISL_867731, EPI_ISL_867795, EPI_ISL_867796, EPI_ISL_867797, EPI_ISL_867800, EPI_ISL_867801, EPI_ISL_867802, EPI_ISL_867804, EPI_ISL_867805, EPI_ISL_867806, EPI_ISL_867807, EPI_ISL_867808, EPI_ISL_867809, EPI_ISL_867810, EPI_ISL_867811, EPI_ISL_867812, EPI_ISL_867815, EPI_ISL_867816, EPI_ISL_867821, EPI_ISL_867823, EPI_ISL_867824, EPI_ISL_867825, EPI_ISL_867828, EPI_ISL_867830, EPI_ISL_867832, EPI_ISL_867833, EPI_ISL_867837, EPI_ISL_867838, EPI_ISL_867839, EPI_ISL_867841, EPI_ISL_867842, EPI_ISL_867843, EPI_ISL_867844, EPI_ISL_867845, EPI_ISL_867847, EPI_ISL_867848, EPI_ISL_867849, EPI_ISL_867850, EPI_ISL_867851, EPI_ISL_867853, EPI_ISL_867855, EPI_ISL_867856, EPI_ISL_867859, EPI_ISL_867860, EPI_ISL_867861, EPI_ISL_867862, EPI_ISL_867863, EPI_ISL_867868, EPI_ISL_867873, EPI_ISL_867875, EPI_ISL_867876, EPI_ISL_867878, EPI_ISL_867879, EPI_ISL_867880, EPI_ISL_867881, EPI_ISL_867883, EPI_ISL_867884, EPI_ISL_867885, EPI_ISL_867889, EPI_ISL_867893, EPI_ISL_867895 | | | |
| see above | Originating lab: Wales Specialist Virology Centre Sequencing lab: Pathogen Genomics Unit | Public Health Wales Microbiology Cardiff Wales Specialist Virology Centre | Catherine Moore, Johnathan Evans, Laura Gifford, Malorie Perry, Simon Cottrell, Angela Marchbank, Alec Birchley, Alexander Adams, Amy Gaskin, Bree Gatica-Wilcox, Jason Coombes, Joel Southgate, Lauren Gilbert, Lee Graham, Nicole Pacchiarini, Sara Kumziene-Summerhayes, Sarah Taylor, Sophie Jones, Sara Rey, Matthew Bull, Joanne Watkins, Sally Corden, Tom Connor |
| EPI_ISL_868142, EPI_ISL_868143, EPI_ISL_868144, EPI_ISL_868145, EPI_ISL_868146, EPI_ISL_868147, EPI_ISL_868161, EPI_ISL_868162, EPI_ISL_868163, EPI_ISL_868164, EPI_ISL_868165, EPI_ISL_868166, EPI_ISL_868167, EPI_ISL_868168, EPI_ISL_868169, EPI_ISL_868170, EPI_ISL_868171, EPI_ISL_868172, EPI_ISL_868173, EPI_ISL_868174, EPI_ISL_868181, EPI_ISL_868182, EPI_ISL_868183, EPI_ISL_868184, EPI_ISL_868185, EPI_ISL_868186, EPI_ISL_868187, EPI_ISL_868188, EPI_ISL_868189, EPI_ISL_868190, EPI_ISL_868191, EPI_ISL_868192, EPI_ISL_868193, EPI_ISL_868194, EPI_ISL_868196, EPI_ISL_868197, EPI_ISL_868198, EPI_ISL_868199, EPI_ISL_868200, EPI_ISL_868201, EPI_ISL_868217, EPI_ISL_868218, EPI_ISL_868332, EPI_ISL_868333, EPI_ISL_868334, EPI_ISL_868335, EPI_ISL_868336, EPI_ISL_868337, EPI_ISL_868338, EPI_ISL_868354, EPI_ISL_868355 | | | |
| see above | Centre for Enzyme Innovation, University of Portsmouth / Translational Research Laboratory, Portsmouth Hospitals NHS Trust | COVID-19 Genomics UK (COG-UK) Consortium | Angela Beckett,Yann Bourgeois,Garry Scarlett,Sharon Glaysher,Scott Elliott,Kelly Bicknell,Robert Impey,Allyson Lloyd,Sarah Wyllie,Ethan Butcher,Anoop Chauhan,Samuel Robson |
| EPI_ISL_868373, EPI_ISL_868381, EPI_ISL_868382, EPI_ISL_868388, EPI_ISL_868395, EPI_ISL_868401, EPI_ISL_868408, EPI_ISL_868409, EPI_ISL_868415, EPI_ISL_868442, EPI_ISL_868464, EPI_ISL_868465, EPI_ISL_868469, EPI_ISL_868480, EPI_ISL_868486, EPI_ISL_868495, EPI_ISL_868508, EPI_ISL_868513, EPI_ISL_868521, EPI_ISL_868524, EPI_ISL_868533, EPI_ISL_868541, EPI_ISL_868560, EPI_ISL_868564, EPI_ISL_868566, EPI_ISL_868588, EPI_ISL_868594, EPI_ISL_868597, EPI_ISL_868599, EPI_ISL_868638, EPI_ISL_868643, EPI_ISL_868651, EPI_ISL_868655, EPI_ISL_868674, EPI_ISL_868681, EPI_ISL_868688, EPI_ISL_868692, EPI_ISL_868701 | | | |
| see above | Virology Department, Sheffield Teaching Hospitals NHS Foundation Trust/Department of Infection, Immunity and Cardiovascular Disease, The Medical School, University of Sheffield | COVID-19 Genomics UK (COG-UK) Consortium | Thushan de Silva, Matthew Parker, Nikki Smith, Adri Angyal, Rebecca Brown, Luke Green, Rachel Tucker, Paul Parsons, Danielle Groves, Katie Johnson, Laura Carrilero, Alex Keeley, Dave Partridge, Matthew Wyles, Benjamin Lindsey, Mehmet Yavuz, Mohammad Raza, Cariad Evans |
| EPI_ISL_869169, EPI_ISL_869172, EPI_ISL_869173, EPI_ISL_869177, EPI_ISL_869178, EPI_ISL_869179, EPI_ISL_869217, EPI_ISL_869218, EPI_ISL_869219, EPI_ISL_869220, EPI_ISL_869221, EPI_ISL_869222, EPI_ISL_869223, EPI_ISL_869224, EPI_ISL_869226, EPI_ISL_869228, EPI_ISL_869229, EPI_ISL_869230, EPI_ISL_869231, EPI_ISL_869232, EPI_ISL_869233, EPI_ISL_869234, EPI_ISL_869235, EPI_ISL_869236, EPI_ISL_869237, EPI_ISL_869238, EPI_ISL_869239 | | | |

| | | | |
|---|---|---|---|
| see above | New Mexico Department of Health Scientific Laboratory | Center for Global Health, University of New Mexico Health Sciences Center | Daryl Domman, Kurt Schwalm, Twila Kunde, Joseph Hicks, Anastacia Griego, Michael Edwards, Darrell Dinwiddie |
| EPI_ISL_871792 | Cerballiance Normandie | CERBA LAB | LAFOREST D, BRIAND G, LEPIGEON, technicien V; COUDRAY J; MARTINS-AMARAL F; COULON C, THOMAS Johan, VITEL R, HAMEL A. |
| EPI_ISL_871821, EPI_ISL_871822, EPI_ISL_871823, EPI_ISL_871824, EPI_ISL_871825, EPI_ISL_871826 | Jessa | Jessa | Brigitte Maes, Bert Cruys |
| EPI_ISL_871840, EPI_ISL_871847, EPI_ISL_871869, EPI_ISL_871873, EPI_ISL_871874, EPI_ISL_871877, EPI_ISL_871878 | Wyoming Public Health Laboratory | Wyoming Public Health Laboratory | Noah Hull, Taylor Fearing, Lynette Gumbleton, Channing Weber, Ashley Norberg, Bailey Bowcutt, and Wanda Manley |
| EPI_ISL_872199 | Hopital | National Reference Center for Viruses of Respiratory Infections, Institut Pasteur, Paris | Marion Barbet, Sylvie Behillil, Méline Bizard, Angela Brisebarre, Camille Capel, Etienne Simon-Lorière, Vincent Enouf, Maud Vanpeene, Sylvie van der Werf,Ramanantsoa CéLine |
| EPI_ISL_872221 | hopital | National Reference Center for Viruses of Respiratory Infections, Institut Pasteur, Paris | Marion Barbet, Sylvie Behillil, Méline Bizard, Angela Brisebarre, Camille Capel, Etienne Simon-Lorière, Vincent Enouf, Maud Vanpeene, Sylvie van der Werf,Combe Patrice |
| EPI_ISL_872228 | Hopital | National Reference Center for Viruses of Respiratory Infections, Institut Pasteur, Paris | Marion Barbet, Sylvie Behillil, Méline Bizard, Angela Brisebarre, Camille Capel, Etienne Simon-Lorière, Vincent Enouf, Maud Vanpeene, Sylvie van der Werf,Irimia Alix |
| EPI_ISL_872229, EPI_ISL_872230 | Hopital | National Reference Center for Viruses of Respiratory Infections, Institut Pasteur, Paris | Marion Barbet, Sylvie Behillil, Méline Bizard, Angela Brisebarre, Camille Capel, Etienne Simon-Lorière, Vincent Enouf, Maud Vanpeene, Sylvie van der Werf,Ramanantsoa CéLine |
| EPI_ISL_872231 | Hopital | National Reference Center for Viruses of Respiratory Infections, Institut Pasteur, Paris | Marion Barbet, Sylvie Behillil, Méline Bizard, Angela Brisebarre, Camille Capel, Etienne Simon-Lorière, Vincent Enouf, Maud Vanpeene, Sylvie van der Werf,Florin CéCile |
| EPI_ISL_872240, EPI_ISL_872241 | hopital | National Reference Center for Viruses of Respiratory Infections, Institut Pasteur, Paris | Marion Barbet, Sylvie Behillil, Méline Bizard, Angela Brisebarre, Camille Capel, Etienne Simon-Lorière, Vincent Enouf, Maud Vanpeene, Sylvie van der Werf,Combe Patrice |
| EPI_ISL_872249, EPI_ISL_872250 | Hopital | National Reference Center for Viruses of Respiratory Infections, Institut Pasteur, Paris | Marion Barbet, Sylvie Behillil, Méline Bizard, Angela Brisebarre, Camille Capel, Etienne Simon-Lorière, Vincent Enouf, Maud Vanpeene, Sylvie van der Werf,Ducancelle Alexandra |
| EPI_ISL_872251 | Hopital | National Reference Center for Viruses of Respiratory Infections, Institut Pasteur, Paris | Marion Barbet, Sylvie Behillil, Méline Bizard, Angela Brisebarre, Camille Capel, Etienne Simon-Lorière, Vincent Enouf, Maud Vanpeene, Sylvie van der Werf,Irimia Alix |
| EPI_ISL_872252 | Hopital | National Reference Center for Viruses of Respiratory Infections, Institut Pasteur, Paris | Marion Barbet, Sylvie Behillil, Méline Bizard, Angela Brisebarre, Camille Capel, Etienne Simon-Lorière, Vincent Enouf, Maud Vanpeene, Sylvie van der Werf,Ramanantsoa CéLine |
| EPI_ISL_872253 | Hopital | National Reference Center for Viruses of Respiratory Infections, Institut Pasteur, Paris | Marion Barbet, Sylvie Behillil, Méline Bizard, Angela Brisebarre, Camille Capel, Etienne Simon-Lorière, Vincent Enouf, Maud Vanpeene, Sylvie van der Werf,Irimia Alix |
| EPI_ISL_872255 | Hopital | National Reference Center for Viruses of Respiratory Infections, Institut Pasteur, Paris | Marion Barbet, Sylvie Behillil, Méline Bizard, Angela Brisebarre, Camille Capel, Etienne Simon-Lorière, Vincent Enouf, Maud Vanpeene, Sylvie van der Werf,Philippi Jean-FrançOis |
| EPI_ISL_872257 | Labo Analyses Med | National Reference Center for Viruses of Respiratory Infections, Institut Pasteur, Paris | Marion Barbet, Sylvie Behillil, Méline Bizard, Angela Brisebarre, Camille Capel, Etienne Simon-Lorière, Vincent Enouf, Maud Vanpeene, Sylvie van der Werf,Bour Jean Baptiste |
| EPI_ISL_872261 | Hopital | National Reference Center for Viruses of Respiratory Infections, Institut Pasteur, Paris | Marion Barbet, Sylvie Behillil, Méline Bizard, Angela Brisebarre, Camille Capel, Etienne Simon-Lorière, Vincent Enouf, Maud Vanpeene, Sylvie van der Werf,Guigon AuréLie |
| EPI_ISL_872275, EPI_ISL_872276, EPI_ISL_872277 | Hopital | National Reference Center for Viruses of Respiratory Infections, Institut Pasteur, Paris | Marion Barbet, Sylvie Behillil, Méline Bizard, Angela Brisebarre, Camille Capel, Etienne Simon-Lorière, Vincent Enouf, Maud Vanpeene, Sylvie van der Werf,Ramanantsoa CéLine |
| EPI_ISL_872280 | Labo Analyses Med | National Reference Center for Viruses of Respiratory Infections, Institut Pasteur, Paris | Marion Barbet, Sylvie Behillil, Méline Bizard, Angela Brisebarre, Camille Capel, Etienne Simon-Lorière, Vincent Enouf, Maud Vanpeene, Sylvie van der Werf,Bour Jean Baptiste |
| EPI_ISL_872283 | Labo Analyses Med | National Reference Center for Viruses of Respiratory Infections, Institut Pasteur, Paris | Marion Barbet, Sylvie Behillil, Méline Bizard, Angela Brisebarre, Camille Capel, Etienne Simon-Lorière, Vincent Enouf, Maud Vanpeene, Sylvie van der Werf,Girard Sophie |
| EPI_ISL_872293, EPI_ISL_872294 | Hopital | National Reference Center for Viruses of Respiratory Infections, Institut Pasteur, Paris | Marion Barbet, Sylvie Behillil, Méline Bizard, Angela Brisebarre, Camille Capel, Etienne Simon-Lorière, Vincent Enouf, Maud Vanpeene, Sylvie van der Werf,Guigon AuréLie |
| EPI_ISL_872300 | Hopital | National Reference Center for Viruses of Respiratory Infections, Institut Pasteur, Paris | Marion Barbet, Sylvie Behillil, Méline Bizard, Angela Brisebarre, Camille Capel, Etienne Simon-Lorière, Vincent Enouf, Maud Vanpeene, Sylvie van der Werf,Florin CéCile |
| EPI_ISL_872320 | Labo Analyses Med | National Reference Center for Viruses of Respiratory Infections, Institut Pasteur, Paris | Marion Barbet, Sylvie Behillil, Méline Bizard, Angela Brisebarre, Camille Capel, Etienne Simon-Lorière, Vincent Enouf, Maud Vanpeene, Sylvie van der Werf |
| EPI_ISL_872326, EPI_ISL_872327, EPI_ISL_872328, EPI_ISL_872329 | hopital | National Reference Center for Viruses of Respiratory Infections, Institut Pasteur, Paris | Marion Barbet, Sylvie Behillil, Méline Bizard, Angela Brisebarre, Camille Capel, Etienne Simon-Lorière, Vincent Enouf, Maud Vanpeene, Sylvie van der Werf,Combe Patrice |
| EPI_ISL_872341, EPI_ISL_872342 | Hopital | National Reference Center for Viruses of Respiratory Infections, Institut Pasteur, Paris | Marion Barbet, Sylvie Behillil, Méline Bizard, Angela Brisebarre, Camille Capel, Etienne Simon-Lorière, Vincent Enouf, Maud Vanpeene, Sylvie van der Werf,Ducancelle Alexandra |
| EPI_ISL_872343 | Hopital | National Reference Center for Viruses of Respiratory Infections, Institut Pasteur, Paris | Marion Barbet, Sylvie Behillil, Méline Bizard, Angela Brisebarre, Camille Capel, Etienne Simon-Lorière, Vincent Enouf, Maud Vanpeene, Sylvie van der Werf,Irimia Alix |
| EPI_ISL_872344, EPI_ISL_872345, EPI_ISL_872346 | Hopital | National Reference Center for Viruses of Respiratory Infections, Institut Pasteur, Paris | Marion Barbet, Sylvie Behillil, Méline Bizard, Angela Brisebarre, Camille Capel, Etienne Simon-Lorière, Vincent Enouf, Maud Vanpeene, Sylvie van der Werf,Ramanantsoa CéLine |
| EPI_ISL_872348 | Hopital | National Reference Center for Viruses of Respiratory Infections, Institut Pasteur, Paris | Marion Barbet, Sylvie Behillil, Méline Bizard, Angela Brisebarre, Camille Capel, Etienne Simon-Lorière, Vincent Enouf, Maud Vanpeene, Sylvie van der Werf,Irimia Alix |
| EPI_ISL_872479, EPI_ISL_872480, EPI_ISL_872481, EPI_ISL_872482, EPI_ISL_872483, EPI_ISL_872484, EPI_ISL_872485, EPI_ISL_872486, EPI_ISL_872487, EPI_ISL_872488, EPI_ISL_872489, EPI_ISL_872490, EPI_ISL_872491, EPI_ISL_872492, EPI_ISL_872493, EPI_ISL_872511 | | | |
| see above | New Mexico Department of Health Scientific Laboratory | Center for Global Health, University of New Mexico Health Sciences Center | Daryl Domman, Kurt Schwalm, Twila Kunde, Joseph Hicks, Anastacia Griego, Michael Edwards, Darrell Dinwiddie |
| EPI_ISL_872579 | Sydney South West Pathology Service (SSWPS) - Royal Prince Alfred Hospital - NSW Health Pathology | NSW Health Pathology - Institute of Clinical Pathology and Medical Research; Westmead Hospital; University of Sydney | CIDM-PH et al. |
| EPI_ISL_872580 | Histopath | NSW Health Pathology - Institute of Clinical Pathology and Medical Research; Westmead Hospital; University of Sydney | CIDM-PH et al. |
| EPI_ISL_872581 | St Vincent's Pathology (SydPath) | NSW Health Pathology - Institute of Clinical Pathology and Medical Research; Westmead Hospital; University of Sydney | CIDM-PH et al. |
| EPI_ISL_872582 | South Eastern Area Laboratory Services (SEALS) | NSW Health Pathology - Institute of Clinical Pathology and Medical Research; Westmead Hospital; University of Sydney | CIDM-PH et al. |
| EPI_ISL_872849, EPI_ISL_872850, | University of Wisconsin-Madison AIDS Vaccine Research | University of Wisconsin-Madison AIDS Vaccine Research | Gage Moreno, Katarina Braun, et al. AIDS Vaccine Research Laboratories |

| | | | |
|---|---|---|---|
| EPI_ISL_872851, EPI_ISL_872852, EPI_ISL_872853 | Laboratories | Laboratories | |
| EPI_ISL_873040 | WHO National Influenza Centre Russian Federation | WHO National Influenza Centre Russian Federation | Andrey Komissarov, Artem Fadeev, Anna Ivanova, Kseniya Komissarova, Dmitry Bazhenov, Mikhail Bakaev, Daria Danilenko, Ksenia Safina, Elena Nabieva, Georgii Bazykin, Dmitry Lioznov |
| EPI_ISL_873110, EPI_ISL_873112, EPI_ISL_873114, EPI_ISL_873115, EPI_ISL_873118, EPI_ISL_873121, EPI_ISL_873123 | University of Michigan Clinical Microbiology Laboratory | Lauring Lab, University of Michigan, Department of Microbiology and Immunology | Valesano |
| EPI_ISL_873166, EPI_ISL_873167, EPI_ISL_873168, EPI_ISL_873169, EPI_ISL_873170, EPI_ISL_873171, EPI_ISL_873172, EPI_ISL_873173, EPI_ISL_873174, EPI_ISL_873175, EPI_ISL_873177, EPI_ISL_873178, EPI_ISL_873179, EPI_ISL_873180, EPI_ISL_873181, EPI_ISL_873182 | | | |
| see above | Microbiology Division, South Carolina Department of Health and Environmental Control (SC DHEC) | Microbiology Division, South Carolina Department of Health and Environmental Control (SC DHEC) | Flores,H., Freeman,J. |
| EPI_ISL_873235, EPI_ISL_873236, EPI_ISL_873237, EPI_ISL_873239, EPI_ISL_873240, EPI_ISL_873242, EPI_ISL_873244, EPI_ISL_873247, EPI_ISL_873248, EPI_ISL_873251, EPI_ISL_873253, EPI_ISL_873255, EPI_ISL_873256, EPI_ISL_873257, EPI_ISL_873259, EPI_ISL_873260, EPI_ISL_873262 | | | |
| see above | M Health Fairview | Minnesota Department of Health, Public Health Laboratory | Alexandra Lorentz, Jacob Garfin, Matt Plumb, and Xiong Wang |
| EPI_ISL_873272 | Vault Health | Minnesota Department of Health, Public Health Laboratory | Alexandra Lorentz, Jacob Garfin, Matt Plumb, and Xiong Wang |
| EPI_ISL_873276 | Hennepin County Medical Center | Minnesota Department of Health, Public Health Laboratory | Alexandra Lorentz, Jacob Garfin, Matt Plumb, and Xiong Wang |
| EPI_ISL_873280, EPI_ISL_873288, EPI_ISL_873289, EPI_ISL_873308, EPI_ISL_873309, EPI_ISL_873312, EPI_ISL_873331, EPI_ISL_873345, EPI_ISL_873352, EPI_ISL_873360, EPI_ISL_873372, EPI_ISL_873377, EPI_ISL_873379, EPI_ISL_873382, EPI_ISL_873383, EPI_ISL_873385, EPI_ISL_873390, EPI_ISL_873395, EPI_ISL_873401, EPI_ISL_873403, EPI_ISL_873407, EPI_ISL_873409, EPI_ISL_873417, EPI_ISL_873419, EPI_ISL_873425, EPI_ISL_873427, EPI_ISL_873447, EPI_ISL_873452, EPI_ISL_873458, EPI_ISL_873463, EPI_ISL_873468, EPI_ISL_873469, EPI_ISL_873483, EPI_ISL_873485, EPI_ISL_873486, EPI_ISL_873488, EPI_ISL_873490, EPI_ISL_873505, EPI_ISL_873517, EPI_ISL_873521, EPI_ISL_873527, EPI_ISL_873528, EPI_ISL_873535, EPI_ISL_873536, EPI_ISL_873538, EPI_ISL_873542, EPI_ISL_873543, EPI_ISL_873546, EPI_ISL_873547, EPI_ISL_873549, EPI_ISL_873554, EPI_ISL_873557, EPI_ISL_873562, EPI_ISL_873563, EPI_ISL_873565, EPI_ISL_873570, EPI_ISL_873575, EPI_ISL_873577, EPI_ISL_873583, EPI_ISL_873589, EPI_ISL_873590, EPI_ISL_873591, EPI_ISL_873595, EPI_ISL_873599, EPI_ISL_873602, EPI_ISL_873618, EPI_ISL_873623, EPI_ISL_873627, EPI_ISL_873656, EPI_ISL_873672, EPI_ISL_873724, EPI_ISL_873726, EPI_ISL_873730, EPI_ISL_873769, EPI_ISL_873786, EPI_ISL_873850, EPI_ISL_873865, EPI_ISL_873875 | | | |
| see above | Lighthouse Lab in Milton Keynes | Wellcome Sanger Institute for the COVID-19 Genomics UK (COG-UK) Consortium | The Lighthouse Lab in Milton Keynes and Alex Alderton, Roberto Amato, Sonia Goncalves, Ewan Harrison, David K. Jackson, Ian Johnston, Dominic Kwiatkowski, Cordelia Langford, John Sillitoe on behalf of the Wellcome Sanger Institute COVID-19 Surveillance Team |
| EPI_ISL_873882 | Lighthouse Lab in Cambridge | Wellcome Sanger Institute for the COVID-19 Genomics UK (COG-UK) Consortium | Rob Howes, The Lighthouse Lab in Cambridge and Alex Alderton, Roberto Amato, Sonia Goncalves, Ewan Harrison, David K. Jackson, Ian Johnston, Dominic Kwiatkowski, Cordelia Langford, John Sillitoe on behalf of the Wellcome Sanger Institute COVID-19 Surveillance Team |
| EPI_ISL_873904, EPI_ISL_873925, EPI_ISL_873935, EPI_ISL_873943, EPI_ISL_873947, EPI_ISL_873954 | Lighthouse Lab in Milton Keynes | Wellcome Sanger Institute for the COVID-19 Genomics UK (COG-UK) Consortium | The Lighthouse Lab in Milton Keynes and Alex Alderton, Roberto Amato, Sonia Goncalves, Ewan Harrison, David K. Jackson, Ian Johnston, Dominic Kwiatkowski, Cordelia Langford, John Sillitoe on behalf of the Wellcome Sanger Institute COVID-19 Surveillance Team |
| EPI_ISL_873962, EPI_ISL_873981, EPI_ISL_873982, EPI_ISL_873986, EPI_ISL_874011, EPI_ISL_874024, EPI_ISL_874070, EPI_ISL_874078, EPI_ISL_874080, EPI_ISL_874081, EPI_ISL_874082, EPI_ISL_874093, EPI_ISL_874098, EPI_ISL_874109, EPI_ISL_874121, EPI_ISL_874145, EPI_ISL_874149, EPI_ISL_874155, EPI_ISL_874159, EPI_ISL_874181, EPI_ISL_874185, EPI_ISL_874196, EPI_ISL_874203, EPI_ISL_874210, EPI_ISL_874218, EPI_ISL_874220, EPI_ISL_874225, EPI_ISL_874227, EPI_ISL_874243, EPI_ISL_874245, EPI_ISL_874249, EPI_ISL_874252, EPI_ISL_874257, EPI_ISL_874259, EPI_ISL_874273, EPI_ISL_874284, EPI_ISL_874287, EPI_ISL_874291, EPI_ISL_874310, EPI_ISL_874502, EPI_ISL_874530, EPI_ISL_874853, EPI_ISL_874966 | | | |
| see above | Lighthouse Lab in Alderley Park | Wellcome Sanger Institute for the COVID-19 Genomics UK (COG-UK) Consortium | Jacquelyn Wynn, Mairead Hyland, The Lighthouse Lab in Alderley Park and Alex Alderton, Roberto Amato, Sonia Goncalves, Ewan Harrison, David K. Jackson, Ian Johnston, Dominic Kwiatkowski, Cordelia Langford, John Sillitoe on behalf of the Wellcome Sanger Institute COVID-19 Surveillance Team |
| EPI_ISL_875342 | Pakistan Institute of Medical Sciences | Department of Virology | Massab Umair, Aamer Ikram, Muhammad Salman, Sana Tamim, Nazish Badar, Adnan Khurshid, Zaira Rahman, Abdul Ahad, Hamza Ahmad |
| EPI_ISL_875343, EPI_ISL_875345 | Pakistan Institute of Medical Sciences | Department of Virology | Massab Umair, Aamer Ikram, Muhammad Salman, Nazish Badar, Sana Tamim, Adnan Khurshid, Zaira Rahman, Abdul Ahad, Hamza Ahmad |
| EPI_ISL_875410, EPI_ISL_875411, EPI_ISL_875412, EPI_ISL_875413, EPI_ISL_875414, EPI_ISL_875415, EPI_ISL_875416, EPI_ISL_875417, EPI_ISL_875418, EPI_ISL_875419, EPI_ISL_875420, EPI_ISL_875421, EPI_ISL_875422, EPI_ISL_875423, EPI_ISL_875424, EPI_ISL_875425, EPI_ISL_875426, EPI_ISL_875427, EPI_ISL_875428, EPI_ISL_875429, EPI_ISL_875430, EPI_ISL_875431, EPI_ISL_875432, EPI_ISL_875433, EPI_ISL_875434, EPI_ISL_875435, EPI_ISL_875436, EPI_ISL_875437, EPI_ISL_875438, EPI_ISL_875439, EPI_ISL_875440, EPI_ISL_875441, EPI_ISL_875442, EPI_ISL_875443, EPI_ISL_875444, EPI_ISL_875445, EPI_ISL_875446, EPI_ISL_875447, EPI_ISL_875448, EPI_ISL_875449, EPI_ISL_875450, EPI_ISL_875451, EPI_ISL_875452, EPI_ISL_875453, EPI_ISL_875454, EPI_ISL_875455, EPI_ISL_875456, EPI_ISL_875457, EPI_ISL_875458, EPI_ISL_875459 | | | |
| see above | National Virus Reference Laboratory | National Virus Reference Laboratory | Michael Carr, Gabriel Gonzalez, Jonathan Dean, Cillian F De Gascun |
| EPI_ISL_875519 | Institute of Virology, Biomedical Research Center of the Slovak Academy of Sciences, Bratislava | Faculty of Natural Sciences, Comenius University, Bratislava | Broa Brejová, Viktória abanová, Kristína Boršová, Viktória Hodorová, Sabina Fumaová Havlíková, Juraj Kopáek, Martina Liková, ubomíra Lukáiková, Martina Neboháová, Monika Sláviková, Tomáš Vina, Jozef Nosek, Boris Klempa |
| EPI_ISL_875529 | Institute of Virology, Biomedical Research Center of the Slovak Academy of Sciences, Bratislava | Faculty of Natural Sciences, Comenius University, Bratislava | Kristína Boršová, Viktória abanová, Broa Brejová, Viktória Hodorová, Sabina Fumaová Havlíková, Juraj Kopáek, Martina Liková, ubomíra Lukáiková, Martina Neboháová, Monika Sláviková, Tomáš Vina, Boris Klempa, Jozef Nosek |
| EPI_ISL_875532 | Institute of Virology, Biomedical Research Center of the Slovak Academy of Sciences, Bratislava | Faculty of Natural Sciences, Comenius University, Bratislava | Broa Brejová, Viktória abanová, Kristína Boršová, Viktória Hodorová, Sabina Fumaová Havlíková, Juraj Kopáek, Martina Liková, ubomíra Lukáiková, Martina Neboháová, Monika Sláviková, Tomáš Vina, Jozef Nosek, Boris Klempa |
| EPI_ISL_875662, EPI_ISL_875663 | University of Michigan Clinical Microbiology Laboratory | Lauring Lab, University of Michigan, Department of Microbiology and Immunology | Valesano |
| EPI_ISL_875671 | CHU Purpan - Laboratoire de Virologie - Institut Fédératif de Biologie | CHU Purpan - Laboratoire de Virologie - Institut Fédératif de Biologie | Latour J., Ranger N., Dubois M., Carcenac R., Harter A., Boyer P., Tremeaux P., Izopet J. |
| EPI_ISL_875685 | National Virus Reference Laboratory | National Virus Reference Laboratory | Michael Carr, Gabriel Gonzalez, Jonathan Dean, Cillian F De Gascun |
| EPI_ISL_876063, EPI_ISL_876077, EPI_ISL_876111, EPI_ISL_876144, EPI_ISL_876155, EPI_ISL_876244 | Massachusetts State Public Health Laboratory | Massachusetts State Public Health Laboratory | Andrew Lang, Timelia Fink, Glen Gallagher, Sandra Smole |
| EPI_ISL_876589 | Blackhawk Genomics | Pathogen Discovery, Respiratory Viruses Branch, Division of Viral Diseases, Centers for Disease Control and Prevention | Ying Tao, Yan Li, Jing Zhang, Krista Queen, Anna Uehara, Peter Cook, Clinton R. Paden, Haibin Wang, Suxiang Tong |
| EPI_ISL_877210, EPI_ISL_877211, EPI_ISL_877212, EPI_ISL_877213 | LabPLUS | Institute of Environmental Science and Research (ESR) | Xiaoyun Ren, Matt Storey, Nikki Freed, Muhammad Faisal, Jing Wang, Hermes Perez, Anja Werno, Antje van der Linden, Arlo Upton, Chris Mansell, David Hammer, Dragana Drinkovic, Gary McAuliffe, Hana Sofia Andersson, James Ussher, Jill Sherwood, Josh Freeman, Julia Howard, Juliet Elvy, Mary DeAlmeida, Matt Blakiston, Matthew Rogers, Max Bloomfield, Michael Addidle, Michelle Balm, Sally Roberts, Sarah Jefferies, Sharmini Muttaiyah, Susan Morpeth, Susan Taylor, Timothy Blackmore, Vani Sathyendran, Veronica Playle, Virginia Hope, Erasmus Smit, Lauren Jelly, Olin Silander, Joep de Ligt |
| EPI_ISL_877215, EPI_ISL_877216, EPI_ISL_877217, EPI_ISL_877218 | Middlemore Hospital | Institute of Environmental Science and Research (ESR) | Xiaoyun Ren, Matt Storey, Nikki Freed, Muhammad Faisal, Jing Wang, Hermes Perez, Anja Werno, Antje van der Linden, Arlo Upton, Chris Mansell, David Hammer, Dragana Drinkovic, Gary McAuliffe, Hana Sofia Andersson, James Ussher, Jill Sherwood, Josh Freeman, Julia Howard, Juliet Elvy, Mary DeAlmeida, Matt Blakiston, Matthew Rogers, Max Bloomfield, Michael Addidle, Michelle Balm, Sally Roberts, Sarah Jefferies, Sharmini Muttaiyah, Susan Morpeth, Susan Taylor, Timothy Blackmore, Vani Sathyendran, Veronica Playle, Virginia Hope, Erasmus Smit, Lauren Jelly, Olin Silander, Joep de Ligt |
| EPI_ISL_877227 | Thai Red Cross Emerging Infectious Diseases Health Science Centre, Chulalongkorn Hospital, Faculty of Medicine, Chulalongkorn University | Thai Red Cross Emerging Infectious Diseases Center and Faculty of Medicine, Chulalongkorn University | Krirk Asavametha, Panpit Suwangool, Sininat Petcharat, Yutthana Joyjinda, Weenassarin Ampoot, Apaporn Rodpan, Opass Putcharoen, Thiravat Hemachudha, Supaporn Wacharapluesadee |
| EPI_ISL_877229, EPI_ISL_877230, EPI_ISL_877231, EPI_ISL_877232, EPI_ISL_877233, EPI_ISL_877234, EPI_ISL_877235 | University College Sedaya International (UCSI University) | Institute for Medical Research, Infectious Disease Research Centre, National Institutes of Health, Ministry of Health Malaysia | Suppiah J, Kamel K, Azizan MA, Sekaran SD, Thayan R |
| EPI_ISL_878833 | Lighthouse Lab in Milton Keynes | Wellcome Sanger Institute for the COVID-19 Genomics UK (COG-UK) Consortium | The Lighthouse Lab in Milton Keynes and Alex Alderton, Roberto Amato, Sonia Goncalves, Ewan Harrison, David K. Jackson, Ian Johnston, Dominic Kwiatkowski, Cordelia Langford, John Sillitoe on behalf of the Wellcome Sanger Institute COVID-19 Surveillance Team |
| EPI_ISL_880040, EPI_ISL_880042, EPI_ISL_880058, EPI_ISL_880066, EPI_ISL_880069, EPI_ISL_880077, EPI_ISL_880085, EPI_ISL_880087, EPI_ISL_880098, EPI_ISL_880109, EPI_ISL_880111, EPI_ISL_880114, EPI_ISL_880142, EPI_ISL_880145, EPI_ISL_880148, EPI_ISL_880153, EPI_ISL_880154, EPI_ISL_880155, | | | |

| | | | |
|---|---|---|---|
| EPI_ISL_880158, EPI_ISL_880160, EPI_ISL_880163, EPI_ISL_880166, EPI_ISL_880167, EPI_ISL_880170 | | | |
| see above | Sharp HealthCare Laboratory | Andersen lab at Scripps Research | SEARCH Alliance San Diego with Aaron Harding, Jacquelyn Berumen, Cathy Woerle, Liam McGinnis, Art Mendoza, Omid Bakhtar |
| EPI_ISL_880179 | Rady's Childrens Hospital | Andersen lab at Scripps Research | SEARCH Alliance San Diego with Nanda Radamchar, David Dimmock, Linda Luo, Christina Clarke, Kathryn Bouic, Teresa Mueller, Denise Malicki |
| EPI_ISL_880601, EPI_ISL_880620, EPI_ISL_880629, EPI_ISL_880637, EPI_ISL_880669, EPI_ISL_880674, EPI_ISL_880682, EPI_ISL_880709, EPI_ISL_880710, EPI_ISL_880717, EPI_ISL_880718, EPI_ISL_880721, EPI_ISL_880746, EPI_ISL_880753, EPI_ISL_880763, EPI_ISL_880774, EPI_ISL_880775, EPI_ISL_880776, EPI_ISL_880779, EPI_ISL_880789, EPI_ISL_880799, EPI_ISL_880812, EPI_ISL_880831, EPI_ISL_880835, EPI_ISL_880853, EPI_ISL_880868, EPI_ISL_880879, EPI_ISL_880915 | | | |
| see above | Lighthouse Lab in Milton Keynes | Wellcome Sanger Institute for the COVID-19 Genomics UK (COG-UK) Consortium | The Lighthouse Lab in Milton Keynes and Alex Alderton, Roberto Amato, Sonia Goncalves, Ewan Harrison, David K. Jackson, Ian Johnston, Dominic Kwiatkowski, Cordelia Langford, John Sillitoe on behalf of the Wellcome Sanger Institute COVID-19 Surveillance Team |
| EPI_ISL_881318, EPI_ISL_881326 | Lighthouse Lab in Alderley Park | Wellcome Sanger Institute for the COVID-19 Genomics UK (COG-UK) Consortium | Jacquelyn Wynn, Mairead Hyland, The Lighthouse Lab in Alderley Park and Alex Alderton, Roberto Amato, Sonia Goncalves, Ewan Harrison, David K. Jackson, Ian Johnston, Dominic Kwiatkowski, Cordelia Langford, John Sillitoe on behalf of the Wellcome Sanger Institute COVID-19 Surveillance Team |
| EPI_ISL_881618, EPI_ISL_881621 | Lighthouse Lab in Milton Keynes | Wellcome Sanger Institute for the COVID-19 Genomics UK (COG-UK) Consortium | The Lighthouse Lab in Milton Keynes and Alex Alderton, Roberto Amato, Sonia Goncalves, Ewan Harrison, David K. Jackson, Ian Johnston, Dominic Kwiatkowski, Cordelia Langford, John Sillitoe on behalf of the Wellcome Sanger Institute COVID-19 Surveillance Team |
| EPI_ISL_881623, EPI_ISL_881624, EPI_ISL_881625, EPI_ISL_881626, EPI_ISL_881627, EPI_ISL_881628, EPI_ISL_881629, EPI_ISL_881630, EPI_ISL_881631, EPI_ISL_881632, EPI_ISL_881633, EPI_ISL_881634, EPI_ISL_881635, EPI_ISL_881636, EPI_ISL_881637, EPI_ISL_881638, EPI_ISL_881639, EPI_ISL_881640, EPI_ISL_881641, EPI_ISL_881642, EPI_ISL_881643, EPI_ISL_881644, EPI_ISL_881645, EPI_ISL_881646, EPI_ISL_881647, EPI_ISL_881648, EPI_ISL_881649, EPI_ISL_881650, EPI_ISL_881651, EPI_ISL_881652, EPI_ISL_881653, EPI_ISL_881654, EPI_ISL_881655, EPI_ISL_881656, EPI_ISL_881657, EPI_ISL_881658, EPI_ISL_881659, EPI_ISL_881660, EPI_ISL_881661, EPI_ISL_881662, EPI_ISL_881663, EPI_ISL_881664, EPI_ISL_881665, EPI_ISL_881666, EPI_ISL_881667, EPI_ISL_881668, EPI_ISL_881669, EPI_ISL_881670, EPI_ISL_881671, EPI_ISL_881672, EPI_ISL_881673, EPI_ISL_881674, EPI_ISL_881675, EPI_ISL_881676, EPI_ISL_881677, EPI_ISL_881678, EPI_ISL_881679, EPI_ISL_881680, EPI_ISL_881681, EPI_ISL_881682, EPI_ISL_881683, EPI_ISL_881684, EPI_ISL_881685, EPI_ISL_881686, EPI_ISL_881687, EPI_ISL_881688, EPI_ISL_881689, EPI_ISL_881690, EPI_ISL_881691, EPI_ISL_881692, EPI_ISL_881693, EPI_ISL_881694, EPI_ISL_881695, EPI_ISL_881696, EPI_ISL_881697, EPI_ISL_881698, EPI_ISL_881699, EPI_ISL_881700, EPI_ISL_881701, EPI_ISL_881702, EPI_ISL_881703, EPI_ISL_881704, EPI_ISL_881705, EPI_ISL_881706, EPI_ISL_881707, EPI_ISL_881708, EPI_ISL_881709, EPI_ISL_881710, EPI_ISL_881711, EPI_ISL_881712, EPI_ISL_881713, EPI_ISL_881714, EPI_ISL_881715, EPI_ISL_881716, EPI_ISL_881717, EPI_ISL_881718, EPI_ISL_881719, EPI_ISL_881720, EPI_ISL_881721, EPI_ISL_881722, EPI_ISL_881723, EPI_ISL_881724, EPI_ISL_881725, EPI_ISL_881726, EPI_ISL_881727, EPI_ISL_881728, EPI_ISL_881729, EPI_ISL_881730, EPI_ISL_881731, EPI_ISL_881732, EPI_ISL_881733, EPI_ISL_881734, EPI_ISL_881735, EPI_ISL_881736, EPI_ISL_881737, EPI_ISL_881738, EPI_ISL_881739, EPI_ISL_881740, EPI_ISL_881741, EPI_ISL_881742, EPI_ISL_881743, EPI_ISL_881744, EPI_ISL_881745, EPI_ISL_881746, EPI_ISL_881747, EPI_ISL_881748, EPI_ISL_881749, EPI_ISL_881750, EPI_ISL_881751, EPI_ISL_881752, EPI_ISL_881753, EPI_ISL_881754, EPI_ISL_881755, EPI_ISL_881756, EPI_ISL_881757, EPI_ISL_881758, EPI_ISL_881759, EPI_ISL_881760, EPI_ISL_881761, EPI_ISL_881762, EPI_ISL_881763, EPI_ISL_881764, EPI_ISL_881765, EPI_ISL_881766, EPI_ISL_881767, EPI_ISL_881768, EPI_ISL_881769, EPI_ISL_881770, EPI_ISL_881771, EPI_ISL_881772, EPI_ISL_881773, EPI_ISL_881774, EPI_ISL_881775, EPI_ISL_881776, EPI_ISL_881777, EPI_ISL_881778, EPI_ISL_881779, EPI_ISL_881780, EPI_ISL_881781, EPI_ISL_881782, EPI_ISL_881783, EPI_ISL_881784, EPI_ISL_881785, EPI_ISL_881786, EPI_ISL_881787, EPI_ISL_881788, EPI_ISL_881789, EPI_ISL_881790, EPI_ISL_881791, EPI_ISL_881792, EPI_ISL_881793, EPI_ISL_881794, EPI_ISL_881795, EPI_ISL_881796, EPI_ISL_881797, EPI_ISL_881798, EPI_ISL_881799, EPI_ISL_881800, EPI_ISL_881801, EPI_ISL_881802, EPI_ISL_881803, EPI_ISL_881804, EPI_ISL_881805, EPI_ISL_881806, EPI_ISL_881807, EPI_ISL_881808, EPI_ISL_881809, EPI_ISL_881810, EPI_ISL_881811, EPI_ISL_881812, EPI_ISL_881813, EPI_ISL_881814, EPI_ISL_881815, EPI_ISL_881816, EPI_ISL_881817, EPI_ISL_881818, EPI_ISL_881819, EPI_ISL_881820, EPI_ISL_881821, EPI_ISL_881822, EPI_ISL_881823, EPI_ISL_881824, EPI_ISL_881825, EPI_ISL_881826, EPI_ISL_881827, EPI_ISL_881828, EPI_ISL_881829, EPI_ISL_881830, EPI_ISL_881831, EPI_ISL_881832, EPI_ISL_881833, EPI_ISL_881834, EPI_ISL_881835, EPI_ISL_881836, EPI_ISL_881837, EPI_ISL_881838, EPI_ISL_881839, EPI_ISL_881840, EPI_ISL_881841, EPI_ISL_881842, EPI_ISL_881843, EPI_ISL_881844, EPI_ISL_881845, EPI_ISL_881846, EPI_ISL_881847, EPI_ISL_881848, EPI_ISL_881849, EPI_ISL_881850, EPI_ISL_881851, EPI_ISL_881852, EPI_ISL_881853, EPI_ISL_881854, EPI_ISL_881855, EPI_ISL_881856, EPI_ISL_881857, EPI_ISL_881858, EPI_ISL_881859, EPI_ISL_881860, EPI_ISL_881861, EPI_ISL_881862, EPI_ISL_881863, EPI_ISL_881864, EPI_ISL_881865, EPI_ISL_881866, EPI_ISL_881867, EPI_ISL_881868, EPI_ISL_881869, EPI_ISL_881870, EPI_ISL_881871, EPI_ISL_881872, EPI_ISL_881873, EPI_ISL_881874, EPI_ISL_881875, EPI_ISL_881876, EPI_ISL_881877, EPI_ISL_881878, EPI_ISL_881879, EPI_ISL_881880, EPI_ISL_881881, EPI_ISL_881882, EPI_ISL_881883, EPI_ISL_881884, EPI_ISL_881885, EPI_ISL_881886, EPI_ISL_881887, EPI_ISL_881888, EPI_ISL_881889, EPI_ISL_881890, EPI_ISL_881891, EPI_ISL_881892, EPI_ISL_881893, EPI_ISL_881894, EPI_ISL_881895, EPI_ISL_881896, EPI_ISL_881897, EPI_ISL_881898, EPI_ISL_881899, EPI_ISL_881900, EPI_ISL_881901, EPI_ISL_881902, EPI_ISL_881903, EPI_ISL_881904, EPI_ISL_881905, EPI_ISL_881906, EPI_ISL_881907, EPI_ISL_881908, EPI_ISL_881909, EPI_ISL_881910, EPI_ISL_881911, EPI_ISL_881912, EPI_ISL_881913, EPI_ISL_881914, EPI_ISL_881915, EPI_ISL_881916, EPI_ISL_881917, EPI_ISL_881918, EPI_ISL_881919, EPI_ISL_881920, EPI_ISL_881921, EPI_ISL_881922, EPI_ISL_881923, EPI_ISL_881924, EPI_ISL_881925, EPI_ISL_881926, EPI_ISL_881927, EPI_ISL_881928, EPI_ISL_881929, EPI_ISL_881930, EPI_ISL_881931, EPI_ISL_881932, EPI_ISL_881933, EPI_ISL_881934, EPI_ISL_881935, EPI_ISL_881936, EPI_ISL_881937, EPI_ISL_881938, EPI_ISL_881939, EPI_ISL_881940, EPI_ISL_881941, EPI_ISL_881942, EPI_ISL_881943, EPI_ISL_881944, EPI_ISL_881945, EPI_ISL_881946, EPI_ISL_881947, EPI_ISL_881948, EPI_ISL_881949, EPI_ISL_881950, EPI_ISL_881951, EPI_ISL_881952, EPI_ISL_881953, EPI_ISL_881954, EPI_ISL_881955, EPI_ISL_881956, EPI_ISL_881957, EPI_ISL_881958, EPI_ISL_881959, EPI_ISL_881960, EPI_ISL_881961, EPI_ISL_881962, EPI_ISL_881963, EPI_ISL_881964, EPI_ISL_881965, EPI_ISL_881966, EPI_ISL_881967, EPI_ISL_881968, EPI_ISL_881969, EPI_ISL_881970, EPI_ISL_881971, EPI_ISL_881972, EPI_ISL_881973, EPI_ISL_881974, EPI_ISL_881975, EPI_ISL_881976, EPI_ISL_881977, EPI_ISL_881978, EPI_ISL_881979, EPI_ISL_881980, EPI_ISL_881981, EPI_ISL_881982, EPI_ISL_881983, EPI_ISL_881984, EPI_ISL_881985, EPI_ISL_881986, EPI_ISL_881987, EPI_ISL_881988, EPI_ISL_881989, EPI_ISL_881990, EPI_ISL_881991, EPI_ISL_881992, EPI_ISL_881993, EPI_ISL_881994, EPI_ISL_881995, EPI_ISL_881996, EPI_ISL_881997, EPI_ISL_881998, EPI_ISL_881999, EPI_ISL_882000, EPI_ISL_882001, EPI_ISL_882002, EPI_ISL_882003, EPI_ISL_882004, EPI_ISL_882005, EPI_ISL_882006, EPI_ISL_882007, EPI_ISL_882008, EPI_ISL_882009, EPI_ISL_882010, EPI_ISL_882011, EPI_ISL_882012, EPI_ISL_882013, EPI_ISL_882014, EPI_ISL_882015, EPI_ISL_882016, EPI_ISL_882017, EPI_ISL_882018, EPI_ISL_882019, EPI_ISL_882020, EPI_ISL_882021, EPI_ISL_882022, EPI_ISL_882023, EPI_ISL_882024, EPI_ISL_882025, EPI_ISL_882026, EPI_ISL_882027, EPI_ISL_882028, EPI_ISL_882029, EPI_ISL_882030, EPI_ISL_882031, EPI_ISL_882032, EPI_ISL_882033, EPI_ISL_882034, EPI_ISL_882035, EPI_ISL_882036, EPI_ISL_882037, EPI_ISL_882038, EPI_ISL_882039, EPI_ISL_882040, EPI_ISL_882041, EPI_ISL_882042, EPI_ISL_882043, EPI_ISL_882044, EPI_ISL_882045, EPI_ISL_882046, EPI_ISL_882047, EPI_ISL_882048, EPI_ISL_882049, EPI_ISL_882050, EPI_ISL_882051, EPI_ISL_882052, EPI_ISL_882053, EPI_ISL_882054, EPI_ISL_882055, EPI_ISL_882056, EPI_ISL_882057, EPI_ISL_882058, EPI_ISL_882059, EPI_ISL_882060, EPI_ISL_882061, EPI_ISL_882062, EPI_ISL_882063, EPI_ISL_882064, EPI_ISL_882065, EPI_ISL_882066, EPI_ISL_882067, EPI_ISL_882068, EPI_ISL_882069, EPI_ISL_882070, EPI_ISL_882071, EPI_ISL_882072, EPI_ISL_882073, EPI_ISL_882074, EPI_ISL_882075, EPI_ISL_882076, EPI_ISL_882077, EPI_ISL_882078, EPI_ISL_882079, EPI_ISL_882080, EPI_ISL_882081, EPI_ISL_882082, EPI_ISL_882083, EPI_ISL_882084, EPI_ISL_882085, EPI_ISL_882086, EPI_ISL_882087, EPI_ISL_882088, EPI_ISL_882089, EPI_ISL_882090, EPI_ISL_882091, EPI_ISL_882092, EPI_ISL_882093, EPI_ISL_882094, EPI_ISL_882095, EPI_ISL_882096, EPI_ISL_882097, EPI_ISL_882098, EPI_ISL_882099, EPI_ISL_882100, EPI_ISL_882101, EPI_ISL_882102, EPI_ISL_882103, EPI_ISL_882104, EPI_ISL_882105, EPI_ISL_882106, EPI_ISL_882107, EPI_ISL_882108, EPI_ISL_882109, EPI_ISL_882110, EPI_ISL_882111, EPI_ISL_882112, EPI_ISL_882113, EPI_ISL_882114, EPI_ISL_882115, EPI_ISL_882116, EPI_ISL_882117, EPI_ISL_882118, EPI_ISL_882119, EPI_ISL_882120, EPI_ISL_882121, EPI_ISL_882122, EPI_ISL_882123, EPI_ISL_882124, EPI_ISL_882125, EPI_ISL_882126, EPI_ISL_882127, EPI_ISL_882128, EPI_ISL_882129, EPI_ISL_882130, EPI_ISL_882131, EPI_ISL_882132, EPI_ISL_882133, EPI_ISL_882134, EPI_ISL_882135, EPI_ISL_882136, EPI_ISL_882137, EPI_ISL_882138, EPI_ISL_882139, EPI_ISL_882140, EPI_ISL_882141, EPI_ISL_882142, EPI_ISL_882143, EPI_ISL_882144, EPI_ISL_882145, EPI_ISL_882146, EPI_ISL_882147, EPI_ISL_882148, EPI_ISL_882149, EPI_ISL_882150, EPI_ISL_882151, EPI_ISL_882152, EPI_ISL_882153, EPI_ISL_882154, EPI_ISL_882155, EPI_ISL_882156, EPI_ISL_882157, EPI_ISL_882158, EPI_ISL_882159, EPI_ISL_882160, EPI_ISL_882161, EPI_ISL_882162, EPI_ISL_882163, EPI_ISL_882164, EPI_ISL_882165, EPI_ISL_882166, EPI_ISL_882167, EPI_ISL_882168, EPI_ISL_882169, EPI_ISL_882170, EPI_ISL_882171, EPI_ISL_882172, EPI_ISL_882173, EPI_ISL_882174, EPI_ISL_882175, EPI_ISL_882176, EPI_ISL_882177, EPI_ISL_882178, EPI_ISL_882179, EPI_ISL_882180, EPI_ISL_882181, EPI_ISL_882182, EPI_ISL_882183, EPI_ISL_882184, EPI_ISL_882185, EPI_ISL_882186, EPI_ISL_882187, EPI_ISL_882188, EPI_ISL_882189, EPI_ISL_882190, EPI_ISL_882191, EPI_ISL_882192, EPI_ISL_882193, EPI_ISL_882194, EPI_ISL_882195, EPI_ISL_882196, EPI_ISL_882197, EPI_ISL_882198, EPI_ISL_882199, EPI_ISL_882200, EPI_ISL_882201, EPI_ISL_882202, EPI_ISL_882203, EPI_ISL_882204, EPI_ISL_882205, EPI_ISL_882206, EPI_ISL_882207, EPI_ISL_882208, EPI_ISL_882209, EPI_ISL_882210, EPI_ISL_882211, EPI_ISL_882212, EPI_ISL_882213, EPI_ISL_882214, EPI_ISL_882215, EPI_ISL_882216, EPI_ISL_882217, EPI_ISL_882218, EPI_ISL_882219, EPI_ISL_882220, EPI_ISL_882221, EPI_ISL_882222, EPI_ISL_882223, EPI_ISL_882224, EPI_ISL_882225, EPI_ISL_882226, EPI_ISL_882227, EPI_ISL_882228, EPI_ISL_882229, EPI_ISL_882230, EPI_ISL_882231, EPI_ISL_882232, EPI_ISL_882233, EPI_ISL_882234, EPI_ISL_882235, EPI_ISL_882236, EPI_ISL_882237, EPI_ISL_882238, EPI_ISL_882239, EPI_ISL_882240, EPI_ISL_882241, EPI_ISL_882242, EPI_ISL_882243, EPI_ISL_882244, EPI_ISL_882245, EPI_ISL_882246, EPI_ISL_882247, EPI_ISL_882248, EPI_ISL_882249, EPI_ISL_882250, EPI_ISL_882251, EPI_ISL_882252, EPI_ISL_882253, EPI_ISL_882254, EPI_ISL_882255, EPI_ISL_882256, EPI_ISL_882257, EPI_ISL_882258, EPI_ISL_882259, EPI_ISL_882260, EPI_ISL_882261, EPI_ISL_882262, EPI_ISL_882263, EPI_ISL_882264, EPI_ISL_882265, EPI_ISL_882266, EPI_ISL_882267, EPI_ISL_882268, EPI_ISL_882269, EPI_ISL_882270, EPI_ISL_882271, EPI_ISL_882272, EPI_ISL_882273, EPI_ISL_882274, EPI_ISL_882275, EPI_ISL_882276, EPI_ISL_882277, EPI_ISL_882278, EPI_ISL_882279, EPI_ISL_882280, EPI_ISL_882281, EPI_ISL_882282, EPI_ISL_882283, EPI_ISL_882284, EPI_ISL_882285, EPI_ISL_882286, EPI_ISL_882287, EPI_ISL_882288, EPI_ISL_882289, EPI_ISL_882290, EPI_ISL_882291, EPI_ISL_882292, EPI_ISL_882293, EPI_ISL_882294, EPI_ISL_882295, EPI_ISL_882296, EPI_ISL_882297, EPI_ISL_882298 | | | |
| see above | Lighthouse Lab in Alderley Park | Wellcome Sanger Institute for the COVID-19 Genomics UK (COG-UK) Consortium | Jacquelyn Wynn, Mairead Hyland, The Lighthouse Lab in Alderley Park and Alex Alderton, Roberto Amato, Sonia Goncalves, Ewan Harrison, David K. Jackson, Ian Johnston, Dominic Kwiatkowski, Cordelia Langford, John Sillitoe on behalf of the Wellcome Sanger Institute COVID-19 Surveillance Team |
| EPI_ISL_882620, EPI_ISL_882624, EPI_ISL_882625, EPI_ISL_882627, EPI_ISL_882631, EPI_ISL_882632 | Hospital Ramón y Cajal | Hospital Ramón y Cajal | José M Gonzalez-Alba, Concepción Rodríguez, Melanie Abreu, Laura Martínez, Val F Lanza, Luz Leticia Olavarrieta, Rafael Cantón, JC Galán |
| EPI_ISL_882666 | UMS de Juquitiba | Instituto Adolfo Lutz, Interdiciplinary Procedures Center, Strategic Laboratory | Claudio Tavares Sacchi, Claudia Regina Gonçalves, Erica Valessa Ramos Gomes, Karoline Rodrigues Campos |
| EPI_ISL_882667 | Hospital de Clinicas Caieiras | Instituto Adolfo Lutz, Interdiciplinary Procedures Center, Strategic Laboratory | Claudio Tavares Sacchi, Claudia Regina Gonçalves, Erica Valessa Ramos Gomes, Karoline Rodrigues Campos |
| EPI_ISL_882668 | UMS de Juquitiba | Instituto Adolfo Lutz, Interdiciplinary Procedures Center, Strategic Laboratory | Claudio Tavares Sacchi, Claudia Regina Gonçalves, Erica Valessa Ramos Gomes, Karoline Rodrigues Campos |
| EPI_ISL_882963, EPI_ISL_882964, EPI_ISL_882965, EPI_ISL_882966, EPI_ISL_882986, EPI_ISL_882988, EPI_ISL_882989, EPI_ISL_882990, EPI_ISL_882991, EPI_ISL_882992, EPI_ISL_882993, EPI_ISL_882994, EPI_ISL_882995, EPI_ISL_882996 | | | |
| see above | MD Laboratories | Los Angeles County PHL | P. Hemarajata et al. |
| EPI_ISL_883012, EPI_ISL_883013, EPI_ISL_883014, EPI_ISL_883016, EPI_ISL_883017 | Maryland Public Health Laboratory | Maryland Public Health Laboratory | Maryland Department of Health Laboratories Administration |
| EPI_ISL_883428, EPI_ISL_883431, EPI_ISL_883432, EPI_ISL_883433, EPI_ISL_883434, EPI_ISL_883445, EPI_ISL_883446, EPI_ISL_883447, EPI_ISL_883471, EPI_ISL_883472, EPI_ISL_883474, EPI_ISL_883475, EPI_ISL_883476, EPI_ISL_883477, EPI_ISL_883478, EPI_ISL_883479, EPI_ISL_883483, EPI_ISL_883484, EPI_ISL_883485, EPI_ISL_883486, EPI_ISL_883491, EPI_ISL_883494, EPI_ISL_883498 | | | |
| see above | NORTHWELL HEALTH LABORATORIES | Wadsworth Center, New York State Department of Health | Kirsten St. George, Daryl M. Lamson, Alexis Russel, Matthew Shudt, Melissa A Leisner, Jonathan Plitnick, Navjot Singh, John Kelly, Erasmus Schneider, Erica Lasek-Nesselquist |
| EPI_ISL_883959 | Labo Analyses Med | National Reference Center for Viruses of Respiratory Infections, Institut Pasteur, Paris | Marion Barbet, Sylvie Behillil, Méline Bizard, Angela Brisebarre, Camille Capel, Etienne Simon-Lorière, Vincent Enouf, Maud Vanpeene, Sylvie van der Werf |
| EPI_ISL_883965, EPI_ISL_883966, EPI_ISL_883971 | Labo Analyses med | National Reference Center for Viruses of Respiratory Infections, Institut Pasteur, Paris | Marion Barbet, Sylvie Behillil, Méline Bizard, Angela Brisebarre, Camille Capel, Etienne Simon-Lorière, Vincent Enouf, Maud Vanpeene, Sylvie van der Werf,Amzalag Jonas |
| EPI_ISL_883977, EPI_ISL_883978, EPI_ISL_883982, EPI_ISL_883983, EPI_ISL_883984 | Hopital | National Reference Center for Viruses of Respiratory Infections, Institut Pasteur, Paris | Marion Barbet, Sylvie Behillil, Méline Bizard, Angela Brisebarre, Camille Capel, Etienne Simon-Lorière, Vincent Enouf, Maud Vanpeene, Sylvie van der Werf,Fourgeaud Jacques |
| EPI_ISL_883997 | Labo Analyses Med | National Reference Center for Viruses of Respiratory | Marion Barbet, Sylvie Behillil, Méline Bizard, Angela Brisebarre, Camille Capel, Etienne Simon-Lorière, Vincent Enouf, Maud Vanpeene, Sylvie van der |

| | | | |
|---|---|---|---|
| | | Infections, Institut Pasteur, Paris | Werf,Selas Labomaine |
| EPI_ISL_884004, EPI_ISL_884005 | Labo Analyses Med | National Reference Center for Viruses of Respiratory Infections, Institut Pasteur, Paris | Marion Barbet, Sylvie Behillil, Méline Bizard, Angela Brisebarre, Camille Capel, Etienne Simon-Lorière, Vincent Enouf, Maud Vanpeene, Sylvie van der Werf,Rousset Dominique |
| EPI_ISL_884006, EPI_ISL_884007 | hopital | National Reference Center for Viruses of Respiratory Infections, Institut Pasteur, Paris | Marion Barbet, Sylvie Behillil, Méline Bizard, Angela Brisebarre, Camille Capel, Etienne Simon-Lorière, Vincent Enouf, Maud Vanpeene, Sylvie van der Werf,Combe Patrice |
| EPI_ISL_884065, EPI_ISL_884066, EPI_ISL_884067, EPI_ISL_884068, EPI_ISL_884069, EPI_ISL_884070, EPI_ISL_884071, EPI_ISL_884072, EPI_ISL_884073, EPI_ISL_884074, EPI_ISL_884075, EPI_ISL_884076, EPI_ISL_884077, EPI_ISL_884078, EPI_ISL_884079 | | | |
| see above | ALBANY MEDICAL CENTER HOSPITAL CLINICAL LABORATORIES | Wadsworth Center, New York State Department of Health | Kirsten St. George, Daryl M. Lamson, Alexis Russel, Matthew Shudt, Melissa A Leisner, Jonathan Plitnick, Navjot Singh, John Kelly, Erasmus Schneider, Erica Lasek-Nesselquist |
| EPI_ISL_884223, EPI_ISL_884224, EPI_ISL_884225, EPI_ISL_884226, EPI_ISL_884242, EPI_ISL_884243, EPI_ISL_884244, EPI_ISL_884245, EPI_ISL_884246, EPI_ISL_884247 | Kansas Health and Environmental Lab | Kansas Health and Environmental Lab | Mike Grose, Paige Drury, Carissa Robertson, Ben Olsen, and Phil Adam |
| EPI_ISL_884789, EPI_ISL_884790, EPI_ISL_884791, EPI_ISL_884792, EPI_ISL_884793, EPI_ISL_884794, EPI_ISL_884795, EPI_ISL_884796, EPI_ISL_884797, EPI_ISL_884798, EPI_ISL_884799, EPI_ISL_884800, EPI_ISL_884801, EPI_ISL_884802, EPI_ISL_884803, EPI_ISL_884804, EPI_ISL_884805, EPI_ISL_884806, EPI_ISL_884807, EPI_ISL_884808, EPI_ISL_884809, EPI_ISL_884810, EPI_ISL_884811, EPI_ISL_884812, EPI_ISL_884813, EPI_ISL_884814, EPI_ISL_884815, EPI_ISL_884816, EPI_ISL_884817, EPI_ISL_884818, EPI_ISL_884819, EPI_ISL_884820, EPI_ISL_884821, EPI_ISL_884822, EPI_ISL_884823, EPI_ISL_884824, EPI_ISL_884825 | | | |
| see above | Respiratory Viruses Branch, Centers for Disease Control and Prevention | Respiratory Viruses Branch, Centers for Disease Control and Prevention | Cook,P.W., Batra,D., Rambo-Martin,B.L., de Feo,E., Antico,J., Tran,C., Tolentino,M., Wickline,S., Gietzen,K., Sickler,B., Liu,J., Allen,E., Febbo,P., Galloway,S., Washington,N.L., White,S., Levan,G., Barret,K.S., Cirulli,E., Bolze,A., Ascencio,A., Rivera-Garcia,C., Cho,R., Nguyen,J., Wang,S., Ramirez,J., Cassens,T., Sandoval,E., Isaksson,M., Lee,W., Becker,D., Laurent,M., Lu,J., Paden,C.R., Tong,S., MacCannell,D. |
| EPI_ISL_886158, EPI_ISL_886160, EPI_ISL_886174, EPI_ISL_886176, EPI_ISL_886177, EPI_ISL_886180, EPI_ISL_886181, EPI_ISL_886184, EPI_ISL_886188, EPI_ISL_886189, EPI_ISL_886191, EPI_ISL_886199, EPI_ISL_886200, EPI_ISL_886201, EPI_ISL_886204, EPI_ISL_886205, EPI_ISL_886207, EPI_ISL_886212, EPI_ISL_886218, EPI_ISL_886220, EPI_ISL_886223, EPI_ISL_886224, EPI_ISL_886233, EPI_ISL_886237, EPI_ISL_886241, EPI_ISL_886243, EPI_ISL_886244, EPI_ISL_886245, EPI_ISL_886254, EPI_ISL_886259, EPI_ISL_886263, EPI_ISL_886265, EPI_ISL_886267, EPI_ISL_886269, EPI_ISL_886271, EPI_ISL_886276, EPI_ISL_886280, EPI_ISL_886281, EPI_ISL_886289, EPI_ISL_886293, EPI_ISL_886297, EPI_ISL_886298, EPI_ISL_886302, EPI_ISL_886303, EPI_ISL_886321, EPI_ISL_886330, EPI_ISL_886331, EPI_ISL_886333, EPI_ISL_886336, EPI_ISL_886337, EPI_ISL_886338, EPI_ISL_886339, EPI_ISL_886341, EPI_ISL_886342, EPI_ISL_886345, EPI_ISL_886348, EPI_ISL_886350, EPI_ISL_886352, EPI_ISL_886353, EPI_ISL_886354, EPI_ISL_886355, EPI_ISL_886358, EPI_ISL_886363, EPI_ISL_886367, EPI_ISL_886369, EPI_ISL_886370, EPI_ISL_886371, EPI_ISL_886372, EPI_ISL_886374, EPI_ISL_886375, EPI_ISL_886376, EPI_ISL_886378, EPI_ISL_886385, EPI_ISL_886388, EPI_ISL_886391, EPI_ISL_886402, EPI_ISL_886403, EPI_ISL_886406, EPI_ISL_886409, EPI_ISL_886411, EPI_ISL_886416, EPI_ISL_886418, EPI_ISL_886423, EPI_ISL_886426, EPI_ISL_886430, EPI_ISL_886431, EPI_ISL_886432, EPI_ISL_886433, EPI_ISL_886434, EPI_ISL_886435, EPI_ISL_886436, EPI_ISL_886438, EPI_ISL_886443, EPI_ISL_886445, EPI_ISL_886447, EPI_ISL_886448, EPI_ISL_886451, EPI_ISL_886463, EPI_ISL_886464, EPI_ISL_886466, EPI_ISL_886467, EPI_ISL_886469, EPI_ISL_886473, EPI_ISL_886479, EPI_ISL_886482, EPI_ISL_886483, EPI_ISL_886487, EPI_ISL_886488, EPI_ISL_886489, EPI_ISL_886495, EPI_ISL_886496, EPI_ISL_886498, EPI_ISL_886499, EPI_ISL_886504, EPI_ISL_886510, EPI_ISL_886511, EPI_ISL_886514, EPI_ISL_886515, EPI_ISL_886518, EPI_ISL_886519, EPI_ISL_886520, EPI_ISL_886525, EPI_ISL_886528, EPI_ISL_886530, EPI_ISL_886534, EPI_ISL_886535, EPI_ISL_886547, EPI_ISL_886553, EPI_ISL_886557, EPI_ISL_886558, EPI_ISL_886564, EPI_ISL_886568, EPI_ISL_886569, EPI_ISL_886571, EPI_ISL_886572, EPI_ISL_886574, EPI_ISL_886575, EPI_ISL_886577, EPI_ISL_886584, EPI_ISL_886587, EPI_ISL_886588, EPI_ISL_886591, EPI_ISL_886594, EPI_ISL_886597, EPI_ISL_886599, EPI_ISL_886602, EPI_ISL_886606, EPI_ISL_886608, EPI_ISL_886609, EPI_ISL_886618, EPI_ISL_886620, EPI_ISL_886622, EPI_ISL_886625, EPI_ISL_886630, EPI_ISL_886632, EPI_ISL_886637, EPI_ISL_886638, EPI_ISL_886645, EPI_ISL_886646, EPI_ISL_886649, EPI_ISL_886653, EPI_ISL_886655, EPI_ISL_886656, EPI_ISL_886657, EPI_ISL_886659, EPI_ISL_886663, EPI_ISL_886668, EPI_ISL_886669, EPI_ISL_886672, EPI_ISL_886684, EPI_ISL_886685, EPI_ISL_886688, EPI_ISL_886699, EPI_ISL_886706, EPI_ISL_886707, EPI_ISL_886710, EPI_ISL_886711, EPI_ISL_886716, EPI_ISL_886719, EPI_ISL_886721, EPI_ISL_886725, EPI_ISL_886726, EPI_ISL_886728, EPI_ISL_886729, EPI_ISL_886730, EPI_ISL_886743, EPI_ISL_886745, EPI_ISL_886746, EPI_ISL_886747, EPI_ISL_886749, EPI_ISL_886751, EPI_ISL_886757, EPI_ISL_886764, EPI_ISL_886767, EPI_ISL_886771, EPI_ISL_886773, EPI_ISL_886774, EPI_ISL_886777, EPI_ISL_886778, EPI_ISL_886779, EPI_ISL_886783, EPI_ISL_886788, EPI_ISL_886794, EPI_ISL_886796, EPI_ISL_886802, EPI_ISL_886807, EPI_ISL_886813, EPI_ISL_886818, EPI_ISL_886824, EPI_ISL_886828, EPI_ISL_886829, EPI_ISL_886835, EPI_ISL_886836, EPI_ISL_886840, EPI_ISL_886842, EPI_ISL_886847, EPI_ISL_886848, EPI_ISL_886849, EPI_ISL_886850, EPI_ISL_886851, EPI_ISL_886852, EPI_ISL_886856, EPI_ISL_886861, EPI_ISL_886867, EPI_ISL_886869, EPI_ISL_886873, EPI_ISL_886874, EPI_ISL_886875, EPI_ISL_886876, EPI_ISL_886883, EPI_ISL_886885, EPI_ISL_886888, EPI_ISL_886894, EPI_ISL_886896, EPI_ISL_886901, EPI_ISL_886902, EPI_ISL_886904, EPI_ISL_886905, EPI_ISL_886906, EPI_ISL_886910, EPI_ISL_886914, EPI_ISL_886915, EPI_ISL_886926, EPI_ISL_886934, EPI_ISL_886935, EPI_ISL_886936, EPI_ISL_886940, EPI_ISL_886941, EPI_ISL_886948, EPI_ISL_886950, EPI_ISL_886952, EPI_ISL_886953, EPI_ISL_886957, EPI_ISL_886958, EPI_ISL_886959, EPI_ISL_886961, EPI_ISL_886964, EPI_ISL_886966, EPI_ISL_886969, EPI_ISL_886970, EPI_ISL_886971, EPI_ISL_886972, EPI_ISL_886974, EPI_ISL_886976, EPI_ISL_886978, EPI_ISL_886981, EPI_ISL_886983, EPI_ISL_886986, EPI_ISL_886988, EPI_ISL_887012, EPI_ISL_887017, EPI_ISL_887018, EPI_ISL_887020, EPI_ISL_887025, EPI_ISL_887028, EPI_ISL_887032, EPI_ISL_887033, EPI_ISL_887037, EPI_ISL_887041, EPI_ISL_887043, EPI_ISL_887048, EPI_ISL_887049, EPI_ISL_887050, EPI_ISL_887052, EPI_ISL_887061, EPI_ISL_887073, EPI_ISL_887082, EPI_ISL_887089, EPI_ISL_887094, EPI_ISL_887095, EPI_ISL_887097, EPI_ISL_887098, EPI_ISL_887099 | | | |
| see above | Labcorp | Genomics and Discovery, Respiratory Viruses Branch, Division of Viral Diseases, Centers for Disease Control and Prevention | Peter W. Cook,Dhwani Batra,Ben L. Rambo-Martin,Summer Galloway,Brian Krueger,Minoo Agarwal,Eyad Almasri,Debbie Boles,Ayla Burns,Nuthawin Charoensri,Oren Cohen,Susan Countryman,Mary Ann Cristobal,Bobbi Croy,Suzanne Dale,Hrushikesh Deshmukh,Amanda Douglas,Vincent Drouillon,Marcia Eisenberg,Howard Engler,Rama Ghatti,Prashant Gupta,Susan Hicks,Jake Humphrey,Lax Iyer,Manoj Jain,Mohan Kolli,Tim Kuphal,Stanley Letovsky,Michael Levandoski,Craig Lukasik,Jonathan Meltzer,Brian Norvell,Mindy Nye,Scott Parker,Christos Petropoulos,John Pruitt,Steven Ragan,Scott Ryan,Mike Sapeta,Jana Schroth,Suresh Babu Selvaraju,Goran Stevovic,Amanda Suchanek,Andrea Throop,Lyndon Tilson,Thomas Urban,Joe Voshell,Kimberly Wagner,Jonathan Williams,Mary Williamson,Qian Zeng,Tricia Zwiefelhofer,Clinton R. Paden,Suxiang Tong,Duncan MacCannell, |
| EPI_ISL_887133, EPI_ISL_887140 | Institute of Medical Microbiology and Hospital Hygiene | Institute of Medical Microbiology and Hospital Hygiene | Prof. Dr. Achim Kaasch, Aljoscha Tersteegen |
| EPI_ISL_888644, EPI_ISL_888645, EPI_ISL_888646, EPI_ISL_888647, EPI_ISL_888648, EPI_ISL_888651, EPI_ISL_888791, EPI_ISL_888792, EPI_ISL_888793, EPI_ISL_888794, EPI_ISL_888795, EPI_ISL_888796, EPI_ISL_888797, EPI_ISL_888798, EPI_ISL_888799, EPI_ISL_888800, EPI_ISL_888801, EPI_ISL_888802 | | | |
| see above | Univeristy of New Mexico Hospital | Center for Global Health, University of New Mexico Health Sciences Center | Daryl Domman, Kurt Schwalm, Justin Bacca, Jon Femling, Darrell Dinwiddie |
| EPI_ISL_888848, EPI_ISL_888849, EPI_ISL_888872, EPI_ISL_888896, EPI_ISL_888901 | Michigan Department of Health and Human Services, Bureau of Laboratories | Michigan Department of Health and Human Services, Bureau of Laboratories | Blankenship HM, Riner D, Soehnlen MK |
| EPI_ISL_889671, EPI_ISL_889672, EPI_ISL_889673, EPI_ISL_889674, EPI_ISL_889675, EPI_ISL_889676, EPI_ISL_889677, EPI_ISL_889678, EPI_ISL_889679, EPI_ISL_889680, EPI_ISL_889681, EPI_ISL_889682, EPI_ISL_889683, EPI_ISL_889684, EPI_ISL_889685 | | | |
| see above | LSUHS Emerging Viral Threat Laboratory | Microbial Genome Sequencing Center | Jeremy P. Kamil, Jennifer L. Carroll, Camille F. Abshire, Maarten Van Diest, Mohammed N.A. Siddiquey, Andrew D. Yurochko, Martin J. Sapp, Rona S. Scott, Christopher G. Kevil, Daniel J. Snyder, Vaughn S. Cooper, John A. Vanchiere |
| EPI_ISL_890195, EPI_ISL_890196, EPI_ISL_890197, EPI_ISL_890198, EPI_ISL_890199, EPI_ISL_890200, EPI_ISL_890201, EPI_ISL_890202, EPI_ISL_890203, EPI_ISL_890204, EPI_ISL_890205, EPI_ISL_890206, EPI_ISL_890207, EPI_ISL_890208, EPI_ISL_890209, EPI_ISL_890210, EPI_ISL_890211, EPI_ISL_890212, EPI_ISL_890213, EPI_ISL_890214, EPI_ISL_890215, EPI_ISL_890216, EPI_ISL_890217, EPI_ISL_890218, EPI_ISL_890219, EPI_ISL_890220, EPI_ISL_890221, EPI_ISL_890222, EPI_ISL_890223, EPI_ISL_890224, EPI_ISL_890225, EPI_ISL_890226, EPI_ISL_890227, EPI_ISL_890228, EPI_ISL_890229, EPI_ISL_890230, EPI_ISL_890231 | | | |
| see above | Cancer Biology Department, National Cancer Institute | Cancer Biology Department, National Cancer Institute | Zekri,A.N., Sedawy,M.G., Ahmed,O.S., Hafez,M.M., Soliman,H.K., Bahnassy,A.A., Elhosieny,F.W., Gad,A.E., Hamdy,M.S., Soliman,M.S., Soliman,L., Abouelhoda,M. |
| EPI_ISL_890344, EPI_ISL_890348 | KU Leuven, Rega Institute, Clinical and Epidemiological Virology | KU Leuven, Rega Institute, Clinical and Epidemiological Virology | Tony Wawina-Bokalanga, Bert Vanmechelen, Joan Marti-Carerras, Piet Maes |
| EPI_ISL_890355, EPI_ISL_890356, EPI_ISL_890357 | Hospital | National Reference Center for Viruses of Respiratory Infections, Institut Pasteur, Paris | Marion Barbet, Sylvie Behillil, Méline Bizard, Angela Brisebarre, Camille Capel, Etienne Simon-Lorière, Vincent Enouf, Maud Vanpeene, Sylvie van der Werf,Combe Patrice |
| EPI_ISL_890960, EPI_ISL_890961, EPI_ISL_890962, EPI_ISL_890963, EPI_ISL_890964, EPI_ISL_890965, EPI_ISL_890966, EPI_ISL_890968 | Seattle Flu Study | Seattle Flu Study | Deborah A. Nickerson, Chris D. Frazar, Jover Lee, Benjamin Pelle, Erica Ryke, Matthew Richardson, Amanda Adler, Elisabeth Brandstetter, Peter D. Han, Kairsten Fay, Misja Ilcisin, Kirsten Lacombe, Thomas R. Sibley, Melissa Truong, Caitlin R. Wolf, Karen Cowgill, Stephanie Schrag, Jeff Duchin, Michael Boeckh, Janet A. Englund, Michael Famulare, Barry R. Lutz, Mark J. Rieder, Lea M. Starita, Matthew Thompson, Helen Y. Chu, Trevor Bedford, Jay Shendure |
| EPI_ISL_891059 | FidaLab | Seattle Flu Study | Deborah A. Nickerson, Chris D. Frazar, Jover Lee, Benjamin Pelle, Erica Ryke, Matthew Richardson, Amanda Adler, Elisabeth Brandstetter, Peter D. Han, Kairsten Fay, Misja Ilcisin, Kirsten Lacombe, Thomas R. Sibley, Melissa Truong, Caitlin R. Wolf, Romesh Gautom, Geoff Melly, Brian Hiatt, Philip Dykema, Scott Lindquist, Michael Boeckh, Janet A. Englund, Michael Famulare, Barry R. Lutz, Mark J. Rieder, Lea M. Starita, Matthew Thompson, Helen Y. Chu, Jay Shendure, Trevor Bedford |
| EPI_ISL_891084, EPI_ISL_891085, EPI_ISL_891086, EPI_ISL_891087, EPI_ISL_891088, EPI_ISL_891089, EPI_ISL_891090, EPI_ISL_891091 | Seattle Flu Study | Seattle Flu Study | Deborah A. Nickerson, Chris D. Frazar, Jover Lee, Benjamin Pelle, Erica Ryke, Matthew Richardson, Amanda Adler, Elisabeth Brandstetter, Peter D. Han, Kairsten Fay, Misja Ilcisin, Kirsten Lacombe, Thomas R. Sibley, Melissa Truong, Caitlin R. Wolf, Karen Cowgill, Stephanie Schrag, Jeff Duchin, Michael Boeckh, Janet A. Englund, Michael Famulare, Barry R. Lutz, Mark J. Rieder, Lea M. Starita, Matthew Thompson, Helen Y. Chu, Trevor Bedford, Jay Shendure |
| EPI_ISL_891094, EPI_ISL_891095, EPI_ISL_891096, EPI_ISL_891097, EPI_ISL_891098, EPI_ISL_891099, EPI_ISL_891100, EPI_ISL_891101, EPI_ISL_891102, EPI_ISL_891103, EPI_ISL_891104, EPI_ISL_891105, EPI_ISL_891106, EPI_ISL_891107, EPI_ISL_891108, EPI_ISL_891109, EPI_ISL_891110, EPI_ISL_891111, EPI_ISL_891112, EPI_ISL_891113, EPI_ISL_891114, EPI_ISL_891115, EPI_ISL_891116, EPI_ISL_891117, EPI_ISL_891118, EPI_ISL_891119, EPI_ISL_891120, EPI_ISL_891121, EPI_ISL_891122, EPI_ISL_891123, EPI_ISL_891124, EPI_ISL_891125, EPI_ISL_891126, EPI_ISL_891127, EPI_ISL_891128, EPI_ISL_891129, EPI_ISL_891130, EPI_ISL_891131, EPI_ISL_891132, EPI_ISL_891133, EPI_ISL_891134, EPI_ISL_891135 | | | |

| | | | |
|---|---|---|---|
| see above | Altius Institute for Biomedical Sciences | Seattle Flu Study | Deborah A. Nickerson, Chris D. Frazar, Jover Lee, Benjamin Pelle, Erica Ryke, Matthew Richardson, Amanda Adler, Elisabeth Brandstetter, Peter D. Han, Kairsten Fay, Misja Ilcisin, Kirsten Lacombe, Thomas R. Sibley, Melissa Truong, Caitlin R. Wolf, Ryan Alexander, Daniel Bates, Rebecca Bruders, Stephanie DeBaun, Clem Green, Muhammad Halimun, Kneshay Harper, Matt Hartman, Andrew Meuser, Alex Nguyen, Truong Nguyen, Sofia Olsson, Sadie Patraw, Hannah Petersen, Tobias Ragoczy, Joshua Richards, Jacob Rodriguez, John Stamatoyannopoulos, Julia Wald, Olivia Waltner, Michael Boeckh, Janet A. Englund, Michael Famulare, Barry R. Lutz, Mark J. Rieder, Lea M. Starita, Matthew Thompson, Helen Y. Chu, Jay Shendure, Trevor Bedford |
| EPI_ISL_891141 | Indiana Animal Disease Diagnostic Laboratory | Carpi Laboratory - Purdue University | Jack Dorman, Ilinca I Ciubotariu, Lev Gorenstein, Abebe A Fola, G Kenitra Hendrix, Rebecca P Wilkes, Giovanna Carpi |
| EPI_ISL_892052, EPI_ISL_892053, EPI_ISL_892055, EPI_ISL_892067, EPI_ISL_892068, EPI_ISL_892069, EPI_ISL_892070, EPI_ISL_892071, EPI_ISL_892072, EPI_ISL_892073, EPI_ISL_892074, EPI_ISL_892075, EPI_ISL_892076, EPI_ISL_892077, EPI_ISL_892078, EPI_ISL_892079, EPI_ISL_892080, EPI_ISL_892081, EPI_ISL_892082, EPI_ISL_892083, EPI_ISL_892084, EPI_ISL_892085, EPI_ISL_892086, EPI_ISL_892087, EPI_ISL_892088, EPI_ISL_892089, EPI_ISL_892090, EPI_ISL_892091, EPI_ISL_892092, EPI_ISL_892093, EPI_ISL_892094, EPI_ISL_892095, EPI_ISL_892096, EPI_ISL_892097, EPI_ISL_892098, EPI_ISL_892099, EPI_ISL_892100, EPI_ISL_892101, EPI_ISL_892102, EPI_ISL_892103, EPI_ISL_892104, EPI_ISL_892105, EPI_ISL_892106, EPI_ISL_892107, EPI_ISL_892108, EPI_ISL_892109, EPI_ISL_892110, EPI_ISL_892111, EPI_ISL_892112, EPI_ISL_892113, EPI_ISL_892114, EPI_ISL_892115, EPI_ISL_892116, EPI_ISL_892117, EPI_ISL_892118, EPI_ISL_892119, EPI_ISL_892120, EPI_ISL_892121, EPI_ISL_892122, EPI_ISL_892123, EPI_ISL_892124, EPI_ISL_892125, EPI_ISL_892206 | | | |
| see above | Lighthouse Lab in Alderley Park | Wellcome Sanger Institute for the COVID-19 Genomics UK (COG-UK) Consortium | Jacquelyn Wynn, Mairead Hyland, The Lighthouse Lab in Alderley Park and Alex Alderton, Roberto Amato, Sonia Goncalves, Ewan Harrison, David K. Jackson, Ian Johnston, Dominic Kwiatkowski, Cordelia Langford, John Sillitoe on behalf of the Wellcome Sanger Institute COVID-19 Surveillance Team |
| EPI_ISL_892249, EPI_ISL_892251, EPI_ISL_892263, EPI_ISL_892264, EPI_ISL_892265 | MD Laboratories | Los Angeles County PHL | P. Hemarajata et al. |
| EPI_ISL_893214 | Indiana Animal Disease Diagnostic Laboratory | Carpi Laboratory - Purdue University | Jack Dorman, Ilinca I Ciubotariu, Lev Gorenstein, Abebe A Fola, G Kenitra Hendrix, Rebecca P Wilkes, Giovanna Carpi |
| EPI_ISL_893764, EPI_ISL_893766 | Institute of Virology, Medical Center, University of Freiburg, Freiburg, Germany | Institute of Virology, Clinial Virus Genomics, Medical Center, University of Freiburg, Freiburg, Germany | Jonas Fuchs, Lisa Kern, Sandra Reuter, Hajo Grundmann, Marcus Panning |
| EPI_ISL_894175, EPI_ISL_894177, EPI_ISL_894182, EPI_ISL_894184, EPI_ISL_894188, EPI_ISL_894190, EPI_ISL_894194, EPI_ISL_894195, EPI_ISL_894199, EPI_ISL_894200, EPI_ISL_894201, EPI_ISL_894202, EPI_ISL_894203, EPI_ISL_894204, EPI_ISL_894205, EPI_ISL_894206, EPI_ISL_894207, EPI_ISL_894208, EPI_ISL_894214, EPI_ISL_894215, EPI_ISL_894217 | | | |
| see above | KU Leuven, Rega Institute, Clinical and Epidemiological Virology | KU Leuven, Rega Institute, Clinical and Epidemiological Virology | Tony Wawina-Bokalanga, Bert Vanmechelen, Joan Marti-Carerras, Piet Maes |
| EPI_ISL_896069 | Labormedizinisches Zentrum Dr Risch | University Hospital Basel, Clinical Bacteriology | Tim Roloff, Madlen Stange, Helena MB Seth-Smith, Alfredo Mari, Karoline Leuzinger, Julia Bielicki, Nadia Wohlwend,Martin Risch, Lorenz Risch, Manuel Battegay, Hans Hirsch, Adrian Egli |
| EPI_ISL_896085 | Viollier AG | University Hospital Basel, Clinical Bacteriology | Tim Roloff, Madlen Stange, Helena MB Seth-Smith, Alfredo Mari, Karoline Leuzinger, Julia Bielicki, Christiane Beckmann, Manuel Battegay, Hans Hirsch, Adrian Egli |
| EPI_ISL_896099 | Labormedizinisches Zentrum Dr Risch | University Hospital Basel, Clinical Bacteriology | Tim Roloff, Madlen Stange, Helena MB Seth-Smith, Alfredo Mari, Karoline Leuzinger, Julia Bielicki, Nadia Wohlwend,Martin Risch, Lorenz Risch, Manuel Battegay, Hans Hirsch, Adrian Egli |
| EPI_ISL_896149, EPI_ISL_896158 | MEPHI, Aix Marseille University | MEPHI, Aix Marseille University | Anthony LEVASSEUR |
| EPI_ISL_896216, EPI_ISL_896227, EPI_ISL_896234, EPI_ISL_896244, EPI_ISL_896245, EPI_ISL_896246 | Columbia University Irving Medical Center | Wadsworth Center, New York State Department of Health | Kirsten St. George, Daryl M. Lamson, Alexis Russel, Matthew Shudt, Melissa A Leisner, Jonathan Plitnick, Navjot Singh, John Kelly, Erasmus Schneider, Erica Lasek-Nesselquist |
| EPI_ISL_896300 | BIO-REFERENCE LABORATORIES | Wadsworth Center, New York State Department of Health | Kirsten St. George, Daryl M. Lamson, Alexis Russel, Matthew Shudt, Melissa A Leisner, Jonathan Plitnick, Navjot Singh, John Kelly, Erasmus Schneider, Erica Lasek-Nesselquist |
| EPI_ISL_896390, EPI_ISL_896417, EPI_ISL_896418, EPI_ISL_896421 | URMC LABS | Wadsworth Center, New York State Department of Health | Kirsten St. George, Daryl M. Lamson, Alexis Russel, Matthew Shudt, Melissa A Leisner, Jonathan Plitnick, Navjot Singh, John Kelly, Erasmus Schneider, Erica Lasek-Nesselquist |
| EPI_ISL_896488 | KU Leuven, Rega Institute, Clinical and Epidemiological Virology | KU Leuven, Rega Institute, Clinical and Epidemiological Virology | Tony Wawina-Bokalanga, Bert Vanmechelen, Joan Marti-Carerras, Piet Maes |
| EPI_ISL_897687, EPI_ISL_897937 | University Hospitals of Geneva, Laboratory of Virology | HUG, Laboratory of Virology and the Health2030 Genome Center | Samuel Cordey, Ana Rita Goncalves, Laurent Kaiser, Lorenzo Cerutti, Henri Pegeot, Melyssa Elies, Deborah Penet, Keith Harshman, Ioannis Xenarios, Emmanouil Dermitzakis |
| EPI_ISL_898009, EPI_ISL_898024, EPI_ISL_898030, EPI_ISL_898031 | KU Leuven, Rega Institute, Clinical and Epidemiological Virology | KU Leuven, Rega Institute, Clinical and Epidemiological Virology | Tony Wawina-Bokalanga, Bert Vanmechelen, Joan Marti-Carerras, Piet Maes |
| EPI_ISL_899001, EPI_ISL_899012, EPI_ISL_899013, EPI_ISL_899014, EPI_ISL_899018, EPI_ISL_899037, EPI_ISL_899040, EPI_ISL_899055, EPI_ISL_899061, EPI_ISL_899072, EPI_ISL_899077, EPI_ISL_899083, EPI_ISL_899089, EPI_ISL_899099, EPI_ISL_899103, EPI_ISL_899104, EPI_ISL_899158, EPI_ISL_899174, EPI_ISL_899250, EPI_ISL_899251, EPI_ISL_899252, EPI_ISL_899253, EPI_ISL_899254, EPI_ISL_899255, EPI_ISL_899256, EPI_ISL_899257, EPI_ISL_899258, EPI_ISL_899259, EPI_ISL_899260, EPI_ISL_899354, EPI_ISL_899355, EPI_ISL_899356, EPI_ISL_899357, EPI_ISL_899358, EPI_ISL_899359, EPI_ISL_899361, EPI_ISL_899362, EPI_ISL_899363, EPI_ISL_899364, EPI_ISL_899365, EPI_ISL_899366, EPI_ISL_899367, EPI_ISL_899368, EPI_ISL_899369, EPI_ISL_899479, EPI_ISL_899480, EPI_ISL_899481, EPI_ISL_899482, EPI_ISL_899483, EPI_ISL_899484, EPI_ISL_899485, EPI_ISL_899486, EPI_ISL_899487, EPI_ISL_899488, EPI_ISL_899489, EPI_ISL_899490, EPI_ISL_899491, EPI_ISL_899492, EPI_ISL_899493, EPI_ISL_899494, EPI_ISL_899495, EPI_ISL_899496, EPI_ISL_899497, EPI_ISL_899555, EPI_ISL_899556, EPI_ISL_899641, EPI_ISL_899642, EPI_ISL_899739, EPI_ISL_899740, EPI_ISL_899741, EPI_ISL_899742, EPI_ISL_899743, EPI_ISL_899744, EPI_ISL_899745, EPI_ISL_899746, EPI_ISL_899747, EPI_ISL_899748, EPI_ISL_899749, EPI_ISL_899794, EPI_ISL_899795, EPI_ISL_899796, EPI_ISL_899797, EPI_ISL_899798, EPI_ISL_899799, EPI_ISL_899846, EPI_ISL_899886, EPI_ISL_899887, EPI_ISL_899888, EPI_ISL_900024, EPI_ISL_900025, EPI_ISL_900026, EPI_ISL_900027, EPI_ISL_900028, EPI_ISL_900029 | | | |
| see above | Viollier AG | Department of Biosystems Science and Engineering, ETH Zürich | Chaoran Chen, Sarah Nadeau, Ivan Topolsky, Emmanouil Dermitzakis, Keith Harshman, Ioannis Xenarios, Henri Pegeot, Lorenzo Cerutti, Deborah Penet, Philipp Jablonski, Lara Fuhrmann, David Dreifuss, Katharina Jahn, Christiane Beckmann, Maurice Redondo, Olivier Kobel, Christoph Noppen, Sophie Seidel, Noemie Santamaria de Souza, Niko Beerenwinkel, Tanja Stadler |
| EPI_ISL_902894, EPI_ISL_902895, EPI_ISL_902896, EPI_ISL_902897, EPI_ISL_902898, EPI_ISL_902899, EPI_ISL_902900, EPI_ISL_902901, EPI_ISL_902902, EPI_ISL_902903, EPI_ISL_902904, EPI_ISL_902905, EPI_ISL_902906, EPI_ISL_902907, EPI_ISL_902908, EPI_ISL_902909, EPI_ISL_902910, EPI_ISL_902911, EPI_ISL_902912, EPI_ISL_902913, EPI_ISL_902914, EPI_ISL_902915 | | | |
| see above | Department of Virology and Immunology, University of Helsinki and Helsinki University Hospital, Huslab Finland | Department of Virology, Faculty of Medicine, University of Helsinki, Helsinki, Finland | Teemu Smura, Ravi Kant, Phuoc Truong, Hussein Alburkat, Hannimari Kallio-Kokko, Jenni Virtanen, Maija Suvanto, Essi Korhonen, Sari Hannula, Harri Kangas, Hanna Liimatainen, Satu Kurkela, Hanna Jarva, Maija Lappalainen, Pekka Ellonen, Olli Vapalahti |
| EPI_ISL_902925 | MD Laboratories | Los Angeles County PHL | P. Hemarajata et al. |
| EPI_ISL_902926 | SC (UCO) Igiene e Sanità Pubblica (funzione integrata con SC Microbiologia e Virologia) e Laboratory of Molecular Virology of the International Centre for Genetic Engineering and Biotechnology (ICGEB) | ARGO Laboratorio Genomica ed Epigenomica | Licastro D, Dal Monego S, Degasperi M, Marcello A, D'Agaro P |
| EPI_ISL_903081, EPI_ISL_903084 | Seattle Flu Study | Seattle Flu Study | Deborah A. Nickerson, Chris D. Frazar, Jover Lee, Benjamin Pelle, Erica Ryke, Matthew Richardson, Amanda Adler, Elisabeth Brandstetter, Peter D. Han, Kairsten Fay, Misja Ilcisin, Kirsten Lacombe, Thomas R. Sibley, Melissa Truong, Caitlin R. Wolf, Michael Boeckh, Janet A. Englund, Michael Famulare, Barry R. Lutz, Mark J. Rieder, Lea M. Starita, Matthew Thompson, Jay Shendure, Trevor Bedford, Helen Y. Chu |
| EPI_ISL_903085, EPI_ISL_903086, EPI_ISL_903204, EPI_ISL_903206, EPI_ISL_903207 | Seattle Flu Study | Seattle Flu Study | Deborah A. Nickerson, Chris D. Frazar, Jover Lee, Benjamin Pelle, Erica Ryke, Matthew Richardson, Amanda Adler, Elisabeth Brandstetter, Peter D. Han, Kairsten Fay, Misja Ilcisin, Kirsten Lacombe, Thomas R. Sibley, Melissa Truong, Caitlin R. Wolf, Karen Cowgill, Stephanie Schrag, Jeff Duchin, Michael Boeckh, Janet A. Englund, Michael Famulare, Barry R. Lutz, Mark J. Rieder, Lea M. Starita, Matthew Thompson, Helen Y. Chu, Trevor Bedford, Jay Shendure |
| EPI_ISL_903357 | Wyoming Public Health Laboratory | Wyoming Public Health Laboratory | Noah Hull, Taylor Fearing, Lynette Gumbleton, Channing Weber, Ashley Norberg, Bailey Bowcutt, and Wanda Manley |
| EPI_ISL_903373, EPI_ISL_903374 | MD Laboratories | Los Angeles County PHL | P. Hemarajata et al. |
| EPI_ISL_903390, EPI_ISL_903391, EPI_ISL_903392, EPI_ISL_903393, EPI_ISL_903395, EPI_ISL_903396, EPI_ISL_903397, EPI_ISL_903399, EPI_ISL_903401, EPI_ISL_903402, EPI_ISL_903403, EPI_ISL_903404, EPI_ISL_903405, EPI_ISL_903407, EPI_ISL_903408, EPI_ISL_903409, EPI_ISL_903410, EPI_ISL_903411, EPI_ISL_903412, EPI_ISL_903413, EPI_ISL_903414, EPI_ISL_903415, EPI_ISL_903416, EPI_ISL_903418, EPI_ISL_903419, EPI_ISL_903420, EPI_ISL_903422, EPI_ISL_903424, EPI_ISL_903426, EPI_ISL_903427, EPI_ISL_903428, EPI_ISL_903429, EPI_ISL_903430, EPI_ISL_903431, EPI_ISL_903432, EPI_ISL_903433, EPI_ISL_903434, EPI_ISL_903435, EPI_ISL_903436, EPI_ISL_903438, EPI_ISL_903439, EPI_ISL_903440, EPI_ISL_903441, EPI_ISL_903442, EPI_ISL_903443, EPI_ISL_903445, EPI_ISL_903446, EPI_ISL_903447, EPI_ISL_903448, EPI_ISL_903449, EPI_ISL_903450, EPI_ISL_903451, EPI_ISL_903452, EPI_ISL_903453, EPI_ISL_903454, EPI_ISL_903455, EPI_ISL_903456, EPI_ISL_903457, EPI_ISL_903458, EPI_ISL_903460, EPI_ISL_903461, EPI_ISL_903462, EPI_ISL_903463, EPI_ISL_903464, EPI_ISL_903465, EPI_ISL_903466, EPI_ISL_903468, EPI_ISL_903469, EPI_ISL_903470, EPI_ISL_903471, EPI_ISL_903472, EPI_ISL_903474, | | | |

| | | | |
|---|---|---|---|
| EPI_ISL_903475, EPI_ISL_903476, EPI_ISL_903477, EPI_ISL_903479, EPI_ISL_903480, EPI_ISL_903481, EPI_ISL_903482, EPI_ISL_903484, EPI_ISL_903485, EPI_ISL_903486, EPI_ISL_903487, EPI_ISL_903488, EPI_ISL_903489, EPI_ISL_903490, EPI_ISL_903491, EPI_ISL_903492, EPI_ISL_903493, EPI_ISL_903494, EPI_ISL_903495, EPI_ISL_903496, EPI_ISL_903497, EPI_ISL_903499, EPI_ISL_903500, EPI_ISL_903501, EPI_ISL_903502, EPI_ISL_903503, EPI_ISL_903505, EPI_ISL_903506, EPI_ISL_903507, EPI_ISL_903508, EPI_ISL_903509, EPI_ISL_903510, EPI_ISL_903511, EPI_ISL_903512, EPI_ISL_903513, EPI_ISL_903514, EPI_ISL_903515, EPI_ISL_903516, EPI_ISL_903517, EPI_ISL_903518, EPI_ISL_903519, EPI_ISL_903520, EPI_ISL_903521, EPI_ISL_903522, EPI_ISL_903523, EPI_ISL_903524, EPI_ISL_903525, EPI_ISL_903552, EPI_ISL_903554, EPI_ISL_903555, EPI_ISL_903556, EPI_ISL_903557, EPI_ISL_903559, EPI_ISL_903560, EPI_ISL_903561, EPI_ISL_903562 | | | |
| see above | Quest Diagnostics | Quest Diagnostics | Rosenthal,S.H., Gerasimova,A., Kagan,R.M., Anderson, B., Hua, M., Liu Y., Bernstein, L.E., Livingston, K.E., Perez, A., Shalhout, D.F., Shlyakhter, I.A., Owen, R., Tanpaiboon, P., Lacbawan, F. |
| EPI_ISL_903979 | Institute of Virology, Biomedical Research Center of the Slovak Academy of Sciences, Bratislava | Faculty of Natural Sciences, Comenius University, Bratislava | Viktória abanová, Kristína Boršová, Broa Brejová, Viktória Hodorová, Sabina Fumaová Havlíková, Juraj Kopáek, Martina Liková, ubomíra Lukáiková, Martina Neboháová, Monika Sláviková, Tomáš Vina, Jozef Nosek, Boris Klempa |
| EPI_ISL_904064, EPI_ISL_904065, EPI_ISL_904066, EPI_ISL_904067 | New Mexico Department of Health Scientific Laboratory | New Mexico Department of Health Scientific Laboratory | Ellie Johnson, Anastacia Griego-Fisher, D'eldra Malone |
| EPI_ISL_904120, EPI_ISL_904121 | LACEN - Laboratório Central de Saúde Pública do Pará | Evandro Chagas Institute | Santos, M.C.; Silva, A.M.; Junior, W.D.C.; Barbagelata, L.S.; Ferreira, J.A.; Sousa, E.M.A.; da Silva, P.S.; Pinheiro, K.C.; L.C.; Sousa Junior, E.C. |
| EPI_ISL_904127, EPI_ISL_904129 | Quest Diagnostics | Quest Diagnostics | Rosenthal,S.H., Gerasimova,A., Kagan,R.M., Anderson, B., Hua, M., Liu Y., Bernstein, L.E., Livingston, K.E., Perez, A., Shalhout, D.F., Shlyakhter, I.A., Owen, R., Tanpaiboon, P., Lacbawan, F. |
| EPI_ISL_904138, EPI_ISL_904141, EPI_ISL_904142, EPI_ISL_904152, EPI_ISL_904157, EPI_ISL_904173, EPI_ISL_904184, EPI_ISL_904204, EPI_ISL_904205, EPI_ISL_904206, EPI_ISL_904232, EPI_ISL_904255, EPI_ISL_904273, EPI_ISL_904274, EPI_ISL_904311, EPI_ISL_904312, EPI_ISL_904313, EPI_ISL_904368, EPI_ISL_904377, EPI_ISL_904565, EPI_ISL_904566, EPI_ISL_904567, EPI_ISL_904568, EPI_ISL_904569, EPI_ISL_904570, EPI_ISL_904571, EPI_ISL_904572, EPI_ISL_904573, EPI_ISL_904574, EPI_ISL_904575, EPI_ISL_904576, EPI_ISL_904577, EPI_ISL_904578 | | | |
| see above | Dutch COVID-19 response team | Erasmus Medical Center | Bas Oude Munnink, Reina Sikkema, David Nieuwenhuijse, Irina Chestakova, Anne van der Linden, Marjan Boter, Emmanuelle Munger, Corine GeurtsvanKessel, Annemiek van der Eijk, Richard Molenkamp, Marion Koopmans, on behalf of the Dutch national COVID-19 response team. |
| EPI_ISL_904627 | Servicio de Microbiología, Laboratori Clínic Metropolitana Nord. Hospital Universitari Germans Trias i Pujol. Institut d'Investigació en Ciències de la Salut Germans Trias i Pujol (IGTP) | IrsiCaixa - Can Ruti CovidSeq | Marc Noguera-Julian, Mariona Parera, Maria Casadellà, Pilar Armengol, Francesc Catala-Moll, Roger Paredes, Bonaventura Clotet Elisa Martró, Verónica Saludes, Anna Not, Ana Pérez, Montserrat Giménez, Ignacio Blanco, Cristina Casañ, Antoni E. Bordoy, Adrián Antuori |
| EPI_ISL_904665, EPI_ISL_904668, EPI_ISL_904678, EPI_ISL_904715, EPI_ISL_904723, EPI_ISL_904725, EPI_ISL_904726, EPI_ISL_904727, EPI_ISL_904764, EPI_ISL_904800, EPI_ISL_904872, EPI_ISL_904974, EPI_ISL_904986, EPI_ISL_904988, EPI_ISL_904990, EPI_ISL_904997, EPI_ISL_904998, EPI_ISL_904999, EPI_ISL_905001, EPI_ISL_905002, EPI_ISL_905050, EPI_ISL_905059, EPI_ISL_905060, EPI_ISL_905071, EPI_ISL_905110, EPI_ISL_905113, EPI_ISL_905116, EPI_ISL_905122, EPI_ISL_905185, EPI_ISL_905186, EPI_ISL_905188, EPI_ISL_905195, EPI_ISL_905196, EPI_ISL_905243, EPI_ISL_905244, EPI_ISL_905254, EPI_ISL_905260, EPI_ISL_905261, EPI_ISL_905264, EPI_ISL_905265, EPI_ISL_905266, EPI_ISL_905274, EPI_ISL_905280, EPI_ISL_905283, EPI_ISL_905291, EPI_ISL_905312, EPI_ISL_905313, EPI_ISL_905314, EPI_ISL_905315, EPI_ISL_905321, EPI_ISL_905322, EPI_ISL_905323, EPI_ISL_905324, EPI_ISL_905327, EPI_ISL_905328, EPI_ISL_905329, EPI_ISL_905333, EPI_ISL_905334, EPI_ISL_905335, EPI_ISL_905336, EPI_ISL_905337, EPI_ISL_905345, EPI_ISL_905350, EPI_ISL_905355, EPI_ISL_905356, EPI_ISL_905357, EPI_ISL_905358, EPI_ISL_905359, EPI_ISL_905577, EPI_ISL_905578, EPI_ISL_905579, EPI_ISL_905580, EPI_ISL_905581, EPI_ISL_905583, EPI_ISL_905584, EPI_ISL_905586, EPI_ISL_905589, EPI_ISL_905590, EPI_ISL_905591, EPI_ISL_905617, EPI_ISL_905618, EPI_ISL_905619, EPI_ISL_905620, EPI_ISL_905621, EPI_ISL_905623, EPI_ISL_905624, EPI_ISL_905625, EPI_ISL_905626, EPI_ISL_905627, EPI_ISL_905628, EPI_ISL_905637, EPI_ISL_905638, EPI_ISL_905639, EPI_ISL_905640, EPI_ISL_905667, EPI_ISL_905668, EPI_ISL_905669, EPI_ISL_905670, EPI_ISL_905677, EPI_ISL_905678, EPI_ISL_905679, EPI_ISL_905680, EPI_ISL_905700, EPI_ISL_905701, EPI_ISL_905702, EPI_ISL_905703, EPI_ISL_905704, EPI_ISL_905705, EPI_ISL_905706, EPI_ISL_905707, EPI_ISL_905708, EPI_ISL_905716, EPI_ISL_905717, EPI_ISL_905723, EPI_ISL_905724 | | | |
| see above | Dutch COVID-19 response team | National Institute for Public Health and the Environment (RIVM) | Adam Meijer, Harry Vennema, Dirk Eggink, Jeroen Cremer, Sharon van den Brink, Bas van der Veer, AnneMarie van den Brandt, Florian Zwagemaker, Dennis Schmitz, Chantal Reusken, on behalf of the national COVID-19 response team |
| EPI_ISL_905765, EPI_ISL_905766, EPI_ISL_905767, EPI_ISL_905768, EPI_ISL_905769, EPI_ISL_905770, EPI_ISL_905771, EPI_ISL_905772 | MD Laboratories | Los Angeles County PHL | P. Hemarajata et al. |
| EPI_ISL_906047, EPI_ISL_906048 | MD Laboratories | Los Angeles County Public Health Laboratory | P. Hemarajata et al. |
| EPI_ISL_906144 | Laboratorio de Salud Publica de Amazonas | Instituto Nacional de Salud- Dirección de Investigación en Salud Pública, Universidad de los Andes- Applied genomics research group, Vicerrectoria de Investigación y Creación, Universidad de los Andes- Systems and Computing Engineering Department | Katherine Laiton-Donato, Diego A. Álvarez-Díaz, Carlos Franco-Muñoz, Mauricio Pacheco-Montealegre, Héctor Alejandro Ruiz-Moreno, Maria T. Herrera-Sepúlveda, Diego Andrés Prada, Jhonnatan Reales-González, Sheryll Corchuelo, Julian Naizaque, Gerardo Santamaría Jorge Duitama, Laura Natalia Gonzalez, Jorge Ivan Diaz, Silvia Restrepo-Restrepo, Magdalena Wiesner, Martha Lucia Ospina Martinez, Marcela Mercado-Reyes |
| EPI_ISL_906307, EPI_ISL_906308, EPI_ISL_906309, EPI_ISL_906310, EPI_ISL_906311, EPI_ISL_906312, EPI_ISL_906313, EPI_ISL_906314, EPI_ISL_906316, EPI_ISL_906317, EPI_ISL_906318, EPI_ISL_906319, EPI_ISL_906320, EPI_ISL_906321, EPI_ISL_906322, EPI_ISL_906323, EPI_ISL_906324, EPI_ISL_906325, EPI_ISL_906326, EPI_ISL_906327, EPI_ISL_906328, EPI_ISL_906329, EPI_ISL_906330, EPI_ISL_906331, EPI_ISL_906332, EPI_ISL_906333, EPI_ISL_906334, EPI_ISL_906335, EPI_ISL_906336, EPI_ISL_906337, EPI_ISL_906338, EPI_ISL_906340, EPI_ISL_906341, EPI_ISL_906343, EPI_ISL_906344, EPI_ISL_906345, EPI_ISL_906346, EPI_ISL_906347, EPI_ISL_906348, EPI_ISL_906349, EPI_ISL_906350, EPI_ISL_906351, EPI_ISL_906352, EPI_ISL_906353, EPI_ISL_906354, EPI_ISL_906355, EPI_ISL_906356, EPI_ISL_906357, EPI_ISL_906358, EPI_ISL_906359, EPI_ISL_906360, EPI_ISL_906361, EPI_ISL_906362, EPI_ISL_906363, EPI_ISL_906364, EPI_ISL_906366, EPI_ISL_906367, EPI_ISL_906368, EPI_ISL_906370, EPI_ISL_906371, EPI_ISL_906372, EPI_ISL_906373, EPI_ISL_906374, EPI_ISL_906375, EPI_ISL_906376, EPI_ISL_906377, EPI_ISL_906378, EPI_ISL_906379, EPI_ISL_906380, EPI_ISL_906381, EPI_ISL_906382, EPI_ISL_906383, EPI_ISL_906384, EPI_ISL_906385, EPI_ISL_906386, EPI_ISL_906387, EPI_ISL_906388, EPI_ISL_906390, EPI_ISL_906391, EPI_ISL_906392, EPI_ISL_906393, EPI_ISL_906395, EPI_ISL_906397, EPI_ISL_906398, EPI_ISL_906399, EPI_ISL_906400, EPI_ISL_906401, EPI_ISL_906402, EPI_ISL_906404, EPI_ISL_906405, EPI_ISL_906406, EPI_ISL_906407, EPI_ISL_906408, EPI_ISL_906409, EPI_ISL_906410, EPI_ISL_906411, EPI_ISL_906412, EPI_ISL_906413, EPI_ISL_906414, EPI_ISL_906415, EPI_ISL_906416, EPI_ISL_906417, EPI_ISL_906418, EPI_ISL_906419, EPI_ISL_906420, EPI_ISL_906421, EPI_ISL_906422, EPI_ISL_906423, EPI_ISL_906433, EPI_ISL_906436, EPI_ISL_906448, EPI_ISL_906454, EPI_ISL_906455, EPI_ISL_906456, EPI_ISL_906457, EPI_ISL_906458, EPI_ISL_906461, EPI_ISL_906462, EPI_ISL_906463 | | | |
| see above | Quest Diagnostics | Quest Diagnostics | Rosenthal,S.H., Gerasimova,A., Kagan,R.M., Anderson, B., Hua, M., Liu Y., Bernstein, L.E., Livingston, K.E., Perez, A., Shalhout, D.F., Shlyakhter, I.A., Owen, R., Tanpaiboon, P., Lacbawan, F. |
| EPI_ISL_906534, EPI_ISL_906560, EPI_ISL_906561 | Laboratorio de Salud Publica de Amazonas | Instituto Nacional de Salud- Dirección de Investigación en Salud Pública, Universidad de los Andes- Applied genomics research group, Vicerrectoria de Investigación y Creación, Universidad de los Andes- Systems and Computing Engineering Department | Katherine Laiton-Donato, Diego A. Álvarez-Díaz, Carlos Franco-Muñoz, Mauricio Pacheco-Montealegre, Héctor Alejandro Ruiz-Moreno, Maria T. Herrera-Sepúlveda, Diego Andrés Prada, Jhonnatan Reales-González, Sheryll Corchuelo, Julian Naizaque, Gerardo Santamaría Jorge Duitama, Laura Natalia Gonzalez, Jorge Ivan Diaz, Silvia Restrepo-Restrepo, Magdalena Wiesner, Martha Lucia Ospina Martinez, Marcela Mercado-Reyes |
| EPI_ISL_906575, EPI_ISL_906576, EPI_ISL_906577, EPI_ISL_906578, EPI_ISL_906579, EPI_ISL_906580, EPI_ISL_906581, EPI_ISL_906582, EPI_ISL_906583, EPI_ISL_906584, EPI_ISL_906585, EPI_ISL_906586, EPI_ISL_906587, EPI_ISL_906588, EPI_ISL_906589 | | | |
| see above | Maine Health and Environmental Testing Laboratory (Maine HETL) | Tewhey Lab, The Jackson Laboratory | Matluk,N., Dewey,H., Iosue,F., Barter,M., Lynch,R., Munger,H. and Tewhey,R. |
| EPI_ISL_906715, EPI_ISL_906717, EPI_ISL_906721, EPI_ISL_906723, EPI_ISL_906724, EPI_ISL_906729, EPI_ISL_906731, EPI_ISL_906733, EPI_ISL_906735, EPI_ISL_906738, EPI_ISL_906748, EPI_ISL_906749, EPI_ISL_906756 | | | |
| see above | Hematology Laboratory, Section of Molecular Diagnostics, University Clinical Centre, Medical University of Gdansk | Laboratory of Recombinant Vaccines | Lukasz Rabalski, Maciej Kosinski, Maciej Grzybek, Adam Sodol, Aneta Szulc, Krzysztof Lewandowski, Ewa Milosz, Marlena Robakowska, Boguslaw Szewczyk, Krystyna Bienkowska-Szewczyk |
| EPI_ISL_906779 | New Mexico Department of Health Scientific Laboratory | New Mexico Department of Health Scientific Laboratory | Ellie Johnson, Anastacia Griego-Fisher, D'eldra Malone |
| EPI_ISL_906805 | Vestfold Hospital, Toensberg Department of Microbiology | Norwegian Institute of Public Health, Department of Virology | Kathrine Stene-Johansen, Kamilla Heddeland Instefjord, Hilde Elshaug, Atiya R Ali,Marie Paulsen Madsen, Rasmus Riis Kopperud, Hilde Vollan, Karoline Bragstad, Olav Hungnes |
| EPI_ISL_906905, EPI_ISL_906906, EPI_ISL_906907, EPI_ISL_906908, EPI_ISL_906909, EPI_ISL_906915, EPI_ISL_906919 | Bureau of Public Health Laboratories, Florida Department of Health (BPHL, FLDOH) | Bureau of Public Health Laboratories, Florida Department of Health (BPHL, FLDOH) | Schmedes,S., Blanton,J. |
| EPI_ISL_906920, EPI_ISL_906921, EPI_ISL_906922, EPI_ISL_906923, EPI_ISL_906925, EPI_ISL_906926, EPI_ISL_906927, EPI_ISL_906928, EPI_ISL_906929, EPI_ISL_906931, EPI_ISL_906932, EPI_ISL_906933, EPI_ISL_906934, EPI_ISL_906935, EPI_ISL_906936, EPI_ISL_906937, EPI_ISL_906938, EPI_ISL_906939, EPI_ISL_906940, EPI_ISL_906941, EPI_ISL_906942, EPI_ISL_906943, EPI_ISL_906944, EPI_ISL_906945, EPI_ISL_906946, EPI_ISL_906947, EPI_ISL_906948, EPI_ISL_906949, EPI_ISL_906950, EPI_ISL_906951, EPI_ISL_906952, EPI_ISL_906953, EPI_ISL_906955, EPI_ISL_906956, EPI_ISL_906957, EPI_ISL_906958, EPI_ISL_906959, EPI_ISL_906960, EPI_ISL_906961, EPI_ISL_906962, EPI_ISL_906963, EPI_ISL_906965, EPI_ISL_906966, EPI_ISL_906967, EPI_ISL_906968, EPI_ISL_906969, EPI_ISL_906970, EPI_ISL_906971, EPI_ISL_906972, EPI_ISL_906973, EPI_ISL_906974, EPI_ISL_906975, EPI_ISL_906976, EPI_ISL_906977, EPI_ISL_906979, EPI_ISL_906980, EPI_ISL_906981, EPI_ISL_906982, EPI_ISL_906983, EPI_ISL_906984, EPI_ISL_906985, EPI_ISL_906986, EPI_ISL_906987, EPI_ISL_906988, EPI_ISL_906989, EPI_ISL_906990, EPI_ISL_906991, EPI_ISL_906992, EPI_ISL_906993, EPI_ISL_906994, EPI_ISL_906995, EPI_ISL_906996, EPI_ISL_906997, EPI_ISL_906998, EPI_ISL_906999, EPI_ISL_907000, EPI_ISL_907001, EPI_ISL_907002, EPI_ISL_907003, EPI_ISL_907004, EPI_ISL_907005, EPI_ISL_907006, EPI_ISL_907007, EPI_ISL_907008, EPI_ISL_907009, EPI_ISL_907010, EPI_ISL_907011, EPI_ISL_907012, EPI_ISL_907013, EPI_ISL_907014, EPI_ISL_907015, EPI_ISL_907016, EPI_ISL_907017, EPI_ISL_907018, EPI_ISL_907019, EPI_ISL_907020, EPI_ISL_907021, EPI_ISL_907022, EPI_ISL_907023, EPI_ISL_907024, EPI_ISL_907025, EPI_ISL_907026, EPI_ISL_907027, EPI_ISL_907028, EPI_ISL_907029, EPI_ISL_907030, EPI_ISL_907031, EPI_ISL_907032, | | | |

| | | | |
|---|---|---|---|
| EPI_ISL_907033, EPI_ISL_907034, EPI_ISL_907035, EPI_ISL_907036, EPI_ISL_907037, EPI_ISL_907038, EPI_ISL_907039, EPI_ISL_907040, EPI_ISL_907041, EPI_ISL_907042, EPI_ISL_907043, EPI_ISL_907044, EPI_ISL_907045, EPI_ISL_907046, EPI_ISL_907047, EPI_ISL_907048, EPI_ISL_907049, EPI_ISL_907050, EPI_ISL_907051, EPI_ISL_907052, EPI_ISL_907053, EPI_ISL_907054, EPI_ISL_907055, EPI_ISL_907056, EPI_ISL_907057, EPI_ISL_907058, EPI_ISL_907059, EPI_ISL_907060, EPI_ISL_907061, EPI_ISL_907062, EPI_ISL_907063, EPI_ISL_907065, EPI_ISL_907067, EPI_ISL_907069, EPI_ISL_907071, EPI_ISL_907072, EPI_ISL_907073 | | | |
| see above | Infectious Diseases, Quest Diagnostics | Infectious Diseases, Quest Diagnostics | Rosenthal,S.H., Gerasimova,A., Kagan,R.M., Anderson,B., Bernstein,L.E., Livingston,K.E., Hua,M., Liu,Y., Shalhout,D.F., Owen,R., Lacbawan,F. |
| EPI_ISL_909595, EPI_ISL_909596, EPI_ISL_909597, EPI_ISL_909598, EPI_ISL_909599, EPI_ISL_909600, EPI_ISL_909601, EPI_ISL_909602, EPI_ISL_909603, EPI_ISL_909604, EPI_ISL_909605, EPI_ISL_909606, EPI_ISL_909607, EPI_ISL_909608, EPI_ISL_909609, EPI_ISL_909610, EPI_ISL_909611, EPI_ISL_909612, EPI_ISL_909613, EPI_ISL_909614, EPI_ISL_909615, EPI_ISL_909616, EPI_ISL_909617, EPI_ISL_909618, EPI_ISL_909619, EPI_ISL_909620, EPI_ISL_909621, EPI_ISL_909622, EPI_ISL_909623, EPI_ISL_909624, EPI_ISL_909625, EPI_ISL_909626, EPI_ISL_909627, EPI_ISL_909628, EPI_ISL_909629, EPI_ISL_909630, EPI_ISL_909631, EPI_ISL_909632, EPI_ISL_909633, EPI_ISL_909634, EPI_ISL_909635, EPI_ISL_909636, EPI_ISL_909637, EPI_ISL_909638, EPI_ISL_909639, EPI_ISL_909640, EPI_ISL_909641, EPI_ISL_909642, EPI_ISL_909643, EPI_ISL_909644, EPI_ISL_909645, EPI_ISL_909646, EPI_ISL_909647, EPI_ISL_909648, EPI_ISL_909649, EPI_ISL_909650, EPI_ISL_909651 | | | |
| see above | Lighthouse Lab in Alderley Park | Wellcome Sanger Institute for the COVID-19 Genomics UK (COG-UK) Consortium | Jacquelyn Wynn, Mairead Hyland, The Lighthouse Lab in Alderley Park and Alex Alderton, Roberto Amato, Sonia Goncalves, Ewan Harrison, David K. Jackson, Ian Johnston, Dominic Kwiatkowski, Cordelia Langford, John Sillitoe on behalf of the Wellcome Sanger Institute COVID-19 Surveillance Team |
| EPI_ISL_909656, EPI_ISL_909660, EPI_ISL_909661, EPI_ISL_909662 | Labo Analyses med | National Reference Center for Viruses of Respiratory Infections, Institut Pasteur, Paris | Marion Barbet, Sylvie Behillil, Méline Bizard, Angela Brisebarre, Camille Capel, Etienne Simon-Lorière, Vincent Enouf, Maud Vanpeene, Sylvie van der Werf,Amzalag Jonas |
| EPI_ISL_909675, EPI_ISL_909676, EPI_ISL_909677, EPI_ISL_909678, EPI_ISL_909679, EPI_ISL_909680, EPI_ISL_909681 | Labo Analyses Med | National Reference Center for Viruses of Respiratory Infections, Institut Pasteur, Paris | Marion Barbet, Sylvie Behillil, Méline Bizard, Angela Brisebarre, Camille Capel, Etienne Simon-Lorière, Vincent Enouf, Maud Vanpeene, Sylvie van der Werf,Le Vicky |
| EPI_ISL_909949, EPI_ISL_909950 | CUSL/UCLouvain COVID testing federal platform | UCLouvain/IREC/MBLG | Jean Ruelle, Lysa Pinsmaye, Benoit Kabamba Mukadi |
| EPI_ISL_910941, EPI_ISL_910943, EPI_ISL_910944, EPI_ISL_910945, EPI_ISL_910946, EPI_ISL_910947, EPI_ISL_910948, EPI_ISL_910949, EPI_ISL_910950, EPI_ISL_910951, EPI_ISL_910952, EPI_ISL_910953, EPI_ISL_911104, EPI_ISL_911105, EPI_ISL_911106, EPI_ISL_911107, EPI_ISL_911108, EPI_ISL_911109, EPI_ISL_911110, EPI_ISL_911111, EPI_ISL_911112, EPI_ISL_911113, EPI_ISL_911114, EPI_ISL_911115, EPI_ISL_911116, EPI_ISL_911117, EPI_ISL_911118, EPI_ISL_911119, EPI_ISL_911120, EPI_ISL_911121, EPI_ISL_911122, EPI_ISL_911123, EPI_ISL_911124, EPI_ISL_911167, EPI_ISL_911169, EPI_ISL_911170, EPI_ISL_911171, EPI_ISL_911172, EPI_ISL_911173 | | | |
| see above | Laboratoire national de sante, Microbiology, Virology | Laboratoire national de sante, Microbiology, Microbial Genomics Platform | Anke Wienecke-Baldacchino, Catherine Ragimbeau,Jessica Tapp, Fatu Djabi, Lise Pignon, Raoul Salmon, Tamir Abdelrahman |
| EPI_ISL_911297, EPI_ISL_911307, EPI_ISL_911316, EPI_ISL_911327, EPI_ISL_911334, EPI_ISL_911337 | Servicio de Microbiología, Hospital Universitario Son Espases | SeqCOVID-SPAIN consortium/IBV(CSIC) | Carla López-Causapé, Jordi Reina, Antonio Oliver and SeqCOVID-SPAIN consortium |
| EPI_ISL_911634 | Texas Department of State Health Services (Texas Department of State Health Services (TXDSHS)) | Texas Department of State Health Services (TXDSHS) | Bonnie Oh, Anita Pokharel, James Daniel Bonser, Myong Koag, Chung Wang, Rachel Lee, Grace Kubin, Rashmi Tuladhar, Mayela Pedrueza, Maliha Rahman, Jenny Zhang |
| EPI_ISL_911639, EPI_ISL_911640 | Texas Department of State Health Services (TXDSHS) | Texas Department of State Health Services (TXDSHS) | Bonnie Oh, Anita Pokharel, James Daniel Bonser, Myong Koag, Chung Wang, Rachel Lee, Grace Kubin, Rashmi Tuladhar, Mayela Pedrueza, Maliha Rahman, Jenny Zhang |
| EPI_ISL_911704, EPI_ISL_911705, EPI_ISL_911706 | Quest Diagnostics | Quest Diagnostics | Rosenthal,S.H., Gerasimova,A., Kagan,R.M., Anderson, B., Hua, M., Liu Y., Bernstein, L.E., Livingston, K.E., Perez, A., Shalhout, D.F., Shlyakhter, I.A., Owen, R., Tanpaiboon, P., Lacbawan, F. |
| EPI_ISL_911794 | Johns Hopkins Hospital Department of Pathology | Johns Hopkins Hospital Department of Pathology | C. Paul Morris, Chun Huai Luo, Adannaya Amadi, Matthew Schwartz, Nicholas Gallagher, Heba H. Mostafa |
| EPI_ISL_911929, EPI_ISL_911930 | Wyoming Public Health Laboratory | Wyoming Public Health Laboratory | Noah Hull, Taylor Fearing, Lynette Gumbleton, Channing Weber, Ashley Norberg, Bailey Bowcutt, and Wanda Manley |
| EPI_ISL_911972, EPI_ISL_911973, EPI_ISL_911974, EPI_ISL_911975, EPI_ISL_911976, EPI_ISL_911977, EPI_ISL_911978, EPI_ISL_911979, EPI_ISL_911980, EPI_ISL_911981, EPI_ISL_911982, EPI_ISL_911983, EPI_ISL_911984, EPI_ISL_911985, EPI_ISL_911986, EPI_ISL_911987, EPI_ISL_911988, EPI_ISL_911989, EPI_ISL_911990, EPI_ISL_911991, EPI_ISL_911992, EPI_ISL_911993, EPI_ISL_911994, EPI_ISL_911995, EPI_ISL_911996, EPI_ISL_911997, EPI_ISL_911998, EPI_ISL_911999, EPI_ISL_912000, EPI_ISL_912001, EPI_ISL_912002, EPI_ISL_912003, EPI_ISL_912004, EPI_ISL_912005 | | | |
| see above | Altius Institute for Biomedical Sciences | Seattle Flu Study | Deborah A. Nickerson, Chris D. Frazar, Jover Lee, Benjamin Pelle, Erica Ryke, Matthew Richardson, Amanda Adler, Elisabeth Brandstetter, Peter D. Han, Kairsten Fay, Misja Ilcisin, Kirsten Lacombe, Thomas R. Sibley, Melissa Truong, Caitlin R. Wolf, Ryan Alexander, Daniel Bates, Rebecca Bruders, Stephanie DeBaun, Clem Green, Muhammad Halimun, Jessica Halow, Kneshay Harper, Matt Hartman, Andrew Meuser, Alex Nguyen, Truong Nguyen, Sofia Olsson, Sadie Patraw, Hannah Petersen, Tobias Ragoczy, Joshua Richards, Jacob Rodriguez, John Stamatoyannopoulos, Julia Wald, Olivia Waltner, Michael Boeckh, Janet A. Englund, Michael Famulare, Barry R. Lutz, Mark J. Rieder, Lea M. Starita, Matthew Thompson, Helen Y. Chu, Jay Shendure, Trevor Bedford |
| EPI_ISL_912047, EPI_ISL_912048, EPI_ISL_912049, EPI_ISL_912050, EPI_ISL_912051, EPI_ISL_912052, EPI_ISL_912053, EPI_ISL_912054, EPI_ISL_912055, EPI_ISL_912056, EPI_ISL_912057, EPI_ISL_912058, EPI_ISL_912059, EPI_ISL_912060, EPI_ISL_912061, EPI_ISL_912062, EPI_ISL_912063, EPI_ISL_912064, EPI_ISL_912065, EPI_ISL_912066, EPI_ISL_912067, EPI_ISL_912068, EPI_ISL_912069, EPI_ISL_912070, EPI_ISL_912071, EPI_ISL_912072, EPI_ISL_912073, EPI_ISL_912074, EPI_ISL_912075, EPI_ISL_912076, EPI_ISL_912077, EPI_ISL_912078 | | | |
| see above | Washington State Department of Health | Seattle Flu Study | Deborah A. Nickerson, Chris D. Frazar, Jover Lee, Benjamin Pelle, Erica Ryke, Matthew Richardson, Amanda Adler, Elisabeth Brandstetter, Peter D. Han, Kairsten Fay, Misja Ilcisin, Kirsten Lacombe, Thomas R. Sibley, Melissa Truong, Caitlin R. Wolf, Romesh Gautom, Geoff Melly, Brian Hiatt, Philip Dykema, Scott Lindquist, Michael Boeckh, Janet A. Englund, Michael Famulare, Barry R. Lutz, Mark J. Rieder, Lea M. Starita, Matthew Thompson, Helen Y. Chu, Jay Shendure, Trevor Bedford |
| EPI_ISL_912195 | Respiratory Viruses Branch, Centers for Disease Control and Prevention | Respiratory Viruses Branch, Centers for Disease Control and Prevention | Cook,P.W., Batra,D., Rambo-Martin,B.L., Galloway,S., Krueger,B.,Agarwal,M., Almasri,E., Boles,D., Burns,A., Charoensri,N.,Cohen,O., Countryman,S., Cristobal,M.A., Croy,B., Dale,S.,Deshmukh,H., Douglas,A., Drouillon,V., Eisenberg,M., Engler,H.,Ghatti,R., Gupta,P., Hicks,S., Humphrey,J., Iyer,L., Jain,M.,Kolli,M., Kuphal,T., Letovsky,S., Levandoski,M., Lukasik,C.,Meltzer,J., Norvell,B., Nye,M., Parker,S., Petropoulos,C.,Pruitt,J., Ragan,S., Ryan,S., Sapeta,M., Schroth,J.,Selvaraju,S.B., Stevovic,G., Suchanek,A., Throop,A., Tilson,L.,Urban,T., Voshell,J., Wagner,K., Williams,J., Williamson,M.,Zeng,Q., Zwiefelhofer,T., Paden,C.R., Tong,S. and MacCannell,D. |
| EPI_ISL_912226 | Lighthouse Lab in Alderley Park | Wellcome Sanger Institute for the COVID-19 Genomics UK (COG-UK) Consortium | Jacquelyn Wynn, Mairead Hyland, The Lighthouse Lab in Alderley Park and Alex Alderton, Roberto Amato, Sonia Goncalves, Ewan Harrison, David K. Jackson, Ian Johnston, Dominic Kwiatkowski, Cordelia Langford, John Sillitoe on behalf of the Wellcome Sanger Institute COVID-19 Surveillance Team (http://www.sanger.ac.uk/covid-team) |
| EPI_ISL_912256 | Charité Universitätsmedizin Berlin, Institut für Virologie/Labor Berlin | Charité Universitätsmedizin Berlin, Institut für Virologie | Victor M Corman, Barbara Mühlemann, Jörn Beheim-Schwarzbach, Tobias Bleicker, Julia Tesch, Talitha Veith, Julia Schneider, Terry Jones, Christian Drosten |
| EPI_ISL_912401 | PathCare | National Health Laboratory Service (NHLS), Tygerberg | Susan Engelbrecht, Jean Maritz, Bronwyn Kleinhans, Houriiyah Tegally, Eduan Wilkindon, Gert van Zyl, Wolfgang Preiser, Tulio de Oliveira |
| EPI_ISL_912414, EPI_ISL_912415, EPI_ISL_912423, EPI_ISL_912425 | KU Leuven, Rega Institute, Clinical and Epidemiological Virology | KU Leuven, Rega Institute, Clinical and Epidemiological Virology | Tony Wawina-Bokalanga, Bert Vanmechelen, Joan Marti-Carerras, Piet Maes |
| EPI_ISL_912547, EPI_ISL_912548 | Labo Analyses Med | National Reference Center for Viruses of Respiratory Infections, Institut Pasteur, Paris | Marion Barbet, Sylvie Behillil, Méline Bizard, Angela Brisebarre, Camille Capel, Etienne Simon-Lorière, Vincent Enouf, Maud Vanpeene, Sylvie van der Werf,Le Vicky |
| EPI_ISL_912560 | HOPITAL SIMONE VEIL | National Reference Center for Viruses of Respiratory Infections, Institut Pasteur, Paris | Marion Barbet, Sylvie Behillil, Méline Bizard, Angela Brisebarre, Camille Capel, Etienne Simon-Lorière, Vincent Enouf, Maud Vanpeene, Sylvie van der Werf,Moreau Farida |
| EPI_ISL_912584, EPI_ISL_912585, EPI_ISL_912586, EPI_ISL_912587, EPI_ISL_912588, EPI_ISL_912589, EPI_ISL_912590, EPI_ISL_912591, EPI_ISL_912592, EPI_ISL_912593, EPI_ISL_912594, EPI_ISL_912595, EPI_ISL_912596 | | | |
| see above | Labo Analyses Med | National Reference Center for Viruses of Respiratory Infections, Institut Pasteur, Paris | Marion Barbet, Sylvie Behillil, Méline Bizard, Angela Brisebarre, Camille Capel, Etienne Simon-Lorière, Vincent Enouf, Maud Vanpeene, Sylvie van der Werf,Le Vicky |
| EPI_ISL_912635, EPI_ISL_912742, EPI_ISL_912743, EPI_ISL_912744, EPI_ISL_912745, EPI_ISL_912746, EPI_ISL_912747 | Hôpital Henri Mondor | Department of Virology, Henri Mondor University Hospital, Assistance Publique Hôpitaux de Paris, Université Paris-Est Créteil, INSERM U955 | Christophe Rodriguez, Slim Fourati, Vanessa Demontant, Guillaume Gricourt, Melissa N'Debi, Alexandre Soulier, Elisabeth Trawinski, Jean-Michel Pawlotsky |
| EPI_ISL_912909, EPI_ISL_912910, EPI_ISL_912911, EPI_ISL_912923, | Hôpital Pitié-Salpêtrière | Department of Virology, Henri Mondor University Hospital, Assistance Publique Hôpitaux de Paris, Université Paris-Est | Christophe Rodriguez, Slim Fourati, Vanessa Demontant, Guillaume Gricourt, Melissa N'Debi, Alexandre Soulier, Elisabeth Trawinski, Jean-Michel Pawlotsky |

| | | | |
|---|---|---|---|
| EPI_ISL_912926, EPI_ISL_912934 | | Créteil, INSERM U955 | |
| EPI_ISL_913039 | Complejo Hospitalario Xeral-Calde | Instituto de Salud Carlos III | Iglesias-Caballero, M. Camarero, S. Sandonís,V. Vázquez, S. Pozo, F. Casas, I. Jiménez, P. Zaballos, A. Monzón, S. Varona, S. Cuesta, I. Alonso, P. |
| EPI_ISL_913060 | Hospital Universitario Virgen de la Arrixaca | Instituto de Salud Carlos III | Iglesias-Caballero, M. Camarero, S. Sandonís,V. Vázquez, S. Pozo, F. Casas, I. Jiménez, P. Zaballos, A. Monzón, S. Varona, S. Cuesta, I. Moreno, A. |
| EPI_ISL_913130, EPI_ISL_913135, EPI_ISL_913143, EPI_ISL_913146, EPI_ISL_913149, EPI_ISL_913158, EPI_ISL_913159, EPI_ISL_913161, EPI_ISL_913164, EPI_ISL_913165, EPI_ISL_913166, EPI_ISL_913167, EPI_ISL_913170, EPI_ISL_913175 | | | |
| see above | University of Michigan Clinical Microbiology Laboratory | Lauring Lab, University of Michigan, Department of Microbiology and Immunology | Valesano |
| EPI_ISL_913258, EPI_ISL_913260, EPI_ISL_913261, EPI_ISL_913262, EPI_ISL_913264, EPI_ISL_913265, EPI_ISL_913270, EPI_ISL_913271, EPI_ISL_913273, EPI_ISL_913307, EPI_ISL_913308, EPI_ISL_913359, EPI_ISL_913360, EPI_ISL_913361, EPI_ISL_913362 | | | |
| see above | Klinisk mikrobiologi | The Public Health Agency of Sweden | Anna-Malin Linde, Maria Lind Karlberg, Carlo Berg, Oskar Karlsson Lindsjo, Sofia Stamouli, Reza Advani, Mattias Haukland, Petra Holmstrom, Noura Walai, Petra Edquist, Mia Brytting, Anna Risberg, Karin Tegmark-Wisell |
| EPI_ISL_913428 | Massachusetts State Public Health Laboratory | Massachusetts State Public Health Laboratory | Andrew Lang, Timelia Fink, Glen Gallagher, Sandra Smole |
| EPI_ISL_913453 | Klinisk mikrobiologi | The Public Health Agency of Sweden | Anna-Malin Linde, Maria Lind Karlberg, Carlo Berg, Oskar Karlsson Lindsjo, Sofia Stamouli, Reza Advani, Mattias Haukland, Petra Holmstrom, Noura Walai, Petra Edquist, Mia Brytting, Anna Risberg, Karin Tegmark-Wisell |
| EPI_ISL_913454 | Dynamic Code AB | The Public Health Agency of Sweden | Anna-Malin Linde, Maria Lind Karlberg, Carlo Berg, Oskar Karlsson Lindsjo, Sofia Stamouli, Reza Advani, Mattias Haukland, Petra Holmstrom, Noura Walai, Petra Edquist, Mia Brytting, Anna Risberg, Karin Tegmark-Wisell |
| EPI_ISL_913467 | Synlab Medilab, Mikrobiologi | The Public Health Agency of Sweden | Anna-Malin Linde, Maria Lind Karlberg, Carlo Berg, Oskar Karlsson Lindsjo, Sofia Stamouli, Reza Advani, Mattias Haukland, Petra Holmstrom, Noura Walai, Petra Edquist, Mia Brytting, Anna Risberg, Karin Tegmark-Wisell |
| EPI_ISL_913468, EPI_ISL_913494 | The Public Health Agency of Sweden | The Public Health Agency of Sweden | Anna-Malin Linde, Maria Lind Karlberg, Carlo Berg, Oskar Karlsson Lindsjo, Sofia Stamouli, Reza Advani, Mattias Haukland, Petra Holmstrom, Noura Walai, Petra Edquist, Mia Brytting, Anna Risberg, Karin Tegmark-Wisell |
| EPI_ISL_913746, EPI_ISL_913747 | DC Public Health Lab/ Dept. of Forensic Sciences | DC Public Health Lab/ Dept. of Forensic Sciences | Scott Nguyen, Elizabeth Zelaya, Connie Maza, Monica Mann, Brittany Hamilton, David Payne, Jocelyn Hauser |
| EPI_ISL_913769, EPI_ISL_913770, EPI_ISL_913772, EPI_ISL_913773, EPI_ISL_913775, EPI_ISL_913776, EPI_ISL_913777, EPI_ISL_913779, EPI_ISL_913780, EPI_ISL_913783, EPI_ISL_913784, EPI_ISL_913786, EPI_ISL_913787, EPI_ISL_913789, EPI_ISL_913790, EPI_ISL_913792, EPI_ISL_913793 | | | |
| see above | KU Leuven, Rega Institute, Clinical and Epidemiological Virology | KU Leuven, Rega Institute, Clinical and Epidemiological Virology | Tony Wawina-Bokalanga, Bert Vanmechelen, Joan Marti-Carerras, Piet Maes |
| EPI_ISL_914002 | Quest Diagnostics Tucker GA | Pathogen Discovery, Respiratory Viruses Branch, Division of Viral Diseases, Centers for Disease Control and Prevention | Ying Tao, Yan Li, Jing Zhang, Krista Queen, Anna Uehara, Peter Cook, Clinton R. Paden, Haibin Wang, Suxiang Tong |
| EPI_ISL_914015, EPI_ISL_914016, EPI_ISL_914017, EPI_ISL_914018, EPI_ISL_914019, EPI_ISL_914020, EPI_ISL_914021, EPI_ISL_914022, EPI_ISL_914023, EPI_ISL_914024 | IL Department of Public Health Chicago Laboratory | Pathogen Discovery, Respiratory Viruses Branch, Division of Viral Diseases, Centers for Disease Control and Prevention | Ying Tao, Yan Li, Jing Zhang, Krista Queen, Anna Uehara, Peter Cook, Clinton R. Paden, Haibin Wang, Suxiang Tong |
| EPI_ISL_914828 | AREA DE SALUD ALAJUELA SUR | Inciensa, Instituto Costarricense de Investigación y Enseñanza en Nutrición y Salud | Francisco Duarte, Hebleen Porras, Claudio Soto-Garita, Estela Cordero, Adriana Godínez, Melany Calderón & Mariel López |
| EPI_ISL_914831 | AREA DE SALUD CARRILLO | Inciensa, Instituto Costarricense de Investigación y Enseñanza en Nutrición y Salud | Francisco Duarte, Hebleen Porras, Claudio Soto-Garita, Estela Cordero, Adriana Godínez, Melany Calderón & Adriana Bermúdez-Espinoza |
| EPI_ISL_914833 | LABORATORIO CLINICO LABIN | Inciensa, Instituto Costarricense de Investigación y Enseñanza en Nutrición y Salud | Francisco Duarte, Hebleen Porras, Claudio Soto-Garita, Estela Cordero, Adriana Godínez, Melany Calderón & Pei Ling Chan Ma |
| EPI_ISL_914834 | HOSPITAL METROPOLITANO | Inciensa, Instituto Costarricense de Investigación y Enseñanza en Nutrición y Salud | Francisco Duarte, Hebleen Porras, Claudio Soto-Garita, Estela Cordero, Adriana Godínez, Melany Calderón & Margarita Lee-Lui |
| EPI_ISL_915198, EPI_ISL_915202, EPI_ISL_915235, EPI_ISL_915236, EPI_ISL_915237, EPI_ISL_915238, EPI_ISL_915239, EPI_ISL_915284, EPI_ISL_915285, EPI_ISL_915286, EPI_ISL_915287, EPI_ISL_915288, EPI_ISL_915289, EPI_ISL_915323, EPI_ISL_915325 | | | |
| see above | Quest Diagnostics | Quest Diagnostics | Rosenthal,S.H., Gerasimova,A., Kagan,R.M., Anderson, B., Hua, M., Liu Y., Bernstein, L.E., Livingston, K.E., Perez, A., Shalhout, D.F., Shlyakhter, I.A., Owen, R., Tanpaiboon, P., Lacbawan, F. |
| EPI_ISL_915435, EPI_ISL_915436, EPI_ISL_915437, EPI_ISL_917944, EPI_ISL_917945, EPI_ISL_917946, EPI_ISL_917947, EPI_ISL_917948, EPI_ISL_917949, EPI_ISL_917950 | MD Laboratories | Los Angeles County PHL | P. Hemarajata et al. |
| EPI_ISL_918181, EPI_ISL_918201, EPI_ISL_918259 | Innovative Genomics Institute, UC Berkeley | Innovative Genomics Institute, UC Berkeley | Stacia Wyman, Haridha Shivram, Phil Frankino, Liana Lareau, Shana McDevitt, Justin Choi |
| EPI_ISL_918264 | SIESP CHIETI-DRIVE IN CHIETI | Istituto Zooprofilattico Sperimentale dell'Abruzzo e Molise "G. Caporale" | Lorusso A, Marcacci M, Di Domenico M, Ancora M, Curini V, Mangone I, Rinaldi A, Scialabba S, Di Pasquale A, Cammà C, Puglia I, Calistri P, Savini G |
| EPI_ISL_918265 | OSPEDALE ATRI | Istituto Zooprofilattico Sperimentale dell'Abruzzo e Molise "G. Caporale" | Lorusso A, Marcacci M, Di Domenico M, Ancora M, Curini V, Mangone I, Rinaldi A, Scialabba S, Di Pasquale A, Cammà C, Puglia I, Calistri P, Savini G |
| EPI_ISL_918270 | SIESP CHIETI - DRIVE IN LANCIANO | Istituto Zooprofilattico Sperimentale dell'Abruzzo e Molise "G. Caporale" | Lorusso A, Marcacci M, Di Domenico M, Ancora M, Curini V, Mangone I, Rinaldi A, Scialabba S, Di Pasquale A, Cammà C, Puglia I, Calistri P, Savini G |
| EPI_ISL_918280, EPI_ISL_918281 | Hospital Universitari Vall d'Hebron - Vall d'Hebron Institut de Rercerca | Hospital Universitari Vall d'Hebron | Cristina Andrés, Maria Piñana, Josep F Abril, Damir Garcia-Cehic, Ariadna Rando, Juliana Esperalba, Maria Gema Codina, Carla Castillo, Maria Carmen Martin, Tomàs Pumarola, Josep Quer, Andrés Antón |
| EPI_ISL_918356 | Institute of Virology, Medical Center, University of Freiburg, Freiburg, Germany | Institute of Virology, Clinial Virus Genomics, Medical Center, University of Freiburg, Freiburg, Germany | Jonas Fuchs, Lisa Kern, Sandra Reuter, Hajo Grundmann, Marcus Panning |
| EPI_ISL_918490 | CUSL/UCLouvain COVID testing federal platform | UCLouvain/IREC/MBLG | Jean Ruelle, Lysa Pinsmaye, Benoit Kabamba Mukadi |
| EPI_ISL_918539 | LACEN - Laboratório Central de Saúde Pública do Ceara | Evandro Chagas Institute | Santos, M.C.; Silva, A.M.; Junior, W.D.C.; Barbagelata, L.S.; Ferreira, J.A.; Sousa, E.M.A.; da Silva, P.S.; Pinheiro, K.C.; L.C.; Sousa Junior, E.C. |
| EPI_ISL_918560, EPI_ISL_918561 | LACEN - Laboratório Central de Saúde Pública do Amapa | Evandro Chagas Institute | Santos, M.C.; Silva, A.M.; Junior, W.D.C.; Barbagelata, L.S.; Ferreira, J.A.; Sousa, E.M.A.; da Silva, P.S.; Pinheiro, K.C.; L.C.; Sousa Junior, E.C. |
| EPI_ISL_918686, EPI_ISL_918687, EPI_ISL_918688, EPI_ISL_918689, EPI_ISL_918690, EPI_ISL_918691, EPI_ISL_918692, EPI_ISL_918693, EPI_ISL_918694, EPI_ISL_918695, EPI_ISL_918696, EPI_ISL_918701, EPI_ISL_918733, EPI_ISL_918734, EPI_ISL_918860, EPI_ISL_918861, EPI_ISL_918884, EPI_ISL_918885, EPI_ISL_918886, EPI_ISL_918887, EPI_ISL_918888, EPI_ISL_918889, EPI_ISL_918890, EPI_ISL_918891, EPI_ISL_918892, EPI_ISL_918893, EPI_ISL_918894, EPI_ISL_918895, EPI_ISL_918896, EPI_ISL_918897, EPI_ISL_918898, EPI_ISL_918899, EPI_ISL_918916, EPI_ISL_918917, EPI_ISL_918918, EPI_ISL_918919, EPI_ISL_918920, EPI_ISL_918921, EPI_ISL_918922, EPI_ISL_918923, EPI_ISL_918924, EPI_ISL_918925, EPI_ISL_918926, EPI_ISL_918927, EPI_ISL_918928, EPI_ISL_918929, EPI_ISL_918930, EPI_ISL_918931, EPI_ISL_918932, EPI_ISL_918933, EPI_ISL_918934, EPI_ISL_918936, EPI_ISL_918937, EPI_ISL_918938, EPI_ISL_918939, EPI_ISL_918973, EPI_ISL_918974, EPI_ISL_918975, EPI_ISL_918976 | | | |
| see above | University of Birmingham | COVID-19 Genomics UK (COG-UK) Consortium | Institute of Microbiology, University of Birmingham: Claire McMurray, Joanne Stockton, Samuel Nicholls, Radoslaw Poplawski, Will Rowe, Josh Quick, Nicholas Loman. University of Birmingham Testing Laboratory: Celina M Whalley, Andrew Bosworth, Charlotte Poxon, Kasun Wanigasooriya, Oliver Pickles, Mike Kidd, Alex Richter, Andrew D Beggs PHE Heartlands Lab: Husam Osman, Andrew Bosworth. Queen Elizabeth Hospital: Anna Casey |
| EPI_ISL_919029, EPI_ISL_919044, EPI_ISL_919046, EPI_ISL_919048, EPI_ISL_919050, EPI_ISL_919052, | Department of Pathology, University of Cambridge | COVID-19 Genomics UK (COG-UK) Consortium | Aminu S. Jahun, Yasmin Chaudhry, Iliana Georgana, Myra Hosmillo, Rhys Izu, Martin D. Curran, Surendra Parmar, Ian Goodfellow |

| | | | |
|---|---|---|---|
| EPI_ISL_919056 | | | |
| EPI_ISL_919190, EPI_ISL_919191, EPI_ISL_919192, EPI_ISL_919193, EPI_ISL_919194, EPI_ISL_919195 | West of Scotland Specialist Virology Centre, NHSGGC / MRC-University of Glasgow Centre for Virus Research | COVID-19 Genomics UK (COG-UK) Consortium | Ana da Silva Filipe, Natasha Johnson, Kathy Smollett, Daniel Mair, Stephen Carmichael, Alice Broos, Lily Tong, Jenna Nichols, Kyriaki Nomikou; Sarah McDonald; Richard Orton, Joseph Hughes, Sreenu Vattipally, David L Robertson; Alasdair MacLean, Rory Gunson; Sharif Shaaban, Matthew Holden; Rachel Blacow, Guy Mollett, Kathy Li, James Shepherd, Antonia Ho, Emma Thomson |
| EPI_ISL_919337 | Virology Department, Royal Infirmary of Edinburgh, NHS Lothian / School of Biological Sciences, University of Edinburgh / Institute of Genetics and Molecular Medicine, University of Edinburgh | COVID-19 Genomics UK (COG-UK) Consortium | McHugh M, Dewar R, Rooke S, Gallagher M, Balcaza C, O'Toole Á, Scher E, Hill V, McCrone JT, Colquhoun R, Yu X, Jackson B, Rambaut A, Williams TC, Templeton K |
| EPI_ISL_919467, EPI_ISL_919468, EPI_ISL_919469, EPI_ISL_919470, EPI_ISL_919471, EPI_ISL_919472, EPI_ISL_919473, EPI_ISL_919474, EPI_ISL_919593 | Liverpool Clinical Laboratories | COVID-19 Genomics UK (COG-UK) Consortium | Sam Haldenby, Anita Lucaci, Steve Paterson, Julian Hiscox, Alistair Darby, M Almsaud, A Alrezaihi, Muhannad Alruwaili, Stuart D Armstrong, Jones Benjamin, Eleanor G Bentley, Anu Chawla, Jordan J Clark, Angela Cowell, Richard Eccles, Isabel García-Dorival, Matthew Gemmell, Alessandro Gerada, PKF Gilmore, Richard Gregory, Ximeng Han, Catherine Hartley, Margaret Hughes, Miren Iturriza-Gomara, James Johnson, L Luu, Jenifer Manson, Charlotte Nelson, Elaine O'Toole, Cassie Olateju, Rebekah Penrice-Randal , Lucille Rainbow, N.P Randle, Trevor Ian Robinson, Parul Sharma, Ghada T Shawli, James P Stewart, Neil Swainston, Ecaterina Vamos, Joanne Watts, Mark Whitehead |
| EPI_ISL_920000, EPI_ISL_920001, EPI_ISL_920002, EPI_ISL_920003, EPI_ISL_920004, EPI_ISL_920005, EPI_ISL_920006, EPI_ISL_920007, EPI_ISL_920008, EPI_ISL_920009, EPI_ISL_920010, EPI_ISL_920011, EPI_ISL_920014, EPI_ISL_920042, EPI_ISL_920050 | | | |
| see above | University College London, Great Ormond Street Hospital for Children NHS Foundation Trust, Imperial College Healthcare NHS Trust | COVID-19 Genomics UK (COG-UK) Consortium | Sergi Castellano, Rachel Williams, Mark Kristiansen, Paola Resende Silva, Sunando Roy, Tony Brooks, Helena Tutill, Paola Niola, Patricia Dyal, Charlotte Williams, Leysa Forrest, Yasmin Panchbhaya, Jacqueline Findlay, Samuel Weeks, Julianne Brown, Kathryn Harris, Paul Randell, James Price, Alison Holmes, Judith Breuer |
| EPI_ISL_920191, EPI_ISL_920241, EPI_ISL_920298, EPI_ISL_920310, EPI_ISL_920329, EPI_ISL_920345, EPI_ISL_920357, EPI_ISL_920458, EPI_ISL_920467 | University College London Hospital | COVID-19 Genomics UK (COG-UK) Consortium | Judith Heaney, Matthew Byott, Catherine Houlihan, Dan Frampton, Stuart Kirk, Moira Spyer and Eleni Nastouli |
| EPI_ISL_920851, EPI_ISL_920860, EPI_ISL_920861, EPI_ISL_920862, EPI_ISL_920863, EPI_ISL_920864, EPI_ISL_920865, EPI_ISL_920866 | University College London, Great Ormond Street Hospital for Children NHS Foundation Trust, Imperial College Healthcare NHS Trust | COVID-19 Genomics UK (COG-UK) Consortium | Sergi Castellano, Rachel Williams, Mark Kristiansen, Paola Resende Silva, Sunando Roy, Tony Brooks, Helena Tutill, Paola Niola, Patricia Dyal, Charlotte Williams, Leysa Forrest, Yasmin Panchbhaya, Jacqueline Findlay, Samuel Weeks, Julianne Brown, Kathryn Harris, Paul Randell, James Price, Alison Holmes, Judith Breuer |
| EPI_ISL_920900 | Department of Pathology, University of Cambridge | COVID-19 Genomics UK (COG-UK) Consortium | Aminu S. Jahun, Yasmin Chaudhry, Iliana Georgana, Myra Hosmillo, Rhys Izu, Martin D. Curran, Surendra Parmar, Ian Goodfellow |
| EPI_ISL_920996, EPI_ISL_920997, EPI_ISL_920998, EPI_ISL_920999, EPI_ISL_921000, EPI_ISL_921001, EPI_ISL_921002, EPI_ISL_921003, EPI_ISL_921004, EPI_ISL_921005, EPI_ISL_921006, EPI_ISL_921008, EPI_ISL_921009, EPI_ISL_921010, EPI_ISL_921018 | | | |
| see above | Regional Virus Laboratory, Belfast Health and Social Care Trust | COVID-19 Genomics UK (COG-UK) Consortium | Conall McCaughey, James McKenna, Tanya Curran, Susan Feeney, Alison Watt, Ciara Cox, Mairead Connor, Zoltan Molnar, David Simpson, Derek Fairley |
| EPI_ISL_921491, EPI_ISL_921492, EPI_ISL_921495, EPI_ISL_921496, EPI_ISL_921497, EPI_ISL_921498, EPI_ISL_921499, EPI_ISL_921500, EPI_ISL_921506, EPI_ISL_921515, EPI_ISL_921517 | | | |
| see above | Northumbria University / South Tees Hospitals NHS Foundation Trust / North Cumbria Integrated Care NHS Foundation Trust / North Tees and Hartlepool NHS Foundation Trust / Newcastle Hospitals NHS Foundation Trust | COVID-19 Genomics UK (COG-UK) Consortium | Darren L Smith,Andrew Nelson,Matthew Bashton,Greg R Young,Joshua Loh,John Allan,Mohammad A Tariq,Giles S Holt,Gary Black,Wen C Yew,Lynn Dover,Paul Baker,Steve Liggett,Sarah Essex,Jane Greenaway,Debra Padgett,Clive Graham,Garren Scott,Edward Barton,Emma Swindells,Brendan Payne,Jennifer Collins,Yusri Taha,Gary Eltringham |
| EPI_ISL_922071, EPI_ISL_922072, EPI_ISL_922074, EPI_ISL_922075 | Lincolnshire Hospitals and DeepSeq Nottingham | COVID-19 Genomics UK (COG-UK) Consortium | Nichola Duckworth, Tim Sloan, Sarah Walsh, Jonathan Ball, Patrick McClure, Joeseph Chappell, Nadine Holmes, Matthew Carlisle, Christopher Moore, Fei Sang, Johnny Debebe, Victoria Wright, Matthew Loose |
| EPI_ISL_922076 | General Hospital #Abdulah Nakas" Sarajevo | Alea Genetic Centre | Rijad Konjhodzic, Dino Pecar, Lana Salihefendic, Sead Jazic |
| EPI_ISL_922077, EPI_ISL_922078, EPI_ISL_922079, EPI_ISL_922080, EPI_ISL_922081, EPI_ISL_922082, EPI_ISL_922083, EPI_ISL_922084, EPI_ISL_922085, EPI_ISL_922086, EPI_ISL_922087, EPI_ISL_922088, EPI_ISL_922089, EPI_ISL_922090, EPI_ISL_922091, EPI_ISL_922092, EPI_ISL_922093, EPI_ISL_922094, EPI_ISL_922095, EPI_ISL_922096, EPI_ISL_922097, EPI_ISL_922098, EPI_ISL_922099, EPI_ISL_922100, EPI_ISL_922102, EPI_ISL_922103, EPI_ISL_922104, EPI_ISL_922107 | | | |
| see above | Lincolnshire Hospitals and DeepSeq Nottingham | COVID-19 Genomics UK (COG-UK) Consortium | Nichola Duckworth, Tim Sloan, Sarah Walsh, Jonathan Ball, Patrick McClure, Joeseph Chappell, Nadine Holmes, Matthew Carlisle, Christopher Moore, Fei Sang, Johnny Debebe, Victoria Wright, Matthew Loose |
| EPI_ISL_922212, EPI_ISL_922213, EPI_ISL_922214, EPI_ISL_922215, EPI_ISL_922216, EPI_ISL_922217, EPI_ISL_922218, EPI_ISL_922219, EPI_ISL_922220, EPI_ISL_922224, EPI_ISL_922225, EPI_ISL_922226, EPI_ISL_922227, EPI_ISL_922228, EPI_ISL_922229, EPI_ISL_922230, EPI_ISL_922237, EPI_ISL_922259, EPI_ISL_922294 | | | |
| see above | Oxford Viromics, NDM, University of Oxford; Oxford University Hospitals; Basingstoke and North Hampshire Hospital | COVID-19 Genomics UK (COG-UK) Consortium | Tanya Golubchik, David Bonsall, George Macintyre, Amy Trebes, Mariateresa de Cesare, Catrin Moore, Alex Mobbs, Anita Justice, Robert Shaw, Monique Andersson, Timothy Peto, Emma Wise, Nathan Moore, Jessica Lynch, Nick Cortes, Matilde Mori, Stephen Kidd, David Buck, John Todd, Christophe Fraser |
| EPI_ISL_922380, EPI_ISL_922558 | Wales Specialist Virology Centre Sequencing lab: Pathogen Genomics Unit | Public Health Wales Microbiology Cardiff Wales Specialist Virology Centre | Catherine Moore, Johnathan Evans, Laura Gifford, Malorie Perry, Simon Cottrell, Angela Marchbank, Alec Birchley, Alexander Adams, Amy Gaskin, Bree Gatica-Wilcox, Jason Coombes, Joel Southgate, Lauren Gilbert, Lee Graham, Nicole Pacchiarini, Sara Kumziene-Summerhayes, Sarah Taylor, Sophie Jones, Sara Rey, Matthew Bull, Joanne Watkins, Sally Corden, Tom Connor |
| EPI_ISL_923262, EPI_ISL_923373, EPI_ISL_923375, EPI_ISL_923376, EPI_ISL_923378, EPI_ISL_923390, EPI_ISL_923392, EPI_ISL_923393, EPI_ISL_923394, EPI_ISL_923395, EPI_ISL_923396, EPI_ISL_923397, EPI_ISL_923398, EPI_ISL_923399, EPI_ISL_923401, EPI_ISL_923402, EPI_ISL_923403, EPI_ISL_923404, EPI_ISL_923405, EPI_ISL_923406, EPI_ISL_923407, EPI_ISL_923408, EPI_ISL_923409, EPI_ISL_923410, EPI_ISL_923411, EPI_ISL_923412, EPI_ISL_923413, EPI_ISL_923539, EPI_ISL_923593 | | | |
| see above | Centre for Enzyme Innovation, University of Portsmouth / Translational Research Laboratory, Portsmouth Hospitals NHS Trust | COVID-19 Genomics UK (COG-UK) Consortium | Angela Beckett,Salman Goudarzi,Christopher Fearn,Kate Cook,Katie Loveson,Sharon Glaysher,Scott Elliott,Samuel Robson |
| EPI_ISL_924270 | Virology Department, Sheffield Teaching Hospitals NHS Foundation Trust/Department of Infection, Immunity and Cardiovascular Disease, The Medical School, University of Sheffield | COVID-19 Genomics UK (COG-UK) Consortium | Thushan de Silva, Matthew Parker, Nikki Smith, Adri Angyal, Rebecca Brown, Luke Green, Rachel Tucker, Paul Parsons, Danielle Groves, Katie Johnson, Laura Carrilero, Alex Keeley, Dave Partridge, Matthew Wyles, Benjamin Lindsey, Mehmet Yavuz, Mohammad Raza, Cariad Evans |
| EPI_ISL_924729, EPI_ISL_924731, EPI_ISL_924746, EPI_ISL_924749, EPI_ISL_924764, EPI_ISL_924770, EPI_ISL_924774, EPI_ISL_924794, EPI_ISL_924795, EPI_ISL_924797, EPI_ISL_924798, EPI_ISL_924800, EPI_ISL_924805, EPI_ISL_924807, EPI_ISL_924808, EPI_ISL_924816, EPI_ISL_924819, EPI_ISL_924827, EPI_ISL_924828, EPI_ISL_924838, EPI_ISL_924839, EPI_ISL_924848, EPI_ISL_924849, EPI_ISL_924850, EPI_ISL_924851, EPI_ISL_924855, EPI_ISL_924863, EPI_ISL_924864, EPI_ISL_924865, EPI_ISL_924867, EPI_ISL_924871, EPI_ISL_924877, EPI_ISL_924879, EPI_ISL_924880, EPI_ISL_924883, EPI_ISL_924886, EPI_ISL_924887, EPI_ISL_924890, EPI_ISL_924891, EPI_ISL_924892, EPI_ISL_924895, EPI_ISL_924899, EPI_ISL_924901, EPI_ISL_924902, EPI_ISL_924904, EPI_ISL_924905, EPI_ISL_924907, EPI_ISL_924908, EPI_ISL_924909, EPI_ISL_924910, EPI_ISL_924911, EPI_ISL_924912, EPI_ISL_924913, EPI_ISL_924914, EPI_ISL_924915, EPI_ISL_924916, EPI_ISL_924917, EPI_ISL_924918, EPI_ISL_924919, EPI_ISL_924921, EPI_ISL_924922, EPI_ISL_924923, EPI_ISL_924924, EPI_ISL_924925, EPI_ISL_924926, EPI_ISL_924927, EPI_ISL_924928, EPI_ISL_924929, EPI_ISL_924930, EPI_ISL_924931, EPI_ISL_924933, EPI_ISL_924935, EPI_ISL_924936, EPI_ISL_924937, EPI_ISL_924938, EPI_ISL_924939, EPI_ISL_924940, EPI_ISL_924941, EPI_ISL_924942, EPI_ISL_924943, EPI_ISL_924944, EPI_ISL_924945, EPI_ISL_924946, EPI_ISL_924947, EPI_ISL_924950, EPI_ISL_924975, EPI_ISL_924980 | | | |
| see above | Bioinformatics and Biostatistics Lab, Advanced Sequencing Facility | COVID-19 Genomics UK (COG-UK) Consortium | Aengus Stewart,Jerome Nicod,Chelsea Sawyer,Laura Cubitt,Harshil Patel,Margaret Crawford |
| EPI_ISL_925263, EPI_ISL_925284, EPI_ISL_925294, EPI_ISL_925302, EPI_ISL_925305 | Wyoming Public Health Laboratory | Wyoming Public Health Laboratory | Noah Hull, Taylor Fearing, Lynette Gumbleton, Channing Weber, Ashley Norberg, Bailey Bowcutt, and Wanda Manley |
| EPI_ISL_925399, EPI_ISL_925400 | Department of Clinical Microbiology | GIGA Medical Genomics | Keith Durkin, Maria Artesi, Sébastien Bontems, Raphaël Boreux, Bouchra Boujemla, Cécile Meex, Pierrette Melin, Marie-Pierre Hayette, Vincent Bours |
| EPI_ISL_925914 | Nucleic Acid Testing, National Reference Laboratory | GIGA Medical Genomics | Yvan Butera, Keith Durkin, Maria Artesi, Bouchra Boujemla, Robert Rutayisire, Patrick Tuyisenge, Esperence Umumararungu, Sébastien Bontems, Marie-Pierre Hayette, Nathalie Renotte, Swaibu Gatare, Jacob Souopgui, Sabin Nsanzimana, Vincent Bours, Léon Mutesa |
| EPI_ISL_930635, EPI_ISL_930637, EPI_ISL_930638, EPI_ISL_930639, EPI_ISL_930640, EPI_ISL_930641, EPI_ISL_930642, EPI_ISL_930643, EPI_ISL_930644, EPI_ISL_930645, EPI_ISL_930646, EPI_ISL_930647, EPI_ISL_930648, EPI_ISL_930651, EPI_ISL_930652, EPI_ISL_930653, EPI_ISL_930654, EPI_ISL_930655, | | | |

| | | | |
|---|---|---|---|
| EPI_ISL_930662, EPI_ISL_930663 | | | |
| see above | Arizona State Public Health Laboratory | Arizona State Public Health Laboratory | Trung Huynh, Jessica Escobar, Katherine Fullerton, Nobuko Fukushima, Stacy White, Linda Getsinger, Victor Waddell |
| EPI_ISL_933663, EPI_ISL_933690, EPI_ISL_933691, EPI_ISL_933692, EPI_ISL_933705 | Instituto de Diagnostico y Referencia Epidemiologicos INDRE_RNLSP | Instituto de Diagnostico y Referencia Epidemiologicos (INDRE) | Claudia Wong-Arambula, Abril Rodriguez-Maldonado, Fabiola Garces-Ayala, Adnan Araiza-Rodriguez, David Fragoso-Fonseca, Sergio Rangel-Guerrero, Mayra Jimenez-Morales, Nancy Munoz-Hernandez, Natividad Cruz-Ortiz, Tatiana Nunez-Garcia, Gisela Barrera-Badillo, Lucia Hernandez-Rivas, Irma Lopez-Martinez, Ernesto Ramirez-Gonzalez. |
| EPI_ISL_934238 | Vilnius university hospital Santaros Klinikos, Center of Laboratory Medicine | Vilnius university hospital Santaros Klinikos, Center of Laboratory Medicine | Ingrida Olendraite, Daniel Naumovas, Rimvydas Norvilas, Dovile Ezerskyte, Justinas Slikas, Gytis Dudas |
| EPI_ISL_934324, EPI_ISL_934340, EPI_ISL_934350, EPI_ISL_934351, EPI_ISL_934352, EPI_ISL_934353 | Klinisk mikrobiologi | The Public Health Agency of Sweden | Anna-Malin Linde, Maria Lind Karlberg, Carlo Berg, Oskar Karlsson Lindsjo, Sofia Stamouli, Reza Advani, Mattias Haukland, Petra Holmstrom, Noura Walai, Petra Edquist, Mia Brytting, Anna Risberg, Karin Tegmark-Wisell |
| EPI_ISL_934370, EPI_ISL_934371, EPI_ISL_934372, EPI_ISL_934373, EPI_ISL_934374 | Synlab Medilab, Mikrobiologi | The Public Health Agency of Sweden | Anna-Malin Linde, Maria Lind Karlberg, Carlo Berg, Oskar Karlsson Lindsjo, Sofia Stamouli, Reza Advani, Mattias Haukland, Petra Holmstrom, Noura Walai, Petra Edquist, Mia Brytting, Anna Risberg, Karin Tegmark-Wisell |
| EPI_ISL_934437, EPI_ISL_934438, EPI_ISL_934439, EPI_ISL_934440, EPI_ISL_934441, EPI_ISL_934442, EPI_ISL_934443, EPI_ISL_934444, EPI_ISL_934445, EPI_ISL_934446, EPI_ISL_934447, EPI_ISL_934448, EPI_ISL_934449, EPI_ISL_934450, EPI_ISL_934451, EPI_ISL_934455, EPI_ISL_934458, EPI_ISL_934461, EPI_ISL_934462, EPI_ISL_934463, EPI_ISL_934464, EPI_ISL_934473, EPI_ISL_934474, EPI_ISL_934475, EPI_ISL_934476, EPI_ISL_934477, EPI_ISL_934478, EPI_ISL_934480, EPI_ISL_934484, EPI_ISL_934488, EPI_ISL_934489, EPI_ISL_934493, EPI_ISL_934494, EPI_ISL_934503, EPI_ISL_934504, EPI_ISL_934505, EPI_ISL_934506, EPI_ISL_934507, EPI_ISL_934508, EPI_ISL_934509 | | | |
| see above | Austrian Agency for Health and Food Safety (AGES) | Bergthaler laboratory, CeMM Research Center for Molecular Medicine of the Austrian Academy of Sciences | Lukas Endler, Anna Schedl, Thomas Penz, Benedikt Agerer, Maelle Le Moing, Michael Schuster, Bekir Erguner, Jan Laine, Martin Senekowitsch, Christoph Bock, Andreas Bergthaler |
| EPI_ISL_934635, EPI_ISL_934645, EPI_ISL_934648, EPI_ISL_934649 | Department of Microbiology, University Innsbruck | Bergthaler laboratory, CeMM Research Center for Molecular Medicine of the Austrian Academy of Sciences | Lukas Endler, Anna Schedl, Thomas Penz, Benedikt Agerer, Maelle Le Moing, Michael Schuster, Bekir Erguner, Jan Laine, Martin Senekowitsch, Christoph Bock, Andreas Bergthaler |
| EPI_ISL_935156, EPI_ISL_935159 | SIESP CHIETI | Istituto Zooprofilattico Sperimentale dell'Abruzzo e Molise "G. Caporale" | Lorusso A, Marcacci M, Di Domenico M, Ancora M, Curini V, Mangone I, Rinaldi A, Scialabba S, Di Pasquale A, Cammà C, Puglia I, Calistri P, Savini G |
| EPI_ISL_935160 | SIESP CHIETI-DRIVE IN ORTONA | Istituto Zooprofilattico Sperimentale dell'Abruzzo e Molise "G. Caporale" | Lorusso A, Marcacci M, Di Domenico M, Ancora M, Curini V, Mangone I, Rinaldi A, Scialabba S, Di Pasquale A, Cammà C, Puglia I, Calistri P, Savini G |
| EPI_ISL_935161, EPI_ISL_935162 | SIESP CHIETI | Istituto Zooprofilattico Sperimentale dell'Abruzzo e Molise "G. Caporale" | Lorusso A, Marcacci M, Di Domenico M, Ancora M, Curini V, Mangone I, Rinaldi A, Scialabba S, Di Pasquale A, Cammà C, Puglia I, Calistri P, Savini G |
| EPI_ISL_935163, EPI_ISL_935164 | SIESP CHIETI-DRIVE IN ORTONA | Istituto Zooprofilattico Sperimentale dell'Abruzzo e Molise "G. Caporale" | Lorusso A, Marcacci M, Di Domenico M, Ancora M, Curini V, Mangone I, Rinaldi A, Scialabba S, Di Pasquale A, Cammà C, Puglia I, Calistri P, Savini G |
| EPI_ISL_935165, EPI_ISL_935166, EPI_ISL_935167, EPI_ISL_935168, EPI_ISL_935169 | SIESP CHIETI | Istituto Zooprofilattico Sperimentale dell'Abruzzo e Molise "G. Caporale" | Lorusso A, Marcacci M, Di Domenico M, Ancora M, Curini V, Mangone I, Rinaldi A, Scialabba S, Di Pasquale A, Cammà C, Puglia I, Calistri P, Savini G |
| EPI_ISL_935206, EPI_ISL_935208, EPI_ISL_935221, EPI_ISL_935264, EPI_ISL_935265, EPI_ISL_935266, EPI_ISL_935267, EPI_ISL_935268, EPI_ISL_935269, EPI_ISL_935270, EPI_ISL_935271, EPI_ISL_935272, EPI_ISL_935273, EPI_ISL_935301, EPI_ISL_935302, EPI_ISL_935303 | | | |
| see above | KU Leuven, Rega Institute, Clinical and Epidemiological Virology | KU Leuven, Rega Institute, Clinical and Epidemiological Virology | Tony Wawina-Bokalanga, Bert Vanmechelen, Joan Marti-Carerras, Piet Maes |
| EPI_ISL_935971, EPI_ISL_935972 | SIESP CHIETI | Istituto Zooprofilattico Sperimentale dell'Abruzzo e Molise "G. Caporale" | Lorusso A, Marcacci M, Di Domenico M, Ancora M, Curini V, Mangone I, Rinaldi A, Scialabba S, Di Pasquale A, Cammà C, Puglia I, Calistri P, Savini G |
| EPI_ISL_935973, EPI_ISL_935975, EPI_ISL_935991 | GLENS FALLS HOSPITAL LABORATORY | Wadsworth Center, New York State Department of Health | Kirsten St. George, Daryl M. Lamson, Alexis Russel, Matthew Shudt, Melissa A Leisner, Jonathan Plitnick, Navjot Singh, John Kelly, Erasmus Schneider, Erica Lasek-Nesselquist |
| EPI_ISL_936002, EPI_ISL_936006, EPI_ISL_936007, EPI_ISL_936008, EPI_ISL_936011, EPI_ISL_936012, EPI_ISL_936013, EPI_ISL_936014, EPI_ISL_936015, EPI_ISL_936016, EPI_ISL_936017, EPI_ISL_936018, EPI_ISL_936019, EPI_ISL_936020, EPI_ISL_936021 | | | |
| see above | SUNY UPSTATE MEDICAL UNIVERSITY | Wadsworth Center, New York State Department of Health | Kirsten St. George, Daryl M. Lamson, Alexis Russel, Matthew Shudt, Melissa A Leisner, Jonathan Plitnick, Navjot Singh, John Kelly, Erasmus Schneider, Erica Lasek-Nesselquist |
| EPI_ISL_936025, EPI_ISL_936027, EPI_ISL_936028, EPI_ISL_936031 | THE MARY IMOGENE BASSETT HOSPITAL | Wadsworth Center, New York State Department of Health | Kirsten St. George, Daryl M. Lamson, Alexis Russel, Matthew Shudt, Melissa A Leisner, Jonathan Plitnick, Navjot Singh, John Kelly, Erasmus Schneider, Erica Lasek-Nesselquist |
| EPI_ISL_936050, EPI_ISL_936051, EPI_ISL_936055, EPI_ISL_936057, EPI_ISL_936058, EPI_ISL_936059 | ADIRONDACK MEDICAL CENTER | Wadsworth Center, New York State Department of Health | Kirsten St. George, Daryl M. Lamson, Alexis Russel, Matthew Shudt, Melissa A Leisner, Jonathan Plitnick, Navjot Singh, John Kelly, Erasmus Schneider, Erica Lasek-Nesselquist |
| EPI_ISL_936138, EPI_ISL_936140, EPI_ISL_936141 | THE MARY IMOGENE BASSETT HOSPITAL | Wadsworth Center, New York State Department of Health | Kirsten St. George, Daryl M. Lamson, Alexis Russel, Matthew Shudt, Melissa A Leisner, Jonathan Plitnick, Navjot Singh, John Kelly, Erasmus Schneider, Erica Lasek-Nesselquist |
| EPI_ISL_936146 | New York Presbyterian Hospital | Wadsworth Center, New York State Department of Health | Kirsten St. George, Daryl M. Lamson, Alexis Russel, Matthew Shudt, Melissa A Leisner, Jonathan Plitnick, Navjot Singh, John Kelly, Erasmus Schneider, Erica Lasek-Nesselquist |
| EPI_ISL_936153 | THE MARY IMOGENE BASSETT HOSPITAL | Wadsworth Center, New York State Department of Health | Kirsten St. George, Daryl M. Lamson, Alexis Russel, Matthew Shudt, Melissa A Leisner, Jonathan Plitnick, Navjot Singh, John Kelly, Erasmus Schneider, Erica Lasek-Nesselquist |
| EPI_ISL_936388, EPI_ISL_936398 | TGen North | TGen North | Jolene Bowers, Megan Folkerts, Chris French, Hayley Yaglom, Ashlyn Pfeiffer, Darrin Lemmer, Dave Engelthaler, The Arizona COVID Genomics Union (ACGU) |
| EPI_ISL_936466 | DPH, Massachusetts State Public Health Lab | DPH, Massachusetts State Public Health Lab | Lang,A.S., Fink,T., Gallagher,G.R., Smole,S.C. |
| EPI_ISL_936967, EPI_ISL_936968, EPI_ISL_936969, EPI_ISL_936970, EPI_ISL_936971, EPI_ISL_936972, EPI_ISL_936973, EPI_ISL_936974, EPI_ISL_936975, EPI_ISL_936976, EPI_ISL_936977, EPI_ISL_936978, EPI_ISL_936979, EPI_ISL_936980, EPI_ISL_936981, EPI_ISL_936982, EPI_ISL_936983, EPI_ISL_936984, EPI_ISL_936985, EPI_ISL_936986, EPI_ISL_936987, EPI_ISL_936988, EPI_ISL_936989, EPI_ISL_936990, EPI_ISL_936991, EPI_ISL_936992, EPI_ISL_936993, EPI_ISL_936994, EPI_ISL_936995, EPI_ISL_936996, EPI_ISL_936997, EPI_ISL_936998, EPI_ISL_936999, EPI_ISL_937000, EPI_ISL_937001, EPI_ISL_937002, EPI_ISL_937003, EPI_ISL_937004, EPI_ISL_937005, EPI_ISL_937006, EPI_ISL_937007, EPI_ISL_937008, EPI_ISL_937009, EPI_ISL_937010, EPI_ISL_937011, EPI_ISL_937012, EPI_ISL_937013, EPI_ISL_937014, EPI_ISL_937016, EPI_ISL_937017, EPI_ISL_937018, EPI_ISL_937019, EPI_ISL_937020, EPI_ISL_937021, EPI_ISL_937022, EPI_ISL_937023 | | | |
| see above | Northwestern Memorial Hospital | Ozer Lab | Ramon Lorenzo-Redondo, Lacy M. Simons, Chad J. Achenbach, Lawrence J. Jennings, Michael G. Ison, Judd F. Hultquist, Egon A. Ozer |
| EPI_ISL_937437 | Maine Health and Environmental Testing Laboratory (Maine HETL) | Tewhey Lab, The Jackson Laboratory | Matluk,N., Dewey,H., Iosue,F., Barter,M., Lynch,R., Munger,H. and Tewhey,R. |
| EPI_ISL_939628 | Laboratory of Virology and Molecular Diagnostics | Institute of Public Health of Republic of North Macedonia Laboratory of Virology and Molecular Diagnostics | Maja Kuzmanovska. Golubinka Boshevska, Elizabeta Janchevska |
| EPI_ISL_939644, EPI_ISL_939649 | Laboratory of Virology and Molecular Diagnostics | Institute of Public Health of Republic of North Macedonia Laboratory of Virology and Molecular Diagnostics | Maja Kuzmanovska, Golubinka Boshevska, Elizabeta Janchevska |
| EPI_ISL_939650 | Laboratory of Virology and Molecular Diagnostics | Institute of Public Health of Republic of North Macedonia Laboratory of Virology and Molecular Diagnostics | Maja Kuzmanovska,Golubinka Boshevska, Elizabeta Janchevska |
| EPI_ISL_939651 | MD Laboratories | Los Angeles County PHL | P. Hemarajata et al. |
| EPI_ISL_939652 | Los Angeles County PHL | Los Angeles County PHL | P. Hemarajata et al. |

| | | | |
|---|---|---|---|
| EPI_ISL_939653 | MD Laboratories | Los Angeles County PHL | P. Hemarajata et al. |
| EPI_ISL_940623 | Hospital Sao Joaquim - Beneficiencia Portuguesa | Instituto Adolfo Lutz, Interdiciplinary Procedures Center, Strategic Laboratory | Claudio Tavares Sacchi, Claudia Regina Gonçalves, Erica Valessa Ramos Gomes, Karoline Rodrigues Campos |
| EPI_ISL_940727 | City of Milwaukee Health Department Laboratory | City of Milwaukee Health Department Laboratory | Sanjib Bhattacharyya |
| EPI_ISL_940766 | Laboratory of Virology and Molecular Diagnostics | Institute of Public Health of Republic of North Macedonia Laboratory of Virology and Molecular Diagnostics | Maja Kuzmanovska, Golubinka Boshevska, Elizabeta Janchevska |
| EPI_ISL_940767 | Heilig Hart Lier | UAntwerp, Laboratory of Medical Microbiology, Campus Drie Eiken S6.26, Universiteitsplein 1, 2610, Wilrijk, Belgium | Basil Britto Xavier, Jasmine Coppens, Marie Le Mercier, Christine Lammens, Veerle Matheeussen, Herman Goossens |
| EPI_ISL_940864 | Vaccines and Infectious Diseases Analytics Research Unit (VIDA) | KRISP, KZN Research Innovation and Sequencing Platform | Baillie Vicky, du Plessis Jeanine, Giandhari Jennifer, Pillay Sureshnee, Naidoo Yeshnee, Tegally Houriiyah, de Oliveira Tulio, Madhi Shabir |
| EPI_ISL_940995 | Hopital | National Reference Center for Viruses of Respiratory Infections, Institut Pasteur, Paris | Marion Barbet, Sylvie Behillil, Méline Bizard, Angela Brisebarre, Camille Capel, Etienne Simon-Lorière, Vincent Enouf, Maud Vanpeene, Sylvie van der Werf |
| EPI_ISL_940997, EPI_ISL_940998 | Labo Analyses Med | National Reference Center for Viruses of Respiratory Infections, Institut Pasteur, Paris | Marion Barbet, Sylvie Behillil, Méline Bizard, Angela Brisebarre, Camille Capel, Etienne Simon-Lorière, Vincent Enouf, Maud Vanpeene, Sylvie van der Werf, Gestin (B) Brieuc |
| EPI_ISL_941004, EPI_ISL_941007, EPI_ISL_941008, EPI_ISL_941013 | Labo Analyses Med | National Reference Center for Viruses of Respiratory Infections, Institut Pasteur, Paris | Marion Barbet, Sylvie Behillil, Méline Bizard, Angela Brisebarre, Camille Capel, Etienne Simon-Lorière, Vincent Enouf, Maud Vanpeene, Sylvie van der Werf, Amzalag Jonas |
| EPI_ISL_941081 | Labo Analyses Med | National Reference Center for Viruses of Respiratory Infections, Institut Pasteur, Paris | Marion Barbet, Sylvie Behillil, Méline Bizard, Angela Brisebarre, Camille Capel, Etienne Simon-Lorière, Vincent Enouf, Maud Vanpeene, Sylvie van der Werf, Merah Kader |
| EPI_ISL_941163 | Laboratory of Virology and Molecular Diagnostics | Institute of Public Health of Republic of North Macedonia Laboratory of Virology and Molecular Diagnostics | Maja Kuzmanovska |
| EPI_ISL_941228 | Laboratorio de Microbiología. Hospital General Universitario de Elda, Alicante | SeqCOVID-SPAIN consortium/IBV(CSIC) | Mª Isabel Gascón Ros, Cristina Torregrosa Hetland, Eva Pastor Boix, Paloma Cascales Ramos and SeqCOVID-SPAIN consortium |
| EPI_ISL_941336 | Ospedale San Camillo De Lellis di Rieti | INMI Lazzaro Spallanzani IRCCS | Cesare E.M. Gruber, Francesco Messina, Martina Rueca, Emanuela Giombini, Barbara Bartolini, Ornella Butera, Stefano Venarubea, Luca Casertano, Antonino Di Caro, Maria R. Capobianchi |
| EPI_ISL_941337 | Ospedale San Camillo De Lellis di Rieti | INMI Lazzaro Spallanzani IRCCS | Ornella Butera, Barbara Bartolini, Cesare E.M. Gruber, Martina Rueca, Francesco Messina, Emanuela Giombini, Stefano Venarubea, Assunta De Luca, Maria R. Capobianchi, Antonino Di Caro |
| EPI_ISL_941376, EPI_ISL_941377, EPI_ISL_941378, EPI_ISL_941379, EPI_ISL_941380, EPI_ISL_941381, EPI_ISL_941385, EPI_ISL_941386, EPI_ISL_941388, EPI_ISL_941414, EPI_ISL_941426, EPI_ISL_941427, EPI_ISL_941455, EPI_ISL_941456, EPI_ISL_941457, EPI_ISL_941458, EPI_ISL_941459, EPI_ISL_941460, EPI_ISL_941461, EPI_ISL_941631, EPI_ISL_941632, EPI_ISL_941634 | | | |
| see above | Instituto Nacional de Saude (INSA) | Instituto Nacional de Saude (INSA) | Borges et al |
| EPI_ISL_941706, EPI_ISL_941771, EPI_ISL_941796 | Instituto Nacional de Saude (INSA) and Instituto Gulbenkian de Ciencia (IGC) | Instituto Nacional de Saude (INSA) and Instituto Gulbenkian de Ciencia (IGC) | Borges et al |
| EPI_ISL_942166 | Wisconsin State Laboratory of Hygiene Communicable Disease Division | Wisconsin State Laboratory of Hygiene Communicable Disease Division | Kelsey R. Florek, Abigail C. Shockey |
| EPI_ISL_942809, EPI_ISL_942810, EPI_ISL_942811, EPI_ISL_942812, EPI_ISL_942813, EPI_ISL_942814, EPI_ISL_942815, EPI_ISL_942816, EPI_ISL_942817, EPI_ISL_942818 | Gundersen Molecular Diagnostics Laboratory | Kabara Cancer Research Institute | Craig S. Richmond, Paraic A. Kenny |
| EPI_ISL_942916, EPI_ISL_942917, EPI_ISL_942918, EPI_ISL_942919, EPI_ISL_942920, EPI_ISL_942921, EPI_ISL_942922, EPI_ISL_942923, EPI_ISL_942924, EPI_ISL_942925, EPI_ISL_942926 | | | |
| see above | Houston Health Dept. | Houston Health Dept. | Ryker Penn, Pamela Brown, Adolpho Lara |
| EPI_ISL_942945 | Gundersen Molecular Diagnostics Laboratory | Kabara Cancer Research Institute | Craig S. Richmond, Paraic A. Kenny |
| EPI_ISL_942977, EPI_ISL_943012, EPI_ISL_943017, EPI_ISL_943018, EPI_ISL_943020, EPI_ISL_943021, EPI_ISL_943066, EPI_ISL_943144, EPI_ISL_943145, EPI_ISL_943150, EPI_ISL_943151, EPI_ISL_943152, EPI_ISL_943193, EPI_ISL_943194, EPI_ISL_943195, EPI_ISL_943196, EPI_ISL_943197, EPI_ISL_943198, EPI_ISL_943199, EPI_ISL_943243, EPI_ISL_943244, EPI_ISL_943266, EPI_ISL_943269, EPI_ISL_943296, EPI_ISL_943317, EPI_ISL_943360, EPI_ISL_943361, EPI_ISL_943376, EPI_ISL_943377, EPI_ISL_943405, EPI_ISL_943406, EPI_ISL_943407, EPI_ISL_943435, EPI_ISL_943449, EPI_ISL_943461, EPI_ISL_943462, EPI_ISL_943463, EPI_ISL_943464, EPI_ISL_943479, EPI_ISL_943480, EPI_ISL_943496, EPI_ISL_943506, EPI_ISL_943513 | | | |
| see above | Dutch COVID-19 response team | National Institute for Public Health and the Environment (RIVM) | Adam Meijer, Harry Vennema, Dirk Eggink, Jeroen Cremer, Sharon van den Brink, Bas van der Veer, AnneMarie van den Brandt, Florian Zwagemaker, Dennis Schmitz, Chantal Reusken, on behalf of the national COVID-19 response team |
| EPI_ISL_943831, EPI_ISL_943832, EPI_ISL_943833, EPI_ISL_943834, EPI_ISL_943835, EPI_ISL_943940 | Utah Public Health Laboratory | Utah Public Health Laboratory | Erin L. Young, Kelly F. Oakeson, Tara Gallagher |
| EPI_ISL_944623, EPI_ISL_944624, EPI_ISL_944625 | Instituto Nacional de Medicina Genomica | Instituto Nacional de Medicina Genomica | Hidalgo-Miranda A, Mendoza-Vargas A, Reyes-Grajeda JP, Cisneros-Villanueva M, Cedro-Tanda A,Peñaloza-Figueroa F, Herrera-Montalvo LA |
| EPI_ISL_944656, EPI_ISL_944657, EPI_ISL_944669, EPI_ISL_944679, EPI_ISL_944680, EPI_ISL_944681, EPI_ISL_944687, EPI_ISL_944688, EPI_ISL_944693, EPI_ISL_944699, EPI_ISL_944707, EPI_ISL_944711 | | | |
| see above | Department of Biochemistry, Cell and Molecular Biology, West African Centre for Cell Biology of Infectious Pathogens (WACCBIP), University of Ghana | Department of Biochemistry, Cell and Molecular Biology, West African Centre for Cell Biology of Infectious Pathogens (WACCBIP), University of Ghana | Morang'a,C.M., Ngoi,J.M., Quansah,E.B., Saiid,S., Amuzu,D.S., Asante,I., Bonney,J.H., Bonney,E., Odoom,J.K., Ndam,N.T., Tei-Maya,F., Adusei-Poku,M., Ofori-Boadu,L., Ampofo,W.K., Amenga-Etego,L.N., Quashie,P., Bediako,Y., Awandare,G.A. |
| EPI_ISL_944787 | Laboratory of Virology and Molecular Diagnostics | Institute of Public Health of Republic of North Macedonia Laboratory of Virology and Molecular Diagnostics | Maja Kuzmanovska, Golubinka Boshevska, Elizabeta Janchevska |
| EPI_ISL_945049, EPI_ISL_945056, EPI_ISL_945061, EPI_ISL_945072, EPI_ISL_945084, EPI_ISL_945095, EPI_ISL_945114, EPI_ISL_945115, EPI_ISL_945122, EPI_ISL_945125, EPI_ISL_945130, EPI_ISL_945142, EPI_ISL_945143, EPI_ISL_945146, EPI_ISL_945147, EPI_ISL_945152, EPI_ISL_945155, EPI_ISL_945163, EPI_ISL_945169, EPI_ISL_945181, EPI_ISL_945190, EPI_ISL_945197, EPI_ISL_945207, EPI_ISL_945208, EPI_ISL_945212, EPI_ISL_945217, EPI_ISL_945221, EPI_ISL_945236, EPI_ISL_945238, EPI_ISL_945241, EPI_ISL_945243, EPI_ISL_945244, EPI_ISL_945248, EPI_ISL_945250, EPI_ISL_945254, EPI_ISL_945275, EPI_ISL_945279, EPI_ISL_945285, EPI_ISL_945288, EPI_ISL_945297, EPI_ISL_945304, EPI_ISL_945311, EPI_ISL_945319, EPI_ISL_945324, EPI_ISL_945332, EPI_ISL_945334, EPI_ISL_945337, EPI_ISL_945345, EPI_ISL_945347, EPI_ISL_945363, EPI_ISL_945364, EPI_ISL_945366 | | | |
| see above | Lighthouse Lab in Cambridge | Wellcome Sanger Institute for the COVID-19 Genomics UK (COG-UK) Consortium | Rob Howes, The Lighthouse Lab in Cambridge and Alex Alderton, Roberto Amato, Sonia Goncalves, Ewan Harrison, David K. Jackson, Ian Johnston, Dominic Kwiatkowski, Cordelia Langford, John Sillitoe on behalf of the Wellcome Sanger Institute COVID-19 Surveillance Team |
| EPI_ISL_945547, EPI_ISL_946913 | Laboratory of Virology and Molecular Diagnostics | Institute of Public Health of Republic of North Macedonia Laboratory of Virology and Molecular Diagnostics | Maja Kuzmanovska, Golubinka Boshevska, Elizabeta Janchevska |
| EPI_ISL_947328 | RS Mitra Keluarga Depok | Eijkman Institute for Molecular Biology, Ministry of Research and Technology/National Agency for Research and Innovation | Willy Agustine, Edison Johar, Hidayat Trimarsanto, Iskandar Adnan, Lydia V. Panggalo, Sukma Oktavianthi, Frilasita A Yudhaputri, Safarina G Malik, Khin Saw Myint, Amin Soebandrio |
| EPI_ISL_947329, EPI_ISL_947330 | RSU Bunda Mulia | Eijkman Institute for Molecular Biology, Ministry of Research and Technology/National Agency for Research and Innovation | Willy Agustine, Edison Johar, Hidayat Trimarsanto, Iskandar Adnan, Lydia V. Panggalo, Sukma Oktavianthi, Frilasita A Yudhaputri, Safarina G Malik, Khin Saw Myint, Amin Soebandrio |
| EPI_ISL_949124 | SC (UCO) Igiene e Sanità Pubblica (funzione integrata con SC Microbiologia e Virologia) e Laboratory of Molecular Virology of the International Centre for Genetic Engineering | ARGO Laboratorio Genomica ed Epigenomica | Licastro D, Dal Monego S, Degasperi M, Marcello A, D'Agaro P |

| | | | |
|---|---|---|---|
| | and Biotechnology (ICGEB) | | |
| EPI_ISL_949192 | National Institute of Health Research and Development | National Institute of Health Research and Development | Ririn Ramadhany; Yuni Rukminiati; Agustiningsih; Kindi Adam; Holy Arif Wibowo; Hana Apsari Pawestri; Subangkit; Kartika Dewi Puspa; Arie Ardiansyah Nugraha; Hartanti Dian Ikawati; Krisna Nur Andriana Pangesti; Ni Ketut Susilarini; Nur Ika Hariastuti; Uly Alfi Nikmah; Mursinah; Asri Febriyani; Reni Herman; Nike Susanti; Herna; Tati Febriyanti; Natalie Laurentia Kipuw; Fauzul Muna; Irene Lorinda Indalao; Aulia Rizki; Nelly Puspandari; Vivi Setiawaty |
| EPI_ISL_949757, EPI_ISL_949759, EPI_ISL_949760, EPI_ISL_949762, EPI_ISL_949763, EPI_ISL_949765, EPI_ISL_949766, EPI_ISL_949772, EPI_ISL_949774 | Barts Health NHS Trust | COVID-19 Genomics UK (COG-UK) Consortium | CUTINO-MOGUEL, Maria-Teresa; HARRINGTON, David; OWOYEMI, Dola; KULASEGARAN-SHYLINI, Raghavendran; BROAD, Claire; KELE, Beatrix |
| EPI_ISL_949789, EPI_ISL_950069, EPI_ISL_950070, EPI_ISL_950072, EPI_ISL_950074, EPI_ISL_950075, EPI_ISL_950077, EPI_ISL_950079, EPI_ISL_950080, EPI_ISL_950083, EPI_ISL_950084, EPI_ISL_950085, EPI_ISL_950097, EPI_ISL_950098, EPI_ISL_950100, EPI_ISL_950102, EPI_ISL_950110, EPI_ISL_950112, EPI_ISL_950115, EPI_ISL_950126, EPI_ISL_950150, EPI_ISL_950169 | | | |
| see above | University College London, Great Ormond Street Hospital for Children NHS Foundation Trust, Imperial College Healthcare NHS Trust | COVID-19 Genomics UK (COG-UK) Consortium | Sergi Castellano, Rachel Williams, Mark Kristiansen, Paola Resende Silva, Sunando Roy, Tony Brooks, Helena Tutill, Paola Niola, Patricia Dyal, Charlotte Williams, Leysa Forrest, Yasmin Panchbhaya, Jacqueline Findlay, Samuel Weeks, Julianne Brown, Kathryn Harris, Paul Randell, James Price, Alison Holmes, Judith Breuer |
| EPI_ISL_950272 | Northumbria University / South Tees Hospitals NHS Foundation Trust / North Cumbria Integrated Care NHS Foundation Trust / North Tees and Hartlepool NHS Foundation Trust / Newcastle Hospitals NHS Foundation Trust | COVID-19 Genomics UK (COG-UK) Consortium | Darren L Smith,Andrew Nelson,Matthew Bashton,Greg R Young,Joshua Loh,John Allan,Mohammad A Tariq,Giles S Holt,Gary Black,Wen C Yew,Lynn Dover,Paul Baker,Steve Liggett,Sarah Essex,Jane Greenaway,Debra Padgett,Clive Graham,Garren Scott,Edward Barton,Emma Swindells,Brendan Payne,Jennifer Collins,Yusri Taha,Gary Eltringham |
| EPI_ISL_950713 | Lincolnshire Hospitals and DeepSeq Nottingham | COVID-19 Genomics UK (COG-UK) Consortium | Nichola Duckworth, Tim Sloan, Sarah Walsh, Jonathan Ball, Patrick McClure, Joeseph Chappell, Nadine Holmes, Matthew Carlisle, Christopher Moore, Fei Sang, Johnny Debebe, Victoria Wright, Matthew Loose |
| EPI_ISL_951231, EPI_ISL_951232, EPI_ISL_951239, EPI_ISL_951245, EPI_ISL_951246, EPI_ISL_951247, EPI_ISL_951248, EPI_ISL_951252, EPI_ISL_951254, EPI_ISL_951255, EPI_ISL_951256, EPI_ISL_951258, EPI_ISL_951265, EPI_ISL_951268, EPI_ISL_951269, EPI_ISL_951270, EPI_ISL_951271, EPI_ISL_951272, EPI_ISL_951273, EPI_ISL_951275, EPI_ISL_951276, EPI_ISL_951277, EPI_ISL_951281, EPI_ISL_951282, EPI_ISL_951283, EPI_ISL_951286, EPI_ISL_951287, EPI_ISL_951288, EPI_ISL_951289, EPI_ISL_951294, EPI_ISL_951299, EPI_ISL_951300, EPI_ISL_951308, EPI_ISL_951309, EPI_ISL_951316, EPI_ISL_951356, EPI_ISL_951364, EPI_ISL_951366, EPI_ISL_951367, EPI_ISL_951389, EPI_ISL_951391, EPI_ISL_951392, EPI_ISL_951393, EPI_ISL_951395, EPI_ISL_951396, EPI_ISL_951406, EPI_ISL_951407, EPI_ISL_951408, EPI_ISL_951409, EPI_ISL_951414, EPI_ISL_951426, EPI_ISL_951427, EPI_ISL_951429 | | | |
| see above | Oxford Viromics, NDM, University of Oxford; Oxford University Hospitals; Basingstoke and North Hampshire Hospital | COVID-19 Genomics UK (COG-UK) Consortium | Tanya Golubchik, David Bonsall, George Macintyre, Amy Trebes, Mariateresa de Cesare, Catrin Moore, Alex Mobbs, Anita Justice, Robert Shaw, Monique Andersson, Timothy Peto, Emma Wise, Nathan Moore, Jessica Lynch, Nick Cortes, Matilde Mori, Stephen Kidd, David Buck, John Todd, Christophe Fraser |
| EPI_ISL_952381, EPI_ISL_952384, EPI_ISL_952388, EPI_ISL_952396, EPI_ISL_952400, EPI_ISL_952402, EPI_ISL_952406, EPI_ISL_952419, EPI_ISL_952425, EPI_ISL_952426, EPI_ISL_952781, EPI_ISL_952782, EPI_ISL_952784, EPI_ISL_952786, EPI_ISL_952787, EPI_ISL_952788, EPI_ISL_952791, EPI_ISL_952800, EPI_ISL_952802, EPI_ISL_952805, EPI_ISL_952806, EPI_ISL_952807, EPI_ISL_952808, EPI_ISL_952809, EPI_ISL_952810, EPI_ISL_952811, EPI_ISL_952812, EPI_ISL_952819 | | | |
| see above | Centre for Enzyme Innovation, University of Portsmouth / Translational Research Laboratory, Portsmouth Hospitals NHS Trust | COVID-19 Genomics UK (COG-UK) Consortium | Angela Beckett,Salman Goudarzi,Christopher Fearn,Kate Cook,Katie Loveson,Sharon Glaysher,Scott Elliott,Samuel Robson |
| EPI_ISL_953177, EPI_ISL_953180, EPI_ISL_953181, EPI_ISL_953182, EPI_ISL_953185, EPI_ISL_953187, EPI_ISL_953193, EPI_ISL_953199, EPI_ISL_953205, EPI_ISL_953207, EPI_ISL_953216, EPI_ISL_953217, EPI_ISL_953219, EPI_ISL_953222, EPI_ISL_953226, EPI_ISL_953230, EPI_ISL_953231, EPI_ISL_953232, EPI_ISL_953233, EPI_ISL_953236, EPI_ISL_953238, EPI_ISL_953239 | | | |
| see above | Bioinformatics and Biostatistics Lab, Advanced Sequencing Facility | COVID-19 Genomics UK (COG-UK) Consortium | Aengus Stewart,Jerome Nicod,Chelsea Sawyer,Laura Cubitt,Harshil Patel,Margaret Crawford |
| EPI_ISL_953521 | University Hospitals of Geneva, Laboratory of Virology | HUG, Laboratory of Virology and the Health2030 Genome Center | Samuel Cordey, Ana Rita Goncalves, Laurent Kaiser, Lorenzo Cerutti, Henri Pegeot, Melyssa Elies, Deborah Penet, Keith Harshman, Ioannis Xenarios, Emmanouil Dermitzakis |
| EPI_ISL_954128 | Labo Analyses Med | National Reference Center for Viruses of Respiratory Infections, Institut Pasteur, Paris | Marion Barbet, Sylvie Behillil, Méline Bizard, Angela Brisebarre, Camille Capel, Etienne Simon-Lorière, Vincent Enouf, Maud Vanpeene, Sylvie van der Werf,Takoudju Eve-Marie |
| EPI_ISL_954223, EPI_ISL_954224, EPI_ISL_954225 | 1.AO Universitaria 'S. Giovanni di Dio e Ruggi D'Aragona, Scuola Medica Salernitana' Hospital / 2.UOC di Virologia e Microbiologia, Università della Campania 'L. Vanvitelli' / 3.AO Universitaria 'Federico II' Napoli Hospital / 4.AORN 'San Giuseppe Moscati' Avellino Hospital / 5.AO 'San Pio - presidio G. Rummo' Benevento Hospital / 6.AO 'Sant'Anna e San Sebastiano' Caserta Hospital / 7.PO 'Maria Santissima Addolorata' Eboli Hospital / 8.Biogem Istituto di Ricerche Genetiche | 1. Genome Research Center for Health (CRGS) / 2. Laboratory of Molecular Medicine and Genomics(LMMGe) / 3. Center for Research in Pure and Applied Mathematics (CRMPA) | Giorgio Giurato, Francesca Rizzo, Alessandro Weisz, Gianluigi Franci, Giovanni Nassa, Pasquale Pagliano, Roberta Tarallo, Elena Alexandrova, Ylenia D'Agostino, Carlo Ferravante, Jessica Lamberti, Viola Melone, Domenico Memoli, Valeria Mirici Cappa, Domenico Palumbo, Giovanni Pecoraro, Assunta Sellitto, Oriana Strianese, Ilaria Terenzi, Giuseppe Fenza, Aniello Gentile, Antonello Saccomanno, Sonia Amabile, Teresa Rocco, Annamaria Salvati, Emilia Vaccaro, Massimiliano Galdiero, Michele Cennamo, Giuseppe Portella, Maria Grazia Foti, Mariarosaria Ingino, Maria Landi, Maurizio Fumi, Vincenzo Rocco, Rita Greco, Vittoria Letizia, Arnolfo Petruzziello, Maddalena Schioppa, Gregorio Goffredi, Francesca Marciano, Michele Caraglia, Alessia Cossu, Marianna Scrima, Edmondo Adorisio, Morena D'Avenia, Michela Iacobellis, Rosanna Piluscio, Giorgio Dirani, Vittorio Sambri, Simona Semprini, Silvia Zanoli, Francesco Curcio, Stefania Marzinotto, Andreina Baj, Fausto Sessa. |
| EPI_ISL_954781 | Hospital General Universitario de Ciudad Real | Instituto de Salud Carlos III | Iglesias-Caballero, M. Camarero, S. Sandonís,V. Vázquez, S. Pozo, F. Casas, I. Jiménez, P. Zaballos, A. Monzón, S. Varona, S. Cuesta, I. Illescas, S. |
| EPI_ISL_954793 | Hospital Universitario Virgen de la Arrixaca | Instituto de Salud Carlos III | Iglesias-Caballero, M. Camarero, S. Sandonís,V. Vázquez, S. Pozo, F. Casas, I. Jiménez, P. Zaballos, A. Monzón, S. Varona, S. Cuesta, I. Moreno, L. |
| EPI_ISL_954796 | HOSPITAL COMARCAL DE MELILLA | Instituto de Salud Carlos III | Iglesias-Caballero, M. Camarero, S. Sandonís,V. Vázquez, S. Pozo, F. Casas, I. Jiménez, P. Zaballos, A. Monzón, S. Varona, S. Cuesta, I. Illescas, S. |
| EPI_ISL_954804 | Complejo Hospitalario de Navarra | Instituto de Salud Carlos III | Iglesias-Caballero, M. Camarero, S. Sandonís,V. Vázquez, S. Pozo, F. Casas, I. Jiménez, P. Zaballos, A. Monzón, S. Varona, S. Cuesta, I. Ezpeleta,C. |
| EPI_ISL_954858, EPI_ISL_954876, EPI_ISL_954919, EPI_ISL_954920, EPI_ISL_954921, EPI_ISL_954922, EPI_ISL_954923, EPI_ISL_954924, EPI_ISL_954925, EPI_ISL_954926, EPI_ISL_954927, EPI_ISL_954928, EPI_ISL_954929, EPI_ISL_954930 | | | |
| see above | Colorado Department of Public Health and Environment | Colorado Department of Puplic Health and Environment | Laura Bankers, Molly C. Hetherington-Rauth, Diana Ir, Shannon Ely, Shannon R. Matzinger, Sarah Elizabeth Totten, Emily A. Travanty |
| EPI_ISL_955277, EPI_ISL_955319, EPI_ISL_955329, EPI_ISL_955334 | GA Department of Public Health Laboratory | Pathogen Discovery, Respiratory Viruses Branch, Division of Viral Diseases, Centers for Disease Control and Prevention | Ying Tao, Jing Zhang, Yan Li, Krista Queen, Anna Uehara, Peter Cook, Clinton R. Paden, Haibin Wang, Suxiang Tong |
| EPI_ISL_956304, EPI_ISL_956305 | ESE INSTITUTO NACIONAL DE CANCEROLOGIA | Instituto Nacional de Salud- Dirección de Investigación en Salud Pública | Katherine Laiton-Donato, Diego A. Álvarez-Díaz, Carlos Franco-Muñoz, Mauricio Pacheco-Montealegre, Hector Alejandro Ruiz-Moreno, Maria T. Herrera-Sepúlveda, Diego Andrés Prada, Jhonnatan Reales-González, Sheryll Corchuelo, Julian Naizaque, Gerardo Santamaría, Magdalena Wiesner, Martha Lucia Ospina Martinez, Marcela Mercado-Reyes |
| EPI_ISL_956313 | Siti Khodijah Hospital | Institute of Tropical Disease, Universitas Airlangga | Jezzy R Dewantari, Rima R Prasetya, Krisnoadi Rahardjo, Aldise M Nastri, Muhammad Hamdan, Gatot Soegiarto, Laksmi Wulandari, Resti Yudhawati, Yasuko Mori, Soetjipto, Kazufumi Shimizu, Maria I Lusida |
| EPI_ISL_959274 | General Hospital - Tetovo | Research Center for Genetic Engineering and Biotechnology "Georgi D. Efremov" , Macedonian Academy of Sciences and Arts | Aleksandar J. Dimovski, Dijana Plasheska-Karanfilska, Predrag Noveski, Gjorgji Bozinovski, Milena Jakimovska |
| EPI_ISL_959285 | General Hospital - Prilep | Research Center for Genetic Engineering and Biotechnology "Georgi D. Efremov" , Macedonian Academy of Sciences and Arts | Aleksandar J. Dimovski, Dijana Plasheska-Karanfilska, Predrag Noveski, Gjorgji Bozinovski, Milena Jakimovska |
| EPI_ISL_959551, EPI_ISL_959552 | CHC Liège | GIGA Medical Genomics | Keith Durkin, Maria Artesi, Sébastien Bontems, Raphaël Boreux, Bouchra Boujemla, Cécile Meex, Pierrette Melin, Marie-Pierre Hayette, Vincent Bours |
| EPI_ISL_959553 | CHR Namur | GIGA Medical Genomics | Keith Durkin, Maria Artesi, Sébastien Bontems, Raphaël Boreux, Bouchra Boujemla, Cécile Meex, Pierrette Melin, Marie-Pierre Hayette, Vincent Bours |
| EPI_ISL_959554 | Vivalia - Clinique Saint-Joseph | GIGA Medical Genomics | Keith Durkin, Maria Artesi, Sébastien Bontems, Raphaël Boreux, Bouchra Boujemla, Cécile Meex, Pierrette Melin, Marie-Pierre Hayette, Vincent Bours |
| EPI_ISL_959650 | General Hospital "Abdulah Nakas" Sarajevo | Alea Genetic Centre | Dino Pecar, Lana Salihefendic, Sead Jazic, Rijad Konjhodzic |
| EPI_ISL_959855, EPI_ISL_959858 | National Virus Reference Laboratory | National Virus Reference Laboratory | Michael Carr, Gabriel Gonzalez, Jonathan Dean, Cillian F De Gascun |

| | | | |
|---|---|---|---|
| EPI_ISL_960324 | University of Wisconsin-Madison AIDS Vaccine Research Laboratories | University of Wisconsin-Madison AIDS Vaccine Research Laboratories | Gage Moreno, Katarina Braun, et al. AIDS Vaccine Research Laboratories |
| EPI_ISL_960404, EPI_ISL_960407, EPI_ISL_960413, EPI_ISL_960419, EPI_ISL_960430, EPI_ISL_960436 | The National Institute of Public Health | State Veterinary Institute Prague | Nagy,A;Vecerova,J;Cernikova,L;Stara,M;Jirincova,H;Trnka,D |
| EPI_ISL_960834, EPI_ISL_960876 | Institute of Medical Microbiology and Hospital Hygiene | Institute of Medical Microbiology and Hospital Hygiene | Prof. Dr. Achim Kaasch, Aljoscha Tersteegen |
| EPI_ISL_961105, EPI_ISL_961106, EPI_ISL_961107, EPI_ISL_961108, EPI_ISL_961109, EPI_ISL_961110, EPI_ISL_961111, EPI_ISL_961112, EPI_ISL_961113, EPI_ISL_961114, EPI_ISL_961115, EPI_ISL_961116, EPI_ISL_961117, EPI_ISL_961118, EPI_ISL_961119, EPI_ISL_961120, EPI_ISL_961121, EPI_ISL_961122, EPI_ISL_961123, EPI_ISL_961124, EPI_ISL_961125, EPI_ISL_961126, EPI_ISL_961131 | | | |
| see above | Delaware Public Health Laboratory | Delaware Public Health Lab | Gregory Hovan |
| EPI_ISL_961475, EPI_ISL_961487, EPI_ISL_961500, EPI_ISL_961501, EPI_ISL_961507, EPI_ISL_961515, EPI_ISL_961516, EPI_ISL_961547 | Michigan Department of Health and Human Services, Bureau of Laboratories | Michigan Department of Health and Human Services, Bureau of Laboratories | Blankenship HM, Riner D, Soehnlen MK |
| EPI_ISL_961556, EPI_ISL_961557, EPI_ISL_961558, EPI_ISL_961559, EPI_ISL_961598, EPI_ISL_961643, EPI_ISL_961644, EPI_ISL_961645, EPI_ISL_961646, EPI_ISL_961647, EPI_ISL_961648 | | | |
| see above | Hôpital Georges L. Dumont | National Microbiology Laboratory (NML) | Anna Majer, Shari Tyson, Grace Seo, Philip Mabon, Elsie Grudeski, Rhiannon Huzarewich, Russell Mandes, Anneliese Landgraff, Jennifer Tanner, Natalie Knox, Morag Graham, Gary Van Domselaar, Richard Garceau, Guillaume Desnoyers, Nathalie Bastien, Yan Li, Timothy Booth, Darian Hole, Madison Chapel, Kirsten Biggar, CanCOGeN's metadata curation team, Public Health Agency of Canada CanCOGeN team |
| EPI_ISL_961881, EPI_ISL_961882 | E. Gulbja laboratorija | Latvian Biomedical Research and Study Centre | Janis Pjalkovskis, Nikita Zrelovs, Monta Ustinova, Ivars Silamikelis, Liga Birzniece, Kaspars Megnis, Vita Rovite, Lauma Freimane, Laila Silamikele, Laura Ansone, Davids Fridmanis, Mikus Gavars, Dmitrijs Perminovs, Jurijs Perevoscikovs, Uga Dumpis, Janis Klovins |
| EPI_ISL_961897, EPI_ISL_961920, EPI_ISL_961925, EPI_ISL_961930, EPI_ISL_961931, EPI_ISL_961932, EPI_ISL_961935, EPI_ISL_961938, EPI_ISL_961960, EPI_ISL_961962, EPI_ISL_961969, EPI_ISL_961971, EPI_ISL_962154, EPI_ISL_962161, EPI_ISL_962162, EPI_ISL_962163, EPI_ISL_962164, EPI_ISL_962165, EPI_ISL_962166, EPI_ISL_962167, EPI_ISL_962168, EPI_ISL_962169, EPI_ISL_962170, EPI_ISL_962172 | | | |
| see above | Illinois Department of Public Health | Gagnon Lab, Southern Illinois University | Keith Gagnon |
| EPI_ISL_962511 | UCLA Clinical Micro Lab | Los Angeles County PHL | P. Hemarajata et al. |
| EPI_ISL_962516 | MD Laboratories | Los Angeles County PHL | P. Hemarajata et al. |
| EPI_ISL_962812 | Robert Garry lab | Andersen lab at Scripps Research | Allison Smither, Gilberto Sabino-Santos, Patricia Snarski, Lilia Melnik, Antoinette Bell, Kaylynn Genemaras, Arnaud Drouin, Dahlene Fusco, Robert Garry with SEARCH Alliance San Diego |
| EPI_ISL_962825 | Victorian Infectious Diseases Reference Laboratory (VIDRL) | VIDRL and MDU-PHL | Caly L., Seemann T., Sait, M.L., Druce J., Sherry, N.L. |
| EPI_ISL_962889 | Laboratory of Virology and Molecular Diagnostics | Institute of Public Health of Republic of North Macedonia Laboratory of Virology and Molecular Diagnostics | Kuzmanovska M., Boshevska G. |
| EPI_ISL_962915, EPI_ISL_962917, EPI_ISL_962922, EPI_ISL_962923 | Servicio de Microbiología, Laboratori Clínic Metropolitana Nord. Hospital Universitari Germans Trias i Pujol. Institut d'Investigació en Ciències de la Salut Germans Trias i Pujol (IGTP) | SeqCOVID-SPAIN consortium/IBV(CSIC) | Elisa Martró, Antoni E. Bordoy, Anna Not, Adrián Antuori, Anabel Fernández, Nona Romaní, Verónica Saludes, Cristina Casañ and SeqCOVID-SPAIN consortium |
| EPI_ISL_962934 | Hospital Universitario de Gran Canaria Dr. Negrín | SeqCOVID-SPAIN consortium/IBV(CSIC) | M. Carmen Pérez González, Francisco J. Chamizo López, Ana Bordes Benítez and SeqCOVID-SPAIN consortium |
| EPI_ISL_964292 | Oslo University Hospital, Department of Medical Microbiology | Norwegian Institute of Public Health, Department of Virology | Kathrine Stene-Johansen, Kamilla Heddeland Instefjord, Hilde Elshaug, Ignacio Garcia Llorente, Serina B Engebretsen, Atiya R Ali, Marie Paulsen Madsen, Rasmus Riis Kopperud, Hilde Vollan, Karoline Bragstad, Olav Hungnes |
| EPI_ISL_964924 | Instituto Nacional de Saude (INS), Mozambique | KRISP, KZN Research Innovation and Sequencing Platform | Nalia Ismael, Nadia Sitoe, Paulo Arnaldo, Nedio Mabunda, Giandhari J, Pillay S, Emmanuel S, Tegally H, Wilkinson E, de Oliveira T |
| EPI_ISL_964951 | Haukeland University Hospital, Dept. of Microbiology | Norwegian Institute of Public Health, Department of Virology | Kathrine Stene-Johansen, Kamilla Heddeland Instefjord, Hilde Elshaug, Ignacio Garcia Llorente, Serina B Engebretsen, Atiya R Ali,Marie Paulsen Madsen, Rasmus Riis Kopperud, Hilde Vollan, Karoline Bragstad, Olav Hungnes |
| EPI_ISL_965131 | Azienda Ospedaliera San Giovanni Addolorata | INMI Lazzaro Spallanzani IRCCS | O Butera, F Messina, CEM Gruber, B Bartolini, E Giombini, M Rueca, M Gaudio, PM Placanica, MR Capobianchi, A Di Caro |
| EPI_ISL_965136 | Azienda Sanitaria Locale Roma 5 | INMI Lazzaro Spallanzani IRCCS | O Butera, F Messina, CEM Gruber, B Bartolini, E Giombini, M Rueca, D Di Fusco, D Cerini, MR Capobianchi, A Di Caro |
| EPI_ISL_965657, EPI_ISL_965690, EPI_ISL_965763, EPI_ISL_965793, EPI_ISL_965806 | Dutch COVID-19 response team | Medical Microbiology, Maastricht University Medical Centre | Jozef Dingemans*, Brian van der Veer*, Erik Beuken, Carmen Reumkens, Lieke van Alphen, Christian Hoebe, Paul Savelkoul |
| EPI_ISL_966782 | Maine HETL | Tewhey Lab, The Jackson Laboratory | Matluk,N., Dewey,H., Iosue,F., Barter,M., Lynch,R., Munger,H. and Tewhey,R. |
| EPI_ISL_967220 | Helix/Illumina | Respiratory Viruses Branch, Division of Viral Diseases, Centers for Disease Control and Prevention | Peter W. Cook,Dakota Howard,Dhwani Batra,Ben L. Rambo-Martin,Eileen de Feo,Jan Antico,Christine Tran,Matthew Tolentino,Shannon Wickline,Kim Gietzen,Brad Sickler,Jingtao Liu,Eric Allen,Phil Febbo,Summer Galloway,Nicole L. Washington,Simon White,Geraint Levan,Kelly Schiabor Barrett,Elizabeth Cirulli,Alexandre Bolze,Ary Ascencio,Charlotte Rivera-Garcia,Ryan Cho,Jason Nguyen,Sherry Wang,Jimmy Ramirez,Tyler Cassens,Efren Sandoval,Magnus Isaksson,William Lee,David Becker,Marc Laurent,James Lu,Clinton R. Paden,Suxiang Tong,Duncan MacCannell, |
| EPI_ISL_967696, EPI_ISL_967743 | State Laboratories Division, Hawaii State Department of Health | State Laboratories Division, Hawaii State Department of Health | Pamela O'Brien, Drew Kuwazaki, Ayana Garnet, Razvan Sultana, Edward Desmond |
| EPI_ISL_967901, EPI_ISL_967906, EPI_ISL_967938, EPI_ISL_967955, EPI_ISL_967974, EPI_ISL_968001, EPI_ISL_968055 | TGen North | TGen North | "Jolene Bowers, Megan Folkerts, Chris French, Hayley Yaglom, Ashlyn Pfeiffer, Darrin Lemmer, Dave Engelthaler, The Arizona COVID Genomics Union (ACGU)" |
| EPI_ISL_970179 | Hospital San Agustin, Avilés | Laboratorio de Virología HUCA | Castelló C, Gómez de Oña J, Boga JA, Rojo S, Alvarez-Arguelles ME, Abreu F, Costales I, Sandoval M, Perez-Martínez Z, Martín-Rodríguez G, Coto E, Melón S |
| EPI_ISL_976941, EPI_ISL_976942, EPI_ISL_976943, EPI_ISL_976944, EPI_ISL_976946, EPI_ISL_976947, EPI_ISL_976948, EPI_ISL_976949, EPI_ISL_976950, EPI_ISL_976951, EPI_ISL_976952, EPI_ISL_976954, EPI_ISL_976955, EPI_ISL_976956, EPI_ISL_976957, EPI_ISL_976958, EPI_ISL_976959, EPI_ISL_976960, EPI_ISL_976962, EPI_ISL_976963, EPI_ISL_976964, EPI_ISL_976965, EPI_ISL_976966, EPI_ISL_976967, EPI_ISL_976968, EPI_ISL_976970, EPI_ISL_976971, EPI_ISL_976972, EPI_ISL_976973, EPI_ISL_976974, EPI_ISL_976975, EPI_ISL_976976, EPI_ISL_976978, EPI_ISL_976979, EPI_ISL_976980, EPI_ISL_976983, EPI_ISL_976984, EPI_ISL_976985, EPI_ISL_976987, EPI_ISL_976988, EPI_ISL_976989, EPI_ISL_976990, EPI_ISL_976991, EPI_ISL_976992, EPI_ISL_976993, EPI_ISL_976995, EPI_ISL_976996, EPI_ISL_976997, EPI_ISL_976998, EPI_ISL_976999, EPI_ISL_977000, EPI_ISL_977002, EPI_ISL_977003, EPI_ISL_977004, EPI_ISL_977005, EPI_ISL_977006, EPI_ISL_977007, EPI_ISL_977009, EPI_ISL_977010, EPI_ISL_977011, EPI_ISL_977012, EPI_ISL_977013, EPI_ISL_977015, EPI_ISL_977016, EPI_ISL_977017, EPI_ISL_977018, EPI_ISL_977019, EPI_ISL_977020, EPI_ISL_977021, EPI_ISL_977022, EPI_ISL_977023 | | | |
| see above | Broad Institute Clinical Research Sequencing Platform | Infectious Disease Program, Broad Institute of Harvard and MIT | Lemieux,J.E., Siddle,K.J., Adams,G., Gladden-Young,A., Lagerborg,K., Rudy,M., DeRuff,K., Carter,A., Normandin,E., Bauer,M., Reilly,S., Tomkins-Tinch,C., Loreth,C., Chaluvadi,S., Birren,B.W., Gallagher,G., Smole,S., Park,D.J., MacInnis,B.L., and Sabeti,P.C. |
| EPI_ISL_977083 | Massachusetts General Hospital | Infectious Disease Program, Broad Institute of Harvard and MIT | Lemieux,J.E., Siddle,K.J., Shaw,B., Adams,G., Pierce,V., Turbett,S., Anahtar,M., Branda,J., Slater,D., Harris,J., Lin,A.E., Gladden-Young,A., Lagerborg,K., Rudy,M., DeRuff,K., Carter,A., Normandin,E., Bauer,M., Reilly,S., Tomkins-Tinch,C., Loreth,C., Chaluvadi,S., Neumann,A., Cusick,C., Chapman,S.B., Gnirke,A., Flowers,K., Cerrato,F., Birren,B.W., Gallagher,G., Smole,S., Park,D.J., MacInnis,B.L., Ryan,E., LaRocque,R., Rosenberg,E. and Sabeti,P.C. |
| EPI_ISL_977155, EPI_ISL_977157 | ULSS 03 Venezia | Istituto Zooprofilattico Sperimentale delle Venezie | Adelaide Milani, Alessia Schivo, Annalisa Salviato, Erika Giorgia Quaranta, Ambra Pastori, Bianca Zecchin, Alice Fusaro, Isabella Monne, Calogero Terregino, Antonia Ricci |
| EPI_ISL_977347 | University of Zambia, School of Veterinary Medicine | UNZAVET and PATH | Mulenga Mwenda-Chimfwembe, Ngonda Saasa, Daniel Bridges |
| EPI_ISL_977541, EPI_ISL_977542, EPI_ISL_977545, EPI_ISL_977546, | Nigeria Centre of Disease Control (NCDC) | African Centre of Excellence for Genomics of Infectious Diseases (ACEGID), Redeemer's University | Olawoye I. B. et al |

| | | | |
|---|---|---|---|
| EPI_ISL_977547, EPI_ISL_977548, EPI_ISL_977552, EPI_ISL_977563, EPI_ISL_977565 | | | |
| EPI_ISL_977652 | Michigan Department of Health and Human Services, Bureau of Laboratories | Michigan Department of Health and Human Services, Bureau of Laboratories | Blankenship HM, Riner D, Soehnlen MK |
| EPI_ISL_977962, EPI_ISL_977963, EPI_ISL_977964, EPI_ISL_977965, EPI_ISL_977966, EPI_ISL_977967, EPI_ISL_977968, EPI_ISL_977969, EPI_ISL_977970, EPI_ISL_977971, EPI_ISL_977972, EPI_ISL_977973, EPI_ISL_977974, EPI_ISL_977975, EPI_ISL_977976, EPI_ISL_977977, EPI_ISL_977978 | | | |
| see above | Chiu Laboratory, University of California, San Francisco | Chiu Laboratory, University of California, San Francisco | Charles Chiu, Xianding (Wayne) Deng, Candace Wang, Venice Servellita, Jill Hacker, Debra Wadford |
| EPI_ISL_978357, EPI_ISL_978366, EPI_ISL_978367, EPI_ISL_978368, EPI_ISL_978369, EPI_ISL_978370 | Arizona State Public Health Laboratory | Arizona State Public Health Laboratory | Trung Huynh, Jessica Escobar, Katherine Fullerton, Nobuko Fukushima, Stacy White, Linda Getsinger, Victor Waddell |
| EPI_ISL_978625, EPI_ISL_978626, EPI_ISL_978627, EPI_ISL_978628, EPI_ISL_978629, EPI_ISL_978630, EPI_ISL_978631, EPI_ISL_978632, EPI_ISL_978633, EPI_ISL_978634, EPI_ISL_978635, EPI_ISL_978636, EPI_ISL_978637, EPI_ISL_978638, EPI_ISL_978639, EPI_ISL_978640, EPI_ISL_978641, EPI_ISL_978642, EPI_ISL_978643, EPI_ISL_978644, EPI_ISL_978645, EPI_ISL_978646, EPI_ISL_978647, EPI_ISL_978648, EPI_ISL_978649, EPI_ISL_978650, EPI_ISL_978651, EPI_ISL_978652, EPI_ISL_978653, EPI_ISL_978654, EPI_ISL_978655, EPI_ISL_978656, EPI_ISL_978657, EPI_ISL_978658, EPI_ISL_978659, EPI_ISL_978660, EPI_ISL_978661, EPI_ISL_978662, EPI_ISL_978663, EPI_ISL_978664, EPI_ISL_978665, EPI_ISL_978666, EPI_ISL_978667, EPI_ISL_978668, EPI_ISL_978669, EPI_ISL_978670, EPI_ISL_978671, EPI_ISL_978672, EPI_ISL_978673, EPI_ISL_978674, EPI_ISL_978675, EPI_ISL_978676, EPI_ISL_978677, EPI_ISL_978678, EPI_ISL_978679, EPI_ISL_978680, EPI_ISL_978681, EPI_ISL_978682, EPI_ISL_978683, EPI_ISL_978684, EPI_ISL_978685, EPI_ISL_978686, EPI_ISL_978687, EPI_ISL_978688, EPI_ISL_978689, EPI_ISL_978690, EPI_ISL_978691, EPI_ISL_978692, EPI_ISL_978693, EPI_ISL_978694, EPI_ISL_978695, EPI_ISL_978696, EPI_ISL_978697, EPI_ISL_978698, EPI_ISL_978699, EPI_ISL_978700, EPI_ISL_978701, EPI_ISL_978702, EPI_ISL_978703, EPI_ISL_978704, EPI_ISL_978705, EPI_ISL_978706, EPI_ISL_978707, EPI_ISL_978708, EPI_ISL_978709, EPI_ISL_978710, EPI_ISL_978711, EPI_ISL_978712, EPI_ISL_978713, EPI_ISL_978714, EPI_ISL_978715, EPI_ISL_978716, EPI_ISL_978717, EPI_ISL_978718, EPI_ISL_978719, EPI_ISL_978720, EPI_ISL_978721, EPI_ISL_978722, EPI_ISL_978723, EPI_ISL_978724, EPI_ISL_978725, EPI_ISL_978726, EPI_ISL_978727, EPI_ISL_978728, EPI_ISL_978729, EPI_ISL_978730, EPI_ISL_978731, EPI_ISL_978732, EPI_ISL_978733, EPI_ISL_978734, EPI_ISL_978735, EPI_ISL_978736, EPI_ISL_978737, EPI_ISL_978738, EPI_ISL_978739, EPI_ISL_978740 | | | |
| see above | Helix/Illumina | Respiratory Viruses Branch, Division of Viral Diseases, Centers for Disease Control and Prevention | Peter W. Cook,Dakota Howard,Dhwani Batra,Ben L. Rambo-Martin,Eileen de Feo,Jan Antico,Christine Tran,Matthew Tolentino,Shannon Wickline,Kim Gietzen,Brad Sickler,Jingtao Liu,Eric Allen,Phil Febbo,Summer Galloway,Nicole L. Washington,Simon White,Geraint Levan,Kelly Schiabor Barrett,Elizabeth Cirulli,Alexandre Bolze,Ary Ascencio,Charlotte Rivera-Garcia,Ryan Cho,Jason Nguyen,Sherry Wang,Jimmy Ramirez,Tyler Cassens,Efren Sandoval,Magnus Isaksson,William Lee,David Becker,Marc Laurent,James Lu,Clinton R. Paden,Suxiang Tong,Duncan MacCannell, |
| EPI_ISL_978973 | Chiu Laboratory, University of California, San Francisco | Chiu Laboratory, University of California, San Francisco | Charles Chiu, Xianding (Wayne) Deng, Candace Wang, Venice Servellita, Jill Hacker, Debra Wadford |
| EPI_ISL_979085, EPI_ISL_979089, EPI_ISL_979092, EPI_ISL_979093, EPI_ISL_979094, EPI_ISL_979095, EPI_ISL_979096, EPI_ISL_979097, EPI_ISL_979098 | Santa Clara County Public Health Laboratory | Chan-Zuckerberg Biohub | CZB Cliahub Consortium |
| EPI_ISL_979343, EPI_ISL_979344, EPI_ISL_979345, EPI_ISL_979346, EPI_ISL_979347, EPI_ISL_979348, EPI_ISL_979349, EPI_ISL_979350, EPI_ISL_979351, EPI_ISL_979352 | Laboratorio Estatal de Salud Pública de Nuevo León | Laboratorio de Infectología Molecular, Departamento de Bioquímica y Medicina Molecular,Facultad de Medicina - Universidad Autónoma de Nuevo León | Kame A. Galán-Huerta, María F. Herrera-Saldivar, Natalia Martínez-Acuña, Sonia A. Lozano-Sepúlveda, Daniel Arellanos-Soto, Ana M. Rivas-Estilla, Samuel Buentello-Wong, Else del Carmen García-García, Gloria A. Jasso-de-la-Peña, Roberto Montes-de-Oca, Consuelo Treviño-Garza, Manuel E. de-la-O-Cavazos |
| EPI_ISL_979379, EPI_ISL_979380, EPI_ISL_979381, EPI_ISL_979382, EPI_ISL_979383, EPI_ISL_979384, EPI_ISL_979385, EPI_ISL_979386, EPI_ISL_979387, EPI_ISL_979388, EPI_ISL_979389, EPI_ISL_979390, EPI_ISL_979391, EPI_ISL_979392, EPI_ISL_979393, EPI_ISL_979394, EPI_ISL_979395, EPI_ISL_979396, EPI_ISL_979397, EPI_ISL_979398, EPI_ISL_979399, EPI_ISL_979400, EPI_ISL_979401, EPI_ISL_979402, EPI_ISL_979403, EPI_ISL_979404, EPI_ISL_979405, EPI_ISL_979406, EPI_ISL_979407, EPI_ISL_979408, EPI_ISL_979409 | | | |
| see above | Houston Health Dept. | Houston Health Dept. | Ryker Penn, Pamela Brown, Adolpho Lara |
| EPI_ISL_981947, EPI_ISL_981948, EPI_ISL_981949, EPI_ISL_981950, EPI_ISL_981951, EPI_ISL_981952 | Microbiology Service, Hospital Universitario Clinico San Cecilio, Granada | Microbiology Service, Hospital Universitario Clinico San Cecilio, Granada | Adolfo de Salazar, Natalia Chueca, Laura Viñuela, Ana Fuentes, Federico García |
| EPI_ISL_981983, EPI_ISL_981989, EPI_ISL_981992, EPI_ISL_981995, EPI_ISL_982000, EPI_ISL_982013, EPI_ISL_982014, EPI_ISL_982015, EPI_ISL_982017, EPI_ISL_982018, EPI_ISL_982032, EPI_ISL_982035, EPI_ISL_982041, EPI_ISL_982046, EPI_ISL_982049, EPI_ISL_982050, EPI_ISL_982060, EPI_ISL_982061, EPI_ISL_982066, EPI_ISL_982073, EPI_ISL_982074, EPI_ISL_982077, EPI_ISL_982079, EPI_ISL_982083, EPI_ISL_982088, EPI_ISL_982089, EPI_ISL_982090, EPI_ISL_982092, EPI_ISL_982100, EPI_ISL_982108, EPI_ISL_982112 | | | |
| see above | TGen North | Sonora Quest Laboratories | "Jolene Bowers, Megan Folkerts, Chris French, Hayley Yaglom, Ashlyn Pfeiffer, Darrin Lemmer, Dave Engelthaler, The Arizona COVID Genomics Union (ACGU)" |
| EPI_ISL_982232 | MEPHI, Aix Marseille University | MEPHI, Aix Marseille University | Anthony LEVASSEUR |
| EPI_ISL_982246, EPI_ISL_982249, EPI_ISL_982250 | Lab voor klinische biologie | Lab voor klinische biologie | Hannelore Hamerlinck, Marija Janevska, Bruno Verhasselt |
| EPI_ISL_982345, EPI_ISL_982353, EPI_ISL_982354, EPI_ISL_982385, EPI_ISL_982386, EPI_ISL_982387, EPI_ISL_982388, EPI_ISL_982392 | M Health Fairview | Minnesota Department of Health, Public Health Laboratory | Alexandra Lorentz, Jacob Garfin, Matt Plumb, and Xiong Wang |
| EPI_ISL_982539 | Landstuhl Regional Medical Center | US Air Force School of Aerospace Medicine | Anthony Fries, Jennifer Meyer, William Gruner, William Buggele, Amanda Javorina, Sarah Purves, Fritz Castillo, Cole Anderson, Clarise Starr, Elizabeth Macias |
| EPI_ISL_982540, EPI_ISL_982569, EPI_ISL_982578, EPI_ISL_982597, EPI_ISL_982600, EPI_ISL_982623 | US Air Force School of Aerospace Medicine | US Air Force School of Aerospace Medicine | Anthony Fries, Jennifer Meyer, William Gruner, William Buggele, Amanda Javorina, Sarah Purves, Clarise Starr, Elizabeth Macias |
| EPI_ISL_982627, EPI_ISL_982628, EPI_ISL_982629, EPI_ISL_982630 | Landstuhl Regional Medical Center | US Air Force School of Aerospace Medicine | Anthony Fries, Jennifer Meyer, William Gruner, William Buggele, Amanda Javorina, Sarah Purves, Fritz Castillo, Cole Anderson, Clarise Starr, Elizabeth Macias |
| EPI_ISL_982634, EPI_ISL_982661, EPI_ISL_982672, EPI_ISL_982673, EPI_ISL_982676, EPI_ISL_982691, EPI_ISL_982692, EPI_ISL_982706, EPI_ISL_982719 | US Air Force School of Aerospace Medicine | US Air Force School of Aerospace Medicine | Anthony Fries, Jennifer Meyer, William Gruner, William Buggele, Amanda Javorina, Sarah Purves, Clarise Starr, Elizabeth Macias |
| EPI_ISL_983064, EPI_ISL_983065 | Gravity Diagnostics | Kentucky State Public Health Lab | Stephanie Lunn, Karim George, Joshua Tobias, William Grooms, Vaneet Arora, Matthew Johnson, Rachel Zinner, Rhonda Lucas |
| EPI_ISL_983119, EPI_ISL_983120, EPI_ISL_983121, EPI_ISL_983122, EPI_ISL_983123, EPI_ISL_983124, EPI_ISL_983125, EPI_ISL_983129 | THE MARY IMOGENE BASSETT HOSPITAL | Wadsworth Center, New York State Department of Health | Kirsten St. George, Daryl M. Lamson, Alexis Russel, Matthew Shudt, Melissa A Leisner, Jonathan Plitnick, Navjot Singh, John Kelly, Erasmus Schneider, Erica Lasek-Nesselquist |
| EPI_ISL_983251, EPI_ISL_983252, EPI_ISL_983253, EPI_ISL_983254, EPI_ISL_983255, EPI_ISL_983256, EPI_ISL_983257, EPI_ISL_983258, EPI_ISL_983259, EPI_ISL_983260, EPI_ISL_983261, EPI_ISL_983262 | | | |
| see above | KALEIDA CENTER FOR LABORATORY MEDICINE | Wadsworth Center, New York State Department of Health | Kirsten St. George, Daryl M. Lamson, Alexis Russel, Matthew Shudt, Melissa A Leisner, Jonathan Plitnick, Navjot Singh, John Kelly, Erasmus Schneider, Erica Lasek-Nesselquist |
| EPI_ISL_983711, EPI_ISL_983811, EPI_ISL_983812 | Colorado Department of Public Health and Environment | Colorado Department of Puplic Health and Environment | Laura Bankers, Molly C. Hetherington-Rauth, Diana Ir, Shannon Ely, Shannon R. Matzinger, Sarah Elizabeth Totten, Emily A. Travanty |
