## Supplementary material for "Detection and characterization of the SARS-CoV-2 lineage B.1.526 in New York": Supp. Table 3 GISAID Acknowledgment part 2: gisaid_hcov-19_acknowledgement_table_2021_02_13_010-10.pdf

We gratefully acknowledge the following Authors from the Originating laboratories responsible for obtaining the specimens, as well as the Submitting laboratories where the genome data were generated and shared via GISAID, on which this research is based.

All Submitters of data may be contacted directly via www.gisaid.org

Authors are sorted alphabetically.

| Accession ID | Originating Laboratory | Submitting Laboratory | Authors |
|---|---|---|---|
| EPI_ISL_803109 | Pathology West - NSW Health Pathology | NSW Health Pathology - Institute of Clinical Pathology and Medical Research; Westmead Hospital; University of Sydney | CIDM-PH et al. |
| EPI_ISL_825550, EPI_ISL_825551, EPI_ISL_825573, EPI_ISL_825574 | Respiratory Virus Unit, National Infection Service, Public Health England | COVID-19 Genomics UK (COG-UK) Consortium | PHE Covid Sequencing Team |
| EPI_ISL_830726 | Wadsworth Center, New York State Department of Health | Wadsworth Center, New York State Department of Health | Kirsten St. George, Daryl M. Lamson, Alexis Russel, Matthew Shudt, Melissa A Leisner, Jonathan Plitnick, Navjot Singh, John Kelly, Erasmus Schneider, Erica Lasek-Nesselquist |
| EPI_ISL_831669 | Institute of Virology, Biomedical Research Center of the Slovak Academy of Sciences, Bratislava | Faculty of Natural Sciences, Comenius University, Bratislava | Kristína Boršová, Viktória abanová, Broa Brejová, Viktória Hodorová, Sabina Fumaová Havlíková, Juraj Kopáek, Martina Liková, ubomíra Lukáiková, Martina Neboháová, Monika Sláviková, Tomáš Vina, Boris Klempa, Jozef Nosek |
| EPI_ISL_831670 | Institute of Virology, Biomedical Research Center of the Slovak Academy of Sciences, Bratislava | Faculty of Natural Sciences, Comenius University, Bratislava | Viktória abanová, Kristína Boršová, Broa Brejová, Viktória Hodorová, Sabina Fumaová Havlíková, Juraj Kopáek, Martina Liková, ubomíra Lukáiková, Martina Neboháová, Monika Sláviková, Tomáš Vina, Jozef Nosek, Boris Klempa |
| EPI_ISL_831671 | Institute of Virology, Biomedical Research Center of the Slovak Academy of Sciences, Bratislava | Faculty of Natural Sciences, Comenius University, Bratislava | Broa Brejová, Viktória abanová, Kristína Boršová, Viktória Hodorová, Sabina Fumaová Havlíková, Juraj Kopáek, Martina Liková, ubomíra Lukáiková, Martina Neboháová, Monika Sláviková, Tomáš Vina, Jozef Nosek, Boris Klempa |
| EPI_ISL_831673 | Institute of Virology, Biomedical Research Center of the Slovak Academy of Sciences, Bratislava | Faculty of Natural Sciences, Comenius University, Bratislava | Kristína Boršová, Viktória abanová, Broa Brejová, Viktória Hodorová, Sabina Fumaová Havlíková, Juraj Kopáek, Martina Liková, ubomíra Lukáiková, Martina Neboháová, Monika Sláviková, Tomáš Vina, Boris Klempa, Jozef Nosek |
| EPI_ISL_832242 | Department of Clinical Microbiology | GIGA Medical Genomics | Keith Durkin, Maria Artesi, Sébastien Bontems, Raphaël Boreux, Bouchra Boujemla, Cécile Meex, Pierrette Melin, Marie-Pierre Hayette, Vincent Bours |
| EPI_ISL_833378, EPI_ISL_833387 | National Public Health Laboratory, National Centre for Infectious Diseases | National Public Health Laboratory, National Centre for Infectious Diseases | Tze Minn Mak, Sophie Octavia, Zhenyang Zhou, Lin Cui, Raymond Tzer Pin Lin |
| EPI_ISL_833434, EPI_ISL_833481, EPI_ISL_833483, EPI_ISL_833484, EPI_ISL_833486 | UZA/UAntwerpen (BIS lab), University Hospital Antwerp, Edegem, Belgium | Laboratory of Medical Microbiology, University of Antwerp, Campus Drie Eiken, S6.26, Universiteitsplein 1, 2610, Wilrijk, Antwerp, Belgium | Basil Britto Xavier, Jasmine Coppens, Christine Lammens, Veerle Matheeussen, Herman Goossens |
| EPI_ISL_837057, EPI_ISL_837182 | Respiratory Virus Unit, National Infection Service, Public Health England | COVID-19 Genomics UK (COG-UK) Consortium | PHE Covid Sequencing Team |
| EPI_ISL_837973, EPI_ISL_837975, EPI_ISL_837978, EPI_ISL_837984, EPI_ISL_837985, EPI_ISL_837990, EPI_ISL_838010, EPI_ISL_838014, EPI_ISL_838019, EPI_ISL_838020, EPI_ISL_838023, EPI_ISL_838024, EPI_ISL_838027, EPI_ISL_838028, EPI_ISL_838029, EPI_ISL_838030, EPI_ISL_838031, EPI_ISL_838033, EPI_ISL_838036, EPI_ISL_838039, EPI_ISL_838040, EPI_ISL_838041, EPI_ISL_838042, EPI_ISL_838043, EPI_ISL_838044, EPI_ISL_838045, EPI_ISL_838046, EPI_ISL_838047 | | | |
| see above | Department of Pathology, University of Cambridge | COVID-19 Genomics UK (COG-UK) Consortium | Aminu S. Jahun, Yasmin Chaudhry, Grant Hall, Iliana Georgana, Myra Hosmillo, Martin D. Curran, Malte Pinckert, Surendra Parmar, Ian Goodfellow |
| EPI_ISL_838997, EPI_ISL_838998, EPI_ISL_839014, EPI_ISL_839016, EPI_ISL_839017, EPI_ISL_839018, EPI_ISL_839019, EPI_ISL_839021, EPI_ISL_839022, EPI_ISL_839023, EPI_ISL_839024, EPI_ISL_839025 | | | |
| see above | University College London, Great Ormond Street Hospital for Children NHS Foundation Trust, Imperial College Healthcare NHS Trust | COVID-19 Genomics UK (COG-UK) Consortium | Sergi Castellano, Rachel Williams, Mark Kristiansen, Paola Resende Silva, Sunando Roy, Tony Brooks, Helena Tutill, Paola Niola, Patricia Dyal, Charlotte Williams, Leysa Forrest, Yasmin Panchbhaya, Jacqueline Findlay, Samuel Weeks, Julianne Brown, Kathryn Harris, Paul Randell, James Price, Alison Holmes, Judith Breuer |
| EPI_ISL_840061 | Queens Medical Centre, Clinical Microbiology Department / DeepSeq Nottingham | COVID-19 Genomics UK (COG-UK) Consortium | Gemma Clark, Wendy Smith, Manjinder Khakh, Vicki M Fleming, Michelle M Lister, Hannah Howson-Wells, Jonathan Ball, Patrick McClure, Joseph Chappell, Theocharis Tsoleridis, Nadine Holmes, Matthew Carlisle, Christopher Moore, Fei Sang, Johnny Debebe, Victoria Wright, Matthew Loose |
| EPI_ISL_841526, EPI_ISL_841543, EPI_ISL_841544, EPI_ISL_841545, EPI_ISL_841546, EPI_ISL_841552, EPI_ISL_841555, EPI_ISL_841556, EPI_ISL_841741, EPI_ISL_841742, EPI_ISL_841743, EPI_ISL_841744, EPI_ISL_841745, EPI_ISL_841746, EPI_ISL_841747, EPI_ISL_841748, EPI_ISL_841749, EPI_ISL_841750, EPI_ISL_841751, EPI_ISL_841752, EPI_ISL_841753 | | | |
| see above | Originating lab: Wales Specialist Virology Centre Sequencing lab: Pathogen Genomics Unit | Public Health Wales Microbiology Cardiff Wales Specialist Virology Centre | Catherine Moore, Johnathan Evans, Laura Gifford, Malorie Perry, Simon Cottrell, Angela Marchbank, Alec Birchley, Alexander Adams, Amy Gaskin, Bree Gatica-Wilcox, Jason Coombes, Joel Southgate, Lauren Gilbert, Lee Graham, Nicole Pacchiarini, Sara Kumziene-Summerhayes, Sarah Taylor, Sophie Jones, Sara Rey, Matthew Bull, Joanne Watkins, Sally Corden, Tom Connor |
| EPI_ISL_842201, EPI_ISL_842210 | Virology Department, Sheffield Teaching Hospitals NHS Foundation Trust/Department of Infection, Immunity and Cardiovascular Disease, The Medical School, University of Sheffield | COVID-19 Genomics UK (COG-UK) Consortium | Thushan de Silva, Matthew Parker, Nikki Smith, Adri Angyal, Rebecca Brown, Luke Green, Rachel Tucker, Paul Parsons, Danielle Groves, Katie Johnson, Laura Carrilero, Alex Keeley, Dave Partridge, Matthew Wyles, Benjamin Lindsey, Mehmet Yavuz, Mohammad Raza, Cariad Evans |
| EPI_ISL_843188 | Maryland Public Health Laboratory | Maryland Public Health Laboratory | Maryland Department of Health Laboratories Administration |
| EPI_ISL_844543, EPI_ISL_844545, EPI_ISL_844552, EPI_ISL_844553, EPI_ISL_844556, EPI_ISL_844557, EPI_ISL_844558, EPI_ISL_844559, EPI_ISL_844560, EPI_ISL_844561, EPI_ISL_844562, EPI_ISL_844563, EPI_ISL_844564, EPI_ISL_844565, EPI_ISL_844566, EPI_ISL_844567, EPI_ISL_844568, EPI_ISL_844569, EPI_ISL_844570, EPI_ISL_844571, EPI_ISL_844572, EPI_ISL_844573, EPI_ISL_844574, EPI_ISL_844575, EPI_ISL_844576, EPI_ISL_844577, EPI_ISL_844635, EPI_ISL_844636, EPI_ISL_844637, EPI_ISL_844638, EPI_ISL_844639, EPI_ISL_844653, EPI_ISL_844654, EPI_ISL_844655, EPI_ISL_844666, EPI_ISL_844667, EPI_ISL_844669, EPI_ISL_844670, EPI_ISL_844676, EPI_ISL_844680, EPI_ISL_844681, EPI_ISL_844693, EPI_ISL_844694, EPI_ISL_844702, EPI_ISL_844706, EPI_ISL_844707, EPI_ISL_844708, EPI_ISL_844716, EPI_ISL_844717, EPI_ISL_844718, EPI_ISL_844730, EPI_ISL_844731, EPI_ISL_844732, EPI_ISL_844733, EPI_ISL_844734, EPI_ISL_844735, EPI_ISL_844736, EPI_ISL_844737, EPI_ISL_844738, EPI_ISL_844747, EPI_ISL_844758, EPI_ISL_844759, EPI_ISL_844763, EPI_ISL_844766, EPI_ISL_844767, EPI_ISL_844768, EPI_ISL_844788, EPI_ISL_844805, EPI_ISL_844807, EPI_ISL_844808, EPI_ISL_844818, EPI_ISL_844821, EPI_ISL_844824, EPI_ISL_844834, EPI_ISL_844836, EPI_ISL_844837, EPI_ISL_844842, EPI_ISL_844843, EPI_ISL_844844, EPI_ISL_844845, EPI_ISL_844846, EPI_ISL_844847, EPI_ISL_844848, EPI_ISL_844849, EPI_ISL_844860, EPI_ISL_844861, EPI_ISL_844862, EPI_ISL_844863, EPI_ISL_844864, EPI_ISL_844865, EPI_ISL_844911, EPI_ISL_844912, EPI_ISL_844931, EPI_ISL_844932, EPI_ISL_844948, EPI_ISL_844951, EPI_ISL_844953, EPI_ISL_844954, EPI_ISL_844955, EPI_ISL_844956, EPI_ISL_844957, EPI_ISL_844958, EPI_ISL_844978, EPI_ISL_844981, EPI_ISL_844988, EPI_ISL_844991, EPI_ISL_844992, EPI_ISL_844993, EPI_ISL_844998, EPI_ISL_844999, EPI_ISL_845000, EPI_ISL_845001, EPI_ISL_845002, EPI_ISL_845003, EPI_ISL_845004, EPI_ISL_845005, EPI_ISL_845006, EPI_ISL_845007, EPI_ISL_845008, EPI_ISL_845009, EPI_ISL_845010, EPI_ISL_845011, EPI_ISL_845012, EPI_ISL_845013, EPI_ISL_845014, EPI_ISL_845015, EPI_ISL_845016, EPI_ISL_845017, EPI_ISL_845018, EPI_ISL_845019, EPI_ISL_845020, EPI_ISL_845021, EPI_ISL_845022, EPI_ISL_845023, EPI_ISL_845024, EPI_ISL_845025, EPI_ISL_845026, EPI_ISL_845027, EPI_ISL_845028, EPI_ISL_845029, EPI_ISL_845030, EPI_ISL_845031, EPI_ISL_845032, EPI_ISL_845033, EPI_ISL_845034, EPI_ISL_845035, EPI_ISL_845036, EPI_ISL_845037, EPI_ISL_845038, EPI_ISL_845039, EPI_ISL_845040, EPI_ISL_845041, EPI_ISL_845042, EPI_ISL_845043, EPI_ISL_845044, EPI_ISL_845045, EPI_ISL_845046, EPI_ISL_845047, EPI_ISL_845048, EPI_ISL_845049, EPI_ISL_845050, EPI_ISL_845051, EPI_ISL_845052, EPI_ISL_845053, EPI_ISL_845054, EPI_ISL_845055, EPI_ISL_845056, EPI_ISL_845057, EPI_ISL_845058, EPI_ISL_845059, EPI_ISL_845060, EPI_ISL_845061, EPI_ISL_845062, EPI_ISL_845063, EPI_ISL_845064, EPI_ISL_845065, EPI_ISL_845066, EPI_ISL_845067, EPI_ISL_845068, EPI_ISL_845069, EPI_ISL_845070, EPI_ISL_845071, EPI_ISL_845072, EPI_ISL_845073, EPI_ISL_845074, EPI_ISL_845075, EPI_ISL_845076, EPI_ISL_845077, EPI_ISL_845078, EPI_ISL_845079, EPI_ISL_845080, EPI_ISL_845275, EPI_ISL_845276, EPI_ISL_845277, EPI_ISL_845278, EPI_ISL_845279, EPI_ISL_845280, EPI_ISL_845377, EPI_ISL_845378, EPI_ISL_845379, EPI_ISL_845380, EPI_ISL_845381, EPI_ISL_845382, EPI_ISL_845383, EPI_ISL_845384, EPI_ISL_845385, EPI_ISL_845386, EPI_ISL_845387, EPI_ISL_845388, EPI_ISL_845389, EPI_ISL_845390, EPI_ISL_845391, EPI_ISL_845392, EPI_ISL_845393, EPI_ISL_845394, EPI_ISL_845395, EPI_ISL_845396, EPI_ISL_845397, EPI_ISL_845398, EPI_ISL_845399, EPI_ISL_845498, EPI_ISL_845499, EPI_ISL_845500 | | | |
| see above | Department of Virus and Microbiological Special Diagnostics, Statens Serum Institut, Copenhagen, Denmark | Albertsen Lab, Department of Chemistry and Bioscience, Aalborg University, Denmark | Danish Covid-19 Genome Consortium |
| EPI_ISL_846544, EPI_ISL_846550, EPI_ISL_846551, EPI_ISL_846558, EPI_ISL_846561, EPI_ISL_846562, EPI_ISL_846563, EPI_ISL_846564, EPI_ISL_846565, EPI_ISL_846566, EPI_ISL_846567, EPI_ISL_846577 | | | |
| see above | Lab voor klinische biologie | Lab voor klinische biologie | Hannelore Hamerlinck, Bruno Verhasselt |
| EPI_ISL_846775, EPI_ISL_846776, EPI_ISL_846777, EPI_ISL_846778, EPI_ISL_846780, EPI_ISL_846782, EPI_ISL_846783, EPI_ISL_846784, EPI_ISL_846785, EPI_ISL_846786, EPI_ISL_846787, EPI_ISL_846788, EPI_ISL_846789, EPI_ISL_846790, EPI_ISL_846791, EPI_ISL_846792, EPI_ISL_846793, EPI_ISL_846794, EPI_ISL_846795, EPI_ISL_846796, EPI_ISL_846797, EPI_ISL_846798, EPI_ISL_846799, EPI_ISL_846800, EPI_ISL_846801, EPI_ISL_846802, EPI_ISL_846803, EPI_ISL_846804, EPI_ISL_846805, EPI_ISL_846806, EPI_ISL_846807, EPI_ISL_846808, EPI_ISL_846809, EPI_ISL_846810, EPI_ISL_846811, EPI_ISL_846821, EPI_ISL_846822, EPI_ISL_846823, EPI_ISL_846824, EPI_ISL_846825, EPI_ISL_846826, EPI_ISL_846827, EPI_ISL_846828, EPI_ISL_846829, EPI_ISL_846830, EPI_ISL_846831, EPI_ISL_846832, EPI_ISL_846833, EPI_ISL_846834, EPI_ISL_846835, EPI_ISL_846836, EPI_ISL_846837, EPI_ISL_846838, EPI_ISL_846839, EPI_ISL_846840, EPI_ISL_846841, EPI_ISL_846842, EPI_ISL_846843, EPI_ISL_846844, EPI_ISL_846845, EPI_ISL_846846, EPI_ISL_846847, EPI_ISL_846848, EPI_ISL_846849, EPI_ISL_846850, EPI_ISL_846854, EPI_ISL_846855, EPI_ISL_846856, EPI_ISL_846857, EPI_ISL_846858, EPI_ISL_846859, EPI_ISL_846862, EPI_ISL_846863, EPI_ISL_846864, EPI_ISL_846865, EPI_ISL_846867, EPI_ISL_846868, EPI_ISL_846869, EPI_ISL_846870, EPI_ISL_846871, EPI_ISL_846872, EPI_ISL_846873, EPI_ISL_846874, EPI_ISL_846875, EPI_ISL_846876, EPI_ISL_846878, EPI_ISL_846879, EPI_ISL_846880, EPI_ISL_846881, EPI_ISL_846882, | | | |

EPI_ISL_846883, EPI_ISL_846885, EPI_ISL_846887, EPI_ISL_846890, EPI_ISL_846891, EPI_ISL_846892, EPI_ISL_846896, EPI_ISL_846897, EPI_ISL_846899, EPI_ISL_846900, EPI_ISL_846901, EPI_ISL_846902, EPI_ISL_846903, EPI_ISL_846905, EPI_ISL_846906, EPI_ISL_846907, EPI_ISL_846908, EPI_ISL_846909, EPI_ISL_846911, EPI_ISL_846912, EPI_ISL_846914, EPI_ISL_846915, EPI_ISL_846916, EPI_ISL_846917, EPI_ISL_846918, EPI_ISL_846919, EPI_ISL_846920, EPI_ISL_846921, EPI_ISL_846922, EPI_ISL_846923, EPI_ISL_846924, EPI_ISL_846925, EPI_ISL_846926, EPI_ISL_846927, EPI_ISL_846928, EPI_ISL_846929, EPI_ISL_846930, EPI_ISL_846931, EPI_ISL_846932, EPI_ISL_846933, EPI_ISL_846934, EPI_ISL_846935, EPI_ISL_846936, EPI_ISL_846937, EPI_ISL_846938, EPI_ISL_846939, EPI_ISL_846940, EPI_ISL_846941, EPI_ISL_846942, EPI_ISL_846943, EPI_ISL_846944, EPI_ISL_846945, EPI_ISL_846946, EPI_ISL_846947, EPI_ISL_846948, EPI_ISL_846949, EPI_ISL_846950, EPI_ISL_846951, EPI_ISL_846952, EPI_ISL_846953, EPI_ISL_846954, EPI_ISL_846955, EPI_ISL_846956, EPI_ISL_846957, EPI_ISL_846958, EPI_ISL_846959, EPI_ISL_846960, EPI_ISL_846961, EPI_ISL_846962, EPI_ISL_846963, EPI_ISL_846964, EPI_ISL_846965, EPI_ISL_846966, EPI_ISL_846967, EPI_ISL_846968, EPI_ISL_846969, EPI_ISL_846970, EPI_ISL_846971, EPI_ISL_846972, EPI_ISL_846973, EPI_ISL_846974, EPI_ISL_846975, EPI_ISL_846976, EPI_ISL_846977, EPI_ISL_846978, EPI_ISL_846979, EPI_ISL_846980, EPI_ISL_846981, EPI_ISL_846982, EPI_ISL_846983, EPI_ISL_846984, EPI_ISL_846985, EPI_ISL_846986, EPI_ISL_846987, EPI_ISL_846988, EPI_ISL_846989, EPI_ISL_846990, EPI_ISL_846991, EPI_ISL_846992, EPI_ISL_846993, EPI_ISL_846994, EPI_ISL_846995, EPI_ISL_846996, EPI_ISL_846997, EPI_ISL_846998, EPI_ISL_846999, EPI_ISL_847000, EPI_ISL_847001, EPI_ISL_847002, EPI_ISL_847003, EPI_ISL_847004, EPI_ISL_847005, EPI_ISL_847006, EPI_ISL_847007, EPI_ISL_847008, EPI_ISL_847009, EPI_ISL_847010, EPI_ISL_847011, EPI_ISL_847012, EPI_ISL_847013, EPI_ISL_847014, EPI_ISL_847015, EPI_ISL_847016, EPI_ISL_847017, EPI_ISL_847018, EPI_ISL_847019, EPI_ISL_847020, EPI_ISL_847021, EPI_ISL_847022, EPI_ISL_847023, EPI_ISL_847024, EPI_ISL_847025, EPI_ISL_847026, EPI_ISL_847027, EPI_ISL_847028, EPI_ISL_847029, EPI_ISL_847030, EPI_ISL_847031, EPI_ISL_847032, EPI_ISL_847033, EPI_ISL_847034, EPI_ISL_847035, EPI_ISL_847036, EPI_ISL_847037, EPI_ISL_847038, EPI_ISL_847039, EPI_ISL_847040, EPI_ISL_847041, EPI_ISL_847042, EPI_ISL_847043, EPI_ISL_847044, EPI_ISL_847045, EPI_ISL_847046, EPI_ISL_847047, EPI_ISL_847048, EPI_ISL_847049, EPI_ISL_847050, EPI_ISL_847051, EPI_ISL_847052, EPI_ISL_847053, EPI_ISL_847054, EPI_ISL_847055, EPI_ISL_847056, EPI_ISL_847057, EPI_ISL_847058, EPI_ISL_847059, EPI_ISL_847060, EPI_ISL_847061, EPI_ISL_847062, EPI_ISL_847063, EPI_ISL_847064, EPI_ISL_847065, EPI_ISL_847066, EPI_ISL_847067, EPI_ISL_847068, EPI_ISL_847069, EPI_ISL_847070, EPI_ISL_847071, EPI_ISL_847072, EPI_ISL_847073, EPI_ISL_847074, EPI_ISL_847075, EPI_ISL_847076, EPI_ISL_847077, EPI_ISL_847078, EPI_ISL_847079, EPI_ISL_847080, EPI_ISL_847081, EPI_ISL_847082, EPI_ISL_847083, EPI_ISL_847084, EPI_ISL_847085, EPI_ISL_847086, EPI_ISL_847087, EPI_ISL_847088, EPI_ISL_847089, EPI_ISL_847090, EPI_ISL_847091, EPI_ISL_847092, EPI_ISL_847093, EPI_ISL_847094, EPI_ISL_847095, EPI_ISL_847096, EPI_ISL_847097, EPI_ISL_847098, EPI_ISL_847099, EPI_ISL_847100, EPI_ISL_847101, EPI_ISL_847102, EPI_ISL_847103, EPI_ISL_847104, EPI_ISL_847105, EPI_ISL_847106, EPI_ISL_847107, EPI_ISL_847108, EPI_ISL_847109, EPI_ISL_847110, EPI_ISL_847111, EPI_ISL_847112, EPI_ISL_847113, EPI_ISL_847114, EPI_ISL_847115, EPI_ISL_847116, EPI_ISL_847117, EPI_ISL_847118, EPI_ISL_847119, EPI_ISL_847120, EPI_ISL_847121, EPI_ISL_847122, EPI_ISL_847123, EPI_ISL_847124, EPI_ISL_847180, EPI_ISL_847181, EPI_ISL_847182, EPI_ISL_847183, EPI_ISL_847184, EPI_ISL_847185, EPI_ISL_847186, EPI_ISL_847187, EPI_ISL_847188, EPI_ISL_847189, EPI_ISL_847190, EPI_ISL_847191, EPI_ISL_847192, EPI_ISL_847193, EPI_ISL_847194, EPI_ISL_847195, EPI_ISL_847196, EPI_ISL_847197, EPI_ISL_847198, EPI_ISL_847206, EPI_ISL_847207, EPI_ISL_847208, EPI_ISL_847209, EPI_ISL_847210, EPI_ISL_847211, EPI_ISL_847212, EPI_ISL_847213, EPI_ISL_847214, EPI_ISL_847215, EPI_ISL_847216, EPI_ISL_847217, EPI_ISL_847218, EPI_ISL_847219, EPI_ISL_847220, EPI_ISL_847221, EPI_ISL_847222, EPI_ISL_847223, EPI_ISL_847224, EPI_ISL_847225, EPI_ISL_847226, EPI_ISL_847227, EPI_ISL_847228, EPI_ISL_847229, EPI_ISL_847230, EPI_ISL_847231, EPI_ISL_847232, EPI_ISL_847238, EPI_ISL_847239, EPI_ISL_847240, EPI_ISL_847241, EPI_ISL_847242, EPI_ISL_847243, EPI_ISL_847250, EPI_ISL_847251, EPI_ISL_847257, EPI_ISL_847258, EPI_ISL_847259, EPI_ISL_847261, EPI_ISL_847265, EPI_ISL_847267, EPI_ISL_847268, EPI_ISL_847269, EPI_ISL_847271, EPI_ISL_847272, EPI_ISL_847273, EPI_ISL_847275, EPI_ISL_847276, EPI_ISL_847277, EPI_ISL_847278, EPI_ISL_847279, EPI_ISL_847281, EPI_ISL_847282, EPI_ISL_847283, EPI_ISL_847284, EPI_ISL_847285, EPI_ISL_847286, EPI_ISL_847287, EPI_ISL_847288, EPI_ISL_847289, EPI_ISL_847290, EPI_ISL_847291, EPI_ISL_847292, EPI_ISL_847293, EPI_ISL_847294, EPI_ISL_847295, EPI_ISL_847296, EPI_ISL_847297, EPI_ISL_847298, EPI_ISL_847299, EPI_ISL_847300, EPI_ISL_847301, EPI_ISL_847302, EPI_ISL_847303, EPI_ISL_847304, EPI_ISL_847305, EPI_ISL_847306, EPI_ISL_847307, EPI_ISL_847308, EPI_ISL_847309, EPI_ISL_847310, EPI_ISL_847311, EPI_ISL_847312, EPI_ISL_847313, EPI_ISL_847314, EPI_ISL_847315, EPI_ISL_847316, EPI_ISL_847317, EPI_ISL_847318, EPI_ISL_847319, EPI_ISL_847321, EPI_ISL_847322, EPI_ISL_847323, EPI_ISL_847324, EPI_ISL_847325, EPI_ISL_847326, EPI_ISL_847327, EPI_ISL_847328, EPI_ISL_847329, EPI_ISL_847330, EPI_ISL_847331, EPI_ISL_847333, EPI_ISL_847334, EPI_ISL_847336, EPI_ISL_847338, EPI_ISL_847339, EPI_ISL_847340, EPI_ISL_847341, EPI_ISL_847342, EPI_ISL_847343, EPI_ISL_847344, EPI_ISL_847345, EPI_ISL_847346, EPI_ISL_847347, EPI_ISL_847350, EPI_ISL_847354, EPI_ISL_847355, EPI_ISL_847356, EPI_ISL_847357, EPI_ISL_847359, EPI_ISL_847360, EPI_ISL_847361, EPI_ISL_847362, EPI_ISL_847363, EPI_ISL_847364, EPI_ISL_847365, EPI_ISL_847367, EPI_ISL_847371, EPI_ISL_847372, EPI_ISL_847373, EPI_ISL_847374, EPI_ISL_847375, EPI_ISL_847376, EPI_ISL_847378, EPI_ISL_847379, EPI_ISL_847380, EPI_ISL_847381, EPI_ISL_847382, EPI_ISL_847383, EPI_ISL_847384, EPI_ISL_847385, EPI_ISL_847386, EPI_ISL_847387, EPI_ISL_847388, EPI_ISL_847389, EPI_ISL_847390, EPI_ISL_847391, EPI_ISL_847392, EPI_ISL_847393, EPI_ISL_847394, EPI_ISL_847395, EPI_ISL_847396, EPI_ISL_847397, EPI_ISL_847398, EPI_ISL_847399, EPI_ISL_847400, EPI_ISL_847401, EPI_ISL_847402, EPI_ISL_847403, EPI_ISL_847404, EPI_ISL_847405, EPI_ISL_847406, EPI_ISL_847407, EPI_ISL_847408, EPI_ISL_847409, EPI_ISL_847410, EPI_ISL_847411, EPI_ISL_847412, EPI_ISL_847413, EPI_ISL_847414, EPI_ISL_847415, EPI_ISL_847416, EPI_ISL_847417, EPI_ISL_847418, EPI_ISL_847419, EPI_ISL_847420, EPI_ISL_847421, EPI_ISL_847422, EPI_ISL_847423, EPI_ISL_847424, EPI_ISL_847425, EPI_ISL_847426, EPI_ISL_847427, EPI_ISL_847428, EPI_ISL_847429, EPI_ISL_847430, EPI_ISL_847431, EPI_ISL_847432, EPI_ISL_847433, EPI_ISL_847434, EPI_ISL_847435, EPI_ISL_847436, EPI_ISL_847437, EPI_ISL_847438, EPI_ISL_847439, EPI_ISL_847440, EPI_ISL_847441, EPI_ISL_847442, EPI_ISL_847443, EPI_ISL_847444, EPI_ISL_847445, EPI_ISL_847446, EPI_ISL_847447, EPI_ISL_847448, EPI_ISL_847449, EPI_ISL_847450, EPI_ISL_847451, EPI_ISL_847452, EPI_ISL_847453, EPI_ISL_847454, EPI_ISL_847455, EPI_ISL_847456, EPI_ISL_847457, EPI_ISL_847458, EPI_ISL_847459, EPI_ISL_847460, EPI_ISL_847461, EPI_ISL_847462, EPI_ISL_847463, EPI_ISL_847464, EPI_ISL_847465, EPI_ISL_847466, EPI_ISL_847467, EPI_ISL_847468, EPI_ISL_847469, EPI_ISL_847470, EPI_ISL_847471, EPI_ISL_847472, EPI_ISL_847473, EPI_ISL_847474, EPI_ISL_847475, EPI_ISL_847476, EPI_ISL_847477, EPI_ISL_847478, EPI_ISL_847479, EPI_ISL_847480, EPI_ISL_847481, EPI_ISL_847482, EPI_ISL_847483, EPI_ISL_847484, EPI_ISL_847485

| | | | |
|---|---|---|---|
| see above | Department of Virus and Microbiological Special Diagnostics, Statens Serum Institut, Copenhagen, Denmark | Albertsen Lab, Department of Chemistry and Bioscience, Aalborg University, Denmark | Danish Covid-19 Genome Consortium |
| EPI_ISL_847564 | California Department of Public Health | Chiu Laboratory, University of California, San Francisco | Charles Chiu, Xianding (Wayne) Deng, Candace Wang, Brian Bushnell, Scot Federman, Jill Hacker, Debra Wadford |
| EPI_ISL_849758, EPI_ISL_849759, EPI_ISL_849760 | unknown | PHV-FSS | Son Nguyen et al. |

EPI_ISL_851101, EPI_ISL_851126, EPI_ISL_851142, EPI_ISL_851180, EPI_ISL_851187, EPI_ISL_851190, EPI_ISL_851196, EPI_ISL_851217, EPI_ISL_851229, EPI_ISL_851236, EPI_ISL_851241, EPI_ISL_851258, EPI_ISL_851329, EPI_ISL_851365, EPI_ISL_851374, EPI_ISL_851674

| | | | |
|---|---|---|---|
| see above | Lighthouse Lab in Glasgow | Wellcome Sanger Institute for the COVID-19 Genomics UK (COG-UK) Consortium | Harper VanSteenhouse, Yumi Kasai, David Gray, Carol Clugston, Anna Dominiczak and Alex Alderton, Roberto Amato, Sonia Goncalves, Ewan Harrison, David K. Jackson, Ian Johnston, Dominic Kwiatkowski, Cordelia Langford, John Sillitoe on behalf of the Wellcome Sanger Institute COVID-19 Surveillance Team |

EPI_ISL_852949, EPI_ISL_852951, EPI_ISL_852956, EPI_ISL_852957, EPI_ISL_852959, EPI_ISL_852960, EPI_ISL_852965, EPI_ISL_852966, EPI_ISL_852967, EPI_ISL_852969, EPI_ISL_852970, EPI_ISL_852971, EPI_ISL_852972, EPI_ISL_852973, EPI_ISL_852975, EPI_ISL_852976, EPI_ISL_852977, EPI_ISL_852978, EPI_ISL_852979, EPI_ISL_852980, EPI_ISL_852981, EPI_ISL_852982, EPI_ISL_852984, EPI_ISL_852985, EPI_ISL_852986, EPI_ISL_852987, EPI_ISL_853017, EPI_ISL_853030, EPI_ISL_853031

| | | | |
|---|---|---|---|
| see above | Hospital General Universitario Gregorio Marañón | SeqCOVID-SPAIN consortium/IBV(CSIC) | Darío García de Viedma, Laura Pérez-Lago, Pedro J Sola-Campoy, Sergio Buenestado-Serrano, Marta Herranz, Victor Manuel de la Cueva, Julia Suárez, Pilar Catalán, Patricia Muñoz and SeqCOVID-SPAIN consortium |

EPI_ISL_853721, EPI_ISL_853741, EPI_ISL_853746, EPI_ISL_853766, EPI_ISL_853807, EPI_ISL_853809, EPI_ISL_853810, EPI_ISL_854176, EPI_ISL_854177, EPI_ISL_854178, EPI_ISL_854180, EPI_ISL_854181, EPI_ISL_854182, EPI_ISL_854183, EPI_ISL_854235

| | | | |
|---|---|---|---|
| see above | Austrian Agency for Health and Food Safety (AGES) | Bergthaler laboratory, CeMM Research Center for Molecular Medicine of the Austrian Academy of Sciences | Lukas Endler, Alexandra Popa, Benedikt Agerer, Jakob-Wendelin Genger, Alexander Lercher, Anna Schedl, Thomas Penz, Michael Schuster, Jan Laine, Martin Senekowitsch, Christoph Bock, Andreas Bergthaler |
| EPI_ISL_855344 | SA Pathology | SA Pathology | Lex Leong, Chuan Kok Lim, Mark Turra, Karin Kassahn, Ivan Bastian, Geoff Higgins |
| EPI_ISL_855605 | Respiratory Virus Unit, National Infection Service, Public Health England | COVID-19 Genomics UK (COG-UK) Consortium | PHE Covid Sequencing Team |

EPI_ISL_855615, EPI_ISL_855616, EPI_ISL_855617, EPI_ISL_855618, EPI_ISL_855619, EPI_ISL_855620, EPI_ISL_855621, EPI_ISL_855622, EPI_ISL_855623, EPI_ISL_855624, EPI_ISL_855625, EPI_ISL_855626, EPI_ISL_855627, EPI_ISL_855628, EPI_ISL_855629, EPI_ISL_855630, EPI_ISL_855631, EPI_ISL_855632, EPI_ISL_855633, EPI_ISL_855634, EPI_ISL_855635, EPI_ISL_855636, EPI_ISL_855637, EPI_ISL_855638, EPI_ISL_855639, EPI_ISL_855640, EPI_ISL_855641, EPI_ISL_855642, EPI_ISL_855643, EPI_ISL_855644, EPI_ISL_855645, EPI_ISL_855646, EPI_ISL_855647, EPI_ISL_855648, EPI_ISL_855649, EPI_ISL_855650, EPI_ISL_855651, EPI_ISL_855652, EPI_ISL_855653, EPI_ISL_855654, EPI_ISL_855655, EPI_ISL_855656, EPI_ISL_855657, EPI_ISL_855658, EPI_ISL_855659, EPI_ISL_855660, EPI_ISL_855661, EPI_ISL_855662, EPI_ISL_855663, EPI_ISL_855664, EPI_ISL_855665, EPI_ISL_855666, EPI_ISL_855667, EPI_ISL_855668, EPI_ISL_855669, EPI_ISL_855670, EPI_ISL_855671, EPI_ISL_855672, EPI_ISL_855673, EPI_ISL_855674, EPI_ISL_855675, EPI_ISL_855676, EPI_ISL_855677, EPI_ISL_855678, EPI_ISL_855679, EPI_ISL_855680, EPI_ISL_855681, EPI_ISL_855682, EPI_ISL_855683, EPI_ISL_855684, EPI_ISL_855685, EPI_ISL_855686, EPI_ISL_855687, EPI_ISL_855688, EPI_ISL_855689, EPI_ISL_855690, EPI_ISL_855691, EPI_ISL_855692, EPI_ISL_855693, EPI_ISL_855694, EPI_ISL_855695, EPI_ISL_855696, EPI_ISL_855697, EPI_ISL_855698, EPI_ISL_855699, EPI_ISL_855700, EPI_ISL_855701, EPI_ISL_855702, EPI_ISL_855703, EPI_ISL_855704, EPI_ISL_855705, EPI_ISL_855706, EPI_ISL_855707, EPI_ISL_855708, EPI_ISL_855709, EPI_ISL_855710, EPI_ISL_855711, EPI_ISL_855712, EPI_ISL_855713, EPI_ISL_855714, EPI_ISL_855715, EPI_ISL_855716, EPI_ISL_855717, EPI_ISL_855718, EPI_ISL_855719, EPI_ISL_855720, EPI_ISL_855721, EPI_ISL_855722, EPI_ISL_855723, EPI_ISL_855724, EPI_ISL_855725, EPI_ISL_855726, EPI_ISL_855727, EPI_ISL_855728, EPI_ISL_855729, EPI_ISL_855730, EPI_ISL_855731, EPI_ISL_855732, EPI_ISL_855733, EPI_ISL_855734, EPI_ISL_855735, EPI_ISL_855736, EPI_ISL_855737, EPI_ISL_855738, EPI_ISL_855739, EPI_ISL_855740, EPI_ISL_855741, EPI_ISL_855742, EPI_ISL_855743, EPI_ISL_855744, EPI_ISL_855745, EPI_ISL_855746, EPI_ISL_855747, EPI_ISL_855748, EPI_ISL_855749, EPI_ISL_855750, EPI_ISL_855751, EPI_ISL_855752, EPI_ISL_855753, EPI_ISL_855754, EPI_ISL_855755, EPI_ISL_855756, EPI_ISL_855757, EPI_ISL_855758, EPI_ISL_855759, EPI_ISL_855760, EPI_ISL_855761, EPI_ISL_855762, EPI_ISL_855763, EPI_ISL_855764, EPI_ISL_855765, EPI_ISL_855766, EPI_ISL_855767, EPI_ISL_855768, EPI_ISL_855769, EPI_ISL_855770, EPI_ISL_855771, EPI_ISL_855772, EPI_ISL_855773, EPI_ISL_855774, EPI_ISL_855775, EPI_ISL_855776, EPI_ISL_855777, EPI_ISL_855778, EPI_ISL_855779, EPI_ISL_855780, EPI_ISL_855781, EPI_ISL_855782, EPI_ISL_855783, EPI_ISL_855784, EPI_ISL_855785, EPI_ISL_855786, EPI_ISL_855787, EPI_ISL_855788, EPI_ISL_855789, EPI_ISL_855790, EPI_ISL_855791, EPI_ISL_855792, EPI_ISL_855793, EPI_ISL_855794, EPI_ISL_855795, EPI_ISL_855796, EPI_ISL_855797, EPI_ISL_855798, EPI_ISL_855799, EPI_ISL_855800, EPI_ISL_855801, EPI_ISL_855802, EPI_ISL_855803, EPI_ISL_855804, EPI_ISL_855805, EPI_ISL_855806, EPI_ISL_855807, EPI_ISL_855808, EPI_ISL_855809, EPI_ISL_855810, EPI_ISL_855811, EPI_ISL_855812, EPI_ISL_855813, EPI_ISL_855814, EPI_ISL_855815, EPI_ISL_855816, EPI_ISL_855817, EPI_ISL_855818, EPI_ISL_855819, EPI_ISL_855820, EPI_ISL_855821, EPI_ISL_855822, EPI_ISL_855823, EPI_ISL_855824, EPI_ISL_855825, EPI_ISL_855826, EPI_ISL_855827, EPI_ISL_855828, EPI_ISL_855829, EPI_ISL_855830, EPI_ISL_855831, EPI_ISL_855832, EPI_ISL_855833, EPI_ISL_855834, EPI_ISL_855835, EPI_ISL_855836, EPI_ISL_855837, EPI_ISL_855838, EPI_ISL_855839, EPI_ISL_855840, EPI_ISL_855841, EPI_ISL_855842, EPI_ISL_855843, EPI_ISL_855844, EPI_ISL_855845, EPI_ISL_855846, EPI_ISL_855847, EPI_ISL_855848, EPI_ISL_855849, EPI_ISL_855850, EPI_ISL_855851, EPI_ISL_855852, EPI_ISL_855853, EPI_ISL_855854, EPI_ISL_855855, EPI_ISL_855856, EPI_ISL_855857, EPI_ISL_855858, EPI_ISL_855859, EPI_ISL_855860, EPI_ISL_855861, EPI_ISL_855862, EPI_ISL_855863, EPI_ISL_855864, EPI_ISL_855865, EPI_ISL_855866, EPI_ISL_855867, EPI_ISL_855868, EPI_ISL_855869, EPI_ISL_855870, EPI_ISL_855871, EPI_ISL_855872, EPI_ISL_855873, EPI_ISL_855874, EPI_ISL_855875, EPI_ISL_855876, EPI_ISL_855877, EPI_ISL_855878, EPI_ISL_855879, EPI_ISL_855880, EPI_ISL_855881, EPI_ISL_855882, EPI_ISL_855883, EPI_ISL_855884, EPI_ISL_855885, EPI_ISL_855886, EPI_ISL_855887, EPI_ISL_855888, EPI_ISL_855889, EPI_ISL_855890, EPI_ISL_855891, EPI_ISL_855892, EPI_ISL_855893, EPI_ISL_855894, EPI_ISL_855895, EPI_ISL_855896, EPI_ISL_855897, EPI_ISL_855898, EPI_ISL_855899, EPI_ISL_855900, EPI_ISL_855901, EPI_ISL_855903, EPI_ISL_855905, EPI_ISL_855907, EPI_ISL_855909, EPI_ISL_855911, EPI_ISL_855913, EPI_ISL_855915, EPI_ISL_855918, EPI_ISL_855920, EPI_ISL_855922, EPI_ISL_855924, EPI_ISL_855926, EPI_ISL_855929, EPI_ISL_855931, EPI_ISL_855933, EPI_ISL_855935, EPI_ISL_855937, EPI_ISL_855939, EPI_ISL_855941, EPI_ISL_855943, EPI_ISL_855946, EPI_ISL_855948, EPI_ISL_855950, EPI_ISL_855952, EPI_ISL_855954, EPI_ISL_855956, EPI_ISL_855958, EPI_ISL_855961, EPI_ISL_855963, EPI_ISL_855965, EPI_ISL_855967, EPI_ISL_855969, EPI_ISL_855970, EPI_ISL_855971, EPI_ISL_855972, EPI_ISL_855973, EPI_ISL_855974, EPI_ISL_855975, EPI_ISL_855976, EPI_ISL_855977, EPI_ISL_855978, EPI_ISL_855979, EPI_ISL_855980, EPI_ISL_855981, EPI_ISL_855982, EPI_ISL_855983, EPI_ISL_855984, EPI_ISL_855985, EPI_ISL_855986, EPI_ISL_855987, EPI_ISL_855988, EPI_ISL_855989, EPI_ISL_855990, EPI_ISL_855991, EPI_ISL_855992, EPI_ISL_855993, EPI_ISL_855994, EPI_ISL_855995, EPI_ISL_855996, EPI_ISL_855997, EPI_ISL_855998, EPI_ISL_855999, EPI_ISL_856000, EPI_ISL_856001, EPI_ISL_856002, EPI_ISL_856003, EPI_ISL_856004, EPI_ISL_856005, EPI_ISL_856006, EPI_ISL_856007, EPI_ISL_856008, EPI_ISL_856009, EPI_ISL_856010, EPI_ISL_856011, EPI_ISL_856012, EPI_ISL_856013, EPI_ISL_856014, EPI_ISL_856015, EPI_ISL_856016, EPI_ISL_856017, EPI_ISL_856018, EPI_ISL_856019, EPI_ISL_856020, EPI_ISL_856021, EPI_ISL_856022, EPI_ISL_856023, EPI_ISL_856024, EPI_ISL_856025, EPI_ISL_856026, EPI_ISL_856027, EPI_ISL_856028, EPI_ISL_856029, EPI_ISL_856030, EPI_ISL_856031, EPI_ISL_856032, EPI_ISL_856033, EPI_ISL_856034, EPI_ISL_856035, EPI_ISL_856036, EPI_ISL_856037, EPI_ISL_856038, EPI_ISL_856039, EPI_ISL_856040, EPI_ISL_856041, EPI_ISL_856042, EPI_ISL_856043, EPI_ISL_856044, EPI_ISL_856045, EPI_ISL_856046, EPI_ISL_856047, EPI_ISL_856048, EPI_ISL_856049, EPI_ISL_856050, EPI_ISL_856051, EPI_ISL_856052, EPI_ISL_856053, EPI_ISL_856054, EPI_ISL_856055, EPI_ISL_856056, EPI_ISL_856057, EPI_ISL_856058, EPI_ISL_856059, EPI_ISL_856060, EPI_ISL_856061, EPI_ISL_856062, EPI_ISL_856063, EPI_ISL_856064, EPI_ISL_856065, EPI_ISL_856066, EPI_ISL_856067, EPI_ISL_856068, EPI_ISL_856069, EPI_ISL_856070, EPI_ISL_856071, EPI_ISL_856072, EPI_ISL_856073, EPI_ISL_856074, EPI_ISL_856075, EPI_ISL_856076, EPI_ISL_856077, EPI_ISL_856078, EPI_ISL_856079, EPI_ISL_856080, EPI_ISL_856081, EPI_ISL_856082, EPI_ISL_856083, EPI_ISL_856084, EPI_ISL_856085, EPI_ISL_856086, EPI_ISL_856087, EPI_ISL_856088, EPI_ISL_856089, EPI_ISL_856090, EPI_ISL_856091, EPI_ISL_856092, EPI_ISL_856093, EPI_ISL_856094, EPI_ISL_856095, EPI_ISL_856096, EPI_ISL_856097, EPI_ISL_856098, EPI_ISL_856099, EPI_ISL_856100, EPI_ISL_856101, EPI_ISL_856102, EPI_ISL_856103, EPI_ISL_856104, EPI_ISL_856105, EPI_ISL_856106, EPI_ISL_856107, EPI_ISL_856108, EPI_ISL_856109, EPI_ISL_856110, EPI_ISL_856111, EPI_ISL_856112, EPI_ISL_856113, EPI_ISL_856114, EPI_ISL_856115, EPI_ISL_856116, EPI_ISL_856117, EPI_ISL_856118, EPI_ISL_856119, EPI_ISL_856120, EPI_ISL_856121, EPI_ISL_856122, EPI_ISL_856123, EPI_ISL_856124, EPI_ISL_856125, EPI_ISL_856126, EPI_ISL_856127, EPI_ISL_856128, EPI_ISL_856129, EPI_ISL_856130, EPI_ISL_856131, EPI_ISL_856132, EPI_ISL_856133, EPI_ISL_856134, EPI_ISL_856135, EPI_ISL_856136, EPI_ISL_856137, EPI_ISL_856138, EPI_ISL_856139, EPI_ISL_856140, EPI_ISL_856141, EPI_ISL_856142, EPI_ISL_856143, EPI_ISL_856144, EPI_ISL_856145, EPI_ISL_856146, EPI_ISL_856147, EPI_ISL_856148, EPI_ISL_856149, EPI_ISL_856150, EPI_ISL_856151, EPI_ISL_856152, EPI_ISL_856153, EPI_ISL_856154, EPI_ISL_856155, EPI_ISL_856156, EPI_ISL_856157, EPI_ISL_856158, EPI_ISL_856159, EPI_ISL_856160, EPI_ISL_856161, EPI_ISL_856162, EPI_ISL_856163, EPI_ISL_856164, EPI_ISL_856165, EPI_ISL_856166, EPI_ISL_856167, EPI_ISL_856168, EPI_ISL_856169, EPI_ISL_856170, EPI_ISL_856171, EPI_ISL_856172, EPI_ISL_856173, EPI_ISL_856174, EPI_ISL_856175, EPI_ISL_856176, EPI_ISL_856177, EPI_ISL_856178, EPI_ISL_856179, EPI_ISL_856180, EPI_ISL_856181, EPI_ISL_856182, EPI_ISL_856183, EPI_ISL_856184, EPI_ISL_856185, EPI_ISL_856186, EPI_ISL_856187, EPI_ISL_856188, EPI_ISL_856189, EPI_ISL_856190, EPI_ISL_856191, EPI_ISL_856192, EPI_ISL_856193, EPI_ISL_856194, EPI_ISL_856195, EPI_ISL_856196, EPI_ISL_856197, EPI_ISL_856198, EPI_ISL_856199, EPI_ISL_856200, EPI_ISL_856201, EPI_ISL_856202, EPI_ISL_856203, EPI_ISL_856204, EPI_ISL_856205, EPI_ISL_856206, EPI_ISL_856207, EPI_ISL_856208, EPI_ISL_856209, EPI_ISL_856210, EPI_ISL_856211, EPI_ISL_856212, EPI_ISL_856213, EPI_ISL_856214, EPI_ISL_856215, EPI_ISL_856216, EPI_ISL_856217, EPI_ISL_856218, EPI_ISL_856219, EPI_ISL_856220, EPI_ISL_856221, EPI_ISL_856222, EPI_ISL_856223, EPI_ISL_856224, EPI_ISL_856225, EPI_ISL_856226, EPI_ISL_856227, EPI_ISL_856228, EPI_ISL_856229, EPI_ISL_856230, EPI_ISL_856231, EPI_ISL_856232, EPI_ISL_856233, EPI_ISL_856234, EPI_ISL_856235, EPI_ISL_856236, EPI_ISL_856237, EPI_ISL_856238, EPI_ISL_856239, EPI_ISL_856240, EPI_ISL_856241, EPI_ISL_856242, EPI_ISL_856243, EPI_ISL_856244, EPI_ISL_856245, EPI_ISL_856246, EPI_ISL_856247, EPI_ISL_856248, EPI_ISL_856249, EPI_ISL_856250, EPI_ISL_856251, EPI_ISL_856252, EPI_ISL_856253, EPI_ISL_856254, EPI_ISL_856255, EPI_ISL_856256, EPI_ISL_856257, EPI_ISL_856258, EPI_ISL_856259, EPI_ISL_856260, EPI_ISL_856261, EPI_ISL_856262,

| | | | |
|---|---|---|---|
| EPI_ISL_856263, EPI_ISL_856264, EPI_ISL_856265, EPI_ISL_856266, EPI_ISL_856267, EPI_ISL_856268, EPI_ISL_856269, EPI_ISL_856270, EPI_ISL_856271, EPI_ISL_856272, EPI_ISL_856273, EPI_ISL_856274, EPI_ISL_856275, EPI_ISL_856276, EPI_ISL_856277, EPI_ISL_856278, EPI_ISL_856279, EPI_ISL_856280, EPI_ISL_856281, EPI_ISL_856282, EPI_ISL_856283, EPI_ISL_856284, EPI_ISL_856285, EPI_ISL_856286, EPI_ISL_856287, EPI_ISL_856288, EPI_ISL_856289, EPI_ISL_856290, EPI_ISL_856291 | | | |
| see above | Department of Virus and Microbiological Special Diagnostics, Statens Serum Institut, Copenhagen, Denmark | Aalborg University | Danish Covid-19 Genome Consortium |
| EPI_ISL_856723, EPI_ISL_856724, EPI_ISL_856725, EPI_ISL_856732, EPI_ISL_856733, EPI_ISL_856734, EPI_ISL_856735, EPI_ISL_856736, EPI_ISL_856737, EPI_ISL_856738, EPI_ISL_856739, EPI_ISL_856740, EPI_ISL_856741, EPI_ISL_856742, EPI_ISL_856743, EPI_ISL_856744, EPI_ISL_856745, EPI_ISL_856746, EPI_ISL_856747, EPI_ISL_856748 | | | |
| see above | Department of Clinical Microbiology | GIGA Medical Genomics | Keith Durkin, Maria Artesi, Sébastien Bontems, Raphaël Boreux, Bouchra Boujemla, Cécile Meex, Pierrette Melin, Marie-Pierre Hayette, Vincent Bours |
| EPI_ISL_856972 | The Institute of Tropical Medicine in Antwerp, Belgium | UAntwerp, Laboratory of Medical Microbiology, Campus Drie Eiken S6.26, Universiteitsplein 1, 2610, Wilrijk, Antwerp, Belgium | Basil Britto Xavier, Jasmine Coppens, Christine Lammens, Veerle Matheeussen, Herman Goossens |
| EPI_ISL_857158 | DOHMH Corona | New York City Public Health Laboratory | Jade Wang, et al. |
| EPI_ISL_857159, EPI_ISL_857160 | DOHMH Morrisania | New York City Public Health Laboratory | Jade Wang, et al. |
| EPI_ISL_857161, EPI_ISL_857162 | DOHMH PHL | New York City Public Health Laboratory | Jade Wang, et al. |
| EPI_ISL_857163 | DOHMH Central Harlem | New York City Public Health Laboratory | Jade Wang, et al. |
| EPI_ISL_857164 | DOHMH Morrisania | New York City Public Health Laboratory | Jade Wang, et al. |
| EPI_ISL_857165 | DOHMH Corona | New York City Public Health Laboratory | Jade Wang, et al. |
| EPI_ISL_857166, EPI_ISL_857167 | DOHMH Fort Greene | New York City Public Health Laboratory | Jade Wang, et al. |
| EPI_ISL_857168 | DOHMH Corona | New York City Public Health Laboratory | Jade Wang, et al. |
| EPI_ISL_857169, EPI_ISL_857170, EPI_ISL_857171 | DOHMH Morrisania | New York City Public Health Laboratory | Jade Wang, et al. |
| EPI_ISL_857172, EPI_ISL_857173 | DOHMH Crown Heights | New York City Public Health Laboratory | Jade Wang, et al. |
| EPI_ISL_857174, EPI_ISL_857175, EPI_ISL_857176, EPI_ISL_857177 | DOHMH Jamaica | New York City Public Health Laboratory | Jade Wang, et al. |
| EPI_ISL_857178 | DOHMH Riverside | New York City Public Health Laboratory | Jade Wang, et al. |
| EPI_ISL_857179 | DOHMH Jamaica | New York City Public Health Laboratory | Jade Wang, et al. |
| EPI_ISL_857180, EPI_ISL_857181, EPI_ISL_857182, EPI_ISL_857183 | DOHMH Central Harlem | New York City Public Health Laboratory | Jade Wang, et al. |
| EPI_ISL_857184 | DOHMH Riverside | New York City Public Health Laboratory | Jade Wang, et al. |
| EPI_ISL_857185 | DOHMH PHL | New York City Public Health Laboratory | Jade Wang, et al. |
| EPI_ISL_857395 | S.M.S.Medical College, Jaipur, Rajasthan | S.M.S.Medical College, Jaipur, Rajasthan | Dr. Bharti Malhotra, Dr. Swati Gautam |
| EPI_ISL_857444, EPI_ISL_857446 | Maine HETL | Tewhey Lab, The Jackson Laboratory | Matluk,N., Dewey,H., Iosue,F., Barter,M., Lynch,R., Munger,H. and Tewhey,R. |
| EPI_ISL_857466, EPI_ISL_857467 | PathWest Laboratory Medicine WA | PathWest Laboratory Medicine WA Microbial Surveillance Unit | PathWest Laboratory Medicine WA Microbial Surveillance Unit |
| EPI_ISL_857543, EPI_ISL_857554, EPI_ISL_857559, EPI_ISL_857564, EPI_ISL_857566, EPI_ISL_857572, EPI_ISL_857581, EPI_ISL_857587, EPI_ISL_857593, EPI_ISL_857595, EPI_ISL_857596, EPI_ISL_857600, EPI_ISL_857602, EPI_ISL_857605, EPI_ISL_857607, EPI_ISL_857611, EPI_ISL_857613, EPI_ISL_857614, EPI_ISL_857616, EPI_ISL_857619, EPI_ISL_857621, EPI_ISL_857622, EPI_ISL_857625, EPI_ISL_857626, EPI_ISL_857631, EPI_ISL_857632, EPI_ISL_857633, EPI_ISL_857635, EPI_ISL_857638, EPI_ISL_857641, EPI_ISL_857646, EPI_ISL_857649, EPI_ISL_857653, EPI_ISL_857659, EPI_ISL_857660, EPI_ISL_857662, EPI_ISL_857675, EPI_ISL_857677, EPI_ISL_857679, EPI_ISL_857687, EPI_ISL_857691, EPI_ISL_857700, EPI_ISL_857705, EPI_ISL_857709, EPI_ISL_857717, EPI_ISL_857719, EPI_ISL_857720, EPI_ISL_857723, EPI_ISL_857728, EPI_ISL_857731, EPI_ISL_857734, EPI_ISL_857736, EPI_ISL_857738, EPI_ISL_857739, EPI_ISL_857740, EPI_ISL_857743, EPI_ISL_857744, EPI_ISL_857749, EPI_ISL_857751, EPI_ISL_857754, EPI_ISL_857756, EPI_ISL_857763, EPI_ISL_857769, EPI_ISL_857773, EPI_ISL_857776, EPI_ISL_857778, EPI_ISL_857780, EPI_ISL_857782, EPI_ISL_857783, EPI_ISL_857786, EPI_ISL_857787, EPI_ISL_857792, EPI_ISL_857794, EPI_ISL_857795, EPI_ISL_857798, EPI_ISL_857799, EPI_ISL_857803, EPI_ISL_857811, EPI_ISL_857820, EPI_ISL_857825, EPI_ISL_857829, EPI_ISL_857835, EPI_ISL_857839, EPI_ISL_857840, EPI_ISL_857845, EPI_ISL_857846, EPI_ISL_857847, EPI_ISL_857853, EPI_ISL_857857, EPI_ISL_857863, EPI_ISL_857866 | | | |
| see above | Lighthouse Lab in Cambridge | Wellcome Sanger Institute for the COVID-19 Genomics UK (COG-UK) Consortium | Rob Howes, The Lighthouse Lab in Cambridge and Alex Alderton, Roberto Amato, Sonia Goncalves, Ewan Harrison, David K. Jackson, Ian Johnston, Dominic Kwiatkowski, Cordelia Langford, John Sillitoe on behalf of the Wellcome Sanger Institute COVID-19 Surveillance Team |
| EPI_ISL_857892, EPI_ISL_857893, EPI_ISL_857895, EPI_ISL_857896, EPI_ISL_857897, EPI_ISL_857898, EPI_ISL_857899, EPI_ISL_857900, EPI_ISL_857901, EPI_ISL_857902, EPI_ISL_857903, EPI_ISL_857904, EPI_ISL_857905, EPI_ISL_857906, EPI_ISL_857907, EPI_ISL_857908, EPI_ISL_857909, EPI_ISL_857910 | | | |
| see above | Lighthouse Lab in Glasgow | Wellcome Sanger Institute for the COVID-19 Genomics UK (COG-UK) Consortium | Harper VanSteenhouse, Yumi Kasai, David Gray, Carol Clugston, Anna Dominiczak and Alex Alderton, Roberto Amato, Sonia Goncalves, Ewan Harrison, David K. Jackson, Ian Johnston, Dominic Kwiatkowski, Cordelia Langford, John Sillitoe on behalf of the Wellcome Sanger Institute COVID-19 Surveillance Team |
| EPI_ISL_860280, EPI_ISL_860282, EPI_ISL_860283 | Nordland Hospital - Bodo, Laboratory Department, Molecular Biology Unit | Norwegian Institute of Public Health, Department of Virology | Kathrine Stene-Johansen, Kamilla Heddeland Instefjord, Hilde Elshaug, Atiya R Ali,Marie Paulsen Madsen, Rasmus Riis Kopperud, Hilde Vollan, Karoline Bragstad, Olav Hungnes |
| EPI_ISL_860720 | Institute for Infectious Diseases, University of Bern, Switzerland | Institute for Infectious Diseases, University of Bern, Switzerland | Michel C Koch, Christian Baumann, Miguel A Terrazos Miani, Cora Sägesser, Pascal Bittel, Stephen L Leib, Peter Keller, Franziska Suter-Riniker, Alban Ramette |
| EPI_ISL_860785, EPI_ISL_860789, EPI_ISL_860790 | Charité Universitätsmedizin Berlin, Institute of Virology, Charitéplatz 1, 10117 Berlin, Germany | Charité Universitätsmedizin Berlin, Institute of Virology, Charitéplatz 1, 10117 Berlin, Germany | Victor M Corman, Julia Schneider, Jörn Beheim-Schwarzbach, Tobias Bleicker, Julia Tesch, Barbara Mühlemann, Talitha Veith, Terry Jones, Christian Drosten |
| EPI_ISL_860850, EPI_ISL_860851 | HIA PERCY, Biologie médicale | National Reference Center for Viruses of Respiratory Infections, Institut Pasteur, Paris | Marion Barbet, Sylvie Behillil, Méline Bizard, Angela Brisebarre, Camille Capel, Etienne Simon-Lorière, Vincent Enouf, Maud Vanpeene, Sylvie van der Werf,Foissaud Vincent |
| EPI_ISL_861004, EPI_ISL_861014, EPI_ISL_861027, EPI_ISL_861043, EPI_ISL_861048, EPI_ISL_861052, EPI_ISL_861082, EPI_ISL_861104 | Johns Hopkins Hospital Department of Pathology | Johns Hopkins Hospital Department of Pathology | C. Paul Morris, Chun Huai Luo, Adannaya Amadi, Nicholas Gallagher, Heba H. Mostafa |
| EPI_ISL_861183, EPI_ISL_861184, EPI_ISL_861199, EPI_ISL_861200, EPI_ISL_861201, EPI_ISL_861202, EPI_ISL_861203, EPI_ISL_861204, EPI_ISL_861206, EPI_ISL_861207, EPI_ISL_861208, EPI_ISL_861209 | | | |
| see above | URMC LABS | Wadsworth Center, New York State Department of Health | Kirsten St. George, Daryl M. Lamson, Alexis Russel, Matthew Shudt, Melissa A Leisner, Jonathan Plitnick, Navjot Singh, John Kelly, Erasmus Schneider, Erica Lasek-Nesselquist |
| EPI_ISL_861332 | DOHMH Central Harlem | New York City Public Health Laboratory | Jade Wang, et al. |
| EPI_ISL_861446 | Clinical Molecular Microbiology Laboratory, UNC Hospitals | Jeremy Wang | Jeremy Wang, Alexander Rubinsteyn, Colleen Rice, Jason Smedberg, Melissa Miller, Corbin Jones, Robert Hagan |
| EPI_ISL_861749 | Tempus | Grubaugh Lab - Yale School of Public Health | Tara Alpert, Joseph Fauver, Anderson Brito, Mallery Breban, Anne Wyllie, Chantal Vogels, Mary Petrone, Annie Watkins, Chaney Kalinich, Isabel Ott, Nathan Grubaugh |
| EPI_ISL_862038 | Johns Hopkins Hospital Department of Pathology | Johns Hopkins Hospital Department of Pathology | C. Paul Morris, Chun Huai Luo, Adannaya Amadi, Matthew Schwartz, Nicholas Gallagher, Heba H. Mostafa |
| EPI_ISL_862129, EPI_ISL_862130, EPI_ISL_862134, EPI_ISL_862135, EPI_ISL_862140, EPI_ISL_862142, EPI_ISL_862147, EPI_ISL_862150, EPI_ISL_862163, EPI_ISL_862166, EPI_ISL_862170, EPI_ISL_862179, EPI_ISL_862180, EPI_ISL_862181, EPI_ISL_862183, EPI_ISL_862187, EPI_ISL_862659, EPI_ISL_862661, EPI_ISL_862662, EPI_ISL_862663, EPI_ISL_862664 | | | |
| see above | Charité Universitätsmedizin Berlin, Institut fur Virologie/Labor | Charité Universitätsmedizin Berlin, Institut für Virologie | Victor M Corman, Barbara Mühlemann, Jörn Beheim-Schwarzbach, Tobias Bleicker, Julia Tesch, Talitha Veith, Julia Schneider, Terry Jones, Christian |

| | | | |
|---|---|---|---|
| | Berlin | | Drosten |
| EPI_ISL_862670, EPI_ISL_862671, EPI_ISL_862672, EPI_ISL_862673, EPI_ISL_862674, EPI_ISL_862675, EPI_ISL_862677, EPI_ISL_862678, EPI_ISL_862679, EPI_ISL_862680, EPI_ISL_862681, EPI_ISL_862682, EPI_ISL_862683, EPI_ISL_862684, EPI_ISL_862685, EPI_ISL_862686, EPI_ISL_862687, EPI_ISL_862688, EPI_ISL_862689, EPI_ISL_862690, EPI_ISL_862691, EPI_ISL_862692, EPI_ISL_862693, EPI_ISL_862694, EPI_ISL_862695, EPI_ISL_862696, EPI_ISL_862697, EPI_ISL_862698, EPI_ISL_862699, EPI_ISL_862700, EPI_ISL_862701, EPI_ISL_862702, EPI_ISL_862703, EPI_ISL_862704, EPI_ISL_862705, EPI_ISL_862706, EPI_ISL_862707, EPI_ISL_862708, EPI_ISL_862709, EPI_ISL_862713 | | | |
| see above | New Mexico Department of Health Scientific Laboratory | New Mexico Department of Health Scientific Laboratory | D'eldra Malone, Ellie Johnson, Anastacia Griego-Fisher |
| EPI_ISL_862889, EPI_ISL_862890, EPI_ISL_862891, EPI_ISL_862892, EPI_ISL_862893, EPI_ISL_862894, EPI_ISL_862895, EPI_ISL_862896, EPI_ISL_862897, EPI_ISL_862898, EPI_ISL_862899, EPI_ISL_862901, EPI_ISL_862902, EPI_ISL_862903, EPI_ISL_862904, EPI_ISL_862905, EPI_ISL_862906, EPI_ISL_862907, EPI_ISL_862908, EPI_ISL_862909, EPI_ISL_862910, EPI_ISL_862911, EPI_ISL_862912, EPI_ISL_862913, EPI_ISL_862914, EPI_ISL_862916, EPI_ISL_862917, EPI_ISL_862918, EPI_ISL_862919, EPI_ISL_862920, EPI_ISL_862921, EPI_ISL_862922, EPI_ISL_862923, EPI_ISL_862924, EPI_ISL_862925, EPI_ISL_862926, EPI_ISL_862927, EPI_ISL_862928, EPI_ISL_862929, EPI_ISL_862930, EPI_ISL_862931, EPI_ISL_862932, EPI_ISL_862933, EPI_ISL_862934, EPI_ISL_862935, EPI_ISL_862936, EPI_ISL_862937, EPI_ISL_862939, EPI_ISL_862940, EPI_ISL_862941, EPI_ISL_862942, EPI_ISL_862943, EPI_ISL_862944, EPI_ISL_862945, EPI_ISL_862946, EPI_ISL_862947, EPI_ISL_862949, EPI_ISL_862950, EPI_ISL_862951, EPI_ISL_862952, EPI_ISL_862953, EPI_ISL_862954, EPI_ISL_862955, EPI_ISL_862956, EPI_ISL_862957, EPI_ISL_862958, EPI_ISL_862959, EPI_ISL_862960, EPI_ISL_862961, EPI_ISL_862962, EPI_ISL_862963, EPI_ISL_862964, EPI_ISL_862965, EPI_ISL_862966, EPI_ISL_862967, EPI_ISL_862968, EPI_ISL_862969, EPI_ISL_862970, EPI_ISL_862971, EPI_ISL_862972, EPI_ISL_862973, EPI_ISL_862974, EPI_ISL_862975, EPI_ISL_862976, EPI_ISL_862977, EPI_ISL_862978, EPI_ISL_862979, EPI_ISL_862980, EPI_ISL_862981, EPI_ISL_862982, EPI_ISL_862984, EPI_ISL_862985, EPI_ISL_862986, EPI_ISL_862987, EPI_ISL_862988, EPI_ISL_862989, EPI_ISL_862991, EPI_ISL_862992, EPI_ISL_862993, EPI_ISL_862994, EPI_ISL_862995, EPI_ISL_862996, EPI_ISL_862997, EPI_ISL_862998, EPI_ISL_862999, EPI_ISL_863000, EPI_ISL_863001, EPI_ISL_863002, EPI_ISL_863003, EPI_ISL_863004, EPI_ISL_863005, EPI_ISL_863006, EPI_ISL_863007, EPI_ISL_863008, EPI_ISL_863009, EPI_ISL_863010, EPI_ISL_863011, EPI_ISL_863012, EPI_ISL_863014, EPI_ISL_863015, EPI_ISL_863016, EPI_ISL_863017, EPI_ISL_863018, EPI_ISL_863019, EPI_ISL_863020, EPI_ISL_863021, EPI_ISL_863022, EPI_ISL_863024, EPI_ISL_863025, EPI_ISL_863026, EPI_ISL_863027, EPI_ISL_863028, EPI_ISL_863029, EPI_ISL_863030, EPI_ISL_863031, EPI_ISL_863032, EPI_ISL_863033, EPI_ISL_863034, EPI_ISL_863035, EPI_ISL_863036, EPI_ISL_863037, EPI_ISL_863038, EPI_ISL_863039, EPI_ISL_863040, EPI_ISL_863041, EPI_ISL_863042, EPI_ISL_863043, EPI_ISL_863044, EPI_ISL_863045, EPI_ISL_863047, EPI_ISL_863048, EPI_ISL_863049, EPI_ISL_863050, EPI_ISL_863051, EPI_ISL_863052, EPI_ISL_863053, EPI_ISL_863054, EPI_ISL_863055, EPI_ISL_863056, EPI_ISL_863057, EPI_ISL_863059, EPI_ISL_863060, EPI_ISL_863062, EPI_ISL_863063, EPI_ISL_863064, EPI_ISL_863065, EPI_ISL_863066, EPI_ISL_863067, EPI_ISL_863068, EPI_ISL_863069, EPI_ISL_863070, EPI_ISL_863071, EPI_ISL_863072, EPI_ISL_863073, EPI_ISL_863075, EPI_ISL_863076, EPI_ISL_863077, EPI_ISL_863078, EPI_ISL_863079, EPI_ISL_863080, EPI_ISL_863081, EPI_ISL_863082, EPI_ISL_863083, EPI_ISL_863085, EPI_ISL_863086, EPI_ISL_863087, EPI_ISL_863088, EPI_ISL_863090, EPI_ISL_863092, EPI_ISL_863093, EPI_ISL_863094, EPI_ISL_863095, EPI_ISL_863096, EPI_ISL_863097, EPI_ISL_863098, EPI_ISL_863099, EPI_ISL_863100, EPI_ISL_863102, EPI_ISL_863104, EPI_ISL_863105, EPI_ISL_863107, EPI_ISL_863108, EPI_ISL_863109, EPI_ISL_863110, EPI_ISL_863111, EPI_ISL_863113, EPI_ISL_863114, EPI_ISL_863115, EPI_ISL_863116, EPI_ISL_863117, EPI_ISL_863118, EPI_ISL_863119, EPI_ISL_863120, EPI_ISL_863121, EPI_ISL_863123, EPI_ISL_863124, EPI_ISL_863125, EPI_ISL_863127, EPI_ISL_863128, EPI_ISL_863129, EPI_ISL_863130, EPI_ISL_863131, EPI_ISL_863132, EPI_ISL_863133, EPI_ISL_863134, EPI_ISL_863135, EPI_ISL_863139, EPI_ISL_863141, EPI_ISL_863142, EPI_ISL_863143, EPI_ISL_863144, EPI_ISL_863145, EPI_ISL_863146, EPI_ISL_863147, EPI_ISL_863148, EPI_ISL_863149, EPI_ISL_863150, EPI_ISL_863152, EPI_ISL_863153, EPI_ISL_863154, EPI_ISL_863155, EPI_ISL_863157, EPI_ISL_863158, EPI_ISL_863159, EPI_ISL_863160, EPI_ISL_863161, EPI_ISL_863163, EPI_ISL_863164, EPI_ISL_863165, EPI_ISL_863166, EPI_ISL_863167, EPI_ISL_863168, EPI_ISL_863169, EPI_ISL_863170, EPI_ISL_863171, EPI_ISL_863172, EPI_ISL_863173, EPI_ISL_863174, EPI_ISL_863175, EPI_ISL_863176, EPI_ISL_863177, EPI_ISL_863178, EPI_ISL_863181, EPI_ISL_863182, EPI_ISL_863183, EPI_ISL_863184, EPI_ISL_863185, EPI_ISL_863186, EPI_ISL_863187, EPI_ISL_863189, EPI_ISL_863190, EPI_ISL_863191, EPI_ISL_863194, EPI_ISL_863195, EPI_ISL_863196, EPI_ISL_863197, EPI_ISL_863198, EPI_ISL_863199, EPI_ISL_863200, EPI_ISL_863201, EPI_ISL_863202, EPI_ISL_863203, EPI_ISL_863204, EPI_ISL_863205, EPI_ISL_863206, EPI_ISL_863207, EPI_ISL_863208, EPI_ISL_863209, EPI_ISL_863211, EPI_ISL_863212, EPI_ISL_863213, EPI_ISL_863214, EPI_ISL_863215, EPI_ISL_863216, EPI_ISL_863217, EPI_ISL_863219, EPI_ISL_863220, EPI_ISL_863221, EPI_ISL_863222, EPI_ISL_863223, EPI_ISL_863224, EPI_ISL_863225, EPI_ISL_863226, EPI_ISL_863227, EPI_ISL_863228, EPI_ISL_863229, EPI_ISL_863230, EPI_ISL_863231, EPI_ISL_863232, EPI_ISL_863233, EPI_ISL_863234, EPI_ISL_863235, EPI_ISL_863236, EPI_ISL_863237, EPI_ISL_863239, EPI_ISL_863240, EPI_ISL_863241, EPI_ISL_863242, EPI_ISL_863243, EPI_ISL_863244, EPI_ISL_863334, EPI_ISL_863396, EPI_ISL_863401, EPI_ISL_863403, EPI_ISL_863424 | | | |
| see above | Lighthouse Lab in Alderley Park | Wellcome Sanger Institute for the COVID-19 Genomics UK (COG-UK) Consortium | Jacquelyn Wynn, Mairead Hyland, The Lighthouse Lab in Alderley Park and Alex Alderton, Roberto Amato, Sonia Goncalves, Ewan Harrison, David K. Jackson, Ian Johnston, Dominic Kwiatkowski, Cordelia Langford, John Sillitoe on behalf of the Wellcome Sanger Institute COVID-19 Surveillance Team |
| EPI_ISL_863431 | Lighthouse Lab in Glasgow | Wellcome Sanger Institute for the COVID-19 Genomics UK (COG-UK) Consortium | Harper VanSteenhouse, Yumi Kasai, David Gray, Carol Clugston, Anna Dominiczak and Alex Alderton, Roberto Amato, Sonia Goncalves, Ewan Harrison, David K. Jackson, Ian Johnston, Dominic Kwiatkowski, Cordelia Langford, John Sillitoe on behalf of the Wellcome Sanger Institute COVID-19 Surveillance Team |
| EPI_ISL_863432, EPI_ISL_863467, EPI_ISL_863514, EPI_ISL_863529 | Lighthouse Lab in Alderley Park | Wellcome Sanger Institute for the COVID-19 Genomics UK (COG-UK) Consortium | Jacquelyn Wynn, Mairead Hyland, The Lighthouse Lab in Alderley Park and Alex Alderton, Roberto Amato, Sonia Goncalves, Ewan Harrison, David K. Jackson, Ian Johnston, Dominic Kwiatkowski, Cordelia Langford, John Sillitoe on behalf of the Wellcome Sanger Institute COVID-19 Surveillance Team |
| EPI_ISL_863531, EPI_ISL_863533, EPI_ISL_863539, EPI_ISL_863540, EPI_ISL_863542, EPI_ISL_863544, EPI_ISL_863546, EPI_ISL_863549, EPI_ISL_863551, EPI_ISL_863552, EPI_ISL_863555, EPI_ISL_863556, EPI_ISL_863557, EPI_ISL_863558, EPI_ISL_863560, EPI_ISL_863565, EPI_ISL_863566, EPI_ISL_863570, EPI_ISL_863572, EPI_ISL_863574, EPI_ISL_863576, EPI_ISL_863578, EPI_ISL_863579, EPI_ISL_863581, EPI_ISL_863582, EPI_ISL_863583, EPI_ISL_863584, EPI_ISL_863586, EPI_ISL_863591, EPI_ISL_863592, EPI_ISL_863593, EPI_ISL_863594, EPI_ISL_863595, EPI_ISL_863603, EPI_ISL_863604, EPI_ISL_863608, EPI_ISL_863609, EPI_ISL_863610, EPI_ISL_863612, EPI_ISL_863614, EPI_ISL_863616, EPI_ISL_863618, EPI_ISL_863619, EPI_ISL_863621, EPI_ISL_863625, EPI_ISL_863626, EPI_ISL_863627, EPI_ISL_863629, EPI_ISL_863630, EPI_ISL_863636, EPI_ISL_863640, EPI_ISL_863641, EPI_ISL_863643, EPI_ISL_863644, EPI_ISL_863648, EPI_ISL_863652, EPI_ISL_863654, EPI_ISL_863656, EPI_ISL_863658, EPI_ISL_863659, EPI_ISL_863661, EPI_ISL_863662, EPI_ISL_863664, EPI_ISL_863665, EPI_ISL_863667, EPI_ISL_863668, EPI_ISL_863671, EPI_ISL_863672, EPI_ISL_863673, EPI_ISL_863674, EPI_ISL_863675, EPI_ISL_863676, EPI_ISL_863680, EPI_ISL_863682, EPI_ISL_863685, EPI_ISL_863687, EPI_ISL_863688, EPI_ISL_863689, EPI_ISL_863690, EPI_ISL_863691, EPI_ISL_863692, EPI_ISL_863694, EPI_ISL_863695, EPI_ISL_863697, EPI_ISL_863698, EPI_ISL_863701, EPI_ISL_863702, EPI_ISL_863704, EPI_ISL_863706, EPI_ISL_863708, EPI_ISL_863710, EPI_ISL_863712, EPI_ISL_863713, EPI_ISL_863715, EPI_ISL_863720, EPI_ISL_863722, EPI_ISL_863727, EPI_ISL_863728, EPI_ISL_863730, EPI_ISL_863731, EPI_ISL_863736, EPI_ISL_863738, EPI_ISL_863740, EPI_ISL_863743, EPI_ISL_863744, EPI_ISL_863748, EPI_ISL_863749, EPI_ISL_863750, EPI_ISL_863751, EPI_ISL_863752, EPI_ISL_863754, EPI_ISL_863755, EPI_ISL_863756, EPI_ISL_863757, EPI_ISL_863758, EPI_ISL_863760, EPI_ISL_863762, EPI_ISL_863763, EPI_ISL_863764, EPI_ISL_863765, EPI_ISL_863767, EPI_ISL_863769, EPI_ISL_863770, EPI_ISL_863771, EPI_ISL_863772, EPI_ISL_863774, EPI_ISL_863777, EPI_ISL_863778, EPI_ISL_863779, EPI_ISL_863780, EPI_ISL_863782, EPI_ISL_863786, EPI_ISL_863787, EPI_ISL_863790, EPI_ISL_863791, EPI_ISL_863794, EPI_ISL_863798, EPI_ISL_863799, EPI_ISL_863800, EPI_ISL_863801, EPI_ISL_863813, EPI_ISL_863816, EPI_ISL_863818, EPI_ISL_863819, EPI_ISL_863820, EPI_ISL_863821, EPI_ISL_863822, EPI_ISL_863823, EPI_ISL_863824, EPI_ISL_863826, EPI_ISL_863827, EPI_ISL_863828, EPI_ISL_863831, EPI_ISL_863832, EPI_ISL_863836, EPI_ISL_863837, EPI_ISL_863839, EPI_ISL_863840, EPI_ISL_863843, EPI_ISL_863844, EPI_ISL_863846, EPI_ISL_863849, EPI_ISL_863852, EPI_ISL_863854, EPI_ISL_863861, EPI_ISL_863862, EPI_ISL_863864, EPI_ISL_863866, EPI_ISL_863868 | | | |
| see above | Lighthouse Lab in Milton Keynes | Wellcome Sanger Institute for the COVID-19 Genomics UK (COG-UK) Consortium | The Lighthouse Lab in Milton Keynes and Alex Alderton, Roberto Amato, Sonia Goncalves, Ewan Harrison, David K. Jackson, Ian Johnston, Dominic Kwiatkowski, Cordelia Langford, John Sillitoe on behalf of the Wellcome Sanger Institute COVID-19 Surveillance Team |
| EPI_ISL_863873, EPI_ISL_863946, EPI_ISL_864004, EPI_ISL_864064, EPI_ISL_864094, EPI_ISL_864142, EPI_ISL_864145, EPI_ISL_864178 | Lighthouse Lab in Alderley Park | Wellcome Sanger Institute for the COVID-19 Genomics UK (COG-UK) Consortium | Jacquelyn Wynn, Mairead Hyland, The Lighthouse Lab in Alderley Park and Alex Alderton, Roberto Amato, Sonia Goncalves, Ewan Harrison, David K. Jackson, Ian Johnston, Dominic Kwiatkowski, Cordelia Langford, John Sillitoe on behalf of the Wellcome Sanger Institute COVID-19 Surveillance Team |
| EPI_ISL_864595, EPI_ISL_864607, EPI_ISL_864612, EPI_ISL_864616, EPI_ISL_864617, EPI_ISL_864618, EPI_ISL_864625, EPI_ISL_864628, EPI_ISL_864629, EPI_ISL_864634, EPI_ISL_864689, EPI_ISL_864690, EPI_ISL_864691, EPI_ISL_864692, EPI_ISL_864693, EPI_ISL_864694, EPI_ISL_864695, EPI_ISL_864696, EPI_ISL_864697, EPI_ISL_864698, EPI_ISL_864699, EPI_ISL_864700, EPI_ISL_864701, EPI_ISL_864702, EPI_ISL_864703, EPI_ISL_864704, EPI_ISL_864705, EPI_ISL_864706, EPI_ISL_864716, EPI_ISL_864725, EPI_ISL_864726, EPI_ISL_864727 | | | |
| see above | University Hospitals of Geneva, Laboratory of Virology | HUG, Laboratory of Virology and the Health2030 Genome Center | Samuel Cordey, Ana Rita Goncalves, Laurent Kaiser, Lorenzo Cerutti, Henri Pegeot, Melyssa Elies, Deborah Penet, Keith Harshman, Ioannis Xenarios, Emmanouil Dermitzakis |
| EPI_ISL_864926 | Department of Pathology, University of Cambridge | COVID-19 Genomics UK (COG-UK) Consortium | Aminu S. Jahun, Yasmin Chaudhry, Grant Hall, Iliana Georgana, Myra Hosmillo, Martin D. Curran, Malte Pinckert, Surendra Parmar, Ian Goodfellow |
| EPI_ISL_865035, EPI_ISL_865036 | West of Scotland Specialist Virology Centre, NHSGGC / MRC-University of Glasgow Centre for Virus Research | COVID-19 Genomics UK (COG-UK) Consortium | Ana da Silva Filipe, Natasha Johnson, Kathy Smollett, Daniel Mair, Stephen Carmichael, Alice Broos, Lily Tong, Jenna Nichols, Kyriaki Nomikou; Sarah McDonald; Richard Orton, Joseph Hughes, Sreenu Vattipally, David L Robertson; Alasdair MacLean, Rory Gunson; Sharif Shaaban, Matthew Holden; Rachel Blacow, Guy Mollett, Kathy Li, James Shepherd, Antonia Ho, Emma Thomson |
| EPI_ISL_865068 | Lighthouse Lab in Glasgow / MRC-University of Glasgow Centre for Virus Research | COVID-19 Genomics UK (COG-UK) Consortium | Ana da Silva Filipe, Natasha Johnson, Kathy Smollett, Daniel Mair, Stephen Carmichael, Alice Broos, Lily Tong, Jenna Nichols, Kyriaki Nomikou; Sarah McDonald; Harper VanSteenhouse, Yumi Kasai, David Gray, Carol Clugston, Anna Dominiczak; Alasdair MacLean, Rory Gunson; Richard Orton, Joseph Hughes, Sreenu Vattipally, David L Robertson; Sharif Shaaban, Matthew Holden; Kathy Li, James Shepherd, Antonia Ho, Emma Thomson |
| EPI_ISL_865140 | Virology Department, Royal Infirmary of Edinburgh, NHS Lothian / School of Biological Sciences, University of Edinburgh / Institute of Genetics and Molecular Medicine, University of Edinburgh | COVID-19 Genomics UK (COG-UK) Consortium | McHugh M, Dewar R, Rooke S, Gallagher M, Balcaza C, O'Toole Á, Scher E, Hill V, McCrone JT, Colquhoun R, Yu X, Jackson B, Rambaut A, Williams TC, Templeton K |
| EPI_ISL_865343, EPI_ISL_865344, EPI_ISL_865345, EPI_ISL_865348, EPI_ISL_865349, EPI_ISL_865351, EPI_ISL_865353, EPI_ISL_865449 | Liverpool Clinical Laboratories | COVID-19 Genomics UK (COG-UK) Consortium | Sam Haldenby, Anita Lucaci, Steve Paterson, Julian Hiscox, Alistair Darby, M Almsaud, A Alrezaihi, Muhannad Alruwaili, Stuart D Armstrong, Jones Benjamin, Eleanor G Bentley, Anu Chawla, Jordan J Clark, Angela Cowell, Richard Eccles, Isabel García-Dorival, Matthew Gemmell, Alessandro Gerada, PKF Gilmore, Richard Gregory, Ximeng Han, Catherine Hartley, Margaret Hughes, Miren Iturriza-Gomara, James Johnson, L Luu, Jenifer Manson, Charlotte Nelson, Elaine O'Toole, Cassie Olateju, Rebekah Penrice-Randal , Lucille Rainbow, N.P Randle, Trevor Ian Robinson, Parul Sharma, Ghada T Shawli, James P Stewart, Neil Swainston, Ecaterina Vamos, Joanne Watts, Mark Whitehead |
| EPI_ISL_865847 | University College London, Great Ormond Street Hospital for Children NHS Foundation Trust, Imperial College Healthcare NHS Trust | COVID-19 Genomics UK (COG-UK) Consortium | Sergi Castellano, Rachel Williams, Mark Kristiansen, Paola Resende Silva, Sunando Roy, Tony Brooks, Helena Tutill, Paola Niola, Patricia Dyal, Charlotte Williams, Leysa Forrest, Yasmin Panchbhaya, Jacqueline Findlay, Samuel Weeks, Julianne Brown, Kathryn Harris, Paul Randell, James Price, Alison Holmes, Judith Breuer |
| EPI_ISL_866103, EPI_ISL_866104, EPI_ISL_866105, EPI_ISL_866117, EPI_ISL_866120, EPI_ISL_866125, EPI_ISL_866126, EPI_ISL_866134, EPI_ISL_866142, EPI_ISL_866143, EPI_ISL_866146, EPI_ISL_866152, EPI_ISL_866154, EPI_ISL_866160, EPI_ISL_866171, EPI_ISL_866174, EPI_ISL_866178, EPI_ISL_866186 | | | |
| see above | University College London Hospital | COVID-19 Genomics UK (COG-UK) Consortium | Judith Heaney, Matthew Byott, Catherine Houlihan, Dan Frampton, Stuart Kirk, Moira Spyer and Eleni Nastouli |
| EPI_ISL_866195, EPI_ISL_866197, EPI_ISL_866199, EPI_ISL_866200, EPI_ISL_866203, EPI_ISL_866204, EPI_ISL_866205, EPI_ISL_866214, EPI_ISL_866216, EPI_ISL_866218, EPI_ISL_866220, EPI_ISL_866224, EPI_ISL_866226, EPI_ISL_866227, EPI_ISL_866233, EPI_ISL_866234, EPI_ISL_866245, EPI_ISL_866246, EPI_ISL_866248, EPI_ISL_866249, EPI_ISL_866252, EPI_ISL_866254 | | | |
| see above | University College London, Great Ormond Street Hospital for Children NHS Foundation Trust, Imperial College Healthcare NHS Trust | COVID-19 Genomics UK (COG-UK) Consortium | Sergi Castellano, Rachel Williams, Mark Kristiansen, Paola Resende Silva, Sunando Roy, Tony Brooks, Helena Tutill, Paola Niola, Patricia Dyal, Charlotte Williams, Leysa Forrest, Yasmin Panchbhaya, Jacqueline Findlay, Samuel Weeks, Julianne Brown, Kathryn Harris, Paul Randell, James Price, Alison Holmes, Judith Breuer |

| | | | |
|---|---|---|---|
| EPI_ISL_866542, EPI_ISL_866543, EPI_ISL_866544, EPI_ISL_866546, EPI_ISL_866547, EPI_ISL_866549, EPI_ISL_866550, EPI_ISL_866551, EPI_ISL_866552, EPI_ISL_866553, EPI_ISL_866554, EPI_ISL_866555, EPI_ISL_866557, EPI_ISL_866558, EPI_ISL_866559, EPI_ISL_866561, EPI_ISL_866562, EPI_ISL_866563, EPI_ISL_866564, EPI_ISL_866565, EPI_ISL_866567, EPI_ISL_866568, EPI_ISL_866570, EPI_ISL_866571, EPI_ISL_866572, EPI_ISL_866574, EPI_ISL_866575, EPI_ISL_866576, EPI_ISL_866603, EPI_ISL_866604, EPI_ISL_866605, EPI_ISL_866606, EPI_ISL_866607, EPI_ISL_866608, EPI_ISL_866609, EPI_ISL_866610, EPI_ISL_866611, EPI_ISL_866612, EPI_ISL_866613, EPI_ISL_866614, EPI_ISL_866615, EPI_ISL_866616, EPI_ISL_866617, EPI_ISL_866620, EPI_ISL_866623, EPI_ISL_866624, EPI_ISL_866625, EPI_ISL_866626, EPI_ISL_866630, EPI_ISL_866632, EPI_ISL_866633, EPI_ISL_866634, EPI_ISL_866636, EPI_ISL_866637, EPI_ISL_866638, EPI_ISL_866639, EPI_ISL_866640, EPI_ISL_866641, EPI_ISL_866644, EPI_ISL_866645, EPI_ISL_866646, EPI_ISL_866647, EPI_ISL_866648, EPI_ISL_866649, EPI_ISL_866650, EPI_ISL_866739, EPI_ISL_866758, EPI_ISL_866759, EPI_ISL_866761, EPI_ISL_866762, EPI_ISL_866765, EPI_ISL_866766, EPI_ISL_866768, EPI_ISL_866769, EPI_ISL_866771, EPI_ISL_866772, EPI_ISL_866773, EPI_ISL_866776, EPI_ISL_866777, EPI_ISL_866778, EPI_ISL_866779, EPI_ISL_866780, EPI_ISL_866783, EPI_ISL_866784, EPI_ISL_866785, EPI_ISL_866786, EPI_ISL_866787, EPI_ISL_866790, EPI_ISL_866791, EPI_ISL_866792, EPI_ISL_866794, EPI_ISL_866795, EPI_ISL_866796, EPI_ISL_866797, EPI_ISL_866798, EPI_ISL_866799, EPI_ISL_866800 | | | |
| see above | Quadram Institute Bioscience | COVID-19 Genomics UK (COG-UK) Consortium | Dave J. Baker, Gemma L. Kay, Alp Aydin, Thanh Le-Viet, Steven Rudder, Ana P. Tedim, Anastasia Kolyva, Maria Diaz, Leonardo de Oliveira Martins, Nabil-Fareed Alikhan, Lizzie Meadows, Rachael Stanley, Ngozi Elumogo, Muhammed Yasir, Nicholas M. Thomson, Alexander J Trotter, Rachel Gilroy, Samuel Bloomfield, Claire Stuart, Andrew Bell, Reenesh Prakash, Samir Dervisevic, Alison E. Mather, John Wain, Mark Webber, Andrew J. Page, Justin O'Grady |
| EPI_ISL_867192, EPI_ISL_867193, EPI_ISL_867194, EPI_ISL_867195, EPI_ISL_867202, EPI_ISL_867215, EPI_ISL_867227, EPI_ISL_867228, EPI_ISL_867229, EPI_ISL_867230, EPI_ISL_867231, EPI_ISL_867232, EPI_ISL_867233, EPI_ISL_867252, EPI_ISL_867253, EPI_ISL_867254, EPI_ISL_867255, EPI_ISL_867256, EPI_ISL_867257, EPI_ISL_867258, EPI_ISL_867261, EPI_ISL_867269, EPI_ISL_867298, EPI_ISL_867299, EPI_ISL_867300, EPI_ISL_867301, EPI_ISL_867353, EPI_ISL_867358, EPI_ISL_867360, EPI_ISL_867362, EPI_ISL_867364, EPI_ISL_867365, EPI_ISL_867366, EPI_ISL_867367, EPI_ISL_867369, EPI_ISL_867370, EPI_ISL_867372, EPI_ISL_867373, EPI_ISL_867374, EPI_ISL_867375, EPI_ISL_867377, EPI_ISL_867378, EPI_ISL_867379, EPI_ISL_867381, EPI_ISL_867382, EPI_ISL_867383, EPI_ISL_867384, EPI_ISL_867385, EPI_ISL_867386, EPI_ISL_867387, EPI_ISL_867388, EPI_ISL_867390, EPI_ISL_867391, EPI_ISL_867392, EPI_ISL_867393, EPI_ISL_867394, EPI_ISL_867395, EPI_ISL_867396, EPI_ISL_867397, EPI_ISL_867398, EPI_ISL_867399, EPI_ISL_867400, EPI_ISL_867401, EPI_ISL_867403, EPI_ISL_867404, EPI_ISL_867405, EPI_ISL_867419, EPI_ISL_867424, EPI_ISL_867430, EPI_ISL_867450, EPI_ISL_867451, EPI_ISL_867459, EPI_ISL_867491, EPI_ISL_867492, EPI_ISL_867494, EPI_ISL_867525, EPI_ISL_867536, EPI_ISL_867538, EPI_ISL_867539, EPI_ISL_867540, EPI_ISL_867541, EPI_ISL_867556, EPI_ISL_867557, EPI_ISL_867561, EPI_ISL_867562, EPI_ISL_867567, EPI_ISL_867572, EPI_ISL_867573, EPI_ISL_867574, EPI_ISL_867575, EPI_ISL_867579, EPI_ISL_867582, EPI_ISL_867584, EPI_ISL_867586, EPI_ISL_867587, EPI_ISL_867588, EPI_ISL_867589, EPI_ISL_867590, EPI_ISL_867704, EPI_ISL_867705, EPI_ISL_867706, EPI_ISL_867716, EPI_ISL_867726, EPI_ISL_867728, EPI_ISL_867737, EPI_ISL_867738, EPI_ISL_867739, EPI_ISL_867740, EPI_ISL_867741, EPI_ISL_867819, EPI_ISL_867834, EPI_ISL_867890, EPI_ISL_867899, EPI_ISL_867900, EPI_ISL_867901, EPI_ISL_867904, EPI_ISL_867906, EPI_ISL_867911, EPI_ISL_867912, EPI_ISL_867913, EPI_ISL_867914, EPI_ISL_867915, EPI_ISL_867916, EPI_ISL_867918, EPI_ISL_867921 | | | |
| see above | Originating lab: Wales Specialist Virology Centre Sequencing lab: Pathogen Genomics Unit | Public Health Wales Microbiology Cardiff Wales Specialist Virology Centre | Catherine Moore, Johnathan Evans, Laura Gifford, Malorie Perry, Simon Cottrell, Angela Marchbank, Alec Birchley, Alexander Adams, Amy Gaskin, Bree Gatica-Wilcox, Jason Coombes, Joel Southgate, Lauren Gilbert, Lee Graham, Nicole Pacchiarini, Sara Kumziene-Summerhayes, Sarah Taylor, Sophie Jones, Sara Rey, Matthew Bull, Joanne Watkins, Sally Corden, Tom Connor |
| EPI_ISL_868141, EPI_ISL_868148, EPI_ISL_868150, EPI_ISL_868152, EPI_ISL_868153, EPI_ISL_868154, EPI_ISL_868177, EPI_ISL_868178, EPI_ISL_868179, EPI_ISL_868203, EPI_ISL_868204 | | | |
| see above | Centre for Enzyme Innovation, University of Portsmouth / Translational Research Laboratory, Portsmouth Hospitals NHS Trust | COVID-19 Genomics UK (COG-UK) Consortium | Angela Beckett,Yann Bourgeois,Garry Scarlett,Sharon Glaysher,Scott Elliott,Kelly Bicknell,Robert Impey,Allyson Lloyd,Sarah Wyllie,Ethan Butcher,Anoop Chauhan,Samuel Robson |
| EPI_ISL_868358, EPI_ISL_868364, EPI_ISL_868386, EPI_ISL_868399, EPI_ISL_868413, EPI_ISL_868428, EPI_ISL_868430, EPI_ISL_868432, EPI_ISL_868434, EPI_ISL_868445, EPI_ISL_868446, EPI_ISL_868448, EPI_ISL_868459, EPI_ISL_868467, EPI_ISL_868471, EPI_ISL_868487, EPI_ISL_868503, EPI_ISL_868522, EPI_ISL_868535, EPI_ISL_868547, EPI_ISL_868552, EPI_ISL_868556, EPI_ISL_868557, EPI_ISL_868573, EPI_ISL_868593, EPI_ISL_868621, EPI_ISL_868631, EPI_ISL_868648, EPI_ISL_868656, EPI_ISL_868660, EPI_ISL_868677, EPI_ISL_868684, EPI_ISL_868693, EPI_ISL_868700, EPI_ISL_868715 | | | |
| see above | Virology Department, Sheffield Teaching Hospitals NHS Foundation Trust/Department of Infection, Immunity and Cardiovascular Disease, The Medical School, University of Sheffield | COVID-19 Genomics UK (COG-UK) Consortium | Thushan de Silva, Matthew Parker, Nikki Smith, Adri Angyal, Rebecca Brown, Luke Green, Rachel Tucker, Paul Parsons, Danielle Groves, Katie Johnson, Laura Carrilero, Alex Keeley, Dave Partridge, Matthew Wyles, Benjamin Lindsey, Mehmet Yavuz, Mohammad Raza, Cariad Evans |
| EPI_ISL_869132 | Charité Universitätsmedizin Berlin, Institut für Virologie/Labor Berlin | Charité Universitätsmedizin Berlin, Institut für Virologie | Victor M Corman, Barbara Mühlemann, Jörn Beheim-Schwarzbach, Tobias Bleicker, Julia Tesch, Talitha Veith, Julia Schneider, Terry Jones, Christian Drosten |
| EPI_ISL_869644, EPI_ISL_869645, EPI_ISL_869646, EPI_ISL_869647, EPI_ISL_869648, EPI_ISL_869649, EPI_ISL_869650, EPI_ISL_869651, EPI_ISL_869652, EPI_ISL_869653, EPI_ISL_869654, EPI_ISL_869655, EPI_ISL_869656, EPI_ISL_869657, EPI_ISL_869658, EPI_ISL_869659, EPI_ISL_869660, EPI_ISL_869661, EPI_ISL_869662, EPI_ISL_869663, EPI_ISL_869664, EPI_ISL_869665, EPI_ISL_869666, EPI_ISL_869667, EPI_ISL_869668, EPI_ISL_869669, EPI_ISL_869670, EPI_ISL_869671, EPI_ISL_869672, EPI_ISL_869673, EPI_ISL_869674, EPI_ISL_869675, EPI_ISL_869676, EPI_ISL_869677, EPI_ISL_869678, EPI_ISL_869679, EPI_ISL_869680, EPI_ISL_869681, EPI_ISL_869682, EPI_ISL_869683, EPI_ISL_869684, EPI_ISL_869685, EPI_ISL_869686, EPI_ISL_869687, EPI_ISL_869688, EPI_ISL_869689, EPI_ISL_869690, EPI_ISL_869691, EPI_ISL_869692, EPI_ISL_869693, EPI_ISL_869694, EPI_ISL_869695, EPI_ISL_869696, EPI_ISL_869697, EPI_ISL_869698, EPI_ISL_869699, EPI_ISL_869700, EPI_ISL_869701, EPI_ISL_869702, EPI_ISL_869703, EPI_ISL_869704, EPI_ISL_869705, EPI_ISL_869706, EPI_ISL_869707, EPI_ISL_869708, EPI_ISL_869709, EPI_ISL_869710, EPI_ISL_869711, EPI_ISL_869712, EPI_ISL_869713, EPI_ISL_869714, EPI_ISL_869715, EPI_ISL_869716, EPI_ISL_869717, EPI_ISL_869718, EPI_ISL_869719, EPI_ISL_869720, EPI_ISL_869721, EPI_ISL_869722, EPI_ISL_869723, EPI_ISL_869724, EPI_ISL_869725, EPI_ISL_869726, EPI_ISL_869727, EPI_ISL_869728, EPI_ISL_869729, EPI_ISL_869730, EPI_ISL_869731, EPI_ISL_869732, EPI_ISL_869733, EPI_ISL_869734, EPI_ISL_869735, EPI_ISL_869736, EPI_ISL_869737, EPI_ISL_869738, EPI_ISL_869739, EPI_ISL_869740, EPI_ISL_869741, EPI_ISL_869742, EPI_ISL_869743, EPI_ISL_869744, EPI_ISL_869745, EPI_ISL_869746, EPI_ISL_869747, EPI_ISL_869748, EPI_ISL_869749, EPI_ISL_869750, EPI_ISL_869751, EPI_ISL_869752, EPI_ISL_869753, EPI_ISL_869754, EPI_ISL_869755, EPI_ISL_869756, EPI_ISL_869757, EPI_ISL_869758, EPI_ISL_869759, EPI_ISL_869760, EPI_ISL_869761, EPI_ISL_869762, EPI_ISL_869763, EPI_ISL_869764, EPI_ISL_869765, EPI_ISL_869766, EPI_ISL_869767, EPI_ISL_869768, EPI_ISL_869769, EPI_ISL_869770, EPI_ISL_869771, EPI_ISL_869772, EPI_ISL_869773, EPI_ISL_869774, EPI_ISL_869775, EPI_ISL_869776, EPI_ISL_869777, EPI_ISL_869778, EPI_ISL_869779, EPI_ISL_869780, EPI_ISL_869781, EPI_ISL_869782, EPI_ISL_869783, EPI_ISL_869784, EPI_ISL_869785, EPI_ISL_869786, EPI_ISL_869787, EPI_ISL_869788, EPI_ISL_869789, EPI_ISL_869790, EPI_ISL_869791, EPI_ISL_869792, EPI_ISL_869793, EPI_ISL_869794, EPI_ISL_869795, EPI_ISL_869796, EPI_ISL_869797, EPI_ISL_869798, EPI_ISL_869799, EPI_ISL_869800, EPI_ISL_869801, EPI_ISL_869802, EPI_ISL_869803, EPI_ISL_869804, EPI_ISL_869805, EPI_ISL_869806, EPI_ISL_869807, EPI_ISL_869808, EPI_ISL_869809, EPI_ISL_869810, EPI_ISL_869811, EPI_ISL_869812, EPI_ISL_869813, EPI_ISL_869814, EPI_ISL_869815, EPI_ISL_869816, EPI_ISL_869817, EPI_ISL_869818, EPI_ISL_869819, EPI_ISL_869820, EPI_ISL_869821, EPI_ISL_869822, EPI_ISL_869823, EPI_ISL_869824, EPI_ISL_869825, EPI_ISL_869826, EPI_ISL_869827, EPI_ISL_869828, EPI_ISL_869829, EPI_ISL_869830, EPI_ISL_869831, EPI_ISL_869832, EPI_ISL_869833, EPI_ISL_869834, EPI_ISL_869835, EPI_ISL_869836, EPI_ISL_869837, EPI_ISL_869838, EPI_ISL_869839, EPI_ISL_869840, EPI_ISL_869841, EPI_ISL_869842, EPI_ISL_869843, EPI_ISL_869844, EPI_ISL_869845, EPI_ISL_869846, EPI_ISL_869847, EPI_ISL_869848, EPI_ISL_869849, EPI_ISL_869850, EPI_ISL_869851, EPI_ISL_869852, EPI_ISL_869853, EPI_ISL_869854, EPI_ISL_869855, EPI_ISL_869856, EPI_ISL_869857, EPI_ISL_869858, EPI_ISL_869859, EPI_ISL_869860, EPI_ISL_869861, EPI_ISL_869862, EPI_ISL_869863, EPI_ISL_869864, EPI_ISL_869865, EPI_ISL_869866, EPI_ISL_869867, EPI_ISL_869868, EPI_ISL_869869, EPI_ISL_869870, EPI_ISL_869871, EPI_ISL_869872, EPI_ISL_869873, EPI_ISL_869874, EPI_ISL_869875, EPI_ISL_869876, EPI_ISL_869877, EPI_ISL_869878, EPI_ISL_869879, EPI_ISL_869880, EPI_ISL_869881, EPI_ISL_869882, EPI_ISL_869883, EPI_ISL_869884, EPI_ISL_869885, EPI_ISL_869886, EPI_ISL_869887, EPI_ISL_869888, EPI_ISL_869889, EPI_ISL_869890, EPI_ISL_869891, EPI_ISL_869892, EPI_ISL_869893, EPI_ISL_869894, EPI_ISL_869895, EPI_ISL_869896, EPI_ISL_869897, EPI_ISL_869898, EPI_ISL_869899, EPI_ISL_869900, EPI_ISL_869901, EPI_ISL_869902, EPI_ISL_869903, EPI_ISL_869904, EPI_ISL_869905, EPI_ISL_869906, EPI_ISL_869907, EPI_ISL_869908, EPI_ISL_869909, EPI_ISL_869910, EPI_ISL_869911, EPI_ISL_869912, EPI_ISL_869913, EPI_ISL_869914, EPI_ISL_869915, EPI_ISL_869916, EPI_ISL_869917, EPI_ISL_869918, EPI_ISL_869919, EPI_ISL_869920, EPI_ISL_869921, EPI_ISL_869922, EPI_ISL_869923, EPI_ISL_869924, EPI_ISL_869925, EPI_ISL_869926, EPI_ISL_869927, EPI_ISL_869928, EPI_ISL_869929, EPI_ISL_869930, EPI_ISL_869931, EPI_ISL_869932, EPI_ISL_869933, EPI_ISL_869934, EPI_ISL_869935, EPI_ISL_869936, EPI_ISL_869937, EPI_ISL_869938, EPI_ISL_869939, EPI_ISL_869940, EPI_ISL_869941, EPI_ISL_869942, EPI_ISL_869943, EPI_ISL_869944, EPI_ISL_869945, EPI_ISL_869946, EPI_ISL_869947, EPI_ISL_869948, EPI_ISL_869949, EPI_ISL_869950, EPI_ISL_869951, EPI_ISL_869952, EPI_ISL_869953, EPI_ISL_869954, EPI_ISL_869955, EPI_ISL_869956, EPI_ISL_869957, EPI_ISL_869958, EPI_ISL_869959, EPI_ISL_869960, EPI_ISL_869961, EPI_ISL_869962, EPI_ISL_869963, EPI_ISL_869964, EPI_ISL_869965, EPI_ISL_869966, EPI_ISL_869967, EPI_ISL_869968, EPI_ISL_869969, EPI_ISL_869970, EPI_ISL_869971, EPI_ISL_869972, EPI_ISL_869973, EPI_ISL_869974, EPI_ISL_869975, EPI_ISL_869976, EPI_ISL_869977, EPI_ISL_869978, EPI_ISL_869979, EPI_ISL_869980, EPI_ISL_869981, EPI_ISL_869982, EPI_ISL_869983, EPI_ISL_869984, EPI_ISL_869985, EPI_ISL_869986, EPI_ISL_869987, EPI_ISL_869988, EPI_ISL_869989, EPI_ISL_869990, EPI_ISL_869991, EPI_ISL_869992, EPI_ISL_869993, EPI_ISL_869994, EPI_ISL_869995, EPI_ISL_869996, EPI_ISL_869997, EPI_ISL_869998, EPI_ISL_869999, EPI_ISL_870000, EPI_ISL_870001, EPI_ISL_870002, EPI_ISL_870003, EPI_ISL_870004, EPI_ISL_870005, EPI_ISL_870006, EPI_ISL_870007, EPI_ISL_870008, EPI_ISL_870009, EPI_ISL_870010, EPI_ISL_870011, EPI_ISL_870012, EPI_ISL_870013, EPI_ISL_870014, EPI_ISL_870015, EPI_ISL_870016, EPI_ISL_870017, EPI_ISL_870018, EPI_ISL_870019, EPI_ISL_870020, EPI_ISL_870021, EPI_ISL_870022, EPI_ISL_870023, EPI_ISL_870024, EPI_ISL_870025, EPI_ISL_870026, EPI_ISL_870027, EPI_ISL_870028, EPI_ISL_870029, EPI_ISL_870030, EPI_ISL_870031, EPI_ISL_870032, EPI_ISL_870033, EPI_ISL_870034, EPI_ISL_870035, EPI_ISL_870036, EPI_ISL_870037, EPI_ISL_870038, EPI_ISL_870039, EPI_ISL_870040, EPI_ISL_870041, EPI_ISL_870042, EPI_ISL_870043, EPI_ISL_870044, EPI_ISL_870045, EPI_ISL_870046, EPI_ISL_870047, EPI_ISL_870048, EPI_ISL_870049, EPI_ISL_870050, EPI_ISL_870051, EPI_ISL_870052, EPI_ISL_870053, EPI_ISL_870054, EPI_ISL_870055, EPI_ISL_870056, EPI_ISL_870057, EPI_ISL_870058, EPI_ISL_870059, EPI_ISL_870060, EPI_ISL_870061, EPI_ISL_870062, EPI_ISL_870063, EPI_ISL_870064, EPI_ISL_870066, EPI_ISL_870067, EPI_ISL_870068, EPI_ISL_870069, EPI_ISL_870070, EPI_ISL_870071, EPI_ISL_870072, EPI_ISL_870073, EPI_ISL_870074, EPI_ISL_870075, EPI_ISL_870076, EPI_ISL_870077, EPI_ISL_870078, EPI_ISL_870079, EPI_ISL_870080, EPI_ISL_870081, EPI_ISL_870082, EPI_ISL_870083, EPI_ISL_870084, EPI_ISL_870085, EPI_ISL_870086, EPI_ISL_870087, EPI_ISL_870088, EPI_ISL_870089, EPI_ISL_870090, EPI_ISL_870091, EPI_ISL_870092, EPI_ISL_870093, EPI_ISL_870094, EPI_ISL_870095, EPI_ISL_870096, EPI_ISL_870097, EPI_ISL_870098, EPI_ISL_870099, EPI_ISL_870100, EPI_ISL_870101, EPI_ISL_870102, EPI_ISL_870103, EPI_ISL_870104, EPI_ISL_870105, EPI_ISL_870106, EPI_ISL_870107, EPI_ISL_870108, EPI_ISL_870109, EPI_ISL_870110, EPI_ISL_870111, EPI_ISL_870112, EPI_ISL_870113, EPI_ISL_870114, EPI_ISL_870115, EPI_ISL_870116, EPI_ISL_870117, EPI_ISL_870118, EPI_ISL_870119, EPI_ISL_870120, EPI_ISL_870121, EPI_ISL_870122, EPI_ISL_870123, EPI_ISL_870124, EPI_ISL_870125, EPI_ISL_870126, EPI_ISL_870127, EPI_ISL_870128, EPI_ISL_870129, EPI_ISL_870130, EPI_ISL_870131, EPI_ISL_870132, EPI_ISL_870133, EPI_ISL_870134, EPI_ISL_870135, EPI_ISL_870136, EPI_ISL_870137, EPI_ISL_870138, EPI_ISL_870139, EPI_ISL_870140, EPI_ISL_870141, EPI_ISL_870142, EPI_ISL_870143, EPI_ISL_870144, EPI_ISL_870145, EPI_ISL_870146, EPI_ISL_870147, EPI_ISL_870148, EPI_ISL_870149, EPI_ISL_870150, EPI_ISL_870151, EPI_ISL_870152, EPI_ISL_870153, EPI_ISL_870154, EPI_ISL_870155, EPI_ISL_870156, EPI_ISL_870157, EPI_ISL_870158, EPI_ISL_870159, EPI_ISL_870160, EPI_ISL_870161, EPI_ISL_870162, EPI_ISL_870163, EPI_ISL_870164, EPI_ISL_870165, EPI_ISL_870166, EPI_ISL_870167, EPI_ISL_870168, EPI_ISL_870169, EPI_ISL_870170, EPI_ISL_870171, EPI_ISL_870172, EPI_ISL_870173, EPI_ISL_870174, EPI_ISL_870175, EPI_ISL_870176, EPI_ISL_870177, EPI_ISL_870178, EPI_ISL_870179, EPI_ISL_870180, EPI_ISL_870181, EPI_ISL_870182, EPI_ISL_870183, EPI_ISL_870184, EPI_ISL_870185, EPI_ISL_870186, EPI_ISL_870187, EPI_ISL_870188, EPI_ISL_870189, EPI_ISL_870190, EPI_ISL_870191, EPI_ISL_870192, EPI_ISL_870193, EPI_ISL_870194, EPI_ISL_870195, EPI_ISL_870196, EPI_ISL_870197, EPI_ISL_870198, EPI_ISL_870199, EPI_ISL_870200, EPI_ISL_870201, EPI_ISL_870202, EPI_ISL_870203, EPI_ISL_870204, EPI_ISL_870205, EPI_ISL_870206, EPI_ISL_870207, EPI_ISL_870208, EPI_ISL_870209, EPI_ISL_870210, EPI_ISL_870211, EPI_ISL_870212, EPI_ISL_870213, EPI_ISL_870214, EPI_ISL_870215, EPI_ISL_870216, EPI_ISL_870217, EPI_ISL_870218, EPI_ISL_870219, EPI_ISL_870220, EPI_ISL_870221, EPI_ISL_870222, EPI_ISL_870223, EPI_ISL_870224, EPI_ISL_870225, EPI_ISL_870226, EPI_ISL_870227, EPI_ISL_870228, EPI_ISL_870229, EPI_ISL_870230, EPI_ISL_870231, EPI_ISL_870232, EPI_ISL_870233, EPI_ISL_870234, EPI_ISL_870235, EPI_ISL_870236, EPI_ISL_870237, EPI_ISL_870238, EPI_ISL_870239, EPI_ISL_870240, EPI_ISL_870241, EPI_ISL_870242, EPI_ISL_870243, EPI_ISL_870244, EPI_ISL_870245, EPI_ISL_870246, EPI_ISL_870247, EPI_ISL_870248, EPI_ISL_870249, EPI_ISL_870250, EPI_ISL_870251, EPI_ISL_870252, EPI_ISL_870253, EPI_ISL_870254, EPI_ISL_870255, EPI_ISL_870256, EPI_ISL_870257, EPI_ISL_870258, EPI_ISL_870259, EPI_ISL_870260, EPI_ISL_870261, EPI_ISL_870262, EPI_ISL_870263, EPI_ISL_870264, EPI_ISL_870265, EPI_ISL_870266, EPI_ISL_870267, EPI_ISL_870268, EPI_ISL_870269, EPI_ISL_870270, EPI_ISL_870271, EPI_ISL_870272, EPI_ISL_870273, EPI_ISL_870274, EPI_ISL_870275, EPI_ISL_870276, EPI_ISL_870277, EPI_ISL_870278, EPI_ISL_870279, EPI_ISL_870280, EPI_ISL_870281, EPI_ISL_870282, EPI_ISL_870283, EPI_ISL_870284, EPI_ISL_870285, EPI_ISL_870286, EPI_ISL_870287, EPI_ISL_870288, EPI_ISL_870289, EPI_ISL_870290, EPI_ISL_870291, EPI_ISL_870292, EPI_ISL_870293, EPI_ISL_870294, EPI_ISL_870295, EPI_ISL_870296, EPI_ISL_870297, EPI_ISL_870298, EPI_ISL_870299, EPI_ISL_870300, EPI_ISL_870301, EPI_ISL_870302, EPI_ISL_870303, EPI_ISL_870304, EPI_ISL_870305, EPI_ISL_870306, EPI_ISL_870307, EPI_ISL_870308, EPI_ISL_870309, EPI_ISL_870310, EPI_ISL_870311, EPI_ISL_870312, EPI_ISL_870313, EPI_ISL_870314, EPI_ISL_870316, EPI_ISL_870318, EPI_ISL_870320 | | | |
| see above | Department of Virus and Microbiological Special Diagnostics, Statens Serum Institut, Copenhagen, Denmark | Aalborg University | Danish Covid-19 Genome Consortium |
| EPI_ISL_870321 | University of Michigan Clinical Microbiology Laboratory | Lauring Lab, University of Michigan, Department of Microbiology and Immunology | Valesano |
| EPI_ISL_870323, EPI_ISL_870324, EPI_ISL_870325, EPI_ISL_870326, EPI_ISL_870327, EPI_ISL_870328, EPI_ISL_870329, EPI_ISL_870330, EPI_ISL_870331, EPI_ISL_870332, EPI_ISL_870333, EPI_ISL_870334, EPI_ISL_870335, EPI_ISL_870336, EPI_ISL_870337, EPI_ISL_870338, EPI_ISL_870339, EPI_ISL_870340, EPI_ISL_870341, EPI_ISL_870342, EPI_ISL_870343, EPI_ISL_870344, EPI_ISL_870345, EPI_ISL_870346, EPI_ISL_870347, EPI_ISL_870348, EPI_ISL_870349, EPI_ISL_870350, EPI_ISL_870351, EPI_ISL_870352, EPI_ISL_870353, EPI_ISL_870354, EPI_ISL_870355, EPI_ISL_870356, EPI_ISL_870357, EPI_ISL_870358, EPI_ISL_870359, EPI_ISL_870360, EPI_ISL_870361, EPI_ISL_870362, EPI_ISL_870363, EPI_ISL_870364, EPI_ISL_870365, EPI_ISL_870366, EPI_ISL_870367, EPI_ISL_870368, EPI_ISL_870369, EPI_ISL_870370, EPI_ISL_870371, EPI_ISL_870372, EPI_ISL_870373, EPI_ISL_870374, EPI_ISL_870375, EPI_ISL_870376, EPI_ISL_870377, EPI_ISL_870378, EPI_ISL_870379, EPI_ISL_870380, EPI_ISL_870381, EPI_ISL_870382, EPI_ISL_870383, EPI_ISL_870384, EPI_ISL_870385, EPI_ISL_870386, EPI_ISL_870387, EPI_ISL_870388, EPI_ISL_870389, EPI_ISL_870390, EPI_ISL_870391, EPI_ISL_870392, EPI_ISL_870393, EPI_ISL_870394, EPI_ISL_870395, EPI_ISL_870396, EPI_ISL_870397, EPI_ISL_870398, EPI_ISL_870399, EPI_ISL_870400, EPI_ISL_870401, EPI_ISL_870402, EPI_ISL_870403, EPI_ISL_870404, EPI_ISL_870405, EPI_ISL_870406, EPI_ISL_870407, EPI_ISL_870408, EPI_ISL_870409, EPI_ISL_870410, EPI_ISL_870411, EPI_ISL_870412, EPI_ISL_870413, EPI_ISL_870414, EPI_ISL_870415, EPI_ISL_870416, EPI_ISL_870417, EPI_ISL_870418, EPI_ISL_870419, EPI_ISL_870420, EPI_ISL_870421, EPI_ISL_870422, EPI_ISL_870423, EPI_ISL_870424, EPI_ISL_870425, EPI_ISL_870426, EPI_ISL_870427, EPI_ISL_870428, EPI_ISL_870429, EPI_ISL_870430, EPI_ISL_870431, EPI_ISL_870432, EPI_ISL_870433, EPI_ISL_870434, EPI_ISL_870435, EPI_ISL_870436, EPI_ISL_870437, EPI_ISL_870438, EPI_ISL_870439, EPI_ISL_870440, EPI_ISL_870441, EPI_ISL_870442, EPI_ISL_870443, EPI_ISL_870444, EPI_ISL_870445, EPI_ISL_870446, EPI_ISL_870447, EPI_ISL_870448, | | | |

| | | | |
|---|---|---|---|
| EPI_ISL_870449, EPI_ISL_870450, EPI_ISL_870451, EPI_ISL_870452, EPI_ISL_870453, EPI_ISL_870454, EPI_ISL_870455, EPI_ISL_870456, EPI_ISL_870457, EPI_ISL_870458, EPI_ISL_870459, EPI_ISL_870460, EPI_ISL_870461, EPI_ISL_870462, EPI_ISL_870463, EPI_ISL_870464, EPI_ISL_870465, EPI_ISL_870466, EPI_ISL_870467, EPI_ISL_870468, EPI_ISL_870469, EPI_ISL_870470, EPI_ISL_870471, EPI_ISL_870472, EPI_ISL_870473, EPI_ISL_870474, EPI_ISL_870475, EPI_ISL_870476, EPI_ISL_870477, EPI_ISL_870478, EPI_ISL_870479, EPI_ISL_870480, EPI_ISL_870481, EPI_ISL_870482, EPI_ISL_870483, EPI_ISL_870484, EPI_ISL_870485, EPI_ISL_870486, EPI_ISL_870487, EPI_ISL_870488, EPI_ISL_870489, EPI_ISL_870490, EPI_ISL_870491, EPI_ISL_870492, EPI_ISL_870493, EPI_ISL_870494, EPI_ISL_870495, EPI_ISL_870496, EPI_ISL_870497, EPI_ISL_870498, EPI_ISL_870499, EPI_ISL_870500, EPI_ISL_870501, EPI_ISL_870502, EPI_ISL_870503, EPI_ISL_870504, EPI_ISL_870505, EPI_ISL_870506, EPI_ISL_870507, EPI_ISL_870508, EPI_ISL_870509, EPI_ISL_870510, EPI_ISL_870511, EPI_ISL_870512, EPI_ISL_870513, EPI_ISL_870514, EPI_ISL_870515, EPI_ISL_870516, EPI_ISL_870517, EPI_ISL_870518, EPI_ISL_870519, EPI_ISL_870520, EPI_ISL_870521, EPI_ISL_870522, EPI_ISL_870523, EPI_ISL_870524, EPI_ISL_870525, EPI_ISL_870526, EPI_ISL_870527, EPI_ISL_870528, EPI_ISL_870529, EPI_ISL_870530, EPI_ISL_870531, EPI_ISL_870532, EPI_ISL_870533, EPI_ISL_870534, EPI_ISL_870535, EPI_ISL_870536, EPI_ISL_870537, EPI_ISL_870538, EPI_ISL_870539, EPI_ISL_870540, EPI_ISL_870541, EPI_ISL_870542, EPI_ISL_870543, EPI_ISL_870544, EPI_ISL_870545, EPI_ISL_870546, EPI_ISL_870547, EPI_ISL_870548, EPI_ISL_870549, EPI_ISL_870550, EPI_ISL_870551, EPI_ISL_870552, EPI_ISL_870553, EPI_ISL_870554, EPI_ISL_870555, EPI_ISL_870556, EPI_ISL_870557, EPI_ISL_870558, EPI_ISL_870559, EPI_ISL_870560, EPI_ISL_870561, EPI_ISL_870562, EPI_ISL_870563, EPI_ISL_870564, EPI_ISL_870565, EPI_ISL_870566, EPI_ISL_870567, EPI_ISL_870568, EPI_ISL_870569, EPI_ISL_870570, EPI_ISL_870571, EPI_ISL_870572, EPI_ISL_870573, EPI_ISL_870574, EPI_ISL_870575, EPI_ISL_870576, EPI_ISL_870577, EPI_ISL_870578, EPI_ISL_870579, EPI_ISL_870580, EPI_ISL_870581, EPI_ISL_870582, EPI_ISL_870583, EPI_ISL_870584, EPI_ISL_870585, EPI_ISL_870586, EPI_ISL_870587, EPI_ISL_870588, EPI_ISL_870589, EPI_ISL_870590, EPI_ISL_870591, EPI_ISL_870592, EPI_ISL_870593, EPI_ISL_870594, EPI_ISL_870595, EPI_ISL_870596, EPI_ISL_870597, EPI_ISL_870598, EPI_ISL_870599, EPI_ISL_870600, EPI_ISL_870601, EPI_ISL_870602, EPI_ISL_870603, EPI_ISL_870604, EPI_ISL_870605, EPI_ISL_870606, EPI_ISL_870607, EPI_ISL_870608, EPI_ISL_870609, EPI_ISL_870610, EPI_ISL_870611, EPI_ISL_870612, EPI_ISL_870613, EPI_ISL_870614, EPI_ISL_870615, EPI_ISL_870616, EPI_ISL_870617, EPI_ISL_870618, EPI_ISL_870619, EPI_ISL_870620, EPI_ISL_870621, EPI_ISL_870622, EPI_ISL_870623, EPI_ISL_870624, EPI_ISL_870625, EPI_ISL_870626, EPI_ISL_870627, EPI_ISL_870628, EPI_ISL_870629, EPI_ISL_870630, EPI_ISL_870631, EPI_ISL_870632, EPI_ISL_870633, EPI_ISL_870634, EPI_ISL_870635, EPI_ISL_870636, EPI_ISL_870637, EPI_ISL_870638, EPI_ISL_870639, EPI_ISL_870640, EPI_ISL_870641, EPI_ISL_870642, EPI_ISL_870643, EPI_ISL_870644, EPI_ISL_870645, EPI_ISL_870646, EPI_ISL_870647, EPI_ISL_870648, EPI_ISL_870649, EPI_ISL_870650, EPI_ISL_870651, EPI_ISL_870652, EPI_ISL_870653, EPI_ISL_870654, EPI_ISL_870655, EPI_ISL_870656, EPI_ISL_870657, EPI_ISL_870658, EPI_ISL_870659, EPI_ISL_870660, EPI_ISL_870661, EPI_ISL_870662, EPI_ISL_870663, EPI_ISL_870664, EPI_ISL_870665, EPI_ISL_870666, EPI_ISL_870667, EPI_ISL_870668, EPI_ISL_870669, EPI_ISL_870670, EPI_ISL_870671, EPI_ISL_870672, EPI_ISL_870673, EPI_ISL_870674, EPI_ISL_870675, EPI_ISL_870676, EPI_ISL_870677, EPI_ISL_870678, EPI_ISL_870679, EPI_ISL_870680, EPI_ISL_870681, EPI_ISL_870682, EPI_ISL_870683, EPI_ISL_870684, EPI_ISL_870685, EPI_ISL_870686, EPI_ISL_870687, EPI_ISL_870688, EPI_ISL_870689, EPI_ISL_870690, EPI_ISL_870691, EPI_ISL_870692, EPI_ISL_870693, EPI_ISL_870694, EPI_ISL_870695, EPI_ISL_870696, EPI_ISL_870697, EPI_ISL_870698, EPI_ISL_870699, EPI_ISL_870700, EPI_ISL_870701, EPI_ISL_870702, EPI_ISL_870703, EPI_ISL_870704, EPI_ISL_870705, EPI_ISL_870706, EPI_ISL_870707, EPI_ISL_870708, EPI_ISL_870709, EPI_ISL_870710, EPI_ISL_870711, EPI_ISL_870712, EPI_ISL_870713, EPI_ISL_870714, EPI_ISL_870715, EPI_ISL_870716, EPI_ISL_870717, EPI_ISL_870718, EPI_ISL_870719, EPI_ISL_870720, EPI_ISL_870721, EPI_ISL_870722, EPI_ISL_870723, EPI_ISL_870724, EPI_ISL_870725, EPI_ISL_870726, EPI_ISL_870727, EPI_ISL_870728, EPI_ISL_870729, EPI_ISL_870730, EPI_ISL_870731, EPI_ISL_870732, EPI_ISL_870733, EPI_ISL_870734, EPI_ISL_870735, EPI_ISL_870736, EPI_ISL_870737, EPI_ISL_870738, EPI_ISL_870739, EPI_ISL_870740, EPI_ISL_870741, EPI_ISL_870742, EPI_ISL_870743, EPI_ISL_870744, EPI_ISL_870745, EPI_ISL_870746, EPI_ISL_870747, EPI_ISL_870748, EPI_ISL_870749, EPI_ISL_870750, EPI_ISL_870751, EPI_ISL_870752, EPI_ISL_870753, EPI_ISL_870754, EPI_ISL_870755, EPI_ISL_870756, EPI_ISL_870757, EPI_ISL_870758, EPI_ISL_870759, EPI_ISL_870760, EPI_ISL_870761, EPI_ISL_870762, EPI_ISL_870763, EPI_ISL_870764, EPI_ISL_870765, EPI_ISL_870766, EPI_ISL_870767, EPI_ISL_870768, EPI_ISL_870769, EPI_ISL_870770, EPI_ISL_870771, EPI_ISL_870772, EPI_ISL_870773, EPI_ISL_870774, EPI_ISL_870775, EPI_ISL_870776, EPI_ISL_870777, EPI_ISL_870778, EPI_ISL_870779, EPI_ISL_870780, EPI_ISL_870781, EPI_ISL_870782, EPI_ISL_870783, EPI_ISL_870784, EPI_ISL_870785, EPI_ISL_870786, EPI_ISL_870787, EPI_ISL_870788, EPI_ISL_870789, EPI_ISL_870790, EPI_ISL_870791, EPI_ISL_870792, EPI_ISL_870793, EPI_ISL_870794, EPI_ISL_870795, EPI_ISL_870796, EPI_ISL_870797, EPI_ISL_870798, EPI_ISL_870799, EPI_ISL_870800, EPI_ISL_870801, EPI_ISL_870802, EPI_ISL_870803, EPI_ISL_870804, EPI_ISL_870805, EPI_ISL_870806, EPI_ISL_870807, EPI_ISL_870808, EPI_ISL_870809, EPI_ISL_870810, EPI_ISL_870811, EPI_ISL_870812, EPI_ISL_870813, EPI_ISL_870814, EPI_ISL_870815, EPI_ISL_870816, EPI_ISL_870817, EPI_ISL_870818, EPI_ISL_870819, EPI_ISL_870820, EPI_ISL_870821, EPI_ISL_870822, EPI_ISL_870823, EPI_ISL_870824, EPI_ISL_870825, EPI_ISL_870826, EPI_ISL_870827, EPI_ISL_870828, EPI_ISL_870829, EPI_ISL_870830, EPI_ISL_870831, EPI_ISL_870832, EPI_ISL_870833, EPI_ISL_870834, EPI_ISL_870835, EPI_ISL_870836, EPI_ISL_870837, EPI_ISL_870838, EPI_ISL_870839, EPI_ISL_870840, EPI_ISL_870841, EPI_ISL_870842, EPI_ISL_870843, EPI_ISL_870844, EPI_ISL_870845, EPI_ISL_870846, EPI_ISL_870847, EPI_ISL_870848, EPI_ISL_870849, EPI_ISL_870850, EPI_ISL_870851, EPI_ISL_870852, EPI_ISL_870853, EPI_ISL_870854, EPI_ISL_870855, EPI_ISL_870856, EPI_ISL_870857, EPI_ISL_870858, EPI_ISL_870859, EPI_ISL_870860, EPI_ISL_870861, EPI_ISL_870862, EPI_ISL_870863, EPI_ISL_870864, EPI_ISL_870865, EPI_ISL_870866, EPI_ISL_870867, EPI_ISL_870868, EPI_ISL_870869, EPI_ISL_870870, EPI_ISL_870871, EPI_ISL_870872, EPI_ISL_870873, EPI_ISL_870874, EPI_ISL_870875, EPI_ISL_870876, EPI_ISL_870877, EPI_ISL_870878, EPI_ISL_870879, EPI_ISL_870880, EPI_ISL_870881, EPI_ISL_870882, EPI_ISL_870883, EPI_ISL_870884, EPI_ISL_870885, EPI_ISL_870886, EPI_ISL_870887, EPI_ISL_870888, EPI_ISL_870889, EPI_ISL_870890, EPI_ISL_870891, EPI_ISL_870892, EPI_ISL_870893, EPI_ISL_870894, EPI_ISL_870895, EPI_ISL_870896, EPI_ISL_870897, EPI_ISL_870898, EPI_ISL_870899, EPI_ISL_870900, EPI_ISL_870901, EPI_ISL_870902, EPI_ISL_870903, EPI_ISL_870904, EPI_ISL_870905, EPI_ISL_870906, EPI_ISL_870907, EPI_ISL_870908, EPI_ISL_870909, EPI_ISL_870910, EPI_ISL_870911, EPI_ISL_870912, EPI_ISL_870913, EPI_ISL_870914, EPI_ISL_870915, EPI_ISL_870916, EPI_ISL_870917, EPI_ISL_870918, EPI_ISL_870919, EPI_ISL_870920, EPI_ISL_870921, EPI_ISL_870922, EPI_ISL_870923, EPI_ISL_870924, EPI_ISL_870925, EPI_ISL_870926, EPI_ISL_870927, EPI_ISL_870928, EPI_ISL_870929, EPI_ISL_870930, EPI_ISL_870931, EPI_ISL_870932, EPI_ISL_870933, EPI_ISL_870934, EPI_ISL_870935, EPI_ISL_870936, EPI_ISL_870937, EPI_ISL_870938, EPI_ISL_870939, EPI_ISL_870940, EPI_ISL_870941, EPI_ISL_870942, EPI_ISL_870943, EPI_ISL_870944, EPI_ISL_870945, EPI_ISL_870946, EPI_ISL_870947, EPI_ISL_870948, EPI_ISL_870949, EPI_ISL_870950, EPI_ISL_870951, EPI_ISL_870952, EPI_ISL_870953, EPI_ISL_870954, EPI_ISL_870955, EPI_ISL_870956, EPI_ISL_870957, EPI_ISL_870958, EPI_ISL_870959, EPI_ISL_870960, EPI_ISL_870961, EPI_ISL_870962, EPI_ISL_870963, EPI_ISL_870964, EPI_ISL_870965, EPI_ISL_870966, EPI_ISL_870967, EPI_ISL_870968, EPI_ISL_870969, EPI_ISL_870970, EPI_ISL_870971, EPI_ISL_870972, EPI_ISL_870973, EPI_ISL_870974, EPI_ISL_870975, EPI_ISL_870976, EPI_ISL_870977, EPI_ISL_870978, EPI_ISL_870979, EPI_ISL_870980, EPI_ISL_870981, EPI_ISL_870982, EPI_ISL_870983, EPI_ISL_870984, EPI_ISL_870985, EPI_ISL_870986, EPI_ISL_870987, EPI_ISL_870988, EPI_ISL_870989, EPI_ISL_870990, EPI_ISL_870991, EPI_ISL_870992, EPI_ISL_870993, EPI_ISL_870994, EPI_ISL_870995, EPI_ISL_870996, EPI_ISL_870997, EPI_ISL_870998, EPI_ISL_870999, EPI_ISL_871000, EPI_ISL_871001, EPI_ISL_871002, EPI_ISL_871003, EPI_ISL_871004, EPI_ISL_871005, EPI_ISL_871006, EPI_ISL_871007, EPI_ISL_871008, EPI_ISL_871009, EPI_ISL_871010, EPI_ISL_871011, EPI_ISL_871012, EPI_ISL_871013, EPI_ISL_871014, EPI_ISL_871015, EPI_ISL_871016, EPI_ISL_871017, EPI_ISL_871018, EPI_ISL_871019, EPI_ISL_871020, EPI_ISL_871021, EPI_ISL_871022, EPI_ISL_871023, EPI_ISL_871024, EPI_ISL_871025, EPI_ISL_871026, EPI_ISL_871027, EPI_ISL_871028, EPI_ISL_871029, EPI_ISL_871030, EPI_ISL_871031, EPI_ISL_871032, EPI_ISL_871033, EPI_ISL_871034, EPI_ISL_871035, EPI_ISL_871036, EPI_ISL_871037, EPI_ISL_871038, EPI_ISL_871039, EPI_ISL_871040, EPI_ISL_871041, EPI_ISL_871042, EPI_ISL_871043, EPI_ISL_871044, EPI_ISL_871045, EPI_ISL_871046, EPI_ISL_871047, EPI_ISL_871048, EPI_ISL_871049, EPI_ISL_871050, EPI_ISL_871051, EPI_ISL_871052, EPI_ISL_871053, EPI_ISL_871054, EPI_ISL_871055, EPI_ISL_871056, EPI_ISL_871057, EPI_ISL_871058, EPI_ISL_871059, EPI_ISL_871060, EPI_ISL_871061, EPI_ISL_871062, EPI_ISL_871063, EPI_ISL_871064, EPI_ISL_871065, EPI_ISL_871066, EPI_ISL_871067, EPI_ISL_871068, EPI_ISL_871069, EPI_ISL_871070, EPI_ISL_871071, EPI_ISL_871072, EPI_ISL_871073, EPI_ISL_871074, EPI_ISL_871075, EPI_ISL_871076, EPI_ISL_871077, EPI_ISL_871078, EPI_ISL_871079, EPI_ISL_871080, EPI_ISL_871081, EPI_ISL_871082, EPI_ISL_871083, EPI_ISL_871084, EPI_ISL_871085, EPI_ISL_871086, EPI_ISL_871087, EPI_ISL_871088, EPI_ISL_871089, EPI_ISL_871090, EPI_ISL_871091, EPI_ISL_871092, EPI_ISL_871093, EPI_ISL_871094, EPI_ISL_871095, EPI_ISL_871096, EPI_ISL_871097, EPI_ISL_871098, EPI_ISL_871099, EPI_ISL_871100, EPI_ISL_871101, EPI_ISL_871102, EPI_ISL_871103, EPI_ISL_871104, EPI_ISL_871105, EPI_ISL_871106, EPI_ISL_871107, EPI_ISL_871108, EPI_ISL_871109, EPI_ISL_871110, EPI_ISL_871111, EPI_ISL_871112, EPI_ISL_871113, EPI_ISL_871114, EPI_ISL_871115, EPI_ISL_871116, EPI_ISL_871117, EPI_ISL_871118, EPI_ISL_871119, EPI_ISL_871120, EPI_ISL_871121, EPI_ISL_871122, EPI_ISL_871123, EPI_ISL_871124, EPI_ISL_871125, EPI_ISL_871126, EPI_ISL_871127, EPI_ISL_871128, EPI_ISL_871129, EPI_ISL_871130, EPI_ISL_871131, EPI_ISL_871132 | | | |
| see above | Department of Virus and Microbiological Special Diagnostics, Statens Serum Institut, Copenhagen, Denmark | Aalborg University | Danish Covid-19 Genome Consortium |
| EPI_ISL_871820 | Jessa | Jessa | Brigitte Maes, Bert Cruys |
| EPI_ISL_872065, EPI_ISL_872066 | AZ Vesale (Tongres) | GIGA Medical Genomics | Keith Durkin, Maria Artesi, Sébastien Bontems, Raphaël Boreux, Bouchra Boujemla, Cécile Meex, Pierrette Melin, Marie-Pierre Hayette, Vincent Bours |
| EPI_ISL_872197, EPI_ISL_872200, EPI_ISL_872202 | hopital | National Reference Center for Viruses of Respiratory Infections, Institut Pasteur, Paris | Marion Barbet, Sylvie Behillil, Méline Bizard, Angela Brisebarre, Camille Capel, Etienne Simon-Lorière, Vincent Enouf, Maud Vanpeene, Sylvie van der Werf,Combe Patrice |
| EPI_ISL_872207 | Labo Analyses Med | National Reference Center for Viruses of Respiratory Infections, Institut Pasteur, Paris | Marion Barbet, Sylvie Behillil, Méline Bizard, Angela Brisebarre, Camille Capel, Etienne Simon-Lorière, Vincent Enouf, Maud Vanpeene, Sylvie van der Werf |
| EPI_ISL_872210 | Hopital | National Reference Center for Viruses of Respiratory Infections, Institut Pasteur, Paris | Marion Barbet, Sylvie Behillil, Méline Bizard, Angela Brisebarre, Camille Capel, Etienne Simon-Lorière, Vincent Enouf, Maud Vanpeene, Sylvie van der Werf,Guigon AuréLie |
| EPI_ISL_872222, EPI_ISL_872223, EPI_ISL_872224, EPI_ISL_872225, EPI_ISL_872226 | hopital | National Reference Center for Viruses of Respiratory Infections, Institut Pasteur, Paris | Marion Barbet, Sylvie Behillil, Méline Bizard, Angela Brisebarre, Camille Capel, Etienne Simon-Lorière, Vincent Enouf, Maud Vanpeene, Sylvie van der Werf,Combe Patrice |
| EPI_ISL_872227 | Hopital | National Reference Center for Viruses of Respiratory Infections, Institut Pasteur, Paris | Marion Barbet, Sylvie Behillil, Méline Bizard, Angela Brisebarre, Camille Capel, Etienne Simon-Lorière, Vincent Enouf, Maud Vanpeene, Sylvie van der Werf |
| EPI_ISL_872232 | hopital | National Reference Center for Viruses of Respiratory Infections, Institut Pasteur, Paris | Marion Barbet, Sylvie Behillil, Méline Bizard, Angela Brisebarre, Camille Capel, Etienne Simon-Lorière, Vincent Enouf, Maud Vanpeene, Sylvie van der Werf,Combe Patrice |
| EPI_ISL_872235 | Labo Analyses Med | National Reference Center for Viruses of Respiratory Infections, Institut Pasteur, Paris | Marion Barbet, Sylvie Behillil, Méline Bizard, Angela Brisebarre, Camille Capel, Etienne Simon-Lorière, Vincent Enouf, Maud Vanpeene, Sylvie van der Werf |
| EPI_ISL_872237 | Hopital | National Reference Center for Viruses of Respiratory Infections, Institut Pasteur, Paris | Marion Barbet, Sylvie Behillil, Méline Bizard, Angela Brisebarre, Camille Capel, Etienne Simon-Lorière, Vincent Enouf, Maud Vanpeene, Sylvie van der Werf,Florin CéCile |
| EPI_ISL_872242, EPI_ISL_872243, EPI_ISL_872244, EPI_ISL_872245, EPI_ISL_872246, EPI_ISL_872247, EPI_ISL_872248 | hopital | National Reference Center for Viruses of Respiratory Infections, Institut Pasteur, Paris | Marion Barbet, Sylvie Behillil, Méline Bizard, Angela Brisebarre, Camille Capel, Etienne Simon-Lorière, Vincent Enouf, Maud Vanpeene, Sylvie van der Werf,Combe Patrice |
| EPI_ISL_872260 | Labo Analyses Med | National Reference Center for Viruses of Respiratory Infections, Institut Pasteur, Paris | Marion Barbet, Sylvie Behillil, Méline Bizard, Angela Brisebarre, Camille Capel, Etienne Simon-Lorière, Vincent Enouf, Maud Vanpeene, Sylvie van der Werf |
| EPI_ISL_872263 | Hopital | National Reference Center for Viruses of Respiratory Infections, Institut Pasteur, Paris | Marion Barbet, Sylvie Behillil, Méline Bizard, Angela Brisebarre, Camille Capel, Etienne Simon-Lorière, Vincent Enouf, Maud Vanpeene, Sylvie van der Werf,Florin CéCile |
| EPI_ISL_872271, EPI_ISL_872272, EPI_ISL_872273 | hopital | National Reference Center for Viruses of Respiratory Infections, Institut Pasteur, Paris | Marion Barbet, Sylvie Behillil, Méline Bizard, Angela Brisebarre, Camille Capel, Etienne Simon-Lorière, Vincent Enouf, Maud Vanpeene, Sylvie van der Werf,Combe Patrice |
| EPI_ISL_872285, EPI_ISL_872286, EPI_ISL_872287, EPI_ISL_872288, EPI_ISL_872289 | Labo Analyses Med | National Reference Center for Viruses of Respiratory Infections, Institut Pasteur, Paris | Marion Barbet, Sylvie Behillil, Méline Bizard, Angela Brisebarre, Camille Capel, Etienne Simon-Lorière, Vincent Enouf, Maud Vanpeene, Sylvie van der Werf |
| EPI_ISL_872295, EPI_ISL_872296, EPI_ISL_872297 | Hopital | National Reference Center for Viruses of Respiratory Infections, Institut Pasteur, Paris | Marion Barbet, Sylvie Behillil, Méline Bizard, Angela Brisebarre, Camille Capel, Etienne Simon-Lorière, Vincent Enouf, Maud Vanpeene, Sylvie van der Werf,Guigon AuréLie |
| EPI_ISL_872299, EPI_ISL_872301 | Hopital | National Reference Center for Viruses of Respiratory Infections, Institut Pasteur, Paris | Marion Barbet, Sylvie Behillil, Méline Bizard, Angela Brisebarre, Camille Capel, Etienne Simon-Lorière, Vincent Enouf, Maud Vanpeene, Sylvie van der Werf,Florin CéCile |
| EPI_ISL_872308 | Hopital | National Reference Center for Viruses of Respiratory Infections, Institut Pasteur, Paris | Marion Barbet, Sylvie Behillil, Méline Bizard, Angela Brisebarre, Camille Capel, Etienne Simon-Lorière, Vincent Enouf, Maud Vanpeene, Sylvie van der Werf,Brichler Ségolène |

| | | | |
|---|---|---|---|
| EPI_ISL_872321, EPI_ISL_872322 | Labo Analyses Med | National Reference Center for Viruses of Respiratory Infections, Institut Pasteur, Paris | Marion Barbet, Sylvie Behillil, Méline Bizard, Angela Brisebarre, Camille Capel, Etienne Simon-Lorière, Vincent Enouf, Maud Vanpeene, Sylvie van der Werf |
| EPI_ISL_872325 | Hopital | National Reference Center for Viruses of Respiratory Infections, Institut Pasteur, Paris | Marion Barbet, Sylvie Behillil, Méline Bizard, Angela Brisebarre, Camille Capel, Etienne Simon-Lorière, Vincent Enouf, Maud Vanpeene, Sylvie van der Werf,Brichler Ségolène |
| EPI_ISL_872330, EPI_ISL_872331, EPI_ISL_872332, EPI_ISL_872333, EPI_ISL_872334, EPI_ISL_872335, EPI_ISL_872336, EPI_ISL_872337, EPI_ISL_872338 | hopital | National Reference Center for Viruses of Respiratory Infections, Institut Pasteur, Paris | Marion Barbet, Sylvie Behillil, Méline Bizard, Angela Brisebarre, Camille Capel, Etienne Simon-Lorière, Vincent Enouf, Maud Vanpeene, Sylvie van der Werf,Combe Patrice |
| EPI_ISL_872478 | Centre for Dengue Research and AICBU, Department of Immunology and Molecular Medicine | Centre for Dengue Research and AICBU, Department of Immunology and Molecular Medicine | Chandima Jeewandara, Deshni Jayathilaka, Dinuka Ariyaratne, Tibutius Thanesh Pramanayagam, Diyanath Ranasinghe, Laksiri Gomes, Gathsaurie Neelika Malavige |
| EPI_ISL_872494, EPI_ISL_872495, EPI_ISL_872496, EPI_ISL_872497, EPI_ISL_872498, EPI_ISL_872499, EPI_ISL_872500, EPI_ISL_872501, EPI_ISL_872502, EPI_ISL_872503, EPI_ISL_872504, EPI_ISL_872505, EPI_ISL_872506, EPI_ISL_872507, EPI_ISL_872508, EPI_ISL_872509, EPI_ISL_872510 | | | |
| see above | New Mexico Department of Health Scientific Laboratory | Center for Global Health, University of New Mexico Health Sciences Center | Daryl Domman, Kurt Schwalm, Twila Kunde, Joseph Hicks, Anastacia Griego, Michael Edwards, Darrell Dinwiddie |
| EPI_ISL_872781, EPI_ISL_872792, EPI_ISL_872794, EPI_ISL_872795, EPI_ISL_872797, EPI_ISL_872798, EPI_ISL_872799, EPI_ISL_872802, EPI_ISL_872803, EPI_ISL_872804, EPI_ISL_872808, EPI_ISL_872809, EPI_ISL_872811, EPI_ISL_872813, EPI_ISL_872815, EPI_ISL_872818, EPI_ISL_872819, EPI_ISL_872821 | | | |
| see above | University of Wisconsin-Madison AIDS Vaccine Research Laboratories | University of Wisconsin-Madison AIDS Vaccine Research Laboratories | Gage Moreno, Katarina Braun, et al. AIDS Vaccine Research Laboratories |
| EPI_ISL_873157, EPI_ISL_873162 | University of Michigan Clinical Microbiology Laboratory | Lauring Lab, University of Michigan, Department of Microbiology and Immunology | Valesano |
| EPI_ISL_873258, EPI_ISL_873261, EPI_ISL_873263, EPI_ISL_873264, EPI_ISL_873265, EPI_ISL_873266, EPI_ISL_873267, EPI_ISL_873268, EPI_ISL_873269 | M Health Fairview | Minnesota Department of Health, Public Health Laboratory | Alexandra Lorentz, Jacob Garfin, Matt Plumb, and Xiong Wang |
| EPI_ISL_873279 | Lighthouse Lab in Milton Keynes | Wellcome Sanger Institute for the COVID-19 Genomics UK (COG-UK) Consortium | The Lighthouse Lab in Milton Keynes and Alex Alderton, Roberto Amato, Sonia Goncalves, Ewan Harrison, David K. Jackson, Ian Johnston, Dominic Kwiatkowski, Cordelia Langford, John Sillitoe on behalf of the Wellcome Sanger Institute COVID-19 Surveillance Team |
| EPI_ISL_873282 | Lighthouse Lab in Cambridge | Wellcome Sanger Institute for the COVID-19 Genomics UK (COG-UK) Consortium | Rob Howes, The Lighthouse Lab in Cambridge and Alex Alderton, Roberto Amato, Sonia Goncalves, Ewan Harrison, David K. Jackson, Ian Johnston, Dominic Kwiatkowski, Cordelia Langford, John Sillitoe on behalf of the Wellcome Sanger Institute COVID-19 Surveillance Team |
| EPI_ISL_873284, EPI_ISL_873285, EPI_ISL_873287, EPI_ISL_873290, EPI_ISL_873293 | Lighthouse Lab in Milton Keynes | Wellcome Sanger Institute for the COVID-19 Genomics UK (COG-UK) Consortium | The Lighthouse Lab in Milton Keynes and Alex Alderton, Roberto Amato, Sonia Goncalves, Ewan Harrison, David K. Jackson, Ian Johnston, Dominic Kwiatkowski, Cordelia Langford, John Sillitoe on behalf of the Wellcome Sanger Institute COVID-19 Surveillance Team |
| EPI_ISL_873295 | Lighthouse Lab in Cambridge | Wellcome Sanger Institute for the COVID-19 Genomics UK (COG-UK) Consortium | Rob Howes, The Lighthouse Lab in Cambridge and Alex Alderton, Roberto Amato, Sonia Goncalves, Ewan Harrison, David K. Jackson, Ian Johnston, Dominic Kwiatkowski, Cordelia Langford, John Sillitoe on behalf of the Wellcome Sanger Institute COVID-19 Surveillance Team |
| EPI_ISL_873296, EPI_ISL_873299, EPI_ISL_873300, EPI_ISL_873301, EPI_ISL_873303, EPI_ISL_873305, EPI_ISL_873307, EPI_ISL_873314, EPI_ISL_873315, EPI_ISL_873317, EPI_ISL_873319, EPI_ISL_873322, EPI_ISL_873323, EPI_ISL_873327, EPI_ISL_873328, EPI_ISL_873329, EPI_ISL_873332, EPI_ISL_873333, EPI_ISL_873335, EPI_ISL_873338, EPI_ISL_873340, EPI_ISL_873344, EPI_ISL_873346, EPI_ISL_873349, EPI_ISL_873351, EPI_ISL_873353, EPI_ISL_873355, EPI_ISL_873356, EPI_ISL_873358, EPI_ISL_873363 | | | |
| see above | Lighthouse Lab in Milton Keynes | Wellcome Sanger Institute for the COVID-19 Genomics UK (COG-UK) Consortium | The Lighthouse Lab in Milton Keynes and Alex Alderton, Roberto Amato, Sonia Goncalves, Ewan Harrison, David K. Jackson, Ian Johnston, Dominic Kwiatkowski, Cordelia Langford, John Sillitoe on behalf of the Wellcome Sanger Institute COVID-19 Surveillance Team |
| EPI_ISL_873365 | Lighthouse Lab in Cambridge | Wellcome Sanger Institute for the COVID-19 Genomics UK (COG-UK) Consortium | Rob Howes, The Lighthouse Lab in Cambridge and Alex Alderton, Roberto Amato, Sonia Goncalves, Ewan Harrison, David K. Jackson, Ian Johnston, Dominic Kwiatkowski, Cordelia Langford, John Sillitoe on behalf of the Wellcome Sanger Institute COVID-19 Surveillance Team |
| EPI_ISL_873366, EPI_ISL_873367 | Lighthouse Lab in Milton Keynes | Wellcome Sanger Institute for the COVID-19 Genomics UK (COG-UK) Consortium | The Lighthouse Lab in Milton Keynes and Alex Alderton, Roberto Amato, Sonia Goncalves, Ewan Harrison, David K. Jackson, Ian Johnston, Dominic Kwiatkowski, Cordelia Langford, John Sillitoe on behalf of the Wellcome Sanger Institute COVID-19 Surveillance Team |
| EPI_ISL_873368 | Lighthouse Lab in Cambridge | Wellcome Sanger Institute for the COVID-19 Genomics UK (COG-UK) Consortium | Rob Howes, The Lighthouse Lab in Cambridge and Alex Alderton, Roberto Amato, Sonia Goncalves, Ewan Harrison, David K. Jackson, Ian Johnston, Dominic Kwiatkowski, Cordelia Langford, John Sillitoe on behalf of the Wellcome Sanger Institute COVID-19 Surveillance Team |
| EPI_ISL_873373, EPI_ISL_873375, EPI_ISL_873380, EPI_ISL_873384, EPI_ISL_873387, EPI_ISL_873391, EPI_ISL_873392, EPI_ISL_873393, EPI_ISL_873394, EPI_ISL_873397, EPI_ISL_873398, EPI_ISL_873400, EPI_ISL_873402, EPI_ISL_873404, EPI_ISL_873405, EPI_ISL_873406, EPI_ISL_873408, EPI_ISL_873410 | | | |
| see above | Lighthouse Lab in Milton Keynes | Wellcome Sanger Institute for the COVID-19 Genomics UK (COG-UK) Consortium | The Lighthouse Lab in Milton Keynes and Alex Alderton, Roberto Amato, Sonia Goncalves, Ewan Harrison, David K. Jackson, Ian Johnston, Dominic Kwiatkowski, Cordelia Langford, John Sillitoe on behalf of the Wellcome Sanger Institute COVID-19 Surveillance Team |
| EPI_ISL_873418 | Lighthouse Lab in Cambridge | Wellcome Sanger Institute for the COVID-19 Genomics UK (COG-UK) Consortium | Rob Howes, The Lighthouse Lab in Cambridge and Alex Alderton, Roberto Amato, Sonia Goncalves, Ewan Harrison, David K. Jackson, Ian Johnston, Dominic Kwiatkowski, Cordelia Langford, John Sillitoe on behalf of the Wellcome Sanger Institute COVID-19 Surveillance Team |
| EPI_ISL_873421, EPI_ISL_873423, EPI_ISL_873424 | Lighthouse Lab in Milton Keynes | Wellcome Sanger Institute for the COVID-19 Genomics UK (COG-UK) Consortium | The Lighthouse Lab in Milton Keynes and Alex Alderton, Roberto Amato, Sonia Goncalves, Ewan Harrison, David K. Jackson, Ian Johnston, Dominic Kwiatkowski, Cordelia Langford, John Sillitoe on behalf of the Wellcome Sanger Institute COVID-19 Surveillance Team |
| EPI_ISL_873432 | Lighthouse Lab in Cambridge | Wellcome Sanger Institute for the COVID-19 Genomics UK (COG-UK) Consortium | Rob Howes, The Lighthouse Lab in Cambridge and Alex Alderton, Roberto Amato, Sonia Goncalves, Ewan Harrison, David K. Jackson, Ian Johnston, Dominic Kwiatkowski, Cordelia Langford, John Sillitoe on behalf of the Wellcome Sanger Institute COVID-19 Surveillance Team |
| EPI_ISL_873434, EPI_ISL_873436, EPI_ISL_873437, EPI_ISL_873440, EPI_ISL_873441, EPI_ISL_873442, EPI_ISL_873446, EPI_ISL_873448, EPI_ISL_873449, EPI_ISL_873453, EPI_ISL_873456, EPI_ISL_873460, EPI_ISL_873461, EPI_ISL_873464, EPI_ISL_873465, EPI_ISL_873467, EPI_ISL_873470, EPI_ISL_873471, EPI_ISL_873472, EPI_ISL_873474, EPI_ISL_873475, EPI_ISL_873477, EPI_ISL_873478, EPI_ISL_873479, EPI_ISL_873480, EPI_ISL_873484 | | | |
| see above | Lighthouse Lab in Milton Keynes | Wellcome Sanger Institute for the COVID-19 Genomics UK (COG-UK) Consortium | The Lighthouse Lab in Milton Keynes and Alex Alderton, Roberto Amato, Sonia Goncalves, Ewan Harrison, David K. Jackson, Ian Johnston, Dominic Kwiatkowski, Cordelia Langford, John Sillitoe on behalf of the Wellcome Sanger Institute COVID-19 Surveillance Team |
| EPI_ISL_873489 | Lighthouse Lab in Cambridge | Wellcome Sanger Institute for the COVID-19 Genomics UK (COG-UK) Consortium | Rob Howes, The Lighthouse Lab in Cambridge and Alex Alderton, Roberto Amato, Sonia Goncalves, Ewan Harrison, David K. Jackson, Ian Johnston, Dominic Kwiatkowski, Cordelia Langford, John Sillitoe on behalf of the Wellcome Sanger Institute COVID-19 Surveillance Team |
| EPI_ISL_873491, EPI_ISL_873498, EPI_ISL_873502, EPI_ISL_873506, EPI_ISL_873507, EPI_ISL_873508, EPI_ISL_873511, EPI_ISL_873512 | Lighthouse Lab in Milton Keynes | Wellcome Sanger Institute for the COVID-19 Genomics UK (COG-UK) Consortium | The Lighthouse Lab in Milton Keynes and Alex Alderton, Roberto Amato, Sonia Goncalves, Ewan Harrison, David K. Jackson, Ian Johnston, Dominic Kwiatkowski, Cordelia Langford, John Sillitoe on behalf of the Wellcome Sanger Institute COVID-19 Surveillance Team |
| EPI_ISL_873513 | Lighthouse Lab in Cambridge | Wellcome Sanger Institute for the COVID-19 Genomics UK (COG-UK) Consortium | Rob Howes, The Lighthouse Lab in Cambridge and Alex Alderton, Roberto Amato, Sonia Goncalves, Ewan Harrison, David K. Jackson, Ian Johnston, Dominic Kwiatkowski, Cordelia Langford, John Sillitoe on behalf of the Wellcome Sanger Institute COVID-19 Surveillance Team |
| EPI_ISL_873514, EPI_ISL_873516, EPI_ISL_873519, EPI_ISL_873522, EPI_ISL_873523, EPI_ISL_873524, EPI_ISL_873525, EPI_ISL_873526, EPI_ISL_873529, EPI_ISL_873530, EPI_ISL_873533, EPI_ISL_873534, EPI_ISL_873539, EPI_ISL_873540, EPI_ISL_873541, EPI_ISL_873544, EPI_ISL_873545, EPI_ISL_873548, EPI_ISL_873550, EPI_ISL_873553, EPI_ISL_873555, EPI_ISL_873556, EPI_ISL_873558, EPI_ISL_873559, EPI_ISL_873560 | | | |
| see above | Lighthouse Lab in Milton Keynes | Wellcome Sanger Institute for the COVID-19 Genomics UK (COG-UK) Consortium | The Lighthouse Lab in Milton Keynes and Alex Alderton, Roberto Amato, Sonia Goncalves, Ewan Harrison, David K. Jackson, Ian Johnston, Dominic Kwiatkowski, Cordelia Langford, John Sillitoe on behalf of the Wellcome Sanger Institute COVID-19 Surveillance Team |
| EPI_ISL_873561 | Lighthouse Lab in Cambridge | Wellcome Sanger Institute for the COVID-19 Genomics UK (COG-UK) Consortium | Rob Howes, The Lighthouse Lab in Cambridge and Alex Alderton, Roberto Amato, Sonia Goncalves, Ewan Harrison, David K. Jackson, Ian Johnston, Dominic Kwiatkowski, Cordelia Langford, John Sillitoe on behalf of the Wellcome Sanger Institute COVID-19 Surveillance Team |
| EPI_ISL_873564, EPI_ISL_873567, EPI_ISL_873568, EPI_ISL_873569, EPI_ISL_873572, EPI_ISL_873578, EPI_ISL_873579, EPI_ISL_873580, EPI_ISL_873581, EPI_ISL_873582, EPI_ISL_873584, EPI_ISL_873586, EPI_ISL_873587, EPI_ISL_873588, EPI_ISL_873592, EPI_ISL_873593, EPI_ISL_873594, EPI_ISL_873597, | | | |

| | | | |
|---|---|---|---|
| EPI_ISL_873598, EPI_ISL_873601, EPI_ISL_873606, EPI_ISL_873608, EPI_ISL_873609, EPI_ISL_873610 | | | |
| see above | Lighthouse Lab in Milton Keynes | Wellcome Sanger Institute for the COVID-19 Genomics UK (COG-UK) Consortium | The Lighthouse Lab in Milton Keynes and Alex Alderton, Roberto Amato, Sonia Goncalves, Ewan Harrison, David K. Jackson, Ian Johnston, Dominic Kwiatkowski, Cordelia Langford, John Sillitoe on behalf of the Wellcome Sanger Institute COVID-19 Surveillance Team |
| EPI_ISL_873611 | Lighthouse Lab in Cambridge | Wellcome Sanger Institute for the COVID-19 Genomics UK (COG-UK) Consortium | Rob Howes, The Lighthouse Lab in Cambridge and Alex Alderton, Roberto Amato, Sonia Goncalves, Ewan Harrison, David K. Jackson, Ian Johnston, Dominic Kwiatkowski, Cordelia Langford, John Sillitoe on behalf of the Wellcome Sanger Institute COVID-19 Surveillance Team |
| EPI_ISL_873612 | Lighthouse Lab in Milton Keynes | Wellcome Sanger Institute for the COVID-19 Genomics UK (COG-UK) Consortium | The Lighthouse Lab in Milton Keynes and Alex Alderton, Roberto Amato, Sonia Goncalves, Ewan Harrison, David K. Jackson, Ian Johnston, Dominic Kwiatkowski, Cordelia Langford, John Sillitoe on behalf of the Wellcome Sanger Institute COVID-19 Surveillance Team |
| EPI_ISL_873613 | Lighthouse Lab in Cambridge | Wellcome Sanger Institute for the COVID-19 Genomics UK (COG-UK) Consortium | Rob Howes, The Lighthouse Lab in Cambridge and Alex Alderton, Roberto Amato, Sonia Goncalves, Ewan Harrison, David K. Jackson, Ian Johnston, Dominic Kwiatkowski, Cordelia Langford, John Sillitoe on behalf of the Wellcome Sanger Institute COVID-19 Surveillance Team |
| EPI_ISL_873615, EPI_ISL_873616, EPI_ISL_873617, EPI_ISL_873619, EPI_ISL_873620, EPI_ISL_873622, EPI_ISL_873626, EPI_ISL_873628, EPI_ISL_873631, EPI_ISL_873637, EPI_ISL_873641, EPI_ISL_873654, EPI_ISL_873657, EPI_ISL_873660, EPI_ISL_873669, EPI_ISL_873704, EPI_ISL_873725, EPI_ISL_873729, EPI_ISL_873734, EPI_ISL_873741, EPI_ISL_873743, EPI_ISL_873744, EPI_ISL_873749 | | | |
| see above | Lighthouse Lab in Milton Keynes | Wellcome Sanger Institute for the COVID-19 Genomics UK (COG-UK) Consortium | The Lighthouse Lab in Milton Keynes and Alex Alderton, Roberto Amato, Sonia Goncalves, Ewan Harrison, David K. Jackson, Ian Johnston, Dominic Kwiatkowski, Cordelia Langford, John Sillitoe on behalf of the Wellcome Sanger Institute COVID-19 Surveillance Team |
| EPI_ISL_873750, EPI_ISL_873752 | Lighthouse Lab in Cambridge | Wellcome Sanger Institute for the COVID-19 Genomics UK (COG-UK) Consortium | Rob Howes, The Lighthouse Lab in Cambridge and Alex Alderton, Roberto Amato, Sonia Goncalves, Ewan Harrison, David K. Jackson, Ian Johnston, Dominic Kwiatkowski, Cordelia Langford, John Sillitoe on behalf of the Wellcome Sanger Institute COVID-19 Surveillance Team |
| EPI_ISL_873760, EPI_ISL_873764, EPI_ISL_873794, EPI_ISL_873806, EPI_ISL_873809, EPI_ISL_873812, EPI_ISL_873817 | Lighthouse Lab in Milton Keynes | Wellcome Sanger Institute for the COVID-19 Genomics UK (COG-UK) Consortium | The Lighthouse Lab in Milton Keynes and Alex Alderton, Roberto Amato, Sonia Goncalves, Ewan Harrison, David K. Jackson, Ian Johnston, Dominic Kwiatkowski, Cordelia Langford, John Sillitoe on behalf of the Wellcome Sanger Institute COVID-19 Surveillance Team |
| EPI_ISL_873825 | Lighthouse Lab in Cambridge | Wellcome Sanger Institute for the COVID-19 Genomics UK (COG-UK) Consortium | Rob Howes, The Lighthouse Lab in Cambridge and Alex Alderton, Roberto Amato, Sonia Goncalves, Ewan Harrison, David K. Jackson, Ian Johnston, Dominic Kwiatkowski, Cordelia Langford, John Sillitoe on behalf of the Wellcome Sanger Institute COVID-19 Surveillance Team |
| EPI_ISL_873833, EPI_ISL_873834, EPI_ISL_873847, EPI_ISL_873857, EPI_ISL_873859, EPI_ISL_873870, EPI_ISL_873873, EPI_ISL_873880, EPI_ISL_873884, EPI_ISL_873887, EPI_ISL_873900, EPI_ISL_873905, EPI_ISL_873910, EPI_ISL_873913, EPI_ISL_873916, EPI_ISL_873923, EPI_ISL_873924, EPI_ISL_873928, EPI_ISL_873934, EPI_ISL_873937, EPI_ISL_873948, EPI_ISL_873960 | | | |
| see above | Lighthouse Lab in Milton Keynes | Wellcome Sanger Institute for the COVID-19 Genomics UK (COG-UK) Consortium | The Lighthouse Lab in Milton Keynes and Alex Alderton, Roberto Amato, Sonia Goncalves, Ewan Harrison, David K. Jackson, Ian Johnston, Dominic Kwiatkowski, Cordelia Langford, John Sillitoe on behalf of the Wellcome Sanger Institute COVID-19 Surveillance Team |
| EPI_ISL_873961, EPI_ISL_873963, EPI_ISL_873966, EPI_ISL_873967, EPI_ISL_873968, EPI_ISL_873969, EPI_ISL_873970, EPI_ISL_873971, EPI_ISL_873972, EPI_ISL_873973, EPI_ISL_873975, EPI_ISL_873977, EPI_ISL_873978, EPI_ISL_873979, EPI_ISL_873980, EPI_ISL_873983, EPI_ISL_873985, EPI_ISL_873988, EPI_ISL_873989, EPI_ISL_873991, EPI_ISL_873992, EPI_ISL_873993, EPI_ISL_873994, EPI_ISL_873995, EPI_ISL_873996, EPI_ISL_873997, EPI_ISL_873998, EPI_ISL_874000, EPI_ISL_874001, EPI_ISL_874002, EPI_ISL_874003, EPI_ISL_874005, EPI_ISL_874006, EPI_ISL_874008, EPI_ISL_874013, EPI_ISL_874015, EPI_ISL_874016, EPI_ISL_874018, EPI_ISL_874019, EPI_ISL_874020, EPI_ISL_874023, EPI_ISL_874025, EPI_ISL_874026, EPI_ISL_874027, EPI_ISL_874028, EPI_ISL_874029, EPI_ISL_874030, EPI_ISL_874031, EPI_ISL_874033, EPI_ISL_874034, EPI_ISL_874035, EPI_ISL_874036, EPI_ISL_874040, EPI_ISL_874041, EPI_ISL_874042, EPI_ISL_874043, EPI_ISL_874044, EPI_ISL_874045, EPI_ISL_874046, EPI_ISL_874048, EPI_ISL_874049, EPI_ISL_874050, EPI_ISL_874051, EPI_ISL_874052, EPI_ISL_874053, EPI_ISL_874054, EPI_ISL_874056, EPI_ISL_874057, EPI_ISL_874058, EPI_ISL_874059, EPI_ISL_874060, EPI_ISL_874063, EPI_ISL_874065, EPI_ISL_874066, EPI_ISL_874067, EPI_ISL_874068, EPI_ISL_874069, EPI_ISL_874072, EPI_ISL_874073, EPI_ISL_874074, EPI_ISL_874075, EPI_ISL_874077, EPI_ISL_874079, EPI_ISL_874083, EPI_ISL_874084, EPI_ISL_874086, EPI_ISL_874087, EPI_ISL_874088, EPI_ISL_874089, EPI_ISL_874091, EPI_ISL_874094, EPI_ISL_874095, EPI_ISL_874099, EPI_ISL_874100, EPI_ISL_874101, EPI_ISL_874102, EPI_ISL_874103, EPI_ISL_874104, EPI_ISL_874105, EPI_ISL_874106, EPI_ISL_874107, EPI_ISL_874108, EPI_ISL_874110, EPI_ISL_874111, EPI_ISL_874112, EPI_ISL_874113, EPI_ISL_874114, EPI_ISL_874116, EPI_ISL_874117, EPI_ISL_874119, EPI_ISL_874120, EPI_ISL_874122, EPI_ISL_874123, EPI_ISL_874124, EPI_ISL_874125, EPI_ISL_874126, EPI_ISL_874127, EPI_ISL_874128, EPI_ISL_874129, EPI_ISL_874131, EPI_ISL_874132, EPI_ISL_874133, EPI_ISL_874134, EPI_ISL_874135, EPI_ISL_874136, EPI_ISL_874137, EPI_ISL_874138, EPI_ISL_874139, EPI_ISL_874140, EPI_ISL_874142, EPI_ISL_874144, EPI_ISL_874146, EPI_ISL_874147, EPI_ISL_874150, EPI_ISL_874151, EPI_ISL_874152, EPI_ISL_874156, EPI_ISL_874158, EPI_ISL_874161, EPI_ISL_874162, EPI_ISL_874166, EPI_ISL_874167, EPI_ISL_874168, EPI_ISL_874171, EPI_ISL_874172, EPI_ISL_874173, EPI_ISL_874174, EPI_ISL_874175, EPI_ISL_874176, EPI_ISL_874177, EPI_ISL_874178, EPI_ISL_874179, EPI_ISL_874182, EPI_ISL_874183, EPI_ISL_874184, EPI_ISL_874186, EPI_ISL_874187, EPI_ISL_874189, EPI_ISL_874190, EPI_ISL_874192, EPI_ISL_874194, EPI_ISL_874200, EPI_ISL_874201, EPI_ISL_874205, EPI_ISL_874206, EPI_ISL_874208, EPI_ISL_874212, EPI_ISL_874213, EPI_ISL_874214, EPI_ISL_874216, EPI_ISL_874217, EPI_ISL_874219, EPI_ISL_874221, EPI_ISL_874222, EPI_ISL_874223, EPI_ISL_874224, EPI_ISL_874226, EPI_ISL_874228, EPI_ISL_874229, EPI_ISL_874230, EPI_ISL_874231, EPI_ISL_874232, EPI_ISL_874233, EPI_ISL_874234, EPI_ISL_874235, EPI_ISL_874236, EPI_ISL_874237, EPI_ISL_874239, EPI_ISL_874241, EPI_ISL_874242, EPI_ISL_874244, EPI_ISL_874247, EPI_ISL_874250, EPI_ISL_874251, EPI_ISL_874253, EPI_ISL_874255, EPI_ISL_874256, EPI_ISL_874260, EPI_ISL_874261, EPI_ISL_874262, EPI_ISL_874264, EPI_ISL_874265, EPI_ISL_874267, EPI_ISL_874269, EPI_ISL_874270, EPI_ISL_874271, EPI_ISL_874272, EPI_ISL_874274, EPI_ISL_874276, EPI_ISL_874277, EPI_ISL_874279, EPI_ISL_874280, EPI_ISL_874282, EPI_ISL_874285, EPI_ISL_874286, EPI_ISL_874288, EPI_ISL_874289, EPI_ISL_874292, EPI_ISL_874294, EPI_ISL_874295, EPI_ISL_874296, EPI_ISL_874298, EPI_ISL_874299, EPI_ISL_874300, EPI_ISL_874302, EPI_ISL_874303, EPI_ISL_874304, EPI_ISL_874305, EPI_ISL_874306, EPI_ISL_874307, EPI_ISL_874308, EPI_ISL_874309, EPI_ISL_874315, EPI_ISL_874379, EPI_ISL_874401, EPI_ISL_874404, EPI_ISL_874443, EPI_ISL_874448, EPI_ISL_874469, EPI_ISL_874482, EPI_ISL_874499, EPI_ISL_874523, EPI_ISL_874527, EPI_ISL_874528, EPI_ISL_874543, EPI_ISL_874558, EPI_ISL_874624, EPI_ISL_874759, EPI_ISL_874816, EPI_ISL_874938, EPI_ISL_874956 | | | |
| see above | Lighthouse Lab in Alderley Park | Wellcome Sanger Institute for the COVID-19 Genomics UK (COG-UK) Consortium | Jacquelyn Wynn, Mairead Hyland, The Lighthouse Lab in Alderley Park and Alex Alderton, Roberto Amato, Sonia Goncalves, Ewan Harrison, David K. Jackson, Ian Johnston, Dominic Kwiatkowski, Cordelia Langford, John Sillitoe on behalf of the Wellcome Sanger Institute COVID-19 Surveillance Team |
| EPI_ISL_875373, EPI_ISL_875374, EPI_ISL_875375, EPI_ISL_875376, EPI_ISL_875377, EPI_ISL_875378, EPI_ISL_875379, EPI_ISL_875380, EPI_ISL_875381, EPI_ISL_875382, EPI_ISL_875383, EPI_ISL_875384, EPI_ISL_875385, EPI_ISL_875386, EPI_ISL_875387, EPI_ISL_875388, EPI_ISL_875389, EPI_ISL_875390, EPI_ISL_875391, EPI_ISL_875392, EPI_ISL_875393, EPI_ISL_875394, EPI_ISL_875395, EPI_ISL_875396, EPI_ISL_875397, EPI_ISL_875398, EPI_ISL_875399, EPI_ISL_875400, EPI_ISL_875401, EPI_ISL_875402, EPI_ISL_875403, EPI_ISL_875404, EPI_ISL_875405, EPI_ISL_875406, EPI_ISL_875407, EPI_ISL_875408, EPI_ISL_875409 | | | |
| see above | National Virus Reference Laboratory | National Virus Reference Laboratory | Michael Carr, Gabriel Gonzalez, Jonathan Dean, Cillian F De Gascun |
| EPI_ISL_875557 | ULSS 2 Treviso | Istituto Zooprofilattico Sperimentale delle Venezie | Adelaide Milani, Alessia Schivo, Annalisa Salviato, Erika Giorgia Quaranta, Ambra Pastori, Bianca Zecchin, Alice Fusaro, Isabella Monne, Calogero Terregino, Antonia Ricci |
| EPI_ISL_875664, EPI_ISL_875665 | University of Michigan Clinical Microbiology Laboratory | Lauring Lab, University of Michigan, Department of Microbiology and Immunology | Valesano |
| EPI_ISL_876050, EPI_ISL_876070, EPI_ISL_876080, EPI_ISL_876096, EPI_ISL_876162, EPI_ISL_876190, EPI_ISL_876239, EPI_ISL_876240, EPI_ISL_876241, EPI_ISL_876242, EPI_ISL_876243 | | | |
| see above | Massachusetts State Public Health Laboratory | Massachusetts State Public Health Laboratory | Andrew Lang, Timelia Fink, Glen Gallagher, Sandra Smole |
| EPI_ISL_876595 | TN Division of Laboratory Services | Pathogen Discovery, Respiratory Viruses Branch, Division of Viral Diseases, Centers for Disease Control and Prevention | Ying Tao, Yan Li, Jing Zhang, Krista Queen, Anna Uehara, Peter Cook, Clinton R. Paden, Haibin Wang, Suxiang Tong |
| EPI_ISL_876741 | Istituto Zooprofilattico Sperimentale della Puglia e della Basilicata | Istituto Zooprofilattico Sperimentale della Puglia e della Basilicata | Parisi A., Bianco A., Capozzi L., Del Sambro L., Manzulli V, Rondinone V., Pace L., Cipolletta D., Galante D. |
| EPI_ISL_877236, EPI_ISL_877329, EPI_ISL_877330 | University College Sedaya International (UCSI University) | Institute for Medical Research, Infectious Disease Research Centre, National Institutes of Health, Ministry of Health Malaysia | Suppiah J, Kamel K, Azizan MA, Sekaran SD, Thayan R |
| EPI_ISL_877913, EPI_ISL_877955 | Lighthouse Lab in Alderley Park | Wellcome Sanger Institute for the COVID-19 Genomics UK (COG-UK) Consortium | Jacquelyn Wynn, Mairead Hyland, The Lighthouse Lab in Alderley Park and Alex Alderton, Roberto Amato, Sonia Goncalves, Ewan Harrison, David K. Jackson, Ian Johnston, Dominic Kwiatkowski, Cordelia Langford, John Sillitoe on behalf of the Wellcome Sanger Institute COVID-19 Surveillance Team |
| EPI_ISL_878769 | Lighthouse Lab in Milton Keynes | Wellcome Sanger Institute for the COVID-19 Genomics UK (COG-UK) Consortium | The Lighthouse Lab in Milton Keynes and Alex Alderton, Roberto Amato, Sonia Goncalves, Ewan Harrison, David K. Jackson, Ian Johnston, Dominic Kwiatkowski, Cordelia Langford, John Sillitoe on behalf of the Wellcome Sanger Institute COVID-19 Surveillance Team |
| EPI_ISL_878844, EPI_ISL_879274 | Lighthouse Lab in Alderley Park | Wellcome Sanger Institute for the COVID-19 Genomics UK (COG-UK) Consortium | Jacquelyn Wynn, Mairead Hyland, The Lighthouse Lab in Alderley Park and Alex Alderton, Roberto Amato, Sonia Goncalves, Ewan Harrison, David K. Jackson, Ian Johnston, Dominic Kwiatkowski, Cordelia Langford, John Sillitoe on behalf of the Wellcome Sanger Institute COVID-19 Surveillance Team |
| EPI_ISL_879820, EPI_ISL_880038 | Lighthouse Lab in Cambridge | Wellcome Sanger Institute for the COVID-19 Genomics UK (COG-UK) Consortium | Rob Howes, The Lighthouse Lab in Cambridge and Alex Alderton, Roberto Amato, Sonia Goncalves, Ewan Harrison, David K. Jackson, Ian Johnston, Dominic Kwiatkowski, Cordelia Langford, John Sillitoe on behalf of the Wellcome Sanger Institute COVID-19 Surveillance Team |
| EPI_ISL_880586, EPI_ISL_880590, EPI_ISL_880591, EPI_ISL_880595, EPI_ISL_880596, EPI_ISL_880599, EPI_ISL_880602, EPI_ISL_880606, EPI_ISL_880608, EPI_ISL_880609, EPI_ISL_880610, EPI_ISL_880611, EPI_ISL_880613, EPI_ISL_880615, EPI_ISL_880616, EPI_ISL_880618, EPI_ISL_880619, EPI_ISL_880623, EPI_ISL_880626, EPI_ISL_880627, EPI_ISL_880639, EPI_ISL_880641, EPI_ISL_880642, EPI_ISL_880647, EPI_ISL_880651, EPI_ISL_880653, EPI_ISL_880654, EPI_ISL_880655, EPI_ISL_880657, EPI_ISL_880660, EPI_ISL_880662, EPI_ISL_880665, EPI_ISL_880667, EPI_ISL_880668, EPI_ISL_880672, EPI_ISL_880673, EPI_ISL_880675, EPI_ISL_880677, EPI_ISL_880679, EPI_ISL_880695, EPI_ISL_880701, EPI_ISL_880705, EPI_ISL_880706, EPI_ISL_880708, EPI_ISL_880713, EPI_ISL_880722, EPI_ISL_880724, EPI_ISL_880725, EPI_ISL_880726, EPI_ISL_880727, EPI_ISL_880729, EPI_ISL_880736, EPI_ISL_880737, EPI_ISL_880738, EPI_ISL_880739, EPI_ISL_880740, EPI_ISL_880745, EPI_ISL_880751, EPI_ISL_880752, EPI_ISL_880755, EPI_ISL_880759, EPI_ISL_880762, EPI_ISL_880766, EPI_ISL_880769, EPI_ISL_880781, EPI_ISL_880785, EPI_ISL_880786, EPI_ISL_880791, EPI_ISL_880792, EPI_ISL_880797, EPI_ISL_880798, EPI_ISL_880800, EPI_ISL_880803, EPI_ISL_880809, EPI_ISL_880813, EPI_ISL_880817, EPI_ISL_880818, EPI_ISL_880819, EPI_ISL_880822, EPI_ISL_880824, EPI_ISL_880826, EPI_ISL_880829, EPI_ISL_880830, EPI_ISL_880833, EPI_ISL_880836, EPI_ISL_880838, EPI_ISL_880839, EPI_ISL_880840, EPI_ISL_880841, EPI_ISL_880843, EPI_ISL_880847, EPI_ISL_880854, EPI_ISL_880856, EPI_ISL_880857, EPI_ISL_880858, EPI_ISL_880862, EPI_ISL_880866, EPI_ISL_880870, EPI_ISL_880878, EPI_ISL_880883, EPI_ISL_880884, EPI_ISL_880886, EPI_ISL_880889, EPI_ISL_880894, EPI_ISL_880895, EPI_ISL_880898, EPI_ISL_880900, EPI_ISL_880901, EPI_ISL_880903, EPI_ISL_880908, EPI_ISL_880909, EPI_ISL_880910, EPI_ISL_880911, EPI_ISL_880914, EPI_ISL_880916, EPI_ISL_880921, EPI_ISL_880925, EPI_ISL_880933, EPI_ISL_880934, EPI_ISL_880935 | | | |

| | | | |
|---|---|---|---|
| see above | Lighthouse Lab in Milton Keynes | Wellcome Sanger Institute for the COVID-19 Genomics UK (COG-UK) Consortium | The Lighthouse Lab in Milton Keynes and Alex Alderton, Roberto Amato, Sonia Goncalves, Ewan Harrison, David K. Jackson, Ian Johnston, Dominic Kwiatkowski, Cordelia Langford, John Sillitoe on behalf of the Wellcome Sanger Institute COVID-19 Surveillance Team |
| EPI_ISL_881277, EPI_ISL_881281, EPI_ISL_881283, EPI_ISL_881289, EPI_ISL_881290, EPI_ISL_881291, EPI_ISL_881293, EPI_ISL_881298, EPI_ISL_881306, EPI_ISL_881307, EPI_ISL_881309, EPI_ISL_881310, EPI_ISL_881315, EPI_ISL_881341, EPI_ISL_881359, EPI_ISL_881386, EPI_ISL_881388, EPI_ISL_881414, EPI_ISL_881429, EPI_ISL_881455, EPI_ISL_881461, EPI_ISL_881487, EPI_ISL_881494, EPI_ISL_881495, EPI_ISL_881509, EPI_ISL_881518, EPI_ISL_881521, EPI_ISL_881522, EPI_ISL_881531, EPI_ISL_881540, EPI_ISL_881543, EPI_ISL_881546, EPI_ISL_881550, EPI_ISL_881567, EPI_ISL_881570, EPI_ISL_881571, EPI_ISL_881576, EPI_ISL_881580, EPI_ISL_881582, EPI_ISL_881586, EPI_ISL_881588, EPI_ISL_881598, EPI_ISL_881600, EPI_ISL_881607, EPI_ISL_881612 | | | |
| see above | Lighthouse Lab in Alderley Park | Wellcome Sanger Institute for the COVID-19 Genomics UK (COG-UK) Consortium | Jacquelyn Wynn, Mairead Hyland, The Lighthouse Lab in Alderley Park and Alex Alderton, Roberto Amato, Sonia Goncalves, Ewan Harrison, David K. Jackson, Ian Johnston, Dominic Kwiatkowski, Cordelia Langford, John Sillitoe on behalf of the Wellcome Sanger Institute COVID-19 Surveillance Team |
| EPI_ISL_881619 | Lighthouse Lab in Cambridge | Wellcome Sanger Institute for the COVID-19 Genomics UK (COG-UK) Consortium | Rob Howes, The Lighthouse Lab in Cambridge and Alex Alderton, Roberto Amato, Sonia Goncalves, Ewan Harrison, David K. Jackson, Ian Johnston, Dominic Kwiatkowski, Cordelia Langford, John Sillitoe on behalf of the Wellcome Sanger Institute COVID-19 Surveillance Team |
| EPI_ISL_882299 | Lighthouse Lab in Alderley Park | Wellcome Sanger Institute for the COVID-19 Genomics UK (COG-UK) Consortium | Jacquelyn Wynn, Mairead Hyland, The Lighthouse Lab in Alderley Park and Alex Alderton, Roberto Amato, Sonia Goncalves, Ewan Harrison, David K. Jackson, Ian Johnston, Dominic Kwiatkowski, Cordelia Langford, John Sillitoe on behalf of the Wellcome Sanger Institute COVID-19 Surveillance Team |
| EPI_ISL_882669 | UPA Vila Santa Catarina | Instituto Adolfo Lutz, Interdiciplinary Procedures Center, Strategic Laboratory | Claudio Tavares Sacchi, Claudia Regina Gonçalves, Erica Valessa Ramos Gomes, Karoline Rodrigues Campos |
| EPI_ISL_882782 | Institute for Urban Disease Control and Prevention | COVID-19 Network Investigations (CONI) Alliance | Kamolthip Atsawawaranunt, Elizabeth Batty, Wasun Chantratita, Thanat Chookajorn, Stefan Fernandez, Angkana Huang, Anthony R. Jones, Khajohn Joonsalak, Chonticha Klungtong, Theerarat Kochakarn, Prayuth Kaewmalang, Amornmas Kongklieng, Namfon Kotanan, Krittikorn Kumpornsin, Duangkamon Loesbanluechai, Wudtichai Manasatienkij, Anek Mungaomklang, Bhakbhoom Panthan, Pukkaporn Parnwijitkul, Ekawat Pasomsub, Vichan Pawun, Kingkan Rakmanee, Insee Sensorn, Janjira Thaipadungpanit, Arporn Wangwiwatsin, Treewat Watthanachockchai |
| EPI_ISL_882979, EPI_ISL_882980, EPI_ISL_882981, EPI_ISL_882982, EPI_ISL_882983, EPI_ISL_882984 | UCLA Clinical Micro Lab | Los Angeles County PHL | P. Hemarajata et al. |
| EPI_ISL_882985 | MD Laboratories | Los Angeles County PHL | P. Hemarajata et al. |
| EPI_ISL_883004, EPI_ISL_883005, EPI_ISL_883011, EPI_ISL_883015, EPI_ISL_883025, EPI_ISL_883026 | Maryland Public Health Laboratory | Maryland Public Health Laboratory | Maryland Department of Health Laboratories Administration |
| EPI_ISL_883426 | NORTHWELL HEALTH LABORATORIES | Wadsworth Center, New York State Department of Health | Kirsten St. George, Daryl M. Lamson, Alexis Russel, Matthew Shudt, Melissa A Leisner, Jonathan Plitnick, Navjot Singh, John Kelly, Erasmus Schneider, Erica Lasek-Nesselquist |
| EPI_ISL_883429 | BOSTON HEART DIAGNOSTICS CORP | Wadsworth Center, New York State Department of Health | Kirsten St. George, Daryl M. Lamson, Alexis Russel, Matthew Shudt, Melissa A Leisner, Jonathan Plitnick, Navjot Singh, John Kelly, Erasmus Schneider, Erica Lasek-Nesselquist |
| EPI_ISL_883435 | NORTHWELL HEALTH LABORATORIES | Wadsworth Center, New York State Department of Health | Kirsten St. George, Daryl M. Lamson, Alexis Russel, Matthew Shudt, Melissa A Leisner, Jonathan Plitnick, Navjot Singh, John Kelly, Erasmus Schneider, Erica Lasek-Nesselquist |
| EPI_ISL_883439 | ADIRONDACK MEDICAL CENTER | Wadsworth Center, New York State Department of Health | Kirsten St. George, Daryl M. Lamson, Alexis Russel, Matthew Shudt, Melissa A Leisner, Jonathan Plitnick, Navjot Singh, John Kelly, Erasmus Schneider, Erica Lasek-Nesselquist |
| EPI_ISL_883480, EPI_ISL_883482, EPI_ISL_883487, EPI_ISL_883488, EPI_ISL_883489, EPI_ISL_883490, EPI_ISL_883492, EPI_ISL_883493, EPI_ISL_883495, EPI_ISL_883496, EPI_ISL_883497 | | | |
| see above | NORTHWELL HEALTH LABORATORIES | Wadsworth Center, New York State Department of Health | Kirsten St. George, Daryl M. Lamson, Alexis Russel, Matthew Shudt, Melissa A Leisner, Jonathan Plitnick, Navjot Singh, John Kelly, Erasmus Schneider, Erica Lasek-Nesselquist |
| EPI_ISL_883960 | Labo Analyses Med | National Reference Center for Viruses of Respiratory Infections, Institut Pasteur, Paris | Marion Barbet, Sylvie Behillil, Méline Bizard, Angela Brisebarre, Camille Capel, Etienne Simon-Lorière, Vincent Enouf, Maud Vanpeene, Sylvie van der Werf,Jacques Alexandra |
| EPI_ISL_884055, EPI_ISL_884056, EPI_ISL_884057, EPI_ISL_884058 | Wadsworth Center, New York State Department of Health | Wadsworth Center, New York State Department of Health | Kirsten St. George, Daryl M. Lamson, Alexis Russel, Matthew Shudt, Melissa A Leisner, Jonathan Plitnick, Navjot Singh, John Kelly, Erasmus Schneider, Erica Lasek-Nesselquist |
| EPI_ISL_884059, EPI_ISL_884060, EPI_ISL_884061, EPI_ISL_884062, EPI_ISL_884063, EPI_ISL_884064 | ALBANY MEDICAL CENTER HOSPITAL CLINICAL LABORATORIES | Wadsworth Center, New York State Department of Health | Kirsten St. George, Daryl M. Lamson, Alexis Russel, Matthew Shudt, Melissa A Leisner, Jonathan Plitnick, Navjot Singh, John Kelly, Erasmus Schneider, Erica Lasek-Nesselquist |
| EPI_ISL_884204, EPI_ISL_884205, EPI_ISL_884207, EPI_ISL_884210 | Wyoming Public Health Laboratory | Wyoming Public Health Laboratory | Noah Hull, Taylor Fearing, Lynette Gumbleton, Channing Weber, Ashley Norberg, Bailey Bowcutt, and Wanda Manley |
| EPI_ISL_884231, EPI_ISL_884232, EPI_ISL_884233, EPI_ISL_884234, EPI_ISL_884235, EPI_ISL_884236, EPI_ISL_884237, EPI_ISL_884238, EPI_ISL_884239, EPI_ISL_884240, EPI_ISL_884241 | | | |
| see above | Kansas Health and Environmental Lab | Kansas Health and Environmental Lab | Mike Grose, Paige Drury, Carissa Robertson, Ben Olsen, and Phil Adam |
| EPI_ISL_884255 | Maryland Public Health Laboratory | Maryland Public Health Laboratory | Maryland Department of Health Laboratories Administration |
| EPI_ISL_884729, EPI_ISL_884730, EPI_ISL_884731, EPI_ISL_884732, EPI_ISL_884733, EPI_ISL_884734, EPI_ISL_884735, EPI_ISL_884736, EPI_ISL_884737, EPI_ISL_884738, EPI_ISL_884739, EPI_ISL_884740, EPI_ISL_884741, EPI_ISL_884742, EPI_ISL_884743, EPI_ISL_884744, EPI_ISL_884745, EPI_ISL_884746, EPI_ISL_884747, EPI_ISL_884748, EPI_ISL_884749, EPI_ISL_884750, EPI_ISL_884751, EPI_ISL_884752, EPI_ISL_884753, EPI_ISL_884754, EPI_ISL_884755, EPI_ISL_884756, EPI_ISL_884757, EPI_ISL_884758, EPI_ISL_884759, EPI_ISL_884760, EPI_ISL_884761, EPI_ISL_884762, EPI_ISL_884763, EPI_ISL_884764, EPI_ISL_884765, EPI_ISL_884766, EPI_ISL_884767, EPI_ISL_884768, EPI_ISL_884769, EPI_ISL_884770, EPI_ISL_884771, EPI_ISL_884772, EPI_ISL_884773, EPI_ISL_884774, EPI_ISL_884775, EPI_ISL_884776, EPI_ISL_884777, EPI_ISL_884778, EPI_ISL_884779, EPI_ISL_884780, EPI_ISL_884781, EPI_ISL_884782, EPI_ISL_884783, EPI_ISL_884784, EPI_ISL_884785, EPI_ISL_884786, EPI_ISL_884787, EPI_ISL_884788 | | | |
| see above | Respiratory Viruses Branch, Centers for Disease Control and Prevention | Respiratory Viruses Branch, Centers for Disease Control and Prevention | Cook,P.W., Batra,D., Rambo-Martin,B.L., de Feo,E., Antico,J., Tran,C., Tolentino,M., Wickline,S., Gietzen,K., Sickler,B., Liu,J., Allen,E., Febbo,P., Galloway,S., Washington,N.L., White,S., Levan,G., Barret,K.S., Cirulli,E., Bolze,A., Ascencio,A., Rivera-Garcia,C., Cho,R., Nguyen,J., Wang,S., Ramirez,J., Cassens,T., Sandoval,E., Isaksson,M., Lee,W., Becker,D., Laurent,M., Lu,J., Paden,C.R., Tong,S., MacCannell,D. |
| EPI_ISL_885240, EPI_ISL_885327, EPI_ISL_885409, EPI_ISL_885514 | Lighthouse Lab in Alderley Park | Wellcome Sanger Institute for the COVID-19 Genomics UK (COG-UK) Consortium | Jacquelyn Wynn, Mairead Hyland, The Lighthouse Lab in Alderley Park and Alex Alderton, Roberto Amato, Sonia Goncalves, Ewan Harrison, David K. Jackson, Ian Johnston, Dominic Kwiatkowski, Cordelia Langford, John Sillitoe on behalf of the Wellcome Sanger Institute COVID-19 Surveillance Team |
| EPI_ISL_886185, EPI_ISL_886186, EPI_ISL_886194, EPI_ISL_886195, EPI_ISL_886213, EPI_ISL_886215, EPI_ISL_886219, EPI_ISL_886221, EPI_ISL_886227, EPI_ISL_886228, EPI_ISL_886235, EPI_ISL_886238, EPI_ISL_886240, EPI_ISL_886249, EPI_ISL_886252, EPI_ISL_886253, EPI_ISL_886255, EPI_ISL_886261, EPI_ISL_886266, EPI_ISL_886274, EPI_ISL_886283, EPI_ISL_886285, EPI_ISL_886286, EPI_ISL_886288, EPI_ISL_886292, EPI_ISL_886294, EPI_ISL_886295, EPI_ISL_886305, EPI_ISL_886306, EPI_ISL_886308, EPI_ISL_886310, EPI_ISL_886312, EPI_ISL_886324, EPI_ISL_886344, EPI_ISL_886359, EPI_ISL_886360, EPI_ISL_886366, EPI_ISL_886380, EPI_ISL_886384, EPI_ISL_886392, EPI_ISL_886393, EPI_ISL_886401, EPI_ISL_886404, EPI_ISL_886405, EPI_ISL_886407, EPI_ISL_886412, EPI_ISL_886414, EPI_ISL_886420, EPI_ISL_886437, EPI_ISL_886460, EPI_ISL_886470, EPI_ISL_886474, EPI_ISL_886476, EPI_ISL_886505, EPI_ISL_886506, EPI_ISL_886523, EPI_ISL_886529, EPI_ISL_886531, EPI_ISL_886562, EPI_ISL_886565, EPI_ISL_886566, EPI_ISL_886567, EPI_ISL_886570, EPI_ISL_886595, EPI_ISL_886617, EPI_ISL_886626, EPI_ISL_886634, EPI_ISL_886635, EPI_ISL_886647, EPI_ISL_886661, EPI_ISL_886671, EPI_ISL_886677, EPI_ISL_886689, EPI_ISL_886690, EPI_ISL_886697, EPI_ISL_886700, EPI_ISL_886703, EPI_ISL_886705, EPI_ISL_886723, EPI_ISL_886734, EPI_ISL_886736, EPI_ISL_886748, EPI_ISL_886750, EPI_ISL_886753, EPI_ISL_886758, EPI_ISL_886759, EPI_ISL_886765, EPI_ISL_886775, EPI_ISL_886776, EPI_ISL_886784, EPI_ISL_886789, EPI_ISL_886799, EPI_ISL_886814, EPI_ISL_886816, EPI_ISL_886817, EPI_ISL_886819, EPI_ISL_886821, EPI_ISL_886822, EPI_ISL_886827, EPI_ISL_886830, EPI_ISL_886833, EPI_ISL_886834, EPI_ISL_886838, EPI_ISL_886839, EPI_ISL_886841, EPI_ISL_886854, EPI_ISL_886855, EPI_ISL_886857, EPI_ISL_886858, EPI_ISL_886864, EPI_ISL_886871, EPI_ISL_886880, EPI_ISL_886886, EPI_ISL_886890, EPI_ISL_886897, EPI_ISL_886900, EPI_ISL_886918, EPI_ISL_886927, EPI_ISL_886933, EPI_ISL_886944, EPI_ISL_886946, EPI_ISL_886967, EPI_ISL_886968, EPI_ISL_886973, EPI_ISL_886980, EPI_ISL_886991, EPI_ISL_886992, EPI_ISL_886993, EPI_ISL_886996, EPI_ISL_887007, EPI_ISL_887011, EPI_ISL_887013, EPI_ISL_887016, EPI_ISL_887022, EPI_ISL_887023, EPI_ISL_887026, EPI_ISL_887046, EPI_ISL_887053, EPI_ISL_887085, EPI_ISL_887086, EPI_ISL_887088, EPI_ISL_887093 | | | |
| see above | Labcorp | Genomics and Discovery, Respiratory Viruses Branch, Division of Viral Diseases, Centers for Disease Control and Prevention | Peter W. Cook,Dhwani Batra,Ben L. Rambo-Martin,Summer Galloway,Brian Krueger,Minoo Agarwal,Eyad Almasri,Debbie Boles,Ayla Burns,Nuthawin Charoensri,Oren Cohen,Susan Countryman,Mary Ann Cristobal,Bobbi Croy,Suzanne Dale,Hrushikesh Deshmukh,Amanda Douglas,Vincent Drouillon,Marcia Eisenberg,Howard Engler,Rama Ghatti,Prashant Gupta,Susan Hicks,Jake Humphrey,Lax Iyer,Manoj Jain,Mohan Kolli,Tim Kuphal,Stanley Letovsky,Michael Levandoski,Craig Lukasik,Jonathan Meltzer,Brian Norvell,Mindy Nye,Scott Parker,Christos Petropoulos,John Pruitt,Steven Ragan,Scott Ryan,Mike Sapeta,Jana Schroth,Suresh Babu Selvaraju,Goran Stevovic,Amanda Suchanek,Andrea Throop,Lyndon Tilson,Thomas Urban,Joe Voshell,Kimberly Wagner,Jonathan Williams,Mary Williamson,Qian Zeng,Tricia Zwiefelhofer,Clinton R. Paden,Suxiang Tong,Duncan MacCannell, |
| EPI_ISL_887121, EPI_ISL_887130, EPI_ISL_887131, EPI_ISL_887141 | Institute of Medical Microbiology and Hospital Hygiene | Institute of Medical Microbiology and Hospital Hygiene | Prof. Dr. Achim Kaasch, Aljoscha Tersteegen |
| EPI_ISL_888597, EPI_ISL_888598, | Wyoming Public Health Laboratory | Wyoming Public Health Laboratory | Noah Hull, Taylor Fearing, Lynette Gumbleton, Channing Weber, Ashley Norberg, Bailey Bowcutt, and Wanda Manley |

| | | | |
|---|---|---|---|
| EPI_ISL_888601 | | | |
| EPI_ISL_888803, EPI_ISL_888804, EPI_ISL_888805, EPI_ISL_888806, EPI_ISL_888807, EPI_ISL_888808, EPI_ISL_888809 | Univeristy of New Mexico Hospital | Center for Global Health, University of New Mexico Health Sciences Center | Daryl Domman, Kurt Schwalm, Justin Bacca, Jon Femling, Darrell Dinwiddie |
| EPI_ISL_888855, EPI_ISL_888897, EPI_ISL_888898, EPI_ISL_888899, EPI_ISL_888900, EPI_ISL_888902, EPI_ISL_888904, EPI_ISL_888905 | Michigan Department of Health and Human Services, Bureau of Laboratories | Michigan Department of Health and Human Services, Bureau of Laboratories | Blankenship HM, Riner D, Soehnlen MK |
| EPI_ISL_889005 | Eijkman Institute for Molecular Biology, Ministry of Research and Technology/National Agency for Research and Innovation | Eijkman Institute for Molecular Biology, Ministry of Research and Technology/National Agency for Research and Innovation | Willy Agustine, Edison Johar, Hidayat Trimarsanto, Iskandar Adnan, Lydia V. Panggalo, Sukma Oktavianthi, Frilasita A Yudhaputri, Safarina G Malik, Khin Saw Myint, Amin Soebandrio |
| EPI_ISL_889006 | RSU Harapan Bunda | Eijkman Institute for Molecular Biology, Ministry of Research and Technology/National Agency for Research and Innovation | Willy Agustine, Edison Johar, Hidayat Trimarsanto, Iskandar Adnan, Lydia V. Panggalo, Sukma Oktavianthi, Frilasita A Yudhaputri, Safarina G Malik, Khin Saw Myint, Amin Soebandrio |
| EPI_ISL_889007 | RSUD Cileungsi | Eijkman Institute for Molecular Biology, Ministry of Research and Technology/National Agency for Research and Innovation | Willy Agustine, Edison Johar, Hidayat Trimarsanto, Iskandar Adnan, Lydia V. Panggalo, Sukma Oktavianthi, Frilasita A Yudhaputri, Safarina G Malik, Khin Saw Myint, Amin Soebandrio |
| EPI_ISL_889686, EPI_ISL_889687, EPI_ISL_889688, EPI_ISL_889689, EPI_ISL_889690, EPI_ISL_889691, EPI_ISL_889692, EPI_ISL_889693, EPI_ISL_889694, EPI_ISL_889695, EPI_ISL_889696, EPI_ISL_889697, EPI_ISL_889698, EPI_ISL_889699, EPI_ISL_889700, EPI_ISL_889701, EPI_ISL_889702, EPI_ISL_889713 | | | |
| see above | LSUHS Emerging Viral Threat Laboratory | Microbial Genome Sequencing Center | Jeremy P. Kamil, Jennifer L. Carroll, Camille F. Abshire, Maarten Van Diest, Mohammed N.A. Siddiquey, Andrew D. Yurochko, Martin J. Sapp, Rona S. Scott, Christopher G. Kevil, Daniel J. Snyder, Vaughn S. Cooper, John A. Vanchiere |
| EPI_ISL_890358 | Hospital | National Reference Center for Viruses of Respiratory Infections, Institut Pasteur, Paris | Marion Barbet, Sylvie Behillil, Méline Bizard, Angela Brisebarre, Camille Capel, Etienne Simon-Lorière, Vincent Enouf, Maud Vanpeene, Sylvie van der Werf,Combe Patrice |
| EPI_ISL_890967, EPI_ISL_890969 | Seattle Flu Study | Seattle Flu Study | Deborah A. Nickerson, Chris D. Frazar, Jover Lee, Benjamin Pelle, Erica Ryke, Matthew Richardson, Amanda Adler, Elisabeth Brandstetter, Peter D. Han, Kairsten Fay, Misja Ilcisin, Kirsten Lacombe, Thomas R. Sibley, Melissa Truong, Caitlin R. Wolf, Michael Boeckh, Janet A. Englund, Michael Famulare, Barry R. Lutz, Mark J. Rieder, Lea M. Starita, Matthew Thompson, Jay Shendure, Trevor Bedford, Helen Y. Chu |
| EPI_ISL_890970, EPI_ISL_890971, EPI_ISL_890972, EPI_ISL_890973, EPI_ISL_890974, EPI_ISL_890975, EPI_ISL_890976, EPI_ISL_890977, EPI_ISL_890978, EPI_ISL_890979, EPI_ISL_890980, EPI_ISL_890981, EPI_ISL_890982, EPI_ISL_890983, EPI_ISL_890984, EPI_ISL_890990, EPI_ISL_890993 | | | |
| see above | Seattle Flu Study | Seattle Flu Study | Deborah A. Nickerson, Chris D. Frazar, Jover Lee, Benjamin Pelle, Erica Ryke, Matthew Richardson, Amanda Adler, Elisabeth Brandstetter, Peter D. Han, Kairsten Fay, Misja Ilcisin, Kirsten Lacombe, Thomas R. Sibley, Melissa Truong, Caitlin R. Wolf, Karen Cowgill, Stephanie Schrag, Jeff Duchin, Michael Boeckh, Janet A. Englund, Michael Famulare, Barry R. Lutz, Mark J. Rieder, Lea M. Starita, Matthew Thompson, Helen Y. Chu, Trevor Bedford, Jay Shendure |
| EPI_ISL_891145 | University of Wisconsin-Madison AIDS Vaccine Research Laboratories | University of Wisconsin-Madison AIDS Vaccine Research Laboratories | Gage Moreno, Katarina Braun, et al. AIDS Vaccine Research Laboratories |
| EPI_ISL_891204 | DPH, Massachusetts State Public Health Lab | DPH, Massachusetts State Public Health Lab | Lang,A.S., Fink,T., Gallagher,G.R., Smole,S.C. |
| EPI_ISL_892050, EPI_ISL_892051, EPI_ISL_892054, EPI_ISL_892057, EPI_ISL_892058, EPI_ISL_892126, EPI_ISL_892179, EPI_ISL_892189, EPI_ISL_892190, EPI_ISL_892191, EPI_ISL_892192, EPI_ISL_892193, EPI_ISL_892194, EPI_ISL_892195, EPI_ISL_892196, EPI_ISL_892197 | | | |
| see above | Lighthouse Lab in Alderley Park | Wellcome Sanger Institute for the COVID-19 Genomics UK (COG-UK) Consortium | Jacquelyn Wynn, Mairead Hyland, The Lighthouse Lab in Alderley Park and Alex Alderton, Roberto Amato, Sonia Goncalves, Ewan Harrison, David K. Jackson, Ian Johnston, Dominic Kwiatkowski, Cordelia Langford, John Sillitoe on behalf of the Wellcome Sanger Institute COVID-19 Surveillance Team |
| EPI_ISL_892250, EPI_ISL_892252, EPI_ISL_892253, EPI_ISL_892254, EPI_ISL_892255, EPI_ISL_892256, EPI_ISL_892257, EPI_ISL_892258, EPI_ISL_892259, EPI_ISL_892260, EPI_ISL_892261, EPI_ISL_892262 | | | |
| see above | MD Laboratories | Los Angeles County PHL | P. Hemarajata et al. |
| EPI_ISL_894189, EPI_ISL_894191, EPI_ISL_894198, EPI_ISL_894210 | KU Leuven, Rega Institute, Clinical and Epidemiological Virology | KU Leuven, Rega Institute, Clinical and Epidemiological Virology | Tony Wawina-Bokalanga, Bert Vanmechelen, Joan Marti-Carerras, Piet Maes |
| EPI_ISL_894224, EPI_ISL_894225, EPI_ISL_894226, EPI_ISL_894236 | CH de Mayotte - Laboratoire de Biologie | National Reference Center for Viruses of Respiratory Infections, Institut Pasteur, Paris | Marion Barbet, Sylvie Behillil, Méline Bizard, Angela Brisebarre, Camille Capel, Etienne Simon-Lorière, Vincent Enouf, Maud Vanpeene, Sylvie van der Werf,Combe Patrice |
| EPI_ISL_894248, EPI_ISL_894249, EPI_ISL_894250, EPI_ISL_894251, EPI_ISL_894252, EPI_ISL_894253, EPI_ISL_894254 | Ministry of Health Turkey | Ministry of Health Turkey | Fatma Bayrakdar, Yasemin Cogun, Süleyman Yalcin, Aye Baak Alta, Gülay Korukluolu |
| EPI_ISL_896086, EPI_ISL_896100, EPI_ISL_896111 | Viollier AG | University Hospital Basel, Clinical Bacteriology | Tim Roloff, Madlen Stange, Helena MB Seth-Smith, Alfredo Mari, Karoline Leuzinger, Julia Bielicki, Christiane Beckmann, Manuel Battegay, Hans Hirsch, Adrian Egli |
| EPI_ISL_896169, EPI_ISL_896212, EPI_ISL_896214, EPI_ISL_896215 | MEPHI, Aix Marseille University | MEPHI, Aix Marseille University | Anthony LEVASSEUR |
| EPI_ISL_896220, EPI_ISL_896228, EPI_ISL_896248, EPI_ISL_896249, EPI_ISL_896250, EPI_ISL_896251, EPI_ISL_896254, EPI_ISL_896255, EPI_ISL_896290 | SUNY UPSTATE MEDICAL UNIVERSITY | Wadsworth Center, New York State Department of Health | Kirsten St. George, Daryl M. Lamson, Alexis Russel, Matthew Shudt, Melissa A Leisner, Jonathan Plitnick, Navjot Singh, John Kelly, Erasmus Schneider, Erica Lasek-Nesselquist |
| EPI_ISL_896385, EPI_ISL_896387, EPI_ISL_896388, EPI_ISL_896396, EPI_ISL_896397, EPI_ISL_896398, EPI_ISL_896399, EPI_ISL_896400, EPI_ISL_896401, EPI_ISL_896402, EPI_ISL_896403, EPI_ISL_896404, EPI_ISL_896405, EPI_ISL_896406, EPI_ISL_896407, EPI_ISL_896408, EPI_ISL_896409, EPI_ISL_896410, EPI_ISL_896411, EPI_ISL_896412, EPI_ISL_896414, EPI_ISL_896415, EPI_ISL_896416, EPI_ISL_896419, EPI_ISL_896420 | | | |
| see above | URMC LABS | Wadsworth Center, New York State Department of Health | Kirsten St. George, Daryl M. Lamson, Alexis Russel, Matthew Shudt, Melissa A Leisner, Jonathan Plitnick, Navjot Singh, John Kelly, Erasmus Schneider, Erica Lasek-Nesselquist |
| EPI_ISL_896456, EPI_ISL_896474, EPI_ISL_896493, EPI_ISL_896494 | KU Leuven, Rega Institute, Clinical and Epidemiological Virology | KU Leuven, Rega Institute, Clinical and Epidemiological Virology | Tony Wawina-Bokalanga, Bert Vanmechelen, Joan Marti-Carerras, Piet Maes |
| EPI_ISL_897605, EPI_ISL_897673, EPI_ISL_897676, EPI_ISL_897734, EPI_ISL_897934, EPI_ISL_897935, EPI_ISL_897936, EPI_ISL_897938, EPI_ISL_897939, EPI_ISL_897940, EPI_ISL_897941 | | | |
| see above | University Hospitals of Geneva, Laboratory of Virology | HUG, Laboratory of Virology and the Health2030 Genome Center | Samuel Cordey, Ana Rita Goncalves, Laurent Kaiser, Lorenzo Cerutti, Henri Pegeot, Melyssa Elies, Deborah Penet, Keith Harshman, Ioannis Xenarios, Emmanouil Dermitzakis |
| EPI_ISL_897988, EPI_ISL_897991, EPI_ISL_897992, EPI_ISL_897994, EPI_ISL_897995, EPI_ISL_897997, EPI_ISL_897999, EPI_ISL_898003, EPI_ISL_898006, EPI_ISL_898017, EPI_ISL_898018, EPI_ISL_898019, EPI_ISL_898020, EPI_ISL_898021, EPI_ISL_898022, EPI_ISL_898032 | | | |
| see above | KU Leuven, Rega Institute, Clinical and Epidemiological Virology | KU Leuven, Rega Institute, Clinical and Epidemiological Virology | Tony Wawina-Bokalanga, Bert Vanmechelen, Joan Marti-Carerras, Piet Maes |
| EPI_ISL_898997, EPI_ISL_898999, EPI_ISL_899002, EPI_ISL_899004 | Viollier AG | Department of Biosystems Science and Engineering, ETH Zürich | Chaoran Chen, Sarah Nadeau, Ivan Topolsky, Emmanouil Dermitzakis, Keith Harshman, Ioannis Xenarios, Henri Pegeot, Lorenzo Cerutti, Deborah Penet, Philipp Jablonski, Lara Fuhrmann, David Dreifuss, Katharina Jahn, Christiane Beckmann, Maurice Redondo, Olivier Kobel, Christoph Noppen, Sophie Seidel, Noemie Santamaria de Souza, Niko Beerenwinkel, Tanja Stadler |
| EPI_ISL_899009 | Viollier AG | Department of Biosystems Science and Engineering, ETH Zürich | Christian Beisel, Sarah Nadeau, Chaoran Chen, Ivan Topolsky, Philipp Jablonski, Lara Fuhrmann, David Dreifuss, Katharina Jahn, Tobias Schär, Ina Nissen, Natascha Santacroce, Elodie Burcklen, Christiane Beckmann, Maurice Redondo, Olivier Kobel, Christoph Noppen, Sophie Seidel, Noemie Santamaria de Souza, Niko Beerenwinkel, Tanja Stadler |

| | | | |
|---|---|---|---|
| EPI_ISL_899010, EPI_ISL_899016, EPI_ISL_899022, EPI_ISL_899024, EPI_ISL_899025, EPI_ISL_899035, EPI_ISL_899039, EPI_ISL_899043, EPI_ISL_899053, EPI_ISL_899056, EPI_ISL_899060, EPI_ISL_899064 | | | |
| see above | Viollier AG | Department of Biosystems Science and Engineering, ETH Zürich | Chaoran Chen, Sarah Nadeau, Ivan Topolsky, Emmanouil Dermitzakis, Keith Harshman, Ioannis Xenarios, Henri Pegeot, Lorenzo Cerutti, Deborah Penet, Philipp Jablonski, Lara Fuhrmann, David Dreifuss, Katharina Jahn, Christiane Beckmann, Maurice Redondo, Olivier Kobel, Christoph Noppen, Sophie Seidel, Noemie Santamaria de Souza, Niko Beerenwinkel, Tanja Stadler |
| EPI_ISL_899070 | Viollier AG | Department of Biosystems Science and Engineering, ETH Zürich | Christian Beisel, Sarah Nadeau, Chaoran Chen, Ivan Topolsky, Philipp Jablonski, Lara Fuhrmann, David Dreifuss, Katharina Jahn, Tobias Schär, Ina Nissen, Natascha Santacroce, Elodie Burcklen, Christiane Beckmann, Maurice Redondo, Olivier Kobel, Christoph Noppen, Sophie Seidel, Noemie Santamaria de Souza, Niko Beerenwinkel, Tanja Stadler |
| EPI_ISL_899073, EPI_ISL_899076, EPI_ISL_899079 | Viollier AG | Department of Biosystems Science and Engineering, ETH Zürich | Chaoran Chen, Sarah Nadeau, Ivan Topolsky, Emmanouil Dermitzakis, Keith Harshman, Ioannis Xenarios, Henri Pegeot, Lorenzo Cerutti, Deborah Penet, Philipp Jablonski, Lara Fuhrmann, David Dreifuss, Katharina Jahn, Christiane Beckmann, Maurice Redondo, Olivier Kobel, Christoph Noppen, Sophie Seidel, Noemie Santamaria de Souza, Niko Beerenwinkel, Tanja Stadler |
| EPI_ISL_899080 | Viollier AG | Department of Biosystems Science and Engineering, ETH Zürich | Christian Beisel, Sarah Nadeau, Chaoran Chen, Ivan Topolsky, Philipp Jablonski, Lara Fuhrmann, David Dreifuss, Katharina Jahn, Tobias Schär, Ina Nissen, Natascha Santacroce, Elodie Burcklen, Christiane Beckmann, Maurice Redondo, Olivier Kobel, Christoph Noppen, Sophie Seidel, Noemie Santamaria de Souza, Niko Beerenwinkel, Tanja Stadler |
| EPI_ISL_899084, EPI_ISL_899086, EPI_ISL_899087, EPI_ISL_899093 | Viollier AG | Department of Biosystems Science and Engineering, ETH Zürich | Chaoran Chen, Sarah Nadeau, Ivan Topolsky, Emmanouil Dermitzakis, Keith Harshman, Ioannis Xenarios, Henri Pegeot, Lorenzo Cerutti, Deborah Penet, Philipp Jablonski, Lara Fuhrmann, David Dreifuss, Katharina Jahn, Christiane Beckmann, Maurice Redondo, Olivier Kobel, Christoph Noppen, Sophie Seidel, Noemie Santamaria de Souza, Niko Beerenwinkel, Tanja Stadler |
| EPI_ISL_899096 | Viollier AG | Department of Biosystems Science and Engineering, ETH Zürich | Christian Beisel, Sarah Nadeau, Chaoran Chen, Ivan Topolsky, Philipp Jablonski, Lara Fuhrmann, David Dreifuss, Katharina Jahn, Tobias Schär, Ina Nissen, Natascha Santacroce, Elodie Burcklen, Christiane Beckmann, Maurice Redondo, Olivier Kobel, Christoph Noppen, Sophie Seidel, Noemie Santamaria de Souza, Niko Beerenwinkel, Tanja Stadler |
| EPI_ISL_899098, EPI_ISL_899100 | Viollier AG | Department of Biosystems Science and Engineering, ETH Zürich | Chaoran Chen, Sarah Nadeau, Ivan Topolsky, Emmanouil Dermitzakis, Keith Harshman, Ioannis Xenarios, Henri Pegeot, Lorenzo Cerutti, Deborah Penet, Philipp Jablonski, Lara Fuhrmann, David Dreifuss, Katharina Jahn, Christiane Beckmann, Maurice Redondo, Olivier Kobel, Christoph Noppen, Sophie Seidel, Noemie Santamaria de Souza, Niko Beerenwinkel, Tanja Stadler |
| EPI_ISL_899117 | Viollier AG | Department of Biosystems Science and Engineering, ETH Zürich | Christian Beisel, Sarah Nadeau, Chaoran Chen, Ivan Topolsky, Philipp Jablonski, Lara Fuhrmann, David Dreifuss, Katharina Jahn, Tobias Schär, Ina Nissen, Natascha Santacroce, Elodie Burcklen, Christiane Beckmann, Maurice Redondo, Olivier Kobel, Christoph Noppen, Sophie Seidel, Noemie Santamaria de Souza, Niko Beerenwinkel, Tanja Stadler |
| EPI_ISL_899124 | Viollier AG | Department of Biosystems Science and Engineering, ETH Zürich | Chaoran Chen, Sarah Nadeau, Ivan Topolsky, Emmanouil Dermitzakis, Keith Harshman, Ioannis Xenarios, Henri Pegeot, Lorenzo Cerutti, Deborah Penet, Philipp Jablonski, Lara Fuhrmann, David Dreifuss, Katharina Jahn, Christiane Beckmann, Maurice Redondo, Olivier Kobel, Christoph Noppen, Sophie Seidel, Noemie Santamaria de Souza, Niko Beerenwinkel, Tanja Stadler |
| EPI_ISL_899125 | Viollier AG | Department of Biosystems Science and Engineering, ETH Zürich | Christian Beisel, Sarah Nadeau, Chaoran Chen, Ivan Topolsky, Philipp Jablonski, Lara Fuhrmann, David Dreifuss, Katharina Jahn, Tobias Schär, Ina Nissen, Natascha Santacroce, Elodie Burcklen, Christiane Beckmann, Maurice Redondo, Olivier Kobel, Christoph Noppen, Sophie Seidel, Noemie Santamaria de Souza, Niko Beerenwinkel, Tanja Stadler |
| EPI_ISL_899153 | Viollier AG | Department of Biosystems Science and Engineering, ETH Zürich | Chaoran Chen, Sarah Nadeau, Ivan Topolsky, Emmanouil Dermitzakis, Keith Harshman, Ioannis Xenarios, Henri Pegeot, Lorenzo Cerutti, Deborah Penet, Philipp Jablonski, Lara Fuhrmann, David Dreifuss, Katharina Jahn, Christiane Beckmann, Maurice Redondo, Olivier Kobel, Christoph Noppen, Sophie Seidel, Noemie Santamaria de Souza, Niko Beerenwinkel, Tanja Stadler |
| EPI_ISL_899154 | Viollier AG | Department of Biosystems Science and Engineering, ETH Zürich | Christian Beisel, Sarah Nadeau, Chaoran Chen, Ivan Topolsky, Philipp Jablonski, Lara Fuhrmann, David Dreifuss, Katharina Jahn, Tobias Schär, Ina Nissen, Natascha Santacroce, Elodie Burcklen, Christiane Beckmann, Maurice Redondo, Olivier Kobel, Christoph Noppen, Sophie Seidel, Noemie Santamaria de Souza, Niko Beerenwinkel, Tanja Stadler |
| EPI_ISL_899159, EPI_ISL_899160, EPI_ISL_899164 | Viollier AG | Department of Biosystems Science and Engineering, ETH Zürich | Chaoran Chen, Sarah Nadeau, Ivan Topolsky, Emmanouil Dermitzakis, Keith Harshman, Ioannis Xenarios, Henri Pegeot, Lorenzo Cerutti, Deborah Penet, Philipp Jablonski, Lara Fuhrmann, David Dreifuss, Katharina Jahn, Christiane Beckmann, Maurice Redondo, Olivier Kobel, Christoph Noppen, Sophie Seidel, Noemie Santamaria de Souza, Niko Beerenwinkel, Tanja Stadler |
| EPI_ISL_899241, EPI_ISL_899243, EPI_ISL_899246 | Viollier AG | Department of Biosystems Science and Engineering, ETH Zürich | Christian Beisel, Sarah Nadeau, Chaoran Chen, Ivan Topolsky, Philipp Jablonski, Lara Fuhrmann, David Dreifuss, Katharina Jahn, Tobias Schär, Ina Nissen, Natascha Santacroce, Elodie Burcklen, Christiane Beckmann, Maurice Redondo, Olivier Kobel, Christoph Noppen, Sophie Seidel, Noemie Santamaria de Souza, Niko Beerenwinkel, Tanja Stadler |
| EPI_ISL_899261, EPI_ISL_899262, EPI_ISL_899263, EPI_ISL_899264, EPI_ISL_899265, EPI_ISL_899266, EPI_ISL_899267 | Viollier AG | Department of Biosystems Science and Engineering, ETH Zürich | Chaoran Chen, Sarah Nadeau, Ivan Topolsky, Emmanouil Dermitzakis, Keith Harshman, Ioannis Xenarios, Henri Pegeot, Lorenzo Cerutti, Deborah Penet, Philipp Jablonski, Lara Fuhrmann, David Dreifuss, Katharina Jahn, Christiane Beckmann, Maurice Redondo, Olivier Kobel, Christoph Noppen, Sophie Seidel, Noemie Santamaria de Souza, Niko Beerenwinkel, Tanja Stadler |
| EPI_ISL_899340, EPI_ISL_899341, EPI_ISL_899346, EPI_ISL_899347, EPI_ISL_899348, EPI_ISL_899349 | Viollier AG | Department of Biosystems Science and Engineering, ETH Zürich | Christian Beisel, Sarah Nadeau, Chaoran Chen, Ivan Topolsky, Philipp Jablonski, Lara Fuhrmann, David Dreifuss, Katharina Jahn, Tobias Schär, Ina Nissen, Natascha Santacroce, Elodie Burcklen, Christiane Beckmann, Maurice Redondo, Olivier Kobel, Christoph Noppen, Sophie Seidel, Noemie Santamaria de Souza, Niko Beerenwinkel, Tanja Stadler |
| EPI_ISL_899360, EPI_ISL_899370, EPI_ISL_899371, EPI_ISL_899372 | Viollier AG | Department of Biosystems Science and Engineering, ETH Zürich | Chaoran Chen, Sarah Nadeau, Ivan Topolsky, Emmanouil Dermitzakis, Keith Harshman, Ioannis Xenarios, Henri Pegeot, Lorenzo Cerutti, Deborah Penet, Philipp Jablonski, Lara Fuhrmann, David Dreifuss, Katharina Jahn, Christiane Beckmann, Maurice Redondo, Olivier Kobel, Christoph Noppen, Sophie Seidel, Noemie Santamaria de Souza, Niko Beerenwinkel, Tanja Stadler |
| EPI_ISL_899456, EPI_ISL_899457, EPI_ISL_899475, EPI_ISL_899476, EPI_ISL_899477 | Viollier AG | Department of Biosystems Science and Engineering, ETH Zürich | Christian Beisel, Sarah Nadeau, Chaoran Chen, Ivan Topolsky, Philipp Jablonski, Lara Fuhrmann, David Dreifuss, Katharina Jahn, Tobias Schär, Ina Nissen, Natascha Santacroce, Elodie Burcklen, Christiane Beckmann, Maurice Redondo, Olivier Kobel, Christoph Noppen, Sophie Seidel, Noemie Santamaria de Souza, Niko Beerenwinkel, Tanja Stadler |
| EPI_ISL_899498, EPI_ISL_899499, EPI_ISL_899500, EPI_ISL_899501, EPI_ISL_899502, EPI_ISL_899503, EPI_ISL_899504, EPI_ISL_899505, EPI_ISL_899506, EPI_ISL_899507, EPI_ISL_899508 | | | |
| see above | Viollier AG | Department of Biosystems Science and Engineering, ETH Zürich | Chaoran Chen, Sarah Nadeau, Ivan Topolsky, Emmanouil Dermitzakis, Keith Harshman, Ioannis Xenarios, Henri Pegeot, Lorenzo Cerutti, Deborah Penet, Philipp Jablonski, Lara Fuhrmann, David Dreifuss, Katharina Jahn, Christiane Beckmann, Maurice Redondo, Olivier Kobel, Christoph Noppen, Sophie Seidel, Noemie Santamaria de Souza, Niko Beerenwinkel, Tanja Stadler |
| EPI_ISL_899530 | Viollier AG | Department of Biosystems Science and Engineering, ETH Zürich | Christian Beisel, Sarah Nadeau, Chaoran Chen, Ivan Topolsky, Philipp Jablonski, Lara Fuhrmann, David Dreifuss, Katharina Jahn, Tobias Schär, Ina Nissen, Natascha Santacroce, Elodie Burcklen, Christiane Beckmann, Maurice Redondo, Olivier Kobel, Christoph Noppen, Sophie Seidel, Noemie Santamaria de Souza, Niko Beerenwinkel, Tanja Stadler |
| EPI_ISL_899532, EPI_ISL_899533, EPI_ISL_899534 | Viollier AG | Department of Biosystems Science and Engineering, ETH Zürich | Chaoran Chen, Sarah Nadeau, Ivan Topolsky, Emmanouil Dermitzakis, Keith Harshman, Ioannis Xenarios, Henri Pegeot, Lorenzo Cerutti, Deborah Penet, Philipp Jablonski, Lara Fuhrmann, David Dreifuss, Katharina Jahn, Christiane Beckmann, Maurice Redondo, Olivier Kobel, Christoph Noppen, Sophie Seidel, Noemie Santamaria de Souza, Niko Beerenwinkel, Tanja Stadler |
| EPI_ISL_899538, EPI_ISL_899544 | Viollier AG | Department of Biosystems Science and Engineering, ETH Zürich | Christian Beisel, Sarah Nadeau, Chaoran Chen, Ivan Topolsky, Philipp Jablonski, Lara Fuhrmann, David Dreifuss, Katharina Jahn, Tobias Schär, Ina Nissen, Natascha Santacroce, Elodie Burcklen, Christiane Beckmann, Maurice Redondo, Olivier Kobel, Christoph Noppen, Sophie Seidel, Noemie Santamaria de Souza, Niko Beerenwinkel, Tanja Stadler |
| EPI_ISL_899560 | Viollier AG | Department of Biosystems Science and Engineering, ETH Zürich | Chaoran Chen, Sarah Nadeau, Ivan Topolsky, Emmanouil Dermitzakis, Keith Harshman, Ioannis Xenarios, Henri Pegeot, Lorenzo Cerutti, Deborah Penet, Philipp Jablonski, Lara Fuhrmann, David Dreifuss, Katharina Jahn, Christiane Beckmann, Maurice Redondo, Olivier Kobel, Christoph Noppen, Sophie Seidel, Noemie Santamaria de Souza, Niko Beerenwinkel, Tanja Stadler |
| EPI_ISL_899589, EPI_ISL_899590 | Viollier AG | Department of Biosystems Science and Engineering, ETH Zürich | Christian Beisel, Sarah Nadeau, Chaoran Chen, Ivan Topolsky, Philipp Jablonski, Lara Fuhrmann, David Dreifuss, Katharina Jahn, Tobias Schär, Ina Nissen, Natascha Santacroce, Elodie Burcklen, Christiane Beckmann, Maurice Redondo, Olivier Kobel, Christoph Noppen, Sophie Seidel, Noemie Santamaria de Souza, Niko Beerenwinkel, Tanja Stadler |

| | | | |
|---|---|---|---|
| EPI_ISL_899593, EPI_ISL_899594, EPI_ISL_899595 | Viollier AG | Department of Biosystems Science and Engineering, ETH Zürich | Chaoran Chen, Sarah Nadeau, Ivan Topolsky, Emmanouil Dermitzakis, Keith Harshman, Ioannis Xenarios, Henri Pegeot, Lorenzo Cerutti, Deborah Penet, Philipp Jablonski, Lara Fuhrmann, David Dreifuss, Katharina Jahn, Christiane Beckmann, Maurice Redondo, Olivier Kobel, Christoph Noppen, Sophie Seidel, Noemie Santamaria de Souza, Niko Beerenwinkel, Tanja Stadler |
| EPI_ISL_899599, EPI_ISL_899602, EPI_ISL_899603, EPI_ISL_899604, EPI_ISL_899605, EPI_ISL_899607, EPI_ISL_899608 | Viollier AG | Department of Biosystems Science and Engineering, ETH Zürich | Christian Beisel, Sarah Nadeau, Chaoran Chen, Ivan Topolsky, Philipp Jablonski, Lara Fuhrmann, David Dreifuss, Katharina Jahn, Tobias Schär, Ina Nissen, Natascha Santacroce, Elodie Burcklen, Christiane Beckmann, Maurice Redondo, Olivier Kobel, Christoph Noppen, Sophie Seidel, Noemie Santamaria de Souza, Niko Beerenwinkel, Tanja Stadler |
| EPI_ISL_899611, EPI_ISL_899643, EPI_ISL_899644, EPI_ISL_899645 | Viollier AG | Department of Biosystems Science and Engineering, ETH Zürich | Chaoran Chen, Sarah Nadeau, Ivan Topolsky, Emmanouil Dermitzakis, Keith Harshman, Ioannis Xenarios, Henri Pegeot, Lorenzo Cerutti, Deborah Penet, Philipp Jablonski, Lara Fuhrmann, David Dreifuss, Katharina Jahn, Christiane Beckmann, Maurice Redondo, Olivier Kobel, Christoph Noppen, Sophie Seidel, Noemie Santamaria de Souza, Niko Beerenwinkel, Tanja Stadler |
| EPI_ISL_899722, EPI_ISL_899723, EPI_ISL_899724, EPI_ISL_899737 | Viollier AG | Department of Biosystems Science and Engineering, ETH Zürich | Christian Beisel, Sarah Nadeau, Chaoran Chen, Ivan Topolsky, Philipp Jablonski, Lara Fuhrmann, David Dreifuss, Katharina Jahn, Tobias Schär, Ina Nissen, Natascha Santacroce, Elodie Burcklen, Christiane Beckmann, Maurice Redondo, Olivier Kobel, Christoph Noppen, Sophie Seidel, Noemie Santamaria de Souza, Niko Beerenwinkel, Tanja Stadler |
| EPI_ISL_899750, EPI_ISL_899751, EPI_ISL_899752, EPI_ISL_899753, EPI_ISL_899754, EPI_ISL_899755 | Viollier AG | Department of Biosystems Science and Engineering, ETH Zürich | Chaoran Chen, Sarah Nadeau, Ivan Topolsky, Emmanouil Dermitzakis, Keith Harshman, Ioannis Xenarios, Henri Pegeot, Lorenzo Cerutti, Deborah Penet, Philipp Jablonski, Lara Fuhrmann, David Dreifuss, Katharina Jahn, Christiane Beckmann, Maurice Redondo, Olivier Kobel, Christoph Noppen, Sophie Seidel, Noemie Santamaria de Souza, Niko Beerenwinkel, Tanja Stadler |
| EPI_ISL_899792 | Viollier AG | Department of Biosystems Science and Engineering, ETH Zürich | Christian Beisel, Sarah Nadeau, Chaoran Chen, Ivan Topolsky, Philipp Jablonski, Lara Fuhrmann, David Dreifuss, Katharina Jahn, Tobias Schär, Ina Nissen, Natascha Santacroce, Elodie Burcklen, Christiane Beckmann, Maurice Redondo, Olivier Kobel, Christoph Noppen, Sophie Seidel, Noemie Santamaria de Souza, Niko Beerenwinkel, Tanja Stadler |
| EPI_ISL_899800, EPI_ISL_899801, EPI_ISL_899802, EPI_ISL_899803, EPI_ISL_899804 | Viollier AG | Department of Biosystems Science and Engineering, ETH Zürich | Chaoran Chen, Sarah Nadeau, Ivan Topolsky, Emmanouil Dermitzakis, Keith Harshman, Ioannis Xenarios, Henri Pegeot, Lorenzo Cerutti, Deborah Penet, Philipp Jablonski, Lara Fuhrmann, David Dreifuss, Katharina Jahn, Christiane Beckmann, Maurice Redondo, Olivier Kobel, Christoph Noppen, Sophie Seidel, Noemie Santamaria de Souza, Niko Beerenwinkel, Tanja Stadler |
| EPI_ISL_899844, EPI_ISL_899848, EPI_ISL_899849, EPI_ISL_899874, EPI_ISL_899912, EPI_ISL_899915 | Viollier AG | Department of Biosystems Science and Engineering, ETH Zürich | Christian Beisel, Sarah Nadeau, Chaoran Chen, Ivan Topolsky, Philipp Jablonski, Lara Fuhrmann, David Dreifuss, Katharina Jahn, Tobias Schär, Ina Nissen, Natascha Santacroce, Elodie Burcklen, Christiane Beckmann, Maurice Redondo, Olivier Kobel, Christoph Noppen, Sophie Seidel, Noemie Santamaria de Souza, Niko Beerenwinkel, Tanja Stadler |
| EPI_ISL_899917, EPI_ISL_899918 | Viollier AG | Department of Biosystems Science and Engineering, ETH Zürich | Chaoran Chen, Sarah Nadeau, Ivan Topolsky, Emmanouil Dermitzakis, Keith Harshman, Ioannis Xenarios, Henri Pegeot, Lorenzo Cerutti, Deborah Penet, Philipp Jablonski, Lara Fuhrmann, David Dreifuss, Katharina Jahn, Christiane Beckmann, Maurice Redondo, Olivier Kobel, Christoph Noppen, Sophie Seidel, Noemie Santamaria de Souza, Niko Beerenwinkel, Tanja Stadler |
| EPI_ISL_900010, EPI_ISL_900011, EPI_ISL_900012, EPI_ISL_900016, EPI_ISL_900017, EPI_ISL_900018 | Viollier AG | Department of Biosystems Science and Engineering, ETH Zürich | Christian Beisel, Sarah Nadeau, Chaoran Chen, Ivan Topolsky, Philipp Jablonski, Lara Fuhrmann, David Dreifuss, Katharina Jahn, Tobias Schär, Ina Nissen, Natascha Santacroce, Elodie Burcklen, Christiane Beckmann, Maurice Redondo, Olivier Kobel, Christoph Noppen, Sophie Seidel, Noemie Santamaria de Souza, Niko Beerenwinkel, Tanja Stadler |
| EPI_ISL_900030, EPI_ISL_900031, EPI_ISL_900032, EPI_ISL_900033, EPI_ISL_900034, EPI_ISL_900035, EPI_ISL_900036 | Viollier AG | Department of Biosystems Science and Engineering, ETH Zürich | Chaoran Chen, Sarah Nadeau, Ivan Topolsky, Emmanouil Dermitzakis, Keith Harshman, Ioannis Xenarios, Henri Pegeot, Lorenzo Cerutti, Deborah Penet, Philipp Jablonski, Lara Fuhrmann, David Dreifuss, Katharina Jahn, Christiane Beckmann, Maurice Redondo, Olivier Kobel, Christoph Noppen, Sophie Seidel, Noemie Santamaria de Souza, Niko Beerenwinkel, Tanja Stadler |
| EPI_ISL_900061 | MEPHI, Aix Marseille University | MEPHI, Aix Marseille University | Anthony LEVASSEUR |
| EPI_ISL_900499 | Althaia. Xarxa Assistencial Universitària de Manresa | IrsiCaixa - Can Ruti CovidSeq | Fundació irsiCaixa. Hospital Universitari Germans Trias i Pujol(HUGTiP), 2a planta, maternal Ctra Canyet s/n, Badalona Gloria Trujillo, Rafel Perez Vidal, Jaume Trape Pujol, Carolina Gonzalez Fernandez, Roger Paredes, Eulalia Grau, Bonaventura Clotet |
| EPI_ISL_902916 | Department of Virology and Immunology, University of Helsinki and Helsinki University Hospital, Huslab Finland | Department of Virology, Faculty of Medicine, University of Helsinki, Helsinki, Finland | Teemu Smura, Ravi Kant, Phuoc Truong, Hussein Alburkat, Hannimari Kallio-Kokko, Jenni Virtanen, Maija Suvanto, Essi Korhonen, Sari Hannula, Harri Kangas, Hanna Liimatainen, Satu Kurkela, Hanna Jarva, Maija Lappalainen, Pekka Ellonen, Olli Vapalahti |
| EPI_ISL_902948 | Maryland Public Health Laboratory | Maryland Public Health Laboratory | Maryland Department of Health Laboratories Administration |
| EPI_ISL_903347 | Ministry of Health Turkey | Ministry of Health Turkey | Fatma Bayrakdar, Yasemin Cogun, Süleyman Yalcin, Aye Baak Alta, Gülay Korukluolu |
| EPI_ISL_903372 | Los Angeles County PHL | Los Angeles County PHL | P. Hemarajata et al. |
| EPI_ISL_904068, EPI_ISL_904069, EPI_ISL_904070 | New Mexico Department of Health Scientific Laboratory | New Mexico Department of Health Scientific Laboratory | Ellie Johnson, Anastacia Griego-Fisher, D'eldra Malone |
| EPI_ISL_904139, EPI_ISL_904156, EPI_ISL_904163, EPI_ISL_904207, EPI_ISL_904208, EPI_ISL_904209, EPI_ISL_904210, EPI_ISL_904211, EPI_ISL_904249, EPI_ISL_904275, EPI_ISL_904276, EPI_ISL_904277, EPI_ISL_904319, EPI_ISL_904369, EPI_ISL_904579, EPI_ISL_904580, EPI_ISL_904581, EPI_ISL_904582, EPI_ISL_904583, EPI_ISL_904584, EPI_ISL_904585, EPI_ISL_904586, EPI_ISL_904587 | | | |
| see above | Dutch COVID-19 response team | Erasmus Medical Center | Bas Oude Munnink, Reina Sikkema, David Nieuwenhuijse, Irina Chestakova, Anne van der Linden, Marjan Boter, Emmanuelle Munger, Corine GeurtsvanKessel, Annemiek van der Eijk, Richard Molenkamp, Marion Koopmans, on behalf of the Dutch national COVID-19 response team. |
| EPI_ISL_904659, EPI_ISL_904673, EPI_ISL_904690, EPI_ISL_904696, EPI_ISL_904728, EPI_ISL_904729, EPI_ISL_904769, EPI_ISL_904786, EPI_ISL_904789, EPI_ISL_904801, EPI_ISL_904802, EPI_ISL_904803, EPI_ISL_904812, EPI_ISL_904826, EPI_ISL_904827, EPI_ISL_904832, EPI_ISL_904840, EPI_ISL_904852, EPI_ISL_904873, EPI_ISL_904874, EPI_ISL_904878, EPI_ISL_904882, EPI_ISL_904883, EPI_ISL_905048, EPI_ISL_905051, EPI_ISL_905052, EPI_ISL_905053, EPI_ISL_905054, EPI_ISL_905055, EPI_ISL_905056, EPI_ISL_905061, EPI_ISL_905063, EPI_ISL_905066, EPI_ISL_905067, EPI_ISL_905076, EPI_ISL_905118, EPI_ISL_905119, EPI_ISL_905176, EPI_ISL_905178, EPI_ISL_905180, EPI_ISL_905192, EPI_ISL_905193, EPI_ISL_905197, EPI_ISL_905204, EPI_ISL_905245, EPI_ISL_905246, EPI_ISL_905247, EPI_ISL_905248, EPI_ISL_905249, EPI_ISL_905250, EPI_ISL_905262, EPI_ISL_905267, EPI_ISL_905281, EPI_ISL_905299, EPI_ISL_905300, EPI_ISL_905301, EPI_ISL_905302, EPI_ISL_905303, EPI_ISL_905304, EPI_ISL_905305, EPI_ISL_905307, EPI_ISL_905308, EPI_ISL_905316, EPI_ISL_905317, EPI_ISL_905318, EPI_ISL_905325, EPI_ISL_905326, EPI_ISL_905548, EPI_ISL_905549, EPI_ISL_905550 | | | |
| see above | Dutch COVID-19 response team | National Institute for Public Health and the Environment (RIVM) | Adam Meijer, Harry Vennema, Dirk Eggink, Jeroen Cremer, Sharon van den Brink, Bas van der Veer, AnneMarie van den Brandt, Florian Zwagemaker, Dennis Schmitz, Chantal Reusken, on behalf of the national COVID-19 response team |
| EPI_ISL_905741, EPI_ISL_905742, EPI_ISL_905743, EPI_ISL_905744, EPI_ISL_905745, EPI_ISL_905746, EPI_ISL_905747 | National Institute of Public Health - National Institute of Hygiene | National Institute of Public Health - National Institute of Hygiene | Wokowicz Tomasz, Zacharczuk Katarzyna |
| EPI_ISL_906045 | OHSU Lab Services Molecular Microbiology Lab | Oregon SARS-CoV-2 Genome Sequencing Center | Brendan L. O'Connell, Sally Grindstaff, Kayla Carter, Ruth V. Nichols, Alec J. Hirsch, Donna Hansel, Guang Fan, Xuan, Qin, Daniel N. Streblow, William B. Messer, Andrew C. Adey, Benjamin N. Bimber, Brian J. O'Roak |
| EPI_ISL_906279, EPI_ISL_906280, EPI_ISL_906281, EPI_ISL_906284 | Nigeria Centre for Disease Control (NCDC) | African Centre of Excellence for Genomics of Infectious Diseases (ACEGID), Redeemer's University | Oluniyi P.E. et al |
| EPI_ISL_906306 | Gorgas memorial Institute For Health Studies | Gorgas memorial Institute For Health Studies | Diaz Y, Franco D, Moreno B, Moreno A, Gondola Y, Saenz L, Abrego L, Chavarria O, Pitti Y, Castillo M, Lopez-Verguez S, Martinez Alexander A. |
| EPI_ISL_906559 | Laboratorio de Salud Publica de Amazonas | Instituto Nacional de Salud- Dirección de Investigación en Salud Pública, Universidad de los Andes- Applied genomics research group, Vicerrectoria de Investigación y Creación, Universidad de los Andes- Systems and Computing Engineering Department | Katherine Laiton-Donato, Diego A. Álvarez-Díaz, Carlos Franco-Muñoz, Mauricio Pacheco-Montealegre, Héctor Alejandro Ruiz-Moreno, Maria T. Herrera-Sepúlveda, Diego Andrés Prada, Jhonnatan Reales-González, Sheryll Corchuelo, Julian Naizaque, Gerardo Santamaría Jorge Duitama, Laura Natalia Gonzalez, Jorge Ivan Diaz, Silvia Restrepo-Restrepo, Magdalena Wiesner, Martha Lucia Ospina Martinez, Marcela Mercado-Reyes |
| EPI_ISL_906590, EPI_ISL_906591, EPI_ISL_906592, EPI_ISL_906593, EPI_ISL_906594, EPI_ISL_906595, EPI_ISL_906596, EPI_ISL_906597, EPI_ISL_906598, EPI_ISL_906599, EPI_ISL_906600, EPI_ISL_906601, EPI_ISL_906602 | | | |
| see above | Maine Health and Environmental Testing Laboratory (Maine HETL) | Tewhey Lab, The Jackson Laboratory | Matluk,N., Dewey,H., Iosue,F., Barter,M., Lynch,R., Munger,H. and Tewhey,R. |
| EPI_ISL_906806, EPI_ISL_906807 | University Hospital of Northern Norway, Department for | Norwegian Institute of Public Health, Department of Virology | Kathrine Stene-Johansen, Kamilla Heddeland Instefjord, Hilde Elshaug, Atiya R Ali,Marie Paulsen Madsen, Rasmus Riis Kopperud, Hilde Vollan, Karoline |

| | | | |
|---|---|---|---|
| | Microbiology and Infectious Disease Control | | Bragstad, Olav Hungnes |
| EPI_ISL_906818 | Akershus University Hospital, Department for Microbiology and Infectious Disease Control | Norwegian Institute of Public Health, Department of Virology | Kathrine Stene-Johansen, Kamilla Heddeland Instefjord, Hilde Elshaug, Atiya R Ali,Marie Paulsen Madsen, Rasmus Riis Kopperud, Hilde Vollan, Karoline Bragstad, Olav Hungnes |
| EPI_ISL_906910, EPI_ISL_906911, EPI_ISL_906912, EPI_ISL_906913, EPI_ISL_906914 | Bureau of Public Health Laboratories, Florida Department of Health (BPHL, FLDOH) | Bureau of Public Health Laboratories, Florida Department of Health (BPHL, FLDOH) | Schmedes,S., Blanton,J. |
| EPI_ISL_907210, EPI_ISL_907750 | Lighthouse Lab in Glasgow | Wellcome Sanger Institute for the COVID-19 Genomics UK (COG-UK) Consortium | Harper VanSteenhouse, Yumi Kasai, David Gray, Carol Clugston, Anna Dominiczak and Alex Alderton, Roberto Amato, Sonia Goncalves, Ewan Harrison, David K. Jackson, Ian Johnston, Dominic Kwiatkowski, Cordelia Langford, John Sillitoe on behalf of the Wellcome Sanger Institute COVID-19 Surveillance Team |
| EPI_ISL_909665 | Labo Analyses Med | National Reference Center for Viruses of Respiratory Infections, Institut Pasteur, Paris | Marion Barbet, Sylvie Behillil, Méline Bizard, Angela Brisebarre, Camille Capel, Etienne Simon-Lorière, Vincent Enouf, Maud Vanpeene, Sylvie van der Werf,Le Vicky |
| EPI_ISL_909835, EPI_ISL_909836, EPI_ISL_909837, EPI_ISL_909838, EPI_ISL_909839, EPI_ISL_909840, EPI_ISL_909842, EPI_ISL_909843, EPI_ISL_909845, EPI_ISL_909846, EPI_ISL_909848, EPI_ISL_909852, EPI_ISL_909853, EPI_ISL_909858, EPI_ISL_909863, EPI_ISL_909868, EPI_ISL_909873, EPI_ISL_909874, EPI_ISL_909897, EPI_ISL_909904, EPI_ISL_909906, EPI_ISL_909907, EPI_ISL_909913, EPI_ISL_909915, EPI_ISL_909916, EPI_ISL_909919, EPI_ISL_909928, EPI_ISL_909931, EPI_ISL_909937, EPI_ISL_909939 | | | |
| see above | National Virus Reference Laboratory | National Virus Reference Laboratory | Michael Carr, Gabriel Gonzalez, Jonathan Dean, Cillian F De Gascun |
| EPI_ISL_910329, EPI_ISL_910334 | ALEA dr Kandic | Alea Genetic Centre | Rijad Konjhodzic, Dino Pecar, Lana Salihefendic |
| EPI_ISL_910954, EPI_ISL_910955, EPI_ISL_910956, EPI_ISL_911125, EPI_ISL_911126, EPI_ISL_911127, EPI_ISL_911128, EPI_ISL_911129, EPI_ISL_911130, EPI_ISL_911131, EPI_ISL_911132, EPI_ISL_911133, EPI_ISL_911135, EPI_ISL_911168 | | | |
| see above | Laboratoire national de sante, Microbiology, Virology | Laboratoire national de sante, Microbiology, Microbial Genomics Platform | Anke Wienecke-Baldacchino, Catherine Ragimbeau,Jessica Tapp, Fatu Djabi, Lise Pignon, Raoul Salmon, Tamir Abdelrahman |
| EPI_ISL_911245, EPI_ISL_911247 | National Virus Reference Laboratory | National Virus Reference Laboratory | Michael Carr, Gabriel Gonzalez, Jonathan Dean, Cillian F De Gascun |
| EPI_ISL_911289, EPI_ISL_911342, EPI_ISL_911350 | Servicio de Microbiología, Hospital Universitario Son Espases | SeqCOVID-SPAIN consortium/IBV(CSIC) | Carla López-Causapé, Jordi Reina, Antonio Oliver and SeqCOVID-SPAIN consortium |
| EPI_ISL_911931, EPI_ISL_911932, EPI_ISL_911933, EPI_ISL_911934, EPI_ISL_911935, EPI_ISL_911936, EPI_ISL_911937, EPI_ISL_911938, EPI_ISL_911939, EPI_ISL_911940, EPI_ISL_911941, EPI_ISL_911942, EPI_ISL_911943, EPI_ISL_911944, EPI_ISL_911945, EPI_ISL_911946, EPI_ISL_911947, EPI_ISL_911948, EPI_ISL_911949, EPI_ISL_911950, EPI_ISL_911951 | | | |
| see above | Wyoming Public Health Laboratory | Wyoming Public Health Laboratory | Noah Hull, Taylor Fearing, Lynette Gumbleton, Channing Weber, Ashley Norberg, Bailey Bowcutt, and Wanda Manley |
| EPI_ISL_912006, EPI_ISL_912007, EPI_ISL_912008, EPI_ISL_912009, EPI_ISL_912010, EPI_ISL_912011, EPI_ISL_912012, EPI_ISL_912013, EPI_ISL_912014, EPI_ISL_912015, EPI_ISL_912016, EPI_ISL_912017, EPI_ISL_912018, EPI_ISL_912019, EPI_ISL_912020, EPI_ISL_912021, EPI_ISL_912022, EPI_ISL_912023, EPI_ISL_912024, EPI_ISL_912025, EPI_ISL_912026, EPI_ISL_912027, EPI_ISL_912028, EPI_ISL_912029, EPI_ISL_912030, EPI_ISL_912031, EPI_ISL_912032, EPI_ISL_912033, EPI_ISL_912034 | | | |
| see above | Altius Institute for Biomedical Sciences | Seattle Flu Study | Deborah A. Nickerson, Chris D. Frazar, Jover Lee, Benjamin Pelle, Erica Ryke, Matthew Richardson, Amanda Adler, Elisabeth Brandstetter, Peter D. Han, Kairsten Fay, Misja Ilcisin, Kirsten Lacombe, Thomas R. Sibley, Melissa Truong, Caitlin R. Wolf, Ryan Alexander, Daniel Bates, Rebecca Bruders, Stephanie DeBaun, Clem Green, Muhammad Halimun, Jessica Halow, Kneshay Harper, Matt Hartman, Andrew Meuser, Alex Nguyen, Truong Nguyen, Sofia Olsson, Sadie Patraw, Hannah Petersen, Tobias Ragoczy, Joshua Richards, Jacob Rodriguez, John Stamatoyannopoulos, Julia Wald, Olivia Waltner, Michael Boeckh, Janet A. Englund, Michael Famulare, Barry R. Lutz, Mark J. Rieder, Lea M. Starita, Matthew Thompson, Helen Y. Chu, Jay Shendure, Trevor Bedford |
| EPI_ISL_912079, EPI_ISL_912080, EPI_ISL_912081, EPI_ISL_912082, EPI_ISL_912083, EPI_ISL_912084, EPI_ISL_912085, EPI_ISL_912086, EPI_ISL_912087, EPI_ISL_912088, EPI_ISL_912089, EPI_ISL_912090, EPI_ISL_912091, EPI_ISL_912092, EPI_ISL_912093, EPI_ISL_912095 | | | |
| see above | Washington State Department of Health | Seattle Flu Study | Deborah A. Nickerson, Chris D. Frazar, Jover Lee, Benjamin Pelle, Erica Ryke, Matthew Richardson, Amanda Adler, Elisabeth Brandstetter, Peter D. Han, Kairsten Fay, Misja Ilcisin, Kirsten Lacombe, Thomas R. Sibley, Melissa Truong, Caitlin R. Wolf, Romesh Gautom, Geoff Melly, Brian Hiatt, Philip Dykema, Scott Lindquist, Michael Boeckh, Janet A. Englund, Michael Famulare, Barry R. Lutz, Mark J. Rieder, Lea M. Starita, Matthew Thompson, Helen Y. Chu, Jay Shendure, Trevor Bedford |
| EPI_ISL_912166 | Natual State Genomics | Grubaugh Lab - Yale School of Public Health | Tara Alpert, Joseph Fauver, Anderson Brito, Mallery Breban, Anne Wyllie, Chantal Vogels, Mary Petrone, Annie Watkins, Chaney Kalinich, Isabel Ott, Nathan Grubaugh |
| EPI_ISL_912248, EPI_ISL_912255 | Charité Universitätsmedizin Berlin, Institut für Virologie/Labor Berlin | Charité Universitätsmedizin Berlin, Institut für Virologie | Victor M Corman, Barbara Mühlemann, Jörn Beheim-Schwarzbach, Tobias Bleicker, Julia Tesch, Talitha Veith, Julia Schneider, Terry Jones, Christian Drosten |
| EPI_ISL_912413, EPI_ISL_912416 | KU Leuven, Rega Institute, Clinical and Epidemiological Virology | KU Leuven, Rega Institute, Clinical and Epidemiological Virology | Tony Wawina-Bokalanga, Bert Vanmechelen, Joan Marti-Carerras, Piet Maes |
| EPI_ISL_912539, EPI_ISL_912580, EPI_ISL_912581 | CENTRE HOSPITALIER LEON BINET | National Reference Center for Viruses of Respiratory Infections, Institut Pasteur, Paris | Marion Barbet, Sylvie Behillil, Méline Bizard, Angela Brisebarre, Camille Capel, Etienne Simon-Lorière, Vincent Enouf, Maud Vanpeene, Sylvie van der Werf,Meziane Ilham |
| EPI_ISL_912617, EPI_ISL_912618, EPI_ISL_912619, EPI_ISL_912620 | Labo Analyses Med | National Reference Center for Viruses of Respiratory Infections, Institut Pasteur, Paris | Marion Barbet, Sylvie Behillil, Méline Bizard, Angela Brisebarre, Camille Capel, Etienne Simon-Lorière, Vincent Enouf, Maud Vanpeene, Sylvie van der Werf,Le Vicky |
| EPI_ISL_912643 | Hôpital Pitié-Salpêtrière | Department of Virology, Henri Mondor University Hospital, Assistance Publique Hôpitaux de Paris, Université Paris-Est Créteil, INSERM U955 | Christophe Rodriguez, Slim Fourati, Vanessa Demontant, Guillaume Gricourt, Melissa N'Debi, Alexandre Soulier, Elisabeth Trawinski, Jean-Michel Pawlotsky |
| EPI_ISL_912648, EPI_ISL_912672, EPI_ISL_912748, EPI_ISL_912749 | Hôpital Henri Mondor | Department of Virology, Henri Mondor University Hospital, Assistance Publique Hôpitaux de Paris, Université Paris-Est Créteil, INSERM U955 | Christophe Rodriguez, Slim Fourati, Vanessa Demontant, Guillaume Gricourt, Melissa N'Debi, Alexandre Soulier, Elisabeth Trawinski, Jean-Michel Pawlotsky |
| EPI_ISL_912789, EPI_ISL_912790, EPI_ISL_912916 | Hôpital Pitié-Salpêtrière | Department of Virology, Henri Mondor University Hospital, Assistance Publique Hôpitaux de Paris, Université Paris-Est Créteil, INSERM U955 | Christophe Rodriguez, Slim Fourati, Vanessa Demontant, Guillaume Gricourt, Melissa N'Debi, Alexandre Soulier, Elisabeth Trawinski, Jean-Michel Pawlotsky |
| EPI_ISL_912958, EPI_ISL_912959 | Hôpital Henri Mondor | Department of Virology, Henri Mondor University Hospital, Assistance Publique Hôpitaux de Paris, Université Paris-Est Créteil, INSERM U955 | Christophe Rodriguez, Slim Fourati, Vanessa Demontant, Guillaume Gricourt, Melissa N'Debi, Alexandre Soulier, Elisabeth Trawinski, Jean-Michel Pawlotsky |
| EPI_ISL_913032 | Hospital San Pedro de Alcántara | Instituto de Salud Carlos III | Iglesias-Caballero, M. Camarero, S. Sandonís,V. Vázquez, S. Pozo, F. Casas, I. Jiménez, P. Zaballos, A. Monzón, S. Varona, S. Cuesta, I. Rodríguez, G. |
| EPI_ISL_913121, EPI_ISL_913125, EPI_ISL_913136, EPI_ISL_913155, EPI_ISL_913162, EPI_ISL_913163, EPI_ISL_913168, EPI_ISL_913169, EPI_ISL_913171, EPI_ISL_913172, EPI_ISL_913173, EPI_ISL_913176, EPI_ISL_913179, EPI_ISL_913181 | | | |
| see above | University of Michigan Clinical Microbiology Laboratory | Lauring Lab, University of Michigan, Department of Microbiology and Immunology | Valesano |
| EPI_ISL_913412, EPI_ISL_913421, EPI_ISL_913429, EPI_ISL_913431 | Massachusetts State Public Health Laboratory | Massachusetts State Public Health Laboratory | Andrew Lang, Timelia Fink, Glen Gallagher, Sandra Smole |
| EPI_ISL_913469 | Klinisk mikrobiologi | The Public Health Agency of Sweden | Anna-Malin Linde, Maria Lind Karlberg, Carlo Berg, Oskar Karlsson Lindsjo, Sofia Stamouli, Reza Advani, Mattias Haukland, Petra Holmstrom, Noura Walai, Petra Edquist, Mia Brytting, Anna Risberg, Karin Tegmark-Wisell |
| EPI_ISL_913587 | Vault Health | Minnesota Department of Health, Public Health Laboratory | Alexandra Lorentz, Jacob Garfin, Matt Plumb, and Xiong Wang |
| EPI_ISL_913757, EPI_ISL_913803, EPI_ISL_913804, EPI_ISL_913806, EPI_ISL_913807, EPI_ISL_913809, EPI_ISL_913860, EPI_ISL_913861, EPI_ISL_913863, EPI_ISL_913864, EPI_ISL_913865 | | | |
| see above | KU Leuven, Rega Institute, Clinical and Epidemiological Virology | KU Leuven, Rega Institute, Clinical and Epidemiological Virology | Tony Wawina-Bokalanga, Bert Vanmechelen, Joan Marti-Carerras, Piet Maes |

| | | | |
|---|---|---|---|
| EPI_ISL_914025, EPI_ISL_914026 | IL Department of Public Health Chicago Laboratory | Pathogen Discovery, Respiratory Viruses Branch, Division of Viral Diseases, Centers for Disease Control and Prevention | Ying Tao, Yan Li, Jing Zhang, Krista Queen, Anna Uehara, Peter Cook, Clinton R. Paden, Haibin Wang, Suxiang Tong |
| EPI_ISL_914039 | United States Air Force School of Aerospace Medicine | U.S. Air Force School of Aerospace Medicine | Anthony Fries, Jennifer Meyer, William Gruner, William Buggele, Amanda Javorina, Sarah Purves, Clarise Starr, Elizabeth Macias |
| EPI_ISL_914829 | TAMIZAJE COMUNITARIO - PASO CANOAS | Inciensa, Instituto Costarricense de Investigación y Enseñanza en Nutrición y Salud | Francisco Duarte, Hebleen Porras, Claudio Soto-Garita, Estela Cordero, Adriana Godínez, Melany Calderón & Mariel López |
| EPI_ISL_915427 | Bundeswehr Institute of Microbiology | Bundeswehr Institute of Microbiology | Markus Antwerpen, Alexandra Rehn, Mathias Walter, Malena Bestehorn-Willmann, Sabine Zange, Enrico Georgi, Roman Wölfel |
| EPI_ISL_916630, EPI_ISL_916637, EPI_ISL_916651, EPI_ISL_916732, EPI_ISL_916746, EPI_ISL_916766, EPI_ISL_916775, EPI_ISL_916779, EPI_ISL_916849, EPI_ISL_916919, EPI_ISL_916934, EPI_ISL_916950, EPI_ISL_917011, EPI_ISL_917060, EPI_ISL_917095, EPI_ISL_917121, EPI_ISL_917792 | | | |
| see above | Lighthouse Lab in Alderley Park | Wellcome Sanger Institute for the COVID-19 Genomics UK (COG-UK) Consortium | Jacquelyn Wynn, Mairead Hyland, The Lighthouse Lab in Alderley Park and Alex Alderton, Roberto Amato, Sonia Goncalves, Ewan Harrison, David K. Jackson, Ian Johnston, Dominic Kwiatkowski, Cordelia Langford, John Sillitoe on behalf of the Wellcome Sanger Institute COVID-19 Surveillance Team |
| EPI_ISL_918203, EPI_ISL_918229, EPI_ISL_918234, EPI_ISL_918256, EPI_ISL_918257, EPI_ISL_918258, EPI_ISL_918262 | Innovative Genomics Institute, UC Berkeley | Innovative Genomics Institute, UC Berkeley | Stacia Wyman, Haridha Shivram, Phil Frankino, Liana Lareau, Shana McDevitt, Justin Choi |
| EPI_ISL_918282, EPI_ISL_918283, EPI_ISL_918284, EPI_ISL_918285, EPI_ISL_918286, EPI_ISL_918287 | Hospital Universitari Vall d'Hebron - Vall d'Hebron Institut de Rercerca | Hospital Universitari Vall d'Hebron | Cristina Andrés, Maria Piñana, Josep F Abril, Damir Garcia-Cehic, Ariadna Rando, Juliana Esperalba, Maria Gema Codina, Carla Castillo, Maria Carmen Martin, Tomàs Pumarola, Josep Quer, Andrés Antón |
| EPI_ISL_918407 | ALEA Dr Kandic | Alea Genetic Centre | Rijad Konjhodzic, Dino Pecar, Lana Salihefendic |
| EPI_ISL_918550 | LACEN - Laboratório Central de Saúde Pública do Para | Evandro Chagas Institute | Santos, M.C.; Silva, A.M.; Junior, W.D.C.; Barbagelata, L.S.; Ferreira, J.A.; Sousa, E.M.A.; da Silva, P.S.; Pinheiro, K.C.; L.C.; Sousa Junior, E.C. |
| EPI_ISL_918859, EPI_ISL_918862, EPI_ISL_918863, EPI_ISL_918864, EPI_ISL_918866, EPI_ISL_918867, EPI_ISL_918868, EPI_ISL_918870, EPI_ISL_918871, EPI_ISL_918872, EPI_ISL_918873, EPI_ISL_918879, EPI_ISL_918880, EPI_ISL_918881, EPI_ISL_918914, EPI_ISL_918915, EPI_ISL_918935 | | | |
| see above | University of Birmingham | COVID-19 Genomics UK (COG-UK) Consortium | Institute of Microbiology, University of Birmingham: Claire McMurray, Joanne Stockton, Samuel Nicholls, Radoslaw Poplawski, Will Rowe, Josh Quick, Nicholas Loman. University of Birmingham Testing Laboratory: Celina M Whalley, Andrew Bosworth, Charlotte Poxon, Kasun Wanigasooriya, Oliver Pickles, Mike Kidd, Alex Richter, Andrew D Beggs PHE Heartlands Lab: Husam Osman, Andrew Bosworth. Queen Elizabeth Hospital: Anna Casey |
| EPI_ISL_919011, EPI_ISL_919012, EPI_ISL_919017, EPI_ISL_919037, EPI_ISL_919039, EPI_ISL_919041, EPI_ISL_919043, EPI_ISL_919045, EPI_ISL_919047, EPI_ISL_919049, EPI_ISL_919051, EPI_ISL_919089, EPI_ISL_919105 | | | |
| see above | Department of Pathology, University of Cambridge | COVID-19 Genomics UK (COG-UK) Consortium | Aminu S. Jahun, Yasmin Chaudhry, Iliana Georgana, Myra Hosmillo, Rhys Izu, Martin D. Curran, Surendra Parmar, Ian Goodfellow |
| EPI_ISL_919197 | West of Scotland Specialist Virology Centre, NHSGGC / MRC-University of Glasgow Centre for Virus Research | COVID-19 Genomics UK (COG-UK) Consortium | Ana da Silva Filipe, Natasha Johnson, Kathy Smollett, Daniel Mair, Stephen Carmichael, Alice Broos, Lily Tong, Jenna Nichols, Kyriaki Nomikou; Sarah McDonald; Richard Orton, Joseph Hughes, Sreenu Vattipally, David L Robertson; Alasdair MacLean, Rory Gunson; Sharif Shaaban, Matthew Holden; Rachel Blacow, Guy Mollett, Kathy Li, James Shepherd, Antonia Ho, Emma Thomson |
| EPI_ISL_919338 | Virology Department, Royal Infirmary of Edinburgh, NHS Lothian / School of Biological Sciences, University of Edinburgh / Institute of Genetics and Molecular Medicine, University of Edinburgh | COVID-19 Genomics UK (COG-UK) Consortium | McHugh M, Dewar R, Rooke S, Gallagher M, Balcaza C, O'Toole Á, Scher E, Hill V, McCrone JT, Colquhoun R, Yu X, Jackson B, Rambaut A, Williams TC, Templeton K |
| EPI_ISL_919475, EPI_ISL_919478, EPI_ISL_919515, EPI_ISL_919517, EPI_ISL_919591, EPI_ISL_919644 | Liverpool Clinical Laboratories | COVID-19 Genomics UK (COG-UK) Consortium | Sam Haldenby, Anita Lucaci, Steve Paterson, Julian Hiscox, Alistair Darby, M Almsaud, A Alrezaihi, Muhannad Alruwaili, Stuart D Armstrong, Jones Benjamin, Eleanor G Bentley, Anu Chawla, Jordan J Clark, Angela Cowell, Richard Eccles, Isabel García-Dorival, Matthew Gemmell, Alessandro Gerada, PKF Gilmore, Richard Gregory, Ximeng Han, Catherine Hartley, Margaret Hughes, Miren Iturriza-Gomara, James Johnson, L Luu, Jenifer Manson, Charlotte Nelson, Elaine O'Toole, Cassie Olateju, Rebekah Penrice-Randal , Lucille Rainbow, N.P Randle, Trevor Ian Robinson, Parul Sharma, Ghada T Shawli, James P Stewart, Neil Swainston, Ecaterina Vamos, Joanne Watts, Mark Whitehead |
| EPI_ISL_919852 | Barts Health NHS Trust | COVID-19 Genomics UK (COG-UK) Consortium | CUTINO-MOGUEL, Maria-Teresa; HARRINGTON, David; OWOYEMI, Dola; KULASEGARAN-SHYLINI, Raghavendran; BROAD, Claire; KELE, Beatrix |
| EPI_ISL_920012, EPI_ISL_920013, EPI_ISL_920015, EPI_ISL_920016, EPI_ISL_920017, EPI_ISL_920018, EPI_ISL_920019, EPI_ISL_920020, EPI_ISL_920021, EPI_ISL_920022, EPI_ISL_920023, EPI_ISL_920024, EPI_ISL_920025, EPI_ISL_920027, EPI_ISL_920028, EPI_ISL_920029, EPI_ISL_920034 | | | |
| see above | University College London, Great Ormond Street Hospital for Children NHS Foundation Trust, Imperial College Healthcare NHS Trust | COVID-19 Genomics UK (COG-UK) Consortium | Sergi Castellano, Rachel Williams, Mark Kristiansen, Paola Resende Silva, Sunando Roy, Tony Brooks, Helena Tutill, Paola Niola, Patricia Dyal, Charlotte Williams, Leysa Forrest, Yasmin Panchbhaya, Jacqueline Findlay, Samuel Weeks, Julianne Brown, Kathryn Harris, Paul Randell, James Price, Alison Holmes, Judith Breuer |
| EPI_ISL_920188, EPI_ISL_920193, EPI_ISL_920299, EPI_ISL_920309, EPI_ISL_920318, EPI_ISL_920327, EPI_ISL_920328, EPI_ISL_920330, EPI_ISL_920337, EPI_ISL_920338, EPI_ISL_920339, EPI_ISL_920353, EPI_ISL_920362, EPI_ISL_920363, EPI_ISL_920371, EPI_ISL_920466, EPI_ISL_920501 | | | |
| see above | University College London Hospital | COVID-19 Genomics UK (COG-UK) Consortium | Judith Heaney, Matthew Byott, Catherine Houlihan, Dan Frampton, Stuart Kirk, Moira Spyer and Eleni Nastouli |
| EPI_ISL_920867, EPI_ISL_920868, EPI_ISL_920869, EPI_ISL_920870, EPI_ISL_920871, EPI_ISL_920872, EPI_ISL_920873, EPI_ISL_920874, EPI_ISL_920875, EPI_ISL_920876, EPI_ISL_920877, EPI_ISL_920878, EPI_ISL_920879, EPI_ISL_920880, EPI_ISL_920881 | | | |
| see above | University College London, Great Ormond Street Hospital for Children NHS Foundation Trust, Imperial College Healthcare NHS Trust | COVID-19 Genomics UK (COG-UK) Consortium | Sergi Castellano, Rachel Williams, Mark Kristiansen, Paola Resende Silva, Sunando Roy, Tony Brooks, Helena Tutill, Paola Niola, Patricia Dyal, Charlotte Williams, Leysa Forrest, Yasmin Panchbhaya, Jacqueline Findlay, Samuel Weeks, Julianne Brown, Kathryn Harris, Paul Randell, James Price, Alison Holmes, Judith Breuer |
| EPI_ISL_920894, EPI_ISL_920895, EPI_ISL_920896, EPI_ISL_920897, EPI_ISL_920898, EPI_ISL_920899 | Department of Pathology, University of Cambridge | COVID-19 Genomics UK (COG-UK) Consortium | Aminu S. Jahun, Yasmin Chaudhry, Iliana Georgana, Myra Hosmillo, Rhys Izu, Martin D. Curran, Surendra Parmar, Ian Goodfellow |
| EPI_ISL_921007 | Regional Virus Laboratory, Belfast Health and Social Care Trust | COVID-19 Genomics UK (COG-UK) Consortium | Conall McCaughey, James McKenna, Tanya Curran, Susan Feeney, Alison Watt, Ciara Cox, Mairead Connor, Zoltan Molnar, David Simpson, Derek Fairley |
| EPI_ISL_921508, EPI_ISL_921509, EPI_ISL_921510, EPI_ISL_921511, EPI_ISL_921512, EPI_ISL_921513, EPI_ISL_921514, EPI_ISL_921516, EPI_ISL_921518, EPI_ISL_921520 | Northumbria University / South Tees Hospitals NHS Foundation Trust / North Cumbria Integrated Care NHS Foundation Trust / North Tees and Hartlepool NHS Foundation Trust / Newcastle Hospitals NHS Foundation Trust | COVID-19 Genomics UK (COG-UK) Consortium | Darren L Smith,Andrew Nelson,Matthew Bashton,Greg R Young,Joshua Loh,John Allan,Mohammad A Tariq,Giles S Holt,Gary Black,Wen C Yew,Lynn Dover,Paul Baker,Steve Liggett,Sarah Essex,Jane Greenaway,Debra Padgett,Clive Graham,Garren Scott,Edward Barton,Emma Swindells,Brendan Payne,Jennifer Collins,Yusri Taha,Gary Eltringham |
| EPI_ISL_922101, EPI_ISL_922105, EPI_ISL_922106, EPI_ISL_922108, EPI_ISL_922109, EPI_ISL_922110, EPI_ISL_922111, EPI_ISL_922112, EPI_ISL_922115, EPI_ISL_922116 | Lincolnshire Hospitals and DeepSeq Nottingham | COVID-19 Genomics UK (COG-UK) Consortium | Nichola Duckworth, Tim Sloan, Sarah Walsh, Jonathan Ball, Patrick McClure, Joeseph Chappell, Nadine Holmes, Matthew Carlisle, Christopher Moore, Fei Sang, Johnny Debebe, Victoria Wright, Matthew Loose |
| EPI_ISL_922206, EPI_ISL_922207, EPI_ISL_922211, EPI_ISL_922223, EPI_ISL_922349 | Oxford Viromics, NDM, University of Oxford; Oxford University Hospitals; Basingstoke and North Hampshire Hospital | COVID-19 Genomics UK (COG-UK) Consortium | Tanya Golubchik, David Bonsall, George Macintyre, Amy Trebes, Mariateresa de Cesare, Catrin Moore, Alex Mobbs, Anita Justice, Robert Shaw, Monique Andersson, Timothy Peto, Emma Wise, Nathan Moore, Jessica Lynch, Nick Cortes, Matilde Mori, Stephen Kidd, David Buck, John Todd, Christophe Fraser |
| EPI_ISL_922361, EPI_ISL_922364, EPI_ISL_922387, EPI_ISL_922388, EPI_ISL_922393, EPI_ISL_922397, EPI_ISL_922588, EPI_ISL_922589, EPI_ISL_922644 | Wales Specialist Virology Centre Sequencing lab: Pathogen Genomics Unit | Public Health Wales Microbiology Cardiff Wales Specialist Virology Centre | Catherine Moore, Johnathan Evans, Laura Gifford, Malorie Perry, Simon Cottrell, Angela Marchbank, Alec Birchley, Alexander Adams, Amy Gaskin, Bree Gatica-Wilcox, Jason Coombes, Joel Southgate, Lauren Gilbert, Lee Graham, Nicole Pacchiarini, Sara Kumziene-Summerhayes, Sarah Taylor, Sophie Jones, Sara Rey, Matthew Bull, Joanne Watkins, Sally Corden, Tom Connor |
| EPI_ISL_923279, EPI_ISL_923284, EPI_ISL_923293, EPI_ISL_923301, EPI_ISL_923305, EPI_ISL_923306, EPI_ISL_923605, EPI_ISL_923611, EPI_ISL_923613, EPI_ISL_923671, EPI_ISL_923678, EPI_ISL_923679 | | | |
| see above | Centre for Enzyme Innovation, University of Portsmouth / | COVID-19 Genomics UK (COG-UK) Consortium | Angela Beckett,Salman Goudarzi,Christopher Fearn,Kate Cook,Katie Loveson,Sharon Glaysher,Scott Elliott,Samuel Robson |

| | | | |
|---|---|---|---|
| | Translational Research Laboratory, Portsmouth Hospitals NHS Trust | | |
| EPI_ISL_924091, EPI_ISL_924182 | Virology Department, Sheffield Teaching Hospitals NHS Foundation Trust/Department of Infection, Immunity and Cardiovascular Disease, The Medical School, University of Sheffield | COVID-19 Genomics UK (COG-UK) Consortium | Thushan de Silva, Matthew Parker, Nikki Smith, Adri Angyal, Rebecca Brown, Luke Green, Rachel Tucker, Paul Parsons, Danielle Groves, Katie Johnson, Laura Carrilero, Alex Keeley, Dave Partridge, Matthew Wyles, Benjamin Lindsey, Mehmet Yavuz, Mohammad Raza, Cariad Evans |
| EPI_ISL_924809, EPI_ISL_924818, EPI_ISL_924832, EPI_ISL_924836, EPI_ISL_924846, EPI_ISL_924860 | Bioinformatics and Biostatistics Lab, Advanced Sequencing Facility | COVID-19 Genomics UK (COG-UK) Consortium | Aengus Stewart,Jerome Nicod,Chelsea Sawyer,Laura Cubitt,Harshil Patel,Margaret Crawford |
| EPI_ISL_925048, EPI_ISL_925049, EPI_ISL_925050, EPI_ISL_925051, EPI_ISL_925052, EPI_ISL_925053, EPI_ISL_925054, EPI_ISL_925055, EPI_ISL_925056, EPI_ISL_925057, EPI_ISL_925058, EPI_ISL_925059, EPI_ISL_925075, EPI_ISL_925076, EPI_ISL_925077 | | | |
| see above | TXDSHS | TXDSHS | Bonnie Oh, Anita Pokharel, James Daniel Bonser, Myong Koag, Chung Wang, Rachel Lee, Grace Kubin, Rashmi Tuladhar, Mayela Pedrueza, Maliha Rahman, Jenny Zhang |
| EPI_ISL_925209, EPI_ISL_925210, EPI_ISL_925211, EPI_ISL_925212, EPI_ISL_925213, EPI_ISL_925214, EPI_ISL_925215, EPI_ISL_925224, EPI_ISL_925225, EPI_ISL_925227, EPI_ISL_925253 | | | |
| see above | New Mexico Department of Health Scientific Laboratory | New Mexico Department of Health Scientific Laboratory | Ellie Johnson, Anastacia Griego-Fisher, D'eldra Malone, Jennifer Benoit |
| EPI_ISL_925258, EPI_ISL_925261, EPI_ISL_925262, EPI_ISL_925277, EPI_ISL_925282, EPI_ISL_925283, EPI_ISL_925306 | Wyoming Public Health Laboratory | Wyoming Public Health Laboratory | Noah Hull, Taylor Fearing, Lynette Gumbleton, Channing Weber, Ashley Norberg, Bailey Bowcutt, and Wanda Manley |
| EPI_ISL_925401 | Department of Clinical Microbiology | GIGA Medical Genomics | Keith Durkin, Maria Artesi, Sébastien Bontems, Raphaël Boreux, Bouchra Boujemla, Cécile Meex, Pierrette Melin, Marie-Pierre Hayette, Vincent Bours |
| EPI_ISL_925882, EPI_ISL_925884, EPI_ISL_925885, EPI_ISL_925886 | Nucleic Acid Testing, National Reference Laboratory | GIGA Medical Genomics | Yvan Butera, Keith Durkin, Maria Artesi, Bouchra Boujemla, Robert Rutayisire, Patrick Tuyisenge, Esperence Umumararungu, Sébastien Bontems, Marie-Pierre Hayette, Nathalie Renotte, Swaibu Gatare, Jacob Souopgui, Sabin Nsanzimana, Vincent Bours, Léon Mutesa |
| EPI_ISL_925950, EPI_ISL_925961, EPI_ISL_925967, EPI_ISL_925974, EPI_ISL_925976, EPI_ISL_926000, EPI_ISL_926008, EPI_ISL_926021, EPI_ISL_926035, EPI_ISL_926038, EPI_ISL_926047, EPI_ISL_926051, EPI_ISL_926063, EPI_ISL_926064, EPI_ISL_926065, EPI_ISL_926096, EPI_ISL_926103, EPI_ISL_926114, EPI_ISL_926122, EPI_ISL_926139, EPI_ISL_926144, EPI_ISL_926150, EPI_ISL_926169, EPI_ISL_926221, EPI_ISL_926226, EPI_ISL_926281, EPI_ISL_926282, EPI_ISL_926299, EPI_ISL_926313, EPI_ISL_926359, EPI_ISL_926380, EPI_ISL_926385, EPI_ISL_926394, EPI_ISL_926395, EPI_ISL_926419, EPI_ISL_926428, EPI_ISL_926436, EPI_ISL_926439, EPI_ISL_926466, EPI_ISL_926482, EPI_ISL_926491, EPI_ISL_926522, EPI_ISL_926532, EPI_ISL_926553, EPI_ISL_926559, EPI_ISL_926569, EPI_ISL_926570, EPI_ISL_926591, EPI_ISL_926598, EPI_ISL_926605, EPI_ISL_926619, EPI_ISL_926621, EPI_ISL_926633, EPI_ISL_926651, EPI_ISL_926663, EPI_ISL_926692, EPI_ISL_926713, EPI_ISL_926714, EPI_ISL_926730, EPI_ISL_926784, EPI_ISL_926786, EPI_ISL_926789, EPI_ISL_926827, EPI_ISL_926830, EPI_ISL_926893, EPI_ISL_926902, EPI_ISL_926927, EPI_ISL_926933, EPI_ISL_926945, EPI_ISL_926979, EPI_ISL_927053, EPI_ISL_927057, EPI_ISL_927058, EPI_ISL_927061, EPI_ISL_927063, EPI_ISL_927107, EPI_ISL_927128, EPI_ISL_927160, EPI_ISL_927176, EPI_ISL_927185, EPI_ISL_927190, EPI_ISL_927197, EPI_ISL_927200, EPI_ISL_927205, EPI_ISL_927207, EPI_ISL_927209, EPI_ISL_927224, EPI_ISL_927246, EPI_ISL_927257, EPI_ISL_927258, EPI_ISL_927259, EPI_ISL_927274, EPI_ISL_927287, EPI_ISL_927291, EPI_ISL_927301, EPI_ISL_927306, EPI_ISL_927309, EPI_ISL_927320, EPI_ISL_927347, EPI_ISL_927367, EPI_ISL_927370, EPI_ISL_927400, EPI_ISL_927418, EPI_ISL_927423, EPI_ISL_927425, EPI_ISL_927463, EPI_ISL_927465, EPI_ISL_927475, EPI_ISL_927485, EPI_ISL_927486, EPI_ISL_927489, EPI_ISL_927492, EPI_ISL_927507, EPI_ISL_927508, EPI_ISL_927510, EPI_ISL_927526, EPI_ISL_927538, EPI_ISL_927551, EPI_ISL_927553, EPI_ISL_927559, EPI_ISL_927570, EPI_ISL_927573, EPI_ISL_927596, EPI_ISL_927613, EPI_ISL_927622, EPI_ISL_927630, EPI_ISL_927659, EPI_ISL_927674, EPI_ISL_927675, EPI_ISL_927682, EPI_ISL_927706, EPI_ISL_927732, EPI_ISL_927739, EPI_ISL_927752, EPI_ISL_927755, EPI_ISL_927770, EPI_ISL_927782, EPI_ISL_927784, EPI_ISL_927797, EPI_ISL_927813, EPI_ISL_927826, EPI_ISL_927846, EPI_ISL_927847, EPI_ISL_927883, EPI_ISL_927901, EPI_ISL_927902, EPI_ISL_927907, EPI_ISL_927911, EPI_ISL_927934, EPI_ISL_927961, EPI_ISL_927962, EPI_ISL_927968, EPI_ISL_927991, EPI_ISL_928001, EPI_ISL_928006, EPI_ISL_928009, EPI_ISL_928014, EPI_ISL_928020, EPI_ISL_928059, EPI_ISL_928063, EPI_ISL_928097, EPI_ISL_928111, EPI_ISL_928120, EPI_ISL_928121, EPI_ISL_928123, EPI_ISL_928147, EPI_ISL_928164, EPI_ISL_928190, EPI_ISL_928202, EPI_ISL_928214, EPI_ISL_928233, EPI_ISL_928241, EPI_ISL_928248, EPI_ISL_928252, EPI_ISL_928264, EPI_ISL_928266, EPI_ISL_928272, EPI_ISL_928276, EPI_ISL_928281, EPI_ISL_928308, EPI_ISL_928323, EPI_ISL_928332, EPI_ISL_928345, EPI_ISL_928359, EPI_ISL_928361, EPI_ISL_928406, EPI_ISL_928409, EPI_ISL_928413, EPI_ISL_928415, EPI_ISL_928447, EPI_ISL_928450, EPI_ISL_928485, EPI_ISL_928489, EPI_ISL_928495, EPI_ISL_928514, EPI_ISL_928541, EPI_ISL_928552, EPI_ISL_928553, EPI_ISL_928557, EPI_ISL_928570, EPI_ISL_928601, EPI_ISL_928604, EPI_ISL_928623, EPI_ISL_928624, EPI_ISL_928626, EPI_ISL_928633, EPI_ISL_928653, EPI_ISL_928654, EPI_ISL_928661, EPI_ISL_928674, EPI_ISL_928677, EPI_ISL_928679, EPI_ISL_928700, EPI_ISL_928717, EPI_ISL_928728, EPI_ISL_928736, EPI_ISL_928738, EPI_ISL_928745, EPI_ISL_928754, EPI_ISL_928774, EPI_ISL_928789, EPI_ISL_928822, EPI_ISL_928882, EPI_ISL_928883, EPI_ISL_928893, EPI_ISL_928905, EPI_ISL_928961, EPI_ISL_929011, EPI_ISL_929043, EPI_ISL_929044, EPI_ISL_929079, EPI_ISL_929085, EPI_ISL_929092, EPI_ISL_929099, EPI_ISL_929122, EPI_ISL_929128, EPI_ISL_929129, EPI_ISL_929130, EPI_ISL_929132, EPI_ISL_929146, EPI_ISL_929149, EPI_ISL_929156, EPI_ISL_929170, EPI_ISL_929176, EPI_ISL_929184, EPI_ISL_929191, EPI_ISL_929196, EPI_ISL_929201, EPI_ISL_929231, EPI_ISL_929247, EPI_ISL_929254, EPI_ISL_929258, EPI_ISL_929282, EPI_ISL_929284, EPI_ISL_929297, EPI_ISL_929313, EPI_ISL_929318, EPI_ISL_929322, EPI_ISL_929329, EPI_ISL_929344, EPI_ISL_929385, EPI_ISL_929398, EPI_ISL_929400, EPI_ISL_929403, EPI_ISL_929421, EPI_ISL_929424, EPI_ISL_929429, EPI_ISL_929456, EPI_ISL_929526, EPI_ISL_929530, EPI_ISL_929561, EPI_ISL_929565, EPI_ISL_929572, EPI_ISL_929589, EPI_ISL_929595, EPI_ISL_929602, EPI_ISL_929611, EPI_ISL_929613, EPI_ISL_929634, EPI_ISL_929648, EPI_ISL_929669, EPI_ISL_929674, EPI_ISL_929716, EPI_ISL_929737, EPI_ISL_929749, EPI_ISL_929765, EPI_ISL_929776, EPI_ISL_929780, EPI_ISL_929784, EPI_ISL_929794, EPI_ISL_929800, EPI_ISL_929837, EPI_ISL_929848, EPI_ISL_929857, EPI_ISL_929867, EPI_ISL_929892, EPI_ISL_929894, EPI_ISL_929914, EPI_ISL_929933, EPI_ISL_929941, EPI_ISL_929942, EPI_ISL_929948, EPI_ISL_930006, EPI_ISL_930035, EPI_ISL_930037, EPI_ISL_930041, EPI_ISL_930050, EPI_ISL_930069, EPI_ISL_930095, EPI_ISL_930108, EPI_ISL_930110, EPI_ISL_930143, EPI_ISL_930156, EPI_ISL_930162, EPI_ISL_930193, EPI_ISL_930194, EPI_ISL_930216, EPI_ISL_930219, EPI_ISL_930225, EPI_ISL_930228, EPI_ISL_930233, EPI_ISL_930237, EPI_ISL_930244, EPI_ISL_930294, EPI_ISL_930299, EPI_ISL_930310, EPI_ISL_930311, EPI_ISL_930337, EPI_ISL_930345, EPI_ISL_930348, EPI_ISL_930351, EPI_ISL_930353, EPI_ISL_930367, EPI_ISL_930391, EPI_ISL_930404, EPI_ISL_930406, EPI_ISL_930433, EPI_ISL_930443, EPI_ISL_930449, EPI_ISL_930467, EPI_ISL_930490, EPI_ISL_930491, EPI_ISL_930505, EPI_ISL_930506, EPI_ISL_930514, EPI_ISL_930529, EPI_ISL_930553, EPI_ISL_930557 | | | |
| see above | Department of Virus and Microbiological Special Diagnostics, Statens Serum Institut, Copenhagen, Denmark | Aalborg University | Danish Covid-19 Genome Consortium |
| EPI_ISL_930656, EPI_ISL_930657, EPI_ISL_930658, EPI_ISL_930659, EPI_ISL_930660, EPI_ISL_930664 | Arizona State Public Health Laboratory | Arizona State Public Health Laboratory | Trung Huynh, Jessica Escobar, Katherine Fullerton, Nobuko Fukushima, Stacy White, Linda Getsinger, Victor Waddell |
| EPI_ISL_933761 | DPHL | Delaware Public Health Lab | Gregory Hovan |
| EPI_ISL_934241 | Vilnius university hospital Santaros Klinikos, Center of Laboratory Medicine | Vilnius university hospital Santaros Klinikos, Center of Laboratory Medicine | Ingrida Olendraite, Daniel Naumovas, Rimvydas Norvilas, Dovile Ezerskyte, Justinas Slikas, Gytis Dudas |
| EPI_ISL_934378 | Klinisk mikrobiologi | The Public Health Agency of Sweden | Anna-Malin Linde, Maria Lind Karlberg, Carlo Berg, Oskar Karlsson Lindsjo, Sofia Stamouli, Reza Advani, Mattias Haukland, Petra Holmstrom, Noura Walai, Petra Edquist, Mia Brytting, Anna Risberg, Karin Tegmark-Wisell |
| EPI_ISL_934452, EPI_ISL_934453, EPI_ISL_934454, EPI_ISL_934457, EPI_ISL_934459, EPI_ISL_934465, EPI_ISL_934466, EPI_ISL_934467, EPI_ISL_934468, EPI_ISL_934469, EPI_ISL_934470, EPI_ISL_934471, EPI_ISL_934472, EPI_ISL_934479, EPI_ISL_934481, EPI_ISL_934482, EPI_ISL_934483, EPI_ISL_934485, EPI_ISL_934487, EPI_ISL_934490, EPI_ISL_934491, EPI_ISL_934492, EPI_ISL_934495, EPI_ISL_934539 | | | |
| see above | Austrian Agency for Health and Food Safety (AGES) | Bergthaler laboratory, CeMM Research Center for Molecular Medicine of the Austrian Academy of Sciences | Lukas Endler, Anna Schedl, Thomas Penz, Benidekt Agerer, Maelle Le Moing, Michael Schuster, Bekir Erguner, Jan Laine, Martin Senekowitsch, Christoph Bock, Andreas Bergthaler |
| EPI_ISL_934589, EPI_ISL_934590, EPI_ISL_934591, EPI_ISL_934592 | Institute of Legal Medicine, Medical University of Innsbruck | Bergthaler laboratory, CeMM Research Center for Molecular Medicine of the Austrian Academy of Sciences | Lukas Endler, Anna Schedl, Thomas Penz, Benedikt Agerer, Maelle Le Moing, Michael Schuster, Bekir Erguner, Jan Laine, Martin Senekowitsch, Christoph Bock, Andreas Bergthaler |
| EPI_ISL_934696 | Institute for Water Quality and Resource Management, Technical University Vienna | Bergthaler laboratory, CeMM Research Center for Molecular Medicine of the Austrian Academy of Sciences | Lukas Endler, Anna Schedl, Thomas Penz, Benedikt Agerer, Maelle Le Moing, Michael Schuster, Bekir Erguner, Jan Laine, Martin Senekowitsch, Christoph Bock, Andreas Bergthaler |
| EPI_ISL_934986, EPI_ISL_934989, EPI_ISL_934995, EPI_ISL_934996, EPI_ISL_935000, EPI_ISL_935001, EPI_ISL_935002, EPI_ISL_935003, EPI_ISL_935004 | ADMED Microbiologie | Genomics and Transcriptomics, Philip Morris International | Reto Lienhard, Marie-Lise Tritten, Emmanuel Guedj, Nicolas Sierro, Rémi Dulize, David Bornand, Mehdi Auberson, Maxime Berthouzoz, Nikolai Ivanov, Manuel Peitsch |
| EPI_ISL_935199, EPI_ISL_935210, EPI_ISL_935225, EPI_ISL_935274, EPI_ISL_935275, EPI_ISL_935276, EPI_ISL_935277, EPI_ISL_935278, EPI_ISL_935279, EPI_ISL_935280, EPI_ISL_935284, EPI_ISL_935285, EPI_ISL_935286, EPI_ISL_935287, EPI_ISL_935288, EPI_ISL_935289, EPI_ISL_935290, EPI_ISL_935291, EPI_ISL_935292, EPI_ISL_935293, EPI_ISL_935294, EPI_ISL_935295, EPI_ISL_935296, EPI_ISL_935297, EPI_ISL_935298, EPI_ISL_935299, EPI_ISL_935300 | | | |
| see above | KU Leuven, Rega Institute, Clinical and Epidemiological Virology | KU Leuven, Rega Institute, Clinical and Epidemiological Virology | Tony Wawina-Bokalanga, Bert Vanmechelen, Joan Marti-Carerras, Piet Maes |
| EPI_ISL_935532 | Labo Analyses Med | National Reference Center for Viruses of Respiratory Infections, Institut Pasteur, Paris | Marion Barbet, Sylvie Behillil, Méline Bizard, Angela Brisebarre, Camille Capel, Etienne Simon-Lorière, Vincent Enouf, Maud Vanpeene, Sylvie van der Werf,Merah Kader |
| EPI_ISL_935562 | Hopital | National Reference Center for Viruses of Respiratory Infections, Institut Pasteur, Paris | Marion Barbet, Sylvie Behillil, Méline Bizard, Angela Brisebarre, Camille Capel, Etienne Simon-Lorière, Vincent Enouf, Maud Vanpeene, Sylvie van der Werf,Fiette |
| EPI_ISL_936003, EPI_ISL_936004, EPI_ISL_936009, EPI_ISL_936010, EPI_ISL_936022 | SUNY UPSTATE MEDICAL UNIVERSITY | Wadsworth Center, New York State Department of Health | Kirsten St. George, Daryl M. Lamson, Alexis Russel, Matthew Shudt, Melissa A Leisner, Jonathan Plitnick, Navjot Singh, John Kelly, Erasmus Schneider, Erica Lasek-Nesselquist |

| | | | |
|---|---|---|---|
| EPI_ISL_936214, EPI_ISL_936215, EPI_ISL_936216, EPI_ISL_936217, EPI_ISL_936218, EPI_ISL_936219, EPI_ISL_936220, EPI_ISL_936221, EPI_ISL_936222, EPI_ISL_936223, EPI_ISL_936224, EPI_ISL_936225, EPI_ISL_936226, EPI_ISL_936227, EPI_ISL_936228, EPI_ISL_936229, EPI_ISL_936230, EPI_ISL_936231, EPI_ISL_936232, EPI_ISL_936233, EPI_ISL_936234, EPI_ISL_936235, EPI_ISL_936236 | | | |
| see above | Wadsworth Center, New York State Department of Health | Wadsworth Center, New York State Department of Health | Kirsten St. George, Daryl M. Lamson, Alexis Russel, Matthew Shudt, Melissa A Leisner, Jonathan Plitnick, Navjot Singh, John Kelly, Erasmus Schneider, Erica Lasek-Nesselquist |
| EPI_ISL_936476 | DPH, Massachusetts State Public Health Lab | DPH, Massachusetts State Public Health Lab | Lang,A.S., Fink,T., Gallagher,G.R., Smole,S.C. |
| EPI_ISL_937356, EPI_ISL_937357, EPI_ISL_937358, EPI_ISL_937359, EPI_ISL_937360, EPI_ISL_937361, EPI_ISL_937363, EPI_ISL_937373, EPI_ISL_937376, EPI_ISL_937377, EPI_ISL_937379, EPI_ISL_937381, EPI_ISL_937382 | | | |
| see above | Utah Public Health Laboratory | Utah Public Health Laboratory | Erin L. Young, Kelly F. Oakeson, Tara Gallagher |
| EPI_ISL_937435, EPI_ISL_937436, EPI_ISL_937445, EPI_ISL_937446, EPI_ISL_937447, EPI_ISL_937448, EPI_ISL_937457, EPI_ISL_937458 | Maine Health and Environmental Testing Laboratory (Maine HETL) | Tewhey Lab, The Jackson Laboratory | Matluk,N., Dewey,H., Iosue,F., Barter,M., Lynch,R., Munger,H. and Tewhey,R. |
| EPI_ISL_937818 | Lighthouse Lab in Alderley Park | Wellcome Sanger Institute for the COVID-19 Genomics UK (COG-UK) Consortium | Jacquelyn Wynn, Mairead Hyland, The Lighthouse Lab in Alderley Park and Alex Alderton, Roberto Amato, Sonia Goncalves, Ewan Harrison, David K. Jackson, Ian Johnston, Dominic Kwiatkowski, Cordelia Langford, John Sillitoe on behalf of the Wellcome Sanger Institute COVID-19 Surveillance Team |
| EPI_ISL_940146 | Laboratory of Virology and Molecular Diagnostics | Institute of Public Health of Republic of North Macedonia Laboratory of Virology and Molecular Diagnostics | Maja Kuzmanovska, Golubinka Boshevska, Elizabeta Janchevska |
| EPI_ISL_940728, EPI_ISL_940729, EPI_ISL_940730, EPI_ISL_940731 | City of Milwaukee Health Department Laboratory | City of Milwaukee Health Department Laboratory | Sanjib Bhattacharyya |
| EPI_ISL_941083, EPI_ISL_941085, EPI_ISL_941090, EPI_ISL_941098, EPI_ISL_941099, EPI_ISL_941102 | Labo Analyses Med | National Reference Center for Viruses of Respiratory Infections, Institut Pasteur, Paris | Marion Barbet, Sylvie Behillil, Méline Bizard, Angela Brisebarre, Camille Capel, Etienne Simon-Lorière, Vincent Enouf, Maud Vanpeene, Sylvie van der Werf, Merah Kader |
| EPI_ISL_941341, EPI_ISL_941342, EPI_ISL_941343, EPI_ISL_941344, EPI_ISL_941345, EPI_ISL_941382, EPI_ISL_941383, EPI_ISL_941384, EPI_ISL_941387, EPI_ISL_941470, EPI_ISL_941471, EPI_ISL_941472, EPI_ISL_941537, EPI_ISL_941538, EPI_ISL_941543, EPI_ISL_941544, EPI_ISL_941545, EPI_ISL_941546, EPI_ISL_941571, EPI_ISL_941572, EPI_ISL_941573 | | | |
| see above | Instituto Nacional de Saude (INSA) | Instituto Nacional de Saude (INSA) | Borges et al |
| EPI_ISL_941647, EPI_ISL_941648, EPI_ISL_941649, EPI_ISL_941659, EPI_ISL_941662, EPI_ISL_941664, EPI_ISL_941665, EPI_ISL_941666, EPI_ISL_941688, EPI_ISL_941689, EPI_ISL_941690, EPI_ISL_941702, EPI_ISL_941703, EPI_ISL_941709, EPI_ISL_941710, EPI_ISL_941711, EPI_ISL_941726, EPI_ISL_941727, EPI_ISL_941728, EPI_ISL_941729, EPI_ISL_941730, EPI_ISL_941731, EPI_ISL_941732, EPI_ISL_941762, EPI_ISL_941765, EPI_ISL_941766, EPI_ISL_941772, EPI_ISL_941773, EPI_ISL_941774, EPI_ISL_941775, EPI_ISL_941797, EPI_ISL_941798, EPI_ISL_941799, EPI_ISL_941808, EPI_ISL_941818, EPI_ISL_941820, EPI_ISL_941821, EPI_ISL_941822, EPI_ISL_941823, EPI_ISL_941824, EPI_ISL_941825, EPI_ISL_941826, EPI_ISL_941827, EPI_ISL_941835, EPI_ISL_941836, EPI_ISL_941873, EPI_ISL_941877, EPI_ISL_941881, EPI_ISL_941882 | | | |
| see above | Instituto Nacional de Saude (INSA) and Instituto Gulbenkian de Ciencia (IGC) | Instituto Nacional de Saude (INSA) and Instituto Gulbenkian de Ciencia (IGC) | Borges et al |
| EPI_ISL_942819, EPI_ISL_942820, EPI_ISL_942821 | Gundersen Molecular Diagnostics Laboratory | Kabara Cancer Research Institute | Craig S. Richmond, Paraic A. Kenny |
| EPI_ISL_943011, EPI_ISL_943300 | Dutch COVID-19 response team | National Institute for Public Health and the Environment (RIVM) | Adam Meijer, Harry Vennema, Dirk Eggink, Jeroen Cremer, Sharon van den Brink, Bas van der Veer, AnneMarie van den Brandt, Florian Zwagemaker, Dennis Schmitz, Chantal Reusken, on behalf of the national COVID-19 response team |
| EPI_ISL_943927, EPI_ISL_943928, EPI_ISL_943933, EPI_ISL_943962 | Utah Public Health Laboratory | Utah Public Health Laboratory | Erin L. Young, Kelly F. Oakeson, Tara Gallagher |
| EPI_ISL_944203, EPI_ISL_944209, EPI_ISL_944210, EPI_ISL_944211, EPI_ISL_944212, EPI_ISL_944219, EPI_ISL_944223, EPI_ISL_944227, EPI_ISL_944322, EPI_ISL_944323, EPI_ISL_944324, EPI_ISL_944325, EPI_ISL_944326, EPI_ISL_944327, EPI_ISL_944328, EPI_ISL_944329, EPI_ISL_944330, EPI_ISL_944331, EPI_ISL_944332, EPI_ISL_944333, EPI_ISL_944334, EPI_ISL_944335, EPI_ISL_944336, EPI_ISL_944337, EPI_ISL_944338, EPI_ISL_944369, EPI_ISL_944370, EPI_ISL_944374, EPI_ISL_944398, EPI_ISL_944404, EPI_ISL_944405, EPI_ISL_944406, EPI_ISL_944411, EPI_ISL_944417, EPI_ISL_944418, EPI_ISL_944419, EPI_ISL_944423, EPI_ISL_944424, EPI_ISL_944430, EPI_ISL_944464, EPI_ISL_944465, EPI_ISL_944466, EPI_ISL_944467, EPI_ISL_944468, EPI_ISL_944469, EPI_ISL_944470, EPI_ISL_944471, EPI_ISL_944472, EPI_ISL_944473, EPI_ISL_944474, EPI_ISL_944475, EPI_ISL_944476, EPI_ISL_944477, EPI_ISL_944478, EPI_ISL_944479, EPI_ISL_944480, EPI_ISL_944481, EPI_ISL_944482, EPI_ISL_944483, EPI_ISL_944484, EPI_ISL_944485, EPI_ISL_944486, EPI_ISL_944487, EPI_ISL_944488, EPI_ISL_944489, EPI_ISL_944490, EPI_ISL_944491, EPI_ISL_944492, EPI_ISL_944493, EPI_ISL_944494, EPI_ISL_944495, EPI_ISL_944496, EPI_ISL_944497, EPI_ISL_944498, EPI_ISL_944499, EPI_ISL_944500, EPI_ISL_944501, EPI_ISL_944502, EPI_ISL_944503, EPI_ISL_944504, EPI_ISL_944505, EPI_ISL_944506, EPI_ISL_944507, EPI_ISL_944508, EPI_ISL_944509, EPI_ISL_944510, EPI_ISL_944511, EPI_ISL_944512, EPI_ISL_944513, EPI_ISL_944514, EPI_ISL_944515 | | | |
| see above | Israel Central Virology laboratory | Israel National Consortium for SARS-CoV-2 sequencing | Neta Zuckerman, Efrat Dahan Bucris, Michal Mandelboim, Dana Bar-Ilan, Oran Erster, Tzvia Mann, Omer Murik, David A. Zeevi, Assaf Rokney, Joseph Jaffe, Eva Nachum, Maya Davidovich Cohen, Ephraim Fass, Gal Zizelski Valenci, Mor Rubinstein, Efrat Rorman, Israel Nissan, Efrat Glick-Saar, Omri Nayshool, Gideon Rechavi, Ella Mendelson, Orna Mor |
| EPI_ISL_944611, EPI_ISL_944612, EPI_ISL_944613, EPI_ISL_944615, EPI_ISL_944616, EPI_ISL_944619, EPI_ISL_944627, EPI_ISL_944628, EPI_ISL_944631, EPI_ISL_944633 | Instituto Nacional de Medicina Genomica | Instituto Nacional de Medicina Genomica | Hidalgo-Miranda A, Mendoza-Vargas A, Reyes-Grajeda JP, Cisneros-Villanueva M, Cedro-Tanda A,Peñaloza-Figueroa F, Herrera-Montalvo LA |
| EPI_ISL_944660, EPI_ISL_944722 | Department of Biochemistry, Cell and Molecular Biology, West African Centre for Cell Biology of Infectious Pathogens (WACCBIP), University of Ghana | Department of Biochemistry, Cell and Molecular Biology, West African Centre for Cell Biology of Infectious Pathogens (WACCBIP), University of Ghana | Morang'a,C.M., Ngoi,J.M., Quansah,E.B., Saiid,S., Amuzu,D.S., Asante,I., Bonney,J.H., Bonney,E., Odoom,J.K., Ndam,N.T., Tei-Maya,F., Adusei-Poku,M., Ofori-Boadu,L., Ampofo,W.K., Amenga-Etego,L.N., Quashie,P., Bediako,Y., Awandare,G.A. |
| EPI_ISL_946701, EPI_ISL_946780 | Lighthouse Lab in Alderley Park | Wellcome Sanger Institute for the COVID-19 Genomics UK (COG-UK) Consortium | Jacquelyn Wynn, Mairead Hyland, The Lighthouse Lab in Alderley Park and Alex Alderton, Roberto Amato, Sonia Goncalves, Ewan Harrison, David K. Jackson, Ian Johnston, Dominic Kwiatkowski, Cordelia Langford, John Sillitoe on behalf of the Wellcome Sanger Institute COVID-19 Surveillance Team |
| EPI_ISL_949093 | Jessa | Jessa | Jessa_cmdLab |
| EPI_ISL_949750, EPI_ISL_949768, EPI_ISL_949776, EPI_ISL_949779, EPI_ISL_949782 | Barts Health NHS Trust | COVID-19 Genomics UK (COG-UK) Consortium | CUTINO-MOGUEL, Maria-Teresa; HARRINGTON, David; OWOYEMI, Dola; KULASEGARAN-SHYLINI, Raghavendran; BROAD, Claire; KELE, Beatrix |
| EPI_ISL_949790, EPI_ISL_950162, EPI_ISL_950163, EPI_ISL_950164 | University College London, Great Ormond Street Hospital for Children NHS Foundation Trust, Imperial College Healthcare NHS Trust | COVID-19 Genomics UK (COG-UK) Consortium | Sergi Castellano, Rachel Williams, Mark Kristiansen, Paola Resende Silva, Sunando Roy, Tony Brooks, Helena Tutill, Paola Niola, Patricia Dyal, Charlotte Williams, Leysa Forrest, Yasmin Panchbhaya, Jacqueline Findlay, Samuel Weeks, Julianne Brown, Kathryn Harris, Paul Randell, James Price, Alison Holmes, Judith Breuer |
| EPI_ISL_950280, EPI_ISL_950282, EPI_ISL_950283, EPI_ISL_950284, EPI_ISL_950285, EPI_ISL_950289, EPI_ISL_950290, EPI_ISL_950291, EPI_ISL_950292 | Northumbria University / South Tees Hospitals NHS Foundation Trust / North Cumbria Integrated Care NHS Foundation Trust / North Tees and Hartlepool NHS Foundation Trust / Newcastle Hospitals NHS Foundation Trust | COVID-19 Genomics UK (COG-UK) Consortium | Darren L Smith,Andrew Nelson,Matthew Bashton,Greg R Young,Joshua Loh,John Allan,Mohammad A Tariq,Giles S Holt,Gary Black,Wen C Yew,Lynn Dover,Paul Baker,Steve Liggett,Sarah Essex,Jane Greenaway,Debra Padgett,Clive Graham,Garren Scott,Edward Barton,Emma Swindells,Brendan Payne,Jennifer Collins,Yusri Taha,Gary Eltringham |
| EPI_ISL_951274, EPI_ISL_951290, EPI_ISL_951291, EPI_ISL_951293, EPI_ISL_951297, EPI_ISL_951304, EPI_ISL_951307, EPI_ISL_951317, EPI_ISL_951362, EPI_ISL_951363, EPI_ISL_951376, EPI_ISL_951377, EPI_ISL_951412, EPI_ISL_951413, EPI_ISL_951415, EPI_ISL_951417, EPI_ISL_951439 | | | |
| see above | Oxford Viromics, NDM, University of Oxford; Oxford University Hospitals; Basingstoke and North Hampshire Hospital | COVID-19 Genomics UK (COG-UK) Consortium | Tanya Golubchik, David Bonsall, George Macintyre, Amy Trebes, Mariateresa de Cesare, Catrin Moore, Alex Mobbs, Anita Justice, Robert Shaw, Monique Andersson, Timothy Peto, Emma Wise, Nathan Moore, Jessica Lynch, Nick Cortes, Matilde Mori, Stephen Kidd, David Buck, John Todd, Christophe Fraser |
| EPI_ISL_952421, EPI_ISL_952434, EPI_ISL_952559, EPI_ISL_952561, EPI_ISL_952563, EPI_ISL_952565, EPI_ISL_952566, EPI_ISL_952567, EPI_ISL_952568, EPI_ISL_952777, EPI_ISL_952790, EPI_ISL_952792, EPI_ISL_952796, EPI_ISL_952797, EPI_ISL_952801, EPI_ISL_952803 | | | |
| see above | Centre for Enzyme Innovation, University of Portsmouth / Translational Research Laboratory, Portsmouth Hospitals NHS Trust | COVID-19 Genomics UK (COG-UK) Consortium | Angela Beckett,Salman Goudarzi,Christopher Fearn,Kate Cook,Katie Loveson,Sharon Glaysher,Scott Elliott,Samuel Robson |

| | | | |
|---|---|---|---|
| EPI_ISL_953493 | University Hospitals of Geneva, Laboratory of Virology | HUG, Laboratory of Virology and the Health2030 Genome Center | Samuel Cordey, Ana Rita Goncalves, Laurent Kaiser, Lorenzo Cerutti, Henri Pegeot, Melyssa Elies, Deborah Penet, Keith Harshman, Ioannis Xenarios, Emmanouil Dermitzakis |
| EPI_ISL_953952 | Outre Mer | National Reference Center for Viruses of Respiratory Infections, Institut Pasteur, Paris | Marion Barbet, Sylvie Behillil, Méline Bizard, Angela Brisebarre, Camille Capel, Etienne Simon-Lorière, Vincent Enouf, Maud Vanpeene, Sylvie van der Werf,Rousset Dominique |
| EPI_ISL_954126 | Hopital | National Reference Center for Viruses of Respiratory Infections, Institut Pasteur, Paris | Marion Barbet, Sylvie Behillil, Méline Bizard, Angela Brisebarre, Camille Capel, Etienne Simon-Lorière, Vincent Enouf, Maud Vanpeene, Sylvie van der Werf,Lesimple BéAtrice |
| EPI_ISL_954127 | Labo Analyses Med | National Reference Center for Viruses of Respiratory Infections, Institut Pasteur, Paris | Marion Barbet, Sylvie Behillil, Méline Bizard, Angela Brisebarre, Camille Capel, Etienne Simon-Lorière, Vincent Enouf, Maud Vanpeene, Sylvie van der Werf,Takoudju Eve-Marie |
| EPI_ISL_954784 | Complejo Hospitalario de Navarra | Instituto de Salud Carlos III | Iglesias-Caballero, M. Camarero, S. Sandonís,V. Vázquez, S. Pozo, F. Casas, I. Jiménez, P. Zaballos, A. Monzón, S. Varona, S. Cuesta, I. Ezpeleta,C. |
| EPI_ISL_954788 | Servicio Murciano de Salud | Instituto de Salud Carlos III | Iglesias-Caballero, M. Camarero, S. Sandonís,V. Vázquez, S. Pozo, F. Casas, I. Jiménez, P. Zaballos, A. Monzón, S. Varona, S. Cuesta, I. Blázquez, A. |
| EPI_ISL_954842, EPI_ISL_954931, EPI_ISL_954932, EPI_ISL_954933, EPI_ISL_954934, EPI_ISL_954935 | Colorado Department of Public Health and Environment | Colorado Department of Puplic Health and Environment | Laura Bankers, Molly C. Hetherington-Rauth, Diana Ir, Shannon Ely, Shannon R. Matzinger, Sarah Elizabeth Totten, Emily A. Travanty |
| EPI_ISL_955227, EPI_ISL_955228 | Indiana Animal Disease Diagnostic Laboratory | Carpi Laboratory - Purdue University | Jack Dorman, Ilinca I Ciubotariu, Lev Gorenstein, Abebe A Fola, G Kenitra Hendrix, Rebecca P Wilkes, Giovanna Carpi |
| EPI_ISL_956299 | Laboratorio de salud publica Arauca | Instituto Nacional de Salud- Dirección de Investigación en Salud Pública | Katherine Laiton-Donato, Diego A. Álvarez-Díaz, Carlos Franco-Muñoz, Mauricio Pacheco-Montealegre, Hector Alejandro Ruiz-Moreno, Maria T. Herrera-Sepúlveda, Diego Andrés Prada, Jhonnatan Reales-González, Sheryll Corchuelo, Julian Naizaque, Gerardo Santamaría, Magdalena Wiesner, Martha Lucia Ospina Martinez, Marcela Mercado-Reyes |
| EPI_ISL_956314 | Paru Hospital | Institute of Tropical Disease, Universitas Airlangga | Rima R Prasetya, Krisnoadi Rahardjo, Aldise M Nastri, Jezzy R Dewantari, Dyah Retno, Gatot Soegiarto, Laksmi Wulandari, Resti Yudhawati, Yasuko Mori, Soetjipto, Kazufumi Shimizu, Maria I Lusida |
| EPI_ISL_956335 | Utah Public Health Laboratory, Utah Public Health Laboratory Infectious Disease submission group | Utah Public Health Laboratory, Utah Public Health Laboratory Infectious Disease submission group | Gallagher,T., Young,E.L., Oakeson,K.F. |
| EPI_ISL_959273 | Lighthouse Lab in Milton Keynes | Wellcome Sanger Institute for the COVID-19 Genomics UK (COG-UK) Consortium | The Lighthouse Lab in Milton Keynes and Alex Alderton, Roberto Amato, Sonia Goncalves, Ewan Harrison, David K. Jackson, Ian Johnston, Dominic Kwiatkowski, Cordelia Langford, John Sillitoe on behalf of the Wellcome Sanger Institute COVID-19 Surveillance Team |
| EPI_ISL_959286 | General Hospital - Strumica | Research Center for Genetic Engineering and Biotechnology "Georgi D. Efremov" , Macedonian Academy of Sciences and Arts | Aleksandar J. Dimovski, Dijana Plasheska-Karanfilska, Predrag Noveski, Gjorgji Bozinovski, Milena Jakimovska |
| EPI_ISL_959288 | General Hospital - Shtip | Research Center for Genetic Engineering and Biotechnology "Georgi D. Efremov" , Macedonian Academy of Sciences and Arts | Aleksandar J. Dimovski, Dijana Plasheska-Karanfilska, Predrag Noveski, Gjorgji Bozinovski, Milena Jakimovska |
| EPI_ISL_959289, EPI_ISL_959291 | General Hospital - Ohrid | Research Center for Genetic Engineering and Biotechnology "Georgi D. Efremov" , Macedonian Academy of Sciences and Arts | Aleksandar J. Dimovski, Dijana Plasheska-Karanfilska, Predrag Noveski, Gjorgji Bozinovski, Milena Jakimovska |
| EPI_ISL_959850 | National Virus Reference Laboratory | National Virus Reference Laboratory | Michael Carr, Gabriel Gonzalez, Jonathan Dean, Cillian F De Gascun |
| EPI_ISL_960405, EPI_ISL_960431, EPI_ISL_960440 | The National Institute of Public Health | State Veterinary Institute Prague | Nagy,A;Vecerova,J;Cernikova,L;Stara,M;Jirincova,H;Trnka,D |
| EPI_ISL_960470, EPI_ISL_960572 | Istituto Zooprofilattico Sperimentale del Mezzogiorno | TIGEM | Patrizia Annunziata, Andrea Ballabio, Valentina Bouche, Davide Cacchiarelli, Pellegrino Cerino, Chiara Colantuono, Maria Concetta Cuomo, Denise Di Concilio, Lucio Di Filippo, Antonio Grimaldi, Antonio Limone, Anna Manfredi, Francesco Panariello, Biancamaria Pierri, Marcello Salvi |
| EPI_ISL_960877, EPI_ISL_960878, EPI_ISL_960879 | Institute of Medical Microbiology and Hospital Hygiene | Institute of Medical Microbiology and Hospital Hygiene | Prof. Dr. Achim Kaasch, Aljoscha Tersteegen |
| EPI_ISL_960996 | AIID | Irish Coronavirus Sequencing Consortium-Teagasc Grange | Matthew McCabe, Aljandro Abner Garcia Leon, Fiona Crispie, Calum Walsh, Michael Carr, John Kenny, Paul Cotter, Patrick Mallon, Gabriel Gonzalez |
| EPI_ISL_961129, EPI_ISL_961130 | Delaware Public Health Laboratory | Delaware Public Health Lab | Gregory Hovan |
| EPI_ISL_961381 | Toronto Invasive Bacterial Diseases Network | McMaster University | Allison McGeer, Patryk Aftanas, Hooman Derakhshani, Angel Li, Kuganya Nirmalarajah, Emily Panousis, Ahmed Draia, Jalees Nasir, Michael Surette, Samira Mubareka, Andrew G. McArthur |
| EPI_ISL_961642 | Hôpital Georges L. Dumont | National Microbiology Laboratory (NML) | Anna Majer, Shari Tyson, Grace Seo, Philip Mabon, Elsie Grudeski, Rhiannon Huzarewich, Russell Mandes, Anneliese Landgraff, Jennifer Tanner, Natalie Knox, Morag Graham, Gary Van Domselaar, Richard Garceau, Guillaume Desnoyers, Nathalie Bastien, Yan Li, Timothy Booth, Darian Hole, Madison Chapel, Kirsten Biggar, CanCOGeN's metadata curation team, Public Health Agency of Canada CanCOGeN team |
| EPI_ISL_961877, EPI_ISL_961878, EPI_ISL_961879, EPI_ISL_961880 | E. Gulbja laboratorija | Latvian Biomedical Research and Study Centre | Janis Pjalkovskis, Nikita Zrelovs, Monta Ustinova, Ivars Silamikelis, Liga Birzniece, Kaspars Megnis, Vita Rovite, Lauma Freimane, Laila Silamikele, Laura Ansone, Davids Fridmanis, Mikus Gavars, Dmitrijs Perminovs, Jurijs Perevoscikovs, Uga Dumpis, Janis Klovins |
| EPI_ISL_961905, EPI_ISL_961906, EPI_ISL_961907, EPI_ISL_961933, EPI_ISL_961940, EPI_ISL_961944, EPI_ISL_961970, EPI_ISL_962010, EPI_ISL_962011, EPI_ISL_962171, EPI_ISL_962173 | | | |
| see above | Illinois Department of Public Health | Gagnon Lab, Southern Illinois University | Keith Gagnon |
| EPI_ISL_962182 | Ohio Department of Health Laboratory | Ohio Department of Health Laboratory | Holmes, Jennifer; Eric Brandt, Keoni Omura, Glen McGillivary, Caitlin McDonnell, Jade Mowery, Stephanie Mccracken, Tyler Payne, Kirtana Ramadugu, Erica Leasure, Brent Lee, Kelsey Florek, Heather Blankenship, Quanta Brown, and Tammy Bannerman |
| EPI_ISL_962206 | Institute for Medical Research, Infectious Disease Research Centre, National Institutes of Health, Ministry of Health Malaysia | Institute for Medical Research, Infectious Disease Research Centre, National Institutes of Health, Ministry of Health Malaysia | Suppiah J, Kamel K, Azizan MA, Thayan R |
| EPI_ISL_962515 | MD Laboratories | Los Angeles County PHL | P. Hemarajata et al. |
| EPI_ISL_962909, EPI_ISL_962910 | Norwegian Institute of Public Health, Department of Virology | Norwegian Institute of Public Health, Department of Virology | Kathrine Stene-Johansen, Kamilla Heddeland Instefjord, Hilde Elshaug, Ignacio Garcia Llorente, Serina B Engebretsen, Atiya R Ali,Marie Paulsen Madsen, Rasmus Riis Kopperud, Hilde Vollan, Karoline Bragstad, Olav Hungnes |
| EPI_ISL_962924, EPI_ISL_962925, EPI_ISL_962926, EPI_ISL_962927 | Servicio de Microbiología, Laboratori Clínic Metropolitana Nord. Hospital Universitari Germans Trias i Pujol. Institut d'Investigació en Ciències de la Salut Germans Trias i Pujol (IGTP) | SeqCOVID-SPAIN consortium/IBV(CSIC) | Elisa Martró, Antoni E. Bordoy, Anna Not, Adrián Antuori, Anabel Fernández, Nona Romaní, Verónica Saludes, Cristina Casañ and SeqCOVID-SPAIN consortium |
| EPI_ISL_962939 | Hospital Universitario de Gran Canaria Dr. Negrín | SeqCOVID-SPAIN consortium/IBV(CSIC) | M. Carmen Pérez González, Francisco J. Chamizo López, Ana Bordes Benítez and SeqCOVID-SPAIN consortium |
| EPI_ISL_963061, EPI_ISL_963296, EPI_ISL_963955 | Lighthouse Lab in Alderley Park | Wellcome Sanger Institute for the COVID-19 Genomics UK (COG-UK) Consortium | Jacquelyn Wynn, Mairead Hyland, The Lighthouse Lab in Alderley Park and Alex Alderton, Roberto Amato, Sonia Goncalves, Ewan Harrison, David K. Jackson, Ian Johnston, Dominic Kwiatkowski, Cordelia Langford, John Sillitoe on behalf of the Wellcome Sanger Institute COVID-19 Surveillance Team |
| EPI_ISL_964278 | Oslo University Hospital, Department of Medical Microbiology | Norwegian Institute of Public Health, Department of Virology | Kathrine Stene-Johansen, Kamilla Heddeland Instefjord, Hilde Elshaug, Ignacio Garcia Llorente, Serina B Engebretsen, Atiya R Ali, Marie Paulsen Madsen, Rasmus Riis Kopperud, Hilde Vollan, Karoline Bragstad, Olav Hungnes |
| EPI_ISL_964882 | General Hospital "Abdulah Nakas" Sarajevo | Alea Genetic Centre | Lana Salihefendic, Dino Pecar, Adis Kandic, Sead Jazic, Rijad Konjhodzic |
| EPI_ISL_965008 | Nordland Hospital - Bodo, Laboratory Department, Molecular Biology Unit | Norwegian Institute of Public Health, Department of Virology | Kathrine Stene-Johansen, Kamilla Heddeland Instefjord, Hilde Elshaug, Ignacio Garcia Llorente, Serina B Engebretsen, Atiya R Ali, Marie Paulsen Madsen, Rasmus Riis Kopperud, Hilde Vollan, Karoline Bragstad, Olav Hungnes |

| | | | |
|---|---|---|---|
| EPI_ISL_965113, EPI_ISL_965121 | Ohio Department of Health Laboratory | Ohio Department of Health Laboratory | Holmes, Jennifer; Eric Brandt, Keoni Omura, Glen McGillivary, Caitlin McDonnell, Jade Mowery, Stephanie Mccracken, Tyler Payne, Kirtana Ramadugu, Erica Leasure, Brent Lee, Kelsey Florek, Heather Blankenship, Quanta Brown, and Tammy Bannerman |
| EPI_ISL_965129 | Sant'Eugenio/CTO ASL Roma 2. | INMI Lazzaro Spallanzani IRCCS | F Messina, C.E.M Gruber, B Bartolini, E Giombini, M Rueca, O Butera, F Bondanini, GC Cocciolillo, C Disegni, A Di Caro, MR Capobianchi |
| EPI_ISL_965130 | Ospedale Sandro Pertini ASL Roma2 | INMI Lazzaro Spallanzani IRCCS | B Bartolini, O Butera, C.E.M Gruber, M Rueca, F Messina, E Giombini, R Longo, V Michela, MR Capobianchi, A Di Caro |
| EPI_ISL_965132 | Azienda Ospedaliera San Giovanni Addolorata | INMI Lazzaro Spallanzani IRCCS | CEM Gruber, B Bartolini, E Giombini, M Rueca, O Butera, F Messina, M Gaudio, PM Placanica, A Di Caro, MR Capobianchi |
| EPI_ISL_965137 | Fondazione Policlinico Universitario "A. Gemelli" IRCCS | INMI Lazzaro Spallanzani IRCCS | E Giombini, M. Rueca, B Bartolini, O Butera, C.E.M Gruber, F Messina, P Cattani,M Sanguinetti, A Di Caro, MR Capobianchi |
| EPI_ISL_965143, EPI_ISL_965149, EPI_ISL_965301, EPI_ISL_965302, EPI_ISL_965813 | Ohio Department of Health Laboratory | Ohio Department of Health Laboratory | Holmes, Jennifer; Eric Brandt, Keoni Omura, Glen McGillivary, Caitlin McDonnell, Jade Mowery, Stephanie Mccracken, Tyler Payne, Kirtana Ramadugu, Erica Leasure, Brent Lee, Kelsey Florek, Heather Blankenship, Quanta Brown, and Tammy Bannerman |
| EPI_ISL_965890, EPI_ISL_965891, EPI_ISL_965892, EPI_ISL_965893, EPI_ISL_965894 | Massachusetts State Public Health Laboratory | Massachusetts State Public Health Laboratory | Andrew Lang, Timelia Fink, Glen Gallagher, Sandra Smole |
| EPI_ISL_965952 | Consejería de Sanidad y Asuntos Sociales de Castilla La Mancha | Instituto de Salud Carlos III | Iglesias-Caballero, M. Sandonís,V. Vázquez, S. Camarero, S. Pozo, F. Casas, I. Jiménez, P. Zaballos, A. Monzón, S. Varona, S. Cuesta, I. Gutiérrez, G. |
| EPI_ISL_966547, EPI_ISL_966548, EPI_ISL_967186, EPI_ISL_967187, EPI_ISL_967188, EPI_ISL_967189, EPI_ISL_967190, EPI_ISL_967191, EPI_ISL_967216, EPI_ISL_967221, EPI_ISL_967222, EPI_ISL_967223, EPI_ISL_967224, EPI_ISL_967225, EPI_ISL_967226, EPI_ISL_967227, EPI_ISL_967228, EPI_ISL_967229, EPI_ISL_967230, EPI_ISL_967231, EPI_ISL_967232, EPI_ISL_967233, EPI_ISL_967234, EPI_ISL_967235, EPI_ISL_967236, EPI_ISL_967237, EPI_ISL_967238, EPI_ISL_967239, EPI_ISL_967240, EPI_ISL_967241, EPI_ISL_967242 | | | |
| see above | Helix/Illumina | Respiratory Viruses Branch, Division of Viral Diseases, Centers for Disease Control and Prevention | Peter W. Cook,Dakota Howard,Dhwani Batra,Ben L. Rambo-Martin,Eileen de Feo,Jan Antico,Christine Tran,Matthew Tolentino,Shannon Wickline,Kim Gietzen,Brad Sickler,Jingtao Liu,Eric Allen,Phil Febbo,Summer Galloway,Nicole L. Washington,Simon White,Geraint Levan,Kelly Schiabor Barrett,Elizabeth Cirulli,Alexandre Bolze,Ary Ascencio,Charlotte Rivera-Garcia,Ryan Cho,Jason Nguyen,Sherry Wang,Jimmy Ramirez,Tyler Cassens,Efren Sandoval,Magnus Isaksson,William Lee,David Becker,Marc Laurent,James Lu,Clinton R. Paden,Suxiang Tong,Duncan MacCannell, |
| EPI_ISL_967683, EPI_ISL_967713, EPI_ISL_967714, EPI_ISL_967744 | State Laboratories Division, Hawaii State Department of Health | State Laboratories Division, Hawaii State Department of Health | Pamela O'Brien, Drew Kuwazaki, Ayana Garnet, Razvan Sultana, Edward Desmond |
| EPI_ISL_967897, EPI_ISL_967913 | TGen North | TGen North | "Jolene Bowers, Megan Folkerts, Chris French, Hayley Yaglom, Ashlyn Pfeiffer, Darrin Lemmer, Dave Engelthaler, The Arizona COVID Genomics Union (ACGU)" |
| EPI_ISL_967916 | TGen North | Sonora Quest Laboratories | "Jolene Bowers, Megan Folkerts, Chris French, Hayley Yaglom, Ashlyn Pfeiffer, Darrin Lemmer, Dave Engelthaler, The Arizona COVID Genomics Union (ACGU)" |
| EPI_ISL_967928, EPI_ISL_967936 | TGen North | TGen North | "Jolene Bowers, Megan Folkerts, Chris French, Hayley Yaglom, Ashlyn Pfeiffer, Darrin Lemmer, Dave Engelthaler, The Arizona COVID Genomics Union (ACGU)" |
| EPI_ISL_967957, EPI_ISL_967964 | TGen North | Sonora Quest Laboratories | "Jolene Bowers, Megan Folkerts, Chris French, Hayley Yaglom, Ashlyn Pfeiffer, Darrin Lemmer, Dave Engelthaler, The Arizona COVID Genomics Union (ACGU)" |
| EPI_ISL_967966, EPI_ISL_967976, EPI_ISL_967977, EPI_ISL_967982, EPI_ISL_967983, EPI_ISL_967990, EPI_ISL_968002, EPI_ISL_968017, EPI_ISL_968020 | TGen North | TGen North | "Jolene Bowers, Megan Folkerts, Chris French, Hayley Yaglom, Ashlyn Pfeiffer, Darrin Lemmer, Dave Engelthaler, The Arizona COVID Genomics Union (ACGU)" |
| EPI_ISL_968060 | TGen North | Sonora Quest Laboratories | "Jolene Bowers, Megan Folkerts, Chris French, Hayley Yaglom, Ashlyn Pfeiffer, Darrin Lemmer, Dave Engelthaler, The Arizona COVID Genomics Union (ACGU)" |
| EPI_ISL_968063 | TGen North | TGen North | "Jolene Bowers, Megan Folkerts, Chris French, Hayley Yaglom, Ashlyn Pfeiffer, Darrin Lemmer, Dave Engelthaler, The Arizona COVID Genomics Union (ACGU)" |
| EPI_ISL_970829, EPI_ISL_970846, EPI_ISL_970936, EPI_ISL_970992, EPI_ISL_971028, EPI_ISL_971030, EPI_ISL_971065, EPI_ISL_971133, EPI_ISL_971145, EPI_ISL_971155, EPI_ISL_971519, EPI_ISL_971586, EPI_ISL_971610, EPI_ISL_971695, EPI_ISL_971699, EPI_ISL_971791, EPI_ISL_971793, EPI_ISL_971911, EPI_ISL_971919, EPI_ISL_971949, EPI_ISL_971977, EPI_ISL_972044, EPI_ISL_972055, EPI_ISL_972091, EPI_ISL_972094, EPI_ISL_972098, EPI_ISL_972109, EPI_ISL_972178, EPI_ISL_972216, EPI_ISL_972218, EPI_ISL_972221, EPI_ISL_972226, EPI_ISL_972240, EPI_ISL_972270, EPI_ISL_972277, EPI_ISL_972290, EPI_ISL_972291, EPI_ISL_972353, EPI_ISL_972382, EPI_ISL_972392, EPI_ISL_972399, EPI_ISL_972413, EPI_ISL_972442, EPI_ISL_972484, EPI_ISL_972496, EPI_ISL_972502, EPI_ISL_972522, EPI_ISL_972554, EPI_ISL_972557, EPI_ISL_972607, EPI_ISL_972633, EPI_ISL_972635, EPI_ISL_972671, EPI_ISL_972688, EPI_ISL_972709, EPI_ISL_972734, EPI_ISL_972875, EPI_ISL_972885, EPI_ISL_972899, EPI_ISL_972913, EPI_ISL_972933, EPI_ISL_972944, EPI_ISL_972950, EPI_ISL_972969, EPI_ISL_973077, EPI_ISL_973138, EPI_ISL_973157, EPI_ISL_973175, EPI_ISL_973180, EPI_ISL_973197, EPI_ISL_973227, EPI_ISL_973239, EPI_ISL_973265, EPI_ISL_973298, EPI_ISL_973339, EPI_ISL_973350, EPI_ISL_973351, EPI_ISL_973379, EPI_ISL_973389, EPI_ISL_973397, EPI_ISL_973407, EPI_ISL_973420, EPI_ISL_973438, EPI_ISL_973535, EPI_ISL_973614, EPI_ISL_973666, EPI_ISL_973676, EPI_ISL_973752 | | | |
| see above | Department of Virus and Microbiological Special Diagnostics, Statens Serum Institut, Copenhagen, Denmark | Aalborg University | Danish Covid-19 Genome Consortium |
| EPI_ISL_974150 | Laboratorio de Virología / Hospital Universitario Central de Asturias (HUCA) | Laboratorio de Virología / Hospital Universitario Central de Asturias (HUCA) | Castelló C, Gómez de Oña J, Boga JA, Rojo S, Alvarez-Arguelles ME, Abreu F, Costales I, Sandoval M, Perez-Martínez Z, Martín-Rodríguez G, Coto E, Melón S |
| EPI_ISL_977117 | Massachusetts General Hospital | Infectious Disease Program, Broad Institute of Harvard and MIT | Lemieux,J.E., Siddle,K.J., Shaw,B., Adams,G., Pierce,V., Turbett,S., Anahtar,M., Branda,J., Slater,D., Harris,J., Lin,A.E., Gladden-Young,A., Lagerborg,K., Rudy,M., DeRuff,K., Carter,A., Normandin,E., Bauer,M., Reilly,S., Tomkins-Tinch,C., Loreth,C., Chaluvadi,S., Neumann,A., Cusick,C., Chapman,S.B., Gnirke,A., Flowers,K., Cerrato,F., Birren,B.W., Gallagher,G., Smole,S., Park,D.J., MacInnis,B.L., Ryan,E., LaRocque,R., Rosenberg,E. and Sabeti,P.C. |
| EPI_ISL_977120 | Flow Health | Infectious Disease Program, Broad Institute of Harvard and MIT | Lemieux,J.E., Siddle,K.J., Adams,G., Gladden-Young,A., Lagerborg,K., Rudy,M., DeRuff,K., Carter,A., Normandin,E., Bauer,M., Reilly,S., Tomkins-Tinch,C., Loreth,C., Chaluvadi,S., Birren,B.W., Gallagher,G., Smole,S., Park,D.J., MacInnis,B.L., and Sabeti,P.C. |
| EPI_ISL_977278, EPI_ISL_977292, EPI_ISL_977346 | University of Zambia, School of Veterinary Medicine | UNZAVET and PATH | Mulenga Mwenda-Chimfwembe, Ngonda Saasa, Daniel Bridges |
| EPI_ISL_977543, EPI_ISL_977551, EPI_ISL_977557, EPI_ISL_977558, EPI_ISL_977559 | Nigeria Centre of Disease Control (NCDC) | African Centre of Excellence for Genomics of Infectious Diseases (ACEGID), Redeemer's University | Olawoye I. B. et al |
| EPI_ISL_977664 | Hopital | National Reference Center for Viruses of Respiratory Infections, Institut Pasteur, Paris | Marion Barbet, Sylvie Behillil, Méline Bizard, Angela Brisebarre, Camille Capel, Etienne Simon-Lorière, Vincent Enouf, Maud Vanpeene, Sylvie van der Werf,Combe Patrice |
| EPI_ISL_978294, EPI_ISL_978295, EPI_ISL_978296, EPI_ISL_978297, EPI_ISL_978298, EPI_ISL_978299, EPI_ISL_978347, EPI_ISL_978348, EPI_ISL_978349 | Texas Department of State Health Services | Texas Department of State Health Services | Bonnie Oh, Anita Pokharel, James Daniel Bonser, Myong Koag, Chung Wang, Rachel Lee, Grace Kubin, Rashmi Tuladhar, Mayela Pedrueza, Maliha Rahman, Jenny Zhang |
| EPI_ISL_978356, EPI_ISL_978363, EPI_ISL_978364, EPI_ISL_978365 | Arizona State Public Health Laboratory | Arizona State Public Health Laboratory | Trung Huynh, Jessica Escobar, Katherine Fullerton, Nobuko Fukushima, Stacy White, Linda Getsinger, Victor Waddell |
| EPI_ISL_978622, EPI_ISL_978623, EPI_ISL_978624 | Helix/Illumina | Respiratory Viruses Branch, Division of Viral Diseases, Centers for Disease Control and Prevention | Peter W. Cook,Dakota Howard,Dhwani Batra,Ben L. Rambo-Martin,Eileen de Feo,Jan Antico,Christine Tran,Matthew Tolentino,Shannon Wickline,Kim Gietzen,Brad Sickler,Jingtao Liu,Eric Allen,Phil Febbo,Summer Galloway,Nicole L. Washington,Simon White,Geraint Levan,Kelly Schiabor Barrett,Elizabeth Cirulli,Alexandre Bolze,Ary Ascencio,Charlotte Rivera-Garcia,Ryan Cho,Jason Nguyen,Sherry Wang,Jimmy Ramirez,Tyler Cassens,Efren Sandoval,Magnus Isaksson,William Lee,David Becker,Marc Laurent,James Lu,Clinton R. Paden,Suxiang Tong,Duncan MacCannell, |
| EPI_ISL_979084, EPI_ISL_979086, EPI_ISL_979087, EPI_ISL_979088, | Santa Clara County Public Health Laboratory | Chan-Zuckerberg Biohub | CZB Cliahub Consortium |

| | | | |
|---|---|---|---|
| EPI_ISL_979090, EPI_ISL_979091 | | | |
| EPI_ISL_979254, EPI_ISL_979255, EPI_ISL_979256, EPI_ISL_979257, EPI_ISL_979258, EPI_ISL_979259 | Institute of Microbiology and Immunology, Faculty of Medicine, University of Ljubljana | Institute of Microbiology and Immunology, Faculty of Medicine, University of Ljubljana | Samo Zakotnik, Tomaž Mark Zorec, Matic Brvar, Doroteja Vlaj, Patricija Pozvek,Špela Pleh, Miša Korva, Mario Poljak, Tatjana Avši - Županc |
| EPI_ISL_979376 | The Jackson Laboratory | The Jackson Laboratory | Lloyd M, Sanderson B, Srivastava A, Maurya R, Renzette N, Omerza G, Kelly K, Li L, Wei C L, Adams M |
| EPI_ISL_981063, EPI_ISL_981064 | Johns Hopkins Hospital Department of Pathology | Johns Hopkins Hospital Department of Pathology | C. Paul Morris, Chun Huai Luo, Adannaya Amadi, Matthew Schwartz, Nicholas Gallagher, Heba H. Mostafa |
| EPI_ISL_981202, EPI_ISL_981305, EPI_ISL_981306, EPI_ISL_981307, EPI_ISL_981368, EPI_ISL_981369 | Ohio Department of Health Laboratory | Ohio Department of Health Laboratory | Holmes, Jennifer; Eric Brandt, Keoni Omura, Glen McGillivary, Caitlin McDonnell, Jade Mowery, Stephanie Mccracken, Tyler Payne, Kirtana Ramadugu, Erica Leasure, Brent Lee, Kelsey Florek, Heather Blankenship, Quanta Brown, and Tammy Bannerman |
| EPI_ISL_981961, EPI_ISL_981962, EPI_ISL_981963 | Microbiology Service, Hospital Universitario Clinico San Cecilio, Granada | Microbiology Service, Hospital Universitario Clinico San Cecilio, Granada | Adolfo de Salazar, Natalia Chueca, Laura Viñuela, Ana Fuentes, Federico García |
| EPI_ISL_981979, EPI_ISL_982024, EPI_ISL_982082, EPI_ISL_982084, EPI_ISL_982094, EPI_ISL_982099, EPI_ISL_982104 | TGen North | Sonora Quest Laboratories | "Jolene Bowers, Megan Folkerts, Chris French, Hayley Yaglom, Ashlyn Pfeiffer, Darrin Lemmer, Dave Engelthaler, The Arizona COVID Genomics Union (ACGU)" |
| EPI_ISL_982247 | Lab voor klinische biologie | Lab voor klinische biologie | Hannelore Hamerlinck, Marija Janevska, Bruno Verhasselt |
| EPI_ISL_982349, EPI_ISL_982352, EPI_ISL_982393, EPI_ISL_982396, EPI_ISL_982399, EPI_ISL_982403, EPI_ISL_982404, EPI_ISL_982405 | M Health Fairview | Minnesota Department of Health, Public Health Laboratory | Alexandra Lorentz, Jacob Garfin, Matt Plumb, and Xiong Wang |
| EPI_ISL_982499, EPI_ISL_982500, EPI_ISL_982501 | ADIRONDACK MEDICAL CENTER | Wadsworth Center, New York State Department of Health | Kirsten St. George, Daryl M. Lamson, Alexis Russel, Matthew Shudt, Melissa A Leisner, Jonathan Plitnick, Navjot Singh, John Kelly, Erasmus Schneider, Erica Lasek-Nesselquist |
| EPI_ISL_982544, EPI_ISL_982546, EPI_ISL_982560, EPI_ISL_982563, EPI_ISL_982568, EPI_ISL_982591, EPI_ISL_982617, EPI_ISL_982626, EPI_ISL_982652, EPI_ISL_982662, EPI_ISL_982663, EPI_ISL_982677, EPI_ISL_982683, EPI_ISL_982731 | | | |
| see above | US Air Force School of Aerospace Medicine | US Air Force School of Aerospace Medicine | Anthony Fries, Jennifer Meyer, William Gruner, William Buggele, Amanda Javorina, Sarah Purves, Clarise Starr, Elizabeth Macias |
| EPI_ISL_983116, EPI_ISL_983117 | SUNY UPSTATE MEDICAL UNIVERSITY | Wadsworth Center, New York State Department of Health | Kirsten St. George, Daryl M. Lamson, Alexis Russel, Matthew Shudt, Melissa A Leisner, Jonathan Plitnick, Navjot Singh, John Kelly, Erasmus Schneider, Erica Lasek-Nesselquist |
| EPI_ISL_983126, EPI_ISL_983127, EPI_ISL_983128 | THE MARY IMOGENE BASSETT HOSPITAL | Wadsworth Center, New York State Department of Health | Kirsten St. George, Daryl M. Lamson, Alexis Russel, Matthew Shudt, Melissa A Leisner, Jonathan Plitnick, Navjot Singh, John Kelly, Erasmus Schneider, Erica Lasek-Nesselquist |
| EPI_ISL_983322 | ASL LATINA PRESIDIO OSPEDALIERO NORD | INMI Lazzaro Spallanzani IRCCS | E Giombini, M. Rueca, B Bartolini, O Butera, C.E.M Gruber, F Messina, A Lucci, S Pignalosa, A Di Caro, MR Capobianchi |
| EPI_ISL_983618, EPI_ISL_983619, EPI_ISL_983620, EPI_ISL_983621, EPI_ISL_983622, EPI_ISL_983623 | Texas Department of State Health Services | Texas Department of State Health Services | Bonnie Oh, Anita Pokharel, James Daniel Bonser, Myong Koag, Chung Wang, Rachel Lee, Grace Kubin, Rashmi Tuladhar, Mayela Pedrueza, Maliha Rahman, Jenny Zhang |
| EPI_ISL_983713, EPI_ISL_983730, EPI_ISL_983805, EPI_ISL_983806, EPI_ISL_983807, EPI_ISL_983808, EPI_ISL_983809, EPI_ISL_983810 | Colorado Department of Public Health and Environment | Colorado Department of Puplic Health and Environment | Laura Bankers, Molly C. Hetherington-Rauth, Diana Ir, Shannon Ely, Shannon R. Matzinger, Sarah Elizabeth Totten, Emily A. Travanty |
