## Supplementary material for "Detection and characterization of the SARS-CoV-2 lineage B.1.526 in New York": Supp. Table 3 GISAID Acknowledgment part 2: gisaid_hcov-19_acknowledgement_table_2021_02_13_010-11.pdf

We gratefully acknowledge the following Authors from the Originating laboratories responsible for obtaining the specimens, as well as the Submitting laboratories where the genome data were generated and shared via GISAID, on which this research is based.

All Submitters of data may be contacted directly via www.gisaid.org

Authors are sorted alphabetically.

| Accession ID | Originating Laboratory | Submitting Laboratory | Authors |
|---|---|---|---|
| EPI_ISL_812359 | Area of Virology, Serology and Virology Division (SAViD), New South Wales Health Pathology Randwick | Virology Research Laboratory; Area of Virology, Serology and Virology Division (SAViD), New South Wales Health Pathology Randwick | Foster, C.; Au, J.; Ruiz Silva, M.; Deveson, I.; Bull, R.; Van Hal, S.; Rawlinson, W. |
| EPI_ISL_831663 | Institute of Virology, Biomedical Research Center of the Slovak Academy of Sciences, Bratislava | Faculty of Natural Sciences, Comenius University, Bratislava | Kristína Boršová, Viktória abanová, Broa Brejová, Viktória Hodorová, Sabina Fumaová Havlíková, Juraj Kopáek, Martina Liková, ubomíra Lukáiková, Martina Neboháová, Monika Sláviková, Tomáš Vina, Boris Klempa, Jozef Nosek |
| EPI_ISL_831664, EPI_ISL_831665 | Institute of Virology, Biomedical Research Center of the Slovak Academy of Sciences, Bratislava | Faculty of Natural Sciences, Comenius University, Bratislava | Broa Brejová, Viktória abanová, Kristína Boršová, Viktória Hodorová, Sabina Fumaová Havlíková, Juraj Kopáek, Martina Liková, ubomíra Lukáiková, Martina Neboháová, Monika Sláviková, Tomáš Vina, Jozef Nosek, Boris Klempa |
| EPI_ISL_831666 | Institute of Virology, Biomedical Research Center of the Slovak Academy of Sciences, Bratislava | Faculty of Natural Sciences, Comenius University, Bratislava | Viktória abanová, Kristína Boršová, Broa Brejová, Viktória Hodorová, Sabina Fumaová Havlíková, Juraj Kopáek, Martina Liková, ubomíra Lukáiková, Martina Neboháová, Monika Sláviková, Tomáš Vina, Jozef Nosek, Boris Klempa |
| EPI_ISL_833373, EPI_ISL_833374, EPI_ISL_833375, EPI_ISL_833376, EPI_ISL_833377, EPI_ISL_833379, EPI_ISL_833380, EPI_ISL_833381, EPI_ISL_833382, EPI_ISL_833383, EPI_ISL_833384, EPI_ISL_833385, EPI_ISL_833386 | | | |
| see above | National Public Health Laboratory, National Centre for Infectious Diseases | National Public Health Laboratory, National Centre for Infectious Diseases | Tze Minn Mak, Sophie Octavia, Zhenyang Zhou, Lin Cui, Raymond Tzer Pin Lin |
| EPI_ISL_833432 | Torrance Memorial Medical Center Microbiology | Los Angeles County PHL | P. Hemarajata et al. |
| EPI_ISL_837058, EPI_ISL_837063, EPI_ISL_837065, EPI_ISL_837073, EPI_ISL_837087, EPI_ISL_837183, EPI_ISL_837196, EPI_ISL_837197, EPI_ISL_837245, EPI_ISL_837246 | Respiratory Virus Unit, National Infection Service, Public Health England | COVID-19 Genomics UK (COG-UK) Consortium | PHE Covid Sequencing Team |
| EPI_ISL_838015, EPI_ISL_838017, EPI_ISL_838018, EPI_ISL_838021, EPI_ISL_838022, EPI_ISL_838025, EPI_ISL_838035, EPI_ISL_838038, EPI_ISL_838048 | Department of Pathology, University of Cambridge | COVID-19 Genomics UK (COG-UK) Consortium | Aminu S. Jahun, Yasmin Chaudhry, Grant Hall, Iliana Georgana, Myra Hosmillo, Martin D. Curran, Malte Pinckert, Surendra Parmar, Ian Goodfellow |
| EPI_ISL_839006, EPI_ISL_839020 | University College London, Great Ormond Street Hospital for Children NHS Foundation Trust, Imperial College Healthcare NHS Trust | COVID-19 Genomics UK (COG-UK) Consortium | Sergi Castellano, Rachel Williams, Mark Kristiansen, Paola Resende Silva, Sunando Roy, Tony Brooks, Helena Tutill, Paola Niola, Patricia Dyal, Charlotte Williams, Leysa Forrest, Yasmin Panchbhaya, Jacqueline Findlay, Samuel Weeks, Julianne Brown, Kathryn Harris, Paul Randell, James Price, Alison Holmes, Judith Breuer |
| EPI_ISL_845617 | KU Leuven, Rega Institute, Clinical and Epidemiological Virology | KU Leuven, Rega Institute, Clinical and Epidemiological Virology | Tony Wawina-Bokalanga, Bert Vanmechelen, Joan Marti-Carerras, Piet Maes |
| EPI_ISL_845655 | Laboratorio de Salud Pública - Secretaría Distrital de Salud | Instituto Nacional de Salud - Dirección de Investigación en Salud Pública | Katherine Laiton-Donato, Diego A. Álvarez-Díaz, Carlos Franco-Muñoz, Mauricio Pacheco-Montealegre, Maria T. Herrera-Sepúlveda, Jonathan Reales, Sheryll Corchuelo, Julian Naizaque, Gerardo Santamaría, Paola Muñoz-Laiton, Diego Andrés Prada, Magdalena Wiesner, Martha Lucia Ospina Martinez, Marcela Mercado-Reyes |
| EPI_ISL_845806 | Sydney South West Pathology Service (SSWPS) - Royal Prince Alfred Hospital - NSW Health Pathology | NSW Health Pathology - Institute of Clinical Pathology and Medical Research; Westmead Hospital; University of Sydney | CIDM-PH et al. |
| EPI_ISL_846553, EPI_ISL_846554, EPI_ISL_846555, EPI_ISL_846556, EPI_ISL_846557, EPI_ISL_846568, EPI_ISL_846569, EPI_ISL_846570, EPI_ISL_846571, EPI_ISL_846572, EPI_ISL_846573, EPI_ISL_846574, EPI_ISL_846578, EPI_ISL_846579, EPI_ISL_846580 | | | |
| see above | Lab voor klinische biologie | Lab voor klinische biologie | Hannelore Hamerlinck, Bruno Verhasselt |
| EPI_ISL_848200, EPI_ISL_848201, EPI_ISL_848202 | Charité Universitätsmedizin Berlin, Institute of Virology, Charitéplatz 1, 10117 Berlin, Germany | Charité Universitätsmedizin Berlin, Institute of Virology, Charitéplatz 1, 10117 Berlin, Germany | Victor M Corman, Julia Schneider, Jörn Beheim-Schwarzbach, Tobias Bleicker, Julia Tesch, Barbara Mühlemann, Talitha Veith, Terry Jones, Christian Drosten |
| EPI_ISL_849761 | unknown | PHV-FSS | Son Nguyen et al. |
| EPI_ISL_852968, EPI_ISL_853006, EPI_ISL_853013, EPI_ISL_853014, EPI_ISL_853015 | Hospital General Universitario Gregorio Marañón | SeqCOVID-SPAIN consortium/IBV(CSIC) | Darío García de Viedma, Laura Pérez-Lago, Pedro J Sola-Campoy, Sergio Buenestado-Serrano, Marta Herranz, Victor Manuel de la Cueva, Julia Suárez, Pilar Catalán, Patricia Muñoz and SeqCOVID-SPAIN consortium |
| EPI_ISL_854594 | Faroese National Reference Laboratory for Fish and Animal Diseases | Faroese National Reference Laboratory for Fish and Animal Diseases | Maria Marjunardóttir Dahl, Petra Elisabeth Petersen, Arnfinnur Kallsberg Junior, Debes Hammershaimb Christiansen |
| EPI_ISL_854754, EPI_ISL_854758 | Microbiological Diagnostic Unit - Public Health Laboratory (MDU-PHL) | MDU-PHL | Seemann T., Sait, M.L., Sherry, N.L. |
| EPI_ISL_854759, EPI_ISL_854762 | Victorian Infectious Diseases Reference Laboratory (VIDRL) | VIDRL and MDU-PHL | Caly L., Seemann T., Sait, M.L., Druce J., Sherry, N.L. |
| EPI_ISL_855603 | Respiratory Virus Unit, National Infection Service, Public Health England | COVID-19 Genomics UK (COG-UK) Consortium | PHE Covid Sequencing Team |
| EPI_ISL_856707, EPI_ISL_856708, EPI_ISL_856709, EPI_ISL_856710 | MFD Veselbas centrs "Dziedniecba" | Latvian Biomedical Research and Study Centre | Ivars Silamielis, Kaspars Megnis, Monta Ustinova, Jnis Pjalkovskis, ikita Zrelovs, Vita Rovte, Iveta Bere, Marija Donaka, Uga Dumpis, Jnis Kloviš |
| EPI_ISL_856726, EPI_ISL_856727, EPI_ISL_856728, EPI_ISL_856729, EPI_ISL_856749, EPI_ISL_856750, EPI_ISL_856751, EPI_ISL_856752, EPI_ISL_856753, EPI_ISL_856754 | Department of Clinical Microbiology | GIGA Medical Genomics | Keith Durkin, Maria Artesi, Sébastien Bontems, Raphaël Boreux, Bouchra Boujemla, Cécile Meex, Pierrette Melin, Marie-Pierre Hayette, Vincent Bours |
| EPI_ISL_857309 | Medlab Pathology | NSW Health Pathology - Institute of Clinical Pathology and Medical Research; Westmead Hospital; University of Sydney | CIDM-PH et al. |
| EPI_ISL_857348 | Caribbean Public Health Agency | Carrington Lab, Department of PreClinical Sciences, Faculty of Medical Sciences, The University of the West Indies | Nikita S. D. Sahadeo, Arianne Brown-Jordan, Vernie Ramkissoon, Sarah Hill, Vernie Ramkissoon, Roshan Parasram, Naresh Nandram, Avery Hinds, Jerome Foster, Stanley Giddings, Karla Georges, Marsha Ivey, Rahul Naidu, Risha Singh, SueMin Nathaniel, Rajini Haraksingh, Jaya Jayaraman, Chinna Chinnadurai, Adesh Ramsubhag, Nuno Faria, Oliver Pybus, Christopher Oura, Gabriel Escobar, Christine V. F. Carrington |
| EPI_ISL_857396, EPI_ISL_857397, EPI_ISL_857398, EPI_ISL_857399, EPI_ISL_857400, EPI_ISL_857401, EPI_ISL_857407, EPI_ISL_857410, EPI_ISL_857412, EPI_ISL_857413, EPI_ISL_857415, EPI_ISL_857416, EPI_ISL_857417, EPI_ISL_857418, EPI_ISL_857419, EPI_ISL_857420, EPI_ISL_857421, EPI_ISL_857422, EPI_ISL_857423, EPI_ISL_857424, EPI_ISL_857425, EPI_ISL_857426, EPI_ISL_857427, EPI_ISL_857428, EPI_ISL_857429, EPI_ISL_857430, EPI_ISL_857431, EPI_ISL_857432, EPI_ISL_857433, EPI_ISL_857434, EPI_ISL_857435, EPI_ISL_857436, EPI_ISL_857437, EPI_ISL_857438, EPI_ISL_857439, EPI_ISL_857440, | | | |

| | | | |
|---|---|---|---|
| EPI_ISL_857441, EPI_ISL_857442, EPI_ISL_857443, EPI_ISL_857447 | | | |
| see above | Maine HETL | Tewhey Lab, The Jackson Laboratory | Matluk,N., Dewey,H., Iosue,F., Barter,M., Lynch,R., Munger,H. and Tewhey,R. |
| EPI_ISL_857468 | PathWest Laboratory Medicine WA | PathWest Laboratory Medicine WA Microbial Surveillance Unit | PathWest Laboratory Medicine WA Microbial Surveillance Unit |
| EPI_ISL_857485 | National Public Health Laboratory, National Centre for Infectious Diseases | National Public Health Laboratory, National Centre for Infectious Diseases | Tze Minn Mak, Sophie Octavia, Zhenyang Zhou, Lin Cui, Raymond Tzer Pin Lin |
| EPI_ISL_857486 | PathWest Laboratory Medicine WA | PathWest Laboratory Medicine WA Microbial Surveillance Unit | PathWest Laboratory Medicine WA Microbial Surveillance Unit |
| EPI_ISL_857855 | Lighthouse Lab in Cambridge | Wellcome Sanger Institute for the COVID-19 Genomics UK (COG-UK) Consortium | Rob Howes, The Lighthouse Lab in Cambridge and Alex Alderton, Roberto Amato, Sonia Goncalves, Ewan Harrison, David K. Jackson, Ian Johnston, Dominic Kwiatkowski, Cordelia Langford, John Sillitoe on behalf of the Wellcome Sanger Institute COVID-19 Surveillance Team |
| EPI_ISL_860262 | Department of Medical Microbiology - section Molde, Molde Hospital | Norwegian Institute of Public Health, Department of Virology | Kathrine Stene-Johansen, Kamilla Heddeland Instefjord, Hilde Elshaug, Atiya R Ali,Marie Paulsen Madsen, Rasmus Riis Kopperud, Hilde Vollan, Karoline Bragstad, Olav Hungnes |
| EPI_ISL_860278 | Haukeland University Hospital, Dept. of Microbiology | Norwegian Institute of Public Health, Department of Virology | Kathrine Stene-Johansen, Kamilla Heddeland Instefjord, Hilde Elshaug, Atiya R Ali,Marie Paulsen Madsen, Rasmus Riis Kopperud, Hilde Vollan, Karoline Bragstad, Olav Hungnes |
| EPI_ISL_860686, EPI_ISL_860695, EPI_ISL_860709 | Respiratory Virus Unit, National Infection Service, Public Health England | COVID-19 Genomics UK (COG-UK) Consortium | PHE Covid Sequencing Team |
| EPI_ISL_860721, EPI_ISL_860722, EPI_ISL_860723, EPI_ISL_860726 | Institute for Infectious Diseases, University of Bern, Switzerland | Institute for Infectious Diseases, University of Bern, Switzerland | Michel C Koch, Christian Baumann, Miguel A Terrazos Miani, Cora Sägesser, Pascal Bittel, Stephen L Leib, Peter Keller, Franziska Suter-Riniker, Alban Ramette |
| EPI_ISL_860954, EPI_ISL_860959, EPI_ISL_860961, EPI_ISL_860963, EPI_ISL_860966, EPI_ISL_860967, EPI_ISL_860968, EPI_ISL_860975, EPI_ISL_860983, EPI_ISL_860986, EPI_ISL_860990, EPI_ISL_860991, EPI_ISL_860993, EPI_ISL_861001, EPI_ISL_861005, EPI_ISL_861006, EPI_ISL_861007, EPI_ISL_861009, EPI_ISL_861012, EPI_ISL_861015, EPI_ISL_861034, EPI_ISL_861038, EPI_ISL_861046, EPI_ISL_861049, EPI_ISL_861053, EPI_ISL_861057, EPI_ISL_861066, EPI_ISL_861069, EPI_ISL_861071, EPI_ISL_861072, EPI_ISL_861073, EPI_ISL_861084, EPI_ISL_861092, EPI_ISL_861093, EPI_ISL_861094 | | | |
| see above | Johns Hopkins Hospital Department of Pathology | Johns Hopkins Hospital Department of Pathology | C. Paul Morris, Chun Huai Luo, Adannaya Amadi, Nicholas Gallagher, Heba H. Mostafa |
| EPI_ISL_861447, EPI_ISL_861448, EPI_ISL_861452, EPI_ISL_861453, EPI_ISL_861455 | Clinical Molecular Microbiology Laboratory, UNC Hospitals | Jeremy Wang | Jeremy Wang, Alexander Rubinsteyn, Colleen Rice, Jason Smedberg, Melissa Miller, Corbin Jones, Robert Hagan |
| EPI_ISL_861507 | Cerballiance Provence | CERBA LAB | Roig JC; Delaunay E; Prots L. |
| EPI_ISL_861710 | Laboratory for Respiratory Viruses, Cantacuzino National Military-Medical Institute for Research and Development | Cantacuzino Institute Virology | Mihaela Lazar, Luiza Ustea, Nicoleta Parashiv, Tim Durfee |
| EPI_ISL_861711, EPI_ISL_861712, EPI_ISL_861713 | Laboratory for Respiratory Viruses, Cantacuzino National Military-Medical Institute for Research and Development | Cantacuzino Institute Virology | Mihaela Lazar, Luiza Ustea, Nicoleta Paraschiv, Tim Durfee |
| EPI_ISL_861735, EPI_ISL_861738, EPI_ISL_861741 | Tempus | Grubaugh Lab - Yale School of Public Health | Tara Alpert, Joseph Fauver, Anderson Brito, Mallery Breban, Anne Wyllie, Chantal Vogels, Mary Petrone, Annie Watkins, Chaney Kalinich, Isabel Ott, Nathan Grubaugh |
| EPI_ISL_861745 | Yale Pathology Lab | Grubaugh Lab - Yale School of Public Health | Tara Alpert, Joseph Fauver, Chen Liu, Pei Hui, Jianhui Wang, Susan Bell and Han Zhou, Anderson Brito, Mallery Breban, Anne Wyllie, Chantal Vogels, Mary Petrone, Chaney Kalinich, Isabel Ott, Arnau Casanovas, Catherine Muenker, Adam Moore, Alice Lu, Maria Tokuyama, Patrick Wong, Peiwen Lu, Saad Omer, Richard Martinello, Allison Nelson, Shelli Farhadian, Akiko Iwasaki, Charlese Dela Cruz, Albert Ko, Nathan Grubaugh |
| EPI_ISL_861757, EPI_ISL_861759, EPI_ISL_861760 | Tempus | Grubaugh Lab - Yale School of Public Health | Tara Alpert, Joseph Fauver, Anderson Brito, Mallery Breban, Anne Wyllie, Chantal Vogels, Mary Petrone, Annie Watkins, Chaney Kalinich, Isabel Ott, Nathan Grubaugh |
| EPI_ISL_861768 | Yale Pathology Lab | Grubaugh Lab - Yale School of Public Health | Tara Alpert, Joseph Fauver, Chen Liu, Pei Hui, Jianhui Wang, Susan Bell and Han Zhou, Anderson Brito, Mallery Breban, Anne Wyllie, Chantal Vogels, Mary Petrone, Chaney Kalinich, Isabel Ott, Arnau Casanovas, Catherine Muenker, Adam Moore, Alice Lu, Maria Tokuyama, Patrick Wong, Peiwen Lu, Saad Omer, Richard Martinello, Allison Nelson, Shelli Farhadian, Akiko Iwasaki, Charlese Dela Cruz, Albert Ko, Nathan Grubaugh |
| EPI_ISL_861769 | Tempus | Grubaugh Lab - Yale School of Public Health | Tara Alpert, Joseph Fauver, Anderson Brito, Mallery Breban, Anne Wyllie, Chantal Vogels, Mary Petrone, Annie Watkins, Chaney Kalinich, Isabel Ott, Nathan Grubaugh |
| EPI_ISL_861770 | Yale Pathology Lab | Grubaugh Lab - Yale School of Public Health | Tara Alpert, Joseph Fauver, Chen Liu, Pei Hui, Jianhui Wang, Susan Bell and Han Zhou, Anderson Brito, Mallery Breban, Anne Wyllie, Chantal Vogels, Mary Petrone, Chaney Kalinich, Isabel Ott, Arnau Casanovas, Catherine Muenker, Adam Moore, Alice Lu, Maria Tokuyama, Patrick Wong, Peiwen Lu, Saad Omer, Richard Martinello, Allison Nelson, Shelli Farhadian, Akiko Iwasaki, Charlese Dela Cruz, Albert Ko, Nathan Grubaugh |
| EPI_ISL_862124, EPI_ISL_862136, EPI_ISL_862139, EPI_ISL_862141, EPI_ISL_862144, EPI_ISL_862146, EPI_ISL_862151, EPI_ISL_862152, EPI_ISL_862164, EPI_ISL_862165, EPI_ISL_862167, EPI_ISL_862168, EPI_ISL_862169, EPI_ISL_862171, EPI_ISL_862176, EPI_ISL_862177, EPI_ISL_862178 | | | |
| see above | Charité Universitätsmedizin Berlin, Institut fur Virologie/Labor Berlin | Charité Universitätsmedizin Berlin, Institut für Virologie | Victor M Corman, Barbara Mühlemann, Jörn Beheim-Schwarzbach, Tobias Bleicker, Julia Tesch, Talitha Veith, Julia Schneider, Terry Jones, Christian Drosten |
| EPI_ISL_862193 | Respiratory Virus Unit, National Infection Service, Public Health England | COVID-19 Genomics UK (COG-UK) Consortium | PHE Covid Sequencing Team |
| EPI_ISL_862718, EPI_ISL_862719 | Centre for Dengue Research and AICBU, Department of Immunology and Molecular Medicine | Centre for Dengue Research and AICBU, Department of Immunology and Molecular Medicine | Chandima Jeewandara, Deshni Jayathilaka, Dinuka Ariyaratne, Diyanath Ranasinghe, Laksiri Gomes, Gathsaurie Neelika Malavige |
| EPI_ISL_862723 | San Diego County Public Health Laboratory | Andersen lab at Scripps Research | SEARCH Alliance San Diego with Tracy Basler, Jovan Shephard, Brett Austin |
| EPI_ISL_862725 | Centre for Dengue Research and AICBU, Department of Immunology and Molecular Medicine | Centre for Dengue Research and AICBU, Department of Immunology and Molecular Medicine | Chandima Jeewandara, Deshni Jayathilaka, Dinuka Ariyaratne, Tibutius Thanesh Pramanayagam, Diyanath Ranasinghe, Laksiri Gomes, Gathsaurie Neelika Malavige |
| EPI_ISL_862833 | National Institute of Infectious Diseases-Prof. Dr. Matei Bals Molecular Diagnostics Laboratory | National Institute of Infectious Diseases-Prof. Dr. Matei Bals Molecular Diagnostics Laboratory | Leontina Banica, Marius Surleac, Corina Casangiu, Petre Milu, Andreea Tudor, Simona Paraschiv, Dan Otelea |
| EPI_ISL_863245, EPI_ISL_863246, EPI_ISL_863247, EPI_ISL_863248, EPI_ISL_863249, EPI_ISL_863250 | Lighthouse Lab in Glasgow | Wellcome Sanger Institute for the COVID-19 Genomics UK (COG-UK) Consortium | Harper VanSteenhouse, Yumi Kasai, David Gray, Carol Clugston, Anna Dominiczak and Alex Alderton, Roberto Amato, Sonia Goncalves, Ewan Harrison, David K. Jackson, Ian Johnston, Dominic Kwiatkowski, Cordelia Langford, John Sillitoe on behalf of the Wellcome Sanger Institute COVID-19 Surveillance Team |
| EPI_ISL_863251 | Lighthouse Lab in Alderley Park | Wellcome Sanger Institute for the COVID-19 Genomics UK (COG-UK) Consortium | Jacquelyn Wynn, Mairead Hyland, The Lighthouse Lab in Alderley Park and Alex Alderton, Roberto Amato, Sonia Goncalves, Ewan Harrison, David K. Jackson, Ian Johnston, Dominic Kwiatkowski, Cordelia Langford, John Sillitoe on behalf of the Wellcome Sanger Institute COVID-19 Surveillance Team |
| EPI_ISL_863252, EPI_ISL_863253 | Lighthouse Lab in Glasgow | Wellcome Sanger Institute for the COVID-19 Genomics UK (COG-UK) Consortium | Harper VanSteenhouse, Yumi Kasai, David Gray, Carol Clugston, Anna Dominiczak and Alex Alderton, Roberto Amato, Sonia Goncalves, Ewan Harrison, David K. Jackson, Ian Johnston, Dominic Kwiatkowski, Cordelia Langford, John Sillitoe on behalf of the Wellcome Sanger Institute COVID-19 Surveillance Team |
| EPI_ISL_863254 | Lighthouse Lab in Alderley Park | Wellcome Sanger Institute for the COVID-19 Genomics UK (COG-UK) Consortium | Jacquelyn Wynn, Mairead Hyland, The Lighthouse Lab in Alderley Park and Alex Alderton, Roberto Amato, Sonia Goncalves, Ewan Harrison, David K. Jackson, Ian Johnston, Dominic Kwiatkowski, Cordelia Langford, John Sillitoe on behalf of the Wellcome Sanger Institute COVID-19 Surveillance Team |
| EPI_ISL_863255, EPI_ISL_863256, EPI_ISL_863257 | Lighthouse Lab in Glasgow | Wellcome Sanger Institute for the COVID-19 Genomics UK (COG-UK) Consortium | Harper VanSteenhouse, Yumi Kasai, David Gray, Carol Clugston, Anna Dominiczak and Alex Alderton, Roberto Amato, Sonia Goncalves, Ewan Harrison, David K. Jackson, Ian Johnston, Dominic Kwiatkowski, Cordelia Langford, John Sillitoe on behalf of the Wellcome Sanger Institute COVID-19 Surveillance Team |
| EPI_ISL_863258 | Lighthouse Lab in Alderley Park | Wellcome Sanger Institute for the COVID-19 Genomics UK (COG-UK) Consortium | Jacquelyn Wynn, Mairead Hyland, The Lighthouse Lab in Alderley Park and Alex Alderton, Roberto Amato, Sonia Goncalves, Ewan Harrison, David K. Jackson, Ian Johnston, Dominic Kwiatkowski, Cordelia Langford, John Sillitoe on behalf of the Wellcome Sanger Institute COVID-19 Surveillance Team |

| | | | |
|---|---|---|---|
| EPI_ISL_863259, EPI_ISL_863260, EPI_ISL_863261, EPI_ISL_863262, EPI_ISL_863263, EPI_ISL_863264, EPI_ISL_863265, EPI_ISL_863266, EPI_ISL_863267 | Lighthouse Lab in Glasgow | Wellcome Sanger Institute for the COVID-19 Genomics UK (COG-UK) Consortium | Harper VanSteenhouse, Yumi Kasai, David Gray, Carol Clugston, Anna Dominiczak and Alex Alderton, Roberto Amato, Sonia Goncalves, Ewan Harrison, David K. Jackson, Ian Johnston, Dominic Kwiatkowski, Cordelia Langford, John Sillitoe on behalf of the Wellcome Sanger Institute COVID-19 Surveillance Team |
| EPI_ISL_863268 | Lighthouse Lab in Alderley Park | Wellcome Sanger Institute for the COVID-19 Genomics UK (COG-UK) Consortium | Jacquelyn Wynn, Mairead Hyland, The Lighthouse Lab in Alderley Park and Alex Alderton, Roberto Amato, Sonia Goncalves, Ewan Harrison, David K. Jackson, Ian Johnston, Dominic Kwiatkowski, Cordelia Langford, John Sillitoe on behalf of the Wellcome Sanger Institute COVID-19 Surveillance Team |
| EPI_ISL_863269 | Lighthouse Lab in Glasgow | Wellcome Sanger Institute for the COVID-19 Genomics UK (COG-UK) Consortium | Harper VanSteenhouse, Yumi Kasai, David Gray, Carol Clugston, Anna Dominiczak and Alex Alderton, Roberto Amato, Sonia Goncalves, Ewan Harrison, David K. Jackson, Ian Johnston, Dominic Kwiatkowski, Cordelia Langford, John Sillitoe on behalf of the Wellcome Sanger Institute COVID-19 Surveillance Team |
| EPI_ISL_863270, EPI_ISL_863271, EPI_ISL_863272 | Lighthouse Lab in Alderley Park | Wellcome Sanger Institute for the COVID-19 Genomics UK (COG-UK) Consortium | Jacquelyn Wynn, Mairead Hyland, The Lighthouse Lab in Alderley Park and Alex Alderton, Roberto Amato, Sonia Goncalves, Ewan Harrison, David K. Jackson, Ian Johnston, Dominic Kwiatkowski, Cordelia Langford, John Sillitoe on behalf of the Wellcome Sanger Institute COVID-19 Surveillance Team |
| EPI_ISL_863273 | Lighthouse Lab in Glasgow | Wellcome Sanger Institute for the COVID-19 Genomics UK (COG-UK) Consortium | Harper VanSteenhouse, Yumi Kasai, David Gray, Carol Clugston, Anna Dominiczak and Alex Alderton, Roberto Amato, Sonia Goncalves, Ewan Harrison, David K. Jackson, Ian Johnston, Dominic Kwiatkowski, Cordelia Langford, John Sillitoe on behalf of the Wellcome Sanger Institute COVID-19 Surveillance Team |
| EPI_ISL_863274, EPI_ISL_863275 | Lighthouse Lab in Alderley Park | Wellcome Sanger Institute for the COVID-19 Genomics UK (COG-UK) Consortium | Jacquelyn Wynn, Mairead Hyland, The Lighthouse Lab in Alderley Park and Alex Alderton, Roberto Amato, Sonia Goncalves, Ewan Harrison, David K. Jackson, Ian Johnston, Dominic Kwiatkowski, Cordelia Langford, John Sillitoe on behalf of the Wellcome Sanger Institute COVID-19 Surveillance Team |
| EPI_ISL_863276, EPI_ISL_863277, EPI_ISL_863278, EPI_ISL_863279 | Lighthouse Lab in Glasgow | Wellcome Sanger Institute for the COVID-19 Genomics UK (COG-UK) Consortium | Harper VanSteenhouse, Yumi Kasai, David Gray, Carol Clugston, Anna Dominiczak and Alex Alderton, Roberto Amato, Sonia Goncalves, Ewan Harrison, David K. Jackson, Ian Johnston, Dominic Kwiatkowski, Cordelia Langford, John Sillitoe on behalf of the Wellcome Sanger Institute COVID-19 Surveillance Team |
| EPI_ISL_863280, EPI_ISL_863281 | Lighthouse Lab in Alderley Park | Wellcome Sanger Institute for the COVID-19 Genomics UK (COG-UK) Consortium | Jacquelyn Wynn, Mairead Hyland, The Lighthouse Lab in Alderley Park and Alex Alderton, Roberto Amato, Sonia Goncalves, Ewan Harrison, David K. Jackson, Ian Johnston, Dominic Kwiatkowski, Cordelia Langford, John Sillitoe on behalf of the Wellcome Sanger Institute COVID-19 Surveillance Team |
| EPI_ISL_863282, EPI_ISL_863283, EPI_ISL_863284, EPI_ISL_863285, EPI_ISL_863286, EPI_ISL_863287, EPI_ISL_863288 | Lighthouse Lab in Glasgow | Wellcome Sanger Institute for the COVID-19 Genomics UK (COG-UK) Consortium | Harper VanSteenhouse, Yumi Kasai, David Gray, Carol Clugston, Anna Dominiczak and Alex Alderton, Roberto Amato, Sonia Goncalves, Ewan Harrison, David K. Jackson, Ian Johnston, Dominic Kwiatkowski, Cordelia Langford, John Sillitoe on behalf of the Wellcome Sanger Institute COVID-19 Surveillance Team |
| EPI_ISL_863289, EPI_ISL_863290 | Lighthouse Lab in Alderley Park | Wellcome Sanger Institute for the COVID-19 Genomics UK (COG-UK) Consortium | Jacquelyn Wynn, Mairead Hyland, The Lighthouse Lab in Alderley Park and Alex Alderton, Roberto Amato, Sonia Goncalves, Ewan Harrison, David K. Jackson, Ian Johnston, Dominic Kwiatkowski, Cordelia Langford, John Sillitoe on behalf of the Wellcome Sanger Institute COVID-19 Surveillance Team |
| EPI_ISL_863291, EPI_ISL_863292, EPI_ISL_863293, EPI_ISL_863294, EPI_ISL_863295, EPI_ISL_863296, EPI_ISL_863297, EPI_ISL_863298 | Lighthouse Lab in Glasgow | Wellcome Sanger Institute for the COVID-19 Genomics UK (COG-UK) Consortium | Harper VanSteenhouse, Yumi Kasai, David Gray, Carol Clugston, Anna Dominiczak and Alex Alderton, Roberto Amato, Sonia Goncalves, Ewan Harrison, David K. Jackson, Ian Johnston, Dominic Kwiatkowski, Cordelia Langford, John Sillitoe on behalf of the Wellcome Sanger Institute COVID-19 Surveillance Team |
| EPI_ISL_863299 | Lighthouse Lab in Alderley Park | Wellcome Sanger Institute for the COVID-19 Genomics UK (COG-UK) Consortium | Jacquelyn Wynn, Mairead Hyland, The Lighthouse Lab in Alderley Park and Alex Alderton, Roberto Amato, Sonia Goncalves, Ewan Harrison, David K. Jackson, Ian Johnston, Dominic Kwiatkowski, Cordelia Langford, John Sillitoe on behalf of the Wellcome Sanger Institute COVID-19 Surveillance Team |
| EPI_ISL_863300, EPI_ISL_863301 | Lighthouse Lab in Glasgow | Wellcome Sanger Institute for the COVID-19 Genomics UK (COG-UK) Consortium | Harper VanSteenhouse, Yumi Kasai, David Gray, Carol Clugston, Anna Dominiczak and Alex Alderton, Roberto Amato, Sonia Goncalves, Ewan Harrison, David K. Jackson, Ian Johnston, Dominic Kwiatkowski, Cordelia Langford, John Sillitoe on behalf of the Wellcome Sanger Institute COVID-19 Surveillance Team |
| EPI_ISL_863302 | Lighthouse Lab in Alderley Park | Wellcome Sanger Institute for the COVID-19 Genomics UK (COG-UK) Consortium | Jacquelyn Wynn, Mairead Hyland, The Lighthouse Lab in Alderley Park and Alex Alderton, Roberto Amato, Sonia Goncalves, Ewan Harrison, David K. Jackson, Ian Johnston, Dominic Kwiatkowski, Cordelia Langford, John Sillitoe on behalf of the Wellcome Sanger Institute COVID-19 Surveillance Team |
| EPI_ISL_863303, EPI_ISL_863304, EPI_ISL_863305 | Lighthouse Lab in Glasgow | Wellcome Sanger Institute for the COVID-19 Genomics UK (COG-UK) Consortium | Harper VanSteenhouse, Yumi Kasai, David Gray, Carol Clugston, Anna Dominiczak and Alex Alderton, Roberto Amato, Sonia Goncalves, Ewan Harrison, David K. Jackson, Ian Johnston, Dominic Kwiatkowski, Cordelia Langford, John Sillitoe on behalf of the Wellcome Sanger Institute COVID-19 Surveillance Team |
| EPI_ISL_863306 | Lighthouse Lab in Alderley Park | Wellcome Sanger Institute for the COVID-19 Genomics UK (COG-UK) Consortium | Jacquelyn Wynn, Mairead Hyland, The Lighthouse Lab in Alderley Park and Alex Alderton, Roberto Amato, Sonia Goncalves, Ewan Harrison, David K. Jackson, Ian Johnston, Dominic Kwiatkowski, Cordelia Langford, John Sillitoe on behalf of the Wellcome Sanger Institute COVID-19 Surveillance Team |
| EPI_ISL_863307, EPI_ISL_863308, EPI_ISL_863309 | Lighthouse Lab in Glasgow | Wellcome Sanger Institute for the COVID-19 Genomics UK (COG-UK) Consortium | Harper VanSteenhouse, Yumi Kasai, David Gray, Carol Clugston, Anna Dominiczak and Alex Alderton, Roberto Amato, Sonia Goncalves, Ewan Harrison, David K. Jackson, Ian Johnston, Dominic Kwiatkowski, Cordelia Langford, John Sillitoe on behalf of the Wellcome Sanger Institute COVID-19 Surveillance Team |
| EPI_ISL_863310, EPI_ISL_863311 | Lighthouse Lab in Alderley Park | Wellcome Sanger Institute for the COVID-19 Genomics UK (COG-UK) Consortium | Jacquelyn Wynn, Mairead Hyland, The Lighthouse Lab in Alderley Park and Alex Alderton, Roberto Amato, Sonia Goncalves, Ewan Harrison, David K. Jackson, Ian Johnston, Dominic Kwiatkowski, Cordelia Langford, John Sillitoe on behalf of the Wellcome Sanger Institute COVID-19 Surveillance Team |
| EPI_ISL_863312, EPI_ISL_863313, EPI_ISL_863314, EPI_ISL_863315, EPI_ISL_863316, EPI_ISL_863317 | Lighthouse Lab in Glasgow | Wellcome Sanger Institute for the COVID-19 Genomics UK (COG-UK) Consortium | Harper VanSteenhouse, Yumi Kasai, David Gray, Carol Clugston, Anna Dominiczak and Alex Alderton, Roberto Amato, Sonia Goncalves, Ewan Harrison, David K. Jackson, Ian Johnston, Dominic Kwiatkowski, Cordelia Langford, John Sillitoe on behalf of the Wellcome Sanger Institute COVID-19 Surveillance Team |
| EPI_ISL_863318, EPI_ISL_863319 | Lighthouse Lab in Alderley Park | Wellcome Sanger Institute for the COVID-19 Genomics UK (COG-UK) Consortium | Jacquelyn Wynn, Mairead Hyland, The Lighthouse Lab in Alderley Park and Alex Alderton, Roberto Amato, Sonia Goncalves, Ewan Harrison, David K. Jackson, Ian Johnston, Dominic Kwiatkowski, Cordelia Langford, John Sillitoe on behalf of the Wellcome Sanger Institute COVID-19 Surveillance Team |
| EPI_ISL_863320, EPI_ISL_863321, EPI_ISL_863322, EPI_ISL_863323, EPI_ISL_863324, EPI_ISL_863325 | Lighthouse Lab in Glasgow | Wellcome Sanger Institute for the COVID-19 Genomics UK (COG-UK) Consortium | Harper VanSteenhouse, Yumi Kasai, David Gray, Carol Clugston, Anna Dominiczak and Alex Alderton, Roberto Amato, Sonia Goncalves, Ewan Harrison, David K. Jackson, Ian Johnston, Dominic Kwiatkowski, Cordelia Langford, John Sillitoe on behalf of the Wellcome Sanger Institute COVID-19 Surveillance Team |
| EPI_ISL_863326 | Lighthouse Lab in Alderley Park | Wellcome Sanger Institute for the COVID-19 Genomics UK (COG-UK) Consortium | Jacquelyn Wynn, Mairead Hyland, The Lighthouse Lab in Alderley Park and Alex Alderton, Roberto Amato, Sonia Goncalves, Ewan Harrison, David K. Jackson, Ian Johnston, Dominic Kwiatkowski, Cordelia Langford, John Sillitoe on behalf of the Wellcome Sanger Institute COVID-19 Surveillance Team |
| EPI_ISL_863327, EPI_ISL_863328, EPI_ISL_863329, EPI_ISL_863330, EPI_ISL_863331, EPI_ISL_863332, EPI_ISL_863333, EPI_ISL_863335, EPI_ISL_863336, EPI_ISL_863337 | Lighthouse Lab in Glasgow | Wellcome Sanger Institute for the COVID-19 Genomics UK (COG-UK) Consortium | Harper VanSteenhouse, Yumi Kasai, David Gray, Carol Clugston, Anna Dominiczak and Alex Alderton, Roberto Amato, Sonia Goncalves, Ewan Harrison, David K. Jackson, Ian Johnston, Dominic Kwiatkowski, Cordelia Langford, John Sillitoe on behalf of the Wellcome Sanger Institute COVID-19 Surveillance Team |
| EPI_ISL_863338 | Lighthouse Lab in Alderley Park | Wellcome Sanger Institute for the COVID-19 Genomics UK (COG-UK) Consortium | Jacquelyn Wynn, Mairead Hyland, The Lighthouse Lab in Alderley Park and Alex Alderton, Roberto Amato, Sonia Goncalves, Ewan Harrison, David K. Jackson, Ian Johnston, Dominic Kwiatkowski, Cordelia Langford, John Sillitoe on behalf of the Wellcome Sanger Institute COVID-19 Surveillance Team |
| EPI_ISL_863339, EPI_ISL_863340, EPI_ISL_863341 | Lighthouse Lab in Glasgow | Wellcome Sanger Institute for the COVID-19 Genomics UK (COG-UK) Consortium | Harper VanSteenhouse, Yumi Kasai, David Gray, Carol Clugston, Anna Dominiczak and Alex Alderton, Roberto Amato, Sonia Goncalves, Ewan Harrison, David K. Jackson, Ian Johnston, Dominic Kwiatkowski, Cordelia Langford, John Sillitoe on behalf of the Wellcome Sanger Institute COVID-19 Surveillance Team |
| EPI_ISL_863342, EPI_ISL_863343 | Lighthouse Lab in Alderley Park | Wellcome Sanger Institute for the COVID-19 Genomics UK (COG-UK) Consortium | Jacquelyn Wynn, Mairead Hyland, The Lighthouse Lab in Alderley Park and Alex Alderton, Roberto Amato, Sonia Goncalves, Ewan Harrison, David K. Jackson, Ian Johnston, Dominic Kwiatkowski, Cordelia Langford, John Sillitoe on behalf of the Wellcome Sanger Institute COVID-19 Surveillance Team |
| EPI_ISL_863344, EPI_ISL_863345 | Lighthouse Lab in Glasgow | Wellcome Sanger Institute for the COVID-19 Genomics UK (COG-UK) Consortium | Harper VanSteenhouse, Yumi Kasai, David Gray, Carol Clugston, Anna Dominiczak and Alex Alderton, Roberto Amato, Sonia Goncalves, Ewan Harrison, David K. Jackson, Ian Johnston, Dominic Kwiatkowski, Cordelia Langford, John Sillitoe on behalf of the Wellcome Sanger Institute COVID-19 Surveillance Team |

| | | | |
|---|---|---|---|
| EPI_ISL_863346 | Lighthouse Lab in Alderley Park | Wellcome Sanger Institute for the COVID-19 Genomics UK (COG-UK) Consortium | Jacquelyn Wynn, Mairead Hyland, The Lighthouse Lab in Alderley Park and Alex Alderton, Roberto Amato, Sonia Goncalves, Ewan Harrison, David K. Jackson, Ian Johnston, Dominic Kwiatkowski, Cordelia Langford, John Sillitoe on behalf of the Wellcome Sanger Institute COVID-19 Surveillance Team |
| EPI_ISL_863347, EPI_ISL_863348, EPI_ISL_863349, EPI_ISL_863350, EPI_ISL_863351, EPI_ISL_863352, EPI_ISL_863353, EPI_ISL_863354, EPI_ISL_863355 | Lighthouse Lab in Glasgow | Wellcome Sanger Institute for the COVID-19 Genomics UK (COG-UK) Consortium | Harper VanSteenhouse, Yumi Kasai, David Gray, Carol Clugston, Anna Dominiczak and Alex Alderton, Roberto Amato, Sonia Goncalves, Ewan Harrison, David K. Jackson, Ian Johnston, Dominic Kwiatkowski, Cordelia Langford, John Sillitoe on behalf of the Wellcome Sanger Institute COVID-19 Surveillance Team |
| EPI_ISL_863356, EPI_ISL_863357 | Lighthouse Lab in Alderley Park | Wellcome Sanger Institute for the COVID-19 Genomics UK (COG-UK) Consortium | Jacquelyn Wynn, Mairead Hyland, The Lighthouse Lab in Alderley Park and Alex Alderton, Roberto Amato, Sonia Goncalves, Ewan Harrison, David K. Jackson, Ian Johnston, Dominic Kwiatkowski, Cordelia Langford, John Sillitoe on behalf of the Wellcome Sanger Institute COVID-19 Surveillance Team |
| EPI_ISL_863358, EPI_ISL_863359 | Lighthouse Lab in Glasgow | Wellcome Sanger Institute for the COVID-19 Genomics UK (COG-UK) Consortium | Harper VanSteenhouse, Yumi Kasai, David Gray, Carol Clugston, Anna Dominiczak and Alex Alderton, Roberto Amato, Sonia Goncalves, Ewan Harrison, David K. Jackson, Ian Johnston, Dominic Kwiatkowski, Cordelia Langford, John Sillitoe on behalf of the Wellcome Sanger Institute COVID-19 Surveillance Team |
| EPI_ISL_863360 | Lighthouse Lab in Alderley Park | Wellcome Sanger Institute for the COVID-19 Genomics UK (COG-UK) Consortium | Jacquelyn Wynn, Mairead Hyland, The Lighthouse Lab in Alderley Park and Alex Alderton, Roberto Amato, Sonia Goncalves, Ewan Harrison, David K. Jackson, Ian Johnston, Dominic Kwiatkowski, Cordelia Langford, John Sillitoe on behalf of the Wellcome Sanger Institute COVID-19 Surveillance Team |
| EPI_ISL_863361 | Lighthouse Lab in Glasgow | Wellcome Sanger Institute for the COVID-19 Genomics UK (COG-UK) Consortium | Harper VanSteenhouse, Yumi Kasai, David Gray, Carol Clugston, Anna Dominiczak and Alex Alderton, Roberto Amato, Sonia Goncalves, Ewan Harrison, David K. Jackson, Ian Johnston, Dominic Kwiatkowski, Cordelia Langford, John Sillitoe on behalf of the Wellcome Sanger Institute COVID-19 Surveillance Team |
| EPI_ISL_863362, EPI_ISL_863363, EPI_ISL_863364 | Lighthouse Lab in Alderley Park | Wellcome Sanger Institute for the COVID-19 Genomics UK (COG-UK) Consortium | Jacquelyn Wynn, Mairead Hyland, The Lighthouse Lab in Alderley Park and Alex Alderton, Roberto Amato, Sonia Goncalves, Ewan Harrison, David K. Jackson, Ian Johnston, Dominic Kwiatkowski, Cordelia Langford, John Sillitoe on behalf of the Wellcome Sanger Institute COVID-19 Surveillance Team |
| EPI_ISL_863365, EPI_ISL_863366, EPI_ISL_863367, EPI_ISL_863368, EPI_ISL_863369, EPI_ISL_863370, EPI_ISL_863371, EPI_ISL_863372, EPI_ISL_863373, EPI_ISL_863374, EPI_ISL_863375, EPI_ISL_863376 | | | |
| see above | Lighthouse Lab in Glasgow | Wellcome Sanger Institute for the COVID-19 Genomics UK (COG-UK) Consortium | Harper VanSteenhouse, Yumi Kasai, David Gray, Carol Clugston, Anna Dominiczak and Alex Alderton, Roberto Amato, Sonia Goncalves, Ewan Harrison, David K. Jackson, Ian Johnston, Dominic Kwiatkowski, Cordelia Langford, John Sillitoe on behalf of the Wellcome Sanger Institute COVID-19 Surveillance Team |
| EPI_ISL_863377 | Lighthouse Lab in Alderley Park | Wellcome Sanger Institute for the COVID-19 Genomics UK (COG-UK) Consortium | Jacquelyn Wynn, Mairead Hyland, The Lighthouse Lab in Alderley Park and Alex Alderton, Roberto Amato, Sonia Goncalves, Ewan Harrison, David K. Jackson, Ian Johnston, Dominic Kwiatkowski, Cordelia Langford, John Sillitoe on behalf of the Wellcome Sanger Institute COVID-19 Surveillance Team |
| EPI_ISL_863378 | Lighthouse Lab in Glasgow | Wellcome Sanger Institute for the COVID-19 Genomics UK (COG-UK) Consortium | Harper VanSteenhouse, Yumi Kasai, David Gray, Carol Clugston, Anna Dominiczak and Alex Alderton, Roberto Amato, Sonia Goncalves, Ewan Harrison, David K. Jackson, Ian Johnston, Dominic Kwiatkowski, Cordelia Langford, John Sillitoe on behalf of the Wellcome Sanger Institute COVID-19 Surveillance Team |
| EPI_ISL_863379 | Lighthouse Lab in Alderley Park | Wellcome Sanger Institute for the COVID-19 Genomics UK (COG-UK) Consortium | Jacquelyn Wynn, Mairead Hyland, The Lighthouse Lab in Alderley Park and Alex Alderton, Roberto Amato, Sonia Goncalves, Ewan Harrison, David K. Jackson, Ian Johnston, Dominic Kwiatkowski, Cordelia Langford, John Sillitoe on behalf of the Wellcome Sanger Institute COVID-19 Surveillance Team |
| EPI_ISL_863380, EPI_ISL_863381, EPI_ISL_863382 | Lighthouse Lab in Glasgow | Wellcome Sanger Institute for the COVID-19 Genomics UK (COG-UK) Consortium | Harper VanSteenhouse, Yumi Kasai, David Gray, Carol Clugston, Anna Dominiczak and Alex Alderton, Roberto Amato, Sonia Goncalves, Ewan Harrison, David K. Jackson, Ian Johnston, Dominic Kwiatkowski, Cordelia Langford, John Sillitoe on behalf of the Wellcome Sanger Institute COVID-19 Surveillance Team |
| EPI_ISL_863383 | Lighthouse Lab in Alderley Park | Wellcome Sanger Institute for the COVID-19 Genomics UK (COG-UK) Consortium | Jacquelyn Wynn, Mairead Hyland, The Lighthouse Lab in Alderley Park and Alex Alderton, Roberto Amato, Sonia Goncalves, Ewan Harrison, David K. Jackson, Ian Johnston, Dominic Kwiatkowski, Cordelia Langford, John Sillitoe on behalf of the Wellcome Sanger Institute COVID-19 Surveillance Team |
| EPI_ISL_863384, EPI_ISL_863385, EPI_ISL_863386 | Lighthouse Lab in Glasgow | Wellcome Sanger Institute for the COVID-19 Genomics UK (COG-UK) Consortium | Harper VanSteenhouse, Yumi Kasai, David Gray, Carol Clugston, Anna Dominiczak and Alex Alderton, Roberto Amato, Sonia Goncalves, Ewan Harrison, David K. Jackson, Ian Johnston, Dominic Kwiatkowski, Cordelia Langford, John Sillitoe on behalf of the Wellcome Sanger Institute COVID-19 Surveillance Team |
| EPI_ISL_863387 | Lighthouse Lab in Alderley Park | Wellcome Sanger Institute for the COVID-19 Genomics UK (COG-UK) Consortium | Jacquelyn Wynn, Mairead Hyland, The Lighthouse Lab in Alderley Park and Alex Alderton, Roberto Amato, Sonia Goncalves, Ewan Harrison, David K. Jackson, Ian Johnston, Dominic Kwiatkowski, Cordelia Langford, John Sillitoe on behalf of the Wellcome Sanger Institute COVID-19 Surveillance Team |
| EPI_ISL_863388, EPI_ISL_863389, EPI_ISL_863390, EPI_ISL_863391, EPI_ISL_863392, EPI_ISL_863393, EPI_ISL_863394 | Lighthouse Lab in Glasgow | Wellcome Sanger Institute for the COVID-19 Genomics UK (COG-UK) Consortium | Harper VanSteenhouse, Yumi Kasai, David Gray, Carol Clugston, Anna Dominiczak and Alex Alderton, Roberto Amato, Sonia Goncalves, Ewan Harrison, David K. Jackson, Ian Johnston, Dominic Kwiatkowski, Cordelia Langford, John Sillitoe on behalf of the Wellcome Sanger Institute COVID-19 Surveillance Team |
| EPI_ISL_863395, EPI_ISL_863397 | Lighthouse Lab in Alderley Park | Wellcome Sanger Institute for the COVID-19 Genomics UK (COG-UK) Consortium | Jacquelyn Wynn, Mairead Hyland, The Lighthouse Lab in Alderley Park and Alex Alderton, Roberto Amato, Sonia Goncalves, Ewan Harrison, David K. Jackson, Ian Johnston, Dominic Kwiatkowski, Cordelia Langford, John Sillitoe on behalf of the Wellcome Sanger Institute COVID-19 Surveillance Team |
| EPI_ISL_863398, EPI_ISL_863399, EPI_ISL_863400, EPI_ISL_863402 | Lighthouse Lab in Glasgow | Wellcome Sanger Institute for the COVID-19 Genomics UK (COG-UK) Consortium | Harper VanSteenhouse, Yumi Kasai, David Gray, Carol Clugston, Anna Dominiczak and Alex Alderton, Roberto Amato, Sonia Goncalves, Ewan Harrison, David K. Jackson, Ian Johnston, Dominic Kwiatkowski, Cordelia Langford, John Sillitoe on behalf of the Wellcome Sanger Institute COVID-19 Surveillance Team |
| EPI_ISL_863404 | Lighthouse Lab in Alderley Park | Wellcome Sanger Institute for the COVID-19 Genomics UK (COG-UK) Consortium | Jacquelyn Wynn, Mairead Hyland, The Lighthouse Lab in Alderley Park and Alex Alderton, Roberto Amato, Sonia Goncalves, Ewan Harrison, David K. Jackson, Ian Johnston, Dominic Kwiatkowski, Cordelia Langford, John Sillitoe on behalf of the Wellcome Sanger Institute COVID-19 Surveillance Team |
| EPI_ISL_863405 | Lighthouse Lab in Glasgow | Wellcome Sanger Institute for the COVID-19 Genomics UK (COG-UK) Consortium | Harper VanSteenhouse, Yumi Kasai, David Gray, Carol Clugston, Anna Dominiczak and Alex Alderton, Roberto Amato, Sonia Goncalves, Ewan Harrison, David K. Jackson, Ian Johnston, Dominic Kwiatkowski, Cordelia Langford, John Sillitoe on behalf of the Wellcome Sanger Institute COVID-19 Surveillance Team |
| EPI_ISL_863406 | Lighthouse Lab in Alderley Park | Wellcome Sanger Institute for the COVID-19 Genomics UK (COG-UK) Consortium | Jacquelyn Wynn, Mairead Hyland, The Lighthouse Lab in Alderley Park and Alex Alderton, Roberto Amato, Sonia Goncalves, Ewan Harrison, David K. Jackson, Ian Johnston, Dominic Kwiatkowski, Cordelia Langford, John Sillitoe on behalf of the Wellcome Sanger Institute COVID-19 Surveillance Team |
| EPI_ISL_863407 | Lighthouse Lab in Glasgow | Wellcome Sanger Institute for the COVID-19 Genomics UK (COG-UK) Consortium | Harper VanSteenhouse, Yumi Kasai, David Gray, Carol Clugston, Anna Dominiczak and Alex Alderton, Roberto Amato, Sonia Goncalves, Ewan Harrison, David K. Jackson, Ian Johnston, Dominic Kwiatkowski, Cordelia Langford, John Sillitoe on behalf of the Wellcome Sanger Institute COVID-19 Surveillance Team |
| EPI_ISL_863408 | Lighthouse Lab in Alderley Park | Wellcome Sanger Institute for the COVID-19 Genomics UK (COG-UK) Consortium | Jacquelyn Wynn, Mairead Hyland, The Lighthouse Lab in Alderley Park and Alex Alderton, Roberto Amato, Sonia Goncalves, Ewan Harrison, David K. Jackson, Ian Johnston, Dominic Kwiatkowski, Cordelia Langford, John Sillitoe on behalf of the Wellcome Sanger Institute COVID-19 Surveillance Team |
| EPI_ISL_863409, EPI_ISL_863410 | Lighthouse Lab in Glasgow | Wellcome Sanger Institute for the COVID-19 Genomics UK (COG-UK) Consortium | Harper VanSteenhouse, Yumi Kasai, David Gray, Carol Clugston, Anna Dominiczak and Alex Alderton, Roberto Amato, Sonia Goncalves, Ewan Harrison, David K. Jackson, Ian Johnston, Dominic Kwiatkowski, Cordelia Langford, John Sillitoe on behalf of the Wellcome Sanger Institute COVID-19 Surveillance Team |
| EPI_ISL_863411 | Lighthouse Lab in Alderley Park | Wellcome Sanger Institute for the COVID-19 Genomics UK (COG-UK) Consortium | Jacquelyn Wynn, Mairead Hyland, The Lighthouse Lab in Alderley Park and Alex Alderton, Roberto Amato, Sonia Goncalves, Ewan Harrison, David K. Jackson, Ian Johnston, Dominic Kwiatkowski, Cordelia Langford, John Sillitoe on behalf of the Wellcome Sanger Institute COVID-19 Surveillance Team |
| EPI_ISL_863412, EPI_ISL_863413 | Lighthouse Lab in Glasgow | Wellcome Sanger Institute for the COVID-19 Genomics UK (COG-UK) Consortium | Harper VanSteenhouse, Yumi Kasai, David Gray, Carol Clugston, Anna Dominiczak and Alex Alderton, Roberto Amato, Sonia Goncalves, Ewan Harrison, David K. Jackson, Ian Johnston, Dominic Kwiatkowski, Cordelia Langford, John Sillitoe on behalf of the Wellcome Sanger Institute COVID-19 Surveillance Team |
| EPI_ISL_863414 | Lighthouse Lab in Alderley Park | Wellcome Sanger Institute for the COVID-19 Genomics UK (COG-UK) Consortium | Jacquelyn Wynn, Mairead Hyland, The Lighthouse Lab in Alderley Park and Alex Alderton, Roberto Amato, Sonia Goncalves, Ewan Harrison, David K. Jackson, Ian Johnston, Dominic Kwiatkowski, Cordelia Langford, John Sillitoe on behalf of the Wellcome Sanger Institute COVID-19 Surveillance Team |
| EPI_ISL_863415 | Lighthouse Lab in Glasgow | Wellcome Sanger Institute for the COVID-19 Genomics UK (COG-UK) Consortium | Harper VanSteenhouse, Yumi Kasai, David Gray, Carol Clugston, Anna Dominiczak and Alex Alderton, Roberto Amato, Sonia Goncalves, Ewan Harrison, David K. Jackson, Ian Johnston, Dominic Kwiatkowski, Cordelia Langford, John Sillitoe on behalf of the Wellcome Sanger Institute COVID-19 Surveillance |

| | | | |
|---|---|---|---|
| | | | Team |
| EPI_ISL_863416 | Lighthouse Lab in Alderley Park | Wellcome Sanger Institute for the COVID-19 Genomics UK (COG-UK) Consortium | Jacquelyn Wynn, Mairead Hyland, The Lighthouse Lab in Alderley Park and Alex Alderton, Roberto Amato, Sonia Goncalves, Ewan Harrison, David K. Jackson, Ian Johnston, Dominic Kwiatkowski, Cordelia Langford, John Sillitoe on behalf of the Wellcome Sanger Institute COVID-19 Surveillance Team |
| EPI_ISL_863417, EPI_ISL_863418, EPI_ISL_863419 | Lighthouse Lab in Glasgow | Wellcome Sanger Institute for the COVID-19 Genomics UK (COG-UK) Consortium | Harper VanSteenhouse, Yumi Kasai, David Gray, Carol Clugston, Anna Dominiczak and Alex Alderton, Roberto Amato, Sonia Goncalves, Ewan Harrison, David K. Jackson, Ian Johnston, Dominic Kwiatkowski, Cordelia Langford, John Sillitoe on behalf of the Wellcome Sanger Institute COVID-19 Surveillance Team |
| EPI_ISL_863420, EPI_ISL_863421 | Lighthouse Lab in Alderley Park | Wellcome Sanger Institute for the COVID-19 Genomics UK (COG-UK) Consortium | Jacquelyn Wynn, Mairead Hyland, The Lighthouse Lab in Alderley Park and Alex Alderton, Roberto Amato, Sonia Goncalves, Ewan Harrison, David K. Jackson, Ian Johnston, Dominic Kwiatkowski, Cordelia Langford, John Sillitoe on behalf of the Wellcome Sanger Institute COVID-19 Surveillance Team |
| EPI_ISL_863422 | Lighthouse Lab in Glasgow | Wellcome Sanger Institute for the COVID-19 Genomics UK (COG-UK) Consortium | Harper VanSteenhouse, Yumi Kasai, David Gray, Carol Clugston, Anna Dominiczak and Alex Alderton, Roberto Amato, Sonia Goncalves, Ewan Harrison, David K. Jackson, Ian Johnston, Dominic Kwiatkowski, Cordelia Langford, John Sillitoe on behalf of the Wellcome Sanger Institute COVID-19 Surveillance Team |
| EPI_ISL_863423 | Lighthouse Lab in Alderley Park | Wellcome Sanger Institute for the COVID-19 Genomics UK (COG-UK) Consortium | Jacquelyn Wynn, Mairead Hyland, The Lighthouse Lab in Alderley Park and Alex Alderton, Roberto Amato, Sonia Goncalves, Ewan Harrison, David K. Jackson, Ian Johnston, Dominic Kwiatkowski, Cordelia Langford, John Sillitoe on behalf of the Wellcome Sanger Institute COVID-19 Surveillance Team |
| EPI_ISL_863425 | Lighthouse Lab in Glasgow | Wellcome Sanger Institute for the COVID-19 Genomics UK (COG-UK) Consortium | Harper VanSteenhouse, Yumi Kasai, David Gray, Carol Clugston, Anna Dominiczak and Alex Alderton, Roberto Amato, Sonia Goncalves, Ewan Harrison, David K. Jackson, Ian Johnston, Dominic Kwiatkowski, Cordelia Langford, John Sillitoe on behalf of the Wellcome Sanger Institute COVID-19 Surveillance Team |
| EPI_ISL_863426 | Lighthouse Lab in Alderley Park | Wellcome Sanger Institute for the COVID-19 Genomics UK (COG-UK) Consortium | Jacquelyn Wynn, Mairead Hyland, The Lighthouse Lab in Alderley Park and Alex Alderton, Roberto Amato, Sonia Goncalves, Ewan Harrison, David K. Jackson, Ian Johnston, Dominic Kwiatkowski, Cordelia Langford, John Sillitoe on behalf of the Wellcome Sanger Institute COVID-19 Surveillance Team |
| EPI_ISL_863427, EPI_ISL_863428, EPI_ISL_863429, EPI_ISL_863430, EPI_ISL_863433 | Lighthouse Lab in Glasgow | Wellcome Sanger Institute for the COVID-19 Genomics UK (COG-UK) Consortium | Harper VanSteenhouse, Yumi Kasai, David Gray, Carol Clugston, Anna Dominiczak and Alex Alderton, Roberto Amato, Sonia Goncalves, Ewan Harrison, David K. Jackson, Ian Johnston, Dominic Kwiatkowski, Cordelia Langford, John Sillitoe on behalf of the Wellcome Sanger Institute COVID-19 Surveillance Team |
| EPI_ISL_863434 | Lighthouse Lab in Alderley Park | Wellcome Sanger Institute for the COVID-19 Genomics UK (COG-UK) Consortium | Jacquelyn Wynn, Mairead Hyland, The Lighthouse Lab in Alderley Park and Alex Alderton, Roberto Amato, Sonia Goncalves, Ewan Harrison, David K. Jackson, Ian Johnston, Dominic Kwiatkowski, Cordelia Langford, John Sillitoe on behalf of the Wellcome Sanger Institute COVID-19 Surveillance Team |
| EPI_ISL_863435, EPI_ISL_863436 | Lighthouse Lab in Glasgow | Wellcome Sanger Institute for the COVID-19 Genomics UK (COG-UK) Consortium | Harper VanSteenhouse, Yumi Kasai, David Gray, Carol Clugston, Anna Dominiczak and Alex Alderton, Roberto Amato, Sonia Goncalves, Ewan Harrison, David K. Jackson, Ian Johnston, Dominic Kwiatkowski, Cordelia Langford, John Sillitoe on behalf of the Wellcome Sanger Institute COVID-19 Surveillance Team |
| EPI_ISL_863437 | Lighthouse Lab in Alderley Park | Wellcome Sanger Institute for the COVID-19 Genomics UK (COG-UK) Consortium | Jacquelyn Wynn, Mairead Hyland, The Lighthouse Lab in Alderley Park and Alex Alderton, Roberto Amato, Sonia Goncalves, Ewan Harrison, David K. Jackson, Ian Johnston, Dominic Kwiatkowski, Cordelia Langford, John Sillitoe on behalf of the Wellcome Sanger Institute COVID-19 Surveillance Team |
| EPI_ISL_863438, EPI_ISL_863439, EPI_ISL_863440, EPI_ISL_863441, EPI_ISL_863442 | Lighthouse Lab in Glasgow | Wellcome Sanger Institute for the COVID-19 Genomics UK (COG-UK) Consortium | Harper VanSteenhouse, Yumi Kasai, David Gray, Carol Clugston, Anna Dominiczak and Alex Alderton, Roberto Amato, Sonia Goncalves, Ewan Harrison, David K. Jackson, Ian Johnston, Dominic Kwiatkowski, Cordelia Langford, John Sillitoe on behalf of the Wellcome Sanger Institute COVID-19 Surveillance Team |
| EPI_ISL_863443 | Lighthouse Lab in Alderley Park | Wellcome Sanger Institute for the COVID-19 Genomics UK (COG-UK) Consortium | Jacquelyn Wynn, Mairead Hyland, The Lighthouse Lab in Alderley Park and Alex Alderton, Roberto Amato, Sonia Goncalves, Ewan Harrison, David K. Jackson, Ian Johnston, Dominic Kwiatkowski, Cordelia Langford, John Sillitoe on behalf of the Wellcome Sanger Institute COVID-19 Surveillance Team |
| EPI_ISL_863444, EPI_ISL_863445 | Lighthouse Lab in Glasgow | Wellcome Sanger Institute for the COVID-19 Genomics UK (COG-UK) Consortium | Harper VanSteenhouse, Yumi Kasai, David Gray, Carol Clugston, Anna Dominiczak and Alex Alderton, Roberto Amato, Sonia Goncalves, Ewan Harrison, David K. Jackson, Ian Johnston, Dominic Kwiatkowski, Cordelia Langford, John Sillitoe on behalf of the Wellcome Sanger Institute COVID-19 Surveillance Team |
| EPI_ISL_863446, EPI_ISL_863447, EPI_ISL_863448 | Lighthouse Lab in Alderley Park | Wellcome Sanger Institute for the COVID-19 Genomics UK (COG-UK) Consortium | Jacquelyn Wynn, Mairead Hyland, The Lighthouse Lab in Alderley Park and Alex Alderton, Roberto Amato, Sonia Goncalves, Ewan Harrison, David K. Jackson, Ian Johnston, Dominic Kwiatkowski, Cordelia Langford, John Sillitoe on behalf of the Wellcome Sanger Institute COVID-19 Surveillance Team |
| EPI_ISL_863449, EPI_ISL_863450, EPI_ISL_863451, EPI_ISL_863452, EPI_ISL_863453 | Lighthouse Lab in Glasgow | Wellcome Sanger Institute for the COVID-19 Genomics UK (COG-UK) Consortium | Harper VanSteenhouse, Yumi Kasai, David Gray, Carol Clugston, Anna Dominiczak and Alex Alderton, Roberto Amato, Sonia Goncalves, Ewan Harrison, David K. Jackson, Ian Johnston, Dominic Kwiatkowski, Cordelia Langford, John Sillitoe on behalf of the Wellcome Sanger Institute COVID-19 Surveillance Team |
| EPI_ISL_863454 | Lighthouse Lab in Alderley Park | Wellcome Sanger Institute for the COVID-19 Genomics UK (COG-UK) Consortium | Jacquelyn Wynn, Mairead Hyland, The Lighthouse Lab in Alderley Park and Alex Alderton, Roberto Amato, Sonia Goncalves, Ewan Harrison, David K. Jackson, Ian Johnston, Dominic Kwiatkowski, Cordelia Langford, John Sillitoe on behalf of the Wellcome Sanger Institute COVID-19 Surveillance Team |
| EPI_ISL_863455, EPI_ISL_863456, EPI_ISL_863457, EPI_ISL_863458, EPI_ISL_863459, EPI_ISL_863460 | Lighthouse Lab in Glasgow | Wellcome Sanger Institute for the COVID-19 Genomics UK (COG-UK) Consortium | Harper VanSteenhouse, Yumi Kasai, David Gray, Carol Clugston, Anna Dominiczak and Alex Alderton, Roberto Amato, Sonia Goncalves, Ewan Harrison, David K. Jackson, Ian Johnston, Dominic Kwiatkowski, Cordelia Langford, John Sillitoe on behalf of the Wellcome Sanger Institute COVID-19 Surveillance Team |
| EPI_ISL_863461 | Lighthouse Lab in Alderley Park | Wellcome Sanger Institute for the COVID-19 Genomics UK (COG-UK) Consortium | Jacquelyn Wynn, Mairead Hyland, The Lighthouse Lab in Alderley Park and Alex Alderton, Roberto Amato, Sonia Goncalves, Ewan Harrison, David K. Jackson, Ian Johnston, Dominic Kwiatkowski, Cordelia Langford, John Sillitoe on behalf of the Wellcome Sanger Institute COVID-19 Surveillance Team |
| EPI_ISL_863462, EPI_ISL_863463, EPI_ISL_863464, EPI_ISL_863465, EPI_ISL_863466, EPI_ISL_863468, EPI_ISL_863469 | Lighthouse Lab in Glasgow | Wellcome Sanger Institute for the COVID-19 Genomics UK (COG-UK) Consortium | Harper VanSteenhouse, Yumi Kasai, David Gray, Carol Clugston, Anna Dominiczak and Alex Alderton, Roberto Amato, Sonia Goncalves, Ewan Harrison, David K. Jackson, Ian Johnston, Dominic Kwiatkowski, Cordelia Langford, John Sillitoe on behalf of the Wellcome Sanger Institute COVID-19 Surveillance Team |
| EPI_ISL_863470 | Lighthouse Lab in Alderley Park | Wellcome Sanger Institute for the COVID-19 Genomics UK (COG-UK) Consortium | Jacquelyn Wynn, Mairead Hyland, The Lighthouse Lab in Alderley Park and Alex Alderton, Roberto Amato, Sonia Goncalves, Ewan Harrison, David K. Jackson, Ian Johnston, Dominic Kwiatkowski, Cordelia Langford, John Sillitoe on behalf of the Wellcome Sanger Institute COVID-19 Surveillance Team |
| EPI_ISL_863471, EPI_ISL_863472, EPI_ISL_863473, EPI_ISL_863474, EPI_ISL_863475, EPI_ISL_863476 | Lighthouse Lab in Glasgow | Wellcome Sanger Institute for the COVID-19 Genomics UK (COG-UK) Consortium | Harper VanSteenhouse, Yumi Kasai, David Gray, Carol Clugston, Anna Dominiczak and Alex Alderton, Roberto Amato, Sonia Goncalves, Ewan Harrison, David K. Jackson, Ian Johnston, Dominic Kwiatkowski, Cordelia Langford, John Sillitoe on behalf of the Wellcome Sanger Institute COVID-19 Surveillance Team |
| EPI_ISL_863477 | Lighthouse Lab in Alderley Park | Wellcome Sanger Institute for the COVID-19 Genomics UK (COG-UK) Consortium | Jacquelyn Wynn, Mairead Hyland, The Lighthouse Lab in Alderley Park and Alex Alderton, Roberto Amato, Sonia Goncalves, Ewan Harrison, David K. Jackson, Ian Johnston, Dominic Kwiatkowski, Cordelia Langford, John Sillitoe on behalf of the Wellcome Sanger Institute COVID-19 Surveillance Team |
| EPI_ISL_863478 | Lighthouse Lab in Glasgow | Wellcome Sanger Institute for the COVID-19 Genomics UK (COG-UK) Consortium | Harper VanSteenhouse, Yumi Kasai, David Gray, Carol Clugston, Anna Dominiczak and Alex Alderton, Roberto Amato, Sonia Goncalves, Ewan Harrison, David K. Jackson, Ian Johnston, Dominic Kwiatkowski, Cordelia Langford, John Sillitoe on behalf of the Wellcome Sanger Institute COVID-19 Surveillance Team |
| EPI_ISL_863479 | Lighthouse Lab in Alderley Park | Wellcome Sanger Institute for the COVID-19 Genomics UK (COG-UK) Consortium | Jacquelyn Wynn, Mairead Hyland, The Lighthouse Lab in Alderley Park and Alex Alderton, Roberto Amato, Sonia Goncalves, Ewan Harrison, David K. Jackson, Ian Johnston, Dominic Kwiatkowski, Cordelia Langford, John Sillitoe on behalf of the Wellcome Sanger Institute COVID-19 Surveillance Team |
| EPI_ISL_863480, EPI_ISL_863481, EPI_ISL_863482, EPI_ISL_863483 | Lighthouse Lab in Glasgow | Wellcome Sanger Institute for the COVID-19 Genomics UK (COG-UK) Consortium | Harper VanSteenhouse, Yumi Kasai, David Gray, Carol Clugston, Anna Dominiczak and Alex Alderton, Roberto Amato, Sonia Goncalves, Ewan Harrison, David K. Jackson, Ian Johnston, Dominic Kwiatkowski, Cordelia Langford, John Sillitoe on behalf of the Wellcome Sanger Institute COVID-19 Surveillance Team |
| EPI_ISL_863484 | Lighthouse Lab in Alderley Park | Wellcome Sanger Institute for the COVID-19 Genomics UK (COG-UK) Consortium | Jacquelyn Wynn, Mairead Hyland, The Lighthouse Lab in Alderley Park and Alex Alderton, Roberto Amato, Sonia Goncalves, Ewan Harrison, David K. Jackson, Ian Johnston, Dominic Kwiatkowski, Cordelia Langford, John Sillitoe on behalf of the Wellcome Sanger Institute COVID-19 Surveillance Team |
| EPI_ISL_863485, EPI_ISL_863486, EPI_ISL_863487, EPI_ISL_863488, EPI_ISL_863489, EPI_ISL_863490 | Lighthouse Lab in Glasgow | Wellcome Sanger Institute for the COVID-19 Genomics UK (COG-UK) Consortium | Harper VanSteenhouse, Yumi Kasai, David Gray, Carol Clugston, Anna Dominiczak and Alex Alderton, Roberto Amato, Sonia Goncalves, Ewan Harrison, David K. Jackson, Ian Johnston, Dominic Kwiatkowski, Cordelia Langford, John Sillitoe on behalf of the Wellcome Sanger Institute COVID-19 Surveillance Team |
| EPI_ISL_863491 | Lighthouse Lab in Alderley Park | Wellcome Sanger Institute for the COVID-19 Genomics UK | Jacquelyn Wynn, Mairead Hyland, The Lighthouse Lab in Alderley Park and Alex Alderton, Roberto Amato, Sonia Goncalves, Ewan Harrison, David K. |

| | | | |
|---|---|---|---|
| | | (COG-UK) Consortium | Jackson, Ian Johnston, Dominic Kwiatkowski, Cordelia Langford, John Sillitoe on behalf of the Wellcome Sanger Institute COVID-19 Surveillance Team |
| EPI_ISL_863492, EPI_ISL_863493, EPI_ISL_863494, EPI_ISL_863495, EPI_ISL_863496, EPI_ISL_863497, EPI_ISL_863498, EPI_ISL_863499, EPI_ISL_863500, EPI_ISL_863501, EPI_ISL_863502 | | | |
| see above | Lighthouse Lab in Glasgow | Wellcome Sanger Institute for the COVID-19 Genomics UK (COG-UK) Consortium | Harper VanSteenhouse, Yumi Kasai, David Gray, Carol Clugston, Anna Dominiczak and Alex Alderton, Roberto Amato, Sonia Goncalves, Ewan Harrison, David K. Jackson, Ian Johnston, Dominic Kwiatkowski, Cordelia Langford, John Sillitoe on behalf of the Wellcome Sanger Institute COVID-19 Surveillance Team |
| EPI_ISL_863503, EPI_ISL_863504 | Lighthouse Lab in Alderley Park | Wellcome Sanger Institute for the COVID-19 Genomics UK (COG-UK) Consortium | Jacquelyn Wynn, Mairead Hyland, The Lighthouse Lab in Alderley Park and Alex Alderton, Roberto Amato, Sonia Goncalves, Ewan Harrison, David K. Jackson, Ian Johnston, Dominic Kwiatkowski, Cordelia Langford, John Sillitoe on behalf of the Wellcome Sanger Institute COVID-19 Surveillance Team |
| EPI_ISL_863505 | Lighthouse Lab in Glasgow | Wellcome Sanger Institute for the COVID-19 Genomics UK (COG-UK) Consortium | Harper VanSteenhouse, Yumi Kasai, David Gray, Carol Clugston, Anna Dominiczak and Alex Alderton, Roberto Amato, Sonia Goncalves, Ewan Harrison, David K. Jackson, Ian Johnston, Dominic Kwiatkowski, Cordelia Langford, John Sillitoe on behalf of the Wellcome Sanger Institute COVID-19 Surveillance Team |
| EPI_ISL_863506 | Lighthouse Lab in Alderley Park | Wellcome Sanger Institute for the COVID-19 Genomics UK (COG-UK) Consortium | Jacquelyn Wynn, Mairead Hyland, The Lighthouse Lab in Alderley Park and Alex Alderton, Roberto Amato, Sonia Goncalves, Ewan Harrison, David K. Jackson, Ian Johnston, Dominic Kwiatkowski, Cordelia Langford, John Sillitoe on behalf of the Wellcome Sanger Institute COVID-19 Surveillance Team |
| EPI_ISL_863507, EPI_ISL_863508, EPI_ISL_863509, EPI_ISL_863510, EPI_ISL_863511, EPI_ISL_863512, EPI_ISL_863513 | Lighthouse Lab in Glasgow | Wellcome Sanger Institute for the COVID-19 Genomics UK (COG-UK) Consortium | Harper VanSteenhouse, Yumi Kasai, David Gray, Carol Clugston, Anna Dominiczak and Alex Alderton, Roberto Amato, Sonia Goncalves, Ewan Harrison, David K. Jackson, Ian Johnston, Dominic Kwiatkowski, Cordelia Langford, John Sillitoe on behalf of the Wellcome Sanger Institute COVID-19 Surveillance Team |
| EPI_ISL_863515, EPI_ISL_863516 | Lighthouse Lab in Alderley Park | Wellcome Sanger Institute for the COVID-19 Genomics UK (COG-UK) Consortium | Jacquelyn Wynn, Mairead Hyland, The Lighthouse Lab in Alderley Park and Alex Alderton, Roberto Amato, Sonia Goncalves, Ewan Harrison, David K. Jackson, Ian Johnston, Dominic Kwiatkowski, Cordelia Langford, John Sillitoe on behalf of the Wellcome Sanger Institute COVID-19 Surveillance Team |
| EPI_ISL_863517, EPI_ISL_863518 | Lighthouse Lab in Glasgow | Wellcome Sanger Institute for the COVID-19 Genomics UK (COG-UK) Consortium | Harper VanSteenhouse, Yumi Kasai, David Gray, Carol Clugston, Anna Dominiczak and Alex Alderton, Roberto Amato, Sonia Goncalves, Ewan Harrison, David K. Jackson, Ian Johnston, Dominic Kwiatkowski, Cordelia Langford, John Sillitoe on behalf of the Wellcome Sanger Institute COVID-19 Surveillance Team |
| EPI_ISL_863519, EPI_ISL_863520 | Lighthouse Lab in Alderley Park | Wellcome Sanger Institute for the COVID-19 Genomics UK (COG-UK) Consortium | Jacquelyn Wynn, Mairead Hyland, The Lighthouse Lab in Alderley Park and Alex Alderton, Roberto Amato, Sonia Goncalves, Ewan Harrison, David K. Jackson, Ian Johnston, Dominic Kwiatkowski, Cordelia Langford, John Sillitoe on behalf of the Wellcome Sanger Institute COVID-19 Surveillance Team |
| EPI_ISL_863521, EPI_ISL_863522, EPI_ISL_863523 | Lighthouse Lab in Glasgow | Wellcome Sanger Institute for the COVID-19 Genomics UK (COG-UK) Consortium | Harper VanSteenhouse, Yumi Kasai, David Gray, Carol Clugston, Anna Dominiczak and Alex Alderton, Roberto Amato, Sonia Goncalves, Ewan Harrison, David K. Jackson, Ian Johnston, Dominic Kwiatkowski, Cordelia Langford, John Sillitoe on behalf of the Wellcome Sanger Institute COVID-19 Surveillance Team |
| EPI_ISL_863524 | Lighthouse Lab in Alderley Park | Wellcome Sanger Institute for the COVID-19 Genomics UK (COG-UK) Consortium | Jacquelyn Wynn, Mairead Hyland, The Lighthouse Lab in Alderley Park and Alex Alderton, Roberto Amato, Sonia Goncalves, Ewan Harrison, David K. Jackson, Ian Johnston, Dominic Kwiatkowski, Cordelia Langford, John Sillitoe on behalf of the Wellcome Sanger Institute COVID-19 Surveillance Team |
| EPI_ISL_863525, EPI_ISL_863526 | Lighthouse Lab in Glasgow | Wellcome Sanger Institute for the COVID-19 Genomics UK (COG-UK) Consortium | Harper VanSteenhouse, Yumi Kasai, David Gray, Carol Clugston, Anna Dominiczak and Alex Alderton, Roberto Amato, Sonia Goncalves, Ewan Harrison, David K. Jackson, Ian Johnston, Dominic Kwiatkowski, Cordelia Langford, John Sillitoe on behalf of the Wellcome Sanger Institute COVID-19 Surveillance Team |
| EPI_ISL_863527 | Lighthouse Lab in Alderley Park | Wellcome Sanger Institute for the COVID-19 Genomics UK (COG-UK) Consortium | Jacquelyn Wynn, Mairead Hyland, The Lighthouse Lab in Alderley Park and Alex Alderton, Roberto Amato, Sonia Goncalves, Ewan Harrison, David K. Jackson, Ian Johnston, Dominic Kwiatkowski, Cordelia Langford, John Sillitoe on behalf of the Wellcome Sanger Institute COVID-19 Surveillance Team |
| EPI_ISL_863528, EPI_ISL_863530 | Lighthouse Lab in Glasgow | Wellcome Sanger Institute for the COVID-19 Genomics UK (COG-UK) Consortium | Harper VanSteenhouse, Yumi Kasai, David Gray, Carol Clugston, Anna Dominiczak and Alex Alderton, Roberto Amato, Sonia Goncalves, Ewan Harrison, David K. Jackson, Ian Johnston, Dominic Kwiatkowski, Cordelia Langford, John Sillitoe on behalf of the Wellcome Sanger Institute COVID-19 Surveillance Team |
| EPI_ISL_863869, EPI_ISL_863870, EPI_ISL_863871, EPI_ISL_863872, EPI_ISL_863874, EPI_ISL_863875, EPI_ISL_863876, EPI_ISL_863877, EPI_ISL_863878, EPI_ISL_863879, EPI_ISL_863880, EPI_ISL_863881, EPI_ISL_863882, EPI_ISL_863883, EPI_ISL_863884, EPI_ISL_863885, EPI_ISL_863886, EPI_ISL_863887, EPI_ISL_863888, EPI_ISL_863889, EPI_ISL_863890, EPI_ISL_863891, EPI_ISL_863892, EPI_ISL_863893, EPI_ISL_863894, EPI_ISL_863895, EPI_ISL_863896, EPI_ISL_863897, EPI_ISL_863898, EPI_ISL_863900, EPI_ISL_863902, EPI_ISL_863903, EPI_ISL_863904, EPI_ISL_863905, EPI_ISL_863906, EPI_ISL_863907, EPI_ISL_863908, EPI_ISL_863909, EPI_ISL_863910, EPI_ISL_863911, EPI_ISL_863914, EPI_ISL_863915, EPI_ISL_863916, EPI_ISL_863917, EPI_ISL_863918, EPI_ISL_863919, EPI_ISL_863920, EPI_ISL_863923, EPI_ISL_863924, EPI_ISL_863925, EPI_ISL_863926, EPI_ISL_863927, EPI_ISL_863928, EPI_ISL_863929, EPI_ISL_863930, EPI_ISL_863931, EPI_ISL_863932, EPI_ISL_863933, EPI_ISL_863934, EPI_ISL_863935, EPI_ISL_863936, EPI_ISL_863937, EPI_ISL_863938, EPI_ISL_863939, EPI_ISL_863940, EPI_ISL_863941, EPI_ISL_863942, EPI_ISL_863943, EPI_ISL_863944, EPI_ISL_863945, EPI_ISL_863947, EPI_ISL_863948, EPI_ISL_863949, EPI_ISL_863950, EPI_ISL_863951, EPI_ISL_863952, EPI_ISL_863953, EPI_ISL_863954, EPI_ISL_863955, EPI_ISL_863957, EPI_ISL_863958, EPI_ISL_863959, EPI_ISL_863960, EPI_ISL_863961, EPI_ISL_863962, EPI_ISL_863963, EPI_ISL_863964, EPI_ISL_863965, EPI_ISL_863966, EPI_ISL_863967, EPI_ISL_863968, EPI_ISL_863970, EPI_ISL_863971, EPI_ISL_863972, EPI_ISL_863973, EPI_ISL_863974, EPI_ISL_863975, EPI_ISL_863976, EPI_ISL_863977, EPI_ISL_863978, EPI_ISL_863979, EPI_ISL_863980, EPI_ISL_863981, EPI_ISL_863982, EPI_ISL_863983, EPI_ISL_863984, EPI_ISL_863985, EPI_ISL_863986, EPI_ISL_863987, EPI_ISL_863988, EPI_ISL_863989, EPI_ISL_863990, EPI_ISL_863991, EPI_ISL_863992, EPI_ISL_863993, EPI_ISL_863994, EPI_ISL_863995, EPI_ISL_863997, EPI_ISL_863998, EPI_ISL_863999, EPI_ISL_864000, EPI_ISL_864001, EPI_ISL_864002, EPI_ISL_864003, EPI_ISL_864006, EPI_ISL_864007, EPI_ISL_864008, EPI_ISL_864010, EPI_ISL_864011, EPI_ISL_864012, EPI_ISL_864013, EPI_ISL_864014, EPI_ISL_864015, EPI_ISL_864016, EPI_ISL_864017, EPI_ISL_864018, EPI_ISL_864019, EPI_ISL_864020, EPI_ISL_864021, EPI_ISL_864022, EPI_ISL_864023, EPI_ISL_864024, EPI_ISL_864025, EPI_ISL_864027, EPI_ISL_864028, EPI_ISL_864029, EPI_ISL_864030, EPI_ISL_864031, EPI_ISL_864032, EPI_ISL_864033, EPI_ISL_864034, EPI_ISL_864035, EPI_ISL_864037, EPI_ISL_864038, EPI_ISL_864040, EPI_ISL_864041, EPI_ISL_864042, EPI_ISL_864044, EPI_ISL_864045, EPI_ISL_864046, EPI_ISL_864047, EPI_ISL_864048, EPI_ISL_864049, EPI_ISL_864050, EPI_ISL_864051, EPI_ISL_864052, EPI_ISL_864053, EPI_ISL_864054, EPI_ISL_864055, EPI_ISL_864056, EPI_ISL_864057, EPI_ISL_864058, EPI_ISL_864059, EPI_ISL_864060, EPI_ISL_864061, EPI_ISL_864062, EPI_ISL_864063, EPI_ISL_864065, EPI_ISL_864066, EPI_ISL_864067, EPI_ISL_864068, EPI_ISL_864069, EPI_ISL_864070, EPI_ISL_864071, EPI_ISL_864072, EPI_ISL_864073, EPI_ISL_864074, EPI_ISL_864075, EPI_ISL_864076, EPI_ISL_864077, EPI_ISL_864078, EPI_ISL_864079, EPI_ISL_864080, EPI_ISL_864081, EPI_ISL_864082, EPI_ISL_864083, EPI_ISL_864084, EPI_ISL_864085, EPI_ISL_864086, EPI_ISL_864087, EPI_ISL_864088, EPI_ISL_864089, EPI_ISL_864090, EPI_ISL_864091, EPI_ISL_864092, EPI_ISL_864093, EPI_ISL_864095, EPI_ISL_864096, EPI_ISL_864097, EPI_ISL_864099, EPI_ISL_864100, EPI_ISL_864101, EPI_ISL_864102, EPI_ISL_864103, EPI_ISL_864104, EPI_ISL_864105, EPI_ISL_864106, EPI_ISL_864107, EPI_ISL_864108, EPI_ISL_864109, EPI_ISL_864111, EPI_ISL_864112, EPI_ISL_864113, EPI_ISL_864114, EPI_ISL_864115, EPI_ISL_864116, EPI_ISL_864117, EPI_ISL_864118, EPI_ISL_864119, EPI_ISL_864120, EPI_ISL_864121, EPI_ISL_864122, EPI_ISL_864123, EPI_ISL_864125, EPI_ISL_864126, EPI_ISL_864127, EPI_ISL_864128, EPI_ISL_864129, EPI_ISL_864130, EPI_ISL_864131, EPI_ISL_864132, EPI_ISL_864134, EPI_ISL_864135, EPI_ISL_864136, EPI_ISL_864137, EPI_ISL_864138, EPI_ISL_864139, EPI_ISL_864140, EPI_ISL_864141, EPI_ISL_864143, EPI_ISL_864144, EPI_ISL_864146, EPI_ISL_864147, EPI_ISL_864148, EPI_ISL_864149, EPI_ISL_864150, EPI_ISL_864151, EPI_ISL_864152, EPI_ISL_864153, EPI_ISL_864154, EPI_ISL_864155, EPI_ISL_864156, EPI_ISL_864157, EPI_ISL_864158, EPI_ISL_864159, EPI_ISL_864160, EPI_ISL_864161, EPI_ISL_864162, EPI_ISL_864163, EPI_ISL_864164, EPI_ISL_864166, EPI_ISL_864167, EPI_ISL_864168, EPI_ISL_864170, EPI_ISL_864171, EPI_ISL_864172, EPI_ISL_864174, EPI_ISL_864177, EPI_ISL_864179, EPI_ISL_864180, EPI_ISL_864181, EPI_ISL_864182, EPI_ISL_864183, EPI_ISL_864184, EPI_ISL_864185, EPI_ISL_864186, EPI_ISL_864187, EPI_ISL_864188, EPI_ISL_864189, EPI_ISL_864190, EPI_ISL_864192, EPI_ISL_864194, EPI_ISL_864195, EPI_ISL_864196, EPI_ISL_864197, EPI_ISL_864198, EPI_ISL_864199, EPI_ISL_864200, EPI_ISL_864201, EPI_ISL_864202 | | | |
| see above | Lighthouse Lab in Alderley Park | Wellcome Sanger Institute for the COVID-19 Genomics UK (COG-UK) Consortium | Jacquelyn Wynn, Mairead Hyland, The Lighthouse Lab in Alderley Park and Alex Alderton, Roberto Amato, Sonia Goncalves, Ewan Harrison, David K. Jackson, Ian Johnston, Dominic Kwiatkowski, Cordelia Langford, John Sillitoe on behalf of the Wellcome Sanger Institute COVID-19 Surveillance Team |
| EPI_ISL_864581, EPI_ISL_864584, EPI_ISL_864586, EPI_ISL_864588, EPI_ISL_864589, EPI_ISL_864590, EPI_ISL_864591, EPI_ISL_864592, EPI_ISL_864593 | CHU Purpan - Laboratoire de Virologie - Institut Fédératif de Biologie | CHU Purpan - Laboratoire de Virologie - Institut Fédératif de Biologie | Latour J., Ranger N., Dubois M., Carcenac R., Harter A., Boyer P., Tremeaux P., Izopet J. |
| EPI_ISL_864598, EPI_ISL_864599, EPI_ISL_864608, EPI_ISL_864609, EPI_ISL_864610, EPI_ISL_864611, EPI_ISL_864619, EPI_ISL_864620, EPI_ISL_864624, EPI_ISL_864630, EPI_ISL_864635, EPI_ISL_864707, EPI_ISL_864708, EPI_ISL_864709, EPI_ISL_864710, EPI_ISL_864711, EPI_ISL_864712, EPI_ISL_864713, EPI_ISL_864714, EPI_ISL_864715, EPI_ISL_864728, EPI_ISL_864729, EPI_ISL_864730, EPI_ISL_864731, EPI_ISL_864732, EPI_ISL_864733 | | | |
| see above | University Hospitals of Geneva, Laboratory of Virology | HUG, Laboratory of Virology and the Health2030 Genome Center | Samuel Cordey, Ana Rita Goncalves, Laurent Kaiser, Lorenzo Cerutti, Henri Pegeot, Melyssa Elies, Deborah Penet, Keith Harshman, Ioannis Xenarios, Emmanouil Dermitzakis |
| EPI_ISL_864896, EPI_ISL_864897, EPI_ISL_864898, EPI_ISL_864899, EPI_ISL_864900, EPI_ISL_864901, EPI_ISL_864902, EPI_ISL_864903, EPI_ISL_864904, EPI_ISL_864905, EPI_ISL_864906, EPI_ISL_864907, EPI_ISL_864908, EPI_ISL_864909, EPI_ISL_864910, EPI_ISL_864911, EPI_ISL_864912, EPI_ISL_864913, EPI_ISL_864914, EPI_ISL_864915, EPI_ISL_864916, EPI_ISL_864919, EPI_ISL_864920, EPI_ISL_864921, EPI_ISL_864922, EPI_ISL_864923, EPI_ISL_864924, EPI_ISL_864933, EPI_ISL_864935, EPI_ISL_864937, EPI_ISL_864938, EPI_ISL_864939, EPI_ISL_864940, EPI_ISL_864941, EPI_ISL_864942, EPI_ISL_864943, EPI_ISL_864944, EPI_ISL_864949, EPI_ISL_864950, EPI_ISL_864951, EPI_ISL_864952, EPI_ISL_864954, EPI_ISL_864957 | | | |
| see above | Department of Pathology, University of Cambridge | COVID-19 Genomics UK (COG-UK) Consortium | Aminu S. Jahun, Yasmin Chaudhry, Grant Hall, Iliana Georgana, Myra Hosmillo, Martin D. Curran, Malte Pinckert, Surendra Parmar, Ian Goodfellow |
| EPI_ISL_865092, EPI_ISL_865093, EPI_ISL_865105, EPI_ISL_865106, EPI_ISL_865107, EPI_ISL_865108, EPI_ISL_865109, EPI_ISL_865110, EPI_ISL_865136, EPI_ISL_865138, EPI_ISL_865139, EPI_ISL_865141, EPI_ISL_865144 | | | |
| see above | Virology Department, Royal Infirmary of Edinburgh, NHS Lothian / School of Biological Sciences, University of Edinburgh / Institute of Genetics and Molecular Medicine, University of Edinburgh | COVID-19 Genomics UK (COG-UK) Consortium | McHugh M, Dewar R, Rooke S, Gallagher M, Balcaza C, O'Toole Á, Scher E, Hill V, McCrone JT, Colquhoun R, Yu X, Jackson B, Rambaut A, Williams TC, Templeton K |
| EPI_ISL_865472, EPI_ISL_865474, EPI_ISL_865475, EPI_ISL_865476 | Liverpool Clinical Laboratories | COVID-19 Genomics UK (COG-UK) Consortium | Sam Haldenby, Anita Lucaci, Steve Paterson, Julian Hiscox, Alistair Darby, M Almsaud, A Alrezaihi, Muhannad Alruwaili, Stuart D Armstrong, Jones Benjamin, Eleanor G Bentley, Anu Chawla, Jordan J Clark, Angela Cowell, Richard Eccles, Isabel García-Dorival, Matthew Gemmell, Alessandro Gerada, |

| | | | |
|---|---|---|---|
| | | | PKF Gilmore, Richard Gregory, Ximeng Han, Catherine Hartley, Margaret Hughes, Miren Iturriza-Gomara, James Johnson, L Luu, Jenifer Manson, Charlotte Nelson, Elaine O'Toole, Cassie Olateju, Rebekah Penrice-Randal , Lucille Rainbow, N.P Randle, Trevor Ian Robinson, Parul Sharma, Ghada T Shawli, James P Stewart, Neil Swainston, Ecaterina Vamos, Joanne Watts, Mark Whitehead |
| EPI_ISL_865485, EPI_ISL_865486, EPI_ISL_865487, EPI_ISL_865585, EPI_ISL_865586, EPI_ISL_865587, EPI_ISL_865588, EPI_ISL_865589, EPI_ISL_865590, EPI_ISL_865591, EPI_ISL_865592, EPI_ISL_865593, EPI_ISL_865594, EPI_ISL_865595, EPI_ISL_865596, EPI_ISL_865599, EPI_ISL_865602, EPI_ISL_865603, EPI_ISL_865604, EPI_ISL_865848, EPI_ISL_865849, EPI_ISL_865850, EPI_ISL_865851, EPI_ISL_865852, EPI_ISL_865864, EPI_ISL_865867, EPI_ISL_865868, EPI_ISL_865869, EPI_ISL_865870, EPI_ISL_865871, EPI_ISL_865872, EPI_ISL_865873, EPI_ISL_865874, EPI_ISL_865875, EPI_ISL_865876, EPI_ISL_865877, EPI_ISL_865878, EPI_ISL_865879, EPI_ISL_865880, EPI_ISL_865881, EPI_ISL_865883, EPI_ISL_865891, EPI_ISL_865895, EPI_ISL_865897, EPI_ISL_865905, EPI_ISL_865906, EPI_ISL_865909, EPI_ISL_865910, EPI_ISL_865911, EPI_ISL_865913, EPI_ISL_865914, EPI_ISL_865915, EPI_ISL_865916, EPI_ISL_865918, EPI_ISL_865919, EPI_ISL_865921, EPI_ISL_865922, EPI_ISL_865923, EPI_ISL_865924, EPI_ISL_865925, EPI_ISL_865929, EPI_ISL_865930, EPI_ISL_865931, EPI_ISL_865932, EPI_ISL_865934, EPI_ISL_865940, EPI_ISL_865941, EPI_ISL_865942, EPI_ISL_865943, EPI_ISL_865944, EPI_ISL_865945, EPI_ISL_865949, EPI_ISL_865950, EPI_ISL_865952, EPI_ISL_865953, EPI_ISL_865954, EPI_ISL_865955, EPI_ISL_865956, EPI_ISL_865957, EPI_ISL_865958, EPI_ISL_865959, EPI_ISL_865960, EPI_ISL_865961, EPI_ISL_865962, EPI_ISL_865963, EPI_ISL_865964, EPI_ISL_865965, EPI_ISL_865966, EPI_ISL_865968, EPI_ISL_866000, EPI_ISL_866012 | | | |
| see above | University College London, Great Ormond Street Hospital for Children NHS Foundation Trust, Imperial College Healthcare NHS Trust | COVID-19 Genomics UK (COG-UK) Consortium | Sergi Castellano, Rachel Williams, Mark Kristiansen, Paola Resende Silva, Sunando Roy, Tony Brooks, Helena Tutill, Paola Niola, Patricia Dyal, Charlotte Williams, Leysa Forrest, Yasmin Panchbhaya, Jacqueline Findlay, Samuel Weeks, Julianne Brown, Kathryn Harris, Paul Randell, James Price, Alison Holmes, Judith Breuer |
| EPI_ISL_866150, EPI_ISL_866153, EPI_ISL_866155, EPI_ISL_866156, EPI_ISL_866158, EPI_ISL_866159, EPI_ISL_866161, EPI_ISL_866163, EPI_ISL_866165, EPI_ISL_866166, EPI_ISL_866167, EPI_ISL_866169, EPI_ISL_866170, EPI_ISL_866172, EPI_ISL_866173, EPI_ISL_866175, EPI_ISL_866176, EPI_ISL_866177, EPI_ISL_866181, EPI_ISL_866182, EPI_ISL_866184, EPI_ISL_866185 | | | |
| see above | University College London Hospital | COVID-19 Genomics UK (COG-UK) Consortium | Judith Heaney, Matthew Byott, Catherine Houlihan, Dan Frampton, Stuart Kirk, Moira Spyer and Eleni Nastouli |
| EPI_ISL_866198, EPI_ISL_866201, EPI_ISL_866215, EPI_ISL_866217, EPI_ISL_866219, EPI_ISL_866221, EPI_ISL_866231, EPI_ISL_866262, EPI_ISL_866263, EPI_ISL_866264, EPI_ISL_866265, EPI_ISL_866266, EPI_ISL_866267, EPI_ISL_866268, EPI_ISL_866269, EPI_ISL_866270, EPI_ISL_866271, EPI_ISL_866272, EPI_ISL_866273, EPI_ISL_866274, EPI_ISL_866275, EPI_ISL_866276, EPI_ISL_866277, EPI_ISL_866278, EPI_ISL_866279, EPI_ISL_866280, EPI_ISL_866281, EPI_ISL_866282, EPI_ISL_866283, EPI_ISL_866284, EPI_ISL_866285, EPI_ISL_866286, EPI_ISL_866287, EPI_ISL_866288, EPI_ISL_866294, EPI_ISL_866295, EPI_ISL_866296, EPI_ISL_866297, EPI_ISL_866299, EPI_ISL_866300, EPI_ISL_866308, EPI_ISL_866309, EPI_ISL_866310, EPI_ISL_866313, EPI_ISL_866314, EPI_ISL_866315, EPI_ISL_866316, EPI_ISL_866317, EPI_ISL_866318, EPI_ISL_866319 | | | |
| see above | University College London, Great Ormond Street Hospital for Children NHS Foundation Trust, Imperial College Healthcare NHS Trust | COVID-19 Genomics UK (COG-UK) Consortium | Sergi Castellano, Rachel Williams, Mark Kristiansen, Paola Resende Silva, Sunando Roy, Tony Brooks, Helena Tutill, Paola Niola, Patricia Dyal, Charlotte Williams, Leysa Forrest, Yasmin Panchbhaya, Jacqueline Findlay, Samuel Weeks, Julianne Brown, Kathryn Harris, Paul Randell, James Price, Alison Holmes, Judith Breuer |
| EPI_ISL_866541, EPI_ISL_866548, EPI_ISL_866556, EPI_ISL_866566, EPI_ISL_866569, EPI_ISL_866748, EPI_ISL_866749, EPI_ISL_866750, EPI_ISL_866751, EPI_ISL_866752, EPI_ISL_866753, EPI_ISL_866754, EPI_ISL_866755, EPI_ISL_866756, EPI_ISL_866757, EPI_ISL_866760, EPI_ISL_866763, EPI_ISL_866764, EPI_ISL_866767, EPI_ISL_866770, EPI_ISL_866774, EPI_ISL_866775, EPI_ISL_866781, EPI_ISL_866788, EPI_ISL_866789, EPI_ISL_866801, EPI_ISL_866835, EPI_ISL_866836, EPI_ISL_866837, EPI_ISL_866838, EPI_ISL_866839, EPI_ISL_866840, EPI_ISL_866841, EPI_ISL_866842, EPI_ISL_866843, EPI_ISL_866844, EPI_ISL_866845, EPI_ISL_866846, EPI_ISL_866847, EPI_ISL_866848, EPI_ISL_866849, EPI_ISL_866850, EPI_ISL_866851, EPI_ISL_866852, EPI_ISL_866853, EPI_ISL_866854, EPI_ISL_866855, EPI_ISL_866856, EPI_ISL_866857, EPI_ISL_866858, EPI_ISL_866859, EPI_ISL_866860, EPI_ISL_866861, EPI_ISL_866862, EPI_ISL_866863, EPI_ISL_866864, EPI_ISL_866865, EPI_ISL_866866, EPI_ISL_866867, EPI_ISL_866868, EPI_ISL_866869, EPI_ISL_866871, EPI_ISL_866872, EPI_ISL_866873, EPI_ISL_866874, EPI_ISL_866875, EPI_ISL_866876, EPI_ISL_866877, EPI_ISL_866878, EPI_ISL_866879, EPI_ISL_866880, EPI_ISL_866881, EPI_ISL_866882, EPI_ISL_866883, EPI_ISL_866884, EPI_ISL_866885, EPI_ISL_866886, EPI_ISL_866887, EPI_ISL_866888 | | | |
| see above | Quadram Institute Bioscience | COVID-19 Genomics UK (COG-UK) Consortium | Dave J. Baker, Gemma L. Kay, Alp Aydin, Thanh Le-Viet, Steven Rudder, Ana P. Tedim, Anastasia Kolyva, Maria Diaz, Leonardo de Oliveira Martins, Nabil-Fareed Alikhan, Lizzie Meadows, Rachael Stanley, Ngozi Elumogo, Muhammed Yasir, Nicholas M. Thomson, Alexander J Trotter, Rachel Gilroy, Samuel Bloomfield, Claire Stuart, Andrew Bell, Reenesh Prakash, Samir Dervisevic, Alison E. Mather, John Wain, Mark Webber, Andrew J. Page, Justin O'Grady |
| EPI_ISL_866921, EPI_ISL_866922, EPI_ISL_866923, EPI_ISL_866924, EPI_ISL_866925, EPI_ISL_866926, EPI_ISL_866927, EPI_ISL_866928, EPI_ISL_866929, EPI_ISL_866930, EPI_ISL_866931, EPI_ISL_866932, EPI_ISL_866933, EPI_ISL_866934, EPI_ISL_866935, EPI_ISL_866936, EPI_ISL_866937, EPI_ISL_866938, EPI_ISL_866939, EPI_ISL_866940, EPI_ISL_866941, EPI_ISL_866942, EPI_ISL_866971, EPI_ISL_866972, EPI_ISL_866973, EPI_ISL_866974, EPI_ISL_866975, EPI_ISL_866976, EPI_ISL_866977, EPI_ISL_866978, EPI_ISL_866979, EPI_ISL_866980, EPI_ISL_866981, EPI_ISL_866982, EPI_ISL_866983, EPI_ISL_866984, EPI_ISL_866986, EPI_ISL_866987, EPI_ISL_866988, EPI_ISL_866989 | | | |
| see above | Queens Medical Centre, Clinical Microbiology Department / DeepSeq Nottingham | COVID-19 Genomics UK (COG-UK) Consortium | Gemma Clark, Wendy Smith, Manjinder Khakh, Vicki M Fleming, Michelle M Lister, Hannah Howson-Wells, Jonathan Ball, Patrick McClure, Joseph Chappell, Theocharis Tsoleridis, Nadine Holmes, Matthew Carlisle, Christopher Moore, Fei Sang, Johnny Debebe, Victoria Wright, Matthew Loose |
| EPI_ISL_867234, EPI_ISL_867235, EPI_ISL_867236, EPI_ISL_867259, EPI_ISL_867260, EPI_ISL_867262, EPI_ISL_867263, EPI_ISL_867264, EPI_ISL_867265, EPI_ISL_867266, EPI_ISL_867267, EPI_ISL_867268, EPI_ISL_867407, EPI_ISL_867408, EPI_ISL_867409, EPI_ISL_867410, EPI_ISL_867411, EPI_ISL_867412, EPI_ISL_867413, EPI_ISL_867414, EPI_ISL_867415, EPI_ISL_867420, EPI_ISL_867421, EPI_ISL_867422, EPI_ISL_867423, EPI_ISL_867425, EPI_ISL_867426, EPI_ISL_867428, EPI_ISL_867436, EPI_ISL_867444, EPI_ISL_867445, EPI_ISL_867446, EPI_ISL_867447, EPI_ISL_867448, EPI_ISL_867449, EPI_ISL_867452, EPI_ISL_867453, EPI_ISL_867454, EPI_ISL_867455, EPI_ISL_867460, EPI_ISL_867462, EPI_ISL_867463, EPI_ISL_867464, EPI_ISL_867465, EPI_ISL_867466, EPI_ISL_867467, EPI_ISL_867468, EPI_ISL_867469, EPI_ISL_867470, EPI_ISL_867471, EPI_ISL_867472, EPI_ISL_867473, EPI_ISL_867474, EPI_ISL_867475, EPI_ISL_867476, EPI_ISL_867477, EPI_ISL_867478, EPI_ISL_867479, EPI_ISL_867480, EPI_ISL_867481, EPI_ISL_867482, EPI_ISL_867483, EPI_ISL_867484, EPI_ISL_867485, EPI_ISL_867487, EPI_ISL_867488, EPI_ISL_867489, EPI_ISL_867495, EPI_ISL_867496, EPI_ISL_867502, EPI_ISL_867549, EPI_ISL_867576, EPI_ISL_867577, EPI_ISL_867585, EPI_ISL_867591, EPI_ISL_867592, EPI_ISL_867594, EPI_ISL_867595, EPI_ISL_867596, EPI_ISL_867597, EPI_ISL_867599, EPI_ISL_867600, EPI_ISL_867601, EPI_ISL_867602, EPI_ISL_867605, EPI_ISL_867607, EPI_ISL_867608, EPI_ISL_867610, EPI_ISL_867611, EPI_ISL_867613, EPI_ISL_867616, EPI_ISL_867617, EPI_ISL_867618, EPI_ISL_867620, EPI_ISL_867621, EPI_ISL_867623, EPI_ISL_867624, EPI_ISL_867625, EPI_ISL_867628, EPI_ISL_867629, EPI_ISL_867630, EPI_ISL_867631, EPI_ISL_867633, EPI_ISL_867634, EPI_ISL_867635, EPI_ISL_867636, EPI_ISL_867637, EPI_ISL_867639, EPI_ISL_867640, EPI_ISL_867641, EPI_ISL_867642, EPI_ISL_867643, EPI_ISL_867644, EPI_ISL_867647, EPI_ISL_867649, EPI_ISL_867650, EPI_ISL_867651, EPI_ISL_867652, EPI_ISL_867653, EPI_ISL_867654, EPI_ISL_867655, EPI_ISL_867656, EPI_ISL_867657, EPI_ISL_867658, EPI_ISL_867659, EPI_ISL_867660, EPI_ISL_867661, EPI_ISL_867662, EPI_ISL_867663, EPI_ISL_867664, EPI_ISL_867665, EPI_ISL_867667, EPI_ISL_867668, EPI_ISL_867670, EPI_ISL_867671, EPI_ISL_867672, EPI_ISL_867673, EPI_ISL_867674, EPI_ISL_867675, EPI_ISL_867676, EPI_ISL_867677, EPI_ISL_867678, EPI_ISL_867679, EPI_ISL_867680, EPI_ISL_867681, EPI_ISL_867682, EPI_ISL_867683, EPI_ISL_867684, EPI_ISL_867686, EPI_ISL_867688, EPI_ISL_867689, EPI_ISL_867690, EPI_ISL_867692, EPI_ISL_867696, EPI_ISL_867697, EPI_ISL_867698, EPI_ISL_867699, EPI_ISL_867700, EPI_ISL_867701, EPI_ISL_867727, EPI_ISL_867732, EPI_ISL_867733, EPI_ISL_867734, EPI_ISL_867735, EPI_ISL_867909, EPI_ISL_867917, EPI_ISL_867922, EPI_ISL_867923, EPI_ISL_867924, EPI_ISL_867926, EPI_ISL_867927, EPI_ISL_867928, EPI_ISL_867929 | | | |
| see above | Originating lab: Wales Specialist Virology Centre Sequencing lab: Pathogen Genomics Unit | Public Health Wales Microbiology Cardiff Wales Specialist Virology Centre | Catherine Moore, Johnathan Evans, Laura Gifford, Malorie Perry, Simon Cottrell, Angela Marchbank, Alec Birchley, Alexander Adams, Amy Gaskin, Bree Gatica-Wilcox, Jason Coombes, Joel Southgate, Lauren Gilbert, Lee Graham, Nicole Pacchiarini, Sara Kumziene-Summerhayes, Sarah Taylor, Sophie Jones, Sara Rey, Matthew Bull, Joanne Watkins, Sally Corden, Tom Connor |
| EPI_ISL_868034, EPI_ISL_868102, EPI_ISL_868155, EPI_ISL_868156, EPI_ISL_868157, EPI_ISL_868158, EPI_ISL_868159, EPI_ISL_868160, EPI_ISL_868175, EPI_ISL_868176, EPI_ISL_868180, EPI_ISL_868195, EPI_ISL_868205, EPI_ISL_868206, EPI_ISL_868207, EPI_ISL_868208, EPI_ISL_868209, EPI_ISL_868210, EPI_ISL_868212, EPI_ISL_868213, EPI_ISL_868214, EPI_ISL_868215, EPI_ISL_868216, EPI_ISL_868219, EPI_ISL_868220, EPI_ISL_868221, EPI_ISL_868222, EPI_ISL_868224, EPI_ISL_868225, EPI_ISL_868238 | | | |
| see above | Centre for Enzyme Innovation, University of Portsmouth / Translational Research Laboratory, Portsmouth Hospitals NHS Trust | COVID-19 Genomics UK (COG-UK) Consortium | Angela Beckett,Yann Bourgeois,Garry Scarlett,Sharon Glaysher,Scott Elliott,Kelly Bicknell,Robert Impey,Allyson Lloyd,Sarah Wyllie,Ethan Butcher,Anoop Chauhan,Samuel Robson |
| EPI_ISL_868361, EPI_ISL_868362, EPI_ISL_868365, EPI_ISL_868367, EPI_ISL_868368, EPI_ISL_868378, EPI_ISL_868379, EPI_ISL_868389, EPI_ISL_868391, EPI_ISL_868398, EPI_ISL_868407, EPI_ISL_868424, EPI_ISL_868425, EPI_ISL_868440, EPI_ISL_868451, EPI_ISL_868476, EPI_ISL_868482, EPI_ISL_868484, EPI_ISL_868492, EPI_ISL_868496, EPI_ISL_868500, EPI_ISL_868523, EPI_ISL_868527, EPI_ISL_868536, EPI_ISL_868578, EPI_ISL_868579, EPI_ISL_868602, EPI_ISL_868618, EPI_ISL_868626, EPI_ISL_868632, EPI_ISL_868639, EPI_ISL_868642, EPI_ISL_868661, EPI_ISL_868662, EPI_ISL_868679, EPI_ISL_868690, EPI_ISL_868696, EPI_ISL_868698, EPI_ISL_868703 | | | |
| see above | Virology Department, Sheffield Teaching Hospitals NHS Foundation Trust/Department of Infection, Immunity and Cardiovascular Disease, The Medical School, University of Sheffield | COVID-19 Genomics UK (COG-UK) Consortium | Thushan de Silva, Matthew Parker, Nikki Smith, Adri Angyal, Rebecca Brown, Luke Green, Rachel Tucker, Paul Parsons, Danielle Groves, Katie Johnson, Laura Carrilero, Alex Keeley, Dave Partridge, Matthew Wyles, Benjamin Lindsey, Mehmet Yavuz, Mohammad Raza, Cariad Evans |
| EPI_ISL_869082 | A. Krumbholz, Labor Dr. Krause und Kollegen MVZ GmbH, Kiel | Charité Universitätsmedizin Berlin, Institut für Virologie | Victor M Corman, Barbara Mühlemann, Jörn Beheim-Schwarzbach, Tobias Bleicker, Julia Tesch, Talitha Veith, Julia Schneider, Terry Jones, Christian Drosten |
| EPI_ISL_869085 | Charité Universitätsmedizin Berlin, Institut für Virologie/Labor Berlin | Charité Universitätsmedizin Berlin, Institut für Virologie | Victor M Corman, Barbara Mühlemann, Jörn Beheim-Schwarzbach, Tobias Bleicker, Julia Tesch, Talitha Veith, Julia Schneider, Terry Jones, Christian Drosten |
| EPI_ISL_869086, EPI_ISL_869087, EPI_ISL_869088, EPI_ISL_869089, EPI_ISL_869090, EPI_ISL_869091, EPI_ISL_869092, EPI_ISL_869093, EPI_ISL_869094, EPI_ISL_869095, EPI_ISL_869096, EPI_ISL_869097, EPI_ISL_869098, EPI_ISL_869099, EPI_ISL_869100, EPI_ISL_869101 | | | |
| see above | A. Krumbholz, Labor Dr. Krause und Kollegen MVZ GmbH, Kiel | Charité Universitätsmedizin Berlin, Institut für Virologie | Victor M Corman, Barbara Mühlemann, Jörn Beheim-Schwarzbach, Tobias Bleicker, Julia Tesch, Talitha Veith, Julia Schneider, Terry Jones, Christian Drosten |
| EPI_ISL_869131, EPI_ISL_869134, EPI_ISL_869136, EPI_ISL_869143 | Charité Universitätsmedizin Berlin, Institut für Virologie/Labor Berlin | Charité Universitätsmedizin Berlin, Institut für Virologie | Victor M Corman, Barbara Mühlemann, Jörn Beheim-Schwarzbach, Tobias Bleicker, Julia Tesch, Talitha Veith, Julia Schneider, Terry Jones, Christian Drosten |
| EPI_ISL_869241 | Suceava County Emergency Hospital | "Stefan cel Mare" University Metagenomics Laboratory | Lobiuc Andrei, Puscaselu Roxana |
| EPI_ISL_872038, EPI_ISL_872039, EPI_ISL_872042, EPI_ISL_872043, EPI_ISL_872049, EPI_ISL_872050 | Department of Clinical Microbiology | GIGA Medical Genomics | Keith Durkin, Maria Artesi, Sébastien Bontems, Raphaël Boreux, Bouchra Boujemla, Cécile Meex, Pierrette Melin, Marie-Pierre Hayette, Vincent Bours |
| EPI_ISL_872067, EPI_ISL_872068 | AZ Vesale (Tongres) | GIGA Medical Genomics | Keith Durkin, Maria Artesi, Sébastien Bontems, Raphaël Boreux, Bouchra Boujemla, Cécile Meex, Pierrette Melin, Marie-Pierre Hayette, Vincent Bours |
| EPI_ISL_872069, EPI_ISL_872070, | Department of Clinical Microbiology | GIGA Medical Genomics | Keith Durkin, Maria Artesi, Sébastien Bontems, Raphaël Boreux, Bouchra Boujemla, Cécile Meex, Pierrette Melin, Marie-Pierre Hayette, Vincent Bours |

| | | | |
|---|---|---|---|
| EPI_ISL_872071, EPI_ISL_872072 | | | |
| EPI_ISL_872152 | CHR Citadelle | GIGA Medical Genomics | Keith Durkin, Maria Artesi, Sébastien Bontems, Raphaël Boreux, Bouchra Boujemla, Cécile Meex, Pierrette Melin, Marie-Pierre Hayette, Vincent Bours |
| EPI_ISL_872176, EPI_ISL_872177, EPI_ISL_872180, EPI_ISL_872181 | Wyoming Public Health Laboratory | Wyoming Public Health Laboratory | Noah Hull, Taylor Fearing, Lynette Gumbleton, Channing Weber, Ashley Norberg, Bailey Bowcutt, and Wanda Manley |
| EPI_ISL_872211 | Hopital | National Reference Center for Viruses of Respiratory Infections, Institut Pasteur, Paris | Marion Barbet, Sylvie Behillil, Méline Bizard, Angela Brisebarre, Camille Capel, Etienne Simon-Lorière, Vincent Enouf, Maud Vanpeene, Sylvie van der Werf,Guigon AuréLie |
| EPI_ISL_872220 | Hopital | National Reference Center for Viruses of Respiratory Infections, Institut Pasteur, Paris | Marion Barbet, Sylvie Behillil, Méline Bizard, Angela Brisebarre, Camille Capel, Etienne Simon-Lorière, Vincent Enouf, Maud Vanpeene, Sylvie van der Werf,Brichler Ségolène |
| EPI_ISL_872262 | Hopital | National Reference Center for Viruses of Respiratory Infections, Institut Pasteur, Paris | Marion Barbet, Sylvie Behillil, Méline Bizard, Angela Brisebarre, Camille Capel, Etienne Simon-Lorière, Vincent Enouf, Maud Vanpeene, Sylvie van der Werf,Florin CéCile |
| EPI_ISL_872269 | Hopital | National Reference Center for Viruses of Respiratory Infections, Institut Pasteur, Paris | Marion Barbet, Sylvie Behillil, Méline Bizard, Angela Brisebarre, Camille Capel, Etienne Simon-Lorière, Vincent Enouf, Maud Vanpeene, Sylvie van der Werf,Foissaud Vincent |
| EPI_ISL_872284 | Labo Analyses Med | National Reference Center for Viruses of Respiratory Infections, Institut Pasteur, Paris | Marion Barbet, Sylvie Behillil, Méline Bizard, Angela Brisebarre, Camille Capel, Etienne Simon-Lorière, Vincent Enouf, Maud Vanpeene, Sylvie van der Werf |
| EPI_ISL_872302, EPI_ISL_872303, EPI_ISL_872304, EPI_ISL_872305 | Hopital | National Reference Center for Viruses of Respiratory Infections, Institut Pasteur, Paris | Marion Barbet, Sylvie Behillil, Méline Bizard, Angela Brisebarre, Camille Capel, Etienne Simon-Lorière, Vincent Enouf, Maud Vanpeene, Sylvie van der Werf,Florin CéCile |
| EPI_ISL_872306, EPI_ISL_872307 | Hopital | National Reference Center for Viruses of Respiratory Infections, Institut Pasteur, Paris | Marion Barbet, Sylvie Behillil, Méline Bizard, Angela Brisebarre, Camille Capel, Etienne Simon-Lorière, Vincent Enouf, Maud Vanpeene, Sylvie van der Werf,Castelain Sandrine |
| EPI_ISL_872324 | Labo Analyses Med | National Reference Center for Viruses of Respiratory Infections, Institut Pasteur, Paris | Marion Barbet, Sylvie Behillil, Méline Bizard, Angela Brisebarre, Camille Capel, Etienne Simon-Lorière, Vincent Enouf, Maud Vanpeene, Sylvie van der Werf |
| EPI_ISL_872394 | Centre for Dengue Research and AICBU, Department of Immunology and Molecular Medicine | Centre for Dengue Research and AICBU, Department of Immunology and Molecular Medicine | Chandima Jeewandara, Deshni Jayathilaka, Dinuka Ariyaratne, Tibutius Thanesh Pramanayagam, Diyanath Ranasinghe, Laksiri Gomes, Gathsaurie Neelika Malavige |
| EPI_ISL_872513, EPI_ISL_872514, EPI_ISL_872515, EPI_ISL_872516, EPI_ISL_872517, EPI_ISL_872518, EPI_ISL_872519, EPI_ISL_872520, EPI_ISL_872521, EPI_ISL_872522, EPI_ISL_872523, EPI_ISL_872524, EPI_ISL_872525, EPI_ISL_872526, EPI_ISL_872527, EPI_ISL_872528, EPI_ISL_872529, EPI_ISL_872530, EPI_ISL_872531, EPI_ISL_872532, EPI_ISL_872533, EPI_ISL_872534, EPI_ISL_872535, EPI_ISL_872536, EPI_ISL_872537, EPI_ISL_872538, EPI_ISL_872539, EPI_ISL_872540, EPI_ISL_872541, EPI_ISL_872542, EPI_ISL_872543, EPI_ISL_872544, EPI_ISL_872545, EPI_ISL_872546, EPI_ISL_872547, EPI_ISL_872548, EPI_ISL_872549, EPI_ISL_872550, EPI_ISL_872551, EPI_ISL_872552, EPI_ISL_872553, EPI_ISL_872554, EPI_ISL_872555, EPI_ISL_872556, EPI_ISL_872562, EPI_ISL_872563 | | | |
| see above | New Mexico Department of Health Scientific Laboratory | Center for Global Health, University of New Mexico Health Sciences Center | Daryl Domman, Kurt Schwalm, Twila Kunde, Joseph Hicks, Anastacia Griego, Michael Edwards, Darrell Dinwiddie |
| EPI_ISL_872588 | Sydney South West Pathology Service (SSWPS) - Royal Prince Alfred Hospital - NSW Health Pathology | NSW Health Pathology - Institute of Clinical Pathology and Medical Research; Westmead Hospital; University of Sydney | CIDM-PH et al. |
| EPI_ISL_872778, EPI_ISL_872779, EPI_ISL_872780, EPI_ISL_872782, EPI_ISL_872783, EPI_ISL_872784, EPI_ISL_872785, EPI_ISL_872786, EPI_ISL_872787, EPI_ISL_872788, EPI_ISL_872789, EPI_ISL_872790, EPI_ISL_872791, EPI_ISL_872793, EPI_ISL_872796, EPI_ISL_872800, EPI_ISL_872801, EPI_ISL_872805, EPI_ISL_872806, EPI_ISL_872807, EPI_ISL_872810, EPI_ISL_872812, EPI_ISL_872814, EPI_ISL_872816, EPI_ISL_872817, EPI_ISL_872820, EPI_ISL_872822, EPI_ISL_872823, EPI_ISL_872824, EPI_ISL_872828, EPI_ISL_872829, EPI_ISL_872833, EPI_ISL_872835, EPI_ISL_872837, EPI_ISL_872839, EPI_ISL_872840, EPI_ISL_872841, EPI_ISL_872842, EPI_ISL_872843, EPI_ISL_872845, EPI_ISL_872847, EPI_ISL_872854, EPI_ISL_872856, EPI_ISL_872858, EPI_ISL_872862, EPI_ISL_872864, EPI_ISL_872867, EPI_ISL_872869, EPI_ISL_872870, EPI_ISL_872872, EPI_ISL_872874, EPI_ISL_872880, EPI_ISL_872882, EPI_ISL_872886, EPI_ISL_872888, EPI_ISL_872890, EPI_ISL_872891, EPI_ISL_872893, EPI_ISL_872894 | | | |
| see above | University of Wisconsin-Madison AIDS Vaccine Research Laboratories | University of Wisconsin-Madison AIDS Vaccine Research Laboratories | Gage Moreno, Katarina Braun, et al. AIDS Vaccine Research Laboratories |
| EPI_ISL_873158, EPI_ISL_873160 | University of Michigan Clinical Microbiology Laboratory | Lauring Lab, University of Michigan, Department of Microbiology and Immunology | Valesano |
| EPI_ISL_873281 | Lighthouse Lab in Milton Keynes | Wellcome Sanger Institute for the COVID-19 Genomics UK (COG-UK) Consortium | The Lighthouse Lab in Milton Keynes and Alex Alderton, Roberto Amato, Sonia Goncalves, Ewan Harrison, David K. Jackson, Ian Johnston, Dominic Kwiatkowski, Cordelia Langford, John Sillitoe on behalf of the Wellcome Sanger Institute COVID-19 Surveillance Team |
| EPI_ISL_873292 | Lighthouse Lab in Cambridge | Wellcome Sanger Institute for the COVID-19 Genomics UK (COG-UK) Consortium | Rob Howes, The Lighthouse Lab in Cambridge and Alex Alderton, Roberto Amato, Sonia Goncalves, Ewan Harrison, David K. Jackson, Ian Johnston, Dominic Kwiatkowski, Cordelia Langford, John Sillitoe on behalf of the Wellcome Sanger Institute COVID-19 Surveillance Team |
| EPI_ISL_873294 | Lighthouse Lab in Milton Keynes | Wellcome Sanger Institute for the COVID-19 Genomics UK (COG-UK) Consortium | The Lighthouse Lab in Milton Keynes and Alex Alderton, Roberto Amato, Sonia Goncalves, Ewan Harrison, David K. Jackson, Ian Johnston, Dominic Kwiatkowski, Cordelia Langford, John Sillitoe on behalf of the Wellcome Sanger Institute COVID-19 Surveillance Team |
| EPI_ISL_873297 | Lighthouse Lab in Cambridge | Wellcome Sanger Institute for the COVID-19 Genomics UK (COG-UK) Consortium | Rob Howes, The Lighthouse Lab in Cambridge and Alex Alderton, Roberto Amato, Sonia Goncalves, Ewan Harrison, David K. Jackson, Ian Johnston, Dominic Kwiatkowski, Cordelia Langford, John Sillitoe on behalf of the Wellcome Sanger Institute COVID-19 Surveillance Team |
| EPI_ISL_873298 | Lighthouse Lab in Milton Keynes | Wellcome Sanger Institute for the COVID-19 Genomics UK (COG-UK) Consortium | The Lighthouse Lab in Milton Keynes and Alex Alderton, Roberto Amato, Sonia Goncalves, Ewan Harrison, David K. Jackson, Ian Johnston, Dominic Kwiatkowski, Cordelia Langford, John Sillitoe on behalf of the Wellcome Sanger Institute COVID-19 Surveillance Team |
| EPI_ISL_873302, EPI_ISL_873304, EPI_ISL_873306, EPI_ISL_873310 | Lighthouse Lab in Cambridge | Wellcome Sanger Institute for the COVID-19 Genomics UK (COG-UK) Consortium | Rob Howes, The Lighthouse Lab in Cambridge and Alex Alderton, Roberto Amato, Sonia Goncalves, Ewan Harrison, David K. Jackson, Ian Johnston, Dominic Kwiatkowski, Cordelia Langford, John Sillitoe on behalf of the Wellcome Sanger Institute COVID-19 Surveillance Team |
| EPI_ISL_873311 | Lighthouse Lab in Milton Keynes | Wellcome Sanger Institute for the COVID-19 Genomics UK (COG-UK) Consortium | The Lighthouse Lab in Milton Keynes and Alex Alderton, Roberto Amato, Sonia Goncalves, Ewan Harrison, David K. Jackson, Ian Johnston, Dominic Kwiatkowski, Cordelia Langford, John Sillitoe on behalf of the Wellcome Sanger Institute COVID-19 Surveillance Team |
| EPI_ISL_873313, EPI_ISL_873316, EPI_ISL_873318, EPI_ISL_873320 | Lighthouse Lab in Cambridge | Wellcome Sanger Institute for the COVID-19 Genomics UK (COG-UK) Consortium | Rob Howes, The Lighthouse Lab in Cambridge and Alex Alderton, Roberto Amato, Sonia Goncalves, Ewan Harrison, David K. Jackson, Ian Johnston, Dominic Kwiatkowski, Cordelia Langford, John Sillitoe on behalf of the Wellcome Sanger Institute COVID-19 Surveillance Team |
| EPI_ISL_873321 | Lighthouse Lab in Milton Keynes | Wellcome Sanger Institute for the COVID-19 Genomics UK (COG-UK) Consortium | The Lighthouse Lab in Milton Keynes and Alex Alderton, Roberto Amato, Sonia Goncalves, Ewan Harrison, David K. Jackson, Ian Johnston, Dominic Kwiatkowski, Cordelia Langford, John Sillitoe on behalf of the Wellcome Sanger Institute COVID-19 Surveillance Team |
| EPI_ISL_873324, EPI_ISL_873325, EPI_ISL_873326 | Lighthouse Lab in Cambridge | Wellcome Sanger Institute for the COVID-19 Genomics UK (COG-UK) Consortium | Rob Howes, The Lighthouse Lab in Cambridge and Alex Alderton, Roberto Amato, Sonia Goncalves, Ewan Harrison, David K. Jackson, Ian Johnston, Dominic Kwiatkowski, Cordelia Langford, John Sillitoe on behalf of the Wellcome Sanger Institute COVID-19 Surveillance Team |
| EPI_ISL_873330 | Lighthouse Lab in Milton Keynes | Wellcome Sanger Institute for the COVID-19 Genomics UK (COG-UK) Consortium | The Lighthouse Lab in Milton Keynes and Alex Alderton, Roberto Amato, Sonia Goncalves, Ewan Harrison, David K. Jackson, Ian Johnston, Dominic Kwiatkowski, Cordelia Langford, John Sillitoe on behalf of the Wellcome Sanger Institute COVID-19 Surveillance Team |
| EPI_ISL_873334 | Lighthouse Lab in Cambridge | Wellcome Sanger Institute for the COVID-19 Genomics UK (COG-UK) Consortium | Rob Howes, The Lighthouse Lab in Cambridge and Alex Alderton, Roberto Amato, Sonia Goncalves, Ewan Harrison, David K. Jackson, Ian Johnston, Dominic Kwiatkowski, Cordelia Langford, John Sillitoe on behalf of the Wellcome Sanger Institute COVID-19 Surveillance Team |
| EPI_ISL_873336 | Lighthouse Lab in Milton Keynes | Wellcome Sanger Institute for the COVID-19 Genomics UK (COG-UK) Consortium | The Lighthouse Lab in Milton Keynes and Alex Alderton, Roberto Amato, Sonia Goncalves, Ewan Harrison, David K. Jackson, Ian Johnston, Dominic Kwiatkowski, Cordelia Langford, John Sillitoe on behalf of the Wellcome Sanger Institute COVID-19 Surveillance Team |
| EPI_ISL_873337, EPI_ISL_873339, EPI_ISL_873341, EPI_ISL_873342, EPI_ISL_873343, EPI_ISL_873350 | Lighthouse Lab in Cambridge | Wellcome Sanger Institute for the COVID-19 Genomics UK (COG-UK) Consortium | Rob Howes, The Lighthouse Lab in Cambridge and Alex Alderton, Roberto Amato, Sonia Goncalves, Ewan Harrison, David K. Jackson, Ian Johnston, Dominic Kwiatkowski, Cordelia Langford, John Sillitoe on behalf of the Wellcome Sanger Institute COVID-19 Surveillance Team |
| EPI_ISL_873354 | Lighthouse Lab in Milton Keynes | Wellcome Sanger Institute for the COVID-19 Genomics UK (COG-UK) Consortium | The Lighthouse Lab in Milton Keynes and Alex Alderton, Roberto Amato, Sonia Goncalves, Ewan Harrison, David K. Jackson, Ian Johnston, Dominic Kwiatkowski, Cordelia Langford, John Sillitoe on behalf of the Wellcome Sanger Institute COVID-19 Surveillance Team |
| EPI_ISL_873357, EPI_ISL_873359 | Lighthouse Lab in Cambridge | Wellcome Sanger Institute for the COVID-19 Genomics UK (COG-UK) Consortium | Rob Howes, The Lighthouse Lab in Cambridge and Alex Alderton, Roberto Amato, Sonia Goncalves, Ewan Harrison, David K. Jackson, Ian Johnston, Dominic Kwiatkowski, Cordelia Langford, John Sillitoe on behalf of the Wellcome Sanger Institute COVID-19 Surveillance Team |
| EPI_ISL_873361 | Lighthouse Lab in Milton Keynes | Wellcome Sanger Institute for the COVID-19 Genomics UK | The Lighthouse Lab in Milton Keynes and Alex Alderton, Roberto Amato, Sonia Goncalves, Ewan Harrison, David K. Jackson, Ian Johnston, Dominic |

| | | | |
|---|---|---|---|
| | | (COG-UK) Consortium | Kwiatkowski, Cordelia Langford, John Sillitoe on behalf of the Wellcome Sanger Institute COVID-19 Surveillance Team |
| EPI_ISL_873362, EPI_ISL_873364, EPI_ISL_873374 | Lighthouse Lab in Cambridge | Wellcome Sanger Institute for the COVID-19 Genomics UK (COG-UK) Consortium | Rob Howes, The Lighthouse Lab in Cambridge and Alex Alderton, Roberto Amato, Sonia Goncalves, Ewan Harrison, David K. Jackson, Ian Johnston, Dominic Kwiatkowski, Cordelia Langford, John Sillitoe on behalf of the Wellcome Sanger Institute COVID-19 Surveillance Team |
| EPI_ISL_873376, EPI_ISL_873378, EPI_ISL_873386, EPI_ISL_873389, EPI_ISL_873396, EPI_ISL_873411 | Lighthouse Lab in Milton Keynes | Wellcome Sanger Institute for the COVID-19 Genomics UK (COG-UK) Consortium | The Lighthouse Lab in Milton Keynes and Alex Alderton, Roberto Amato, Sonia Goncalves, Ewan Harrison, David K. Jackson, Ian Johnston, Dominic Kwiatkowski, Cordelia Langford, John Sillitoe on behalf of the Wellcome Sanger Institute COVID-19 Surveillance Team |
| EPI_ISL_873412 | Lighthouse Lab in Cambridge | Wellcome Sanger Institute for the COVID-19 Genomics UK (COG-UK) Consortium | Rob Howes, The Lighthouse Lab in Cambridge and Alex Alderton, Roberto Amato, Sonia Goncalves, Ewan Harrison, David K. Jackson, Ian Johnston, Dominic Kwiatkowski, Cordelia Langford, John Sillitoe on behalf of the Wellcome Sanger Institute COVID-19 Surveillance Team |
| EPI_ISL_873413, EPI_ISL_873414, EPI_ISL_873415 | Lighthouse Lab in Milton Keynes | Wellcome Sanger Institute for the COVID-19 Genomics UK (COG-UK) Consortium | The Lighthouse Lab in Milton Keynes and Alex Alderton, Roberto Amato, Sonia Goncalves, Ewan Harrison, David K. Jackson, Ian Johnston, Dominic Kwiatkowski, Cordelia Langford, John Sillitoe on behalf of the Wellcome Sanger Institute COVID-19 Surveillance Team |
| EPI_ISL_873416 | Lighthouse Lab in Cambridge | Wellcome Sanger Institute for the COVID-19 Genomics UK (COG-UK) Consortium | Rob Howes, The Lighthouse Lab in Cambridge and Alex Alderton, Roberto Amato, Sonia Goncalves, Ewan Harrison, David K. Jackson, Ian Johnston, Dominic Kwiatkowski, Cordelia Langford, John Sillitoe on behalf of the Wellcome Sanger Institute COVID-19 Surveillance Team |
| EPI_ISL_873420 | Lighthouse Lab in Milton Keynes | Wellcome Sanger Institute for the COVID-19 Genomics UK (COG-UK) Consortium | The Lighthouse Lab in Milton Keynes and Alex Alderton, Roberto Amato, Sonia Goncalves, Ewan Harrison, David K. Jackson, Ian Johnston, Dominic Kwiatkowski, Cordelia Langford, John Sillitoe on behalf of the Wellcome Sanger Institute COVID-19 Surveillance Team |
| EPI_ISL_873422 | Lighthouse Lab in Cambridge | Wellcome Sanger Institute for the COVID-19 Genomics UK (COG-UK) Consortium | Rob Howes, The Lighthouse Lab in Cambridge and Alex Alderton, Roberto Amato, Sonia Goncalves, Ewan Harrison, David K. Jackson, Ian Johnston, Dominic Kwiatkowski, Cordelia Langford, John Sillitoe on behalf of the Wellcome Sanger Institute COVID-19 Surveillance Team |
| EPI_ISL_873429, EPI_ISL_873430 | Lighthouse Lab in Milton Keynes | Wellcome Sanger Institute for the COVID-19 Genomics UK (COG-UK) Consortium | The Lighthouse Lab in Milton Keynes and Alex Alderton, Roberto Amato, Sonia Goncalves, Ewan Harrison, David K. Jackson, Ian Johnston, Dominic Kwiatkowski, Cordelia Langford, John Sillitoe on behalf of the Wellcome Sanger Institute COVID-19 Surveillance Team |
| EPI_ISL_873431, EPI_ISL_873433 | Lighthouse Lab in Cambridge | Wellcome Sanger Institute for the COVID-19 Genomics UK (COG-UK) Consortium | Rob Howes, The Lighthouse Lab in Cambridge and Alex Alderton, Roberto Amato, Sonia Goncalves, Ewan Harrison, David K. Jackson, Ian Johnston, Dominic Kwiatkowski, Cordelia Langford, John Sillitoe on behalf of the Wellcome Sanger Institute COVID-19 Surveillance Team |
| EPI_ISL_873435, EPI_ISL_873438, EPI_ISL_873439 | Lighthouse Lab in Milton Keynes | Wellcome Sanger Institute for the COVID-19 Genomics UK (COG-UK) Consortium | The Lighthouse Lab in Milton Keynes and Alex Alderton, Roberto Amato, Sonia Goncalves, Ewan Harrison, David K. Jackson, Ian Johnston, Dominic Kwiatkowski, Cordelia Langford, John Sillitoe on behalf of the Wellcome Sanger Institute COVID-19 Surveillance Team |
| EPI_ISL_873443 | Lighthouse Lab in Cambridge | Wellcome Sanger Institute for the COVID-19 Genomics UK (COG-UK) Consortium | Rob Howes, The Lighthouse Lab in Cambridge and Alex Alderton, Roberto Amato, Sonia Goncalves, Ewan Harrison, David K. Jackson, Ian Johnston, Dominic Kwiatkowski, Cordelia Langford, John Sillitoe on behalf of the Wellcome Sanger Institute COVID-19 Surveillance Team |
| EPI_ISL_873444, EPI_ISL_873445, EPI_ISL_873450 | Lighthouse Lab in Milton Keynes | Wellcome Sanger Institute for the COVID-19 Genomics UK (COG-UK) Consortium | The Lighthouse Lab in Milton Keynes and Alex Alderton, Roberto Amato, Sonia Goncalves, Ewan Harrison, David K. Jackson, Ian Johnston, Dominic Kwiatkowski, Cordelia Langford, John Sillitoe on behalf of the Wellcome Sanger Institute COVID-19 Surveillance Team |
| EPI_ISL_873454 | Lighthouse Lab in Cambridge | Wellcome Sanger Institute for the COVID-19 Genomics UK (COG-UK) Consortium | Rob Howes, The Lighthouse Lab in Cambridge and Alex Alderton, Roberto Amato, Sonia Goncalves, Ewan Harrison, David K. Jackson, Ian Johnston, Dominic Kwiatkowski, Cordelia Langford, John Sillitoe on behalf of the Wellcome Sanger Institute COVID-19 Surveillance Team |
| EPI_ISL_873455 | Lighthouse Lab in Milton Keynes | Wellcome Sanger Institute for the COVID-19 Genomics UK (COG-UK) Consortium | The Lighthouse Lab in Milton Keynes and Alex Alderton, Roberto Amato, Sonia Goncalves, Ewan Harrison, David K. Jackson, Ian Johnston, Dominic Kwiatkowski, Cordelia Langford, John Sillitoe on behalf of the Wellcome Sanger Institute COVID-19 Surveillance Team |
| EPI_ISL_873457, EPI_ISL_873459 | Lighthouse Lab in Cambridge | Wellcome Sanger Institute for the COVID-19 Genomics UK (COG-UK) Consortium | Rob Howes, The Lighthouse Lab in Cambridge and Alex Alderton, Roberto Amato, Sonia Goncalves, Ewan Harrison, David K. Jackson, Ian Johnston, Dominic Kwiatkowski, Cordelia Langford, John Sillitoe on behalf of the Wellcome Sanger Institute COVID-19 Surveillance Team |
| EPI_ISL_873462, EPI_ISL_873466, EPI_ISL_873473 | Lighthouse Lab in Milton Keynes | Wellcome Sanger Institute for the COVID-19 Genomics UK (COG-UK) Consortium | The Lighthouse Lab in Milton Keynes and Alex Alderton, Roberto Amato, Sonia Goncalves, Ewan Harrison, David K. Jackson, Ian Johnston, Dominic Kwiatkowski, Cordelia Langford, John Sillitoe on behalf of the Wellcome Sanger Institute COVID-19 Surveillance Team |
| EPI_ISL_873476, EPI_ISL_873481 | Lighthouse Lab in Cambridge | Wellcome Sanger Institute for the COVID-19 Genomics UK (COG-UK) Consortium | Rob Howes, The Lighthouse Lab in Cambridge and Alex Alderton, Roberto Amato, Sonia Goncalves, Ewan Harrison, David K. Jackson, Ian Johnston, Dominic Kwiatkowski, Cordelia Langford, John Sillitoe on behalf of the Wellcome Sanger Institute COVID-19 Surveillance Team |
| EPI_ISL_873487, EPI_ISL_873493 | Lighthouse Lab in Milton Keynes | Wellcome Sanger Institute for the COVID-19 Genomics UK (COG-UK) Consortium | The Lighthouse Lab in Milton Keynes and Alex Alderton, Roberto Amato, Sonia Goncalves, Ewan Harrison, David K. Jackson, Ian Johnston, Dominic Kwiatkowski, Cordelia Langford, John Sillitoe on behalf of the Wellcome Sanger Institute COVID-19 Surveillance Team |
| EPI_ISL_873494, EPI_ISL_873495 | Lighthouse Lab in Cambridge | Wellcome Sanger Institute for the COVID-19 Genomics UK (COG-UK) Consortium | Rob Howes, The Lighthouse Lab in Cambridge and Alex Alderton, Roberto Amato, Sonia Goncalves, Ewan Harrison, David K. Jackson, Ian Johnston, Dominic Kwiatkowski, Cordelia Langford, John Sillitoe on behalf of the Wellcome Sanger Institute COVID-19 Surveillance Team |
| EPI_ISL_873496 | Lighthouse Lab in Milton Keynes | Wellcome Sanger Institute for the COVID-19 Genomics UK (COG-UK) Consortium | The Lighthouse Lab in Milton Keynes and Alex Alderton, Roberto Amato, Sonia Goncalves, Ewan Harrison, David K. Jackson, Ian Johnston, Dominic Kwiatkowski, Cordelia Langford, John Sillitoe on behalf of the Wellcome Sanger Institute COVID-19 Surveillance Team |
| EPI_ISL_873497, EPI_ISL_873499, EPI_ISL_873500 | Lighthouse Lab in Cambridge | Wellcome Sanger Institute for the COVID-19 Genomics UK (COG-UK) Consortium | Rob Howes, The Lighthouse Lab in Cambridge and Alex Alderton, Roberto Amato, Sonia Goncalves, Ewan Harrison, David K. Jackson, Ian Johnston, Dominic Kwiatkowski, Cordelia Langford, John Sillitoe on behalf of the Wellcome Sanger Institute COVID-19 Surveillance Team |
| EPI_ISL_873503, EPI_ISL_873504, EPI_ISL_873509 | Lighthouse Lab in Milton Keynes | Wellcome Sanger Institute for the COVID-19 Genomics UK (COG-UK) Consortium | The Lighthouse Lab in Milton Keynes and Alex Alderton, Roberto Amato, Sonia Goncalves, Ewan Harrison, David K. Jackson, Ian Johnston, Dominic Kwiatkowski, Cordelia Langford, John Sillitoe on behalf of the Wellcome Sanger Institute COVID-19 Surveillance Team |
| EPI_ISL_873510 | Lighthouse Lab in Cambridge | Wellcome Sanger Institute for the COVID-19 Genomics UK (COG-UK) Consortium | Rob Howes, The Lighthouse Lab in Cambridge and Alex Alderton, Roberto Amato, Sonia Goncalves, Ewan Harrison, David K. Jackson, Ian Johnston, Dominic Kwiatkowski, Cordelia Langford, John Sillitoe on behalf of the Wellcome Sanger Institute COVID-19 Surveillance Team |
| EPI_ISL_873515, EPI_ISL_873520, EPI_ISL_873531 | Lighthouse Lab in Milton Keynes | Wellcome Sanger Institute for the COVID-19 Genomics UK (COG-UK) Consortium | The Lighthouse Lab in Milton Keynes and Alex Alderton, Roberto Amato, Sonia Goncalves, Ewan Harrison, David K. Jackson, Ian Johnston, Dominic Kwiatkowski, Cordelia Langford, John Sillitoe on behalf of the Wellcome Sanger Institute COVID-19 Surveillance Team |
| EPI_ISL_873532 | Lighthouse Lab in Cambridge | Wellcome Sanger Institute for the COVID-19 Genomics UK (COG-UK) Consortium | Rob Howes, The Lighthouse Lab in Cambridge and Alex Alderton, Roberto Amato, Sonia Goncalves, Ewan Harrison, David K. Jackson, Ian Johnston, Dominic Kwiatkowski, Cordelia Langford, John Sillitoe on behalf of the Wellcome Sanger Institute COVID-19 Surveillance Team |
| EPI_ISL_873551 | Lighthouse Lab in Milton Keynes | Wellcome Sanger Institute for the COVID-19 Genomics UK (COG-UK) Consortium | The Lighthouse Lab in Milton Keynes and Alex Alderton, Roberto Amato, Sonia Goncalves, Ewan Harrison, David K. Jackson, Ian Johnston, Dominic Kwiatkowski, Cordelia Langford, John Sillitoe on behalf of the Wellcome Sanger Institute COVID-19 Surveillance Team |
| EPI_ISL_873552, EPI_ISL_873566, EPI_ISL_873571, EPI_ISL_873573, EPI_ISL_873576 | Lighthouse Lab in Cambridge | Wellcome Sanger Institute for the COVID-19 Genomics UK (COG-UK) Consortium | Rob Howes, The Lighthouse Lab in Cambridge and Alex Alderton, Roberto Amato, Sonia Goncalves, Ewan Harrison, David K. Jackson, Ian Johnston, Dominic Kwiatkowski, Cordelia Langford, John Sillitoe on behalf of the Wellcome Sanger Institute COVID-19 Surveillance Team |
| EPI_ISL_873585, EPI_ISL_873596, EPI_ISL_873600, EPI_ISL_873603, EPI_ISL_873604 | Lighthouse Lab in Milton Keynes | Wellcome Sanger Institute for the COVID-19 Genomics UK (COG-UK) Consortium | The Lighthouse Lab in Milton Keynes and Alex Alderton, Roberto Amato, Sonia Goncalves, Ewan Harrison, David K. Jackson, Ian Johnston, Dominic Kwiatkowski, Cordelia Langford, John Sillitoe on behalf of the Wellcome Sanger Institute COVID-19 Surveillance Team |
| EPI_ISL_873605 | Lighthouse Lab in Cambridge | Wellcome Sanger Institute for the COVID-19 Genomics UK (COG-UK) Consortium | Rob Howes, The Lighthouse Lab in Cambridge and Alex Alderton, Roberto Amato, Sonia Goncalves, Ewan Harrison, David K. Jackson, Ian Johnston, Dominic Kwiatkowski, Cordelia Langford, John Sillitoe on behalf of the Wellcome Sanger Institute COVID-19 Surveillance Team |
| EPI_ISL_873607 | Lighthouse Lab in Milton Keynes | Wellcome Sanger Institute for the COVID-19 Genomics UK (COG-UK) Consortium | The Lighthouse Lab in Milton Keynes and Alex Alderton, Roberto Amato, Sonia Goncalves, Ewan Harrison, David K. Jackson, Ian Johnston, Dominic Kwiatkowski, Cordelia Langford, John Sillitoe on behalf of the Wellcome Sanger Institute COVID-19 Surveillance Team |
| EPI_ISL_873614, EPI_ISL_873621, EPI_ISL_873629, EPI_ISL_873632, EPI_ISL_873633, EPI_ISL_873634, EPI_ISL_873635, EPI_ISL_873636, EPI_ISL_873638, EPI_ISL_873639, EPI_ISL_873640, EPI_ISL_873642, EPI_ISL_873643, EPI_ISL_873644 | | | |
| see above | Lighthouse Lab in Cambridge | Wellcome Sanger Institute for the COVID-19 Genomics UK (COG-UK) Consortium | Rob Howes, The Lighthouse Lab in Cambridge and Alex Alderton, Roberto Amato, Sonia Goncalves, Ewan Harrison, David K. Jackson, Ian Johnston, Dominic Kwiatkowski, Cordelia Langford, John Sillitoe on behalf of the Wellcome Sanger Institute COVID-19 Surveillance Team |
| EPI_ISL_873645 | Lighthouse Lab in Milton Keynes | Wellcome Sanger Institute for the COVID-19 Genomics UK (COG-UK) Consortium | The Lighthouse Lab in Milton Keynes and Alex Alderton, Roberto Amato, Sonia Goncalves, Ewan Harrison, David K. Jackson, Ian Johnston, Dominic Kwiatkowski, Cordelia Langford, John Sillitoe on behalf of the Wellcome Sanger Institute COVID-19 Surveillance Team |
| EPI_ISL_873646, EPI_ISL_873647, EPI_ISL_873648, EPI_ISL_873649, | Lighthouse Lab in Cambridge | Wellcome Sanger Institute for the COVID-19 Genomics UK (COG-UK) Consortium | Rob Howes, The Lighthouse Lab in Cambridge and Alex Alderton, Roberto Amato, Sonia Goncalves, Ewan Harrison, David K. Jackson, Ian Johnston, Dominic Kwiatkowski, Cordelia Langford, John Sillitoe on behalf of the Wellcome Sanger Institute COVID-19 Surveillance Team |

| | | | |
|---|---|---|---|
| EPI_ISL_873650, EPI_ISL_873651 | | | |
| EPI_ISL_873652 | Lighthouse Lab in Milton Keynes | Wellcome Sanger Institute for the COVID-19 Genomics UK (COG-UK) Consortium | The Lighthouse Lab in Milton Keynes and Alex Alderton, Roberto Amato, Sonia Goncalves, Ewan Harrison, David K. Jackson, Ian Johnston, Dominic Kwiatkowski, Cordelia Langford, John Sillitoe on behalf of the Wellcome Sanger Institute COVID-19 Surveillance Team |
| EPI_ISL_873653, EPI_ISL_873655, EPI_ISL_873658, EPI_ISL_873659, EPI_ISL_873661, EPI_ISL_873662, EPI_ISL_873663, EPI_ISL_873664, EPI_ISL_873665, EPI_ISL_873666, EPI_ISL_873667, EPI_ISL_873668, EPI_ISL_873670, EPI_ISL_873671, EPI_ISL_873673, EPI_ISL_873675, EPI_ISL_873676, EPI_ISL_873677, EPI_ISL_873678 | | | |
| see above | Lighthouse Lab in Cambridge | Wellcome Sanger Institute for the COVID-19 Genomics UK (COG-UK) Consortium | Rob Howes, The Lighthouse Lab in Cambridge and Alex Alderton, Roberto Amato, Sonia Goncalves, Ewan Harrison, David K. Jackson, Ian Johnston, Dominic Kwiatkowski, Cordelia Langford, John Sillitoe on behalf of the Wellcome Sanger Institute COVID-19 Surveillance Team |
| EPI_ISL_873679 | Lighthouse Lab in Milton Keynes | Wellcome Sanger Institute for the COVID-19 Genomics UK (COG-UK) Consortium | The Lighthouse Lab in Milton Keynes and Alex Alderton, Roberto Amato, Sonia Goncalves, Ewan Harrison, David K. Jackson, Ian Johnston, Dominic Kwiatkowski, Cordelia Langford, John Sillitoe on behalf of the Wellcome Sanger Institute COVID-19 Surveillance Team |
| EPI_ISL_873680, EPI_ISL_873681, EPI_ISL_873682, EPI_ISL_873683 | Lighthouse Lab in Cambridge | Wellcome Sanger Institute for the COVID-19 Genomics UK (COG-UK) Consortium | Rob Howes, The Lighthouse Lab in Cambridge and Alex Alderton, Roberto Amato, Sonia Goncalves, Ewan Harrison, David K. Jackson, Ian Johnston, Dominic Kwiatkowski, Cordelia Langford, John Sillitoe on behalf of the Wellcome Sanger Institute COVID-19 Surveillance Team |
| EPI_ISL_873684 | Lighthouse Lab in Milton Keynes | Wellcome Sanger Institute for the COVID-19 Genomics UK (COG-UK) Consortium | The Lighthouse Lab in Milton Keynes and Alex Alderton, Roberto Amato, Sonia Goncalves, Ewan Harrison, David K. Jackson, Ian Johnston, Dominic Kwiatkowski, Cordelia Langford, John Sillitoe on behalf of the Wellcome Sanger Institute COVID-19 Surveillance Team |
| EPI_ISL_873685, EPI_ISL_873686, EPI_ISL_873687, EPI_ISL_873688, EPI_ISL_873689, EPI_ISL_873690, EPI_ISL_873691, EPI_ISL_873692, EPI_ISL_873693, EPI_ISL_873694, EPI_ISL_873695, EPI_ISL_873696, EPI_ISL_873697, EPI_ISL_873698, EPI_ISL_873699, EPI_ISL_873700, EPI_ISL_873701, EPI_ISL_873703, EPI_ISL_873705, EPI_ISL_873706, EPI_ISL_873707, EPI_ISL_873708, EPI_ISL_873709, EPI_ISL_873710, EPI_ISL_873711, EPI_ISL_873712, EPI_ISL_873713, EPI_ISL_873714, EPI_ISL_873715 | | | |
| see above | Lighthouse Lab in Cambridge | Wellcome Sanger Institute for the COVID-19 Genomics UK (COG-UK) Consortium | Rob Howes, The Lighthouse Lab in Cambridge and Alex Alderton, Roberto Amato, Sonia Goncalves, Ewan Harrison, David K. Jackson, Ian Johnston, Dominic Kwiatkowski, Cordelia Langford, John Sillitoe on behalf of the Wellcome Sanger Institute COVID-19 Surveillance Team |
| EPI_ISL_873716 | Lighthouse Lab in Milton Keynes | Wellcome Sanger Institute for the COVID-19 Genomics UK (COG-UK) Consortium | The Lighthouse Lab in Milton Keynes and Alex Alderton, Roberto Amato, Sonia Goncalves, Ewan Harrison, David K. Jackson, Ian Johnston, Dominic Kwiatkowski, Cordelia Langford, John Sillitoe on behalf of the Wellcome Sanger Institute COVID-19 Surveillance Team |
| EPI_ISL_873717, EPI_ISL_873718, EPI_ISL_873719, EPI_ISL_873720, EPI_ISL_873721, EPI_ISL_873722, EPI_ISL_873723, EPI_ISL_873727, EPI_ISL_873728, EPI_ISL_873731, EPI_ISL_873732, EPI_ISL_873733, EPI_ISL_873735, EPI_ISL_873736, EPI_ISL_873738, EPI_ISL_873739, EPI_ISL_873740, EPI_ISL_873742, EPI_ISL_873745, EPI_ISL_873747 | | | |
| see above | Lighthouse Lab in Cambridge | Wellcome Sanger Institute for the COVID-19 Genomics UK (COG-UK) Consortium | Rob Howes, The Lighthouse Lab in Cambridge and Alex Alderton, Roberto Amato, Sonia Goncalves, Ewan Harrison, David K. Jackson, Ian Johnston, Dominic Kwiatkowski, Cordelia Langford, John Sillitoe on behalf of the Wellcome Sanger Institute COVID-19 Surveillance Team |
| EPI_ISL_873748 | Lighthouse Lab in Milton Keynes | Wellcome Sanger Institute for the COVID-19 Genomics UK (COG-UK) Consortium | The Lighthouse Lab in Milton Keynes and Alex Alderton, Roberto Amato, Sonia Goncalves, Ewan Harrison, David K. Jackson, Ian Johnston, Dominic Kwiatkowski, Cordelia Langford, John Sillitoe on behalf of the Wellcome Sanger Institute COVID-19 Surveillance Team |
| EPI_ISL_873751, EPI_ISL_873753, EPI_ISL_873754 | Lighthouse Lab in Cambridge | Wellcome Sanger Institute for the COVID-19 Genomics UK (COG-UK) Consortium | Rob Howes, The Lighthouse Lab in Cambridge and Alex Alderton, Roberto Amato, Sonia Goncalves, Ewan Harrison, David K. Jackson, Ian Johnston, Dominic Kwiatkowski, Cordelia Langford, John Sillitoe on behalf of the Wellcome Sanger Institute COVID-19 Surveillance Team |
| EPI_ISL_873755 | Lighthouse Lab in Milton Keynes | Wellcome Sanger Institute for the COVID-19 Genomics UK (COG-UK) Consortium | The Lighthouse Lab in Milton Keynes and Alex Alderton, Roberto Amato, Sonia Goncalves, Ewan Harrison, David K. Jackson, Ian Johnston, Dominic Kwiatkowski, Cordelia Langford, John Sillitoe on behalf of the Wellcome Sanger Institute COVID-19 Surveillance Team |
| EPI_ISL_873756, EPI_ISL_873757, EPI_ISL_873758, EPI_ISL_873759, EPI_ISL_873761, EPI_ISL_873762, EPI_ISL_873763, EPI_ISL_873765, EPI_ISL_873766, EPI_ISL_873767, EPI_ISL_873768, EPI_ISL_873770, EPI_ISL_873771, EPI_ISL_873772, EPI_ISL_873773, EPI_ISL_873774, EPI_ISL_873775, EPI_ISL_873776, EPI_ISL_873777, EPI_ISL_873778, EPI_ISL_873779, EPI_ISL_873780, EPI_ISL_873781, EPI_ISL_873782 | | | |
| see above | Lighthouse Lab in Cambridge | Wellcome Sanger Institute for the COVID-19 Genomics UK (COG-UK) Consortium | Rob Howes, The Lighthouse Lab in Cambridge and Alex Alderton, Roberto Amato, Sonia Goncalves, Ewan Harrison, David K. Jackson, Ian Johnston, Dominic Kwiatkowski, Cordelia Langford, John Sillitoe on behalf of the Wellcome Sanger Institute COVID-19 Surveillance Team |
| EPI_ISL_873783 | Lighthouse Lab in Milton Keynes | Wellcome Sanger Institute for the COVID-19 Genomics UK (COG-UK) Consortium | The Lighthouse Lab in Milton Keynes and Alex Alderton, Roberto Amato, Sonia Goncalves, Ewan Harrison, David K. Jackson, Ian Johnston, Dominic Kwiatkowski, Cordelia Langford, John Sillitoe on behalf of the Wellcome Sanger Institute COVID-19 Surveillance Team |
| EPI_ISL_873784, EPI_ISL_873785, EPI_ISL_873787, EPI_ISL_873788, EPI_ISL_873789, EPI_ISL_873790, EPI_ISL_873791, EPI_ISL_873792, EPI_ISL_873793, EPI_ISL_873795, EPI_ISL_873796 | | | |
| see above | Lighthouse Lab in Cambridge | Wellcome Sanger Institute for the COVID-19 Genomics UK (COG-UK) Consortium | Rob Howes, The Lighthouse Lab in Cambridge and Alex Alderton, Roberto Amato, Sonia Goncalves, Ewan Harrison, David K. Jackson, Ian Johnston, Dominic Kwiatkowski, Cordelia Langford, John Sillitoe on behalf of the Wellcome Sanger Institute COVID-19 Surveillance Team |
| EPI_ISL_873797 | Lighthouse Lab in Milton Keynes | Wellcome Sanger Institute for the COVID-19 Genomics UK (COG-UK) Consortium | The Lighthouse Lab in Milton Keynes and Alex Alderton, Roberto Amato, Sonia Goncalves, Ewan Harrison, David K. Jackson, Ian Johnston, Dominic Kwiatkowski, Cordelia Langford, John Sillitoe on behalf of the Wellcome Sanger Institute COVID-19 Surveillance Team |
| EPI_ISL_873798, EPI_ISL_873799, EPI_ISL_873800, EPI_ISL_873801, EPI_ISL_873802, EPI_ISL_873803, EPI_ISL_873804, EPI_ISL_873805, EPI_ISL_873807, EPI_ISL_873808, EPI_ISL_873810, EPI_ISL_873811, EPI_ISL_873813, EPI_ISL_873815, EPI_ISL_873816, EPI_ISL_873818, EPI_ISL_873819, EPI_ISL_873821 | | | |
| see above | Lighthouse Lab in Cambridge | Wellcome Sanger Institute for the COVID-19 Genomics UK (COG-UK) Consortium | Rob Howes, The Lighthouse Lab in Cambridge and Alex Alderton, Roberto Amato, Sonia Goncalves, Ewan Harrison, David K. Jackson, Ian Johnston, Dominic Kwiatkowski, Cordelia Langford, John Sillitoe on behalf of the Wellcome Sanger Institute COVID-19 Surveillance Team |
| EPI_ISL_873822 | Lighthouse Lab in Milton Keynes | Wellcome Sanger Institute for the COVID-19 Genomics UK (COG-UK) Consortium | The Lighthouse Lab in Milton Keynes and Alex Alderton, Roberto Amato, Sonia Goncalves, Ewan Harrison, David K. Jackson, Ian Johnston, Dominic Kwiatkowski, Cordelia Langford, John Sillitoe on behalf of the Wellcome Sanger Institute COVID-19 Surveillance Team |
| EPI_ISL_873823, EPI_ISL_873824, EPI_ISL_873826 | Lighthouse Lab in Cambridge | Wellcome Sanger Institute for the COVID-19 Genomics UK (COG-UK) Consortium | Rob Howes, The Lighthouse Lab in Cambridge and Alex Alderton, Roberto Amato, Sonia Goncalves, Ewan Harrison, David K. Jackson, Ian Johnston, Dominic Kwiatkowski, Cordelia Langford, John Sillitoe on behalf of the Wellcome Sanger Institute COVID-19 Surveillance Team |
| EPI_ISL_873827 | Lighthouse Lab in Milton Keynes | Wellcome Sanger Institute for the COVID-19 Genomics UK (COG-UK) Consortium | The Lighthouse Lab in Milton Keynes and Alex Alderton, Roberto Amato, Sonia Goncalves, Ewan Harrison, David K. Jackson, Ian Johnston, Dominic Kwiatkowski, Cordelia Langford, John Sillitoe on behalf of the Wellcome Sanger Institute COVID-19 Surveillance Team |
| EPI_ISL_873828 | Lighthouse Lab in Cambridge | Wellcome Sanger Institute for the COVID-19 Genomics UK (COG-UK) Consortium | Rob Howes, The Lighthouse Lab in Cambridge and Alex Alderton, Roberto Amato, Sonia Goncalves, Ewan Harrison, David K. Jackson, Ian Johnston, Dominic Kwiatkowski, Cordelia Langford, John Sillitoe on behalf of the Wellcome Sanger Institute COVID-19 Surveillance Team |
| EPI_ISL_873829 | Lighthouse Lab in Milton Keynes | Wellcome Sanger Institute for the COVID-19 Genomics UK (COG-UK) Consortium | The Lighthouse Lab in Milton Keynes and Alex Alderton, Roberto Amato, Sonia Goncalves, Ewan Harrison, David K. Jackson, Ian Johnston, Dominic Kwiatkowski, Cordelia Langford, John Sillitoe on behalf of the Wellcome Sanger Institute COVID-19 Surveillance Team |
| EPI_ISL_873830, EPI_ISL_873831, EPI_ISL_873832, EPI_ISL_873835, EPI_ISL_873836, EPI_ISL_873837, EPI_ISL_873838 | Lighthouse Lab in Cambridge | Wellcome Sanger Institute for the COVID-19 Genomics UK (COG-UK) Consortium | Rob Howes, The Lighthouse Lab in Cambridge and Alex Alderton, Roberto Amato, Sonia Goncalves, Ewan Harrison, David K. Jackson, Ian Johnston, Dominic Kwiatkowski, Cordelia Langford, John Sillitoe on behalf of the Wellcome Sanger Institute COVID-19 Surveillance Team |
| EPI_ISL_873840 | Lighthouse Lab in Milton Keynes | Wellcome Sanger Institute for the COVID-19 Genomics UK (COG-UK) Consortium | The Lighthouse Lab in Milton Keynes and Alex Alderton, Roberto Amato, Sonia Goncalves, Ewan Harrison, David K. Jackson, Ian Johnston, Dominic Kwiatkowski, Cordelia Langford, John Sillitoe on behalf of the Wellcome Sanger Institute COVID-19 Surveillance Team |
| EPI_ISL_873841 | Lighthouse Lab in Cambridge | Wellcome Sanger Institute for the COVID-19 Genomics UK (COG-UK) Consortium | Rob Howes, The Lighthouse Lab in Cambridge and Alex Alderton, Roberto Amato, Sonia Goncalves, Ewan Harrison, David K. Jackson, Ian Johnston, Dominic Kwiatkowski, Cordelia Langford, John Sillitoe on behalf of the Wellcome Sanger Institute COVID-19 Surveillance Team |
| EPI_ISL_873842 | Lighthouse Lab in Milton Keynes | Wellcome Sanger Institute for the COVID-19 Genomics UK (COG-UK) Consortium | The Lighthouse Lab in Milton Keynes and Alex Alderton, Roberto Amato, Sonia Goncalves, Ewan Harrison, David K. Jackson, Ian Johnston, Dominic Kwiatkowski, Cordelia Langford, John Sillitoe on behalf of the Wellcome Sanger Institute COVID-19 Surveillance Team |
| EPI_ISL_873843, EPI_ISL_873844, EPI_ISL_873845, EPI_ISL_873846 | Lighthouse Lab in Cambridge | Wellcome Sanger Institute for the COVID-19 Genomics UK (COG-UK) Consortium | Rob Howes, The Lighthouse Lab in Cambridge and Alex Alderton, Roberto Amato, Sonia Goncalves, Ewan Harrison, David K. Jackson, Ian Johnston, Dominic Kwiatkowski, Cordelia Langford, John Sillitoe on behalf of the Wellcome Sanger Institute COVID-19 Surveillance Team |
| EPI_ISL_873848 | Lighthouse Lab in Milton Keynes | Wellcome Sanger Institute for the COVID-19 Genomics UK (COG-UK) Consortium | The Lighthouse Lab in Milton Keynes and Alex Alderton, Roberto Amato, Sonia Goncalves, Ewan Harrison, David K. Jackson, Ian Johnston, Dominic Kwiatkowski, Cordelia Langford, John Sillitoe on behalf of the Wellcome Sanger Institute COVID-19 Surveillance Team |
| EPI_ISL_873849, EPI_ISL_873851, EPI_ISL_873852 | Lighthouse Lab in Cambridge | Wellcome Sanger Institute for the COVID-19 Genomics UK (COG-UK) Consortium | Rob Howes, The Lighthouse Lab in Cambridge and Alex Alderton, Roberto Amato, Sonia Goncalves, Ewan Harrison, David K. Jackson, Ian Johnston, Dominic Kwiatkowski, Cordelia Langford, John Sillitoe on behalf of the Wellcome Sanger Institute COVID-19 Surveillance Team |
| EPI_ISL_873853 | Lighthouse Lab in Milton Keynes | Wellcome Sanger Institute for the COVID-19 Genomics UK | The Lighthouse Lab in Milton Keynes and Alex Alderton, Roberto Amato, Sonia Goncalves, Ewan Harrison, David K. Jackson, Ian Johnston, Dominic |

| | | | |
|---|---|---|---|
| | | (COG-UK) Consortium | Kwiatkowski, Cordelia Langford, John Sillitoe on behalf of the Wellcome Sanger Institute COVID-19 Surveillance Team |
| EPI_ISL_873854, EPI_ISL_873855, EPI_ISL_873856, EPI_ISL_873858 | Lighthouse Lab in Cambridge | Wellcome Sanger Institute for the COVID-19 Genomics UK (COG-UK) Consortium | Rob Howes, The Lighthouse Lab in Cambridge and Alex Alderton, Roberto Amato, Sonia Goncalves, Ewan Harrison, David K. Jackson, Ian Johnston, Dominic Kwiatkowski, Cordelia Langford, John Sillitoe on behalf of the Wellcome Sanger Institute COVID-19 Surveillance Team |
| EPI_ISL_873860 | Lighthouse Lab in Milton Keynes | Wellcome Sanger Institute for the COVID-19 Genomics UK (COG-UK) Consortium | The Lighthouse Lab in Milton Keynes and Alex Alderton, Roberto Amato, Sonia Goncalves, Ewan Harrison, David K. Jackson, Ian Johnston, Dominic Kwiatkowski, Cordelia Langford, John Sillitoe on behalf of the Wellcome Sanger Institute COVID-19 Surveillance Team |
| EPI_ISL_873861 | Lighthouse Lab in Cambridge | Wellcome Sanger Institute for the COVID-19 Genomics UK (COG-UK) Consortium | Rob Howes, The Lighthouse Lab in Cambridge and Alex Alderton, Roberto Amato, Sonia Goncalves, Ewan Harrison, David K. Jackson, Ian Johnston, Dominic Kwiatkowski, Cordelia Langford, John Sillitoe on behalf of the Wellcome Sanger Institute COVID-19 Surveillance Team |
| EPI_ISL_873862 | Lighthouse Lab in Milton Keynes | Wellcome Sanger Institute for the COVID-19 Genomics UK (COG-UK) Consortium | The Lighthouse Lab in Milton Keynes and Alex Alderton, Roberto Amato, Sonia Goncalves, Ewan Harrison, David K. Jackson, Ian Johnston, Dominic Kwiatkowski, Cordelia Langford, John Sillitoe on behalf of the Wellcome Sanger Institute COVID-19 Surveillance Team |
| EPI_ISL_873863, EPI_ISL_873864, EPI_ISL_873866, EPI_ISL_873867, EPI_ISL_873868, EPI_ISL_873869, EPI_ISL_873871, EPI_ISL_873872, EPI_ISL_873874, EPI_ISL_873877, EPI_ISL_873878, EPI_ISL_873879 | | | |
| see above | Lighthouse Lab in Cambridge | Wellcome Sanger Institute for the COVID-19 Genomics UK (COG-UK) Consortium | Rob Howes, The Lighthouse Lab in Cambridge and Alex Alderton, Roberto Amato, Sonia Goncalves, Ewan Harrison, David K. Jackson, Ian Johnston, Dominic Kwiatkowski, Cordelia Langford, John Sillitoe on behalf of the Wellcome Sanger Institute COVID-19 Surveillance Team |
| EPI_ISL_873881 | Lighthouse Lab in Milton Keynes | Wellcome Sanger Institute for the COVID-19 Genomics UK (COG-UK) Consortium | The Lighthouse Lab in Milton Keynes and Alex Alderton, Roberto Amato, Sonia Goncalves, Ewan Harrison, David K. Jackson, Ian Johnston, Dominic Kwiatkowski, Cordelia Langford, John Sillitoe on behalf of the Wellcome Sanger Institute COVID-19 Surveillance Team |
| EPI_ISL_873883, EPI_ISL_873885 | Lighthouse Lab in Cambridge | Wellcome Sanger Institute for the COVID-19 Genomics UK (COG-UK) Consortium | Rob Howes, The Lighthouse Lab in Cambridge and Alex Alderton, Roberto Amato, Sonia Goncalves, Ewan Harrison, David K. Jackson, Ian Johnston, Dominic Kwiatkowski, Cordelia Langford, John Sillitoe on behalf of the Wellcome Sanger Institute COVID-19 Surveillance Team |
| EPI_ISL_873886 | Lighthouse Lab in Milton Keynes | Wellcome Sanger Institute for the COVID-19 Genomics UK (COG-UK) Consortium | The Lighthouse Lab in Milton Keynes and Alex Alderton, Roberto Amato, Sonia Goncalves, Ewan Harrison, David K. Jackson, Ian Johnston, Dominic Kwiatkowski, Cordelia Langford, John Sillitoe on behalf of the Wellcome Sanger Institute COVID-19 Surveillance Team |
| EPI_ISL_873888, EPI_ISL_873889, EPI_ISL_873890, EPI_ISL_873891, EPI_ISL_873892, EPI_ISL_873893, EPI_ISL_873894, EPI_ISL_873895, EPI_ISL_873896, EPI_ISL_873897, EPI_ISL_873898, EPI_ISL_873899, EPI_ISL_873901, EPI_ISL_873902, EPI_ISL_873906, EPI_ISL_873907, EPI_ISL_873908, EPI_ISL_873909, EPI_ISL_873911, EPI_ISL_873912, EPI_ISL_873914, EPI_ISL_873915, EPI_ISL_873917, EPI_ISL_873918, EPI_ISL_873919, EPI_ISL_873920 | | | |
| see above | Lighthouse Lab in Cambridge | Wellcome Sanger Institute for the COVID-19 Genomics UK (COG-UK) Consortium | Rob Howes, The Lighthouse Lab in Cambridge and Alex Alderton, Roberto Amato, Sonia Goncalves, Ewan Harrison, David K. Jackson, Ian Johnston, Dominic Kwiatkowski, Cordelia Langford, John Sillitoe on behalf of the Wellcome Sanger Institute COVID-19 Surveillance Team |
| EPI_ISL_873921 | Lighthouse Lab in Milton Keynes | Wellcome Sanger Institute for the COVID-19 Genomics UK (COG-UK) Consortium | The Lighthouse Lab in Milton Keynes and Alex Alderton, Roberto Amato, Sonia Goncalves, Ewan Harrison, David K. Jackson, Ian Johnston, Dominic Kwiatkowski, Cordelia Langford, John Sillitoe on behalf of the Wellcome Sanger Institute COVID-19 Surveillance Team |
| EPI_ISL_873922 | Lighthouse Lab in Cambridge | Wellcome Sanger Institute for the COVID-19 Genomics UK (COG-UK) Consortium | Rob Howes, The Lighthouse Lab in Cambridge and Alex Alderton, Roberto Amato, Sonia Goncalves, Ewan Harrison, David K. Jackson, Ian Johnston, Dominic Kwiatkowski, Cordelia Langford, John Sillitoe on behalf of the Wellcome Sanger Institute COVID-19 Surveillance Team |
| EPI_ISL_873926 | Lighthouse Lab in Milton Keynes | Wellcome Sanger Institute for the COVID-19 Genomics UK (COG-UK) Consortium | The Lighthouse Lab in Milton Keynes and Alex Alderton, Roberto Amato, Sonia Goncalves, Ewan Harrison, David K. Jackson, Ian Johnston, Dominic Kwiatkowski, Cordelia Langford, John Sillitoe on behalf of the Wellcome Sanger Institute COVID-19 Surveillance Team |
| EPI_ISL_873927, EPI_ISL_873929, EPI_ISL_873930, EPI_ISL_873931, EPI_ISL_873932, EPI_ISL_873933, EPI_ISL_873936, EPI_ISL_873938, EPI_ISL_873939, EPI_ISL_873940, EPI_ISL_873941 | | | |
| see above | Lighthouse Lab in Cambridge | Wellcome Sanger Institute for the COVID-19 Genomics UK (COG-UK) Consortium | Rob Howes, The Lighthouse Lab in Cambridge and Alex Alderton, Roberto Amato, Sonia Goncalves, Ewan Harrison, David K. Jackson, Ian Johnston, Dominic Kwiatkowski, Cordelia Langford, John Sillitoe on behalf of the Wellcome Sanger Institute COVID-19 Surveillance Team |
| EPI_ISL_873942 | Lighthouse Lab in Milton Keynes | Wellcome Sanger Institute for the COVID-19 Genomics UK (COG-UK) Consortium | The Lighthouse Lab in Milton Keynes and Alex Alderton, Roberto Amato, Sonia Goncalves, Ewan Harrison, David K. Jackson, Ian Johnston, Dominic Kwiatkowski, Cordelia Langford, John Sillitoe on behalf of the Wellcome Sanger Institute COVID-19 Surveillance Team |
| EPI_ISL_873944, EPI_ISL_873945, EPI_ISL_873946, EPI_ISL_873949 | Lighthouse Lab in Cambridge | Wellcome Sanger Institute for the COVID-19 Genomics UK (COG-UK) Consortium | Rob Howes, The Lighthouse Lab in Cambridge and Alex Alderton, Roberto Amato, Sonia Goncalves, Ewan Harrison, David K. Jackson, Ian Johnston, Dominic Kwiatkowski, Cordelia Langford, John Sillitoe on behalf of the Wellcome Sanger Institute COVID-19 Surveillance Team |
| EPI_ISL_873950 | Lighthouse Lab in Milton Keynes | Wellcome Sanger Institute for the COVID-19 Genomics UK (COG-UK) Consortium | The Lighthouse Lab in Milton Keynes and Alex Alderton, Roberto Amato, Sonia Goncalves, Ewan Harrison, David K. Jackson, Ian Johnston, Dominic Kwiatkowski, Cordelia Langford, John Sillitoe on behalf of the Wellcome Sanger Institute COVID-19 Surveillance Team |
| EPI_ISL_873951, EPI_ISL_873952, EPI_ISL_873953, EPI_ISL_873955 | Lighthouse Lab in Cambridge | Wellcome Sanger Institute for the COVID-19 Genomics UK (COG-UK) Consortium | Rob Howes, The Lighthouse Lab in Cambridge and Alex Alderton, Roberto Amato, Sonia Goncalves, Ewan Harrison, David K. Jackson, Ian Johnston, Dominic Kwiatkowski, Cordelia Langford, John Sillitoe on behalf of the Wellcome Sanger Institute COVID-19 Surveillance Team |
| EPI_ISL_873956 | Lighthouse Lab in Milton Keynes | Wellcome Sanger Institute for the COVID-19 Genomics UK (COG-UK) Consortium | The Lighthouse Lab in Milton Keynes and Alex Alderton, Roberto Amato, Sonia Goncalves, Ewan Harrison, David K. Jackson, Ian Johnston, Dominic Kwiatkowski, Cordelia Langford, John Sillitoe on behalf of the Wellcome Sanger Institute COVID-19 Surveillance Team |
| EPI_ISL_873957, EPI_ISL_873958, EPI_ISL_873959 | Lighthouse Lab in Cambridge | Wellcome Sanger Institute for the COVID-19 Genomics UK (COG-UK) Consortium | Rob Howes, The Lighthouse Lab in Cambridge and Alex Alderton, Roberto Amato, Sonia Goncalves, Ewan Harrison, David K. Jackson, Ian Johnston, Dominic Kwiatkowski, Cordelia Langford, John Sillitoe on behalf of the Wellcome Sanger Institute COVID-19 Surveillance Team |
| EPI_ISL_873964, EPI_ISL_873965, EPI_ISL_873974, EPI_ISL_873976, EPI_ISL_873984, EPI_ISL_873987, EPI_ISL_873990, EPI_ISL_873999, EPI_ISL_874004, EPI_ISL_874007, EPI_ISL_874009, EPI_ISL_874010, EPI_ISL_874012, EPI_ISL_874014, EPI_ISL_874017, EPI_ISL_874021, EPI_ISL_874022, EPI_ISL_874032, EPI_ISL_874037, EPI_ISL_874038, EPI_ISL_874039, EPI_ISL_874047, EPI_ISL_874055, EPI_ISL_874061, EPI_ISL_874062, EPI_ISL_874064, EPI_ISL_874071, EPI_ISL_874076, EPI_ISL_874085, EPI_ISL_874090, EPI_ISL_874092, EPI_ISL_874096, EPI_ISL_874097, EPI_ISL_874115, EPI_ISL_874118, EPI_ISL_874130, EPI_ISL_874141, EPI_ISL_874143, EPI_ISL_874148, EPI_ISL_874153, EPI_ISL_874154, EPI_ISL_874157, EPI_ISL_874160, EPI_ISL_874163, EPI_ISL_874164, EPI_ISL_874165, EPI_ISL_874169, EPI_ISL_874180, EPI_ISL_874188, EPI_ISL_874191, EPI_ISL_874193, EPI_ISL_874195, EPI_ISL_874197, EPI_ISL_874198, EPI_ISL_874199, EPI_ISL_874202, EPI_ISL_874204, EPI_ISL_874207, EPI_ISL_874209, EPI_ISL_874211, EPI_ISL_874215, EPI_ISL_874238, EPI_ISL_874240, EPI_ISL_874246, EPI_ISL_874248, EPI_ISL_874254, EPI_ISL_874258, EPI_ISL_874263, EPI_ISL_874266, EPI_ISL_874268, EPI_ISL_874275, EPI_ISL_874278, EPI_ISL_874283, EPI_ISL_874290, EPI_ISL_874293, EPI_ISL_874297, EPI_ISL_874301, EPI_ISL_874311, EPI_ISL_874312, EPI_ISL_874313, EPI_ISL_874314, EPI_ISL_874316, EPI_ISL_874317, EPI_ISL_874318, EPI_ISL_874319, EPI_ISL_874320, EPI_ISL_874321, EPI_ISL_874322, EPI_ISL_874323, EPI_ISL_874324, EPI_ISL_874325, EPI_ISL_874326, EPI_ISL_874327, EPI_ISL_874328, EPI_ISL_874329, EPI_ISL_874330, EPI_ISL_874331, EPI_ISL_874332, EPI_ISL_874333, EPI_ISL_874334, EPI_ISL_874335, EPI_ISL_874336, EPI_ISL_874337, EPI_ISL_874338, EPI_ISL_874339, EPI_ISL_874340, EPI_ISL_874341, EPI_ISL_874342, EPI_ISL_874343, EPI_ISL_874344, EPI_ISL_874345, EPI_ISL_874346, EPI_ISL_874347, EPI_ISL_874348, EPI_ISL_874349, EPI_ISL_874350, EPI_ISL_874351, EPI_ISL_874352, EPI_ISL_874353, EPI_ISL_874354, EPI_ISL_874355, EPI_ISL_874356, EPI_ISL_874357, EPI_ISL_874358, EPI_ISL_874359, EPI_ISL_874360, EPI_ISL_874361, EPI_ISL_874362, EPI_ISL_874363, EPI_ISL_874364, EPI_ISL_874365, EPI_ISL_874366, EPI_ISL_874367, EPI_ISL_874368, EPI_ISL_874369, EPI_ISL_874370, EPI_ISL_874371, EPI_ISL_874372, EPI_ISL_874373, EPI_ISL_874374, EPI_ISL_874375, EPI_ISL_874376, EPI_ISL_874377, EPI_ISL_874378, EPI_ISL_874380, EPI_ISL_874381, EPI_ISL_874382, EPI_ISL_874383, EPI_ISL_874384, EPI_ISL_874385, EPI_ISL_874386, EPI_ISL_874387, EPI_ISL_874388, EPI_ISL_874389, EPI_ISL_874390, EPI_ISL_874391, EPI_ISL_874392, EPI_ISL_874393, EPI_ISL_874394, EPI_ISL_874395, EPI_ISL_874396, EPI_ISL_874397, EPI_ISL_874398, EPI_ISL_874399, EPI_ISL_874400, EPI_ISL_874402, EPI_ISL_874403, EPI_ISL_874405, EPI_ISL_874406, EPI_ISL_874407, EPI_ISL_874408, EPI_ISL_874409, EPI_ISL_874410, EPI_ISL_874411, EPI_ISL_874412, EPI_ISL_874413, EPI_ISL_874414, EPI_ISL_874415, EPI_ISL_874416, EPI_ISL_874417, EPI_ISL_874418, EPI_ISL_874419, EPI_ISL_874420, EPI_ISL_874421, EPI_ISL_874422, EPI_ISL_874423, EPI_ISL_874424, EPI_ISL_874425, EPI_ISL_874426, EPI_ISL_874427, EPI_ISL_874428, EPI_ISL_874429, EPI_ISL_874430, EPI_ISL_874431, EPI_ISL_874432, EPI_ISL_874433, EPI_ISL_874434, EPI_ISL_874435, EPI_ISL_874436, EPI_ISL_874437, EPI_ISL_874438, EPI_ISL_874439, EPI_ISL_874440, EPI_ISL_874441, EPI_ISL_874442, EPI_ISL_874444, EPI_ISL_874445, EPI_ISL_874446, EPI_ISL_874447, EPI_ISL_874449, EPI_ISL_874450, EPI_ISL_874451, EPI_ISL_874452, EPI_ISL_874453, EPI_ISL_874454, EPI_ISL_874455, EPI_ISL_874456, EPI_ISL_874457, EPI_ISL_874458, EPI_ISL_874459, EPI_ISL_874460, EPI_ISL_874461, EPI_ISL_874462, EPI_ISL_874463, EPI_ISL_874464, EPI_ISL_874465, EPI_ISL_874466, EPI_ISL_874467, EPI_ISL_874468, EPI_ISL_874470, EPI_ISL_874471, EPI_ISL_874472, EPI_ISL_874473, EPI_ISL_874474, EPI_ISL_874475, EPI_ISL_874476, EPI_ISL_874477, EPI_ISL_874478, EPI_ISL_874479, EPI_ISL_874480, EPI_ISL_874481, EPI_ISL_874483, EPI_ISL_874484, EPI_ISL_874485, EPI_ISL_874486, EPI_ISL_874487, EPI_ISL_874488, EPI_ISL_874489, EPI_ISL_874490, EPI_ISL_874491, EPI_ISL_874492, EPI_ISL_874493, EPI_ISL_874494, EPI_ISL_874495, EPI_ISL_874496, EPI_ISL_874497, EPI_ISL_874498, EPI_ISL_874500, EPI_ISL_874501, EPI_ISL_874503, EPI_ISL_874504, EPI_ISL_874505, EPI_ISL_874506, EPI_ISL_874507, EPI_ISL_874508, EPI_ISL_874509, EPI_ISL_874510, EPI_ISL_874511, EPI_ISL_874512, EPI_ISL_874513, EPI_ISL_874514, EPI_ISL_874515, EPI_ISL_874516, EPI_ISL_874517, EPI_ISL_874518, EPI_ISL_874519, EPI_ISL_874520, EPI_ISL_874521, EPI_ISL_874522, EPI_ISL_874524, EPI_ISL_874525, EPI_ISL_874526, EPI_ISL_874529, EPI_ISL_874531, EPI_ISL_874532, EPI_ISL_874533, EPI_ISL_874534, EPI_ISL_874535, EPI_ISL_874536, EPI_ISL_874537, EPI_ISL_874538, EPI_ISL_874539, EPI_ISL_874540, EPI_ISL_874541, EPI_ISL_874542, EPI_ISL_874544, EPI_ISL_874545, EPI_ISL_874546, EPI_ISL_874547, EPI_ISL_874548, EPI_ISL_874549, EPI_ISL_874550, EPI_ISL_874551, EPI_ISL_874552, EPI_ISL_874553, EPI_ISL_874554, EPI_ISL_874555, EPI_ISL_874556, EPI_ISL_874557, EPI_ISL_874559, EPI_ISL_874560, EPI_ISL_874561, EPI_ISL_874562, EPI_ISL_874563, EPI_ISL_874564, EPI_ISL_874565, EPI_ISL_874566, EPI_ISL_874567, EPI_ISL_874568, EPI_ISL_874569, EPI_ISL_874570, EPI_ISL_874571, EPI_ISL_874572, EPI_ISL_874573, EPI_ISL_874574, EPI_ISL_874575, EPI_ISL_874576, EPI_ISL_874577, EPI_ISL_874578, EPI_ISL_874579, EPI_ISL_874580, EPI_ISL_874581, EPI_ISL_874582, EPI_ISL_874583, EPI_ISL_874584, EPI_ISL_874585, EPI_ISL_874586, EPI_ISL_874587, EPI_ISL_874588, EPI_ISL_874589, EPI_ISL_874590, EPI_ISL_874591, EPI_ISL_874592, EPI_ISL_874593, EPI_ISL_874594, EPI_ISL_874595, EPI_ISL_874596, EPI_ISL_874597, EPI_ISL_874598, EPI_ISL_874599, EPI_ISL_874600, EPI_ISL_874601, EPI_ISL_874602, EPI_ISL_874603, EPI_ISL_874604, EPI_ISL_874605, EPI_ISL_874606, EPI_ISL_874607, EPI_ISL_874608, EPI_ISL_874609, EPI_ISL_874610, EPI_ISL_874611, EPI_ISL_874612, EPI_ISL_874613, EPI_ISL_874614, EPI_ISL_874615, EPI_ISL_874616, EPI_ISL_874617, EPI_ISL_874618, EPI_ISL_874619, EPI_ISL_874620, EPI_ISL_874621, EPI_ISL_874622, EPI_ISL_874623, EPI_ISL_874625, EPI_ISL_874626, EPI_ISL_874627, EPI_ISL_874628, EPI_ISL_874629, EPI_ISL_874630, EPI_ISL_874631, EPI_ISL_874632, EPI_ISL_874633, EPI_ISL_874634, EPI_ISL_874635, EPI_ISL_874636, EPI_ISL_874637, EPI_ISL_874638, EPI_ISL_874639, EPI_ISL_874640, EPI_ISL_874641, EPI_ISL_874642, EPI_ISL_874643, EPI_ISL_874644, EPI_ISL_874645, EPI_ISL_874646, EPI_ISL_874647, EPI_ISL_874648, EPI_ISL_874649, EPI_ISL_874650, EPI_ISL_874651, EPI_ISL_874652, EPI_ISL_874653, EPI_ISL_874654, EPI_ISL_874655 | | | |
| see above | Lighthouse Lab in Alderley Park | Wellcome Sanger Institute for the COVID-19 Genomics UK (COG-UK) Consortium | Jacquelyn Wynn, Mairead Hyland, The Lighthouse Lab in Alderley Park and Alex Alderton, Roberto Amato, Sonia Goncalves, Ewan Harrison, David K. Jackson, Ian Johnston, Dominic Kwiatkowski, Cordelia Langford, John Sillitoe on behalf of the Wellcome Sanger Institute COVID-19 Surveillance Team |
| EPI_ISL_874656 | Lighthouse Lab in Glasgow | Wellcome Sanger Institute for the COVID-19 Genomics UK (COG-UK) Consortium | Harper VanSteenhouse, Yumi Kasai, David Gray, Carol Clugston, Anna Dominiczak and Alex Alderton, Roberto Amato, Sonia Goncalves, Ewan Harrison, David K. Jackson, Ian Johnston, Dominic Kwiatkowski, Cordelia Langford, John Sillitoe on behalf of the Wellcome Sanger Institute COVID-19 Surveillance Team |
| EPI_ISL_874657, EPI_ISL_874658, EPI_ISL_874659, EPI_ISL_874660, EPI_ISL_874661, EPI_ISL_874662, EPI_ISL_874663, EPI_ISL_874664, EPI_ISL_874665, EPI_ISL_874666, EPI_ISL_874667, EPI_ISL_874668, EPI_ISL_874669, EPI_ISL_874670, EPI_ISL_874671, EPI_ISL_874672, EPI_ISL_874673, EPI_ISL_874674, EPI_ISL_874675 | | | |
| see above | Lighthouse Lab in Alderley Park | Wellcome Sanger Institute for the COVID-19 Genomics UK | Jacquelyn Wynn, Mairead Hyland, The Lighthouse Lab in Alderley Park and Alex Alderton, Roberto Amato, Sonia Goncalves, Ewan Harrison, David K. |

| | | | |
|---|---|---|---|
| | | (COG-UK) Consortium | Jackson, Ian Johnston, Dominic Kwiatkowski, Cordelia Langford, John Sillitoe on behalf of the Wellcome Sanger Institute COVID-19 Surveillance Team |
| EPI_ISL_874676 | Lighthouse Lab in Glasgow | Wellcome Sanger Institute for the COVID-19 Genomics UK (COG-UK) Consortium | Harper VanSteenhouse, Yumi Kasai, David Gray, Carol Clugston, Anna Dominiczak and Alex Alderton, Roberto Amato, Sonia Goncalves, Ewan Harrison, David K. Jackson, Ian Johnston, Dominic Kwiatkowski, Cordelia Langford, John Sillitoe on behalf of the Wellcome Sanger Institute COVID-19 Surveillance Team |
| EPI_ISL_874677, EPI_ISL_874678, EPI_ISL_874679, EPI_ISL_874680, EPI_ISL_874681, EPI_ISL_874682, EPI_ISL_874683, EPI_ISL_874684, EPI_ISL_874685, EPI_ISL_874686, EPI_ISL_874687, EPI_ISL_874688, EPI_ISL_874689, EPI_ISL_874690, EPI_ISL_874691, EPI_ISL_874692, EPI_ISL_874693, EPI_ISL_874694, EPI_ISL_874695, EPI_ISL_874696, EPI_ISL_874697, EPI_ISL_874698, EPI_ISL_874699, EPI_ISL_874700, EPI_ISL_874701, EPI_ISL_874702, EPI_ISL_874703, EPI_ISL_874704, EPI_ISL_874705, EPI_ISL_874706, EPI_ISL_874707, EPI_ISL_874708, EPI_ISL_874709, EPI_ISL_874710, EPI_ISL_874711, EPI_ISL_874712, EPI_ISL_874713, EPI_ISL_874714, EPI_ISL_874715, EPI_ISL_874716, EPI_ISL_874717, EPI_ISL_874718, EPI_ISL_874719, EPI_ISL_874720, EPI_ISL_874721, EPI_ISL_874722, EPI_ISL_874723, EPI_ISL_874724, EPI_ISL_874725, EPI_ISL_874726, EPI_ISL_874727, EPI_ISL_874728, EPI_ISL_874729, EPI_ISL_874730, EPI_ISL_874731, EPI_ISL_874732 | | | |
| see above | Lighthouse Lab in Alderley Park | Wellcome Sanger Institute for the COVID-19 Genomics UK (COG-UK) Consortium | Jacquelyn Wynn, Mairead Hyland, The Lighthouse Lab in Alderley Park and Alex Alderton, Roberto Amato, Sonia Goncalves, Ewan Harrison, David K. Jackson, Ian Johnston, Dominic Kwiatkowski, Cordelia Langford, John Sillitoe on behalf of the Wellcome Sanger Institute COVID-19 Surveillance Team |
| EPI_ISL_874733 | Lighthouse Lab in Glasgow | Wellcome Sanger Institute for the COVID-19 Genomics UK (COG-UK) Consortium | Harper VanSteenhouse, Yumi Kasai, David Gray, Carol Clugston, Anna Dominiczak and Alex Alderton, Roberto Amato, Sonia Goncalves, Ewan Harrison, David K. Jackson, Ian Johnston, Dominic Kwiatkowski, Cordelia Langford, John Sillitoe on behalf of the Wellcome Sanger Institute COVID-19 Surveillance Team |
| EPI_ISL_874734, EPI_ISL_874735 | Lighthouse Lab in Alderley Park | Wellcome Sanger Institute for the COVID-19 Genomics UK (COG-UK) Consortium | Jacquelyn Wynn, Mairead Hyland, The Lighthouse Lab in Alderley Park and Alex Alderton, Roberto Amato, Sonia Goncalves, Ewan Harrison, David K. Jackson, Ian Johnston, Dominic Kwiatkowski, Cordelia Langford, John Sillitoe on behalf of the Wellcome Sanger Institute COVID-19 Surveillance Team |
| EPI_ISL_874736, EPI_ISL_874737 | Lighthouse Lab in Glasgow | Wellcome Sanger Institute for the COVID-19 Genomics UK (COG-UK) Consortium | Harper VanSteenhouse, Yumi Kasai, David Gray, Carol Clugston, Anna Dominiczak and Alex Alderton, Roberto Amato, Sonia Goncalves, Ewan Harrison, David K. Jackson, Ian Johnston, Dominic Kwiatkowski, Cordelia Langford, John Sillitoe on behalf of the Wellcome Sanger Institute COVID-19 Surveillance Team |
| EPI_ISL_874738 | Lighthouse Lab in Alderley Park | Wellcome Sanger Institute for the COVID-19 Genomics UK (COG-UK) Consortium | Jacquelyn Wynn, Mairead Hyland, The Lighthouse Lab in Alderley Park and Alex Alderton, Roberto Amato, Sonia Goncalves, Ewan Harrison, David K. Jackson, Ian Johnston, Dominic Kwiatkowski, Cordelia Langford, John Sillitoe on behalf of the Wellcome Sanger Institute COVID-19 Surveillance Team |
| EPI_ISL_874739 | Lighthouse Lab in Glasgow | Wellcome Sanger Institute for the COVID-19 Genomics UK (COG-UK) Consortium | Harper VanSteenhouse, Yumi Kasai, David Gray, Carol Clugston, Anna Dominiczak and Alex Alderton, Roberto Amato, Sonia Goncalves, Ewan Harrison, David K. Jackson, Ian Johnston, Dominic Kwiatkowski, Cordelia Langford, John Sillitoe on behalf of the Wellcome Sanger Institute COVID-19 Surveillance Team |
| EPI_ISL_874740, EPI_ISL_874741, EPI_ISL_874742, EPI_ISL_874743, EPI_ISL_874744, EPI_ISL_874745, EPI_ISL_874746, EPI_ISL_874747, EPI_ISL_874748, EPI_ISL_874749, EPI_ISL_874750, EPI_ISL_874751, EPI_ISL_874752 | | | |
| see above | Lighthouse Lab in Alderley Park | Wellcome Sanger Institute for the COVID-19 Genomics UK (COG-UK) Consortium | Jacquelyn Wynn, Mairead Hyland, The Lighthouse Lab in Alderley Park and Alex Alderton, Roberto Amato, Sonia Goncalves, Ewan Harrison, David K. Jackson, Ian Johnston, Dominic Kwiatkowski, Cordelia Langford, John Sillitoe on behalf of the Wellcome Sanger Institute COVID-19 Surveillance Team |
| EPI_ISL_874753 | Lighthouse Lab in Glasgow | Wellcome Sanger Institute for the COVID-19 Genomics UK (COG-UK) Consortium | Harper VanSteenhouse, Yumi Kasai, David Gray, Carol Clugston, Anna Dominiczak and Alex Alderton, Roberto Amato, Sonia Goncalves, Ewan Harrison, David K. Jackson, Ian Johnston, Dominic Kwiatkowski, Cordelia Langford, John Sillitoe on behalf of the Wellcome Sanger Institute COVID-19 Surveillance Team |
| EPI_ISL_874754, EPI_ISL_874755, EPI_ISL_874756, EPI_ISL_874757, EPI_ISL_874758, EPI_ISL_874760, EPI_ISL_874761, EPI_ISL_874762, EPI_ISL_874763, EPI_ISL_874764, EPI_ISL_874765, EPI_ISL_874766, EPI_ISL_874767, EPI_ISL_874768, EPI_ISL_874769, EPI_ISL_874770, EPI_ISL_874771, EPI_ISL_874772, EPI_ISL_874773, EPI_ISL_874774 | | | |
| see above | Lighthouse Lab in Alderley Park | Wellcome Sanger Institute for the COVID-19 Genomics UK (COG-UK) Consortium | Jacquelyn Wynn, Mairead Hyland, The Lighthouse Lab in Alderley Park and Alex Alderton, Roberto Amato, Sonia Goncalves, Ewan Harrison, David K. Jackson, Ian Johnston, Dominic Kwiatkowski, Cordelia Langford, John Sillitoe on behalf of the Wellcome Sanger Institute COVID-19 Surveillance Team |
| EPI_ISL_874775 | Lighthouse Lab in Glasgow | Wellcome Sanger Institute for the COVID-19 Genomics UK (COG-UK) Consortium | Harper VanSteenhouse, Yumi Kasai, David Gray, Carol Clugston, Anna Dominiczak and Alex Alderton, Roberto Amato, Sonia Goncalves, Ewan Harrison, David K. Jackson, Ian Johnston, Dominic Kwiatkowski, Cordelia Langford, John Sillitoe on behalf of the Wellcome Sanger Institute COVID-19 Surveillance Team |
| EPI_ISL_874776, EPI_ISL_874777, EPI_ISL_874778, EPI_ISL_874779, EPI_ISL_874780, EPI_ISL_874781, EPI_ISL_874782, EPI_ISL_874783, EPI_ISL_874784, EPI_ISL_874785, EPI_ISL_874786, EPI_ISL_874787, EPI_ISL_874788, EPI_ISL_874789, EPI_ISL_874790, EPI_ISL_874791, EPI_ISL_874792, EPI_ISL_874793, EPI_ISL_874794, EPI_ISL_874795, EPI_ISL_874796, EPI_ISL_874797 | | | |
| see above | Lighthouse Lab in Alderley Park | Wellcome Sanger Institute for the COVID-19 Genomics UK (COG-UK) Consortium | Jacquelyn Wynn, Mairead Hyland, The Lighthouse Lab in Alderley Park and Alex Alderton, Roberto Amato, Sonia Goncalves, Ewan Harrison, David K. Jackson, Ian Johnston, Dominic Kwiatkowski, Cordelia Langford, John Sillitoe on behalf of the Wellcome Sanger Institute COVID-19 Surveillance Team |
| EPI_ISL_874798 | Lighthouse Lab in Glasgow | Wellcome Sanger Institute for the COVID-19 Genomics UK (COG-UK) Consortium | Harper VanSteenhouse, Yumi Kasai, David Gray, Carol Clugston, Anna Dominiczak and Alex Alderton, Roberto Amato, Sonia Goncalves, Ewan Harrison, David K. Jackson, Ian Johnston, Dominic Kwiatkowski, Cordelia Langford, John Sillitoe on behalf of the Wellcome Sanger Institute COVID-19 Surveillance Team |
| EPI_ISL_874799, EPI_ISL_874800, EPI_ISL_874801, EPI_ISL_874802, EPI_ISL_874803, EPI_ISL_874804, EPI_ISL_874805 | Lighthouse Lab in Alderley Park | Wellcome Sanger Institute for the COVID-19 Genomics UK (COG-UK) Consortium | Jacquelyn Wynn, Mairead Hyland, The Lighthouse Lab in Alderley Park and Alex Alderton, Roberto Amato, Sonia Goncalves, Ewan Harrison, David K. Jackson, Ian Johnston, Dominic Kwiatkowski, Cordelia Langford, John Sillitoe on behalf of the Wellcome Sanger Institute COVID-19 Surveillance Team |
| EPI_ISL_874806 | Lighthouse Lab in Glasgow | Wellcome Sanger Institute for the COVID-19 Genomics UK (COG-UK) Consortium | Harper VanSteenhouse, Yumi Kasai, David Gray, Carol Clugston, Anna Dominiczak and Alex Alderton, Roberto Amato, Sonia Goncalves, Ewan Harrison, David K. Jackson, Ian Johnston, Dominic Kwiatkowski, Cordelia Langford, John Sillitoe on behalf of the Wellcome Sanger Institute COVID-19 Surveillance Team |
| EPI_ISL_874807 | Lighthouse Lab in Alderley Park | Wellcome Sanger Institute for the COVID-19 Genomics UK (COG-UK) Consortium | Jacquelyn Wynn, Mairead Hyland, The Lighthouse Lab in Alderley Park and Alex Alderton, Roberto Amato, Sonia Goncalves, Ewan Harrison, David K. Jackson, Ian Johnston, Dominic Kwiatkowski, Cordelia Langford, John Sillitoe on behalf of the Wellcome Sanger Institute COVID-19 Surveillance Team |
| EPI_ISL_874808 | Lighthouse Lab in Glasgow | Wellcome Sanger Institute for the COVID-19 Genomics UK (COG-UK) Consortium | Harper VanSteenhouse, Yumi Kasai, David Gray, Carol Clugston, Anna Dominiczak and Alex Alderton, Roberto Amato, Sonia Goncalves, Ewan Harrison, David K. Jackson, Ian Johnston, Dominic Kwiatkowski, Cordelia Langford, John Sillitoe on behalf of the Wellcome Sanger Institute COVID-19 Surveillance Team |
| EPI_ISL_874809, EPI_ISL_874810, EPI_ISL_874811, EPI_ISL_874812, EPI_ISL_874813, EPI_ISL_874814, EPI_ISL_874815, EPI_ISL_874817, EPI_ISL_874818, EPI_ISL_874819, EPI_ISL_874820, EPI_ISL_874821, EPI_ISL_874822, EPI_ISL_874823, EPI_ISL_874824, EPI_ISL_874825, EPI_ISL_874826, EPI_ISL_874827, EPI_ISL_874828, EPI_ISL_874829, EPI_ISL_874830, EPI_ISL_874831, EPI_ISL_874832, EPI_ISL_874833, EPI_ISL_874834, EPI_ISL_874835, EPI_ISL_874836, EPI_ISL_874837, EPI_ISL_874838, EPI_ISL_874839, EPI_ISL_874840, EPI_ISL_874841, EPI_ISL_874842, EPI_ISL_874843, EPI_ISL_874844, EPI_ISL_874845, EPI_ISL_874846, EPI_ISL_874847, EPI_ISL_874848, EPI_ISL_874849, EPI_ISL_874850, EPI_ISL_874851, EPI_ISL_874852, EPI_ISL_874854, EPI_ISL_874855, EPI_ISL_874856, EPI_ISL_874857, EPI_ISL_874858, EPI_ISL_874859, EPI_ISL_874860, EPI_ISL_874861, EPI_ISL_874862 | | | |
| see above | Lighthouse Lab in Alderley Park | Wellcome Sanger Institute for the COVID-19 Genomics UK (COG-UK) Consortium | Jacquelyn Wynn, Mairead Hyland, The Lighthouse Lab in Alderley Park and Alex Alderton, Roberto Amato, Sonia Goncalves, Ewan Harrison, David K. Jackson, Ian Johnston, Dominic Kwiatkowski, Cordelia Langford, John Sillitoe on behalf of the Wellcome Sanger Institute COVID-19 Surveillance Team |
| EPI_ISL_874863 | Lighthouse Lab in Glasgow | Wellcome Sanger Institute for the COVID-19 Genomics UK (COG-UK) Consortium | Harper VanSteenhouse, Yumi Kasai, David Gray, Carol Clugston, Anna Dominiczak and Alex Alderton, Roberto Amato, Sonia Goncalves, Ewan Harrison, David K. Jackson, Ian Johnston, Dominic Kwiatkowski, Cordelia Langford, John Sillitoe on behalf of the Wellcome Sanger Institute COVID-19 Surveillance Team |
| EPI_ISL_874864, EPI_ISL_874865, EPI_ISL_874866, EPI_ISL_874867, EPI_ISL_874868, EPI_ISL_874869, EPI_ISL_874870, EPI_ISL_874871, EPI_ISL_874872, EPI_ISL_874873, EPI_ISL_874874, EPI_ISL_874875, EPI_ISL_874876, EPI_ISL_874877, EPI_ISL_874878, EPI_ISL_874879, EPI_ISL_874880, EPI_ISL_874881, EPI_ISL_874882, EPI_ISL_874883, EPI_ISL_874884, EPI_ISL_874885, EPI_ISL_874886, EPI_ISL_874887, EPI_ISL_874888, EPI_ISL_874889, EPI_ISL_874890, EPI_ISL_874891, EPI_ISL_874892, EPI_ISL_874893, EPI_ISL_874894, EPI_ISL_874895 | | | |
| see above | Lighthouse Lab in Alderley Park | Wellcome Sanger Institute for the COVID-19 Genomics UK (COG-UK) Consortium | Jacquelyn Wynn, Mairead Hyland, The Lighthouse Lab in Alderley Park and Alex Alderton, Roberto Amato, Sonia Goncalves, Ewan Harrison, David K. Jackson, Ian Johnston, Dominic Kwiatkowski, Cordelia Langford, John Sillitoe on behalf of the Wellcome Sanger Institute COVID-19 Surveillance Team |
| EPI_ISL_874896 | Lighthouse Lab in Glasgow | Wellcome Sanger Institute for the COVID-19 Genomics UK (COG-UK) Consortium | Harper VanSteenhouse, Yumi Kasai, David Gray, Carol Clugston, Anna Dominiczak and Alex Alderton, Roberto Amato, Sonia Goncalves, Ewan Harrison, David K. Jackson, Ian Johnston, Dominic Kwiatkowski, Cordelia Langford, John Sillitoe on behalf of the Wellcome Sanger Institute COVID-19 Surveillance Team |
| EPI_ISL_874897 | Lighthouse Lab in Alderley Park | Wellcome Sanger Institute for the COVID-19 Genomics UK (COG-UK) Consortium | Jacquelyn Wynn, Mairead Hyland, The Lighthouse Lab in Alderley Park and Alex Alderton, Roberto Amato, Sonia Goncalves, Ewan Harrison, David K. Jackson, Ian Johnston, Dominic Kwiatkowski, Cordelia Langford, John Sillitoe on behalf of the Wellcome Sanger Institute COVID-19 Surveillance Team |
| EPI_ISL_874898 | Lighthouse Lab in Glasgow | Wellcome Sanger Institute for the COVID-19 Genomics UK | Harper VanSteenhouse, Yumi Kasai, David Gray, Carol Clugston, Anna Dominiczak and Alex Alderton, Roberto Amato, Sonia Goncalves, Ewan Harrison, |

| | | | |
|---|---|---|---|
| | | (COG-UK) Consortium | David K. Jackson, Ian Johnston, Dominic Kwiatkowski, Cordelia Langford, John Sillitoe on behalf of the Wellcome Sanger Institute COVID-19 Surveillance Team |
| EPI_ISL_874899, EPI_ISL_874900, EPI_ISL_874901, EPI_ISL_874902, EPI_ISL_874903, EPI_ISL_874904 | Lighthouse Lab in Alderley Park | Wellcome Sanger Institute for the COVID-19 Genomics UK (COG-UK) Consortium | Jacquelyn Wynn, Mairead Hyland, The Lighthouse Lab in Alderley Park and Alex Alderton, Roberto Amato, Sonia Goncalves, Ewan Harrison, David K. Jackson, Ian Johnston, Dominic Kwiatkowski, Cordelia Langford, John Sillitoe on behalf of the Wellcome Sanger Institute COVID-19 Surveillance Team |
| EPI_ISL_874905 | Lighthouse Lab in Glasgow | Wellcome Sanger Institute for the COVID-19 Genomics UK (COG-UK) Consortium | Harper VanSteenhouse, Yumi Kasai, David Gray, Carol Clugston, Anna Dominiczak and Alex Alderton, Roberto Amato, Sonia Goncalves, Ewan Harrison, David K. Jackson, Ian Johnston, Dominic Kwiatkowski, Cordelia Langford, John Sillitoe on behalf of the Wellcome Sanger Institute COVID-19 Surveillance Team |
| EPI_ISL_874906, EPI_ISL_874907, EPI_ISL_874908, EPI_ISL_874909, EPI_ISL_874910, EPI_ISL_874911, EPI_ISL_874912, EPI_ISL_874913, EPI_ISL_874914, EPI_ISL_874915, EPI_ISL_874916, EPI_ISL_874917, EPI_ISL_874918, EPI_ISL_874919, EPI_ISL_874920, EPI_ISL_874921 | | | |
| see above | Lighthouse Lab in Alderley Park | Wellcome Sanger Institute for the COVID-19 Genomics UK (COG-UK) Consortium | Jacquelyn Wynn, Mairead Hyland, The Lighthouse Lab in Alderley Park and Alex Alderton, Roberto Amato, Sonia Goncalves, Ewan Harrison, David K. Jackson, Ian Johnston, Dominic Kwiatkowski, Cordelia Langford, John Sillitoe on behalf of the Wellcome Sanger Institute COVID-19 Surveillance Team |
| EPI_ISL_874922 | Lighthouse Lab in Glasgow | Wellcome Sanger Institute for the COVID-19 Genomics UK (COG-UK) Consortium | Harper VanSteenhouse, Yumi Kasai, David Gray, Carol Clugston, Anna Dominiczak and Alex Alderton, Roberto Amato, Sonia Goncalves, Ewan Harrison, David K. Jackson, Ian Johnston, Dominic Kwiatkowski, Cordelia Langford, John Sillitoe on behalf of the Wellcome Sanger Institute COVID-19 Surveillance Team |
| EPI_ISL_874923, EPI_ISL_874924, EPI_ISL_874925, EPI_ISL_874926, EPI_ISL_874927, EPI_ISL_874928, EPI_ISL_874929, EPI_ISL_874930, EPI_ISL_874931, EPI_ISL_874932, EPI_ISL_874933, EPI_ISL_874934 | | | |
| see above | Lighthouse Lab in Alderley Park | Wellcome Sanger Institute for the COVID-19 Genomics UK (COG-UK) Consortium | Jacquelyn Wynn, Mairead Hyland, The Lighthouse Lab in Alderley Park and Alex Alderton, Roberto Amato, Sonia Goncalves, Ewan Harrison, David K. Jackson, Ian Johnston, Dominic Kwiatkowski, Cordelia Langford, John Sillitoe on behalf of the Wellcome Sanger Institute COVID-19 Surveillance Team |
| EPI_ISL_874935 | Lighthouse Lab in Glasgow | Wellcome Sanger Institute for the COVID-19 Genomics UK (COG-UK) Consortium | Harper VanSteenhouse, Yumi Kasai, David Gray, Carol Clugston, Anna Dominiczak and Alex Alderton, Roberto Amato, Sonia Goncalves, Ewan Harrison, David K. Jackson, Ian Johnston, Dominic Kwiatkowski, Cordelia Langford, John Sillitoe on behalf of the Wellcome Sanger Institute COVID-19 Surveillance Team |
| EPI_ISL_874936, EPI_ISL_874937, EPI_ISL_874939, EPI_ISL_874940, EPI_ISL_874941, EPI_ISL_874942 | Lighthouse Lab in Alderley Park | Wellcome Sanger Institute for the COVID-19 Genomics UK (COG-UK) Consortium | Jacquelyn Wynn, Mairead Hyland, The Lighthouse Lab in Alderley Park and Alex Alderton, Roberto Amato, Sonia Goncalves, Ewan Harrison, David K. Jackson, Ian Johnston, Dominic Kwiatkowski, Cordelia Langford, John Sillitoe on behalf of the Wellcome Sanger Institute COVID-19 Surveillance Team |
| EPI_ISL_874943 | Lighthouse Lab in Glasgow | Wellcome Sanger Institute for the COVID-19 Genomics UK (COG-UK) Consortium | Harper VanSteenhouse, Yumi Kasai, David Gray, Carol Clugston, Anna Dominiczak and Alex Alderton, Roberto Amato, Sonia Goncalves, Ewan Harrison, David K. Jackson, Ian Johnston, Dominic Kwiatkowski, Cordelia Langford, John Sillitoe on behalf of the Wellcome Sanger Institute COVID-19 Surveillance Team |
| EPI_ISL_874944, EPI_ISL_874945, EPI_ISL_874946, EPI_ISL_874947, EPI_ISL_874948, EPI_ISL_874949 | Lighthouse Lab in Alderley Park | Wellcome Sanger Institute for the COVID-19 Genomics UK (COG-UK) Consortium | Jacquelyn Wynn, Mairead Hyland, The Lighthouse Lab in Alderley Park and Alex Alderton, Roberto Amato, Sonia Goncalves, Ewan Harrison, David K. Jackson, Ian Johnston, Dominic Kwiatkowski, Cordelia Langford, John Sillitoe on behalf of the Wellcome Sanger Institute COVID-19 Surveillance Team |
| EPI_ISL_874950 | Lighthouse Lab in Glasgow | Wellcome Sanger Institute for the COVID-19 Genomics UK (COG-UK) Consortium | Harper VanSteenhouse, Yumi Kasai, David Gray, Carol Clugston, Anna Dominiczak and Alex Alderton, Roberto Amato, Sonia Goncalves, Ewan Harrison, David K. Jackson, Ian Johnston, Dominic Kwiatkowski, Cordelia Langford, John Sillitoe on behalf of the Wellcome Sanger Institute COVID-19 Surveillance Team |
| EPI_ISL_874951, EPI_ISL_874952, EPI_ISL_874953, EPI_ISL_874954, EPI_ISL_874955, EPI_ISL_874957, EPI_ISL_874958, EPI_ISL_874959, EPI_ISL_874960, EPI_ISL_874961, EPI_ISL_874962, EPI_ISL_874963, EPI_ISL_874964 | | | |
| see above | Lighthouse Lab in Alderley Park | Wellcome Sanger Institute for the COVID-19 Genomics UK (COG-UK) Consortium | Jacquelyn Wynn, Mairead Hyland, The Lighthouse Lab in Alderley Park and Alex Alderton, Roberto Amato, Sonia Goncalves, Ewan Harrison, David K. Jackson, Ian Johnston, Dominic Kwiatkowski, Cordelia Langford, John Sillitoe on behalf of the Wellcome Sanger Institute COVID-19 Surveillance Team |
| EPI_ISL_874965, EPI_ISL_874967 | Lighthouse Lab in Glasgow | Wellcome Sanger Institute for the COVID-19 Genomics UK (COG-UK) Consortium | Harper VanSteenhouse, Yumi Kasai, David Gray, Carol Clugston, Anna Dominiczak and Alex Alderton, Roberto Amato, Sonia Goncalves, Ewan Harrison, David K. Jackson, Ian Johnston, Dominic Kwiatkowski, Cordelia Langford, John Sillitoe on behalf of the Wellcome Sanger Institute COVID-19 Surveillance Team |
| EPI_ISL_874968, EPI_ISL_874969, EPI_ISL_874970, EPI_ISL_874971, EPI_ISL_874972, EPI_ISL_874973, EPI_ISL_874974, EPI_ISL_874975, EPI_ISL_874976, EPI_ISL_874977, EPI_ISL_874978, EPI_ISL_874979, EPI_ISL_874980, EPI_ISL_874981, EPI_ISL_874982, EPI_ISL_874983, EPI_ISL_874984, EPI_ISL_874985, EPI_ISL_874986, EPI_ISL_874987, EPI_ISL_874988, EPI_ISL_874989 | | | |
| see above | Lighthouse Lab in Alderley Park | Wellcome Sanger Institute for the COVID-19 Genomics UK (COG-UK) Consortium | Jacquelyn Wynn, Mairead Hyland, The Lighthouse Lab in Alderley Park and Alex Alderton, Roberto Amato, Sonia Goncalves, Ewan Harrison, David K. Jackson, Ian Johnston, Dominic Kwiatkowski, Cordelia Langford, John Sillitoe on behalf of the Wellcome Sanger Institute COVID-19 Surveillance Team |
| EPI_ISL_875353, EPI_ISL_875354, EPI_ISL_875355, EPI_ISL_875356, EPI_ISL_875357, EPI_ISL_875358, EPI_ISL_875359, EPI_ISL_875360, EPI_ISL_875361, EPI_ISL_875362, EPI_ISL_875363, EPI_ISL_875364, EPI_ISL_875365, EPI_ISL_875366, EPI_ISL_875367, EPI_ISL_875368, EPI_ISL_875369, EPI_ISL_875370, EPI_ISL_875371, EPI_ISL_875372 | | | |
| see above | National Virus Reference Laboratory | National Virus Reference Laboratory | Michael Carr, Gabriel Gonzalez, Jonathan Dean, Cillian F De Gascun |
| EPI_ISL_875539 | Institute of Virology, Biomedical Research Center of the Slovak Academy of Sciences, Bratislava | Faculty of Natural Sciences, Comenius University, Bratislava | Kristína Boršová, Viktória abanová, Broa Brejová, Viktória Hodorová, Sabina Fumaová Havlíková, Juraj Kopáek, Martina Liková, ubomíra Lukáiková, Martina Neboháová, Monika Sláviková, Tomáš Vina, Boris Klempa, Jozef Nosek |
| EPI_ISL_875558 | Microbiologia e Virologia | Istituto Zooprofilattico Sperimentale delle Venezie | Adelaide Milani, Alessia Schivo, Annalisa Salviato, Erika Giorgia Quaranta, Ambra Pastori, Bianca Zecchin, Alice Fusaro, Isabella Monne, Calogero Terregino, Antonia Ricci |
| EPI_ISL_875666, EPI_ISL_875667, EPI_ISL_875668 | CHU Purpan - Laboratoire de Virologie - Institut Fédératif de Biologie | CHU Purpan - Laboratoire de Virologie - Institut Fédératif de Biologie | Latour J., Ranger N., Dubois M., Carcenac R., Harter A., Boyer P., Tremeaux P., Izopet J. |
| EPI_ISL_876065, EPI_ISL_876087, EPI_ISL_876095, EPI_ISL_876099, EPI_ISL_876124, EPI_ISL_876202, EPI_ISL_876215, EPI_ISL_876234, EPI_ISL_876235, EPI_ISL_876236, EPI_ISL_876237, EPI_ISL_876238 | | | |
| see above | Massachusetts State Public Health Laboratory | Massachusetts State Public Health Laboratory | Andrew Lang, Timelia Fink, Glen Gallagher, Sandra Smole |
| EPI_ISL_877044, EPI_ISL_877045, EPI_ISL_877051, EPI_ISL_877052, EPI_ISL_877053, EPI_ISL_877054, EPI_ISL_877055, EPI_ISL_877056, EPI_ISL_877057, EPI_ISL_877058 | Quest Diagnostics | Quest Diagnostics | Rosenthal,S.H., Gerasimova,A., Kagan,R.M., Anderson, B., Hua, M., Liu Y., Bernstein, L.E., Livingston, K.E., Perez, A., Shalhout, D.F., Shlyakhter, I.A., Owen, R., Tanpaiboon, P., Lacbawan, F. |
| EPI_ISL_877133, EPI_ISL_877134, EPI_ISL_877135, EPI_ISL_877136, EPI_ISL_877137, EPI_ISL_877138, EPI_ISL_877139, EPI_ISL_877140, EPI_ISL_877141, EPI_ISL_877142, EPI_ISL_877143, EPI_ISL_877144, EPI_ISL_877145, EPI_ISL_877153 | | | |
| see above | Univeristy of New Mexico Hospital | Center for Global Health, University of New Mexico Health Sciences Center | Daryl Domman, Kurt Schwalm, Justin Bacca, Jon Femling, Darrell Dinwiddie |
| EPI_ISL_877219 | Canterbury Health Laboratories | Institute of Environmental Science and Research (ESR) | Xiaoyun Ren, Matt Storey, Nikki Freed, Muhammad Faisal, Jing Wang, Hermes Perez, Anja Werno, Antje van der Linden, Arlo Upton, Chris Mansell, David Hammer, Dragana Drinkovic, Gary McAuliffe, Hana Sofia Andersson, James Ussher, Jill Sherwood, Josh Freeman, Julia Howard, Juliet Elvy, Mary DeAlmeida, Matt Blakiston, Matthew Rogers, Max Bloomfield, Michael Addidle, Michelle Balm, Sally Roberts, Sarah Jefferies, Sharmini Muttaiyah, Susan Morpeth, Susan Taylor, Timothy Blackmore, Vani Sathyendran, Veronica Playle, Virginia Hope, Erasmus Smit, Lauren Jelly, Olin Silander, Joep de Ligt |
| EPI_ISL_877221, EPI_ISL_877222, EPI_ISL_877223, EPI_ISL_877224, EPI_ISL_877225, EPI_ISL_877226 | LabPLUS | Institute of Environmental Science and Research (ESR) | Xiaoyun Ren, Matt Storey, Nikki Freed, Muhammad Faisal, Jing Wang, Hermes Perez, Anja Werno, Antje van der Linden, Arlo Upton, Chris Mansell, David Hammer, Dragana Drinkovic, Gary McAuliffe, Hana Sofia Andersson, James Ussher, Jill Sherwood, Josh Freeman, Julia Howard, Juliet Elvy, Mary DeAlmeida, Matt Blakiston, Matthew Rogers, Max Bloomfield, Michael Addidle, Michelle Balm, Sally Roberts, Sarah Jefferies, Sharmini Muttaiyah, Susan Morpeth, Susan Taylor, Timothy Blackmore, Vani Sathyendran, Veronica Playle, Virginia Hope, Erasmus Smit, Lauren Jelly, Olin Silander, Joep de Ligt |
| EPI_ISL_877551 | Institute of Microbiology, Universidad San Francisco de Quito | Institute of Microbiology, Universidad San Francisco de Quito | Belén Prado-Vivar, Sully Márquez, Juan José Guadalupe, Monica Becerra-Wong, Bernardo Gutiérrez, Marcos DiStefano, Verónica Barragán, Patricio Rojas-Silva, Gabriel Trueba, Michelle Grunauer, Paúl Cárdenas |
| EPI_ISL_877771, EPI_ISL_877807, EPI_ISL_877821, EPI_ISL_877828, EPI_ISL_877844, EPI_ISL_877903, EPI_ISL_877940, EPI_ISL_877972, EPI_ISL_878089, EPI_ISL_878118, EPI_ISL_878124, EPI_ISL_878126, EPI_ISL_878163, EPI_ISL_878173, EPI_ISL_878176, EPI_ISL_878179, EPI_ISL_878192, EPI_ISL_878208, | | | |

| | | | |
|---|---|---|---|
| EPI_ISL_878249, EPI_ISL_878252, EPI_ISL_878265, EPI_ISL_878277, EPI_ISL_878285, EPI_ISL_878323, EPI_ISL_878337, EPI_ISL_878339, EPI_ISL_878420, EPI_ISL_878422, EPI_ISL_878449, EPI_ISL_878454, EPI_ISL_878456, EPI_ISL_878458, EPI_ISL_878466, EPI_ISL_878481, EPI_ISL_878484, EPI_ISL_878486, EPI_ISL_878505, EPI_ISL_878507, EPI_ISL_878512, EPI_ISL_878530, EPI_ISL_878542, EPI_ISL_878556, EPI_ISL_878570, EPI_ISL_878572, EPI_ISL_878586, EPI_ISL_878598, EPI_ISL_878609, EPI_ISL_878614, EPI_ISL_878618, EPI_ISL_878652, EPI_ISL_878654, EPI_ISL_878681, EPI_ISL_878683, EPI_ISL_878688, EPI_ISL_878689, EPI_ISL_878701, EPI_ISL_878704, EPI_ISL_878706, EPI_ISL_878712, EPI_ISL_878730 | | | |
| see above | Lighthouse Lab in Alderley Park | Wellcome Sanger Institute for the COVID-19 Genomics UK (COG-UK) Consortium | Jacquelyn Wynn, Mairead Hyland, The Lighthouse Lab in Alderley Park and Alex Alderton, Roberto Amato, Sonia Goncalves, Ewan Harrison, David K. Jackson, Ian Johnston, Dominic Kwiatkowski, Cordelia Langford, John Sillitoe on behalf of the Wellcome Sanger Institute COVID-19 Surveillance Team |
| EPI_ISL_878751, EPI_ISL_878756, EPI_ISL_878759, EPI_ISL_878772, EPI_ISL_878775, EPI_ISL_878778, EPI_ISL_878780, EPI_ISL_878782, EPI_ISL_878792, EPI_ISL_878793, EPI_ISL_878798, EPI_ISL_878800, EPI_ISL_878806, EPI_ISL_878807, EPI_ISL_878814 | | | |
| see above | Lighthouse Lab in Milton Keynes | Wellcome Sanger Institute for the COVID-19 Genomics UK (COG-UK) Consortium | The Lighthouse Lab in Milton Keynes and Alex Alderton, Roberto Amato, Sonia Goncalves, Ewan Harrison, David K. Jackson, Ian Johnston, Dominic Kwiatkowski, Cordelia Langford, John Sillitoe on behalf of the Wellcome Sanger Institute COVID-19 Surveillance Team |
| EPI_ISL_878819 | Lighthouse Lab in Alderley Park | Wellcome Sanger Institute for the COVID-19 Genomics UK (COG-UK) Consortium | Jacquelyn Wynn, Mairead Hyland, The Lighthouse Lab in Alderley Park and Alex Alderton, Roberto Amato, Sonia Goncalves, Ewan Harrison, David K. Jackson, Ian Johnston, Dominic Kwiatkowski, Cordelia Langford, John Sillitoe on behalf of the Wellcome Sanger Institute COVID-19 Surveillance Team |
| EPI_ISL_878824, EPI_ISL_878842, EPI_ISL_878845, EPI_ISL_878846, EPI_ISL_878848, EPI_ISL_878851, EPI_ISL_878852, EPI_ISL_878854, EPI_ISL_878858, EPI_ISL_878860, EPI_ISL_878861, EPI_ISL_878874 | | | |
| see above | Lighthouse Lab in Milton Keynes | Wellcome Sanger Institute for the COVID-19 Genomics UK (COG-UK) Consortium | The Lighthouse Lab in Milton Keynes and Alex Alderton, Roberto Amato, Sonia Goncalves, Ewan Harrison, David K. Jackson, Ian Johnston, Dominic Kwiatkowski, Cordelia Langford, John Sillitoe on behalf of the Wellcome Sanger Institute COVID-19 Surveillance Team |
| EPI_ISL_878876 | Lighthouse Lab in Alderley Park | Wellcome Sanger Institute for the COVID-19 Genomics UK (COG-UK) Consortium | Jacquelyn Wynn, Mairead Hyland, The Lighthouse Lab in Alderley Park and Alex Alderton, Roberto Amato, Sonia Goncalves, Ewan Harrison, David K. Jackson, Ian Johnston, Dominic Kwiatkowski, Cordelia Langford, John Sillitoe on behalf of the Wellcome Sanger Institute COVID-19 Surveillance Team |
| EPI_ISL_878879, EPI_ISL_878880, EPI_ISL_878882, EPI_ISL_878884, EPI_ISL_878892, EPI_ISL_878896, EPI_ISL_878902, EPI_ISL_878906 | Lighthouse Lab in Milton Keynes | Wellcome Sanger Institute for the COVID-19 Genomics UK (COG-UK) Consortium | The Lighthouse Lab in Milton Keynes and Alex Alderton, Roberto Amato, Sonia Goncalves, Ewan Harrison, David K. Jackson, Ian Johnston, Dominic Kwiatkowski, Cordelia Langford, John Sillitoe on behalf of the Wellcome Sanger Institute COVID-19 Surveillance Team |
| EPI_ISL_878913 | Lighthouse Lab in Alderley Park | Wellcome Sanger Institute for the COVID-19 Genomics UK (COG-UK) Consortium | Jacquelyn Wynn, Mairead Hyland, The Lighthouse Lab in Alderley Park and Alex Alderton, Roberto Amato, Sonia Goncalves, Ewan Harrison, David K. Jackson, Ian Johnston, Dominic Kwiatkowski, Cordelia Langford, John Sillitoe on behalf of the Wellcome Sanger Institute COVID-19 Surveillance Team |
| EPI_ISL_878918, EPI_ISL_878919, EPI_ISL_878934, EPI_ISL_878941, EPI_ISL_878942, EPI_ISL_878943, EPI_ISL_878944, EPI_ISL_878945, EPI_ISL_878951, EPI_ISL_878952, EPI_ISL_878960, EPI_ISL_878962, EPI_ISL_878967, EPI_ISL_878971, EPI_ISL_878972, EPI_ISL_878982, EPI_ISL_878990 | | | |
| see above | Lighthouse Lab in Milton Keynes | Wellcome Sanger Institute for the COVID-19 Genomics UK (COG-UK) Consortium | The Lighthouse Lab in Milton Keynes and Alex Alderton, Roberto Amato, Sonia Goncalves, Ewan Harrison, David K. Jackson, Ian Johnston, Dominic Kwiatkowski, Cordelia Langford, John Sillitoe on behalf of the Wellcome Sanger Institute COVID-19 Surveillance Team |
| EPI_ISL_878993 | Lighthouse Lab in Alderley Park | Wellcome Sanger Institute for the COVID-19 Genomics UK (COG-UK) Consortium | Jacquelyn Wynn, Mairead Hyland, The Lighthouse Lab in Alderley Park and Alex Alderton, Roberto Amato, Sonia Goncalves, Ewan Harrison, David K. Jackson, Ian Johnston, Dominic Kwiatkowski, Cordelia Langford, John Sillitoe on behalf of the Wellcome Sanger Institute COVID-19 Surveillance Team |
| EPI_ISL_879001, EPI_ISL_879009, EPI_ISL_879013, EPI_ISL_879014, EPI_ISL_879016, EPI_ISL_879019, EPI_ISL_879021, EPI_ISL_879023, EPI_ISL_879025, EPI_ISL_879028, EPI_ISL_879032, EPI_ISL_879034, EPI_ISL_879037, EPI_ISL_879038 | | | |
| see above | Lighthouse Lab in Milton Keynes | Wellcome Sanger Institute for the COVID-19 Genomics UK (COG-UK) Consortium | The Lighthouse Lab in Milton Keynes and Alex Alderton, Roberto Amato, Sonia Goncalves, Ewan Harrison, David K. Jackson, Ian Johnston, Dominic Kwiatkowski, Cordelia Langford, John Sillitoe on behalf of the Wellcome Sanger Institute COVID-19 Surveillance Team |
| EPI_ISL_879040 | Lighthouse Lab in Alderley Park | Wellcome Sanger Institute for the COVID-19 Genomics UK (COG-UK) Consortium | Jacquelyn Wynn, Mairead Hyland, The Lighthouse Lab in Alderley Park and Alex Alderton, Roberto Amato, Sonia Goncalves, Ewan Harrison, David K. Jackson, Ian Johnston, Dominic Kwiatkowski, Cordelia Langford, John Sillitoe on behalf of the Wellcome Sanger Institute COVID-19 Surveillance Team |
| EPI_ISL_879042, EPI_ISL_879043, EPI_ISL_879044, EPI_ISL_879047, EPI_ISL_879049, EPI_ISL_879050, EPI_ISL_879052, EPI_ISL_879055 | Lighthouse Lab in Milton Keynes | Wellcome Sanger Institute for the COVID-19 Genomics UK (COG-UK) Consortium | The Lighthouse Lab in Milton Keynes and Alex Alderton, Roberto Amato, Sonia Goncalves, Ewan Harrison, David K. Jackson, Ian Johnston, Dominic Kwiatkowski, Cordelia Langford, John Sillitoe on behalf of the Wellcome Sanger Institute COVID-19 Surveillance Team |
| EPI_ISL_879056 | Lighthouse Lab in Alderley Park | Wellcome Sanger Institute for the COVID-19 Genomics UK (COG-UK) Consortium | Jacquelyn Wynn, Mairead Hyland, The Lighthouse Lab in Alderley Park and Alex Alderton, Roberto Amato, Sonia Goncalves, Ewan Harrison, David K. Jackson, Ian Johnston, Dominic Kwiatkowski, Cordelia Langford, John Sillitoe on behalf of the Wellcome Sanger Institute COVID-19 Surveillance Team |
| EPI_ISL_879057, EPI_ISL_879063, EPI_ISL_879064, EPI_ISL_879065 | Lighthouse Lab in Milton Keynes | Wellcome Sanger Institute for the COVID-19 Genomics UK (COG-UK) Consortium | The Lighthouse Lab in Milton Keynes and Alex Alderton, Roberto Amato, Sonia Goncalves, Ewan Harrison, David K. Jackson, Ian Johnston, Dominic Kwiatkowski, Cordelia Langford, John Sillitoe on behalf of the Wellcome Sanger Institute COVID-19 Surveillance Team |
| EPI_ISL_879073 | Lighthouse Lab in Alderley Park | Wellcome Sanger Institute for the COVID-19 Genomics UK (COG-UK) Consortium | Jacquelyn Wynn, Mairead Hyland, The Lighthouse Lab in Alderley Park and Alex Alderton, Roberto Amato, Sonia Goncalves, Ewan Harrison, David K. Jackson, Ian Johnston, Dominic Kwiatkowski, Cordelia Langford, John Sillitoe on behalf of the Wellcome Sanger Institute COVID-19 Surveillance Team |
| EPI_ISL_879075, EPI_ISL_879086, EPI_ISL_879087, EPI_ISL_879089, EPI_ISL_879092, EPI_ISL_879093, EPI_ISL_879094, EPI_ISL_879096 | Lighthouse Lab in Milton Keynes | Wellcome Sanger Institute for the COVID-19 Genomics UK (COG-UK) Consortium | The Lighthouse Lab in Milton Keynes and Alex Alderton, Roberto Amato, Sonia Goncalves, Ewan Harrison, David K. Jackson, Ian Johnston, Dominic Kwiatkowski, Cordelia Langford, John Sillitoe on behalf of the Wellcome Sanger Institute COVID-19 Surveillance Team |
| EPI_ISL_879116, EPI_ISL_879117, EPI_ISL_879120, EPI_ISL_879138, EPI_ISL_879154, EPI_ISL_879185, EPI_ISL_879193, EPI_ISL_879220, EPI_ISL_879235, EPI_ISL_879253, EPI_ISL_879278, EPI_ISL_879290, EPI_ISL_879314, EPI_ISL_879322, EPI_ISL_879331, EPI_ISL_879338, EPI_ISL_879344, EPI_ISL_879363, EPI_ISL_879377 | | | |
| see above | Lighthouse Lab in Alderley Park | Wellcome Sanger Institute for the COVID-19 Genomics UK (COG-UK) Consortium | Jacquelyn Wynn, Mairead Hyland, The Lighthouse Lab in Alderley Park and Alex Alderton, Roberto Amato, Sonia Goncalves, Ewan Harrison, David K. Jackson, Ian Johnston, Dominic Kwiatkowski, Cordelia Langford, John Sillitoe on behalf of the Wellcome Sanger Institute COVID-19 Surveillance Team |
| EPI_ISL_879378, EPI_ISL_879379, EPI_ISL_879380 | Lighthouse Lab in Milton Keynes | Wellcome Sanger Institute for the COVID-19 Genomics UK (COG-UK) Consortium | The Lighthouse Lab in Milton Keynes and Alex Alderton, Roberto Amato, Sonia Goncalves, Ewan Harrison, David K. Jackson, Ian Johnston, Dominic Kwiatkowski, Cordelia Langford, John Sillitoe on behalf of the Wellcome Sanger Institute COVID-19 Surveillance Team |
| EPI_ISL_879384 | Lighthouse Lab in Alderley Park | Wellcome Sanger Institute for the COVID-19 Genomics UK (COG-UK) Consortium | Jacquelyn Wynn, Mairead Hyland, The Lighthouse Lab in Alderley Park and Alex Alderton, Roberto Amato, Sonia Goncalves, Ewan Harrison, David K. Jackson, Ian Johnston, Dominic Kwiatkowski, Cordelia Langford, John Sillitoe on behalf of the Wellcome Sanger Institute COVID-19 Surveillance Team |
| EPI_ISL_879388, EPI_ISL_879395, EPI_ISL_879396, EPI_ISL_879397, EPI_ISL_879399, EPI_ISL_879404, EPI_ISL_879409, EPI_ISL_879412, EPI_ISL_879413, EPI_ISL_879424, EPI_ISL_879425, EPI_ISL_879426, EPI_ISL_879429 | | | |
| see above | Lighthouse Lab in Milton Keynes | Wellcome Sanger Institute for the COVID-19 Genomics UK (COG-UK) Consortium | The Lighthouse Lab in Milton Keynes and Alex Alderton, Roberto Amato, Sonia Goncalves, Ewan Harrison, David K. Jackson, Ian Johnston, Dominic Kwiatkowski, Cordelia Langford, John Sillitoe on behalf of the Wellcome Sanger Institute COVID-19 Surveillance Team |
| EPI_ISL_879434 | Lighthouse Lab in Alderley Park | Wellcome Sanger Institute for the COVID-19 Genomics UK (COG-UK) Consortium | Jacquelyn Wynn, Mairead Hyland, The Lighthouse Lab in Alderley Park and Alex Alderton, Roberto Amato, Sonia Goncalves, Ewan Harrison, David K. Jackson, Ian Johnston, Dominic Kwiatkowski, Cordelia Langford, John Sillitoe on behalf of the Wellcome Sanger Institute COVID-19 Surveillance Team |
| EPI_ISL_879442, EPI_ISL_879448, EPI_ISL_879449, EPI_ISL_879450 | Lighthouse Lab in Milton Keynes | Wellcome Sanger Institute for the COVID-19 Genomics UK (COG-UK) Consortium | The Lighthouse Lab in Milton Keynes and Alex Alderton, Roberto Amato, Sonia Goncalves, Ewan Harrison, David K. Jackson, Ian Johnston, Dominic Kwiatkowski, Cordelia Langford, John Sillitoe on behalf of the Wellcome Sanger Institute COVID-19 Surveillance Team |
| EPI_ISL_879452 | Lighthouse Lab in Alderley Park | Wellcome Sanger Institute for the COVID-19 Genomics UK (COG-UK) Consortium | Jacquelyn Wynn, Mairead Hyland, The Lighthouse Lab in Alderley Park and Alex Alderton, Roberto Amato, Sonia Goncalves, Ewan Harrison, David K. Jackson, Ian Johnston, Dominic Kwiatkowski, Cordelia Langford, John Sillitoe on behalf of the Wellcome Sanger Institute COVID-19 Surveillance Team |
| EPI_ISL_879455, EPI_ISL_879457, EPI_ISL_879458, EPI_ISL_879459, EPI_ISL_879461 | Lighthouse Lab in Milton Keynes | Wellcome Sanger Institute for the COVID-19 Genomics UK (COG-UK) Consortium | The Lighthouse Lab in Milton Keynes and Alex Alderton, Roberto Amato, Sonia Goncalves, Ewan Harrison, David K. Jackson, Ian Johnston, Dominic Kwiatkowski, Cordelia Langford, John Sillitoe on behalf of the Wellcome Sanger Institute COVID-19 Surveillance Team |
| EPI_ISL_879462 | Lighthouse Lab in Alderley Park | Wellcome Sanger Institute for the COVID-19 Genomics UK (COG-UK) Consortium | Jacquelyn Wynn, Mairead Hyland, The Lighthouse Lab in Alderley Park and Alex Alderton, Roberto Amato, Sonia Goncalves, Ewan Harrison, David K. Jackson, Ian Johnston, Dominic Kwiatkowski, Cordelia Langford, John Sillitoe on behalf of the Wellcome Sanger Institute COVID-19 Surveillance Team |
| EPI_ISL_879464 | Lighthouse Lab in Milton Keynes | Wellcome Sanger Institute for the COVID-19 Genomics UK (COG-UK) Consortium | The Lighthouse Lab in Milton Keynes and Alex Alderton, Roberto Amato, Sonia Goncalves, Ewan Harrison, David K. Jackson, Ian Johnston, Dominic Kwiatkowski, Cordelia Langford, John Sillitoe on behalf of the Wellcome Sanger Institute COVID-19 Surveillance Team |
| EPI_ISL_879466 | Lighthouse Lab in Alderley Park | Wellcome Sanger Institute for the COVID-19 Genomics UK (COG-UK) Consortium | Jacquelyn Wynn, Mairead Hyland, The Lighthouse Lab in Alderley Park and Alex Alderton, Roberto Amato, Sonia Goncalves, Ewan Harrison, David K. Jackson, Ian Johnston, Dominic Kwiatkowski, Cordelia Langford, John Sillitoe on behalf of the Wellcome Sanger Institute COVID-19 Surveillance Team |

| | | | |
|---|---|---|---|
| EPI_ISL_879467, EPI_ISL_879471 | Lighthouse Lab in Milton Keynes | Wellcome Sanger Institute for the COVID-19 Genomics UK (COG-UK) Consortium | The Lighthouse Lab in Milton Keynes and Alex Alderton, Roberto Amato, Sonia Goncalves, Ewan Harrison, David K. Jackson, Ian Johnston, Dominic Kwiatkowski, Cordelia Langford, John Sillitoe on behalf of the Wellcome Sanger Institute COVID-19 Surveillance Team |
| EPI_ISL_879473, EPI_ISL_879475 | Lighthouse Lab in Alderley Park | Wellcome Sanger Institute for the COVID-19 Genomics UK (COG-UK) Consortium | Jacquelyn Wynn, Mairead Hyland, The Lighthouse Lab in Alderley Park and Alex Alderton, Roberto Amato, Sonia Goncalves, Ewan Harrison, David K. Jackson, Ian Johnston, Dominic Kwiatkowski, Cordelia Langford, John Sillitoe on behalf of the Wellcome Sanger Institute COVID-19 Surveillance Team |
| EPI_ISL_879477, EPI_ISL_879479 | Lighthouse Lab in Milton Keynes | Wellcome Sanger Institute for the COVID-19 Genomics UK (COG-UK) Consortium | The Lighthouse Lab in Milton Keynes and Alex Alderton, Roberto Amato, Sonia Goncalves, Ewan Harrison, David K. Jackson, Ian Johnston, Dominic Kwiatkowski, Cordelia Langford, John Sillitoe on behalf of the Wellcome Sanger Institute COVID-19 Surveillance Team |
| EPI_ISL_879486, EPI_ISL_879487 | Lighthouse Lab in Alderley Park | Wellcome Sanger Institute for the COVID-19 Genomics UK (COG-UK) Consortium | Jacquelyn Wynn, Mairead Hyland, The Lighthouse Lab in Alderley Park and Alex Alderton, Roberto Amato, Sonia Goncalves, Ewan Harrison, David K. Jackson, Ian Johnston, Dominic Kwiatkowski, Cordelia Langford, John Sillitoe on behalf of the Wellcome Sanger Institute COVID-19 Surveillance Team |
| EPI_ISL_879492, EPI_ISL_879495, EPI_ISL_879499, EPI_ISL_879502 | Lighthouse Lab in Milton Keynes | Wellcome Sanger Institute for the COVID-19 Genomics UK (COG-UK) Consortium | The Lighthouse Lab in Milton Keynes and Alex Alderton, Roberto Amato, Sonia Goncalves, Ewan Harrison, David K. Jackson, Ian Johnston, Dominic Kwiatkowski, Cordelia Langford, John Sillitoe on behalf of the Wellcome Sanger Institute COVID-19 Surveillance Team |
| EPI_ISL_879504, EPI_ISL_879510 | Lighthouse Lab in Alderley Park | Wellcome Sanger Institute for the COVID-19 Genomics UK (COG-UK) Consortium | Jacquelyn Wynn, Mairead Hyland, The Lighthouse Lab in Alderley Park and Alex Alderton, Roberto Amato, Sonia Goncalves, Ewan Harrison, David K. Jackson, Ian Johnston, Dominic Kwiatkowski, Cordelia Langford, John Sillitoe on behalf of the Wellcome Sanger Institute COVID-19 Surveillance Team |
| EPI_ISL_879514 | Lighthouse Lab in Milton Keynes | Wellcome Sanger Institute for the COVID-19 Genomics UK (COG-UK) Consortium | The Lighthouse Lab in Milton Keynes and Alex Alderton, Roberto Amato, Sonia Goncalves, Ewan Harrison, David K. Jackson, Ian Johnston, Dominic Kwiatkowski, Cordelia Langford, John Sillitoe on behalf of the Wellcome Sanger Institute COVID-19 Surveillance Team |
| EPI_ISL_879519, EPI_ISL_879520 | Lighthouse Lab in Alderley Park | Wellcome Sanger Institute for the COVID-19 Genomics UK (COG-UK) Consortium | Jacquelyn Wynn, Mairead Hyland, The Lighthouse Lab in Alderley Park and Alex Alderton, Roberto Amato, Sonia Goncalves, Ewan Harrison, David K. Jackson, Ian Johnston, Dominic Kwiatkowski, Cordelia Langford, John Sillitoe on behalf of the Wellcome Sanger Institute COVID-19 Surveillance Team |
| EPI_ISL_879526, EPI_ISL_879532, EPI_ISL_879538, EPI_ISL_879539, EPI_ISL_879540 | Lighthouse Lab in Milton Keynes | Wellcome Sanger Institute for the COVID-19 Genomics UK (COG-UK) Consortium | The Lighthouse Lab in Milton Keynes and Alex Alderton, Roberto Amato, Sonia Goncalves, Ewan Harrison, David K. Jackson, Ian Johnston, Dominic Kwiatkowski, Cordelia Langford, John Sillitoe on behalf of the Wellcome Sanger Institute COVID-19 Surveillance Team |
| EPI_ISL_879546, EPI_ISL_879547 | Lighthouse Lab in Alderley Park | Wellcome Sanger Institute for the COVID-19 Genomics UK (COG-UK) Consortium | Jacquelyn Wynn, Mairead Hyland, The Lighthouse Lab in Alderley Park and Alex Alderton, Roberto Amato, Sonia Goncalves, Ewan Harrison, David K. Jackson, Ian Johnston, Dominic Kwiatkowski, Cordelia Langford, John Sillitoe on behalf of the Wellcome Sanger Institute COVID-19 Surveillance Team |
| EPI_ISL_879548, EPI_ISL_879549 | Lighthouse Lab in Milton Keynes | Wellcome Sanger Institute for the COVID-19 Genomics UK (COG-UK) Consortium | The Lighthouse Lab in Milton Keynes and Alex Alderton, Roberto Amato, Sonia Goncalves, Ewan Harrison, David K. Jackson, Ian Johnston, Dominic Kwiatkowski, Cordelia Langford, John Sillitoe on behalf of the Wellcome Sanger Institute COVID-19 Surveillance Team |
| EPI_ISL_879550 | Lighthouse Lab in Alderley Park | Wellcome Sanger Institute for the COVID-19 Genomics UK (COG-UK) Consortium | Jacquelyn Wynn, Mairead Hyland, The Lighthouse Lab in Alderley Park and Alex Alderton, Roberto Amato, Sonia Goncalves, Ewan Harrison, David K. Jackson, Ian Johnston, Dominic Kwiatkowski, Cordelia Langford, John Sillitoe on behalf of the Wellcome Sanger Institute COVID-19 Surveillance Team |
| EPI_ISL_879552, EPI_ISL_879553, EPI_ISL_879554 | Lighthouse Lab in Milton Keynes | Wellcome Sanger Institute for the COVID-19 Genomics UK (COG-UK) Consortium | The Lighthouse Lab in Milton Keynes and Alex Alderton, Roberto Amato, Sonia Goncalves, Ewan Harrison, David K. Jackson, Ian Johnston, Dominic Kwiatkowski, Cordelia Langford, John Sillitoe on behalf of the Wellcome Sanger Institute COVID-19 Surveillance Team |
| EPI_ISL_879556 | Lighthouse Lab in Alderley Park | Wellcome Sanger Institute for the COVID-19 Genomics UK (COG-UK) Consortium | Jacquelyn Wynn, Mairead Hyland, The Lighthouse Lab in Alderley Park and Alex Alderton, Roberto Amato, Sonia Goncalves, Ewan Harrison, David K. Jackson, Ian Johnston, Dominic Kwiatkowski, Cordelia Langford, John Sillitoe on behalf of the Wellcome Sanger Institute COVID-19 Surveillance Team |
| EPI_ISL_879560, EPI_ISL_879561 | Lighthouse Lab in Milton Keynes | Wellcome Sanger Institute for the COVID-19 Genomics UK (COG-UK) Consortium | The Lighthouse Lab in Milton Keynes and Alex Alderton, Roberto Amato, Sonia Goncalves, Ewan Harrison, David K. Jackson, Ian Johnston, Dominic Kwiatkowski, Cordelia Langford, John Sillitoe on behalf of the Wellcome Sanger Institute COVID-19 Surveillance Team |
| EPI_ISL_879565 | Lighthouse Lab in Alderley Park | Wellcome Sanger Institute for the COVID-19 Genomics UK (COG-UK) Consortium | Jacquelyn Wynn, Mairead Hyland, The Lighthouse Lab in Alderley Park and Alex Alderton, Roberto Amato, Sonia Goncalves, Ewan Harrison, David K. Jackson, Ian Johnston, Dominic Kwiatkowski, Cordelia Langford, John Sillitoe on behalf of the Wellcome Sanger Institute COVID-19 Surveillance Team |
| EPI_ISL_879566, EPI_ISL_879571, EPI_ISL_879572, EPI_ISL_879575, EPI_ISL_879576, EPI_ISL_879581, EPI_ISL_879585, EPI_ISL_879586, EPI_ISL_879591, EPI_ISL_879592, EPI_ISL_879594, EPI_ISL_879597, EPI_ISL_879600, EPI_ISL_879601, EPI_ISL_879608, EPI_ISL_879611, EPI_ISL_879613 | | | |
| see above | Lighthouse Lab in Milton Keynes | Wellcome Sanger Institute for the COVID-19 Genomics UK (COG-UK) Consortium | The Lighthouse Lab in Milton Keynes and Alex Alderton, Roberto Amato, Sonia Goncalves, Ewan Harrison, David K. Jackson, Ian Johnston, Dominic Kwiatkowski, Cordelia Langford, John Sillitoe on behalf of the Wellcome Sanger Institute COVID-19 Surveillance Team |
| EPI_ISL_879617 | Lighthouse Lab in Alderley Park | Wellcome Sanger Institute for the COVID-19 Genomics UK (COG-UK) Consortium | Jacquelyn Wynn, Mairead Hyland, The Lighthouse Lab in Alderley Park and Alex Alderton, Roberto Amato, Sonia Goncalves, Ewan Harrison, David K. Jackson, Ian Johnston, Dominic Kwiatkowski, Cordelia Langford, John Sillitoe on behalf of the Wellcome Sanger Institute COVID-19 Surveillance Team |
| EPI_ISL_879618, EPI_ISL_879621, EPI_ISL_879623, EPI_ISL_879628 | Lighthouse Lab in Milton Keynes | Wellcome Sanger Institute for the COVID-19 Genomics UK (COG-UK) Consortium | The Lighthouse Lab in Milton Keynes and Alex Alderton, Roberto Amato, Sonia Goncalves, Ewan Harrison, David K. Jackson, Ian Johnston, Dominic Kwiatkowski, Cordelia Langford, John Sillitoe on behalf of the Wellcome Sanger Institute COVID-19 Surveillance Team |
| EPI_ISL_879629, EPI_ISL_879631 | Lighthouse Lab in Alderley Park | Wellcome Sanger Institute for the COVID-19 Genomics UK (COG-UK) Consortium | Jacquelyn Wynn, Mairead Hyland, The Lighthouse Lab in Alderley Park and Alex Alderton, Roberto Amato, Sonia Goncalves, Ewan Harrison, David K. Jackson, Ian Johnston, Dominic Kwiatkowski, Cordelia Langford, John Sillitoe on behalf of the Wellcome Sanger Institute COVID-19 Surveillance Team |
| EPI_ISL_879642, EPI_ISL_879643 | Lighthouse Lab in Milton Keynes | Wellcome Sanger Institute for the COVID-19 Genomics UK (COG-UK) Consortium | The Lighthouse Lab in Milton Keynes and Alex Alderton, Roberto Amato, Sonia Goncalves, Ewan Harrison, David K. Jackson, Ian Johnston, Dominic Kwiatkowski, Cordelia Langford, John Sillitoe on behalf of the Wellcome Sanger Institute COVID-19 Surveillance Team |
| EPI_ISL_879646 | Lighthouse Lab in Alderley Park | Wellcome Sanger Institute for the COVID-19 Genomics UK (COG-UK) Consortium | Jacquelyn Wynn, Mairead Hyland, The Lighthouse Lab in Alderley Park and Alex Alderton, Roberto Amato, Sonia Goncalves, Ewan Harrison, David K. Jackson, Ian Johnston, Dominic Kwiatkowski, Cordelia Langford, John Sillitoe on behalf of the Wellcome Sanger Institute COVID-19 Surveillance Team |
| EPI_ISL_879653, EPI_ISL_879654, EPI_ISL_879655, EPI_ISL_879656, EPI_ISL_879663, EPI_ISL_879664, EPI_ISL_879675, EPI_ISL_879678, EPI_ISL_879679, EPI_ISL_879681, EPI_ISL_879686, EPI_ISL_879689 | | | |
| see above | Lighthouse Lab in Milton Keynes | Wellcome Sanger Institute for the COVID-19 Genomics UK (COG-UK) Consortium | The Lighthouse Lab in Milton Keynes and Alex Alderton, Roberto Amato, Sonia Goncalves, Ewan Harrison, David K. Jackson, Ian Johnston, Dominic Kwiatkowski, Cordelia Langford, John Sillitoe on behalf of the Wellcome Sanger Institute COVID-19 Surveillance Team |
| EPI_ISL_879692, EPI_ISL_879693, EPI_ISL_879700 | Lighthouse Lab in Alderley Park | Wellcome Sanger Institute for the COVID-19 Genomics UK (COG-UK) Consortium | Jacquelyn Wynn, Mairead Hyland, The Lighthouse Lab in Alderley Park and Alex Alderton, Roberto Amato, Sonia Goncalves, Ewan Harrison, David K. Jackson, Ian Johnston, Dominic Kwiatkowski, Cordelia Langford, John Sillitoe on behalf of the Wellcome Sanger Institute COVID-19 Surveillance Team |
| EPI_ISL_879703, EPI_ISL_879707, EPI_ISL_879710, EPI_ISL_879711, EPI_ISL_879712, EPI_ISL_879713 | Lighthouse Lab in Milton Keynes | Wellcome Sanger Institute for the COVID-19 Genomics UK (COG-UK) Consortium | The Lighthouse Lab in Milton Keynes and Alex Alderton, Roberto Amato, Sonia Goncalves, Ewan Harrison, David K. Jackson, Ian Johnston, Dominic Kwiatkowski, Cordelia Langford, John Sillitoe on behalf of the Wellcome Sanger Institute COVID-19 Surveillance Team |
| EPI_ISL_879714 | Lighthouse Lab in Alderley Park | Wellcome Sanger Institute for the COVID-19 Genomics UK (COG-UK) Consortium | Jacquelyn Wynn, Mairead Hyland, The Lighthouse Lab in Alderley Park and Alex Alderton, Roberto Amato, Sonia Goncalves, Ewan Harrison, David K. Jackson, Ian Johnston, Dominic Kwiatkowski, Cordelia Langford, John Sillitoe on behalf of the Wellcome Sanger Institute COVID-19 Surveillance Team |
| EPI_ISL_879715, EPI_ISL_879717 | Lighthouse Lab in Milton Keynes | Wellcome Sanger Institute for the COVID-19 Genomics UK (COG-UK) Consortium | The Lighthouse Lab in Milton Keynes and Alex Alderton, Roberto Amato, Sonia Goncalves, Ewan Harrison, David K. Jackson, Ian Johnston, Dominic Kwiatkowski, Cordelia Langford, John Sillitoe on behalf of the Wellcome Sanger Institute COVID-19 Surveillance Team |
| EPI_ISL_879719 | Lighthouse Lab in Alderley Park | Wellcome Sanger Institute for the COVID-19 Genomics UK (COG-UK) Consortium | Jacquelyn Wynn, Mairead Hyland, The Lighthouse Lab in Alderley Park and Alex Alderton, Roberto Amato, Sonia Goncalves, Ewan Harrison, David K. Jackson, Ian Johnston, Dominic Kwiatkowski, Cordelia Langford, John Sillitoe on behalf of the Wellcome Sanger Institute COVID-19 Surveillance Team |
| EPI_ISL_879720 | Lighthouse Lab in Cambridge | Wellcome Sanger Institute for the COVID-19 Genomics UK (COG-UK) Consortium | Rob Howes, The Lighthouse Lab in Cambridge and Alex Alderton, Roberto Amato, Sonia Goncalves, Ewan Harrison, David K. Jackson, Ian Johnston, Dominic Kwiatkowski, Cordelia Langford, John Sillitoe on behalf of the Wellcome Sanger Institute COVID-19 Surveillance Team |
| EPI_ISL_879721 | Lighthouse Lab in Alderley Park | Wellcome Sanger Institute for the COVID-19 Genomics UK (COG-UK) Consortium | Jacquelyn Wynn, Mairead Hyland, The Lighthouse Lab in Alderley Park and Alex Alderton, Roberto Amato, Sonia Goncalves, Ewan Harrison, David K. Jackson, Ian Johnston, Dominic Kwiatkowski, Cordelia Langford, John Sillitoe on behalf of the Wellcome Sanger Institute COVID-19 Surveillance Team |
| EPI_ISL_879723, EPI_ISL_879724, EPI_ISL_879725 | Lighthouse Lab in Cambridge | Wellcome Sanger Institute for the COVID-19 Genomics UK (COG-UK) Consortium | Rob Howes, The Lighthouse Lab in Cambridge and Alex Alderton, Roberto Amato, Sonia Goncalves, Ewan Harrison, David K. Jackson, Ian Johnston, Dominic Kwiatkowski, Cordelia Langford, John Sillitoe on behalf of the Wellcome Sanger Institute COVID-19 Surveillance Team |
| EPI_ISL_879726 | Lighthouse Lab in Alderley Park | Wellcome Sanger Institute for the COVID-19 Genomics UK (COG-UK) Consortium | Jacquelyn Wynn, Mairead Hyland, The Lighthouse Lab in Alderley Park and Alex Alderton, Roberto Amato, Sonia Goncalves, Ewan Harrison, David K. Jackson, Ian Johnston, Dominic Kwiatkowski, Cordelia Langford, John Sillitoe on behalf of the Wellcome Sanger Institute COVID-19 Surveillance Team |
| EPI_ISL_879727 | Lighthouse Lab in Cambridge | Wellcome Sanger Institute for the COVID-19 Genomics UK (COG-UK) Consortium | Rob Howes, The Lighthouse Lab in Cambridge and Alex Alderton, Roberto Amato, Sonia Goncalves, Ewan Harrison, David K. Jackson, Ian Johnston, Dominic Kwiatkowski, Cordelia Langford, John Sillitoe on behalf of the Wellcome Sanger Institute COVID-19 Surveillance Team |

| | | | |
|---|---|---|---|
| EPI_ISL_879728 | Lighthouse Lab in Alderley Park | Wellcome Sanger Institute for the COVID-19 Genomics UK (COG-UK) Consortium | Jacquelyn Wynn, Mairead Hyland, The Lighthouse Lab in Alderley Park and Alex Alderton, Roberto Amato, Sonia Goncalves, Ewan Harrison, David K. Jackson, Ian Johnston, Dominic Kwiatkowski, Cordelia Langford, John Sillitoe on behalf of the Wellcome Sanger Institute COVID-19 Surveillance Team |
| EPI_ISL_879729, EPI_ISL_879730, EPI_ISL_879731, EPI_ISL_879732, EPI_ISL_879734 | Lighthouse Lab in Cambridge | Wellcome Sanger Institute for the COVID-19 Genomics UK (COG-UK) Consortium | Rob Howes, The Lighthouse Lab in Cambridge and Alex Alderton, Roberto Amato, Sonia Goncalves, Ewan Harrison, David K. Jackson, Ian Johnston, Dominic Kwiatkowski, Cordelia Langford, John Sillitoe on behalf of the Wellcome Sanger Institute COVID-19 Surveillance Team |
| EPI_ISL_879736, EPI_ISL_879737 | Lighthouse Lab in Alderley Park | Wellcome Sanger Institute for the COVID-19 Genomics UK (COG-UK) Consortium | Jacquelyn Wynn, Mairead Hyland, The Lighthouse Lab in Alderley Park and Alex Alderton, Roberto Amato, Sonia Goncalves, Ewan Harrison, David K. Jackson, Ian Johnston, Dominic Kwiatkowski, Cordelia Langford, John Sillitoe on behalf of the Wellcome Sanger Institute COVID-19 Surveillance Team |
| EPI_ISL_879738, EPI_ISL_879739 | Lighthouse Lab in Cambridge | Wellcome Sanger Institute for the COVID-19 Genomics UK (COG-UK) Consortium | Rob Howes, The Lighthouse Lab in Cambridge and Alex Alderton, Roberto Amato, Sonia Goncalves, Ewan Harrison, David K. Jackson, Ian Johnston, Dominic Kwiatkowski, Cordelia Langford, John Sillitoe on behalf of the Wellcome Sanger Institute COVID-19 Surveillance Team |
| EPI_ISL_879740 | Lighthouse Lab in Alderley Park | Wellcome Sanger Institute for the COVID-19 Genomics UK (COG-UK) Consortium | Jacquelyn Wynn, Mairead Hyland, The Lighthouse Lab in Alderley Park and Alex Alderton, Roberto Amato, Sonia Goncalves, Ewan Harrison, David K. Jackson, Ian Johnston, Dominic Kwiatkowski, Cordelia Langford, John Sillitoe on behalf of the Wellcome Sanger Institute COVID-19 Surveillance Team |
| EPI_ISL_879741 | Lighthouse Lab in Cambridge | Wellcome Sanger Institute for the COVID-19 Genomics UK (COG-UK) Consortium | Rob Howes, The Lighthouse Lab in Cambridge and Alex Alderton, Roberto Amato, Sonia Goncalves, Ewan Harrison, David K. Jackson, Ian Johnston, Dominic Kwiatkowski, Cordelia Langford, John Sillitoe on behalf of the Wellcome Sanger Institute COVID-19 Surveillance Team |
| EPI_ISL_879742, EPI_ISL_879744, EPI_ISL_879747, EPI_ISL_879748 | Lighthouse Lab in Alderley Park | Wellcome Sanger Institute for the COVID-19 Genomics UK (COG-UK) Consortium | Jacquelyn Wynn, Mairead Hyland, The Lighthouse Lab in Alderley Park and Alex Alderton, Roberto Amato, Sonia Goncalves, Ewan Harrison, David K. Jackson, Ian Johnston, Dominic Kwiatkowski, Cordelia Langford, John Sillitoe on behalf of the Wellcome Sanger Institute COVID-19 Surveillance Team |
| EPI_ISL_879749 | Lighthouse Lab in Cambridge | Wellcome Sanger Institute for the COVID-19 Genomics UK (COG-UK) Consortium | Rob Howes, The Lighthouse Lab in Cambridge and Alex Alderton, Roberto Amato, Sonia Goncalves, Ewan Harrison, David K. Jackson, Ian Johnston, Dominic Kwiatkowski, Cordelia Langford, John Sillitoe on behalf of the Wellcome Sanger Institute COVID-19 Surveillance Team |
| EPI_ISL_879750, EPI_ISL_879751, EPI_ISL_879752, EPI_ISL_879753 | Lighthouse Lab in Alderley Park | Wellcome Sanger Institute for the COVID-19 Genomics UK (COG-UK) Consortium | Jacquelyn Wynn, Mairead Hyland, The Lighthouse Lab in Alderley Park and Alex Alderton, Roberto Amato, Sonia Goncalves, Ewan Harrison, David K. Jackson, Ian Johnston, Dominic Kwiatkowski, Cordelia Langford, John Sillitoe on behalf of the Wellcome Sanger Institute COVID-19 Surveillance Team |
| EPI_ISL_879754 | Lighthouse Lab in Cambridge | Wellcome Sanger Institute for the COVID-19 Genomics UK (COG-UK) Consortium | Rob Howes, The Lighthouse Lab in Cambridge and Alex Alderton, Roberto Amato, Sonia Goncalves, Ewan Harrison, David K. Jackson, Ian Johnston, Dominic Kwiatkowski, Cordelia Langford, John Sillitoe on behalf of the Wellcome Sanger Institute COVID-19 Surveillance Team |
| EPI_ISL_879755, EPI_ISL_879756, EPI_ISL_879757, EPI_ISL_879758, EPI_ISL_879759, EPI_ISL_879760, EPI_ISL_879761, EPI_ISL_879762 | Lighthouse Lab in Alderley Park | Wellcome Sanger Institute for the COVID-19 Genomics UK (COG-UK) Consortium | Jacquelyn Wynn, Mairead Hyland, The Lighthouse Lab in Alderley Park and Alex Alderton, Roberto Amato, Sonia Goncalves, Ewan Harrison, David K. Jackson, Ian Johnston, Dominic Kwiatkowski, Cordelia Langford, John Sillitoe on behalf of the Wellcome Sanger Institute COVID-19 Surveillance Team |
| EPI_ISL_879763, EPI_ISL_879764 | Lighthouse Lab in Cambridge | Wellcome Sanger Institute for the COVID-19 Genomics UK (COG-UK) Consortium | Rob Howes, The Lighthouse Lab in Cambridge and Alex Alderton, Roberto Amato, Sonia Goncalves, Ewan Harrison, David K. Jackson, Ian Johnston, Dominic Kwiatkowski, Cordelia Langford, John Sillitoe on behalf of the Wellcome Sanger Institute COVID-19 Surveillance Team |
| EPI_ISL_879765, EPI_ISL_879766, EPI_ISL_879767 | Lighthouse Lab in Alderley Park | Wellcome Sanger Institute for the COVID-19 Genomics UK (COG-UK) Consortium | Jacquelyn Wynn, Mairead Hyland, The Lighthouse Lab in Alderley Park and Alex Alderton, Roberto Amato, Sonia Goncalves, Ewan Harrison, David K. Jackson, Ian Johnston, Dominic Kwiatkowski, Cordelia Langford, John Sillitoe on behalf of the Wellcome Sanger Institute COVID-19 Surveillance Team |
| EPI_ISL_879768, EPI_ISL_879769, EPI_ISL_879771, EPI_ISL_879772, EPI_ISL_879773, EPI_ISL_879774 | Lighthouse Lab in Cambridge | Wellcome Sanger Institute for the COVID-19 Genomics UK (COG-UK) Consortium | Rob Howes, The Lighthouse Lab in Cambridge and Alex Alderton, Roberto Amato, Sonia Goncalves, Ewan Harrison, David K. Jackson, Ian Johnston, Dominic Kwiatkowski, Cordelia Langford, John Sillitoe on behalf of the Wellcome Sanger Institute COVID-19 Surveillance Team |
| EPI_ISL_879775 | Lighthouse Lab in Alderley Park | Wellcome Sanger Institute for the COVID-19 Genomics UK (COG-UK) Consortium | Jacquelyn Wynn, Mairead Hyland, The Lighthouse Lab in Alderley Park and Alex Alderton, Roberto Amato, Sonia Goncalves, Ewan Harrison, David K. Jackson, Ian Johnston, Dominic Kwiatkowski, Cordelia Langford, John Sillitoe on behalf of the Wellcome Sanger Institute COVID-19 Surveillance Team |
| EPI_ISL_879776, EPI_ISL_879777 | Lighthouse Lab in Cambridge | Wellcome Sanger Institute for the COVID-19 Genomics UK (COG-UK) Consortium | Rob Howes, The Lighthouse Lab in Cambridge and Alex Alderton, Roberto Amato, Sonia Goncalves, Ewan Harrison, David K. Jackson, Ian Johnston, Dominic Kwiatkowski, Cordelia Langford, John Sillitoe on behalf of the Wellcome Sanger Institute COVID-19 Surveillance Team |
| EPI_ISL_879778 | Lighthouse Lab in Alderley Park | Wellcome Sanger Institute for the COVID-19 Genomics UK (COG-UK) Consortium | Jacquelyn Wynn, Mairead Hyland, The Lighthouse Lab in Alderley Park and Alex Alderton, Roberto Amato, Sonia Goncalves, Ewan Harrison, David K. Jackson, Ian Johnston, Dominic Kwiatkowski, Cordelia Langford, John Sillitoe on behalf of the Wellcome Sanger Institute COVID-19 Surveillance Team |
| EPI_ISL_879779 | Lighthouse Lab in Cambridge | Wellcome Sanger Institute for the COVID-19 Genomics UK (COG-UK) Consortium | Rob Howes, The Lighthouse Lab in Cambridge and Alex Alderton, Roberto Amato, Sonia Goncalves, Ewan Harrison, David K. Jackson, Ian Johnston, Dominic Kwiatkowski, Cordelia Langford, John Sillitoe on behalf of the Wellcome Sanger Institute COVID-19 Surveillance Team |
| EPI_ISL_879780 | Lighthouse Lab in Alderley Park | Wellcome Sanger Institute for the COVID-19 Genomics UK (COG-UK) Consortium | Jacquelyn Wynn, Mairead Hyland, The Lighthouse Lab in Alderley Park and Alex Alderton, Roberto Amato, Sonia Goncalves, Ewan Harrison, David K. Jackson, Ian Johnston, Dominic Kwiatkowski, Cordelia Langford, John Sillitoe on behalf of the Wellcome Sanger Institute COVID-19 Surveillance Team |
| EPI_ISL_879781 | Lighthouse Lab in Cambridge | Wellcome Sanger Institute for the COVID-19 Genomics UK (COG-UK) Consortium | Rob Howes, The Lighthouse Lab in Cambridge and Alex Alderton, Roberto Amato, Sonia Goncalves, Ewan Harrison, David K. Jackson, Ian Johnston, Dominic Kwiatkowski, Cordelia Langford, John Sillitoe on behalf of the Wellcome Sanger Institute COVID-19 Surveillance Team |
| EPI_ISL_879783, EPI_ISL_879784, EPI_ISL_879785, EPI_ISL_879787 | Lighthouse Lab in Alderley Park | Wellcome Sanger Institute for the COVID-19 Genomics UK (COG-UK) Consortium | Jacquelyn Wynn, Mairead Hyland, The Lighthouse Lab in Alderley Park and Alex Alderton, Roberto Amato, Sonia Goncalves, Ewan Harrison, David K. Jackson, Ian Johnston, Dominic Kwiatkowski, Cordelia Langford, John Sillitoe on behalf of the Wellcome Sanger Institute COVID-19 Surveillance Team |
| EPI_ISL_879788 | Lighthouse Lab in Cambridge | Wellcome Sanger Institute for the COVID-19 Genomics UK (COG-UK) Consortium | Rob Howes, The Lighthouse Lab in Cambridge and Alex Alderton, Roberto Amato, Sonia Goncalves, Ewan Harrison, David K. Jackson, Ian Johnston, Dominic Kwiatkowski, Cordelia Langford, John Sillitoe on behalf of the Wellcome Sanger Institute COVID-19 Surveillance Team |
| EPI_ISL_879789 | Lighthouse Lab in Alderley Park | Wellcome Sanger Institute for the COVID-19 Genomics UK (COG-UK) Consortium | Jacquelyn Wynn, Mairead Hyland, The Lighthouse Lab in Alderley Park and Alex Alderton, Roberto Amato, Sonia Goncalves, Ewan Harrison, David K. Jackson, Ian Johnston, Dominic Kwiatkowski, Cordelia Langford, John Sillitoe on behalf of the Wellcome Sanger Institute COVID-19 Surveillance Team |
| EPI_ISL_879790 | Lighthouse Lab in Cambridge | Wellcome Sanger Institute for the COVID-19 Genomics UK (COG-UK) Consortium | Rob Howes, The Lighthouse Lab in Cambridge and Alex Alderton, Roberto Amato, Sonia Goncalves, Ewan Harrison, David K. Jackson, Ian Johnston, Dominic Kwiatkowski, Cordelia Langford, John Sillitoe on behalf of the Wellcome Sanger Institute COVID-19 Surveillance Team |
| EPI_ISL_879791, EPI_ISL_879792, EPI_ISL_879793, EPI_ISL_879794 | Lighthouse Lab in Alderley Park | Wellcome Sanger Institute for the COVID-19 Genomics UK (COG-UK) Consortium | Jacquelyn Wynn, Mairead Hyland, The Lighthouse Lab in Alderley Park and Alex Alderton, Roberto Amato, Sonia Goncalves, Ewan Harrison, David K. Jackson, Ian Johnston, Dominic Kwiatkowski, Cordelia Langford, John Sillitoe on behalf of the Wellcome Sanger Institute COVID-19 Surveillance Team |
| EPI_ISL_879795, EPI_ISL_879797, EPI_ISL_879798 | Lighthouse Lab in Cambridge | Wellcome Sanger Institute for the COVID-19 Genomics UK (COG-UK) Consortium | Rob Howes, The Lighthouse Lab in Cambridge and Alex Alderton, Roberto Amato, Sonia Goncalves, Ewan Harrison, David K. Jackson, Ian Johnston, Dominic Kwiatkowski, Cordelia Langford, John Sillitoe on behalf of the Wellcome Sanger Institute COVID-19 Surveillance Team |
| EPI_ISL_879800, EPI_ISL_879801, EPI_ISL_879802, EPI_ISL_879803 | Lighthouse Lab in Alderley Park | Wellcome Sanger Institute for the COVID-19 Genomics UK (COG-UK) Consortium | Jacquelyn Wynn, Mairead Hyland, The Lighthouse Lab in Alderley Park and Alex Alderton, Roberto Amato, Sonia Goncalves, Ewan Harrison, David K. Jackson, Ian Johnston, Dominic Kwiatkowski, Cordelia Langford, John Sillitoe on behalf of the Wellcome Sanger Institute COVID-19 Surveillance Team |
| EPI_ISL_879806 | Lighthouse Lab in Cambridge | Wellcome Sanger Institute for the COVID-19 Genomics UK (COG-UK) Consortium | Rob Howes, The Lighthouse Lab in Cambridge and Alex Alderton, Roberto Amato, Sonia Goncalves, Ewan Harrison, David K. Jackson, Ian Johnston, Dominic Kwiatkowski, Cordelia Langford, John Sillitoe on behalf of the Wellcome Sanger Institute COVID-19 Surveillance Team |
| EPI_ISL_879808 | Lighthouse Lab in Alderley Park | Wellcome Sanger Institute for the COVID-19 Genomics UK (COG-UK) Consortium | Jacquelyn Wynn, Mairead Hyland, The Lighthouse Lab in Alderley Park and Alex Alderton, Roberto Amato, Sonia Goncalves, Ewan Harrison, David K. Jackson, Ian Johnston, Dominic Kwiatkowski, Cordelia Langford, John Sillitoe on behalf of the Wellcome Sanger Institute COVID-19 Surveillance Team |
| EPI_ISL_879809 | Lighthouse Lab in Cambridge | Wellcome Sanger Institute for the COVID-19 Genomics UK (COG-UK) Consortium | Rob Howes, The Lighthouse Lab in Cambridge and Alex Alderton, Roberto Amato, Sonia Goncalves, Ewan Harrison, David K. Jackson, Ian Johnston, Dominic Kwiatkowski, Cordelia Langford, John Sillitoe on behalf of the Wellcome Sanger Institute COVID-19 Surveillance Team |
| EPI_ISL_879810, EPI_ISL_879811, EPI_ISL_879813, EPI_ISL_879814 | Lighthouse Lab in Alderley Park | Wellcome Sanger Institute for the COVID-19 Genomics UK (COG-UK) Consortium | Jacquelyn Wynn, Mairead Hyland, The Lighthouse Lab in Alderley Park and Alex Alderton, Roberto Amato, Sonia Goncalves, Ewan Harrison, David K. Jackson, Ian Johnston, Dominic Kwiatkowski, Cordelia Langford, John Sillitoe on behalf of the Wellcome Sanger Institute COVID-19 Surveillance Team |
| EPI_ISL_879816 | Lighthouse Lab in Cambridge | Wellcome Sanger Institute for the COVID-19 Genomics UK (COG-UK) Consortium | Rob Howes, The Lighthouse Lab in Cambridge and Alex Alderton, Roberto Amato, Sonia Goncalves, Ewan Harrison, David K. Jackson, Ian Johnston, Dominic Kwiatkowski, Cordelia Langford, John Sillitoe on behalf of the Wellcome Sanger Institute COVID-19 Surveillance Team |
| EPI_ISL_879817 | Lighthouse Lab in Alderley Park | Wellcome Sanger Institute for the COVID-19 Genomics UK (COG-UK) Consortium | Jacquelyn Wynn, Mairead Hyland, The Lighthouse Lab in Alderley Park and Alex Alderton, Roberto Amato, Sonia Goncalves, Ewan Harrison, David K. Jackson, Ian Johnston, Dominic Kwiatkowski, Cordelia Langford, John Sillitoe on behalf of the Wellcome Sanger Institute COVID-19 Surveillance Team |
| EPI_ISL_879818 | Lighthouse Lab in Cambridge | Wellcome Sanger Institute for the COVID-19 Genomics UK | Rob Howes, The Lighthouse Lab in Cambridge and Alex Alderton, Roberto Amato, Sonia Goncalves, Ewan Harrison, David K. Jackson, Ian Johnston, |

| | | | |
|---|---|---|---|
| | | (COG-UK) Consortium | Dominic Kwiatkowski, Cordelia Langford, John Sillitoe on behalf of the Wellcome Sanger Institute COVID-19 Surveillance Team |
| EPI_ISL_879819, EPI_ISL_879821, EPI_ISL_879822 | Lighthouse Lab in Alderley Park | Wellcome Sanger Institute for the COVID-19 Genomics UK (COG-UK) Consortium | Jacquelyn Wynn, Mairead Hyland, The Lighthouse Lab in Alderley Park and Alex Alderton, Roberto Amato, Sonia Goncalves, Ewan Harrison, David K. Jackson, Ian Johnston, Dominic Kwiatkowski, Cordelia Langford, John Sillitoe on behalf of the Wellcome Sanger Institute COVID-19 Surveillance Team |
| EPI_ISL_879823 | Lighthouse Lab in Cambridge | Wellcome Sanger Institute for the COVID-19 Genomics UK (COG-UK) Consortium | Rob Howes, The Lighthouse Lab in Cambridge and Alex Alderton, Roberto Amato, Sonia Goncalves, Ewan Harrison, David K. Jackson, Ian Johnston, Dominic Kwiatkowski, Cordelia Langford, John Sillitoe on behalf of the Wellcome Sanger Institute COVID-19 Surveillance Team |
| EPI_ISL_879825, EPI_ISL_879826, EPI_ISL_879827 | Lighthouse Lab in Alderley Park | Wellcome Sanger Institute for the COVID-19 Genomics UK (COG-UK) Consortium | Jacquelyn Wynn, Mairead Hyland, The Lighthouse Lab in Alderley Park and Alex Alderton, Roberto Amato, Sonia Goncalves, Ewan Harrison, David K. Jackson, Ian Johnston, Dominic Kwiatkowski, Cordelia Langford, John Sillitoe on behalf of the Wellcome Sanger Institute COVID-19 Surveillance Team |
| EPI_ISL_879828 | Lighthouse Lab in Cambridge | Wellcome Sanger Institute for the COVID-19 Genomics UK (COG-UK) Consortium | Rob Howes, The Lighthouse Lab in Cambridge and Alex Alderton, Roberto Amato, Sonia Goncalves, Ewan Harrison, David K. Jackson, Ian Johnston, Dominic Kwiatkowski, Cordelia Langford, John Sillitoe on behalf of the Wellcome Sanger Institute COVID-19 Surveillance Team |
| EPI_ISL_879829, EPI_ISL_879831 | Lighthouse Lab in Alderley Park | Wellcome Sanger Institute for the COVID-19 Genomics UK (COG-UK) Consortium | Jacquelyn Wynn, Mairead Hyland, The Lighthouse Lab in Alderley Park and Alex Alderton, Roberto Amato, Sonia Goncalves, Ewan Harrison, David K. Jackson, Ian Johnston, Dominic Kwiatkowski, Cordelia Langford, John Sillitoe on behalf of the Wellcome Sanger Institute COVID-19 Surveillance Team |
| EPI_ISL_879832, EPI_ISL_879833, EPI_ISL_879834, EPI_ISL_879835 | Lighthouse Lab in Cambridge | Wellcome Sanger Institute for the COVID-19 Genomics UK (COG-UK) Consortium | Rob Howes, The Lighthouse Lab in Cambridge and Alex Alderton, Roberto Amato, Sonia Goncalves, Ewan Harrison, David K. Jackson, Ian Johnston, Dominic Kwiatkowski, Cordelia Langford, John Sillitoe on behalf of the Wellcome Sanger Institute COVID-19 Surveillance Team |
| EPI_ISL_879836, EPI_ISL_879837, EPI_ISL_879838 | Lighthouse Lab in Alderley Park | Wellcome Sanger Institute for the COVID-19 Genomics UK (COG-UK) Consortium | Jacquelyn Wynn, Mairead Hyland, The Lighthouse Lab in Alderley Park and Alex Alderton, Roberto Amato, Sonia Goncalves, Ewan Harrison, David K. Jackson, Ian Johnston, Dominic Kwiatkowski, Cordelia Langford, John Sillitoe on behalf of the Wellcome Sanger Institute COVID-19 Surveillance Team |
| EPI_ISL_879839 | Lighthouse Lab in Cambridge | Wellcome Sanger Institute for the COVID-19 Genomics UK (COG-UK) Consortium | Rob Howes, The Lighthouse Lab in Cambridge and Alex Alderton, Roberto Amato, Sonia Goncalves, Ewan Harrison, David K. Jackson, Ian Johnston, Dominic Kwiatkowski, Cordelia Langford, John Sillitoe on behalf of the Wellcome Sanger Institute COVID-19 Surveillance Team |
| EPI_ISL_879841, EPI_ISL_879842, EPI_ISL_879843, EPI_ISL_879844 | Lighthouse Lab in Alderley Park | Wellcome Sanger Institute for the COVID-19 Genomics UK (COG-UK) Consortium | Jacquelyn Wynn, Mairead Hyland, The Lighthouse Lab in Alderley Park and Alex Alderton, Roberto Amato, Sonia Goncalves, Ewan Harrison, David K. Jackson, Ian Johnston, Dominic Kwiatkowski, Cordelia Langford, John Sillitoe on behalf of the Wellcome Sanger Institute COVID-19 Surveillance Team |
| EPI_ISL_879845 | Lighthouse Lab in Cambridge | Wellcome Sanger Institute for the COVID-19 Genomics UK (COG-UK) Consortium | Rob Howes, The Lighthouse Lab in Cambridge and Alex Alderton, Roberto Amato, Sonia Goncalves, Ewan Harrison, David K. Jackson, Ian Johnston, Dominic Kwiatkowski, Cordelia Langford, John Sillitoe on behalf of the Wellcome Sanger Institute COVID-19 Surveillance Team |
| EPI_ISL_879846 | Lighthouse Lab in Alderley Park | Wellcome Sanger Institute for the COVID-19 Genomics UK (COG-UK) Consortium | Jacquelyn Wynn, Mairead Hyland, The Lighthouse Lab in Alderley Park and Alex Alderton, Roberto Amato, Sonia Goncalves, Ewan Harrison, David K. Jackson, Ian Johnston, Dominic Kwiatkowski, Cordelia Langford, John Sillitoe on behalf of the Wellcome Sanger Institute COVID-19 Surveillance Team |
| EPI_ISL_879848 | Lighthouse Lab in Cambridge | Wellcome Sanger Institute for the COVID-19 Genomics UK (COG-UK) Consortium | Rob Howes, The Lighthouse Lab in Cambridge and Alex Alderton, Roberto Amato, Sonia Goncalves, Ewan Harrison, David K. Jackson, Ian Johnston, Dominic Kwiatkowski, Cordelia Langford, John Sillitoe on behalf of the Wellcome Sanger Institute COVID-19 Surveillance Team |
| EPI_ISL_879849 | Lighthouse Lab in Alderley Park | Wellcome Sanger Institute for the COVID-19 Genomics UK (COG-UK) Consortium | Jacquelyn Wynn, Mairead Hyland, The Lighthouse Lab in Alderley Park and Alex Alderton, Roberto Amato, Sonia Goncalves, Ewan Harrison, David K. Jackson, Ian Johnston, Dominic Kwiatkowski, Cordelia Langford, John Sillitoe on behalf of the Wellcome Sanger Institute COVID-19 Surveillance Team |
| EPI_ISL_879850 | Lighthouse Lab in Cambridge | Wellcome Sanger Institute for the COVID-19 Genomics UK (COG-UK) Consortium | Rob Howes, The Lighthouse Lab in Cambridge and Alex Alderton, Roberto Amato, Sonia Goncalves, Ewan Harrison, David K. Jackson, Ian Johnston, Dominic Kwiatkowski, Cordelia Langford, John Sillitoe on behalf of the Wellcome Sanger Institute COVID-19 Surveillance Team |
| EPI_ISL_879851 | Lighthouse Lab in Alderley Park | Wellcome Sanger Institute for the COVID-19 Genomics UK (COG-UK) Consortium | Jacquelyn Wynn, Mairead Hyland, The Lighthouse Lab in Alderley Park and Alex Alderton, Roberto Amato, Sonia Goncalves, Ewan Harrison, David K. Jackson, Ian Johnston, Dominic Kwiatkowski, Cordelia Langford, John Sillitoe on behalf of the Wellcome Sanger Institute COVID-19 Surveillance Team |
| EPI_ISL_879852 | Lighthouse Lab in Cambridge | Wellcome Sanger Institute for the COVID-19 Genomics UK (COG-UK) Consortium | Rob Howes, The Lighthouse Lab in Cambridge and Alex Alderton, Roberto Amato, Sonia Goncalves, Ewan Harrison, David K. Jackson, Ian Johnston, Dominic Kwiatkowski, Cordelia Langford, John Sillitoe on behalf of the Wellcome Sanger Institute COVID-19 Surveillance Team |
| EPI_ISL_879854, EPI_ISL_879855, EPI_ISL_879856, EPI_ISL_879858, EPI_ISL_879859 | Lighthouse Lab in Alderley Park | Wellcome Sanger Institute for the COVID-19 Genomics UK (COG-UK) Consortium | Jacquelyn Wynn, Mairead Hyland, The Lighthouse Lab in Alderley Park and Alex Alderton, Roberto Amato, Sonia Goncalves, Ewan Harrison, David K. Jackson, Ian Johnston, Dominic Kwiatkowski, Cordelia Langford, John Sillitoe on behalf of the Wellcome Sanger Institute COVID-19 Surveillance Team |
| EPI_ISL_879860, EPI_ISL_879861, EPI_ISL_879862 | Lighthouse Lab in Cambridge | Wellcome Sanger Institute for the COVID-19 Genomics UK (COG-UK) Consortium | Rob Howes, The Lighthouse Lab in Cambridge and Alex Alderton, Roberto Amato, Sonia Goncalves, Ewan Harrison, David K. Jackson, Ian Johnston, Dominic Kwiatkowski, Cordelia Langford, John Sillitoe on behalf of the Wellcome Sanger Institute COVID-19 Surveillance Team |
| EPI_ISL_879863 | Lighthouse Lab in Alderley Park | Wellcome Sanger Institute for the COVID-19 Genomics UK (COG-UK) Consortium | Jacquelyn Wynn, Mairead Hyland, The Lighthouse Lab in Alderley Park and Alex Alderton, Roberto Amato, Sonia Goncalves, Ewan Harrison, David K. Jackson, Ian Johnston, Dominic Kwiatkowski, Cordelia Langford, John Sillitoe on behalf of the Wellcome Sanger Institute COVID-19 Surveillance Team |
| EPI_ISL_879864 | Lighthouse Lab in Cambridge | Wellcome Sanger Institute for the COVID-19 Genomics UK (COG-UK) Consortium | Rob Howes, The Lighthouse Lab in Cambridge and Alex Alderton, Roberto Amato, Sonia Goncalves, Ewan Harrison, David K. Jackson, Ian Johnston, Dominic Kwiatkowski, Cordelia Langford, John Sillitoe on behalf of the Wellcome Sanger Institute COVID-19 Surveillance Team |
| EPI_ISL_879865, EPI_ISL_879866 | Lighthouse Lab in Alderley Park | Wellcome Sanger Institute for the COVID-19 Genomics UK (COG-UK) Consortium | Jacquelyn Wynn, Mairead Hyland, The Lighthouse Lab in Alderley Park and Alex Alderton, Roberto Amato, Sonia Goncalves, Ewan Harrison, David K. Jackson, Ian Johnston, Dominic Kwiatkowski, Cordelia Langford, John Sillitoe on behalf of the Wellcome Sanger Institute COVID-19 Surveillance Team |
| EPI_ISL_879867 | Lighthouse Lab in Cambridge | Wellcome Sanger Institute for the COVID-19 Genomics UK (COG-UK) Consortium | Rob Howes, The Lighthouse Lab in Cambridge and Alex Alderton, Roberto Amato, Sonia Goncalves, Ewan Harrison, David K. Jackson, Ian Johnston, Dominic Kwiatkowski, Cordelia Langford, John Sillitoe on behalf of the Wellcome Sanger Institute COVID-19 Surveillance Team |
| EPI_ISL_879868, EPI_ISL_879869, EPI_ISL_879870, EPI_ISL_879871, EPI_ISL_879872, EPI_ISL_879873 | Lighthouse Lab in Alderley Park | Wellcome Sanger Institute for the COVID-19 Genomics UK (COG-UK) Consortium | Jacquelyn Wynn, Mairead Hyland, The Lighthouse Lab in Alderley Park and Alex Alderton, Roberto Amato, Sonia Goncalves, Ewan Harrison, David K. Jackson, Ian Johnston, Dominic Kwiatkowski, Cordelia Langford, John Sillitoe on behalf of the Wellcome Sanger Institute COVID-19 Surveillance Team |
| EPI_ISL_879874 | Lighthouse Lab in Cambridge | Wellcome Sanger Institute for the COVID-19 Genomics UK (COG-UK) Consortium | Rob Howes, The Lighthouse Lab in Cambridge and Alex Alderton, Roberto Amato, Sonia Goncalves, Ewan Harrison, David K. Jackson, Ian Johnston, Dominic Kwiatkowski, Cordelia Langford, John Sillitoe on behalf of the Wellcome Sanger Institute COVID-19 Surveillance Team |
| EPI_ISL_879875 | Lighthouse Lab in Alderley Park | Wellcome Sanger Institute for the COVID-19 Genomics UK (COG-UK) Consortium | Jacquelyn Wynn, Mairead Hyland, The Lighthouse Lab in Alderley Park and Alex Alderton, Roberto Amato, Sonia Goncalves, Ewan Harrison, David K. Jackson, Ian Johnston, Dominic Kwiatkowski, Cordelia Langford, John Sillitoe on behalf of the Wellcome Sanger Institute COVID-19 Surveillance Team |
| EPI_ISL_879876, EPI_ISL_879877 | Lighthouse Lab in Cambridge | Wellcome Sanger Institute for the COVID-19 Genomics UK (COG-UK) Consortium | Rob Howes, The Lighthouse Lab in Cambridge and Alex Alderton, Roberto Amato, Sonia Goncalves, Ewan Harrison, David K. Jackson, Ian Johnston, Dominic Kwiatkowski, Cordelia Langford, John Sillitoe on behalf of the Wellcome Sanger Institute COVID-19 Surveillance Team |
| EPI_ISL_879878, EPI_ISL_879879 | Lighthouse Lab in Alderley Park | Wellcome Sanger Institute for the COVID-19 Genomics UK (COG-UK) Consortium | Jacquelyn Wynn, Mairead Hyland, The Lighthouse Lab in Alderley Park and Alex Alderton, Roberto Amato, Sonia Goncalves, Ewan Harrison, David K. Jackson, Ian Johnston, Dominic Kwiatkowski, Cordelia Langford, John Sillitoe on behalf of the Wellcome Sanger Institute COVID-19 Surveillance Team |
| EPI_ISL_879880 | Lighthouse Lab in Cambridge | Wellcome Sanger Institute for the COVID-19 Genomics UK (COG-UK) Consortium | Rob Howes, The Lighthouse Lab in Cambridge and Alex Alderton, Roberto Amato, Sonia Goncalves, Ewan Harrison, David K. Jackson, Ian Johnston, Dominic Kwiatkowski, Cordelia Langford, John Sillitoe on behalf of the Wellcome Sanger Institute COVID-19 Surveillance Team |
| EPI_ISL_879882, EPI_ISL_879884, EPI_ISL_879885, EPI_ISL_879886, EPI_ISL_879887, EPI_ISL_879888 | Lighthouse Lab in Alderley Park | Wellcome Sanger Institute for the COVID-19 Genomics UK (COG-UK) Consortium | Jacquelyn Wynn, Mairead Hyland, The Lighthouse Lab in Alderley Park and Alex Alderton, Roberto Amato, Sonia Goncalves, Ewan Harrison, David K. Jackson, Ian Johnston, Dominic Kwiatkowski, Cordelia Langford, John Sillitoe on behalf of the Wellcome Sanger Institute COVID-19 Surveillance Team |
| EPI_ISL_879889, EPI_ISL_879890, EPI_ISL_879891, EPI_ISL_879892 | Lighthouse Lab in Cambridge | Wellcome Sanger Institute for the COVID-19 Genomics UK (COG-UK) Consortium | Rob Howes, The Lighthouse Lab in Cambridge and Alex Alderton, Roberto Amato, Sonia Goncalves, Ewan Harrison, David K. Jackson, Ian Johnston, Dominic Kwiatkowski, Cordelia Langford, John Sillitoe on behalf of the Wellcome Sanger Institute COVID-19 Surveillance Team |
| EPI_ISL_879894, EPI_ISL_879895 | Lighthouse Lab in Alderley Park | Wellcome Sanger Institute for the COVID-19 Genomics UK (COG-UK) Consortium | Jacquelyn Wynn, Mairead Hyland, The Lighthouse Lab in Alderley Park and Alex Alderton, Roberto Amato, Sonia Goncalves, Ewan Harrison, David K. Jackson, Ian Johnston, Dominic Kwiatkowski, Cordelia Langford, John Sillitoe on behalf of the Wellcome Sanger Institute COVID-19 Surveillance Team |
| EPI_ISL_879896 | Lighthouse Lab in Cambridge | Wellcome Sanger Institute for the COVID-19 Genomics UK (COG-UK) Consortium | Rob Howes, The Lighthouse Lab in Cambridge and Alex Alderton, Roberto Amato, Sonia Goncalves, Ewan Harrison, David K. Jackson, Ian Johnston, Dominic Kwiatkowski, Cordelia Langford, John Sillitoe on behalf of the Wellcome Sanger Institute COVID-19 Surveillance Team |
| EPI_ISL_879897, EPI_ISL_879898, EPI_ISL_879901, EPI_ISL_879903 | Lighthouse Lab in Alderley Park | Wellcome Sanger Institute for the COVID-19 Genomics UK (COG-UK) Consortium | Jacquelyn Wynn, Mairead Hyland, The Lighthouse Lab in Alderley Park and Alex Alderton, Roberto Amato, Sonia Goncalves, Ewan Harrison, David K. Jackson, Ian Johnston, Dominic Kwiatkowski, Cordelia Langford, John Sillitoe on behalf of the Wellcome Sanger Institute COVID-19 Surveillance Team |
| EPI_ISL_879904 | Lighthouse Lab in Cambridge | Wellcome Sanger Institute for the COVID-19 Genomics UK | Rob Howes, The Lighthouse Lab in Cambridge and Alex Alderton, Roberto Amato, Sonia Goncalves, Ewan Harrison, David K. Jackson, Ian Johnston, |

| | | | |
|---|---|---|---|
| | | (COG-UK) Consortium | Dominic Kwiatkowski, Cordelia Langford, John Sillitoe on behalf of the Wellcome Sanger Institute COVID-19 Surveillance Team |
| EPI_ISL_879905, EPI_ISL_879906, EPI_ISL_879907 | Lighthouse Lab in Alderley Park | Wellcome Sanger Institute for the COVID-19 Genomics UK (COG-UK) Consortium | Jacquelyn Wynn, Mairead Hyland, The Lighthouse Lab in Alderley Park and Alex Alderton, Roberto Amato, Sonia Goncalves, Ewan Harrison, David K. Jackson, Ian Johnston, Dominic Kwiatkowski, Cordelia Langford, John Sillitoe on behalf of the Wellcome Sanger Institute COVID-19 Surveillance Team |
| EPI_ISL_879908, EPI_ISL_879909, EPI_ISL_879911 | Lighthouse Lab in Cambridge | Wellcome Sanger Institute for the COVID-19 Genomics UK (COG-UK) Consortium | Rob Howes, The Lighthouse Lab in Cambridge and Alex Alderton, Roberto Amato, Sonia Goncalves, Ewan Harrison, David K. Jackson, Ian Johnston, Dominic Kwiatkowski, Cordelia Langford, John Sillitoe on behalf of the Wellcome Sanger Institute COVID-19 Surveillance Team |
| EPI_ISL_879912, EPI_ISL_879913, EPI_ISL_879914, EPI_ISL_879915, EPI_ISL_879916, EPI_ISL_879918, EPI_ISL_879919 | Lighthouse Lab in Alderley Park | Wellcome Sanger Institute for the COVID-19 Genomics UK (COG-UK) Consortium | Jacquelyn Wynn, Mairead Hyland, The Lighthouse Lab in Alderley Park and Alex Alderton, Roberto Amato, Sonia Goncalves, Ewan Harrison, David K. Jackson, Ian Johnston, Dominic Kwiatkowski, Cordelia Langford, John Sillitoe on behalf of the Wellcome Sanger Institute COVID-19 Surveillance Team |
| EPI_ISL_879920, EPI_ISL_879921 | Lighthouse Lab in Cambridge | Wellcome Sanger Institute for the COVID-19 Genomics UK (COG-UK) Consortium | Rob Howes, The Lighthouse Lab in Cambridge and Alex Alderton, Roberto Amato, Sonia Goncalves, Ewan Harrison, David K. Jackson, Ian Johnston, Dominic Kwiatkowski, Cordelia Langford, John Sillitoe on behalf of the Wellcome Sanger Institute COVID-19 Surveillance Team |
| EPI_ISL_879922, EPI_ISL_879923, EPI_ISL_879925, EPI_ISL_879927 | Lighthouse Lab in Alderley Park | Wellcome Sanger Institute for the COVID-19 Genomics UK (COG-UK) Consortium | Jacquelyn Wynn, Mairead Hyland, The Lighthouse Lab in Alderley Park and Alex Alderton, Roberto Amato, Sonia Goncalves, Ewan Harrison, David K. Jackson, Ian Johnston, Dominic Kwiatkowski, Cordelia Langford, John Sillitoe on behalf of the Wellcome Sanger Institute COVID-19 Surveillance Team |
| EPI_ISL_879928, EPI_ISL_879930 | Lighthouse Lab in Cambridge | Wellcome Sanger Institute for the COVID-19 Genomics UK (COG-UK) Consortium | Rob Howes, The Lighthouse Lab in Cambridge and Alex Alderton, Roberto Amato, Sonia Goncalves, Ewan Harrison, David K. Jackson, Ian Johnston, Dominic Kwiatkowski, Cordelia Langford, John Sillitoe on behalf of the Wellcome Sanger Institute COVID-19 Surveillance Team |
| EPI_ISL_879931, EPI_ISL_879933 | Lighthouse Lab in Alderley Park | Wellcome Sanger Institute for the COVID-19 Genomics UK (COG-UK) Consortium | Jacquelyn Wynn, Mairead Hyland, The Lighthouse Lab in Alderley Park and Alex Alderton, Roberto Amato, Sonia Goncalves, Ewan Harrison, David K. Jackson, Ian Johnston, Dominic Kwiatkowski, Cordelia Langford, John Sillitoe on behalf of the Wellcome Sanger Institute COVID-19 Surveillance Team |
| EPI_ISL_879935 | Lighthouse Lab in Cambridge | Wellcome Sanger Institute for the COVID-19 Genomics UK (COG-UK) Consortium | Rob Howes, The Lighthouse Lab in Cambridge and Alex Alderton, Roberto Amato, Sonia Goncalves, Ewan Harrison, David K. Jackson, Ian Johnston, Dominic Kwiatkowski, Cordelia Langford, John Sillitoe on behalf of the Wellcome Sanger Institute COVID-19 Surveillance Team |
| EPI_ISL_879936, EPI_ISL_879938 | Lighthouse Lab in Alderley Park | Wellcome Sanger Institute for the COVID-19 Genomics UK (COG-UK) Consortium | Jacquelyn Wynn, Mairead Hyland, The Lighthouse Lab in Alderley Park and Alex Alderton, Roberto Amato, Sonia Goncalves, Ewan Harrison, David K. Jackson, Ian Johnston, Dominic Kwiatkowski, Cordelia Langford, John Sillitoe on behalf of the Wellcome Sanger Institute COVID-19 Surveillance Team |
| EPI_ISL_879940, EPI_ISL_879941, EPI_ISL_879944, EPI_ISL_879946 | Lighthouse Lab in Cambridge | Wellcome Sanger Institute for the COVID-19 Genomics UK (COG-UK) Consortium | Rob Howes, The Lighthouse Lab in Cambridge and Alex Alderton, Roberto Amato, Sonia Goncalves, Ewan Harrison, David K. Jackson, Ian Johnston, Dominic Kwiatkowski, Cordelia Langford, John Sillitoe on behalf of the Wellcome Sanger Institute COVID-19 Surveillance Team |
| EPI_ISL_879948, EPI_ISL_879949 | Lighthouse Lab in Alderley Park | Wellcome Sanger Institute for the COVID-19 Genomics UK (COG-UK) Consortium | Jacquelyn Wynn, Mairead Hyland, The Lighthouse Lab in Alderley Park and Alex Alderton, Roberto Amato, Sonia Goncalves, Ewan Harrison, David K. Jackson, Ian Johnston, Dominic Kwiatkowski, Cordelia Langford, John Sillitoe on behalf of the Wellcome Sanger Institute COVID-19 Surveillance Team |
| EPI_ISL_879954 | Lighthouse Lab in Cambridge | Wellcome Sanger Institute for the COVID-19 Genomics UK (COG-UK) Consortium | Rob Howes, The Lighthouse Lab in Cambridge and Alex Alderton, Roberto Amato, Sonia Goncalves, Ewan Harrison, David K. Jackson, Ian Johnston, Dominic Kwiatkowski, Cordelia Langford, John Sillitoe on behalf of the Wellcome Sanger Institute COVID-19 Surveillance Team |
| EPI_ISL_879955, EPI_ISL_879957 | Lighthouse Lab in Alderley Park | Wellcome Sanger Institute for the COVID-19 Genomics UK (COG-UK) Consortium | Jacquelyn Wynn, Mairead Hyland, The Lighthouse Lab in Alderley Park and Alex Alderton, Roberto Amato, Sonia Goncalves, Ewan Harrison, David K. Jackson, Ian Johnston, Dominic Kwiatkowski, Cordelia Langford, John Sillitoe on behalf of the Wellcome Sanger Institute COVID-19 Surveillance Team |
| EPI_ISL_879958, EPI_ISL_879960, EPI_ISL_879962, EPI_ISL_879963 | Lighthouse Lab in Cambridge | Wellcome Sanger Institute for the COVID-19 Genomics UK (COG-UK) Consortium | Rob Howes, The Lighthouse Lab in Cambridge and Alex Alderton, Roberto Amato, Sonia Goncalves, Ewan Harrison, David K. Jackson, Ian Johnston, Dominic Kwiatkowski, Cordelia Langford, John Sillitoe on behalf of the Wellcome Sanger Institute COVID-19 Surveillance Team |
| EPI_ISL_879965, EPI_ISL_879966, EPI_ISL_879968, EPI_ISL_879970, EPI_ISL_879971 | Lighthouse Lab in Alderley Park | Wellcome Sanger Institute for the COVID-19 Genomics UK (COG-UK) Consortium | Jacquelyn Wynn, Mairead Hyland, The Lighthouse Lab in Alderley Park and Alex Alderton, Roberto Amato, Sonia Goncalves, Ewan Harrison, David K. Jackson, Ian Johnston, Dominic Kwiatkowski, Cordelia Langford, John Sillitoe on behalf of the Wellcome Sanger Institute COVID-19 Surveillance Team |
| EPI_ISL_879973 | Lighthouse Lab in Cambridge | Wellcome Sanger Institute for the COVID-19 Genomics UK (COG-UK) Consortium | Rob Howes, The Lighthouse Lab in Cambridge and Alex Alderton, Roberto Amato, Sonia Goncalves, Ewan Harrison, David K. Jackson, Ian Johnston, Dominic Kwiatkowski, Cordelia Langford, John Sillitoe on behalf of the Wellcome Sanger Institute COVID-19 Surveillance Team |
| EPI_ISL_879974, EPI_ISL_879976, EPI_ISL_879978, EPI_ISL_879979, EPI_ISL_879981, EPI_ISL_879984, EPI_ISL_879986, EPI_ISL_879988 | Lighthouse Lab in Alderley Park | Wellcome Sanger Institute for the COVID-19 Genomics UK (COG-UK) Consortium | Jacquelyn Wynn, Mairead Hyland, The Lighthouse Lab in Alderley Park and Alex Alderton, Roberto Amato, Sonia Goncalves, Ewan Harrison, David K. Jackson, Ian Johnston, Dominic Kwiatkowski, Cordelia Langford, John Sillitoe on behalf of the Wellcome Sanger Institute COVID-19 Surveillance Team |
| EPI_ISL_879989 | Lighthouse Lab in Cambridge | Wellcome Sanger Institute for the COVID-19 Genomics UK (COG-UK) Consortium | Rob Howes, The Lighthouse Lab in Cambridge and Alex Alderton, Roberto Amato, Sonia Goncalves, Ewan Harrison, David K. Jackson, Ian Johnston, Dominic Kwiatkowski, Cordelia Langford, John Sillitoe on behalf of the Wellcome Sanger Institute COVID-19 Surveillance Team |
| EPI_ISL_879991 | Lighthouse Lab in Alderley Park | Wellcome Sanger Institute for the COVID-19 Genomics UK (COG-UK) Consortium | Jacquelyn Wynn, Mairead Hyland, The Lighthouse Lab in Alderley Park and Alex Alderton, Roberto Amato, Sonia Goncalves, Ewan Harrison, David K. Jackson, Ian Johnston, Dominic Kwiatkowski, Cordelia Langford, John Sillitoe on behalf of the Wellcome Sanger Institute COVID-19 Surveillance Team |
| EPI_ISL_879994, EPI_ISL_879996 | Lighthouse Lab in Cambridge | Wellcome Sanger Institute for the COVID-19 Genomics UK (COG-UK) Consortium | Rob Howes, The Lighthouse Lab in Cambridge and Alex Alderton, Roberto Amato, Sonia Goncalves, Ewan Harrison, David K. Jackson, Ian Johnston, Dominic Kwiatkowski, Cordelia Langford, John Sillitoe on behalf of the Wellcome Sanger Institute COVID-19 Surveillance Team |
| EPI_ISL_879997, EPI_ISL_879999, EPI_ISL_880002 | Lighthouse Lab in Alderley Park | Wellcome Sanger Institute for the COVID-19 Genomics UK (COG-UK) Consortium | Jacquelyn Wynn, Mairead Hyland, The Lighthouse Lab in Alderley Park and Alex Alderton, Roberto Amato, Sonia Goncalves, Ewan Harrison, David K. Jackson, Ian Johnston, Dominic Kwiatkowski, Cordelia Langford, John Sillitoe on behalf of the Wellcome Sanger Institute COVID-19 Surveillance Team |
| EPI_ISL_880004, EPI_ISL_880005 | Lighthouse Lab in Cambridge | Wellcome Sanger Institute for the COVID-19 Genomics UK (COG-UK) Consortium | Rob Howes, The Lighthouse Lab in Cambridge and Alex Alderton, Roberto Amato, Sonia Goncalves, Ewan Harrison, David K. Jackson, Ian Johnston, Dominic Kwiatkowski, Cordelia Langford, John Sillitoe on behalf of the Wellcome Sanger Institute COVID-19 Surveillance Team |
| EPI_ISL_880007, EPI_ISL_880008, EPI_ISL_880010 | Lighthouse Lab in Alderley Park | Wellcome Sanger Institute for the COVID-19 Genomics UK (COG-UK) Consortium | Jacquelyn Wynn, Mairead Hyland, The Lighthouse Lab in Alderley Park and Alex Alderton, Roberto Amato, Sonia Goncalves, Ewan Harrison, David K. Jackson, Ian Johnston, Dominic Kwiatkowski, Cordelia Langford, John Sillitoe on behalf of the Wellcome Sanger Institute COVID-19 Surveillance Team |
| EPI_ISL_880012, EPI_ISL_880013, EPI_ISL_880015 | Lighthouse Lab in Cambridge | Wellcome Sanger Institute for the COVID-19 Genomics UK (COG-UK) Consortium | Rob Howes, The Lighthouse Lab in Cambridge and Alex Alderton, Roberto Amato, Sonia Goncalves, Ewan Harrison, David K. Jackson, Ian Johnston, Dominic Kwiatkowski, Cordelia Langford, John Sillitoe on behalf of the Wellcome Sanger Institute COVID-19 Surveillance Team |
| EPI_ISL_880016, EPI_ISL_880018 | Lighthouse Lab in Alderley Park | Wellcome Sanger Institute for the COVID-19 Genomics UK (COG-UK) Consortium | Jacquelyn Wynn, Mairead Hyland, The Lighthouse Lab in Alderley Park and Alex Alderton, Roberto Amato, Sonia Goncalves, Ewan Harrison, David K. Jackson, Ian Johnston, Dominic Kwiatkowski, Cordelia Langford, John Sillitoe on behalf of the Wellcome Sanger Institute COVID-19 Surveillance Team |
| EPI_ISL_880020 | Lighthouse Lab in Cambridge | Wellcome Sanger Institute for the COVID-19 Genomics UK (COG-UK) Consortium | Rob Howes, The Lighthouse Lab in Cambridge and Alex Alderton, Roberto Amato, Sonia Goncalves, Ewan Harrison, David K. Jackson, Ian Johnston, Dominic Kwiatkowski, Cordelia Langford, John Sillitoe on behalf of the Wellcome Sanger Institute COVID-19 Surveillance Team |
| EPI_ISL_880021, EPI_ISL_880023 | Lighthouse Lab in Alderley Park | Wellcome Sanger Institute for the COVID-19 Genomics UK (COG-UK) Consortium | Jacquelyn Wynn, Mairead Hyland, The Lighthouse Lab in Alderley Park and Alex Alderton, Roberto Amato, Sonia Goncalves, Ewan Harrison, David K. Jackson, Ian Johnston, Dominic Kwiatkowski, Cordelia Langford, John Sillitoe on behalf of the Wellcome Sanger Institute COVID-19 Surveillance Team |
| EPI_ISL_880025 | Lighthouse Lab in Cambridge | Wellcome Sanger Institute for the COVID-19 Genomics UK (COG-UK) Consortium | Rob Howes, The Lighthouse Lab in Cambridge and Alex Alderton, Roberto Amato, Sonia Goncalves, Ewan Harrison, David K. Jackson, Ian Johnston, Dominic Kwiatkowski, Cordelia Langford, John Sillitoe on behalf of the Wellcome Sanger Institute COVID-19 Surveillance Team |
| EPI_ISL_880028, EPI_ISL_880030, EPI_ISL_880031 | Lighthouse Lab in Alderley Park | Wellcome Sanger Institute for the COVID-19 Genomics UK (COG-UK) Consortium | Jacquelyn Wynn, Mairead Hyland, The Lighthouse Lab in Alderley Park and Alex Alderton, Roberto Amato, Sonia Goncalves, Ewan Harrison, David K. Jackson, Ian Johnston, Dominic Kwiatkowski, Cordelia Langford, John Sillitoe on behalf of the Wellcome Sanger Institute COVID-19 Surveillance Team |
| EPI_ISL_880033, EPI_ISL_880035 | Lighthouse Lab in Cambridge | Wellcome Sanger Institute for the COVID-19 Genomics UK (COG-UK) Consortium | Rob Howes, The Lighthouse Lab in Cambridge and Alex Alderton, Roberto Amato, Sonia Goncalves, Ewan Harrison, David K. Jackson, Ian Johnston, Dominic Kwiatkowski, Cordelia Langford, John Sillitoe on behalf of the Wellcome Sanger Institute COVID-19 Surveillance Team |
| EPI_ISL_880036 | Lighthouse Lab in Alderley Park | Wellcome Sanger Institute for the COVID-19 Genomics UK (COG-UK) Consortium | Jacquelyn Wynn, Mairead Hyland, The Lighthouse Lab in Alderley Park and Alex Alderton, Roberto Amato, Sonia Goncalves, Ewan Harrison, David K. Jackson, Ian Johnston, Dominic Kwiatkowski, Cordelia Langford, John Sillitoe on behalf of the Wellcome Sanger Institute COVID-19 Surveillance Team |
| EPI_ISL_880039 | Lighthouse Lab in Cambridge | Wellcome Sanger Institute for the COVID-19 Genomics UK (COG-UK) Consortium | Rob Howes, The Lighthouse Lab in Cambridge and Alex Alderton, Roberto Amato, Sonia Goncalves, Ewan Harrison, David K. Jackson, Ian Johnston, Dominic Kwiatkowski, Cordelia Langford, John Sillitoe on behalf of the Wellcome Sanger Institute COVID-19 Surveillance Team |
| EPI_ISL_880041, EPI_ISL_880043 | Lighthouse Lab in Alderley Park | Wellcome Sanger Institute for the COVID-19 Genomics UK | Jacquelyn Wynn, Mairead Hyland, The Lighthouse Lab in Alderley Park and Alex Alderton, Roberto Amato, Sonia Goncalves, Ewan Harrison, David K. |

| | | | |
|---|---|---|---|
| | | (COG-UK) Consortium | Jackson, Ian Johnston, Dominic Kwiatkowski, Cordelia Langford, John Sillitoe on behalf of the Wellcome Sanger Institute COVID-19 Surveillance Team |
| EPI_ISL_880044, EPI_ISL_880046, EPI_ISL_880047, EPI_ISL_880049 | Lighthouse Lab in Cambridge | Wellcome Sanger Institute for the COVID-19 Genomics UK (COG-UK) Consortium | Rob Howes, The Lighthouse Lab in Cambridge and Alex Alderton, Roberto Amato, Sonia Goncalves, Ewan Harrison, David K. Jackson, Ian Johnston, Dominic Kwiatkowski, Cordelia Langford, John Sillitoe on behalf of the Wellcome Sanger Institute COVID-19 Surveillance Team |
| EPI_ISL_880052 | Lighthouse Lab in Alderley Park | Wellcome Sanger Institute for the COVID-19 Genomics UK (COG-UK) Consortium | Jacquelyn Wynn, Mairead Hyland, The Lighthouse Lab in Alderley Park and Alex Alderton, Roberto Amato, Sonia Goncalves, Ewan Harrison, David K. Jackson, Ian Johnston, Dominic Kwiatkowski, Cordelia Langford, John Sillitoe on behalf of the Wellcome Sanger Institute COVID-19 Surveillance Team |
| EPI_ISL_880054 | Lighthouse Lab in Cambridge | Wellcome Sanger Institute for the COVID-19 Genomics UK (COG-UK) Consortium | Rob Howes, The Lighthouse Lab in Cambridge and Alex Alderton, Roberto Amato, Sonia Goncalves, Ewan Harrison, David K. Jackson, Ian Johnston, Dominic Kwiatkowski, Cordelia Langford, John Sillitoe on behalf of the Wellcome Sanger Institute COVID-19 Surveillance Team |
| EPI_ISL_880055, EPI_ISL_880057 | Lighthouse Lab in Alderley Park | Wellcome Sanger Institute for the COVID-19 Genomics UK (COG-UK) Consortium | Jacquelyn Wynn, Mairead Hyland, The Lighthouse Lab in Alderley Park and Alex Alderton, Roberto Amato, Sonia Goncalves, Ewan Harrison, David K. Jackson, Ian Johnston, Dominic Kwiatkowski, Cordelia Langford, John Sillitoe on behalf of the Wellcome Sanger Institute COVID-19 Surveillance Team |
| EPI_ISL_880059, EPI_ISL_880060 | Lighthouse Lab in Cambridge | Wellcome Sanger Institute for the COVID-19 Genomics UK (COG-UK) Consortium | Rob Howes, The Lighthouse Lab in Cambridge and Alex Alderton, Roberto Amato, Sonia Goncalves, Ewan Harrison, David K. Jackson, Ian Johnston, Dominic Kwiatkowski, Cordelia Langford, John Sillitoe on behalf of the Wellcome Sanger Institute COVID-19 Surveillance Team |
| EPI_ISL_880062, EPI_ISL_880064 | Lighthouse Lab in Alderley Park | Wellcome Sanger Institute for the COVID-19 Genomics UK (COG-UK) Consortium | Jacquelyn Wynn, Mairead Hyland, The Lighthouse Lab in Alderley Park and Alex Alderton, Roberto Amato, Sonia Goncalves, Ewan Harrison, David K. Jackson, Ian Johnston, Dominic Kwiatkowski, Cordelia Langford, John Sillitoe on behalf of the Wellcome Sanger Institute COVID-19 Surveillance Team |
| EPI_ISL_880065 | Lighthouse Lab in Cambridge | Wellcome Sanger Institute for the COVID-19 Genomics UK (COG-UK) Consortium | Rob Howes, The Lighthouse Lab in Cambridge and Alex Alderton, Roberto Amato, Sonia Goncalves, Ewan Harrison, David K. Jackson, Ian Johnston, Dominic Kwiatkowski, Cordelia Langford, John Sillitoe on behalf of the Wellcome Sanger Institute COVID-19 Surveillance Team |
| EPI_ISL_880067 | Lighthouse Lab in Alderley Park | Wellcome Sanger Institute for the COVID-19 Genomics UK (COG-UK) Consortium | Jacquelyn Wynn, Mairead Hyland, The Lighthouse Lab in Alderley Park and Alex Alderton, Roberto Amato, Sonia Goncalves, Ewan Harrison, David K. Jackson, Ian Johnston, Dominic Kwiatkowski, Cordelia Langford, John Sillitoe on behalf of the Wellcome Sanger Institute COVID-19 Surveillance Team |
| EPI_ISL_880068, EPI_ISL_880070 | Lighthouse Lab in Cambridge | Wellcome Sanger Institute for the COVID-19 Genomics UK (COG-UK) Consortium | Rob Howes, The Lighthouse Lab in Cambridge and Alex Alderton, Roberto Amato, Sonia Goncalves, Ewan Harrison, David K. Jackson, Ian Johnston, Dominic Kwiatkowski, Cordelia Langford, John Sillitoe on behalf of the Wellcome Sanger Institute COVID-19 Surveillance Team |
| EPI_ISL_880072 | Lighthouse Lab in Alderley Park | Wellcome Sanger Institute for the COVID-19 Genomics UK (COG-UK) Consortium | Jacquelyn Wynn, Mairead Hyland, The Lighthouse Lab in Alderley Park and Alex Alderton, Roberto Amato, Sonia Goncalves, Ewan Harrison, David K. Jackson, Ian Johnston, Dominic Kwiatkowski, Cordelia Langford, John Sillitoe on behalf of the Wellcome Sanger Institute COVID-19 Surveillance Team |
| EPI_ISL_880075, EPI_ISL_880076, EPI_ISL_880078 | Lighthouse Lab in Cambridge | Wellcome Sanger Institute for the COVID-19 Genomics UK (COG-UK) Consortium | Rob Howes, The Lighthouse Lab in Cambridge and Alex Alderton, Roberto Amato, Sonia Goncalves, Ewan Harrison, David K. Jackson, Ian Johnston, Dominic Kwiatkowski, Cordelia Langford, John Sillitoe on behalf of the Wellcome Sanger Institute COVID-19 Surveillance Team |
| EPI_ISL_880083, EPI_ISL_880084, EPI_ISL_880086, EPI_ISL_880088, EPI_ISL_880089, EPI_ISL_880093, EPI_ISL_880094, EPI_ISL_880096, EPI_ISL_880097, EPI_ISL_880099, EPI_ISL_880102, EPI_ISL_880104, EPI_ISL_880105, EPI_ISL_880107, EPI_ISL_880108, EPI_ISL_880110, EPI_ISL_880115, EPI_ISL_880116, EPI_ISL_880118, EPI_ISL_880121, EPI_ISL_880127, EPI_ISL_880129, EPI_ISL_880131, EPI_ISL_880132, EPI_ISL_880134, EPI_ISL_880137, EPI_ISL_880139, EPI_ISL_880210, EPI_ISL_880211, EPI_ISL_880213, EPI_ISL_880214, EPI_ISL_880217, EPI_ISL_880219, EPI_ISL_880221, EPI_ISL_880224, EPI_ISL_880226, EPI_ISL_880227, EPI_ISL_880229, EPI_ISL_880230, EPI_ISL_880232, EPI_ISL_880234, EPI_ISL_880235, EPI_ISL_880237, EPI_ISL_880238, EPI_ISL_880240, EPI_ISL_880242, EPI_ISL_880243, EPI_ISL_880245, EPI_ISL_880247, EPI_ISL_880248, EPI_ISL_880250, EPI_ISL_880252, EPI_ISL_880254, EPI_ISL_880255, EPI_ISL_880257, EPI_ISL_880262, EPI_ISL_880267, EPI_ISL_880269, EPI_ISL_880270, EPI_ISL_880272, EPI_ISL_880273, EPI_ISL_880275, EPI_ISL_880276, EPI_ISL_880278, EPI_ISL_880280, EPI_ISL_880281, EPI_ISL_880283, EPI_ISL_880285, EPI_ISL_880289, EPI_ISL_880293, EPI_ISL_880295, EPI_ISL_880296, EPI_ISL_880299, EPI_ISL_880302, EPI_ISL_880304, EPI_ISL_880307, EPI_ISL_880309, EPI_ISL_880311, EPI_ISL_880312, EPI_ISL_880314, EPI_ISL_880315, EPI_ISL_880317, EPI_ISL_880318, EPI_ISL_880320, EPI_ISL_880322, EPI_ISL_880323, EPI_ISL_880325, EPI_ISL_880327, EPI_ISL_880328, EPI_ISL_880330, EPI_ISL_880331, EPI_ISL_880333, EPI_ISL_880334, EPI_ISL_880336, EPI_ISL_880338, EPI_ISL_880339, EPI_ISL_880341, EPI_ISL_880342, EPI_ISL_880344, EPI_ISL_880345, EPI_ISL_880346, EPI_ISL_880347, EPI_ISL_880349, EPI_ISL_880350, EPI_ISL_880351, EPI_ISL_880353, EPI_ISL_880354, EPI_ISL_880355, EPI_ISL_880357, EPI_ISL_880358, EPI_ISL_880359, EPI_ISL_880360, EPI_ISL_880362, EPI_ISL_880363, EPI_ISL_880364, EPI_ISL_880365, EPI_ISL_880367, EPI_ISL_880368, EPI_ISL_880369, EPI_ISL_880370, EPI_ISL_880371, EPI_ISL_880372, EPI_ISL_880373, EPI_ISL_880374, EPI_ISL_880377, EPI_ISL_880378, EPI_ISL_880380, EPI_ISL_880381, EPI_ISL_880382, EPI_ISL_880383, EPI_ISL_880384, EPI_ISL_880385, EPI_ISL_880386, EPI_ISL_880388, EPI_ISL_880389, EPI_ISL_880391, EPI_ISL_880393, EPI_ISL_880394, EPI_ISL_880395, EPI_ISL_880396, EPI_ISL_880397, EPI_ISL_880398, EPI_ISL_880399, EPI_ISL_880401, EPI_ISL_880403, EPI_ISL_880404, EPI_ISL_880405, EPI_ISL_880406, EPI_ISL_880407, EPI_ISL_880408, EPI_ISL_880409, EPI_ISL_880410, EPI_ISL_880411, EPI_ISL_880412, EPI_ISL_880413, EPI_ISL_880414, EPI_ISL_880415, EPI_ISL_880416, EPI_ISL_880419, EPI_ISL_880420, EPI_ISL_880421, EPI_ISL_880422, EPI_ISL_880423, EPI_ISL_880424, EPI_ISL_880425, EPI_ISL_880428, EPI_ISL_880429, EPI_ISL_880430, EPI_ISL_880431, EPI_ISL_880432, EPI_ISL_880434, EPI_ISL_880437, EPI_ISL_880438, EPI_ISL_880439, EPI_ISL_880440, EPI_ISL_880441, EPI_ISL_880442, EPI_ISL_880444, EPI_ISL_880445, EPI_ISL_880447, EPI_ISL_880448, EPI_ISL_880449, EPI_ISL_880450, EPI_ISL_880451, EPI_ISL_880452, EPI_ISL_880453, EPI_ISL_880454, EPI_ISL_880455, EPI_ISL_880457, EPI_ISL_880459, EPI_ISL_880460, EPI_ISL_880461, EPI_ISL_880462, EPI_ISL_880463, EPI_ISL_880465, EPI_ISL_880466, EPI_ISL_880468, EPI_ISL_880469, EPI_ISL_880470, EPI_ISL_880471, EPI_ISL_880472, EPI_ISL_880473, EPI_ISL_880474, EPI_ISL_880476, EPI_ISL_880478, EPI_ISL_880479, EPI_ISL_880480, EPI_ISL_880481, EPI_ISL_880483, EPI_ISL_880484, EPI_ISL_880485, EPI_ISL_880486, EPI_ISL_880487, EPI_ISL_880488, EPI_ISL_880489, EPI_ISL_880490, EPI_ISL_880491, EPI_ISL_880492, EPI_ISL_880494, EPI_ISL_880495, EPI_ISL_880496, EPI_ISL_880497, EPI_ISL_880499, EPI_ISL_880500, EPI_ISL_880501, EPI_ISL_880503, EPI_ISL_880504, EPI_ISL_880505, EPI_ISL_880506, EPI_ISL_880507, EPI_ISL_880510, EPI_ISL_880511, EPI_ISL_880512, EPI_ISL_880513, EPI_ISL_880514, EPI_ISL_880515, EPI_ISL_880516, EPI_ISL_880517, EPI_ISL_880520, EPI_ISL_880521, EPI_ISL_880522, EPI_ISL_880523, EPI_ISL_880524, EPI_ISL_880525, EPI_ISL_880526, EPI_ISL_880527, EPI_ISL_880528, EPI_ISL_880530, EPI_ISL_880532, EPI_ISL_880533, EPI_ISL_880534, EPI_ISL_880535, EPI_ISL_880536, EPI_ISL_880537, EPI_ISL_880538, EPI_ISL_880539, EPI_ISL_880540, EPI_ISL_880541, EPI_ISL_880542, EPI_ISL_880543, EPI_ISL_880544, EPI_ISL_880545, EPI_ISL_880546, EPI_ISL_880547, EPI_ISL_880548, EPI_ISL_880549, EPI_ISL_880550, EPI_ISL_880551, EPI_ISL_880552, EPI_ISL_880553, EPI_ISL_880554, EPI_ISL_880555, EPI_ISL_880556, EPI_ISL_880557, EPI_ISL_880558, EPI_ISL_880559, EPI_ISL_880560, EPI_ISL_880561, EPI_ISL_880562, EPI_ISL_880563, EPI_ISL_880564, EPI_ISL_880565, EPI_ISL_880566, EPI_ISL_880567, EPI_ISL_880568, EPI_ISL_880569, EPI_ISL_880570, EPI_ISL_880571, EPI_ISL_880572, EPI_ISL_880573, EPI_ISL_880574, EPI_ISL_880576, EPI_ISL_880577, EPI_ISL_880578, EPI_ISL_880579, EPI_ISL_880580, EPI_ISL_880581, EPI_ISL_880582 | | | |
| see above | Lighthouse Lab in Alderley Park | Wellcome Sanger Institute for the COVID-19 Genomics UK (COG-UK) Consortium | Jacquelyn Wynn, Mairead Hyland, The Lighthouse Lab in Alderley Park and Alex Alderton, Roberto Amato, Sonia Goncalves, Ewan Harrison, David K. Jackson, Ian Johnston, Dominic Kwiatkowski, Cordelia Langford, John Sillitoe on behalf of the Wellcome Sanger Institute COVID-19 Surveillance Team |
| EPI_ISL_880584, EPI_ISL_880585, EPI_ISL_880587, EPI_ISL_880588, EPI_ISL_880589, EPI_ISL_880592, EPI_ISL_880593, EPI_ISL_880594, EPI_ISL_880597, EPI_ISL_880598, EPI_ISL_880600, EPI_ISL_880603, EPI_ISL_880604, EPI_ISL_880605, EPI_ISL_880607, EPI_ISL_880612, EPI_ISL_880614, EPI_ISL_880617, EPI_ISL_880621, EPI_ISL_880622, EPI_ISL_880624, EPI_ISL_880625, EPI_ISL_880628, EPI_ISL_880630, EPI_ISL_880631, EPI_ISL_880632, EPI_ISL_880633, EPI_ISL_880634, EPI_ISL_880635, EPI_ISL_880636, EPI_ISL_880638, EPI_ISL_880640, EPI_ISL_880643, EPI_ISL_880644, EPI_ISL_880645, EPI_ISL_880646, EPI_ISL_880648, EPI_ISL_880649, EPI_ISL_880650, EPI_ISL_880652, EPI_ISL_880656, EPI_ISL_880658, EPI_ISL_880659, EPI_ISL_880661, EPI_ISL_880663, EPI_ISL_880664, EPI_ISL_880666, EPI_ISL_880670, EPI_ISL_880671, EPI_ISL_880676, EPI_ISL_880678, EPI_ISL_880680, EPI_ISL_880681, EPI_ISL_880683, EPI_ISL_880684, EPI_ISL_880685, EPI_ISL_880686, EPI_ISL_880687, EPI_ISL_880688, EPI_ISL_880689, EPI_ISL_880690, EPI_ISL_880691, EPI_ISL_880692, EPI_ISL_880693, EPI_ISL_880694, EPI_ISL_880696, EPI_ISL_880697, EPI_ISL_880698, EPI_ISL_880699, EPI_ISL_880700, EPI_ISL_880702, EPI_ISL_880703, EPI_ISL_880704, EPI_ISL_880707, EPI_ISL_880711, EPI_ISL_880712, EPI_ISL_880714, EPI_ISL_880715, EPI_ISL_880716, EPI_ISL_880719, EPI_ISL_880720, EPI_ISL_880723, EPI_ISL_880728, EPI_ISL_880730, EPI_ISL_880731, EPI_ISL_880732, EPI_ISL_880733, EPI_ISL_880734, EPI_ISL_880735, EPI_ISL_880741, EPI_ISL_880742, EPI_ISL_880743, EPI_ISL_880744, EPI_ISL_880747, EPI_ISL_880748, EPI_ISL_880749, EPI_ISL_880750, EPI_ISL_880754, EPI_ISL_880756, EPI_ISL_880757, EPI_ISL_880758, EPI_ISL_880760, EPI_ISL_880761, EPI_ISL_880764, EPI_ISL_880765, EPI_ISL_880767, EPI_ISL_880768, EPI_ISL_880770, EPI_ISL_880771, EPI_ISL_880772, EPI_ISL_880773, EPI_ISL_880777, EPI_ISL_880778, EPI_ISL_880780, EPI_ISL_880782, EPI_ISL_880783, EPI_ISL_880784, EPI_ISL_880787, EPI_ISL_880788, EPI_ISL_880790, EPI_ISL_880793, EPI_ISL_880794, EPI_ISL_880795, EPI_ISL_880796, EPI_ISL_880801, EPI_ISL_880802, EPI_ISL_880804, EPI_ISL_880805, EPI_ISL_880806, EPI_ISL_880807, EPI_ISL_880808, EPI_ISL_880810, EPI_ISL_880811, EPI_ISL_880814, EPI_ISL_880815, EPI_ISL_880816, EPI_ISL_880820, EPI_ISL_880821, EPI_ISL_880823, EPI_ISL_880825, EPI_ISL_880827, EPI_ISL_880828, EPI_ISL_880832, EPI_ISL_880834, EPI_ISL_880837, EPI_ISL_880842, EPI_ISL_880844, EPI_ISL_880845, EPI_ISL_880846, EPI_ISL_880848, EPI_ISL_880849, EPI_ISL_880850, EPI_ISL_880851, EPI_ISL_880852, EPI_ISL_880855, EPI_ISL_880859, EPI_ISL_880860, EPI_ISL_880861, EPI_ISL_880863, EPI_ISL_880864, EPI_ISL_880865, EPI_ISL_880867, EPI_ISL_880869, EPI_ISL_880871, EPI_ISL_880872, EPI_ISL_880873, EPI_ISL_880874, EPI_ISL_880875, EPI_ISL_880876, EPI_ISL_880877, EPI_ISL_880880, EPI_ISL_880881, EPI_ISL_880882, EPI_ISL_880885, EPI_ISL_880887, EPI_ISL_880888, EPI_ISL_880890, EPI_ISL_880891, EPI_ISL_880892, EPI_ISL_880893, EPI_ISL_880896, EPI_ISL_880897, EPI_ISL_880899, EPI_ISL_880902, EPI_ISL_880904, EPI_ISL_880905, EPI_ISL_880906, EPI_ISL_880907, EPI_ISL_880912, EPI_ISL_880913, EPI_ISL_880917, EPI_ISL_880918, EPI_ISL_880919, EPI_ISL_880920, EPI_ISL_880922, EPI_ISL_880923, EPI_ISL_880924, EPI_ISL_880926, EPI_ISL_880927, EPI_ISL_880928, EPI_ISL_880929, EPI_ISL_880930, EPI_ISL_880931, EPI_ISL_880932 | | | |
| see above | Lighthouse Lab in Milton Keynes | Wellcome Sanger Institute for the COVID-19 Genomics UK (COG-UK) Consortium | The Lighthouse Lab in Milton Keynes and Alex Alderton, Roberto Amato, Sonia Goncalves, Ewan Harrison, David K. Jackson, Ian Johnston, Dominic Kwiatkowski, Cordelia Langford, John Sillitoe on behalf of the Wellcome Sanger Institute COVID-19 Surveillance Team |
| EPI_ISL_880936, EPI_ISL_880937, EPI_ISL_880938, EPI_ISL_880939, EPI_ISL_880940, EPI_ISL_880941, EPI_ISL_880942, EPI_ISL_880943, EPI_ISL_880944, EPI_ISL_880945, EPI_ISL_880946, EPI_ISL_880947, EPI_ISL_880948, EPI_ISL_880949, EPI_ISL_880950, EPI_ISL_880951, EPI_ISL_880952, EPI_ISL_880953, EPI_ISL_880954, EPI_ISL_880955, EPI_ISL_880956, EPI_ISL_880957, EPI_ISL_880958, EPI_ISL_880959, EPI_ISL_880960, EPI_ISL_880961, EPI_ISL_880962, EPI_ISL_880963, EPI_ISL_880964, EPI_ISL_880965, EPI_ISL_880966, EPI_ISL_880967, EPI_ISL_880968, EPI_ISL_880969, EPI_ISL_880970, EPI_ISL_880971, EPI_ISL_880972, EPI_ISL_880973, EPI_ISL_880974, EPI_ISL_880975, EPI_ISL_880976, EPI_ISL_880977, EPI_ISL_880978, EPI_ISL_880979, EPI_ISL_880980, EPI_ISL_880981, EPI_ISL_880982, EPI_ISL_880983, EPI_ISL_880984, EPI_ISL_880985, EPI_ISL_880986, EPI_ISL_880987, EPI_ISL_880988, EPI_ISL_880989, EPI_ISL_880990, EPI_ISL_880991, EPI_ISL_880992, EPI_ISL_880993, EPI_ISL_880994, EPI_ISL_880995, EPI_ISL_880996, EPI_ISL_880997, EPI_ISL_880998, EPI_ISL_880999, EPI_ISL_881000, EPI_ISL_881001, EPI_ISL_881002, EPI_ISL_881003, EPI_ISL_881004, EPI_ISL_881005, EPI_ISL_881006, EPI_ISL_881007, EPI_ISL_881008, EPI_ISL_881009, EPI_ISL_881010, EPI_ISL_881011, EPI_ISL_881012, EPI_ISL_881013, EPI_ISL_881014, EPI_ISL_881015, EPI_ISL_881016, EPI_ISL_881017, EPI_ISL_881018, EPI_ISL_881019, EPI_ISL_881020, EPI_ISL_881021, EPI_ISL_881022, EPI_ISL_881023, EPI_ISL_881024, EPI_ISL_881025, EPI_ISL_881026, EPI_ISL_881027, EPI_ISL_881028, EPI_ISL_881029, EPI_ISL_881030, EPI_ISL_881031, EPI_ISL_881032, EPI_ISL_881033, EPI_ISL_881034, EPI_ISL_881035, EPI_ISL_881036, EPI_ISL_881037, EPI_ISL_881038, EPI_ISL_881039, EPI_ISL_881040, EPI_ISL_881041, EPI_ISL_881042, EPI_ISL_881043, EPI_ISL_881044, EPI_ISL_881045, EPI_ISL_881046, EPI_ISL_881047, EPI_ISL_881048, EPI_ISL_881049, EPI_ISL_881050, EPI_ISL_881051, EPI_ISL_881052, EPI_ISL_881053, EPI_ISL_881054, EPI_ISL_881055, EPI_ISL_881056, EPI_ISL_881057, EPI_ISL_881058, EPI_ISL_881059, EPI_ISL_881060, EPI_ISL_881061, EPI_ISL_881062, EPI_ISL_881063, EPI_ISL_881064, EPI_ISL_881065, EPI_ISL_881066, EPI_ISL_881067, EPI_ISL_881068, EPI_ISL_881069, EPI_ISL_881070, EPI_ISL_881071, EPI_ISL_881072, EPI_ISL_881073, EPI_ISL_881074, EPI_ISL_881075, EPI_ISL_881076, EPI_ISL_881077, EPI_ISL_881078, EPI_ISL_881079, EPI_ISL_881080, EPI_ISL_881081, EPI_ISL_881082, EPI_ISL_881083, EPI_ISL_881084, EPI_ISL_881085, EPI_ISL_881086, EPI_ISL_881087, EPI_ISL_881088, EPI_ISL_881089, EPI_ISL_881090, EPI_ISL_881091, EPI_ISL_881092, EPI_ISL_881093, EPI_ISL_881094, EPI_ISL_881095, EPI_ISL_881096, EPI_ISL_881097, EPI_ISL_881098, EPI_ISL_881099, EPI_ISL_881100, EPI_ISL_881101, EPI_ISL_881102, EPI_ISL_881103, EPI_ISL_881104, EPI_ISL_881105, EPI_ISL_881106, EPI_ISL_881107, EPI_ISL_881108, EPI_ISL_881109, EPI_ISL_881110, EPI_ISL_881111, EPI_ISL_881112, EPI_ISL_881113, EPI_ISL_881114, EPI_ISL_881115, EPI_ISL_881116, EPI_ISL_881117, EPI_ISL_881118, EPI_ISL_881119, EPI_ISL_881120, EPI_ISL_881121, EPI_ISL_881122, EPI_ISL_881123, EPI_ISL_881124, EPI_ISL_881125, EPI_ISL_881126, EPI_ISL_881127, EPI_ISL_881128, EPI_ISL_881129, EPI_ISL_881130, EPI_ISL_881131, EPI_ISL_881132, EPI_ISL_881133, EPI_ISL_881134, EPI_ISL_881135, EPI_ISL_881136, EPI_ISL_881137, EPI_ISL_881138, EPI_ISL_881139, EPI_ISL_881140, EPI_ISL_881141, EPI_ISL_881142, EPI_ISL_881143, EPI_ISL_881144, EPI_ISL_881145, EPI_ISL_881146, EPI_ISL_881147, EPI_ISL_881148, EPI_ISL_881149, EPI_ISL_881150, EPI_ISL_881151, EPI_ISL_881152, EPI_ISL_881153, EPI_ISL_881154, EPI_ISL_881155, EPI_ISL_881156, EPI_ISL_881157, EPI_ISL_881158, EPI_ISL_881159, EPI_ISL_881160, EPI_ISL_881161, EPI_ISL_881162, EPI_ISL_881163, EPI_ISL_881164, EPI_ISL_881165, EPI_ISL_881166, EPI_ISL_881167, EPI_ISL_881168, EPI_ISL_881169, EPI_ISL_881170, EPI_ISL_881171, EPI_ISL_881172, EPI_ISL_881173, EPI_ISL_881174, EPI_ISL_881175, EPI_ISL_881176, EPI_ISL_881177, EPI_ISL_881178, EPI_ISL_881179, EPI_ISL_881180, EPI_ISL_881181, EPI_ISL_881182, EPI_ISL_881183, EPI_ISL_881184, EPI_ISL_881185, EPI_ISL_881186, EPI_ISL_881187, EPI_ISL_881188, EPI_ISL_881189, EPI_ISL_881190, EPI_ISL_881191, EPI_ISL_881192, EPI_ISL_881193, EPI_ISL_881194, EPI_ISL_881195, EPI_ISL_881196, EPI_ISL_881197, EPI_ISL_881198, EPI_ISL_881199, EPI_ISL_881200, EPI_ISL_881201, EPI_ISL_881202, EPI_ISL_881203, EPI_ISL_881204, EPI_ISL_881205, EPI_ISL_881206, EPI_ISL_881207, EPI_ISL_881208, EPI_ISL_881209, EPI_ISL_881210, EPI_ISL_881211, EPI_ISL_881212, EPI_ISL_881213, EPI_ISL_881214, EPI_ISL_881215, EPI_ISL_881216, EPI_ISL_881217, EPI_ISL_881218, EPI_ISL_881219, EPI_ISL_881220, EPI_ISL_881221, EPI_ISL_881222, EPI_ISL_881223, EPI_ISL_881224, EPI_ISL_881225, EPI_ISL_881226, EPI_ISL_881227, EPI_ISL_881228, EPI_ISL_881229, EPI_ISL_881230, EPI_ISL_881231, EPI_ISL_881232, EPI_ISL_881233, EPI_ISL_881234, EPI_ISL_881235, EPI_ISL_881236, EPI_ISL_881237, EPI_ISL_881238, EPI_ISL_881239, EPI_ISL_881240, EPI_ISL_881241, EPI_ISL_881242, EPI_ISL_881243, EPI_ISL_881244, EPI_ISL_881245, EPI_ISL_881246, EPI_ISL_881247, EPI_ISL_881248, EPI_ISL_881249, EPI_ISL_881250, EPI_ISL_881251, EPI_ISL_881252, EPI_ISL_881253, EPI_ISL_881254, EPI_ISL_881255, EPI_ISL_881256, EPI_ISL_881257, EPI_ISL_881258, EPI_ISL_881259, EPI_ISL_881260, EPI_ISL_881261, EPI_ISL_881262, EPI_ISL_881263, EPI_ISL_881264, EPI_ISL_881265, EPI_ISL_881266, EPI_ISL_881267, EPI_ISL_881268, EPI_ISL_881269, EPI_ISL_881270, EPI_ISL_881271, EPI_ISL_881272, EPI_ISL_881273, EPI_ISL_881274, EPI_ISL_881275, EPI_ISL_881276, EPI_ISL_881278, EPI_ISL_881279, EPI_ISL_881280, EPI_ISL_881282, EPI_ISL_881284, EPI_ISL_881285, EPI_ISL_881286, EPI_ISL_881287, EPI_ISL_881288, EPI_ISL_881292, EPI_ISL_881294, EPI_ISL_881295, EPI_ISL_881296, EPI_ISL_881297, EPI_ISL_881299, EPI_ISL_881300, EPI_ISL_881301, EPI_ISL_881302, EPI_ISL_881303, EPI_ISL_881304, EPI_ISL_881305, EPI_ISL_881308, EPI_ISL_881311, EPI_ISL_881312, EPI_ISL_881313, EPI_ISL_881314, EPI_ISL_881316, EPI_ISL_881317, EPI_ISL_881319, EPI_ISL_881320, EPI_ISL_881321, EPI_ISL_881322, EPI_ISL_881323, EPI_ISL_881324, EPI_ISL_881325, EPI_ISL_881327, EPI_ISL_881328, EPI_ISL_881329, EPI_ISL_881330, EPI_ISL_881331, EPI_ISL_881332, EPI_ISL_881333, EPI_ISL_881334, EPI_ISL_881335, EPI_ISL_881336, EPI_ISL_881337, EPI_ISL_881338, EPI_ISL_881339, EPI_ISL_881340, EPI_ISL_881342, EPI_ISL_881343, EPI_ISL_881344, EPI_ISL_881345, EPI_ISL_881346, EPI_ISL_881347, EPI_ISL_881348, EPI_ISL_881349, EPI_ISL_881350, EPI_ISL_881351, EPI_ISL_881352, EPI_ISL_881353, EPI_ISL_881354, EPI_ISL_881355, EPI_ISL_881356, EPI_ISL_881357, EPI_ISL_881358, EPI_ISL_881360, EPI_ISL_881361, EPI_ISL_881362, EPI_ISL_881363, EPI_ISL_881364, EPI_ISL_881365, EPI_ISL_881366, | | | |

| | | | |
|---|---|---|---|
| EPI_ISL_881367, EPI_ISL_881368, EPI_ISL_881369, EPI_ISL_881370, EPI_ISL_881371, EPI_ISL_881372, EPI_ISL_881373, EPI_ISL_881374, EPI_ISL_881375, EPI_ISL_881376, EPI_ISL_881377, EPI_ISL_881378, EPI_ISL_881379, EPI_ISL_881380, EPI_ISL_881381, EPI_ISL_881382, EPI_ISL_881383, EPI_ISL_881384, EPI_ISL_881385, EPI_ISL_881387, EPI_ISL_881389, EPI_ISL_881390, EPI_ISL_881391, EPI_ISL_881392, EPI_ISL_881393, EPI_ISL_881394, EPI_ISL_881395, EPI_ISL_881396, EPI_ISL_881397, EPI_ISL_881398, EPI_ISL_881399, EPI_ISL_881400, EPI_ISL_881401, EPI_ISL_881402, EPI_ISL_881403, EPI_ISL_881404, EPI_ISL_881405, EPI_ISL_881406, EPI_ISL_881407, EPI_ISL_881408, EPI_ISL_881409, EPI_ISL_881410, EPI_ISL_881411, EPI_ISL_881412, EPI_ISL_881413, EPI_ISL_881415, EPI_ISL_881416, EPI_ISL_881417, EPI_ISL_881418, EPI_ISL_881419, EPI_ISL_881420, EPI_ISL_881421, EPI_ISL_881422, EPI_ISL_881423, EPI_ISL_881424, EPI_ISL_881425, EPI_ISL_881426, EPI_ISL_881427, EPI_ISL_881428, EPI_ISL_881430, EPI_ISL_881431, EPI_ISL_881432, EPI_ISL_881433, EPI_ISL_881434, EPI_ISL_881435, EPI_ISL_881436, EPI_ISL_881437, EPI_ISL_881438, EPI_ISL_881439, EPI_ISL_881440, EPI_ISL_881441, EPI_ISL_881442, EPI_ISL_881443, EPI_ISL_881444, EPI_ISL_881445, EPI_ISL_881446, EPI_ISL_881447, EPI_ISL_881448, EPI_ISL_881449, EPI_ISL_881450, EPI_ISL_881451, EPI_ISL_881452, EPI_ISL_881453, EPI_ISL_881454, EPI_ISL_881456, EPI_ISL_881457, EPI_ISL_881458, EPI_ISL_881459, EPI_ISL_881460, EPI_ISL_881462, EPI_ISL_881463, EPI_ISL_881464, EPI_ISL_881465, EPI_ISL_881466, EPI_ISL_881467, EPI_ISL_881468, EPI_ISL_881469, EPI_ISL_881470, EPI_ISL_881471, EPI_ISL_881472, EPI_ISL_881473, EPI_ISL_881474, EPI_ISL_881475, EPI_ISL_881476, EPI_ISL_881477, EPI_ISL_881478, EPI_ISL_881479, EPI_ISL_881480, EPI_ISL_881481, EPI_ISL_881482, EPI_ISL_881483, EPI_ISL_881484, EPI_ISL_881485, EPI_ISL_881486, EPI_ISL_881488, EPI_ISL_881489, EPI_ISL_881490, EPI_ISL_881491, EPI_ISL_881492, EPI_ISL_881493, EPI_ISL_881496, EPI_ISL_881497, EPI_ISL_881498, EPI_ISL_881499, EPI_ISL_881500, EPI_ISL_881501, EPI_ISL_881502, EPI_ISL_881503, EPI_ISL_881504, EPI_ISL_881505, EPI_ISL_881506, EPI_ISL_881507, EPI_ISL_881508, EPI_ISL_881510, EPI_ISL_881511, EPI_ISL_881512, EPI_ISL_881513, EPI_ISL_881514, EPI_ISL_881515, EPI_ISL_881516, EPI_ISL_881517, EPI_ISL_881519, EPI_ISL_881520, EPI_ISL_881523, EPI_ISL_881524, EPI_ISL_881525, EPI_ISL_881526, EPI_ISL_881527, EPI_ISL_881528, EPI_ISL_881529, EPI_ISL_881530, EPI_ISL_881532, EPI_ISL_881533, EPI_ISL_881534, EPI_ISL_881535, EPI_ISL_881536, EPI_ISL_881537, EPI_ISL_881538, EPI_ISL_881539, EPI_ISL_881541, EPI_ISL_881542, EPI_ISL_881544, EPI_ISL_881545, EPI_ISL_881547, EPI_ISL_881548, EPI_ISL_881549, EPI_ISL_881551, EPI_ISL_881552, EPI_ISL_881553, EPI_ISL_881554, EPI_ISL_881555, EPI_ISL_881556, EPI_ISL_881557, EPI_ISL_881558, EPI_ISL_881559, EPI_ISL_881560, EPI_ISL_881561, EPI_ISL_881562, EPI_ISL_881563, EPI_ISL_881564, EPI_ISL_881565, EPI_ISL_881566, EPI_ISL_881568, EPI_ISL_881569, EPI_ISL_881572, EPI_ISL_881573, EPI_ISL_881574, EPI_ISL_881575, EPI_ISL_881577, EPI_ISL_881578, EPI_ISL_881579, EPI_ISL_881581, EPI_ISL_881583, EPI_ISL_881584, EPI_ISL_881585, EPI_ISL_881587, EPI_ISL_881589, EPI_ISL_881590, EPI_ISL_881591, EPI_ISL_881592, EPI_ISL_881593, EPI_ISL_881594, EPI_ISL_881595, EPI_ISL_881596, EPI_ISL_881597, EPI_ISL_881599, EPI_ISL_881601, EPI_ISL_881602, EPI_ISL_881603, EPI_ISL_881604, EPI_ISL_881605, EPI_ISL_881606, EPI_ISL_881608, EPI_ISL_881609, EPI_ISL_881610, EPI_ISL_881611, EPI_ISL_881613, EPI_ISL_881614, EPI_ISL_881615, EPI_ISL_881616, EPI_ISL_881617 | | | |
| see above | Lighthouse Lab in Alderley Park | Wellcome Sanger Institute for the COVID-19 Genomics UK (COG-UK) Consortium | Jacquelyn Wynn, Mairead Hyland, The Lighthouse Lab in Alderley Park and Alex Alderton, Roberto Amato, Sonia Goncalves, Ewan Harrison, David K. Jackson, Ian Johnston, Dominic Kwiatkowski, Cordelia Langford, John Sillitoe on behalf of the Wellcome Sanger Institute COVID-19 Surveillance Team |
| EPI_ISL_881620, EPI_ISL_881622 | Lighthouse Lab in Cambridge | Wellcome Sanger Institute for the COVID-19 Genomics UK (COG-UK) Consortium | Rob Howes, The Lighthouse Lab in Cambridge and Alex Alderton, Roberto Amato, Sonia Goncalves, Ewan Harrison, David K. Jackson, Ian Johnston, Dominic Kwiatkowski, Cordelia Langford, John Sillitoe on behalf of the Wellcome Sanger Institute COVID-19 Surveillance Team |
| EPI_ISL_882300, EPI_ISL_882605, EPI_ISL_882606, EPI_ISL_882607, EPI_ISL_882608 | Lighthouse Lab in Alderley Park | Wellcome Sanger Institute for the COVID-19 Genomics UK (COG-UK) Consortium | Jacquelyn Wynn, Mairead Hyland, The Lighthouse Lab in Alderley Park and Alex Alderton, Roberto Amato, Sonia Goncalves, Ewan Harrison, David K. Jackson, Ian Johnston, Dominic Kwiatkowski, Cordelia Langford, John Sillitoe on behalf of the Wellcome Sanger Institute COVID-19 Surveillance Team |
| EPI_ISL_882609 | Lighthouse Lab in Glasgow | Wellcome Sanger Institute for the COVID-19 Genomics UK (COG-UK) Consortium | Harper VanSteenhouse, Yumi Kasai, David Gray, Carol Clugston, Anna Dominiczak and Alex Alderton, Roberto Amato, Sonia Goncalves, Ewan Harrison, David K. Jackson, Ian Johnston, Dominic Kwiatkowski, Cordelia Langford, John Sillitoe on behalf of the Wellcome Sanger Institute COVID-19 Surveillance Team |
| EPI_ISL_882670 | Hospital Samaritano Paulista | Instituto Adolfo Lutz, Interdiciplinary Procedures Center, Strategic Laboratory | Claudio Tavares Sacchi, Claudia Regina Gonçalves, Erica Valessa Ramos Gomes, Karoline Rodrigues Campos |
| EPI_ISL_882671, EPI_ISL_882672 | Hospital Municipal Dr. Guido Guida | Instituto Adolfo Lutz, Interdiciplinary Procedures Center, Strategic Laboratory | Claudio Tavares Sacchi, Claudia Regina Gonçalves, Erica Valessa Ramos Gomes, Karoline Rodrigues Campos |
| EPI_ISL_882776 | Institute for Urban Disease Control and Prevention | COVID-19 Network Investigations (CONI) Alliance | Kamolthip Atsawawaranunt, Elizabeth Batty, Wasun Chantratita, Thanat Chookajorn, Stefan Fernandez, Angkana Huang, Anthony R. Jones, Khajohn Joonsalak, Chonticha Klungtong, Theerarat Kochakarn, Prayuth Kaewmalang, Amornmas Kongklieng, Namfon Kotanan, Krittikorn Kumpornsin, Duangkamon Loesbanluechai, Wudtichai Manasatienkij, Anek Mungaomklang, Bhakbhoom Panthan, Pukkaporn Parnwijitkul, Ekawat Pasomsub, Vichan Pawun, Kingkan Rakmanee, Insee Sensorn, Janjira Thaipadungpanit, Arporn Wangwiwatsin, Treewat Watthanachockchai |
| EPI_ISL_882922 | Sant'Eugenio/CTO ASL Roma 2 | INMI Lazzaro Spallanzani IRCCS | E Giombini, M Rueca, B Bartolini, O Butera, C.E.M Gruber, F Messina, F Bondanini, GC Cocciolillo, C Disegni, MR Capobianchi, A Di Caro |
| EPI_ISL_882933 | Sant'Eugenio/CTO ASL Roma 2 | INMI Lazzaro Spallanzani IRCCS | E Giombini, M. Rueca, B Bartolini, O Butera, C.E.M Gruber, F Bondanini, GC Cocciolillo, C Disegni, F Messina, MR Capobianchi, A Di Caro |
| EPI_ISL_882934 | Sant'Eugenio/CTO ASL Roma 2 | INMI Lazzaro Spallanzani IRCCS | B Bartolini, O Butera, C.E.M Gruber, M Rueca, F Messina, E Giombini, F Bondanini, GC Cocciolillo, C Disegni, MR Capobianchi, A Di Caro |
| EPI_ISL_882935 | Sant'Eugenio/CTO ASL Roma 2 | INMI Lazzaro Spallanzani IRCCS | F Messina, O Butera, E Giombini, M Rueca, B Bartolini, C.E.M Gruber, F Bondanini, C Disegni, GC Cocciolillo, MR Capobianchi, A Di Caro |
| EPI_ISL_882936 | Sant'Eugenio/CTO ASL Roma 2 | INMI Lazzaro Spallanzani IRCCS | E Giombini, M Rueca, B Bartolini, O Butera, C.E.M Gruber, F Messina, F Bondanini, GC Cocciolillo, C Disegni, MR Capobianchi, A Di Caro |
| EPI_ISL_882937 | Azienda Ospedaliera San Giovanni Addolorata | INMI Lazzaro Spallanzani IRCCS | F Messina, O Butera, E Giombini, M Rueca, B Bartolini, C.E.M Gruber, M Gaudio, PM Placanica, MR Capobianchi, A Di Caro |
| EPI_ISL_882938 | Azienda Ospedaliera San Giovanni Addolorata | INMI Lazzaro Spallanzani IRCCS | C.E.M Gruber, B Bartolini, E Giombini, M Rueca, O Butera, F Messina, PM Placanica, M Gaudio, A Di Caro, MR Capobianchi |
| EPI_ISL_882939 | Azienda Ospedaliera San Giovanni Addolorata | INMI Lazzaro Spallanzani IRCCS | B Bartolini, O Butera, C.E.M Gruber, M Rueca, F Messina, E Giombini, M Gaudio, PM Placanica, MR Capobianchi, A Di Caro |
| EPI_ISL_882940 | Ospedale San Camillo De Lellis Rieti | INMI Lazzaro Spallanzani IRCCS | M Rueca, E Giombini, C.E.M Gruber, B Bartolini, O Butera, S Venarubea, A De Luca, L Casertano, F Messina, A Di Caro, MR Capobianchi |
| EPI_ISL_882941 | Ospedale San Camillo De Lellis Rieti | INMI Lazzaro Spallanzani IRCCS | C.E.M Gruber, B Bartolini, E Giombini, M Rueca, O Butera, F Messina, S Venarubea, L Casertano, A De Luca, A Di Caro, MR Capobianchi |
| EPI_ISL_882942 | Ospedale San Camillo De Lellis Rieti | INMI Lazzaro Spallanzani IRCCS | M Rueca, E Giombini, C.E.M Gruber, B Bartolini, O Butera, F Messina, S Venarubea, A De Luca, L Casertano, A Di Caro, MR Capobianchi |
| EPI_ISL_882967, EPI_ISL_882968, EPI_ISL_882969, EPI_ISL_882970, EPI_ISL_882971, EPI_ISL_882972, EPI_ISL_882973, EPI_ISL_882974, EPI_ISL_882975, EPI_ISL_882976, EPI_ISL_882977, EPI_ISL_882978 | | | |
| see above | UCLA Clinical Micro Lab | Los Angeles County PHL | P. Hemarajata et al. |
| EPI_ISL_882999, EPI_ISL_883000, EPI_ISL_883001, EPI_ISL_883002, EPI_ISL_883018, EPI_ISL_883019, EPI_ISL_883020, EPI_ISL_883021, EPI_ISL_883022, EPI_ISL_883023, EPI_ISL_883024, EPI_ISL_883027, EPI_ISL_883028, EPI_ISL_883029, EPI_ISL_883030, EPI_ISL_883031, EPI_ISL_883032 | | | |
| see above | Maryland Public Health Laboratory | Maryland Public Health Laboratory | Maryland Department of Health Laboratories Administration |
| EPI_ISL_883309 | DOHMH Central Harlem | New York City Public Health Laboratory | Jade Wang, et al. |
| EPI_ISL_883310 | DOHMH Jamaica | New York City Public Health Laboratory | Jade Wang, et al. |
| EPI_ISL_883312, EPI_ISL_883313 | DOHMH Corona | New York City Public Health Laboratory | Jade Wang, et al. |
| EPI_ISL_883314 | DOHMH Jamaica | New York City Public Health Laboratory | Jade Wang, et al. |
| EPI_ISL_883315, EPI_ISL_883316 | OCME Office Of Chief Medical Examiner | New York City Public Health Laboratory | Jade Wang, et al. |
| EPI_ISL_883318, EPI_ISL_883319, EPI_ISL_883320, EPI_ISL_883321, EPI_ISL_883322, EPI_ISL_883323 | DOHMH Jamaica | New York City Public Health Laboratory | Jade Wang, et al. |
| EPI_ISL_883326 | Department of Homeless Services | New York City Public Health Laboratory | Jade Wang, et al. |
| EPI_ISL_883327 | DOHMH Central Harlem | New York City Public Health Laboratory | Jade Wang, et al. |
| EPI_ISL_883328 | OCME Office Of Chief Medical Examiner | New York City Public Health Laboratory | Jade Wang, et al. |
| EPI_ISL_883329 | DOHMH Central Harlem | New York City Public Health Laboratory | Jade Wang, et al. |
| EPI_ISL_883333, EPI_ISL_883334 | DOHMH Riverside | New York City Public Health Laboratory | Jade Wang, et al. |
| EPI_ISL_883335 | DOHMH Chelsea | New York City Public Health Laboratory | Jade Wang, et al. |
| EPI_ISL_883336 | OCME Office Of Chief Medical Examiner | New York City Public Health Laboratory | Jade Wang, et al. |
| EPI_ISL_883338 | DOHMH Morrisania | New York City Public Health Laboratory | Jade Wang, et al. |
| EPI_ISL_883339 | DOHMH Chelsea | New York City Public Health Laboratory | Jade Wang, et al. |
| EPI_ISL_883340, EPI_ISL_883341, EPI_ISL_883342, EPI_ISL_883343, EPI_ISL_883346 | DOHMH PHL | New York City Public Health Laboratory | Jade Wang, et al. |
| EPI_ISL_883348, EPI_ISL_883354, EPI_ISL_883355, EPI_ISL_883356 | DOHMH Corona | New York City Public Health Laboratory | Jade Wang, et al. |

| | | | |
|---|---|---|---|
| EPI_ISL_883357, EPI_ISL_883358, EPI_ISL_883359, EPI_ISL_883360 | DOHMH Crown Heights | New York City Public Health Laboratory | Jade Wang, et al. |
| EPI_ISL_883361, EPI_ISL_883362, EPI_ISL_883363 | DOHMH PHL | New York City Public Health Laboratory | Jade Wang, et al. |
| EPI_ISL_883364 | DOHMH Crown Heights | New York City Public Health Laboratory | Jade Wang, et al. |
| EPI_ISL_883365, EPI_ISL_883366, EPI_ISL_883367 | DOHMH Morrisania | New York City Public Health Laboratory | Jade Wang, et al. |
| EPI_ISL_883368, EPI_ISL_883369 | DOHMH Chelsea | New York City Public Health Laboratory | Jade Wang, et al. |
| EPI_ISL_883370 | DOHMH Crown Heights | New York City Public Health Laboratory | Jade Wang, et al. |
| EPI_ISL_883371, EPI_ISL_883372 | DOHMH Central Harlem | New York City Public Health Laboratory | Jade Wang, et al. |
| EPI_ISL_883373, EPI_ISL_883374, EPI_ISL_883375 | DOHMH Jamaica | New York City Public Health Laboratory | Jade Wang, et al. |
| EPI_ISL_883376, EPI_ISL_883377, EPI_ISL_883378, EPI_ISL_883379 | DOHMH Central Harlem | New York City Public Health Laboratory | Jade Wang, et al. |
| EPI_ISL_883380, EPI_ISL_883381, EPI_ISL_883382 | DOHMH PHL | New York City Public Health Laboratory | Jade Wang, et al. |
| EPI_ISL_883390, EPI_ISL_883391, EPI_ISL_883392, EPI_ISL_883393, EPI_ISL_883394 | OCME Office Of Chief Medical Examiner | New York City Public Health Laboratory | Jade Wang, et al. |
| EPI_ISL_883395, EPI_ISL_883396 | Department of Homeless Services | New York City Public Health Laboratory | Jade Wang, et al. |
| EPI_ISL_883397, EPI_ISL_883398, EPI_ISL_883399, EPI_ISL_883400, EPI_ISL_883401, EPI_ISL_883402 | OCME Office Of Chief Medical Examiner | New York City Public Health Laboratory | Jade Wang, et al. |
| EPI_ISL_883409 | DOHMH Central Harlem | New York City Public Health Laboratory | Jade Wang, et al. |
| EPI_ISL_883423 | DOHMH Jamaica | New York City Public Health Laboratory | Jade Wang, et al. |
| EPI_ISL_883425 | BOSTON HEART DIAGNOSTICS CORP | Wadsworth Center, New York State Department of Health | Kirsten St. George, Daryl M. Lamson, Alexis Russel, Matthew Shudt, Melissa A Leisner, Jonathan Plitnick, Navjot Singh, John Kelly, Erasmus Schneider, Erica Lasek-Nesselquist |
| EPI_ISL_883436, EPI_ISL_883437, EPI_ISL_883438, EPI_ISL_883440, EPI_ISL_883441, EPI_ISL_883442, EPI_ISL_883443, EPI_ISL_883444, EPI_ISL_883473 | ADIRONDACK MEDICAL CENTER | Wadsworth Center, New York State Department of Health | Kirsten St. George, Daryl M. Lamson, Alexis Russel, Matthew Shudt, Melissa A Leisner, Jonathan Plitnick, Navjot Singh, John Kelly, Erasmus Schneider, Erica Lasek-Nesselquist |
| EPI_ISL_884021, EPI_ISL_884022, EPI_ISL_884023, EPI_ISL_884024, EPI_ISL_884025, EPI_ISL_884026, EPI_ISL_884027, EPI_ISL_884028, EPI_ISL_884029, EPI_ISL_884030, EPI_ISL_884031 | | | |
| see above | ALBANY MEDICAL CENTER HOSPITAL CLINICAL LABORATORIES | Wadsworth Center, New York State Department of Health | Kirsten St. George, Daryl M. Lamson, Alexis Russel, Matthew Shudt, Melissa A Leisner, Jonathan Plitnick, Navjot Singh, John Kelly, Erasmus Schneider, Erica Lasek-Nesselquist |
| EPI_ISL_884032, EPI_ISL_884033, EPI_ISL_884034, EPI_ISL_884035, EPI_ISL_884036, EPI_ISL_884037, EPI_ISL_884038, EPI_ISL_884039, EPI_ISL_884040, EPI_ISL_884041 | Wadsworth Center, New York State Department of Health | Wadsworth Center, New York State Department of Health | Kirsten St. George, Daryl M. Lamson, Alexis Russel, Matthew Shudt, Melissa A Leisner, Jonathan Plitnick, Navjot Singh, John Kelly, Erasmus Schneider, Erica Lasek-Nesselquist |
| EPI_ISL_884042, EPI_ISL_884043, EPI_ISL_884044, EPI_ISL_884045, EPI_ISL_884046 | ALBANY MEDICAL CENTER HOSPITAL CLINICAL LABORATORIES | Wadsworth Center, New York State Department of Health | Kirsten St. George, Daryl M. Lamson, Alexis Russel, Matthew Shudt, Melissa A Leisner, Jonathan Plitnick, Navjot Singh, John Kelly, Erasmus Schneider, Erica Lasek-Nesselquist |
| EPI_ISL_884047 | Wadsworth Center, New York State Department of Health | Wadsworth Center, New York State Department of Health | Kirsten St. George, Daryl M. Lamson, Alexis Russel, Matthew Shudt, Melissa A Leisner, Jonathan Plitnick, Navjot Singh, John Kelly, Erasmus Schneider, Erica Lasek-Nesselquist |
| EPI_ISL_884048, EPI_ISL_884049, EPI_ISL_884050, EPI_ISL_884051, EPI_ISL_884052, EPI_ISL_884053 | ALBANY MEDICAL CENTER HOSPITAL CLINICAL LABORATORIES | Wadsworth Center, New York State Department of Health | Kirsten St. George, Daryl M. Lamson, Alexis Russel, Matthew Shudt, Melissa A Leisner, Jonathan Plitnick, Navjot Singh, John Kelly, Erasmus Schneider, Erica Lasek-Nesselquist |
| EPI_ISL_884054 | TEMPUS LABS INC | Wadsworth Center, New York State Department of Health | Kirsten St. George, Daryl M. Lamson, Alexis Russel, Matthew Shudt, Melissa A Leisner, Jonathan Plitnick, Navjot Singh, John Kelly, Erasmus Schneider, Erica Lasek-Nesselquist |
| EPI_ISL_884206, EPI_ISL_884212, EPI_ISL_884213 | Wyoming Public Health Laboratory | Wyoming Public Health Laboratory | Noah Hull, Taylor Fearing, Lynette Gumbleton, Channing Weber, Ashley Norberg, Bailey Bowcutt, and Wanda Manley |
| EPI_ISL_884227, EPI_ISL_884228, EPI_ISL_884229, EPI_ISL_884230 | Kansas Health and Environmental Lab | Kansas Health and Environmental Lab | Mike Grose, Paige Drury, Carissa Robertson, Ben Olsen, and Phil Adam |
| EPI_ISL_884256, EPI_ISL_884257, EPI_ISL_884258, EPI_ISL_884259 | Maryland Public Health Laboratory | Maryland Public Health Laboratory | Maryland Department of Health Laboratories Administration |
| EPI_ISL_884590, EPI_ISL_884591, EPI_ISL_884592, EPI_ISL_884593, EPI_ISL_884594, EPI_ISL_884595, EPI_ISL_884596, EPI_ISL_884597, EPI_ISL_884598, EPI_ISL_884599, EPI_ISL_884600, EPI_ISL_884601, EPI_ISL_884602, EPI_ISL_884603, EPI_ISL_884604, EPI_ISL_884605, EPI_ISL_884606, EPI_ISL_884607, EPI_ISL_884608, EPI_ISL_884609, EPI_ISL_884610, EPI_ISL_884611, EPI_ISL_884612, EPI_ISL_884613, EPI_ISL_884614, EPI_ISL_884615, EPI_ISL_884616, EPI_ISL_884617, EPI_ISL_884618, EPI_ISL_884619, EPI_ISL_884620, EPI_ISL_884621, EPI_ISL_884622, EPI_ISL_884623, EPI_ISL_884624, EPI_ISL_884625, EPI_ISL_884626, EPI_ISL_884627, EPI_ISL_884628, EPI_ISL_884629, EPI_ISL_884630, EPI_ISL_884631, EPI_ISL_884632, EPI_ISL_884633, EPI_ISL_884634, EPI_ISL_884635, EPI_ISL_884636, EPI_ISL_884637, EPI_ISL_884638, EPI_ISL_884639, EPI_ISL_884640, EPI_ISL_884641, EPI_ISL_884642, EPI_ISL_884643, EPI_ISL_884644, EPI_ISL_884645, EPI_ISL_884646, EPI_ISL_884647, EPI_ISL_884648, EPI_ISL_884649, EPI_ISL_884650, EPI_ISL_884651, EPI_ISL_884652, EPI_ISL_884653, EPI_ISL_884654, EPI_ISL_884655, EPI_ISL_884656, EPI_ISL_884657, EPI_ISL_884658, EPI_ISL_884659, EPI_ISL_884660, EPI_ISL_884661, EPI_ISL_884662, EPI_ISL_884663, EPI_ISL_884664, EPI_ISL_884665, EPI_ISL_884666, EPI_ISL_884667, EPI_ISL_884668, EPI_ISL_884669, EPI_ISL_884670, EPI_ISL_884671, EPI_ISL_884672, EPI_ISL_884673, EPI_ISL_884674, EPI_ISL_884675, EPI_ISL_884676, EPI_ISL_884677, EPI_ISL_884678, EPI_ISL_884679, EPI_ISL_884680, EPI_ISL_884681, EPI_ISL_884682, EPI_ISL_884683, EPI_ISL_884684, EPI_ISL_884685, EPI_ISL_884686, EPI_ISL_884687, EPI_ISL_884688, EPI_ISL_884689, EPI_ISL_884690, EPI_ISL_884691, EPI_ISL_884692, EPI_ISL_884693, EPI_ISL_884694, EPI_ISL_884695, EPI_ISL_884696, EPI_ISL_884697, EPI_ISL_884698, EPI_ISL_884699, EPI_ISL_884700, EPI_ISL_884701, EPI_ISL_884702, EPI_ISL_884703, EPI_ISL_884704, EPI_ISL_884705, EPI_ISL_884706, EPI_ISL_884707, EPI_ISL_884708, EPI_ISL_884709, EPI_ISL_884710, EPI_ISL_884711, EPI_ISL_884712, EPI_ISL_884713, EPI_ISL_884714, EPI_ISL_884715, EPI_ISL_884716, EPI_ISL_884717, EPI_ISL_884718, EPI_ISL_884719, EPI_ISL_884720, EPI_ISL_884721, EPI_ISL_884722, EPI_ISL_884723, EPI_ISL_884724, EPI_ISL_884725, EPI_ISL_884726, EPI_ISL_884727, EPI_ISL_884728 | | | |
| see above | Respiratory Viruses Branch, Centers for Disease Control and Prevention | Respiratory Viruses Branch, Centers for Disease Control and Prevention | Cook,P.W., Batra,D., Rambo-Martin,B.L., de Feo,E., Antico,J., Tran,C., Tolentino,M., Wickline,S., Gietzen,K., Sickler,B., Liu,J., Allen,E., Febbo,P., Galloway,S., Washington,N.L., White,S., Levan,G., Barret,K.S., Cirulli,E., Bolze,A., Ascencio,A., Rivera-Garcia,C., Cho,R., Nguyen,J., Wang,S., Ramirez,J., Cassens,T., Sandoval,E., Isaksson,M., Lee,W., Becker,D., Laurent,M., Lu,J., Paden,C.R., Tong,S., MacCannell,D. |
| EPI_ISL_885146 | Genomic Laboratory (GLAB) (Conjoint lab of Health Directorate of Istanbul and Istanbul Technical University) | Genomic Laboratory (GLAB), Istanbul Technical University | Ilker Karacan, Tugba Kizilboga Akgun, Payam Zolfagharian, Nisan Denizce Can, Pari Sharifli, Levent Doganay, Gizem Dinler Doganay |
| EPI_ISL_885183, EPI_ISL_885204, EPI_ISL_885211, EPI_ISL_885244, EPI_ISL_885291, EPI_ISL_885321, EPI_ISL_885349, EPI_ISL_885359, EPI_ISL_885392, EPI_ISL_885411, EPI_ISL_885434, EPI_ISL_885503, EPI_ISL_885580, EPI_ISL_885589, EPI_ISL_885813, EPI_ISL_885880, EPI_ISL_885889, EPI_ISL_885916, EPI_ISL_886016 | | | |
| see above | Lighthouse Lab in Alderley Park | Wellcome Sanger Institute for the COVID-19 Genomics UK | Jacquelyn Wynn, Mairead Hyland, The Lighthouse Lab in Alderley Park and Alex Alderton, Roberto Amato, Sonia Goncalves, Ewan Harrison, David K. |

| | | | |
|---|---|---|---|
| | | (COG-UK) Consortium | Jackson, Ian Johnston, Dominic Kwiatkowski, Cordelia Langford, John Sillitoe on behalf of the Wellcome Sanger Institute COVID-19 Surveillance Team |
| EPI_ISL_886169, EPI_ISL_886172, EPI_ISL_886214, EPI_ISL_886246, EPI_ISL_886247, EPI_ISL_886248, EPI_ISL_886270, EPI_ISL_886318, EPI_ISL_886322, EPI_ISL_886347, EPI_ISL_886351, EPI_ISL_886362, EPI_ISL_886364, EPI_ISL_886365, EPI_ISL_886424, EPI_ISL_886442, EPI_ISL_886452, EPI_ISL_886465, EPI_ISL_886471, EPI_ISL_886494, EPI_ISL_886522, EPI_ISL_886545, EPI_ISL_886546, EPI_ISL_886554, EPI_ISL_886580, EPI_ISL_886583, EPI_ISL_886592, EPI_ISL_886616, EPI_ISL_886633, EPI_ISL_886636, EPI_ISL_886650, EPI_ISL_886662, EPI_ISL_886666, EPI_ISL_886701, EPI_ISL_886720, EPI_ISL_886735, EPI_ISL_886761, EPI_ISL_886790, EPI_ISL_886808, EPI_ISL_886837, EPI_ISL_886853, EPI_ISL_886860, EPI_ISL_886866, EPI_ISL_886870, EPI_ISL_886895, EPI_ISL_886898, EPI_ISL_886907, EPI_ISL_886963, EPI_ISL_886965, EPI_ISL_886975, EPI_ISL_886990, EPI_ISL_886997, EPI_ISL_887006, EPI_ISL_887067, EPI_ISL_887087, EPI_ISL_887091 | | | |
| see above | Labcorp | Genomics and Discovery, Respiratory Viruses Branch, Division of Viral Diseases, Centers for Disease Control and Prevention | Peter W. Cook,Dhwani Batra,Ben L. Rambo-Martin,Summer Galloway,Brian Krueger,Minoo Agarwal,Eyad Almasri,Debbie Boles,Ayla Burns,Nuthawin Charoensri,Oren Cohen,Susan Countryman,Mary Ann Cristobal,Bobbi Croy,Suzanne Dale,Hrushikesh Deshmukh,Amanda Douglas,Vincent Drouillon,Marcia Eisenberg,Howard Engler,Rama Ghatti,Prashant Gupta,Susan Hicks,Jake Humphrey,Lax Iyer,Manoj Jain,Mohan Kolli,Tim Kuphal,Stanley Letovsky,Michael Levandoski,Craig Lukasik,Jonathan Meltzer,Brian Norvell,Mindy Nye,Scott Parker,Christos Petropoulos,John Pruitt,Steven Ragan,Scott Ryan,Mike Sapeta,Jana Schroth,Suresh Babu Selvaraju,Goran Stevovic,Amanda Suchanek,Andrea Throop,Lyndon Tilson,Thomas Urban,Joe Voshell,Kimberly Wagner,Jonathan Williams,Mary Williamson,Qian Zeng,Tricia Zwiefelhofer,Clinton R. Paden,Suxiang Tong,Duncan MacCannell, |
| EPI_ISL_887136, EPI_ISL_887137, EPI_ISL_887144 | Institute of Medical Microbiology and Hospital Hygiene | Institute of Medical Microbiology and Hospital Hygiene | Prof. Dr. Achim Kaasch, Aljoscha Tersteegen |
| EPI_ISL_887508, EPI_ISL_887509, EPI_ISL_887510, EPI_ISL_887511, EPI_ISL_887513, EPI_ISL_887522, EPI_ISL_887523, EPI_ISL_887529, EPI_ISL_887530, EPI_ISL_887532, EPI_ISL_887533, EPI_ISL_887538, EPI_ISL_887544, EPI_ISL_887550, EPI_ISL_887552, EPI_ISL_887558, EPI_ISL_887559, EPI_ISL_887560, EPI_ISL_887563, EPI_ISL_887567, EPI_ISL_887568, EPI_ISL_887577, EPI_ISL_887580, EPI_ISL_887581, EPI_ISL_887582, EPI_ISL_887583 | | | |
| see above | Johns Hopkins Hospital Department of Pathology | Johns Hopkins Hospital Department of Pathology | C. Paul Morris, Chun Huai Luo, Adannaya Amadi, Matthew Schwartz, Nicholas Gallagher, Heba H. Mostafa |
| EPI_ISL_888599, EPI_ISL_888600, EPI_ISL_888602, EPI_ISL_888603, EPI_ISL_888604, EPI_ISL_888605, EPI_ISL_888606, EPI_ISL_888607, EPI_ISL_888608, EPI_ISL_888609, EPI_ISL_888610, EPI_ISL_888611, EPI_ISL_888612, EPI_ISL_888613, EPI_ISL_888614, EPI_ISL_888615, EPI_ISL_888616, EPI_ISL_888617, EPI_ISL_888618, EPI_ISL_888619, EPI_ISL_888620, EPI_ISL_888621, EPI_ISL_888622 | | | |
| see above | Wyoming Public Health Laboratory | Wyoming Public Health Laboratory | Noah Hull, Taylor Fearing, Lynette Gumbleton, Channing Weber, Ashley Norberg, Bailey Bowcutt, and Wanda Manley |
| EPI_ISL_888667, EPI_ISL_888668, EPI_ISL_888669, EPI_ISL_888670 | University of Michigan Clinical Microbiology Laboratory | Lauring Lab, University of Michigan, Department of Microbiology and Immunology | Valesano |
| EPI_ISL_888810, EPI_ISL_888811, EPI_ISL_888812, EPI_ISL_888813, EPI_ISL_888814, EPI_ISL_888815, EPI_ISL_888816, EPI_ISL_888817 | Univeristy of New Mexico Hospital | Center for Global Health, University of New Mexico Health Sciences Center | Daryl Domman, Kurt Schwalm, Justin Bacca, Jon Femling, Darrell Dinwiddie |
| EPI_ISL_888830, EPI_ISL_888831 | National Virus Reference Laboratory | National Virus Reference Laboratory | Michael Carr, Gabriel Gonzalez, Jonathan Dean, Cillian F De Gascun |
| EPI_ISL_888851, EPI_ISL_888856, EPI_ISL_888903 | Michigan Department of Health and Human Services, Bureau of Laboratories | Michigan Department of Health and Human Services, Bureau of Laboratories | Blankenship HM, Riner D, Soehnlen MK |
| EPI_ISL_888908, EPI_ISL_888910, EPI_ISL_888911, EPI_ISL_888914, EPI_ISL_888934, EPI_ISL_888936, EPI_ISL_888941, EPI_ISL_888942, EPI_ISL_888943, EPI_ISL_888944, EPI_ISL_888949, EPI_ISL_888959 | | | |
| see above | Wyoming Public Health Laboratory | Wyoming Public Health Laboratory | Noah Hull, Taylor Fearing, Lynette Gumbleton, Channing Weber, Ashley Norberg, Bailey Bowcutt, and Wanda Manley |
| EPI_ISL_889703, EPI_ISL_889704, EPI_ISL_889705, EPI_ISL_889706, EPI_ISL_889707, EPI_ISL_889708, EPI_ISL_889709, EPI_ISL_889710, EPI_ISL_889711, EPI_ISL_889712, EPI_ISL_889714, EPI_ISL_889715, EPI_ISL_889716, EPI_ISL_889717, EPI_ISL_889718, EPI_ISL_889719, EPI_ISL_889720, EPI_ISL_889721, EPI_ISL_889722, EPI_ISL_889723, EPI_ISL_889724, EPI_ISL_889725, EPI_ISL_889726, EPI_ISL_889727, EPI_ISL_889728, EPI_ISL_889729, EPI_ISL_889730, EPI_ISL_889731, EPI_ISL_889732, EPI_ISL_889733, EPI_ISL_889734, EPI_ISL_889735, EPI_ISL_889736, EPI_ISL_889737, EPI_ISL_889738, EPI_ISL_889739, EPI_ISL_889740, EPI_ISL_889741, EPI_ISL_889742, EPI_ISL_889745 | | | |
| see above | LSUHS Emerging Viral Threat Laboratory | Microbial Genome Sequencing Center | Jeremy P. Kamil, Jennifer L. Carroll, Camille F. Abshire, Maarten Van Diest, Mohammed N.A. Siddiquey, Andrew D. Yurochko, Martin J. Sapp, Rona S. Scott, Christopher G. Kevil, Daniel J. Snyder, Vaughn S. Cooper, John A. Vanchiere |
| EPI_ISL_890232 | School of Pharmacy, Shenandoah University | School of Pharmacy, Shenandoah University | Adams,S.M., Harralson,A.F., Kidd,R.S., Sawyer,G.W. |
| EPI_ISL_890365, EPI_ISL_890366 | LSUHS Emerging Viral Threat Laboratory | Microbial Genome Sequencing Center | Jeremy P. Kamil, Jennifer L. Carroll, Camille F. Abshire, Maarten Van Diest, Mohammed N.A. Siddiquey, Andrew D. Yurochko, Martin J. Sapp, Rona S. Scott, Christopher G. Kevil, Daniel J. Snyder, Vaughn S. Cooper, John A. Vanchiere |
| EPI_ISL_890985, EPI_ISL_890986, EPI_ISL_890987, EPI_ISL_890988 | Seattle Flu Study | Seattle Flu Study | Deborah A. Nickerson, Chris D. Frazar, Jover Lee, Benjamin Pelle, Erica Ryke, Matthew Richardson, Amanda Adler, Elisabeth Brandstetter, Peter D. Han, Kairsten Fay, Misja Ilcisin, Kirsten Lacombe, Thomas R. Sibley, Melissa Truong, Caitlin R. Wolf, Michael Boeckh, Janet A. Englund, Michael Famulare, Barry R. Lutz, Mark J. Rieder, Lea M. Starita, Matthew Thompson, Jay Shendure, Trevor Bedford, Helen Y. Chu |
| EPI_ISL_890989 | Seattle Flu Study | Seattle Flu Study | Deborah A. Nickerson, Chris D. Frazar, Jover Lee, Benjamin Pelle, Erica Ryke, Matthew Richardson, Amanda Adler, Elisabeth Brandstetter, Peter D. Han, Kairsten Fay, Misja Ilcisin, Kirsten Lacombe, Thomas R. Sibley, Melissa Truong, Caitlin R. Wolf, Karen Cowgill, Stephanie Schrag, Jeff Duchin, Michael Boeckh, Janet A. Englund, Michael Famulare, Barry R. Lutz, Mark J. Rieder, Lea M. Starita, Matthew Thompson, Helen Y. Chu, Trevor Bedford, Jay Shendure |
| EPI_ISL_890991, EPI_ISL_890992, EPI_ISL_890994, EPI_ISL_890995, EPI_ISL_890996, EPI_ISL_890997 | Seattle Flu Study | Seattle Flu Study | Deborah A. Nickerson, Chris D. Frazar, Jover Lee, Benjamin Pelle, Erica Ryke, Matthew Richardson, Amanda Adler, Elisabeth Brandstetter, Peter D. Han, Kairsten Fay, Misja Ilcisin, Kirsten Lacombe, Thomas R. Sibley, Melissa Truong, Caitlin R. Wolf, Michael Boeckh, Janet A. Englund, Michael Famulare, Barry R. Lutz, Mark J. Rieder, Lea M. Starita, Matthew Thompson, Jay Shendure, Trevor Bedford, Helen Y. Chu |
| EPI_ISL_891058 | Seattle Flu Study | Seattle Flu Study | Deborah A. Nickerson, Chris D. Frazar, Jover Lee, Benjamin Pelle, Erica Ryke, Matthew Richardson, Amanda Adler, Elisabeth Brandstetter, Peter D. Han, Kairsten Fay, Misja Ilcisin, Kirsten Lacombe, Thomas R. Sibley, Melissa Truong, Caitlin R. Wolf, Karen Cowgill, Stephanie Schrag, Jeff Duchin, Michael Boeckh, Janet A. Englund, Michael Famulare, Barry R. Lutz, Mark J. Rieder, Lea M. Starita, Matthew Thompson, Helen Y. Chu, Trevor Bedford, Jay Shendure |
| EPI_ISL_891060, EPI_ISL_891061, EPI_ISL_891062, EPI_ISL_891063 | Seattle Flu Study | Seattle Flu Study | Deborah A. Nickerson, Chris D. Frazar, Jover Lee, Benjamin Pelle, Erica Ryke, Matthew Richardson, Amanda Adler, Elisabeth Brandstetter, Peter D. Han, Kairsten Fay, Misja Ilcisin, Kirsten Lacombe, Thomas R. Sibley, Melissa Truong, Caitlin R. Wolf, Michael Boeckh, Janet A. Englund, Michael Famulare, Barry R. Lutz, Mark J. Rieder, Lea M. Starita, Matthew Thompson, Jay Shendure, Trevor Bedford, Helen Y. Chu |
| EPI_ISL_891064, EPI_ISL_891065, EPI_ISL_891066, EPI_ISL_891067, EPI_ISL_891068, EPI_ISL_891069 | Seattle Flu Study | Seattle Flu Study | Deborah A. Nickerson, Chris D. Frazar, Jover Lee, Benjamin Pelle, Erica Ryke, Matthew Richardson, Amanda Adler, Elisabeth Brandstetter, Peter D. Han, Kairsten Fay, Misja Ilcisin, Kirsten Lacombe, Thomas R. Sibley, Melissa Truong, Caitlin R. Wolf, Karen Cowgill, Stephanie Schrag, Jeff Duchin, Michael Boeckh, Janet A. Englund, Michael Famulare, Barry R. Lutz, Mark J. Rieder, Lea M. Starita, Matthew Thompson, Helen Y. Chu, Trevor Bedford, Jay Shendure |
| EPI_ISL_891144 | Indiana Animal Disease Diagnostic Laboratory | Carpi Laboratory - Purdue University | Jack Dorman, Ilinca I Ciubotariu, Lev Gorenstein, Abebe A Fola, G Kenitra Hendrix, Rebecca P Wilkes, Giovanna Carpi |
| EPI_ISL_891146, EPI_ISL_891147, EPI_ISL_891148, EPI_ISL_891149, EPI_ISL_891150 | University of Wisconsin-Madison AIDS Vaccine Research Laboratories | University of Wisconsin-Madison AIDS Vaccine Research Laboratories | Gage Moreno, Katarina Braun, et al. AIDS Vaccine Research Laboratories |
| EPI_ISL_891206 | DPH, Massachusetts State Public Health Lab | DPH, Massachusetts State Public Health Lab | Lang,A.S., Fink,T., Gallagher,G.R., Smole,S.C. |
| EPI_ISL_891252 | Singapore General Hospital | Department of Microbiology | Nurdyana Abdul Rahman, Kun Lee Lim, Chenhao Li, Sui Sin Goh, Kenneth Xin Long Chan, Kian Sing Chan, Lynette Oon, Kern Rei Chng, Niranjan Nagarajan, Karrie Ko |
| EPI_ISL_891903, EPI_ISL_891909, EPI_ISL_891918, EPI_ISL_891928, EPI_ISL_891945, EPI_ISL_891958, EPI_ISL_891967, EPI_ISL_891972, EPI_ISL_891973, EPI_ISL_891974, EPI_ISL_891975, EPI_ISL_891976, EPI_ISL_891977, EPI_ISL_891978, EPI_ISL_891979, EPI_ISL_891980, EPI_ISL_891981, EPI_ISL_891982, EPI_ISL_891983, EPI_ISL_891984, EPI_ISL_891985, EPI_ISL_891986, EPI_ISL_891987, EPI_ISL_891988, EPI_ISL_891989, EPI_ISL_891990, EPI_ISL_891991, EPI_ISL_891992, EPI_ISL_891993, EPI_ISL_891994, EPI_ISL_891995, EPI_ISL_891996, EPI_ISL_891997, EPI_ISL_891998, EPI_ISL_891999, EPI_ISL_892000, EPI_ISL_892001, EPI_ISL_892003, EPI_ISL_892004, EPI_ISL_892007, EPI_ISL_892008, EPI_ISL_892013, EPI_ISL_892014, EPI_ISL_892015, EPI_ISL_892016, EPI_ISL_892017, EPI_ISL_892018, EPI_ISL_892019, EPI_ISL_892020, EPI_ISL_892021, EPI_ISL_892022, EPI_ISL_892023, EPI_ISL_892024, EPI_ISL_892025, EPI_ISL_892026, EPI_ISL_892027, EPI_ISL_892028, EPI_ISL_892029, EPI_ISL_892030, EPI_ISL_892031, EPI_ISL_892032, EPI_ISL_892033, EPI_ISL_892034, EPI_ISL_892035, EPI_ISL_892036, EPI_ISL_892037, EPI_ISL_892038, EPI_ISL_892039, EPI_ISL_892040, EPI_ISL_892041, EPI_ISL_892042, EPI_ISL_892043, EPI_ISL_892044, EPI_ISL_892045, EPI_ISL_892046, EPI_ISL_892047, EPI_ISL_892048, EPI_ISL_892049, EPI_ISL_892056, EPI_ISL_892127, EPI_ISL_892128, EPI_ISL_892129, EPI_ISL_892130, EPI_ISL_892131, EPI_ISL_892132, EPI_ISL_892133, EPI_ISL_892134, EPI_ISL_892135, EPI_ISL_892136, EPI_ISL_892137, | | | |

| | | | |
|---|---|---|---|
| EPI_ISL_892138, EPI_ISL_892139, EPI_ISL_892140, EPI_ISL_892141, EPI_ISL_892142, EPI_ISL_892143, EPI_ISL_892144, EPI_ISL_892145, EPI_ISL_892146, EPI_ISL_892147, EPI_ISL_892148, EPI_ISL_892149, EPI_ISL_892150, EPI_ISL_892151, EPI_ISL_892152, EPI_ISL_892153, EPI_ISL_892154, EPI_ISL_892155, EPI_ISL_892156, EPI_ISL_892157, EPI_ISL_892158, EPI_ISL_892159, EPI_ISL_892160, EPI_ISL_892161, EPI_ISL_892162, EPI_ISL_892163, EPI_ISL_892164, EPI_ISL_892165, EPI_ISL_892166, EPI_ISL_892167, EPI_ISL_892168, EPI_ISL_892169, EPI_ISL_892170, EPI_ISL_892171, EPI_ISL_892172, EPI_ISL_892173, EPI_ISL_892174, EPI_ISL_892175, EPI_ISL_892176, EPI_ISL_892177, EPI_ISL_892178, EPI_ISL_892180, EPI_ISL_892181, EPI_ISL_892182, EPI_ISL_892183, EPI_ISL_892184, EPI_ISL_892185, EPI_ISL_892186, EPI_ISL_892187, EPI_ISL_892188, EPI_ISL_892198, EPI_ISL_892199, EPI_ISL_892200 | | | |
| see above | Lighthouse Lab in Alderley Park | Wellcome Sanger Institute for the COVID-19 Genomics UK (COG-UK) Consortium | Jacquelyn Wynn, Mairead Hyland, The Lighthouse Lab in Alderley Park and Alex Alderton, Roberto Amato, Sonia Goncalves, Ewan Harrison, David K. Jackson, Ian Johnston, Dominic Kwiatkowski, Cordelia Langford, John Sillitoe on behalf of the Wellcome Sanger Institute COVID-19 Surveillance Team |
| EPI_ISL_892201 | Lighthouse Lab in Glasgow | Wellcome Sanger Institute for the COVID-19 Genomics UK (COG-UK) Consortium | Harper VanSteenhouse, Yumi Kasai, David Gray, Carol Clugston, Anna Dominiczak and Alex Alderton, Roberto Amato, Sonia Goncalves, Ewan Harrison, David K. Jackson, Ian Johnston, Dominic Kwiatkowski, Cordelia Langford, John Sillitoe on behalf of the Wellcome Sanger Institute COVID-19 Surveillance Team |
| EPI_ISL_892202, EPI_ISL_892203, EPI_ISL_892204, EPI_ISL_892205 | Lighthouse Lab in Alderley Park | Wellcome Sanger Institute for the COVID-19 Genomics UK (COG-UK) Consortium | Jacquelyn Wynn, Mairead Hyland, The Lighthouse Lab in Alderley Park and Alex Alderton, Roberto Amato, Sonia Goncalves, Ewan Harrison, David K. Jackson, Ian Johnston, Dominic Kwiatkowski, Cordelia Langford, John Sillitoe on behalf of the Wellcome Sanger Institute COVID-19 Surveillance Team |
| EPI_ISL_893781 | Institute of Virology, Medical Center, University of Freiburg, Freiburg, Germany | Institute of Virology, Clinial Virus Genomics, Medical Center, University of Freiburg, Freiburg, Germany | Jonas Fuchs, Lisa Kern, Sandra Reuter, Hajo Grundmann, Marcus Panning |
| EPI_ISL_894220, EPI_ISL_894221, EPI_ISL_894222, EPI_ISL_894227, EPI_ISL_894228, EPI_ISL_894229, EPI_ISL_894230, EPI_ISL_894231 | CH de Mayotte - Laboratoire de Biologie | National Reference Center for Viruses of Respiratory Infections, Institut Pasteur, Paris | Marion Barbet, Sylvie Behillil, Méline Bizard, Angela Brisebarre, Camille Capel, Etienne Simon-Lorière, Vincent Enouf, Maud Vanpeene, Sylvie van der Werf,Combe Patrice |
| EPI_ISL_894255, EPI_ISL_894256, EPI_ISL_894257, EPI_ISL_894258, EPI_ISL_894259, EPI_ISL_894260, EPI_ISL_894261, EPI_ISL_894262, EPI_ISL_894263, EPI_ISL_894264, EPI_ISL_894265, EPI_ISL_894266, EPI_ISL_894267, EPI_ISL_894268, EPI_ISL_894269, EPI_ISL_894270, EPI_ISL_894271, EPI_ISL_894272, EPI_ISL_894273, EPI_ISL_894274, EPI_ISL_894275, EPI_ISL_894276 | | | |
| see above | Ministry of Health Turkey | Ministry of Health Turkey | Fatma Bayrakdar, Yasemin Cogun, Süleyman Yalcin, Aye Baak Alta, Gülay Korukluolu |
| EPI_ISL_896070 | Rothen Medizinische Laboratorien AG | University Hospital Basel, Clinical Bacteriology | Tim Roloff, Madlen Stange, Helena MB Seth-Smith, Alfredo Mari, Karoline Leuzinger, Julia Bielicki, Ingrid Steffen, Manuel Battegay, Hans Hirsch, Adrian Egli |
| EPI_ISL_896211, EPI_ISL_896213 | MEPHI, Aix Marseille University | MEPHI, Aix Marseille University | Anthony LEVASSEUR |
| EPI_ISL_896219, EPI_ISL_896221, EPI_ISL_896222, EPI_ISL_896223, EPI_ISL_896225, EPI_ISL_896230, EPI_ISL_896235, EPI_ISL_896247, EPI_ISL_896252, EPI_ISL_896253, EPI_ISL_896256, EPI_ISL_896257, EPI_ISL_896258, EPI_ISL_896259, EPI_ISL_896260, EPI_ISL_896261, EPI_ISL_896262, EPI_ISL_896263, EPI_ISL_896264, EPI_ISL_896265, EPI_ISL_896266, EPI_ISL_896267, EPI_ISL_896268, EPI_ISL_896269, EPI_ISL_896270, EPI_ISL_896271, EPI_ISL_896272, EPI_ISL_896273, EPI_ISL_896274, EPI_ISL_896275, EPI_ISL_896276, EPI_ISL_896277, EPI_ISL_896279, EPI_ISL_896280, EPI_ISL_896283, EPI_ISL_896284, EPI_ISL_896285, EPI_ISL_896287, EPI_ISL_896288, EPI_ISL_896292, EPI_ISL_896293 | | | |
| see above | SUNY UPSTATE MEDICAL UNIVERSITY | Wadsworth Center, New York State Department of Health | Kirsten St. George, Daryl M. Lamson, Alexis Russel, Matthew Shudt, Melissa A Leisner, Jonathan Plitnick, Navjot Singh, John Kelly, Erasmus Schneider, Erica Lasek-Nesselquist |
| EPI_ISL_896380 | Suceava County Emergency Hospital | "Stefan cel Mare" University Metagenomics Lab | Lobiuc Andrei, Gheorghita Roxana |
| EPI_ISL_896413 | URMC LABS | Wadsworth Center, New York State Department of Health | Kirsten St. George, Daryl M. Lamson, Alexis Russel, Matthew Shudt, Melissa A Leisner, Jonathan Plitnick, Navjot Singh, John Kelly, Erasmus Schneider, Erica Lasek-Nesselquist |
| EPI_ISL_896451, EPI_ISL_896452, EPI_ISL_896453, EPI_ISL_896455, EPI_ISL_896457, EPI_ISL_896461, EPI_ISL_896462, EPI_ISL_896463, EPI_ISL_896464, EPI_ISL_896465, EPI_ISL_896469, EPI_ISL_896470, EPI_ISL_896471, EPI_ISL_896472, EPI_ISL_896475 | | | |
| see above | KU Leuven, Rega Institute, Clinical and Epidemiological Virology | KU Leuven, Rega Institute, Clinical and Epidemiological Virology | Tony Wawina-Bokalanga, Bert Vanmechelen, Joan Marti-Carerras, Piet Maes |
| EPI_ISL_897598, EPI_ISL_897603, EPI_ISL_897604, EPI_ISL_897611, EPI_ISL_897616, EPI_ISL_897617, EPI_ISL_897618, EPI_ISL_897621, EPI_ISL_897622, EPI_ISL_897623, EPI_ISL_897624, EPI_ISL_897625, EPI_ISL_897670, EPI_ISL_897674, EPI_ISL_897677, EPI_ISL_897680, EPI_ISL_897681, EPI_ISL_897682, EPI_ISL_897685, EPI_ISL_897688, EPI_ISL_897689, EPI_ISL_897692, EPI_ISL_897693, EPI_ISL_897694, EPI_ISL_897697, EPI_ISL_897699, EPI_ISL_897700, EPI_ISL_897704, EPI_ISL_897705, EPI_ISL_897708, EPI_ISL_897711, EPI_ISL_897713, EPI_ISL_897714, EPI_ISL_897716, EPI_ISL_897721, EPI_ISL_897722, EPI_ISL_897723, EPI_ISL_897724, EPI_ISL_897725, EPI_ISL_897726, EPI_ISL_897727, EPI_ISL_897728, EPI_ISL_897729, EPI_ISL_897730, EPI_ISL_897731, EPI_ISL_897732, EPI_ISL_897733, EPI_ISL_897735, EPI_ISL_897736, EPI_ISL_897737, EPI_ISL_897738, EPI_ISL_897739, EPI_ISL_897740, EPI_ISL_897741, EPI_ISL_897742, EPI_ISL_897743, EPI_ISL_897744, EPI_ISL_897745, EPI_ISL_897746, EPI_ISL_897747, EPI_ISL_897748, EPI_ISL_897749, EPI_ISL_897750, EPI_ISL_897751, EPI_ISL_897752, EPI_ISL_897753, EPI_ISL_897754, EPI_ISL_897755, EPI_ISL_897756, EPI_ISL_897757, EPI_ISL_897758, EPI_ISL_897759, EPI_ISL_897760, EPI_ISL_897761, EPI_ISL_897762, EPI_ISL_897763, EPI_ISL_897764, EPI_ISL_897765, EPI_ISL_897766, EPI_ISL_897767, EPI_ISL_897768, EPI_ISL_897769, EPI_ISL_897770, EPI_ISL_897903, EPI_ISL_897904, EPI_ISL_897906, EPI_ISL_897907, EPI_ISL_897908, EPI_ISL_897909, EPI_ISL_897910, EPI_ISL_897911, EPI_ISL_897912, EPI_ISL_897913, EPI_ISL_897914, EPI_ISL_897915, EPI_ISL_897916, EPI_ISL_897917, EPI_ISL_897918, EPI_ISL_897919, EPI_ISL_897942, EPI_ISL_897943, EPI_ISL_897944, EPI_ISL_897945, EPI_ISL_897946, EPI_ISL_897947, EPI_ISL_897948, EPI_ISL_897949, EPI_ISL_897950, EPI_ISL_897951, EPI_ISL_897952, EPI_ISL_897954, EPI_ISL_897955, EPI_ISL_897956, EPI_ISL_897957, EPI_ISL_897958, EPI_ISL_897959, EPI_ISL_897960, EPI_ISL_897961, EPI_ISL_897962, EPI_ISL_897963, EPI_ISL_897964, EPI_ISL_897965, EPI_ISL_897980, EPI_ISL_897981, EPI_ISL_897982, EPI_ISL_897983, EPI_ISL_897984, EPI_ISL_897985 | | | |
| see above | University Hospitals of Geneva, Laboratory of Virology | HUG, Laboratory of Virology and the Health2030 Genome Center | Samuel Cordey, Ana Rita Goncalves, Laurent Kaiser, Lorenzo Cerutti, Henri Pegeot, Melyssa Elies, Deborah Penet, Keith Harshman, Ioannis Xenarios, Emmanouil Dermitzakis |
| EPI_ISL_897986, EPI_ISL_897989, EPI_ISL_897990, EPI_ISL_897993, EPI_ISL_897996, EPI_ISL_897998, EPI_ISL_898000, EPI_ISL_898001, EPI_ISL_898002, EPI_ISL_898004, EPI_ISL_898005, EPI_ISL_898007, EPI_ISL_898015, EPI_ISL_898016, EPI_ISL_898023 | | | |
| see above | KU Leuven, Rega Institute, Clinical and Epidemiological Virology | KU Leuven, Rega Institute, Clinical and Epidemiological Virology | Tony Wawina-Bokalanga, Bert Vanmechelen, Joan Marti-Carerras, Piet Maes |
| EPI_ISL_898993, EPI_ISL_898994, EPI_ISL_898995, EPI_ISL_898996 | Viollier AG | Department of Biosystems Science and Engineering, ETH Zürich | Christian Beisel, Sarah Nadeau, Chaoran Chen, Ivan Topolsky, Philipp Jablonski, Lara Fuhrmann, David Dreifuss, Katharina Jahn, Tobias Schär, Ina Nissen, Natascha Santacroce, Elodie Burcklen, Christiane Beckmann, Maurice Redondo, Olivier Kobel, Christoph Noppen, Sophie Seidel, Noemie Santamaria de Souza, Niko Beerenwinkel, Tanja Stadler |
| EPI_ISL_898998, EPI_ISL_899000, EPI_ISL_899005, EPI_ISL_899007, EPI_ISL_899011 | Viollier AG | Department of Biosystems Science and Engineering, ETH Zürich | Chaoran Chen, Sarah Nadeau, Ivan Topolsky, Emmanouil Dermitzakis, Keith Harshman, Ioannis Xenarios, Henri Pegeot, Lorenzo Cerutti, Deborah Penet, Philipp Jablonski, Lara Fuhrmann, David Dreifuss, Katharina Jahn, Christiane Beckmann, Maurice Redondo, Olivier Kobel, Christoph Noppen, Sophie Seidel, Noemie Santamaria de Souza, Niko Beerenwinkel, Tanja Stadler |
| EPI_ISL_899023 | Viollier AG | Department of Biosystems Science and Engineering, ETH Zürich | Christian Beisel, Sarah Nadeau, Chaoran Chen, Ivan Topolsky, Philipp Jablonski, Lara Fuhrmann, David Dreifuss, Katharina Jahn, Tobias Schär, Ina Nissen, Natascha Santacroce, Elodie Burcklen, Christiane Beckmann, Maurice Redondo, Olivier Kobel, Christoph Noppen, Sophie Seidel, Noemie Santamaria de Souza, Niko Beerenwinkel, Tanja Stadler |
| EPI_ISL_899026, EPI_ISL_899028, EPI_ISL_899029, EPI_ISL_899030 | Viollier AG | Department of Biosystems Science and Engineering, ETH Zürich | Chaoran Chen, Sarah Nadeau, Ivan Topolsky, Emmanouil Dermitzakis, Keith Harshman, Ioannis Xenarios, Henri Pegeot, Lorenzo Cerutti, Deborah Penet, Philipp Jablonski, Lara Fuhrmann, David Dreifuss, Katharina Jahn, Christiane Beckmann, Maurice Redondo, Olivier Kobel, Christoph Noppen, Sophie Seidel, Noemie Santamaria de Souza, Niko Beerenwinkel, Tanja Stadler |
| EPI_ISL_899031 | Viollier AG | Department of Biosystems Science and Engineering, ETH Zürich | Christian Beisel, Sarah Nadeau, Chaoran Chen, Ivan Topolsky, Philipp Jablonski, Lara Fuhrmann, David Dreifuss, Katharina Jahn, Tobias Schär, Ina Nissen, Natascha Santacroce, Elodie Burcklen, Christiane Beckmann, Maurice Redondo, Olivier Kobel, Christoph Noppen, Sophie Seidel, Noemie Santamaria de Souza, Niko Beerenwinkel, Tanja Stadler |
| EPI_ISL_899032, EPI_ISL_899034, EPI_ISL_899036, EPI_ISL_899038, EPI_ISL_899041, EPI_ISL_899042, EPI_ISL_899044 | Viollier AG | Department of Biosystems Science and Engineering, ETH Zürich | Chaoran Chen, Sarah Nadeau, Ivan Topolsky, Emmanouil Dermitzakis, Keith Harshman, Ioannis Xenarios, Henri Pegeot, Lorenzo Cerutti, Deborah Penet, Philipp Jablonski, Lara Fuhrmann, David Dreifuss, Katharina Jahn, Christiane Beckmann, Maurice Redondo, Olivier Kobel, Christoph Noppen, Sophie Seidel, Noemie Santamaria de Souza, Niko Beerenwinkel, Tanja Stadler |
| EPI_ISL_899052 | Viollier AG | Department of Biosystems Science and Engineering, ETH Zürich | Christian Beisel, Sarah Nadeau, Chaoran Chen, Ivan Topolsky, Philipp Jablonski, Lara Fuhrmann, David Dreifuss, Katharina Jahn, Tobias Schär, Ina Nissen, Natascha Santacroce, Elodie Burcklen, Christiane Beckmann, Maurice Redondo, Olivier Kobel, Christoph Noppen, Sophie Seidel, Noemie Santamaria de Souza, Niko Beerenwinkel, Tanja Stadler |
| EPI_ISL_899054, EPI_ISL_899057, EPI_ISL_899058, EPI_ISL_899059 | Viollier AG | Department of Biosystems Science and Engineering, ETH Zürich | Chaoran Chen, Sarah Nadeau, Ivan Topolsky, Emmanouil Dermitzakis, Keith Harshman, Ioannis Xenarios, Henri Pegeot, Lorenzo Cerutti, Deborah Penet, Philipp Jablonski, Lara Fuhrmann, David Dreifuss, Katharina Jahn, Christiane Beckmann, Maurice Redondo, Olivier Kobel, Christoph Noppen, Sophie Seidel, Noemie Santamaria de Souza, Niko Beerenwinkel, Tanja Stadler |
| EPI_ISL_899069 | Viollier AG | Department of Biosystems Science and Engineering, ETH Zürich | Christian Beisel, Sarah Nadeau, Chaoran Chen, Ivan Topolsky, Philipp Jablonski, Lara Fuhrmann, David Dreifuss, Katharina Jahn, Tobias Schär, Ina Nissen, Natascha Santacroce, Elodie Burcklen, Christiane Beckmann, Maurice Redondo, Olivier Kobel, Christoph Noppen, Sophie Seidel, Noemie |

| | | | |
|---|---|---|---|
| | | | Santamaria de Souza, Niko Beerenwinkel, Tanja Stadler |
| EPI_ISL_899071, EPI_ISL_899074, EPI_ISL_899075, EPI_ISL_899078, EPI_ISL_899081, EPI_ISL_899082 | Viollier AG | Department of Biosystems Science and Engineering, ETH Zürich | Chaoran Chen, Sarah Nadeau, Ivan Topolsky, Emmanouil Dermitzakis, Keith Harshman, Ioannis Xenarios, Henri Pegeot, Lorenzo Cerutti, Deborah Penet, Philipp Jablonski, Lara Fuhrmann, David Dreifuss, Katharina Jahn, Christiane Beckmann, Maurice Redondo, Olivier Kobel, Christoph Noppen, Sophie Seidel, Noemie Santamaria de Souza, Niko Beerenwinkel, Tanja Stadler |
| EPI_ISL_899090 | Viollier AG | Department of Biosystems Science and Engineering, ETH Zürich | Christian Beisel, Sarah Nadeau, Chaoran Chen, Ivan Topolsky, Philipp Jablonski, Lara Fuhrmann, David Dreifuss, Katharina Jahn, Tobias Schär, Ina Nissen, Natascha Santacroce, Elodie Burcklen, Christiane Beckmann, Maurice Redondo, Olivier Kobel, Christoph Noppen, Sophie Seidel, Noemie Santamaria de Souza, Niko Beerenwinkel, Tanja Stadler |
| EPI_ISL_899094, EPI_ISL_899106 | Viollier AG | Department of Biosystems Science and Engineering, ETH Zürich | Chaoran Chen, Sarah Nadeau, Ivan Topolsky, Emmanouil Dermitzakis, Keith Harshman, Ioannis Xenarios, Henri Pegeot, Lorenzo Cerutti, Deborah Penet, Philipp Jablonski, Lara Fuhrmann, David Dreifuss, Katharina Jahn, Christiane Beckmann, Maurice Redondo, Olivier Kobel, Christoph Noppen, Sophie Seidel, Noemie Santamaria de Souza, Niko Beerenwinkel, Tanja Stadler |
| EPI_ISL_899123 | Viollier AG | Department of Biosystems Science and Engineering, ETH Zürich | Christian Beisel, Sarah Nadeau, Chaoran Chen, Ivan Topolsky, Philipp Jablonski, Lara Fuhrmann, David Dreifuss, Katharina Jahn, Tobias Schär, Ina Nissen, Natascha Santacroce, Elodie Burcklen, Christiane Beckmann, Maurice Redondo, Olivier Kobel, Christoph Noppen, Sophie Seidel, Noemie Santamaria de Souza, Niko Beerenwinkel, Tanja Stadler |
| EPI_ISL_899126, EPI_ISL_899132, EPI_ISL_899133 | Viollier AG | Department of Biosystems Science and Engineering, ETH Zürich | Chaoran Chen, Sarah Nadeau, Ivan Topolsky, Emmanouil Dermitzakis, Keith Harshman, Ioannis Xenarios, Henri Pegeot, Lorenzo Cerutti, Deborah Penet, Philipp Jablonski, Lara Fuhrmann, David Dreifuss, Katharina Jahn, Christiane Beckmann, Maurice Redondo, Olivier Kobel, Christoph Noppen, Sophie Seidel, Noemie Santamaria de Souza, Niko Beerenwinkel, Tanja Stadler |
| EPI_ISL_899137, EPI_ISL_899138, EPI_ISL_899151, EPI_ISL_899152, EPI_ISL_899167 | Viollier AG | Department of Biosystems Science and Engineering, ETH Zürich | Christian Beisel, Sarah Nadeau, Chaoran Chen, Ivan Topolsky, Philipp Jablonski, Lara Fuhrmann, David Dreifuss, Katharina Jahn, Tobias Schär, Ina Nissen, Natascha Santacroce, Elodie Burcklen, Christiane Beckmann, Maurice Redondo, Olivier Kobel, Christoph Noppen, Sophie Seidel, Noemie Santamaria de Souza, Niko Beerenwinkel, Tanja Stadler |
| EPI_ISL_899169 | Viollier AG | Department of Biosystems Science and Engineering, ETH Zürich | Chaoran Chen, Sarah Nadeau, Ivan Topolsky, Emmanouil Dermitzakis, Keith Harshman, Ioannis Xenarios, Henri Pegeot, Lorenzo Cerutti, Deborah Penet, Philipp Jablonski, Lara Fuhrmann, David Dreifuss, Katharina Jahn, Christiane Beckmann, Maurice Redondo, Olivier Kobel, Christoph Noppen, Sophie Seidel, Noemie Santamaria de Souza, Niko Beerenwinkel, Tanja Stadler |
| EPI_ISL_899176, EPI_ISL_899242, EPI_ISL_899244, EPI_ISL_899245, EPI_ISL_899247, EPI_ISL_899248, EPI_ISL_899249 | Viollier AG | Department of Biosystems Science and Engineering, ETH Zürich | Christian Beisel, Sarah Nadeau, Chaoran Chen, Ivan Topolsky, Philipp Jablonski, Lara Fuhrmann, David Dreifuss, Katharina Jahn, Tobias Schär, Ina Nissen, Natascha Santacroce, Elodie Burcklen, Christiane Beckmann, Maurice Redondo, Olivier Kobel, Christoph Noppen, Sophie Seidel, Noemie Santamaria de Souza, Niko Beerenwinkel, Tanja Stadler |
| EPI_ISL_899268, EPI_ISL_899269, EPI_ISL_899270, EPI_ISL_899271, EPI_ISL_899272, EPI_ISL_899273, EPI_ISL_899274, EPI_ISL_899275, EPI_ISL_899276, EPI_ISL_899277 | Viollier AG | Department of Biosystems Science and Engineering, ETH Zürich | Chaoran Chen, Sarah Nadeau, Ivan Topolsky, Emmanouil Dermitzakis, Keith Harshman, Ioannis Xenarios, Henri Pegeot, Lorenzo Cerutti, Deborah Penet, Philipp Jablonski, Lara Fuhrmann, David Dreifuss, Katharina Jahn, Christiane Beckmann, Maurice Redondo, Olivier Kobel, Christoph Noppen, Sophie Seidel, Noemie Santamaria de Souza, Niko Beerenwinkel, Tanja Stadler |
| EPI_ISL_899342, EPI_ISL_899343, EPI_ISL_899344, EPI_ISL_899345, EPI_ISL_899350, EPI_ISL_899351, EPI_ISL_899352, EPI_ISL_899353 | Viollier AG | Department of Biosystems Science and Engineering, ETH Zürich | Christian Beisel, Sarah Nadeau, Chaoran Chen, Ivan Topolsky, Philipp Jablonski, Lara Fuhrmann, David Dreifuss, Katharina Jahn, Tobias Schär, Ina Nissen, Natascha Santacroce, Elodie Burcklen, Christiane Beckmann, Maurice Redondo, Olivier Kobel, Christoph Noppen, Sophie Seidel, Noemie Santamaria de Souza, Niko Beerenwinkel, Tanja Stadler |
| EPI_ISL_899373, EPI_ISL_899374, EPI_ISL_899375, EPI_ISL_899376, EPI_ISL_899377, EPI_ISL_899378, EPI_ISL_899379, EPI_ISL_899380, EPI_ISL_899381, EPI_ISL_899382, EPI_ISL_899383, EPI_ISL_899384 | | | |
| see above | Viollier AG | Department of Biosystems Science and Engineering, ETH Zürich | Chaoran Chen, Sarah Nadeau, Ivan Topolsky, Emmanouil Dermitzakis, Keith Harshman, Ioannis Xenarios, Henri Pegeot, Lorenzo Cerutti, Deborah Penet, Philipp Jablonski, Lara Fuhrmann, David Dreifuss, Katharina Jahn, Christiane Beckmann, Maurice Redondo, Olivier Kobel, Christoph Noppen, Sophie Seidel, Noemie Santamaria de Souza, Niko Beerenwinkel, Tanja Stadler |
| EPI_ISL_899458, EPI_ISL_899459, EPI_ISL_899460, EPI_ISL_899461, EPI_ISL_899462, EPI_ISL_899463, EPI_ISL_899464, EPI_ISL_899465, EPI_ISL_899466, EPI_ISL_899467, EPI_ISL_899468, EPI_ISL_899469, EPI_ISL_899470, EPI_ISL_899471, EPI_ISL_899472, EPI_ISL_899473, EPI_ISL_899474, EPI_ISL_899478 | | | |
| see above | Viollier AG | Department of Biosystems Science and Engineering, ETH Zürich | Christian Beisel, Sarah Nadeau, Chaoran Chen, Ivan Topolsky, Philipp Jablonski, Lara Fuhrmann, David Dreifuss, Katharina Jahn, Tobias Schär, Ina Nissen, Natascha Santacroce, Elodie Burcklen, Christiane Beckmann, Maurice Redondo, Olivier Kobel, Christoph Noppen, Sophie Seidel, Noemie Santamaria de Souza, Niko Beerenwinkel, Tanja Stadler |
| EPI_ISL_899509, EPI_ISL_899510, EPI_ISL_899511, EPI_ISL_899512, EPI_ISL_899513, EPI_ISL_899514 | Viollier AG | Department of Biosystems Science and Engineering, ETH Zürich | Chaoran Chen, Sarah Nadeau, Ivan Topolsky, Emmanouil Dermitzakis, Keith Harshman, Ioannis Xenarios, Henri Pegeot, Lorenzo Cerutti, Deborah Penet, Philipp Jablonski, Lara Fuhrmann, David Dreifuss, Katharina Jahn, Christiane Beckmann, Maurice Redondo, Olivier Kobel, Christoph Noppen, Sophie Seidel, Noemie Santamaria de Souza, Niko Beerenwinkel, Tanja Stadler |
| EPI_ISL_899531 | Viollier AG | Department of Biosystems Science and Engineering, ETH Zürich | Christian Beisel, Sarah Nadeau, Chaoran Chen, Ivan Topolsky, Philipp Jablonski, Lara Fuhrmann, David Dreifuss, Katharina Jahn, Tobias Schär, Ina Nissen, Natascha Santacroce, Elodie Burcklen, Christiane Beckmann, Maurice Redondo, Olivier Kobel, Christoph Noppen, Sophie Seidel, Noemie Santamaria de Souza, Niko Beerenwinkel, Tanja Stadler |
| EPI_ISL_899535, EPI_ISL_899536 | Viollier AG | Department of Biosystems Science and Engineering, ETH Zürich | Chaoran Chen, Sarah Nadeau, Ivan Topolsky, Emmanouil Dermitzakis, Keith Harshman, Ioannis Xenarios, Henri Pegeot, Lorenzo Cerutti, Deborah Penet, Philipp Jablonski, Lara Fuhrmann, David Dreifuss, Katharina Jahn, Christiane Beckmann, Maurice Redondo, Olivier Kobel, Christoph Noppen, Sophie Seidel, Noemie Santamaria de Souza, Niko Beerenwinkel, Tanja Stadler |
| EPI_ISL_899542, EPI_ISL_899543, EPI_ISL_899545 | Viollier AG | Department of Biosystems Science and Engineering, ETH Zürich | Christian Beisel, Sarah Nadeau, Chaoran Chen, Ivan Topolsky, Philipp Jablonski, Lara Fuhrmann, David Dreifuss, Katharina Jahn, Tobias Schär, Ina Nissen, Natascha Santacroce, Elodie Burcklen, Christiane Beckmann, Maurice Redondo, Olivier Kobel, Christoph Noppen, Sophie Seidel, Noemie Santamaria de Souza, Niko Beerenwinkel, Tanja Stadler |
| EPI_ISL_899546 | Viollier AG | Department of Biosystems Science and Engineering, ETH Zürich | Chaoran Chen, Sarah Nadeau, Ivan Topolsky, Emmanouil Dermitzakis, Keith Harshman, Ioannis Xenarios, Henri Pegeot, Lorenzo Cerutti, Deborah Penet, Philipp Jablonski, Lara Fuhrmann, David Dreifuss, Katharina Jahn, Christiane Beckmann, Maurice Redondo, Olivier Kobel, Christoph Noppen, Sophie Seidel, Noemie Santamaria de Souza, Niko Beerenwinkel, Tanja Stadler |
| EPI_ISL_899554 | Viollier AG | Department of Biosystems Science and Engineering, ETH Zürich | Christian Beisel, Sarah Nadeau, Chaoran Chen, Ivan Topolsky, Philipp Jablonski, Lara Fuhrmann, David Dreifuss, Katharina Jahn, Tobias Schär, Ina Nissen, Natascha Santacroce, Elodie Burcklen, Christiane Beckmann, Maurice Redondo, Olivier Kobel, Christoph Noppen, Sophie Seidel, Noemie Santamaria de Souza, Niko Beerenwinkel, Tanja Stadler |
| EPI_ISL_899557, EPI_ISL_899558, EPI_ISL_899561 | Viollier AG | Department of Biosystems Science and Engineering, ETH Zürich | Chaoran Chen, Sarah Nadeau, Ivan Topolsky, Emmanouil Dermitzakis, Keith Harshman, Ioannis Xenarios, Henri Pegeot, Lorenzo Cerutti, Deborah Penet, Philipp Jablonski, Lara Fuhrmann, David Dreifuss, Katharina Jahn, Christiane Beckmann, Maurice Redondo, Olivier Kobel, Christoph Noppen, Sophie Seidel, Noemie Santamaria de Souza, Niko Beerenwinkel, Tanja Stadler |
| EPI_ISL_899571, EPI_ISL_899572, EPI_ISL_899573 | Viollier AG | Department of Biosystems Science and Engineering, ETH Zürich | Christian Beisel, Sarah Nadeau, Chaoran Chen, Ivan Topolsky, Philipp Jablonski, Lara Fuhrmann, David Dreifuss, Katharina Jahn, Tobias Schär, Ina Nissen, Natascha Santacroce, Elodie Burcklen, Christiane Beckmann, Maurice Redondo, Olivier Kobel, Christoph Noppen, Sophie Seidel, Noemie Santamaria de Souza, Niko Beerenwinkel, Tanja Stadler |
| EPI_ISL_899574, EPI_ISL_899575 | Viollier AG | Department of Biosystems Science and Engineering, ETH Zürich | Chaoran Chen, Sarah Nadeau, Ivan Topolsky, Emmanouil Dermitzakis, Keith Harshman, Ioannis Xenarios, Henri Pegeot, Lorenzo Cerutti, Deborah Penet, Philipp Jablonski, Lara Fuhrmann, David Dreifuss, Katharina Jahn, Christiane Beckmann, Maurice Redondo, Olivier Kobel, Christoph Noppen, Sophie Seidel, Noemie Santamaria de Souza, Niko Beerenwinkel, Tanja Stadler |
| EPI_ISL_899583, EPI_ISL_899584, EPI_ISL_899585, EPI_ISL_899586, EPI_ISL_899587, EPI_ISL_899588, EPI_ISL_899591, EPI_ISL_899592 | Viollier AG | Department of Biosystems Science and Engineering, ETH Zürich | Christian Beisel, Sarah Nadeau, Chaoran Chen, Ivan Topolsky, Philipp Jablonski, Lara Fuhrmann, David Dreifuss, Katharina Jahn, Tobias Schär, Ina Nissen, Natascha Santacroce, Elodie Burcklen, Christiane Beckmann, Maurice Redondo, Olivier Kobel, Christoph Noppen, Sophie Seidel, Noemie Santamaria de Souza, Niko Beerenwinkel, Tanja Stadler |
| EPI_ISL_899596 | Viollier AG | Department of Biosystems Science and Engineering, ETH | Chaoran Chen, Sarah Nadeau, Ivan Topolsky, Emmanouil Dermitzakis, Keith Harshman, Ioannis Xenarios, Henri Pegeot, Lorenzo Cerutti, Deborah Penet, |

| | | | |
|---|---|---|---|
| | | Zürich | Philipp Jablonski, Lara Fuhrmann, David Dreifuss, Katharina Jahn, Christiane Beckmann, Maurice Redondo, Olivier Kobel, Christoph Noppen, Sophie Seidel, Noemie Santamaria de Souza, Niko Beerenwinkel, Tanja Stadler |
| EPI_ISL_899600, EPI_ISL_899601, EPI_ISL_899606, EPI_ISL_899609, EPI_ISL_899610 | Viollier AG | Department of Biosystems Science and Engineering, ETH Zürich | Christian Beisel, Sarah Nadeau, Chaoran Chen, Ivan Topolsky, Philipp Jablonski, Lara Fuhrmann, David Dreifuss, Katharina Jahn, Tobias Schär, Ina Nissen, Natascha Santacroce, Elodie Burcklen, Christiane Beckmann, Maurice Redondo, Olivier Kobel, Christoph Noppen, Sophie Seidel, Noemie Santamaria de Souza, Niko Beerenwinkel, Tanja Stadler |
| EPI_ISL_899612, EPI_ISL_899613 | Viollier AG | Department of Biosystems Science and Engineering, ETH Zürich | Chaoran Chen, Sarah Nadeau, Ivan Topolsky, Emmanouil Dermitzakis, Keith Harshman, Ioannis Xenarios, Henri Pegeot, Lorenzo Cerutti, Deborah Penet, Philipp Jablonski, Lara Fuhrmann, David Dreifuss, Katharina Jahn, Christiane Beckmann, Maurice Redondo, Olivier Kobel, Christoph Noppen, Sophie Seidel, Noemie Santamaria de Souza, Niko Beerenwinkel, Tanja Stadler |
| EPI_ISL_899640 | Viollier AG | Department of Biosystems Science and Engineering, ETH Zürich | Christian Beisel, Sarah Nadeau, Chaoran Chen, Ivan Topolsky, Philipp Jablonski, Lara Fuhrmann, David Dreifuss, Katharina Jahn, Tobias Schär, Ina Nissen, Natascha Santacroce, Elodie Burcklen, Christiane Beckmann, Maurice Redondo, Olivier Kobel, Christoph Noppen, Sophie Seidel, Noemie Santamaria de Souza, Niko Beerenwinkel, Tanja Stadler |
| EPI_ISL_899646 | Viollier AG | Department of Biosystems Science and Engineering, ETH Zürich | Chaoran Chen, Sarah Nadeau, Ivan Topolsky, Emmanouil Dermitzakis, Keith Harshman, Ioannis Xenarios, Henri Pegeot, Lorenzo Cerutti, Deborah Penet, Philipp Jablonski, Lara Fuhrmann, David Dreifuss, Katharina Jahn, Christiane Beckmann, Maurice Redondo, Olivier Kobel, Christoph Noppen, Sophie Seidel, Noemie Santamaria de Souza, Niko Beerenwinkel, Tanja Stadler |
| EPI_ISL_899721, EPI_ISL_899725, EPI_ISL_899726, EPI_ISL_899727, EPI_ISL_899728, EPI_ISL_899729, EPI_ISL_899730, EPI_ISL_899731, EPI_ISL_899732, EPI_ISL_899733, EPI_ISL_899734, EPI_ISL_899735, EPI_ISL_899736, EPI_ISL_899738 | | | |
| see above | Viollier AG | Department of Biosystems Science and Engineering, ETH Zürich | Christian Beisel, Sarah Nadeau, Chaoran Chen, Ivan Topolsky, Philipp Jablonski, Lara Fuhrmann, David Dreifuss, Katharina Jahn, Tobias Schär, Ina Nissen, Natascha Santacroce, Elodie Burcklen, Christiane Beckmann, Maurice Redondo, Olivier Kobel, Christoph Noppen, Sophie Seidel, Noemie Santamaria de Souza, Niko Beerenwinkel, Tanja Stadler |
| EPI_ISL_899756, EPI_ISL_899757, EPI_ISL_899758 | Viollier AG | Department of Biosystems Science and Engineering, ETH Zürich | Chaoran Chen, Sarah Nadeau, Ivan Topolsky, Emmanouil Dermitzakis, Keith Harshman, Ioannis Xenarios, Henri Pegeot, Lorenzo Cerutti, Deborah Penet, Philipp Jablonski, Lara Fuhrmann, David Dreifuss, Katharina Jahn, Christiane Beckmann, Maurice Redondo, Olivier Kobel, Christoph Noppen, Sophie Seidel, Noemie Santamaria de Souza, Niko Beerenwinkel, Tanja Stadler |
| EPI_ISL_899787, EPI_ISL_899788, EPI_ISL_899789, EPI_ISL_899790, EPI_ISL_899791, EPI_ISL_899793 | Viollier AG | Department of Biosystems Science and Engineering, ETH Zürich | Christian Beisel, Sarah Nadeau, Chaoran Chen, Ivan Topolsky, Philipp Jablonski, Lara Fuhrmann, David Dreifuss, Katharina Jahn, Tobias Schär, Ina Nissen, Natascha Santacroce, Elodie Burcklen, Christiane Beckmann, Maurice Redondo, Olivier Kobel, Christoph Noppen, Sophie Seidel, Noemie Santamaria de Souza, Niko Beerenwinkel, Tanja Stadler |
| EPI_ISL_899805, EPI_ISL_899806, EPI_ISL_899807, EPI_ISL_899808, EPI_ISL_899809 | Viollier AG | Department of Biosystems Science and Engineering, ETH Zürich | Chaoran Chen, Sarah Nadeau, Ivan Topolsky, Emmanouil Dermitzakis, Keith Harshman, Ioannis Xenarios, Henri Pegeot, Lorenzo Cerutti, Deborah Penet, Philipp Jablonski, Lara Fuhrmann, David Dreifuss, Katharina Jahn, Christiane Beckmann, Maurice Redondo, Olivier Kobel, Christoph Noppen, Sophie Seidel, Noemie Santamaria de Souza, Niko Beerenwinkel, Tanja Stadler |
| EPI_ISL_899843, EPI_ISL_899845 | Viollier AG | Department of Biosystems Science and Engineering, ETH Zürich | Christian Beisel, Sarah Nadeau, Chaoran Chen, Ivan Topolsky, Philipp Jablonski, Lara Fuhrmann, David Dreifuss, Katharina Jahn, Tobias Schär, Ina Nissen, Natascha Santacroce, Elodie Burcklen, Christiane Beckmann, Maurice Redondo, Olivier Kobel, Christoph Noppen, Sophie Seidel, Noemie Santamaria de Souza, Niko Beerenwinkel, Tanja Stadler |
| EPI_ISL_899847 | Viollier AG | Department of Biosystems Science and Engineering, ETH Zürich | Chaoran Chen, Sarah Nadeau, Ivan Topolsky, Emmanouil Dermitzakis, Keith Harshman, Ioannis Xenarios, Henri Pegeot, Lorenzo Cerutti, Deborah Penet, Philipp Jablonski, Lara Fuhrmann, David Dreifuss, Katharina Jahn, Christiane Beckmann, Maurice Redondo, Olivier Kobel, Christoph Noppen, Sophie Seidel, Noemie Santamaria de Souza, Niko Beerenwinkel, Tanja Stadler |
| EPI_ISL_899850 | Viollier AG | Department of Biosystems Science and Engineering, ETH Zürich | Christian Beisel, Sarah Nadeau, Chaoran Chen, Ivan Topolsky, Philipp Jablonski, Lara Fuhrmann, David Dreifuss, Katharina Jahn, Tobias Schär, Ina Nissen, Natascha Santacroce, Elodie Burcklen, Christiane Beckmann, Maurice Redondo, Olivier Kobel, Christoph Noppen, Sophie Seidel, Noemie Santamaria de Souza, Niko Beerenwinkel, Tanja Stadler |
| EPI_ISL_899851, EPI_ISL_899852, EPI_ISL_899853 | Viollier AG | Department of Biosystems Science and Engineering, ETH Zürich | Chaoran Chen, Sarah Nadeau, Ivan Topolsky, Emmanouil Dermitzakis, Keith Harshman, Ioannis Xenarios, Henri Pegeot, Lorenzo Cerutti, Deborah Penet, Philipp Jablonski, Lara Fuhrmann, David Dreifuss, Katharina Jahn, Christiane Beckmann, Maurice Redondo, Olivier Kobel, Christoph Noppen, Sophie Seidel, Noemie Santamaria de Souza, Niko Beerenwinkel, Tanja Stadler |
| EPI_ISL_899854, EPI_ISL_899855, EPI_ISL_899856, EPI_ISL_899857, EPI_ISL_899858, EPI_ISL_899859, EPI_ISL_899860, EPI_ISL_899861, EPI_ISL_899864, EPI_ISL_899865, EPI_ISL_899866, EPI_ISL_899867, EPI_ISL_899868, EPI_ISL_899869, EPI_ISL_899870, EPI_ISL_899871, EPI_ISL_899872, EPI_ISL_899873, EPI_ISL_899885 | | | |
| see above | Viollier AG | Department of Biosystems Science and Engineering, ETH Zürich | Christian Beisel, Sarah Nadeau, Chaoran Chen, Ivan Topolsky, Philipp Jablonski, Lara Fuhrmann, David Dreifuss, Katharina Jahn, Tobias Schär, Ina Nissen, Natascha Santacroce, Elodie Burcklen, Christiane Beckmann, Maurice Redondo, Olivier Kobel, Christoph Noppen, Sophie Seidel, Noemie Santamaria de Souza, Niko Beerenwinkel, Tanja Stadler |
| EPI_ISL_899889, EPI_ISL_899890, EPI_ISL_899891, EPI_ISL_899892, EPI_ISL_899893, EPI_ISL_899894, EPI_ISL_899895, EPI_ISL_899896, EPI_ISL_899897, EPI_ISL_899898, EPI_ISL_899899, EPI_ISL_899900, EPI_ISL_899901, EPI_ISL_899902, EPI_ISL_899903, EPI_ISL_899904, EPI_ISL_899905, EPI_ISL_899906 | | | |
| see above | Viollier AG | Department of Biosystems Science and Engineering, ETH Zürich | Chaoran Chen, Sarah Nadeau, Ivan Topolsky, Emmanouil Dermitzakis, Keith Harshman, Ioannis Xenarios, Henri Pegeot, Lorenzo Cerutti, Deborah Penet, Philipp Jablonski, Lara Fuhrmann, David Dreifuss, Katharina Jahn, Christiane Beckmann, Maurice Redondo, Olivier Kobel, Christoph Noppen, Sophie Seidel, Noemie Santamaria de Souza, Niko Beerenwinkel, Tanja Stadler |
| EPI_ISL_899913, EPI_ISL_899914, EPI_ISL_899916 | Viollier AG | Department of Biosystems Science and Engineering, ETH Zürich | Christian Beisel, Sarah Nadeau, Chaoran Chen, Ivan Topolsky, Philipp Jablonski, Lara Fuhrmann, David Dreifuss, Katharina Jahn, Tobias Schär, Ina Nissen, Natascha Santacroce, Elodie Burcklen, Christiane Beckmann, Maurice Redondo, Olivier Kobel, Christoph Noppen, Sophie Seidel, Noemie Santamaria de Souza, Niko Beerenwinkel, Tanja Stadler |
| EPI_ISL_899919, EPI_ISL_899920, EPI_ISL_899922 | Viollier AG | Department of Biosystems Science and Engineering, ETH Zürich | Chaoran Chen, Sarah Nadeau, Ivan Topolsky, Emmanouil Dermitzakis, Keith Harshman, Ioannis Xenarios, Henri Pegeot, Lorenzo Cerutti, Deborah Penet, Philipp Jablonski, Lara Fuhrmann, David Dreifuss, Katharina Jahn, Christiane Beckmann, Maurice Redondo, Olivier Kobel, Christoph Noppen, Sophie Seidel, Noemie Santamaria de Souza, Niko Beerenwinkel, Tanja Stadler |
| EPI_ISL_900008, EPI_ISL_900009, EPI_ISL_900013, EPI_ISL_900014, EPI_ISL_900015, EPI_ISL_900019, EPI_ISL_900020, EPI_ISL_900021, EPI_ISL_900022 | Viollier AG | Department of Biosystems Science and Engineering, ETH Zürich | Christian Beisel, Sarah Nadeau, Chaoran Chen, Ivan Topolsky, Philipp Jablonski, Lara Fuhrmann, David Dreifuss, Katharina Jahn, Tobias Schär, Ina Nissen, Natascha Santacroce, Elodie Burcklen, Christiane Beckmann, Maurice Redondo, Olivier Kobel, Christoph Noppen, Sophie Seidel, Noemie Santamaria de Souza, Niko Beerenwinkel, Tanja Stadler |
| EPI_ISL_900037, EPI_ISL_900038, EPI_ISL_900039, EPI_ISL_900040, EPI_ISL_900041, EPI_ISL_900042, EPI_ISL_900043, EPI_ISL_900044, EPI_ISL_900045, EPI_ISL_900046, EPI_ISL_900047, EPI_ISL_900048, EPI_ISL_900049, EPI_ISL_900050, EPI_ISL_900051, EPI_ISL_900052, EPI_ISL_900053 | | | |
| see above | Viollier AG | Department of Biosystems Science and Engineering, ETH Zürich | Chaoran Chen, Sarah Nadeau, Ivan Topolsky, Emmanouil Dermitzakis, Keith Harshman, Ioannis Xenarios, Henri Pegeot, Lorenzo Cerutti, Deborah Penet, Philipp Jablonski, Lara Fuhrmann, David Dreifuss, Katharina Jahn, Christiane Beckmann, Maurice Redondo, Olivier Kobel, Christoph Noppen, Sophie Seidel, Noemie Santamaria de Souza, Niko Beerenwinkel, Tanja Stadler |
| EPI_ISL_900492, EPI_ISL_900496, EPI_ISL_900501 | Althaia. Xarxa Assistencial Universitària de Manresa | IrsiCaixa - Can Ruti CovidSeq | Fundació irsiCaixa. Hospital Universitari Germans Trias i Pujol(HUGTiP), 2a planta, maternal Ctra Canyet s/n, Badalona Gloria Trujillo, Rafel Perez Vidal, Jaume Trape Pujol, Carolina Gonzalez Fernandez, Roger Paredes, Eulalia Grau, Bonaventura Clotet |
| EPI_ISL_902750 | Ospedale "F. Spaziani" Frosinone | INMI Lazzaro Spallanzani IRCCS | F Messina, C.E.M Gruber, B Bartolini, E Giombini, M Rueca, O Butera, C. Gargiulo, C Sias, A Di Caro, MR Capobianchi |
| EPI_ISL_902751 | Ospedale Sandro Pertini ASL Roma2 | INMI Lazzaro Spallanzani IRCCS | B Bartolini, O Butera, C.E.M Gruber, M Rueca, F Messina, E Giombini, G Cappiello, MC Cava, MR Capobianchi, A Di Caro |
| EPI_ISL_902752 | Ospedale Sandro Pertini ASL Roma2 | INMI Lazzaro Spallanzani IRCCS | O Butera, F Messina, C.E.M Gruber, B Bartolini, E Giombini, M Rueca, S Romano, MC Cava, A Di Caro, MR Capobianchi |
| EPI_ISL_902753 | Azienda Ospedaliera San Camillo Forlanini | INMI Lazzaro Spallanzani IRCCS | C.E.M Gruber, B Bartolini, E Giombini, M Rueca, O Butera, F Messina, D.Gallone, G Parisi, A Di Caro, MR Capobianchi |
| EPI_ISL_902757 | San Camillo De Lellis Rieti Hospital | INMI Lazzaro Spallanzani IRCCS | C.E.M Gruber, B Bartolini, E Giombini, M Rueca, O Butera, F Messina, S Venarubea, A De Luca, A Di Caro, MR Capobianchi |
| EPI_ISL_902927, EPI_ISL_902928, EPI_ISL_902929, EPI_ISL_902930, EPI_ISL_902931, EPI_ISL_902932, EPI_ISL_902935, EPI_ISL_902936, EPI_ISL_902937, EPI_ISL_902938, EPI_ISL_902939, EPI_ISL_902940, EPI_ISL_902941, EPI_ISL_902943, EPI_ISL_902944, EPI_ISL_902945, EPI_ISL_902946, EPI_ISL_902947 | | | |
| see above | Maryland Public Health Laboratory | Maryland Public Health Laboratory | Maryland Department of Health Laboratories Administration |

| | | | |
|---|---|---|---|
| EPI_ISL_903348, EPI_ISL_903358 | Wyoming Public Health Laboratory | Wyoming Public Health Laboratory | Noah Hull, Taylor Fearing, Lynette Gumbleton, Channing Weber, Ashley Norberg, Bailey Bowcutt, and Wanda Manley |
| EPI_ISL_903371 | Los Angeles County PHL | Los Angeles County PHL | P. Hemarajata et al. |
| EPI_ISL_904118 | New Mexico Department of Health Scientific Laboratory | New Mexico Department of Health Scientific Laboratory | Ellie Johnson, Anastacia Griego-Fisher, D'eldra Malone |
| EPI_ISL_904123, EPI_ISL_904124 | National Institute of Public Health - National Institute of Hygiene | National Institute of Public Health - National Institute of Hygiene | Wokowicz Tomasz, Zacharczuk Katarzyna |
| EPI_ISL_904135, EPI_ISL_904212, EPI_ISL_904213, EPI_ISL_904214, EPI_ISL_904215, EPI_ISL_904216, EPI_ISL_904247, EPI_ISL_904314, EPI_ISL_904374, EPI_ISL_904378, EPI_ISL_904379, EPI_ISL_904380, EPI_ISL_904588, EPI_ISL_904589, EPI_ISL_904590, EPI_ISL_904591, EPI_ISL_904592, EPI_ISL_904593, EPI_ISL_904594, EPI_ISL_904595, EPI_ISL_904596, EPI_ISL_904597, EPI_ISL_904598, EPI_ISL_904599, EPI_ISL_904600, EPI_ISL_904601, EPI_ISL_904602 | | | |
| see above | Dutch COVID-19 response team | Erasmus Medical Center | Bas Oude Munnink, Reina Sikkema, David Nieuwenhuijse, Irina Chestakova, Anne van der Linden, Marjan Boter, Emmanuelle Munger, Corine GeurtsvanKessel, Annemiek van der Eijk, Richard Molenkamp, Marion Koopmans, on behalf of the Dutch national COVID-19 response team. |
| EPI_ISL_904660, EPI_ISL_904682, EPI_ISL_904683, EPI_ISL_904684, EPI_ISL_904686, EPI_ISL_904687, EPI_ISL_904688, EPI_ISL_904691, EPI_ISL_904692, EPI_ISL_904693, EPI_ISL_904694, EPI_ISL_904697, EPI_ISL_904698, EPI_ISL_904699, EPI_ISL_904710, EPI_ISL_904714, EPI_ISL_904719, EPI_ISL_904720, EPI_ISL_904722, EPI_ISL_904724, EPI_ISL_904730, EPI_ISL_904731, EPI_ISL_904732, EPI_ISL_904740, EPI_ISL_904741, EPI_ISL_904744, EPI_ISL_904752, EPI_ISL_904753, EPI_ISL_904754, EPI_ISL_904770, EPI_ISL_904771, EPI_ISL_904778, EPI_ISL_904779, EPI_ISL_904782, EPI_ISL_904783, EPI_ISL_904787, EPI_ISL_904788, EPI_ISL_904791, EPI_ISL_904792, EPI_ISL_904794, EPI_ISL_904799, EPI_ISL_904804, EPI_ISL_904805, EPI_ISL_904806, EPI_ISL_904807, EPI_ISL_904810, EPI_ISL_904814, EPI_ISL_904815, EPI_ISL_904816, EPI_ISL_904817, EPI_ISL_904818, EPI_ISL_904819, EPI_ISL_904820, EPI_ISL_904821, EPI_ISL_904822, EPI_ISL_904823, EPI_ISL_904824, EPI_ISL_904825, EPI_ISL_904828, EPI_ISL_904829, EPI_ISL_904833, EPI_ISL_904834, EPI_ISL_904835, EPI_ISL_904841, EPI_ISL_904843, EPI_ISL_904844, EPI_ISL_904845, EPI_ISL_904851, EPI_ISL_904853, EPI_ISL_904854, EPI_ISL_904855, EPI_ISL_904856, EPI_ISL_904857, EPI_ISL_904858, EPI_ISL_904859, EPI_ISL_904866, EPI_ISL_904867, EPI_ISL_904869, EPI_ISL_904870, EPI_ISL_904875, EPI_ISL_904876, EPI_ISL_904877, EPI_ISL_904879, EPI_ISL_904880, EPI_ISL_904894, EPI_ISL_904906, EPI_ISL_904907, EPI_ISL_904908, EPI_ISL_904909, EPI_ISL_904910, EPI_ISL_904911, EPI_ISL_904912, EPI_ISL_904913, EPI_ISL_904925, EPI_ISL_904926, EPI_ISL_904958, EPI_ISL_904959, EPI_ISL_904960, EPI_ISL_904961, EPI_ISL_904962, EPI_ISL_904963, EPI_ISL_904964, EPI_ISL_904965, EPI_ISL_904966, EPI_ISL_904967, EPI_ISL_904968, EPI_ISL_904969, EPI_ISL_904970, EPI_ISL_904971, EPI_ISL_904976, EPI_ISL_904977, EPI_ISL_904978, EPI_ISL_904979, EPI_ISL_904980, EPI_ISL_904981, EPI_ISL_904995, EPI_ISL_905017, EPI_ISL_905031, EPI_ISL_905032, EPI_ISL_905057, EPI_ISL_905058, EPI_ISL_905062, EPI_ISL_905064, EPI_ISL_905065, EPI_ISL_905068, EPI_ISL_905112, EPI_ISL_905114, EPI_ISL_905115, EPI_ISL_905117, EPI_ISL_905171, EPI_ISL_905177, EPI_ISL_905191, EPI_ISL_905194, EPI_ISL_905239, EPI_ISL_905240, EPI_ISL_905241, EPI_ISL_905242, EPI_ISL_905251, EPI_ISL_905252, EPI_ISL_905253, EPI_ISL_905255, EPI_ISL_905256, EPI_ISL_905257, EPI_ISL_905258, EPI_ISL_905259, EPI_ISL_905306, EPI_ISL_905309, EPI_ISL_905310, EPI_ISL_905608, EPI_ISL_905611, EPI_ISL_905612, EPI_ISL_905613, EPI_ISL_905614, EPI_ISL_905660, EPI_ISL_905694, EPI_ISL_905695, EPI_ISL_905696, EPI_ISL_905721, EPI_ISL_905722, EPI_ISL_905726, EPI_ISL_905727, EPI_ISL_905730 | | | |
| see above | Dutch COVID-19 response team | National Institute for Public Health and the Environment (RIVM) | Adam Meijer, Harry Vennema, Dirk Eggink, Jeroen Cremer, Sharon van den Brink, Bas van der Veer, AnneMarie van den Brandt, Florian Zwagemaker, Dennis Schmitz, Chantal Reusken, on behalf of the national COVID-19 response team |
| EPI_ISL_905739, EPI_ISL_905748, EPI_ISL_905749, EPI_ISL_905750, EPI_ISL_905751, EPI_ISL_905754, EPI_ISL_905755 | National Institute of Public Health - National Institute of Hygiene | National Institute of Public Health - National Institute of Hygiene | Wokowicz Tomasz, Zacharczuk Katarzyna |
| EPI_ISL_906149, EPI_ISL_906151, EPI_ISL_906152, EPI_ISL_906154, EPI_ISL_906155, EPI_ISL_906156, EPI_ISL_906157, EPI_ISL_906160, EPI_ISL_906162, EPI_ISL_906163, EPI_ISL_906165, EPI_ISL_906166, EPI_ISL_906168, EPI_ISL_906170, EPI_ISL_906171, EPI_ISL_906173, EPI_ISL_906175, EPI_ISL_906177, EPI_ISL_906179, EPI_ISL_906180, EPI_ISL_906183, EPI_ISL_906184, EPI_ISL_906186, EPI_ISL_906188 | | | |
| see above | University of Wisconsin-Madison AIDS Vaccine Research Laboratories | University of Wisconsin-Madison AIDS Vaccine Research Laboratories | Gage Moreno, Katarina Braun, et al. AIDS Vaccine Research Laboratories |
| EPI_ISL_906282, EPI_ISL_906283 | Nigeria Centre for Disease Control (NCDC) | African Centre of Excellence for Genomics of Infectious Diseases (ACEGID), Redeemer's University | Oluniyi P.E. et al |
| EPI_ISL_906530 | Laboratorio de salud publica de Bogota | Instituto Nacional de Salud- Dirección de Investigación en Salud Pública, Universidad de los Andes- Applied genomics research group, Vicerrectoria de Investigación y Creación, Universidad de los Andes- Systems and Computing Engineering Department | Katherine Laiton-Donato, Diego A. Álvarez-Díaz, Carlos Franco-Muñoz, Mauricio Pacheco-Montealegre, Héctor Alejandro Ruiz-Moreno, Maria T. Herrera-Sepúlveda, Diego Andrés Prada, Jhonnatan Reales-González, Sheryll Corchuelo, Julian Naizaque, Gerardo Santamaría Jorge Duitama, Laura Natalia Gonzalez, Jorge Ivan Diaz, Silvia Restrepo-Restrepo, Magdalena Wiesner, Martha Lucia Ospina Martinez, Marcela Mercado-Reyes |
| EPI_ISL_906531 | Laboratorio de salud publica del Valle del Cauca | Instituto Nacional de Salud- Dirección de Investigación en Salud Pública, Universidad de los Andes- Applied genomics research group, Vicerrectoria de Investigación y Creación, Universidad de los Andes- Systems and Computing Engineering Department | Katherine Laiton-Donato, Diego A. Álvarez-Díaz, Carlos Franco-Muñoz, Mauricio Pacheco-Montealegre, Héctor Alejandro Ruiz-Moreno, Maria T. Herrera-Sepúlveda, Diego Andrés Prada, Jhonnatan Reales-González, Sheryll Corchuelo, Julian Naizaque, Gerardo Santamaría Jorge Duitama, Laura Natalia Gonzalez, Jorge Ivan Diaz, Silvia Restrepo-Restrepo, Magdalena Wiesner, Martha Lucia Ospina Martinez, Marcela Mercado-Reyes |
| EPI_ISL_906532 | Hospital San Jose de Maicao | Instituto Nacional de Salud- Dirección de Investigación en Salud Pública, Universidad de los Andes- Applied genomics research group, Vicerrectoria de Investigación y Creación, Universidad de los Andes- Systems and Computing Engineering Department | Katherine Laiton-Donato, Diego A. Álvarez-Díaz, Carlos Franco-Muñoz, Mauricio Pacheco-Montealegre, Héctor Alejandro Ruiz-Moreno, Maria T. Herrera-Sepúlveda, Diego Andrés Prada, Jhonnatan Reales-González, Sheryll Corchuelo, Julian Naizaque, Gerardo Santamaría Jorge Duitama, Laura Natalia Gonzalez, Jorge Ivan Diaz, Silvia Restrepo-Restrepo, Magdalena Wiesner, Martha Lucia Ospina Martinez, Marcela Mercado-Reyes |
| EPI_ISL_906552 | Laboratorio Angel Diagnostica | Instituto Nacional de Salud- Dirección de Investigación en Salud Pública, Universidad de los Andes- Applied genomics research group, Vicerrectoria de Investigación y Creación, Universidad de los Andes- Systems and Computing Engineering Department | Katherine Laiton-Donato, Diego A. Álvarez-Díaz, Carlos Franco-Muñoz, Mauricio Pacheco-Montealegre, Héctor Alejandro Ruiz-Moreno, Maria T. Herrera-Sepúlveda, Diego Andrés Prada, Jhonnatan Reales-González, Sheryll Corchuelo, Julian Naizaque, Gerardo Santamaría Jorge Duitama, Laura Natalia Gonzalez, Jorge Ivan Diaz, Silvia Restrepo-Restrepo, Magdalena Wiesner, Martha Lucia Ospina Martinez, Marcela Mercado-Reyes |
| EPI_ISL_906556 | DIRECCION DEPARTAMENTAL DE SALUD DE GUAJIRA | Instituto Nacional de Salud- Dirección de Investigación en Salud Pública, Universidad de los Andes- Applied genomics research group, Vicerrectoria de Investigación y Creación, Universidad de los Andes- Systems and Computing Engineering Department | Katherine Laiton-Donato, Diego A. Álvarez-Díaz, Carlos Franco-Muñoz, Mauricio Pacheco-Montealegre, Héctor Alejandro Ruiz-Moreno, Maria T. Herrera-Sepúlveda, Diego Andrés Prada, Jhonnatan Reales-González, Sheryll Corchuelo, Julian Naizaque, Gerardo Santamaría Jorge Duitama, Laura Natalia Gonzalez, Jorge Ivan Diaz, Silvia Restrepo-Restrepo, Magdalena Wiesner, Martha Lucia Ospina Martinez, Marcela Mercado-Reyes |
| EPI_ISL_906557 | Laboratorio de Salud Publica de Amazonas | Instituto Nacional de Salud- Dirección de Investigación en Salud Pública, Universidad de los Andes- Applied genomics research group, Vicerrectoria de Investigación y Creación, Universidad de los Andes- Systems and Computing Engineering Department | Katherine Laiton-Donato, Diego A. Álvarez-Díaz, Carlos Franco-Muñoz, Mauricio Pacheco-Montealegre, Héctor Alejandro Ruiz-Moreno, Maria T. Herrera-Sepúlveda, Diego Andrés Prada, Jhonnatan Reales-González, Sheryll Corchuelo, Julian Naizaque, Gerardo Santamaría Jorge Duitama, Laura Natalia Gonzalez, Jorge Ivan Diaz, Silvia Restrepo-Restrepo, Magdalena Wiesner, Martha Lucia Ospina Martinez, Marcela Mercado-Reyes |
| EPI_ISL_906603, EPI_ISL_906604, EPI_ISL_906605, EPI_ISL_906606, EPI_ISL_906607, EPI_ISL_906608, EPI_ISL_906609, EPI_ISL_906610, EPI_ISL_906611, EPI_ISL_906612, EPI_ISL_906613, EPI_ISL_906614, EPI_ISL_906615, EPI_ISL_906616, EPI_ISL_906617, EPI_ISL_906618, EPI_ISL_906619, EPI_ISL_906620, EPI_ISL_906621, EPI_ISL_906622, EPI_ISL_906623, EPI_ISL_906624, EPI_ISL_906625, EPI_ISL_906626, EPI_ISL_906627, EPI_ISL_906628, EPI_ISL_906629, EPI_ISL_906630, EPI_ISL_906631, EPI_ISL_906632, EPI_ISL_906633, EPI_ISL_906634, EPI_ISL_906635, EPI_ISL_906636, EPI_ISL_906637, EPI_ISL_906638, EPI_ISL_906639, EPI_ISL_906640, EPI_ISL_906641, EPI_ISL_906642, EPI_ISL_906643, EPI_ISL_906644, EPI_ISL_906645, EPI_ISL_906646, EPI_ISL_906647, EPI_ISL_906648, EPI_ISL_906649, EPI_ISL_906650, EPI_ISL_906651 | | | |
| see above | Maine Health and Environmental Testing Laboratory (Maine HETL) | Tewhey Lab, The Jackson Laboratory | Matluk,N., Dewey,H., Iosue,F., Barter,M., Lynch,R., Munger,H. and Tewhey,R. |
| EPI_ISL_906720, EPI_ISL_906727, EPI_ISL_906730, EPI_ISL_906736 | Laboratory of Molecular Biology, Diagnostyka sp. z o.o. | Laboratory of Recombinant Vaccines | Lukasz Rabalski, Maciej Kosinski, Anna Piotrowska-Mietelska,Izabela Szczygielska, Boguslaw Szewczyk, Krystyna Bienkowska-Szewczyk |
| EPI_ISL_906793, EPI_ISL_906796 | Innlandet Hospital Trust, Division Lillehammer, Department for Medical Microbiology | Norwegian Institute of Public Health, Department of Virology | Kathrine Stene-Johansen, Kamilla Heddeland Instefjord, Hilde Elshaug, Atiya R Ali,Marie Paulsen Madsen, Rasmus Riis Kopperud, Hilde Vollan, Karoline Bragstad, Olav Hungnes |
| EPI_ISL_906808, EPI_ISL_906809, EPI_ISL_906810, EPI_ISL_906811 | University Hospital of Northern Norway, Department for Microbiology and Infectious Disease Control | Norwegian Institute of Public Health, Department of Virology | Kathrine Stene-Johansen, Kamilla Heddeland Instefjord, Hilde Elshaug, Atiya R Ali,Marie Paulsen Madsen, Rasmus Riis Kopperud, Hilde Vollan, Karoline Bragstad, Olav Hungnes |
| EPI_ISL_906812 | Innlandet Hospital Trust, Division Lillehammer, Department for Medical Microbiology | Norwegian Institute of Public Health, Department of Virology | Kathrine Stene-Johansen, Kamilla Heddeland Instefjord, Hilde Elshaug, Atiya R Ali,Marie Paulsen Madsen, Rasmus Riis Kopperud, Hilde Vollan, Karoline Bragstad, Olav Hungnes |
| EPI_ISL_906817, EPI_ISL_906819, EPI_ISL_906820 | Akershus University Hospital, Department for Microbiology and Infectious Disease Control | Norwegian Institute of Public Health, Department of Virology | Kathrine Stene-Johansen, Kamilla Heddeland Instefjord, Hilde Elshaug, Atiya R Ali,Marie Paulsen Madsen, Rasmus Riis Kopperud, Hilde Vollan, Karoline Bragstad, Olav Hungnes |

| | | | |
|---|---|---|---|
| EPI_ISL_906821 | Oslo University Hospital, Department of Medical Microbiology | Norwegian Institute of Public Health, Department of Virology | Kathrine Stene-Johansen, Kamilla Heddeland Instefjord, Hilde Elshaug, Atiya R Ali,Marie Paulsen Madsen, Rasmus Riis Kopperud, Hilde Vollan, Karoline Bragstad, Olav Hungnes |
| EPI_ISL_906825 | Department of Medical Microbiology - section Molde, Molde Hospital | Norwegian Institute of Public Health, Department of Virology | Kathrine Stene-Johansen, Kamilla Heddeland Instefjord, Hilde Elshaug, Atiya R Ali,Marie Paulsen Madsen, Rasmus Riis Kopperud, Hilde Vollan, Karoline Bragstad, Olav Hungnes |
| EPI_ISL_906916, EPI_ISL_906917 | Bureau of Public Health Laboratories, Florida Department of Health (BPHL, FLDOH) | Bureau of Public Health Laboratories, Florida Department of Health (BPHL, FLDOH) | Schmedes,S., Blanton,J. |
| EPI_ISL_908934, EPI_ISL_908966, EPI_ISL_908969, EPI_ISL_908971, EPI_ISL_908972, EPI_ISL_908973, EPI_ISL_908975, EPI_ISL_908980, EPI_ISL_908981, EPI_ISL_909000, EPI_ISL_909001, EPI_ISL_909033, EPI_ISL_909045, EPI_ISL_909074, EPI_ISL_909098, EPI_ISL_909112, EPI_ISL_909115, EPI_ISL_909140, EPI_ISL_909142, EPI_ISL_909151, EPI_ISL_909164, EPI_ISL_909168, EPI_ISL_909175, EPI_ISL_909188, EPI_ISL_909190, EPI_ISL_909225, EPI_ISL_909239, EPI_ISL_909249, EPI_ISL_909253 | | | |
| see above | Lighthouse Lab in Milton Keynes | Wellcome Sanger Institute for the COVID-19 Genomics UK (COG-UK) Consortium | The Lighthouse Lab in Milton Keynes and Alex Alderton, Roberto Amato, Sonia Goncalves, Ewan Harrison, David K. Jackson, Ian Johnston, Dominic Kwiatkowski, Cordelia Langford, John Sillitoe on behalf of the Wellcome Sanger Institute COVID-19 Surveillance Team |
| EPI_ISL_909657, EPI_ISL_909659 | Labo Analyses med | National Reference Center for Viruses of Respiratory Infections, Institut Pasteur, Paris | Marion Barbet, Sylvie Behillil, Méline Bizard, Angela Brisebarre, Camille Capel, Etienne Simon-Lorière, Vincent Enouf, Maud Vanpeene, Sylvie van der Werf,Amzalag Jonas |
| EPI_ISL_909666 | Hopital Avicenne. Laboratoire central de bactériologie virologie hygiène | National Reference Center for Viruses of Respiratory Infections, Institut Pasteur, Paris | Marion Barbet, Sylvie Behillil, Méline Bizard, Angela Brisebarre, Camille Capel, Etienne Simon-Lorière, Vincent Enouf, Maud Vanpeene, Sylvie van der Werf,Brichler Ségolène |
| EPI_ISL_909667, EPI_ISL_909668 | Labo Analyses Med | National Reference Center for Viruses of Respiratory Infections, Institut Pasteur, Paris | Marion Barbet, Sylvie Behillil, Méline Bizard, Angela Brisebarre, Camille Capel, Etienne Simon-Lorière, Vincent Enouf, Maud Vanpeene, Sylvie van der Werf,Le Vicky |
| EPI_ISL_909674 | CH de Mayotte - Laboratoire de Biologie | National Reference Center for Viruses of Respiratory Infections, Institut Pasteur, Paris | Marion Barbet, Sylvie Behillil, Méline Bizard, Angela Brisebarre, Camille Capel, Etienne Simon-Lorière, Vincent Enouf, Maud Vanpeene, Sylvie van der Werf,Combe Patrice |
| EPI_ISL_909682, EPI_ISL_909683 | Hopital Avicenne. Laboratoire central de bactériologie virologie hygiène | National Reference Center for Viruses of Respiratory Infections, Institut Pasteur, Paris | Marion Barbet, Sylvie Behillil, Méline Bizard, Angela Brisebarre, Camille Capel, Etienne Simon-Lorière, Vincent Enouf, Maud Vanpeene, Sylvie van der Werf,Brichler Ségolène |
| EPI_ISL_909687, EPI_ISL_909697 | CH de Mayotte - Laboratoire de Biologie | National Reference Center for Viruses of Respiratory Infections, Institut Pasteur, Paris | Marion Barbet, Sylvie Behillil, Méline Bizard, Angela Brisebarre, Camille Capel, Etienne Simon-Lorière, Vincent Enouf, Maud Vanpeene, Sylvie van der Werf,Combe Patrice |
| EPI_ISL_909702 | CHU de Nantes - Hôtel Dieu. Laboratoire de Virologie | National Reference Center for Viruses of Respiratory Infections, Institut Pasteur, Paris | Marion Barbet, Sylvie Behillil, Méline Bizard, Angela Brisebarre, Camille Capel, Etienne Simon-Lorière, Vincent Enouf, Maud Vanpeene, Sylvie van der Werf,Bressollette CéLine |
| EPI_ISL_909707 | Labo Analyses Med | National Reference Center for Viruses of Respiratory Infections, Institut Pasteur, Paris | Marion Barbet, Sylvie Behillil, Méline Bizard, Angela Brisebarre, Camille Capel, Etienne Simon-Lorière, Vincent Enouf, Maud Vanpeene, Sylvie van der Werf |
| EPI_ISL_909708, EPI_ISL_909709, EPI_ISL_909714, EPI_ISL_909723 | CH de Mayotte - Laboratoire de Biologie | National Reference Center for Viruses of Respiratory Infections, Institut Pasteur, Paris | Marion Barbet, Sylvie Behillil, Méline Bizard, Angela Brisebarre, Camille Capel, Etienne Simon-Lorière, Vincent Enouf, Maud Vanpeene, Sylvie van der Werf,Combe Patrice |
| EPI_ISL_909724, EPI_ISL_909725, EPI_ISL_909726, EPI_ISL_909727, EPI_ISL_909729, EPI_ISL_909730, EPI_ISL_909731 | Labo Analyses Med | National Reference Center for Viruses of Respiratory Infections, Institut Pasteur, Paris | Marion Barbet, Sylvie Behillil, Méline Bizard, Angela Brisebarre, Camille Capel, Etienne Simon-Lorière, Vincent Enouf, Maud Vanpeene, Sylvie van der Werf,Nizard Patrice |
| EPI_ISL_909733, EPI_ISL_909734, EPI_ISL_909735 | Labo Analyses Med | National Reference Center for Viruses of Respiratory Infections, Institut Pasteur, Paris | Marion Barbet, Sylvie Behillil, Méline Bizard, Angela Brisebarre, Camille Capel, Etienne Simon-Lorière, Vincent Enouf, Maud Vanpeene, Sylvie van der Werf |
| EPI_ISL_909747, EPI_ISL_909752, EPI_ISL_909753, EPI_ISL_909754, EPI_ISL_909758, EPI_ISL_909759, EPI_ISL_909760, EPI_ISL_909771 | Charité Universitätsmedizin Berlin, Institut für Virologie/Labor Berlin | Charité Universitätsmedizin Berlin, Institut für Virologie | Victor M Corman, Barbara Mühlemann, Jörn Beheim-Schwarzbach, Tobias Bleicker, Julia Tesch, Talitha Veith, Julia Schneider, Terry Jones, Christian Drosten |
| EPI_ISL_909775, EPI_ISL_909816, EPI_ISL_909817, EPI_ISL_909818, EPI_ISL_909827, EPI_ISL_909828, EPI_ISL_909829, EPI_ISL_909830, EPI_ISL_909831, EPI_ISL_909833, EPI_ISL_909834, EPI_ISL_909841, EPI_ISL_909844, EPI_ISL_909847, EPI_ISL_909851, EPI_ISL_909854, EPI_ISL_909855, EPI_ISL_909856, EPI_ISL_909857, EPI_ISL_909859, EPI_ISL_909860, EPI_ISL_909861, EPI_ISL_909862, EPI_ISL_909864, EPI_ISL_909865, EPI_ISL_909866, EPI_ISL_909867, EPI_ISL_909869, EPI_ISL_909870, EPI_ISL_909871, EPI_ISL_909872, EPI_ISL_909875, EPI_ISL_909903, EPI_ISL_909905, EPI_ISL_909908, EPI_ISL_909909, EPI_ISL_909910, EPI_ISL_909911, EPI_ISL_909912, EPI_ISL_909914, EPI_ISL_909918, EPI_ISL_909920, EPI_ISL_909921, EPI_ISL_909923, EPI_ISL_909924, EPI_ISL_909929, EPI_ISL_909933, EPI_ISL_909936, EPI_ISL_909938, EPI_ISL_909940, EPI_ISL_909942, EPI_ISL_909943, EPI_ISL_909944 | | | |
| see above | National Virus Reference Laboratory | National Virus Reference Laboratory | Michael Carr, Gabriel Gonzalez, Jonathan Dean, Cillian F De Gascun |
| EPI_ISL_911134, EPI_ISL_911136, EPI_ISL_911137, EPI_ISL_911138, EPI_ISL_911139, EPI_ISL_911140, EPI_ISL_911141, EPI_ISL_911142, EPI_ISL_911143, EPI_ISL_911144, EPI_ISL_911145, EPI_ISL_911146, EPI_ISL_911147, EPI_ISL_911148, EPI_ISL_911149, EPI_ISL_911150, EPI_ISL_911151, EPI_ISL_911152, EPI_ISL_911153, EPI_ISL_911154, EPI_ISL_911155, EPI_ISL_911156, EPI_ISL_911157, EPI_ISL_911158, EPI_ISL_911159, EPI_ISL_911160, EPI_ISL_911161, EPI_ISL_911162, EPI_ISL_911163, EPI_ISL_911164, EPI_ISL_911165, EPI_ISL_911166, EPI_ISL_911232, EPI_ISL_911234 | | | |
| see above | Laboratoire national de sante, Microbiology, Virology | Laboratoire national de sante, Microbiology, Microbial Genomics Platform | Anke Wienecke-Baldacchino, Catherine Ragimbeau,Jessica Tapp, Fatu Djabi, Lise Pignon, Raoul Salmon, Tamir Abdelrahman |
| EPI_ISL_911242, EPI_ISL_911243, EPI_ISL_911244, EPI_ISL_911246 | National Virus Reference Laboratory | National Virus Reference Laboratory | Michael Carr, Gabriel Gonzalez, Jonathan Dean, Cillian F De Gascun |
| EPI_ISL_911295, EPI_ISL_911305, EPI_ISL_911315, EPI_ISL_911325, EPI_ISL_911332, EPI_ISL_911340, EPI_ISL_911349 | Servicio de Microbiología, Hospital Universitario Son Espases | SeqCOVID-SPAIN consortium/IBV(CSIC) | Carla López-Causapé, Jordi Reina, Antonio Oliver and SeqCOVID-SPAIN consortium |
| EPI_ISL_911383, EPI_ISL_911384, EPI_ISL_911385 | University of Michigan Clinical Microbiology Laboratory | Lauring Lab, University of Michigan, Department of Microbiology and Immunology | Valesano |
| EPI_ISL_911580 | Clinical Molecular Microbiology Laboratory, UNC Hospitals | Jeremy Wang | Jeremy Wang, Alexander Rubinsteyn, Colleen Rice, Jason Smedberg, Melissa Miller, Corbin Jones, Robert Hagan |
| EPI_ISL_911746, EPI_ISL_911747 | Toronto Invasive Bacterial Diseases Network | McMaster University | Allison McGeer, Patryk Aftanas, Hooman Derakhshani, Angel Li, Kuganya Nirmalarajah, Emily Panousis, Ahmed Draia, Jalees Nasir, Michael Surette, Samira Mubareka, Andrew G. McArthur |
| EPI_ISL_911767, EPI_ISL_911774, EPI_ISL_911792, EPI_ISL_911793, EPI_ISL_911796, EPI_ISL_911797, EPI_ISL_911798, EPI_ISL_911902, EPI_ISL_911903, EPI_ISL_911907, EPI_ISL_911908, EPI_ISL_911909, EPI_ISL_911910 | | | |
| see above | Johns Hopkins Hospital Department of Pathology | Johns Hopkins Hospital Department of Pathology | C. Paul Morris, Chun Huai Luo, Adannaya Amadi, Matthew Schwartz, Nicholas Gallagher, Heba H. Mostafa |
| EPI_ISL_911918, EPI_ISL_911919, EPI_ISL_911920, EPI_ISL_911921, EPI_ISL_911922 | SRL (Sonic Reference Laboratory - Sonic Healthcare) | Pathogen Discovery, Respiratory Viruses Branch, Division of Viral Diseases, Centers for Disease Control and Prevention | Ying Tao, Yan Li, Jing Zhang, Krista Queen, Anna Uehara, Peter Cook, Clinton R. Paden, Haibin Wang, Suxiang Tong |
| EPI_ISL_911952, EPI_ISL_911953, EPI_ISL_911954, EPI_ISL_911955, EPI_ISL_911956, EPI_ISL_911957, EPI_ISL_911958, EPI_ISL_911959, EPI_ISL_911960, EPI_ISL_911961, EPI_ISL_911962, EPI_ISL_911963, EPI_ISL_911964, EPI_ISL_911965 | | | |
| see above | Wyoming Public Health Laboratory | Wyoming Public Health Laboratory | Noah Hull, Taylor Fearing, Lynette Gumbleton, Channing Weber, Ashley Norberg, Bailey Bowcutt, and Wanda Manley |
| EPI_ISL_912094, EPI_ISL_912096, EPI_ISL_912097, EPI_ISL_912098, EPI_ISL_912099, EPI_ISL_912100, EPI_ISL_912101, EPI_ISL_912102, EPI_ISL_912103, EPI_ISL_912104, EPI_ISL_912105, EPI_ISL_912106, EPI_ISL_912107, EPI_ISL_912108 | | | |
| see above | Washington State Department of Health | Seattle Flu Study | Deborah A. Nickerson, Chris D. Frazar, Jover Lee, Benjamin Pelle, Erica Ryke, Matthew Richardson, Amanda Adler, Elisabeth Brandstetter, Peter D. Han, Kairsten Fay, Misja Ilcisin, Kirsten Lacombe, Thomas R. Sibley, Melissa Truong, Caitlin R. Wolf, Romesh Gautom, Geoff Melly, Brian Hiatt, Philip Dykema, Scott Lindquist, Michael Boeckh, Janet A. Englund, Michael Famulare, Barry R. Lutz, Mark J. Rieder, Lea M. Starita, Matthew Thompson, Helen Y. Chu, Jay Shendure, Trevor Bedford |
| EPI_ISL_912227, EPI_ISL_912228, EPI_ISL_912229, EPI_ISL_912230, EPI_ISL_912231, EPI_ISL_912232, EPI_ISL_912233, EPI_ISL_912234, EPI_ISL_912235, EPI_ISL_912247, EPI_ISL_912249 | | | |
| see above | Charité Universitätsmedizin Berlin, Institut für Virologie/Labor | Charité Universitätsmedizin Berlin, Institut für Virologie | Victor M Corman, Barbara Mühlemann, Jörn Beheim-Schwarzbach, Tobias Bleicker, Julia Tesch, Talitha Veith, Julia Schneider, Terry Jones, Christian |

| | | | |
|---|---|---|---|
| | Berlin | | Drosten |
| EPI_ISL_912274, EPI_ISL_912275, EPI_ISL_912277, EPI_ISL_912278, EPI_ISL_912281, EPI_ISL_912284, EPI_ISL_912286, EPI_ISL_912290, EPI_ISL_912303, EPI_ISL_912310, EPI_ISL_912314 | | | |
| see above | Hospital General Universitario Gregorio Marañón | SeqCOVID-SPAIN consortium / IBV (CSIC) | Darío García de Viedma, Laura Pérez-Lago, Pedro J Sola-Campoy, Sergio Buenestado-Serrano, Marta Herranz, Victor Manuel de la Cueva, Julia Suárez, Pilar Catalán, Patricia Muñoz and SeqCOVID-SPAIN consortium |
| EPI_ISL_912316 | Hospital General Universitario Gregorio Marañón | SeqCOVID-SPAIN consortium / IBV (CSIC) | Sergio Buenestado Serrano, Pedro J. Sola Campoy, Laura Pérez-Lago, Pilar Catalán, Arturo fraile Torres, Andrés Von Wernitz, Carmen del Arco, Patricia Muñoz, Laura Cardeñoso, Darío García de Viedma and SeqCOVID-SPAIN consortium |
| EPI_ISL_912320, EPI_ISL_912323, EPI_ISL_912328, EPI_ISL_912329 | Hospital General Universitario Gregorio Marañón | SeqCOVID-SPAIN consortium / IBV (CSIC) | Darío García de Viedma, Laura Pérez-Lago, Pedro J Sola-Campoy, Sergio Buenestado-Serrano, Marta Herranz, Victor Manuel de la Cueva, Julia Suárez, Pilar Catalán, Patricia Muñoz and SeqCOVID-SPAIN consortium |
| EPI_ISL_912337, EPI_ISL_912338 | Hospital General Universitario Gregorio Marañón | SeqCOVID-SPAIN consortium / IBV (CSIC) | Sergio Buenestado Serrano, Pedro J. Sola Campoy, Laura Pérez-Lago, Pilar Catalán, Arturo fraile Torres, Andrés Von Wernitz, Carmen del Arco, Patricia Muñoz, Laura Cardeñoso, Darío García de Viedma and SeqCOVID-SPAIN consortium |
| EPI_ISL_912340, EPI_ISL_912343, EPI_ISL_912344, EPI_ISL_912348, EPI_ISL_912350 | Hospital General Universitario Gregorio Marañón | SeqCOVID-SPAIN consortium / IBV (CSIC) | Darío García de Viedma, Laura Pérez-Lago, Pedro J Sola-Campoy, Sergio Buenestado-Serrano, Marta Herranz, Victor Manuel de la Cueva, Julia Suárez, Pilar Catalán, Patricia Muñoz and SeqCOVID-SPAIN consortium |
| EPI_ISL_912402 | CH de Mayotte - Laboratoire de Biologie | National Reference Center for Viruses of Respiratory Infections, Institut Pasteur, Paris | Marion Barbet, Sylvie Behillil, Méline Bizard, Angela Brisebarre, Camille Capel, Etienne Simon-Lorière, Vincent Enouf, Maud Vanpeene, Sylvie van der Werf,Combe Patrice |
| EPI_ISL_912424, EPI_ISL_912428, EPI_ISL_912430, EPI_ISL_912431, EPI_ISL_912432, EPI_ISL_912433, EPI_ISL_912434, EPI_ISL_912435, EPI_ISL_912436, EPI_ISL_912437, EPI_ISL_912438, EPI_ISL_912439, EPI_ISL_912440, EPI_ISL_912441, EPI_ISL_912442, EPI_ISL_912443, EPI_ISL_912444, EPI_ISL_912445, EPI_ISL_912446, EPI_ISL_912447, EPI_ISL_912448 | | | |
| see above | KU Leuven, Rega Institute, Clinical and Epidemiological Virology | KU Leuven, Rega Institute, Clinical and Epidemiological Virology | Tony Wawina-Bokalanga, Bert Vanmechelen, Joan Marti-Carerras, Piet Maes |
| EPI_ISL_912554, EPI_ISL_912555, EPI_ISL_912556 | C.H.R. d'Orléans - Hôpital de la Source | National Reference Center for Viruses of Respiratory Infections, Institut Pasteur, Paris | Marion Barbet, Sylvie Behillil, Méline Bizard, Angela Brisebarre, Camille Capel, Etienne Simon-Lorière, Vincent Enouf, Maud Vanpeene, Sylvie van der Werf,Guinard Jérome |
| EPI_ISL_912559 | Hopital Avicenne Laboratoire central de bactériologie virologie hygiène | National Reference Center for Viruses of Respiratory Infections, Institut Pasteur, Paris | Marion Barbet, Sylvie Behillil, Méline Bizard, Angela Brisebarre, Camille Capel, Etienne Simon-Lorière, Vincent Enouf, Maud Vanpeene, Sylvie van der Werf,Brichler Ségolène |
| EPI_ISL_912563, EPI_ISL_912564 | Laboratoire Central - American Hospital of Paris | National Reference Center for Viruses of Respiratory Infections, Institut Pasteur, Paris | Marion Barbet, Sylvie Behillil, Méline Bizard, Angela Brisebarre, Camille Capel, Etienne Simon-Lorière, Vincent Enouf, Maud Vanpeene, Sylvie van der Werf,Ramirez José-Manuel |
| EPI_ISL_912566 | Hopital | National Reference Center for Viruses of Respiratory Infections, Institut Pasteur, Paris | Marion Barbet, Sylvie Behillil, Méline Bizard, Angela Brisebarre, Camille Capel, Etienne Simon-Lorière, Vincent Enouf, Maud Vanpeene, Sylvie van der Werf |
| EPI_ISL_912569 | Labo Analyses Med | National Reference Center for Viruses of Respiratory Infections, Institut Pasteur, Paris | Marion Barbet, Sylvie Behillil, Méline Bizard, Angela Brisebarre, Camille Capel, Etienne Simon-Lorière, Vincent Enouf, Maud Vanpeene, Sylvie van der Werf |
| EPI_ISL_912573 | C.H.R. d'Orléans - Hôpital de la Source | National Reference Center for Viruses of Respiratory Infections, Institut Pasteur, Paris | Marion Barbet, Sylvie Behillil, Méline Bizard, Angela Brisebarre, Camille Capel, Etienne Simon-Lorière, Vincent Enouf, Maud Vanpeene, Sylvie van der Werf,Guinard Jérome |
| EPI_ISL_912574 | Laboratoire Central - American Hospital of Paris | National Reference Center for Viruses of Respiratory Infections, Institut Pasteur, Paris | Marion Barbet, Sylvie Behillil, Méline Bizard, Angela Brisebarre, Camille Capel, Etienne Simon-Lorière, Vincent Enouf, Maud Vanpeene, Sylvie van der Werf,Ramirez José-Manuel |
| EPI_ISL_912575, EPI_ISL_912576 | Laboratoire de Virologie du CHU de Lille Bâtiment Paul Boulanger | National Reference Center for Viruses of Respiratory Infections, Institut Pasteur, Paris | Marion Barbet, Sylvie Behillil, Méline Bizard, Angela Brisebarre, Camille Capel, Etienne Simon-Lorière, Vincent Enouf, Maud Vanpeene, Sylvie van der Werf,Guigon Aurélie |
| EPI_ISL_912582, EPI_ISL_912583 | CENTRE HOSPITALIER LEON BINET | National Reference Center for Viruses of Respiratory Infections, Institut Pasteur, Paris | Marion Barbet, Sylvie Behillil, Méline Bizard, Angela Brisebarre, Camille Capel, Etienne Simon-Lorière, Vincent Enouf, Maud Vanpeene, Sylvie van der Werf,Meziane Ilham |
| EPI_ISL_912599 | HOPITAL SIMONE VEIL | National Reference Center for Viruses of Respiratory Infections, Institut Pasteur, Paris | Marion Barbet, Sylvie Behillil, Méline Bizard, Angela Brisebarre, Camille Capel, Etienne Simon-Lorière, Vincent Enouf, Maud Vanpeene, Sylvie van der Werf,Moreau Farida |
| EPI_ISL_912600 | Labo Analyses Med | National Reference Center for Viruses of Respiratory Infections, Institut Pasteur, Paris | Marion Barbet, Sylvie Behillil, Méline Bizard, Angela Brisebarre, Camille Capel, Etienne Simon-Lorière, Vincent Enouf, Maud Vanpeene, Sylvie van der Werf,Gaumery Bertrand |
| EPI_ISL_912602 | Laboratoire de Virologie du CHU de Lille Bâtiment Paul Boulanger | National Reference Center for Viruses of Respiratory Infections, Institut Pasteur, Paris | Marion Barbet, Sylvie Behillil, Méline Bizard, Angela Brisebarre, Camille Capel, Etienne Simon-Lorière, Vincent Enouf, Maud Vanpeene, Sylvie van der Werf,Guigon Aurélie |
| EPI_ISL_912604 | Labo Analyses Med | National Reference Center for Viruses of Respiratory Infections, Institut Pasteur, Paris | Marion Barbet, Sylvie Behillil, Méline Bizard, Angela Brisebarre, Camille Capel, Etienne Simon-Lorière, Vincent Enouf, Maud Vanpeene, Sylvie van der Werf |
| EPI_ISL_912607, EPI_ISL_912608, EPI_ISL_912609, EPI_ISL_912610, EPI_ISL_912611 | Labo Analyses Med | National Reference Center for Viruses of Respiratory Infections, Institut Pasteur, Paris | Marion Barbet, Sylvie Behillil, Méline Bizard, Angela Brisebarre, Camille Capel, Etienne Simon-Lorière, Vincent Enouf, Maud Vanpeene, Sylvie van der Werf,Holstein Anne |
| EPI_ISL_912612, EPI_ISL_912613 | Centre Hospitalier de Compiègne Laboratoire Compiègne | National Reference Center for Viruses of Respiratory Infections, Institut Pasteur, Paris | Marion Barbet, Sylvie Behillil, Méline Bizard, Angela Brisebarre, Camille Capel, Etienne Simon-Lorière, Vincent Enouf, Maud Vanpeene, Sylvie van der Werf,Romand Sylvie |
| EPI_ISL_912622 | Labo Analyses Med | National Reference Center for Viruses of Respiratory Infections, Institut Pasteur, Paris | Marion Barbet, Sylvie Behillil, Méline Bizard, Angela Brisebarre, Camille Capel, Etienne Simon-Lorière, Vincent Enouf, Maud Vanpeene, Sylvie van der Werf |
| EPI_ISL_912642 | Hôpital Henri Mondor | Department of Virology, Henri Mondor University Hospital, Assistance Publique Hôpitaux de Paris, Université Paris-Est Créteil, INSERM U955 | Christophe Rodriguez, Slim Fourati, Vanessa Demontant, Guillaume Gricourt, Melissa N'Debi, Alexandre Soulier, Elisabeth Trawinski, Jean-Michel Pawlotsky |
| EPI_ISL_912660 | Hôpital Pitié-Salpêtrière | Department of Virology, Henri Mondor University Hospital, Assistance Publique Hôpitaux de Paris, Université Paris-Est Créteil, INSERM U955 | Christophe Rodriguez, Slim Fourati, Vanessa Demontant, Guillaume Gricourt, Melissa N'Debi, Alexandre Soulier, Elisabeth Trawinski, Jean-Michel Pawlotsky |
| EPI_ISL_912695, EPI_ISL_912713, EPI_ISL_912734 | Hôpital Henri Mondor | Department of Virology, Henri Mondor University Hospital, Assistance Publique Hôpitaux de Paris, Université Paris-Est Créteil, INSERM U955 | Christophe Rodriguez, Slim Fourati, Vanessa Demontant, Guillaume Gricourt, Melissa N'Debi, Alexandre Soulier, Elisabeth Trawinski, Jean-Michel Pawlotsky |
| EPI_ISL_912791, EPI_ISL_912792 | Hôpital Pitié-Salpêtrière | Department of Virology, Henri Mondor University Hospital, Assistance Publique Hôpitaux de Paris, Université Paris-Est Créteil, INSERM U955 | Christophe Rodriguez, Slim Fourati, Vanessa Demontant, Guillaume Gricourt, Melissa N'Debi, Alexandre Soulier, Elisabeth Trawinski, Jean-Michel Pawlotsky |
| EPI_ISL_912812, EPI_ISL_912814, EPI_ISL_912818, EPI_ISL_912821, EPI_ISL_912822, EPI_ISL_912823, EPI_ISL_912827, EPI_ISL_912828, EPI_ISL_912830, EPI_ISL_912837, EPI_ISL_912838, EPI_ISL_912840, EPI_ISL_912841, EPI_ISL_912844, EPI_ISL_912845, EPI_ISL_912851, EPI_ISL_912852, EPI_ISL_912853, EPI_ISL_912854, EPI_ISL_912855, EPI_ISL_912856, EPI_ISL_912857, EPI_ISL_912858, EPI_ISL_912859, EPI_ISL_912860, EPI_ISL_912861, EPI_ISL_912862, EPI_ISL_912899, EPI_ISL_912908 | | | |
| see above | Hôpital Henri Mondor | Department of Virology, Henri Mondor University Hospital, Assistance Publique Hôpitaux de Paris, Université Paris-Est Créteil, INSERM U955 | Christophe Rodriguez, Slim Fourati, Vanessa Demontant, Guillaume Gricourt, Melissa N'Debi, Alexandre Soulier, Elisabeth Trawinski, Jean-Michel Pawlotsky |
| EPI_ISL_912912, EPI_ISL_912913, EPI_ISL_912914, EPI_ISL_912915, EPI_ISL_912917, EPI_ISL_912918 | Hôpital Pitié-Salpêtrière | Department of Virology, Henri Mondor University Hospital, Assistance Publique Hôpitaux de Paris, Université Paris-Est Créteil, INSERM U955 | Christophe Rodriguez, Slim Fourati, Vanessa Demontant, Guillaume Gricourt, Melissa N'Debi, Alexandre Soulier, Elisabeth Trawinski, Jean-Michel Pawlotsky |

| | | | |
|---|---|---|---|
| EPI_ISL_912953, EPI_ISL_912954, EPI_ISL_912955, EPI_ISL_912956, EPI_ISL_912957 | Hôpital Henri Mondor | Department of Virology, Henri Mondor University Hospital, Assistance Publique Hôpitaux de Paris, Université Paris-Est Créteil, INSERM U955 | Christophe Rodriguez, Slim Fourati, Vanessa Demontant, Guillaume Gricourt, Melissa N'Debi, Alexandre Soulier, Elisabeth Trawinski, Jean-Michel Pawlotsky |
| EPI_ISL_913028, EPI_ISL_913030, EPI_ISL_913031, EPI_ISL_913036, EPI_ISL_913037 | Hospital los Arcos | Instituto de Salud Carlos III | Iglesias-Caballero, M. Camarero, S. Sandonís,V. Vázquez, S. Pozo, F. Casas, I. Jiménez, P. Zaballos, A. Monzón, S. Varona, S. Cuesta, I. Cámara, M. |
| EPI_ISL_913057 | Hospital San Pedro de Alcántara | Instituto de Salud Carlos III | Iglesias-Caballero, M. Camarero, S. Sandonís,V. Vázquez, S. Pozo, F. Casas, I. Jiménez, P. Zaballos, A. Monzón, S. Varona, S. Cuesta, I. Rodríguez, G. |
| EPI_ISL_913061, EPI_ISL_913062 | Hospital Nuestra Señora de Sonsoles | Instituto de Salud Carlos III | Iglesias-Caballero, M. Camarero, S. Sandonís,V. Vázquez, S. Pozo, F. Casas, I. Jiménez, P. Zaballos, A. Monzón, S. Varona, S. Cuesta, I. Sánchez, R. |
| EPI_ISL_913114 | CHU Purpan - Laboratoire de Virologie - Institut Fédératif de Biologie | CHU Purpan - Laboratoire de Virologie - Institut Fédératif de Biologie | Latour J., Ranger N., Dubois M., Carcenac R., Harter A., Boyer P., Tremeaux P., Izopet J. |
| EPI_ISL_913116, EPI_ISL_913118, EPI_ISL_913119, EPI_ISL_913120, EPI_ISL_913126, EPI_ISL_913128, EPI_ISL_913129, EPI_ISL_913131, EPI_ISL_913133, EPI_ISL_913137, EPI_ISL_913138, EPI_ISL_913139, EPI_ISL_913141, EPI_ISL_913142, EPI_ISL_913144, EPI_ISL_913150, EPI_ISL_913174, EPI_ISL_913177, EPI_ISL_913178, EPI_ISL_913180, EPI_ISL_913182, EPI_ISL_913183, EPI_ISL_913184, EPI_ISL_913185, EPI_ISL_913186, EPI_ISL_913187, EPI_ISL_913188, EPI_ISL_913189, EPI_ISL_913190, EPI_ISL_913191, EPI_ISL_913192, EPI_ISL_913193, EPI_ISL_913194, EPI_ISL_913195, EPI_ISL_913196, EPI_ISL_913197, EPI_ISL_913198, EPI_ISL_913199, EPI_ISL_913200 | | | |
| see above | University of Michigan Clinical Microbiology Laboratory | Lauring Lab, University of Michigan, Department of Microbiology and Immunology | Valesano |
| EPI_ISL_913266 | Unilabs AB Skovde | The Public Health Agency of Sweden | Anna-Malin Linde, Maria Lind Karlberg, Carlo Berg, Oskar Karlsson Lindsjo, Sofia Stamouli, Reza Advani, Mattias Haukland, Petra Holmstrom, Noura Walai, Petra Edquist, Mia Brytting, Anna Risberg, Karin Tegmark-Wisell |
| EPI_ISL_913373 | Klinisk mikrobiologi | The Public Health Agency of Sweden | Anna-Malin Linde, Maria Lind Karlberg, Carlo Berg, Oskar Karlsson Lindsjo, Sofia Stamouli, Reza Advani, Mattias Haukland, Petra Holmstrom, Noura Walai, Petra Edquist, Mia Brytting, Anna Risberg, Karin Tegmark-Wisell |
| EPI_ISL_913394, EPI_ISL_913396, EPI_ISL_913401, EPI_ISL_913404, EPI_ISL_913405, EPI_ISL_913406, EPI_ISL_913409, EPI_ISL_913411, EPI_ISL_913420 | Massachusetts State Public Health Laboratory | Massachusetts State Public Health Laboratory | Andrew Lang, Timelia Fink, Glen Gallagher, Sandra Smole |
| EPI_ISL_913455 | Unilabs AB Skovde | The Public Health Agency of Sweden | Anna-Malin Linde, Maria Lind Karlberg, Carlo Berg, Oskar Karlsson Lindsjo, Sofia Stamouli, Reza Advani, Mattias Haukland, Petra Holmstrom, Noura Walai, Petra Edquist, Mia Brytting, Anna Risberg, Karin Tegmark-Wisell |
| EPI_ISL_913456 | Synlab Medilab, Mikrobiologi | The Public Health Agency of Sweden | Anna-Malin Linde, Maria Lind Karlberg, Carlo Berg, Oskar Karlsson Lindsjo, Sofia Stamouli, Reza Advani, Mattias Haukland, Petra Holmstrom, Noura Walai, Petra Edquist, Mia Brytting, Anna Risberg, Karin Tegmark-Wisell |
| EPI_ISL_913470 | Klinisk mikrobiologi | The Public Health Agency of Sweden | Anna-Malin Linde, Maria Lind Karlberg, Carlo Berg, Oskar Karlsson Lindsjo, Sofia Stamouli, Reza Advani, Mattias Haukland, Petra Holmstrom, Noura Walai, Petra Edquist, Mia Brytting, Anna Risberg, Karin Tegmark-Wisell |
| EPI_ISL_913514, EPI_ISL_913515, EPI_ISL_913516 | University of Michigan Clinical Microbiology Laboratory | Lauring Lab, University of Michigan, Department of Microbiology and Immunology | Valesano |
| EPI_ISL_913584 | M Health Fairview | Minnesota Department of Health, Public Health Laboratory | Alexandra Lorentz, Jacob Garfin, Matt Plumb, and Xiong Wang |
| EPI_ISL_913586, EPI_ISL_913588, EPI_ISL_913589, EPI_ISL_913590, EPI_ISL_913591 | Vault Health | Minnesota Department of Health, Public Health Laboratory | Alexandra Lorentz, Jacob Garfin, Matt Plumb, and Xiong Wang |
| EPI_ISL_913616 | Michigan Department of Health and Human Services, Bureau of Laboratories | Michigan Department of Health and Human Services, Bureau of Laboratories | Blankenship HM, Riner D, Soehnlen MK |
| EPI_ISL_913667 | LA Office of Public Health Laboratories | Centers for Disease Control and Prevention Division of Viral Diseases, Pathogen Discovery | Ying Tao, Yan Li, Jing Zhang, Krista Queen, Anna Uehara, Peter Cook, Clinton R. Paden, Haibin Wang, Suxiang Tong |
| EPI_ISL_913748, EPI_ISL_913749, EPI_ISL_913750, EPI_ISL_913751, EPI_ISL_913752, EPI_ISL_913753, EPI_ISL_913754, EPI_ISL_913755, EPI_ISL_913756, EPI_ISL_913758, EPI_ISL_913759, EPI_ISL_913760, EPI_ISL_913761, EPI_ISL_913762, EPI_ISL_913763, EPI_ISL_913764, EPI_ISL_913765, EPI_ISL_913768, EPI_ISL_913794, EPI_ISL_913796, EPI_ISL_913797, EPI_ISL_913799, EPI_ISL_913800, EPI_ISL_913810, EPI_ISL_913812, EPI_ISL_913813, EPI_ISL_913814, EPI_ISL_913816, EPI_ISL_913817, EPI_ISL_913819, EPI_ISL_913820, EPI_ISL_913821, EPI_ISL_913823, EPI_ISL_913824, EPI_ISL_913826, EPI_ISL_913827, EPI_ISL_913828, EPI_ISL_913830, EPI_ISL_913831, EPI_ISL_913833, EPI_ISL_913834, EPI_ISL_913835, EPI_ISL_913837, EPI_ISL_913838, EPI_ISL_913840, EPI_ISL_913841, EPI_ISL_913843, EPI_ISL_913844, EPI_ISL_913845, EPI_ISL_913847, EPI_ISL_913848, EPI_ISL_913850, EPI_ISL_913851, EPI_ISL_913853, EPI_ISL_913854, EPI_ISL_913855, EPI_ISL_913857, EPI_ISL_913858, EPI_ISL_913867, EPI_ISL_913868, EPI_ISL_913869, EPI_ISL_913871, EPI_ISL_913872, EPI_ISL_913874, EPI_ISL_913875, EPI_ISL_913876, EPI_ISL_913878 | | | |
| see above | KU Leuven, Rega Institute, Clinical and Epidemiological Virology | KU Leuven, Rega Institute, Clinical and Epidemiological Virology | Tony Wawina-Bokalanga, Bert Vanmechelen, Joan Marti-Carerras, Piet Maes |
| EPI_ISL_913985, EPI_ISL_913986, EPI_ISL_913987, EPI_ISL_913988, EPI_ISL_913989 | TX DSHS, Lab Services Section MC 1947 | Pathogen Discovery, Respiratory Viruses Branch, Division of Viral Diseases, Centers for Disease Control and Prevention | Ying Tao, Yan Li, Jing Zhang, Krista Queen, Anna Uehara, Peter Cook, Clinton R. Paden, Haibin Wang, Suxiang Tong |
| EPI_ISL_913992, EPI_ISL_913993, EPI_ISL_913994 | TN Division of Laboratory Services | Pathogen Discovery, Respiratory Viruses Branch, Division of Viral Diseases, Centers for Disease Control and Prevention | Ying Tao, Yan Li, Jing Zhang, Krista Queen, Anna Uehara, Peter Cook, Clinton R. Paden, Haibin Wang, Suxiang Tong |
| EPI_ISL_914027, EPI_ISL_914028, EPI_ISL_914029 | IL Department of Public Health Chicago Laboratory | Pathogen Discovery, Respiratory Viruses Branch, Division of Viral Diseases, Centers for Disease Control and Prevention | Ying Tao, Yan Li, Jing Zhang, Krista Queen, Anna Uehara, Peter Cook, Clinton R. Paden, Haibin Wang, Suxiang Tong |
| EPI_ISL_914738, EPI_ISL_914742, EPI_ISL_914752, EPI_ISL_914754, EPI_ISL_914772 | Wyoming Public Health Laboratory | Wyoming Public Health Laboratory | Noah Hull, Taylor Fearing, Lynette Gumbleton, Channing Weber, Ashley Norberg, Bailey Bowcutt, and Wanda Manley |
| EPI_ISL_914832 | LABORATORIO CLINICO LABIN | Inciensa, Instituto Costarricense de Investigación y Enseñanza en Nutrición y Salud | Francisco Duarte, Hebleen Porras, Claudio Soto-Garita, Estela Cordero, Adriana Godínez, Melany Calderón & Pei Ling Chan Ma |
| EPI_ISL_914835 | AREA DE SALUD PARAISO-CERVANTES | Inciensa, Instituto Costarricense de Investigación y Enseñanza en Nutrición y Salud | Francisco Duarte, Hebleen Porras, Claudio Soto-Garita, Estela Cordero, Adriana Godínez, Melany Calderón & Mariel López |
| EPI_ISL_914836 | HOSPITAL MEXICO | Inciensa, Instituto Costarricense de Investigación y Enseñanza en Nutrición y Salud | Francisco Duarte, Hebleen Porras, Claudio Soto-Garita, Estela Cordero, Adriana Godínez, Melany Calderón & Mariel López |
| EPI_ISL_914837, EPI_ISL_914838, EPI_ISL_914839 | HOSPITAL MEXICO | Inciensa, Instituto Costarricense de Investigación y Enseñanza en Nutrición y Salud | Francisco Duarte, Hebleen Porras, Claudio Soto-Garita, Estela Cordero, Adriana Godínez, Melany Calderón & Teresita Somogyi |
| EPI_ISL_914866, EPI_ISL_914867 | Kansas Health and Environmental Lab | Kansas Health and Environmental Lab | Mike Grose, Paige Drury, Carissa Robertson, Ben Olsen, and Phil Adam |
| EPI_ISL_914890, EPI_ISL_914891, EPI_ISL_914892, EPI_ISL_914893, EPI_ISL_914894, EPI_ISL_914895, EPI_ISL_914896, EPI_ISL_914897, EPI_ISL_914898, EPI_ISL_914899, EPI_ISL_914900, EPI_ISL_914901, EPI_ISL_914902, EPI_ISL_914903, EPI_ISL_914904, EPI_ISL_914905, EPI_ISL_914906, EPI_ISL_914907, EPI_ISL_914908, EPI_ISL_914909, EPI_ISL_914910, EPI_ISL_914911, EPI_ISL_914912, EPI_ISL_914913, EPI_ISL_914915, EPI_ISL_914916, EPI_ISL_914917, EPI_ISL_914918, EPI_ISL_914919, EPI_ISL_914920, EPI_ISL_914921, EPI_ISL_914922, EPI_ISL_914923, EPI_ISL_914924, EPI_ISL_914925, EPI_ISL_914926, EPI_ISL_914927, EPI_ISL_914928, EPI_ISL_914929, EPI_ISL_914930, EPI_ISL_914931, EPI_ISL_914932, EPI_ISL_914933, EPI_ISL_914934, EPI_ISL_914935, EPI_ISL_914936, EPI_ISL_914937, EPI_ISL_914938, EPI_ISL_914939, EPI_ISL_914940, EPI_ISL_914941, EPI_ISL_914942, EPI_ISL_914943, EPI_ISL_914945, EPI_ISL_914946, EPI_ISL_914947, EPI_ISL_914948, EPI_ISL_914949, EPI_ISL_914950, EPI_ISL_914951, EPI_ISL_914952, EPI_ISL_914953, EPI_ISL_914954, EPI_ISL_914955, EPI_ISL_914956, EPI_ISL_914957, EPI_ISL_914958, EPI_ISL_914959, EPI_ISL_914960, EPI_ISL_914961, EPI_ISL_914962, EPI_ISL_914963, EPI_ISL_914964, EPI_ISL_914965, EPI_ISL_914966, EPI_ISL_914967, EPI_ISL_914968, EPI_ISL_914969, EPI_ISL_915195, EPI_ISL_915196, EPI_ISL_915197, EPI_ISL_915199, EPI_ISL_915200, EPI_ISL_915201, EPI_ISL_915203, EPI_ISL_915204, EPI_ISL_915205, EPI_ISL_915208, EPI_ISL_915209, EPI_ISL_915210, EPI_ISL_915214, EPI_ISL_915216, EPI_ISL_915217, EPI_ISL_915218, EPI_ISL_915219, EPI_ISL_915220, EPI_ISL_915221, EPI_ISL_915222, EPI_ISL_915223, EPI_ISL_915225, EPI_ISL_915229, EPI_ISL_915233, EPI_ISL_915240, EPI_ISL_915241, EPI_ISL_915242, EPI_ISL_915243, EPI_ISL_915244, EPI_ISL_915245, | | | |

| | | | |
|---|---|---|---|
| EPI_ISL_915246, EPI_ISL_915247, EPI_ISL_915248, EPI_ISL_915249, EPI_ISL_915250, EPI_ISL_915251, EPI_ISL_915252, EPI_ISL_915253, EPI_ISL_915254, EPI_ISL_915255, EPI_ISL_915256, EPI_ISL_915257, EPI_ISL_915258, EPI_ISL_915259, EPI_ISL_915260, EPI_ISL_915261, EPI_ISL_915262, EPI_ISL_915263, EPI_ISL_915265, EPI_ISL_915266, EPI_ISL_915275, EPI_ISL_915277, EPI_ISL_915290, EPI_ISL_915291, EPI_ISL_915292, EPI_ISL_915293, EPI_ISL_915294, EPI_ISL_915295, EPI_ISL_915296, EPI_ISL_915297, EPI_ISL_915298, EPI_ISL_915299, EPI_ISL_915300, EPI_ISL_915301, EPI_ISL_915303, EPI_ISL_915304, EPI_ISL_915305, EPI_ISL_915307, EPI_ISL_915309, EPI_ISL_915315, EPI_ISL_915317, EPI_ISL_915327, EPI_ISL_915342, EPI_ISL_915343, EPI_ISL_915345, EPI_ISL_915350, EPI_ISL_915352, EPI_ISL_915354 | | | |
| see above | Quest Diagnostics | Quest Diagnostics | Rosenthal,S.H., Gerasimova,A., Kagan,R.M., Anderson, B., Hua, M., Liu Y., Bernstein, L.E., Livingston, K.E., Perez, A., Shalhout, D.F., Shlyakhter, I.A., Owen, R., Tanpaiboon, P., Lacbawan, F. |
| EPI_ISL_916622, EPI_ISL_916623, EPI_ISL_916624 | Lighthouse Lab in Alderley Park | Wellcome Sanger Institute for the COVID-19 Genomics UK (COG-UK) Consortium | Jacquelyn Wynn, Mairead Hyland, The Lighthouse Lab in Alderley Park and Alex Alderton, Roberto Amato, Sonia Goncalves, Ewan Harrison, David K. Jackson, Ian Johnston, Dominic Kwiatkowski, Cordelia Langford, John Sillitoe on behalf of the Wellcome Sanger Institute COVID-19 Surveillance Team |
| EPI_ISL_916625, EPI_ISL_916626 | Lighthouse Lab in Glasgow | Wellcome Sanger Institute for the COVID-19 Genomics UK (COG-UK) Consortium | Harper VanSteenhouse, Yumi Kasai, David Gray, Carol Clugston, Anna Dominiczak and Alex Alderton, Roberto Amato, Sonia Goncalves, Ewan Harrison, David K. Jackson, Ian Johnston, Dominic Kwiatkowski, Cordelia Langford, John Sillitoe on behalf of the Wellcome Sanger Institute COVID-19 Surveillance Team |
| EPI_ISL_916627, EPI_ISL_916628, EPI_ISL_916629, EPI_ISL_916631, EPI_ISL_916632, EPI_ISL_916634, EPI_ISL_916635, EPI_ISL_916636, EPI_ISL_916638, EPI_ISL_916639, EPI_ISL_916640, EPI_ISL_916641, EPI_ISL_916642, EPI_ISL_916643, EPI_ISL_916644 | | | |
| see above | Lighthouse Lab in Alderley Park | Wellcome Sanger Institute for the COVID-19 Genomics UK (COG-UK) Consortium | Jacquelyn Wynn, Mairead Hyland, The Lighthouse Lab in Alderley Park and Alex Alderton, Roberto Amato, Sonia Goncalves, Ewan Harrison, David K. Jackson, Ian Johnston, Dominic Kwiatkowski, Cordelia Langford, John Sillitoe on behalf of the Wellcome Sanger Institute COVID-19 Surveillance Team |
| EPI_ISL_916645 | Lighthouse Lab in Glasgow | Wellcome Sanger Institute for the COVID-19 Genomics UK (COG-UK) Consortium | Harper VanSteenhouse, Yumi Kasai, David Gray, Carol Clugston, Anna Dominiczak and Alex Alderton, Roberto Amato, Sonia Goncalves, Ewan Harrison, David K. Jackson, Ian Johnston, Dominic Kwiatkowski, Cordelia Langford, John Sillitoe on behalf of the Wellcome Sanger Institute COVID-19 Surveillance Team |
| EPI_ISL_916646, EPI_ISL_916647 | Lighthouse Lab in Alderley Park | Wellcome Sanger Institute for the COVID-19 Genomics UK (COG-UK) Consortium | Jacquelyn Wynn, Mairead Hyland, The Lighthouse Lab in Alderley Park and Alex Alderton, Roberto Amato, Sonia Goncalves, Ewan Harrison, David K. Jackson, Ian Johnston, Dominic Kwiatkowski, Cordelia Langford, John Sillitoe on behalf of the Wellcome Sanger Institute COVID-19 Surveillance Team |
| EPI_ISL_916648 | Lighthouse Lab in Glasgow | Wellcome Sanger Institute for the COVID-19 Genomics UK (COG-UK) Consortium | Harper VanSteenhouse, Yumi Kasai, David Gray, Carol Clugston, Anna Dominiczak and Alex Alderton, Roberto Amato, Sonia Goncalves, Ewan Harrison, David K. Jackson, Ian Johnston, Dominic Kwiatkowski, Cordelia Langford, John Sillitoe on behalf of the Wellcome Sanger Institute COVID-19 Surveillance Team |
| EPI_ISL_916650, EPI_ISL_916652, EPI_ISL_916653, EPI_ISL_916654, EPI_ISL_916655, EPI_ISL_916656, EPI_ISL_916657, EPI_ISL_916658, EPI_ISL_916659, EPI_ISL_916660 | Lighthouse Lab in Alderley Park | Wellcome Sanger Institute for the COVID-19 Genomics UK (COG-UK) Consortium | Jacquelyn Wynn, Mairead Hyland, The Lighthouse Lab in Alderley Park and Alex Alderton, Roberto Amato, Sonia Goncalves, Ewan Harrison, David K. Jackson, Ian Johnston, Dominic Kwiatkowski, Cordelia Langford, John Sillitoe on behalf of the Wellcome Sanger Institute COVID-19 Surveillance Team |
| EPI_ISL_916662 | Lighthouse Lab in Glasgow | Wellcome Sanger Institute for the COVID-19 Genomics UK (COG-UK) Consortium | Harper VanSteenhouse, Yumi Kasai, David Gray, Carol Clugston, Anna Dominiczak and Alex Alderton, Roberto Amato, Sonia Goncalves, Ewan Harrison, David K. Jackson, Ian Johnston, Dominic Kwiatkowski, Cordelia Langford, John Sillitoe on behalf of the Wellcome Sanger Institute COVID-19 Surveillance Team |
| EPI_ISL_916663, EPI_ISL_916664, EPI_ISL_916665, EPI_ISL_916666, EPI_ISL_916667 | Lighthouse Lab in Alderley Park | Wellcome Sanger Institute for the COVID-19 Genomics UK (COG-UK) Consortium | Jacquelyn Wynn, Mairead Hyland, The Lighthouse Lab in Alderley Park and Alex Alderton, Roberto Amato, Sonia Goncalves, Ewan Harrison, David K. Jackson, Ian Johnston, Dominic Kwiatkowski, Cordelia Langford, John Sillitoe on behalf of the Wellcome Sanger Institute COVID-19 Surveillance Team |
| EPI_ISL_916668 | Lighthouse Lab in Glasgow | Wellcome Sanger Institute for the COVID-19 Genomics UK (COG-UK) Consortium | Harper VanSteenhouse, Yumi Kasai, David Gray, Carol Clugston, Anna Dominiczak and Alex Alderton, Roberto Amato, Sonia Goncalves, Ewan Harrison, David K. Jackson, Ian Johnston, Dominic Kwiatkowski, Cordelia Langford, John Sillitoe on behalf of the Wellcome Sanger Institute COVID-19 Surveillance Team |
| EPI_ISL_916669, EPI_ISL_916670, EPI_ISL_916671, EPI_ISL_916672, EPI_ISL_916673, EPI_ISL_916674, EPI_ISL_916676, EPI_ISL_916677, EPI_ISL_916678, EPI_ISL_916679, EPI_ISL_916680, EPI_ISL_916681, EPI_ISL_916683 | | | |
| see above | Lighthouse Lab in Alderley Park | Wellcome Sanger Institute for the COVID-19 Genomics UK (COG-UK) Consortium | Jacquelyn Wynn, Mairead Hyland, The Lighthouse Lab in Alderley Park and Alex Alderton, Roberto Amato, Sonia Goncalves, Ewan Harrison, David K. Jackson, Ian Johnston, Dominic Kwiatkowski, Cordelia Langford, John Sillitoe on behalf of the Wellcome Sanger Institute COVID-19 Surveillance Team |
| EPI_ISL_916684 | Lighthouse Lab in Glasgow | Wellcome Sanger Institute for the COVID-19 Genomics UK (COG-UK) Consortium | Harper VanSteenhouse, Yumi Kasai, David Gray, Carol Clugston, Anna Dominiczak and Alex Alderton, Roberto Amato, Sonia Goncalves, Ewan Harrison, David K. Jackson, Ian Johnston, Dominic Kwiatkowski, Cordelia Langford, John Sillitoe on behalf of the Wellcome Sanger Institute COVID-19 Surveillance Team |
| EPI_ISL_916685, EPI_ISL_916686, EPI_ISL_916687, EPI_ISL_916689, EPI_ISL_916690, EPI_ISL_916691, EPI_ISL_916692, EPI_ISL_916693, EPI_ISL_916695, EPI_ISL_916697, EPI_ISL_916698, EPI_ISL_916699, EPI_ISL_916700, EPI_ISL_916701, EPI_ISL_916702, EPI_ISL_916703, EPI_ISL_916706, EPI_ISL_916707, EPI_ISL_916708, EPI_ISL_916709, EPI_ISL_916710 | | | |
| see above | Lighthouse Lab in Alderley Park | Wellcome Sanger Institute for the COVID-19 Genomics UK (COG-UK) Consortium | Jacquelyn Wynn, Mairead Hyland, The Lighthouse Lab in Alderley Park and Alex Alderton, Roberto Amato, Sonia Goncalves, Ewan Harrison, David K. Jackson, Ian Johnston, Dominic Kwiatkowski, Cordelia Langford, John Sillitoe on behalf of the Wellcome Sanger Institute COVID-19 Surveillance Team |
| EPI_ISL_916711 | Lighthouse Lab in Glasgow | Wellcome Sanger Institute for the COVID-19 Genomics UK (COG-UK) Consortium | Harper VanSteenhouse, Yumi Kasai, David Gray, Carol Clugston, Anna Dominiczak and Alex Alderton, Roberto Amato, Sonia Goncalves, Ewan Harrison, David K. Jackson, Ian Johnston, Dominic Kwiatkowski, Cordelia Langford, John Sillitoe on behalf of the Wellcome Sanger Institute COVID-19 Surveillance Team |
| EPI_ISL_916712, EPI_ISL_916714, EPI_ISL_916715 | Lighthouse Lab in Alderley Park | Wellcome Sanger Institute for the COVID-19 Genomics UK (COG-UK) Consortium | Jacquelyn Wynn, Mairead Hyland, The Lighthouse Lab in Alderley Park and Alex Alderton, Roberto Amato, Sonia Goncalves, Ewan Harrison, David K. Jackson, Ian Johnston, Dominic Kwiatkowski, Cordelia Langford, John Sillitoe on behalf of the Wellcome Sanger Institute COVID-19 Surveillance Team |
| EPI_ISL_916716 | Lighthouse Lab in Glasgow | Wellcome Sanger Institute for the COVID-19 Genomics UK (COG-UK) Consortium | Harper VanSteenhouse, Yumi Kasai, David Gray, Carol Clugston, Anna Dominiczak and Alex Alderton, Roberto Amato, Sonia Goncalves, Ewan Harrison, David K. Jackson, Ian Johnston, Dominic Kwiatkowski, Cordelia Langford, John Sillitoe on behalf of the Wellcome Sanger Institute COVID-19 Surveillance Team |
| EPI_ISL_916717, EPI_ISL_916718, EPI_ISL_916719, EPI_ISL_916720, EPI_ISL_916721, EPI_ISL_916722, EPI_ISL_916723, EPI_ISL_916724, EPI_ISL_916725, EPI_ISL_916726, EPI_ISL_916727, EPI_ISL_916729, EPI_ISL_916731, EPI_ISL_916733, EPI_ISL_916734, EPI_ISL_916735 | | | |
| see above | Lighthouse Lab in Alderley Park | Wellcome Sanger Institute for the COVID-19 Genomics UK (COG-UK) Consortium | Jacquelyn Wynn, Mairead Hyland, The Lighthouse Lab in Alderley Park and Alex Alderton, Roberto Amato, Sonia Goncalves, Ewan Harrison, David K. Jackson, Ian Johnston, Dominic Kwiatkowski, Cordelia Langford, John Sillitoe on behalf of the Wellcome Sanger Institute COVID-19 Surveillance Team |
| EPI_ISL_916736 | Lighthouse Lab in Glasgow | Wellcome Sanger Institute for the COVID-19 Genomics UK (COG-UK) Consortium | Harper VanSteenhouse, Yumi Kasai, David Gray, Carol Clugston, Anna Dominiczak and Alex Alderton, Roberto Amato, Sonia Goncalves, Ewan Harrison, David K. Jackson, Ian Johnston, Dominic Kwiatkowski, Cordelia Langford, John Sillitoe on behalf of the Wellcome Sanger Institute COVID-19 Surveillance Team |
| EPI_ISL_916737, EPI_ISL_916739, EPI_ISL_916740, EPI_ISL_916741, EPI_ISL_916742, EPI_ISL_916743, EPI_ISL_916744, EPI_ISL_916745, EPI_ISL_916747, EPI_ISL_916748, EPI_ISL_916749, EPI_ISL_916750, EPI_ISL_916751, EPI_ISL_916752, EPI_ISL_916753, EPI_ISL_916754, EPI_ISL_916755, EPI_ISL_916756, EPI_ISL_916757 | | | |
| see above | Lighthouse Lab in Alderley Park | Wellcome Sanger Institute for the COVID-19 Genomics UK (COG-UK) Consortium | Jacquelyn Wynn, Mairead Hyland, The Lighthouse Lab in Alderley Park and Alex Alderton, Roberto Amato, Sonia Goncalves, Ewan Harrison, David K. Jackson, Ian Johnston, Dominic Kwiatkowski, Cordelia Langford, John Sillitoe on behalf of the Wellcome Sanger Institute COVID-19 Surveillance Team |
| EPI_ISL_916758 | Lighthouse Lab in Glasgow | Wellcome Sanger Institute for the COVID-19 Genomics UK (COG-UK) Consortium | Harper VanSteenhouse, Yumi Kasai, David Gray, Carol Clugston, Anna Dominiczak and Alex Alderton, Roberto Amato, Sonia Goncalves, Ewan Harrison, David K. Jackson, Ian Johnston, Dominic Kwiatkowski, Cordelia Langford, John Sillitoe on behalf of the Wellcome Sanger Institute COVID-19 Surveillance Team |
| EPI_ISL_916759, EPI_ISL_916760 | Lighthouse Lab in Alderley Park | Wellcome Sanger Institute for the COVID-19 Genomics UK (COG-UK) Consortium | Jacquelyn Wynn, Mairead Hyland, The Lighthouse Lab in Alderley Park and Alex Alderton, Roberto Amato, Sonia Goncalves, Ewan Harrison, David K. Jackson, Ian Johnston, Dominic Kwiatkowski, Cordelia Langford, John Sillitoe on behalf of the Wellcome Sanger Institute COVID-19 Surveillance Team |
| EPI_ISL_916761 | Lighthouse Lab in Glasgow | Wellcome Sanger Institute for the COVID-19 Genomics UK (COG-UK) Consortium | Harper VanSteenhouse, Yumi Kasai, David Gray, Carol Clugston, Anna Dominiczak and Alex Alderton, Roberto Amato, Sonia Goncalves, Ewan Harrison, David K. Jackson, Ian Johnston, Dominic Kwiatkowski, Cordelia Langford, John Sillitoe on behalf of the Wellcome Sanger Institute COVID-19 Surveillance Team |
| EPI_ISL_916762, EPI_ISL_916763, EPI_ISL_916764, EPI_ISL_916765, EPI_ISL_916767, EPI_ISL_916768, EPI_ISL_916769, EPI_ISL_916770, EPI_ISL_916772, EPI_ISL_916773, EPI_ISL_916774, EPI_ISL_916776, EPI_ISL_916777, EPI_ISL_916778, EPI_ISL_916780, EPI_ISL_916781, EPI_ISL_916783, EPI_ISL_916785, EPI_ISL_916786, EPI_ISL_916787, EPI_ISL_916788, EPI_ISL_916789, EPI_ISL_916790, EPI_ISL_916791, EPI_ISL_916792 | | | |

| | | | |
|---|---|---|---|
| see above | Lighthouse Lab in Alderley Park | Wellcome Sanger Institute for the COVID-19 Genomics UK (COG-UK) Consortium | Jacquelyn Wynn, Mairead Hyland, The Lighthouse Lab in Alderley Park and Alex Alderton, Roberto Amato, Sonia Goncalves, Ewan Harrison, David K. Jackson, Ian Johnston, Dominic Kwiatkowski, Cordelia Langford, John Sillitoe on behalf of the Wellcome Sanger Institute COVID-19 Surveillance Team |
| EPI_ISL_916793 | Lighthouse Lab in Glasgow | Wellcome Sanger Institute for the COVID-19 Genomics UK (COG-UK) Consortium | Harper VanSteenhouse, Yumi Kasai, David Gray, Carol Clugston, Anna Dominiczak and Alex Alderton, Roberto Amato, Sonia Goncalves, Ewan Harrison, David K. Jackson, Ian Johnston, Dominic Kwiatkowski, Cordelia Langford, John Sillitoe on behalf of the Wellcome Sanger Institute COVID-19 Surveillance Team |
| EPI_ISL_916794, EPI_ISL_916795, EPI_ISL_916796, EPI_ISL_916797, EPI_ISL_916798, EPI_ISL_916799, EPI_ISL_916801, EPI_ISL_916803, EPI_ISL_916804, EPI_ISL_916805, EPI_ISL_916806, EPI_ISL_916807, EPI_ISL_916808, EPI_ISL_916811, EPI_ISL_916813, EPI_ISL_916814, EPI_ISL_916816, EPI_ISL_916817, EPI_ISL_916818, EPI_ISL_916819 | | | |
| see above | Lighthouse Lab in Alderley Park | Wellcome Sanger Institute for the COVID-19 Genomics UK (COG-UK) Consortium | Jacquelyn Wynn, Mairead Hyland, The Lighthouse Lab in Alderley Park and Alex Alderton, Roberto Amato, Sonia Goncalves, Ewan Harrison, David K. Jackson, Ian Johnston, Dominic Kwiatkowski, Cordelia Langford, John Sillitoe on behalf of the Wellcome Sanger Institute COVID-19 Surveillance Team |
| EPI_ISL_916820 | Lighthouse Lab in Glasgow | Wellcome Sanger Institute for the COVID-19 Genomics UK (COG-UK) Consortium | Harper VanSteenhouse, Yumi Kasai, David Gray, Carol Clugston, Anna Dominiczak and Alex Alderton, Roberto Amato, Sonia Goncalves, Ewan Harrison, David K. Jackson, Ian Johnston, Dominic Kwiatkowski, Cordelia Langford, John Sillitoe on behalf of the Wellcome Sanger Institute COVID-19 Surveillance Team |
| EPI_ISL_916822, EPI_ISL_916823, EPI_ISL_916824, EPI_ISL_916826, EPI_ISL_916827, EPI_ISL_916828, EPI_ISL_916829, EPI_ISL_916830, EPI_ISL_916831, EPI_ISL_916832, EPI_ISL_916835, EPI_ISL_916836, EPI_ISL_916837, EPI_ISL_916839 | | | |
| see above | Lighthouse Lab in Alderley Park | Wellcome Sanger Institute for the COVID-19 Genomics UK (COG-UK) Consortium | Jacquelyn Wynn, Mairead Hyland, The Lighthouse Lab in Alderley Park and Alex Alderton, Roberto Amato, Sonia Goncalves, Ewan Harrison, David K. Jackson, Ian Johnston, Dominic Kwiatkowski, Cordelia Langford, John Sillitoe on behalf of the Wellcome Sanger Institute COVID-19 Surveillance Team |
| EPI_ISL_916840 | Lighthouse Lab in Glasgow | Wellcome Sanger Institute for the COVID-19 Genomics UK (COG-UK) Consortium | Harper VanSteenhouse, Yumi Kasai, David Gray, Carol Clugston, Anna Dominiczak and Alex Alderton, Roberto Amato, Sonia Goncalves, Ewan Harrison, David K. Jackson, Ian Johnston, Dominic Kwiatkowski, Cordelia Langford, John Sillitoe on behalf of the Wellcome Sanger Institute COVID-19 Surveillance Team |
| EPI_ISL_916841 | Lighthouse Lab in Alderley Park | Wellcome Sanger Institute for the COVID-19 Genomics UK (COG-UK) Consortium | Jacquelyn Wynn, Mairead Hyland, The Lighthouse Lab in Alderley Park and Alex Alderton, Roberto Amato, Sonia Goncalves, Ewan Harrison, David K. Jackson, Ian Johnston, Dominic Kwiatkowski, Cordelia Langford, John Sillitoe on behalf of the Wellcome Sanger Institute COVID-19 Surveillance Team |
| EPI_ISL_916842 | Lighthouse Lab in Glasgow | Wellcome Sanger Institute for the COVID-19 Genomics UK (COG-UK) Consortium | Harper VanSteenhouse, Yumi Kasai, David Gray, Carol Clugston, Anna Dominiczak and Alex Alderton, Roberto Amato, Sonia Goncalves, Ewan Harrison, David K. Jackson, Ian Johnston, Dominic Kwiatkowski, Cordelia Langford, John Sillitoe on behalf of the Wellcome Sanger Institute COVID-19 Surveillance Team |
| EPI_ISL_916843, EPI_ISL_916844, EPI_ISL_916846, EPI_ISL_916848, EPI_ISL_916851, EPI_ISL_916852, EPI_ISL_916853, EPI_ISL_916854 | Lighthouse Lab in Alderley Park | Wellcome Sanger Institute for the COVID-19 Genomics UK (COG-UK) Consortium | Jacquelyn Wynn, Mairead Hyland, The Lighthouse Lab in Alderley Park and Alex Alderton, Roberto Amato, Sonia Goncalves, Ewan Harrison, David K. Jackson, Ian Johnston, Dominic Kwiatkowski, Cordelia Langford, John Sillitoe on behalf of the Wellcome Sanger Institute COVID-19 Surveillance Team |
| EPI_ISL_916855 | Lighthouse Lab in Glasgow | Wellcome Sanger Institute for the COVID-19 Genomics UK (COG-UK) Consortium | Harper VanSteenhouse, Yumi Kasai, David Gray, Carol Clugston, Anna Dominiczak and Alex Alderton, Roberto Amato, Sonia Goncalves, Ewan Harrison, David K. Jackson, Ian Johnston, Dominic Kwiatkowski, Cordelia Langford, John Sillitoe on behalf of the Wellcome Sanger Institute COVID-19 Surveillance Team |
| EPI_ISL_916856, EPI_ISL_916857, EPI_ISL_916858, EPI_ISL_916859, EPI_ISL_916860, EPI_ISL_916861, EPI_ISL_916862, EPI_ISL_916863, EPI_ISL_916864, EPI_ISL_916865, EPI_ISL_916866, EPI_ISL_916867, EPI_ISL_916868 | | | |
| see above | Lighthouse Lab in Alderley Park | Wellcome Sanger Institute for the COVID-19 Genomics UK (COG-UK) Consortium | Jacquelyn Wynn, Mairead Hyland, The Lighthouse Lab in Alderley Park and Alex Alderton, Roberto Amato, Sonia Goncalves, Ewan Harrison, David K. Jackson, Ian Johnston, Dominic Kwiatkowski, Cordelia Langford, John Sillitoe on behalf of the Wellcome Sanger Institute COVID-19 Surveillance Team |
| EPI_ISL_916869 | Lighthouse Lab in Glasgow | Wellcome Sanger Institute for the COVID-19 Genomics UK (COG-UK) Consortium | Harper VanSteenhouse, Yumi Kasai, David Gray, Carol Clugston, Anna Dominiczak and Alex Alderton, Roberto Amato, Sonia Goncalves, Ewan Harrison, David K. Jackson, Ian Johnston, Dominic Kwiatkowski, Cordelia Langford, John Sillitoe on behalf of the Wellcome Sanger Institute COVID-19 Surveillance Team |
| EPI_ISL_916870, EPI_ISL_916871, EPI_ISL_916872, EPI_ISL_916873, EPI_ISL_916874, EPI_ISL_916875, EPI_ISL_916876, EPI_ISL_916877, EPI_ISL_916878 | Lighthouse Lab in Alderley Park | Wellcome Sanger Institute for the COVID-19 Genomics UK (COG-UK) Consortium | Jacquelyn Wynn, Mairead Hyland, The Lighthouse Lab in Alderley Park and Alex Alderton, Roberto Amato, Sonia Goncalves, Ewan Harrison, David K. Jackson, Ian Johnston, Dominic Kwiatkowski, Cordelia Langford, John Sillitoe on behalf of the Wellcome Sanger Institute COVID-19 Surveillance Team |
| EPI_ISL_916879 | Lighthouse Lab in Glasgow | Wellcome Sanger Institute for the COVID-19 Genomics UK (COG-UK) Consortium | Harper VanSteenhouse, Yumi Kasai, David Gray, Carol Clugston, Anna Dominiczak and Alex Alderton, Roberto Amato, Sonia Goncalves, Ewan Harrison, David K. Jackson, Ian Johnston, Dominic Kwiatkowski, Cordelia Langford, John Sillitoe on behalf of the Wellcome Sanger Institute COVID-19 Surveillance Team |
| EPI_ISL_916880, EPI_ISL_916881, EPI_ISL_916883, EPI_ISL_916884, EPI_ISL_916885, EPI_ISL_916886, EPI_ISL_916888, EPI_ISL_916889, EPI_ISL_916890 | Lighthouse Lab in Alderley Park | Wellcome Sanger Institute for the COVID-19 Genomics UK (COG-UK) Consortium | Jacquelyn Wynn, Mairead Hyland, The Lighthouse Lab in Alderley Park and Alex Alderton, Roberto Amato, Sonia Goncalves, Ewan Harrison, David K. Jackson, Ian Johnston, Dominic Kwiatkowski, Cordelia Langford, John Sillitoe on behalf of the Wellcome Sanger Institute COVID-19 Surveillance Team |
| EPI_ISL_916891 | Lighthouse Lab in Glasgow | Wellcome Sanger Institute for the COVID-19 Genomics UK (COG-UK) Consortium | Harper VanSteenhouse, Yumi Kasai, David Gray, Carol Clugston, Anna Dominiczak and Alex Alderton, Roberto Amato, Sonia Goncalves, Ewan Harrison, David K. Jackson, Ian Johnston, Dominic Kwiatkowski, Cordelia Langford, John Sillitoe on behalf of the Wellcome Sanger Institute COVID-19 Surveillance Team |
| EPI_ISL_916892, EPI_ISL_916893 | Lighthouse Lab in Alderley Park | Wellcome Sanger Institute for the COVID-19 Genomics UK (COG-UK) Consortium | Jacquelyn Wynn, Mairead Hyland, The Lighthouse Lab in Alderley Park and Alex Alderton, Roberto Amato, Sonia Goncalves, Ewan Harrison, David K. Jackson, Ian Johnston, Dominic Kwiatkowski, Cordelia Langford, John Sillitoe on behalf of the Wellcome Sanger Institute COVID-19 Surveillance Team |
| EPI_ISL_916894 | Lighthouse Lab in Glasgow | Wellcome Sanger Institute for the COVID-19 Genomics UK (COG-UK) Consortium | Harper VanSteenhouse, Yumi Kasai, David Gray, Carol Clugston, Anna Dominiczak and Alex Alderton, Roberto Amato, Sonia Goncalves, Ewan Harrison, David K. Jackson, Ian Johnston, Dominic Kwiatkowski, Cordelia Langford, John Sillitoe on behalf of the Wellcome Sanger Institute COVID-19 Surveillance Team |
| EPI_ISL_916895, EPI_ISL_916897 | Lighthouse Lab in Alderley Park | Wellcome Sanger Institute for the COVID-19 Genomics UK (COG-UK) Consortium | Jacquelyn Wynn, Mairead Hyland, The Lighthouse Lab in Alderley Park and Alex Alderton, Roberto Amato, Sonia Goncalves, Ewan Harrison, David K. Jackson, Ian Johnston, Dominic Kwiatkowski, Cordelia Langford, John Sillitoe on behalf of the Wellcome Sanger Institute COVID-19 Surveillance Team |
| EPI_ISL_916899, EPI_ISL_916900 | Lighthouse Lab in Glasgow | Wellcome Sanger Institute for the COVID-19 Genomics UK (COG-UK) Consortium | Harper VanSteenhouse, Yumi Kasai, David Gray, Carol Clugston, Anna Dominiczak and Alex Alderton, Roberto Amato, Sonia Goncalves, Ewan Harrison, David K. Jackson, Ian Johnston, Dominic Kwiatkowski, Cordelia Langford, John Sillitoe on behalf of the Wellcome Sanger Institute COVID-19 Surveillance Team |
| EPI_ISL_916901, EPI_ISL_916902, EPI_ISL_916903, EPI_ISL_916904, EPI_ISL_916906, EPI_ISL_916910, EPI_ISL_916913, EPI_ISL_916914, EPI_ISL_916915, EPI_ISL_916917, EPI_ISL_916920, EPI_ISL_916921, EPI_ISL_916922, EPI_ISL_916924, EPI_ISL_916925, EPI_ISL_916926, EPI_ISL_916927, EPI_ISL_916928, EPI_ISL_916929, EPI_ISL_916930, EPI_ISL_916931, EPI_ISL_916932, EPI_ISL_916933, EPI_ISL_916935, EPI_ISL_916936, EPI_ISL_916937, EPI_ISL_916938, EPI_ISL_916939, EPI_ISL_916941 | | | |
| see above | Lighthouse Lab in Alderley Park | Wellcome Sanger Institute for the COVID-19 Genomics UK (COG-UK) Consortium | Jacquelyn Wynn, Mairead Hyland, The Lighthouse Lab in Alderley Park and Alex Alderton, Roberto Amato, Sonia Goncalves, Ewan Harrison, David K. Jackson, Ian Johnston, Dominic Kwiatkowski, Cordelia Langford, John Sillitoe on behalf of the Wellcome Sanger Institute COVID-19 Surveillance Team |
| EPI_ISL_916942 | Lighthouse Lab in Glasgow | Wellcome Sanger Institute for the COVID-19 Genomics UK (COG-UK) Consortium | Harper VanSteenhouse, Yumi Kasai, David Gray, Carol Clugston, Anna Dominiczak and Alex Alderton, Roberto Amato, Sonia Goncalves, Ewan Harrison, David K. Jackson, Ian Johnston, Dominic Kwiatkowski, Cordelia Langford, John Sillitoe on behalf of the Wellcome Sanger Institute COVID-19 Surveillance Team |
| EPI_ISL_916943, EPI_ISL_916944, EPI_ISL_916947 | Lighthouse Lab in Alderley Park | Wellcome Sanger Institute for the COVID-19 Genomics UK (COG-UK) Consortium | Jacquelyn Wynn, Mairead Hyland, The Lighthouse Lab in Alderley Park and Alex Alderton, Roberto Amato, Sonia Goncalves, Ewan Harrison, David K. Jackson, Ian Johnston, Dominic Kwiatkowski, Cordelia Langford, John Sillitoe on behalf of the Wellcome Sanger Institute COVID-19 Surveillance Team |
| EPI_ISL_916951 | Lighthouse Lab in Glasgow | Wellcome Sanger Institute for the COVID-19 Genomics UK (COG-UK) Consortium | Harper VanSteenhouse, Yumi Kasai, David Gray, Carol Clugston, Anna Dominiczak and Alex Alderton, Roberto Amato, Sonia Goncalves, Ewan Harrison, David K. Jackson, Ian Johnston, Dominic Kwiatkowski, Cordelia Langford, John Sillitoe on behalf of the Wellcome Sanger Institute COVID-19 Surveillance Team |

| | | | |
|---|---|---|---|
| EPI_ISL_916952, EPI_ISL_916955, EPI_ISL_916957, EPI_ISL_916958, EPI_ISL_916959, EPI_ISL_916960, EPI_ISL_916962, EPI_ISL_916963, EPI_ISL_916964, EPI_ISL_916965, EPI_ISL_916966 | | | |
| see above | Lighthouse Lab in Alderley Park | Wellcome Sanger Institute for the COVID-19 Genomics UK (COG-UK) Consortium | Jacquelyn Wynn, Mairead Hyland, The Lighthouse Lab in Alderley Park and Alex Alderton, Roberto Amato, Sonia Goncalves, Ewan Harrison, David K. Jackson, Ian Johnston, Dominic Kwiatkowski, Cordelia Langford, John Sillitoe on behalf of the Wellcome Sanger Institute COVID-19 Surveillance Team |
| EPI_ISL_916967 | Lighthouse Lab in Glasgow | Wellcome Sanger Institute for the COVID-19 Genomics UK (COG-UK) Consortium | Harper VanSteenhouse, Yumi Kasai, David Gray, Carol Clugston, Anna Dominiczak and Alex Alderton, Roberto Amato, Sonia Goncalves, Ewan Harrison, David K. Jackson, Ian Johnston, Dominic Kwiatkowski, Cordelia Langford, John Sillitoe on behalf of the Wellcome Sanger Institute COVID-19 Surveillance Team |
| EPI_ISL_916969, EPI_ISL_916971, EPI_ISL_916972, EPI_ISL_916973, EPI_ISL_916975, EPI_ISL_916978 | Lighthouse Lab in Alderley Park | Wellcome Sanger Institute for the COVID-19 Genomics UK (COG-UK) Consortium | Jacquelyn Wynn, Mairead Hyland, The Lighthouse Lab in Alderley Park and Alex Alderton, Roberto Amato, Sonia Goncalves, Ewan Harrison, David K. Jackson, Ian Johnston, Dominic Kwiatkowski, Cordelia Langford, John Sillitoe on behalf of the Wellcome Sanger Institute COVID-19 Surveillance Team |
| EPI_ISL_916979 | Lighthouse Lab in Glasgow | Wellcome Sanger Institute for the COVID-19 Genomics UK (COG-UK) Consortium | Harper VanSteenhouse, Yumi Kasai, David Gray, Carol Clugston, Anna Dominiczak and Alex Alderton, Roberto Amato, Sonia Goncalves, Ewan Harrison, David K. Jackson, Ian Johnston, Dominic Kwiatkowski, Cordelia Langford, John Sillitoe on behalf of the Wellcome Sanger Institute COVID-19 Surveillance Team |
| EPI_ISL_916980, EPI_ISL_916982, EPI_ISL_916983, EPI_ISL_916986, EPI_ISL_916988, EPI_ISL_916989, EPI_ISL_916990, EPI_ISL_916992, EPI_ISL_916993, EPI_ISL_916994, EPI_ISL_916995, EPI_ISL_916996, EPI_ISL_916997 | | | |
| see above | Lighthouse Lab in Alderley Park | Wellcome Sanger Institute for the COVID-19 Genomics UK (COG-UK) Consortium | Jacquelyn Wynn, Mairead Hyland, The Lighthouse Lab in Alderley Park and Alex Alderton, Roberto Amato, Sonia Goncalves, Ewan Harrison, David K. Jackson, Ian Johnston, Dominic Kwiatkowski, Cordelia Langford, John Sillitoe on behalf of the Wellcome Sanger Institute COVID-19 Surveillance Team |
| EPI_ISL_916998 | Lighthouse Lab in Glasgow | Wellcome Sanger Institute for the COVID-19 Genomics UK (COG-UK) Consortium | Harper VanSteenhouse, Yumi Kasai, David Gray, Carol Clugston, Anna Dominiczak and Alex Alderton, Roberto Amato, Sonia Goncalves, Ewan Harrison, David K. Jackson, Ian Johnston, Dominic Kwiatkowski, Cordelia Langford, John Sillitoe on behalf of the Wellcome Sanger Institute COVID-19 Surveillance Team |
| EPI_ISL_917000, EPI_ISL_917001 | Lighthouse Lab in Alderley Park | Wellcome Sanger Institute for the COVID-19 Genomics UK (COG-UK) Consortium | Jacquelyn Wynn, Mairead Hyland, The Lighthouse Lab in Alderley Park and Alex Alderton, Roberto Amato, Sonia Goncalves, Ewan Harrison, David K. Jackson, Ian Johnston, Dominic Kwiatkowski, Cordelia Langford, John Sillitoe on behalf of the Wellcome Sanger Institute COVID-19 Surveillance Team |
| EPI_ISL_917002 | Lighthouse Lab in Glasgow | Wellcome Sanger Institute for the COVID-19 Genomics UK (COG-UK) Consortium | Harper VanSteenhouse, Yumi Kasai, David Gray, Carol Clugston, Anna Dominiczak and Alex Alderton, Roberto Amato, Sonia Goncalves, Ewan Harrison, David K. Jackson, Ian Johnston, Dominic Kwiatkowski, Cordelia Langford, John Sillitoe on behalf of the Wellcome Sanger Institute COVID-19 Surveillance Team |
| EPI_ISL_917003, EPI_ISL_917004, EPI_ISL_917005, EPI_ISL_917006, EPI_ISL_917007, EPI_ISL_917008, EPI_ISL_917010, EPI_ISL_917015, EPI_ISL_917016, EPI_ISL_917017, EPI_ISL_917018, EPI_ISL_917020, EPI_ISL_917023, EPI_ISL_917027, EPI_ISL_917028, EPI_ISL_917030, EPI_ISL_917033, EPI_ISL_917034, EPI_ISL_917035, EPI_ISL_917037, EPI_ISL_917039, EPI_ISL_917040, EPI_ISL_917041, EPI_ISL_917043, EPI_ISL_917044, EPI_ISL_917045, EPI_ISL_917046, EPI_ISL_917048 | | | |
| see above | Lighthouse Lab in Alderley Park | Wellcome Sanger Institute for the COVID-19 Genomics UK (COG-UK) Consortium | Jacquelyn Wynn, Mairead Hyland, The Lighthouse Lab in Alderley Park and Alex Alderton, Roberto Amato, Sonia Goncalves, Ewan Harrison, David K. Jackson, Ian Johnston, Dominic Kwiatkowski, Cordelia Langford, John Sillitoe on behalf of the Wellcome Sanger Institute COVID-19 Surveillance Team |
| EPI_ISL_917049 | Lighthouse Lab in Glasgow | Wellcome Sanger Institute for the COVID-19 Genomics UK (COG-UK) Consortium | Harper VanSteenhouse, Yumi Kasai, David Gray, Carol Clugston, Anna Dominiczak and Alex Alderton, Roberto Amato, Sonia Goncalves, Ewan Harrison, David K. Jackson, Ian Johnston, Dominic Kwiatkowski, Cordelia Langford, John Sillitoe on behalf of the Wellcome Sanger Institute COVID-19 Surveillance Team |
| EPI_ISL_917050, EPI_ISL_917051, EPI_ISL_917052 | Lighthouse Lab in Alderley Park | Wellcome Sanger Institute for the COVID-19 Genomics UK (COG-UK) Consortium | Jacquelyn Wynn, Mairead Hyland, The Lighthouse Lab in Alderley Park and Alex Alderton, Roberto Amato, Sonia Goncalves, Ewan Harrison, David K. Jackson, Ian Johnston, Dominic Kwiatkowski, Cordelia Langford, John Sillitoe on behalf of the Wellcome Sanger Institute COVID-19 Surveillance Team |
| EPI_ISL_917053 | Lighthouse Lab in Glasgow | Wellcome Sanger Institute for the COVID-19 Genomics UK (COG-UK) Consortium | Harper VanSteenhouse, Yumi Kasai, David Gray, Carol Clugston, Anna Dominiczak and Alex Alderton, Roberto Amato, Sonia Goncalves, Ewan Harrison, David K. Jackson, Ian Johnston, Dominic Kwiatkowski, Cordelia Langford, John Sillitoe on behalf of the Wellcome Sanger Institute COVID-19 Surveillance Team |
| EPI_ISL_917056, EPI_ISL_917057, EPI_ISL_917058, EPI_ISL_917059, EPI_ISL_917061, EPI_ISL_917062, EPI_ISL_917064, EPI_ISL_917065, EPI_ISL_917066, EPI_ISL_917068, EPI_ISL_917069, EPI_ISL_917072, EPI_ISL_917073 | | | |
| see above | Lighthouse Lab in Alderley Park | Wellcome Sanger Institute for the COVID-19 Genomics UK (COG-UK) Consortium | Jacquelyn Wynn, Mairead Hyland, The Lighthouse Lab in Alderley Park and Alex Alderton, Roberto Amato, Sonia Goncalves, Ewan Harrison, David K. Jackson, Ian Johnston, Dominic Kwiatkowski, Cordelia Langford, John Sillitoe on behalf of the Wellcome Sanger Institute COVID-19 Surveillance Team |
| EPI_ISL_917076 | Lighthouse Lab in Glasgow | Wellcome Sanger Institute for the COVID-19 Genomics UK (COG-UK) Consortium | Harper VanSteenhouse, Yumi Kasai, David Gray, Carol Clugston, Anna Dominiczak and Alex Alderton, Roberto Amato, Sonia Goncalves, Ewan Harrison, David K. Jackson, Ian Johnston, Dominic Kwiatkowski, Cordelia Langford, John Sillitoe on behalf of the Wellcome Sanger Institute COVID-19 Surveillance Team |
| EPI_ISL_917078, EPI_ISL_917080, EPI_ISL_917084, EPI_ISL_917085, EPI_ISL_917087, EPI_ISL_917089, EPI_ISL_917090, EPI_ISL_917092, EPI_ISL_917094, EPI_ISL_917097, EPI_ISL_917098, EPI_ISL_917100, EPI_ISL_917101, EPI_ISL_917102, EPI_ISL_917103 | | | |
| see above | Lighthouse Lab in Alderley Park | Wellcome Sanger Institute for the COVID-19 Genomics UK (COG-UK) Consortium | Jacquelyn Wynn, Mairead Hyland, The Lighthouse Lab in Alderley Park and Alex Alderton, Roberto Amato, Sonia Goncalves, Ewan Harrison, David K. Jackson, Ian Johnston, Dominic Kwiatkowski, Cordelia Langford, John Sillitoe on behalf of the Wellcome Sanger Institute COVID-19 Surveillance Team |
| EPI_ISL_917104 | Lighthouse Lab in Glasgow | Wellcome Sanger Institute for the COVID-19 Genomics UK (COG-UK) Consortium | Harper VanSteenhouse, Yumi Kasai, David Gray, Carol Clugston, Anna Dominiczak and Alex Alderton, Roberto Amato, Sonia Goncalves, Ewan Harrison, David K. Jackson, Ian Johnston, Dominic Kwiatkowski, Cordelia Langford, John Sillitoe on behalf of the Wellcome Sanger Institute COVID-19 Surveillance Team |
| EPI_ISL_917106, EPI_ISL_917108, EPI_ISL_917109, EPI_ISL_917111 | Lighthouse Lab in Alderley Park | Wellcome Sanger Institute for the COVID-19 Genomics UK (COG-UK) Consortium | Jacquelyn Wynn, Mairead Hyland, The Lighthouse Lab in Alderley Park and Alex Alderton, Roberto Amato, Sonia Goncalves, Ewan Harrison, David K. Jackson, Ian Johnston, Dominic Kwiatkowski, Cordelia Langford, John Sillitoe on behalf of the Wellcome Sanger Institute COVID-19 Surveillance Team |
| EPI_ISL_917112 | Lighthouse Lab in Glasgow | Wellcome Sanger Institute for the COVID-19 Genomics UK (COG-UK) Consortium | Harper VanSteenhouse, Yumi Kasai, David Gray, Carol Clugston, Anna Dominiczak and Alex Alderton, Roberto Amato, Sonia Goncalves, Ewan Harrison, David K. Jackson, Ian Johnston, Dominic Kwiatkowski, Cordelia Langford, John Sillitoe on behalf of the Wellcome Sanger Institute COVID-19 Surveillance Team |
| EPI_ISL_917113 | Lighthouse Lab in Alderley Park | Wellcome Sanger Institute for the COVID-19 Genomics UK (COG-UK) Consortium | Jacquelyn Wynn, Mairead Hyland, The Lighthouse Lab in Alderley Park and Alex Alderton, Roberto Amato, Sonia Goncalves, Ewan Harrison, David K. Jackson, Ian Johnston, Dominic Kwiatkowski, Cordelia Langford, John Sillitoe on behalf of the Wellcome Sanger Institute COVID-19 Surveillance Team |
| EPI_ISL_917114 | Lighthouse Lab in Glasgow | Wellcome Sanger Institute for the COVID-19 Genomics UK (COG-UK) Consortium | Harper VanSteenhouse, Yumi Kasai, David Gray, Carol Clugston, Anna Dominiczak and Alex Alderton, Roberto Amato, Sonia Goncalves, Ewan Harrison, David K. Jackson, Ian Johnston, Dominic Kwiatkowski, Cordelia Langford, John Sillitoe on behalf of the Wellcome Sanger Institute COVID-19 Surveillance Team |
| EPI_ISL_917115, EPI_ISL_917117, EPI_ISL_917118, EPI_ISL_917119, EPI_ISL_917122, EPI_ISL_917123, EPI_ISL_917124, EPI_ISL_917126, EPI_ISL_917127, EPI_ISL_917130, EPI_ISL_917131, EPI_ISL_917132, EPI_ISL_917134, EPI_ISL_917135, EPI_ISL_917136, EPI_ISL_917137, EPI_ISL_917138, EPI_ISL_917140, EPI_ISL_917142 | | | |
| see above | Lighthouse Lab in Alderley Park | Wellcome Sanger Institute for the COVID-19 Genomics UK (COG-UK) Consortium | Jacquelyn Wynn, Mairead Hyland, The Lighthouse Lab in Alderley Park and Alex Alderton, Roberto Amato, Sonia Goncalves, Ewan Harrison, David K. Jackson, Ian Johnston, Dominic Kwiatkowski, Cordelia Langford, John Sillitoe on behalf of the Wellcome Sanger Institute COVID-19 Surveillance Team |
| EPI_ISL_917144 | Lighthouse Lab in Glasgow | Wellcome Sanger Institute for the COVID-19 Genomics UK (COG-UK) Consortium | Harper VanSteenhouse, Yumi Kasai, David Gray, Carol Clugston, Anna Dominiczak and Alex Alderton, Roberto Amato, Sonia Goncalves, Ewan Harrison, David K. Jackson, Ian Johnston, Dominic Kwiatkowski, Cordelia Langford, John Sillitoe on behalf of the Wellcome Sanger Institute COVID-19 Surveillance Team |
| EPI_ISL_917146, EPI_ISL_917147, EPI_ISL_917150, EPI_ISL_917152, EPI_ISL_917153, EPI_ISL_917154, EPI_ISL_917155, EPI_ISL_917157, EPI_ISL_917158 | Lighthouse Lab in Alderley Park | Wellcome Sanger Institute for the COVID-19 Genomics UK (COG-UK) Consortium | Jacquelyn Wynn, Mairead Hyland, The Lighthouse Lab in Alderley Park and Alex Alderton, Roberto Amato, Sonia Goncalves, Ewan Harrison, David K. Jackson, Ian Johnston, Dominic Kwiatkowski, Cordelia Langford, John Sillitoe on behalf of the Wellcome Sanger Institute COVID-19 Surveillance Team |
| EPI_ISL_917161 | Lighthouse Lab in Glasgow | Wellcome Sanger Institute for the COVID-19 Genomics UK (COG-UK) Consortium | Harper VanSteenhouse, Yumi Kasai, David Gray, Carol Clugston, Anna Dominiczak and Alex Alderton, Roberto Amato, Sonia Goncalves, Ewan Harrison, David K. Jackson, Ian Johnston, Dominic Kwiatkowski, Cordelia Langford, John Sillitoe on behalf of the Wellcome Sanger Institute COVID-19 Surveillance Team |

| | | | |
|---|---|---|---|
| EPI_ISL_917162, EPI_ISL_917164, EPI_ISL_917165 | Lighthouse Lab in Alderley Park | Wellcome Sanger Institute for the COVID-19 Genomics UK (COG-UK) Consortium | Jacquelyn Wynn, Mairead Hyland, The Lighthouse Lab in Alderley Park and Alex Alderton, Roberto Amato, Sonia Goncalves, Ewan Harrison, David K. Jackson, Ian Johnston, Dominic Kwiatkowski, Cordelia Langford, John Sillitoe on behalf of the Wellcome Sanger Institute COVID-19 Surveillance Team |
| EPI_ISL_917166 | Lighthouse Lab in Glasgow | Wellcome Sanger Institute for the COVID-19 Genomics UK (COG-UK) Consortium | Harper VanSteenhouse, Yumi Kasai, David Gray, Carol Clugston, Anna Dominiczak and Alex Alderton, Roberto Amato, Sonia Goncalves, Ewan Harrison, David K. Jackson, Ian Johnston, Dominic Kwiatkowski, Cordelia Langford, John Sillitoe on behalf of the Wellcome Sanger Institute COVID-19 Surveillance Team |
| EPI_ISL_917167, EPI_ISL_918061 | Lighthouse Lab in Alderley Park | Wellcome Sanger Institute for the COVID-19 Genomics UK (COG-UK) Consortium | Jacquelyn Wynn, Mairead Hyland, The Lighthouse Lab in Alderley Park and Alex Alderton, Roberto Amato, Sonia Goncalves, Ewan Harrison, David K. Jackson, Ian Johnston, Dominic Kwiatkowski, Cordelia Langford, John Sillitoe on behalf of the Wellcome Sanger Institute COVID-19 Surveillance Team |
| EPI_ISL_918062, EPI_ISL_918063, EPI_ISL_918064, EPI_ISL_918065, EPI_ISL_918066, EPI_ISL_918067, EPI_ISL_918068, EPI_ISL_918069, EPI_ISL_918070, EPI_ISL_918071, EPI_ISL_918072, EPI_ISL_918073, EPI_ISL_918074, EPI_ISL_918075, EPI_ISL_918076, EPI_ISL_918077, EPI_ISL_918078, EPI_ISL_918079, EPI_ISL_918080, EPI_ISL_918081, EPI_ISL_918082, EPI_ISL_918083, EPI_ISL_918084, EPI_ISL_918085, EPI_ISL_918086, EPI_ISL_918087, EPI_ISL_918088, EPI_ISL_918089, EPI_ISL_918090, EPI_ISL_918091, EPI_ISL_918092, EPI_ISL_918093, EPI_ISL_918094, EPI_ISL_918095, EPI_ISL_918096, EPI_ISL_918097, EPI_ISL_918098, EPI_ISL_918099, EPI_ISL_918100, EPI_ISL_918101, EPI_ISL_918102, EPI_ISL_918103, EPI_ISL_918104, EPI_ISL_918105, EPI_ISL_918106, EPI_ISL_918107, EPI_ISL_918108, EPI_ISL_918109, EPI_ISL_918110, EPI_ISL_918111, EPI_ISL_918112, EPI_ISL_918113, EPI_ISL_918114, EPI_ISL_918115, EPI_ISL_918116, EPI_ISL_918117, EPI_ISL_918118, EPI_ISL_918119, EPI_ISL_918120, EPI_ISL_918121, EPI_ISL_918122, EPI_ISL_918123, EPI_ISL_918124 | | | |
| see above | Lighthouse Lab in Glasgow | Wellcome Sanger Institute for the COVID-19 Genomics UK (COG-UK) Consortium | Harper VanSteenhouse, Yumi Kasai, David Gray, Carol Clugston, Anna Dominiczak and Alex Alderton, Roberto Amato, Sonia Goncalves, Ewan Harrison, David K. Jackson, Ian Johnston, Dominic Kwiatkowski, Cordelia Langford, John Sillitoe on behalf of the Wellcome Sanger Institute COVID-19 Surveillance Team |
| EPI_ISL_918165 | UMR190-Unité des virus emergents | UMR190-Unité des virus emergents | Cecile Baronti |
| EPI_ISL_918179, EPI_ISL_918182, EPI_ISL_918188, EPI_ISL_918190, EPI_ISL_918195, EPI_ISL_918196, EPI_ISL_918198, EPI_ISL_918199, EPI_ISL_918204, EPI_ISL_918226, EPI_ISL_918231, EPI_ISL_918232, EPI_ISL_918235, EPI_ISL_918236, EPI_ISL_918238, EPI_ISL_918245, EPI_ISL_918246, EPI_ISL_918247, EPI_ISL_918252, EPI_ISL_918253, EPI_ISL_918260 | | | |
| see above | Innovative Genomics Institute, UC Berkeley | Innovative Genomics Institute, UC Berkeley | Stacia Wyman, Haridha Shivram, Phil Frankino, Liana Lareau, Shana McDevitt, Justin Choi |
| EPI_ISL_918288 | Hospital Universitari Vall d'Hebron - Vall d'Hebron Institut de Rercerca | Hospital Universitari Vall d'Hebron | Cristina Andrés, Maria Piñana, Josep F Abril, Damir Garcia-Cehic, Ariadna Rando, Juliana Esperalba, Maria Gema Codina, Carla Castillo, Maria Carmen Martin, Tomàs Pumarola, Josep Quer, Andrés Antón |
| EPI_ISL_918543, EPI_ISL_918544 | LACEN - Laboratório Central de Saúde Pública do Ceara | Evandro Chagas Institute | Santos, M.C.; Silva, A.M.; Junior, W.D.C.; Barbagelata, L.S.; Ferreira, J.A.; Sousa, E.M.A.; da Silva, P.S.; Pinheiro, K.C.; L.C.; Sousa Junior, E.C. |
| EPI_ISL_918764, EPI_ISL_918765, EPI_ISL_918766, EPI_ISL_918767, EPI_ISL_918768, EPI_ISL_918769, EPI_ISL_918770, EPI_ISL_918771, EPI_ISL_918772, EPI_ISL_918773, EPI_ISL_918774, EPI_ISL_918776, EPI_ISL_918777, EPI_ISL_918778, EPI_ISL_918780, EPI_ISL_918781, EPI_ISL_918782, EPI_ISL_918783, EPI_ISL_918784, EPI_ISL_918785, EPI_ISL_918786, EPI_ISL_918787, EPI_ISL_918788, EPI_ISL_918789, EPI_ISL_918790, EPI_ISL_918791, EPI_ISL_918792, EPI_ISL_918793, EPI_ISL_918794, EPI_ISL_918795, EPI_ISL_918796, EPI_ISL_918797, EPI_ISL_918798, EPI_ISL_918799, EPI_ISL_918800, EPI_ISL_918801, EPI_ISL_918802, EPI_ISL_918803, EPI_ISL_918804, EPI_ISL_918805, EPI_ISL_918806, EPI_ISL_918807, EPI_ISL_918808, EPI_ISL_918809, EPI_ISL_918810, EPI_ISL_918811, EPI_ISL_918812, EPI_ISL_918813, EPI_ISL_918814, EPI_ISL_918815, EPI_ISL_918816, EPI_ISL_918817, EPI_ISL_918818, EPI_ISL_918819, EPI_ISL_918820, EPI_ISL_918821, EPI_ISL_918822, EPI_ISL_918823, EPI_ISL_918824, EPI_ISL_918825, EPI_ISL_918826, EPI_ISL_918827, EPI_ISL_918842, EPI_ISL_918843, EPI_ISL_918844, EPI_ISL_918845, EPI_ISL_918856, EPI_ISL_918869, EPI_ISL_918874, EPI_ISL_918875, EPI_ISL_918876, EPI_ISL_918877, EPI_ISL_918878, EPI_ISL_918882, EPI_ISL_918912, EPI_ISL_918913, EPI_ISL_918977, EPI_ISL_918978, EPI_ISL_918979, EPI_ISL_918980, EPI_ISL_918981, EPI_ISL_918982, EPI_ISL_918983 | | | |
| see above | University of Birmingham | COVID-19 Genomics UK (COG-UK) Consortium | Institute of Microbiology, University of Birmingham: Claire McMurray, Joanne Stockton, Samuel Nicholls, Radoslaw Poplawski, Will Rowe, Josh Quick, Nicholas Loman. University of Birmingham Testing Laboratory: Celina M Whalley, Andrew Bosworth, Charlotte Poxon, Kasun Wanigasooriya, Oliver Pickles, Mike Kidd, Alex Richter, Andrew D Beggs PHE Heartlands Lab: Husam Osman, Andrew Bosworth. Queen Elizabeth Hospital: Anna Casey |
| EPI_ISL_919057, EPI_ISL_919059, EPI_ISL_919060, EPI_ISL_919061, EPI_ISL_919064, EPI_ISL_919065, EPI_ISL_919066, EPI_ISL_919068, EPI_ISL_919069, EPI_ISL_919072, EPI_ISL_919076, EPI_ISL_919078, EPI_ISL_919083, EPI_ISL_919085, EPI_ISL_919086, EPI_ISL_919088, EPI_ISL_919090, EPI_ISL_919094, EPI_ISL_919096, EPI_ISL_919098, EPI_ISL_919100 | | | |
| see above | Department of Pathology, University of Cambridge | COVID-19 Genomics UK (COG-UK) Consortium | Aminu S. Jahun, Yasmin Chaudhry, Iliana Georgana, Myra Hosmillo, Rhys Izu, Martin D. Curran, Surendra Parmar, Ian Goodfellow |
| EPI_ISL_919198, EPI_ISL_919199, EPI_ISL_919200, EPI_ISL_919201, EPI_ISL_919202, EPI_ISL_919203, EPI_ISL_919204 | West of Scotland Specialist Virology Centre, NHSGGC / MRC-University of Glasgow Centre for Virus Research | COVID-19 Genomics UK (COG-UK) Consortium | Ana da Silva Filipe, Natasha Johnson, Kathy Smollett, Daniel Mair, Stephen Carmichael, Alice Broos, Lily Tong, Jenna Nichols, Kyriaki Nomikou; Sarah McDonald; Richard Orton, Joseph Hughes, Sreenu Vattipally, David L Robertson; Alasdair MacLean, Rory Gunson; Sharif Shaaban, Matthew Holden; Rachel Blacow, Guy Mollett, Kathy Li, James Shepherd, Antonia Ho, Emma Thomson |
| EPI_ISL_919339, EPI_ISL_919340, EPI_ISL_919341 | Virology Department, Royal Infirmary of Edinburgh, NHS Lothian / School of Biological Sciences, University of Edinburgh / Institute of Genetics and Molecular Medicine, University of Edinburgh | COVID-19 Genomics UK (COG-UK) Consortium | McHugh M, Dewar R, Rooke S, Gallagher M, Balcaza C, O'Toole Á, Scher E, Hill V, McCrone JT, Colquhoun R, Yu X, Jackson B, Rambaut A, Williams TC, Templeton K |
| EPI_ISL_919479, EPI_ISL_919480, EPI_ISL_919481, EPI_ISL_919482, EPI_ISL_919483, EPI_ISL_919484, EPI_ISL_919485, EPI_ISL_919486, EPI_ISL_919487, EPI_ISL_919516, EPI_ISL_919524, EPI_ISL_919526, EPI_ISL_919527, EPI_ISL_919528, EPI_ISL_919539, EPI_ISL_919540, EPI_ISL_919546, EPI_ISL_919553, EPI_ISL_919555, EPI_ISL_919560, EPI_ISL_919566, EPI_ISL_919567, EPI_ISL_919569, EPI_ISL_919580, EPI_ISL_919583, EPI_ISL_919584, EPI_ISL_919586, EPI_ISL_919592, EPI_ISL_919596, EPI_ISL_919597, EPI_ISL_919598, EPI_ISL_919599, EPI_ISL_919600, EPI_ISL_919601, EPI_ISL_919602, EPI_ISL_919632, EPI_ISL_919634, EPI_ISL_919637, EPI_ISL_919641, EPI_ISL_919642, EPI_ISL_919643, EPI_ISL_919645, EPI_ISL_919646, EPI_ISL_919651, EPI_ISL_919660 | | | |
| see above | Liverpool Clinical Laboratories | COVID-19 Genomics UK (COG-UK) Consortium | Sam Haldenby, Anita Lucaci, Steve Paterson, Julian Hiscox, Alistair Darby, M Almsaud, A Alrezaihi, Muhannad Alruwaili, Stuart D Armstrong, Jones Benjamin, Eleanor G Bentley, Anu Chawla, Jordan J Clark, Angela Cowell, Richard Eccles, Isabel García-Dorival, Matthew Gemmell, Alessandro Gerada, PKF Gilmore, Richard Gregory, Ximeng Han, Catherine Hartley, Margaret Hughes, Miren Iturriza-Gomara, James Johnson, L Luu, Jenifer Manson, Charlotte Nelson, Elaine O'Toole, Cassie Olateju, Rebekah Penrice-Randal , Lucille Rainbow, N.P Randle, Trevor Ian Robinson, Parul Sharma, Ghada T Shawli, James P Stewart, Neil Swainston, Ecaterina Vamos, Joanne Watts, Mark Whitehead |
| EPI_ISL_919853 | Barts Health NHS Trust | COVID-19 Genomics UK (COG-UK) Consortium | CUTINO-MOGUEL, Maria-Teresa; HARRINGTON, David; OWOYEMI, Dola; KULASEGARAN-SHYLINI, Raghavendran; BROAD, Claire; KELE, Beatrix |
| EPI_ISL_920030, EPI_ISL_920031, EPI_ISL_920032, EPI_ISL_920033, EPI_ISL_920035, EPI_ISL_920036, EPI_ISL_920037, EPI_ISL_920038, EPI_ISL_920039, EPI_ISL_920043, EPI_ISL_920044, EPI_ISL_920045 | | | |
| see above | University College London, Great Ormond Street Hospital for Children NHS Foundation Trust, Imperial College Healthcare NHS Trust | COVID-19 Genomics UK (COG-UK) Consortium | Sergi Castellano, Rachel Williams, Mark Kristiansen, Paola Resende Silva, Sunando Roy, Tony Brooks, Helena Tutill, Paola Niola, Patricia Dyal, Charlotte Williams, Leysa Forrest, Yasmin Panchbhaya, Jacqueline Findlay, Samuel Weeks, Julianne Brown, Kathryn Harris, Paul Randell, James Price, Alison Holmes, Judith Breuer |
| EPI_ISL_920174, EPI_ISL_920175, EPI_ISL_920177, EPI_ISL_920180, EPI_ISL_920181, EPI_ISL_920182, EPI_ISL_920183, EPI_ISL_920184, EPI_ISL_920186, EPI_ISL_920190, EPI_ISL_920194, EPI_ISL_920196, EPI_ISL_920197, EPI_ISL_920198, EPI_ISL_920200, EPI_ISL_920201, EPI_ISL_920202, EPI_ISL_920203, EPI_ISL_920205, EPI_ISL_920206, EPI_ISL_920207, EPI_ISL_920208, EPI_ISL_920210, EPI_ISL_920211, EPI_ISL_920212, EPI_ISL_920215, EPI_ISL_920217, EPI_ISL_920219, EPI_ISL_920222, EPI_ISL_920224, EPI_ISL_920225, EPI_ISL_920226, EPI_ISL_920227, EPI_ISL_920228, EPI_ISL_920229, EPI_ISL_920230, EPI_ISL_920232, EPI_ISL_920233, EPI_ISL_920237, EPI_ISL_920238, EPI_ISL_920245, EPI_ISL_920246, EPI_ISL_920253, EPI_ISL_920254, EPI_ISL_920263, EPI_ISL_920264, EPI_ISL_920265, EPI_ISL_920273, EPI_ISL_920274, EPI_ISL_920275, EPI_ISL_920285, EPI_ISL_920286, EPI_ISL_920294, EPI_ISL_920295, EPI_ISL_920296, EPI_ISL_920305, EPI_ISL_920306, EPI_ISL_920315, EPI_ISL_920360 | | | |
| see above | University College London Hospital | COVID-19 Genomics UK (COG-UK) Consortium | Judith Heaney, Matthew Byott, Catherine Houlihan, Dan Frampton, Stuart Kirk, Moira Spyer and Eleni Nastouli |
| EPI_ISL_920882, EPI_ISL_920883, EPI_ISL_920884, EPI_ISL_920885, EPI_ISL_920886, EPI_ISL_920887, EPI_ISL_920888, EPI_ISL_920889, EPI_ISL_920890, EPI_ISL_920891, EPI_ISL_920892 | | | |
| see above | University College London, Great Ormond Street Hospital for Children NHS Foundation Trust, Imperial College Healthcare NHS Trust | COVID-19 Genomics UK (COG-UK) Consortium | Sergi Castellano, Rachel Williams, Mark Kristiansen, Paola Resende Silva, Sunando Roy, Tony Brooks, Helena Tutill, Paola Niola, Patricia Dyal, Charlotte Williams, Leysa Forrest, Yasmin Panchbhaya, Jacqueline Findlay, Samuel Weeks, Julianne Brown, Kathryn Harris, Paul Randell, James Price, Alison Holmes, Judith Breuer |
| EPI_ISL_920901, EPI_ISL_920902, EPI_ISL_920903, EPI_ISL_920904, EPI_ISL_920905, EPI_ISL_920906, EPI_ISL_920907, EPI_ISL_920908, EPI_ISL_920909, EPI_ISL_920911, EPI_ISL_920912, EPI_ISL_920913, EPI_ISL_920914, EPI_ISL_920915, EPI_ISL_920916, EPI_ISL_920917, EPI_ISL_920918, EPI_ISL_920919, EPI_ISL_920920, EPI_ISL_920921, EPI_ISL_920922, EPI_ISL_920923, EPI_ISL_920925, EPI_ISL_920926, EPI_ISL_920927, EPI_ISL_920928, EPI_ISL_920929, EPI_ISL_920930, EPI_ISL_920931, EPI_ISL_920932, EPI_ISL_920933, EPI_ISL_920935, EPI_ISL_920936, EPI_ISL_920937, EPI_ISL_920938, EPI_ISL_920939, EPI_ISL_920940, EPI_ISL_920941, EPI_ISL_920942, EPI_ISL_920944, EPI_ISL_920945, EPI_ISL_920946, EPI_ISL_920947, EPI_ISL_920948, EPI_ISL_920949, EPI_ISL_920950, EPI_ISL_920951, EPI_ISL_920952, EPI_ISL_920953, EPI_ISL_920954, EPI_ISL_920955, EPI_ISL_920956, EPI_ISL_920957, EPI_ISL_920958, EPI_ISL_920959, EPI_ISL_920960, EPI_ISL_920961, EPI_ISL_920962, EPI_ISL_920963, EPI_ISL_920964, EPI_ISL_920965, EPI_ISL_920966, EPI_ISL_920967, EPI_ISL_920968, EPI_ISL_920969, EPI_ISL_920970, EPI_ISL_920971, EPI_ISL_920972, EPI_ISL_920973, EPI_ISL_920974 | | | |
| see above | Department of Pathology, University of Cambridge | COVID-19 Genomics UK (COG-UK) Consortium | Aminu S. Jahun, Yasmin Chaudhry, Iliana Georgana, Myra Hosmillo, Rhys Izu, Martin D. Curran, Surendra Parmar, Ian Goodfellow |
| EPI_ISL_920976, EPI_ISL_920977, EPI_ISL_920978, EPI_ISL_920979, EPI_ISL_920981, EPI_ISL_920982, EPI_ISL_920983, EPI_ISL_920984, EPI_ISL_920985, EPI_ISL_920986, EPI_ISL_920987, EPI_ISL_920988, EPI_ISL_920989, EPI_ISL_920990, EPI_ISL_920991, EPI_ISL_920992, EPI_ISL_920993, EPI_ISL_920994, EPI_ISL_920995, EPI_ISL_921011, EPI_ISL_921012, EPI_ISL_921013, EPI_ISL_921014, EPI_ISL_921015, EPI_ISL_921016, EPI_ISL_921017 | | | |
| see above | Regional Virus Laboratory, Belfast Health and Social Care Trust | COVID-19 Genomics UK (COG-UK) Consortium | Conall McCaughey, James McKenna, Tanya Curran, Susan Feeney, Alison Watt, Ciara Cox, Mairead Connor, Zoltan Molnar, David Simpson, Derek Fairley |
| EPI_ISL_921428, EPI_ISL_921486, EPI_ISL_921487, EPI_ISL_921502, EPI_ISL_921521, EPI_ISL_921523, EPI_ISL_921524, EPI_ISL_921525, EPI_ISL_921526, EPI_ISL_921527, EPI_ISL_921528, EPI_ISL_921529, EPI_ISL_921531, EPI_ISL_921532, EPI_ISL_921533, EPI_ISL_921534, EPI_ISL_921535, EPI_ISL_921536, EPI_ISL_921537, EPI_ISL_921538, EPI_ISL_921539, EPI_ISL_921540, EPI_ISL_921542, EPI_ISL_921543, EPI_ISL_921544, EPI_ISL_921545, EPI_ISL_921547, EPI_ISL_921548, EPI_ISL_921549, EPI_ISL_921550, EPI_ISL_921551, EPI_ISL_921552, EPI_ISL_921553, EPI_ISL_921554, EPI_ISL_921555, EPI_ISL_921556, | | | |

| | | | |
|---|---|---|---|
| EPI_ISL_921557, EPI_ISL_921559, EPI_ISL_921562, EPI_ISL_921567, EPI_ISL_921568, EPI_ISL_921570, EPI_ISL_921571, EPI_ISL_921573, EPI_ISL_921574 | | | |
| see above | Northumbria University / South Tees Hospitals NHS Foundation Trust / North Cumbria Integrated Care NHS Foundation Trust / North Tees and Hartlepool NHS Foundation Trust / Newcastle Hospitals NHS Foundation Trust | COVID-19 Genomics UK (COG-UK) Consortium | Darren L Smith,Andrew Nelson,Matthew Bashton,Greg R Young,Joshua Loh,John Allan,Mohammad A Tariq,Giles S Holt,Gary Black,Wen C Yew,Lynn Dover,Paul Baker,Steve Liggett,Sarah Essex,Jane Greenaway,Debra Padgett,Clive Graham,Garren Scott,Edward Barton,Emma Swindells,Brendan Payne,Jennifer Collins,Yusri Taha,Gary Eltringham |
| EPI_ISL_922113, EPI_ISL_922114, EPI_ISL_922117, EPI_ISL_922118, EPI_ISL_922119, EPI_ISL_922120, EPI_ISL_922121, EPI_ISL_922122, EPI_ISL_922123, EPI_ISL_922124, EPI_ISL_922125, EPI_ISL_922126, EPI_ISL_922127, EPI_ISL_922128, EPI_ISL_922129, EPI_ISL_922130, EPI_ISL_922131, EPI_ISL_922132, EPI_ISL_922133, EPI_ISL_922134 | | | |
| see above | Lincolnshire Hospitals and DeepSeq Nottingham | COVID-19 Genomics UK (COG-UK) Consortium | Nichola Duckworth, Tim Sloan, Sarah Walsh, Jonathan Ball, Patrick McClure, Joeseph Chappell, Nadine Holmes, Matthew Carlisle, Christopher Moore, Fei Sang, Johnny Debebe, Victoria Wright, Matthew Loose |
| EPI_ISL_922231, EPI_ISL_922232, EPI_ISL_922233, EPI_ISL_922234, EPI_ISL_922238, EPI_ISL_922239, EPI_ISL_922240 | Oxford Viromics, NDM, University of Oxford; Oxford University Hospitals; Basingstoke and North Hampshire Hospital | COVID-19 Genomics UK (COG-UK) Consortium | Tanya Golubchik, David Bonsall, George Macintyre, Amy Trebes, Mariateresa de Cesare, Catrin Moore, Alex Mobbs, Anita Justice, Robert Shaw, Monique Andersson, Timothy Peto, Emma Wise, Nathan Moore, Jessica Lynch, Nick Cortes, Matilde Mori, Stephen Kidd, David Buck, John Todd, Christophe Fraser |
| EPI_ISL_922360, EPI_ISL_922362, EPI_ISL_922363, EPI_ISL_922365, EPI_ISL_922384, EPI_ISL_922392, EPI_ISL_922395, EPI_ISL_922396, EPI_ISL_922398, EPI_ISL_922399, EPI_ISL_922401, EPI_ISL_922402, EPI_ISL_922403, EPI_ISL_922404, EPI_ISL_922406, EPI_ISL_922407, EPI_ISL_922408, EPI_ISL_922409, EPI_ISL_922410, EPI_ISL_922411, EPI_ISL_922413, EPI_ISL_922415, EPI_ISL_922416, EPI_ISL_922417, EPI_ISL_922418, EPI_ISL_922419, EPI_ISL_922420, EPI_ISL_922421, EPI_ISL_922423, EPI_ISL_922425, EPI_ISL_922428, EPI_ISL_922430, EPI_ISL_922432, EPI_ISL_922433, EPI_ISL_922434, EPI_ISL_922435, EPI_ISL_922440, EPI_ISL_922442, EPI_ISL_922444, EPI_ISL_922445, EPI_ISL_922446, EPI_ISL_922448, EPI_ISL_922452, EPI_ISL_922455, EPI_ISL_922457, EPI_ISL_922458, EPI_ISL_922459, EPI_ISL_922460, EPI_ISL_922462, EPI_ISL_922463, EPI_ISL_922464, EPI_ISL_922465, EPI_ISL_922466, EPI_ISL_922467, EPI_ISL_922468, EPI_ISL_922469, EPI_ISL_922470, EPI_ISL_922473, EPI_ISL_922474, EPI_ISL_922481, EPI_ISL_922553, EPI_ISL_922563, EPI_ISL_922565, EPI_ISL_922566, EPI_ISL_922567, EPI_ISL_922568, EPI_ISL_922570, EPI_ISL_922571, EPI_ISL_922572, EPI_ISL_922574, EPI_ISL_922580, EPI_ISL_922581, EPI_ISL_922582, EPI_ISL_922590, EPI_ISL_922591, EPI_ISL_922592, EPI_ISL_922593, EPI_ISL_922594, EPI_ISL_922595, EPI_ISL_922596, EPI_ISL_922597, EPI_ISL_922598, EPI_ISL_922599, EPI_ISL_922601, EPI_ISL_922633, EPI_ISL_922634, EPI_ISL_922643, EPI_ISL_922645, EPI_ISL_922646, EPI_ISL_922647, EPI_ISL_922648, EPI_ISL_922649, EPI_ISL_922650, EPI_ISL_922651, EPI_ISL_922652, EPI_ISL_922653, EPI_ISL_922654, EPI_ISL_922655, EPI_ISL_922659, EPI_ISL_922660, EPI_ISL_922663, EPI_ISL_922664, EPI_ISL_922665, EPI_ISL_922666, EPI_ISL_922667, EPI_ISL_922668, EPI_ISL_922669, EPI_ISL_922672, EPI_ISL_922673, EPI_ISL_922674, EPI_ISL_922675, EPI_ISL_922676, EPI_ISL_922677, EPI_ISL_922678, EPI_ISL_922679, EPI_ISL_922680, EPI_ISL_922681, EPI_ISL_922682, EPI_ISL_922683, EPI_ISL_922684, EPI_ISL_922685, EPI_ISL_922686, EPI_ISL_922687, EPI_ISL_922688, EPI_ISL_922689, EPI_ISL_922690, EPI_ISL_922693, EPI_ISL_922695, EPI_ISL_922696, EPI_ISL_922698, EPI_ISL_922701, EPI_ISL_922702, EPI_ISL_922705, EPI_ISL_922706, EPI_ISL_922707, EPI_ISL_922710, EPI_ISL_922895, EPI_ISL_922903, EPI_ISL_922904, EPI_ISL_922905, EPI_ISL_922913, EPI_ISL_922923, EPI_ISL_922925, EPI_ISL_922926, EPI_ISL_922927, EPI_ISL_922928, EPI_ISL_922929, EPI_ISL_922930, EPI_ISL_922931, EPI_ISL_922965, EPI_ISL_922970, EPI_ISL_922974, EPI_ISL_922975, EPI_ISL_922976, EPI_ISL_922977, EPI_ISL_922978, EPI_ISL_922979, EPI_ISL_922980, EPI_ISL_922981, EPI_ISL_922982, EPI_ISL_922983, EPI_ISL_922984, EPI_ISL_922985, EPI_ISL_922986, EPI_ISL_922987, EPI_ISL_922988, EPI_ISL_922989, EPI_ISL_922990, EPI_ISL_922991, EPI_ISL_922992, EPI_ISL_922993, EPI_ISL_922994, EPI_ISL_922995, EPI_ISL_922996, EPI_ISL_922997, EPI_ISL_922998, EPI_ISL_922999, EPI_ISL_923000, EPI_ISL_923001, EPI_ISL_923002, EPI_ISL_923003, EPI_ISL_923004, EPI_ISL_923005, EPI_ISL_923006, EPI_ISL_923007, EPI_ISL_923008, EPI_ISL_923009, EPI_ISL_923010, EPI_ISL_923011, EPI_ISL_923013, EPI_ISL_923014, EPI_ISL_923015 | | | |
| see above | Wales Specialist Virology Centre Sequencing lab: Pathogen Genomics Unit | Public Health Wales Microbiology Cardiff Wales Specialist Virology Centre | Catherine Moore, Johnathan Evans, Laura Gifford, Malorie Perry, Simon Cottrell, Angela Marchbank, Alec Birchley, Alexander Adams, Amy Gaskin, Bree Gatica-Wilcox, Jason Coombes, Joel Southgate, Lauren Gilbert, Lee Graham, Nicole Pacchiarini, Sara Kumziene-Summerhayes, Sarah Taylor, Sophie Jones, Sara Rey, Matthew Bull, Joanne Watkins, Sally Corden, Tom Connor |
| EPI_ISL_923276, EPI_ISL_923280, EPI_ISL_923281, EPI_ISL_923287, EPI_ISL_923290, EPI_ISL_923291, EPI_ISL_923292, EPI_ISL_923294, EPI_ISL_923298, EPI_ISL_923302, EPI_ISL_923303, EPI_ISL_923304, EPI_ISL_923307, EPI_ISL_923308, EPI_ISL_923309, EPI_ISL_923310, EPI_ISL_923311, EPI_ISL_923312, EPI_ISL_923313, EPI_ISL_923314, EPI_ISL_923317, EPI_ISL_923355, EPI_ISL_923356, EPI_ISL_923359, EPI_ISL_923361, EPI_ISL_923364, EPI_ISL_923365, EPI_ISL_923371, EPI_ISL_923414, EPI_ISL_923416, EPI_ISL_923419, EPI_ISL_923438, EPI_ISL_923589, EPI_ISL_923590, EPI_ISL_923594, EPI_ISL_923628, EPI_ISL_923660, EPI_ISL_923664, EPI_ISL_923665, EPI_ISL_923666, EPI_ISL_923667, EPI_ISL_923668, EPI_ISL_923669, EPI_ISL_923670, EPI_ISL_923672, EPI_ISL_923673, EPI_ISL_923674, EPI_ISL_923675, EPI_ISL_923676, EPI_ISL_923677, EPI_ISL_923681, EPI_ISL_923683 | | | |
| see above | Centre for Enzyme Innovation, University of Portsmouth / Translational Research Laboratory, Portsmouth Hospitals NHS Trust | COVID-19 Genomics UK (COG-UK) Consortium | Angela Beckett,Salman Goudarzi,Christopher Fearn,Kate Cook,Katie Loveson,Sharon Glaysher,Scott Elliott,Samuel Robson |
| EPI_ISL_923685, EPI_ISL_923692, EPI_ISL_923695, EPI_ISL_923712, EPI_ISL_923713, EPI_ISL_923714, EPI_ISL_923715, EPI_ISL_923726, EPI_ISL_923738, EPI_ISL_923742, EPI_ISL_923748, EPI_ISL_923752, EPI_ISL_923767, EPI_ISL_923768, EPI_ISL_923776, EPI_ISL_923778, EPI_ISL_923798, EPI_ISL_923804, EPI_ISL_923810, EPI_ISL_923864, EPI_ISL_923916, EPI_ISL_923924, EPI_ISL_923925, EPI_ISL_923939, EPI_ISL_923950, EPI_ISL_923951, EPI_ISL_923952, EPI_ISL_923959, EPI_ISL_923962, EPI_ISL_923982, EPI_ISL_923983, EPI_ISL_923990, EPI_ISL_923996, EPI_ISL_924001, EPI_ISL_924009, EPI_ISL_924014, EPI_ISL_924026, EPI_ISL_924038, EPI_ISL_924041, EPI_ISL_924043, EPI_ISL_924047, EPI_ISL_924048, EPI_ISL_924051, EPI_ISL_924053, EPI_ISL_924067, EPI_ISL_924070, EPI_ISL_924076 | | | |
| see above | Department of Pathology, University of Cambridge | COVID-19 Genomics UK (COG-UK) Consortium | Aminu S. Jahun, Yasmin Chaudhry, Iliana Georgana, Myra Hosmillo, Rhys Izu, Martin D. Curran, Surendra Parmar, Ian Goodfellow |
| EPI_ISL_924083, EPI_ISL_924145, EPI_ISL_924150, EPI_ISL_924177, EPI_ISL_924181, EPI_ISL_924220, EPI_ISL_924224, EPI_ISL_924230, EPI_ISL_924233, EPI_ISL_924239, EPI_ISL_924242, EPI_ISL_924244, EPI_ISL_924247, EPI_ISL_924278, EPI_ISL_924303, EPI_ISL_924325, EPI_ISL_924326, EPI_ISL_924399 | | | |
| see above | Virology Department, Sheffield Teaching Hospitals NHS Foundation Trust/Department of Infection, Immunity and Cardiovascular Disease, The Medical School, University of Sheffield | COVID-19 Genomics UK (COG-UK) Consortium | Thushan de Silva, Matthew Parker, Nikki Smith, Adri Angyal, Rebecca Brown, Luke Green, Rachel Tucker, Paul Parsons, Danielle Groves, Katie Johnson, Laura Carrilero, Alex Keeley, Dave Partridge, Matthew Wyles, Benjamin Lindsey, Mehmet Yavuz, Mohammad Raza, Cariad Evans |
| EPI_ISL_924796, EPI_ISL_924801, EPI_ISL_924802, EPI_ISL_924803, EPI_ISL_924804, EPI_ISL_924806, EPI_ISL_924810, EPI_ISL_924811, EPI_ISL_924812, EPI_ISL_924813, EPI_ISL_924814, EPI_ISL_924815, EPI_ISL_924817, EPI_ISL_924820, EPI_ISL_924821, EPI_ISL_924822, EPI_ISL_924823, EPI_ISL_924824, EPI_ISL_924825, EPI_ISL_924826, EPI_ISL_924830, EPI_ISL_924834, EPI_ISL_924841, EPI_ISL_924842, EPI_ISL_924845, EPI_ISL_924857, EPI_ISL_924859, EPI_ISL_924948, EPI_ISL_924949, EPI_ISL_924952, EPI_ISL_924953, EPI_ISL_924954, EPI_ISL_924956, EPI_ISL_924957, EPI_ISL_924958, EPI_ISL_924959, EPI_ISL_924960, EPI_ISL_924961, EPI_ISL_924962, EPI_ISL_924963, EPI_ISL_924964, EPI_ISL_924965, EPI_ISL_924966, EPI_ISL_924967, EPI_ISL_924968, EPI_ISL_924969, EPI_ISL_924970, EPI_ISL_924971, EPI_ISL_924972, EPI_ISL_924973, EPI_ISL_924974, EPI_ISL_924976, EPI_ISL_924977, EPI_ISL_924978, EPI_ISL_924979, EPI_ISL_924981, EPI_ISL_924982, EPI_ISL_924983, EPI_ISL_924984, EPI_ISL_924985, EPI_ISL_924986, EPI_ISL_924988, EPI_ISL_924989, EPI_ISL_924990, EPI_ISL_924991, EPI_ISL_924992, EPI_ISL_924993, EPI_ISL_924994, EPI_ISL_924995, EPI_ISL_924996, EPI_ISL_924997, EPI_ISL_924998, EPI_ISL_924999, EPI_ISL_925000, EPI_ISL_925001, EPI_ISL_925002, EPI_ISL_925003, EPI_ISL_925004, EPI_ISL_925005, EPI_ISL_925006, EPI_ISL_925007, EPI_ISL_925008, EPI_ISL_925009, EPI_ISL_925010, EPI_ISL_925011, EPI_ISL_925012, EPI_ISL_925013 | | | |
| see above | Bioinformatics and Biostatistics Lab, Advanced Sequencing Facility | COVID-19 Genomics UK (COG-UK) Consortium | Aengus Stewart,Jerome Nicod,Chelsea Sawyer,Laura Cubitt,Harshil Patel,Margaret Crawford |
| EPI_ISL_925078, EPI_ISL_925079, EPI_ISL_925080 | TXDSHS | TXDSHS | Bonnie Oh, Anita Pokharel, James Daniel Bonser, Myong Koag, Chung Wang, Rachel Lee, Grace Kubin, Rashmi Tuladhar, Mayela Pedrueza, Maliha Rahman, Jenny Zhang |
| EPI_ISL_925137, EPI_ISL_925138, EPI_ISL_925139, EPI_ISL_925140 | Quest Diagnostics | Quest Diagnostics | Rosenthal,S.H., Gerasimova,A., Kagan,R.M., Anderson, B., Hua, M., Liu Y., Bernstein, L.E., Livingston, K.E., Perez, A., Shalhout, D.F., Shlyakhter, I.A., Owen, R., Tanpaiboon, P., Lacbawan, F. |
| EPI_ISL_925216, EPI_ISL_925217, EPI_ISL_925218, EPI_ISL_925219, EPI_ISL_925220, EPI_ISL_925221, EPI_ISL_925222, EPI_ISL_925223, EPI_ISL_925226, EPI_ISL_925228, EPI_ISL_925229, EPI_ISL_925230, EPI_ISL_925231, EPI_ISL_925232, EPI_ISL_925233, EPI_ISL_925234, EPI_ISL_925235, EPI_ISL_925236, EPI_ISL_925237, EPI_ISL_925238, EPI_ISL_925239, EPI_ISL_925240, EPI_ISL_925241, EPI_ISL_925242, EPI_ISL_925243, EPI_ISL_925244, EPI_ISL_925245, EPI_ISL_925246, EPI_ISL_925247, EPI_ISL_925248, EPI_ISL_925249, EPI_ISL_925250, EPI_ISL_925251, EPI_ISL_925252, EPI_ISL_925254, EPI_ISL_925255, EPI_ISL_925256, EPI_ISL_925257 | | | |
| see above | New Mexico Department of Health Scientific Laboratory | New Mexico Department of Health Scientific Laboratory | Ellie Johnson, Anastacia Griego-Fisher, D'eldra Malone, Jennifer Benoit |
| EPI_ISL_925266, EPI_ISL_925268, EPI_ISL_925270, EPI_ISL_925271, EPI_ISL_925274, EPI_ISL_925276, EPI_ISL_925288, EPI_ISL_925292, EPI_ISL_925297, EPI_ISL_925298, EPI_ISL_925299 | | | |
| see above | Wyoming Public Health Laboratory | Wyoming Public Health Laboratory | Noah Hull, Taylor Fearing, Lynette Gumbleton, Channing Weber, Ashley Norberg, Bailey Bowcutt, and Wanda Manley |
| EPI_ISL_925402, EPI_ISL_925403, EPI_ISL_925404, EPI_ISL_925405 | Department of Clinical Microbiology | GIGA Medical Genomics | Keith Durkin, Maria Artesi, Sébastien Bontems, Raphaël Boreux, Bouchra Boujemla, Cécile Meex, Pierrette Melin, Marie-Pierre Hayette, Vincent Bours |
| EPI_ISL_925888, EPI_ISL_925889, EPI_ISL_925890, EPI_ISL_925891, EPI_ISL_925892, EPI_ISL_925909, EPI_ISL_925910 | Nucleic Acid Testing, National Reference Laboratory | GIGA Medical Genomics | Yvan Butera, Keith Durkin, Maria Artesi, Bouchra Boujemla, Robert Rutayisire, Patrick Tuyisenge, Esperence Umumararungu, Sébastien Bontems, Marie-Pierre Hayette, Nathalie Renotte, Swaibu Gatare, Jacob Souopgui, Sabin Nsanzimana, Vincent Bours, Léon Mutesa |
| EPI_ISL_930784 | Hematology Laboratory, Section of Molecular Diagnostics, University Clinical Centre, Medical University of Gdansk | Laboratory of Recombinant Vaccines | Lukasz Rabalski, Maciej Kosinski, Maciej Grzybek, Adam Sodol, Aneta Szulc, Krzysztof Lewandowski, Ewa Milosz, Marlena Robakowska, Boguslaw Szewczyk, Krystyna Bienkowska-Szewczyk |
| EPI_ISL_930785 | Laboratory of Molecular Biology, Diagnostyka sp. z o.o. | Laboratory of Recombinant Vaccines | Lukasz Rabalski, Maciej Kosinski, Anna Piotrowska-Mietelska,Izabela Szczygielska, Boguslaw Szewczyk, Krystyna Bienkowska-Szewczyk |
| EPI_ISL_931584, EPI_ISL_933718, EPI_ISL_933721, EPI_ISL_933734, EPI_ISL_933740 | Servicio de Microbiología Hospital Ramón y Cajal | Servicio de Microbiología Hospital Ramón y Cajal | José M Gonzalez-Alba, Concepción Rodríguez, Melanie Abreu, Laura Martínez, Val F Lanza, Luz Leticia Olavarrieta, Rafael Cantón, JC Galán |
| EPI_ISL_933760 | DPHL | Delaware Public Health Lab | Gregory Hovan |
| EPI_ISL_933768, EPI_ISL_933770 | Servicio de Microbiología Hospital Ramón y Cajal | Servicio de Microbiología Hospital Ramón y Cajal | José M Gonzalez-Alba, Concepción Rodríguez, Melanie Abreu, Laura Martínez, Val F Lanza, Luz Leticia Olavarrieta, Rafael Cantón, JC Galán |

| | | | |
|---|---|---|---|
| EPI_ISL_933791 | PathWest Laboratory Medicine WA | PathWest Laboratory Medicine WA Microbial Surveillance Unit | PathWest Laboratory Medicine WA Microbial Surveillance Unit |
| EPI_ISL_934234, EPI_ISL_934235, EPI_ISL_934236, EPI_ISL_934237, EPI_ISL_934239, EPI_ISL_934240, EPI_ISL_934242, EPI_ISL_934243, EPI_ISL_934244, EPI_ISL_934245, EPI_ISL_934246, EPI_ISL_934247, EPI_ISL_934248, EPI_ISL_934249, EPI_ISL_934250, EPI_ISL_934251, EPI_ISL_934252, EPI_ISL_934253, EPI_ISL_934254, EPI_ISL_934255, EPI_ISL_934256, EPI_ISL_934257, EPI_ISL_934258, EPI_ISL_934259, EPI_ISL_934260, EPI_ISL_934261, EPI_ISL_934262, EPI_ISL_934263, EPI_ISL_934264, EPI_ISL_934265, EPI_ISL_934266, EPI_ISL_934267, EPI_ISL_934268, EPI_ISL_934269, EPI_ISL_934270, EPI_ISL_934271, EPI_ISL_934272, EPI_ISL_934273, EPI_ISL_934274, EPI_ISL_934275, EPI_ISL_934276, EPI_ISL_934277, EPI_ISL_934278, EPI_ISL_934279 | | | |
| see above | Vilnius university hospital Santaros Klinikos, Center of Laboratory Medicine | Vilnius university hospital Santaros Klinikos, Center of Laboratory Medicine | Ingrida Olendraite, Daniel Naumovas, Rimvydas Norvilas, Dovile Ezerskyte, Justinas Slikas, Gytis Dudas |
| EPI_ISL_934315 | Narhalsan Fjallbacka VC | The Public Health Agency of Sweden | Anna-Malin Linde, Maria Lind Karlberg, Carlo Berg, Oskar Karlsson Lindsjo, Sofia Stamouli, Reza Advani, Mattias Haukland, Petra Holmstrom, Noura Walai, Petra Edquist, Mia Brytting, Anna Risberg, Karin Tegmark-Wisell |
| EPI_ISL_934375, EPI_ISL_934376, EPI_ISL_934379, EPI_ISL_934380 | Klinisk mikrobiologi | The Public Health Agency of Sweden | Anna-Malin Linde, Maria Lind Karlberg, Carlo Berg, Oskar Karlsson Lindsjo, Sofia Stamouli, Reza Advani, Mattias Haukland, Petra Holmstrom, Noura Walai, Petra Edquist, Mia Brytting, Anna Risberg, Karin Tegmark-Wisell |
| EPI_ISL_934399, EPI_ISL_934401, EPI_ISL_934402, EPI_ISL_934403, EPI_ISL_934404 | Synlab Medilab, Mikrobiologi | The Public Health Agency of Sweden | Anna-Malin Linde, Maria Lind Karlberg, Carlo Berg, Oskar Karlsson Lindsjo, Sofia Stamouli, Reza Advani, Mattias Haukland, Petra Holmstrom, Noura Walai, Petra Edquist, Mia Brytting, Anna Risberg, Karin Tegmark-Wisell |
| EPI_ISL_934408, EPI_ISL_934409 | Klinisk Mikrobiologi | The Public Health Agency of Sweden | Anna-Malin Linde, Maria Lind Karlberg, Carlo Berg, Oskar Karlsson Lindsjo, Sofia Stamouli, Reza Advani, Mattias Haukland, Petra Holmstrom, Noura Walai, Petra Edquist, Mia Brytting, Anna Risberg, Karin Tegmark-Wisell |
| EPI_ISL_934410, EPI_ISL_934411, EPI_ISL_934412 | Synlab Medilab, Mikrobiologi | The Public Health Agency of Sweden | Anna-Malin Linde, Maria Lind Karlberg, Carlo Berg, Oskar Karlsson Lindsjo, Sofia Stamouli, Reza Advani, Mattias Haukland, Petra Holmstrom, Noura Walai, Petra Edquist, Mia Brytting, Anna Risberg, Karin Tegmark-Wisell |
| EPI_ISL_934548, EPI_ISL_934549, EPI_ISL_934550, EPI_ISL_934551, EPI_ISL_934552, EPI_ISL_934553, EPI_ISL_934554, EPI_ISL_934555, EPI_ISL_934556, EPI_ISL_934557, EPI_ISL_934558, EPI_ISL_934559, EPI_ISL_934560, EPI_ISL_934561, EPI_ISL_934562, EPI_ISL_934563, EPI_ISL_934564, EPI_ISL_934565, EPI_ISL_934566, EPI_ISL_934567, EPI_ISL_934568 | | | |
| see above | Department of Laboratory Medicine, Division of Clinical Virology, University of Medicine, Vienna | Bergthaler laboratory, CeMM Research Center for Molecular Medicine of the Austrian Academy of Sciences | Lukas Endler, Anna Schedl, Thomas Penz, Benedikt Agerer, Maelle Le Moing, Michael Schuster, Bekir Erguner, Jan Laine, Martin Senekowitsch, Christoph Bock, Andreas Bergthaler |
| EPI_ISL_934651 | Institute for Water Quality and Resource Management, Technical University Vienna | Bergthaler laboratory, CeMM Research Center for Molecular Medicine of the Austrian Academy of Sciences | Lukas Endler, Anna Schedl, Thomas Penz, Benedikt Agerer, Maelle Le Moing, Michael Schuster, Bekir Erguner, Jan Laine, Martin Senekowitsch, Christoph Bock, Andreas Bergthaler |
| EPI_ISL_934652, EPI_ISL_934653, EPI_ISL_934654 | Klinikum Wels-Grieskirchen | Bergthaler laboratory, CeMM Research Center for Molecular Medicine of the Austrian Academy of Sciences | Lukas Endler, Anna Schedl, Thomas Penz, Benedikt Agerer, Maelle Le Moing, Michael Schuster, Bekir Erguner, Jan Laine, Martin Senekowitsch, Christoph Bock, Andreas Bergthaler |
| EPI_ISL_934682, EPI_ISL_934683, EPI_ISL_934684, EPI_ISL_934685, EPI_ISL_934686, EPI_ISL_934687, EPI_ISL_934688, EPI_ISL_934689, EPI_ISL_934690, EPI_ISL_934691, EPI_ISL_934692, EPI_ISL_934693, EPI_ISL_934694, EPI_ISL_934695 | | | |
| see above | Department of Laboratory Medicine, Division of Clinical Virology, University of Medicine, Vienna | Bergthaler laboratory, CeMM Research Center for Molecular Medicine of the Austrian Academy of Sciences | Lukas Endler, Anna Schedl, Thomas Penz, Benedikt Agerer, Maelle Le Moing, Michael Schuster, Bekir Erguner, Jan Laine, Martin Senekowitsch, Christoph Bock, Andreas Bergthaler |
| EPI_ISL_934980, EPI_ISL_934990, EPI_ISL_934997, EPI_ISL_935005, EPI_ISL_935006, EPI_ISL_935007, EPI_ISL_935008, EPI_ISL_935009, EPI_ISL_935010 | ADMED Microbiologie | Genomics and Transcriptomics, Philip Morris International | Reto Lienhard, Marie-Lise Tritten, Emmanuel Guedj, Nicolas Sierro, Rémi Dulize, David Bornand, Mehdi Auberson, Maxime Berthouzoz, Nikolai Ivanov, Manuel Peitsch |
| EPI_ISL_935195, EPI_ISL_935212, EPI_ISL_935213, EPI_ISL_935222, EPI_ISL_935237, EPI_ISL_935238, EPI_ISL_935239 | KU Leuven, Rega Institute, Clinical and Epidemiological Virology | KU Leuven, Rega Institute, Clinical and Epidemiological Virology | Tony Wawina-Bokalanga, Bert Vanmechelen, Joan Marti-Carerras, Piet Maes |
| EPI_ISL_935527, EPI_ISL_935528, EPI_ISL_935529, EPI_ISL_935530, EPI_ISL_935531 | Labo Analyses Med | National Reference Center for Viruses of Respiratory Infections, Institut Pasteur, Paris | Marion Barbet, Sylvie Behillil, Méline Bizard, Angela Brisebarre, Camille Capel, Etienne Simon-Lorière, Vincent Enouf, Maud Vanpeene, Sylvie van der Werf,Merah Kader |
| EPI_ISL_935537 | Labo Analyses Med | National Reference Center for Viruses of Respiratory Infections, Institut Pasteur, Paris | Marion Barbet, Sylvie Behillil, Méline Bizard, Angela Brisebarre, Camille Capel, Etienne Simon-Lorière, Vincent Enouf, Maud Vanpeene, Sylvie van der Werf,Bonnaudet GéRaldine |
| EPI_ISL_935547 | Labo Analyses Med | National Reference Center for Viruses of Respiratory Infections, Institut Pasteur, Paris | Marion Barbet, Sylvie Behillil, Méline Bizard, Angela Brisebarre, Camille Capel, Etienne Simon-Lorière, Vincent Enouf, Maud Vanpeene, Sylvie van der Werf,Besson J |
| EPI_ISL_935548 | Labo Analyses Med | National Reference Center for Viruses of Respiratory Infections, Institut Pasteur, Paris | Marion Barbet, Sylvie Behillil, Méline Bizard, Angela Brisebarre, Camille Capel, Etienne Simon-Lorière, Vincent Enouf, Maud Vanpeene, Sylvie van der Werf,Pasquier |
| EPI_ISL_935551, EPI_ISL_935552, EPI_ISL_935553, EPI_ISL_935554, EPI_ISL_935555 | Hopital | National Reference Center for Viruses of Respiratory Infections, Institut Pasteur, Paris | Marion Barbet, Sylvie Behillil, Méline Bizard, Angela Brisebarre, Camille Capel, Etienne Simon-Lorière, Vincent Enouf, Maud Vanpeene, Sylvie van der Werf,Goudeau Alain |
| EPI_ISL_935567, EPI_ISL_935568 | Hopital | National Reference Center for Viruses of Respiratory Infections, Institut Pasteur, Paris | Marion Barbet, Sylvie Behillil, Méline Bizard, Angela Brisebarre, Camille Capel, Etienne Simon-Lorière, Vincent Enouf, Maud Vanpeene, Sylvie van der Werf,Bret Laurent |
| EPI_ISL_935581 | Hopital | National Reference Center for Viruses of Respiratory Infections, Institut Pasteur, Paris | Marion Barbet, Sylvie Behillil, Méline Bizard, Angela Brisebarre, Camille Capel, Etienne Simon-Lorière, Vincent Enouf, Maud Vanpeene, Sylvie van der Werf,Brichler SéGolèNe |
| EPI_ISL_935644 | Labo Analyses Med | National Reference Center for Viruses of Respiratory Infections, Institut Pasteur, Paris | Marion Barbet, Sylvie Behillil, Méline Bizard, Angela Brisebarre, Camille Capel, Etienne Simon-Lorière, Vincent Enouf, Maud Vanpeene, Sylvie van der Werf |
| EPI_ISL_936005 | SUNY UPSTATE MEDICAL UNIVERSITY | Wadsworth Center, New York State Department of Health | Kirsten St. George, Daryl M. Lamson, Alexis Russel, Matthew Shudt, Melissa A Leisner, Jonathan Plitnick, Navjot Singh, John Kelly, Erasmus Schneider, Erica Lasek-Nesselquist |
| EPI_ISL_936026, EPI_ISL_936034, EPI_ISL_936035 | THE MARY IMOGENE BASSETT HOSPITAL | Wadsworth Center, New York State Department of Health | Kirsten St. George, Daryl M. Lamson, Alexis Russel, Matthew Shudt, Melissa A Leisner, Jonathan Plitnick, Navjot Singh, John Kelly, Erasmus Schneider, Erica Lasek-Nesselquist |
| EPI_ISL_936237, EPI_ISL_936238, EPI_ISL_936239, EPI_ISL_936240, EPI_ISL_936241 | Wadsworth Center, New York State Department of Health | Wadsworth Center, New York State Department of Health | Kirsten St. George, Daryl M. Lamson, Alexis Russel, Matthew Shudt, Melissa A Leisner, Jonathan Plitnick, Navjot Singh, John Kelly, Erasmus Schneider, Erica Lasek-Nesselquist |
| EPI_ISL_936249, EPI_ISL_936260, EPI_ISL_936261, EPI_ISL_936264, EPI_ISL_936266, EPI_ISL_936267, EPI_ISL_936269, EPI_ISL_936272, EPI_ISL_936273, EPI_ISL_936279, EPI_ISL_936285, EPI_ISL_936288, EPI_ISL_936289, EPI_ISL_936292, EPI_ISL_936296 | | | |
| see above | MONTEFIORE MEDICAL CENTER LABORATORIES | Wadsworth Center, New York State Department of Health | Kirsten St. George, Daryl M. Lamson, Alexis Russel, Matthew Shudt, Melissa A Leisner, Jonathan Plitnick, Navjot Singh, John Kelly, Erasmus Schneider, Erica Lasek-Nesselquist |
| EPI_ISL_936464, EPI_ISL_936465, EPI_ISL_936468, EPI_ISL_936479 | DPH, Massachusetts State Public Health Lab | DPH, Massachusetts State Public Health Lab | Lang,A.S., Fink,T., Gallagher,G.R., Smole,S.C. |
| EPI_ISL_937122, EPI_ISL_937123, EPI_ISL_937124, EPI_ISL_937125 | Quest Diagnostics | Quest Diagnostics | Rosenthal,S.H., Gerasimova,A., Kagan,R.M., Anderson, B., Hua, M., Liu Y., Bernstein, L.E., Livingston, K.E., Perez, A., Shalhout, D.F., Shlyakhter, I.A., Owen, R., Tanpaiboon, P., Lacbawan, F. |
| EPI_ISL_937362, EPI_ISL_937364, EPI_ISL_937365, EPI_ISL_937366, EPI_ISL_937367, EPI_ISL_937368, EPI_ISL_937369, EPI_ISL_937370, EPI_ISL_937371, EPI_ISL_937372, EPI_ISL_937374, EPI_ISL_937375, EPI_ISL_937378, EPI_ISL_937380 | | | |
| see above | Utah Public Health Laboratory | Utah Public Health Laboratory | Erin L. Young, Kelly F. Oakeson, Tara Gallagher |

| | | | |
|---|---|---|---|
| EPI_ISL_937408 | Maine Health and Environmental Testing Laboratory (Maine HETL) | Tewhey Lab, The Jackson Laboratory | Matluk,N., Dewey,H., Iosue,F., Barter,M., Lynch,R., Munger,H. and Tewhey,R. |
| EPI_ISL_940068, EPI_ISL_940071, EPI_ISL_940072, EPI_ISL_940073 | University Hospitals of Geneva, Laboratory of Virology | HUG, Laboratory of Virology and the Health2030 Genome Center | Samuel Cordey, Ana Rita Goncalves, Laurent Kaiser, Lorenzo Cerutti, Henri Pegeot, Melyssa Elies, Deborah Penet, Keith Harshman, Ioannis Xenarios, Emmanouil Dermitzakis |
| EPI_ISL_940157 | Laboratory of Virology and Molecular Diagnostics | Institute of Public Health of Republic of North Macedonia Laboratory of Virology and Molecular Diagnostics | Maja Kuzmanovska, Golubinka Bosehvska, Elizabeta Janchevska |
| EPI_ISL_940620, EPI_ISL_940621 | Hospital Sao Joaquim - Beneficiencia Portuguesa | Instituto Adolfo Lutz, Interdiciplinary Procedures Center, Strategic Laboratory | Claudio Tavares Sacchi, Claudia Regina Gonçalves, Erica Valessa Ramos Gomes, Karoline Rodrigues Campos |
| EPI_ISL_940732 | City of Milwaukee Health Department Laboratory | City of Milwaukee Health Department Laboratory | Sanjib Bhattacharyya |
| EPI_ISL_940749 | University of Bari Biomedical Sciences and Human Oncology | University of Bari Biomedical Sciences and Human Oncology | Chironna M., Sallustio A., Loconsole D., Accogli M. |
| EPI_ISL_940868, EPI_ISL_940871, EPI_ISL_940875, EPI_ISL_940888, EPI_ISL_940889 | Vaccines and Infectious Diseases Analytics Research Unit (VIDA) | KRISP, KZN Research Innovation and Sequencing Platform | Baillie Vicky, du Plessis Jeanine, Giandhari Jennifer, Pillay Sureshnee, Naidoo Yeshnee, Tegally Houriiyah, de Oliveira Tulio, Madhi Shabir |
| EPI_ISL_940999 | Labo Analyses Med | National Reference Center for Viruses of Respiratory Infections, Institut Pasteur, Paris | Marion Barbet, Sylvie Behillil, Méline Bizard, Angela Brisebarre, Camille Capel, Etienne Simon-Lorière, Vincent Enouf, Maud Vanpeene, Sylvie van der Werf, Gestin (B) Brieuc |
| EPI_ISL_941000 | Hopital | National Reference Center for Viruses of Respiratory Infections, Institut Pasteur, Paris | Marion Barbet, Sylvie Behillil, Méline Bizard, Angela Brisebarre, Camille Capel, Etienne Simon-Lorière, Vincent Enouf, Maud Vanpeene, Sylvie van der Werf, Cron Sophie |
| EPI_ISL_941079, EPI_ISL_941080, EPI_ISL_941082, EPI_ISL_941084, EPI_ISL_941086, EPI_ISL_941087, EPI_ISL_941088, EPI_ISL_941091, EPI_ISL_941093, EPI_ISL_941095, EPI_ISL_941096, EPI_ISL_941097, EPI_ISL_941100, EPI_ISL_941101, EPI_ISL_941103 | | | |
| see above | Labo Analyses Med | National Reference Center for Viruses of Respiratory Infections, Institut Pasteur, Paris | Marion Barbet, Sylvie Behillil, Méline Bizard, Angela Brisebarre, Camille Capel, Etienne Simon-Lorière, Vincent Enouf, Maud Vanpeene, Sylvie van der Werf, Merah Kader |
| EPI_ISL_941165, EPI_ISL_941166, EPI_ISL_941167, EPI_ISL_941168, EPI_ISL_941169, EPI_ISL_941170, EPI_ISL_941171 | Hospital Universitari i Politècnic La Fe de València | SeqCOVID-SPAIN consortium/IBV(CSIC) | María Dolores Gómez Ruiz, Eva González Barberá, Ana Gil Brusola, Salvador Giner Almaraz, José Luis López Hontangas and SeqCOVID-SPAIN consortium |
| EPI_ISL_941229, EPI_ISL_941230, EPI_ISL_941231, EPI_ISL_941232 | Laboratorio de Microbiología. Hospital General Universitario de Elda, Alicante | SeqCOVID-SPAIN consortium/IBV(CSIC) | Mª Isabel Gascón Ros, Cristina Torregrosa Hetland, Eva Pastor Boix, Paloma Cascales Ramos and SeqCOVID-SPAIN consortium |
| EPI_ISL_941293 | Nigeria Centre for Disease Control (NCDC) | African Centre of Excellence for Genomics of Infectious Diseases (ACEGID), Redeemer's University | Oluniyi P.E. et al |
| EPI_ISL_941338 | INMI Lazzaro Spallanzani IRCCS | INMI Lazzaro Spallanzani IRCCS | Emanuela Giombini, Ornella Butera, Cesare E.M. Gruber, Martina Rueca, Barbara Bartolini, Francesco Messina, Giandomenico Russo, Daniele Castiglia , Stefania Madonna, Antonino Di Caro, Maria R. Capobianchi |
| EPI_ISL_941346, EPI_ISL_941347, EPI_ISL_941348, EPI_ISL_941349, EPI_ISL_941350, EPI_ISL_941351, EPI_ISL_941352, EPI_ISL_941353, EPI_ISL_941354, EPI_ISL_941355, EPI_ISL_941356, EPI_ISL_941357, EPI_ISL_941358, EPI_ISL_941359, EPI_ISL_941389, EPI_ISL_941390, EPI_ISL_941391, EPI_ISL_941394, EPI_ISL_941395, EPI_ISL_941396, EPI_ISL_941397, EPI_ISL_941398, EPI_ISL_941399, EPI_ISL_941400, EPI_ISL_941401, EPI_ISL_941403, EPI_ISL_941404, EPI_ISL_941411, EPI_ISL_941412, EPI_ISL_941415, EPI_ISL_941416, EPI_ISL_941417, EPI_ISL_941418, EPI_ISL_941419, EPI_ISL_941420, EPI_ISL_941421, EPI_ISL_941422, EPI_ISL_941428, EPI_ISL_941429, EPI_ISL_941433, EPI_ISL_941434, EPI_ISL_941435, EPI_ISL_941436, EPI_ISL_941462, EPI_ISL_941463, EPI_ISL_941464, EPI_ISL_941465, EPI_ISL_941466, EPI_ISL_941467, EPI_ISL_941468, EPI_ISL_941469, EPI_ISL_941473, EPI_ISL_941474, EPI_ISL_941475, EPI_ISL_941476, EPI_ISL_941477, EPI_ISL_941478, EPI_ISL_941479, EPI_ISL_941480, EPI_ISL_941481, EPI_ISL_941482, EPI_ISL_941483, EPI_ISL_941484, EPI_ISL_941485, EPI_ISL_941508, EPI_ISL_941510, EPI_ISL_941511, EPI_ISL_941512, EPI_ISL_941513, EPI_ISL_941515, EPI_ISL_941517, EPI_ISL_941520, EPI_ISL_941521, EPI_ISL_941522, EPI_ISL_941527, EPI_ISL_941528, EPI_ISL_941529, EPI_ISL_941530, EPI_ISL_941531, EPI_ISL_941532, EPI_ISL_941533, EPI_ISL_941534, EPI_ISL_941535, EPI_ISL_941539, EPI_ISL_941540, EPI_ISL_941541, EPI_ISL_941542, EPI_ISL_941547, EPI_ISL_941548, EPI_ISL_941549, EPI_ISL_941550, EPI_ISL_941551, EPI_ISL_941552, EPI_ISL_941574, EPI_ISL_941575, EPI_ISL_941576, EPI_ISL_941577, EPI_ISL_941578, EPI_ISL_941579, EPI_ISL_941580, EPI_ISL_941581, EPI_ISL_941582, EPI_ISL_941583, EPI_ISL_941584, EPI_ISL_941585, EPI_ISL_941586, EPI_ISL_941587, EPI_ISL_941588, EPI_ISL_941589, EPI_ISL_941590, EPI_ISL_941591, EPI_ISL_941592, EPI_ISL_941593, EPI_ISL_941594, EPI_ISL_941595, EPI_ISL_941603, EPI_ISL_941633, EPI_ISL_941636, EPI_ISL_941637, EPI_ISL_941638, EPI_ISL_941640, EPI_ISL_941641, EPI_ISL_941642 | | | |
| see above | Instituto Nacional de Saude (INSA) | Instituto Nacional de Saude (INSA) | Borges et al |
| EPI_ISL_941650, EPI_ISL_941651, EPI_ISL_941652, EPI_ISL_941663, EPI_ISL_941667, EPI_ISL_941670, EPI_ISL_941671, EPI_ISL_941672, EPI_ISL_941673, EPI_ISL_941674, EPI_ISL_941675, EPI_ISL_941676, EPI_ISL_941677, EPI_ISL_941678, EPI_ISL_941680, EPI_ISL_941681, EPI_ISL_941682, EPI_ISL_941683, EPI_ISL_941685, EPI_ISL_941686, EPI_ISL_941687, EPI_ISL_941691, EPI_ISL_941692, EPI_ISL_941693, EPI_ISL_941694, EPI_ISL_941695, EPI_ISL_941704, EPI_ISL_941705, EPI_ISL_941707, EPI_ISL_941712, EPI_ISL_941713, EPI_ISL_941714, EPI_ISL_941715, EPI_ISL_941716, EPI_ISL_941717, EPI_ISL_941718, EPI_ISL_941719, EPI_ISL_941733, EPI_ISL_941734, EPI_ISL_941735, EPI_ISL_941736, EPI_ISL_941737, EPI_ISL_941738, EPI_ISL_941739, EPI_ISL_941740, EPI_ISL_941741, EPI_ISL_941742, EPI_ISL_941743, EPI_ISL_941744, EPI_ISL_941745, EPI_ISL_941746, EPI_ISL_941747, EPI_ISL_941748, EPI_ISL_941749, EPI_ISL_941750, EPI_ISL_941751, EPI_ISL_941752, EPI_ISL_941753, EPI_ISL_941754, EPI_ISL_941755, EPI_ISL_941759, EPI_ISL_941760, EPI_ISL_941761, EPI_ISL_941767, EPI_ISL_941768, EPI_ISL_941769, EPI_ISL_941770, EPI_ISL_941776, EPI_ISL_941777, EPI_ISL_941778, EPI_ISL_941779, EPI_ISL_941780, EPI_ISL_941781, EPI_ISL_941782, EPI_ISL_941783, EPI_ISL_941784, EPI_ISL_941785, EPI_ISL_941786, EPI_ISL_941787, EPI_ISL_941788, EPI_ISL_941800, EPI_ISL_941801, EPI_ISL_941802, EPI_ISL_941809, EPI_ISL_941810, EPI_ISL_941811, EPI_ISL_941812, EPI_ISL_941813, EPI_ISL_941819, EPI_ISL_941828, EPI_ISL_941829, EPI_ISL_941830, EPI_ISL_941831, EPI_ISL_941837, EPI_ISL_941838, EPI_ISL_941839, EPI_ISL_941840, EPI_ISL_941841, EPI_ISL_941846, EPI_ISL_941860, EPI_ISL_941862, EPI_ISL_941863, EPI_ISL_941864, EPI_ISL_941865, EPI_ISL_941866, EPI_ISL_941868, EPI_ISL_941869, EPI_ISL_941870, EPI_ISL_941871, EPI_ISL_941874, EPI_ISL_941875, EPI_ISL_941878, EPI_ISL_941883, EPI_ISL_941884, EPI_ISL_941890, EPI_ISL_941891, EPI_ISL_941892, EPI_ISL_941893, EPI_ISL_941894, EPI_ISL_941895, EPI_ISL_941896, EPI_ISL_941897 | | | |
| see above | Instituto Nacional de Saude (INSA) and Instituto Gulbenkian de Ciencia (IGC) | Instituto Nacional de Saude (INSA) and Instituto Gulbenkian de Ciencia (IGC) | Borges et al |
| EPI_ISL_942822, EPI_ISL_942823, EPI_ISL_942824, EPI_ISL_942825, EPI_ISL_942826, EPI_ISL_942827, EPI_ISL_942828, EPI_ISL_942829, EPI_ISL_942830, EPI_ISL_942831, EPI_ISL_942832, EPI_ISL_942833, EPI_ISL_942834 | | | |
| see above | Gundersen Molecular Diagnostics Laboratory | Kabara Cancer Research Institute | Craig S. Richmond, Paraic A. Kenny |
| EPI_ISL_942928 | Instituto de Diagnostico y Referencia Epidemiologicos INDRE_RNLSP | Instituto de Diagnostico y Referencia Epidemiologicos (INDRE) | Claudia Wong-Arambula, Abril Rodriguez-Maldonado, Fabiola Garces-Ayala, Natividad Cruz-Ortiz, Tatiana Nunez-Garcia, Gisela Barrera-Badillo, Lucia Hernandez-Rivas, Irma Lopez-Martinez, Ernesto Ramirez-Gonzalez. |
| EPI_ISL_943025, EPI_ISL_943053, EPI_ISL_943065, EPI_ISL_943075, EPI_ISL_943076, EPI_ISL_943077, EPI_ISL_943078, EPI_ISL_943086, EPI_ISL_943087, EPI_ISL_943088, EPI_ISL_943089, EPI_ISL_943090, EPI_ISL_943091, EPI_ISL_943117, EPI_ISL_943174, EPI_ISL_943275, EPI_ISL_943280, EPI_ISL_943284, EPI_ISL_943287, EPI_ISL_943288, EPI_ISL_943331, EPI_ISL_943335, EPI_ISL_943336, EPI_ISL_943341, EPI_ISL_943346, EPI_ISL_943353, EPI_ISL_943371, EPI_ISL_943381, EPI_ISL_943507, EPI_ISL_943523 | | | |
| see above | Dutch COVID-19 response team | National Institute for Public Health and the Environment (RIVM) | Adam Meijer, Harry Vennema, Dirk Eggink, Jeroen Cremer, Sharon van den Brink, Bas van der Veer, AnneMarie van den Brandt, Florian Zwagemaker, Dennis Schmitz, Chantal Reusken, on behalf of the national COVID-19 response team |
| EPI_ISL_943805, EPI_ISL_943809 | Servizo de Microbioloxía. Complexo Hospitalario Universitario de Santiago de Compostela | Servizo de Microbioloxía. Complexo Hospitalario Universitario de Santiago de Compostela | Antonio Aguilera, Gema Barbeito, Amparo Coira, José Costa, Rocio Trastoy, María Luisa Pérez del Molino. |
| EPI_ISL_943810, EPI_ISL_943811, EPI_ISL_943841, EPI_ISL_943842, EPI_ISL_943843, EPI_ISL_943844, EPI_ISL_943845, EPI_ISL_943846, EPI_ISL_943847, EPI_ISL_943868, EPI_ISL_943917, EPI_ISL_943918, EPI_ISL_943919, EPI_ISL_943920, EPI_ISL_943921, EPI_ISL_943925, EPI_ISL_943926, EPI_ISL_943935, EPI_ISL_943945, EPI_ISL_943948, EPI_ISL_943953, EPI_ISL_943954, EPI_ISL_943959, EPI_ISL_943960, EPI_ISL_943961, EPI_ISL_943963, EPI_ISL_943965 | | | |
| see above | Utah Public Health Laboratory | Utah Public Health Laboratory | Erin L. Young, Kelly F. Oakeson, Tara Gallagher |
| EPI_ISL_944189, EPI_ISL_944190, EPI_ISL_944191, EPI_ISL_944192, EPI_ISL_944194, EPI_ISL_944195, EPI_ISL_944199, EPI_ISL_944200, EPI_ISL_944201, EPI_ISL_944206, EPI_ISL_944207, EPI_ISL_944214, EPI_ISL_944215, EPI_ISL_944217, EPI_ISL_944218, EPI_ISL_944220, EPI_ISL_944221, EPI_ISL_944224, EPI_ISL_944228, EPI_ISL_944230, EPI_ISL_944231, EPI_ISL_944242, EPI_ISL_944247, EPI_ISL_944248, EPI_ISL_944251, EPI_ISL_944252, EPI_ISL_944253, EPI_ISL_944254, EPI_ISL_944255, EPI_ISL_944339, EPI_ISL_944340, EPI_ISL_944341, EPI_ISL_944342, EPI_ISL_944343, EPI_ISL_944344, EPI_ISL_944345, EPI_ISL_944346, EPI_ISL_944347, EPI_ISL_944348, EPI_ISL_944349, EPI_ISL_944350, EPI_ISL_944351, EPI_ISL_944352, EPI_ISL_944353, EPI_ISL_944354, EPI_ISL_944355, EPI_ISL_944356, EPI_ISL_944357, EPI_ISL_944358, EPI_ISL_944359, EPI_ISL_944360, EPI_ISL_944361, EPI_ISL_944362, EPI_ISL_944363, EPI_ISL_944364, EPI_ISL_944365, EPI_ISL_944367, EPI_ISL_944368, EPI_ISL_944372, EPI_ISL_944375, EPI_ISL_944376, EPI_ISL_944377, EPI_ISL_944384, EPI_ISL_944385, EPI_ISL_944386, EPI_ISL_944390, EPI_ISL_944391, EPI_ISL_944392, EPI_ISL_944393, EPI_ISL_944394, EPI_ISL_944395, EPI_ISL_944397, EPI_ISL_944401, EPI_ISL_944402, EPI_ISL_944407, EPI_ISL_944408, EPI_ISL_944409, EPI_ISL_944410, EPI_ISL_944420, EPI_ISL_944425, EPI_ISL_944426, EPI_ISL_944427, EPI_ISL_944428, EPI_ISL_944516, EPI_ISL_944517, EPI_ISL_944518, EPI_ISL_944519, EPI_ISL_944520, EPI_ISL_944521, EPI_ISL_944522, EPI_ISL_944523, EPI_ISL_944524, EPI_ISL_944525, EPI_ISL_944526, EPI_ISL_944527, EPI_ISL_944528, EPI_ISL_944529, EPI_ISL_944530, EPI_ISL_944531, EPI_ISL_944532, EPI_ISL_944533, EPI_ISL_944534, EPI_ISL_944535, EPI_ISL_944536, EPI_ISL_944537, EPI_ISL_944538, EPI_ISL_944539, EPI_ISL_944540, EPI_ISL_944541, EPI_ISL_944542, EPI_ISL_944543, EPI_ISL_944544, EPI_ISL_944545, EPI_ISL_944546, EPI_ISL_944547, EPI_ISL_944548, EPI_ISL_944549, EPI_ISL_944550, EPI_ISL_944551, EPI_ISL_944552, EPI_ISL_944553, EPI_ISL_944554, EPI_ISL_944555, EPI_ISL_944556, EPI_ISL_944557, EPI_ISL_944558, EPI_ISL_944559, EPI_ISL_944560, EPI_ISL_944561, EPI_ISL_944562, EPI_ISL_944563, EPI_ISL_944564, EPI_ISL_944565 | | | |
| see above | Israel Central Virology laboratory | Israel National Consortium for SARS-CoV-2 sequencing | Neta Zuckerman, Efrat Dahan Bucris, Michal Mandelboim, Dana Bar-Ilan, Oran Erster, Tzvia Mann, Omer Murik, David A. Zeevi, Assaf Rokney, Joseph Jaffe, Eva Nachum, Maya Davidovich Cohen, Ephraim Fass, Gal Zizelski Valenci, Mor Rubinstein, Efrat Rorman, Israel Nissan, Efrat Glick-Saar, Omri Nayshool, Gideon Rechavi, Ella Mendelson, Orna Mor |
| EPI_ISL_944617, EPI_ISL_944618, EPI_ISL_944620, EPI_ISL_944621, | Instituto Nacional de Medicina Genomica | Instituto Nacional de Medicina Genomica | Hidalgo-Miranda A, Mendoza-Vargas A, Reyes-Grajeda JP, Cisneros-Villanueva M, Cedro-Tanda A,Peñaloza-Figueroa F, Herrera-Montalvo LA |

| | | | |
|---|---|---|---|
| EPI_ISL_944622, EPI_ISL_944626, EPI_ISL_944629, EPI_ISL_944630, EPI_ISL_944632 | | | |
| EPI_ISL_944672, EPI_ISL_944674, EPI_ISL_944676, EPI_ISL_944682, EPI_ISL_944685, EPI_ISL_944686, EPI_ISL_944690, EPI_ISL_944697, EPI_ISL_944698, EPI_ISL_944706, EPI_ISL_944708, EPI_ISL_944714, EPI_ISL_944734 | | | |
| see above | Department of Biochemistry, Cell and Molecular Biology, West African Centre for Cell Biology of Infectious Pathogens (WACCBIP), University of Ghana | Department of Biochemistry, Cell and Molecular Biology, West African Centre for Cell Biology of Infectious Pathogens (WACCBIP), University of Ghana | Morang'a,C.M., Ngoi,J.M., Quansah,E.B., Saiid,S., Amuzu,D.S., Asante,I., Bonney,J.H., Bonney,E., Odoom,J.K., Ndam,N.T., Tei-Maya,F., Adusei-Poku,M., Ofori-Boadu,L., Ampofo,W.K., Amenga-Etego,L.N., Quashie,P., Bediako,Y., Awandare,G.A. |
| EPI_ISL_944738 | unknown | Public Health Virology-Forensic and Scientific Services (PHV-FSS) | Son Nguyen et al. |
| EPI_ISL_949094 | Jessa | Jessa | Jessa_cmdLab |
| EPI_ISL_949318 | Department of Pathology, University of Cambridge | COVID-19 Genomics UK (COG-UK) Consortium | Aminu S. Jahun, Yasmin Chaudhry, Iliana Georgana, Myra Hosmillo, Rhys Izu, Martin D. Curran, Surendra Parmar, Ian Goodfellow |
| EPI_ISL_949640, EPI_ISL_949643, EPI_ISL_949644 | Virology Department, Royal Infirmary of Edinburgh, NHS Lothian / School of Biological Sciences, University of Edinburgh / Institute of Genetics and Molecular Medicine, University of Edinburgh | COVID-19 Genomics UK (COG-UK) Consortium | McHugh M, Dewar R, Rooke S, Gallagher M, Balcaza C, O'Toole Á, Scher E, Hill V, McCrone JT, Colquhoun R, Yu X, Jackson B, Rambaut A, Williams TC, Templeton K |
| EPI_ISL_949749, EPI_ISL_949777, EPI_ISL_949778, EPI_ISL_949780, EPI_ISL_949783 | Barts Health NHS Trust | COVID-19 Genomics UK (COG-UK) Consortium | CUTINO-MOGUEL, Maria-Teresa; HARRINGTON, David; OWOYEMI, Dola; KULASEGARAN-SHYLINI, Raghavendran; BROAD, Claire; KELE, Beatrix |
| EPI_ISL_950211, EPI_ISL_950212, EPI_ISL_950213, EPI_ISL_950214, EPI_ISL_950215, EPI_ISL_950216, EPI_ISL_950217, EPI_ISL_950218, EPI_ISL_950219, EPI_ISL_950220, EPI_ISL_950221, EPI_ISL_950229, EPI_ISL_950230, EPI_ISL_950231 | | | |
| see above | University College London, Great Ormond Street Hospital for Children NHS Foundation Trust, Imperial College Healthcare NHS Trust | COVID-19 Genomics UK (COG-UK) Consortium | Sergi Castellano, Rachel Williams, Mark Kristiansen, Paola Resende Silva, Sunando Roy, Tony Brooks, Helena Tutill, Paola Niola, Patricia Dyal, Charlotte Williams, Leysa Forrest, Yasmin Panchbhaya, Jacqueline Findlay, Samuel Weeks, Julianne Brown, Kathryn Harris, Paul Randell, James Price, Alison Holmes, Judith Breuer |
| EPI_ISL_950304, EPI_ISL_950306, EPI_ISL_950307, EPI_ISL_950308, EPI_ISL_950309, EPI_ISL_950310, EPI_ISL_950311, EPI_ISL_950312, EPI_ISL_950317, EPI_ISL_950324, EPI_ISL_950325, EPI_ISL_950330, EPI_ISL_950331, EPI_ISL_950332, EPI_ISL_950333, EPI_ISL_950335, EPI_ISL_950336 | | | |
| see above | Northumbria University / South Tees Hospitals NHS Foundation Trust / North Cumbria Integrated Care NHS Foundation Trust / North Tees and Hartlepool NHS Foundation Trust / Newcastle Hospitals NHS Foundation Trust | COVID-19 Genomics UK (COG-UK) Consortium | Darren L Smith,Andrew Nelson,Matthew Bashton,Greg R Young,Joshua Loh,John Allan,Mohammad A Tariq,Giles S Holt,Gary Black,Wen C Yew,Lynn Dover,Paul Baker,Steve Liggett,Sarah Essex,Jane Greenaway,Debra Padgett,Clive Graham,Garren Scott,Edward Barton,Emma Swindells,Brendan Payne,Jennifer Collins,Yusri Taha,Gary Eltringham |
| EPI_ISL_950714 | Lincolnshire Hospitals and DeepSeq Nottingham | COVID-19 Genomics UK (COG-UK) Consortium | Nichola Duckworth, Tim Sloan, Sarah Walsh, Jonathan Ball, Patrick McClure, Joeseph Chappell, Nadine Holmes, Matthew Carlisle, Christopher Moore, Fei Sang, Johnny Debebe, Victoria Wright, Matthew Loose |
| EPI_ISL_951305, EPI_ISL_951311, EPI_ISL_951313, EPI_ISL_951318, EPI_ISL_951374, EPI_ISL_951375, EPI_ISL_951378, EPI_ISL_951379, EPI_ISL_951380, EPI_ISL_951383, EPI_ISL_951387, EPI_ISL_951388, EPI_ISL_951416, EPI_ISL_951419, EPI_ISL_951420, EPI_ISL_951421, EPI_ISL_951423, EPI_ISL_951424, EPI_ISL_951425, EPI_ISL_951428, EPI_ISL_951435, EPI_ISL_951436, EPI_ISL_951437, EPI_ISL_951438, EPI_ISL_951442, EPI_ISL_951444, EPI_ISL_951445, EPI_ISL_951446, EPI_ISL_951447, EPI_ISL_951450, EPI_ISL_951451, EPI_ISL_951452, EPI_ISL_951453, EPI_ISL_951456, EPI_ISL_951457 | | | |
| see above | Oxford Viromics, NDM, University of Oxford; Oxford University Hospitals; Basingstoke and North Hampshire Hospital | COVID-19 Genomics UK (COG-UK) Consortium | Tanya Golubchik, David Bonsall, George Macintyre, Amy Trebes, Mariateresa de Cesare, Catrin Moore, Alex Mobbs, Anita Justice, Robert Shaw, Monique Andersson, Timothy Peto, Emma Wise, Nathan Moore, Jessica Lynch, Nick Cortes, Matilde Mori, Stephen Kidd, David Buck, John Todd, Christophe Fraser |
| EPI_ISL_951629, EPI_ISL_951816 | Originating lab: Wales Specialist Virology Centre Sequencing lab: Pathogen Genomics Unit | Public Health Wales Microbiology Cardiff Wales Specialist Virology Centre | Catherine Moore, Johnathan Evans, Laura Gifford, Malorie Perry, Simon Cottrell, Angela Marchbank, Alec Birchley, Alexander Adams, Amy Gaskin, Bree Gatica-Wilcox, Jason Coombes, Joel Southgate, Lauren Gilbert, Lee Graham, Nicole Pacchiarini, Sara Kumziene-Summerhayes, Sarah Taylor, Sophie Jones, Sara Rey, Matthew Bull, Joanne Watkins, Sally Corden, Tom Connor |
| EPI_ISL_952377, EPI_ISL_952379, EPI_ISL_952382, EPI_ISL_952383, EPI_ISL_952385, EPI_ISL_952392, EPI_ISL_952394, EPI_ISL_952397, EPI_ISL_952398, EPI_ISL_952409, EPI_ISL_952411, EPI_ISL_952414, EPI_ISL_952418, EPI_ISL_952422, EPI_ISL_952424, EPI_ISL_952427, EPI_ISL_952429, EPI_ISL_952433, EPI_ISL_952471, EPI_ISL_952472, EPI_ISL_952473, EPI_ISL_952474, EPI_ISL_952475, EPI_ISL_952476, EPI_ISL_952526, EPI_ISL_952569, EPI_ISL_952570, EPI_ISL_952571, EPI_ISL_952572, EPI_ISL_952573, EPI_ISL_952574, EPI_ISL_952575, EPI_ISL_952576, EPI_ISL_952578, EPI_ISL_952580, EPI_ISL_952581, EPI_ISL_952582, EPI_ISL_952583, EPI_ISL_952584, EPI_ISL_952585, EPI_ISL_952586, EPI_ISL_952587, EPI_ISL_952588, EPI_ISL_952589, EPI_ISL_952590, EPI_ISL_952591, EPI_ISL_952592, EPI_ISL_952594, EPI_ISL_952595, EPI_ISL_952597, EPI_ISL_952598, EPI_ISL_952599, EPI_ISL_952600, EPI_ISL_952778, EPI_ISL_952779, EPI_ISL_952780, EPI_ISL_952789, EPI_ISL_952793, EPI_ISL_952794, EPI_ISL_952795, EPI_ISL_952798, EPI_ISL_952799, EPI_ISL_952804 | | | |
| see above | Centre for Enzyme Innovation, University of Portsmouth / Translational Research Laboratory, Portsmouth Hospitals NHS Trust | COVID-19 Genomics UK (COG-UK) Consortium | Angela Beckett,Salman Goudarzi,Christopher Fearn,Kate Cook,Katie Loveson,Sharon Glaysher,Scott Elliott,Samuel Robson |
| EPI_ISL_952899, EPI_ISL_952908, EPI_ISL_952909, EPI_ISL_952912 | Department of Pathology, University of Cambridge | COVID-19 Genomics UK (COG-UK) Consortium | Aminu S. Jahun, Yasmin Chaudhry, Iliana Georgana, Myra Hosmillo, Rhys Izu, Martin D. Curran, Surendra Parmar, Ian Goodfellow |
| EPI_ISL_953024, EPI_ISL_953025, EPI_ISL_953026, EPI_ISL_953027, EPI_ISL_953028, EPI_ISL_953029, EPI_ISL_953030, EPI_ISL_953031, EPI_ISL_953032, EPI_ISL_953033, EPI_ISL_953034, EPI_ISL_953035, EPI_ISL_953036, EPI_ISL_953037, EPI_ISL_953038, EPI_ISL_953039, EPI_ISL_953040, EPI_ISL_953041, EPI_ISL_953042, EPI_ISL_953043, EPI_ISL_953044, EPI_ISL_953045, EPI_ISL_953046, EPI_ISL_953047, EPI_ISL_953048, EPI_ISL_953049, EPI_ISL_953050, EPI_ISL_953051, EPI_ISL_953052, EPI_ISL_953053, EPI_ISL_953054, EPI_ISL_953055, EPI_ISL_953056, EPI_ISL_953057, EPI_ISL_953058, EPI_ISL_953059, EPI_ISL_953060, EPI_ISL_953061, EPI_ISL_953062, EPI_ISL_953063, EPI_ISL_953064, EPI_ISL_953066, EPI_ISL_953067, EPI_ISL_953068, EPI_ISL_953069, EPI_ISL_953070, EPI_ISL_953071, EPI_ISL_953072, EPI_ISL_953073, EPI_ISL_953074, EPI_ISL_953075 | | | |
| see above | Bioinformatics and Biostatistics Lab, Advanced Sequencing Facility | COVID-19 Genomics UK (COG-UK) Consortium | Aengus Stewart,Jerome Nicod,Chelsea Sawyer,Laura Cubitt,Harshil Patel,Margaret Crawford |
| EPI_ISL_953522, EPI_ISL_953666, EPI_ISL_953667 | University Hospitals of Geneva, Laboratory of Virology | HUG, Laboratory of Virology and the Health2030 Genome Center | Samuel Cordey, Ana Rita Goncalves, Laurent Kaiser, Lorenzo Cerutti, Henri Pegeot, Melyssa Elies, Deborah Penet, Keith Harshman, Ioannis Xenarios, Emmanouil Dermitzakis |
| EPI_ISL_953940, EPI_ISL_953941 | Hopital | National Reference Center for Viruses of Respiratory Infections, Institut Pasteur, Paris | Marion Barbet, Sylvie Behillil, Méline Bizard, Angela Brisebarre, Camille Capel, Etienne Simon-Lorière, Vincent Enouf, Maud Vanpeene, Sylvie van der Werf,Lagathu GisèLe |
| EPI_ISL_953954 | Outre Mer | National Reference Center for Viruses of Respiratory Infections, Institut Pasteur, Paris | Marion Barbet, Sylvie Behillil, Méline Bizard, Angela Brisebarre, Camille Capel, Etienne Simon-Lorière, Vincent Enouf, Maud Vanpeene, Sylvie van der Werf,Rousset Dominique |
| EPI_ISL_953994 | Labo Analyses Med | National Reference Center for Viruses of Respiratory Infections, Institut Pasteur, Paris | Marion Barbet, Sylvie Behillil, Méline Bizard, Angela Brisebarre, Camille Capel, Etienne Simon-Lorière, Vincent Enouf, Maud Vanpeene, Sylvie van der Werf,Amzalag Jonas |
| EPI_ISL_953995, EPI_ISL_953996 | Hopital | National Reference Center for Viruses of Respiratory Infections, Institut Pasteur, Paris | Marion Barbet, Sylvie Behillil, Méline Bizard, Angela Brisebarre, Camille Capel, Etienne Simon-Lorière, Vincent Enouf, Maud Vanpeene, Sylvie van der Werf,Lagathu GisèLe |
| EPI_ISL_954007, EPI_ISL_954022, EPI_ISL_954023, EPI_ISL_954024, EPI_ISL_954025, EPI_ISL_954031 | Labo Analyses Med | National Reference Center for Viruses of Respiratory Infections, Institut Pasteur, Paris | Marion Barbet, Sylvie Behillil, Méline Bizard, Angela Brisebarre, Camille Capel, Etienne Simon-Lorière, Vincent Enouf, Maud Vanpeene, Sylvie van der Werf,Amzalag Jonas |
| EPI_ISL_954035 | Outre Mer | National Reference Center for Viruses of Respiratory Infections, Institut Pasteur, Paris | Marion Barbet, Sylvie Behillil, Méline Bizard, Angela Brisebarre, Camille Capel, Etienne Simon-Lorière, Vincent Enouf, Maud Vanpeene, Sylvie van der Werf,Rousset Dominique |
| EPI_ISL_954119 | Labo Analyses Med | National Reference Center for Viruses of Respiratory Infections, Institut Pasteur, Paris | Marion Barbet, Sylvie Behillil, Méline Bizard, Angela Brisebarre, Camille Capel, Etienne Simon-Lorière, Vincent Enouf, Maud Vanpeene, Sylvie van der Werf,Takoudju Eve-Marie |
| EPI_ISL_954120 | Hopital | National Reference Center for Viruses of Respiratory Infections, Institut Pasteur, Paris | Marion Barbet, Sylvie Behillil, Méline Bizard, Angela Brisebarre, Camille Capel, Etienne Simon-Lorière, Vincent Enouf, Maud Vanpeene, Sylvie van der Werf,Lagathu GisèLe |
| EPI_ISL_954121 | Labo Analyses Med | National Reference Center for Viruses of Respiratory Infections, Institut Pasteur, Paris | Marion Barbet, Sylvie Behillil, Méline Bizard, Angela Brisebarre, Camille Capel, Etienne Simon-Lorière, Vincent Enouf, Maud Vanpeene, Sylvie van der Werf,Takoudju Eve-Marie |
| EPI_ISL_954122 | Hopital | National Reference Center for Viruses of Respiratory | Marion Barbet, Sylvie Behillil, Méline Bizard, Angela Brisebarre, Camille Capel, Etienne Simon-Lorière, Vincent Enouf, Maud Vanpeene, Sylvie van der |

| | | | |
|---|---|---|---|
| | | Infections, Institut Pasteur, Paris | Werf,Lagathu GisèLe |
| EPI_ISL_954123, EPI_ISL_954124 | Labo Analyses Med | National Reference Center for Viruses of Respiratory Infections, Institut Pasteur, Paris | Marion Barbet, Sylvie Behillil, Méline Bizard, Angela Brisebarre, Camille Capel, Etienne Simon-Lorière, Vincent Enouf, Maud Vanpeene, Sylvie van der Werf,Takoudju Eve-Marie |
| EPI_ISL_954125 | Outre Mer | National Reference Center for Viruses of Respiratory Infections, Institut Pasteur, Paris | Marion Barbet, Sylvie Behillil, Méline Bizard, Angela Brisebarre, Camille Capel, Etienne Simon-Lorière, Vincent Enouf, Maud Vanpeene, Sylvie van der Werf,Rousset Dominique |
| EPI_ISL_954165, EPI_ISL_954166, EPI_ISL_954167, EPI_ISL_954168, EPI_ISL_954169, EPI_ISL_954170, EPI_ISL_954171, EPI_ISL_954175, EPI_ISL_954176, EPI_ISL_954177, EPI_ISL_954178 | | | |
| see above | Hospital Universitari Vall d'Hebron - Vall d'Hebron Institut de Rercerca | Hospital Universitari Vall d'Hebron | Cristina Andrés, Maria Piñana, Josep F Abril, Damir Garcia-Cehic, Ariadna Rando, Juliana Esperalba, Maria Gema Codina, Carla Castillo, Maria Carmen Martin, Tomàs Pumarola, Josep Quer, Andrés Antón |
| EPI_ISL_954750 | Diagen | Diagen | Koliada O |
| EPI_ISL_954754 | City of Milwaukee Health Department Laboratory | City of Milwaukee Health Department Laboratory | Sanjib Bhattacharyya |
| EPI_ISL_954780 | COMPLEJO HOSPITALARIO XERAL-CALDE | Instituto de Salud Carlos III | Iglesias-Caballero, M. Camarero, S. Sandonís,V. Vázquez, S. Pozo, F. Casas, I. Jiménez, P. Zaballos, A. Monzón, S. Varona, S. Cuesta, I. Alonso, P. |
| EPI_ISL_954786 | Complejo Hospitalario de Navarra | Instituto de Salud Carlos III | Iglesias-Caballero, M. Camarero, S. Sandonís,V. Vázquez, S. Pozo, F. Casas, I. Jiménez, P. Zaballos, A. Monzón, S. Varona, S. Cuesta, I. Ezpeleta,C. |
| EPI_ISL_954798 | Hospital Comarcal de Melilla | Instituto de Salud Carlos III | Iglesias-Caballero, M. Camarero, S. Sandonís,V. Vázquez, S. Pozo, F. Casas, I. Jiménez, P. Zaballos, A. Monzón, S. Varona, S. Cuesta, I. Roman, S. |
| EPI_ISL_954806 | Gerencia de Asistencia Sanitaria de Soria | Instituto de Salud Carlos III | Iglesias-Caballero, M. Camarero, S. Sandonís,V. Vázquez, S. Pozo, F. Casas, I. Jiménez, P. Zaballos, A. Monzón, S. Varona, S. Cuesta, I. Aldea, C. |
| EPI_ISL_954807, EPI_ISL_954808 | Complejo Hospitalario Universitario La Coruña | Instituto de Salud Carlos III | Iglesias-Caballero, M. Camarero, S. Sandonís,V. Vázquez, S. Pozo, F. Casas, I. Jiménez, P. Zaballos, A. Monzón, S. Varona, S. Cuesta, I. Peña, F. |
| EPI_ISL_954811, EPI_ISL_954813 | HOSPITAL UNIVERSITARIO DE GUADALAJARA | Instituto de Salud Carlos III | Iglesias-Caballero, M. Camarero, S. Sandonís,V. Vázquez, S. Pozo, F. Casas, I. Jiménez, P. Zaballos, A. Monzón, S. Varona, S. Cuesta, I. González, A. |
| EPI_ISL_954818, EPI_ISL_954830, EPI_ISL_954840, EPI_ISL_954845, EPI_ISL_954854, EPI_ISL_954879, EPI_ISL_954936, EPI_ISL_954937, EPI_ISL_954938, EPI_ISL_954939, EPI_ISL_954940, EPI_ISL_954941, EPI_ISL_954942 | | | |
| see above | Colorado Department of Public Health and Environment | Colorado Department of Puplic Health and Environment | Laura Bankers, Molly C. Hetherington-Rauth, Diana Ir, Shannon Ely, Shannon R. Matzinger, Sarah Elizabeth Totten, Emily A. Travanty |
| EPI_ISL_954943 | Laboratory of Virology and Molecular Diagnostics | Institute of Public Health of Republic of North Macedonia Laboratory of Virology and Molecular Diagnostics | M. Kuzmanovska, G. Boshevska |
| EPI_ISL_954944, EPI_ISL_954945, EPI_ISL_954946, EPI_ISL_954947, EPI_ISL_954948, EPI_ISL_954949, EPI_ISL_954950 | Colorado Department of Public Health and Environment | Colorado Department of Puplic Health and Environment | Laura Bankers, Molly C. Hetherington-Rauth, Diana Ir, Shannon Ely, Shannon R. Matzinger, Sarah Elizabeth Totten, Emily A. Travanty |
| EPI_ISL_955129 | Innovative Genomics Institute, UC Berkeley | Innovative Genomics Institute, UC Berkeley | Stacia Wyman, Haridha Shivram, Phil Frankino, Liana Lareau, Shana McDevitt, Justin Choi |
| EPI_ISL_955153, EPI_ISL_955154 | Genomic Laboratory (GLAB) (Conjoint lab of Health Directorate of Istanbul and Istanbul Technical University) | Genomic Laboratory (GLAB), Istanbul Technical University | Ilker Karacan, Tugba Kizilboga Akgun, Payam Zolfagharian, Nisan Denizce Can, Pari Sharifli, Levent Doganay, Gizem Dinler Doganay |
| EPI_ISL_955182 | Labo Analyses Med | National Reference Center for Viruses of Respiratory Infections, Institut Pasteur, Paris | Marion Barbet, Sylvie Behillil, Méline Bizard, Angela Brisebarre, Camille Capel, Etienne Simon-Lorière, Vincent Enouf, Maud Vanpeene, Sylvie van der Werf |
| EPI_ISL_955212, EPI_ISL_955216, EPI_ISL_955217, EPI_ISL_955226 | Indiana Animal Disease Diagnostic Laboratory | Carpi Laboratory - Purdue University | Jack Dorman, Ilinca I Ciubotariu, Lev Gorenstein, Abebe A Fola, G Kenitra Hendrix, Rebecca P Wilkes, Giovanna Carpi |
| EPI_ISL_955230 | Mary Eliza Mahoney HC | NJ Public Health and Environmental Laboratories | Lindsey Bodnar, Shiv Verma, Dana Woell, Byeong Jeong |
| EPI_ISL_955323 | American Esoteric Laboratory | Pathogen Discovery, Respiratory Viruses Branch, Division of Viral Diseases, Centers for Disease Control and Prevention | Ying Tao, Jing Zhang, Yan Li, Krista Queen, Anna Uehara, Peter Cook, Clinton R. Paden, Haibin Wang, Suxiang Tong |
| EPI_ISL_955426 | Alameda County Public Health Lab | Chan-Zuckerberg Biohub | CZB Cliahub Consortium |
| EPI_ISL_955779, EPI_ISL_955781, EPI_ISL_955786, EPI_ISL_955789, EPI_ISL_955790 | University of Michigan Clinical Microbiology Laboratory | Lauring Lab, University of Michigan, Department of Microbiology and Immunology | Valesano |
| EPI_ISL_956282, EPI_ISL_956288, EPI_ISL_956289, EPI_ISL_956290 | Instituto Nacional de Salud- Dirección de Redes de Laboratorios de Salud Pública | Instituto Nacional de Salud- Dirección de Investigación en Salud Pública | Katherine Laiton-Donato, Diego A. Álvarez-Díaz, Carlos Franco-Muñoz, Mauricio Pacheco-Montealegre, Hector Alejandro Ruiz-Moreno, Maria T. Herrera-Sepúlveda, Diego Andrés Prada, Jhonnatan Reales-González, Sheryll Corchuelo, Julian Naizaque, Gerardo Santamaría, Magdalena Wiesner, Martha Lucia Ospina Martinez, Marcela Mercado-Reyes |
| EPI_ISL_956298 | Laboratorio de salud publica del Valle del Cauca | Instituto Nacional de Salud- Dirección de Investigación en Salud Pública | Katherine Laiton-Donato, Diego A. Álvarez-Díaz, Carlos Franco-Muñoz, Mauricio Pacheco-Montealegre, Hector Alejandro Ruiz-Moreno, Maria T. Herrera-Sepúlveda, Diego Andrés Prada, Jhonnatan Reales-González, Sheryll Corchuelo, Julian Naizaque, Gerardo Santamaría, Magdalena Wiesner, Martha Lucia Ospina Martinez, Marcela Mercado-Reyes |
| EPI_ISL_956413, EPI_ISL_956416, EPI_ISL_956420, EPI_ISL_959287 | Laboratory of Virology and Molecular Diagnostics | Institute of Public Health of Republic of North Macedonia Laboratory of Virology and Molecular Diagnostics | Maja Kuzmanovska, Golubinka Boshevska |
| EPI_ISL_959290, EPI_ISL_959375, EPI_ISL_959376 | Laboratory of Virology and Molecular Diagnostics | Institute of Public Health of Republic of North Macedonia Laboratory of Virology and Molecular Diagnostics | Kuzmanovska M., Boshevska G. |
| EPI_ISL_959426 | Servicio de Microbiología, Hospital Universitario Son Espases | SeqCOVID-SPAIN consortium/IBV(CSIC) | Carla López-Causapé, Jordi Reina, Antonio Oliver and SeqCOVID-SPAIN consortium |
| EPI_ISL_959544 | Laboratory of Virology and Molecular Diagnostics | Institute of Public Health of Republic of North Macedonia Laboratory of Virology and Molecular Diagnostics | Kuzmanovska M., Boshevska G. |
| EPI_ISL_959555, EPI_ISL_959556, EPI_ISL_959557, EPI_ISL_959558 | Synlab | GIGA Medical Genomics | Keith Durkin, Maria Artesi, Sébastien Bontems, Raphaël Boreux, Bouchra Boujemla, Cécile Meex, Pierrette Melin, Marie-Pierre Hayette, Vincent Bours |
| EPI_ISL_959559, EPI_ISL_959560 | Vivalia - Clinique Saint-Joseph | GIGA Medical Genomics | Keith Durkin, Maria Artesi, Sébastien Bontems, Raphaël Boreux, Bouchra Boujemla, Cécile Meex, Pierrette Melin, Marie-Pierre Hayette, Vincent Bours |
| EPI_ISL_959561 | University of Liège COVID-19 testing center | GIGA Medical Genomics | Keith Durkin, Maria Artesi, Sébastien Bontems, Raphaël Boreux, Bouchra Boujemla, Cécile Meex, Pierrette Melin, Marie-Pierre Hayette, Vincent Bours |
| EPI_ISL_959562 | Vivalia - Clinique Saint-Joseph | GIGA Medical Genomics | Keith Durkin, Maria Artesi, Sébastien Bontems, Raphaël Boreux, Bouchra Boujemla, Cécile Meex, Pierrette Melin, Marie-Pierre Hayette, Vincent Bours |
| EPI_ISL_959840, EPI_ISL_959841, EPI_ISL_959842, EPI_ISL_959843, EPI_ISL_959844, EPI_ISL_959845, EPI_ISL_959846, EPI_ISL_959847, EPI_ISL_959848 | National Virus Reference Laboratory | National Virus Reference Laboratory | Michael Carr, Gabriel Gonzalez, Jonathan Dean, Cillian F De Gascun |
| EPI_ISL_960223 | FidaLab | Seattle Flu Study | Deborah A. Nickerson, Chris D. Frazar, Jover Lee, Benjamin Pelle, Erica Ryke, Matthew Richardson, Amanda Adler, Elisabeth Brandstetter, Peter D. Han, Kairsten Fay, Misja Ilcisin, Kirsten Lacombe, Thomas R. Sibley, Melissa Truong, Caitlin R. Wolf, Romesh Gautom, Geoff Melly, Brian Hiatt, Philip Dykema, Scott Lindquist, Michael Boeckh, Janet A. Englund, Michael Famulare, Barry R. Lutz, Mark J. Rieder, Lea M. Starita, Matthew Thompson, Helen Y. Chu, Jay Shendure, Trevor Bedford |
| EPI_ISL_960415 | The National Institute of Public Health | State Veterinary Institute Prague | Nagy,A;Vecerova,J;Cernikova,L;Stara,M;Jirincova,H;Trnka,D |
| EPI_ISL_960461, EPI_ISL_960573 | Istituto Zooprofilattico Sperimentale del Mezzogiorno | TIGEM | Patrizia Annunziata, Andrea Ballabio, Valentina Bouche, Davide Cacchiarelli, Pellegrino Cerino, Chiara Colantuono, Maria Concetta Cuomo, Denise Di Concilio, Lucio Di Filippo, Antonio Grimaldi, Antonio Limone, Anna Manfredi, Francesco Panariello, Biancamaria Pierri, Marcello Salvi |

| | | | |
|---|---|---|---|
| EPI_ISL_960989, EPI_ISL_960990, EPI_ISL_960992, EPI_ISL_960993, EPI_ISL_960995, EPI_ISL_960997, EPI_ISL_960998, EPI_ISL_961001, EPI_ISL_961003, EPI_ISL_961004, EPI_ISL_961005, EPI_ISL_961006, EPI_ISL_961007, EPI_ISL_961008 | | | |
| see above | AIID | Irish Coronavirus Sequencing Consortium-Teagasc Grange | Matthew McCabe, Aljandro Abner Garcia Leon, Fiona Crispie, Calum Walsh, Michael Carr, John Kenny, Paul Cotter, Patrick Mallon, Gabriel Gonzalez |
| EPI_ISL_961466 | Laboratorios Lister | Instituto de Diagnostico y Referencia Epidemiologicos (INDRE) | Claudia Wong-Arambula, Abril Rodriguez-Maldonado, Fabiola Garces-Ayala, Natividad Cruz-Ortiz, Tatiana Nunez-Garcia, Gisela Barrera-Badillo, Lucia Hernandez-Rivas, Irma Lopez-Martinez, Ernesto Ramirez-Gonzalez. |
| EPI_ISL_961597, EPI_ISL_961609, EPI_ISL_961637, EPI_ISL_961638, EPI_ISL_961639, EPI_ISL_961640, EPI_ISL_961641 | Hôpital Georges L. Dumont | National Microbiology Laboratory (NML) | Anna Majer, Shari Tyson, Grace Seo, Philip Mabon, Elsie Grudeski, Rhiannon Huzarewich, Russell Mandes, Anneliese Landgraff, Jennifer Tanner, Natalie Knox, Morag Graham, Gary Van Domselaar, Richard Garceau, Guillaume Desnoyers, Nathalie Bastien, Yan Li, Timothy Booth, Darian Hole, Madison Chapel, Kirsten Biggar, CanCOGeN's metadata curation team, Public Health Agency of Canada CanCOGeN team |
| EPI_ISL_961875 | E. Gulbja laboratorija | Latvian Biomedical Research and Study Centre | Janis Pjalkovskis, Nikita Zrelovs, Monta Ustinova, Ivars Silamikelis, Liga Birzniece, Kaspars Megnis, Vita Rovite, Lauma Freimane, Laila Silamikele, Laura Ansone, Davids Fridmanis, Mikus Gavars, Dmitrijs Perminovs, Jurijs Perevoscikovs, Uga Dumpis, Janis Klovins |
| EPI_ISL_961876 | Latvijas Infektologijas Centrs | Latvian Biomedical Research and Study Centre | Janis Pjalkovskis, Nikita Zrelovs, Monta Ustinova, Ivars Silamikelis, Liga Birzniece, Kaspars Megnis, Vita Rovite, Lauma Freimane, Laila Silamikele, Laura Ansone, Davids Fridmanis,Reinis Zeltmatis, Diana Dusacka, Jurijs Perevoscikovs, Uga Dumpis, Janis Klovins |
| EPI_ISL_961899, EPI_ISL_961926, EPI_ISL_961934, EPI_ISL_961941, EPI_ISL_961942, EPI_ISL_961949, EPI_ISL_961950, EPI_ISL_961957, EPI_ISL_961974, EPI_ISL_961981, EPI_ISL_961982, EPI_ISL_961991, EPI_ISL_962014, EPI_ISL_962174, EPI_ISL_962175 | | | |
| see above | Illinois Department of Public Health | Gagnon Lab, Southern Illinois University | Keith Gagnon |
| EPI_ISL_962368 | Seattle Flu Study | Seattle Flu Study | Deborah A. Nickerson, Chris D. Frazar, Jover Lee, Benjamin Pelle, Erica Ryke, Matthew Richardson, Amanda Adler, Elisabeth Brandstetter, Peter D. Han, Kairsten Fay, Misja Ilcisin, Kirsten Lacombe, Thomas R. Sibley, Melissa Truong, Caitlin R. Wolf, Karen Cowgill, Stephanie Schrag, Jeff Duchin, Michael Boeckh, Janet A. Englund, Michael Famulare, Barry R. Lutz, Mark J. Rieder, Lea M. Starita, Matthew Thompson, Helen Y. Chu, Trevor Bedford, Jay Shendure |
| EPI_ISL_962379, EPI_ISL_962380, EPI_ISL_962381, EPI_ISL_962382, EPI_ISL_962383, EPI_ISL_962384, EPI_ISL_962385, EPI_ISL_962386, EPI_ISL_962387, EPI_ISL_962388 | Washington State Department of Health | Seattle Flu Study | Deborah A. Nickerson, Chris D. Frazar, Jover Lee, Benjamin Pelle, Erica Ryke, Matthew Richardson, Amanda Adler, Elisabeth Brandstetter, Peter D. Han, Kairsten Fay, Misja Ilcisin, Kirsten Lacombe, Thomas R. Sibley, Melissa Truong, Caitlin R. Wolf, Romesh Gautom, Geoff Melly, Brian Hiatt, Philip Dykema, Scott Lindquist, Michael Boeckh, Janet A. Englund, Michael Famulare, Barry R. Lutz, Mark J. Rieder, Lea M. Starita, Matthew Thompson, Helen Y. Chu, Jay Shendure, Trevor Bedford |
| EPI_ISL_962512 | UCLA Clinical Micro Lab | Los Angeles County PHL | P. Hemarajata et al. |
| EPI_ISL_962602, EPI_ISL_962603, EPI_ISL_962604, EPI_ISL_962605, EPI_ISL_962606, EPI_ISL_962607, EPI_ISL_962608 | Sharp HealthCare Laboratory | Andersen lab at Scripps Research | SEARCH Alliance San Diego with Aaron Harding, Jacquelyn Berumen, Cathy Woerle, Liam McGinnis, Art Mendoza, Omid Bakhtar |
| EPI_ISL_962619, EPI_ISL_962620, EPI_ISL_962624, EPI_ISL_962626, EPI_ISL_962627, EPI_ISL_962632, EPI_ISL_962633, EPI_ISL_962636, EPI_ISL_962643, EPI_ISL_962645, EPI_ISL_962648, EPI_ISL_962649 | | | |
| see above | Scripps Medical Laboratory | Andersen lab at Scripps Research | SEARCH Alliance San Diego with Michael Quigley, Ellen Stefanski, Ian Mchardy |
| EPI_ISL_962650, EPI_ISL_962651, EPI_ISL_962652, EPI_ISL_962653, EPI_ISL_962654, EPI_ISL_962655, EPI_ISL_962656, EPI_ISL_962657, EPI_ISL_962658, EPI_ISL_962659, EPI_ISL_962660, EPI_ISL_962661, EPI_ISL_962662, EPI_ISL_962663, EPI_ISL_962664, EPI_ISL_962665, EPI_ISL_962666, EPI_ISL_962667, EPI_ISL_962668, EPI_ISL_962669, EPI_ISL_962670, EPI_ISL_962671, EPI_ISL_962672, EPI_ISL_962673, EPI_ISL_962674, EPI_ISL_962675, EPI_ISL_962676, EPI_ISL_962677, EPI_ISL_962678, EPI_ISL_962679, EPI_ISL_962680, EPI_ISL_962681, EPI_ISL_962682, EPI_ISL_962683, EPI_ISL_962684, EPI_ISL_962685, EPI_ISL_962686, EPI_ISL_962687, EPI_ISL_962688, EPI_ISL_962689, EPI_ISL_962690, EPI_ISL_962691, EPI_ISL_962692, EPI_ISL_962693, EPI_ISL_962694, EPI_ISL_962695, EPI_ISL_962696, EPI_ISL_962697, EPI_ISL_962698, EPI_ISL_962699, EPI_ISL_962700, EPI_ISL_962701, EPI_ISL_962702, EPI_ISL_962703, EPI_ISL_962704, EPI_ISL_962705, EPI_ISL_962706, EPI_ISL_962707, EPI_ISL_962708, EPI_ISL_962709, EPI_ISL_962719, EPI_ISL_962727, EPI_ISL_962730, EPI_ISL_962739, EPI_ISL_962744, EPI_ISL_962745, EPI_ISL_962754, EPI_ISL_962756, EPI_ISL_962758, EPI_ISL_962759, EPI_ISL_962765 | | | |
| see above | Sharp HealthCare Laboratory | Andersen lab at Scripps Research | SEARCH Alliance San Diego with Aaron Harding, Jacquelyn Berumen, Cathy Woerle, Liam McGinnis, Art Mendoza, Omid Bakhtar |
| EPI_ISL_962896 | Norwegian Institute of Public Health, Department of Virology | Norwegian Institute of Public Health, Department of Virology | Kathrine Stene-Johansen, Kamilla Heddeland Instefjord, Hilde Elshaug, Ignacio Garcia Llorente, Serina B Engebretsen, Atiya R Ali,Marie Paulsen Madsen, Rasmus Riis Kopperud, Hilde Vollan, Karoline Bragstad, Olav Hungnes |
| EPI_ISL_962906 | Ostfold Hospital Trust - Kalnes, Centre for Laboratory Medicine, Section for gene technology and infection serology | Norwegian Institute of Public Health, Department of Virology | Kathrine Stene-Johansen, Kamilla Heddeland Instefjord, Hilde Elshaug, Ignacio Garcia Llorente, Serina B Engebretsen, Atiya R Ali,Marie Paulsen Madsen, Rasmus Riis Kopperud, Hilde Vollan, Karoline Bragstad, Olav Hungnes |
| EPI_ISL_962945, EPI_ISL_962946 | Hospital Universitario de Gran Canaria Dr. Negrín | SeqCOVID-SPAIN consortium/IBV(CSIC) | M. Carmen Pérez González, Francisco J. Chamizo López, Ana Bordes Benítez and SeqCOVID-SPAIN consortium |
| EPI_ISL_963321 | Lighthouse Lab in Alderley Park | Wellcome Sanger Institute for the COVID-19 Genomics UK (COG-UK) Consortium | Jacquelyn Wynn, Mairead Hyland, The Lighthouse Lab in Alderley Park and Alex Alderton, Roberto Amato, Sonia Goncalves, Ewan Harrison, David K. Jackson, Ian Johnston, Dominic Kwiatkowski, Cordelia Langford, John Sillitoe on behalf of the Wellcome Sanger Institute COVID-19 Surveillance Team |
| EPI_ISL_964257 | Akershus University Hospital, Department for Microbiology and Infectious Disease Control | Norwegian Institute of Public Health, Department of Virology | Kathrine Stene-Johansen, Kamilla Heddeland Instefjord, Hilde Elshaug, Ignacio Garcia Llorente, Serina B Engebretsen, Atiya R Ali, Marie Paulsen Madsen, Rasmus Riis Kopperud, Hilde Vollan, Karoline Bragstad, Olav Hungnes |
| EPI_ISL_964918, EPI_ISL_964921, EPI_ISL_964932, EPI_ISL_964933, EPI_ISL_964940, EPI_ISL_964941 | Instituto Nacional de Saude (INS), Mozambique | KRISP, KZN Research Innovation and Sequencing Platform | Nalia Ismael, Nadia Sitoe, Paulo Arnaldo, Nedio Mabunda, Giandhari J, Pillay S, Emmanuel S, Tegally H, Wilkinson E, de Oliveira T |
| EPI_ISL_964952 | Ostfold Hospital Trust - Kalnes, Centre for Laboratory Medicine, Section for gene technology and infection serology | Norwegian Institute of Public Health, Department of Virology | Kathrine Stene-Johansen, Kamilla Heddeland Instefjord, Hilde Elshaug, Ignacio Garcia Llorente, Serina B Engebretsen, Atiya R Ali,Marie Paulsen Madsen, Rasmus Riis Kopperud, Hilde Vollan, Karoline Bragstad, Olav Hungnes |
| EPI_ISL_964953, EPI_ISL_964954 | Akershus University Hospital, Department for Microbiology and Infectious Disease Control | Norwegian Institute of Public Health, Department of Virology | Kathrine Stene-Johansen, Kamilla Heddeland Instefjord, Hilde Elshaug, Ignacio Garcia Llorente, Serina B Engebretsen, Atiya R Ali,Marie Paulsen Madsen, Rasmus Riis Kopperud, Hilde Vollan, Karoline Bragstad, Olav Hungnes |
| EPI_ISL_964955, EPI_ISL_964960 | Oslo University Hospital, Department of Medical Microbiology | Norwegian Institute of Public Health, Department of Virology | Kathrine Stene-Johansen, Kamilla Heddeland Instefjord, Hilde Elshaug, Ignacio Garcia Llorente, Serina B Engebretsen, Atiya R Ali,Marie Paulsen Madsen, Rasmus Riis Kopperud, Hilde Vollan, Karoline Bragstad, Olav Hungnes |
| EPI_ISL_964974 | Ostfold Hospital Trust - Kalnes, Centre for Laboratory Medicine, Section for gene technology and infection serology | Norwegian Institute of Public Health, Department of Virology | Kathrine Stene-Johansen, Kamilla Heddeland Instefjord, Hilde Elshaug, Ignacio Garcia Llorente, Serina B Engebretsen, Atiya R Ali,Marie Paulsen Madsen, Rasmus Riis Kopperud, Hilde Vollan, Karoline Bragstad, Olav Hungnes |
| EPI_ISL_964980 | Oslo University Hospital, Department of Medical Microbiology | Norwegian Institute of Public Health, Department of Virology | Kathrine Stene-Johansen, Kamilla Heddeland Instefjord, Hilde Elshaug, Ignacio Garcia Llorente, Serina B Engebretsen, Atiya R Ali,Marie Paulsen Madsen, Rasmus Riis Kopperud, Hilde Vollan, Karoline Bragstad, Olav Hungnes |
| EPI_ISL_965009, EPI_ISL_965010 | Nordland Hospital - Bodo, Laboratory Department, Molecular Biology Unit | Norwegian Institute of Public Health, Department of Virology | Kathrine Stene-Johansen, Kamilla Heddeland Instefjord, Hilde Elshaug, Ignacio Garcia Llorente, Serina B Engebretsen, Atiya R Ali, Marie Paulsen Madsen, Rasmus Riis Kopperud, Hilde Vollan, Karoline Bragstad, Olav Hungnes |
| EPI_ISL_965133 | Azienda Ospedaliera San Camillo Forlanini | INMI Lazzaro Spallanzani IRCCS | B Bartolini, O Butera, C.E.M Gruber, M Rueca, F Messina, E Giombini, F Basile, A D'Agostino, G Parisi, MR Capobianchi, A Di Caro |
| EPI_ISL_965134 | San Gallicano Dermatological Institute I.F.O. | INMI Lazzaro Spallanzani IRCCS | M Rueca, O Butera, F Messina, CEM Gruber, B Bartolini, E Giombini, F Pimpinelli, F Ensoli, A Mastrofrancesco, A Di Caro, MR Capobianchi |
| EPI_ISL_965135 | Presidio Ospedaliero G. B. Grassi ASL RM3 | INMI Lazzaro Spallanzani IRCCS | F Messina, C.E.M Gruber, B Bartolini, E Giombini, M Rueca, O Butera, F Tabacco, E Ristori, A Di Caro, MR Capobianchi |
| EPI_ISL_965139 | Fondazione Policlinico Universitario "A. Gemelli" IRCCS | INMI Lazzaro Spallanzani IRCCS | B Bartolini, O Butera, C.E.M Gruber, M Rueca, F Messina, E Giombini, P Cattani,M Sanguinetti, MR Capobianchi, A Di Caro |
| EPI_ISL_965559, EPI_ISL_965780 | Dutch COVID-19 response team | Medical Microbiology, Maastricht University Medical Centre | Jozef Dingemans*, Brian van der Veer*, Erik Beuken, Carmen Reumkens, Lieke van Alphen, Christian Hoebe, Paul Savelkoul |
| EPI_ISL_965840, EPI_ISL_965841, EPI_ISL_965842, EPI_ISL_965843, EPI_ISL_965844, EPI_ISL_965845, EPI_ISL_965846, EPI_ISL_965853, EPI_ISL_965854 | GA Department of Public Health | GA Department of Public Health | Stacy Reeves, Jonathan Edwards, Cynthia Dixey, Tonia Parrott |
| EPI_ISL_965934 | Hospital General Universitario de Ciudad Real | Instituto de Salud Carlos III | Sandonís,V. Vázquez, S. Iglesias-Caballero, M. Camarero, S. Pozo, F. Casas, I. Jiménez, P. Zaballos, A. Monzón, S. Varona, S. Cuesta, I. Illescas, S. |

| | | | |
|---|---|---|---|
| EPI_ISL_965938 | Consejería de Sanidad y Asuntos Sociales de Castilla La Mancha | Instituto de Salud Carlos III | Sandonís,V. Vázquez, S. Iglesias-Caballero, M. Camarero, S. Pozo, F. Casas, I. Jiménez, P. Zaballos, A. Monzón, S. Varona, S. Cuesta, I. Gutiérrez, G. |
| EPI_ISL_965959 | Hospital General Universitario de Ciudad Real | Instituto de Salud Carlos III | Iglesias-Caballero, M. Sandonís,V. Vázquez, S. Camarero, S. Pozo, F. Casas, I. Jiménez, P. Zaballos, A. Monzón, S. Varona, S. Cuesta, I. Illescas, S. |
| EPI_ISL_965974 | Vancouver Clinic | Seattle Flu Study | Deborah A. Nickerson, Chris D. Frazar, Jover Lee, Benjamin Pelle, Erica Ryke, Matthew Richardson, Amanda Adler, Elisabeth Brandstetter, Peter D. Han, Kairsten Fay, Misja Ilcisin, Kirsten Lacombe, Thomas R. Sibley, Melissa Truong, Caitlin R. Wolf, Romesh Gautom, Geoff Melly, Brian Hiatt, Philip Dykema, Scott Lindquist, Michael Boeckh, Janet A. Englund, Michael Famulare, Barry R. Lutz, Mark J. Rieder, Lea M. Starita, Matthew Thompson, Helen Y. Chu, Jay Shendure, Trevor Bedford |
| EPI_ISL_966312 | Toronto Invasive Bacterial Diseases Network | McMaster University | Allison McGeer, Patryk Aftanas, Hooman Derakhshani, Angel Li, Kuganya Nirmalarajah, Emily Panousis, Ahmed Draia, Jalees Nasir, Michael Surette, Samira Mubareka, Andrew G. McArthur |
| EPI_ISL_966532, EPI_ISL_966534, EPI_ISL_966535, EPI_ISL_966536, EPI_ISL_966537, EPI_ISL_966538, EPI_ISL_966539, EPI_ISL_966540, EPI_ISL_966541, EPI_ISL_966542, EPI_ISL_966543, EPI_ISL_966544, EPI_ISL_966545, EPI_ISL_966546, EPI_ISL_967169, EPI_ISL_967171, EPI_ISL_967174, EPI_ISL_967175, EPI_ISL_967179, EPI_ISL_967180, EPI_ISL_967181, EPI_ISL_967183, EPI_ISL_967184, EPI_ISL_967192, EPI_ISL_967193, EPI_ISL_967194, EPI_ISL_967195, EPI_ISL_967196, EPI_ISL_967197, EPI_ISL_967198, EPI_ISL_967199, EPI_ISL_967200, EPI_ISL_967201, EPI_ISL_967213, EPI_ISL_967214, EPI_ISL_967218, EPI_ISL_967243, EPI_ISL_967244, EPI_ISL_967245, EPI_ISL_967246, EPI_ISL_967247, EPI_ISL_967248, EPI_ISL_967249, EPI_ISL_967250, EPI_ISL_967251, EPI_ISL_967252, EPI_ISL_967253, EPI_ISL_967254, EPI_ISL_967255, EPI_ISL_967256, EPI_ISL_967257, EPI_ISL_967258, EPI_ISL_967259, EPI_ISL_967260, EPI_ISL_967261, EPI_ISL_967262, EPI_ISL_967263, EPI_ISL_967264, EPI_ISL_967265, EPI_ISL_967266, EPI_ISL_967267, EPI_ISL_967268, EPI_ISL_967269, EPI_ISL_967270, EPI_ISL_967271, EPI_ISL_967272, EPI_ISL_967273, EPI_ISL_967274, EPI_ISL_967275, EPI_ISL_967276, EPI_ISL_967277, EPI_ISL_967278, EPI_ISL_967279, EPI_ISL_967280, EPI_ISL_967281, EPI_ISL_967282, EPI_ISL_967283, EPI_ISL_967284, EPI_ISL_967285, EPI_ISL_967286, EPI_ISL_967287, EPI_ISL_967288, EPI_ISL_967289, EPI_ISL_967290, EPI_ISL_967291, EPI_ISL_967292, EPI_ISL_967293, EPI_ISL_967294, EPI_ISL_967295, EPI_ISL_967296, EPI_ISL_967297, EPI_ISL_967298, EPI_ISL_967299, EPI_ISL_967300, EPI_ISL_967301, EPI_ISL_967302, EPI_ISL_967303, EPI_ISL_967304, EPI_ISL_967305, EPI_ISL_967306, EPI_ISL_967307, EPI_ISL_967308, EPI_ISL_967309, EPI_ISL_967310, EPI_ISL_967311, EPI_ISL_967312, EPI_ISL_967313, EPI_ISL_967314, EPI_ISL_967315, EPI_ISL_967316, EPI_ISL_967317, EPI_ISL_967318, EPI_ISL_967319, EPI_ISL_967320, EPI_ISL_967321, EPI_ISL_967322, EPI_ISL_967323, EPI_ISL_967324, EPI_ISL_967325, EPI_ISL_967326 | | | |
| see above | Helix/Illumina | Respiratory Viruses Branch, Division of Viral Diseases, Centers for Disease Control and Prevention | Peter W. Cook,Dakota Howard,Dhwani Batra,Ben L. Rambo-Martin,Eileen de Feo,Jan Antico,Christine Tran,Matthew Tolentino,Shannon Wickline,Kim Gietzen,Brad Sickler,Jingtao Liu,Eric Allen,Phil Febbo,Summer Galloway,Nicole L. Washington,Simon White,Geraint Levan,Kelly Schiabor Barrett,Elizabeth Cirulli,Alexandre Bolze,Ary Ascencio,Charlotte Rivera-Garcia,Ryan Cho,Jason Nguyen,Sherry Wang,Jimmy Ramirez,Tyler Cassens,Efren Sandoval,Magnus Isaksson,William Lee,David Becker,Marc Laurent,James Lu,Clinton R. Paden,Suxiang Tong,Duncan MacCannell, |
| EPI_ISL_967697, EPI_ISL_967702, EPI_ISL_967703, EPI_ISL_967745 | State Laboratories Division, Hawaii State Department of Health | State Laboratories Division, Hawaii State Department of Health | Pamela O'Brien, Drew Kuwazaki, Ayana Garnet, Razvan Sultana, Edward Desmond |
| EPI_ISL_967843, EPI_ISL_967844, EPI_ISL_967845, EPI_ISL_967846, EPI_ISL_967847, EPI_ISL_967848, EPI_ISL_967849, EPI_ISL_967850, EPI_ISL_967851, EPI_ISL_967852, EPI_ISL_967853, EPI_ISL_967854, EPI_ISL_967855, EPI_ISL_967856, EPI_ISL_967857, EPI_ISL_967858, EPI_ISL_967859, EPI_ISL_967860, EPI_ISL_967861, EPI_ISL_967862, EPI_ISL_967863, EPI_ISL_967864, EPI_ISL_967865, EPI_ISL_967866, EPI_ISL_967867, EPI_ISL_967868, EPI_ISL_967869, EPI_ISL_967870, EPI_ISL_967871, EPI_ISL_967872, EPI_ISL_967873, EPI_ISL_967874, EPI_ISL_967875, EPI_ISL_967876, EPI_ISL_967877, EPI_ISL_967878, EPI_ISL_967879, EPI_ISL_967880, EPI_ISL_967881 | | | |
| see above | Helix/Illumina | Respiratory Viruses Branch, Division of Viral Diseases, Centers for Disease Control and Prevention | Peter W. Cook,Dakota Howard,Dhwani Batra,Ben L. Rambo-Martin,Eileen de Feo,Jan Antico,Christine Tran,Matthew Tolentino,Shannon Wickline,Kim Gietzen,Brad Sickler,Jingtao Liu,Eric Allen,Phil Febbo,Summer Galloway,Nicole L. Washington,Simon White,Geraint Levan,Kelly Schiabor Barrett,Elizabeth Cirulli,Alexandre Bolze,Ary Ascencio,Charlotte Rivera-Garcia,Ryan Cho,Jason Nguyen,Sherry Wang,Jimmy Ramirez,Tyler Cassens,Efren Sandoval,Magnus Isaksson,William Lee,David Becker,Marc Laurent,James Lu,Clinton R. Paden,Suxiang Tong,Duncan MacCannell, |
| EPI_ISL_967885, EPI_ISL_967886, EPI_ISL_967888, EPI_ISL_967889 | TGen North | TGen North | "Jolene Bowers, Megan Folkerts, Chris French, Hayley Yaglom, Ashlyn Pfeiffer, Darrin Lemmer, Dave Engelthaler, The Arizona COVID Genomics Union (ACGU)" |
| EPI_ISL_967890, EPI_ISL_967893 | TGen North | Sonora Quest Laboratories | "Jolene Bowers, Megan Folkerts, Chris French, Hayley Yaglom, Ashlyn Pfeiffer, Darrin Lemmer, Dave Engelthaler, The Arizona COVID Genomics Union (ACGU)" |
| EPI_ISL_967894 | TGen North | TGen North | "Jolene Bowers, Megan Folkerts, Chris French, Hayley Yaglom, Ashlyn Pfeiffer, Darrin Lemmer, Dave Engelthaler, The Arizona COVID Genomics Union (ACGU)" |
| EPI_ISL_967895, EPI_ISL_967896, EPI_ISL_967898 | TGen North | Sonora Quest Laboratories | "Jolene Bowers, Megan Folkerts, Chris French, Hayley Yaglom, Ashlyn Pfeiffer, Darrin Lemmer, Dave Engelthaler, The Arizona COVID Genomics Union (ACGU)" |
| EPI_ISL_967902 | TGen North | TGen North | "Jolene Bowers, Megan Folkerts, Chris French, Hayley Yaglom, Ashlyn Pfeiffer, Darrin Lemmer, Dave Engelthaler, The Arizona COVID Genomics Union (ACGU)" |
| EPI_ISL_967904 | TGen North | Sonora Quest Laboratories | "Jolene Bowers, Megan Folkerts, Chris French, Hayley Yaglom, Ashlyn Pfeiffer, Darrin Lemmer, Dave Engelthaler, The Arizona COVID Genomics Union (ACGU)" |
| EPI_ISL_967905, EPI_ISL_967908 | TGen North | TGen North | "Jolene Bowers, Megan Folkerts, Chris French, Hayley Yaglom, Ashlyn Pfeiffer, Darrin Lemmer, Dave Engelthaler, The Arizona COVID Genomics Union (ACGU)" |
| EPI_ISL_967909 | TGen North | Sonora Quest Laboratories | "Jolene Bowers, Megan Folkerts, Chris French, Hayley Yaglom, Ashlyn Pfeiffer, Darrin Lemmer, Dave Engelthaler, The Arizona COVID Genomics Union (ACGU)" |
| EPI_ISL_967910, EPI_ISL_967911, EPI_ISL_967912, EPI_ISL_967914 | TGen North | TGen North | "Jolene Bowers, Megan Folkerts, Chris French, Hayley Yaglom, Ashlyn Pfeiffer, Darrin Lemmer, Dave Engelthaler, The Arizona COVID Genomics Union (ACGU)" |
| EPI_ISL_967917, EPI_ISL_967918 | TGen North | Sonora Quest Laboratories | "Jolene Bowers, Megan Folkerts, Chris French, Hayley Yaglom, Ashlyn Pfeiffer, Darrin Lemmer, Dave Engelthaler, The Arizona COVID Genomics Union (ACGU)" |
| EPI_ISL_967919, EPI_ISL_967921 | TGen North | TGen North | "Jolene Bowers, Megan Folkerts, Chris French, Hayley Yaglom, Ashlyn Pfeiffer, Darrin Lemmer, Dave Engelthaler, The Arizona COVID Genomics Union (ACGU)" |
| EPI_ISL_967922 | TGen North | Sonora Quest Laboratories | "Jolene Bowers, Megan Folkerts, Chris French, Hayley Yaglom, Ashlyn Pfeiffer, Darrin Lemmer, Dave Engelthaler, The Arizona COVID Genomics Union (ACGU)" |
| EPI_ISL_967923 | TGen North | TGen North | "Jolene Bowers, Megan Folkerts, Chris French, Hayley Yaglom, Ashlyn Pfeiffer, Darrin Lemmer, Dave Engelthaler, The Arizona COVID Genomics Union (ACGU)" |
| EPI_ISL_967924 | TGen North | Sonora Quest Laboratories | "Jolene Bowers, Megan Folkerts, Chris French, Hayley Yaglom, Ashlyn Pfeiffer, Darrin Lemmer, Dave Engelthaler, The Arizona COVID Genomics Union (ACGU)" |
| EPI_ISL_967925 | TGen North | TGen North | "Jolene Bowers, Megan Folkerts, Chris French, Hayley Yaglom, Ashlyn Pfeiffer, Darrin Lemmer, Dave Engelthaler, The Arizona COVID Genomics Union (ACGU)" |
| EPI_ISL_967927 | TGen North | Sonora Quest Laboratories | "Jolene Bowers, Megan Folkerts, Chris French, Hayley Yaglom, Ashlyn Pfeiffer, Darrin Lemmer, Dave Engelthaler, The Arizona COVID Genomics Union (ACGU)" |
| EPI_ISL_967930, EPI_ISL_967931, EPI_ISL_967935, EPI_ISL_967937, EPI_ISL_967939, EPI_ISL_967941, EPI_ISL_967944, EPI_ISL_967945, EPI_ISL_967947, EPI_ISL_967948, EPI_ISL_967949, EPI_ISL_967950, EPI_ISL_967952, EPI_ISL_967953, EPI_ISL_967956, EPI_ISL_967959, EPI_ISL_967960, EPI_ISL_967961 | | | |
| see above | TGen North | TGen North | "Jolene Bowers, Megan Folkerts, Chris French, Hayley Yaglom, Ashlyn Pfeiffer, Darrin Lemmer, Dave Engelthaler, The Arizona COVID Genomics Union (ACGU)" |
| EPI_ISL_967962, EPI_ISL_967963 | TGen North | Sonora Quest Laboratories | "Jolene Bowers, Megan Folkerts, Chris French, Hayley Yaglom, Ashlyn Pfeiffer, Darrin Lemmer, Dave Engelthaler, The Arizona COVID Genomics Union (ACGU)" |
| EPI_ISL_967970, EPI_ISL_967971, EPI_ISL_967973, EPI_ISL_967975 | TGen North | TGen North | "Jolene Bowers, Megan Folkerts, Chris French, Hayley Yaglom, Ashlyn Pfeiffer, Darrin Lemmer, Dave Engelthaler, The Arizona COVID Genomics Union (ACGU)" |
| EPI_ISL_967985 | TGen North | Sonora Quest Laboratories | "Jolene Bowers, Megan Folkerts, Chris French, Hayley Yaglom, Ashlyn Pfeiffer, Darrin Lemmer, Dave Engelthaler, The Arizona COVID Genomics Union (ACGU)" |
| EPI_ISL_967986, EPI_ISL_967987, EPI_ISL_967988 | TGen North | TGen North | "Jolene Bowers, Megan Folkerts, Chris French, Hayley Yaglom, Ashlyn Pfeiffer, Darrin Lemmer, Dave Engelthaler, The Arizona COVID Genomics Union (ACGU)" |

| | | | |
|---|---|---|---|
| EPI_ISL_967991 | TGen North | Sonora Quest Laboratories | "Jolene Bowers, Megan Folkerts, Chris French, Hayley Yaglom, Ashlyn Pfeiffer, Darrin Lemmer, Dave Engelthaler, The Arizona COVID Genomics Union (ACGU)" |
| EPI_ISL_967992 | TGen North | TGen North | "Jolene Bowers, Megan Folkerts, Chris French, Hayley Yaglom, Ashlyn Pfeiffer, Darrin Lemmer, Dave Engelthaler, The Arizona COVID Genomics Union (ACGU)" |
| EPI_ISL_967995 | TGen North | Sonora Quest Laboratories | "Jolene Bowers, Megan Folkerts, Chris French, Hayley Yaglom, Ashlyn Pfeiffer, Darrin Lemmer, Dave Engelthaler, The Arizona COVID Genomics Union (ACGU)" |
| EPI_ISL_967997, EPI_ISL_967999, EPI_ISL_968003, EPI_ISL_968005, EPI_ISL_968007 | TGen North | TGen North | "Jolene Bowers, Megan Folkerts, Chris French, Hayley Yaglom, Ashlyn Pfeiffer, Darrin Lemmer, Dave Engelthaler, The Arizona COVID Genomics Union (ACGU)" |
| EPI_ISL_968008 | TGen North | Sonora Quest Laboratories | "Jolene Bowers, Megan Folkerts, Chris French, Hayley Yaglom, Ashlyn Pfeiffer, Darrin Lemmer, Dave Engelthaler, The Arizona COVID Genomics Union (ACGU)" |
| EPI_ISL_968009, EPI_ISL_968011, EPI_ISL_968012, EPI_ISL_968013, EPI_ISL_968015, EPI_ISL_968016 | TGen North | TGen North | "Jolene Bowers, Megan Folkerts, Chris French, Hayley Yaglom, Ashlyn Pfeiffer, Darrin Lemmer, Dave Engelthaler, The Arizona COVID Genomics Union (ACGU)" |
| EPI_ISL_968025 | TGen North | Sonora Quest Laboratories | "Jolene Bowers, Megan Folkerts, Chris French, Hayley Yaglom, Ashlyn Pfeiffer, Darrin Lemmer, Dave Engelthaler, The Arizona COVID Genomics Union (ACGU)" |
| EPI_ISL_968032, EPI_ISL_968034 | TGen North | TGen North | "Jolene Bowers, Megan Folkerts, Chris French, Hayley Yaglom, Ashlyn Pfeiffer, Darrin Lemmer, Dave Engelthaler, The Arizona COVID Genomics Union (ACGU)" |
| EPI_ISL_968035 | TGen North | Sonora Quest Laboratories | "Jolene Bowers, Megan Folkerts, Chris French, Hayley Yaglom, Ashlyn Pfeiffer, Darrin Lemmer, Dave Engelthaler, The Arizona COVID Genomics Union (ACGU)" |
| EPI_ISL_968037, EPI_ISL_968038 | TGen North | TGen North | "Jolene Bowers, Megan Folkerts, Chris French, Hayley Yaglom, Ashlyn Pfeiffer, Darrin Lemmer, Dave Engelthaler, The Arizona COVID Genomics Union (ACGU)" |
| EPI_ISL_968039 | TGen North | Sonora Quest Laboratories | "Jolene Bowers, Megan Folkerts, Chris French, Hayley Yaglom, Ashlyn Pfeiffer, Darrin Lemmer, Dave Engelthaler, The Arizona COVID Genomics Union (ACGU)" |
| EPI_ISL_968041 | TGen North | TGen North | "Jolene Bowers, Megan Folkerts, Chris French, Hayley Yaglom, Ashlyn Pfeiffer, Darrin Lemmer, Dave Engelthaler, The Arizona COVID Genomics Union (ACGU)" |
| EPI_ISL_968043 | TGen North | Sonora Quest Laboratories | "Jolene Bowers, Megan Folkerts, Chris French, Hayley Yaglom, Ashlyn Pfeiffer, Darrin Lemmer, Dave Engelthaler, The Arizona COVID Genomics Union (ACGU)" |
| EPI_ISL_968044, EPI_ISL_968045 | TGen North | TGen North | "Jolene Bowers, Megan Folkerts, Chris French, Hayley Yaglom, Ashlyn Pfeiffer, Darrin Lemmer, Dave Engelthaler, The Arizona COVID Genomics Union (ACGU)" |
| EPI_ISL_968046, EPI_ISL_968049 | TGen North | Sonora Quest Laboratories | "Jolene Bowers, Megan Folkerts, Chris French, Hayley Yaglom, Ashlyn Pfeiffer, Darrin Lemmer, Dave Engelthaler, The Arizona COVID Genomics Union (ACGU)" |
| EPI_ISL_968050, EPI_ISL_968051, EPI_ISL_968053 | TGen North | TGen North | "Jolene Bowers, Megan Folkerts, Chris French, Hayley Yaglom, Ashlyn Pfeiffer, Darrin Lemmer, Dave Engelthaler, The Arizona COVID Genomics Union (ACGU)" |
| EPI_ISL_968058, EPI_ISL_968059, EPI_ISL_968061 | TGen North | Sonora Quest Laboratories | "Jolene Bowers, Megan Folkerts, Chris French, Hayley Yaglom, Ashlyn Pfeiffer, Darrin Lemmer, Dave Engelthaler, The Arizona COVID Genomics Union (ACGU)" |
| EPI_ISL_968064 | TGen North | TGen North | "Jolene Bowers, Megan Folkerts, Chris French, Hayley Yaglom, Ashlyn Pfeiffer, Darrin Lemmer, Dave Engelthaler, The Arizona COVID Genomics Union (ACGU)" |
| EPI_ISL_968066 | TGen North | Sonora Quest Laboratories | "Jolene Bowers, Megan Folkerts, Chris French, Hayley Yaglom, Ashlyn Pfeiffer, Darrin Lemmer, Dave Engelthaler, The Arizona COVID Genomics Union (ACGU)" |
| EPI_ISL_968067, EPI_ISL_968068 | TGen North | TGen North | "Jolene Bowers, Megan Folkerts, Chris French, Hayley Yaglom, Ashlyn Pfeiffer, Darrin Lemmer, Dave Engelthaler, The Arizona COVID Genomics Union (ACGU)" |
| EPI_ISL_968079 | Monterey County Public Health Laboratory | Monterey County Public Health Laboratory | Monterey County Public Health Laboratory |
| EPI_ISL_977035, EPI_ISL_977036, EPI_ISL_977037, EPI_ISL_977038, EPI_ISL_977039, EPI_ISL_977040, EPI_ISL_977041, EPI_ISL_977042, EPI_ISL_977043, EPI_ISL_977044, EPI_ISL_977045, EPI_ISL_977046, EPI_ISL_977047, EPI_ISL_977048, EPI_ISL_977049, EPI_ISL_977050, EPI_ISL_977051, EPI_ISL_977052, EPI_ISL_977053, EPI_ISL_977054, EPI_ISL_977055, EPI_ISL_977056, EPI_ISL_977057, EPI_ISL_977058, EPI_ISL_977059, EPI_ISL_977060, EPI_ISL_977061, EPI_ISL_977062, EPI_ISL_977063, EPI_ISL_977064, EPI_ISL_977073, EPI_ISL_977074, EPI_ISL_977075 | | | |
| see above | Rhode Island Department of Health | Infectious Disease Program, Broad Institute of Harvard and MIT | Lemieux,J.E., Siddle,K.J., Huard,R., King,E., Azevedo,K., Miller,A., Adams,G., Gladden-Young,A., Lagerborg,K., Rudy,M., DeRuff,K., Carter,A., Normandin,E., Bauer,M., Reilly,S., Tomkins-Tinch,C., Loreth,C., Chaluvadi,S., Birren,B.W., Gallagher,G., Smole,S., Park,D.J., MacInnis,B.L., and Sabeti,P.C. |
| EPI_ISL_977101, EPI_ISL_977112, EPI_ISL_977113, EPI_ISL_977114, EPI_ISL_977118, EPI_ISL_977119 | Massachusetts General Hospital | Infectious Disease Program, Broad Institute of Harvard and MIT | Lemieux,J.E., Siddle,K.J., Shaw,B., Adams,G., Pierce,V., Turbett,S., Anahtar,M., Branda,J., Slater,D., Harris,J., Lin,A.E., Gladden-Young,A., Lagerborg,K., Rudy,M., DeRuff,K., Carter,A., Normandin,E., Bauer,M., Reilly,S., Tomkins-Tinch,C., Loreth,C., Chaluvadi,S., Neumann,A., Cusick,C., Chapman,S.B., Gnirke,A., Flowers,K., Cerrato,F., Birren,B.W., Gallagher,G., Smole,S., Park,D.J., MacInnis,B.L., Ryan,E., LaRocque,R., Rosenberg,E. and Sabeti,P.C. |
| EPI_ISL_977121, EPI_ISL_977122, EPI_ISL_977123, EPI_ISL_977124, EPI_ISL_977125, EPI_ISL_977126, EPI_ISL_977127, EPI_ISL_977128, EPI_ISL_977129 | Flow Health | Infectious Disease Program, Broad Institute of Harvard and MIT | Lemieux,J.E., Siddle,K.J., Adams,G., Gladden-Young,A., Lagerborg,K., Rudy,M., DeRuff,K., Carter,A., Normandin,E., Bauer,M., Reilly,S., Tomkins-Tinch,C., Loreth,C., Chaluvadi,S., Birren,B.W., Gallagher,G., Smole,S., Park,D.J., MacInnis,B.L., and Sabeti,P.C. |
| EPI_ISL_977160, EPI_ISL_977246 | Microbiologia e Virologia | Istituto Zooprofilattico Sperimentale delle Venezie | Adelaide Milani, Alessia Schivo, Annalisa Salviato, Erika Giorgia Quaranta, Ambra Pastori, Bianca Zecchin, Alice Fusaro, Isabella Monne, Calogero Terregino, Antonia Ricci |
| EPI_ISL_977348, EPI_ISL_977349 | University of Zambia, School of Veterinary Medicine | UNZAVET and PATH | Mulenga Mwenda-Chimfwembe, Ngonda Saasa, Daniel Bridges |
| EPI_ISL_977540, EPI_ISL_977550, EPI_ISL_977553, EPI_ISL_977554 | Nigeria Centre of Disease Control (NCDC) | African Centre of Excellence for Genomics of Infectious Diseases (ACEGID), Redeemer's University | Olawoye I. B. et al |
| EPI_ISL_977581, EPI_ISL_977584, EPI_ISL_977594 | Caribbean Public Health Agency | Carrington Lab, Department of PreClinical Sciences | Nikita S. D. Sahadeo, Arianne Brown-Jordan, Vernie Ramkissoon, Sarah Hill, Naresh Nandram, Avery Hinds, Dr. Sharon Belmar-George, Jerome Foster, Stanley Giddings, Karla Georges, Marsha Ivey, Rahul Naidu, Risha Singh, SueMin Nathaniel, Rajini Haraksingh, Jaya Jayaraman, Chinna Chinnadurai, Adesh Ramsubhag, Nuno Faria, Oliver Pybus, Christopher Oura, Gabriel Escobar, Christine V. F. Carrington |
| EPI_ISL_977940, EPI_ISL_977941, EPI_ISL_977942, EPI_ISL_977943, EPI_ISL_977944, EPI_ISL_977945, EPI_ISL_977946, EPI_ISL_977947, EPI_ISL_977948, EPI_ISL_977949, EPI_ISL_977950, EPI_ISL_977951, EPI_ISL_977952, EPI_ISL_977953, EPI_ISL_977954, EPI_ISL_977955, EPI_ISL_977956, EPI_ISL_977957, EPI_ISL_977958, EPI_ISL_977959, EPI_ISL_977960, EPI_ISL_977961, EPI_ISL_977980 | | | |
| see above | Chiu Laboratory, University of California, San Francisco | Chiu Laboratory, University of California, San Francisco | Charles Chiu, Xianding (Wayne) Deng, Candace Wang, Venice Servellita, Jill Hacker, Debra Wadford |
| EPI_ISL_978219, EPI_ISL_978220, EPI_ISL_978221 | Virginia Division of Consolidated Laboratory Services | Virginia Division of Consolidated Laboratory Services | Virginia DCLS |
| EPI_ISL_978282, EPI_ISL_978283, EPI_ISL_978284, EPI_ISL_978285, EPI_ISL_978286, EPI_ISL_978287, EPI_ISL_978288, EPI_ISL_978289, EPI_ISL_978290, EPI_ISL_978291, EPI_ISL_978292, EPI_ISL_978293 | | | |
| see above | Texas Department of State Health Services | Texas Department of State Health Services | Bonnie Oh, Anita Pokharel, James Daniel Bonser, Myong Koag, Chung Wang, Rachel Lee, Grace Kubin, Rashmi Tuladhar, Mayela Pedrueza, Maliha |

| | | | |
|---|---|---|---|
| | | | Rahman, Jenny Zhang |
| EPI_ISL_978353, EPI_ISL_978354, EPI_ISL_978355, EPI_ISL_978358, EPI_ISL_978359, EPI_ISL_978360, EPI_ISL_978361, EPI_ISL_978362, EPI_ISL_978396 | Arizona State Public Health Laboratory | Arizona State Public Health Laboratory | Trung Huynh, Jessica Escobar, Katherine Fullerton, Nobuko Fukushima, Stacy White, Linda Getsinger, Victor Waddell |
| EPI_ISL_978811, EPI_ISL_978926, EPI_ISL_978927, EPI_ISL_978928 | Centre for Dengue Research and AICBU, Department of Immunology and Molecular Medicine | Centre for Dengue Research and AICBU, Department of Immunology and Molecular Medicine | Chandima Jeewandara, Deshni Jayathilaka, Dinuka Ariyaratne, Tibutius Thanesh Pramanayagam, Diyanath Ranasinghe, Laksiri Gomes, Gathsaurie Neelika Malavige |
| EPI_ISL_979056, EPI_ISL_979057, EPI_ISL_979058, EPI_ISL_979059, EPI_ISL_979061, EPI_ISL_979062, EPI_ISL_979063, EPI_ISL_979064, EPI_ISL_979065, EPI_ISL_979066, EPI_ISL_979067, EPI_ISL_979068, EPI_ISL_979069, EPI_ISL_979070, EPI_ISL_979071, EPI_ISL_979072, EPI_ISL_979073, EPI_ISL_979074, EPI_ISL_979075, EPI_ISL_979076, EPI_ISL_979077, EPI_ISL_979078, EPI_ISL_979079, EPI_ISL_979080, EPI_ISL_979081, EPI_ISL_979082, EPI_ISL_979083, EPI_ISL_979099, EPI_ISL_979100, EPI_ISL_979101, EPI_ISL_979102, EPI_ISL_979108, EPI_ISL_979109, EPI_ISL_979111, EPI_ISL_979113, EPI_ISL_979114, EPI_ISL_979123, EPI_ISL_979127, EPI_ISL_979128 | | | |
| see above | Santa Clara County Public Health Laboratory | Chan-Zuckerberg Biohub | CZB Cliahub Consortium |
| EPI_ISL_979160, EPI_ISL_979161 | County of San Luis Obispo Public Health Laboratory | Chan-Zuckerberg Biohub | CZB Cliahub Consortium |
| EPI_ISL_979260, EPI_ISL_979261, EPI_ISL_979262, EPI_ISL_979263, EPI_ISL_979264, EPI_ISL_979265, EPI_ISL_979266, EPI_ISL_979267 | Institute of Microbiology and Immunology, Faculty of Medicine, University of Ljubljana | Institute of Microbiology and Immunology, Faculty of Medicine, University of Ljubljana | Samo Zakotnik, Tomaž Mark Zorec, Matic Brvar, Doroteja Vlaj, Patricija Pozvek,Špela Pleh, Miša Korva, Mario Poljak, Tatjana Avši - Županc |
| EPI_ISL_979321, EPI_ISL_979323 | Cadham Provincial laboratory | National Microbiology Laboratory (NML) | Anna Majer, Shari Tyson, Grace Seo, Philip Mabon, Elsie Grudeski, Rhiannon Huzarewich, Russell Mandes, Anneliese Landgraff, Jennifer Tanner, Natalie Knox, Morag Graham, Gary Van Domselaar, Paul Van Caeseele, Jared Bullard, David Alexander, Kerry Dust, Nathalie Bastien, Yan Li, Timothy Booth, Darian Hole, Madison Chapel, Kirsten Biggar, CanCOGeN's metadata curation team, Public Health Agency of Canada CanCOGeN team |
| EPI_ISL_979368, EPI_ISL_979370, EPI_ISL_979371, EPI_ISL_979373, EPI_ISL_979374, EPI_ISL_979375, EPI_ISL_979377 | The Jackson Laboratory | The Jackson Laboratory | Lloyd M, Sanderson B, Srivastava A, Maurya R, Renzette N, Omerza G, Kelly K, Li L, Wei C L, Adams M |
| EPI_ISL_980845, EPI_ISL_980852, EPI_ISL_980854, EPI_ISL_980855, EPI_ISL_980859, EPI_ISL_980897, EPI_ISL_980898, EPI_ISL_980899, EPI_ISL_980902, EPI_ISL_980916, EPI_ISL_980920, EPI_ISL_980921, EPI_ISL_980922, EPI_ISL_980935, EPI_ISL_980936, EPI_ISL_980937, EPI_ISL_980977, EPI_ISL_980978, EPI_ISL_980979, EPI_ISL_980980, EPI_ISL_980981 | | | |
| see above | Innovative Genomics Institute, UC Berkeley | Innovative Genomics Institute, UC Berkeley | Stacia Wyman, Haridha Shivram, Phil Frankino, Liana Lareau |
| EPI_ISL_981071 | Johns Hopkins Hospital Department of Pathology | Johns Hopkins Hospital Department of Pathology | C. Paul Morris, Chun Huai Luo, Adannaya Amadi, Matthew Schwartz, Nicholas Gallagher, Heba H. Mostafa |
| EPI_ISL_981271, EPI_ISL_981273, EPI_ISL_981274, EPI_ISL_981275, EPI_ISL_981276, EPI_ISL_981277, EPI_ISL_981278, EPI_ISL_981279, EPI_ISL_981280, EPI_ISL_981281, EPI_ISL_981282, EPI_ISL_981283, EPI_ISL_981284, EPI_ISL_981285, EPI_ISL_981286, EPI_ISL_981287, EPI_ISL_981288, EPI_ISL_981289, EPI_ISL_981290, EPI_ISL_981291, EPI_ISL_981292, EPI_ISL_981293, EPI_ISL_981294, EPI_ISL_981296 | | | |
| see above | Hospital Universitari de Bellvitge | Hospital Universitari Vall d'Hebron | Cristina Andrés, Maria Piñana, Josep F Abril, Damir Garcia-Cehic, Ariadna Rando, Juliana Esperalba, Maria Gema Codina, Carla Castillo, Maria Carmen Martin, Tomàs Pumarola, Josep Quer, Andrés Antón |
| EPI_ISL_981297, EPI_ISL_981300, EPI_ISL_981304 | Hospital Universitari Vall d'Hebron | Hospital Universitari Vall d'Hebron | Cristina Andrés, Maria Piñana, Josep F Abril, Damir Garcia-Cehic, Ariadna Rando, Juliana Esperalba, Maria Gema Codina, Carla Castillo, Maria Carmen Martin, Tomàs Pumarola, Josep Quer, Andrés Antón |
| EPI_ISL_981964, EPI_ISL_981965, EPI_ISL_981966, EPI_ISL_981969 | Microbiology Service, Hospital Universitario Clinico San Cecilio, Granada | Microbiology Service, Hospital Universitario Clinico San Cecilio, Granada | Adolfo de Salazar, Natalia Chueca, Laura Viñuela, Ana Fuentes, Federico García |
| EPI_ISL_981981, EPI_ISL_982001 | TGen North | TGen North | "Jolene Bowers, Megan Folkerts, Chris French, Hayley Yaglom, Ashlyn Pfeiffer, Darrin Lemmer, Dave Engelthaler, The Arizona COVID Genomics Union (ACGU)" |
| EPI_ISL_982045, EPI_ISL_982064, EPI_ISL_982067 | TGen North | Sonora Quest Laboratories | "Jolene Bowers, Megan Folkerts, Chris French, Hayley Yaglom, Ashlyn Pfeiffer, Darrin Lemmer, Dave Engelthaler, The Arizona COVID Genomics Union (ACGU)" |
| EPI_ISL_982241, EPI_ISL_982245, EPI_ISL_982248, EPI_ISL_982283 | Lab voor klinische biologie | Lab voor klinische biologie | Hannelore Hamerlinck, Marija Janevska, Bruno Verhasselt |
| EPI_ISL_982344, EPI_ISL_982346, EPI_ISL_982347, EPI_ISL_982389, EPI_ISL_982390, EPI_ISL_982391, EPI_ISL_982394, EPI_ISL_982397, EPI_ISL_982398, EPI_ISL_982400, EPI_ISL_982401, EPI_ISL_982402, EPI_ISL_982406, EPI_ISL_982407, EPI_ISL_982408, EPI_ISL_982409, EPI_ISL_982410, EPI_ISL_982411, EPI_ISL_982412, EPI_ISL_982413, EPI_ISL_982414, EPI_ISL_982415, EPI_ISL_982416, EPI_ISL_982417, EPI_ISL_982418, EPI_ISL_982419 | | | |
| see above | M Health Fairview | Minnesota Department of Health, Public Health Laboratory | Alexandra Lorentz, Jacob Garfin, Matt Plumb, and Xiong Wang |
| EPI_ISL_982502, EPI_ISL_982503, EPI_ISL_982504 | ADIRONDACK MEDICAL CENTER | Wadsworth Center, New York State Department of Health | Kirsten St. George, Daryl M. Lamson, Alexis Russel, Matthew Shudt, Melissa A Leisner, Jonathan Plitnick, Navjot Singh, John Kelly, Erasmus Schneider, Erica Lasek-Nesselquist |
| EPI_ISL_982529, EPI_ISL_982535, EPI_ISL_982537, EPI_ISL_982545, EPI_ISL_982547, EPI_ISL_982548, EPI_ISL_982550, EPI_ISL_982551, EPI_ISL_982552, EPI_ISL_982555, EPI_ISL_982557, EPI_ISL_982558, EPI_ISL_982561, EPI_ISL_982567, EPI_ISL_982573, EPI_ISL_982576, EPI_ISL_982583, EPI_ISL_982589, EPI_ISL_982596, EPI_ISL_982603, EPI_ISL_982609, EPI_ISL_982615, EPI_ISL_982624, EPI_ISL_982635, EPI_ISL_982637, EPI_ISL_982641, EPI_ISL_982642, EPI_ISL_982647, EPI_ISL_982664, EPI_ISL_982665, EPI_ISL_982666, EPI_ISL_982667, EPI_ISL_982668, EPI_ISL_982670, EPI_ISL_982674, EPI_ISL_982684, EPI_ISL_982693, EPI_ISL_982695, EPI_ISL_982707, EPI_ISL_982708, EPI_ISL_982709, EPI_ISL_982720, EPI_ISL_982721, EPI_ISL_982722, EPI_ISL_982723, EPI_ISL_982724, EPI_ISL_982725, EPI_ISL_982726, EPI_ISL_982727, EPI_ISL_982732, EPI_ISL_982733, EPI_ISL_982738, EPI_ISL_982742, EPI_ISL_982746, EPI_ISL_982758, EPI_ISL_982760 | | | |
| see above | US Air Force School of Aerospace Medicine | US Air Force School of Aerospace Medicine | Anthony Fries, Jennifer Meyer, William Gruner, William Buggele, Amanda Javorina, Sarah Purves, Clarise Starr, Elizabeth Macias |
| EPI_ISL_983063, EPI_ISL_983086, EPI_ISL_983087 | Gravity Diagnostics | Kentucky State Public Health Lab | Stephanie Lunn, Karim George, Joshua Tobias, William Grooms, Vaneet Arora, Matthew Johnson, Rachel Zinner, Rhonda Lucas |
| EPI_ISL_983118 | SUNY UPSTATE MEDICAL UNIVERSITY | Wadsworth Center, New York State Department of Health | Kirsten St. George, Daryl M. Lamson, Alexis Russel, Matthew Shudt, Melissa A Leisner, Jonathan Plitnick, Navjot Singh, John Kelly, Erasmus Schneider, Erica Lasek-Nesselquist |
| EPI_ISL_983130, EPI_ISL_983131 | THE MARY IMOGENE BASSETT HOSPITAL | Wadsworth Center, New York State Department of Health | Kirsten St. George, Daryl M. Lamson, Alexis Russel, Matthew Shudt, Melissa A Leisner, Jonathan Plitnick, Navjot Singh, John Kelly, Erasmus Schneider, Erica Lasek-Nesselquist |
| EPI_ISL_983321 | Eurofins Genoma Group s.r.l. | INMI Lazzaro Spallanzani IRCCS | O Butera, F Messina, CEM Gruber, B Bartolini, E Giombini, M Rueca, F Spinella, F Fiorentino, MR Capobianchi, A Di Caro |
| EPI_ISL_983616, EPI_ISL_983617 | Texas Department of State Health Services | Texas Department of State Health Services | Bonnie Oh, Anita Pokharel, James Daniel Bonser, Myong Koag, Chung Wang, Rachel Lee, Grace Kubin, Rashmi Tuladhar, Mayela Pedrueza, Maliha Rahman, Jenny Zhang |
| EPI_ISL_983698 | Vault Health | Minnesota Department of Health, Public Health Laboratory | Alexandra Lorentz, Jacob Garfin, Matt Plumb, and Xiong Wang |
| EPI_ISL_983710, EPI_ISL_983717, EPI_ISL_983801, EPI_ISL_983802, EPI_ISL_983803, EPI_ISL_983804 | Colorado Department of Public Health and Environment | Colorado Department of Puplic Health and Environment | Laura Bankers, Molly C. Hetherington-Rauth, Diana Ir, Shannon Ely, Shannon R. Matzinger, Sarah Elizabeth Totten, Emily A. Travanty |
