## Supplementary material for "Detection and characterization of the SARS-CoV-2 lineage B.1.526 in New York": Supp. Table 3 GISAID Acknowledgment part 2: gisaid_hcov-19_acknowledgement_table_2021_02_13_010-12.pdf

We gratefully acknowledge the following Authors from the Originating laboratories responsible for obtaining the specimens, as well as the Submitting laboratories where the genome data were generated and shared via GISAID, on which this research is based.

All Submitters of data may be contacted directly via www.gisaid.org

Authors are sorted alphabetically.

| Accession ID | Originating Laboratory | Submitting Laboratory | Authors |
|---|---|---|---|
| EPI_ISL_845802, EPI_ISL_845803, EPI_ISL_845804 | Medlab Pathology | NSW Health Pathology - Institute of Clinical Pathology and Medical Research; Westmead Hospital; University of Sydney | CIDM-PH et al. |
| EPI_ISL_845805 | Sydney South West Pathology Service (SSWPS) - Royal Prince Alfred Hospital - NSW Health Pathology | NSW Health Pathology - Institute of Clinical Pathology and Medical Research; Westmead Hospital; University of Sydney | CIDM-PH et al. |
| EPI_ISL_846774 | Respiratory Virus Unit, National Infection Service, Public Health England | COVID-19 Genomics UK (COG-UK) Consortium | PHE Covid Sequencing Team |
| EPI_ISL_849762 | unknown | PHV-FSS | Son Nguyen et al. |
| EPI_ISL_852014 | Lighthouse Lab in Milton Keynes | Wellcome Sanger Institute for the COVID-19 Genomics UK (COG-UK) Consortium | The Lighthouse Lab in Milton Keynes and Alex Alderton, Roberto Amato, Sonia Goncalves, Ewan Harrison, David K. Jackson, Ian Johnston, Dominic Kwiatkowski, Cordelia Langford, John Sillitoe on behalf of the Wellcome Sanger Institute COVID-19 Surveillance Team |
| EPI_ISL_855604 | Respiratory Virus Unit, National Infection Service, Public Health England | COVID-19 Genomics UK (COG-UK) Consortium | PHE Covid Sequencing Team |
| EPI_ISL_857310, EPI_ISL_857311, EPI_ISL_857312, EPI_ISL_857313 | South Eastern Area Laboratory Services (SEALS) | NSW Health Pathology - Institute of Clinical Pathology and Medical Research; Westmead Hospital; University of Sydney | CIDM-PH et al. |
| EPI_ISL_857408, EPI_ISL_857409, EPI_ISL_857411, EPI_ISL_857414 | Maine HETL | Tewhey Lab, The Jackson Laboratory | Matluk,N., Dewey,H., Iosue,F., Barter,M., Lynch,R., Munger,H. and Tewhey,R. |
| EPI_ISL_857469, EPI_ISL_857470, EPI_ISL_857471, EPI_ISL_857472, EPI_ISL_857474, EPI_ISL_857475, EPI_ISL_857476, EPI_ISL_857477, EPI_ISL_857479, EPI_ISL_857480, EPI_ISL_857481, EPI_ISL_857482, EPI_ISL_857483, EPI_ISL_857484 | | | |
| see above | National Public Health Laboratory, National Centre for Infectious Diseases | National Public Health Laboratory, National Centre for Infectious Diseases | Tze Minn Mak, Sophie Octavia, Zhenyang Zhou, Lin Cui, Raymond Tzer Pin Lin |
| EPI_ISL_857487, EPI_ISL_857488 | PathWest Laboratory Medicine WA | PathWest Laboratory Medicine WA Microbial Surveillance Unit | PathWest Laboratory Medicine WA Microbial Surveillance Unit |
| EPI_ISL_860691, EPI_ISL_860692, EPI_ISL_860693, EPI_ISL_860694, EPI_ISL_860696, EPI_ISL_860697, EPI_ISL_860698, EPI_ISL_860699, EPI_ISL_860700, EPI_ISL_860701, EPI_ISL_860702, EPI_ISL_860704, EPI_ISL_860705, EPI_ISL_860706, EPI_ISL_860707, EPI_ISL_860708, EPI_ISL_860783 | | | |
| see above | Respiratory Virus Unit, National Infection Service, Public Health England | COVID-19 Genomics UK (COG-UK) Consortium | PHE Covid Sequencing Team |
| EPI_ISL_861003, EPI_ISL_861010, EPI_ISL_861011, EPI_ISL_861013, EPI_ISL_861036, EPI_ISL_861054, EPI_ISL_861063, EPI_ISL_861067, EPI_ISL_861075, EPI_ISL_861080, EPI_ISL_861085, EPI_ISL_861089, EPI_ISL_861090, EPI_ISL_861091, EPI_ISL_861095, EPI_ISL_861096 | | | |
| see above | Johns Hopkins Hospital Department of Pathology | Johns Hopkins Hospital Department of Pathology | C. Paul Morris, Chun Huai Luo, Adannaya Amadi, Nicholas Gallagher, Heba H. Mostafa |
| EPI_ISL_861443, EPI_ISL_861444, EPI_ISL_861449, EPI_ISL_861450, EPI_ISL_861451, EPI_ISL_861454 | Clinical Molecular Microbiology Laboratory, UNC Hospitals | Jeremy Wang | Jeremy Wang, Alexander Rubinsteyn, Colleen Rice, Jason Smedberg, Melissa Miller, Corbin Jones, Robert Hagan |
| EPI_ISL_861730, EPI_ISL_861731 | UW Virology Lab | UW Virology Lab | Pavitra Roychoudhury, Hong Xie, Lasata Shrestha, Michelle Lin, Meei-Li Huang, Keith R Jerome, Alexander Greninger |
| EPI_ISL_861737, EPI_ISL_861746, EPI_ISL_861751, EPI_ISL_861754, EPI_ISL_861756, EPI_ISL_861772 | Tempus | Grubaugh Lab - Yale School of Public Health | Tara Alpert, Joseph Fauver, Anderson Brito, Mallery Breban, Anne Wyllie, Chantal Vogels, Mary Petrone, Annie Watkins, Chaney Kalinich, Isabel Ott, Nathan Grubaugh |
| EPI_ISL_862191, EPI_ISL_862192, EPI_ISL_862259, EPI_ISL_862260, EPI_ISL_862261, EPI_ISL_862262, EPI_ISL_862263, EPI_ISL_862264 | Respiratory Virus Unit, National Infection Service, Public Health England | COVID-19 Genomics UK (COG-UK) Consortium | PHE Covid Sequencing Team |
| EPI_ISL_862834 | National Institute of Infectious Diseases-Prof. Dr. Matei Bals Molecular Diagnostics Laboratory | National Institute of Infectious Diseases-Prof. Dr. Matei Bals Molecular Diagnostics Laboratory | Leontina Banica, Marius Surleac, Corina Casangiu, Petre Milu, Andreea Tudor, Simona Paraschiv, Dan Otelea |
| EPI_ISL_864583, EPI_ISL_864585, EPI_ISL_864587 | CHU Purpan - Laboratoire de Virologie - Institut Fédératif de Biologie | CHU Purpan - Laboratoire de Virologie - Institut Fédératif de Biologie | Latour J., Ranger N., Dubois M., Carcenac R., Harter A., Boyer P., Tremeaux P., Izopet J. |
| EPI_ISL_864927, EPI_ISL_864928, EPI_ISL_864929, EPI_ISL_864930, EPI_ISL_864931, EPI_ISL_864932, EPI_ISL_864934, EPI_ISL_864936, EPI_ISL_864945, EPI_ISL_864946, EPI_ISL_864953, EPI_ISL_864955, EPI_ISL_864956, EPI_ISL_864958, EPI_ISL_864959 | | | |
| see above | Department of Pathology, University of Cambridge | COVID-19 Genomics UK (COG-UK) Consortium | Aminu S. Jahun, Yasmin Chaudhry, Grant Hall, Iliana Georgana, Myra Hosmillo, Martin D. Curran, Malte Pinckert, Surendra Parmar, Ian Goodfellow |
| EPI_ISL_865163 | University of Exeter | COVID-19 Genomics UK (COG-UK) Consortium | Ben Temperton,Aaron Jeffries,Michelle Michelsen,Joanna Warwick-Dugdale,Audrey Farbos,Robyn Manley,Stephen Michell,Jane Masoli |
| EPI_ISL_865467, EPI_ISL_865468, EPI_ISL_865469, EPI_ISL_865470, EPI_ISL_865471, EPI_ISL_865477 | Liverpool Clinical Laboratories | COVID-19 Genomics UK (COG-UK) Consortium | Sam Haldenby, Anita Lucaci, Steve Paterson, Julian Hiscox, Alistair Darby, M Almsaud, A Alrezaihi, Muhannad Alruwaili, Stuart D Armstrong, Jones Benjamin, Eleanor G Bentley, Anu Chawla, Jordan J Clark, Angela Cowell, Richard Eccles, Isabel García-Dorival, Matthew Gemmell, Alessandro Gerada, PKF Gilmore, Richard Gregory, Ximeng Han, Catherine Hartley, Margaret Hughes, Miren Iturriza-Gomara, James Johnson, L Luu, Jenifer Manson, Charlotte Nelson, Elaine O'Toole, Cassie Olateju, Rebekah Penrice-Randal , Lucille Rainbow, N.P Randle, Trevor Ian Robinson, Parul Sharma, Ghada T Shawli, James P Stewart, Neil Swainston, Ecaterina Vamos, Joanne Watts, Mark Whitehead |
| EPI_ISL_865597, EPI_ISL_865598, EPI_ISL_865601, EPI_ISL_865605, EPI_ISL_865606, EPI_ISL_865607, EPI_ISL_865608, EPI_ISL_865609, EPI_ISL_865610, EPI_ISL_865611, EPI_ISL_865612, EPI_ISL_865613, EPI_ISL_865619, EPI_ISL_865620, EPI_ISL_865621, EPI_ISL_865622, EPI_ISL_865624, EPI_ISL_865625, EPI_ISL_865626, EPI_ISL_865627, EPI_ISL_865628, EPI_ISL_865629, EPI_ISL_865632, EPI_ISL_865633, EPI_ISL_865635, EPI_ISL_865636, EPI_ISL_865637, EPI_ISL_865641, EPI_ISL_865642, EPI_ISL_865643, EPI_ISL_865644, EPI_ISL_865645, EPI_ISL_865646, EPI_ISL_865647, EPI_ISL_865649, EPI_ISL_865650, EPI_ISL_865651, EPI_ISL_865652, EPI_ISL_865653, EPI_ISL_865654, EPI_ISL_865655, EPI_ISL_865656, EPI_ISL_865657, EPI_ISL_865853, EPI_ISL_865855, EPI_ISL_865856, EPI_ISL_865857, EPI_ISL_865858, EPI_ISL_865859, EPI_ISL_865860, EPI_ISL_865861, EPI_ISL_865862, EPI_ISL_865863, EPI_ISL_865865, EPI_ISL_865866, EPI_ISL_865882, EPI_ISL_865884, EPI_ISL_865885, EPI_ISL_865886, EPI_ISL_865887, EPI_ISL_865888, EPI_ISL_865889, EPI_ISL_865890, EPI_ISL_865902, EPI_ISL_865904, EPI_ISL_865907, EPI_ISL_865908, EPI_ISL_865917, EPI_ISL_865920, EPI_ISL_865926, EPI_ISL_865927, EPI_ISL_865928, EPI_ISL_865935, EPI_ISL_865936, EPI_ISL_865937, EPI_ISL_865938, EPI_ISL_865939, EPI_ISL_865946, EPI_ISL_865947, EPI_ISL_865948, EPI_ISL_865951, EPI_ISL_865967, EPI_ISL_865969, EPI_ISL_865970, EPI_ISL_865975, EPI_ISL_865976, EPI_ISL_865977, EPI_ISL_865978, EPI_ISL_865980, EPI_ISL_865985, EPI_ISL_865986, EPI_ISL_865987, EPI_ISL_865993, EPI_ISL_865994, EPI_ISL_865995, EPI_ISL_865996, EPI_ISL_866001, EPI_ISL_866002, EPI_ISL_866003, EPI_ISL_866007, EPI_ISL_866008, EPI_ISL_866009, EPI_ISL_866011, EPI_ISL_866013, EPI_ISL_866014, EPI_ISL_866015, EPI_ISL_866020, EPI_ISL_866290, EPI_ISL_866291, EPI_ISL_866292, EPI_ISL_866293, EPI_ISL_866298, EPI_ISL_866301, EPI_ISL_866302, EPI_ISL_866303, EPI_ISL_866304, EPI_ISL_866305, EPI_ISL_866306, EPI_ISL_866307, EPI_ISL_866311, EPI_ISL_866312, EPI_ISL_866320, EPI_ISL_866321, EPI_ISL_866322, EPI_ISL_866323, EPI_ISL_866324, EPI_ISL_866326, EPI_ISL_866327, EPI_ISL_866329, EPI_ISL_866330, EPI_ISL_866331, EPI_ISL_866332, EPI_ISL_866333 | | | |
| see above | University College London, Great Ormond Street Hospital for Children NHS Foundation Trust, Imperial College Healthcare NHS Trust | COVID-19 Genomics UK (COG-UK) Consortium | Sergi Castellano, Rachel Williams, Mark Kristiansen, Paola Resende Silva, Sunando Roy, Tony Brooks, Helena Tutill, Paola Niola, Patricia Dyal, Charlotte Williams, Leysa Forrest, Yasmin Panchbhaya, Jacqueline Findlay, Samuel Weeks, Julianne Brown, Kathryn Harris, Paul Randell, James Price, Alison Holmes, Judith Breuer |
| EPI_ISL_866432 | Respiratory Virus Unit, National Infection Service, Public Health England | COVID-19 Genomics UK (COG-UK) Consortium | PHE Covid Sequencing Team |
| EPI_ISL_866943, EPI_ISL_866944, EPI_ISL_866945, EPI_ISL_866946, EPI_ISL_866947, EPI_ISL_866948, EPI_ISL_866949, EPI_ISL_866950, EPI_ISL_866951, EPI_ISL_866952, EPI_ISL_866953, EPI_ISL_866954, EPI_ISL_866955, EPI_ISL_866956, EPI_ISL_866957, EPI_ISL_866958, EPI_ISL_866959, EPI_ISL_866960, | | | |

| | | | |
|---|---|---|---|
| EPI_ISL_866961 | | | |
| see above | Queens Medical Centre, Clinical Microbiology Department / DeepSeq Nottingham | COVID-19 Genomics UK (COG-UK) Consortium | Gemma Clark, Wendy Smith, Manjinder Khakh, Vicki M Fleming, Michelle M Lister, Hannah Howson-Wells, Jonathan Ball, Patrick McClure, Joseph Chappell, Theocharis Tsoleridis, Nadine Holmes, Matthew Carlisle, Christopher Moore, Fei Sang, Johnny Debebe, Victoria Wright, Matthew Loose |
| EPI_ISL_867993, EPI_ISL_867999, EPI_ISL_868226, EPI_ISL_868228, EPI_ISL_868229, EPI_ISL_868230, EPI_ISL_868231, EPI_ISL_868232, EPI_ISL_868233, EPI_ISL_868234, EPI_ISL_868235, EPI_ISL_868236, EPI_ISL_868237, EPI_ISL_868240, EPI_ISL_868241, EPI_ISL_868242, EPI_ISL_868243, EPI_ISL_868244, EPI_ISL_868245, EPI_ISL_868246, EPI_ISL_868247, EPI_ISL_868248, EPI_ISL_868249, EPI_ISL_868250, EPI_ISL_868252, EPI_ISL_868253, EPI_ISL_868254, EPI_ISL_868255, EPI_ISL_868256 | | | |
| see above | Centre for Enzyme Innovation, University of Portsmouth / Translational Research Laboratory, Portsmouth Hospitals NHS Trust | COVID-19 Genomics UK (COG-UK) Consortium | Angela Beckett,Yann Bourgeois,Garry Scarlett,Sharon Glaysher,Scott Elliott,Kelly Bicknell,Robert Impey,Allyson Lloyd,Sarah Wyllie,Ethan Butcher,Anoop Chauhan,Samuel Robson |
| EPI_ISL_868363, EPI_ISL_868366, EPI_ISL_868371, EPI_ISL_868385, EPI_ISL_868387, EPI_ISL_868397, EPI_ISL_868417, EPI_ISL_868421, EPI_ISL_868426, EPI_ISL_868437, EPI_ISL_868439, EPI_ISL_868447, EPI_ISL_868456, EPI_ISL_868458, EPI_ISL_868461, EPI_ISL_868463, EPI_ISL_868489, EPI_ISL_868493, EPI_ISL_868497, EPI_ISL_868501, EPI_ISL_868504, EPI_ISL_868509, EPI_ISL_868510, EPI_ISL_868517, EPI_ISL_868529, EPI_ISL_868532, EPI_ISL_868544, EPI_ISL_868546, EPI_ISL_868562, EPI_ISL_868563, EPI_ISL_868568, EPI_ISL_868569, EPI_ISL_868580, EPI_ISL_868581, EPI_ISL_868582, EPI_ISL_868584, EPI_ISL_868591, EPI_ISL_868592, EPI_ISL_868595, EPI_ISL_868596, EPI_ISL_868611, EPI_ISL_868623, EPI_ISL_868634, EPI_ISL_868646, EPI_ISL_868652, EPI_ISL_868657, EPI_ISL_868666, EPI_ISL_868667, EPI_ISL_868668, EPI_ISL_868687, EPI_ISL_868689, EPI_ISL_868691, EPI_ISL_868695, EPI_ISL_868697, EPI_ISL_868705, EPI_ISL_868706, EPI_ISL_868707, EPI_ISL_868708, EPI_ISL_868714 | | | |
| see above | Virology Department, Sheffield Teaching Hospitals NHS Foundation Trust/Department of Infection, Immunity and Cardiovascular Disease, The Medical School, University of Sheffield | COVID-19 Genomics UK (COG-UK) Consortium | Thushan de Silva, Matthew Parker, Nikki Smith, Adri Angyal, Rebecca Brown, Luke Green, Rachel Tucker, Paul Parsons, Danielle Groves, Katie Johnson, Laura Carrilero, Alex Keeley, Dave Partridge, Matthew Wyles, Benjamin Lindsey, Mehmet Yavuz, Mohammad Raza, Cariad Evans |
| EPI_ISL_869084, EPI_ISL_869102, EPI_ISL_869103, EPI_ISL_869104, EPI_ISL_869105, EPI_ISL_869106, EPI_ISL_869107, EPI_ISL_869108, EPI_ISL_869109, EPI_ISL_869110, EPI_ISL_869111, EPI_ISL_869112, EPI_ISL_869113, EPI_ISL_869114, EPI_ISL_869115, EPI_ISL_869116, EPI_ISL_869117 | | | |
| see above | A. Krumbholz, Labor Dr. Krause und Kollegen MVZ GmbH, Kiel | Charité Universitätsmedizin Berlin, Institut für Virologie | Victor M Corman, Barbara Mühlemann, Jörn Beheim-Schwarzbach, Tobias Bleicker, Julia Tesch, Talitha Veith, Julia Schneider, Terry Jones, Christian Drosten |
| EPI_ISL_869119 | Charité Universitätsmedizin Berlin, Institut für Virologie/Labor Berlin | Charité Universitätsmedizin Berlin, Institut für Virologie | Victor M Corman, Barbara Mühlemann, Jörn Beheim-Schwarzbach, Tobias Bleicker, Julia Tesch, Talitha Veith, Julia Schneider, Terry Jones, Christian Drosten |
| EPI_ISL_869120, EPI_ISL_869121, EPI_ISL_869122, EPI_ISL_869123, EPI_ISL_869137, EPI_ISL_869138, EPI_ISL_869139, EPI_ISL_869140, EPI_ISL_869141, EPI_ISL_869142 | A. Krumbholz, Labor Dr. Krause und Kollegen MVZ GmbH, Kiel | Charité Universitätsmedizin Berlin, Institut für Virologie | Victor M Corman, Barbara Mühlemann, Jörn Beheim-Schwarzbach, Tobias Bleicker, Julia Tesch, Talitha Veith, Julia Schneider, Terry Jones, Christian Drosten |
| EPI_ISL_871842, EPI_ISL_871843, EPI_ISL_871850, EPI_ISL_871851, EPI_ISL_871852, EPI_ISL_871853, EPI_ISL_871854, EPI_ISL_871855, EPI_ISL_871856, EPI_ISL_871857, EPI_ISL_871858, EPI_ISL_871859, EPI_ISL_871860, EPI_ISL_871861, EPI_ISL_871862, EPI_ISL_871863, EPI_ISL_871864, EPI_ISL_871865, EPI_ISL_871866, EPI_ISL_871867, EPI_ISL_871868, EPI_ISL_871970, EPI_ISL_871971, EPI_ISL_872000 | | | |
| see above | Wyoming Public Health Laboratory | Wyoming Public Health Laboratory | Noah Hull, Taylor Fearing, Lynette Gumbleton, Channing Weber, Ashley Norberg, Bailey Bowcutt, and Wanda Manley |
| EPI_ISL_872044, EPI_ISL_872045, EPI_ISL_872046, EPI_ISL_872047, EPI_ISL_872048 | Department of Clinical Microbiology | GIGA Medical Genomics | Keith Durkin, Maria Artesi, Sébastien Bontems, Raphaël Boreux, Bouchra Boujemla, Cécile Meex, Pierrette Melin, Marie-Pierre Hayette, Vincent Bours |
| EPI_ISL_872051 | CHC | GIGA Medical Genomics | Keith Durkin, Maria Artesi, Sébastien Bontems, Raphaël Boreux, Bouchra Boujemla, Cécile Meex, Pierrette Melin, Marie-Pierre Hayette, Vincent Bours |
| EPI_ISL_872073, EPI_ISL_872074, EPI_ISL_872075, EPI_ISL_872076, EPI_ISL_872077, EPI_ISL_872078, EPI_ISL_872079, EPI_ISL_872080 | Department of Clinical Microbiology | GIGA Medical Genomics | Keith Durkin, Maria Artesi, Sébastien Bontems, Raphaël Boreux, Bouchra Boujemla, Cécile Meex, Pierrette Melin, Marie-Pierre Hayette, Vincent Bours |
| EPI_ISL_872151 | CHR Citadelle | GIGA Medical Genomics | Keith Durkin, Maria Artesi, Sébastien Bontems, Raphaël Boreux, Bouchra Boujemla, Cécile Meex, Pierrette Melin, Marie-Pierre Hayette, Vincent Bours |
| EPI_ISL_872156, EPI_ISL_872157, EPI_ISL_872158, EPI_ISL_872159, EPI_ISL_872160, EPI_ISL_872161, EPI_ISL_872162, EPI_ISL_872163, EPI_ISL_872165, EPI_ISL_872166, EPI_ISL_872168, EPI_ISL_872169, EPI_ISL_872173, EPI_ISL_872174, EPI_ISL_872175, EPI_ISL_872178, EPI_ISL_872179, EPI_ISL_872182, EPI_ISL_872183, EPI_ISL_872184, EPI_ISL_872185, EPI_ISL_872186, EPI_ISL_872187 | | | |
| see above | Wyoming Public Health Laboratory | Wyoming Public Health Laboratory | Noah Hull, Taylor Fearing, Lynette Gumbleton, Channing Weber, Ashley Norberg, Bailey Bowcutt, and Wanda Manley |
| EPI_ISL_872191 | Conjunto Hospitalar do Mandaqui de Sao Paulo | Instituto Adolfo Lutz, Interdiciplinary Procedures Center, Strategic Laboratory | Claudio Tavares Sacchi, Claudia Regina Gonçalves, Erica Valessa Ramos Gomes, Karoline Rodrigues Campos, Katia Correa de Oliveira Santos, Ana Lucia de Carvalho Avelino, Fabiana Cristina Pereira dos Santos |
| EPI_ISL_872557, EPI_ISL_872558, EPI_ISL_872559, EPI_ISL_872560, EPI_ISL_872561, EPI_ISL_872564, EPI_ISL_872565, EPI_ISL_872566 | New Mexico Department of Health Scientific Laboratory | Center for Global Health, University of New Mexico Health Sciences Center | Daryl Domman, Kurt Schwalm, Twila Kunde, Joseph Hicks, Anastacia Griego, Michael Edwards, Darrell Dinwiddie |
| EPI_ISL_872567 | Centre for Dengue Research and AICBU, Department of Immunology and Molecular Medicine | Centre for Dengue Research and AICBU, Department of Immunology and Molecular Medicine | Chandima Jeewandara, Deshni Jayathilaka, Dinuka Ariyaratne, Tibutius Thanesh Pramanayagam, Diyanath Ranasinghe, Laksiri Gomes, Gathsaurie Neelika Malavige |
| EPI_ISL_873270, EPI_ISL_873271 | Minnesota Department of Health, Public Health Laboratory | Minnesota Department of Health, Public Health Laboratory | Alexandra Lorentz, Jacob Garfin, Matt Plumb, and Xiong Wang |
| EPI_ISL_873283, EPI_ISL_873286, EPI_ISL_873291, EPI_ISL_873347, EPI_ISL_873348, EPI_ISL_873369, EPI_ISL_873370, EPI_ISL_873371, EPI_ISL_873381, EPI_ISL_873388, EPI_ISL_873399, EPI_ISL_873426, EPI_ISL_873428, EPI_ISL_873451, EPI_ISL_873482, EPI_ISL_873492, EPI_ISL_873501, EPI_ISL_873518, EPI_ISL_873537, EPI_ISL_873574, EPI_ISL_873625, EPI_ISL_873630, EPI_ISL_873674, EPI_ISL_873702, EPI_ISL_873737, EPI_ISL_873814, EPI_ISL_873820, EPI_ISL_873839, EPI_ISL_873876, EPI_ISL_873903 | | | |
| see above | Lighthouse Lab in Cambridge | Wellcome Sanger Institute for the COVID-19 Genomics UK (COG-UK) Consortium | Rob Howes, The Lighthouse Lab in Cambridge and Alex Alderton, Roberto Amato, Sonia Goncalves, Ewan Harrison, David K. Jackson, Ian Johnston, Dominic Kwiatkowski, Cordelia Langford, John Sillitoe on behalf of the Wellcome Sanger Institute COVID-19 Surveillance Team |
| EPI_ISL_875344 | Charité Universitätsmedizin Berlin, Institute of Virology, Charitéplatz 1, 10117 Berlin, Germany | Charité Universitätsmedizin Berlin, Institute of Virology, Charitéplatz 1, 10117 Berlin, Germany | Victor M Corman, Julia Schneider, Jörn Beheim-Schwarzbach, Tobias Bleicker, Barbara Mühlemann, Talitha Veith, Terry Jones, Christian Drosten |
| EPI_ISL_875349, EPI_ISL_875350, EPI_ISL_875351, EPI_ISL_875352 | National Virus Reference Laboratory | National Virus Reference Laboratory | Michael Carr, Gabriel Gonzalez, Jonathan Dean, Cillian F De Gascun |
| EPI_ISL_875669, EPI_ISL_875670 | CHU Purpan - Laboratoire de Virologie - Institut Fédératif de Biologie | CHU Purpan - Laboratoire de Virologie - Institut Fédératif de Biologie | Latour J., Ranger N., Dubois M., Carcenac R., Harter A., Boyer P., Tremeaux P., Izopet J. |
| EPI_ISL_875689 | Hospital do Servidor Publico | Instituto Adolfo Lutz, Interdiciplinary Procedures Center, Strategic Laboratory | Claudio Tavares Sacchi, Claudia Regina Gonçalves, Erica Valessa Ramos Gomes, Karoline Rodrigues Campos |
| EPI_ISL_876231, EPI_ISL_876232, EPI_ISL_876233 | Massachusetts State Public Health Laboratory | Massachusetts State Public Health Laboratory | Andrew Lang, Timelia Fink, Glen Gallagher, Sandra Smole |
| EPI_ISL_877037, EPI_ISL_877038, EPI_ISL_877039, EPI_ISL_877040, EPI_ISL_877041, EPI_ISL_877042, EPI_ISL_877043, EPI_ISL_877046, EPI_ISL_877047, EPI_ISL_877048, EPI_ISL_877049, EPI_ISL_877050, EPI_ISL_877059, EPI_ISL_877060, EPI_ISL_877061, EPI_ISL_877062, EPI_ISL_877063, EPI_ISL_877064, EPI_ISL_877065, EPI_ISL_877066, EPI_ISL_877067, EPI_ISL_877068, EPI_ISL_877069, EPI_ISL_877070, EPI_ISL_877071, EPI_ISL_877072, EPI_ISL_877073, EPI_ISL_877074, EPI_ISL_877075, EPI_ISL_877076, EPI_ISL_877077, EPI_ISL_877078, EPI_ISL_877079, EPI_ISL_877080, EPI_ISL_877081, EPI_ISL_877082, EPI_ISL_877083, EPI_ISL_877084, EPI_ISL_877085, EPI_ISL_877086, EPI_ISL_877087, EPI_ISL_877088, EPI_ISL_877089, EPI_ISL_877090, EPI_ISL_877091, EPI_ISL_877092, EPI_ISL_877093, EPI_ISL_877094, EPI_ISL_877095, EPI_ISL_877096, EPI_ISL_877097, EPI_ISL_877098, EPI_ISL_877099, EPI_ISL_877100, EPI_ISL_877101, EPI_ISL_877102, EPI_ISL_877103, EPI_ISL_877104, EPI_ISL_877105, EPI_ISL_877106, EPI_ISL_877107, EPI_ISL_877108, EPI_ISL_877109, EPI_ISL_877110, EPI_ISL_877111, EPI_ISL_877112, EPI_ISL_877113, EPI_ISL_877114, EPI_ISL_877115, EPI_ISL_877116, EPI_ISL_877117, EPI_ISL_877118 | | | |
| see above | Quest Diagnostics | Quest Diagnostics | Rosenthal,S.H., Gerasimova,A., Kagan,R.M., Anderson, B., Hua, M., Liu Y., Bernstein, L.E., Livingston, K.E., Perez, A., Shalhout, D.F., Shlyakhter, I.A., Owen, R., Tanpaiboon, P., Lacbawan, F. |
| EPI_ISL_877146, EPI_ISL_877147, EPI_ISL_877148, EPI_ISL_877149, EPI_ISL_877150, EPI_ISL_877151, EPI_ISL_877152, EPI_ISL_877154, EPI_ISL_877164, EPI_ISL_877165, EPI_ISL_877166, EPI_ISL_877167, EPI_ISL_877168, EPI_ISL_877169, EPI_ISL_877170, EPI_ISL_877171, EPI_ISL_877172, EPI_ISL_877173, EPI_ISL_877174, EPI_ISL_877175, EPI_ISL_877176, EPI_ISL_877177, EPI_ISL_877178, EPI_ISL_877179, EPI_ISL_877180 | | | |

| | | | |
|---|---|---|---|
| see above | Univeristy of New Mexico Hospital | Center for Global Health, University of New Mexico Health Sciences Center | Daryl Domman, Kurt Schwalm, Justin Bacca, Jon Femling, Darrell Dinwiddie |
| EPI_ISL_877220 | Canterbury Health Laboratories | Institute of Environmental Science and Research (ESR) | Xiaoyun Ren, Matt Storey, Nikki Freed, Muhammad Faisal, Jing Wang, Hermes Perez, Anja Werno, Antje van der Linden, Arlo Upton, Chris Mansell, David Hammer, Dragana Drinkovic, Gary McAuliffe, Hana Sofia Andersson, James Ussher, Jill Sherwood, Josh Freeman, Julia Howard, Juliet Elvy, Mary DeAlmeida, Matt Blakiston, Matthew Rogers, Max Bloomfield, Michael Addidle, Michelle Balm, Sally Roberts, Sarah Jefferies, Sharmini Muttaiyah, Susan Morpeth, Susan Taylor, Timothy Blackmore, Vani Sathyendran, Veronica Playle, Virginia Hope, Erasmus Smit, Lauren Jelly, Olin Silander, Joep de Ligt |
| EPI_ISL_877552, EPI_ISL_877553, EPI_ISL_877554, EPI_ISL_877555 | Institute of Microbiology, Universidad San Francisco de Quito | Institute of Microbiology, Universidad San Francisco de Quito | Belén Prado-Vivar, Sully Márquez, Juan José Guadalupe, Monica Becerra-Wong, Bernardo Gutiérrez, Kyllen Briones, Edmundo Encalada, Ninfa Hernandez, Francisco Cordova, Verónica Barragán, Patricio Rojas-Silva, Gabriel Trueba, Michelle Grunauer, Paúl Cárdenas |
| EPI_ISL_877556, EPI_ISL_877557, EPI_ISL_877558, EPI_ISL_877559, EPI_ISL_877560 | Institute of Microbiology, Universidad San Francisco de Quito | Institute of Microbiology, Universidad San Francisco de Quito | Belén Prado-Vivar, Sully Márquez, Juan José Guadalupe, Monica Becerra-Wong, Bernardo Gutiérrez, Eulalia Pazmiño, Katalina Pacheco, Verónica Barragán, Patricio Rojas-Silva, Gabriel Trueba, Michelle Grunauer, Paúl Cárdenas |
| EPI_ISL_877564, EPI_ISL_877565, EPI_ISL_877566, EPI_ISL_877567, EPI_ISL_877568, EPI_ISL_877569 | Victorian Infectious Diseases Reference Laboratory (VIDRL) | VIDRL and MDU-PHL | Caly L., Seemann T., Sait, M.L., Druce J., Sherry, N.L. |
| EPI_ISL_877571, EPI_ISL_877572, EPI_ISL_877573, EPI_ISL_877574, EPI_ISL_877575 | Microbiological Diagnostic Unit - Public Health Laboratory (MDU-PHL) | MDU-PHL | Seemann T., Sait, M.L., Sherry, N.L. |
| EPI_ISL_877766 | Thai Red Cross Emerging Infectious Diseases Health Science Centre, Chulalongkorn Hospital, Faculty of Medicine, Chulalongkorn University | Thai Red Cross Emerging Infectious Diseases Center and Faculty of Medicine, Chulalongkorn University | Rome Buathong, Sopon Iamsirithaworn, Sininat Petcharat, Yutthana Joyjinda, Weenassarin Ampoot, Apaporn Rodpan, Opass Putcharoen, Thiravat Hemachudha, Supaporn Wacharapluesadee |
| EPI_ISL_877770, EPI_ISL_877772, EPI_ISL_877773, EPI_ISL_877774, EPI_ISL_877775, EPI_ISL_877776, EPI_ISL_877777, EPI_ISL_877778, EPI_ISL_877779, EPI_ISL_877780, EPI_ISL_877781, EPI_ISL_877782, EPI_ISL_877783, EPI_ISL_877784, EPI_ISL_877785, EPI_ISL_877786, EPI_ISL_877787, EPI_ISL_877788, EPI_ISL_877789, EPI_ISL_877790, EPI_ISL_877791, EPI_ISL_877792, EPI_ISL_877793, EPI_ISL_877794, EPI_ISL_877795, EPI_ISL_877796, EPI_ISL_877797, EPI_ISL_877798, EPI_ISL_877799, EPI_ISL_877800, EPI_ISL_877801, EPI_ISL_877802, EPI_ISL_877803, EPI_ISL_877804, EPI_ISL_877805, EPI_ISL_877806, EPI_ISL_877808, EPI_ISL_877809, EPI_ISL_877810, EPI_ISL_877811, EPI_ISL_877812, EPI_ISL_877813, EPI_ISL_877814, EPI_ISL_877815, EPI_ISL_877816, EPI_ISL_877817, EPI_ISL_877818, EPI_ISL_877819, EPI_ISL_877820, EPI_ISL_877822, EPI_ISL_877823, EPI_ISL_877824, EPI_ISL_877825, EPI_ISL_877826, EPI_ISL_877827, EPI_ISL_877829, EPI_ISL_877830, EPI_ISL_877831, EPI_ISL_877832, EPI_ISL_877833, EPI_ISL_877834, EPI_ISL_877835, EPI_ISL_877836, EPI_ISL_877837, EPI_ISL_877838, EPI_ISL_877839, EPI_ISL_877840, EPI_ISL_877841, EPI_ISL_877842, EPI_ISL_877843, EPI_ISL_877845, EPI_ISL_877846, EPI_ISL_877847, EPI_ISL_877848, EPI_ISL_877849, EPI_ISL_877850, EPI_ISL_877851, EPI_ISL_877852, EPI_ISL_877853, EPI_ISL_877854, EPI_ISL_877855, EPI_ISL_877856, EPI_ISL_877857, EPI_ISL_877858, EPI_ISL_877859, EPI_ISL_877860, EPI_ISL_877861, EPI_ISL_877862, EPI_ISL_877863, EPI_ISL_877864, EPI_ISL_877865, EPI_ISL_877866, EPI_ISL_877867, EPI_ISL_877868, EPI_ISL_877869, EPI_ISL_877870, EPI_ISL_877871, EPI_ISL_877872, EPI_ISL_877873, EPI_ISL_877874, EPI_ISL_877875, EPI_ISL_877876, EPI_ISL_877877, EPI_ISL_877878, EPI_ISL_877879, EPI_ISL_877880, EPI_ISL_877881, EPI_ISL_877882, EPI_ISL_877883, EPI_ISL_877884, EPI_ISL_877885, EPI_ISL_877886, EPI_ISL_877887, EPI_ISL_877888, EPI_ISL_877889, EPI_ISL_877890, EPI_ISL_877891, EPI_ISL_877892, EPI_ISL_877893, EPI_ISL_877894, EPI_ISL_877895, EPI_ISL_877896, EPI_ISL_877897, EPI_ISL_877898, EPI_ISL_877899, EPI_ISL_877900, EPI_ISL_877901, EPI_ISL_877902, EPI_ISL_877904, EPI_ISL_877905, EPI_ISL_877906, EPI_ISL_877907, EPI_ISL_877908, EPI_ISL_877909, EPI_ISL_877910, EPI_ISL_877911, EPI_ISL_877912, EPI_ISL_877914, EPI_ISL_877915, EPI_ISL_877916, EPI_ISL_877917, EPI_ISL_877918, EPI_ISL_877919, EPI_ISL_877920, EPI_ISL_877921, EPI_ISL_877922, EPI_ISL_877923, EPI_ISL_877924, EPI_ISL_877925, EPI_ISL_877926, EPI_ISL_877927, EPI_ISL_877928, EPI_ISL_877929, EPI_ISL_877930, EPI_ISL_877931, EPI_ISL_877932, EPI_ISL_877933, EPI_ISL_877934, EPI_ISL_877935, EPI_ISL_877936, EPI_ISL_877937, EPI_ISL_877938, EPI_ISL_877939, EPI_ISL_877941, EPI_ISL_877942, EPI_ISL_877943, EPI_ISL_877944, EPI_ISL_877945, EPI_ISL_877946, EPI_ISL_877947, EPI_ISL_877948, EPI_ISL_877949, EPI_ISL_877950, EPI_ISL_877951, EPI_ISL_877952, EPI_ISL_877953, EPI_ISL_877954, EPI_ISL_877956, EPI_ISL_877957, EPI_ISL_877958, EPI_ISL_877959, EPI_ISL_877960, EPI_ISL_877961, EPI_ISL_877962, EPI_ISL_877963, EPI_ISL_877964, EPI_ISL_877965, EPI_ISL_877966, EPI_ISL_877967, EPI_ISL_877968, EPI_ISL_877969, EPI_ISL_877970, EPI_ISL_877971, EPI_ISL_877973, EPI_ISL_877974, EPI_ISL_877975, EPI_ISL_877976, EPI_ISL_877977, EPI_ISL_877978, EPI_ISL_877979, EPI_ISL_877980, EPI_ISL_877981, EPI_ISL_877982, EPI_ISL_877983, EPI_ISL_877984, EPI_ISL_877985, EPI_ISL_877986, EPI_ISL_877987, EPI_ISL_877988, EPI_ISL_877989, EPI_ISL_877990, EPI_ISL_877991, EPI_ISL_877992, EPI_ISL_877993, EPI_ISL_877994, EPI_ISL_877995, EPI_ISL_877996, EPI_ISL_877997, EPI_ISL_877998, EPI_ISL_877999, EPI_ISL_878000, EPI_ISL_878001, EPI_ISL_878002, EPI_ISL_878003, EPI_ISL_878004, EPI_ISL_878005, EPI_ISL_878006, EPI_ISL_878007, EPI_ISL_878008, EPI_ISL_878009, EPI_ISL_878010, EPI_ISL_878011, EPI_ISL_878012, EPI_ISL_878013, EPI_ISL_878014, EPI_ISL_878015, EPI_ISL_878016, EPI_ISL_878017, EPI_ISL_878018, EPI_ISL_878019, EPI_ISL_878020, EPI_ISL_878021, EPI_ISL_878022, EPI_ISL_878023, EPI_ISL_878024, EPI_ISL_878025, EPI_ISL_878026, EPI_ISL_878027, EPI_ISL_878028, EPI_ISL_878029, EPI_ISL_878030, EPI_ISL_878031, EPI_ISL_878032, EPI_ISL_878033, EPI_ISL_878034, EPI_ISL_878035, EPI_ISL_878036, EPI_ISL_878037, EPI_ISL_878038, EPI_ISL_878039, EPI_ISL_878040, EPI_ISL_878041, EPI_ISL_878042, EPI_ISL_878043, EPI_ISL_878044, EPI_ISL_878045, EPI_ISL_878046, EPI_ISL_878047, EPI_ISL_878048, EPI_ISL_878049, EPI_ISL_878050, EPI_ISL_878051, EPI_ISL_878052, EPI_ISL_878053, EPI_ISL_878054, EPI_ISL_878055, EPI_ISL_878056, EPI_ISL_878057, EPI_ISL_878058, EPI_ISL_878059, EPI_ISL_878060, EPI_ISL_878061, EPI_ISL_878062, EPI_ISL_878063, EPI_ISL_878064, EPI_ISL_878065, EPI_ISL_878066, EPI_ISL_878067, EPI_ISL_878068, EPI_ISL_878069, EPI_ISL_878070, EPI_ISL_878071, EPI_ISL_878072, EPI_ISL_878073, EPI_ISL_878074, EPI_ISL_878075, EPI_ISL_878076, EPI_ISL_878077, EPI_ISL_878078, EPI_ISL_878079, EPI_ISL_878080, EPI_ISL_878081, EPI_ISL_878082, EPI_ISL_878083, EPI_ISL_878084, EPI_ISL_878085, EPI_ISL_878086, EPI_ISL_878087, EPI_ISL_878088, EPI_ISL_878090, EPI_ISL_878091, EPI_ISL_878092, EPI_ISL_878093, EPI_ISL_878094, EPI_ISL_878095, EPI_ISL_878096, EPI_ISL_878097, EPI_ISL_878098, EPI_ISL_878099, EPI_ISL_878100, EPI_ISL_878101, EPI_ISL_878102, EPI_ISL_878103, EPI_ISL_878104, EPI_ISL_878105, EPI_ISL_878106, EPI_ISL_878107, EPI_ISL_878108, EPI_ISL_878109, EPI_ISL_878110, EPI_ISL_878111, EPI_ISL_878112, EPI_ISL_878113, EPI_ISL_878114, EPI_ISL_878115, EPI_ISL_878116, EPI_ISL_878117, EPI_ISL_878119, EPI_ISL_878120, EPI_ISL_878121, EPI_ISL_878122, EPI_ISL_878123, EPI_ISL_878125, EPI_ISL_878127, EPI_ISL_878128, EPI_ISL_878129, EPI_ISL_878130, EPI_ISL_878131, EPI_ISL_878132, EPI_ISL_878133, EPI_ISL_878134, EPI_ISL_878135, EPI_ISL_878136, EPI_ISL_878137, EPI_ISL_878138, EPI_ISL_878139, EPI_ISL_878140, EPI_ISL_878141, EPI_ISL_878142, EPI_ISL_878143, EPI_ISL_878144, EPI_ISL_878145, EPI_ISL_878146, EPI_ISL_878147, EPI_ISL_878148, EPI_ISL_878149, EPI_ISL_878150, EPI_ISL_878151, EPI_ISL_878153, EPI_ISL_878154, EPI_ISL_878156, EPI_ISL_878158, EPI_ISL_878160, EPI_ISL_878161, EPI_ISL_878164, EPI_ISL_878166, EPI_ISL_878168, EPI_ISL_878169, EPI_ISL_878171, EPI_ISL_878174, EPI_ISL_878177, EPI_ISL_878181, EPI_ISL_878182, EPI_ISL_878184, EPI_ISL_878185, EPI_ISL_878187, EPI_ISL_878188, EPI_ISL_878190, EPI_ISL_878193, EPI_ISL_878195, EPI_ISL_878196, EPI_ISL_878198, EPI_ISL_878200, EPI_ISL_878201, EPI_ISL_878203, EPI_ISL_878204, EPI_ISL_878206, EPI_ISL_878210, EPI_ISL_878211, EPI_ISL_878213, EPI_ISL_878214, EPI_ISL_878216, EPI_ISL_878218, EPI_ISL_878219, EPI_ISL_878220, EPI_ISL_878222, EPI_ISL_878224, EPI_ISL_878225, EPI_ISL_878227, EPI_ISL_878228, EPI_ISL_878230, EPI_ISL_878231, EPI_ISL_878233, EPI_ISL_878235, EPI_ISL_878236, EPI_ISL_878238, EPI_ISL_878240, EPI_ISL_878241, EPI_ISL_878243, EPI_ISL_878244, EPI_ISL_878246, EPI_ISL_878247, EPI_ISL_878251, EPI_ISL_878254, EPI_ISL_878255, EPI_ISL_878257, EPI_ISL_878259, EPI_ISL_878261, EPI_ISL_878262, EPI_ISL_878264, EPI_ISL_878267, EPI_ISL_878269, EPI_ISL_878270, EPI_ISL_878272, EPI_ISL_878273, EPI_ISL_878275, EPI_ISL_878278, EPI_ISL_878280, EPI_ISL_878282, EPI_ISL_878283, EPI_ISL_878286, EPI_ISL_878288, EPI_ISL_878289, EPI_ISL_878291, EPI_ISL_878293, EPI_ISL_878294, EPI_ISL_878296, EPI_ISL_878297, EPI_ISL_878299, EPI_ISL_878301, EPI_ISL_878302, EPI_ISL_878304, EPI_ISL_878306, EPI_ISL_878307, EPI_ISL_878309, EPI_ISL_878310, EPI_ISL_878312, EPI_ISL_878313, EPI_ISL_878315, EPI_ISL_878317, EPI_ISL_878318, EPI_ISL_878320, EPI_ISL_878321, EPI_ISL_878324, EPI_ISL_878326, EPI_ISL_878328, EPI_ISL_878329, EPI_ISL_878331, EPI_ISL_878332, EPI_ISL_878334, EPI_ISL_878335, EPI_ISL_878340, EPI_ISL_878409, EPI_ISL_878410, EPI_ISL_878412, EPI_ISL_878414, EPI_ISL_878415, EPI_ISL_878417, EPI_ISL_878419, EPI_ISL_878424, EPI_ISL_878425, EPI_ISL_878427, EPI_ISL_878428, EPI_ISL_878430, EPI_ISL_878432, EPI_ISL_878433, EPI_ISL_878435, EPI_ISL_878436, EPI_ISL_878438, EPI_ISL_878439, EPI_ISL_878441, EPI_ISL_878442, EPI_ISL_878444, EPI_ISL_878446, EPI_ISL_878447, EPI_ISL_878451, EPI_ISL_878453, EPI_ISL_878459, EPI_ISL_878461, EPI_ISL_878462, EPI_ISL_878464, EPI_ISL_878467, EPI_ISL_878469, EPI_ISL_878471, EPI_ISL_878473, EPI_ISL_878475, EPI_ISL_878476, EPI_ISL_878478, EPI_ISL_878479, EPI_ISL_878482, EPI_ISL_878487, EPI_ISL_878489, EPI_ISL_878490, EPI_ISL_878493, EPI_ISL_878494, EPI_ISL_878496, EPI_ISL_878497, EPI_ISL_878499, EPI_ISL_878501, EPI_ISL_878502, EPI_ISL_878504, EPI_ISL_878508, EPI_ISL_878510, EPI_ISL_878513, EPI_ISL_878515, EPI_ISL_878517, EPI_ISL_878518, EPI_ISL_878520, EPI_ISL_878521, EPI_ISL_878523, EPI_ISL_878525, EPI_ISL_878527, EPI_ISL_878528, EPI_ISL_878532, EPI_ISL_878533, EPI_ISL_878535, EPI_ISL_878536, EPI_ISL_878538, EPI_ISL_878540, EPI_ISL_878543, EPI_ISL_878545, EPI_ISL_878546, EPI_ISL_878548, EPI_ISL_878550, EPI_ISL_878551, EPI_ISL_878553, EPI_ISL_878555, EPI_ISL_878558, EPI_ISL_878559, EPI_ISL_878561, EPI_ISL_878562, EPI_ISL_878564, EPI_ISL_878566, EPI_ISL_878567, EPI_ISL_878569, EPI_ISL_878573, EPI_ISL_878575, EPI_ISL_878577, EPI_ISL_878578, EPI_ISL_878580, EPI_ISL_878582, EPI_ISL_878583, EPI_ISL_878585, EPI_ISL_878588, EPI_ISL_878590, EPI_ISL_878591, EPI_ISL_878593, EPI_ISL_878594, EPI_ISL_878596, EPI_ISL_878599, EPI_ISL_878601, EPI_ISL_878602, EPI_ISL_878604, EPI_ISL_878606, EPI_ISL_878607, EPI_ISL_878610, EPI_ISL_878612, EPI_ISL_878615, EPI_ISL_878617, EPI_ISL_878620, EPI_ISL_878621, EPI_ISL_878623, EPI_ISL_878625, EPI_ISL_878626, EPI_ISL_878628, EPI_ISL_878630, EPI_ISL_878631, EPI_ISL_878633, EPI_ISL_878634, EPI_ISL_878636, EPI_ISL_878638, EPI_ISL_878639, EPI_ISL_878641, EPI_ISL_878642, EPI_ISL_878644, EPI_ISL_878646, EPI_ISL_878647, EPI_ISL_878649, EPI_ISL_878650, EPI_ISL_878655, EPI_ISL_878657, EPI_ISL_878659, EPI_ISL_878660, EPI_ISL_878662, EPI_ISL_878663, EPI_ISL_878665, EPI_ISL_878667, EPI_ISL_878668, EPI_ISL_878670, EPI_ISL_878672, EPI_ISL_878673, EPI_ISL_878675, EPI_ISL_878676, EPI_ISL_878678, EPI_ISL_878680, EPI_ISL_878685, EPI_ISL_878686, EPI_ISL_878691, EPI_ISL_878692, EPI_ISL_878694, EPI_ISL_878696, EPI_ISL_878698, EPI_ISL_878699, EPI_ISL_878703, EPI_ISL_878708, EPI_ISL_878709, EPI_ISL_878711, EPI_ISL_878714, EPI_ISL_878715, EPI_ISL_878717, EPI_ISL_878718, EPI_ISL_878720, EPI_ISL_878722, EPI_ISL_878723, EPI_ISL_878725, EPI_ISL_878727 | | | |
| see above | Lighthouse Lab in Alderley Park | Wellcome Sanger Institute for the COVID-19 Genomics UK (COG-UK) Consortium | Jacquelyn Wynn, Mairead Hyland, The Lighthouse Lab in Alderley Park and Alex Alderton, Roberto Amato, Sonia Goncalves, Ewan Harrison, David K. Jackson, Ian Johnston, Dominic Kwiatkowski, Cordelia Langford, John Sillitoe on behalf of the Wellcome Sanger Institute COVID-19 Surveillance Team |
| EPI_ISL_878728, EPI_ISL_878731, EPI_ISL_878733, EPI_ISL_878735, EPI_ISL_878737, EPI_ISL_878738, EPI_ISL_878740 | Lighthouse Lab in Cambridge | Wellcome Sanger Institute for the COVID-19 Genomics UK (COG-UK) Consortium | Rob Howes, The Lighthouse Lab in Cambridge and Alex Alderton, Roberto Amato, Sonia Goncalves, Ewan Harrison, David K. Jackson, Ian Johnston, Dominic Kwiatkowski, Cordelia Langford, John Sillitoe on behalf of the Wellcome Sanger Institute COVID-19 Surveillance Team |
| EPI_ISL_878741 | Lighthouse Lab in Alderley Park | Wellcome Sanger Institute for the COVID-19 Genomics UK (COG-UK) Consortium | Jacquelyn Wynn, Mairead Hyland, The Lighthouse Lab in Alderley Park and Alex Alderton, Roberto Amato, Sonia Goncalves, Ewan Harrison, David K. Jackson, Ian Johnston, Dominic Kwiatkowski, Cordelia Langford, John Sillitoe on behalf of the Wellcome Sanger Institute COVID-19 Surveillance Team |
| EPI_ISL_878743, EPI_ISL_878745 | Lighthouse Lab in Cambridge | Wellcome Sanger Institute for the COVID-19 Genomics UK (COG-UK) Consortium | Rob Howes, The Lighthouse Lab in Cambridge and Alex Alderton, Roberto Amato, Sonia Goncalves, Ewan Harrison, David K. Jackson, Ian Johnston, Dominic Kwiatkowski, Cordelia Langford, John Sillitoe on behalf of the Wellcome Sanger Institute COVID-19 Surveillance Team |
| EPI_ISL_878746 | Lighthouse Lab in Alderley Park | Wellcome Sanger Institute for the COVID-19 Genomics UK (COG-UK) Consortium | Jacquelyn Wynn, Mairead Hyland, The Lighthouse Lab in Alderley Park and Alex Alderton, Roberto Amato, Sonia Goncalves, Ewan Harrison, David K. Jackson, Ian Johnston, Dominic Kwiatkowski, Cordelia Langford, John Sillitoe on behalf of the Wellcome Sanger Institute COVID-19 Surveillance Team |
| EPI_ISL_878748, EPI_ISL_878749, EPI_ISL_878753 | Lighthouse Lab in Cambridge | Wellcome Sanger Institute for the COVID-19 Genomics UK (COG-UK) Consortium | Rob Howes, The Lighthouse Lab in Cambridge and Alex Alderton, Roberto Amato, Sonia Goncalves, Ewan Harrison, David K. Jackson, Ian Johnston, Dominic Kwiatkowski, Cordelia Langford, John Sillitoe on behalf of the Wellcome Sanger Institute COVID-19 Surveillance Team |
| EPI_ISL_878754 | Lighthouse Lab in Alderley Park | Wellcome Sanger Institute for the COVID-19 Genomics UK (COG-UK) Consortium | Jacquelyn Wynn, Mairead Hyland, The Lighthouse Lab in Alderley Park and Alex Alderton, Roberto Amato, Sonia Goncalves, Ewan Harrison, David K. Jackson, Ian Johnston, Dominic Kwiatkowski, Cordelia Langford, John Sillitoe on behalf of the Wellcome Sanger Institute COVID-19 Surveillance Team |
| EPI_ISL_878758, EPI_ISL_878761, EPI_ISL_878762, EPI_ISL_878764, EPI_ISL_878766, EPI_ISL_878767, EPI_ISL_878770, EPI_ISL_878774 | Lighthouse Lab in Cambridge | Wellcome Sanger Institute for the COVID-19 Genomics UK (COG-UK) Consortium | Rob Howes, The Lighthouse Lab in Cambridge and Alex Alderton, Roberto Amato, Sonia Goncalves, Ewan Harrison, David K. Jackson, Ian Johnston, Dominic Kwiatkowski, Cordelia Langford, John Sillitoe on behalf of the Wellcome Sanger Institute COVID-19 Surveillance Team |
| EPI_ISL_878777 | Lighthouse Lab in Alderley Park | Wellcome Sanger Institute for the COVID-19 Genomics UK (COG-UK) Consortium | Jacquelyn Wynn, Mairead Hyland, The Lighthouse Lab in Alderley Park and Alex Alderton, Roberto Amato, Sonia Goncalves, Ewan Harrison, David K. Jackson, Ian Johnston, Dominic Kwiatkowski, Cordelia Langford, John Sillitoe on behalf of the Wellcome Sanger Institute COVID-19 Surveillance Team |
| EPI_ISL_878783 | Lighthouse Lab in Cambridge | Wellcome Sanger Institute for the COVID-19 Genomics UK | Rob Howes, The Lighthouse Lab in Cambridge and Alex Alderton, Roberto Amato, Sonia Goncalves, Ewan Harrison, David K. Jackson, Ian Johnston, |

| | | | |
|---|---|---|---|
| | | (COG-UK) Consortium | Dominic Kwiatkowski, Cordelia Langford, John Sillitoe on behalf of the Wellcome Sanger Institute COVID-19 Surveillance Team |
| EPI_ISL_878785 | Lighthouse Lab in Alderley Park | Wellcome Sanger Institute for the COVID-19 Genomics UK (COG-UK) Consortium | Jacquelyn Wynn, Mairead Hyland, The Lighthouse Lab in Alderley Park and Alex Alderton, Roberto Amato, Sonia Goncalves, Ewan Harrison, David K. Jackson, Ian Johnston, Dominic Kwiatkowski, Cordelia Langford, John Sillitoe on behalf of the Wellcome Sanger Institute COVID-19 Surveillance Team |
| EPI_ISL_878787, EPI_ISL_878788 | Lighthouse Lab in Cambridge | Wellcome Sanger Institute for the COVID-19 Genomics UK (COG-UK) Consortium | Rob Howes, The Lighthouse Lab in Cambridge and Alex Alderton, Roberto Amato, Sonia Goncalves, Ewan Harrison, David K. Jackson, Ian Johnston, Dominic Kwiatkowski, Cordelia Langford, John Sillitoe on behalf of the Wellcome Sanger Institute COVID-19 Surveillance Team |
| EPI_ISL_878790 | Lighthouse Lab in Alderley Park | Wellcome Sanger Institute for the COVID-19 Genomics UK (COG-UK) Consortium | Jacquelyn Wynn, Mairead Hyland, The Lighthouse Lab in Alderley Park and Alex Alderton, Roberto Amato, Sonia Goncalves, Ewan Harrison, David K. Jackson, Ian Johnston, Dominic Kwiatkowski, Cordelia Langford, John Sillitoe on behalf of the Wellcome Sanger Institute COVID-19 Surveillance Team |
| EPI_ISL_878795, EPI_ISL_878796, EPI_ISL_878799, EPI_ISL_878801, EPI_ISL_878802, EPI_ISL_878803 | Lighthouse Lab in Cambridge | Wellcome Sanger Institute for the COVID-19 Genomics UK (COG-UK) Consortium | Rob Howes, The Lighthouse Lab in Cambridge and Alex Alderton, Roberto Amato, Sonia Goncalves, Ewan Harrison, David K. Jackson, Ian Johnston, Dominic Kwiatkowski, Cordelia Langford, John Sillitoe on behalf of the Wellcome Sanger Institute COVID-19 Surveillance Team |
| EPI_ISL_878804, EPI_ISL_878805, EPI_ISL_878808, EPI_ISL_878809 | Lighthouse Lab in Alderley Park | Wellcome Sanger Institute for the COVID-19 Genomics UK (COG-UK) Consortium | Jacquelyn Wynn, Mairead Hyland, The Lighthouse Lab in Alderley Park and Alex Alderton, Roberto Amato, Sonia Goncalves, Ewan Harrison, David K. Jackson, Ian Johnston, Dominic Kwiatkowski, Cordelia Langford, John Sillitoe on behalf of the Wellcome Sanger Institute COVID-19 Surveillance Team |
| EPI_ISL_878810, EPI_ISL_878811 | Lighthouse Lab in Cambridge | Wellcome Sanger Institute for the COVID-19 Genomics UK (COG-UK) Consortium | Rob Howes, The Lighthouse Lab in Cambridge and Alex Alderton, Roberto Amato, Sonia Goncalves, Ewan Harrison, David K. Jackson, Ian Johnston, Dominic Kwiatkowski, Cordelia Langford, John Sillitoe on behalf of the Wellcome Sanger Institute COVID-19 Surveillance Team |
| EPI_ISL_878812 | Lighthouse Lab in Alderley Park | Wellcome Sanger Institute for the COVID-19 Genomics UK (COG-UK) Consortium | Jacquelyn Wynn, Mairead Hyland, The Lighthouse Lab in Alderley Park and Alex Alderton, Roberto Amato, Sonia Goncalves, Ewan Harrison, David K. Jackson, Ian Johnston, Dominic Kwiatkowski, Cordelia Langford, John Sillitoe on behalf of the Wellcome Sanger Institute COVID-19 Surveillance Team |
| EPI_ISL_878813, EPI_ISL_878815, EPI_ISL_878816 | Lighthouse Lab in Cambridge | Wellcome Sanger Institute for the COVID-19 Genomics UK (COG-UK) Consortium | Rob Howes, The Lighthouse Lab in Cambridge and Alex Alderton, Roberto Amato, Sonia Goncalves, Ewan Harrison, David K. Jackson, Ian Johnston, Dominic Kwiatkowski, Cordelia Langford, John Sillitoe on behalf of the Wellcome Sanger Institute COVID-19 Surveillance Team |
| EPI_ISL_878817 | Lighthouse Lab in Alderley Park | Wellcome Sanger Institute for the COVID-19 Genomics UK (COG-UK) Consortium | Jacquelyn Wynn, Mairead Hyland, The Lighthouse Lab in Alderley Park and Alex Alderton, Roberto Amato, Sonia Goncalves, Ewan Harrison, David K. Jackson, Ian Johnston, Dominic Kwiatkowski, Cordelia Langford, John Sillitoe on behalf of the Wellcome Sanger Institute COVID-19 Surveillance Team |
| EPI_ISL_878818 | Lighthouse Lab in Cambridge | Wellcome Sanger Institute for the COVID-19 Genomics UK (COG-UK) Consortium | Rob Howes, The Lighthouse Lab in Cambridge and Alex Alderton, Roberto Amato, Sonia Goncalves, Ewan Harrison, David K. Jackson, Ian Johnston, Dominic Kwiatkowski, Cordelia Langford, John Sillitoe on behalf of the Wellcome Sanger Institute COVID-19 Surveillance Team |
| EPI_ISL_878820 | Lighthouse Lab in Alderley Park | Wellcome Sanger Institute for the COVID-19 Genomics UK (COG-UK) Consortium | Jacquelyn Wynn, Mairead Hyland, The Lighthouse Lab in Alderley Park and Alex Alderton, Roberto Amato, Sonia Goncalves, Ewan Harrison, David K. Jackson, Ian Johnston, Dominic Kwiatkowski, Cordelia Langford, John Sillitoe on behalf of the Wellcome Sanger Institute COVID-19 Surveillance Team |
| EPI_ISL_878821 | Lighthouse Lab in Cambridge | Wellcome Sanger Institute for the COVID-19 Genomics UK (COG-UK) Consortium | Rob Howes, The Lighthouse Lab in Cambridge and Alex Alderton, Roberto Amato, Sonia Goncalves, Ewan Harrison, David K. Jackson, Ian Johnston, Dominic Kwiatkowski, Cordelia Langford, John Sillitoe on behalf of the Wellcome Sanger Institute COVID-19 Surveillance Team |
| EPI_ISL_878822 | Lighthouse Lab in Alderley Park | Wellcome Sanger Institute for the COVID-19 Genomics UK (COG-UK) Consortium | Jacquelyn Wynn, Mairead Hyland, The Lighthouse Lab in Alderley Park and Alex Alderton, Roberto Amato, Sonia Goncalves, Ewan Harrison, David K. Jackson, Ian Johnston, Dominic Kwiatkowski, Cordelia Langford, John Sillitoe on behalf of the Wellcome Sanger Institute COVID-19 Surveillance Team |
| EPI_ISL_878823 | Lighthouse Lab in Cambridge | Wellcome Sanger Institute for the COVID-19 Genomics UK (COG-UK) Consortium | Rob Howes, The Lighthouse Lab in Cambridge and Alex Alderton, Roberto Amato, Sonia Goncalves, Ewan Harrison, David K. Jackson, Ian Johnston, Dominic Kwiatkowski, Cordelia Langford, John Sillitoe on behalf of the Wellcome Sanger Institute COVID-19 Surveillance Team |
| EPI_ISL_878825 | Lighthouse Lab in Alderley Park | Wellcome Sanger Institute for the COVID-19 Genomics UK (COG-UK) Consortium | Jacquelyn Wynn, Mairead Hyland, The Lighthouse Lab in Alderley Park and Alex Alderton, Roberto Amato, Sonia Goncalves, Ewan Harrison, David K. Jackson, Ian Johnston, Dominic Kwiatkowski, Cordelia Langford, John Sillitoe on behalf of the Wellcome Sanger Institute COVID-19 Surveillance Team |
| EPI_ISL_878826, EPI_ISL_878827, EPI_ISL_878828, EPI_ISL_878829, EPI_ISL_878830, EPI_ISL_878831, EPI_ISL_878832, EPI_ISL_878834 | Lighthouse Lab in Cambridge | Wellcome Sanger Institute for the COVID-19 Genomics UK (COG-UK) Consortium | Rob Howes, The Lighthouse Lab in Cambridge and Alex Alderton, Roberto Amato, Sonia Goncalves, Ewan Harrison, David K. Jackson, Ian Johnston, Dominic Kwiatkowski, Cordelia Langford, John Sillitoe on behalf of the Wellcome Sanger Institute COVID-19 Surveillance Team |
| EPI_ISL_878835, EPI_ISL_878836 | Lighthouse Lab in Alderley Park | Wellcome Sanger Institute for the COVID-19 Genomics UK (COG-UK) Consortium | Jacquelyn Wynn, Mairead Hyland, The Lighthouse Lab in Alderley Park and Alex Alderton, Roberto Amato, Sonia Goncalves, Ewan Harrison, David K. Jackson, Ian Johnston, Dominic Kwiatkowski, Cordelia Langford, John Sillitoe on behalf of the Wellcome Sanger Institute COVID-19 Surveillance Team |
| EPI_ISL_878837, EPI_ISL_878838, EPI_ISL_878839 | Lighthouse Lab in Cambridge | Wellcome Sanger Institute for the COVID-19 Genomics UK (COG-UK) Consortium | Rob Howes, The Lighthouse Lab in Cambridge and Alex Alderton, Roberto Amato, Sonia Goncalves, Ewan Harrison, David K. Jackson, Ian Johnston, Dominic Kwiatkowski, Cordelia Langford, John Sillitoe on behalf of the Wellcome Sanger Institute COVID-19 Surveillance Team |
| EPI_ISL_878840 | Lighthouse Lab in Alderley Park | Wellcome Sanger Institute for the COVID-19 Genomics UK (COG-UK) Consortium | Jacquelyn Wynn, Mairead Hyland, The Lighthouse Lab in Alderley Park and Alex Alderton, Roberto Amato, Sonia Goncalves, Ewan Harrison, David K. Jackson, Ian Johnston, Dominic Kwiatkowski, Cordelia Langford, John Sillitoe on behalf of the Wellcome Sanger Institute COVID-19 Surveillance Team |
| EPI_ISL_878841 | Lighthouse Lab in Cambridge | Wellcome Sanger Institute for the COVID-19 Genomics UK (COG-UK) Consortium | Rob Howes, The Lighthouse Lab in Cambridge and Alex Alderton, Roberto Amato, Sonia Goncalves, Ewan Harrison, David K. Jackson, Ian Johnston, Dominic Kwiatkowski, Cordelia Langford, John Sillitoe on behalf of the Wellcome Sanger Institute COVID-19 Surveillance Team |
| EPI_ISL_878843 | Lighthouse Lab in Alderley Park | Wellcome Sanger Institute for the COVID-19 Genomics UK (COG-UK) Consortium | Jacquelyn Wynn, Mairead Hyland, The Lighthouse Lab in Alderley Park and Alex Alderton, Roberto Amato, Sonia Goncalves, Ewan Harrison, David K. Jackson, Ian Johnston, Dominic Kwiatkowski, Cordelia Langford, John Sillitoe on behalf of the Wellcome Sanger Institute COVID-19 Surveillance Team |
| EPI_ISL_878847 | Lighthouse Lab in Cambridge | Wellcome Sanger Institute for the COVID-19 Genomics UK (COG-UK) Consortium | Rob Howes, The Lighthouse Lab in Cambridge and Alex Alderton, Roberto Amato, Sonia Goncalves, Ewan Harrison, David K. Jackson, Ian Johnston, Dominic Kwiatkowski, Cordelia Langford, John Sillitoe on behalf of the Wellcome Sanger Institute COVID-19 Surveillance Team |
| EPI_ISL_878849 | Lighthouse Lab in Alderley Park | Wellcome Sanger Institute for the COVID-19 Genomics UK (COG-UK) Consortium | Jacquelyn Wynn, Mairead Hyland, The Lighthouse Lab in Alderley Park and Alex Alderton, Roberto Amato, Sonia Goncalves, Ewan Harrison, David K. Jackson, Ian Johnston, Dominic Kwiatkowski, Cordelia Langford, John Sillitoe on behalf of the Wellcome Sanger Institute COVID-19 Surveillance Team |
| EPI_ISL_878850 | Lighthouse Lab in Cambridge | Wellcome Sanger Institute for the COVID-19 Genomics UK (COG-UK) Consortium | Rob Howes, The Lighthouse Lab in Cambridge and Alex Alderton, Roberto Amato, Sonia Goncalves, Ewan Harrison, David K. Jackson, Ian Johnston, Dominic Kwiatkowski, Cordelia Langford, John Sillitoe on behalf of the Wellcome Sanger Institute COVID-19 Surveillance Team |
| EPI_ISL_878853 | Lighthouse Lab in Alderley Park | Wellcome Sanger Institute for the COVID-19 Genomics UK (COG-UK) Consortium | Jacquelyn Wynn, Mairead Hyland, The Lighthouse Lab in Alderley Park and Alex Alderton, Roberto Amato, Sonia Goncalves, Ewan Harrison, David K. Jackson, Ian Johnston, Dominic Kwiatkowski, Cordelia Langford, John Sillitoe on behalf of the Wellcome Sanger Institute COVID-19 Surveillance Team |
| EPI_ISL_878855 | Lighthouse Lab in Cambridge | Wellcome Sanger Institute for the COVID-19 Genomics UK (COG-UK) Consortium | Rob Howes, The Lighthouse Lab in Cambridge and Alex Alderton, Roberto Amato, Sonia Goncalves, Ewan Harrison, David K. Jackson, Ian Johnston, Dominic Kwiatkowski, Cordelia Langford, John Sillitoe on behalf of the Wellcome Sanger Institute COVID-19 Surveillance Team |
| EPI_ISL_878856 | Lighthouse Lab in Alderley Park | Wellcome Sanger Institute for the COVID-19 Genomics UK (COG-UK) Consortium | Jacquelyn Wynn, Mairead Hyland, The Lighthouse Lab in Alderley Park and Alex Alderton, Roberto Amato, Sonia Goncalves, Ewan Harrison, David K. Jackson, Ian Johnston, Dominic Kwiatkowski, Cordelia Langford, John Sillitoe on behalf of the Wellcome Sanger Institute COVID-19 Surveillance Team |
| EPI_ISL_878857 | Lighthouse Lab in Cambridge | Wellcome Sanger Institute for the COVID-19 Genomics UK (COG-UK) Consortium | Rob Howes, The Lighthouse Lab in Cambridge and Alex Alderton, Roberto Amato, Sonia Goncalves, Ewan Harrison, David K. Jackson, Ian Johnston, Dominic Kwiatkowski, Cordelia Langford, John Sillitoe on behalf of the Wellcome Sanger Institute COVID-19 Surveillance Team |
| EPI_ISL_878859 | Lighthouse Lab in Alderley Park | Wellcome Sanger Institute for the COVID-19 Genomics UK (COG-UK) Consortium | Jacquelyn Wynn, Mairead Hyland, The Lighthouse Lab in Alderley Park and Alex Alderton, Roberto Amato, Sonia Goncalves, Ewan Harrison, David K. Jackson, Ian Johnston, Dominic Kwiatkowski, Cordelia Langford, John Sillitoe on behalf of the Wellcome Sanger Institute COVID-19 Surveillance Team |
| EPI_ISL_878862, EPI_ISL_878863, EPI_ISL_878864 | Lighthouse Lab in Cambridge | Wellcome Sanger Institute for the COVID-19 Genomics UK (COG-UK) Consortium | Rob Howes, The Lighthouse Lab in Cambridge and Alex Alderton, Roberto Amato, Sonia Goncalves, Ewan Harrison, David K. Jackson, Ian Johnston, Dominic Kwiatkowski, Cordelia Langford, John Sillitoe on behalf of the Wellcome Sanger Institute COVID-19 Surveillance Team |
| EPI_ISL_878865, EPI_ISL_878866 | Lighthouse Lab in Alderley Park | Wellcome Sanger Institute for the COVID-19 Genomics UK (COG-UK) Consortium | Jacquelyn Wynn, Mairead Hyland, The Lighthouse Lab in Alderley Park and Alex Alderton, Roberto Amato, Sonia Goncalves, Ewan Harrison, David K. Jackson, Ian Johnston, Dominic Kwiatkowski, Cordelia Langford, John Sillitoe on behalf of the Wellcome Sanger Institute COVID-19 Surveillance Team |
| EPI_ISL_878867 | Lighthouse Lab in Cambridge | Wellcome Sanger Institute for the COVID-19 Genomics UK (COG-UK) Consortium | Rob Howes, The Lighthouse Lab in Cambridge and Alex Alderton, Roberto Amato, Sonia Goncalves, Ewan Harrison, David K. Jackson, Ian Johnston, Dominic Kwiatkowski, Cordelia Langford, John Sillitoe on behalf of the Wellcome Sanger Institute COVID-19 Surveillance Team |
| EPI_ISL_878868 | Lighthouse Lab in Alderley Park | Wellcome Sanger Institute for the COVID-19 Genomics UK (COG-UK) Consortium | Jacquelyn Wynn, Mairead Hyland, The Lighthouse Lab in Alderley Park and Alex Alderton, Roberto Amato, Sonia Goncalves, Ewan Harrison, David K. Jackson, Ian Johnston, Dominic Kwiatkowski, Cordelia Langford, John Sillitoe on behalf of the Wellcome Sanger Institute COVID-19 Surveillance Team |
| EPI_ISL_878869, EPI_ISL_878870, | Lighthouse Lab in Cambridge | Wellcome Sanger Institute for the COVID-19 Genomics UK | Rob Howes, The Lighthouse Lab in Cambridge and Alex Alderton, Roberto Amato, Sonia Goncalves, Ewan Harrison, David K. Jackson, Ian Johnston, |

| | | | |
|---|---|---|---|
| EPI_ISL_878871, EPI_ISL_878872, EPI_ISL_878873 | | (COG-UK) Consortium | Dominic Kwiatkowski, Cordelia Langford, John Sillitoe on behalf of the Wellcome Sanger Institute COVID-19 Surveillance Team |
| EPI_ISL_878875 | Lighthouse Lab in Alderley Park | Wellcome Sanger Institute for the COVID-19 Genomics UK (COG-UK) Consortium | Jacquelyn Wynn, Mairead Hyland, The Lighthouse Lab in Alderley Park and Alex Alderton, Roberto Amato, Sonia Goncalves, Ewan Harrison, David K. Jackson, Ian Johnston, Dominic Kwiatkowski, Cordelia Langford, John Sillitoe on behalf of the Wellcome Sanger Institute COVID-19 Surveillance Team |
| EPI_ISL_878877 | Lighthouse Lab in Cambridge | Wellcome Sanger Institute for the COVID-19 Genomics UK (COG-UK) Consortium | Rob Howes, The Lighthouse Lab in Cambridge and Alex Alderton, Roberto Amato, Sonia Goncalves, Ewan Harrison, David K. Jackson, Ian Johnston, Dominic Kwiatkowski, Cordelia Langford, John Sillitoe on behalf of the Wellcome Sanger Institute COVID-19 Surveillance Team |
| EPI_ISL_878878 | Lighthouse Lab in Alderley Park | Wellcome Sanger Institute for the COVID-19 Genomics UK (COG-UK) Consortium | Jacquelyn Wynn, Mairead Hyland, The Lighthouse Lab in Alderley Park and Alex Alderton, Roberto Amato, Sonia Goncalves, Ewan Harrison, David K. Jackson, Ian Johnston, Dominic Kwiatkowski, Cordelia Langford, John Sillitoe on behalf of the Wellcome Sanger Institute COVID-19 Surveillance Team |
| EPI_ISL_878881 | Lighthouse Lab in Cambridge | Wellcome Sanger Institute for the COVID-19 Genomics UK (COG-UK) Consortium | Rob Howes, The Lighthouse Lab in Cambridge and Alex Alderton, Roberto Amato, Sonia Goncalves, Ewan Harrison, David K. Jackson, Ian Johnston, Dominic Kwiatkowski, Cordelia Langford, John Sillitoe on behalf of the Wellcome Sanger Institute COVID-19 Surveillance Team |
| EPI_ISL_878883 | Lighthouse Lab in Alderley Park | Wellcome Sanger Institute for the COVID-19 Genomics UK (COG-UK) Consortium | Jacquelyn Wynn, Mairead Hyland, The Lighthouse Lab in Alderley Park and Alex Alderton, Roberto Amato, Sonia Goncalves, Ewan Harrison, David K. Jackson, Ian Johnston, Dominic Kwiatkowski, Cordelia Langford, John Sillitoe on behalf of the Wellcome Sanger Institute COVID-19 Surveillance Team |
| EPI_ISL_878885 | Lighthouse Lab in Cambridge | Wellcome Sanger Institute for the COVID-19 Genomics UK (COG-UK) Consortium | Rob Howes, The Lighthouse Lab in Cambridge and Alex Alderton, Roberto Amato, Sonia Goncalves, Ewan Harrison, David K. Jackson, Ian Johnston, Dominic Kwiatkowski, Cordelia Langford, John Sillitoe on behalf of the Wellcome Sanger Institute COVID-19 Surveillance Team |
| EPI_ISL_878886, EPI_ISL_878887, EPI_ISL_878888 | Lighthouse Lab in Alderley Park | Wellcome Sanger Institute for the COVID-19 Genomics UK (COG-UK) Consortium | Jacquelyn Wynn, Mairead Hyland, The Lighthouse Lab in Alderley Park and Alex Alderton, Roberto Amato, Sonia Goncalves, Ewan Harrison, David K. Jackson, Ian Johnston, Dominic Kwiatkowski, Cordelia Langford, John Sillitoe on behalf of the Wellcome Sanger Institute COVID-19 Surveillance Team |
| EPI_ISL_878889, EPI_ISL_878890 | Lighthouse Lab in Cambridge | Wellcome Sanger Institute for the COVID-19 Genomics UK (COG-UK) Consortium | Rob Howes, The Lighthouse Lab in Cambridge and Alex Alderton, Roberto Amato, Sonia Goncalves, Ewan Harrison, David K. Jackson, Ian Johnston, Dominic Kwiatkowski, Cordelia Langford, John Sillitoe on behalf of the Wellcome Sanger Institute COVID-19 Surveillance Team |
| EPI_ISL_878891 | Lighthouse Lab in Alderley Park | Wellcome Sanger Institute for the COVID-19 Genomics UK (COG-UK) Consortium | Jacquelyn Wynn, Mairead Hyland, The Lighthouse Lab in Alderley Park and Alex Alderton, Roberto Amato, Sonia Goncalves, Ewan Harrison, David K. Jackson, Ian Johnston, Dominic Kwiatkowski, Cordelia Langford, John Sillitoe on behalf of the Wellcome Sanger Institute COVID-19 Surveillance Team |
| EPI_ISL_878893, EPI_ISL_878894, EPI_ISL_878895, EPI_ISL_878897, EPI_ISL_878898, EPI_ISL_878899, EPI_ISL_878900, EPI_ISL_878901, EPI_ISL_878903, EPI_ISL_878904, EPI_ISL_878905, EPI_ISL_878907, EPI_ISL_878908 | | | |
| see above | Lighthouse Lab in Cambridge | Wellcome Sanger Institute for the COVID-19 Genomics UK (COG-UK) Consortium | Rob Howes, The Lighthouse Lab in Cambridge and Alex Alderton, Roberto Amato, Sonia Goncalves, Ewan Harrison, David K. Jackson, Ian Johnston, Dominic Kwiatkowski, Cordelia Langford, John Sillitoe on behalf of the Wellcome Sanger Institute COVID-19 Surveillance Team |
| EPI_ISL_878909 | Lighthouse Lab in Alderley Park | Wellcome Sanger Institute for the COVID-19 Genomics UK (COG-UK) Consortium | Jacquelyn Wynn, Mairead Hyland, The Lighthouse Lab in Alderley Park and Alex Alderton, Roberto Amato, Sonia Goncalves, Ewan Harrison, David K. Jackson, Ian Johnston, Dominic Kwiatkowski, Cordelia Langford, John Sillitoe on behalf of the Wellcome Sanger Institute COVID-19 Surveillance Team |
| EPI_ISL_878910, EPI_ISL_878911, EPI_ISL_878912 | Lighthouse Lab in Cambridge | Wellcome Sanger Institute for the COVID-19 Genomics UK (COG-UK) Consortium | Rob Howes, The Lighthouse Lab in Cambridge and Alex Alderton, Roberto Amato, Sonia Goncalves, Ewan Harrison, David K. Jackson, Ian Johnston, Dominic Kwiatkowski, Cordelia Langford, John Sillitoe on behalf of the Wellcome Sanger Institute COVID-19 Surveillance Team |
| EPI_ISL_878914, EPI_ISL_878915 | Lighthouse Lab in Alderley Park | Wellcome Sanger Institute for the COVID-19 Genomics UK (COG-UK) Consortium | Jacquelyn Wynn, Mairead Hyland, The Lighthouse Lab in Alderley Park and Alex Alderton, Roberto Amato, Sonia Goncalves, Ewan Harrison, David K. Jackson, Ian Johnston, Dominic Kwiatkowski, Cordelia Langford, John Sillitoe on behalf of the Wellcome Sanger Institute COVID-19 Surveillance Team |
| EPI_ISL_878916 | Lighthouse Lab in Cambridge | Wellcome Sanger Institute for the COVID-19 Genomics UK (COG-UK) Consortium | Rob Howes, The Lighthouse Lab in Cambridge and Alex Alderton, Roberto Amato, Sonia Goncalves, Ewan Harrison, David K. Jackson, Ian Johnston, Dominic Kwiatkowski, Cordelia Langford, John Sillitoe on behalf of the Wellcome Sanger Institute COVID-19 Surveillance Team |
| EPI_ISL_878917, EPI_ISL_878920 | Lighthouse Lab in Alderley Park | Wellcome Sanger Institute for the COVID-19 Genomics UK (COG-UK) Consortium | Jacquelyn Wynn, Mairead Hyland, The Lighthouse Lab in Alderley Park and Alex Alderton, Roberto Amato, Sonia Goncalves, Ewan Harrison, David K. Jackson, Ian Johnston, Dominic Kwiatkowski, Cordelia Langford, John Sillitoe on behalf of the Wellcome Sanger Institute COVID-19 Surveillance Team |
| EPI_ISL_878921, EPI_ISL_878922, EPI_ISL_878923, EPI_ISL_878924 | Lighthouse Lab in Cambridge | Wellcome Sanger Institute for the COVID-19 Genomics UK (COG-UK) Consortium | Rob Howes, The Lighthouse Lab in Cambridge and Alex Alderton, Roberto Amato, Sonia Goncalves, Ewan Harrison, David K. Jackson, Ian Johnston, Dominic Kwiatkowski, Cordelia Langford, John Sillitoe on behalf of the Wellcome Sanger Institute COVID-19 Surveillance Team |
| EPI_ISL_878925, EPI_ISL_878926, EPI_ISL_878927 | Lighthouse Lab in Alderley Park | Wellcome Sanger Institute for the COVID-19 Genomics UK (COG-UK) Consortium | Jacquelyn Wynn, Mairead Hyland, The Lighthouse Lab in Alderley Park and Alex Alderton, Roberto Amato, Sonia Goncalves, Ewan Harrison, David K. Jackson, Ian Johnston, Dominic Kwiatkowski, Cordelia Langford, John Sillitoe on behalf of the Wellcome Sanger Institute COVID-19 Surveillance Team |
| EPI_ISL_878928 | Lighthouse Lab in Cambridge | Wellcome Sanger Institute for the COVID-19 Genomics UK (COG-UK) Consortium | Rob Howes, The Lighthouse Lab in Cambridge and Alex Alderton, Roberto Amato, Sonia Goncalves, Ewan Harrison, David K. Jackson, Ian Johnston, Dominic Kwiatkowski, Cordelia Langford, John Sillitoe on behalf of the Wellcome Sanger Institute COVID-19 Surveillance Team |
| EPI_ISL_878929 | Lighthouse Lab in Alderley Park | Wellcome Sanger Institute for the COVID-19 Genomics UK (COG-UK) Consortium | Jacquelyn Wynn, Mairead Hyland, The Lighthouse Lab in Alderley Park and Alex Alderton, Roberto Amato, Sonia Goncalves, Ewan Harrison, David K. Jackson, Ian Johnston, Dominic Kwiatkowski, Cordelia Langford, John Sillitoe on behalf of the Wellcome Sanger Institute COVID-19 Surveillance Team |
| EPI_ISL_878930, EPI_ISL_878931 | Lighthouse Lab in Cambridge | Wellcome Sanger Institute for the COVID-19 Genomics UK (COG-UK) Consortium | Rob Howes, The Lighthouse Lab in Cambridge and Alex Alderton, Roberto Amato, Sonia Goncalves, Ewan Harrison, David K. Jackson, Ian Johnston, Dominic Kwiatkowski, Cordelia Langford, John Sillitoe on behalf of the Wellcome Sanger Institute COVID-19 Surveillance Team |
| EPI_ISL_878932 | Lighthouse Lab in Alderley Park | Wellcome Sanger Institute for the COVID-19 Genomics UK (COG-UK) Consortium | Jacquelyn Wynn, Mairead Hyland, The Lighthouse Lab in Alderley Park and Alex Alderton, Roberto Amato, Sonia Goncalves, Ewan Harrison, David K. Jackson, Ian Johnston, Dominic Kwiatkowski, Cordelia Langford, John Sillitoe on behalf of the Wellcome Sanger Institute COVID-19 Surveillance Team |
| EPI_ISL_878933, EPI_ISL_878935 | Lighthouse Lab in Cambridge | Wellcome Sanger Institute for the COVID-19 Genomics UK (COG-UK) Consortium | Rob Howes, The Lighthouse Lab in Cambridge and Alex Alderton, Roberto Amato, Sonia Goncalves, Ewan Harrison, David K. Jackson, Ian Johnston, Dominic Kwiatkowski, Cordelia Langford, John Sillitoe on behalf of the Wellcome Sanger Institute COVID-19 Surveillance Team |
| EPI_ISL_878936 | Lighthouse Lab in Alderley Park | Wellcome Sanger Institute for the COVID-19 Genomics UK (COG-UK) Consortium | Jacquelyn Wynn, Mairead Hyland, The Lighthouse Lab in Alderley Park and Alex Alderton, Roberto Amato, Sonia Goncalves, Ewan Harrison, David K. Jackson, Ian Johnston, Dominic Kwiatkowski, Cordelia Langford, John Sillitoe on behalf of the Wellcome Sanger Institute COVID-19 Surveillance Team |
| EPI_ISL_878937 | Lighthouse Lab in Cambridge | Wellcome Sanger Institute for the COVID-19 Genomics UK (COG-UK) Consortium | Rob Howes, The Lighthouse Lab in Cambridge and Alex Alderton, Roberto Amato, Sonia Goncalves, Ewan Harrison, David K. Jackson, Ian Johnston, Dominic Kwiatkowski, Cordelia Langford, John Sillitoe on behalf of the Wellcome Sanger Institute COVID-19 Surveillance Team |
| EPI_ISL_878938 | Lighthouse Lab in Alderley Park | Wellcome Sanger Institute for the COVID-19 Genomics UK (COG-UK) Consortium | Jacquelyn Wynn, Mairead Hyland, The Lighthouse Lab in Alderley Park and Alex Alderton, Roberto Amato, Sonia Goncalves, Ewan Harrison, David K. Jackson, Ian Johnston, Dominic Kwiatkowski, Cordelia Langford, John Sillitoe on behalf of the Wellcome Sanger Institute COVID-19 Surveillance Team |
| EPI_ISL_878939, EPI_ISL_878940 | Lighthouse Lab in Cambridge | Wellcome Sanger Institute for the COVID-19 Genomics UK (COG-UK) Consortium | Rob Howes, The Lighthouse Lab in Cambridge and Alex Alderton, Roberto Amato, Sonia Goncalves, Ewan Harrison, David K. Jackson, Ian Johnston, Dominic Kwiatkowski, Cordelia Langford, John Sillitoe on behalf of the Wellcome Sanger Institute COVID-19 Surveillance Team |
| EPI_ISL_878946, EPI_ISL_878947 | Lighthouse Lab in Alderley Park | Wellcome Sanger Institute for the COVID-19 Genomics UK (COG-UK) Consortium | Jacquelyn Wynn, Mairead Hyland, The Lighthouse Lab in Alderley Park and Alex Alderton, Roberto Amato, Sonia Goncalves, Ewan Harrison, David K. Jackson, Ian Johnston, Dominic Kwiatkowski, Cordelia Langford, John Sillitoe on behalf of the Wellcome Sanger Institute COVID-19 Surveillance Team |
| EPI_ISL_878948, EPI_ISL_878949, EPI_ISL_878950, EPI_ISL_878953, EPI_ISL_878954 | Lighthouse Lab in Cambridge | Wellcome Sanger Institute for the COVID-19 Genomics UK (COG-UK) Consortium | Rob Howes, The Lighthouse Lab in Cambridge and Alex Alderton, Roberto Amato, Sonia Goncalves, Ewan Harrison, David K. Jackson, Ian Johnston, Dominic Kwiatkowski, Cordelia Langford, John Sillitoe on behalf of the Wellcome Sanger Institute COVID-19 Surveillance Team |
| EPI_ISL_878955 | Lighthouse Lab in Alderley Park | Wellcome Sanger Institute for the COVID-19 Genomics UK (COG-UK) Consortium | Jacquelyn Wynn, Mairead Hyland, The Lighthouse Lab in Alderley Park and Alex Alderton, Roberto Amato, Sonia Goncalves, Ewan Harrison, David K. Jackson, Ian Johnston, Dominic Kwiatkowski, Cordelia Langford, John Sillitoe on behalf of the Wellcome Sanger Institute COVID-19 Surveillance Team |
| EPI_ISL_878956, EPI_ISL_878957 | Lighthouse Lab in Cambridge | Wellcome Sanger Institute for the COVID-19 Genomics UK (COG-UK) Consortium | Rob Howes, The Lighthouse Lab in Cambridge and Alex Alderton, Roberto Amato, Sonia Goncalves, Ewan Harrison, David K. Jackson, Ian Johnston, Dominic Kwiatkowski, Cordelia Langford, John Sillitoe on behalf of the Wellcome Sanger Institute COVID-19 Surveillance Team |
| EPI_ISL_878958, EPI_ISL_878959, EPI_ISL_878961 | Lighthouse Lab in Alderley Park | Wellcome Sanger Institute for the COVID-19 Genomics UK (COG-UK) Consortium | Jacquelyn Wynn, Mairead Hyland, The Lighthouse Lab in Alderley Park and Alex Alderton, Roberto Amato, Sonia Goncalves, Ewan Harrison, David K. Jackson, Ian Johnston, Dominic Kwiatkowski, Cordelia Langford, John Sillitoe on behalf of the Wellcome Sanger Institute COVID-19 Surveillance Team |
| EPI_ISL_878963, EPI_ISL_878964 | Lighthouse Lab in Cambridge | Wellcome Sanger Institute for the COVID-19 Genomics UK (COG-UK) Consortium | Rob Howes, The Lighthouse Lab in Cambridge and Alex Alderton, Roberto Amato, Sonia Goncalves, Ewan Harrison, David K. Jackson, Ian Johnston, Dominic Kwiatkowski, Cordelia Langford, John Sillitoe on behalf of the Wellcome Sanger Institute COVID-19 Surveillance Team |
| EPI_ISL_878965, EPI_ISL_878966 | Lighthouse Lab in Alderley Park | Wellcome Sanger Institute for the COVID-19 Genomics UK (COG-UK) Consortium | Jacquelyn Wynn, Mairead Hyland, The Lighthouse Lab in Alderley Park and Alex Alderton, Roberto Amato, Sonia Goncalves, Ewan Harrison, David K. Jackson, Ian Johnston, Dominic Kwiatkowski, Cordelia Langford, John Sillitoe on behalf of the Wellcome Sanger Institute COVID-19 Surveillance Team |

| | | | |
|---|---|---|---|
| EPI_ISL_878968, EPI_ISL_878969 | Lighthouse Lab in Cambridge | Wellcome Sanger Institute for the COVID-19 Genomics UK (COG-UK) Consortium | Rob Howes, The Lighthouse Lab in Cambridge and Alex Alderton, Roberto Amato, Sonia Goncalves, Ewan Harrison, David K. Jackson, Ian Johnston, Dominic Kwiatkowski, Cordelia Langford, John Sillitoe on behalf of the Wellcome Sanger Institute COVID-19 Surveillance Team |
| EPI_ISL_878970 | Lighthouse Lab in Alderley Park | Wellcome Sanger Institute for the COVID-19 Genomics UK (COG-UK) Consortium | Jacquelyn Wynn, Mairead Hyland, The Lighthouse Lab in Alderley Park and Alex Alderton, Roberto Amato, Sonia Goncalves, Ewan Harrison, David K. Jackson, Ian Johnston, Dominic Kwiatkowski, Cordelia Langford, John Sillitoe on behalf of the Wellcome Sanger Institute COVID-19 Surveillance Team |
| EPI_ISL_878973, EPI_ISL_878974, EPI_ISL_878975, EPI_ISL_878976, EPI_ISL_878977, EPI_ISL_878978 | Lighthouse Lab in Cambridge | Wellcome Sanger Institute for the COVID-19 Genomics UK (COG-UK) Consortium | Rob Howes, The Lighthouse Lab in Cambridge and Alex Alderton, Roberto Amato, Sonia Goncalves, Ewan Harrison, David K. Jackson, Ian Johnston, Dominic Kwiatkowski, Cordelia Langford, John Sillitoe on behalf of the Wellcome Sanger Institute COVID-19 Surveillance Team |
| EPI_ISL_878979 | Lighthouse Lab in Alderley Park | Wellcome Sanger Institute for the COVID-19 Genomics UK (COG-UK) Consortium | Jacquelyn Wynn, Mairead Hyland, The Lighthouse Lab in Alderley Park and Alex Alderton, Roberto Amato, Sonia Goncalves, Ewan Harrison, David K. Jackson, Ian Johnston, Dominic Kwiatkowski, Cordelia Langford, John Sillitoe on behalf of the Wellcome Sanger Institute COVID-19 Surveillance Team |
| EPI_ISL_878980 | Lighthouse Lab in Cambridge | Wellcome Sanger Institute for the COVID-19 Genomics UK (COG-UK) Consortium | Rob Howes, The Lighthouse Lab in Cambridge and Alex Alderton, Roberto Amato, Sonia Goncalves, Ewan Harrison, David K. Jackson, Ian Johnston, Dominic Kwiatkowski, Cordelia Langford, John Sillitoe on behalf of the Wellcome Sanger Institute COVID-19 Surveillance Team |
| EPI_ISL_878981, EPI_ISL_878983 | Lighthouse Lab in Alderley Park | Wellcome Sanger Institute for the COVID-19 Genomics UK (COG-UK) Consortium | Jacquelyn Wynn, Mairead Hyland, The Lighthouse Lab in Alderley Park and Alex Alderton, Roberto Amato, Sonia Goncalves, Ewan Harrison, David K. Jackson, Ian Johnston, Dominic Kwiatkowski, Cordelia Langford, John Sillitoe on behalf of the Wellcome Sanger Institute COVID-19 Surveillance Team |
| EPI_ISL_878984 | Lighthouse Lab in Cambridge | Wellcome Sanger Institute for the COVID-19 Genomics UK (COG-UK) Consortium | Rob Howes, The Lighthouse Lab in Cambridge and Alex Alderton, Roberto Amato, Sonia Goncalves, Ewan Harrison, David K. Jackson, Ian Johnston, Dominic Kwiatkowski, Cordelia Langford, John Sillitoe on behalf of the Wellcome Sanger Institute COVID-19 Surveillance Team |
| EPI_ISL_878985 | Lighthouse Lab in Alderley Park | Wellcome Sanger Institute for the COVID-19 Genomics UK (COG-UK) Consortium | Jacquelyn Wynn, Mairead Hyland, The Lighthouse Lab in Alderley Park and Alex Alderton, Roberto Amato, Sonia Goncalves, Ewan Harrison, David K. Jackson, Ian Johnston, Dominic Kwiatkowski, Cordelia Langford, John Sillitoe on behalf of the Wellcome Sanger Institute COVID-19 Surveillance Team |
| EPI_ISL_878986, EPI_ISL_878987, EPI_ISL_878988, EPI_ISL_878989, EPI_ISL_878991 | Lighthouse Lab in Cambridge | Wellcome Sanger Institute for the COVID-19 Genomics UK (COG-UK) Consortium | Rob Howes, The Lighthouse Lab in Cambridge and Alex Alderton, Roberto Amato, Sonia Goncalves, Ewan Harrison, David K. Jackson, Ian Johnston, Dominic Kwiatkowski, Cordelia Langford, John Sillitoe on behalf of the Wellcome Sanger Institute COVID-19 Surveillance Team |
| EPI_ISL_878992, EPI_ISL_878994 | Lighthouse Lab in Alderley Park | Wellcome Sanger Institute for the COVID-19 Genomics UK (COG-UK) Consortium | Jacquelyn Wynn, Mairead Hyland, The Lighthouse Lab in Alderley Park and Alex Alderton, Roberto Amato, Sonia Goncalves, Ewan Harrison, David K. Jackson, Ian Johnston, Dominic Kwiatkowski, Cordelia Langford, John Sillitoe on behalf of the Wellcome Sanger Institute COVID-19 Surveillance Team |
| EPI_ISL_878995, EPI_ISL_878996, EPI_ISL_878997, EPI_ISL_878998 | Lighthouse Lab in Cambridge | Wellcome Sanger Institute for the COVID-19 Genomics UK (COG-UK) Consortium | Rob Howes, The Lighthouse Lab in Cambridge and Alex Alderton, Roberto Amato, Sonia Goncalves, Ewan Harrison, David K. Jackson, Ian Johnston, Dominic Kwiatkowski, Cordelia Langford, John Sillitoe on behalf of the Wellcome Sanger Institute COVID-19 Surveillance Team |
| EPI_ISL_878999, EPI_ISL_879000 | Lighthouse Lab in Alderley Park | Wellcome Sanger Institute for the COVID-19 Genomics UK (COG-UK) Consortium | Jacquelyn Wynn, Mairead Hyland, The Lighthouse Lab in Alderley Park and Alex Alderton, Roberto Amato, Sonia Goncalves, Ewan Harrison, David K. Jackson, Ian Johnston, Dominic Kwiatkowski, Cordelia Langford, John Sillitoe on behalf of the Wellcome Sanger Institute COVID-19 Surveillance Team |
| EPI_ISL_879002 | Lighthouse Lab in Cambridge | Wellcome Sanger Institute for the COVID-19 Genomics UK (COG-UK) Consortium | Rob Howes, The Lighthouse Lab in Cambridge and Alex Alderton, Roberto Amato, Sonia Goncalves, Ewan Harrison, David K. Jackson, Ian Johnston, Dominic Kwiatkowski, Cordelia Langford, John Sillitoe on behalf of the Wellcome Sanger Institute COVID-19 Surveillance Team |
| EPI_ISL_879003 | Lighthouse Lab in Alderley Park | Wellcome Sanger Institute for the COVID-19 Genomics UK (COG-UK) Consortium | Jacquelyn Wynn, Mairead Hyland, The Lighthouse Lab in Alderley Park and Alex Alderton, Roberto Amato, Sonia Goncalves, Ewan Harrison, David K. Jackson, Ian Johnston, Dominic Kwiatkowski, Cordelia Langford, John Sillitoe on behalf of the Wellcome Sanger Institute COVID-19 Surveillance Team |
| EPI_ISL_879004, EPI_ISL_879005, EPI_ISL_879006 | Lighthouse Lab in Cambridge | Wellcome Sanger Institute for the COVID-19 Genomics UK (COG-UK) Consortium | Rob Howes, The Lighthouse Lab in Cambridge and Alex Alderton, Roberto Amato, Sonia Goncalves, Ewan Harrison, David K. Jackson, Ian Johnston, Dominic Kwiatkowski, Cordelia Langford, John Sillitoe on behalf of the Wellcome Sanger Institute COVID-19 Surveillance Team |
| EPI_ISL_879007, EPI_ISL_879008 | Lighthouse Lab in Alderley Park | Wellcome Sanger Institute for the COVID-19 Genomics UK (COG-UK) Consortium | Jacquelyn Wynn, Mairead Hyland, The Lighthouse Lab in Alderley Park and Alex Alderton, Roberto Amato, Sonia Goncalves, Ewan Harrison, David K. Jackson, Ian Johnston, Dominic Kwiatkowski, Cordelia Langford, John Sillitoe on behalf of the Wellcome Sanger Institute COVID-19 Surveillance Team |
| EPI_ISL_879010, EPI_ISL_879011, EPI_ISL_879012 | Lighthouse Lab in Cambridge | Wellcome Sanger Institute for the COVID-19 Genomics UK (COG-UK) Consortium | Rob Howes, The Lighthouse Lab in Cambridge and Alex Alderton, Roberto Amato, Sonia Goncalves, Ewan Harrison, David K. Jackson, Ian Johnston, Dominic Kwiatkowski, Cordelia Langford, John Sillitoe on behalf of the Wellcome Sanger Institute COVID-19 Surveillance Team |
| EPI_ISL_879015 | Lighthouse Lab in Alderley Park | Wellcome Sanger Institute for the COVID-19 Genomics UK (COG-UK) Consortium | Jacquelyn Wynn, Mairead Hyland, The Lighthouse Lab in Alderley Park and Alex Alderton, Roberto Amato, Sonia Goncalves, Ewan Harrison, David K. Jackson, Ian Johnston, Dominic Kwiatkowski, Cordelia Langford, John Sillitoe on behalf of the Wellcome Sanger Institute COVID-19 Surveillance Team |
| EPI_ISL_879017 | Lighthouse Lab in Cambridge | Wellcome Sanger Institute for the COVID-19 Genomics UK (COG-UK) Consortium | Rob Howes, The Lighthouse Lab in Cambridge and Alex Alderton, Roberto Amato, Sonia Goncalves, Ewan Harrison, David K. Jackson, Ian Johnston, Dominic Kwiatkowski, Cordelia Langford, John Sillitoe on behalf of the Wellcome Sanger Institute COVID-19 Surveillance Team |
| EPI_ISL_879018 | Lighthouse Lab in Alderley Park | Wellcome Sanger Institute for the COVID-19 Genomics UK (COG-UK) Consortium | Jacquelyn Wynn, Mairead Hyland, The Lighthouse Lab in Alderley Park and Alex Alderton, Roberto Amato, Sonia Goncalves, Ewan Harrison, David K. Jackson, Ian Johnston, Dominic Kwiatkowski, Cordelia Langford, John Sillitoe on behalf of the Wellcome Sanger Institute COVID-19 Surveillance Team |
| EPI_ISL_879020, EPI_ISL_879022, EPI_ISL_879024 | Lighthouse Lab in Cambridge | Wellcome Sanger Institute for the COVID-19 Genomics UK (COG-UK) Consortium | Rob Howes, The Lighthouse Lab in Cambridge and Alex Alderton, Roberto Amato, Sonia Goncalves, Ewan Harrison, David K. Jackson, Ian Johnston, Dominic Kwiatkowski, Cordelia Langford, John Sillitoe on behalf of the Wellcome Sanger Institute COVID-19 Surveillance Team |
| EPI_ISL_879026 | Lighthouse Lab in Alderley Park | Wellcome Sanger Institute for the COVID-19 Genomics UK (COG-UK) Consortium | Jacquelyn Wynn, Mairead Hyland, The Lighthouse Lab in Alderley Park and Alex Alderton, Roberto Amato, Sonia Goncalves, Ewan Harrison, David K. Jackson, Ian Johnston, Dominic Kwiatkowski, Cordelia Langford, John Sillitoe on behalf of the Wellcome Sanger Institute COVID-19 Surveillance Team |
| EPI_ISL_879027, EPI_ISL_879029, EPI_ISL_879030, EPI_ISL_879031 | Lighthouse Lab in Cambridge | Wellcome Sanger Institute for the COVID-19 Genomics UK (COG-UK) Consortium | Rob Howes, The Lighthouse Lab in Cambridge and Alex Alderton, Roberto Amato, Sonia Goncalves, Ewan Harrison, David K. Jackson, Ian Johnston, Dominic Kwiatkowski, Cordelia Langford, John Sillitoe on behalf of the Wellcome Sanger Institute COVID-19 Surveillance Team |
| EPI_ISL_879033 | Lighthouse Lab in Alderley Park | Wellcome Sanger Institute for the COVID-19 Genomics UK (COG-UK) Consortium | Jacquelyn Wynn, Mairead Hyland, The Lighthouse Lab in Alderley Park and Alex Alderton, Roberto Amato, Sonia Goncalves, Ewan Harrison, David K. Jackson, Ian Johnston, Dominic Kwiatkowski, Cordelia Langford, John Sillitoe on behalf of the Wellcome Sanger Institute COVID-19 Surveillance Team |
| EPI_ISL_879035, EPI_ISL_879036, EPI_ISL_879039, EPI_ISL_879041, EPI_ISL_879045, EPI_ISL_879046, EPI_ISL_879048, EPI_ISL_879051, EPI_ISL_879053, EPI_ISL_879054, EPI_ISL_879058, EPI_ISL_879059, EPI_ISL_879060 | | | |
| see above | Lighthouse Lab in Cambridge | Wellcome Sanger Institute for the COVID-19 Genomics UK (COG-UK) Consortium | Rob Howes, The Lighthouse Lab in Cambridge and Alex Alderton, Roberto Amato, Sonia Goncalves, Ewan Harrison, David K. Jackson, Ian Johnston, Dominic Kwiatkowski, Cordelia Langford, John Sillitoe on behalf of the Wellcome Sanger Institute COVID-19 Surveillance Team |
| EPI_ISL_879061 | Lighthouse Lab in Alderley Park | Wellcome Sanger Institute for the COVID-19 Genomics UK (COG-UK) Consortium | Jacquelyn Wynn, Mairead Hyland, The Lighthouse Lab in Alderley Park and Alex Alderton, Roberto Amato, Sonia Goncalves, Ewan Harrison, David K. Jackson, Ian Johnston, Dominic Kwiatkowski, Cordelia Langford, John Sillitoe on behalf of the Wellcome Sanger Institute COVID-19 Surveillance Team |
| EPI_ISL_879062 | Lighthouse Lab in Cambridge | Wellcome Sanger Institute for the COVID-19 Genomics UK (COG-UK) Consortium | Rob Howes, The Lighthouse Lab in Cambridge and Alex Alderton, Roberto Amato, Sonia Goncalves, Ewan Harrison, David K. Jackson, Ian Johnston, Dominic Kwiatkowski, Cordelia Langford, John Sillitoe on behalf of the Wellcome Sanger Institute COVID-19 Surveillance Team |
| EPI_ISL_879066 | Lighthouse Lab in Alderley Park | Wellcome Sanger Institute for the COVID-19 Genomics UK (COG-UK) Consortium | Jacquelyn Wynn, Mairead Hyland, The Lighthouse Lab in Alderley Park and Alex Alderton, Roberto Amato, Sonia Goncalves, Ewan Harrison, David K. Jackson, Ian Johnston, Dominic Kwiatkowski, Cordelia Langford, John Sillitoe on behalf of the Wellcome Sanger Institute COVID-19 Surveillance Team |
| EPI_ISL_879067, EPI_ISL_879068, EPI_ISL_879069 | Lighthouse Lab in Cambridge | Wellcome Sanger Institute for the COVID-19 Genomics UK (COG-UK) Consortium | Rob Howes, The Lighthouse Lab in Cambridge and Alex Alderton, Roberto Amato, Sonia Goncalves, Ewan Harrison, David K. Jackson, Ian Johnston, Dominic Kwiatkowski, Cordelia Langford, John Sillitoe on behalf of the Wellcome Sanger Institute COVID-19 Surveillance Team |
| EPI_ISL_879070 | Lighthouse Lab in Alderley Park | Wellcome Sanger Institute for the COVID-19 Genomics UK (COG-UK) Consortium | Jacquelyn Wynn, Mairead Hyland, The Lighthouse Lab in Alderley Park and Alex Alderton, Roberto Amato, Sonia Goncalves, Ewan Harrison, David K. Jackson, Ian Johnston, Dominic Kwiatkowski, Cordelia Langford, John Sillitoe on behalf of the Wellcome Sanger Institute COVID-19 Surveillance Team |
| EPI_ISL_879071 | Lighthouse Lab in Cambridge | Wellcome Sanger Institute for the COVID-19 Genomics UK (COG-UK) Consortium | Rob Howes, The Lighthouse Lab in Cambridge and Alex Alderton, Roberto Amato, Sonia Goncalves, Ewan Harrison, David K. Jackson, Ian Johnston, Dominic Kwiatkowski, Cordelia Langford, John Sillitoe on behalf of the Wellcome Sanger Institute COVID-19 Surveillance Team |
| EPI_ISL_879072 | Lighthouse Lab in Alderley Park | Wellcome Sanger Institute for the COVID-19 Genomics UK (COG-UK) Consortium | Jacquelyn Wynn, Mairead Hyland, The Lighthouse Lab in Alderley Park and Alex Alderton, Roberto Amato, Sonia Goncalves, Ewan Harrison, David K. Jackson, Ian Johnston, Dominic Kwiatkowski, Cordelia Langford, John Sillitoe on behalf of the Wellcome Sanger Institute COVID-19 Surveillance Team |
| EPI_ISL_879074, EPI_ISL_879076 | Lighthouse Lab in Cambridge | Wellcome Sanger Institute for the COVID-19 Genomics UK (COG-UK) Consortium | Rob Howes, The Lighthouse Lab in Cambridge and Alex Alderton, Roberto Amato, Sonia Goncalves, Ewan Harrison, David K. Jackson, Ian Johnston, Dominic Kwiatkowski, Cordelia Langford, John Sillitoe on behalf of the Wellcome Sanger Institute COVID-19 Surveillance Team |
| EPI_ISL_879077 | Lighthouse Lab in Alderley Park | Wellcome Sanger Institute for the COVID-19 Genomics UK (COG-UK) Consortium | Jacquelyn Wynn, Mairead Hyland, The Lighthouse Lab in Alderley Park and Alex Alderton, Roberto Amato, Sonia Goncalves, Ewan Harrison, David K. Jackson, Ian Johnston, Dominic Kwiatkowski, Cordelia Langford, John Sillitoe on behalf of the Wellcome Sanger Institute COVID-19 Surveillance Team |

| | | | |
|---|---|---|---|
| EPI_ISL_879078 | Lighthouse Lab in Cambridge | Wellcome Sanger Institute for the COVID-19 Genomics UK (COG-UK) Consortium | Rob Howes, The Lighthouse Lab in Cambridge and Alex Alderton, Roberto Amato, Sonia Goncalves, Ewan Harrison, David K. Jackson, Ian Johnston, Dominic Kwiatkowski, Cordelia Langford, John Sillitoe on behalf of the Wellcome Sanger Institute COVID-19 Surveillance Team |
| EPI_ISL_879079 | Lighthouse Lab in Alderley Park | Wellcome Sanger Institute for the COVID-19 Genomics UK (COG-UK) Consortium | Jacquelyn Wynn, Mairead Hyland, The Lighthouse Lab in Alderley Park and Alex Alderton, Roberto Amato, Sonia Goncalves, Ewan Harrison, David K. Jackson, Ian Johnston, Dominic Kwiatkowski, Cordelia Langford, John Sillitoe on behalf of the Wellcome Sanger Institute COVID-19 Surveillance Team |
| EPI_ISL_879080 | Lighthouse Lab in Cambridge | Wellcome Sanger Institute for the COVID-19 Genomics UK (COG-UK) Consortium | Rob Howes, The Lighthouse Lab in Cambridge and Alex Alderton, Roberto Amato, Sonia Goncalves, Ewan Harrison, David K. Jackson, Ian Johnston, Dominic Kwiatkowski, Cordelia Langford, John Sillitoe on behalf of the Wellcome Sanger Institute COVID-19 Surveillance Team |
| EPI_ISL_879081 | Lighthouse Lab in Alderley Park | Wellcome Sanger Institute for the COVID-19 Genomics UK (COG-UK) Consortium | Jacquelyn Wynn, Mairead Hyland, The Lighthouse Lab in Alderley Park and Alex Alderton, Roberto Amato, Sonia Goncalves, Ewan Harrison, David K. Jackson, Ian Johnston, Dominic Kwiatkowski, Cordelia Langford, John Sillitoe on behalf of the Wellcome Sanger Institute COVID-19 Surveillance Team |
| EPI_ISL_879082 | Lighthouse Lab in Cambridge | Wellcome Sanger Institute for the COVID-19 Genomics UK (COG-UK) Consortium | Rob Howes, The Lighthouse Lab in Cambridge and Alex Alderton, Roberto Amato, Sonia Goncalves, Ewan Harrison, David K. Jackson, Ian Johnston, Dominic Kwiatkowski, Cordelia Langford, John Sillitoe on behalf of the Wellcome Sanger Institute COVID-19 Surveillance Team |
| EPI_ISL_879083 | Lighthouse Lab in Alderley Park | Wellcome Sanger Institute for the COVID-19 Genomics UK (COG-UK) Consortium | Jacquelyn Wynn, Mairead Hyland, The Lighthouse Lab in Alderley Park and Alex Alderton, Roberto Amato, Sonia Goncalves, Ewan Harrison, David K. Jackson, Ian Johnston, Dominic Kwiatkowski, Cordelia Langford, John Sillitoe on behalf of the Wellcome Sanger Institute COVID-19 Surveillance Team |
| EPI_ISL_879084, EPI_ISL_879085 | Lighthouse Lab in Cambridge | Wellcome Sanger Institute for the COVID-19 Genomics UK (COG-UK) Consortium | Rob Howes, The Lighthouse Lab in Cambridge and Alex Alderton, Roberto Amato, Sonia Goncalves, Ewan Harrison, David K. Jackson, Ian Johnston, Dominic Kwiatkowski, Cordelia Langford, John Sillitoe on behalf of the Wellcome Sanger Institute COVID-19 Surveillance Team |
| EPI_ISL_879088 | Lighthouse Lab in Alderley Park | Wellcome Sanger Institute for the COVID-19 Genomics UK (COG-UK) Consortium | Jacquelyn Wynn, Mairead Hyland, The Lighthouse Lab in Alderley Park and Alex Alderton, Roberto Amato, Sonia Goncalves, Ewan Harrison, David K. Jackson, Ian Johnston, Dominic Kwiatkowski, Cordelia Langford, John Sillitoe on behalf of the Wellcome Sanger Institute COVID-19 Surveillance Team |
| EPI_ISL_879090 | Lighthouse Lab in Cambridge | Wellcome Sanger Institute for the COVID-19 Genomics UK (COG-UK) Consortium | Rob Howes, The Lighthouse Lab in Cambridge and Alex Alderton, Roberto Amato, Sonia Goncalves, Ewan Harrison, David K. Jackson, Ian Johnston, Dominic Kwiatkowski, Cordelia Langford, John Sillitoe on behalf of the Wellcome Sanger Institute COVID-19 Surveillance Team |
| EPI_ISL_879091 | Lighthouse Lab in Alderley Park | Wellcome Sanger Institute for the COVID-19 Genomics UK (COG-UK) Consortium | Jacquelyn Wynn, Mairead Hyland, The Lighthouse Lab in Alderley Park and Alex Alderton, Roberto Amato, Sonia Goncalves, Ewan Harrison, David K. Jackson, Ian Johnston, Dominic Kwiatkowski, Cordelia Langford, John Sillitoe on behalf of the Wellcome Sanger Institute COVID-19 Surveillance Team |
| EPI_ISL_879095 | Lighthouse Lab in Cambridge | Wellcome Sanger Institute for the COVID-19 Genomics UK (COG-UK) Consortium | Rob Howes, The Lighthouse Lab in Cambridge and Alex Alderton, Roberto Amato, Sonia Goncalves, Ewan Harrison, David K. Jackson, Ian Johnston, Dominic Kwiatkowski, Cordelia Langford, John Sillitoe on behalf of the Wellcome Sanger Institute COVID-19 Surveillance Team |
| EPI_ISL_879097, EPI_ISL_879098, EPI_ISL_879099, EPI_ISL_879100, EPI_ISL_879101 | Lighthouse Lab in Alderley Park | Wellcome Sanger Institute for the COVID-19 Genomics UK (COG-UK) Consortium | Jacquelyn Wynn, Mairead Hyland, The Lighthouse Lab in Alderley Park and Alex Alderton, Roberto Amato, Sonia Goncalves, Ewan Harrison, David K. Jackson, Ian Johnston, Dominic Kwiatkowski, Cordelia Langford, John Sillitoe on behalf of the Wellcome Sanger Institute COVID-19 Surveillance Team |
| EPI_ISL_879102 | Lighthouse Lab in Cambridge | Wellcome Sanger Institute for the COVID-19 Genomics UK (COG-UK) Consortium | Rob Howes, The Lighthouse Lab in Cambridge and Alex Alderton, Roberto Amato, Sonia Goncalves, Ewan Harrison, David K. Jackson, Ian Johnston, Dominic Kwiatkowski, Cordelia Langford, John Sillitoe on behalf of the Wellcome Sanger Institute COVID-19 Surveillance Team |
| EPI_ISL_879103 | Lighthouse Lab in Alderley Park | Wellcome Sanger Institute for the COVID-19 Genomics UK (COG-UK) Consortium | Jacquelyn Wynn, Mairead Hyland, The Lighthouse Lab in Alderley Park and Alex Alderton, Roberto Amato, Sonia Goncalves, Ewan Harrison, David K. Jackson, Ian Johnston, Dominic Kwiatkowski, Cordelia Langford, John Sillitoe on behalf of the Wellcome Sanger Institute COVID-19 Surveillance Team |
| EPI_ISL_879104, EPI_ISL_879105, EPI_ISL_879106, EPI_ISL_879107 | Lighthouse Lab in Cambridge | Wellcome Sanger Institute for the COVID-19 Genomics UK (COG-UK) Consortium | Rob Howes, The Lighthouse Lab in Cambridge and Alex Alderton, Roberto Amato, Sonia Goncalves, Ewan Harrison, David K. Jackson, Ian Johnston, Dominic Kwiatkowski, Cordelia Langford, John Sillitoe on behalf of the Wellcome Sanger Institute COVID-19 Surveillance Team |
| EPI_ISL_879108, EPI_ISL_879109, EPI_ISL_879110, EPI_ISL_879111 | Lighthouse Lab in Alderley Park | Wellcome Sanger Institute for the COVID-19 Genomics UK (COG-UK) Consortium | Jacquelyn Wynn, Mairead Hyland, The Lighthouse Lab in Alderley Park and Alex Alderton, Roberto Amato, Sonia Goncalves, Ewan Harrison, David K. Jackson, Ian Johnston, Dominic Kwiatkowski, Cordelia Langford, John Sillitoe on behalf of the Wellcome Sanger Institute COVID-19 Surveillance Team |
| EPI_ISL_879112 | Lighthouse Lab in Cambridge | Wellcome Sanger Institute for the COVID-19 Genomics UK (COG-UK) Consortium | Rob Howes, The Lighthouse Lab in Cambridge and Alex Alderton, Roberto Amato, Sonia Goncalves, Ewan Harrison, David K. Jackson, Ian Johnston, Dominic Kwiatkowski, Cordelia Langford, John Sillitoe on behalf of the Wellcome Sanger Institute COVID-19 Surveillance Team |
| EPI_ISL_879113, EPI_ISL_879114 | Lighthouse Lab in Alderley Park | Wellcome Sanger Institute for the COVID-19 Genomics UK (COG-UK) Consortium | Jacquelyn Wynn, Mairead Hyland, The Lighthouse Lab in Alderley Park and Alex Alderton, Roberto Amato, Sonia Goncalves, Ewan Harrison, David K. Jackson, Ian Johnston, Dominic Kwiatkowski, Cordelia Langford, John Sillitoe on behalf of the Wellcome Sanger Institute COVID-19 Surveillance Team |
| EPI_ISL_879115 | Lighthouse Lab in Cambridge | Wellcome Sanger Institute for the COVID-19 Genomics UK (COG-UK) Consortium | Rob Howes, The Lighthouse Lab in Cambridge and Alex Alderton, Roberto Amato, Sonia Goncalves, Ewan Harrison, David K. Jackson, Ian Johnston, Dominic Kwiatkowski, Cordelia Langford, John Sillitoe on behalf of the Wellcome Sanger Institute COVID-19 Surveillance Team |
| EPI_ISL_879118 | Lighthouse Lab in Alderley Park | Wellcome Sanger Institute for the COVID-19 Genomics UK (COG-UK) Consortium | Jacquelyn Wynn, Mairead Hyland, The Lighthouse Lab in Alderley Park and Alex Alderton, Roberto Amato, Sonia Goncalves, Ewan Harrison, David K. Jackson, Ian Johnston, Dominic Kwiatkowski, Cordelia Langford, John Sillitoe on behalf of the Wellcome Sanger Institute COVID-19 Surveillance Team |
| EPI_ISL_879119 | Lighthouse Lab in Cambridge | Wellcome Sanger Institute for the COVID-19 Genomics UK (COG-UK) Consortium | Rob Howes, The Lighthouse Lab in Cambridge and Alex Alderton, Roberto Amato, Sonia Goncalves, Ewan Harrison, David K. Jackson, Ian Johnston, Dominic Kwiatkowski, Cordelia Langford, John Sillitoe on behalf of the Wellcome Sanger Institute COVID-19 Surveillance Team |
| EPI_ISL_879121 | Lighthouse Lab in Alderley Park | Wellcome Sanger Institute for the COVID-19 Genomics UK (COG-UK) Consortium | Jacquelyn Wynn, Mairead Hyland, The Lighthouse Lab in Alderley Park and Alex Alderton, Roberto Amato, Sonia Goncalves, Ewan Harrison, David K. Jackson, Ian Johnston, Dominic Kwiatkowski, Cordelia Langford, John Sillitoe on behalf of the Wellcome Sanger Institute COVID-19 Surveillance Team |
| EPI_ISL_879122, EPI_ISL_879123 | Lighthouse Lab in Cambridge | Wellcome Sanger Institute for the COVID-19 Genomics UK (COG-UK) Consortium | Rob Howes, The Lighthouse Lab in Cambridge and Alex Alderton, Roberto Amato, Sonia Goncalves, Ewan Harrison, David K. Jackson, Ian Johnston, Dominic Kwiatkowski, Cordelia Langford, John Sillitoe on behalf of the Wellcome Sanger Institute COVID-19 Surveillance Team |
| EPI_ISL_879124, EPI_ISL_879125, EPI_ISL_879126 | Lighthouse Lab in Alderley Park | Wellcome Sanger Institute for the COVID-19 Genomics UK (COG-UK) Consortium | Jacquelyn Wynn, Mairead Hyland, The Lighthouse Lab in Alderley Park and Alex Alderton, Roberto Amato, Sonia Goncalves, Ewan Harrison, David K. Jackson, Ian Johnston, Dominic Kwiatkowski, Cordelia Langford, John Sillitoe on behalf of the Wellcome Sanger Institute COVID-19 Surveillance Team |
| EPI_ISL_879127 | Lighthouse Lab in Cambridge | Wellcome Sanger Institute for the COVID-19 Genomics UK (COG-UK) Consortium | Rob Howes, The Lighthouse Lab in Cambridge and Alex Alderton, Roberto Amato, Sonia Goncalves, Ewan Harrison, David K. Jackson, Ian Johnston, Dominic Kwiatkowski, Cordelia Langford, John Sillitoe on behalf of the Wellcome Sanger Institute COVID-19 Surveillance Team |
| EPI_ISL_879128, EPI_ISL_879129, EPI_ISL_879130, EPI_ISL_879131, EPI_ISL_879132 | Lighthouse Lab in Alderley Park | Wellcome Sanger Institute for the COVID-19 Genomics UK (COG-UK) Consortium | Jacquelyn Wynn, Mairead Hyland, The Lighthouse Lab in Alderley Park and Alex Alderton, Roberto Amato, Sonia Goncalves, Ewan Harrison, David K. Jackson, Ian Johnston, Dominic Kwiatkowski, Cordelia Langford, John Sillitoe on behalf of the Wellcome Sanger Institute COVID-19 Surveillance Team |
| EPI_ISL_879133 | Lighthouse Lab in Cambridge | Wellcome Sanger Institute for the COVID-19 Genomics UK (COG-UK) Consortium | Rob Howes, The Lighthouse Lab in Cambridge and Alex Alderton, Roberto Amato, Sonia Goncalves, Ewan Harrison, David K. Jackson, Ian Johnston, Dominic Kwiatkowski, Cordelia Langford, John Sillitoe on behalf of the Wellcome Sanger Institute COVID-19 Surveillance Team |
| EPI_ISL_879134 | Lighthouse Lab in Alderley Park | Wellcome Sanger Institute for the COVID-19 Genomics UK (COG-UK) Consortium | Jacquelyn Wynn, Mairead Hyland, The Lighthouse Lab in Alderley Park and Alex Alderton, Roberto Amato, Sonia Goncalves, Ewan Harrison, David K. Jackson, Ian Johnston, Dominic Kwiatkowski, Cordelia Langford, John Sillitoe on behalf of the Wellcome Sanger Institute COVID-19 Surveillance Team |
| EPI_ISL_879135, EPI_ISL_879136 | Lighthouse Lab in Cambridge | Wellcome Sanger Institute for the COVID-19 Genomics UK (COG-UK) Consortium | Rob Howes, The Lighthouse Lab in Cambridge and Alex Alderton, Roberto Amato, Sonia Goncalves, Ewan Harrison, David K. Jackson, Ian Johnston, Dominic Kwiatkowski, Cordelia Langford, John Sillitoe on behalf of the Wellcome Sanger Institute COVID-19 Surveillance Team |
| EPI_ISL_879137 | Lighthouse Lab in Alderley Park | Wellcome Sanger Institute for the COVID-19 Genomics UK (COG-UK) Consortium | Jacquelyn Wynn, Mairead Hyland, The Lighthouse Lab in Alderley Park and Alex Alderton, Roberto Amato, Sonia Goncalves, Ewan Harrison, David K. Jackson, Ian Johnston, Dominic Kwiatkowski, Cordelia Langford, John Sillitoe on behalf of the Wellcome Sanger Institute COVID-19 Surveillance Team |
| EPI_ISL_879139 | Lighthouse Lab in Cambridge | Wellcome Sanger Institute for the COVID-19 Genomics UK (COG-UK) Consortium | Rob Howes, The Lighthouse Lab in Cambridge and Alex Alderton, Roberto Amato, Sonia Goncalves, Ewan Harrison, David K. Jackson, Ian Johnston, Dominic Kwiatkowski, Cordelia Langford, John Sillitoe on behalf of the Wellcome Sanger Institute COVID-19 Surveillance Team |
| EPI_ISL_879140 | Lighthouse Lab in Alderley Park | Wellcome Sanger Institute for the COVID-19 Genomics UK (COG-UK) Consortium | Jacquelyn Wynn, Mairead Hyland, The Lighthouse Lab in Alderley Park and Alex Alderton, Roberto Amato, Sonia Goncalves, Ewan Harrison, David K. Jackson, Ian Johnston, Dominic Kwiatkowski, Cordelia Langford, John Sillitoe on behalf of the Wellcome Sanger Institute COVID-19 Surveillance Team |
| EPI_ISL_879141, EPI_ISL_879142, EPI_ISL_879143 | Lighthouse Lab in Cambridge | Wellcome Sanger Institute for the COVID-19 Genomics UK (COG-UK) Consortium | Rob Howes, The Lighthouse Lab in Cambridge and Alex Alderton, Roberto Amato, Sonia Goncalves, Ewan Harrison, David K. Jackson, Ian Johnston, Dominic Kwiatkowski, Cordelia Langford, John Sillitoe on behalf of the Wellcome Sanger Institute COVID-19 Surveillance Team |
| EPI_ISL_879144, EPI_ISL_879145, EPI_ISL_879147 | Lighthouse Lab in Alderley Park | Wellcome Sanger Institute for the COVID-19 Genomics UK (COG-UK) Consortium | Jacquelyn Wynn, Mairead Hyland, The Lighthouse Lab in Alderley Park and Alex Alderton, Roberto Amato, Sonia Goncalves, Ewan Harrison, David K. Jackson, Ian Johnston, Dominic Kwiatkowski, Cordelia Langford, John Sillitoe on behalf of the Wellcome Sanger Institute COVID-19 Surveillance Team |

| | | | |
|---|---|---|---|
| EPI_ISL_879148 | Lighthouse Lab in Cambridge | Wellcome Sanger Institute for the COVID-19 Genomics UK (COG-UK) Consortium | Rob Howes, The Lighthouse Lab in Cambridge and Alex Alderton, Roberto Amato, Sonia Goncalves, Ewan Harrison, David K. Jackson, Ian Johnston, Dominic Kwiatkowski, Cordelia Langford, John Sillitoe on behalf of the Wellcome Sanger Institute COVID-19 Surveillance Team |
| EPI_ISL_879149 | Lighthouse Lab in Alderley Park | Wellcome Sanger Institute for the COVID-19 Genomics UK (COG-UK) Consortium | Jacquelyn Wynn, Mairead Hyland, The Lighthouse Lab in Alderley Park and Alex Alderton, Roberto Amato, Sonia Goncalves, Ewan Harrison, David K. Jackson, Ian Johnston, Dominic Kwiatkowski, Cordelia Langford, John Sillitoe on behalf of the Wellcome Sanger Institute COVID-19 Surveillance Team |
| EPI_ISL_879150 | Lighthouse Lab in Cambridge | Wellcome Sanger Institute for the COVID-19 Genomics UK (COG-UK) Consortium | Rob Howes, The Lighthouse Lab in Cambridge and Alex Alderton, Roberto Amato, Sonia Goncalves, Ewan Harrison, David K. Jackson, Ian Johnston, Dominic Kwiatkowski, Cordelia Langford, John Sillitoe on behalf of the Wellcome Sanger Institute COVID-19 Surveillance Team |
| EPI_ISL_879151, EPI_ISL_879152 | Lighthouse Lab in Alderley Park | Wellcome Sanger Institute for the COVID-19 Genomics UK (COG-UK) Consortium | Jacquelyn Wynn, Mairead Hyland, The Lighthouse Lab in Alderley Park and Alex Alderton, Roberto Amato, Sonia Goncalves, Ewan Harrison, David K. Jackson, Ian Johnston, Dominic Kwiatkowski, Cordelia Langford, John Sillitoe on behalf of the Wellcome Sanger Institute COVID-19 Surveillance Team |
| EPI_ISL_879153, EPI_ISL_879155, EPI_ISL_879156 | Lighthouse Lab in Cambridge | Wellcome Sanger Institute for the COVID-19 Genomics UK (COG-UK) Consortium | Rob Howes, The Lighthouse Lab in Cambridge and Alex Alderton, Roberto Amato, Sonia Goncalves, Ewan Harrison, David K. Jackson, Ian Johnston, Dominic Kwiatkowski, Cordelia Langford, John Sillitoe on behalf of the Wellcome Sanger Institute COVID-19 Surveillance Team |
| EPI_ISL_879157 | Lighthouse Lab in Alderley Park | Wellcome Sanger Institute for the COVID-19 Genomics UK (COG-UK) Consortium | Jacquelyn Wynn, Mairead Hyland, The Lighthouse Lab in Alderley Park and Alex Alderton, Roberto Amato, Sonia Goncalves, Ewan Harrison, David K. Jackson, Ian Johnston, Dominic Kwiatkowski, Cordelia Langford, John Sillitoe on behalf of the Wellcome Sanger Institute COVID-19 Surveillance Team |
| EPI_ISL_879158 | Lighthouse Lab in Cambridge | Wellcome Sanger Institute for the COVID-19 Genomics UK (COG-UK) Consortium | Rob Howes, The Lighthouse Lab in Cambridge and Alex Alderton, Roberto Amato, Sonia Goncalves, Ewan Harrison, David K. Jackson, Ian Johnston, Dominic Kwiatkowski, Cordelia Langford, John Sillitoe on behalf of the Wellcome Sanger Institute COVID-19 Surveillance Team |
| EPI_ISL_879159, EPI_ISL_879160, EPI_ISL_879161, EPI_ISL_879162 | Lighthouse Lab in Alderley Park | Wellcome Sanger Institute for the COVID-19 Genomics UK (COG-UK) Consortium | Jacquelyn Wynn, Mairead Hyland, The Lighthouse Lab in Alderley Park and Alex Alderton, Roberto Amato, Sonia Goncalves, Ewan Harrison, David K. Jackson, Ian Johnston, Dominic Kwiatkowski, Cordelia Langford, John Sillitoe on behalf of the Wellcome Sanger Institute COVID-19 Surveillance Team |
| EPI_ISL_879163 | Lighthouse Lab in Cambridge | Wellcome Sanger Institute for the COVID-19 Genomics UK (COG-UK) Consortium | Rob Howes, The Lighthouse Lab in Cambridge and Alex Alderton, Roberto Amato, Sonia Goncalves, Ewan Harrison, David K. Jackson, Ian Johnston, Dominic Kwiatkowski, Cordelia Langford, John Sillitoe on behalf of the Wellcome Sanger Institute COVID-19 Surveillance Team |
| EPI_ISL_879164 | Lighthouse Lab in Alderley Park | Wellcome Sanger Institute for the COVID-19 Genomics UK (COG-UK) Consortium | Jacquelyn Wynn, Mairead Hyland, The Lighthouse Lab in Alderley Park and Alex Alderton, Roberto Amato, Sonia Goncalves, Ewan Harrison, David K. Jackson, Ian Johnston, Dominic Kwiatkowski, Cordelia Langford, John Sillitoe on behalf of the Wellcome Sanger Institute COVID-19 Surveillance Team |
| EPI_ISL_879165, EPI_ISL_879166 | Lighthouse Lab in Cambridge | Wellcome Sanger Institute for the COVID-19 Genomics UK (COG-UK) Consortium | Rob Howes, The Lighthouse Lab in Cambridge and Alex Alderton, Roberto Amato, Sonia Goncalves, Ewan Harrison, David K. Jackson, Ian Johnston, Dominic Kwiatkowski, Cordelia Langford, John Sillitoe on behalf of the Wellcome Sanger Institute COVID-19 Surveillance Team |
| EPI_ISL_879167 | Lighthouse Lab in Alderley Park | Wellcome Sanger Institute for the COVID-19 Genomics UK (COG-UK) Consortium | Jacquelyn Wynn, Mairead Hyland, The Lighthouse Lab in Alderley Park and Alex Alderton, Roberto Amato, Sonia Goncalves, Ewan Harrison, David K. Jackson, Ian Johnston, Dominic Kwiatkowski, Cordelia Langford, John Sillitoe on behalf of the Wellcome Sanger Institute COVID-19 Surveillance Team |
| EPI_ISL_879168 | Lighthouse Lab in Cambridge | Wellcome Sanger Institute for the COVID-19 Genomics UK (COG-UK) Consortium | Rob Howes, The Lighthouse Lab in Cambridge and Alex Alderton, Roberto Amato, Sonia Goncalves, Ewan Harrison, David K. Jackson, Ian Johnston, Dominic Kwiatkowski, Cordelia Langford, John Sillitoe on behalf of the Wellcome Sanger Institute COVID-19 Surveillance Team |
| EPI_ISL_879169 | Lighthouse Lab in Alderley Park | Wellcome Sanger Institute for the COVID-19 Genomics UK (COG-UK) Consortium | Jacquelyn Wynn, Mairead Hyland, The Lighthouse Lab in Alderley Park and Alex Alderton, Roberto Amato, Sonia Goncalves, Ewan Harrison, David K. Jackson, Ian Johnston, Dominic Kwiatkowski, Cordelia Langford, John Sillitoe on behalf of the Wellcome Sanger Institute COVID-19 Surveillance Team |
| EPI_ISL_879170 | Lighthouse Lab in Cambridge | Wellcome Sanger Institute for the COVID-19 Genomics UK (COG-UK) Consortium | Rob Howes, The Lighthouse Lab in Cambridge and Alex Alderton, Roberto Amato, Sonia Goncalves, Ewan Harrison, David K. Jackson, Ian Johnston, Dominic Kwiatkowski, Cordelia Langford, John Sillitoe on behalf of the Wellcome Sanger Institute COVID-19 Surveillance Team |
| EPI_ISL_879171 | Lighthouse Lab in Alderley Park | Wellcome Sanger Institute for the COVID-19 Genomics UK (COG-UK) Consortium | Jacquelyn Wynn, Mairead Hyland, The Lighthouse Lab in Alderley Park and Alex Alderton, Roberto Amato, Sonia Goncalves, Ewan Harrison, David K. Jackson, Ian Johnston, Dominic Kwiatkowski, Cordelia Langford, John Sillitoe on behalf of the Wellcome Sanger Institute COVID-19 Surveillance Team |
| EPI_ISL_879172 | Lighthouse Lab in Cambridge | Wellcome Sanger Institute for the COVID-19 Genomics UK (COG-UK) Consortium | Rob Howes, The Lighthouse Lab in Cambridge and Alex Alderton, Roberto Amato, Sonia Goncalves, Ewan Harrison, David K. Jackson, Ian Johnston, Dominic Kwiatkowski, Cordelia Langford, John Sillitoe on behalf of the Wellcome Sanger Institute COVID-19 Surveillance Team |
| EPI_ISL_879173 | Lighthouse Lab in Alderley Park | Wellcome Sanger Institute for the COVID-19 Genomics UK (COG-UK) Consortium | Jacquelyn Wynn, Mairead Hyland, The Lighthouse Lab in Alderley Park and Alex Alderton, Roberto Amato, Sonia Goncalves, Ewan Harrison, David K. Jackson, Ian Johnston, Dominic Kwiatkowski, Cordelia Langford, John Sillitoe on behalf of the Wellcome Sanger Institute COVID-19 Surveillance Team |
| EPI_ISL_879174, EPI_ISL_879175, EPI_ISL_879176, EPI_ISL_879177, EPI_ISL_879178, EPI_ISL_879179, EPI_ISL_879180 | Lighthouse Lab in Cambridge | Wellcome Sanger Institute for the COVID-19 Genomics UK (COG-UK) Consortium | Rob Howes, The Lighthouse Lab in Cambridge and Alex Alderton, Roberto Amato, Sonia Goncalves, Ewan Harrison, David K. Jackson, Ian Johnston, Dominic Kwiatkowski, Cordelia Langford, John Sillitoe on behalf of the Wellcome Sanger Institute COVID-19 Surveillance Team |
| EPI_ISL_879181 | Lighthouse Lab in Alderley Park | Wellcome Sanger Institute for the COVID-19 Genomics UK (COG-UK) Consortium | Jacquelyn Wynn, Mairead Hyland, The Lighthouse Lab in Alderley Park and Alex Alderton, Roberto Amato, Sonia Goncalves, Ewan Harrison, David K. Jackson, Ian Johnston, Dominic Kwiatkowski, Cordelia Langford, John Sillitoe on behalf of the Wellcome Sanger Institute COVID-19 Surveillance Team |
| EPI_ISL_879182, EPI_ISL_879183, EPI_ISL_879184, EPI_ISL_879186 | Lighthouse Lab in Cambridge | Wellcome Sanger Institute for the COVID-19 Genomics UK (COG-UK) Consortium | Rob Howes, The Lighthouse Lab in Cambridge and Alex Alderton, Roberto Amato, Sonia Goncalves, Ewan Harrison, David K. Jackson, Ian Johnston, Dominic Kwiatkowski, Cordelia Langford, John Sillitoe on behalf of the Wellcome Sanger Institute COVID-19 Surveillance Team |
| EPI_ISL_879187, EPI_ISL_879188 | Lighthouse Lab in Alderley Park | Wellcome Sanger Institute for the COVID-19 Genomics UK (COG-UK) Consortium | Jacquelyn Wynn, Mairead Hyland, The Lighthouse Lab in Alderley Park and Alex Alderton, Roberto Amato, Sonia Goncalves, Ewan Harrison, David K. Jackson, Ian Johnston, Dominic Kwiatkowski, Cordelia Langford, John Sillitoe on behalf of the Wellcome Sanger Institute COVID-19 Surveillance Team |
| EPI_ISL_879189, EPI_ISL_879190 | Lighthouse Lab in Cambridge | Wellcome Sanger Institute for the COVID-19 Genomics UK (COG-UK) Consortium | Rob Howes, The Lighthouse Lab in Cambridge and Alex Alderton, Roberto Amato, Sonia Goncalves, Ewan Harrison, David K. Jackson, Ian Johnston, Dominic Kwiatkowski, Cordelia Langford, John Sillitoe on behalf of the Wellcome Sanger Institute COVID-19 Surveillance Team |
| EPI_ISL_879191 | Lighthouse Lab in Alderley Park | Wellcome Sanger Institute for the COVID-19 Genomics UK (COG-UK) Consortium | Jacquelyn Wynn, Mairead Hyland, The Lighthouse Lab in Alderley Park and Alex Alderton, Roberto Amato, Sonia Goncalves, Ewan Harrison, David K. Jackson, Ian Johnston, Dominic Kwiatkowski, Cordelia Langford, John Sillitoe on behalf of the Wellcome Sanger Institute COVID-19 Surveillance Team |
| EPI_ISL_879192 | Lighthouse Lab in Cambridge | Wellcome Sanger Institute for the COVID-19 Genomics UK (COG-UK) Consortium | Rob Howes, The Lighthouse Lab in Cambridge and Alex Alderton, Roberto Amato, Sonia Goncalves, Ewan Harrison, David K. Jackson, Ian Johnston, Dominic Kwiatkowski, Cordelia Langford, John Sillitoe on behalf of the Wellcome Sanger Institute COVID-19 Surveillance Team |
| EPI_ISL_879194, EPI_ISL_879195 | Lighthouse Lab in Alderley Park | Wellcome Sanger Institute for the COVID-19 Genomics UK (COG-UK) Consortium | Jacquelyn Wynn, Mairead Hyland, The Lighthouse Lab in Alderley Park and Alex Alderton, Roberto Amato, Sonia Goncalves, Ewan Harrison, David K. Jackson, Ian Johnston, Dominic Kwiatkowski, Cordelia Langford, John Sillitoe on behalf of the Wellcome Sanger Institute COVID-19 Surveillance Team |
| EPI_ISL_879196, EPI_ISL_879197, EPI_ISL_879198 | Lighthouse Lab in Cambridge | Wellcome Sanger Institute for the COVID-19 Genomics UK (COG-UK) Consortium | Rob Howes, The Lighthouse Lab in Cambridge and Alex Alderton, Roberto Amato, Sonia Goncalves, Ewan Harrison, David K. Jackson, Ian Johnston, Dominic Kwiatkowski, Cordelia Langford, John Sillitoe on behalf of the Wellcome Sanger Institute COVID-19 Surveillance Team |
| EPI_ISL_879199 | Lighthouse Lab in Alderley Park | Wellcome Sanger Institute for the COVID-19 Genomics UK (COG-UK) Consortium | Jacquelyn Wynn, Mairead Hyland, The Lighthouse Lab in Alderley Park and Alex Alderton, Roberto Amato, Sonia Goncalves, Ewan Harrison, David K. Jackson, Ian Johnston, Dominic Kwiatkowski, Cordelia Langford, John Sillitoe on behalf of the Wellcome Sanger Institute COVID-19 Surveillance Team |
| EPI_ISL_879200 | Lighthouse Lab in Cambridge | Wellcome Sanger Institute for the COVID-19 Genomics UK (COG-UK) Consortium | Rob Howes, The Lighthouse Lab in Cambridge and Alex Alderton, Roberto Amato, Sonia Goncalves, Ewan Harrison, David K. Jackson, Ian Johnston, Dominic Kwiatkowski, Cordelia Langford, John Sillitoe on behalf of the Wellcome Sanger Institute COVID-19 Surveillance Team |
| EPI_ISL_879201, EPI_ISL_879202, EPI_ISL_879203 | Lighthouse Lab in Alderley Park | Wellcome Sanger Institute for the COVID-19 Genomics UK (COG-UK) Consortium | Jacquelyn Wynn, Mairead Hyland, The Lighthouse Lab in Alderley Park and Alex Alderton, Roberto Amato, Sonia Goncalves, Ewan Harrison, David K. Jackson, Ian Johnston, Dominic Kwiatkowski, Cordelia Langford, John Sillitoe on behalf of the Wellcome Sanger Institute COVID-19 Surveillance Team |
| EPI_ISL_879204, EPI_ISL_879205 | Lighthouse Lab in Cambridge | Wellcome Sanger Institute for the COVID-19 Genomics UK (COG-UK) Consortium | Rob Howes, The Lighthouse Lab in Cambridge and Alex Alderton, Roberto Amato, Sonia Goncalves, Ewan Harrison, David K. Jackson, Ian Johnston, Dominic Kwiatkowski, Cordelia Langford, John Sillitoe on behalf of the Wellcome Sanger Institute COVID-19 Surveillance Team |
| EPI_ISL_879206, EPI_ISL_879207 | Lighthouse Lab in Alderley Park | Wellcome Sanger Institute for the COVID-19 Genomics UK (COG-UK) Consortium | Jacquelyn Wynn, Mairead Hyland, The Lighthouse Lab in Alderley Park and Alex Alderton, Roberto Amato, Sonia Goncalves, Ewan Harrison, David K. Jackson, Ian Johnston, Dominic Kwiatkowski, Cordelia Langford, John Sillitoe on behalf of the Wellcome Sanger Institute COVID-19 Surveillance Team |
| EPI_ISL_879208, EPI_ISL_879209, EPI_ISL_879210, EPI_ISL_879211 | Lighthouse Lab in Cambridge | Wellcome Sanger Institute for the COVID-19 Genomics UK (COG-UK) Consortium | Rob Howes, The Lighthouse Lab in Cambridge and Alex Alderton, Roberto Amato, Sonia Goncalves, Ewan Harrison, David K. Jackson, Ian Johnston, Dominic Kwiatkowski, Cordelia Langford, John Sillitoe on behalf of the Wellcome Sanger Institute COVID-19 Surveillance Team |
| EPI_ISL_879212 | Lighthouse Lab in Alderley Park | Wellcome Sanger Institute for the COVID-19 Genomics UK (COG-UK) Consortium | Jacquelyn Wynn, Mairead Hyland, The Lighthouse Lab in Alderley Park and Alex Alderton, Roberto Amato, Sonia Goncalves, Ewan Harrison, David K. Jackson, Ian Johnston, Dominic Kwiatkowski, Cordelia Langford, John Sillitoe on behalf of the Wellcome Sanger Institute COVID-19 Surveillance Team |
| EPI_ISL_879213 | Lighthouse Lab in Cambridge | Wellcome Sanger Institute for the COVID-19 Genomics UK | Rob Howes, The Lighthouse Lab in Cambridge and Alex Alderton, Roberto Amato, Sonia Goncalves, Ewan Harrison, David K. Jackson, Ian Johnston, |

| | | | |
|---|---|---|---|
| | | (COG-UK) Consortium | Dominic Kwiatkowski, Cordelia Langford, John Sillitoe on behalf of the Wellcome Sanger Institute COVID-19 Surveillance Team |
| EPI_ISL_879214 | Lighthouse Lab in Alderley Park | Wellcome Sanger Institute for the COVID-19 Genomics UK (COG-UK) Consortium | Jacquelyn Wynn, Mairead Hyland, The Lighthouse Lab in Alderley Park and Alex Alderton, Roberto Amato, Sonia Goncalves, Ewan Harrison, David K. Jackson, Ian Johnston, Dominic Kwiatkowski, Cordelia Langford, John Sillitoe on behalf of the Wellcome Sanger Institute COVID-19 Surveillance Team |
| EPI_ISL_879215, EPI_ISL_879216, EPI_ISL_879217 | Lighthouse Lab in Cambridge | Wellcome Sanger Institute for the COVID-19 Genomics UK (COG-UK) Consortium | Rob Howes, The Lighthouse Lab in Cambridge and Alex Alderton, Roberto Amato, Sonia Goncalves, Ewan Harrison, David K. Jackson, Ian Johnston, Dominic Kwiatkowski, Cordelia Langford, John Sillitoe on behalf of the Wellcome Sanger Institute COVID-19 Surveillance Team |
| EPI_ISL_879218, EPI_ISL_879219 | Lighthouse Lab in Alderley Park | Wellcome Sanger Institute for the COVID-19 Genomics UK (COG-UK) Consortium | Jacquelyn Wynn, Mairead Hyland, The Lighthouse Lab in Alderley Park and Alex Alderton, Roberto Amato, Sonia Goncalves, Ewan Harrison, David K. Jackson, Ian Johnston, Dominic Kwiatkowski, Cordelia Langford, John Sillitoe on behalf of the Wellcome Sanger Institute COVID-19 Surveillance Team |
| EPI_ISL_879221, EPI_ISL_879222, EPI_ISL_879223 | Lighthouse Lab in Cambridge | Wellcome Sanger Institute for the COVID-19 Genomics UK (COG-UK) Consortium | Rob Howes, The Lighthouse Lab in Cambridge and Alex Alderton, Roberto Amato, Sonia Goncalves, Ewan Harrison, David K. Jackson, Ian Johnston, Dominic Kwiatkowski, Cordelia Langford, John Sillitoe on behalf of the Wellcome Sanger Institute COVID-19 Surveillance Team |
| EPI_ISL_879224, EPI_ISL_879225, EPI_ISL_879226, EPI_ISL_879227 | Lighthouse Lab in Alderley Park | Wellcome Sanger Institute for the COVID-19 Genomics UK (COG-UK) Consortium | Jacquelyn Wynn, Mairead Hyland, The Lighthouse Lab in Alderley Park and Alex Alderton, Roberto Amato, Sonia Goncalves, Ewan Harrison, David K. Jackson, Ian Johnston, Dominic Kwiatkowski, Cordelia Langford, John Sillitoe on behalf of the Wellcome Sanger Institute COVID-19 Surveillance Team |
| EPI_ISL_879228 | Lighthouse Lab in Cambridge | Wellcome Sanger Institute for the COVID-19 Genomics UK (COG-UK) Consortium | Rob Howes, The Lighthouse Lab in Cambridge and Alex Alderton, Roberto Amato, Sonia Goncalves, Ewan Harrison, David K. Jackson, Ian Johnston, Dominic Kwiatkowski, Cordelia Langford, John Sillitoe on behalf of the Wellcome Sanger Institute COVID-19 Surveillance Team |
| EPI_ISL_879229, EPI_ISL_879230, EPI_ISL_879231 | Lighthouse Lab in Alderley Park | Wellcome Sanger Institute for the COVID-19 Genomics UK (COG-UK) Consortium | Jacquelyn Wynn, Mairead Hyland, The Lighthouse Lab in Alderley Park and Alex Alderton, Roberto Amato, Sonia Goncalves, Ewan Harrison, David K. Jackson, Ian Johnston, Dominic Kwiatkowski, Cordelia Langford, John Sillitoe on behalf of the Wellcome Sanger Institute COVID-19 Surveillance Team |
| EPI_ISL_879232, EPI_ISL_879233 | Lighthouse Lab in Cambridge | Wellcome Sanger Institute for the COVID-19 Genomics UK (COG-UK) Consortium | Rob Howes, The Lighthouse Lab in Cambridge and Alex Alderton, Roberto Amato, Sonia Goncalves, Ewan Harrison, David K. Jackson, Ian Johnston, Dominic Kwiatkowski, Cordelia Langford, John Sillitoe on behalf of the Wellcome Sanger Institute COVID-19 Surveillance Team |
| EPI_ISL_879234 | Lighthouse Lab in Alderley Park | Wellcome Sanger Institute for the COVID-19 Genomics UK (COG-UK) Consortium | Jacquelyn Wynn, Mairead Hyland, The Lighthouse Lab in Alderley Park and Alex Alderton, Roberto Amato, Sonia Goncalves, Ewan Harrison, David K. Jackson, Ian Johnston, Dominic Kwiatkowski, Cordelia Langford, John Sillitoe on behalf of the Wellcome Sanger Institute COVID-19 Surveillance Team |
| EPI_ISL_879236 | Lighthouse Lab in Cambridge | Wellcome Sanger Institute for the COVID-19 Genomics UK (COG-UK) Consortium | Rob Howes, The Lighthouse Lab in Cambridge and Alex Alderton, Roberto Amato, Sonia Goncalves, Ewan Harrison, David K. Jackson, Ian Johnston, Dominic Kwiatkowski, Cordelia Langford, John Sillitoe on behalf of the Wellcome Sanger Institute COVID-19 Surveillance Team |
| EPI_ISL_879237 | Lighthouse Lab in Alderley Park | Wellcome Sanger Institute for the COVID-19 Genomics UK (COG-UK) Consortium | Jacquelyn Wynn, Mairead Hyland, The Lighthouse Lab in Alderley Park and Alex Alderton, Roberto Amato, Sonia Goncalves, Ewan Harrison, David K. Jackson, Ian Johnston, Dominic Kwiatkowski, Cordelia Langford, John Sillitoe on behalf of the Wellcome Sanger Institute COVID-19 Surveillance Team |
| EPI_ISL_879238 | Lighthouse Lab in Cambridge | Wellcome Sanger Institute for the COVID-19 Genomics UK (COG-UK) Consortium | Rob Howes, The Lighthouse Lab in Cambridge and Alex Alderton, Roberto Amato, Sonia Goncalves, Ewan Harrison, David K. Jackson, Ian Johnston, Dominic Kwiatkowski, Cordelia Langford, John Sillitoe on behalf of the Wellcome Sanger Institute COVID-19 Surveillance Team |
| EPI_ISL_879239 | Lighthouse Lab in Alderley Park | Wellcome Sanger Institute for the COVID-19 Genomics UK (COG-UK) Consortium | Jacquelyn Wynn, Mairead Hyland, The Lighthouse Lab in Alderley Park and Alex Alderton, Roberto Amato, Sonia Goncalves, Ewan Harrison, David K. Jackson, Ian Johnston, Dominic Kwiatkowski, Cordelia Langford, John Sillitoe on behalf of the Wellcome Sanger Institute COVID-19 Surveillance Team |
| EPI_ISL_879240 | Lighthouse Lab in Cambridge | Wellcome Sanger Institute for the COVID-19 Genomics UK (COG-UK) Consortium | Rob Howes, The Lighthouse Lab in Cambridge and Alex Alderton, Roberto Amato, Sonia Goncalves, Ewan Harrison, David K. Jackson, Ian Johnston, Dominic Kwiatkowski, Cordelia Langford, John Sillitoe on behalf of the Wellcome Sanger Institute COVID-19 Surveillance Team |
| EPI_ISL_879241 | Lighthouse Lab in Alderley Park | Wellcome Sanger Institute for the COVID-19 Genomics UK (COG-UK) Consortium | Jacquelyn Wynn, Mairead Hyland, The Lighthouse Lab in Alderley Park and Alex Alderton, Roberto Amato, Sonia Goncalves, Ewan Harrison, David K. Jackson, Ian Johnston, Dominic Kwiatkowski, Cordelia Langford, John Sillitoe on behalf of the Wellcome Sanger Institute COVID-19 Surveillance Team |
| EPI_ISL_879242 | Lighthouse Lab in Cambridge | Wellcome Sanger Institute for the COVID-19 Genomics UK (COG-UK) Consortium | Rob Howes, The Lighthouse Lab in Cambridge and Alex Alderton, Roberto Amato, Sonia Goncalves, Ewan Harrison, David K. Jackson, Ian Johnston, Dominic Kwiatkowski, Cordelia Langford, John Sillitoe on behalf of the Wellcome Sanger Institute COVID-19 Surveillance Team |
| EPI_ISL_879243, EPI_ISL_879244 | Lighthouse Lab in Alderley Park | Wellcome Sanger Institute for the COVID-19 Genomics UK (COG-UK) Consortium | Jacquelyn Wynn, Mairead Hyland, The Lighthouse Lab in Alderley Park and Alex Alderton, Roberto Amato, Sonia Goncalves, Ewan Harrison, David K. Jackson, Ian Johnston, Dominic Kwiatkowski, Cordelia Langford, John Sillitoe on behalf of the Wellcome Sanger Institute COVID-19 Surveillance Team |
| EPI_ISL_879245, EPI_ISL_879246 | Lighthouse Lab in Cambridge | Wellcome Sanger Institute for the COVID-19 Genomics UK (COG-UK) Consortium | Rob Howes, The Lighthouse Lab in Cambridge and Alex Alderton, Roberto Amato, Sonia Goncalves, Ewan Harrison, David K. Jackson, Ian Johnston, Dominic Kwiatkowski, Cordelia Langford, John Sillitoe on behalf of the Wellcome Sanger Institute COVID-19 Surveillance Team |
| EPI_ISL_879247, EPI_ISL_879248, EPI_ISL_879249, EPI_ISL_879250 | Lighthouse Lab in Alderley Park | Wellcome Sanger Institute for the COVID-19 Genomics UK (COG-UK) Consortium | Jacquelyn Wynn, Mairead Hyland, The Lighthouse Lab in Alderley Park and Alex Alderton, Roberto Amato, Sonia Goncalves, Ewan Harrison, David K. Jackson, Ian Johnston, Dominic Kwiatkowski, Cordelia Langford, John Sillitoe on behalf of the Wellcome Sanger Institute COVID-19 Surveillance Team |
| EPI_ISL_879251 | Lighthouse Lab in Cambridge | Wellcome Sanger Institute for the COVID-19 Genomics UK (COG-UK) Consortium | Rob Howes, The Lighthouse Lab in Cambridge and Alex Alderton, Roberto Amato, Sonia Goncalves, Ewan Harrison, David K. Jackson, Ian Johnston, Dominic Kwiatkowski, Cordelia Langford, John Sillitoe on behalf of the Wellcome Sanger Institute COVID-19 Surveillance Team |
| EPI_ISL_879252, EPI_ISL_879254 | Lighthouse Lab in Alderley Park | Wellcome Sanger Institute for the COVID-19 Genomics UK (COG-UK) Consortium | Jacquelyn Wynn, Mairead Hyland, The Lighthouse Lab in Alderley Park and Alex Alderton, Roberto Amato, Sonia Goncalves, Ewan Harrison, David K. Jackson, Ian Johnston, Dominic Kwiatkowski, Cordelia Langford, John Sillitoe on behalf of the Wellcome Sanger Institute COVID-19 Surveillance Team |
| EPI_ISL_879255 | Lighthouse Lab in Cambridge | Wellcome Sanger Institute for the COVID-19 Genomics UK (COG-UK) Consortium | Rob Howes, The Lighthouse Lab in Cambridge and Alex Alderton, Roberto Amato, Sonia Goncalves, Ewan Harrison, David K. Jackson, Ian Johnston, Dominic Kwiatkowski, Cordelia Langford, John Sillitoe on behalf of the Wellcome Sanger Institute COVID-19 Surveillance Team |
| EPI_ISL_879256 | Lighthouse Lab in Alderley Park | Wellcome Sanger Institute for the COVID-19 Genomics UK (COG-UK) Consortium | Jacquelyn Wynn, Mairead Hyland, The Lighthouse Lab in Alderley Park and Alex Alderton, Roberto Amato, Sonia Goncalves, Ewan Harrison, David K. Jackson, Ian Johnston, Dominic Kwiatkowski, Cordelia Langford, John Sillitoe on behalf of the Wellcome Sanger Institute COVID-19 Surveillance Team |
| EPI_ISL_879257 | Lighthouse Lab in Cambridge | Wellcome Sanger Institute for the COVID-19 Genomics UK (COG-UK) Consortium | Rob Howes, The Lighthouse Lab in Cambridge and Alex Alderton, Roberto Amato, Sonia Goncalves, Ewan Harrison, David K. Jackson, Ian Johnston, Dominic Kwiatkowski, Cordelia Langford, John Sillitoe on behalf of the Wellcome Sanger Institute COVID-19 Surveillance Team |
| EPI_ISL_879258, EPI_ISL_879259 | Lighthouse Lab in Alderley Park | Wellcome Sanger Institute for the COVID-19 Genomics UK (COG-UK) Consortium | Jacquelyn Wynn, Mairead Hyland, The Lighthouse Lab in Alderley Park and Alex Alderton, Roberto Amato, Sonia Goncalves, Ewan Harrison, David K. Jackson, Ian Johnston, Dominic Kwiatkowski, Cordelia Langford, John Sillitoe on behalf of the Wellcome Sanger Institute COVID-19 Surveillance Team |
| EPI_ISL_879260, EPI_ISL_879261 | Lighthouse Lab in Cambridge | Wellcome Sanger Institute for the COVID-19 Genomics UK (COG-UK) Consortium | Rob Howes, The Lighthouse Lab in Cambridge and Alex Alderton, Roberto Amato, Sonia Goncalves, Ewan Harrison, David K. Jackson, Ian Johnston, Dominic Kwiatkowski, Cordelia Langford, John Sillitoe on behalf of the Wellcome Sanger Institute COVID-19 Surveillance Team |
| EPI_ISL_879262, EPI_ISL_879263 | Lighthouse Lab in Alderley Park | Wellcome Sanger Institute for the COVID-19 Genomics UK (COG-UK) Consortium | Jacquelyn Wynn, Mairead Hyland, The Lighthouse Lab in Alderley Park and Alex Alderton, Roberto Amato, Sonia Goncalves, Ewan Harrison, David K. Jackson, Ian Johnston, Dominic Kwiatkowski, Cordelia Langford, John Sillitoe on behalf of the Wellcome Sanger Institute COVID-19 Surveillance Team |
| EPI_ISL_879264 | Lighthouse Lab in Cambridge | Wellcome Sanger Institute for the COVID-19 Genomics UK (COG-UK) Consortium | Rob Howes, The Lighthouse Lab in Cambridge and Alex Alderton, Roberto Amato, Sonia Goncalves, Ewan Harrison, David K. Jackson, Ian Johnston, Dominic Kwiatkowski, Cordelia Langford, John Sillitoe on behalf of the Wellcome Sanger Institute COVID-19 Surveillance Team |
| EPI_ISL_879265, EPI_ISL_879266 | Lighthouse Lab in Alderley Park | Wellcome Sanger Institute for the COVID-19 Genomics UK (COG-UK) Consortium | Jacquelyn Wynn, Mairead Hyland, The Lighthouse Lab in Alderley Park and Alex Alderton, Roberto Amato, Sonia Goncalves, Ewan Harrison, David K. Jackson, Ian Johnston, Dominic Kwiatkowski, Cordelia Langford, John Sillitoe on behalf of the Wellcome Sanger Institute COVID-19 Surveillance Team |
| EPI_ISL_879267 | Lighthouse Lab in Cambridge | Wellcome Sanger Institute for the COVID-19 Genomics UK (COG-UK) Consortium | Rob Howes, The Lighthouse Lab in Cambridge and Alex Alderton, Roberto Amato, Sonia Goncalves, Ewan Harrison, David K. Jackson, Ian Johnston, Dominic Kwiatkowski, Cordelia Langford, John Sillitoe on behalf of the Wellcome Sanger Institute COVID-19 Surveillance Team |
| EPI_ISL_879268 | Lighthouse Lab in Alderley Park | Wellcome Sanger Institute for the COVID-19 Genomics UK (COG-UK) Consortium | Jacquelyn Wynn, Mairead Hyland, The Lighthouse Lab in Alderley Park and Alex Alderton, Roberto Amato, Sonia Goncalves, Ewan Harrison, David K. Jackson, Ian Johnston, Dominic Kwiatkowski, Cordelia Langford, John Sillitoe on behalf of the Wellcome Sanger Institute COVID-19 Surveillance Team |
| EPI_ISL_879269 | Lighthouse Lab in Cambridge | Wellcome Sanger Institute for the COVID-19 Genomics UK (COG-UK) Consortium | Rob Howes, The Lighthouse Lab in Cambridge and Alex Alderton, Roberto Amato, Sonia Goncalves, Ewan Harrison, David K. Jackson, Ian Johnston, Dominic Kwiatkowski, Cordelia Langford, John Sillitoe on behalf of the Wellcome Sanger Institute COVID-19 Surveillance Team |
| EPI_ISL_879270, EPI_ISL_879271 | Lighthouse Lab in Alderley Park | Wellcome Sanger Institute for the COVID-19 Genomics UK (COG-UK) Consortium | Jacquelyn Wynn, Mairead Hyland, The Lighthouse Lab in Alderley Park and Alex Alderton, Roberto Amato, Sonia Goncalves, Ewan Harrison, David K. Jackson, Ian Johnston, Dominic Kwiatkowski, Cordelia Langford, John Sillitoe on behalf of the Wellcome Sanger Institute COVID-19 Surveillance Team |
| EPI_ISL_879272 | Lighthouse Lab in Cambridge | Wellcome Sanger Institute for the COVID-19 Genomics UK (COG-UK) Consortium | Rob Howes, The Lighthouse Lab in Cambridge and Alex Alderton, Roberto Amato, Sonia Goncalves, Ewan Harrison, David K. Jackson, Ian Johnston, Dominic Kwiatkowski, Cordelia Langford, John Sillitoe on behalf of the Wellcome Sanger Institute COVID-19 Surveillance Team |
| EPI_ISL_879273 | Lighthouse Lab in Alderley Park | Wellcome Sanger Institute for the COVID-19 Genomics UK | Jacquelyn Wynn, Mairead Hyland, The Lighthouse Lab in Alderley Park and Alex Alderton, Roberto Amato, Sonia Goncalves, Ewan Harrison, David K. |

| | | | |
|---|---|---|---|
| | | (COG-UK) Consortium | Jackson, Ian Johnston, Dominic Kwiatkowski, Cordelia Langford, John Sillitoe on behalf of the Wellcome Sanger Institute COVID-19 Surveillance Team |
| EPI_ISL_879275 | Lighthouse Lab in Cambridge | Wellcome Sanger Institute for the COVID-19 Genomics UK (COG-UK) Consortium | Rob Howes, The Lighthouse Lab in Cambridge and Alex Alderton, Roberto Amato, Sonia Goncalves, Ewan Harrison, David K. Jackson, Ian Johnston, Dominic Kwiatkowski, Cordelia Langford, John Sillitoe on behalf of the Wellcome Sanger Institute COVID-19 Surveillance Team |
| EPI_ISL_879276, EPI_ISL_879277, EPI_ISL_879279 | Lighthouse Lab in Alderley Park | Wellcome Sanger Institute for the COVID-19 Genomics UK (COG-UK) Consortium | Jacquelyn Wynn, Mairead Hyland, The Lighthouse Lab in Alderley Park and Alex Alderton, Roberto Amato, Sonia Goncalves, Ewan Harrison, David K. Jackson, Ian Johnston, Dominic Kwiatkowski, Cordelia Langford, John Sillitoe on behalf of the Wellcome Sanger Institute COVID-19 Surveillance Team |
| EPI_ISL_879280, EPI_ISL_879281, EPI_ISL_879282, EPI_ISL_879283 | Lighthouse Lab in Cambridge | Wellcome Sanger Institute for the COVID-19 Genomics UK (COG-UK) Consortium | Rob Howes, The Lighthouse Lab in Cambridge and Alex Alderton, Roberto Amato, Sonia Goncalves, Ewan Harrison, David K. Jackson, Ian Johnston, Dominic Kwiatkowski, Cordelia Langford, John Sillitoe on behalf of the Wellcome Sanger Institute COVID-19 Surveillance Team |
| EPI_ISL_879284 | Lighthouse Lab in Alderley Park | Wellcome Sanger Institute for the COVID-19 Genomics UK (COG-UK) Consortium | Jacquelyn Wynn, Mairead Hyland, The Lighthouse Lab in Alderley Park and Alex Alderton, Roberto Amato, Sonia Goncalves, Ewan Harrison, David K. Jackson, Ian Johnston, Dominic Kwiatkowski, Cordelia Langford, John Sillitoe on behalf of the Wellcome Sanger Institute COVID-19 Surveillance Team |
| EPI_ISL_879285 | Lighthouse Lab in Cambridge | Wellcome Sanger Institute for the COVID-19 Genomics UK (COG-UK) Consortium | Rob Howes, The Lighthouse Lab in Cambridge and Alex Alderton, Roberto Amato, Sonia Goncalves, Ewan Harrison, David K. Jackson, Ian Johnston, Dominic Kwiatkowski, Cordelia Langford, John Sillitoe on behalf of the Wellcome Sanger Institute COVID-19 Surveillance Team |
| EPI_ISL_879286 | Lighthouse Lab in Alderley Park | Wellcome Sanger Institute for the COVID-19 Genomics UK (COG-UK) Consortium | Jacquelyn Wynn, Mairead Hyland, The Lighthouse Lab in Alderley Park and Alex Alderton, Roberto Amato, Sonia Goncalves, Ewan Harrison, David K. Jackson, Ian Johnston, Dominic Kwiatkowski, Cordelia Langford, John Sillitoe on behalf of the Wellcome Sanger Institute COVID-19 Surveillance Team |
| EPI_ISL_879287, EPI_ISL_879288, EPI_ISL_879289, EPI_ISL_879291, EPI_ISL_879292, EPI_ISL_879293, EPI_ISL_879294, EPI_ISL_879295 | Lighthouse Lab in Cambridge | Wellcome Sanger Institute for the COVID-19 Genomics UK (COG-UK) Consortium | Rob Howes, The Lighthouse Lab in Cambridge and Alex Alderton, Roberto Amato, Sonia Goncalves, Ewan Harrison, David K. Jackson, Ian Johnston, Dominic Kwiatkowski, Cordelia Langford, John Sillitoe on behalf of the Wellcome Sanger Institute COVID-19 Surveillance Team |
| EPI_ISL_879296, EPI_ISL_879297, EPI_ISL_879298, EPI_ISL_879299, EPI_ISL_879300 | Lighthouse Lab in Alderley Park | Wellcome Sanger Institute for the COVID-19 Genomics UK (COG-UK) Consortium | Jacquelyn Wynn, Mairead Hyland, The Lighthouse Lab in Alderley Park and Alex Alderton, Roberto Amato, Sonia Goncalves, Ewan Harrison, David K. Jackson, Ian Johnston, Dominic Kwiatkowski, Cordelia Langford, John Sillitoe on behalf of the Wellcome Sanger Institute COVID-19 Surveillance Team |
| EPI_ISL_879301 | Lighthouse Lab in Cambridge | Wellcome Sanger Institute for the COVID-19 Genomics UK (COG-UK) Consortium | Rob Howes, The Lighthouse Lab in Cambridge and Alex Alderton, Roberto Amato, Sonia Goncalves, Ewan Harrison, David K. Jackson, Ian Johnston, Dominic Kwiatkowski, Cordelia Langford, John Sillitoe on behalf of the Wellcome Sanger Institute COVID-19 Surveillance Team |
| EPI_ISL_879302, EPI_ISL_879303 | Lighthouse Lab in Alderley Park | Wellcome Sanger Institute for the COVID-19 Genomics UK (COG-UK) Consortium | Jacquelyn Wynn, Mairead Hyland, The Lighthouse Lab in Alderley Park and Alex Alderton, Roberto Amato, Sonia Goncalves, Ewan Harrison, David K. Jackson, Ian Johnston, Dominic Kwiatkowski, Cordelia Langford, John Sillitoe on behalf of the Wellcome Sanger Institute COVID-19 Surveillance Team |
| EPI_ISL_879304 | Lighthouse Lab in Cambridge | Wellcome Sanger Institute for the COVID-19 Genomics UK (COG-UK) Consortium | Rob Howes, The Lighthouse Lab in Cambridge and Alex Alderton, Roberto Amato, Sonia Goncalves, Ewan Harrison, David K. Jackson, Ian Johnston, Dominic Kwiatkowski, Cordelia Langford, John Sillitoe on behalf of the Wellcome Sanger Institute COVID-19 Surveillance Team |
| EPI_ISL_879305, EPI_ISL_879306, EPI_ISL_879307, EPI_ISL_879308 | Lighthouse Lab in Alderley Park | Wellcome Sanger Institute for the COVID-19 Genomics UK (COG-UK) Consortium | Jacquelyn Wynn, Mairead Hyland, The Lighthouse Lab in Alderley Park and Alex Alderton, Roberto Amato, Sonia Goncalves, Ewan Harrison, David K. Jackson, Ian Johnston, Dominic Kwiatkowski, Cordelia Langford, John Sillitoe on behalf of the Wellcome Sanger Institute COVID-19 Surveillance Team |
| EPI_ISL_879309, EPI_ISL_879310, EPI_ISL_879311, EPI_ISL_879312 | Lighthouse Lab in Cambridge | Wellcome Sanger Institute for the COVID-19 Genomics UK (COG-UK) Consortium | Rob Howes, The Lighthouse Lab in Cambridge and Alex Alderton, Roberto Amato, Sonia Goncalves, Ewan Harrison, David K. Jackson, Ian Johnston, Dominic Kwiatkowski, Cordelia Langford, John Sillitoe on behalf of the Wellcome Sanger Institute COVID-19 Surveillance Team |
| EPI_ISL_879313 | Lighthouse Lab in Alderley Park | Wellcome Sanger Institute for the COVID-19 Genomics UK (COG-UK) Consortium | Jacquelyn Wynn, Mairead Hyland, The Lighthouse Lab in Alderley Park and Alex Alderton, Roberto Amato, Sonia Goncalves, Ewan Harrison, David K. Jackson, Ian Johnston, Dominic Kwiatkowski, Cordelia Langford, John Sillitoe on behalf of the Wellcome Sanger Institute COVID-19 Surveillance Team |
| EPI_ISL_879315 | Lighthouse Lab in Cambridge | Wellcome Sanger Institute for the COVID-19 Genomics UK (COG-UK) Consortium | Rob Howes, The Lighthouse Lab in Cambridge and Alex Alderton, Roberto Amato, Sonia Goncalves, Ewan Harrison, David K. Jackson, Ian Johnston, Dominic Kwiatkowski, Cordelia Langford, John Sillitoe on behalf of the Wellcome Sanger Institute COVID-19 Surveillance Team |
| EPI_ISL_879316 | Lighthouse Lab in Alderley Park | Wellcome Sanger Institute for the COVID-19 Genomics UK (COG-UK) Consortium | Jacquelyn Wynn, Mairead Hyland, The Lighthouse Lab in Alderley Park and Alex Alderton, Roberto Amato, Sonia Goncalves, Ewan Harrison, David K. Jackson, Ian Johnston, Dominic Kwiatkowski, Cordelia Langford, John Sillitoe on behalf of the Wellcome Sanger Institute COVID-19 Surveillance Team |
| EPI_ISL_879317 | Lighthouse Lab in Cambridge | Wellcome Sanger Institute for the COVID-19 Genomics UK (COG-UK) Consortium | Rob Howes, The Lighthouse Lab in Cambridge and Alex Alderton, Roberto Amato, Sonia Goncalves, Ewan Harrison, David K. Jackson, Ian Johnston, Dominic Kwiatkowski, Cordelia Langford, John Sillitoe on behalf of the Wellcome Sanger Institute COVID-19 Surveillance Team |
| EPI_ISL_879318 | Lighthouse Lab in Alderley Park | Wellcome Sanger Institute for the COVID-19 Genomics UK (COG-UK) Consortium | Jacquelyn Wynn, Mairead Hyland, The Lighthouse Lab in Alderley Park and Alex Alderton, Roberto Amato, Sonia Goncalves, Ewan Harrison, David K. Jackson, Ian Johnston, Dominic Kwiatkowski, Cordelia Langford, John Sillitoe on behalf of the Wellcome Sanger Institute COVID-19 Surveillance Team |
| EPI_ISL_879319, EPI_ISL_879320, EPI_ISL_879321 | Lighthouse Lab in Cambridge | Wellcome Sanger Institute for the COVID-19 Genomics UK (COG-UK) Consortium | Rob Howes, The Lighthouse Lab in Cambridge and Alex Alderton, Roberto Amato, Sonia Goncalves, Ewan Harrison, David K. Jackson, Ian Johnston, Dominic Kwiatkowski, Cordelia Langford, John Sillitoe on behalf of the Wellcome Sanger Institute COVID-19 Surveillance Team |
| EPI_ISL_879323, EPI_ISL_879324 | Lighthouse Lab in Alderley Park | Wellcome Sanger Institute for the COVID-19 Genomics UK (COG-UK) Consortium | Jacquelyn Wynn, Mairead Hyland, The Lighthouse Lab in Alderley Park and Alex Alderton, Roberto Amato, Sonia Goncalves, Ewan Harrison, David K. Jackson, Ian Johnston, Dominic Kwiatkowski, Cordelia Langford, John Sillitoe on behalf of the Wellcome Sanger Institute COVID-19 Surveillance Team |
| EPI_ISL_879325, EPI_ISL_879326, EPI_ISL_879327, EPI_ISL_879328 | Lighthouse Lab in Cambridge | Wellcome Sanger Institute for the COVID-19 Genomics UK (COG-UK) Consortium | Rob Howes, The Lighthouse Lab in Cambridge and Alex Alderton, Roberto Amato, Sonia Goncalves, Ewan Harrison, David K. Jackson, Ian Johnston, Dominic Kwiatkowski, Cordelia Langford, John Sillitoe on behalf of the Wellcome Sanger Institute COVID-19 Surveillance Team |
| EPI_ISL_879329 | Lighthouse Lab in Alderley Park | Wellcome Sanger Institute for the COVID-19 Genomics UK (COG-UK) Consortium | Jacquelyn Wynn, Mairead Hyland, The Lighthouse Lab in Alderley Park and Alex Alderton, Roberto Amato, Sonia Goncalves, Ewan Harrison, David K. Jackson, Ian Johnston, Dominic Kwiatkowski, Cordelia Langford, John Sillitoe on behalf of the Wellcome Sanger Institute COVID-19 Surveillance Team |
| EPI_ISL_879330, EPI_ISL_879332 | Lighthouse Lab in Cambridge | Wellcome Sanger Institute for the COVID-19 Genomics UK (COG-UK) Consortium | Rob Howes, The Lighthouse Lab in Cambridge and Alex Alderton, Roberto Amato, Sonia Goncalves, Ewan Harrison, David K. Jackson, Ian Johnston, Dominic Kwiatkowski, Cordelia Langford, John Sillitoe on behalf of the Wellcome Sanger Institute COVID-19 Surveillance Team |
| EPI_ISL_879333 | Lighthouse Lab in Alderley Park | Wellcome Sanger Institute for the COVID-19 Genomics UK (COG-UK) Consortium | Jacquelyn Wynn, Mairead Hyland, The Lighthouse Lab in Alderley Park and Alex Alderton, Roberto Amato, Sonia Goncalves, Ewan Harrison, David K. Jackson, Ian Johnston, Dominic Kwiatkowski, Cordelia Langford, John Sillitoe on behalf of the Wellcome Sanger Institute COVID-19 Surveillance Team |
| EPI_ISL_879334, EPI_ISL_879335, EPI_ISL_879336, EPI_ISL_879337 | Lighthouse Lab in Cambridge | Wellcome Sanger Institute for the COVID-19 Genomics UK (COG-UK) Consortium | Rob Howes, The Lighthouse Lab in Cambridge and Alex Alderton, Roberto Amato, Sonia Goncalves, Ewan Harrison, David K. Jackson, Ian Johnston, Dominic Kwiatkowski, Cordelia Langford, John Sillitoe on behalf of the Wellcome Sanger Institute COVID-19 Surveillance Team |
| EPI_ISL_879339, EPI_ISL_879340, EPI_ISL_879341, EPI_ISL_879342, EPI_ISL_879343, EPI_ISL_879345, EPI_ISL_879346, EPI_ISL_879347, EPI_ISL_879348 | Lighthouse Lab in Alderley Park | Wellcome Sanger Institute for the COVID-19 Genomics UK (COG-UK) Consortium | Jacquelyn Wynn, Mairead Hyland, The Lighthouse Lab in Alderley Park and Alex Alderton, Roberto Amato, Sonia Goncalves, Ewan Harrison, David K. Jackson, Ian Johnston, Dominic Kwiatkowski, Cordelia Langford, John Sillitoe on behalf of the Wellcome Sanger Institute COVID-19 Surveillance Team |
| EPI_ISL_879349, EPI_ISL_879350 | Lighthouse Lab in Cambridge | Wellcome Sanger Institute for the COVID-19 Genomics UK (COG-UK) Consortium | Rob Howes, The Lighthouse Lab in Cambridge and Alex Alderton, Roberto Amato, Sonia Goncalves, Ewan Harrison, David K. Jackson, Ian Johnston, Dominic Kwiatkowski, Cordelia Langford, John Sillitoe on behalf of the Wellcome Sanger Institute COVID-19 Surveillance Team |
| EPI_ISL_879351 | Lighthouse Lab in Alderley Park | Wellcome Sanger Institute for the COVID-19 Genomics UK (COG-UK) Consortium | Jacquelyn Wynn, Mairead Hyland, The Lighthouse Lab in Alderley Park and Alex Alderton, Roberto Amato, Sonia Goncalves, Ewan Harrison, David K. Jackson, Ian Johnston, Dominic Kwiatkowski, Cordelia Langford, John Sillitoe on behalf of the Wellcome Sanger Institute COVID-19 Surveillance Team |
| EPI_ISL_879352, EPI_ISL_879353 | Lighthouse Lab in Cambridge | Wellcome Sanger Institute for the COVID-19 Genomics UK (COG-UK) Consortium | Rob Howes, The Lighthouse Lab in Cambridge and Alex Alderton, Roberto Amato, Sonia Goncalves, Ewan Harrison, David K. Jackson, Ian Johnston, Dominic Kwiatkowski, Cordelia Langford, John Sillitoe on behalf of the Wellcome Sanger Institute COVID-19 Surveillance Team |
| EPI_ISL_879354 | Lighthouse Lab in Alderley Park | Wellcome Sanger Institute for the COVID-19 Genomics UK (COG-UK) Consortium | Jacquelyn Wynn, Mairead Hyland, The Lighthouse Lab in Alderley Park and Alex Alderton, Roberto Amato, Sonia Goncalves, Ewan Harrison, David K. Jackson, Ian Johnston, Dominic Kwiatkowski, Cordelia Langford, John Sillitoe on behalf of the Wellcome Sanger Institute COVID-19 Surveillance Team |
| EPI_ISL_879355 | Lighthouse Lab in Cambridge | Wellcome Sanger Institute for the COVID-19 Genomics UK (COG-UK) Consortium | Rob Howes, The Lighthouse Lab in Cambridge and Alex Alderton, Roberto Amato, Sonia Goncalves, Ewan Harrison, David K. Jackson, Ian Johnston, Dominic Kwiatkowski, Cordelia Langford, John Sillitoe on behalf of the Wellcome Sanger Institute COVID-19 Surveillance Team |
| EPI_ISL_879356, EPI_ISL_879357, EPI_ISL_879358, EPI_ISL_879359, EPI_ISL_879360, EPI_ISL_879361, | Lighthouse Lab in Alderley Park | Wellcome Sanger Institute for the COVID-19 Genomics UK (COG-UK) Consortium | Jacquelyn Wynn, Mairead Hyland, The Lighthouse Lab in Alderley Park and Alex Alderton, Roberto Amato, Sonia Goncalves, Ewan Harrison, David K. Jackson, Ian Johnston, Dominic Kwiatkowski, Cordelia Langford, John Sillitoe on behalf of the Wellcome Sanger Institute COVID-19 Surveillance Team |

| | | | |
|---|---|---|---|
| EPI_ISL_879362 | | | |
| EPI_ISL_879364, EPI_ISL_879365, EPI_ISL_879366 | Lighthouse Lab in Cambridge | Wellcome Sanger Institute for the COVID-19 Genomics UK (COG-UK) Consortium | Rob Howes, The Lighthouse Lab in Cambridge and Alex Alderton, Roberto Amato, Sonia Goncalves, Ewan Harrison, David K. Jackson, Ian Johnston, Dominic Kwiatkowski, Cordelia Langford, John Sillitoe on behalf of the Wellcome Sanger Institute COVID-19 Surveillance Team |
| EPI_ISL_879367 | Lighthouse Lab in Alderley Park | Wellcome Sanger Institute for the COVID-19 Genomics UK (COG-UK) Consortium | Jacquelyn Wynn, Mairead Hyland, The Lighthouse Lab in Alderley Park and Alex Alderton, Roberto Amato, Sonia Goncalves, Ewan Harrison, David K. Jackson, Ian Johnston, Dominic Kwiatkowski, Cordelia Langford, John Sillitoe on behalf of the Wellcome Sanger Institute COVID-19 Surveillance Team |
| EPI_ISL_879368, EPI_ISL_879369, EPI_ISL_879370 | Lighthouse Lab in Cambridge | Wellcome Sanger Institute for the COVID-19 Genomics UK (COG-UK) Consortium | Rob Howes, The Lighthouse Lab in Cambridge and Alex Alderton, Roberto Amato, Sonia Goncalves, Ewan Harrison, David K. Jackson, Ian Johnston, Dominic Kwiatkowski, Cordelia Langford, John Sillitoe on behalf of the Wellcome Sanger Institute COVID-19 Surveillance Team |
| EPI_ISL_879371 | Lighthouse Lab in Alderley Park | Wellcome Sanger Institute for the COVID-19 Genomics UK (COG-UK) Consortium | Jacquelyn Wynn, Mairead Hyland, The Lighthouse Lab in Alderley Park and Alex Alderton, Roberto Amato, Sonia Goncalves, Ewan Harrison, David K. Jackson, Ian Johnston, Dominic Kwiatkowski, Cordelia Langford, John Sillitoe on behalf of the Wellcome Sanger Institute COVID-19 Surveillance Team |
| EPI_ISL_879372, EPI_ISL_879373 | Lighthouse Lab in Cambridge | Wellcome Sanger Institute for the COVID-19 Genomics UK (COG-UK) Consortium | Rob Howes, The Lighthouse Lab in Cambridge and Alex Alderton, Roberto Amato, Sonia Goncalves, Ewan Harrison, David K. Jackson, Ian Johnston, Dominic Kwiatkowski, Cordelia Langford, John Sillitoe on behalf of the Wellcome Sanger Institute COVID-19 Surveillance Team |
| EPI_ISL_879374, EPI_ISL_879375 | Lighthouse Lab in Alderley Park | Wellcome Sanger Institute for the COVID-19 Genomics UK (COG-UK) Consortium | Jacquelyn Wynn, Mairead Hyland, The Lighthouse Lab in Alderley Park and Alex Alderton, Roberto Amato, Sonia Goncalves, Ewan Harrison, David K. Jackson, Ian Johnston, Dominic Kwiatkowski, Cordelia Langford, John Sillitoe on behalf of the Wellcome Sanger Institute COVID-19 Surveillance Team |
| EPI_ISL_879376, EPI_ISL_879381 | Lighthouse Lab in Cambridge | Wellcome Sanger Institute for the COVID-19 Genomics UK (COG-UK) Consortium | Rob Howes, The Lighthouse Lab in Cambridge and Alex Alderton, Roberto Amato, Sonia Goncalves, Ewan Harrison, David K. Jackson, Ian Johnston, Dominic Kwiatkowski, Cordelia Langford, John Sillitoe on behalf of the Wellcome Sanger Institute COVID-19 Surveillance Team |
| EPI_ISL_879382 | Lighthouse Lab in Alderley Park | Wellcome Sanger Institute for the COVID-19 Genomics UK (COG-UK) Consortium | Jacquelyn Wynn, Mairead Hyland, The Lighthouse Lab in Alderley Park and Alex Alderton, Roberto Amato, Sonia Goncalves, Ewan Harrison, David K. Jackson, Ian Johnston, Dominic Kwiatkowski, Cordelia Langford, John Sillitoe on behalf of the Wellcome Sanger Institute COVID-19 Surveillance Team |
| EPI_ISL_879383, EPI_ISL_879385 | Lighthouse Lab in Cambridge | Wellcome Sanger Institute for the COVID-19 Genomics UK (COG-UK) Consortium | Rob Howes, The Lighthouse Lab in Cambridge and Alex Alderton, Roberto Amato, Sonia Goncalves, Ewan Harrison, David K. Jackson, Ian Johnston, Dominic Kwiatkowski, Cordelia Langford, John Sillitoe on behalf of the Wellcome Sanger Institute COVID-19 Surveillance Team |
| EPI_ISL_879386 | Lighthouse Lab in Alderley Park | Wellcome Sanger Institute for the COVID-19 Genomics UK (COG-UK) Consortium | Jacquelyn Wynn, Mairead Hyland, The Lighthouse Lab in Alderley Park and Alex Alderton, Roberto Amato, Sonia Goncalves, Ewan Harrison, David K. Jackson, Ian Johnston, Dominic Kwiatkowski, Cordelia Langford, John Sillitoe on behalf of the Wellcome Sanger Institute COVID-19 Surveillance Team |
| EPI_ISL_879387, EPI_ISL_879389, EPI_ISL_879390, EPI_ISL_879391 | Lighthouse Lab in Cambridge | Wellcome Sanger Institute for the COVID-19 Genomics UK (COG-UK) Consortium | Rob Howes, The Lighthouse Lab in Cambridge and Alex Alderton, Roberto Amato, Sonia Goncalves, Ewan Harrison, David K. Jackson, Ian Johnston, Dominic Kwiatkowski, Cordelia Langford, John Sillitoe on behalf of the Wellcome Sanger Institute COVID-19 Surveillance Team |
| EPI_ISL_879392, EPI_ISL_879393 | Lighthouse Lab in Alderley Park | Wellcome Sanger Institute for the COVID-19 Genomics UK (COG-UK) Consortium | Jacquelyn Wynn, Mairead Hyland, The Lighthouse Lab in Alderley Park and Alex Alderton, Roberto Amato, Sonia Goncalves, Ewan Harrison, David K. Jackson, Ian Johnston, Dominic Kwiatkowski, Cordelia Langford, John Sillitoe on behalf of the Wellcome Sanger Institute COVID-19 Surveillance Team |
| EPI_ISL_879394, EPI_ISL_879398, EPI_ISL_879400, EPI_ISL_879401, EPI_ISL_879402, EPI_ISL_879403, EPI_ISL_879405, EPI_ISL_879406, EPI_ISL_879407, EPI_ISL_879408, EPI_ISL_879410, EPI_ISL_879411, EPI_ISL_879414 | | | |
| see above | Lighthouse Lab in Cambridge | Wellcome Sanger Institute for the COVID-19 Genomics UK (COG-UK) Consortium | Rob Howes, The Lighthouse Lab in Cambridge and Alex Alderton, Roberto Amato, Sonia Goncalves, Ewan Harrison, David K. Jackson, Ian Johnston, Dominic Kwiatkowski, Cordelia Langford, John Sillitoe on behalf of the Wellcome Sanger Institute COVID-19 Surveillance Team |
| EPI_ISL_879415, EPI_ISL_879416, EPI_ISL_879417, EPI_ISL_879418 | Lighthouse Lab in Alderley Park | Wellcome Sanger Institute for the COVID-19 Genomics UK (COG-UK) Consortium | Jacquelyn Wynn, Mairead Hyland, The Lighthouse Lab in Alderley Park and Alex Alderton, Roberto Amato, Sonia Goncalves, Ewan Harrison, David K. Jackson, Ian Johnston, Dominic Kwiatkowski, Cordelia Langford, John Sillitoe on behalf of the Wellcome Sanger Institute COVID-19 Surveillance Team |
| EPI_ISL_879419 | Lighthouse Lab in Cambridge | Wellcome Sanger Institute for the COVID-19 Genomics UK (COG-UK) Consortium | Rob Howes, The Lighthouse Lab in Cambridge and Alex Alderton, Roberto Amato, Sonia Goncalves, Ewan Harrison, David K. Jackson, Ian Johnston, Dominic Kwiatkowski, Cordelia Langford, John Sillitoe on behalf of the Wellcome Sanger Institute COVID-19 Surveillance Team |
| EPI_ISL_879420 | Lighthouse Lab in Alderley Park | Wellcome Sanger Institute for the COVID-19 Genomics UK (COG-UK) Consortium | Jacquelyn Wynn, Mairead Hyland, The Lighthouse Lab in Alderley Park and Alex Alderton, Roberto Amato, Sonia Goncalves, Ewan Harrison, David K. Jackson, Ian Johnston, Dominic Kwiatkowski, Cordelia Langford, John Sillitoe on behalf of the Wellcome Sanger Institute COVID-19 Surveillance Team |
| EPI_ISL_879421, EPI_ISL_879422, EPI_ISL_879423 | Lighthouse Lab in Cambridge | Wellcome Sanger Institute for the COVID-19 Genomics UK (COG-UK) Consortium | Rob Howes, The Lighthouse Lab in Cambridge and Alex Alderton, Roberto Amato, Sonia Goncalves, Ewan Harrison, David K. Jackson, Ian Johnston, Dominic Kwiatkowski, Cordelia Langford, John Sillitoe on behalf of the Wellcome Sanger Institute COVID-19 Surveillance Team |
| EPI_ISL_879427 | Lighthouse Lab in Alderley Park | Wellcome Sanger Institute for the COVID-19 Genomics UK (COG-UK) Consortium | Jacquelyn Wynn, Mairead Hyland, The Lighthouse Lab in Alderley Park and Alex Alderton, Roberto Amato, Sonia Goncalves, Ewan Harrison, David K. Jackson, Ian Johnston, Dominic Kwiatkowski, Cordelia Langford, John Sillitoe on behalf of the Wellcome Sanger Institute COVID-19 Surveillance Team |
| EPI_ISL_879428, EPI_ISL_879430, EPI_ISL_879431, EPI_ISL_879432 | Lighthouse Lab in Cambridge | Wellcome Sanger Institute for the COVID-19 Genomics UK (COG-UK) Consortium | Rob Howes, The Lighthouse Lab in Cambridge and Alex Alderton, Roberto Amato, Sonia Goncalves, Ewan Harrison, David K. Jackson, Ian Johnston, Dominic Kwiatkowski, Cordelia Langford, John Sillitoe on behalf of the Wellcome Sanger Institute COVID-19 Surveillance Team |
| EPI_ISL_879433 | Lighthouse Lab in Alderley Park | Wellcome Sanger Institute for the COVID-19 Genomics UK (COG-UK) Consortium | Jacquelyn Wynn, Mairead Hyland, The Lighthouse Lab in Alderley Park and Alex Alderton, Roberto Amato, Sonia Goncalves, Ewan Harrison, David K. Jackson, Ian Johnston, Dominic Kwiatkowski, Cordelia Langford, John Sillitoe on behalf of the Wellcome Sanger Institute COVID-19 Surveillance Team |
| EPI_ISL_879435, EPI_ISL_879436, EPI_ISL_879437, EPI_ISL_879438, EPI_ISL_879439, EPI_ISL_879440, EPI_ISL_879441, EPI_ISL_879443 | Lighthouse Lab in Cambridge | Wellcome Sanger Institute for the COVID-19 Genomics UK (COG-UK) Consortium | Rob Howes, The Lighthouse Lab in Cambridge and Alex Alderton, Roberto Amato, Sonia Goncalves, Ewan Harrison, David K. Jackson, Ian Johnston, Dominic Kwiatkowski, Cordelia Langford, John Sillitoe on behalf of the Wellcome Sanger Institute COVID-19 Surveillance Team |
| EPI_ISL_879444 | Lighthouse Lab in Alderley Park | Wellcome Sanger Institute for the COVID-19 Genomics UK (COG-UK) Consortium | Jacquelyn Wynn, Mairead Hyland, The Lighthouse Lab in Alderley Park and Alex Alderton, Roberto Amato, Sonia Goncalves, Ewan Harrison, David K. Jackson, Ian Johnston, Dominic Kwiatkowski, Cordelia Langford, John Sillitoe on behalf of the Wellcome Sanger Institute COVID-19 Surveillance Team |
| EPI_ISL_879445 | Lighthouse Lab in Cambridge | Wellcome Sanger Institute for the COVID-19 Genomics UK (COG-UK) Consortium | Rob Howes, The Lighthouse Lab in Cambridge and Alex Alderton, Roberto Amato, Sonia Goncalves, Ewan Harrison, David K. Jackson, Ian Johnston, Dominic Kwiatkowski, Cordelia Langford, John Sillitoe on behalf of the Wellcome Sanger Institute COVID-19 Surveillance Team |
| EPI_ISL_879446 | Lighthouse Lab in Alderley Park | Wellcome Sanger Institute for the COVID-19 Genomics UK (COG-UK) Consortium | Jacquelyn Wynn, Mairead Hyland, The Lighthouse Lab in Alderley Park and Alex Alderton, Roberto Amato, Sonia Goncalves, Ewan Harrison, David K. Jackson, Ian Johnston, Dominic Kwiatkowski, Cordelia Langford, John Sillitoe on behalf of the Wellcome Sanger Institute COVID-19 Surveillance Team |
| EPI_ISL_879447 | Lighthouse Lab in Cambridge | Wellcome Sanger Institute for the COVID-19 Genomics UK (COG-UK) Consortium | Rob Howes, The Lighthouse Lab in Cambridge and Alex Alderton, Roberto Amato, Sonia Goncalves, Ewan Harrison, David K. Jackson, Ian Johnston, Dominic Kwiatkowski, Cordelia Langford, John Sillitoe on behalf of the Wellcome Sanger Institute COVID-19 Surveillance Team |
| EPI_ISL_879451 | Lighthouse Lab in Alderley Park | Wellcome Sanger Institute for the COVID-19 Genomics UK (COG-UK) Consortium | Jacquelyn Wynn, Mairead Hyland, The Lighthouse Lab in Alderley Park and Alex Alderton, Roberto Amato, Sonia Goncalves, Ewan Harrison, David K. Jackson, Ian Johnston, Dominic Kwiatkowski, Cordelia Langford, John Sillitoe on behalf of the Wellcome Sanger Institute COVID-19 Surveillance Team |
| EPI_ISL_879453, EPI_ISL_879454 | Lighthouse Lab in Cambridge | Wellcome Sanger Institute for the COVID-19 Genomics UK (COG-UK) Consortium | Rob Howes, The Lighthouse Lab in Cambridge and Alex Alderton, Roberto Amato, Sonia Goncalves, Ewan Harrison, David K. Jackson, Ian Johnston, Dominic Kwiatkowski, Cordelia Langford, John Sillitoe on behalf of the Wellcome Sanger Institute COVID-19 Surveillance Team |
| EPI_ISL_879456, EPI_ISL_879460 | Lighthouse Lab in Alderley Park | Wellcome Sanger Institute for the COVID-19 Genomics UK (COG-UK) Consortium | Jacquelyn Wynn, Mairead Hyland, The Lighthouse Lab in Alderley Park and Alex Alderton, Roberto Amato, Sonia Goncalves, Ewan Harrison, David K. Jackson, Ian Johnston, Dominic Kwiatkowski, Cordelia Langford, John Sillitoe on behalf of the Wellcome Sanger Institute COVID-19 Surveillance Team |
| EPI_ISL_879463, EPI_ISL_879465, EPI_ISL_879468, EPI_ISL_879469, EPI_ISL_879470, EPI_ISL_879472, EPI_ISL_879474 | Lighthouse Lab in Cambridge | Wellcome Sanger Institute for the COVID-19 Genomics UK (COG-UK) Consortium | Rob Howes, The Lighthouse Lab in Cambridge and Alex Alderton, Roberto Amato, Sonia Goncalves, Ewan Harrison, David K. Jackson, Ian Johnston, Dominic Kwiatkowski, Cordelia Langford, John Sillitoe on behalf of the Wellcome Sanger Institute COVID-19 Surveillance Team |
| EPI_ISL_879476 | Lighthouse Lab in Alderley Park | Wellcome Sanger Institute for the COVID-19 Genomics UK (COG-UK) Consortium | Jacquelyn Wynn, Mairead Hyland, The Lighthouse Lab in Alderley Park and Alex Alderton, Roberto Amato, Sonia Goncalves, Ewan Harrison, David K. Jackson, Ian Johnston, Dominic Kwiatkowski, Cordelia Langford, John Sillitoe on behalf of the Wellcome Sanger Institute COVID-19 Surveillance Team |
| EPI_ISL_879478 | Lighthouse Lab in Cambridge | Wellcome Sanger Institute for the COVID-19 Genomics UK (COG-UK) Consortium | Rob Howes, The Lighthouse Lab in Cambridge and Alex Alderton, Roberto Amato, Sonia Goncalves, Ewan Harrison, David K. Jackson, Ian Johnston, Dominic Kwiatkowski, Cordelia Langford, John Sillitoe on behalf of the Wellcome Sanger Institute COVID-19 Surveillance Team |
| EPI_ISL_879480 | Lighthouse Lab in Alderley Park | Wellcome Sanger Institute for the COVID-19 Genomics UK (COG-UK) Consortium | Jacquelyn Wynn, Mairead Hyland, The Lighthouse Lab in Alderley Park and Alex Alderton, Roberto Amato, Sonia Goncalves, Ewan Harrison, David K. Jackson, Ian Johnston, Dominic Kwiatkowski, Cordelia Langford, John Sillitoe on behalf of the Wellcome Sanger Institute COVID-19 Surveillance Team |
| EPI_ISL_879481, EPI_ISL_879482 | Lighthouse Lab in Cambridge | Wellcome Sanger Institute for the COVID-19 Genomics UK | Rob Howes, The Lighthouse Lab in Cambridge and Alex Alderton, Roberto Amato, Sonia Goncalves, Ewan Harrison, David K. Jackson, Ian Johnston, |

| | | | |
|---|---|---|---|
| | | (COG-UK) Consortium | Dominic Kwiatkowski, Cordelia Langford, John Sillitoe on behalf of the Wellcome Sanger Institute COVID-19 Surveillance Team |
| EPI_ISL_879483 | Lighthouse Lab in Alderley Park | Wellcome Sanger Institute for the COVID-19 Genomics UK (COG-UK) Consortium | Jacquelyn Wynn, Mairead Hyland, The Lighthouse Lab in Alderley Park and Alex Alderton, Roberto Amato, Sonia Goncalves, Ewan Harrison, David K. Jackson, Ian Johnston, Dominic Kwiatkowski, Cordelia Langford, John Sillitoe on behalf of the Wellcome Sanger Institute COVID-19 Surveillance Team |
| EPI_ISL_879484, EPI_ISL_879485, EPI_ISL_879488, EPI_ISL_879489, EPI_ISL_879490, EPI_ISL_879491 | Lighthouse Lab in Cambridge | Wellcome Sanger Institute for the COVID-19 Genomics UK (COG-UK) Consortium | Rob Howes, The Lighthouse Lab in Cambridge and Alex Alderton, Roberto Amato, Sonia Goncalves, Ewan Harrison, David K. Jackson, Ian Johnston, Dominic Kwiatkowski, Cordelia Langford, John Sillitoe on behalf of the Wellcome Sanger Institute COVID-19 Surveillance Team |
| EPI_ISL_879493 | Lighthouse Lab in Alderley Park | Wellcome Sanger Institute for the COVID-19 Genomics UK (COG-UK) Consortium | Jacquelyn Wynn, Mairead Hyland, The Lighthouse Lab in Alderley Park and Alex Alderton, Roberto Amato, Sonia Goncalves, Ewan Harrison, David K. Jackson, Ian Johnston, Dominic Kwiatkowski, Cordelia Langford, John Sillitoe on behalf of the Wellcome Sanger Institute COVID-19 Surveillance Team |
| EPI_ISL_879494, EPI_ISL_879496 | Lighthouse Lab in Cambridge | Wellcome Sanger Institute for the COVID-19 Genomics UK (COG-UK) Consortium | Rob Howes, The Lighthouse Lab in Cambridge and Alex Alderton, Roberto Amato, Sonia Goncalves, Ewan Harrison, David K. Jackson, Ian Johnston, Dominic Kwiatkowski, Cordelia Langford, John Sillitoe on behalf of the Wellcome Sanger Institute COVID-19 Surveillance Team |
| EPI_ISL_879497 | Lighthouse Lab in Alderley Park | Wellcome Sanger Institute for the COVID-19 Genomics UK (COG-UK) Consortium | Jacquelyn Wynn, Mairead Hyland, The Lighthouse Lab in Alderley Park and Alex Alderton, Roberto Amato, Sonia Goncalves, Ewan Harrison, David K. Jackson, Ian Johnston, Dominic Kwiatkowski, Cordelia Langford, John Sillitoe on behalf of the Wellcome Sanger Institute COVID-19 Surveillance Team |
| EPI_ISL_879498, EPI_ISL_879500 | Lighthouse Lab in Cambridge | Wellcome Sanger Institute for the COVID-19 Genomics UK (COG-UK) Consortium | Rob Howes, The Lighthouse Lab in Cambridge and Alex Alderton, Roberto Amato, Sonia Goncalves, Ewan Harrison, David K. Jackson, Ian Johnston, Dominic Kwiatkowski, Cordelia Langford, John Sillitoe on behalf of the Wellcome Sanger Institute COVID-19 Surveillance Team |
| EPI_ISL_879501 | Lighthouse Lab in Alderley Park | Wellcome Sanger Institute for the COVID-19 Genomics UK (COG-UK) Consortium | Jacquelyn Wynn, Mairead Hyland, The Lighthouse Lab in Alderley Park and Alex Alderton, Roberto Amato, Sonia Goncalves, Ewan Harrison, David K. Jackson, Ian Johnston, Dominic Kwiatkowski, Cordelia Langford, John Sillitoe on behalf of the Wellcome Sanger Institute COVID-19 Surveillance Team |
| EPI_ISL_879503 | Lighthouse Lab in Cambridge | Wellcome Sanger Institute for the COVID-19 Genomics UK (COG-UK) Consortium | Rob Howes, The Lighthouse Lab in Cambridge and Alex Alderton, Roberto Amato, Sonia Goncalves, Ewan Harrison, David K. Jackson, Ian Johnston, Dominic Kwiatkowski, Cordelia Langford, John Sillitoe on behalf of the Wellcome Sanger Institute COVID-19 Surveillance Team |
| EPI_ISL_879505 | Lighthouse Lab in Alderley Park | Wellcome Sanger Institute for the COVID-19 Genomics UK (COG-UK) Consortium | Jacquelyn Wynn, Mairead Hyland, The Lighthouse Lab in Alderley Park and Alex Alderton, Roberto Amato, Sonia Goncalves, Ewan Harrison, David K. Jackson, Ian Johnston, Dominic Kwiatkowski, Cordelia Langford, John Sillitoe on behalf of the Wellcome Sanger Institute COVID-19 Surveillance Team |
| EPI_ISL_879506, EPI_ISL_879507, EPI_ISL_879508, EPI_ISL_879509, EPI_ISL_879511, EPI_ISL_879512, EPI_ISL_879513 | Lighthouse Lab in Cambridge | Wellcome Sanger Institute for the COVID-19 Genomics UK (COG-UK) Consortium | Rob Howes, The Lighthouse Lab in Cambridge and Alex Alderton, Roberto Amato, Sonia Goncalves, Ewan Harrison, David K. Jackson, Ian Johnston, Dominic Kwiatkowski, Cordelia Langford, John Sillitoe on behalf of the Wellcome Sanger Institute COVID-19 Surveillance Team |
| EPI_ISL_879515 | Lighthouse Lab in Alderley Park | Wellcome Sanger Institute for the COVID-19 Genomics UK (COG-UK) Consortium | Jacquelyn Wynn, Mairead Hyland, The Lighthouse Lab in Alderley Park and Alex Alderton, Roberto Amato, Sonia Goncalves, Ewan Harrison, David K. Jackson, Ian Johnston, Dominic Kwiatkowski, Cordelia Langford, John Sillitoe on behalf of the Wellcome Sanger Institute COVID-19 Surveillance Team |
| EPI_ISL_879516, EPI_ISL_879517 | Lighthouse Lab in Cambridge | Wellcome Sanger Institute for the COVID-19 Genomics UK (COG-UK) Consortium | Rob Howes, The Lighthouse Lab in Cambridge and Alex Alderton, Roberto Amato, Sonia Goncalves, Ewan Harrison, David K. Jackson, Ian Johnston, Dominic Kwiatkowski, Cordelia Langford, John Sillitoe on behalf of the Wellcome Sanger Institute COVID-19 Surveillance Team |
| EPI_ISL_879518 | Lighthouse Lab in Alderley Park | Wellcome Sanger Institute for the COVID-19 Genomics UK (COG-UK) Consortium | Jacquelyn Wynn, Mairead Hyland, The Lighthouse Lab in Alderley Park and Alex Alderton, Roberto Amato, Sonia Goncalves, Ewan Harrison, David K. Jackson, Ian Johnston, Dominic Kwiatkowski, Cordelia Langford, John Sillitoe on behalf of the Wellcome Sanger Institute COVID-19 Surveillance Team |
| EPI_ISL_879521 | Lighthouse Lab in Cambridge | Wellcome Sanger Institute for the COVID-19 Genomics UK (COG-UK) Consortium | Rob Howes, The Lighthouse Lab in Cambridge and Alex Alderton, Roberto Amato, Sonia Goncalves, Ewan Harrison, David K. Jackson, Ian Johnston, Dominic Kwiatkowski, Cordelia Langford, John Sillitoe on behalf of the Wellcome Sanger Institute COVID-19 Surveillance Team |
| EPI_ISL_879522 | Lighthouse Lab in Alderley Park | Wellcome Sanger Institute for the COVID-19 Genomics UK (COG-UK) Consortium | Jacquelyn Wynn, Mairead Hyland, The Lighthouse Lab in Alderley Park and Alex Alderton, Roberto Amato, Sonia Goncalves, Ewan Harrison, David K. Jackson, Ian Johnston, Dominic Kwiatkowski, Cordelia Langford, John Sillitoe on behalf of the Wellcome Sanger Institute COVID-19 Surveillance Team |
| EPI_ISL_879523 | Lighthouse Lab in Cambridge | Wellcome Sanger Institute for the COVID-19 Genomics UK (COG-UK) Consortium | Rob Howes, The Lighthouse Lab in Cambridge and Alex Alderton, Roberto Amato, Sonia Goncalves, Ewan Harrison, David K. Jackson, Ian Johnston, Dominic Kwiatkowski, Cordelia Langford, John Sillitoe on behalf of the Wellcome Sanger Institute COVID-19 Surveillance Team |
| EPI_ISL_879524 | Lighthouse Lab in Alderley Park | Wellcome Sanger Institute for the COVID-19 Genomics UK (COG-UK) Consortium | Jacquelyn Wynn, Mairead Hyland, The Lighthouse Lab in Alderley Park and Alex Alderton, Roberto Amato, Sonia Goncalves, Ewan Harrison, David K. Jackson, Ian Johnston, Dominic Kwiatkowski, Cordelia Langford, John Sillitoe on behalf of the Wellcome Sanger Institute COVID-19 Surveillance Team |
| EPI_ISL_879525, EPI_ISL_879527, EPI_ISL_879528, EPI_ISL_879529, EPI_ISL_879530, EPI_ISL_879531, EPI_ISL_879533 | Lighthouse Lab in Cambridge | Wellcome Sanger Institute for the COVID-19 Genomics UK (COG-UK) Consortium | Rob Howes, The Lighthouse Lab in Cambridge and Alex Alderton, Roberto Amato, Sonia Goncalves, Ewan Harrison, David K. Jackson, Ian Johnston, Dominic Kwiatkowski, Cordelia Langford, John Sillitoe on behalf of the Wellcome Sanger Institute COVID-19 Surveillance Team |
| EPI_ISL_879534, EPI_ISL_879535 | Lighthouse Lab in Alderley Park | Wellcome Sanger Institute for the COVID-19 Genomics UK (COG-UK) Consortium | Jacquelyn Wynn, Mairead Hyland, The Lighthouse Lab in Alderley Park and Alex Alderton, Roberto Amato, Sonia Goncalves, Ewan Harrison, David K. Jackson, Ian Johnston, Dominic Kwiatkowski, Cordelia Langford, John Sillitoe on behalf of the Wellcome Sanger Institute COVID-19 Surveillance Team |
| EPI_ISL_879536, EPI_ISL_879537, EPI_ISL_879541 | Lighthouse Lab in Cambridge | Wellcome Sanger Institute for the COVID-19 Genomics UK (COG-UK) Consortium | Rob Howes, The Lighthouse Lab in Cambridge and Alex Alderton, Roberto Amato, Sonia Goncalves, Ewan Harrison, David K. Jackson, Ian Johnston, Dominic Kwiatkowski, Cordelia Langford, John Sillitoe on behalf of the Wellcome Sanger Institute COVID-19 Surveillance Team |
| EPI_ISL_879542 | Lighthouse Lab in Alderley Park | Wellcome Sanger Institute for the COVID-19 Genomics UK (COG-UK) Consortium | Jacquelyn Wynn, Mairead Hyland, The Lighthouse Lab in Alderley Park and Alex Alderton, Roberto Amato, Sonia Goncalves, Ewan Harrison, David K. Jackson, Ian Johnston, Dominic Kwiatkowski, Cordelia Langford, John Sillitoe on behalf of the Wellcome Sanger Institute COVID-19 Surveillance Team |
| EPI_ISL_879543, EPI_ISL_879544, EPI_ISL_879545, EPI_ISL_879551, EPI_ISL_879555, EPI_ISL_879557, EPI_ISL_879558, EPI_ISL_879559, EPI_ISL_879562 | Lighthouse Lab in Cambridge | Wellcome Sanger Institute for the COVID-19 Genomics UK (COG-UK) Consortium | Rob Howes, The Lighthouse Lab in Cambridge and Alex Alderton, Roberto Amato, Sonia Goncalves, Ewan Harrison, David K. Jackson, Ian Johnston, Dominic Kwiatkowski, Cordelia Langford, John Sillitoe on behalf of the Wellcome Sanger Institute COVID-19 Surveillance Team |
| EPI_ISL_879563, EPI_ISL_879564 | Lighthouse Lab in Alderley Park | Wellcome Sanger Institute for the COVID-19 Genomics UK (COG-UK) Consortium | Jacquelyn Wynn, Mairead Hyland, The Lighthouse Lab in Alderley Park and Alex Alderton, Roberto Amato, Sonia Goncalves, Ewan Harrison, David K. Jackson, Ian Johnston, Dominic Kwiatkowski, Cordelia Langford, John Sillitoe on behalf of the Wellcome Sanger Institute COVID-19 Surveillance Team |
| EPI_ISL_879567, EPI_ISL_879568, EPI_ISL_879569, EPI_ISL_879570, EPI_ISL_879573, EPI_ISL_879574, EPI_ISL_879577 | Lighthouse Lab in Cambridge | Wellcome Sanger Institute for the COVID-19 Genomics UK (COG-UK) Consortium | Rob Howes, The Lighthouse Lab in Cambridge and Alex Alderton, Roberto Amato, Sonia Goncalves, Ewan Harrison, David K. Jackson, Ian Johnston, Dominic Kwiatkowski, Cordelia Langford, John Sillitoe on behalf of the Wellcome Sanger Institute COVID-19 Surveillance Team |
| EPI_ISL_879578 | Lighthouse Lab in Alderley Park | Wellcome Sanger Institute for the COVID-19 Genomics UK (COG-UK) Consortium | Jacquelyn Wynn, Mairead Hyland, The Lighthouse Lab in Alderley Park and Alex Alderton, Roberto Amato, Sonia Goncalves, Ewan Harrison, David K. Jackson, Ian Johnston, Dominic Kwiatkowski, Cordelia Langford, John Sillitoe on behalf of the Wellcome Sanger Institute COVID-19 Surveillance Team |
| EPI_ISL_879579, EPI_ISL_879580 | Lighthouse Lab in Cambridge | Wellcome Sanger Institute for the COVID-19 Genomics UK (COG-UK) Consortium | Rob Howes, The Lighthouse Lab in Cambridge and Alex Alderton, Roberto Amato, Sonia Goncalves, Ewan Harrison, David K. Jackson, Ian Johnston, Dominic Kwiatkowski, Cordelia Langford, John Sillitoe on behalf of the Wellcome Sanger Institute COVID-19 Surveillance Team |
| EPI_ISL_879582 | Lighthouse Lab in Alderley Park | Wellcome Sanger Institute for the COVID-19 Genomics UK (COG-UK) Consortium | Jacquelyn Wynn, Mairead Hyland, The Lighthouse Lab in Alderley Park and Alex Alderton, Roberto Amato, Sonia Goncalves, Ewan Harrison, David K. Jackson, Ian Johnston, Dominic Kwiatkowski, Cordelia Langford, John Sillitoe on behalf of the Wellcome Sanger Institute COVID-19 Surveillance Team |
| EPI_ISL_879583, EPI_ISL_879584, EPI_ISL_879587, EPI_ISL_879588 | Lighthouse Lab in Cambridge | Wellcome Sanger Institute for the COVID-19 Genomics UK (COG-UK) Consortium | Rob Howes, The Lighthouse Lab in Cambridge and Alex Alderton, Roberto Amato, Sonia Goncalves, Ewan Harrison, David K. Jackson, Ian Johnston, Dominic Kwiatkowski, Cordelia Langford, John Sillitoe on behalf of the Wellcome Sanger Institute COVID-19 Surveillance Team |
| EPI_ISL_879589 | Lighthouse Lab in Alderley Park | Wellcome Sanger Institute for the COVID-19 Genomics UK (COG-UK) Consortium | Jacquelyn Wynn, Mairead Hyland, The Lighthouse Lab in Alderley Park and Alex Alderton, Roberto Amato, Sonia Goncalves, Ewan Harrison, David K. Jackson, Ian Johnston, Dominic Kwiatkowski, Cordelia Langford, John Sillitoe on behalf of the Wellcome Sanger Institute COVID-19 Surveillance Team |
| EPI_ISL_879590 | Lighthouse Lab in Cambridge | Wellcome Sanger Institute for the COVID-19 Genomics UK (COG-UK) Consortium | Rob Howes, The Lighthouse Lab in Cambridge and Alex Alderton, Roberto Amato, Sonia Goncalves, Ewan Harrison, David K. Jackson, Ian Johnston, Dominic Kwiatkowski, Cordelia Langford, John Sillitoe on behalf of the Wellcome Sanger Institute COVID-19 Surveillance Team |
| EPI_ISL_879593 | Lighthouse Lab in Alderley Park | Wellcome Sanger Institute for the COVID-19 Genomics UK | Jacquelyn Wynn, Mairead Hyland, The Lighthouse Lab in Alderley Park and Alex Alderton, Roberto Amato, Sonia Goncalves, Ewan Harrison, David K. |

| | | | |
|---|---|---|---|
| | | (COG-UK) Consortium | Jackson, Ian Johnston, Dominic Kwiatkowski, Cordelia Langford, John Sillitoe on behalf of the Wellcome Sanger Institute COVID-19 Surveillance Team |
| EPI_ISL_879595 | Lighthouse Lab in Cambridge | Wellcome Sanger Institute for the COVID-19 Genomics UK (COG-UK) Consortium | Rob Howes, The Lighthouse Lab in Cambridge and Alex Alderton, Roberto Amato, Sonia Goncalves, Ewan Harrison, David K. Jackson, Ian Johnston, Dominic Kwiatkowski, Cordelia Langford, John Sillitoe on behalf of the Wellcome Sanger Institute COVID-19 Surveillance Team |
| EPI_ISL_879596 | Lighthouse Lab in Alderley Park | Wellcome Sanger Institute for the COVID-19 Genomics UK (COG-UK) Consortium | Jacquelyn Wynn, Mairead Hyland, The Lighthouse Lab in Alderley Park and Alex Alderton, Roberto Amato, Sonia Goncalves, Ewan Harrison, David K. Jackson, Ian Johnston, Dominic Kwiatkowski, Cordelia Langford, John Sillitoe on behalf of the Wellcome Sanger Institute COVID-19 Surveillance Team |
| EPI_ISL_879598, EPI_ISL_879599 | Lighthouse Lab in Cambridge | Wellcome Sanger Institute for the COVID-19 Genomics UK (COG-UK) Consortium | Rob Howes, The Lighthouse Lab in Cambridge and Alex Alderton, Roberto Amato, Sonia Goncalves, Ewan Harrison, David K. Jackson, Ian Johnston, Dominic Kwiatkowski, Cordelia Langford, John Sillitoe on behalf of the Wellcome Sanger Institute COVID-19 Surveillance Team |
| EPI_ISL_879602, EPI_ISL_879603, EPI_ISL_879604 | Lighthouse Lab in Alderley Park | Wellcome Sanger Institute for the COVID-19 Genomics UK (COG-UK) Consortium | Jacquelyn Wynn, Mairead Hyland, The Lighthouse Lab in Alderley Park and Alex Alderton, Roberto Amato, Sonia Goncalves, Ewan Harrison, David K. Jackson, Ian Johnston, Dominic Kwiatkowski, Cordelia Langford, John Sillitoe on behalf of the Wellcome Sanger Institute COVID-19 Surveillance Team |
| EPI_ISL_879605, EPI_ISL_879606, EPI_ISL_879607, EPI_ISL_879609, EPI_ISL_879610, EPI_ISL_879612 | Lighthouse Lab in Cambridge | Wellcome Sanger Institute for the COVID-19 Genomics UK (COG-UK) Consortium | Rob Howes, The Lighthouse Lab in Cambridge and Alex Alderton, Roberto Amato, Sonia Goncalves, Ewan Harrison, David K. Jackson, Ian Johnston, Dominic Kwiatkowski, Cordelia Langford, John Sillitoe on behalf of the Wellcome Sanger Institute COVID-19 Surveillance Team |
| EPI_ISL_879614 | Lighthouse Lab in Alderley Park | Wellcome Sanger Institute for the COVID-19 Genomics UK (COG-UK) Consortium | Jacquelyn Wynn, Mairead Hyland, The Lighthouse Lab in Alderley Park and Alex Alderton, Roberto Amato, Sonia Goncalves, Ewan Harrison, David K. Jackson, Ian Johnston, Dominic Kwiatkowski, Cordelia Langford, John Sillitoe on behalf of the Wellcome Sanger Institute COVID-19 Surveillance Team |
| EPI_ISL_879615 | Lighthouse Lab in Cambridge | Wellcome Sanger Institute for the COVID-19 Genomics UK (COG-UK) Consortium | Rob Howes, The Lighthouse Lab in Cambridge and Alex Alderton, Roberto Amato, Sonia Goncalves, Ewan Harrison, David K. Jackson, Ian Johnston, Dominic Kwiatkowski, Cordelia Langford, John Sillitoe on behalf of the Wellcome Sanger Institute COVID-19 Surveillance Team |
| EPI_ISL_879616, EPI_ISL_879619 | Lighthouse Lab in Alderley Park | Wellcome Sanger Institute for the COVID-19 Genomics UK (COG-UK) Consortium | Jacquelyn Wynn, Mairead Hyland, The Lighthouse Lab in Alderley Park and Alex Alderton, Roberto Amato, Sonia Goncalves, Ewan Harrison, David K. Jackson, Ian Johnston, Dominic Kwiatkowski, Cordelia Langford, John Sillitoe on behalf of the Wellcome Sanger Institute COVID-19 Surveillance Team |
| EPI_ISL_879620, EPI_ISL_879622, EPI_ISL_879624, EPI_ISL_879625, EPI_ISL_879626 | Lighthouse Lab in Cambridge | Wellcome Sanger Institute for the COVID-19 Genomics UK (COG-UK) Consortium | Rob Howes, The Lighthouse Lab in Cambridge and Alex Alderton, Roberto Amato, Sonia Goncalves, Ewan Harrison, David K. Jackson, Ian Johnston, Dominic Kwiatkowski, Cordelia Langford, John Sillitoe on behalf of the Wellcome Sanger Institute COVID-19 Surveillance Team |
| EPI_ISL_879627 | Lighthouse Lab in Alderley Park | Wellcome Sanger Institute for the COVID-19 Genomics UK (COG-UK) Consortium | Jacquelyn Wynn, Mairead Hyland, The Lighthouse Lab in Alderley Park and Alex Alderton, Roberto Amato, Sonia Goncalves, Ewan Harrison, David K. Jackson, Ian Johnston, Dominic Kwiatkowski, Cordelia Langford, John Sillitoe on behalf of the Wellcome Sanger Institute COVID-19 Surveillance Team |
| EPI_ISL_879630, EPI_ISL_879632, EPI_ISL_879633, EPI_ISL_879634, EPI_ISL_879635, EPI_ISL_879636, EPI_ISL_879637, EPI_ISL_879638, EPI_ISL_879639, EPI_ISL_879640 | Lighthouse Lab in Cambridge | Wellcome Sanger Institute for the COVID-19 Genomics UK (COG-UK) Consortium | Rob Howes, The Lighthouse Lab in Cambridge and Alex Alderton, Roberto Amato, Sonia Goncalves, Ewan Harrison, David K. Jackson, Ian Johnston, Dominic Kwiatkowski, Cordelia Langford, John Sillitoe on behalf of the Wellcome Sanger Institute COVID-19 Surveillance Team |
| EPI_ISL_879641 | Lighthouse Lab in Alderley Park | Wellcome Sanger Institute for the COVID-19 Genomics UK (COG-UK) Consortium | Jacquelyn Wynn, Mairead Hyland, The Lighthouse Lab in Alderley Park and Alex Alderton, Roberto Amato, Sonia Goncalves, Ewan Harrison, David K. Jackson, Ian Johnston, Dominic Kwiatkowski, Cordelia Langford, John Sillitoe on behalf of the Wellcome Sanger Institute COVID-19 Surveillance Team |
| EPI_ISL_879644, EPI_ISL_879645 | Lighthouse Lab in Cambridge | Wellcome Sanger Institute for the COVID-19 Genomics UK (COG-UK) Consortium | Rob Howes, The Lighthouse Lab in Cambridge and Alex Alderton, Roberto Amato, Sonia Goncalves, Ewan Harrison, David K. Jackson, Ian Johnston, Dominic Kwiatkowski, Cordelia Langford, John Sillitoe on behalf of the Wellcome Sanger Institute COVID-19 Surveillance Team |
| EPI_ISL_879647 | Lighthouse Lab in Alderley Park | Wellcome Sanger Institute for the COVID-19 Genomics UK (COG-UK) Consortium | Jacquelyn Wynn, Mairead Hyland, The Lighthouse Lab in Alderley Park and Alex Alderton, Roberto Amato, Sonia Goncalves, Ewan Harrison, David K. Jackson, Ian Johnston, Dominic Kwiatkowski, Cordelia Langford, John Sillitoe on behalf of the Wellcome Sanger Institute COVID-19 Surveillance Team |
| EPI_ISL_879648 | Lighthouse Lab in Cambridge | Wellcome Sanger Institute for the COVID-19 Genomics UK (COG-UK) Consortium | Rob Howes, The Lighthouse Lab in Cambridge and Alex Alderton, Roberto Amato, Sonia Goncalves, Ewan Harrison, David K. Jackson, Ian Johnston, Dominic Kwiatkowski, Cordelia Langford, John Sillitoe on behalf of the Wellcome Sanger Institute COVID-19 Surveillance Team |
| EPI_ISL_879649 | Lighthouse Lab in Alderley Park | Wellcome Sanger Institute for the COVID-19 Genomics UK (COG-UK) Consortium | Jacquelyn Wynn, Mairead Hyland, The Lighthouse Lab in Alderley Park and Alex Alderton, Roberto Amato, Sonia Goncalves, Ewan Harrison, David K. Jackson, Ian Johnston, Dominic Kwiatkowski, Cordelia Langford, John Sillitoe on behalf of the Wellcome Sanger Institute COVID-19 Surveillance Team |
| EPI_ISL_879650, EPI_ISL_879651, EPI_ISL_879652, EPI_ISL_879657, EPI_ISL_879658, EPI_ISL_879659, EPI_ISL_879660, EPI_ISL_879661, EPI_ISL_879662 | Lighthouse Lab in Cambridge | Wellcome Sanger Institute for the COVID-19 Genomics UK (COG-UK) Consortium | Rob Howes, The Lighthouse Lab in Cambridge and Alex Alderton, Roberto Amato, Sonia Goncalves, Ewan Harrison, David K. Jackson, Ian Johnston, Dominic Kwiatkowski, Cordelia Langford, John Sillitoe on behalf of the Wellcome Sanger Institute COVID-19 Surveillance Team |
| EPI_ISL_879665, EPI_ISL_879666 | Lighthouse Lab in Alderley Park | Wellcome Sanger Institute for the COVID-19 Genomics UK (COG-UK) Consortium | Jacquelyn Wynn, Mairead Hyland, The Lighthouse Lab in Alderley Park and Alex Alderton, Roberto Amato, Sonia Goncalves, Ewan Harrison, David K. Jackson, Ian Johnston, Dominic Kwiatkowski, Cordelia Langford, John Sillitoe on behalf of the Wellcome Sanger Institute COVID-19 Surveillance Team |
| EPI_ISL_879667, EPI_ISL_879668, EPI_ISL_879669, EPI_ISL_879670, EPI_ISL_879671 | Lighthouse Lab in Cambridge | Wellcome Sanger Institute for the COVID-19 Genomics UK (COG-UK) Consortium | Rob Howes, The Lighthouse Lab in Cambridge and Alex Alderton, Roberto Amato, Sonia Goncalves, Ewan Harrison, David K. Jackson, Ian Johnston, Dominic Kwiatkowski, Cordelia Langford, John Sillitoe on behalf of the Wellcome Sanger Institute COVID-19 Surveillance Team |
| EPI_ISL_879672 | Lighthouse Lab in Alderley Park | Wellcome Sanger Institute for the COVID-19 Genomics UK (COG-UK) Consortium | Jacquelyn Wynn, Mairead Hyland, The Lighthouse Lab in Alderley Park and Alex Alderton, Roberto Amato, Sonia Goncalves, Ewan Harrison, David K. Jackson, Ian Johnston, Dominic Kwiatkowski, Cordelia Langford, John Sillitoe on behalf of the Wellcome Sanger Institute COVID-19 Surveillance Team |
| EPI_ISL_879673 | Lighthouse Lab in Cambridge | Wellcome Sanger Institute for the COVID-19 Genomics UK (COG-UK) Consortium | Rob Howes, The Lighthouse Lab in Cambridge and Alex Alderton, Roberto Amato, Sonia Goncalves, Ewan Harrison, David K. Jackson, Ian Johnston, Dominic Kwiatkowski, Cordelia Langford, John Sillitoe on behalf of the Wellcome Sanger Institute COVID-19 Surveillance Team |
| EPI_ISL_879674 | Lighthouse Lab in Alderley Park | Wellcome Sanger Institute for the COVID-19 Genomics UK (COG-UK) Consortium | Jacquelyn Wynn, Mairead Hyland, The Lighthouse Lab in Alderley Park and Alex Alderton, Roberto Amato, Sonia Goncalves, Ewan Harrison, David K. Jackson, Ian Johnston, Dominic Kwiatkowski, Cordelia Langford, John Sillitoe on behalf of the Wellcome Sanger Institute COVID-19 Surveillance Team |
| EPI_ISL_879676, EPI_ISL_879677, EPI_ISL_879680 | Lighthouse Lab in Cambridge | Wellcome Sanger Institute for the COVID-19 Genomics UK (COG-UK) Consortium | Rob Howes, The Lighthouse Lab in Cambridge and Alex Alderton, Roberto Amato, Sonia Goncalves, Ewan Harrison, David K. Jackson, Ian Johnston, Dominic Kwiatkowski, Cordelia Langford, John Sillitoe on behalf of the Wellcome Sanger Institute COVID-19 Surveillance Team |
| EPI_ISL_879682, EPI_ISL_879683 | Lighthouse Lab in Alderley Park | Wellcome Sanger Institute for the COVID-19 Genomics UK (COG-UK) Consortium | Jacquelyn Wynn, Mairead Hyland, The Lighthouse Lab in Alderley Park and Alex Alderton, Roberto Amato, Sonia Goncalves, Ewan Harrison, David K. Jackson, Ian Johnston, Dominic Kwiatkowski, Cordelia Langford, John Sillitoe on behalf of the Wellcome Sanger Institute COVID-19 Surveillance Team |
| EPI_ISL_879684, EPI_ISL_879685 | Lighthouse Lab in Cambridge | Wellcome Sanger Institute for the COVID-19 Genomics UK (COG-UK) Consortium | Rob Howes, The Lighthouse Lab in Cambridge and Alex Alderton, Roberto Amato, Sonia Goncalves, Ewan Harrison, David K. Jackson, Ian Johnston, Dominic Kwiatkowski, Cordelia Langford, John Sillitoe on behalf of the Wellcome Sanger Institute COVID-19 Surveillance Team |
| EPI_ISL_879687 | Lighthouse Lab in Alderley Park | Wellcome Sanger Institute for the COVID-19 Genomics UK (COG-UK) Consortium | Jacquelyn Wynn, Mairead Hyland, The Lighthouse Lab in Alderley Park and Alex Alderton, Roberto Amato, Sonia Goncalves, Ewan Harrison, David K. Jackson, Ian Johnston, Dominic Kwiatkowski, Cordelia Langford, John Sillitoe on behalf of the Wellcome Sanger Institute COVID-19 Surveillance Team |
| EPI_ISL_879688, EPI_ISL_879690, EPI_ISL_879691, EPI_ISL_879694 | Lighthouse Lab in Cambridge | Wellcome Sanger Institute for the COVID-19 Genomics UK (COG-UK) Consortium | Rob Howes, The Lighthouse Lab in Cambridge and Alex Alderton, Roberto Amato, Sonia Goncalves, Ewan Harrison, David K. Jackson, Ian Johnston, Dominic Kwiatkowski, Cordelia Langford, John Sillitoe on behalf of the Wellcome Sanger Institute COVID-19 Surveillance Team |
| EPI_ISL_879695 | Lighthouse Lab in Alderley Park | Wellcome Sanger Institute for the COVID-19 Genomics UK (COG-UK) Consortium | Jacquelyn Wynn, Mairead Hyland, The Lighthouse Lab in Alderley Park and Alex Alderton, Roberto Amato, Sonia Goncalves, Ewan Harrison, David K. Jackson, Ian Johnston, Dominic Kwiatkowski, Cordelia Langford, John Sillitoe on behalf of the Wellcome Sanger Institute COVID-19 Surveillance Team |
| EPI_ISL_879696, EPI_ISL_879697, EPI_ISL_879698, EPI_ISL_879699, EPI_ISL_879701 | Lighthouse Lab in Cambridge | Wellcome Sanger Institute for the COVID-19 Genomics UK (COG-UK) Consortium | Rob Howes, The Lighthouse Lab in Cambridge and Alex Alderton, Roberto Amato, Sonia Goncalves, Ewan Harrison, David K. Jackson, Ian Johnston, Dominic Kwiatkowski, Cordelia Langford, John Sillitoe on behalf of the Wellcome Sanger Institute COVID-19 Surveillance Team |
| EPI_ISL_879702 | Lighthouse Lab in Alderley Park | Wellcome Sanger Institute for the COVID-19 Genomics UK (COG-UK) Consortium | Jacquelyn Wynn, Mairead Hyland, The Lighthouse Lab in Alderley Park and Alex Alderton, Roberto Amato, Sonia Goncalves, Ewan Harrison, David K. Jackson, Ian Johnston, Dominic Kwiatkowski, Cordelia Langford, John Sillitoe on behalf of the Wellcome Sanger Institute COVID-19 Surveillance Team |
| EPI_ISL_879704, EPI_ISL_879705, | Lighthouse Lab in Cambridge | Wellcome Sanger Institute for the COVID-19 Genomics UK | Rob Howes, The Lighthouse Lab in Cambridge and Alex Alderton, Roberto Amato, Sonia Goncalves, Ewan Harrison, David K. Jackson, Ian Johnston, |

| | | | |
|---|---|---|---|
| EPI_ISL_879706, EPI_ISL_879708, EPI_ISL_879709, EPI_ISL_879716 | | (COG-UK) Consortium | Dominic Kwiatkowski, Cordelia Langford, John Sillitoe on behalf of the Wellcome Sanger Institute COVID-19 Surveillance Team |
| EPI_ISL_879718 | Lighthouse Lab in Alderley Park | Wellcome Sanger Institute for the COVID-19 Genomics UK (COG-UK) Consortium | Jacquelyn Wynn, Mairead Hyland, The Lighthouse Lab in Alderley Park and Alex Alderton, Roberto Amato, Sonia Goncalves, Ewan Harrison, David K. Jackson, Ian Johnston, Dominic Kwiatkowski, Cordelia Langford, John Sillitoe on behalf of the Wellcome Sanger Institute COVID-19 Surveillance Team |
| EPI_ISL_879722, EPI_ISL_879733, EPI_ISL_879735, EPI_ISL_879743, EPI_ISL_879745, EPI_ISL_879746, EPI_ISL_879770, EPI_ISL_879782, EPI_ISL_879786, EPI_ISL_879796, EPI_ISL_879799, EPI_ISL_879804, EPI_ISL_879805, EPI_ISL_879807, EPI_ISL_879812, EPI_ISL_879815, EPI_ISL_879824, EPI_ISL_879830, EPI_ISL_879840, EPI_ISL_879847, EPI_ISL_879853, EPI_ISL_879857, EPI_ISL_879881, EPI_ISL_879883, EPI_ISL_879893, EPI_ISL_879899, EPI_ISL_879900, EPI_ISL_879902, EPI_ISL_879910, EPI_ISL_879917, EPI_ISL_879943, EPI_ISL_879951, EPI_ISL_879952, EPI_ISL_879983, EPI_ISL_879992, EPI_ISL_880001, EPI_ISL_880026, EPI_ISL_880051, EPI_ISL_880073, EPI_ISL_880080, EPI_ISL_880081 | | | |
| see above | Lighthouse Lab in Cambridge | Wellcome Sanger Institute for the COVID-19 Genomics UK (COG-UK) Consortium | Rob Howes, The Lighthouse Lab in Cambridge and Alex Alderton, Roberto Amato, Sonia Goncalves, Ewan Harrison, David K. Jackson, Ian Johnston, Dominic Kwiatkowski, Cordelia Langford, John Sillitoe on behalf of the Wellcome Sanger Institute COVID-19 Surveillance Team |
| EPI_ISL_880091, EPI_ISL_880100, EPI_ISL_880112, EPI_ISL_880113, EPI_ISL_880120, EPI_ISL_880123, EPI_ISL_880124, EPI_ISL_880126, EPI_ISL_880136, EPI_ISL_880216, EPI_ISL_880222, EPI_ISL_880258, EPI_ISL_880260, EPI_ISL_880264, EPI_ISL_880265, EPI_ISL_880286, EPI_ISL_880288, EPI_ISL_880291, EPI_ISL_880298, EPI_ISL_880301, EPI_ISL_880306, EPI_ISL_880332, EPI_ISL_880335, EPI_ISL_880337, EPI_ISL_880340, EPI_ISL_880343, EPI_ISL_880348, EPI_ISL_880352, EPI_ISL_880356, EPI_ISL_880361, EPI_ISL_880366, EPI_ISL_880375, EPI_ISL_880376, EPI_ISL_880379, EPI_ISL_880387, EPI_ISL_880390, EPI_ISL_880392, EPI_ISL_880400, EPI_ISL_880402, EPI_ISL_880417, EPI_ISL_880418, EPI_ISL_880426, EPI_ISL_880427, EPI_ISL_880433, EPI_ISL_880435, EPI_ISL_880436, EPI_ISL_880443, EPI_ISL_880446, EPI_ISL_880456, EPI_ISL_880458, EPI_ISL_880464, EPI_ISL_880467, EPI_ISL_880475, EPI_ISL_880477, EPI_ISL_880482, EPI_ISL_880493, EPI_ISL_880498, EPI_ISL_880502, EPI_ISL_880508, EPI_ISL_880509, EPI_ISL_880518, EPI_ISL_880519, EPI_ISL_880529, EPI_ISL_880531, EPI_ISL_880575, EPI_ISL_880583 | | | |
| see above | Lighthouse Lab in Alderley Park | Wellcome Sanger Institute for the COVID-19 Genomics UK (COG-UK) Consortium | Jacquelyn Wynn, Mairead Hyland, The Lighthouse Lab in Alderley Park and Alex Alderton, Roberto Amato, Sonia Goncalves, Ewan Harrison, David K. Jackson, Ian Johnston, Dominic Kwiatkowski, Cordelia Langford, John Sillitoe on behalf of the Wellcome Sanger Institute COVID-19 Surveillance Team |
| EPI_ISL_882673 | Hospital Municipal Dr. Guido Guida | Instituto Adolfo Lutz, Interdiciplinary Procedures Center, Strategic Laboratory | Claudio Tavares Sacchi, Claudia Regina Gonçalves, Erica Valessa Ramos Gomes, Karoline Rodrigues Campos |
| EPI_ISL_883307, EPI_ISL_883308, EPI_ISL_883311, EPI_ISL_883317 | Pandemic Response Laboratory | New York City Public Health Laboratory | Jade Wang, et al. |
| EPI_ISL_883325, EPI_ISL_883330 | OCME Office Of Chief Medical Examiner | New York City Public Health Laboratory | Jade Wang, et al. |
| EPI_ISL_883331 | DOHMH Morrisania | New York City Public Health Laboratory | Jade Wang, et al. |
| EPI_ISL_883337 | DOHMH Crown Heights | New York City Public Health Laboratory | Jade Wang, et al. |
| EPI_ISL_883344, EPI_ISL_883345 | OCME Office Of Chief Medical Examiner | New York City Public Health Laboratory | Jade Wang, et al. |
| EPI_ISL_883347 | DOHMH Morrisania | New York City Public Health Laboratory | Jade Wang, et al. |
| EPI_ISL_883349 | DOHMH Jamaica | New York City Public Health Laboratory | Jade Wang, et al. |
| EPI_ISL_883383, EPI_ISL_883384, EPI_ISL_883385, EPI_ISL_883386, EPI_ISL_883387, EPI_ISL_883388, EPI_ISL_883389 | Pandemic Response Laboratory | New York City Public Health Laboratory | Jade Wang, et al. |
| EPI_ISL_883403 | Department of Homeless Services | New York City Public Health Laboratory | Jade Wang, et al. |
| EPI_ISL_883404, EPI_ISL_883405, EPI_ISL_883406, EPI_ISL_883407, EPI_ISL_883408 | OCME Office Of Chief Medical Examiner | New York City Public Health Laboratory | Jade Wang, et al. |
| EPI_ISL_883410, EPI_ISL_883411 | DOHMH Corona | New York City Public Health Laboratory | Jade Wang, et al. |
| EPI_ISL_883412 | DOHMH Crown Heights | New York City Public Health Laboratory | Jade Wang, et al. |
| EPI_ISL_883413, EPI_ISL_883414, EPI_ISL_883415 | DOHMH Central Harlem | New York City Public Health Laboratory | Jade Wang, et al. |
| EPI_ISL_883416, EPI_ISL_883417 | DOHMH Fort Greene | New York City Public Health Laboratory | Jade Wang, et al. |
| EPI_ISL_883418, EPI_ISL_883419, EPI_ISL_883420, EPI_ISL_883421, EPI_ISL_883422 | DOHMH Jamaica | New York City Public Health Laboratory | Jade Wang, et al. |
| EPI_ISL_883424 | DOHMH PHL | New York City Public Health Laboratory | Jade Wang, et al. |
| EPI_ISL_883430 | BOSTON HEART DIAGNOSTICS CORP | Wadsworth Center, New York State Department of Health | Kirsten St. George, Daryl M. Lamson, Alexis Russel, Matthew Shudt, Melissa A Leisner, Jonathan Plitnick, Navjot Singh, John Kelly, Erasmus Schneider, Erica Lasek-Nesselquist |
| EPI_ISL_884008, EPI_ISL_884009, EPI_ISL_884010, EPI_ISL_884011, EPI_ISL_884012 | ALBANY MEDICAL CENTER HOSPITAL CLINICAL LABORATORIES | Wadsworth Center, New York State Department of Health | Kirsten St. George, Daryl M. Lamson, Alexis Russel, Matthew Shudt, Melissa A Leisner, Jonathan Plitnick, Navjot Singh, John Kelly, Erasmus Schneider, Erica Lasek-Nesselquist |
| EPI_ISL_884013, EPI_ISL_884014, EPI_ISL_884015, EPI_ISL_884016 | Wadsworth Center, New York State Department of Health | Wadsworth Center, New York State Department of Health | Kirsten St. George, Daryl M. Lamson, Alexis Russel, Matthew Shudt, Melissa A Leisner, Jonathan Plitnick, Navjot Singh, John Kelly, Erasmus Schneider, Erica Lasek-Nesselquist |
| EPI_ISL_884017, EPI_ISL_884018, EPI_ISL_884019, EPI_ISL_884020 | ALBANY MEDICAL CENTER HOSPITAL CLINICAL LABORATORIES | Wadsworth Center, New York State Department of Health | Kirsten St. George, Daryl M. Lamson, Alexis Russel, Matthew Shudt, Melissa A Leisner, Jonathan Plitnick, Navjot Singh, John Kelly, Erasmus Schneider, Erica Lasek-Nesselquist |
| EPI_ISL_884194, EPI_ISL_884195 | Wyoming Public Health Laboratory | Wyoming Public Health Laboratory | Noah Hull, Taylor Fearing, Lynette Gumbleton, Channing Weber, Ashley Norberg, Bailey Bowcutt, and Wanda Manley |
| EPI_ISL_884580, EPI_ISL_884581, EPI_ISL_884582, EPI_ISL_884583, EPI_ISL_884584, EPI_ISL_884585, EPI_ISL_884586, EPI_ISL_884587, EPI_ISL_884588, EPI_ISL_884589 | Respiratory Viruses Branch, Centers for Disease Control and Prevention | Respiratory Viruses Branch, Centers for Disease Control and Prevention | Cook,P.W., Batra,D., Rambo-Martin,B.L., de Feo,E., Antico,J., Tran,C., Tolentino,M., Wickline,S., Gietzen,K., Sickler,B., Liu,J., Allen,E., Febbo,P., Galloway,S., Washington,N.L., White,S., Levan,G., Barret,K.S., Cirulli,E., Bolze,A., Ascencio,A., Rivera-Garcia,C., Cho,R., Nguyen,J., Wang,S., Ramirez,J., Cassens,T., Sandoval,E., Isaksson,M., Lee,W., Becker,D., Laurent,M., Lu,J., Paden,C.R., Tong,S., MacCannell,D. |
| EPI_ISL_885155, EPI_ISL_885156, EPI_ISL_885157, EPI_ISL_885158, EPI_ISL_885159, EPI_ISL_885160, EPI_ISL_885161, EPI_ISL_885162, EPI_ISL_885163, EPI_ISL_885164, EPI_ISL_885165, EPI_ISL_885166, EPI_ISL_885167, EPI_ISL_885168, EPI_ISL_885169, EPI_ISL_885170, EPI_ISL_885171, EPI_ISL_885172, EPI_ISL_885173, EPI_ISL_885174, EPI_ISL_885175, EPI_ISL_885176, EPI_ISL_885177, EPI_ISL_885178, EPI_ISL_885179, EPI_ISL_885180, EPI_ISL_885181, EPI_ISL_885182, EPI_ISL_885184, EPI_ISL_885185, EPI_ISL_885186, EPI_ISL_885187, EPI_ISL_885188, EPI_ISL_885189, EPI_ISL_885190, EPI_ISL_885191, EPI_ISL_885192, EPI_ISL_885193, EPI_ISL_885194, EPI_ISL_885195, EPI_ISL_885196, EPI_ISL_885197, EPI_ISL_885198, EPI_ISL_885199, EPI_ISL_885200, EPI_ISL_885201, EPI_ISL_885202, EPI_ISL_885203, EPI_ISL_885205, EPI_ISL_885206, EPI_ISL_885207, EPI_ISL_885208, EPI_ISL_885209, EPI_ISL_885210, EPI_ISL_885212, EPI_ISL_885213, EPI_ISL_885214, EPI_ISL_885215, EPI_ISL_885216, EPI_ISL_885217, EPI_ISL_885218, EPI_ISL_885219, EPI_ISL_885220, EPI_ISL_885221, EPI_ISL_885222, EPI_ISL_885223, EPI_ISL_885224, EPI_ISL_885225, EPI_ISL_885226, EPI_ISL_885227, EPI_ISL_885228, EPI_ISL_885229, EPI_ISL_885230, EPI_ISL_885231, EPI_ISL_885232, EPI_ISL_885233, EPI_ISL_885234, EPI_ISL_885235, EPI_ISL_885236, EPI_ISL_885237, EPI_ISL_885238, EPI_ISL_885239, EPI_ISL_885241, EPI_ISL_885242, EPI_ISL_885243, EPI_ISL_885245, EPI_ISL_885246, EPI_ISL_885247, EPI_ISL_885248, EPI_ISL_885249, EPI_ISL_885250, EPI_ISL_885251, EPI_ISL_885252, EPI_ISL_885253, EPI_ISL_885254, EPI_ISL_885255, EPI_ISL_885256, EPI_ISL_885257, EPI_ISL_885258, EPI_ISL_885259, EPI_ISL_885260, EPI_ISL_885261, EPI_ISL_885262, EPI_ISL_885263, EPI_ISL_885264, EPI_ISL_885265, EPI_ISL_885266, EPI_ISL_885267, EPI_ISL_885268, EPI_ISL_885269, EPI_ISL_885270, EPI_ISL_885271, EPI_ISL_885272, EPI_ISL_885273, EPI_ISL_885274, EPI_ISL_885275, EPI_ISL_885276, EPI_ISL_885277, EPI_ISL_885278, EPI_ISL_885279, EPI_ISL_885280, EPI_ISL_885281, EPI_ISL_885282, EPI_ISL_885283, EPI_ISL_885284, EPI_ISL_885285, EPI_ISL_885286, EPI_ISL_885287, EPI_ISL_885288, EPI_ISL_885289, EPI_ISL_885290, EPI_ISL_885292, EPI_ISL_885293, EPI_ISL_885294, EPI_ISL_885295, EPI_ISL_885296, EPI_ISL_885297, EPI_ISL_885298, EPI_ISL_885299, EPI_ISL_885300, EPI_ISL_885301, EPI_ISL_885302, EPI_ISL_885303, EPI_ISL_885304, EPI_ISL_885305, EPI_ISL_885306, EPI_ISL_885307, EPI_ISL_885308, EPI_ISL_885309, EPI_ISL_885310, EPI_ISL_885311, EPI_ISL_885312, EPI_ISL_885313, EPI_ISL_885314, EPI_ISL_885315, EPI_ISL_885316, EPI_ISL_885317, EPI_ISL_885318, EPI_ISL_885319, EPI_ISL_885320, EPI_ISL_885322, EPI_ISL_885323, EPI_ISL_885324, EPI_ISL_885325, EPI_ISL_885326, EPI_ISL_885328, EPI_ISL_885329, EPI_ISL_885330, EPI_ISL_885331, EPI_ISL_885332, EPI_ISL_885333, EPI_ISL_885334, EPI_ISL_885335, EPI_ISL_885336, EPI_ISL_885337, EPI_ISL_885338, EPI_ISL_885339, EPI_ISL_885340, EPI_ISL_885341, EPI_ISL_885342, EPI_ISL_885343, EPI_ISL_885344, EPI_ISL_885345, EPI_ISL_885346, EPI_ISL_885347, EPI_ISL_885348, EPI_ISL_885350, EPI_ISL_885351, EPI_ISL_885352, EPI_ISL_885353, EPI_ISL_885354, EPI_ISL_885355, EPI_ISL_885356, EPI_ISL_885357, EPI_ISL_885358, EPI_ISL_885360, EPI_ISL_885361, EPI_ISL_885362, EPI_ISL_885363, EPI_ISL_885364, EPI_ISL_885365, EPI_ISL_885366, EPI_ISL_885367, EPI_ISL_885368, EPI_ISL_885369, EPI_ISL_885370, EPI_ISL_885371, EPI_ISL_885372, EPI_ISL_885373, EPI_ISL_885374, EPI_ISL_885375, EPI_ISL_885376, EPI_ISL_885377, EPI_ISL_885378, EPI_ISL_885379, EPI_ISL_885380, EPI_ISL_885381, EPI_ISL_885382, EPI_ISL_885383, EPI_ISL_885384, EPI_ISL_885385, EPI_ISL_885386, EPI_ISL_885387, EPI_ISL_885388, EPI_ISL_885389, EPI_ISL_885390, EPI_ISL_885391, EPI_ISL_885393, EPI_ISL_885394, EPI_ISL_885395, EPI_ISL_885396, EPI_ISL_885397, EPI_ISL_885398, EPI_ISL_885399, EPI_ISL_885400, EPI_ISL_885401, EPI_ISL_885402, EPI_ISL_885403, EPI_ISL_885404, EPI_ISL_885405, EPI_ISL_885406, EPI_ISL_885407, EPI_ISL_885408, EPI_ISL_885410, EPI_ISL_885412, EPI_ISL_885413, EPI_ISL_885414, EPI_ISL_885415, EPI_ISL_885416, EPI_ISL_885417, EPI_ISL_885418, EPI_ISL_885419, EPI_ISL_885420, EPI_ISL_885421, EPI_ISL_885422, EPI_ISL_885423, EPI_ISL_885424, EPI_ISL_885425, EPI_ISL_885426, EPI_ISL_885427, EPI_ISL_885428, EPI_ISL_885429, EPI_ISL_885430, EPI_ISL_885431, EPI_ISL_885432, EPI_ISL_885433, EPI_ISL_885435, EPI_ISL_885436, EPI_ISL_885437, EPI_ISL_885438, EPI_ISL_885439, EPI_ISL_885440, EPI_ISL_885441, EPI_ISL_885442, EPI_ISL_885443, EPI_ISL_885444, EPI_ISL_885445, EPI_ISL_885446, EPI_ISL_885447, EPI_ISL_885448, EPI_ISL_885449, EPI_ISL_885450, EPI_ISL_885451, EPI_ISL_885452, EPI_ISL_885453, EPI_ISL_885454, EPI_ISL_885455, EPI_ISL_885456, | | | |

| | | | |
|---|---|---|---|
| EPI_ISL_885457, EPI_ISL_885458, EPI_ISL_885459, EPI_ISL_885460, EPI_ISL_885461, EPI_ISL_885462, EPI_ISL_885463, EPI_ISL_885464, EPI_ISL_885465, EPI_ISL_885466, EPI_ISL_885467, EPI_ISL_885468, EPI_ISL_885469, EPI_ISL_885470, EPI_ISL_885471, EPI_ISL_885472, EPI_ISL_885473, EPI_ISL_885474, EPI_ISL_885475, EPI_ISL_885476, EPI_ISL_885477, EPI_ISL_885478, EPI_ISL_885479, EPI_ISL_885480, EPI_ISL_885481, EPI_ISL_885482, EPI_ISL_885483, EPI_ISL_885484, EPI_ISL_885485, EPI_ISL_885486, EPI_ISL_885487, EPI_ISL_885488, EPI_ISL_885489, EPI_ISL_885490, EPI_ISL_885493, EPI_ISL_885495, EPI_ISL_885498, EPI_ISL_885499, EPI_ISL_885500, EPI_ISL_885501, EPI_ISL_885505, EPI_ISL_885506, EPI_ISL_885509, EPI_ISL_885510, EPI_ISL_885513, EPI_ISL_885515, EPI_ISL_885518, EPI_ISL_885521, EPI_ISL_885522, EPI_ISL_885523, EPI_ISL_885525, EPI_ISL_885527, EPI_ISL_885529, EPI_ISL_885530, EPI_ISL_885531, EPI_ISL_885532, EPI_ISL_885533, EPI_ISL_885536, EPI_ISL_885537, EPI_ISL_885539, EPI_ISL_885540, EPI_ISL_885541, EPI_ISL_885542, EPI_ISL_885543, EPI_ISL_885544, EPI_ISL_885547, EPI_ISL_885548, EPI_ISL_885549, EPI_ISL_885552, EPI_ISL_885553, EPI_ISL_885554, EPI_ISL_885555, EPI_ISL_885556, EPI_ISL_885557, EPI_ISL_885558, EPI_ISL_885559, EPI_ISL_885560, EPI_ISL_885561, EPI_ISL_885565, EPI_ISL_885566, EPI_ISL_885568, EPI_ISL_885569, EPI_ISL_885570, EPI_ISL_885571, EPI_ISL_885572, EPI_ISL_885573, EPI_ISL_885576, EPI_ISL_885577, EPI_ISL_885578, EPI_ISL_885581, EPI_ISL_885583, EPI_ISL_885584, EPI_ISL_885586, EPI_ISL_885587, EPI_ISL_885590, EPI_ISL_885591, EPI_ISL_885592, EPI_ISL_885594, EPI_ISL_885595, EPI_ISL_885596, EPI_ISL_885597, EPI_ISL_885598, EPI_ISL_885599, EPI_ISL_885600, EPI_ISL_885601, EPI_ISL_885603, EPI_ISL_885605, EPI_ISL_885606, EPI_ISL_885610, EPI_ISL_885611, EPI_ISL_885612, EPI_ISL_885613, EPI_ISL_885615, EPI_ISL_885616, EPI_ISL_885617, EPI_ISL_885618, EPI_ISL_885619, EPI_ISL_885620, EPI_ISL_885622, EPI_ISL_885623, EPI_ISL_885624, EPI_ISL_885629, EPI_ISL_885630, EPI_ISL_885631, EPI_ISL_885632, EPI_ISL_885633, EPI_ISL_885635, EPI_ISL_885637, EPI_ISL_885638, EPI_ISL_885640, EPI_ISL_885641, EPI_ISL_885642, EPI_ISL_885643, EPI_ISL_885644, EPI_ISL_885645, EPI_ISL_885646, EPI_ISL_885648, EPI_ISL_885649, EPI_ISL_885650, EPI_ISL_885651, EPI_ISL_885652, EPI_ISL_885653, EPI_ISL_885654, EPI_ISL_885655, EPI_ISL_885656, EPI_ISL_885657, EPI_ISL_885659, EPI_ISL_885661, EPI_ISL_885662, EPI_ISL_885663, EPI_ISL_885664, EPI_ISL_885667, EPI_ISL_885668, EPI_ISL_885671, EPI_ISL_885673, EPI_ISL_885674, EPI_ISL_885676, EPI_ISL_885677, EPI_ISL_885679, EPI_ISL_885680, EPI_ISL_885682, EPI_ISL_885683, EPI_ISL_885686, EPI_ISL_885690, EPI_ISL_885691, EPI_ISL_885694, EPI_ISL_885695, EPI_ISL_885697, EPI_ISL_885698, EPI_ISL_885700, EPI_ISL_885701, EPI_ISL_885702, EPI_ISL_885703, EPI_ISL_885705, EPI_ISL_885706, EPI_ISL_885707, EPI_ISL_885708, EPI_ISL_885709, EPI_ISL_885711, EPI_ISL_885716, EPI_ISL_885717, EPI_ISL_885718, EPI_ISL_885719, EPI_ISL_885721, EPI_ISL_885723, EPI_ISL_885725, EPI_ISL_885726, EPI_ISL_885727, EPI_ISL_885728, EPI_ISL_885729, EPI_ISL_885730, EPI_ISL_885731, EPI_ISL_885732, EPI_ISL_885735, EPI_ISL_885736, EPI_ISL_885739, EPI_ISL_885740, EPI_ISL_885741, EPI_ISL_885744, EPI_ISL_885746, EPI_ISL_885747, EPI_ISL_885749, EPI_ISL_885751, EPI_ISL_885752, EPI_ISL_885753, EPI_ISL_885754, EPI_ISL_885757, EPI_ISL_885758, EPI_ISL_885759, EPI_ISL_885760, EPI_ISL_885762, EPI_ISL_885763, EPI_ISL_885766, EPI_ISL_885767, EPI_ISL_885769, EPI_ISL_885771, EPI_ISL_885772, EPI_ISL_885774, EPI_ISL_885777, EPI_ISL_885778, EPI_ISL_885781, EPI_ISL_885782, EPI_ISL_885783, EPI_ISL_885784, EPI_ISL_885786, EPI_ISL_885787, EPI_ISL_885789, EPI_ISL_885790, EPI_ISL_885791, EPI_ISL_885796, EPI_ISL_885797, EPI_ISL_885798, EPI_ISL_885799, EPI_ISL_885801, EPI_ISL_885802, EPI_ISL_885803, EPI_ISL_885805, EPI_ISL_885806, EPI_ISL_885808, EPI_ISL_885809, EPI_ISL_885811, EPI_ISL_885814, EPI_ISL_885816, EPI_ISL_885817, EPI_ISL_885818, EPI_ISL_885819, EPI_ISL_885821, EPI_ISL_885823, EPI_ISL_885827, EPI_ISL_885828, EPI_ISL_885833, EPI_ISL_885835, EPI_ISL_885838, EPI_ISL_885843, EPI_ISL_885844, EPI_ISL_885846, EPI_ISL_885848, EPI_ISL_885855, EPI_ISL_885856, EPI_ISL_885860, EPI_ISL_885864, EPI_ISL_885867, EPI_ISL_885868, EPI_ISL_885870, EPI_ISL_885871, EPI_ISL_885874, EPI_ISL_885875, EPI_ISL_885879, EPI_ISL_885883, EPI_ISL_885885, EPI_ISL_885888, EPI_ISL_885894, EPI_ISL_885898, EPI_ISL_885899, EPI_ISL_885902, EPI_ISL_885906, EPI_ISL_885910, EPI_ISL_885912, EPI_ISL_885915, EPI_ISL_885917, EPI_ISL_885918, EPI_ISL_885919, EPI_ISL_885924, EPI_ISL_885926, EPI_ISL_885927, EPI_ISL_885928, EPI_ISL_885929, EPI_ISL_885931, EPI_ISL_885934, EPI_ISL_885935, EPI_ISL_885937, EPI_ISL_885939, EPI_ISL_885946, EPI_ISL_885951, EPI_ISL_885954, EPI_ISL_885958, EPI_ISL_885959, EPI_ISL_885961, EPI_ISL_885962, EPI_ISL_885963, EPI_ISL_885966, EPI_ISL_885967, EPI_ISL_885970, EPI_ISL_885974, EPI_ISL_885976, EPI_ISL_885980, EPI_ISL_885982, EPI_ISL_885983, EPI_ISL_885984, EPI_ISL_885987, EPI_ISL_885988, EPI_ISL_885990, EPI_ISL_885992, EPI_ISL_885993, EPI_ISL_885995, EPI_ISL_885996, EPI_ISL_885997, EPI_ISL_885998, EPI_ISL_885999, EPI_ISL_886001, EPI_ISL_886002, EPI_ISL_886004, EPI_ISL_886005, EPI_ISL_886006, EPI_ISL_886009, EPI_ISL_886014, EPI_ISL_886015, EPI_ISL_886018, EPI_ISL_886020, EPI_ISL_886021, EPI_ISL_886027, EPI_ISL_886029, EPI_ISL_886032, EPI_ISL_886037, EPI_ISL_886038, EPI_ISL_886039, EPI_ISL_886040, EPI_ISL_886043, EPI_ISL_886044, EPI_ISL_886046, EPI_ISL_886047, EPI_ISL_886049, EPI_ISL_886050, EPI_ISL_886053, EPI_ISL_886056, EPI_ISL_886057, EPI_ISL_886060, EPI_ISL_886063, EPI_ISL_886064, EPI_ISL_886066, EPI_ISL_886067, EPI_ISL_886070, EPI_ISL_886071, EPI_ISL_886074, EPI_ISL_886075, EPI_ISL_886076, EPI_ISL_886077, EPI_ISL_886078, EPI_ISL_886081, EPI_ISL_886083, EPI_ISL_886089, EPI_ISL_886090, EPI_ISL_886091, EPI_ISL_886096, EPI_ISL_886097, EPI_ISL_886098, EPI_ISL_886099, EPI_ISL_886101, EPI_ISL_886103, EPI_ISL_886106, EPI_ISL_886110, EPI_ISL_886112, EPI_ISL_886116, EPI_ISL_886117, EPI_ISL_886118, EPI_ISL_886119, EPI_ISL_886120, EPI_ISL_886121, EPI_ISL_886127, EPI_ISL_886129, EPI_ISL_886132, EPI_ISL_886133, EPI_ISL_886134, EPI_ISL_886136, EPI_ISL_886137, EPI_ISL_886140, EPI_ISL_886141, EPI_ISL_886142, EPI_ISL_886143, EPI_ISL_886144, EPI_ISL_886145, EPI_ISL_886146, EPI_ISL_886148, EPI_ISL_886149, EPI_ISL_886150, EPI_ISL_886154 | | | |
| see above | Lighthouse Lab in Alderley Park | Wellcome Sanger Institute for the COVID-19 Genomics UK (COG-UK) Consortium | Jacquelyn Wynn, Mairead Hyland, The Lighthouse Lab in Alderley Park and Alex Alderton, Roberto Amato, Sonia Goncalves, Ewan Harrison, David K. Jackson, Ian Johnston, Dominic Kwiatkowski, Cordelia Langford, John Sillitoe on behalf of the Wellcome Sanger Institute COVID-19 Surveillance Team |
| EPI_ISL_887116, EPI_ISL_887119, EPI_ISL_887122, EPI_ISL_887128, EPI_ISL_887138, EPI_ISL_887139 | Institute of Medical Microbiology and Hospital Hygiene | Institute of Medical Microbiology and Hospital Hygiene | Prof. Dr. Achim Kaasch, Aljoscha Tersteegen |
| EPI_ISL_887512, EPI_ISL_887516, EPI_ISL_887517, EPI_ISL_887518, EPI_ISL_887524, EPI_ISL_887528, EPI_ISL_887539, EPI_ISL_887540, EPI_ISL_887541, EPI_ISL_887542, EPI_ISL_887543, EPI_ISL_887545, EPI_ISL_887546, EPI_ISL_887547, EPI_ISL_887548, EPI_ISL_887549, EPI_ISL_887551, EPI_ISL_887553, EPI_ISL_887554, EPI_ISL_887555, EPI_ISL_887556, EPI_ISL_887557, EPI_ISL_887569, EPI_ISL_887570, EPI_ISL_887571, EPI_ISL_887572, EPI_ISL_887573, EPI_ISL_887574, EPI_ISL_887575, EPI_ISL_887576, EPI_ISL_887578, EPI_ISL_887579 | | | |
| see above | Johns Hopkins Hospital Department of Pathology | Johns Hopkins Hospital Department of Pathology | C. Paul Morris, Chun Huai Luo, Adannaya Amadi, Matthew Schwartz, Nicholas Gallagher, Heba H. Mostafa |
| EPI_ISL_888850, EPI_ISL_888906 | Michigan Department of Health and Human Services, Bureau of Laboratories | Michigan Department of Health and Human Services, Bureau of Laboratories | Blankenship HM, Riner D, Soehnlen MK |
| EPI_ISL_888909, EPI_ISL_888919, EPI_ISL_888930, EPI_ISL_888945, EPI_ISL_888946 | Wyoming Public Health Laboratory | Wyoming Public Health Laboratory | Noah Hull, Taylor Fearing, Lynette Gumbleton, Channing Weber, Ashley Norberg, Bailey Bowcutt, and Wanda Manley |
| EPI_ISL_889743, EPI_ISL_889744, EPI_ISL_889746, EPI_ISL_889747, EPI_ISL_889748, EPI_ISL_889749, EPI_ISL_889750, EPI_ISL_889751, EPI_ISL_889752, EPI_ISL_889753, EPI_ISL_889754, EPI_ISL_889755, EPI_ISL_889756, EPI_ISL_889757, EPI_ISL_889758, EPI_ISL_889759, EPI_ISL_889760, EPI_ISL_889761 | | | |
| see above | LSUHS Emerging Viral Threat Laboratory | Microbial Genome Sequencing Center | Jeremy P. Kamil, Jennifer L. Carroll, Camille F. Abshire, Maarten Van Diest, Mohammed N.A. Siddiquey, Andrew D. Yurochko, Martin J. Sapp, Rona S. Scott, Christopher G. Kevil, Daniel J. Snyder, Vaughn S. Cooper, John A. Vanchiere |
| EPI_ISL_891070, EPI_ISL_891071, EPI_ISL_891072 | Seattle Flu Study | Seattle Flu Study | Deborah A. Nickerson, Chris D. Frazar, Jover Lee, Benjamin Pelle, Erica Ryke, Matthew Richardson, Amanda Adler, Elisabeth Brandstetter, Peter D. Han, Kairsten Fay, Misja Ilcisin, Kirsten Lacombe, Thomas R. Sibley, Melissa Truong, Caitlin R. Wolf, Michael Boeckh, Janet A. Englund, Michael Famulare, Barry R. Lutz, Mark J. Rieder, Lea M. Starita, Matthew Thompson, Jay Shendure, Trevor Bedford, Helen Y. Chu |
| EPI_ISL_891073, EPI_ISL_891074 | Seattle Flu Study | Seattle Flu Study | Deborah A. Nickerson, Chris D. Frazar, Jover Lee, Benjamin Pelle, Erica Ryke, Matthew Richardson, Amanda Adler, Elisabeth Brandstetter, Peter D. Han, Kairsten Fay, Misja Ilcisin, Kirsten Lacombe, Thomas R. Sibley, Melissa Truong, Caitlin R. Wolf, Karen Cowgill, Stephanie Schrag, Jeff Duchin, Michael Boeckh, Janet A. Englund, Michael Famulare, Barry R. Lutz, Mark J. Rieder, Lea M. Starita, Matthew Thompson, Helen Y. Chu, Trevor Bedford, Jay Shendure |
| EPI_ISL_891075, EPI_ISL_891076 | Seattle Flu Study | Seattle Flu Study | Deborah A. Nickerson, Chris D. Frazar, Jover Lee, Benjamin Pelle, Erica Ryke, Matthew Richardson, Amanda Adler, Elisabeth Brandstetter, Peter D. Han, Kairsten Fay, Misja Ilcisin, Kirsten Lacombe, Thomas R. Sibley, Melissa Truong, Caitlin R. Wolf, Michael Boeckh, Janet A. Englund, Michael Famulare, Barry R. Lutz, Mark J. Rieder, Lea M. Starita, Matthew Thompson, Jay Shendure, Trevor Bedford, Helen Y. Chu |
| EPI_ISL_891077, EPI_ISL_891078, EPI_ISL_891079, EPI_ISL_891080, EPI_ISL_891081 | Seattle Flu Study | Seattle Flu Study | Deborah A. Nickerson, Chris D. Frazar, Jover Lee, Benjamin Pelle, Erica Ryke, Matthew Richardson, Amanda Adler, Elisabeth Brandstetter, Peter D. Han, Kairsten Fay, Misja Ilcisin, Kirsten Lacombe, Thomas R. Sibley, Melissa Truong, Caitlin R. Wolf, Karen Cowgill, Stephanie Schrag, Jeff Duchin, Michael Boeckh, Janet A. Englund, Michael Famulare, Barry R. Lutz, Mark J. Rieder, Lea M. Starita, Matthew Thompson, Helen Y. Chu, Trevor Bedford, Jay Shendure |
| EPI_ISL_891082, EPI_ISL_891083 | Seattle Flu Study | Seattle Flu Study | Deborah A. Nickerson, Chris D. Frazar, Jover Lee, Benjamin Pelle, Erica Ryke, Matthew Richardson, Amanda Adler, Elisabeth Brandstetter, Peter D. Han, Kairsten Fay, Misja Ilcisin, Kirsten Lacombe, Thomas R. Sibley, Melissa Truong, Caitlin R. Wolf, Michael Boeckh, Janet A. Englund, Michael Famulare, Barry R. Lutz, Mark J. Rieder, Lea M. Starita, Matthew Thompson, Jay Shendure, Trevor Bedford, Helen Y. Chu |
| EPI_ISL_891092 | Seattle Flu Study | Seattle Flu Study | Deborah A. Nickerson, Chris D. Frazar, Jover Lee, Benjamin Pelle, Erica Ryke, Matthew Richardson, Amanda Adler, Elisabeth Brandstetter, Peter D. Han, Kairsten Fay, Misja Ilcisin, Kirsten Lacombe, Thomas R. Sibley, Melissa Truong, Caitlin R. Wolf, Karen Cowgill, Stephanie Schrag, Jeff Duchin, Michael Boeckh, Janet A. Englund, Michael Famulare, Barry R. Lutz, Mark J. Rieder, Lea M. Starita, Matthew Thompson, Helen Y. Chu, Trevor Bedford, Jay Shendure |
| EPI_ISL_891274, EPI_ISL_891277, EPI_ISL_891287, EPI_ISL_891291, EPI_ISL_891302, EPI_ISL_891303, EPI_ISL_891321, EPI_ISL_891336, EPI_ISL_891341, EPI_ISL_891343, EPI_ISL_891348, EPI_ISL_891362, EPI_ISL_891368, EPI_ISL_891381, EPI_ISL_891390, EPI_ISL_891407, EPI_ISL_891408, EPI_ISL_891414, EPI_ISL_891418, EPI_ISL_891425, EPI_ISL_891428, EPI_ISL_891429, EPI_ISL_891432, EPI_ISL_891443 | | | |
| see above | Lighthouse Lab in Cambridge | Wellcome Sanger Institute for the COVID-19 Genomics UK (COG-UK) Consortium | Rob Howes, The Lighthouse Lab in Cambridge and Alex Alderton, Roberto Amato, Sonia Goncalves, Ewan Harrison, David K. Jackson, Ian Johnston, Dominic Kwiatkowski, Cordelia Langford, John Sillitoe on behalf of the Wellcome Sanger Institute COVID-19 Surveillance Team |
| EPI_ISL_891445 | Lighthouse Lab in Glasgow | Wellcome Sanger Institute for the COVID-19 Genomics UK (COG-UK) Consortium | Harper VanSteenhouse, Yumi Kasai, David Gray, Carol Clugston, Anna Dominiczak and Alex Alderton, Roberto Amato, Sonia Goncalves, Ewan Harrison, David K. Jackson, Ian Johnston, Dominic Kwiatkowski, Cordelia Langford, John Sillitoe on behalf of the Wellcome Sanger Institute COVID-19 Surveillance Team |
| EPI_ISL_891449 | Lighthouse Lab in Cambridge | Wellcome Sanger Institute for the COVID-19 Genomics UK (COG-UK) Consortium | Rob Howes, The Lighthouse Lab in Cambridge and Alex Alderton, Roberto Amato, Sonia Goncalves, Ewan Harrison, David K. Jackson, Ian Johnston, Dominic Kwiatkowski, Cordelia Langford, John Sillitoe on behalf of the Wellcome Sanger Institute COVID-19 Surveillance Team |
| EPI_ISL_891458 | Lighthouse Lab in Alderley Park | Wellcome Sanger Institute for the COVID-19 Genomics UK (COG-UK) Consortium | Jacquelyn Wynn, Mairead Hyland, The Lighthouse Lab in Alderley Park and Alex Alderton, Roberto Amato, Sonia Goncalves, Ewan Harrison, David K. Jackson, Ian Johnston, Dominic Kwiatkowski, Cordelia Langford, John Sillitoe on behalf of the Wellcome Sanger Institute COVID-19 Surveillance Team |
| EPI_ISL_891468, EPI_ISL_891471, EPI_ISL_891474, EPI_ISL_891482 | Lighthouse Lab in Cambridge | Wellcome Sanger Institute for the COVID-19 Genomics UK (COG-UK) Consortium | Rob Howes, The Lighthouse Lab in Cambridge and Alex Alderton, Roberto Amato, Sonia Goncalves, Ewan Harrison, David K. Jackson, Ian Johnston, Dominic Kwiatkowski, Cordelia Langford, John Sillitoe on behalf of the Wellcome Sanger Institute COVID-19 Surveillance Team |
| EPI_ISL_891507 | Lighthouse Lab in Alderley Park | Wellcome Sanger Institute for the COVID-19 Genomics UK (COG-UK) Consortium | Jacquelyn Wynn, Mairead Hyland, The Lighthouse Lab in Alderley Park and Alex Alderton, Roberto Amato, Sonia Goncalves, Ewan Harrison, David K. Jackson, Ian Johnston, Dominic Kwiatkowski, Cordelia Langford, John Sillitoe on behalf of the Wellcome Sanger Institute COVID-19 Surveillance Team |
| EPI_ISL_891523, EPI_ISL_891524, EPI_ISL_891533, EPI_ISL_891546, | Lighthouse Lab in Cambridge | Wellcome Sanger Institute for the COVID-19 Genomics UK (COG-UK) Consortium | Rob Howes, The Lighthouse Lab in Cambridge and Alex Alderton, Roberto Amato, Sonia Goncalves, Ewan Harrison, David K. Jackson, Ian Johnston, Dominic Kwiatkowski, Cordelia Langford, John Sillitoe on behalf of the Wellcome Sanger Institute COVID-19 Surveillance Team |

| | | | |
|---|---|---|---|
| EPI_ISL_891552, EPI_ISL_891573, EPI_ISL_891576, EPI_ISL_891655 | | | |
| EPI_ISL_891700 | Lighthouse Lab in Glasgow | Wellcome Sanger Institute for the COVID-19 Genomics UK (COG-UK) Consortium | Harper VanSteenhouse, Yumi Kasai, David Gray, Carol Clugston, Anna Dominiczak and Alex Alderton, Roberto Amato, Sonia Goncalves, Ewan Harrison, David K. Jackson, Ian Johnston, Dominic Kwiatkowski, Cordelia Langford, John Sillitoe on behalf of the Wellcome Sanger Institute COVID-19 Surveillance Team |
| EPI_ISL_891709, EPI_ISL_891755, EPI_ISL_891799 | Lighthouse Lab in Cambridge | Wellcome Sanger Institute for the COVID-19 Genomics UK (COG-UK) Consortium | Rob Howes, The Lighthouse Lab in Cambridge and Alex Alderton, Roberto Amato, Sonia Goncalves, Ewan Harrison, David K. Jackson, Ian Johnston, Dominic Kwiatkowski, Cordelia Langford, John Sillitoe on behalf of the Wellcome Sanger Institute COVID-19 Surveillance Team |
| EPI_ISL_891822 | Lighthouse Lab in Alderley Park | Wellcome Sanger Institute for the COVID-19 Genomics UK (COG-UK) Consortium | Jacquelyn Wynn, Mairead Hyland, The Lighthouse Lab in Alderley Park and Alex Alderton, Roberto Amato, Sonia Goncalves, Ewan Harrison, David K. Jackson, Ian Johnston, Dominic Kwiatkowski, Cordelia Langford, John Sillitoe on behalf of the Wellcome Sanger Institute COVID-19 Surveillance Team |
| EPI_ISL_891832 | Lighthouse Lab in Cambridge | Wellcome Sanger Institute for the COVID-19 Genomics UK (COG-UK) Consortium | Rob Howes, The Lighthouse Lab in Cambridge and Alex Alderton, Roberto Amato, Sonia Goncalves, Ewan Harrison, David K. Jackson, Ian Johnston, Dominic Kwiatkowski, Cordelia Langford, John Sillitoe on behalf of the Wellcome Sanger Institute COVID-19 Surveillance Team |
| EPI_ISL_891850, EPI_ISL_891860, EPI_ISL_891867 | Lighthouse Lab in Alderley Park | Wellcome Sanger Institute for the COVID-19 Genomics UK (COG-UK) Consortium | Jacquelyn Wynn, Mairead Hyland, The Lighthouse Lab in Alderley Park and Alex Alderton, Roberto Amato, Sonia Goncalves, Ewan Harrison, David K. Jackson, Ian Johnston, Dominic Kwiatkowski, Cordelia Langford, John Sillitoe on behalf of the Wellcome Sanger Institute COVID-19 Surveillance Team |
| EPI_ISL_891882, EPI_ISL_891896 | Lighthouse Lab in Cambridge | Wellcome Sanger Institute for the COVID-19 Genomics UK (COG-UK) Consortium | Rob Howes, The Lighthouse Lab in Cambridge and Alex Alderton, Roberto Amato, Sonia Goncalves, Ewan Harrison, David K. Jackson, Ian Johnston, Dominic Kwiatkowski, Cordelia Langford, John Sillitoe on behalf of the Wellcome Sanger Institute COVID-19 Surveillance Team |
| EPI_ISL_891899, EPI_ISL_891900, EPI_ISL_891901, EPI_ISL_891902, EPI_ISL_891904, EPI_ISL_891905, EPI_ISL_891906, EPI_ISL_891907, EPI_ISL_891908, EPI_ISL_891910, EPI_ISL_891911, EPI_ISL_891912, EPI_ISL_891913, EPI_ISL_891914, EPI_ISL_891915, EPI_ISL_891916, EPI_ISL_891917, EPI_ISL_891919, EPI_ISL_891920, EPI_ISL_891921, EPI_ISL_891922, EPI_ISL_891923, EPI_ISL_891924, EPI_ISL_891925, EPI_ISL_891926, EPI_ISL_891927, EPI_ISL_891929, EPI_ISL_891930, EPI_ISL_891931, EPI_ISL_891932, EPI_ISL_891933, EPI_ISL_891934, EPI_ISL_891935, EPI_ISL_891936, EPI_ISL_891937, EPI_ISL_891938, EPI_ISL_891939, EPI_ISL_891940, EPI_ISL_891941, EPI_ISL_891942, EPI_ISL_891943, EPI_ISL_891944, EPI_ISL_891946, EPI_ISL_891947, EPI_ISL_891948, EPI_ISL_891949, EPI_ISL_891950, EPI_ISL_891951, EPI_ISL_891952, EPI_ISL_891953, EPI_ISL_891954, EPI_ISL_891955, EPI_ISL_891956, EPI_ISL_891957, EPI_ISL_891959, EPI_ISL_891960, EPI_ISL_891961, EPI_ISL_891962, EPI_ISL_891963, EPI_ISL_891964, EPI_ISL_891965, EPI_ISL_891966, EPI_ISL_891968, EPI_ISL_891969, EPI_ISL_891970, EPI_ISL_891971, EPI_ISL_892002, EPI_ISL_892005, EPI_ISL_892006, EPI_ISL_892009, EPI_ISL_892010, EPI_ISL_892011, EPI_ISL_892012 | | | |
| see above | Lighthouse Lab in Alderley Park | Wellcome Sanger Institute for the COVID-19 Genomics UK (COG-UK) Consortium | Jacquelyn Wynn, Mairead Hyland, The Lighthouse Lab in Alderley Park and Alex Alderton, Roberto Amato, Sonia Goncalves, Ewan Harrison, David K. Jackson, Ian Johnston, Dominic Kwiatkowski, Cordelia Langford, John Sillitoe on behalf of the Wellcome Sanger Institute COVID-19 Surveillance Team |
| EPI_ISL_896109 | Abklärungs- und _Teststation BL | University Hospital Basel, Clinical Bacteriology | Tim Roloff, Madlen Stange, Helena MB Seth-Smith, Alfredo Mari, Karoline Leuzinger, Julia Bielicki, Samuel Erny, Thomas Goetz, Juerg Sommer, Manuel Battegay, Hans Hirsch, Adrian Egli |
| EPI_ISL_896218, EPI_ISL_896224, EPI_ISL_896229 | SUNY UPSTATE MEDICAL UNIVERSITY | Wadsworth Center, New York State Department of Health | Kirsten St. George, Daryl M. Lamson, Alexis Russel, Matthew Shudt, Melissa A Leisner, Jonathan Plitnick, Navjot Singh, John Kelly, Erasmus Schneider, Erica Lasek-Nesselquist |
| EPI_ISL_896236, EPI_ISL_896237, EPI_ISL_896238, EPI_ISL_896239 | Columbia University Irving Medical Center | Wadsworth Center, New York State Department of Health | Kirsten St. George, Daryl M. Lamson, Alexis Russel, Matthew Shudt, Melissa A Leisner, Jonathan Plitnick, Navjot Singh, John Kelly, Erasmus Schneider, Erica Lasek-Nesselquist |
| EPI_ISL_896278, EPI_ISL_896281, EPI_ISL_896282, EPI_ISL_896286, EPI_ISL_896289, EPI_ISL_896291 | SUNY UPSTATE MEDICAL UNIVERSITY | Wadsworth Center, New York State Department of Health | Kirsten St. George, Daryl M. Lamson, Alexis Russel, Matthew Shudt, Melissa A Leisner, Jonathan Plitnick, Navjot Singh, John Kelly, Erasmus Schneider, Erica Lasek-Nesselquist |
| EPI_ISL_896307, EPI_ISL_896308, EPI_ISL_896309 | Wadsworth Center, New York State Department of Health | Wadsworth Center, New York State Department of Health | Kirsten St. George, Daryl M. Lamson, Alexis Russel, Matthew Shudt, Melissa A Leisner, Jonathan Plitnick, Navjot Singh, John Kelly, Erasmus Schneider, Erica Lasek-Nesselquist |
| EPI_ISL_896381 | Suceava County Emergency Hospital | "Stefan cel Mare" University Metagenomics Lab | Lobiuc Andrei, Gheorghita Roxana |
| EPI_ISL_896447, EPI_ISL_896448, EPI_ISL_896449, EPI_ISL_896450, EPI_ISL_896454, EPI_ISL_896458, EPI_ISL_896459, EPI_ISL_896460, EPI_ISL_896466, EPI_ISL_896467, EPI_ISL_896468, EPI_ISL_896473, EPI_ISL_896476, EPI_ISL_896477, EPI_ISL_896478, EPI_ISL_896479, EPI_ISL_896480, EPI_ISL_896481, EPI_ISL_896482, EPI_ISL_896483, EPI_ISL_896484, EPI_ISL_896485, EPI_ISL_896486, EPI_ISL_896489, EPI_ISL_896490, EPI_ISL_896491, EPI_ISL_896492 | | | |
| see above | KU Leuven, Rega Institute, Clinical and Epidemiological Virology | KU Leuven, Rega Institute, Clinical and Epidemiological Virology | Tony Wawina-Bokalanga, Bert Vanmechelen, Joan Marti-Carerras, Piet Maes |
| EPI_ISL_896503, EPI_ISL_896514, EPI_ISL_896516, EPI_ISL_896517, EPI_ISL_896518, EPI_ISL_896519 | URMC LABS | Wadsworth Center, New York State Department of Health | Kirsten St. George, Daryl M. Lamson, Alexis Russel, Matthew Shudt, Melissa A Leisner, Jonathan Plitnick, Navjot Singh, John Kelly, Erasmus Schneider, Erica Lasek-Nesselquist |
| EPI_ISL_897599, EPI_ISL_897600, EPI_ISL_897601, EPI_ISL_897602, EPI_ISL_897607, EPI_ISL_897608, EPI_ISL_897609, EPI_ISL_897610, EPI_ISL_897612, EPI_ISL_897613, EPI_ISL_897614, EPI_ISL_897615, EPI_ISL_897619, EPI_ISL_897620, EPI_ISL_897626, EPI_ISL_897627, EPI_ISL_897628, EPI_ISL_897629, EPI_ISL_897630, EPI_ISL_897631, EPI_ISL_897632, EPI_ISL_897633, EPI_ISL_897634, EPI_ISL_897635, EPI_ISL_897636, EPI_ISL_897637, EPI_ISL_897638, EPI_ISL_897639, EPI_ISL_897640, EPI_ISL_897641, EPI_ISL_897642, EPI_ISL_897643, EPI_ISL_897644, EPI_ISL_897645, EPI_ISL_897646, EPI_ISL_897678, EPI_ISL_897683, EPI_ISL_897686, EPI_ISL_897696, EPI_ISL_897698, EPI_ISL_897701, EPI_ISL_897702, EPI_ISL_897703, EPI_ISL_897709, EPI_ISL_897710, EPI_ISL_897715, EPI_ISL_897720, EPI_ISL_897771, EPI_ISL_897772, EPI_ISL_897773, EPI_ISL_897774, EPI_ISL_897775, EPI_ISL_897776, EPI_ISL_897777, EPI_ISL_897778, EPI_ISL_897779, EPI_ISL_897780, EPI_ISL_897781, EPI_ISL_897782, EPI_ISL_897783, EPI_ISL_897784, EPI_ISL_897785, EPI_ISL_897786, EPI_ISL_897787, EPI_ISL_897788, EPI_ISL_897789, EPI_ISL_897790, EPI_ISL_897791, EPI_ISL_897792, EPI_ISL_897793, EPI_ISL_897794, EPI_ISL_897795, EPI_ISL_897796, EPI_ISL_897797, EPI_ISL_897798, EPI_ISL_897799, EPI_ISL_897800, EPI_ISL_897801, EPI_ISL_897802, EPI_ISL_897803, EPI_ISL_897804, EPI_ISL_897805, EPI_ISL_897806, EPI_ISL_897807, EPI_ISL_897808, EPI_ISL_897809, EPI_ISL_897810, EPI_ISL_897811, EPI_ISL_897812, EPI_ISL_897813, EPI_ISL_897814, EPI_ISL_897815, EPI_ISL_897816, EPI_ISL_897817, EPI_ISL_897818, EPI_ISL_897819, EPI_ISL_897820, EPI_ISL_897821, EPI_ISL_897822, EPI_ISL_897823, EPI_ISL_897824, EPI_ISL_897825, EPI_ISL_897826, EPI_ISL_897827, EPI_ISL_897828, EPI_ISL_897829, EPI_ISL_897830, EPI_ISL_897831, EPI_ISL_897832, EPI_ISL_897833, EPI_ISL_897834, EPI_ISL_897835, EPI_ISL_897836, EPI_ISL_897837, EPI_ISL_897838, EPI_ISL_897839, EPI_ISL_897840, EPI_ISL_897841, EPI_ISL_897842, EPI_ISL_897843, EPI_ISL_897844, EPI_ISL_897845, EPI_ISL_897859, EPI_ISL_897905, EPI_ISL_897920, EPI_ISL_897921, EPI_ISL_897922, EPI_ISL_897923, EPI_ISL_897929, EPI_ISL_897930, EPI_ISL_897953, EPI_ISL_897967, EPI_ISL_897968, EPI_ISL_897969, EPI_ISL_897970, EPI_ISL_897971, EPI_ISL_897972, EPI_ISL_897973, EPI_ISL_897974, EPI_ISL_897975, EPI_ISL_897976, EPI_ISL_897977, EPI_ISL_897978, EPI_ISL_897979 | | | |
| see above | University Hospitals of Geneva, Laboratory of Virology | HUG, Laboratory of Virology and the Health2030 Genome Center | Samuel Cordey, Ana Rita Goncalves, Laurent Kaiser, Lorenzo Cerutti, Henri Pegeot, Melyssa Elies, Deborah Penet, Keith Harshman, Ioannis Xenarios, Emmanouil Dermitzakis |
| EPI_ISL_900491, EPI_ISL_900493, EPI_ISL_900494, EPI_ISL_900495, EPI_ISL_900497, EPI_ISL_900498, EPI_ISL_900500 | Althaia. Xarxa Assistencial Universitària de Manresa | IrsiCaixa - Can Ruti CovidSeq | Fundació irsiCaixa. Hospital Universitari Germans Trias i Pujol(HUGTiP), 2a planta, maternal Ctra Canyet s/n, Badalona Gloria Trujillo, Rafel Perez Vidal, Jaume Trape Pujol, Carolina Gonzalez Fernandez, Roger Paredes, Eulalia Grau, Bonaventura Clotet |
| EPI_ISL_902754 | University Hospital Sant'Andrea-Sapienza | INMI Lazzaro Spallanzani IRCCS | B Bartolini, E Giombini, M Rueca, O Butera, F Messina, C.E.M Gruber, M Simmaco, I Santino, A Di Caro, MR Capobianchi |
| EPI_ISL_902755 | IRCCS San Raffaele | INMI Lazzaro Spallanzani IRCCS | E Giombini, M Rueca, O Butera, F Messina, C.E.M Gruber, B Bartolini, D Russo, D Limongi, MR Capobianchi, A Di Caro |
| EPI_ISL_902942 | Maryland Public Health Laboratory | Maryland Public Health Laboratory | Maryland Department of Health Laboratories Administration |
| EPI_ISL_903366 | University Medical Center Hamburg Eppendorf | Heinrich Pette Institute, Leibniz Institute for Experimental Virology | Alexis Robitaille, Thomas Günther, Johannes Knobloch, Martin Aepfelbacher, Nicole Fischer, Adam Grundhoff |
| EPI_ISL_904071, EPI_ISL_904072, EPI_ISL_904073, EPI_ISL_904074, EPI_ISL_904075, EPI_ISL_904076, EPI_ISL_904077, EPI_ISL_904078, EPI_ISL_904079, EPI_ISL_904080, EPI_ISL_904081, EPI_ISL_904082, EPI_ISL_904083, EPI_ISL_904084, EPI_ISL_904085 | | | |
| see above | New Mexico Department of Health Scientific Laboratory | New Mexico Department of Health Scientific Laboratory | Ellie Johnson, Anastacia Griego-Fisher, D'eldra Malone |
| EPI_ISL_904217, EPI_ISL_904218, EPI_ISL_904254, EPI_ISL_904603, EPI_ISL_904604, EPI_ISL_904605, EPI_ISL_904606 | Dutch COVID-19 response team | Erasmus Medical Center | Bas Oude Munnink, Reina Sikkema, David Nieuwenhuijse, Irina Chestakova, Anne van der Linden, Marjan Boter, Emmanuelle Munger, Corine GeurtsvanKessel, Annemiek van der Eijk, Richard Molenkamp, Marion Koopmans, on behalf of the Dutch national COVID-19 response team. |
| EPI_ISL_904661, EPI_ISL_904662, EPI_ISL_904663, EPI_ISL_904675, EPI_ISL_904676, EPI_ISL_904679, EPI_ISL_904680, EPI_ISL_904681, EPI_ISL_904685, EPI_ISL_904713, EPI_ISL_904734, EPI_ISL_904739, EPI_ISL_904745, EPI_ISL_904758, EPI_ISL_904763, EPI_ISL_904772, EPI_ISL_904773, EPI_ISL_904774, EPI_ISL_904777, EPI_ISL_904780, EPI_ISL_904790, EPI_ISL_904793, EPI_ISL_904808, EPI_ISL_904809, EPI_ISL_904811, EPI_ISL_904813, EPI_ISL_904830, EPI_ISL_904831, EPI_ISL_904836, EPI_ISL_904837, EPI_ISL_904838, EPI_ISL_904839, EPI_ISL_904842, EPI_ISL_904868, EPI_ISL_904881, EPI_ISL_904884, EPI_ISL_905179, EPI_ISL_905184, EPI_ISL_905189, EPI_ISL_905198, EPI_ISL_905199, EPI_ISL_905205, EPI_ISL_905206, EPI_ISL_905207, EPI_ISL_905208, EPI_ISL_905209, EPI_ISL_905210, EPI_ISL_905234, EPI_ISL_905236, EPI_ISL_905292, EPI_ISL_905293, EPI_ISL_905311, EPI_ISL_905319, EPI_ISL_905320, EPI_ISL_905330, EPI_ISL_905331, EPI_ISL_905332, EPI_ISL_905338, EPI_ISL_905341, EPI_ISL_905342, EPI_ISL_905343, EPI_ISL_905344, EPI_ISL_905346, EPI_ISL_905347, EPI_ISL_905348 | | | |
| see above | Dutch COVID-19 response team | National Institute for Public Health and the Environment (RIVM) | Adam Meijer, Harry Vennema, Dirk Eggink, Jeroen Cremer, Sharon van den Brink, Bas van der Veer, AnneMarie van den Brandt, Florian Zwagemaker, Dennis Schmitz, Chantal Reusken, on behalf of the national COVID-19 response team |
| EPI_ISL_905788, EPI_ISL_905789, | UCLA Clinical Micro Lab | Los Angeles County PHL | P. Hemarajata et al. |

| | | | |
|---|---|---|---|
| EPI_ISL_905790 | | | |
| EPI_ISL_906068, EPI_ISL_906069 | Instituto Adolfo Lutz - Regional de Campinas | Instituto Adolfo Lutz, Interdiciplinary Procedures Center, Strategic Laboratory | Claudio Tavares Sacchi, Claudia Regina Gonçalves, Erica Valessa Ramos Gomes, Karoline Rodrigues Campos |
| EPI_ISL_906190, EPI_ISL_906191, EPI_ISL_906192, EPI_ISL_906193, EPI_ISL_906194, EPI_ISL_906195, EPI_ISL_906196, EPI_ISL_906197, EPI_ISL_906198, EPI_ISL_906199, EPI_ISL_906201, EPI_ISL_906202, EPI_ISL_906203, EPI_ISL_906204, EPI_ISL_906206, EPI_ISL_906208, EPI_ISL_906209, EPI_ISL_906211, EPI_ISL_906213, EPI_ISL_906214, EPI_ISL_906216, EPI_ISL_906217, EPI_ISL_906219, EPI_ISL_906227, EPI_ISL_906229, EPI_ISL_906233, EPI_ISL_906238, EPI_ISL_906240, EPI_ISL_906241, EPI_ISL_906242, EPI_ISL_906246, EPI_ISL_906249, EPI_ISL_906266, EPI_ISL_906268, EPI_ISL_906269, EPI_ISL_906270, EPI_ISL_906271, EPI_ISL_906272 | | | |
| see above | University of Wisconsin-Madison AIDS Vaccine Research Laboratories | University of Wisconsin-Madison AIDS Vaccine Research Laboratories | Gage Moreno, Katarina Braun, et al. AIDS Vaccine Research Laboratories |
| EPI_ISL_906558 | Laboratorio de Salud Publica de Amazonas | Instituto Nacional de Salud- Dirección de Investigación en Salud Pública, Universidad de los Andes- Applied genomics research group, Vicerrectoria de Investigación y Creación, Universidad de los Andes- Systems and Computing Engineering Department | Katherine Laiton-Donato, Diego A. Álvarez-Díaz, Carlos Franco-Muñoz, Mauricio Pacheco-Montealegre, Héctor Alejandro Ruiz-Moreno, Maria T. Herrera-Sepúlveda, Diego Andrés Prada, Jhonnatan Reales-González, Sheryll Corchuelo, Julian Naizaque, Gerardo Santamaría Jorge Duitama, Laura Natalia Gonzalez, Jorge Ivan Diaz, Silvia Restrepo-Restrepo, Magdalena Wiesner, Martha Lucia Ospina Martinez, Marcela Mercado-Reyes |
| EPI_ISL_906652, EPI_ISL_906653, EPI_ISL_906654 | Maine Health and Environmental Testing Laboratory (Maine HETL) | Tewhey Lab, The Jackson Laboratory | Matluk,N., Dewey,H., Iosue,F., Barter,M., Lynch,R., Munger,H. and Tewhey,R. |
| EPI_ISL_906794, EPI_ISL_906795 | Norwegian Institute of Public Health, Department of Virology | Norwegian Institute of Public Health, Department of Virology | Kathrine Stene-Johansen, Kamilla Heddeland Instefjord, Hilde Elshaug, Atiya R Ali,Marie Paulsen Madsen, Rasmus Riis Kopperud, Hilde Vollan, Karoline Bragstad, Olav Hungnes |
| EPI_ISL_906797 | Dept. of Medical Microbiology, Stavanger University Hospital, Helse Stavanger HF | Norwegian Institute of Public Health, Department of Virology | Kathrine Stene-Johansen, Kamilla Heddeland Instefjord, Hilde Elshaug, Atiya R Ali,Marie Paulsen Madsen, Rasmus Riis Kopperud, Hilde Vollan, Karoline Bragstad, Olav Hungnes |
| EPI_ISL_906815, EPI_ISL_906816 | Medical Microbiology Unit, Department for Laboratory Medicine, Drammen Hospital, Vestre Viken Health Trust, | Norwegian Institute of Public Health, Department of Virology | Kathrine Stene-Johansen, Kamilla Heddeland Instefjord, Hilde Elshaug, Atiya R Ali,Marie Paulsen Madsen, Rasmus Riis Kopperud, Hilde Vollan, Karoline Bragstad, Olav Hungnes |
| EPI_ISL_907725, EPI_ISL_907775, EPI_ISL_907779, EPI_ISL_907783, EPI_ISL_907841, EPI_ISL_907884, EPI_ISL_907909, EPI_ISL_907933, EPI_ISL_907978, EPI_ISL_908001 | Lighthouse Lab in Alderley Park | Wellcome Sanger Institute for the COVID-19 Genomics UK (COG-UK) Consortium | Jacquelyn Wynn, Mairead Hyland, The Lighthouse Lab in Alderley Park and Alex Alderton, Roberto Amato, Sonia Goncalves, Ewan Harrison, David K. Jackson, Ian Johnston, Dominic Kwiatkowski, Cordelia Langford, John Sillitoe on behalf of the Wellcome Sanger Institute COVID-19 Surveillance Team |
| EPI_ISL_908035 | Lighthouse Lab in Cambridge | Wellcome Sanger Institute for the COVID-19 Genomics UK (COG-UK) Consortium | Rob Howes, The Lighthouse Lab in Cambridge and Alex Alderton, Roberto Amato, Sonia Goncalves, Ewan Harrison, David K. Jackson, Ian Johnston, Dominic Kwiatkowski, Cordelia Langford, John Sillitoe on behalf of the Wellcome Sanger Institute COVID-19 Surveillance Team |
| EPI_ISL_908036 | Lighthouse Lab in Glasgow | Wellcome Sanger Institute for the COVID-19 Genomics UK (COG-UK) Consortium | Harper VanSteenhouse, Yumi Kasai, David Gray, Carol Clugston, Anna Dominiczak and Alex Alderton, Roberto Amato, Sonia Goncalves, Ewan Harrison, David K. Jackson, Ian Johnston, Dominic Kwiatkowski, Cordelia Langford, John Sillitoe on behalf of the Wellcome Sanger Institute COVID-19 Surveillance Team |
| EPI_ISL_908037, EPI_ISL_908038, EPI_ISL_908039, EPI_ISL_908040, EPI_ISL_908041, EPI_ISL_908042, EPI_ISL_908043, EPI_ISL_908044, EPI_ISL_908045, EPI_ISL_908046, EPI_ISL_908047, EPI_ISL_908048, EPI_ISL_908049, EPI_ISL_908050 | | | |
| see above | Lighthouse Lab in Cambridge | Wellcome Sanger Institute for the COVID-19 Genomics UK (COG-UK) Consortium | Rob Howes, The Lighthouse Lab in Cambridge and Alex Alderton, Roberto Amato, Sonia Goncalves, Ewan Harrison, David K. Jackson, Ian Johnston, Dominic Kwiatkowski, Cordelia Langford, John Sillitoe on behalf of the Wellcome Sanger Institute COVID-19 Surveillance Team |
| EPI_ISL_908051 | Lighthouse Lab in Glasgow | Wellcome Sanger Institute for the COVID-19 Genomics UK (COG-UK) Consortium | Harper VanSteenhouse, Yumi Kasai, David Gray, Carol Clugston, Anna Dominiczak and Alex Alderton, Roberto Amato, Sonia Goncalves, Ewan Harrison, David K. Jackson, Ian Johnston, Dominic Kwiatkowski, Cordelia Langford, John Sillitoe on behalf of the Wellcome Sanger Institute COVID-19 Surveillance Team |
| EPI_ISL_908052, EPI_ISL_908053 | Lighthouse Lab in Cambridge | Wellcome Sanger Institute for the COVID-19 Genomics UK (COG-UK) Consortium | Rob Howes, The Lighthouse Lab in Cambridge and Alex Alderton, Roberto Amato, Sonia Goncalves, Ewan Harrison, David K. Jackson, Ian Johnston, Dominic Kwiatkowski, Cordelia Langford, John Sillitoe on behalf of the Wellcome Sanger Institute COVID-19 Surveillance Team |
| EPI_ISL_908054 | Lighthouse Lab in Glasgow | Wellcome Sanger Institute for the COVID-19 Genomics UK (COG-UK) Consortium | Harper VanSteenhouse, Yumi Kasai, David Gray, Carol Clugston, Anna Dominiczak and Alex Alderton, Roberto Amato, Sonia Goncalves, Ewan Harrison, David K. Jackson, Ian Johnston, Dominic Kwiatkowski, Cordelia Langford, John Sillitoe on behalf of the Wellcome Sanger Institute COVID-19 Surveillance Team |
| EPI_ISL_908055 | Lighthouse Lab in Cambridge | Wellcome Sanger Institute for the COVID-19 Genomics UK (COG-UK) Consortium | Rob Howes, The Lighthouse Lab in Cambridge and Alex Alderton, Roberto Amato, Sonia Goncalves, Ewan Harrison, David K. Jackson, Ian Johnston, Dominic Kwiatkowski, Cordelia Langford, John Sillitoe on behalf of the Wellcome Sanger Institute COVID-19 Surveillance Team |
| EPI_ISL_908056 | Lighthouse Lab in Glasgow | Wellcome Sanger Institute for the COVID-19 Genomics UK (COG-UK) Consortium | Harper VanSteenhouse, Yumi Kasai, David Gray, Carol Clugston, Anna Dominiczak and Alex Alderton, Roberto Amato, Sonia Goncalves, Ewan Harrison, David K. Jackson, Ian Johnston, Dominic Kwiatkowski, Cordelia Langford, John Sillitoe on behalf of the Wellcome Sanger Institute COVID-19 Surveillance Team |
| EPI_ISL_908057, EPI_ISL_908058 | Lighthouse Lab in Cambridge | Wellcome Sanger Institute for the COVID-19 Genomics UK (COG-UK) Consortium | Rob Howes, The Lighthouse Lab in Cambridge and Alex Alderton, Roberto Amato, Sonia Goncalves, Ewan Harrison, David K. Jackson, Ian Johnston, Dominic Kwiatkowski, Cordelia Langford, John Sillitoe on behalf of the Wellcome Sanger Institute COVID-19 Surveillance Team |
| EPI_ISL_908059, EPI_ISL_908060, EPI_ISL_908061 | Lighthouse Lab in Glasgow | Wellcome Sanger Institute for the COVID-19 Genomics UK (COG-UK) Consortium | Harper VanSteenhouse, Yumi Kasai, David Gray, Carol Clugston, Anna Dominiczak and Alex Alderton, Roberto Amato, Sonia Goncalves, Ewan Harrison, David K. Jackson, Ian Johnston, Dominic Kwiatkowski, Cordelia Langford, John Sillitoe on behalf of the Wellcome Sanger Institute COVID-19 Surveillance Team |
| EPI_ISL_908062, EPI_ISL_908063 | Lighthouse Lab in Cambridge | Wellcome Sanger Institute for the COVID-19 Genomics UK (COG-UK) Consortium | Rob Howes, The Lighthouse Lab in Cambridge and Alex Alderton, Roberto Amato, Sonia Goncalves, Ewan Harrison, David K. Jackson, Ian Johnston, Dominic Kwiatkowski, Cordelia Langford, John Sillitoe on behalf of the Wellcome Sanger Institute COVID-19 Surveillance Team |
| EPI_ISL_908064 | Lighthouse Lab in Glasgow | Wellcome Sanger Institute for the COVID-19 Genomics UK (COG-UK) Consortium | Harper VanSteenhouse, Yumi Kasai, David Gray, Carol Clugston, Anna Dominiczak and Alex Alderton, Roberto Amato, Sonia Goncalves, Ewan Harrison, David K. Jackson, Ian Johnston, Dominic Kwiatkowski, Cordelia Langford, John Sillitoe on behalf of the Wellcome Sanger Institute COVID-19 Surveillance Team |
| EPI_ISL_908065, EPI_ISL_908066, EPI_ISL_908067, EPI_ISL_908068, EPI_ISL_908069 | Lighthouse Lab in Cambridge | Wellcome Sanger Institute for the COVID-19 Genomics UK (COG-UK) Consortium | Rob Howes, The Lighthouse Lab in Cambridge and Alex Alderton, Roberto Amato, Sonia Goncalves, Ewan Harrison, David K. Jackson, Ian Johnston, Dominic Kwiatkowski, Cordelia Langford, John Sillitoe on behalf of the Wellcome Sanger Institute COVID-19 Surveillance Team |
| EPI_ISL_908070 | Lighthouse Lab in Glasgow | Wellcome Sanger Institute for the COVID-19 Genomics UK (COG-UK) Consortium | Harper VanSteenhouse, Yumi Kasai, David Gray, Carol Clugston, Anna Dominiczak and Alex Alderton, Roberto Amato, Sonia Goncalves, Ewan Harrison, David K. Jackson, Ian Johnston, Dominic Kwiatkowski, Cordelia Langford, John Sillitoe on behalf of the Wellcome Sanger Institute COVID-19 Surveillance Team |
| EPI_ISL_908071 | Lighthouse Lab in Cambridge | Wellcome Sanger Institute for the COVID-19 Genomics UK (COG-UK) Consortium | Rob Howes, The Lighthouse Lab in Cambridge and Alex Alderton, Roberto Amato, Sonia Goncalves, Ewan Harrison, David K. Jackson, Ian Johnston, Dominic Kwiatkowski, Cordelia Langford, John Sillitoe on behalf of the Wellcome Sanger Institute COVID-19 Surveillance Team |
| EPI_ISL_908072, EPI_ISL_908073, EPI_ISL_908074, EPI_ISL_908075 | Lighthouse Lab in Glasgow | Wellcome Sanger Institute for the COVID-19 Genomics UK (COG-UK) Consortium | Harper VanSteenhouse, Yumi Kasai, David Gray, Carol Clugston, Anna Dominiczak and Alex Alderton, Roberto Amato, Sonia Goncalves, Ewan Harrison, David K. Jackson, Ian Johnston, Dominic Kwiatkowski, Cordelia Langford, John Sillitoe on behalf of the Wellcome Sanger Institute COVID-19 Surveillance Team |
| EPI_ISL_908076, EPI_ISL_908077, EPI_ISL_908078, EPI_ISL_908079, EPI_ISL_908080 | Lighthouse Lab in Cambridge | Wellcome Sanger Institute for the COVID-19 Genomics UK (COG-UK) Consortium | Rob Howes, The Lighthouse Lab in Cambridge and Alex Alderton, Roberto Amato, Sonia Goncalves, Ewan Harrison, David K. Jackson, Ian Johnston, Dominic Kwiatkowski, Cordelia Langford, John Sillitoe on behalf of the Wellcome Sanger Institute COVID-19 Surveillance Team |
| EPI_ISL_908081 | Lighthouse Lab in Glasgow | Wellcome Sanger Institute for the COVID-19 Genomics UK (COG-UK) Consortium | Harper VanSteenhouse, Yumi Kasai, David Gray, Carol Clugston, Anna Dominiczak and Alex Alderton, Roberto Amato, Sonia Goncalves, Ewan Harrison, David K. Jackson, Ian Johnston, Dominic Kwiatkowski, Cordelia Langford, John Sillitoe on behalf of the Wellcome Sanger Institute COVID-19 Surveillance |

| | | | |
|---|---|---|---|
| | | | Team |
| EPI_ISL_908082, EPI_ISL_908083 | Lighthouse Lab in Cambridge | Wellcome Sanger Institute for the COVID-19 Genomics UK (COG-UK) Consortium | Rob Howes, The Lighthouse Lab in Cambridge and Alex Alderton, Roberto Amato, Sonia Goncalves, Ewan Harrison, David K. Jackson, Ian Johnston, Dominic Kwiatkowski, Cordelia Langford, John Sillitoe on behalf of the Wellcome Sanger Institute COVID-19 Surveillance Team |
| EPI_ISL_908084, EPI_ISL_908085, EPI_ISL_908086, EPI_ISL_908087 | Lighthouse Lab in Glasgow | Wellcome Sanger Institute for the COVID-19 Genomics UK (COG-UK) Consortium | Harper VanSteenhouse, Yumi Kasai, David Gray, Carol Clugston, Anna Dominiczak and Alex Alderton, Roberto Amato, Sonia Goncalves, Ewan Harrison, David K. Jackson, Ian Johnston, Dominic Kwiatkowski, Cordelia Langford, John Sillitoe on behalf of the Wellcome Sanger Institute COVID-19 Surveillance Team |
| EPI_ISL_908088 | Lighthouse Lab in Cambridge | Wellcome Sanger Institute for the COVID-19 Genomics UK (COG-UK) Consortium | Rob Howes, The Lighthouse Lab in Cambridge and Alex Alderton, Roberto Amato, Sonia Goncalves, Ewan Harrison, David K. Jackson, Ian Johnston, Dominic Kwiatkowski, Cordelia Langford, John Sillitoe on behalf of the Wellcome Sanger Institute COVID-19 Surveillance Team |
| EPI_ISL_908089 | Lighthouse Lab in Glasgow | Wellcome Sanger Institute for the COVID-19 Genomics UK (COG-UK) Consortium | Harper VanSteenhouse, Yumi Kasai, David Gray, Carol Clugston, Anna Dominiczak and Alex Alderton, Roberto Amato, Sonia Goncalves, Ewan Harrison, David K. Jackson, Ian Johnston, Dominic Kwiatkowski, Cordelia Langford, John Sillitoe on behalf of the Wellcome Sanger Institute COVID-19 Surveillance Team |
| EPI_ISL_908090, EPI_ISL_908091 | Lighthouse Lab in Cambridge | Wellcome Sanger Institute for the COVID-19 Genomics UK (COG-UK) Consortium | Rob Howes, The Lighthouse Lab in Cambridge and Alex Alderton, Roberto Amato, Sonia Goncalves, Ewan Harrison, David K. Jackson, Ian Johnston, Dominic Kwiatkowski, Cordelia Langford, John Sillitoe on behalf of the Wellcome Sanger Institute COVID-19 Surveillance Team |
| EPI_ISL_908092 | Lighthouse Lab in Glasgow | Wellcome Sanger Institute for the COVID-19 Genomics UK (COG-UK) Consortium | Harper VanSteenhouse, Yumi Kasai, David Gray, Carol Clugston, Anna Dominiczak and Alex Alderton, Roberto Amato, Sonia Goncalves, Ewan Harrison, David K. Jackson, Ian Johnston, Dominic Kwiatkowski, Cordelia Langford, John Sillitoe on behalf of the Wellcome Sanger Institute COVID-19 Surveillance Team |
| EPI_ISL_908093 | Lighthouse Lab in Cambridge | Wellcome Sanger Institute for the COVID-19 Genomics UK (COG-UK) Consortium | Rob Howes, The Lighthouse Lab in Cambridge and Alex Alderton, Roberto Amato, Sonia Goncalves, Ewan Harrison, David K. Jackson, Ian Johnston, Dominic Kwiatkowski, Cordelia Langford, John Sillitoe on behalf of the Wellcome Sanger Institute COVID-19 Surveillance Team |
| EPI_ISL_908094 | Lighthouse Lab in Glasgow | Wellcome Sanger Institute for the COVID-19 Genomics UK (COG-UK) Consortium | Harper VanSteenhouse, Yumi Kasai, David Gray, Carol Clugston, Anna Dominiczak and Alex Alderton, Roberto Amato, Sonia Goncalves, Ewan Harrison, David K. Jackson, Ian Johnston, Dominic Kwiatkowski, Cordelia Langford, John Sillitoe on behalf of the Wellcome Sanger Institute COVID-19 Surveillance Team |
| EPI_ISL_908095 | Lighthouse Lab in Cambridge | Wellcome Sanger Institute for the COVID-19 Genomics UK (COG-UK) Consortium | Rob Howes, The Lighthouse Lab in Cambridge and Alex Alderton, Roberto Amato, Sonia Goncalves, Ewan Harrison, David K. Jackson, Ian Johnston, Dominic Kwiatkowski, Cordelia Langford, John Sillitoe on behalf of the Wellcome Sanger Institute COVID-19 Surveillance Team |
| EPI_ISL_908096, EPI_ISL_908097 | Lighthouse Lab in Glasgow | Wellcome Sanger Institute for the COVID-19 Genomics UK (COG-UK) Consortium | Harper VanSteenhouse, Yumi Kasai, David Gray, Carol Clugston, Anna Dominiczak and Alex Alderton, Roberto Amato, Sonia Goncalves, Ewan Harrison, David K. Jackson, Ian Johnston, Dominic Kwiatkowski, Cordelia Langford, John Sillitoe on behalf of the Wellcome Sanger Institute COVID-19 Surveillance Team |
| EPI_ISL_908098, EPI_ISL_908099, EPI_ISL_908100, EPI_ISL_908101, EPI_ISL_908102, EPI_ISL_908103 | Lighthouse Lab in Cambridge | Wellcome Sanger Institute for the COVID-19 Genomics UK (COG-UK) Consortium | Rob Howes, The Lighthouse Lab in Cambridge and Alex Alderton, Roberto Amato, Sonia Goncalves, Ewan Harrison, David K. Jackson, Ian Johnston, Dominic Kwiatkowski, Cordelia Langford, John Sillitoe on behalf of the Wellcome Sanger Institute COVID-19 Surveillance Team |
| EPI_ISL_908104 | Lighthouse Lab in Glasgow | Wellcome Sanger Institute for the COVID-19 Genomics UK (COG-UK) Consortium | Harper VanSteenhouse, Yumi Kasai, David Gray, Carol Clugston, Anna Dominiczak and Alex Alderton, Roberto Amato, Sonia Goncalves, Ewan Harrison, David K. Jackson, Ian Johnston, Dominic Kwiatkowski, Cordelia Langford, John Sillitoe on behalf of the Wellcome Sanger Institute COVID-19 Surveillance Team |
| EPI_ISL_908105, EPI_ISL_908106, EPI_ISL_908107, EPI_ISL_908108, EPI_ISL_908109, EPI_ISL_908110, EPI_ISL_908111, EPI_ISL_908112 | Lighthouse Lab in Cambridge | Wellcome Sanger Institute for the COVID-19 Genomics UK (COG-UK) Consortium | Rob Howes, The Lighthouse Lab in Cambridge and Alex Alderton, Roberto Amato, Sonia Goncalves, Ewan Harrison, David K. Jackson, Ian Johnston, Dominic Kwiatkowski, Cordelia Langford, John Sillitoe on behalf of the Wellcome Sanger Institute COVID-19 Surveillance Team |
| EPI_ISL_908113 | Lighthouse Lab in Glasgow | Wellcome Sanger Institute for the COVID-19 Genomics UK (COG-UK) Consortium | Harper VanSteenhouse, Yumi Kasai, David Gray, Carol Clugston, Anna Dominiczak and Alex Alderton, Roberto Amato, Sonia Goncalves, Ewan Harrison, David K. Jackson, Ian Johnston, Dominic Kwiatkowski, Cordelia Langford, John Sillitoe on behalf of the Wellcome Sanger Institute COVID-19 Surveillance Team |
| EPI_ISL_908114, EPI_ISL_908115 | Lighthouse Lab in Cambridge | Wellcome Sanger Institute for the COVID-19 Genomics UK (COG-UK) Consortium | Rob Howes, The Lighthouse Lab in Cambridge and Alex Alderton, Roberto Amato, Sonia Goncalves, Ewan Harrison, David K. Jackson, Ian Johnston, Dominic Kwiatkowski, Cordelia Langford, John Sillitoe on behalf of the Wellcome Sanger Institute COVID-19 Surveillance Team |
| EPI_ISL_908116 | Lighthouse Lab in Glasgow | Wellcome Sanger Institute for the COVID-19 Genomics UK (COG-UK) Consortium | Harper VanSteenhouse, Yumi Kasai, David Gray, Carol Clugston, Anna Dominiczak and Alex Alderton, Roberto Amato, Sonia Goncalves, Ewan Harrison, David K. Jackson, Ian Johnston, Dominic Kwiatkowski, Cordelia Langford, John Sillitoe on behalf of the Wellcome Sanger Institute COVID-19 Surveillance Team |
| EPI_ISL_908117, EPI_ISL_908118 | Lighthouse Lab in Cambridge | Wellcome Sanger Institute for the COVID-19 Genomics UK (COG-UK) Consortium | Rob Howes, The Lighthouse Lab in Cambridge and Alex Alderton, Roberto Amato, Sonia Goncalves, Ewan Harrison, David K. Jackson, Ian Johnston, Dominic Kwiatkowski, Cordelia Langford, John Sillitoe on behalf of the Wellcome Sanger Institute COVID-19 Surveillance Team |
| EPI_ISL_908119, EPI_ISL_908120 | Lighthouse Lab in Glasgow | Wellcome Sanger Institute for the COVID-19 Genomics UK (COG-UK) Consortium | Harper VanSteenhouse, Yumi Kasai, David Gray, Carol Clugston, Anna Dominiczak and Alex Alderton, Roberto Amato, Sonia Goncalves, Ewan Harrison, David K. Jackson, Ian Johnston, Dominic Kwiatkowski, Cordelia Langford, John Sillitoe on behalf of the Wellcome Sanger Institute COVID-19 Surveillance Team |
| EPI_ISL_908121 | Lighthouse Lab in Cambridge | Wellcome Sanger Institute for the COVID-19 Genomics UK (COG-UK) Consortium | Rob Howes, The Lighthouse Lab in Cambridge and Alex Alderton, Roberto Amato, Sonia Goncalves, Ewan Harrison, David K. Jackson, Ian Johnston, Dominic Kwiatkowski, Cordelia Langford, John Sillitoe on behalf of the Wellcome Sanger Institute COVID-19 Surveillance Team |
| EPI_ISL_908122 | Lighthouse Lab in Glasgow | Wellcome Sanger Institute for the COVID-19 Genomics UK (COG-UK) Consortium | Harper VanSteenhouse, Yumi Kasai, David Gray, Carol Clugston, Anna Dominiczak and Alex Alderton, Roberto Amato, Sonia Goncalves, Ewan Harrison, David K. Jackson, Ian Johnston, Dominic Kwiatkowski, Cordelia Langford, John Sillitoe on behalf of the Wellcome Sanger Institute COVID-19 Surveillance Team |
| EPI_ISL_908123, EPI_ISL_908124, EPI_ISL_908125 | Lighthouse Lab in Cambridge | Wellcome Sanger Institute for the COVID-19 Genomics UK (COG-UK) Consortium | Rob Howes, The Lighthouse Lab in Cambridge and Alex Alderton, Roberto Amato, Sonia Goncalves, Ewan Harrison, David K. Jackson, Ian Johnston, Dominic Kwiatkowski, Cordelia Langford, John Sillitoe on behalf of the Wellcome Sanger Institute COVID-19 Surveillance Team |
| EPI_ISL_908126 | Lighthouse Lab in Glasgow | Wellcome Sanger Institute for the COVID-19 Genomics UK (COG-UK) Consortium | Harper VanSteenhouse, Yumi Kasai, David Gray, Carol Clugston, Anna Dominiczak and Alex Alderton, Roberto Amato, Sonia Goncalves, Ewan Harrison, David K. Jackson, Ian Johnston, Dominic Kwiatkowski, Cordelia Langford, John Sillitoe on behalf of the Wellcome Sanger Institute COVID-19 Surveillance Team |
| EPI_ISL_908127, EPI_ISL_908128, EPI_ISL_908129, EPI_ISL_908130 | Lighthouse Lab in Cambridge | Wellcome Sanger Institute for the COVID-19 Genomics UK (COG-UK) Consortium | Rob Howes, The Lighthouse Lab in Cambridge and Alex Alderton, Roberto Amato, Sonia Goncalves, Ewan Harrison, David K. Jackson, Ian Johnston, Dominic Kwiatkowski, Cordelia Langford, John Sillitoe on behalf of the Wellcome Sanger Institute COVID-19 Surveillance Team |
| EPI_ISL_908131 | Lighthouse Lab in Glasgow | Wellcome Sanger Institute for the COVID-19 Genomics UK (COG-UK) Consortium | Harper VanSteenhouse, Yumi Kasai, David Gray, Carol Clugston, Anna Dominiczak and Alex Alderton, Roberto Amato, Sonia Goncalves, Ewan Harrison, David K. Jackson, Ian Johnston, Dominic Kwiatkowski, Cordelia Langford, John Sillitoe on behalf of the Wellcome Sanger Institute COVID-19 Surveillance Team |
| EPI_ISL_908132, EPI_ISL_908133, EPI_ISL_908134, EPI_ISL_908135, EPI_ISL_908136, EPI_ISL_908137, EPI_ISL_908138 | Lighthouse Lab in Cambridge | Wellcome Sanger Institute for the COVID-19 Genomics UK (COG-UK) Consortium | Rob Howes, The Lighthouse Lab in Cambridge and Alex Alderton, Roberto Amato, Sonia Goncalves, Ewan Harrison, David K. Jackson, Ian Johnston, Dominic Kwiatkowski, Cordelia Langford, John Sillitoe on behalf of the Wellcome Sanger Institute COVID-19 Surveillance Team |
| EPI_ISL_908139 | Lighthouse Lab in Glasgow | Wellcome Sanger Institute for the COVID-19 Genomics UK (COG-UK) Consortium | Harper VanSteenhouse, Yumi Kasai, David Gray, Carol Clugston, Anna Dominiczak and Alex Alderton, Roberto Amato, Sonia Goncalves, Ewan Harrison, David K. Jackson, Ian Johnston, Dominic Kwiatkowski, Cordelia Langford, John Sillitoe on behalf of the Wellcome Sanger Institute COVID-19 Surveillance Team |
| EPI_ISL_908140, EPI_ISL_908141, EPI_ISL_908142, EPI_ISL_908143, EPI_ISL_908144, EPI_ISL_908145, EPI_ISL_908146, EPI_ISL_908147, EPI_ISL_908148, EPI_ISL_908149, EPI_ISL_908150 | | | |
| see above | Lighthouse Lab in Cambridge | Wellcome Sanger Institute for the COVID-19 Genomics UK | Rob Howes, The Lighthouse Lab in Cambridge and Alex Alderton, Roberto Amato, Sonia Goncalves, Ewan Harrison, David K. Jackson, Ian Johnston, |

| | | | |
|---|---|---|---|
| | | (COG-UK) Consortium | Dominic Kwiatkowski, Cordelia Langford, John Sillitoe on behalf of the Wellcome Sanger Institute COVID-19 Surveillance Team |
| EPI_ISL_908151, EPI_ISL_908152 | Lighthouse Lab in Glasgow | Wellcome Sanger Institute for the COVID-19 Genomics UK (COG-UK) Consortium | Harper VanSteenhouse, Yumi Kasai, David Gray, Carol Clugston, Anna Dominiczak and Alex Alderton, Roberto Amato, Sonia Goncalves, Ewan Harrison, David K. Jackson, Ian Johnston, Dominic Kwiatkowski, Cordelia Langford, John Sillitoe on behalf of the Wellcome Sanger Institute COVID-19 Surveillance Team |
| EPI_ISL_908153, EPI_ISL_908154, EPI_ISL_908155, EPI_ISL_908156, EPI_ISL_908157, EPI_ISL_908158, EPI_ISL_908159, EPI_ISL_908160, EPI_ISL_908161, EPI_ISL_908162, EPI_ISL_908163, EPI_ISL_908164, EPI_ISL_908165, EPI_ISL_908166, EPI_ISL_908167 | | | |
| see above | Lighthouse Lab in Cambridge | Wellcome Sanger Institute for the COVID-19 Genomics UK (COG-UK) Consortium | Rob Howes, The Lighthouse Lab in Cambridge and Alex Alderton, Roberto Amato, Sonia Goncalves, Ewan Harrison, David K. Jackson, Ian Johnston, Dominic Kwiatkowski, Cordelia Langford, John Sillitoe on behalf of the Wellcome Sanger Institute COVID-19 Surveillance Team |
| EPI_ISL_908168 | Lighthouse Lab in Glasgow | Wellcome Sanger Institute for the COVID-19 Genomics UK (COG-UK) Consortium | Harper VanSteenhouse, Yumi Kasai, David Gray, Carol Clugston, Anna Dominiczak and Alex Alderton, Roberto Amato, Sonia Goncalves, Ewan Harrison, David K. Jackson, Ian Johnston, Dominic Kwiatkowski, Cordelia Langford, John Sillitoe on behalf of the Wellcome Sanger Institute COVID-19 Surveillance Team |
| EPI_ISL_908169, EPI_ISL_908170 | Lighthouse Lab in Cambridge | Wellcome Sanger Institute for the COVID-19 Genomics UK (COG-UK) Consortium | Rob Howes, The Lighthouse Lab in Cambridge and Alex Alderton, Roberto Amato, Sonia Goncalves, Ewan Harrison, David K. Jackson, Ian Johnston, Dominic Kwiatkowski, Cordelia Langford, John Sillitoe on behalf of the Wellcome Sanger Institute COVID-19 Surveillance Team |
| EPI_ISL_908171 | Lighthouse Lab in Glasgow | Wellcome Sanger Institute for the COVID-19 Genomics UK (COG-UK) Consortium | Harper VanSteenhouse, Yumi Kasai, David Gray, Carol Clugston, Anna Dominiczak and Alex Alderton, Roberto Amato, Sonia Goncalves, Ewan Harrison, David K. Jackson, Ian Johnston, Dominic Kwiatkowski, Cordelia Langford, John Sillitoe on behalf of the Wellcome Sanger Institute COVID-19 Surveillance Team |
| EPI_ISL_908172, EPI_ISL_908173 | Lighthouse Lab in Cambridge | Wellcome Sanger Institute for the COVID-19 Genomics UK (COG-UK) Consortium | Rob Howes, The Lighthouse Lab in Cambridge and Alex Alderton, Roberto Amato, Sonia Goncalves, Ewan Harrison, David K. Jackson, Ian Johnston, Dominic Kwiatkowski, Cordelia Langford, John Sillitoe on behalf of the Wellcome Sanger Institute COVID-19 Surveillance Team |
| EPI_ISL_908174 | Lighthouse Lab in Glasgow | Wellcome Sanger Institute for the COVID-19 Genomics UK (COG-UK) Consortium | Harper VanSteenhouse, Yumi Kasai, David Gray, Carol Clugston, Anna Dominiczak and Alex Alderton, Roberto Amato, Sonia Goncalves, Ewan Harrison, David K. Jackson, Ian Johnston, Dominic Kwiatkowski, Cordelia Langford, John Sillitoe on behalf of the Wellcome Sanger Institute COVID-19 Surveillance Team |
| EPI_ISL_908175, EPI_ISL_908176 | Lighthouse Lab in Cambridge | Wellcome Sanger Institute for the COVID-19 Genomics UK (COG-UK) Consortium | Rob Howes, The Lighthouse Lab in Cambridge and Alex Alderton, Roberto Amato, Sonia Goncalves, Ewan Harrison, David K. Jackson, Ian Johnston, Dominic Kwiatkowski, Cordelia Langford, John Sillitoe on behalf of the Wellcome Sanger Institute COVID-19 Surveillance Team |
| EPI_ISL_908177 | Lighthouse Lab in Glasgow | Wellcome Sanger Institute for the COVID-19 Genomics UK (COG-UK) Consortium | Harper VanSteenhouse, Yumi Kasai, David Gray, Carol Clugston, Anna Dominiczak and Alex Alderton, Roberto Amato, Sonia Goncalves, Ewan Harrison, David K. Jackson, Ian Johnston, Dominic Kwiatkowski, Cordelia Langford, John Sillitoe on behalf of the Wellcome Sanger Institute COVID-19 Surveillance Team |
| EPI_ISL_908178 | Lighthouse Lab in Cambridge | Wellcome Sanger Institute for the COVID-19 Genomics UK (COG-UK) Consortium | Rob Howes, The Lighthouse Lab in Cambridge and Alex Alderton, Roberto Amato, Sonia Goncalves, Ewan Harrison, David K. Jackson, Ian Johnston, Dominic Kwiatkowski, Cordelia Langford, John Sillitoe on behalf of the Wellcome Sanger Institute COVID-19 Surveillance Team |
| EPI_ISL_908179 | Lighthouse Lab in Glasgow | Wellcome Sanger Institute for the COVID-19 Genomics UK (COG-UK) Consortium | Harper VanSteenhouse, Yumi Kasai, David Gray, Carol Clugston, Anna Dominiczak and Alex Alderton, Roberto Amato, Sonia Goncalves, Ewan Harrison, David K. Jackson, Ian Johnston, Dominic Kwiatkowski, Cordelia Langford, John Sillitoe on behalf of the Wellcome Sanger Institute COVID-19 Surveillance Team |
| EPI_ISL_908180 | Lighthouse Lab in Cambridge | Wellcome Sanger Institute for the COVID-19 Genomics UK (COG-UK) Consortium | Rob Howes, The Lighthouse Lab in Cambridge and Alex Alderton, Roberto Amato, Sonia Goncalves, Ewan Harrison, David K. Jackson, Ian Johnston, Dominic Kwiatkowski, Cordelia Langford, John Sillitoe on behalf of the Wellcome Sanger Institute COVID-19 Surveillance Team |
| EPI_ISL_908181 | Lighthouse Lab in Glasgow | Wellcome Sanger Institute for the COVID-19 Genomics UK (COG-UK) Consortium | Harper VanSteenhouse, Yumi Kasai, David Gray, Carol Clugston, Anna Dominiczak and Alex Alderton, Roberto Amato, Sonia Goncalves, Ewan Harrison, David K. Jackson, Ian Johnston, Dominic Kwiatkowski, Cordelia Langford, John Sillitoe on behalf of the Wellcome Sanger Institute COVID-19 Surveillance Team |
| EPI_ISL_908182 | Lighthouse Lab in Cambridge | Wellcome Sanger Institute for the COVID-19 Genomics UK (COG-UK) Consortium | Rob Howes, The Lighthouse Lab in Cambridge and Alex Alderton, Roberto Amato, Sonia Goncalves, Ewan Harrison, David K. Jackson, Ian Johnston, Dominic Kwiatkowski, Cordelia Langford, John Sillitoe on behalf of the Wellcome Sanger Institute COVID-19 Surveillance Team |
| EPI_ISL_908183 | Lighthouse Lab in Glasgow | Wellcome Sanger Institute for the COVID-19 Genomics UK (COG-UK) Consortium | Harper VanSteenhouse, Yumi Kasai, David Gray, Carol Clugston, Anna Dominiczak and Alex Alderton, Roberto Amato, Sonia Goncalves, Ewan Harrison, David K. Jackson, Ian Johnston, Dominic Kwiatkowski, Cordelia Langford, John Sillitoe on behalf of the Wellcome Sanger Institute COVID-19 Surveillance Team |
| EPI_ISL_908184, EPI_ISL_908185, EPI_ISL_908186 | Lighthouse Lab in Cambridge | Wellcome Sanger Institute for the COVID-19 Genomics UK (COG-UK) Consortium | Rob Howes, The Lighthouse Lab in Cambridge and Alex Alderton, Roberto Amato, Sonia Goncalves, Ewan Harrison, David K. Jackson, Ian Johnston, Dominic Kwiatkowski, Cordelia Langford, John Sillitoe on behalf of the Wellcome Sanger Institute COVID-19 Surveillance Team |
| EPI_ISL_908187 | Lighthouse Lab in Glasgow | Wellcome Sanger Institute for the COVID-19 Genomics UK (COG-UK) Consortium | Harper VanSteenhouse, Yumi Kasai, David Gray, Carol Clugston, Anna Dominiczak and Alex Alderton, Roberto Amato, Sonia Goncalves, Ewan Harrison, David K. Jackson, Ian Johnston, Dominic Kwiatkowski, Cordelia Langford, John Sillitoe on behalf of the Wellcome Sanger Institute COVID-19 Surveillance Team |
| EPI_ISL_908188, EPI_ISL_908189, EPI_ISL_908190, EPI_ISL_908191, EPI_ISL_908192, EPI_ISL_908193 | Lighthouse Lab in Cambridge | Wellcome Sanger Institute for the COVID-19 Genomics UK (COG-UK) Consortium | Rob Howes, The Lighthouse Lab in Cambridge and Alex Alderton, Roberto Amato, Sonia Goncalves, Ewan Harrison, David K. Jackson, Ian Johnston, Dominic Kwiatkowski, Cordelia Langford, John Sillitoe on behalf of the Wellcome Sanger Institute COVID-19 Surveillance Team |
| EPI_ISL_908194 | Lighthouse Lab in Glasgow | Wellcome Sanger Institute for the COVID-19 Genomics UK (COG-UK) Consortium | Harper VanSteenhouse, Yumi Kasai, David Gray, Carol Clugston, Anna Dominiczak and Alex Alderton, Roberto Amato, Sonia Goncalves, Ewan Harrison, David K. Jackson, Ian Johnston, Dominic Kwiatkowski, Cordelia Langford, John Sillitoe on behalf of the Wellcome Sanger Institute COVID-19 Surveillance Team |
| EPI_ISL_908195, EPI_ISL_908196, EPI_ISL_908197, EPI_ISL_908198, EPI_ISL_908199 | Lighthouse Lab in Cambridge | Wellcome Sanger Institute for the COVID-19 Genomics UK (COG-UK) Consortium | Rob Howes, The Lighthouse Lab in Cambridge and Alex Alderton, Roberto Amato, Sonia Goncalves, Ewan Harrison, David K. Jackson, Ian Johnston, Dominic Kwiatkowski, Cordelia Langford, John Sillitoe on behalf of the Wellcome Sanger Institute COVID-19 Surveillance Team |
| EPI_ISL_908200 | Lighthouse Lab in Glasgow | Wellcome Sanger Institute for the COVID-19 Genomics UK (COG-UK) Consortium | Harper VanSteenhouse, Yumi Kasai, David Gray, Carol Clugston, Anna Dominiczak and Alex Alderton, Roberto Amato, Sonia Goncalves, Ewan Harrison, David K. Jackson, Ian Johnston, Dominic Kwiatkowski, Cordelia Langford, John Sillitoe on behalf of the Wellcome Sanger Institute COVID-19 Surveillance Team |
| EPI_ISL_908201, EPI_ISL_908202, EPI_ISL_908203, EPI_ISL_908204, EPI_ISL_908205 | Lighthouse Lab in Cambridge | Wellcome Sanger Institute for the COVID-19 Genomics UK (COG-UK) Consortium | Rob Howes, The Lighthouse Lab in Cambridge and Alex Alderton, Roberto Amato, Sonia Goncalves, Ewan Harrison, David K. Jackson, Ian Johnston, Dominic Kwiatkowski, Cordelia Langford, John Sillitoe on behalf of the Wellcome Sanger Institute COVID-19 Surveillance Team |
| EPI_ISL_908206 | Lighthouse Lab in Glasgow | Wellcome Sanger Institute for the COVID-19 Genomics UK (COG-UK) Consortium | Harper VanSteenhouse, Yumi Kasai, David Gray, Carol Clugston, Anna Dominiczak and Alex Alderton, Roberto Amato, Sonia Goncalves, Ewan Harrison, David K. Jackson, Ian Johnston, Dominic Kwiatkowski, Cordelia Langford, John Sillitoe on behalf of the Wellcome Sanger Institute COVID-19 Surveillance Team |
| EPI_ISL_908207 | Lighthouse Lab in Cambridge | Wellcome Sanger Institute for the COVID-19 Genomics UK (COG-UK) Consortium | Rob Howes, The Lighthouse Lab in Cambridge and Alex Alderton, Roberto Amato, Sonia Goncalves, Ewan Harrison, David K. Jackson, Ian Johnston, Dominic Kwiatkowski, Cordelia Langford, John Sillitoe on behalf of the Wellcome Sanger Institute COVID-19 Surveillance Team |
| EPI_ISL_908208 | Lighthouse Lab in Glasgow | Wellcome Sanger Institute for the COVID-19 Genomics UK (COG-UK) Consortium | Harper VanSteenhouse, Yumi Kasai, David Gray, Carol Clugston, Anna Dominiczak and Alex Alderton, Roberto Amato, Sonia Goncalves, Ewan Harrison, David K. Jackson, Ian Johnston, Dominic Kwiatkowski, Cordelia Langford, John Sillitoe on behalf of the Wellcome Sanger Institute COVID-19 Surveillance Team |
| EPI_ISL_908209, EPI_ISL_908210, EPI_ISL_908211, EPI_ISL_908212, EPI_ISL_908213 | Lighthouse Lab in Cambridge | Wellcome Sanger Institute for the COVID-19 Genomics UK (COG-UK) Consortium | Rob Howes, The Lighthouse Lab in Cambridge and Alex Alderton, Roberto Amato, Sonia Goncalves, Ewan Harrison, David K. Jackson, Ian Johnston, Dominic Kwiatkowski, Cordelia Langford, John Sillitoe on behalf of the Wellcome Sanger Institute COVID-19 Surveillance Team |
| EPI_ISL_908214 | Lighthouse Lab in Glasgow | Wellcome Sanger Institute for the COVID-19 Genomics UK (COG-UK) Consortium | Harper VanSteenhouse, Yumi Kasai, David Gray, Carol Clugston, Anna Dominiczak and Alex Alderton, Roberto Amato, Sonia Goncalves, Ewan Harrison, David K. Jackson, Ian Johnston, Dominic Kwiatkowski, Cordelia Langford, John Sillitoe on behalf of the Wellcome Sanger Institute COVID-19 Surveillance |

| | | | |
|---|---|---|---|
| | | | Team |
| EPI_ISL_908215, EPI_ISL_908216, EPI_ISL_908217 | Lighthouse Lab in Cambridge | Wellcome Sanger Institute for the COVID-19 Genomics UK (COG-UK) Consortium | Rob Howes, The Lighthouse Lab in Cambridge and Alex Alderton, Roberto Amato, Sonia Goncalves, Ewan Harrison, David K. Jackson, Ian Johnston, Dominic Kwiatkowski, Cordelia Langford, John Sillitoe on behalf of the Wellcome Sanger Institute COVID-19 Surveillance Team |
| EPI_ISL_908218 | Lighthouse Lab in Glasgow | Wellcome Sanger Institute for the COVID-19 Genomics UK (COG-UK) Consortium | Harper VanSteenhouse, Yumi Kasai, David Gray, Carol Clugston, Anna Dominiczak and Alex Alderton, Roberto Amato, Sonia Goncalves, Ewan Harrison, David K. Jackson, Ian Johnston, Dominic Kwiatkowski, Cordelia Langford, John Sillitoe on behalf of the Wellcome Sanger Institute COVID-19 Surveillance Team |
| EPI_ISL_908219, EPI_ISL_908220, EPI_ISL_908221, EPI_ISL_908222, EPI_ISL_908223, EPI_ISL_908224, EPI_ISL_908225, EPI_ISL_908226, EPI_ISL_908227, EPI_ISL_908228 | Lighthouse Lab in Cambridge | Wellcome Sanger Institute for the COVID-19 Genomics UK (COG-UK) Consortium | Rob Howes, The Lighthouse Lab in Cambridge and Alex Alderton, Roberto Amato, Sonia Goncalves, Ewan Harrison, David K. Jackson, Ian Johnston, Dominic Kwiatkowski, Cordelia Langford, John Sillitoe on behalf of the Wellcome Sanger Institute COVID-19 Surveillance Team |
| EPI_ISL_908229 | Lighthouse Lab in Glasgow | Wellcome Sanger Institute for the COVID-19 Genomics UK (COG-UK) Consortium | Harper VanSteenhouse, Yumi Kasai, David Gray, Carol Clugston, Anna Dominiczak and Alex Alderton, Roberto Amato, Sonia Goncalves, Ewan Harrison, David K. Jackson, Ian Johnston, Dominic Kwiatkowski, Cordelia Langford, John Sillitoe on behalf of the Wellcome Sanger Institute COVID-19 Surveillance Team |
| EPI_ISL_908230 | Lighthouse Lab in Cambridge | Wellcome Sanger Institute for the COVID-19 Genomics UK (COG-UK) Consortium | Rob Howes, The Lighthouse Lab in Cambridge and Alex Alderton, Roberto Amato, Sonia Goncalves, Ewan Harrison, David K. Jackson, Ian Johnston, Dominic Kwiatkowski, Cordelia Langford, John Sillitoe on behalf of the Wellcome Sanger Institute COVID-19 Surveillance Team |
| EPI_ISL_908231 | Lighthouse Lab in Glasgow | Wellcome Sanger Institute for the COVID-19 Genomics UK (COG-UK) Consortium | Harper VanSteenhouse, Yumi Kasai, David Gray, Carol Clugston, Anna Dominiczak and Alex Alderton, Roberto Amato, Sonia Goncalves, Ewan Harrison, David K. Jackson, Ian Johnston, Dominic Kwiatkowski, Cordelia Langford, John Sillitoe on behalf of the Wellcome Sanger Institute COVID-19 Surveillance Team |
| EPI_ISL_908232, EPI_ISL_908233, EPI_ISL_908234, EPI_ISL_908235, EPI_ISL_908236 | Lighthouse Lab in Cambridge | Wellcome Sanger Institute for the COVID-19 Genomics UK (COG-UK) Consortium | Rob Howes, The Lighthouse Lab in Cambridge and Alex Alderton, Roberto Amato, Sonia Goncalves, Ewan Harrison, David K. Jackson, Ian Johnston, Dominic Kwiatkowski, Cordelia Langford, John Sillitoe on behalf of the Wellcome Sanger Institute COVID-19 Surveillance Team |
| EPI_ISL_908237, EPI_ISL_908238 | Lighthouse Lab in Glasgow | Wellcome Sanger Institute for the COVID-19 Genomics UK (COG-UK) Consortium | Harper VanSteenhouse, Yumi Kasai, David Gray, Carol Clugston, Anna Dominiczak and Alex Alderton, Roberto Amato, Sonia Goncalves, Ewan Harrison, David K. Jackson, Ian Johnston, Dominic Kwiatkowski, Cordelia Langford, John Sillitoe on behalf of the Wellcome Sanger Institute COVID-19 Surveillance Team |
| EPI_ISL_908239, EPI_ISL_908240 | Lighthouse Lab in Cambridge | Wellcome Sanger Institute for the COVID-19 Genomics UK (COG-UK) Consortium | Rob Howes, The Lighthouse Lab in Cambridge and Alex Alderton, Roberto Amato, Sonia Goncalves, Ewan Harrison, David K. Jackson, Ian Johnston, Dominic Kwiatkowski, Cordelia Langford, John Sillitoe on behalf of the Wellcome Sanger Institute COVID-19 Surveillance Team |
| EPI_ISL_908241, EPI_ISL_908242, EPI_ISL_908243 | Lighthouse Lab in Glasgow | Wellcome Sanger Institute for the COVID-19 Genomics UK (COG-UK) Consortium | Harper VanSteenhouse, Yumi Kasai, David Gray, Carol Clugston, Anna Dominiczak and Alex Alderton, Roberto Amato, Sonia Goncalves, Ewan Harrison, David K. Jackson, Ian Johnston, Dominic Kwiatkowski, Cordelia Langford, John Sillitoe on behalf of the Wellcome Sanger Institute COVID-19 Surveillance Team |
| EPI_ISL_908244, EPI_ISL_908245, EPI_ISL_908246, EPI_ISL_908247, EPI_ISL_908248, EPI_ISL_908249, EPI_ISL_908250, EPI_ISL_908251 | Lighthouse Lab in Cambridge | Wellcome Sanger Institute for the COVID-19 Genomics UK (COG-UK) Consortium | Rob Howes, The Lighthouse Lab in Cambridge and Alex Alderton, Roberto Amato, Sonia Goncalves, Ewan Harrison, David K. Jackson, Ian Johnston, Dominic Kwiatkowski, Cordelia Langford, John Sillitoe on behalf of the Wellcome Sanger Institute COVID-19 Surveillance Team |
| EPI_ISL_908252 | Lighthouse Lab in Glasgow | Wellcome Sanger Institute for the COVID-19 Genomics UK (COG-UK) Consortium | Harper VanSteenhouse, Yumi Kasai, David Gray, Carol Clugston, Anna Dominiczak and Alex Alderton, Roberto Amato, Sonia Goncalves, Ewan Harrison, David K. Jackson, Ian Johnston, Dominic Kwiatkowski, Cordelia Langford, John Sillitoe on behalf of the Wellcome Sanger Institute COVID-19 Surveillance Team |
| EPI_ISL_908253 | Lighthouse Lab in Cambridge | Wellcome Sanger Institute for the COVID-19 Genomics UK (COG-UK) Consortium | Rob Howes, The Lighthouse Lab in Cambridge and Alex Alderton, Roberto Amato, Sonia Goncalves, Ewan Harrison, David K. Jackson, Ian Johnston, Dominic Kwiatkowski, Cordelia Langford, John Sillitoe on behalf of the Wellcome Sanger Institute COVID-19 Surveillance Team |
| EPI_ISL_908254 | Lighthouse Lab in Glasgow | Wellcome Sanger Institute for the COVID-19 Genomics UK (COG-UK) Consortium | Harper VanSteenhouse, Yumi Kasai, David Gray, Carol Clugston, Anna Dominiczak and Alex Alderton, Roberto Amato, Sonia Goncalves, Ewan Harrison, David K. Jackson, Ian Johnston, Dominic Kwiatkowski, Cordelia Langford, John Sillitoe on behalf of the Wellcome Sanger Institute COVID-19 Surveillance Team |
| EPI_ISL_908255 | Lighthouse Lab in Cambridge | Wellcome Sanger Institute for the COVID-19 Genomics UK (COG-UK) Consortium | Rob Howes, The Lighthouse Lab in Cambridge and Alex Alderton, Roberto Amato, Sonia Goncalves, Ewan Harrison, David K. Jackson, Ian Johnston, Dominic Kwiatkowski, Cordelia Langford, John Sillitoe on behalf of the Wellcome Sanger Institute COVID-19 Surveillance Team |
| EPI_ISL_908256 | Lighthouse Lab in Glasgow | Wellcome Sanger Institute for the COVID-19 Genomics UK (COG-UK) Consortium | Harper VanSteenhouse, Yumi Kasai, David Gray, Carol Clugston, Anna Dominiczak and Alex Alderton, Roberto Amato, Sonia Goncalves, Ewan Harrison, David K. Jackson, Ian Johnston, Dominic Kwiatkowski, Cordelia Langford, John Sillitoe on behalf of the Wellcome Sanger Institute COVID-19 Surveillance Team |
| EPI_ISL_908257, EPI_ISL_908258, EPI_ISL_908259 | Lighthouse Lab in Cambridge | Wellcome Sanger Institute for the COVID-19 Genomics UK (COG-UK) Consortium | Rob Howes, The Lighthouse Lab in Cambridge and Alex Alderton, Roberto Amato, Sonia Goncalves, Ewan Harrison, David K. Jackson, Ian Johnston, Dominic Kwiatkowski, Cordelia Langford, John Sillitoe on behalf of the Wellcome Sanger Institute COVID-19 Surveillance Team |
| EPI_ISL_908260, EPI_ISL_908261, EPI_ISL_908262, EPI_ISL_908263 | Lighthouse Lab in Glasgow | Wellcome Sanger Institute for the COVID-19 Genomics UK (COG-UK) Consortium | Harper VanSteenhouse, Yumi Kasai, David Gray, Carol Clugston, Anna Dominiczak and Alex Alderton, Roberto Amato, Sonia Goncalves, Ewan Harrison, David K. Jackson, Ian Johnston, Dominic Kwiatkowski, Cordelia Langford, John Sillitoe on behalf of the Wellcome Sanger Institute COVID-19 Surveillance Team |
| EPI_ISL_908264 | Lighthouse Lab in Cambridge | Wellcome Sanger Institute for the COVID-19 Genomics UK (COG-UK) Consortium | Rob Howes, The Lighthouse Lab in Cambridge and Alex Alderton, Roberto Amato, Sonia Goncalves, Ewan Harrison, David K. Jackson, Ian Johnston, Dominic Kwiatkowski, Cordelia Langford, John Sillitoe on behalf of the Wellcome Sanger Institute COVID-19 Surveillance Team |
| EPI_ISL_908265, EPI_ISL_908266 | Lighthouse Lab in Glasgow | Wellcome Sanger Institute for the COVID-19 Genomics UK (COG-UK) Consortium | Harper VanSteenhouse, Yumi Kasai, David Gray, Carol Clugston, Anna Dominiczak and Alex Alderton, Roberto Amato, Sonia Goncalves, Ewan Harrison, David K. Jackson, Ian Johnston, Dominic Kwiatkowski, Cordelia Langford, John Sillitoe on behalf of the Wellcome Sanger Institute COVID-19 Surveillance Team |
| EPI_ISL_908267 | Lighthouse Lab in Cambridge | Wellcome Sanger Institute for the COVID-19 Genomics UK (COG-UK) Consortium | Rob Howes, The Lighthouse Lab in Cambridge and Alex Alderton, Roberto Amato, Sonia Goncalves, Ewan Harrison, David K. Jackson, Ian Johnston, Dominic Kwiatkowski, Cordelia Langford, John Sillitoe on behalf of the Wellcome Sanger Institute COVID-19 Surveillance Team |
| EPI_ISL_908376, EPI_ISL_908465, EPI_ISL_908474, EPI_ISL_908513, EPI_ISL_908570, EPI_ISL_908592 | Lighthouse Lab in Alderley Park | Wellcome Sanger Institute for the COVID-19 Genomics UK (COG-UK) Consortium | Jacquelyn Wynn, Mairead Hyland, The Lighthouse Lab in Alderley Park and Alex Alderton, Roberto Amato, Sonia Goncalves, Ewan Harrison, David K. Jackson, Ian Johnston, Dominic Kwiatkowski, Cordelia Langford, John Sillitoe on behalf of the Wellcome Sanger Institute COVID-19 Surveillance Team |
| EPI_ISL_908932, EPI_ISL_908936, EPI_ISL_908937, EPI_ISL_908938, EPI_ISL_908939, EPI_ISL_908940, EPI_ISL_908941, EPI_ISL_908943, EPI_ISL_908944, EPI_ISL_908947, EPI_ISL_908948, EPI_ISL_908950, EPI_ISL_908954, EPI_ISL_908956, EPI_ISL_908960, EPI_ISL_908964, EPI_ISL_908970, EPI_ISL_908976 | | | |
| see above | Lighthouse Lab in Milton Keynes | Wellcome Sanger Institute for the COVID-19 Genomics UK (COG-UK) Consortium | The Lighthouse Lab in Milton Keynes and Alex Alderton, Roberto Amato, Sonia Goncalves, Ewan Harrison, David K. Jackson, Ian Johnston, Dominic Kwiatkowski, Cordelia Langford, John Sillitoe on behalf of the Wellcome Sanger Institute COVID-19 Surveillance Team |
| EPI_ISL_908982 | Lighthouse Lab in Glasgow | Wellcome Sanger Institute for the COVID-19 Genomics UK (COG-UK) Consortium | Harper VanSteenhouse, Yumi Kasai, David Gray, Carol Clugston, Anna Dominiczak and Alex Alderton, Roberto Amato, Sonia Goncalves, Ewan Harrison, David K. Jackson, Ian Johnston, Dominic Kwiatkowski, Cordelia Langford, John Sillitoe on behalf of the Wellcome Sanger Institute COVID-19 Surveillance Team |
| EPI_ISL_908984, EPI_ISL_908987, EPI_ISL_908988, EPI_ISL_908994, EPI_ISL_908995, EPI_ISL_908996, EPI_ISL_908997, EPI_ISL_909003, EPI_ISL_909012, EPI_ISL_909013, EPI_ISL_909014, EPI_ISL_909016, EPI_ISL_909018, EPI_ISL_909020, EPI_ISL_909022, EPI_ISL_909023, EPI_ISL_909025, EPI_ISL_909026, EPI_ISL_909027, EPI_ISL_909028, EPI_ISL_909031, EPI_ISL_909032, EPI_ISL_909037, EPI_ISL_909039, EPI_ISL_909041, EPI_ISL_909042, EPI_ISL_909043, EPI_ISL_909044, EPI_ISL_909051, EPI_ISL_909054, EPI_ISL_909055, EPI_ISL_909059, EPI_ISL_909062, EPI_ISL_909064, EPI_ISL_909065, EPI_ISL_909070, EPI_ISL_909073, EPI_ISL_909076, EPI_ISL_909077, EPI_ISL_909078, EPI_ISL_909082, EPI_ISL_909084, EPI_ISL_909091, EPI_ISL_909092, EPI_ISL_909099, EPI_ISL_909100, EPI_ISL_909104, EPI_ISL_909107, EPI_ISL_909108, EPI_ISL_909110, EPI_ISL_909120, EPI_ISL_909123, EPI_ISL_909124, EPI_ISL_909125, EPI_ISL_909126, EPI_ISL_909127, EPI_ISL_909129, EPI_ISL_909130, EPI_ISL_909132, EPI_ISL_909133, EPI_ISL_909134, EPI_ISL_909135, EPI_ISL_909136, EPI_ISL_909137, EPI_ISL_909138, EPI_ISL_909139, EPI_ISL_909141, EPI_ISL_909143, EPI_ISL_909146, EPI_ISL_909147, EPI_ISL_909149, EPI_ISL_909152, | | | |

| | | | |
|---|---|---|---|
| EPI_ISL_909153, EPI_ISL_909154, EPI_ISL_909159, EPI_ISL_909160, EPI_ISL_909161, EPI_ISL_909163, EPI_ISL_909166, EPI_ISL_909167, EPI_ISL_909172, EPI_ISL_909174, EPI_ISL_909179, EPI_ISL_909181, EPI_ISL_909183, EPI_ISL_909185, EPI_ISL_909186, EPI_ISL_909191, EPI_ISL_909195, EPI_ISL_909196, EPI_ISL_909197, EPI_ISL_909198, EPI_ISL_909199, EPI_ISL_909201, EPI_ISL_909205, EPI_ISL_909207, EPI_ISL_909209, EPI_ISL_909212, EPI_ISL_909213, EPI_ISL_909214, EPI_ISL_909215, EPI_ISL_909216, EPI_ISL_909221, EPI_ISL_909222, EPI_ISL_909224, EPI_ISL_909226, EPI_ISL_909228, EPI_ISL_909229, EPI_ISL_909231, EPI_ISL_909232, EPI_ISL_909241, EPI_ISL_909244, EPI_ISL_909245, EPI_ISL_909246, EPI_ISL_909247, EPI_ISL_909248, EPI_ISL_909250, EPI_ISL_909252, EPI_ISL_909256, EPI_ISL_909258, EPI_ISL_909261, EPI_ISL_909264, EPI_ISL_909265, EPI_ISL_909266, EPI_ISL_909267, EPI_ISL_909268 | | | |
| see above | Lighthouse Lab in Milton Keynes | Wellcome Sanger Institute for the COVID-19 Genomics UK (COG-UK) Consortium | The Lighthouse Lab in Milton Keynes and Alex Alderton, Roberto Amato, Sonia Goncalves, Ewan Harrison, David K. Jackson, Ian Johnston, Dominic Kwiatkowski, Cordelia Langford, John Sillitoe on behalf of the Wellcome Sanger Institute COVID-19 Surveillance Team |
| EPI_ISL_909273 | Lighthouse Lab in Alderley Park | Wellcome Sanger Institute for the COVID-19 Genomics UK (COG-UK) Consortium | Jacquelyn Wynn, Mairead Hyland, The Lighthouse Lab in Alderley Park and Alex Alderton, Roberto Amato, Sonia Goncalves, Ewan Harrison, David K. Jackson, Ian Johnston, Dominic Kwiatkowski, Cordelia Langford, John Sillitoe on behalf of the Wellcome Sanger Institute COVID-19 Surveillance Team |
| EPI_ISL_909274, EPI_ISL_909279, EPI_ISL_909280 | Lighthouse Lab in Milton Keynes | Wellcome Sanger Institute for the COVID-19 Genomics UK (COG-UK) Consortium | The Lighthouse Lab in Milton Keynes and Alex Alderton, Roberto Amato, Sonia Goncalves, Ewan Harrison, David K. Jackson, Ian Johnston, Dominic Kwiatkowski, Cordelia Langford, John Sillitoe on behalf of the Wellcome Sanger Institute COVID-19 Surveillance Team |
| EPI_ISL_909281 | Lighthouse Lab in Alderley Park | Wellcome Sanger Institute for the COVID-19 Genomics UK (COG-UK) Consortium | Jacquelyn Wynn, Mairead Hyland, The Lighthouse Lab in Alderley Park and Alex Alderton, Roberto Amato, Sonia Goncalves, Ewan Harrison, David K. Jackson, Ian Johnston, Dominic Kwiatkowski, Cordelia Langford, John Sillitoe on behalf of the Wellcome Sanger Institute COVID-19 Surveillance Team |
| EPI_ISL_909282, EPI_ISL_909286, EPI_ISL_909288, EPI_ISL_909290, EPI_ISL_909295, EPI_ISL_909299, EPI_ISL_909300, EPI_ISL_909305, EPI_ISL_909307, EPI_ISL_909308 | Lighthouse Lab in Milton Keynes | Wellcome Sanger Institute for the COVID-19 Genomics UK (COG-UK) Consortium | The Lighthouse Lab in Milton Keynes and Alex Alderton, Roberto Amato, Sonia Goncalves, Ewan Harrison, David K. Jackson, Ian Johnston, Dominic Kwiatkowski, Cordelia Langford, John Sillitoe on behalf of the Wellcome Sanger Institute COVID-19 Surveillance Team |
| EPI_ISL_909311 | Lighthouse Lab in Alderley Park | Wellcome Sanger Institute for the COVID-19 Genomics UK (COG-UK) Consortium | Jacquelyn Wynn, Mairead Hyland, The Lighthouse Lab in Alderley Park and Alex Alderton, Roberto Amato, Sonia Goncalves, Ewan Harrison, David K. Jackson, Ian Johnston, Dominic Kwiatkowski, Cordelia Langford, John Sillitoe on behalf of the Wellcome Sanger Institute COVID-19 Surveillance Team |
| EPI_ISL_909313, EPI_ISL_909315 | Lighthouse Lab in Milton Keynes | Wellcome Sanger Institute for the COVID-19 Genomics UK (COG-UK) Consortium | The Lighthouse Lab in Milton Keynes and Alex Alderton, Roberto Amato, Sonia Goncalves, Ewan Harrison, David K. Jackson, Ian Johnston, Dominic Kwiatkowski, Cordelia Langford, John Sillitoe on behalf of the Wellcome Sanger Institute COVID-19 Surveillance Team |
| EPI_ISL_909318 | Lighthouse Lab in Alderley Park | Wellcome Sanger Institute for the COVID-19 Genomics UK (COG-UK) Consortium | Jacquelyn Wynn, Mairead Hyland, The Lighthouse Lab in Alderley Park and Alex Alderton, Roberto Amato, Sonia Goncalves, Ewan Harrison, David K. Jackson, Ian Johnston, Dominic Kwiatkowski, Cordelia Langford, John Sillitoe on behalf of the Wellcome Sanger Institute COVID-19 Surveillance Team |
| EPI_ISL_909319, EPI_ISL_909320, EPI_ISL_909323, EPI_ISL_909324 | Lighthouse Lab in Milton Keynes | Wellcome Sanger Institute for the COVID-19 Genomics UK (COG-UK) Consortium | The Lighthouse Lab in Milton Keynes and Alex Alderton, Roberto Amato, Sonia Goncalves, Ewan Harrison, David K. Jackson, Ian Johnston, Dominic Kwiatkowski, Cordelia Langford, John Sillitoe on behalf of the Wellcome Sanger Institute COVID-19 Surveillance Team |
| EPI_ISL_909328 | Lighthouse Lab in Alderley Park | Wellcome Sanger Institute for the COVID-19 Genomics UK (COG-UK) Consortium | Jacquelyn Wynn, Mairead Hyland, The Lighthouse Lab in Alderley Park and Alex Alderton, Roberto Amato, Sonia Goncalves, Ewan Harrison, David K. Jackson, Ian Johnston, Dominic Kwiatkowski, Cordelia Langford, John Sillitoe on behalf of the Wellcome Sanger Institute COVID-19 Surveillance Team |
| EPI_ISL_909330, EPI_ISL_909332, EPI_ISL_909336, EPI_ISL_909337 | Lighthouse Lab in Milton Keynes | Wellcome Sanger Institute for the COVID-19 Genomics UK (COG-UK) Consortium | The Lighthouse Lab in Milton Keynes and Alex Alderton, Roberto Amato, Sonia Goncalves, Ewan Harrison, David K. Jackson, Ian Johnston, Dominic Kwiatkowski, Cordelia Langford, John Sillitoe on behalf of the Wellcome Sanger Institute COVID-19 Surveillance Team |
| EPI_ISL_909344 | Lighthouse Lab in Alderley Park | Wellcome Sanger Institute for the COVID-19 Genomics UK (COG-UK) Consortium | Jacquelyn Wynn, Mairead Hyland, The Lighthouse Lab in Alderley Park and Alex Alderton, Roberto Amato, Sonia Goncalves, Ewan Harrison, David K. Jackson, Ian Johnston, Dominic Kwiatkowski, Cordelia Langford, John Sillitoe on behalf of the Wellcome Sanger Institute COVID-19 Surveillance Team |
| EPI_ISL_909346, EPI_ISL_909347, EPI_ISL_909357, EPI_ISL_909360, EPI_ISL_909362, EPI_ISL_909363 | Lighthouse Lab in Milton Keynes | Wellcome Sanger Institute for the COVID-19 Genomics UK (COG-UK) Consortium | The Lighthouse Lab in Milton Keynes and Alex Alderton, Roberto Amato, Sonia Goncalves, Ewan Harrison, David K. Jackson, Ian Johnston, Dominic Kwiatkowski, Cordelia Langford, John Sillitoe on behalf of the Wellcome Sanger Institute COVID-19 Surveillance Team |
| EPI_ISL_909364, EPI_ISL_909373 | Lighthouse Lab in Alderley Park | Wellcome Sanger Institute for the COVID-19 Genomics UK (COG-UK) Consortium | Jacquelyn Wynn, Mairead Hyland, The Lighthouse Lab in Alderley Park and Alex Alderton, Roberto Amato, Sonia Goncalves, Ewan Harrison, David K. Jackson, Ian Johnston, Dominic Kwiatkowski, Cordelia Langford, John Sillitoe on behalf of the Wellcome Sanger Institute COVID-19 Surveillance Team |
| EPI_ISL_909377, EPI_ISL_909378, EPI_ISL_909384, EPI_ISL_909395, EPI_ISL_909398, EPI_ISL_909399, EPI_ISL_909400 | Lighthouse Lab in Milton Keynes | Wellcome Sanger Institute for the COVID-19 Genomics UK (COG-UK) Consortium | The Lighthouse Lab in Milton Keynes and Alex Alderton, Roberto Amato, Sonia Goncalves, Ewan Harrison, David K. Jackson, Ian Johnston, Dominic Kwiatkowski, Cordelia Langford, John Sillitoe on behalf of the Wellcome Sanger Institute COVID-19 Surveillance Team |
| EPI_ISL_909401 | Lighthouse Lab in Alderley Park | Wellcome Sanger Institute for the COVID-19 Genomics UK (COG-UK) Consortium | Jacquelyn Wynn, Mairead Hyland, The Lighthouse Lab in Alderley Park and Alex Alderton, Roberto Amato, Sonia Goncalves, Ewan Harrison, David K. Jackson, Ian Johnston, Dominic Kwiatkowski, Cordelia Langford, John Sillitoe on behalf of the Wellcome Sanger Institute COVID-19 Surveillance Team |
| EPI_ISL_909402, EPI_ISL_909404 | Lighthouse Lab in Milton Keynes | Wellcome Sanger Institute for the COVID-19 Genomics UK (COG-UK) Consortium | The Lighthouse Lab in Milton Keynes and Alex Alderton, Roberto Amato, Sonia Goncalves, Ewan Harrison, David K. Jackson, Ian Johnston, Dominic Kwiatkowski, Cordelia Langford, John Sillitoe on behalf of the Wellcome Sanger Institute COVID-19 Surveillance Team |
| EPI_ISL_909414 | Lighthouse Lab in Alderley Park | Wellcome Sanger Institute for the COVID-19 Genomics UK (COG-UK) Consortium | Jacquelyn Wynn, Mairead Hyland, The Lighthouse Lab in Alderley Park and Alex Alderton, Roberto Amato, Sonia Goncalves, Ewan Harrison, David K. Jackson, Ian Johnston, Dominic Kwiatkowski, Cordelia Langford, John Sillitoe on behalf of the Wellcome Sanger Institute COVID-19 Surveillance Team |
| EPI_ISL_909419, EPI_ISL_909420, EPI_ISL_909423 | Lighthouse Lab in Milton Keynes | Wellcome Sanger Institute for the COVID-19 Genomics UK (COG-UK) Consortium | The Lighthouse Lab in Milton Keynes and Alex Alderton, Roberto Amato, Sonia Goncalves, Ewan Harrison, David K. Jackson, Ian Johnston, Dominic Kwiatkowski, Cordelia Langford, John Sillitoe on behalf of the Wellcome Sanger Institute COVID-19 Surveillance Team |
| EPI_ISL_909425, EPI_ISL_909426 | Lighthouse Lab in Alderley Park | Wellcome Sanger Institute for the COVID-19 Genomics UK (COG-UK) Consortium | Jacquelyn Wynn, Mairead Hyland, The Lighthouse Lab in Alderley Park and Alex Alderton, Roberto Amato, Sonia Goncalves, Ewan Harrison, David K. Jackson, Ian Johnston, Dominic Kwiatkowski, Cordelia Langford, John Sillitoe on behalf of the Wellcome Sanger Institute COVID-19 Surveillance Team |
| EPI_ISL_909430, EPI_ISL_909432, EPI_ISL_909433, EPI_ISL_909434, EPI_ISL_909436, EPI_ISL_909437, EPI_ISL_909439, EPI_ISL_909441, EPI_ISL_909445 | Lighthouse Lab in Milton Keynes | Wellcome Sanger Institute for the COVID-19 Genomics UK (COG-UK) Consortium | The Lighthouse Lab in Milton Keynes and Alex Alderton, Roberto Amato, Sonia Goncalves, Ewan Harrison, David K. Jackson, Ian Johnston, Dominic Kwiatkowski, Cordelia Langford, John Sillitoe on behalf of the Wellcome Sanger Institute COVID-19 Surveillance Team |
| EPI_ISL_909447 | Lighthouse Lab in Alderley Park | Wellcome Sanger Institute for the COVID-19 Genomics UK (COG-UK) Consortium | Jacquelyn Wynn, Mairead Hyland, The Lighthouse Lab in Alderley Park and Alex Alderton, Roberto Amato, Sonia Goncalves, Ewan Harrison, David K. Jackson, Ian Johnston, Dominic Kwiatkowski, Cordelia Langford, John Sillitoe on behalf of the Wellcome Sanger Institute COVID-19 Surveillance Team |
| EPI_ISL_909448, EPI_ISL_909449, EPI_ISL_909451, EPI_ISL_909454, EPI_ISL_909455, EPI_ISL_909458, EPI_ISL_909459, EPI_ISL_909461, EPI_ISL_909462, EPI_ISL_909466 | Lighthouse Lab in Milton Keynes | Wellcome Sanger Institute for the COVID-19 Genomics UK (COG-UK) Consortium | The Lighthouse Lab in Milton Keynes and Alex Alderton, Roberto Amato, Sonia Goncalves, Ewan Harrison, David K. Jackson, Ian Johnston, Dominic Kwiatkowski, Cordelia Langford, John Sillitoe on behalf of the Wellcome Sanger Institute COVID-19 Surveillance Team |
| EPI_ISL_909467 | Lighthouse Lab in Alderley Park | Wellcome Sanger Institute for the COVID-19 Genomics UK (COG-UK) Consortium | Jacquelyn Wynn, Mairead Hyland, The Lighthouse Lab in Alderley Park and Alex Alderton, Roberto Amato, Sonia Goncalves, Ewan Harrison, David K. Jackson, Ian Johnston, Dominic Kwiatkowski, Cordelia Langford, John Sillitoe on behalf of the Wellcome Sanger Institute COVID-19 Surveillance Team |
| EPI_ISL_909469, EPI_ISL_909470, EPI_ISL_909472, EPI_ISL_909473, EPI_ISL_909477, EPI_ISL_909483, EPI_ISL_909485, EPI_ISL_909486 | Lighthouse Lab in Milton Keynes | Wellcome Sanger Institute for the COVID-19 Genomics UK (COG-UK) Consortium | The Lighthouse Lab in Milton Keynes and Alex Alderton, Roberto Amato, Sonia Goncalves, Ewan Harrison, David K. Jackson, Ian Johnston, Dominic Kwiatkowski, Cordelia Langford, John Sillitoe on behalf of the Wellcome Sanger Institute COVID-19 Surveillance Team |
| EPI_ISL_909488 | Lighthouse Lab in Alderley Park | Wellcome Sanger Institute for the COVID-19 Genomics UK (COG-UK) Consortium | Jacquelyn Wynn, Mairead Hyland, The Lighthouse Lab in Alderley Park and Alex Alderton, Roberto Amato, Sonia Goncalves, Ewan Harrison, David K. Jackson, Ian Johnston, Dominic Kwiatkowski, Cordelia Langford, John Sillitoe on behalf of the Wellcome Sanger Institute COVID-19 Surveillance Team |
| EPI_ISL_909489, EPI_ISL_909491, EPI_ISL_909494, EPI_ISL_909495, EPI_ISL_909498, EPI_ISL_909499, EPI_ISL_909500, EPI_ISL_909501, EPI_ISL_909502, EPI_ISL_909507, EPI_ISL_909509, EPI_ISL_909516, EPI_ISL_909517, EPI_ISL_909519, EPI_ISL_909520, EPI_ISL_909521, EPI_ISL_909523, EPI_ISL_909527, EPI_ISL_909530, EPI_ISL_909535 | | | |
| see above | Lighthouse Lab in Milton Keynes | Wellcome Sanger Institute for the COVID-19 Genomics UK (COG-UK) Consortium | The Lighthouse Lab in Milton Keynes and Alex Alderton, Roberto Amato, Sonia Goncalves, Ewan Harrison, David K. Jackson, Ian Johnston, Dominic Kwiatkowski, Cordelia Langford, John Sillitoe on behalf of the Wellcome Sanger Institute COVID-19 Surveillance Team |

| | | | |
|---|---|---|---|
| EPI_ISL_909536 | Lighthouse Lab in Alderley Park | Wellcome Sanger Institute for the COVID-19 Genomics UK (COG-UK) Consortium | Jacquelyn Wynn, Mairead Hyland, The Lighthouse Lab in Alderley Park and Alex Alderton, Roberto Amato, Sonia Goncalves, Ewan Harrison, David K. Jackson, Ian Johnston, Dominic Kwiatkowski, Cordelia Langford, John Sillitoe on behalf of the Wellcome Sanger Institute COVID-19 Surveillance Team |
| EPI_ISL_909542, EPI_ISL_909543, EPI_ISL_909547, EPI_ISL_909548, EPI_ISL_909550, EPI_ISL_909551, EPI_ISL_909555, EPI_ISL_909558, EPI_ISL_909559, EPI_ISL_909563, EPI_ISL_909565, EPI_ISL_909567, EPI_ISL_909574, EPI_ISL_909579, EPI_ISL_909585, EPI_ISL_909587, EPI_ISL_909591, EPI_ISL_909592 | | | |
| see above | Lighthouse Lab in Milton Keynes | Wellcome Sanger Institute for the COVID-19 Genomics UK (COG-UK) Consortium | The Lighthouse Lab in Milton Keynes and Alex Alderton, Roberto Amato, Sonia Goncalves, Ewan Harrison, David K. Jackson, Ian Johnston, Dominic Kwiatkowski, Cordelia Langford, John Sillitoe on behalf of the Wellcome Sanger Institute COVID-19 Surveillance Team |
| EPI_ISL_909594 | Lighthouse Lab in Alderley Park | Wellcome Sanger Institute for the COVID-19 Genomics UK (COG-UK) Consortium | Jacquelyn Wynn, Mairead Hyland, The Lighthouse Lab in Alderley Park and Alex Alderton, Roberto Amato, Sonia Goncalves, Ewan Harrison, David K. Jackson, Ian Johnston, Dominic Kwiatkowski, Cordelia Langford, John Sillitoe on behalf of the Wellcome Sanger Institute COVID-19 Surveillance Team |
| EPI_ISL_909655, EPI_ISL_909658, EPI_ISL_909663, EPI_ISL_909664, EPI_ISL_909669, EPI_ISL_909670, EPI_ISL_909671, EPI_ISL_909672, EPI_ISL_909673 | Labo Analyses med | National Reference Center for Viruses of Respiratory Infections, Institut Pasteur, Paris | Marion Barbet, Sylvie Behillil, Méline Bizard, Angela Brisebarre, Camille Capel, Etienne Simon-Lorière, Vincent Enouf, Maud Vanpeene, Sylvie van der Werf,Amzalag Jonas |
| EPI_ISL_909688, EPI_ISL_909689, EPI_ISL_909690, EPI_ISL_909691, EPI_ISL_909692, EPI_ISL_909693, EPI_ISL_909694, EPI_ISL_909695, EPI_ISL_909696, EPI_ISL_909698, EPI_ISL_909699, EPI_ISL_909700, EPI_ISL_909701, EPI_ISL_909710, EPI_ISL_909711, EPI_ISL_909712, EPI_ISL_909713, EPI_ISL_909715, EPI_ISL_909716, EPI_ISL_909717, EPI_ISL_909718, EPI_ISL_909719, EPI_ISL_909720, EPI_ISL_909721, EPI_ISL_909722 | | | |
| see above | CH de Mayotte - Laboratoire de Biologie | National Reference Center for Viruses of Respiratory Infections, Institut Pasteur, Paris | Marion Barbet, Sylvie Behillil, Méline Bizard, Angela Brisebarre, Camille Capel, Etienne Simon-Lorière, Vincent Enouf, Maud Vanpeene, Sylvie van der Werf,Combe Patrice |
| EPI_ISL_909755, EPI_ISL_909756 | Charité Universitätsmedizin Berlin, Institut für Virologie/Labor Berlin | Charité Universitätsmedizin Berlin, Institut für Virologie | Victor M Corman, Barbara Mühlemann, Jörn Beheim-Schwarzbach, Tobias Bleicker, Julia Tesch, Talitha Veith, Julia Schneider, Terry Jones, Christian Drosten |
| EPI_ISL_909832, EPI_ISL_909850, EPI_ISL_909917, EPI_ISL_909932 | National Virus Reference Laboratory | National Virus Reference Laboratory | Michael Carr, Gabriel Gonzalez, Jonathan Dean, Cillian F De Gascun |
| EPI_ISL_911175, EPI_ISL_911176, EPI_ISL_911177, EPI_ISL_911178, EPI_ISL_911179, EPI_ISL_911180, EPI_ISL_911181, EPI_ISL_911182, EPI_ISL_911183, EPI_ISL_911184, EPI_ISL_911185, EPI_ISL_911186, EPI_ISL_911187, EPI_ISL_911188, EPI_ISL_911189, EPI_ISL_911190, EPI_ISL_911191, EPI_ISL_911192, EPI_ISL_911193, EPI_ISL_911194, EPI_ISL_911195, EPI_ISL_911196, EPI_ISL_911197, EPI_ISL_911198, EPI_ISL_911199, EPI_ISL_911200, EPI_ISL_911201, EPI_ISL_911202, EPI_ISL_911203, EPI_ISL_911204, EPI_ISL_911205, EPI_ISL_911206, EPI_ISL_911207, EPI_ISL_911208 | | | |
| see above | Laboratoire national de sante, Microbiology, Virology | Laboratoire national de sante, Microbiology, Microbial Genomics Platform | Anke Wienecke-Baldacchino, Catherine Ragimbeau,Jessica Tapp, Fatu Djabi, Lise Pignon, Raoul Salmon, Tamir Abdelrahman |
| EPI_ISL_911285, EPI_ISL_911287, EPI_ISL_911288, EPI_ISL_911291, EPI_ISL_911293, EPI_ISL_911294, EPI_ISL_911296, EPI_ISL_911301, EPI_ISL_911303, EPI_ISL_911304, EPI_ISL_911306, EPI_ISL_911312, EPI_ISL_911313, EPI_ISL_911322, EPI_ISL_911323, EPI_ISL_911326, EPI_ISL_911328, EPI_ISL_911330, EPI_ISL_911333, EPI_ISL_911338, EPI_ISL_911339, EPI_ISL_911347 | | | |
| see above | Servicio de Microbiología, Hospital Universitario Son Espases | SeqCOVID-SPAIN consortium/IBV(CSIC) | Carla López-Causapé, Jordi Reina, Antonio Oliver and SeqCOVID-SPAIN consortium |
| EPI_ISL_911599 | Clinical Molecular Microbiology Laboratory, UNC Hospitals | Jeremy Wang | Jeremy Wang, Alexander Rubinsteyn, Colleen Rice, Jason Smedberg, Melissa Miller, Corbin Jones, Robert Hagan |
| EPI_ISL_911759, EPI_ISL_911760, EPI_ISL_911761, EPI_ISL_911769, EPI_ISL_911770, EPI_ISL_911771, EPI_ISL_911773, EPI_ISL_911783, EPI_ISL_911823, EPI_ISL_911904 | Johns Hopkins Hospital Department of Pathology | Johns Hopkins Hospital Department of Pathology | C. Paul Morris, Chun Huai Luo, Adannaya Amadi, Matthew Schwartz, Nicholas Gallagher, Heba H. Mostafa |
| EPI_ISL_912160 | Yale Clinical Virology Lab | Grubaugh Lab - Yale School of Public Health | Tara Alpert, Joseph Fauver, Anderson Brito, Mallery Breban, Anne Wyllie, Chantal Vogels, Mary Petrone, Annie Watkins, Chaney Kalinich, Isabel Ott, Nathan Grubaugh |
| EPI_ISL_912161 | Tempus | Grubaugh Lab - Yale School of Public Health | Tara Alpert, Joseph Fauver, Anderson Brito, Mallery Breban, Anne Wyllie, Chantal Vogels, Mary Petrone, Annie Watkins, Chaney Kalinich, Isabel Ott, Nathan Grubaugh |
| EPI_ISL_912168 | Yale Pathology Lab | Grubaugh Lab - Yale School of Public Health | Tara Alpert, Joseph Fauver, Chen Liu, Pei Hui, Jianhui Wang, Susan Bell and Han Zhou, Anderson Brito, Mallery Breban, Anne Wyllie, Chantal Vogels, Mary Petrone, Chaney Kalinich, Isabel Ott, Arnau Casanovas, Catherine Muenker, Adam Moore, Alice Lu, Maria Tokuyama, Patrick Wong, Peiwen Lu, Saad Omer, Richard Martinello, Allison Nelson, Shelli Farhadian, Akiko Iwasaki, Charlese Dela Cruz, Albert Ko, Nathan Grubaugh |
| EPI_ISL_912224, EPI_ISL_912225 | Lighthouse Lab in Milton Keynes | Wellcome Sanger Institute for the COVID-19 Genomics UK (COG-UK) Consortium | The Lighthouse Lab in Milton Keynes and Alex Alderton, Roberto Amato, Sonia Goncalves, Ewan Harrison, David K. Jackson, Ian Johnston, Dominic Kwiatkowski, Cordelia Langford, John Sillitoe on behalf of the Wellcome Sanger Institute COVID-19 Surveillance Team (http://www.sanger.ac.uk/covid-team) |
| EPI_ISL_912236, EPI_ISL_912237, EPI_ISL_912238, EPI_ISL_912239, EPI_ISL_912240, EPI_ISL_912241, EPI_ISL_912242, EPI_ISL_912243, EPI_ISL_912244 | Charité Universitätsmedizin Berlin, Institut für Virologie/Labor Berlin | Charité Universitätsmedizin Berlin, Institut für Virologie | Victor M Corman, Barbara Mühlemann, Jörn Beheim-Schwarzbach, Tobias Bleicker, Julia Tesch, Talitha Veith, Julia Schneider, Terry Jones, Christian Drosten |
| EPI_ISL_912268, EPI_ISL_912271, EPI_ISL_912282, EPI_ISL_912285, EPI_ISL_912287, EPI_ISL_912289, EPI_ISL_912291, EPI_ISL_912292, EPI_ISL_912294, EPI_ISL_912295, EPI_ISL_912296, EPI_ISL_912300, EPI_ISL_912301, EPI_ISL_912302, EPI_ISL_912304, EPI_ISL_912307, EPI_ISL_912308, EPI_ISL_912322, EPI_ISL_912326, EPI_ISL_912327, EPI_ISL_912334, EPI_ISL_912341, EPI_ISL_912345, EPI_ISL_912346, EPI_ISL_912347 | | | |
| see above | Hospital General Universitario Gregorio Marañón | SeqCOVID-SPAIN consortium / IBV (CSIC) | Darío García de Viedma, Laura Pérez-Lago, Pedro J Sola-Campoy, Sergio Buenestado-Serrano, Marta Herranz, Victor Manuel de la Cueva, Julia Suárez, Pilar Catalán, Patricia Muñoz and SeqCOVID-SPAIN consortium |
| EPI_ISL_912403 | CH de Mayotte - Laboratoire de Biologie | National Reference Center for Viruses of Respiratory Infections, Institut Pasteur, Paris | Marion Barbet, Sylvie Behillil, Méline Bizard, Angela Brisebarre, Camille Capel, Etienne Simon-Lorière, Vincent Enouf, Maud Vanpeene, Sylvie van der Werf,Combe Patrice |
| EPI_ISL_912545, EPI_ISL_912549 | Labo Analyses Med | National Reference Center for Viruses of Respiratory Infections, Institut Pasteur, Paris | Marion Barbet, Sylvie Behillil, Méline Bizard, Angela Brisebarre, Camille Capel, Etienne Simon-Lorière, Vincent Enouf, Maud Vanpeene, Sylvie van der Werf |
| EPI_ISL_912557, EPI_ISL_912558 | Centre Hospitalier de Gisors Route de Rouen | National Reference Center for Viruses of Respiratory Infections, Institut Pasteur, Paris | Marion Barbet, Sylvie Behillil, Méline Bizard, Angela Brisebarre, Camille Capel, Etienne Simon-Lorière, Vincent Enouf, Maud Vanpeene, Sylvie van der Werf |
| EPI_ISL_912561, EPI_ISL_912562 | HOPITAL SIMONE VEIL | National Reference Center for Viruses of Respiratory Infections, Institut Pasteur, Paris | Marion Barbet, Sylvie Behillil, Méline Bizard, Angela Brisebarre, Camille Capel, Etienne Simon-Lorière, Vincent Enouf, Maud Vanpeene, Sylvie van der Werf,Moreau Farida |
| EPI_ISL_912565, EPI_ISL_912568, EPI_ISL_912577, EPI_ISL_912578 | Laboratoire de Virologie du CHU de Lille Bâtiment Paul Boulanger | National Reference Center for Viruses of Respiratory Infections, Institut Pasteur, Paris | Marion Barbet, Sylvie Behillil, Méline Bizard, Angela Brisebarre, Camille Capel, Etienne Simon-Lorière, Vincent Enouf, Maud Vanpeene, Sylvie van der Werf,Guigon Aurélie |
| EPI_ISL_912598 | Hopital Avicenne Laboratoire central de bactériologie virologie hygiène | National Reference Center for Viruses of Respiratory Infections, Institut Pasteur, Paris | Marion Barbet, Sylvie Behillil, Méline Bizard, Angela Brisebarre, Camille Capel, Etienne Simon-Lorière, Vincent Enouf, Maud Vanpeene, Sylvie van der Werf,Brichler Ségolène |
| EPI_ISL_912601, EPI_ISL_912603, EPI_ISL_912605 | Labo Analyses Med | National Reference Center for Viruses of Respiratory Infections, Institut Pasteur, Paris | Marion Barbet, Sylvie Behillil, Méline Bizard, Angela Brisebarre, Camille Capel, Etienne Simon-Lorière, Vincent Enouf, Maud Vanpeene, Sylvie van der Werf |
| EPI_ISL_912614 | CENTRE HOSPITALIER JACQUES LACARIN LABORATOIRE | National Reference Center for Viruses of Respiratory Infections, Institut Pasteur, Paris | Marion Barbet, Sylvie Behillil, Méline Bizard, Angela Brisebarre, Camille Capel, Etienne Simon-Lorière, Vincent Enouf, Maud Vanpeene, Sylvie van der Werf |
| EPI_ISL_912621 | HOPITAL YVES LE FOLL Laboratoire de Biologie Médicale | National Reference Center for Viruses of Respiratory Infections, Institut Pasteur, Paris | Marion Barbet, Sylvie Behillil, Méline Bizard, Angela Brisebarre, Camille Capel, Etienne Simon-Lorière, Vincent Enouf, Maud Vanpeene, Sylvie van der Werf,Chrevet Laurent |
| EPI_ISL_912623 | Labo Analyses Med | National Reference Center for Viruses of Respiratory Infections, Institut Pasteur, Paris | Marion Barbet, Sylvie Behillil, Méline Bizard, Angela Brisebarre, Camille Capel, Etienne Simon-Lorière, Vincent Enouf, Maud Vanpeene, Sylvie van der Werf |

| | | | |
|---|---|---|---|
| EPI_ISL_912636 | Hôpital Henri Mondor | Department of Virology, Henri Mondor University Hospital, Assistance Publique Hôpitaux de Paris, Université Paris-Est Créteil, INSERM U955 | Christophe Rodriguez, Slim Fourati, Vanessa Demontant, Guillaume Gricourt, Melissa N'Debi, Alexandre Soulier, Elisabeth Trawinski, Jean-Michel Pawlotsky |
| EPI_ISL_912637 | Hôpital Pitié-Salpêtrière | Department of Virology, Henri Mondor University Hospital, Assistance Publique Hôpitaux de Paris, Université Paris-Est Créteil, INSERM U955 | Christophe Rodriguez, Slim Fourati, Vanessa Demontant, Guillaume Gricourt, Melissa N'Debi, Alexandre Soulier, Elisabeth Trawinski, Jean-Michel Pawlotsky |
| EPI_ISL_912638 | Hôpital Henri Mondor | Department of Virology, Henri Mondor University Hospital, Assistance Publique Hôpitaux de Paris, Université Paris-Est Créteil, INSERM U955 | Christophe Rodriguez, Slim Fourati, Vanessa Demontant, Guillaume Gricourt, Melissa N'Debi, Alexandre Soulier, Elisabeth Trawinski, Jean-Michel Pawlotsky |
| EPI_ISL_912644 | Hôpital Pitié-Salpêtrière | Department of Virology, Henri Mondor University Hospital, Assistance Publique Hôpitaux de Paris, Université Paris-Est Créteil, INSERM U955 | Christophe Rodriguez, Slim Fourati, Vanessa Demontant, Guillaume Gricourt, Melissa N'Debi, Alexandre Soulier, Elisabeth Trawinski, Jean-Michel Pawlotsky |
| EPI_ISL_912653, EPI_ISL_912657, EPI_ISL_912659 | Hôpital Henri Mondor | Department of Virology, Henri Mondor University Hospital, Assistance Publique Hôpitaux de Paris, Université Paris-Est Créteil, INSERM U955 | Christophe Rodriguez, Slim Fourati, Vanessa Demontant, Guillaume Gricourt, Melissa N'Debi, Alexandre Soulier, Elisabeth Trawinski, Jean-Michel Pawlotsky |
| EPI_ISL_912662 | Hôpital Pitié-Salpêtrière | Department of Virology, Henri Mondor University Hospital, Assistance Publique Hôpitaux de Paris, Université Paris-Est Créteil, INSERM U955 | Christophe Rodriguez, Slim Fourati, Vanessa Demontant, Guillaume Gricourt, Melissa N'Debi, Alexandre Soulier, Elisabeth Trawinski, Jean-Michel Pawlotsky |
| EPI_ISL_912663 | Hôpital Henri Mondor | Department of Virology, Henri Mondor University Hospital, Assistance Publique Hôpitaux de Paris, Université Paris-Est Créteil, INSERM U955 | Christophe Rodriguez, Slim Fourati, Vanessa Demontant, Guillaume Gricourt, Melissa N'Debi, Alexandre Soulier, Elisabeth Trawinski, Jean-Michel Pawlotsky |
| EPI_ISL_912667 | Hôpital Pitié-Salpêtrière | Department of Virology, Henri Mondor University Hospital, Assistance Publique Hôpitaux de Paris, Université Paris-Est Créteil, INSERM U955 | Christophe Rodriguez, Slim Fourati, Vanessa Demontant, Guillaume Gricourt, Melissa N'Debi, Alexandre Soulier, Elisabeth Trawinski, Jean-Michel Pawlotsky |
| EPI_ISL_912668, EPI_ISL_912671, EPI_ISL_912694, EPI_ISL_912776, EPI_ISL_912777, EPI_ISL_912778, EPI_ISL_912779, EPI_ISL_912780 | Hôpital Henri Mondor | Department of Virology, Henri Mondor University Hospital, Assistance Publique Hôpitaux de Paris, Université Paris-Est Créteil, INSERM U955 | Christophe Rodriguez, Slim Fourati, Vanessa Demontant, Guillaume Gricourt, Melissa N'Debi, Alexandre Soulier, Elisabeth Trawinski, Jean-Michel Pawlotsky |
| EPI_ISL_912788, EPI_ISL_912793, EPI_ISL_912794 | Hôpital Pitié-Salpêtrière | Department of Virology, Henri Mondor University Hospital, Assistance Publique Hôpitaux de Paris, Université Paris-Est Créteil, INSERM U955 | Christophe Rodriguez, Slim Fourati, Vanessa Demontant, Guillaume Gricourt, Melissa N'Debi, Alexandre Soulier, Elisabeth Trawinski, Jean-Michel Pawlotsky |
| EPI_ISL_912804, EPI_ISL_912805, EPI_ISL_912806, EPI_ISL_912807, EPI_ISL_912808, EPI_ISL_912809, EPI_ISL_912810, EPI_ISL_912863 | Hôpital Henri Mondor | Department of Virology, Henri Mondor University Hospital, Assistance Publique Hôpitaux de Paris, Université Paris-Est Créteil, INSERM U955 | Christophe Rodriguez, Slim Fourati, Vanessa Demontant, Guillaume Gricourt, Melissa N'Debi, Alexandre Soulier, Elisabeth Trawinski, Jean-Michel Pawlotsky |
| EPI_ISL_912891 | CH.INTERCOMMUNAL DE CRETEIL | Department of Virology, Henri Mondor University Hospital, Assistance Publique Hôpitaux de Paris, Université Paris-Est Créteil, INSERM U955 | Christophe Rodriguez, Slim Fourati, Vanessa Demontant, Guillaume Gricourt, Melissa N'Debi, Alexandre Soulier, Elisabeth Trawinski, Jean-Michel Pawlotsky |
| EPI_ISL_912897, EPI_ISL_912898 | Hôpital Henri Mondor | Department of Virology, Henri Mondor University Hospital, Assistance Publique Hôpitaux de Paris, Université Paris-Est Créteil, INSERM U955 | Christophe Rodriguez, Slim Fourati, Vanessa Demontant, Guillaume Gricourt, Melissa N'Debi, Alexandre Soulier, Elisabeth Trawinski, Jean-Michel Pawlotsky |
| EPI_ISL_912907 | CH.INTERCOMMUNAL DE CRETEIL | Department of Virology, Henri Mondor University Hospital, Assistance Publique Hôpitaux de Paris, Université Paris-Est Créteil, INSERM U955 | Christophe Rodriguez, Slim Fourati, Vanessa Demontant, Guillaume Gricourt, Melissa N'Debi, Alexandre Soulier, Elisabeth Trawinski, Jean-Michel Pawlotsky |
| EPI_ISL_912933, EPI_ISL_912936, EPI_ISL_912940 | Hôpital Pitié-Salpêtrière | Department of Virology, Henri Mondor University Hospital, Assistance Publique Hôpitaux de Paris, Université Paris-Est Créteil, INSERM U955 | Christophe Rodriguez, Slim Fourati, Vanessa Demontant, Guillaume Gricourt, Melissa N'Debi, Alexandre Soulier, Elisabeth Trawinski, Jean-Michel Pawlotsky |
| EPI_ISL_912942, EPI_ISL_912944, EPI_ISL_912945, EPI_ISL_912946, EPI_ISL_912947, EPI_ISL_912948, EPI_ISL_912949, EPI_ISL_912950, EPI_ISL_912951, EPI_ISL_912952, EPI_ISL_912960 | | | |
| see above | Hôpital Henri Mondor | Department of Virology, Henri Mondor University Hospital, Assistance Publique Hôpitaux de Paris, Université Paris-Est Créteil, INSERM U955 | Christophe Rodriguez, Slim Fourati, Vanessa Demontant, Guillaume Gricourt, Melissa N'Debi, Alexandre Soulier, Elisabeth Trawinski, Jean-Michel Pawlotsky |
| EPI_ISL_912961 | CH.INTERCOMMUNAL DE CRETEIL | Department of Virology, Henri Mondor University Hospital, Assistance Publique Hôpitaux de Paris, Université Paris-Est Créteil, INSERM U955 | Christophe Rodriguez, Slim Fourati, Vanessa Demontant, Guillaume Gricourt, Melissa N'Debi, Alexandre Soulier, Elisabeth Trawinski, Jean-Michel Pawlotsky |
| EPI_ISL_913025, EPI_ISL_913026, EPI_ISL_913027, EPI_ISL_913029 | Hospital los Arcos | Instituto de Salud Carlos III | Iglesias-Caballero, M. Camarero, S. Sandonís,V. Vázquez, S. Pozo, F. Casas, I. Jiménez, P. Zaballos, A. Monzón, S. Varona, S. Cuesta, I. Cámara, M. |
| EPI_ISL_913283 | Klinisk mikrobiologi | The Public Health Agency of Sweden | Anna-Malin Linde, Maria Lind Karlberg, Carlo Berg, Oskar Karlsson Lindsjo, Sofia Stamouli, Reza Advani, Mattias Haukland, Petra Holmstrom, Noura Walai, Petra Edquist, Mia Brytting, Anna Risberg, Karin Tegmark-Wisell |
| EPI_ISL_913393, EPI_ISL_913397 | Massachusetts State Public Health Laboratory | Massachusetts State Public Health Laboratory | Andrew Lang, Timelia Fink, Glen Gallagher, Sandra Smole |
| EPI_ISL_913446 | Laboratorio Genzano - ASL RM 6 | INMI Lazzaro Spallanzani IRCCS | Emanuela Giombini, Martina Rueca, Barbara Bartolini, Ornella Butera, Cesare E.M. Gruber, Francesco Messina, Grazia Tramini, Emanuela Conti, Antonino Di Caro, Maria R. Capobianchi |
| EPI_ISL_913447 | Laboratorio Genzano - ASL RM 6 | INMI Lazzaro Spallanzani IRCCS | Francesco Messina, Emanuela Giombini, Ornella Butera, Cesare EM Gruber, Martina Rueca, Barbara Bartolini, Grazia Tramini, Emanuela Conti, Maria R Capobianchi, Antonino Di Caro |
| EPI_ISL_913457, EPI_ISL_913458 | Klinisk mikrobiologi | The Public Health Agency of Sweden | Anna-Malin Linde, Maria Lind Karlberg, Carlo Berg, Oskar Karlsson Lindsjo, Sofia Stamouli, Reza Advani, Mattias Haukland, Petra Holmstrom, Noura Walai, Petra Edquist, Mia Brytting, Anna Risberg, Karin Tegmark-Wisell |
| EPI_ISL_913594, EPI_ISL_913595, EPI_ISL_913596, EPI_ISL_913597, EPI_ISL_913598, EPI_ISL_913599, EPI_ISL_913600, EPI_ISL_913601, EPI_ISL_913602, EPI_ISL_913603, EPI_ISL_913604, EPI_ISL_913605 | | | |
| see above | AZDelta | AZDelta | Geert Martens; Dieter De Smet |
| EPI_ISL_913607 | Hopital Avicenne Laboratoire central de bactériologie virologie hygiène | National Reference Center for Viruses of Respiratory Infections, Institut Pasteur, Paris | Marion Barbet, Sylvie Behillil, Méline Bizard, Angela Brisebarre, Camille Capel, Etienne Simon-Lorière, Vincent Enouf, Maud Vanpeene, Sylvie van der Werf,Brichler Ségolène |
| EPI_ISL_913608, EPI_ISL_913609, EPI_ISL_913610 | Laboratoire de Virologie du CHU de Lille Bâtiment Paul Boulanger | National Reference Center for Viruses of Respiratory Infections, Institut Pasteur, Paris | Marion Barbet, Sylvie Behillil, Méline Bizard, Angela Brisebarre, Camille Capel, Etienne Simon-Lorière, Vincent Enouf, Maud Vanpeene, Sylvie van der Werf,Guigon Aurélie |
| EPI_ISL_913615 | Michigan Department of Health and Human Services, Bureau of Laboratories | Michigan Department of Health and Human Services, Bureau of Laboratories | Blankenship HM, Riner D, Soehnlen MK |

| | | | |
|---|---|---|---|
| EPI_ISL_913767 | KU Leuven, Rega Institute, Clinical and Epidemiological Virology | KU Leuven, Rega Institute, Clinical and Epidemiological Virology | Tony Wawina-Bokalanga, Bert Vanmechelen, Joan Marti-Carerras, Piet Maes |
| EPI_ISL_913991, EPI_ISL_913995, EPI_ISL_913996 | IL Dept. of Public Health Springfield Laboratory | Pathogen Discovery, Respiratory Viruses Branch, Division of Viral Diseases, Centers for Disease Control and Prevention | Ying Tao, Yan Li, Jing Zhang, Krista Queen, Anna Uehara, Peter Cook, Clinton R. Paden, Haibin Wang, Suxiang Tong |
| EPI_ISL_914005 | Clinical Pathology Lab | Pathogen Discovery, Respiratory Viruses Branch, Division of Viral Diseases, Centers for Disease Control and Prevention | Ying Tao, Yan Li, Jing Zhang, Krista Queen, Anna Uehara, Peter Cook, Clinton R. Paden, Haibin Wang, Suxiang Tong |
| EPI_ISL_914035 | IN State Department of Health Laboratory Service | Pathogen Discovery, Respiratory Viruses Branch, Division of Viral Diseases, Centers for Disease Control and Prevention | Ying Tao, Yan Li, Jing Zhang, Krista Queen, Anna Uehara, Peter Cook, Clinton R. Paden, Haibin Wang, Suxiang Tong |
| EPI_ISL_914642 | Santa Clara County Public Health Laboratory | Santa Clara County Public Health Laboratory | Santa Clara County Public Health Department |
| EPI_ISL_914745, EPI_ISL_914755, EPI_ISL_914765, EPI_ISL_914773, EPI_ISL_914776 | Wyoming Public Health Laboratory | Wyoming Public Health Laboratory | Noah Hull, Taylor Fearing, Lynette Gumbleton, Channing Weber, Ashley Norberg, Bailey Bowcutt, and Wanda Manley |
| EPI_ISL_914914, EPI_ISL_914944, EPI_ISL_915193, EPI_ISL_915194, EPI_ISL_915206, EPI_ISL_915207, EPI_ISL_915211, EPI_ISL_915212, EPI_ISL_915213, EPI_ISL_915215, EPI_ISL_915224, EPI_ISL_915226, EPI_ISL_915227, EPI_ISL_915228, EPI_ISL_915230, EPI_ISL_915231, EPI_ISL_915232, EPI_ISL_915234, EPI_ISL_915264, EPI_ISL_915267, EPI_ISL_915268, EPI_ISL_915269, EPI_ISL_915270, EPI_ISL_915271, EPI_ISL_915272, EPI_ISL_915273, EPI_ISL_915274, EPI_ISL_915276, EPI_ISL_915278, EPI_ISL_915279, EPI_ISL_915280, EPI_ISL_915281, EPI_ISL_915282, EPI_ISL_915283, EPI_ISL_915302, EPI_ISL_915306, EPI_ISL_915308, EPI_ISL_915310, EPI_ISL_915311, EPI_ISL_915312, EPI_ISL_915313, EPI_ISL_915314, EPI_ISL_915316, EPI_ISL_915318, EPI_ISL_915319, EPI_ISL_915320, EPI_ISL_915321, EPI_ISL_915322, EPI_ISL_915324, EPI_ISL_915328, EPI_ISL_915329, EPI_ISL_915330, EPI_ISL_915331, EPI_ISL_915332, EPI_ISL_915335, EPI_ISL_915336, EPI_ISL_915337, EPI_ISL_915338, EPI_ISL_915339, EPI_ISL_915340, EPI_ISL_915344, EPI_ISL_915346, EPI_ISL_915347, EPI_ISL_915348, EPI_ISL_915349, EPI_ISL_915351, EPI_ISL_915353, EPI_ISL_915355, EPI_ISL_915356, EPI_ISL_915357 | | | |
| see above | Quest Diagnostics | Quest Diagnostics | Rosenthal,S.H., Gerasimova,A., Kagan,R.M., Anderson, B., Hua, M., Liu Y., Bernstein, L.E., Livingston, K.E., Perez, A., Shalhout, D.F., Shlyakhter, I.A., Owen, R., Tanpaiboon, P., Lacbawan, F. |
| EPI_ISL_915421 | MRCG at LSHTM Genomics lab | MRCG at LSHTM Genomics lab | Abdul Karim sesay, Abdoulie Kanteh, Jarra Manneh, Mariama Kujabi, Bakary Sanyang |
| EPI_ISL_916633, EPI_ISL_916649, EPI_ISL_916661, EPI_ISL_916675, EPI_ISL_916682, EPI_ISL_916688, EPI_ISL_916694, EPI_ISL_916696, EPI_ISL_916704, EPI_ISL_916705, EPI_ISL_916713, EPI_ISL_916728, EPI_ISL_916730, EPI_ISL_916738, EPI_ISL_916771, EPI_ISL_916782, EPI_ISL_916784, EPI_ISL_916800, EPI_ISL_916802, EPI_ISL_916809, EPI_ISL_916810, EPI_ISL_916812, EPI_ISL_916815, EPI_ISL_916821, EPI_ISL_916825, EPI_ISL_916833, EPI_ISL_916834, EPI_ISL_916838, EPI_ISL_916845, EPI_ISL_916847, EPI_ISL_916850, EPI_ISL_916882, EPI_ISL_916887, EPI_ISL_916896, EPI_ISL_916898, EPI_ISL_916905, EPI_ISL_916907, EPI_ISL_916908, EPI_ISL_916909, EPI_ISL_916911, EPI_ISL_916912, EPI_ISL_916916, EPI_ISL_916918, EPI_ISL_916923, EPI_ISL_916940, EPI_ISL_916945, EPI_ISL_916946, EPI_ISL_916948, EPI_ISL_916949, EPI_ISL_916953, EPI_ISL_916954, EPI_ISL_916956, EPI_ISL_916961, EPI_ISL_916968, EPI_ISL_916970, EPI_ISL_916974, EPI_ISL_916976, EPI_ISL_916977, EPI_ISL_916981, EPI_ISL_916984, EPI_ISL_916985, EPI_ISL_916987, EPI_ISL_916991, EPI_ISL_916999, EPI_ISL_917009, EPI_ISL_917012, EPI_ISL_917013, EPI_ISL_917014, EPI_ISL_917019, EPI_ISL_917021, EPI_ISL_917022, EPI_ISL_917024, EPI_ISL_917025, EPI_ISL_917026, EPI_ISL_917029, EPI_ISL_917031, EPI_ISL_917032, EPI_ISL_917036, EPI_ISL_917038, EPI_ISL_917042, EPI_ISL_917047, EPI_ISL_917054, EPI_ISL_917055, EPI_ISL_917063, EPI_ISL_917067, EPI_ISL_917070, EPI_ISL_917071, EPI_ISL_917074, EPI_ISL_917075, EPI_ISL_917077, EPI_ISL_917079, EPI_ISL_917081, EPI_ISL_917082, EPI_ISL_917083, EPI_ISL_917086, EPI_ISL_917088, EPI_ISL_917091, EPI_ISL_917093, EPI_ISL_917096, EPI_ISL_917099, EPI_ISL_917105, EPI_ISL_917107, EPI_ISL_917110, EPI_ISL_917116, EPI_ISL_917120, EPI_ISL_917125, EPI_ISL_917128, EPI_ISL_917129, EPI_ISL_917133, EPI_ISL_917139, EPI_ISL_917141, EPI_ISL_917143, EPI_ISL_917145, EPI_ISL_917148, EPI_ISL_917149, EPI_ISL_917151, EPI_ISL_917156, EPI_ISL_917159, EPI_ISL_917160, EPI_ISL_917163, EPI_ISL_917849, EPI_ISL_917850, EPI_ISL_917851, EPI_ISL_917852, EPI_ISL_917853, EPI_ISL_917854, EPI_ISL_917855, EPI_ISL_917856, EPI_ISL_917857, EPI_ISL_917858, EPI_ISL_917859, EPI_ISL_917860, EPI_ISL_917861, EPI_ISL_917862, EPI_ISL_917863, EPI_ISL_917864, EPI_ISL_917865, EPI_ISL_917866, EPI_ISL_917867, EPI_ISL_917868, EPI_ISL_917869, EPI_ISL_917870, EPI_ISL_917871, EPI_ISL_917872, EPI_ISL_917873, EPI_ISL_917874, EPI_ISL_917875, EPI_ISL_917876, EPI_ISL_917877, EPI_ISL_917878, EPI_ISL_917879, EPI_ISL_917880, EPI_ISL_917881, EPI_ISL_917882, EPI_ISL_917883, EPI_ISL_917884, EPI_ISL_917885, EPI_ISL_917886, EPI_ISL_917887, EPI_ISL_917888, EPI_ISL_917889, EPI_ISL_917890, EPI_ISL_917891, EPI_ISL_917892, EPI_ISL_917893, EPI_ISL_917894, EPI_ISL_917895, EPI_ISL_917896, EPI_ISL_917897, EPI_ISL_917898, EPI_ISL_917899, EPI_ISL_917900, EPI_ISL_917901, EPI_ISL_917902, EPI_ISL_917903, EPI_ISL_917904, EPI_ISL_917905, EPI_ISL_917906, EPI_ISL_917907, EPI_ISL_917908, EPI_ISL_917909, EPI_ISL_917910, EPI_ISL_917911, EPI_ISL_917912, EPI_ISL_917913, EPI_ISL_917914, EPI_ISL_917915, EPI_ISL_917916, EPI_ISL_917917, EPI_ISL_917918, EPI_ISL_917919, EPI_ISL_917920, EPI_ISL_917921, EPI_ISL_917922, EPI_ISL_917923, EPI_ISL_917924, EPI_ISL_917925, EPI_ISL_917926, EPI_ISL_917927, EPI_ISL_917928, EPI_ISL_917929, EPI_ISL_917930, EPI_ISL_917931, EPI_ISL_917932, EPI_ISL_917933, EPI_ISL_917934, EPI_ISL_917935, EPI_ISL_917937, EPI_ISL_917939, EPI_ISL_917940, EPI_ISL_917942, EPI_ISL_917956, EPI_ISL_917957, EPI_ISL_917958, EPI_ISL_917959, EPI_ISL_917960, EPI_ISL_917961, EPI_ISL_917962, EPI_ISL_917963, EPI_ISL_917964, EPI_ISL_917965, EPI_ISL_917966, EPI_ISL_917967, EPI_ISL_917968, EPI_ISL_917969, EPI_ISL_917970, EPI_ISL_917971, EPI_ISL_917972, EPI_ISL_917973, EPI_ISL_917974, EPI_ISL_917975, EPI_ISL_917976, EPI_ISL_917977, EPI_ISL_917978, EPI_ISL_917979, EPI_ISL_917980, EPI_ISL_917981, EPI_ISL_917982, EPI_ISL_917983, EPI_ISL_917984 | | | |
| see above | Lighthouse Lab in Glasgow | Wellcome Sanger Institute for the COVID-19 Genomics UK (COG-UK) Consortium | Harper VanSteenhouse, Yumi Kasai, David Gray, Carol Clugston, Anna Dominiczak and Alex Alderton, Roberto Amato, Sonia Goncalves, Ewan Harrison, David K. Jackson, Ian Johnston, Dominic Kwiatkowski, Cordelia Langford, John Sillitoe on behalf of the Wellcome Sanger Institute COVID-19 Surveillance Team |
| EPI_ISL_918060 | Lighthouse Lab in Alderley Park | Wellcome Sanger Institute for the COVID-19 Genomics UK (COG-UK) Consortium | Jacquelyn Wynn, Mairead Hyland, The Lighthouse Lab in Alderley Park and Alex Alderton, Roberto Amato, Sonia Goncalves, Ewan Harrison, David K. Jackson, Ian Johnston, Dominic Kwiatkowski, Cordelia Langford, John Sillitoe on behalf of the Wellcome Sanger Institute COVID-19 Surveillance Team |
| EPI_ISL_918230 | Innovative Genomics Institute, UC Berkeley | Innovative Genomics Institute, UC Berkeley | Stacia Wyman, Haridha Shivram, Phil Frankino, Liana Lareau, Shana McDevitt, Justin Choi |
| EPI_ISL_918289, EPI_ISL_918290 | Hospital Universitari Vall d'Hebron - Vall d'Hebron Institut de Rercerca | Hospital Universitari Vall d'Hebron | Cristina Andrés, Maria Piñana, Josep F Abril, Damir Garcia-Cehic, Ariadna Rando, Juliana Esperalba, Maria Gema Codina, Carla Castillo, Maria Carmen Martin, Tomàs Pumarola, Josep Quer, Andrés Antón |
| EPI_ISL_918493 | Florida Bureau of Public Health Laboratories | Florida Bureau of Public Health Laboratories | Sarah Schmedes, Jason Blanton |
| EPI_ISL_918986, EPI_ISL_918987, EPI_ISL_918988, EPI_ISL_918989, EPI_ISL_918994, EPI_ISL_918995, EPI_ISL_918996 | Department of Pathology, University of Cambridge | COVID-19 Genomics UK (COG-UK) Consortium | Aminu S. Jahun, Yasmin Chaudhry, Iliana Georgana, Myra Hosmillo, Rhys Izu, Martin D. Curran, Surendra Parmar, Ian Goodfellow |
| EPI_ISL_919205, EPI_ISL_919206, EPI_ISL_919207, EPI_ISL_919208, EPI_ISL_919209, EPI_ISL_919210, EPI_ISL_919211, EPI_ISL_919212, EPI_ISL_919213, EPI_ISL_919214, EPI_ISL_919215, EPI_ISL_919216, EPI_ISL_919217, EPI_ISL_919218, EPI_ISL_919219, EPI_ISL_919220, EPI_ISL_919221, EPI_ISL_919222, EPI_ISL_919223, EPI_ISL_919224, EPI_ISL_919225, EPI_ISL_919226, EPI_ISL_919227, EPI_ISL_919228, EPI_ISL_919229, EPI_ISL_919230 | | | |
| see above | West of Scotland Specialist Virology Centre, NHSGGC / MRC-University of Glasgow Centre for Virus Research | COVID-19 Genomics UK (COG-UK) Consortium | Ana da Silva Filipe, Natasha Johnson, Kathy Smollett, Daniel Mair, Stephen Carmichael, Alice Broos, Lily Tong, Jenna Nichols, Kyriaki Nomikou; Sarah McDonald; Richard Orton, Joseph Hughes, Sreenu Vattipally, David L Robertson; Alasdair MacLean, Rory Gunson; Sharif Shaaban, Matthew Holden; Rachel Blacow, Guy Mollett, Kathy Li, James Shepherd, Antonia Ho, Emma Thomson |
| EPI_ISL_919342, EPI_ISL_919345 | Virology Department, Royal Infirmary of Edinburgh, NHS Lothian / School of Biological Sciences, University of Edinburgh / Institute of Genetics and Molecular Medicine, University of Edinburgh | COVID-19 Genomics UK (COG-UK) Consortium | McHugh M, Dewar R, Rooke S, Gallagher M, Balcaza C, O'Toole Á, Scher E, Hill V, McCrone JT, Colquhoun R, Yu X, Jackson B, Rambaut A, Williams TC, Templeton K |
| EPI_ISL_919488, EPI_ISL_919495, EPI_ISL_919500, EPI_ISL_919502, EPI_ISL_919503, EPI_ISL_919504, EPI_ISL_919505, EPI_ISL_919507, EPI_ISL_919508, EPI_ISL_919509, EPI_ISL_919510, EPI_ISL_919529, EPI_ISL_919530, EPI_ISL_919531, EPI_ISL_919532, EPI_ISL_919533, EPI_ISL_919535, EPI_ISL_919536, EPI_ISL_919537, EPI_ISL_919538, EPI_ISL_919541, EPI_ISL_919542, EPI_ISL_919544, EPI_ISL_919545, EPI_ISL_919547, EPI_ISL_919548, EPI_ISL_919549, EPI_ISL_919550, EPI_ISL_919551, EPI_ISL_919552, EPI_ISL_919554, EPI_ISL_919556, EPI_ISL_919557, EPI_ISL_919558, EPI_ISL_919559, EPI_ISL_919561, EPI_ISL_919562, EPI_ISL_919563, EPI_ISL_919564, EPI_ISL_919565, EPI_ISL_919568, EPI_ISL_919570, EPI_ISL_919571, EPI_ISL_919572, EPI_ISL_919573, EPI_ISL_919575, EPI_ISL_919576, EPI_ISL_919577, EPI_ISL_919578, EPI_ISL_919581, EPI_ISL_919585, EPI_ISL_919587, EPI_ISL_919589, EPI_ISL_919590, EPI_ISL_919614, EPI_ISL_919647, EPI_ISL_919648, EPI_ISL_919649, EPI_ISL_919650, EPI_ISL_919652, EPI_ISL_919653, EPI_ISL_919654, EPI_ISL_919655, EPI_ISL_919656, EPI_ISL_919657, EPI_ISL_919659, EPI_ISL_919661, EPI_ISL_919662 | | | |
| see above | Liverpool Clinical Laboratories | COVID-19 Genomics UK (COG-UK) Consortium | Sam Haldenby, Anita Lucaci, Steve Paterson, Julian Hiscox, Alistair Darby, M Almsaud, A Alrezaihi, Muhannad Alruwaili, Stuart D Armstrong, Jones Benjamin, Eleanor G Bentley, Anu Chawla, Jordan J Clark, Angela Cowell, Richard Eccles, Isabel García-Dorival, Matthew Gemmell, Alessandro Gerada, PKF Gilmore, Richard Gregory, Ximeng Han, Catherine Hartley, Margaret Hughes, Miren Iturriza-Gomara, James Johnson, L Luu, Jenifer Manson, Charlotte Nelson, Elaine O'Toole, Cassie Olateju, Rebekah Penrice-Randal , Lucille Rainbow, N.P Randle, Trevor Ian Robinson, Parul Sharma, Ghada T Shawli, James P Stewart, Neil Swainston, Ecaterina Vamos, Joanne Watts, Mark Whitehead |
| EPI_ISL_919854 | Barts Health NHS Trust | COVID-19 Genomics UK (COG-UK) Consortium | CUTINO-MOGUEL, Maria-Teresa; HARRINGTON, David; OWOYEMI, Dola; KULASEGARAN-SHYLINI, Raghavendran; BROAD, Claire; KELE, Beatrix |
| EPI_ISL_920052, EPI_ISL_920053, EPI_ISL_920054, EPI_ISL_920055, EPI_ISL_920056, EPI_ISL_920057, EPI_ISL_920058, EPI_ISL_920059, EPI_ISL_920060, EPI_ISL_920079, EPI_ISL_920080 | | | |
| see above | University College London, Great Ormond Street Hospital for Children NHS Foundation Trust, Imperial College Healthcare NHS Trust | COVID-19 Genomics UK (COG-UK) Consortium | Sergi Castellano, Rachel Williams, Mark Kristiansen, Paola Resende Silva, Sunando Roy, Tony Brooks, Helena Tutill, Paola Niola, Patricia Dyal, Charlotte Williams, Leysa Forrest, Yasmin Panchbhaya, Jacqueline Findlay, Samuel Weeks, Julianne Brown, Kathryn Harris, Paul Randell, James Price, Alison Holmes, Judith Breuer |
| EPI_ISL_920195, EPI_ISL_920214, EPI_ISL_920218, EPI_ISL_920231, EPI_ISL_920234, EPI_ISL_920235, EPI_ISL_920236, EPI_ISL_920239, EPI_ISL_920244, EPI_ISL_920247, EPI_ISL_920252, EPI_ISL_920255, EPI_ISL_920261, EPI_ISL_920262, EPI_ISL_920266, EPI_ISL_920271, EPI_ISL_920272, EPI_ISL_920276, EPI_ISL_920283, EPI_ISL_920284, EPI_ISL_920287, EPI_ISL_920289, EPI_ISL_920293, EPI_ISL_920300, EPI_ISL_920307, EPI_ISL_920308, EPI_ISL_920312, EPI_ISL_920313, EPI_ISL_920314, EPI_ISL_920316, EPI_ISL_920317, EPI_ISL_920323, EPI_ISL_920324, EPI_ISL_920325, EPI_ISL_920326, EPI_ISL_920332, EPI_ISL_920333, EPI_ISL_920334, EPI_ISL_920335, EPI_ISL_920336, EPI_ISL_920340, EPI_ISL_920341, EPI_ISL_920342, EPI_ISL_920343, EPI_ISL_920347, EPI_ISL_920348, EPI_ISL_920349, EPI_ISL_920350, EPI_ISL_920351, EPI_ISL_920352, EPI_ISL_920358, EPI_ISL_920359, EPI_ISL_920361, EPI_ISL_920366, EPI_ISL_920367, EPI_ISL_920368, EPI_ISL_920369, EPI_ISL_920374, EPI_ISL_920375, EPI_ISL_920400, EPI_ISL_920409, EPI_ISL_920456, EPI_ISL_920464, EPI_ISL_920481, EPI_ISL_920489, EPI_ISL_920490, EPI_ISL_920497, EPI_ISL_920498, EPI_ISL_920507, EPI_ISL_920514, EPI_ISL_920670, EPI_ISL_920683, EPI_ISL_920684, EPI_ISL_920691, EPI_ISL_920700, EPI_ISL_920705, EPI_ISL_920714, EPI_ISL_920738, EPI_ISL_920747, EPI_ISL_920748, EPI_ISL_920755, EPI_ISL_920770, EPI_ISL_920775 | | | |
| see above | University College London Hospital | COVID-19 Genomics UK (COG-UK) Consortium | Judith Heaney, Matthew Byott, Catherine Houlihan, Dan Frampton, Stuart Kirk, Moira Spyer and Eleni Nastouli |

| | | | |
|---|---|---|---|
| EPI_ISL_920782, EPI_ISL_920784, EPI_ISL_920785, EPI_ISL_920786, EPI_ISL_920787, EPI_ISL_920792, EPI_ISL_920793, EPI_ISL_920796 | University College London, Great Ormond Street Hospital for Children NHS Foundation Trust, Imperial College Healthcare NHS Trust | COVID-19 Genomics UK (COG-UK) Consortium | Sergi Castellano, Rachel Williams, Mark Kristiansen, Paola Resende Silva, Sunando Roy, Tony Brooks, Helena Tutill, Paola Niola, Patricia Dyal, Charlotte Williams, Leysa Forrest, Yasmin Panchbhaya, Jacqueline Findlay, Samuel Weeks, Julianne Brown, Kathryn Harris, Paul Randell, James Price, Alison Holmes, Judith Breuer |
| EPI_ISL_921563, EPI_ISL_921564, EPI_ISL_921578, EPI_ISL_921579, EPI_ISL_921580, EPI_ISL_921582, EPI_ISL_921583, EPI_ISL_921584, EPI_ISL_921585, EPI_ISL_921586, EPI_ISL_921587, EPI_ISL_921589, EPI_ISL_921591, EPI_ISL_921596, EPI_ISL_921597, EPI_ISL_921598, EPI_ISL_921599, EPI_ISL_921600, EPI_ISL_921602, EPI_ISL_921604, EPI_ISL_921610 | | | |
| see above | Northumbria University / South Tees Hospitals NHS Foundation Trust / North Cumbria Integrated Care NHS Foundation Trust / North Tees and Hartlepool NHS Foundation Trust / Newcastle Hospitals NHS Foundation Trust | COVID-19 Genomics UK (COG-UK) Consortium | Darren L Smith,Andrew Nelson,Matthew Bashton,Greg R Young,Joshua Loh,John Allan,Mohammad A Tariq,Giles S Holt,Gary Black,Wen C Yew,Lynn Dover,Paul Baker,Steve Liggett,Sarah Essex,Jane Greenaway,Debra Padgett,Clive Graham,Garren Scott,Edward Barton,Emma Swindells,Brendan Payne,Jennifer Collins,Yusri Taha,Gary Eltringham |
| EPI_ISL_922135, EPI_ISL_922136, EPI_ISL_922137, EPI_ISL_922138, EPI_ISL_922139, EPI_ISL_922140, EPI_ISL_922141, EPI_ISL_922142, EPI_ISL_922143, EPI_ISL_922144, EPI_ISL_922145, EPI_ISL_922146, EPI_ISL_922147, EPI_ISL_922148, EPI_ISL_922149, EPI_ISL_922150, EPI_ISL_922151, EPI_ISL_922152, EPI_ISL_922153, EPI_ISL_922154, EPI_ISL_922155, EPI_ISL_922156, EPI_ISL_922157, EPI_ISL_922158, EPI_ISL_922159, EPI_ISL_922160, EPI_ISL_922161, EPI_ISL_922162, EPI_ISL_922163 | | | |
| see above | Lincolnshire Hospitals and DeepSeq Nottingham | COVID-19 Genomics UK (COG-UK) Consortium | Nichola Duckworth, Tim Sloan, Sarah Walsh, Jonathan Ball, Patrick McClure, Joeseph Chappell, Nadine Holmes, Matthew Carlisle, Christopher Moore, Fei Sang, Johnny Debebe, Victoria Wright, Matthew Loose |
| EPI_ISL_922353, EPI_ISL_922356, EPI_ISL_922357, EPI_ISL_922358, EPI_ISL_922359, EPI_ISL_922367, EPI_ISL_922368, EPI_ISL_922369, EPI_ISL_922370, EPI_ISL_922371, EPI_ISL_922372, EPI_ISL_922373, EPI_ISL_922436, EPI_ISL_922437, EPI_ISL_922439, EPI_ISL_922443, EPI_ISL_922483, EPI_ISL_922485, EPI_ISL_922488, EPI_ISL_922491, EPI_ISL_922494, EPI_ISL_922495, EPI_ISL_922497, EPI_ISL_922498, EPI_ISL_922499, EPI_ISL_922500, EPI_ISL_922502, EPI_ISL_922504, EPI_ISL_922505, EPI_ISL_922507, EPI_ISL_922510, EPI_ISL_922511, EPI_ISL_922512, EPI_ISL_922513, EPI_ISL_922514, EPI_ISL_922515, EPI_ISL_922516, EPI_ISL_922518, EPI_ISL_922520, EPI_ISL_922521, EPI_ISL_922522, EPI_ISL_922530, EPI_ISL_922536, EPI_ISL_922537, EPI_ISL_922538, EPI_ISL_922542, EPI_ISL_922546, EPI_ISL_922552, EPI_ISL_922555, EPI_ISL_922556, EPI_ISL_922559, EPI_ISL_922560, EPI_ISL_922561, EPI_ISL_922564, EPI_ISL_922575, EPI_ISL_922576, EPI_ISL_922577, EPI_ISL_922585, EPI_ISL_922586, EPI_ISL_922587, EPI_ISL_922600, EPI_ISL_922602, EPI_ISL_922603, EPI_ISL_922604, EPI_ISL_922605, EPI_ISL_922606, EPI_ISL_922607, EPI_ISL_922608, EPI_ISL_922609, EPI_ISL_922610, EPI_ISL_922611, EPI_ISL_922616, EPI_ISL_922618, EPI_ISL_922626, EPI_ISL_922627, EPI_ISL_922628, EPI_ISL_922629, EPI_ISL_922656, EPI_ISL_922692, EPI_ISL_922697, EPI_ISL_922699, EPI_ISL_922700, EPI_ISL_922703, EPI_ISL_922704, EPI_ISL_922708, EPI_ISL_922709, EPI_ISL_922712, EPI_ISL_922713, EPI_ISL_922714, EPI_ISL_922716, EPI_ISL_922717, EPI_ISL_922719, EPI_ISL_922721, EPI_ISL_922722, EPI_ISL_922723, EPI_ISL_922724, EPI_ISL_922726, EPI_ISL_922727, EPI_ISL_922728, EPI_ISL_922731, EPI_ISL_922732, EPI_ISL_922733, EPI_ISL_922734, EPI_ISL_922735, EPI_ISL_922737, EPI_ISL_922739, EPI_ISL_922740, EPI_ISL_922741, EPI_ISL_922743, EPI_ISL_922744, EPI_ISL_922746, EPI_ISL_922747, EPI_ISL_922748, EPI_ISL_922749, EPI_ISL_922751, EPI_ISL_922752, EPI_ISL_922753, EPI_ISL_922754, EPI_ISL_922755, EPI_ISL_922756, EPI_ISL_922757, EPI_ISL_922758, EPI_ISL_922759, EPI_ISL_922760, EPI_ISL_922761, EPI_ISL_922762, EPI_ISL_922763, EPI_ISL_922764, EPI_ISL_922765, EPI_ISL_922766, EPI_ISL_922767, EPI_ISL_922768, EPI_ISL_922770, EPI_ISL_922771, EPI_ISL_922772, EPI_ISL_922773, EPI_ISL_922774, EPI_ISL_922775, EPI_ISL_922776, EPI_ISL_922777, EPI_ISL_922779, EPI_ISL_922781, EPI_ISL_922782, EPI_ISL_922783, EPI_ISL_922784, EPI_ISL_922785, EPI_ISL_922786, EPI_ISL_922787, EPI_ISL_922788, EPI_ISL_922789, EPI_ISL_922790, EPI_ISL_922791, EPI_ISL_922792, EPI_ISL_922793, EPI_ISL_922794, EPI_ISL_922796, EPI_ISL_922798, EPI_ISL_922800, EPI_ISL_922806, EPI_ISL_922810, EPI_ISL_922811, EPI_ISL_922812, EPI_ISL_922813, EPI_ISL_922814, EPI_ISL_922815, EPI_ISL_922816, EPI_ISL_922817, EPI_ISL_922819, EPI_ISL_922820, EPI_ISL_922821, EPI_ISL_922822, EPI_ISL_922823, EPI_ISL_922824, EPI_ISL_922875, EPI_ISL_922906, EPI_ISL_922907, EPI_ISL_922908, EPI_ISL_922909, EPI_ISL_922910, EPI_ISL_922911, EPI_ISL_922912, EPI_ISL_922914, EPI_ISL_922915, EPI_ISL_922932, EPI_ISL_922933, EPI_ISL_922934, EPI_ISL_922935, EPI_ISL_922956, EPI_ISL_922966, EPI_ISL_923016, EPI_ISL_923017, EPI_ISL_923018, EPI_ISL_923019, EPI_ISL_923020, EPI_ISL_923021, EPI_ISL_923022, EPI_ISL_923023, EPI_ISL_923025, EPI_ISL_923026, EPI_ISL_923027, EPI_ISL_923028, EPI_ISL_923045, EPI_ISL_923046, EPI_ISL_923049, EPI_ISL_923050 | | | |
| see above | Wales Specialist Virology Centre Sequencing lab: Pathogen Genomics Unit | Public Health Wales Microbiology Cardiff Wales Specialist Virology Centre | Catherine Moore, Johnathan Evans, Laura Gifford, Malorie Perry, Simon Cottrell, Angela Marchbank, Alec Birchley, Alexander Adams, Amy Gaskin, Bree Gatica-Wilcox, Jason Coombes, Joel Southgate, Lauren Gilbert, Lee Graham, Nicole Pacchiarini, Sara Kumziene-Summerhayes, Sarah Taylor, Sophie Jones, Sara Rey, Matthew Bull, Joanne Watkins, Sally Corden, Tom Connor |
| EPI_ISL_923315, EPI_ISL_923316, EPI_ISL_923318, EPI_ISL_923319, EPI_ISL_923357, EPI_ISL_923360, EPI_ISL_923362, EPI_ISL_923363, EPI_ISL_923366, EPI_ISL_923367, EPI_ISL_923368, EPI_ISL_923369, EPI_ISL_923370, EPI_ISL_923415, EPI_ISL_923417, EPI_ISL_923418, EPI_ISL_923420, EPI_ISL_923421, EPI_ISL_923591, EPI_ISL_923592, EPI_ISL_923595, EPI_ISL_923596, EPI_ISL_923597, EPI_ISL_923680, EPI_ISL_923682 | | | |
| see above | Centre for Enzyme Innovation, University of Portsmouth / Translational Research Laboratory, Portsmouth Hospitals NHS Trust | COVID-19 Genomics UK (COG-UK) Consortium | Angela Beckett,Salman Goudarzi,Christopher Fearn,Kate Cook,Katie Loveson,Sharon Glaysher,Scott Elliott,Samuel Robson |
| EPI_ISL_923686, EPI_ISL_923687, EPI_ISL_923688, EPI_ISL_923689, EPI_ISL_923691, EPI_ISL_923693, EPI_ISL_923694, EPI_ISL_923696, EPI_ISL_923697, EPI_ISL_923698, EPI_ISL_923699, EPI_ISL_923700, EPI_ISL_923701, EPI_ISL_923702, EPI_ISL_923703, EPI_ISL_923704, EPI_ISL_923705, EPI_ISL_923706, EPI_ISL_923707, EPI_ISL_923708, EPI_ISL_923709, EPI_ISL_923710, EPI_ISL_923711, EPI_ISL_923717, EPI_ISL_923719, EPI_ISL_923720, EPI_ISL_923721, EPI_ISL_923722, EPI_ISL_923723, EPI_ISL_923724, EPI_ISL_923725, EPI_ISL_923727, EPI_ISL_923728, EPI_ISL_923729, EPI_ISL_923730, EPI_ISL_923731, EPI_ISL_923732, EPI_ISL_923733, EPI_ISL_923734, EPI_ISL_923735, EPI_ISL_923736, EPI_ISL_923737, EPI_ISL_923739, EPI_ISL_923740, EPI_ISL_923741, EPI_ISL_923743, EPI_ISL_923744, EPI_ISL_923745, EPI_ISL_923746, EPI_ISL_923747, EPI_ISL_923749, EPI_ISL_923750, EPI_ISL_923751, EPI_ISL_923753, EPI_ISL_923754, EPI_ISL_923755, EPI_ISL_923756, EPI_ISL_923757, EPI_ISL_923759, EPI_ISL_923761, EPI_ISL_923762, EPI_ISL_923763, EPI_ISL_923764, EPI_ISL_923765, EPI_ISL_923766, EPI_ISL_923769, EPI_ISL_923770, EPI_ISL_923771, EPI_ISL_923772, EPI_ISL_923773, EPI_ISL_923774, EPI_ISL_923775, EPI_ISL_923779, EPI_ISL_923782, EPI_ISL_923783, EPI_ISL_923784, EPI_ISL_923785, EPI_ISL_923786, EPI_ISL_923787, EPI_ISL_923788, EPI_ISL_923789, EPI_ISL_923790, EPI_ISL_923791, EPI_ISL_923792, EPI_ISL_923793, EPI_ISL_923794, EPI_ISL_923795, EPI_ISL_923796, EPI_ISL_923797, EPI_ISL_923799, EPI_ISL_923800, EPI_ISL_923802, EPI_ISL_923803, EPI_ISL_923805, EPI_ISL_923807, EPI_ISL_923808, EPI_ISL_923811, EPI_ISL_923812, EPI_ISL_923813, EPI_ISL_923814, EPI_ISL_923815, EPI_ISL_923816, EPI_ISL_923818, EPI_ISL_923820, EPI_ISL_923821, EPI_ISL_923822, EPI_ISL_923824, EPI_ISL_923825, EPI_ISL_923826, EPI_ISL_923828, EPI_ISL_923829, EPI_ISL_923830, EPI_ISL_923831, EPI_ISL_923832, EPI_ISL_923833, EPI_ISL_923834, EPI_ISL_923835, EPI_ISL_923836, EPI_ISL_923837, EPI_ISL_923841, EPI_ISL_923844, EPI_ISL_923845, EPI_ISL_923847, EPI_ISL_923848, EPI_ISL_923849, EPI_ISL_923850, EPI_ISL_923851, EPI_ISL_923853, EPI_ISL_923854, EPI_ISL_923856, EPI_ISL_923857, EPI_ISL_923858, EPI_ISL_923859, EPI_ISL_923860, EPI_ISL_923861, EPI_ISL_923862, EPI_ISL_923863, EPI_ISL_923865, EPI_ISL_923866, EPI_ISL_923867, EPI_ISL_923868, EPI_ISL_923869, EPI_ISL_923870, EPI_ISL_923871, EPI_ISL_923872, EPI_ISL_923873, EPI_ISL_923874, EPI_ISL_923876, EPI_ISL_923878, EPI_ISL_923879, EPI_ISL_923880, EPI_ISL_923881, EPI_ISL_923883, EPI_ISL_923884, EPI_ISL_923885, EPI_ISL_923886, EPI_ISL_923887, EPI_ISL_923888, EPI_ISL_923890, EPI_ISL_923891, EPI_ISL_923892, EPI_ISL_923893, EPI_ISL_923894, EPI_ISL_923895, EPI_ISL_923896, EPI_ISL_923897, EPI_ISL_923898, EPI_ISL_923900, EPI_ISL_923901, EPI_ISL_923904, EPI_ISL_923905, EPI_ISL_923906, EPI_ISL_923907, EPI_ISL_923908, EPI_ISL_923909, EPI_ISL_923910, EPI_ISL_923911, EPI_ISL_923912, EPI_ISL_923913, EPI_ISL_923914, EPI_ISL_923915, EPI_ISL_923917, EPI_ISL_923919, EPI_ISL_923920, EPI_ISL_923921, EPI_ISL_923922, EPI_ISL_923923, EPI_ISL_923926, EPI_ISL_923927, EPI_ISL_923928, EPI_ISL_923929, EPI_ISL_923930, EPI_ISL_923931, EPI_ISL_923932, EPI_ISL_923933, EPI_ISL_923934, EPI_ISL_923935, EPI_ISL_923936, EPI_ISL_923937, EPI_ISL_923938, EPI_ISL_923940, EPI_ISL_923941, EPI_ISL_923942, EPI_ISL_923943, EPI_ISL_923944, EPI_ISL_923945, EPI_ISL_923946, EPI_ISL_923947, EPI_ISL_923948, EPI_ISL_923949, EPI_ISL_923954, EPI_ISL_923955, EPI_ISL_923956, EPI_ISL_923957, EPI_ISL_923958, EPI_ISL_923961, EPI_ISL_923964, EPI_ISL_923965, EPI_ISL_923966, EPI_ISL_923967, EPI_ISL_923968, EPI_ISL_923969, EPI_ISL_923970, EPI_ISL_923971, EPI_ISL_923972, EPI_ISL_923973, EPI_ISL_923974, EPI_ISL_923975, EPI_ISL_923976, EPI_ISL_923977, EPI_ISL_923978, EPI_ISL_923979, EPI_ISL_923980, EPI_ISL_923981, EPI_ISL_923984, EPI_ISL_923985, EPI_ISL_923986, EPI_ISL_923989, EPI_ISL_923991, EPI_ISL_923992, EPI_ISL_923993, EPI_ISL_923994, EPI_ISL_923997, EPI_ISL_923999, EPI_ISL_924002, EPI_ISL_924003, EPI_ISL_924004, EPI_ISL_924005, EPI_ISL_924006, EPI_ISL_924008, EPI_ISL_924011, EPI_ISL_924012, EPI_ISL_924013, EPI_ISL_924015, EPI_ISL_924016, EPI_ISL_924017, EPI_ISL_924018, EPI_ISL_924019, EPI_ISL_924020, EPI_ISL_924021, EPI_ISL_924022, EPI_ISL_924023, EPI_ISL_924024, EPI_ISL_924025, EPI_ISL_924027, EPI_ISL_924028, EPI_ISL_924029, EPI_ISL_924030, EPI_ISL_924031, EPI_ISL_924032, EPI_ISL_924033, EPI_ISL_924034, EPI_ISL_924035, EPI_ISL_924036, EPI_ISL_924039, EPI_ISL_924042, EPI_ISL_924044, EPI_ISL_924045, EPI_ISL_924046, EPI_ISL_924049, EPI_ISL_924050, EPI_ISL_924052, EPI_ISL_924054, EPI_ISL_924055, EPI_ISL_924056, EPI_ISL_924057, EPI_ISL_924058, EPI_ISL_924059, EPI_ISL_924060, EPI_ISL_924062, EPI_ISL_924063, EPI_ISL_924064, EPI_ISL_924065, EPI_ISL_924066, EPI_ISL_924068, EPI_ISL_924069, EPI_ISL_924071, EPI_ISL_924072, EPI_ISL_924073, EPI_ISL_924074, EPI_ISL_924075 | | | |
| see above | Department of Pathology, University of Cambridge | COVID-19 Genomics UK (COG-UK) Consortium | Aminu S. Jahun, Yasmin Chaudhry, Iliana Georgana, Myra Hosmillo, Rhys Izu, Martin D. Curran, Surendra Parmar, Ian Goodfellow |
| EPI_ISL_924096, EPI_ISL_924121, EPI_ISL_924199, EPI_ISL_924218, EPI_ISL_924295, EPI_ISL_924319, EPI_ISL_924324, EPI_ISL_924354, EPI_ISL_924408 | Virology Department, Sheffield Teaching Hospitals NHS Foundation Trust/Department of Infection, Immunity and Cardiovascular Disease, The Medical School, University of Sheffield | COVID-19 Genomics UK (COG-UK) Consortium | Thushan de Silva, Matthew Parker, Nikki Smith, Adri Angyal, Rebecca Brown, Luke Green, Rachel Tucker, Paul Parsons, Danielle Groves, Katie Johnson, Laura Carrilero, Alex Keeley, Dave Partridge, Matthew Wyles, Benjamin Lindsey, Mehmet Yavuz, Mohammad Raza, Cariad Evans |
| EPI_ISL_924858, EPI_ISL_924869, EPI_ISL_924873, EPI_ISL_924882, EPI_ISL_924884 | Bioinformatics and Biostatistics Lab, Advanced Sequencing Facility | COVID-19 Genomics UK (COG-UK) Consortium | Aengus Stewart,Jerome Nicod,Chelsea Sawyer,Laura Cubitt,Harshil Patel,Margaret Crawford |
| EPI_ISL_925208 | Baylor Scott & White-Temple | Baylor Scott & White-Temple | Ari Rao, Linden Morales, Kimberly Walker, Marcus Volz, Shelby Hendrickson |
| EPI_ISL_925278 | Wyoming Public Health Laboratory | Wyoming Public Health Laboratory | Noah Hull, Taylor Fearing, Lynette Gumbleton, Channing Weber, Ashley Norberg, Bailey Bowcutt, and Wanda Manley |
| EPI_ISL_930588, EPI_ISL_930597 | Synlab | GIGA Medical Genomics | Keith Durkin, Maria Artesi, Sébastien Bontems, Raphaël Boreux, Bouchra Boujemla, Cécile Meex, Pierrette Melin, Marie-Pierre Hayette, Vincent Bours |
| EPI_ISL_930792 | Monterey County Public Health Laboratory | Monterey County Public Health Laboratory | C. Anaya et al., |
| EPI_ISL_930794, EPI_ISL_930796 | Monterey County Public Health Laboratory | Monterey County Public Health Laboratory | C. Anaya et al. |
| EPI_ISL_930797 | Monterey County Public Health Laboratory | Monterey County Public Health Laboratory | C. Anaya et al., |
| EPI_ISL_930800 | Monterey County Public Health Laboratory | Monterey County Public Health Laboratory | C. Anaya et al. |
| EPI_ISL_930804 | Monterey County Public Health Laboratory | Monterey County Public Health Laboratory | C. Anaya et al., |
| EPI_ISL_930805 | Monterey County Public Health Laboratory | Monterey County Public Health Laboratory | C. Anaya et al. |
| EPI_ISL_930830 | Monterey County Public Health Laboratory | Monterey County Public Health Laboratory | C. Anaya et al., |
| EPI_ISL_931451, EPI_ISL_931452, EPI_ISL_931453, EPI_ISL_931454 | Maryland Public Health Laboratory (MD PHL) | Maryland Public Health Laboratory (MD PHL) | Maryland Department of Health Laboratories Administration |
| EPI_ISL_933723 | Servicio de Microbiología Hospital Ramón y Cajal | Servicio de Microbiología Hospital Ramón y Cajal | José M Gonzalez-Alba, Concepción Rodríguez, Melanie Abreu, Laura Martínez, Val F Lanza, Luz Leticia Olavarrieta, Rafael Cantón, JC Galán |
| EPI_ISL_933724 | Botswana Harvard HIV Reference Laboratory | Botswana Harvard HIV Reference Laboratory | Sikhulile Moyo, Dorcas Maruapula, Wonderful T. Choga, Botshelo Radibe, Boitumelo Zuze, David Lawrence, Roger Shapiro, Shahin Lockman, Mosepele Mosepele, Joseph Makhema, Simani Gaseitsiwe |

| | | | |
|---|---|---|---|
| EPI_ISL_933725, EPI_ISL_933727, EPI_ISL_933728, EPI_ISL_933729 | Servicio de Microbiología Hospital Ramón y Cajal | Servicio de Microbiología Hospital Ramón y Cajal | José M Gonzalez-Alba, Concepción Rodríguez, Melanie Abreu, Laura Martínez, Val F Lanza, Luz Leticia Olavarrieta, Rafael Cantón, JC Galán |
| EPI_ISL_933739, EPI_ISL_933741, EPI_ISL_933742, EPI_ISL_933743, EPI_ISL_933744, EPI_ISL_933745, EPI_ISL_933746, EPI_ISL_933747, EPI_ISL_933748, EPI_ISL_933749, EPI_ISL_933750, EPI_ISL_933751, EPI_ISL_933752, EPI_ISL_933753, EPI_ISL_933754, EPI_ISL_933755, EPI_ISL_933756, EPI_ISL_933757, EPI_ISL_933758, EPI_ISL_933759, EPI_ISL_933764, EPI_ISL_933765 | | | |
| see above | DPHL | Delaware Public Health Lab | Gregory Hovan |
| EPI_ISL_933771, EPI_ISL_933773, EPI_ISL_933774 | Servicio de Microbiología Hospital Ramón y Cajal | Servicio de Microbiología Hospital Ramón y Cajal | José M Gonzalez-Alba, Concepción Rodríguez, Melanie Abreu, Laura Martínez, Val F Lanza, Luz Leticia Olavarrieta, Rafael Cantón, JC Galán |
| EPI_ISL_934280, EPI_ISL_934281, EPI_ISL_934282, EPI_ISL_934283, EPI_ISL_934284, EPI_ISL_934285, EPI_ISL_934286, EPI_ISL_934287, EPI_ISL_934288, EPI_ISL_934289, EPI_ISL_934290, EPI_ISL_934291, EPI_ISL_934292, EPI_ISL_934293, EPI_ISL_934294, EPI_ISL_934295, EPI_ISL_934296, EPI_ISL_934297, EPI_ISL_934298, EPI_ISL_934299, EPI_ISL_934300, EPI_ISL_934301, EPI_ISL_934302, EPI_ISL_934303, EPI_ISL_934304, EPI_ISL_934305, EPI_ISL_934306, EPI_ISL_934307 | | | |
| see above | Vilnius university hospital Santaros Klinikos, Center of Laboratory Medicine | Vilnius university hospital Santaros Klinikos, Center of Laboratory Medicine | Ingrida Olendraite, Daniel Naumovas, Rimvydas Norvilas, Dovile Ezerskyte, Justinas Slikas, Gytis Dudas |
| EPI_ISL_934381 | Klinisk mikrobiologi | The Public Health Agency of Sweden | Anna-Malin Linde, Maria Lind Karlberg, Carlo Berg, Oskar Karlsson Lindsjo, Sofia Stamouli, Reza Advani, Mattias Haukland, Petra Holmstrom, Noura Walai, Petra Edquist, Mia Brytting, Anna Risberg, Karin Tegmark-Wisell |
| EPI_ISL_934397, EPI_ISL_934398, EPI_ISL_934400 | Synlab Medilab, Mikrobiologi | The Public Health Agency of Sweden | Anna-Malin Linde, Maria Lind Karlberg, Carlo Berg, Oskar Karlsson Lindsjo, Sofia Stamouli, Reza Advani, Mattias Haukland, Petra Holmstrom, Noura Walai, Petra Edquist, Mia Brytting, Anna Risberg, Karin Tegmark-Wisell |
| EPI_ISL_934406 | The Public Health Agency of Sweden | The Public Health Agency of Sweden | Anna-Malin Linde, Maria Lind Karlberg, Carlo Berg, Oskar Karlsson Lindsjo, Sofia Stamouli, Reza Advani, Mattias Haukland, Petra Holmstrom, Noura Walai, Petra Edquist, Mia Brytting, Anna Risberg, Karin Tegmark-Wisell |
| EPI_ISL_934655, EPI_ISL_934656, EPI_ISL_934657, EPI_ISL_934658, EPI_ISL_934659, EPI_ISL_934660, EPI_ISL_934661, EPI_ISL_934662, EPI_ISL_934663, EPI_ISL_934664, EPI_ISL_934665, EPI_ISL_934666, EPI_ISL_934667, EPI_ISL_934668, EPI_ISL_934669, EPI_ISL_934670, EPI_ISL_934671, EPI_ISL_934672, EPI_ISL_934673, EPI_ISL_934674, EPI_ISL_934675, EPI_ISL_934676, EPI_ISL_934677, EPI_ISL_934678, EPI_ISL_934679, EPI_ISL_934680, EPI_ISL_934681 | | | |
| see above | Department of Laboratory Medicine, Division of Clinical Virology, University of Medicine, Vienna | Bergthaler laboratory, CeMM Research Center for Molecular Medicine of the Austrian Academy of Sciences | Lukas Endler, Anna Schedl, Thomas Penz, Benedikt Agerer, Maelle Le Moing, Michael Schuster, Bekir Erguner, Jan Laine, Martin Senekowitsch, Christoph Bock, Andreas Bergthaler |
| EPI_ISL_934974 | Novabio | CNR Virus des Infections Respiratoires - France SUD | Antonin Bal, Gregory Destras, Gwendolyne Burfin, Hadrien Règue, Quentin Semanas, Martine Valette, Bruno Lina, Laurence Josset |
| EPI_ISL_934978, EPI_ISL_934981, EPI_ISL_934982, EPI_ISL_934983, EPI_ISL_934991, EPI_ISL_935011 | ADMED Microbiologie | Genomics and Transcriptomics, Philip Morris International | Reto Lienhard, Marie-Lise Tritten, Emmanuel Guedj, Nicolas Sierro, Rémi Dulize, David Bornand, Mehdi Auberson, Maxime Berthouzoz, Nikolai Ivanov, Manuel Peitsch |
| EPI_ISL_935040 | Botswana Harvard HIV Reference Laboratory | Botswana Harvard HIV Reference Laboratory | Sikhulile Moyo, Wonderful T. Choga, Dorcas Maruapula, Botshelo Radibe, Boitumelo Zuze, David Lawrence, Roger Shapiro, Shahin Lockman, Mosepele Mosepele, Joseph, Makhema, Simani Gaseitsiwe |
| EPI_ISL_935042 | Botswana Harvard HIV Reference Laboratory | Botswana Harvard HIV Reference Laboratory | Sikhulile Moyo, Dorcas Maruapula, Wonderful Choga, Botshelo Radibe, Boitumelo Zuze, David Lawrence, Roger Shapiro, Shahin Lockman, Mosepele Mosepele, Joseph Makhema, Simani Gaseitsiwe |
| EPI_ISL_935045 | Botswana Harvard HIV Reference Laboratory | Botswana Harvard HIV Reference Laboratory | Sikhulile Moyo, Dorcas Maruapula, Wonderful T. Choga, Botshelo Radibe, Boitumelo Zuze, David Lawrence, Roger Shapiro, Shahin Lockman, Mosepele Mosepele, Joseph Makhema, Simani Gaseitsiwe |
| EPI_ISL_935196, EPI_ISL_935197, EPI_ISL_935200, EPI_ISL_935202, EPI_ISL_935203, EPI_ISL_935204, EPI_ISL_935205 | KU Leuven, Rega Institute, Clinical and Epidemiological Virology | KU Leuven, Rega Institute, Clinical and Epidemiological Virology | Tony Wawina-Bokalanga, Bert Vanmechelen, Joan Marti-Carerras, Piet Maes |
| EPI_ISL_935538 | Labo Analyses Med | National Reference Center for Viruses of Respiratory Infections, Institut Pasteur, Paris | Marion Barbet, Sylvie Behillil, Méline Bizard, Angela Brisebarre, Camille Capel, Etienne Simon-Lorière, Vincent Enouf, Maud Vanpeene, Sylvie van der Werf,Bonnaudet GéRaldine |
| EPI_ISL_935539 | Labo Analyses Med | National Reference Center for Viruses of Respiratory Infections, Institut Pasteur, Paris | Marion Barbet, Sylvie Behillil, Méline Bizard, Angela Brisebarre, Camille Capel, Etienne Simon-Lorière, Vincent Enouf, Maud Vanpeene, Sylvie van der Werf,Tomasi Florent |
| EPI_ISL_935550 | Labo Analyses Med | National Reference Center for Viruses of Respiratory Infections, Institut Pasteur, Paris | Marion Barbet, Sylvie Behillil, Méline Bizard, Angela Brisebarre, Camille Capel, Etienne Simon-Lorière, Vincent Enouf, Maud Vanpeene, Sylvie van der Werf,Lefaure Brieuc |
| EPI_ISL_935556, EPI_ISL_935557 | Hopital | National Reference Center for Viruses of Respiratory Infections, Institut Pasteur, Paris | Marion Barbet, Sylvie Behillil, Méline Bizard, Angela Brisebarre, Camille Capel, Etienne Simon-Lorière, Vincent Enouf, Maud Vanpeene, Sylvie van der Werf,Goudeau Alain |
| EPI_ISL_935564 | Labo Analyses Med | National Reference Center for Viruses of Respiratory Infections, Institut Pasteur, Paris | Marion Barbet, Sylvie Behillil, Méline Bizard, Angela Brisebarre, Camille Capel, Etienne Simon-Lorière, Vincent Enouf, Maud Vanpeene, Sylvie van der Werf |
| EPI_ISL_935565, EPI_ISL_935566 | Hopital | National Reference Center for Viruses of Respiratory Infections, Institut Pasteur, Paris | Marion Barbet, Sylvie Behillil, Méline Bizard, Angela Brisebarre, Camille Capel, Etienne Simon-Lorière, Vincent Enouf, Maud Vanpeene, Sylvie van der Werf,Bret Laurent |
| EPI_ISL_935570, EPI_ISL_935571, EPI_ISL_935572, EPI_ISL_935573, EPI_ISL_935574, EPI_ISL_935575 | Hopital | National Reference Center for Viruses of Respiratory Infections, Institut Pasteur, Paris | Marion Barbet, Sylvie Behillil, Méline Bizard, Angela Brisebarre, Camille Capel, Etienne Simon-Lorière, Vincent Enouf, Maud Vanpeene, Sylvie van der Werf,Castelain Sandrine |
| EPI_ISL_935582 | Hopital | National Reference Center for Viruses of Respiratory Infections, Institut Pasteur, Paris | Marion Barbet, Sylvie Behillil, Méline Bizard, Angela Brisebarre, Camille Capel, Etienne Simon-Lorière, Vincent Enouf, Maud Vanpeene, Sylvie van der Werf,Blazejewski C. |
| EPI_ISL_935583, EPI_ISL_935584, EPI_ISL_935585 | Hopital | National Reference Center for Viruses of Respiratory Infections, Institut Pasteur, Paris | Marion Barbet, Sylvie Behillil, Méline Bizard, Angela Brisebarre, Camille Capel, Etienne Simon-Lorière, Vincent Enouf, Maud Vanpeene, Sylvie van der Werf,Fourgeaud Jacques |
| EPI_ISL_935587, EPI_ISL_935588, EPI_ISL_935589, EPI_ISL_935595, EPI_ISL_935596, EPI_ISL_935597, EPI_ISL_935598, EPI_ISL_935600, EPI_ISL_935601, EPI_ISL_935621, EPI_ISL_935622, EPI_ISL_935623, EPI_ISL_935624, EPI_ISL_935625, EPI_ISL_935626, EPI_ISL_935627, EPI_ISL_935628 | | | |
| see above | Labo Analyses med | National Reference Center for Viruses of Respiratory Infections, Institut Pasteur, Paris | Marion Barbet, Sylvie Behillil, Méline Bizard, Angela Brisebarre, Camille Capel, Etienne Simon-Lorière, Vincent Enouf, Maud Vanpeene, Sylvie van der Werf,Amzalag Jonas |
| EPI_ISL_936242, EPI_ISL_936243, EPI_ISL_936244, EPI_ISL_936245, EPI_ISL_936246, EPI_ISL_936247 | Wadsworth Center, New York State Department of Health | Wadsworth Center, New York State Department of Health | Kirsten St. George, Daryl M. Lamson, Alexis Russel, Matthew Shudt, Melissa A Leisner, Jonathan Plitnick, Navjot Singh, John Kelly, Erasmus Schneider, Erica Lasek-Nesselquist |
| EPI_ISL_936248, EPI_ISL_936250, EPI_ISL_936251, EPI_ISL_936253, EPI_ISL_936259, EPI_ISL_936265, EPI_ISL_936271, EPI_ISL_936275, EPI_ISL_936281, EPI_ISL_936282, EPI_ISL_936286, EPI_ISL_936287, EPI_ISL_936290, EPI_ISL_936297, EPI_ISL_936299 | | | |
| see above | MONTEFIORE MEDICAL CENTER LABORATORIES | Wadsworth Center, New York State Department of Health | Kirsten St. George, Daryl M. Lamson, Alexis Russel, Matthew Shudt, Melissa A Leisner, Jonathan Plitnick, Navjot Singh, John Kelly, Erasmus Schneider, Erica Lasek-Nesselquist |
| EPI_ISL_936467, EPI_ISL_936472 | DPH, Massachusetts State Public Health Lab | DPH, Massachusetts State Public Health Lab | Lang,A.S., Fink,T., Gallagher,G.R., Smole,S.C. |
| EPI_ISL_937121 | Quest Diagnostics | Quest Diagnostics | Rosenthal,S.H., Gerasimova,A., Kagan,R.M., Anderson, B., Hua, M., Liu Y., Bernstein, L.E., Livingston, K.E., Perez, A., Shalhout, D.F., Shlyakhter, I.A., Owen, R., Tanpaiboon, P., Lacbawan, F. |
| EPI_ISL_937154, EPI_ISL_937155, EPI_ISL_937156 | DOHMH PHL | New York City Public Health Laboratory | Jade Wang, et al. |
| EPI_ISL_937186, EPI_ISL_937187, EPI_ISL_937189, EPI_ISL_937190 | OCME Office Of Chief Medical Examiner | New York City Public Health Laboratory | Jade Wang, et al. |

| | | | |
|---|---|---|---|
| EPI_ISL_937266, EPI_ISL_937269, EPI_ISL_937272 | NYC HH Elmhurst Hospital Medical Center | New York City Public Health Laboratory | Jade Wang, et al. |
| EPI_ISL_937392, EPI_ISL_937394, EPI_ISL_937395, EPI_ISL_937396, EPI_ISL_937397, EPI_ISL_937398, EPI_ISL_937399, EPI_ISL_937400, EPI_ISL_937401, EPI_ISL_937402, EPI_ISL_937403, EPI_ISL_937404, EPI_ISL_937405, EPI_ISL_937406, EPI_ISL_937407, EPI_ISL_937409, EPI_ISL_937410, EPI_ISL_937411, EPI_ISL_937412, EPI_ISL_937413 | | | |
| see above | Maine Health and Environmental Testing Laboratory (Maine HETL) | Tewhey Lab, The Jackson Laboratory | Matluk,N., Dewey,H., Iosue,F., Barter,M., Lynch,R., Munger,H. and Tewhey,R. |
| EPI_ISL_939608 | Lighthouse Lab in Alderley Park | Wellcome Sanger Institute for the COVID-19 Genomics UK (COG-UK) Consortium | Jacquelyn Wynn, Mairead Hyland, The Lighthouse Lab in Alderley Park and Alex Alderton, Roberto Amato, Sonia Goncalves, Ewan Harrison, David K. Jackson, Ian Johnston, Dominic Kwiatkowski, Cordelia Langford, John Sillitoe on behalf of the Wellcome Sanger Institute COVID-19 Surveillance Team |
| EPI_ISL_939636, EPI_ISL_939640, EPI_ISL_939641 | MEPHI, Aix Marseille University | MEPHI, Aix Marseille University | Anthony LEVASSEUR |
| EPI_ISL_940074 | University Hospitals of Geneva, Laboratory of Virology | HUG, Laboratory of Virology and the Health2030 Genome Center | Samuel Cordey, Ana Rita Goncalves, Laurent Kaiser, Lorenzo Cerutti, Henri Pegeot, Melyssa Elies, Deborah Penet, Keith Harshman, Ioannis Xenarios, Emmanouil Dermitzakis |
| EPI_ISL_940612 | Hospital de Campanha para Enfrentamento do Coronavirus - Goiania | Instituto Adolfo Lutz, Interdiciplinary Procedures Center, Strategic Laboratory | Claudio Tavares Sacchi, Claudia Regina Gonçalves, Erica Valessa Ramos Gomes, Karoline Rodrigues Campos |
| EPI_ISL_940619, EPI_ISL_940622, EPI_ISL_940624 | Hospital Sao Joaquim - Beneficiencia Portuguesa | Instituto Adolfo Lutz, Interdiciplinary Procedures Center, Strategic Laboratory | Claudio Tavares Sacchi, Claudia Regina Gonçalves, Erica Valessa Ramos Gomes, Karoline Rodrigues Campos |
| EPI_ISL_940772 | Platform BIS UZA/UAntwerpen | UAntwerp, Laboratory of Medical Microbiology, Campus Drie Eiken S6.26, Universiteitsplein 1, 2610, Wilrijk, Belgium | Basil Britto Xavier, Jasmine Coppens, Marie Le Mercier, Christine Lammens, Veerle Matheeussen, Herman Goossens |
| EPI_ISL_940890 | Vaccines and Infectious Diseases Analytics Research Unit (VIDA) | KRISP, KZN Research Innovation and Sequencing Platform | Baillie Vicky, du Plessis Jeanine, Giandhari Jennifer, Pillay Sureshnee, Naidoo Yeshnee, Tegally Houriiyah, de Oliveira Tulio, Madhi Shabir |
| EPI_ISL_941001, EPI_ISL_941002, EPI_ISL_941003 | Hopital | National Reference Center for Viruses of Respiratory Infections, Institut Pasteur, Paris | Marion Barbet, Sylvie Behillil, Méline Bizard, Angela Brisebarre, Camille Capel, Etienne Simon-Lorière, Vincent Enouf, Maud Vanpeene, Sylvie van der Werf, Cron Sophie |
| EPI_ISL_941028, EPI_ISL_941029, EPI_ISL_941030, EPI_ISL_941031, EPI_ISL_941032, EPI_ISL_941033, EPI_ISL_941034, EPI_ISL_941038, EPI_ISL_941039, EPI_ISL_941042, EPI_ISL_941043, EPI_ISL_941044, EPI_ISL_941045, EPI_ISL_941046, EPI_ISL_941047, EPI_ISL_941048, EPI_ISL_941049, EPI_ISL_941050, EPI_ISL_941051, EPI_ISL_941052, EPI_ISL_941053, EPI_ISL_941054, EPI_ISL_941055, EPI_ISL_941056, EPI_ISL_941057, EPI_ISL_941059, EPI_ISL_941060, EPI_ISL_941061, EPI_ISL_941062, EPI_ISL_941064, EPI_ISL_941066, EPI_ISL_941067, EPI_ISL_941068, EPI_ISL_941069, EPI_ISL_941070, EPI_ISL_941072, EPI_ISL_941073, EPI_ISL_941074, EPI_ISL_941075, EPI_ISL_941076, EPI_ISL_941077, EPI_ISL_941078 | | | |
| see above | Labo Analyses Med | National Reference Center for Viruses of Respiratory Infections, Institut Pasteur, Paris | Marion Barbet, Sylvie Behillil, Méline Bizard, Angela Brisebarre, Camille Capel, Etienne Simon-Lorière, Vincent Enouf, Maud Vanpeene, Sylvie van der Werf, Merah Kader |
| EPI_ISL_941212, EPI_ISL_941213, EPI_ISL_941214, EPI_ISL_941215, EPI_ISL_941216, EPI_ISL_941217, EPI_ISL_941218, EPI_ISL_941219 | Servicio de Microbiología. Hospital Clínico Universitario de Valencia | SeqCOVID-SPAIN consortium/IBV(CSIC) | David Navarro Ortega, Eliseo Albert Vicent, Ignacio Torres and SeqCOVID-SPAIN consortium |
| EPI_ISL_941294, EPI_ISL_941295 | Nigeria Centre for Disease Control (NCDC) | African Centre of Excellence for Genomics of Infectious Diseases (ACEGID), Redeemer's University | Oluniyi P.E. et al |
| EPI_ISL_941360, EPI_ISL_941361, EPI_ISL_941362, EPI_ISL_941363, EPI_ISL_941364, EPI_ISL_941365, EPI_ISL_941366, EPI_ISL_941367, EPI_ISL_941368, EPI_ISL_941392, EPI_ISL_941393, EPI_ISL_941402, EPI_ISL_941405, EPI_ISL_941406, EPI_ISL_941407, EPI_ISL_941408, EPI_ISL_941409, EPI_ISL_941410, EPI_ISL_941413, EPI_ISL_941423, EPI_ISL_941424, EPI_ISL_941425, EPI_ISL_941430, EPI_ISL_941431, EPI_ISL_941432, EPI_ISL_941486, EPI_ISL_941487, EPI_ISL_941488, EPI_ISL_941489, EPI_ISL_941490, EPI_ISL_941491, EPI_ISL_941492, EPI_ISL_941493, EPI_ISL_941494, EPI_ISL_941495, EPI_ISL_941496, EPI_ISL_941497, EPI_ISL_941498, EPI_ISL_941499, EPI_ISL_941500, EPI_ISL_941501, EPI_ISL_941502, EPI_ISL_941503, EPI_ISL_941504, EPI_ISL_941505, EPI_ISL_941506, EPI_ISL_941507, EPI_ISL_941509, EPI_ISL_941514, EPI_ISL_941516, EPI_ISL_941518, EPI_ISL_941519, EPI_ISL_941523, EPI_ISL_941524, EPI_ISL_941525, EPI_ISL_941526, EPI_ISL_941536, EPI_ISL_941553, EPI_ISL_941554, EPI_ISL_941555, EPI_ISL_941556, EPI_ISL_941557, EPI_ISL_941558, EPI_ISL_941559, EPI_ISL_941560, EPI_ISL_941561, EPI_ISL_941562, EPI_ISL_941563, EPI_ISL_941564, EPI_ISL_941565, EPI_ISL_941566, EPI_ISL_941567, EPI_ISL_941568, EPI_ISL_941569, EPI_ISL_941570, EPI_ISL_941596, EPI_ISL_941597, EPI_ISL_941598, EPI_ISL_941599, EPI_ISL_941604, EPI_ISL_941605, EPI_ISL_941606, EPI_ISL_941607, EPI_ISL_941608, EPI_ISL_941609, EPI_ISL_941635, EPI_ISL_941639, EPI_ISL_941643, EPI_ISL_941644, EPI_ISL_941645, EPI_ISL_941646 | | | |
| see above | Instituto Nacional de Saude (INSA) | Instituto Nacional de Saude (INSA) | Borges et al |
| EPI_ISL_941653, EPI_ISL_941654, EPI_ISL_941655, EPI_ISL_941656, EPI_ISL_941657, EPI_ISL_941658, EPI_ISL_941660, EPI_ISL_941661, EPI_ISL_941668, EPI_ISL_941669, EPI_ISL_941679, EPI_ISL_941684, EPI_ISL_941696, EPI_ISL_941697, EPI_ISL_941698, EPI_ISL_941699, EPI_ISL_941700, EPI_ISL_941701, EPI_ISL_941708, EPI_ISL_941720, EPI_ISL_941721, EPI_ISL_941722, EPI_ISL_941723, EPI_ISL_941724, EPI_ISL_941725, EPI_ISL_941756, EPI_ISL_941757, EPI_ISL_941758, EPI_ISL_941763, EPI_ISL_941764, EPI_ISL_941789, EPI_ISL_941790, EPI_ISL_941791, EPI_ISL_941792, EPI_ISL_941793, EPI_ISL_941794, EPI_ISL_941795, EPI_ISL_941803, EPI_ISL_941804, EPI_ISL_941805, EPI_ISL_941806, EPI_ISL_941807, EPI_ISL_941814, EPI_ISL_941815, EPI_ISL_941816, EPI_ISL_941817, EPI_ISL_941832, EPI_ISL_941833, EPI_ISL_941834, EPI_ISL_941843, EPI_ISL_941844, EPI_ISL_941847, EPI_ISL_941848, EPI_ISL_941851, EPI_ISL_941861, EPI_ISL_941872, EPI_ISL_941876, EPI_ISL_941879, EPI_ISL_941880, EPI_ISL_941885, EPI_ISL_941886, EPI_ISL_941887, EPI_ISL_941888, EPI_ISL_941889, EPI_ISL_941898, EPI_ISL_941899, EPI_ISL_941900 | | | |
| see above | Instituto Nacional de Saude (INSA) and Instituto Gulbenkian de Ciencia (IGC) | Instituto Nacional de Saude (INSA) and Instituto Gulbenkian de Ciencia (IGC) | Borges et al |
| EPI_ISL_942835, EPI_ISL_942836, EPI_ISL_942837, EPI_ISL_942838, EPI_ISL_942839, EPI_ISL_942840, EPI_ISL_942841, EPI_ISL_942842, EPI_ISL_942843, EPI_ISL_942844, EPI_ISL_942845, EPI_ISL_942846, EPI_ISL_942847, EPI_ISL_942848, EPI_ISL_942849, EPI_ISL_942850 | | | |
| see above | Gundersen Molecular Diagnostics Laboratory | Kabara Cancer Research Institute | Craig S. Richmond, Paraic A. Kenny |
| EPI_ISL_942978, EPI_ISL_942994, EPI_ISL_942995, EPI_ISL_943024, EPI_ISL_943059, EPI_ISL_943079, EPI_ISL_943080, EPI_ISL_943092, EPI_ISL_943093, EPI_ISL_943094, EPI_ISL_943095, EPI_ISL_943096, EPI_ISL_943097, EPI_ISL_943098, EPI_ISL_943103, EPI_ISL_943114, EPI_ISL_943115, EPI_ISL_943116, EPI_ISL_943124, EPI_ISL_943125, EPI_ISL_943137, EPI_ISL_943138, EPI_ISL_943139, EPI_ISL_943164, EPI_ISL_943165, EPI_ISL_943166, EPI_ISL_943167, EPI_ISL_943231, EPI_ISL_943272, EPI_ISL_943276, EPI_ISL_943281, EPI_ISL_943289, EPI_ISL_943293, EPI_ISL_943301, EPI_ISL_943304, EPI_ISL_943307, EPI_ISL_943308, EPI_ISL_943309, EPI_ISL_943310, EPI_ISL_943322, EPI_ISL_943330, EPI_ISL_943338, EPI_ISL_943342, EPI_ISL_943349, EPI_ISL_943354, EPI_ISL_943384, EPI_ISL_943426, EPI_ISL_943429, EPI_ISL_943441, EPI_ISL_943442, EPI_ISL_943482, EPI_ISL_943516, EPI_ISL_943521, EPI_ISL_943524, EPI_ISL_943527, EPI_ISL_943528 | | | |
| see above | Dutch COVID-19 response team | National Institute for Public Health and the Environment (RIVM) | Adam Meijer, Harry Vennema, Dirk Eggink, Jeroen Cremer, Sharon van den Brink, Bas van der Veer, AnneMarie van den Brandt, Florian Zwagemaker, Dennis Schmitz, Chantal Reusken, on behalf of the national COVID-19 response team |
| EPI_ISL_943550 | Baylor Scott & White-Temple | Baylor Scott & White-Temple | Ari Rao, Linden Morales, Kimberly Walker, Marcus Volz, Shelby Hendrickson |
| EPI_ISL_943813, EPI_ISL_943848, EPI_ISL_943849, EPI_ISL_943850, EPI_ISL_943851, EPI_ISL_943852, EPI_ISL_943853, EPI_ISL_943854, EPI_ISL_943855, EPI_ISL_943856, EPI_ISL_943857, EPI_ISL_943858, EPI_ISL_943859, EPI_ISL_943860, EPI_ISL_943861, EPI_ISL_943862, EPI_ISL_943863, EPI_ISL_943864, EPI_ISL_943865, EPI_ISL_943866, EPI_ISL_943867, EPI_ISL_943869, EPI_ISL_943870, EPI_ISL_943871, EPI_ISL_943872, EPI_ISL_943873, EPI_ISL_943874, EPI_ISL_943875, EPI_ISL_943876, EPI_ISL_943877, EPI_ISL_943878, EPI_ISL_943879, EPI_ISL_943880, EPI_ISL_943881, EPI_ISL_943882, EPI_ISL_943883, EPI_ISL_943884, EPI_ISL_943885, EPI_ISL_943886, EPI_ISL_943887, EPI_ISL_943888, EPI_ISL_943889, EPI_ISL_943890, EPI_ISL_943891, EPI_ISL_943892, EPI_ISL_943893, EPI_ISL_943894, EPI_ISL_943895, EPI_ISL_943896, EPI_ISL_943897, EPI_ISL_943898, EPI_ISL_943899, EPI_ISL_943900, EPI_ISL_943901, EPI_ISL_943902, EPI_ISL_943903, EPI_ISL_943904, EPI_ISL_943905, EPI_ISL_943906, EPI_ISL_943907, EPI_ISL_943908, EPI_ISL_943909, EPI_ISL_943910, EPI_ISL_943911, EPI_ISL_943912, EPI_ISL_943913, EPI_ISL_943914, EPI_ISL_943915, EPI_ISL_943916, EPI_ISL_943922, EPI_ISL_943923, EPI_ISL_943924, EPI_ISL_943929, EPI_ISL_943932, EPI_ISL_943936, EPI_ISL_943938, EPI_ISL_943939, EPI_ISL_943941, EPI_ISL_943942, EPI_ISL_943943, EPI_ISL_943944, EPI_ISL_943946, EPI_ISL_943947, EPI_ISL_943951, EPI_ISL_943952, EPI_ISL_943955, EPI_ISL_943956, EPI_ISL_943957, EPI_ISL_943958, EPI_ISL_943964, EPI_ISL_943966 | | | |
| see above | Utah Public Health Laboratory | Utah Public Health Laboratory | Erin L. Young, Kelly F. Oakeson, Tara Gallagher |
| EPI_ISL_943992 | Botswana Harvard HIV Reference Laboratory | Botswana Harvard HIV Reference Laboratory | Sikhulile Moyo, Wonderful T. Choga, Dorcas Maruapula, Botshelo Radibe, Boitumelo Zuze, David Lawrence, Roger Shapiro, Shahin Lockman, Mosepele Mosepele, Joseph, Makhema, Simani Gaseitsiwe |
| EPI_ISL_944229, EPI_ISL_944366, EPI_ISL_944380, EPI_ISL_944387, EPI_ISL_944412, EPI_ISL_944566, EPI_ISL_944567, EPI_ISL_944568, EPI_ISL_944569, EPI_ISL_944570 | Israel Central Virology laboratory | Israel National Consortium for SARS-CoV-2 sequencing | Neta Zuckerman, Efrat Dahan Bucris, Michal Mandelboim, Dana Bar-Ilan, Oran Erster, Tzvia Mann, Omer Murik, David A. Zeevi, Assaf Rokney, Joseph Jaffe, Eva Nachum, Maya Davidovich Cohen, Ephraim Fass, Gal Zizelski Valenci, Mor Rubinstein, Efrat Rorman, Israel Nissan, Efrat Glick-Saar, Omri Nayshool, Gideon Rechavi, Ella Mendelson, Orna Mor |
| EPI_ISL_944739 | unknown | Public Health Virology-Forensic and Scientific Services (PHV-FSS) | Son Nguyen et al. |
| EPI_ISL_948967, EPI_ISL_948968, EPI_ISL_948969, EPI_ISL_948970, EPI_ISL_948971, EPI_ISL_948972, EPI_ISL_948974, EPI_ISL_948976, EPI_ISL_948977, EPI_ISL_948979, EPI_ISL_948980, EPI_ISL_948982, EPI_ISL_948983, EPI_ISL_948985 | | | |
| see above | Lighthouse Lab in Glasgow | Wellcome Sanger Institute for the COVID-19 Genomics UK | Harper VanSteenhouse, Yumi Kasai, David Gray, Carol Clugston, Anna Dominiczak and Alex Alderton, Roberto Amato, Sonia Goncalves, Ewan Harrison, |

| | | | |
|---|---|---|---|
| | | (COG-UK) Consortium | David K. Jackson, Ian Johnston, Dominic Kwiatkowski, Cordelia Langford, John Sillitoe on behalf of the Wellcome Sanger Institute COVID-19 Surveillance Team |
| EPI_ISL_948986 | Jessa | Jessa | Jessa_cmdLab |
| EPI_ISL_948987, EPI_ISL_948988, EPI_ISL_948990, EPI_ISL_948992, EPI_ISL_948993, EPI_ISL_948995, EPI_ISL_948996, EPI_ISL_948998, EPI_ISL_948999, EPI_ISL_949001, EPI_ISL_949002, EPI_ISL_949004, EPI_ISL_949006, EPI_ISL_949007, EPI_ISL_949009, EPI_ISL_949010, EPI_ISL_949012, EPI_ISL_949014, EPI_ISL_949015, EPI_ISL_949017, EPI_ISL_949018, EPI_ISL_949020, EPI_ISL_949021, EPI_ISL_949023, EPI_ISL_949025, EPI_ISL_949026, EPI_ISL_949027, EPI_ISL_949028, EPI_ISL_949029, EPI_ISL_949030, EPI_ISL_949031, EPI_ISL_949032, EPI_ISL_949033, EPI_ISL_949034, EPI_ISL_949035, EPI_ISL_949036, EPI_ISL_949037, EPI_ISL_949038, EPI_ISL_949039, EPI_ISL_949040, EPI_ISL_949041, EPI_ISL_949042, EPI_ISL_949043, EPI_ISL_949044, EPI_ISL_949045, EPI_ISL_949046, EPI_ISL_949047, EPI_ISL_949048, EPI_ISL_949049, EPI_ISL_949050, EPI_ISL_949051, EPI_ISL_949052, EPI_ISL_949053, EPI_ISL_949054, EPI_ISL_949055, EPI_ISL_949056, EPI_ISL_949057, EPI_ISL_949058, EPI_ISL_949059, EPI_ISL_949060, EPI_ISL_949061, EPI_ISL_949062, EPI_ISL_949063, EPI_ISL_949064, EPI_ISL_949065, EPI_ISL_949066 | | | |
| see above | Lighthouse Lab in Glasgow | Wellcome Sanger Institute for the COVID-19 Genomics UK (COG-UK) Consortium | Harper VanSteenhouse, Yumi Kasai, David Gray, Carol Clugston, Anna Dominiczak and Alex Alderton, Roberto Amato, Sonia Goncalves, Ewan Harrison, David K. Jackson, Ian Johnston, Dominic Kwiatkowski, Cordelia Langford, John Sillitoe on behalf of the Wellcome Sanger Institute COVID-19 Surveillance Team |
| EPI_ISL_949255 | Jessa | Jessa | Jessa_cmdLab |
| EPI_ISL_949256, EPI_ISL_949258, EPI_ISL_949259, EPI_ISL_949260, EPI_ISL_949261, EPI_ISL_949262, EPI_ISL_949263, EPI_ISL_949265, EPI_ISL_949266, EPI_ISL_949267, EPI_ISL_949268, EPI_ISL_949269, EPI_ISL_949270, EPI_ISL_949272, EPI_ISL_949273, EPI_ISL_949274, EPI_ISL_949275, EPI_ISL_949276, EPI_ISL_949277, EPI_ISL_949278, EPI_ISL_949279, EPI_ISL_949280, EPI_ISL_949281, EPI_ISL_949282, EPI_ISL_949284, EPI_ISL_949285, EPI_ISL_949286, EPI_ISL_949287, EPI_ISL_949288, EPI_ISL_949289, EPI_ISL_949291, EPI_ISL_949293, EPI_ISL_949294, EPI_ISL_949295, EPI_ISL_949296, EPI_ISL_949297, EPI_ISL_949298, EPI_ISL_949299, EPI_ISL_949300, EPI_ISL_949301, EPI_ISL_949302, EPI_ISL_949303, EPI_ISL_949304, EPI_ISL_949305, EPI_ISL_949306, EPI_ISL_949307, EPI_ISL_949308, EPI_ISL_949309, EPI_ISL_949311, EPI_ISL_949312, EPI_ISL_949313, EPI_ISL_949314, EPI_ISL_949315, EPI_ISL_949316, EPI_ISL_949317, EPI_ISL_949319, EPI_ISL_949320, EPI_ISL_949321, EPI_ISL_949322, EPI_ISL_949323, EPI_ISL_949324, EPI_ISL_949325, EPI_ISL_949326 | | | |
| see above | Department of Pathology, University of Cambridge | COVID-19 Genomics UK (COG-UK) Consortium | Aminu S. Jahun, Yasmin Chaudhry, Iliana Georgana, Myra Hosmillo, Rhys Izu, Martin D. Curran, Surendra Parmar, Ian Goodfellow |
| EPI_ISL_949407, EPI_ISL_949417, EPI_ISL_949418, EPI_ISL_949419, EPI_ISL_949420, EPI_ISL_949421, EPI_ISL_949422, EPI_ISL_949423, EPI_ISL_949424, EPI_ISL_949425 | University of Birmingham | COVID-19 Genomics UK (COG-UK) Consortium | Institute of Microbiology, University of Birmingham: Claire McMurray, Joanne Stockton, Samuel Nicholls, Radoslaw Poplawski, Will Rowe, Josh Quick, Nicholas Loman. University of Birmingham Testing Laboratory: Celina M Whalley, Andrew Bosworth, Charlotte Poxon, Kasun Wanigasooriya, Oliver Pickles, Mike Kidd, Alex Richter, Andrew D Beggs PHE Heartlands Lab: Husam Osman, Andrew Bosworth. Queen Elizabeth Hospital: Anna Casey |
| EPI_ISL_949492, EPI_ISL_949494, EPI_ISL_949496, EPI_ISL_949498, EPI_ISL_949504, EPI_ISL_949506 | Department of Pathology, University of Cambridge | COVID-19 Genomics UK (COG-UK) Consortium | Aminu S. Jahun, Yasmin Chaudhry, Iliana Georgana, Myra Hosmillo, Rhys Izu, Martin D. Curran, Surendra Parmar, Ian Goodfellow |
| EPI_ISL_949637, EPI_ISL_949638, EPI_ISL_949641 | Virology Department, Royal Infirmary of Edinburgh, NHS Lothian / School of Biological Sciences, University of Edinburgh / Institute of Genetics and Molecular Medicine, University of Edinburgh | COVID-19 Genomics UK (COG-UK) Consortium | McHugh M, Dewar R, Rooke S, Gallagher M, Balcaza C, O'Toole Á, Scher E, Hill V, McCrone JT, Colquhoun R, Yu X, Jackson B, Rambaut A, Williams TC, Templeton K |
| EPI_ISL_949905, EPI_ISL_950156, EPI_ISL_950157, EPI_ISL_950159, EPI_ISL_950160, EPI_ISL_950161, EPI_ISL_950222, EPI_ISL_950223, EPI_ISL_950224, EPI_ISL_950225, EPI_ISL_950226, EPI_ISL_950227, EPI_ISL_950228 | | | |
| see above | University College London, Great Ormond Street Hospital for Children NHS Foundation Trust, Imperial College Healthcare NHS Trust | COVID-19 Genomics UK (COG-UK) Consortium | Sergi Castellano, Rachel Williams, Mark Kristiansen, Paola Resende Silva, Sunando Roy, Tony Brooks, Helena Tutill, Paola Niola, Patricia Dyal, Charlotte Williams, Leysa Forrest, Yasmin Panchbhaya, Jacqueline Findlay, Samuel Weeks, Julianne Brown, Kathryn Harris, Paul Randell, James Price, Alison Holmes, Judith Breuer |
| EPI_ISL_950715 | Lincolnshire Hospitals and DeepSeq Nottingham | COVID-19 Genomics UK (COG-UK) Consortium | Nichola Duckworth, Tim Sloan, Sarah Walsh, Jonathan Ball, Patrick McClure, Joeseph Chappell, Nadine Holmes, Matthew Carlisle, Christopher Moore, Fei Sang, Johnny Debebe, Victoria Wright, Matthew Loose |
| EPI_ISL_951359, EPI_ISL_951361, EPI_ISL_951368, EPI_ISL_951372, EPI_ISL_951373, EPI_ISL_951449, EPI_ISL_951458, EPI_ISL_951459, EPI_ISL_951461, EPI_ISL_951462, EPI_ISL_951463, EPI_ISL_951464, EPI_ISL_951469, EPI_ISL_951475, EPI_ISL_951477, EPI_ISL_951478 | | | |
| see above | Oxford Viromics, NDM, University of Oxford; Oxford University Hospitals; Basingstoke and North Hampshire Hospital | COVID-19 Genomics UK (COG-UK) Consortium | Tanya Golubchik, David Bonsall, George Macintyre, Amy Trebes, Mariateresa de Cesare, Catrin Moore, Alex Mobbs, Anita Justice, Robert Shaw, Monique Andersson, Timothy Peto, Emma Wise, Nathan Moore, Jessica Lynch, Nick Cortes, Matilde Mori, Stephen Kidd, David Buck, John Todd, Christophe Fraser |
| EPI_ISL_951819 | Originating lab: Wales Specialist Virology Centre Sequencing lab: Pathogen Genomics Unit | Public Health Wales Microbiology Cardiff Wales Specialist Virology Centre | Catherine Moore, Johnathan Evans, Laura Gifford, Malorie Perry, Simon Cottrell, Angela Marchbank, Alec Birchley, Alexander Adams, Amy Gaskin, Bree Gatica-Wilcox, Jason Coombes, Joel Southgate, Lauren Gilbert, Lee Graham, Nicole Pacchiarini, Sara Kumziene-Summerhayes, Sarah Taylor, Sophie Jones, Sara Rey, Matthew Bull, Joanne Watkins, Sally Corden, Tom Connor |
| EPI_ISL_952378, EPI_ISL_952386, EPI_ISL_952387, EPI_ISL_952389, EPI_ISL_952393, EPI_ISL_952554, EPI_ISL_952601, EPI_ISL_952602, EPI_ISL_952624, EPI_ISL_952783, EPI_ISL_952821, EPI_ISL_952833, EPI_ISL_952834, EPI_ISL_952835, EPI_ISL_952854 | | | |
| see above | Centre for Enzyme Innovation, University of Portsmouth / Translational Research Laboratory, Portsmouth Hospitals NHS Trust | COVID-19 Genomics UK (COG-UK) Consortium | Angela Beckett,Salman Goudarzi,Christopher Fearn,Kate Cook,Katie Loveson,Sharon Glaysher,Scott Elliott,Samuel Robson |
| EPI_ISL_952858, EPI_ISL_952859, EPI_ISL_952860, EPI_ISL_952862, EPI_ISL_952863, EPI_ISL_952865, EPI_ISL_952866, EPI_ISL_952867, EPI_ISL_952868, EPI_ISL_952871, EPI_ISL_952874, EPI_ISL_952876, EPI_ISL_952877, EPI_ISL_952878, EPI_ISL_952879, EPI_ISL_952880, EPI_ISL_952881, EPI_ISL_952882, EPI_ISL_952883, EPI_ISL_952884, EPI_ISL_952885, EPI_ISL_952886, EPI_ISL_952888, EPI_ISL_952889, EPI_ISL_952890, EPI_ISL_952891, EPI_ISL_952892, EPI_ISL_952893, EPI_ISL_952894, EPI_ISL_952896, EPI_ISL_952897, EPI_ISL_952900, EPI_ISL_952901, EPI_ISL_952902, EPI_ISL_952903, EPI_ISL_952904, EPI_ISL_952905, EPI_ISL_952906, EPI_ISL_952907, EPI_ISL_952910, EPI_ISL_952913, EPI_ISL_952914, EPI_ISL_952916, EPI_ISL_952917, EPI_ISL_952919, EPI_ISL_952920, EPI_ISL_952923, EPI_ISL_952924, EPI_ISL_952925, EPI_ISL_952926, EPI_ISL_952929, EPI_ISL_952932, EPI_ISL_952933, EPI_ISL_952934, EPI_ISL_952935, EPI_ISL_952937, EPI_ISL_952938, EPI_ISL_952939, EPI_ISL_952940, EPI_ISL_952941 | | | |
| see above | Department of Pathology, University of Cambridge | COVID-19 Genomics UK (COG-UK) Consortium | Aminu S. Jahun, Yasmin Chaudhry, Iliana Georgana, Myra Hosmillo, Rhys Izu, Martin D. Curran, Surendra Parmar, Ian Goodfellow |
| EPI_ISL_953065, EPI_ISL_953076, EPI_ISL_953077, EPI_ISL_953078, EPI_ISL_953079, EPI_ISL_953080, EPI_ISL_953081, EPI_ISL_953082, EPI_ISL_953083, EPI_ISL_953085, EPI_ISL_953086, EPI_ISL_953087, EPI_ISL_953088, EPI_ISL_953089, EPI_ISL_953090, EPI_ISL_953091, EPI_ISL_953092, EPI_ISL_953093, EPI_ISL_953094, EPI_ISL_953095, EPI_ISL_953096, EPI_ISL_953097, EPI_ISL_953098, EPI_ISL_953099, EPI_ISL_953100, EPI_ISL_953101, EPI_ISL_953102, EPI_ISL_953103, EPI_ISL_953104, EPI_ISL_953105, EPI_ISL_953106, EPI_ISL_953107, EPI_ISL_953108, EPI_ISL_953109, EPI_ISL_953111, EPI_ISL_953113, EPI_ISL_953114, EPI_ISL_953115, EPI_ISL_953116, EPI_ISL_953117 | | | |
| see above | Bioinformatics and Biostatistics Lab, Advanced Sequencing Facility | COVID-19 Genomics UK (COG-UK) Consortium | Aengus Stewart,Jerome Nicod,Chelsea Sawyer,Laura Cubitt,Harshil Patel,Margaret Crawford |
| EPI_ISL_953498, EPI_ISL_953668, EPI_ISL_953669, EPI_ISL_953670 | University Hospitals of Geneva, Laboratory of Virology | HUG, Laboratory of Virology and the Health2030 Genome Center | Samuel Cordey, Ana Rita Goncalves, Laurent Kaiser, Lorenzo Cerutti, Henri Pegeot, Melyssa Elies, Deborah Penet, Keith Harshman, Ioannis Xenarios, Emmanouil Dermitzakis |
| EPI_ISL_953949, EPI_ISL_953950 | Hopital | National Reference Center for Viruses of Respiratory Infections, Institut Pasteur, Paris | Marion Barbet, Sylvie Behillil, Méline Bizard, Angela Brisebarre, Camille Capel, Etienne Simon-Lorière, Vincent Enouf, Maud Vanpeene, Sylvie van der Werf,Cady Anne |
| EPI_ISL_953957, EPI_ISL_953991, EPI_ISL_953992, EPI_ISL_953993 | Hopital | National Reference Center for Viruses of Respiratory Infections, Institut Pasteur, Paris | Marion Barbet, Sylvie Behillil, Méline Bizard, Angela Brisebarre, Camille Capel, Etienne Simon-Lorière, Vincent Enouf, Maud Vanpeene, Sylvie van der Werf,Lagathu GisèLe |
| EPI_ISL_954112 | Hopital | National Reference Center for Viruses of Respiratory Infections, Institut Pasteur, Paris | Marion Barbet, Sylvie Behillil, Méline Bizard, Angela Brisebarre, Camille Capel, Etienne Simon-Lorière, Vincent Enouf, Maud Vanpeene, Sylvie van der Werf,Lesimple BéAtrice |
| EPI_ISL_954113 | Hopital | National Reference Center for Viruses of Respiratory Infections, Institut Pasteur, Paris | Marion Barbet, Sylvie Behillil, Méline Bizard, Angela Brisebarre, Camille Capel, Etienne Simon-Lorière, Vincent Enouf, Maud Vanpeene, Sylvie van der Werf,Lagathu GisèLe |
| EPI_ISL_954114 | Labo Analyses Med | National Reference Center for Viruses of Respiratory Infections, Institut Pasteur, Paris | Marion Barbet, Sylvie Behillil, Méline Bizard, Angela Brisebarre, Camille Capel, Etienne Simon-Lorière, Vincent Enouf, Maud Vanpeene, Sylvie van der Werf,Selas Labomaine |
| EPI_ISL_954115 | Hopital | National Reference Center for Viruses of Respiratory Infections, Institut Pasteur, Paris | Marion Barbet, Sylvie Behillil, Méline Bizard, Angela Brisebarre, Camille Capel, Etienne Simon-Lorière, Vincent Enouf, Maud Vanpeene, Sylvie van der Werf,Lagathu GisèLe |
| EPI_ISL_954116, EPI_ISL_954117, EPI_ISL_954118 | Outre Mer | National Reference Center for Viruses of Respiratory Infections, Institut Pasteur, Paris | Marion Barbet, Sylvie Behillil, Méline Bizard, Angela Brisebarre, Camille Capel, Etienne Simon-Lorière, Vincent Enouf, Maud Vanpeene, Sylvie van der Werf,Rousset Dominique |
| EPI_ISL_954787, EPI_ISL_954790 | Hospital J.M. Morales Meseguer | Instituto de Salud Carlos III | Iglesias-Caballero, M. Camarero, S. Sandonís,V. Vázquez, S. Pozo, F. Casas, I. Jiménez, P. Zaballos, A. Monzón, S. Varona, S. Cuesta, I. Navarro, D. |
| EPI_ISL_954792 | HOSPITAL COMARCAL DE MELILLA | Instituto de Salud Carlos III | Iglesias-Caballero, M. Camarero, S. Sandonís,V. Vázquez, S. Pozo, F. Casas, I. Jiménez, P. Zaballos, A. Monzón, S. Varona, S. Cuesta, I. Roman, S. |

| | | | |
|---|---|---|---|
| EPI_ISL_954795 | HOSPITAL GENERAL RIO CARRION | Instituto de Salud Carlos III | Iglesias-Caballero, M. Camarero, S. Sandonís,V. Vázquez, S. Pozo, F. Casas, I. Jiménez, P. Zaballos, A. Monzón, S. Varona, S. Cuesta, I. García, A. |
| EPI_ISL_954812 | Hospital Universitario Virgen de la Arrixaca | Instituto de Salud Carlos III | Iglesias-Caballero, M. Camarero, S. Sandonís,V. Vázquez, S. Pozo, F. Casas, I. Jiménez, P. Zaballos, A. Monzón, S. Varona, S. Cuesta, I. Moreno, L. |
| EPI_ISL_954819, EPI_ISL_954822, EPI_ISL_954823, EPI_ISL_954824, EPI_ISL_954826, EPI_ISL_954827, EPI_ISL_954833, EPI_ISL_954834, EPI_ISL_954836, EPI_ISL_954837, EPI_ISL_954839, EPI_ISL_954841, EPI_ISL_954843, EPI_ISL_954849, EPI_ISL_954850, EPI_ISL_954851, EPI_ISL_954853, EPI_ISL_954859, EPI_ISL_954860, EPI_ISL_954861, EPI_ISL_954866, EPI_ISL_954870, EPI_ISL_954874, EPI_ISL_954880, EPI_ISL_954882, EPI_ISL_954888, EPI_ISL_954889, EPI_ISL_954890, EPI_ISL_954891, EPI_ISL_954951, EPI_ISL_954952, EPI_ISL_954953, EPI_ISL_954954, EPI_ISL_954955, EPI_ISL_954956, EPI_ISL_954957, EPI_ISL_954958, EPI_ISL_954959, EPI_ISL_954960, EPI_ISL_954961, EPI_ISL_954962, EPI_ISL_954963, EPI_ISL_954964, EPI_ISL_954965, EPI_ISL_954966, EPI_ISL_954967, EPI_ISL_954968, EPI_ISL_954969, EPI_ISL_954970, EPI_ISL_954971, EPI_ISL_954972, EPI_ISL_954973, EPI_ISL_954974, EPI_ISL_954975, EPI_ISL_954976, EPI_ISL_954977, EPI_ISL_954978, EPI_ISL_954979, EPI_ISL_954980, EPI_ISL_954981, EPI_ISL_954982, EPI_ISL_954983, EPI_ISL_954984, EPI_ISL_954985, EPI_ISL_954986, EPI_ISL_954987, EPI_ISL_954988, EPI_ISL_954989, EPI_ISL_954990, EPI_ISL_954991, EPI_ISL_954992, EPI_ISL_954993, EPI_ISL_954994, EPI_ISL_954995, EPI_ISL_954996, EPI_ISL_954997, EPI_ISL_954998, EPI_ISL_955131, EPI_ISL_955132, EPI_ISL_955133, EPI_ISL_955134, EPI_ISL_955135 | | | |
| see above | Colorado Department of Public Health and Environment | Colorado Department of Puplic Health and Environment | Laura Bankers, Molly C. Hetherington-Rauth, Diana Ir, Shannon Ely, Shannon R. Matzinger, Sarah Elizabeth Totten, Emily A. Travanty |
| EPI_ISL_955174 | Hopital | National Reference Center for Viruses of Respiratory Infections, Institut Pasteur, Paris | Marion Barbet, Sylvie Behillil, Méline Bizard, Angela Brisebarre, Camille Capel, Etienne Simon-Lorière, Vincent Enouf, Maud Vanpeene, Sylvie van der Werf,Goudeau Alain |
| EPI_ISL_955175 | Hopital | National Reference Center for Viruses of Respiratory Infections, Institut Pasteur, Paris | Marion Barbet, Sylvie Behillil, Méline Bizard, Angela Brisebarre, Camille Capel, Etienne Simon-Lorière, Vincent Enouf, Maud Vanpeene, Sylvie van der Werf |
| EPI_ISL_955186 | University of Sarajevo, Veterinary Faculty, Laboratory for Molecular Diagnostic and Research Laboratory | University of Sarajevo, Veterinary Faculty, Laboratory for Molecular Diagnostic and Research Laboratory | Goleti Š., Goleti T., Ali-Šeho A., Softi A., Terzi I., Jaži A., Nicevi M., Hodži A., Šabi E. |
| EPI_ISL_955187 | University of Sarajevo, Veterinary Faculty, Laboratory for Molecular Diagnostic and Research Laboratory | University of Sarajevo, Veterinary Faculty, Laboratory for Molecular Diagnostic and Research Laboratory | Goleti Š., Goleti T., Ali-Šeho A., Softi A., Hodži A., Terzi I., Šabi E., Jaži A., Nicevi M. |
| EPI_ISL_955273 | American Esoteric Laboratory | Pathogen Discovery, Respiratory Viruses Branch, Division of Viral Diseases, Centers for Disease Control and Prevention | Ying Tao, Jing Zhang, Yan Li, Krista Queen, Anna Uehara, Peter Cook, Clinton R. Paden, Haibin Wang, Suxiang Tong |
| EPI_ISL_955391 | Alameda County Public Health Lab | Chan-Zuckerberg Biohub | CZB Cliahub Consortium |
| EPI_ISL_955778, EPI_ISL_955780, EPI_ISL_955783, EPI_ISL_955784, EPI_ISL_955788, EPI_ISL_955791, EPI_ISL_955792, EPI_ISL_955793, EPI_ISL_955802, EPI_ISL_955840, EPI_ISL_955841, EPI_ISL_955842, EPI_ISL_955843, EPI_ISL_955844 | | | |
| see above | University of Michigan Clinical Microbiology Laboratory | Lauring Lab, University of Michigan, Department of Microbiology and Immunology | Valesano |
| EPI_ISL_956283 | Instituto Nacional de Salud- Dirección de Redes de Laboratorios de Salud Pública | Instituto Nacional de Salud- Dirección de Investigación en Salud Pública | Katherine Laiton-Donato, Diego A. Álvarez-Díaz, Carlos Franco-Muñoz, Mauricio Pacheco-Montealegre, Hector Alejandro Ruiz-Moreno, Maria T. Herrera-Sepúlveda, Diego Andrés Prada, Jhonnatan Reales-González, Sheryll Corchuelo, Julian Naizaque, Gerardo Santamaría, Magdalena Wiesner, Martha Lucia Ospina Martinez, Marcela Mercado-Reyes |
| EPI_ISL_956303 | LABORATORIO ANALIZAR SYNLAB | Instituto Nacional de Salud- Dirección de Investigación en Salud Pública | Katherine Laiton-Donato, Diego A. Álvarez-Díaz, Carlos Franco-Muñoz, Mauricio Pacheco-Montealegre, Hector Alejandro Ruiz-Moreno, Maria T. Herrera-Sepúlveda, Diego Andrés Prada, Jhonnatan Reales-González, Sheryll Corchuelo, Julian Naizaque, Gerardo Santamaría, Magdalena Wiesner, Martha Lucia Ospina Martinez, Marcela Mercado-Reyes |
| EPI_ISL_956334, EPI_ISL_956336 | Utah Public Health Laboratory, Utah Public Health Laboratory Infectious Disease submission group | Utah Public Health Laboratory, Utah Public Health Laboratory Infectious Disease submission group | Gallagher,T., Young,E.L., Oakeson,K.F. |
| EPI_ISL_959444, EPI_ISL_959468, EPI_ISL_959469, EPI_ISL_959470, EPI_ISL_959471 | Servicio de Microbiología, Hospital Universitario Son Espases | SeqCOVID-SPAIN consortium/IBV(CSIC) | Carla López-Causapé, Jordi Reina, Antonio Oliver and SeqCOVID-SPAIN consortium |
| EPI_ISL_959542 | University of Michigan Clinical Microbiology Laboratory | Lauring Lab, University of Michigan, Department of Microbiology and Immunology | Valesano |
| EPI_ISL_959546 | Laboratory of Virology and Molecular Diagnostics | Institute of Public Health of Republic of North Macedonia Laboratory of Virology and Molecular Diagnostics | Kuzmanovska M., Boshevska G. |
| EPI_ISL_959563 | University of Liège COVID-19 testing center | GIGA Medical Genomics | Keith Durkin, Maria Artesi, Sébastien Bontems, Raphaël Boreux, Bouchra Boujemla, Cécile Meex, Pierrette Melin, Marie-Pierre Hayette, Vincent Bours |
| EPI_ISL_959564, EPI_ISL_959565 | Vivalia - Clinique Saint-Joseph | GIGA Medical Genomics | Keith Durkin, Maria Artesi, Sébastien Bontems, Raphaël Boreux, Bouchra Boujemla, Cécile Meex, Pierrette Melin, Marie-Pierre Hayette, Vincent Bours |
| EPI_ISL_959566 | University of Liège COVID-19 testing center | GIGA Medical Genomics | Keith Durkin, Maria Artesi, Sébastien Bontems, Raphaël Boreux, Bouchra Boujemla, Cécile Meex, Pierrette Melin, Marie-Pierre Hayette, Vincent Bours |
| EPI_ISL_959839, EPI_ISL_959856 | National Virus Reference Laboratory | National Virus Reference Laboratory | Michael Carr, Gabriel Gonzalez, Jonathan Dean, Cillian F De Gascun |
| EPI_ISL_960226 | Botswana Harvard HIV Reference Laboratory | Botswana Harvard HIV Reference Laboratory | Sikhulile Moyo,Dorcas Maruapula, Wonderful T. Choga, Botshelo Radibe, Boitumelo Zuze, David Lawrence, Roger Shapiro, Shahin Lockman, Mosepele Mosepele, Joseph Makhema, Simani Gaseitsiwe |
| EPI_ISL_960303 | Botswana Harvard HIV Reference Laboratory | Botswana Harvard HIV Reference Laboratory | Sikhulile Moyo, Dorcas Maruapula, Wonderful Choga, Botshelo Radibe, Boitumelo Zuze, David Lawrence, Roger Shapiro, Shahin Lockman, Mosepele Mosepele, Joseph Makhema, Simani Gaseitsiwe |
| EPI_ISL_960314, EPI_ISL_960315, EPI_ISL_960316, EPI_ISL_960317 | University of Wisconsin-Madison AIDS Vaccine Research Laboratories | University of Wisconsin-Madison AIDS Vaccine Research Laboratories | Gage Moreno, Katarina Braun, et al. AIDS Vaccine Research Laboratories |
| EPI_ISL_960406, EPI_ISL_960420, EPI_ISL_960421, EPI_ISL_960428 | The National Institute of Public Health | State Veterinary Institute Prague | Nagy,A;Vecerova,J;Cernikova,L;Stara,M;Jirincova,H;Trnka,D |
| EPI_ISL_960986, EPI_ISL_960987, EPI_ISL_960988, EPI_ISL_960991, EPI_ISL_960994, EPI_ISL_960999, EPI_ISL_961000, EPI_ISL_961002 | AIID | Irish Coronavirus Sequencing Consortium-Teagasc Grange | Matthew McCabe, Aljandro Abner Garcia Leon, Fiona Crispie, Calum Walsh, Michael Carr, John Kenny, Paul Cotter, Patrick Mallon, Gabriel Gonzalez |
| EPI_ISL_961134, EPI_ISL_961166 | Texas Department of State Health Services | Texas Department of State Health Services | Bonnie Oh, Anita Pokharel, James Daniel Bonser, Myong Koag, Chung Wang, Rachel Lee, Grace Kubin, Rashmi Tuladhar, Mayela Pedrueza, Maliha Rahman, Jenny Zhang |
| EPI_ISL_961228, EPI_ISL_961229 | Hospital General Universitario de Alicante - Instituto de Investigación Sanitaria y Biomédica de Alicante | SeqCOVID-SPAIN consortium/IBV(CSIC) | Maripaz Ventero Martín, Carmen Molina Pardines and SeqCOVID-SPAIN consortium |
| EPI_ISL_961471, EPI_ISL_961484, EPI_ISL_961494, EPI_ISL_961497, EPI_ISL_961502, EPI_ISL_961503, EPI_ISL_961511, EPI_ISL_961520, EPI_ISL_961526, EPI_ISL_961544, EPI_ISL_961545, EPI_ISL_961546 | | | |
| see above | Michigan Department of Health and Human Services, Bureau of Laboratories | Michigan Department of Health and Human Services, Bureau of Laboratories | Blankenship HM, Riner D, Soehnlen MK |
| EPI_ISL_961591, EPI_ISL_961592, EPI_ISL_961593, EPI_ISL_961594, EPI_ISL_961595, EPI_ISL_961596, EPI_ISL_961630, EPI_ISL_961631, EPI_ISL_961632, EPI_ISL_961633, EPI_ISL_961634, EPI_ISL_961635, EPI_ISL_961636 | | | |
| see above | Hôpital Georges L. Dumont | National Microbiology Laboratory (NML) | Anna Majer, Shari Tyson, Grace Seo, Philip Mabon, Elsie Grudeski, Rhiannon Huzarewich, Russell Mandes, Anneliese Landgraff, Jennifer Tanner, Natalie Knox, Morag Graham, Gary Van Domselaar, Richard Garceau, Guillaume Desnoyers, Nathalie Bastien, Yan Li, Timothy Booth, Darian Hole, Madison Chapel, Kirsten Biggar, CanCOGeN's metadata curation team, Public Health Agency of Canada CanCOGeN team |
| EPI_ISL_961873, EPI_ISL_961874 | E. Gulbja laboratorija | Latvian Biomedical Research and Study Centre | Janis Pjalkovskis, Nikita Zrelovs, Monta Ustinova, Ivars Silamikelis, Liga Birzniece, Kaspars Megnis, Vita Rovite, Lauma Freimane, Laila Silamikele, Laura Ansone, Davids Fridmanis, Mikus Gavars, Dmitrijs Perminovs, Jurijs Perevoscikovs, Uga Dumpis, Janis Klovins |
| EPI_ISL_961972, EPI_ISL_962176 | Illinois Department of Public Health | Gagnon Lab, Southern Illinois University | Keith Gagnon |
| EPI_ISL_962203, EPI_ISL_962204, EPI_ISL_962207 | Institute for Medical Research, Infectious Disease Research Centre, National Institutes of Health, Ministry of Health Malaysia | Institute for Medical Research, Infectious Disease Research Centre, National Institutes of Health, Ministry of Health Malaysia | Suppiah J, Kamel K, Azizan MA, Thayan R |

| | | | |
|---|---|---|---|
| EPI_ISL_962278, EPI_ISL_962279, EPI_ISL_962280, EPI_ISL_962281, EPI_ISL_962282, EPI_ISL_962283, EPI_ISL_962284, EPI_ISL_962287 | Seattle Flu Study | Seattle Flu Study | Deborah A. Nickerson, Chris D. Frazar, Jover Lee, Benjamin Pelle, Erica Ryke, Matthew Richardson, Amanda Adler, Elisabeth Brandstetter, Peter D. Han, Kairsten Fay, Misja Ilcisin, Kirsten Lacombe, Thomas R. Sibley, Melissa Truong, Caitlin R. Wolf, Karen Cowgill, Stephanie Schrag, Jeff Duchin, Michael Boeckh, Janet A. Englund, Michael Famulare, Barry R. Lutz, Mark J. Rieder, Lea M. Starita, Matthew Thompson, Helen Y. Chu, Trevor Bedford, Jay Shendure |
| EPI_ISL_962389, EPI_ISL_962390, EPI_ISL_962391, EPI_ISL_962392, EPI_ISL_962393, EPI_ISL_962394, EPI_ISL_962395, EPI_ISL_962396, EPI_ISL_962397, EPI_ISL_962398, EPI_ISL_962399, EPI_ISL_962400, EPI_ISL_962401, EPI_ISL_962402 | | | |
| see above | Washington State Department of Health | Seattle Flu Study | Deborah A. Nickerson, Chris D. Frazar, Jover Lee, Benjamin Pelle, Erica Ryke, Matthew Richardson, Amanda Adler, Elisabeth Brandstetter, Peter D. Han, Kairsten Fay, Misja Ilcisin, Kirsten Lacombe, Thomas R. Sibley, Melissa Truong, Caitlin R. Wolf, Romesh Gautom, Geoff Melly, Brian Hiatt, Philip Dykema, Scott Lindquist, Michael Boeckh, Janet A. Englund, Michael Famulare, Barry R. Lutz, Mark J. Rieder, Lea M. Starita, Matthew Thompson, Helen Y. Chu, Jay Shendure, Trevor Bedford |
| EPI_ISL_962525, EPI_ISL_962526 | Institute for Medical Research, Infectious Disease Research Centre, National Institutes of Health, Ministry of Health Malaysia | Institute for Medical Research, Infectious Disease Research Centre, National Institutes of Health, Ministry of Health Malaysia | Suppiah J, Kamel K, Azizan MA, Thayan R |
| EPI_ISL_962621, EPI_ISL_962622, EPI_ISL_962623, EPI_ISL_962625, EPI_ISL_962628, EPI_ISL_962630, EPI_ISL_962631, EPI_ISL_962634, EPI_ISL_962635, EPI_ISL_962637, EPI_ISL_962638, EPI_ISL_962639, EPI_ISL_962640, EPI_ISL_962642, EPI_ISL_962644, EPI_ISL_962646, EPI_ISL_962647 | | | |
| see above | Scripps Medical Laboratory | Andersen lab at Scripps Research | SEARCH Alliance San Diego with Michael Quigley, Ellen Stefanski, Ian Mchardy |
| EPI_ISL_962710, EPI_ISL_962711, EPI_ISL_962712, EPI_ISL_962713, EPI_ISL_962714, EPI_ISL_962715, EPI_ISL_962716, EPI_ISL_962717, EPI_ISL_962718, EPI_ISL_962720, EPI_ISL_962721, EPI_ISL_962722, EPI_ISL_962723, EPI_ISL_962724, EPI_ISL_962725, EPI_ISL_962726, EPI_ISL_962728, EPI_ISL_962729, EPI_ISL_962731, EPI_ISL_962732, EPI_ISL_962733, EPI_ISL_962734, EPI_ISL_962735, EPI_ISL_962736, EPI_ISL_962737, EPI_ISL_962738, EPI_ISL_962740, EPI_ISL_962741, EPI_ISL_962742, EPI_ISL_962743, EPI_ISL_962746, EPI_ISL_962747, EPI_ISL_962748, EPI_ISL_962749, EPI_ISL_962750, EPI_ISL_962751, EPI_ISL_962752, EPI_ISL_962753, EPI_ISL_962755, EPI_ISL_962757, EPI_ISL_962760, EPI_ISL_962761, EPI_ISL_962762, EPI_ISL_962763, EPI_ISL_962764 | | | |
| see above | Sharp HealthCare Laboratory | Andersen lab at Scripps Research | SEARCH Alliance San Diego with Aaron Harding, Jacquelyn Berumen, Cathy Woerle, Liam McGinnis, Art Mendoza, Omid Bakhtar |
| EPI_ISL_962813 | Robert Garry lab | Andersen lab at Scripps Research | Allison Smither, Gilberto Sabino-Santos, Patricia Snarski, Lilia Melnik, Antoinette Bell, Kaylynn Genemaras, Arnaud Drouin, Dahlene Fusco, Robert Garry with SEARCH Alliance San Diego |
| EPI_ISL_962887, EPI_ISL_962890 | Laboratory of Virology and Molecular Diagnostics | Institute of Public Health of Republic of North Macedonia Laboratory of Virology and Molecular Diagnostics | Kuzmanovska M., Boshevska G. |
| EPI_ISL_962891 | Laboratory of Virology and Molecular Diagnostics | Laboratory of Virology and Molecular Diagnostics, Institute of Public Health of Republic of North Macedonia | Kuzmanovska M., Boshevska G. |
| EPI_ISL_962893, EPI_ISL_962894 | Laboratory of Virology and Molecular Diagnostics | Institute of Public Health of Republic of North Macedonia Laboratory of Virology and Molecular Diagnostics | Kuzmanovska M., Boshevska G. |
| EPI_ISL_962944, EPI_ISL_962954, EPI_ISL_962955 | Hospital Universitario de Gran Canaria Dr. Negrín | SeqCOVID-SPAIN consortium/IBV(CSIC) | M. Carmen Pérez González, Francisco J. Chamizo López, Ana Bordes Benítez and SeqCOVID-SPAIN consortium |
| EPI_ISL_964272, EPI_ISL_964275 | Oslo University Hospital, Department of Medical Microbiology | Norwegian Institute of Public Health, Department of Virology | Kathrine Stene-Johansen, Kamilla Heddeland Instefjord, Hilde Elshaug, Ignacio Garcia Llorente, Serina B Engebretsen, Atiya R Ali, Marie Paulsen Madsen, Rasmus Riis Kopperud, Hilde Vollan, Karoline Bragstad, Olav Hungnes |
| EPI_ISL_964881, EPI_ISL_964883 | Laboratory of Virology and Molecular Diagnostics | Institute of Public Health of Republic of North Macedonia Laboratory of Virology and Molecular Diagnostics | Kuzmanovska M., Boshevska G. |
| EPI_ISL_964919, EPI_ISL_964920, EPI_ISL_964922, EPI_ISL_964923, EPI_ISL_964925, EPI_ISL_964926, EPI_ISL_964927, EPI_ISL_964928, EPI_ISL_964929, EPI_ISL_964930, EPI_ISL_964931, EPI_ISL_964934, EPI_ISL_964935, EPI_ISL_964936, EPI_ISL_964937, EPI_ISL_964939, EPI_ISL_964942, EPI_ISL_964943, EPI_ISL_964944, EPI_ISL_964945, EPI_ISL_964946, EPI_ISL_964947 | | | |
| see above | Instituto Nacional de Saude (INS), Mozambique | KRISP, KZN Research Innovation and Sequencing Platform | Nalia Ismael, Nadia Sitoe, Paulo Arnaldo, Nedio Mabunda, Giandhari J, Pillay S, Emmanuel S, Tegally H, Wilkinson E, de Oliveira T |
| EPI_ISL_964995 | Foerde Hospital, Department of Microbiology | Norwegian Institute of Public Health, Department of Virology | Kathrine Stene-Johansen, Kamilla Heddeland Instefjord, Hilde Elshaug, Ignacio Garcia Llorente, Serina B Engebretsen, Atiya R Ali, Marie Paulsen Madsen, Rasmus Riis Kopperud, Hilde Vollan, Karoline Bragstad, Olav Hungnes |
| EPI_ISL_965021, EPI_ISL_965022, EPI_ISL_965023, EPI_ISL_965024 | Florida Bureau of Public Health Laboratories | Florida Bureau of Public Health Laboratories | Sarah Schmedes, Jason Blanton |
| EPI_ISL_965101, EPI_ISL_965102, EPI_ISL_965103, EPI_ISL_965104, EPI_ISL_965105, EPI_ISL_965106, EPI_ISL_965107, EPI_ISL_965108, EPI_ISL_965109, EPI_ISL_965110, EPI_ISL_965111, EPI_ISL_965112 | | | |
| see above | Wyoming Public Health Laboratory | Wyoming Public Health Laboratory | Noah Hull, Taylor Fearing, Lynette Gumbleton, Channing Weber, Ashley Norberg, Bailey Bowcutt, and Wanda Manley |
| EPI_ISL_965127 | INMI Lazzaro Spallanzani IRCCS | INMI Lazzaro Spallanzani IRCCS | CEM Gruber, B Bartolini, E Giombini, M Rueca, O Butera, F Messina, A Di Caro, MR Capobianchi |
| EPI_ISL_965550, EPI_ISL_965738, EPI_ISL_965745, EPI_ISL_965746 | Dutch COVID-19 response team | Medical Microbiology, Maastricht University Medical Centre | Jozef Dingemans*, Brian van der Veer*, Erik Beuken, Carmen Reumkens, Lieke van Alphen, Christian Hoebe, Paul Savelkoul |
| EPI_ISL_965847, EPI_ISL_965848, EPI_ISL_965849, EPI_ISL_965850, EPI_ISL_965851, EPI_ISL_965852 | GA Department of Public Health | GA Department of Public Health | Stacy Reeves, Jonathan Edwards, Cynthia Dixey, Tonia Parrott |
| EPI_ISL_965863 | Massachusetts State Public Health Laboratory | Massachusetts State Public Health Laboratory | Andrew Lang, Timelia Fink, Glen Gallagher, Sandra Smole |
| EPI_ISL_965912 | Servicio Murciano de Salud | Instituto de Salud Carlos III | Vázquez, S. Iglesias-Caballero, M. Sandonís,V. Camarero, S. Pozo, F. Casas, I. Jiménez, P. Zaballos, A. Monzón, S. Varona, S. Cuesta, I. Blázquez, A. |
| EPI_ISL_965921 | Consejería de Sanidad y Asuntos Sociales de Castilla La Mancha | Instituto de Salud Carlos III | Vázquez, S. Iglesias-Caballero, M. Sandonís,V. Camarero, S. Pozo, F. Casas, I. Jiménez, P. Zaballos, A. Monzón, S. Varona, S. Cuesta, I. Gutiérrez, G. |
| EPI_ISL_965922, EPI_ISL_965925 | Servicio Murciano de Salud | Instituto de Salud Carlos III | Vázquez, S. Iglesias-Caballero, M. Sandonís,V. Camarero, S. Pozo, F. Casas, I. Jiménez, P. Zaballos, A. Monzón, S. Varona, S. Cuesta, I. Blázquez, A. |
| EPI_ISL_965927, EPI_ISL_965931 | Servicio Murciano de Salud | Instituto de Salud Carlos III | Sandonís,V. Vázquez, S. Iglesias-Caballero, M. Camarero, S. Pozo, F. Casas, I. Jiménez, P. Zaballos, A. Monzón, S. Varona, S. Cuesta, I. Blázquez, A. |
| EPI_ISL_965953 | Servicio Murciano de Salud | Instituto de Salud Carlos III | Iglesias-Caballero, M. Sandonís,V. Vázquez, S. Camarero, S. Pozo, F. Casas, I. Jiménez, P. Zaballos, A. Monzón, S. Varona, S. Cuesta, I. Blázquez, A. |
| EPI_ISL_966500, EPI_ISL_966501, EPI_ISL_966502, EPI_ISL_966503, EPI_ISL_966504, EPI_ISL_966505, EPI_ISL_966506, EPI_ISL_966507, EPI_ISL_966508, EPI_ISL_966509, EPI_ISL_966510, EPI_ISL_966511, EPI_ISL_966512, EPI_ISL_966514, EPI_ISL_966515, EPI_ISL_966516, EPI_ISL_966517, EPI_ISL_966518, EPI_ISL_966519, EPI_ISL_966520, EPI_ISL_966521, EPI_ISL_966522, EPI_ISL_966523, EPI_ISL_966524, EPI_ISL_966525, EPI_ISL_966526, EPI_ISL_966527, EPI_ISL_966528, EPI_ISL_966529, EPI_ISL_966530, EPI_ISL_966531, EPI_ISL_966577, EPI_ISL_966578, EPI_ISL_966579, EPI_ISL_966580, EPI_ISL_966581, EPI_ISL_966582, EPI_ISL_966583, EPI_ISL_966584, EPI_ISL_966585, EPI_ISL_966586, EPI_ISL_966587, EPI_ISL_966588, EPI_ISL_966589, EPI_ISL_966590, EPI_ISL_966600, EPI_ISL_966601, EPI_ISL_966604, EPI_ISL_967170, EPI_ISL_967172, EPI_ISL_967173, EPI_ISL_967176, EPI_ISL_967177, EPI_ISL_967178, EPI_ISL_967185, EPI_ISL_967202, EPI_ISL_967203, EPI_ISL_967204, EPI_ISL_967205, EPI_ISL_967206, EPI_ISL_967207, EPI_ISL_967208, EPI_ISL_967209, EPI_ISL_967210, EPI_ISL_967211, EPI_ISL_967212, EPI_ISL_967215, EPI_ISL_967217, EPI_ISL_967219, EPI_ISL_967327, EPI_ISL_967328, EPI_ISL_967329, EPI_ISL_967330, EPI_ISL_967331, EPI_ISL_967332, EPI_ISL_967333, EPI_ISL_967334, EPI_ISL_967335, EPI_ISL_967336, EPI_ISL_967337, EPI_ISL_967338, EPI_ISL_967339, EPI_ISL_967340, EPI_ISL_967341, EPI_ISL_967342, EPI_ISL_967343, EPI_ISL_967344, EPI_ISL_967345, EPI_ISL_967346, EPI_ISL_967347, EPI_ISL_967348, EPI_ISL_967349, EPI_ISL_967350, EPI_ISL_967351, EPI_ISL_967352, EPI_ISL_967353, EPI_ISL_967354, EPI_ISL_967355, EPI_ISL_967356, EPI_ISL_967357, EPI_ISL_967358, EPI_ISL_967359, EPI_ISL_967360, EPI_ISL_967361, EPI_ISL_967362, EPI_ISL_967363, EPI_ISL_967364, EPI_ISL_967365, EPI_ISL_967366, EPI_ISL_967367, EPI_ISL_967368, EPI_ISL_967369, EPI_ISL_967370, EPI_ISL_967371, EPI_ISL_967372, EPI_ISL_967373, EPI_ISL_967374, EPI_ISL_967375, EPI_ISL_967376, EPI_ISL_967377, EPI_ISL_967378, EPI_ISL_967379, EPI_ISL_967380, EPI_ISL_967381, EPI_ISL_967382, EPI_ISL_967383, EPI_ISL_967384, EPI_ISL_967385, EPI_ISL_967386, EPI_ISL_967387, EPI_ISL_967388, EPI_ISL_967389, EPI_ISL_967390, EPI_ISL_967391, EPI_ISL_967392, EPI_ISL_967393, EPI_ISL_967394, EPI_ISL_967395, EPI_ISL_967396, EPI_ISL_967397, EPI_ISL_967398, EPI_ISL_967399, EPI_ISL_967400, EPI_ISL_967401, EPI_ISL_967402, EPI_ISL_967403, EPI_ISL_967404, EPI_ISL_967405, EPI_ISL_967406, EPI_ISL_967407, EPI_ISL_967408, EPI_ISL_967409, EPI_ISL_967410, EPI_ISL_967411, EPI_ISL_967412, EPI_ISL_967413, EPI_ISL_967414, EPI_ISL_967415, EPI_ISL_967416, EPI_ISL_967417, EPI_ISL_967418, EPI_ISL_967419, EPI_ISL_967420, EPI_ISL_967421, EPI_ISL_967422, EPI_ISL_967423, EPI_ISL_967424 | | | |
| see above | Helix/Illumina | Respiratory Viruses Branch, Division of Viral Diseases, Centers for Disease Control and Prevention | Peter W. Cook,Dakota Howard,Dhwani Batra,Ben L. Rambo-Martin,Eileen de Feo,Jan Antico,Christine Tran,Matthew Tolentino,Shannon Wickline,Kim Gietzen,Brad Sickler,Jingtao Liu,Eric Allen,Phil Febbo,Summer Galloway,Nicole L. Washington,Simon White,Geraint Levan,Kelly Schiabor Barrett,Elizabeth Cirulli,Alexandre Bolze,Ary Ascencio,Charlotte Rivera-Garcia,Ryan Cho,Jason Nguyen,Sherry Wang,Jimmy Ramirez,Tyler Cassens,Efren Sandoval,Magnus Isaksson,William Lee,David Becker,Marc Laurent,James Lu,Clinton R. Paden,Suxiang Tong,Duncan MacCannell, |
| EPI_ISL_967704, EPI_ISL_967715, EPI_ISL_967720, EPI_ISL_967721, EPI_ISL_967746, EPI_ISL_967757 | State Laboratories Division, Hawaii State Department of Health | State Laboratories Division, Hawaii State Department of Health | Pamela O'Brien, Drew Kuwazaki, Ayana Garnet, Razvan Sultana, Edward Desmond |

| | | | |
|---|---|---|---|
| EPI_ISL_967781, EPI_ISL_967782, EPI_ISL_967783, EPI_ISL_967784, EPI_ISL_967785, EPI_ISL_967786, EPI_ISL_967787, EPI_ISL_967788, EPI_ISL_967789, EPI_ISL_967790, EPI_ISL_967791, EPI_ISL_967792, EPI_ISL_967793, EPI_ISL_967794, EPI_ISL_967795, EPI_ISL_967796, EPI_ISL_967797, EPI_ISL_967798, EPI_ISL_967799, EPI_ISL_967800, EPI_ISL_967801, EPI_ISL_967802, EPI_ISL_967803, EPI_ISL_967804, EPI_ISL_967805, EPI_ISL_967806, EPI_ISL_967807, EPI_ISL_967808, EPI_ISL_967809, EPI_ISL_967810, EPI_ISL_967811, EPI_ISL_967812, EPI_ISL_967813, EPI_ISL_967814, EPI_ISL_967815, EPI_ISL_967816, EPI_ISL_967817, EPI_ISL_967818, EPI_ISL_967819, EPI_ISL_967820, EPI_ISL_967821, EPI_ISL_967822, EPI_ISL_967823, EPI_ISL_967824, EPI_ISL_967825, EPI_ISL_967826, EPI_ISL_967827, EPI_ISL_967828, EPI_ISL_967829, EPI_ISL_967830, EPI_ISL_967831, EPI_ISL_967832, EPI_ISL_967833, EPI_ISL_967834, EPI_ISL_967835, EPI_ISL_967836, EPI_ISL_967837, EPI_ISL_967838, EPI_ISL_967839, EPI_ISL_967840, EPI_ISL_967841, EPI_ISL_967842 | | | |
| see above | Helix/Illumina | Respiratory Viruses Branch, Division of Viral Diseases, Centers for Disease Control and Prevention | Peter W. Cook,Dakota Howard,Dhwani Batra,Ben L. Rambo-Martin,Eileen de Feo,Jan Antico,Christine Tran,Matthew Tolentino,Shannon Wickline,Kim Gietzen,Brad Sickler,Jingtao Liu,Eric Allen,Phil Febbo,Summer Galloway,Nicole L. Washington,Simon White,Geraint Levan,Kelly Schiabor Barrett,Elizabeth Cirulli,Alexandre Bolze,Ary Ascencio,Charlotte Rivera-Garcia,Ryan Cho,Jason Nguyen,Sherry Wang,Jimmy Ramirez,Tyler Cassens,Efren Sandoval,Magnus Isaksson,William Lee,David Becker,Marc Laurent,James Lu,Clinton R. Paden,Suxiang Tong,Duncan MacCannell, |
| EPI_ISL_967887 | TGen North | TGen North | "Jolene Bowers, Megan Folkerts, Chris French, Hayley Yaglom, Ashlyn Pfeiffer, Darrin Lemmer, Dave Engelthaler, The Arizona COVID Genomics Union (ACGU)" |
| EPI_ISL_967891, EPI_ISL_967899, EPI_ISL_967900, EPI_ISL_967907, EPI_ISL_967926, EPI_ISL_967932, EPI_ISL_967933, EPI_ISL_967934, EPI_ISL_967942, EPI_ISL_967943, EPI_ISL_967965, EPI_ISL_967969, EPI_ISL_967978 | | | |
| see above | TGen North | Sonora Quest Laboratories | "Jolene Bowers, Megan Folkerts, Chris French, Hayley Yaglom, Ashlyn Pfeiffer, Darrin Lemmer, Dave Engelthaler, The Arizona COVID Genomics Union (ACGU)" |
| EPI_ISL_967979 | TGen North | TGen North | "Jolene Bowers, Megan Folkerts, Chris French, Hayley Yaglom, Ashlyn Pfeiffer, Darrin Lemmer, Dave Engelthaler, The Arizona COVID Genomics Union (ACGU)" |
| EPI_ISL_967980, EPI_ISL_967989, EPI_ISL_967993, EPI_ISL_967994, EPI_ISL_968000, EPI_ISL_968004, EPI_ISL_968014, EPI_ISL_968023, EPI_ISL_968024, EPI_ISL_968028, EPI_ISL_968031, EPI_ISL_968033, EPI_ISL_968036, EPI_ISL_968052, EPI_ISL_968056 | | | |
| see above | TGen North | Sonora Quest Laboratories | "Jolene Bowers, Megan Folkerts, Chris French, Hayley Yaglom, Ashlyn Pfeiffer, Darrin Lemmer, Dave Engelthaler, The Arizona COVID Genomics Union (ACGU)" |
| EPI_ISL_968057 | TGen North | TGen North | "Jolene Bowers, Megan Folkerts, Chris French, Hayley Yaglom, Ashlyn Pfeiffer, Darrin Lemmer, Dave Engelthaler, The Arizona COVID Genomics Union (ACGU)" |
| EPI_ISL_968062 | TGen North | Sonora Quest Laboratories | "Jolene Bowers, Megan Folkerts, Chris French, Hayley Yaglom, Ashlyn Pfeiffer, Darrin Lemmer, Dave Engelthaler, The Arizona COVID Genomics Union (ACGU)" |
| EPI_ISL_968065 | TGen North | TGen North | "Jolene Bowers, Megan Folkerts, Chris French, Hayley Yaglom, Ashlyn Pfeiffer, Darrin Lemmer, Dave Engelthaler, The Arizona COVID Genomics Union (ACGU)" |
| EPI_ISL_968998, EPI_ISL_968999, EPI_ISL_969003, EPI_ISL_969050, EPI_ISL_969051 | KEMRI-Wellcome Trust Research Programme/KEMRI-CGMR-C Kilifi | KEMRI-Wellcome Trust Research Programme/KEMRI-CGMR-C Kilifi | Githinji et al |
| EPI_ISL_977102, EPI_ISL_977103, EPI_ISL_977104, EPI_ISL_977105, EPI_ISL_977106, EPI_ISL_977107, EPI_ISL_977108, EPI_ISL_977109, EPI_ISL_977110, EPI_ISL_977111, EPI_ISL_977115, EPI_ISL_977116 | | | |
| see above | Massachusetts General Hospital | Infectious Disease Program, Broad Institute of Harvard and MIT | Lemieux,J.E., Siddle,K.J., Shaw,B., Adams,G., Pierce,V., Turbett,S., Anahtar,M., Branda,J., Slater,D., Harris,J., Lin,A.E., Gladden-Young,A., Lagerborg,K., Rudy,M., DeRuff,K., Carter,A., Normandin,E., Bauer,M., Reilly,S., Tomkins-Tinch,C., Loreth,C., Chaluvadi,S., Neumann,A., Cusick,C., Chapman,S.B., Gnirke,A., Flowers,K., Cerrato,F., Birren,B.W., Gallagher,G., Smole,S., Park,D.J., MacInnis,B.L., Ryan,E., LaRocque,R., Rosenberg,E. and Sabeti,P.C. |
| EPI_ISL_977131, EPI_ISL_977132, EPI_ISL_977138, EPI_ISL_977139, EPI_ISL_977140, EPI_ISL_977141, EPI_ISL_977142, EPI_ISL_977143 | Flow Health | Infectious Disease Program, Broad Institute of Harvard and MIT | Lemieux,J.E., Siddle,K.J., Adams,G., Gladden-Young,A., Lagerborg,K., Rudy,M., DeRuff,K., Carter,A., Normandin,E., Bauer,M., Reilly,S., Tomkins-Tinch,C., Loreth,C., Chaluvadi,S., Birren,B.W., Gallagher,G., Smole,S., Park,D.J., MacInnis,B.L., and Sabeti,P.C. |
| EPI_ISL_977583 | Caribbean Public Health Agency | Carrington Lab, Department of PreClinical Sciences | Nikita S. D. Sahadeo, Arianne Brown-Jordan, Vernie Ramkissoon, Sarah Hill, Naresh Nandram, Avery Hinds, Dr. Sharon Belmar-George, Jerome Foster, Stanley Giddings, Karla Georges, Marsha Ivey, Rahul Naidu, Risha Singh, SueMin Nathaniel, Rajini Haraksingh, Jaya Jayaraman, Chinna Chinnadurai, Adesh Ramsubhag, Nuno Faria, Oliver Pybus, Christopher Oura, Gabriel Escobar, Christine V. F. Carrington |
| EPI_ISL_977597 | Servicio Murciano de Salud | Instituto de Salud Carlos III | Vázquez, S. Iglesias-Caballero, M. Sandonís,V. Camarero, S. Pozo, F. Casas, I. Jiménez, P. Zaballos, A. Monzón, S. Varona, S. Cuesta, I. Blázquez, A. |
| EPI_ISL_977654 | Michigan Department of Health and Human Services, Bureau of Laboratories | Michigan Department of Health and Human Services, Bureau of Laboratories | Blankenship HM, Riner D, Soehnlen MK |
| EPI_ISL_977666 | NYU Langone Health | Departments of Pathology and Medicine, New York University School of Medicine | Adriana Heguy, Dacia Dimartino, Emily Guzman, Christian Marier, Peter Meyn, Sitharam Ramaswami, Gael Westby, Paul Zappile, Yutong Zhang, Paolo Cotzia, Guiqing Wang |
| EPI_ISL_977918, EPI_ISL_977919, EPI_ISL_977920, EPI_ISL_977921, EPI_ISL_977922, EPI_ISL_977923, EPI_ISL_977924, EPI_ISL_977925, EPI_ISL_977926, EPI_ISL_977927, EPI_ISL_977928, EPI_ISL_977929, EPI_ISL_977930, EPI_ISL_977931, EPI_ISL_977932, EPI_ISL_977933, EPI_ISL_977934, EPI_ISL_977935, EPI_ISL_977936, EPI_ISL_977937, EPI_ISL_977938, EPI_ISL_977939, EPI_ISL_977979 | | | |
| see above | Chiu Laboratory, University of California, San Francisco | Chiu Laboratory, University of California, San Francisco | Charles Chiu, Xianding (Wayne) Deng, Candace Wang, Venice Servellita, Jill Hacker, Debra Wadford |
| EPI_ISL_978222, EPI_ISL_978223, EPI_ISL_978224, EPI_ISL_978225 | Virginia Division of Consolidated Laboratory Services | Virginia Division of Consolidated Laboratory Services | Virginia DCLS |
| EPI_ISL_978280, EPI_ISL_978281 | Texas Department of State Health Services | Texas Department of State Health Services | Bonnie Oh, Anita Pokharel, James Daniel Bonser, Myong Koag, Chung Wang, Rachel Lee, Grace Kubin, Rashmi Tuladhar, Mayela Pedrueza, Maliha Rahman, Jenny Zhang |
| EPI_ISL_978391, EPI_ISL_978392, EPI_ISL_978393, EPI_ISL_978394, EPI_ISL_978395, EPI_ISL_978462, EPI_ISL_978463, EPI_ISL_978464, EPI_ISL_978465, EPI_ISL_978466, EPI_ISL_978467 | | | |
| see above | Arizona State Public Health Laboratory | Arizona State Public Health Laboratory | Trung Huynh, Jessica Escobar, Katherine Fullerton, Nobuko Fukushima, Stacy White, Linda Getsinger, Victor Waddell |
| EPI_ISL_978867, EPI_ISL_978930 | Centre for Dengue Research and AICBU, Department of Immunology and Molecular Medicine | Centre for Dengue Research and AICBU, Department of Immunology and Molecular Medicine | Chandima Jeewandara, Deshni Jayathilaka, Dinuka Ariyaratne, Tibutius Thanesh Pramanayagam, Diyanath Ranasinghe, Laksiri Gomes, Gathsaurie Neelika Malavige |
| EPI_ISL_979103, EPI_ISL_979104, EPI_ISL_979105, EPI_ISL_979106, EPI_ISL_979107, EPI_ISL_979110, EPI_ISL_979112, EPI_ISL_979115, EPI_ISL_979116, EPI_ISL_979117, EPI_ISL_979118, EPI_ISL_979119, EPI_ISL_979120, EPI_ISL_979121, EPI_ISL_979122, EPI_ISL_979124, EPI_ISL_979125, EPI_ISL_979126, EPI_ISL_979129, EPI_ISL_979130, EPI_ISL_979131, EPI_ISL_979132, EPI_ISL_979135 | | | |
| see above | Santa Clara County Public Health Laboratory | Chan-Zuckerberg Biohub | CZB Cliahub Consortium |
| EPI_ISL_979251, EPI_ISL_979268 | Institute of Microbiology and Immunology, Faculty of Medicine, University of Ljubljana | Institute of Microbiology and Immunology, Faculty of Medicine, University of Ljubljana | Samo Zakotnik, Tomaž Mark Zorec, Matic Brvar, Doroteja Vlaj, Patricija Pozvek,Špela Pleh, Miša Korva, Mario Poljak, Tatjana Avši - Županc |
| EPI_ISL_979315 | Cadham Provincial laboratory | National Microbiology Laboratory (NML) | Anna Majer, Shari Tyson, Grace Seo, Philip Mabon, Elsie Grudeski, Rhiannon Huzarewich, Russell Mandes, Anneliese Landgraff, Jennifer Tanner, Natalie Knox, Morag Graham, Gary Van Domselaar, Paul Van Caeseele, Jared Bullard, David Alexander, Kerry Dust, Nathalie Bastien, Yan Li, Timothy Booth, Darian Hole, Madison Chapel, Kirsten Biggar, CanCOGeN's metadata curation team, Public Health Agency of Canada CanCOGeN team |
| EPI_ISL_979453 | University of Iowa Hospitals and Clinics | State Hygienic Laboratory at the University of Iowa | Valerie Reeb, Wes Hottel, Alankar Kampoowale |
| EPI_ISL_980905, EPI_ISL_980933, EPI_ISL_980938, EPI_ISL_980939, EPI_ISL_980982, EPI_ISL_980983 | Innovative Genomics Institute, UC Berkeley | Innovative Genomics Institute, UC Berkeley | Stacia Wyman, Haridha Shivram, Phil Frankino, Liana Lareau |
| EPI_ISL_981371 | AZ Klina | AZ Klina | Dr. C. Vael |
| EPI_ISL_981967, EPI_ISL_981970, EPI_ISL_981974 | Microbiology Service, Hospital Universitario Clinico San Cecilio, Granada | Microbiology Service, Hospital Universitario Clinico San Cecilio, Granada | Adolfo de Salazar, Natalia Chueca, Laura Viñuela, Ana Fuentes, Federico García |
| EPI_ISL_982081 | TGen North | Sonora Quest Laboratories | "Jolene Bowers, Megan Folkerts, Chris French, Hayley Yaglom, Ashlyn Pfeiffer, Darrin Lemmer, Dave Engelthaler, The Arizona COVID Genomics Union (ACGU)" |

| | | | |
|---|---|---|---|
| EPI_ISL_982259, EPI_ISL_982268, EPI_ISL_982269 | Lab voor klinische biologie | Lab voor klinische biologie | Hannelore Hamerlinck, Marija Janevska, Bruno Verhasselt |
| EPI_ISL_982420, EPI_ISL_982421, EPI_ISL_982422, EPI_ISL_982423, EPI_ISL_982424 | M Health Fairview | Minnesota Department of Health, Public Health Laboratory | Alexandra Lorentz, Jacob Garfin, Matt Plumb, and Xiong Wang |
| EPI_ISL_982538, EPI_ISL_982553, EPI_ISL_982579, EPI_ISL_982611, EPI_ISL_982625, EPI_ISL_982643, EPI_ISL_982675, EPI_ISL_982679, EPI_ISL_982680, EPI_ISL_982730, EPI_ISL_982734, EPI_ISL_982735, EPI_ISL_982747, EPI_ISL_982761 | | | |
| see above | US Air Force School of Aerospace Medicine | US Air Force School of Aerospace Medicine | Anthony Fries, Jennifer Meyer, William Gruner, William Buggele, Amanda Javorina, Sarah Purves, Clarise Starr, Elizabeth Macias |
| EPI_ISL_982860 | Botswana Harvard HIV Reference Laboratory | Botswana Harvard HIV Reference Laboratory | Sikhulile Moyo, Dorcas Maruapula, Wonderful T. Choga, Botshelo Radibe, Boitumelo Zuze, David Lawrence, Roger Shapiro, Shahin Lockman, Mosepele Mosepele, Joseph Makhema, Simani Gaseitsiwe |
| EPI_ISL_983083 | Gravity Diagnostics | Kentucky State Public Health Lab | Stephanie Lunn, Karim George, Joshua Tobias, William Grooms, Vaneet Arora, Matthew Johnson, Rachel Zinner, Rhonda Lucas |
