## Supplementary material for "Detection and characterization of the SARS-CoV-2 lineage B.1.526 in New York": Supp. Table 3 GISAID Acknowledgment part 2: gisaid_hcov-19_acknowledgement_table_2021_02_13_010-13.pdf

We gratefully acknowledge the following Authors from the Originating laboratories responsible for obtaining the specimens, as well as the Submitting laboratories where the genome data were generated and shared via GISAID, on which this research is based.

All Submitters of data may be contacted directly via www.gisaid.org

Authors are sorted alphabetically.

| Accession ID | Originating Laboratory | Submitting Laboratory | Authors |
|---|---|---|---|
| EPI_ISL_857473 | National Public Health Laboratory, National Centre for Infectious Diseases | National Public Health Laboratory, National Centre for Infectious Diseases | Tze Minn Mak, Sophie Octavia, Zhenyang Zhou, Lin Cui, Raymond Tzer Pin Lin |
| EPI_ISL_860727, EPI_ISL_860728 | University of Michigan Clinical Microbiology Laboratory | Lauring Lab, University of Michigan, Department of Microbiology and Immunology | Valesano |
| EPI_ISL_861035, EPI_ISL_861055, EPI_ISL_861070 | Johns Hopkins Hospital Department of Pathology | Johns Hopkins Hospital Department of Pathology | C. Paul Morris, Chun Huai Luo, Adannaya Amadi, Nicholas Gallagher, Heba H. Mostafa |
| EPI_ISL_861488 | Cerballiance Côte d'Azur | CERBA LAB | Prots.L; Pichon R; Barrieu-Moussat S. |
| EPI_ISL_861489 | Cerballiance Côte d'Azur | CERBA LAB | Adjtoutah Z; Mangiarotti T; Prots L. |
| EPI_ISL_861490 | Cerballiance Côte d'Azur | CERBA LAB | De La Chapelle A; Prots.L; |
| EPI_ISL_861491 | Cerballiance Côte d'Azur | CERBA LAB | Chobert F; Prots.L |
| EPI_ISL_861492 | Cerballiance Côte d'Azur | CERBA LAB | Barrieu-Moussat S;Adoul O;Boudjahem Z; Roman E; Chaudet L; Martin L; Aankour N; Jaugin P; Gastaud S; Prots L. |
| EPI_ISL_861493 | Cerballiance Côte d'Azur | CERBA LAB | Marius M; Barsanti M; Maltese S; Francois J; Sinsoulieu T; Van Der Rhyn I; Morello V; Marcadal M; Gandillet A; Prots L. |
| EPI_ISL_861494 | Cerballiance Côte d'Azur | CERBA LAB | Prots L. Raimondi V. |
| EPI_ISL_861495 | Cerballiance Côte d'Azur | CERBA LAB | Pichon R. Quintle S; Sanza O; Mangiarotti T; Viltard B; Prots L. |
| EPI_ISL_861497 | Cerballiance Centre Est | CERBA LAB | Chanard C; Gazzano V; Munier C; Prots L. |
| EPI_ISL_861500, EPI_ISL_861501, EPI_ISL_861502 | Cerballiance Centre Est | CERBA LAB | Chanard E; Gazzano V; Munier C; Prots L. |
| EPI_ISL_861736, EPI_ISL_861767 | Yale Pathology Lab | Grubaugh Lab - Yale School of Public Health | Tara Alpert, Joseph Fauver, Chen Liu, Pei Hui, Jianhui Wang, Susan Bell and Han Zhou, Anderson Brito, Mallery Breban, Anne Wyllie, Chantal Vogels, Mary Petrone, Chaney Kalinich, Isabel Ott, Arnau Casanovas, Catherine Muenker, Adam Moore, Alice Lu, Maria Tokuyama, Patrick Wong, Peiwen Lu, Saad Omer, Richard Martinello, Allison Nelson, Shelli Farhadian, Akiko Iwasaki, Charlese Dela Cruz, Albert Ko, Nathan Grubaugh |
| EPI_ISL_862048 | Respiratory Virus Unit, National Infection Service, Public Health England | COVID-19 Genomics UK (COG-UK) Consortium | PHE Covid Sequencing Team |
| EPI_ISL_862049 | Department of Microbiology, National Influenza Reference Laboratory for Northern Greece | Department of Microbiology, National Influenza Reference Laboratory for Northern Greece | Anna Papa, Styliani Pappa |
| EPI_ISL_862188, EPI_ISL_862189, EPI_ISL_862194, EPI_ISL_862195, EPI_ISL_862196, EPI_ISL_862197, EPI_ISL_862198, EPI_ISL_862199, EPI_ISL_862200, EPI_ISL_862201, EPI_ISL_862202, EPI_ISL_862203, EPI_ISL_862204, EPI_ISL_862205, EPI_ISL_862206, EPI_ISL_862207, EPI_ISL_862208, EPI_ISL_862209, EPI_ISL_862210, EPI_ISL_862211, EPI_ISL_862212, EPI_ISL_862213, EPI_ISL_862214, EPI_ISL_862215, EPI_ISL_862216, EPI_ISL_862217, EPI_ISL_862218, EPI_ISL_862219, EPI_ISL_862220, EPI_ISL_862221, EPI_ISL_862222, EPI_ISL_862223, EPI_ISL_862224, EPI_ISL_862225, EPI_ISL_862226, EPI_ISL_862227, EPI_ISL_862228, EPI_ISL_862229, EPI_ISL_862230, EPI_ISL_862231, EPI_ISL_862232, EPI_ISL_862233, EPI_ISL_862234, EPI_ISL_862235, EPI_ISL_862236, EPI_ISL_862237, EPI_ISL_862238, EPI_ISL_862239, EPI_ISL_862240, EPI_ISL_862241, EPI_ISL_862242, EPI_ISL_862243, EPI_ISL_862244, EPI_ISL_862245, EPI_ISL_862246, EPI_ISL_862247, EPI_ISL_862248, EPI_ISL_862249, EPI_ISL_862250, EPI_ISL_862252, EPI_ISL_862253, EPI_ISL_862254, EPI_ISL_862255, EPI_ISL_862256, EPI_ISL_862257, EPI_ISL_862258, EPI_ISL_862265, EPI_ISL_862266 | | | |
| see above | Respiratory Virus Unit, National Infection Service, Public Health England | COVID-19 Genomics UK (COG-UK) Consortium | PHE Covid Sequencing Team |
| EPI_ISL_862716, EPI_ISL_862717, EPI_ISL_862720, EPI_ISL_862721 | Centre for Dengue Research and AICBU, Department of Immunology and Molecular Medicine | Centre for Dengue Research and AICBU, Department of Immunology and Molecular Medicine | Chandima Jeewandara, Deshni Jayathilaka, Dinuka Ariyaratne, Diyanath Ranasinghe, Laksiri Gomes, Gathsaurie Neelika Malavige |
| EPI_ISL_862724, EPI_ISL_862726, EPI_ISL_862727 | Centre for Dengue Research and AICBU, Department of Immunology and Molecular Medicine | Centre for Dengue Research and AICBU, Department of Immunology and Molecular Medicine | Chandima Jeewandara, Deshni Jayathilaka, Dinuka Ariyaratne, Tibutius Thanesh Pramanayagam, Diyanath Ranasinghe, Laksiri Gomes, Gathsaurie Neelika Malavige |
| EPI_ISL_862816, EPI_ISL_862817, EPI_ISL_862818, EPI_ISL_862820, EPI_ISL_862821, EPI_ISL_862822, EPI_ISL_862823, EPI_ISL_862824, EPI_ISL_862826, EPI_ISL_862827, EPI_ISL_862828, EPI_ISL_862829, EPI_ISL_862830 | | | |
| see above | National Public Health Laboratory, National Centre for Infectious Diseases | National Public Health Laboratory, National Centre for Infectious Diseases | Tze Minn Mak, Zhenyang Zhou, Lin Cui, Raymond Tzer Pin Lin |
| EPI_ISL_862835, EPI_ISL_862836 | National Institute of Infectious Diseases-Prof. Dr. Matei Bals Molecular Diagnostics Laboratory | National Institute of Infectious Diseases-Prof. Dr. Matei Bals Molecular Diagnostics Laboratory | Leontina Banica, Marius Surleac, Corina Casangiu, Petre Milu, Andreea Tudor, Simona Paraschiv, Dan Otelea |
| EPI_ISL_864579 | Institute of Medical Microbiology and Hospital Hygiene | Institute of Medical Microbiology and Hospital Hygiene | Prof. Dr. Achim Kaasch, Aljoscha Tersteegen |
| EPI_ISL_865631, EPI_ISL_865634, EPI_ISL_865638, EPI_ISL_865640, EPI_ISL_865648, EPI_ISL_865658, EPI_ISL_865659, EPI_ISL_865660, EPI_ISL_865661, EPI_ISL_865663, EPI_ISL_865664, EPI_ISL_865665, EPI_ISL_865666, EPI_ISL_865667, EPI_ISL_865668, EPI_ISL_865671, EPI_ISL_865672, EPI_ISL_865673, EPI_ISL_865674, EPI_ISL_865675, EPI_ISL_865676, EPI_ISL_865677, EPI_ISL_865678, EPI_ISL_865679, EPI_ISL_865680, EPI_ISL_865681, EPI_ISL_865971, EPI_ISL_865972, EPI_ISL_865973, EPI_ISL_865974, EPI_ISL_865981, EPI_ISL_865982, EPI_ISL_865983, EPI_ISL_865984, EPI_ISL_865988, EPI_ISL_865989, EPI_ISL_865990, EPI_ISL_865992, EPI_ISL_865997, EPI_ISL_865998, EPI_ISL_865999, EPI_ISL_866004, EPI_ISL_866005, EPI_ISL_866006, EPI_ISL_866010, EPI_ISL_866016, EPI_ISL_866018, EPI_ISL_866022, EPI_ISL_866023, EPI_ISL_866025, EPI_ISL_866026, EPI_ISL_866029, EPI_ISL_866038, EPI_ISL_866040, EPI_ISL_866043, EPI_ISL_866044 | | | |
| see above | University College London, Great Ormond Street Hospital for Children NHS Foundation Trust, Imperial College Healthcare NHS Trust | COVID-19 Genomics UK (COG-UK) Consortium | Sergi Castellano, Rachel Williams, Mark Kristiansen, Paola Resende Silva, Sunando Roy, Tony Brooks, Helena Tutill, Paola Niola, Patricia Dyal, Charlotte Williams, Leysa Forrest, Yasmin Panchbhaya, Jacqueline Findlay, Samuel Weeks, Julianne Brown, Kathryn Harris, Paul Randell, James Price, Alison Holmes, Judith Breuer |
| EPI_ISL_866619, EPI_ISL_866793 | Respiratory Virus Unit, National Infection Service, Public Health England | COVID-19 Genomics UK (COG-UK) Consortium | PHE Covid Sequencing Team |
| EPI_ISL_866962, EPI_ISL_866963, EPI_ISL_866964, EPI_ISL_866965, EPI_ISL_866966, EPI_ISL_866967, EPI_ISL_866968, EPI_ISL_866969, EPI_ISL_866970 | Queens Medical Centre, Clinical Microbiology Department / DeepSeq Nottingham | COVID-19 Genomics UK (COG-UK) Consortium | Gemma Clark, Wendy Smith, Manjinder Khakh, Vicki M Fleming, Michelle M Lister, Hannah Howson-Wells, Jonathan Ball, Patrick McClure, Joseph Chappell, Theocharis Tsoleridis, Nadine Holmes, Matthew Carlisle, Christopher Moore, Fei Sang, Johnny Debebe, Victoria Wright, Matthew Loose |
| EPI_ISL_866985, EPI_ISL_867156 | Respiratory Virus Unit, National Infection Service, Public Health England | COVID-19 Genomics UK (COG-UK) Consortium | PHE Covid Sequencing Team |
| EPI_ISL_868257 | Centre for Enzyme Innovation, University of Portsmouth / Translational Research Laboratory, Portsmouth Hospitals NHS Trust | COVID-19 Genomics UK (COG-UK) Consortium | Angela Beckett,Yann Bourgeois,Garry Scarlett,Sharon Glaysher,Scott Elliott,Kelly Bicknell,Robert Impey,Allyson Lloyd,Sarah Wyllie,Ethan Butcher,Anoop Chauhan,Samuel Robson |
| EPI_ISL_868369, EPI_ISL_868374, EPI_ISL_868375, EPI_ISL_868376, EPI_ISL_868380, EPI_ISL_868383, EPI_ISL_868384, EPI_ISL_868394, EPI_ISL_868396, EPI_ISL_868404, EPI_ISL_868405, EPI_ISL_868410, EPI_ISL_868412, EPI_ISL_868435, EPI_ISL_868457, EPI_ISL_868460, EPI_ISL_868462, EPI_ISL_868473, | | | |

| | | | |
|---|---|---|---|
| EPI_ISL_868477, EPI_ISL_868490, EPI_ISL_868494, EPI_ISL_868506, EPI_ISL_868519, EPI_ISL_868528, EPI_ISL_868550, EPI_ISL_868551, EPI_ISL_868553, EPI_ISL_868555, EPI_ISL_868571, EPI_ISL_868583, EPI_ISL_868585, EPI_ISL_868587, EPI_ISL_868600, EPI_ISL_868604, EPI_ISL_868605, EPI_ISL_868610, EPI_ISL_868616, EPI_ISL_868619, EPI_ISL_868624, EPI_ISL_868633, EPI_ISL_868636, EPI_ISL_868637, EPI_ISL_868640, EPI_ISL_868641, EPI_ISL_868650, EPI_ISL_868672, EPI_ISL_868676, EPI_ISL_868682, EPI_ISL_868685, EPI_ISL_868694, EPI_ISL_868709, EPI_ISL_868711, EPI_ISL_868712 | | | |
| see above | Virology Department, Sheffield Teaching Hospitals NHS Foundation Trust/Department of Infection, Immunity and Cardiovascular Disease, The Medical School, University of Sheffield | COVID-19 Genomics UK (COG-UK) Consortium | Thushan de Silva, Matthew Parker, Nikki Smith, Adri Angyal, Rebecca Brown, Luke Green, Rachel Tucker, Paul Parsons, Danielle Groves, Katie Johnson, Laura Carrilero, Alex Keeley, Dave Partridge, Matthew Wyles, Benjamin Lindsey, Mehmet Yavuz, Mohammad Raza, Cariad Evans |
| EPI_ISL_869135 | Charité Universitätsmedizin Berlin, Institut für Virologie/Labor Berlin | Charité Universitätsmedizin Berlin, Institut für Virologie | Victor M Corman, Barbara Mühlemann, Jörn Beheim-Schwarzbach, Tobias Bleicker, Julia Tesch, Talitha Veith, Julia Schneider, Terry Jones, Christian Drosten |
| EPI_ISL_869265, EPI_ISL_869266, EPI_ISL_869267, EPI_ISL_869268, EPI_ISL_869269, EPI_ISL_869270, EPI_ISL_869271, EPI_ISL_869272, EPI_ISL_869273, EPI_ISL_869274, EPI_ISL_869275, EPI_ISL_869276, EPI_ISL_869277, EPI_ISL_869278, EPI_ISL_869279, EPI_ISL_869280, EPI_ISL_869281, EPI_ISL_869282, EPI_ISL_869283, EPI_ISL_869284, EPI_ISL_869285, EPI_ISL_869286, EPI_ISL_869287, EPI_ISL_869288, EPI_ISL_869289, EPI_ISL_869290, EPI_ISL_869291, EPI_ISL_869292, EPI_ISL_869293, EPI_ISL_869294, EPI_ISL_869295, EPI_ISL_869296, EPI_ISL_869297, EPI_ISL_869298, EPI_ISL_869299, EPI_ISL_869300, EPI_ISL_869301, EPI_ISL_869302, EPI_ISL_869303, EPI_ISL_869304, EPI_ISL_869305, EPI_ISL_869306, EPI_ISL_869307, EPI_ISL_869308, EPI_ISL_869309, EPI_ISL_869310, EPI_ISL_869311, EPI_ISL_869312, EPI_ISL_869313, EPI_ISL_869314, EPI_ISL_869315, EPI_ISL_869316, EPI_ISL_869317, EPI_ISL_869318, EPI_ISL_869319, EPI_ISL_869320, EPI_ISL_869321, EPI_ISL_869322, EPI_ISL_869323, EPI_ISL_869324, EPI_ISL_869325, EPI_ISL_869326, EPI_ISL_869327, EPI_ISL_869328, EPI_ISL_869329, EPI_ISL_869330, EPI_ISL_869331, EPI_ISL_869332, EPI_ISL_869333, EPI_ISL_869334, EPI_ISL_869335, EPI_ISL_869336, EPI_ISL_869337, EPI_ISL_869338, EPI_ISL_869339, EPI_ISL_869340, EPI_ISL_869341, EPI_ISL_869342, EPI_ISL_869343, EPI_ISL_869344, EPI_ISL_869345, EPI_ISL_869346, EPI_ISL_869347, EPI_ISL_869348, EPI_ISL_869349, EPI_ISL_869350, EPI_ISL_869351, EPI_ISL_869352, EPI_ISL_869353, EPI_ISL_869354, EPI_ISL_869355, EPI_ISL_869356, EPI_ISL_869357, EPI_ISL_869358, EPI_ISL_869359, EPI_ISL_869360, EPI_ISL_869361, EPI_ISL_869362, EPI_ISL_869363, EPI_ISL_869364, EPI_ISL_869365, EPI_ISL_869366, EPI_ISL_869367, EPI_ISL_869368, EPI_ISL_869369, EPI_ISL_869370, EPI_ISL_869371, EPI_ISL_869372, EPI_ISL_869373, EPI_ISL_869374, EPI_ISL_869375, EPI_ISL_869376, EPI_ISL_869377, EPI_ISL_869378, EPI_ISL_869379, EPI_ISL_869380, EPI_ISL_869381, EPI_ISL_869382, EPI_ISL_869383, EPI_ISL_869384, EPI_ISL_869385, EPI_ISL_869386, EPI_ISL_869387, EPI_ISL_869388, EPI_ISL_869389, EPI_ISL_869390, EPI_ISL_869391, EPI_ISL_869392, EPI_ISL_869393, EPI_ISL_869394, EPI_ISL_869395, EPI_ISL_869396, EPI_ISL_869397, EPI_ISL_869398, EPI_ISL_869399, EPI_ISL_869400, EPI_ISL_869401, EPI_ISL_869402, EPI_ISL_869403, EPI_ISL_869404, EPI_ISL_869405, EPI_ISL_869406, EPI_ISL_869407, EPI_ISL_869408, EPI_ISL_869409, EPI_ISL_869410, EPI_ISL_869411, EPI_ISL_869412, EPI_ISL_869413, EPI_ISL_869414, EPI_ISL_869415, EPI_ISL_869416, EPI_ISL_869417, EPI_ISL_869418, EPI_ISL_869419, EPI_ISL_869420, EPI_ISL_869421, EPI_ISL_869422, EPI_ISL_869423, EPI_ISL_869424, EPI_ISL_869425, EPI_ISL_869426, EPI_ISL_869427, EPI_ISL_869428, EPI_ISL_869429, EPI_ISL_869430, EPI_ISL_869431, EPI_ISL_869432, EPI_ISL_869433, EPI_ISL_869434, EPI_ISL_869435, EPI_ISL_869436, EPI_ISL_869437, EPI_ISL_869438, EPI_ISL_869439, EPI_ISL_869440, EPI_ISL_869441, EPI_ISL_869442, EPI_ISL_869443, EPI_ISL_869444, EPI_ISL_869445, EPI_ISL_869446, EPI_ISL_869447, EPI_ISL_869448, EPI_ISL_869449, EPI_ISL_869450, EPI_ISL_869451, EPI_ISL_869452, EPI_ISL_869453, EPI_ISL_869454, EPI_ISL_869455, EPI_ISL_869456, EPI_ISL_869457, EPI_ISL_869458, EPI_ISL_869459, EPI_ISL_869460, EPI_ISL_869461, EPI_ISL_869462, EPI_ISL_869463, EPI_ISL_869464, EPI_ISL_869465, EPI_ISL_869466, EPI_ISL_869467, EPI_ISL_869468, EPI_ISL_869469, EPI_ISL_869470, EPI_ISL_869471, EPI_ISL_869472, EPI_ISL_869473, EPI_ISL_869474, EPI_ISL_869475, EPI_ISL_869476, EPI_ISL_869477, EPI_ISL_869478, EPI_ISL_869479, EPI_ISL_869480, EPI_ISL_869481, EPI_ISL_869482, EPI_ISL_869483, EPI_ISL_869484, EPI_ISL_869485, EPI_ISL_869486, EPI_ISL_869487, EPI_ISL_869488, EPI_ISL_869489, EPI_ISL_869490, EPI_ISL_869491, EPI_ISL_869492, EPI_ISL_869493, EPI_ISL_869494, EPI_ISL_869495, EPI_ISL_869496, EPI_ISL_869497, EPI_ISL_869498, EPI_ISL_869499, EPI_ISL_869500, EPI_ISL_869501, EPI_ISL_869502, EPI_ISL_869503, EPI_ISL_869504, EPI_ISL_869505, EPI_ISL_869506, EPI_ISL_869507, EPI_ISL_869508, EPI_ISL_869509, EPI_ISL_869510, EPI_ISL_869511, EPI_ISL_869512, EPI_ISL_869513, EPI_ISL_869514, EPI_ISL_869515, EPI_ISL_869516, EPI_ISL_869517, EPI_ISL_869518, EPI_ISL_869519, EPI_ISL_869520, EPI_ISL_869521, EPI_ISL_869522, EPI_ISL_869523, EPI_ISL_869524, EPI_ISL_869525, EPI_ISL_869526, EPI_ISL_869527, EPI_ISL_869528, EPI_ISL_869529, EPI_ISL_869530, EPI_ISL_869531, EPI_ISL_869532, EPI_ISL_869533, EPI_ISL_869534, EPI_ISL_869535, EPI_ISL_869536, EPI_ISL_869537, EPI_ISL_869538, EPI_ISL_869539, EPI_ISL_869540, EPI_ISL_869541, EPI_ISL_869542, EPI_ISL_869543, EPI_ISL_869544, EPI_ISL_869545, EPI_ISL_869546, EPI_ISL_869547, EPI_ISL_869548, EPI_ISL_869549, EPI_ISL_869550, EPI_ISL_869551, EPI_ISL_869552, EPI_ISL_869553, EPI_ISL_869554, EPI_ISL_869555, EPI_ISL_869556, EPI_ISL_869557, EPI_ISL_869558, EPI_ISL_869559, EPI_ISL_869560, EPI_ISL_869561, EPI_ISL_869562, EPI_ISL_869563, EPI_ISL_869564, EPI_ISL_869565, EPI_ISL_869566, EPI_ISL_869567, EPI_ISL_869568, EPI_ISL_869569, EPI_ISL_869570, EPI_ISL_869571, EPI_ISL_869572, EPI_ISL_869573, EPI_ISL_869574, EPI_ISL_869575, EPI_ISL_869576, EPI_ISL_869577, EPI_ISL_869578, EPI_ISL_869579, EPI_ISL_869580, EPI_ISL_869581, EPI_ISL_869582, EPI_ISL_869583, EPI_ISL_869584, EPI_ISL_869585, EPI_ISL_869586, EPI_ISL_869587, EPI_ISL_869588, EPI_ISL_869589, EPI_ISL_869590, EPI_ISL_869591, EPI_ISL_869592, EPI_ISL_869593, EPI_ISL_869594, EPI_ISL_869595, EPI_ISL_869596, EPI_ISL_869597, EPI_ISL_869598, EPI_ISL_869599, EPI_ISL_869600, EPI_ISL_869601, EPI_ISL_869602, EPI_ISL_869603, EPI_ISL_869604, EPI_ISL_869605, EPI_ISL_869606, EPI_ISL_869607, EPI_ISL_869608, EPI_ISL_869609, EPI_ISL_869610, EPI_ISL_869611, EPI_ISL_869612, EPI_ISL_869613, EPI_ISL_869614, EPI_ISL_869615, EPI_ISL_869616, EPI_ISL_869617, EPI_ISL_869618, EPI_ISL_869619, EPI_ISL_869620, EPI_ISL_869621, EPI_ISL_869622, EPI_ISL_869623, EPI_ISL_869624, EPI_ISL_869625, EPI_ISL_869626, EPI_ISL_869627, EPI_ISL_869628, EPI_ISL_869629, EPI_ISL_869630, EPI_ISL_869631, EPI_ISL_869632, EPI_ISL_869633, EPI_ISL_869634, EPI_ISL_869635, EPI_ISL_869636, EPI_ISL_869637, EPI_ISL_869638, EPI_ISL_869639, EPI_ISL_869640, EPI_ISL_869641, EPI_ISL_869642, EPI_ISL_869643 | | | |
| see above | Department of Virus and Microbiological Special Diagnostics, Statens Serum Institut, Copenhagen, Denmark | Aalborg University | Danish Covid-19 Genome Consortium |
| EPI_ISL_870315, EPI_ISL_870317, EPI_ISL_870319 | University of Michigan Clinical Microbiology Laboratory | Lauring Lab, University of Michigan, Department of Microbiology and Immunology | Valesano |
| EPI_ISL_871872 | Botswana Harvard HIV Reference Laboratory | Botswana Harvard HIV Reference Laboratory | Sikhulile Moyo, Dorcas Maruapula, Wonderful Choga, Botshelo Radibe, Boitumelo Zuze, David Lawrence, Roger Shapiro, Shahin Lockman, Mosepele Mosepele, Joseph, Makhema, Simani Gaseitsiwe |
| EPI_ISL_872052, EPI_ISL_872053, EPI_ISL_872054, EPI_ISL_872055, EPI_ISL_872056 | CHR Citadelle | GIGA Medical Genomics | Keith Durkin, Maria Artesi, Sébastien Bontems, Raphaël Boreux, Bouchra Boujemla, Cécile Meex, Pierrette Melin, Marie-Pierre Hayette, Vincent Bours |
| EPI_ISL_872057, EPI_ISL_872081, EPI_ISL_872082, EPI_ISL_872083, EPI_ISL_872084, EPI_ISL_872085, EPI_ISL_872086, EPI_ISL_872087, EPI_ISL_872105, EPI_ISL_872106, EPI_ISL_872107, EPI_ISL_872108 | | | |
| see above | Department of Clinical Microbiology | GIGA Medical Genomics | Keith Durkin, Maria Artesi, Sébastien Bontems, Raphaël Boreux, Bouchra Boujemla, Cécile Meex, Pierrette Melin, Marie-Pierre Hayette, Vincent Bours |
| EPI_ISL_872110, EPI_ISL_872111 | CHC | GIGA Medical Genomics | Keith Durkin, Maria Artesi, Sébastien Bontems, Raphaël Boreux, Bouchra Boujemla, Cécile Meex, Pierrette Melin, Marie-Pierre Hayette, Vincent Bours |
| EPI_ISL_872112, EPI_ISL_872115, EPI_ISL_872116, EPI_ISL_872122 | Department of Clinical Microbiology | GIGA Medical Genomics | Keith Durkin, Maria Artesi, Sébastien Bontems, Raphaël Boreux, Bouchra Boujemla, Cécile Meex, Pierrette Melin, Marie-Pierre Hayette, Vincent Bours |
| EPI_ISL_872123, EPI_ISL_872124, EPI_ISL_872125, EPI_ISL_872126 | CHR Citadelle | GIGA Medical Genomics | Keith Durkin, Maria Artesi, Sébastien Bontems, Raphaël Boreux, Bouchra Boujemla, Cécile Meex, Pierrette Melin, Marie-Pierre Hayette, Vincent Bours |
| EPI_ISL_872127, EPI_ISL_872128, EPI_ISL_872129, EPI_ISL_872130, EPI_ISL_872135, EPI_ISL_872136, EPI_ISL_872137, EPI_ISL_872138, EPI_ISL_872139, EPI_ISL_872140, EPI_ISL_872141, EPI_ISL_872142, EPI_ISL_872143, EPI_ISL_872144, EPI_ISL_872145 | | | |
| see above | Department of Clinical Microbiology | GIGA Medical Genomics | Keith Durkin, Maria Artesi, Sébastien Bontems, Raphaël Boreux, Bouchra Boujemla, Cécile Meex, Pierrette Melin, Marie-Pierre Hayette, Vincent Bours |
| EPI_ISL_872146, EPI_ISL_872147 | CHC | GIGA Medical Genomics | Keith Durkin, Maria Artesi, Sébastien Bontems, Raphaël Boreux, Bouchra Boujemla, Cécile Meex, Pierrette Melin, Marie-Pierre Hayette, Vincent Bours |
| EPI_ISL_872148, EPI_ISL_872149 | Department of Clinical Microbiology | GIGA Medical Genomics | Keith Durkin, Maria Artesi, Sébastien Bontems, Raphaël Boreux, Bouchra Boujemla, Cécile Meex, Pierrette Melin, Marie-Pierre Hayette, Vincent Bours |
| EPI_ISL_872150 | CHR Citadelle | GIGA Medical Genomics | Keith Durkin, Maria Artesi, Sébastien Bontems, Raphaël Boreux, Bouchra Boujemla, Cécile Meex, Pierrette Melin, Marie-Pierre Hayette, Vincent Bours |
| EPI_ISL_872164, EPI_ISL_872167, EPI_ISL_872170, EPI_ISL_872171, EPI_ISL_872172 | Wyoming Public Health Laboratory | Wyoming Public Health Laboratory | Noah Hull, Taylor Fearing, Lynette Gumbleton, Channing Weber, Ashley Norberg, Bailey Bowcutt, and Wanda Manley |
| EPI_ISL_872192 | Conjunto Hospitalar do Mandaqui de Sao Paulo | Instituto Adolfo Lutz, Interdiciplinary Procedures Center, Strategic Laboratory | Claudio Tavares Sacchi, Claudia Regina Gonçalves, Erica Valessa Ramos Gomes, Karoline Rodrigues Campos, Katia Correa de Oliveira Santos, Ana Lucia de Carvalho Avelino, Fabiana Cristina Pereira dos Santos |
| EPI_ISL_872568 | Centre for Dengue Research and AICBU, Department of Immunology and Molecular Medicine | Centre for Dengue Research and AICBU, Department of Immunology and Molecular Medicine | Chandima Jeewandara, Deshni Jayathilaka, Dinuka Ariyaratne, Tibutius Thanesh Pramanayagam, Diyanath Ranasinghe, Laksiri Gomes, Gathsaurie Neelika Malavige |
| EPI_ISL_872583, EPI_ISL_872584 | South Eastern Area Laboratory Services (SEALS) | NSW Health Pathology - Institute of Clinical Pathology and Medical Research; Westmead Hospital; University of Sydney | CIDM-PH et al. |
| EPI_ISL_872586 | Sydney South West Pathology Service (SSWPS) - Royal Prince Alfred Hospital - NSW Health Pathology | NSW Health Pathology - Institute of Clinical Pathology and Medical Research; Westmead Hospital; University of Sydney | CIDM-PH et al. |
| EPI_ISL_875566, EPI_ISL_875567, EPI_ISL_875568 | SIESP L'AQUILA | Istituto Zooprofilattico Sperimentale dell'Abruzzo e Molise "G. Caporale" | Lorusso A, Marcacci M, Di Domenico M, Ancora M, Curini V, Mangone I, Rinaldi A, Scialabba S, Di Pasquale A, Cammà C, Puglia I, Calistri P, Savini G |
| EPI_ISL_875672, EPI_ISL_875673 | CHU Purpan - Laboratoire de Virologie - Institut Fédératif de Biologie | CHU Purpan - Laboratoire de Virologie - Institut Fédératif de Biologie | Latour J., Ranger N., Dubois M., Carcenac R., Harter A., Boyer P., Tremeaux P., Izopet J. |
| EPI_ISL_876061, EPI_ISL_876118, EPI_ISL_876136, EPI_ISL_876143, EPI_ISL_876153, EPI_ISL_876161, EPI_ISL_876218, EPI_ISL_876219, EPI_ISL_876220, EPI_ISL_876221, EPI_ISL_876222, EPI_ISL_876223, EPI_ISL_876224, EPI_ISL_876225, EPI_ISL_876226, EPI_ISL_876227, EPI_ISL_876228, EPI_ISL_876229, EPI_ISL_876230 | | | |
| see above | Massachusetts State Public Health Laboratory | Massachusetts State Public Health Laboratory | Andrew Lang, Timelia Fink, Glen Gallagher, Sandra Smole |
| EPI_ISL_877155, EPI_ISL_877156, EPI_ISL_877157, EPI_ISL_877158, EPI_ISL_877159, EPI_ISL_877160, EPI_ISL_877161, EPI_ISL_877162, EPI_ISL_877163, EPI_ISL_877181, EPI_ISL_877182, EPI_ISL_877183, EPI_ISL_877184, EPI_ISL_877185 | | | |
| see above | Univeristy of New Mexico Hospital | Center for Global Health, University of New Mexico Health Sciences Center | Daryl Domman, Kurt Schwalm, Justin Bacca, Jon Femling, Darrell Dinwiddie |
| EPI_ISL_877454 | National laboratory of health, environment and food Kranj | Institute of Microbiology and Immunology, Faculty of Medicine, | Samo Zakotnik, Tomaž Mark Zorec, Matic Brvar, Miša Korva, Mario Poljak, Tatjana Avši - Županc |

| | | | |
|---|---|---|---|
| | | University of Ljubljana | |
| EPI_ISL_877570 | Microbiological Diagnostic Unit - Public Health Laboratory (MDU-PHL) | MDU-PHL | Seemann T., Sait, M.L., Sherry, N.L. |
| EPI_ISL_877576, EPI_ISL_877578 | Victorian Infectious Diseases Reference Laboratory (VIDRL) | VIDRL and MDU-PHL | Caly L., Seemann T., Sait, M.L., Druce J., Sherry, N.L. |
| EPI_ISL_879146 | Lighthouse Lab in Alderley Park | Wellcome Sanger Institute for the COVID-19 Genomics UK (COG-UK) Consortium | Jacquelyn Wynn, Mairead Hyland, The Lighthouse Lab in Alderley Park and Alex Alderton, Roberto Amato, Sonia Goncalves, Ewan Harrison, David K. Jackson, Ian Johnston, Dominic Kwiatkowski, Cordelia Langford, John Sillitoe on behalf of the Wellcome Sanger Institute COVID-19 Surveillance Team |
| EPI_ISL_882997, EPI_ISL_882998 | Motion Picture Television Fund Hospital Lab | Los Angeles County PHL | P. Hemarajata et al. |
| EPI_ISL_884182, EPI_ISL_884183, EPI_ISL_884184, EPI_ISL_884185, EPI_ISL_884186, EPI_ISL_884187, EPI_ISL_884188, EPI_ISL_884190, EPI_ISL_884191, EPI_ISL_884193 | Wyoming Public Health Laboratory | Wyoming Public Health Laboratory | Noah Hull, Taylor Fearing, Lynette Gumbleton, Channing Weber, Ashley Norberg, Bailey Bowcutt, and Wanda Manley |
| EPI_ISL_885152 | National Institute of Infectious Diseases-Prof. Dr. Matei Bals Molecular Diagnostics Laboratory | National Institute of Infectious Diseases-Prof. Dr. Matei Bals Molecular Diagnostics Laboratory | Leontina Banica, Marius Surleac, Corina Casangiu, Petre Milu, Andreea Tudor, Simona Paraschiv, Dan Otelea |
| EPI_ISL_885491, EPI_ISL_885492, EPI_ISL_885494, EPI_ISL_885496, EPI_ISL_885497, EPI_ISL_885502, EPI_ISL_885504, EPI_ISL_885507, EPI_ISL_885508, EPI_ISL_885511, EPI_ISL_885512, EPI_ISL_885516, EPI_ISL_885517, EPI_ISL_885519, EPI_ISL_885520, EPI_ISL_885524, EPI_ISL_885526, EPI_ISL_885528, EPI_ISL_885534, EPI_ISL_885535, EPI_ISL_885538, EPI_ISL_885545, EPI_ISL_885546, EPI_ISL_885550, EPI_ISL_885551, EPI_ISL_885562, EPI_ISL_885563, EPI_ISL_885564, EPI_ISL_885567, EPI_ISL_885574, EPI_ISL_885575, EPI_ISL_885579, EPI_ISL_885582, EPI_ISL_885585, EPI_ISL_885588, EPI_ISL_885593, EPI_ISL_885602, EPI_ISL_885604, EPI_ISL_885607, EPI_ISL_885608, EPI_ISL_885609, EPI_ISL_885614, EPI_ISL_885621, EPI_ISL_885625, EPI_ISL_885626, EPI_ISL_885627, EPI_ISL_885628, EPI_ISL_885634, EPI_ISL_885636, EPI_ISL_885639, EPI_ISL_885647, EPI_ISL_885658, EPI_ISL_885660, EPI_ISL_885665, EPI_ISL_885666, EPI_ISL_885669, EPI_ISL_885670, EPI_ISL_885672, EPI_ISL_885675, EPI_ISL_885678, EPI_ISL_885681, EPI_ISL_885684, EPI_ISL_885685, EPI_ISL_885687, EPI_ISL_885688, EPI_ISL_885689, EPI_ISL_885692, EPI_ISL_885693, EPI_ISL_885696, EPI_ISL_885699, EPI_ISL_885704, EPI_ISL_885710, EPI_ISL_885712, EPI_ISL_885713, EPI_ISL_885714, EPI_ISL_885715, EPI_ISL_885720, EPI_ISL_885722, EPI_ISL_885724, EPI_ISL_885733, EPI_ISL_885734, EPI_ISL_885737, EPI_ISL_885738, EPI_ISL_885742, EPI_ISL_885743, EPI_ISL_885745, EPI_ISL_885748, EPI_ISL_885750, EPI_ISL_885755, EPI_ISL_885756, EPI_ISL_885761, EPI_ISL_885764, EPI_ISL_885765, EPI_ISL_885768, EPI_ISL_885770, EPI_ISL_885773, EPI_ISL_885775, EPI_ISL_885776, EPI_ISL_885779, EPI_ISL_885780, EPI_ISL_885785, EPI_ISL_885788, EPI_ISL_885792, EPI_ISL_885793, EPI_ISL_885794, EPI_ISL_885795, EPI_ISL_885800, EPI_ISL_885804, EPI_ISL_885807, EPI_ISL_885810, EPI_ISL_885812, EPI_ISL_885815, EPI_ISL_885820, EPI_ISL_885822, EPI_ISL_885824, EPI_ISL_885825, EPI_ISL_885826, EPI_ISL_885829, EPI_ISL_885830, EPI_ISL_885831, EPI_ISL_885832, EPI_ISL_885834, EPI_ISL_885836, EPI_ISL_885837, EPI_ISL_885839, EPI_ISL_885840, EPI_ISL_885841, EPI_ISL_885842, EPI_ISL_885845, EPI_ISL_885847, EPI_ISL_885849, EPI_ISL_885850, EPI_ISL_885851, EPI_ISL_885852, EPI_ISL_885853, EPI_ISL_885854, EPI_ISL_885857, EPI_ISL_885858, EPI_ISL_885859, EPI_ISL_885861, EPI_ISL_885862, EPI_ISL_885863, EPI_ISL_885865, EPI_ISL_885866, EPI_ISL_885869, EPI_ISL_885872, EPI_ISL_885873, EPI_ISL_885876, EPI_ISL_885877, EPI_ISL_885878, EPI_ISL_885881, EPI_ISL_885882, EPI_ISL_885884, EPI_ISL_885886, EPI_ISL_885887, EPI_ISL_885890, EPI_ISL_885891, EPI_ISL_885892, EPI_ISL_885893, EPI_ISL_885895, EPI_ISL_885896, EPI_ISL_885897, EPI_ISL_885900, EPI_ISL_885901, EPI_ISL_885903, EPI_ISL_885904, EPI_ISL_885905, EPI_ISL_885907, EPI_ISL_885908, EPI_ISL_885909, EPI_ISL_885911, EPI_ISL_885913, EPI_ISL_885914, EPI_ISL_885920, EPI_ISL_885921, EPI_ISL_885922, EPI_ISL_885923, EPI_ISL_885925, EPI_ISL_885930, EPI_ISL_885932, EPI_ISL_885933, EPI_ISL_885936, EPI_ISL_885938, EPI_ISL_885940, EPI_ISL_885941, EPI_ISL_885942, EPI_ISL_885943, EPI_ISL_885944, EPI_ISL_885945, EPI_ISL_885947, EPI_ISL_885948, EPI_ISL_885949, EPI_ISL_885950, EPI_ISL_885952, EPI_ISL_885953, EPI_ISL_885955, EPI_ISL_885956, EPI_ISL_885957, EPI_ISL_885960, EPI_ISL_885964, EPI_ISL_885965, EPI_ISL_885968, EPI_ISL_885969, EPI_ISL_885971, EPI_ISL_885972, EPI_ISL_885973, EPI_ISL_885975, EPI_ISL_885977, EPI_ISL_885978, EPI_ISL_885979, EPI_ISL_885981, EPI_ISL_885985, EPI_ISL_885986, EPI_ISL_885989, EPI_ISL_885991, EPI_ISL_885994, EPI_ISL_886000, EPI_ISL_886003, EPI_ISL_886007, EPI_ISL_886008, EPI_ISL_886010, EPI_ISL_886011, EPI_ISL_886012, EPI_ISL_886013, EPI_ISL_886017, EPI_ISL_886019, EPI_ISL_886022, EPI_ISL_886023, EPI_ISL_886024, EPI_ISL_886025, EPI_ISL_886026, EPI_ISL_886028, EPI_ISL_886030, EPI_ISL_886031, EPI_ISL_886033, EPI_ISL_886034, EPI_ISL_886035, EPI_ISL_886036, EPI_ISL_886041, EPI_ISL_886042, EPI_ISL_886045, EPI_ISL_886048, EPI_ISL_886051, EPI_ISL_886052, EPI_ISL_886054, EPI_ISL_886055, EPI_ISL_886058, EPI_ISL_886059, EPI_ISL_886061, EPI_ISL_886062, EPI_ISL_886065, EPI_ISL_886068, EPI_ISL_886069, EPI_ISL_886072, EPI_ISL_886073, EPI_ISL_886079, EPI_ISL_886080, EPI_ISL_886082, EPI_ISL_886084, EPI_ISL_886085, EPI_ISL_886086, EPI_ISL_886087, EPI_ISL_886088, EPI_ISL_886092, EPI_ISL_886093, EPI_ISL_886094, EPI_ISL_886095, EPI_ISL_886100, EPI_ISL_886102, EPI_ISL_886104, EPI_ISL_886105, EPI_ISL_886107, EPI_ISL_886108, EPI_ISL_886109, EPI_ISL_886111, EPI_ISL_886113, EPI_ISL_886114, EPI_ISL_886115, EPI_ISL_886122, EPI_ISL_886123, EPI_ISL_886124, EPI_ISL_886125, EPI_ISL_886126, EPI_ISL_886128, EPI_ISL_886130, EPI_ISL_886131, EPI_ISL_886135, EPI_ISL_886138, EPI_ISL_886139, EPI_ISL_886147, EPI_ISL_886151, EPI_ISL_886152, EPI_ISL_886153, EPI_ISL_886155, EPI_ISL_886156 | | | |
| see above | Lighthouse Lab in Alderley Park | Wellcome Sanger Institute for the COVID-19 Genomics UK (COG-UK) Consortium | Jacquelyn Wynn, Mairead Hyland, The Lighthouse Lab in Alderley Park and Alex Alderton, Roberto Amato, Sonia Goncalves, Ewan Harrison, David K. Jackson, Ian Johnston, Dominic Kwiatkowski, Cordelia Langford, John Sillitoe on behalf of the Wellcome Sanger Institute COVID-19 Surveillance Team |
| EPI_ISL_887120, EPI_ISL_887124, EPI_ISL_887125, EPI_ISL_887132 | Institute of Medical Microbiology and Hospital Hygiene | Institute of Medical Microbiology and Hospital Hygiene | Prof. Dr. Achim Kaasch, Aljoscha Tersteegen |
| EPI_ISL_888663, EPI_ISL_888664, EPI_ISL_888665 | University of Michigan Clinical Microbiology Laboratory | Lauring Lab, University of Michigan, Department of Microbiology and Immunology | Valesano |
| EPI_ISL_888673 | Montana Public Health Laboratory | Wyoming Public Health Laboratory | Noah Hull, Joy Ritter, Taylor Fearing, Lynette Gumbleton, Channing Weber, Ashley Norberg, Bailey Bowcutt, Wanda Manley, Deborah Gibson |
| EPI_ISL_888824, EPI_ISL_888825 | GZA Sint-Augustinus Hospital | UAntwerp, Laboratory of Medical Microbiology, Campus Drie Eiken S6.26, Universiteitsplein 1, 2610, Wilrijk, Antwerp, Belgium | Basil Britto Xavier, Jasmine Coppens, Christine Lammens, Veerle Matheeussen, Herman Goossens |
| EPI_ISL_888907, EPI_ISL_888912, EPI_ISL_888913, EPI_ISL_888915, EPI_ISL_888916, EPI_ISL_888917, EPI_ISL_888918, EPI_ISL_888920, EPI_ISL_888921, EPI_ISL_888922, EPI_ISL_888923, EPI_ISL_888924, EPI_ISL_888925, EPI_ISL_888926, EPI_ISL_888927, EPI_ISL_888928, EPI_ISL_888929, EPI_ISL_888931, EPI_ISL_888932, EPI_ISL_888933, EPI_ISL_888965, EPI_ISL_888967, EPI_ISL_888969, EPI_ISL_888971, EPI_ISL_888972, EPI_ISL_888973, EPI_ISL_888974, EPI_ISL_888975, EPI_ISL_888976, EPI_ISL_888977 | | | |
| see above | Wyoming Public Health Laboratory | Wyoming Public Health Laboratory | Noah Hull, Taylor Fearing, Lynette Gumbleton, Channing Weber, Ashley Norberg, Bailey Bowcutt, and Wanda Manley |
| EPI_ISL_889762, EPI_ISL_889763, EPI_ISL_889764, EPI_ISL_889765, EPI_ISL_889766, EPI_ISL_889767, EPI_ISL_889768, EPI_ISL_889769, EPI_ISL_889770, EPI_ISL_889771, EPI_ISL_889772 | | | |
| see above | LSUHS Emerging Viral Threat Laboratory | Microbial Genome Sequencing Center | Jeremy P. Kamil, Jennifer L. Carroll, Camille F. Abshire, Maarten Van Diest, Mohammed N.A. Siddiquey, Andrew D. Yurochko, Martin J. Sapp, Rona S. Scott, Christopher G. Kevil, Daniel J. Snyder, Vaughn S. Cooper, John A. Vanchiere |
| EPI_ISL_889773, EPI_ISL_889774, EPI_ISL_889775 | LSUHS Emerging Viral Threat Laboratory | Microbial Genome Sequencing Center | Mohammed N.A. Siddiquey, Jennifer L. Carroll, Camille F. Abshire, Maarten Van Diest, Andrew D. Yurochko, Martin J. Sapp, Rona S. Scott, Christopher G. Kevil, Daniel J. Snyder, Vaughn S. Cooper, John A. Vanchiere, Jeremy P. Kamil, |
| EPI_ISL_889776, EPI_ISL_889777, EPI_ISL_889778, EPI_ISL_889779, EPI_ISL_889780, EPI_ISL_889781, EPI_ISL_889782, EPI_ISL_889783, EPI_ISL_889784, EPI_ISL_889785 | LSUHS Emerging Viral Threat Laboratory | Microbial Genome Sequencing Center | Maarten Van Diest, Jennifer L. Carroll, Camille F. Abshire, Mohammed N.A. Siddiquey, Andrew D. Yurochko, Martin J. Sapp, Rona S. Scott, Christopher G. Kevil, Daniel J. Snyder, Vaughn S. Cooper, John A. Vanchiere, Jeremy P. Kamil |
| EPI_ISL_889786, EPI_ISL_889787, EPI_ISL_889788, EPI_ISL_889789, EPI_ISL_889790, EPI_ISL_889791 | LSUHS Emerging Viral Threat Laboratory | Microbial Genome Sequencing Center | Jennifer L. Carroll, Rona S. Scott, Camille F. Abshire, Maarten Van Diest, Mohammed N.A. Siddiquey, Andrew D. Yurochko, Martin J. Sapp, Christopher G. Kevil, Daniel J. Snyder, Vaughn S. Cooper, John A. Vanchiere, Jeremy P. Kamil, |
| EPI_ISL_889792, EPI_ISL_889793, EPI_ISL_889794, EPI_ISL_889795, EPI_ISL_889796, EPI_ISL_889797, EPI_ISL_889798, EPI_ISL_889799, EPI_ISL_889800, EPI_ISL_889801 | LSUHS Emerging Viral Threat Laboratory | Microbial Genome Sequencing Center | Rona S. Scott, Jennifer L. Carroll, Camille F. Abshire, Maarten Van Diest, Mohammed N.A. Siddiquey, Andrew D. Yurochko, Martin J. Sapp, Christopher G. Kevil, Daniel J. Snyder, Vaughn S. Cooper, John A. Vanchiere, Jeremy P. Kamil, |
| EPI_ISL_890233 | School of Pharmacy, Shenandoah University | School of Pharmacy, Shenandoah University | Adams,S.M., Harralson,A.F., Kidd,R.S., Sawyer,G.W. |
| EPI_ISL_890367, EPI_ISL_890368, EPI_ISL_890369, EPI_ISL_890370 | LSUHS Emerging Viral Threat Laboratory | Microbial Genome Sequencing Center | Jennifer L. Carroll, Rona S. Scott, Camille F. Abshire, Maarten Van Diest, Mohammed N.A. Siddiquey, Andrew D. Yurochko, Martin J. Sapp, Christopher G. Kevil, Daniel J. Snyder, Vaughn S. Cooper, John A. Vanchiere, Jeremy P. Kamil, |
| EPI_ISL_891093 | Seattle Flu Study | Seattle Flu Study | Deborah A. Nickerson, Chris D. Frazar, Jover Lee, Benjamin Pelle, Erica Ryke, Matthew Richardson, Amanda Adler, Elisabeth Brandstetter, Peter D. Han, Kairsten Fay, Misja Ilcisin, Kirsten Lacombe, Thomas R. Sibley, Melissa Truong, Caitlin R. Wolf, Michael Boeckh, Janet A. Englund, Michael Famulare, Barry R. Lutz, Mark J. Rieder, Lea M. Starita, Matthew Thompson, Jay Shendure, Trevor Bedford, Helen Y. Chu |
| EPI_ISL_891205 | DPH, Massachusetts State Public Health Lab | DPH, Massachusetts State Public Health Lab | Lang,A.S., Fink,T., Gallagher,G.R., Smole,S.C. |
| EPI_ISL_891253 | Singapore General Hospital | Department of Microbiology | Nurdyana Abdul Rahman, Kun Lee Lim, Chenhao Li, Sui Sin Goh, Kenneth Xin Long Chan, Kian Sing Chan, Lynette Oon, Kern Rei Chng, Niranjan Nagarajan, Karrie Ko |
| EPI_ISL_891273, EPI_ISL_891275, EPI_ISL_891276 | Lighthouse Lab in Cambridge | Wellcome Sanger Institute for the COVID-19 Genomics UK (COG-UK) Consortium | Rob Howes, The Lighthouse Lab in Cambridge and Alex Alderton, Roberto Amato, Sonia Goncalves, Ewan Harrison, David K. Jackson, Ian Johnston, Dominic Kwiatkowski, Cordelia Langford, John Sillitoe on behalf of the Wellcome Sanger Institute COVID-19 Surveillance Team |
| EPI_ISL_891278, EPI_ISL_891280 | Lighthouse Lab in Alderley Park | Wellcome Sanger Institute for the COVID-19 Genomics UK | Jacquelyn Wynn, Mairead Hyland, The Lighthouse Lab in Alderley Park and Alex Alderton, Roberto Amato, Sonia Goncalves, Ewan Harrison, David K. |

| | | | |
|---|---|---|---|
| | | (COG-UK) Consortium | Jackson, Ian Johnston, Dominic Kwiatkowski, Cordelia Langford, John Sillitoe on behalf of the Wellcome Sanger Institute COVID-19 Surveillance Team |
| EPI_ISL_891281 | Lighthouse Lab in Glasgow | Wellcome Sanger Institute for the COVID-19 Genomics UK (COG-UK) Consortium | Harper VanSteenhouse, Yumi Kasai, David Gray, Carol Clugston, Anna Dominiczak and Alex Alderton, Roberto Amato, Sonia Goncalves, Ewan Harrison, David K. Jackson, Ian Johnston, Dominic Kwiatkowski, Cordelia Langford, John Sillitoe on behalf of the Wellcome Sanger Institute COVID-19 Surveillance Team |
| EPI_ISL_891282 | Lighthouse Lab in Cambridge | Wellcome Sanger Institute for the COVID-19 Genomics UK (COG-UK) Consortium | Rob Howes, The Lighthouse Lab in Cambridge and Alex Alderton, Roberto Amato, Sonia Goncalves, Ewan Harrison, David K. Jackson, Ian Johnston, Dominic Kwiatkowski, Cordelia Langford, John Sillitoe on behalf of the Wellcome Sanger Institute COVID-19 Surveillance Team |
| EPI_ISL_891283 | Lighthouse Lab in Glasgow | Wellcome Sanger Institute for the COVID-19 Genomics UK (COG-UK) Consortium | Harper VanSteenhouse, Yumi Kasai, David Gray, Carol Clugston, Anna Dominiczak and Alex Alderton, Roberto Amato, Sonia Goncalves, Ewan Harrison, David K. Jackson, Ian Johnston, Dominic Kwiatkowski, Cordelia Langford, John Sillitoe on behalf of the Wellcome Sanger Institute COVID-19 Surveillance Team |
| EPI_ISL_891284 | Lighthouse Lab in Cambridge | Wellcome Sanger Institute for the COVID-19 Genomics UK (COG-UK) Consortium | Rob Howes, The Lighthouse Lab in Cambridge and Alex Alderton, Roberto Amato, Sonia Goncalves, Ewan Harrison, David K. Jackson, Ian Johnston, Dominic Kwiatkowski, Cordelia Langford, John Sillitoe on behalf of the Wellcome Sanger Institute COVID-19 Surveillance Team |
| EPI_ISL_891285 | Lighthouse Lab in Alderley Park | Wellcome Sanger Institute for the COVID-19 Genomics UK (COG-UK) Consortium | Jacquelyn Wynn, Mairead Hyland, The Lighthouse Lab in Alderley Park and Alex Alderton, Roberto Amato, Sonia Goncalves, Ewan Harrison, David K. Jackson, Ian Johnston, Dominic Kwiatkowski, Cordelia Langford, John Sillitoe on behalf of the Wellcome Sanger Institute COVID-19 Surveillance Team |
| EPI_ISL_891286, EPI_ISL_891288 | Lighthouse Lab in Glasgow | Wellcome Sanger Institute for the COVID-19 Genomics UK (COG-UK) Consortium | Harper VanSteenhouse, Yumi Kasai, David Gray, Carol Clugston, Anna Dominiczak and Alex Alderton, Roberto Amato, Sonia Goncalves, Ewan Harrison, David K. Jackson, Ian Johnston, Dominic Kwiatkowski, Cordelia Langford, John Sillitoe on behalf of the Wellcome Sanger Institute COVID-19 Surveillance Team |
| EPI_ISL_891289, EPI_ISL_891290, EPI_ISL_891292, EPI_ISL_891293, EPI_ISL_891294 | Lighthouse Lab in Cambridge | Wellcome Sanger Institute for the COVID-19 Genomics UK (COG-UK) Consortium | Rob Howes, The Lighthouse Lab in Cambridge and Alex Alderton, Roberto Amato, Sonia Goncalves, Ewan Harrison, David K. Jackson, Ian Johnston, Dominic Kwiatkowski, Cordelia Langford, John Sillitoe on behalf of the Wellcome Sanger Institute COVID-19 Surveillance Team |
| EPI_ISL_891295 | Lighthouse Lab in Alderley Park | Wellcome Sanger Institute for the COVID-19 Genomics UK (COG-UK) Consortium | Jacquelyn Wynn, Mairead Hyland, The Lighthouse Lab in Alderley Park and Alex Alderton, Roberto Amato, Sonia Goncalves, Ewan Harrison, David K. Jackson, Ian Johnston, Dominic Kwiatkowski, Cordelia Langford, John Sillitoe on behalf of the Wellcome Sanger Institute COVID-19 Surveillance Team |
| EPI_ISL_891296 | Lighthouse Lab in Cambridge | Wellcome Sanger Institute for the COVID-19 Genomics UK (COG-UK) Consortium | Rob Howes, The Lighthouse Lab in Cambridge and Alex Alderton, Roberto Amato, Sonia Goncalves, Ewan Harrison, David K. Jackson, Ian Johnston, Dominic Kwiatkowski, Cordelia Langford, John Sillitoe on behalf of the Wellcome Sanger Institute COVID-19 Surveillance Team |
| EPI_ISL_891297, EPI_ISL_891298 | Lighthouse Lab in Glasgow | Wellcome Sanger Institute for the COVID-19 Genomics UK (COG-UK) Consortium | Harper VanSteenhouse, Yumi Kasai, David Gray, Carol Clugston, Anna Dominiczak and Alex Alderton, Roberto Amato, Sonia Goncalves, Ewan Harrison, David K. Jackson, Ian Johnston, Dominic Kwiatkowski, Cordelia Langford, John Sillitoe on behalf of the Wellcome Sanger Institute COVID-19 Surveillance Team |
| EPI_ISL_891299 | Lighthouse Lab in Alderley Park | Wellcome Sanger Institute for the COVID-19 Genomics UK (COG-UK) Consortium | Jacquelyn Wynn, Mairead Hyland, The Lighthouse Lab in Alderley Park and Alex Alderton, Roberto Amato, Sonia Goncalves, Ewan Harrison, David K. Jackson, Ian Johnston, Dominic Kwiatkowski, Cordelia Langford, John Sillitoe on behalf of the Wellcome Sanger Institute COVID-19 Surveillance Team |
| EPI_ISL_891300, EPI_ISL_891301 | Lighthouse Lab in Cambridge | Wellcome Sanger Institute for the COVID-19 Genomics UK (COG-UK) Consortium | Rob Howes, The Lighthouse Lab in Cambridge and Alex Alderton, Roberto Amato, Sonia Goncalves, Ewan Harrison, David K. Jackson, Ian Johnston, Dominic Kwiatkowski, Cordelia Langford, John Sillitoe on behalf of the Wellcome Sanger Institute COVID-19 Surveillance Team |
| EPI_ISL_891304 | Lighthouse Lab in Glasgow | Wellcome Sanger Institute for the COVID-19 Genomics UK (COG-UK) Consortium | Harper VanSteenhouse, Yumi Kasai, David Gray, Carol Clugston, Anna Dominiczak and Alex Alderton, Roberto Amato, Sonia Goncalves, Ewan Harrison, David K. Jackson, Ian Johnston, Dominic Kwiatkowski, Cordelia Langford, John Sillitoe on behalf of the Wellcome Sanger Institute COVID-19 Surveillance Team |
| EPI_ISL_891305 | Lighthouse Lab in Alderley Park | Wellcome Sanger Institute for the COVID-19 Genomics UK (COG-UK) Consortium | Jacquelyn Wynn, Mairead Hyland, The Lighthouse Lab in Alderley Park and Alex Alderton, Roberto Amato, Sonia Goncalves, Ewan Harrison, David K. Jackson, Ian Johnston, Dominic Kwiatkowski, Cordelia Langford, John Sillitoe on behalf of the Wellcome Sanger Institute COVID-19 Surveillance Team |
| EPI_ISL_891306, EPI_ISL_891307, EPI_ISL_891308, EPI_ISL_891309, EPI_ISL_891310 | Lighthouse Lab in Cambridge | Wellcome Sanger Institute for the COVID-19 Genomics UK (COG-UK) Consortium | Rob Howes, The Lighthouse Lab in Cambridge and Alex Alderton, Roberto Amato, Sonia Goncalves, Ewan Harrison, David K. Jackson, Ian Johnston, Dominic Kwiatkowski, Cordelia Langford, John Sillitoe on behalf of the Wellcome Sanger Institute COVID-19 Surveillance Team |
| EPI_ISL_891311, EPI_ISL_891312, EPI_ISL_891313 | Lighthouse Lab in Alderley Park | Wellcome Sanger Institute for the COVID-19 Genomics UK (COG-UK) Consortium | Jacquelyn Wynn, Mairead Hyland, The Lighthouse Lab in Alderley Park and Alex Alderton, Roberto Amato, Sonia Goncalves, Ewan Harrison, David K. Jackson, Ian Johnston, Dominic Kwiatkowski, Cordelia Langford, John Sillitoe on behalf of the Wellcome Sanger Institute COVID-19 Surveillance Team |
| EPI_ISL_891314, EPI_ISL_891316 | Lighthouse Lab in Cambridge | Wellcome Sanger Institute for the COVID-19 Genomics UK (COG-UK) Consortium | Rob Howes, The Lighthouse Lab in Cambridge and Alex Alderton, Roberto Amato, Sonia Goncalves, Ewan Harrison, David K. Jackson, Ian Johnston, Dominic Kwiatkowski, Cordelia Langford, John Sillitoe on behalf of the Wellcome Sanger Institute COVID-19 Surveillance Team |
| EPI_ISL_891317, EPI_ISL_891318, EPI_ISL_891319 | Lighthouse Lab in Alderley Park | Wellcome Sanger Institute for the COVID-19 Genomics UK (COG-UK) Consortium | Jacquelyn Wynn, Mairead Hyland, The Lighthouse Lab in Alderley Park and Alex Alderton, Roberto Amato, Sonia Goncalves, Ewan Harrison, David K. Jackson, Ian Johnston, Dominic Kwiatkowski, Cordelia Langford, John Sillitoe on behalf of the Wellcome Sanger Institute COVID-19 Surveillance Team |
| EPI_ISL_891320 | Lighthouse Lab in Glasgow | Wellcome Sanger Institute for the COVID-19 Genomics UK (COG-UK) Consortium | Harper VanSteenhouse, Yumi Kasai, David Gray, Carol Clugston, Anna Dominiczak and Alex Alderton, Roberto Amato, Sonia Goncalves, Ewan Harrison, David K. Jackson, Ian Johnston, Dominic Kwiatkowski, Cordelia Langford, John Sillitoe on behalf of the Wellcome Sanger Institute COVID-19 Surveillance Team |
| EPI_ISL_891322 | Lighthouse Lab in Cambridge | Wellcome Sanger Institute for the COVID-19 Genomics UK (COG-UK) Consortium | Rob Howes, The Lighthouse Lab in Cambridge and Alex Alderton, Roberto Amato, Sonia Goncalves, Ewan Harrison, David K. Jackson, Ian Johnston, Dominic Kwiatkowski, Cordelia Langford, John Sillitoe on behalf of the Wellcome Sanger Institute COVID-19 Surveillance Team |
| EPI_ISL_891323, EPI_ISL_891324 | Lighthouse Lab in Alderley Park | Wellcome Sanger Institute for the COVID-19 Genomics UK (COG-UK) Consortium | Jacquelyn Wynn, Mairead Hyland, The Lighthouse Lab in Alderley Park and Alex Alderton, Roberto Amato, Sonia Goncalves, Ewan Harrison, David K. Jackson, Ian Johnston, Dominic Kwiatkowski, Cordelia Langford, John Sillitoe on behalf of the Wellcome Sanger Institute COVID-19 Surveillance Team |
| EPI_ISL_891325 | Lighthouse Lab in Cambridge | Wellcome Sanger Institute for the COVID-19 Genomics UK (COG-UK) Consortium | Rob Howes, The Lighthouse Lab in Cambridge and Alex Alderton, Roberto Amato, Sonia Goncalves, Ewan Harrison, David K. Jackson, Ian Johnston, Dominic Kwiatkowski, Cordelia Langford, John Sillitoe on behalf of the Wellcome Sanger Institute COVID-19 Surveillance Team |
| EPI_ISL_891326 | Lighthouse Lab in Alderley Park | Wellcome Sanger Institute for the COVID-19 Genomics UK (COG-UK) Consortium | Jacquelyn Wynn, Mairead Hyland, The Lighthouse Lab in Alderley Park and Alex Alderton, Roberto Amato, Sonia Goncalves, Ewan Harrison, David K. Jackson, Ian Johnston, Dominic Kwiatkowski, Cordelia Langford, John Sillitoe on behalf of the Wellcome Sanger Institute COVID-19 Surveillance Team |
| EPI_ISL_891327 | Lighthouse Lab in Glasgow | Wellcome Sanger Institute for the COVID-19 Genomics UK (COG-UK) Consortium | Harper VanSteenhouse, Yumi Kasai, David Gray, Carol Clugston, Anna Dominiczak and Alex Alderton, Roberto Amato, Sonia Goncalves, Ewan Harrison, David K. Jackson, Ian Johnston, Dominic Kwiatkowski, Cordelia Langford, John Sillitoe on behalf of the Wellcome Sanger Institute COVID-19 Surveillance Team |
| EPI_ISL_891328, EPI_ISL_891329 | Lighthouse Lab in Cambridge | Wellcome Sanger Institute for the COVID-19 Genomics UK (COG-UK) Consortium | Rob Howes, The Lighthouse Lab in Cambridge and Alex Alderton, Roberto Amato, Sonia Goncalves, Ewan Harrison, David K. Jackson, Ian Johnston, Dominic Kwiatkowski, Cordelia Langford, John Sillitoe on behalf of the Wellcome Sanger Institute COVID-19 Surveillance Team |
| EPI_ISL_891330 | Lighthouse Lab in Glasgow | Wellcome Sanger Institute for the COVID-19 Genomics UK (COG-UK) Consortium | Harper VanSteenhouse, Yumi Kasai, David Gray, Carol Clugston, Anna Dominiczak and Alex Alderton, Roberto Amato, Sonia Goncalves, Ewan Harrison, David K. Jackson, Ian Johnston, Dominic Kwiatkowski, Cordelia Langford, John Sillitoe on behalf of the Wellcome Sanger Institute COVID-19 Surveillance Team |
| EPI_ISL_891331, EPI_ISL_891332, EPI_ISL_891333 | Lighthouse Lab in Cambridge | Wellcome Sanger Institute for the COVID-19 Genomics UK (COG-UK) Consortium | Rob Howes, The Lighthouse Lab in Cambridge and Alex Alderton, Roberto Amato, Sonia Goncalves, Ewan Harrison, David K. Jackson, Ian Johnston, Dominic Kwiatkowski, Cordelia Langford, John Sillitoe on behalf of the Wellcome Sanger Institute COVID-19 Surveillance Team |
| EPI_ISL_891334 | Lighthouse Lab in Alderley Park | Wellcome Sanger Institute for the COVID-19 Genomics UK (COG-UK) Consortium | Jacquelyn Wynn, Mairead Hyland, The Lighthouse Lab in Alderley Park and Alex Alderton, Roberto Amato, Sonia Goncalves, Ewan Harrison, David K. Jackson, Ian Johnston, Dominic Kwiatkowski, Cordelia Langford, John Sillitoe on behalf of the Wellcome Sanger Institute COVID-19 Surveillance Team |
| EPI_ISL_891335 | Lighthouse Lab in Glasgow | Wellcome Sanger Institute for the COVID-19 Genomics UK (COG-UK) Consortium | Harper VanSteenhouse, Yumi Kasai, David Gray, Carol Clugston, Anna Dominiczak and Alex Alderton, Roberto Amato, Sonia Goncalves, Ewan Harrison, David K. Jackson, Ian Johnston, Dominic Kwiatkowski, Cordelia Langford, John Sillitoe on behalf of the Wellcome Sanger Institute COVID-19 Surveillance Team |
| EPI_ISL_891337 | Lighthouse Lab in Alderley Park | Wellcome Sanger Institute for the COVID-19 Genomics UK (COG-UK) Consortium | Jacquelyn Wynn, Mairead Hyland, The Lighthouse Lab in Alderley Park and Alex Alderton, Roberto Amato, Sonia Goncalves, Ewan Harrison, David K. Jackson, Ian Johnston, Dominic Kwiatkowski, Cordelia Langford, John Sillitoe on behalf of the Wellcome Sanger Institute COVID-19 Surveillance Team |

| | | | |
|---|---|---|---|
| EPI_ISL_891338 | Lighthouse Lab in Glasgow | Wellcome Sanger Institute for the COVID-19 Genomics UK (COG-UK) Consortium | Harper VanSteenhouse, Yumi Kasai, David Gray, Carol Clugston, Anna Dominiczak and Alex Alderton, Roberto Amato, Sonia Goncalves, Ewan Harrison, David K. Jackson, Ian Johnston, Dominic Kwiatkowski, Cordelia Langford, John Sillitoe on behalf of the Wellcome Sanger Institute COVID-19 Surveillance Team |
| EPI_ISL_891339 | Lighthouse Lab in Cambridge | Wellcome Sanger Institute for the COVID-19 Genomics UK (COG-UK) Consortium | Rob Howes, The Lighthouse Lab in Cambridge and Alex Alderton, Roberto Amato, Sonia Goncalves, Ewan Harrison, David K. Jackson, Ian Johnston, Dominic Kwiatkowski, Cordelia Langford, John Sillitoe on behalf of the Wellcome Sanger Institute COVID-19 Surveillance Team |
| EPI_ISL_891340, EPI_ISL_891342, EPI_ISL_891344 | Lighthouse Lab in Alderley Park | Wellcome Sanger Institute for the COVID-19 Genomics UK (COG-UK) Consortium | Jacquelyn Wynn, Mairead Hyland, The Lighthouse Lab in Alderley Park and Alex Alderton, Roberto Amato, Sonia Goncalves, Ewan Harrison, David K. Jackson, Ian Johnston, Dominic Kwiatkowski, Cordelia Langford, John Sillitoe on behalf of the Wellcome Sanger Institute COVID-19 Surveillance Team |
| EPI_ISL_891345 | Lighthouse Lab in Glasgow | Wellcome Sanger Institute for the COVID-19 Genomics UK (COG-UK) Consortium | Harper VanSteenhouse, Yumi Kasai, David Gray, Carol Clugston, Anna Dominiczak and Alex Alderton, Roberto Amato, Sonia Goncalves, Ewan Harrison, David K. Jackson, Ian Johnston, Dominic Kwiatkowski, Cordelia Langford, John Sillitoe on behalf of the Wellcome Sanger Institute COVID-19 Surveillance Team |
| EPI_ISL_891346, EPI_ISL_891347 | Lighthouse Lab in Cambridge | Wellcome Sanger Institute for the COVID-19 Genomics UK (COG-UK) Consortium | Rob Howes, The Lighthouse Lab in Cambridge and Alex Alderton, Roberto Amato, Sonia Goncalves, Ewan Harrison, David K. Jackson, Ian Johnston, Dominic Kwiatkowski, Cordelia Langford, John Sillitoe on behalf of the Wellcome Sanger Institute COVID-19 Surveillance Team |
| EPI_ISL_891349 | Lighthouse Lab in Alderley Park | Wellcome Sanger Institute for the COVID-19 Genomics UK (COG-UK) Consortium | Jacquelyn Wynn, Mairead Hyland, The Lighthouse Lab in Alderley Park and Alex Alderton, Roberto Amato, Sonia Goncalves, Ewan Harrison, David K. Jackson, Ian Johnston, Dominic Kwiatkowski, Cordelia Langford, John Sillitoe on behalf of the Wellcome Sanger Institute COVID-19 Surveillance Team |
| EPI_ISL_891350, EPI_ISL_891351 | Lighthouse Lab in Cambridge | Wellcome Sanger Institute for the COVID-19 Genomics UK (COG-UK) Consortium | Rob Howes, The Lighthouse Lab in Cambridge and Alex Alderton, Roberto Amato, Sonia Goncalves, Ewan Harrison, David K. Jackson, Ian Johnston, Dominic Kwiatkowski, Cordelia Langford, John Sillitoe on behalf of the Wellcome Sanger Institute COVID-19 Surveillance Team |
| EPI_ISL_891352 | Lighthouse Lab in Alderley Park | Wellcome Sanger Institute for the COVID-19 Genomics UK (COG-UK) Consortium | Jacquelyn Wynn, Mairead Hyland, The Lighthouse Lab in Alderley Park and Alex Alderton, Roberto Amato, Sonia Goncalves, Ewan Harrison, David K. Jackson, Ian Johnston, Dominic Kwiatkowski, Cordelia Langford, John Sillitoe on behalf of the Wellcome Sanger Institute COVID-19 Surveillance Team |
| EPI_ISL_891354, EPI_ISL_891355, EPI_ISL_891356 | Lighthouse Lab in Cambridge | Wellcome Sanger Institute for the COVID-19 Genomics UK (COG-UK) Consortium | Rob Howes, The Lighthouse Lab in Cambridge and Alex Alderton, Roberto Amato, Sonia Goncalves, Ewan Harrison, David K. Jackson, Ian Johnston, Dominic Kwiatkowski, Cordelia Langford, John Sillitoe on behalf of the Wellcome Sanger Institute COVID-19 Surveillance Team |
| EPI_ISL_891357 | Lighthouse Lab in Alderley Park | Wellcome Sanger Institute for the COVID-19 Genomics UK (COG-UK) Consortium | Jacquelyn Wynn, Mairead Hyland, The Lighthouse Lab in Alderley Park and Alex Alderton, Roberto Amato, Sonia Goncalves, Ewan Harrison, David K. Jackson, Ian Johnston, Dominic Kwiatkowski, Cordelia Langford, John Sillitoe on behalf of the Wellcome Sanger Institute COVID-19 Surveillance Team |
| EPI_ISL_891359 | Lighthouse Lab in Cambridge | Wellcome Sanger Institute for the COVID-19 Genomics UK (COG-UK) Consortium | Rob Howes, The Lighthouse Lab in Cambridge and Alex Alderton, Roberto Amato, Sonia Goncalves, Ewan Harrison, David K. Jackson, Ian Johnston, Dominic Kwiatkowski, Cordelia Langford, John Sillitoe on behalf of the Wellcome Sanger Institute COVID-19 Surveillance Team |
| EPI_ISL_891360 | Lighthouse Lab in Alderley Park | Wellcome Sanger Institute for the COVID-19 Genomics UK (COG-UK) Consortium | Jacquelyn Wynn, Mairead Hyland, The Lighthouse Lab in Alderley Park and Alex Alderton, Roberto Amato, Sonia Goncalves, Ewan Harrison, David K. Jackson, Ian Johnston, Dominic Kwiatkowski, Cordelia Langford, John Sillitoe on behalf of the Wellcome Sanger Institute COVID-19 Surveillance Team |
| EPI_ISL_891361 | Lighthouse Lab in Glasgow | Wellcome Sanger Institute for the COVID-19 Genomics UK (COG-UK) Consortium | Harper VanSteenhouse, Yumi Kasai, David Gray, Carol Clugston, Anna Dominiczak and Alex Alderton, Roberto Amato, Sonia Goncalves, Ewan Harrison, David K. Jackson, Ian Johnston, Dominic Kwiatkowski, Cordelia Langford, John Sillitoe on behalf of the Wellcome Sanger Institute COVID-19 Surveillance Team |
| EPI_ISL_891363 | Lighthouse Lab in Cambridge | Wellcome Sanger Institute for the COVID-19 Genomics UK (COG-UK) Consortium | Rob Howes, The Lighthouse Lab in Cambridge and Alex Alderton, Roberto Amato, Sonia Goncalves, Ewan Harrison, David K. Jackson, Ian Johnston, Dominic Kwiatkowski, Cordelia Langford, John Sillitoe on behalf of the Wellcome Sanger Institute COVID-19 Surveillance Team |
| EPI_ISL_891364 | Lighthouse Lab in Alderley Park | Wellcome Sanger Institute for the COVID-19 Genomics UK (COG-UK) Consortium | Jacquelyn Wynn, Mairead Hyland, The Lighthouse Lab in Alderley Park and Alex Alderton, Roberto Amato, Sonia Goncalves, Ewan Harrison, David K. Jackson, Ian Johnston, Dominic Kwiatkowski, Cordelia Langford, John Sillitoe on behalf of the Wellcome Sanger Institute COVID-19 Surveillance Team |
| EPI_ISL_891366, EPI_ISL_891367, EPI_ISL_891369, EPI_ISL_891370 | Lighthouse Lab in Cambridge | Wellcome Sanger Institute for the COVID-19 Genomics UK (COG-UK) Consortium | Rob Howes, The Lighthouse Lab in Cambridge and Alex Alderton, Roberto Amato, Sonia Goncalves, Ewan Harrison, David K. Jackson, Ian Johnston, Dominic Kwiatkowski, Cordelia Langford, John Sillitoe on behalf of the Wellcome Sanger Institute COVID-19 Surveillance Team |
| EPI_ISL_891371 | Lighthouse Lab in Alderley Park | Wellcome Sanger Institute for the COVID-19 Genomics UK (COG-UK) Consortium | Jacquelyn Wynn, Mairead Hyland, The Lighthouse Lab in Alderley Park and Alex Alderton, Roberto Amato, Sonia Goncalves, Ewan Harrison, David K. Jackson, Ian Johnston, Dominic Kwiatkowski, Cordelia Langford, John Sillitoe on behalf of the Wellcome Sanger Institute COVID-19 Surveillance Team |
| EPI_ISL_891372 | Lighthouse Lab in Glasgow | Wellcome Sanger Institute for the COVID-19 Genomics UK (COG-UK) Consortium | Harper VanSteenhouse, Yumi Kasai, David Gray, Carol Clugston, Anna Dominiczak and Alex Alderton, Roberto Amato, Sonia Goncalves, Ewan Harrison, David K. Jackson, Ian Johnston, Dominic Kwiatkowski, Cordelia Langford, John Sillitoe on behalf of the Wellcome Sanger Institute COVID-19 Surveillance Team |
| EPI_ISL_891373 | Lighthouse Lab in Cambridge | Wellcome Sanger Institute for the COVID-19 Genomics UK (COG-UK) Consortium | Rob Howes, The Lighthouse Lab in Cambridge and Alex Alderton, Roberto Amato, Sonia Goncalves, Ewan Harrison, David K. Jackson, Ian Johnston, Dominic Kwiatkowski, Cordelia Langford, John Sillitoe on behalf of the Wellcome Sanger Institute COVID-19 Surveillance Team |
| EPI_ISL_891374, EPI_ISL_891376, EPI_ISL_891377, EPI_ISL_891378 | Lighthouse Lab in Alderley Park | Wellcome Sanger Institute for the COVID-19 Genomics UK (COG-UK) Consortium | Jacquelyn Wynn, Mairead Hyland, The Lighthouse Lab in Alderley Park and Alex Alderton, Roberto Amato, Sonia Goncalves, Ewan Harrison, David K. Jackson, Ian Johnston, Dominic Kwiatkowski, Cordelia Langford, John Sillitoe on behalf of the Wellcome Sanger Institute COVID-19 Surveillance Team |
| EPI_ISL_891379 | Lighthouse Lab in Cambridge | Wellcome Sanger Institute for the COVID-19 Genomics UK (COG-UK) Consortium | Rob Howes, The Lighthouse Lab in Cambridge and Alex Alderton, Roberto Amato, Sonia Goncalves, Ewan Harrison, David K. Jackson, Ian Johnston, Dominic Kwiatkowski, Cordelia Langford, John Sillitoe on behalf of the Wellcome Sanger Institute COVID-19 Surveillance Team |
| EPI_ISL_891380 | Lighthouse Lab in Glasgow | Wellcome Sanger Institute for the COVID-19 Genomics UK (COG-UK) Consortium | Harper VanSteenhouse, Yumi Kasai, David Gray, Carol Clugston, Anna Dominiczak and Alex Alderton, Roberto Amato, Sonia Goncalves, Ewan Harrison, David K. Jackson, Ian Johnston, Dominic Kwiatkowski, Cordelia Langford, John Sillitoe on behalf of the Wellcome Sanger Institute COVID-19 Surveillance Team |
| EPI_ISL_891383, EPI_ISL_891384, EPI_ISL_891385, EPI_ISL_891386 | Lighthouse Lab in Alderley Park | Wellcome Sanger Institute for the COVID-19 Genomics UK (COG-UK) Consortium | Jacquelyn Wynn, Mairead Hyland, The Lighthouse Lab in Alderley Park and Alex Alderton, Roberto Amato, Sonia Goncalves, Ewan Harrison, David K. Jackson, Ian Johnston, Dominic Kwiatkowski, Cordelia Langford, John Sillitoe on behalf of the Wellcome Sanger Institute COVID-19 Surveillance Team |
| EPI_ISL_891387, EPI_ISL_891388, EPI_ISL_891389 | Lighthouse Lab in Cambridge | Wellcome Sanger Institute for the COVID-19 Genomics UK (COG-UK) Consortium | Rob Howes, The Lighthouse Lab in Cambridge and Alex Alderton, Roberto Amato, Sonia Goncalves, Ewan Harrison, David K. Jackson, Ian Johnston, Dominic Kwiatkowski, Cordelia Langford, John Sillitoe on behalf of the Wellcome Sanger Institute COVID-19 Surveillance Team |
| EPI_ISL_891391 | Lighthouse Lab in Glasgow | Wellcome Sanger Institute for the COVID-19 Genomics UK (COG-UK) Consortium | Harper VanSteenhouse, Yumi Kasai, David Gray, Carol Clugston, Anna Dominiczak and Alex Alderton, Roberto Amato, Sonia Goncalves, Ewan Harrison, David K. Jackson, Ian Johnston, Dominic Kwiatkowski, Cordelia Langford, John Sillitoe on behalf of the Wellcome Sanger Institute COVID-19 Surveillance Team |
| EPI_ISL_891392 | Lighthouse Lab in Alderley Park | Wellcome Sanger Institute for the COVID-19 Genomics UK (COG-UK) Consortium | Jacquelyn Wynn, Mairead Hyland, The Lighthouse Lab in Alderley Park and Alex Alderton, Roberto Amato, Sonia Goncalves, Ewan Harrison, David K. Jackson, Ian Johnston, Dominic Kwiatkowski, Cordelia Langford, John Sillitoe on behalf of the Wellcome Sanger Institute COVID-19 Surveillance Team |
| EPI_ISL_891393 | Lighthouse Lab in Glasgow | Wellcome Sanger Institute for the COVID-19 Genomics UK (COG-UK) Consortium | Harper VanSteenhouse, Yumi Kasai, David Gray, Carol Clugston, Anna Dominiczak and Alex Alderton, Roberto Amato, Sonia Goncalves, Ewan Harrison, David K. Jackson, Ian Johnston, Dominic Kwiatkowski, Cordelia Langford, John Sillitoe on behalf of the Wellcome Sanger Institute COVID-19 Surveillance Team |
| EPI_ISL_891394, EPI_ISL_891395 | Lighthouse Lab in Cambridge | Wellcome Sanger Institute for the COVID-19 Genomics UK (COG-UK) Consortium | Rob Howes, The Lighthouse Lab in Cambridge and Alex Alderton, Roberto Amato, Sonia Goncalves, Ewan Harrison, David K. Jackson, Ian Johnston, Dominic Kwiatkowski, Cordelia Langford, John Sillitoe on behalf of the Wellcome Sanger Institute COVID-19 Surveillance Team |
| EPI_ISL_891396 | Lighthouse Lab in Glasgow | Wellcome Sanger Institute for the COVID-19 Genomics UK (COG-UK) Consortium | Harper VanSteenhouse, Yumi Kasai, David Gray, Carol Clugston, Anna Dominiczak and Alex Alderton, Roberto Amato, Sonia Goncalves, Ewan Harrison, David K. Jackson, Ian Johnston, Dominic Kwiatkowski, Cordelia Langford, John Sillitoe on behalf of the Wellcome Sanger Institute COVID-19 Surveillance Team |
| EPI_ISL_891397, EPI_ISL_891398, EPI_ISL_891399 | Lighthouse Lab in Cambridge | Wellcome Sanger Institute for the COVID-19 Genomics UK (COG-UK) Consortium | Rob Howes, The Lighthouse Lab in Cambridge and Alex Alderton, Roberto Amato, Sonia Goncalves, Ewan Harrison, David K. Jackson, Ian Johnston, Dominic Kwiatkowski, Cordelia Langford, John Sillitoe on behalf of the Wellcome Sanger Institute COVID-19 Surveillance Team |
| EPI_ISL_891400, EPI_ISL_891401, EPI_ISL_891403, EPI_ISL_891404 | Lighthouse Lab in Alderley Park | Wellcome Sanger Institute for the COVID-19 Genomics UK (COG-UK) Consortium | Jacquelyn Wynn, Mairead Hyland, The Lighthouse Lab in Alderley Park and Alex Alderton, Roberto Amato, Sonia Goncalves, Ewan Harrison, David K. Jackson, Ian Johnston, Dominic Kwiatkowski, Cordelia Langford, John Sillitoe on behalf of the Wellcome Sanger Institute COVID-19 Surveillance Team |
| EPI_ISL_891406, EPI_ISL_891409 | Lighthouse Lab in Cambridge | Wellcome Sanger Institute for the COVID-19 Genomics UK (COG-UK) Consortium | Rob Howes, The Lighthouse Lab in Cambridge and Alex Alderton, Roberto Amato, Sonia Goncalves, Ewan Harrison, David K. Jackson, Ian Johnston, Dominic Kwiatkowski, Cordelia Langford, John Sillitoe on behalf of the Wellcome Sanger Institute COVID-19 Surveillance Team |

| | | | |
|---|---|---|---|
| EPI_ISL_891410 | Lighthouse Lab in Glasgow | Wellcome Sanger Institute for the COVID-19 Genomics UK (COG-UK) Consortium | Harper VanSteenhouse, Yumi Kasai, David Gray, Carol Clugston, Anna Dominiczak and Alex Alderton, Roberto Amato, Sonia Goncalves, Ewan Harrison, David K. Jackson, Ian Johnston, Dominic Kwiatkowski, Cordelia Langford, John Sillitoe on behalf of the Wellcome Sanger Institute COVID-19 Surveillance Team |
| EPI_ISL_891411 | Lighthouse Lab in Alderley Park | Wellcome Sanger Institute for the COVID-19 Genomics UK (COG-UK) Consortium | Jacquelyn Wynn, Mairead Hyland, The Lighthouse Lab in Alderley Park and Alex Alderton, Roberto Amato, Sonia Goncalves, Ewan Harrison, David K. Jackson, Ian Johnston, Dominic Kwiatkowski, Cordelia Langford, John Sillitoe on behalf of the Wellcome Sanger Institute COVID-19 Surveillance Team |
| EPI_ISL_891412, EPI_ISL_891413 | Lighthouse Lab in Cambridge | Wellcome Sanger Institute for the COVID-19 Genomics UK (COG-UK) Consortium | Rob Howes, The Lighthouse Lab in Cambridge and Alex Alderton, Roberto Amato, Sonia Goncalves, Ewan Harrison, David K. Jackson, Ian Johnston, Dominic Kwiatkowski, Cordelia Langford, John Sillitoe on behalf of the Wellcome Sanger Institute COVID-19 Surveillance Team |
| EPI_ISL_891415 | Lighthouse Lab in Glasgow | Wellcome Sanger Institute for the COVID-19 Genomics UK (COG-UK) Consortium | Harper VanSteenhouse, Yumi Kasai, David Gray, Carol Clugston, Anna Dominiczak and Alex Alderton, Roberto Amato, Sonia Goncalves, Ewan Harrison, David K. Jackson, Ian Johnston, Dominic Kwiatkowski, Cordelia Langford, John Sillitoe on behalf of the Wellcome Sanger Institute COVID-19 Surveillance Team |
| EPI_ISL_891416, EPI_ISL_891417, EPI_ISL_891419 | Lighthouse Lab in Alderley Park | Wellcome Sanger Institute for the COVID-19 Genomics UK (COG-UK) Consortium | Jacquelyn Wynn, Mairead Hyland, The Lighthouse Lab in Alderley Park and Alex Alderton, Roberto Amato, Sonia Goncalves, Ewan Harrison, David K. Jackson, Ian Johnston, Dominic Kwiatkowski, Cordelia Langford, John Sillitoe on behalf of the Wellcome Sanger Institute COVID-19 Surveillance Team |
| EPI_ISL_891420 | Lighthouse Lab in Glasgow | Wellcome Sanger Institute for the COVID-19 Genomics UK (COG-UK) Consortium | Harper VanSteenhouse, Yumi Kasai, David Gray, Carol Clugston, Anna Dominiczak and Alex Alderton, Roberto Amato, Sonia Goncalves, Ewan Harrison, David K. Jackson, Ian Johnston, Dominic Kwiatkowski, Cordelia Langford, John Sillitoe on behalf of the Wellcome Sanger Institute COVID-19 Surveillance Team |
| EPI_ISL_891421 | Lighthouse Lab in Alderley Park | Wellcome Sanger Institute for the COVID-19 Genomics UK (COG-UK) Consortium | Jacquelyn Wynn, Mairead Hyland, The Lighthouse Lab in Alderley Park and Alex Alderton, Roberto Amato, Sonia Goncalves, Ewan Harrison, David K. Jackson, Ian Johnston, Dominic Kwiatkowski, Cordelia Langford, John Sillitoe on behalf of the Wellcome Sanger Institute COVID-19 Surveillance Team |
| EPI_ISL_891422 | Lighthouse Lab in Cambridge | Wellcome Sanger Institute for the COVID-19 Genomics UK (COG-UK) Consortium | Rob Howes, The Lighthouse Lab in Cambridge and Alex Alderton, Roberto Amato, Sonia Goncalves, Ewan Harrison, David K. Jackson, Ian Johnston, Dominic Kwiatkowski, Cordelia Langford, John Sillitoe on behalf of the Wellcome Sanger Institute COVID-19 Surveillance Team |
| EPI_ISL_891423 | Lighthouse Lab in Alderley Park | Wellcome Sanger Institute for the COVID-19 Genomics UK (COG-UK) Consortium | Jacquelyn Wynn, Mairead Hyland, The Lighthouse Lab in Alderley Park and Alex Alderton, Roberto Amato, Sonia Goncalves, Ewan Harrison, David K. Jackson, Ian Johnston, Dominic Kwiatkowski, Cordelia Langford, John Sillitoe on behalf of the Wellcome Sanger Institute COVID-19 Surveillance Team |
| EPI_ISL_891426 | Lighthouse Lab in Glasgow | Wellcome Sanger Institute for the COVID-19 Genomics UK (COG-UK) Consortium | Harper VanSteenhouse, Yumi Kasai, David Gray, Carol Clugston, Anna Dominiczak and Alex Alderton, Roberto Amato, Sonia Goncalves, Ewan Harrison, David K. Jackson, Ian Johnston, Dominic Kwiatkowski, Cordelia Langford, John Sillitoe on behalf of the Wellcome Sanger Institute COVID-19 Surveillance Team |
| EPI_ISL_891427, EPI_ISL_891431, EPI_ISL_891433, EPI_ISL_891434 | Lighthouse Lab in Cambridge | Wellcome Sanger Institute for the COVID-19 Genomics UK (COG-UK) Consortium | Rob Howes, The Lighthouse Lab in Cambridge and Alex Alderton, Roberto Amato, Sonia Goncalves, Ewan Harrison, David K. Jackson, Ian Johnston, Dominic Kwiatkowski, Cordelia Langford, John Sillitoe on behalf of the Wellcome Sanger Institute COVID-19 Surveillance Team |
| EPI_ISL_891435 | Lighthouse Lab in Glasgow | Wellcome Sanger Institute for the COVID-19 Genomics UK (COG-UK) Consortium | Harper VanSteenhouse, Yumi Kasai, David Gray, Carol Clugston, Anna Dominiczak and Alex Alderton, Roberto Amato, Sonia Goncalves, Ewan Harrison, David K. Jackson, Ian Johnston, Dominic Kwiatkowski, Cordelia Langford, John Sillitoe on behalf of the Wellcome Sanger Institute COVID-19 Surveillance Team |
| EPI_ISL_891436, EPI_ISL_891437 | Lighthouse Lab in Cambridge | Wellcome Sanger Institute for the COVID-19 Genomics UK (COG-UK) Consortium | Rob Howes, The Lighthouse Lab in Cambridge and Alex Alderton, Roberto Amato, Sonia Goncalves, Ewan Harrison, David K. Jackson, Ian Johnston, Dominic Kwiatkowski, Cordelia Langford, John Sillitoe on behalf of the Wellcome Sanger Institute COVID-19 Surveillance Team |
| EPI_ISL_891438 | Lighthouse Lab in Alderley Park | Wellcome Sanger Institute for the COVID-19 Genomics UK (COG-UK) Consortium | Jacquelyn Wynn, Mairead Hyland, The Lighthouse Lab in Alderley Park and Alex Alderton, Roberto Amato, Sonia Goncalves, Ewan Harrison, David K. Jackson, Ian Johnston, Dominic Kwiatkowski, Cordelia Langford, John Sillitoe on behalf of the Wellcome Sanger Institute COVID-19 Surveillance Team |
| EPI_ISL_891439, EPI_ISL_891441, EPI_ISL_891442 | Lighthouse Lab in Cambridge | Wellcome Sanger Institute for the COVID-19 Genomics UK (COG-UK) Consortium | Rob Howes, The Lighthouse Lab in Cambridge and Alex Alderton, Roberto Amato, Sonia Goncalves, Ewan Harrison, David K. Jackson, Ian Johnston, Dominic Kwiatkowski, Cordelia Langford, John Sillitoe on behalf of the Wellcome Sanger Institute COVID-19 Surveillance Team |
| EPI_ISL_891444 | Lighthouse Lab in Alderley Park | Wellcome Sanger Institute for the COVID-19 Genomics UK (COG-UK) Consortium | Jacquelyn Wynn, Mairead Hyland, The Lighthouse Lab in Alderley Park and Alex Alderton, Roberto Amato, Sonia Goncalves, Ewan Harrison, David K. Jackson, Ian Johnston, Dominic Kwiatkowski, Cordelia Langford, John Sillitoe on behalf of the Wellcome Sanger Institute COVID-19 Surveillance Team |
| EPI_ISL_891446 | Lighthouse Lab in Glasgow | Wellcome Sanger Institute for the COVID-19 Genomics UK (COG-UK) Consortium | Harper VanSteenhouse, Yumi Kasai, David Gray, Carol Clugston, Anna Dominiczak and Alex Alderton, Roberto Amato, Sonia Goncalves, Ewan Harrison, David K. Jackson, Ian Johnston, Dominic Kwiatkowski, Cordelia Langford, John Sillitoe on behalf of the Wellcome Sanger Institute COVID-19 Surveillance Team |
| EPI_ISL_891447 | Lighthouse Lab in Cambridge | Wellcome Sanger Institute for the COVID-19 Genomics UK (COG-UK) Consortium | Rob Howes, The Lighthouse Lab in Cambridge and Alex Alderton, Roberto Amato, Sonia Goncalves, Ewan Harrison, David K. Jackson, Ian Johnston, Dominic Kwiatkowski, Cordelia Langford, John Sillitoe on behalf of the Wellcome Sanger Institute COVID-19 Surveillance Team |
| EPI_ISL_891448 | Lighthouse Lab in Alderley Park | Wellcome Sanger Institute for the COVID-19 Genomics UK (COG-UK) Consortium | Jacquelyn Wynn, Mairead Hyland, The Lighthouse Lab in Alderley Park and Alex Alderton, Roberto Amato, Sonia Goncalves, Ewan Harrison, David K. Jackson, Ian Johnston, Dominic Kwiatkowski, Cordelia Langford, John Sillitoe on behalf of the Wellcome Sanger Institute COVID-19 Surveillance Team |
| EPI_ISL_891450 | Lighthouse Lab in Cambridge | Wellcome Sanger Institute for the COVID-19 Genomics UK (COG-UK) Consortium | Rob Howes, The Lighthouse Lab in Cambridge and Alex Alderton, Roberto Amato, Sonia Goncalves, Ewan Harrison, David K. Jackson, Ian Johnston, Dominic Kwiatkowski, Cordelia Langford, John Sillitoe on behalf of the Wellcome Sanger Institute COVID-19 Surveillance Team |
| EPI_ISL_891451 | Lighthouse Lab in Alderley Park | Wellcome Sanger Institute for the COVID-19 Genomics UK (COG-UK) Consortium | Jacquelyn Wynn, Mairead Hyland, The Lighthouse Lab in Alderley Park and Alex Alderton, Roberto Amato, Sonia Goncalves, Ewan Harrison, David K. Jackson, Ian Johnston, Dominic Kwiatkowski, Cordelia Langford, John Sillitoe on behalf of the Wellcome Sanger Institute COVID-19 Surveillance Team |
| EPI_ISL_891452 | Lighthouse Lab in Cambridge | Wellcome Sanger Institute for the COVID-19 Genomics UK (COG-UK) Consortium | Rob Howes, The Lighthouse Lab in Cambridge and Alex Alderton, Roberto Amato, Sonia Goncalves, Ewan Harrison, David K. Jackson, Ian Johnston, Dominic Kwiatkowski, Cordelia Langford, John Sillitoe on behalf of the Wellcome Sanger Institute COVID-19 Surveillance Team |
| EPI_ISL_891453 | Lighthouse Lab in Glasgow | Wellcome Sanger Institute for the COVID-19 Genomics UK (COG-UK) Consortium | Harper VanSteenhouse, Yumi Kasai, David Gray, Carol Clugston, Anna Dominiczak and Alex Alderton, Roberto Amato, Sonia Goncalves, Ewan Harrison, David K. Jackson, Ian Johnston, Dominic Kwiatkowski, Cordelia Langford, John Sillitoe on behalf of the Wellcome Sanger Institute COVID-19 Surveillance Team |
| EPI_ISL_891454, EPI_ISL_891455 | Lighthouse Lab in Alderley Park | Wellcome Sanger Institute for the COVID-19 Genomics UK (COG-UK) Consortium | Jacquelyn Wynn, Mairead Hyland, The Lighthouse Lab in Alderley Park and Alex Alderton, Roberto Amato, Sonia Goncalves, Ewan Harrison, David K. Jackson, Ian Johnston, Dominic Kwiatkowski, Cordelia Langford, John Sillitoe on behalf of the Wellcome Sanger Institute COVID-19 Surveillance Team |
| EPI_ISL_891457 | Lighthouse Lab in Cambridge | Wellcome Sanger Institute for the COVID-19 Genomics UK (COG-UK) Consortium | Rob Howes, The Lighthouse Lab in Cambridge and Alex Alderton, Roberto Amato, Sonia Goncalves, Ewan Harrison, David K. Jackson, Ian Johnston, Dominic Kwiatkowski, Cordelia Langford, John Sillitoe on behalf of the Wellcome Sanger Institute COVID-19 Surveillance Team |
| EPI_ISL_891459, EPI_ISL_891460 | Lighthouse Lab in Alderley Park | Wellcome Sanger Institute for the COVID-19 Genomics UK (COG-UK) Consortium | Jacquelyn Wynn, Mairead Hyland, The Lighthouse Lab in Alderley Park and Alex Alderton, Roberto Amato, Sonia Goncalves, Ewan Harrison, David K. Jackson, Ian Johnston, Dominic Kwiatkowski, Cordelia Langford, John Sillitoe on behalf of the Wellcome Sanger Institute COVID-19 Surveillance Team |
| EPI_ISL_891461 | Lighthouse Lab in Cambridge | Wellcome Sanger Institute for the COVID-19 Genomics UK (COG-UK) Consortium | Rob Howes, The Lighthouse Lab in Cambridge and Alex Alderton, Roberto Amato, Sonia Goncalves, Ewan Harrison, David K. Jackson, Ian Johnston, Dominic Kwiatkowski, Cordelia Langford, John Sillitoe on behalf of the Wellcome Sanger Institute COVID-19 Surveillance Team |
| EPI_ISL_891462 | Lighthouse Lab in Glasgow | Wellcome Sanger Institute for the COVID-19 Genomics UK (COG-UK) Consortium | Harper VanSteenhouse, Yumi Kasai, David Gray, Carol Clugston, Anna Dominiczak and Alex Alderton, Roberto Amato, Sonia Goncalves, Ewan Harrison, David K. Jackson, Ian Johnston, Dominic Kwiatkowski, Cordelia Langford, John Sillitoe on behalf of the Wellcome Sanger Institute COVID-19 Surveillance Team |
| EPI_ISL_891463 | Lighthouse Lab in Alderley Park | Wellcome Sanger Institute for the COVID-19 Genomics UK (COG-UK) Consortium | Jacquelyn Wynn, Mairead Hyland, The Lighthouse Lab in Alderley Park and Alex Alderton, Roberto Amato, Sonia Goncalves, Ewan Harrison, David K. Jackson, Ian Johnston, Dominic Kwiatkowski, Cordelia Langford, John Sillitoe on behalf of the Wellcome Sanger Institute COVID-19 Surveillance Team |
| EPI_ISL_891464, EPI_ISL_891465 | Lighthouse Lab in Glasgow | Wellcome Sanger Institute for the COVID-19 Genomics UK (COG-UK) Consortium | Harper VanSteenhouse, Yumi Kasai, David Gray, Carol Clugston, Anna Dominiczak and Alex Alderton, Roberto Amato, Sonia Goncalves, Ewan Harrison, David K. Jackson, Ian Johnston, Dominic Kwiatkowski, Cordelia Langford, John Sillitoe on behalf of the Wellcome Sanger Institute COVID-19 Surveillance Team |
| EPI_ISL_891466, EPI_ISL_891467, EPI_ISL_891470, EPI_ISL_891472 | Lighthouse Lab in Cambridge | Wellcome Sanger Institute for the COVID-19 Genomics UK (COG-UK) Consortium | Rob Howes, The Lighthouse Lab in Cambridge and Alex Alderton, Roberto Amato, Sonia Goncalves, Ewan Harrison, David K. Jackson, Ian Johnston, Dominic Kwiatkowski, Cordelia Langford, John Sillitoe on behalf of the Wellcome Sanger Institute COVID-19 Surveillance Team |
| EPI_ISL_891473 | Lighthouse Lab in Glasgow | Wellcome Sanger Institute for the COVID-19 Genomics UK | Harper VanSteenhouse, Yumi Kasai, David Gray, Carol Clugston, Anna Dominiczak and Alex Alderton, Roberto Amato, Sonia Goncalves, Ewan Harrison, |

| | | | |
|---|---|---|---|
| | | (COG-UK) Consortium | David K. Jackson, Ian Johnston, Dominic Kwiatkowski, Cordelia Langford, John Sillitoe on behalf of the Wellcome Sanger Institute COVID-19 Surveillance Team |
| EPI_ISL_891475, EPI_ISL_891476, EPI_ISL_891477 | Lighthouse Lab in Cambridge | Wellcome Sanger Institute for the COVID-19 Genomics UK (COG-UK) Consortium | Rob Howes, The Lighthouse Lab in Cambridge and Alex Alderton, Roberto Amato, Sonia Goncalves, Ewan Harrison, David K. Jackson, Ian Johnston, Dominic Kwiatkowski, Cordelia Langford, John Sillitoe on behalf of the Wellcome Sanger Institute COVID-19 Surveillance Team |
| EPI_ISL_891478 | Lighthouse Lab in Alderley Park | Wellcome Sanger Institute for the COVID-19 Genomics UK (COG-UK) Consortium | Jacquelyn Wynn, Mairead Hyland, The Lighthouse Lab in Alderley Park and Alex Alderton, Roberto Amato, Sonia Goncalves, Ewan Harrison, David K. Jackson, Ian Johnston, Dominic Kwiatkowski, Cordelia Langford, John Sillitoe on behalf of the Wellcome Sanger Institute COVID-19 Surveillance Team |
| EPI_ISL_891480, EPI_ISL_891481, EPI_ISL_891483 | Lighthouse Lab in Cambridge | Wellcome Sanger Institute for the COVID-19 Genomics UK (COG-UK) Consortium | Rob Howes, The Lighthouse Lab in Cambridge and Alex Alderton, Roberto Amato, Sonia Goncalves, Ewan Harrison, David K. Jackson, Ian Johnston, Dominic Kwiatkowski, Cordelia Langford, John Sillitoe on behalf of the Wellcome Sanger Institute COVID-19 Surveillance Team |
| EPI_ISL_891484, EPI_ISL_891485 | Lighthouse Lab in Alderley Park | Wellcome Sanger Institute for the COVID-19 Genomics UK (COG-UK) Consortium | Jacquelyn Wynn, Mairead Hyland, The Lighthouse Lab in Alderley Park and Alex Alderton, Roberto Amato, Sonia Goncalves, Ewan Harrison, David K. Jackson, Ian Johnston, Dominic Kwiatkowski, Cordelia Langford, John Sillitoe on behalf of the Wellcome Sanger Institute COVID-19 Surveillance Team |
| EPI_ISL_891486, EPI_ISL_891488 | Lighthouse Lab in Cambridge | Wellcome Sanger Institute for the COVID-19 Genomics UK (COG-UK) Consortium | Rob Howes, The Lighthouse Lab in Cambridge and Alex Alderton, Roberto Amato, Sonia Goncalves, Ewan Harrison, David K. Jackson, Ian Johnston, Dominic Kwiatkowski, Cordelia Langford, John Sillitoe on behalf of the Wellcome Sanger Institute COVID-19 Surveillance Team |
| EPI_ISL_891489, EPI_ISL_891490 | Lighthouse Lab in Alderley Park | Wellcome Sanger Institute for the COVID-19 Genomics UK (COG-UK) Consortium | Jacquelyn Wynn, Mairead Hyland, The Lighthouse Lab in Alderley Park and Alex Alderton, Roberto Amato, Sonia Goncalves, Ewan Harrison, David K. Jackson, Ian Johnston, Dominic Kwiatkowski, Cordelia Langford, John Sillitoe on behalf of the Wellcome Sanger Institute COVID-19 Surveillance Team |
| EPI_ISL_891491 | Lighthouse Lab in Glasgow | Wellcome Sanger Institute for the COVID-19 Genomics UK (COG-UK) Consortium | Harper VanSteenhouse, Yumi Kasai, David Gray, Carol Clugston, Anna Dominiczak and Alex Alderton, Roberto Amato, Sonia Goncalves, Ewan Harrison, David K. Jackson, Ian Johnston, Dominic Kwiatkowski, Cordelia Langford, John Sillitoe on behalf of the Wellcome Sanger Institute COVID-19 Surveillance Team |
| EPI_ISL_891492, EPI_ISL_891493, EPI_ISL_891494 | Lighthouse Lab in Cambridge | Wellcome Sanger Institute for the COVID-19 Genomics UK (COG-UK) Consortium | Rob Howes, The Lighthouse Lab in Cambridge and Alex Alderton, Roberto Amato, Sonia Goncalves, Ewan Harrison, David K. Jackson, Ian Johnston, Dominic Kwiatkowski, Cordelia Langford, John Sillitoe on behalf of the Wellcome Sanger Institute COVID-19 Surveillance Team |
| EPI_ISL_891495 | Lighthouse Lab in Glasgow | Wellcome Sanger Institute for the COVID-19 Genomics UK (COG-UK) Consortium | Harper VanSteenhouse, Yumi Kasai, David Gray, Carol Clugston, Anna Dominiczak and Alex Alderton, Roberto Amato, Sonia Goncalves, Ewan Harrison, David K. Jackson, Ian Johnston, Dominic Kwiatkowski, Cordelia Langford, John Sillitoe on behalf of the Wellcome Sanger Institute COVID-19 Surveillance Team |
| EPI_ISL_891496 | Lighthouse Lab in Alderley Park | Wellcome Sanger Institute for the COVID-19 Genomics UK (COG-UK) Consortium | Jacquelyn Wynn, Mairead Hyland, The Lighthouse Lab in Alderley Park and Alex Alderton, Roberto Amato, Sonia Goncalves, Ewan Harrison, David K. Jackson, Ian Johnston, Dominic Kwiatkowski, Cordelia Langford, John Sillitoe on behalf of the Wellcome Sanger Institute COVID-19 Surveillance Team |
| EPI_ISL_891497 | Lighthouse Lab in Glasgow | Wellcome Sanger Institute for the COVID-19 Genomics UK (COG-UK) Consortium | Harper VanSteenhouse, Yumi Kasai, David Gray, Carol Clugston, Anna Dominiczak and Alex Alderton, Roberto Amato, Sonia Goncalves, Ewan Harrison, David K. Jackson, Ian Johnston, Dominic Kwiatkowski, Cordelia Langford, John Sillitoe on behalf of the Wellcome Sanger Institute COVID-19 Surveillance Team |
| EPI_ISL_891498 | Lighthouse Lab in Cambridge | Wellcome Sanger Institute for the COVID-19 Genomics UK (COG-UK) Consortium | Rob Howes, The Lighthouse Lab in Cambridge and Alex Alderton, Roberto Amato, Sonia Goncalves, Ewan Harrison, David K. Jackson, Ian Johnston, Dominic Kwiatkowski, Cordelia Langford, John Sillitoe on behalf of the Wellcome Sanger Institute COVID-19 Surveillance Team |
| EPI_ISL_891499 | Lighthouse Lab in Alderley Park | Wellcome Sanger Institute for the COVID-19 Genomics UK (COG-UK) Consortium | Jacquelyn Wynn, Mairead Hyland, The Lighthouse Lab in Alderley Park and Alex Alderton, Roberto Amato, Sonia Goncalves, Ewan Harrison, David K. Jackson, Ian Johnston, Dominic Kwiatkowski, Cordelia Langford, John Sillitoe on behalf of the Wellcome Sanger Institute COVID-19 Surveillance Team |
| EPI_ISL_891500, EPI_ISL_891501 | Lighthouse Lab in Cambridge | Wellcome Sanger Institute for the COVID-19 Genomics UK (COG-UK) Consortium | Rob Howes, The Lighthouse Lab in Cambridge and Alex Alderton, Roberto Amato, Sonia Goncalves, Ewan Harrison, David K. Jackson, Ian Johnston, Dominic Kwiatkowski, Cordelia Langford, John Sillitoe on behalf of the Wellcome Sanger Institute COVID-19 Surveillance Team |
| EPI_ISL_891502 | Lighthouse Lab in Glasgow | Wellcome Sanger Institute for the COVID-19 Genomics UK (COG-UK) Consortium | Harper VanSteenhouse, Yumi Kasai, David Gray, Carol Clugston, Anna Dominiczak and Alex Alderton, Roberto Amato, Sonia Goncalves, Ewan Harrison, David K. Jackson, Ian Johnston, Dominic Kwiatkowski, Cordelia Langford, John Sillitoe on behalf of the Wellcome Sanger Institute COVID-19 Surveillance Team |
| EPI_ISL_891503 | Lighthouse Lab in Alderley Park | Wellcome Sanger Institute for the COVID-19 Genomics UK (COG-UK) Consortium | Jacquelyn Wynn, Mairead Hyland, The Lighthouse Lab in Alderley Park and Alex Alderton, Roberto Amato, Sonia Goncalves, Ewan Harrison, David K. Jackson, Ian Johnston, Dominic Kwiatkowski, Cordelia Langford, John Sillitoe on behalf of the Wellcome Sanger Institute COVID-19 Surveillance Team |
| EPI_ISL_891504 | Lighthouse Lab in Cambridge | Wellcome Sanger Institute for the COVID-19 Genomics UK (COG-UK) Consortium | Rob Howes, The Lighthouse Lab in Cambridge and Alex Alderton, Roberto Amato, Sonia Goncalves, Ewan Harrison, David K. Jackson, Ian Johnston, Dominic Kwiatkowski, Cordelia Langford, John Sillitoe on behalf of the Wellcome Sanger Institute COVID-19 Surveillance Team |
| EPI_ISL_891505 | Lighthouse Lab in Glasgow | Wellcome Sanger Institute for the COVID-19 Genomics UK (COG-UK) Consortium | Harper VanSteenhouse, Yumi Kasai, David Gray, Carol Clugston, Anna Dominiczak and Alex Alderton, Roberto Amato, Sonia Goncalves, Ewan Harrison, David K. Jackson, Ian Johnston, Dominic Kwiatkowski, Cordelia Langford, John Sillitoe on behalf of the Wellcome Sanger Institute COVID-19 Surveillance Team |
| EPI_ISL_891506 | Lighthouse Lab in Cambridge | Wellcome Sanger Institute for the COVID-19 Genomics UK (COG-UK) Consortium | Rob Howes, The Lighthouse Lab in Cambridge and Alex Alderton, Roberto Amato, Sonia Goncalves, Ewan Harrison, David K. Jackson, Ian Johnston, Dominic Kwiatkowski, Cordelia Langford, John Sillitoe on behalf of the Wellcome Sanger Institute COVID-19 Surveillance Team |
| EPI_ISL_891508, EPI_ISL_891509 | Lighthouse Lab in Glasgow | Wellcome Sanger Institute for the COVID-19 Genomics UK (COG-UK) Consortium | Harper VanSteenhouse, Yumi Kasai, David Gray, Carol Clugston, Anna Dominiczak and Alex Alderton, Roberto Amato, Sonia Goncalves, Ewan Harrison, David K. Jackson, Ian Johnston, Dominic Kwiatkowski, Cordelia Langford, John Sillitoe on behalf of the Wellcome Sanger Institute COVID-19 Surveillance Team |
| EPI_ISL_891510 | Lighthouse Lab in Cambridge | Wellcome Sanger Institute for the COVID-19 Genomics UK (COG-UK) Consortium | Rob Howes, The Lighthouse Lab in Cambridge and Alex Alderton, Roberto Amato, Sonia Goncalves, Ewan Harrison, David K. Jackson, Ian Johnston, Dominic Kwiatkowski, Cordelia Langford, John Sillitoe on behalf of the Wellcome Sanger Institute COVID-19 Surveillance Team |
| EPI_ISL_891511 | Lighthouse Lab in Alderley Park | Wellcome Sanger Institute for the COVID-19 Genomics UK (COG-UK) Consortium | Jacquelyn Wynn, Mairead Hyland, The Lighthouse Lab in Alderley Park and Alex Alderton, Roberto Amato, Sonia Goncalves, Ewan Harrison, David K. Jackson, Ian Johnston, Dominic Kwiatkowski, Cordelia Langford, John Sillitoe on behalf of the Wellcome Sanger Institute COVID-19 Surveillance Team |
| EPI_ISL_891513 | Lighthouse Lab in Cambridge | Wellcome Sanger Institute for the COVID-19 Genomics UK (COG-UK) Consortium | Rob Howes, The Lighthouse Lab in Cambridge and Alex Alderton, Roberto Amato, Sonia Goncalves, Ewan Harrison, David K. Jackson, Ian Johnston, Dominic Kwiatkowski, Cordelia Langford, John Sillitoe on behalf of the Wellcome Sanger Institute COVID-19 Surveillance Team |
| EPI_ISL_891514 | Lighthouse Lab in Alderley Park | Wellcome Sanger Institute for the COVID-19 Genomics UK (COG-UK) Consortium | Jacquelyn Wynn, Mairead Hyland, The Lighthouse Lab in Alderley Park and Alex Alderton, Roberto Amato, Sonia Goncalves, Ewan Harrison, David K. Jackson, Ian Johnston, Dominic Kwiatkowski, Cordelia Langford, John Sillitoe on behalf of the Wellcome Sanger Institute COVID-19 Surveillance Team |
| EPI_ISL_891515 | Lighthouse Lab in Cambridge | Wellcome Sanger Institute for the COVID-19 Genomics UK (COG-UK) Consortium | Rob Howes, The Lighthouse Lab in Cambridge and Alex Alderton, Roberto Amato, Sonia Goncalves, Ewan Harrison, David K. Jackson, Ian Johnston, Dominic Kwiatkowski, Cordelia Langford, John Sillitoe on behalf of the Wellcome Sanger Institute COVID-19 Surveillance Team |
| EPI_ISL_891516 | Lighthouse Lab in Glasgow | Wellcome Sanger Institute for the COVID-19 Genomics UK (COG-UK) Consortium | Harper VanSteenhouse, Yumi Kasai, David Gray, Carol Clugston, Anna Dominiczak and Alex Alderton, Roberto Amato, Sonia Goncalves, Ewan Harrison, David K. Jackson, Ian Johnston, Dominic Kwiatkowski, Cordelia Langford, John Sillitoe on behalf of the Wellcome Sanger Institute COVID-19 Surveillance Team |
| EPI_ISL_891517, EPI_ISL_891518, EPI_ISL_891519 | Lighthouse Lab in Cambridge | Wellcome Sanger Institute for the COVID-19 Genomics UK (COG-UK) Consortium | Rob Howes, The Lighthouse Lab in Cambridge and Alex Alderton, Roberto Amato, Sonia Goncalves, Ewan Harrison, David K. Jackson, Ian Johnston, Dominic Kwiatkowski, Cordelia Langford, John Sillitoe on behalf of the Wellcome Sanger Institute COVID-19 Surveillance Team |
| EPI_ISL_891520 | Lighthouse Lab in Alderley Park | Wellcome Sanger Institute for the COVID-19 Genomics UK (COG-UK) Consortium | Jacquelyn Wynn, Mairead Hyland, The Lighthouse Lab in Alderley Park and Alex Alderton, Roberto Amato, Sonia Goncalves, Ewan Harrison, David K. Jackson, Ian Johnston, Dominic Kwiatkowski, Cordelia Langford, John Sillitoe on behalf of the Wellcome Sanger Institute COVID-19 Surveillance Team |
| EPI_ISL_891521 | Lighthouse Lab in Cambridge | Wellcome Sanger Institute for the COVID-19 Genomics UK (COG-UK) Consortium | Rob Howes, The Lighthouse Lab in Cambridge and Alex Alderton, Roberto Amato, Sonia Goncalves, Ewan Harrison, David K. Jackson, Ian Johnston, Dominic Kwiatkowski, Cordelia Langford, John Sillitoe on behalf of the Wellcome Sanger Institute COVID-19 Surveillance Team |
| EPI_ISL_891522, EPI_ISL_891525 | Lighthouse Lab in Alderley Park | Wellcome Sanger Institute for the COVID-19 Genomics UK (COG-UK) Consortium | Jacquelyn Wynn, Mairead Hyland, The Lighthouse Lab in Alderley Park and Alex Alderton, Roberto Amato, Sonia Goncalves, Ewan Harrison, David K. Jackson, Ian Johnston, Dominic Kwiatkowski, Cordelia Langford, John Sillitoe on behalf of the Wellcome Sanger Institute COVID-19 Surveillance Team |
| EPI_ISL_891526 | Lighthouse Lab in Cambridge | Wellcome Sanger Institute for the COVID-19 Genomics UK (COG-UK) Consortium | Rob Howes, The Lighthouse Lab in Cambridge and Alex Alderton, Roberto Amato, Sonia Goncalves, Ewan Harrison, David K. Jackson, Ian Johnston, Dominic Kwiatkowski, Cordelia Langford, John Sillitoe on behalf of the Wellcome Sanger Institute COVID-19 Surveillance Team |
| EPI_ISL_891527, EPI_ISL_891528, | Lighthouse Lab in Alderley Park | Wellcome Sanger Institute for the COVID-19 Genomics UK | Jacquelyn Wynn, Mairead Hyland, The Lighthouse Lab in Alderley Park and Alex Alderton, Roberto Amato, Sonia Goncalves, Ewan Harrison, David K. |

| | | | |
|---|---|---|---|
| EPI_ISL_891529 | | (COG-UK) Consortium | Jackson, Ian Johnston, Dominic Kwiatkowski, Cordelia Langford, John Sillitoe on behalf of the Wellcome Sanger Institute COVID-19 Surveillance Team |
| EPI_ISL_891530 | Lighthouse Lab in Cambridge | Wellcome Sanger Institute for the COVID-19 Genomics UK (COG-UK) Consortium | Rob Howes, The Lighthouse Lab in Cambridge and Alex Alderton, Roberto Amato, Sonia Goncalves, Ewan Harrison, David K. Jackson, Ian Johnston, Dominic Kwiatkowski, Cordelia Langford, John Sillitoe on behalf of the Wellcome Sanger Institute COVID-19 Surveillance Team |
| EPI_ISL_891531 | Lighthouse Lab in Alderley Park | Wellcome Sanger Institute for the COVID-19 Genomics UK (COG-UK) Consortium | Jacquelyn Wynn, Mairead Hyland, The Lighthouse Lab in Alderley Park and Alex Alderton, Roberto Amato, Sonia Goncalves, Ewan Harrison, David K. Jackson, Ian Johnston, Dominic Kwiatkowski, Cordelia Langford, John Sillitoe on behalf of the Wellcome Sanger Institute COVID-19 Surveillance Team |
| EPI_ISL_891534, EPI_ISL_891535 | Lighthouse Lab in Cambridge | Wellcome Sanger Institute for the COVID-19 Genomics UK (COG-UK) Consortium | Rob Howes, The Lighthouse Lab in Cambridge and Alex Alderton, Roberto Amato, Sonia Goncalves, Ewan Harrison, David K. Jackson, Ian Johnston, Dominic Kwiatkowski, Cordelia Langford, John Sillitoe on behalf of the Wellcome Sanger Institute COVID-19 Surveillance Team |
| EPI_ISL_891536 | Lighthouse Lab in Alderley Park | Wellcome Sanger Institute for the COVID-19 Genomics UK (COG-UK) Consortium | Jacquelyn Wynn, Mairead Hyland, The Lighthouse Lab in Alderley Park and Alex Alderton, Roberto Amato, Sonia Goncalves, Ewan Harrison, David K. Jackson, Ian Johnston, Dominic Kwiatkowski, Cordelia Langford, John Sillitoe on behalf of the Wellcome Sanger Institute COVID-19 Surveillance Team |
| EPI_ISL_891537 | Lighthouse Lab in Cambridge | Wellcome Sanger Institute for the COVID-19 Genomics UK (COG-UK) Consortium | Rob Howes, The Lighthouse Lab in Cambridge and Alex Alderton, Roberto Amato, Sonia Goncalves, Ewan Harrison, David K. Jackson, Ian Johnston, Dominic Kwiatkowski, Cordelia Langford, John Sillitoe on behalf of the Wellcome Sanger Institute COVID-19 Surveillance Team |
| EPI_ISL_891539 | Lighthouse Lab in Glasgow | Wellcome Sanger Institute for the COVID-19 Genomics UK (COG-UK) Consortium | Harper VanSteenhouse, Yumi Kasai, David Gray, Carol Clugston, Anna Dominiczak and Alex Alderton, Roberto Amato, Sonia Goncalves, Ewan Harrison, David K. Jackson, Ian Johnston, Dominic Kwiatkowski, Cordelia Langford, John Sillitoe on behalf of the Wellcome Sanger Institute COVID-19 Surveillance Team |
| EPI_ISL_891540, EPI_ISL_891541, EPI_ISL_891543 | Lighthouse Lab in Cambridge | Wellcome Sanger Institute for the COVID-19 Genomics UK (COG-UK) Consortium | Rob Howes, The Lighthouse Lab in Cambridge and Alex Alderton, Roberto Amato, Sonia Goncalves, Ewan Harrison, David K. Jackson, Ian Johnston, Dominic Kwiatkowski, Cordelia Langford, John Sillitoe on behalf of the Wellcome Sanger Institute COVID-19 Surveillance Team |
| EPI_ISL_891544, EPI_ISL_891545, EPI_ISL_891547, EPI_ISL_891548 | Lighthouse Lab in Alderley Park | Wellcome Sanger Institute for the COVID-19 Genomics UK (COG-UK) Consortium | Jacquelyn Wynn, Mairead Hyland, The Lighthouse Lab in Alderley Park and Alex Alderton, Roberto Amato, Sonia Goncalves, Ewan Harrison, David K. Jackson, Ian Johnston, Dominic Kwiatkowski, Cordelia Langford, John Sillitoe on behalf of the Wellcome Sanger Institute COVID-19 Surveillance Team |
| EPI_ISL_891549 | Lighthouse Lab in Glasgow | Wellcome Sanger Institute for the COVID-19 Genomics UK (COG-UK) Consortium | Harper VanSteenhouse, Yumi Kasai, David Gray, Carol Clugston, Anna Dominiczak and Alex Alderton, Roberto Amato, Sonia Goncalves, Ewan Harrison, David K. Jackson, Ian Johnston, Dominic Kwiatkowski, Cordelia Langford, John Sillitoe on behalf of the Wellcome Sanger Institute COVID-19 Surveillance Team |
| EPI_ISL_891550, EPI_ISL_891551, EPI_ISL_891553, EPI_ISL_891554 | Lighthouse Lab in Cambridge | Wellcome Sanger Institute for the COVID-19 Genomics UK (COG-UK) Consortium | Rob Howes, The Lighthouse Lab in Cambridge and Alex Alderton, Roberto Amato, Sonia Goncalves, Ewan Harrison, David K. Jackson, Ian Johnston, Dominic Kwiatkowski, Cordelia Langford, John Sillitoe on behalf of the Wellcome Sanger Institute COVID-19 Surveillance Team |
| EPI_ISL_891555 | Lighthouse Lab in Alderley Park | Wellcome Sanger Institute for the COVID-19 Genomics UK (COG-UK) Consortium | Jacquelyn Wynn, Mairead Hyland, The Lighthouse Lab in Alderley Park and Alex Alderton, Roberto Amato, Sonia Goncalves, Ewan Harrison, David K. Jackson, Ian Johnston, Dominic Kwiatkowski, Cordelia Langford, John Sillitoe on behalf of the Wellcome Sanger Institute COVID-19 Surveillance Team |
| EPI_ISL_891556, EPI_ISL_891557, EPI_ISL_891558 | Lighthouse Lab in Cambridge | Wellcome Sanger Institute for the COVID-19 Genomics UK (COG-UK) Consortium | Rob Howes, The Lighthouse Lab in Cambridge and Alex Alderton, Roberto Amato, Sonia Goncalves, Ewan Harrison, David K. Jackson, Ian Johnston, Dominic Kwiatkowski, Cordelia Langford, John Sillitoe on behalf of the Wellcome Sanger Institute COVID-19 Surveillance Team |
| EPI_ISL_891560 | Lighthouse Lab in Alderley Park | Wellcome Sanger Institute for the COVID-19 Genomics UK (COG-UK) Consortium | Jacquelyn Wynn, Mairead Hyland, The Lighthouse Lab in Alderley Park and Alex Alderton, Roberto Amato, Sonia Goncalves, Ewan Harrison, David K. Jackson, Ian Johnston, Dominic Kwiatkowski, Cordelia Langford, John Sillitoe on behalf of the Wellcome Sanger Institute COVID-19 Surveillance Team |
| EPI_ISL_891561 | Lighthouse Lab in Cambridge | Wellcome Sanger Institute for the COVID-19 Genomics UK (COG-UK) Consortium | Rob Howes, The Lighthouse Lab in Cambridge and Alex Alderton, Roberto Amato, Sonia Goncalves, Ewan Harrison, David K. Jackson, Ian Johnston, Dominic Kwiatkowski, Cordelia Langford, John Sillitoe on behalf of the Wellcome Sanger Institute COVID-19 Surveillance Team |
| EPI_ISL_891562 | Lighthouse Lab in Glasgow | Wellcome Sanger Institute for the COVID-19 Genomics UK (COG-UK) Consortium | Harper VanSteenhouse, Yumi Kasai, David Gray, Carol Clugston, Anna Dominiczak and Alex Alderton, Roberto Amato, Sonia Goncalves, Ewan Harrison, David K. Jackson, Ian Johnston, Dominic Kwiatkowski, Cordelia Langford, John Sillitoe on behalf of the Wellcome Sanger Institute COVID-19 Surveillance Team |
| EPI_ISL_891563, EPI_ISL_891564 | Lighthouse Lab in Alderley Park | Wellcome Sanger Institute for the COVID-19 Genomics UK (COG-UK) Consortium | Jacquelyn Wynn, Mairead Hyland, The Lighthouse Lab in Alderley Park and Alex Alderton, Roberto Amato, Sonia Goncalves, Ewan Harrison, David K. Jackson, Ian Johnston, Dominic Kwiatkowski, Cordelia Langford, John Sillitoe on behalf of the Wellcome Sanger Institute COVID-19 Surveillance Team |
| EPI_ISL_891565, EPI_ISL_891566, EPI_ISL_891567 | Lighthouse Lab in Cambridge | Wellcome Sanger Institute for the COVID-19 Genomics UK (COG-UK) Consortium | Rob Howes, The Lighthouse Lab in Cambridge and Alex Alderton, Roberto Amato, Sonia Goncalves, Ewan Harrison, David K. Jackson, Ian Johnston, Dominic Kwiatkowski, Cordelia Langford, John Sillitoe on behalf of the Wellcome Sanger Institute COVID-19 Surveillance Team |
| EPI_ISL_891568 | Lighthouse Lab in Glasgow | Wellcome Sanger Institute for the COVID-19 Genomics UK (COG-UK) Consortium | Harper VanSteenhouse, Yumi Kasai, David Gray, Carol Clugston, Anna Dominiczak and Alex Alderton, Roberto Amato, Sonia Goncalves, Ewan Harrison, David K. Jackson, Ian Johnston, Dominic Kwiatkowski, Cordelia Langford, John Sillitoe on behalf of the Wellcome Sanger Institute COVID-19 Surveillance Team |
| EPI_ISL_891569 | Lighthouse Lab in Alderley Park | Wellcome Sanger Institute for the COVID-19 Genomics UK (COG-UK) Consortium | Jacquelyn Wynn, Mairead Hyland, The Lighthouse Lab in Alderley Park and Alex Alderton, Roberto Amato, Sonia Goncalves, Ewan Harrison, David K. Jackson, Ian Johnston, Dominic Kwiatkowski, Cordelia Langford, John Sillitoe on behalf of the Wellcome Sanger Institute COVID-19 Surveillance Team |
| EPI_ISL_891570 | Lighthouse Lab in Cambridge | Wellcome Sanger Institute for the COVID-19 Genomics UK (COG-UK) Consortium | Rob Howes, The Lighthouse Lab in Cambridge and Alex Alderton, Roberto Amato, Sonia Goncalves, Ewan Harrison, David K. Jackson, Ian Johnston, Dominic Kwiatkowski, Cordelia Langford, John Sillitoe on behalf of the Wellcome Sanger Institute COVID-19 Surveillance Team |
| EPI_ISL_891571 | Lighthouse Lab in Glasgow | Wellcome Sanger Institute for the COVID-19 Genomics UK (COG-UK) Consortium | Harper VanSteenhouse, Yumi Kasai, David Gray, Carol Clugston, Anna Dominiczak and Alex Alderton, Roberto Amato, Sonia Goncalves, Ewan Harrison, David K. Jackson, Ian Johnston, Dominic Kwiatkowski, Cordelia Langford, John Sillitoe on behalf of the Wellcome Sanger Institute COVID-19 Surveillance Team |
| EPI_ISL_891572 | Lighthouse Lab in Alderley Park | Wellcome Sanger Institute for the COVID-19 Genomics UK (COG-UK) Consortium | Jacquelyn Wynn, Mairead Hyland, The Lighthouse Lab in Alderley Park and Alex Alderton, Roberto Amato, Sonia Goncalves, Ewan Harrison, David K. Jackson, Ian Johnston, Dominic Kwiatkowski, Cordelia Langford, John Sillitoe on behalf of the Wellcome Sanger Institute COVID-19 Surveillance Team |
| EPI_ISL_891574 | Lighthouse Lab in Glasgow | Wellcome Sanger Institute for the COVID-19 Genomics UK (COG-UK) Consortium | Harper VanSteenhouse, Yumi Kasai, David Gray, Carol Clugston, Anna Dominiczak and Alex Alderton, Roberto Amato, Sonia Goncalves, Ewan Harrison, David K. Jackson, Ian Johnston, Dominic Kwiatkowski, Cordelia Langford, John Sillitoe on behalf of the Wellcome Sanger Institute COVID-19 Surveillance Team |
| EPI_ISL_891575, EPI_ISL_891577, EPI_ISL_891578, EPI_ISL_891579, EPI_ISL_891580, EPI_ISL_891581 | Lighthouse Lab in Cambridge | Wellcome Sanger Institute for the COVID-19 Genomics UK (COG-UK) Consortium | Rob Howes, The Lighthouse Lab in Cambridge and Alex Alderton, Roberto Amato, Sonia Goncalves, Ewan Harrison, David K. Jackson, Ian Johnston, Dominic Kwiatkowski, Cordelia Langford, John Sillitoe on behalf of the Wellcome Sanger Institute COVID-19 Surveillance Team |
| EPI_ISL_891582 | Lighthouse Lab in Alderley Park | Wellcome Sanger Institute for the COVID-19 Genomics UK (COG-UK) Consortium | Jacquelyn Wynn, Mairead Hyland, The Lighthouse Lab in Alderley Park and Alex Alderton, Roberto Amato, Sonia Goncalves, Ewan Harrison, David K. Jackson, Ian Johnston, Dominic Kwiatkowski, Cordelia Langford, John Sillitoe on behalf of the Wellcome Sanger Institute COVID-19 Surveillance Team |
| EPI_ISL_891583, EPI_ISL_891584 | Lighthouse Lab in Cambridge | Wellcome Sanger Institute for the COVID-19 Genomics UK (COG-UK) Consortium | Rob Howes, The Lighthouse Lab in Cambridge and Alex Alderton, Roberto Amato, Sonia Goncalves, Ewan Harrison, David K. Jackson, Ian Johnston, Dominic Kwiatkowski, Cordelia Langford, John Sillitoe on behalf of the Wellcome Sanger Institute COVID-19 Surveillance Team |
| EPI_ISL_891585 | Lighthouse Lab in Glasgow | Wellcome Sanger Institute for the COVID-19 Genomics UK (COG-UK) Consortium | Harper VanSteenhouse, Yumi Kasai, David Gray, Carol Clugston, Anna Dominiczak and Alex Alderton, Roberto Amato, Sonia Goncalves, Ewan Harrison, David K. Jackson, Ian Johnston, Dominic Kwiatkowski, Cordelia Langford, John Sillitoe on behalf of the Wellcome Sanger Institute COVID-19 Surveillance Team |
| EPI_ISL_891586 | Lighthouse Lab in Alderley Park | Wellcome Sanger Institute for the COVID-19 Genomics UK (COG-UK) Consortium | Jacquelyn Wynn, Mairead Hyland, The Lighthouse Lab in Alderley Park and Alex Alderton, Roberto Amato, Sonia Goncalves, Ewan Harrison, David K. Jackson, Ian Johnston, Dominic Kwiatkowski, Cordelia Langford, John Sillitoe on behalf of the Wellcome Sanger Institute COVID-19 Surveillance Team |
| EPI_ISL_891587, EPI_ISL_891588 | Lighthouse Lab in Glasgow | Wellcome Sanger Institute for the COVID-19 Genomics UK (COG-UK) Consortium | Harper VanSteenhouse, Yumi Kasai, David Gray, Carol Clugston, Anna Dominiczak and Alex Alderton, Roberto Amato, Sonia Goncalves, Ewan Harrison, David K. Jackson, Ian Johnston, Dominic Kwiatkowski, Cordelia Langford, John Sillitoe on behalf of the Wellcome Sanger Institute COVID-19 Surveillance Team |
| EPI_ISL_891589, EPI_ISL_891590, EPI_ISL_891591, EPI_ISL_891592, EPI_ISL_891593, EPI_ISL_891594, EPI_ISL_891595 | Lighthouse Lab in Cambridge | Wellcome Sanger Institute for the COVID-19 Genomics UK (COG-UK) Consortium | Rob Howes, The Lighthouse Lab in Cambridge and Alex Alderton, Roberto Amato, Sonia Goncalves, Ewan Harrison, David K. Jackson, Ian Johnston, Dominic Kwiatkowski, Cordelia Langford, John Sillitoe on behalf of the Wellcome Sanger Institute COVID-19 Surveillance Team |

| | | | |
|---|---|---|---|
| EPI_ISL_891596 | Lighthouse Lab in Alderley Park | Wellcome Sanger Institute for the COVID-19 Genomics UK (COG-UK) Consortium | Jacquelyn Wynn, Mairead Hyland, The Lighthouse Lab in Alderley Park and Alex Alderton, Roberto Amato, Sonia Goncalves, Ewan Harrison, David K. Jackson, Ian Johnston, Dominic Kwiatkowski, Cordelia Langford, John Sillitoe on behalf of the Wellcome Sanger Institute COVID-19 Surveillance Team |
| EPI_ISL_891597, EPI_ISL_891598 | Lighthouse Lab in Cambridge | Wellcome Sanger Institute for the COVID-19 Genomics UK (COG-UK) Consortium | Rob Howes, The Lighthouse Lab in Cambridge and Alex Alderton, Roberto Amato, Sonia Goncalves, Ewan Harrison, David K. Jackson, Ian Johnston, Dominic Kwiatkowski, Cordelia Langford, John Sillitoe on behalf of the Wellcome Sanger Institute COVID-19 Surveillance Team |
| EPI_ISL_891599, EPI_ISL_891600 | Lighthouse Lab in Alderley Park | Wellcome Sanger Institute for the COVID-19 Genomics UK (COG-UK) Consortium | Jacquelyn Wynn, Mairead Hyland, The Lighthouse Lab in Alderley Park and Alex Alderton, Roberto Amato, Sonia Goncalves, Ewan Harrison, David K. Jackson, Ian Johnston, Dominic Kwiatkowski, Cordelia Langford, John Sillitoe on behalf of the Wellcome Sanger Institute COVID-19 Surveillance Team |
| EPI_ISL_891601 | Lighthouse Lab in Cambridge | Wellcome Sanger Institute for the COVID-19 Genomics UK (COG-UK) Consortium | Rob Howes, The Lighthouse Lab in Cambridge and Alex Alderton, Roberto Amato, Sonia Goncalves, Ewan Harrison, David K. Jackson, Ian Johnston, Dominic Kwiatkowski, Cordelia Langford, John Sillitoe on behalf of the Wellcome Sanger Institute COVID-19 Surveillance Team |
| EPI_ISL_891602 | Lighthouse Lab in Alderley Park | Wellcome Sanger Institute for the COVID-19 Genomics UK (COG-UK) Consortium | Jacquelyn Wynn, Mairead Hyland, The Lighthouse Lab in Alderley Park and Alex Alderton, Roberto Amato, Sonia Goncalves, Ewan Harrison, David K. Jackson, Ian Johnston, Dominic Kwiatkowski, Cordelia Langford, John Sillitoe on behalf of the Wellcome Sanger Institute COVID-19 Surveillance Team |
| EPI_ISL_891604, EPI_ISL_891605 | Lighthouse Lab in Glasgow | Wellcome Sanger Institute for the COVID-19 Genomics UK (COG-UK) Consortium | Harper VanSteenhouse, Yumi Kasai, David Gray, Carol Clugston, Anna Dominiczak and Alex Alderton, Roberto Amato, Sonia Goncalves, Ewan Harrison, David K. Jackson, Ian Johnston, Dominic Kwiatkowski, Cordelia Langford, John Sillitoe on behalf of the Wellcome Sanger Institute COVID-19 Surveillance Team |
| EPI_ISL_891606 | Lighthouse Lab in Alderley Park | Wellcome Sanger Institute for the COVID-19 Genomics UK (COG-UK) Consortium | Jacquelyn Wynn, Mairead Hyland, The Lighthouse Lab in Alderley Park and Alex Alderton, Roberto Amato, Sonia Goncalves, Ewan Harrison, David K. Jackson, Ian Johnston, Dominic Kwiatkowski, Cordelia Langford, John Sillitoe on behalf of the Wellcome Sanger Institute COVID-19 Surveillance Team |
| EPI_ISL_891607, EPI_ISL_891608 | Lighthouse Lab in Cambridge | Wellcome Sanger Institute for the COVID-19 Genomics UK (COG-UK) Consortium | Rob Howes, The Lighthouse Lab in Cambridge and Alex Alderton, Roberto Amato, Sonia Goncalves, Ewan Harrison, David K. Jackson, Ian Johnston, Dominic Kwiatkowski, Cordelia Langford, John Sillitoe on behalf of the Wellcome Sanger Institute COVID-19 Surveillance Team |
| EPI_ISL_891610, EPI_ISL_891612 | Lighthouse Lab in Glasgow | Wellcome Sanger Institute for the COVID-19 Genomics UK (COG-UK) Consortium | Harper VanSteenhouse, Yumi Kasai, David Gray, Carol Clugston, Anna Dominiczak and Alex Alderton, Roberto Amato, Sonia Goncalves, Ewan Harrison, David K. Jackson, Ian Johnston, Dominic Kwiatkowski, Cordelia Langford, John Sillitoe on behalf of the Wellcome Sanger Institute COVID-19 Surveillance Team |
| EPI_ISL_891614 | Lighthouse Lab in Cambridge | Wellcome Sanger Institute for the COVID-19 Genomics UK (COG-UK) Consortium | Rob Howes, The Lighthouse Lab in Cambridge and Alex Alderton, Roberto Amato, Sonia Goncalves, Ewan Harrison, David K. Jackson, Ian Johnston, Dominic Kwiatkowski, Cordelia Langford, John Sillitoe on behalf of the Wellcome Sanger Institute COVID-19 Surveillance Team |
| EPI_ISL_891615, EPI_ISL_891616 | Lighthouse Lab in Alderley Park | Wellcome Sanger Institute for the COVID-19 Genomics UK (COG-UK) Consortium | Jacquelyn Wynn, Mairead Hyland, The Lighthouse Lab in Alderley Park and Alex Alderton, Roberto Amato, Sonia Goncalves, Ewan Harrison, David K. Jackson, Ian Johnston, Dominic Kwiatkowski, Cordelia Langford, John Sillitoe on behalf of the Wellcome Sanger Institute COVID-19 Surveillance Team |
| EPI_ISL_891617 | Lighthouse Lab in Cambridge | Wellcome Sanger Institute for the COVID-19 Genomics UK (COG-UK) Consortium | Rob Howes, The Lighthouse Lab in Cambridge and Alex Alderton, Roberto Amato, Sonia Goncalves, Ewan Harrison, David K. Jackson, Ian Johnston, Dominic Kwiatkowski, Cordelia Langford, John Sillitoe on behalf of the Wellcome Sanger Institute COVID-19 Surveillance Team |
| EPI_ISL_891618 | Lighthouse Lab in Glasgow | Wellcome Sanger Institute for the COVID-19 Genomics UK (COG-UK) Consortium | Harper VanSteenhouse, Yumi Kasai, David Gray, Carol Clugston, Anna Dominiczak and Alex Alderton, Roberto Amato, Sonia Goncalves, Ewan Harrison, David K. Jackson, Ian Johnston, Dominic Kwiatkowski, Cordelia Langford, John Sillitoe on behalf of the Wellcome Sanger Institute COVID-19 Surveillance Team |
| EPI_ISL_891619, EPI_ISL_891620, EPI_ISL_891621 | Lighthouse Lab in Cambridge | Wellcome Sanger Institute for the COVID-19 Genomics UK (COG-UK) Consortium | Rob Howes, The Lighthouse Lab in Cambridge and Alex Alderton, Roberto Amato, Sonia Goncalves, Ewan Harrison, David K. Jackson, Ian Johnston, Dominic Kwiatkowski, Cordelia Langford, John Sillitoe on behalf of the Wellcome Sanger Institute COVID-19 Surveillance Team |
| EPI_ISL_891623 | Lighthouse Lab in Alderley Park | Wellcome Sanger Institute for the COVID-19 Genomics UK (COG-UK) Consortium | Jacquelyn Wynn, Mairead Hyland, The Lighthouse Lab in Alderley Park and Alex Alderton, Roberto Amato, Sonia Goncalves, Ewan Harrison, David K. Jackson, Ian Johnston, Dominic Kwiatkowski, Cordelia Langford, John Sillitoe on behalf of the Wellcome Sanger Institute COVID-19 Surveillance Team |
| EPI_ISL_891624 | Lighthouse Lab in Glasgow | Wellcome Sanger Institute for the COVID-19 Genomics UK (COG-UK) Consortium | Harper VanSteenhouse, Yumi Kasai, David Gray, Carol Clugston, Anna Dominiczak and Alex Alderton, Roberto Amato, Sonia Goncalves, Ewan Harrison, David K. Jackson, Ian Johnston, Dominic Kwiatkowski, Cordelia Langford, John Sillitoe on behalf of the Wellcome Sanger Institute COVID-19 Surveillance Team |
| EPI_ISL_891625 | Lighthouse Lab in Alderley Park | Wellcome Sanger Institute for the COVID-19 Genomics UK (COG-UK) Consortium | Jacquelyn Wynn, Mairead Hyland, The Lighthouse Lab in Alderley Park and Alex Alderton, Roberto Amato, Sonia Goncalves, Ewan Harrison, David K. Jackson, Ian Johnston, Dominic Kwiatkowski, Cordelia Langford, John Sillitoe on behalf of the Wellcome Sanger Institute COVID-19 Surveillance Team |
| EPI_ISL_891626 | Lighthouse Lab in Glasgow | Wellcome Sanger Institute for the COVID-19 Genomics UK (COG-UK) Consortium | Harper VanSteenhouse, Yumi Kasai, David Gray, Carol Clugston, Anna Dominiczak and Alex Alderton, Roberto Amato, Sonia Goncalves, Ewan Harrison, David K. Jackson, Ian Johnston, Dominic Kwiatkowski, Cordelia Langford, John Sillitoe on behalf of the Wellcome Sanger Institute COVID-19 Surveillance Team |
| EPI_ISL_891627 | Lighthouse Lab in Cambridge | Wellcome Sanger Institute for the COVID-19 Genomics UK (COG-UK) Consortium | Rob Howes, The Lighthouse Lab in Cambridge and Alex Alderton, Roberto Amato, Sonia Goncalves, Ewan Harrison, David K. Jackson, Ian Johnston, Dominic Kwiatkowski, Cordelia Langford, John Sillitoe on behalf of the Wellcome Sanger Institute COVID-19 Surveillance Team |
| EPI_ISL_891628 | Lighthouse Lab in Glasgow | Wellcome Sanger Institute for the COVID-19 Genomics UK (COG-UK) Consortium | Harper VanSteenhouse, Yumi Kasai, David Gray, Carol Clugston, Anna Dominiczak and Alex Alderton, Roberto Amato, Sonia Goncalves, Ewan Harrison, David K. Jackson, Ian Johnston, Dominic Kwiatkowski, Cordelia Langford, John Sillitoe on behalf of the Wellcome Sanger Institute COVID-19 Surveillance Team |
| EPI_ISL_891629, EPI_ISL_891630 | Lighthouse Lab in Alderley Park | Wellcome Sanger Institute for the COVID-19 Genomics UK (COG-UK) Consortium | Jacquelyn Wynn, Mairead Hyland, The Lighthouse Lab in Alderley Park and Alex Alderton, Roberto Amato, Sonia Goncalves, Ewan Harrison, David K. Jackson, Ian Johnston, Dominic Kwiatkowski, Cordelia Langford, John Sillitoe on behalf of the Wellcome Sanger Institute COVID-19 Surveillance Team |
| EPI_ISL_891631, EPI_ISL_891632 | Lighthouse Lab in Cambridge | Wellcome Sanger Institute for the COVID-19 Genomics UK (COG-UK) Consortium | Rob Howes, The Lighthouse Lab in Cambridge and Alex Alderton, Roberto Amato, Sonia Goncalves, Ewan Harrison, David K. Jackson, Ian Johnston, Dominic Kwiatkowski, Cordelia Langford, John Sillitoe on behalf of the Wellcome Sanger Institute COVID-19 Surveillance Team |
| EPI_ISL_891633, EPI_ISL_891634 | Lighthouse Lab in Alderley Park | Wellcome Sanger Institute for the COVID-19 Genomics UK (COG-UK) Consortium | Jacquelyn Wynn, Mairead Hyland, The Lighthouse Lab in Alderley Park and Alex Alderton, Roberto Amato, Sonia Goncalves, Ewan Harrison, David K. Jackson, Ian Johnston, Dominic Kwiatkowski, Cordelia Langford, John Sillitoe on behalf of the Wellcome Sanger Institute COVID-19 Surveillance Team |
| EPI_ISL_891635 | Lighthouse Lab in Cambridge | Wellcome Sanger Institute for the COVID-19 Genomics UK (COG-UK) Consortium | Rob Howes, The Lighthouse Lab in Cambridge and Alex Alderton, Roberto Amato, Sonia Goncalves, Ewan Harrison, David K. Jackson, Ian Johnston, Dominic Kwiatkowski, Cordelia Langford, John Sillitoe on behalf of the Wellcome Sanger Institute COVID-19 Surveillance Team |
| EPI_ISL_891636 | Lighthouse Lab in Glasgow | Wellcome Sanger Institute for the COVID-19 Genomics UK (COG-UK) Consortium | Harper VanSteenhouse, Yumi Kasai, David Gray, Carol Clugston, Anna Dominiczak and Alex Alderton, Roberto Amato, Sonia Goncalves, Ewan Harrison, David K. Jackson, Ian Johnston, Dominic Kwiatkowski, Cordelia Langford, John Sillitoe on behalf of the Wellcome Sanger Institute COVID-19 Surveillance Team |
| EPI_ISL_891637, EPI_ISL_891638 | Lighthouse Lab in Cambridge | Wellcome Sanger Institute for the COVID-19 Genomics UK (COG-UK) Consortium | Rob Howes, The Lighthouse Lab in Cambridge and Alex Alderton, Roberto Amato, Sonia Goncalves, Ewan Harrison, David K. Jackson, Ian Johnston, Dominic Kwiatkowski, Cordelia Langford, John Sillitoe on behalf of the Wellcome Sanger Institute COVID-19 Surveillance Team |
| EPI_ISL_891640 | Lighthouse Lab in Alderley Park | Wellcome Sanger Institute for the COVID-19 Genomics UK (COG-UK) Consortium | Jacquelyn Wynn, Mairead Hyland, The Lighthouse Lab in Alderley Park and Alex Alderton, Roberto Amato, Sonia Goncalves, Ewan Harrison, David K. Jackson, Ian Johnston, Dominic Kwiatkowski, Cordelia Langford, John Sillitoe on behalf of the Wellcome Sanger Institute COVID-19 Surveillance Team |
| EPI_ISL_891641, EPI_ISL_891642 | Lighthouse Lab in Cambridge | Wellcome Sanger Institute for the COVID-19 Genomics UK (COG-UK) Consortium | Rob Howes, The Lighthouse Lab in Cambridge and Alex Alderton, Roberto Amato, Sonia Goncalves, Ewan Harrison, David K. Jackson, Ian Johnston, Dominic Kwiatkowski, Cordelia Langford, John Sillitoe on behalf of the Wellcome Sanger Institute COVID-19 Surveillance Team |
| EPI_ISL_891643 | Lighthouse Lab in Alderley Park | Wellcome Sanger Institute for the COVID-19 Genomics UK (COG-UK) Consortium | Jacquelyn Wynn, Mairead Hyland, The Lighthouse Lab in Alderley Park and Alex Alderton, Roberto Amato, Sonia Goncalves, Ewan Harrison, David K. Jackson, Ian Johnston, Dominic Kwiatkowski, Cordelia Langford, John Sillitoe on behalf of the Wellcome Sanger Institute COVID-19 Surveillance Team |
| EPI_ISL_891644, EPI_ISL_891645, EPI_ISL_891646 | Lighthouse Lab in Cambridge | Wellcome Sanger Institute for the COVID-19 Genomics UK (COG-UK) Consortium | Rob Howes, The Lighthouse Lab in Cambridge and Alex Alderton, Roberto Amato, Sonia Goncalves, Ewan Harrison, David K. Jackson, Ian Johnston, Dominic Kwiatkowski, Cordelia Langford, John Sillitoe on behalf of the Wellcome Sanger Institute COVID-19 Surveillance Team |
| EPI_ISL_891647 | Lighthouse Lab in Alderley Park | Wellcome Sanger Institute for the COVID-19 Genomics UK (COG-UK) Consortium | Jacquelyn Wynn, Mairead Hyland, The Lighthouse Lab in Alderley Park and Alex Alderton, Roberto Amato, Sonia Goncalves, Ewan Harrison, David K. Jackson, Ian Johnston, Dominic Kwiatkowski, Cordelia Langford, John Sillitoe on behalf of the Wellcome Sanger Institute COVID-19 Surveillance Team |
| EPI_ISL_891648, EPI_ISL_891649 | Lighthouse Lab in Cambridge | Wellcome Sanger Institute for the COVID-19 Genomics UK (COG-UK) Consortium | Rob Howes, The Lighthouse Lab in Cambridge and Alex Alderton, Roberto Amato, Sonia Goncalves, Ewan Harrison, David K. Jackson, Ian Johnston, Dominic Kwiatkowski, Cordelia Langford, John Sillitoe on behalf of the Wellcome Sanger Institute COVID-19 Surveillance Team |
| EPI_ISL_891650 | Lighthouse Lab in Alderley Park | Wellcome Sanger Institute for the COVID-19 Genomics UK | Jacquelyn Wynn, Mairead Hyland, The Lighthouse Lab in Alderley Park and Alex Alderton, Roberto Amato, Sonia Goncalves, Ewan Harrison, David K. |

| | | | |
|---|---|---|---|
| | | (COG-UK) Consortium | Jackson, Ian Johnston, Dominic Kwiatkowski, Cordelia Langford, John Sillitoe on behalf of the Wellcome Sanger Institute COVID-19 Surveillance Team |
| EPI_ISL_891651, EPI_ISL_891652, EPI_ISL_891653, EPI_ISL_891654 | Lighthouse Lab in Cambridge | Wellcome Sanger Institute for the COVID-19 Genomics UK (COG-UK) Consortium | Rob Howes, The Lighthouse Lab in Cambridge and Alex Alderton, Roberto Amato, Sonia Goncalves, Ewan Harrison, David K. Jackson, Ian Johnston, Dominic Kwiatkowski, Cordelia Langford, John Sillitoe on behalf of the Wellcome Sanger Institute COVID-19 Surveillance Team |
| EPI_ISL_891656 | Lighthouse Lab in Glasgow | Wellcome Sanger Institute for the COVID-19 Genomics UK (COG-UK) Consortium | Harper VanSteenhouse, Yumi Kasai, David Gray, Carol Clugston, Anna Dominiczak and Alex Alderton, Roberto Amato, Sonia Goncalves, Ewan Harrison, David K. Jackson, Ian Johnston, Dominic Kwiatkowski, Cordelia Langford, John Sillitoe on behalf of the Wellcome Sanger Institute COVID-19 Surveillance Team |
| EPI_ISL_891657 | Lighthouse Lab in Cambridge | Wellcome Sanger Institute for the COVID-19 Genomics UK (COG-UK) Consortium | Rob Howes, The Lighthouse Lab in Cambridge and Alex Alderton, Roberto Amato, Sonia Goncalves, Ewan Harrison, David K. Jackson, Ian Johnston, Dominic Kwiatkowski, Cordelia Langford, John Sillitoe on behalf of the Wellcome Sanger Institute COVID-19 Surveillance Team |
| EPI_ISL_891658 | Lighthouse Lab in Glasgow | Wellcome Sanger Institute for the COVID-19 Genomics UK (COG-UK) Consortium | Harper VanSteenhouse, Yumi Kasai, David Gray, Carol Clugston, Anna Dominiczak and Alex Alderton, Roberto Amato, Sonia Goncalves, Ewan Harrison, David K. Jackson, Ian Johnston, Dominic Kwiatkowski, Cordelia Langford, John Sillitoe on behalf of the Wellcome Sanger Institute COVID-19 Surveillance Team |
| EPI_ISL_891659 | Lighthouse Lab in Alderley Park | Wellcome Sanger Institute for the COVID-19 Genomics UK (COG-UK) Consortium | Jacquelyn Wynn, Mairead Hyland, The Lighthouse Lab in Alderley Park and Alex Alderton, Roberto Amato, Sonia Goncalves, Ewan Harrison, David K. Jackson, Ian Johnston, Dominic Kwiatkowski, Cordelia Langford, John Sillitoe on behalf of the Wellcome Sanger Institute COVID-19 Surveillance Team |
| EPI_ISL_891660 | Lighthouse Lab in Cambridge | Wellcome Sanger Institute for the COVID-19 Genomics UK (COG-UK) Consortium | Rob Howes, The Lighthouse Lab in Cambridge and Alex Alderton, Roberto Amato, Sonia Goncalves, Ewan Harrison, David K. Jackson, Ian Johnston, Dominic Kwiatkowski, Cordelia Langford, John Sillitoe on behalf of the Wellcome Sanger Institute COVID-19 Surveillance Team |
| EPI_ISL_891661, EPI_ISL_891662 | Lighthouse Lab in Alderley Park | Wellcome Sanger Institute for the COVID-19 Genomics UK (COG-UK) Consortium | Jacquelyn Wynn, Mairead Hyland, The Lighthouse Lab in Alderley Park and Alex Alderton, Roberto Amato, Sonia Goncalves, Ewan Harrison, David K. Jackson, Ian Johnston, Dominic Kwiatkowski, Cordelia Langford, John Sillitoe on behalf of the Wellcome Sanger Institute COVID-19 Surveillance Team |
| EPI_ISL_891663, EPI_ISL_891665, EPI_ISL_891666, EPI_ISL_891667, EPI_ISL_891669 | Lighthouse Lab in Cambridge | Wellcome Sanger Institute for the COVID-19 Genomics UK (COG-UK) Consortium | Rob Howes, The Lighthouse Lab in Cambridge and Alex Alderton, Roberto Amato, Sonia Goncalves, Ewan Harrison, David K. Jackson, Ian Johnston, Dominic Kwiatkowski, Cordelia Langford, John Sillitoe on behalf of the Wellcome Sanger Institute COVID-19 Surveillance Team |
| EPI_ISL_891670 | Lighthouse Lab in Alderley Park | Wellcome Sanger Institute for the COVID-19 Genomics UK (COG-UK) Consortium | Jacquelyn Wynn, Mairead Hyland, The Lighthouse Lab in Alderley Park and Alex Alderton, Roberto Amato, Sonia Goncalves, Ewan Harrison, David K. Jackson, Ian Johnston, Dominic Kwiatkowski, Cordelia Langford, John Sillitoe on behalf of the Wellcome Sanger Institute COVID-19 Surveillance Team |
| EPI_ISL_891671 | Lighthouse Lab in Cambridge | Wellcome Sanger Institute for the COVID-19 Genomics UK (COG-UK) Consortium | Rob Howes, The Lighthouse Lab in Cambridge and Alex Alderton, Roberto Amato, Sonia Goncalves, Ewan Harrison, David K. Jackson, Ian Johnston, Dominic Kwiatkowski, Cordelia Langford, John Sillitoe on behalf of the Wellcome Sanger Institute COVID-19 Surveillance Team |
| EPI_ISL_891672, EPI_ISL_891673 | Lighthouse Lab in Alderley Park | Wellcome Sanger Institute for the COVID-19 Genomics UK (COG-UK) Consortium | Jacquelyn Wynn, Mairead Hyland, The Lighthouse Lab in Alderley Park and Alex Alderton, Roberto Amato, Sonia Goncalves, Ewan Harrison, David K. Jackson, Ian Johnston, Dominic Kwiatkowski, Cordelia Langford, John Sillitoe on behalf of the Wellcome Sanger Institute COVID-19 Surveillance Team |
| EPI_ISL_891674, EPI_ISL_891675, EPI_ISL_891676 | Lighthouse Lab in Cambridge | Wellcome Sanger Institute for the COVID-19 Genomics UK (COG-UK) Consortium | Rob Howes, The Lighthouse Lab in Cambridge and Alex Alderton, Roberto Amato, Sonia Goncalves, Ewan Harrison, David K. Jackson, Ian Johnston, Dominic Kwiatkowski, Cordelia Langford, John Sillitoe on behalf of the Wellcome Sanger Institute COVID-19 Surveillance Team |
| EPI_ISL_891677 | Lighthouse Lab in Alderley Park | Wellcome Sanger Institute for the COVID-19 Genomics UK (COG-UK) Consortium | Jacquelyn Wynn, Mairead Hyland, The Lighthouse Lab in Alderley Park and Alex Alderton, Roberto Amato, Sonia Goncalves, Ewan Harrison, David K. Jackson, Ian Johnston, Dominic Kwiatkowski, Cordelia Langford, John Sillitoe on behalf of the Wellcome Sanger Institute COVID-19 Surveillance Team |
| EPI_ISL_891678, EPI_ISL_891679 | Lighthouse Lab in Cambridge | Wellcome Sanger Institute for the COVID-19 Genomics UK (COG-UK) Consortium | Rob Howes, The Lighthouse Lab in Cambridge and Alex Alderton, Roberto Amato, Sonia Goncalves, Ewan Harrison, David K. Jackson, Ian Johnston, Dominic Kwiatkowski, Cordelia Langford, John Sillitoe on behalf of the Wellcome Sanger Institute COVID-19 Surveillance Team |
| EPI_ISL_891680, EPI_ISL_891681 | Lighthouse Lab in Alderley Park | Wellcome Sanger Institute for the COVID-19 Genomics UK (COG-UK) Consortium | Jacquelyn Wynn, Mairead Hyland, The Lighthouse Lab in Alderley Park and Alex Alderton, Roberto Amato, Sonia Goncalves, Ewan Harrison, David K. Jackson, Ian Johnston, Dominic Kwiatkowski, Cordelia Langford, John Sillitoe on behalf of the Wellcome Sanger Institute COVID-19 Surveillance Team |
| EPI_ISL_891682 | Lighthouse Lab in Cambridge | Wellcome Sanger Institute for the COVID-19 Genomics UK (COG-UK) Consortium | Rob Howes, The Lighthouse Lab in Cambridge and Alex Alderton, Roberto Amato, Sonia Goncalves, Ewan Harrison, David K. Jackson, Ian Johnston, Dominic Kwiatkowski, Cordelia Langford, John Sillitoe on behalf of the Wellcome Sanger Institute COVID-19 Surveillance Team |
| EPI_ISL_891683 | Lighthouse Lab in Alderley Park | Wellcome Sanger Institute for the COVID-19 Genomics UK (COG-UK) Consortium | Jacquelyn Wynn, Mairead Hyland, The Lighthouse Lab in Alderley Park and Alex Alderton, Roberto Amato, Sonia Goncalves, Ewan Harrison, David K. Jackson, Ian Johnston, Dominic Kwiatkowski, Cordelia Langford, John Sillitoe on behalf of the Wellcome Sanger Institute COVID-19 Surveillance Team |
| EPI_ISL_891684 | Lighthouse Lab in Cambridge | Wellcome Sanger Institute for the COVID-19 Genomics UK (COG-UK) Consortium | Rob Howes, The Lighthouse Lab in Cambridge and Alex Alderton, Roberto Amato, Sonia Goncalves, Ewan Harrison, David K. Jackson, Ian Johnston, Dominic Kwiatkowski, Cordelia Langford, John Sillitoe on behalf of the Wellcome Sanger Institute COVID-19 Surveillance Team |
| EPI_ISL_891685 | Lighthouse Lab in Glasgow | Wellcome Sanger Institute for the COVID-19 Genomics UK (COG-UK) Consortium | Harper VanSteenhouse, Yumi Kasai, David Gray, Carol Clugston, Anna Dominiczak and Alex Alderton, Roberto Amato, Sonia Goncalves, Ewan Harrison, David K. Jackson, Ian Johnston, Dominic Kwiatkowski, Cordelia Langford, John Sillitoe on behalf of the Wellcome Sanger Institute COVID-19 Surveillance Team |
| EPI_ISL_891686 | Lighthouse Lab in Cambridge | Wellcome Sanger Institute for the COVID-19 Genomics UK (COG-UK) Consortium | Rob Howes, The Lighthouse Lab in Cambridge and Alex Alderton, Roberto Amato, Sonia Goncalves, Ewan Harrison, David K. Jackson, Ian Johnston, Dominic Kwiatkowski, Cordelia Langford, John Sillitoe on behalf of the Wellcome Sanger Institute COVID-19 Surveillance Team |
| EPI_ISL_891687 | Lighthouse Lab in Alderley Park | Wellcome Sanger Institute for the COVID-19 Genomics UK (COG-UK) Consortium | Jacquelyn Wynn, Mairead Hyland, The Lighthouse Lab in Alderley Park and Alex Alderton, Roberto Amato, Sonia Goncalves, Ewan Harrison, David K. Jackson, Ian Johnston, Dominic Kwiatkowski, Cordelia Langford, John Sillitoe on behalf of the Wellcome Sanger Institute COVID-19 Surveillance Team |
| EPI_ISL_891688 | Lighthouse Lab in Cambridge | Wellcome Sanger Institute for the COVID-19 Genomics UK (COG-UK) Consortium | Rob Howes, The Lighthouse Lab in Cambridge and Alex Alderton, Roberto Amato, Sonia Goncalves, Ewan Harrison, David K. Jackson, Ian Johnston, Dominic Kwiatkowski, Cordelia Langford, John Sillitoe on behalf of the Wellcome Sanger Institute COVID-19 Surveillance Team |
| EPI_ISL_891689 | Lighthouse Lab in Glasgow | Wellcome Sanger Institute for the COVID-19 Genomics UK (COG-UK) Consortium | Harper VanSteenhouse, Yumi Kasai, David Gray, Carol Clugston, Anna Dominiczak and Alex Alderton, Roberto Amato, Sonia Goncalves, Ewan Harrison, David K. Jackson, Ian Johnston, Dominic Kwiatkowski, Cordelia Langford, John Sillitoe on behalf of the Wellcome Sanger Institute COVID-19 Surveillance Team |
| EPI_ISL_891690, EPI_ISL_891691, EPI_ISL_891692, EPI_ISL_891693, EPI_ISL_891695, EPI_ISL_891696, EPI_ISL_891697, EPI_ISL_891698, EPI_ISL_891699, EPI_ISL_891701, EPI_ISL_891702, EPI_ISL_891703 | | | |
| see above | Lighthouse Lab in Cambridge | Wellcome Sanger Institute for the COVID-19 Genomics UK (COG-UK) Consortium | Rob Howes, The Lighthouse Lab in Cambridge and Alex Alderton, Roberto Amato, Sonia Goncalves, Ewan Harrison, David K. Jackson, Ian Johnston, Dominic Kwiatkowski, Cordelia Langford, John Sillitoe on behalf of the Wellcome Sanger Institute COVID-19 Surveillance Team |
| EPI_ISL_891704 | Lighthouse Lab in Alderley Park | Wellcome Sanger Institute for the COVID-19 Genomics UK (COG-UK) Consortium | Jacquelyn Wynn, Mairead Hyland, The Lighthouse Lab in Alderley Park and Alex Alderton, Roberto Amato, Sonia Goncalves, Ewan Harrison, David K. Jackson, Ian Johnston, Dominic Kwiatkowski, Cordelia Langford, John Sillitoe on behalf of the Wellcome Sanger Institute COVID-19 Surveillance Team |
| EPI_ISL_891705 | Lighthouse Lab in Glasgow | Wellcome Sanger Institute for the COVID-19 Genomics UK (COG-UK) Consortium | Harper VanSteenhouse, Yumi Kasai, David Gray, Carol Clugston, Anna Dominiczak and Alex Alderton, Roberto Amato, Sonia Goncalves, Ewan Harrison, David K. Jackson, Ian Johnston, Dominic Kwiatkowski, Cordelia Langford, John Sillitoe on behalf of the Wellcome Sanger Institute COVID-19 Surveillance Team |
| EPI_ISL_891706, EPI_ISL_891707, EPI_ISL_891708 | Lighthouse Lab in Cambridge | Wellcome Sanger Institute for the COVID-19 Genomics UK (COG-UK) Consortium | Rob Howes, The Lighthouse Lab in Cambridge and Alex Alderton, Roberto Amato, Sonia Goncalves, Ewan Harrison, David K. Jackson, Ian Johnston, Dominic Kwiatkowski, Cordelia Langford, John Sillitoe on behalf of the Wellcome Sanger Institute COVID-19 Surveillance Team |
| EPI_ISL_891710 | Lighthouse Lab in Alderley Park | Wellcome Sanger Institute for the COVID-19 Genomics UK (COG-UK) Consortium | Jacquelyn Wynn, Mairead Hyland, The Lighthouse Lab in Alderley Park and Alex Alderton, Roberto Amato, Sonia Goncalves, Ewan Harrison, David K. Jackson, Ian Johnston, Dominic Kwiatkowski, Cordelia Langford, John Sillitoe on behalf of the Wellcome Sanger Institute COVID-19 Surveillance Team |
| EPI_ISL_891712 | Lighthouse Lab in Cambridge | Wellcome Sanger Institute for the COVID-19 Genomics UK (COG-UK) Consortium | Rob Howes, The Lighthouse Lab in Cambridge and Alex Alderton, Roberto Amato, Sonia Goncalves, Ewan Harrison, David K. Jackson, Ian Johnston, Dominic Kwiatkowski, Cordelia Langford, John Sillitoe on behalf of the Wellcome Sanger Institute COVID-19 Surveillance Team |
| EPI_ISL_891713, EPI_ISL_891714 | Lighthouse Lab in Alderley Park | Wellcome Sanger Institute for the COVID-19 Genomics UK (COG-UK) Consortium | Jacquelyn Wynn, Mairead Hyland, The Lighthouse Lab in Alderley Park and Alex Alderton, Roberto Amato, Sonia Goncalves, Ewan Harrison, David K. Jackson, Ian Johnston, Dominic Kwiatkowski, Cordelia Langford, John Sillitoe on behalf of the Wellcome Sanger Institute COVID-19 Surveillance Team |
| EPI_ISL_891715, EPI_ISL_891716 | Lighthouse Lab in Cambridge | Wellcome Sanger Institute for the COVID-19 Genomics UK (COG-UK) Consortium | Rob Howes, The Lighthouse Lab in Cambridge and Alex Alderton, Roberto Amato, Sonia Goncalves, Ewan Harrison, David K. Jackson, Ian Johnston, Dominic Kwiatkowski, Cordelia Langford, John Sillitoe on behalf of the Wellcome Sanger Institute COVID-19 Surveillance Team |
| EPI_ISL_891717 | Lighthouse Lab in Glasgow | Wellcome Sanger Institute for the COVID-19 Genomics UK | Harper VanSteenhouse, Yumi Kasai, David Gray, Carol Clugston, Anna Dominiczak and Alex Alderton, Roberto Amato, Sonia Goncalves, Ewan Harrison, |

| | | | |
|---|---|---|---|
| | | (COG-UK) Consortium | David K. Jackson, Ian Johnston, Dominic Kwiatkowski, Cordelia Langford, John Sillitoe on behalf of the Wellcome Sanger Institute COVID-19 Surveillance Team |
| EPI_ISL_891718, EPI_ISL_891719 | Lighthouse Lab in Alderley Park | Wellcome Sanger Institute for the COVID-19 Genomics UK (COG-UK) Consortium | Jacquelyn Wynn, Mairead Hyland, The Lighthouse Lab in Alderley Park and Alex Alderton, Roberto Amato, Sonia Goncalves, Ewan Harrison, David K. Jackson, Ian Johnston, Dominic Kwiatkowski, Cordelia Langford, John Sillitoe on behalf of the Wellcome Sanger Institute COVID-19 Surveillance Team |
| EPI_ISL_891720 | Lighthouse Lab in Cambridge | Wellcome Sanger Institute for the COVID-19 Genomics UK (COG-UK) Consortium | Rob Howes, The Lighthouse Lab in Cambridge and Alex Alderton, Roberto Amato, Sonia Goncalves, Ewan Harrison, David K. Jackson, Ian Johnston, Dominic Kwiatkowski, Cordelia Langford, John Sillitoe on behalf of the Wellcome Sanger Institute COVID-19 Surveillance Team |
| EPI_ISL_891721, EPI_ISL_891722 | Lighthouse Lab in Glasgow | Wellcome Sanger Institute for the COVID-19 Genomics UK (COG-UK) Consortium | Harper VanSteenhouse, Yumi Kasai, David Gray, Carol Clugston, Anna Dominiczak and Alex Alderton, Roberto Amato, Sonia Goncalves, Ewan Harrison, David K. Jackson, Ian Johnston, Dominic Kwiatkowski, Cordelia Langford, John Sillitoe on behalf of the Wellcome Sanger Institute COVID-19 Surveillance Team |
| EPI_ISL_891723, EPI_ISL_891724 | Lighthouse Lab in Alderley Park | Wellcome Sanger Institute for the COVID-19 Genomics UK (COG-UK) Consortium | Jacquelyn Wynn, Mairead Hyland, The Lighthouse Lab in Alderley Park and Alex Alderton, Roberto Amato, Sonia Goncalves, Ewan Harrison, David K. Jackson, Ian Johnston, Dominic Kwiatkowski, Cordelia Langford, John Sillitoe on behalf of the Wellcome Sanger Institute COVID-19 Surveillance Team |
| EPI_ISL_891725 | Lighthouse Lab in Glasgow | Wellcome Sanger Institute for the COVID-19 Genomics UK (COG-UK) Consortium | Harper VanSteenhouse, Yumi Kasai, David Gray, Carol Clugston, Anna Dominiczak and Alex Alderton, Roberto Amato, Sonia Goncalves, Ewan Harrison, David K. Jackson, Ian Johnston, Dominic Kwiatkowski, Cordelia Langford, John Sillitoe on behalf of the Wellcome Sanger Institute COVID-19 Surveillance Team |
| EPI_ISL_891726 | Lighthouse Lab in Alderley Park | Wellcome Sanger Institute for the COVID-19 Genomics UK (COG-UK) Consortium | Jacquelyn Wynn, Mairead Hyland, The Lighthouse Lab in Alderley Park and Alex Alderton, Roberto Amato, Sonia Goncalves, Ewan Harrison, David K. Jackson, Ian Johnston, Dominic Kwiatkowski, Cordelia Langford, John Sillitoe on behalf of the Wellcome Sanger Institute COVID-19 Surveillance Team |
| EPI_ISL_891727 | Lighthouse Lab in Glasgow | Wellcome Sanger Institute for the COVID-19 Genomics UK (COG-UK) Consortium | Harper VanSteenhouse, Yumi Kasai, David Gray, Carol Clugston, Anna Dominiczak and Alex Alderton, Roberto Amato, Sonia Goncalves, Ewan Harrison, David K. Jackson, Ian Johnston, Dominic Kwiatkowski, Cordelia Langford, John Sillitoe on behalf of the Wellcome Sanger Institute COVID-19 Surveillance Team |
| EPI_ISL_891728 | Lighthouse Lab in Cambridge | Wellcome Sanger Institute for the COVID-19 Genomics UK (COG-UK) Consortium | Rob Howes, The Lighthouse Lab in Cambridge and Alex Alderton, Roberto Amato, Sonia Goncalves, Ewan Harrison, David K. Jackson, Ian Johnston, Dominic Kwiatkowski, Cordelia Langford, John Sillitoe on behalf of the Wellcome Sanger Institute COVID-19 Surveillance Team |
| EPI_ISL_891729 | Lighthouse Lab in Alderley Park | Wellcome Sanger Institute for the COVID-19 Genomics UK (COG-UK) Consortium | Jacquelyn Wynn, Mairead Hyland, The Lighthouse Lab in Alderley Park and Alex Alderton, Roberto Amato, Sonia Goncalves, Ewan Harrison, David K. Jackson, Ian Johnston, Dominic Kwiatkowski, Cordelia Langford, John Sillitoe on behalf of the Wellcome Sanger Institute COVID-19 Surveillance Team |
| EPI_ISL_891730, EPI_ISL_891731, EPI_ISL_891732, EPI_ISL_891733, EPI_ISL_891734 | Lighthouse Lab in Cambridge | Wellcome Sanger Institute for the COVID-19 Genomics UK (COG-UK) Consortium | Rob Howes, The Lighthouse Lab in Cambridge and Alex Alderton, Roberto Amato, Sonia Goncalves, Ewan Harrison, David K. Jackson, Ian Johnston, Dominic Kwiatkowski, Cordelia Langford, John Sillitoe on behalf of the Wellcome Sanger Institute COVID-19 Surveillance Team |
| EPI_ISL_891735, EPI_ISL_891736 | Lighthouse Lab in Alderley Park | Wellcome Sanger Institute for the COVID-19 Genomics UK (COG-UK) Consortium | Jacquelyn Wynn, Mairead Hyland, The Lighthouse Lab in Alderley Park and Alex Alderton, Roberto Amato, Sonia Goncalves, Ewan Harrison, David K. Jackson, Ian Johnston, Dominic Kwiatkowski, Cordelia Langford, John Sillitoe on behalf of the Wellcome Sanger Institute COVID-19 Surveillance Team |
| EPI_ISL_891737 | Lighthouse Lab in Cambridge | Wellcome Sanger Institute for the COVID-19 Genomics UK (COG-UK) Consortium | Rob Howes, The Lighthouse Lab in Cambridge and Alex Alderton, Roberto Amato, Sonia Goncalves, Ewan Harrison, David K. Jackson, Ian Johnston, Dominic Kwiatkowski, Cordelia Langford, John Sillitoe on behalf of the Wellcome Sanger Institute COVID-19 Surveillance Team |
| EPI_ISL_891738 | Lighthouse Lab in Alderley Park | Wellcome Sanger Institute for the COVID-19 Genomics UK (COG-UK) Consortium | Jacquelyn Wynn, Mairead Hyland, The Lighthouse Lab in Alderley Park and Alex Alderton, Roberto Amato, Sonia Goncalves, Ewan Harrison, David K. Jackson, Ian Johnston, Dominic Kwiatkowski, Cordelia Langford, John Sillitoe on behalf of the Wellcome Sanger Institute COVID-19 Surveillance Team |
| EPI_ISL_891739 | Lighthouse Lab in Cambridge | Wellcome Sanger Institute for the COVID-19 Genomics UK (COG-UK) Consortium | Rob Howes, The Lighthouse Lab in Cambridge and Alex Alderton, Roberto Amato, Sonia Goncalves, Ewan Harrison, David K. Jackson, Ian Johnston, Dominic Kwiatkowski, Cordelia Langford, John Sillitoe on behalf of the Wellcome Sanger Institute COVID-19 Surveillance Team |
| EPI_ISL_891740, EPI_ISL_891741 | Lighthouse Lab in Glasgow | Wellcome Sanger Institute for the COVID-19 Genomics UK (COG-UK) Consortium | Harper VanSteenhouse, Yumi Kasai, David Gray, Carol Clugston, Anna Dominiczak and Alex Alderton, Roberto Amato, Sonia Goncalves, Ewan Harrison, David K. Jackson, Ian Johnston, Dominic Kwiatkowski, Cordelia Langford, John Sillitoe on behalf of the Wellcome Sanger Institute COVID-19 Surveillance Team |
| EPI_ISL_891742 | Lighthouse Lab in Alderley Park | Wellcome Sanger Institute for the COVID-19 Genomics UK (COG-UK) Consortium | Jacquelyn Wynn, Mairead Hyland, The Lighthouse Lab in Alderley Park and Alex Alderton, Roberto Amato, Sonia Goncalves, Ewan Harrison, David K. Jackson, Ian Johnston, Dominic Kwiatkowski, Cordelia Langford, John Sillitoe on behalf of the Wellcome Sanger Institute COVID-19 Surveillance Team |
| EPI_ISL_891743, EPI_ISL_891744, EPI_ISL_891745 | Lighthouse Lab in Cambridge | Wellcome Sanger Institute for the COVID-19 Genomics UK (COG-UK) Consortium | Rob Howes, The Lighthouse Lab in Cambridge and Alex Alderton, Roberto Amato, Sonia Goncalves, Ewan Harrison, David K. Jackson, Ian Johnston, Dominic Kwiatkowski, Cordelia Langford, John Sillitoe on behalf of the Wellcome Sanger Institute COVID-19 Surveillance Team |
| EPI_ISL_891746 | Lighthouse Lab in Alderley Park | Wellcome Sanger Institute for the COVID-19 Genomics UK (COG-UK) Consortium | Jacquelyn Wynn, Mairead Hyland, The Lighthouse Lab in Alderley Park and Alex Alderton, Roberto Amato, Sonia Goncalves, Ewan Harrison, David K. Jackson, Ian Johnston, Dominic Kwiatkowski, Cordelia Langford, John Sillitoe on behalf of the Wellcome Sanger Institute COVID-19 Surveillance Team |
| EPI_ISL_891748, EPI_ISL_891750 | Lighthouse Lab in Cambridge | Wellcome Sanger Institute for the COVID-19 Genomics UK (COG-UK) Consortium | Rob Howes, The Lighthouse Lab in Cambridge and Alex Alderton, Roberto Amato, Sonia Goncalves, Ewan Harrison, David K. Jackson, Ian Johnston, Dominic Kwiatkowski, Cordelia Langford, John Sillitoe on behalf of the Wellcome Sanger Institute COVID-19 Surveillance Team |
| EPI_ISL_891751 | Lighthouse Lab in Alderley Park | Wellcome Sanger Institute for the COVID-19 Genomics UK (COG-UK) Consortium | Jacquelyn Wynn, Mairead Hyland, The Lighthouse Lab in Alderley Park and Alex Alderton, Roberto Amato, Sonia Goncalves, Ewan Harrison, David K. Jackson, Ian Johnston, Dominic Kwiatkowski, Cordelia Langford, John Sillitoe on behalf of the Wellcome Sanger Institute COVID-19 Surveillance Team |
| EPI_ISL_891752 | Lighthouse Lab in Cambridge | Wellcome Sanger Institute for the COVID-19 Genomics UK (COG-UK) Consortium | Rob Howes, The Lighthouse Lab in Cambridge and Alex Alderton, Roberto Amato, Sonia Goncalves, Ewan Harrison, David K. Jackson, Ian Johnston, Dominic Kwiatkowski, Cordelia Langford, John Sillitoe on behalf of the Wellcome Sanger Institute COVID-19 Surveillance Team |
| EPI_ISL_891753 | Lighthouse Lab in Alderley Park | Wellcome Sanger Institute for the COVID-19 Genomics UK (COG-UK) Consortium | Jacquelyn Wynn, Mairead Hyland, The Lighthouse Lab in Alderley Park and Alex Alderton, Roberto Amato, Sonia Goncalves, Ewan Harrison, David K. Jackson, Ian Johnston, Dominic Kwiatkowski, Cordelia Langford, John Sillitoe on behalf of the Wellcome Sanger Institute COVID-19 Surveillance Team |
| EPI_ISL_891754, EPI_ISL_891756 | Lighthouse Lab in Cambridge | Wellcome Sanger Institute for the COVID-19 Genomics UK (COG-UK) Consortium | Rob Howes, The Lighthouse Lab in Cambridge and Alex Alderton, Roberto Amato, Sonia Goncalves, Ewan Harrison, David K. Jackson, Ian Johnston, Dominic Kwiatkowski, Cordelia Langford, John Sillitoe on behalf of the Wellcome Sanger Institute COVID-19 Surveillance Team |
| EPI_ISL_891757 | Lighthouse Lab in Alderley Park | Wellcome Sanger Institute for the COVID-19 Genomics UK (COG-UK) Consortium | Jacquelyn Wynn, Mairead Hyland, The Lighthouse Lab in Alderley Park and Alex Alderton, Roberto Amato, Sonia Goncalves, Ewan Harrison, David K. Jackson, Ian Johnston, Dominic Kwiatkowski, Cordelia Langford, John Sillitoe on behalf of the Wellcome Sanger Institute COVID-19 Surveillance Team |
| EPI_ISL_891758, EPI_ISL_891759 | Lighthouse Lab in Cambridge | Wellcome Sanger Institute for the COVID-19 Genomics UK (COG-UK) Consortium | Rob Howes, The Lighthouse Lab in Cambridge and Alex Alderton, Roberto Amato, Sonia Goncalves, Ewan Harrison, David K. Jackson, Ian Johnston, Dominic Kwiatkowski, Cordelia Langford, John Sillitoe on behalf of the Wellcome Sanger Institute COVID-19 Surveillance Team |
| EPI_ISL_891760 | Lighthouse Lab in Alderley Park | Wellcome Sanger Institute for the COVID-19 Genomics UK (COG-UK) Consortium | Jacquelyn Wynn, Mairead Hyland, The Lighthouse Lab in Alderley Park and Alex Alderton, Roberto Amato, Sonia Goncalves, Ewan Harrison, David K. Jackson, Ian Johnston, Dominic Kwiatkowski, Cordelia Langford, John Sillitoe on behalf of the Wellcome Sanger Institute COVID-19 Surveillance Team |
| EPI_ISL_891761 | Lighthouse Lab in Glasgow | Wellcome Sanger Institute for the COVID-19 Genomics UK (COG-UK) Consortium | Harper VanSteenhouse, Yumi Kasai, David Gray, Carol Clugston, Anna Dominiczak and Alex Alderton, Roberto Amato, Sonia Goncalves, Ewan Harrison, David K. Jackson, Ian Johnston, Dominic Kwiatkowski, Cordelia Langford, John Sillitoe on behalf of the Wellcome Sanger Institute COVID-19 Surveillance Team |
| EPI_ISL_891762 | Lighthouse Lab in Alderley Park | Wellcome Sanger Institute for the COVID-19 Genomics UK (COG-UK) Consortium | Jacquelyn Wynn, Mairead Hyland, The Lighthouse Lab in Alderley Park and Alex Alderton, Roberto Amato, Sonia Goncalves, Ewan Harrison, David K. Jackson, Ian Johnston, Dominic Kwiatkowski, Cordelia Langford, John Sillitoe on behalf of the Wellcome Sanger Institute COVID-19 Surveillance Team |
| EPI_ISL_891763, EPI_ISL_891764 | Lighthouse Lab in Cambridge | Wellcome Sanger Institute for the COVID-19 Genomics UK (COG-UK) Consortium | Rob Howes, The Lighthouse Lab in Cambridge and Alex Alderton, Roberto Amato, Sonia Goncalves, Ewan Harrison, David K. Jackson, Ian Johnston, Dominic Kwiatkowski, Cordelia Langford, John Sillitoe on behalf of the Wellcome Sanger Institute COVID-19 Surveillance Team |
| EPI_ISL_891765, EPI_ISL_891766 | Lighthouse Lab in Alderley Park | Wellcome Sanger Institute for the COVID-19 Genomics UK (COG-UK) Consortium | Jacquelyn Wynn, Mairead Hyland, The Lighthouse Lab in Alderley Park and Alex Alderton, Roberto Amato, Sonia Goncalves, Ewan Harrison, David K. Jackson, Ian Johnston, Dominic Kwiatkowski, Cordelia Langford, John Sillitoe on behalf of the Wellcome Sanger Institute COVID-19 Surveillance Team |
| EPI_ISL_891767, EPI_ISL_891768, EPI_ISL_891770 | Lighthouse Lab in Cambridge | Wellcome Sanger Institute for the COVID-19 Genomics UK (COG-UK) Consortium | Rob Howes, The Lighthouse Lab in Cambridge and Alex Alderton, Roberto Amato, Sonia Goncalves, Ewan Harrison, David K. Jackson, Ian Johnston, Dominic Kwiatkowski, Cordelia Langford, John Sillitoe on behalf of the Wellcome Sanger Institute COVID-19 Surveillance Team |
| EPI_ISL_891771, EPI_ISL_891772 | Lighthouse Lab in Glasgow | Wellcome Sanger Institute for the COVID-19 Genomics UK (COG-UK) Consortium | Harper VanSteenhouse, Yumi Kasai, David Gray, Carol Clugston, Anna Dominiczak and Alex Alderton, Roberto Amato, Sonia Goncalves, Ewan Harrison, David K. Jackson, Ian Johnston, Dominic Kwiatkowski, Cordelia Langford, John Sillitoe on behalf of the Wellcome Sanger Institute COVID-19 Surveillance |

| | | | |
|---|---|---|---|
| | | | Team |
| EPI_ISL_891773, EPI_ISL_891774, EPI_ISL_891775, EPI_ISL_891776, EPI_ISL_891777 | Lighthouse Lab in Alderley Park | Wellcome Sanger Institute for the COVID-19 Genomics UK (COG-UK) Consortium | Jacquelyn Wynn, Mairead Hyland, The Lighthouse Lab in Alderley Park and Alex Alderton, Roberto Amato, Sonia Goncalves, Ewan Harrison, David K. Jackson, Ian Johnston, Dominic Kwiatkowski, Cordelia Langford, John Sillitoe on behalf of the Wellcome Sanger Institute COVID-19 Surveillance Team |
| EPI_ISL_891778, EPI_ISL_891779 | Lighthouse Lab in Cambridge | Wellcome Sanger Institute for the COVID-19 Genomics UK (COG-UK) Consortium | Rob Howes, The Lighthouse Lab in Cambridge and Alex Alderton, Roberto Amato, Sonia Goncalves, Ewan Harrison, David K. Jackson, Ian Johnston, Dominic Kwiatkowski, Cordelia Langford, John Sillitoe on behalf of the Wellcome Sanger Institute COVID-19 Surveillance Team |
| EPI_ISL_891780 | Lighthouse Lab in Alderley Park | Wellcome Sanger Institute for the COVID-19 Genomics UK (COG-UK) Consortium | Jacquelyn Wynn, Mairead Hyland, The Lighthouse Lab in Alderley Park and Alex Alderton, Roberto Amato, Sonia Goncalves, Ewan Harrison, David K. Jackson, Ian Johnston, Dominic Kwiatkowski, Cordelia Langford, John Sillitoe on behalf of the Wellcome Sanger Institute COVID-19 Surveillance Team |
| EPI_ISL_891781, EPI_ISL_891782, EPI_ISL_891783, EPI_ISL_891784, EPI_ISL_891785 | Lighthouse Lab in Cambridge | Wellcome Sanger Institute for the COVID-19 Genomics UK (COG-UK) Consortium | Rob Howes, The Lighthouse Lab in Cambridge and Alex Alderton, Roberto Amato, Sonia Goncalves, Ewan Harrison, David K. Jackson, Ian Johnston, Dominic Kwiatkowski, Cordelia Langford, John Sillitoe on behalf of the Wellcome Sanger Institute COVID-19 Surveillance Team |
| EPI_ISL_891786, EPI_ISL_891787, EPI_ISL_891788, EPI_ISL_891789 | Lighthouse Lab in Alderley Park | Wellcome Sanger Institute for the COVID-19 Genomics UK (COG-UK) Consortium | Jacquelyn Wynn, Mairead Hyland, The Lighthouse Lab in Alderley Park and Alex Alderton, Roberto Amato, Sonia Goncalves, Ewan Harrison, David K. Jackson, Ian Johnston, Dominic Kwiatkowski, Cordelia Langford, John Sillitoe on behalf of the Wellcome Sanger Institute COVID-19 Surveillance Team |
| EPI_ISL_891790, EPI_ISL_891791 | Lighthouse Lab in Glasgow | Wellcome Sanger Institute for the COVID-19 Genomics UK (COG-UK) Consortium | Harper VanSteenhouse, Yumi Kasai, David Gray, Carol Clugston, Anna Dominiczak and Alex Alderton, Roberto Amato, Sonia Goncalves, Ewan Harrison, David K. Jackson, Ian Johnston, Dominic Kwiatkowski, Cordelia Langford, John Sillitoe on behalf of the Wellcome Sanger Institute COVID-19 Surveillance Team |
| EPI_ISL_891792 | Lighthouse Lab in Cambridge | Wellcome Sanger Institute for the COVID-19 Genomics UK (COG-UK) Consortium | Rob Howes, The Lighthouse Lab in Cambridge and Alex Alderton, Roberto Amato, Sonia Goncalves, Ewan Harrison, David K. Jackson, Ian Johnston, Dominic Kwiatkowski, Cordelia Langford, John Sillitoe on behalf of the Wellcome Sanger Institute COVID-19 Surveillance Team |
| EPI_ISL_891793 | Lighthouse Lab in Glasgow | Wellcome Sanger Institute for the COVID-19 Genomics UK (COG-UK) Consortium | Harper VanSteenhouse, Yumi Kasai, David Gray, Carol Clugston, Anna Dominiczak and Alex Alderton, Roberto Amato, Sonia Goncalves, Ewan Harrison, David K. Jackson, Ian Johnston, Dominic Kwiatkowski, Cordelia Langford, John Sillitoe on behalf of the Wellcome Sanger Institute COVID-19 Surveillance Team |
| EPI_ISL_891794 | Lighthouse Lab in Alderley Park | Wellcome Sanger Institute for the COVID-19 Genomics UK (COG-UK) Consortium | Jacquelyn Wynn, Mairead Hyland, The Lighthouse Lab in Alderley Park and Alex Alderton, Roberto Amato, Sonia Goncalves, Ewan Harrison, David K. Jackson, Ian Johnston, Dominic Kwiatkowski, Cordelia Langford, John Sillitoe on behalf of the Wellcome Sanger Institute COVID-19 Surveillance Team |
| EPI_ISL_891795, EPI_ISL_891796, EPI_ISL_891797, EPI_ISL_891798, EPI_ISL_891800, EPI_ISL_891801 | Lighthouse Lab in Cambridge | Wellcome Sanger Institute for the COVID-19 Genomics UK (COG-UK) Consortium | Rob Howes, The Lighthouse Lab in Cambridge and Alex Alderton, Roberto Amato, Sonia Goncalves, Ewan Harrison, David K. Jackson, Ian Johnston, Dominic Kwiatkowski, Cordelia Langford, John Sillitoe on behalf of the Wellcome Sanger Institute COVID-19 Surveillance Team |
| EPI_ISL_891802 | Lighthouse Lab in Glasgow | Wellcome Sanger Institute for the COVID-19 Genomics UK (COG-UK) Consortium | Harper VanSteenhouse, Yumi Kasai, David Gray, Carol Clugston, Anna Dominiczak and Alex Alderton, Roberto Amato, Sonia Goncalves, Ewan Harrison, David K. Jackson, Ian Johnston, Dominic Kwiatkowski, Cordelia Langford, John Sillitoe on behalf of the Wellcome Sanger Institute COVID-19 Surveillance Team |
| EPI_ISL_891803 | Lighthouse Lab in Cambridge | Wellcome Sanger Institute for the COVID-19 Genomics UK (COG-UK) Consortium | Rob Howes, The Lighthouse Lab in Cambridge and Alex Alderton, Roberto Amato, Sonia Goncalves, Ewan Harrison, David K. Jackson, Ian Johnston, Dominic Kwiatkowski, Cordelia Langford, John Sillitoe on behalf of the Wellcome Sanger Institute COVID-19 Surveillance Team |
| EPI_ISL_891804 | Lighthouse Lab in Alderley Park | Wellcome Sanger Institute for the COVID-19 Genomics UK (COG-UK) Consortium | Jacquelyn Wynn, Mairead Hyland, The Lighthouse Lab in Alderley Park and Alex Alderton, Roberto Amato, Sonia Goncalves, Ewan Harrison, David K. Jackson, Ian Johnston, Dominic Kwiatkowski, Cordelia Langford, John Sillitoe on behalf of the Wellcome Sanger Institute COVID-19 Surveillance Team |
| EPI_ISL_891805, EPI_ISL_891806 | Lighthouse Lab in Cambridge | Wellcome Sanger Institute for the COVID-19 Genomics UK (COG-UK) Consortium | Rob Howes, The Lighthouse Lab in Cambridge and Alex Alderton, Roberto Amato, Sonia Goncalves, Ewan Harrison, David K. Jackson, Ian Johnston, Dominic Kwiatkowski, Cordelia Langford, John Sillitoe on behalf of the Wellcome Sanger Institute COVID-19 Surveillance Team |
| EPI_ISL_891807 | Lighthouse Lab in Alderley Park | Wellcome Sanger Institute for the COVID-19 Genomics UK (COG-UK) Consortium | Jacquelyn Wynn, Mairead Hyland, The Lighthouse Lab in Alderley Park and Alex Alderton, Roberto Amato, Sonia Goncalves, Ewan Harrison, David K. Jackson, Ian Johnston, Dominic Kwiatkowski, Cordelia Langford, John Sillitoe on behalf of the Wellcome Sanger Institute COVID-19 Surveillance Team |
| EPI_ISL_891808 | Lighthouse Lab in Cambridge | Wellcome Sanger Institute for the COVID-19 Genomics UK (COG-UK) Consortium | Rob Howes, The Lighthouse Lab in Cambridge and Alex Alderton, Roberto Amato, Sonia Goncalves, Ewan Harrison, David K. Jackson, Ian Johnston, Dominic Kwiatkowski, Cordelia Langford, John Sillitoe on behalf of the Wellcome Sanger Institute COVID-19 Surveillance Team |
| EPI_ISL_891809 | Lighthouse Lab in Glasgow | Wellcome Sanger Institute for the COVID-19 Genomics UK (COG-UK) Consortium | Harper VanSteenhouse, Yumi Kasai, David Gray, Carol Clugston, Anna Dominiczak and Alex Alderton, Roberto Amato, Sonia Goncalves, Ewan Harrison, David K. Jackson, Ian Johnston, Dominic Kwiatkowski, Cordelia Langford, John Sillitoe on behalf of the Wellcome Sanger Institute COVID-19 Surveillance Team |
| EPI_ISL_891810 | Lighthouse Lab in Cambridge | Wellcome Sanger Institute for the COVID-19 Genomics UK (COG-UK) Consortium | Rob Howes, The Lighthouse Lab in Cambridge and Alex Alderton, Roberto Amato, Sonia Goncalves, Ewan Harrison, David K. Jackson, Ian Johnston, Dominic Kwiatkowski, Cordelia Langford, John Sillitoe on behalf of the Wellcome Sanger Institute COVID-19 Surveillance Team |
| EPI_ISL_891811 | Lighthouse Lab in Alderley Park | Wellcome Sanger Institute for the COVID-19 Genomics UK (COG-UK) Consortium | Jacquelyn Wynn, Mairead Hyland, The Lighthouse Lab in Alderley Park and Alex Alderton, Roberto Amato, Sonia Goncalves, Ewan Harrison, David K. Jackson, Ian Johnston, Dominic Kwiatkowski, Cordelia Langford, John Sillitoe on behalf of the Wellcome Sanger Institute COVID-19 Surveillance Team |
| EPI_ISL_891812 | Lighthouse Lab in Cambridge | Wellcome Sanger Institute for the COVID-19 Genomics UK (COG-UK) Consortium | Rob Howes, The Lighthouse Lab in Cambridge and Alex Alderton, Roberto Amato, Sonia Goncalves, Ewan Harrison, David K. Jackson, Ian Johnston, Dominic Kwiatkowski, Cordelia Langford, John Sillitoe on behalf of the Wellcome Sanger Institute COVID-19 Surveillance Team |
| EPI_ISL_891813 | Lighthouse Lab in Alderley Park | Wellcome Sanger Institute for the COVID-19 Genomics UK (COG-UK) Consortium | Jacquelyn Wynn, Mairead Hyland, The Lighthouse Lab in Alderley Park and Alex Alderton, Roberto Amato, Sonia Goncalves, Ewan Harrison, David K. Jackson, Ian Johnston, Dominic Kwiatkowski, Cordelia Langford, John Sillitoe on behalf of the Wellcome Sanger Institute COVID-19 Surveillance Team |
| EPI_ISL_891814, EPI_ISL_891815, EPI_ISL_891816, EPI_ISL_891817, EPI_ISL_891818, EPI_ISL_891820 | Lighthouse Lab in Cambridge | Wellcome Sanger Institute for the COVID-19 Genomics UK (COG-UK) Consortium | Rob Howes, The Lighthouse Lab in Cambridge and Alex Alderton, Roberto Amato, Sonia Goncalves, Ewan Harrison, David K. Jackson, Ian Johnston, Dominic Kwiatkowski, Cordelia Langford, John Sillitoe on behalf of the Wellcome Sanger Institute COVID-19 Surveillance Team |
| EPI_ISL_891824 | Lighthouse Lab in Alderley Park | Wellcome Sanger Institute for the COVID-19 Genomics UK (COG-UK) Consortium | Jacquelyn Wynn, Mairead Hyland, The Lighthouse Lab in Alderley Park and Alex Alderton, Roberto Amato, Sonia Goncalves, Ewan Harrison, David K. Jackson, Ian Johnston, Dominic Kwiatkowski, Cordelia Langford, John Sillitoe on behalf of the Wellcome Sanger Institute COVID-19 Surveillance Team |
| EPI_ISL_891825, EPI_ISL_891827 | Lighthouse Lab in Cambridge | Wellcome Sanger Institute for the COVID-19 Genomics UK (COG-UK) Consortium | Rob Howes, The Lighthouse Lab in Cambridge and Alex Alderton, Roberto Amato, Sonia Goncalves, Ewan Harrison, David K. Jackson, Ian Johnston, Dominic Kwiatkowski, Cordelia Langford, John Sillitoe on behalf of the Wellcome Sanger Institute COVID-19 Surveillance Team |
| EPI_ISL_891828 | Lighthouse Lab in Glasgow | Wellcome Sanger Institute for the COVID-19 Genomics UK (COG-UK) Consortium | Harper VanSteenhouse, Yumi Kasai, David Gray, Carol Clugston, Anna Dominiczak and Alex Alderton, Roberto Amato, Sonia Goncalves, Ewan Harrison, David K. Jackson, Ian Johnston, Dominic Kwiatkowski, Cordelia Langford, John Sillitoe on behalf of the Wellcome Sanger Institute COVID-19 Surveillance Team |
| EPI_ISL_891829, EPI_ISL_891830, EPI_ISL_891831 | Lighthouse Lab in Cambridge | Wellcome Sanger Institute for the COVID-19 Genomics UK (COG-UK) Consortium | Rob Howes, The Lighthouse Lab in Cambridge and Alex Alderton, Roberto Amato, Sonia Goncalves, Ewan Harrison, David K. Jackson, Ian Johnston, Dominic Kwiatkowski, Cordelia Langford, John Sillitoe on behalf of the Wellcome Sanger Institute COVID-19 Surveillance Team |
| EPI_ISL_891833 | Lighthouse Lab in Alderley Park | Wellcome Sanger Institute for the COVID-19 Genomics UK (COG-UK) Consortium | Jacquelyn Wynn, Mairead Hyland, The Lighthouse Lab in Alderley Park and Alex Alderton, Roberto Amato, Sonia Goncalves, Ewan Harrison, David K. Jackson, Ian Johnston, Dominic Kwiatkowski, Cordelia Langford, John Sillitoe on behalf of the Wellcome Sanger Institute COVID-19 Surveillance Team |
| EPI_ISL_891835 | Lighthouse Lab in Cambridge | Wellcome Sanger Institute for the COVID-19 Genomics UK (COG-UK) Consortium | Rob Howes, The Lighthouse Lab in Cambridge and Alex Alderton, Roberto Amato, Sonia Goncalves, Ewan Harrison, David K. Jackson, Ian Johnston, Dominic Kwiatkowski, Cordelia Langford, John Sillitoe on behalf of the Wellcome Sanger Institute COVID-19 Surveillance Team |
| EPI_ISL_891836 | Lighthouse Lab in Glasgow | Wellcome Sanger Institute for the COVID-19 Genomics UK (COG-UK) Consortium | Harper VanSteenhouse, Yumi Kasai, David Gray, Carol Clugston, Anna Dominiczak and Alex Alderton, Roberto Amato, Sonia Goncalves, Ewan Harrison, David K. Jackson, Ian Johnston, Dominic Kwiatkowski, Cordelia Langford, John Sillitoe on behalf of the Wellcome Sanger Institute COVID-19 Surveillance Team |
| EPI_ISL_891837 | Lighthouse Lab in Cambridge | Wellcome Sanger Institute for the COVID-19 Genomics UK (COG-UK) Consortium | Rob Howes, The Lighthouse Lab in Cambridge and Alex Alderton, Roberto Amato, Sonia Goncalves, Ewan Harrison, David K. Jackson, Ian Johnston, Dominic Kwiatkowski, Cordelia Langford, John Sillitoe on behalf of the Wellcome Sanger Institute COVID-19 Surveillance Team |
| EPI_ISL_891838, EPI_ISL_891839 | Lighthouse Lab in Alderley Park | Wellcome Sanger Institute for the COVID-19 Genomics UK | Jacquelyn Wynn, Mairead Hyland, The Lighthouse Lab in Alderley Park and Alex Alderton, Roberto Amato, Sonia Goncalves, Ewan Harrison, David K. |

| | | | |
|---|---|---|---|
| | | (COG-UK) Consortium | Jackson, Ian Johnston, Dominic Kwiatkowski, Cordelia Langford, John Sillitoe on behalf of the Wellcome Sanger Institute COVID-19 Surveillance Team |
| EPI_ISL_891840, EPI_ISL_891841, EPI_ISL_891842 | Lighthouse Lab in Cambridge | Wellcome Sanger Institute for the COVID-19 Genomics UK (COG-UK) Consortium | Rob Howes, The Lighthouse Lab in Cambridge and Alex Alderton, Roberto Amato, Sonia Goncalves, Ewan Harrison, David K. Jackson, Ian Johnston, Dominic Kwiatkowski, Cordelia Langford, John Sillitoe on behalf of the Wellcome Sanger Institute COVID-19 Surveillance Team |
| EPI_ISL_891843 | Lighthouse Lab in Alderley Park | Wellcome Sanger Institute for the COVID-19 Genomics UK (COG-UK) Consortium | Jacquelyn Wynn, Mairead Hyland, The Lighthouse Lab in Alderley Park and Alex Alderton, Roberto Amato, Sonia Goncalves, Ewan Harrison, David K. Jackson, Ian Johnston, Dominic Kwiatkowski, Cordelia Langford, John Sillitoe on behalf of the Wellcome Sanger Institute COVID-19 Surveillance Team |
| EPI_ISL_891844 | Lighthouse Lab in Cambridge | Wellcome Sanger Institute for the COVID-19 Genomics UK (COG-UK) Consortium | Rob Howes, The Lighthouse Lab in Cambridge and Alex Alderton, Roberto Amato, Sonia Goncalves, Ewan Harrison, David K. Jackson, Ian Johnston, Dominic Kwiatkowski, Cordelia Langford, John Sillitoe on behalf of the Wellcome Sanger Institute COVID-19 Surveillance Team |
| EPI_ISL_891846 | Lighthouse Lab in Glasgow | Wellcome Sanger Institute for the COVID-19 Genomics UK (COG-UK) Consortium | Harper VanSteenhouse, Yumi Kasai, David Gray, Carol Clugston, Anna Dominiczak and Alex Alderton, Roberto Amato, Sonia Goncalves, Ewan Harrison, David K. Jackson, Ian Johnston, Dominic Kwiatkowski, Cordelia Langford, John Sillitoe on behalf of the Wellcome Sanger Institute COVID-19 Surveillance Team |
| EPI_ISL_891847, EPI_ISL_891848 | Lighthouse Lab in Alderley Park | Wellcome Sanger Institute for the COVID-19 Genomics UK (COG-UK) Consortium | Jacquelyn Wynn, Mairead Hyland, The Lighthouse Lab in Alderley Park and Alex Alderton, Roberto Amato, Sonia Goncalves, Ewan Harrison, David K. Jackson, Ian Johnston, Dominic Kwiatkowski, Cordelia Langford, John Sillitoe on behalf of the Wellcome Sanger Institute COVID-19 Surveillance Team |
| EPI_ISL_891849 | Lighthouse Lab in Cambridge | Wellcome Sanger Institute for the COVID-19 Genomics UK (COG-UK) Consortium | Rob Howes, The Lighthouse Lab in Cambridge and Alex Alderton, Roberto Amato, Sonia Goncalves, Ewan Harrison, David K. Jackson, Ian Johnston, Dominic Kwiatkowski, Cordelia Langford, John Sillitoe on behalf of the Wellcome Sanger Institute COVID-19 Surveillance Team |
| EPI_ISL_891851, EPI_ISL_891852, EPI_ISL_891853 | Lighthouse Lab in Glasgow | Wellcome Sanger Institute for the COVID-19 Genomics UK (COG-UK) Consortium | Harper VanSteenhouse, Yumi Kasai, David Gray, Carol Clugston, Anna Dominiczak and Alex Alderton, Roberto Amato, Sonia Goncalves, Ewan Harrison, David K. Jackson, Ian Johnston, Dominic Kwiatkowski, Cordelia Langford, John Sillitoe on behalf of the Wellcome Sanger Institute COVID-19 Surveillance Team |
| EPI_ISL_891854 | Lighthouse Lab in Cambridge | Wellcome Sanger Institute for the COVID-19 Genomics UK (COG-UK) Consortium | Rob Howes, The Lighthouse Lab in Cambridge and Alex Alderton, Roberto Amato, Sonia Goncalves, Ewan Harrison, David K. Jackson, Ian Johnston, Dominic Kwiatkowski, Cordelia Langford, John Sillitoe on behalf of the Wellcome Sanger Institute COVID-19 Surveillance Team |
| EPI_ISL_891855 | Lighthouse Lab in Glasgow | Wellcome Sanger Institute for the COVID-19 Genomics UK (COG-UK) Consortium | Harper VanSteenhouse, Yumi Kasai, David Gray, Carol Clugston, Anna Dominiczak and Alex Alderton, Roberto Amato, Sonia Goncalves, Ewan Harrison, David K. Jackson, Ian Johnston, Dominic Kwiatkowski, Cordelia Langford, John Sillitoe on behalf of the Wellcome Sanger Institute COVID-19 Surveillance Team |
| EPI_ISL_891856 | Lighthouse Lab in Alderley Park | Wellcome Sanger Institute for the COVID-19 Genomics UK (COG-UK) Consortium | Jacquelyn Wynn, Mairead Hyland, The Lighthouse Lab in Alderley Park and Alex Alderton, Roberto Amato, Sonia Goncalves, Ewan Harrison, David K. Jackson, Ian Johnston, Dominic Kwiatkowski, Cordelia Langford, John Sillitoe on behalf of the Wellcome Sanger Institute COVID-19 Surveillance Team |
| EPI_ISL_891857 | Lighthouse Lab in Glasgow | Wellcome Sanger Institute for the COVID-19 Genomics UK (COG-UK) Consortium | Harper VanSteenhouse, Yumi Kasai, David Gray, Carol Clugston, Anna Dominiczak and Alex Alderton, Roberto Amato, Sonia Goncalves, Ewan Harrison, David K. Jackson, Ian Johnston, Dominic Kwiatkowski, Cordelia Langford, John Sillitoe on behalf of the Wellcome Sanger Institute COVID-19 Surveillance Team |
| EPI_ISL_891858 | Lighthouse Lab in Alderley Park | Wellcome Sanger Institute for the COVID-19 Genomics UK (COG-UK) Consortium | Jacquelyn Wynn, Mairead Hyland, The Lighthouse Lab in Alderley Park and Alex Alderton, Roberto Amato, Sonia Goncalves, Ewan Harrison, David K. Jackson, Ian Johnston, Dominic Kwiatkowski, Cordelia Langford, John Sillitoe on behalf of the Wellcome Sanger Institute COVID-19 Surveillance Team |
| EPI_ISL_891859 | Lighthouse Lab in Cambridge | Wellcome Sanger Institute for the COVID-19 Genomics UK (COG-UK) Consortium | Rob Howes, The Lighthouse Lab in Cambridge and Alex Alderton, Roberto Amato, Sonia Goncalves, Ewan Harrison, David K. Jackson, Ian Johnston, Dominic Kwiatkowski, Cordelia Langford, John Sillitoe on behalf of the Wellcome Sanger Institute COVID-19 Surveillance Team |
| EPI_ISL_891861 | Lighthouse Lab in Glasgow | Wellcome Sanger Institute for the COVID-19 Genomics UK (COG-UK) Consortium | Harper VanSteenhouse, Yumi Kasai, David Gray, Carol Clugston, Anna Dominiczak and Alex Alderton, Roberto Amato, Sonia Goncalves, Ewan Harrison, David K. Jackson, Ian Johnston, Dominic Kwiatkowski, Cordelia Langford, John Sillitoe on behalf of the Wellcome Sanger Institute COVID-19 Surveillance Team |
| EPI_ISL_891862 | Lighthouse Lab in Cambridge | Wellcome Sanger Institute for the COVID-19 Genomics UK (COG-UK) Consortium | Rob Howes, The Lighthouse Lab in Cambridge and Alex Alderton, Roberto Amato, Sonia Goncalves, Ewan Harrison, David K. Jackson, Ian Johnston, Dominic Kwiatkowski, Cordelia Langford, John Sillitoe on behalf of the Wellcome Sanger Institute COVID-19 Surveillance Team |
| EPI_ISL_891863, EPI_ISL_891864 | Lighthouse Lab in Alderley Park | Wellcome Sanger Institute for the COVID-19 Genomics UK (COG-UK) Consortium | Jacquelyn Wynn, Mairead Hyland, The Lighthouse Lab in Alderley Park and Alex Alderton, Roberto Amato, Sonia Goncalves, Ewan Harrison, David K. Jackson, Ian Johnston, Dominic Kwiatkowski, Cordelia Langford, John Sillitoe on behalf of the Wellcome Sanger Institute COVID-19 Surveillance Team |
| EPI_ISL_891865 | Lighthouse Lab in Glasgow | Wellcome Sanger Institute for the COVID-19 Genomics UK (COG-UK) Consortium | Harper VanSteenhouse, Yumi Kasai, David Gray, Carol Clugston, Anna Dominiczak and Alex Alderton, Roberto Amato, Sonia Goncalves, Ewan Harrison, David K. Jackson, Ian Johnston, Dominic Kwiatkowski, Cordelia Langford, John Sillitoe on behalf of the Wellcome Sanger Institute COVID-19 Surveillance Team |
| EPI_ISL_891866 | Lighthouse Lab in Alderley Park | Wellcome Sanger Institute for the COVID-19 Genomics UK (COG-UK) Consortium | Jacquelyn Wynn, Mairead Hyland, The Lighthouse Lab in Alderley Park and Alex Alderton, Roberto Amato, Sonia Goncalves, Ewan Harrison, David K. Jackson, Ian Johnston, Dominic Kwiatkowski, Cordelia Langford, John Sillitoe on behalf of the Wellcome Sanger Institute COVID-19 Surveillance Team |
| EPI_ISL_891868 | Lighthouse Lab in Cambridge | Wellcome Sanger Institute for the COVID-19 Genomics UK (COG-UK) Consortium | Rob Howes, The Lighthouse Lab in Cambridge and Alex Alderton, Roberto Amato, Sonia Goncalves, Ewan Harrison, David K. Jackson, Ian Johnston, Dominic Kwiatkowski, Cordelia Langford, John Sillitoe on behalf of the Wellcome Sanger Institute COVID-19 Surveillance Team |
| EPI_ISL_891869 | Lighthouse Lab in Glasgow | Wellcome Sanger Institute for the COVID-19 Genomics UK (COG-UK) Consortium | Harper VanSteenhouse, Yumi Kasai, David Gray, Carol Clugston, Anna Dominiczak and Alex Alderton, Roberto Amato, Sonia Goncalves, Ewan Harrison, David K. Jackson, Ian Johnston, Dominic Kwiatkowski, Cordelia Langford, John Sillitoe on behalf of the Wellcome Sanger Institute COVID-19 Surveillance Team |
| EPI_ISL_891870 | Lighthouse Lab in Alderley Park | Wellcome Sanger Institute for the COVID-19 Genomics UK (COG-UK) Consortium | Jacquelyn Wynn, Mairead Hyland, The Lighthouse Lab in Alderley Park and Alex Alderton, Roberto Amato, Sonia Goncalves, Ewan Harrison, David K. Jackson, Ian Johnston, Dominic Kwiatkowski, Cordelia Langford, John Sillitoe on behalf of the Wellcome Sanger Institute COVID-19 Surveillance Team |
| EPI_ISL_891871, EPI_ISL_891872 | Lighthouse Lab in Cambridge | Wellcome Sanger Institute for the COVID-19 Genomics UK (COG-UK) Consortium | Rob Howes, The Lighthouse Lab in Cambridge and Alex Alderton, Roberto Amato, Sonia Goncalves, Ewan Harrison, David K. Jackson, Ian Johnston, Dominic Kwiatkowski, Cordelia Langford, John Sillitoe on behalf of the Wellcome Sanger Institute COVID-19 Surveillance Team |
| EPI_ISL_891873 | Lighthouse Lab in Glasgow | Wellcome Sanger Institute for the COVID-19 Genomics UK (COG-UK) Consortium | Harper VanSteenhouse, Yumi Kasai, David Gray, Carol Clugston, Anna Dominiczak and Alex Alderton, Roberto Amato, Sonia Goncalves, Ewan Harrison, David K. Jackson, Ian Johnston, Dominic Kwiatkowski, Cordelia Langford, John Sillitoe on behalf of the Wellcome Sanger Institute COVID-19 Surveillance Team |
| EPI_ISL_891874 | Lighthouse Lab in Cambridge | Wellcome Sanger Institute for the COVID-19 Genomics UK (COG-UK) Consortium | Rob Howes, The Lighthouse Lab in Cambridge and Alex Alderton, Roberto Amato, Sonia Goncalves, Ewan Harrison, David K. Jackson, Ian Johnston, Dominic Kwiatkowski, Cordelia Langford, John Sillitoe on behalf of the Wellcome Sanger Institute COVID-19 Surveillance Team |
| EPI_ISL_891875 | Lighthouse Lab in Glasgow | Wellcome Sanger Institute for the COVID-19 Genomics UK (COG-UK) Consortium | Harper VanSteenhouse, Yumi Kasai, David Gray, Carol Clugston, Anna Dominiczak and Alex Alderton, Roberto Amato, Sonia Goncalves, Ewan Harrison, David K. Jackson, Ian Johnston, Dominic Kwiatkowski, Cordelia Langford, John Sillitoe on behalf of the Wellcome Sanger Institute COVID-19 Surveillance Team |
| EPI_ISL_891876, EPI_ISL_891877 | Lighthouse Lab in Cambridge | Wellcome Sanger Institute for the COVID-19 Genomics UK (COG-UK) Consortium | Rob Howes, The Lighthouse Lab in Cambridge and Alex Alderton, Roberto Amato, Sonia Goncalves, Ewan Harrison, David K. Jackson, Ian Johnston, Dominic Kwiatkowski, Cordelia Langford, John Sillitoe on behalf of the Wellcome Sanger Institute COVID-19 Surveillance Team |
| EPI_ISL_891878 | Lighthouse Lab in Alderley Park | Wellcome Sanger Institute for the COVID-19 Genomics UK (COG-UK) Consortium | Jacquelyn Wynn, Mairead Hyland, The Lighthouse Lab in Alderley Park and Alex Alderton, Roberto Amato, Sonia Goncalves, Ewan Harrison, David K. Jackson, Ian Johnston, Dominic Kwiatkowski, Cordelia Langford, John Sillitoe on behalf of the Wellcome Sanger Institute COVID-19 Surveillance Team |
| EPI_ISL_891879, EPI_ISL_891880 | Lighthouse Lab in Cambridge | Wellcome Sanger Institute for the COVID-19 Genomics UK (COG-UK) Consortium | Rob Howes, The Lighthouse Lab in Cambridge and Alex Alderton, Roberto Amato, Sonia Goncalves, Ewan Harrison, David K. Jackson, Ian Johnston, Dominic Kwiatkowski, Cordelia Langford, John Sillitoe on behalf of the Wellcome Sanger Institute COVID-19 Surveillance Team |
| EPI_ISL_891881 | Lighthouse Lab in Glasgow | Wellcome Sanger Institute for the COVID-19 Genomics UK (COG-UK) Consortium | Harper VanSteenhouse, Yumi Kasai, David Gray, Carol Clugston, Anna Dominiczak and Alex Alderton, Roberto Amato, Sonia Goncalves, Ewan Harrison, David K. Jackson, Ian Johnston, Dominic Kwiatkowski, Cordelia Langford, John Sillitoe on behalf of the Wellcome Sanger Institute COVID-19 Surveillance Team |
| EPI_ISL_891883 | Lighthouse Lab in Cambridge | Wellcome Sanger Institute for the COVID-19 Genomics UK (COG-UK) Consortium | Rob Howes, The Lighthouse Lab in Cambridge and Alex Alderton, Roberto Amato, Sonia Goncalves, Ewan Harrison, David K. Jackson, Ian Johnston, Dominic Kwiatkowski, Cordelia Langford, John Sillitoe on behalf of the Wellcome Sanger Institute COVID-19 Surveillance Team |
| EPI_ISL_891884, EPI_ISL_891885 | Lighthouse Lab in Glasgow | Wellcome Sanger Institute for the COVID-19 Genomics UK | Harper VanSteenhouse, Yumi Kasai, David Gray, Carol Clugston, Anna Dominiczak and Alex Alderton, Roberto Amato, Sonia Goncalves, Ewan Harrison, |

| | | | |
|---|---|---|---|
| | | (COG-UK) Consortium | David K. Jackson, Ian Johnston, Dominic Kwiatkowski, Cordelia Langford, John Sillitoe on behalf of the Wellcome Sanger Institute COVID-19 Surveillance Team |
| EPI_ISL_891886 | Lighthouse Lab in Cambridge | Wellcome Sanger Institute for the COVID-19 Genomics UK (COG-UK) Consortium | Rob Howes, The Lighthouse Lab in Cambridge and Alex Alderton, Roberto Amato, Sonia Goncalves, Ewan Harrison, David K. Jackson, Ian Johnston, Dominic Kwiatkowski, Cordelia Langford, John Sillitoe on behalf of the Wellcome Sanger Institute COVID-19 Surveillance Team |
| EPI_ISL_891887 | Lighthouse Lab in Alderley Park | Wellcome Sanger Institute for the COVID-19 Genomics UK (COG-UK) Consortium | Jacquelyn Wynn, Mairead Hyland, The Lighthouse Lab in Alderley Park and Alex Alderton, Roberto Amato, Sonia Goncalves, Ewan Harrison, David K. Jackson, Ian Johnston, Dominic Kwiatkowski, Cordelia Langford, John Sillitoe on behalf of the Wellcome Sanger Institute COVID-19 Surveillance Team |
| EPI_ISL_891888, EPI_ISL_891889, EPI_ISL_891890, EPI_ISL_891891, EPI_ISL_891892, EPI_ISL_891893, EPI_ISL_891894 | Lighthouse Lab in Cambridge | Wellcome Sanger Institute for the COVID-19 Genomics UK (COG-UK) Consortium | Rob Howes, The Lighthouse Lab in Cambridge and Alex Alderton, Roberto Amato, Sonia Goncalves, Ewan Harrison, David K. Jackson, Ian Johnston, Dominic Kwiatkowski, Cordelia Langford, John Sillitoe on behalf of the Wellcome Sanger Institute COVID-19 Surveillance Team |
| EPI_ISL_891895 | Lighthouse Lab in Alderley Park | Wellcome Sanger Institute for the COVID-19 Genomics UK (COG-UK) Consortium | Jacquelyn Wynn, Mairead Hyland, The Lighthouse Lab in Alderley Park and Alex Alderton, Roberto Amato, Sonia Goncalves, Ewan Harrison, David K. Jackson, Ian Johnston, Dominic Kwiatkowski, Cordelia Langford, John Sillitoe on behalf of the Wellcome Sanger Institute COVID-19 Surveillance Team |
| EPI_ISL_891897, EPI_ISL_891898 | Lighthouse Lab in Cambridge | Wellcome Sanger Institute for the COVID-19 Genomics UK (COG-UK) Consortium | Rob Howes, The Lighthouse Lab in Cambridge and Alex Alderton, Roberto Amato, Sonia Goncalves, Ewan Harrison, David K. Jackson, Ian Johnston, Dominic Kwiatkowski, Cordelia Langford, John Sillitoe on behalf of the Wellcome Sanger Institute COVID-19 Surveillance Team |
| EPI_ISL_896217 | Columbia University Irving Medical Center | Wadsworth Center, New York State Department of Health | Kirsten St. George, Daryl M. Lamson, Alexis Russel, Matthew Shudt, Melissa A Leisner, Jonathan Plitnick, Navjot Singh, John Kelly, Erasmus Schneider, Erica Lasek-Nesselquist |
| EPI_ISL_896226 | TEMPUS LABS INC | Wadsworth Center, New York State Department of Health | Kirsten St. George, Daryl M. Lamson, Alexis Russel, Matthew Shudt, Melissa A Leisner, Jonathan Plitnick, Navjot Singh, John Kelly, Erasmus Schneider, Erica Lasek-Nesselquist |
| EPI_ISL_896231, EPI_ISL_896232, EPI_ISL_896233, EPI_ISL_896240, EPI_ISL_896241, EPI_ISL_896242, EPI_ISL_896243 | Columbia University Irving Medical Center | Wadsworth Center, New York State Department of Health | Kirsten St. George, Daryl M. Lamson, Alexis Russel, Matthew Shudt, Melissa A Leisner, Jonathan Plitnick, Navjot Singh, John Kelly, Erasmus Schneider, Erica Lasek-Nesselquist |
| EPI_ISL_896495, EPI_ISL_896496, EPI_ISL_896497, EPI_ISL_896502, EPI_ISL_896508, EPI_ISL_896509, EPI_ISL_896510, EPI_ISL_896511, EPI_ISL_896512, EPI_ISL_896513, EPI_ISL_896515, EPI_ISL_896563, EPI_ISL_896564, EPI_ISL_896565, EPI_ISL_896566, EPI_ISL_896567, EPI_ISL_896568, EPI_ISL_896569 | | | |
| see above | URMC LABS | Wadsworth Center, New York State Department of Health | Kirsten St. George, Daryl M. Lamson, Alexis Russel, Matthew Shudt, Melissa A Leisner, Jonathan Plitnick, Navjot Singh, John Kelly, Erasmus Schneider, Erica Lasek-Nesselquist |
| EPI_ISL_897606, EPI_ISL_897647, EPI_ISL_897648, EPI_ISL_897649, EPI_ISL_897650, EPI_ISL_897651, EPI_ISL_897652, EPI_ISL_897653, EPI_ISL_897654, EPI_ISL_897655, EPI_ISL_897656, EPI_ISL_897657, EPI_ISL_897658, EPI_ISL_897659, EPI_ISL_897660, EPI_ISL_897661, EPI_ISL_897662, EPI_ISL_897663, EPI_ISL_897664, EPI_ISL_897665, EPI_ISL_897666, EPI_ISL_897667, EPI_ISL_897668, EPI_ISL_897669, EPI_ISL_897671, EPI_ISL_897672, EPI_ISL_897675, EPI_ISL_897679, EPI_ISL_897684, EPI_ISL_897690, EPI_ISL_897691, EPI_ISL_897695, EPI_ISL_897706, EPI_ISL_897707, EPI_ISL_897712, EPI_ISL_897717, EPI_ISL_897718, EPI_ISL_897719, EPI_ISL_897846, EPI_ISL_897847, EPI_ISL_897848, EPI_ISL_897849, EPI_ISL_897850, EPI_ISL_897851, EPI_ISL_897852, EPI_ISL_897853, EPI_ISL_897854, EPI_ISL_897855, EPI_ISL_897856, EPI_ISL_897857, EPI_ISL_897858, EPI_ISL_897860, EPI_ISL_897861, EPI_ISL_897862, EPI_ISL_897863, EPI_ISL_897864, EPI_ISL_897865, EPI_ISL_897866, EPI_ISL_897867, EPI_ISL_897868, EPI_ISL_897869, EPI_ISL_897870, EPI_ISL_897871, EPI_ISL_897872, EPI_ISL_897873, EPI_ISL_897874, EPI_ISL_897875, EPI_ISL_897876, EPI_ISL_897877, EPI_ISL_897878, EPI_ISL_897879, EPI_ISL_897880, EPI_ISL_897881, EPI_ISL_897882, EPI_ISL_897883, EPI_ISL_897884, EPI_ISL_897885, EPI_ISL_897886, EPI_ISL_897887, EPI_ISL_897888, EPI_ISL_897889, EPI_ISL_897890, EPI_ISL_897891, EPI_ISL_897892, EPI_ISL_897893, EPI_ISL_897894, EPI_ISL_897895, EPI_ISL_897896, EPI_ISL_897897, EPI_ISL_897898, EPI_ISL_897899, EPI_ISL_897900, EPI_ISL_897901, EPI_ISL_897902, EPI_ISL_897924, EPI_ISL_897925, EPI_ISL_897926, EPI_ISL_897927, EPI_ISL_897928, EPI_ISL_897931, EPI_ISL_897932, EPI_ISL_897933, EPI_ISL_897966 | | | |
| see above | University Hospitals of Geneva, Laboratory of Virology | HUG, Laboratory of Virology and the Health2030 Genome Center | Samuel Cordey, Ana Rita Goncalves, Laurent Kaiser, Lorenzo Cerutti, Henri Pegeot, Melyssa Elies, Deborah Penet, Keith Harshman, Ioannis Xenarios, Emmanouil Dermitzakis |
| EPI_ISL_902730, EPI_ISL_902731, EPI_ISL_902733 | Santa Clara County Public Health Laboratory | Santa Clara County Public Health Laboratory | Santa Clara County Public Health Department |
| EPI_ISL_902756 | Fondazione Policlinico Universitario "A. Gemelli" IRCCS | INMI Lazzaro Spallanzani IRCCS | M Rueca, O Butera, F Messina, C.E.M Gruber, B Bartolini, E Giombini, P Cattani, M Sanguinetti, MR Capobianchi, A Di Caro |
| EPI_ISL_903208 | University of Michigan Clinical Microbiology Laboratory | Lauring Lab, University of Michigan, Department of Microbiology and Immunology | Valesano |
| EPI_ISL_903364, EPI_ISL_903365 | University Medical Center Hamburg Eppendorf | Heinrich Pette Institute, Leibniz Institute for Experimental Virology | Alexis Robitaille, Thomas Günther, Johannes Knobloch, Martin Aepfelbacher, Nicole Fischer, Adam Grundhoff |
| EPI_ISL_904086, EPI_ISL_904087, EPI_ISL_904088, EPI_ISL_904089, EPI_ISL_904090, EPI_ISL_904091, EPI_ISL_904092, EPI_ISL_904093, EPI_ISL_904094, EPI_ISL_904095, EPI_ISL_904096, EPI_ISL_904097, EPI_ISL_904098, EPI_ISL_904099, EPI_ISL_904100, EPI_ISL_904101, EPI_ISL_904102, EPI_ISL_904103, EPI_ISL_904104, EPI_ISL_904105, EPI_ISL_904106, EPI_ISL_904107, EPI_ISL_904108, EPI_ISL_904109, EPI_ISL_904110, EPI_ISL_904111, EPI_ISL_904112, EPI_ISL_904113, EPI_ISL_904114, EPI_ISL_904115, EPI_ISL_904116 | | | |
| see above | New Mexico Department of Health Scientific Laboratory | New Mexico Department of Health Scientific Laboratory | Ellie Johnson, Anastacia Griego-Fisher, D'eldra Malone |
| EPI_ISL_904179, EPI_ISL_904222, EPI_ISL_904607, EPI_ISL_904608, EPI_ISL_904609 | Dutch COVID-19 response team | Erasmus Medical Center | Bas Oude Munnink, Reina Sikkema, David Nieuwenhuijse, Irina Chestakova, Anne van der Linden, Marjan Boter, Emmanuelle Munger, Corine GeurtsvanKessel, Annemiek van der Eijk, Richard Molenkamp, Marion Koopmans, on behalf of the Dutch national COVID-19 response team. |
| EPI_ISL_904669, EPI_ISL_904670, EPI_ISL_904671, EPI_ISL_904672, EPI_ISL_904674, EPI_ISL_904677, EPI_ISL_904689, EPI_ISL_904716, EPI_ISL_904721, EPI_ISL_904733, EPI_ISL_904756, EPI_ISL_904759, EPI_ISL_904775, EPI_ISL_904846, EPI_ISL_904847, EPI_ISL_904848, EPI_ISL_904849, EPI_ISL_904871, EPI_ISL_904893, EPI_ISL_904914, EPI_ISL_904915, EPI_ISL_904916, EPI_ISL_904917, EPI_ISL_904918, EPI_ISL_904919, EPI_ISL_904920, EPI_ISL_904921, EPI_ISL_904922, EPI_ISL_904923, EPI_ISL_904924 | | | |
| see above | Dutch COVID-19 response team | National Institute for Public Health and the Environment (RIVM) | Adam Meijer, Harry Vennema, Dirk Eggink, Jeroen Cremer, Sharon van den Brink, Bas van der Veer, AnneMarie van den Brandt, Florian Zwagemaker, Dennis Schmitz, Chantal Reusken, on behalf of the national COVID-19 response team |
| EPI_ISL_905764 | Zakad Diagnostyki Laboratoryjnej i Mikrobiologii, Wojewódzki Szpital Podkarpacki w Kronie | National Institute of Public Health - National Institute of Hygiene | Wokowicz Tomasz, Zacharczuk Katarzyna |
| EPI_ISL_905773, EPI_ISL_905774, EPI_ISL_905775, EPI_ISL_905776, EPI_ISL_905777, EPI_ISL_905778, EPI_ISL_905779, EPI_ISL_905780, EPI_ISL_905781, EPI_ISL_905782, EPI_ISL_905783, EPI_ISL_905784, EPI_ISL_905785, EPI_ISL_905786, EPI_ISL_905787, EPI_ISL_905791 | | | |
| see above | UCLA Clinical Micro Lab | Los Angeles County PHL | P. Hemarajata et al. |
| EPI_ISL_906053 | Tilia Laboratories s.r.o. | Tilia Laboratories s.r.o. | Sona Pekova, MD, PhD. |
| EPI_ISL_906070 | UPA Dr. Akira Tada | Instituto Adolfo Lutz, Interdiciplinary Procedures Center, Strategic Laboratory | Claudio Tavares Sacchi, Claudia Regina Gonçalves, Erica Valessa Ramos Gomes, Karoline Rodrigues Campos |
| EPI_ISL_906071 | LACEN-PI DR. Costa Alvarenga | Instituto Adolfo Lutz, Interdiciplinary Procedures Center, Strategic Laboratory | Claudio Tavares Sacchi, Claudia Regina Gonçalves, Erica Valessa Ramos Gomes, Karoline Rodrigues Campos |
| EPI_ISL_906072 | UPA Dr. Akira Tada | Instituto Adolfo Lutz, Interdiciplinary Procedures Center, Strategic Laboratory | Claudio Tavares Sacchi, Claudia Regina Gonçalves, Erica Valessa Ramos Gomes, Karoline Rodrigues Campos |
| EPI_ISL_906073 | Hospital Vila Lobos | Instituto Adolfo Lutz, Interdiciplinary Procedures Center, Strategic Laboratory | Claudio Tavares Sacchi, Claudia Regina Gonçalves, Erica Valessa Ramos Gomes, Karoline Rodrigues Campos |
| EPI_ISL_906074 | Searom Laboratorio Diagnostico | Instituto Adolfo Lutz, Interdiciplinary Procedures Center, Strategic Laboratory | Claudio Tavares Sacchi, Claudia Regina Gonçalves, Erica Valessa Ramos Gomes, Karoline Rodrigues Campos |
| EPI_ISL_906075 | Hospital Geral de Vila Penteado Dr Jose Pangella Sao Paulo | Instituto Adolfo Lutz, Interdiciplinary Procedures Center, Strategic Laboratory | Claudio Tavares Sacchi, Claudia Regina Gonçalves, Erica Valessa Ramos Gomes, Karoline Rodrigues Campos |
| EPI_ISL_906076, EPI_ISL_906077 | Hospital Sao Luiz Sao Caetano | Instituto Adolfo Lutz, Interdiciplinary Procedures Center, Strategic Laboratory | Claudio Tavares Sacchi, Claudia Regina Gonçalves, Erica Valessa Ramos Gomes, Karoline Rodrigues Campos |
| EPI_ISL_906131 | National laboratory of health, environment and food Kranj | Institute of Microbiology and Immunology, Faculty of Medicine, University of Ljubljana | Samo Zakotnik, Tomaž Mark Zorec, Matic Brvar, Miša Korva, Mario Poljak, Tatjana Avši - Županc |

| | | | |
|---|---|---|---|
| EPI_ISL_906200, EPI_ISL_906205, EPI_ISL_906207, EPI_ISL_906210, EPI_ISL_906221, EPI_ISL_906223, EPI_ISL_906224, EPI_ISL_906228, EPI_ISL_906231, EPI_ISL_906232, EPI_ISL_906234, EPI_ISL_906235, EPI_ISL_906237, EPI_ISL_906239, EPI_ISL_906243, EPI_ISL_906244, EPI_ISL_906245, EPI_ISL_906247, EPI_ISL_906251, EPI_ISL_906253, EPI_ISL_906265, EPI_ISL_906267 | | | |
| see above | University of Wisconsin-Madison AIDS Vaccine Research Laboratories | University of Wisconsin-Madison AIDS Vaccine Research Laboratories | Gage Moreno, Katarina Braun, et al. AIDS Vaccine Research Laboratories |
| EPI_ISL_906538 | Laboratorio de Virologia-Instituto Nacional de Salud | Instituto Nacional de Salud- Dirección de Investigación en Salud Pública, Universidad de los Andes- Applied genomics research group, Vicerrectoria de Investigación y Creación, Universidad de los Andes- Systems and Computing Engineering Department | Katherine Laiton-Donato, Diego A. Álvarez-Díaz, Carlos Franco-Muñoz, Mauricio Pacheco-Montealegre, Héctor Alejandro Ruiz-Moreno, Maria T. Herrera-Sepúlveda, Diego Andrés Prada, Jhonnatan Reales-González, Sheryll Corchuelo, Julian Naizaque, Gerardo Santamaría Jorge Duitama, Laura Natalia Gonzalez, Jorge Ivan Diaz, Silvia Restrepo-Restrepo, Magdalena Wiesner, Martha Lucia Ospina Martinez, Marcela Mercado-Reyes |
| EPI_ISL_906655, EPI_ISL_906656, EPI_ISL_906657, EPI_ISL_906658, EPI_ISL_906659, EPI_ISL_906660, EPI_ISL_906661, EPI_ISL_906662, EPI_ISL_906663, EPI_ISL_906664, EPI_ISL_906665, EPI_ISL_906666, EPI_ISL_906667, EPI_ISL_906668 | | | |
| see above | Maine Health and Environmental Testing Laboratory (Maine HETL) | Tewhey Lab, The Jackson Laboratory | Matluk,N., Dewey,H., Iosue,F., Barter,M., Lynch,R., Munger,H. and Tewhey,R. |
| EPI_ISL_906804 | Norwegian Institute of Public Health, Department of Virology | Norwegian Institute of Public Health, Department of Virology | Kathrine Stene-Johansen, Kamilla Heddeland Instefjord, Hilde Elshaug, Atiya R Ali,Marie Paulsen Madsen, Rasmus Riis Kopperud, Hilde Vollan, Karoline Bragstad, Olav Hungnes |
| EPI_ISL_906813, EPI_ISL_906814 | Medical Microbiology Unit, Department for Laboratory Medicine, Drammen Hospital, Vestre Viken Health Trust, | Norwegian Institute of Public Health, Department of Virology | Kathrine Stene-Johansen, Kamilla Heddeland Instefjord, Hilde Elshaug, Atiya R Ali,Marie Paulsen Madsen, Rasmus Riis Kopperud, Hilde Vollan, Karoline Bragstad, Olav Hungnes |
| EPI_ISL_907119, EPI_ISL_907133, EPI_ISL_907136, EPI_ISL_907139, EPI_ISL_907144, EPI_ISL_907148, EPI_ISL_907152, EPI_ISL_907155, EPI_ISL_907160, EPI_ISL_907164, EPI_ISL_907167, EPI_ISL_907173, EPI_ISL_907179, EPI_ISL_907182, EPI_ISL_907186, EPI_ISL_907187, EPI_ISL_907190, EPI_ISL_907193, EPI_ISL_907195, EPI_ISL_907200, EPI_ISL_907206, EPI_ISL_907209, EPI_ISL_907212, EPI_ISL_907214, EPI_ISL_907219, EPI_ISL_907221, EPI_ISL_907223, EPI_ISL_907224, EPI_ISL_907225, EPI_ISL_907238, EPI_ISL_907241, EPI_ISL_907250, EPI_ISL_907254, EPI_ISL_907256, EPI_ISL_907260, EPI_ISL_907263, EPI_ISL_907264, EPI_ISL_907267, EPI_ISL_907269, EPI_ISL_907279, EPI_ISL_907284, EPI_ISL_907287, EPI_ISL_907288, EPI_ISL_907292, EPI_ISL_907300, EPI_ISL_907304, EPI_ISL_907308, EPI_ISL_907315, EPI_ISL_907318, EPI_ISL_907321, EPI_ISL_907322, EPI_ISL_907323, EPI_ISL_907325, EPI_ISL_907328, EPI_ISL_907336, EPI_ISL_907341, EPI_ISL_907347, EPI_ISL_907351, EPI_ISL_907352, EPI_ISL_907353, EPI_ISL_907355, EPI_ISL_907362, EPI_ISL_907363, EPI_ISL_907367 | | | |
| see above | Lighthouse Lab in Cambridge | Wellcome Sanger Institute for the COVID-19 Genomics UK (COG-UK) Consortium | Rob Howes, The Lighthouse Lab in Cambridge and Alex Alderton, Roberto Amato, Sonia Goncalves, Ewan Harrison, David K. Jackson, Ian Johnston, Dominic Kwiatkowski, Cordelia Langford, John Sillitoe on behalf of the Wellcome Sanger Institute COVID-19 Surveillance Team |
| EPI_ISL_907479, EPI_ISL_907492, EPI_ISL_907636, EPI_ISL_907700, EPI_ISL_907701, EPI_ISL_907704, EPI_ISL_907709, EPI_ISL_907710, EPI_ISL_907711, EPI_ISL_907712, EPI_ISL_907713, EPI_ISL_907714, EPI_ISL_907715, EPI_ISL_907717, EPI_ISL_907718, EPI_ISL_907719, EPI_ISL_907720, EPI_ISL_907724, EPI_ISL_907726, EPI_ISL_907727, EPI_ISL_907728, EPI_ISL_907729, EPI_ISL_907730, EPI_ISL_907732, EPI_ISL_907733, EPI_ISL_907734, EPI_ISL_907737, EPI_ISL_907739, EPI_ISL_907740, EPI_ISL_907741, EPI_ISL_907742, EPI_ISL_907743, EPI_ISL_907744, EPI_ISL_907746, EPI_ISL_907748, EPI_ISL_907751, EPI_ISL_907752, EPI_ISL_907753, EPI_ISL_907754, EPI_ISL_907758, EPI_ISL_907759, EPI_ISL_907761, EPI_ISL_907762, EPI_ISL_907764, EPI_ISL_907766, EPI_ISL_907768, EPI_ISL_907769, EPI_ISL_907770, EPI_ISL_907774, EPI_ISL_907777, EPI_ISL_907780, EPI_ISL_907781, EPI_ISL_907782, EPI_ISL_907784, EPI_ISL_907785 | | | |
| see above | Lighthouse Lab in Alderley Park | Wellcome Sanger Institute for the COVID-19 Genomics UK (COG-UK) Consortium | Jacquelyn Wynn, Mairead Hyland, The Lighthouse Lab in Alderley Park and Alex Alderton, Roberto Amato, Sonia Goncalves, Ewan Harrison, David K. Jackson, Ian Johnston, Dominic Kwiatkowski, Cordelia Langford, John Sillitoe on behalf of the Wellcome Sanger Institute COVID-19 Surveillance Team |
| EPI_ISL_907788 | Lighthouse Lab in Glasgow | Wellcome Sanger Institute for the COVID-19 Genomics UK (COG-UK) Consortium | Harper VanSteenhouse, Yumi Kasai, David Gray, Carol Clugston, Anna Dominiczak and Alex Alderton, Roberto Amato, Sonia Goncalves, Ewan Harrison, David K. Jackson, Ian Johnston, Dominic Kwiatkowski, Cordelia Langford, John Sillitoe on behalf of the Wellcome Sanger Institute COVID-19 Surveillance Team |
| EPI_ISL_907790, EPI_ISL_907791, EPI_ISL_907792, EPI_ISL_907798, EPI_ISL_907799, EPI_ISL_907803, EPI_ISL_907804, EPI_ISL_907806, EPI_ISL_907807, EPI_ISL_907808, EPI_ISL_907810, EPI_ISL_907815, EPI_ISL_907816, EPI_ISL_907818, EPI_ISL_907819, EPI_ISL_907822, EPI_ISL_907823, EPI_ISL_907825, EPI_ISL_907827, EPI_ISL_907828, EPI_ISL_907829, EPI_ISL_907830, EPI_ISL_907831, EPI_ISL_907836, EPI_ISL_907839, EPI_ISL_907840, EPI_ISL_907842, EPI_ISL_907843, EPI_ISL_907844, EPI_ISL_907845, EPI_ISL_907846, EPI_ISL_907848, EPI_ISL_907849, EPI_ISL_907850, EPI_ISL_907853, EPI_ISL_907856, EPI_ISL_907857, EPI_ISL_907858, EPI_ISL_907859, EPI_ISL_907861, EPI_ISL_907862, EPI_ISL_907864, EPI_ISL_907866, EPI_ISL_907869, EPI_ISL_907870, EPI_ISL_907871, EPI_ISL_907873, EPI_ISL_907876, EPI_ISL_907880, EPI_ISL_907883, EPI_ISL_907885, EPI_ISL_907886, EPI_ISL_907887, EPI_ISL_907888, EPI_ISL_907892, EPI_ISL_907896, EPI_ISL_907897, EPI_ISL_907900, EPI_ISL_907901, EPI_ISL_907904, EPI_ISL_907905, EPI_ISL_907906, EPI_ISL_907907, EPI_ISL_907908, EPI_ISL_907910, EPI_ISL_907913, EPI_ISL_907916, EPI_ISL_907917, EPI_ISL_907918, EPI_ISL_907919, EPI_ISL_907920, EPI_ISL_907922, EPI_ISL_907924, EPI_ISL_907927, EPI_ISL_907928, EPI_ISL_907929, EPI_ISL_907930, EPI_ISL_907932, EPI_ISL_907935, EPI_ISL_907937, EPI_ISL_907938, EPI_ISL_907939, EPI_ISL_907941, EPI_ISL_907942, EPI_ISL_907947, EPI_ISL_907950, EPI_ISL_907951, EPI_ISL_907952, EPI_ISL_907954, EPI_ISL_907955, EPI_ISL_907957, EPI_ISL_907958, EPI_ISL_907959, EPI_ISL_907961, EPI_ISL_907962, EPI_ISL_907963, EPI_ISL_907964, EPI_ISL_907965, EPI_ISL_907966, EPI_ISL_907968, EPI_ISL_907971, EPI_ISL_907977, EPI_ISL_907980, EPI_ISL_907983, EPI_ISL_907985, EPI_ISL_907986, EPI_ISL_907990, EPI_ISL_907991, EPI_ISL_907992, EPI_ISL_907993, EPI_ISL_907996, EPI_ISL_907997, EPI_ISL_907999, EPI_ISL_908000, EPI_ISL_908004, EPI_ISL_908005, EPI_ISL_908006, EPI_ISL_908008, EPI_ISL_908012, EPI_ISL_908014, EPI_ISL_908015, EPI_ISL_908018, EPI_ISL_908019, EPI_ISL_908020, EPI_ISL_908023, EPI_ISL_908024, EPI_ISL_908029, EPI_ISL_908031, EPI_ISL_908032, EPI_ISL_908268, EPI_ISL_908271, EPI_ISL_908272, EPI_ISL_908273, EPI_ISL_908276, EPI_ISL_908278, EPI_ISL_908280, EPI_ISL_908281, EPI_ISL_908283, EPI_ISL_908286, EPI_ISL_908287, EPI_ISL_908288, EPI_ISL_908294, EPI_ISL_908297 | | | |
| see above | Lighthouse Lab in Alderley Park | Wellcome Sanger Institute for the COVID-19 Genomics UK (COG-UK) Consortium | Jacquelyn Wynn, Mairead Hyland, The Lighthouse Lab in Alderley Park and Alex Alderton, Roberto Amato, Sonia Goncalves, Ewan Harrison, David K. Jackson, Ian Johnston, Dominic Kwiatkowski, Cordelia Langford, John Sillitoe on behalf of the Wellcome Sanger Institute COVID-19 Surveillance Team |
| EPI_ISL_908298 | Lighthouse Lab in Glasgow | Wellcome Sanger Institute for the COVID-19 Genomics UK (COG-UK) Consortium | Harper VanSteenhouse, Yumi Kasai, David Gray, Carol Clugston, Anna Dominiczak and Alex Alderton, Roberto Amato, Sonia Goncalves, Ewan Harrison, David K. Jackson, Ian Johnston, Dominic Kwiatkowski, Cordelia Langford, John Sillitoe on behalf of the Wellcome Sanger Institute COVID-19 Surveillance Team |
| EPI_ISL_908299, EPI_ISL_908300, EPI_ISL_908304, EPI_ISL_908305, EPI_ISL_908311, EPI_ISL_908312, EPI_ISL_908313, EPI_ISL_908314, EPI_ISL_908318, EPI_ISL_908319, EPI_ISL_908320, EPI_ISL_908324, EPI_ISL_908325, EPI_ISL_908327, EPI_ISL_908328, EPI_ISL_908330, EPI_ISL_908331, EPI_ISL_908332, EPI_ISL_908336, EPI_ISL_908339, EPI_ISL_908342, EPI_ISL_908345, EPI_ISL_908346, EPI_ISL_908347, EPI_ISL_908349, EPI_ISL_908350, EPI_ISL_908352, EPI_ISL_908354, EPI_ISL_908355, EPI_ISL_908356, EPI_ISL_908358, EPI_ISL_908359, EPI_ISL_908361, EPI_ISL_908362, EPI_ISL_908363, EPI_ISL_908364, EPI_ISL_908368, EPI_ISL_908370, EPI_ISL_908371, EPI_ISL_908373, EPI_ISL_908374, EPI_ISL_908375, EPI_ISL_908377, EPI_ISL_908378, EPI_ISL_908379, EPI_ISL_908381, EPI_ISL_908382, EPI_ISL_908386, EPI_ISL_908388, EPI_ISL_908390, EPI_ISL_908391, EPI_ISL_908392, EPI_ISL_908395, EPI_ISL_908396, EPI_ISL_908398, EPI_ISL_908399, EPI_ISL_908400, EPI_ISL_908404, EPI_ISL_908406, EPI_ISL_908409, EPI_ISL_908410, EPI_ISL_908414, EPI_ISL_908417, EPI_ISL_908418 | | | |
| see above | Lighthouse Lab in Alderley Park | Wellcome Sanger Institute for the COVID-19 Genomics UK (COG-UK) Consortium | Jacquelyn Wynn, Mairead Hyland, The Lighthouse Lab in Alderley Park and Alex Alderton, Roberto Amato, Sonia Goncalves, Ewan Harrison, David K. Jackson, Ian Johnston, Dominic Kwiatkowski, Cordelia Langford, John Sillitoe on behalf of the Wellcome Sanger Institute COVID-19 Surveillance Team |
| EPI_ISL_908419 | Lighthouse Lab in Glasgow | Wellcome Sanger Institute for the COVID-19 Genomics UK (COG-UK) Consortium | Harper VanSteenhouse, Yumi Kasai, David Gray, Carol Clugston, Anna Dominiczak and Alex Alderton, Roberto Amato, Sonia Goncalves, Ewan Harrison, David K. Jackson, Ian Johnston, Dominic Kwiatkowski, Cordelia Langford, John Sillitoe on behalf of the Wellcome Sanger Institute COVID-19 Surveillance Team |
| EPI_ISL_908420, EPI_ISL_908423, EPI_ISL_908424, EPI_ISL_908425, EPI_ISL_908426, EPI_ISL_908427, EPI_ISL_908428, EPI_ISL_908430, EPI_ISL_908432, EPI_ISL_908434, EPI_ISL_908436, EPI_ISL_908437, EPI_ISL_908438, EPI_ISL_908440, EPI_ISL_908441, EPI_ISL_908445, EPI_ISL_908448, EPI_ISL_908449, EPI_ISL_908450, EPI_ISL_908454, EPI_ISL_908457, EPI_ISL_908458, EPI_ISL_908459, EPI_ISL_908460, EPI_ISL_908461, EPI_ISL_908463, EPI_ISL_908464, EPI_ISL_908466, EPI_ISL_908468, EPI_ISL_908469, EPI_ISL_908472, EPI_ISL_908473, EPI_ISL_908475, EPI_ISL_908476, EPI_ISL_908477, EPI_ISL_908480, EPI_ISL_908482, EPI_ISL_908483, EPI_ISL_908484, EPI_ISL_908485, EPI_ISL_908487, EPI_ISL_908490, EPI_ISL_908491, EPI_ISL_908492, EPI_ISL_908493, EPI_ISL_908497, EPI_ISL_908498, EPI_ISL_908499, EPI_ISL_908501, EPI_ISL_908502, EPI_ISL_908503, EPI_ISL_908504, EPI_ISL_908505, EPI_ISL_908510, EPI_ISL_908514, EPI_ISL_908515, EPI_ISL_908517, EPI_ISL_908523, EPI_ISL_908531, EPI_ISL_908532, EPI_ISL_908533, EPI_ISL_908537, EPI_ISL_908539, EPI_ISL_908540, EPI_ISL_908541, EPI_ISL_908543, EPI_ISL_908544, EPI_ISL_908545, EPI_ISL_908548, EPI_ISL_908549, EPI_ISL_908552, EPI_ISL_908554, EPI_ISL_908555, EPI_ISL_908556, EPI_ISL_908562, EPI_ISL_908563, EPI_ISL_908565, EPI_ISL_908566, EPI_ISL_908567, EPI_ISL_908569, EPI_ISL_908572, EPI_ISL_908574, EPI_ISL_908575, EPI_ISL_908577, EPI_ISL_908578, EPI_ISL_908579, EPI_ISL_908583, EPI_ISL_908584, EPI_ISL_908586, EPI_ISL_908587, EPI_ISL_908588, EPI_ISL_908593, EPI_ISL_908594 | | | |
| see above | Lighthouse Lab in Alderley Park | Wellcome Sanger Institute for the COVID-19 Genomics UK (COG-UK) Consortium | Jacquelyn Wynn, Mairead Hyland, The Lighthouse Lab in Alderley Park and Alex Alderton, Roberto Amato, Sonia Goncalves, Ewan Harrison, David K. Jackson, Ian Johnston, Dominic Kwiatkowski, Cordelia Langford, John Sillitoe on behalf of the Wellcome Sanger Institute COVID-19 Surveillance Team |
| EPI_ISL_908597, EPI_ISL_908598, EPI_ISL_908599, EPI_ISL_908600, EPI_ISL_908601, EPI_ISL_908602, EPI_ISL_908603, EPI_ISL_908604, EPI_ISL_908605, EPI_ISL_908606, EPI_ISL_908607, EPI_ISL_908608, EPI_ISL_908609, EPI_ISL_908610, EPI_ISL_908611, EPI_ISL_908612, EPI_ISL_908613, EPI_ISL_908614, EPI_ISL_908615, EPI_ISL_908616, EPI_ISL_908617, EPI_ISL_908618, EPI_ISL_908619, EPI_ISL_908620, EPI_ISL_908621, EPI_ISL_908622, EPI_ISL_908623, EPI_ISL_908624, EPI_ISL_908625, EPI_ISL_908626, EPI_ISL_908627, EPI_ISL_908628, EPI_ISL_908629, EPI_ISL_908630, EPI_ISL_908631, EPI_ISL_908632, EPI_ISL_908633, EPI_ISL_908634, EPI_ISL_908635, EPI_ISL_908636, EPI_ISL_908637, EPI_ISL_908638, EPI_ISL_908639, EPI_ISL_908640, EPI_ISL_908641, EPI_ISL_908642, EPI_ISL_908643, EPI_ISL_908644, EPI_ISL_908645, EPI_ISL_908646, EPI_ISL_908647, EPI_ISL_908648, EPI_ISL_908649, EPI_ISL_908650, EPI_ISL_908651, EPI_ISL_908652, EPI_ISL_908653, EPI_ISL_908654, EPI_ISL_908655, EPI_ISL_908656, EPI_ISL_908657, EPI_ISL_908658, EPI_ISL_908659, EPI_ISL_908660, EPI_ISL_908661, EPI_ISL_908662, EPI_ISL_908663, EPI_ISL_908664, EPI_ISL_908665, EPI_ISL_908666, EPI_ISL_908667, EPI_ISL_908668, EPI_ISL_908669, EPI_ISL_908670, EPI_ISL_908671, EPI_ISL_908672, EPI_ISL_908673, EPI_ISL_908674, EPI_ISL_908675, EPI_ISL_908676, EPI_ISL_908677, EPI_ISL_908678, EPI_ISL_908679, EPI_ISL_908680, EPI_ISL_908681, EPI_ISL_908682, EPI_ISL_908683, EPI_ISL_908684, EPI_ISL_908685, EPI_ISL_908686, EPI_ISL_908687, EPI_ISL_908688, EPI_ISL_908689, EPI_ISL_908690, EPI_ISL_908691, EPI_ISL_908692, EPI_ISL_908693, EPI_ISL_908694, EPI_ISL_908695, EPI_ISL_908696, EPI_ISL_908697, EPI_ISL_908698, EPI_ISL_908699, EPI_ISL_908700, EPI_ISL_908701, EPI_ISL_908702, EPI_ISL_908703, EPI_ISL_908704, EPI_ISL_908705, EPI_ISL_908706, EPI_ISL_908707, EPI_ISL_908708, EPI_ISL_908709, EPI_ISL_908710, EPI_ISL_908711, EPI_ISL_908712, EPI_ISL_908713, EPI_ISL_908714, EPI_ISL_908715, EPI_ISL_908716, EPI_ISL_908717, EPI_ISL_908718, EPI_ISL_908719, EPI_ISL_908720, EPI_ISL_908721, EPI_ISL_908722, EPI_ISL_908723, EPI_ISL_908724, EPI_ISL_908725, EPI_ISL_908726, EPI_ISL_908727, EPI_ISL_908728, EPI_ISL_908729, EPI_ISL_908730, EPI_ISL_908731, EPI_ISL_908732, EPI_ISL_908733, EPI_ISL_908734, EPI_ISL_908735, EPI_ISL_908736, EPI_ISL_908737, EPI_ISL_908738, EPI_ISL_908739, EPI_ISL_908740, EPI_ISL_908741, EPI_ISL_908742, EPI_ISL_908743, EPI_ISL_908744, EPI_ISL_908745, EPI_ISL_908746, EPI_ISL_908747, EPI_ISL_908748, EPI_ISL_908749, EPI_ISL_908750, EPI_ISL_908751, EPI_ISL_908752, EPI_ISL_908753, EPI_ISL_908754, EPI_ISL_908755, EPI_ISL_908756, EPI_ISL_908757, EPI_ISL_908758, EPI_ISL_908759, EPI_ISL_908760, EPI_ISL_908762, EPI_ISL_908763, EPI_ISL_908764, EPI_ISL_908765, EPI_ISL_908766, EPI_ISL_908767, EPI_ISL_908768, EPI_ISL_908769, EPI_ISL_908770, EPI_ISL_908771, EPI_ISL_908772, EPI_ISL_908773, EPI_ISL_908774, EPI_ISL_908775, EPI_ISL_908776, EPI_ISL_908777, EPI_ISL_908778, EPI_ISL_908779, EPI_ISL_908780, EPI_ISL_908781, EPI_ISL_908782, EPI_ISL_908783, EPI_ISL_908784, EPI_ISL_908785, EPI_ISL_908786, EPI_ISL_908787, EPI_ISL_908788, EPI_ISL_908789, EPI_ISL_908790, EPI_ISL_908791, EPI_ISL_908792, EPI_ISL_908793, EPI_ISL_908794, EPI_ISL_908795, EPI_ISL_908796, EPI_ISL_908797, EPI_ISL_908798, EPI_ISL_908799, EPI_ISL_908800, EPI_ISL_908801, EPI_ISL_908802, EPI_ISL_908803, EPI_ISL_908804, EPI_ISL_908805, EPI_ISL_908806, EPI_ISL_908807, EPI_ISL_908808, EPI_ISL_908809, EPI_ISL_908810, EPI_ISL_908811, EPI_ISL_908812, EPI_ISL_908813, EPI_ISL_908814, EPI_ISL_908815, EPI_ISL_908816, EPI_ISL_908817, EPI_ISL_908818, EPI_ISL_908819, EPI_ISL_908820, EPI_ISL_908821, EPI_ISL_908822, EPI_ISL_908823, EPI_ISL_908824, EPI_ISL_908825, EPI_ISL_908826, EPI_ISL_908827, EPI_ISL_908828, EPI_ISL_908829, EPI_ISL_908830, EPI_ISL_908831, EPI_ISL_908832, EPI_ISL_908833, EPI_ISL_908834, EPI_ISL_908835, EPI_ISL_908836, EPI_ISL_908837, EPI_ISL_908838, EPI_ISL_908839, EPI_ISL_908840, EPI_ISL_908841, EPI_ISL_908842, EPI_ISL_908843, EPI_ISL_908844, EPI_ISL_908845, EPI_ISL_908846, EPI_ISL_908847, EPI_ISL_908848, EPI_ISL_908849, EPI_ISL_908850, EPI_ISL_908851, EPI_ISL_908852, EPI_ISL_908853, EPI_ISL_908854, EPI_ISL_908855, EPI_ISL_908856, EPI_ISL_908857, EPI_ISL_908858, EPI_ISL_908859, EPI_ISL_908860, EPI_ISL_908861, EPI_ISL_908862, EPI_ISL_908863, EPI_ISL_908864, EPI_ISL_908865, EPI_ISL_908866, EPI_ISL_908867, EPI_ISL_908868, EPI_ISL_908869, EPI_ISL_908870, EPI_ISL_908871, EPI_ISL_908872, EPI_ISL_908873, EPI_ISL_908874, EPI_ISL_908875, EPI_ISL_908876, EPI_ISL_908877, EPI_ISL_908878, EPI_ISL_908879, EPI_ISL_908880, EPI_ISL_908881, EPI_ISL_908882, EPI_ISL_908883, EPI_ISL_908884, EPI_ISL_908885, | | | |

EPI_ISL_908886, EPI_ISL_908887, EPI_ISL_908888, EPI_ISL_908889, EPI_ISL_908890, EPI_ISL_908891, EPI_ISL_908892, EPI_ISL_908893, EPI_ISL_908894, EPI_ISL_908895, EPI_ISL_908896, EPI_ISL_908897, EPI_ISL_908898, EPI_ISL_908899, EPI_ISL_908900, EPI_ISL_908901, EPI_ISL_908902, EPI_ISL_908903, EPI_ISL_908904, EPI_ISL_908905, EPI_ISL_908906, EPI_ISL_908907, EPI_ISL_908908, EPI_ISL_908909, EPI_ISL_908910, EPI_ISL_908911, EPI_ISL_908912, EPI_ISL_908913, EPI_ISL_908914, EPI_ISL_908915, EPI_ISL_908916, EPI_ISL_908917, EPI_ISL_908918, EPI_ISL_908919, EPI_ISL_908920, EPI_ISL_908921, EPI_ISL_908922, EPI_ISL_908923, EPI_ISL_908924, EPI_ISL_908925, EPI_ISL_908926, EPI_ISL_908927, EPI_ISL_908928, EPI_ISL_908929, EPI_ISL_908930, EPI_ISL_908931, EPI_ISL_908933, EPI_ISL_908935, EPI_ISL_908942

| | | | |
|---|---|---|---|
| see above | Lighthouse Lab in Cambridge | Wellcome Sanger Institute for the COVID-19 Genomics UK (COG-UK) Consortium | Rob Howes, The Lighthouse Lab in Cambridge and Alex Alderton, Roberto Amato, Sonia Goncalves, Ewan Harrison, David K. Jackson, Ian Johnston, Dominic Kwiatkowski, Cordelia Langford, John Sillitoe on behalf of the Wellcome Sanger Institute COVID-19 Surveillance Team |
| EPI_ISL_908945 | Lighthouse Lab in Glasgow | Wellcome Sanger Institute for the COVID-19 Genomics UK (COG-UK) Consortium | Harper VanSteenhouse, Yumi Kasai, David Gray, Carol Clugston, Anna Dominiczak and Alex Alderton, Roberto Amato, Sonia Goncalves, Ewan Harrison, David K. Jackson, Ian Johnston, Dominic Kwiatkowski, Cordelia Langford, John Sillitoe on behalf of the Wellcome Sanger Institute COVID-19 Surveillance Team |
| EPI_ISL_908946, EPI_ISL_908949, EPI_ISL_908951, EPI_ISL_908952 | Lighthouse Lab in Cambridge | Wellcome Sanger Institute for the COVID-19 Genomics UK (COG-UK) Consortium | Rob Howes, The Lighthouse Lab in Cambridge and Alex Alderton, Roberto Amato, Sonia Goncalves, Ewan Harrison, David K. Jackson, Ian Johnston, Dominic Kwiatkowski, Cordelia Langford, John Sillitoe on behalf of the Wellcome Sanger Institute COVID-19 Surveillance Team |
| EPI_ISL_908953 | Lighthouse Lab in Glasgow | Wellcome Sanger Institute for the COVID-19 Genomics UK (COG-UK) Consortium | Harper VanSteenhouse, Yumi Kasai, David Gray, Carol Clugston, Anna Dominiczak and Alex Alderton, Roberto Amato, Sonia Goncalves, Ewan Harrison, David K. Jackson, Ian Johnston, Dominic Kwiatkowski, Cordelia Langford, John Sillitoe on behalf of the Wellcome Sanger Institute COVID-19 Surveillance Team |
| EPI_ISL_908955, EPI_ISL_908957 | Lighthouse Lab in Cambridge | Wellcome Sanger Institute for the COVID-19 Genomics UK (COG-UK) Consortium | Rob Howes, The Lighthouse Lab in Cambridge and Alex Alderton, Roberto Amato, Sonia Goncalves, Ewan Harrison, David K. Jackson, Ian Johnston, Dominic Kwiatkowski, Cordelia Langford, John Sillitoe on behalf of the Wellcome Sanger Institute COVID-19 Surveillance Team |
| EPI_ISL_908958 | Lighthouse Lab in Glasgow | Wellcome Sanger Institute for the COVID-19 Genomics UK (COG-UK) Consortium | Harper VanSteenhouse, Yumi Kasai, David Gray, Carol Clugston, Anna Dominiczak and Alex Alderton, Roberto Amato, Sonia Goncalves, Ewan Harrison, David K. Jackson, Ian Johnston, Dominic Kwiatkowski, Cordelia Langford, John Sillitoe on behalf of the Wellcome Sanger Institute COVID-19 Surveillance Team |
| EPI_ISL_908959, EPI_ISL_908961 | Lighthouse Lab in Cambridge | Wellcome Sanger Institute for the COVID-19 Genomics UK (COG-UK) Consortium | Rob Howes, The Lighthouse Lab in Cambridge and Alex Alderton, Roberto Amato, Sonia Goncalves, Ewan Harrison, David K. Jackson, Ian Johnston, Dominic Kwiatkowski, Cordelia Langford, John Sillitoe on behalf of the Wellcome Sanger Institute COVID-19 Surveillance Team |
| EPI_ISL_908962 | Lighthouse Lab in Glasgow | Wellcome Sanger Institute for the COVID-19 Genomics UK (COG-UK) Consortium | Harper VanSteenhouse, Yumi Kasai, David Gray, Carol Clugston, Anna Dominiczak and Alex Alderton, Roberto Amato, Sonia Goncalves, Ewan Harrison, David K. Jackson, Ian Johnston, Dominic Kwiatkowski, Cordelia Langford, John Sillitoe on behalf of the Wellcome Sanger Institute COVID-19 Surveillance Team |
| EPI_ISL_908963 | Lighthouse Lab in Cambridge | Wellcome Sanger Institute for the COVID-19 Genomics UK (COG-UK) Consortium | Rob Howes, The Lighthouse Lab in Cambridge and Alex Alderton, Roberto Amato, Sonia Goncalves, Ewan Harrison, David K. Jackson, Ian Johnston, Dominic Kwiatkowski, Cordelia Langford, John Sillitoe on behalf of the Wellcome Sanger Institute COVID-19 Surveillance Team |
| EPI_ISL_908965 | Lighthouse Lab in Glasgow | Wellcome Sanger Institute for the COVID-19 Genomics UK (COG-UK) Consortium | Harper VanSteenhouse, Yumi Kasai, David Gray, Carol Clugston, Anna Dominiczak and Alex Alderton, Roberto Amato, Sonia Goncalves, Ewan Harrison, David K. Jackson, Ian Johnston, Dominic Kwiatkowski, Cordelia Langford, John Sillitoe on behalf of the Wellcome Sanger Institute COVID-19 Surveillance Team |
| EPI_ISL_908967, EPI_ISL_908968 | Lighthouse Lab in Cambridge | Wellcome Sanger Institute for the COVID-19 Genomics UK (COG-UK) Consortium | Rob Howes, The Lighthouse Lab in Cambridge and Alex Alderton, Roberto Amato, Sonia Goncalves, Ewan Harrison, David K. Jackson, Ian Johnston, Dominic Kwiatkowski, Cordelia Langford, John Sillitoe on behalf of the Wellcome Sanger Institute COVID-19 Surveillance Team |
| EPI_ISL_908974 | Lighthouse Lab in Glasgow | Wellcome Sanger Institute for the COVID-19 Genomics UK (COG-UK) Consortium | Harper VanSteenhouse, Yumi Kasai, David Gray, Carol Clugston, Anna Dominiczak and Alex Alderton, Roberto Amato, Sonia Goncalves, Ewan Harrison, David K. Jackson, Ian Johnston, Dominic Kwiatkowski, Cordelia Langford, John Sillitoe on behalf of the Wellcome Sanger Institute COVID-19 Surveillance Team |
| EPI_ISL_908977, EPI_ISL_908978, EPI_ISL_908979, EPI_ISL_908983, EPI_ISL_908985 | Lighthouse Lab in Cambridge | Wellcome Sanger Institute for the COVID-19 Genomics UK (COG-UK) Consortium | Rob Howes, The Lighthouse Lab in Cambridge and Alex Alderton, Roberto Amato, Sonia Goncalves, Ewan Harrison, David K. Jackson, Ian Johnston, Dominic Kwiatkowski, Cordelia Langford, John Sillitoe on behalf of the Wellcome Sanger Institute COVID-19 Surveillance Team |
| EPI_ISL_908986, EPI_ISL_908989 | Lighthouse Lab in Glasgow | Wellcome Sanger Institute for the COVID-19 Genomics UK (COG-UK) Consortium | Harper VanSteenhouse, Yumi Kasai, David Gray, Carol Clugston, Anna Dominiczak and Alex Alderton, Roberto Amato, Sonia Goncalves, Ewan Harrison, David K. Jackson, Ian Johnston, Dominic Kwiatkowski, Cordelia Langford, John Sillitoe on behalf of the Wellcome Sanger Institute COVID-19 Surveillance Team |
| EPI_ISL_908990, EPI_ISL_908991, EPI_ISL_908992 | Lighthouse Lab in Cambridge | Wellcome Sanger Institute for the COVID-19 Genomics UK (COG-UK) Consortium | Rob Howes, The Lighthouse Lab in Cambridge and Alex Alderton, Roberto Amato, Sonia Goncalves, Ewan Harrison, David K. Jackson, Ian Johnston, Dominic Kwiatkowski, Cordelia Langford, John Sillitoe on behalf of the Wellcome Sanger Institute COVID-19 Surveillance Team |
| EPI_ISL_908993 | Lighthouse Lab in Glasgow | Wellcome Sanger Institute for the COVID-19 Genomics UK (COG-UK) Consortium | Harper VanSteenhouse, Yumi Kasai, David Gray, Carol Clugston, Anna Dominiczak and Alex Alderton, Roberto Amato, Sonia Goncalves, Ewan Harrison, David K. Jackson, Ian Johnston, Dominic Kwiatkowski, Cordelia Langford, John Sillitoe on behalf of the Wellcome Sanger Institute COVID-19 Surveillance Team |
| EPI_ISL_908998, EPI_ISL_908999 | Lighthouse Lab in Cambridge | Wellcome Sanger Institute for the COVID-19 Genomics UK (COG-UK) Consortium | Rob Howes, The Lighthouse Lab in Cambridge and Alex Alderton, Roberto Amato, Sonia Goncalves, Ewan Harrison, David K. Jackson, Ian Johnston, Dominic Kwiatkowski, Cordelia Langford, John Sillitoe on behalf of the Wellcome Sanger Institute COVID-19 Surveillance Team |
| EPI_ISL_909002, EPI_ISL_909004, EPI_ISL_909005 | Lighthouse Lab in Glasgow | Wellcome Sanger Institute for the COVID-19 Genomics UK (COG-UK) Consortium | Harper VanSteenhouse, Yumi Kasai, David Gray, Carol Clugston, Anna Dominiczak and Alex Alderton, Roberto Amato, Sonia Goncalves, Ewan Harrison, David K. Jackson, Ian Johnston, Dominic Kwiatkowski, Cordelia Langford, John Sillitoe on behalf of the Wellcome Sanger Institute COVID-19 Surveillance Team |
| EPI_ISL_909006, EPI_ISL_909007 | Lighthouse Lab in Cambridge | Wellcome Sanger Institute for the COVID-19 Genomics UK (COG-UK) Consortium | Rob Howes, The Lighthouse Lab in Cambridge and Alex Alderton, Roberto Amato, Sonia Goncalves, Ewan Harrison, David K. Jackson, Ian Johnston, Dominic Kwiatkowski, Cordelia Langford, John Sillitoe on behalf of the Wellcome Sanger Institute COVID-19 Surveillance Team |
| EPI_ISL_909008 | Lighthouse Lab in Glasgow | Wellcome Sanger Institute for the COVID-19 Genomics UK (COG-UK) Consortium | Harper VanSteenhouse, Yumi Kasai, David Gray, Carol Clugston, Anna Dominiczak and Alex Alderton, Roberto Amato, Sonia Goncalves, Ewan Harrison, David K. Jackson, Ian Johnston, Dominic Kwiatkowski, Cordelia Langford, John Sillitoe on behalf of the Wellcome Sanger Institute COVID-19 Surveillance Team |
| EPI_ISL_909009 | Lighthouse Lab in Cambridge | Wellcome Sanger Institute for the COVID-19 Genomics UK (COG-UK) Consortium | Rob Howes, The Lighthouse Lab in Cambridge and Alex Alderton, Roberto Amato, Sonia Goncalves, Ewan Harrison, David K. Jackson, Ian Johnston, Dominic Kwiatkowski, Cordelia Langford, John Sillitoe on behalf of the Wellcome Sanger Institute COVID-19 Surveillance Team |
| EPI_ISL_909010 | Lighthouse Lab in Glasgow | Wellcome Sanger Institute for the COVID-19 Genomics UK (COG-UK) Consortium | Harper VanSteenhouse, Yumi Kasai, David Gray, Carol Clugston, Anna Dominiczak and Alex Alderton, Roberto Amato, Sonia Goncalves, Ewan Harrison, David K. Jackson, Ian Johnston, Dominic Kwiatkowski, Cordelia Langford, John Sillitoe on behalf of the Wellcome Sanger Institute COVID-19 Surveillance Team |
| EPI_ISL_909011, EPI_ISL_909015 | Lighthouse Lab in Cambridge | Wellcome Sanger Institute for the COVID-19 Genomics UK (COG-UK) Consortium | Rob Howes, The Lighthouse Lab in Cambridge and Alex Alderton, Roberto Amato, Sonia Goncalves, Ewan Harrison, David K. Jackson, Ian Johnston, Dominic Kwiatkowski, Cordelia Langford, John Sillitoe on behalf of the Wellcome Sanger Institute COVID-19 Surveillance Team |
| EPI_ISL_909017 | Lighthouse Lab in Glasgow | Wellcome Sanger Institute for the COVID-19 Genomics UK (COG-UK) Consortium | Harper VanSteenhouse, Yumi Kasai, David Gray, Carol Clugston, Anna Dominiczak and Alex Alderton, Roberto Amato, Sonia Goncalves, Ewan Harrison, David K. Jackson, Ian Johnston, Dominic Kwiatkowski, Cordelia Langford, John Sillitoe on behalf of the Wellcome Sanger Institute COVID-19 Surveillance Team |
| EPI_ISL_909019 | Lighthouse Lab in Cambridge | Wellcome Sanger Institute for the COVID-19 Genomics UK (COG-UK) Consortium | Rob Howes, The Lighthouse Lab in Cambridge and Alex Alderton, Roberto Amato, Sonia Goncalves, Ewan Harrison, David K. Jackson, Ian Johnston, Dominic Kwiatkowski, Cordelia Langford, John Sillitoe on behalf of the Wellcome Sanger Institute COVID-19 Surveillance Team |
| EPI_ISL_909021, EPI_ISL_909024, EPI_ISL_909029, EPI_ISL_909030, EPI_ISL_909034 | Lighthouse Lab in Glasgow | Wellcome Sanger Institute for the COVID-19 Genomics UK (COG-UK) Consortium | Harper VanSteenhouse, Yumi Kasai, David Gray, Carol Clugston, Anna Dominiczak and Alex Alderton, Roberto Amato, Sonia Goncalves, Ewan Harrison, David K. Jackson, Ian Johnston, Dominic Kwiatkowski, Cordelia Langford, John Sillitoe on behalf of the Wellcome Sanger Institute COVID-19 Surveillance Team |
| EPI_ISL_909035, EPI_ISL_909036 | Lighthouse Lab in Cambridge | Wellcome Sanger Institute for the COVID-19 Genomics UK (COG-UK) Consortium | Rob Howes, The Lighthouse Lab in Cambridge and Alex Alderton, Roberto Amato, Sonia Goncalves, Ewan Harrison, David K. Jackson, Ian Johnston, Dominic Kwiatkowski, Cordelia Langford, John Sillitoe on behalf of the Wellcome Sanger Institute COVID-19 Surveillance Team |
| EPI_ISL_909038 | Lighthouse Lab in Glasgow | Wellcome Sanger Institute for the COVID-19 Genomics UK (COG-UK) Consortium | Harper VanSteenhouse, Yumi Kasai, David Gray, Carol Clugston, Anna Dominiczak and Alex Alderton, Roberto Amato, Sonia Goncalves, Ewan Harrison, David K. Jackson, Ian Johnston, Dominic Kwiatkowski, Cordelia Langford, John Sillitoe on behalf of the Wellcome Sanger Institute COVID-19 Surveillance |

| | | | |
|---|---|---|---|
| | | | Team |
| EPI_ISL_909040, EPI_ISL_909046, EPI_ISL_909047, EPI_ISL_909048, EPI_ISL_909049, EPI_ISL_909050, EPI_ISL_909052 | Lighthouse Lab in Cambridge | Wellcome Sanger Institute for the COVID-19 Genomics UK (COG-UK) Consortium | Rob Howes, The Lighthouse Lab in Cambridge and Alex Alderton, Roberto Amato, Sonia Goncalves, Ewan Harrison, David K. Jackson, Ian Johnston, Dominic Kwiatkowski, Cordelia Langford, John Sillitoe on behalf of the Wellcome Sanger Institute COVID-19 Surveillance Team |
| EPI_ISL_909053, EPI_ISL_909056, EPI_ISL_909057, EPI_ISL_909058 | Lighthouse Lab in Glasgow | Wellcome Sanger Institute for the COVID-19 Genomics UK (COG-UK) Consortium | Harper VanSteenhouse, Yumi Kasai, David Gray, Carol Clugston, Anna Dominiczak and Alex Alderton, Roberto Amato, Sonia Goncalves, Ewan Harrison, David K. Jackson, Ian Johnston, Dominic Kwiatkowski, Cordelia Langford, John Sillitoe on behalf of the Wellcome Sanger Institute COVID-19 Surveillance Team |
| EPI_ISL_909060 | Lighthouse Lab in Cambridge | Wellcome Sanger Institute for the COVID-19 Genomics UK (COG-UK) Consortium | Rob Howes, The Lighthouse Lab in Cambridge and Alex Alderton, Roberto Amato, Sonia Goncalves, Ewan Harrison, David K. Jackson, Ian Johnston, Dominic Kwiatkowski, Cordelia Langford, John Sillitoe on behalf of the Wellcome Sanger Institute COVID-19 Surveillance Team |
| EPI_ISL_909061, EPI_ISL_909063 | Lighthouse Lab in Glasgow | Wellcome Sanger Institute for the COVID-19 Genomics UK (COG-UK) Consortium | Harper VanSteenhouse, Yumi Kasai, David Gray, Carol Clugston, Anna Dominiczak and Alex Alderton, Roberto Amato, Sonia Goncalves, Ewan Harrison, David K. Jackson, Ian Johnston, Dominic Kwiatkowski, Cordelia Langford, John Sillitoe on behalf of the Wellcome Sanger Institute COVID-19 Surveillance Team |
| EPI_ISL_909066, EPI_ISL_909067 | Lighthouse Lab in Cambridge | Wellcome Sanger Institute for the COVID-19 Genomics UK (COG-UK) Consortium | Rob Howes, The Lighthouse Lab in Cambridge and Alex Alderton, Roberto Amato, Sonia Goncalves, Ewan Harrison, David K. Jackson, Ian Johnston, Dominic Kwiatkowski, Cordelia Langford, John Sillitoe on behalf of the Wellcome Sanger Institute COVID-19 Surveillance Team |
| EPI_ISL_909068, EPI_ISL_909069 | Lighthouse Lab in Glasgow | Wellcome Sanger Institute for the COVID-19 Genomics UK (COG-UK) Consortium | Harper VanSteenhouse, Yumi Kasai, David Gray, Carol Clugston, Anna Dominiczak and Alex Alderton, Roberto Amato, Sonia Goncalves, Ewan Harrison, David K. Jackson, Ian Johnston, Dominic Kwiatkowski, Cordelia Langford, John Sillitoe on behalf of the Wellcome Sanger Institute COVID-19 Surveillance Team |
| EPI_ISL_909071, EPI_ISL_909072, EPI_ISL_909075, EPI_ISL_909079, EPI_ISL_909080, EPI_ISL_909081 | Lighthouse Lab in Cambridge | Wellcome Sanger Institute for the COVID-19 Genomics UK (COG-UK) Consortium | Rob Howes, The Lighthouse Lab in Cambridge and Alex Alderton, Roberto Amato, Sonia Goncalves, Ewan Harrison, David K. Jackson, Ian Johnston, Dominic Kwiatkowski, Cordelia Langford, John Sillitoe on behalf of the Wellcome Sanger Institute COVID-19 Surveillance Team |
| EPI_ISL_909083, EPI_ISL_909085, EPI_ISL_909086, EPI_ISL_909087 | Lighthouse Lab in Glasgow | Wellcome Sanger Institute for the COVID-19 Genomics UK (COG-UK) Consortium | Harper VanSteenhouse, Yumi Kasai, David Gray, Carol Clugston, Anna Dominiczak and Alex Alderton, Roberto Amato, Sonia Goncalves, Ewan Harrison, David K. Jackson, Ian Johnston, Dominic Kwiatkowski, Cordelia Langford, John Sillitoe on behalf of the Wellcome Sanger Institute COVID-19 Surveillance Team |
| EPI_ISL_909088 | Lighthouse Lab in Cambridge | Wellcome Sanger Institute for the COVID-19 Genomics UK (COG-UK) Consortium | Rob Howes, The Lighthouse Lab in Cambridge and Alex Alderton, Roberto Amato, Sonia Goncalves, Ewan Harrison, David K. Jackson, Ian Johnston, Dominic Kwiatkowski, Cordelia Langford, John Sillitoe on behalf of the Wellcome Sanger Institute COVID-19 Surveillance Team |
| EPI_ISL_909089, EPI_ISL_909090 | Lighthouse Lab in Glasgow | Wellcome Sanger Institute for the COVID-19 Genomics UK (COG-UK) Consortium | Harper VanSteenhouse, Yumi Kasai, David Gray, Carol Clugston, Anna Dominiczak and Alex Alderton, Roberto Amato, Sonia Goncalves, Ewan Harrison, David K. Jackson, Ian Johnston, Dominic Kwiatkowski, Cordelia Langford, John Sillitoe on behalf of the Wellcome Sanger Institute COVID-19 Surveillance Team |
| EPI_ISL_909093 | Lighthouse Lab in Cambridge | Wellcome Sanger Institute for the COVID-19 Genomics UK (COG-UK) Consortium | Rob Howes, The Lighthouse Lab in Cambridge and Alex Alderton, Roberto Amato, Sonia Goncalves, Ewan Harrison, David K. Jackson, Ian Johnston, Dominic Kwiatkowski, Cordelia Langford, John Sillitoe on behalf of the Wellcome Sanger Institute COVID-19 Surveillance Team |
| EPI_ISL_909094 | Lighthouse Lab in Glasgow | Wellcome Sanger Institute for the COVID-19 Genomics UK (COG-UK) Consortium | Harper VanSteenhouse, Yumi Kasai, David Gray, Carol Clugston, Anna Dominiczak and Alex Alderton, Roberto Amato, Sonia Goncalves, Ewan Harrison, David K. Jackson, Ian Johnston, Dominic Kwiatkowski, Cordelia Langford, John Sillitoe on behalf of the Wellcome Sanger Institute COVID-19 Surveillance Team |
| EPI_ISL_909095, EPI_ISL_909096, EPI_ISL_909097 | Lighthouse Lab in Cambridge | Wellcome Sanger Institute for the COVID-19 Genomics UK (COG-UK) Consortium | Rob Howes, The Lighthouse Lab in Cambridge and Alex Alderton, Roberto Amato, Sonia Goncalves, Ewan Harrison, David K. Jackson, Ian Johnston, Dominic Kwiatkowski, Cordelia Langford, John Sillitoe on behalf of the Wellcome Sanger Institute COVID-19 Surveillance Team |
| EPI_ISL_909101 | Lighthouse Lab in Glasgow | Wellcome Sanger Institute for the COVID-19 Genomics UK (COG-UK) Consortium | Harper VanSteenhouse, Yumi Kasai, David Gray, Carol Clugston, Anna Dominiczak and Alex Alderton, Roberto Amato, Sonia Goncalves, Ewan Harrison, David K. Jackson, Ian Johnston, Dominic Kwiatkowski, Cordelia Langford, John Sillitoe on behalf of the Wellcome Sanger Institute COVID-19 Surveillance Team |
| EPI_ISL_909102, EPI_ISL_909103, EPI_ISL_909105 | Lighthouse Lab in Cambridge | Wellcome Sanger Institute for the COVID-19 Genomics UK (COG-UK) Consortium | Rob Howes, The Lighthouse Lab in Cambridge and Alex Alderton, Roberto Amato, Sonia Goncalves, Ewan Harrison, David K. Jackson, Ian Johnston, Dominic Kwiatkowski, Cordelia Langford, John Sillitoe on behalf of the Wellcome Sanger Institute COVID-19 Surveillance Team |
| EPI_ISL_909106 | Lighthouse Lab in Glasgow | Wellcome Sanger Institute for the COVID-19 Genomics UK (COG-UK) Consortium | Harper VanSteenhouse, Yumi Kasai, David Gray, Carol Clugston, Anna Dominiczak and Alex Alderton, Roberto Amato, Sonia Goncalves, Ewan Harrison, David K. Jackson, Ian Johnston, Dominic Kwiatkowski, Cordelia Langford, John Sillitoe on behalf of the Wellcome Sanger Institute COVID-19 Surveillance Team |
| EPI_ISL_909109 | Lighthouse Lab in Cambridge | Wellcome Sanger Institute for the COVID-19 Genomics UK (COG-UK) Consortium | Rob Howes, The Lighthouse Lab in Cambridge and Alex Alderton, Roberto Amato, Sonia Goncalves, Ewan Harrison, David K. Jackson, Ian Johnston, Dominic Kwiatkowski, Cordelia Langford, John Sillitoe on behalf of the Wellcome Sanger Institute COVID-19 Surveillance Team |
| EPI_ISL_909111, EPI_ISL_909113, EPI_ISL_909114, EPI_ISL_909116, EPI_ISL_909117, EPI_ISL_909118 | Lighthouse Lab in Glasgow | Wellcome Sanger Institute for the COVID-19 Genomics UK (COG-UK) Consortium | Harper VanSteenhouse, Yumi Kasai, David Gray, Carol Clugston, Anna Dominiczak and Alex Alderton, Roberto Amato, Sonia Goncalves, Ewan Harrison, David K. Jackson, Ian Johnston, Dominic Kwiatkowski, Cordelia Langford, John Sillitoe on behalf of the Wellcome Sanger Institute COVID-19 Surveillance Team |
| EPI_ISL_909119 | Lighthouse Lab in Cambridge | Wellcome Sanger Institute for the COVID-19 Genomics UK (COG-UK) Consortium | Rob Howes, The Lighthouse Lab in Cambridge and Alex Alderton, Roberto Amato, Sonia Goncalves, Ewan Harrison, David K. Jackson, Ian Johnston, Dominic Kwiatkowski, Cordelia Langford, John Sillitoe on behalf of the Wellcome Sanger Institute COVID-19 Surveillance Team |
| EPI_ISL_909121 | Lighthouse Lab in Glasgow | Wellcome Sanger Institute for the COVID-19 Genomics UK (COG-UK) Consortium | Harper VanSteenhouse, Yumi Kasai, David Gray, Carol Clugston, Anna Dominiczak and Alex Alderton, Roberto Amato, Sonia Goncalves, Ewan Harrison, David K. Jackson, Ian Johnston, Dominic Kwiatkowski, Cordelia Langford, John Sillitoe on behalf of the Wellcome Sanger Institute COVID-19 Surveillance Team |
| EPI_ISL_909122, EPI_ISL_909128 | Lighthouse Lab in Cambridge | Wellcome Sanger Institute for the COVID-19 Genomics UK (COG-UK) Consortium | Rob Howes, The Lighthouse Lab in Cambridge and Alex Alderton, Roberto Amato, Sonia Goncalves, Ewan Harrison, David K. Jackson, Ian Johnston, Dominic Kwiatkowski, Cordelia Langford, John Sillitoe on behalf of the Wellcome Sanger Institute COVID-19 Surveillance Team |
| EPI_ISL_909131 | Lighthouse Lab in Glasgow | Wellcome Sanger Institute for the COVID-19 Genomics UK (COG-UK) Consortium | Harper VanSteenhouse, Yumi Kasai, David Gray, Carol Clugston, Anna Dominiczak and Alex Alderton, Roberto Amato, Sonia Goncalves, Ewan Harrison, David K. Jackson, Ian Johnston, Dominic Kwiatkowski, Cordelia Langford, John Sillitoe on behalf of the Wellcome Sanger Institute COVID-19 Surveillance Team |
| EPI_ISL_909144, EPI_ISL_909145, EPI_ISL_909148 | Lighthouse Lab in Cambridge | Wellcome Sanger Institute for the COVID-19 Genomics UK (COG-UK) Consortium | Rob Howes, The Lighthouse Lab in Cambridge and Alex Alderton, Roberto Amato, Sonia Goncalves, Ewan Harrison, David K. Jackson, Ian Johnston, Dominic Kwiatkowski, Cordelia Langford, John Sillitoe on behalf of the Wellcome Sanger Institute COVID-19 Surveillance Team |
| EPI_ISL_909150 | Lighthouse Lab in Glasgow | Wellcome Sanger Institute for the COVID-19 Genomics UK (COG-UK) Consortium | Harper VanSteenhouse, Yumi Kasai, David Gray, Carol Clugston, Anna Dominiczak and Alex Alderton, Roberto Amato, Sonia Goncalves, Ewan Harrison, David K. Jackson, Ian Johnston, Dominic Kwiatkowski, Cordelia Langford, John Sillitoe on behalf of the Wellcome Sanger Institute COVID-19 Surveillance Team |
| EPI_ISL_909155, EPI_ISL_909156, EPI_ISL_909157 | Lighthouse Lab in Cambridge | Wellcome Sanger Institute for the COVID-19 Genomics UK (COG-UK) Consortium | Rob Howes, The Lighthouse Lab in Cambridge and Alex Alderton, Roberto Amato, Sonia Goncalves, Ewan Harrison, David K. Jackson, Ian Johnston, Dominic Kwiatkowski, Cordelia Langford, John Sillitoe on behalf of the Wellcome Sanger Institute COVID-19 Surveillance Team |
| EPI_ISL_909158, EPI_ISL_909162, EPI_ISL_909165, EPI_ISL_909169, EPI_ISL_909170, EPI_ISL_909171 | Lighthouse Lab in Glasgow | Wellcome Sanger Institute for the COVID-19 Genomics UK (COG-UK) Consortium | Harper VanSteenhouse, Yumi Kasai, David Gray, Carol Clugston, Anna Dominiczak and Alex Alderton, Roberto Amato, Sonia Goncalves, Ewan Harrison, David K. Jackson, Ian Johnston, Dominic Kwiatkowski, Cordelia Langford, John Sillitoe on behalf of the Wellcome Sanger Institute COVID-19 Surveillance Team |
| EPI_ISL_909173 | Lighthouse Lab in Cambridge | Wellcome Sanger Institute for the COVID-19 Genomics UK (COG-UK) Consortium | Rob Howes, The Lighthouse Lab in Cambridge and Alex Alderton, Roberto Amato, Sonia Goncalves, Ewan Harrison, David K. Jackson, Ian Johnston, Dominic Kwiatkowski, Cordelia Langford, John Sillitoe on behalf of the Wellcome Sanger Institute COVID-19 Surveillance Team |
| EPI_ISL_909176, EPI_ISL_909177 | Lighthouse Lab in Glasgow | Wellcome Sanger Institute for the COVID-19 Genomics UK (COG-UK) Consortium | Harper VanSteenhouse, Yumi Kasai, David Gray, Carol Clugston, Anna Dominiczak and Alex Alderton, Roberto Amato, Sonia Goncalves, Ewan Harrison, David K. Jackson, Ian Johnston, Dominic Kwiatkowski, Cordelia Langford, John Sillitoe on behalf of the Wellcome Sanger Institute COVID-19 Surveillance |

| | | | |
|---|---|---|---|
| | | | Team |
| EPI_ISL_909178, EPI_ISL_909180, EPI_ISL_909182 | Lighthouse Lab in Cambridge | Wellcome Sanger Institute for the COVID-19 Genomics UK (COG-UK) Consortium | Rob Howes, The Lighthouse Lab in Cambridge and Alex Alderton, Roberto Amato, Sonia Goncalves, Ewan Harrison, David K. Jackson, Ian Johnston, Dominic Kwiatkowski, Cordelia Langford, John Sillitoe on behalf of the Wellcome Sanger Institute COVID-19 Surveillance Team |
| EPI_ISL_909184 | Lighthouse Lab in Glasgow | Wellcome Sanger Institute for the COVID-19 Genomics UK (COG-UK) Consortium | Harper VanSteenhouse, Yumi Kasai, David Gray, Carol Clugston, Anna Dominiczak and Alex Alderton, Roberto Amato, Sonia Goncalves, Ewan Harrison, David K. Jackson, Ian Johnston, Dominic Kwiatkowski, Cordelia Langford, John Sillitoe on behalf of the Wellcome Sanger Institute COVID-19 Surveillance Team |
| EPI_ISL_909187, EPI_ISL_909189, EPI_ISL_909192, EPI_ISL_909193 | Lighthouse Lab in Cambridge | Wellcome Sanger Institute for the COVID-19 Genomics UK (COG-UK) Consortium | Rob Howes, The Lighthouse Lab in Cambridge and Alex Alderton, Roberto Amato, Sonia Goncalves, Ewan Harrison, David K. Jackson, Ian Johnston, Dominic Kwiatkowski, Cordelia Langford, John Sillitoe on behalf of the Wellcome Sanger Institute COVID-19 Surveillance Team |
| EPI_ISL_909194, EPI_ISL_909200 | Lighthouse Lab in Glasgow | Wellcome Sanger Institute for the COVID-19 Genomics UK (COG-UK) Consortium | Harper VanSteenhouse, Yumi Kasai, David Gray, Carol Clugston, Anna Dominiczak and Alex Alderton, Roberto Amato, Sonia Goncalves, Ewan Harrison, David K. Jackson, Ian Johnston, Dominic Kwiatkowski, Cordelia Langford, John Sillitoe on behalf of the Wellcome Sanger Institute COVID-19 Surveillance Team |
| EPI_ISL_909202, EPI_ISL_909203, EPI_ISL_909204 | Lighthouse Lab in Cambridge | Wellcome Sanger Institute for the COVID-19 Genomics UK (COG-UK) Consortium | Rob Howes, The Lighthouse Lab in Cambridge and Alex Alderton, Roberto Amato, Sonia Goncalves, Ewan Harrison, David K. Jackson, Ian Johnston, Dominic Kwiatkowski, Cordelia Langford, John Sillitoe on behalf of the Wellcome Sanger Institute COVID-19 Surveillance Team |
| EPI_ISL_909206, EPI_ISL_909208, EPI_ISL_909210, EPI_ISL_909211 | Lighthouse Lab in Glasgow | Wellcome Sanger Institute for the COVID-19 Genomics UK (COG-UK) Consortium | Harper VanSteenhouse, Yumi Kasai, David Gray, Carol Clugston, Anna Dominiczak and Alex Alderton, Roberto Amato, Sonia Goncalves, Ewan Harrison, David K. Jackson, Ian Johnston, Dominic Kwiatkowski, Cordelia Langford, John Sillitoe on behalf of the Wellcome Sanger Institute COVID-19 Surveillance Team |
| EPI_ISL_909217 | Lighthouse Lab in Cambridge | Wellcome Sanger Institute for the COVID-19 Genomics UK (COG-UK) Consortium | Rob Howes, The Lighthouse Lab in Cambridge and Alex Alderton, Roberto Amato, Sonia Goncalves, Ewan Harrison, David K. Jackson, Ian Johnston, Dominic Kwiatkowski, Cordelia Langford, John Sillitoe on behalf of the Wellcome Sanger Institute COVID-19 Surveillance Team |
| EPI_ISL_909218, EPI_ISL_909219 | Lighthouse Lab in Glasgow | Wellcome Sanger Institute for the COVID-19 Genomics UK (COG-UK) Consortium | Harper VanSteenhouse, Yumi Kasai, David Gray, Carol Clugston, Anna Dominiczak and Alex Alderton, Roberto Amato, Sonia Goncalves, Ewan Harrison, David K. Jackson, Ian Johnston, Dominic Kwiatkowski, Cordelia Langford, John Sillitoe on behalf of the Wellcome Sanger Institute COVID-19 Surveillance Team |
| EPI_ISL_909220, EPI_ISL_909223 | Lighthouse Lab in Cambridge | Wellcome Sanger Institute for the COVID-19 Genomics UK (COG-UK) Consortium | Rob Howes, The Lighthouse Lab in Cambridge and Alex Alderton, Roberto Amato, Sonia Goncalves, Ewan Harrison, David K. Jackson, Ian Johnston, Dominic Kwiatkowski, Cordelia Langford, John Sillitoe on behalf of the Wellcome Sanger Institute COVID-19 Surveillance Team |
| EPI_ISL_909227 | Lighthouse Lab in Glasgow | Wellcome Sanger Institute for the COVID-19 Genomics UK (COG-UK) Consortium | Harper VanSteenhouse, Yumi Kasai, David Gray, Carol Clugston, Anna Dominiczak and Alex Alderton, Roberto Amato, Sonia Goncalves, Ewan Harrison, David K. Jackson, Ian Johnston, Dominic Kwiatkowski, Cordelia Langford, John Sillitoe on behalf of the Wellcome Sanger Institute COVID-19 Surveillance Team |
| EPI_ISL_909230 | Lighthouse Lab in Cambridge | Wellcome Sanger Institute for the COVID-19 Genomics UK (COG-UK) Consortium | Rob Howes, The Lighthouse Lab in Cambridge and Alex Alderton, Roberto Amato, Sonia Goncalves, Ewan Harrison, David K. Jackson, Ian Johnston, Dominic Kwiatkowski, Cordelia Langford, John Sillitoe on behalf of the Wellcome Sanger Institute COVID-19 Surveillance Team |
| EPI_ISL_909233 | Lighthouse Lab in Glasgow | Wellcome Sanger Institute for the COVID-19 Genomics UK (COG-UK) Consortium | Harper VanSteenhouse, Yumi Kasai, David Gray, Carol Clugston, Anna Dominiczak and Alex Alderton, Roberto Amato, Sonia Goncalves, Ewan Harrison, David K. Jackson, Ian Johnston, Dominic Kwiatkowski, Cordelia Langford, John Sillitoe on behalf of the Wellcome Sanger Institute COVID-19 Surveillance Team |
| EPI_ISL_909234 | Lighthouse Lab in Cambridge | Wellcome Sanger Institute for the COVID-19 Genomics UK (COG-UK) Consortium | Rob Howes, The Lighthouse Lab in Cambridge and Alex Alderton, Roberto Amato, Sonia Goncalves, Ewan Harrison, David K. Jackson, Ian Johnston, Dominic Kwiatkowski, Cordelia Langford, John Sillitoe on behalf of the Wellcome Sanger Institute COVID-19 Surveillance Team |
| EPI_ISL_909235, EPI_ISL_909236, EPI_ISL_909237 | Lighthouse Lab in Glasgow | Wellcome Sanger Institute for the COVID-19 Genomics UK (COG-UK) Consortium | Harper VanSteenhouse, Yumi Kasai, David Gray, Carol Clugston, Anna Dominiczak and Alex Alderton, Roberto Amato, Sonia Goncalves, Ewan Harrison, David K. Jackson, Ian Johnston, Dominic Kwiatkowski, Cordelia Langford, John Sillitoe on behalf of the Wellcome Sanger Institute COVID-19 Surveillance Team |
| EPI_ISL_909238, EPI_ISL_909240 | Lighthouse Lab in Cambridge | Wellcome Sanger Institute for the COVID-19 Genomics UK (COG-UK) Consortium | Rob Howes, The Lighthouse Lab in Cambridge and Alex Alderton, Roberto Amato, Sonia Goncalves, Ewan Harrison, David K. Jackson, Ian Johnston, Dominic Kwiatkowski, Cordelia Langford, John Sillitoe on behalf of the Wellcome Sanger Institute COVID-19 Surveillance Team |
| EPI_ISL_909242 | Lighthouse Lab in Glasgow | Wellcome Sanger Institute for the COVID-19 Genomics UK (COG-UK) Consortium | Harper VanSteenhouse, Yumi Kasai, David Gray, Carol Clugston, Anna Dominiczak and Alex Alderton, Roberto Amato, Sonia Goncalves, Ewan Harrison, David K. Jackson, Ian Johnston, Dominic Kwiatkowski, Cordelia Langford, John Sillitoe on behalf of the Wellcome Sanger Institute COVID-19 Surveillance Team |
| EPI_ISL_909243, EPI_ISL_909251, EPI_ISL_909254 | Lighthouse Lab in Cambridge | Wellcome Sanger Institute for the COVID-19 Genomics UK (COG-UK) Consortium | Rob Howes, The Lighthouse Lab in Cambridge and Alex Alderton, Roberto Amato, Sonia Goncalves, Ewan Harrison, David K. Jackson, Ian Johnston, Dominic Kwiatkowski, Cordelia Langford, John Sillitoe on behalf of the Wellcome Sanger Institute COVID-19 Surveillance Team |
| EPI_ISL_909255, EPI_ISL_909257 | Lighthouse Lab in Glasgow | Wellcome Sanger Institute for the COVID-19 Genomics UK (COG-UK) Consortium | Harper VanSteenhouse, Yumi Kasai, David Gray, Carol Clugston, Anna Dominiczak and Alex Alderton, Roberto Amato, Sonia Goncalves, Ewan Harrison, David K. Jackson, Ian Johnston, Dominic Kwiatkowski, Cordelia Langford, John Sillitoe on behalf of the Wellcome Sanger Institute COVID-19 Surveillance Team |
| EPI_ISL_909259 | Lighthouse Lab in Milton Keynes | Wellcome Sanger Institute for the COVID-19 Genomics UK (COG-UK) Consortium | The Lighthouse Lab in Milton Keynes and Alex Alderton, Roberto Amato, Sonia Goncalves, Ewan Harrison, David K. Jackson, Ian Johnston, Dominic Kwiatkowski, Cordelia Langford, John Sillitoe on behalf of the Wellcome Sanger Institute COVID-19 Surveillance Team |
| EPI_ISL_909260 | Lighthouse Lab in Alderley Park | Wellcome Sanger Institute for the COVID-19 Genomics UK (COG-UK) Consortium | Jacquelyn Wynn, Mairead Hyland, The Lighthouse Lab in Alderley Park and Alex Alderton, Roberto Amato, Sonia Goncalves, Ewan Harrison, David K. Jackson, Ian Johnston, Dominic Kwiatkowski, Cordelia Langford, John Sillitoe on behalf of the Wellcome Sanger Institute COVID-19 Surveillance Team |
| EPI_ISL_909263, EPI_ISL_909269 | Lighthouse Lab in Milton Keynes | Wellcome Sanger Institute for the COVID-19 Genomics UK (COG-UK) Consortium | The Lighthouse Lab in Milton Keynes and Alex Alderton, Roberto Amato, Sonia Goncalves, Ewan Harrison, David K. Jackson, Ian Johnston, Dominic Kwiatkowski, Cordelia Langford, John Sillitoe on behalf of the Wellcome Sanger Institute COVID-19 Surveillance Team |
| EPI_ISL_909270 | Lighthouse Lab in Alderley Park | Wellcome Sanger Institute for the COVID-19 Genomics UK (COG-UK) Consortium | Jacquelyn Wynn, Mairead Hyland, The Lighthouse Lab in Alderley Park and Alex Alderton, Roberto Amato, Sonia Goncalves, Ewan Harrison, David K. Jackson, Ian Johnston, Dominic Kwiatkowski, Cordelia Langford, John Sillitoe on behalf of the Wellcome Sanger Institute COVID-19 Surveillance Team |
| EPI_ISL_909271 | Lighthouse Lab in Milton Keynes | Wellcome Sanger Institute for the COVID-19 Genomics UK (COG-UK) Consortium | The Lighthouse Lab in Milton Keynes and Alex Alderton, Roberto Amato, Sonia Goncalves, Ewan Harrison, David K. Jackson, Ian Johnston, Dominic Kwiatkowski, Cordelia Langford, John Sillitoe on behalf of the Wellcome Sanger Institute COVID-19 Surveillance Team |
| EPI_ISL_909275 | Lighthouse Lab in Alderley Park | Wellcome Sanger Institute for the COVID-19 Genomics UK (COG-UK) Consortium | Jacquelyn Wynn, Mairead Hyland, The Lighthouse Lab in Alderley Park and Alex Alderton, Roberto Amato, Sonia Goncalves, Ewan Harrison, David K. Jackson, Ian Johnston, Dominic Kwiatkowski, Cordelia Langford, John Sillitoe on behalf of the Wellcome Sanger Institute COVID-19 Surveillance Team |
| EPI_ISL_909276, EPI_ISL_909277, EPI_ISL_909284 | Lighthouse Lab in Milton Keynes | Wellcome Sanger Institute for the COVID-19 Genomics UK (COG-UK) Consortium | The Lighthouse Lab in Milton Keynes and Alex Alderton, Roberto Amato, Sonia Goncalves, Ewan Harrison, David K. Jackson, Ian Johnston, Dominic Kwiatkowski, Cordelia Langford, John Sillitoe on behalf of the Wellcome Sanger Institute COVID-19 Surveillance Team |
| EPI_ISL_909285 | Lighthouse Lab in Alderley Park | Wellcome Sanger Institute for the COVID-19 Genomics UK (COG-UK) Consortium | Jacquelyn Wynn, Mairead Hyland, The Lighthouse Lab in Alderley Park and Alex Alderton, Roberto Amato, Sonia Goncalves, Ewan Harrison, David K. Jackson, Ian Johnston, Dominic Kwiatkowski, Cordelia Langford, John Sillitoe on behalf of the Wellcome Sanger Institute COVID-19 Surveillance Team |
| EPI_ISL_909287, EPI_ISL_909291 | Lighthouse Lab in Milton Keynes | Wellcome Sanger Institute for the COVID-19 Genomics UK (COG-UK) Consortium | The Lighthouse Lab in Milton Keynes and Alex Alderton, Roberto Amato, Sonia Goncalves, Ewan Harrison, David K. Jackson, Ian Johnston, Dominic Kwiatkowski, Cordelia Langford, John Sillitoe on behalf of the Wellcome Sanger Institute COVID-19 Surveillance Team |
| EPI_ISL_909292 | Lighthouse Lab in Alderley Park | Wellcome Sanger Institute for the COVID-19 Genomics UK (COG-UK) Consortium | Jacquelyn Wynn, Mairead Hyland, The Lighthouse Lab in Alderley Park and Alex Alderton, Roberto Amato, Sonia Goncalves, Ewan Harrison, David K. Jackson, Ian Johnston, Dominic Kwiatkowski, Cordelia Langford, John Sillitoe on behalf of the Wellcome Sanger Institute COVID-19 Surveillance Team |
| EPI_ISL_909293 | Lighthouse Lab in Milton Keynes | Wellcome Sanger Institute for the COVID-19 Genomics UK (COG-UK) Consortium | The Lighthouse Lab in Milton Keynes and Alex Alderton, Roberto Amato, Sonia Goncalves, Ewan Harrison, David K. Jackson, Ian Johnston, Dominic Kwiatkowski, Cordelia Langford, John Sillitoe on behalf of the Wellcome Sanger Institute COVID-19 Surveillance Team |
| EPI_ISL_909294, EPI_ISL_909297 | Lighthouse Lab in Alderley Park | Wellcome Sanger Institute for the COVID-19 Genomics UK (COG-UK) Consortium | Jacquelyn Wynn, Mairead Hyland, The Lighthouse Lab in Alderley Park and Alex Alderton, Roberto Amato, Sonia Goncalves, Ewan Harrison, David K. Jackson, Ian Johnston, Dominic Kwiatkowski, Cordelia Langford, John Sillitoe on behalf of the Wellcome Sanger Institute COVID-19 Surveillance Team |
| EPI_ISL_909301 | Lighthouse Lab in Milton Keynes | Wellcome Sanger Institute for the COVID-19 Genomics UK (COG-UK) Consortium | The Lighthouse Lab in Milton Keynes and Alex Alderton, Roberto Amato, Sonia Goncalves, Ewan Harrison, David K. Jackson, Ian Johnston, Dominic Kwiatkowski, Cordelia Langford, John Sillitoe on behalf of the Wellcome Sanger Institute COVID-19 Surveillance Team |

| | | | |
|---|---|---|---|
| EPI_ISL_909302, EPI_ISL_909309 | Lighthouse Lab in Alderley Park | Wellcome Sanger Institute for the COVID-19 Genomics UK (COG-UK) Consortium | Jacquelyn Wynn, Mairead Hyland, The Lighthouse Lab in Alderley Park and Alex Alderton, Roberto Amato, Sonia Goncalves, Ewan Harrison, David K. Jackson, Ian Johnston, Dominic Kwiatkowski, Cordelia Langford, John Sillitoe on behalf of the Wellcome Sanger Institute COVID-19 Surveillance Team |
| EPI_ISL_909310 | Lighthouse Lab in Milton Keynes | Wellcome Sanger Institute for the COVID-19 Genomics UK (COG-UK) Consortium | The Lighthouse Lab in Milton Keynes and Alex Alderton, Roberto Amato, Sonia Goncalves, Ewan Harrison, David K. Jackson, Ian Johnston, Dominic Kwiatkowski, Cordelia Langford, John Sillitoe on behalf of the Wellcome Sanger Institute COVID-19 Surveillance Team |
| EPI_ISL_909312, EPI_ISL_909314, EPI_ISL_909316, EPI_ISL_909321, EPI_ISL_909322, EPI_ISL_909326 | Lighthouse Lab in Alderley Park | Wellcome Sanger Institute for the COVID-19 Genomics UK (COG-UK) Consortium | Jacquelyn Wynn, Mairead Hyland, The Lighthouse Lab in Alderley Park and Alex Alderton, Roberto Amato, Sonia Goncalves, Ewan Harrison, David K. Jackson, Ian Johnston, Dominic Kwiatkowski, Cordelia Langford, John Sillitoe on behalf of the Wellcome Sanger Institute COVID-19 Surveillance Team |
| EPI_ISL_909327 | Lighthouse Lab in Glasgow | Wellcome Sanger Institute for the COVID-19 Genomics UK (COG-UK) Consortium | Harper VanSteenhouse, Yumi Kasai, David Gray, Carol Clugston, Anna Dominiczak and Alex Alderton, Roberto Amato, Sonia Goncalves, Ewan Harrison, David K. Jackson, Ian Johnston, Dominic Kwiatkowski, Cordelia Langford, John Sillitoe on behalf of the Wellcome Sanger Institute COVID-19 Surveillance Team |
| EPI_ISL_909329, EPI_ISL_909331 | Lighthouse Lab in Milton Keynes | Wellcome Sanger Institute for the COVID-19 Genomics UK (COG-UK) Consortium | The Lighthouse Lab in Milton Keynes and Alex Alderton, Roberto Amato, Sonia Goncalves, Ewan Harrison, David K. Jackson, Ian Johnston, Dominic Kwiatkowski, Cordelia Langford, John Sillitoe on behalf of the Wellcome Sanger Institute COVID-19 Surveillance Team |
| EPI_ISL_909334 | Lighthouse Lab in Alderley Park | Wellcome Sanger Institute for the COVID-19 Genomics UK (COG-UK) Consortium | Jacquelyn Wynn, Mairead Hyland, The Lighthouse Lab in Alderley Park and Alex Alderton, Roberto Amato, Sonia Goncalves, Ewan Harrison, David K. Jackson, Ian Johnston, Dominic Kwiatkowski, Cordelia Langford, John Sillitoe on behalf of the Wellcome Sanger Institute COVID-19 Surveillance Team |
| EPI_ISL_909335, EPI_ISL_909338 | Lighthouse Lab in Milton Keynes | Wellcome Sanger Institute for the COVID-19 Genomics UK (COG-UK) Consortium | The Lighthouse Lab in Milton Keynes and Alex Alderton, Roberto Amato, Sonia Goncalves, Ewan Harrison, David K. Jackson, Ian Johnston, Dominic Kwiatkowski, Cordelia Langford, John Sillitoe on behalf of the Wellcome Sanger Institute COVID-19 Surveillance Team |
| EPI_ISL_909339 | Lighthouse Lab in Alderley Park | Wellcome Sanger Institute for the COVID-19 Genomics UK (COG-UK) Consortium | Jacquelyn Wynn, Mairead Hyland, The Lighthouse Lab in Alderley Park and Alex Alderton, Roberto Amato, Sonia Goncalves, Ewan Harrison, David K. Jackson, Ian Johnston, Dominic Kwiatkowski, Cordelia Langford, John Sillitoe on behalf of the Wellcome Sanger Institute COVID-19 Surveillance Team |
| EPI_ISL_909340 | Lighthouse Lab in Milton Keynes | Wellcome Sanger Institute for the COVID-19 Genomics UK (COG-UK) Consortium | The Lighthouse Lab in Milton Keynes and Alex Alderton, Roberto Amato, Sonia Goncalves, Ewan Harrison, David K. Jackson, Ian Johnston, Dominic Kwiatkowski, Cordelia Langford, John Sillitoe on behalf of the Wellcome Sanger Institute COVID-19 Surveillance Team |
| EPI_ISL_909341, EPI_ISL_909343 | Lighthouse Lab in Alderley Park | Wellcome Sanger Institute for the COVID-19 Genomics UK (COG-UK) Consortium | Jacquelyn Wynn, Mairead Hyland, The Lighthouse Lab in Alderley Park and Alex Alderton, Roberto Amato, Sonia Goncalves, Ewan Harrison, David K. Jackson, Ian Johnston, Dominic Kwiatkowski, Cordelia Langford, John Sillitoe on behalf of the Wellcome Sanger Institute COVID-19 Surveillance Team |
| EPI_ISL_909345, EPI_ISL_909350 | Lighthouse Lab in Milton Keynes | Wellcome Sanger Institute for the COVID-19 Genomics UK (COG-UK) Consortium | The Lighthouse Lab in Milton Keynes and Alex Alderton, Roberto Amato, Sonia Goncalves, Ewan Harrison, David K. Jackson, Ian Johnston, Dominic Kwiatkowski, Cordelia Langford, John Sillitoe on behalf of the Wellcome Sanger Institute COVID-19 Surveillance Team |
| EPI_ISL_909351 | Lighthouse Lab in Glasgow | Wellcome Sanger Institute for the COVID-19 Genomics UK (COG-UK) Consortium | Harper VanSteenhouse, Yumi Kasai, David Gray, Carol Clugston, Anna Dominiczak and Alex Alderton, Roberto Amato, Sonia Goncalves, Ewan Harrison, David K. Jackson, Ian Johnston, Dominic Kwiatkowski, Cordelia Langford, John Sillitoe on behalf of the Wellcome Sanger Institute COVID-19 Surveillance Team |
| EPI_ISL_909352, EPI_ISL_909354, EPI_ISL_909355 | Lighthouse Lab in Milton Keynes | Wellcome Sanger Institute for the COVID-19 Genomics UK (COG-UK) Consortium | The Lighthouse Lab in Milton Keynes and Alex Alderton, Roberto Amato, Sonia Goncalves, Ewan Harrison, David K. Jackson, Ian Johnston, Dominic Kwiatkowski, Cordelia Langford, John Sillitoe on behalf of the Wellcome Sanger Institute COVID-19 Surveillance Team |
| EPI_ISL_909356 | Lighthouse Lab in Glasgow | Wellcome Sanger Institute for the COVID-19 Genomics UK (COG-UK) Consortium | Harper VanSteenhouse, Yumi Kasai, David Gray, Carol Clugston, Anna Dominiczak and Alex Alderton, Roberto Amato, Sonia Goncalves, Ewan Harrison, David K. Jackson, Ian Johnston, Dominic Kwiatkowski, Cordelia Langford, John Sillitoe on behalf of the Wellcome Sanger Institute COVID-19 Surveillance Team |
| EPI_ISL_909358, EPI_ISL_909359 | Lighthouse Lab in Alderley Park | Wellcome Sanger Institute for the COVID-19 Genomics UK (COG-UK) Consortium | Jacquelyn Wynn, Mairead Hyland, The Lighthouse Lab in Alderley Park and Alex Alderton, Roberto Amato, Sonia Goncalves, Ewan Harrison, David K. Jackson, Ian Johnston, Dominic Kwiatkowski, Cordelia Langford, John Sillitoe on behalf of the Wellcome Sanger Institute COVID-19 Surveillance Team |
| EPI_ISL_909365 | Lighthouse Lab in Milton Keynes | Wellcome Sanger Institute for the COVID-19 Genomics UK (COG-UK) Consortium | The Lighthouse Lab in Milton Keynes and Alex Alderton, Roberto Amato, Sonia Goncalves, Ewan Harrison, David K. Jackson, Ian Johnston, Dominic Kwiatkowski, Cordelia Langford, John Sillitoe on behalf of the Wellcome Sanger Institute COVID-19 Surveillance Team |
| EPI_ISL_909366, EPI_ISL_909367, EPI_ISL_909368 | Lighthouse Lab in Alderley Park | Wellcome Sanger Institute for the COVID-19 Genomics UK (COG-UK) Consortium | Jacquelyn Wynn, Mairead Hyland, The Lighthouse Lab in Alderley Park and Alex Alderton, Roberto Amato, Sonia Goncalves, Ewan Harrison, David K. Jackson, Ian Johnston, Dominic Kwiatkowski, Cordelia Langford, John Sillitoe on behalf of the Wellcome Sanger Institute COVID-19 Surveillance Team |
| EPI_ISL_909369 | Lighthouse Lab in Glasgow | Wellcome Sanger Institute for the COVID-19 Genomics UK (COG-UK) Consortium | Harper VanSteenhouse, Yumi Kasai, David Gray, Carol Clugston, Anna Dominiczak and Alex Alderton, Roberto Amato, Sonia Goncalves, Ewan Harrison, David K. Jackson, Ian Johnston, Dominic Kwiatkowski, Cordelia Langford, John Sillitoe on behalf of the Wellcome Sanger Institute COVID-19 Surveillance Team |
| EPI_ISL_909370, EPI_ISL_909372 | Lighthouse Lab in Alderley Park | Wellcome Sanger Institute for the COVID-19 Genomics UK (COG-UK) Consortium | Jacquelyn Wynn, Mairead Hyland, The Lighthouse Lab in Alderley Park and Alex Alderton, Roberto Amato, Sonia Goncalves, Ewan Harrison, David K. Jackson, Ian Johnston, Dominic Kwiatkowski, Cordelia Langford, John Sillitoe on behalf of the Wellcome Sanger Institute COVID-19 Surveillance Team |
| EPI_ISL_909375 | Lighthouse Lab in Glasgow | Wellcome Sanger Institute for the COVID-19 Genomics UK (COG-UK) Consortium | Harper VanSteenhouse, Yumi Kasai, David Gray, Carol Clugston, Anna Dominiczak and Alex Alderton, Roberto Amato, Sonia Goncalves, Ewan Harrison, David K. Jackson, Ian Johnston, Dominic Kwiatkowski, Cordelia Langford, John Sillitoe on behalf of the Wellcome Sanger Institute COVID-19 Surveillance Team |
| EPI_ISL_909380, EPI_ISL_909381 | Lighthouse Lab in Alderley Park | Wellcome Sanger Institute for the COVID-19 Genomics UK (COG-UK) Consortium | Jacquelyn Wynn, Mairead Hyland, The Lighthouse Lab in Alderley Park and Alex Alderton, Roberto Amato, Sonia Goncalves, Ewan Harrison, David K. Jackson, Ian Johnston, Dominic Kwiatkowski, Cordelia Langford, John Sillitoe on behalf of the Wellcome Sanger Institute COVID-19 Surveillance Team |
| EPI_ISL_909382 | Lighthouse Lab in Glasgow | Wellcome Sanger Institute for the COVID-19 Genomics UK (COG-UK) Consortium | Harper VanSteenhouse, Yumi Kasai, David Gray, Carol Clugston, Anna Dominiczak and Alex Alderton, Roberto Amato, Sonia Goncalves, Ewan Harrison, David K. Jackson, Ian Johnston, Dominic Kwiatkowski, Cordelia Langford, John Sillitoe on behalf of the Wellcome Sanger Institute COVID-19 Surveillance Team |
| EPI_ISL_909383 | Lighthouse Lab in Milton Keynes | Wellcome Sanger Institute for the COVID-19 Genomics UK (COG-UK) Consortium | The Lighthouse Lab in Milton Keynes and Alex Alderton, Roberto Amato, Sonia Goncalves, Ewan Harrison, David K. Jackson, Ian Johnston, Dominic Kwiatkowski, Cordelia Langford, John Sillitoe on behalf of the Wellcome Sanger Institute COVID-19 Surveillance Team |
| EPI_ISL_909385 | Lighthouse Lab in Alderley Park | Wellcome Sanger Institute for the COVID-19 Genomics UK (COG-UK) Consortium | Jacquelyn Wynn, Mairead Hyland, The Lighthouse Lab in Alderley Park and Alex Alderton, Roberto Amato, Sonia Goncalves, Ewan Harrison, David K. Jackson, Ian Johnston, Dominic Kwiatkowski, Cordelia Langford, John Sillitoe on behalf of the Wellcome Sanger Institute COVID-19 Surveillance Team |
| EPI_ISL_909386 | Lighthouse Lab in Milton Keynes | Wellcome Sanger Institute for the COVID-19 Genomics UK (COG-UK) Consortium | The Lighthouse Lab in Milton Keynes and Alex Alderton, Roberto Amato, Sonia Goncalves, Ewan Harrison, David K. Jackson, Ian Johnston, Dominic Kwiatkowski, Cordelia Langford, John Sillitoe on behalf of the Wellcome Sanger Institute COVID-19 Surveillance Team |
| EPI_ISL_909388, EPI_ISL_909389 | Lighthouse Lab in Alderley Park | Wellcome Sanger Institute for the COVID-19 Genomics UK (COG-UK) Consortium | Jacquelyn Wynn, Mairead Hyland, The Lighthouse Lab in Alderley Park and Alex Alderton, Roberto Amato, Sonia Goncalves, Ewan Harrison, David K. Jackson, Ian Johnston, Dominic Kwiatkowski, Cordelia Langford, John Sillitoe on behalf of the Wellcome Sanger Institute COVID-19 Surveillance Team |
| EPI_ISL_909390 | Lighthouse Lab in Glasgow | Wellcome Sanger Institute for the COVID-19 Genomics UK (COG-UK) Consortium | Harper VanSteenhouse, Yumi Kasai, David Gray, Carol Clugston, Anna Dominiczak and Alex Alderton, Roberto Amato, Sonia Goncalves, Ewan Harrison, David K. Jackson, Ian Johnston, Dominic Kwiatkowski, Cordelia Langford, John Sillitoe on behalf of the Wellcome Sanger Institute COVID-19 Surveillance Team |
| EPI_ISL_909394, EPI_ISL_909396 | Lighthouse Lab in Milton Keynes | Wellcome Sanger Institute for the COVID-19 Genomics UK (COG-UK) Consortium | The Lighthouse Lab in Milton Keynes and Alex Alderton, Roberto Amato, Sonia Goncalves, Ewan Harrison, David K. Jackson, Ian Johnston, Dominic Kwiatkowski, Cordelia Langford, John Sillitoe on behalf of the Wellcome Sanger Institute COVID-19 Surveillance Team |
| EPI_ISL_909397 | Lighthouse Lab in Alderley Park | Wellcome Sanger Institute for the COVID-19 Genomics UK (COG-UK) Consortium | Jacquelyn Wynn, Mairead Hyland, The Lighthouse Lab in Alderley Park and Alex Alderton, Roberto Amato, Sonia Goncalves, Ewan Harrison, David K. Jackson, Ian Johnston, Dominic Kwiatkowski, Cordelia Langford, John Sillitoe on behalf of the Wellcome Sanger Institute COVID-19 Surveillance Team |
| EPI_ISL_909405 | Lighthouse Lab in Glasgow | Wellcome Sanger Institute for the COVID-19 Genomics UK (COG-UK) Consortium | Harper VanSteenhouse, Yumi Kasai, David Gray, Carol Clugston, Anna Dominiczak and Alex Alderton, Roberto Amato, Sonia Goncalves, Ewan Harrison, David K. Jackson, Ian Johnston, Dominic Kwiatkowski, Cordelia Langford, John Sillitoe on behalf of the Wellcome Sanger Institute COVID-19 Surveillance Team |
| EPI_ISL_909409 | Lighthouse Lab in Alderley Park | Wellcome Sanger Institute for the COVID-19 Genomics UK (COG-UK) Consortium | Jacquelyn Wynn, Mairead Hyland, The Lighthouse Lab in Alderley Park and Alex Alderton, Roberto Amato, Sonia Goncalves, Ewan Harrison, David K. Jackson, Ian Johnston, Dominic Kwiatkowski, Cordelia Langford, John Sillitoe on behalf of the Wellcome Sanger Institute COVID-19 Surveillance Team |
| EPI_ISL_909411, EPI_ISL_909412, | Lighthouse Lab in Milton Keynes | Wellcome Sanger Institute for the COVID-19 Genomics UK | The Lighthouse Lab in Milton Keynes and Alex Alderton, Roberto Amato, Sonia Goncalves, Ewan Harrison, David K. Jackson, Ian Johnston, Dominic |

| | | | |
|---|---|---|---|
| EPI_ISL_909413, EPI_ISL_909415 | | (COG-UK) Consortium | Kwiatkowski, Cordelia Langford, John Sillitoe on behalf of the Wellcome Sanger Institute COVID-19 Surveillance Team |
| EPI_ISL_909416, EPI_ISL_909417 | Lighthouse Lab in Alderley Park | Wellcome Sanger Institute for the COVID-19 Genomics UK (COG-UK) Consortium | Jacquelyn Wynn, Mairead Hyland, The Lighthouse Lab in Alderley Park and Alex Alderton, Roberto Amato, Sonia Goncalves, Ewan Harrison, David K. Jackson, Ian Johnston, Dominic Kwiatkowski, Cordelia Langford, John Sillitoe on behalf of the Wellcome Sanger Institute COVID-19 Surveillance Team |
| EPI_ISL_909418 | Lighthouse Lab in Milton Keynes | Wellcome Sanger Institute for the COVID-19 Genomics UK (COG-UK) Consortium | The Lighthouse Lab in Milton Keynes and Alex Alderton, Roberto Amato, Sonia Goncalves, Ewan Harrison, David K. Jackson, Ian Johnston, Dominic Kwiatkowski, Cordelia Langford, John Sillitoe on behalf of the Wellcome Sanger Institute COVID-19 Surveillance Team |
| EPI_ISL_909421, EPI_ISL_909424 | Lighthouse Lab in Alderley Park | Wellcome Sanger Institute for the COVID-19 Genomics UK (COG-UK) Consortium | Jacquelyn Wynn, Mairead Hyland, The Lighthouse Lab in Alderley Park and Alex Alderton, Roberto Amato, Sonia Goncalves, Ewan Harrison, David K. Jackson, Ian Johnston, Dominic Kwiatkowski, Cordelia Langford, John Sillitoe on behalf of the Wellcome Sanger Institute COVID-19 Surveillance Team |
| EPI_ISL_909427, EPI_ISL_909431 | Lighthouse Lab in Milton Keynes | Wellcome Sanger Institute for the COVID-19 Genomics UK (COG-UK) Consortium | The Lighthouse Lab in Milton Keynes and Alex Alderton, Roberto Amato, Sonia Goncalves, Ewan Harrison, David K. Jackson, Ian Johnston, Dominic Kwiatkowski, Cordelia Langford, John Sillitoe on behalf of the Wellcome Sanger Institute COVID-19 Surveillance Team |
| EPI_ISL_909435 | Lighthouse Lab in Glasgow | Wellcome Sanger Institute for the COVID-19 Genomics UK (COG-UK) Consortium | Harper VanSteenhouse, Yumi Kasai, David Gray, Carol Clugston, Anna Dominiczak and Alex Alderton, Roberto Amato, Sonia Goncalves, Ewan Harrison, David K. Jackson, Ian Johnston, Dominic Kwiatkowski, Cordelia Langford, John Sillitoe on behalf of the Wellcome Sanger Institute COVID-19 Surveillance Team |
| EPI_ISL_909440 | Lighthouse Lab in Alderley Park | Wellcome Sanger Institute for the COVID-19 Genomics UK (COG-UK) Consortium | Jacquelyn Wynn, Mairead Hyland, The Lighthouse Lab in Alderley Park and Alex Alderton, Roberto Amato, Sonia Goncalves, Ewan Harrison, David K. Jackson, Ian Johnston, Dominic Kwiatkowski, Cordelia Langford, John Sillitoe on behalf of the Wellcome Sanger Institute COVID-19 Surveillance Team |
| EPI_ISL_909442, EPI_ISL_909443, EPI_ISL_909444 | Lighthouse Lab in Milton Keynes | Wellcome Sanger Institute for the COVID-19 Genomics UK (COG-UK) Consortium | The Lighthouse Lab in Milton Keynes and Alex Alderton, Roberto Amato, Sonia Goncalves, Ewan Harrison, David K. Jackson, Ian Johnston, Dominic Kwiatkowski, Cordelia Langford, John Sillitoe on behalf of the Wellcome Sanger Institute COVID-19 Surveillance Team |
| EPI_ISL_909446, EPI_ISL_909450 | Lighthouse Lab in Alderley Park | Wellcome Sanger Institute for the COVID-19 Genomics UK (COG-UK) Consortium | Jacquelyn Wynn, Mairead Hyland, The Lighthouse Lab in Alderley Park and Alex Alderton, Roberto Amato, Sonia Goncalves, Ewan Harrison, David K. Jackson, Ian Johnston, Dominic Kwiatkowski, Cordelia Langford, John Sillitoe on behalf of the Wellcome Sanger Institute COVID-19 Surveillance Team |
| EPI_ISL_909452 | Lighthouse Lab in Milton Keynes | Wellcome Sanger Institute for the COVID-19 Genomics UK (COG-UK) Consortium | The Lighthouse Lab in Milton Keynes and Alex Alderton, Roberto Amato, Sonia Goncalves, Ewan Harrison, David K. Jackson, Ian Johnston, Dominic Kwiatkowski, Cordelia Langford, John Sillitoe on behalf of the Wellcome Sanger Institute COVID-19 Surveillance Team |
| EPI_ISL_909453, EPI_ISL_909456, EPI_ISL_909457 | Lighthouse Lab in Alderley Park | Wellcome Sanger Institute for the COVID-19 Genomics UK (COG-UK) Consortium | Jacquelyn Wynn, Mairead Hyland, The Lighthouse Lab in Alderley Park and Alex Alderton, Roberto Amato, Sonia Goncalves, Ewan Harrison, David K. Jackson, Ian Johnston, Dominic Kwiatkowski, Cordelia Langford, John Sillitoe on behalf of the Wellcome Sanger Institute COVID-19 Surveillance Team |
| EPI_ISL_909460, EPI_ISL_909463, EPI_ISL_909464 | Lighthouse Lab in Milton Keynes | Wellcome Sanger Institute for the COVID-19 Genomics UK (COG-UK) Consortium | The Lighthouse Lab in Milton Keynes and Alex Alderton, Roberto Amato, Sonia Goncalves, Ewan Harrison, David K. Jackson, Ian Johnston, Dominic Kwiatkowski, Cordelia Langford, John Sillitoe on behalf of the Wellcome Sanger Institute COVID-19 Surveillance Team |
| EPI_ISL_909465, EPI_ISL_909471 | Lighthouse Lab in Alderley Park | Wellcome Sanger Institute for the COVID-19 Genomics UK (COG-UK) Consortium | Jacquelyn Wynn, Mairead Hyland, The Lighthouse Lab in Alderley Park and Alex Alderton, Roberto Amato, Sonia Goncalves, Ewan Harrison, David K. Jackson, Ian Johnston, Dominic Kwiatkowski, Cordelia Langford, John Sillitoe on behalf of the Wellcome Sanger Institute COVID-19 Surveillance Team |
| EPI_ISL_909474, EPI_ISL_909475 | Lighthouse Lab in Milton Keynes | Wellcome Sanger Institute for the COVID-19 Genomics UK (COG-UK) Consortium | The Lighthouse Lab in Milton Keynes and Alex Alderton, Roberto Amato, Sonia Goncalves, Ewan Harrison, David K. Jackson, Ian Johnston, Dominic Kwiatkowski, Cordelia Langford, John Sillitoe on behalf of the Wellcome Sanger Institute COVID-19 Surveillance Team |
| EPI_ISL_909476, EPI_ISL_909478 | Lighthouse Lab in Alderley Park | Wellcome Sanger Institute for the COVID-19 Genomics UK (COG-UK) Consortium | Jacquelyn Wynn, Mairead Hyland, The Lighthouse Lab in Alderley Park and Alex Alderton, Roberto Amato, Sonia Goncalves, Ewan Harrison, David K. Jackson, Ian Johnston, Dominic Kwiatkowski, Cordelia Langford, John Sillitoe on behalf of the Wellcome Sanger Institute COVID-19 Surveillance Team |
| EPI_ISL_909479, EPI_ISL_909480 | Lighthouse Lab in Milton Keynes | Wellcome Sanger Institute for the COVID-19 Genomics UK (COG-UK) Consortium | The Lighthouse Lab in Milton Keynes and Alex Alderton, Roberto Amato, Sonia Goncalves, Ewan Harrison, David K. Jackson, Ian Johnston, Dominic Kwiatkowski, Cordelia Langford, John Sillitoe on behalf of the Wellcome Sanger Institute COVID-19 Surveillance Team |
| EPI_ISL_909482 | Lighthouse Lab in Alderley Park | Wellcome Sanger Institute for the COVID-19 Genomics UK (COG-UK) Consortium | Jacquelyn Wynn, Mairead Hyland, The Lighthouse Lab in Alderley Park and Alex Alderton, Roberto Amato, Sonia Goncalves, Ewan Harrison, David K. Jackson, Ian Johnston, Dominic Kwiatkowski, Cordelia Langford, John Sillitoe on behalf of the Wellcome Sanger Institute COVID-19 Surveillance Team |
| EPI_ISL_909484 | Lighthouse Lab in Milton Keynes | Wellcome Sanger Institute for the COVID-19 Genomics UK (COG-UK) Consortium | The Lighthouse Lab in Milton Keynes and Alex Alderton, Roberto Amato, Sonia Goncalves, Ewan Harrison, David K. Jackson, Ian Johnston, Dominic Kwiatkowski, Cordelia Langford, John Sillitoe on behalf of the Wellcome Sanger Institute COVID-19 Surveillance Team |
| EPI_ISL_909487 | Lighthouse Lab in Alderley Park | Wellcome Sanger Institute for the COVID-19 Genomics UK (COG-UK) Consortium | Jacquelyn Wynn, Mairead Hyland, The Lighthouse Lab in Alderley Park and Alex Alderton, Roberto Amato, Sonia Goncalves, Ewan Harrison, David K. Jackson, Ian Johnston, Dominic Kwiatkowski, Cordelia Langford, John Sillitoe on behalf of the Wellcome Sanger Institute COVID-19 Surveillance Team |
| EPI_ISL_909492, EPI_ISL_909493 | Lighthouse Lab in Milton Keynes | Wellcome Sanger Institute for the COVID-19 Genomics UK (COG-UK) Consortium | The Lighthouse Lab in Milton Keynes and Alex Alderton, Roberto Amato, Sonia Goncalves, Ewan Harrison, David K. Jackson, Ian Johnston, Dominic Kwiatkowski, Cordelia Langford, John Sillitoe on behalf of the Wellcome Sanger Institute COVID-19 Surveillance Team |
| EPI_ISL_909497 | Lighthouse Lab in Glasgow | Wellcome Sanger Institute for the COVID-19 Genomics UK (COG-UK) Consortium | Harper VanSteenhouse, Yumi Kasai, David Gray, Carol Clugston, Anna Dominiczak and Alex Alderton, Roberto Amato, Sonia Goncalves, Ewan Harrison, David K. Jackson, Ian Johnston, Dominic Kwiatkowski, Cordelia Langford, John Sillitoe on behalf of the Wellcome Sanger Institute COVID-19 Surveillance Team |
| EPI_ISL_909503, EPI_ISL_909504 | Lighthouse Lab in Alderley Park | Wellcome Sanger Institute for the COVID-19 Genomics UK (COG-UK) Consortium | Jacquelyn Wynn, Mairead Hyland, The Lighthouse Lab in Alderley Park and Alex Alderton, Roberto Amato, Sonia Goncalves, Ewan Harrison, David K. Jackson, Ian Johnston, Dominic Kwiatkowski, Cordelia Langford, John Sillitoe on behalf of the Wellcome Sanger Institute COVID-19 Surveillance Team |
| EPI_ISL_909505, EPI_ISL_909506 | Lighthouse Lab in Milton Keynes | Wellcome Sanger Institute for the COVID-19 Genomics UK (COG-UK) Consortium | The Lighthouse Lab in Milton Keynes and Alex Alderton, Roberto Amato, Sonia Goncalves, Ewan Harrison, David K. Jackson, Ian Johnston, Dominic Kwiatkowski, Cordelia Langford, John Sillitoe on behalf of the Wellcome Sanger Institute COVID-19 Surveillance Team |
| EPI_ISL_909508 | Lighthouse Lab in Glasgow | Wellcome Sanger Institute for the COVID-19 Genomics UK (COG-UK) Consortium | Harper VanSteenhouse, Yumi Kasai, David Gray, Carol Clugston, Anna Dominiczak and Alex Alderton, Roberto Amato, Sonia Goncalves, Ewan Harrison, David K. Jackson, Ian Johnston, Dominic Kwiatkowski, Cordelia Langford, John Sillitoe on behalf of the Wellcome Sanger Institute COVID-19 Surveillance Team |
| EPI_ISL_909510, EPI_ISL_909511 | Lighthouse Lab in Milton Keynes | Wellcome Sanger Institute for the COVID-19 Genomics UK (COG-UK) Consortium | The Lighthouse Lab in Milton Keynes and Alex Alderton, Roberto Amato, Sonia Goncalves, Ewan Harrison, David K. Jackson, Ian Johnston, Dominic Kwiatkowski, Cordelia Langford, John Sillitoe on behalf of the Wellcome Sanger Institute COVID-19 Surveillance Team |
| EPI_ISL_909513 | Lighthouse Lab in Alderley Park | Wellcome Sanger Institute for the COVID-19 Genomics UK (COG-UK) Consortium | Jacquelyn Wynn, Mairead Hyland, The Lighthouse Lab in Alderley Park and Alex Alderton, Roberto Amato, Sonia Goncalves, Ewan Harrison, David K. Jackson, Ian Johnston, Dominic Kwiatkowski, Cordelia Langford, John Sillitoe on behalf of the Wellcome Sanger Institute COVID-19 Surveillance Team |
| EPI_ISL_909514, EPI_ISL_909515 | Lighthouse Lab in Milton Keynes | Wellcome Sanger Institute for the COVID-19 Genomics UK (COG-UK) Consortium | The Lighthouse Lab in Milton Keynes and Alex Alderton, Roberto Amato, Sonia Goncalves, Ewan Harrison, David K. Jackson, Ian Johnston, Dominic Kwiatkowski, Cordelia Langford, John Sillitoe on behalf of the Wellcome Sanger Institute COVID-19 Surveillance Team |
| EPI_ISL_909518, EPI_ISL_909526 | Lighthouse Lab in Alderley Park | Wellcome Sanger Institute for the COVID-19 Genomics UK (COG-UK) Consortium | Jacquelyn Wynn, Mairead Hyland, The Lighthouse Lab in Alderley Park and Alex Alderton, Roberto Amato, Sonia Goncalves, Ewan Harrison, David K. Jackson, Ian Johnston, Dominic Kwiatkowski, Cordelia Langford, John Sillitoe on behalf of the Wellcome Sanger Institute COVID-19 Surveillance Team |
| EPI_ISL_909529 | Lighthouse Lab in Milton Keynes | Wellcome Sanger Institute for the COVID-19 Genomics UK (COG-UK) Consortium | The Lighthouse Lab in Milton Keynes and Alex Alderton, Roberto Amato, Sonia Goncalves, Ewan Harrison, David K. Jackson, Ian Johnston, Dominic Kwiatkowski, Cordelia Langford, John Sillitoe on behalf of the Wellcome Sanger Institute COVID-19 Surveillance Team |
| EPI_ISL_909531 | Lighthouse Lab in Alderley Park | Wellcome Sanger Institute for the COVID-19 Genomics UK (COG-UK) Consortium | Jacquelyn Wynn, Mairead Hyland, The Lighthouse Lab in Alderley Park and Alex Alderton, Roberto Amato, Sonia Goncalves, Ewan Harrison, David K. Jackson, Ian Johnston, Dominic Kwiatkowski, Cordelia Langford, John Sillitoe on behalf of the Wellcome Sanger Institute COVID-19 Surveillance Team |
| EPI_ISL_909533 | Lighthouse Lab in Milton Keynes | Wellcome Sanger Institute for the COVID-19 Genomics UK (COG-UK) Consortium | The Lighthouse Lab in Milton Keynes and Alex Alderton, Roberto Amato, Sonia Goncalves, Ewan Harrison, David K. Jackson, Ian Johnston, Dominic Kwiatkowski, Cordelia Langford, John Sillitoe on behalf of the Wellcome Sanger Institute COVID-19 Surveillance Team |
| EPI_ISL_909538 | Lighthouse Lab in Glasgow | Wellcome Sanger Institute for the COVID-19 Genomics UK (COG-UK) Consortium | Harper VanSteenhouse, Yumi Kasai, David Gray, Carol Clugston, Anna Dominiczak and Alex Alderton, Roberto Amato, Sonia Goncalves, Ewan Harrison, David K. Jackson, Ian Johnston, Dominic Kwiatkowski, Cordelia Langford, John Sillitoe on behalf of the Wellcome Sanger Institute COVID-19 Surveillance Team |
| EPI_ISL_909539 | Lighthouse Lab in Alderley Park | Wellcome Sanger Institute for the COVID-19 Genomics UK (COG-UK) Consortium | Jacquelyn Wynn, Mairead Hyland, The Lighthouse Lab in Alderley Park and Alex Alderton, Roberto Amato, Sonia Goncalves, Ewan Harrison, David K. Jackson, Ian Johnston, Dominic Kwiatkowski, Cordelia Langford, John Sillitoe on behalf of the Wellcome Sanger Institute COVID-19 Surveillance Team |
| EPI_ISL_909540, EPI_ISL_909541 | Lighthouse Lab in Milton Keynes | Wellcome Sanger Institute for the COVID-19 Genomics UK (COG-UK) Consortium | The Lighthouse Lab in Milton Keynes and Alex Alderton, Roberto Amato, Sonia Goncalves, Ewan Harrison, David K. Jackson, Ian Johnston, Dominic Kwiatkowski, Cordelia Langford, John Sillitoe on behalf of the Wellcome Sanger Institute COVID-19 Surveillance Team |

| | | | |
|---|---|---|---|
| EPI_ISL_909545, EPI_ISL_909546 | Lighthouse Lab in Alderley Park | Wellcome Sanger Institute for the COVID-19 Genomics UK (COG-UK) Consortium | Jacquelyn Wynn, Mairead Hyland, The Lighthouse Lab in Alderley Park and Alex Alderton, Roberto Amato, Sonia Goncalves, Ewan Harrison, David K. Jackson, Ian Johnston, Dominic Kwiatkowski, Cordelia Langford, John Sillitoe on behalf of the Wellcome Sanger Institute COVID-19 Surveillance Team |
| EPI_ISL_909549 | Lighthouse Lab in Glasgow | Wellcome Sanger Institute for the COVID-19 Genomics UK (COG-UK) Consortium | Harper VanSteenhouse, Yumi Kasai, David Gray, Carol Clugston, Anna Dominiczak and Alex Alderton, Roberto Amato, Sonia Goncalves, Ewan Harrison, David K. Jackson, Ian Johnston, Dominic Kwiatkowski, Cordelia Langford, John Sillitoe on behalf of the Wellcome Sanger Institute COVID-19 Surveillance Team |
| EPI_ISL_909552, EPI_ISL_909553 | Lighthouse Lab in Alderley Park | Wellcome Sanger Institute for the COVID-19 Genomics UK (COG-UK) Consortium | Jacquelyn Wynn, Mairead Hyland, The Lighthouse Lab in Alderley Park and Alex Alderton, Roberto Amato, Sonia Goncalves, Ewan Harrison, David K. Jackson, Ian Johnston, Dominic Kwiatkowski, Cordelia Langford, John Sillitoe on behalf of the Wellcome Sanger Institute COVID-19 Surveillance Team |
| EPI_ISL_909556, EPI_ISL_909560 | Lighthouse Lab in Milton Keynes | Wellcome Sanger Institute for the COVID-19 Genomics UK (COG-UK) Consortium | The Lighthouse Lab in Milton Keynes and Alex Alderton, Roberto Amato, Sonia Goncalves, Ewan Harrison, David K. Jackson, Ian Johnston, Dominic Kwiatkowski, Cordelia Langford, John Sillitoe on behalf of the Wellcome Sanger Institute COVID-19 Surveillance Team |
| EPI_ISL_909561 | Lighthouse Lab in Alderley Park | Wellcome Sanger Institute for the COVID-19 Genomics UK (COG-UK) Consortium | Jacquelyn Wynn, Mairead Hyland, The Lighthouse Lab in Alderley Park and Alex Alderton, Roberto Amato, Sonia Goncalves, Ewan Harrison, David K. Jackson, Ian Johnston, Dominic Kwiatkowski, Cordelia Langford, John Sillitoe on behalf of the Wellcome Sanger Institute COVID-19 Surveillance Team |
| EPI_ISL_909562 | Lighthouse Lab in Milton Keynes | Wellcome Sanger Institute for the COVID-19 Genomics UK (COG-UK) Consortium | The Lighthouse Lab in Milton Keynes and Alex Alderton, Roberto Amato, Sonia Goncalves, Ewan Harrison, David K. Jackson, Ian Johnston, Dominic Kwiatkowski, Cordelia Langford, John Sillitoe on behalf of the Wellcome Sanger Institute COVID-19 Surveillance Team |
| EPI_ISL_909564 | Lighthouse Lab in Alderley Park | Wellcome Sanger Institute for the COVID-19 Genomics UK (COG-UK) Consortium | Jacquelyn Wynn, Mairead Hyland, The Lighthouse Lab in Alderley Park and Alex Alderton, Roberto Amato, Sonia Goncalves, Ewan Harrison, David K. Jackson, Ian Johnston, Dominic Kwiatkowski, Cordelia Langford, John Sillitoe on behalf of the Wellcome Sanger Institute COVID-19 Surveillance Team |
| EPI_ISL_909566 | Lighthouse Lab in Milton Keynes | Wellcome Sanger Institute for the COVID-19 Genomics UK (COG-UK) Consortium | The Lighthouse Lab in Milton Keynes and Alex Alderton, Roberto Amato, Sonia Goncalves, Ewan Harrison, David K. Jackson, Ian Johnston, Dominic Kwiatkowski, Cordelia Langford, John Sillitoe on behalf of the Wellcome Sanger Institute COVID-19 Surveillance Team |
| EPI_ISL_909568 | Lighthouse Lab in Glasgow | Wellcome Sanger Institute for the COVID-19 Genomics UK (COG-UK) Consortium | Harper VanSteenhouse, Yumi Kasai, David Gray, Carol Clugston, Anna Dominiczak and Alex Alderton, Roberto Amato, Sonia Goncalves, Ewan Harrison, David K. Jackson, Ian Johnston, Dominic Kwiatkowski, Cordelia Langford, John Sillitoe on behalf of the Wellcome Sanger Institute COVID-19 Surveillance Team |
| EPI_ISL_909569 | Lighthouse Lab in Milton Keynes | Wellcome Sanger Institute for the COVID-19 Genomics UK (COG-UK) Consortium | The Lighthouse Lab in Milton Keynes and Alex Alderton, Roberto Amato, Sonia Goncalves, Ewan Harrison, David K. Jackson, Ian Johnston, Dominic Kwiatkowski, Cordelia Langford, John Sillitoe on behalf of the Wellcome Sanger Institute COVID-19 Surveillance Team |
| EPI_ISL_909571, EPI_ISL_909572, EPI_ISL_909573, EPI_ISL_909575 | Lighthouse Lab in Alderley Park | Wellcome Sanger Institute for the COVID-19 Genomics UK (COG-UK) Consortium | Jacquelyn Wynn, Mairead Hyland, The Lighthouse Lab in Alderley Park and Alex Alderton, Roberto Amato, Sonia Goncalves, Ewan Harrison, David K. Jackson, Ian Johnston, Dominic Kwiatkowski, Cordelia Langford, John Sillitoe on behalf of the Wellcome Sanger Institute COVID-19 Surveillance Team |
| EPI_ISL_909576 | Lighthouse Lab in Milton Keynes | Wellcome Sanger Institute for the COVID-19 Genomics UK (COG-UK) Consortium | The Lighthouse Lab in Milton Keynes and Alex Alderton, Roberto Amato, Sonia Goncalves, Ewan Harrison, David K. Jackson, Ian Johnston, Dominic Kwiatkowski, Cordelia Langford, John Sillitoe on behalf of the Wellcome Sanger Institute COVID-19 Surveillance Team |
| EPI_ISL_909577, EPI_ISL_909578, EPI_ISL_909580, EPI_ISL_909581 | Lighthouse Lab in Alderley Park | Wellcome Sanger Institute for the COVID-19 Genomics UK (COG-UK) Consortium | Jacquelyn Wynn, Mairead Hyland, The Lighthouse Lab in Alderley Park and Alex Alderton, Roberto Amato, Sonia Goncalves, Ewan Harrison, David K. Jackson, Ian Johnston, Dominic Kwiatkowski, Cordelia Langford, John Sillitoe on behalf of the Wellcome Sanger Institute COVID-19 Surveillance Team |
| EPI_ISL_909584 | Lighthouse Lab in Milton Keynes | Wellcome Sanger Institute for the COVID-19 Genomics UK (COG-UK) Consortium | The Lighthouse Lab in Milton Keynes and Alex Alderton, Roberto Amato, Sonia Goncalves, Ewan Harrison, David K. Jackson, Ian Johnston, Dominic Kwiatkowski, Cordelia Langford, John Sillitoe on behalf of the Wellcome Sanger Institute COVID-19 Surveillance Team |
| EPI_ISL_909586 | Lighthouse Lab in Alderley Park | Wellcome Sanger Institute for the COVID-19 Genomics UK (COG-UK) Consortium | Jacquelyn Wynn, Mairead Hyland, The Lighthouse Lab in Alderley Park and Alex Alderton, Roberto Amato, Sonia Goncalves, Ewan Harrison, David K. Jackson, Ian Johnston, Dominic Kwiatkowski, Cordelia Langford, John Sillitoe on behalf of the Wellcome Sanger Institute COVID-19 Surveillance Team |
| EPI_ISL_909588, EPI_ISL_909590 | Lighthouse Lab in Glasgow | Wellcome Sanger Institute for the COVID-19 Genomics UK (COG-UK) Consortium | Harper VanSteenhouse, Yumi Kasai, David Gray, Carol Clugston, Anna Dominiczak and Alex Alderton, Roberto Amato, Sonia Goncalves, Ewan Harrison, David K. Jackson, Ian Johnston, Dominic Kwiatkowski, Cordelia Langford, John Sillitoe on behalf of the Wellcome Sanger Institute COVID-19 Surveillance Team |
| EPI_ISL_909745, EPI_ISL_909746, EPI_ISL_909748, EPI_ISL_909749 | A. Krumbholz, Labor Dr. Krause und Kollegen MVZ GmbH, Kiel | Charité Universitätsmedizin Berlin, Institut für Virologie | Victor M Corman, Barbara Mühlemann, Jörn Beheim-Schwarzbach, Tobias Bleicker, Julia Tesch, Talitha Veith, Julia Schneider, Terry Jones, Christian Drosten |
| EPI_ISL_909750, EPI_ISL_909757 | Charité Universitätsmedizin Berlin, Institut für Virologie/Labor Berlin | Charité Universitätsmedizin Berlin, Institut für Virologie | Victor M Corman, Barbara Mühlemann, Jörn Beheim-Schwarzbach, Tobias Bleicker, Julia Tesch, Talitha Veith, Julia Schneider, Terry Jones, Christian Drosten |
| EPI_ISL_909773, EPI_ISL_909782, EPI_ISL_909791, EPI_ISL_909792, EPI_ISL_909793, EPI_ISL_909795, EPI_ISL_909796, EPI_ISL_909800, EPI_ISL_909805, EPI_ISL_909807, EPI_ISL_909808, EPI_ISL_909810, EPI_ISL_909812, EPI_ISL_909823, EPI_ISL_909824, EPI_ISL_909825, EPI_ISL_909876, EPI_ISL_909880, EPI_ISL_909882, EPI_ISL_909885, EPI_ISL_909891, EPI_ISL_909892, EPI_ISL_909893, EPI_ISL_909894, EPI_ISL_909902, EPI_ISL_909922, EPI_ISL_909926, EPI_ISL_909927, EPI_ISL_909930, EPI_ISL_909935 | | | |
| see above | National Virus Reference Laboratory | National Virus Reference Laboratory | Michael Carr, Gabriel Gonzalez, Jonathan Dean, Cillian F De Gascun |
| EPI_ISL_911209, EPI_ISL_911210, EPI_ISL_911211, EPI_ISL_911212, EPI_ISL_911213, EPI_ISL_911214, EPI_ISL_911215, EPI_ISL_911216, EPI_ISL_911217, EPI_ISL_911218, EPI_ISL_911219, EPI_ISL_911220, EPI_ISL_911221, EPI_ISL_911222, EPI_ISL_911223, EPI_ISL_911224, EPI_ISL_911225, EPI_ISL_911226, EPI_ISL_911227, EPI_ISL_911228, EPI_ISL_911229, EPI_ISL_911230, EPI_ISL_911231 | | | |
| see above | Laboratoire national de sante, Microbiology, Virology | Laboratoire national de sante, Microbiology, Microbial Genomics Platform | Anke Wienecke-Baldacchino, Catherine Ragimbeau,Jessica Tapp, Fatu Djabi, Lise Pignon, Raoul Salmon, Tamir Abdelrahman |
| EPI_ISL_911314, EPI_ISL_911324, EPI_ISL_911341 | Servicio de Microbiología, Hospital Universitario Son Espases | SeqCOVID-SPAIN consortium/IBV(CSIC) | Carla López-Causapé, Jordi Reina, Antonio Oliver and SeqCOVID-SPAIN consortium |
| EPI_ISL_911386, EPI_ISL_911387, EPI_ISL_911388, EPI_ISL_911389, EPI_ISL_911430, EPI_ISL_911431, EPI_ISL_911432, EPI_ISL_911433, EPI_ISL_911434, EPI_ISL_911435, EPI_ISL_911436, EPI_ISL_911437, EPI_ISL_911438, EPI_ISL_911439, EPI_ISL_911440, EPI_ISL_911441, EPI_ISL_911442 | | | |
| see above | AZDelta | AZDelta | Geert Martens; Dieter De Smet |
| EPI_ISL_911525 | Microbiology and Virology Unit, Florence Careggi University Hospital | Microbiology and Virology Unit, Florence Careggi University Hospital | Vincenzo Di Pilato, Marco Coppi, Fabio Morecchiato, Noemi Aiezza, Ilaria Baccani, Alberto Antonelli, Emanuele Gori, Gian Maria Rossolini |
| EPI_ISL_911582, EPI_ISL_911584, EPI_ISL_911585, EPI_ISL_911587, EPI_ISL_911588, EPI_ISL_911593, EPI_ISL_911595, EPI_ISL_911597, EPI_ISL_911598, EPI_ISL_911602, EPI_ISL_911674 | | | |
| see above | Clinical Molecular Microbiology Laboratory, UNC Hospitals | Jeremy Wang | Jeremy Wang, Alexander Rubinsteyn, Colleen Rice, Jason Smedberg, Melissa Miller, Corbin Jones, Robert Hagan |
| EPI_ISL_911748, EPI_ISL_911749 | Toronto Invasive Bacterial Diseases Network | McMaster University | Allison McGeer, Patryk Aftanas, Hooman Derakhshani, Angel Li, Kuganya Nirmalarajah, Emily Panousis, Ahmed Draia, Jalees Nasir, Michael Surette, Samira Mubareka, Andrew G. McArthur |
| EPI_ISL_911755, EPI_ISL_911757, EPI_ISL_911758, EPI_ISL_911762, EPI_ISL_911763, EPI_ISL_911764, EPI_ISL_911765, EPI_ISL_911766, EPI_ISL_911768, EPI_ISL_911772, EPI_ISL_911775, EPI_ISL_911776, EPI_ISL_911777, EPI_ISL_911778, EPI_ISL_911779, EPI_ISL_911780, EPI_ISL_911781, EPI_ISL_911782, EPI_ISL_911791, EPI_ISL_911865, EPI_ISL_911900, EPI_ISL_911901, EPI_ISL_911905, EPI_ISL_911906 | | | |
| see above | Johns Hopkins Hospital Department of Pathology | Johns Hopkins Hospital Department of Pathology | C. Paul Morris, Chun Huai Luo, Adannaya Amadi, Matthew Schwartz, Nicholas Gallagher, Heba H. Mostafa |
| EPI_ISL_912109, EPI_ISL_912110, EPI_ISL_912111, EPI_ISL_912112, EPI_ISL_912113, EPI_ISL_912114, EPI_ISL_912115, EPI_ISL_912116, EPI_ISL_912117, EPI_ISL_912118, EPI_ISL_912119, EPI_ISL_912120, EPI_ISL_912121, EPI_ISL_912122, EPI_ISL_912123, EPI_ISL_912124, EPI_ISL_912125, EPI_ISL_912126, EPI_ISL_912127, EPI_ISL_912128, EPI_ISL_912129, EPI_ISL_912130, EPI_ISL_912131, EPI_ISL_912132, EPI_ISL_912133, EPI_ISL_912134, EPI_ISL_912135, EPI_ISL_912136, EPI_ISL_912137, EPI_ISL_912138, EPI_ISL_912139, EPI_ISL_912140, EPI_ISL_912141, EPI_ISL_912142, EPI_ISL_912143, EPI_ISL_912144, EPI_ISL_912145, EPI_ISL_912146, EPI_ISL_912147, EPI_ISL_912148, EPI_ISL_912149, EPI_ISL_912150, EPI_ISL_912151, EPI_ISL_912152, EPI_ISL_912153, EPI_ISL_912154, EPI_ISL_912155, EPI_ISL_912156, EPI_ISL_912157, EPI_ISL_912158, EPI_ISL_912159 | | | |
| see above | Altius Institute for Biomedical Sciences | Seattle Flu Study | Deborah A. Nickerson, Chris D. Frazar, Jover Lee, Benjamin Pelle, Erica Ryke, Matthew Richardson, Amanda Adler, Elisabeth Brandstetter, Peter D. Han, Kairsten Fay, Misja Ilcisin, Kirsten Lacombe, Thomas R. Sibley, Melissa Truong, Caitlin R. Wolf, Ryan Alexander, Daniel Bates, Rebecca Bruders, Stephanie DeBaun, Clem Green, Muhammad Halimun, Jessica Halow, Kneshay Harper, Matt Hartman, Andrew Meuser, Alex Nguyen, Truong Nguyen, Sofia Olsson, Sadie Patraw, Hannah Petersen, Tobias Ragoczy, Joshua Richards, Jacob Rodriguez, John Stamatoyannopoulos, Julia Wald, Olivia Waltner, Michael Boeckh, Janet A. Englund, Michael Famulare, Barry R. Lutz, Mark J. Rieder, Lea M. Starita, Matthew Thompson, Helen Y. Chu, Jay Shendure, Trevor Bedford |
| EPI_ISL_912171, EPI_ISL_912172, EPI_ISL_912173, EPI_ISL_912174 | Tempus | Grubaugh Lab - Yale School of Public Health | Tara Alpert, Joseph Fauver, Anderson Brito, Mallery Breban, Anne Wyllie, Chantal Vogels, Mary Petrone, Annie Watkins, Chaney Kalinich, Isabel Ott, Nathan Grubaugh |
| EPI_ISL_912214 | Lighthouse Lab in Cambridge | Wellcome Sanger Institute for the COVID-19 Genomics UK | Rob Howes, The Lighthouse Lab in Cambridge and Alex Alderton, Roberto Amato, Sonia Goncalves, Ewan Harrison, David K. Jackson, Ian Johnston, |

| | | | |
|---|---|---|---|
| | | (COG-UK) Consortium | Dominic Kwiatkowski, Cordelia Langford, John Sillitoe on behalf of the Wellcome Sanger Institute COVID-19 Surveillance Team (http://www.sanger.ac.uk/covid-team) |
| EPI_ISL_912220, EPI_ISL_912221, EPI_ISL_912222, EPI_ISL_912223 | Lighthouse Lab in Alderley Park | Wellcome Sanger Institute for the COVID-19 Genomics UK (COG-UK) Consortium | Jacquelyn Wynn, Mairead Hyland, The Lighthouse Lab in Alderley Park and Alex Alderton, Roberto Amato, Sonia Goncalves, Ewan Harrison, David K. Jackson, Ian Johnston, Dominic Kwiatkowski, Cordelia Langford, John Sillitoe on behalf of the Wellcome Sanger Institute COVID-19 Surveillance Team (http://www.sanger.ac.uk/covid-team) |
| EPI_ISL_912245, EPI_ISL_912246, EPI_ISL_912250, EPI_ISL_912251, EPI_ISL_912252, EPI_ISL_912253, EPI_ISL_912254, EPI_ISL_912257, EPI_ISL_912258, EPI_ISL_912260, EPI_ISL_912263, EPI_ISL_912264 | | | |
| see above | Charité Universitätsmedizin Berlin, Institut für Virologie/Labor Berlin | Charité Universitätsmedizin Berlin, Institut für Virologie | Victor M Corman, Barbara Mühlemann, Jörn Beheim-Schwarzbach, Tobias Bleicker, Julia Tesch, Talitha Veith, Julia Schneider, Terry Jones, Christian Drosten |
| EPI_ISL_912579 | Service Biologie -CH de Versailles | National Reference Center for Viruses of Respiratory Infections, Institut Pasteur, Paris | Marion Barbet, Sylvie Behillil, Méline Bizard, Angela Brisebarre, Camille Capel, Etienne Simon-Lorière, Vincent Enouf, Maud Vanpeene, Sylvie van der Werf,Marque Juillet StéPhanie |
| EPI_ISL_912639 | Hôpital Henri Mondor | Department of Virology, Henri Mondor University Hospital, Assistance Publique Hôpitaux de Paris, Université Paris-Est Créteil, INSERM U955 | Christophe Rodriguez, Slim Fourati, Vanessa Demontant, Guillaume Gricourt, Melissa N'Debi, Alexandre Soulier, Elisabeth Trawinski, Jean-Michel Pawlotsky |
| EPI_ISL_912649, EPI_ISL_912655 | Hôpital Pitié-Salpêtrière | Department of Virology, Henri Mondor University Hospital, Assistance Publique Hôpitaux de Paris, Université Paris-Est Créteil, INSERM U955 | Christophe Rodriguez, Slim Fourati, Vanessa Demontant, Guillaume Gricourt, Melissa N'Debi, Alexandre Soulier, Elisabeth Trawinski, Jean-Michel Pawlotsky |
| EPI_ISL_912665, EPI_ISL_912666, EPI_ISL_912696, EPI_ISL_912697, EPI_ISL_912698, EPI_ISL_912699, EPI_ISL_912700, EPI_ISL_912703, EPI_ISL_912704, EPI_ISL_912716, EPI_ISL_912717 | | | |
| see above | Hôpital Henri Mondor | Department of Virology, Henri Mondor University Hospital, Assistance Publique Hôpitaux de Paris, Université Paris-Est Créteil, INSERM U955 | Christophe Rodriguez, Slim Fourati, Vanessa Demontant, Guillaume Gricourt, Melissa N'Debi, Alexandre Soulier, Elisabeth Trawinski, Jean-Michel Pawlotsky |
| EPI_ISL_912719 | Hôpital Pitié-Salpêtrière | Department of Virology, Henri Mondor University Hospital, Assistance Publique Hôpitaux de Paris, Université Paris-Est Créteil, INSERM U955 | Christophe Rodriguez, Slim Fourati, Vanessa Demontant, Guillaume Gricourt, Melissa N'Debi, Alexandre Soulier, Elisabeth Trawinski, Jean-Michel Pawlotsky |
| EPI_ISL_912750, EPI_ISL_912751, EPI_ISL_912752, EPI_ISL_912753, EPI_ISL_912754, EPI_ISL_912755, EPI_ISL_912756, EPI_ISL_912757, EPI_ISL_912758, EPI_ISL_912759, EPI_ISL_912760, EPI_ISL_912761 | | | |
| see above | Hôpital Henri Mondor | Department of Virology, Henri Mondor University Hospital, Assistance Publique Hôpitaux de Paris, Université Paris-Est Créteil, INSERM U955 | Christophe Rodriguez, Slim Fourati, Vanessa Demontant, Guillaume Gricourt, Melissa N'Debi, Alexandre Soulier, Elisabeth Trawinski, Jean-Michel Pawlotsky |
| EPI_ISL_912786, EPI_ISL_912787 | Hôpital Pitié-Salpêtrière | Department of Virology, Henri Mondor University Hospital, Assistance Publique Hôpitaux de Paris, Université Paris-Est Créteil, INSERM U955 | Christophe Rodriguez, Slim Fourati, Vanessa Demontant, Guillaume Gricourt, Melissa N'Debi, Alexandre Soulier, Elisabeth Trawinski, Jean-Michel Pawlotsky |
| EPI_ISL_913402, EPI_ISL_913403, EPI_ISL_913413, EPI_ISL_913419, EPI_ISL_913422, EPI_ISL_913423, EPI_ISL_913432, EPI_ISL_913433 | Massachusetts State Public Health Laboratory | Massachusetts State Public Health Laboratory | Andrew Lang, Timelia Fink, Glen Gallagher, Sandra Smole |
| EPI_ISL_913445 | Institute for Infectious Diseases, University of Bern, Switzerland | Institute for Infectious Diseases, University of Bern, Switzerland | Michel C Koch, Christian Baumann, Miguel A Terrazos Miani, Cora Sägesser, Pascal Bittel, Stephen L Leib, Peter Keller, Franziska Suter-Riniker, Alban Ramette |
| EPI_ISL_913496, EPI_ISL_913497, EPI_ISL_913498, EPI_ISL_913499, EPI_ISL_913500, EPI_ISL_913501, EPI_ISL_913502, EPI_ISL_913503, EPI_ISL_913504, EPI_ISL_913505, EPI_ISL_913506, EPI_ISL_913507, EPI_ISL_913508, EPI_ISL_913509, EPI_ISL_913510, EPI_ISL_913511, EPI_ISL_913512, EPI_ISL_913513 | | | |
| see above | AZDelta | AZDelta | Geert Martens; Dieter De Smet |
| EPI_ISL_913617, EPI_ISL_913618, EPI_ISL_913619, EPI_ISL_913620 | Michigan Department of Health and Human Services, Bureau of Laboratories | Michigan Department of Health and Human Services, Bureau of Laboratories | Blankenship HM, Riner D, Soehnlen MK |
| EPI_ISL_914003, EPI_ISL_914004, EPI_ISL_914006, EPI_ISL_914007, EPI_ISL_914008, EPI_ISL_914009 | Clinical Pathology Lab | Pathogen Discovery, Respiratory Viruses Branch, Division of Viral Diseases, Centers for Disease Control and Prevention | Ying Tao, Yan Li, Jing Zhang, Krista Queen, Anna Uehara, Peter Cook, Clinton R. Paden, Haibin Wang, Suxiang Tong |
| EPI_ISL_914640 | Santa Clara County Public Health Laboratory | Santa Clara County Public Health Laboratory | Santa Clara County Public Health Department |
| EPI_ISL_914702, EPI_ISL_914703, EPI_ISL_914704, EPI_ISL_914705, EPI_ISL_914706, EPI_ISL_914707, EPI_ISL_914708, EPI_ISL_914709, EPI_ISL_914710, EPI_ISL_914711, EPI_ISL_914712, EPI_ISL_914713, EPI_ISL_914714, EPI_ISL_914716, EPI_ISL_914717, EPI_ISL_914718, EPI_ISL_914719, EPI_ISL_914720, EPI_ISL_914721, EPI_ISL_914722, EPI_ISL_914723, EPI_ISL_914724 | | | |
| see above | Maryland Public Health Laboratory (MD PHL) | Maryland Public Health Laboratory (MD PHL) | Maryland Department of Health Laboratories Administration |
| EPI_ISL_914725, EPI_ISL_914726, EPI_ISL_914727, EPI_ISL_914728, EPI_ISL_914730, EPI_ISL_914731, EPI_ISL_914732, EPI_ISL_914733, EPI_ISL_914734, EPI_ISL_914735, EPI_ISL_914736, EPI_ISL_914737, EPI_ISL_914739, EPI_ISL_914740, EPI_ISL_914743, EPI_ISL_914744, EPI_ISL_914748, EPI_ISL_914749, EPI_ISL_914750, EPI_ISL_914751, EPI_ISL_914753, EPI_ISL_914779, EPI_ISL_914781, EPI_ISL_914784, EPI_ISL_914785, EPI_ISL_914786, EPI_ISL_914787, EPI_ISL_914788, EPI_ISL_914789, EPI_ISL_914792 | | | |
| see above | Wyoming Public Health Laboratory | Wyoming Public Health Laboratory | Noah Hull, Taylor Fearing, Lynette Gumbleton, Channing Weber, Ashley Norberg, Bailey Bowcutt, and Wanda Manley |
| EPI_ISL_914840 | HOSPITAL DR. RAUL BLANCO CERVANTES | Inciensa, Instituto Costarricense de Investigación y Enseñanza en Nutrición y Salud | Francisco Duarte, Hebleen Porras, Claudio Soto-Garita, Estela Cordero, Adriana Godínez, Melany Calderón & Rebeca Ruiz-Sánchez |
| EPI_ISL_916191, EPI_ISL_916192, EPI_ISL_916196, EPI_ISL_916210, EPI_ISL_916211, EPI_ISL_916212, EPI_ISL_916229, EPI_ISL_916231, EPI_ISL_916237, EPI_ISL_916242, EPI_ISL_916243, EPI_ISL_916247, EPI_ISL_916249, EPI_ISL_916252, EPI_ISL_916260, EPI_ISL_916262, EPI_ISL_916269, EPI_ISL_916270, EPI_ISL_916271, EPI_ISL_916274, EPI_ISL_916277, EPI_ISL_916280, EPI_ISL_916292, EPI_ISL_916297, EPI_ISL_916304, EPI_ISL_916313, EPI_ISL_916331, EPI_ISL_916334, EPI_ISL_916346, EPI_ISL_916348, EPI_ISL_916354, EPI_ISL_916355, EPI_ISL_916362, EPI_ISL_916363, EPI_ISL_916376, EPI_ISL_916382, EPI_ISL_916391, EPI_ISL_916392, EPI_ISL_916414, EPI_ISL_916416, EPI_ISL_916420, EPI_ISL_916427, EPI_ISL_916435, EPI_ISL_916444, EPI_ISL_916445, EPI_ISL_916450, EPI_ISL_916461, EPI_ISL_916470, EPI_ISL_916475, EPI_ISL_916495, EPI_ISL_916496, EPI_ISL_916497 | | | |
| see above | Lighthouse Lab in Cambridge | Wellcome Sanger Institute for the COVID-19 Genomics UK (COG-UK) Consortium | Rob Howes, The Lighthouse Lab in Cambridge and Alex Alderton, Roberto Amato, Sonia Goncalves, Ewan Harrison, David K. Jackson, Ian Johnston, Dominic Kwiatkowski, Cordelia Langford, John Sillitoe on behalf of the Wellcome Sanger Institute COVID-19 Surveillance Team |
| EPI_ISL_917214, EPI_ISL_917501 | Lighthouse Lab in Alderley Park | Wellcome Sanger Institute for the COVID-19 Genomics UK (COG-UK) Consortium | Jacquelyn Wynn, Mairead Hyland, The Lighthouse Lab in Alderley Park and Alex Alderton, Roberto Amato, Sonia Goncalves, Ewan Harrison, David K. Jackson, Ian Johnston, Dominic Kwiatkowski, Cordelia Langford, John Sillitoe on behalf of the Wellcome Sanger Institute COVID-19 Surveillance Team |
| EPI_ISL_917951, EPI_ISL_917952, EPI_ISL_917953 | Los Angeles County PHL | Los Angeles County PHL | P. Hemarajata et al. |
| EPI_ISL_918020, EPI_ISL_918021, EPI_ISL_918022, EPI_ISL_918023, EPI_ISL_918024, EPI_ISL_918025, EPI_ISL_918026, EPI_ISL_918027, EPI_ISL_918028, EPI_ISL_918029, EPI_ISL_918030, EPI_ISL_918031, EPI_ISL_918032, EPI_ISL_918033, EPI_ISL_918034, EPI_ISL_918035, EPI_ISL_918036, EPI_ISL_918037, EPI_ISL_918038 | | | |
| see above | Lighthouse Lab in Glasgow | Wellcome Sanger Institute for the COVID-19 Genomics UK (COG-UK) Consortium | Harper VanSteenhouse, Yumi Kasai, David Gray, Carol Clugston, Anna Dominiczak and Alex Alderton, Roberto Amato, Sonia Goncalves, Ewan Harrison, David K. Jackson, Ian Johnston, Dominic Kwiatkowski, Cordelia Langford, John Sillitoe on behalf of the Wellcome Sanger Institute COVID-19 Surveillance Team |
| EPI_ISL_918191 | Innovative Genomics Institute, UC Berkeley | Innovative Genomics Institute, UC Berkeley | Stacia Wyman, Haridha Shivram, Phil Frankino, Liana Lareau, Shana McDevitt, Justin Choi |
| EPI_ISL_918292, EPI_ISL_918293, EPI_ISL_918298, EPI_ISL_918299, EPI_ISL_918300, EPI_ISL_918301, EPI_ISL_918303, EPI_ISL_918304, EPI_ISL_918305, EPI_ISL_918306, EPI_ISL_918307, EPI_ISL_918308, EPI_ISL_918309, EPI_ISL_918310, EPI_ISL_918311, EPI_ISL_918312, EPI_ISL_918314, EPI_ISL_918318, EPI_ISL_918319, EPI_ISL_918321, EPI_ISL_918322, EPI_ISL_918323, EPI_ISL_918325, EPI_ISL_918326, EPI_ISL_918330, EPI_ISL_918332, EPI_ISL_918333, EPI_ISL_918334, EPI_ISL_918335, EPI_ISL_918337, EPI_ISL_918338 | | | |
| see above | Hospital Universitari Vall d'Hebron - Vall d'Hebron Institut de Rercerca | Hospital Universitari Vall d'Hebron | Cristina Andrés, Maria Piñana, Josep F Abril, Damir Garcia-Cehic, Ariadna Rando, Juliana Esperalba, Maria Gema Codina, Carla Castillo, Maria Carmen Martin, Tomàs Pumarola, Josep Quer, Andrés Antón |

| | | | |
|---|---|---|---|
| EPI_ISL_918495 | Florida Bureau of Public Health Laboratories | Florida Bureau of Public Health Laboratories | Sarah Schmedes, Jason Blanton |
| EPI_ISL_918985, EPI_ISL_918990, EPI_ISL_918991, EPI_ISL_918992, EPI_ISL_918993, EPI_ISL_918998, EPI_ISL_918999, EPI_ISL_919000, EPI_ISL_919001, EPI_ISL_919002, EPI_ISL_919003, EPI_ISL_919004, EPI_ISL_919006, EPI_ISL_919008, EPI_ISL_919009, EPI_ISL_919010, EPI_ISL_919013, EPI_ISL_919014, EPI_ISL_919015, EPI_ISL_919016, EPI_ISL_919115, EPI_ISL_919148 | | | |
| see above | Department of Pathology, University of Cambridge | COVID-19 Genomics UK (COG-UK) Consortium | Aminu S. Jahun, Yasmin Chaudhry, Iliana Georgana, Myra Hosmillo, Rhys Izu, Martin D. Curran, Surendra Parmar, Ian Goodfellow |
| EPI_ISL_919231, EPI_ISL_919277, EPI_ISL_919278, EPI_ISL_919279, EPI_ISL_919280 | West of Scotland Specialist Virology Centre, NHSGGC / MRC-University of Glasgow Centre for Virus Research | COVID-19 Genomics UK (COG-UK) Consortium | Ana da Silva Filipe, Natasha Johnson, Kathy Smollett, Daniel Mair, Stephen Carmichael, Alice Broos, Lily Tong, Jenna Nichols, Kyriaki Nomikou; Sarah McDonald; Richard Orton, Joseph Hughes, Sreenu Vattipally, David L Robertson; Alasdair MacLean, Rory Gunson; Sharif Shaaban, Matthew Holden; Rachel Blacow, Guy Mollett, Kathy Li, James Shepherd, Antonia Ho, Emma Thomson |
| EPI_ISL_919346 | Virology Department, Royal Infirmary of Edinburgh, NHS Lothian / School of Biological Sciences, University of Edinburgh / Institute of Genetics and Molecular Medicine, University of Edinburgh | COVID-19 Genomics UK (COG-UK) Consortium | McHugh M, Dewar R, Rooke S, Gallagher M, Balcaza C, O'Toole Á, Scher E, Hill V, McCrone JT, Colquhoun R, Yu X, Jackson B, Rambaut A, Williams TC, Templeton K |
| EPI_ISL_919489, EPI_ISL_919490, EPI_ISL_919491, EPI_ISL_919492, EPI_ISL_919493, EPI_ISL_919494, EPI_ISL_919496, EPI_ISL_919497, EPI_ISL_919498, EPI_ISL_919499, EPI_ISL_919501, EPI_ISL_919512, EPI_ISL_919514, EPI_ISL_919518, EPI_ISL_919525, EPI_ISL_919543, EPI_ISL_919574, EPI_ISL_919579, EPI_ISL_919582, EPI_ISL_919588, EPI_ISL_919604, EPI_ISL_919605, EPI_ISL_919606, EPI_ISL_919607, EPI_ISL_919608, EPI_ISL_919609, EPI_ISL_919611, EPI_ISL_919612, EPI_ISL_919613, EPI_ISL_919615, EPI_ISL_919617, EPI_ISL_919618, EPI_ISL_919619, EPI_ISL_919635, EPI_ISL_919638, EPI_ISL_919639, EPI_ISL_919691 | | | |
| see above | Liverpool Clinical Laboratories | COVID-19 Genomics UK (COG-UK) Consortium | Sam Haldenby, Anita Lucaci, Steve Paterson, Julian Hiscox, Alistair Darby, M Almsaud, A Alrezaihi, Muhannad Alruwaili, Stuart D Armstrong, Jones Benjamin, Eleanor G Bentley, Anu Chawla, Jordan J Clark, Angela Cowell, Richard Eccles, Isabel García-Dorival, Matthew Gemmell, Alessandro Gerada, PKF Gilmore, Richard Gregory, Ximeng Han, Catherine Hartley, Margaret Hughes, Miren Iturriza-Gomara, James Johnson, L Luu, Jenifer Manson, Charlotte Nelson, Elaine O'Toole, Cassie Olateju, Rebekah Penrice-Randal , Lucille Rainbow, N.P Randle, Trevor Ian Robinson, Parul Sharma, Ghada T Shawli, James P Stewart, Neil Swainston, Ecaterina Vamos, Joanne Watts, Mark Whitehead |
| EPI_ISL_919855 | Barts Health NHS Trust | COVID-19 Genomics UK (COG-UK) Consortium | CUTINO-MOGUEL, Maria-Teresa; HARRINGTON, David; OWOYEMI, Dola; KULASEGARAN-SHYLINI, Raghavendran; BROAD, Claire; KELE, Beatrix |
| EPI_ISL_920061, EPI_ISL_920062, EPI_ISL_920063, EPI_ISL_920064, EPI_ISL_920066, EPI_ISL_920068, EPI_ISL_920071, EPI_ISL_920072 | University College London, Great Ormond Street Hospital for Children NHS Foundation Trust, Imperial College Healthcare NHS Trust | COVID-19 Genomics UK (COG-UK) Consortium | Sergi Castellano, Rachel Williams, Mark Kristiansen, Paola Resende Silva, Sunando Roy, Tony Brooks, Helena Tutill, Paola Niola, Patricia Dyal, Charlotte Williams, Leysa Forrest, Yasmin Panchbhaya, Jacqueline Findlay, Samuel Weeks, Julianne Brown, Kathryn Harris, Paul Randell, James Price, Alison Holmes, Judith Breuer |
| EPI_ISL_920379, EPI_ISL_920380, EPI_ISL_920381, EPI_ISL_920383, EPI_ISL_920384, EPI_ISL_920385, EPI_ISL_920386, EPI_ISL_920387, EPI_ISL_920388, EPI_ISL_920389, EPI_ISL_920390, EPI_ISL_920391, EPI_ISL_920392, EPI_ISL_920393, EPI_ISL_920394, EPI_ISL_920395, EPI_ISL_920396, EPI_ISL_920397, EPI_ISL_920398, EPI_ISL_920399, EPI_ISL_920401, EPI_ISL_920403, EPI_ISL_920404, EPI_ISL_920405, EPI_ISL_920406, EPI_ISL_920407, EPI_ISL_920408, EPI_ISL_920411, EPI_ISL_920412, EPI_ISL_920414, EPI_ISL_920415, EPI_ISL_920416, EPI_ISL_920418, EPI_ISL_920419, EPI_ISL_920420, EPI_ISL_920421, EPI_ISL_920422, EPI_ISL_920423, EPI_ISL_920425, EPI_ISL_920426, EPI_ISL_920428, EPI_ISL_920429, EPI_ISL_920430, EPI_ISL_920431, EPI_ISL_920432, EPI_ISL_920433, EPI_ISL_920434, EPI_ISL_920435, EPI_ISL_920436, EPI_ISL_920438, EPI_ISL_920439, EPI_ISL_920441, EPI_ISL_920442, EPI_ISL_920443, EPI_ISL_920444, EPI_ISL_920445, EPI_ISL_920446, EPI_ISL_920448, EPI_ISL_920449, EPI_ISL_920450, EPI_ISL_920451, EPI_ISL_920452, EPI_ISL_920453, EPI_ISL_920627, EPI_ISL_920739 | | | |
| see above | University College London Hospital | COVID-19 Genomics UK (COG-UK) Consortium | Judith Heaney, Matthew Byott, Catherine Houlihan, Dan Frampton, Stuart Kirk, Moira Spyer and Eleni Nastouli |
| EPI_ISL_920783, EPI_ISL_920788, EPI_ISL_920789, EPI_ISL_920790, EPI_ISL_920791, EPI_ISL_920794, EPI_ISL_920795, EPI_ISL_920797, EPI_ISL_920798, EPI_ISL_920799, EPI_ISL_920800, EPI_ISL_920801, EPI_ISL_920802, EPI_ISL_920803, EPI_ISL_920804, EPI_ISL_920805, EPI_ISL_920806, EPI_ISL_920807, EPI_ISL_920809, EPI_ISL_920811, EPI_ISL_920812, EPI_ISL_920813, EPI_ISL_920816, EPI_ISL_920817, EPI_ISL_920831, EPI_ISL_920832 | | | |
| see above | University College London, Great Ormond Street Hospital for Children NHS Foundation Trust, Imperial College Healthcare NHS Trust | COVID-19 Genomics UK (COG-UK) Consortium | Sergi Castellano, Rachel Williams, Mark Kristiansen, Paola Resende Silva, Sunando Roy, Tony Brooks, Helena Tutill, Paola Niola, Patricia Dyal, Charlotte Williams, Leysa Forrest, Yasmin Panchbhaya, Jacqueline Findlay, Samuel Weeks, Julianne Brown, Kathryn Harris, Paul Randell, James Price, Alison Holmes, Judith Breuer |
| EPI_ISL_921608, EPI_ISL_921609, EPI_ISL_921617, EPI_ISL_921618, EPI_ISL_921620, EPI_ISL_921621, EPI_ISL_921622, EPI_ISL_921623, EPI_ISL_921624, EPI_ISL_921625, EPI_ISL_921626, EPI_ISL_921627, EPI_ISL_921628, EPI_ISL_921629, EPI_ISL_921630 | | | |
| see above | Northumbria University / South Tees Hospitals NHS Foundation Trust / North Cumbria Integrated Care NHS Foundation Trust / North Tees and Hartlepool NHS Foundation Trust / Newcastle Hospitals NHS Foundation Trust | COVID-19 Genomics UK (COG-UK) Consortium | Darren L Smith,Andrew Nelson,Matthew Bashton,Greg R Young,Joshua Loh,John Allan,Mohammad A Tariq,Giles S Holt,Gary Black,Wen C Yew,Lynn Dover,Paul Baker,Steve Liggett,Sarah Essex,Jane Greenaway,Debra Padgett,Clive Graham,Garren Scott,Edward Barton,Emma Swindells,Brendan Payne,Jennifer Collins,Yusri Taha,Gary Eltringham |
| EPI_ISL_921854, EPI_ISL_921855, EPI_ISL_921856, EPI_ISL_921860, EPI_ISL_921862, EPI_ISL_921869, EPI_ISL_921871, EPI_ISL_921873, EPI_ISL_921878, EPI_ISL_921880, EPI_ISL_921881, EPI_ISL_921882, EPI_ISL_921883, EPI_ISL_921884, EPI_ISL_921885, EPI_ISL_921886, EPI_ISL_921887, EPI_ISL_921888, EPI_ISL_921889, EPI_ISL_921890, EPI_ISL_921891, EPI_ISL_921892, EPI_ISL_921893, EPI_ISL_921894, EPI_ISL_921896, EPI_ISL_921897, EPI_ISL_921898, EPI_ISL_921899, EPI_ISL_921900, EPI_ISL_921901, EPI_ISL_921902, EPI_ISL_921903, EPI_ISL_921905, EPI_ISL_921906, EPI_ISL_921907, EPI_ISL_921908, EPI_ISL_921911, EPI_ISL_921915, EPI_ISL_921916, EPI_ISL_921918, EPI_ISL_921919, EPI_ISL_921920, EPI_ISL_921921, EPI_ISL_921922, EPI_ISL_921924, EPI_ISL_921926, EPI_ISL_921927, EPI_ISL_921928, EPI_ISL_921929, EPI_ISL_921930, EPI_ISL_921932, EPI_ISL_921936, EPI_ISL_921937, EPI_ISL_921939, EPI_ISL_921941, EPI_ISL_921943, EPI_ISL_921944, EPI_ISL_921945, EPI_ISL_921946, EPI_ISL_921949, EPI_ISL_921950, EPI_ISL_921951, EPI_ISL_921954, EPI_ISL_921955, EPI_ISL_921956, EPI_ISL_921957, EPI_ISL_921958, EPI_ISL_921959, EPI_ISL_921960, EPI_ISL_921961, EPI_ISL_921962, EPI_ISL_921963, EPI_ISL_921964, EPI_ISL_921965, EPI_ISL_921967, EPI_ISL_921970, EPI_ISL_921971, EPI_ISL_921972, EPI_ISL_921973, EPI_ISL_921974, EPI_ISL_921975, EPI_ISL_921976, EPI_ISL_921977, EPI_ISL_921979, EPI_ISL_921980 | | | |
| see above | Quadram Institute Bioscience | COVID-19 Genomics UK (COG-UK) Consortium | Dave J. Baker, Gemma L. Kay, Alp Aydin, Thanh Le-Viet, Steven Rudder, Ana P. Tedim, Anastasia Kolyva, Maria Diaz, Leonardo de Oliveira Martins, Nabil-Fareed Alikhan, Lizzie Meadows, Rachael Stanley, Ngozi Elumogo, Muhammed Yasir, Nicholas M. Thomson, Alexander J Trotter, Rachel Gilroy, Samuel Bloomfield, Claire Stuart, Andrew Bell, Reenesh Prakash, Samir Dervisevic, Alison E. Mather, John Wain, Mark Webber, Andrew J. Page, Justin O'Grady |
| EPI_ISL_921981, EPI_ISL_921982, EPI_ISL_921983, EPI_ISL_921984, EPI_ISL_921985, EPI_ISL_921986, EPI_ISL_921987, EPI_ISL_921988, EPI_ISL_921989, EPI_ISL_921990, EPI_ISL_921991, EPI_ISL_921992, EPI_ISL_921993, EPI_ISL_921994, EPI_ISL_921995, EPI_ISL_921996, EPI_ISL_921997, EPI_ISL_921998 | | | |
| see above | Queens Medical Centre, Clinical Microbiology Department / DeepSeq Nottingham | COVID-19 Genomics UK (COG-UK) Consortium | Gemma Clark, Wendy Smith, Manjinder Khakh, Vicki M Fleming, Michelle M Lister, Hannah Howson-Wells, Jonathan Ball, Patrick McClure, Joseph Chappell, Theocharis Tsoleridis, Nadine Holmes, Matthew Carlisle, Christopher Moore, Fei Sang, Johnny Debebe, Victoria Wright, Matthew Loose |
| EPI_ISL_922164, EPI_ISL_922165, EPI_ISL_922166, EPI_ISL_922167, EPI_ISL_922168 | Lincolnshire Hospitals and DeepSeq Nottingham | COVID-19 Genomics UK (COG-UK) Consortium | Nichola Duckworth, Tim Sloan, Sarah Walsh, Jonathan Ball, Patrick McClure, Joeseph Chappell, Nadine Holmes, Matthew Carlisle, Christopher Moore, Fei Sang, Johnny Debebe, Victoria Wright, Matthew Loose |
| EPI_ISL_922545, EPI_ISL_922557, EPI_ISL_922562, EPI_ISL_922569, EPI_ISL_922578, EPI_ISL_922579, EPI_ISL_922583, EPI_ISL_922584, EPI_ISL_922612, EPI_ISL_922613, EPI_ISL_922614, EPI_ISL_922615, EPI_ISL_922617, EPI_ISL_922619, EPI_ISL_922620, EPI_ISL_922621, EPI_ISL_922622, EPI_ISL_922623, EPI_ISL_922624, EPI_ISL_922625, EPI_ISL_922630, EPI_ISL_922631, EPI_ISL_922632, EPI_ISL_922635, EPI_ISL_922636, EPI_ISL_922736, EPI_ISL_922742, EPI_ISL_922769, EPI_ISL_922778, EPI_ISL_922795, EPI_ISL_922799, EPI_ISL_922804, EPI_ISL_922808, EPI_ISL_922826, EPI_ISL_922827, EPI_ISL_922828, EPI_ISL_922829, EPI_ISL_922830, EPI_ISL_922831, EPI_ISL_922832, EPI_ISL_922833, EPI_ISL_922834, EPI_ISL_922839, EPI_ISL_922840, EPI_ISL_922842, EPI_ISL_922843, EPI_ISL_922844, EPI_ISL_922845, EPI_ISL_922847, EPI_ISL_922848, EPI_ISL_922850, EPI_ISL_922851, EPI_ISL_922854, EPI_ISL_922857, EPI_ISL_922859, EPI_ISL_922860, EPI_ISL_922861, EPI_ISL_922863, EPI_ISL_922864, EPI_ISL_922865, EPI_ISL_922866, EPI_ISL_922867, EPI_ISL_922868, EPI_ISL_922869, EPI_ISL_922870, EPI_ISL_922871, EPI_ISL_922872, EPI_ISL_922873, EPI_ISL_922874, EPI_ISL_922876, EPI_ISL_922877, EPI_ISL_922878, EPI_ISL_922879, EPI_ISL_922880, EPI_ISL_922881, EPI_ISL_922885, EPI_ISL_922886, EPI_ISL_922888, EPI_ISL_922889, EPI_ISL_922890, EPI_ISL_922896, EPI_ISL_922897, EPI_ISL_922898, EPI_ISL_922899, EPI_ISL_922900, EPI_ISL_922916, EPI_ISL_922917, EPI_ISL_922918, EPI_ISL_922919, EPI_ISL_922920, EPI_ISL_922921, EPI_ISL_922922, EPI_ISL_922936, EPI_ISL_922937, EPI_ISL_922939, EPI_ISL_922962, EPI_ISL_922963, EPI_ISL_922971, EPI_ISL_923048, EPI_ISL_923052, EPI_ISL_923053, EPI_ISL_923054, EPI_ISL_923055, EPI_ISL_923056, EPI_ISL_923057, EPI_ISL_923058, EPI_ISL_923059, EPI_ISL_923060, EPI_ISL_923061, EPI_ISL_923062, EPI_ISL_923063, EPI_ISL_923064, EPI_ISL_923065, EPI_ISL_923066, EPI_ISL_923067, EPI_ISL_923068, EPI_ISL_923069, EPI_ISL_923070, EPI_ISL_923071, EPI_ISL_923073, EPI_ISL_923074, EPI_ISL_923075, EPI_ISL_923076, EPI_ISL_923084, EPI_ISL_923088, EPI_ISL_923089, EPI_ISL_923091, EPI_ISL_923092, EPI_ISL_923093, EPI_ISL_923095, EPI_ISL_923096, EPI_ISL_923097, EPI_ISL_923098, EPI_ISL_923099, EPI_ISL_923100, EPI_ISL_923103, EPI_ISL_923104, EPI_ISL_923105, EPI_ISL_923106, EPI_ISL_923107, EPI_ISL_923108, EPI_ISL_923109, EPI_ISL_923111, EPI_ISL_923112, EPI_ISL_923115, EPI_ISL_923116, EPI_ISL_923118, EPI_ISL_923121, EPI_ISL_923123, EPI_ISL_923125, EPI_ISL_923126, EPI_ISL_923127, EPI_ISL_923129, EPI_ISL_923130, EPI_ISL_923131, EPI_ISL_923133, EPI_ISL_923134, EPI_ISL_923135, EPI_ISL_923137, EPI_ISL_923138, EPI_ISL_923139, EPI_ISL_923140, EPI_ISL_923141, EPI_ISL_923144, EPI_ISL_923145, EPI_ISL_923148, EPI_ISL_923149, EPI_ISL_923150, EPI_ISL_923151, EPI_ISL_923152, EPI_ISL_923154, EPI_ISL_923157, EPI_ISL_923161, EPI_ISL_923163, EPI_ISL_923164, EPI_ISL_923165, EPI_ISL_923166, EPI_ISL_923167 | | | |
| see above | Wales Specialist Virology Centre Sequencing lab: Pathogen Genomics Unit | Public Health Wales Microbiology Cardiff Wales Specialist Virology Centre | Catherine Moore, Johnathan Evans, Laura Gifford, Malorie Perry, Simon Cottrell, Angela Marchbank, Alec Birchley, Alexander Adams, Amy Gaskin, Bree Gatica-Wilcox, Jason Coombes, Joel Southgate, Lauren Gilbert, Lee Graham, Nicole Pacchiarini, Sara Kumziene-Summerhayes, Sarah Taylor, Sophie Jones, Sara Rey, Matthew Bull, Joanne Watkins, Sally Corden, Tom Connor |
| EPI_ISL_923320, EPI_ISL_923321, EPI_ISL_923322, EPI_ISL_923323, EPI_ISL_923324, EPI_ISL_923325, EPI_ISL_923326, EPI_ISL_923327, EPI_ISL_923329, EPI_ISL_923331, EPI_ISL_923332, EPI_ISL_923333, EPI_ISL_923335, EPI_ISL_923336, EPI_ISL_923337, EPI_ISL_923338, EPI_ISL_923339, EPI_ISL_923340, EPI_ISL_923341, EPI_ISL_923342, EPI_ISL_923343, EPI_ISL_923344, EPI_ISL_923346, EPI_ISL_923347, EPI_ISL_923348, EPI_ISL_923350, EPI_ISL_923352, EPI_ISL_923353, EPI_ISL_923354, EPI_ISL_923436, EPI_ISL_923598 | | | |
| see above | Centre for Enzyme Innovation, University of Portsmouth / Translational Research Laboratory, Portsmouth Hospitals NHS Trust | COVID-19 Genomics UK (COG-UK) Consortium | Angela Beckett,Salman Goudarzi,Christopher Fearn,Kate Cook,Katie Loveson,Sharon Glaysher,Scott Elliott,Samuel Robson |
| EPI_ISL_924098, EPI_ISL_924102, EPI_ISL_924106, EPI_ISL_924114, EPI_ISL_924123, EPI_ISL_924159, EPI_ISL_924169, EPI_ISL_924170, EPI_ISL_924174, EPI_ISL_924189, EPI_ISL_924204, EPI_ISL_924206, EPI_ISL_924207, EPI_ISL_924208, EPI_ISL_924253, EPI_ISL_924255, EPI_ISL_924256, EPI_ISL_924266, EPI_ISL_924284, EPI_ISL_924291, EPI_ISL_924299, EPI_ISL_924307, EPI_ISL_924310, EPI_ISL_924311, EPI_ISL_924312, EPI_ISL_924314, EPI_ISL_924337, EPI_ISL_924343, EPI_ISL_924345, EPI_ISL_924348, EPI_ISL_924358, EPI_ISL_924360, EPI_ISL_924363, EPI_ISL_924364, EPI_ISL_924380, EPI_ISL_924386, EPI_ISL_924388, EPI_ISL_924394, EPI_ISL_924413 | | | |

| | | | |
|---|---|---|---|
| see above | Virology Department, Sheffield Teaching Hospitals NHS Foundation Trust/Department of Infection, Immunity and Cardiovascular Disease, The Medical School, University of Sheffield | COVID-19 Genomics UK (COG-UK) Consortium | Thushan de Silva, Matthew Parker, Nikki Smith, Adri Angyal, Rebecca Brown, Luke Green, Rachel Tucker, Paul Parsons, Danielle Groves, Katie Johnson, Laura Carrilero, Alex Keeley, Dave Partridge, Matthew Wyles, Benjamin Lindsey, Mehmet Yavuz, Mohammad Raza, Cariad Evans |
| EPI_ISL_924652, EPI_ISL_924654, EPI_ISL_924655, EPI_ISL_924656, EPI_ISL_924658, EPI_ISL_924659, EPI_ISL_924660, EPI_ISL_924662, EPI_ISL_924663, EPI_ISL_924665, EPI_ISL_924670, EPI_ISL_924673, EPI_ISL_924678, EPI_ISL_924679, EPI_ISL_924680, EPI_ISL_924681, EPI_ISL_924682, EPI_ISL_924686, EPI_ISL_924687, EPI_ISL_924703, EPI_ISL_924710, EPI_ISL_924711, EPI_ISL_924829, EPI_ISL_924831, EPI_ISL_924833, EPI_ISL_924835, EPI_ISL_924837, EPI_ISL_924840, EPI_ISL_924843, EPI_ISL_924844, EPI_ISL_924847, EPI_ISL_924852, EPI_ISL_924854, EPI_ISL_924856, EPI_ISL_924861, EPI_ISL_924862, EPI_ISL_924866, EPI_ISL_924868, EPI_ISL_924870, EPI_ISL_924872, EPI_ISL_924874, EPI_ISL_924875, EPI_ISL_924878, EPI_ISL_924881, EPI_ISL_924885, EPI_ISL_924888, EPI_ISL_924889 | | | |
| see above | Bioinformatics and Biostatistics Lab, Advanced Sequencing Facility | COVID-19 Genomics UK (COG-UK) Consortium | Aengus Stewart,Jerome Nicod,Chelsea Sawyer,Laura Cubitt,Harshil Patel,Margaret Crawford |
| EPI_ISL_925173 | Laboratorio de Referencia Nacional de Enteropatógenos. Instituto Nacional de Salud del Perú | Laboratorio de Referencia Nacional de Enteropatógenos. Instituto Nacional de Salud del Perú | Ronnie Gavilan Chavez, Junior Caro Castro, Willi Quino Sifuentes, Veronica Hurtado Vela, Iris Silva Molina, Fiorella Orellana Peralta, |
| EPI_ISL_925917, EPI_ISL_925919, EPI_ISL_925920, EPI_ISL_925921, EPI_ISL_925922, EPI_ISL_925925, EPI_ISL_925926, EPI_ISL_925927, EPI_ISL_925928, EPI_ISL_925930, EPI_ISL_925931, EPI_ISL_925932, EPI_ISL_925934, EPI_ISL_925935, EPI_ISL_925936, EPI_ISL_925938, EPI_ISL_925941, EPI_ISL_925943, EPI_ISL_925945, EPI_ISL_925946, EPI_ISL_925947, EPI_ISL_925948, EPI_ISL_925951, EPI_ISL_925954, EPI_ISL_925956, EPI_ISL_925957, EPI_ISL_925959, EPI_ISL_925960, EPI_ISL_925962, EPI_ISL_925963, EPI_ISL_925964, EPI_ISL_925966, EPI_ISL_925972, EPI_ISL_925973, EPI_ISL_925975, EPI_ISL_925977, EPI_ISL_925978, EPI_ISL_925980, EPI_ISL_925981, EPI_ISL_925982, EPI_ISL_925984, EPI_ISL_925986, EPI_ISL_925989, EPI_ISL_925991, EPI_ISL_925993, EPI_ISL_925994, EPI_ISL_925995, EPI_ISL_925996, EPI_ISL_925997, EPI_ISL_925998, EPI_ISL_925999, EPI_ISL_926001, EPI_ISL_926004, EPI_ISL_926006, EPI_ISL_926009, EPI_ISL_926013, EPI_ISL_926014, EPI_ISL_926016, EPI_ISL_926019, EPI_ISL_926022, EPI_ISL_926023, EPI_ISL_926026, EPI_ISL_926027, EPI_ISL_926028, EPI_ISL_926032, EPI_ISL_926033, EPI_ISL_926034, EPI_ISL_926036, EPI_ISL_926039, EPI_ISL_926040, EPI_ISL_926042, EPI_ISL_926049, EPI_ISL_926052, EPI_ISL_926054, EPI_ISL_926056, EPI_ISL_926057, EPI_ISL_926058, EPI_ISL_926060, EPI_ISL_926061, EPI_ISL_926062, EPI_ISL_926066, EPI_ISL_926067, EPI_ISL_926068, EPI_ISL_926069, EPI_ISL_926072, EPI_ISL_926074, EPI_ISL_926076, EPI_ISL_926077, EPI_ISL_926078, EPI_ISL_926080, EPI_ISL_926081, EPI_ISL_926084, EPI_ISL_926085, EPI_ISL_926087, EPI_ISL_926089, EPI_ISL_926091, EPI_ISL_926092, EPI_ISL_926094, EPI_ISL_926097, EPI_ISL_926098, EPI_ISL_926100, EPI_ISL_926101, EPI_ISL_926102, EPI_ISL_926105, EPI_ISL_926106, EPI_ISL_926108, EPI_ISL_926109, EPI_ISL_926110, EPI_ISL_926111, EPI_ISL_926112, EPI_ISL_926115, EPI_ISL_926116, EPI_ISL_926119, EPI_ISL_926121, EPI_ISL_926123, EPI_ISL_926125, EPI_ISL_926127, EPI_ISL_926130, EPI_ISL_926131, EPI_ISL_926132, EPI_ISL_926133, EPI_ISL_926135, EPI_ISL_926136, EPI_ISL_926138, EPI_ISL_926141, EPI_ISL_926143, EPI_ISL_926145, EPI_ISL_926146, EPI_ISL_926148, EPI_ISL_926149, EPI_ISL_926152, EPI_ISL_926154, EPI_ISL_926155, EPI_ISL_926159, EPI_ISL_926160, EPI_ISL_926161, EPI_ISL_926165, EPI_ISL_926167, EPI_ISL_926168, EPI_ISL_926170, EPI_ISL_926171, EPI_ISL_926172, EPI_ISL_926173, EPI_ISL_926175, EPI_ISL_926176, EPI_ISL_926179, EPI_ISL_926180, EPI_ISL_926181, EPI_ISL_926182, EPI_ISL_926184, EPI_ISL_926185, EPI_ISL_926186, EPI_ISL_926188, EPI_ISL_926189, EPI_ISL_926190, EPI_ISL_926191, EPI_ISL_926194, EPI_ISL_926195, EPI_ISL_926196, EPI_ISL_926197, EPI_ISL_926198, EPI_ISL_926200, EPI_ISL_926202, EPI_ISL_926203, EPI_ISL_926204, EPI_ISL_926205, EPI_ISL_926206, EPI_ISL_926207, EPI_ISL_926208, EPI_ISL_926212, EPI_ISL_926213, EPI_ISL_926214, EPI_ISL_926215, EPI_ISL_926217, EPI_ISL_926219, EPI_ISL_926220, EPI_ISL_926223, EPI_ISL_926228, EPI_ISL_926229, EPI_ISL_926230, EPI_ISL_926231, EPI_ISL_926232, EPI_ISL_926234, EPI_ISL_926236, EPI_ISL_926239, EPI_ISL_926240, EPI_ISL_926241, EPI_ISL_926242, EPI_ISL_926244, EPI_ISL_926246, EPI_ISL_926248, EPI_ISL_926249, EPI_ISL_926250, EPI_ISL_926251, EPI_ISL_926253, EPI_ISL_926254, EPI_ISL_926256, EPI_ISL_926258, EPI_ISL_926260, EPI_ISL_926262, EPI_ISL_926263, EPI_ISL_926264, EPI_ISL_926266, EPI_ISL_926267, EPI_ISL_926268, EPI_ISL_926269, EPI_ISL_926271, EPI_ISL_926273, EPI_ISL_926276, EPI_ISL_926277, EPI_ISL_926278, EPI_ISL_926279, EPI_ISL_926280, EPI_ISL_926284, EPI_ISL_926285, EPI_ISL_926287, EPI_ISL_926288, EPI_ISL_926290, EPI_ISL_926293, EPI_ISL_926294, EPI_ISL_926296, EPI_ISL_926298, EPI_ISL_926300, EPI_ISL_926301, EPI_ISL_926302, EPI_ISL_926305, EPI_ISL_926306, EPI_ISL_926307, EPI_ISL_926309, EPI_ISL_926310, EPI_ISL_926312, EPI_ISL_926314, EPI_ISL_926317, EPI_ISL_926318, EPI_ISL_926319, EPI_ISL_926322, EPI_ISL_926325, EPI_ISL_926326, EPI_ISL_926329, EPI_ISL_926331, EPI_ISL_926332, EPI_ISL_926333, EPI_ISL_926335, EPI_ISL_926336, EPI_ISL_926338, EPI_ISL_926340, EPI_ISL_926341, EPI_ISL_926343, EPI_ISL_926344, EPI_ISL_926349, EPI_ISL_926351, EPI_ISL_926352, EPI_ISL_926353, EPI_ISL_926354, EPI_ISL_926355, EPI_ISL_926356, EPI_ISL_926358, EPI_ISL_926362, EPI_ISL_926363, EPI_ISL_926366, EPI_ISL_926368, EPI_ISL_926369, EPI_ISL_926370, EPI_ISL_926371, EPI_ISL_926373, EPI_ISL_926375, EPI_ISL_926376, EPI_ISL_926377, EPI_ISL_926378, EPI_ISL_926379, EPI_ISL_926382, EPI_ISL_926383, EPI_ISL_926384, EPI_ISL_926387, EPI_ISL_926391, EPI_ISL_926392, EPI_ISL_926393, EPI_ISL_926400, EPI_ISL_926401, EPI_ISL_926403, EPI_ISL_926405, EPI_ISL_926408, EPI_ISL_926409, EPI_ISL_926411, EPI_ISL_926412, EPI_ISL_926413, EPI_ISL_926414, EPI_ISL_926415, EPI_ISL_926417, EPI_ISL_926418, EPI_ISL_926421, EPI_ISL_926424, EPI_ISL_926425, EPI_ISL_926426, EPI_ISL_926427, EPI_ISL_926429, EPI_ISL_926430, EPI_ISL_926431, EPI_ISL_926433, EPI_ISL_926434, EPI_ISL_926443, EPI_ISL_926444, EPI_ISL_926445, EPI_ISL_926447, EPI_ISL_926448, EPI_ISL_926449, EPI_ISL_926450, EPI_ISL_926453, EPI_ISL_926454, EPI_ISL_926456, EPI_ISL_926457, EPI_ISL_926458, EPI_ISL_926460, EPI_ISL_926462, EPI_ISL_926463, EPI_ISL_926464, EPI_ISL_926467, EPI_ISL_926468, EPI_ISL_926470, EPI_ISL_926471, EPI_ISL_926472, EPI_ISL_926473, EPI_ISL_926474, EPI_ISL_926475, EPI_ISL_926477, EPI_ISL_926480, EPI_ISL_926481, EPI_ISL_926486, EPI_ISL_926488, EPI_ISL_926490, EPI_ISL_926493, EPI_ISL_926495, EPI_ISL_926496, EPI_ISL_926497, EPI_ISL_926498, EPI_ISL_926500, EPI_ISL_926501, EPI_ISL_926502, EPI_ISL_926503, EPI_ISL_926506, EPI_ISL_926507, EPI_ISL_926508, EPI_ISL_926509, EPI_ISL_926510, EPI_ISL_926511, EPI_ISL_926513, EPI_ISL_926514, EPI_ISL_926516, EPI_ISL_926518, EPI_ISL_926519, EPI_ISL_926523, EPI_ISL_926524, EPI_ISL_926526, EPI_ISL_926527, EPI_ISL_926529, EPI_ISL_926530, EPI_ISL_926531, EPI_ISL_926533, EPI_ISL_926534, EPI_ISL_926540, EPI_ISL_926542, EPI_ISL_926543, EPI_ISL_926544, EPI_ISL_926545, EPI_ISL_926548, EPI_ISL_926550, EPI_ISL_926554, EPI_ISL_926556, EPI_ISL_926557, EPI_ISL_926561, EPI_ISL_926562, EPI_ISL_926563, EPI_ISL_926565, EPI_ISL_926568, EPI_ISL_926571, EPI_ISL_926572, EPI_ISL_926574, EPI_ISL_926577, EPI_ISL_926578, EPI_ISL_926579, EPI_ISL_926580, EPI_ISL_926581, EPI_ISL_926582, EPI_ISL_926583, EPI_ISL_926584, EPI_ISL_926585, EPI_ISL_926589, EPI_ISL_926590, EPI_ISL_926592, EPI_ISL_926595, EPI_ISL_926597, EPI_ISL_926599, EPI_ISL_926600, EPI_ISL_926602, EPI_ISL_926604, EPI_ISL_926606, EPI_ISL_926608, EPI_ISL_926610, EPI_ISL_926611, EPI_ISL_926612, EPI_ISL_926613, EPI_ISL_926614, EPI_ISL_926618, EPI_ISL_926623, EPI_ISL_926624, EPI_ISL_926625, EPI_ISL_926626, EPI_ISL_926629, EPI_ISL_926630, EPI_ISL_926634, EPI_ISL_926635, EPI_ISL_926636, EPI_ISL_926637, EPI_ISL_926638, EPI_ISL_926639, EPI_ISL_926640, EPI_ISL_926641, EPI_ISL_926644, EPI_ISL_926645, EPI_ISL_926646, EPI_ISL_926647, EPI_ISL_926650, EPI_ISL_926652, EPI_ISL_926653, EPI_ISL_926654, EPI_ISL_926657, EPI_ISL_926658, EPI_ISL_926659, EPI_ISL_926660, EPI_ISL_926662, EPI_ISL_926664, EPI_ISL_926665, EPI_ISL_926669, EPI_ISL_926670, EPI_ISL_926672, EPI_ISL_926673, EPI_ISL_926675, EPI_ISL_926677, EPI_ISL_926678, EPI_ISL_926680, EPI_ISL_926682, EPI_ISL_926683, EPI_ISL_926685, EPI_ISL_926686, EPI_ISL_926688, EPI_ISL_926689, EPI_ISL_926690, EPI_ISL_926691, EPI_ISL_926694, EPI_ISL_926695, EPI_ISL_926696, EPI_ISL_926699, EPI_ISL_926700, EPI_ISL_926701, EPI_ISL_926702, EPI_ISL_926703, EPI_ISL_926704, EPI_ISL_926705, EPI_ISL_926706, EPI_ISL_926707, EPI_ISL_926709, EPI_ISL_926711, EPI_ISL_926712, EPI_ISL_926715, EPI_ISL_926718, EPI_ISL_926719, EPI_ISL_926720, EPI_ISL_926723, EPI_ISL_926724, EPI_ISL_926725, EPI_ISL_926726, EPI_ISL_926728, EPI_ISL_926729, EPI_ISL_926731, EPI_ISL_926732, EPI_ISL_926733, EPI_ISL_926734, EPI_ISL_926736, EPI_ISL_926737, EPI_ISL_926738, EPI_ISL_926739, EPI_ISL_926740, EPI_ISL_926741, EPI_ISL_926742, EPI_ISL_926743, EPI_ISL_926744, EPI_ISL_926745, EPI_ISL_926746, EPI_ISL_926747, EPI_ISL_926748, EPI_ISL_926749, EPI_ISL_926750, EPI_ISL_926753, EPI_ISL_926754, EPI_ISL_926755, EPI_ISL_926756, EPI_ISL_926759, EPI_ISL_926761, EPI_ISL_926762, EPI_ISL_926764, EPI_ISL_926766, EPI_ISL_926769, EPI_ISL_926771, EPI_ISL_926772, EPI_ISL_926774, EPI_ISL_926777, EPI_ISL_926779, EPI_ISL_926780, EPI_ISL_926781, EPI_ISL_926782, EPI_ISL_926783, EPI_ISL_926785, EPI_ISL_926788, EPI_ISL_926790, EPI_ISL_926792, EPI_ISL_926796, EPI_ISL_926798, EPI_ISL_926802, EPI_ISL_926803, EPI_ISL_926804, EPI_ISL_926807, EPI_ISL_926808, EPI_ISL_926812, EPI_ISL_926814, EPI_ISL_926815, EPI_ISL_926816, EPI_ISL_926819, EPI_ISL_926822, EPI_ISL_926823, EPI_ISL_926824, EPI_ISL_926828, EPI_ISL_926829, EPI_ISL_926831, EPI_ISL_926832, EPI_ISL_926834, EPI_ISL_926835, EPI_ISL_926837, EPI_ISL_926839, EPI_ISL_926840, EPI_ISL_926841, EPI_ISL_926842, EPI_ISL_926843, EPI_ISL_926845, EPI_ISL_926846, EPI_ISL_926847, EPI_ISL_926849, EPI_ISL_926850, EPI_ISL_926854, EPI_ISL_926856, EPI_ISL_926857, EPI_ISL_926858, EPI_ISL_926860, EPI_ISL_926861, EPI_ISL_926863, EPI_ISL_926864, EPI_ISL_926867, EPI_ISL_926868, EPI_ISL_926869, EPI_ISL_926871, EPI_ISL_926874, EPI_ISL_926875, EPI_ISL_926876, EPI_ISL_926877, EPI_ISL_926879, EPI_ISL_926883, EPI_ISL_926884, EPI_ISL_926886, EPI_ISL_926887, EPI_ISL_926888, EPI_ISL_926889, EPI_ISL_926890, EPI_ISL_926891, EPI_ISL_926892, EPI_ISL_926894, EPI_ISL_926895, EPI_ISL_926896, EPI_ISL_926897, EPI_ISL_926898, EPI_ISL_926901, EPI_ISL_926904, EPI_ISL_926905, EPI_ISL_926906, EPI_ISL_926907, EPI_ISL_926908, EPI_ISL_926910, EPI_ISL_926912, EPI_ISL_926913, EPI_ISL_926918, EPI_ISL_926919, EPI_ISL_926920, EPI_ISL_926921, EPI_ISL_926922, EPI_ISL_926924, EPI_ISL_926925, EPI_ISL_926926, EPI_ISL_926928, EPI_ISL_926929, EPI_ISL_926930, EPI_ISL_926935, EPI_ISL_926938, EPI_ISL_926939, EPI_ISL_926943, EPI_ISL_926944, EPI_ISL_926946, EPI_ISL_926947, EPI_ISL_926948, EPI_ISL_926950, EPI_ISL_926951, EPI_ISL_926952, EPI_ISL_926953, EPI_ISL_926954, EPI_ISL_926956, EPI_ISL_926957, EPI_ISL_926958, EPI_ISL_926961, EPI_ISL_926963, EPI_ISL_926966, EPI_ISL_926969, EPI_ISL_926971, EPI_ISL_926974, EPI_ISL_926975, EPI_ISL_926977, EPI_ISL_926978, EPI_ISL_926983, EPI_ISL_926984, EPI_ISL_926985, EPI_ISL_926986, EPI_ISL_926988, EPI_ISL_926991, EPI_ISL_926992, EPI_ISL_926993, EPI_ISL_926994, EPI_ISL_926996, EPI_ISL_926997, EPI_ISL_926998, EPI_ISL_927000, EPI_ISL_927001, EPI_ISL_927005, EPI_ISL_927006, EPI_ISL_927007, EPI_ISL_927008, EPI_ISL_927009, EPI_ISL_927010, EPI_ISL_927012, EPI_ISL_927013, EPI_ISL_927015, EPI_ISL_927016, EPI_ISL_927017, EPI_ISL_927019, EPI_ISL_927021, EPI_ISL_927022, EPI_ISL_927023, EPI_ISL_927026, EPI_ISL_927027, EPI_ISL_927029, EPI_ISL_927030, EPI_ISL_927031, EPI_ISL_927033, EPI_ISL_927034, EPI_ISL_927035, EPI_ISL_927037, EPI_ISL_927038, EPI_ISL_927040, EPI_ISL_927041, EPI_ISL_927042, EPI_ISL_927043, EPI_ISL_927044, EPI_ISL_927045, EPI_ISL_927046, EPI_ISL_927048, EPI_ISL_927049, EPI_ISL_927050, EPI_ISL_927051, EPI_ISL_927052, EPI_ISL_927054, EPI_ISL_927055, EPI_ISL_927059, EPI_ISL_927062, EPI_ISL_927064, EPI_ISL_927065, EPI_ISL_927067, EPI_ISL_927068, EPI_ISL_927071, EPI_ISL_927073, EPI_ISL_927074, EPI_ISL_927076, EPI_ISL_927077, EPI_ISL_927078, EPI_ISL_927080, EPI_ISL_927083, EPI_ISL_927085, EPI_ISL_927086, EPI_ISL_927087, EPI_ISL_927088, EPI_ISL_927090, EPI_ISL_927093, EPI_ISL_927094, EPI_ISL_927096, EPI_ISL_927098, EPI_ISL_927102, EPI_ISL_927104, EPI_ISL_927105, EPI_ISL_927108, EPI_ISL_927109, EPI_ISL_927110, EPI_ISL_927111, EPI_ISL_927112, EPI_ISL_927113, EPI_ISL_927115, EPI_ISL_927116, EPI_ISL_927120, EPI_ISL_927121, EPI_ISL_927124, EPI_ISL_927126, EPI_ISL_927129, EPI_ISL_927130, EPI_ISL_927131, EPI_ISL_927135, EPI_ISL_927136, EPI_ISL_927137, EPI_ISL_927138, EPI_ISL_927139, EPI_ISL_927142, EPI_ISL_927143, EPI_ISL_927146, EPI_ISL_927147, EPI_ISL_927150, EPI_ISL_927152, EPI_ISL_927154, EPI_ISL_927155, EPI_ISL_927156, EPI_ISL_927158, EPI_ISL_927159, EPI_ISL_927163, EPI_ISL_927165, EPI_ISL_927166, EPI_ISL_927169, EPI_ISL_927170, EPI_ISL_927171, EPI_ISL_927172, EPI_ISL_927173, EPI_ISL_927175, EPI_ISL_927178, EPI_ISL_927179, EPI_ISL_927180, EPI_ISL_927183, EPI_ISL_927187, EPI_ISL_927188, EPI_ISL_927191, EPI_ISL_927193, EPI_ISL_927195, EPI_ISL_927198, EPI_ISL_927201, EPI_ISL_927202, EPI_ISL_927204, EPI_ISL_927206, EPI_ISL_927208, EPI_ISL_927210, EPI_ISL_927212, EPI_ISL_927214, EPI_ISL_927217, EPI_ISL_927218, EPI_ISL_927219, EPI_ISL_927220, EPI_ISL_927221, EPI_ISL_927225, EPI_ISL_927226, EPI_ISL_927227, EPI_ISL_927229, EPI_ISL_927230, EPI_ISL_927231, EPI_ISL_927232, EPI_ISL_927233, EPI_ISL_927234, EPI_ISL_927235, EPI_ISL_927237, EPI_ISL_927238, EPI_ISL_927242, EPI_ISL_927244, EPI_ISL_927245, EPI_ISL_927250, EPI_ISL_927251, EPI_ISL_927253, EPI_ISL_927254, EPI_ISL_927256, EPI_ISL_927260, EPI_ISL_927264, EPI_ISL_927265, EPI_ISL_927267, EPI_ISL_927268, EPI_ISL_927269, EPI_ISL_927271, EPI_ISL_927272, EPI_ISL_927273, EPI_ISL_927275, EPI_ISL_927279, EPI_ISL_927280, EPI_ISL_927281, EPI_ISL_927282, EPI_ISL_927285, EPI_ISL_927286, EPI_ISL_927289, EPI_ISL_927290, EPI_ISL_927292, EPI_ISL_927294, EPI_ISL_927296, EPI_ISL_927297, EPI_ISL_927303, EPI_ISL_927304, EPI_ISL_927305, EPI_ISL_927307, EPI_ISL_927308, EPI_ISL_927310, EPI_ISL_927312, EPI_ISL_927313, EPI_ISL_927315, EPI_ISL_927316, EPI_ISL_927317, EPI_ISL_927318, EPI_ISL_927322, EPI_ISL_927323, EPI_ISL_927324, EPI_ISL_927326, EPI_ISL_927328, EPI_ISL_927329, EPI_ISL_927330, EPI_ISL_927331, EPI_ISL_927332, EPI_ISL_927333, EPI_ISL_927334, EPI_ISL_927335, EPI_ISL_927336, EPI_ISL_927339, EPI_ISL_927340, EPI_ISL_927343, EPI_ISL_927344, EPI_ISL_927349, EPI_ISL_927350, EPI_ISL_927354, EPI_ISL_927355, EPI_ISL_927357, EPI_ISL_927358, EPI_ISL_927359, EPI_ISL_927360, EPI_ISL_927361, EPI_ISL_927362, EPI_ISL_927363, EPI_ISL_927365, EPI_ISL_927368, EPI_ISL_927369, EPI_ISL_927372, EPI_ISL_927373, EPI_ISL_927374, EPI_ISL_927376, EPI_ISL_927377, EPI_ISL_927378, EPI_ISL_927379, EPI_ISL_927380, EPI_ISL_927381, EPI_ISL_927382, EPI_ISL_927383, EPI_ISL_927386, EPI_ISL_927387, EPI_ISL_927388, EPI_ISL_927390, EPI_ISL_927391, EPI_ISL_927392, EPI_ISL_927393, EPI_ISL_927394, EPI_ISL_927395, EPI_ISL_927396, EPI_ISL_927401, EPI_ISL_927403, EPI_ISL_927404, EPI_ISL_927405, EPI_ISL_927406, EPI_ISL_927407, EPI_ISL_927409, EPI_ISL_927410, EPI_ISL_927412, EPI_ISL_927413, EPI_ISL_927414, EPI_ISL_927415, EPI_ISL_927416, EPI_ISL_927417, EPI_ISL_927419, EPI_ISL_927421, EPI_ISL_927426, EPI_ISL_927427, EPI_ISL_927429, EPI_ISL_927430, EPI_ISL_927434, EPI_ISL_927435, EPI_ISL_927436, EPI_ISL_927437, EPI_ISL_927438, EPI_ISL_927442, EPI_ISL_927443, EPI_ISL_927445, EPI_ISL_927446, EPI_ISL_927448, EPI_ISL_927449, EPI_ISL_927450, EPI_ISL_927451, EPI_ISL_927452, EPI_ISL_927453, EPI_ISL_927454, EPI_ISL_927457, EPI_ISL_927458, EPI_ISL_927462, EPI_ISL_927466, EPI_ISL_927467, EPI_ISL_927468, EPI_ISL_927469, EPI_ISL_927471, EPI_ISL_927477, EPI_ISL_927478, EPI_ISL_927481, EPI_ISL_927482, EPI_ISL_927487, EPI_ISL_927488, EPI_ISL_927490, EPI_ISL_927494, EPI_ISL_927495, EPI_ISL_927496, EPI_ISL_927497, EPI_ISL_927499, EPI_ISL_927500, EPI_ISL_927503, EPI_ISL_927506, EPI_ISL_927509, EPI_ISL_927511, EPI_ISL_927512, EPI_ISL_927513, EPI_ISL_927514, EPI_ISL_927515, EPI_ISL_927516, EPI_ISL_927517, EPI_ISL_927518, EPI_ISL_927520, EPI_ISL_927521, EPI_ISL_927523, EPI_ISL_927524, EPI_ISL_927527, EPI_ISL_927528, EPI_ISL_927529, EPI_ISL_927530, EPI_ISL_927531, EPI_ISL_927532, EPI_ISL_927533, EPI_ISL_927535, EPI_ISL_927537, EPI_ISL_927539, EPI_ISL_927540, EPI_ISL_927541, EPI_ISL_927542, EPI_ISL_927545, EPI_ISL_927547, EPI_ISL_927548, EPI_ISL_927549, EPI_ISL_927552, EPI_ISL_927556, EPI_ISL_927557, EPI_ISL_927560, EPI_ISL_927561, EPI_ISL_927562, EPI_ISL_927563, EPI_ISL_927564, EPI_ISL_927567, EPI_ISL_927572, EPI_ISL_927574, EPI_ISL_927575, EPI_ISL_927576, EPI_ISL_927579, EPI_ISL_927581, EPI_ISL_927582, EPI_ISL_927583, EPI_ISL_927584, EPI_ISL_927585, EPI_ISL_927586, EPI_ISL_927588, EPI_ISL_927590, EPI_ISL_927591, EPI_ISL_927592, EPI_ISL_927594, EPI_ISL_927597, EPI_ISL_927604, EPI_ISL_927605, EPI_ISL_927606, EPI_ISL_927607, EPI_ISL_927609, EPI_ISL_927611, EPI_ISL_927618, EPI_ISL_927619, EPI_ISL_927620, EPI_ISL_927625, EPI_ISL_927629, EPI_ISL_927631, EPI_ISL_927632, EPI_ISL_927635, EPI_ISL_927638, EPI_ISL_927640, EPI_ISL_927642, EPI_ISL_927645, EPI_ISL_927646, EPI_ISL_927647, EPI_ISL_927649, EPI_ISL_927650, EPI_ISL_927654, EPI_ISL_927655, EPI_ISL_927656, EPI_ISL_927661, EPI_ISL_927663, EPI_ISL_927664, EPI_ISL_927666, EPI_ISL_927667, EPI_ISL_927670, EPI_ISL_927672, EPI_ISL_927676, EPI_ISL_927677, EPI_ISL_927678, EPI_ISL_927680, EPI_ISL_927681, EPI_ISL_927684, EPI_ISL_927686, EPI_ISL_927687, EPI_ISL_927688, EPI_ISL_927690, EPI_ISL_927691, EPI_ISL_927692, EPI_ISL_927693, EPI_ISL_927694, EPI_ISL_927696, EPI_ISL_927697, EPI_ISL_927699, EPI_ISL_927700, EPI_ISL_927701, EPI_ISL_927702, EPI_ISL_927704, EPI_ISL_927710, EPI_ISL_927712, EPI_ISL_927713, EPI_ISL_927714, EPI_ISL_927715, EPI_ISL_927717, EPI_ISL_927718, EPI_ISL_927719, EPI_ISL_927720, EPI_ISL_927722, EPI_ISL_927724, EPI_ISL_927725, EPI_ISL_927728, EPI_ISL_927729, EPI_ISL_927731, EPI_ISL_927734, EPI_ISL_927735, EPI_ISL_927736, EPI_ISL_927737, EPI_ISL_927738, EPI_ISL_927741, EPI_ISL_927742, EPI_ISL_927743, EPI_ISL_927745, EPI_ISL_927746, EPI_ISL_927747, EPI_ISL_927749, EPI_ISL_927750, EPI_ISL_927753, EPI_ISL_927754, EPI_ISL_927757, EPI_ISL_927759, EPI_ISL_927760, EPI_ISL_927761, EPI_ISL_927762, EPI_ISL_927764, EPI_ISL_927765, EPI_ISL_927766, EPI_ISL_927767, EPI_ISL_927768, EPI_ISL_927772, EPI_ISL_927773, EPI_ISL_927775, EPI_ISL_927776, EPI_ISL_927777, EPI_ISL_927779, EPI_ISL_927780, EPI_ISL_927783, EPI_ISL_927785, EPI_ISL_927786, EPI_ISL_927790, EPI_ISL_927791, EPI_ISL_927793, EPI_ISL_927795, EPI_ISL_927796, EPI_ISL_927798, EPI_ISL_927799, EPI_ISL_927800, EPI_ISL_927801, EPI_ISL_927802, EPI_ISL_927803, EPI_ISL_927804, EPI_ISL_927805, EPI_ISL_927806, EPI_ISL_927807, EPI_ISL_927809, EPI_ISL_927810, EPI_ISL_927811, EPI_ISL_927812, EPI_ISL_927814, EPI_ISL_927815, EPI_ISL_927816, EPI_ISL_927817, EPI_ISL_927819, EPI_ISL_927821, EPI_ISL_927823, EPI_ISL_927825, EPI_ISL_927827, EPI_ISL_927828, EPI_ISL_927829, EPI_ISL_927834, EPI_ISL_927835, EPI_ISL_927836, EPI_ISL_927837, EPI_ISL_927838, EPI_ISL_927841, EPI_ISL_927842, EPI_ISL_927844, EPI_ISL_927848, EPI_ISL_927850, EPI_ISL_927851, EPI_ISL_927852, EPI_ISL_927853, EPI_ISL_927854, EPI_ISL_927857, EPI_ISL_927865, EPI_ISL_927867, EPI_ISL_927870, EPI_ISL_927874, EPI_ISL_927875, EPI_ISL_927884, EPI_ISL_927885, EPI_ISL_927886, EPI_ISL_927889, EPI_ISL_927890, EPI_ISL_927892, EPI_ISL_927893, EPI_ISL_927894, EPI_ISL_927898, EPI_ISL_927899, EPI_ISL_927900, EPI_ISL_927903, EPI_ISL_927906, EPI_ISL_927908, EPI_ISL_927909, EPI_ISL_927910, EPI_ISL_927912, EPI_ISL_927913, EPI_ISL_927914, EPI_ISL_927915, EPI_ISL_927916, EPI_ISL_927917, EPI_ISL_927920, EPI_ISL_927921, EPI_ISL_927922, EPI_ISL_927923, EPI_ISL_927924, EPI_ISL_927929, EPI_ISL_927931, EPI_ISL_927935, EPI_ISL_927936, EPI_ISL_927937, EPI_ISL_927938, EPI_ISL_927939, EPI_ISL_927940, EPI_ISL_927941, EPI_ISL_927942, EPI_ISL_927944, EPI_ISL_927947, EPI_ISL_927948, EPI_ISL_927949, EPI_ISL_927950, EPI_ISL_927951, EPI_ISL_927954, EPI_ISL_927960, EPI_ISL_927963, EPI_ISL_927965, EPI_ISL_927966, EPI_ISL_927970, EPI_ISL_927971, EPI_ISL_927973, EPI_ISL_927974, EPI_ISL_927975, EPI_ISL_927977, EPI_ISL_927983, EPI_ISL_927985, EPI_ISL_927986, EPI_ISL_927989, EPI_ISL_927990, EPI_ISL_927992, EPI_ISL_927993, EPI_ISL_927994, EPI_ISL_927995, EPI_ISL_927997, EPI_ISL_927999, EPI_ISL_928000, EPI_ISL_928002, EPI_ISL_928004, EPI_ISL_928007, EPI_ISL_928008, EPI_ISL_928010, EPI_ISL_928011, EPI_ISL_928013, EPI_ISL_928015, EPI_ISL_928016, EPI_ISL_928017, EPI_ISL_928018, EPI_ISL_928019, EPI_ISL_928021, EPI_ISL_928023, EPI_ISL_928024, EPI_ISL_928027, EPI_ISL_928028, EPI_ISL_928029, EPI_ISL_928030, EPI_ISL_928031, EPI_ISL_928033, EPI_ISL_928034, EPI_ISL_928035, EPI_ISL_928036, EPI_ISL_928037, EPI_ISL_928039, EPI_ISL_928040, EPI_ISL_928041, EPI_ISL_928042, EPI_ISL_928045, EPI_ISL_928046, EPI_ISL_928050, EPI_ISL_928052, EPI_ISL_928053, EPI_ISL_928054, EPI_ISL_928055, EPI_ISL_928056, EPI_ISL_928057, EPI_ISL_928058, EPI_ISL_928060, EPI_ISL_928065, EPI_ISL_928069, EPI_ISL_928075, EPI_ISL_928077, EPI_ISL_928078, EPI_ISL_928079, EPI_ISL_928082, EPI_ISL_928083, EPI_ISL_928084, EPI_ISL_928086, EPI_ISL_928088, EPI_ISL_928090, EPI_ISL_928091, EPI_ISL_928093, EPI_ISL_928094, EPI_ISL_928095, EPI_ISL_928098, EPI_ISL_928099, EPI_ISL_928101, EPI_ISL_928103, EPI_ISL_928104, EPI_ISL_928107, EPI_ISL_928108, EPI_ISL_928110, EPI_ISL_928114, EPI_ISL_928115, EPI_ISL_928116, EPI_ISL_928117, EPI_ISL_928119, EPI_ISL_928122, EPI_ISL_928124, EPI_ISL_928126, EPI_ISL_928127, EPI_ISL_928128, EPI_ISL_928129, EPI_ISL_928130, EPI_ISL_928132, EPI_ISL_928133, EPI_ISL_928134, EPI_ISL_928138, EPI_ISL_928139, EPI_ISL_928140, EPI_ISL_928142, EPI_ISL_928143, EPI_ISL_928144, EPI_ISL_928145, EPI_ISL_928146, EPI_ISL_928151, EPI_ISL_928152, EPI_ISL_928153, EPI_ISL_928156, EPI_ISL_928158, EPI_ISL_928159, EPI_ISL_928160, EPI_ISL_928161, EPI_ISL_928162, EPI_ISL_928165, EPI_ISL_928168, EPI_ISL_928169, EPI_ISL_928172, EPI_ISL_928173, EPI_ISL_928174, EPI_ISL_928176, EPI_ISL_928177, EPI_ISL_928178, EPI_ISL_928180, EPI_ISL_928181, EPI_ISL_928182, EPI_ISL_928183, EPI_ISL_928184, EPI_ISL_928186, EPI_ISL_928187, EPI_ISL_928188, EPI_ISL_928189, EPI_ISL_928193, EPI_ISL_928194, EPI_ISL_928195, EPI_ISL_928196, EPI_ISL_928198, EPI_ISL_928199, EPI_ISL_928201, EPI_ISL_928203, EPI_ISL_928204, EPI_ISL_928205, EPI_ISL_928206, EPI_ISL_928207, EPI_ISL_928209, | | | |

EPI_ISL_928215, EPI_ISL_928216, EPI_ISL_928219, EPI_ISL_928220, EPI_ISL_928222, EPI_ISL_928223, EPI_ISL_928226, EPI_ISL_928227, EPI_ISL_928228, EPI_ISL_928232, EPI_ISL_928234, EPI_ISL_928235, EPI_ISL_928238, EPI_ISL_928242, EPI_ISL_928243, EPI_ISL_928244, EPI_ISL_928245, EPI_ISL_928246, EPI_ISL_928253, EPI_ISL_928254, EPI_ISL_928255, EPI_ISL_928256, EPI_ISL_928257, EPI_ISL_928260, EPI_ISL_928265, EPI_ISL_928267, EPI_ISL_928268, EPI_ISL_928269, EPI_ISL_928270, EPI_ISL_928271, EPI_ISL_928275, EPI_ISL_928277, EPI_ISL_928278, EPI_ISL_928279, EPI_ISL_928280, EPI_ISL_928283, EPI_ISL_928284, EPI_ISL_928286, EPI_ISL_928287, EPI_ISL_928288, EPI_ISL_928289, EPI_ISL_928290, EPI_ISL_928291, EPI_ISL_928292, EPI_ISL_928293, EPI_ISL_928294, EPI_ISL_928295, EPI_ISL_928296, EPI_ISL_928300, EPI_ISL_928301, EPI_ISL_928302, EPI_ISL_928303, EPI_ISL_928304, EPI_ISL_928306, EPI_ISL_928309, EPI_ISL_928310, EPI_ISL_928311, EPI_ISL_928312, EPI_ISL_928313, EPI_ISL_928314, EPI_ISL_928317, EPI_ISL_928318, EPI_ISL_928319, EPI_ISL_928321, EPI_ISL_928324, EPI_ISL_928325, EPI_ISL_928326, EPI_ISL_928328, EPI_ISL_928329, EPI_ISL_928330, EPI_ISL_928331, EPI_ISL_928334, EPI_ISL_928336, EPI_ISL_928337, EPI_ISL_928338, EPI_ISL_928339, EPI_ISL_928341, EPI_ISL_928342, EPI_ISL_928346, EPI_ISL_928347, EPI_ISL_928348, EPI_ISL_928349, EPI_ISL_928350, EPI_ISL_928351, EPI_ISL_928352, EPI_ISL_928353, EPI_ISL_928354, EPI_ISL_928357, EPI_ISL_928358, EPI_ISL_928360, EPI_ISL_928364, EPI_ISL_928366, EPI_ISL_928368, EPI_ISL_928369, EPI_ISL_928370, EPI_ISL_928371, EPI_ISL_928373, EPI_ISL_928374, EPI_ISL_928375, EPI_ISL_928376, EPI_ISL_928377, EPI_ISL_928378, EPI_ISL_928379, EPI_ISL_928381, EPI_ISL_928382, EPI_ISL_928384, EPI_ISL_928385, EPI_ISL_928387, EPI_ISL_928390, EPI_ISL_928391, EPI_ISL_928393, EPI_ISL_928394, EPI_ISL_928396, EPI_ISL_928398, EPI_ISL_928399, EPI_ISL_928400, EPI_ISL_928401, EPI_ISL_928402, EPI_ISL_928405, EPI_ISL_928407, EPI_ISL_928408, EPI_ISL_928410, EPI_ISL_928411, EPI_ISL_928414, EPI_ISL_928416, EPI_ISL_928418, EPI_ISL_928419, EPI_ISL_928421, EPI_ISL_928423, EPI_ISL_928429, EPI_ISL_928433, EPI_ISL_928435, EPI_ISL_928436, EPI_ISL_928437, EPI_ISL_928438, EPI_ISL_928440, EPI_ISL_928448, EPI_ISL_928449, EPI_ISL_928451, EPI_ISL_928452, EPI_ISL_928453, EPI_ISL_928457, EPI_ISL_928458, EPI_ISL_928460, EPI_ISL_928461, EPI_ISL_928463, EPI_ISL_928465, EPI_ISL_928469, EPI_ISL_928471, EPI_ISL_928472, EPI_ISL_928473, EPI_ISL_928474, EPI_ISL_928475, EPI_ISL_928476, EPI_ISL_928477, EPI_ISL_928478, EPI_ISL_928479, EPI_ISL_928482, EPI_ISL_928484, EPI_ISL_928486, EPI_ISL_928487, EPI_ISL_928488, EPI_ISL_928492, EPI_ISL_928496, EPI_ISL_928497, EPI_ISL_928498, EPI_ISL_928500, EPI_ISL_928501, EPI_ISL_928502, EPI_ISL_928505, EPI_ISL_928506, EPI_ISL_928507, EPI_ISL_928508, EPI_ISL_928510, EPI_ISL_928511, EPI_ISL_928512, EPI_ISL_928515, EPI_ISL_928516, EPI_ISL_928517, EPI_ISL_928518, EPI_ISL_928519, EPI_ISL_928520, EPI_ISL_928521, EPI_ISL_928526, EPI_ISL_928530, EPI_ISL_928531, EPI_ISL_928533, EPI_ISL_928534, EPI_ISL_928535, EPI_ISL_928536, EPI_ISL_928538, EPI_ISL_928540, EPI_ISL_928542, EPI_ISL_928543, EPI_ISL_928544, EPI_ISL_928545, EPI_ISL_928547, EPI_ISL_928549, EPI_ISL_928551, EPI_ISL_928554, EPI_ISL_928555, EPI_ISL_928556, EPI_ISL_928558, EPI_ISL_928559, EPI_ISL_928562, EPI_ISL_928564, EPI_ISL_928566, EPI_ISL_928567, EPI_ISL_928571, EPI_ISL_928572, EPI_ISL_928573, EPI_ISL_928574, EPI_ISL_928575, EPI_ISL_928576, EPI_ISL_928577, EPI_ISL_928578, EPI_ISL_928579, EPI_ISL_928582, EPI_ISL_928583, EPI_ISL_928584, EPI_ISL_928589, EPI_ISL_928590, EPI_ISL_928591, EPI_ISL_928592, EPI_ISL_928593, EPI_ISL_928594, EPI_ISL_928596, EPI_ISL_928597, EPI_ISL_928598, EPI_ISL_928599, EPI_ISL_928602, EPI_ISL_928603, EPI_ISL_928605, EPI_ISL_928607, EPI_ISL_928608, EPI_ISL_928610, EPI_ISL_928611, EPI_ISL_928612, EPI_ISL_928613, EPI_ISL_928615, EPI_ISL_928616, EPI_ISL_928618, EPI_ISL_928619, EPI_ISL_928620, EPI_ISL_928622, EPI_ISL_928627, EPI_ISL_928632, EPI_ISL_928634, EPI_ISL_928635, EPI_ISL_928637, EPI_ISL_928639, EPI_ISL_928640, EPI_ISL_928641, EPI_ISL_928642, EPI_ISL_928643, EPI_ISL_928644, EPI_ISL_928645, EPI_ISL_928647, EPI_ISL_928649, EPI_ISL_928656, EPI_ISL_928658, EPI_ISL_928659, EPI_ISL_928662, EPI_ISL_928663, EPI_ISL_928666, EPI_ISL_928667, EPI_ISL_928668, EPI_ISL_928669, EPI_ISL_928671, EPI_ISL_928672, EPI_ISL_928673, EPI_ISL_928680, EPI_ISL_928681, EPI_ISL_928684, EPI_ISL_928685, EPI_ISL_928686, EPI_ISL_928691, EPI_ISL_928692, EPI_ISL_928693, EPI_ISL_928694, EPI_ISL_928695, EPI_ISL_928697, EPI_ISL_928698, EPI_ISL_928701, EPI_ISL_928702, EPI_ISL_928703, EPI_ISL_928704, EPI_ISL_928707, EPI_ISL_928709, EPI_ISL_928710, EPI_ISL_928712, EPI_ISL_928713, EPI_ISL_928715, EPI_ISL_928721, EPI_ISL_928724, EPI_ISL_928725, EPI_ISL_928726, EPI_ISL_928729, EPI_ISL_928731, EPI_ISL_928732, EPI_ISL_928733, EPI_ISL_928734, EPI_ISL_928737, EPI_ISL_928739, EPI_ISL_928740, EPI_ISL_928741, EPI_ISL_928742, EPI_ISL_928744, EPI_ISL_928746, EPI_ISL_928747, EPI_ISL_928748, EPI_ISL_928749, EPI_ISL_928752, EPI_ISL_928753, EPI_ISL_928756, EPI_ISL_928758, EPI_ISL_928759, EPI_ISL_928760, EPI_ISL_928761, EPI_ISL_928762, EPI_ISL_928763, EPI_ISL_928764, EPI_ISL_928765, EPI_ISL_928766, EPI_ISL_928767, EPI_ISL_928770, EPI_ISL_928771, EPI_ISL_928773, EPI_ISL_928775, EPI_ISL_928776, EPI_ISL_928779, EPI_ISL_928781, EPI_ISL_928782, EPI_ISL_928783, EPI_ISL_928784, EPI_ISL_928785, EPI_ISL_928787, EPI_ISL_928790, EPI_ISL_928792, EPI_ISL_928793, EPI_ISL_928795, EPI_ISL_928798, EPI_ISL_928799, EPI_ISL_928800, EPI_ISL_928801, EPI_ISL_928802, EPI_ISL_928803, EPI_ISL_928805, EPI_ISL_928806, EPI_ISL_928807, EPI_ISL_928809, EPI_ISL_928810, EPI_ISL_928811, EPI_ISL_928813, EPI_ISL_928815, EPI_ISL_928818, EPI_ISL_928819, EPI_ISL_928824, EPI_ISL_928825, EPI_ISL_928826, EPI_ISL_928827, EPI_ISL_928828, EPI_ISL_928829, EPI_ISL_928830, EPI_ISL_928831, EPI_ISL_928832, EPI_ISL_928835, EPI_ISL_928836, EPI_ISL_928837, EPI_ISL_928838, EPI_ISL_928839, EPI_ISL_928840, EPI_ISL_928841, EPI_ISL_928844, EPI_ISL_928845, EPI_ISL_928846, EPI_ISL_928847, EPI_ISL_928848, EPI_ISL_928851, EPI_ISL_928852, EPI_ISL_928854, EPI_ISL_928855, EPI_ISL_928856, EPI_ISL_928857, EPI_ISL_928858, EPI_ISL_928862, EPI_ISL_928865, EPI_ISL_928866, EPI_ISL_928868, EPI_ISL_928870, EPI_ISL_928871, EPI_ISL_928873, EPI_ISL_928874, EPI_ISL_928875, EPI_ISL_928877, EPI_ISL_928879, EPI_ISL_928880, EPI_ISL_928887, EPI_ISL_928889, EPI_ISL_928890, EPI_ISL_928891, EPI_ISL_928896, EPI_ISL_928897, EPI_ISL_928898, EPI_ISL_928900, EPI_ISL_928901, EPI_ISL_928902, EPI_ISL_928903, EPI_ISL_928904, EPI_ISL_928906, EPI_ISL_928907, EPI_ISL_928908, EPI_ISL_928910, EPI_ISL_928911, EPI_ISL_928912, EPI_ISL_928913, EPI_ISL_928914, EPI_ISL_928917, EPI_ISL_928920, EPI_ISL_928921, EPI_ISL_928922, EPI_ISL_928923, EPI_ISL_928925, EPI_ISL_928927, EPI_ISL_928928, EPI_ISL_928929, EPI_ISL_928930, EPI_ISL_928931, EPI_ISL_928933, EPI_ISL_928936, EPI_ISL_928937, EPI_ISL_928938, EPI_ISL_928941, EPI_ISL_928945, EPI_ISL_928947, EPI_ISL_928948, EPI_ISL_928949, EPI_ISL_928950, EPI_ISL_928952, EPI_ISL_928953, EPI_ISL_928955, EPI_ISL_928956, EPI_ISL_928957, EPI_ISL_928958, EPI_ISL_928959, EPI_ISL_928960, EPI_ISL_928962, EPI_ISL_928963, EPI_ISL_928964, EPI_ISL_928968, EPI_ISL_928969, EPI_ISL_928970, EPI_ISL_928971, EPI_ISL_928972, EPI_ISL_928973, EPI_ISL_928974, EPI_ISL_928975, EPI_ISL_928977, EPI_ISL_928979, EPI_ISL_928981, EPI_ISL_928982, EPI_ISL_928986, EPI_ISL_928990, EPI_ISL_928991, EPI_ISL_928992, EPI_ISL_928993, EPI_ISL_928996, EPI_ISL_928997, EPI_ISL_928998, EPI_ISL_928999, EPI_ISL_929000, EPI_ISL_929001, EPI_ISL_929002, EPI_ISL_929003, EPI_ISL_929004, EPI_ISL_929007, EPI_ISL_929008, EPI_ISL_929009, EPI_ISL_929012, EPI_ISL_929013, EPI_ISL_929014, EPI_ISL_929016, EPI_ISL_929017, EPI_ISL_929019, EPI_ISL_929021, EPI_ISL_929023, EPI_ISL_929024, EPI_ISL_929025, EPI_ISL_929028, EPI_ISL_929029, EPI_ISL_929031, EPI_ISL_929032, EPI_ISL_929033, EPI_ISL_929036, EPI_ISL_929037, EPI_ISL_929038, EPI_ISL_929041, EPI_ISL_929042, EPI_ISL_929045, EPI_ISL_929048, EPI_ISL_929049, EPI_ISL_929050, EPI_ISL_929051, EPI_ISL_929052, EPI_ISL_929053, EPI_ISL_929054, EPI_ISL_929059, EPI_ISL_929060, EPI_ISL_929065, EPI_ISL_929067, EPI_ISL_929068, EPI_ISL_929070, EPI_ISL_929071, EPI_ISL_929072, EPI_ISL_929073, EPI_ISL_929074, EPI_ISL_929077, EPI_ISL_929078, EPI_ISL_929080, EPI_ISL_929081, EPI_ISL_929084, EPI_ISL_929087, EPI_ISL_929089, EPI_ISL_929091, EPI_ISL_929094, EPI_ISL_929095, EPI_ISL_929098, EPI_ISL_929100, EPI_ISL_929101, EPI_ISL_929102, EPI_ISL_929103, EPI_ISL_929104, EPI_ISL_929105, EPI_ISL_929106, EPI_ISL_929107, EPI_ISL_929108, EPI_ISL_929109, EPI_ISL_929110, EPI_ISL_929111, EPI_ISL_929113, EPI_ISL_929114, EPI_ISL_929115, EPI_ISL_929117, EPI_ISL_929118, EPI_ISL_929121, EPI_ISL_929123, EPI_ISL_929125, EPI_ISL_929126, EPI_ISL_929127, EPI_ISL_929131, EPI_ISL_929133, EPI_ISL_929134, EPI_ISL_929135, EPI_ISL_929137, EPI_ISL_929138, EPI_ISL_929139, EPI_ISL_929143, EPI_ISL_929144, EPI_ISL_929148, EPI_ISL_929150, EPI_ISL_929151, EPI_ISL_929152, EPI_ISL_929154, EPI_ISL_929155, EPI_ISL_929160, EPI_ISL_929161, EPI_ISL_929163, EPI_ISL_929164, EPI_ISL_929167, EPI_ISL_929172, EPI_ISL_929174, EPI_ISL_929177, EPI_ISL_929178, EPI_ISL_929179, EPI_ISL_929181, EPI_ISL_929183, EPI_ISL_929186, EPI_ISL_929187, EPI_ISL_929189, EPI_ISL_929190, EPI_ISL_929192, EPI_ISL_929193, EPI_ISL_929195, EPI_ISL_929204, EPI_ISL_929206, EPI_ISL_929208, EPI_ISL_929209, EPI_ISL_929210, EPI_ISL_929211, EPI_ISL_929212, EPI_ISL_929216, EPI_ISL_929217, EPI_ISL_929218, EPI_ISL_929220, EPI_ISL_929221, EPI_ISL_929225, EPI_ISL_929232, EPI_ISL_929233, EPI_ISL_929234, EPI_ISL_929236, EPI_ISL_929237, EPI_ISL_929238, EPI_ISL_929239, EPI_ISL_929240, EPI_ISL_929242, EPI_ISL_929244, EPI_ISL_929245, EPI_ISL_929246, EPI_ISL_929248, EPI_ISL_929249, EPI_ISL_929251, EPI_ISL_929252, EPI_ISL_929255, EPI_ISL_929256, EPI_ISL_929257, EPI_ISL_929259, EPI_ISL_929260, EPI_ISL_929263, EPI_ISL_929264, EPI_ISL_929266, EPI_ISL_929267, EPI_ISL_929271, EPI_ISL_929272, EPI_ISL_929273, EPI_ISL_929274, EPI_ISL_929275, EPI_ISL_929276, EPI_ISL_929277, EPI_ISL_929278, EPI_ISL_929283, EPI_ISL_929285, EPI_ISL_929287, EPI_ISL_929288, EPI_ISL_929289, EPI_ISL_929290, EPI_ISL_929293, EPI_ISL_929294, EPI_ISL_929295, EPI_ISL_929296, EPI_ISL_929298, EPI_ISL_929300, EPI_ISL_929301, EPI_ISL_929302, EPI_ISL_929303, EPI_ISL_929304, EPI_ISL_929305, EPI_ISL_929306, EPI_ISL_929310, EPI_ISL_929311, EPI_ISL_929312, EPI_ISL_929315, EPI_ISL_929316, EPI_ISL_929320, EPI_ISL_929321, EPI_ISL_929324, EPI_ISL_929325, EPI_ISL_929327, EPI_ISL_929328, EPI_ISL_929330, EPI_ISL_929333, EPI_ISL_929335, EPI_ISL_929336, EPI_ISL_929337, EPI_ISL_929341, EPI_ISL_929342, EPI_ISL_929343, EPI_ISL_929345, EPI_ISL_929346, EPI_ISL_929347, EPI_ISL_929348, EPI_ISL_929350, EPI_ISL_929351, EPI_ISL_929352, EPI_ISL_929354, EPI_ISL_929358, EPI_ISL_929359, EPI_ISL_929361, EPI_ISL_929363, EPI_ISL_929364, EPI_ISL_929365, EPI_ISL_929367, EPI_ISL_929369, EPI_ISL_929373, EPI_ISL_929374, EPI_ISL_929375, EPI_ISL_929376, EPI_ISL_929377, EPI_ISL_929378, EPI_ISL_929379, EPI_ISL_929380, EPI_ISL_929381, EPI_ISL_929382, EPI_ISL_929384, EPI_ISL_929386, EPI_ISL_929392, EPI_ISL_929393, EPI_ISL_929394, EPI_ISL_929395, EPI_ISL_929396, EPI_ISL_929397, EPI_ISL_929399, EPI_ISL_929401, EPI_ISL_929402, EPI_ISL_929404, EPI_ISL_929405, EPI_ISL_929406, EPI_ISL_929407, EPI_ISL_929408, EPI_ISL_929410, EPI_ISL_929411, EPI_ISL_929412, EPI_ISL_929413, EPI_ISL_929414, EPI_ISL_929416, EPI_ISL_929417, EPI_ISL_929419, EPI_ISL_929420, EPI_ISL_929423, EPI_ISL_929426, EPI_ISL_929427, EPI_ISL_929430, EPI_ISL_929431, EPI_ISL_929432, EPI_ISL_929434, EPI_ISL_929436, EPI_ISL_929437, EPI_ISL_929439, EPI_ISL_929440, EPI_ISL_929441, EPI_ISL_929442, EPI_ISL_929443, EPI_ISL_929445, EPI_ISL_929447, EPI_ISL_929448, EPI_ISL_929453, EPI_ISL_929454, EPI_ISL_929455, EPI_ISL_929457, EPI_ISL_929459, EPI_ISL_929460, EPI_ISL_929465, EPI_ISL_929470, EPI_ISL_929471, EPI_ISL_929472, EPI_ISL_929473, EPI_ISL_929474, EPI_ISL_929475, EPI_ISL_929478, EPI_ISL_929479, EPI_ISL_929480, EPI_ISL_929481, EPI_ISL_929482, EPI_ISL_929483, EPI_ISL_929484, EPI_ISL_929485, EPI_ISL_929487, EPI_ISL_929489, EPI_ISL_929490, EPI_ISL_929491, EPI_ISL_929492, EPI_ISL_929493, EPI_ISL_929496, EPI_ISL_929498, EPI_ISL_929499, EPI_ISL_929500, EPI_ISL_929502, EPI_ISL_929503, EPI_ISL_929504, EPI_ISL_929505, EPI_ISL_929506, EPI_ISL_929507, EPI_ISL_929508, EPI_ISL_929509, EPI_ISL_929510, EPI_ISL_929511, EPI_ISL_929512, EPI_ISL_929514, EPI_ISL_929516, EPI_ISL_929517, EPI_ISL_929520, EPI_ISL_929521, EPI_ISL_929522, EPI_ISL_929525, EPI_ISL_929528, EPI_ISL_929529, EPI_ISL_929532, EPI_ISL_929536, EPI_ISL_929537, EPI_ISL_929539, EPI_ISL_929541, EPI_ISL_929542, EPI_ISL_929543, EPI_ISL_929544, EPI_ISL_929545, EPI_ISL_929548, EPI_ISL_929549, EPI_ISL_929550, EPI_ISL_929551, EPI_ISL_929553, EPI_ISL_929554, EPI_ISL_929555, EPI_ISL_929556, EPI_ISL_929557, EPI_ISL_929558, EPI_ISL_929562, EPI_ISL_929563, EPI_ISL_929564, EPI_ISL_929566, EPI_ISL_929567, EPI_ISL_929569, EPI_ISL_929571, EPI_ISL_929573, EPI_ISL_929574, EPI_ISL_929576, EPI_ISL_929577, EPI_ISL_929580, EPI_ISL_929581, EPI_ISL_929582, EPI_ISL_929584, EPI_ISL_929585, EPI_ISL_929586, EPI_ISL_929587, EPI_ISL_929590, EPI_ISL_929591, EPI_ISL_929592, EPI_ISL_929593, EPI_ISL_929594, EPI_ISL_929596, EPI_ISL_929597, EPI_ISL_929599, EPI_ISL_929604, EPI_ISL_929606, EPI_ISL_929615, EPI_ISL_929616, EPI_ISL_929620, EPI_ISL_929621, EPI_ISL_929622, EPI_ISL_929625, EPI_ISL_929626, EPI_ISL_929629, EPI_ISL_929630, EPI_ISL_929632, EPI_ISL_929633, EPI_ISL_929635, EPI_ISL_929636, EPI_ISL_929640, EPI_ISL_929641, EPI_ISL_929642, EPI_ISL_929644, EPI_ISL_929646, EPI_ISL_929649, EPI_ISL_929650, EPI_ISL_929652, EPI_ISL_929654, EPI_ISL_929656, EPI_ISL_929663, EPI_ISL_929664, EPI_ISL_929665, EPI_ISL_929666, EPI_ISL_929667, EPI_ISL_929668, EPI_ISL_929670, EPI_ISL_929671, EPI_ISL_929675, EPI_ISL_929679, EPI_ISL_929680, EPI_ISL_929681, EPI_ISL_929682, EPI_ISL_929684, EPI_ISL_929690, EPI_ISL_929691, EPI_ISL_929692, EPI_ISL_929696, EPI_ISL_929697, EPI_ISL_929698, EPI_ISL_929699, EPI_ISL_929700, EPI_ISL_929703, EPI_ISL_929704, EPI_ISL_929705, EPI_ISL_929707, EPI_ISL_929708, EPI_ISL_929710, EPI_ISL_929713, EPI_ISL_929714, EPI_ISL_929715, EPI_ISL_929717, EPI_ISL_929718, EPI_ISL_929719, EPI_ISL_929721, EPI_ISL_929724, EPI_ISL_929727, EPI_ISL_929728, EPI_ISL_929729, EPI_ISL_929731, EPI_ISL_929732, EPI_ISL_929733, EPI_ISL_929736, EPI_ISL_929739, EPI_ISL_929742, EPI_ISL_929743, EPI_ISL_929747, EPI_ISL_929751, EPI_ISL_929752, EPI_ISL_929753, EPI_ISL_929754, EPI_ISL_929759, EPI_ISL_929760, EPI_ISL_929761, EPI_ISL_929764, EPI_ISL_929766, EPI_ISL_929767, EPI_ISL_929768, EPI_ISL_929769, EPI_ISL_929774, EPI_ISL_929775, EPI_ISL_929777, EPI_ISL_929778, EPI_ISL_929779, EPI_ISL_929785, EPI_ISL_929786, EPI_ISL_929787, EPI_ISL_929789, EPI_ISL_929791, EPI_ISL_929792, EPI_ISL_929793, EPI_ISL_929795, EPI_ISL_929798, EPI_ISL_929799, EPI_ISL_929801, EPI_ISL_929804, EPI_ISL_929805, EPI_ISL_929806, EPI_ISL_929809, EPI_ISL_929812, EPI_ISL_929813, EPI_ISL_929814, EPI_ISL_929815, EPI_ISL_929817, EPI_ISL_929818, EPI_ISL_929819, EPI_ISL_929820, EPI_ISL_929821, EPI_ISL_929822, EPI_ISL_929824, EPI_ISL_929825, EPI_ISL_929826, EPI_ISL_929828, EPI_ISL_929831, EPI_ISL_929834, EPI_ISL_929835, EPI_ISL_929836, EPI_ISL_929838, EPI_ISL_929839, EPI_ISL_929841, EPI_ISL_929842, EPI_ISL_929843, EPI_ISL_929844, EPI_ISL_929846, EPI_ISL_929847, EPI_ISL_929849, EPI_ISL_929850, EPI_ISL_929851, EPI_ISL_929852, EPI_ISL_929853, EPI_ISL_929855, EPI_ISL_929858, EPI_ISL_929859, EPI_ISL_929862, EPI_ISL_929863, EPI_ISL_929865, EPI_ISL_929869, EPI_ISL_929871, EPI_ISL_929874, EPI_ISL_929875, EPI_ISL_929876, EPI_ISL_929877, EPI_ISL_929878, EPI_ISL_929879, EPI_ISL_929883, EPI_ISL_929884, EPI_ISL_929885, EPI_ISL_929888, EPI_ISL_929891, EPI_ISL_929893, EPI_ISL_929895, EPI_ISL_929896, EPI_ISL_929897, EPI_ISL_929898, EPI_ISL_929901, EPI_ISL_929902, EPI_ISL_929903, EPI_ISL_929904, EPI_ISL_929905, EPI_ISL_929906, EPI_ISL_929909, EPI_ISL_929912, EPI_ISL_929915, EPI_ISL_929918, EPI_ISL_929920, EPI_ISL_929922, EPI_ISL_929924, EPI_ISL_929925, EPI_ISL_929927, EPI_ISL_929930, EPI_ISL_929932, EPI_ISL_929934, EPI_ISL_929935, EPI_ISL_929938, EPI_ISL_929943, EPI_ISL_929946, EPI_ISL_929947, EPI_ISL_929949, EPI_ISL_929950, EPI_ISL_929952, EPI_ISL_929953, EPI_ISL_929955, EPI_ISL_929959, EPI_ISL_929961, EPI_ISL_929962, EPI_ISL_929963, EPI_ISL_929965, EPI_ISL_929967, EPI_ISL_929968, EPI_ISL_929969, EPI_ISL_929970, EPI_ISL_929971, EPI_ISL_929973, EPI_ISL_929974, EPI_ISL_929975, EPI_ISL_929977, EPI_ISL_929978, EPI_ISL_929979, EPI_ISL_929981, EPI_ISL_929982, EPI_ISL_929983, EPI_ISL_929984, EPI_ISL_929985, EPI_ISL_929986, EPI_ISL_929989, EPI_ISL_929990, EPI_ISL_929991, EPI_ISL_929992, EPI_ISL_929996, EPI_ISL_929997, EPI_ISL_929999, EPI_ISL_930000, EPI_ISL_930002, EPI_ISL_930005, EPI_ISL_930009, EPI_ISL_930012, EPI_ISL_930013, EPI_ISL_930014, EPI_ISL_930015, EPI_ISL_930018, EPI_ISL_930020, EPI_ISL_930022, EPI_ISL_930023, EPI_ISL_930024, EPI_ISL_930025, EPI_ISL_930026, EPI_ISL_930028, EPI_ISL_930032, EPI_ISL_930033, EPI_ISL_930036, EPI_ISL_930038, EPI_ISL_930039, EPI_ISL_930042, EPI_ISL_930046, EPI_ISL_930047, EPI_ISL_930048, EPI_ISL_930051, EPI_ISL_930052, EPI_ISL_930055, EPI_ISL_930056, EPI_ISL_930057, EPI_ISL_930060, EPI_ISL_930061, EPI_ISL_930066, EPI_ISL_930067, EPI_ISL_930070, EPI_ISL_930071, EPI_ISL_930072, EPI_ISL_930073, EPI_ISL_930076, EPI_ISL_930077, EPI_ISL_930078, EPI_ISL_930084, EPI_ISL_930085, EPI_ISL_930086, EPI_ISL_930087, EPI_ISL_930089, EPI_ISL_930091, EPI_ISL_930092, EPI_ISL_930094, EPI_ISL_930097, EPI_ISL_930098, EPI_ISL_930099, EPI_ISL_930101, EPI_ISL_930102, EPI_ISL_930103, EPI_ISL_930106, EPI_ISL_930113, EPI_ISL_930114, EPI_ISL_930116, EPI_ISL_930118, EPI_ISL_930119, EPI_ISL_930120, EPI_ISL_930121, EPI_ISL_930122, EPI_ISL_930124, EPI_ISL_930125, EPI_ISL_930127, EPI_ISL_930130, EPI_ISL_930131, EPI_ISL_930132, EPI_ISL_930133, EPI_ISL_930136, EPI_ISL_930139, EPI_ISL_930141, EPI_ISL_930142, EPI_ISL_930144, EPI_ISL_930145, EPI_ISL_930146, EPI_ISL_930147, EPI_ISL_930148, EPI_ISL_930149, EPI_ISL_930150, EPI_ISL_930152, EPI_ISL_930155, EPI_ISL_930159, EPI_ISL_930163, EPI_ISL_930164, EPI_ISL_930165, EPI_ISL_930167, EPI_ISL_930168, EPI_ISL_930169, EPI_ISL_930171, EPI_ISL_930175, EPI_ISL_930176, EPI_ISL_930177, EPI_ISL_930179, EPI_ISL_930182, EPI_ISL_930183, EPI_ISL_930190, EPI_ISL_930191, EPI_ISL_930192, EPI_ISL_930195, EPI_ISL_930196, EPI_ISL_930197, EPI_ISL_930198, EPI_ISL_930199, EPI_ISL_930201, EPI_ISL_930203, EPI_ISL_930204, EPI_ISL_930205, EPI_ISL_930207, EPI_ISL_930209, EPI_ISL_930213, EPI_ISL_930214, EPI_ISL_930217, EPI_ISL_930218, EPI_ISL_930226, EPI_ISL_930229, EPI_ISL_930230, EPI_ISL_930231, EPI_ISL_930232, EPI_ISL_930234, EPI_ISL_930236, EPI_ISL_930240, EPI_ISL_930241, EPI_ISL_930242, EPI_ISL_930243, EPI_ISL_930245, EPI_ISL_930246, EPI_ISL_930248, EPI_ISL_930250, EPI_ISL_930251, EPI_ISL_930256, EPI_ISL_930260, EPI_ISL_930261, EPI_ISL_930262, EPI_ISL_930264, EPI_ISL_930266, EPI_ISL_930267, EPI_ISL_930269, EPI_ISL_930271, EPI_ISL_930276, EPI_ISL_930277, EPI_ISL_930278, EPI_ISL_930279, EPI_ISL_930281, EPI_ISL_930282, EPI_ISL_930283, EPI_ISL_930287, EPI_ISL_930291, EPI_ISL_930293, EPI_ISL_930295, EPI_ISL_930297, EPI_ISL_930298, EPI_ISL_930301, EPI_ISL_930302, EPI_ISL_930304, EPI_ISL_930305, EPI_ISL_930306, EPI_ISL_930308, EPI_ISL_930309, EPI_ISL_930312, EPI_ISL_930313, EPI_ISL_930316, EPI_ISL_930317, EPI_ISL_930318, EPI_ISL_930323, EPI_ISL_930325, EPI_ISL_930326, EPI_ISL_930328, EPI_ISL_930329, EPI_ISL_930330, EPI_ISL_930331, EPI_ISL_930332, EPI_ISL_930336, EPI_ISL_930339, EPI_ISL_930340, EPI_ISL_930344, EPI_ISL_930346, EPI_ISL_930349, EPI_ISL_930350, EPI_ISL_930352, EPI_ISL_930354, EPI_ISL_930355, EPI_ISL_930357, EPI_ISL_930358, EPI_ISL_930359, EPI_ISL_930361, EPI_ISL_930362, EPI_ISL_930363, EPI_ISL_930364, EPI_ISL_930365, EPI_ISL_930366, EPI_ISL_930369, EPI_ISL_930370, EPI_ISL_930373, EPI_ISL_930374, EPI_ISL_930377, EPI_ISL_930378, EPI_ISL_930379, EPI_ISL_930380, EPI_ISL_930381, EPI_ISL_930382, EPI_ISL_930383, EPI_ISL_930384, EPI_ISL_930386, EPI_ISL_930388, EPI_ISL_930389, EPI_ISL_930393, EPI_ISL_930395, EPI_ISL_930397, EPI_ISL_930398, EPI_ISL_930400, EPI_ISL_930401, EPI_ISL_930402, EPI_ISL_930405, EPI_ISL_930408, EPI_ISL_930411, EPI_ISL_930413, EPI_ISL_930415, EPI_ISL_930416, EPI_ISL_930417, EPI_ISL_930418, EPI_ISL_930420, EPI_ISL_930424, EPI_ISL_930425, EPI_ISL_930426, EPI_ISL_930427, EPI_ISL_930428, EPI_ISL_930429, EPI_ISL_930435, EPI_ISL_930436, EPI_ISL_930437, EPI_ISL_930438, EPI_ISL_930439, EPI_ISL_930440, EPI_ISL_930441, EPI_ISL_930445, EPI_ISL_930446, EPI_ISL_930447, EPI_ISL_930448, EPI_ISL_930451, EPI_ISL_930456, EPI_ISL_930459, EPI_ISL_930462, EPI_ISL_930463, EPI_ISL_930466, EPI_ISL_930468, EPI_ISL_930469, EPI_ISL_930470, EPI_ISL_930471, EPI_ISL_930472, EPI_ISL_930473, EPI_ISL_930474, EPI_ISL_930475, EPI_ISL_930477, EPI_ISL_930479, EPI_ISL_930480, EPI_ISL_930481, EPI_ISL_930482, EPI_ISL_930483, EPI_ISL_930484, EPI_ISL_930487, EPI_ISL_930489, EPI_ISL_930492, EPI_ISL_930493, EPI_ISL_930495, EPI_ISL_930498, EPI_ISL_930500, EPI_ISL_930501, EPI_ISL_930502, EPI_ISL_930507, EPI_ISL_930508, EPI_ISL_930512, EPI_ISL_930515, EPI_ISL_930517, EPI_ISL_930520, EPI_ISL_930521, EPI_ISL_930525, EPI_ISL_930528, EPI_ISL_930530, EPI_ISL_930531, EPI_ISL_930532, EPI_ISL_930533, EPI_ISL_930534, EPI_ISL_930535, EPI_ISL_930536, EPI_ISL_930537, EPI_ISL_930538, EPI_ISL_930539, EPI_ISL_930540, EPI_ISL_930541, EPI_ISL_930542, EPI_ISL_930543, EPI_ISL_930544, EPI_ISL_930545, EPI_ISL_930548, EPI_ISL_930549, EPI_ISL_930550, EPI_ISL_930551, EPI_ISL_930555, EPI_ISL_930556, EPI_ISL_930558, EPI_ISL_930560, EPI_ISL_930561, EPI_ISL_930564, EPI_ISL_930565

| | | | |
|---|---|---|---|
| see above | Department of Virus and Microbiological Special Diagnostics, Statens Serum Institut, Copenhagen, Denmark | Aalborg University | Danish Covid-19 Genome Consortium |

EPI_ISL_930579, EPI_ISL_930585, EPI_ISL_930586, EPI_ISL_930587, EPI_ISL_930589, EPI_ISL_930590, EPI_ISL_930591, EPI_ISL_930592, EPI_ISL_930593, EPI_ISL_930598, EPI_ISL_930599, EPI_ISL_930600, EPI_ISL_930601, EPI_ISL_930602

| | | | |
|---|---|---|---|
| see above | Synlab | GIGA Medical Genomics | Keith Durkin, Maria Artesi, Sébastien Bontems, Raphaël Boreux, Bouchra Boujemla, Cécile Meex, Pierrette Melin, Marie-Pierre Hayette, Vincent Bours |
| EPI_ISL_930788, EPI_ISL_930789, EPI_ISL_930791, EPI_ISL_930833, EPI_ISL_930845, EPI_ISL_930847, EPI_ISL_930848, EPI_ISL_930851 | Monterey County Public Health Laboratory | Monterey County Public Health Laboratory | C. Anaya et al., |
| EPI_ISL_930852 | Monterey County Public Health Laboratory | Monterey County Public Health Laboratory | C. Anaya et al. |

| | | | |
|---|---|---|---|
| EPI_ISL_931455, EPI_ISL_931456, EPI_ISL_931457, EPI_ISL_931458, EPI_ISL_931459, EPI_ISL_931487 | Maryland Public Health Laboratory (MD PHL) | Maryland Public Health Laboratory (MD PHL) | Maryland Department of Health Laboratories Administration |
| EPI_ISL_932257, EPI_ISL_932263, EPI_ISL_932276, EPI_ISL_932294, EPI_ISL_932319, EPI_ISL_932329, EPI_ISL_932358, EPI_ISL_932366, EPI_ISL_932372, EPI_ISL_932380, EPI_ISL_932391, EPI_ISL_932395, EPI_ISL_932403, EPI_ISL_932417, EPI_ISL_932456, EPI_ISL_932505, EPI_ISL_932507, EPI_ISL_932542, EPI_ISL_932563, EPI_ISL_932575, EPI_ISL_932627, EPI_ISL_932631, EPI_ISL_932756, EPI_ISL_932793, EPI_ISL_932874, EPI_ISL_932886, EPI_ISL_932895, EPI_ISL_932913, EPI_ISL_932974, EPI_ISL_933022, EPI_ISL_933061 | | | |
| see above | Lighthouse Lab in Milton Keynes | Wellcome Sanger Institute for the COVID-19 Genomics UK (COG-UK) Consortium | The Lighthouse Lab in Milton Keynes and Alex Alderton, Roberto Amato, Sonia Goncalves, Ewan Harrison, David K. Jackson, Ian Johnston, Dominic Kwiatkowski, Cordelia Langford, John Sillitoe on behalf of the Wellcome Sanger Institute COVID-19 Surveillance Team |
| EPI_ISL_933113, EPI_ISL_933123, EPI_ISL_933124, EPI_ISL_933127, EPI_ISL_933130, EPI_ISL_933134, EPI_ISL_933136, EPI_ISL_933141, EPI_ISL_933143, EPI_ISL_933149, EPI_ISL_933150, EPI_ISL_933154, EPI_ISL_933158, EPI_ISL_933166, EPI_ISL_933167, EPI_ISL_933170, EPI_ISL_933171, EPI_ISL_933181, EPI_ISL_933187, EPI_ISL_933188, EPI_ISL_933196, EPI_ISL_933199, EPI_ISL_933200, EPI_ISL_933207, EPI_ISL_933213, EPI_ISL_933214, EPI_ISL_933215, EPI_ISL_933221, EPI_ISL_933226, EPI_ISL_933227, EPI_ISL_933228, EPI_ISL_933233, EPI_ISL_933239, EPI_ISL_933243, EPI_ISL_933256, EPI_ISL_933259, EPI_ISL_933265, EPI_ISL_933266, EPI_ISL_933271, EPI_ISL_933272, EPI_ISL_933273, EPI_ISL_933277, EPI_ISL_933282, EPI_ISL_933283, EPI_ISL_933289, EPI_ISL_933293, EPI_ISL_933298, EPI_ISL_933303, EPI_ISL_933321, EPI_ISL_933325, EPI_ISL_933326, EPI_ISL_933336, EPI_ISL_933340, EPI_ISL_933341, EPI_ISL_933343, EPI_ISL_933344, EPI_ISL_933345, EPI_ISL_933346, EPI_ISL_933348, EPI_ISL_933350, EPI_ISL_933354, EPI_ISL_933358, EPI_ISL_933359, EPI_ISL_933364, EPI_ISL_933371, EPI_ISL_933375, EPI_ISL_933382, EPI_ISL_933386, EPI_ISL_933387, EPI_ISL_933388, EPI_ISL_933393, EPI_ISL_933396, EPI_ISL_933397, EPI_ISL_933400, EPI_ISL_933401, EPI_ISL_933410, EPI_ISL_933412, EPI_ISL_933414, EPI_ISL_933417 | | | |
| see above | Lighthouse Lab in Cambridge | Wellcome Sanger Institute for the COVID-19 Genomics UK (COG-UK) Consortium | Rob Howes, The Lighthouse Lab in Cambridge and Alex Alderton, Roberto Amato, Sonia Goncalves, Ewan Harrison, David K. Jackson, Ian Johnston, Dominic Kwiatkowski, Cordelia Langford, John Sillitoe on behalf of the Wellcome Sanger Institute COVID-19 Surveillance Team |
| EPI_ISL_933730, EPI_ISL_933731, EPI_ISL_933733 | Servicio de Microbiología Hospital Ramón y Cajal | Servicio de Microbiología Hospital Ramón y Cajal | José M Gonzalez-Alba, Concepción Rodríguez, Melanie Abreu, Laura Martínez, Val F Lanza, Luz Leticia Olavarrieta, Rafael Cantón, JC Galán |
| EPI_ISL_933736, EPI_ISL_933737, EPI_ISL_933738, EPI_ISL_933762, EPI_ISL_933763 | DPHL | Delaware Public Health Lab | Gregory Hovan |
| EPI_ISL_933767 | Servicio de Microbiología Hospital Ramón y Cajal | Servicio de Microbiología Hospital Ramón y Cajal | José M Gonzalez-Alba, Concepción Rodríguez, Melanie Abreu, Laura Martínez, Val F Lanza, Luz Leticia Olavarrieta, Rafael Cantón, JC Galán |
| EPI_ISL_933790 | PathWest Laboratory Medicine WA | PathWest Laboratory Medicine WA Microbial Surveillance Unit | PathWest Laboratory Medicine WA Microbial Surveillance Unit |
| EPI_ISL_934308, EPI_ISL_934309, EPI_ISL_934310, EPI_ISL_934311, EPI_ISL_934312, EPI_ISL_934313 | Vilnius university hospital Santaros Klinikos, Center of Laboratory Medicine | Vilnius university hospital Santaros Klinikos, Center of Laboratory Medicine | Ingrida Olendraite, Daniel Naumovas, Rimvydas Norvilas, Dovile Ezerskyte, Justinas Slikas, Gytis Dudas |
| EPI_ISL_934382, EPI_ISL_934383, EPI_ISL_934387 | Klinisk mikrobiologi | The Public Health Agency of Sweden | Anna-Malin Linde, Maria Lind Karlberg, Carlo Berg, Oskar Karlsson Lindsjo, Sofia Stamouli, Reza Advani, Mattias Haukland, Petra Holmstrom, Noura Walai, Petra Edquist, Mia Brytting, Anna Risberg, Karin Tegmark-Wisell |
| EPI_ISL_934391, EPI_ISL_934392, EPI_ISL_934393, EPI_ISL_934394, EPI_ISL_934396 | Klinisk Mikrobiologi | The Public Health Agency of Sweden | Anna-Malin Linde, Maria Lind Karlberg, Carlo Berg, Oskar Karlsson Lindsjo, Sofia Stamouli, Reza Advani, Mattias Haukland, Petra Holmstrom, Noura Walai, Petra Edquist, Mia Brytting, Anna Risberg, Karin Tegmark-Wisell |
| EPI_ISL_934979, EPI_ISL_934985, EPI_ISL_934988, EPI_ISL_934992 | ADMED Microbiologie | Genomics and Transcriptomics, Philip Morris International | Reto Lienhard, Marie-Lise Tritten, Emmanuel Guedj, Nicolas Sierro, Rémi Dulize, David Bornand, Mehdi Auberson, Maxime Berthouzoz, Nikolai Ivanov, Manuel Peitsch |
| EPI_ISL_935048, EPI_ISL_935049, EPI_ISL_935050, EPI_ISL_935051, EPI_ISL_935052, EPI_ISL_935053, EPI_ISL_935054, EPI_ISL_935055, EPI_ISL_935056, EPI_ISL_935057, EPI_ISL_935058, EPI_ISL_935059, EPI_ISL_935060, EPI_ISL_935061, EPI_ISL_935062, EPI_ISL_935063, EPI_ISL_935064, EPI_ISL_935065, EPI_ISL_935066, EPI_ISL_935067, EPI_ISL_935068, EPI_ISL_935069 | | | |
| see above | Ministry of Health Turkey | Ministry of Health Turkey | Fatma Bayrakdar, Yasemin Cogun, Süleyman Yalcin, Aye Baak Alta, Gülay Korukluolu |
| EPI_ISL_935187, EPI_ISL_935188, EPI_ISL_935189, EPI_ISL_935190, EPI_ISL_935191, EPI_ISL_935192, EPI_ISL_935193, EPI_ISL_935194, EPI_ISL_935198, EPI_ISL_935201, EPI_ISL_935227, EPI_ISL_935228, EPI_ISL_935229, EPI_ISL_935230, EPI_ISL_935231, EPI_ISL_935232, EPI_ISL_935233, EPI_ISL_935234, EPI_ISL_935235, EPI_ISL_935236, EPI_ISL_935244, EPI_ISL_935245, EPI_ISL_935246, EPI_ISL_935247, EPI_ISL_935248, EPI_ISL_935249, EPI_ISL_935250, EPI_ISL_935251, EPI_ISL_935252, EPI_ISL_935253, EPI_ISL_935254, EPI_ISL_935255, EPI_ISL_935256, EPI_ISL_935257, EPI_ISL_935258, EPI_ISL_935259, EPI_ISL_935260, EPI_ISL_935261, EPI_ISL_935262, EPI_ISL_935263, EPI_ISL_935281, EPI_ISL_935282, EPI_ISL_935283, EPI_ISL_935326, EPI_ISL_935329 | | | |
| see above | KU Leuven, Rega Institute, Clinical and Epidemiological Virology | KU Leuven, Rega Institute, Clinical and Epidemiological Virology | Tony Wawina-Bokalanga, Bert Vanmechelen, Joan Marti-Carerras, Piet Maes |
| EPI_ISL_935533 | Labo Analyses Med | National Reference Center for Viruses of Respiratory Infections, Institut Pasteur, Paris | Marion Barbet, Sylvie Behillil, Méline Bizard, Angela Brisebarre, Camille Capel, Etienne Simon-Lorière, Vincent Enouf, Maud Vanpeene, Sylvie van der Werf,Jacques Alexandra |
| EPI_ISL_935534, EPI_ISL_935535 | Labo Analyses Med | National Reference Center for Viruses of Respiratory Infections, Institut Pasteur, Paris | Marion Barbet, Sylvie Behillil, Méline Bizard, Angela Brisebarre, Camille Capel, Etienne Simon-Lorière, Vincent Enouf, Maud Vanpeene, Sylvie van der Werf,Garel FréDéRick |
| EPI_ISL_935536 | Labo Analyses Med | National Reference Center for Viruses of Respiratory Infections, Institut Pasteur, Paris | Marion Barbet, Sylvie Behillil, Méline Bizard, Angela Brisebarre, Camille Capel, Etienne Simon-Lorière, Vincent Enouf, Maud Vanpeene, Sylvie van der Werf,Wittersheim Pascal |
| EPI_ISL_935546 | Labo Analyses Med | National Reference Center for Viruses of Respiratory Infections, Institut Pasteur, Paris | Marion Barbet, Sylvie Behillil, Méline Bizard, Angela Brisebarre, Camille Capel, Etienne Simon-Lorière, Vincent Enouf, Maud Vanpeene, Sylvie van der Werf,Besson J |
| EPI_ISL_935549 | Labo Analyses Med | National Reference Center for Viruses of Respiratory Infections, Institut Pasteur, Paris | Marion Barbet, Sylvie Behillil, Méline Bizard, Angela Brisebarre, Camille Capel, Etienne Simon-Lorière, Vincent Enouf, Maud Vanpeene, Sylvie van der Werf |
| EPI_ISL_935558, EPI_ISL_935559 | Hopital | National Reference Center for Viruses of Respiratory Infections, Institut Pasteur, Paris | Marion Barbet, Sylvie Behillil, Méline Bizard, Angela Brisebarre, Camille Capel, Etienne Simon-Lorière, Vincent Enouf, Maud Vanpeene, Sylvie van der Werf |
| EPI_ISL_935561 | Hopital | National Reference Center for Viruses of Respiratory Infections, Institut Pasteur, Paris | Marion Barbet, Sylvie Behillil, Méline Bizard, Angela Brisebarre, Camille Capel, Etienne Simon-Lorière, Vincent Enouf, Maud Vanpeene, Sylvie van der Werf,Gaschet Anne |
| EPI_ISL_935563 | Labo Analyses Med | National Reference Center for Viruses of Respiratory Infections, Institut Pasteur, Paris | Marion Barbet, Sylvie Behillil, Méline Bizard, Angela Brisebarre, Camille Capel, Etienne Simon-Lorière, Vincent Enouf, Maud Vanpeene, Sylvie van der Werf |
| EPI_ISL_935569 | Hopital | National Reference Center for Viruses of Respiratory Infections, Institut Pasteur, Paris | Marion Barbet, Sylvie Behillil, Méline Bizard, Angela Brisebarre, Camille Capel, Etienne Simon-Lorière, Vincent Enouf, Maud Vanpeene, Sylvie van der Werf,Bret Laurent |
| EPI_ISL_935576 | Labo Analyses Med | National Reference Center for Viruses of Respiratory Infections, Institut Pasteur, Paris | Marion Barbet, Sylvie Behillil, Méline Bizard, Angela Brisebarre, Camille Capel, Etienne Simon-Lorière, Vincent Enouf, Maud Vanpeene, Sylvie van der Werf,Breton-Cazaux JéRôMe |
| EPI_ISL_935577, EPI_ISL_935578, EPI_ISL_935579, EPI_ISL_935580 | Hopital | National Reference Center for Viruses of Respiratory Infections, Institut Pasteur, Paris | Marion Barbet, Sylvie Behillil, Méline Bizard, Angela Brisebarre, Camille Capel, Etienne Simon-Lorière, Vincent Enouf, Maud Vanpeene, Sylvie van der Werf,Brichler SéGolèNe |
| EPI_ISL_935586 | Hopital | National Reference Center for Viruses of Respiratory Infections, Institut Pasteur, Paris | Marion Barbet, Sylvie Behillil, Méline Bizard, Angela Brisebarre, Camille Capel, Etienne Simon-Lorière, Vincent Enouf, Maud Vanpeene, Sylvie van der Werf,Fourgeaud Jacques |
| EPI_ISL_935590 | Labo Analyses Med | National Reference Center for Viruses of Respiratory Infections, Institut Pasteur, Paris | Marion Barbet, Sylvie Behillil, Méline Bizard, Angela Brisebarre, Camille Capel, Etienne Simon-Lorière, Vincent Enouf, Maud Vanpeene, Sylvie van der Werf,Amzalag Jonas |
| EPI_ISL_935591, EPI_ISL_935592, EPI_ISL_935593, EPI_ISL_935594, EPI_ISL_935599, EPI_ISL_935604 | Labo Analyses med | National Reference Center for Viruses of Respiratory Infections, Institut Pasteur, Paris | Marion Barbet, Sylvie Behillil, Méline Bizard, Angela Brisebarre, Camille Capel, Etienne Simon-Lorière, Vincent Enouf, Maud Vanpeene, Sylvie van der Werf,Amzalag Jonas |
| EPI_ISL_935605, EPI_ISL_935606 | Labo Analyses Med | National Reference Center for Viruses of Respiratory Infections, Institut Pasteur, Paris | Marion Barbet, Sylvie Behillil, Méline Bizard, Angela Brisebarre, Camille Capel, Etienne Simon-Lorière, Vincent Enouf, Maud Vanpeene, Sylvie van der Werf,Amzalag Jonas |
| EPI_ISL_935607, EPI_ISL_935608, EPI_ISL_935609, EPI_ISL_935610, EPI_ISL_935611, EPI_ISL_935612, EPI_ISL_935613, EPI_ISL_935614, EPI_ISL_935615, EPI_ISL_935616, EPI_ISL_935617, EPI_ISL_935618, EPI_ISL_935619, EPI_ISL_935620, EPI_ISL_935629, EPI_ISL_935630, EPI_ISL_935631, EPI_ISL_935632, | | | |

| | | | |
|---|---|---|---|
| EPI_ISL_935633, EPI_ISL_935634, EPI_ISL_935635, EPI_ISL_935636 | | | |
| see above | Labo Analyses med | National Reference Center for Viruses of Respiratory Infections, Institut Pasteur, Paris | Marion Barbet, Sylvie Behillil, Méline Bizard, Angela Brisebarre, Camille Capel, Etienne Simon-Lorière, Vincent Enouf, Maud Vanpeene, Sylvie van der Werf,Amzalag Jonas |
| EPI_ISL_935637 | Labo Analyses Med | National Reference Center for Viruses of Respiratory Infections, Institut Pasteur, Paris | Marion Barbet, Sylvie Behillil, Méline Bizard, Angela Brisebarre, Camille Capel, Etienne Simon-Lorière, Vincent Enouf, Maud Vanpeene, Sylvie van der Werf,Jacques Alexandra |
| EPI_ISL_935639 | Hopital | National Reference Center for Viruses of Respiratory Infections, Institut Pasteur, Paris | Marion Barbet, Sylvie Behillil, Méline Bizard, Angela Brisebarre, Camille Capel, Etienne Simon-Lorière, Vincent Enouf, Maud Vanpeene, Sylvie van der Werf,Breuil J. |
| EPI_ISL_935642, EPI_ISL_935645, EPI_ISL_935646, EPI_ISL_935647, EPI_ISL_935648, EPI_ISL_935649 | Labo Analyses Med | National Reference Center for Viruses of Respiratory Infections, Institut Pasteur, Paris | Marion Barbet, Sylvie Behillil, Méline Bizard, Angela Brisebarre, Camille Capel, Etienne Simon-Lorière, Vincent Enouf, Maud Vanpeene, Sylvie van der Werf |
| EPI_ISL_936252, EPI_ISL_936254, EPI_ISL_936255, EPI_ISL_936256, EPI_ISL_936257, EPI_ISL_936262, EPI_ISL_936263, EPI_ISL_936268, EPI_ISL_936270, EPI_ISL_936274, EPI_ISL_936276, EPI_ISL_936277, EPI_ISL_936278, EPI_ISL_936280, EPI_ISL_936283, EPI_ISL_936284, EPI_ISL_936291, EPI_ISL_936294, EPI_ISL_936295, EPI_ISL_936298 | | | |
| see above | MONTEFIORE MEDICAL CENTER LABORATORIES | Wadsworth Center, New York State Department of Health | Kirsten St. George, Daryl M. Lamson, Alexis Russel, Matthew Shudt, Melissa A Leisner, Jonathan Plitnick, Navjot Singh, John Kelly, Erasmus Schneider, Erica Lasek-Nesselquist |
| EPI_ISL_936478, EPI_ISL_936480, EPI_ISL_936481 | DPH, Massachusetts State Public Health Lab | DPH, Massachusetts State Public Health Lab | Lang,A.S., Fink,T., Gallagher,G.R., Smole,S.C. |
| EPI_ISL_937126, EPI_ISL_937127, EPI_ISL_937128 | DOHMH Riverside | New York City Public Health Laboratory | Jade Wang, et al. |
| EPI_ISL_937129, EPI_ISL_937130, EPI_ISL_937131 | DOHMH Morrisania | New York City Public Health Laboratory | Jade Wang, et al. |
| EPI_ISL_937132, EPI_ISL_937133 | DOHMH Central Harlem | New York City Public Health Laboratory | Jade Wang, et al. |
| EPI_ISL_937134, EPI_ISL_937135 | DOHMH Fort Greene | New York City Public Health Laboratory | Jade Wang, et al. |
| EPI_ISL_937136, EPI_ISL_937137 | DOHMH Chelsea | New York City Public Health Laboratory | Jade Wang, et al. |
| EPI_ISL_937138 | DOHMH Crown Heights | New York City Public Health Laboratory | Jade Wang, et al. |
| EPI_ISL_937139, EPI_ISL_937140, EPI_ISL_937141, EPI_ISL_937142, EPI_ISL_937143, EPI_ISL_937144 | DOHMH Corona | New York City Public Health Laboratory | Jade Wang, et al. |
| EPI_ISL_937145, EPI_ISL_937146, EPI_ISL_937147, EPI_ISL_937148, EPI_ISL_937149, EPI_ISL_937150, EPI_ISL_937151 | DOHMH Jamaica | New York City Public Health Laboratory | Jade Wang, et al. |
| EPI_ISL_937152, EPI_ISL_937153 | DOHMH PHL | New York City Public Health Laboratory | Jade Wang, et al. |
| EPI_ISL_937167 | DOHMH Central Harlem | New York City Public Health Laboratory | Jade Wang, et al. |
| EPI_ISL_937168 | DOHMH Crown Heights | New York City Public Health Laboratory | Jade Wang, et al. |
| EPI_ISL_937188 | OCME Office Of Chief Medical Examiner | New York City Public Health Laboratory | Jade Wang, et al. |
| EPI_ISL_937191 | Department of Homeless Services | New York City Public Health Laboratory | Jade Wang, et al. |
| EPI_ISL_937193, EPI_ISL_937194 | OCME Office Of Chief Medical Examiner | New York City Public Health Laboratory | Jade Wang, et al. |
| EPI_ISL_937383, EPI_ISL_937384, EPI_ISL_937388, EPI_ISL_937389, EPI_ISL_937390, EPI_ISL_937391, EPI_ISL_937414, EPI_ISL_937415, EPI_ISL_937416, EPI_ISL_937417, EPI_ISL_937418, EPI_ISL_937419, EPI_ISL_937420, EPI_ISL_937428, EPI_ISL_937429, EPI_ISL_937430, EPI_ISL_937431, EPI_ISL_937432, EPI_ISL_937433, EPI_ISL_937434, EPI_ISL_937438, EPI_ISL_937439, EPI_ISL_937440, EPI_ISL_937441, EPI_ISL_937442, EPI_ISL_937443, EPI_ISL_937444, EPI_ISL_937449, EPI_ISL_937450, EPI_ISL_937451, EPI_ISL_937452, EPI_ISL_937453, EPI_ISL_937454, EPI_ISL_937455, EPI_ISL_937456 | | | |
| see above | Maine Health and Environmental Testing Laboratory (Maine HETL) | Tewhey Lab, The Jackson Laboratory | Matluk,N., Dewey,H., Iosue,F., Barter,M., Lynch,R., Munger,H. and Tewhey,R. |
| EPI_ISL_938057, EPI_ISL_938120, EPI_ISL_938151 | Lighthouse Lab in Milton Keynes | Wellcome Sanger Institute for the COVID-19 Genomics UK (COG-UK) Consortium | The Lighthouse Lab in Milton Keynes and Alex Alderton, Roberto Amato, Sonia Goncalves, Ewan Harrison, David K. Jackson, Ian Johnston, Dominic Kwiatkowski, Cordelia Langford, John Sillitoe on behalf of the Wellcome Sanger Institute COVID-19 Surveillance Team |
| EPI_ISL_939630, EPI_ISL_939631, EPI_ISL_939632, EPI_ISL_939633, EPI_ISL_939634, EPI_ISL_939635, EPI_ISL_939637, EPI_ISL_939638, EPI_ISL_939639 | MEPHI, Aix Marseille University | MEPHI, Aix Marseille University | Anthony LEVASSEUR |
| EPI_ISL_940069, EPI_ISL_940070 | University Hospitals of Geneva, Laboratory of Virology | HUG, Laboratory of Virology and the Health2030 Genome Center | Samuel Cordey, Ana Rita Goncalves, Laurent Kaiser, Lorenzo Cerutti, Henri Pegeot, Melyssa Elies, Deborah Penet, Keith Harshman, Ioannis Xenarios, Emmanouil Dermitzakis |
| EPI_ISL_940564 | Servizo de Microbioloxía. Complexo Hospitalario Universitario de Santiago de Compostela | Servizo de Microbioloxía. Complexo Hospitalario Universitario de Santiago de Compostela | Antonio Aguilera, Gema Barbeito, Amparo Coira, Rocio Trastoy, María Luisa Pérez del Molino |
| EPI_ISL_940565 | University of Bari Biomedical Sciences and Human Oncology | University of Bari Biomedical Sciences and Human Oncology | Chironna M., Sallustio A., Loconsole D., Accogli M. |
| EPI_ISL_940609 | Hospital e Maternidade Celso Pierro | Instituto Adolfo Lutz, Interdiciplinary Procedures Center, Strategic Laboratory | Claudio Tavares Sacchi, Claudia Regina Gonçalves, Erica Valessa Ramos Gomes, Karoline Rodrigues Campos |
| EPI_ISL_940613, EPI_ISL_940614, EPI_ISL_940615, EPI_ISL_940616, EPI_ISL_940617, EPI_ISL_940618 | LACEN-PI DR. Costa Alvarenga | Instituto Adolfo Lutz, Interdiciplinary Procedures Center, Strategic Laboratory | Claudio Tavares Sacchi, Claudia Regina Gonçalves, Erica Valessa Ramos Gomes, Karoline Rodrigues Campos |
| EPI_ISL_940625 | Hospital Sao Joaquim - Beneficiencia Portuguesa | Instituto Adolfo Lutz, Interdiciplinary Procedures Center, Strategic Laboratory | Claudio Tavares Sacchi, Claudia Regina Gonçalves, Erica Valessa Ramos Gomes, Karoline Rodrigues Campos |
| EPI_ISL_940631, EPI_ISL_940632 | University of Bari Biomedical Sciences and Human Oncology | University of Bari Biomedical Sciences and Human Oncology | Chironna M., Sallustio A., Loconsole D., Accogli M. |
| EPI_ISL_940644, EPI_ISL_940645, EPI_ISL_940646, EPI_ISL_940647, EPI_ISL_940648, EPI_ISL_940649, EPI_ISL_940650, EPI_ISL_940651, EPI_ISL_940652, EPI_ISL_940653 | Ministry of Health Turkey | Ministry of Health Turkey | Fatma Bayrakdar, Yasemin Cogun, Süleyman Yalcin, Aye Baak Alta, Gülay Korukluolu |
| EPI_ISL_940683 | Servizo de Microbioloxía. Complexo Hospitalario Universitario de Santiago de Compostela | Servizo de Microbioloxía. Complexo Hospitalario Universitario de Santiago de Compostela | Antonio Aguilera, Gema Barbeito, Amparo Coira, Rocio Trastoy, María Luisa Pérez del Molino |
| EPI_ISL_940712 | Ministry of Health Turkey | Ministry of Health Turkey | Fatma Bayrakdar, Yasemin Cogun, Süleyman Yalcin, Aye Baak Alta, Gülay Korukluolu |
| EPI_ISL_940757 | GZA Sint-Augustinus Hospital | UAntwerp, Laboratory of Medical Microbiology, Campus Drie | Basil Britto Xavier, Jasmine Coppens, Marie Le Mercier, Christine Lammens, Veerle Matheeussen, Herman Goossens |

| | | | |
|---|---|---|---|
| | | Eiken S6.26, Universiteitsplein 1, 2610, Wilrijk, Belgium | |
| EPI_ISL_940762 | Botswana Harvard HIV Reference Laboratory | Botswana Harvard HIV Reference Laboratory | Sikhulile Moyo, Wonderful Choga, Dorcas Maruapula, Botshelo Radibe, Boitumelo Zuze, David Lawrence, Roger Shapiro, Shahin Lockman, Mosepele Mosepele, Joseph, Makhema, Simani Gaseitsiwe |
| EPI_ISL_940763 | ZNA Middelheim | UAntwerp, Laboratory of Medical Microbiology, Campus Drie Eiken S6.26, Universiteitsplein 1, 2610, Wilrijk, Belgium | Basil Britto Xavier, Jasmine Coppens, Marie Le Mercier, Christine Lammens, Veerle Matheeussen, Herman Goossens |
| EPI_ISL_940784, EPI_ISL_940786, EPI_ISL_940788, EPI_ISL_940789, EPI_ISL_940790, EPI_ISL_940791 | City of Milwaukee Health Department Laboratory | City of Milwaukee Health Department Laboratory | Sanjib Bhattacharyya |
| EPI_ISL_940843, EPI_ISL_940847, EPI_ISL_940849 | Platform BIS UZA/UAntwerpen | UAntwerp, Laboratory of Medical Microbiology | Basil Britto Xavier, Jasmine Coppens, Marie Le Mercier, Christine Lammens, Veerle Matheeussen, Herman Goossens |
| EPI_ISL_940891 | Vaccines and Infectious Diseases Analytics Research Unit (VIDA) | KRISP, KZN Research Innovation and Sequencing Platform | Baillie Vicky, du Plessis Jeanine, Giandhari Jennifer, Pillay Sureshnee, Naidoo Yeshnee, Tegally Houriiyah, de Oliveira Tulio, Madhi Shabir |
| EPI_ISL_940893 | Platform BIS UZA/UAntwerpen | UAntwerp, Laboratory of Medical Microbiology, Campus Drie Eiken S6.26, Universiteitsplein 1, 2610, Wilrijk, Belgium | Basil Britto Xavier, Jasmine Coppens, Marie Le Mercier, Christine Lammens, Veerle Matheeussen, Herman Goossens |
| EPI_ISL_940996 | Sentinelles Idf | National Reference Center for Viruses of Respiratory Infections, Institut Pasteur, Paris | Marion Barbet, Sylvie Behillil, Méline Bizard, Angela Brisebarre, Camille Capel, Etienne Simon-Lorière, Vincent Enouf, Maud Vanpeene, Sylvie van der Werf, Garrigues Anne |
| EPI_ISL_941026 | Labo Analyses Med | National Reference Center for Viruses of Respiratory Infections, Institut Pasteur, Paris | Marion Barbet, Sylvie Behillil, Méline Bizard, Angela Brisebarre, Camille Capel, Etienne Simon-Lorière, Vincent Enouf, Maud Vanpeene, Sylvie van der Werf, Griscelli Franck |
| EPI_ISL_941027 | Hopital | National Reference Center for Viruses of Respiratory Infections, Institut Pasteur, Paris | Marion Barbet, Sylvie Behillil, Méline Bizard, Angela Brisebarre, Camille Capel, Etienne Simon-Lorière, Vincent Enouf, Maud Vanpeene, Sylvie van der Werf, Marque Juillet StéPhanie |
| EPI_ISL_941035, EPI_ISL_941036, EPI_ISL_941037, EPI_ISL_941040, EPI_ISL_941041, EPI_ISL_941058, EPI_ISL_941063, EPI_ISL_941065, EPI_ISL_941071 | Labo Analyses Med | National Reference Center for Viruses of Respiratory Infections, Institut Pasteur, Paris | Marion Barbet, Sylvie Behillil, Méline Bizard, Angela Brisebarre, Camille Capel, Etienne Simon-Lorière, Vincent Enouf, Maud Vanpeene, Sylvie van der Werf, Merah Kader |
| EPI_ISL_941296, EPI_ISL_941297 | Nigeria Centre for Disease Control (NCDC) | African Centre of Excellence for Genomics of Infectious Diseases (ACEGID), Redeemer's University | Oluniyi P.E. et al |
| EPI_ISL_941437, EPI_ISL_941438, EPI_ISL_941439, EPI_ISL_941440, EPI_ISL_941441, EPI_ISL_941610, EPI_ISL_941611 | Instituto Nacional de Saude (INSA) | Instituto Nacional de Saude (INSA) | Borges et al |
| EPI_ISL_941842, EPI_ISL_941845, EPI_ISL_941849, EPI_ISL_941850, EPI_ISL_941852, EPI_ISL_941853, EPI_ISL_941854, EPI_ISL_941855, EPI_ISL_941856, EPI_ISL_941857, EPI_ISL_941858, EPI_ISL_941859, EPI_ISL_941867 | | | |
| see above | Instituto Nacional de Saude (INSA) and Instituto Gulbenkian de Ciencia (IGC) | Instituto Nacional de Saude (INSA) and Instituto Gulbenkian de Ciencia (IGC) | Borges et al |
| EPI_ISL_942376, EPI_ISL_942406 | Viollier AG | University Hospital Basel, Clinical Bacteriology | Tim Roloff, Madlen Stange, Helena MB Seth-Smith, Alfredo Mari, Karoline Leuzinger, Julia Bielicki, Christiane Beckmann, Manuel Battegay, Hans Hirsch, Adrian Egli |
| EPI_ISL_942515 | University Hospital Basel, Clinical Virology | University Hospital Basel, Clinical Bacteriology | Tim Roloff, Madlen Stange, Helena MB Seth-Smith, Alfredo Mari, Karoline Leuzinger, Julia Bielicki, Manuel Battegay, Hans Hirsch, Adrian Egli |
| EPI_ISL_942851, EPI_ISL_942852, EPI_ISL_942853, EPI_ISL_942854, EPI_ISL_942855, EPI_ISL_942856, EPI_ISL_942857, EPI_ISL_942858, EPI_ISL_942859, EPI_ISL_942860, EPI_ISL_942861 | | | |
| see above | Gundersen Molecular Diagnostics Laboratory | Kabara Cancer Research Institute | Craig S. Richmond, Paraic A. Kenny |
| EPI_ISL_942927 | LESP Nuevo Leon/Grupo de Diagnostico ARIES | Instituto de Diagnostico y Referencia Epidemiologicos (INDRE) | Claudia Wong-Arambula, Abril Rodriguez-Maldonado, Fabiola Garces-Ayala, Natividad Cruz-Ortiz, Tatiana Nunez-Garcia, Gisela Barrera-Badillo, Lucia Hernandez-Rivas, Irma Lopez-Martinez, Ernesto Ramirez-Gonzalez. |
| EPI_ISL_942946 | Gundersen Molecular Diagnostics Laboratory | Kabara Cancer Research Institute | Craig S. Richmond, Paraic A. Kenny |
| EPI_ISL_942972, EPI_ISL_942973, EPI_ISL_942974 | Servizo de Microbioloxía. Complexo Hospitalario Universitario de Santiago de Compostela | Servizo de Microbioloxía. Complexo Hospitalario Universitario de Santiago de Compostela | Antonio Aguilera, Gema Barbeito, Amparo Coira, José Costa, Rocio Trastoy, María Luisa Pérez del Molino. |
| EPI_ISL_942998, EPI_ISL_943027, EPI_ISL_943042, EPI_ISL_943043, EPI_ISL_943048, EPI_ISL_943060, EPI_ISL_943061, EPI_ISL_943062, EPI_ISL_943081, EPI_ISL_943083, EPI_ISL_943140, EPI_ISL_943146, EPI_ISL_943168, EPI_ISL_943169, EPI_ISL_943170, EPI_ISL_943171, EPI_ISL_943172, EPI_ISL_943175, EPI_ISL_943178, EPI_ISL_943179, EPI_ISL_943180, EPI_ISL_943237, EPI_ISL_943238, EPI_ISL_943239, EPI_ISL_943259, EPI_ISL_943261, EPI_ISL_943273, EPI_ISL_943285, EPI_ISL_943311, EPI_ISL_943312, EPI_ISL_943315, EPI_ISL_943323, EPI_ISL_943324, EPI_ISL_943325, EPI_ISL_943352, EPI_ISL_943378, EPI_ISL_943385, EPI_ISL_943388, EPI_ISL_943389, EPI_ISL_943391, EPI_ISL_943392, EPI_ISL_943440, EPI_ISL_943470, EPI_ISL_943471, EPI_ISL_943472, EPI_ISL_943476, EPI_ISL_943487, EPI_ISL_943488, EPI_ISL_943517, EPI_ISL_943546 | | | |
| see above | Dutch COVID-19 response team | National Institute for Public Health and the Environment (RIVM) | Adam Meijer, Harry Vennema, Dirk Eggink, Jeroen Cremer, Sharon van den Brink, Bas van der Veer, AnneMarie van den Brandt, Florian Zwagemaker, Dennis Schmitz, Chantal Reusken, on behalf of the national COVID-19 response team |
| EPI_ISL_943549 | National Institute of Laboratory Medicine and Referral Center | Genomic Research Lab, BCSIR | Md. Murshed Hasan Sarkar, Mohammad Samir Uzzaman,Eshrar Osman, Md. Ahashan Habib, Shahina Akter, Tanjina Akhtar Banu,Abu Sayeed Mohammad Mahmud,Barna Goswami,Iffat Jahan, Md. Saddam Hossain, Tasnim Nafisa, Md. Maruf Ahmed Molla, Mahmuda Yeasmin, Asish Kumar Ghosh, Arifa Akram, A. K. M. Shamsuzzaman,Md. Salim Khan |
| EPI_ISL_943554 | National Institute of Laboratory Medicine and Referral Center | Genomic Research Lab, BCSIR | Barna Goswami,Mohammad Samir Uzzaman, Eshrar Osman, Md. Ahashan Habib,Shahina Akter, Tanjina Akhter Banu, Abu Sayeed Mohammad Mahmud, Md. Murshed Hasan Sarkar,Iffat Jahan, Md. Saddam Hossain, Tasnim Nafisa, Md. Maruf Ahmed Molla, Mahmuda Yeasmin, Asish Kumar Ghosh,Arifa Akram, A. K. M. Shamsuzzaman, Md. Salim Khan |
| EPI_ISL_943556, EPI_ISL_943558 | Servizo de Microbioloxía. Complexo Hospitalario Universitario de Santiago de Compostela | Servizo de Microbioloxía. Complexo Hospitalario Universitario de Santiago de Compostela | Antonio Aguilera, Gema Barbeito, Amparo Coira, José Costa, Rocio Trastoy, María Luisa Pérez del Molino. |
| EPI_ISL_943559 | National Institute of Laboratory Medicine and Referral Center | Genomic Research Lab, BCSIR | Shahina Akter, Mohammad Samir Uzzaman, Eshrar Osman, Md. Ahashan Habib,Tanjina Akhtar Banu, Abu Sayeed Mohammad Mahmud, Md. Murshed Hasan Sarkar, Barna Goswami, Iffat Jahan, Md. Saddam Hossain, Tasnim Nafisa, Md. Maruf Ahmed Molla, Mahmuda Yeasmin, Asish Kumar Ghosh, Arifa Akram, A. K. M. Shamsuzzaman, Md. Salim Khan |
| EPI_ISL_943560, EPI_ISL_943562 | Servizo de Microbioloxía. Complexo Hospitalario Universitario de Santiago de Compostela | Servizo de Microbioloxía. Complexo Hospitalario Universitario de Santiago de Compostela | Antonio Aguilera, Gema Barbeito, Amparo Coira, José Costa, Rocio Trastoy, María Luisa Pérez del Molino. |
| EPI_ISL_943563 | National Institute of Laboratory Medicine and Referral Center | Genomic Research Lab, BCSIR | Md. Saddam Hossain,Mohammad Samir Uzzaman, Eshrar Osman, Md. Ahashan Habib,Shahina Akter,Tanjina Akhter Banu, Abu Sayeed Mohammad Mahmud, Md. Murshed Hasan Sarkar, Barna Goswami, Iffat Jahan, Tasnim Nafisa, Md. Maruf Ahmed Molla, Mahmuda Yeasmin, Asish Kumar Ghosh, Arifa Akram, A. K. M. Shamsuzzaman, Md. Salim Khan |
| EPI_ISL_943572 | National Institute of Laboratory Medicine and Referral Center | Genomic Research Lab, BCSIR | Md. Maruf Ahmed Molla,Mohammad Samir Uzzaman, Eshrar Osman, Md. Ahashan Habib, Shahina Akter,Tanjina Akhtar Banu, Abu Sayeed Mohammad Mahmud, Md. Murshed Hasan Sarkar, Barna Goswami, Iffat Jahan, Md. Saddam Hossain, Tasnim Nafisa,Mahmuda Yeasmin, Asish Kumar Ghosh,Arifa Akram, A. K. M. Shamsuzzaman,Md. Salim Khan |
| EPI_ISL_943814, EPI_ISL_943815, EPI_ISL_943816, EPI_ISL_943817, EPI_ISL_943930, EPI_ISL_943934, EPI_ISL_943937 | Utah Public Health Laboratory | Utah Public Health Laboratory | Erin L. Young, Kelly F. Oakeson, Tara Gallagher |

| | | | |
|---|---|---|---|
| EPI_ISL_943990 | LACEN do Estado de Goias | Instituto Adolfo Lutz, Interdiciplinary Procedures Center, Strategic Laboratory | Claudio Tavares Sacchi, Claudia Regina Gonçalves, Erica Valessa Ramos Gomes, Karoline Rodrigues Campos |
| EPI_ISL_944590 | Botswana Harvard HIV Reference Laboratory | Botswana Harvard HIV Reference Laboratory | Sikhulile Moyo, Wonderful T. Choga, Dorcas Maruapula, Botshelo Radibe, Boitumelo Zuze, David Lawrence, Roger Shapiro, Shahin Lockman, Mosepele Mosepele, Joseph Makhema, Simani Gaseitsiwe |
| EPI_ISL_944740, EPI_ISL_944741, EPI_ISL_944742, EPI_ISL_944743 | unknown | Public Health Virology-Forensic and Scientific Services (PHV-FSS) | Son Nguyen et al. |
| EPI_ISL_944782 | Botswana Harvard HIV Reference Laboratory | Botswana Harvard HIV Reference Laboratory | Sikhulile Moyo, Wonderful T. Choga, Dorcas Maruapula, Botshelo Radibe, Boitumelo Zuze, David Lawrence, Roger Shapiro, Shahin Lockman, Mosepele Mosepele, Joseph Makhema, Simani Gaseitsiwe |
| EPI_ISL_948973, EPI_ISL_948975, EPI_ISL_948978, EPI_ISL_948981, EPI_ISL_948984, EPI_ISL_948989, EPI_ISL_948991, EPI_ISL_948994, EPI_ISL_948997, EPI_ISL_949000, EPI_ISL_949003, EPI_ISL_949005, EPI_ISL_949008, EPI_ISL_949011, EPI_ISL_949013, EPI_ISL_949016, EPI_ISL_949019, EPI_ISL_949022, EPI_ISL_949024, EPI_ISL_949069, EPI_ISL_949070, EPI_ISL_949095, EPI_ISL_949096, EPI_ISL_949097, EPI_ISL_949098, EPI_ISL_949099, EPI_ISL_949100, EPI_ISL_949101, EPI_ISL_949102, EPI_ISL_949103, EPI_ISL_949104, EPI_ISL_949105 | | | |
| see above | Jessa | Jessa | Jessa_cmdLab |
| EPI_ISL_949182 | University of Bari Biomedical Sciences and Human Oncology | University of Bari Biomedical Sciences and Human Oncology | Chironna M., Sallustio A., Loconsole D., Accogli M. |
| EPI_ISL_949254 | Jessa | Jessa | Jessa_cmdLab |
| EPI_ISL_949408, EPI_ISL_949426 | University of Birmingham | COVID-19 Genomics UK (COG-UK) Consortium | Institute of Microbiology, University of Birmingham: Claire McMurray, Joanne Stockton, Samuel Nicholls, Radoslaw Poplawski, Will Rowe, Josh Quick, Nicholas Loman. University of Birmingham Testing Laboratory: Celina M Whalley, Andrew Bosworth, Charlotte Poxon, Kasun Wanigasooriya, Oliver Pickles, Mike Kidd, Alex Richter, Andrew D Beggs PHE Heartlands Lab: Husam Osman, Andrew Bosworth. Queen Elizabeth Hospital: Anna Casey |
| EPI_ISL_949428, EPI_ISL_949442, EPI_ISL_949443, EPI_ISL_949444, EPI_ISL_949448, EPI_ISL_949478, EPI_ISL_949480, EPI_ISL_949481, EPI_ISL_949482, EPI_ISL_949483, EPI_ISL_949484, EPI_ISL_949486, EPI_ISL_949488, EPI_ISL_949489, EPI_ISL_949491, EPI_ISL_949493, EPI_ISL_949495, EPI_ISL_949497, EPI_ISL_949499, EPI_ISL_949501, EPI_ISL_949503, EPI_ISL_949505, EPI_ISL_949507, EPI_ISL_949509, EPI_ISL_949511, EPI_ISL_949513, EPI_ISL_949515, EPI_ISL_949517 | | | |
| see above | Department of Pathology, University of Cambridge | COVID-19 Genomics UK (COG-UK) Consortium | Aminu S. Jahun, Yasmin Chaudhry, Iliana Georgana, Myra Hosmillo, Rhys Izu, Martin D. Curran, Surendra Parmar, Ian Goodfellow |
| EPI_ISL_949612 | West of Scotland Specialist Virology Centre, NHSGGC / MRC-University of Glasgow Centre for Virus Research | COVID-19 Genomics UK (COG-UK) Consortium | Ana da Silva Filipe, Natasha Johnson, Kathy Smollett, Daniel Mair, Stephen Carmichael, Alice Broos, Lily Tong, Jenna Nichols, Kyriaki Nomikou; Sarah McDonald; Richard Orton, Joseph Hughes, Sreenu Vattipally, David L Robertson; Alasdair MacLean, Rory Gunson; Sharif Shaaban, Matthew Holden; Rachel Blacow, Guy Mollett, Kathy Li, James Shepherd, Antonia Ho, Emma Thomson |
| EPI_ISL_949639 | Virology Department, Royal Infirmary of Edinburgh, NHS Lothian / School of Biological Sciences, University of Edinburgh / Institute of Genetics and Molecular Medicine, University of Edinburgh | COVID-19 Genomics UK (COG-UK) Consortium | McHugh M, Dewar R, Rooke S, Gallagher M, Balcaza C, O'Toole Á, Scher E, Hill V, McCrone JT, Colquhoun R, Yu X, Jackson B, Rambaut A, Williams TC, Templeton K |
| EPI_ISL_949791, EPI_ISL_949979 | University College London, Great Ormond Street Hospital for Children NHS Foundation Trust, Imperial College Healthcare NHS Trust | COVID-19 Genomics UK (COG-UK) Consortium | Sergi Castellano, Rachel Williams, Mark Kristiansen, Paola Resende Silva, Sunando Roy, Tony Brooks, Helena Tutill, Paola Niola, Patricia Dyal, Charlotte Williams, Leysa Forrest, Yasmin Panchbhaya, Jacqueline Findlay, Samuel Weeks, Julianne Brown, Kathryn Harris, Paul Randell, James Price, Alison Holmes, Judith Breuer |
| EPI_ISL_950339, EPI_ISL_950340, EPI_ISL_950343, EPI_ISL_950344, EPI_ISL_950345 | Northumbria University / South Tees Hospitals NHS Foundation Trust / North Cumbria Integrated Care NHS Foundation Trust / North Tees and Hartlepool NHS Foundation Trust / Newcastle Hospitals NHS Foundation Trust | COVID-19 Genomics UK (COG-UK) Consortium | Darren L Smith,Andrew Nelson,Matthew Bashton,Greg R Young,Joshua Loh,John Allan,Mohammad A Tariq,Giles S Holt,Gary Black,Wen C Yew,Lynn Dover,Paul Baker,Steve Liggett,Sarah Essex,Jane Greenaway,Debra Padgett,Clive Graham,Garren Scott,Edward Barton,Emma Swindells,Brendan Payne,Jennifer Collins,Yusri Taha,Gary Eltringham |
| EPI_ISL_950563, EPI_ISL_950564, EPI_ISL_950565, EPI_ISL_950567, EPI_ISL_950568, EPI_ISL_950569, EPI_ISL_950570, EPI_ISL_950571 | Quadram Institute Bioscience | COVID-19 Genomics UK (COG-UK) Consortium | Dave J. Baker, Gemma L. Kay, Alp Aydin, Thanh Le-Viet, Steven Rudder, Ana P. Tedim, Anastasia Kolyva, Maria Diaz, Leonardo de Oliveira Martins, Nabil-Fareed Alikhan, Lizzie Meadows, Rachael Stanley, Ngozi Elumogo, Muhammed Yasir, Nicholas M. Thomson, Alexander J Trotter, Rachel Gilroy, Samuel Bloomfield, Claire Stuart, Andrew Bell, Reenesh Prakash, Samir Dervisevic, Alison E. Mather, John Wain, Mark Webber, Andrew J. Page, Justin O'Grady |
| EPI_ISL_950716 | Lincolnshire Hospitals and DeepSeq Nottingham | COVID-19 Genomics UK (COG-UK) Consortium | Nichola Duckworth, Tim Sloan, Sarah Walsh, Jonathan Ball, Patrick McClure, Joeseph Chappell, Nadine Holmes, Matthew Carlisle, Christopher Moore, Fei Sang, Johnny Debebe, Victoria Wright, Matthew Loose |
| EPI_ISL_951360, EPI_ISL_951369, EPI_ISL_951370, EPI_ISL_951371, EPI_ISL_951466, EPI_ISL_951468, EPI_ISL_951470, EPI_ISL_951471, EPI_ISL_951476, EPI_ISL_951479, EPI_ISL_951480, EPI_ISL_951481, EPI_ISL_951482, EPI_ISL_951484, EPI_ISL_951485, EPI_ISL_951489 | | | |
| see above | Oxford Viromics, NDM, University of Oxford; Oxford University Hospitals; Basingstoke and North Hampshire Hospital | COVID-19 Genomics UK (COG-UK) Consortium | Tanya Golubchik, David Bonsall, George Macintyre, Amy Trebes, Mariateresa de Cesare, Catrin Moore, Alex Mobbs, Anita Justice, Robert Shaw, Monique Andersson, Timothy Peto, Emma Wise, Nathan Moore, Jessica Lynch, Nick Cortes, Matilde Mori, Stephen Kidd, David Buck, John Todd, Christophe Fraser |
| EPI_ISL_951590, EPI_ISL_951591, EPI_ISL_951592, EPI_ISL_951593, EPI_ISL_951632, EPI_ISL_951633, EPI_ISL_951634, EPI_ISL_951635, EPI_ISL_951637, EPI_ISL_951648, EPI_ISL_951740, EPI_ISL_951756, EPI_ISL_952187, EPI_ISL_952188, EPI_ISL_952189, EPI_ISL_952190 | | | |
| see above | Originating lab: Wales Specialist Virology Centre Sequencing lab: Pathogen Genomics Unit | Public Health Wales Microbiology Cardiff Wales Specialist Virology Centre | Catherine Moore, Johnathan Evans, Laura Gifford, Malorie Perry, Simon Cottrell, Angela Marchbank, Alec Birchley, Alexander Adams, Amy Gaskin, Bree Gatica-Wilcox, Jason Coombes, Joel Southgate, Lauren Gilbert, Lee Graham, Nicole Pacchiarini, Sara Kumziene-Summerhayes, Sarah Taylor, Sophie Jones, Sara Rey, Matthew Bull, Joanne Watkins, Sally Corden, Tom Connor |
| EPI_ISL_952405, EPI_ISL_952410, EPI_ISL_952412, EPI_ISL_952416, EPI_ISL_952432, EPI_ISL_952489, EPI_ISL_952490, EPI_ISL_952521, EPI_ISL_952522, EPI_ISL_952528, EPI_ISL_952529, EPI_ISL_952538, EPI_ISL_952673, EPI_ISL_952675, EPI_ISL_952677, EPI_ISL_952680, EPI_ISL_952689, EPI_ISL_952820, EPI_ISL_952830, EPI_ISL_952831, EPI_ISL_952832, EPI_ISL_952842, EPI_ISL_952844, EPI_ISL_952845, EPI_ISL_952846, EPI_ISL_952847, EPI_ISL_952848, EPI_ISL_952849, EPI_ISL_952850, EPI_ISL_952852, EPI_ISL_952853 | | | |
| see above | Centre for Enzyme Innovation, University of Portsmouth / Translational Research Laboratory, Portsmouth Hospitals NHS Trust | COVID-19 Genomics UK (COG-UK) Consortium | Angela Beckett,Salman Goudarzi,Christopher Fearn,Kate Cook,Katie Loveson,Sharon Glaysher,Scott Elliott,Samuel Robson |
| EPI_ISL_953504, EPI_ISL_953515, EPI_ISL_953523, EPI_ISL_953524, EPI_ISL_953525, EPI_ISL_953599, EPI_ISL_953600, EPI_ISL_953601, EPI_ISL_953626, EPI_ISL_953652, EPI_ISL_953662, EPI_ISL_953671, EPI_ISL_953672, EPI_ISL_953673, EPI_ISL_953674, EPI_ISL_953675, EPI_ISL_953676, EPI_ISL_953677, EPI_ISL_953678, EPI_ISL_953679, EPI_ISL_953680, EPI_ISL_953681, EPI_ISL_953682, EPI_ISL_953683, EPI_ISL_953684, EPI_ISL_953685, EPI_ISL_953686, EPI_ISL_953687, EPI_ISL_953688, EPI_ISL_953689, EPI_ISL_953690, EPI_ISL_953691, EPI_ISL_953692, EPI_ISL_953693, EPI_ISL_953694, EPI_ISL_953695, EPI_ISL_953696, EPI_ISL_953697, EPI_ISL_953698, EPI_ISL_953699, EPI_ISL_953700, EPI_ISL_953701, EPI_ISL_953702, EPI_ISL_953703, EPI_ISL_953704, EPI_ISL_953705, EPI_ISL_953706 | | | |
| see above | University Hospitals of Geneva, Laboratory of Virology | HUG, Laboratory of Virology and the Health2030 Genome Center | Samuel Cordey, Ana Rita Goncalves, Laurent Kaiser, Lorenzo Cerutti, Henri Pegeot, Melyssa Elies, Deborah Penet, Keith Harshman, Ioannis Xenarios, Emmanouil Dermitzakis |
| EPI_ISL_953946, EPI_ISL_953947, EPI_ISL_953948 | Hopital | National Reference Center for Viruses of Respiratory Infections, Institut Pasteur, Paris | Marion Barbet, Sylvie Behillil, Méline Bizard, Angela Brisebarre, Camille Capel, Etienne Simon-Lorière, Vincent Enouf, Maud Vanpeene, Sylvie van der Werf,Cady Anne |
| EPI_ISL_953986 | Labo Analyses Med | National Reference Center for Viruses of Respiratory Infections, Institut Pasteur, Paris | Marion Barbet, Sylvie Behillil, Méline Bizard, Angela Brisebarre, Camille Capel, Etienne Simon-Lorière, Vincent Enouf, Maud Vanpeene, Sylvie van der Werf |
| EPI_ISL_953987 | Hopital | National Reference Center for Viruses of Respiratory Infections, Institut Pasteur, Paris | Marion Barbet, Sylvie Behillil, Méline Bizard, Angela Brisebarre, Camille Capel, Etienne Simon-Lorière, Vincent Enouf, Maud Vanpeene, Sylvie van der Werf,Lagathu GisèLe |
| EPI_ISL_953988 | Labo Analyses Med | National Reference Center for Viruses of Respiratory Infections, Institut Pasteur, Paris | Marion Barbet, Sylvie Behillil, Méline Bizard, Angela Brisebarre, Camille Capel, Etienne Simon-Lorière, Vincent Enouf, Maud Vanpeene, Sylvie van der Werf,Selas Labomaine |
| EPI_ISL_953989 | Hopital | National Reference Center for Viruses of Respiratory Infections, Institut Pasteur, Paris | Marion Barbet, Sylvie Behillil, Méline Bizard, Angela Brisebarre, Camille Capel, Etienne Simon-Lorière, Vincent Enouf, Maud Vanpeene, Sylvie van der Werf,Lesimple BéAtrice |
| EPI_ISL_953990 | Hopital | National Reference Center for Viruses of Respiratory Infections, Institut Pasteur, Paris | Marion Barbet, Sylvie Behillil, Méline Bizard, Angela Brisebarre, Camille Capel, Etienne Simon-Lorière, Vincent Enouf, Maud Vanpeene, Sylvie van der Werf,Eloy Clarence |
| EPI_ISL_954020 | Hopital | National Reference Center for Viruses of Respiratory Infections, Institut Pasteur, Paris | Marion Barbet, Sylvie Behillil, Méline Bizard, Angela Brisebarre, Camille Capel, Etienne Simon-Lorière, Vincent Enouf, Maud Vanpeene, Sylvie van der Werf,Breuil J. |
| EPI_ISL_954021 | hopital | National Reference Center for Viruses of Respiratory Infections, Institut Pasteur, Paris | Marion Barbet, Sylvie Behillil, Méline Bizard, Angela Brisebarre, Camille Capel, Etienne Simon-Lorière, Vincent Enouf, Maud Vanpeene, Sylvie van der Werf,Isabelle Joly |

| | | | |
|---|---|---|---|
| EPI_ISL_954091 | Labo Analyses Med | National Reference Center for Viruses of Respiratory Infections, Institut Pasteur, Paris | Marion Barbet, Sylvie Behillil, Méline Bizard, Angela Brisebarre, Camille Capel, Etienne Simon-Lorière, Vincent Enouf, Maud Vanpeene, Sylvie van der Werf,Besson J |
| EPI_ISL_954092 | Labo Analyses Med | National Reference Center for Viruses of Respiratory Infections, Institut Pasteur, Paris | Marion Barbet, Sylvie Behillil, Méline Bizard, Angela Brisebarre, Camille Capel, Etienne Simon-Lorière, Vincent Enouf, Maud Vanpeene, Sylvie van der Werf,Dlab Dieppe |
| EPI_ISL_954093 | Labo Analyses Med | National Reference Center for Viruses of Respiratory Infections, Institut Pasteur, Paris | Marion Barbet, Sylvie Behillil, Méline Bizard, Angela Brisebarre, Camille Capel, Etienne Simon-Lorière, Vincent Enouf, Maud Vanpeene, Sylvie van der Werf,Holstein Anne |
| EPI_ISL_954094, EPI_ISL_954095, EPI_ISL_954096, EPI_ISL_954097, EPI_ISL_954098, EPI_ISL_954099, EPI_ISL_954100, EPI_ISL_954101, EPI_ISL_954102 | Outre Mer | National Reference Center for Viruses of Respiratory Infections, Institut Pasteur, Paris | Marion Barbet, Sylvie Behillil, Méline Bizard, Angela Brisebarre, Camille Capel, Etienne Simon-Lorière, Vincent Enouf, Maud Vanpeene, Sylvie van der Werf,Rousset Dominique |
| EPI_ISL_954103, EPI_ISL_954104 | Hopital | National Reference Center for Viruses of Respiratory Infections, Institut Pasteur, Paris | Marion Barbet, Sylvie Behillil, Méline Bizard, Angela Brisebarre, Camille Capel, Etienne Simon-Lorière, Vincent Enouf, Maud Vanpeene, Sylvie van der Werf,Raulin Olivia |
| EPI_ISL_954105 | Labo Analyses Med | National Reference Center for Viruses of Respiratory Infections, Institut Pasteur, Paris | Marion Barbet, Sylvie Behillil, Méline Bizard, Angela Brisebarre, Camille Capel, Etienne Simon-Lorière, Vincent Enouf, Maud Vanpeene, Sylvie van der Werf,Takoudju Eve-Marie |
| EPI_ISL_954106, EPI_ISL_954107 | Hopital | National Reference Center for Viruses of Respiratory Infections, Institut Pasteur, Paris | Marion Barbet, Sylvie Behillil, Méline Bizard, Angela Brisebarre, Camille Capel, Etienne Simon-Lorière, Vincent Enouf, Maud Vanpeene, Sylvie van der Werf,Lesimple BéAtrice |
| EPI_ISL_954108, EPI_ISL_954109, EPI_ISL_954110, EPI_ISL_954111 | Outre Mer | National Reference Center for Viruses of Respiratory Infections, Institut Pasteur, Paris | Marion Barbet, Sylvie Behillil, Méline Bizard, Angela Brisebarre, Camille Capel, Etienne Simon-Lorière, Vincent Enouf, Maud Vanpeene, Sylvie van der Werf,Rousset Dominique |
| EPI_ISL_954142, EPI_ISL_954156, EPI_ISL_954157, EPI_ISL_954158, EPI_ISL_954159, EPI_ISL_954160, EPI_ISL_954161, EPI_ISL_954162, EPI_ISL_954163, EPI_ISL_954164, EPI_ISL_954173, EPI_ISL_954174, EPI_ISL_954182 | | | |
| see above | Hospital Universitari Vall d'Hebron - Vall d'Hebron Institut de Rercerca | Hospital Universitari Vall d'Hebron | Cristina Andrés, Maria Piñana, Josep F Abril, Damir Garcia-Cehic, Ariadna Rando, Juliana Esperalba, Maria Gema Codina, Carla Castillo, Maria Carmen Martin, Tomàs Pumarola, Josep Quer, Andrés Antón |
| EPI_ISL_954753 | City of Milwaukee Health Department Laboratory | City of Milwaukee Health Department Laboratory | Sanjib Bhattacharyya |
| EPI_ISL_954783 | Hospital Universitario Virgen de la Arrixaca | Instituto de Salud Carlos III | Iglesias-Caballero, M. Camarero, S. Sandonís,V. Vázquez, S. Pozo, F. Casas, I. Jiménez, P. Zaballos, A. Monzón, S. Varona, S. Cuesta, I. Moreno, L. |
| EPI_ISL_954785 | Complejo Asistencial Universitario de Salamanca | Instituto de Salud Carlos III | Iglesias-Caballero, M. Camarero, S. Sandonís,V. Vázquez, S. Pozo, F. Casas, I. Jiménez, P. Zaballos, A. Monzón, S. Varona, S. Cuesta, I. Ávila,A. |
| EPI_ISL_954797, EPI_ISL_954801, EPI_ISL_954803 | Hospital Los Arcos | Instituto de Salud Carlos III | Iglesias-Caballero, M. Camarero, S. Sandonís,V. Vázquez, S. Pozo, F. Casas, I. Jiménez, P. Zaballos, A. Monzón, S. Varona, S. Cuesta, I. Camara, M. |
| EPI_ISL_954821, EPI_ISL_954828, EPI_ISL_954847, EPI_ISL_954855, EPI_ISL_954863, EPI_ISL_954868, EPI_ISL_954873, EPI_ISL_954875, EPI_ISL_954999, EPI_ISL_955000, EPI_ISL_955001, EPI_ISL_955002, EPI_ISL_955003, EPI_ISL_955004, EPI_ISL_955005, EPI_ISL_955006, EPI_ISL_955007, EPI_ISL_955008, EPI_ISL_955009, EPI_ISL_955010, EPI_ISL_955011, EPI_ISL_955012, EPI_ISL_955013, EPI_ISL_955014, EPI_ISL_955015, EPI_ISL_955016, EPI_ISL_955017, EPI_ISL_955018, EPI_ISL_955019, EPI_ISL_955020, EPI_ISL_955021, EPI_ISL_955022, EPI_ISL_955023, EPI_ISL_955024, EPI_ISL_955025, EPI_ISL_955026, EPI_ISL_955027, EPI_ISL_955028, EPI_ISL_955029, EPI_ISL_955030, EPI_ISL_955031, EPI_ISL_955032, EPI_ISL_955033, EPI_ISL_955034, EPI_ISL_955035, EPI_ISL_955036, EPI_ISL_955037, EPI_ISL_955038, EPI_ISL_955039, EPI_ISL_955040 | | | |
| see above | Colorado Department of Public Health and Environment | Colorado Department of Puplic Health and Environment | Laura Bankers, Molly C. Hetherington-Rauth, Diana Ir, Shannon Ely, Shannon R. Matzinger, Sarah Elizabeth Totten, Emily A. Travanty |
| EPI_ISL_955064, EPI_ISL_955075, EPI_ISL_955100 | Hopital | National Reference Center for Viruses of Respiratory Infections, Institut Pasteur, Paris | Marion Barbet, Sylvie Behillil, Méline Bizard, Angela Brisebarre, Camille Capel, Etienne Simon-Lorière, Vincent Enouf, Maud Vanpeene, Sylvie van der Werf,Guinoiseau Thibault |
| EPI_ISL_955103 | National Institute of Laboratory Medicine and Referral Center | Genomic Research Lab, BCSIR | Md. Ahashan Habib,Mohammad Samir Uzzaman, Eshrar Osman, Shahina Akter,Tanjina Akhtar Banu, Abu Sayeed Mohammad Mahmud, Md. Murshed Hasan Sarkar, Barna Goswami, Iffat Jahan, Md. Saddam Hossain, Tasnim Nafisa, Md. Maruf Ahmed Molla, Mahmuda Yeasmin, Asish Kumar Ghosh, Arifa Akram, A. K. M. Shamsuzzaman, Md. Salim Khan |
| EPI_ISL_955120, EPI_ISL_955121, EPI_ISL_955122, EPI_ISL_955123, EPI_ISL_955124, EPI_ISL_955125 | Maryland Public Health Laboratory | Maryland Public Health Laboratory | Maryland Department of Health Laboratories Administration |
| EPI_ISL_955143, EPI_ISL_955145 | Platform BIS UZA/UAntwerpen | UAntwerp, Laboratory of Medical Microbiology | Basil Britto Xavier, Jasmine Coppens, Marie Le Mercier, Christine Lammens, Veerle Matheeussen, Herman Goossens |
| EPI_ISL_955176 | Hopital | National Reference Center for Viruses of Respiratory Infections, Institut Pasteur, Paris | Marion Barbet, Sylvie Behillil, Méline Bizard, Angela Brisebarre, Camille Capel, Etienne Simon-Lorière, Vincent Enouf, Maud Vanpeene, Sylvie van der Werf,Brichler SéGolèNe |
| EPI_ISL_955177 | Hopital | National Reference Center for Viruses of Respiratory Infections, Institut Pasteur, Paris | Marion Barbet, Sylvie Behillil, Méline Bizard, Angela Brisebarre, Camille Capel, Etienne Simon-Lorière, Vincent Enouf, Maud Vanpeene, Sylvie van der Werf,Fourgeaud Jacques |
| EPI_ISL_955178, EPI_ISL_955179 | Labo Analyses med | National Reference Center for Viruses of Respiratory Infections, Institut Pasteur, Paris | Marion Barbet, Sylvie Behillil, Méline Bizard, Angela Brisebarre, Camille Capel, Etienne Simon-Lorière, Vincent Enouf, Maud Vanpeene, Sylvie van der Werf,Amzalag Jonas |
| EPI_ISL_955180, EPI_ISL_955181 | Labo Analyses Med | National Reference Center for Viruses of Respiratory Infections, Institut Pasteur, Paris | Marion Barbet, Sylvie Behillil, Méline Bizard, Angela Brisebarre, Camille Capel, Etienne Simon-Lorière, Vincent Enouf, Maud Vanpeene, Sylvie van der Werf,Amzalag Jonas |
| EPI_ISL_955183 | Labo Analyses Med | National Reference Center for Viruses of Respiratory Infections, Institut Pasteur, Paris | Marion Barbet, Sylvie Behillil, Méline Bizard, Angela Brisebarre, Camille Capel, Etienne Simon-Lorière, Vincent Enouf, Maud Vanpeene, Sylvie van der Werf,Le Berre (D) David |
| EPI_ISL_955224, EPI_ISL_955225 | Indiana Animal Disease Diagnostic Laboratory | Carpi Laboratory - Purdue University | Jack Dorman, Ilinca I Ciubotariu, Lev Gorenstein, Abebe A Fola, G Kenitra Hendrix, Rebecca P Wilkes, Giovanna Carpi |
| EPI_ISL_955782, EPI_ISL_955785, EPI_ISL_955787, EPI_ISL_955794, EPI_ISL_955795, EPI_ISL_955796, EPI_ISL_955797, EPI_ISL_955798, EPI_ISL_955799, EPI_ISL_955800, EPI_ISL_955801, EPI_ISL_955803, EPI_ISL_955804, EPI_ISL_955805, EPI_ISL_955806, EPI_ISL_955807, EPI_ISL_955808, EPI_ISL_955809, EPI_ISL_955812, EPI_ISL_955813, EPI_ISL_955814, EPI_ISL_955837, EPI_ISL_955838, EPI_ISL_955839, EPI_ISL_955845, EPI_ISL_955846, EPI_ISL_955847, EPI_ISL_955848, EPI_ISL_955849, EPI_ISL_955850, EPI_ISL_955851 | | | |
| see above | University of Michigan Clinical Microbiology Laboratory | Lauring Lab, University of Michigan, Department of Microbiology and Immunology | Valesano |
| EPI_ISL_956284, EPI_ISL_956285, EPI_ISL_956286, EPI_ISL_956294 | Instituto Nacional de Salud- Dirección de Redes de Laboratorios de Salud Pública | Instituto Nacional de Salud- Dirección de Investigación en Salud Pública | Katherine Laiton-Donato, Diego A. Álvarez-Díaz, Carlos Franco-Muñoz, Mauricio Pacheco-Montealegre, Hector Alejandro Ruiz-Moreno, Maria T. Herrera-Sepúlveda, Diego Andrés Prada, Jhonnatan Reales-González, Sheryll Corchuelo, Julian Naizaque, Gerardo Santamaría, Magdalena Wiesner, Martha Lucia Ospina Martinez, Marcela Mercado-Reyes |
| EPI_ISL_956329 | Laboratory Medicine | Department of Laboratory Medicine, Lin-Kou Chang Gung Memorial Hospital, Taoyuan, Taiwan | Kuo-Chien Tsao, Yu-Nong Gong, Shu-Li Yang, Yi-Chun Liu, Chung-Guei Huang, Mei-Jen Hsiao, Po-Wei Huang, Cheng-Ta Yang, Cheng-Hsun Chiu, Peng-Nien Huang, Kuo-Ming Lee, Guang-Wu Chen, Shin-Ru Shih |
| EPI_ISL_956333 | Utah Public Health Laboratory, Utah Public Health Laboratory Infectious Disease submission group | Utah Public Health Laboratory, Utah Public Health Laboratory Infectious Disease submission group | Gallagher,T., Young,E.L., Oakeson,K.F. |
| EPI_ISL_959309 | National Institute of Laboratory Medicine and Referral Center | Genomic Research Lab, BCSIR | Md. Maruf Ahmed Molla, Mohammad Samir Uzzaman, Eshrar Osman, Md. Ahashan Habib, Shahina Akter,Tanjina Akhtar Banu, Abu Sayeed Mohammad Mahmud, Md. Murshed Hasan Sarkar, Barna Goswami, Iffat Jahan, Md. Saddam Hossain, Tasnim Nafisa, Mahmuda Yeasmin, Asish Kumar Ghosh, Arifa Akram, A. K. M. Shamsuzzaman, Md. Salim Khan |
| EPI_ISL_959311, EPI_ISL_959312, EPI_ISL_959313, EPI_ISL_959316, EPI_ISL_959317, EPI_ISL_959318, EPI_ISL_959319, EPI_ISL_959320, EPI_ISL_959321, EPI_ISL_959322, EPI_ISL_959323, EPI_ISL_959324, EPI_ISL_959325, EPI_ISL_959326, EPI_ISL_959327, EPI_ISL_959328, EPI_ISL_959329, EPI_ISL_959330, EPI_ISL_959331, EPI_ISL_959332, EPI_ISL_959334 | | | |
| see above | Servicio de Microbiología, Laboratori Clínic Metropolitana Nord. Hospital Universitari Germans Trias i Pujol. Institut d'Investigació en Ciències de la Salut Germans Trias i Pujol (IGTP) | IrsiCaixa - Can Ruti CovidSeq | Marc Noguera-Julian, Mariona Parera, Maria Casadellà, Pilar Armengol, Francesc Catala-Moll, Roger Paredes, Bonaventura Clotet Elisa Martró, Verónica Saludes, Anna Not, Ana Pérez, Montserrat Giménez, Ignacio Blanco, Cristina Casañ, Antoni E. Bordoy, Adrián Antuori |

| | | | |
|---|---|---|---|
| EPI_ISL_959371 | National Institute of Laboratory Medicine and Referral Center | Genomic Research Lab, BCSIR | Shahina Akter, Mohammad Samir Uzzaman, Eshrar Osman, Md. Ahashan Habib, Tanjina Akhtar Banu, Abu Sayeed Mohammad Mahmud, Md. Murshed Hasan Sarkar, Barna Goswami, Iffat Jahan, Md. Saddam Hossain, Tasnim Nafisa, Md. Maruf Ahmed Molla, Mahmuda Yeasmin, Asish Kumar Ghosh, Arifa Akram, A. K. M. Shamsuzzaman, Md. Salim Khan |
| EPI_ISL_959372 | National Institute of Laboratory Medicine and Referral Center | Genomic Research Lab, BCSIR | Tanjina Akhtar Banu, Mohammad Samir Uzzaman, Eshrar Osman, Md. Ahashan Habib, Shahina Akter, Abu Sayeed Mohammad Mahmud, Md. Murshed Hasan Sarkar, Barna Goswami, Iffat Jahan, Md. Saddam Hossain, Tasnim Nafisa, Md. Maruf Ahmed Molla, Mahmuda Yeasmin, Asish Kumar Ghosh, Arifa Akram, A. K. M. Shamsuzzaman, Md. Salim Khan |
| EPI_ISL_959424, EPI_ISL_959427, EPI_ISL_959428, EPI_ISL_959430, EPI_ISL_959431, EPI_ISL_959439, EPI_ISL_959440, EPI_ISL_959446, EPI_ISL_959463, EPI_ISL_959464, EPI_ISL_959465, EPI_ISL_959466, EPI_ISL_959467 | | | |
| see above | Servicio de Microbiología, Hospital Universitario Son Espases | SeqCOVID-SPAIN consortium/IBV(CSIC) | Carla López-Causapé, Jordi Reina, Antonio Oliver and SeqCOVID-SPAIN consortium |
| EPI_ISL_959538 | National Institute of Laboratory Medicine and Referral Center | Genomic Research Lab, BCSIR | Barna Goswami, Mohammad Samir Uzzaman, Eshrar Osman, Md. Ahashan Habib, Shahina Akter, Tanjina Akhtar Banu, Abu Sayeed Mohammad Mahmud, Md. Murshed Hasan Sarkar, Iffat Jahan, Md. Saddam Hossain, Tasnim Nafisa, Md. Maruf Ahmed Molla, Mahmuda Yeasmin, Asish Kumar Ghosh, Arifa Akram, A. K. M. Shamsuzzaman, Md. Salim Khan |
| EPI_ISL_959539, EPI_ISL_959540 | University of Michigan Clinical Microbiology Laboratory | Lauring Lab, University of Michigan, Department of Microbiology and Immunology | Valesano |
| EPI_ISL_959545 | National Institute of Laboratory Medicine and Referral Center | Genomic Research Lab, BCSIR | Md. Saddam Hossain, Mohammad Samir Uzzaman, Eshrar Osman, Md. Ahashan Habib, Shahina Akter, Tanjina Akhtar Banu, Abu Sayeed Mohammad Mahmud, Md. Murshed Hasan Sarkar, Barna Goswami, Iffat Jahan, Tasnim Nafisa, Md. Maruf Ahmed Molla, Mahmuda Yeasmin, Asish Kumar Ghosh, Arifa Akram, A. K. M.Shamsuzzaman, Md. Salim Khan |
| EPI_ISL_959567, EPI_ISL_959568 | Vivalia - Clinique Saint-Joseph | GIGA Medical Genomics | Keith Durkin, Maria Artesi, Sébastien Bontems, Raphaël Boreux, Bouchra Boujemla, Cécile Meex, Pierrette Melin, Marie-Pierre Hayette, Vincent Bours |
| EPI_ISL_959569, EPI_ISL_959570, EPI_ISL_959571, EPI_ISL_959572, EPI_ISL_959573, EPI_ISL_959574, EPI_ISL_959575, EPI_ISL_959576, EPI_ISL_959577, EPI_ISL_959578, EPI_ISL_959579, EPI_ISL_959580, EPI_ISL_959581, EPI_ISL_959582, EPI_ISL_959583, EPI_ISL_959584 | | | |
| see above | Labo Luc Olivier | GIGA Medical Genomics | Keith Durkin, Maria Artesi, Sébastien Bontems, Raphaël Boreux, Bouchra Boujemla, Cécile Meex, Pierrette Melin, Marie-Pierre Hayette, Vincent Bours |
| EPI_ISL_959774, EPI_ISL_959775, EPI_ISL_959776, EPI_ISL_959777, EPI_ISL_959778, EPI_ISL_959779, EPI_ISL_959780, EPI_ISL_959781, EPI_ISL_959782, EPI_ISL_959783, EPI_ISL_959784, EPI_ISL_959785, EPI_ISL_959786, EPI_ISL_959787, EPI_ISL_959788, EPI_ISL_959789, EPI_ISL_959790, EPI_ISL_959791, EPI_ISL_959792, EPI_ISL_959793, EPI_ISL_959794, EPI_ISL_959795, EPI_ISL_959796, EPI_ISL_959797, EPI_ISL_959798, EPI_ISL_959799, EPI_ISL_959800, EPI_ISL_959801, EPI_ISL_959802, EPI_ISL_959803, EPI_ISL_959804, EPI_ISL_959805, EPI_ISL_959806, EPI_ISL_959807, EPI_ISL_959808, EPI_ISL_959809, EPI_ISL_959810, EPI_ISL_959811, EPI_ISL_959812, EPI_ISL_959813, EPI_ISL_959814, EPI_ISL_959815, EPI_ISL_959816, EPI_ISL_959817, EPI_ISL_959818, EPI_ISL_959819, EPI_ISL_959820, EPI_ISL_959821, EPI_ISL_959822, EPI_ISL_959823, EPI_ISL_959824, EPI_ISL_959825, EPI_ISL_959826, EPI_ISL_959827, EPI_ISL_959828, EPI_ISL_959829, EPI_ISL_959830, EPI_ISL_959831, EPI_ISL_959832, EPI_ISL_959833, EPI_ISL_959834, EPI_ISL_959835, EPI_ISL_959836, EPI_ISL_959837, EPI_ISL_959838, EPI_ISL_959854, EPI_ISL_959873 | | | |
| see above | National Virus Reference Laboratory | National Virus Reference Laboratory | Michael Carr, Gabriel Gonzalez, Jonathan Dean, Cillian F De Gascun |
| EPI_ISL_960312 | Botswana Harvard HIV Reference Laboratory | Botswana Harvard HIV Reference Laboratory | Sikhulile Moyo, Wonderful T. Choga, Dorcas Maruapula, Botshelo Radibe, Boitumelo Zuze, David Lawrence, Roger Shapiro, Shahin Lockman, Mosepele Mosepele, Joseph Makhema, Simani Gaseitsiwe |
| EPI_ISL_960422, EPI_ISL_960427, EPI_ISL_960429 | The National Institute of Public Health | State Veterinary Institute Prague | Nagy,A;Vecerova,J;Cernikova,L;Stara,M;Jirincova,H;Trnka,D |
| EPI_ISL_960855, EPI_ISL_960860, EPI_ISL_960862, EPI_ISL_960869, EPI_ISL_960870, EPI_ISL_960871, EPI_ISL_960880 | Institute of Medical Microbiology and Hospital Hygiene | Institute of Medical Microbiology and Hospital Hygiene | Prof. Dr. Achim Kaasch, Aljoscha Tersteegen |
| EPI_ISL_961052, EPI_ISL_961053, EPI_ISL_961054 | SIESP DIPARTIMENTO DI PREVENZIONE CHIETI | Istituto Zooprofilattico Sperimentale dell'Abruzzo e Molise "G. Caporale" | Lorusso A, Marcacci M, Di Domenico M, Ancora M, Curini V, Mangone I, Rinaldi A, Scialabba S, Di Pasquale A, Cammà C, Puglia I, Calistri P, Savini G |
| EPI_ISL_961055, EPI_ISL_961056 | SIESP CHIETI - DRIVE IN ORTONA | Istituto Zooprofilattico Sperimentale dell'Abruzzo e Molise "G. Caporale" | Lorusso A, Marcacci M, Di Domenico M, Ancora M, Curini V, Mangone I, Rinaldi A, Scialabba S, Di Pasquale A, Cammà C, Puglia I, Calistri P, Savini G |
| EPI_ISL_961057, EPI_ISL_961058, EPI_ISL_961059, EPI_ISL_961060 | SIESP CHIETI - DRIVE IN CHIETI | Istituto Zooprofilattico Sperimentale dell'Abruzzo e Molise "G. Caporale" | Lorusso A, Marcacci M, Di Domenico M, Ancora M, Curini V, Mangone I, Rinaldi A, Scialabba S, Di Pasquale A, Cammà C, Puglia I, Calistri P, Savini G |
| EPI_ISL_961061 | SIESP CHIETI - DRIVE IN ORTONA | Istituto Zooprofilattico Sperimentale dell'Abruzzo e Molise "G. Caporale" | Lorusso A, Marcacci M, Di Domenico M, Ancora M, Curini V, Mangone I, Rinaldi A, Scialabba S, Di Pasquale A, Cammà C, Puglia I, Calistri P, Savini G |
| EPI_ISL_961062 | SIESP CHIETI - DRIVE IN CHIETI | Istituto Zooprofilattico Sperimentale dell'Abruzzo e Molise "G. Caporale" | Lorusso A, Marcacci M, Di Domenico M, Ancora M, Curini V, Mangone I, Rinaldi A, Scialabba S, Di Pasquale A, Cammà C, Puglia I, Calistri P, Savini G |
| EPI_ISL_961063 | SIESP DIPARTIMENTO DI PREVENZIONE CHIETI | Istituto Zooprofilattico Sperimentale dell'Abruzzo e Molise "G. Caporale" | Lorusso A, Marcacci M, Di Domenico M, Ancora M, Curini V, Mangone I, Rinaldi A, Scialabba S, Di Pasquale A, Cammà C, Puglia I, Calistri P, Savini G |
| EPI_ISL_961064, EPI_ISL_961065 | SIESP CHIETI - DRIVE IN ORTONA | Istituto Zooprofilattico Sperimentale dell'Abruzzo e Molise "G. Caporale" | Lorusso A, Marcacci M, Di Domenico M, Ancora M, Curini V, Mangone I, Rinaldi A, Scialabba S, Di Pasquale A, Cammà C, Puglia I, Calistri P, Savini G |
| EPI_ISL_961084 | SIESP TERAMO | Istituto Zooprofilattico Sperimentale dell'Abruzzo e Molise "G. Caporale" | Lorusso A, Marcacci M, Di Domenico M, Ancora M, Curini V, Mangone I, Rinaldi A, Scialabba S, Di Pasquale A, Cammà C, Puglia I, Calistri P, Savini G |
| EPI_ISL_961085 | SIESP CHIETI - DRIVE IN ORTONA | Istituto Zooprofilattico Sperimentale dell'Abruzzo e Molise "G. Caporale" | Lorusso A, Marcacci M, Di Domenico M, Ancora M, Curini V, Mangone I, Rinaldi A, Scialabba S, Di Pasquale A, Cammà C, Puglia I, Calistri P, Savini G |
| EPI_ISL_961086 | SIESP CHIETI - DRIVE IN CHIETI | Istituto Zooprofilattico Sperimentale dell'Abruzzo e Molise "G. Caporale" | Lorusso A, Marcacci M, Di Domenico M, Ancora M, Curini V, Mangone I, Rinaldi A, Scialabba S, Di Pasquale A, Cammà C, Puglia I, Calistri P, Savini G |
| EPI_ISL_961087, EPI_ISL_961088, EPI_ISL_961089 | SIESP DIPARTIMENTO DI PREVENZIONE CHIETI | Istituto Zooprofilattico Sperimentale dell'Abruzzo e Molise "G. Caporale" | Lorusso A, Marcacci M, Di Domenico M, Ancora M, Curini V, Mangone I, Rinaldi A, Scialabba S, Di Pasquale A, Cammà C, Puglia I, Calistri P, Savini G |
| EPI_ISL_961090, EPI_ISL_961091, EPI_ISL_961092 | SIESP CHIETI - DRIVE IN CHIETI | Istituto Zooprofilattico Sperimentale dell'Abruzzo e Molise "G. Caporale" | Lorusso A, Marcacci M, Di Domenico M, Ancora M, Curini V, Mangone I, Rinaldi A, Scialabba S, Di Pasquale A, Cammà C, Puglia I, Calistri P, Savini G |
| EPI_ISL_961093 | Ospedale SS Annunziata | Istituto Zooprofilattico Sperimentale dell'Abruzzo e Molise "G. Caporale" | Lorusso A, Marcacci M, Di Domenico M, Ancora M, Curini V, Mangone I, Rinaldi A, Scialabba S, Di Pasquale A, Cammà C, Puglia I, Calistri P, Savini G |
| EPI_ISL_961094, EPI_ISL_961095 | DIP. PREV. AVEZZANO SERVIZIO DI IGIENE EPIDEMIOLOGIAE SANITA' PUBBLICA | Istituto Zooprofilattico Sperimentale dell'Abruzzo e Molise "G. Caporale" | Lorusso A, Marcacci M, Di Domenico M, Ancora M, Curini V, Mangone I, Rinaldi A, Scialabba S, Di Pasquale A, Cammà C, Puglia I, Calistri P, Savini G |
| EPI_ISL_961171, EPI_ISL_961175 | Texas Department of State Health Services | Texas Department of State Health Services | Bonnie Oh, Anita Pokharel, James Daniel Bonser, Myong Koag, Chung Wang, Rachel Lee, Grace Kubin, Rashmi Tuladhar, Mayela Pedrueza, Maliha Rahman, Jenny Zhang |
| EPI_ISL_961215, EPI_ISL_961217, EPI_ISL_961220, EPI_ISL_961221, EPI_ISL_961222, EPI_ISL_961224, EPI_ISL_961227, EPI_ISL_961234, EPI_ISL_961238 | Hospital General Universitario de Alicante - Instituto de Investigación Sanitaria y Biomédica de Alicante | SeqCOVID-SPAIN consortium/IBV(CSIC) | Maripaz Ventero Martín, Carmen Molina Pardines and SeqCOVID-SPAIN consortium |
| EPI_ISL_961368, EPI_ISL_961369, EPI_ISL_961371, EPI_ISL_961372, EPI_ISL_961373, EPI_ISL_961374, EPI_ISL_961375, EPI_ISL_961376, EPI_ISL_961377, EPI_ISL_961378, EPI_ISL_961382, EPI_ISL_961383, EPI_ISL_961384, EPI_ISL_961385, EPI_ISL_961386, EPI_ISL_961387, EPI_ISL_961388, EPI_ISL_961389, EPI_ISL_961390, EPI_ISL_961391, EPI_ISL_961392, EPI_ISL_961393, EPI_ISL_961394, EPI_ISL_961395, EPI_ISL_961396, EPI_ISL_961397, EPI_ISL_961398, EPI_ISL_961399, EPI_ISL_961400, EPI_ISL_961401, EPI_ISL_961402, EPI_ISL_961403, EPI_ISL_961404, EPI_ISL_961405, EPI_ISL_961406, EPI_ISL_961407, EPI_ISL_961408, EPI_ISL_961409, EPI_ISL_961410, EPI_ISL_961411, EPI_ISL_961412, EPI_ISL_961413, EPI_ISL_961414, EPI_ISL_961415, EPI_ISL_961416, EPI_ISL_961417, EPI_ISL_961418, EPI_ISL_961419, EPI_ISL_961420, EPI_ISL_961421, EPI_ISL_961422, EPI_ISL_961423, EPI_ISL_961424, EPI_ISL_961425 | | | |
| see above | Toronto Invasive Bacterial Diseases Network | McMaster University | Allison McGeer, Patryk Aftanas, Hooman Derakhshani, Angel Li, Kuganya Nirmalarajah, Emily Panousis, Ahmed Draia, Jalees Nasir, Michael Surette, |

| | | | |
|---|---|---|---|
| | | | Samira Mubareka, Andrew G. McArthur |
| EPI_ISL_961467, EPI_ISL_961468, EPI_ISL_961469 | LESP Nuevo Leon/Grupo de Diagnostico ARIES | Instituto de Diagnostico y Referencia Epidemiologicos (INDRE) | Claudia Wong-Arambula, Abril Rodriguez-Maldonado, Fabiola Garces-Ayala, Natividad Cruz-Ortiz, Tatiana Nunez-Garcia, Gisela Barrera-Badillo, Lucia Hernandez-Rivas, Irma Lopez-Martinez, Ernesto Ramirez-Gonzalez. |
| EPI_ISL_961472, EPI_ISL_961504, EPI_ISL_961505, EPI_ISL_961517, EPI_ISL_961523, EPI_ISL_961539, EPI_ISL_961540, EPI_ISL_961541, EPI_ISL_961542, EPI_ISL_961543 | Michigan Department of Health and Human Services, Bureau of Laboratories | Michigan Department of Health and Human Services, Bureau of Laboratories | Blankenship HM, Riner D, Soehnlen MK |
| EPI_ISL_961581, EPI_ISL_961582, EPI_ISL_961588, EPI_ISL_961589, EPI_ISL_961590, EPI_ISL_961620, EPI_ISL_961621, EPI_ISL_961622, EPI_ISL_961623, EPI_ISL_961624, EPI_ISL_961625, EPI_ISL_961626, EPI_ISL_961627, EPI_ISL_961628, EPI_ISL_961629 | | | |
| see above | Hôpital Georges L. Dumont | National Microbiology Laboratory (NML) | Anna Majer, Shari Tyson, Grace Seo, Philip Mabon, Elsie Grudeski, Rhiannon Huzarewich, Russell Mandes, Anneliese Landgraff, Jennifer Tanner, Natalie Knox, Morag Graham, Gary Van Domselaar, Richard Garceau, Guillaume Desnoyers, Nathalie Bastien, Yan Li, Timothy Booth, Darian Hole, Madison Chapel, Kirsten Biggar, CanCOGeN's metadata curation team, Public Health Agency of Canada CanCOGeN team |
| EPI_ISL_961802, EPI_ISL_961804 | E. Gulbja laboratorija | Latvian Biomedical Research and Study Centre | Janis Pjalkovskis, Nikita Zrelovs, Monta Ustinova, Ivars Silamikelis, Liga Birzniece, Kaspars Megnis, Vita Rovite, Lauma Freimane, Laila Silamikele, Laura Ansone, Davids Fridmanis, Mikus Gavars, Dmitrijs Perminovs, Jurijs Perevoscikovs, Uga Dumpis, Janis Klovins |
| EPI_ISL_961806 | Latvijas Infektologijas Centrs | Latvian Biomedical Research and Study Centre | Janis Pjalkovskis, Nikita Zrelovs, Monta Ustinova, Ivars Silamikelis, Liga Birzniece, Kaspars Megnis, Vita Rovite, Lauma Freimane, Laila Silamikele, Laura Ansone, Davids Fridmanis,Reinis Zeltmatis, Diana Dusacka, Jurijs Perevoscikovs, Uga Dumpis, Janis Klovins |
| EPI_ISL_961853, EPI_ISL_961854, EPI_ISL_961855, EPI_ISL_961856, EPI_ISL_961857, EPI_ISL_961858, EPI_ISL_961859, EPI_ISL_961860, EPI_ISL_961861 | E. Gulbja laboratorija | Latvian Biomedical Research and Study Centre | Janis Pjalkovskis, Nikita Zrelovs, Monta Ustinova, Ivars Silamikelis, Liga Birzniece, Kaspars Megnis, Vita Rovite, Lauma Freimane, Laila Silamikele, Laura Ansone, Davids Fridmanis, Mikus Gavars, Dmitrijs Perminovs, Jurijs Perevoscikovs, Uga Dumpis, Janis Klovins |
| EPI_ISL_961862, EPI_ISL_961863 | Latvijas Infektologijas Centrs | Latvian Biomedical Research and Study Centre | Janis Pjalkovskis, Nikita Zrelovs, Monta Ustinova, Ivars Silamikelis, Liga Birzniece, Kaspars Megnis, Vita Rovite, Lauma Freimane, Laila Silamikele, Laura Ansone, Davids Fridmanis,Reinis Zeltmatis, Diana Dusacka, Jurijs Perevoscikovs, Uga Dumpis, Janis Klovins |
| EPI_ISL_961864, EPI_ISL_961865 | E. Gulbja laboratorija | Latvian Biomedical Research and Study Centre | Janis Pjalkovskis, Nikita Zrelovs, Monta Ustinova, Ivars Silamikelis, Liga Birzniece, Kaspars Megnis, Vita Rovite, Lauma Freimane, Laila Silamikele, Laura Ansone, Davids Fridmanis, Mikus Gavars, Dmitrijs Perminovs, Jurijs Perevoscikovs, Uga Dumpis, Janis Klovins |
| EPI_ISL_961866 | Latvijas Infektologijas Centrs | Latvian Biomedical Research and Study Centre | Janis Pjalkovskis, Nikita Zrelovs, Monta Ustinova, Ivars Silamikelis, Liga Birzniece, Kaspars Megnis, Vita Rovite, Lauma Freimane, Laila Silamikele, Laura Ansone, Davids Fridmanis,Reinis Zeltmatis, Diana Dusacka, Jurijs Perevoscikovs, Uga Dumpis, Janis Klovins |
| EPI_ISL_961867, EPI_ISL_961868 | E. Gulbja laboratorija | Latvian Biomedical Research and Study Centre | Janis Pjalkovskis, Nikita Zrelovs, Monta Ustinova, Ivars Silamikelis, Liga Birzniece, Kaspars Megnis, Vita Rovite, Lauma Freimane, Laila Silamikele, Laura Ansone, Davids Fridmanis, Mikus Gavars, Dmitrijs Perminovs, Jurijs Perevoscikovs, Uga Dumpis, Janis Klovins |
| EPI_ISL_961869, EPI_ISL_961870 | Latvijas Infektologijas Centrs | Latvian Biomedical Research and Study Centre | Janis Pjalkovskis, Nikita Zrelovs, Monta Ustinova, Ivars Silamikelis, Liga Birzniece, Kaspars Megnis, Vita Rovite, Lauma Freimane, Laila Silamikele, Laura Ansone, Davids Fridmanis,Reinis Zeltmatis, Diana Dusacka, Jurijs Perevoscikovs, Uga Dumpis, Janis Klovins |
| EPI_ISL_961871, EPI_ISL_961872 | E. Gulbja laboratorija | Latvian Biomedical Research and Study Centre | Janis Pjalkovskis, Nikita Zrelovs, Monta Ustinova, Ivars Silamikelis, Liga Birzniece, Kaspars Megnis, Vita Rovite, Lauma Freimane, Laila Silamikele, Laura Ansone, Davids Fridmanis, Mikus Gavars, Dmitrijs Perminovs, Jurijs Perevoscikovs, Uga Dumpis, Janis Klovins |
| EPI_ISL_962183 | Toronto Invasive Bacterial Diseases Network | McMaster University | Allison McGeer, Patryk Aftanas, Hooman Derakhshani, Angel Li, Kuganya Nirmalarajah, Emily Panousis, Ahmed Draia, Jalees Nasir, Michael Surette, Samira Mubareka, Andrew G. McArthur |
| EPI_ISL_962205 | Institute for Medical Research, Infectious Disease Research Centre, National Institutes of Health, Ministry of Health Malaysia | Institute for Medical Research, Infectious Disease Research Centre, National Institutes of Health, Ministry of Health Malaysia | Suppiah J, Kamel K, Azizan MA, Thayan R |
| EPI_ISL_962264, EPI_ISL_962265, EPI_ISL_962266, EPI_ISL_962267, EPI_ISL_962268, EPI_ISL_962269 | Seattle Flu Study | Seattle Flu Study | Deborah A. Nickerson, Chris D. Frazar, Jover Lee, Benjamin Pelle, Erica Ryke, Matthew Richardson, Amanda Adler, Elisabeth Brandstetter, Peter D. Han, Kairsten Fay, Misja Ilcisin, Kirsten Lacombe, Thomas R. Sibley, Melissa Truong, Caitlin R. Wolf, Michael Boeckh, Janet A. Englund, Michael Famulare, Barry R. Lutz, Mark J. Rieder, Lea M. Starita, Matthew Thompson, Jay Shendure, Trevor Bedford, Helen Y. Chu |
| EPI_ISL_962285, EPI_ISL_962286, EPI_ISL_962288, EPI_ISL_962289, EPI_ISL_962290, EPI_ISL_962292, EPI_ISL_962293, EPI_ISL_962294, EPI_ISL_962295, EPI_ISL_962296, EPI_ISL_962297, EPI_ISL_962298, EPI_ISL_962299 | | | |
| see above | Seattle Flu Study | Seattle Flu Study | Deborah A. Nickerson, Chris D. Frazar, Jover Lee, Benjamin Pelle, Erica Ryke, Matthew Richardson, Amanda Adler, Elisabeth Brandstetter, Peter D. Han, Kairsten Fay, Misja Ilcisin, Kirsten Lacombe, Thomas R. Sibley, Melissa Truong, Caitlin R. Wolf, Karen Cowgill, Stephanie Schrag, Jeff Duchin, Michael Boeckh, Janet A. Englund, Michael Famulare, Barry R. Lutz, Mark J. Rieder, Lea M. Starita, Matthew Thompson, Helen Y. Chu, Trevor Bedford, Jay Shendure |
| EPI_ISL_962305, EPI_ISL_962306, EPI_ISL_962307, EPI_ISL_962308, EPI_ISL_962309, EPI_ISL_962310, EPI_ISL_962311, EPI_ISL_962312, EPI_ISL_962313, EPI_ISL_962314, EPI_ISL_962315, EPI_ISL_962316, EPI_ISL_962317, EPI_ISL_962318, EPI_ISL_962319, EPI_ISL_962320, EPI_ISL_962321, EPI_ISL_962322, EPI_ISL_962323, EPI_ISL_962324, EPI_ISL_962325, EPI_ISL_962326, EPI_ISL_962327, EPI_ISL_962328, EPI_ISL_962329, EPI_ISL_962330, EPI_ISL_962331, EPI_ISL_962332, EPI_ISL_962333, EPI_ISL_962334, EPI_ISL_962335, EPI_ISL_962336, EPI_ISL_962337, EPI_ISL_962338, EPI_ISL_962339, EPI_ISL_962340, EPI_ISL_962341, EPI_ISL_962342, EPI_ISL_962343, EPI_ISL_962344, EPI_ISL_962345, EPI_ISL_962346, EPI_ISL_962347, EPI_ISL_962348, EPI_ISL_962349, EPI_ISL_962350, EPI_ISL_962351, EPI_ISL_962352, EPI_ISL_962353, EPI_ISL_962354, EPI_ISL_962355, EPI_ISL_962356, EPI_ISL_962357, EPI_ISL_962358, EPI_ISL_962359, EPI_ISL_962360, EPI_ISL_962361, EPI_ISL_962362, EPI_ISL_962363, EPI_ISL_962364, EPI_ISL_962365, EPI_ISL_962366, EPI_ISL_962367 | | | |
| see above | Altius Institute for Biomedical Sciences | Seattle Flu Study | Deborah A. Nickerson, Chris D. Frazar, Jover Lee, Benjamin Pelle, Erica Ryke, Matthew Richardson, Amanda Adler, Elisabeth Brandstetter, Peter D. Han, Kairsten Fay, Misja Ilcisin, Kirsten Lacombe, Thomas R. Sibley, Melissa Truong, Caitlin R. Wolf, Ryan Alexander, Daniel Bates, Rebecca Bruders, Stephanie DeBaun, Clem Green, Muhammad Halimun, Jessica Halow, Kneshay Harper, Matt Hartman, Andrew Meuser, Alex Nguyen, Truong Nguyen, Sofia Olsson, Sadie Patraw, Hannah Petersen, Tobias Ragoczy, Joshua Richards, Jacob Rodriguez, John Stamatoyannopoulos, Julia Wald, Olivia Waltner, Michael Boeckh, Janet A. Englund, Michael Famulare, Barry R. Lutz, Mark J. Rieder, Lea M. Starita, Matthew Thompson, Helen Y. Chu, Jay Shendure, Trevor Bedford |
| EPI_ISL_962369 | Seattle Flu Study | Seattle Flu Study | Deborah A. Nickerson, Chris D. Frazar, Jover Lee, Benjamin Pelle, Erica Ryke, Matthew Richardson, Amanda Adler, Elisabeth Brandstetter, Peter D. Han, Kairsten Fay, Misja Ilcisin, Kirsten Lacombe, Thomas R. Sibley, Melissa Truong, Caitlin R. Wolf, Karen Cowgill, Stephanie Schrag, Jeff Duchin, Michael Boeckh, Janet A. Englund, Michael Famulare, Barry R. Lutz, Mark J. Rieder, Lea M. Starita, Matthew Thompson, Helen Y. Chu, Trevor Bedford, Jay Shendure |
| EPI_ISL_962403, EPI_ISL_962404, EPI_ISL_962405, EPI_ISL_962406, EPI_ISL_962407, EPI_ISL_962408, EPI_ISL_962409, EPI_ISL_962410, EPI_ISL_962411, EPI_ISL_962412, EPI_ISL_962413, EPI_ISL_962414, EPI_ISL_962415, EPI_ISL_962416, EPI_ISL_962418, EPI_ISL_962419, EPI_ISL_962420 | | | |
| see above | Washington State Department of Health | Seattle Flu Study | Deborah A. Nickerson, Chris D. Frazar, Jover Lee, Benjamin Pelle, Erica Ryke, Matthew Richardson, Amanda Adler, Elisabeth Brandstetter, Peter D. Han, Kairsten Fay, Misja Ilcisin, Kirsten Lacombe, Thomas R. Sibley, Melissa Truong, Caitlin R. Wolf, Romesh Gautom, Geoff Melly, Brian Hiatt, Philip Dykema, Scott Lindquist, Michael Boeckh, Janet A. Englund, Michael Famulare, Barry R. Lutz, Mark J. Rieder, Lea M. Starita, Matthew Thompson, Helen Y. Chu, Jay Shendure, Trevor Bedford |
| EPI_ISL_962521 | UCLA Clinical Micro Lab | Los Angeles County PHL | P. Hemarajata et al. |
| EPI_ISL_962527 | Institute for Medical Research, Infectious Disease Research Centre, National Institutes of Health, Ministry of Health Malaysia | Institute for Medical Research, Infectious Disease Research Centre, National Institutes of Health, Ministry of Health Malaysia | Suppiah J, Kamel K, Azizan MA, Thayan R |
| EPI_ISL_962618, EPI_ISL_962629, EPI_ISL_962641 | Scripps Medical Laboratory | Andersen lab at Scripps Research | SEARCH Alliance San Diego with Michael Quigley, Ellen Stefanski, Ian Mchardy |
| EPI_ISL_962814 | Robert Garry lab | Andersen lab at Scripps Research | Allison Smither, Gilberto Sabino-Santos, Patricia Snarski, Lilia Melnik, Antoinette Bell, Kaylynn Genemaras, Arnaud Drouin, Dahlene Fusco, Robert Garry with SEARCH Alliance San Diego |

| | | | |
|---|---|---|---|
| EPI_ISL_962827 | Microbiological Diagnostic Unit - Public Health Laboratory (MDU-PHL) | MDU-PHL | Seemann T., Sait, M.L., Sherry, N.L. |
| EPI_ISL_962847 | MD Laboratories | Los Angeles County PHL | P. Hemarajata et al. |
| EPI_ISL_962897 | Akershus University Hospital, Department for Microbiology and Infectious Disease Control | Norwegian Institute of Public Health, Department of Virology | Kathrine Stene-Johansen, Kamilla Heddeland Instefjord, Hilde Elshaug, Ignacio Garcia Llorente, Serina B Engebretsen, Atiya R Ali,Marie Paulsen Madsen, Rasmus Riis Kopperud, Hilde Vollan, Karoline Bragstad, Olav Hungnes |
| EPI_ISL_962901, EPI_ISL_962902, EPI_ISL_962903 | Furst Medical Laboratory | Norwegian Institute of Public Health, Department of Virology | Kathrine Stene-Johansen, Kamilla Heddeland Instefjord, Hilde Elshaug, Ignacio Garcia Llorente, Serina B Engebretsen, Atiya R Ali,Marie Paulsen Madsen, Rasmus Riis Kopperud, Hilde Vollan, Karoline Bragstad, Olav Hungnes |
| EPI_ISL_962908 | Medical Microbiology Unit, Department for Laboratory Medicine, Drammen Hospital, Vestre Viken Health Trust, | Norwegian Institute of Public Health, Department of Virology | Kathrine Stene-Johansen, Kamilla Heddeland Instefjord, Hilde Elshaug, Ignacio Garcia Llorente, Serina B Engebretsen, Atiya R Ali,Marie Paulsen Madsen, Rasmus Riis Kopperud, Hilde Vollan, Karoline Bragstad, Olav Hungnes |
| EPI_ISL_962928 | Servicio de Microbiología, Laboratori Clínic Metropolitana Nord. Hospital Universitari Germans Trias i Pujol. Institut d'Investigació en Ciències de la Salut Germans Trias i Pujol (IGTP) | SeqCOVID-SPAIN consortium/IBV(CSIC) | Elisa Martró, Antoni E. Bordoy, Anna Not, Adrián Antuori, Anabel Fernández, Nona Romaní, Verónica Saludes, Cristina Casañ and SeqCOVID-SPAIN consortium |
| EPI_ISL_962947, EPI_ISL_962948, EPI_ISL_962949, EPI_ISL_962950, EPI_ISL_962951, EPI_ISL_962952, EPI_ISL_962953, EPI_ISL_962956, EPI_ISL_962957 | Hospital Universitario de Gran Canaria Dr. Negrín | SeqCOVID-SPAIN consortium/IBV(CSIC) | M. Carmen Pérez González, Francisco J. Chamizo López, Ana Bordes Benítez and SeqCOVID-SPAIN consortium |
| EPI_ISL_964295 | Oslo University Hospital, Department of Medical Microbiology | Norwegian Institute of Public Health, Department of Virology | Kathrine Stene-Johansen, Kamilla Heddeland Instefjord, Hilde Elshaug, Ignacio Garcia Llorente, Serina B Engebretsen, Atiya R Ali, Marie Paulsen Madsen, Rasmus Riis Kopperud, Hilde Vollan, Karoline Bragstad, Olav Hungnes |
| EPI_ISL_964880 | National Institute of Laboratory Medicine and Referral Center | Sher-e-Bangla Nagar, Agargaon, Dhaka-1207, Bangladesh. | Shahina Akter, Mohammad Samir Uzzaman, Eshrar Osman, Md. Ahashan Habib, Tanjina Akhtar Banu, Abu Sayeed Mohammad Mahmud, Md. Murshed Hasan Sarkar, Barna Goswami, Iffat Jahan, Md. Saddam Hossain, Tasnim Nafisa, Md. Maruf Ahmed Molla, Mahmuda Yeasmin, Asish Kumar Ghosh, Arifa Akram, A. K. M.Shamsuzzaman, Md. Salim Khan |
| EPI_ISL_964886 | National Institute of Laboratory Medicine and Referral Center | Genomic Research Lab, BCSIR | Tasnim Nafisa, Mohammad Samir Uzzaman, Eshrar Osman, Md. Ahashan Habib, Shahina Akter,Tanjina Akhtar Banu, Abu Sayeed Mohammad Mahmud, Md. Murshed Hasan Sarkar, Barna Goswami, Iffat Jahan, Md. Saddam Hossain, Md. Maruf Ahmed Molla, Mahmuda Yeasmin, Asish Kumar Ghosh, Arifa Akram, A. K. M.Shamsuzzaman, Md. Salim Khan |
| EPI_ISL_964950 | Laboratory of Virology and Molecular Diagnostics | Institute of Public Health of Republic of North Macedonia Laboratory of Virology and Molecular Diagnostics | Kuzmanovska M., Boshevska G. |
| EPI_ISL_964958 | Akershus University Hospital, Department for Microbiology and Infectious Disease Control | Norwegian Institute of Public Health, Department of Virology | Kathrine Stene-Johansen, Kamilla Heddeland Instefjord, Hilde Elshaug, Ignacio Garcia Llorente, Serina B Engebretsen, Atiya R Ali,Marie Paulsen Madsen, Rasmus Riis Kopperud, Hilde Vollan, Karoline Bragstad, Olav Hungnes |
| EPI_ISL_964961 | Dept. of Medical Microbiology, Stavanger University Hospital, Helse Stavanger HF | Norwegian Institute of Public Health, Department of Virology | Kathrine Stene-Johansen, Kamilla Heddeland Instefjord, Hilde Elshaug, Ignacio Garcia Llorente, Serina B Engebretsen, Atiya R Ali,Marie Paulsen Madsen, Rasmus Riis Kopperud, Hilde Vollan, Karoline Bragstad, Olav Hungnes |
| EPI_ISL_964969 | University Hospital of Northern Norway, Department for Microbiology and Infectious Disease Control | Norwegian Institute of Public Health, Department of Virology | Kathrine Stene-Johansen, Kamilla Heddeland Instefjord, Hilde Elshaug, Ignacio Garcia Llorente, Serina B Engebretsen, Atiya R Ali,Marie Paulsen Madsen, Rasmus Riis Kopperud, Hilde Vollan, Karoline Bragstad, Olav Hungnes |
| EPI_ISL_964973 | Ostfold Hospital Trust - Kalnes, Centre for Laboratory Medicine, Section for gene technology and infection serology | Norwegian Institute of Public Health, Department of Virology | Kathrine Stene-Johansen, Kamilla Heddeland Instefjord, Hilde Elshaug, Ignacio Garcia Llorente, Serina B Engebretsen, Atiya R Ali,Marie Paulsen Madsen, Rasmus Riis Kopperud, Hilde Vollan, Karoline Bragstad, Olav Hungnes |
| EPI_ISL_964981, EPI_ISL_964982 | Department of Medical Microbiology - section Molde, Molde Hospital | Norwegian Institute of Public Health, Department of Virology | Kathrine Stene-Johansen, Kamilla Heddeland Instefjord, Hilde Elshaug, Ignacio Garcia Llorente, Serina B Engebretsen, Atiya R Ali,Marie Paulsen Madsen, Rasmus Riis Kopperud, Hilde Vollan, Karoline Bragstad, Olav Hungnes |
| EPI_ISL_965005, EPI_ISL_965006, EPI_ISL_965007 | Dept. of Medical Microbiology, Stavanger University Hospital, Helse Stavanger HF | Norwegian Institute of Public Health, Department of Virology | Kathrine Stene-Johansen, Kamilla Heddeland Instefjord, Hilde Elshaug, Ignacio Garcia Llorente, Serina B Engebretsen, Atiya R Ali, Marie Paulsen Madsen, Rasmus Riis Kopperud, Hilde Vollan, Karoline Bragstad, Olav Hungnes |
| EPI_ISL_965015 | Medical Microbiology Unit, Department for Laboratory Medicine, Drammen Hospital, Vestre Viken Health Trust, | Norwegian Institute of Public Health, Department of Virology | Kathrine Stene-Johansen, Kamilla Heddeland Instefjord, Hilde Elshaug, Ignacio Garcia Llorente, Serina B Engebretsen, Atiya R Ali, Marie Paulsen Madsen, Rasmus Riis Kopperud, Hilde Vollan, Karoline Bragstad, Olav Hungnes |
| EPI_ISL_965020 | Florida Bureau of Public Health Laboratories | Florida Bureau of Public Health Laboratories | Sarah Schmedes, Jason Blanton |
| EPI_ISL_965086, EPI_ISL_965087, EPI_ISL_965088, EPI_ISL_965089, EPI_ISL_965090, EPI_ISL_965091, EPI_ISL_965092, EPI_ISL_965093, EPI_ISL_965094, EPI_ISL_965095, EPI_ISL_965096, EPI_ISL_965097, EPI_ISL_965098, EPI_ISL_965099, EPI_ISL_965100 | | | |
| see above | Wyoming Public Health Laboratory | Wyoming Public Health Laboratory | Noah Hull, Taylor Fearing, Lynette Gumbleton, Channing Weber, Ashley Norberg, Bailey Bowcutt, and Wanda Manley |
| EPI_ISL_965128 | INMI Lazzaro Spallanzani IRCCS | INMI Lazzaro Spallanzani IRCCS | M. Rueca, E Giombini, B Bartolini, O Butera, C.E.M Gruber, F Messina, A Di Caro, MR Capobianchi |
| EPI_ISL_965140 | UOC "Microbiologia e Virologia" AOU "Policlinico Umberto I", Roma; Dipartimento di Medicina Molecolare Sapienza Università di Roma | INMI Lazzaro Spallanzani IRCCS | F Messina, C.E.M Gruber, B Bartolini, E Giombini, M Rueca, O Butera, G Antonelli, O Turriziani, MR Capobianchi, A Di Caro |
| EPI_ISL_965700 | Dutch COVID-19 response team | Medical Microbiology, Maastricht University Medical Centre | Jozef Dingemans*, Brian van der Veer*, Erik Beuken, Carmen Reumkens, Lieke van Alphen, Christian Hoebe, Paul Savelkoul |
| EPI_ISL_965909 | Botswana Harvard HIV Reference Laboratory | Botswana Harvard HIV Reference Laboratory | Sikhulile Moyo, Wonderful T. Choga, Dorcas Maruapula, Botshelo Radibe, Boitumelo Zuze, David Lawrence, Roger Shapiro, Shahin Lockman, Mosepele Mosepele, Joseph Makhema, Simani Gaseitsiwe |
| EPI_ISL_965914 | Hospital General Universitario de Ciudad Real | Instituto de Salud Carlos III | Vázquez, S. Iglesias-Caballero, M. Sandonís,V. Camarero, S. Pozo, F. Casas, I. Jiménez, P. Zaballos, A. Monzón, S. Varona, S. Cuesta, I. Illescas, S. |
| EPI_ISL_965920, EPI_ISL_965924 | Servicio Murciano de Salud | Instituto de Salud Carlos III | Vázquez, S. Iglesias-Caballero, M. Sandonís,V. Camarero, S. Pozo, F. Casas, I. Jiménez, P. Zaballos, A. Monzón, S. Varona, S. Cuesta, I. Blázquez, A. |
| EPI_ISL_965926 | Hospital General Universitario de Ciudad Real | Instituto de Salud Carlos III | Sandonís,V. Vázquez, S. Iglesias-Caballero, M. Camarero, S. Pozo, F. Casas, I. Jiménez, P. Zaballos, A. Monzón, S. Varona, S. Cuesta, I. Illescas, S. |
| EPI_ISL_965946 | Hospital General Universitario de Ciudad Real | Instituto de Salud Carlos III | Iglesias-Caballero, M. Sandonís,V. Vázquez, S. Camarero, S. Pozo, F. Casas, I. Jiménez, P. Zaballos, A. Monzón, S. Varona, S. Cuesta, I. Illescas, S. |
| EPI_ISL_965954 | Servicio Murciano de Salud | Instituto de Salud Carlos III | Iglesias-Caballero, M. Sandonís,V. Vázquez, S. Camarero, S. Pozo, F. Casas, I. Jiménez, P. Zaballos, A. Monzón, S. Varona, S. Cuesta, I. Blázquez, A. |
| EPI_ISL_966497, EPI_ISL_966498, EPI_ISL_966499, EPI_ISL_966553, EPI_ISL_966554, EPI_ISL_966556, EPI_ISL_966557, EPI_ISL_966558, EPI_ISL_966559, EPI_ISL_966564, EPI_ISL_966566, EPI_ISL_966567, EPI_ISL_966568, EPI_ISL_966569, EPI_ISL_966570, EPI_ISL_966571, EPI_ISL_966572, EPI_ISL_966573, EPI_ISL_966574, EPI_ISL_966575, EPI_ISL_966576, EPI_ISL_966592, EPI_ISL_966593, EPI_ISL_966594, EPI_ISL_966595, EPI_ISL_966596, EPI_ISL_966597, EPI_ISL_966598, EPI_ISL_966599, EPI_ISL_966602, EPI_ISL_966608, EPI_ISL_966618, EPI_ISL_966752, EPI_ISL_966753, EPI_ISL_966754, EPI_ISL_966755, EPI_ISL_967182, EPI_ISL_967425, EPI_ISL_967426, EPI_ISL_967427, EPI_ISL_967428, EPI_ISL_967429, EPI_ISL_967430, EPI_ISL_967456, EPI_ISL_967457, EPI_ISL_967458, EPI_ISL_967459, EPI_ISL_967460, EPI_ISL_967461, EPI_ISL_967462, EPI_ISL_967463, EPI_ISL_967464, EPI_ISL_967465, EPI_ISL_967466, EPI_ISL_967467, EPI_ISL_967468, EPI_ISL_967469, EPI_ISL_967470, EPI_ISL_967471, EPI_ISL_967472, EPI_ISL_967473, EPI_ISL_967474, EPI_ISL_967475, EPI_ISL_967476, EPI_ISL_967477, EPI_ISL_967478, EPI_ISL_967479, EPI_ISL_967480, EPI_ISL_967481, EPI_ISL_967482, EPI_ISL_967483, EPI_ISL_967484, EPI_ISL_967485, EPI_ISL_967486, EPI_ISL_967487, EPI_ISL_967488, EPI_ISL_967489, EPI_ISL_967490, EPI_ISL_967491, EPI_ISL_967492, EPI_ISL_967493, EPI_ISL_967494, EPI_ISL_967495, EPI_ISL_967496, EPI_ISL_967497, EPI_ISL_967498, EPI_ISL_967499, EPI_ISL_967500, EPI_ISL_967501, EPI_ISL_967502, EPI_ISL_967503, EPI_ISL_967504, EPI_ISL_967505, EPI_ISL_967506, EPI_ISL_967507, EPI_ISL_967508, EPI_ISL_967509, EPI_ISL_967510, EPI_ISL_967511, EPI_ISL_967512, EPI_ISL_967513, EPI_ISL_967514, EPI_ISL_967515, EPI_ISL_967516, EPI_ISL_967517, EPI_ISL_967518, EPI_ISL_967519, EPI_ISL_967520, EPI_ISL_967521, EPI_ISL_967522, EPI_ISL_967524, EPI_ISL_967531, EPI_ISL_967537, EPI_ISL_967547, EPI_ISL_967554, EPI_ISL_967567, EPI_ISL_967576, EPI_ISL_967583, EPI_ISL_967590, EPI_ISL_967599, EPI_ISL_967609, EPI_ISL_967617, EPI_ISL_967629, EPI_ISL_967642, EPI_ISL_967655, EPI_ISL_967662, EPI_ISL_967671, EPI_ISL_967678, EPI_ISL_967684, EPI_ISL_967693, EPI_ISL_967700, EPI_ISL_967710, EPI_ISL_967719, EPI_ISL_967726, EPI_ISL_967736, EPI_ISL_967748, EPI_ISL_967753 | | | |
| see above | Helix/Illumina | Respiratory Viruses Branch, Division of Viral Diseases, Centers for Disease Control and Prevention | Peter W. Cook,Dakota Howard,Dhwani Batra,Ben L. Rambo-Martin,Eileen de Feo,Jan Antico,Christine Tran,Matthew Tolentino,Shannon Wickline,Kim Gietzen,Brad Sickler,Jingtao Liu,Eric Allen,Phil Febbo,Summer Galloway,Nicole L. Washington,Simon White,Geraint Levan,Kelly Schiabor Barrett,Elizabeth Cirulli,Alexandre Bolze,Ary Ascencio,Charlotte Rivera-Garcia,Ryan Cho,Jason Nguyen,Sherry Wang,Jimmy Ramirez,Tyler Cassens,Efren Sandoval,Magnus Isaksson,William Lee,David Becker,Marc Laurent,James Lu,Clinton R. Paden,Suxiang Tong,Duncan MacCannell, |
| EPI_ISL_967755 | State Laboratories Division, Hawaii State Department of Health | State Laboratories Division, Hawaii State Department of Health | Pamela O'Brien, Drew Kuwazaki, Ayana Garnet, Razvan Sultana, Edward Desmond |

| | | | |
|---|---|---|---|
| EPI_ISL_967761, EPI_ISL_967767 | Helix/Illumina | Respiratory Viruses Branch, Division of Viral Diseases, Centers for Disease Control and Prevention | Peter W. Cook,Dakota Howard,Dhwani Batra,Ben L. Rambo-Martin,Eileen de Feo,Jan Antico,Christine Tran,Matthew Tolentino,Shannon Wickline,Kim Gietzen,Brad Sickler,Jingtao Liu,Eric Allen,Phil Febbo,Summer Galloway,Nicole L. Washington,Simon White,Geraint Levan,Kelly Schiabor Barrett,Elizabeth Cirulli,Alexandre Bolze,Ary Ascencio,Charlotte Rivera-Garcia,Ryan Cho,Jason Nguyen,Sherry Wang,Jimmy Ramirez,Tyler Cassens,Efren Sandoval,Magnus Isaksson,William Lee,David Becker,Marc Laurent,James Lu,Clinton R. Paden,Suxiang Tong,Duncan MacCannell, |
| EPI_ISL_967768, EPI_ISL_967769 | State Laboratories Division, Hawaii State Department of Health | State Laboratories Division, Hawaii State Department of Health | Pamela O'Brien, Drew Kuwazaki, Ayana Garnet, Razvan Sultana, Edward Desmond |
| EPI_ISL_967772, EPI_ISL_967773, EPI_ISL_967774, EPI_ISL_967775, EPI_ISL_967776, EPI_ISL_967777, EPI_ISL_967778, EPI_ISL_967779, EPI_ISL_967780 | Helix/Illumina | Respiratory Viruses Branch, Division of Viral Diseases, Centers for Disease Control and Prevention | Peter W. Cook,Dakota Howard,Dhwani Batra,Ben L. Rambo-Martin,Eileen de Feo,Jan Antico,Christine Tran,Matthew Tolentino,Shannon Wickline,Kim Gietzen,Brad Sickler,Jingtao Liu,Eric Allen,Phil Febbo,Summer Galloway,Nicole L. Washington,Simon White,Geraint Levan,Kelly Schiabor Barrett,Elizabeth Cirulli,Alexandre Bolze,Ary Ascencio,Charlotte Rivera-Garcia,Ryan Cho,Jason Nguyen,Sherry Wang,Jimmy Ramirez,Tyler Cassens,Efren Sandoval,Magnus Isaksson,William Lee,David Becker,Marc Laurent,James Lu,Clinton R. Paden,Suxiang Tong,Duncan MacCannell, |
| EPI_ISL_968069, EPI_ISL_968070, EPI_ISL_968071, EPI_ISL_968072, EPI_ISL_968073, EPI_ISL_968074, EPI_ISL_968075, EPI_ISL_968076 | Monterey County Public Health Laboratory | Monterey County Public Health Laboratory | Monterey County Public Health Laboratory |
| EPI_ISL_968252 | Botswana Harvard HIV Reference Laboratory | Botswana Harvard HIV Reference Laboratory | Sikhulile Moyo, Dorcas Maruapula, Wonderful Choga, Botshelo Radibe, Boitumelo Zuze, David Lawrence, Roger Shapiro, Shahin Lockman, Mosepele Mosepele, Joseph Makhema, Simani Gaseitsiwe |
| EPI_ISL_969001, EPI_ISL_969002, EPI_ISL_969023, EPI_ISL_969052 | KEMRI-Wellcome Trust Research Programme/KEMRI-CGMR-C Kilifi | KEMRI-Wellcome Trust Research Programme/KEMRI-CGMR-C Kilifi | Githinji et al |
| EPI_ISL_970827, EPI_ISL_970849, EPI_ISL_970867, EPI_ISL_970907, EPI_ISL_970961, EPI_ISL_970966, EPI_ISL_971266, EPI_ISL_971346, EPI_ISL_971349, EPI_ISL_971391, EPI_ISL_971436, EPI_ISL_971473, EPI_ISL_971478, EPI_ISL_971560, EPI_ISL_971566, EPI_ISL_971613, EPI_ISL_971690, EPI_ISL_971719, EPI_ISL_971839, EPI_ISL_972072, EPI_ISL_972137, EPI_ISL_972139, EPI_ISL_972202, EPI_ISL_972225, EPI_ISL_972232, EPI_ISL_972251, EPI_ISL_972273, EPI_ISL_972275, EPI_ISL_972297, EPI_ISL_972352, EPI_ISL_972363, EPI_ISL_972369, EPI_ISL_972371, EPI_ISL_972418, EPI_ISL_972456, EPI_ISL_972514, EPI_ISL_972575, EPI_ISL_972608, EPI_ISL_972618, EPI_ISL_972722, EPI_ISL_972750, EPI_ISL_972761, EPI_ISL_972778, EPI_ISL_972793, EPI_ISL_972835, EPI_ISL_972874, EPI_ISL_972884, EPI_ISL_973028, EPI_ISL_973035, EPI_ISL_973078, EPI_ISL_973128, EPI_ISL_973131, EPI_ISL_973263, EPI_ISL_973271, EPI_ISL_973310, EPI_ISL_973325, EPI_ISL_973390, EPI_ISL_973492, EPI_ISL_973523, EPI_ISL_973568, EPI_ISL_973591, EPI_ISL_973672, EPI_ISL_973804 | | | |
| see above | Department of Virus and Microbiological Special Diagnostics, Statens Serum Institut, Copenhagen, Denmark | Aalborg University | Danish Covid-19 Genome Consortium |
| EPI_ISL_977065, EPI_ISL_977066, EPI_ISL_977067, EPI_ISL_977069, EPI_ISL_977070, EPI_ISL_977071, EPI_ISL_977072, EPI_ISL_977076, EPI_ISL_977077 | Rhode Island Department of Health | Infectious Disease Program, Broad Institute of Harvard and MIT | Lemieux,J.E., Siddle,K.J., Huard,R., King,E., Azevedo,K., Miller,A., Adams,G., Gladden-Young,A., Lagerborg,K., Rudy,M., DeRuff,K., Carter,A., Normandin,E., Bauer,M., Reilly,S., Tomkins-Tinch,C., Loreth,C., Chaluvadi,S., Birren,B.W., Gallagher,G., Smole,S., Park,D.J., MacInnis,B.L., and Sabeti,P.C. |
| EPI_ISL_977133, EPI_ISL_977134, EPI_ISL_977135, EPI_ISL_977136, EPI_ISL_977137 | Flow Health | Infectious Disease Program, Broad Institute of Harvard and MIT | Lemieux,J.E., Siddle,K.J., Adams,G., Gladden-Young,A., Lagerborg,K., Rudy,M., DeRuff,K., Carter,A., Normandin,E., Bauer,M., Reilly,S., Tomkins-Tinch,C., Loreth,C., Chaluvadi,S., Birren,B.W., Gallagher,G., Smole,S., Park,D.J., MacInnis,B.L., and Sabeti,P.C. |
| EPI_ISL_977270, EPI_ISL_977334, EPI_ISL_977350, EPI_ISL_977351, EPI_ISL_977468 | University of Zambia, School of Veterinary Medicine | UNZAVET and PATH | Mulenga Mwenda-Chimfwembe, Ngonda Saasa, Daniel Bridges |
| EPI_ISL_977538 | Caribbean Public Health Agency | Carrington Lab, Department of PreClinical Sciences | Nikita S. D. Sahadeo, Arianne Brown-Jordan, Vernie Ramkissoon, Sarah Hill, Naresh Nandram, Dr. Sharon Belmar-George Avery Hinds, Jerome Foster, Stanley Giddings, Karla Georges, Marsha Ivey, Rahul Naidu, Risha Singh, SueMin Nathaniel, Rajini Haraksingh, Jaya Jayaraman, Chinna Chinnadurai, Adesh Ramsubhag, Nuno Faria, Oliver Pybus, Christopher Oura, Gabriel Escobar, Christine V. F. Carrington |
| EPI_ISL_977539 | Caribbean Public Health Agency | Carrington Lab, Department of PreClinical Sciences | Nikita S. D. Sahadeo, Arianne Brown-Jordan, Vernie Ramkissoon, Sarah Hill, Naresh Nandram, Avery Hinds, Dr. Sharon Belmar-George, Jerome Foster, Stanley Giddings, Karla Georges, Marsha Ivey, Rahul Naidu, Risha Singh, SueMin Nathaniel, Rajini Haraksingh, Jaya Jayaraman, Chinna Chinnadurai, Adesh Ramsubhag, Nuno Faria, Oliver Pybus, Christopher Oura, Gabriel Escobar, Christine V. F. Carrington |
| EPI_ISL_978158, EPI_ISL_978159 | Chiu Laboratory, University of California, San Francisco | Chiu Laboratory, University of California, San Francisco | Charles Chiu, Xianding (Wayne) Deng, Candace Wang, Venice Servellita, Jill Hacker, Debra Wadford |
| EPI_ISL_978226 | Virginia Division of Consolidated Laboratory Services | Virginia Division of Consolidated Laboratory Services | Virginia DCLS |
| EPI_ISL_978274, EPI_ISL_978275, EPI_ISL_978276, EPI_ISL_978277, EPI_ISL_978278, EPI_ISL_978279, EPI_ISL_978350, EPI_ISL_978351, EPI_ISL_978352 | Texas Department of State Health Services | Texas Department of State Health Services | Bonnie Oh, Anita Pokharel, James Daniel Bonser, Myong Koag, Chung Wang, Rachel Lee, Grace Kubin, Rashmi Tuladhar, Mayela Pedrueza, Maliha Rahman, Jenny Zhang |
| EPI_ISL_978383, EPI_ISL_978384, EPI_ISL_978385, EPI_ISL_978386, EPI_ISL_978387, EPI_ISL_978388, EPI_ISL_978389, EPI_ISL_978390, EPI_ISL_978446, EPI_ISL_978447, EPI_ISL_978448, EPI_ISL_978449, EPI_ISL_978450, EPI_ISL_978451, EPI_ISL_978452, EPI_ISL_978453, EPI_ISL_978454, EPI_ISL_978455, EPI_ISL_978456, EPI_ISL_978457, EPI_ISL_978458, EPI_ISL_978459, EPI_ISL_978460, EPI_ISL_978461, EPI_ISL_978468, EPI_ISL_978469 | | | |
| see above | Arizona State Public Health Laboratory | Arizona State Public Health Laboratory | Trung Huynh, Jessica Escobar, Katherine Fullerton, Nobuko Fukushima, Stacy White, Linda Getsinger, Victor Waddell |
| EPI_ISL_978880, EPI_ISL_978881, EPI_ISL_978882, EPI_ISL_978883, EPI_ISL_978884, EPI_ISL_978885, EPI_ISL_978932, EPI_ISL_978934, EPI_ISL_978937 | Centre for Dengue Research and AICBU, Department of Immunology and Molecular Medicine | Centre for Dengue Research and AICBU, Department of Immunology and Molecular Medicine | Chandima Jeewandara, Deshni Jayathilaka, Dinuka Ariyaratne, Tibutius Thanesh Pramanayagam, Diyanath Ranasinghe, Laksiri Gomes, Gathsaurie Neelika Malavige |
| EPI_ISL_979133, EPI_ISL_979134, EPI_ISL_979143, EPI_ISL_979144 | Santa Clara County Public Health Laboratory | Chan-Zuckerberg Biohub | CZB Cliahub Consortium |
| EPI_ISL_979252 | Institute of Microbiology and Immunology, Faculty of Medicine, University of Ljubljana | Institute of Microbiology and Immunology, Faculty of Medicine, University of Ljubljana | Samo Zakotnik, Tomaž Mark Zorec, Matic Brvar, Doroteja Vlaj, Patricija Pozvek,Špela Pleh, Miša Korva, Mario Poljak, Tatjana Avši - Županc |
| EPI_ISL_979312, EPI_ISL_979324 | Cadham Provincial laboratory | National Microbiology Laboratory (NML) | Anna Majer, Shari Tyson, Grace Seo, Philip Mabon, Elsie Grudeski, Rhiannon Huzarewich, Russell Mandes, Anneliese Landgraff, Jennifer Tanner, Natalie Knox, Morag Graham, Gary Van Domselaar, Paul Van Caeseele, Jared Bullard, David Alexander, Kerry Dust, Nathalie Bastien, Yan Li, Timothy Booth, Darian Hole, Madison Chapel, Kirsten Biggar, CanCOGeN's metadata curation team, Public Health Agency of Canada CanCOGeN team |
| EPI_ISL_979372 | The Jackson Laboratory | The Jackson Laboratory | Lloyd M, Sanderson B, Srivastava A, Maurya R, Renzette N, Omerza G, Kelly K, Li L, Wei C L, Adams M |
| EPI_ISL_979638, EPI_ISL_979639, EPI_ISL_979640, EPI_ISL_979641, EPI_ISL_979642 | Santa Clara County Public Health Laboratory | Chan-Zuckerberg Biohub | CZB Cliahub Consortium |
| EPI_ISL_980907, EPI_ISL_980908, EPI_ISL_980914, EPI_ISL_980940 | Innovative Genomics Institute, UC Berkeley | Innovative Genomics Institute, UC Berkeley | Stacia Wyman, Haridha Shivram, Phil Frankino, Liana Lareau |
| EPI_ISL_981059, EPI_ISL_981065, EPI_ISL_981066, EPI_ISL_981067, EPI_ISL_981072 | Johns Hopkins Hospital Department of Pathology | Johns Hopkins Hospital Department of Pathology | C. Paul Morris, Chun Huai Luo, Adannaya Amadi, Matthew Schwartz, Nicholas Gallagher, Heba H. Mostafa |
| EPI_ISL_981270, EPI_ISL_981272, | Hospital Universitari de Bellvitge | Hospital Universitari Vall d'Hebron | Cristina Andrés, Maria Piñana, Josep F Abril, Damir Garcia-Cehic, Ariadna Rando, Juliana Esperalba, Maria Gema Codina, Carla Castillo, Maria Carmen |

| | | | |
|---|---|---|---|
| EPI_ISL_981295 | | | Martin, Tomàs Pumarola, Josep Quer, Andrés Antón |
| EPI_ISL_981298, EPI_ISL_981299, EPI_ISL_981301, EPI_ISL_981302, EPI_ISL_981303 | Hospital Universitari Vall d'Hebron | Hospital Universitari Vall d'Hebron | Cristina Andrés, Maria Piñana, Josep F Abril, Damir Garcia-Cehic, Ariadna Rando, Juliana Esperalba, Maria Gema Codina, Carla Castillo, Maria Carmen Martin, Tomàs Pumarola, Josep Quer, Andrés Antón |
| EPI_ISL_981373 | AZ Klina | AZ Klina | Dr. C. Vael |
| EPI_ISL_981384, EPI_ISL_981386 | Botswana Harvard HIV Reference Laboratory | Botswana Harvard HIV Reference Laboratory | Sikhulile Moyo, Dorcas Maruapula, Wonderful Choga, Botshelo Radibe, Boitumelo Zuze, David Lawrence, Roger Shapiro, Shahin Lockman, Mosepele Mosepele, Joseph Makhema, Simani Gaseitsiwe |
| EPI_ISL_981387 | IAL Regional de Bauru | Instituto Adolfo Lutz, Interdiciplinary Procedures Center, Strategic Laboratory | Claudio Tavares Sacchi, Claudia Regina Gonçalves, Erica Valessa Ramos Gomes, Karoline Rodrigues Campos |
| EPI_ISL_981548 | Botswana Harvard HIV Reference Laboratory | Botswana Harvard HIV Reference Laboratory | Sikhulile Moyo, Dorcas Maruapula, Wonderful T. Choga, Botshelo Radibe, Boitumelo Zuze, David Lawrence, Roger Shapiro, Shahin Lockman, Mosepele Mosepele, Joseph Makhema, Simani Gaseitsiwe |
| EPI_ISL_981971, EPI_ISL_981972, EPI_ISL_981973 | Microbiology Service, Hospital Universitario Clinico San Cecilio, Granada | Microbiology Service, Hospital Universitario Clinico San Cecilio, Granada | Adolfo de Salazar, Natalia Chueca, Laura Viñuela, Ana Fuentes, Federico García |
| EPI_ISL_982048, EPI_ISL_982057, EPI_ISL_982097 | TGen North | TGen North | "Jolene Bowers, Megan Folkerts, Chris French, Hayley Yaglom, Ashlyn Pfeiffer, Darrin Lemmer, Dave Engelthaler, The Arizona COVID Genomics Union (ACGU)" |
| EPI_ISL_982244, EPI_ISL_982273, EPI_ISL_982285 | Lab voor klinische biologie | Lab voor klinische biologie | Hannelore Hamerlinck, Marija Janevska, Bruno Verhasselt |
| EPI_ISL_982395 | M Health Fairview | Minnesota Department of Health, Public Health Laboratory | Alexandra Lorentz, Jacob Garfin, Matt Plumb, and Xiong Wang |
| EPI_ISL_982505 | National Institute of Laboratory Medicine and Referral Center | Genomic Research Lab, BCSIR | Iffat Jahan, Mohammad Samir Uzzaman, Eshrar Osman, Md. Ahashan Habib, Shahina Akter, Tanjina Akhtar Banu, Abu Sayeed Mohammad Mahmud, Md. Murshed Hasan Sarkar, Barna Goswami, Md. Saddam Hossain, Tasnim Nafisa, Md. Maruf Ahmed Molla, Mahmuda Yeasmin, Asish Kumar Ghosh, Arifa Akram, A. K. M.Shamsuzzaman, Md. Salim Khan |
| EPI_ISL_982781, EPI_ISL_982782, EPI_ISL_982784, EPI_ISL_982785, EPI_ISL_982786, EPI_ISL_982788, EPI_ISL_982789, EPI_ISL_982790, EPI_ISL_982791, EPI_ISL_982792, EPI_ISL_982793, EPI_ISL_982794, EPI_ISL_982795, EPI_ISL_982796, EPI_ISL_982797, EPI_ISL_982798, EPI_ISL_982799, EPI_ISL_982800, EPI_ISL_982801, EPI_ISL_982802, EPI_ISL_982803, EPI_ISL_982804, EPI_ISL_982805, EPI_ISL_982806, EPI_ISL_982807, EPI_ISL_982808, EPI_ISL_982809, EPI_ISL_982810, EPI_ISL_982812, EPI_ISL_982813, EPI_ISL_982814, EPI_ISL_982816, EPI_ISL_982819, EPI_ISL_982820, EPI_ISL_982821, EPI_ISL_982822, EPI_ISL_982823, EPI_ISL_982824, EPI_ISL_982825, EPI_ISL_982826, EPI_ISL_982827, EPI_ISL_982828, EPI_ISL_982829, EPI_ISL_982830, EPI_ISL_982832, EPI_ISL_982833, EPI_ISL_982834, EPI_ISL_982835, EPI_ISL_982836, EPI_ISL_982837, EPI_ISL_982838, EPI_ISL_982840 | | | |
| see above | University Health Network/Mount Sinai Hospital Department of Microbiology | Ontario Institute for Cancer Research | Marie-Ming Aynaud, Javier Hernandez, Seda Barutcu, Kin Chan, Jessica Bourke, Marc Mazzulli, Tony Mazzulli, Laurence Pelletier, Jeff Wrana, Aimee Paterson, Angel Liu, Allison McGeer, Patryk Aftanas, Kuganya Nirmalarajah, Samira Mubareka, Ilinca Lungu, Cassandra Bergwerff, Lubaina Kothari, Bernard Lam, Paul Krzyzanowski, Michael Laszloffy, Lawrence E. Heisler, Richard de Borja, Jared T. Simpson |
| EPI_ISL_982841 | National Institute of Laboratory Medicine and Referral Center | Genomic Research Lab, BCSIR | Md. Saddam Hossain, Mohammad Samir Uzzaman, Eshrar Osman, Md. Ahashan Habib, Shahina Akter, Tanjina Akhtar Banu, Abu Sayeed Mohammad Mahmud, Md. Murshed Hasan Sarkar, Barna Goswami, Iffat Jahan, Tasnim Nafisa, Md. Maruf Ahmed Molla, Mahmuda Yeasmin, Asish Kumar Ghosh, Arifa Akram, A. K. M.Shamsuzzaman, Md. Salim Khan |
| EPI_ISL_982885, EPI_ISL_982886, EPI_ISL_982892, EPI_ISL_982893, EPI_ISL_982924, EPI_ISL_982925 | MEPHI Aix Marseille University (AMU) | MEPHI Aix Marseille University (AMU) | Anthony LEVASSEUR |
| EPI_ISL_982937, EPI_ISL_982965, EPI_ISL_982968, EPI_ISL_982971, EPI_ISL_982975, EPI_ISL_982989, EPI_ISL_982997, EPI_ISL_982998, EPI_ISL_983003, EPI_ISL_983004, EPI_ISL_983006, EPI_ISL_983007, EPI_ISL_983009, EPI_ISL_983010, EPI_ISL_983012, EPI_ISL_983013, EPI_ISL_983014, EPI_ISL_983015, EPI_ISL_983016, EPI_ISL_983017, EPI_ISL_983018, EPI_ISL_983019, EPI_ISL_983020, EPI_ISL_983023, EPI_ISL_983025, EPI_ISL_983026, EPI_ISL_983029, EPI_ISL_983033, EPI_ISL_983034, EPI_ISL_983035, EPI_ISL_983036, EPI_ISL_983037, EPI_ISL_983038, EPI_ISL_983040, EPI_ISL_983041, EPI_ISL_983042, EPI_ISL_983043, EPI_ISL_983046, EPI_ISL_983047, EPI_ISL_983048, EPI_ISL_983050, EPI_ISL_983052, EPI_ISL_983056, EPI_ISL_983057, EPI_ISL_983058, EPI_ISL_983059, EPI_ISL_983060, EPI_ISL_983062 | | | |
| see above | University Health Network/Mount Sinai Hospital Department of Microbiology | Ontario Institute for Cancer Research | Marie-Ming Aynaud, Javier Hernandez, Seda Barutcu, Kin Chan, Jessica Bourke, Marc Mazzulli, Tony Mazzulli, Laurence Pelletier, Jeff Wrana, Aimee Paterson, Angel Liu, Allison McGeer, Patryk Aftanas, Kuganya Nirmalarajah, Samira Mubareka, Ilinca Lungu, Cassandra Bergwerff, Lubaina Kothari, Bernard Lam, Paul Krzyzanowski, Michael Laszloffy, Lawrence E. Heisler, Richard de Borja, Jared T. Simpson |
| EPI_ISL_983082 | Kentucky State Public Health Lab | Kentucky State Public Health Lab | Stephanie Lunn, Karim George, Joshua Tobias, William Grooms, Vaneet Arora, Matthew Johnson, Rachel Zinner, Rhonda Lucas |
| EPI_ISL_983089, EPI_ISL_983090, EPI_ISL_983091 | Gravity Diagnostics | Kentucky State Public Health Lab | Stephanie Lunn, Karim George, Joshua Tobias, William Grooms, Vaneet Arora, Matthew Johnson, Rachel Zinner, Rhonda Lucas |
| EPI_ISL_983092, EPI_ISL_983095 | Kentucky State Public Health Lab | Kentucky State Public Health Lab | Stephanie Lunn, Karim George, Joshua Tobias, William Grooms, Vaneet Arora, Matthew Johnson, Rachel Zinner, Rhonda Lucas |
| EPI_ISL_983171, EPI_ISL_983172, EPI_ISL_983173, EPI_ISL_983174, EPI_ISL_983175, EPI_ISL_983176, EPI_ISL_983177, EPI_ISL_983178, EPI_ISL_983179, EPI_ISL_983180, EPI_ISL_983181, EPI_ISL_983182, EPI_ISL_983183, EPI_ISL_983184, EPI_ISL_983185, EPI_ISL_983186, EPI_ISL_983187, EPI_ISL_983188, EPI_ISL_983189, EPI_ISL_983190, EPI_ISL_983191, EPI_ISL_983192, EPI_ISL_983193, EPI_ISL_983194, EPI_ISL_983195, EPI_ISL_983196, EPI_ISL_983197, EPI_ISL_983198, EPI_ISL_983199, EPI_ISL_983200, EPI_ISL_983201, EPI_ISL_983202, EPI_ISL_983203, EPI_ISL_983204, EPI_ISL_983205, EPI_ISL_983206, EPI_ISL_983207, EPI_ISL_983208, EPI_ISL_983209, EPI_ISL_983210, EPI_ISL_983211, EPI_ISL_983212, EPI_ISL_983213, EPI_ISL_983214, EPI_ISL_983215, EPI_ISL_983216, EPI_ISL_983217, EPI_ISL_983218, EPI_ISL_983219, EPI_ISL_983220, EPI_ISL_983221, EPI_ISL_983222, EPI_ISL_983223, EPI_ISL_983224, EPI_ISL_983225, EPI_ISL_983226, EPI_ISL_983227, EPI_ISL_983228, EPI_ISL_983229, EPI_ISL_983230, EPI_ISL_983231, EPI_ISL_983232, EPI_ISL_983233, EPI_ISL_983234, EPI_ISL_983235, EPI_ISL_983236, EPI_ISL_983237, EPI_ISL_983238, EPI_ISL_983239 | | | |
| see above | University Health Network/Mount Sinai Hospital Department of Microbiology | Ontario Institute for Cancer Research | Marie-Ming Aynaud, Javier Hernandez, Seda Barutcu, Kin Chan, Jessica Bourke, Marc Mazzulli, Tony Mazzulli, Laurence Pelletier, Jeff Wrana, Aimee Paterson, Angel Liu, Allison McGeer, Patryk Aftanas, Kuganya Nirmalarajah, Samira Mubareka, Ilinca Lungu, Cassandra Bergwerff, Lubaina Kothari, Bernard Lam, Paul Krzyzanowski, Michael Laszloffy, Lawrence E. Heisler, Richard de Borja, Jared T. Simpson |
| EPI_ISL_983372, EPI_ISL_983373 | Utah Public Health Laboratory | Utah Public Health Laboratory | Erin L. Young, Kelly F. Oakeson, Tara Gallagher |
| EPI_ISL_983500 | National Institute of Laboratory Medicine and Referral Center | Genomic Research Lab, BCSIR | Iffat Jahan, Mohammad Samir Uzzaman, Eshrar Osman, Md. Ahashan Habib, Shahina Akter, Tanjina Akhtar Banu, Abu Sayeed Mohammad Mahmud, Md. Murshed Hasan Sarkar, Barna Goswami, Md. Saddam Hossain, Tasnim Nafisa, Md. Maruf Ahmed Molla, Mahmuda Yeasmin, Asish Kumar Ghosh, Arifa Akram, A. K. M. Shamsuzzaman, Md. Salim Khan |
| EPI_ISL_983614, EPI_ISL_983615 | Texas Department of State Health Services | Texas Department of State Health Services | Bonnie Oh, Anita Pokharel, James Daniel Bonser, Myong Koag, Chung Wang, Rachel Lee, Grace Kubin, Rashmi Tuladhar, Mayela Pedrueza, Maliha Rahman, Jenny Zhang |
| EPI_ISL_983799, EPI_ISL_983800 | Colorado Department of Public Health and Environment | Colorado Department of Puplic Health and Environment | Laura Bankers, Molly C. Hetherington-Rauth, Diana Ir, Shannon Ely, Shannon R. Matzinger, Sarah Elizabeth Totten, Emily A. Travanty |
