## Supplementary material for "Detection and characterization of the SARS-CoV-2 lineage B.1.526 in New York": Supp. Table 3 GISAID Acknowledgment part 2: gisaid_hcov-19_acknowledgement_table_2021_02_13_010.pdf

We gratefully acknowledge the following Authors from the Originating laboratories responsible for obtaining the specimens, as well as the Submitting laboratories where the genome data were generated and shared via GISAID, on which this research is based.

All Submitters of data may be contacted directly via www.gisaid.org

Authors are sorted alphabetically.

| Accession ID | Originating Laboratory | Submitting Laboratory | Authors |
|---|---|---|---|
| EPI_ISL_710129 | Pathology North - Royal North Shore Hospital - NSW Health Pathology | CIDM-PH et al. | CIDM-PH et al. |
| EPI_ISL_710513 | Area of Virology, Serology and Virology Division (SAViD), New South Wales Health Pathology Randwick | Area of Virology, Serology and Virology Division (SAViD), New South Wales Health Pathology Randwick | Rawlinson, W., Deveson, I., Bull, R. |
| EPI_ISL_727861, EPI_ISL_727874, EPI_ISL_727877, EPI_ISL_727879, EPI_ISL_727880, EPI_ISL_727882, EPI_ISL_727887, EPI_ISL_727890, EPI_ISL_727896, EPI_ISL_727897, EPI_ISL_727901, EPI_ISL_727909, EPI_ISL_727911, EPI_ISL_727915, EPI_ISL_727927, EPI_ISL_727932, EPI_ISL_727942, EPI_ISL_727948, EPI_ISL_727949, EPI_ISL_727950, EPI_ISL_727954, EPI_ISL_727955, EPI_ISL_727956, EPI_ISL_727960, EPI_ISL_727961, EPI_ISL_727963, EPI_ISL_727973, EPI_ISL_727979, EPI_ISL_727980 | | | |
| see above | Virology Department, Sheffield Teaching Hospitals NHS Foundation Trust/Department of Infection, Immunity and Cardiovascular Disease, The Medical School, University of Sheffield | COVID-19 Genomics UK (COG-UK) Consortium | Thushan de Silva, Matthew Parker, Nikki Smith, Adri Angyal, Rebecca Brown, Luke Green, Rachel Tucker, Paul Parsons, Danielle Groves, Katie Johnson, Laura Carrilero, Alex Keeley, Dave Partridge, Matthew Wyles, Benjamin Lindsey, Mehmet Yavuz, Mohammad Raza, Cariad Evans |
| EPI_ISL_731347 | Lighthouse Lab in Glasgow | Wellcome Sanger Institute for the COVID-19 Genomics UK (COG-UK) Consortium | Harper VanSteenhouse, Yumi Kasai, David Gray, Carol Clugston, Anna Dominiczak and Alex Alderton, Roberto Amato, Sonia Goncalves, Ewan Harrison, David K. Jackson, Ian Johnston, Dominic Kwiatkowski, Cordelia Langford, John Sillitoe on behalf of the Wellcome Sanger Institute COVID-19 Surveillance Team |
| EPI_ISL_733498 | Central Virology Laboratory, Israel Ministry of Health | Central Virology Laboratory, Israel Ministry of Health | Neta S. Zuckerman, Efrat Dahan Bucris, Oran Erster, Ella Mendelson, Michal Mandelboim, Orna Mor |
| EPI_ISL_733570 | POK OI HOSPITAL | Hong Kong Department of Health | Alan K.L. Tsang, Peter C.W. Yip, Edman T.K. Lam, Rickjason C.W. Chan, Dominic N.C. Tsang |
| EPI_ISL_735386 | National Virus Reference Laboratory | National Virus Reference Laboratory | Michael Carr, Gabriel Gonzalez, Jonathan Dean, Daniel Hare, Cillian F De Gascun |
| EPI_ISL_735491 | Virology Laboratory, Ospedali Riuniti, Ancona | Virology Laboratory, Ospedali Riuniti, Ancona | Stefano Menzo, Roberta Longo, Laura Di Sante, Sara Caucci, Patrizia Bagnarelli |
| EPI_ISL_737237, EPI_ISL_737238, EPI_ISL_737239, EPI_ISL_737240, EPI_ISL_737241, EPI_ISL_737242, EPI_ISL_737254, EPI_ISL_737255, EPI_ISL_737256, EPI_ISL_737257, EPI_ISL_737258, EPI_ISL_737259, EPI_ISL_737260, EPI_ISL_737261 | | | |
| see above | Los Angeles County PHL | Los Angeles County PHL | P. Hemarajata et al. |
| EPI_ISL_737345, EPI_ISL_737346, EPI_ISL_737347, EPI_ISL_737348, EPI_ISL_737349, EPI_ISL_737350, EPI_ISL_737351, EPI_ISL_737352, EPI_ISL_737353, EPI_ISL_737354, EPI_ISL_737355, EPI_ISL_737356, EPI_ISL_737357, EPI_ISL_737358, EPI_ISL_737359, EPI_ISL_737360, EPI_ISL_737361, EPI_ISL_737362, EPI_ISL_737363, EPI_ISL_737364, EPI_ISL_737365, EPI_ISL_737366, EPI_ISL_737367, EPI_ISL_737368, EPI_ISL_737369, EPI_ISL_737370, EPI_ISL_737371, EPI_ISL_737372, EPI_ISL_737373, EPI_ISL_737374, EPI_ISL_737375, EPI_ISL_737376, EPI_ISL_737377, EPI_ISL_737378, EPI_ISL_737379, EPI_ISL_737380, EPI_ISL_737381, EPI_ISL_737382, EPI_ISL_737383 | | | |
| see above | Department of Clinical Microbiology | GIGA Medical Genomics | Keith Durkin, Maria Artesi, Sébastien Bontems, Raphaël Boreux, Bouchra Boujemla, Cécile Meex, Pierrette Melin, Marie-Pierre Hayette, Vincent Bours |
| EPI_ISL_737397, EPI_ISL_737398, EPI_ISL_737401, EPI_ISL_737402, EPI_ISL_737403, EPI_ISL_737409, EPI_ISL_737411, EPI_ISL_737414, EPI_ISL_737415, EPI_ISL_737417, EPI_ISL_737419, EPI_ISL_737421, EPI_ISL_737423, EPI_ISL_737429, EPI_ISL_737433, EPI_ISL_737434, EPI_ISL_737439, EPI_ISL_737440, EPI_ISL_737447, EPI_ISL_737449, EPI_ISL_737456, EPI_ISL_737458, EPI_ISL_737459, EPI_ISL_737464, EPI_ISL_737469, EPI_ISL_737471, EPI_ISL_737480, EPI_ISL_737481, EPI_ISL_737482, EPI_ISL_737483, EPI_ISL_737484, EPI_ISL_737492, EPI_ISL_737497, EPI_ISL_737498, EPI_ISL_737500, EPI_ISL_737505, EPI_ISL_737512, EPI_ISL_737513, EPI_ISL_737515, EPI_ISL_737518, EPI_ISL_737525, EPI_ISL_737527, EPI_ISL_737530, EPI_ISL_737533, EPI_ISL_737535, EPI_ISL_737538, EPI_ISL_737547, EPI_ISL_737548, EPI_ISL_737550, EPI_ISL_737560, EPI_ISL_737568, EPI_ISL_737569, EPI_ISL_737575, EPI_ISL_737576, EPI_ISL_737578, EPI_ISL_737581, EPI_ISL_737582, EPI_ISL_737583, EPI_ISL_737585, EPI_ISL_737589, EPI_ISL_737591, EPI_ISL_737600, EPI_ISL_737603, EPI_ISL_737610, EPI_ISL_737612, EPI_ISL_737613, EPI_ISL_737614, EPI_ISL_737615, EPI_ISL_737619, EPI_ISL_737620, EPI_ISL_737623, EPI_ISL_737625, EPI_ISL_737626, EPI_ISL_737631, EPI_ISL_737632, EPI_ISL_737633, EPI_ISL_737634, EPI_ISL_737635, EPI_ISL_737644, EPI_ISL_737649, EPI_ISL_737650, EPI_ISL_737651, EPI_ISL_737653, EPI_ISL_737654, EPI_ISL_737655, EPI_ISL_737662, EPI_ISL_737663, EPI_ISL_737664, EPI_ISL_737665, EPI_ISL_737709, EPI_ISL_737746, EPI_ISL_737770, EPI_ISL_737771, EPI_ISL_737782, EPI_ISL_737785, EPI_ISL_737786, EPI_ISL_737787, EPI_ISL_737788, EPI_ISL_737789, EPI_ISL_737790, EPI_ISL_737792, EPI_ISL_737793, EPI_ISL_737794, EPI_ISL_737798, EPI_ISL_737799, EPI_ISL_737814, EPI_ISL_737817, EPI_ISL_737818, EPI_ISL_737819, EPI_ISL_737820, EPI_ISL_737821, EPI_ISL_737840, EPI_ISL_737841, EPI_ISL_737842, EPI_ISL_737844, EPI_ISL_737847, EPI_ISL_737901, EPI_ISL_737910, EPI_ISL_737911, EPI_ISL_737912, EPI_ISL_737914, EPI_ISL_737915, EPI_ISL_737916 | | | |
| see above | Viollier AG | Department of Biosystems Science and Engineering, ETH Zürich | Chaoran Chen, Sarah Nadeau, Catharine Aquino, Ivan Topolsky, Philipp Jablonski, Lara Fuhrmann, David Dreifuss, Katharina Jahn, Andreia Cabral de Gouvea, Maria Domenica Moccia, Simon Grüter, Timothy Sykes, Lennart Opitz, Griffin White, Laura Neff, Doris Popovic, Andrea Patrignani, Jay Tracy, Ralph Schlapbach, Christiane Beckmann, Maurice Redondo, Olivier Kobel, Christoph Noppen, Sophie Seidel, Noemie Santamaria de Souza, Niko Beerenwinkel, Tanja Stadler |
| EPI_ISL_738044 | SIESP CHIETI - DRIVE IN LANCIANO | Istituto Zooprofilattico Sperimentale dell'Abruzzo e Molise "G. Caporale" | Lorusso A, Marcacci M, Di Domenico M, Ancora M, Curini V, Mangone I, Rinaldi A, Di Pasquale A, Cammà C, Puglia I, Savini G |
| EPI_ISL_738045 | SIESP DIPARTIMENTO DI PREVENZIONE CHIETI | Istituto Zooprofilattico Sperimentale dell'Abruzzo e Molise "G. Caporale" | Lorusso A, Marcacci M, Di Domenico M, Ancora M, Curini V, Mangone I, Rinaldi A, Di Pasquale A, Cammà C, Puglia I, Savini G |
| EPI_ISL_738046, EPI_ISL_738047, EPI_ISL_738048 | SIESP CHIETI - DRIVE IN ORTONA | Istituto Zooprofilattico Sperimentale dell'Abruzzo e Molise "G. Caporale" | Lorusso A, Marcacci M, Di Domenico M, Ancora M, Curini V, Mangone I, Rinaldi A, Di Pasquale A, Cammà C, Puglia I, Savini G |
| EPI_ISL_738099, EPI_ISL_738100, EPI_ISL_738101, EPI_ISL_738102, EPI_ISL_738103, EPI_ISL_738104, EPI_ISL_738105 | Instituto Nacional de Saude (INSA) | Instituto Nacional de Saude (INSA) | Borges et al |
| EPI_ISL_738135 | Department of Virology and Immunology, University of Helsinki and Helsinki University Hospital, Huslab Finland | Department of Virology, Faculty of Medicine, University of Helsinki, Helsinki, Finland | Teemu Smura, Olli Vapalahti, Maija Lappalainen, Satu Kurkela |
| EPI_ISL_738146 | Instituto Nacional de Saude (INSA) | Instituto Nacional de Saude (INSA) | Borges et al |
| EPI_ISL_738316, EPI_ISL_738353, EPI_ISL_738354, EPI_ISL_738356, EPI_ISL_738358, EPI_ISL_738359 | Utah Public Health Laboratory | Utah Public Health Laboratory | Erin Young, Kelly Oakeson, Tara Gallagher |
| EPI_ISL_739712, EPI_ISL_739714, EPI_ISL_739850, EPI_ISL_739910, EPI_ISL_739937, EPI_ISL_739943, EPI_ISL_739971, EPI_ISL_740039, EPI_ISL_740056, EPI_ISL_740061, EPI_ISL_740077, EPI_ISL_740111, EPI_ISL_740205, EPI_ISL_740214, EPI_ISL_740237, EPI_ISL_740267, EPI_ISL_740271, EPI_ISL_740274, EPI_ISL_740330, EPI_ISL_740375, EPI_ISL_740387, EPI_ISL_740424, EPI_ISL_740435, EPI_ISL_740503, EPI_ISL_740546 | | | |
| see above | Laboratoire national de santé, Microbiology, Virology | Laboratoire national de santé, Microbiology, Microbial Genomics Platform | Anke Wienecke-Baldacchino, Catherine Ragimbeau,Jessica Tapp, Fatu Djabi, Lise Pignon, Raoul Salmon, Tamir Abdelrahman |
| EPI_ISL_740866, EPI_ISL_740875, EPI_ISL_740876, EPI_ISL_740877, EPI_ISL_740878 | Pathology North - Royal North Shore Hospital - NSW Health Pathology | NSW Health Pathology - Institute of Clinical Pathology and Medical Research; Westmead Hospital; University of Sydney | CIDM-PH et al. |
| EPI_ISL_740879, EPI_ISL_740880 | Histopath | NSW Health Pathology - Institute of Clinical Pathology and Medical Research; Westmead Hospital; University of Sydney | CIDM-PH et al. |
| EPI_ISL_740883, EPI_ISL_740884, EPI_ISL_740885, EPI_ISL_740886, EPI_ISL_740887, EPI_ISL_740889 | Pathology North - Royal North Shore Hospital - NSW Health Pathology | NSW Health Pathology - Institute of Clinical Pathology and Medical Research; Westmead Hospital; University of Sydney | CIDM-PH et al. |

| | | | |
|---|---|---|---|
| EPI_ISL_740891 | Pathology North - Hunter - NSW Health Pathology | NSW Health Pathology - Institute of Clinical Pathology and Medical Research; Westmead Hospital; University of Sydney | CIDM-PH et al. |
| EPI_ISL_740892, EPI_ISL_740893 | Laverty Pathology | NSW Health Pathology - Institute of Clinical Pathology and Medical Research; Westmead Hospital; University of Sydney | CIDM-PH et al. |
| EPI_ISL_741337, EPI_ISL_741338, EPI_ISL_741339, EPI_ISL_741341, EPI_ISL_741342, EPI_ISL_741343, EPI_ISL_741344, EPI_ISL_741345 | University College London, Great Ormond Street Hospital for Children NHS Foundation Trust, Imperial College Healthcare NHS Trust | COVID-19 Genomics UK (COG-UK) Consortium | Sergi Castellano, Rachel Williams, Mark Kristiansen, Paola Resende Silva, Sunando Roy, Tony Brooks, Helena Tutill, Paola Niola, Patricia Dyal, Charlotte Williams, Leysa Forrest, Yasmin Panchbhaya, Jacqueline Findlay, Samuel Weeks, Julianne Brown, Kathryn Harris, Paul Randell, James Price, Alison Holmes, Judith Breuer |
| EPI_ISL_741486, EPI_ISL_741487, EPI_ISL_741488, EPI_ISL_741489, EPI_ISL_741490, EPI_ISL_741491, EPI_ISL_741493, EPI_ISL_741494, EPI_ISL_741497, EPI_ISL_741499, EPI_ISL_741503, EPI_ISL_741504, EPI_ISL_741563, EPI_ISL_741565, EPI_ISL_741566, EPI_ISL_741570, EPI_ISL_741572, EPI_ISL_741574, EPI_ISL_741575, EPI_ISL_741576, EPI_ISL_741577, EPI_ISL_741578, EPI_ISL_741579, EPI_ISL_741581, EPI_ISL_741582 | | | |
| see above | Quadram Institute Bioscience | COVID-19 Genomics UK (COG-UK) Consortium | Dave J. Baker, Gemma L. Kay, Alp Aydin, Thanh Le-Viet, Steven Rudder, Ana P. Tedim, Anastasia Kolyva, Maria Diaz, Leonardo de Oliveira Martins, Nabil-Fareed Alikhan, Lizzie Meadows, Rachael Stanley, Ngozi Elumogo, Muhammed Yasir, Nicholas M. Thomson, Alexander J Trotter, Rachel Gilroy, Samuel Bloomfield, Claire Stuart, Andrew Bell, Reenesh Prakash, Samir Dervisevic, Alison E. Mather, John Wain, Mark Webber, Andrew J. Page, Justin O'Grady |
| EPI_ISL_741631, EPI_ISL_741632, EPI_ISL_741633, EPI_ISL_741634, EPI_ISL_741635, EPI_ISL_741636, EPI_ISL_741637, EPI_ISL_741638, EPI_ISL_741639, EPI_ISL_741640, EPI_ISL_741641, EPI_ISL_741642, EPI_ISL_741643, EPI_ISL_741644, EPI_ISL_741645, EPI_ISL_741646, EPI_ISL_741647, EPI_ISL_741648, EPI_ISL_741649, EPI_ISL_741650 | | | |
| see above | Queens Medical Centre, Clinical Microbiology Department / DeepSeq Nottingham | COVID-19 Genomics UK (COG-UK) Consortium | Gemma Clark, Wendy Smith, Manjinder Khakh, Vicki M Fleming, Michelle M Lister, Hannah Howson-Wells, Jonathan Ball, Patrick McClure, Joseph Chappell, Theocharis Tsoleridis, Nadine Holmes, Matthew Carlisle, Christopher Moore, Fei Sang, Johnny Debebe, Victoria Wright, Matthew Loose |
| EPI_ISL_741942, EPI_ISL_741948, EPI_ISL_741950, EPI_ISL_741951, EPI_ISL_741954, EPI_ISL_741956, EPI_ISL_741960, EPI_ISL_741962, EPI_ISL_741965, EPI_ISL_741969, EPI_ISL_741971, EPI_ISL_741977, EPI_ISL_741983, EPI_ISL_741984, EPI_ISL_741989, EPI_ISL_741990, EPI_ISL_741991, EPI_ISL_742000, EPI_ISL_742004, EPI_ISL_742010, EPI_ISL_742015, EPI_ISL_742018, EPI_ISL_742025, EPI_ISL_742030, EPI_ISL_742034, EPI_ISL_742041, EPI_ISL_742046, EPI_ISL_742048, EPI_ISL_742054, EPI_ISL_742067, EPI_ISL_742074, EPI_ISL_742079, EPI_ISL_742080, EPI_ISL_742086, EPI_ISL_742090, EPI_ISL_742092, EPI_ISL_742094, EPI_ISL_742109, EPI_ISL_742112, EPI_ISL_742113, EPI_ISL_742115, EPI_ISL_742116 | | | |
| see above | Virology Department, Sheffield Teaching Hospitals NHS Foundation Trust/Department of Infection, Immunity and Cardiovascular Disease, The Medical School, University of Sheffield | COVID-19 Genomics UK (COG-UK) Consortium | Thushan de Silva, Matthew Parker, Nikki Smith, Adri Angyal, Rebecca Brown, Luke Green, Rachel Tucker, Paul Parsons, Danielle Groves, Katie Johnson, Laura Carrilero, Alex Keeley, Dave Partridge, Matthew Wyles, Benjamin Lindsey, Mehmet Yavuz, Mohammad Raza, Cariad Evans |
| EPI_ISL_742203, EPI_ISL_742204, EPI_ISL_742205, EPI_ISL_742206, EPI_ISL_742207, EPI_ISL_742208, EPI_ISL_742209, EPI_ISL_742210, EPI_ISL_742211, EPI_ISL_742212, EPI_ISL_742213, EPI_ISL_742214, EPI_ISL_742215, EPI_ISL_742216, EPI_ISL_742217, EPI_ISL_742218, EPI_ISL_742219, EPI_ISL_742220, EPI_ISL_742221, EPI_ISL_742222, EPI_ISL_742223, EPI_ISL_742224, EPI_ISL_742225, EPI_ISL_742226, EPI_ISL_742227, EPI_ISL_742228, EPI_ISL_742229, EPI_ISL_742230, EPI_ISL_742231, EPI_ISL_742232, EPI_ISL_742233, EPI_ISL_742234, EPI_ISL_742235, EPI_ISL_742236, EPI_ISL_742237, EPI_ISL_742238, EPI_ISL_742239, EPI_ISL_742240, EPI_ISL_742241, EPI_ISL_742242 | | | |
| see above | Virology Department, Royal Infirmary of Edinburgh, NHS Lothian / School of Biological Sciences, University of Edinburgh / Institute of Genetics and Molecular Medicine, University of Edinburgh | COVID-19 Genomics UK (COG-UK) Consortium | McHugh M, Dewar R, Rooke S, Gallagher M, Balcaza C, O'Toole Á, Scher E, Hill V, McCrone JT, Colquhoun R, Yu X, Jackson B, Rambaut A, Williams TC, Templeton K |
| EPI_ISL_743878, EPI_ISL_743928, EPI_ISL_743929, EPI_ISL_743930, EPI_ISL_743931, EPI_ISL_743932, EPI_ISL_743933, EPI_ISL_743934, EPI_ISL_743935, EPI_ISL_743936, EPI_ISL_744067, EPI_ISL_744134 | | | |
| see above | Wales Specialist Virology Centre Sequencing lab: Pathogen Genomics Unit | COVID-19 Genomics UK (COG-UK) Consortium | Catherine Moore, Johnathan Evans, Laura Gifford, Malorie Perry, Simon Cottrell, Angela Marchbank, Alec Birchley, Alexander Adams, Amy Gaskin, Bree Gatica-Wilcox, Jason Coombes, Joel Southgate, Lauren Gilbert, Lee Graham, Nicole Pacchiarini, Sara Kumziene-Summerhayes, Sarah Taylor, Sophie Jones, Sara Rey, Matthew Bull, Joanne Watkins, Sally Corden, Tom Connor |
| EPI_ISL_744197, EPI_ISL_744203, EPI_ISL_744211, EPI_ISL_744264, EPI_ISL_744280, EPI_ISL_744294, EPI_ISL_744395, EPI_ISL_744396, EPI_ISL_744400, EPI_ISL_744435, EPI_ISL_744483, EPI_ISL_744557, EPI_ISL_744605, EPI_ISL_744676, EPI_ISL_744734, EPI_ISL_744756, EPI_ISL_744780, EPI_ISL_744792, EPI_ISL_744809, EPI_ISL_744879, EPI_ISL_744893 | | | |
| see above | Laboratoire national de santé, Microbiology, Virology | Laboratoire national de santé, Microbiology, Microbial Genomics Platform | Anke Wienecke-Baldacchino, Catherine Ragimbeau,Jessica Tapp, Fatu Djabi, Lise Pignon, Raoul Salmon, Tamir Abdelrahman |
| EPI_ISL_745078, EPI_ISL_745079, EPI_ISL_745080, EPI_ISL_745081, EPI_ISL_745083, EPI_ISL_745084, EPI_ISL_745085, EPI_ISL_745086 | Israel Central Virology laboratory | Israel Central Virology laboratory | Neta Zuckerman, Efrat Dahan Bucris, Oran Erster, Michal Mandelboim, Orna Mor, Ella Mendelson |
| EPI_ISL_745091, EPI_ISL_745092 | Australian Clinical Labs | CIDM-PH, Westmead Hospital | CIDM-PH et al. |
| EPI_ISL_745093, EPI_ISL_745094 | Histopath | CIDM-PH, Westmead Hospital | CIDM-PH et al. |
| EPI_ISL_745095 | Laverty Pathology | CIDM-PH, Westmead Hospital | CIDM-PH et al. |
| EPI_ISL_745096 | Pathology North - Hunter - NSW Health Pathology | CIDM-PH, Westmead Hospital | CIDM-PH et al. |
| EPI_ISL_745097, EPI_ISL_745098, EPI_ISL_745099, EPI_ISL_745100, EPI_ISL_745101, EPI_ISL_745102, EPI_ISL_745103, EPI_ISL_745104, EPI_ISL_745105, EPI_ISL_745106, EPI_ISL_745107 | | | |
| see above | Pathology North - Royal North Shore Hospital - NSW Health Pathology | CIDM-PH, Westmead Hospital | CIDM-PH et al. |
| EPI_ISL_745108 | St Vincent's Pathology (SydPath) | CIDM-PH, Westmead Hospital | CIDM-PH et al. |
| EPI_ISL_745311, EPI_ISL_745333, EPI_ISL_745334, EPI_ISL_745390, EPI_ISL_745391, EPI_ISL_745392 | CNR Virus des Infections Respiratoires - France SUD | CNR Virus des Infections Respiratoires - France SUD | Antonin Bal, Gregory Destras, Claudia Gonzalez, Gwendolyne Burfin, Quentin Semanas, Martine Valette, Bruno Lina, Laurence Josset |
| EPI_ISL_746830 | National Institute for Infectious Diseases, INMI, "L. Spallanzani" IRCCS | National Institute for Infectious Diseases, INMI, "L. Spallanzani" IRCCS | E Giombini, M Rueca, B Bartolini, C.E.M Gruber, F Messina, A Di Caro, MR Capobianchi |
| EPI_ISL_746831 | National Institute for Infectious Diseases, INMI, "L. Spallanzani" IRCCS | National Institute for Infectious Diseases, INMI, "L. Spallanzani" IRCCS | C.E.M Gruber, F Messina, M Rueca, B Bartolini, E Giombini, MR Capobianchi, A Di Caro |
| EPI_ISL_746832 | National Institute for Infectious Diseases, INMI, "L. Spallanzani" IRCCS | National Institute for Infectious Diseases, INMI, "L. Spallanzani" IRCCS | E Giombini, C.E.M Gruber, M Rueca, B Bartolini, F Messina, A Di Caro, MR Capobianchi |
| EPI_ISL_746833 | National Institute for Infectious Diseases, INMI, "L. Spallanzani" IRCCS | National Institute for Infectious Diseases, INMI, "L. Spallanzani" IRCCS | F Messina, M Rueca, B Bartolini, C.E.M Gruber, E Giombini, MR Capobianchi, A Di Caro |
| EPI_ISL_746834 | National Institute for Infectious Diseases, INMI, "L. Spallanzani" IRCCS | National Institute for Infectious Diseases, INMI, "L. Spallanzani" IRCCS | B Bartolini, M Rueca, C.E.M Gruber, F Messina, E Giombini, A Di Caro, MR Capobianchi |
| EPI_ISL_747521 | Dutch COVID-19 response team | National Institute for Public Health and the Environment (RIVM) | Adam Meijer, Harry Vennema, Jeroen Cremer, Sharon van den Brink, Bas van der Veer, AnneMarie van den Brandt, Florian Zwagemaker, Dennis Schmitz, Chantal Reusken, on behalf of the national COVID-19 response team |
| EPI_ISL_751193 | University Hospital Zürich | Institute of Medical Virology, University of Zurich | Stefan Schmutz, Kevin Steiner, Verena Kufner, Maryam Zaheri, Gabriela Ziltener, Jürg Böni, Michael Huber, Alexandra Trkola |
| EPI_ISL_752600 | Utah Public Health Laboratory, Utah Public Health Laboratory Infectious Disease submission group | Utah Public Health Laboratory, Utah Public Health Laboratory Infectious Disease submission group | Gallagher,T., Young,E., Oakeson,K. |
| EPI_ISL_753675, EPI_ISL_753676, EPI_ISL_753677, EPI_ISL_753678, EPI_ISL_753679, EPI_ISL_753680, EPI_ISL_753681, EPI_ISL_753682, EPI_ISL_753683, EPI_ISL_753684, EPI_ISL_753685, EPI_ISL_753686, EPI_ISL_753687, EPI_ISL_753688, EPI_ISL_753690, EPI_ISL_753692 | | | |

| | | | |
|---|---|---|---|
| see above | Clinical virology Laboratory, Children's Hospital Los Angeles | Center for Personalized Medicine, Children's Hospital Los Angeles | Gai et al |
| EPI_ISL_754080, EPI_ISL_754083, EPI_ISL_754084, EPI_ISL_754086, EPI_ISL_754088, EPI_ISL_754089, EPI_ISL_754090, EPI_ISL_754091, EPI_ISL_754092, EPI_ISL_754094 | National Public Health Laboratory, National Centre for Infectious Diseases | National Public Health Laboratory, National Centre for Infectious Diseases | Tze Minn Mak, Sophie Octavia, Zhenyang Zhou, Lin Cui, Raymond Tzer Pin Lin |
| EPI_ISL_754126, EPI_ISL_754127, EPI_ISL_754128, EPI_ISL_754129 | USC Clinical Lab | Los Angeles County PHL | P. Hemarajata et al. |
| EPI_ISL_754131 | Los Angeles County PHL | Los Angeles County PHL | P. Hemarajata et al. |
| EPI_ISL_754133, EPI_ISL_754134 | USC Clinical Lab | Los Angeles County PHL | P. Hemarajata et al. |
| EPI_ISL_754268, EPI_ISL_754273, EPI_ISL_754278, EPI_ISL_754284, EPI_ISL_754313, EPI_ISL_754324, EPI_ISL_754325, EPI_ISL_754326, EPI_ISL_754327, EPI_ISL_754328, EPI_ISL_754363, EPI_ISL_754365, EPI_ISL_754366, EPI_ISL_754368, EPI_ISL_754369, EPI_ISL_754370, EPI_ISL_754377, EPI_ISL_754379, EPI_ISL_754381, EPI_ISL_754383, EPI_ISL_754384, EPI_ISL_754385, EPI_ISL_754386, EPI_ISL_754387 | | | |
| see above | Respiratory Virus Unit, National Infection Service, Public Health England | COVID-19 Genomics UK (COG-UK) Consortium | PHE Covid Sequencing Team |
| EPI_ISL_754619, EPI_ISL_754648, EPI_ISL_754649, EPI_ISL_754650, EPI_ISL_754651, EPI_ISL_754652, EPI_ISL_754653, EPI_ISL_754654, EPI_ISL_754655, EPI_ISL_754656 | University of Wisconsin-Madison AIDS Vaccine Research Laboratories | University of Wisconsin-Madison AIDS Vaccine Research Laboratories | Gage Moreno, Katarina Braun, et al. AIDS Vaccine Research Laboratories |
| EPI_ISL_754843, EPI_ISL_754844, EPI_ISL_754845, EPI_ISL_754846, EPI_ISL_754849, EPI_ISL_754850, EPI_ISL_754851, EPI_ISL_754852 | Laboratoire de Virologie Hôpital Robert Debré | National Reference Center for Viruses of Respiratory Infections, Institut Pasteur, Paris | Marion Barbet, Sylvie Behillil, Méline Bizard, Angela Brisebarre, Camille Capel, Etienne Simon-Lorière, Vincent Enouf, Maud Vanpeene, Sylvie van der Werf, Andreoletti Laurent |
| EPI_ISL_754853, EPI_ISL_754854, EPI_ISL_754855 | CHU - Hôpital Cavale Blanche - Labo. de Virologie | National Reference Center for Viruses of Respiratory Infections, Institut Pasteur, Paris | Marion Barbet, Sylvie Behillil, Méline Bizard, Angela Brisebarre, Camille Capel, Etienne Simon-Lorière, Vincent Enouf, Maud Vanpeene, Sylvie van der Werf, Pilorge léa |
| EPI_ISL_755181, EPI_ISL_755182, EPI_ISL_755184, EPI_ISL_755185, EPI_ISL_755186, EPI_ISL_755187, EPI_ISL_755188, EPI_ISL_755189, EPI_ISL_755194 | UCSD EXCITE lab | Andersen lab at Scripps Research | SEARCH Alliance San Diego |
| EPI_ISL_755511, EPI_ISL_755512, EPI_ISL_755513, EPI_ISL_755514, EPI_ISL_755515, EPI_ISL_755516, EPI_ISL_755517, EPI_ISL_755518, EPI_ISL_755519, EPI_ISL_755520, EPI_ISL_755521, EPI_ISL_755522, EPI_ISL_755523, EPI_ISL_755524, EPI_ISL_755525, EPI_ISL_755526, EPI_ISL_755527, EPI_ISL_755528, EPI_ISL_755529, EPI_ISL_755530, EPI_ISL_755531, EPI_ISL_755532, EPI_ISL_755533, EPI_ISL_755534, EPI_ISL_755535, EPI_ISL_755536, EPI_ISL_755537, EPI_ISL_755538, EPI_ISL_755539, EPI_ISL_755540, EPI_ISL_755541, EPI_ISL_755542, EPI_ISL_755543, EPI_ISL_755544, EPI_ISL_755545, EPI_ISL_755546, EPI_ISL_755547, EPI_ISL_755548, EPI_ISL_755549, EPI_ISL_755553, EPI_ISL_755560 | | | |
| see above | Maine Health and Environmental Testing Laboratory | Tewhey Lab, The Jackson Laboratory | Matluk,N., Dewey,H., Iosue,F., Barter,M., Lynch,R., Munger,H. and Tewhey,R. |
| EPI_ISL_755616 | Helix/Illumina | Genomics and Discovery, Respiratory Viruses Branch, Division of Viral Diseases, Centers for Disease Control and Prevention | Peter W. Cook, Dhwani Batra, Eileen de Feo, Jan Antico, Christine Tran, Matthew Tolentino, Shannon Wickline, Kim Gietzen, Brad Sickler, Jingtao Liu, Eric Allen, Phil Febbo, Summer Galloway, Nicole L. Washington, Simon White, Geraint Levan, Kelly Schiabor Barrett, Elizabeth Cirulli, Alexandre Bolze, Ary Ascencio, Charlotte Rivera-Garcia, Ryan Cho, Jason Nguyen, Sherry Wang, Jimmy Ramirez, Tyler Cassens, Efren Sandoval, Magnus Isaksson, William Lee, David Becker, Marc Laurent, James Lu, Clinton R. Paden, Suxiang Tong, Duncan MacCannell |
| EPI_ISL_755620 | Canterbury Health Laboratories | Institute of Environmental Science and Research (ESR) | Xiaoyun Ren, Matt Storey, Nikki Freed, Muhammad Faisal, Jing Wang, Hermes Perez, Anja Werno, Antje van der Linden, Arlo Upton, Chris Mansell, David Hammer, Dragana Drinkovic, Gary McAuliffe, Hana Sofia Andersson, James Ussher, Jill Sherwood, Josh Freeman, Julia Howard, Juliet Elvy, Mary DeAlmeida, Matt Blakiston, Matthew Rogers, Max Bloomfield, Michael Addidle, Michelle Balm, Sally Roberts, Sarah Jefferies, Sharmini Muttaiyah, Susan Morpeth, Susan Taylor, Timothy Blackmore, Vani Sathyendran, Veronica Playle, Virginia Hope, Erasmus Smit, Lauren Jelly, Olin Silander, Joep de Ligt |
| EPI_ISL_755625 | Middlemore Hospital | Institute of Environmental Science and Research (ESR) | Xiaoyun Ren, Matt Storey, Nikki Freed, Muhammad Faisal, Jing Wang, Hermes Perez, Anja Werno, Antje van der Linden, Arlo Upton, Chris Mansell, David Hammer, Dragana Drinkovic, Gary McAuliffe, Hana Sofia Andersson, James Ussher, Jill Sherwood, Josh Freeman, Julia Howard, Juliet Elvy, Mary DeAlmeida, Matt Blakiston, Matthew Rogers, Max Bloomfield, Michael Addidle, Michelle Balm, Sally Roberts, Sarah Jefferies, Sharmini Muttaiyah, Susan Morpeth, Susan Taylor, Timothy Blackmore, Vani Sathyendran, Veronica Playle, Virginia Hope, Erasmus Smit, Lauren Jelly, Olin Silander, Joep de Ligt |
| EPI_ISL_755628 | LabPLUS | Institute of Environmental Science and Research (ESR) | Xiaoyun Ren, Matt Storey, Nikki Freed, Muhammad Faisal, Jing Wang, Hermes Perez, Anja Werno, Antje van der Linden, Arlo Upton, Chris Mansell, David Hammer, Dragana Drinkovic, Gary McAuliffe, Hana Sofia Andersson, James Ussher, Jill Sherwood, Josh Freeman, Julia Howard, Juliet Elvy, Mary DeAlmeida, Matt Blakiston, Matthew Rogers, Max Bloomfield, Michael Addidle, Michelle Balm, Sally Roberts, Sarah Jefferies, Sharmini Muttaiyah, Susan Morpeth, Susan Taylor, Timothy Blackmore, Vani Sathyendran, Veronica Playle, Virginia Hope, Erasmus Smit, Lauren Jelly, Olin Silander, Joep de Ligt |
| EPI_ISL_755804 | Toronto Invasive Bacterial Diseases Network | McMaster University | Allison McGeer, Patryk Aftanas, Hooman Derakhshani, Angel Li, Kuganya Nirmalarajah, Emily Panousis, Ahmed Draia, Jalees Nasir, Michael Surette, Samira Mubareka, Andrew G. McArthur |
| EPI_ISL_756135, EPI_ISL_756136, EPI_ISL_756137, EPI_ISL_756138, EPI_ISL_756139, EPI_ISL_756140, EPI_ISL_756141, EPI_ISL_756142, EPI_ISL_756143, EPI_ISL_756144, EPI_ISL_756145, EPI_ISL_756146, EPI_ISL_756147, EPI_ISL_756148, EPI_ISL_756149, EPI_ISL_756150, EPI_ISL_756151, EPI_ISL_756152, EPI_ISL_756153, EPI_ISL_756154, EPI_ISL_756155, EPI_ISL_756156, EPI_ISL_756157, EPI_ISL_756158, EPI_ISL_756159, EPI_ISL_756160, EPI_ISL_756161, EPI_ISL_756162, EPI_ISL_756163 | | | |
| see above | Respiratory Virus Unit, National Infection Service, Public Health England | COVID-19 Genomics UK (COG-UK) Consortium | PHE Covid Sequencing Team |
| EPI_ISL_756228, EPI_ISL_756229, EPI_ISL_756230, EPI_ISL_756234, EPI_ISL_756235, EPI_ISL_756236, EPI_ISL_756237, EPI_ISL_756238, EPI_ISL_756239, EPI_ISL_756240, EPI_ISL_756295 | | | |
| see above | UW Virology Lab | UW Virology Lab | Pavitra Roychoudhury, Hong Xie, Lasata Shrestha, Meei-Li Huang, Keith R Jerome, Alexander Greninger |
| EPI_ISL_756310 | The Caribbean Public Health Agency | Carrington Lab, Department of PreClinical Sciences, Faculty of Medical Sciences, The University of the West Indies | Nikita S. D. Sahadeo, Arianne Brown-Jordan, Sarah Hill, Vernie Ramkissoon, Roshan Parasram, Naresh Nandram, Avery Hinds, Jerome Foster, Stanley Giddings, Karla Georges, Marsha Ivey, Rahul Naidu, Risha Singh, SueMin Nathaniel, Rajini Haraksingh, Jaya Jayaraman, Chinna Chinnadurai, Adesh Ramsubhag, Nuno Faria, Oliver Pybus, Christopher Oura, Gabriel Escobar, Christine V. F. Carrington |
| EPI_ISL_756378 | Lighthouse Lab in Alderley Park | Wellcome Sanger Institute for the COVID-19 Genomics UK (COG-UK) Consortium | Jacquelyn Wynn, Mairead Hyland, The Lighthouse Lab in Alderley Park and Alex Alderton, Roberto Amato, Sonia Goncalves, Ewan Harrison, David K. Jackson, Ian Johnston, Dominic Kwiatkowski, Cordelia Langford, John Sillitoe on behalf of the Wellcome Sanger Institute COVID-19 Surveillance Team |
| EPI_ISL_756379, EPI_ISL_756380 | Lighthouse Lab in Cambridge | Wellcome Sanger Institute for the COVID-19 Genomics UK (COG-UK) Consortium | Rob Howes, The Lighthouse Lab in Cambridge and Alex Alderton, Roberto Amato, Sonia Goncalves, Ewan Harrison, David K. Jackson, Ian Johnston, Dominic Kwiatkowski, Cordelia Langford, John Sillitoe on behalf of the Wellcome Sanger Institute COVID-19 Surveillance Team |
| EPI_ISL_756381, EPI_ISL_756382, EPI_ISL_756383 | Lighthouse Lab in Alderley Park | Wellcome Sanger Institute for the COVID-19 Genomics UK (COG-UK) Consortium | Jacquelyn Wynn, Mairead Hyland, The Lighthouse Lab in Alderley Park and Alex Alderton, Roberto Amato, Sonia Goncalves, Ewan Harrison, David K. Jackson, Ian Johnston, Dominic Kwiatkowski, Cordelia Langford, John Sillitoe on behalf of the Wellcome Sanger Institute COVID-19 Surveillance Team |
| EPI_ISL_756384 | Lighthouse Lab in Cambridge | Wellcome Sanger Institute for the COVID-19 Genomics UK (COG-UK) Consortium | Rob Howes, The Lighthouse Lab in Cambridge and Alex Alderton, Roberto Amato, Sonia Goncalves, Ewan Harrison, David K. Jackson, Ian Johnston, Dominic Kwiatkowski, Cordelia Langford, John Sillitoe on behalf of the Wellcome Sanger Institute COVID-19 Surveillance Team |
| EPI_ISL_756385 | Lighthouse Lab in Alderley Park | Wellcome Sanger Institute for the COVID-19 Genomics UK (COG-UK) Consortium | Jacquelyn Wynn, Mairead Hyland, The Lighthouse Lab in Alderley Park and Alex Alderton, Roberto Amato, Sonia Goncalves, Ewan Harrison, David K. Jackson, Ian Johnston, Dominic Kwiatkowski, Cordelia Langford, John Sillitoe on behalf of the Wellcome Sanger Institute COVID-19 Surveillance Team |
| EPI_ISL_756386 | Lighthouse Lab in Cambridge | Wellcome Sanger Institute for the COVID-19 Genomics UK | Rob Howes, The Lighthouse Lab in Cambridge and Alex Alderton, Roberto Amato, Sonia Goncalves, Ewan Harrison, David K. Jackson, Ian Johnston, |

| | | | |
|---|---|---|---|
| | | (COG-UK) Consortium | Dominic Kwiatkowski, Cordelia Langford, John Sillitoe on behalf of the Wellcome Sanger Institute COVID-19 Surveillance Team |
| EPI_ISL_756387, EPI_ISL_756388 | Lighthouse Lab in Alderley Park | Wellcome Sanger Institute for the COVID-19 Genomics UK (COG-UK) Consortium | Jacquelyn Wynn, Mairead Hyland, The Lighthouse Lab in Alderley Park and Alex Alderton, Roberto Amato, Sonia Goncalves, Ewan Harrison, David K. Jackson, Ian Johnston, Dominic Kwiatkowski, Cordelia Langford, John Sillitoe on behalf of the Wellcome Sanger Institute COVID-19 Surveillance Team |
| EPI_ISL_756389 | Lighthouse Lab in Cambridge | Wellcome Sanger Institute for the COVID-19 Genomics UK (COG-UK) Consortium | Rob Howes, The Lighthouse Lab in Cambridge and Alex Alderton, Roberto Amato, Sonia Goncalves, Ewan Harrison, David K. Jackson, Ian Johnston, Dominic Kwiatkowski, Cordelia Langford, John Sillitoe on behalf of the Wellcome Sanger Institute COVID-19 Surveillance Team |
| EPI_ISL_756390, EPI_ISL_756391 | Lighthouse Lab in Alderley Park | Wellcome Sanger Institute for the COVID-19 Genomics UK (COG-UK) Consortium | Jacquelyn Wynn, Mairead Hyland, The Lighthouse Lab in Alderley Park and Alex Alderton, Roberto Amato, Sonia Goncalves, Ewan Harrison, David K. Jackson, Ian Johnston, Dominic Kwiatkowski, Cordelia Langford, John Sillitoe on behalf of the Wellcome Sanger Institute COVID-19 Surveillance Team |
| EPI_ISL_756392, EPI_ISL_756393 | Lighthouse Lab in Cambridge | Wellcome Sanger Institute for the COVID-19 Genomics UK (COG-UK) Consortium | Rob Howes, The Lighthouse Lab in Cambridge and Alex Alderton, Roberto Amato, Sonia Goncalves, Ewan Harrison, David K. Jackson, Ian Johnston, Dominic Kwiatkowski, Cordelia Langford, John Sillitoe on behalf of the Wellcome Sanger Institute COVID-19 Surveillance Team |
| EPI_ISL_756394 | Lighthouse Lab in Alderley Park | Wellcome Sanger Institute for the COVID-19 Genomics UK (COG-UK) Consortium | Jacquelyn Wynn, Mairead Hyland, The Lighthouse Lab in Alderley Park and Alex Alderton, Roberto Amato, Sonia Goncalves, Ewan Harrison, David K. Jackson, Ian Johnston, Dominic Kwiatkowski, Cordelia Langford, John Sillitoe on behalf of the Wellcome Sanger Institute COVID-19 Surveillance Team |
| EPI_ISL_756395 | Lighthouse Lab in Cambridge | Wellcome Sanger Institute for the COVID-19 Genomics UK (COG-UK) Consortium | Rob Howes, The Lighthouse Lab in Cambridge and Alex Alderton, Roberto Amato, Sonia Goncalves, Ewan Harrison, David K. Jackson, Ian Johnston, Dominic Kwiatkowski, Cordelia Langford, John Sillitoe on behalf of the Wellcome Sanger Institute COVID-19 Surveillance Team |
| EPI_ISL_756396, EPI_ISL_756397, EPI_ISL_756398, EPI_ISL_756399, EPI_ISL_756400, EPI_ISL_756402 | Lighthouse Lab in Alderley Park | Wellcome Sanger Institute for the COVID-19 Genomics UK (COG-UK) Consortium | Jacquelyn Wynn, Mairead Hyland, The Lighthouse Lab in Alderley Park and Alex Alderton, Roberto Amato, Sonia Goncalves, Ewan Harrison, David K. Jackson, Ian Johnston, Dominic Kwiatkowski, Cordelia Langford, John Sillitoe on behalf of the Wellcome Sanger Institute COVID-19 Surveillance Team |
| EPI_ISL_756403 | Lighthouse Lab in Cambridge | Wellcome Sanger Institute for the COVID-19 Genomics UK (COG-UK) Consortium | Rob Howes, The Lighthouse Lab in Cambridge and Alex Alderton, Roberto Amato, Sonia Goncalves, Ewan Harrison, David K. Jackson, Ian Johnston, Dominic Kwiatkowski, Cordelia Langford, John Sillitoe on behalf of the Wellcome Sanger Institute COVID-19 Surveillance Team |
| EPI_ISL_756404, EPI_ISL_756405, EPI_ISL_756406, EPI_ISL_756407, EPI_ISL_756408 | Lighthouse Lab in Alderley Park | Wellcome Sanger Institute for the COVID-19 Genomics UK (COG-UK) Consortium | Jacquelyn Wynn, Mairead Hyland, The Lighthouse Lab in Alderley Park and Alex Alderton, Roberto Amato, Sonia Goncalves, Ewan Harrison, David K. Jackson, Ian Johnston, Dominic Kwiatkowski, Cordelia Langford, John Sillitoe on behalf of the Wellcome Sanger Institute COVID-19 Surveillance Team |
| EPI_ISL_756409 | Lighthouse Lab in Cambridge | Wellcome Sanger Institute for the COVID-19 Genomics UK (COG-UK) Consortium | Rob Howes, The Lighthouse Lab in Cambridge and Alex Alderton, Roberto Amato, Sonia Goncalves, Ewan Harrison, David K. Jackson, Ian Johnston, Dominic Kwiatkowski, Cordelia Langford, John Sillitoe on behalf of the Wellcome Sanger Institute COVID-19 Surveillance Team |
| EPI_ISL_756410, EPI_ISL_756411, EPI_ISL_756412 | Lighthouse Lab in Alderley Park | Wellcome Sanger Institute for the COVID-19 Genomics UK (COG-UK) Consortium | Jacquelyn Wynn, Mairead Hyland, The Lighthouse Lab in Alderley Park and Alex Alderton, Roberto Amato, Sonia Goncalves, Ewan Harrison, David K. Jackson, Ian Johnston, Dominic Kwiatkowski, Cordelia Langford, John Sillitoe on behalf of the Wellcome Sanger Institute COVID-19 Surveillance Team |
| EPI_ISL_756413 | Lighthouse Lab in Cambridge | Wellcome Sanger Institute for the COVID-19 Genomics UK (COG-UK) Consortium | Rob Howes, The Lighthouse Lab in Cambridge and Alex Alderton, Roberto Amato, Sonia Goncalves, Ewan Harrison, David K. Jackson, Ian Johnston, Dominic Kwiatkowski, Cordelia Langford, John Sillitoe on behalf of the Wellcome Sanger Institute COVID-19 Surveillance Team |
| EPI_ISL_756414 | Lighthouse Lab in Alderley Park | Wellcome Sanger Institute for the COVID-19 Genomics UK (COG-UK) Consortium | Jacquelyn Wynn, Mairead Hyland, The Lighthouse Lab in Alderley Park and Alex Alderton, Roberto Amato, Sonia Goncalves, Ewan Harrison, David K. Jackson, Ian Johnston, Dominic Kwiatkowski, Cordelia Langford, John Sillitoe on behalf of the Wellcome Sanger Institute COVID-19 Surveillance Team |
| EPI_ISL_756415, EPI_ISL_756416, EPI_ISL_756417, EPI_ISL_756418, EPI_ISL_756419, EPI_ISL_756420 | Lighthouse Lab in Cambridge | Wellcome Sanger Institute for the COVID-19 Genomics UK (COG-UK) Consortium | Rob Howes, The Lighthouse Lab in Cambridge and Alex Alderton, Roberto Amato, Sonia Goncalves, Ewan Harrison, David K. Jackson, Ian Johnston, Dominic Kwiatkowski, Cordelia Langford, John Sillitoe on behalf of the Wellcome Sanger Institute COVID-19 Surveillance Team |
| EPI_ISL_756421 | Lighthouse Lab in Alderley Park | Wellcome Sanger Institute for the COVID-19 Genomics UK (COG-UK) Consortium | Jacquelyn Wynn, Mairead Hyland, The Lighthouse Lab in Alderley Park and Alex Alderton, Roberto Amato, Sonia Goncalves, Ewan Harrison, David K. Jackson, Ian Johnston, Dominic Kwiatkowski, Cordelia Langford, John Sillitoe on behalf of the Wellcome Sanger Institute COVID-19 Surveillance Team |
| EPI_ISL_756422 | Lighthouse Lab in Cambridge | Wellcome Sanger Institute for the COVID-19 Genomics UK (COG-UK) Consortium | Rob Howes, The Lighthouse Lab in Cambridge and Alex Alderton, Roberto Amato, Sonia Goncalves, Ewan Harrison, David K. Jackson, Ian Johnston, Dominic Kwiatkowski, Cordelia Langford, John Sillitoe on behalf of the Wellcome Sanger Institute COVID-19 Surveillance Team |
| EPI_ISL_756423 | Lighthouse Lab in Alderley Park | Wellcome Sanger Institute for the COVID-19 Genomics UK (COG-UK) Consortium | Jacquelyn Wynn, Mairead Hyland, The Lighthouse Lab in Alderley Park and Alex Alderton, Roberto Amato, Sonia Goncalves, Ewan Harrison, David K. Jackson, Ian Johnston, Dominic Kwiatkowski, Cordelia Langford, John Sillitoe on behalf of the Wellcome Sanger Institute COVID-19 Surveillance Team |
| EPI_ISL_756424 | Lighthouse Lab in Cambridge | Wellcome Sanger Institute for the COVID-19 Genomics UK (COG-UK) Consortium | Rob Howes, The Lighthouse Lab in Cambridge and Alex Alderton, Roberto Amato, Sonia Goncalves, Ewan Harrison, David K. Jackson, Ian Johnston, Dominic Kwiatkowski, Cordelia Langford, John Sillitoe on behalf of the Wellcome Sanger Institute COVID-19 Surveillance Team |
| EPI_ISL_756425 | Lighthouse Lab in Alderley Park | Wellcome Sanger Institute for the COVID-19 Genomics UK (COG-UK) Consortium | Jacquelyn Wynn, Mairead Hyland, The Lighthouse Lab in Alderley Park and Alex Alderton, Roberto Amato, Sonia Goncalves, Ewan Harrison, David K. Jackson, Ian Johnston, Dominic Kwiatkowski, Cordelia Langford, John Sillitoe on behalf of the Wellcome Sanger Institute COVID-19 Surveillance Team |
| EPI_ISL_756426 | Lighthouse Lab in Cambridge | Wellcome Sanger Institute for the COVID-19 Genomics UK (COG-UK) Consortium | Rob Howes, The Lighthouse Lab in Cambridge and Alex Alderton, Roberto Amato, Sonia Goncalves, Ewan Harrison, David K. Jackson, Ian Johnston, Dominic Kwiatkowski, Cordelia Langford, John Sillitoe on behalf of the Wellcome Sanger Institute COVID-19 Surveillance Team |
| EPI_ISL_756427, EPI_ISL_756428 | Lighthouse Lab in Alderley Park | Wellcome Sanger Institute for the COVID-19 Genomics UK (COG-UK) Consortium | Jacquelyn Wynn, Mairead Hyland, The Lighthouse Lab in Alderley Park and Alex Alderton, Roberto Amato, Sonia Goncalves, Ewan Harrison, David K. Jackson, Ian Johnston, Dominic Kwiatkowski, Cordelia Langford, John Sillitoe on behalf of the Wellcome Sanger Institute COVID-19 Surveillance Team |
| EPI_ISL_756429 | Lighthouse Lab in Cambridge | Wellcome Sanger Institute for the COVID-19 Genomics UK (COG-UK) Consortium | Rob Howes, The Lighthouse Lab in Cambridge and Alex Alderton, Roberto Amato, Sonia Goncalves, Ewan Harrison, David K. Jackson, Ian Johnston, Dominic Kwiatkowski, Cordelia Langford, John Sillitoe on behalf of the Wellcome Sanger Institute COVID-19 Surveillance Team |
| EPI_ISL_756430 | Lighthouse Lab in Alderley Park | Wellcome Sanger Institute for the COVID-19 Genomics UK (COG-UK) Consortium | Jacquelyn Wynn, Mairead Hyland, The Lighthouse Lab in Alderley Park and Alex Alderton, Roberto Amato, Sonia Goncalves, Ewan Harrison, David K. Jackson, Ian Johnston, Dominic Kwiatkowski, Cordelia Langford, John Sillitoe on behalf of the Wellcome Sanger Institute COVID-19 Surveillance Team |
| EPI_ISL_756431, EPI_ISL_756432 | Lighthouse Lab in Cambridge | Wellcome Sanger Institute for the COVID-19 Genomics UK (COG-UK) Consortium | Rob Howes, The Lighthouse Lab in Cambridge and Alex Alderton, Roberto Amato, Sonia Goncalves, Ewan Harrison, David K. Jackson, Ian Johnston, Dominic Kwiatkowski, Cordelia Langford, John Sillitoe on behalf of the Wellcome Sanger Institute COVID-19 Surveillance Team |
| EPI_ISL_756433 | Lighthouse Lab in Alderley Park | Wellcome Sanger Institute for the COVID-19 Genomics UK (COG-UK) Consortium | Jacquelyn Wynn, Mairead Hyland, The Lighthouse Lab in Alderley Park and Alex Alderton, Roberto Amato, Sonia Goncalves, Ewan Harrison, David K. Jackson, Ian Johnston, Dominic Kwiatkowski, Cordelia Langford, John Sillitoe on behalf of the Wellcome Sanger Institute COVID-19 Surveillance Team |
| EPI_ISL_756434 | Lighthouse Lab in Cambridge | Wellcome Sanger Institute for the COVID-19 Genomics UK (COG-UK) Consortium | Rob Howes, The Lighthouse Lab in Cambridge and Alex Alderton, Roberto Amato, Sonia Goncalves, Ewan Harrison, David K. Jackson, Ian Johnston, Dominic Kwiatkowski, Cordelia Langford, John Sillitoe on behalf of the Wellcome Sanger Institute COVID-19 Surveillance Team |
| EPI_ISL_756435, EPI_ISL_756436, EPI_ISL_756437 | Lighthouse Lab in Alderley Park | Wellcome Sanger Institute for the COVID-19 Genomics UK (COG-UK) Consortium | Jacquelyn Wynn, Mairead Hyland, The Lighthouse Lab in Alderley Park and Alex Alderton, Roberto Amato, Sonia Goncalves, Ewan Harrison, David K. Jackson, Ian Johnston, Dominic Kwiatkowski, Cordelia Langford, John Sillitoe on behalf of the Wellcome Sanger Institute COVID-19 Surveillance Team |
| EPI_ISL_756438 | Lighthouse Lab in Cambridge | Wellcome Sanger Institute for the COVID-19 Genomics UK (COG-UK) Consortium | Rob Howes, The Lighthouse Lab in Cambridge and Alex Alderton, Roberto Amato, Sonia Goncalves, Ewan Harrison, David K. Jackson, Ian Johnston, Dominic Kwiatkowski, Cordelia Langford, John Sillitoe on behalf of the Wellcome Sanger Institute COVID-19 Surveillance Team |
| EPI_ISL_756439, EPI_ISL_756440, EPI_ISL_756441 | Lighthouse Lab in Alderley Park | Wellcome Sanger Institute for the COVID-19 Genomics UK (COG-UK) Consortium | Jacquelyn Wynn, Mairead Hyland, The Lighthouse Lab in Alderley Park and Alex Alderton, Roberto Amato, Sonia Goncalves, Ewan Harrison, David K. Jackson, Ian Johnston, Dominic Kwiatkowski, Cordelia Langford, John Sillitoe on behalf of the Wellcome Sanger Institute COVID-19 Surveillance Team |
| EPI_ISL_756442 | Lighthouse Lab in Cambridge | Wellcome Sanger Institute for the COVID-19 Genomics UK (COG-UK) Consortium | Rob Howes, The Lighthouse Lab in Cambridge and Alex Alderton, Roberto Amato, Sonia Goncalves, Ewan Harrison, David K. Jackson, Ian Johnston, Dominic Kwiatkowski, Cordelia Langford, John Sillitoe on behalf of the Wellcome Sanger Institute COVID-19 Surveillance Team |
| EPI_ISL_756443 | Lighthouse Lab in Alderley Park | Wellcome Sanger Institute for the COVID-19 Genomics UK (COG-UK) Consortium | Jacquelyn Wynn, Mairead Hyland, The Lighthouse Lab in Alderley Park and Alex Alderton, Roberto Amato, Sonia Goncalves, Ewan Harrison, David K. Jackson, Ian Johnston, Dominic Kwiatkowski, Cordelia Langford, John Sillitoe on behalf of the Wellcome Sanger Institute COVID-19 Surveillance Team |
| EPI_ISL_756444 | Lighthouse Lab in Cambridge | Wellcome Sanger Institute for the COVID-19 Genomics UK (COG-UK) Consortium | Rob Howes, The Lighthouse Lab in Cambridge and Alex Alderton, Roberto Amato, Sonia Goncalves, Ewan Harrison, David K. Jackson, Ian Johnston, Dominic Kwiatkowski, Cordelia Langford, John Sillitoe on behalf of the Wellcome Sanger Institute COVID-19 Surveillance Team |
| EPI_ISL_756445 | Lighthouse Lab in Alderley Park | Wellcome Sanger Institute for the COVID-19 Genomics UK (COG-UK) Consortium | Jacquelyn Wynn, Mairead Hyland, The Lighthouse Lab in Alderley Park and Alex Alderton, Roberto Amato, Sonia Goncalves, Ewan Harrison, David K. Jackson, Ian Johnston, Dominic Kwiatkowski, Cordelia Langford, John Sillitoe on behalf of the Wellcome Sanger Institute COVID-19 Surveillance Team |
| EPI_ISL_756446 | Lighthouse Lab in Cambridge | Wellcome Sanger Institute for the COVID-19 Genomics UK | Rob Howes, The Lighthouse Lab in Cambridge and Alex Alderton, Roberto Amato, Sonia Goncalves, Ewan Harrison, David K. Jackson, Ian Johnston, |

| | | | |
|---|---|---|---|
| | | (COG-UK) Consortium | Dominic Kwiatkowski, Cordelia Langford, John Sillitoe on behalf of the Wellcome Sanger Institute COVID-19 Surveillance Team |
| EPI_ISL_756447 | Lighthouse Lab in Alderley Park | Wellcome Sanger Institute for the COVID-19 Genomics UK (COG-UK) Consortium | Jacquelyn Wynn, Mairead Hyland, The Lighthouse Lab in Alderley Park and Alex Alderton, Roberto Amato, Sonia Goncalves, Ewan Harrison, David K. Jackson, Ian Johnston, Dominic Kwiatkowski, Cordelia Langford, John Sillitoe on behalf of the Wellcome Sanger Institute COVID-19 Surveillance Team |
| EPI_ISL_756448 | Lighthouse Lab in Cambridge | Wellcome Sanger Institute for the COVID-19 Genomics UK (COG-UK) Consortium | Rob Howes, The Lighthouse Lab in Cambridge and Alex Alderton, Roberto Amato, Sonia Goncalves, Ewan Harrison, David K. Jackson, Ian Johnston, Dominic Kwiatkowski, Cordelia Langford, John Sillitoe on behalf of the Wellcome Sanger Institute COVID-19 Surveillance Team |
| EPI_ISL_756449, EPI_ISL_756450, EPI_ISL_756451, EPI_ISL_756452 | Lighthouse Lab in Alderley Park | Wellcome Sanger Institute for the COVID-19 Genomics UK (COG-UK) Consortium | Jacquelyn Wynn, Mairead Hyland, The Lighthouse Lab in Alderley Park and Alex Alderton, Roberto Amato, Sonia Goncalves, Ewan Harrison, David K. Jackson, Ian Johnston, Dominic Kwiatkowski, Cordelia Langford, John Sillitoe on behalf of the Wellcome Sanger Institute COVID-19 Surveillance Team |
| EPI_ISL_756453, EPI_ISL_756454 | Lighthouse Lab in Cambridge | Wellcome Sanger Institute for the COVID-19 Genomics UK (COG-UK) Consortium | Rob Howes, The Lighthouse Lab in Cambridge and Alex Alderton, Roberto Amato, Sonia Goncalves, Ewan Harrison, David K. Jackson, Ian Johnston, Dominic Kwiatkowski, Cordelia Langford, John Sillitoe on behalf of the Wellcome Sanger Institute COVID-19 Surveillance Team |
| EPI_ISL_756455 | Lighthouse Lab in Alderley Park | Wellcome Sanger Institute for the COVID-19 Genomics UK (COG-UK) Consortium | Jacquelyn Wynn, Mairead Hyland, The Lighthouse Lab in Alderley Park and Alex Alderton, Roberto Amato, Sonia Goncalves, Ewan Harrison, David K. Jackson, Ian Johnston, Dominic Kwiatkowski, Cordelia Langford, John Sillitoe on behalf of the Wellcome Sanger Institute COVID-19 Surveillance Team |
| EPI_ISL_756456 | Lighthouse Lab in Cambridge | Wellcome Sanger Institute for the COVID-19 Genomics UK (COG-UK) Consortium | Rob Howes, The Lighthouse Lab in Cambridge and Alex Alderton, Roberto Amato, Sonia Goncalves, Ewan Harrison, David K. Jackson, Ian Johnston, Dominic Kwiatkowski, Cordelia Langford, John Sillitoe on behalf of the Wellcome Sanger Institute COVID-19 Surveillance Team |
| EPI_ISL_756457, EPI_ISL_756458, EPI_ISL_756459 | Lighthouse Lab in Alderley Park | Wellcome Sanger Institute for the COVID-19 Genomics UK (COG-UK) Consortium | Jacquelyn Wynn, Mairead Hyland, The Lighthouse Lab in Alderley Park and Alex Alderton, Roberto Amato, Sonia Goncalves, Ewan Harrison, David K. Jackson, Ian Johnston, Dominic Kwiatkowski, Cordelia Langford, John Sillitoe on behalf of the Wellcome Sanger Institute COVID-19 Surveillance Team |
| EPI_ISL_756460 | Lighthouse Lab in Cambridge | Wellcome Sanger Institute for the COVID-19 Genomics UK (COG-UK) Consortium | Rob Howes, The Lighthouse Lab in Cambridge and Alex Alderton, Roberto Amato, Sonia Goncalves, Ewan Harrison, David K. Jackson, Ian Johnston, Dominic Kwiatkowski, Cordelia Langford, John Sillitoe on behalf of the Wellcome Sanger Institute COVID-19 Surveillance Team |
| EPI_ISL_756461, EPI_ISL_756462, EPI_ISL_756463, EPI_ISL_756464, EPI_ISL_756465, EPI_ISL_756466 | Lighthouse Lab in Alderley Park | Wellcome Sanger Institute for the COVID-19 Genomics UK (COG-UK) Consortium | Jacquelyn Wynn, Mairead Hyland, The Lighthouse Lab in Alderley Park and Alex Alderton, Roberto Amato, Sonia Goncalves, Ewan Harrison, David K. Jackson, Ian Johnston, Dominic Kwiatkowski, Cordelia Langford, John Sillitoe on behalf of the Wellcome Sanger Institute COVID-19 Surveillance Team |
| EPI_ISL_756467 | Lighthouse Lab in Cambridge | Wellcome Sanger Institute for the COVID-19 Genomics UK (COG-UK) Consortium | Rob Howes, The Lighthouse Lab in Cambridge and Alex Alderton, Roberto Amato, Sonia Goncalves, Ewan Harrison, David K. Jackson, Ian Johnston, Dominic Kwiatkowski, Cordelia Langford, John Sillitoe on behalf of the Wellcome Sanger Institute COVID-19 Surveillance Team |
| EPI_ISL_756468 | Lighthouse Lab in Alderley Park | Wellcome Sanger Institute for the COVID-19 Genomics UK (COG-UK) Consortium | Jacquelyn Wynn, Mairead Hyland, The Lighthouse Lab in Alderley Park and Alex Alderton, Roberto Amato, Sonia Goncalves, Ewan Harrison, David K. Jackson, Ian Johnston, Dominic Kwiatkowski, Cordelia Langford, John Sillitoe on behalf of the Wellcome Sanger Institute COVID-19 Surveillance Team |
| EPI_ISL_756469 | Lighthouse Lab in Cambridge | Wellcome Sanger Institute for the COVID-19 Genomics UK (COG-UK) Consortium | Rob Howes, The Lighthouse Lab in Cambridge and Alex Alderton, Roberto Amato, Sonia Goncalves, Ewan Harrison, David K. Jackson, Ian Johnston, Dominic Kwiatkowski, Cordelia Langford, John Sillitoe on behalf of the Wellcome Sanger Institute COVID-19 Surveillance Team |
| EPI_ISL_756471, EPI_ISL_756472, EPI_ISL_756473 | Lighthouse Lab in Alderley Park | Wellcome Sanger Institute for the COVID-19 Genomics UK (COG-UK) Consortium | Jacquelyn Wynn, Mairead Hyland, The Lighthouse Lab in Alderley Park and Alex Alderton, Roberto Amato, Sonia Goncalves, Ewan Harrison, David K. Jackson, Ian Johnston, Dominic Kwiatkowski, Cordelia Langford, John Sillitoe on behalf of the Wellcome Sanger Institute COVID-19 Surveillance Team |
| EPI_ISL_756474, EPI_ISL_756475 | Lighthouse Lab in Cambridge | Wellcome Sanger Institute for the COVID-19 Genomics UK (COG-UK) Consortium | Rob Howes, The Lighthouse Lab in Cambridge and Alex Alderton, Roberto Amato, Sonia Goncalves, Ewan Harrison, David K. Jackson, Ian Johnston, Dominic Kwiatkowski, Cordelia Langford, John Sillitoe on behalf of the Wellcome Sanger Institute COVID-19 Surveillance Team |
| EPI_ISL_756476, EPI_ISL_756477 | Lighthouse Lab in Alderley Park | Wellcome Sanger Institute for the COVID-19 Genomics UK (COG-UK) Consortium | Jacquelyn Wynn, Mairead Hyland, The Lighthouse Lab in Alderley Park and Alex Alderton, Roberto Amato, Sonia Goncalves, Ewan Harrison, David K. Jackson, Ian Johnston, Dominic Kwiatkowski, Cordelia Langford, John Sillitoe on behalf of the Wellcome Sanger Institute COVID-19 Surveillance Team |
| EPI_ISL_756478 | Lighthouse Lab in Cambridge | Wellcome Sanger Institute for the COVID-19 Genomics UK (COG-UK) Consortium | Rob Howes, The Lighthouse Lab in Cambridge and Alex Alderton, Roberto Amato, Sonia Goncalves, Ewan Harrison, David K. Jackson, Ian Johnston, Dominic Kwiatkowski, Cordelia Langford, John Sillitoe on behalf of the Wellcome Sanger Institute COVID-19 Surveillance Team |
| EPI_ISL_756479 | Lighthouse Lab in Alderley Park | Wellcome Sanger Institute for the COVID-19 Genomics UK (COG-UK) Consortium | Jacquelyn Wynn, Mairead Hyland, The Lighthouse Lab in Alderley Park and Alex Alderton, Roberto Amato, Sonia Goncalves, Ewan Harrison, David K. Jackson, Ian Johnston, Dominic Kwiatkowski, Cordelia Langford, John Sillitoe on behalf of the Wellcome Sanger Institute COVID-19 Surveillance Team |
| EPI_ISL_756480, EPI_ISL_756481 | Lighthouse Lab in Cambridge | Wellcome Sanger Institute for the COVID-19 Genomics UK (COG-UK) Consortium | Rob Howes, The Lighthouse Lab in Cambridge and Alex Alderton, Roberto Amato, Sonia Goncalves, Ewan Harrison, David K. Jackson, Ian Johnston, Dominic Kwiatkowski, Cordelia Langford, John Sillitoe on behalf of the Wellcome Sanger Institute COVID-19 Surveillance Team |
| EPI_ISL_756482, EPI_ISL_756483 | Lighthouse Lab in Alderley Park | Wellcome Sanger Institute for the COVID-19 Genomics UK (COG-UK) Consortium | Jacquelyn Wynn, Mairead Hyland, The Lighthouse Lab in Alderley Park and Alex Alderton, Roberto Amato, Sonia Goncalves, Ewan Harrison, David K. Jackson, Ian Johnston, Dominic Kwiatkowski, Cordelia Langford, John Sillitoe on behalf of the Wellcome Sanger Institute COVID-19 Surveillance Team |
| EPI_ISL_756484, EPI_ISL_756485, EPI_ISL_756486, EPI_ISL_756487 | Lighthouse Lab in Cambridge | Wellcome Sanger Institute for the COVID-19 Genomics UK (COG-UK) Consortium | Rob Howes, The Lighthouse Lab in Cambridge and Alex Alderton, Roberto Amato, Sonia Goncalves, Ewan Harrison, David K. Jackson, Ian Johnston, Dominic Kwiatkowski, Cordelia Langford, John Sillitoe on behalf of the Wellcome Sanger Institute COVID-19 Surveillance Team |
| EPI_ISL_756488, EPI_ISL_756489 | Lighthouse Lab in Alderley Park | Wellcome Sanger Institute for the COVID-19 Genomics UK (COG-UK) Consortium | Jacquelyn Wynn, Mairead Hyland, The Lighthouse Lab in Alderley Park and Alex Alderton, Roberto Amato, Sonia Goncalves, Ewan Harrison, David K. Jackson, Ian Johnston, Dominic Kwiatkowski, Cordelia Langford, John Sillitoe on behalf of the Wellcome Sanger Institute COVID-19 Surveillance Team |
| EPI_ISL_756490, EPI_ISL_756491, EPI_ISL_756492 | Lighthouse Lab in Cambridge | Wellcome Sanger Institute for the COVID-19 Genomics UK (COG-UK) Consortium | Rob Howes, The Lighthouse Lab in Cambridge and Alex Alderton, Roberto Amato, Sonia Goncalves, Ewan Harrison, David K. Jackson, Ian Johnston, Dominic Kwiatkowski, Cordelia Langford, John Sillitoe on behalf of the Wellcome Sanger Institute COVID-19 Surveillance Team |
| EPI_ISL_756493, EPI_ISL_756494, EPI_ISL_756495 | Lighthouse Lab in Alderley Park | Wellcome Sanger Institute for the COVID-19 Genomics UK (COG-UK) Consortium | Jacquelyn Wynn, Mairead Hyland, The Lighthouse Lab in Alderley Park and Alex Alderton, Roberto Amato, Sonia Goncalves, Ewan Harrison, David K. Jackson, Ian Johnston, Dominic Kwiatkowski, Cordelia Langford, John Sillitoe on behalf of the Wellcome Sanger Institute COVID-19 Surveillance Team |
| EPI_ISL_756496, EPI_ISL_756497, EPI_ISL_756498, EPI_ISL_756499, EPI_ISL_756500, EPI_ISL_756501, EPI_ISL_756502 | Lighthouse Lab in Cambridge | Wellcome Sanger Institute for the COVID-19 Genomics UK (COG-UK) Consortium | Rob Howes, The Lighthouse Lab in Cambridge and Alex Alderton, Roberto Amato, Sonia Goncalves, Ewan Harrison, David K. Jackson, Ian Johnston, Dominic Kwiatkowski, Cordelia Langford, John Sillitoe on behalf of the Wellcome Sanger Institute COVID-19 Surveillance Team |
| EPI_ISL_756503, EPI_ISL_756504 | Lighthouse Lab in Alderley Park | Wellcome Sanger Institute for the COVID-19 Genomics UK (COG-UK) Consortium | Jacquelyn Wynn, Mairead Hyland, The Lighthouse Lab in Alderley Park and Alex Alderton, Roberto Amato, Sonia Goncalves, Ewan Harrison, David K. Jackson, Ian Johnston, Dominic Kwiatkowski, Cordelia Langford, John Sillitoe on behalf of the Wellcome Sanger Institute COVID-19 Surveillance Team |
| EPI_ISL_756505, EPI_ISL_756506, EPI_ISL_756507, EPI_ISL_756508 | Lighthouse Lab in Cambridge | Wellcome Sanger Institute for the COVID-19 Genomics UK (COG-UK) Consortium | Rob Howes, The Lighthouse Lab in Cambridge and Alex Alderton, Roberto Amato, Sonia Goncalves, Ewan Harrison, David K. Jackson, Ian Johnston, Dominic Kwiatkowski, Cordelia Langford, John Sillitoe on behalf of the Wellcome Sanger Institute COVID-19 Surveillance Team |
| EPI_ISL_756509, EPI_ISL_756510 | Lighthouse Lab in Alderley Park | Wellcome Sanger Institute for the COVID-19 Genomics UK (COG-UK) Consortium | Jacquelyn Wynn, Mairead Hyland, The Lighthouse Lab in Alderley Park and Alex Alderton, Roberto Amato, Sonia Goncalves, Ewan Harrison, David K. Jackson, Ian Johnston, Dominic Kwiatkowski, Cordelia Langford, John Sillitoe on behalf of the Wellcome Sanger Institute COVID-19 Surveillance Team |
| EPI_ISL_756511, EPI_ISL_756512, EPI_ISL_756513 | Lighthouse Lab in Cambridge | Wellcome Sanger Institute for the COVID-19 Genomics UK (COG-UK) Consortium | Rob Howes, The Lighthouse Lab in Cambridge and Alex Alderton, Roberto Amato, Sonia Goncalves, Ewan Harrison, David K. Jackson, Ian Johnston, Dominic Kwiatkowski, Cordelia Langford, John Sillitoe on behalf of the Wellcome Sanger Institute COVID-19 Surveillance Team |
| EPI_ISL_756514, EPI_ISL_756515, EPI_ISL_756516 | Lighthouse Lab in Alderley Park | Wellcome Sanger Institute for the COVID-19 Genomics UK (COG-UK) Consortium | Jacquelyn Wynn, Mairead Hyland, The Lighthouse Lab in Alderley Park and Alex Alderton, Roberto Amato, Sonia Goncalves, Ewan Harrison, David K. Jackson, Ian Johnston, Dominic Kwiatkowski, Cordelia Langford, John Sillitoe on behalf of the Wellcome Sanger Institute COVID-19 Surveillance Team |
| EPI_ISL_756517, EPI_ISL_756518, EPI_ISL_756519 | Lighthouse Lab in Cambridge | Wellcome Sanger Institute for the COVID-19 Genomics UK (COG-UK) Consortium | Rob Howes, The Lighthouse Lab in Cambridge and Alex Alderton, Roberto Amato, Sonia Goncalves, Ewan Harrison, David K. Jackson, Ian Johnston, Dominic Kwiatkowski, Cordelia Langford, John Sillitoe on behalf of the Wellcome Sanger Institute COVID-19 Surveillance Team |
| EPI_ISL_756520 | Lighthouse Lab in Alderley Park | Wellcome Sanger Institute for the COVID-19 Genomics UK (COG-UK) Consortium | Jacquelyn Wynn, Mairead Hyland, The Lighthouse Lab in Alderley Park and Alex Alderton, Roberto Amato, Sonia Goncalves, Ewan Harrison, David K. Jackson, Ian Johnston, Dominic Kwiatkowski, Cordelia Langford, John Sillitoe on behalf of the Wellcome Sanger Institute COVID-19 Surveillance Team |
| EPI_ISL_756521, EPI_ISL_756522 | Lighthouse Lab in Cambridge | Wellcome Sanger Institute for the COVID-19 Genomics UK (COG-UK) Consortium | Rob Howes, The Lighthouse Lab in Cambridge and Alex Alderton, Roberto Amato, Sonia Goncalves, Ewan Harrison, David K. Jackson, Ian Johnston, Dominic Kwiatkowski, Cordelia Langford, John Sillitoe on behalf of the Wellcome Sanger Institute COVID-19 Surveillance Team |
| EPI_ISL_756523, EPI_ISL_756524 | Lighthouse Lab in Alderley Park | Wellcome Sanger Institute for the COVID-19 Genomics UK (COG-UK) Consortium | Jacquelyn Wynn, Mairead Hyland, The Lighthouse Lab in Alderley Park and Alex Alderton, Roberto Amato, Sonia Goncalves, Ewan Harrison, David K. Jackson, Ian Johnston, Dominic Kwiatkowski, Cordelia Langford, John Sillitoe on behalf of the Wellcome Sanger Institute COVID-19 Surveillance Team |
| EPI_ISL_756525 | Lighthouse Lab in Cambridge | Wellcome Sanger Institute for the COVID-19 Genomics UK | Rob Howes, The Lighthouse Lab in Cambridge and Alex Alderton, Roberto Amato, Sonia Goncalves, Ewan Harrison, David K. Jackson, Ian Johnston, |

| | | | |
|---|---|---|---|
| | | (COG-UK) Consortium | Dominic Kwiatkowski, Cordelia Langford, John Sillitoe on behalf of the Wellcome Sanger Institute COVID-19 Surveillance Team |
| EPI_ISL_756526, EPI_ISL_756527 | Lighthouse Lab in Alderley Park | Wellcome Sanger Institute for the COVID-19 Genomics UK (COG-UK) Consortium | Jacquelyn Wynn, Mairead Hyland, The Lighthouse Lab in Alderley Park and Alex Alderton, Roberto Amato, Sonia Goncalves, Ewan Harrison, David K. Jackson, Ian Johnston, Dominic Kwiatkowski, Cordelia Langford, John Sillitoe on behalf of the Wellcome Sanger Institute COVID-19 Surveillance Team |
| EPI_ISL_756528, EPI_ISL_756529 | Lighthouse Lab in Cambridge | Wellcome Sanger Institute for the COVID-19 Genomics UK (COG-UK) Consortium | Rob Howes, The Lighthouse Lab in Cambridge and Alex Alderton, Roberto Amato, Sonia Goncalves, Ewan Harrison, David K. Jackson, Ian Johnston, Dominic Kwiatkowski, Cordelia Langford, John Sillitoe on behalf of the Wellcome Sanger Institute COVID-19 Surveillance Team |
| EPI_ISL_756530, EPI_ISL_756531, EPI_ISL_756532, EPI_ISL_756533 | Lighthouse Lab in Alderley Park | Wellcome Sanger Institute for the COVID-19 Genomics UK (COG-UK) Consortium | Jacquelyn Wynn, Mairead Hyland, The Lighthouse Lab in Alderley Park and Alex Alderton, Roberto Amato, Sonia Goncalves, Ewan Harrison, David K. Jackson, Ian Johnston, Dominic Kwiatkowski, Cordelia Langford, John Sillitoe on behalf of the Wellcome Sanger Institute COVID-19 Surveillance Team |
| EPI_ISL_756534, EPI_ISL_756535 | Lighthouse Lab in Cambridge | Wellcome Sanger Institute for the COVID-19 Genomics UK (COG-UK) Consortium | Rob Howes, The Lighthouse Lab in Cambridge and Alex Alderton, Roberto Amato, Sonia Goncalves, Ewan Harrison, David K. Jackson, Ian Johnston, Dominic Kwiatkowski, Cordelia Langford, John Sillitoe on behalf of the Wellcome Sanger Institute COVID-19 Surveillance Team |
| EPI_ISL_756536, EPI_ISL_756537, EPI_ISL_756538, EPI_ISL_756539 | Lighthouse Lab in Alderley Park | Wellcome Sanger Institute for the COVID-19 Genomics UK (COG-UK) Consortium | Jacquelyn Wynn, Mairead Hyland, The Lighthouse Lab in Alderley Park and Alex Alderton, Roberto Amato, Sonia Goncalves, Ewan Harrison, David K. Jackson, Ian Johnston, Dominic Kwiatkowski, Cordelia Langford, John Sillitoe on behalf of the Wellcome Sanger Institute COVID-19 Surveillance Team |
| EPI_ISL_756540 | Lighthouse Lab in Cambridge | Wellcome Sanger Institute for the COVID-19 Genomics UK (COG-UK) Consortium | Rob Howes, The Lighthouse Lab in Cambridge and Alex Alderton, Roberto Amato, Sonia Goncalves, Ewan Harrison, David K. Jackson, Ian Johnston, Dominic Kwiatkowski, Cordelia Langford, John Sillitoe on behalf of the Wellcome Sanger Institute COVID-19 Surveillance Team |
| EPI_ISL_756541 | Lighthouse Lab in Alderley Park | Wellcome Sanger Institute for the COVID-19 Genomics UK (COG-UK) Consortium | Jacquelyn Wynn, Mairead Hyland, The Lighthouse Lab in Alderley Park and Alex Alderton, Roberto Amato, Sonia Goncalves, Ewan Harrison, David K. Jackson, Ian Johnston, Dominic Kwiatkowski, Cordelia Langford, John Sillitoe on behalf of the Wellcome Sanger Institute COVID-19 Surveillance Team |
| EPI_ISL_756542, EPI_ISL_756543 | Lighthouse Lab in Cambridge | Wellcome Sanger Institute for the COVID-19 Genomics UK (COG-UK) Consortium | Rob Howes, The Lighthouse Lab in Cambridge and Alex Alderton, Roberto Amato, Sonia Goncalves, Ewan Harrison, David K. Jackson, Ian Johnston, Dominic Kwiatkowski, Cordelia Langford, John Sillitoe on behalf of the Wellcome Sanger Institute COVID-19 Surveillance Team |
| EPI_ISL_756544 | Lighthouse Lab in Alderley Park | Wellcome Sanger Institute for the COVID-19 Genomics UK (COG-UK) Consortium | Jacquelyn Wynn, Mairead Hyland, The Lighthouse Lab in Alderley Park and Alex Alderton, Roberto Amato, Sonia Goncalves, Ewan Harrison, David K. Jackson, Ian Johnston, Dominic Kwiatkowski, Cordelia Langford, John Sillitoe on behalf of the Wellcome Sanger Institute COVID-19 Surveillance Team |
| EPI_ISL_756545 | Lighthouse Lab in Cambridge | Wellcome Sanger Institute for the COVID-19 Genomics UK (COG-UK) Consortium | Rob Howes, The Lighthouse Lab in Cambridge and Alex Alderton, Roberto Amato, Sonia Goncalves, Ewan Harrison, David K. Jackson, Ian Johnston, Dominic Kwiatkowski, Cordelia Langford, John Sillitoe on behalf of the Wellcome Sanger Institute COVID-19 Surveillance Team |
| EPI_ISL_756546, EPI_ISL_756547, EPI_ISL_756548 | Lighthouse Lab in Alderley Park | Wellcome Sanger Institute for the COVID-19 Genomics UK (COG-UK) Consortium | Jacquelyn Wynn, Mairead Hyland, The Lighthouse Lab in Alderley Park and Alex Alderton, Roberto Amato, Sonia Goncalves, Ewan Harrison, David K. Jackson, Ian Johnston, Dominic Kwiatkowski, Cordelia Langford, John Sillitoe on behalf of the Wellcome Sanger Institute COVID-19 Surveillance Team |
| EPI_ISL_756549 | Lighthouse Lab in Cambridge | Wellcome Sanger Institute for the COVID-19 Genomics UK (COG-UK) Consortium | Rob Howes, The Lighthouse Lab in Cambridge and Alex Alderton, Roberto Amato, Sonia Goncalves, Ewan Harrison, David K. Jackson, Ian Johnston, Dominic Kwiatkowski, Cordelia Langford, John Sillitoe on behalf of the Wellcome Sanger Institute COVID-19 Surveillance Team |
| EPI_ISL_756550, EPI_ISL_756551, EPI_ISL_756552 | Lighthouse Lab in Alderley Park | Wellcome Sanger Institute for the COVID-19 Genomics UK (COG-UK) Consortium | Jacquelyn Wynn, Mairead Hyland, The Lighthouse Lab in Alderley Park and Alex Alderton, Roberto Amato, Sonia Goncalves, Ewan Harrison, David K. Jackson, Ian Johnston, Dominic Kwiatkowski, Cordelia Langford, John Sillitoe on behalf of the Wellcome Sanger Institute COVID-19 Surveillance Team |
| EPI_ISL_756553, EPI_ISL_756554 | Lighthouse Lab in Cambridge | Wellcome Sanger Institute for the COVID-19 Genomics UK (COG-UK) Consortium | Rob Howes, The Lighthouse Lab in Cambridge and Alex Alderton, Roberto Amato, Sonia Goncalves, Ewan Harrison, David K. Jackson, Ian Johnston, Dominic Kwiatkowski, Cordelia Langford, John Sillitoe on behalf of the Wellcome Sanger Institute COVID-19 Surveillance Team |
| EPI_ISL_756555, EPI_ISL_756556, EPI_ISL_756557 | Lighthouse Lab in Alderley Park | Wellcome Sanger Institute for the COVID-19 Genomics UK (COG-UK) Consortium | Jacquelyn Wynn, Mairead Hyland, The Lighthouse Lab in Alderley Park and Alex Alderton, Roberto Amato, Sonia Goncalves, Ewan Harrison, David K. Jackson, Ian Johnston, Dominic Kwiatkowski, Cordelia Langford, John Sillitoe on behalf of the Wellcome Sanger Institute COVID-19 Surveillance Team |
| EPI_ISL_756558, EPI_ISL_756559 | Lighthouse Lab in Cambridge | Wellcome Sanger Institute for the COVID-19 Genomics UK (COG-UK) Consortium | Rob Howes, The Lighthouse Lab in Cambridge and Alex Alderton, Roberto Amato, Sonia Goncalves, Ewan Harrison, David K. Jackson, Ian Johnston, Dominic Kwiatkowski, Cordelia Langford, John Sillitoe on behalf of the Wellcome Sanger Institute COVID-19 Surveillance Team |
| EPI_ISL_756560, EPI_ISL_756561, EPI_ISL_756562, EPI_ISL_756563 | Lighthouse Lab in Alderley Park | Wellcome Sanger Institute for the COVID-19 Genomics UK (COG-UK) Consortium | Jacquelyn Wynn, Mairead Hyland, The Lighthouse Lab in Alderley Park and Alex Alderton, Roberto Amato, Sonia Goncalves, Ewan Harrison, David K. Jackson, Ian Johnston, Dominic Kwiatkowski, Cordelia Langford, John Sillitoe on behalf of the Wellcome Sanger Institute COVID-19 Surveillance Team |
| EPI_ISL_756564 | Lighthouse Lab in Cambridge | Wellcome Sanger Institute for the COVID-19 Genomics UK (COG-UK) Consortium | Rob Howes, The Lighthouse Lab in Cambridge and Alex Alderton, Roberto Amato, Sonia Goncalves, Ewan Harrison, David K. Jackson, Ian Johnston, Dominic Kwiatkowski, Cordelia Langford, John Sillitoe on behalf of the Wellcome Sanger Institute COVID-19 Surveillance Team |
| EPI_ISL_756565, EPI_ISL_756566 | Lighthouse Lab in Alderley Park | Wellcome Sanger Institute for the COVID-19 Genomics UK (COG-UK) Consortium | Jacquelyn Wynn, Mairead Hyland, The Lighthouse Lab in Alderley Park and Alex Alderton, Roberto Amato, Sonia Goncalves, Ewan Harrison, David K. Jackson, Ian Johnston, Dominic Kwiatkowski, Cordelia Langford, John Sillitoe on behalf of the Wellcome Sanger Institute COVID-19 Surveillance Team |
| EPI_ISL_756567, EPI_ISL_756568 | Lighthouse Lab in Cambridge | Wellcome Sanger Institute for the COVID-19 Genomics UK (COG-UK) Consortium | Rob Howes, The Lighthouse Lab in Cambridge and Alex Alderton, Roberto Amato, Sonia Goncalves, Ewan Harrison, David K. Jackson, Ian Johnston, Dominic Kwiatkowski, Cordelia Langford, John Sillitoe on behalf of the Wellcome Sanger Institute COVID-19 Surveillance Team |
| EPI_ISL_756569 | Lighthouse Lab in Alderley Park | Wellcome Sanger Institute for the COVID-19 Genomics UK (COG-UK) Consortium | Jacquelyn Wynn, Mairead Hyland, The Lighthouse Lab in Alderley Park and Alex Alderton, Roberto Amato, Sonia Goncalves, Ewan Harrison, David K. Jackson, Ian Johnston, Dominic Kwiatkowski, Cordelia Langford, John Sillitoe on behalf of the Wellcome Sanger Institute COVID-19 Surveillance Team |
| EPI_ISL_756570 | Lighthouse Lab in Cambridge | Wellcome Sanger Institute for the COVID-19 Genomics UK (COG-UK) Consortium | Rob Howes, The Lighthouse Lab in Cambridge and Alex Alderton, Roberto Amato, Sonia Goncalves, Ewan Harrison, David K. Jackson, Ian Johnston, Dominic Kwiatkowski, Cordelia Langford, John Sillitoe on behalf of the Wellcome Sanger Institute COVID-19 Surveillance Team |
| EPI_ISL_756571 | Lighthouse Lab in Alderley Park | Wellcome Sanger Institute for the COVID-19 Genomics UK (COG-UK) Consortium | Jacquelyn Wynn, Mairead Hyland, The Lighthouse Lab in Alderley Park and Alex Alderton, Roberto Amato, Sonia Goncalves, Ewan Harrison, David K. Jackson, Ian Johnston, Dominic Kwiatkowski, Cordelia Langford, John Sillitoe on behalf of the Wellcome Sanger Institute COVID-19 Surveillance Team |
| EPI_ISL_756572 | Lighthouse Lab in Cambridge | Wellcome Sanger Institute for the COVID-19 Genomics UK (COG-UK) Consortium | Rob Howes, The Lighthouse Lab in Cambridge and Alex Alderton, Roberto Amato, Sonia Goncalves, Ewan Harrison, David K. Jackson, Ian Johnston, Dominic Kwiatkowski, Cordelia Langford, John Sillitoe on behalf of the Wellcome Sanger Institute COVID-19 Surveillance Team |
| EPI_ISL_756573, EPI_ISL_756574, EPI_ISL_756575 | Lighthouse Lab in Alderley Park | Wellcome Sanger Institute for the COVID-19 Genomics UK (COG-UK) Consortium | Jacquelyn Wynn, Mairead Hyland, The Lighthouse Lab in Alderley Park and Alex Alderton, Roberto Amato, Sonia Goncalves, Ewan Harrison, David K. Jackson, Ian Johnston, Dominic Kwiatkowski, Cordelia Langford, John Sillitoe on behalf of the Wellcome Sanger Institute COVID-19 Surveillance Team |
| EPI_ISL_756576, EPI_ISL_756577 | Lighthouse Lab in Cambridge | Wellcome Sanger Institute for the COVID-19 Genomics UK (COG-UK) Consortium | Rob Howes, The Lighthouse Lab in Cambridge and Alex Alderton, Roberto Amato, Sonia Goncalves, Ewan Harrison, David K. Jackson, Ian Johnston, Dominic Kwiatkowski, Cordelia Langford, John Sillitoe on behalf of the Wellcome Sanger Institute COVID-19 Surveillance Team |
| EPI_ISL_756578 | Lighthouse Lab in Alderley Park | Wellcome Sanger Institute for the COVID-19 Genomics UK (COG-UK) Consortium | Jacquelyn Wynn, Mairead Hyland, The Lighthouse Lab in Alderley Park and Alex Alderton, Roberto Amato, Sonia Goncalves, Ewan Harrison, David K. Jackson, Ian Johnston, Dominic Kwiatkowski, Cordelia Langford, John Sillitoe on behalf of the Wellcome Sanger Institute COVID-19 Surveillance Team |
| EPI_ISL_756579 | Lighthouse Lab in Cambridge | Wellcome Sanger Institute for the COVID-19 Genomics UK (COG-UK) Consortium | Rob Howes, The Lighthouse Lab in Cambridge and Alex Alderton, Roberto Amato, Sonia Goncalves, Ewan Harrison, David K. Jackson, Ian Johnston, Dominic Kwiatkowski, Cordelia Langford, John Sillitoe on behalf of the Wellcome Sanger Institute COVID-19 Surveillance Team |
| EPI_ISL_756580, EPI_ISL_756581 | Lighthouse Lab in Alderley Park | Wellcome Sanger Institute for the COVID-19 Genomics UK (COG-UK) Consortium | Jacquelyn Wynn, Mairead Hyland, The Lighthouse Lab in Alderley Park and Alex Alderton, Roberto Amato, Sonia Goncalves, Ewan Harrison, David K. Jackson, Ian Johnston, Dominic Kwiatkowski, Cordelia Langford, John Sillitoe on behalf of the Wellcome Sanger Institute COVID-19 Surveillance Team |
| EPI_ISL_756582 | Lighthouse Lab in Cambridge | Wellcome Sanger Institute for the COVID-19 Genomics UK (COG-UK) Consortium | Rob Howes, The Lighthouse Lab in Cambridge and Alex Alderton, Roberto Amato, Sonia Goncalves, Ewan Harrison, David K. Jackson, Ian Johnston, Dominic Kwiatkowski, Cordelia Langford, John Sillitoe on behalf of the Wellcome Sanger Institute COVID-19 Surveillance Team |
| EPI_ISL_756583 | Lighthouse Lab in Alderley Park | Wellcome Sanger Institute for the COVID-19 Genomics UK (COG-UK) Consortium | Jacquelyn Wynn, Mairead Hyland, The Lighthouse Lab in Alderley Park and Alex Alderton, Roberto Amato, Sonia Goncalves, Ewan Harrison, David K. Jackson, Ian Johnston, Dominic Kwiatkowski, Cordelia Langford, John Sillitoe on behalf of the Wellcome Sanger Institute COVID-19 Surveillance Team |
| EPI_ISL_756584, EPI_ISL_756585, EPI_ISL_756586 | Lighthouse Lab in Cambridge | Wellcome Sanger Institute for the COVID-19 Genomics UK (COG-UK) Consortium | Rob Howes, The Lighthouse Lab in Cambridge and Alex Alderton, Roberto Amato, Sonia Goncalves, Ewan Harrison, David K. Jackson, Ian Johnston, Dominic Kwiatkowski, Cordelia Langford, John Sillitoe on behalf of the Wellcome Sanger Institute COVID-19 Surveillance Team |
| EPI_ISL_756587 | Lighthouse Lab in Alderley Park | Wellcome Sanger Institute for the COVID-19 Genomics UK (COG-UK) Consortium | Jacquelyn Wynn, Mairead Hyland, The Lighthouse Lab in Alderley Park and Alex Alderton, Roberto Amato, Sonia Goncalves, Ewan Harrison, David K. Jackson, Ian Johnston, Dominic Kwiatkowski, Cordelia Langford, John Sillitoe on behalf of the Wellcome Sanger Institute COVID-19 Surveillance Team |
| EPI_ISL_756588, EPI_ISL_756589, EPI_ISL_756590, EPI_ISL_756591 | Lighthouse Lab in Cambridge | Wellcome Sanger Institute for the COVID-19 Genomics UK (COG-UK) Consortium | Rob Howes, The Lighthouse Lab in Cambridge and Alex Alderton, Roberto Amato, Sonia Goncalves, Ewan Harrison, David K. Jackson, Ian Johnston, Dominic Kwiatkowski, Cordelia Langford, John Sillitoe on behalf of the Wellcome Sanger Institute COVID-19 Surveillance Team |
| EPI_ISL_756592, EPI_ISL_756593 | Lighthouse Lab in Alderley Park | Wellcome Sanger Institute for the COVID-19 Genomics UK | Jacquelyn Wynn, Mairead Hyland, The Lighthouse Lab in Alderley Park and Alex Alderton, Roberto Amato, Sonia Goncalves, Ewan Harrison, David K. |

| | | | |
|---|---|---|---|
| | | (COG-UK) Consortium | Jackson, Ian Johnston, Dominic Kwiatkowski, Cordelia Langford, John Sillitoe on behalf of the Wellcome Sanger Institute COVID-19 Surveillance Team |
| EPI_ISL_756594, EPI_ISL_756595 | Lighthouse Lab in Cambridge | Wellcome Sanger Institute for the COVID-19 Genomics UK (COG-UK) Consortium | Rob Howes, The Lighthouse Lab in Cambridge and Alex Alderton, Roberto Amato, Sonia Goncalves, Ewan Harrison, David K. Jackson, Ian Johnston, Dominic Kwiatkowski, Cordelia Langford, John Sillitoe on behalf of the Wellcome Sanger Institute COVID-19 Surveillance Team |
| EPI_ISL_756596 | Lighthouse Lab in Alderley Park | Wellcome Sanger Institute for the COVID-19 Genomics UK (COG-UK) Consortium | Jacquelyn Wynn, Mairead Hyland, The Lighthouse Lab in Alderley Park and Alex Alderton, Roberto Amato, Sonia Goncalves, Ewan Harrison, David K. Jackson, Ian Johnston, Dominic Kwiatkowski, Cordelia Langford, John Sillitoe on behalf of the Wellcome Sanger Institute COVID-19 Surveillance Team |
| EPI_ISL_756597 | Lighthouse Lab in Cambridge | Wellcome Sanger Institute for the COVID-19 Genomics UK (COG-UK) Consortium | Rob Howes, The Lighthouse Lab in Cambridge and Alex Alderton, Roberto Amato, Sonia Goncalves, Ewan Harrison, David K. Jackson, Ian Johnston, Dominic Kwiatkowski, Cordelia Langford, John Sillitoe on behalf of the Wellcome Sanger Institute COVID-19 Surveillance Team |
| EPI_ISL_756598, EPI_ISL_756599, EPI_ISL_756600 | Lighthouse Lab in Alderley Park | Wellcome Sanger Institute for the COVID-19 Genomics UK (COG-UK) Consortium | Jacquelyn Wynn, Mairead Hyland, The Lighthouse Lab in Alderley Park and Alex Alderton, Roberto Amato, Sonia Goncalves, Ewan Harrison, David K. Jackson, Ian Johnston, Dominic Kwiatkowski, Cordelia Langford, John Sillitoe on behalf of the Wellcome Sanger Institute COVID-19 Surveillance Team |
| EPI_ISL_756601 | Lighthouse Lab in Cambridge | Wellcome Sanger Institute for the COVID-19 Genomics UK (COG-UK) Consortium | Rob Howes, The Lighthouse Lab in Cambridge and Alex Alderton, Roberto Amato, Sonia Goncalves, Ewan Harrison, David K. Jackson, Ian Johnston, Dominic Kwiatkowski, Cordelia Langford, John Sillitoe on behalf of the Wellcome Sanger Institute COVID-19 Surveillance Team |
| EPI_ISL_756602, EPI_ISL_756603 | Lighthouse Lab in Alderley Park | Wellcome Sanger Institute for the COVID-19 Genomics UK (COG-UK) Consortium | Jacquelyn Wynn, Mairead Hyland, The Lighthouse Lab in Alderley Park and Alex Alderton, Roberto Amato, Sonia Goncalves, Ewan Harrison, David K. Jackson, Ian Johnston, Dominic Kwiatkowski, Cordelia Langford, John Sillitoe on behalf of the Wellcome Sanger Institute COVID-19 Surveillance Team |
| EPI_ISL_756604, EPI_ISL_756605, EPI_ISL_756606, EPI_ISL_756607, EPI_ISL_756608 | Lighthouse Lab in Cambridge | Wellcome Sanger Institute for the COVID-19 Genomics UK (COG-UK) Consortium | Rob Howes, The Lighthouse Lab in Cambridge and Alex Alderton, Roberto Amato, Sonia Goncalves, Ewan Harrison, David K. Jackson, Ian Johnston, Dominic Kwiatkowski, Cordelia Langford, John Sillitoe on behalf of the Wellcome Sanger Institute COVID-19 Surveillance Team |
| EPI_ISL_756609 | Lighthouse Lab in Alderley Park | Wellcome Sanger Institute for the COVID-19 Genomics UK (COG-UK) Consortium | Jacquelyn Wynn, Mairead Hyland, The Lighthouse Lab in Alderley Park and Alex Alderton, Roberto Amato, Sonia Goncalves, Ewan Harrison, David K. Jackson, Ian Johnston, Dominic Kwiatkowski, Cordelia Langford, John Sillitoe on behalf of the Wellcome Sanger Institute COVID-19 Surveillance Team |
| EPI_ISL_756610, EPI_ISL_756611 | Lighthouse Lab in Cambridge | Wellcome Sanger Institute for the COVID-19 Genomics UK (COG-UK) Consortium | Rob Howes, The Lighthouse Lab in Cambridge and Alex Alderton, Roberto Amato, Sonia Goncalves, Ewan Harrison, David K. Jackson, Ian Johnston, Dominic Kwiatkowski, Cordelia Langford, John Sillitoe on behalf of the Wellcome Sanger Institute COVID-19 Surveillance Team |
| EPI_ISL_756612 | Lighthouse Lab in Alderley Park | Wellcome Sanger Institute for the COVID-19 Genomics UK (COG-UK) Consortium | Jacquelyn Wynn, Mairead Hyland, The Lighthouse Lab in Alderley Park and Alex Alderton, Roberto Amato, Sonia Goncalves, Ewan Harrison, David K. Jackson, Ian Johnston, Dominic Kwiatkowski, Cordelia Langford, John Sillitoe on behalf of the Wellcome Sanger Institute COVID-19 Surveillance Team |
| EPI_ISL_756613, EPI_ISL_756614, EPI_ISL_756615 | Lighthouse Lab in Cambridge | Wellcome Sanger Institute for the COVID-19 Genomics UK (COG-UK) Consortium | Rob Howes, The Lighthouse Lab in Cambridge and Alex Alderton, Roberto Amato, Sonia Goncalves, Ewan Harrison, David K. Jackson, Ian Johnston, Dominic Kwiatkowski, Cordelia Langford, John Sillitoe on behalf of the Wellcome Sanger Institute COVID-19 Surveillance Team |
| EPI_ISL_756616 | Lighthouse Lab in Alderley Park | Wellcome Sanger Institute for the COVID-19 Genomics UK (COG-UK) Consortium | Jacquelyn Wynn, Mairead Hyland, The Lighthouse Lab in Alderley Park and Alex Alderton, Roberto Amato, Sonia Goncalves, Ewan Harrison, David K. Jackson, Ian Johnston, Dominic Kwiatkowski, Cordelia Langford, John Sillitoe on behalf of the Wellcome Sanger Institute COVID-19 Surveillance Team |
| EPI_ISL_756617 | Lighthouse Lab in Cambridge | Wellcome Sanger Institute for the COVID-19 Genomics UK (COG-UK) Consortium | Rob Howes, The Lighthouse Lab in Cambridge and Alex Alderton, Roberto Amato, Sonia Goncalves, Ewan Harrison, David K. Jackson, Ian Johnston, Dominic Kwiatkowski, Cordelia Langford, John Sillitoe on behalf of the Wellcome Sanger Institute COVID-19 Surveillance Team |
| EPI_ISL_756618, EPI_ISL_756619, EPI_ISL_756620 | Lighthouse Lab in Alderley Park | Wellcome Sanger Institute for the COVID-19 Genomics UK (COG-UK) Consortium | Jacquelyn Wynn, Mairead Hyland, The Lighthouse Lab in Alderley Park and Alex Alderton, Roberto Amato, Sonia Goncalves, Ewan Harrison, David K. Jackson, Ian Johnston, Dominic Kwiatkowski, Cordelia Langford, John Sillitoe on behalf of the Wellcome Sanger Institute COVID-19 Surveillance Team |
| EPI_ISL_756621, EPI_ISL_756622, EPI_ISL_756623 | Lighthouse Lab in Cambridge | Wellcome Sanger Institute for the COVID-19 Genomics UK (COG-UK) Consortium | Rob Howes, The Lighthouse Lab in Cambridge and Alex Alderton, Roberto Amato, Sonia Goncalves, Ewan Harrison, David K. Jackson, Ian Johnston, Dominic Kwiatkowski, Cordelia Langford, John Sillitoe on behalf of the Wellcome Sanger Institute COVID-19 Surveillance Team |
| EPI_ISL_756624 | Lighthouse Lab in Alderley Park | Wellcome Sanger Institute for the COVID-19 Genomics UK (COG-UK) Consortium | Jacquelyn Wynn, Mairead Hyland, The Lighthouse Lab in Alderley Park and Alex Alderton, Roberto Amato, Sonia Goncalves, Ewan Harrison, David K. Jackson, Ian Johnston, Dominic Kwiatkowski, Cordelia Langford, John Sillitoe on behalf of the Wellcome Sanger Institute COVID-19 Surveillance Team |
| EPI_ISL_756625 | Lighthouse Lab in Cambridge | Wellcome Sanger Institute for the COVID-19 Genomics UK (COG-UK) Consortium | Rob Howes, The Lighthouse Lab in Cambridge and Alex Alderton, Roberto Amato, Sonia Goncalves, Ewan Harrison, David K. Jackson, Ian Johnston, Dominic Kwiatkowski, Cordelia Langford, John Sillitoe on behalf of the Wellcome Sanger Institute COVID-19 Surveillance Team |
| EPI_ISL_756626 | Lighthouse Lab in Alderley Park | Wellcome Sanger Institute for the COVID-19 Genomics UK (COG-UK) Consortium | Jacquelyn Wynn, Mairead Hyland, The Lighthouse Lab in Alderley Park and Alex Alderton, Roberto Amato, Sonia Goncalves, Ewan Harrison, David K. Jackson, Ian Johnston, Dominic Kwiatkowski, Cordelia Langford, John Sillitoe on behalf of the Wellcome Sanger Institute COVID-19 Surveillance Team |
| EPI_ISL_756627, EPI_ISL_756628, EPI_ISL_756629, EPI_ISL_756630, EPI_ISL_756631 | Lighthouse Lab in Cambridge | Wellcome Sanger Institute for the COVID-19 Genomics UK (COG-UK) Consortium | Rob Howes, The Lighthouse Lab in Cambridge and Alex Alderton, Roberto Amato, Sonia Goncalves, Ewan Harrison, David K. Jackson, Ian Johnston, Dominic Kwiatkowski, Cordelia Langford, John Sillitoe on behalf of the Wellcome Sanger Institute COVID-19 Surveillance Team |
| EPI_ISL_756632 | Lighthouse Lab in Alderley Park | Wellcome Sanger Institute for the COVID-19 Genomics UK (COG-UK) Consortium | Jacquelyn Wynn, Mairead Hyland, The Lighthouse Lab in Alderley Park and Alex Alderton, Roberto Amato, Sonia Goncalves, Ewan Harrison, David K. Jackson, Ian Johnston, Dominic Kwiatkowski, Cordelia Langford, John Sillitoe on behalf of the Wellcome Sanger Institute COVID-19 Surveillance Team |
| EPI_ISL_756633 | Lighthouse Lab in Cambridge | Wellcome Sanger Institute for the COVID-19 Genomics UK (COG-UK) Consortium | Rob Howes, The Lighthouse Lab in Cambridge and Alex Alderton, Roberto Amato, Sonia Goncalves, Ewan Harrison, David K. Jackson, Ian Johnston, Dominic Kwiatkowski, Cordelia Langford, John Sillitoe on behalf of the Wellcome Sanger Institute COVID-19 Surveillance Team |
| EPI_ISL_756634, EPI_ISL_756635, EPI_ISL_756636 | Lighthouse Lab in Alderley Park | Wellcome Sanger Institute for the COVID-19 Genomics UK (COG-UK) Consortium | Jacquelyn Wynn, Mairead Hyland, The Lighthouse Lab in Alderley Park and Alex Alderton, Roberto Amato, Sonia Goncalves, Ewan Harrison, David K. Jackson, Ian Johnston, Dominic Kwiatkowski, Cordelia Langford, John Sillitoe on behalf of the Wellcome Sanger Institute COVID-19 Surveillance Team |
| EPI_ISL_756637 | Lighthouse Lab in Cambridge | Wellcome Sanger Institute for the COVID-19 Genomics UK (COG-UK) Consortium | Rob Howes, The Lighthouse Lab in Cambridge and Alex Alderton, Roberto Amato, Sonia Goncalves, Ewan Harrison, David K. Jackson, Ian Johnston, Dominic Kwiatkowski, Cordelia Langford, John Sillitoe on behalf of the Wellcome Sanger Institute COVID-19 Surveillance Team |
| EPI_ISL_756638, EPI_ISL_756639, EPI_ISL_756640, EPI_ISL_756641 | Lighthouse Lab in Alderley Park | Wellcome Sanger Institute for the COVID-19 Genomics UK (COG-UK) Consortium | Jacquelyn Wynn, Mairead Hyland, The Lighthouse Lab in Alderley Park and Alex Alderton, Roberto Amato, Sonia Goncalves, Ewan Harrison, David K. Jackson, Ian Johnston, Dominic Kwiatkowski, Cordelia Langford, John Sillitoe on behalf of the Wellcome Sanger Institute COVID-19 Surveillance Team |
| EPI_ISL_756642, EPI_ISL_756643 | Lighthouse Lab in Cambridge | Wellcome Sanger Institute for the COVID-19 Genomics UK (COG-UK) Consortium | Rob Howes, The Lighthouse Lab in Cambridge and Alex Alderton, Roberto Amato, Sonia Goncalves, Ewan Harrison, David K. Jackson, Ian Johnston, Dominic Kwiatkowski, Cordelia Langford, John Sillitoe on behalf of the Wellcome Sanger Institute COVID-19 Surveillance Team |
| EPI_ISL_756644, EPI_ISL_756645, EPI_ISL_756646, EPI_ISL_756647, EPI_ISL_756648 | Lighthouse Lab in Alderley Park | Wellcome Sanger Institute for the COVID-19 Genomics UK (COG-UK) Consortium | Jacquelyn Wynn, Mairead Hyland, The Lighthouse Lab in Alderley Park and Alex Alderton, Roberto Amato, Sonia Goncalves, Ewan Harrison, David K. Jackson, Ian Johnston, Dominic Kwiatkowski, Cordelia Langford, John Sillitoe on behalf of the Wellcome Sanger Institute COVID-19 Surveillance Team |
| EPI_ISL_756649 | Lighthouse Lab in Cambridge | Wellcome Sanger Institute for the COVID-19 Genomics UK (COG-UK) Consortium | Rob Howes, The Lighthouse Lab in Cambridge and Alex Alderton, Roberto Amato, Sonia Goncalves, Ewan Harrison, David K. Jackson, Ian Johnston, Dominic Kwiatkowski, Cordelia Langford, John Sillitoe on behalf of the Wellcome Sanger Institute COVID-19 Surveillance Team |
| EPI_ISL_756650, EPI_ISL_756651, EPI_ISL_756652 | Lighthouse Lab in Alderley Park | Wellcome Sanger Institute for the COVID-19 Genomics UK (COG-UK) Consortium | Jacquelyn Wynn, Mairead Hyland, The Lighthouse Lab in Alderley Park and Alex Alderton, Roberto Amato, Sonia Goncalves, Ewan Harrison, David K. Jackson, Ian Johnston, Dominic Kwiatkowski, Cordelia Langford, John Sillitoe on behalf of the Wellcome Sanger Institute COVID-19 Surveillance Team |
| EPI_ISL_756653 | Lighthouse Lab in Cambridge | Wellcome Sanger Institute for the COVID-19 Genomics UK (COG-UK) Consortium | Rob Howes, The Lighthouse Lab in Cambridge and Alex Alderton, Roberto Amato, Sonia Goncalves, Ewan Harrison, David K. Jackson, Ian Johnston, Dominic Kwiatkowski, Cordelia Langford, John Sillitoe on behalf of the Wellcome Sanger Institute COVID-19 Surveillance Team |
| EPI_ISL_756654 | Lighthouse Lab in Alderley Park | Wellcome Sanger Institute for the COVID-19 Genomics UK (COG-UK) Consortium | Jacquelyn Wynn, Mairead Hyland, The Lighthouse Lab in Alderley Park and Alex Alderton, Roberto Amato, Sonia Goncalves, Ewan Harrison, David K. Jackson, Ian Johnston, Dominic Kwiatkowski, Cordelia Langford, John Sillitoe on behalf of the Wellcome Sanger Institute COVID-19 Surveillance Team |
| EPI_ISL_756655 | Lighthouse Lab in Cambridge | Wellcome Sanger Institute for the COVID-19 Genomics UK (COG-UK) Consortium | Rob Howes, The Lighthouse Lab in Cambridge and Alex Alderton, Roberto Amato, Sonia Goncalves, Ewan Harrison, David K. Jackson, Ian Johnston, Dominic Kwiatkowski, Cordelia Langford, John Sillitoe on behalf of the Wellcome Sanger Institute COVID-19 Surveillance Team |
| EPI_ISL_756656, EPI_ISL_756657, EPI_ISL_756658 | Lighthouse Lab in Alderley Park | Wellcome Sanger Institute for the COVID-19 Genomics UK (COG-UK) Consortium | Jacquelyn Wynn, Mairead Hyland, The Lighthouse Lab in Alderley Park and Alex Alderton, Roberto Amato, Sonia Goncalves, Ewan Harrison, David K. Jackson, Ian Johnston, Dominic Kwiatkowski, Cordelia Langford, John Sillitoe on behalf of the Wellcome Sanger Institute COVID-19 Surveillance Team |
| EPI_ISL_756659, EPI_ISL_756660 | Lighthouse Lab in Cambridge | Wellcome Sanger Institute for the COVID-19 Genomics UK (COG-UK) Consortium | Rob Howes, The Lighthouse Lab in Cambridge and Alex Alderton, Roberto Amato, Sonia Goncalves, Ewan Harrison, David K. Jackson, Ian Johnston, Dominic Kwiatkowski, Cordelia Langford, John Sillitoe on behalf of the Wellcome Sanger Institute COVID-19 Surveillance Team |
| EPI_ISL_756661, EPI_ISL_756662, | Lighthouse Lab in Alderley Park | Wellcome Sanger Institute for the COVID-19 Genomics UK | Jacquelyn Wynn, Mairead Hyland, The Lighthouse Lab in Alderley Park and Alex Alderton, Roberto Amato, Sonia Goncalves, Ewan Harrison, David K. |

| | | | |
|---|---|---|---|
| EPI_ISL_756663, EPI_ISL_756664 | | (COG-UK) Consortium | Jackson, Ian Johnston, Dominic Kwiatkowski, Cordelia Langford, John Sillitoe on behalf of the Wellcome Sanger Institute COVID-19 Surveillance Team |
| EPI_ISL_756665, EPI_ISL_756666, EPI_ISL_756667, EPI_ISL_756668 | Lighthouse Lab in Cambridge | Wellcome Sanger Institute for the COVID-19 Genomics UK (COG-UK) Consortium | Rob Howes, The Lighthouse Lab in Cambridge and Alex Alderton, Roberto Amato, Sonia Goncalves, Ewan Harrison, David K. Jackson, Ian Johnston, Dominic Kwiatkowski, Cordelia Langford, John Sillitoe on behalf of the Wellcome Sanger Institute COVID-19 Surveillance Team |
| EPI_ISL_756669, EPI_ISL_756670 | Lighthouse Lab in Alderley Park | Wellcome Sanger Institute for the COVID-19 Genomics UK (COG-UK) Consortium | Jacquelyn Wynn, Mairead Hyland, The Lighthouse Lab in Alderley Park and Alex Alderton, Roberto Amato, Sonia Goncalves, Ewan Harrison, David K. Jackson, Ian Johnston, Dominic Kwiatkowski, Cordelia Langford, John Sillitoe on behalf of the Wellcome Sanger Institute COVID-19 Surveillance Team |
| EPI_ISL_756671, EPI_ISL_756672 | Lighthouse Lab in Cambridge | Wellcome Sanger Institute for the COVID-19 Genomics UK (COG-UK) Consortium | Rob Howes, The Lighthouse Lab in Cambridge and Alex Alderton, Roberto Amato, Sonia Goncalves, Ewan Harrison, David K. Jackson, Ian Johnston, Dominic Kwiatkowski, Cordelia Langford, John Sillitoe on behalf of the Wellcome Sanger Institute COVID-19 Surveillance Team |
| EPI_ISL_756673 | Lighthouse Lab in Alderley Park | Wellcome Sanger Institute for the COVID-19 Genomics UK (COG-UK) Consortium | Jacquelyn Wynn, Mairead Hyland, The Lighthouse Lab in Alderley Park and Alex Alderton, Roberto Amato, Sonia Goncalves, Ewan Harrison, David K. Jackson, Ian Johnston, Dominic Kwiatkowski, Cordelia Langford, John Sillitoe on behalf of the Wellcome Sanger Institute COVID-19 Surveillance Team |
| EPI_ISL_756674, EPI_ISL_756675, EPI_ISL_756676, EPI_ISL_756677, EPI_ISL_756678, EPI_ISL_756679, EPI_ISL_756680 | Lighthouse Lab in Cambridge | Wellcome Sanger Institute for the COVID-19 Genomics UK (COG-UK) Consortium | Rob Howes, The Lighthouse Lab in Cambridge and Alex Alderton, Roberto Amato, Sonia Goncalves, Ewan Harrison, David K. Jackson, Ian Johnston, Dominic Kwiatkowski, Cordelia Langford, John Sillitoe on behalf of the Wellcome Sanger Institute COVID-19 Surveillance Team |
| EPI_ISL_756681, EPI_ISL_756682, EPI_ISL_756683, EPI_ISL_756684, EPI_ISL_756685, EPI_ISL_756686, EPI_ISL_756687, EPI_ISL_756688 | Lighthouse Lab in Alderley Park | Wellcome Sanger Institute for the COVID-19 Genomics UK (COG-UK) Consortium | Jacquelyn Wynn, Mairead Hyland, The Lighthouse Lab in Alderley Park and Alex Alderton, Roberto Amato, Sonia Goncalves, Ewan Harrison, David K. Jackson, Ian Johnston, Dominic Kwiatkowski, Cordelia Langford, John Sillitoe on behalf of the Wellcome Sanger Institute COVID-19 Surveillance Team |
| EPI_ISL_756690, EPI_ISL_756695, EPI_ISL_756696, EPI_ISL_756702, EPI_ISL_756705, EPI_ISL_756706, EPI_ISL_756709, EPI_ISL_756711, EPI_ISL_756716, EPI_ISL_756723, EPI_ISL_756729, EPI_ISL_756737, EPI_ISL_756744, EPI_ISL_756745, EPI_ISL_756748, EPI_ISL_756750, EPI_ISL_756761, EPI_ISL_756773, EPI_ISL_756775, EPI_ISL_756780, EPI_ISL_756784, EPI_ISL_756789, EPI_ISL_756795, EPI_ISL_756800, EPI_ISL_756803, EPI_ISL_756805, EPI_ISL_756824, EPI_ISL_756835, EPI_ISL_756840, EPI_ISL_756849, EPI_ISL_756850, EPI_ISL_756852, EPI_ISL_756861, EPI_ISL_756867, EPI_ISL_756877, EPI_ISL_756878, EPI_ISL_756879, EPI_ISL_756881, EPI_ISL_756885, EPI_ISL_756886, EPI_ISL_756888, EPI_ISL_756889, EPI_ISL_756893, EPI_ISL_756894, EPI_ISL_756895, EPI_ISL_756897, EPI_ISL_756900, EPI_ISL_756909, EPI_ISL_756910, EPI_ISL_756912, EPI_ISL_756918, EPI_ISL_756920, EPI_ISL_756923, EPI_ISL_756928, EPI_ISL_756931, EPI_ISL_756935, EPI_ISL_756940, EPI_ISL_756947, EPI_ISL_756951, EPI_ISL_756952, EPI_ISL_756953, EPI_ISL_756955, EPI_ISL_756959, EPI_ISL_756969, EPI_ISL_756970, EPI_ISL_756971, EPI_ISL_756973, EPI_ISL_756977, EPI_ISL_756982, EPI_ISL_756998, EPI_ISL_757005, EPI_ISL_757007, EPI_ISL_757008, EPI_ISL_757009, EPI_ISL_757011, EPI_ISL_757012, EPI_ISL_757017, EPI_ISL_757018, EPI_ISL_757021, EPI_ISL_757027, EPI_ISL_757028, EPI_ISL_757032 | | | |
| see above | Lighthouse Lab in Glasgow | Wellcome Sanger Institute for the COVID-19 Genomics UK (COG-UK) Consortium | Harper VanSteenhouse, Yumi Kasai, David Gray, Carol Clugston, Anna Dominiczak and Alex Alderton, Roberto Amato, Sonia Goncalves, Ewan Harrison, David K. Jackson, Ian Johnston, Dominic Kwiatkowski, Cordelia Langford, John Sillitoe on behalf of the Wellcome Sanger Institute COVID-19 Surveillance Team |
| EPI_ISL_759726, EPI_ISL_759727, EPI_ISL_759739, EPI_ISL_759740, EPI_ISL_759742 | University of Wisconsin-Madison AIDS Vaccine Research Laboratories | University of Wisconsin-Madison AIDS Vaccine Research Laboratories | Gage Moreno, Katarina Braun, et al. AIDS Vaccine Research Laboratories |
| EPI_ISL_760276, EPI_ISL_760284, EPI_ISL_760305, EPI_ISL_760326, EPI_ISL_760333, EPI_ISL_760344, EPI_ISL_760350, EPI_ISL_760421, EPI_ISL_760450, EPI_ISL_760495, EPI_ISL_760498, EPI_ISL_760549, EPI_ISL_760587, EPI_ISL_760637, EPI_ISL_760644 | | | |
| see above | Lighthouse Lab in Alderley Park | Wellcome Sanger Institute for the COVID-19 Genomics UK (COG-UK) Consortium | Jacquelyn Wynn, Mairead Hyland, The Lighthouse Lab in Alderley Park and Alex Alderton, Roberto Amato, Sonia Goncalves, Ewan Harrison, David K. Jackson, Ian Johnston, Dominic Kwiatkowski, Cordelia Langford, John Sillitoe on behalf of the Wellcome Sanger Institute COVID-19 Surveillance Team |
| EPI_ISL_760657 | Lighthouse Lab in Glasgow | Wellcome Sanger Institute for the COVID-19 Genomics UK (COG-UK) Consortium | Harper VanSteenhouse, Yumi Kasai, David Gray, Carol Clugston, Anna Dominiczak and Alex Alderton, Roberto Amato, Sonia Goncalves, Ewan Harrison, David K. Jackson, Ian Johnston, Dominic Kwiatkowski, Cordelia Langford, John Sillitoe on behalf of the Wellcome Sanger Institute COVID-19 Surveillance Team |
| EPI_ISL_760698 | Lighthouse Lab in Milton Keynes | Wellcome Sanger Institute for the COVID-19 Genomics UK (COG-UK) Consortium | The Lighthouse Lab in Milton Keynes and Alex Alderton, Roberto Amato, Sonia Goncalves, Ewan Harrison, David K. Jackson, Ian Johnston, Dominic Kwiatkowski, Cordelia Langford, John Sillitoe on behalf of the Wellcome Sanger Institute COVID-19 Surveillance Team |
| EPI_ISL_760724, EPI_ISL_760728 | Lighthouse Lab in Alderley Park | Wellcome Sanger Institute for the COVID-19 Genomics UK (COG-UK) Consortium | Jacquelyn Wynn, Mairead Hyland, The Lighthouse Lab in Alderley Park and Alex Alderton, Roberto Amato, Sonia Goncalves, Ewan Harrison, David K. Jackson, Ian Johnston, Dominic Kwiatkowski, Cordelia Langford, John Sillitoe on behalf of the Wellcome Sanger Institute COVID-19 Surveillance Team |
| EPI_ISL_760740 | Lighthouse Lab in Glasgow | Wellcome Sanger Institute for the COVID-19 Genomics UK (COG-UK) Consortium | Harper VanSteenhouse, Yumi Kasai, David Gray, Carol Clugston, Anna Dominiczak and Alex Alderton, Roberto Amato, Sonia Goncalves, Ewan Harrison, David K. Jackson, Ian Johnston, Dominic Kwiatkowski, Cordelia Langford, John Sillitoe on behalf of the Wellcome Sanger Institute COVID-19 Surveillance Team |
| EPI_ISL_760752, EPI_ISL_760759 | Lighthouse Lab in Alderley Park | Wellcome Sanger Institute for the COVID-19 Genomics UK (COG-UK) Consortium | Jacquelyn Wynn, Mairead Hyland, The Lighthouse Lab in Alderley Park and Alex Alderton, Roberto Amato, Sonia Goncalves, Ewan Harrison, David K. Jackson, Ian Johnston, Dominic Kwiatkowski, Cordelia Langford, John Sillitoe on behalf of the Wellcome Sanger Institute COVID-19 Surveillance Team |
| EPI_ISL_760782 | Lighthouse Lab in Glasgow | Wellcome Sanger Institute for the COVID-19 Genomics UK (COG-UK) Consortium | Harper VanSteenhouse, Yumi Kasai, David Gray, Carol Clugston, Anna Dominiczak and Alex Alderton, Roberto Amato, Sonia Goncalves, Ewan Harrison, David K. Jackson, Ian Johnston, Dominic Kwiatkowski, Cordelia Langford, John Sillitoe on behalf of the Wellcome Sanger Institute COVID-19 Surveillance Team |
| EPI_ISL_760799, EPI_ISL_760852, EPI_ISL_760857, EPI_ISL_760861, EPI_ISL_760862, EPI_ISL_760864, EPI_ISL_760868, EPI_ISL_760887, EPI_ISL_760906, EPI_ISL_760928, EPI_ISL_760968, EPI_ISL_761025, EPI_ISL_761048, EPI_ISL_761052, EPI_ISL_761064, EPI_ISL_761065, EPI_ISL_761069, EPI_ISL_761081, EPI_ISL_761083, EPI_ISL_761086, EPI_ISL_761087, EPI_ISL_761089, EPI_ISL_761092, EPI_ISL_761093, EPI_ISL_761099, EPI_ISL_761100, EPI_ISL_761105, EPI_ISL_761107, EPI_ISL_761112, EPI_ISL_761115, EPI_ISL_761117, EPI_ISL_761119, EPI_ISL_761122, EPI_ISL_761123, EPI_ISL_761126, EPI_ISL_761129, EPI_ISL_761130, EPI_ISL_761132, EPI_ISL_761136, EPI_ISL_761139, EPI_ISL_761141, EPI_ISL_761144, EPI_ISL_761146, EPI_ISL_761149, EPI_ISL_761151, EPI_ISL_761155, EPI_ISL_761159, EPI_ISL_761160, EPI_ISL_761161, EPI_ISL_761171, EPI_ISL_761173, EPI_ISL_761174, EPI_ISL_761175, EPI_ISL_761178, EPI_ISL_761179, EPI_ISL_761183, EPI_ISL_761184, EPI_ISL_761190, EPI_ISL_761192, EPI_ISL_761196, EPI_ISL_761199, EPI_ISL_761200, EPI_ISL_761202, EPI_ISL_761203, EPI_ISL_761205, EPI_ISL_761206, EPI_ISL_761208, EPI_ISL_761211, EPI_ISL_761213, EPI_ISL_761214, EPI_ISL_761215, EPI_ISL_761217, EPI_ISL_761218, EPI_ISL_761219, EPI_ISL_761222, EPI_ISL_761224, EPI_ISL_761232, EPI_ISL_761233, EPI_ISL_761234, EPI_ISL_761237, EPI_ISL_761242, EPI_ISL_761245, EPI_ISL_761246, EPI_ISL_761249, EPI_ISL_761250, EPI_ISL_761251, EPI_ISL_761254, EPI_ISL_761260, EPI_ISL_761262, EPI_ISL_761263, EPI_ISL_761265, EPI_ISL_761267, EPI_ISL_761285, EPI_ISL_761286, EPI_ISL_761288, EPI_ISL_761291, EPI_ISL_761294, EPI_ISL_761299, EPI_ISL_761300, EPI_ISL_761304, EPI_ISL_761305, EPI_ISL_761306, EPI_ISL_761309, EPI_ISL_761310, EPI_ISL_761314, EPI_ISL_761318, EPI_ISL_761319, EPI_ISL_761321, EPI_ISL_761322, EPI_ISL_761323, EPI_ISL_761332, EPI_ISL_761333, EPI_ISL_761335, EPI_ISL_761337, EPI_ISL_761341, EPI_ISL_761342, EPI_ISL_761346, EPI_ISL_761347, EPI_ISL_761354 | | | |
| see above | Lighthouse Lab in Alderley Park | Wellcome Sanger Institute for the COVID-19 Genomics UK (COG-UK) Consortium | Jacquelyn Wynn, Mairead Hyland, The Lighthouse Lab in Alderley Park and Alex Alderton, Roberto Amato, Sonia Goncalves, Ewan Harrison, David K. Jackson, Ian Johnston, Dominic Kwiatkowski, Cordelia Langford, John Sillitoe on behalf of the Wellcome Sanger Institute COVID-19 Surveillance Team |
| EPI_ISL_761357, EPI_ISL_761358, EPI_ISL_761359, EPI_ISL_761360, EPI_ISL_761361, EPI_ISL_761362, EPI_ISL_761363, EPI_ISL_761364, EPI_ISL_761365, EPI_ISL_761366, EPI_ISL_761367, EPI_ISL_761368, EPI_ISL_761369, EPI_ISL_761370, EPI_ISL_761371, EPI_ISL_761372, EPI_ISL_761373, EPI_ISL_761374, EPI_ISL_761375, EPI_ISL_761376, EPI_ISL_761377, EPI_ISL_761378, EPI_ISL_761379, EPI_ISL_761380, EPI_ISL_761381, EPI_ISL_761382, EPI_ISL_761383, EPI_ISL_761384, EPI_ISL_761385, EPI_ISL_761386, EPI_ISL_761387, EPI_ISL_761388, EPI_ISL_761389, EPI_ISL_761390, EPI_ISL_761391, EPI_ISL_761392, EPI_ISL_761393, EPI_ISL_761394, EPI_ISL_761395, EPI_ISL_761396, EPI_ISL_761397, EPI_ISL_761398, EPI_ISL_761399, EPI_ISL_761400, EPI_ISL_761401, EPI_ISL_761402, EPI_ISL_761403, EPI_ISL_761404, EPI_ISL_761405, EPI_ISL_761406, EPI_ISL_761407, EPI_ISL_761408, EPI_ISL_761409, EPI_ISL_761410, EPI_ISL_761411, EPI_ISL_761412, EPI_ISL_761413, EPI_ISL_761414, EPI_ISL_761415, EPI_ISL_761416, EPI_ISL_761417, EPI_ISL_761418, EPI_ISL_761419, EPI_ISL_761420, EPI_ISL_761421, EPI_ISL_761422, EPI_ISL_761423, EPI_ISL_761424, EPI_ISL_761425, EPI_ISL_761426, EPI_ISL_761427, EPI_ISL_761428, EPI_ISL_761429, EPI_ISL_761430, EPI_ISL_761431, EPI_ISL_761432, EPI_ISL_761433, EPI_ISL_761434, EPI_ISL_761435, EPI_ISL_761436, EPI_ISL_761437, EPI_ISL_761438, EPI_ISL_761439, EPI_ISL_761440, EPI_ISL_761441, EPI_ISL_761442, EPI_ISL_761443, EPI_ISL_761444, EPI_ISL_761445, EPI_ISL_761446, EPI_ISL_761447, EPI_ISL_761448, EPI_ISL_761449, EPI_ISL_761450, EPI_ISL_761451, EPI_ISL_761452, EPI_ISL_761453, EPI_ISL_761454, EPI_ISL_761455, EPI_ISL_761456, EPI_ISL_761457, EPI_ISL_761458, EPI_ISL_761459, EPI_ISL_761460, EPI_ISL_761461, EPI_ISL_761462, EPI_ISL_761463, EPI_ISL_761464, EPI_ISL_761465, EPI_ISL_761466, EPI_ISL_761467, EPI_ISL_761468, EPI_ISL_761469, EPI_ISL_761470, EPI_ISL_761471, EPI_ISL_761472, EPI_ISL_761473, EPI_ISL_761474, EPI_ISL_761475, EPI_ISL_761476, EPI_ISL_761477, EPI_ISL_761478, EPI_ISL_761479, EPI_ISL_761480, EPI_ISL_761481, EPI_ISL_761482, EPI_ISL_761483, EPI_ISL_761484, EPI_ISL_761485, EPI_ISL_761486, EPI_ISL_761487, EPI_ISL_761488, EPI_ISL_761489, EPI_ISL_761490, EPI_ISL_761491, EPI_ISL_761492, EPI_ISL_761493, EPI_ISL_761494, EPI_ISL_761495, EPI_ISL_761496, EPI_ISL_761497, EPI_ISL_761498, EPI_ISL_761499, EPI_ISL_761500, EPI_ISL_761501, EPI_ISL_761502, EPI_ISL_761503, EPI_ISL_761504, EPI_ISL_761505, EPI_ISL_761506, EPI_ISL_761507, EPI_ISL_761508, EPI_ISL_761509, EPI_ISL_761510, EPI_ISL_761511, EPI_ISL_761512, EPI_ISL_761513, EPI_ISL_761514, EPI_ISL_761515, EPI_ISL_761516, EPI_ISL_761517, EPI_ISL_761518, EPI_ISL_761519, EPI_ISL_761520, EPI_ISL_761521, EPI_ISL_761522, EPI_ISL_761523, EPI_ISL_761524, EPI_ISL_761525, EPI_ISL_761526, EPI_ISL_761527, EPI_ISL_761528, EPI_ISL_761529, EPI_ISL_761530, EPI_ISL_761531, EPI_ISL_761532, EPI_ISL_761533, EPI_ISL_761534, EPI_ISL_761535, EPI_ISL_761536, EPI_ISL_761537, EPI_ISL_761538, EPI_ISL_761539, EPI_ISL_761540, EPI_ISL_761541, EPI_ISL_761542, EPI_ISL_761543, EPI_ISL_761544, EPI_ISL_761545, EPI_ISL_761546, EPI_ISL_761547, EPI_ISL_761548, EPI_ISL_761549, EPI_ISL_761550, EPI_ISL_761551, EPI_ISL_761552, EPI_ISL_761553, EPI_ISL_761554, EPI_ISL_761555, EPI_ISL_761556, EPI_ISL_761557, EPI_ISL_761558, EPI_ISL_761559, EPI_ISL_761560, EPI_ISL_761561, EPI_ISL_761562, EPI_ISL_761563, EPI_ISL_761564, EPI_ISL_761565, EPI_ISL_761566, EPI_ISL_761567, EPI_ISL_761568, EPI_ISL_761569, EPI_ISL_761570, EPI_ISL_761571, EPI_ISL_761572, EPI_ISL_761573, EPI_ISL_761574, EPI_ISL_761575, EPI_ISL_761576, EPI_ISL_761577, EPI_ISL_761578, EPI_ISL_761579, EPI_ISL_761580, EPI_ISL_761581, EPI_ISL_761582, EPI_ISL_761583, EPI_ISL_761584, EPI_ISL_761585, EPI_ISL_761586, EPI_ISL_761587, EPI_ISL_761588, EPI_ISL_761589, EPI_ISL_761590, EPI_ISL_761591, EPI_ISL_761592, EPI_ISL_761593, EPI_ISL_761594, EPI_ISL_761595, EPI_ISL_761596, EPI_ISL_761597, EPI_ISL_761598, EPI_ISL_761599, EPI_ISL_761600, EPI_ISL_761601, EPI_ISL_761602, EPI_ISL_761603, EPI_ISL_761604, EPI_ISL_761605, EPI_ISL_761606, EPI_ISL_761607, EPI_ISL_761608, EPI_ISL_761609, EPI_ISL_761610, EPI_ISL_761611, EPI_ISL_761612, EPI_ISL_761613, EPI_ISL_761614, EPI_ISL_761615, EPI_ISL_761617, EPI_ISL_761618, EPI_ISL_761619, EPI_ISL_761620, EPI_ISL_761621, EPI_ISL_761622, EPI_ISL_761623, EPI_ISL_761624, EPI_ISL_761625, EPI_ISL_761626, EPI_ISL_761627, EPI_ISL_761628, EPI_ISL_761629, EPI_ISL_761630, EPI_ISL_761631, EPI_ISL_761632, EPI_ISL_761633, EPI_ISL_761634, EPI_ISL_761635, EPI_ISL_761636, EPI_ISL_761637, EPI_ISL_761638, EPI_ISL_761639, EPI_ISL_761640, EPI_ISL_761641, EPI_ISL_761642, EPI_ISL_761643, EPI_ISL_761644, EPI_ISL_761645, EPI_ISL_761646, EPI_ISL_761647, EPI_ISL_761648, EPI_ISL_761649, EPI_ISL_761650, EPI_ISL_761651, EPI_ISL_761652, EPI_ISL_761653, EPI_ISL_761654, EPI_ISL_761655, EPI_ISL_761656, EPI_ISL_761657, EPI_ISL_761658, EPI_ISL_761659, EPI_ISL_761660, EPI_ISL_761661, EPI_ISL_761662, EPI_ISL_761663, EPI_ISL_761664 | | | |
| see above | Lighthouse Lab in Milton Keynes | Wellcome Sanger Institute for the COVID-19 Genomics UK (COG-UK) Consortium | The Lighthouse Lab in Milton Keynes and Alex Alderton, Roberto Amato, Sonia Goncalves, Ewan Harrison, David K. Jackson, Ian Johnston, Dominic Kwiatkowski, Cordelia Langford, John Sillitoe on behalf of the Wellcome Sanger Institute COVID-19 Surveillance Team |
| EPI_ISL_761665, EPI_ISL_761666, EPI_ISL_761667, EPI_ISL_761668, EPI_ISL_761669, EPI_ISL_761672, EPI_ISL_761675, EPI_ISL_761676, EPI_ISL_761677, EPI_ISL_761679, EPI_ISL_761680, EPI_ISL_761681, EPI_ISL_761684, EPI_ISL_761685, EPI_ISL_761686, EPI_ISL_761694, EPI_ISL_761695, EPI_ISL_761696, | | | |

EPI_ISL_761697, EPI_ISL_761698, EPI_ISL_761699, EPI_ISL_761702, EPI_ISL_761704, EPI_ISL_761706, EPI_ISL_761707, EPI_ISL_761711, EPI_ISL_761712, EPI_ISL_761714, EPI_ISL_761715, EPI_ISL_761716, EPI_ISL_761717, EPI_ISL_761719, EPI_ISL_761720, EPI_ISL_761721, EPI_ISL_761723, EPI_ISL_761724, EPI_ISL_761728, EPI_ISL_761729, EPI_ISL_761730, EPI_ISL_761731, EPI_ISL_761732, EPI_ISL_761736, EPI_ISL_761737, EPI_ISL_761738, EPI_ISL_761739, EPI_ISL_761740, EPI_ISL_761741, EPI_ISL_761742, EPI_ISL_761744, EPI_ISL_761745, EPI_ISL_761746, EPI_ISL_761748, EPI_ISL_761749, EPI_ISL_761751, EPI_ISL_761754, EPI_ISL_761756, EPI_ISL_761757, EPI_ISL_761760, EPI_ISL_761761, EPI_ISL_761763, EPI_ISL_761764, EPI_ISL_761765, EPI_ISL_761766, EPI_ISL_761767, EPI_ISL_761768, EPI_ISL_761769, EPI_ISL_761770, EPI_ISL_761771, EPI_ISL_761773, EPI_ISL_761774, EPI_ISL_761776, EPI_ISL_761777, EPI_ISL_761778, EPI_ISL_761780, EPI_ISL_761781, EPI_ISL_761782, EPI_ISL_761783, EPI_ISL_761785, EPI_ISL_761786, EPI_ISL_761788, EPI_ISL_761789, EPI_ISL_761790, EPI_ISL_761792, EPI_ISL_761794, EPI_ISL_761796, EPI_ISL_761800, EPI_ISL_761801, EPI_ISL_761802, EPI_ISL_761803, EPI_ISL_761804, EPI_ISL_761806, EPI_ISL_761807, EPI_ISL_761808, EPI_ISL_761809, EPI_ISL_761810, EPI_ISL_761813, EPI_ISL_761814, EPI_ISL_761815, EPI_ISL_761816, EPI_ISL_761818, EPI_ISL_761820, EPI_ISL_761821, EPI_ISL_761822, EPI_ISL_761823, EPI_ISL_761825, EPI_ISL_761827, EPI_ISL_761828, EPI_ISL_761831, EPI_ISL_761833, EPI_ISL_761835, EPI_ISL_761836, EPI_ISL_761837, EPI_ISL_761838, EPI_ISL_761839, EPI_ISL_761840, EPI_ISL_761841, EPI_ISL_761842, EPI_ISL_761845, EPI_ISL_761846, EPI_ISL_761847, EPI_ISL_761848, EPI_ISL_761849, EPI_ISL_761851, EPI_ISL_761852, EPI_ISL_761855, EPI_ISL_761859, EPI_ISL_761860, EPI_ISL_761862, EPI_ISL_761863, EPI_ISL_761865, EPI_ISL_761867, EPI_ISL_761868, EPI_ISL_761870, EPI_ISL_761872, EPI_ISL_761873, EPI_ISL_761875, EPI_ISL_761876, EPI_ISL_761877, EPI_ISL_761878, EPI_ISL_761879, EPI_ISL_761880, EPI_ISL_761884, EPI_ISL_761887, EPI_ISL_761888, EPI_ISL_761889, EPI_ISL_761890, EPI_ISL_761893, EPI_ISL_761894, EPI_ISL_761895, EPI_ISL_761896, EPI_ISL_761897, EPI_ISL_761898, EPI_ISL_761900, EPI_ISL_761901, EPI_ISL_761902, EPI_ISL_761903, EPI_ISL_761904, EPI_ISL_761905, EPI_ISL_761906, EPI_ISL_761908, EPI_ISL_761909, EPI_ISL_761910, EPI_ISL_761911, EPI_ISL_761912, EPI_ISL_761913, EPI_ISL_761914, EPI_ISL_761917, EPI_ISL_761918, EPI_ISL_761919, EPI_ISL_761920

| | | | |
|---|---|---|---|
| see above | Lighthouse Lab in Alderley Park | Wellcome Sanger Institute for the COVID-19 Genomics UK (COG-UK) Consortium | Jacquelyn Wynn, Mairead Hyland, The Lighthouse Lab in Alderley Park and Alex Alderton, Roberto Amato, Sonia Goncalves, Ewan Harrison, David K. Jackson, Ian Johnston, Dominic Kwiatkowski, Cordelia Langford, John Sillitoe on behalf of the Wellcome Sanger Institute COVID-19 Surveillance Team |
| EPI_ISL_761921 | Lighthouse Lab in Glasgow | Wellcome Sanger Institute for the COVID-19 Genomics UK (COG-UK) Consortium | Harper VanSteenhouse, Yumi Kasai, David Gray, Carol Clugston, Anna Dominiczak and Alex Alderton, Roberto Amato, Sonia Goncalves, Ewan Harrison, David K. Jackson, Ian Johnston, Dominic Kwiatkowski, Cordelia Langford, John Sillitoe on behalf of the Wellcome Sanger Institute COVID-19 Surveillance Team |
| EPI_ISL_761922 | Lighthouse Lab in Cambridge | Wellcome Sanger Institute for the COVID-19 Genomics UK (COG-UK) Consortium | Rob Howes, The Lighthouse Lab in Cambridge and Alex Alderton, Roberto Amato, Sonia Goncalves, Ewan Harrison, David K. Jackson, Ian Johnston, Dominic Kwiatkowski, Cordelia Langford, John Sillitoe on behalf of the Wellcome Sanger Institute COVID-19 Surveillance Team |
| EPI_ISL_761923, EPI_ISL_761924, EPI_ISL_761925, EPI_ISL_761926 | Lighthouse Lab in Glasgow | Wellcome Sanger Institute for the COVID-19 Genomics UK (COG-UK) Consortium | Harper VanSteenhouse, Yumi Kasai, David Gray, Carol Clugston, Anna Dominiczak and Alex Alderton, Roberto Amato, Sonia Goncalves, Ewan Harrison, David K. Jackson, Ian Johnston, Dominic Kwiatkowski, Cordelia Langford, John Sillitoe on behalf of the Wellcome Sanger Institute COVID-19 Surveillance Team |
| EPI_ISL_761927 | Lighthouse Lab in Cambridge | Wellcome Sanger Institute for the COVID-19 Genomics UK (COG-UK) Consortium | Rob Howes, The Lighthouse Lab in Cambridge and Alex Alderton, Roberto Amato, Sonia Goncalves, Ewan Harrison, David K. Jackson, Ian Johnston, Dominic Kwiatkowski, Cordelia Langford, John Sillitoe on behalf of the Wellcome Sanger Institute COVID-19 Surveillance Team |
| EPI_ISL_761928 | Lighthouse Lab in Glasgow | Wellcome Sanger Institute for the COVID-19 Genomics UK (COG-UK) Consortium | Harper VanSteenhouse, Yumi Kasai, David Gray, Carol Clugston, Anna Dominiczak and Alex Alderton, Roberto Amato, Sonia Goncalves, Ewan Harrison, David K. Jackson, Ian Johnston, Dominic Kwiatkowski, Cordelia Langford, John Sillitoe on behalf of the Wellcome Sanger Institute COVID-19 Surveillance Team |
| EPI_ISL_761929 | Lighthouse Lab in Cambridge | Wellcome Sanger Institute for the COVID-19 Genomics UK (COG-UK) Consortium | Rob Howes, The Lighthouse Lab in Cambridge and Alex Alderton, Roberto Amato, Sonia Goncalves, Ewan Harrison, David K. Jackson, Ian Johnston, Dominic Kwiatkowski, Cordelia Langford, John Sillitoe on behalf of the Wellcome Sanger Institute COVID-19 Surveillance Team |
| EPI_ISL_761930, EPI_ISL_761931, EPI_ISL_761932, EPI_ISL_761933, EPI_ISL_761934 | Lighthouse Lab in Glasgow | Wellcome Sanger Institute for the COVID-19 Genomics UK (COG-UK) Consortium | Harper VanSteenhouse, Yumi Kasai, David Gray, Carol Clugston, Anna Dominiczak and Alex Alderton, Roberto Amato, Sonia Goncalves, Ewan Harrison, David K. Jackson, Ian Johnston, Dominic Kwiatkowski, Cordelia Langford, John Sillitoe on behalf of the Wellcome Sanger Institute COVID-19 Surveillance Team |
| EPI_ISL_761935 | Lighthouse Lab in Cambridge | Wellcome Sanger Institute for the COVID-19 Genomics UK (COG-UK) Consortium | Rob Howes, The Lighthouse Lab in Cambridge and Alex Alderton, Roberto Amato, Sonia Goncalves, Ewan Harrison, David K. Jackson, Ian Johnston, Dominic Kwiatkowski, Cordelia Langford, John Sillitoe on behalf of the Wellcome Sanger Institute COVID-19 Surveillance Team |
| EPI_ISL_761936 | Lighthouse Lab in Glasgow | Wellcome Sanger Institute for the COVID-19 Genomics UK (COG-UK) Consortium | Harper VanSteenhouse, Yumi Kasai, David Gray, Carol Clugston, Anna Dominiczak and Alex Alderton, Roberto Amato, Sonia Goncalves, Ewan Harrison, David K. Jackson, Ian Johnston, Dominic Kwiatkowski, Cordelia Langford, John Sillitoe on behalf of the Wellcome Sanger Institute COVID-19 Surveillance Team |
| EPI_ISL_761937 | Lighthouse Lab in Cambridge | Wellcome Sanger Institute for the COVID-19 Genomics UK (COG-UK) Consortium | Rob Howes, The Lighthouse Lab in Cambridge and Alex Alderton, Roberto Amato, Sonia Goncalves, Ewan Harrison, David K. Jackson, Ian Johnston, Dominic Kwiatkowski, Cordelia Langford, John Sillitoe on behalf of the Wellcome Sanger Institute COVID-19 Surveillance Team |
| EPI_ISL_761938 | Lighthouse Lab in Glasgow | Wellcome Sanger Institute for the COVID-19 Genomics UK (COG-UK) Consortium | Harper VanSteenhouse, Yumi Kasai, David Gray, Carol Clugston, Anna Dominiczak and Alex Alderton, Roberto Amato, Sonia Goncalves, Ewan Harrison, David K. Jackson, Ian Johnston, Dominic Kwiatkowski, Cordelia Langford, John Sillitoe on behalf of the Wellcome Sanger Institute COVID-19 Surveillance Team |
| EPI_ISL_761939 | Lighthouse Lab in Cambridge | Wellcome Sanger Institute for the COVID-19 Genomics UK (COG-UK) Consortium | Rob Howes, The Lighthouse Lab in Cambridge and Alex Alderton, Roberto Amato, Sonia Goncalves, Ewan Harrison, David K. Jackson, Ian Johnston, Dominic Kwiatkowski, Cordelia Langford, John Sillitoe on behalf of the Wellcome Sanger Institute COVID-19 Surveillance Team |
| EPI_ISL_761940, EPI_ISL_761941, EPI_ISL_761942, EPI_ISL_761943, EPI_ISL_761944, EPI_ISL_761945 | Lighthouse Lab in Glasgow | Wellcome Sanger Institute for the COVID-19 Genomics UK (COG-UK) Consortium | Harper VanSteenhouse, Yumi Kasai, David Gray, Carol Clugston, Anna Dominiczak and Alex Alderton, Roberto Amato, Sonia Goncalves, Ewan Harrison, David K. Jackson, Ian Johnston, Dominic Kwiatkowski, Cordelia Langford, John Sillitoe on behalf of the Wellcome Sanger Institute COVID-19 Surveillance Team |
| EPI_ISL_761946 | Lighthouse Lab in Cambridge | Wellcome Sanger Institute for the COVID-19 Genomics UK (COG-UK) Consortium | Rob Howes, The Lighthouse Lab in Cambridge and Alex Alderton, Roberto Amato, Sonia Goncalves, Ewan Harrison, David K. Jackson, Ian Johnston, Dominic Kwiatkowski, Cordelia Langford, John Sillitoe on behalf of the Wellcome Sanger Institute COVID-19 Surveillance Team |
| EPI_ISL_761947, EPI_ISL_761948, EPI_ISL_761949, EPI_ISL_761950, EPI_ISL_761951 | Lighthouse Lab in Glasgow | Wellcome Sanger Institute for the COVID-19 Genomics UK (COG-UK) Consortium | Harper VanSteenhouse, Yumi Kasai, David Gray, Carol Clugston, Anna Dominiczak and Alex Alderton, Roberto Amato, Sonia Goncalves, Ewan Harrison, David K. Jackson, Ian Johnston, Dominic Kwiatkowski, Cordelia Langford, John Sillitoe on behalf of the Wellcome Sanger Institute COVID-19 Surveillance Team |
| EPI_ISL_761952, EPI_ISL_761953 | Lighthouse Lab in Cambridge | Wellcome Sanger Institute for the COVID-19 Genomics UK (COG-UK) Consortium | Rob Howes, The Lighthouse Lab in Cambridge and Alex Alderton, Roberto Amato, Sonia Goncalves, Ewan Harrison, David K. Jackson, Ian Johnston, Dominic Kwiatkowski, Cordelia Langford, John Sillitoe on behalf of the Wellcome Sanger Institute COVID-19 Surveillance Team |
| EPI_ISL_761954 | Lighthouse Lab in Glasgow | Wellcome Sanger Institute for the COVID-19 Genomics UK (COG-UK) Consortium | Harper VanSteenhouse, Yumi Kasai, David Gray, Carol Clugston, Anna Dominiczak and Alex Alderton, Roberto Amato, Sonia Goncalves, Ewan Harrison, David K. Jackson, Ian Johnston, Dominic Kwiatkowski, Cordelia Langford, John Sillitoe on behalf of the Wellcome Sanger Institute COVID-19 Surveillance Team |
| EPI_ISL_761955, EPI_ISL_761956 | Lighthouse Lab in Cambridge | Wellcome Sanger Institute for the COVID-19 Genomics UK (COG-UK) Consortium | Rob Howes, The Lighthouse Lab in Cambridge and Alex Alderton, Roberto Amato, Sonia Goncalves, Ewan Harrison, David K. Jackson, Ian Johnston, Dominic Kwiatkowski, Cordelia Langford, John Sillitoe on behalf of the Wellcome Sanger Institute COVID-19 Surveillance Team |
| EPI_ISL_761957, EPI_ISL_761958, EPI_ISL_761959, EPI_ISL_761960 | Lighthouse Lab in Glasgow | Wellcome Sanger Institute for the COVID-19 Genomics UK (COG-UK) Consortium | Harper VanSteenhouse, Yumi Kasai, David Gray, Carol Clugston, Anna Dominiczak and Alex Alderton, Roberto Amato, Sonia Goncalves, Ewan Harrison, David K. Jackson, Ian Johnston, Dominic Kwiatkowski, Cordelia Langford, John Sillitoe on behalf of the Wellcome Sanger Institute COVID-19 Surveillance Team |
| EPI_ISL_761961 | Lighthouse Lab in Cambridge | Wellcome Sanger Institute for the COVID-19 Genomics UK (COG-UK) Consortium | Rob Howes, The Lighthouse Lab in Cambridge and Alex Alderton, Roberto Amato, Sonia Goncalves, Ewan Harrison, David K. Jackson, Ian Johnston, Dominic Kwiatkowski, Cordelia Langford, John Sillitoe on behalf of the Wellcome Sanger Institute COVID-19 Surveillance Team |
| EPI_ISL_761962, EPI_ISL_761963, EPI_ISL_761964, EPI_ISL_761965, EPI_ISL_761966 | Lighthouse Lab in Glasgow | Wellcome Sanger Institute for the COVID-19 Genomics UK (COG-UK) Consortium | Harper VanSteenhouse, Yumi Kasai, David Gray, Carol Clugston, Anna Dominiczak and Alex Alderton, Roberto Amato, Sonia Goncalves, Ewan Harrison, David K. Jackson, Ian Johnston, Dominic Kwiatkowski, Cordelia Langford, John Sillitoe on behalf of the Wellcome Sanger Institute COVID-19 Surveillance Team |
| EPI_ISL_761967, EPI_ISL_761968 | Lighthouse Lab in Cambridge | Wellcome Sanger Institute for the COVID-19 Genomics UK (COG-UK) Consortium | Rob Howes, The Lighthouse Lab in Cambridge and Alex Alderton, Roberto Amato, Sonia Goncalves, Ewan Harrison, David K. Jackson, Ian Johnston, Dominic Kwiatkowski, Cordelia Langford, John Sillitoe on behalf of the Wellcome Sanger Institute COVID-19 Surveillance Team |
| EPI_ISL_761969 | Lighthouse Lab in Glasgow | Wellcome Sanger Institute for the COVID-19 Genomics UK (COG-UK) Consortium | Harper VanSteenhouse, Yumi Kasai, David Gray, Carol Clugston, Anna Dominiczak and Alex Alderton, Roberto Amato, Sonia Goncalves, Ewan Harrison, David K. Jackson, Ian Johnston, Dominic Kwiatkowski, Cordelia Langford, John Sillitoe on behalf of the Wellcome Sanger Institute COVID-19 Surveillance Team |
| EPI_ISL_761970 | Lighthouse Lab in Cambridge | Wellcome Sanger Institute for the COVID-19 Genomics UK (COG-UK) Consortium | Rob Howes, The Lighthouse Lab in Cambridge and Alex Alderton, Roberto Amato, Sonia Goncalves, Ewan Harrison, David K. Jackson, Ian Johnston, Dominic Kwiatkowski, Cordelia Langford, John Sillitoe on behalf of the Wellcome Sanger Institute COVID-19 Surveillance Team |
| EPI_ISL_761971, EPI_ISL_761972, EPI_ISL_761973, EPI_ISL_761974, EPI_ISL_761975, EPI_ISL_761976, | Lighthouse Lab in Glasgow | Wellcome Sanger Institute for the COVID-19 Genomics UK (COG-UK) Consortium | Harper VanSteenhouse, Yumi Kasai, David Gray, Carol Clugston, Anna Dominiczak and Alex Alderton, Roberto Amato, Sonia Goncalves, Ewan Harrison, David K. Jackson, Ian Johnston, Dominic Kwiatkowski, Cordelia Langford, John Sillitoe on behalf of the Wellcome Sanger Institute COVID-19 Surveillance Team |

| | | | |
|---|---|---|---|
| EPI_ISL_761977, EPI_ISL_761978 | | | |
| EPI_ISL_761979 | Lighthouse Lab in Cambridge | Wellcome Sanger Institute for the COVID-19 Genomics UK (COG-UK) Consortium | Rob Howes, The Lighthouse Lab in Cambridge and Alex Alderton, Roberto Amato, Sonia Goncalves, Ewan Harrison, David K. Jackson, Ian Johnston, Dominic Kwiatkowski, Cordelia Langford, John Sillitoe on behalf of the Wellcome Sanger Institute COVID-19 Surveillance Team |
| EPI_ISL_761980 | Lighthouse Lab in Glasgow | Wellcome Sanger Institute for the COVID-19 Genomics UK (COG-UK) Consortium | Harper VanSteenhouse, Yumi Kasai, David Gray, Carol Clugston, Anna Dominiczak and Alex Alderton, Roberto Amato, Sonia Goncalves, Ewan Harrison, David K. Jackson, Ian Johnston, Dominic Kwiatkowski, Cordelia Langford, John Sillitoe on behalf of the Wellcome Sanger Institute COVID-19 Surveillance Team |
| EPI_ISL_761981 | Lighthouse Lab in Cambridge | Wellcome Sanger Institute for the COVID-19 Genomics UK (COG-UK) Consortium | Rob Howes, The Lighthouse Lab in Cambridge and Alex Alderton, Roberto Amato, Sonia Goncalves, Ewan Harrison, David K. Jackson, Ian Johnston, Dominic Kwiatkowski, Cordelia Langford, John Sillitoe on behalf of the Wellcome Sanger Institute COVID-19 Surveillance Team |
| EPI_ISL_761982, EPI_ISL_761983, EPI_ISL_761984, EPI_ISL_761986, EPI_ISL_761987, EPI_ISL_761988, EPI_ISL_761989, EPI_ISL_761990, EPI_ISL_761991, EPI_ISL_761992, EPI_ISL_761993 | | | |
| see above | Lighthouse Lab in Glasgow | Wellcome Sanger Institute for the COVID-19 Genomics UK (COG-UK) Consortium | Harper VanSteenhouse, Yumi Kasai, David Gray, Carol Clugston, Anna Dominiczak and Alex Alderton, Roberto Amato, Sonia Goncalves, Ewan Harrison, David K. Jackson, Ian Johnston, Dominic Kwiatkowski, Cordelia Langford, John Sillitoe on behalf of the Wellcome Sanger Institute COVID-19 Surveillance Team |
| EPI_ISL_761994, EPI_ISL_761995 | Lighthouse Lab in Cambridge | Wellcome Sanger Institute for the COVID-19 Genomics UK (COG-UK) Consortium | Rob Howes, The Lighthouse Lab in Cambridge and Alex Alderton, Roberto Amato, Sonia Goncalves, Ewan Harrison, David K. Jackson, Ian Johnston, Dominic Kwiatkowski, Cordelia Langford, John Sillitoe on behalf of the Wellcome Sanger Institute COVID-19 Surveillance Team |
| EPI_ISL_761996 | Lighthouse Lab in Glasgow | Wellcome Sanger Institute for the COVID-19 Genomics UK (COG-UK) Consortium | Harper VanSteenhouse, Yumi Kasai, David Gray, Carol Clugston, Anna Dominiczak and Alex Alderton, Roberto Amato, Sonia Goncalves, Ewan Harrison, David K. Jackson, Ian Johnston, Dominic Kwiatkowski, Cordelia Langford, John Sillitoe on behalf of the Wellcome Sanger Institute COVID-19 Surveillance Team |
| EPI_ISL_761997, EPI_ISL_761999 | Lighthouse Lab in Cambridge | Wellcome Sanger Institute for the COVID-19 Genomics UK (COG-UK) Consortium | Rob Howes, The Lighthouse Lab in Cambridge and Alex Alderton, Roberto Amato, Sonia Goncalves, Ewan Harrison, David K. Jackson, Ian Johnston, Dominic Kwiatkowski, Cordelia Langford, John Sillitoe on behalf of the Wellcome Sanger Institute COVID-19 Surveillance Team |
| EPI_ISL_762000, EPI_ISL_762001, EPI_ISL_762002, EPI_ISL_762003, EPI_ISL_762004, EPI_ISL_762005, EPI_ISL_762006 | Lighthouse Lab in Glasgow | Wellcome Sanger Institute for the COVID-19 Genomics UK (COG-UK) Consortium | Harper VanSteenhouse, Yumi Kasai, David Gray, Carol Clugston, Anna Dominiczak and Alex Alderton, Roberto Amato, Sonia Goncalves, Ewan Harrison, David K. Jackson, Ian Johnston, Dominic Kwiatkowski, Cordelia Langford, John Sillitoe on behalf of the Wellcome Sanger Institute COVID-19 Surveillance Team |
| EPI_ISL_762007, EPI_ISL_762008, EPI_ISL_762009 | Lighthouse Lab in Cambridge | Wellcome Sanger Institute for the COVID-19 Genomics UK (COG-UK) Consortium | Rob Howes, The Lighthouse Lab in Cambridge and Alex Alderton, Roberto Amato, Sonia Goncalves, Ewan Harrison, David K. Jackson, Ian Johnston, Dominic Kwiatkowski, Cordelia Langford, John Sillitoe on behalf of the Wellcome Sanger Institute COVID-19 Surveillance Team |
| EPI_ISL_762010, EPI_ISL_762011, EPI_ISL_762012 | Lighthouse Lab in Glasgow | Wellcome Sanger Institute for the COVID-19 Genomics UK (COG-UK) Consortium | Harper VanSteenhouse, Yumi Kasai, David Gray, Carol Clugston, Anna Dominiczak and Alex Alderton, Roberto Amato, Sonia Goncalves, Ewan Harrison, David K. Jackson, Ian Johnston, Dominic Kwiatkowski, Cordelia Langford, John Sillitoe on behalf of the Wellcome Sanger Institute COVID-19 Surveillance Team |
| EPI_ISL_762014, EPI_ISL_762015, EPI_ISL_762016 | Lighthouse Lab in Cambridge | Wellcome Sanger Institute for the COVID-19 Genomics UK (COG-UK) Consortium | Rob Howes, The Lighthouse Lab in Cambridge and Alex Alderton, Roberto Amato, Sonia Goncalves, Ewan Harrison, David K. Jackson, Ian Johnston, Dominic Kwiatkowski, Cordelia Langford, John Sillitoe on behalf of the Wellcome Sanger Institute COVID-19 Surveillance Team |
| EPI_ISL_762017 | Lighthouse Lab in Glasgow | Wellcome Sanger Institute for the COVID-19 Genomics UK (COG-UK) Consortium | Harper VanSteenhouse, Yumi Kasai, David Gray, Carol Clugston, Anna Dominiczak and Alex Alderton, Roberto Amato, Sonia Goncalves, Ewan Harrison, David K. Jackson, Ian Johnston, Dominic Kwiatkowski, Cordelia Langford, John Sillitoe on behalf of the Wellcome Sanger Institute COVID-19 Surveillance Team |
| EPI_ISL_762018, EPI_ISL_762019 | Lighthouse Lab in Cambridge | Wellcome Sanger Institute for the COVID-19 Genomics UK (COG-UK) Consortium | Rob Howes, The Lighthouse Lab in Cambridge and Alex Alderton, Roberto Amato, Sonia Goncalves, Ewan Harrison, David K. Jackson, Ian Johnston, Dominic Kwiatkowski, Cordelia Langford, John Sillitoe on behalf of the Wellcome Sanger Institute COVID-19 Surveillance Team |
| EPI_ISL_762020, EPI_ISL_762021, EPI_ISL_762022 | Lighthouse Lab in Glasgow | Wellcome Sanger Institute for the COVID-19 Genomics UK (COG-UK) Consortium | Harper VanSteenhouse, Yumi Kasai, David Gray, Carol Clugston, Anna Dominiczak and Alex Alderton, Roberto Amato, Sonia Goncalves, Ewan Harrison, David K. Jackson, Ian Johnston, Dominic Kwiatkowski, Cordelia Langford, John Sillitoe on behalf of the Wellcome Sanger Institute COVID-19 Surveillance Team |
| EPI_ISL_762023, EPI_ISL_762024, EPI_ISL_762025, EPI_ISL_762026, EPI_ISL_762027, EPI_ISL_762028, EPI_ISL_762029, EPI_ISL_762030 | Lighthouse Lab in Cambridge | Wellcome Sanger Institute for the COVID-19 Genomics UK (COG-UK) Consortium | Rob Howes, The Lighthouse Lab in Cambridge and Alex Alderton, Roberto Amato, Sonia Goncalves, Ewan Harrison, David K. Jackson, Ian Johnston, Dominic Kwiatkowski, Cordelia Langford, John Sillitoe on behalf of the Wellcome Sanger Institute COVID-19 Surveillance Team |
| EPI_ISL_762031, EPI_ISL_762032, EPI_ISL_762033, EPI_ISL_762034, EPI_ISL_762035 | Lighthouse Lab in Glasgow | Wellcome Sanger Institute for the COVID-19 Genomics UK (COG-UK) Consortium | Harper VanSteenhouse, Yumi Kasai, David Gray, Carol Clugston, Anna Dominiczak and Alex Alderton, Roberto Amato, Sonia Goncalves, Ewan Harrison, David K. Jackson, Ian Johnston, Dominic Kwiatkowski, Cordelia Langford, John Sillitoe on behalf of the Wellcome Sanger Institute COVID-19 Surveillance Team |
| EPI_ISL_762036, EPI_ISL_762037, EPI_ISL_762038 | Lighthouse Lab in Cambridge | Wellcome Sanger Institute for the COVID-19 Genomics UK (COG-UK) Consortium | Rob Howes, The Lighthouse Lab in Cambridge and Alex Alderton, Roberto Amato, Sonia Goncalves, Ewan Harrison, David K. Jackson, Ian Johnston, Dominic Kwiatkowski, Cordelia Langford, John Sillitoe on behalf of the Wellcome Sanger Institute COVID-19 Surveillance Team |
| EPI_ISL_762039, EPI_ISL_762040 | Lighthouse Lab in Glasgow | Wellcome Sanger Institute for the COVID-19 Genomics UK (COG-UK) Consortium | Harper VanSteenhouse, Yumi Kasai, David Gray, Carol Clugston, Anna Dominiczak and Alex Alderton, Roberto Amato, Sonia Goncalves, Ewan Harrison, David K. Jackson, Ian Johnston, Dominic Kwiatkowski, Cordelia Langford, John Sillitoe on behalf of the Wellcome Sanger Institute COVID-19 Surveillance Team |
| EPI_ISL_762041, EPI_ISL_762042, EPI_ISL_762043 | Lighthouse Lab in Cambridge | Wellcome Sanger Institute for the COVID-19 Genomics UK (COG-UK) Consortium | Rob Howes, The Lighthouse Lab in Cambridge and Alex Alderton, Roberto Amato, Sonia Goncalves, Ewan Harrison, David K. Jackson, Ian Johnston, Dominic Kwiatkowski, Cordelia Langford, John Sillitoe on behalf of the Wellcome Sanger Institute COVID-19 Surveillance Team |
| EPI_ISL_762044 | Lighthouse Lab in Glasgow | Wellcome Sanger Institute for the COVID-19 Genomics UK (COG-UK) Consortium | Harper VanSteenhouse, Yumi Kasai, David Gray, Carol Clugston, Anna Dominiczak and Alex Alderton, Roberto Amato, Sonia Goncalves, Ewan Harrison, David K. Jackson, Ian Johnston, Dominic Kwiatkowski, Cordelia Langford, John Sillitoe on behalf of the Wellcome Sanger Institute COVID-19 Surveillance Team |
| EPI_ISL_762045 | Lighthouse Lab in Cambridge | Wellcome Sanger Institute for the COVID-19 Genomics UK (COG-UK) Consortium | Rob Howes, The Lighthouse Lab in Cambridge and Alex Alderton, Roberto Amato, Sonia Goncalves, Ewan Harrison, David K. Jackson, Ian Johnston, Dominic Kwiatkowski, Cordelia Langford, John Sillitoe on behalf of the Wellcome Sanger Institute COVID-19 Surveillance Team |
| EPI_ISL_762046, EPI_ISL_762047, EPI_ISL_762048, EPI_ISL_762049 | Lighthouse Lab in Glasgow | Wellcome Sanger Institute for the COVID-19 Genomics UK (COG-UK) Consortium | Harper VanSteenhouse, Yumi Kasai, David Gray, Carol Clugston, Anna Dominiczak and Alex Alderton, Roberto Amato, Sonia Goncalves, Ewan Harrison, David K. Jackson, Ian Johnston, Dominic Kwiatkowski, Cordelia Langford, John Sillitoe on behalf of the Wellcome Sanger Institute COVID-19 Surveillance Team |
| EPI_ISL_762050 | Lighthouse Lab in Cambridge | Wellcome Sanger Institute for the COVID-19 Genomics UK (COG-UK) Consortium | Rob Howes, The Lighthouse Lab in Cambridge and Alex Alderton, Roberto Amato, Sonia Goncalves, Ewan Harrison, David K. Jackson, Ian Johnston, Dominic Kwiatkowski, Cordelia Langford, John Sillitoe on behalf of the Wellcome Sanger Institute COVID-19 Surveillance Team |
| EPI_ISL_762051, EPI_ISL_762052, EPI_ISL_762053, EPI_ISL_762054, EPI_ISL_762055, EPI_ISL_762056 | Lighthouse Lab in Glasgow | Wellcome Sanger Institute for the COVID-19 Genomics UK (COG-UK) Consortium | Harper VanSteenhouse, Yumi Kasai, David Gray, Carol Clugston, Anna Dominiczak and Alex Alderton, Roberto Amato, Sonia Goncalves, Ewan Harrison, David K. Jackson, Ian Johnston, Dominic Kwiatkowski, Cordelia Langford, John Sillitoe on behalf of the Wellcome Sanger Institute COVID-19 Surveillance Team |
| EPI_ISL_762057 | Lighthouse Lab in Cambridge | Wellcome Sanger Institute for the COVID-19 Genomics UK (COG-UK) Consortium | Rob Howes, The Lighthouse Lab in Cambridge and Alex Alderton, Roberto Amato, Sonia Goncalves, Ewan Harrison, David K. Jackson, Ian Johnston, Dominic Kwiatkowski, Cordelia Langford, John Sillitoe on behalf of the Wellcome Sanger Institute COVID-19 Surveillance Team |
| EPI_ISL_762058 | Lighthouse Lab in Glasgow | Wellcome Sanger Institute for the COVID-19 Genomics UK (COG-UK) Consortium | Harper VanSteenhouse, Yumi Kasai, David Gray, Carol Clugston, Anna Dominiczak and Alex Alderton, Roberto Amato, Sonia Goncalves, Ewan Harrison, David K. Jackson, Ian Johnston, Dominic Kwiatkowski, Cordelia Langford, John Sillitoe on behalf of the Wellcome Sanger Institute COVID-19 Surveillance Team |
| EPI_ISL_762059, EPI_ISL_762060, EPI_ISL_762061, EPI_ISL_762063, EPI_ISL_762064, EPI_ISL_762065, | Lighthouse Lab in Cambridge | Wellcome Sanger Institute for the COVID-19 Genomics UK (COG-UK) Consortium | Rob Howes, The Lighthouse Lab in Cambridge and Alex Alderton, Roberto Amato, Sonia Goncalves, Ewan Harrison, David K. Jackson, Ian Johnston, Dominic Kwiatkowski, Cordelia Langford, John Sillitoe on behalf of the Wellcome Sanger Institute COVID-19 Surveillance Team |

| | | | |
|---|---|---|---|
| EPI_ISL_762066, EPI_ISL_762067 | | | |
| EPI_ISL_762068 | Lighthouse Lab in Glasgow | Wellcome Sanger Institute for the COVID-19 Genomics UK (COG-UK) Consortium | Harper VanSteenhouse, Yumi Kasai, David Gray, Carol Clugston, Anna Dominiczak and Alex Alderton, Roberto Amato, Sonia Goncalves, Ewan Harrison, David K. Jackson, Ian Johnston, Dominic Kwiatkowski, Cordelia Langford, John Sillitoe on behalf of the Wellcome Sanger Institute COVID-19 Surveillance Team |
| EPI_ISL_762069 | Lighthouse Lab in Cambridge | Wellcome Sanger Institute for the COVID-19 Genomics UK (COG-UK) Consortium | Rob Howes, The Lighthouse Lab in Cambridge and Alex Alderton, Roberto Amato, Sonia Goncalves, Ewan Harrison, David K. Jackson, Ian Johnston, Dominic Kwiatkowski, Cordelia Langford, John Sillitoe on behalf of the Wellcome Sanger Institute COVID-19 Surveillance Team |
| EPI_ISL_762070, EPI_ISL_762071 | Lighthouse Lab in Glasgow | Wellcome Sanger Institute for the COVID-19 Genomics UK (COG-UK) Consortium | Harper VanSteenhouse, Yumi Kasai, David Gray, Carol Clugston, Anna Dominiczak and Alex Alderton, Roberto Amato, Sonia Goncalves, Ewan Harrison, David K. Jackson, Ian Johnston, Dominic Kwiatkowski, Cordelia Langford, John Sillitoe on behalf of the Wellcome Sanger Institute COVID-19 Surveillance Team |
| EPI_ISL_762072, EPI_ISL_762073, EPI_ISL_762074, EPI_ISL_762075 | Lighthouse Lab in Cambridge | Wellcome Sanger Institute for the COVID-19 Genomics UK (COG-UK) Consortium | Rob Howes, The Lighthouse Lab in Cambridge and Alex Alderton, Roberto Amato, Sonia Goncalves, Ewan Harrison, David K. Jackson, Ian Johnston, Dominic Kwiatkowski, Cordelia Langford, John Sillitoe on behalf of the Wellcome Sanger Institute COVID-19 Surveillance Team |
| EPI_ISL_762076, EPI_ISL_762077 | Lighthouse Lab in Glasgow | Wellcome Sanger Institute for the COVID-19 Genomics UK (COG-UK) Consortium | Harper VanSteenhouse, Yumi Kasai, David Gray, Carol Clugston, Anna Dominiczak and Alex Alderton, Roberto Amato, Sonia Goncalves, Ewan Harrison, David K. Jackson, Ian Johnston, Dominic Kwiatkowski, Cordelia Langford, John Sillitoe on behalf of the Wellcome Sanger Institute COVID-19 Surveillance Team |
| EPI_ISL_762078 | Lighthouse Lab in Cambridge | Wellcome Sanger Institute for the COVID-19 Genomics UK (COG-UK) Consortium | Rob Howes, The Lighthouse Lab in Cambridge and Alex Alderton, Roberto Amato, Sonia Goncalves, Ewan Harrison, David K. Jackson, Ian Johnston, Dominic Kwiatkowski, Cordelia Langford, John Sillitoe on behalf of the Wellcome Sanger Institute COVID-19 Surveillance Team |
| EPI_ISL_762079, EPI_ISL_762080, EPI_ISL_762081, EPI_ISL_762082, EPI_ISL_762084 | Lighthouse Lab in Glasgow | Wellcome Sanger Institute for the COVID-19 Genomics UK (COG-UK) Consortium | Harper VanSteenhouse, Yumi Kasai, David Gray, Carol Clugston, Anna Dominiczak and Alex Alderton, Roberto Amato, Sonia Goncalves, Ewan Harrison, David K. Jackson, Ian Johnston, Dominic Kwiatkowski, Cordelia Langford, John Sillitoe on behalf of the Wellcome Sanger Institute COVID-19 Surveillance Team |
| EPI_ISL_762085 | Lighthouse Lab in Cambridge | Wellcome Sanger Institute for the COVID-19 Genomics UK (COG-UK) Consortium | Rob Howes, The Lighthouse Lab in Cambridge and Alex Alderton, Roberto Amato, Sonia Goncalves, Ewan Harrison, David K. Jackson, Ian Johnston, Dominic Kwiatkowski, Cordelia Langford, John Sillitoe on behalf of the Wellcome Sanger Institute COVID-19 Surveillance Team |
| EPI_ISL_762086 | Lighthouse Lab in Glasgow | Wellcome Sanger Institute for the COVID-19 Genomics UK (COG-UK) Consortium | Harper VanSteenhouse, Yumi Kasai, David Gray, Carol Clugston, Anna Dominiczak and Alex Alderton, Roberto Amato, Sonia Goncalves, Ewan Harrison, David K. Jackson, Ian Johnston, Dominic Kwiatkowski, Cordelia Langford, John Sillitoe on behalf of the Wellcome Sanger Institute COVID-19 Surveillance Team |
| EPI_ISL_762087 | Lighthouse Lab in Cambridge | Wellcome Sanger Institute for the COVID-19 Genomics UK (COG-UK) Consortium | Rob Howes, The Lighthouse Lab in Cambridge and Alex Alderton, Roberto Amato, Sonia Goncalves, Ewan Harrison, David K. Jackson, Ian Johnston, Dominic Kwiatkowski, Cordelia Langford, John Sillitoe on behalf of the Wellcome Sanger Institute COVID-19 Surveillance Team |
| EPI_ISL_762088 | Lighthouse Lab in Glasgow | Wellcome Sanger Institute for the COVID-19 Genomics UK (COG-UK) Consortium | Harper VanSteenhouse, Yumi Kasai, David Gray, Carol Clugston, Anna Dominiczak and Alex Alderton, Roberto Amato, Sonia Goncalves, Ewan Harrison, David K. Jackson, Ian Johnston, Dominic Kwiatkowski, Cordelia Langford, John Sillitoe on behalf of the Wellcome Sanger Institute COVID-19 Surveillance Team |
| EPI_ISL_762089, EPI_ISL_762090, EPI_ISL_762091 | Lighthouse Lab in Cambridge | Wellcome Sanger Institute for the COVID-19 Genomics UK (COG-UK) Consortium | Rob Howes, The Lighthouse Lab in Cambridge and Alex Alderton, Roberto Amato, Sonia Goncalves, Ewan Harrison, David K. Jackson, Ian Johnston, Dominic Kwiatkowski, Cordelia Langford, John Sillitoe on behalf of the Wellcome Sanger Institute COVID-19 Surveillance Team |
| EPI_ISL_762092, EPI_ISL_762093, EPI_ISL_762094, EPI_ISL_762095 | Lighthouse Lab in Glasgow | Wellcome Sanger Institute for the COVID-19 Genomics UK (COG-UK) Consortium | Harper VanSteenhouse, Yumi Kasai, David Gray, Carol Clugston, Anna Dominiczak and Alex Alderton, Roberto Amato, Sonia Goncalves, Ewan Harrison, David K. Jackson, Ian Johnston, Dominic Kwiatkowski, Cordelia Langford, John Sillitoe on behalf of the Wellcome Sanger Institute COVID-19 Surveillance Team |
| EPI_ISL_762096, EPI_ISL_762097, EPI_ISL_762098 | Lighthouse Lab in Cambridge | Wellcome Sanger Institute for the COVID-19 Genomics UK (COG-UK) Consortium | Rob Howes, The Lighthouse Lab in Cambridge and Alex Alderton, Roberto Amato, Sonia Goncalves, Ewan Harrison, David K. Jackson, Ian Johnston, Dominic Kwiatkowski, Cordelia Langford, John Sillitoe on behalf of the Wellcome Sanger Institute COVID-19 Surveillance Team |
| EPI_ISL_762099, EPI_ISL_762100, EPI_ISL_762101, EPI_ISL_762102 | Lighthouse Lab in Glasgow | Wellcome Sanger Institute for the COVID-19 Genomics UK (COG-UK) Consortium | Harper VanSteenhouse, Yumi Kasai, David Gray, Carol Clugston, Anna Dominiczak and Alex Alderton, Roberto Amato, Sonia Goncalves, Ewan Harrison, David K. Jackson, Ian Johnston, Dominic Kwiatkowski, Cordelia Langford, John Sillitoe on behalf of the Wellcome Sanger Institute COVID-19 Surveillance Team |
| EPI_ISL_762103 | Lighthouse Lab in Cambridge | Wellcome Sanger Institute for the COVID-19 Genomics UK (COG-UK) Consortium | Rob Howes, The Lighthouse Lab in Cambridge and Alex Alderton, Roberto Amato, Sonia Goncalves, Ewan Harrison, David K. Jackson, Ian Johnston, Dominic Kwiatkowski, Cordelia Langford, John Sillitoe on behalf of the Wellcome Sanger Institute COVID-19 Surveillance Team |
| EPI_ISL_762104, EPI_ISL_762105, EPI_ISL_762106, EPI_ISL_762107, EPI_ISL_762108 | Lighthouse Lab in Glasgow | Wellcome Sanger Institute for the COVID-19 Genomics UK (COG-UK) Consortium | Harper VanSteenhouse, Yumi Kasai, David Gray, Carol Clugston, Anna Dominiczak and Alex Alderton, Roberto Amato, Sonia Goncalves, Ewan Harrison, David K. Jackson, Ian Johnston, Dominic Kwiatkowski, Cordelia Langford, John Sillitoe on behalf of the Wellcome Sanger Institute COVID-19 Surveillance Team |
| EPI_ISL_762109 | Lighthouse Lab in Cambridge | Wellcome Sanger Institute for the COVID-19 Genomics UK (COG-UK) Consortium | Rob Howes, The Lighthouse Lab in Cambridge and Alex Alderton, Roberto Amato, Sonia Goncalves, Ewan Harrison, David K. Jackson, Ian Johnston, Dominic Kwiatkowski, Cordelia Langford, John Sillitoe on behalf of the Wellcome Sanger Institute COVID-19 Surveillance Team |
| EPI_ISL_762111 | Lighthouse Lab in Glasgow | Wellcome Sanger Institute for the COVID-19 Genomics UK (COG-UK) Consortium | Harper VanSteenhouse, Yumi Kasai, David Gray, Carol Clugston, Anna Dominiczak and Alex Alderton, Roberto Amato, Sonia Goncalves, Ewan Harrison, David K. Jackson, Ian Johnston, Dominic Kwiatkowski, Cordelia Langford, John Sillitoe on behalf of the Wellcome Sanger Institute COVID-19 Surveillance Team |
| EPI_ISL_762112, EPI_ISL_762113, EPI_ISL_762114 | Lighthouse Lab in Cambridge | Wellcome Sanger Institute for the COVID-19 Genomics UK (COG-UK) Consortium | Rob Howes, The Lighthouse Lab in Cambridge and Alex Alderton, Roberto Amato, Sonia Goncalves, Ewan Harrison, David K. Jackson, Ian Johnston, Dominic Kwiatkowski, Cordelia Langford, John Sillitoe on behalf of the Wellcome Sanger Institute COVID-19 Surveillance Team |
| EPI_ISL_762116 | Lighthouse Lab in Glasgow | Wellcome Sanger Institute for the COVID-19 Genomics UK (COG-UK) Consortium | Harper VanSteenhouse, Yumi Kasai, David Gray, Carol Clugston, Anna Dominiczak and Alex Alderton, Roberto Amato, Sonia Goncalves, Ewan Harrison, David K. Jackson, Ian Johnston, Dominic Kwiatkowski, Cordelia Langford, John Sillitoe on behalf of the Wellcome Sanger Institute COVID-19 Surveillance Team |
| EPI_ISL_762117, EPI_ISL_762118 | Lighthouse Lab in Cambridge | Wellcome Sanger Institute for the COVID-19 Genomics UK (COG-UK) Consortium | Rob Howes, The Lighthouse Lab in Cambridge and Alex Alderton, Roberto Amato, Sonia Goncalves, Ewan Harrison, David K. Jackson, Ian Johnston, Dominic Kwiatkowski, Cordelia Langford, John Sillitoe on behalf of the Wellcome Sanger Institute COVID-19 Surveillance Team |
| EPI_ISL_762119, EPI_ISL_762120, EPI_ISL_762121 | Lighthouse Lab in Glasgow | Wellcome Sanger Institute for the COVID-19 Genomics UK (COG-UK) Consortium | Harper VanSteenhouse, Yumi Kasai, David Gray, Carol Clugston, Anna Dominiczak and Alex Alderton, Roberto Amato, Sonia Goncalves, Ewan Harrison, David K. Jackson, Ian Johnston, Dominic Kwiatkowski, Cordelia Langford, John Sillitoe on behalf of the Wellcome Sanger Institute COVID-19 Surveillance Team |
| EPI_ISL_762122 | Lighthouse Lab in Cambridge | Wellcome Sanger Institute for the COVID-19 Genomics UK (COG-UK) Consortium | Rob Howes, The Lighthouse Lab in Cambridge and Alex Alderton, Roberto Amato, Sonia Goncalves, Ewan Harrison, David K. Jackson, Ian Johnston, Dominic Kwiatkowski, Cordelia Langford, John Sillitoe on behalf of the Wellcome Sanger Institute COVID-19 Surveillance Team |
| EPI_ISL_762123 | Lighthouse Lab in Glasgow | Wellcome Sanger Institute for the COVID-19 Genomics UK (COG-UK) Consortium | Harper VanSteenhouse, Yumi Kasai, David Gray, Carol Clugston, Anna Dominiczak and Alex Alderton, Roberto Amato, Sonia Goncalves, Ewan Harrison, David K. Jackson, Ian Johnston, Dominic Kwiatkowski, Cordelia Langford, John Sillitoe on behalf of the Wellcome Sanger Institute COVID-19 Surveillance Team |
| EPI_ISL_762124, EPI_ISL_762125, EPI_ISL_762126 | Lighthouse Lab in Cambridge | Wellcome Sanger Institute for the COVID-19 Genomics UK (COG-UK) Consortium | Rob Howes, The Lighthouse Lab in Cambridge and Alex Alderton, Roberto Amato, Sonia Goncalves, Ewan Harrison, David K. Jackson, Ian Johnston, Dominic Kwiatkowski, Cordelia Langford, John Sillitoe on behalf of the Wellcome Sanger Institute COVID-19 Surveillance Team |
| EPI_ISL_762127, EPI_ISL_762128, EPI_ISL_762129, EPI_ISL_762130 | Lighthouse Lab in Glasgow | Wellcome Sanger Institute for the COVID-19 Genomics UK (COG-UK) Consortium | Harper VanSteenhouse, Yumi Kasai, David Gray, Carol Clugston, Anna Dominiczak and Alex Alderton, Roberto Amato, Sonia Goncalves, Ewan Harrison, David K. Jackson, Ian Johnston, Dominic Kwiatkowski, Cordelia Langford, John Sillitoe on behalf of the Wellcome Sanger Institute COVID-19 Surveillance Team |
| EPI_ISL_762131 | Lighthouse Lab in Cambridge | Wellcome Sanger Institute for the COVID-19 Genomics UK (COG-UK) Consortium | Rob Howes, The Lighthouse Lab in Cambridge and Alex Alderton, Roberto Amato, Sonia Goncalves, Ewan Harrison, David K. Jackson, Ian Johnston, Dominic Kwiatkowski, Cordelia Langford, John Sillitoe on behalf of the Wellcome Sanger Institute COVID-19 Surveillance Team |
| EPI_ISL_762132 | Lighthouse Lab in Glasgow | Wellcome Sanger Institute for the COVID-19 Genomics UK (COG-UK) Consortium | Harper VanSteenhouse, Yumi Kasai, David Gray, Carol Clugston, Anna Dominiczak and Alex Alderton, Roberto Amato, Sonia Goncalves, Ewan Harrison, David K. Jackson, Ian Johnston, Dominic Kwiatkowski, Cordelia Langford, John Sillitoe on behalf of the Wellcome Sanger Institute COVID-19 Surveillance |

| | | | |
|---|---|---|---|
| | | | Team |
| EPI_ISL_762133 | Lighthouse Lab in Cambridge | Wellcome Sanger Institute for the COVID-19 Genomics UK (COG-UK) Consortium | Rob Howes, The Lighthouse Lab in Cambridge and Alex Alderton, Roberto Amato, Sonia Goncalves, Ewan Harrison, David K. Jackson, Ian Johnston, Dominic Kwiatkowski, Cordelia Langford, John Sillitoe on behalf of the Wellcome Sanger Institute COVID-19 Surveillance Team |
| EPI_ISL_762134 | Lighthouse Lab in Glasgow | Wellcome Sanger Institute for the COVID-19 Genomics UK (COG-UK) Consortium | Harper VanSteenhouse, Yumi Kasai, David Gray, Carol Clugston, Anna Dominiczak and Alex Alderton, Roberto Amato, Sonia Goncalves, Ewan Harrison, David K. Jackson, Ian Johnston, Dominic Kwiatkowski, Cordelia Langford, John Sillitoe on behalf of the Wellcome Sanger Institute COVID-19 Surveillance Team |
| EPI_ISL_762135, EPI_ISL_762136, EPI_ISL_762137, EPI_ISL_762138, EPI_ISL_762139, EPI_ISL_762140 | Lighthouse Lab in Cambridge | Wellcome Sanger Institute for the COVID-19 Genomics UK (COG-UK) Consortium | Rob Howes, The Lighthouse Lab in Cambridge and Alex Alderton, Roberto Amato, Sonia Goncalves, Ewan Harrison, David K. Jackson, Ian Johnston, Dominic Kwiatkowski, Cordelia Langford, John Sillitoe on behalf of the Wellcome Sanger Institute COVID-19 Surveillance Team |
| EPI_ISL_762141 | Lighthouse Lab in Glasgow | Wellcome Sanger Institute for the COVID-19 Genomics UK (COG-UK) Consortium | Harper VanSteenhouse, Yumi Kasai, David Gray, Carol Clugston, Anna Dominiczak and Alex Alderton, Roberto Amato, Sonia Goncalves, Ewan Harrison, David K. Jackson, Ian Johnston, Dominic Kwiatkowski, Cordelia Langford, John Sillitoe on behalf of the Wellcome Sanger Institute COVID-19 Surveillance Team |
| EPI_ISL_762142 | Lighthouse Lab in Cambridge | Wellcome Sanger Institute for the COVID-19 Genomics UK (COG-UK) Consortium | Rob Howes, The Lighthouse Lab in Cambridge and Alex Alderton, Roberto Amato, Sonia Goncalves, Ewan Harrison, David K. Jackson, Ian Johnston, Dominic Kwiatkowski, Cordelia Langford, John Sillitoe on behalf of the Wellcome Sanger Institute COVID-19 Surveillance Team |
| EPI_ISL_762143, EPI_ISL_762144, EPI_ISL_762145 | Lighthouse Lab in Glasgow | Wellcome Sanger Institute for the COVID-19 Genomics UK (COG-UK) Consortium | Harper VanSteenhouse, Yumi Kasai, David Gray, Carol Clugston, Anna Dominiczak and Alex Alderton, Roberto Amato, Sonia Goncalves, Ewan Harrison, David K. Jackson, Ian Johnston, Dominic Kwiatkowski, Cordelia Langford, John Sillitoe on behalf of the Wellcome Sanger Institute COVID-19 Surveillance Team |
| EPI_ISL_762146, EPI_ISL_762147, EPI_ISL_762148 | Lighthouse Lab in Cambridge | Wellcome Sanger Institute for the COVID-19 Genomics UK (COG-UK) Consortium | Rob Howes, The Lighthouse Lab in Cambridge and Alex Alderton, Roberto Amato, Sonia Goncalves, Ewan Harrison, David K. Jackson, Ian Johnston, Dominic Kwiatkowski, Cordelia Langford, John Sillitoe on behalf of the Wellcome Sanger Institute COVID-19 Surveillance Team |
| EPI_ISL_762149, EPI_ISL_762150, EPI_ISL_762151 | Lighthouse Lab in Glasgow | Wellcome Sanger Institute for the COVID-19 Genomics UK (COG-UK) Consortium | Harper VanSteenhouse, Yumi Kasai, David Gray, Carol Clugston, Anna Dominiczak and Alex Alderton, Roberto Amato, Sonia Goncalves, Ewan Harrison, David K. Jackson, Ian Johnston, Dominic Kwiatkowski, Cordelia Langford, John Sillitoe on behalf of the Wellcome Sanger Institute COVID-19 Surveillance Team |
| EPI_ISL_762152, EPI_ISL_762153 | Lighthouse Lab in Cambridge | Wellcome Sanger Institute for the COVID-19 Genomics UK (COG-UK) Consortium | Rob Howes, The Lighthouse Lab in Cambridge and Alex Alderton, Roberto Amato, Sonia Goncalves, Ewan Harrison, David K. Jackson, Ian Johnston, Dominic Kwiatkowski, Cordelia Langford, John Sillitoe on behalf of the Wellcome Sanger Institute COVID-19 Surveillance Team |
| EPI_ISL_762154 | Lighthouse Lab in Glasgow | Wellcome Sanger Institute for the COVID-19 Genomics UK (COG-UK) Consortium | Harper VanSteenhouse, Yumi Kasai, David Gray, Carol Clugston, Anna Dominiczak and Alex Alderton, Roberto Amato, Sonia Goncalves, Ewan Harrison, David K. Jackson, Ian Johnston, Dominic Kwiatkowski, Cordelia Langford, John Sillitoe on behalf of the Wellcome Sanger Institute COVID-19 Surveillance Team |
| EPI_ISL_762155 | Lighthouse Lab in Cambridge | Wellcome Sanger Institute for the COVID-19 Genomics UK (COG-UK) Consortium | Rob Howes, The Lighthouse Lab in Cambridge and Alex Alderton, Roberto Amato, Sonia Goncalves, Ewan Harrison, David K. Jackson, Ian Johnston, Dominic Kwiatkowski, Cordelia Langford, John Sillitoe on behalf of the Wellcome Sanger Institute COVID-19 Surveillance Team |
| EPI_ISL_762157, EPI_ISL_762158, EPI_ISL_762159, EPI_ISL_762160, EPI_ISL_762161, EPI_ISL_762162 | Lighthouse Lab in Glasgow | Wellcome Sanger Institute for the COVID-19 Genomics UK (COG-UK) Consortium | Harper VanSteenhouse, Yumi Kasai, David Gray, Carol Clugston, Anna Dominiczak and Alex Alderton, Roberto Amato, Sonia Goncalves, Ewan Harrison, David K. Jackson, Ian Johnston, Dominic Kwiatkowski, Cordelia Langford, John Sillitoe on behalf of the Wellcome Sanger Institute COVID-19 Surveillance Team |
| EPI_ISL_762163 | Lighthouse Lab in Cambridge | Wellcome Sanger Institute for the COVID-19 Genomics UK (COG-UK) Consortium | Rob Howes, The Lighthouse Lab in Cambridge and Alex Alderton, Roberto Amato, Sonia Goncalves, Ewan Harrison, David K. Jackson, Ian Johnston, Dominic Kwiatkowski, Cordelia Langford, John Sillitoe on behalf of the Wellcome Sanger Institute COVID-19 Surveillance Team |
| EPI_ISL_762164, EPI_ISL_762165 | Lighthouse Lab in Glasgow | Wellcome Sanger Institute for the COVID-19 Genomics UK (COG-UK) Consortium | Harper VanSteenhouse, Yumi Kasai, David Gray, Carol Clugston, Anna Dominiczak and Alex Alderton, Roberto Amato, Sonia Goncalves, Ewan Harrison, David K. Jackson, Ian Johnston, Dominic Kwiatkowski, Cordelia Langford, John Sillitoe on behalf of the Wellcome Sanger Institute COVID-19 Surveillance Team |
| EPI_ISL_762166 | Lighthouse Lab in Cambridge | Wellcome Sanger Institute for the COVID-19 Genomics UK (COG-UK) Consortium | Rob Howes, The Lighthouse Lab in Cambridge and Alex Alderton, Roberto Amato, Sonia Goncalves, Ewan Harrison, David K. Jackson, Ian Johnston, Dominic Kwiatkowski, Cordelia Langford, John Sillitoe on behalf of the Wellcome Sanger Institute COVID-19 Surveillance Team |
| EPI_ISL_762167, EPI_ISL_762168, EPI_ISL_762170, EPI_ISL_762172 | Lighthouse Lab in Glasgow | Wellcome Sanger Institute for the COVID-19 Genomics UK (COG-UK) Consortium | Harper VanSteenhouse, Yumi Kasai, David Gray, Carol Clugston, Anna Dominiczak and Alex Alderton, Roberto Amato, Sonia Goncalves, Ewan Harrison, David K. Jackson, Ian Johnston, Dominic Kwiatkowski, Cordelia Langford, John Sillitoe on behalf of the Wellcome Sanger Institute COVID-19 Surveillance Team |
| EPI_ISL_762173 | Lighthouse Lab in Cambridge | Wellcome Sanger Institute for the COVID-19 Genomics UK (COG-UK) Consortium | Rob Howes, The Lighthouse Lab in Cambridge and Alex Alderton, Roberto Amato, Sonia Goncalves, Ewan Harrison, David K. Jackson, Ian Johnston, Dominic Kwiatkowski, Cordelia Langford, John Sillitoe on behalf of the Wellcome Sanger Institute COVID-19 Surveillance Team |
| EPI_ISL_762174 | Lighthouse Lab in Glasgow | Wellcome Sanger Institute for the COVID-19 Genomics UK (COG-UK) Consortium | Harper VanSteenhouse, Yumi Kasai, David Gray, Carol Clugston, Anna Dominiczak and Alex Alderton, Roberto Amato, Sonia Goncalves, Ewan Harrison, David K. Jackson, Ian Johnston, Dominic Kwiatkowski, Cordelia Langford, John Sillitoe on behalf of the Wellcome Sanger Institute COVID-19 Surveillance Team |
| EPI_ISL_762175 | Lighthouse Lab in Cambridge | Wellcome Sanger Institute for the COVID-19 Genomics UK (COG-UK) Consortium | Rob Howes, The Lighthouse Lab in Cambridge and Alex Alderton, Roberto Amato, Sonia Goncalves, Ewan Harrison, David K. Jackson, Ian Johnston, Dominic Kwiatkowski, Cordelia Langford, John Sillitoe on behalf of the Wellcome Sanger Institute COVID-19 Surveillance Team |
| EPI_ISL_762176, EPI_ISL_762177 | Lighthouse Lab in Glasgow | Wellcome Sanger Institute for the COVID-19 Genomics UK (COG-UK) Consortium | Harper VanSteenhouse, Yumi Kasai, David Gray, Carol Clugston, Anna Dominiczak and Alex Alderton, Roberto Amato, Sonia Goncalves, Ewan Harrison, David K. Jackson, Ian Johnston, Dominic Kwiatkowski, Cordelia Langford, John Sillitoe on behalf of the Wellcome Sanger Institute COVID-19 Surveillance Team |
| EPI_ISL_762178 | Lighthouse Lab in Cambridge | Wellcome Sanger Institute for the COVID-19 Genomics UK (COG-UK) Consortium | Rob Howes, The Lighthouse Lab in Cambridge and Alex Alderton, Roberto Amato, Sonia Goncalves, Ewan Harrison, David K. Jackson, Ian Johnston, Dominic Kwiatkowski, Cordelia Langford, John Sillitoe on behalf of the Wellcome Sanger Institute COVID-19 Surveillance Team |
| EPI_ISL_762179, EPI_ISL_762180 | Lighthouse Lab in Glasgow | Wellcome Sanger Institute for the COVID-19 Genomics UK (COG-UK) Consortium | Harper VanSteenhouse, Yumi Kasai, David Gray, Carol Clugston, Anna Dominiczak and Alex Alderton, Roberto Amato, Sonia Goncalves, Ewan Harrison, David K. Jackson, Ian Johnston, Dominic Kwiatkowski, Cordelia Langford, John Sillitoe on behalf of the Wellcome Sanger Institute COVID-19 Surveillance Team |
| EPI_ISL_762181 | Lighthouse Lab in Cambridge | Wellcome Sanger Institute for the COVID-19 Genomics UK (COG-UK) Consortium | Rob Howes, The Lighthouse Lab in Cambridge and Alex Alderton, Roberto Amato, Sonia Goncalves, Ewan Harrison, David K. Jackson, Ian Johnston, Dominic Kwiatkowski, Cordelia Langford, John Sillitoe on behalf of the Wellcome Sanger Institute COVID-19 Surveillance Team |
| EPI_ISL_762182 | Lighthouse Lab in Glasgow | Wellcome Sanger Institute for the COVID-19 Genomics UK (COG-UK) Consortium | Harper VanSteenhouse, Yumi Kasai, David Gray, Carol Clugston, Anna Dominiczak and Alex Alderton, Roberto Amato, Sonia Goncalves, Ewan Harrison, David K. Jackson, Ian Johnston, Dominic Kwiatkowski, Cordelia Langford, John Sillitoe on behalf of the Wellcome Sanger Institute COVID-19 Surveillance Team |
| EPI_ISL_762183 | Lighthouse Lab in Cambridge | Wellcome Sanger Institute for the COVID-19 Genomics UK (COG-UK) Consortium | Rob Howes, The Lighthouse Lab in Cambridge and Alex Alderton, Roberto Amato, Sonia Goncalves, Ewan Harrison, David K. Jackson, Ian Johnston, Dominic Kwiatkowski, Cordelia Langford, John Sillitoe on behalf of the Wellcome Sanger Institute COVID-19 Surveillance Team |
| EPI_ISL_762184, EPI_ISL_762185 | Lighthouse Lab in Glasgow | Wellcome Sanger Institute for the COVID-19 Genomics UK (COG-UK) Consortium | Harper VanSteenhouse, Yumi Kasai, David Gray, Carol Clugston, Anna Dominiczak and Alex Alderton, Roberto Amato, Sonia Goncalves, Ewan Harrison, David K. Jackson, Ian Johnston, Dominic Kwiatkowski, Cordelia Langford, John Sillitoe on behalf of the Wellcome Sanger Institute COVID-19 Surveillance Team |
| EPI_ISL_762186, EPI_ISL_762187 | Lighthouse Lab in Cambridge | Wellcome Sanger Institute for the COVID-19 Genomics UK (COG-UK) Consortium | Rob Howes, The Lighthouse Lab in Cambridge and Alex Alderton, Roberto Amato, Sonia Goncalves, Ewan Harrison, David K. Jackson, Ian Johnston, Dominic Kwiatkowski, Cordelia Langford, John Sillitoe on behalf of the Wellcome Sanger Institute COVID-19 Surveillance Team |
| EPI_ISL_762188 | Lighthouse Lab in Glasgow | Wellcome Sanger Institute for the COVID-19 Genomics UK (COG-UK) Consortium | Harper VanSteenhouse, Yumi Kasai, David Gray, Carol Clugston, Anna Dominiczak and Alex Alderton, Roberto Amato, Sonia Goncalves, Ewan Harrison, David K. Jackson, Ian Johnston, Dominic Kwiatkowski, Cordelia Langford, John Sillitoe on behalf of the Wellcome Sanger Institute COVID-19 Surveillance Team |
| EPI_ISL_762189 | Lighthouse Lab in Cambridge | Wellcome Sanger Institute for the COVID-19 Genomics UK | Rob Howes, The Lighthouse Lab in Cambridge and Alex Alderton, Roberto Amato, Sonia Goncalves, Ewan Harrison, David K. Jackson, Ian Johnston, |

| | | | |
|---|---|---|---|
| | | (COG-UK) Consortium | Dominic Kwiatkowski, Cordelia Langford, John Sillitoe on behalf of the Wellcome Sanger Institute COVID-19 Surveillance Team |
| EPI_ISL_762190, EPI_ISL_762191 | Lighthouse Lab in Glasgow | Wellcome Sanger Institute for the COVID-19 Genomics UK (COG-UK) Consortium | Harper VanSteenhouse, Yumi Kasai, David Gray, Carol Clugston, Anna Dominiczak and Alex Alderton, Roberto Amato, Sonia Goncalves, Ewan Harrison, David K. Jackson, Ian Johnston, Dominic Kwiatkowski, Cordelia Langford, John Sillitoe on behalf of the Wellcome Sanger Institute COVID-19 Surveillance Team |
| EPI_ISL_762192, EPI_ISL_762193 | Lighthouse Lab in Cambridge | Wellcome Sanger Institute for the COVID-19 Genomics UK (COG-UK) Consortium | Rob Howes, The Lighthouse Lab in Cambridge and Alex Alderton, Roberto Amato, Sonia Goncalves, Ewan Harrison, David K. Jackson, Ian Johnston, Dominic Kwiatkowski, Cordelia Langford, John Sillitoe on behalf of the Wellcome Sanger Institute COVID-19 Surveillance Team |
| EPI_ISL_762194 | Lighthouse Lab in Glasgow | Wellcome Sanger Institute for the COVID-19 Genomics UK (COG-UK) Consortium | Harper VanSteenhouse, Yumi Kasai, David Gray, Carol Clugston, Anna Dominiczak and Alex Alderton, Roberto Amato, Sonia Goncalves, Ewan Harrison, David K. Jackson, Ian Johnston, Dominic Kwiatkowski, Cordelia Langford, John Sillitoe on behalf of the Wellcome Sanger Institute COVID-19 Surveillance Team |
| EPI_ISL_762195 | Lighthouse Lab in Cambridge | Wellcome Sanger Institute for the COVID-19 Genomics UK (COG-UK) Consortium | Rob Howes, The Lighthouse Lab in Cambridge and Alex Alderton, Roberto Amato, Sonia Goncalves, Ewan Harrison, David K. Jackson, Ian Johnston, Dominic Kwiatkowski, Cordelia Langford, John Sillitoe on behalf of the Wellcome Sanger Institute COVID-19 Surveillance Team |
| EPI_ISL_762196, EPI_ISL_762197 | Lighthouse Lab in Glasgow | Wellcome Sanger Institute for the COVID-19 Genomics UK (COG-UK) Consortium | Harper VanSteenhouse, Yumi Kasai, David Gray, Carol Clugston, Anna Dominiczak and Alex Alderton, Roberto Amato, Sonia Goncalves, Ewan Harrison, David K. Jackson, Ian Johnston, Dominic Kwiatkowski, Cordelia Langford, John Sillitoe on behalf of the Wellcome Sanger Institute COVID-19 Surveillance Team |
| EPI_ISL_762199, EPI_ISL_762200, EPI_ISL_762201, EPI_ISL_762202, EPI_ISL_762203, EPI_ISL_762206, EPI_ISL_762207, EPI_ISL_762209, EPI_ISL_762210, EPI_ISL_762211, EPI_ISL_762212, EPI_ISL_762213, EPI_ISL_762214, EPI_ISL_762215, EPI_ISL_762216, EPI_ISL_762217, EPI_ISL_762218, EPI_ISL_762219, EPI_ISL_762220, EPI_ISL_762221, EPI_ISL_762222, EPI_ISL_762223, EPI_ISL_762224, EPI_ISL_762225, EPI_ISL_762226, EPI_ISL_762228, EPI_ISL_762229, EPI_ISL_762230, EPI_ISL_762232, EPI_ISL_762233, EPI_ISL_762234, EPI_ISL_762235, EPI_ISL_762236, EPI_ISL_762239, EPI_ISL_762240, EPI_ISL_762241, EPI_ISL_762242, EPI_ISL_762243, EPI_ISL_762245, EPI_ISL_762246, EPI_ISL_762247, EPI_ISL_762248, EPI_ISL_762249, EPI_ISL_762250, EPI_ISL_762251, EPI_ISL_762252, EPI_ISL_762253, EPI_ISL_762255, EPI_ISL_762256, EPI_ISL_762257, EPI_ISL_762259, EPI_ISL_762260, EPI_ISL_762261, EPI_ISL_762262, EPI_ISL_762263, EPI_ISL_762264, EPI_ISL_762265, EPI_ISL_762266, EPI_ISL_762267, EPI_ISL_762268, EPI_ISL_762269, EPI_ISL_762270, EPI_ISL_762272, EPI_ISL_762274, EPI_ISL_762275, EPI_ISL_762276, EPI_ISL_762277, EPI_ISL_762279, EPI_ISL_762280, EPI_ISL_762282, EPI_ISL_762283, EPI_ISL_762284, EPI_ISL_762286, EPI_ISL_762288, EPI_ISL_762289, EPI_ISL_762290, EPI_ISL_762291, EPI_ISL_762292, EPI_ISL_762294, EPI_ISL_762295, EPI_ISL_762296, EPI_ISL_762297, EPI_ISL_762298, EPI_ISL_762299, EPI_ISL_762300, EPI_ISL_762301, EPI_ISL_762302, EPI_ISL_762303, EPI_ISL_762304, EPI_ISL_762306, EPI_ISL_762307, EPI_ISL_762308, EPI_ISL_762310, EPI_ISL_762311, EPI_ISL_762312, EPI_ISL_762313, EPI_ISL_762314, EPI_ISL_762315, EPI_ISL_762316, EPI_ISL_762317, EPI_ISL_762318, EPI_ISL_762319, EPI_ISL_762320, EPI_ISL_762321, EPI_ISL_762322, EPI_ISL_762323, EPI_ISL_762325, EPI_ISL_762326, EPI_ISL_762327, EPI_ISL_762329, EPI_ISL_762330, EPI_ISL_762331, EPI_ISL_762332, EPI_ISL_762334, EPI_ISL_762335, EPI_ISL_762336, EPI_ISL_762337, EPI_ISL_762338, EPI_ISL_762339, EPI_ISL_762341, EPI_ISL_762342, EPI_ISL_762343, EPI_ISL_762344, EPI_ISL_762345, EPI_ISL_762346, EPI_ISL_762347, EPI_ISL_762348, EPI_ISL_762349, EPI_ISL_762350, EPI_ISL_762352, EPI_ISL_762353, EPI_ISL_762354, EPI_ISL_762355, EPI_ISL_762356, EPI_ISL_762357, EPI_ISL_762358, EPI_ISL_762359, EPI_ISL_762360, EPI_ISL_762361, EPI_ISL_762362, EPI_ISL_762363, EPI_ISL_762364, EPI_ISL_762365, EPI_ISL_762366, EPI_ISL_762367, EPI_ISL_762368, EPI_ISL_762369, EPI_ISL_762370, EPI_ISL_762371, EPI_ISL_762372, EPI_ISL_762373, EPI_ISL_762374, EPI_ISL_762375, EPI_ISL_762376, EPI_ISL_762379, EPI_ISL_762380, EPI_ISL_762382, EPI_ISL_762383, EPI_ISL_762384, EPI_ISL_762385, EPI_ISL_762386, EPI_ISL_762387, EPI_ISL_762388, EPI_ISL_762389, EPI_ISL_762390, EPI_ISL_762391, EPI_ISL_762393, EPI_ISL_762394, EPI_ISL_762396, EPI_ISL_762397, EPI_ISL_762398, EPI_ISL_762401, EPI_ISL_762402, EPI_ISL_762403, EPI_ISL_762404, EPI_ISL_762405, EPI_ISL_762406, EPI_ISL_762407, EPI_ISL_762409, EPI_ISL_762410, EPI_ISL_762411, EPI_ISL_762412, EPI_ISL_762413, EPI_ISL_762414, EPI_ISL_762415, EPI_ISL_762416, EPI_ISL_762417, EPI_ISL_762418, EPI_ISL_762420, EPI_ISL_762421, EPI_ISL_762422, EPI_ISL_762424, EPI_ISL_762425, EPI_ISL_762426, EPI_ISL_762427, EPI_ISL_762428, EPI_ISL_762429, EPI_ISL_762430, EPI_ISL_762431, EPI_ISL_762432, EPI_ISL_762433, EPI_ISL_762435, EPI_ISL_762437, EPI_ISL_762439, EPI_ISL_762441, EPI_ISL_762442, EPI_ISL_762443, EPI_ISL_762444, EPI_ISL_762445, EPI_ISL_762446, EPI_ISL_762447, EPI_ISL_762448, EPI_ISL_762449, EPI_ISL_762450, EPI_ISL_762451, EPI_ISL_762452, EPI_ISL_762453, EPI_ISL_762454, EPI_ISL_762455, EPI_ISL_762457, EPI_ISL_762458, EPI_ISL_762459, EPI_ISL_762460, EPI_ISL_762461, EPI_ISL_762464, EPI_ISL_762465, EPI_ISL_762466, EPI_ISL_762467, EPI_ISL_762468, EPI_ISL_762469, EPI_ISL_762470, EPI_ISL_762471, EPI_ISL_762472, EPI_ISL_762473, EPI_ISL_762474, EPI_ISL_762475, EPI_ISL_762476, EPI_ISL_762477, EPI_ISL_762478, EPI_ISL_762479, EPI_ISL_762481, EPI_ISL_762482, EPI_ISL_762483, EPI_ISL_762484, EPI_ISL_762485, EPI_ISL_762486, EPI_ISL_762487, EPI_ISL_762488, EPI_ISL_762489, EPI_ISL_762490, EPI_ISL_762491, EPI_ISL_762492, EPI_ISL_762495, EPI_ISL_762496, EPI_ISL_762497, EPI_ISL_762499, EPI_ISL_762500, EPI_ISL_762501, EPI_ISL_762503, EPI_ISL_762504, EPI_ISL_762506, EPI_ISL_762507, EPI_ISL_762508, EPI_ISL_762510, EPI_ISL_762511, EPI_ISL_762512, EPI_ISL_762513, EPI_ISL_762514, EPI_ISL_762515, EPI_ISL_762517, EPI_ISL_762518, EPI_ISL_762521, EPI_ISL_762522, EPI_ISL_762524, EPI_ISL_762525, EPI_ISL_762526, EPI_ISL_762527, EPI_ISL_762528, EPI_ISL_762529, EPI_ISL_762530, EPI_ISL_762531, EPI_ISL_762532, EPI_ISL_762533, EPI_ISL_762534, EPI_ISL_762535, EPI_ISL_762536, EPI_ISL_762537, EPI_ISL_762538, EPI_ISL_762539, EPI_ISL_762540, EPI_ISL_762541, EPI_ISL_762542, EPI_ISL_762543, EPI_ISL_762544, EPI_ISL_762545 | | | |
| see above | Lighthouse Lab in Milton Keynes | Wellcome Sanger Institute for the COVID-19 Genomics UK (COG-UK) Consortium | The Lighthouse Lab in Milton Keynes and Alex Alderton, Roberto Amato, Sonia Goncalves, Ewan Harrison, David K. Jackson, Ian Johnston, Dominic Kwiatkowski, Cordelia Langford, John Sillitoe on behalf of the Wellcome Sanger Institute COVID-19 Surveillance Team |
| EPI_ISL_762546 | Lighthouse Lab in Cambridge | Wellcome Sanger Institute for the COVID-19 Genomics UK (COG-UK) Consortium | Rob Howes, The Lighthouse Lab in Cambridge and Alex Alderton, Roberto Amato, Sonia Goncalves, Ewan Harrison, David K. Jackson, Ian Johnston, Dominic Kwiatkowski, Cordelia Langford, John Sillitoe on behalf of the Wellcome Sanger Institute COVID-19 Surveillance Team |
| EPI_ISL_762547 | Lighthouse Lab in Alderley Park | Wellcome Sanger Institute for the COVID-19 Genomics UK (COG-UK) Consortium | Jacquelyn Wynn, Mairead Hyland, The Lighthouse Lab in Alderley Park and Alex Alderton, Roberto Amato, Sonia Goncalves, Ewan Harrison, David K. Jackson, Ian Johnston, Dominic Kwiatkowski, Cordelia Langford, John Sillitoe on behalf of the Wellcome Sanger Institute COVID-19 Surveillance Team |
| EPI_ISL_762548 | Lighthouse Lab in Glasgow | Wellcome Sanger Institute for the COVID-19 Genomics UK (COG-UK) Consortium | Harper VanSteenhouse, Yumi Kasai, David Gray, Carol Clugston, Anna Dominiczak and Alex Alderton, Roberto Amato, Sonia Goncalves, Ewan Harrison, David K. Jackson, Ian Johnston, Dominic Kwiatkowski, Cordelia Langford, John Sillitoe on behalf of the Wellcome Sanger Institute COVID-19 Surveillance Team |
| EPI_ISL_762549, EPI_ISL_762550, EPI_ISL_762551, EPI_ISL_762552, EPI_ISL_762553 | Lighthouse Lab in Cambridge | Wellcome Sanger Institute for the COVID-19 Genomics UK (COG-UK) Consortium | Rob Howes, The Lighthouse Lab in Cambridge and Alex Alderton, Roberto Amato, Sonia Goncalves, Ewan Harrison, David K. Jackson, Ian Johnston, Dominic Kwiatkowski, Cordelia Langford, John Sillitoe on behalf of the Wellcome Sanger Institute COVID-19 Surveillance Team |
| EPI_ISL_762554 | Lighthouse Lab in Alderley Park | Wellcome Sanger Institute for the COVID-19 Genomics UK (COG-UK) Consortium | Jacquelyn Wynn, Mairead Hyland, The Lighthouse Lab in Alderley Park and Alex Alderton, Roberto Amato, Sonia Goncalves, Ewan Harrison, David K. Jackson, Ian Johnston, Dominic Kwiatkowski, Cordelia Langford, John Sillitoe on behalf of the Wellcome Sanger Institute COVID-19 Surveillance Team |
| EPI_ISL_762555, EPI_ISL_762556 | Lighthouse Lab in Glasgow | Wellcome Sanger Institute for the COVID-19 Genomics UK (COG-UK) Consortium | Harper VanSteenhouse, Yumi Kasai, David Gray, Carol Clugston, Anna Dominiczak and Alex Alderton, Roberto Amato, Sonia Goncalves, Ewan Harrison, David K. Jackson, Ian Johnston, Dominic Kwiatkowski, Cordelia Langford, John Sillitoe on behalf of the Wellcome Sanger Institute COVID-19 Surveillance Team |
| EPI_ISL_762557, EPI_ISL_762558 | Lighthouse Lab in Alderley Park | Wellcome Sanger Institute for the COVID-19 Genomics UK (COG-UK) Consortium | Jacquelyn Wynn, Mairead Hyland, The Lighthouse Lab in Alderley Park and Alex Alderton, Roberto Amato, Sonia Goncalves, Ewan Harrison, David K. Jackson, Ian Johnston, Dominic Kwiatkowski, Cordelia Langford, John Sillitoe on behalf of the Wellcome Sanger Institute COVID-19 Surveillance Team |
| EPI_ISL_762559 | Lighthouse Lab in Cambridge | Wellcome Sanger Institute for the COVID-19 Genomics UK (COG-UK) Consortium | Rob Howes, The Lighthouse Lab in Cambridge and Alex Alderton, Roberto Amato, Sonia Goncalves, Ewan Harrison, David K. Jackson, Ian Johnston, Dominic Kwiatkowski, Cordelia Langford, John Sillitoe on behalf of the Wellcome Sanger Institute COVID-19 Surveillance Team |
| EPI_ISL_762560, EPI_ISL_762561 | Lighthouse Lab in Glasgow | Wellcome Sanger Institute for the COVID-19 Genomics UK (COG-UK) Consortium | Harper VanSteenhouse, Yumi Kasai, David Gray, Carol Clugston, Anna Dominiczak and Alex Alderton, Roberto Amato, Sonia Goncalves, Ewan Harrison, David K. Jackson, Ian Johnston, Dominic Kwiatkowski, Cordelia Langford, John Sillitoe on behalf of the Wellcome Sanger Institute COVID-19 Surveillance Team |
| EPI_ISL_762562 | Lighthouse Lab in Alderley Park | Wellcome Sanger Institute for the COVID-19 Genomics UK (COG-UK) Consortium | Jacquelyn Wynn, Mairead Hyland, The Lighthouse Lab in Alderley Park and Alex Alderton, Roberto Amato, Sonia Goncalves, Ewan Harrison, David K. Jackson, Ian Johnston, Dominic Kwiatkowski, Cordelia Langford, John Sillitoe on behalf of the Wellcome Sanger Institute COVID-19 Surveillance Team |
| EPI_ISL_762563 | Lighthouse Lab in Cambridge | Wellcome Sanger Institute for the COVID-19 Genomics UK (COG-UK) Consortium | Rob Howes, The Lighthouse Lab in Cambridge and Alex Alderton, Roberto Amato, Sonia Goncalves, Ewan Harrison, David K. Jackson, Ian Johnston, Dominic Kwiatkowski, Cordelia Langford, John Sillitoe on behalf of the Wellcome Sanger Institute COVID-19 Surveillance Team |
| EPI_ISL_762564 | Lighthouse Lab in Alderley Park | Wellcome Sanger Institute for the COVID-19 Genomics UK (COG-UK) Consortium | Jacquelyn Wynn, Mairead Hyland, The Lighthouse Lab in Alderley Park and Alex Alderton, Roberto Amato, Sonia Goncalves, Ewan Harrison, David K. Jackson, Ian Johnston, Dominic Kwiatkowski, Cordelia Langford, John Sillitoe on behalf of the Wellcome Sanger Institute COVID-19 Surveillance Team |
| EPI_ISL_762565, EPI_ISL_762566 | Lighthouse Lab in Cambridge | Wellcome Sanger Institute for the COVID-19 Genomics UK (COG-UK) Consortium | Rob Howes, The Lighthouse Lab in Cambridge and Alex Alderton, Roberto Amato, Sonia Goncalves, Ewan Harrison, David K. Jackson, Ian Johnston, Dominic Kwiatkowski, Cordelia Langford, John Sillitoe on behalf of the Wellcome Sanger Institute COVID-19 Surveillance Team |
| EPI_ISL_762567 | Lighthouse Lab in Glasgow | Wellcome Sanger Institute for the COVID-19 Genomics UK (COG-UK) Consortium | Harper VanSteenhouse, Yumi Kasai, David Gray, Carol Clugston, Anna Dominiczak and Alex Alderton, Roberto Amato, Sonia Goncalves, Ewan Harrison, David K. Jackson, Ian Johnston, Dominic Kwiatkowski, Cordelia Langford, John Sillitoe on behalf of the Wellcome Sanger Institute COVID-19 Surveillance Team |
| EPI_ISL_762568, EPI_ISL_762569, EPI_ISL_762570, EPI_ISL_762571, EPI_ISL_762572 | Lighthouse Lab in Cambridge | Wellcome Sanger Institute for the COVID-19 Genomics UK (COG-UK) Consortium | Rob Howes, The Lighthouse Lab in Cambridge and Alex Alderton, Roberto Amato, Sonia Goncalves, Ewan Harrison, David K. Jackson, Ian Johnston, Dominic Kwiatkowski, Cordelia Langford, John Sillitoe on behalf of the Wellcome Sanger Institute COVID-19 Surveillance Team |
| EPI_ISL_762573, EPI_ISL_762574 | Lighthouse Lab in Glasgow | Wellcome Sanger Institute for the COVID-19 Genomics UK (COG-UK) Consortium | Harper VanSteenhouse, Yumi Kasai, David Gray, Carol Clugston, Anna Dominiczak and Alex Alderton, Roberto Amato, Sonia Goncalves, Ewan Harrison, David K. Jackson, Ian Johnston, Dominic Kwiatkowski, Cordelia Langford, John Sillitoe on behalf of the Wellcome Sanger Institute COVID-19 Surveillance Team |
| EPI_ISL_762575 | Lighthouse Lab in Cambridge | Wellcome Sanger Institute for the COVID-19 Genomics UK (COG-UK) Consortium | Rob Howes, The Lighthouse Lab in Cambridge and Alex Alderton, Roberto Amato, Sonia Goncalves, Ewan Harrison, David K. Jackson, Ian Johnston, Dominic Kwiatkowski, Cordelia Langford, John Sillitoe on behalf of the Wellcome Sanger Institute COVID-19 Surveillance Team |
| EPI_ISL_762576 | Lighthouse Lab in Glasgow | Wellcome Sanger Institute for the COVID-19 Genomics UK | Harper VanSteenhouse, Yumi Kasai, David Gray, Carol Clugston, Anna Dominiczak and Alex Alderton, Roberto Amato, Sonia Goncalves, Ewan Harrison, |

| | | | |
|---|---|---|---|
| | | (COG-UK) Consortium | David K. Jackson, Ian Johnston, Dominic Kwiatkowski, Cordelia Langford, John Sillitoe on behalf of the Wellcome Sanger Institute COVID-19 Surveillance Team |
| EPI_ISL_762577, EPI_ISL_762578, EPI_ISL_762579, EPI_ISL_762580 | Lighthouse Lab in Cambridge | Wellcome Sanger Institute for the COVID-19 Genomics UK (COG-UK) Consortium | Rob Howes, The Lighthouse Lab in Cambridge and Alex Alderton, Roberto Amato, Sonia Goncalves, Ewan Harrison, David K. Jackson, Ian Johnston, Dominic Kwiatkowski, Cordelia Langford, John Sillitoe on behalf of the Wellcome Sanger Institute COVID-19 Surveillance Team |
| EPI_ISL_762581 | Lighthouse Lab in Alderley Park | Wellcome Sanger Institute for the COVID-19 Genomics UK (COG-UK) Consortium | Jacquelyn Wynn, Mairead Hyland, The Lighthouse Lab in Alderley Park and Alex Alderton, Roberto Amato, Sonia Goncalves, Ewan Harrison, David K. Jackson, Ian Johnston, Dominic Kwiatkowski, Cordelia Langford, John Sillitoe on behalf of the Wellcome Sanger Institute COVID-19 Surveillance Team |
| EPI_ISL_762582 | Lighthouse Lab in Cambridge | Wellcome Sanger Institute for the COVID-19 Genomics UK (COG-UK) Consortium | Rob Howes, The Lighthouse Lab in Cambridge and Alex Alderton, Roberto Amato, Sonia Goncalves, Ewan Harrison, David K. Jackson, Ian Johnston, Dominic Kwiatkowski, Cordelia Langford, John Sillitoe on behalf of the Wellcome Sanger Institute COVID-19 Surveillance Team |
| EPI_ISL_762583 | Lighthouse Lab in Glasgow | Wellcome Sanger Institute for the COVID-19 Genomics UK (COG-UK) Consortium | Harper VanSteenhouse, Yumi Kasai, David Gray, Carol Clugston, Anna Dominiczak and Alex Alderton, Roberto Amato, Sonia Goncalves, Ewan Harrison, David K. Jackson, Ian Johnston, Dominic Kwiatkowski, Cordelia Langford, John Sillitoe on behalf of the Wellcome Sanger Institute COVID-19 Surveillance Team |
| EPI_ISL_762584, EPI_ISL_762585 | Lighthouse Lab in Cambridge | Wellcome Sanger Institute for the COVID-19 Genomics UK (COG-UK) Consortium | Rob Howes, The Lighthouse Lab in Cambridge and Alex Alderton, Roberto Amato, Sonia Goncalves, Ewan Harrison, David K. Jackson, Ian Johnston, Dominic Kwiatkowski, Cordelia Langford, John Sillitoe on behalf of the Wellcome Sanger Institute COVID-19 Surveillance Team |
| EPI_ISL_762586 | Lighthouse Lab in Glasgow | Wellcome Sanger Institute for the COVID-19 Genomics UK (COG-UK) Consortium | Harper VanSteenhouse, Yumi Kasai, David Gray, Carol Clugston, Anna Dominiczak and Alex Alderton, Roberto Amato, Sonia Goncalves, Ewan Harrison, David K. Jackson, Ian Johnston, Dominic Kwiatkowski, Cordelia Langford, John Sillitoe on behalf of the Wellcome Sanger Institute COVID-19 Surveillance Team |
| EPI_ISL_762587 | Lighthouse Lab in Cambridge | Wellcome Sanger Institute for the COVID-19 Genomics UK (COG-UK) Consortium | Rob Howes, The Lighthouse Lab in Cambridge and Alex Alderton, Roberto Amato, Sonia Goncalves, Ewan Harrison, David K. Jackson, Ian Johnston, Dominic Kwiatkowski, Cordelia Langford, John Sillitoe on behalf of the Wellcome Sanger Institute COVID-19 Surveillance Team |
| EPI_ISL_762588 | Lighthouse Lab in Alderley Park | Wellcome Sanger Institute for the COVID-19 Genomics UK (COG-UK) Consortium | Jacquelyn Wynn, Mairead Hyland, The Lighthouse Lab in Alderley Park and Alex Alderton, Roberto Amato, Sonia Goncalves, Ewan Harrison, David K. Jackson, Ian Johnston, Dominic Kwiatkowski, Cordelia Langford, John Sillitoe on behalf of the Wellcome Sanger Institute COVID-19 Surveillance Team |
| EPI_ISL_762589 | Lighthouse Lab in Glasgow | Wellcome Sanger Institute for the COVID-19 Genomics UK (COG-UK) Consortium | Harper VanSteenhouse, Yumi Kasai, David Gray, Carol Clugston, Anna Dominiczak and Alex Alderton, Roberto Amato, Sonia Goncalves, Ewan Harrison, David K. Jackson, Ian Johnston, Dominic Kwiatkowski, Cordelia Langford, John Sillitoe on behalf of the Wellcome Sanger Institute COVID-19 Surveillance Team |
| EPI_ISL_762590, EPI_ISL_762591, EPI_ISL_762592, EPI_ISL_762593, EPI_ISL_762594, EPI_ISL_762595, EPI_ISL_762596 | Lighthouse Lab in Cambridge | Wellcome Sanger Institute for the COVID-19 Genomics UK (COG-UK) Consortium | Rob Howes, The Lighthouse Lab in Cambridge and Alex Alderton, Roberto Amato, Sonia Goncalves, Ewan Harrison, David K. Jackson, Ian Johnston, Dominic Kwiatkowski, Cordelia Langford, John Sillitoe on behalf of the Wellcome Sanger Institute COVID-19 Surveillance Team |
| EPI_ISL_762597 | Lighthouse Lab in Glasgow | Wellcome Sanger Institute for the COVID-19 Genomics UK (COG-UK) Consortium | Harper VanSteenhouse, Yumi Kasai, David Gray, Carol Clugston, Anna Dominiczak and Alex Alderton, Roberto Amato, Sonia Goncalves, Ewan Harrison, David K. Jackson, Ian Johnston, Dominic Kwiatkowski, Cordelia Langford, John Sillitoe on behalf of the Wellcome Sanger Institute COVID-19 Surveillance Team |
| EPI_ISL_762598 | Lighthouse Lab in Cambridge | Wellcome Sanger Institute for the COVID-19 Genomics UK (COG-UK) Consortium | Rob Howes, The Lighthouse Lab in Cambridge and Alex Alderton, Roberto Amato, Sonia Goncalves, Ewan Harrison, David K. Jackson, Ian Johnston, Dominic Kwiatkowski, Cordelia Langford, John Sillitoe on behalf of the Wellcome Sanger Institute COVID-19 Surveillance Team |
| EPI_ISL_762599 | Lighthouse Lab in Alderley Park | Wellcome Sanger Institute for the COVID-19 Genomics UK (COG-UK) Consortium | Jacquelyn Wynn, Mairead Hyland, The Lighthouse Lab in Alderley Park and Alex Alderton, Roberto Amato, Sonia Goncalves, Ewan Harrison, David K. Jackson, Ian Johnston, Dominic Kwiatkowski, Cordelia Langford, John Sillitoe on behalf of the Wellcome Sanger Institute COVID-19 Surveillance Team |
| EPI_ISL_762600, EPI_ISL_762601 | Lighthouse Lab in Cambridge | Wellcome Sanger Institute for the COVID-19 Genomics UK (COG-UK) Consortium | Rob Howes, The Lighthouse Lab in Cambridge and Alex Alderton, Roberto Amato, Sonia Goncalves, Ewan Harrison, David K. Jackson, Ian Johnston, Dominic Kwiatkowski, Cordelia Langford, John Sillitoe on behalf of the Wellcome Sanger Institute COVID-19 Surveillance Team |
| EPI_ISL_762602 | Lighthouse Lab in Glasgow | Wellcome Sanger Institute for the COVID-19 Genomics UK (COG-UK) Consortium | Harper VanSteenhouse, Yumi Kasai, David Gray, Carol Clugston, Anna Dominiczak and Alex Alderton, Roberto Amato, Sonia Goncalves, Ewan Harrison, David K. Jackson, Ian Johnston, Dominic Kwiatkowski, Cordelia Langford, John Sillitoe on behalf of the Wellcome Sanger Institute COVID-19 Surveillance Team |
| EPI_ISL_762603 | Lighthouse Lab in Cambridge | Wellcome Sanger Institute for the COVID-19 Genomics UK (COG-UK) Consortium | Rob Howes, The Lighthouse Lab in Cambridge and Alex Alderton, Roberto Amato, Sonia Goncalves, Ewan Harrison, David K. Jackson, Ian Johnston, Dominic Kwiatkowski, Cordelia Langford, John Sillitoe on behalf of the Wellcome Sanger Institute COVID-19 Surveillance Team |
| EPI_ISL_762604 | Lighthouse Lab in Glasgow | Wellcome Sanger Institute for the COVID-19 Genomics UK (COG-UK) Consortium | Harper VanSteenhouse, Yumi Kasai, David Gray, Carol Clugston, Anna Dominiczak and Alex Alderton, Roberto Amato, Sonia Goncalves, Ewan Harrison, David K. Jackson, Ian Johnston, Dominic Kwiatkowski, Cordelia Langford, John Sillitoe on behalf of the Wellcome Sanger Institute COVID-19 Surveillance Team |
| EPI_ISL_762605, EPI_ISL_762606 | Lighthouse Lab in Alderley Park | Wellcome Sanger Institute for the COVID-19 Genomics UK (COG-UK) Consortium | Jacquelyn Wynn, Mairead Hyland, The Lighthouse Lab in Alderley Park and Alex Alderton, Roberto Amato, Sonia Goncalves, Ewan Harrison, David K. Jackson, Ian Johnston, Dominic Kwiatkowski, Cordelia Langford, John Sillitoe on behalf of the Wellcome Sanger Institute COVID-19 Surveillance Team |
| EPI_ISL_762607, EPI_ISL_762608 | Lighthouse Lab in Cambridge | Wellcome Sanger Institute for the COVID-19 Genomics UK (COG-UK) Consortium | Rob Howes, The Lighthouse Lab in Cambridge and Alex Alderton, Roberto Amato, Sonia Goncalves, Ewan Harrison, David K. Jackson, Ian Johnston, Dominic Kwiatkowski, Cordelia Langford, John Sillitoe on behalf of the Wellcome Sanger Institute COVID-19 Surveillance Team |
| EPI_ISL_762609 | Lighthouse Lab in Glasgow | Wellcome Sanger Institute for the COVID-19 Genomics UK (COG-UK) Consortium | Harper VanSteenhouse, Yumi Kasai, David Gray, Carol Clugston, Anna Dominiczak and Alex Alderton, Roberto Amato, Sonia Goncalves, Ewan Harrison, David K. Jackson, Ian Johnston, Dominic Kwiatkowski, Cordelia Langford, John Sillitoe on behalf of the Wellcome Sanger Institute COVID-19 Surveillance Team |
| EPI_ISL_762610 | Lighthouse Lab in Alderley Park | Wellcome Sanger Institute for the COVID-19 Genomics UK (COG-UK) Consortium | Jacquelyn Wynn, Mairead Hyland, The Lighthouse Lab in Alderley Park and Alex Alderton, Roberto Amato, Sonia Goncalves, Ewan Harrison, David K. Jackson, Ian Johnston, Dominic Kwiatkowski, Cordelia Langford, John Sillitoe on behalf of the Wellcome Sanger Institute COVID-19 Surveillance Team |
| EPI_ISL_762611, EPI_ISL_762612, EPI_ISL_762613 | Lighthouse Lab in Cambridge | Wellcome Sanger Institute for the COVID-19 Genomics UK (COG-UK) Consortium | Rob Howes, The Lighthouse Lab in Cambridge and Alex Alderton, Roberto Amato, Sonia Goncalves, Ewan Harrison, David K. Jackson, Ian Johnston, Dominic Kwiatkowski, Cordelia Langford, John Sillitoe on behalf of the Wellcome Sanger Institute COVID-19 Surveillance Team |
| EPI_ISL_762614 | Lighthouse Lab in Glasgow | Wellcome Sanger Institute for the COVID-19 Genomics UK (COG-UK) Consortium | Harper VanSteenhouse, Yumi Kasai, David Gray, Carol Clugston, Anna Dominiczak and Alex Alderton, Roberto Amato, Sonia Goncalves, Ewan Harrison, David K. Jackson, Ian Johnston, Dominic Kwiatkowski, Cordelia Langford, John Sillitoe on behalf of the Wellcome Sanger Institute COVID-19 Surveillance Team |
| EPI_ISL_762615, EPI_ISL_762616 | Lighthouse Lab in Cambridge | Wellcome Sanger Institute for the COVID-19 Genomics UK (COG-UK) Consortium | Rob Howes, The Lighthouse Lab in Cambridge and Alex Alderton, Roberto Amato, Sonia Goncalves, Ewan Harrison, David K. Jackson, Ian Johnston, Dominic Kwiatkowski, Cordelia Langford, John Sillitoe on behalf of the Wellcome Sanger Institute COVID-19 Surveillance Team |
| EPI_ISL_762617 | Lighthouse Lab in Alderley Park | Wellcome Sanger Institute for the COVID-19 Genomics UK (COG-UK) Consortium | Jacquelyn Wynn, Mairead Hyland, The Lighthouse Lab in Alderley Park and Alex Alderton, Roberto Amato, Sonia Goncalves, Ewan Harrison, David K. Jackson, Ian Johnston, Dominic Kwiatkowski, Cordelia Langford, John Sillitoe on behalf of the Wellcome Sanger Institute COVID-19 Surveillance Team |
| EPI_ISL_762618 | Lighthouse Lab in Glasgow | Wellcome Sanger Institute for the COVID-19 Genomics UK (COG-UK) Consortium | Harper VanSteenhouse, Yumi Kasai, David Gray, Carol Clugston, Anna Dominiczak and Alex Alderton, Roberto Amato, Sonia Goncalves, Ewan Harrison, David K. Jackson, Ian Johnston, Dominic Kwiatkowski, Cordelia Langford, John Sillitoe on behalf of the Wellcome Sanger Institute COVID-19 Surveillance Team |
| EPI_ISL_762619 | Lighthouse Lab in Cambridge | Wellcome Sanger Institute for the COVID-19 Genomics UK (COG-UK) Consortium | Rob Howes, The Lighthouse Lab in Cambridge and Alex Alderton, Roberto Amato, Sonia Goncalves, Ewan Harrison, David K. Jackson, Ian Johnston, Dominic Kwiatkowski, Cordelia Langford, John Sillitoe on behalf of the Wellcome Sanger Institute COVID-19 Surveillance Team |
| EPI_ISL_762620 | Lighthouse Lab in Alderley Park | Wellcome Sanger Institute for the COVID-19 Genomics UK (COG-UK) Consortium | Jacquelyn Wynn, Mairead Hyland, The Lighthouse Lab in Alderley Park and Alex Alderton, Roberto Amato, Sonia Goncalves, Ewan Harrison, David K. Jackson, Ian Johnston, Dominic Kwiatkowski, Cordelia Langford, John Sillitoe on behalf of the Wellcome Sanger Institute COVID-19 Surveillance Team |
| EPI_ISL_762621 | Lighthouse Lab in Cambridge | Wellcome Sanger Institute for the COVID-19 Genomics UK (COG-UK) Consortium | Rob Howes, The Lighthouse Lab in Cambridge and Alex Alderton, Roberto Amato, Sonia Goncalves, Ewan Harrison, David K. Jackson, Ian Johnston, Dominic Kwiatkowski, Cordelia Langford, John Sillitoe on behalf of the Wellcome Sanger Institute COVID-19 Surveillance Team |
| EPI_ISL_762622 | Lighthouse Lab in Glasgow | Wellcome Sanger Institute for the COVID-19 Genomics UK (COG-UK) Consortium | Harper VanSteenhouse, Yumi Kasai, David Gray, Carol Clugston, Anna Dominiczak and Alex Alderton, Roberto Amato, Sonia Goncalves, Ewan Harrison, David K. Jackson, Ian Johnston, Dominic Kwiatkowski, Cordelia Langford, John Sillitoe on behalf of the Wellcome Sanger Institute COVID-19 Surveillance |

| | | | |
|---|---|---|---|
| | | | Team |
| EPI_ISL_762623, EPI_ISL_762624 | Lighthouse Lab in Cambridge | Wellcome Sanger Institute for the COVID-19 Genomics UK (COG-UK) Consortium | Rob Howes, The Lighthouse Lab in Cambridge and Alex Alderton, Roberto Amato, Sonia Goncalves, Ewan Harrison, David K. Jackson, Ian Johnston, Dominic Kwiatkowski, Cordelia Langford, John Sillitoe on behalf of the Wellcome Sanger Institute COVID-19 Surveillance Team |
| EPI_ISL_762625 | Lighthouse Lab in Alderley Park | Wellcome Sanger Institute for the COVID-19 Genomics UK (COG-UK) Consortium | Jacquelyn Wynn, Mairead Hyland, The Lighthouse Lab in Alderley Park and Alex Alderton, Roberto Amato, Sonia Goncalves, Ewan Harrison, David K. Jackson, Ian Johnston, Dominic Kwiatkowski, Cordelia Langford, John Sillitoe on behalf of the Wellcome Sanger Institute COVID-19 Surveillance Team |
| EPI_ISL_762626, EPI_ISL_762627, EPI_ISL_762628 | Lighthouse Lab in Cambridge | Wellcome Sanger Institute for the COVID-19 Genomics UK (COG-UK) Consortium | Rob Howes, The Lighthouse Lab in Cambridge and Alex Alderton, Roberto Amato, Sonia Goncalves, Ewan Harrison, David K. Jackson, Ian Johnston, Dominic Kwiatkowski, Cordelia Langford, John Sillitoe on behalf of the Wellcome Sanger Institute COVID-19 Surveillance Team |
| EPI_ISL_762629 | Lighthouse Lab in Alderley Park | Wellcome Sanger Institute for the COVID-19 Genomics UK (COG-UK) Consortium | Jacquelyn Wynn, Mairead Hyland, The Lighthouse Lab in Alderley Park and Alex Alderton, Roberto Amato, Sonia Goncalves, Ewan Harrison, David K. Jackson, Ian Johnston, Dominic Kwiatkowski, Cordelia Langford, John Sillitoe on behalf of the Wellcome Sanger Institute COVID-19 Surveillance Team |
| EPI_ISL_762630, EPI_ISL_762631 | Lighthouse Lab in Glasgow | Wellcome Sanger Institute for the COVID-19 Genomics UK (COG-UK) Consortium | Harper VanSteenhouse, Yumi Kasai, David Gray, Carol Clugston, Anna Dominiczak and Alex Alderton, Roberto Amato, Sonia Goncalves, Ewan Harrison, David K. Jackson, Ian Johnston, Dominic Kwiatkowski, Cordelia Langford, John Sillitoe on behalf of the Wellcome Sanger Institute COVID-19 Surveillance Team |
| EPI_ISL_762632 | Lighthouse Lab in Cambridge | Wellcome Sanger Institute for the COVID-19 Genomics UK (COG-UK) Consortium | Rob Howes, The Lighthouse Lab in Cambridge and Alex Alderton, Roberto Amato, Sonia Goncalves, Ewan Harrison, David K. Jackson, Ian Johnston, Dominic Kwiatkowski, Cordelia Langford, John Sillitoe on behalf of the Wellcome Sanger Institute COVID-19 Surveillance Team |
| EPI_ISL_762633 | Lighthouse Lab in Glasgow | Wellcome Sanger Institute for the COVID-19 Genomics UK (COG-UK) Consortium | Harper VanSteenhouse, Yumi Kasai, David Gray, Carol Clugston, Anna Dominiczak and Alex Alderton, Roberto Amato, Sonia Goncalves, Ewan Harrison, David K. Jackson, Ian Johnston, Dominic Kwiatkowski, Cordelia Langford, John Sillitoe on behalf of the Wellcome Sanger Institute COVID-19 Surveillance Team |
| EPI_ISL_762634 | Lighthouse Lab in Alderley Park | Wellcome Sanger Institute for the COVID-19 Genomics UK (COG-UK) Consortium | Jacquelyn Wynn, Mairead Hyland, The Lighthouse Lab in Alderley Park and Alex Alderton, Roberto Amato, Sonia Goncalves, Ewan Harrison, David K. Jackson, Ian Johnston, Dominic Kwiatkowski, Cordelia Langford, John Sillitoe on behalf of the Wellcome Sanger Institute COVID-19 Surveillance Team |
| EPI_ISL_762635, EPI_ISL_762636, EPI_ISL_762637 | Lighthouse Lab in Cambridge | Wellcome Sanger Institute for the COVID-19 Genomics UK (COG-UK) Consortium | Rob Howes, The Lighthouse Lab in Cambridge and Alex Alderton, Roberto Amato, Sonia Goncalves, Ewan Harrison, David K. Jackson, Ian Johnston, Dominic Kwiatkowski, Cordelia Langford, John Sillitoe on behalf of the Wellcome Sanger Institute COVID-19 Surveillance Team |
| EPI_ISL_762638 | Lighthouse Lab in Alderley Park | Wellcome Sanger Institute for the COVID-19 Genomics UK (COG-UK) Consortium | Jacquelyn Wynn, Mairead Hyland, The Lighthouse Lab in Alderley Park and Alex Alderton, Roberto Amato, Sonia Goncalves, Ewan Harrison, David K. Jackson, Ian Johnston, Dominic Kwiatkowski, Cordelia Langford, John Sillitoe on behalf of the Wellcome Sanger Institute COVID-19 Surveillance Team |
| EPI_ISL_762639 | Lighthouse Lab in Glasgow | Wellcome Sanger Institute for the COVID-19 Genomics UK (COG-UK) Consortium | Harper VanSteenhouse, Yumi Kasai, David Gray, Carol Clugston, Anna Dominiczak and Alex Alderton, Roberto Amato, Sonia Goncalves, Ewan Harrison, David K. Jackson, Ian Johnston, Dominic Kwiatkowski, Cordelia Langford, John Sillitoe on behalf of the Wellcome Sanger Institute COVID-19 Surveillance Team |
| EPI_ISL_762640, EPI_ISL_762641 | Lighthouse Lab in Cambridge | Wellcome Sanger Institute for the COVID-19 Genomics UK (COG-UK) Consortium | Rob Howes, The Lighthouse Lab in Cambridge and Alex Alderton, Roberto Amato, Sonia Goncalves, Ewan Harrison, David K. Jackson, Ian Johnston, Dominic Kwiatkowski, Cordelia Langford, John Sillitoe on behalf of the Wellcome Sanger Institute COVID-19 Surveillance Team |
| EPI_ISL_762642, EPI_ISL_762643 | Lighthouse Lab in Alderley Park | Wellcome Sanger Institute for the COVID-19 Genomics UK (COG-UK) Consortium | Jacquelyn Wynn, Mairead Hyland, The Lighthouse Lab in Alderley Park and Alex Alderton, Roberto Amato, Sonia Goncalves, Ewan Harrison, David K. Jackson, Ian Johnston, Dominic Kwiatkowski, Cordelia Langford, John Sillitoe on behalf of the Wellcome Sanger Institute COVID-19 Surveillance Team |
| EPI_ISL_762644 | Lighthouse Lab in Glasgow | Wellcome Sanger Institute for the COVID-19 Genomics UK (COG-UK) Consortium | Harper VanSteenhouse, Yumi Kasai, David Gray, Carol Clugston, Anna Dominiczak and Alex Alderton, Roberto Amato, Sonia Goncalves, Ewan Harrison, David K. Jackson, Ian Johnston, Dominic Kwiatkowski, Cordelia Langford, John Sillitoe on behalf of the Wellcome Sanger Institute COVID-19 Surveillance Team |
| EPI_ISL_762645 | Lighthouse Lab in Cambridge | Wellcome Sanger Institute for the COVID-19 Genomics UK (COG-UK) Consortium | Rob Howes, The Lighthouse Lab in Cambridge and Alex Alderton, Roberto Amato, Sonia Goncalves, Ewan Harrison, David K. Jackson, Ian Johnston, Dominic Kwiatkowski, Cordelia Langford, John Sillitoe on behalf of the Wellcome Sanger Institute COVID-19 Surveillance Team |
| EPI_ISL_762646 | Lighthouse Lab in Glasgow | Wellcome Sanger Institute for the COVID-19 Genomics UK (COG-UK) Consortium | Harper VanSteenhouse, Yumi Kasai, David Gray, Carol Clugston, Anna Dominiczak and Alex Alderton, Roberto Amato, Sonia Goncalves, Ewan Harrison, David K. Jackson, Ian Johnston, Dominic Kwiatkowski, Cordelia Langford, John Sillitoe on behalf of the Wellcome Sanger Institute COVID-19 Surveillance Team |
| EPI_ISL_762647 | Lighthouse Lab in Alderley Park | Wellcome Sanger Institute for the COVID-19 Genomics UK (COG-UK) Consortium | Jacquelyn Wynn, Mairead Hyland, The Lighthouse Lab in Alderley Park and Alex Alderton, Roberto Amato, Sonia Goncalves, Ewan Harrison, David K. Jackson, Ian Johnston, Dominic Kwiatkowski, Cordelia Langford, John Sillitoe on behalf of the Wellcome Sanger Institute COVID-19 Surveillance Team |
| EPI_ISL_762648 | Lighthouse Lab in Glasgow | Wellcome Sanger Institute for the COVID-19 Genomics UK (COG-UK) Consortium | Harper VanSteenhouse, Yumi Kasai, David Gray, Carol Clugston, Anna Dominiczak and Alex Alderton, Roberto Amato, Sonia Goncalves, Ewan Harrison, David K. Jackson, Ian Johnston, Dominic Kwiatkowski, Cordelia Langford, John Sillitoe on behalf of the Wellcome Sanger Institute COVID-19 Surveillance Team |
| EPI_ISL_762649, EPI_ISL_762650, EPI_ISL_762651 | Lighthouse Lab in Cambridge | Wellcome Sanger Institute for the COVID-19 Genomics UK (COG-UK) Consortium | Rob Howes, The Lighthouse Lab in Cambridge and Alex Alderton, Roberto Amato, Sonia Goncalves, Ewan Harrison, David K. Jackson, Ian Johnston, Dominic Kwiatkowski, Cordelia Langford, John Sillitoe on behalf of the Wellcome Sanger Institute COVID-19 Surveillance Team |
| EPI_ISL_762652, EPI_ISL_762653, EPI_ISL_762654 | Lighthouse Lab in Glasgow | Wellcome Sanger Institute for the COVID-19 Genomics UK (COG-UK) Consortium | Harper VanSteenhouse, Yumi Kasai, David Gray, Carol Clugston, Anna Dominiczak and Alex Alderton, Roberto Amato, Sonia Goncalves, Ewan Harrison, David K. Jackson, Ian Johnston, Dominic Kwiatkowski, Cordelia Langford, John Sillitoe on behalf of the Wellcome Sanger Institute COVID-19 Surveillance Team |
| EPI_ISL_762655 | Lighthouse Lab in Alderley Park | Wellcome Sanger Institute for the COVID-19 Genomics UK (COG-UK) Consortium | Jacquelyn Wynn, Mairead Hyland, The Lighthouse Lab in Alderley Park and Alex Alderton, Roberto Amato, Sonia Goncalves, Ewan Harrison, David K. Jackson, Ian Johnston, Dominic Kwiatkowski, Cordelia Langford, John Sillitoe on behalf of the Wellcome Sanger Institute COVID-19 Surveillance Team |
| EPI_ISL_762656, EPI_ISL_762657 | Lighthouse Lab in Glasgow | Wellcome Sanger Institute for the COVID-19 Genomics UK (COG-UK) Consortium | Harper VanSteenhouse, Yumi Kasai, David Gray, Carol Clugston, Anna Dominiczak and Alex Alderton, Roberto Amato, Sonia Goncalves, Ewan Harrison, David K. Jackson, Ian Johnston, Dominic Kwiatkowski, Cordelia Langford, John Sillitoe on behalf of the Wellcome Sanger Institute COVID-19 Surveillance Team |
| EPI_ISL_762658 | Lighthouse Lab in Cambridge | Wellcome Sanger Institute for the COVID-19 Genomics UK (COG-UK) Consortium | Rob Howes, The Lighthouse Lab in Cambridge and Alex Alderton, Roberto Amato, Sonia Goncalves, Ewan Harrison, David K. Jackson, Ian Johnston, Dominic Kwiatkowski, Cordelia Langford, John Sillitoe on behalf of the Wellcome Sanger Institute COVID-19 Surveillance Team |
| EPI_ISL_762659 | Lighthouse Lab in Alderley Park | Wellcome Sanger Institute for the COVID-19 Genomics UK (COG-UK) Consortium | Jacquelyn Wynn, Mairead Hyland, The Lighthouse Lab in Alderley Park and Alex Alderton, Roberto Amato, Sonia Goncalves, Ewan Harrison, David K. Jackson, Ian Johnston, Dominic Kwiatkowski, Cordelia Langford, John Sillitoe on behalf of the Wellcome Sanger Institute COVID-19 Surveillance Team |
| EPI_ISL_762660 | Lighthouse Lab in Cambridge | Wellcome Sanger Institute for the COVID-19 Genomics UK (COG-UK) Consortium | Rob Howes, The Lighthouse Lab in Cambridge and Alex Alderton, Roberto Amato, Sonia Goncalves, Ewan Harrison, David K. Jackson, Ian Johnston, Dominic Kwiatkowski, Cordelia Langford, John Sillitoe on behalf of the Wellcome Sanger Institute COVID-19 Surveillance Team |
| EPI_ISL_762661, EPI_ISL_762662, EPI_ISL_762663 | Lighthouse Lab in Glasgow | Wellcome Sanger Institute for the COVID-19 Genomics UK (COG-UK) Consortium | Harper VanSteenhouse, Yumi Kasai, David Gray, Carol Clugston, Anna Dominiczak and Alex Alderton, Roberto Amato, Sonia Goncalves, Ewan Harrison, David K. Jackson, Ian Johnston, Dominic Kwiatkowski, Cordelia Langford, John Sillitoe on behalf of the Wellcome Sanger Institute COVID-19 Surveillance Team |
| EPI_ISL_762664, EPI_ISL_762665 | Lighthouse Lab in Cambridge | Wellcome Sanger Institute for the COVID-19 Genomics UK (COG-UK) Consortium | Rob Howes, The Lighthouse Lab in Cambridge and Alex Alderton, Roberto Amato, Sonia Goncalves, Ewan Harrison, David K. Jackson, Ian Johnston, Dominic Kwiatkowski, Cordelia Langford, John Sillitoe on behalf of the Wellcome Sanger Institute COVID-19 Surveillance Team |
| EPI_ISL_762666 | Lighthouse Lab in Alderley Park | Wellcome Sanger Institute for the COVID-19 Genomics UK (COG-UK) Consortium | Jacquelyn Wynn, Mairead Hyland, The Lighthouse Lab in Alderley Park and Alex Alderton, Roberto Amato, Sonia Goncalves, Ewan Harrison, David K. Jackson, Ian Johnston, Dominic Kwiatkowski, Cordelia Langford, John Sillitoe on behalf of the Wellcome Sanger Institute COVID-19 Surveillance Team |
| EPI_ISL_762667, EPI_ISL_762668 | Lighthouse Lab in Cambridge | Wellcome Sanger Institute for the COVID-19 Genomics UK (COG-UK) Consortium | Rob Howes, The Lighthouse Lab in Cambridge and Alex Alderton, Roberto Amato, Sonia Goncalves, Ewan Harrison, David K. Jackson, Ian Johnston, Dominic Kwiatkowski, Cordelia Langford, John Sillitoe on behalf of the Wellcome Sanger Institute COVID-19 Surveillance Team |
| EPI_ISL_762669 | Lighthouse Lab in Glasgow | Wellcome Sanger Institute for the COVID-19 Genomics UK (COG-UK) Consortium | Harper VanSteenhouse, Yumi Kasai, David Gray, Carol Clugston, Anna Dominiczak and Alex Alderton, Roberto Amato, Sonia Goncalves, Ewan Harrison, David K. Jackson, Ian Johnston, Dominic Kwiatkowski, Cordelia Langford, John Sillitoe on behalf of the Wellcome Sanger Institute COVID-19 Surveillance Team |
| EPI_ISL_762670 | Lighthouse Lab in Cambridge | Wellcome Sanger Institute for the COVID-19 Genomics UK | Rob Howes, The Lighthouse Lab in Cambridge and Alex Alderton, Roberto Amato, Sonia Goncalves, Ewan Harrison, David K. Jackson, Ian Johnston, |

| | | | |
|---|---|---|---|
| | | (COG-UK) Consortium | Dominic Kwiatkowski, Cordelia Langford, John Sillitoe on behalf of the Wellcome Sanger Institute COVID-19 Surveillance Team |
| EPI_ISL_762671, EPI_ISL_762672 | Lighthouse Lab in Alderley Park | Wellcome Sanger Institute for the COVID-19 Genomics UK (COG-UK) Consortium | Jacquelyn Wynn, Mairead Hyland, The Lighthouse Lab in Alderley Park and Alex Alderton, Roberto Amato, Sonia Goncalves, Ewan Harrison, David K. Jackson, Ian Johnston, Dominic Kwiatkowski, Cordelia Langford, John Sillitoe on behalf of the Wellcome Sanger Institute COVID-19 Surveillance Team |
| EPI_ISL_762673 | Lighthouse Lab in Cambridge | Wellcome Sanger Institute for the COVID-19 Genomics UK (COG-UK) Consortium | Rob Howes, The Lighthouse Lab in Cambridge and Alex Alderton, Roberto Amato, Sonia Goncalves, Ewan Harrison, David K. Jackson, Ian Johnston, Dominic Kwiatkowski, Cordelia Langford, John Sillitoe on behalf of the Wellcome Sanger Institute COVID-19 Surveillance Team |
| EPI_ISL_762674 | Lighthouse Lab in Alderley Park | Wellcome Sanger Institute for the COVID-19 Genomics UK (COG-UK) Consortium | Jacquelyn Wynn, Mairead Hyland, The Lighthouse Lab in Alderley Park and Alex Alderton, Roberto Amato, Sonia Goncalves, Ewan Harrison, David K. Jackson, Ian Johnston, Dominic Kwiatkowski, Cordelia Langford, John Sillitoe on behalf of the Wellcome Sanger Institute COVID-19 Surveillance Team |
| EPI_ISL_762675 | Lighthouse Lab in Cambridge | Wellcome Sanger Institute for the COVID-19 Genomics UK (COG-UK) Consortium | Rob Howes, The Lighthouse Lab in Cambridge and Alex Alderton, Roberto Amato, Sonia Goncalves, Ewan Harrison, David K. Jackson, Ian Johnston, Dominic Kwiatkowski, Cordelia Langford, John Sillitoe on behalf of the Wellcome Sanger Institute COVID-19 Surveillance Team |
| EPI_ISL_762676 | Lighthouse Lab in Glasgow | Wellcome Sanger Institute for the COVID-19 Genomics UK (COG-UK) Consortium | Harper VanSteenhouse, Yumi Kasai, David Gray, Carol Clugston, Anna Dominiczak and Alex Alderton, Roberto Amato, Sonia Goncalves, Ewan Harrison, David K. Jackson, Ian Johnston, Dominic Kwiatkowski, Cordelia Langford, John Sillitoe on behalf of the Wellcome Sanger Institute COVID-19 Surveillance Team |
| EPI_ISL_762677 | Lighthouse Lab in Cambridge | Wellcome Sanger Institute for the COVID-19 Genomics UK (COG-UK) Consortium | Rob Howes, The Lighthouse Lab in Cambridge and Alex Alderton, Roberto Amato, Sonia Goncalves, Ewan Harrison, David K. Jackson, Ian Johnston, Dominic Kwiatkowski, Cordelia Langford, John Sillitoe on behalf of the Wellcome Sanger Institute COVID-19 Surveillance Team |
| EPI_ISL_762678 | Lighthouse Lab in Glasgow | Wellcome Sanger Institute for the COVID-19 Genomics UK (COG-UK) Consortium | Harper VanSteenhouse, Yumi Kasai, David Gray, Carol Clugston, Anna Dominiczak and Alex Alderton, Roberto Amato, Sonia Goncalves, Ewan Harrison, David K. Jackson, Ian Johnston, Dominic Kwiatkowski, Cordelia Langford, John Sillitoe on behalf of the Wellcome Sanger Institute COVID-19 Surveillance Team |
| EPI_ISL_762679, EPI_ISL_762680, EPI_ISL_762681, EPI_ISL_762682 | Lighthouse Lab in Cambridge | Wellcome Sanger Institute for the COVID-19 Genomics UK (COG-UK) Consortium | Rob Howes, The Lighthouse Lab in Cambridge and Alex Alderton, Roberto Amato, Sonia Goncalves, Ewan Harrison, David K. Jackson, Ian Johnston, Dominic Kwiatkowski, Cordelia Langford, John Sillitoe on behalf of the Wellcome Sanger Institute COVID-19 Surveillance Team |
| EPI_ISL_762683 | Lighthouse Lab in Alderley Park | Wellcome Sanger Institute for the COVID-19 Genomics UK (COG-UK) Consortium | Jacquelyn Wynn, Mairead Hyland, The Lighthouse Lab in Alderley Park and Alex Alderton, Roberto Amato, Sonia Goncalves, Ewan Harrison, David K. Jackson, Ian Johnston, Dominic Kwiatkowski, Cordelia Langford, John Sillitoe on behalf of the Wellcome Sanger Institute COVID-19 Surveillance Team |
| EPI_ISL_762684, EPI_ISL_762685 | Lighthouse Lab in Cambridge | Wellcome Sanger Institute for the COVID-19 Genomics UK (COG-UK) Consortium | Rob Howes, The Lighthouse Lab in Cambridge and Alex Alderton, Roberto Amato, Sonia Goncalves, Ewan Harrison, David K. Jackson, Ian Johnston, Dominic Kwiatkowski, Cordelia Langford, John Sillitoe on behalf of the Wellcome Sanger Institute COVID-19 Surveillance Team |
| EPI_ISL_762686 | Lighthouse Lab in Alderley Park | Wellcome Sanger Institute for the COVID-19 Genomics UK (COG-UK) Consortium | Jacquelyn Wynn, Mairead Hyland, The Lighthouse Lab in Alderley Park and Alex Alderton, Roberto Amato, Sonia Goncalves, Ewan Harrison, David K. Jackson, Ian Johnston, Dominic Kwiatkowski, Cordelia Langford, John Sillitoe on behalf of the Wellcome Sanger Institute COVID-19 Surveillance Team |
| EPI_ISL_762687, EPI_ISL_762688, EPI_ISL_762689 | Lighthouse Lab in Cambridge | Wellcome Sanger Institute for the COVID-19 Genomics UK (COG-UK) Consortium | Rob Howes, The Lighthouse Lab in Cambridge and Alex Alderton, Roberto Amato, Sonia Goncalves, Ewan Harrison, David K. Jackson, Ian Johnston, Dominic Kwiatkowski, Cordelia Langford, John Sillitoe on behalf of the Wellcome Sanger Institute COVID-19 Surveillance Team |
| EPI_ISL_762690 | Lighthouse Lab in Glasgow | Wellcome Sanger Institute for the COVID-19 Genomics UK (COG-UK) Consortium | Harper VanSteenhouse, Yumi Kasai, David Gray, Carol Clugston, Anna Dominiczak and Alex Alderton, Roberto Amato, Sonia Goncalves, Ewan Harrison, David K. Jackson, Ian Johnston, Dominic Kwiatkowski, Cordelia Langford, John Sillitoe on behalf of the Wellcome Sanger Institute COVID-19 Surveillance Team |
| EPI_ISL_762691, EPI_ISL_762692, EPI_ISL_762693 | Lighthouse Lab in Alderley Park | Wellcome Sanger Institute for the COVID-19 Genomics UK (COG-UK) Consortium | Jacquelyn Wynn, Mairead Hyland, The Lighthouse Lab in Alderley Park and Alex Alderton, Roberto Amato, Sonia Goncalves, Ewan Harrison, David K. Jackson, Ian Johnston, Dominic Kwiatkowski, Cordelia Langford, John Sillitoe on behalf of the Wellcome Sanger Institute COVID-19 Surveillance Team |
| EPI_ISL_762694 | Lighthouse Lab in Glasgow | Wellcome Sanger Institute for the COVID-19 Genomics UK (COG-UK) Consortium | Harper VanSteenhouse, Yumi Kasai, David Gray, Carol Clugston, Anna Dominiczak and Alex Alderton, Roberto Amato, Sonia Goncalves, Ewan Harrison, David K. Jackson, Ian Johnston, Dominic Kwiatkowski, Cordelia Langford, John Sillitoe on behalf of the Wellcome Sanger Institute COVID-19 Surveillance Team |
| EPI_ISL_762695, EPI_ISL_762696, EPI_ISL_762697 | Lighthouse Lab in Cambridge | Wellcome Sanger Institute for the COVID-19 Genomics UK (COG-UK) Consortium | Rob Howes, The Lighthouse Lab in Cambridge and Alex Alderton, Roberto Amato, Sonia Goncalves, Ewan Harrison, David K. Jackson, Ian Johnston, Dominic Kwiatkowski, Cordelia Langford, John Sillitoe on behalf of the Wellcome Sanger Institute COVID-19 Surveillance Team |
| EPI_ISL_762698 | Lighthouse Lab in Glasgow | Wellcome Sanger Institute for the COVID-19 Genomics UK (COG-UK) Consortium | Harper VanSteenhouse, Yumi Kasai, David Gray, Carol Clugston, Anna Dominiczak and Alex Alderton, Roberto Amato, Sonia Goncalves, Ewan Harrison, David K. Jackson, Ian Johnston, Dominic Kwiatkowski, Cordelia Langford, John Sillitoe on behalf of the Wellcome Sanger Institute COVID-19 Surveillance Team |
| EPI_ISL_762699, EPI_ISL_762700 | Lighthouse Lab in Cambridge | Wellcome Sanger Institute for the COVID-19 Genomics UK (COG-UK) Consortium | Rob Howes, The Lighthouse Lab in Cambridge and Alex Alderton, Roberto Amato, Sonia Goncalves, Ewan Harrison, David K. Jackson, Ian Johnston, Dominic Kwiatkowski, Cordelia Langford, John Sillitoe on behalf of the Wellcome Sanger Institute COVID-19 Surveillance Team |
| EPI_ISL_762701 | Lighthouse Lab in Glasgow | Wellcome Sanger Institute for the COVID-19 Genomics UK (COG-UK) Consortium | Harper VanSteenhouse, Yumi Kasai, David Gray, Carol Clugston, Anna Dominiczak and Alex Alderton, Roberto Amato, Sonia Goncalves, Ewan Harrison, David K. Jackson, Ian Johnston, Dominic Kwiatkowski, Cordelia Langford, John Sillitoe on behalf of the Wellcome Sanger Institute COVID-19 Surveillance Team |
| EPI_ISL_762702, EPI_ISL_762703 | Lighthouse Lab in Alderley Park | Wellcome Sanger Institute for the COVID-19 Genomics UK (COG-UK) Consortium | Jacquelyn Wynn, Mairead Hyland, The Lighthouse Lab in Alderley Park and Alex Alderton, Roberto Amato, Sonia Goncalves, Ewan Harrison, David K. Jackson, Ian Johnston, Dominic Kwiatkowski, Cordelia Langford, John Sillitoe on behalf of the Wellcome Sanger Institute COVID-19 Surveillance Team |
| EPI_ISL_762704 | Lighthouse Lab in Glasgow | Wellcome Sanger Institute for the COVID-19 Genomics UK (COG-UK) Consortium | Harper VanSteenhouse, Yumi Kasai, David Gray, Carol Clugston, Anna Dominiczak and Alex Alderton, Roberto Amato, Sonia Goncalves, Ewan Harrison, David K. Jackson, Ian Johnston, Dominic Kwiatkowski, Cordelia Langford, John Sillitoe on behalf of the Wellcome Sanger Institute COVID-19 Surveillance Team |
| EPI_ISL_762705 | Lighthouse Lab in Cambridge | Wellcome Sanger Institute for the COVID-19 Genomics UK (COG-UK) Consortium | Rob Howes, The Lighthouse Lab in Cambridge and Alex Alderton, Roberto Amato, Sonia Goncalves, Ewan Harrison, David K. Jackson, Ian Johnston, Dominic Kwiatkowski, Cordelia Langford, John Sillitoe on behalf of the Wellcome Sanger Institute COVID-19 Surveillance Team |
| EPI_ISL_762706, EPI_ISL_762707 | Lighthouse Lab in Alderley Park | Wellcome Sanger Institute for the COVID-19 Genomics UK (COG-UK) Consortium | Jacquelyn Wynn, Mairead Hyland, The Lighthouse Lab in Alderley Park and Alex Alderton, Roberto Amato, Sonia Goncalves, Ewan Harrison, David K. Jackson, Ian Johnston, Dominic Kwiatkowski, Cordelia Langford, John Sillitoe on behalf of the Wellcome Sanger Institute COVID-19 Surveillance Team |
| EPI_ISL_762708 | Lighthouse Lab in Glasgow | Wellcome Sanger Institute for the COVID-19 Genomics UK (COG-UK) Consortium | Harper VanSteenhouse, Yumi Kasai, David Gray, Carol Clugston, Anna Dominiczak and Alex Alderton, Roberto Amato, Sonia Goncalves, Ewan Harrison, David K. Jackson, Ian Johnston, Dominic Kwiatkowski, Cordelia Langford, John Sillitoe on behalf of the Wellcome Sanger Institute COVID-19 Surveillance Team |
| EPI_ISL_762709, EPI_ISL_762710 | Lighthouse Lab in Alderley Park | Wellcome Sanger Institute for the COVID-19 Genomics UK (COG-UK) Consortium | Jacquelyn Wynn, Mairead Hyland, The Lighthouse Lab in Alderley Park and Alex Alderton, Roberto Amato, Sonia Goncalves, Ewan Harrison, David K. Jackson, Ian Johnston, Dominic Kwiatkowski, Cordelia Langford, John Sillitoe on behalf of the Wellcome Sanger Institute COVID-19 Surveillance Team |
| EPI_ISL_762711 | Lighthouse Lab in Glasgow | Wellcome Sanger Institute for the COVID-19 Genomics UK (COG-UK) Consortium | Harper VanSteenhouse, Yumi Kasai, David Gray, Carol Clugston, Anna Dominiczak and Alex Alderton, Roberto Amato, Sonia Goncalves, Ewan Harrison, David K. Jackson, Ian Johnston, Dominic Kwiatkowski, Cordelia Langford, John Sillitoe on behalf of the Wellcome Sanger Institute COVID-19 Surveillance Team |
| EPI_ISL_762712 | Lighthouse Lab in Alderley Park | Wellcome Sanger Institute for the COVID-19 Genomics UK (COG-UK) Consortium | Jacquelyn Wynn, Mairead Hyland, The Lighthouse Lab in Alderley Park and Alex Alderton, Roberto Amato, Sonia Goncalves, Ewan Harrison, David K. Jackson, Ian Johnston, Dominic Kwiatkowski, Cordelia Langford, John Sillitoe on behalf of the Wellcome Sanger Institute COVID-19 Surveillance Team |
| EPI_ISL_762713 | Lighthouse Lab in Cambridge | Wellcome Sanger Institute for the COVID-19 Genomics UK (COG-UK) Consortium | Rob Howes, The Lighthouse Lab in Cambridge and Alex Alderton, Roberto Amato, Sonia Goncalves, Ewan Harrison, David K. Jackson, Ian Johnston, Dominic Kwiatkowski, Cordelia Langford, John Sillitoe on behalf of the Wellcome Sanger Institute COVID-19 Surveillance Team |
| EPI_ISL_762714 | Lighthouse Lab in Alderley Park | Wellcome Sanger Institute for the COVID-19 Genomics UK (COG-UK) Consortium | Jacquelyn Wynn, Mairead Hyland, The Lighthouse Lab in Alderley Park and Alex Alderton, Roberto Amato, Sonia Goncalves, Ewan Harrison, David K. Jackson, Ian Johnston, Dominic Kwiatkowski, Cordelia Langford, John Sillitoe on behalf of the Wellcome Sanger Institute COVID-19 Surveillance Team |
| EPI_ISL_762715, EPI_ISL_762716, EPI_ISL_762717, EPI_ISL_762718, EPI_ISL_762719 | Lighthouse Lab in Cambridge | Wellcome Sanger Institute for the COVID-19 Genomics UK (COG-UK) Consortium | Rob Howes, The Lighthouse Lab in Cambridge and Alex Alderton, Roberto Amato, Sonia Goncalves, Ewan Harrison, David K. Jackson, Ian Johnston, Dominic Kwiatkowski, Cordelia Langford, John Sillitoe on behalf of the Wellcome Sanger Institute COVID-19 Surveillance Team |
| EPI_ISL_762720 | Lighthouse Lab in Alderley Park | Wellcome Sanger Institute for the COVID-19 Genomics UK | Jacquelyn Wynn, Mairead Hyland, The Lighthouse Lab in Alderley Park and Alex Alderton, Roberto Amato, Sonia Goncalves, Ewan Harrison, David K. |

| | | | |
|---|---|---|---|
| | | (COG-UK) Consortium | Jackson, Ian Johnston, Dominic Kwiatkowski, Cordelia Langford, John Sillitoe on behalf of the Wellcome Sanger Institute COVID-19 Surveillance Team |
| EPI_ISL_762721 | Lighthouse Lab in Cambridge | Wellcome Sanger Institute for the COVID-19 Genomics UK (COG-UK) Consortium | Rob Howes, The Lighthouse Lab in Cambridge and Alex Alderton, Roberto Amato, Sonia Goncalves, Ewan Harrison, David K. Jackson, Ian Johnston, Dominic Kwiatkowski, Cordelia Langford, John Sillitoe on behalf of the Wellcome Sanger Institute COVID-19 Surveillance Team |
| EPI_ISL_762722 | Lighthouse Lab in Alderley Park | Wellcome Sanger Institute for the COVID-19 Genomics UK (COG-UK) Consortium | Jacquelyn Wynn, Mairead Hyland, The Lighthouse Lab in Alderley Park and Alex Alderton, Roberto Amato, Sonia Goncalves, Ewan Harrison, David K. Jackson, Ian Johnston, Dominic Kwiatkowski, Cordelia Langford, John Sillitoe on behalf of the Wellcome Sanger Institute COVID-19 Surveillance Team |
| EPI_ISL_762723 | Lighthouse Lab in Cambridge | Wellcome Sanger Institute for the COVID-19 Genomics UK (COG-UK) Consortium | Rob Howes, The Lighthouse Lab in Cambridge and Alex Alderton, Roberto Amato, Sonia Goncalves, Ewan Harrison, David K. Jackson, Ian Johnston, Dominic Kwiatkowski, Cordelia Langford, John Sillitoe on behalf of the Wellcome Sanger Institute COVID-19 Surveillance Team |
| EPI_ISL_762724 | Lighthouse Lab in Alderley Park | Wellcome Sanger Institute for the COVID-19 Genomics UK (COG-UK) Consortium | Jacquelyn Wynn, Mairead Hyland, The Lighthouse Lab in Alderley Park and Alex Alderton, Roberto Amato, Sonia Goncalves, Ewan Harrison, David K. Jackson, Ian Johnston, Dominic Kwiatkowski, Cordelia Langford, John Sillitoe on behalf of the Wellcome Sanger Institute COVID-19 Surveillance Team |
| EPI_ISL_762725 | Lighthouse Lab in Cambridge | Wellcome Sanger Institute for the COVID-19 Genomics UK (COG-UK) Consortium | Rob Howes, The Lighthouse Lab in Cambridge and Alex Alderton, Roberto Amato, Sonia Goncalves, Ewan Harrison, David K. Jackson, Ian Johnston, Dominic Kwiatkowski, Cordelia Langford, John Sillitoe on behalf of the Wellcome Sanger Institute COVID-19 Surveillance Team |
| EPI_ISL_762726 | Lighthouse Lab in Glasgow | Wellcome Sanger Institute for the COVID-19 Genomics UK (COG-UK) Consortium | Harper VanSteenhouse, Yumi Kasai, David Gray, Carol Clugston, Anna Dominiczak and Alex Alderton, Roberto Amato, Sonia Goncalves, Ewan Harrison, David K. Jackson, Ian Johnston, Dominic Kwiatkowski, Cordelia Langford, John Sillitoe on behalf of the Wellcome Sanger Institute COVID-19 Surveillance Team |
| EPI_ISL_762727 | Lighthouse Lab in Cambridge | Wellcome Sanger Institute for the COVID-19 Genomics UK (COG-UK) Consortium | Rob Howes, The Lighthouse Lab in Cambridge and Alex Alderton, Roberto Amato, Sonia Goncalves, Ewan Harrison, David K. Jackson, Ian Johnston, Dominic Kwiatkowski, Cordelia Langford, John Sillitoe on behalf of the Wellcome Sanger Institute COVID-19 Surveillance Team |
| EPI_ISL_762728 | Lighthouse Lab in Glasgow | Wellcome Sanger Institute for the COVID-19 Genomics UK (COG-UK) Consortium | Harper VanSteenhouse, Yumi Kasai, David Gray, Carol Clugston, Anna Dominiczak and Alex Alderton, Roberto Amato, Sonia Goncalves, Ewan Harrison, David K. Jackson, Ian Johnston, Dominic Kwiatkowski, Cordelia Langford, John Sillitoe on behalf of the Wellcome Sanger Institute COVID-19 Surveillance Team |
| EPI_ISL_762729 | Lighthouse Lab in Cambridge | Wellcome Sanger Institute for the COVID-19 Genomics UK (COG-UK) Consortium | Rob Howes, The Lighthouse Lab in Cambridge and Alex Alderton, Roberto Amato, Sonia Goncalves, Ewan Harrison, David K. Jackson, Ian Johnston, Dominic Kwiatkowski, Cordelia Langford, John Sillitoe on behalf of the Wellcome Sanger Institute COVID-19 Surveillance Team |
| EPI_ISL_762730 | Lighthouse Lab in Alderley Park | Wellcome Sanger Institute for the COVID-19 Genomics UK (COG-UK) Consortium | Jacquelyn Wynn, Mairead Hyland, The Lighthouse Lab in Alderley Park and Alex Alderton, Roberto Amato, Sonia Goncalves, Ewan Harrison, David K. Jackson, Ian Johnston, Dominic Kwiatkowski, Cordelia Langford, John Sillitoe on behalf of the Wellcome Sanger Institute COVID-19 Surveillance Team |
| EPI_ISL_762731, EPI_ISL_762732 | Lighthouse Lab in Glasgow | Wellcome Sanger Institute for the COVID-19 Genomics UK (COG-UK) Consortium | Harper VanSteenhouse, Yumi Kasai, David Gray, Carol Clugston, Anna Dominiczak and Alex Alderton, Roberto Amato, Sonia Goncalves, Ewan Harrison, David K. Jackson, Ian Johnston, Dominic Kwiatkowski, Cordelia Langford, John Sillitoe on behalf of the Wellcome Sanger Institute COVID-19 Surveillance Team |
| EPI_ISL_762733 | Lighthouse Lab in Alderley Park | Wellcome Sanger Institute for the COVID-19 Genomics UK (COG-UK) Consortium | Jacquelyn Wynn, Mairead Hyland, The Lighthouse Lab in Alderley Park and Alex Alderton, Roberto Amato, Sonia Goncalves, Ewan Harrison, David K. Jackson, Ian Johnston, Dominic Kwiatkowski, Cordelia Langford, John Sillitoe on behalf of the Wellcome Sanger Institute COVID-19 Surveillance Team |
| EPI_ISL_762734 | Lighthouse Lab in Glasgow | Wellcome Sanger Institute for the COVID-19 Genomics UK (COG-UK) Consortium | Harper VanSteenhouse, Yumi Kasai, David Gray, Carol Clugston, Anna Dominiczak and Alex Alderton, Roberto Amato, Sonia Goncalves, Ewan Harrison, David K. Jackson, Ian Johnston, Dominic Kwiatkowski, Cordelia Langford, John Sillitoe on behalf of the Wellcome Sanger Institute COVID-19 Surveillance Team |
| EPI_ISL_762735 | Lighthouse Lab in Alderley Park | Wellcome Sanger Institute for the COVID-19 Genomics UK (COG-UK) Consortium | Jacquelyn Wynn, Mairead Hyland, The Lighthouse Lab in Alderley Park and Alex Alderton, Roberto Amato, Sonia Goncalves, Ewan Harrison, David K. Jackson, Ian Johnston, Dominic Kwiatkowski, Cordelia Langford, John Sillitoe on behalf of the Wellcome Sanger Institute COVID-19 Surveillance Team |
| EPI_ISL_762736, EPI_ISL_762737, EPI_ISL_762738 | Lighthouse Lab in Cambridge | Wellcome Sanger Institute for the COVID-19 Genomics UK (COG-UK) Consortium | Rob Howes, The Lighthouse Lab in Cambridge and Alex Alderton, Roberto Amato, Sonia Goncalves, Ewan Harrison, David K. Jackson, Ian Johnston, Dominic Kwiatkowski, Cordelia Langford, John Sillitoe on behalf of the Wellcome Sanger Institute COVID-19 Surveillance Team |
| EPI_ISL_762739 | Lighthouse Lab in Glasgow | Wellcome Sanger Institute for the COVID-19 Genomics UK (COG-UK) Consortium | Harper VanSteenhouse, Yumi Kasai, David Gray, Carol Clugston, Anna Dominiczak and Alex Alderton, Roberto Amato, Sonia Goncalves, Ewan Harrison, David K. Jackson, Ian Johnston, Dominic Kwiatkowski, Cordelia Langford, John Sillitoe on behalf of the Wellcome Sanger Institute COVID-19 Surveillance Team |
| EPI_ISL_762740, EPI_ISL_762741, EPI_ISL_762742 | Lighthouse Lab in Cambridge | Wellcome Sanger Institute for the COVID-19 Genomics UK (COG-UK) Consortium | Rob Howes, The Lighthouse Lab in Cambridge and Alex Alderton, Roberto Amato, Sonia Goncalves, Ewan Harrison, David K. Jackson, Ian Johnston, Dominic Kwiatkowski, Cordelia Langford, John Sillitoe on behalf of the Wellcome Sanger Institute COVID-19 Surveillance Team |
| EPI_ISL_762743, EPI_ISL_762744 | Lighthouse Lab in Glasgow | Wellcome Sanger Institute for the COVID-19 Genomics UK (COG-UK) Consortium | Harper VanSteenhouse, Yumi Kasai, David Gray, Carol Clugston, Anna Dominiczak and Alex Alderton, Roberto Amato, Sonia Goncalves, Ewan Harrison, David K. Jackson, Ian Johnston, Dominic Kwiatkowski, Cordelia Langford, John Sillitoe on behalf of the Wellcome Sanger Institute COVID-19 Surveillance Team |
| EPI_ISL_762745 | Lighthouse Lab in Cambridge | Wellcome Sanger Institute for the COVID-19 Genomics UK (COG-UK) Consortium | Rob Howes, The Lighthouse Lab in Cambridge and Alex Alderton, Roberto Amato, Sonia Goncalves, Ewan Harrison, David K. Jackson, Ian Johnston, Dominic Kwiatkowski, Cordelia Langford, John Sillitoe on behalf of the Wellcome Sanger Institute COVID-19 Surveillance Team |
| EPI_ISL_762746 | Lighthouse Lab in Glasgow | Wellcome Sanger Institute for the COVID-19 Genomics UK (COG-UK) Consortium | Harper VanSteenhouse, Yumi Kasai, David Gray, Carol Clugston, Anna Dominiczak and Alex Alderton, Roberto Amato, Sonia Goncalves, Ewan Harrison, David K. Jackson, Ian Johnston, Dominic Kwiatkowski, Cordelia Langford, John Sillitoe on behalf of the Wellcome Sanger Institute COVID-19 Surveillance Team |
| EPI_ISL_762747, EPI_ISL_762748 | Lighthouse Lab in Cambridge | Wellcome Sanger Institute for the COVID-19 Genomics UK (COG-UK) Consortium | Rob Howes, The Lighthouse Lab in Cambridge and Alex Alderton, Roberto Amato, Sonia Goncalves, Ewan Harrison, David K. Jackson, Ian Johnston, Dominic Kwiatkowski, Cordelia Langford, John Sillitoe on behalf of the Wellcome Sanger Institute COVID-19 Surveillance Team |
| EPI_ISL_762749 | Lighthouse Lab in Glasgow | Wellcome Sanger Institute for the COVID-19 Genomics UK (COG-UK) Consortium | Harper VanSteenhouse, Yumi Kasai, David Gray, Carol Clugston, Anna Dominiczak and Alex Alderton, Roberto Amato, Sonia Goncalves, Ewan Harrison, David K. Jackson, Ian Johnston, Dominic Kwiatkowski, Cordelia Langford, John Sillitoe on behalf of the Wellcome Sanger Institute COVID-19 Surveillance Team |
| EPI_ISL_762750 | Lighthouse Lab in Alderley Park | Wellcome Sanger Institute for the COVID-19 Genomics UK (COG-UK) Consortium | Jacquelyn Wynn, Mairead Hyland, The Lighthouse Lab in Alderley Park and Alex Alderton, Roberto Amato, Sonia Goncalves, Ewan Harrison, David K. Jackson, Ian Johnston, Dominic Kwiatkowski, Cordelia Langford, John Sillitoe on behalf of the Wellcome Sanger Institute COVID-19 Surveillance Team |
| EPI_ISL_762751, EPI_ISL_762752, EPI_ISL_762753 | Lighthouse Lab in Cambridge | Wellcome Sanger Institute for the COVID-19 Genomics UK (COG-UK) Consortium | Rob Howes, The Lighthouse Lab in Cambridge and Alex Alderton, Roberto Amato, Sonia Goncalves, Ewan Harrison, David K. Jackson, Ian Johnston, Dominic Kwiatkowski, Cordelia Langford, John Sillitoe on behalf of the Wellcome Sanger Institute COVID-19 Surveillance Team |
| EPI_ISL_762754 | Lighthouse Lab in Alderley Park | Wellcome Sanger Institute for the COVID-19 Genomics UK (COG-UK) Consortium | Jacquelyn Wynn, Mairead Hyland, The Lighthouse Lab in Alderley Park and Alex Alderton, Roberto Amato, Sonia Goncalves, Ewan Harrison, David K. Jackson, Ian Johnston, Dominic Kwiatkowski, Cordelia Langford, John Sillitoe on behalf of the Wellcome Sanger Institute COVID-19 Surveillance Team |
| EPI_ISL_762755 | Lighthouse Lab in Cambridge | Wellcome Sanger Institute for the COVID-19 Genomics UK (COG-UK) Consortium | Rob Howes, The Lighthouse Lab in Cambridge and Alex Alderton, Roberto Amato, Sonia Goncalves, Ewan Harrison, David K. Jackson, Ian Johnston, Dominic Kwiatkowski, Cordelia Langford, John Sillitoe on behalf of the Wellcome Sanger Institute COVID-19 Surveillance Team |
| EPI_ISL_762756 | Lighthouse Lab in Alderley Park | Wellcome Sanger Institute for the COVID-19 Genomics UK (COG-UK) Consortium | Jacquelyn Wynn, Mairead Hyland, The Lighthouse Lab in Alderley Park and Alex Alderton, Roberto Amato, Sonia Goncalves, Ewan Harrison, David K. Jackson, Ian Johnston, Dominic Kwiatkowski, Cordelia Langford, John Sillitoe on behalf of the Wellcome Sanger Institute COVID-19 Surveillance Team |
| EPI_ISL_762757 | Lighthouse Lab in Cambridge | Wellcome Sanger Institute for the COVID-19 Genomics UK (COG-UK) Consortium | Rob Howes, The Lighthouse Lab in Cambridge and Alex Alderton, Roberto Amato, Sonia Goncalves, Ewan Harrison, David K. Jackson, Ian Johnston, Dominic Kwiatkowski, Cordelia Langford, John Sillitoe on behalf of the Wellcome Sanger Institute COVID-19 Surveillance Team |
| EPI_ISL_762758, EPI_ISL_762759, EPI_ISL_762760, EPI_ISL_762761 | Lighthouse Lab in Glasgow | Wellcome Sanger Institute for the COVID-19 Genomics UK (COG-UK) Consortium | Harper VanSteenhouse, Yumi Kasai, David Gray, Carol Clugston, Anna Dominiczak and Alex Alderton, Roberto Amato, Sonia Goncalves, Ewan Harrison, David K. Jackson, Ian Johnston, Dominic Kwiatkowski, Cordelia Langford, John Sillitoe on behalf of the Wellcome Sanger Institute COVID-19 Surveillance Team |
| EPI_ISL_762762 | Lighthouse Lab in Cambridge | Wellcome Sanger Institute for the COVID-19 Genomics UK (COG-UK) Consortium | Rob Howes, The Lighthouse Lab in Cambridge and Alex Alderton, Roberto Amato, Sonia Goncalves, Ewan Harrison, David K. Jackson, Ian Johnston, Dominic Kwiatkowski, Cordelia Langford, John Sillitoe on behalf of the Wellcome Sanger Institute COVID-19 Surveillance Team |
| EPI_ISL_762763, EPI_ISL_762764 | Lighthouse Lab in Alderley Park | Wellcome Sanger Institute for the COVID-19 Genomics UK (COG-UK) Consortium | Jacquelyn Wynn, Mairead Hyland, The Lighthouse Lab in Alderley Park and Alex Alderton, Roberto Amato, Sonia Goncalves, Ewan Harrison, David K. Jackson, Ian Johnston, Dominic Kwiatkowski, Cordelia Langford, John Sillitoe on behalf of the Wellcome Sanger Institute COVID-19 Surveillance Team |

| | | | |
|---|---|---|---|
| EPI_ISL_762765 | Lighthouse Lab in Cambridge | Wellcome Sanger Institute for the COVID-19 Genomics UK (COG-UK) Consortium | Rob Howes, The Lighthouse Lab in Cambridge and Alex Alderton, Roberto Amato, Sonia Goncalves, Ewan Harrison, David K. Jackson, Ian Johnston, Dominic Kwiatkowski, Cordelia Langford, John Sillitoe on behalf of the Wellcome Sanger Institute COVID-19 Surveillance Team |
| EPI_ISL_762766 | Lighthouse Lab in Glasgow | Wellcome Sanger Institute for the COVID-19 Genomics UK (COG-UK) Consortium | Harper VanSteenhouse, Yumi Kasai, David Gray, Carol Clugston, Anna Dominiczak and Alex Alderton, Roberto Amato, Sonia Goncalves, Ewan Harrison, David K. Jackson, Ian Johnston, Dominic Kwiatkowski, Cordelia Langford, John Sillitoe on behalf of the Wellcome Sanger Institute COVID-19 Surveillance Team |
| EPI_ISL_762767, EPI_ISL_762768, EPI_ISL_762769 | Lighthouse Lab in Cambridge | Wellcome Sanger Institute for the COVID-19 Genomics UK (COG-UK) Consortium | Rob Howes, The Lighthouse Lab in Cambridge and Alex Alderton, Roberto Amato, Sonia Goncalves, Ewan Harrison, David K. Jackson, Ian Johnston, Dominic Kwiatkowski, Cordelia Langford, John Sillitoe on behalf of the Wellcome Sanger Institute COVID-19 Surveillance Team |
| EPI_ISL_762770 | Lighthouse Lab in Glasgow | Wellcome Sanger Institute for the COVID-19 Genomics UK (COG-UK) Consortium | Harper VanSteenhouse, Yumi Kasai, David Gray, Carol Clugston, Anna Dominiczak and Alex Alderton, Roberto Amato, Sonia Goncalves, Ewan Harrison, David K. Jackson, Ian Johnston, Dominic Kwiatkowski, Cordelia Langford, John Sillitoe on behalf of the Wellcome Sanger Institute COVID-19 Surveillance Team |
| EPI_ISL_762771 | Lighthouse Lab in Alderley Park | Wellcome Sanger Institute for the COVID-19 Genomics UK (COG-UK) Consortium | Jacquelyn Wynn, Mairead Hyland, The Lighthouse Lab in Alderley Park and Alex Alderton, Roberto Amato, Sonia Goncalves, Ewan Harrison, David K. Jackson, Ian Johnston, Dominic Kwiatkowski, Cordelia Langford, John Sillitoe on behalf of the Wellcome Sanger Institute COVID-19 Surveillance Team |
| EPI_ISL_762772 | Lighthouse Lab in Cambridge | Wellcome Sanger Institute for the COVID-19 Genomics UK (COG-UK) Consortium | Rob Howes, The Lighthouse Lab in Cambridge and Alex Alderton, Roberto Amato, Sonia Goncalves, Ewan Harrison, David K. Jackson, Ian Johnston, Dominic Kwiatkowski, Cordelia Langford, John Sillitoe on behalf of the Wellcome Sanger Institute COVID-19 Surveillance Team |
| EPI_ISL_762773 | Lighthouse Lab in Glasgow | Wellcome Sanger Institute for the COVID-19 Genomics UK (COG-UK) Consortium | Harper VanSteenhouse, Yumi Kasai, David Gray, Carol Clugston, Anna Dominiczak and Alex Alderton, Roberto Amato, Sonia Goncalves, Ewan Harrison, David K. Jackson, Ian Johnston, Dominic Kwiatkowski, Cordelia Langford, John Sillitoe on behalf of the Wellcome Sanger Institute COVID-19 Surveillance Team |
| EPI_ISL_762774 | Lighthouse Lab in Cambridge | Wellcome Sanger Institute for the COVID-19 Genomics UK (COG-UK) Consortium | Rob Howes, The Lighthouse Lab in Cambridge and Alex Alderton, Roberto Amato, Sonia Goncalves, Ewan Harrison, David K. Jackson, Ian Johnston, Dominic Kwiatkowski, Cordelia Langford, John Sillitoe on behalf of the Wellcome Sanger Institute COVID-19 Surveillance Team |
| EPI_ISL_762775, EPI_ISL_762776 | Lighthouse Lab in Glasgow | Wellcome Sanger Institute for the COVID-19 Genomics UK (COG-UK) Consortium | Harper VanSteenhouse, Yumi Kasai, David Gray, Carol Clugston, Anna Dominiczak and Alex Alderton, Roberto Amato, Sonia Goncalves, Ewan Harrison, David K. Jackson, Ian Johnston, Dominic Kwiatkowski, Cordelia Langford, John Sillitoe on behalf of the Wellcome Sanger Institute COVID-19 Surveillance Team |
| EPI_ISL_762777 | Lighthouse Lab in Alderley Park | Wellcome Sanger Institute for the COVID-19 Genomics UK (COG-UK) Consortium | Jacquelyn Wynn, Mairead Hyland, The Lighthouse Lab in Alderley Park and Alex Alderton, Roberto Amato, Sonia Goncalves, Ewan Harrison, David K. Jackson, Ian Johnston, Dominic Kwiatkowski, Cordelia Langford, John Sillitoe on behalf of the Wellcome Sanger Institute COVID-19 Surveillance Team |
| EPI_ISL_762778 | Lighthouse Lab in Cambridge | Wellcome Sanger Institute for the COVID-19 Genomics UK (COG-UK) Consortium | Rob Howes, The Lighthouse Lab in Cambridge and Alex Alderton, Roberto Amato, Sonia Goncalves, Ewan Harrison, David K. Jackson, Ian Johnston, Dominic Kwiatkowski, Cordelia Langford, John Sillitoe on behalf of the Wellcome Sanger Institute COVID-19 Surveillance Team |
| EPI_ISL_762779 | Lighthouse Lab in Glasgow | Wellcome Sanger Institute for the COVID-19 Genomics UK (COG-UK) Consortium | Harper VanSteenhouse, Yumi Kasai, David Gray, Carol Clugston, Anna Dominiczak and Alex Alderton, Roberto Amato, Sonia Goncalves, Ewan Harrison, David K. Jackson, Ian Johnston, Dominic Kwiatkowski, Cordelia Langford, John Sillitoe on behalf of the Wellcome Sanger Institute COVID-19 Surveillance Team |
| EPI_ISL_762780, EPI_ISL_762781, EPI_ISL_762782, EPI_ISL_762783, EPI_ISL_762784 | Lighthouse Lab in Cambridge | Wellcome Sanger Institute for the COVID-19 Genomics UK (COG-UK) Consortium | Rob Howes, The Lighthouse Lab in Cambridge and Alex Alderton, Roberto Amato, Sonia Goncalves, Ewan Harrison, David K. Jackson, Ian Johnston, Dominic Kwiatkowski, Cordelia Langford, John Sillitoe on behalf of the Wellcome Sanger Institute COVID-19 Surveillance Team |
| EPI_ISL_762785 | Lighthouse Lab in Glasgow | Wellcome Sanger Institute for the COVID-19 Genomics UK (COG-UK) Consortium | Harper VanSteenhouse, Yumi Kasai, David Gray, Carol Clugston, Anna Dominiczak and Alex Alderton, Roberto Amato, Sonia Goncalves, Ewan Harrison, David K. Jackson, Ian Johnston, Dominic Kwiatkowski, Cordelia Langford, John Sillitoe on behalf of the Wellcome Sanger Institute COVID-19 Surveillance Team |
| EPI_ISL_762786 | Lighthouse Lab in Alderley Park | Wellcome Sanger Institute for the COVID-19 Genomics UK (COG-UK) Consortium | Jacquelyn Wynn, Mairead Hyland, The Lighthouse Lab in Alderley Park and Alex Alderton, Roberto Amato, Sonia Goncalves, Ewan Harrison, David K. Jackson, Ian Johnston, Dominic Kwiatkowski, Cordelia Langford, John Sillitoe on behalf of the Wellcome Sanger Institute COVID-19 Surveillance Team |
| EPI_ISL_762787, EPI_ISL_762788 | Lighthouse Lab in Cambridge | Wellcome Sanger Institute for the COVID-19 Genomics UK (COG-UK) Consortium | Rob Howes, The Lighthouse Lab in Cambridge and Alex Alderton, Roberto Amato, Sonia Goncalves, Ewan Harrison, David K. Jackson, Ian Johnston, Dominic Kwiatkowski, Cordelia Langford, John Sillitoe on behalf of the Wellcome Sanger Institute COVID-19 Surveillance Team |
| EPI_ISL_762789 | Lighthouse Lab in Alderley Park | Wellcome Sanger Institute for the COVID-19 Genomics UK (COG-UK) Consortium | Jacquelyn Wynn, Mairead Hyland, The Lighthouse Lab in Alderley Park and Alex Alderton, Roberto Amato, Sonia Goncalves, Ewan Harrison, David K. Jackson, Ian Johnston, Dominic Kwiatkowski, Cordelia Langford, John Sillitoe on behalf of the Wellcome Sanger Institute COVID-19 Surveillance Team |
| EPI_ISL_762790 | Lighthouse Lab in Cambridge | Wellcome Sanger Institute for the COVID-19 Genomics UK (COG-UK) Consortium | Rob Howes, The Lighthouse Lab in Cambridge and Alex Alderton, Roberto Amato, Sonia Goncalves, Ewan Harrison, David K. Jackson, Ian Johnston, Dominic Kwiatkowski, Cordelia Langford, John Sillitoe on behalf of the Wellcome Sanger Institute COVID-19 Surveillance Team |
| EPI_ISL_762791 | Lighthouse Lab in Alderley Park | Wellcome Sanger Institute for the COVID-19 Genomics UK (COG-UK) Consortium | Jacquelyn Wynn, Mairead Hyland, The Lighthouse Lab in Alderley Park and Alex Alderton, Roberto Amato, Sonia Goncalves, Ewan Harrison, David K. Jackson, Ian Johnston, Dominic Kwiatkowski, Cordelia Langford, John Sillitoe on behalf of the Wellcome Sanger Institute COVID-19 Surveillance Team |
| EPI_ISL_762792 | Lighthouse Lab in Cambridge | Wellcome Sanger Institute for the COVID-19 Genomics UK (COG-UK) Consortium | Rob Howes, The Lighthouse Lab in Cambridge and Alex Alderton, Roberto Amato, Sonia Goncalves, Ewan Harrison, David K. Jackson, Ian Johnston, Dominic Kwiatkowski, Cordelia Langford, John Sillitoe on behalf of the Wellcome Sanger Institute COVID-19 Surveillance Team |
| EPI_ISL_762793 | Lighthouse Lab in Alderley Park | Wellcome Sanger Institute for the COVID-19 Genomics UK (COG-UK) Consortium | Jacquelyn Wynn, Mairead Hyland, The Lighthouse Lab in Alderley Park and Alex Alderton, Roberto Amato, Sonia Goncalves, Ewan Harrison, David K. Jackson, Ian Johnston, Dominic Kwiatkowski, Cordelia Langford, John Sillitoe on behalf of the Wellcome Sanger Institute COVID-19 Surveillance Team |
| EPI_ISL_762794 | Lighthouse Lab in Glasgow | Wellcome Sanger Institute for the COVID-19 Genomics UK (COG-UK) Consortium | Harper VanSteenhouse, Yumi Kasai, David Gray, Carol Clugston, Anna Dominiczak and Alex Alderton, Roberto Amato, Sonia Goncalves, Ewan Harrison, David K. Jackson, Ian Johnston, Dominic Kwiatkowski, Cordelia Langford, John Sillitoe on behalf of the Wellcome Sanger Institute COVID-19 Surveillance Team |
| EPI_ISL_762795 | Lighthouse Lab in Cambridge | Wellcome Sanger Institute for the COVID-19 Genomics UK (COG-UK) Consortium | Rob Howes, The Lighthouse Lab in Cambridge and Alex Alderton, Roberto Amato, Sonia Goncalves, Ewan Harrison, David K. Jackson, Ian Johnston, Dominic Kwiatkowski, Cordelia Langford, John Sillitoe on behalf of the Wellcome Sanger Institute COVID-19 Surveillance Team |
| EPI_ISL_762796 | Lighthouse Lab in Alderley Park | Wellcome Sanger Institute for the COVID-19 Genomics UK (COG-UK) Consortium | Jacquelyn Wynn, Mairead Hyland, The Lighthouse Lab in Alderley Park and Alex Alderton, Roberto Amato, Sonia Goncalves, Ewan Harrison, David K. Jackson, Ian Johnston, Dominic Kwiatkowski, Cordelia Langford, John Sillitoe on behalf of the Wellcome Sanger Institute COVID-19 Surveillance Team |
| EPI_ISL_762797 | Lighthouse Lab in Cambridge | Wellcome Sanger Institute for the COVID-19 Genomics UK (COG-UK) Consortium | Rob Howes, The Lighthouse Lab in Cambridge and Alex Alderton, Roberto Amato, Sonia Goncalves, Ewan Harrison, David K. Jackson, Ian Johnston, Dominic Kwiatkowski, Cordelia Langford, John Sillitoe on behalf of the Wellcome Sanger Institute COVID-19 Surveillance Team |
| EPI_ISL_762798, EPI_ISL_762799 | Lighthouse Lab in Alderley Park | Wellcome Sanger Institute for the COVID-19 Genomics UK (COG-UK) Consortium | Jacquelyn Wynn, Mairead Hyland, The Lighthouse Lab in Alderley Park and Alex Alderton, Roberto Amato, Sonia Goncalves, Ewan Harrison, David K. Jackson, Ian Johnston, Dominic Kwiatkowski, Cordelia Langford, John Sillitoe on behalf of the Wellcome Sanger Institute COVID-19 Surveillance Team |
| EPI_ISL_762800 | Lighthouse Lab in Cambridge | Wellcome Sanger Institute for the COVID-19 Genomics UK (COG-UK) Consortium | Rob Howes, The Lighthouse Lab in Cambridge and Alex Alderton, Roberto Amato, Sonia Goncalves, Ewan Harrison, David K. Jackson, Ian Johnston, Dominic Kwiatkowski, Cordelia Langford, John Sillitoe on behalf of the Wellcome Sanger Institute COVID-19 Surveillance Team |
| EPI_ISL_762801, EPI_ISL_762802 | Lighthouse Lab in Glasgow | Wellcome Sanger Institute for the COVID-19 Genomics UK (COG-UK) Consortium | Harper VanSteenhouse, Yumi Kasai, David Gray, Carol Clugston, Anna Dominiczak and Alex Alderton, Roberto Amato, Sonia Goncalves, Ewan Harrison, David K. Jackson, Ian Johnston, Dominic Kwiatkowski, Cordelia Langford, John Sillitoe on behalf of the Wellcome Sanger Institute COVID-19 Surveillance Team |
| EPI_ISL_762803, EPI_ISL_762804, EPI_ISL_762805, EPI_ISL_762806, EPI_ISL_762807, EPI_ISL_762808 | Lighthouse Lab in Cambridge | Wellcome Sanger Institute for the COVID-19 Genomics UK (COG-UK) Consortium | Rob Howes, The Lighthouse Lab in Cambridge and Alex Alderton, Roberto Amato, Sonia Goncalves, Ewan Harrison, David K. Jackson, Ian Johnston, Dominic Kwiatkowski, Cordelia Langford, John Sillitoe on behalf of the Wellcome Sanger Institute COVID-19 Surveillance Team |
| EPI_ISL_762809 | Lighthouse Lab in Glasgow | Wellcome Sanger Institute for the COVID-19 Genomics UK (COG-UK) Consortium | Harper VanSteenhouse, Yumi Kasai, David Gray, Carol Clugston, Anna Dominiczak and Alex Alderton, Roberto Amato, Sonia Goncalves, Ewan Harrison, David K. Jackson, Ian Johnston, Dominic Kwiatkowski, Cordelia Langford, John Sillitoe on behalf of the Wellcome Sanger Institute COVID-19 Surveillance Team |
| EPI_ISL_762810, EPI_ISL_762811 | Lighthouse Lab in Cambridge | Wellcome Sanger Institute for the COVID-19 Genomics UK (COG-UK) Consortium | Rob Howes, The Lighthouse Lab in Cambridge and Alex Alderton, Roberto Amato, Sonia Goncalves, Ewan Harrison, David K. Jackson, Ian Johnston, Dominic Kwiatkowski, Cordelia Langford, John Sillitoe on behalf of the Wellcome Sanger Institute COVID-19 Surveillance Team |

| | | | |
|---|---|---|---|
| EPI_ISL_762812 | Lighthouse Lab in Alderley Park | Wellcome Sanger Institute for the COVID-19 Genomics UK (COG-UK) Consortium | Jacquelyn Wynn, Mairead Hyland, The Lighthouse Lab in Alderley Park and Alex Alderton, Roberto Amato, Sonia Goncalves, Ewan Harrison, David K. Jackson, Ian Johnston, Dominic Kwiatkowski, Cordelia Langford, John Sillitoe on behalf of the Wellcome Sanger Institute COVID-19 Surveillance Team |
| EPI_ISL_762813, EPI_ISL_762814 | Lighthouse Lab in Glasgow | Wellcome Sanger Institute for the COVID-19 Genomics UK (COG-UK) Consortium | Harper VanSteenhouse, Yumi Kasai, David Gray, Carol Clugston, Anna Dominiczak and Alex Alderton, Roberto Amato, Sonia Goncalves, Ewan Harrison, David K. Jackson, Ian Johnston, Dominic Kwiatkowski, Cordelia Langford, John Sillitoe on behalf of the Wellcome Sanger Institute COVID-19 Surveillance Team |
| EPI_ISL_762815 | Lighthouse Lab in Alderley Park | Wellcome Sanger Institute for the COVID-19 Genomics UK (COG-UK) Consortium | Jacquelyn Wynn, Mairead Hyland, The Lighthouse Lab in Alderley Park and Alex Alderton, Roberto Amato, Sonia Goncalves, Ewan Harrison, David K. Jackson, Ian Johnston, Dominic Kwiatkowski, Cordelia Langford, John Sillitoe on behalf of the Wellcome Sanger Institute COVID-19 Surveillance Team |
| EPI_ISL_762816 | Lighthouse Lab in Glasgow | Wellcome Sanger Institute for the COVID-19 Genomics UK (COG-UK) Consortium | Harper VanSteenhouse, Yumi Kasai, David Gray, Carol Clugston, Anna Dominiczak and Alex Alderton, Roberto Amato, Sonia Goncalves, Ewan Harrison, David K. Jackson, Ian Johnston, Dominic Kwiatkowski, Cordelia Langford, John Sillitoe on behalf of the Wellcome Sanger Institute COVID-19 Surveillance Team |
| EPI_ISL_762817, EPI_ISL_762818 | Lighthouse Lab in Cambridge | Wellcome Sanger Institute for the COVID-19 Genomics UK (COG-UK) Consortium | Rob Howes, The Lighthouse Lab in Cambridge and Alex Alderton, Roberto Amato, Sonia Goncalves, Ewan Harrison, David K. Jackson, Ian Johnston, Dominic Kwiatkowski, Cordelia Langford, John Sillitoe on behalf of the Wellcome Sanger Institute COVID-19 Surveillance Team |
| EPI_ISL_762819 | Lighthouse Lab in Alderley Park | Wellcome Sanger Institute for the COVID-19 Genomics UK (COG-UK) Consortium | Jacquelyn Wynn, Mairead Hyland, The Lighthouse Lab in Alderley Park and Alex Alderton, Roberto Amato, Sonia Goncalves, Ewan Harrison, David K. Jackson, Ian Johnston, Dominic Kwiatkowski, Cordelia Langford, John Sillitoe on behalf of the Wellcome Sanger Institute COVID-19 Surveillance Team |
| EPI_ISL_762820 | Lighthouse Lab in Cambridge | Wellcome Sanger Institute for the COVID-19 Genomics UK (COG-UK) Consortium | Rob Howes, The Lighthouse Lab in Cambridge and Alex Alderton, Roberto Amato, Sonia Goncalves, Ewan Harrison, David K. Jackson, Ian Johnston, Dominic Kwiatkowski, Cordelia Langford, John Sillitoe on behalf of the Wellcome Sanger Institute COVID-19 Surveillance Team |
| EPI_ISL_762821 | Lighthouse Lab in Alderley Park | Wellcome Sanger Institute for the COVID-19 Genomics UK (COG-UK) Consortium | Jacquelyn Wynn, Mairead Hyland, The Lighthouse Lab in Alderley Park and Alex Alderton, Roberto Amato, Sonia Goncalves, Ewan Harrison, David K. Jackson, Ian Johnston, Dominic Kwiatkowski, Cordelia Langford, John Sillitoe on behalf of the Wellcome Sanger Institute COVID-19 Surveillance Team |
| EPI_ISL_762822 | Lighthouse Lab in Cambridge | Wellcome Sanger Institute for the COVID-19 Genomics UK (COG-UK) Consortium | Rob Howes, The Lighthouse Lab in Cambridge and Alex Alderton, Roberto Amato, Sonia Goncalves, Ewan Harrison, David K. Jackson, Ian Johnston, Dominic Kwiatkowski, Cordelia Langford, John Sillitoe on behalf of the Wellcome Sanger Institute COVID-19 Surveillance Team |
| EPI_ISL_762823 | Lighthouse Lab in Alderley Park | Wellcome Sanger Institute for the COVID-19 Genomics UK (COG-UK) Consortium | Jacquelyn Wynn, Mairead Hyland, The Lighthouse Lab in Alderley Park and Alex Alderton, Roberto Amato, Sonia Goncalves, Ewan Harrison, David K. Jackson, Ian Johnston, Dominic Kwiatkowski, Cordelia Langford, John Sillitoe on behalf of the Wellcome Sanger Institute COVID-19 Surveillance Team |
| EPI_ISL_762824 | Lighthouse Lab in Glasgow | Wellcome Sanger Institute for the COVID-19 Genomics UK (COG-UK) Consortium | Harper VanSteenhouse, Yumi Kasai, David Gray, Carol Clugston, Anna Dominiczak and Alex Alderton, Roberto Amato, Sonia Goncalves, Ewan Harrison, David K. Jackson, Ian Johnston, Dominic Kwiatkowski, Cordelia Langford, John Sillitoe on behalf of the Wellcome Sanger Institute COVID-19 Surveillance Team |
| EPI_ISL_762825 | Lighthouse Lab in Cambridge | Wellcome Sanger Institute for the COVID-19 Genomics UK (COG-UK) Consortium | Rob Howes, The Lighthouse Lab in Cambridge and Alex Alderton, Roberto Amato, Sonia Goncalves, Ewan Harrison, David K. Jackson, Ian Johnston, Dominic Kwiatkowski, Cordelia Langford, John Sillitoe on behalf of the Wellcome Sanger Institute COVID-19 Surveillance Team |
| EPI_ISL_762826 | Lighthouse Lab in Glasgow | Wellcome Sanger Institute for the COVID-19 Genomics UK (COG-UK) Consortium | Harper VanSteenhouse, Yumi Kasai, David Gray, Carol Clugston, Anna Dominiczak and Alex Alderton, Roberto Amato, Sonia Goncalves, Ewan Harrison, David K. Jackson, Ian Johnston, Dominic Kwiatkowski, Cordelia Langford, John Sillitoe on behalf of the Wellcome Sanger Institute COVID-19 Surveillance Team |
| EPI_ISL_762827 | Lighthouse Lab in Cambridge | Wellcome Sanger Institute for the COVID-19 Genomics UK (COG-UK) Consortium | Rob Howes, The Lighthouse Lab in Cambridge and Alex Alderton, Roberto Amato, Sonia Goncalves, Ewan Harrison, David K. Jackson, Ian Johnston, Dominic Kwiatkowski, Cordelia Langford, John Sillitoe on behalf of the Wellcome Sanger Institute COVID-19 Surveillance Team |
| EPI_ISL_762828 | Lighthouse Lab in Alderley Park | Wellcome Sanger Institute for the COVID-19 Genomics UK (COG-UK) Consortium | Jacquelyn Wynn, Mairead Hyland, The Lighthouse Lab in Alderley Park and Alex Alderton, Roberto Amato, Sonia Goncalves, Ewan Harrison, David K. Jackson, Ian Johnston, Dominic Kwiatkowski, Cordelia Langford, John Sillitoe on behalf of the Wellcome Sanger Institute COVID-19 Surveillance Team |
| EPI_ISL_762829 | Lighthouse Lab in Cambridge | Wellcome Sanger Institute for the COVID-19 Genomics UK (COG-UK) Consortium | Rob Howes, The Lighthouse Lab in Cambridge and Alex Alderton, Roberto Amato, Sonia Goncalves, Ewan Harrison, David K. Jackson, Ian Johnston, Dominic Kwiatkowski, Cordelia Langford, John Sillitoe on behalf of the Wellcome Sanger Institute COVID-19 Surveillance Team |
| EPI_ISL_762830 | Lighthouse Lab in Alderley Park | Wellcome Sanger Institute for the COVID-19 Genomics UK (COG-UK) Consortium | Jacquelyn Wynn, Mairead Hyland, The Lighthouse Lab in Alderley Park and Alex Alderton, Roberto Amato, Sonia Goncalves, Ewan Harrison, David K. Jackson, Ian Johnston, Dominic Kwiatkowski, Cordelia Langford, John Sillitoe on behalf of the Wellcome Sanger Institute COVID-19 Surveillance Team |
| EPI_ISL_762831 | Lighthouse Lab in Cambridge | Wellcome Sanger Institute for the COVID-19 Genomics UK (COG-UK) Consortium | Rob Howes, The Lighthouse Lab in Cambridge and Alex Alderton, Roberto Amato, Sonia Goncalves, Ewan Harrison, David K. Jackson, Ian Johnston, Dominic Kwiatkowski, Cordelia Langford, John Sillitoe on behalf of the Wellcome Sanger Institute COVID-19 Surveillance Team |
| EPI_ISL_762832 | Lighthouse Lab in Alderley Park | Wellcome Sanger Institute for the COVID-19 Genomics UK (COG-UK) Consortium | Jacquelyn Wynn, Mairead Hyland, The Lighthouse Lab in Alderley Park and Alex Alderton, Roberto Amato, Sonia Goncalves, Ewan Harrison, David K. Jackson, Ian Johnston, Dominic Kwiatkowski, Cordelia Langford, John Sillitoe on behalf of the Wellcome Sanger Institute COVID-19 Surveillance Team |
| EPI_ISL_762833, EPI_ISL_762834 | Lighthouse Lab in Cambridge | Wellcome Sanger Institute for the COVID-19 Genomics UK (COG-UK) Consortium | Rob Howes, The Lighthouse Lab in Cambridge and Alex Alderton, Roberto Amato, Sonia Goncalves, Ewan Harrison, David K. Jackson, Ian Johnston, Dominic Kwiatkowski, Cordelia Langford, John Sillitoe on behalf of the Wellcome Sanger Institute COVID-19 Surveillance Team |
| EPI_ISL_762835 | Lighthouse Lab in Alderley Park | Wellcome Sanger Institute for the COVID-19 Genomics UK (COG-UK) Consortium | Jacquelyn Wynn, Mairead Hyland, The Lighthouse Lab in Alderley Park and Alex Alderton, Roberto Amato, Sonia Goncalves, Ewan Harrison, David K. Jackson, Ian Johnston, Dominic Kwiatkowski, Cordelia Langford, John Sillitoe on behalf of the Wellcome Sanger Institute COVID-19 Surveillance Team |
| EPI_ISL_762836 | Lighthouse Lab in Cambridge | Wellcome Sanger Institute for the COVID-19 Genomics UK (COG-UK) Consortium | Rob Howes, The Lighthouse Lab in Cambridge and Alex Alderton, Roberto Amato, Sonia Goncalves, Ewan Harrison, David K. Jackson, Ian Johnston, Dominic Kwiatkowski, Cordelia Langford, John Sillitoe on behalf of the Wellcome Sanger Institute COVID-19 Surveillance Team |
| EPI_ISL_762837 | Lighthouse Lab in Glasgow | Wellcome Sanger Institute for the COVID-19 Genomics UK (COG-UK) Consortium | Harper VanSteenhouse, Yumi Kasai, David Gray, Carol Clugston, Anna Dominiczak and Alex Alderton, Roberto Amato, Sonia Goncalves, Ewan Harrison, David K. Jackson, Ian Johnston, Dominic Kwiatkowski, Cordelia Langford, John Sillitoe on behalf of the Wellcome Sanger Institute COVID-19 Surveillance Team |
| EPI_ISL_762838 | Lighthouse Lab in Cambridge | Wellcome Sanger Institute for the COVID-19 Genomics UK (COG-UK) Consortium | Rob Howes, The Lighthouse Lab in Cambridge and Alex Alderton, Roberto Amato, Sonia Goncalves, Ewan Harrison, David K. Jackson, Ian Johnston, Dominic Kwiatkowski, Cordelia Langford, John Sillitoe on behalf of the Wellcome Sanger Institute COVID-19 Surveillance Team |
| EPI_ISL_762839 | Lighthouse Lab in Alderley Park | Wellcome Sanger Institute for the COVID-19 Genomics UK (COG-UK) Consortium | Jacquelyn Wynn, Mairead Hyland, The Lighthouse Lab in Alderley Park and Alex Alderton, Roberto Amato, Sonia Goncalves, Ewan Harrison, David K. Jackson, Ian Johnston, Dominic Kwiatkowski, Cordelia Langford, John Sillitoe on behalf of the Wellcome Sanger Institute COVID-19 Surveillance Team |
| EPI_ISL_762840 | Lighthouse Lab in Cambridge | Wellcome Sanger Institute for the COVID-19 Genomics UK (COG-UK) Consortium | Rob Howes, The Lighthouse Lab in Cambridge and Alex Alderton, Roberto Amato, Sonia Goncalves, Ewan Harrison, David K. Jackson, Ian Johnston, Dominic Kwiatkowski, Cordelia Langford, John Sillitoe on behalf of the Wellcome Sanger Institute COVID-19 Surveillance Team |
| EPI_ISL_762841 | Lighthouse Lab in Alderley Park | Wellcome Sanger Institute for the COVID-19 Genomics UK (COG-UK) Consortium | Jacquelyn Wynn, Mairead Hyland, The Lighthouse Lab in Alderley Park and Alex Alderton, Roberto Amato, Sonia Goncalves, Ewan Harrison, David K. Jackson, Ian Johnston, Dominic Kwiatkowski, Cordelia Langford, John Sillitoe on behalf of the Wellcome Sanger Institute COVID-19 Surveillance Team |
| EPI_ISL_762842 | Lighthouse Lab in Cambridge | Wellcome Sanger Institute for the COVID-19 Genomics UK (COG-UK) Consortium | Rob Howes, The Lighthouse Lab in Cambridge and Alex Alderton, Roberto Amato, Sonia Goncalves, Ewan Harrison, David K. Jackson, Ian Johnston, Dominic Kwiatkowski, Cordelia Langford, John Sillitoe on behalf of the Wellcome Sanger Institute COVID-19 Surveillance Team |
| EPI_ISL_762843, EPI_ISL_762844, EPI_ISL_762845 | Lighthouse Lab in Alderley Park | Wellcome Sanger Institute for the COVID-19 Genomics UK (COG-UK) Consortium | Jacquelyn Wynn, Mairead Hyland, The Lighthouse Lab in Alderley Park and Alex Alderton, Roberto Amato, Sonia Goncalves, Ewan Harrison, David K. Jackson, Ian Johnston, Dominic Kwiatkowski, Cordelia Langford, John Sillitoe on behalf of the Wellcome Sanger Institute COVID-19 Surveillance Team |
| EPI_ISL_762846 | Lighthouse Lab in Glasgow | Wellcome Sanger Institute for the COVID-19 Genomics UK (COG-UK) Consortium | Harper VanSteenhouse, Yumi Kasai, David Gray, Carol Clugston, Anna Dominiczak and Alex Alderton, Roberto Amato, Sonia Goncalves, Ewan Harrison, David K. Jackson, Ian Johnston, Dominic Kwiatkowski, Cordelia Langford, John Sillitoe on behalf of the Wellcome Sanger Institute COVID-19 Surveillance Team |
| EPI_ISL_762847 | Lighthouse Lab in Alderley Park | Wellcome Sanger Institute for the COVID-19 Genomics UK (COG-UK) Consortium | Jacquelyn Wynn, Mairead Hyland, The Lighthouse Lab in Alderley Park and Alex Alderton, Roberto Amato, Sonia Goncalves, Ewan Harrison, David K. Jackson, Ian Johnston, Dominic Kwiatkowski, Cordelia Langford, John Sillitoe on behalf of the Wellcome Sanger Institute COVID-19 Surveillance Team |
| EPI_ISL_762848 | Lighthouse Lab in Cambridge | Wellcome Sanger Institute for the COVID-19 Genomics UK (COG-UK) Consortium | Rob Howes, The Lighthouse Lab in Cambridge and Alex Alderton, Roberto Amato, Sonia Goncalves, Ewan Harrison, David K. Jackson, Ian Johnston, Dominic Kwiatkowski, Cordelia Langford, John Sillitoe on behalf of the Wellcome Sanger Institute COVID-19 Surveillance Team |
| EPI_ISL_762849 | Lighthouse Lab in Alderley Park | Wellcome Sanger Institute for the COVID-19 Genomics UK | Jacquelyn Wynn, Mairead Hyland, The Lighthouse Lab in Alderley Park and Alex Alderton, Roberto Amato, Sonia Goncalves, Ewan Harrison, David K. |

| | | | |
|---|---|---|---|
| | | (COG-UK) Consortium | Jackson, Ian Johnston, Dominic Kwiatkowski, Cordelia Langford, John Sillitoe on behalf of the Wellcome Sanger Institute COVID-19 Surveillance Team |
| EPI_ISL_762850, EPI_ISL_762851 | Lighthouse Lab in Glasgow | Wellcome Sanger Institute for the COVID-19 Genomics UK (COG-UK) Consortium | Harper VanSteenhouse, Yumi Kasai, David Gray, Carol Clugston, Anna Dominiczak and Alex Alderton, Roberto Amato, Sonia Goncalves, Ewan Harrison, David K. Jackson, Ian Johnston, Dominic Kwiatkowski, Cordelia Langford, John Sillitoe on behalf of the Wellcome Sanger Institute COVID-19 Surveillance Team |
| EPI_ISL_762852 | Lighthouse Lab in Cambridge | Wellcome Sanger Institute for the COVID-19 Genomics UK (COG-UK) Consortium | Rob Howes, The Lighthouse Lab in Cambridge and Alex Alderton, Roberto Amato, Sonia Goncalves, Ewan Harrison, David K. Jackson, Ian Johnston, Dominic Kwiatkowski, Cordelia Langford, John Sillitoe on behalf of the Wellcome Sanger Institute COVID-19 Surveillance Team |
| EPI_ISL_762853, EPI_ISL_762854 | Lighthouse Lab in Glasgow | Wellcome Sanger Institute for the COVID-19 Genomics UK (COG-UK) Consortium | Harper VanSteenhouse, Yumi Kasai, David Gray, Carol Clugston, Anna Dominiczak and Alex Alderton, Roberto Amato, Sonia Goncalves, Ewan Harrison, David K. Jackson, Ian Johnston, Dominic Kwiatkowski, Cordelia Langford, John Sillitoe on behalf of the Wellcome Sanger Institute COVID-19 Surveillance Team |
| EPI_ISL_762855 | Lighthouse Lab in Cambridge | Wellcome Sanger Institute for the COVID-19 Genomics UK (COG-UK) Consortium | Rob Howes, The Lighthouse Lab in Cambridge and Alex Alderton, Roberto Amato, Sonia Goncalves, Ewan Harrison, David K. Jackson, Ian Johnston, Dominic Kwiatkowski, Cordelia Langford, John Sillitoe on behalf of the Wellcome Sanger Institute COVID-19 Surveillance Team |
| EPI_ISL_762856 | Lighthouse Lab in Glasgow | Wellcome Sanger Institute for the COVID-19 Genomics UK (COG-UK) Consortium | Harper VanSteenhouse, Yumi Kasai, David Gray, Carol Clugston, Anna Dominiczak and Alex Alderton, Roberto Amato, Sonia Goncalves, Ewan Harrison, David K. Jackson, Ian Johnston, Dominic Kwiatkowski, Cordelia Langford, John Sillitoe on behalf of the Wellcome Sanger Institute COVID-19 Surveillance Team |
| EPI_ISL_762857, EPI_ISL_762858 | Lighthouse Lab in Cambridge | Wellcome Sanger Institute for the COVID-19 Genomics UK (COG-UK) Consortium | Rob Howes, The Lighthouse Lab in Cambridge and Alex Alderton, Roberto Amato, Sonia Goncalves, Ewan Harrison, David K. Jackson, Ian Johnston, Dominic Kwiatkowski, Cordelia Langford, John Sillitoe on behalf of the Wellcome Sanger Institute COVID-19 Surveillance Team |
| EPI_ISL_762859, EPI_ISL_762860, EPI_ISL_762861 | Lighthouse Lab in Glasgow | Wellcome Sanger Institute for the COVID-19 Genomics UK (COG-UK) Consortium | Harper VanSteenhouse, Yumi Kasai, David Gray, Carol Clugston, Anna Dominiczak and Alex Alderton, Roberto Amato, Sonia Goncalves, Ewan Harrison, David K. Jackson, Ian Johnston, Dominic Kwiatkowski, Cordelia Langford, John Sillitoe on behalf of the Wellcome Sanger Institute COVID-19 Surveillance Team |
| EPI_ISL_762862 | Lighthouse Lab in Alderley Park | Wellcome Sanger Institute for the COVID-19 Genomics UK (COG-UK) Consortium | Jacquelyn Wynn, Mairead Hyland, The Lighthouse Lab in Alderley Park and Alex Alderton, Roberto Amato, Sonia Goncalves, Ewan Harrison, David K. Jackson, Ian Johnston, Dominic Kwiatkowski, Cordelia Langford, John Sillitoe on behalf of the Wellcome Sanger Institute COVID-19 Surveillance Team |
| EPI_ISL_762863, EPI_ISL_762865, EPI_ISL_762866, EPI_ISL_762868 | Lighthouse Lab in Glasgow | Wellcome Sanger Institute for the COVID-19 Genomics UK (COG-UK) Consortium | Harper VanSteenhouse, Yumi Kasai, David Gray, Carol Clugston, Anna Dominiczak and Alex Alderton, Roberto Amato, Sonia Goncalves, Ewan Harrison, David K. Jackson, Ian Johnston, Dominic Kwiatkowski, Cordelia Langford, John Sillitoe on behalf of the Wellcome Sanger Institute COVID-19 Surveillance Team |
| EPI_ISL_762869 | Lighthouse Lab in Alderley Park | Wellcome Sanger Institute for the COVID-19 Genomics UK (COG-UK) Consortium | Jacquelyn Wynn, Mairead Hyland, The Lighthouse Lab in Alderley Park and Alex Alderton, Roberto Amato, Sonia Goncalves, Ewan Harrison, David K. Jackson, Ian Johnston, Dominic Kwiatkowski, Cordelia Langford, John Sillitoe on behalf of the Wellcome Sanger Institute COVID-19 Surveillance Team |
| EPI_ISL_762870, EPI_ISL_762871, EPI_ISL_762872, EPI_ISL_762873, EPI_ISL_762874, EPI_ISL_762876, EPI_ISL_762877 | Lighthouse Lab in Glasgow | Wellcome Sanger Institute for the COVID-19 Genomics UK (COG-UK) Consortium | Harper VanSteenhouse, Yumi Kasai, David Gray, Carol Clugston, Anna Dominiczak and Alex Alderton, Roberto Amato, Sonia Goncalves, Ewan Harrison, David K. Jackson, Ian Johnston, Dominic Kwiatkowski, Cordelia Langford, John Sillitoe on behalf of the Wellcome Sanger Institute COVID-19 Surveillance Team |
| EPI_ISL_762878, EPI_ISL_762879 | Lighthouse Lab in Alderley Park | Wellcome Sanger Institute for the COVID-19 Genomics UK (COG-UK) Consortium | Jacquelyn Wynn, Mairead Hyland, The Lighthouse Lab in Alderley Park and Alex Alderton, Roberto Amato, Sonia Goncalves, Ewan Harrison, David K. Jackson, Ian Johnston, Dominic Kwiatkowski, Cordelia Langford, John Sillitoe on behalf of the Wellcome Sanger Institute COVID-19 Surveillance Team |
| EPI_ISL_762881, EPI_ISL_762882 | Lighthouse Lab in Glasgow | Wellcome Sanger Institute for the COVID-19 Genomics UK (COG-UK) Consortium | Harper VanSteenhouse, Yumi Kasai, David Gray, Carol Clugston, Anna Dominiczak and Alex Alderton, Roberto Amato, Sonia Goncalves, Ewan Harrison, David K. Jackson, Ian Johnston, Dominic Kwiatkowski, Cordelia Langford, John Sillitoe on behalf of the Wellcome Sanger Institute COVID-19 Surveillance Team |
| EPI_ISL_762883 | Lighthouse Lab in Alderley Park | Wellcome Sanger Institute for the COVID-19 Genomics UK (COG-UK) Consortium | Jacquelyn Wynn, Mairead Hyland, The Lighthouse Lab in Alderley Park and Alex Alderton, Roberto Amato, Sonia Goncalves, Ewan Harrison, David K. Jackson, Ian Johnston, Dominic Kwiatkowski, Cordelia Langford, John Sillitoe on behalf of the Wellcome Sanger Institute COVID-19 Surveillance Team |
| EPI_ISL_762885 | Lighthouse Lab in Glasgow | Wellcome Sanger Institute for the COVID-19 Genomics UK (COG-UK) Consortium | Harper VanSteenhouse, Yumi Kasai, David Gray, Carol Clugston, Anna Dominiczak and Alex Alderton, Roberto Amato, Sonia Goncalves, Ewan Harrison, David K. Jackson, Ian Johnston, Dominic Kwiatkowski, Cordelia Langford, John Sillitoe on behalf of the Wellcome Sanger Institute COVID-19 Surveillance Team |
| EPI_ISL_762886, EPI_ISL_762887 | Lighthouse Lab in Alderley Park | Wellcome Sanger Institute for the COVID-19 Genomics UK (COG-UK) Consortium | Jacquelyn Wynn, Mairead Hyland, The Lighthouse Lab in Alderley Park and Alex Alderton, Roberto Amato, Sonia Goncalves, Ewan Harrison, David K. Jackson, Ian Johnston, Dominic Kwiatkowski, Cordelia Langford, John Sillitoe on behalf of the Wellcome Sanger Institute COVID-19 Surveillance Team |
| EPI_ISL_762888, EPI_ISL_762889 | Lighthouse Lab in Glasgow | Wellcome Sanger Institute for the COVID-19 Genomics UK (COG-UK) Consortium | Harper VanSteenhouse, Yumi Kasai, David Gray, Carol Clugston, Anna Dominiczak and Alex Alderton, Roberto Amato, Sonia Goncalves, Ewan Harrison, David K. Jackson, Ian Johnston, Dominic Kwiatkowski, Cordelia Langford, John Sillitoe on behalf of the Wellcome Sanger Institute COVID-19 Surveillance Team |
| EPI_ISL_762890 | Lighthouse Lab in Alderley Park | Wellcome Sanger Institute for the COVID-19 Genomics UK (COG-UK) Consortium | Jacquelyn Wynn, Mairead Hyland, The Lighthouse Lab in Alderley Park and Alex Alderton, Roberto Amato, Sonia Goncalves, Ewan Harrison, David K. Jackson, Ian Johnston, Dominic Kwiatkowski, Cordelia Langford, John Sillitoe on behalf of the Wellcome Sanger Institute COVID-19 Surveillance Team |
| EPI_ISL_762892, EPI_ISL_762893 | Lighthouse Lab in Glasgow | Wellcome Sanger Institute for the COVID-19 Genomics UK (COG-UK) Consortium | Harper VanSteenhouse, Yumi Kasai, David Gray, Carol Clugston, Anna Dominiczak and Alex Alderton, Roberto Amato, Sonia Goncalves, Ewan Harrison, David K. Jackson, Ian Johnston, Dominic Kwiatkowski, Cordelia Langford, John Sillitoe on behalf of the Wellcome Sanger Institute COVID-19 Surveillance Team |
| EPI_ISL_762894 | Lighthouse Lab in Alderley Park | Wellcome Sanger Institute for the COVID-19 Genomics UK (COG-UK) Consortium | Jacquelyn Wynn, Mairead Hyland, The Lighthouse Lab in Alderley Park and Alex Alderton, Roberto Amato, Sonia Goncalves, Ewan Harrison, David K. Jackson, Ian Johnston, Dominic Kwiatkowski, Cordelia Langford, John Sillitoe on behalf of the Wellcome Sanger Institute COVID-19 Surveillance Team |
| EPI_ISL_762897, EPI_ISL_762898, EPI_ISL_762899 | Lighthouse Lab in Glasgow | Wellcome Sanger Institute for the COVID-19 Genomics UK (COG-UK) Consortium | Harper VanSteenhouse, Yumi Kasai, David Gray, Carol Clugston, Anna Dominiczak and Alex Alderton, Roberto Amato, Sonia Goncalves, Ewan Harrison, David K. Jackson, Ian Johnston, Dominic Kwiatkowski, Cordelia Langford, John Sillitoe on behalf of the Wellcome Sanger Institute COVID-19 Surveillance Team |
| EPI_ISL_762900, EPI_ISL_762901, EPI_ISL_762902 | Lighthouse Lab in Alderley Park | Wellcome Sanger Institute for the COVID-19 Genomics UK (COG-UK) Consortium | Jacquelyn Wynn, Mairead Hyland, The Lighthouse Lab in Alderley Park and Alex Alderton, Roberto Amato, Sonia Goncalves, Ewan Harrison, David K. Jackson, Ian Johnston, Dominic Kwiatkowski, Cordelia Langford, John Sillitoe on behalf of the Wellcome Sanger Institute COVID-19 Surveillance Team |
| EPI_ISL_762904, EPI_ISL_762905 | Lighthouse Lab in Glasgow | Wellcome Sanger Institute for the COVID-19 Genomics UK (COG-UK) Consortium | Harper VanSteenhouse, Yumi Kasai, David Gray, Carol Clugston, Anna Dominiczak and Alex Alderton, Roberto Amato, Sonia Goncalves, Ewan Harrison, David K. Jackson, Ian Johnston, Dominic Kwiatkowski, Cordelia Langford, John Sillitoe on behalf of the Wellcome Sanger Institute COVID-19 Surveillance Team |
| EPI_ISL_762906 | Lighthouse Lab in Alderley Park | Wellcome Sanger Institute for the COVID-19 Genomics UK (COG-UK) Consortium | Jacquelyn Wynn, Mairead Hyland, The Lighthouse Lab in Alderley Park and Alex Alderton, Roberto Amato, Sonia Goncalves, Ewan Harrison, David K. Jackson, Ian Johnston, Dominic Kwiatkowski, Cordelia Langford, John Sillitoe on behalf of the Wellcome Sanger Institute COVID-19 Surveillance Team |
| EPI_ISL_762907, EPI_ISL_762909 | Lighthouse Lab in Glasgow | Wellcome Sanger Institute for the COVID-19 Genomics UK (COG-UK) Consortium | Harper VanSteenhouse, Yumi Kasai, David Gray, Carol Clugston, Anna Dominiczak and Alex Alderton, Roberto Amato, Sonia Goncalves, Ewan Harrison, David K. Jackson, Ian Johnston, Dominic Kwiatkowski, Cordelia Langford, John Sillitoe on behalf of the Wellcome Sanger Institute COVID-19 Surveillance Team |
| EPI_ISL_762910, EPI_ISL_762911, EPI_ISL_762913 | Lighthouse Lab in Alderley Park | Wellcome Sanger Institute for the COVID-19 Genomics UK (COG-UK) Consortium | Jacquelyn Wynn, Mairead Hyland, The Lighthouse Lab in Alderley Park and Alex Alderton, Roberto Amato, Sonia Goncalves, Ewan Harrison, David K. Jackson, Ian Johnston, Dominic Kwiatkowski, Cordelia Langford, John Sillitoe on behalf of the Wellcome Sanger Institute COVID-19 Surveillance Team |
| EPI_ISL_762915 | Lighthouse Lab in Glasgow | Wellcome Sanger Institute for the COVID-19 Genomics UK (COG-UK) Consortium | Harper VanSteenhouse, Yumi Kasai, David Gray, Carol Clugston, Anna Dominiczak and Alex Alderton, Roberto Amato, Sonia Goncalves, Ewan Harrison, David K. Jackson, Ian Johnston, Dominic Kwiatkowski, Cordelia Langford, John Sillitoe on behalf of the Wellcome Sanger Institute COVID-19 Surveillance Team |
| EPI_ISL_762916 | Lighthouse Lab in Alderley Park | Wellcome Sanger Institute for the COVID-19 Genomics UK (COG-UK) Consortium | Jacquelyn Wynn, Mairead Hyland, The Lighthouse Lab in Alderley Park and Alex Alderton, Roberto Amato, Sonia Goncalves, Ewan Harrison, David K. Jackson, Ian Johnston, Dominic Kwiatkowski, Cordelia Langford, John Sillitoe on behalf of the Wellcome Sanger Institute COVID-19 Surveillance Team |
| EPI_ISL_762918 | Lighthouse Lab in Glasgow | Wellcome Sanger Institute for the COVID-19 Genomics UK | Harper VanSteenhouse, Yumi Kasai, David Gray, Carol Clugston, Anna Dominiczak and Alex Alderton, Roberto Amato, Sonia Goncalves, Ewan Harrison, |

| | | | |
|---|---|---|---|
| | | (COG-UK) Consortium | David K. Jackson, Ian Johnston, Dominic Kwiatkowski, Cordelia Langford, John Sillitoe on behalf of the Wellcome Sanger Institute COVID-19 Surveillance Team |
| EPI_ISL_762919, EPI_ISL_762921, EPI_ISL_762922 | Lighthouse Lab in Alderley Park | Wellcome Sanger Institute for the COVID-19 Genomics UK (COG-UK) Consortium | Jacquelyn Wynn, Mairead Hyland, The Lighthouse Lab in Alderley Park and Alex Alderton, Roberto Amato, Sonia Goncalves, Ewan Harrison, David K. Jackson, Ian Johnston, Dominic Kwiatkowski, Cordelia Langford, John Sillitoe on behalf of the Wellcome Sanger Institute COVID-19 Surveillance Team |
| EPI_ISL_762923, EPI_ISL_762924, EPI_ISL_762925 | Lighthouse Lab in Glasgow | Wellcome Sanger Institute for the COVID-19 Genomics UK (COG-UK) Consortium | Harper VanSteenhouse, Yumi Kasai, David Gray, Carol Clugston, Anna Dominiczak and Alex Alderton, Roberto Amato, Sonia Goncalves, Ewan Harrison, David K. Jackson, Ian Johnston, Dominic Kwiatkowski, Cordelia Langford, John Sillitoe on behalf of the Wellcome Sanger Institute COVID-19 Surveillance Team |
| EPI_ISL_762926 | Lighthouse Lab in Alderley Park | Wellcome Sanger Institute for the COVID-19 Genomics UK (COG-UK) Consortium | Jacquelyn Wynn, Mairead Hyland, The Lighthouse Lab in Alderley Park and Alex Alderton, Roberto Amato, Sonia Goncalves, Ewan Harrison, David K. Jackson, Ian Johnston, Dominic Kwiatkowski, Cordelia Langford, John Sillitoe on behalf of the Wellcome Sanger Institute COVID-19 Surveillance Team |
| EPI_ISL_762927, EPI_ISL_762928, EPI_ISL_762929, EPI_ISL_762930 | Lighthouse Lab in Glasgow | Wellcome Sanger Institute for the COVID-19 Genomics UK (COG-UK) Consortium | Harper VanSteenhouse, Yumi Kasai, David Gray, Carol Clugston, Anna Dominiczak and Alex Alderton, Roberto Amato, Sonia Goncalves, Ewan Harrison, David K. Jackson, Ian Johnston, Dominic Kwiatkowski, Cordelia Langford, John Sillitoe on behalf of the Wellcome Sanger Institute COVID-19 Surveillance Team |
| EPI_ISL_762931, EPI_ISL_762932, EPI_ISL_762933 | Lighthouse Lab in Alderley Park | Wellcome Sanger Institute for the COVID-19 Genomics UK (COG-UK) Consortium | Jacquelyn Wynn, Mairead Hyland, The Lighthouse Lab in Alderley Park and Alex Alderton, Roberto Amato, Sonia Goncalves, Ewan Harrison, David K. Jackson, Ian Johnston, Dominic Kwiatkowski, Cordelia Langford, John Sillitoe on behalf of the Wellcome Sanger Institute COVID-19 Surveillance Team |
| EPI_ISL_762934 | Lighthouse Lab in Glasgow | Wellcome Sanger Institute for the COVID-19 Genomics UK (COG-UK) Consortium | Harper VanSteenhouse, Yumi Kasai, David Gray, Carol Clugston, Anna Dominiczak and Alex Alderton, Roberto Amato, Sonia Goncalves, Ewan Harrison, David K. Jackson, Ian Johnston, Dominic Kwiatkowski, Cordelia Langford, John Sillitoe on behalf of the Wellcome Sanger Institute COVID-19 Surveillance Team |
| EPI_ISL_762935 | Lighthouse Lab in Alderley Park | Wellcome Sanger Institute for the COVID-19 Genomics UK (COG-UK) Consortium | Jacquelyn Wynn, Mairead Hyland, The Lighthouse Lab in Alderley Park and Alex Alderton, Roberto Amato, Sonia Goncalves, Ewan Harrison, David K. Jackson, Ian Johnston, Dominic Kwiatkowski, Cordelia Langford, John Sillitoe on behalf of the Wellcome Sanger Institute COVID-19 Surveillance Team |
| EPI_ISL_762936 | Lighthouse Lab in Glasgow | Wellcome Sanger Institute for the COVID-19 Genomics UK (COG-UK) Consortium | Harper VanSteenhouse, Yumi Kasai, David Gray, Carol Clugston, Anna Dominiczak and Alex Alderton, Roberto Amato, Sonia Goncalves, Ewan Harrison, David K. Jackson, Ian Johnston, Dominic Kwiatkowski, Cordelia Langford, John Sillitoe on behalf of the Wellcome Sanger Institute COVID-19 Surveillance Team |
| EPI_ISL_762938, EPI_ISL_762939 | Lighthouse Lab in Alderley Park | Wellcome Sanger Institute for the COVID-19 Genomics UK (COG-UK) Consortium | Jacquelyn Wynn, Mairead Hyland, The Lighthouse Lab in Alderley Park and Alex Alderton, Roberto Amato, Sonia Goncalves, Ewan Harrison, David K. Jackson, Ian Johnston, Dominic Kwiatkowski, Cordelia Langford, John Sillitoe on behalf of the Wellcome Sanger Institute COVID-19 Surveillance Team |
| EPI_ISL_762940, EPI_ISL_762941, EPI_ISL_762942, EPI_ISL_762944, EPI_ISL_762946 | Lighthouse Lab in Glasgow | Wellcome Sanger Institute for the COVID-19 Genomics UK (COG-UK) Consortium | Harper VanSteenhouse, Yumi Kasai, David Gray, Carol Clugston, Anna Dominiczak and Alex Alderton, Roberto Amato, Sonia Goncalves, Ewan Harrison, David K. Jackson, Ian Johnston, Dominic Kwiatkowski, Cordelia Langford, John Sillitoe on behalf of the Wellcome Sanger Institute COVID-19 Surveillance Team |
| EPI_ISL_762947 | Lighthouse Lab in Alderley Park | Wellcome Sanger Institute for the COVID-19 Genomics UK (COG-UK) Consortium | Jacquelyn Wynn, Mairead Hyland, The Lighthouse Lab in Alderley Park and Alex Alderton, Roberto Amato, Sonia Goncalves, Ewan Harrison, David K. Jackson, Ian Johnston, Dominic Kwiatkowski, Cordelia Langford, John Sillitoe on behalf of the Wellcome Sanger Institute COVID-19 Surveillance Team |
| EPI_ISL_762948 | Lighthouse Lab in Glasgow | Wellcome Sanger Institute for the COVID-19 Genomics UK (COG-UK) Consortium | Harper VanSteenhouse, Yumi Kasai, David Gray, Carol Clugston, Anna Dominiczak and Alex Alderton, Roberto Amato, Sonia Goncalves, Ewan Harrison, David K. Jackson, Ian Johnston, Dominic Kwiatkowski, Cordelia Langford, John Sillitoe on behalf of the Wellcome Sanger Institute COVID-19 Surveillance Team |
| EPI_ISL_762949 | Lighthouse Lab in Alderley Park | Wellcome Sanger Institute for the COVID-19 Genomics UK (COG-UK) Consortium | Jacquelyn Wynn, Mairead Hyland, The Lighthouse Lab in Alderley Park and Alex Alderton, Roberto Amato, Sonia Goncalves, Ewan Harrison, David K. Jackson, Ian Johnston, Dominic Kwiatkowski, Cordelia Langford, John Sillitoe on behalf of the Wellcome Sanger Institute COVID-19 Surveillance Team |
| EPI_ISL_762950 | Lighthouse Lab in Glasgow | Wellcome Sanger Institute for the COVID-19 Genomics UK (COG-UK) Consortium | Harper VanSteenhouse, Yumi Kasai, David Gray, Carol Clugston, Anna Dominiczak and Alex Alderton, Roberto Amato, Sonia Goncalves, Ewan Harrison, David K. Jackson, Ian Johnston, Dominic Kwiatkowski, Cordelia Langford, John Sillitoe on behalf of the Wellcome Sanger Institute COVID-19 Surveillance Team |
| EPI_ISL_762951, EPI_ISL_762952 | Lighthouse Lab in Alderley Park | Wellcome Sanger Institute for the COVID-19 Genomics UK (COG-UK) Consortium | Jacquelyn Wynn, Mairead Hyland, The Lighthouse Lab in Alderley Park and Alex Alderton, Roberto Amato, Sonia Goncalves, Ewan Harrison, David K. Jackson, Ian Johnston, Dominic Kwiatkowski, Cordelia Langford, John Sillitoe on behalf of the Wellcome Sanger Institute COVID-19 Surveillance Team |
| EPI_ISL_762953, EPI_ISL_762954 | Lighthouse Lab in Glasgow | Wellcome Sanger Institute for the COVID-19 Genomics UK (COG-UK) Consortium | Harper VanSteenhouse, Yumi Kasai, David Gray, Carol Clugston, Anna Dominiczak and Alex Alderton, Roberto Amato, Sonia Goncalves, Ewan Harrison, David K. Jackson, Ian Johnston, Dominic Kwiatkowski, Cordelia Langford, John Sillitoe on behalf of the Wellcome Sanger Institute COVID-19 Surveillance Team |
| EPI_ISL_762955 | Lighthouse Lab in Alderley Park | Wellcome Sanger Institute for the COVID-19 Genomics UK (COG-UK) Consortium | Jacquelyn Wynn, Mairead Hyland, The Lighthouse Lab in Alderley Park and Alex Alderton, Roberto Amato, Sonia Goncalves, Ewan Harrison, David K. Jackson, Ian Johnston, Dominic Kwiatkowski, Cordelia Langford, John Sillitoe on behalf of the Wellcome Sanger Institute COVID-19 Surveillance Team |
| EPI_ISL_762956 | Lighthouse Lab in Glasgow | Wellcome Sanger Institute for the COVID-19 Genomics UK (COG-UK) Consortium | Harper VanSteenhouse, Yumi Kasai, David Gray, Carol Clugston, Anna Dominiczak and Alex Alderton, Roberto Amato, Sonia Goncalves, Ewan Harrison, David K. Jackson, Ian Johnston, Dominic Kwiatkowski, Cordelia Langford, John Sillitoe on behalf of the Wellcome Sanger Institute COVID-19 Surveillance Team |
| EPI_ISL_762957 | Lighthouse Lab in Alderley Park | Wellcome Sanger Institute for the COVID-19 Genomics UK (COG-UK) Consortium | Jacquelyn Wynn, Mairead Hyland, The Lighthouse Lab in Alderley Park and Alex Alderton, Roberto Amato, Sonia Goncalves, Ewan Harrison, David K. Jackson, Ian Johnston, Dominic Kwiatkowski, Cordelia Langford, John Sillitoe on behalf of the Wellcome Sanger Institute COVID-19 Surveillance Team |
| EPI_ISL_762958, EPI_ISL_762959, EPI_ISL_762960, EPI_ISL_762961, EPI_ISL_762962 | Lighthouse Lab in Glasgow | Wellcome Sanger Institute for the COVID-19 Genomics UK (COG-UK) Consortium | Harper VanSteenhouse, Yumi Kasai, David Gray, Carol Clugston, Anna Dominiczak and Alex Alderton, Roberto Amato, Sonia Goncalves, Ewan Harrison, David K. Jackson, Ian Johnston, Dominic Kwiatkowski, Cordelia Langford, John Sillitoe on behalf of the Wellcome Sanger Institute COVID-19 Surveillance Team |
| EPI_ISL_762964 | Lighthouse Lab in Alderley Park | Wellcome Sanger Institute for the COVID-19 Genomics UK (COG-UK) Consortium | Jacquelyn Wynn, Mairead Hyland, The Lighthouse Lab in Alderley Park and Alex Alderton, Roberto Amato, Sonia Goncalves, Ewan Harrison, David K. Jackson, Ian Johnston, Dominic Kwiatkowski, Cordelia Langford, John Sillitoe on behalf of the Wellcome Sanger Institute COVID-19 Surveillance Team |
| EPI_ISL_762969 | Lighthouse Lab in Glasgow | Wellcome Sanger Institute for the COVID-19 Genomics UK (COG-UK) Consortium | Harper VanSteenhouse, Yumi Kasai, David Gray, Carol Clugston, Anna Dominiczak and Alex Alderton, Roberto Amato, Sonia Goncalves, Ewan Harrison, David K. Jackson, Ian Johnston, Dominic Kwiatkowski, Cordelia Langford, John Sillitoe on behalf of the Wellcome Sanger Institute COVID-19 Surveillance Team |
| EPI_ISL_762970, EPI_ISL_762971 | Lighthouse Lab in Alderley Park | Wellcome Sanger Institute for the COVID-19 Genomics UK (COG-UK) Consortium | Jacquelyn Wynn, Mairead Hyland, The Lighthouse Lab in Alderley Park and Alex Alderton, Roberto Amato, Sonia Goncalves, Ewan Harrison, David K. Jackson, Ian Johnston, Dominic Kwiatkowski, Cordelia Langford, John Sillitoe on behalf of the Wellcome Sanger Institute COVID-19 Surveillance Team |
| EPI_ISL_762972 | Lighthouse Lab in Glasgow | Wellcome Sanger Institute for the COVID-19 Genomics UK (COG-UK) Consortium | Harper VanSteenhouse, Yumi Kasai, David Gray, Carol Clugston, Anna Dominiczak and Alex Alderton, Roberto Amato, Sonia Goncalves, Ewan Harrison, David K. Jackson, Ian Johnston, Dominic Kwiatkowski, Cordelia Langford, John Sillitoe on behalf of the Wellcome Sanger Institute COVID-19 Surveillance Team |
| EPI_ISL_762973, EPI_ISL_762974, EPI_ISL_762975, EPI_ISL_762977 | Lighthouse Lab in Alderley Park | Wellcome Sanger Institute for the COVID-19 Genomics UK (COG-UK) Consortium | Jacquelyn Wynn, Mairead Hyland, The Lighthouse Lab in Alderley Park and Alex Alderton, Roberto Amato, Sonia Goncalves, Ewan Harrison, David K. Jackson, Ian Johnston, Dominic Kwiatkowski, Cordelia Langford, John Sillitoe on behalf of the Wellcome Sanger Institute COVID-19 Surveillance Team |
| EPI_ISL_762980, EPI_ISL_762982, EPI_ISL_762983, EPI_ISL_762984, EPI_ISL_762985, EPI_ISL_762987 | Lighthouse Lab in Glasgow | Wellcome Sanger Institute for the COVID-19 Genomics UK (COG-UK) Consortium | Harper VanSteenhouse, Yumi Kasai, David Gray, Carol Clugston, Anna Dominiczak and Alex Alderton, Roberto Amato, Sonia Goncalves, Ewan Harrison, David K. Jackson, Ian Johnston, Dominic Kwiatkowski, Cordelia Langford, John Sillitoe on behalf of the Wellcome Sanger Institute COVID-19 Surveillance Team |
| EPI_ISL_762988, EPI_ISL_762989 | Lighthouse Lab in Alderley Park | Wellcome Sanger Institute for the COVID-19 Genomics UK (COG-UK) Consortium | Jacquelyn Wynn, Mairead Hyland, The Lighthouse Lab in Alderley Park and Alex Alderton, Roberto Amato, Sonia Goncalves, Ewan Harrison, David K. Jackson, Ian Johnston, Dominic Kwiatkowski, Cordelia Langford, John Sillitoe on behalf of the Wellcome Sanger Institute COVID-19 Surveillance Team |
| EPI_ISL_762995, EPI_ISL_762996 | Respiratory Virus Unit, National Infection Service, Public Health England | COVID-19 Genomics UK (COG-UK) Consortium | PHE Covid Sequencing Team |
| EPI_ISL_763000, EPI_ISL_763010, EPI_ISL_763011, EPI_ISL_763053, | Unit 17: Influenza & Other Respiratory Viruses, German National Influenza Center | Project group Epidemiology of Highly Pathogenic Microorganisms, Robert Koch-Institute | Ariane Düx, Andreas Sachse, Grit Schubert, Sébastien Calvignac-Spencer, Fabian Leendertz, Thorsten Wolff, Ralf Dürrwald, Djin-Ye Oh, Marianne Wedde |

| | | | |
|---|---|---|---|
| EPI_ISL_763054 | | | |
| EPI_ISL_763203, EPI_ISL_763204, EPI_ISL_763205, EPI_ISL_763207, EPI_ISL_763208, EPI_ISL_763209, EPI_ISL_763210, EPI_ISL_763247, EPI_ISL_763286, EPI_ISL_763288, EPI_ISL_763291, EPI_ISL_763292, EPI_ISL_763349, EPI_ISL_763354 | | | |
| see above | Dutch COVID-19 response team | Erasmus Medical Center | Bas Oude Munnink, Reina Sikkema, David Nieuwenhuijse, Irina Chestakova, Anne van der Linden, Marjan Boter, Emmanuelle Munger, Corine GeurtsvanKessel, Annemiek van der Eijk, Richard Molenkamp, Marion Koopmans, on behalf of the Dutch national COVID-19 response team. |
| EPI_ISL_763365 | Virology Department, Sheffield Teaching Hospitals NHS Foundation Trust/Department of Infection, Immunity and Cardiovascular Disease, The Medical School, University of Sheffield | COVID-19 Genomics UK (COG-UK) Consortium | Thushan de Silva, Matthew Parker, Nikki Smith, Adri Angyal, Rebecca Brown, Luke Green, Rachel Tucker, Paul Parsons, Danielle Groves, Katie Johnson, Laura Carrilero, Alex Keeley, Dave Partridge, Matthew Wyles, Benjamin Lindsey, Mehmet Yavuz, Mohammad Raza, Cariad Evans |
| EPI_ISL_763371 | Wales Specialist Virology Centre Sequencing lab: Pathogen Genomics Unit | COVID-19 Genomics UK (COG-UK) Consortium | Catherine Moore, Johnathan Evans, Laura Gifford, Malorie Perry, Simon Cottrell, Angela Marchbank, Alec Birchley, Alexander Adams, Amy Gaskin, Bree Gatica-Wilcox, Jason Coombes, Joel Southgate, Lauren Gilbert, Lee Graham, Nicole Pacchiarini, Sara Kumziene-Summerhayes, Sarah Taylor, Sophie Jones, Sara Rey, Matthew Bull, Joanne Watkins, Sally Corden, Tom Connor |
| EPI_ISL_763376 | University of Exeter | COVID-19 Genomics UK (COG-UK) Consortium | Ben Temperton,Aaron Jeffries,Michelle Michelsen,Joanna Warwick-Dugdale,Audrey Farbos,Robyn Manley,Stephen Michell,Jane Masoli |
| EPI_ISL_763377 | University College London, Great Ormond Street Hospital for Children NHS Foundation Trust, Imperial College Healthcare NHS Trust | COVID-19 Genomics UK (COG-UK) Consortium | Sergi Castellano, Rachel Williams, Mark Kristiansen, Paola Resende Silva, Sunando Roy, Tony Brooks, Helena Tutill, Paola Niola, Patricia Dyal, Charlotte Williams, Leysa Forrest, Yasmin Panchbhaya, Jacqueline Findlay, Samuel Weeks, Julianne Brown, Kathryn Harris, Paul Randell, James Price, Alison Holmes, Judith Breuer |
| EPI_ISL_763378 | Wales Specialist Virology Centre Sequencing lab: Pathogen Genomics Unit | COVID-19 Genomics UK (COG-UK) Consortium | Catherine Moore, Johnathan Evans, Laura Gifford, Malorie Perry, Simon Cottrell, Angela Marchbank, Alec Birchley, Alexander Adams, Amy Gaskin, Bree Gatica-Wilcox, Jason Coombes, Joel Southgate, Lauren Gilbert, Lee Graham, Nicole Pacchiarini, Sara Kumziene-Summerhayes, Sarah Taylor, Sophie Jones, Sara Rey, Matthew Bull, Joanne Watkins, Sally Corden, Tom Connor |
| EPI_ISL_763387 | University of Exeter | COVID-19 Genomics UK (COG-UK) Consortium | Ben Temperton,Aaron Jeffries,Michelle Michelsen,Joanna Warwick-Dugdale,Audrey Farbos,Robyn Manley,Stephen Michell,Jane Masoli |
| EPI_ISL_763397, EPI_ISL_763398 | University College London, Great Ormond Street Hospital for Children NHS Foundation Trust, Imperial College Healthcare NHS Trust | COVID-19 Genomics UK (COG-UK) Consortium | Sergi Castellano, Rachel Williams, Mark Kristiansen, Paola Resende Silva, Sunando Roy, Tony Brooks, Helena Tutill, Paola Niola, Patricia Dyal, Charlotte Williams, Leysa Forrest, Yasmin Panchbhaya, Jacqueline Findlay, Samuel Weeks, Julianne Brown, Kathryn Harris, Paul Randell, James Price, Alison Holmes, Judith Breuer |
| EPI_ISL_763402, EPI_ISL_763403 | Virology Department, Sheffield Teaching Hospitals NHS Foundation Trust/Department of Infection, Immunity and Cardiovascular Disease, The Medical School, University of Sheffield | COVID-19 Genomics UK (COG-UK) Consortium | Thushan de Silva, Matthew Parker, Nikki Smith, Adri Angyal, Rebecca Brown, Luke Green, Rachel Tucker, Paul Parsons, Danielle Groves, Katie Johnson, Laura Carrilero, Alex Keeley, Dave Partridge, Matthew Wyles, Benjamin Lindsey, Mehmet Yavuz, Mohammad Raza, Cariad Evans |
| EPI_ISL_763404 | University College London, Great Ormond Street Hospital for Children NHS Foundation Trust, Imperial College Healthcare NHS Trust | COVID-19 Genomics UK (COG-UK) Consortium | Sergi Castellano, Rachel Williams, Mark Kristiansen, Paola Resende Silva, Sunando Roy, Tony Brooks, Helena Tutill, Paola Niola, Patricia Dyal, Charlotte Williams, Leysa Forrest, Yasmin Panchbhaya, Jacqueline Findlay, Samuel Weeks, Julianne Brown, Kathryn Harris, Paul Randell, James Price, Alison Holmes, Judith Breuer |
| EPI_ISL_763410, EPI_ISL_763411 | Wales Specialist Virology Centre Sequencing lab: Pathogen Genomics Unit | COVID-19 Genomics UK (COG-UK) Consortium | Catherine Moore, Johnathan Evans, Laura Gifford, Malorie Perry, Simon Cottrell, Angela Marchbank, Alec Birchley, Alexander Adams, Amy Gaskin, Bree Gatica-Wilcox, Jason Coombes, Joel Southgate, Lauren Gilbert, Lee Graham, Nicole Pacchiarini, Sara Kumziene-Summerhayes, Sarah Taylor, Sophie Jones, Sara Rey, Matthew Bull, Joanne Watkins, Sally Corden, Tom Connor |
| EPI_ISL_763412 | University of Exeter | COVID-19 Genomics UK (COG-UK) Consortium | Ben Temperton,Aaron Jeffries,Michelle Michelsen,Joanna Warwick-Dugdale,Audrey Farbos,Robyn Manley,Stephen Michell,Jane Masoli |
| EPI_ISL_763428 | Virology Department, Sheffield Teaching Hospitals NHS Foundation Trust/Department of Infection, Immunity and Cardiovascular Disease, The Medical School, University of Sheffield | COVID-19 Genomics UK (COG-UK) Consortium | Thushan de Silva, Matthew Parker, Nikki Smith, Adri Angyal, Rebecca Brown, Luke Green, Rachel Tucker, Paul Parsons, Danielle Groves, Katie Johnson, Laura Carrilero, Alex Keeley, Dave Partridge, Matthew Wyles, Benjamin Lindsey, Mehmet Yavuz, Mohammad Raza, Cariad Evans |
| EPI_ISL_763431, EPI_ISL_763434, EPI_ISL_763435, EPI_ISL_763436, EPI_ISL_763437 | University of Exeter | COVID-19 Genomics UK (COG-UK) Consortium | Ben Temperton,Aaron Jeffries,Michelle Michelsen,Joanna Warwick-Dugdale,Audrey Farbos,Robyn Manley,Stephen Michell,Jane Masoli |
| EPI_ISL_763442 | University College London, Great Ormond Street Hospital for Children NHS Foundation Trust, Imperial College Healthcare NHS Trust | COVID-19 Genomics UK (COG-UK) Consortium | Sergi Castellano, Rachel Williams, Mark Kristiansen, Paola Resende Silva, Sunando Roy, Tony Brooks, Helena Tutill, Paola Niola, Patricia Dyal, Charlotte Williams, Leysa Forrest, Yasmin Panchbhaya, Jacqueline Findlay, Samuel Weeks, Julianne Brown, Kathryn Harris, Paul Randell, James Price, Alison Holmes, Judith Breuer |
| EPI_ISL_763443, EPI_ISL_763445 | Wales Specialist Virology Centre Sequencing lab: Pathogen Genomics Unit | COVID-19 Genomics UK (COG-UK) Consortium | Catherine Moore, Johnathan Evans, Laura Gifford, Malorie Perry, Simon Cottrell, Angela Marchbank, Alec Birchley, Alexander Adams, Amy Gaskin, Bree Gatica-Wilcox, Jason Coombes, Joel Southgate, Lauren Gilbert, Lee Graham, Nicole Pacchiarini, Sara Kumziene-Summerhayes, Sarah Taylor, Sophie Jones, Sara Rey, Matthew Bull, Joanne Watkins, Sally Corden, Tom Connor |
| EPI_ISL_763451 | Centre for Enzyme Innovation, University of Portsmouth / Translational Research Laboratory, Portsmouth Hospitals NHS Trust | COVID-19 Genomics UK (COG-UK) Consortium | Angela Beckett,Yann Bourgeois,Garry Scarlett,Sharon Glaysher,Scott Elliott,Kelly Bicknell,Robert Impey,Allyson Lloyd,Sarah Wyllie,Ethan Butcher,Anoop Chauhan,Samuel Robson |
| EPI_ISL_763461 | University of Exeter | COVID-19 Genomics UK (COG-UK) Consortium | Ben Temperton,Aaron Jeffries,Michelle Michelsen,Joanna Warwick-Dugdale,Audrey Farbos,Robyn Manley,Stephen Michell,Jane Masoli |
| EPI_ISL_763468 | University College London, Great Ormond Street Hospital for Children NHS Foundation Trust, Imperial College Healthcare NHS Trust | COVID-19 Genomics UK (COG-UK) Consortium | Sergi Castellano, Rachel Williams, Mark Kristiansen, Paola Resende Silva, Sunando Roy, Tony Brooks, Helena Tutill, Paola Niola, Patricia Dyal, Charlotte Williams, Leysa Forrest, Yasmin Panchbhaya, Jacqueline Findlay, Samuel Weeks, Julianne Brown, Kathryn Harris, Paul Randell, James Price, Alison Holmes, Judith Breuer |
| EPI_ISL_763473, EPI_ISL_763474 | Wales Specialist Virology Centre Sequencing lab: Pathogen Genomics Unit | COVID-19 Genomics UK (COG-UK) Consortium | Catherine Moore, Johnathan Evans, Laura Gifford, Malorie Perry, Simon Cottrell, Angela Marchbank, Alec Birchley, Alexander Adams, Amy Gaskin, Bree Gatica-Wilcox, Jason Coombes, Joel Southgate, Lauren Gilbert, Lee Graham, Nicole Pacchiarini, Sara Kumziene-Summerhayes, Sarah Taylor, Sophie Jones, Sara Rey, Matthew Bull, Joanne Watkins, Sally Corden, Tom Connor |
| EPI_ISL_763483 | Quadram Institute Bioscience | COVID-19 Genomics UK (COG-UK) Consortium | Dave J. Baker, Gemma L. Kay, Alp Aydin, Thanh Le-Viet, Steven Rudder, Ana P. Tedim, Anastasia Kolyva, Maria Diaz, Leonardo de Oliveira Martins, Nabil-Fareed Alikhan, Lizzie Meadows, Rachael Stanley, Ngozi Elumogo, Muhammed Yasir, Nicholas M. Thomson, Alexander J Trotter, Rachel Gilroy, Samuel Bloomfield, Claire Stuart, Andrew Bell, Reenesh Prakash, Samir Dervisevic, Alison E. Mather, John Wain, Mark Webber, Andrew J. Page, Justin O'Grady |
| EPI_ISL_763486 | University College London, Great Ormond Street Hospital for Children NHS Foundation Trust, Imperial College Healthcare NHS Trust | COVID-19 Genomics UK (COG-UK) Consortium | Sergi Castellano, Rachel Williams, Mark Kristiansen, Paola Resende Silva, Sunando Roy, Tony Brooks, Helena Tutill, Paola Niola, Patricia Dyal, Charlotte Williams, Leysa Forrest, Yasmin Panchbhaya, Jacqueline Findlay, Samuel Weeks, Julianne Brown, Kathryn Harris, Paul Randell, James Price, Alison Holmes, Judith Breuer |
| EPI_ISL_763488 | Wales Specialist Virology Centre Sequencing lab: Pathogen Genomics Unit | COVID-19 Genomics UK (COG-UK) Consortium | Catherine Moore, Johnathan Evans, Laura Gifford, Malorie Perry, Simon Cottrell, Angela Marchbank, Alec Birchley, Alexander Adams, Amy Gaskin, Bree Gatica-Wilcox, Jason Coombes, Joel Southgate, Lauren Gilbert, Lee Graham, Nicole Pacchiarini, Sara Kumziene-Summerhayes, Sarah Taylor, Sophie Jones, Sara Rey, Matthew Bull, Joanne Watkins, Sally Corden, Tom Connor |
| EPI_ISL_763491 | University College London, Great Ormond Street Hospital for Children NHS Foundation Trust, Imperial College Healthcare NHS Trust | COVID-19 Genomics UK (COG-UK) Consortium | Sergi Castellano, Rachel Williams, Mark Kristiansen, Paola Resende Silva, Sunando Roy, Tony Brooks, Helena Tutill, Paola Niola, Patricia Dyal, Charlotte Williams, Leysa Forrest, Yasmin Panchbhaya, Jacqueline Findlay, Samuel Weeks, Julianne Brown, Kathryn Harris, Paul Randell, James Price, Alison Holmes, Judith Breuer |
| EPI_ISL_763494 | Wales Specialist Virology Centre Sequencing lab: Pathogen Genomics Unit | COVID-19 Genomics UK (COG-UK) Consortium | Catherine Moore, Johnathan Evans, Laura Gifford, Malorie Perry, Simon Cottrell, Angela Marchbank, Alec Birchley, Alexander Adams, Amy Gaskin, Bree Gatica-Wilcox, Jason Coombes, Joel Southgate, Lauren Gilbert, Lee Graham, Nicole Pacchiarini, Sara Kumziene-Summerhayes, Sarah Taylor, Sophie Jones, Sara Rey, Matthew Bull, Joanne Watkins, Sally Corden, Tom Connor |
| EPI_ISL_763495 | Centre for Enzyme Innovation, University of Portsmouth / Translational Research Laboratory, Portsmouth Hospitals NHS | COVID-19 Genomics UK (COG-UK) Consortium | Angela Beckett,Yann Bourgeois,Garry Scarlett,Sharon Glaysher,Scott Elliott,Kelly Bicknell,Robert Impey,Allyson Lloyd,Sarah Wyllie,Ethan Butcher,Anoop Chauhan,Samuel Robson |

| | | | |
|---|---|---|---|
| | Trust | | |
| EPI_ISL_763500 | University College London, Great Ormond Street Hospital for Children NHS Foundation Trust, Imperial College Healthcare NHS Trust | COVID-19 Genomics UK (COG-UK) Consortium | Sergi Castellano, Rachel Williams, Mark Kristiansen, Paola Resende Silva, Sunando Roy, Tony Brooks, Helena Tutill, Paola Niola, Patricia Dyal, Charlotte Williams, Leysa Forrest, Yasmin Panchbhaya, Jacqueline Findlay, Samuel Weeks, Julianne Brown, Kathryn Harris, Paul Randell, James Price, Alison Holmes, Judith Breuer |
| EPI_ISL_763509 | Wales Specialist Virology Centre Sequencing lab: Pathogen Genomics Unit | COVID-19 Genomics UK (COG-UK) Consortium | Catherine Moore, Johnathan Evans, Laura Gifford, Malorie Perry, Simon Cottrell, Angela Marchbank, Alec Birchley, Alexander Adams, Amy Gaskin, Bree Gatica-Wilcox, Jason Coombes, Joel Southgate, Lauren Gilbert, Lee Graham, Nicole Pacchiarini, Sara Kumziene-Summerhayes, Sarah Taylor, Sophie Jones, Sara Rey, Matthew Bull, Joanne Watkins, Sally Corden, Tom Connor |
| EPI_ISL_763510 | University College London, Great Ormond Street Hospital for Children NHS Foundation Trust, Imperial College Healthcare NHS Trust | COVID-19 Genomics UK (COG-UK) Consortium | Sergi Castellano, Rachel Williams, Mark Kristiansen, Paola Resende Silva, Sunando Roy, Tony Brooks, Helena Tutill, Paola Niola, Patricia Dyal, Charlotte Williams, Leysa Forrest, Yasmin Panchbhaya, Jacqueline Findlay, Samuel Weeks, Julianne Brown, Kathryn Harris, Paul Randell, James Price, Alison Holmes, Judith Breuer |
| EPI_ISL_763525 | Wales Specialist Virology Centre Sequencing lab: Pathogen Genomics Unit | COVID-19 Genomics UK (COG-UK) Consortium | Catherine Moore, Johnathan Evans, Laura Gifford, Malorie Perry, Simon Cottrell, Angela Marchbank, Alec Birchley, Alexander Adams, Amy Gaskin, Bree Gatica-Wilcox, Jason Coombes, Joel Southgate, Lauren Gilbert, Lee Graham, Nicole Pacchiarini, Sara Kumziene-Summerhayes, Sarah Taylor, Sophie Jones, Sara Rey, Matthew Bull, Joanne Watkins, Sally Corden, Tom Connor |
| EPI_ISL_763533, EPI_ISL_763534, EPI_ISL_763535, EPI_ISL_763536 | University of Exeter | COVID-19 Genomics UK (COG-UK) Consortium | Ben Temperton,Aaron Jeffries,Michelle Michelsen,Joanna Warwick-Dugdale,Audrey Farbos,Robyn Manley,Stephen Michell,Jane Masoli |
| EPI_ISL_763538 | Wales Specialist Virology Centre Sequencing lab: Pathogen Genomics Unit | COVID-19 Genomics UK (COG-UK) Consortium | Catherine Moore, Johnathan Evans, Laura Gifford, Malorie Perry, Simon Cottrell, Angela Marchbank, Alec Birchley, Alexander Adams, Amy Gaskin, Bree Gatica-Wilcox, Jason Coombes, Joel Southgate, Lauren Gilbert, Lee Graham, Nicole Pacchiarini, Sara Kumziene-Summerhayes, Sarah Taylor, Sophie Jones, Sara Rey, Matthew Bull, Joanne Watkins, Sally Corden, Tom Connor |
| EPI_ISL_763540, EPI_ISL_763545, EPI_ISL_763551, EPI_ISL_763552 | University College London, Great Ormond Street Hospital for Children NHS Foundation Trust, Imperial College Healthcare NHS Trust | COVID-19 Genomics UK (COG-UK) Consortium | Sergi Castellano, Rachel Williams, Mark Kristiansen, Paola Resende Silva, Sunando Roy, Tony Brooks, Helena Tutill, Paola Niola, Patricia Dyal, Charlotte Williams, Leysa Forrest, Yasmin Panchbhaya, Jacqueline Findlay, Samuel Weeks, Julianne Brown, Kathryn Harris, Paul Randell, James Price, Alison Holmes, Judith Breuer |
| EPI_ISL_763557 | Wales Specialist Virology Centre Sequencing lab: Pathogen Genomics Unit | COVID-19 Genomics UK (COG-UK) Consortium | Catherine Moore, Johnathan Evans, Laura Gifford, Malorie Perry, Simon Cottrell, Angela Marchbank, Alec Birchley, Alexander Adams, Amy Gaskin, Bree Gatica-Wilcox, Jason Coombes, Joel Southgate, Lauren Gilbert, Lee Graham, Nicole Pacchiarini, Sara Kumziene-Summerhayes, Sarah Taylor, Sophie Jones, Sara Rey, Matthew Bull, Joanne Watkins, Sally Corden, Tom Connor |
| EPI_ISL_763558, EPI_ISL_763559 | University of Exeter | COVID-19 Genomics UK (COG-UK) Consortium | Ben Temperton,Aaron Jeffries,Michelle Michelsen,Joanna Warwick-Dugdale,Audrey Farbos,Robyn Manley,Stephen Michell,Jane Masoli |
| EPI_ISL_763577, EPI_ISL_763579, EPI_ISL_763581 | Wales Specialist Virology Centre Sequencing lab: Pathogen Genomics Unit | COVID-19 Genomics UK (COG-UK) Consortium | Catherine Moore, Johnathan Evans, Laura Gifford, Malorie Perry, Simon Cottrell, Angela Marchbank, Alec Birchley, Alexander Adams, Amy Gaskin, Bree Gatica-Wilcox, Jason Coombes, Joel Southgate, Lauren Gilbert, Lee Graham, Nicole Pacchiarini, Sara Kumziene-Summerhayes, Sarah Taylor, Sophie Jones, Sara Rey, Matthew Bull, Joanne Watkins, Sally Corden, Tom Connor |
| EPI_ISL_763582 | University College London, Great Ormond Street Hospital for Children NHS Foundation Trust, Imperial College Healthcare NHS Trust | COVID-19 Genomics UK (COG-UK) Consortium | Sergi Castellano, Rachel Williams, Mark Kristiansen, Paola Resende Silva, Sunando Roy, Tony Brooks, Helena Tutill, Paola Niola, Patricia Dyal, Charlotte Williams, Leysa Forrest, Yasmin Panchbhaya, Jacqueline Findlay, Samuel Weeks, Julianne Brown, Kathryn Harris, Paul Randell, James Price, Alison Holmes, Judith Breuer |
| EPI_ISL_763585 | Wales Specialist Virology Centre Sequencing lab: Pathogen Genomics Unit | COVID-19 Genomics UK (COG-UK) Consortium | Catherine Moore, Johnathan Evans, Laura Gifford, Malorie Perry, Simon Cottrell, Angela Marchbank, Alec Birchley, Alexander Adams, Amy Gaskin, Bree Gatica-Wilcox, Jason Coombes, Joel Southgate, Lauren Gilbert, Lee Graham, Nicole Pacchiarini, Sara Kumziene-Summerhayes, Sarah Taylor, Sophie Jones, Sara Rey, Matthew Bull, Joanne Watkins, Sally Corden, Tom Connor |
| EPI_ISL_763588 | Virology Department, Sheffield Teaching Hospitals NHS Foundation Trust/Department of Infection, Immunity and Cardiovascular Disease, The Medical School, University of Sheffield | COVID-19 Genomics UK (COG-UK) Consortium | Thushan de Silva, Matthew Parker, Nikki Smith, Adri Angyal, Rebecca Brown, Luke Green, Rachel Tucker, Paul Parsons, Danielle Groves, Katie Johnson, Laura Carrilero, Alex Keeley, Dave Partridge, Matthew Wyles, Benjamin Lindsey, Mehmet Yavuz, Mohammad Raza, Cariad Evans |
| EPI_ISL_763589 | Wales Specialist Virology Centre Sequencing lab: Pathogen Genomics Unit | COVID-19 Genomics UK (COG-UK) Consortium | Catherine Moore, Johnathan Evans, Laura Gifford, Malorie Perry, Simon Cottrell, Angela Marchbank, Alec Birchley, Alexander Adams, Amy Gaskin, Bree Gatica-Wilcox, Jason Coombes, Joel Southgate, Lauren Gilbert, Lee Graham, Nicole Pacchiarini, Sara Kumziene-Summerhayes, Sarah Taylor, Sophie Jones, Sara Rey, Matthew Bull, Joanne Watkins, Sally Corden, Tom Connor |
| EPI_ISL_763595 | Centre for Enzyme Innovation, University of Portsmouth / Translational Research Laboratory, Portsmouth Hospitals NHS Trust | COVID-19 Genomics UK (COG-UK) Consortium | Angela Beckett,Yann Bourgeois,Garry Scarlett,Sharon Glaysher,Scott Elliott,Kelly Bicknell,Robert Impey,Allyson Lloyd,Sarah Wyllie,Ethan Butcher,Anoop Chauhan,Samuel Robson |
| EPI_ISL_763597, EPI_ISL_763599 | Wales Specialist Virology Centre Sequencing lab: Pathogen Genomics Unit | COVID-19 Genomics UK (COG-UK) Consortium | Catherine Moore, Johnathan Evans, Laura Gifford, Malorie Perry, Simon Cottrell, Angela Marchbank, Alec Birchley, Alexander Adams, Amy Gaskin, Bree Gatica-Wilcox, Jason Coombes, Joel Southgate, Lauren Gilbert, Lee Graham, Nicole Pacchiarini, Sara Kumziene-Summerhayes, Sarah Taylor, Sophie Jones, Sara Rey, Matthew Bull, Joanne Watkins, Sally Corden, Tom Connor |
| EPI_ISL_763609, EPI_ISL_763612, EPI_ISL_763624 | Centre for Enzyme Innovation, University of Portsmouth / Translational Research Laboratory, Portsmouth Hospitals NHS Trust | COVID-19 Genomics UK (COG-UK) Consortium | Angela Beckett,Yann Bourgeois,Garry Scarlett,Sharon Glaysher,Scott Elliott,Kelly Bicknell,Robert Impey,Allyson Lloyd,Sarah Wyllie,Ethan Butcher,Anoop Chauhan,Samuel Robson |
| EPI_ISL_763628 | University College London, Great Ormond Street Hospital for Children NHS Foundation Trust, Imperial College Healthcare NHS Trust | COVID-19 Genomics UK (COG-UK) Consortium | Sergi Castellano, Rachel Williams, Mark Kristiansen, Paola Resende Silva, Sunando Roy, Tony Brooks, Helena Tutill, Paola Niola, Patricia Dyal, Charlotte Williams, Leysa Forrest, Yasmin Panchbhaya, Jacqueline Findlay, Samuel Weeks, Julianne Brown, Kathryn Harris, Paul Randell, James Price, Alison Holmes, Judith Breuer |
| EPI_ISL_763633 | Wales Specialist Virology Centre Sequencing lab: Pathogen Genomics Unit | COVID-19 Genomics UK (COG-UK) Consortium | Catherine Moore, Johnathan Evans, Laura Gifford, Malorie Perry, Simon Cottrell, Angela Marchbank, Alec Birchley, Alexander Adams, Amy Gaskin, Bree Gatica-Wilcox, Jason Coombes, Joel Southgate, Lauren Gilbert, Lee Graham, Nicole Pacchiarini, Sara Kumziene-Summerhayes, Sarah Taylor, Sophie Jones, Sara Rey, Matthew Bull, Joanne Watkins, Sally Corden, Tom Connor |
| EPI_ISL_763635, EPI_ISL_763639 | Virology Department, Sheffield Teaching Hospitals NHS Foundation Trust/Department of Infection, Immunity and Cardiovascular Disease, The Medical School, University of Sheffield | COVID-19 Genomics UK (COG-UK) Consortium | Thushan de Silva, Matthew Parker, Nikki Smith, Adri Angyal, Rebecca Brown, Luke Green, Rachel Tucker, Paul Parsons, Danielle Groves, Katie Johnson, Laura Carrilero, Alex Keeley, Dave Partridge, Matthew Wyles, Benjamin Lindsey, Mehmet Yavuz, Mohammad Raza, Cariad Evans |
| EPI_ISL_763640, EPI_ISL_763643, EPI_ISL_763644 | University College London, Great Ormond Street Hospital for Children NHS Foundation Trust, Imperial College Healthcare NHS Trust | COVID-19 Genomics UK (COG-UK) Consortium | Sergi Castellano, Rachel Williams, Mark Kristiansen, Paola Resende Silva, Sunando Roy, Tony Brooks, Helena Tutill, Paola Niola, Patricia Dyal, Charlotte Williams, Leysa Forrest, Yasmin Panchbhaya, Jacqueline Findlay, Samuel Weeks, Julianne Brown, Kathryn Harris, Paul Randell, James Price, Alison Holmes, Judith Breuer |
| EPI_ISL_763645 | Virology Department, Sheffield Teaching Hospitals NHS Foundation Trust/Department of Infection, Immunity and Cardiovascular Disease, The Medical School, University of Sheffield | COVID-19 Genomics UK (COG-UK) Consortium | Thushan de Silva, Matthew Parker, Nikki Smith, Adri Angyal, Rebecca Brown, Luke Green, Rachel Tucker, Paul Parsons, Danielle Groves, Katie Johnson, Laura Carrilero, Alex Keeley, Dave Partridge, Matthew Wyles, Benjamin Lindsey, Mehmet Yavuz, Mohammad Raza, Cariad Evans |
| EPI_ISL_763647 | Virology Department, Royal Infirmary of Edinburgh, NHS Lothian / School of Biological Sciences, University of Edinburgh / Institute of Genetics and Molecular Medicine, University of Edinburgh | COVID-19 Genomics UK (COG-UK) Consortium | McHugh M, Dewar R, Rooke S, Gallagher M, Balcaza C, O'Toole Á, Scher E, Hill V, McCrone JT, Colquhoun R, Yu X, Jackson B, Rambaut A, Williams TC, Templeton K |
| EPI_ISL_763653, EPI_ISL_763654 | Virology Department, Sheffield Teaching Hospitals NHS Foundation Trust/Department of Infection, Immunity and Cardiovascular Disease, The Medical School, University of Sheffield | COVID-19 Genomics UK (COG-UK) Consortium | Thushan de Silva, Matthew Parker, Nikki Smith, Adri Angyal, Rebecca Brown, Luke Green, Rachel Tucker, Paul Parsons, Danielle Groves, Katie Johnson, Laura Carrilero, Alex Keeley, Dave Partridge, Matthew Wyles, Benjamin Lindsey, Mehmet Yavuz, Mohammad Raza, Cariad Evans |

| | | | |
|---|---|---|---|
| EPI_ISL_763657 | Wales Specialist Virology Centre Sequencing lab: Pathogen Genomics Unit | COVID-19 Genomics UK (COG-UK) Consortium | Catherine Moore, Johnathan Evans, Laura Gifford, Malorie Perry, Simon Cottrell, Angela Marchbank, Alec Birchley, Alexander Adams, Amy Gaskin, Bree Gatica-Wilcox, Jason Coombes, Joel Southgate, Lauren Gilbert, Lee Graham, Nicole Pacchiarini, Sara Kumziene-Summerhayes, Sarah Taylor, Sophie Jones, Sara Rey, Matthew Bull, Joanne Watkins, Sally Corden, Tom Connor |
| EPI_ISL_763659 | Centre for Enzyme Innovation, University of Portsmouth / Translational Research Laboratory, Portsmouth Hospitals NHS Trust | COVID-19 Genomics UK (COG-UK) Consortium | Angela Beckett,Yann Bourgeois,Garry Scarlett,Sharon Glaysher,Scott Elliott,Kelly Bicknell,Robert Impey,Allyson Lloyd,Sarah Wyllie,Ethan Butcher,Anoop Chauhan,Samuel Robson |
| EPI_ISL_763670 | University of Exeter | COVID-19 Genomics UK (COG-UK) Consortium | Ben Temperton,Aaron Jeffries,Michelle Michelsen,Joanna Warwick-Dugdale,Audrey Farbos,Robyn Manley,Stephen Michell,Jane Masoli |
| EPI_ISL_763671 | Wales Specialist Virology Centre Sequencing lab: Pathogen Genomics Unit | COVID-19 Genomics UK (COG-UK) Consortium | Catherine Moore, Johnathan Evans, Laura Gifford, Malorie Perry, Simon Cottrell, Angela Marchbank, Alec Birchley, Alexander Adams, Amy Gaskin, Bree Gatica-Wilcox, Jason Coombes, Joel Southgate, Lauren Gilbert, Lee Graham, Nicole Pacchiarini, Sara Kumziene-Summerhayes, Sarah Taylor, Sophie Jones, Sara Rey, Matthew Bull, Joanne Watkins, Sally Corden, Tom Connor |
| EPI_ISL_763672 | Centre for Enzyme Innovation, University of Portsmouth / Translational Research Laboratory, Portsmouth Hospitals NHS Trust | COVID-19 Genomics UK (COG-UK) Consortium | Angela Beckett,Yann Bourgeois,Garry Scarlett,Sharon Glaysher,Scott Elliott,Kelly Bicknell,Robert Impey,Allyson Lloyd,Sarah Wyllie,Ethan Butcher,Anoop Chauhan,Samuel Robson |
| EPI_ISL_763680 | Wales Specialist Virology Centre Sequencing lab: Pathogen Genomics Unit | COVID-19 Genomics UK (COG-UK) Consortium | Catherine Moore, Johnathan Evans, Laura Gifford, Malorie Perry, Simon Cottrell, Angela Marchbank, Alec Birchley, Alexander Adams, Amy Gaskin, Bree Gatica-Wilcox, Jason Coombes, Joel Southgate, Lauren Gilbert, Lee Graham, Nicole Pacchiarini, Sara Kumziene-Summerhayes, Sarah Taylor, Sophie Jones, Sara Rey, Matthew Bull, Joanne Watkins, Sally Corden, Tom Connor |
| EPI_ISL_763682 | University College London, Great Ormond Street Hospital for Children NHS Foundation Trust, Imperial College Healthcare NHS Trust | COVID-19 Genomics UK (COG-UK) Consortium | Sergi Castellano, Rachel Williams, Mark Kristiansen, Paola Resende Silva, Sunando Roy, Tony Brooks, Helena Tutill, Paola Niola, Patricia Dyal, Charlotte Williams, Leysa Forrest, Yasmin Panchbhaya, Jacqueline Findlay, Samuel Weeks, Julianne Brown, Kathryn Harris, Paul Randell, James Price, Alison Holmes, Judith Breuer |
| EPI_ISL_763683 | Quadram Institute Bioscience | COVID-19 Genomics UK (COG-UK) Consortium | Dave J. Baker, Gemma L. Kay, Alp Aydin, Thanh Le-Viet, Steven Rudder, Ana P. Tedim, Anastasia Kolyva, Maria Diaz, Leonardo de Oliveira Martins, Nabil-Fareed Alikhan, Lizzie Meadows, Rachael Stanley, Ngozi Elumogo, Muhammed Yasir, Nicholas M. Thomson, Alexander J Trotter, Rachel Gilroy, Samuel Bloomfield, Claire Stuart, Andrew Bell, Reenesh Prakash, Samir Dervisevic, Alison E. Mather, John Wain, Mark Webber, Andrew J. Page, Justin O'Grady |
| EPI_ISL_763687 | University College London, Great Ormond Street Hospital for Children NHS Foundation Trust, Imperial College Healthcare NHS Trust | COVID-19 Genomics UK (COG-UK) Consortium | Sergi Castellano, Rachel Williams, Mark Kristiansen, Paola Resende Silva, Sunando Roy, Tony Brooks, Helena Tutill, Paola Niola, Patricia Dyal, Charlotte Williams, Leysa Forrest, Yasmin Panchbhaya, Jacqueline Findlay, Samuel Weeks, Julianne Brown, Kathryn Harris, Paul Randell, James Price, Alison Holmes, Judith Breuer |
| EPI_ISL_763689 | Quadram Institute Bioscience | COVID-19 Genomics UK (COG-UK) Consortium | Dave J. Baker, Gemma L. Kay, Alp Aydin, Thanh Le-Viet, Steven Rudder, Ana P. Tedim, Anastasia Kolyva, Maria Diaz, Leonardo de Oliveira Martins, Nabil-Fareed Alikhan, Lizzie Meadows, Rachael Stanley, Ngozi Elumogo, Muhammed Yasir, Nicholas M. Thomson, Alexander J Trotter, Rachel Gilroy, Samuel Bloomfield, Claire Stuart, Andrew Bell, Reenesh Prakash, Samir Dervisevic, Alison E. Mather, John Wain, Mark Webber, Andrew J. Page, Justin O'Grady |
| EPI_ISL_763690 | Centre for Enzyme Innovation, University of Portsmouth / Translational Research Laboratory, Portsmouth Hospitals NHS Trust | COVID-19 Genomics UK (COG-UK) Consortium | Angela Beckett,Yann Bourgeois,Garry Scarlett,Sharon Glaysher,Scott Elliott,Kelly Bicknell,Robert Impey,Allyson Lloyd,Sarah Wyllie,Ethan Butcher,Anoop Chauhan,Samuel Robson |
| EPI_ISL_763692 | Wales Specialist Virology Centre Sequencing lab: Pathogen Genomics Unit | COVID-19 Genomics UK (COG-UK) Consortium | Catherine Moore, Johnathan Evans, Laura Gifford, Malorie Perry, Simon Cottrell, Angela Marchbank, Alec Birchley, Alexander Adams, Amy Gaskin, Bree Gatica-Wilcox, Jason Coombes, Joel Southgate, Lauren Gilbert, Lee Graham, Nicole Pacchiarini, Sara Kumziene-Summerhayes, Sarah Taylor, Sophie Jones, Sara Rey, Matthew Bull, Joanne Watkins, Sally Corden, Tom Connor |
| EPI_ISL_763713, EPI_ISL_763714, EPI_ISL_763715, EPI_ISL_763716, EPI_ISL_763717 | University of Exeter | COVID-19 Genomics UK (COG-UK) Consortium | Ben Temperton,Aaron Jeffries,Michelle Michelsen,Joanna Warwick-Dugdale,Audrey Farbos,Robyn Manley,Stephen Michell,Jane Masoli |
| EPI_ISL_763718 | Wales Specialist Virology Centre Sequencing lab: Pathogen Genomics Unit | COVID-19 Genomics UK (COG-UK) Consortium | Catherine Moore, Johnathan Evans, Laura Gifford, Malorie Perry, Simon Cottrell, Angela Marchbank, Alec Birchley, Alexander Adams, Amy Gaskin, Bree Gatica-Wilcox, Jason Coombes, Joel Southgate, Lauren Gilbert, Lee Graham, Nicole Pacchiarini, Sara Kumziene-Summerhayes, Sarah Taylor, Sophie Jones, Sara Rey, Matthew Bull, Joanne Watkins, Sally Corden, Tom Connor |
| EPI_ISL_763720 | University College London, Great Ormond Street Hospital for Children NHS Foundation Trust, Imperial College Healthcare NHS Trust | COVID-19 Genomics UK (COG-UK) Consortium | Sergi Castellano, Rachel Williams, Mark Kristiansen, Paola Resende Silva, Sunando Roy, Tony Brooks, Helena Tutill, Paola Niola, Patricia Dyal, Charlotte Williams, Leysa Forrest, Yasmin Panchbhaya, Jacqueline Findlay, Samuel Weeks, Julianne Brown, Kathryn Harris, Paul Randell, James Price, Alison Holmes, Judith Breuer |
| EPI_ISL_763722 | Wales Specialist Virology Centre Sequencing lab: Pathogen Genomics Unit | COVID-19 Genomics UK (COG-UK) Consortium | Catherine Moore, Johnathan Evans, Laura Gifford, Malorie Perry, Simon Cottrell, Angela Marchbank, Alec Birchley, Alexander Adams, Amy Gaskin, Bree Gatica-Wilcox, Jason Coombes, Joel Southgate, Lauren Gilbert, Lee Graham, Nicole Pacchiarini, Sara Kumziene-Summerhayes, Sarah Taylor, Sophie Jones, Sara Rey, Matthew Bull, Joanne Watkins, Sally Corden, Tom Connor |
| EPI_ISL_763728, EPI_ISL_763730 | University College London, Great Ormond Street Hospital for Children NHS Foundation Trust, Imperial College Healthcare NHS Trust | COVID-19 Genomics UK (COG-UK) Consortium | Sergi Castellano, Rachel Williams, Mark Kristiansen, Paola Resende Silva, Sunando Roy, Tony Brooks, Helena Tutill, Paola Niola, Patricia Dyal, Charlotte Williams, Leysa Forrest, Yasmin Panchbhaya, Jacqueline Findlay, Samuel Weeks, Julianne Brown, Kathryn Harris, Paul Randell, James Price, Alison Holmes, Judith Breuer |
| EPI_ISL_763735 | Wales Specialist Virology Centre Sequencing lab: Pathogen Genomics Unit | COVID-19 Genomics UK (COG-UK) Consortium | Catherine Moore, Johnathan Evans, Laura Gifford, Malorie Perry, Simon Cottrell, Angela Marchbank, Alec Birchley, Alexander Adams, Amy Gaskin, Bree Gatica-Wilcox, Jason Coombes, Joel Southgate, Lauren Gilbert, Lee Graham, Nicole Pacchiarini, Sara Kumziene-Summerhayes, Sarah Taylor, Sophie Jones, Sara Rey, Matthew Bull, Joanne Watkins, Sally Corden, Tom Connor |
| EPI_ISL_763745 | University College London, Great Ormond Street Hospital for Children NHS Foundation Trust, Imperial College Healthcare NHS Trust | COVID-19 Genomics UK (COG-UK) Consortium | Sergi Castellano, Rachel Williams, Mark Kristiansen, Paola Resende Silva, Sunando Roy, Tony Brooks, Helena Tutill, Paola Niola, Patricia Dyal, Charlotte Williams, Leysa Forrest, Yasmin Panchbhaya, Jacqueline Findlay, Samuel Weeks, Julianne Brown, Kathryn Harris, Paul Randell, James Price, Alison Holmes, Judith Breuer |
| EPI_ISL_763748, EPI_ISL_763752, EPI_ISL_763753 | Wales Specialist Virology Centre Sequencing lab: Pathogen Genomics Unit | COVID-19 Genomics UK (COG-UK) Consortium | Catherine Moore, Johnathan Evans, Laura Gifford, Malorie Perry, Simon Cottrell, Angela Marchbank, Alec Birchley, Alexander Adams, Amy Gaskin, Bree Gatica-Wilcox, Jason Coombes, Joel Southgate, Lauren Gilbert, Lee Graham, Nicole Pacchiarini, Sara Kumziene-Summerhayes, Sarah Taylor, Sophie Jones, Sara Rey, Matthew Bull, Joanne Watkins, Sally Corden, Tom Connor |
| EPI_ISL_763755 | Centre for Enzyme Innovation, University of Portsmouth / Translational Research Laboratory, Portsmouth Hospitals NHS Trust | COVID-19 Genomics UK (COG-UK) Consortium | Angela Beckett,Yann Bourgeois,Garry Scarlett,Sharon Glaysher,Scott Elliott,Kelly Bicknell,Robert Impey,Allyson Lloyd,Sarah Wyllie,Ethan Butcher,Anoop Chauhan,Samuel Robson |
| EPI_ISL_763756 | University College London, Great Ormond Street Hospital for Children NHS Foundation Trust, Imperial College Healthcare NHS Trust | COVID-19 Genomics UK (COG-UK) Consortium | Sergi Castellano, Rachel Williams, Mark Kristiansen, Paola Resende Silva, Sunando Roy, Tony Brooks, Helena Tutill, Paola Niola, Patricia Dyal, Charlotte Williams, Leysa Forrest, Yasmin Panchbhaya, Jacqueline Findlay, Samuel Weeks, Julianne Brown, Kathryn Harris, Paul Randell, James Price, Alison Holmes, Judith Breuer |
| EPI_ISL_763758, EPI_ISL_763759, EPI_ISL_763761, EPI_ISL_763765, EPI_ISL_763766 | Wales Specialist Virology Centre Sequencing lab: Pathogen Genomics Unit | COVID-19 Genomics UK (COG-UK) Consortium | Catherine Moore, Johnathan Evans, Laura Gifford, Malorie Perry, Simon Cottrell, Angela Marchbank, Alec Birchley, Alexander Adams, Amy Gaskin, Bree Gatica-Wilcox, Jason Coombes, Joel Southgate, Lauren Gilbert, Lee Graham, Nicole Pacchiarini, Sara Kumziene-Summerhayes, Sarah Taylor, Sophie Jones, Sara Rey, Matthew Bull, Joanne Watkins, Sally Corden, Tom Connor |
| EPI_ISL_763767, EPI_ISL_763772 | University College London, Great Ormond Street Hospital for Children NHS Foundation Trust, Imperial College Healthcare NHS Trust | COVID-19 Genomics UK (COG-UK) Consortium | Sergi Castellano, Rachel Williams, Mark Kristiansen, Paola Resende Silva, Sunando Roy, Tony Brooks, Helena Tutill, Paola Niola, Patricia Dyal, Charlotte Williams, Leysa Forrest, Yasmin Panchbhaya, Jacqueline Findlay, Samuel Weeks, Julianne Brown, Kathryn Harris, Paul Randell, James Price, Alison Holmes, Judith Breuer |
| EPI_ISL_763773, EPI_ISL_763774, EPI_ISL_763775, EPI_ISL_763779 | Wales Specialist Virology Centre Sequencing lab: Pathogen Genomics Unit | COVID-19 Genomics UK (COG-UK) Consortium | Catherine Moore, Johnathan Evans, Laura Gifford, Malorie Perry, Simon Cottrell, Angela Marchbank, Alec Birchley, Alexander Adams, Amy Gaskin, Bree Gatica-Wilcox, Jason Coombes, Joel Southgate, Lauren Gilbert, Lee Graham, Nicole Pacchiarini, Sara Kumziene-Summerhayes, Sarah Taylor, Sophie Jones, Sara Rey, Matthew Bull, Joanne Watkins, Sally Corden, Tom Connor |

| | | | |
|---|---|---|---|
| EPI_ISL_763781 | Quadram Institute Bioscience | COVID-19 Genomics UK (COG-UK) Consortium | Dave J. Baker, Gemma L. Kay, Alp Aydin, Thanh Le-Viet, Steven Rudder, Ana P. Tedim, Anastasia Kolyva, Maria Diaz, Leonardo de Oliveira Martins, Nabil-Fareed Alikhan, Lizzie Meadows, Rachael Stanley, Ngozi Elumogo, Muhammed Yasir, Nicholas M. Thomson, Alexander J Trotter, Rachel Gilroy, Samuel Bloomfield, Claire Stuart, Andrew Bell, Reenesh Prakash, Samir Dervisevic, Alison E. Mather, John Wain, Mark Webber, Andrew J. Page, Justin O'Grady |
| EPI_ISL_763783 | Wales Specialist Virology Centre Sequencing lab: Pathogen Genomics Unit | COVID-19 Genomics UK (COG-UK) Consortium | Catherine Moore, Johnathan Evans, Laura Gifford, Malorie Perry, Simon Cottrell, Angela Marchbank, Alec Birchley, Alexander Adams, Amy Gaskin, Bree Gatica-Wilcox, Jason Coombes, Joel Southgate, Lauren Gilbert, Lee Graham, Nicole Pacchiarini, Sara Kumziene-Summerhayes, Sarah Taylor, Sophie Jones, Sara Rey, Matthew Bull, Joanne Watkins, Sally Corden, Tom Connor |
| EPI_ISL_763784 | University College London, Great Ormond Street Hospital for Children NHS Foundation Trust, Imperial College Healthcare NHS Trust | COVID-19 Genomics UK (COG-UK) Consortium | Sergi Castellano, Rachel Williams, Mark Kristiansen, Paola Resende Silva, Sunando Roy, Tony Brooks, Helena Tutill, Paola Niola, Patricia Dyal, Charlotte Williams, Leysa Forrest, Yasmin Panchbhaya, Jacqueline Findlay, Samuel Weeks, Julianne Brown, Kathryn Harris, Paul Randell, James Price, Alison Holmes, Judith Breuer |
| EPI_ISL_763791, EPI_ISL_763799, EPI_ISL_763804 | Centre for Enzyme Innovation, University of Portsmouth / Translational Research Laboratory, Portsmouth Hospitals NHS Trust | COVID-19 Genomics UK (COG-UK) Consortium | Angela Beckett,Yann Bourgeois,Garry Scarlett,Sharon Glaysher,Scott Elliott,Kelly Bicknell,Robert Impey,Allyson Lloyd,Sarah Wyllie,Ethan Butcher,Anoop Chauhan,Samuel Robson |
| EPI_ISL_763809, EPI_ISL_763810, EPI_ISL_763811, EPI_ISL_763813 | University College London, Great Ormond Street Hospital for Children NHS Foundation Trust, Imperial College Healthcare NHS Trust | COVID-19 Genomics UK (COG-UK) Consortium | Sergi Castellano, Rachel Williams, Mark Kristiansen, Paola Resende Silva, Sunando Roy, Tony Brooks, Helena Tutill, Paola Niola, Patricia Dyal, Charlotte Williams, Leysa Forrest, Yasmin Panchbhaya, Jacqueline Findlay, Samuel Weeks, Julianne Brown, Kathryn Harris, Paul Randell, James Price, Alison Holmes, Judith Breuer |
| EPI_ISL_763814 | Wales Specialist Virology Centre Sequencing lab: Pathogen Genomics Unit | COVID-19 Genomics UK (COG-UK) Consortium | Catherine Moore, Johnathan Evans, Laura Gifford, Malorie Perry, Simon Cottrell, Angela Marchbank, Alec Birchley, Alexander Adams, Amy Gaskin, Bree Gatica-Wilcox, Jason Coombes, Joel Southgate, Lauren Gilbert, Lee Graham, Nicole Pacchiarini, Sara Kumziene-Summerhayes, Sarah Taylor, Sophie Jones, Sara Rey, Matthew Bull, Joanne Watkins, Sally Corden, Tom Connor |
| EPI_ISL_763818 | University of Exeter | COVID-19 Genomics UK (COG-UK) Consortium | Ben Temperton,Aaron Jeffries,Michelle Michelsen,Joanna Warwick-Dugdale,Audrey Farbos,Robyn Manley,Stephen Michell,Jane Masoli |
| EPI_ISL_763822 | Wales Specialist Virology Centre Sequencing lab: Pathogen Genomics Unit | COVID-19 Genomics UK (COG-UK) Consortium | Catherine Moore, Johnathan Evans, Laura Gifford, Malorie Perry, Simon Cottrell, Angela Marchbank, Alec Birchley, Alexander Adams, Amy Gaskin, Bree Gatica-Wilcox, Jason Coombes, Joel Southgate, Lauren Gilbert, Lee Graham, Nicole Pacchiarini, Sara Kumziene-Summerhayes, Sarah Taylor, Sophie Jones, Sara Rey, Matthew Bull, Joanne Watkins, Sally Corden, Tom Connor |
| EPI_ISL_763825 | Centre for Enzyme Innovation, University of Portsmouth / Translational Research Laboratory, Portsmouth Hospitals NHS Trust | COVID-19 Genomics UK (COG-UK) Consortium | Angela Beckett,Yann Bourgeois,Garry Scarlett,Sharon Glaysher,Scott Elliott,Kelly Bicknell,Robert Impey,Allyson Lloyd,Sarah Wyllie,Ethan Butcher,Anoop Chauhan,Samuel Robson |
| EPI_ISL_763826 | University College London, Great Ormond Street Hospital for Children NHS Foundation Trust, Imperial College Healthcare NHS Trust | COVID-19 Genomics UK (COG-UK) Consortium | Sergi Castellano, Rachel Williams, Mark Kristiansen, Paola Resende Silva, Sunando Roy, Tony Brooks, Helena Tutill, Paola Niola, Patricia Dyal, Charlotte Williams, Leysa Forrest, Yasmin Panchbhaya, Jacqueline Findlay, Samuel Weeks, Julianne Brown, Kathryn Harris, Paul Randell, James Price, Alison Holmes, Judith Breuer |
| EPI_ISL_763835 | Centre for Enzyme Innovation, University of Portsmouth / Translational Research Laboratory, Portsmouth Hospitals NHS Trust | COVID-19 Genomics UK (COG-UK) Consortium | Angela Beckett,Yann Bourgeois,Garry Scarlett,Sharon Glaysher,Scott Elliott,Kelly Bicknell,Robert Impey,Allyson Lloyd,Sarah Wyllie,Ethan Butcher,Anoop Chauhan,Samuel Robson |
| EPI_ISL_763845, EPI_ISL_763846 | Oxford Viromics, NDM, University of Oxford; Oxford University Hospitals; Basingstoke and North Hampshire Hospital | COVID-19 Genomics UK (COG-UK) Consortium | Tanya Golubchik, David Bonsall, George Macintyre, Amy Trebes, Mariateresa de Cesare, Catrin Moore, Alex Mobbs, Anita Justice, Robert Shaw, Monique Andersson, Timothy Peto, Emma Wise, Nathan Moore, Jessica Lynch, Nick Cortes, Matilde Mori, Stephen Kidd, David Buck, John Todd, Christophe Fraser |
| EPI_ISL_763864 | Wales Specialist Virology Centre Sequencing lab: Pathogen Genomics Unit | COVID-19 Genomics UK (COG-UK) Consortium | Catherine Moore, Johnathan Evans, Laura Gifford, Malorie Perry, Simon Cottrell, Angela Marchbank, Alec Birchley, Alexander Adams, Amy Gaskin, Bree Gatica-Wilcox, Jason Coombes, Joel Southgate, Lauren Gilbert, Lee Graham, Nicole Pacchiarini, Sara Kumziene-Summerhayes, Sarah Taylor, Sophie Jones, Sara Rey, Matthew Bull, Joanne Watkins, Sally Corden, Tom Connor |
| EPI_ISL_763865 | Centre for Enzyme Innovation, University of Portsmouth / Translational Research Laboratory, Portsmouth Hospitals NHS Trust | COVID-19 Genomics UK (COG-UK) Consortium | Angela Beckett,Yann Bourgeois,Garry Scarlett,Sharon Glaysher,Scott Elliott,Kelly Bicknell,Robert Impey,Allyson Lloyd,Sarah Wyllie,Ethan Butcher,Anoop Chauhan,Samuel Robson |
| EPI_ISL_763869 | Virology Department, Sheffield Teaching Hospitals NHS Foundation Trust/Department of Infection, Immunity and Cardiovascular Disease, The Medical School, University of Sheffield | COVID-19 Genomics UK (COG-UK) Consortium | Thushan de Silva, Matthew Parker, Nikki Smith, Adri Angyal, Rebecca Brown, Luke Green, Rachel Tucker, Paul Parsons, Danielle Groves, Katie Johnson, Laura Carrilero, Alex Keeley, Dave Partridge, Matthew Wyles, Benjamin Lindsey, Mehmet Yavuz, Mohammad Raza, Cariad Evans |
| EPI_ISL_763893, EPI_ISL_763894, EPI_ISL_763895, EPI_ISL_763896, EPI_ISL_763897, EPI_ISL_763898, EPI_ISL_763899, EPI_ISL_763900, EPI_ISL_763901, EPI_ISL_763905, EPI_ISL_763906 | | | |
| see above | Virology Department, Royal Infirmary of Edinburgh, NHS Lothian / School of Biological Sciences, University of Edinburgh / Institute of Genetics and Molecular Medicine, University of Edinburgh | COVID-19 Genomics UK (COG-UK) Consortium | McHugh M, Dewar R, Rooke S, Gallagher M, Balcaza C, O'Toole Á, Scher E, Hill V, McCrone JT, Colquhoun R, Yu X, Jackson B, Rambaut A, Williams TC, Templeton K |
| EPI_ISL_763908, EPI_ISL_763909, EPI_ISL_763910, EPI_ISL_763912, EPI_ISL_763913, EPI_ISL_763914, EPI_ISL_763915, EPI_ISL_763916, EPI_ISL_763918, EPI_ISL_763919, EPI_ISL_763920, EPI_ISL_763921, EPI_ISL_763922, EPI_ISL_763923 | | | |
| see above | Quadram Institute Bioscience | COVID-19 Genomics UK (COG-UK) Consortium | Dave J. Baker, Gemma L. Kay, Alp Aydin, Thanh Le-Viet, Steven Rudder, Ana P. Tedim, Anastasia Kolyva, Maria Diaz, Leonardo de Oliveira Martins, Nabil-Fareed Alikhan, Lizzie Meadows, Rachael Stanley, Ngozi Elumogo, Muhammed Yasir, Nicholas M. Thomson, Alexander J Trotter, Rachel Gilroy, Samuel Bloomfield, Claire Stuart, Andrew Bell, Reenesh Prakash, Samir Dervisevic, Alison E. Mather, John Wain, Mark Webber, Andrew J. Page, Justin O'Grady |
| EPI_ISL_763975, EPI_ISL_763983, EPI_ISL_763984, EPI_ISL_763985, EPI_ISL_763987, EPI_ISL_763988, EPI_ISL_763991, EPI_ISL_763992, EPI_ISL_763993, EPI_ISL_763994, EPI_ISL_763995, EPI_ISL_763996, EPI_ISL_763997, EPI_ISL_763998, EPI_ISL_763999, EPI_ISL_764000, EPI_ISL_764001, EPI_ISL_764002, EPI_ISL_764003, EPI_ISL_764004, EPI_ISL_764005 | | | |
| see above | Wales Specialist Virology Centre Sequencing lab: Pathogen Genomics Unit | COVID-19 Genomics UK (COG-UK) Consortium | Catherine Moore, Johnathan Evans, Laura Gifford, Malorie Perry, Simon Cottrell, Angela Marchbank, Alec Birchley, Alexander Adams, Amy Gaskin, Bree Gatica-Wilcox, Jason Coombes, Joel Southgate, Lauren Gilbert, Lee Graham, Nicole Pacchiarini, Sara Kumziene-Summerhayes, Sarah Taylor, Sophie Jones, Sara Rey, Matthew Bull, Joanne Watkins, Sally Corden, Tom Connor |
| EPI_ISL_764013, EPI_ISL_764020, EPI_ISL_764021, EPI_ISL_764022, EPI_ISL_764023, EPI_ISL_764024 | Centre for Enzyme Innovation, University of Portsmouth / Translational Research Laboratory, Portsmouth Hospitals NHS Trust | COVID-19 Genomics UK (COG-UK) Consortium | Angela Beckett,Yann Bourgeois,Garry Scarlett,Sharon Glaysher,Scott Elliott,Kelly Bicknell,Robert Impey,Allyson Lloyd,Sarah Wyllie,Ethan Butcher,Anoop Chauhan,Samuel Robson |
| EPI_ISL_764033 | Virology Department, Sheffield Teaching Hospitals NHS Foundation Trust/Department of Infection, Immunity and Cardiovascular Disease, The Medical School, University of Sheffield | COVID-19 Genomics UK (COG-UK) Consortium | Thushan de Silva, Matthew Parker, Nikki Smith, Adri Angyal, Rebecca Brown, Luke Green, Rachel Tucker, Paul Parsons, Danielle Groves, Katie Johnson, Laura Carrilero, Alex Keeley, Dave Partridge, Matthew Wyles, Benjamin Lindsey, Mehmet Yavuz, Mohammad Raza, Cariad Evans |
| EPI_ISL_764034 | Wales Specialist Virology Centre Sequencing lab: Pathogen Genomics Unit | COVID-19 Genomics UK (COG-UK) Consortium | Catherine Moore, Johnathan Evans, Laura Gifford, Malorie Perry, Simon Cottrell, Angela Marchbank, Alec Birchley, Alexander Adams, Amy Gaskin, Bree Gatica-Wilcox, Jason Coombes, Joel Southgate, Lauren Gilbert, Lee Graham, Nicole Pacchiarini, Sara Kumziene-Summerhayes, Sarah Taylor, Sophie Jones, Sara Rey, Matthew Bull, Joanne Watkins, Sally Corden, Tom Connor |
| EPI_ISL_764037, EPI_ISL_764038, EPI_ISL_764039, EPI_ISL_764040, EPI_ISL_764041, EPI_ISL_764043, EPI_ISL_764044, EPI_ISL_764045, EPI_ISL_764047, EPI_ISL_764048, EPI_ISL_764053, EPI_ISL_764054, EPI_ISL_764055, EPI_ISL_764056, EPI_ISL_764057, EPI_ISL_764058, EPI_ISL_764059, EPI_ISL_764060 | | | |
| see above | University College London, Great Ormond Street Hospital for Children NHS Foundation Trust, Imperial College Healthcare NHS Trust | COVID-19 Genomics UK (COG-UK) Consortium | Sergi Castellano, Rachel Williams, Mark Kristiansen, Paola Resende Silva, Sunando Roy, Tony Brooks, Helena Tutill, Paola Niola, Patricia Dyal, Charlotte Williams, Leysa Forrest, Yasmin Panchbhaya, Jacqueline Findlay, Samuel Weeks, Julianne Brown, Kathryn Harris, Paul Randell, James Price, Alison Holmes, Judith Breuer |
| EPI_ISL_764063, EPI_ISL_764065, | Wales Specialist Virology Centre Sequencing lab: Pathogen | COVID-19 Genomics UK (COG-UK) Consortium | Catherine Moore, Johnathan Evans, Laura Gifford, Malorie Perry, Simon Cottrell, Angela Marchbank, Alec Birchley, Alexander Adams, Amy Gaskin, Bree |

| | | | |
|---|---|---|---|
| EPI_ISL_764080 | Genomics Unit | | Gatica-Wilcox, Jason Coombes, Joel Southgate, Lauren Gilbert, Lee Graham, Nicole Pacchiarini, Sara Kumziene-Summerhayes, Sarah Taylor, Sophie Jones, Sara Rey, Matthew Bull, Joanne Watkins, Sally Corden, Tom Connor |
| EPI_ISL_764083 | Virology Department, Royal Infirmary of Edinburgh, NHS Lothian / School of Biological Sciences, University of Edinburgh / Institute of Genetics and Molecular Medicine, University of Edinburgh | COVID-19 Genomics UK (COG-UK) Consortium | McHugh M, Dewar R, Rooke S, Gallagher M, Balcaza C, O'Toole Á, Scher E, Hill V, McCrone JT, Colquhoun R, Yu X, Jackson B, Rambaut A, Williams TC, Templeton K |
| EPI_ISL_764121, EPI_ISL_764122, EPI_ISL_764123 | Centre for Enzyme Innovation, University of Portsmouth / Translational Research Laboratory, Portsmouth Hospitals NHS Trust | COVID-19 Genomics UK (COG-UK) Consortium | Angela Beckett,Yann Bourgeois,Garry Scarlett,Sharon Glaysher,Scott Elliott,Kelly Bicknell,Robert Impey,Allyson Lloyd,Sarah Wyllie,Ethan Butcher,Anoop Chauhan,Samuel Robson |
| EPI_ISL_764128 | University College London, Great Ormond Street Hospital for Children NHS Foundation Trust, Imperial College Healthcare NHS Trust | COVID-19 Genomics UK (COG-UK) Consortium | Sergi Castellano, Rachel Williams, Mark Kristiansen, Paola Resende Silva, Sunando Roy, Tony Brooks, Helena Tutill, Paola Niola, Patricia Dyal, Charlotte Williams, Leysa Forrest, Yasmin Panchbhaya, Jacqueline Findlay, Samuel Weeks, Julianne Brown, Kathryn Harris, Paul Randell, James Price, Alison Holmes, Judith Breuer |
| EPI_ISL_764129 | University of Exeter | COVID-19 Genomics UK (COG-UK) Consortium | Ben Temperton,Aaron Jeffries,Michelle Michelsen,Joanna Warwick-Dugdale,Audrey Farbos,Robyn Manley,Stephen Michell,Jane Masoli |
| EPI_ISL_764131 | University College London, Great Ormond Street Hospital for Children NHS Foundation Trust, Imperial College Healthcare NHS Trust | COVID-19 Genomics UK (COG-UK) Consortium | Sergi Castellano, Rachel Williams, Mark Kristiansen, Paola Resende Silva, Sunando Roy, Tony Brooks, Helena Tutill, Paola Niola, Patricia Dyal, Charlotte Williams, Leysa Forrest, Yasmin Panchbhaya, Jacqueline Findlay, Samuel Weeks, Julianne Brown, Kathryn Harris, Paul Randell, James Price, Alison Holmes, Judith Breuer |
| EPI_ISL_764136 | Centre for Enzyme Innovation, University of Portsmouth / Translational Research Laboratory, Portsmouth Hospitals NHS Trust | COVID-19 Genomics UK (COG-UK) Consortium | Angela Beckett,Yann Bourgeois,Garry Scarlett,Sharon Glaysher,Scott Elliott,Kelly Bicknell,Robert Impey,Allyson Lloyd,Sarah Wyllie,Ethan Butcher,Anoop Chauhan,Samuel Robson |
| EPI_ISL_764139 | University of Exeter | COVID-19 Genomics UK (COG-UK) Consortium | Ben Temperton,Aaron Jeffries,Michelle Michelsen,Joanna Warwick-Dugdale,Audrey Farbos,Robyn Manley,Stephen Michell,Jane Masoli |
| EPI_ISL_764142 | Wales Specialist Virology Centre Sequencing lab: Pathogen Genomics Unit | COVID-19 Genomics UK (COG-UK) Consortium | Catherine Moore, Johnathan Evans, Laura Gifford, Malorie Perry, Simon Cottrell, Angela Marchbank, Alec Birchley, Alexander Adams, Amy Gaskin, Bree Gatica-Wilcox, Jason Coombes, Joel Southgate, Lauren Gilbert, Lee Graham, Nicole Pacchiarini, Sara Kumziene-Summerhayes, Sarah Taylor, Sophie Jones, Sara Rey, Matthew Bull, Joanne Watkins, Sally Corden, Tom Connor |
| EPI_ISL_764146, EPI_ISL_764160 | University College London, Great Ormond Street Hospital for Children NHS Foundation Trust, Imperial College Healthcare NHS Trust | COVID-19 Genomics UK (COG-UK) Consortium | Sergi Castellano, Rachel Williams, Mark Kristiansen, Paola Resende Silva, Sunando Roy, Tony Brooks, Helena Tutill, Paola Niola, Patricia Dyal, Charlotte Williams, Leysa Forrest, Yasmin Panchbhaya, Jacqueline Findlay, Samuel Weeks, Julianne Brown, Kathryn Harris, Paul Randell, James Price, Alison Holmes, Judith Breuer |
| EPI_ISL_764168 | Wales Specialist Virology Centre Sequencing lab: Pathogen Genomics Unit | COVID-19 Genomics UK (COG-UK) Consortium | Catherine Moore, Johnathan Evans, Laura Gifford, Malorie Perry, Simon Cottrell, Angela Marchbank, Alec Birchley, Alexander Adams, Amy Gaskin, Bree Gatica-Wilcox, Jason Coombes, Joel Southgate, Lauren Gilbert, Lee Graham, Nicole Pacchiarini, Sara Kumziene-Summerhayes, Sarah Taylor, Sophie Jones, Sara Rey, Matthew Bull, Joanne Watkins, Sally Corden, Tom Connor |
| EPI_ISL_764172 | Centre for Enzyme Innovation, University of Portsmouth / Translational Research Laboratory, Portsmouth Hospitals NHS Trust | COVID-19 Genomics UK (COG-UK) Consortium | Angela Beckett,Yann Bourgeois,Garry Scarlett,Sharon Glaysher,Scott Elliott,Kelly Bicknell,Robert Impey,Allyson Lloyd,Sarah Wyllie,Ethan Butcher,Anoop Chauhan,Samuel Robson |
| EPI_ISL_764185 | University of Exeter | COVID-19 Genomics UK (COG-UK) Consortium | Ben Temperton,Aaron Jeffries,Michelle Michelsen,Joanna Warwick-Dugdale,Audrey Farbos,Robyn Manley,Stephen Michell,Jane Masoli |
| EPI_ISL_764194 | Quadram Institute Bioscience | COVID-19 Genomics UK (COG-UK) Consortium | Dave J. Baker, Gemma L. Kay, Alp Aydin, Thanh Le-Viet, Steven Rudder, Ana P. Tedim, Anastasia Kolyva, Maria Diaz, Leonardo de Oliveira Martins, Nabil-Fareed Alikhan, Lizzie Meadows, Rachael Stanley, Ngozi Elumogo, Muhammed Yasir, Nicholas M. Thomson, Alexander J Trotter, Rachel Gilroy, Samuel Bloomfield, Claire Stuart, Andrew Bell, Reenesh Prakash, Samir Dervisevic, Alison E. Mather, John Wain, Mark Webber, Andrew J. Page, Justin O'Grady |
| EPI_ISL_764196 | University of Exeter | COVID-19 Genomics UK (COG-UK) Consortium | Ben Temperton,Aaron Jeffries,Michelle Michelsen,Joanna Warwick-Dugdale,Audrey Farbos,Robyn Manley,Stephen Michell,Jane Masoli |
| EPI_ISL_764199 | Wales Specialist Virology Centre Sequencing lab: Pathogen Genomics Unit | COVID-19 Genomics UK (COG-UK) Consortium | Catherine Moore, Johnathan Evans, Laura Gifford, Malorie Perry, Simon Cottrell, Angela Marchbank, Alec Birchley, Alexander Adams, Amy Gaskin, Bree Gatica-Wilcox, Jason Coombes, Joel Southgate, Lauren Gilbert, Lee Graham, Nicole Pacchiarini, Sara Kumziene-Summerhayes, Sarah Taylor, Sophie Jones, Sara Rey, Matthew Bull, Joanne Watkins, Sally Corden, Tom Connor |
| EPI_ISL_764205 | Quadram Institute Bioscience | COVID-19 Genomics UK (COG-UK) Consortium | Dave J. Baker, Gemma L. Kay, Alp Aydin, Thanh Le-Viet, Steven Rudder, Ana P. Tedim, Anastasia Kolyva, Maria Diaz, Leonardo de Oliveira Martins, Nabil-Fareed Alikhan, Lizzie Meadows, Rachael Stanley, Ngozi Elumogo, Muhammed Yasir, Nicholas M. Thomson, Alexander J Trotter, Rachel Gilroy, Samuel Bloomfield, Claire Stuart, Andrew Bell, Reenesh Prakash, Samir Dervisevic, Alison E. Mather, John Wain, Mark Webber, Andrew J. Page, Justin O'Grady |
| EPI_ISL_764212 | Wales Specialist Virology Centre Sequencing lab: Pathogen Genomics Unit | COVID-19 Genomics UK (COG-UK) Consortium | Catherine Moore, Johnathan Evans, Laura Gifford, Malorie Perry, Simon Cottrell, Angela Marchbank, Alec Birchley, Alexander Adams, Amy Gaskin, Bree Gatica-Wilcox, Jason Coombes, Joel Southgate, Lauren Gilbert, Lee Graham, Nicole Pacchiarini, Sara Kumziene-Summerhayes, Sarah Taylor, Sophie Jones, Sara Rey, Matthew Bull, Joanne Watkins, Sally Corden, Tom Connor |
| EPI_ISL_764214, EPI_ISL_764215 | University of Exeter | COVID-19 Genomics UK (COG-UK) Consortium | Ben Temperton,Aaron Jeffries,Michelle Michelsen,Joanna Warwick-Dugdale,Audrey Farbos,Robyn Manley,Stephen Michell,Jane Masoli |
| EPI_ISL_764222 | Wales Specialist Virology Centre Sequencing lab: Pathogen Genomics Unit | COVID-19 Genomics UK (COG-UK) Consortium | Catherine Moore, Johnathan Evans, Laura Gifford, Malorie Perry, Simon Cottrell, Angela Marchbank, Alec Birchley, Alexander Adams, Amy Gaskin, Bree Gatica-Wilcox, Jason Coombes, Joel Southgate, Lauren Gilbert, Lee Graham, Nicole Pacchiarini, Sara Kumziene-Summerhayes, Sarah Taylor, Sophie Jones, Sara Rey, Matthew Bull, Joanne Watkins, Sally Corden, Tom Connor |
| EPI_ISL_764225 | University of Exeter | COVID-19 Genomics UK (COG-UK) Consortium | Ben Temperton,Aaron Jeffries,Michelle Michelsen,Joanna Warwick-Dugdale,Audrey Farbos,Robyn Manley,Stephen Michell,Jane Masoli |
| EPI_ISL_764226 | Virology Department, Sheffield Teaching Hospitals NHS Foundation Trust/Department of Infection, Immunity and Cardiovascular Disease, The Medical School, University of Sheffield | COVID-19 Genomics UK (COG-UK) Consortium | Thushan de Silva, Matthew Parker, Nikki Smith, Adri Angyal, Rebecca Brown, Luke Green, Rachel Tucker, Paul Parsons, Danielle Groves, Katie Johnson, Laura Carrilero, Alex Keeley, Dave Partridge, Matthew Wyles, Benjamin Lindsey, Mehmet Yavuz, Mohammad Raza, Cariad Evans |
| EPI_ISL_764228 | University College London, Great Ormond Street Hospital for Children NHS Foundation Trust, Imperial College Healthcare NHS Trust | COVID-19 Genomics UK (COG-UK) Consortium | Sergi Castellano, Rachel Williams, Mark Kristiansen, Paola Resende Silva, Sunando Roy, Tony Brooks, Helena Tutill, Paola Niola, Patricia Dyal, Charlotte Williams, Leysa Forrest, Yasmin Panchbhaya, Jacqueline Findlay, Samuel Weeks, Julianne Brown, Kathryn Harris, Paul Randell, James Price, Alison Holmes, Judith Breuer |
| EPI_ISL_764237 | Quadram Institute Bioscience | COVID-19 Genomics UK (COG-UK) Consortium | Dave J. Baker, Gemma L. Kay, Alp Aydin, Thanh Le-Viet, Steven Rudder, Ana P. Tedim, Anastasia Kolyva, Maria Diaz, Leonardo de Oliveira Martins, Nabil-Fareed Alikhan, Lizzie Meadows, Rachael Stanley, Ngozi Elumogo, Muhammed Yasir, Nicholas M. Thomson, Alexander J Trotter, Rachel Gilroy, Samuel Bloomfield, Claire Stuart, Andrew Bell, Reenesh Prakash, Samir Dervisevic, Alison E. Mather, John Wain, Mark Webber, Andrew J. Page, Justin O'Grady |
| EPI_ISL_764241 | University College London, Great Ormond Street Hospital for Children NHS Foundation Trust, Imperial College Healthcare NHS Trust | COVID-19 Genomics UK (COG-UK) Consortium | Sergi Castellano, Rachel Williams, Mark Kristiansen, Paola Resende Silva, Sunando Roy, Tony Brooks, Helena Tutill, Paola Niola, Patricia Dyal, Charlotte Williams, Leysa Forrest, Yasmin Panchbhaya, Jacqueline Findlay, Samuel Weeks, Julianne Brown, Kathryn Harris, Paul Randell, James Price, Alison Holmes, Judith Breuer |
| EPI_ISL_764244, EPI_ISL_764246, EPI_ISL_764248, EPI_ISL_764249, EPI_ISL_764251, EPI_ISL_764252, EPI_ISL_764253, EPI_ISL_764256, EPI_ISL_764257, EPI_ISL_764259, EPI_ISL_764260, EPI_ISL_764268 | | | |
| see above | Wales Specialist Virology Centre Sequencing lab: Pathogen Genomics Unit | COVID-19 Genomics UK (COG-UK) Consortium | Catherine Moore, Johnathan Evans, Laura Gifford, Malorie Perry, Simon Cottrell, Angela Marchbank, Alec Birchley, Alexander Adams, Amy Gaskin, Bree Gatica-Wilcox, Jason Coombes, Joel Southgate, Lauren Gilbert, Lee Graham, Nicole Pacchiarini, Sara Kumziene-Summerhayes, Sarah Taylor, Sophie Jones, Sara Rey, Matthew Bull, Joanne Watkins, Sally Corden, Tom Connor |
| EPI_ISL_764270 | Quadram Institute Bioscience | COVID-19 Genomics UK (COG-UK) Consortium | Dave J. Baker, Gemma L. Kay, Alp Aydin, Thanh Le-Viet, Steven Rudder, Ana P. Tedim, Anastasia Kolyva, Maria Diaz, Leonardo de Oliveira Martins, Nabil-Fareed Alikhan, Lizzie Meadows, Rachael Stanley, Ngozi Elumogo, Muhammed Yasir, Nicholas M. Thomson, Alexander J Trotter, Rachel Gilroy, Samuel Bloomfield, Claire Stuart, Andrew Bell, Reenesh Prakash, Samir Dervisevic, Alison E. Mather, John Wain, Mark Webber, Andrew J. Page, Justin |

| | | | |
|---|---|---|---|
| | | | O'Grady |
| EPI_ISL_764273, EPI_ISL_764274, EPI_ISL_764277, EPI_ISL_764283 | Wales Specialist Virology Centre Sequencing lab: Pathogen Genomics Unit | COVID-19 Genomics UK (COG-UK) Consortium | Catherine Moore, Johnathan Evans, Laura Gifford, Malorie Perry, Simon Cottrell, Angela Marchbank, Alec Birchley, Alexander Adams, Amy Gaskin, Bree Gatica-Wilcox, Jason Coombes, Joel Southgate, Lauren Gilbert, Lee Graham, Nicole Pacchiarini, Sara Kumziene-Summerhayes, Sarah Taylor, Sophie Jones, Sara Rey, Matthew Bull, Joanne Watkins, Sally Corden, Tom Connor |
| EPI_ISL_764378, EPI_ISL_764379, EPI_ISL_764380, EPI_ISL_764381, EPI_ISL_764382 | University of Exeter | COVID-19 Genomics UK (COG-UK) Consortium | Ben Temperton,Aaron Jeffries,Michelle Michelsen,Joanna Warwick-Dugdale,Audrey Farbos,Robyn Manley,Stephen Michell,Jane Masoli |
| EPI_ISL_764383, EPI_ISL_764384 | University College London, Great Ormond Street Hospital for Children NHS Foundation Trust, Imperial College Healthcare NHS Trust | COVID-19 Genomics UK (COG-UK) Consortium | Sergi Castellano, Rachel Williams, Mark Kristiansen, Paola Resende Silva, Sunando Roy, Tony Brooks, Helena Tutill, Paola Niola, Patricia Dyal, Charlotte Williams, Leysa Forrest, Yasmin Panchbhaya, Jacqueline Findlay, Samuel Weeks, Julianne Brown, Kathryn Harris, Paul Randell, James Price, Alison Holmes, Judith Breuer |
| EPI_ISL_764445, EPI_ISL_764446, EPI_ISL_764447, EPI_ISL_764448, EPI_ISL_764450, EPI_ISL_764452, EPI_ISL_764454, EPI_ISL_764455, EPI_ISL_764456, EPI_ISL_764458, EPI_ISL_764459, EPI_ISL_764460, EPI_ISL_764462, EPI_ISL_764463, EPI_ISL_764464, EPI_ISL_764466, EPI_ISL_764469, EPI_ISL_764470, EPI_ISL_764471, EPI_ISL_764472, EPI_ISL_764473, EPI_ISL_764474, EPI_ISL_764475, EPI_ISL_764478, EPI_ISL_764479, EPI_ISL_764480, EPI_ISL_764481, EPI_ISL_764484 | | | |
| see above | Quadram Institute Bioscience | COVID-19 Genomics UK (COG-UK) Consortium | Dave J. Baker, Gemma L. Kay, Alp Aydin, Thanh Le-Viet, Steven Rudder, Ana P. Tedim, Anastasia Kolyva, Maria Diaz, Leonardo de Oliveira Martins, Nabil-Fareed Alikhan, Lizzie Meadows, Rachael Stanley, Ngozi Elumogo, Muhammed Yasir, Nicholas M. Thomson, Alexander J Trotter, Rachel Gilroy, Samuel Bloomfield, Claire Stuart, Andrew Bell, Reenesh Prakash, Samir Dervisevic, Alison E. Mather, John Wain, Mark Webber, Andrew J. Page, Justin O'Grady |
| EPI_ISL_764558, EPI_ISL_764560, EPI_ISL_764562, EPI_ISL_764563, EPI_ISL_764566, EPI_ISL_764571 | Oxford Viromics, NDM, University of Oxford; Oxford University Hospitals; Basingstoke and North Hampshire Hospital | COVID-19 Genomics UK (COG-UK) Consortium | Tanya Golubchik, David Bonsall, George Macintyre, Amy Trebes, Mariateresa de Cesare, Catrin Moore, Alex Mobbs, Anita Justice, Robert Shaw, Monique Andersson, Timothy Peto, Emma Wise, Nathan Moore, Jessica Lynch, Nick Cortes, Matilde Mori, Stephen Kidd, David Buck, John Todd, Christophe Fraser |
| EPI_ISL_764578, EPI_ISL_764579, EPI_ISL_764580, EPI_ISL_764588, EPI_ISL_764589, EPI_ISL_764590, EPI_ISL_764591, EPI_ISL_764592, EPI_ISL_764593, EPI_ISL_764594, EPI_ISL_764595, EPI_ISL_764596, EPI_ISL_764597, EPI_ISL_764659, EPI_ISL_764662, EPI_ISL_764663, EPI_ISL_764664, EPI_ISL_764665, EPI_ISL_764666, EPI_ISL_764667, EPI_ISL_764668, EPI_ISL_764669, EPI_ISL_764670, EPI_ISL_764671, EPI_ISL_764672, EPI_ISL_764673, EPI_ISL_764674, EPI_ISL_764675, EPI_ISL_764676, EPI_ISL_764677, EPI_ISL_764680, EPI_ISL_764681, EPI_ISL_764682, EPI_ISL_764683, EPI_ISL_764684, EPI_ISL_764685, EPI_ISL_764686, EPI_ISL_764687, EPI_ISL_764688, EPI_ISL_764689, EPI_ISL_764690, EPI_ISL_764691, EPI_ISL_764692, EPI_ISL_764693, EPI_ISL_764694, EPI_ISL_764695, EPI_ISL_764696, EPI_ISL_764697, EPI_ISL_764698, EPI_ISL_764699, EPI_ISL_764700, EPI_ISL_764701, EPI_ISL_764702, EPI_ISL_764703, EPI_ISL_764704, EPI_ISL_764705, EPI_ISL_764706, EPI_ISL_764707, EPI_ISL_764708, EPI_ISL_764709, EPI_ISL_764710, EPI_ISL_764711, EPI_ISL_764712, EPI_ISL_764713, EPI_ISL_764714, EPI_ISL_764715, EPI_ISL_764716, EPI_ISL_764717, EPI_ISL_764718, EPI_ISL_764719, EPI_ISL_764720, EPI_ISL_764721, EPI_ISL_764722, EPI_ISL_764723, EPI_ISL_764724, EPI_ISL_764725, EPI_ISL_764726, EPI_ISL_764727, EPI_ISL_764728, EPI_ISL_764729, EPI_ISL_764730, EPI_ISL_764731, EPI_ISL_764732, EPI_ISL_764733, EPI_ISL_764734, EPI_ISL_764735, EPI_ISL_764736, EPI_ISL_764737, EPI_ISL_764738, EPI_ISL_764744, EPI_ISL_764745, EPI_ISL_764765, EPI_ISL_764872, EPI_ISL_764873, EPI_ISL_764876, EPI_ISL_764877, EPI_ISL_764878, EPI_ISL_764880, EPI_ISL_764882, EPI_ISL_764885, EPI_ISL_764888, EPI_ISL_764889, EPI_ISL_764890, EPI_ISL_764891, EPI_ISL_764892, EPI_ISL_764893, EPI_ISL_764894, EPI_ISL_764895, EPI_ISL_764896, EPI_ISL_764908, EPI_ISL_764909, EPI_ISL_764910, EPI_ISL_764911, EPI_ISL_764912, EPI_ISL_764913, EPI_ISL_764914, EPI_ISL_764915, EPI_ISL_764916, EPI_ISL_764917, EPI_ISL_764918, EPI_ISL_764919, EPI_ISL_764920, EPI_ISL_764921, EPI_ISL_764922, EPI_ISL_764923, EPI_ISL_764924, EPI_ISL_764925, EPI_ISL_764926, EPI_ISL_764927, EPI_ISL_764928, EPI_ISL_764929, EPI_ISL_764930, EPI_ISL_764931, EPI_ISL_764932, EPI_ISL_764933, EPI_ISL_764934, EPI_ISL_764935, EPI_ISL_764936, EPI_ISL_764937, EPI_ISL_764938, EPI_ISL_764947, EPI_ISL_764948, EPI_ISL_764949, EPI_ISL_764950, EPI_ISL_764951, EPI_ISL_764989, EPI_ISL_764990, EPI_ISL_764996, EPI_ISL_764998, EPI_ISL_764999, EPI_ISL_765000, EPI_ISL_765001, EPI_ISL_765003, EPI_ISL_765004, EPI_ISL_765005, EPI_ISL_765006, EPI_ISL_765007, EPI_ISL_765008, EPI_ISL_765009, EPI_ISL_765010, EPI_ISL_765011, EPI_ISL_765012, EPI_ISL_765013, EPI_ISL_765014, EPI_ISL_765015, EPI_ISL_765016, EPI_ISL_765017, EPI_ISL_765018, EPI_ISL_765019, EPI_ISL_765020, EPI_ISL_765021, EPI_ISL_765022, EPI_ISL_765023, EPI_ISL_765024, EPI_ISL_765025, EPI_ISL_765026, EPI_ISL_765027, EPI_ISL_765028, EPI_ISL_765029, EPI_ISL_765030, EPI_ISL_765031, EPI_ISL_765032, EPI_ISL_765033, EPI_ISL_765034, EPI_ISL_765035, EPI_ISL_765036, EPI_ISL_765037, EPI_ISL_765038, EPI_ISL_765039, EPI_ISL_765040, EPI_ISL_765041, EPI_ISL_765042, EPI_ISL_765043, EPI_ISL_765044, EPI_ISL_765045, EPI_ISL_765046, EPI_ISL_765047, EPI_ISL_765048, EPI_ISL_765049, EPI_ISL_765050, EPI_ISL_765051, EPI_ISL_765052, EPI_ISL_765053, EPI_ISL_765054, EPI_ISL_765055, EPI_ISL_765056, EPI_ISL_765057, EPI_ISL_765058, EPI_ISL_765059, EPI_ISL_765060, EPI_ISL_765061, EPI_ISL_765062, EPI_ISL_765063, EPI_ISL_765064, EPI_ISL_765065, EPI_ISL_765066, EPI_ISL_765067, EPI_ISL_765068, EPI_ISL_765069, EPI_ISL_765070, EPI_ISL_765071, EPI_ISL_765072, EPI_ISL_765073, EPI_ISL_765074, EPI_ISL_765075, EPI_ISL_765076, EPI_ISL_765077, EPI_ISL_765078, EPI_ISL_765079, EPI_ISL_765080, EPI_ISL_765081, EPI_ISL_765082, EPI_ISL_765083, EPI_ISL_765084, EPI_ISL_765085, EPI_ISL_765086, EPI_ISL_765087, EPI_ISL_765088, EPI_ISL_765089, EPI_ISL_765090, EPI_ISL_765091, EPI_ISL_765092, EPI_ISL_765093, EPI_ISL_765094, EPI_ISL_765095, EPI_ISL_765096, EPI_ISL_765097, EPI_ISL_765098, EPI_ISL_765099, EPI_ISL_765100, EPI_ISL_765101, EPI_ISL_765102, EPI_ISL_765103, EPI_ISL_765104, EPI_ISL_765105, EPI_ISL_765106, EPI_ISL_765107, EPI_ISL_765108, EPI_ISL_765109, EPI_ISL_765110, EPI_ISL_765111 | | | |
| see above | Wales Specialist Virology Centre Sequencing lab: Pathogen Genomics Unit | COVID-19 Genomics UK (COG-UK) Consortium | Catherine Moore, Johnathan Evans, Laura Gifford, Malorie Perry, Simon Cottrell, Angela Marchbank, Alec Birchley, Alexander Adams, Amy Gaskin, Bree Gatica-Wilcox, Jason Coombes, Joel Southgate, Lauren Gilbert, Lee Graham, Nicole Pacchiarini, Sara Kumziene-Summerhayes, Sarah Taylor, Sophie Jones, Sara Rey, Matthew Bull, Joanne Watkins, Sally Corden, Tom Connor |
| EPI_ISL_765131, EPI_ISL_765132, EPI_ISL_765133, EPI_ISL_765134, EPI_ISL_765135, EPI_ISL_765136, EPI_ISL_765137, EPI_ISL_765138, EPI_ISL_765145 | Centre for Enzyme Innovation, University of Portsmouth / Translational Research Laboratory, Portsmouth Hospitals NHS Trust | COVID-19 Genomics UK (COG-UK) Consortium | Angela Beckett,Yann Bourgeois,Garry Scarlett,Sharon Glaysher,Scott Elliott,Kelly Bicknell,Robert Impey,Allyson Lloyd,Sarah Wyllie,Ethan Butcher,Anoop Chauhan,Samuel Robson |
| EPI_ISL_765152, EPI_ISL_765157, EPI_ISL_765162, EPI_ISL_765166, EPI_ISL_765175, EPI_ISL_765180, EPI_ISL_765183, EPI_ISL_765193, EPI_ISL_765194, EPI_ISL_765198, EPI_ISL_765199 | | | |
| see above | Virology Department, Sheffield Teaching Hospitals NHS Foundation Trust/Department of Infection, Immunity and Cardiovascular Disease, The Medical School, University of Sheffield | COVID-19 Genomics UK (COG-UK) Consortium | Thushan de Silva, Matthew Parker, Nikki Smith, Adri Angyal, Rebecca Brown, Luke Green, Rachel Tucker, Paul Parsons, Danielle Groves, Katie Johnson, Laura Carrilero, Alex Keeley, Dave Partridge, Matthew Wyles, Benjamin Lindsey, Mehmet Yavuz, Mohammad Raza, Cariad Evans |
| EPI_ISL_765479 | Laboratoire de Virologie, Hôpital Robert Debré | National Reference Center for Viruses of Respiratory Infections, Institut Pasteur, Paris | Marion Barbet, Sylvie Behillil, Méline Bizard, Angela Brisebarre, Camille Capel, Etienne Simon-Lorière, Vincent Enouf, Maud Vanpeene, Sylvie van der Werf, Andreoletti Laurent |
| EPI_ISL_765483 | SARATOGA HOSPITAL LABORATORY | Wadsworth Center, New York State Department.of Health | Kirsten St. George, Daryl M. Lamson, Alexis Russel, Matthew Shudt, Melissa A Leisner, Jonathan Plitnick, Navjot Singh, John Kelly, Sara Griesemer, Erasmus Schneider, Erica Lasek-Nesselquist |
| EPI_ISL_765491, EPI_ISL_765494, EPI_ISL_765495 | MONTEFIORE MEDICAL CENTER LABORATORIES | Wadsworth Center, New York State Department.of Health | Kirsten St. George, Daryl M. Lamson, Alexis Russel, Matthew Shudt, Melissa A Leisner, Jonathan Plitnick, Navjot Singh, John Kelly, Sara Griesemer, Erasmus Schneider, Erica Lasek-Nesselquist |
| EPI_ISL_765524, EPI_ISL_765525 | SARATOGA HOSPITAL LABORATORY | Wadsworth Center, New York State Department.of Health | Kirsten St. George, Daryl M. Lamson, Alexis Russel, Matthew Shudt, Melissa A Leisner, Jonathan Plitnick, Navjot Singh, John Kelly, Sara Griesemer, Erasmus Schneider, Erica Lasek-Nesselquist |
| EPI_ISL_765531, EPI_ISL_765532, EPI_ISL_765533, EPI_ISL_765534, EPI_ISL_765535, EPI_ISL_765539, EPI_ISL_765540, EPI_ISL_765542, EPI_ISL_765544, EPI_ISL_765547, EPI_ISL_765549, EPI_ISL_765550, EPI_ISL_765551, EPI_ISL_765553, EPI_ISL_765555, EPI_ISL_765556, EPI_ISL_765557, EPI_ISL_765559, EPI_ISL_765560, EPI_ISL_765561, EPI_ISL_765562, EPI_ISL_765566 | | | |
| see above | MONTEFIORE MEDICAL CENTER LABORATORIES | Wadsworth Center, New York State Department.of Health | Kirsten St. George, Daryl M. Lamson, Alexis Russel, Matthew Shudt, Melissa A Leisner, Jonathan Plitnick, Navjot Singh, John Kelly, Sara Griesemer, Erasmus Schneider, Erica Lasek-Nesselquist |
| EPI_ISL_765979, EPI_ISL_765980, EPI_ISL_765981, EPI_ISL_765982, EPI_ISL_765983, EPI_ISL_765984, EPI_ISL_765985 | Wyoming Public Health Laboratory | Wyoming Public Health Laboratory | Noah Hull, Taylor Fearing, Channing Weber, Ashley Norberg, Bailey Bowcutt, and Wanda Manley |
| EPI_ISL_765998 | Los Angeles County PHL | Los Angeles County PHL | P. Hemarajata et al. |
| EPI_ISL_766025 | Department of Virology and Immunology, University of Helsinki and Helsinki University Hospital, Huslab Finland | Department of Virology, Faculty of Medicine, University of Helsinki, Helsinki, Finland | Teemu Smura, Olli Vapalahti, Maija Lappalainen, Satu Kurkela |
| EPI_ISL_766060, EPI_ISL_766062, EPI_ISL_766065, EPI_ISL_766075, EPI_ISL_766082, EPI_ISL_766085, EPI_ISL_766086, EPI_ISL_766088, EPI_ISL_766089, EPI_ISL_766093, EPI_ISL_766099, EPI_ISL_766100, EPI_ISL_766104, EPI_ISL_766108, EPI_ISL_766111, EPI_ISL_766143, EPI_ISL_766145, EPI_ISL_766146, EPI_ISL_766147, EPI_ISL_766148, EPI_ISL_766154, EPI_ISL_766155, EPI_ISL_766156, EPI_ISL_766157, EPI_ISL_766159, EPI_ISL_766160, EPI_ISL_766161, EPI_ISL_766162, EPI_ISL_766163, EPI_ISL_766164, EPI_ISL_766165, EPI_ISL_766166, EPI_ISL_766167, EPI_ISL_766168, EPI_ISL_766169, EPI_ISL_766170, EPI_ISL_766171, EPI_ISL_766172, EPI_ISL_766173, EPI_ISL_766174, EPI_ISL_766175, EPI_ISL_766176, EPI_ISL_766177, EPI_ISL_766178, EPI_ISL_766179, EPI_ISL_766180, EPI_ISL_766181, EPI_ISL_766182, EPI_ISL_766183, EPI_ISL_766184, EPI_ISL_766185, EPI_ISL_766186, EPI_ISL_766187, EPI_ISL_766188, EPI_ISL_766189, EPI_ISL_766190, EPI_ISL_766191, EPI_ISL_766192, EPI_ISL_766193, EPI_ISL_766195, EPI_ISL_766205, EPI_ISL_766215, EPI_ISL_766228, EPI_ISL_766229, EPI_ISL_766234, EPI_ISL_766236, EPI_ISL_766243, EPI_ISL_766244, EPI_ISL_766251, EPI_ISL_766252, EPI_ISL_766253, EPI_ISL_766254, EPI_ISL_766257, EPI_ISL_766398, EPI_ISL_766402, EPI_ISL_766404, EPI_ISL_766406, EPI_ISL_766408, EPI_ISL_766413, EPI_ISL_766414, EPI_ISL_766416, EPI_ISL_766417, EPI_ISL_766418, EPI_ISL_766419, EPI_ISL_766420, EPI_ISL_766422, EPI_ISL_766423, EPI_ISL_766424, EPI_ISL_766426, EPI_ISL_766428, EPI_ISL_766429, EPI_ISL_766430, EPI_ISL_766431, EPI_ISL_766432, EPI_ISL_766433, EPI_ISL_766434, EPI_ISL_766435, EPI_ISL_766436, EPI_ISL_766440, EPI_ISL_766441, EPI_ISL_766442, EPI_ISL_766443, EPI_ISL_766444, EPI_ISL_766445, EPI_ISL_766446, EPI_ISL_766447, EPI_ISL_766448, EPI_ISL_766449, EPI_ISL_766450, EPI_ISL_766451, EPI_ISL_766452, EPI_ISL_766453, EPI_ISL_766454, EPI_ISL_766455, EPI_ISL_766456, EPI_ISL_766457, EPI_ISL_766458, EPI_ISL_766459, EPI_ISL_766460, EPI_ISL_766461, EPI_ISL_766462, EPI_ISL_766464, EPI_ISL_766465, EPI_ISL_766466, EPI_ISL_766467, EPI_ISL_766468, EPI_ISL_766469, EPI_ISL_766470, EPI_ISL_766471, EPI_ISL_766472, EPI_ISL_766473, EPI_ISL_766474, EPI_ISL_766475, EPI_ISL_766476, EPI_ISL_766477, EPI_ISL_766478, EPI_ISL_766479, EPI_ISL_766480, EPI_ISL_766481, EPI_ISL_766482, EPI_ISL_766483, EPI_ISL_766484, EPI_ISL_766485, EPI_ISL_766486, EPI_ISL_766487, EPI_ISL_766488, EPI_ISL_766489, EPI_ISL_766490, EPI_ISL_766491, EPI_ISL_766492, EPI_ISL_766493, EPI_ISL_766494, EPI_ISL_766495, EPI_ISL_766496, EPI_ISL_766497, EPI_ISL_766498, EPI_ISL_766499, EPI_ISL_766500, EPI_ISL_766501, EPI_ISL_766502, EPI_ISL_766517, EPI_ISL_766520, EPI_ISL_766553, EPI_ISL_766554, EPI_ISL_766560, EPI_ISL_766565 | | | |

| | | | |
|---|---|---|---|
| see above | Respiratory Virus Unit, National Infection Service, Public Health England | COVID-19 Genomics UK (COG-UK) Consortium | PHE Covid Sequencing Team |
| EPI_ISL_766633 | Klinisk mikrobiologi | The Public Health Agency of Sweden | Department of Microbiology, The Public Health Agency of Sweden |
| EPI_ISL_766713 | Klinisk Mikrobiologi | The Public Health Agency of Sweden | Department of Microbiology, The Public Health Agency of Sweden |
| EPI_ISL_766775, EPI_ISL_766776, EPI_ISL_766777, EPI_ISL_766778, EPI_ISL_766779, EPI_ISL_766780, EPI_ISL_766781, EPI_ISL_766782, EPI_ISL_766783, EPI_ISL_766784, EPI_ISL_766785, EPI_ISL_766786, EPI_ISL_766787, EPI_ISL_766788, EPI_ISL_766789, EPI_ISL_766790 | | | |
| see above | Wyoming Public Health Laboratory | Wyoming Public Health Laboratory | Noah Hull, Taylor Fearing, Channing Weber, Ashley Norberg, Bailey Bowcutt, and Wanda Manley |
| EPI_ISL_766860 | Respiratory Virus Unit, National Infection Service, Public Health England | COVID-19 Genomics UK (COG-UK) Consortium | PHE Covid Sequencing Team |
| EPI_ISL_766898, EPI_ISL_766909, EPI_ISL_766910 | New Mexico Department of Health Scientific Laboratory | New Mexico Department of Health Scientific Laboratory | D'eldra Malone, Ellie Johnson, Anastacia Griego-Fisher |
| EPI_ISL_766988 | Delaware Public Health Laboratory | Delaware Public Health Laboratory | Gregory Hovan |
| EPI_ISL_767089, EPI_ISL_767093, EPI_ISL_767094 | Lighthouse Lab in Alderley Park | Wellcome Sanger Institute for the COVID-19 Genomics UK (COG-UK) Consortium | Jacquelyn Wynn, Mairead Hyland, The Lighthouse Lab in Alderley Park and Alex Alderton, Roberto Amato, Sonia Goncalves, Ewan Harrison, David K. Jackson, Ian Johnston, Dominic Kwiatkowski, Cordelia Langford, John Sillitoe on behalf of the Wellcome Sanger Institute COVID-19 Surveillance Team |
| EPI_ISL_767098, EPI_ISL_767099, EPI_ISL_767100, EPI_ISL_767101 | Lighthouse Lab in Milton Keynes | Wellcome Sanger Institute for the COVID-19 Genomics UK (COG-UK) Consortium | The Lighthouse Lab in Milton Keynes and Alex Alderton, Roberto Amato, Sonia Goncalves, Ewan Harrison, David K. Jackson, Ian Johnston, Dominic Kwiatkowski, Cordelia Langford, John Sillitoe on behalf of the Wellcome Sanger Institute COVID-19 Surveillance Team |
| EPI_ISL_767102, EPI_ISL_767103, EPI_ISL_767104 | Lighthouse Lab in Alderley Park | Wellcome Sanger Institute for the COVID-19 Genomics UK (COG-UK) Consortium | Jacquelyn Wynn, Mairead Hyland, The Lighthouse Lab in Alderley Park and Alex Alderton, Roberto Amato, Sonia Goncalves, Ewan Harrison, David K. Jackson, Ian Johnston, Dominic Kwiatkowski, Cordelia Langford, John Sillitoe on behalf of the Wellcome Sanger Institute COVID-19 Surveillance Team |
| EPI_ISL_767107, EPI_ISL_767108, EPI_ISL_767110, EPI_ISL_767111, EPI_ISL_767112 | Lighthouse Lab in Milton Keynes | Wellcome Sanger Institute for the COVID-19 Genomics UK (COG-UK) Consortium | The Lighthouse Lab in Milton Keynes and Alex Alderton, Roberto Amato, Sonia Goncalves, Ewan Harrison, David K. Jackson, Ian Johnston, Dominic Kwiatkowski, Cordelia Langford, John Sillitoe on behalf of the Wellcome Sanger Institute COVID-19 Surveillance Team |
| EPI_ISL_767113, EPI_ISL_767114 | Lighthouse Lab in Glasgow | Wellcome Sanger Institute for the COVID-19 Genomics UK (COG-UK) Consortium | Harper VanSteenhouse, Yumi Kasai, David Gray, Carol Clugston, Anna Dominiczak and Alex Alderton, Roberto Amato, Sonia Goncalves, Ewan Harrison, David K. Jackson, Ian Johnston, Dominic Kwiatkowski, Cordelia Langford, John Sillitoe on behalf of the Wellcome Sanger Institute COVID-19 Surveillance Team |
| EPI_ISL_767115 | Lighthouse Lab in Cambridge | Wellcome Sanger Institute for the COVID-19 Genomics UK (COG-UK) Consortium | Rob Howes, The Lighthouse Lab in Cambridge and Alex Alderton, Roberto Amato, Sonia Goncalves, Ewan Harrison, David K. Jackson, Ian Johnston, Dominic Kwiatkowski, Cordelia Langford, John Sillitoe on behalf of the Wellcome Sanger Institute COVID-19 Surveillance Team |
| EPI_ISL_767116 | Lighthouse Lab in Alderley Park | Wellcome Sanger Institute for the COVID-19 Genomics UK (COG-UK) Consortium | Jacquelyn Wynn, Mairead Hyland, The Lighthouse Lab in Alderley Park and Alex Alderton, Roberto Amato, Sonia Goncalves, Ewan Harrison, David K. Jackson, Ian Johnston, Dominic Kwiatkowski, Cordelia Langford, John Sillitoe on behalf of the Wellcome Sanger Institute COVID-19 Surveillance Team |
| EPI_ISL_767117, EPI_ISL_767118 | Lighthouse Lab in Glasgow | Wellcome Sanger Institute for the COVID-19 Genomics UK (COG-UK) Consortium | Harper VanSteenhouse, Yumi Kasai, David Gray, Carol Clugston, Anna Dominiczak and Alex Alderton, Roberto Amato, Sonia Goncalves, Ewan Harrison, David K. Jackson, Ian Johnston, Dominic Kwiatkowski, Cordelia Langford, John Sillitoe on behalf of the Wellcome Sanger Institute COVID-19 Surveillance Team |
| EPI_ISL_767119 | Lighthouse Lab in Alderley Park | Wellcome Sanger Institute for the COVID-19 Genomics UK (COG-UK) Consortium | Jacquelyn Wynn, Mairead Hyland, The Lighthouse Lab in Alderley Park and Alex Alderton, Roberto Amato, Sonia Goncalves, Ewan Harrison, David K. Jackson, Ian Johnston, Dominic Kwiatkowski, Cordelia Langford, John Sillitoe on behalf of the Wellcome Sanger Institute COVID-19 Surveillance Team |
| EPI_ISL_767479, EPI_ISL_767480, EPI_ISL_767481, EPI_ISL_767482, EPI_ISL_767483, EPI_ISL_767484, EPI_ISL_767485, EPI_ISL_767486, EPI_ISL_767487, EPI_ISL_767488, EPI_ISL_767489, EPI_ISL_767490, EPI_ISL_767491, EPI_ISL_767492, EPI_ISL_767493, EPI_ISL_767494 | | | |
| see above | Wadsworth Center, New York State Department.of Health | Wadsworth Center, New York State Department.of Health | Kirsten St. George, Daryl M. Lamson, Alexis Russel, Matthew Shudt, Melissa A Leisner, Jonathan Plitnick, Navjot Singh, John Kelly, Sara Griesemer, Erasmus Schneider, Erica Lasek-Nesselquist |
| EPI_ISL_767648, EPI_ISL_767665, EPI_ISL_767666, EPI_ISL_767668, EPI_ISL_767669, EPI_ISL_767673 | BIO-REFERENCE LABORATORIES | Wadsworth Center, New York State Department.of Health | Kirsten St. George, Daryl M. Lamson, Alexis Russel, Matthew Shudt, Melissa A Leisner, Jonathan Plitnick, Navjot Singh, John Kelly, Sara Griesemer, Erasmus Schneider, Erica Lasek-Nesselquist |
| EPI_ISL_767868 | Pathology North - Royal North Shore Hospital - NSW Health Pathology | NSW Health Pathology - Institute of Clinical Pathology and Medical Research; Westmead Hospital; University of Sydney | CIDM-PH et al. |
| EPI_ISL_767869 | Laverty Pathology | NSW Health Pathology - Institute of Clinical Pathology and Medical Research; Westmead Hospital; University of Sydney | CIDM-PH et al. |
| EPI_ISL_767871 | Pathology North - Royal North Shore Hospital - NSW Health Pathology | NSW Health Pathology - Institute of Clinical Pathology and Medical Research; Westmead Hospital; University of Sydney | CIDM-PH et al. |
| EPI_ISL_767872, EPI_ISL_767873, EPI_ISL_767874, EPI_ISL_767875 | Histopath | NSW Health Pathology - Institute of Clinical Pathology and Medical Research; Westmead Hospital; University of Sydney | CIDM-PH et al. |
| EPI_ISL_767878 | Laverty Pathology | NSW Health Pathology - Institute of Clinical Pathology and Medical Research; Westmead Hospital; University of Sydney | CIDM-PH et al. |
| EPI_ISL_767884 | Australian Clinical Labs (formerly Healthscope Pathology) | NSW Health Pathology - Institute of Clinical Pathology and Medical Research; Westmead Hospital; University of Sydney | CIDM-PH et al. |
| EPI_ISL_767887, EPI_ISL_767888, EPI_ISL_767891, EPI_ISL_767892 | South Eastern Area Laboratory Services (SEALS) | NSW Health Pathology - Institute of Clinical Pathology and Medical Research; Westmead Hospital; University of Sydney | CIDM-PH et al. |
| EPI_ISL_767944, EPI_ISL_767961, EPI_ISL_767964, EPI_ISL_767966, EPI_ISL_767979, EPI_ISL_767989, EPI_ISL_767990, EPI_ISL_767994, EPI_ISL_767999, EPI_ISL_768003, EPI_ISL_768006, EPI_ISL_768012, EPI_ISL_768020, EPI_ISL_768023, EPI_ISL_768037, EPI_ISL_768041, EPI_ISL_768050, EPI_ISL_768054, EPI_ISL_768060, EPI_ISL_768061, EPI_ISL_768069, EPI_ISL_768072, EPI_ISL_768082, EPI_ISL_768087, EPI_ISL_768101, EPI_ISL_768106, EPI_ISL_768118, EPI_ISL_768121, EPI_ISL_768123, EPI_ISL_768140, EPI_ISL_768142, EPI_ISL_768144, EPI_ISL_768146, EPI_ISL_768149, EPI_ISL_768152, EPI_ISL_768154, EPI_ISL_768157, EPI_ISL_768160, EPI_ISL_768162, EPI_ISL_768164, EPI_ISL_768165, EPI_ISL_768173, EPI_ISL_768174, EPI_ISL_768176, EPI_ISL_768180, EPI_ISL_768181, EPI_ISL_768183, EPI_ISL_768236, EPI_ISL_768244, EPI_ISL_768246, EPI_ISL_768247, EPI_ISL_768248, EPI_ISL_768250, EPI_ISL_768251, EPI_ISL_768252, EPI_ISL_768253, EPI_ISL_768254, EPI_ISL_768256, EPI_ISL_768258, EPI_ISL_768260, EPI_ISL_768261, EPI_ISL_768265, EPI_ISL_768270, EPI_ISL_768276, EPI_ISL_768277, EPI_ISL_768279, EPI_ISL_768281, EPI_ISL_768283, EPI_ISL_768285, EPI_ISL_768288, EPI_ISL_768291, EPI_ISL_768292, EPI_ISL_768296, EPI_ISL_768297, EPI_ISL_768306, EPI_ISL_768313, EPI_ISL_768317, EPI_ISL_768319, EPI_ISL_768321 | | | |
| see above | Viollier AG | Department of Biosystems Science and Engineering, ETH Zürich | Chaoran Chen, Sarah Nadeau, Catharine Aquino, Ivan Topolsky, Philipp Jablonski, Lara Fuhrmann, David Dreifuss, Katharina Jahn, Andreia Cabral de Gouvea, Maria Domenica Moccia, Simon Grüter, Timothy Sykes, Lennart Opitz, Griffin White, Laura Neff, Doris Popovic, Andrea Patrignani, Jay Tracy, Ralph Schlapbach, Christiane Beckmann, Maurice Redondo, Olivier Kobel, Christoph Noppen, Sophie Seidel, Noemie Santamaria de Souza, Niko Beerenwinkel, Tanja Stadler |
| EPI_ISL_768496 | LSUHS Emerging Viral Threat Laboratory | Microbial Genome Sequencing Center | Jeremy P. Kamil, Jennifer L. Carroll, Camille F. Abshire, Maarten Van Diest, Andrew D. Yurochko, Martin J. Sapp, Rona S. Scott, Christopher G. Kevil, Daniel J. Snyder, Vaughn S. Cooper, John A. Vanchiere |
| EPI_ISL_768497, EPI_ISL_768499, EPI_ISL_768500, EPI_ISL_768501, EPI_ISL_768502, EPI_ISL_768503, EPI_ISL_768504, EPI_ISL_768505, EPI_ISL_768506, EPI_ISL_768507, EPI_ISL_768508 | | | |
| see above | LSUHS Emerging Viral Threat Laboratory | Microbial Genome Sequencing Center | Jennifer L. Carroll, Jeremy P. Kamil, Camille F. Abshire, Maarten Van Diest, Andrew D. Yurochko, Martin J. Sapp, Rona S. Scott, Christopher G. Kevil, Daniel J. Snyder, Vaughn S. Cooper, John A. Vanchiere |
| EPI_ISL_768509, EPI_ISL_768510, EPI_ISL_768511, EPI_ISL_768512, EPI_ISL_768513, EPI_ISL_768514, EPI_ISL_768515, EPI_ISL_768516, EPI_ISL_768517, EPI_ISL_768518, EPI_ISL_768519, EPI_ISL_768520 | | | |
| see above | LSUHS Emerging Viral Threat Laboratory | Microbial Genome Sequencing Center | Maarten Van Diest, Jeremy P. Kamil, Jennifer L. Carroll, Camille F. Abshire, Andrew D. Yurochko, Martin J. Sapp, Rona S. Scott, Christopher G. Kevil, Daniel J. Snyder, Vaughn S. Cooper, John A. Vanchiere |
| EPI_ISL_768536, EPI_ISL_768539, EPI_ISL_768544 | Regional Medical Sciences Center 5 Samut Songkhram | National Institute of Health, Department of Medical Sciences, Ministry of Public Health, Thailand | Pilailuk Okada; Siripaporn Phuygun; Sittiporn Parnmen; Ratana Tacharoenmuang; Pakorn Piromtong; Natchaya Khiadsang; Thanutsapa Thanadachakul; Warawan Wongboot; sirikanda wimol; Sunthareeya Waicharoen; |

| | | | |
|---|---|---|---|
| EPI_ISL_768840 | Laboratoire Biolife | Laboratoire de Biotechnologie | Mouna Ouadghiri, Tarik Aanniz, Mohammed Walid Chemao Elfihri, Mohamed Chenaoui, Hanae Dakka, Afaf Alaoui, Otmane Touzani, Amina Benouda, Bouchra Belfquih, Lahcen belyamani, Saaid Amzazi and Azeddine Ibrahimi |
| EPI_ISL_768841, EPI_ISL_768843 | Lighthouse Lab in Alderley Park | Wellcome Sanger Institute for the COVID-19 Genomics UK (COG-UK) Consortium | Jacquelyn Wynn, Mairead Hyland, The Lighthouse Lab in Alderley Park and Alex Alderton, Roberto Amato, Sonia Goncalves, Ewan Harrison, David K. Jackson, Ian Johnston, Dominic Kwiatkowski, Cordelia Langford, John Sillitoe on behalf of the Wellcome Sanger Institute COVID-19 Surveillance Team |
| EPI_ISL_768845 | Lighthouse Lab in Cambridge | Wellcome Sanger Institute for the COVID-19 Genomics UK (COG-UK) Consortium | Rob Howes, The Lighthouse Lab in Cambridge and Alex Alderton, Roberto Amato, Sonia Goncalves, Ewan Harrison, David K. Jackson, Ian Johnston, Dominic Kwiatkowski, Cordelia Langford, John Sillitoe on behalf of the Wellcome Sanger Institute COVID-19 Surveillance Team |
| EPI_ISL_768846, EPI_ISL_768847, EPI_ISL_768848, EPI_ISL_768849, EPI_ISL_768850, EPI_ISL_768851, EPI_ISL_768852, EPI_ISL_768853, EPI_ISL_768854, EPI_ISL_768855, EPI_ISL_768856, EPI_ISL_768857, EPI_ISL_768858, EPI_ISL_768859, EPI_ISL_768860, EPI_ISL_768861, EPI_ISL_768862, EPI_ISL_768863, EPI_ISL_768864, EPI_ISL_768865, EPI_ISL_768866, EPI_ISL_768867, EPI_ISL_768868, EPI_ISL_768869, EPI_ISL_768870, EPI_ISL_768871, EPI_ISL_768872, EPI_ISL_768873, EPI_ISL_768874, EPI_ISL_768875, EPI_ISL_768876, EPI_ISL_768877, EPI_ISL_768878, EPI_ISL_768879, EPI_ISL_768880, EPI_ISL_768881, EPI_ISL_768882, EPI_ISL_768883, EPI_ISL_768884, EPI_ISL_768885, EPI_ISL_768886, EPI_ISL_768887, EPI_ISL_768888, EPI_ISL_768889, EPI_ISL_768890, EPI_ISL_768891, EPI_ISL_768892, EPI_ISL_768893, EPI_ISL_768894, EPI_ISL_768895, EPI_ISL_768896, EPI_ISL_768897 | | | |
| see above | Lighthouse Lab in Milton Keynes | Wellcome Sanger Institute for the COVID-19 Genomics UK (COG-UK) Consortium | The Lighthouse Lab in Milton Keynes and Alex Alderton, Roberto Amato, Sonia Goncalves, Ewan Harrison, David K. Jackson, Ian Johnston, Dominic Kwiatkowski, Cordelia Langford, John Sillitoe on behalf of the Wellcome Sanger Institute COVID-19 Surveillance Team |
| EPI_ISL_768900 | Lighthouse Lab in Alderley Park | Wellcome Sanger Institute for the COVID-19 Genomics UK (COG-UK) Consortium | Jacquelyn Wynn, Mairead Hyland, The Lighthouse Lab in Alderley Park and Alex Alderton, Roberto Amato, Sonia Goncalves, Ewan Harrison, David K. Jackson, Ian Johnston, Dominic Kwiatkowski, Cordelia Langford, John Sillitoe on behalf of the Wellcome Sanger Institute COVID-19 Surveillance Team |
| EPI_ISL_768901, EPI_ISL_768911, EPI_ISL_768916, EPI_ISL_768919, EPI_ISL_768925, EPI_ISL_768928, EPI_ISL_768931, EPI_ISL_768932, EPI_ISL_768936, EPI_ISL_768937, EPI_ISL_768939, EPI_ISL_768945, EPI_ISL_768946, EPI_ISL_768948, EPI_ISL_768949, EPI_ISL_768952, EPI_ISL_768953, EPI_ISL_768957, EPI_ISL_768959, EPI_ISL_768963, EPI_ISL_768965, EPI_ISL_768966, EPI_ISL_768972, EPI_ISL_768976, EPI_ISL_768977, EPI_ISL_768992, EPI_ISL_768993, EPI_ISL_769002, EPI_ISL_769005, EPI_ISL_769016, EPI_ISL_769017, EPI_ISL_769027, EPI_ISL_769029, EPI_ISL_769033, EPI_ISL_769036, EPI_ISL_769038, EPI_ISL_769043, EPI_ISL_769045, EPI_ISL_769049, EPI_ISL_769057, EPI_ISL_769060, EPI_ISL_769062, EPI_ISL_769076, EPI_ISL_769080, EPI_ISL_769084, EPI_ISL_769087, EPI_ISL_769089, EPI_ISL_769093, EPI_ISL_769094, EPI_ISL_769098, EPI_ISL_769105, EPI_ISL_769109, EPI_ISL_769110, EPI_ISL_769117, EPI_ISL_769119, EPI_ISL_769122, EPI_ISL_769138, EPI_ISL_769141, EPI_ISL_769152, EPI_ISL_769157, EPI_ISL_769168, EPI_ISL_769169, EPI_ISL_769171, EPI_ISL_769173, EPI_ISL_769176, EPI_ISL_769193, EPI_ISL_769194, EPI_ISL_769195, EPI_ISL_769196, EPI_ISL_769198, EPI_ISL_769206, EPI_ISL_769207, EPI_ISL_769211, EPI_ISL_769213, EPI_ISL_769215, EPI_ISL_769218, EPI_ISL_769219 | | | |
| see above | Lighthouse Lab in Cambridge | Wellcome Sanger Institute for the COVID-19 Genomics UK (COG-UK) Consortium | Rob Howes, The Lighthouse Lab in Cambridge and Alex Alderton, Roberto Amato, Sonia Goncalves, Ewan Harrison, David K. Jackson, Ian Johnston, Dominic Kwiatkowski, Cordelia Langford, John Sillitoe on behalf of the Wellcome Sanger Institute COVID-19 Surveillance Team |
| EPI_ISL_769873 | Respiratory Virus Unit, National Infection Service, Public Health England | COVID-19 Genomics UK (COG-UK) Consortium | PHE Covid Sequencing Team |
| EPI_ISL_769920 | Albany Medical Center Hospital Clinical Laboratories | Wadsworth Center, New York State Department.of Health | Kirsten St. George, Daryl M. Lamson, Alexis Russel, Matthew Shudt, Melissa A Leisner, Jonathan Plitnick, Navjot Singh, John Kelly, Sara Griesemer, Erasmus Schneider, Erica Lasek-Nesselquist |
| EPI_ISL_770471, EPI_ISL_770472, EPI_ISL_770474 | National Health laboratory | Botswana Institute for Technology Research and Innovation | Kefentse Arnold Tumedi, Madisa Mine, Dineo Emang Tshiamo. Gape Nyepetsi, Thongbotho Mphoyakgosi, Malebogo Kebabonye, Maitshwarelo Ignatius Matsheka |
| EPI_ISL_770500, EPI_ISL_770501, EPI_ISL_770502, EPI_ISL_770503, EPI_ISL_770504 | Lithuanian University of Health Sciences Hospital, Department of Laboratory Medicine | Lithuanian University of Health Sciences, Molecular cardiology lab. | Lukas Zemaitis, Ingrida Olendrait, Arnoldas Pautienius, Kamile Tamusauskaite, Dovydas Gecys, Laura Pareckaite, Vaiva Lesauskaite, Astra Vitkauskiene |
| EPI_ISL_775222 | Laboratoire Biolife | Laboratoire de Biotechnologie | Mouna Ouadghiri, Tarik Aanniz, Mohammed Walid Chemao Elfihri, Mohamed Chenaoui, Hanae Dakka, Afaf Alaoui, Otmane Touzani, Amina Benouda, Bouchra Belfquih, Lahcen belyamani, Saaid Amzazi and Azeddine Ibrahimi |
| EPI_ISL_775269, EPI_ISL_775275 | Akershus University Hospital, Department for Microbiology and Infectious Disease Control | Norwegian Institute of Public Health, Department of Virology | Kathrine Stene-Johansen, Kamilla Heddeland Instefjord, Hilde Elshaug, Atiya R Ali,Marie Paulsen Madsen, Rasmus Riis Kopperud, Hilde Vollan, Karoline Bragstad, Olav Hungnes |
| EPI_ISL_775278, EPI_ISL_775424, EPI_ISL_775425, EPI_ISL_775426 | Oslo University Hospital, Department of Medical Microbiology | Norwegian Institute of Public Health, Department of Virology | Kathrine Stene-Johansen, Kamilla Heddeland Instefjord, Hilde Elshaug, Atiya R Ali,Marie Paulsen Madsen, Rasmus Riis Kopperud, Hilde Vollan, Karoline Bragstad, Olav Hungnes |
| EPI_ISL_775438, EPI_ISL_775439 | Nordland Hospital - Bodo, Laboratory Department, Molecular Biology Unit | Norwegian Institute of Public Health, Department of Virology | Kathrine Stene-Johansen, Kamilla Heddeland Instefjord, Hilde Elshaug, Atiya R Ali,Marie Paulsen Madsen, Rasmus Riis Kopperud, Hilde Vollan, Karoline Bragstad, Olav Hungnes |
| EPI_ISL_775458 | University Hospital of Northern Norway, Department for Microbiology and Infectious Disease Control | Norwegian Institute of Public Health, Department of Virology | Kathrine Stene-Johansen, Kamilla Heddeland Instefjord, Hilde Elshaug, Atiya R Ali,Marie Paulsen Madsen, Rasmus Riis Kopperud, Hilde Vollan, Karoline Bragstad, Olav Hungnes |
| EPI_ISL_775528, EPI_ISL_775530 | Akershus University Hospital, Department for Microbiology and Infectious Disease Control | Norwegian Institute of Public Health, Department of Virology | Kathrine Stene-Johansen, Kamilla Heddeland Instefjord, Hilde Elshaug, Atiya R Ali,Marie Paulsen Madsen, Rasmus Riis Kopperud, Hilde Vollan, Karoline Bragstad, Olav Hungnes |
| EPI_ISL_775546, EPI_ISL_775547, EPI_ISL_775548 | Klinisk Mikrobiologi | The Public Health Agency of Sweden | Department of Microbiology, The Public Health Agency of Sweden |
| EPI_ISL_775593 | TATAA Biocenter | The Public Health Agency of Sweden | Department of Microbiology, The Public Health Agency of Sweden |
| EPI_ISL_776731 | UW Virology Lab | UW Virology Lab | Pavitra Roychoudhury, Hong Xie, Lasata Shrestha, Meei-Li Huang, Keith R Jerome, Alexander Greninger |
| EPI_ISL_777016, EPI_ISL_777019, EPI_ISL_777021, EPI_ISL_777029, EPI_ISL_777031, EPI_ISL_777037, EPI_ISL_777038, EPI_ISL_777040, EPI_ISL_777047, EPI_ISL_777059, EPI_ISL_777061, EPI_ISL_777062, EPI_ISL_777066, EPI_ISL_777068, EPI_ISL_777073, EPI_ISL_777082, EPI_ISL_777089, EPI_ISL_777099, EPI_ISL_777100, EPI_ISL_777112, EPI_ISL_777117, EPI_ISL_777118, EPI_ISL_777120, EPI_ISL_777121, EPI_ISL_777122, EPI_ISL_777124, EPI_ISL_777127, EPI_ISL_777130, EPI_ISL_777131, EPI_ISL_777140, EPI_ISL_777146, EPI_ISL_777151, EPI_ISL_777152, EPI_ISL_777154, EPI_ISL_777160, EPI_ISL_777161, EPI_ISL_777162, EPI_ISL_777164, EPI_ISL_777176, EPI_ISL_777180, EPI_ISL_777189, EPI_ISL_777191, EPI_ISL_777205, EPI_ISL_777208, EPI_ISL_777209, EPI_ISL_777215, EPI_ISL_777219, EPI_ISL_777222, EPI_ISL_777223, EPI_ISL_777225, EPI_ISL_777227, EPI_ISL_777230, EPI_ISL_777232, EPI_ISL_777233, EPI_ISL_777235, EPI_ISL_777236, EPI_ISL_777238, EPI_ISL_777241, EPI_ISL_777242, EPI_ISL_777246, EPI_ISL_777252, EPI_ISL_777268, EPI_ISL_777283, EPI_ISL_777298, EPI_ISL_777300, EPI_ISL_777301, EPI_ISL_777302, EPI_ISL_777305, EPI_ISL_777306, EPI_ISL_777316, EPI_ISL_777321, EPI_ISL_777324, EPI_ISL_777325 | | | |
| see above | Lighthouse Lab in Glasgow | Wellcome Sanger Institute for the COVID-19 Genomics UK (COG-UK) Consortium | Harper VanSteenhouse, Yumi Kasai, David Gray, Carol Clugston, Anna Dominiczak and Alex Alderton, Roberto Amato, Sonia Goncalves, Ewan Harrison, David K. Jackson, Ian Johnston, Dominic Kwiatkowski, Cordelia Langford, John Sillitoe on behalf of the Wellcome Sanger Institute COVID-19 Surveillance Team |
| EPI_ISL_777328, EPI_ISL_777329, EPI_ISL_777330, EPI_ISL_777331, EPI_ISL_777332, EPI_ISL_777334 | Lighthouse Lab in Cambridge | Wellcome Sanger Institute for the COVID-19 Genomics UK (COG-UK) Consortium | Rob Howes, The Lighthouse Lab in Cambridge and Alex Alderton, Roberto Amato, Sonia Goncalves, Ewan Harrison, David K. Jackson, Ian Johnston, Dominic Kwiatkowski, Cordelia Langford, John Sillitoe on behalf of the Wellcome Sanger Institute COVID-19 Surveillance Team |
| EPI_ISL_777335, EPI_ISL_777337 | Lighthouse Lab in Glasgow | Wellcome Sanger Institute for the COVID-19 Genomics UK (COG-UK) Consortium | Harper VanSteenhouse, Yumi Kasai, David Gray, Carol Clugston, Anna Dominiczak and Alex Alderton, Roberto Amato, Sonia Goncalves, Ewan Harrison, David K. Jackson, Ian Johnston, Dominic Kwiatkowski, Cordelia Langford, John Sillitoe on behalf of the Wellcome Sanger Institute COVID-19 Surveillance Team |
| EPI_ISL_777339 | Lighthouse Lab in Cambridge | Wellcome Sanger Institute for the COVID-19 Genomics UK (COG-UK) Consortium | Rob Howes, The Lighthouse Lab in Cambridge and Alex Alderton, Roberto Amato, Sonia Goncalves, Ewan Harrison, David K. Jackson, Ian Johnston, Dominic Kwiatkowski, Cordelia Langford, John Sillitoe on behalf of the Wellcome Sanger Institute COVID-19 Surveillance Team |
| EPI_ISL_777340, EPI_ISL_777341 | Lighthouse Lab in Glasgow | Wellcome Sanger Institute for the COVID-19 Genomics UK (COG-UK) Consortium | Harper VanSteenhouse, Yumi Kasai, David Gray, Carol Clugston, Anna Dominiczak and Alex Alderton, Roberto Amato, Sonia Goncalves, Ewan Harrison, David K. Jackson, Ian Johnston, Dominic Kwiatkowski, Cordelia Langford, John Sillitoe on behalf of the Wellcome Sanger Institute COVID-19 Surveillance Team |
| EPI_ISL_777342, EPI_ISL_777343, EPI_ISL_777344, EPI_ISL_777345 | Lighthouse Lab in Cambridge | Wellcome Sanger Institute for the COVID-19 Genomics UK (COG-UK) Consortium | Rob Howes, The Lighthouse Lab in Cambridge and Alex Alderton, Roberto Amato, Sonia Goncalves, Ewan Harrison, David K. Jackson, Ian Johnston, Dominic Kwiatkowski, Cordelia Langford, John Sillitoe on behalf of the Wellcome Sanger Institute COVID-19 Surveillance Team |
| EPI_ISL_777346 | Lighthouse Lab in Glasgow | Wellcome Sanger Institute for the COVID-19 Genomics UK (COG-UK) Consortium | Harper VanSteenhouse, Yumi Kasai, David Gray, Carol Clugston, Anna Dominiczak and Alex Alderton, Roberto Amato, Sonia Goncalves, Ewan Harrison, David K. Jackson, Ian Johnston, Dominic Kwiatkowski, Cordelia Langford, John Sillitoe on behalf of the Wellcome Sanger Institute COVID-19 Surveillance Team |
| EPI_ISL_777347, EPI_ISL_777348, EPI_ISL_777350, EPI_ISL_777351, EPI_ISL_777352 | Lighthouse Lab in Cambridge | Wellcome Sanger Institute for the COVID-19 Genomics UK (COG-UK) Consortium | Rob Howes, The Lighthouse Lab in Cambridge and Alex Alderton, Roberto Amato, Sonia Goncalves, Ewan Harrison, David K. Jackson, Ian Johnston, Dominic Kwiatkowski, Cordelia Langford, John Sillitoe on behalf of the Wellcome Sanger Institute COVID-19 Surveillance Team |

| | | | |
|---|---|---|---|
| EPI_ISL_777353 | Lighthouse Lab in Glasgow | Wellcome Sanger Institute for the COVID-19 Genomics UK (COG-UK) Consortium | Harper VanSteenhouse, Yumi Kasai, David Gray, Carol Clugston, Anna Dominiczak and Alex Alderton, Roberto Amato, Sonia Goncalves, Ewan Harrison, David K. Jackson, Ian Johnston, Dominic Kwiatkowski, Cordelia Langford, John Sillitoe on behalf of the Wellcome Sanger Institute COVID-19 Surveillance Team |
| EPI_ISL_777354, EPI_ISL_777355 | Lighthouse Lab in Cambridge | Wellcome Sanger Institute for the COVID-19 Genomics UK (COG-UK) Consortium | Rob Howes, The Lighthouse Lab in Cambridge and Alex Alderton, Roberto Amato, Sonia Goncalves, Ewan Harrison, David K. Jackson, Ian Johnston, Dominic Kwiatkowski, Cordelia Langford, John Sillitoe on behalf of the Wellcome Sanger Institute COVID-19 Surveillance Team |
| EPI_ISL_777356 | Lighthouse Lab in Glasgow | Wellcome Sanger Institute for the COVID-19 Genomics UK (COG-UK) Consortium | Harper VanSteenhouse, Yumi Kasai, David Gray, Carol Clugston, Anna Dominiczak and Alex Alderton, Roberto Amato, Sonia Goncalves, Ewan Harrison, David K. Jackson, Ian Johnston, Dominic Kwiatkowski, Cordelia Langford, John Sillitoe on behalf of the Wellcome Sanger Institute COVID-19 Surveillance Team |
| EPI_ISL_777357 | Lighthouse Lab in Cambridge | Wellcome Sanger Institute for the COVID-19 Genomics UK (COG-UK) Consortium | Rob Howes, The Lighthouse Lab in Cambridge and Alex Alderton, Roberto Amato, Sonia Goncalves, Ewan Harrison, David K. Jackson, Ian Johnston, Dominic Kwiatkowski, Cordelia Langford, John Sillitoe on behalf of the Wellcome Sanger Institute COVID-19 Surveillance Team |
| EPI_ISL_777358 | Lighthouse Lab in Glasgow | Wellcome Sanger Institute for the COVID-19 Genomics UK (COG-UK) Consortium | Harper VanSteenhouse, Yumi Kasai, David Gray, Carol Clugston, Anna Dominiczak and Alex Alderton, Roberto Amato, Sonia Goncalves, Ewan Harrison, David K. Jackson, Ian Johnston, Dominic Kwiatkowski, Cordelia Langford, John Sillitoe on behalf of the Wellcome Sanger Institute COVID-19 Surveillance Team |
| EPI_ISL_777359, EPI_ISL_777360 | Lighthouse Lab in Cambridge | Wellcome Sanger Institute for the COVID-19 Genomics UK (COG-UK) Consortium | Rob Howes, The Lighthouse Lab in Cambridge and Alex Alderton, Roberto Amato, Sonia Goncalves, Ewan Harrison, David K. Jackson, Ian Johnston, Dominic Kwiatkowski, Cordelia Langford, John Sillitoe on behalf of the Wellcome Sanger Institute COVID-19 Surveillance Team |
| EPI_ISL_777361 | Lighthouse Lab in Glasgow | Wellcome Sanger Institute for the COVID-19 Genomics UK (COG-UK) Consortium | Harper VanSteenhouse, Yumi Kasai, David Gray, Carol Clugston, Anna Dominiczak and Alex Alderton, Roberto Amato, Sonia Goncalves, Ewan Harrison, David K. Jackson, Ian Johnston, Dominic Kwiatkowski, Cordelia Langford, John Sillitoe on behalf of the Wellcome Sanger Institute COVID-19 Surveillance Team |
| EPI_ISL_777362 | Lighthouse Lab in Cambridge | Wellcome Sanger Institute for the COVID-19 Genomics UK (COG-UK) Consortium | Rob Howes, The Lighthouse Lab in Cambridge and Alex Alderton, Roberto Amato, Sonia Goncalves, Ewan Harrison, David K. Jackson, Ian Johnston, Dominic Kwiatkowski, Cordelia Langford, John Sillitoe on behalf of the Wellcome Sanger Institute COVID-19 Surveillance Team |
| EPI_ISL_777363, EPI_ISL_777364 | Lighthouse Lab in Glasgow | Wellcome Sanger Institute for the COVID-19 Genomics UK (COG-UK) Consortium | Harper VanSteenhouse, Yumi Kasai, David Gray, Carol Clugston, Anna Dominiczak and Alex Alderton, Roberto Amato, Sonia Goncalves, Ewan Harrison, David K. Jackson, Ian Johnston, Dominic Kwiatkowski, Cordelia Langford, John Sillitoe on behalf of the Wellcome Sanger Institute COVID-19 Surveillance Team |
| EPI_ISL_777365, EPI_ISL_777366, EPI_ISL_777367, EPI_ISL_777368 | Lighthouse Lab in Cambridge | Wellcome Sanger Institute for the COVID-19 Genomics UK (COG-UK) Consortium | Rob Howes, The Lighthouse Lab in Cambridge and Alex Alderton, Roberto Amato, Sonia Goncalves, Ewan Harrison, David K. Jackson, Ian Johnston, Dominic Kwiatkowski, Cordelia Langford, John Sillitoe on behalf of the Wellcome Sanger Institute COVID-19 Surveillance Team |
| EPI_ISL_777369 | Lighthouse Lab in Glasgow | Wellcome Sanger Institute for the COVID-19 Genomics UK (COG-UK) Consortium | Harper VanSteenhouse, Yumi Kasai, David Gray, Carol Clugston, Anna Dominiczak and Alex Alderton, Roberto Amato, Sonia Goncalves, Ewan Harrison, David K. Jackson, Ian Johnston, Dominic Kwiatkowski, Cordelia Langford, John Sillitoe on behalf of the Wellcome Sanger Institute COVID-19 Surveillance Team |
| EPI_ISL_777370, EPI_ISL_777371 | Lighthouse Lab in Cambridge | Wellcome Sanger Institute for the COVID-19 Genomics UK (COG-UK) Consortium | Rob Howes, The Lighthouse Lab in Cambridge and Alex Alderton, Roberto Amato, Sonia Goncalves, Ewan Harrison, David K. Jackson, Ian Johnston, Dominic Kwiatkowski, Cordelia Langford, John Sillitoe on behalf of the Wellcome Sanger Institute COVID-19 Surveillance Team |
| EPI_ISL_777372, EPI_ISL_777373 | Lighthouse Lab in Glasgow | Wellcome Sanger Institute for the COVID-19 Genomics UK (COG-UK) Consortium | Harper VanSteenhouse, Yumi Kasai, David Gray, Carol Clugston, Anna Dominiczak and Alex Alderton, Roberto Amato, Sonia Goncalves, Ewan Harrison, David K. Jackson, Ian Johnston, Dominic Kwiatkowski, Cordelia Langford, John Sillitoe on behalf of the Wellcome Sanger Institute COVID-19 Surveillance Team |
| EPI_ISL_777374 | Lighthouse Lab in Cambridge | Wellcome Sanger Institute for the COVID-19 Genomics UK (COG-UK) Consortium | Rob Howes, The Lighthouse Lab in Cambridge and Alex Alderton, Roberto Amato, Sonia Goncalves, Ewan Harrison, David K. Jackson, Ian Johnston, Dominic Kwiatkowski, Cordelia Langford, John Sillitoe on behalf of the Wellcome Sanger Institute COVID-19 Surveillance Team |
| EPI_ISL_777376, EPI_ISL_777377 | Lighthouse Lab in Glasgow | Wellcome Sanger Institute for the COVID-19 Genomics UK (COG-UK) Consortium | Harper VanSteenhouse, Yumi Kasai, David Gray, Carol Clugston, Anna Dominiczak and Alex Alderton, Roberto Amato, Sonia Goncalves, Ewan Harrison, David K. Jackson, Ian Johnston, Dominic Kwiatkowski, Cordelia Langford, John Sillitoe on behalf of the Wellcome Sanger Institute COVID-19 Surveillance Team |
| EPI_ISL_777378, EPI_ISL_777379, EPI_ISL_777381 | Lighthouse Lab in Cambridge | Wellcome Sanger Institute for the COVID-19 Genomics UK (COG-UK) Consortium | Rob Howes, The Lighthouse Lab in Cambridge and Alex Alderton, Roberto Amato, Sonia Goncalves, Ewan Harrison, David K. Jackson, Ian Johnston, Dominic Kwiatkowski, Cordelia Langford, John Sillitoe on behalf of the Wellcome Sanger Institute COVID-19 Surveillance Team |
| EPI_ISL_777382, EPI_ISL_777383 | Lighthouse Lab in Glasgow | Wellcome Sanger Institute for the COVID-19 Genomics UK (COG-UK) Consortium | Harper VanSteenhouse, Yumi Kasai, David Gray, Carol Clugston, Anna Dominiczak and Alex Alderton, Roberto Amato, Sonia Goncalves, Ewan Harrison, David K. Jackson, Ian Johnston, Dominic Kwiatkowski, Cordelia Langford, John Sillitoe on behalf of the Wellcome Sanger Institute COVID-19 Surveillance Team |
| EPI_ISL_777384 | Lighthouse Lab in Cambridge | Wellcome Sanger Institute for the COVID-19 Genomics UK (COG-UK) Consortium | Rob Howes, The Lighthouse Lab in Cambridge and Alex Alderton, Roberto Amato, Sonia Goncalves, Ewan Harrison, David K. Jackson, Ian Johnston, Dominic Kwiatkowski, Cordelia Langford, John Sillitoe on behalf of the Wellcome Sanger Institute COVID-19 Surveillance Team |
| EPI_ISL_777385 | Lighthouse Lab in Glasgow | Wellcome Sanger Institute for the COVID-19 Genomics UK (COG-UK) Consortium | Harper VanSteenhouse, Yumi Kasai, David Gray, Carol Clugston, Anna Dominiczak and Alex Alderton, Roberto Amato, Sonia Goncalves, Ewan Harrison, David K. Jackson, Ian Johnston, Dominic Kwiatkowski, Cordelia Langford, John Sillitoe on behalf of the Wellcome Sanger Institute COVID-19 Surveillance Team |
| EPI_ISL_777387, EPI_ISL_777388, EPI_ISL_777390, EPI_ISL_777391 | Lighthouse Lab in Cambridge | Wellcome Sanger Institute for the COVID-19 Genomics UK (COG-UK) Consortium | Rob Howes, The Lighthouse Lab in Cambridge and Alex Alderton, Roberto Amato, Sonia Goncalves, Ewan Harrison, David K. Jackson, Ian Johnston, Dominic Kwiatkowski, Cordelia Langford, John Sillitoe on behalf of the Wellcome Sanger Institute COVID-19 Surveillance Team |
| EPI_ISL_777392, EPI_ISL_777393, EPI_ISL_777395 | Lighthouse Lab in Glasgow | Wellcome Sanger Institute for the COVID-19 Genomics UK (COG-UK) Consortium | Harper VanSteenhouse, Yumi Kasai, David Gray, Carol Clugston, Anna Dominiczak and Alex Alderton, Roberto Amato, Sonia Goncalves, Ewan Harrison, David K. Jackson, Ian Johnston, Dominic Kwiatkowski, Cordelia Langford, John Sillitoe on behalf of the Wellcome Sanger Institute COVID-19 Surveillance Team |
| EPI_ISL_777396 | Lighthouse Lab in Cambridge | Wellcome Sanger Institute for the COVID-19 Genomics UK (COG-UK) Consortium | Rob Howes, The Lighthouse Lab in Cambridge and Alex Alderton, Roberto Amato, Sonia Goncalves, Ewan Harrison, David K. Jackson, Ian Johnston, Dominic Kwiatkowski, Cordelia Langford, John Sillitoe on behalf of the Wellcome Sanger Institute COVID-19 Surveillance Team |
| EPI_ISL_777397 | Lighthouse Lab in Glasgow | Wellcome Sanger Institute for the COVID-19 Genomics UK (COG-UK) Consortium | Harper VanSteenhouse, Yumi Kasai, David Gray, Carol Clugston, Anna Dominiczak and Alex Alderton, Roberto Amato, Sonia Goncalves, Ewan Harrison, David K. Jackson, Ian Johnston, Dominic Kwiatkowski, Cordelia Langford, John Sillitoe on behalf of the Wellcome Sanger Institute COVID-19 Surveillance Team |
| EPI_ISL_777398 | Lighthouse Lab in Cambridge | Wellcome Sanger Institute for the COVID-19 Genomics UK (COG-UK) Consortium | Rob Howes, The Lighthouse Lab in Cambridge and Alex Alderton, Roberto Amato, Sonia Goncalves, Ewan Harrison, David K. Jackson, Ian Johnston, Dominic Kwiatkowski, Cordelia Langford, John Sillitoe on behalf of the Wellcome Sanger Institute COVID-19 Surveillance Team |
| EPI_ISL_777399 | Lighthouse Lab in Glasgow | Wellcome Sanger Institute for the COVID-19 Genomics UK (COG-UK) Consortium | Harper VanSteenhouse, Yumi Kasai, David Gray, Carol Clugston, Anna Dominiczak and Alex Alderton, Roberto Amato, Sonia Goncalves, Ewan Harrison, David K. Jackson, Ian Johnston, Dominic Kwiatkowski, Cordelia Langford, John Sillitoe on behalf of the Wellcome Sanger Institute COVID-19 Surveillance Team |
| EPI_ISL_777400, EPI_ISL_777401 | Lighthouse Lab in Cambridge | Wellcome Sanger Institute for the COVID-19 Genomics UK (COG-UK) Consortium | Rob Howes, The Lighthouse Lab in Cambridge and Alex Alderton, Roberto Amato, Sonia Goncalves, Ewan Harrison, David K. Jackson, Ian Johnston, Dominic Kwiatkowski, Cordelia Langford, John Sillitoe on behalf of the Wellcome Sanger Institute COVID-19 Surveillance Team |
| EPI_ISL_777402 | Lighthouse Lab in Glasgow | Wellcome Sanger Institute for the COVID-19 Genomics UK (COG-UK) Consortium | Harper VanSteenhouse, Yumi Kasai, David Gray, Carol Clugston, Anna Dominiczak and Alex Alderton, Roberto Amato, Sonia Goncalves, Ewan Harrison, David K. Jackson, Ian Johnston, Dominic Kwiatkowski, Cordelia Langford, John Sillitoe on behalf of the Wellcome Sanger Institute COVID-19 Surveillance Team |
| EPI_ISL_777403, EPI_ISL_777404, EPI_ISL_777405, EPI_ISL_777406 | Lighthouse Lab in Cambridge | Wellcome Sanger Institute for the COVID-19 Genomics UK (COG-UK) Consortium | Rob Howes, The Lighthouse Lab in Cambridge and Alex Alderton, Roberto Amato, Sonia Goncalves, Ewan Harrison, David K. Jackson, Ian Johnston, Dominic Kwiatkowski, Cordelia Langford, John Sillitoe on behalf of the Wellcome Sanger Institute COVID-19 Surveillance Team |
| EPI_ISL_777407 | Lighthouse Lab in Glasgow | Wellcome Sanger Institute for the COVID-19 Genomics UK (COG-UK) Consortium | Harper VanSteenhouse, Yumi Kasai, David Gray, Carol Clugston, Anna Dominiczak and Alex Alderton, Roberto Amato, Sonia Goncalves, Ewan Harrison, David K. Jackson, Ian Johnston, Dominic Kwiatkowski, Cordelia Langford, John Sillitoe on behalf of the Wellcome Sanger Institute COVID-19 Surveillance Team |

| | | | |
|---|---|---|---|
| EPI_ISL_777408, EPI_ISL_777409 | Lighthouse Lab in Cambridge | Wellcome Sanger Institute for the COVID-19 Genomics UK (COG-UK) Consortium | Rob Howes, The Lighthouse Lab in Cambridge and Alex Alderton, Roberto Amato, Sonia Goncalves, Ewan Harrison, David K. Jackson, Ian Johnston, Dominic Kwiatkowski, Cordelia Langford, John Sillitoe on behalf of the Wellcome Sanger Institute COVID-19 Surveillance Team |
| EPI_ISL_777410 | Lighthouse Lab in Glasgow | Wellcome Sanger Institute for the COVID-19 Genomics UK (COG-UK) Consortium | Harper VanSteenhouse, Yumi Kasai, David Gray, Carol Clugston, Anna Dominiczak and Alex Alderton, Roberto Amato, Sonia Goncalves, Ewan Harrison, David K. Jackson, Ian Johnston, Dominic Kwiatkowski, Cordelia Langford, John Sillitoe on behalf of the Wellcome Sanger Institute COVID-19 Surveillance Team |
| EPI_ISL_777413, EPI_ISL_777414, EPI_ISL_777415 | Lighthouse Lab in Cambridge | Wellcome Sanger Institute for the COVID-19 Genomics UK (COG-UK) Consortium | Rob Howes, The Lighthouse Lab in Cambridge and Alex Alderton, Roberto Amato, Sonia Goncalves, Ewan Harrison, David K. Jackson, Ian Johnston, Dominic Kwiatkowski, Cordelia Langford, John Sillitoe on behalf of the Wellcome Sanger Institute COVID-19 Surveillance Team |
| EPI_ISL_777417 | Lighthouse Lab in Glasgow | Wellcome Sanger Institute for the COVID-19 Genomics UK (COG-UK) Consortium | Harper VanSteenhouse, Yumi Kasai, David Gray, Carol Clugston, Anna Dominiczak and Alex Alderton, Roberto Amato, Sonia Goncalves, Ewan Harrison, David K. Jackson, Ian Johnston, Dominic Kwiatkowski, Cordelia Langford, John Sillitoe on behalf of the Wellcome Sanger Institute COVID-19 Surveillance Team |
| EPI_ISL_777418, EPI_ISL_777419 | Lighthouse Lab in Cambridge | Wellcome Sanger Institute for the COVID-19 Genomics UK (COG-UK) Consortium | Rob Howes, The Lighthouse Lab in Cambridge and Alex Alderton, Roberto Amato, Sonia Goncalves, Ewan Harrison, David K. Jackson, Ian Johnston, Dominic Kwiatkowski, Cordelia Langford, John Sillitoe on behalf of the Wellcome Sanger Institute COVID-19 Surveillance Team |
| EPI_ISL_777420 | Lighthouse Lab in Glasgow | Wellcome Sanger Institute for the COVID-19 Genomics UK (COG-UK) Consortium | Harper VanSteenhouse, Yumi Kasai, David Gray, Carol Clugston, Anna Dominiczak and Alex Alderton, Roberto Amato, Sonia Goncalves, Ewan Harrison, David K. Jackson, Ian Johnston, Dominic Kwiatkowski, Cordelia Langford, John Sillitoe on behalf of the Wellcome Sanger Institute COVID-19 Surveillance Team |
| EPI_ISL_777422 | Lighthouse Lab in Cambridge | Wellcome Sanger Institute for the COVID-19 Genomics UK (COG-UK) Consortium | Rob Howes, The Lighthouse Lab in Cambridge and Alex Alderton, Roberto Amato, Sonia Goncalves, Ewan Harrison, David K. Jackson, Ian Johnston, Dominic Kwiatkowski, Cordelia Langford, John Sillitoe on behalf of the Wellcome Sanger Institute COVID-19 Surveillance Team |
| EPI_ISL_777423 | Lighthouse Lab in Glasgow | Wellcome Sanger Institute for the COVID-19 Genomics UK (COG-UK) Consortium | Harper VanSteenhouse, Yumi Kasai, David Gray, Carol Clugston, Anna Dominiczak and Alex Alderton, Roberto Amato, Sonia Goncalves, Ewan Harrison, David K. Jackson, Ian Johnston, Dominic Kwiatkowski, Cordelia Langford, John Sillitoe on behalf of the Wellcome Sanger Institute COVID-19 Surveillance Team |
| EPI_ISL_777426, EPI_ISL_777428, EPI_ISL_777429, EPI_ISL_777431 | Lighthouse Lab in Cambridge | Wellcome Sanger Institute for the COVID-19 Genomics UK (COG-UK) Consortium | Rob Howes, The Lighthouse Lab in Cambridge and Alex Alderton, Roberto Amato, Sonia Goncalves, Ewan Harrison, David K. Jackson, Ian Johnston, Dominic Kwiatkowski, Cordelia Langford, John Sillitoe on behalf of the Wellcome Sanger Institute COVID-19 Surveillance Team |
| EPI_ISL_777432 | Lighthouse Lab in Glasgow | Wellcome Sanger Institute for the COVID-19 Genomics UK (COG-UK) Consortium | Harper VanSteenhouse, Yumi Kasai, David Gray, Carol Clugston, Anna Dominiczak and Alex Alderton, Roberto Amato, Sonia Goncalves, Ewan Harrison, David K. Jackson, Ian Johnston, Dominic Kwiatkowski, Cordelia Langford, John Sillitoe on behalf of the Wellcome Sanger Institute COVID-19 Surveillance Team |
| EPI_ISL_777433, EPI_ISL_777434 | Lighthouse Lab in Cambridge | Wellcome Sanger Institute for the COVID-19 Genomics UK (COG-UK) Consortium | Rob Howes, The Lighthouse Lab in Cambridge and Alex Alderton, Roberto Amato, Sonia Goncalves, Ewan Harrison, David K. Jackson, Ian Johnston, Dominic Kwiatkowski, Cordelia Langford, John Sillitoe on behalf of the Wellcome Sanger Institute COVID-19 Surveillance Team |
| EPI_ISL_777436, EPI_ISL_777438 | Lighthouse Lab in Glasgow | Wellcome Sanger Institute for the COVID-19 Genomics UK (COG-UK) Consortium | Harper VanSteenhouse, Yumi Kasai, David Gray, Carol Clugston, Anna Dominiczak and Alex Alderton, Roberto Amato, Sonia Goncalves, Ewan Harrison, David K. Jackson, Ian Johnston, Dominic Kwiatkowski, Cordelia Langford, John Sillitoe on behalf of the Wellcome Sanger Institute COVID-19 Surveillance Team |
| EPI_ISL_777439, EPI_ISL_777441, EPI_ISL_777442 | Lighthouse Lab in Cambridge | Wellcome Sanger Institute for the COVID-19 Genomics UK (COG-UK) Consortium | Rob Howes, The Lighthouse Lab in Cambridge and Alex Alderton, Roberto Amato, Sonia Goncalves, Ewan Harrison, David K. Jackson, Ian Johnston, Dominic Kwiatkowski, Cordelia Langford, John Sillitoe on behalf of the Wellcome Sanger Institute COVID-19 Surveillance Team |
| EPI_ISL_777443, EPI_ISL_777444 | Lighthouse Lab in Glasgow | Wellcome Sanger Institute for the COVID-19 Genomics UK (COG-UK) Consortium | Harper VanSteenhouse, Yumi Kasai, David Gray, Carol Clugston, Anna Dominiczak and Alex Alderton, Roberto Amato, Sonia Goncalves, Ewan Harrison, David K. Jackson, Ian Johnston, Dominic Kwiatkowski, Cordelia Langford, John Sillitoe on behalf of the Wellcome Sanger Institute COVID-19 Surveillance Team |
| EPI_ISL_777445, EPI_ISL_777446 | Lighthouse Lab in Cambridge | Wellcome Sanger Institute for the COVID-19 Genomics UK (COG-UK) Consortium | Rob Howes, The Lighthouse Lab in Cambridge and Alex Alderton, Roberto Amato, Sonia Goncalves, Ewan Harrison, David K. Jackson, Ian Johnston, Dominic Kwiatkowski, Cordelia Langford, John Sillitoe on behalf of the Wellcome Sanger Institute COVID-19 Surveillance Team |
| EPI_ISL_777447, EPI_ISL_777449 | Lighthouse Lab in Glasgow | Wellcome Sanger Institute for the COVID-19 Genomics UK (COG-UK) Consortium | Harper VanSteenhouse, Yumi Kasai, David Gray, Carol Clugston, Anna Dominiczak and Alex Alderton, Roberto Amato, Sonia Goncalves, Ewan Harrison, David K. Jackson, Ian Johnston, Dominic Kwiatkowski, Cordelia Langford, John Sillitoe on behalf of the Wellcome Sanger Institute COVID-19 Surveillance Team |
| EPI_ISL_777450, EPI_ISL_777452, EPI_ISL_777453, EPI_ISL_777454, EPI_ISL_777456, EPI_ISL_777457 | Lighthouse Lab in Cambridge | Wellcome Sanger Institute for the COVID-19 Genomics UK (COG-UK) Consortium | Rob Howes, The Lighthouse Lab in Cambridge and Alex Alderton, Roberto Amato, Sonia Goncalves, Ewan Harrison, David K. Jackson, Ian Johnston, Dominic Kwiatkowski, Cordelia Langford, John Sillitoe on behalf of the Wellcome Sanger Institute COVID-19 Surveillance Team |
| EPI_ISL_777459 | Lighthouse Lab in Glasgow | Wellcome Sanger Institute for the COVID-19 Genomics UK (COG-UK) Consortium | Harper VanSteenhouse, Yumi Kasai, David Gray, Carol Clugston, Anna Dominiczak and Alex Alderton, Roberto Amato, Sonia Goncalves, Ewan Harrison, David K. Jackson, Ian Johnston, Dominic Kwiatkowski, Cordelia Langford, John Sillitoe on behalf of the Wellcome Sanger Institute COVID-19 Surveillance Team |
| EPI_ISL_777460 | Lighthouse Lab in Cambridge | Wellcome Sanger Institute for the COVID-19 Genomics UK (COG-UK) Consortium | Rob Howes, The Lighthouse Lab in Cambridge and Alex Alderton, Roberto Amato, Sonia Goncalves, Ewan Harrison, David K. Jackson, Ian Johnston, Dominic Kwiatkowski, Cordelia Langford, John Sillitoe on behalf of the Wellcome Sanger Institute COVID-19 Surveillance Team |
| EPI_ISL_777461, EPI_ISL_777462 | Lighthouse Lab in Glasgow | Wellcome Sanger Institute for the COVID-19 Genomics UK (COG-UK) Consortium | Harper VanSteenhouse, Yumi Kasai, David Gray, Carol Clugston, Anna Dominiczak and Alex Alderton, Roberto Amato, Sonia Goncalves, Ewan Harrison, David K. Jackson, Ian Johnston, Dominic Kwiatkowski, Cordelia Langford, John Sillitoe on behalf of the Wellcome Sanger Institute COVID-19 Surveillance Team |
| EPI_ISL_777463, EPI_ISL_777464 | Lighthouse Lab in Cambridge | Wellcome Sanger Institute for the COVID-19 Genomics UK (COG-UK) Consortium | Rob Howes, The Lighthouse Lab in Cambridge and Alex Alderton, Roberto Amato, Sonia Goncalves, Ewan Harrison, David K. Jackson, Ian Johnston, Dominic Kwiatkowski, Cordelia Langford, John Sillitoe on behalf of the Wellcome Sanger Institute COVID-19 Surveillance Team |
| EPI_ISL_777466 | Lighthouse Lab in Glasgow | Wellcome Sanger Institute for the COVID-19 Genomics UK (COG-UK) Consortium | Harper VanSteenhouse, Yumi Kasai, David Gray, Carol Clugston, Anna Dominiczak and Alex Alderton, Roberto Amato, Sonia Goncalves, Ewan Harrison, David K. Jackson, Ian Johnston, Dominic Kwiatkowski, Cordelia Langford, John Sillitoe on behalf of the Wellcome Sanger Institute COVID-19 Surveillance Team |
| EPI_ISL_777468, EPI_ISL_777469 | Lighthouse Lab in Cambridge | Wellcome Sanger Institute for the COVID-19 Genomics UK (COG-UK) Consortium | Rob Howes, The Lighthouse Lab in Cambridge and Alex Alderton, Roberto Amato, Sonia Goncalves, Ewan Harrison, David K. Jackson, Ian Johnston, Dominic Kwiatkowski, Cordelia Langford, John Sillitoe on behalf of the Wellcome Sanger Institute COVID-19 Surveillance Team |
| EPI_ISL_777471 | Lighthouse Lab in Glasgow | Wellcome Sanger Institute for the COVID-19 Genomics UK (COG-UK) Consortium | Harper VanSteenhouse, Yumi Kasai, David Gray, Carol Clugston, Anna Dominiczak and Alex Alderton, Roberto Amato, Sonia Goncalves, Ewan Harrison, David K. Jackson, Ian Johnston, Dominic Kwiatkowski, Cordelia Langford, John Sillitoe on behalf of the Wellcome Sanger Institute COVID-19 Surveillance Team |
| EPI_ISL_777472, EPI_ISL_777473 | Lighthouse Lab in Cambridge | Wellcome Sanger Institute for the COVID-19 Genomics UK (COG-UK) Consortium | Rob Howes, The Lighthouse Lab in Cambridge and Alex Alderton, Roberto Amato, Sonia Goncalves, Ewan Harrison, David K. Jackson, Ian Johnston, Dominic Kwiatkowski, Cordelia Langford, John Sillitoe on behalf of the Wellcome Sanger Institute COVID-19 Surveillance Team |
| EPI_ISL_777474, EPI_ISL_777475 | Lighthouse Lab in Glasgow | Wellcome Sanger Institute for the COVID-19 Genomics UK (COG-UK) Consortium | Harper VanSteenhouse, Yumi Kasai, David Gray, Carol Clugston, Anna Dominiczak and Alex Alderton, Roberto Amato, Sonia Goncalves, Ewan Harrison, David K. Jackson, Ian Johnston, Dominic Kwiatkowski, Cordelia Langford, John Sillitoe on behalf of the Wellcome Sanger Institute COVID-19 Surveillance Team |
| EPI_ISL_777476, EPI_ISL_777478, EPI_ISL_777480, EPI_ISL_777482, EPI_ISL_777483 | Lighthouse Lab in Cambridge | Wellcome Sanger Institute for the COVID-19 Genomics UK (COG-UK) Consortium | Rob Howes, The Lighthouse Lab in Cambridge and Alex Alderton, Roberto Amato, Sonia Goncalves, Ewan Harrison, David K. Jackson, Ian Johnston, Dominic Kwiatkowski, Cordelia Langford, John Sillitoe on behalf of the Wellcome Sanger Institute COVID-19 Surveillance Team |
| EPI_ISL_777484 | Lighthouse Lab in Glasgow | Wellcome Sanger Institute for the COVID-19 Genomics UK (COG-UK) Consortium | Harper VanSteenhouse, Yumi Kasai, David Gray, Carol Clugston, Anna Dominiczak and Alex Alderton, Roberto Amato, Sonia Goncalves, Ewan Harrison, David K. Jackson, Ian Johnston, Dominic Kwiatkowski, Cordelia Langford, John Sillitoe on behalf of the Wellcome Sanger Institute COVID-19 Surveillance Team |
| EPI_ISL_777485, EPI_ISL_777486 | Lighthouse Lab in Cambridge | Wellcome Sanger Institute for the COVID-19 Genomics UK | Rob Howes, The Lighthouse Lab in Cambridge and Alex Alderton, Roberto Amato, Sonia Goncalves, Ewan Harrison, David K. Jackson, Ian Johnston, |

| | | | |
|---|---|---|---|
| | | (COG-UK) Consortium | Dominic Kwiatkowski, Cordelia Langford, John Sillitoe on behalf of the Wellcome Sanger Institute COVID-19 Surveillance Team |
| EPI_ISL_777487 | Lighthouse Lab in Glasgow | Wellcome Sanger Institute for the COVID-19 Genomics UK (COG-UK) Consortium | Harper VanSteenhouse, Yumi Kasai, David Gray, Carol Clugston, Anna Dominiczak and Alex Alderton, Roberto Amato, Sonia Goncalves, Ewan Harrison, David K. Jackson, Ian Johnston, Dominic Kwiatkowski, Cordelia Langford, John Sillitoe on behalf of the Wellcome Sanger Institute COVID-19 Surveillance Team |
| EPI_ISL_777488, EPI_ISL_777492 | Lighthouse Lab in Cambridge | Wellcome Sanger Institute for the COVID-19 Genomics UK (COG-UK) Consortium | Rob Howes, The Lighthouse Lab in Cambridge and Alex Alderton, Roberto Amato, Sonia Goncalves, Ewan Harrison, David K. Jackson, Ian Johnston, Dominic Kwiatkowski, Cordelia Langford, John Sillitoe on behalf of the Wellcome Sanger Institute COVID-19 Surveillance Team |
| EPI_ISL_777493 | Lighthouse Lab in Glasgow | Wellcome Sanger Institute for the COVID-19 Genomics UK (COG-UK) Consortium | Harper VanSteenhouse, Yumi Kasai, David Gray, Carol Clugston, Anna Dominiczak and Alex Alderton, Roberto Amato, Sonia Goncalves, Ewan Harrison, David K. Jackson, Ian Johnston, Dominic Kwiatkowski, Cordelia Langford, John Sillitoe on behalf of the Wellcome Sanger Institute COVID-19 Surveillance Team |
| EPI_ISL_777494 | Lighthouse Lab in Cambridge | Wellcome Sanger Institute for the COVID-19 Genomics UK (COG-UK) Consortium | Rob Howes, The Lighthouse Lab in Cambridge and Alex Alderton, Roberto Amato, Sonia Goncalves, Ewan Harrison, David K. Jackson, Ian Johnston, Dominic Kwiatkowski, Cordelia Langford, John Sillitoe on behalf of the Wellcome Sanger Institute COVID-19 Surveillance Team |
| EPI_ISL_777496, EPI_ISL_777497, EPI_ISL_777498 | Lighthouse Lab in Glasgow | Wellcome Sanger Institute for the COVID-19 Genomics UK (COG-UK) Consortium | Harper VanSteenhouse, Yumi Kasai, David Gray, Carol Clugston, Anna Dominiczak and Alex Alderton, Roberto Amato, Sonia Goncalves, Ewan Harrison, David K. Jackson, Ian Johnston, Dominic Kwiatkowski, Cordelia Langford, John Sillitoe on behalf of the Wellcome Sanger Institute COVID-19 Surveillance Team |
| EPI_ISL_777499, EPI_ISL_777500, EPI_ISL_777501, EPI_ISL_777503 | Lighthouse Lab in Cambridge | Wellcome Sanger Institute for the COVID-19 Genomics UK (COG-UK) Consortium | Rob Howes, The Lighthouse Lab in Cambridge and Alex Alderton, Roberto Amato, Sonia Goncalves, Ewan Harrison, David K. Jackson, Ian Johnston, Dominic Kwiatkowski, Cordelia Langford, John Sillitoe on behalf of the Wellcome Sanger Institute COVID-19 Surveillance Team |
| EPI_ISL_777504, EPI_ISL_777505, EPI_ISL_777506 | Lighthouse Lab in Glasgow | Wellcome Sanger Institute for the COVID-19 Genomics UK (COG-UK) Consortium | Harper VanSteenhouse, Yumi Kasai, David Gray, Carol Clugston, Anna Dominiczak and Alex Alderton, Roberto Amato, Sonia Goncalves, Ewan Harrison, David K. Jackson, Ian Johnston, Dominic Kwiatkowski, Cordelia Langford, John Sillitoe on behalf of the Wellcome Sanger Institute COVID-19 Surveillance Team |
| EPI_ISL_777507, EPI_ISL_777508 | Lighthouse Lab in Cambridge | Wellcome Sanger Institute for the COVID-19 Genomics UK (COG-UK) Consortium | Rob Howes, The Lighthouse Lab in Cambridge and Alex Alderton, Roberto Amato, Sonia Goncalves, Ewan Harrison, David K. Jackson, Ian Johnston, Dominic Kwiatkowski, Cordelia Langford, John Sillitoe on behalf of the Wellcome Sanger Institute COVID-19 Surveillance Team |
| EPI_ISL_777510 | Lighthouse Lab in Glasgow | Wellcome Sanger Institute for the COVID-19 Genomics UK (COG-UK) Consortium | Harper VanSteenhouse, Yumi Kasai, David Gray, Carol Clugston, Anna Dominiczak and Alex Alderton, Roberto Amato, Sonia Goncalves, Ewan Harrison, David K. Jackson, Ian Johnston, Dominic Kwiatkowski, Cordelia Langford, John Sillitoe on behalf of the Wellcome Sanger Institute COVID-19 Surveillance Team |
| EPI_ISL_777511, EPI_ISL_777512, EPI_ISL_777513 | Lighthouse Lab in Cambridge | Wellcome Sanger Institute for the COVID-19 Genomics UK (COG-UK) Consortium | Rob Howes, The Lighthouse Lab in Cambridge and Alex Alderton, Roberto Amato, Sonia Goncalves, Ewan Harrison, David K. Jackson, Ian Johnston, Dominic Kwiatkowski, Cordelia Langford, John Sillitoe on behalf of the Wellcome Sanger Institute COVID-19 Surveillance Team |
| EPI_ISL_777514 | Lighthouse Lab in Glasgow | Wellcome Sanger Institute for the COVID-19 Genomics UK (COG-UK) Consortium | Harper VanSteenhouse, Yumi Kasai, David Gray, Carol Clugston, Anna Dominiczak and Alex Alderton, Roberto Amato, Sonia Goncalves, Ewan Harrison, David K. Jackson, Ian Johnston, Dominic Kwiatkowski, Cordelia Langford, John Sillitoe on behalf of the Wellcome Sanger Institute COVID-19 Surveillance Team |
| EPI_ISL_777516, EPI_ISL_777517 | Lighthouse Lab in Cambridge | Wellcome Sanger Institute for the COVID-19 Genomics UK (COG-UK) Consortium | Rob Howes, The Lighthouse Lab in Cambridge and Alex Alderton, Roberto Amato, Sonia Goncalves, Ewan Harrison, David K. Jackson, Ian Johnston, Dominic Kwiatkowski, Cordelia Langford, John Sillitoe on behalf of the Wellcome Sanger Institute COVID-19 Surveillance Team |
| EPI_ISL_777518 | Lighthouse Lab in Glasgow | Wellcome Sanger Institute for the COVID-19 Genomics UK (COG-UK) Consortium | Harper VanSteenhouse, Yumi Kasai, David Gray, Carol Clugston, Anna Dominiczak and Alex Alderton, Roberto Amato, Sonia Goncalves, Ewan Harrison, David K. Jackson, Ian Johnston, Dominic Kwiatkowski, Cordelia Langford, John Sillitoe on behalf of the Wellcome Sanger Institute COVID-19 Surveillance Team |
| EPI_ISL_777519, EPI_ISL_777521 | Lighthouse Lab in Cambridge | Wellcome Sanger Institute for the COVID-19 Genomics UK (COG-UK) Consortium | Rob Howes, The Lighthouse Lab in Cambridge and Alex Alderton, Roberto Amato, Sonia Goncalves, Ewan Harrison, David K. Jackson, Ian Johnston, Dominic Kwiatkowski, Cordelia Langford, John Sillitoe on behalf of the Wellcome Sanger Institute COVID-19 Surveillance Team |
| EPI_ISL_777523, EPI_ISL_777526 | Lighthouse Lab in Glasgow | Wellcome Sanger Institute for the COVID-19 Genomics UK (COG-UK) Consortium | Harper VanSteenhouse, Yumi Kasai, David Gray, Carol Clugston, Anna Dominiczak and Alex Alderton, Roberto Amato, Sonia Goncalves, Ewan Harrison, David K. Jackson, Ian Johnston, Dominic Kwiatkowski, Cordelia Langford, John Sillitoe on behalf of the Wellcome Sanger Institute COVID-19 Surveillance Team |
| EPI_ISL_777527, EPI_ISL_777528 | Lighthouse Lab in Cambridge | Wellcome Sanger Institute for the COVID-19 Genomics UK (COG-UK) Consortium | Rob Howes, The Lighthouse Lab in Cambridge and Alex Alderton, Roberto Amato, Sonia Goncalves, Ewan Harrison, David K. Jackson, Ian Johnston, Dominic Kwiatkowski, Cordelia Langford, John Sillitoe on behalf of the Wellcome Sanger Institute COVID-19 Surveillance Team |
| EPI_ISL_777531, EPI_ISL_777533 | Lighthouse Lab in Glasgow | Wellcome Sanger Institute for the COVID-19 Genomics UK (COG-UK) Consortium | Harper VanSteenhouse, Yumi Kasai, David Gray, Carol Clugston, Anna Dominiczak and Alex Alderton, Roberto Amato, Sonia Goncalves, Ewan Harrison, David K. Jackson, Ian Johnston, Dominic Kwiatkowski, Cordelia Langford, John Sillitoe on behalf of the Wellcome Sanger Institute COVID-19 Surveillance Team |
| EPI_ISL_777536, EPI_ISL_777540, EPI_ISL_777541, EPI_ISL_777542, EPI_ISL_777544, EPI_ISL_777545, EPI_ISL_777546, EPI_ISL_777547, EPI_ISL_777549, EPI_ISL_777551, EPI_ISL_777553 | | | |
| see above | Lighthouse Lab in Cambridge | Wellcome Sanger Institute for the COVID-19 Genomics UK (COG-UK) Consortium | Rob Howes, The Lighthouse Lab in Cambridge and Alex Alderton, Roberto Amato, Sonia Goncalves, Ewan Harrison, David K. Jackson, Ian Johnston, Dominic Kwiatkowski, Cordelia Langford, John Sillitoe on behalf of the Wellcome Sanger Institute COVID-19 Surveillance Team |
| EPI_ISL_777554 | Lighthouse Lab in Glasgow | Wellcome Sanger Institute for the COVID-19 Genomics UK (COG-UK) Consortium | Harper VanSteenhouse, Yumi Kasai, David Gray, Carol Clugston, Anna Dominiczak and Alex Alderton, Roberto Amato, Sonia Goncalves, Ewan Harrison, David K. Jackson, Ian Johnston, Dominic Kwiatkowski, Cordelia Langford, John Sillitoe on behalf of the Wellcome Sanger Institute COVID-19 Surveillance Team |
| EPI_ISL_777555, EPI_ISL_777556 | Lighthouse Lab in Cambridge | Wellcome Sanger Institute for the COVID-19 Genomics UK (COG-UK) Consortium | Rob Howes, The Lighthouse Lab in Cambridge and Alex Alderton, Roberto Amato, Sonia Goncalves, Ewan Harrison, David K. Jackson, Ian Johnston, Dominic Kwiatkowski, Cordelia Langford, John Sillitoe on behalf of the Wellcome Sanger Institute COVID-19 Surveillance Team |
| EPI_ISL_777557 | Lighthouse Lab in Glasgow | Wellcome Sanger Institute for the COVID-19 Genomics UK (COG-UK) Consortium | Harper VanSteenhouse, Yumi Kasai, David Gray, Carol Clugston, Anna Dominiczak and Alex Alderton, Roberto Amato, Sonia Goncalves, Ewan Harrison, David K. Jackson, Ian Johnston, Dominic Kwiatkowski, Cordelia Langford, John Sillitoe on behalf of the Wellcome Sanger Institute COVID-19 Surveillance Team |
| EPI_ISL_777559 | Lighthouse Lab in Cambridge | Wellcome Sanger Institute for the COVID-19 Genomics UK (COG-UK) Consortium | Rob Howes, The Lighthouse Lab in Cambridge and Alex Alderton, Roberto Amato, Sonia Goncalves, Ewan Harrison, David K. Jackson, Ian Johnston, Dominic Kwiatkowski, Cordelia Langford, John Sillitoe on behalf of the Wellcome Sanger Institute COVID-19 Surveillance Team |
| EPI_ISL_777560, EPI_ISL_777562 | Lighthouse Lab in Glasgow | Wellcome Sanger Institute for the COVID-19 Genomics UK (COG-UK) Consortium | Harper VanSteenhouse, Yumi Kasai, David Gray, Carol Clugston, Anna Dominiczak and Alex Alderton, Roberto Amato, Sonia Goncalves, Ewan Harrison, David K. Jackson, Ian Johnston, Dominic Kwiatkowski, Cordelia Langford, John Sillitoe on behalf of the Wellcome Sanger Institute COVID-19 Surveillance Team |
| EPI_ISL_777564, EPI_ISL_777565, EPI_ISL_777566 | Lighthouse Lab in Cambridge | Wellcome Sanger Institute for the COVID-19 Genomics UK (COG-UK) Consortium | Rob Howes, The Lighthouse Lab in Cambridge and Alex Alderton, Roberto Amato, Sonia Goncalves, Ewan Harrison, David K. Jackson, Ian Johnston, Dominic Kwiatkowski, Cordelia Langford, John Sillitoe on behalf of the Wellcome Sanger Institute COVID-19 Surveillance Team |
| EPI_ISL_777569 | Lighthouse Lab in Glasgow | Wellcome Sanger Institute for the COVID-19 Genomics UK (COG-UK) Consortium | Harper VanSteenhouse, Yumi Kasai, David Gray, Carol Clugston, Anna Dominiczak and Alex Alderton, Roberto Amato, Sonia Goncalves, Ewan Harrison, David K. Jackson, Ian Johnston, Dominic Kwiatkowski, Cordelia Langford, John Sillitoe on behalf of the Wellcome Sanger Institute COVID-19 Surveillance Team |
| EPI_ISL_777570, EPI_ISL_777571 | Lighthouse Lab in Cambridge | Wellcome Sanger Institute for the COVID-19 Genomics UK (COG-UK) Consortium | Rob Howes, The Lighthouse Lab in Cambridge and Alex Alderton, Roberto Amato, Sonia Goncalves, Ewan Harrison, David K. Jackson, Ian Johnston, Dominic Kwiatkowski, Cordelia Langford, John Sillitoe on behalf of the Wellcome Sanger Institute COVID-19 Surveillance Team |
| EPI_ISL_777574 | Lighthouse Lab in Glasgow | Wellcome Sanger Institute for the COVID-19 Genomics UK (COG-UK) Consortium | Harper VanSteenhouse, Yumi Kasai, David Gray, Carol Clugston, Anna Dominiczak and Alex Alderton, Roberto Amato, Sonia Goncalves, Ewan Harrison, David K. Jackson, Ian Johnston, Dominic Kwiatkowski, Cordelia Langford, John Sillitoe on behalf of the Wellcome Sanger Institute COVID-19 Surveillance Team |
| EPI_ISL_777575, EPI_ISL_777577, EPI_ISL_777578 | Lighthouse Lab in Cambridge | Wellcome Sanger Institute for the COVID-19 Genomics UK (COG-UK) Consortium | Rob Howes, The Lighthouse Lab in Cambridge and Alex Alderton, Roberto Amato, Sonia Goncalves, Ewan Harrison, David K. Jackson, Ian Johnston, Dominic Kwiatkowski, Cordelia Langford, John Sillitoe on behalf of the Wellcome Sanger Institute COVID-19 Surveillance Team |
| EPI_ISL_777579 | Lighthouse Lab in Glasgow | Wellcome Sanger Institute for the COVID-19 Genomics UK | Harper VanSteenhouse, Yumi Kasai, David Gray, Carol Clugston, Anna Dominiczak and Alex Alderton, Roberto Amato, Sonia Goncalves, Ewan Harrison, |

| | | | |
|---|---|---|---|
| | | (COG-UK) Consortium | David K. Jackson, Ian Johnston, Dominic Kwiatkowski, Cordelia Langford, John Sillitoe on behalf of the Wellcome Sanger Institute COVID-19 Surveillance Team |
| EPI_ISL_777581 | Lighthouse Lab in Cambridge | Wellcome Sanger Institute for the COVID-19 Genomics UK (COG-UK) Consortium | Rob Howes, The Lighthouse Lab in Cambridge and Alex Alderton, Roberto Amato, Sonia Goncalves, Ewan Harrison, David K. Jackson, Ian Johnston, Dominic Kwiatkowski, Cordelia Langford, John Sillitoe on behalf of the Wellcome Sanger Institute COVID-19 Surveillance Team |
| EPI_ISL_777583 | Lighthouse Lab in Glasgow | Wellcome Sanger Institute for the COVID-19 Genomics UK (COG-UK) Consortium | Harper VanSteenhouse, Yumi Kasai, David Gray, Carol Clugston, Anna Dominiczak and Alex Alderton, Roberto Amato, Sonia Goncalves, Ewan Harrison, David K. Jackson, Ian Johnston, Dominic Kwiatkowski, Cordelia Langford, John Sillitoe on behalf of the Wellcome Sanger Institute COVID-19 Surveillance Team |
| EPI_ISL_777585, EPI_ISL_777586, EPI_ISL_777588, EPI_ISL_777590, EPI_ISL_777594, EPI_ISL_777596, EPI_ISL_777598, EPI_ISL_777599 | Lighthouse Lab in Cambridge | Wellcome Sanger Institute for the COVID-19 Genomics UK (COG-UK) Consortium | Rob Howes, The Lighthouse Lab in Cambridge and Alex Alderton, Roberto Amato, Sonia Goncalves, Ewan Harrison, David K. Jackson, Ian Johnston, Dominic Kwiatkowski, Cordelia Langford, John Sillitoe on behalf of the Wellcome Sanger Institute COVID-19 Surveillance Team |
| EPI_ISL_777600, EPI_ISL_777601 | Lighthouse Lab in Glasgow | Wellcome Sanger Institute for the COVID-19 Genomics UK (COG-UK) Consortium | Harper VanSteenhouse, Yumi Kasai, David Gray, Carol Clugston, Anna Dominiczak and Alex Alderton, Roberto Amato, Sonia Goncalves, Ewan Harrison, David K. Jackson, Ian Johnston, Dominic Kwiatkowski, Cordelia Langford, John Sillitoe on behalf of the Wellcome Sanger Institute COVID-19 Surveillance Team |
| EPI_ISL_777602, EPI_ISL_777603, EPI_ISL_777604, EPI_ISL_777606, EPI_ISL_777607, EPI_ISL_777608, EPI_ISL_777609 | Lighthouse Lab in Cambridge | Wellcome Sanger Institute for the COVID-19 Genomics UK (COG-UK) Consortium | Rob Howes, The Lighthouse Lab in Cambridge and Alex Alderton, Roberto Amato, Sonia Goncalves, Ewan Harrison, David K. Jackson, Ian Johnston, Dominic Kwiatkowski, Cordelia Langford, John Sillitoe on behalf of the Wellcome Sanger Institute COVID-19 Surveillance Team |
| EPI_ISL_777610 | Lighthouse Lab in Glasgow | Wellcome Sanger Institute for the COVID-19 Genomics UK (COG-UK) Consortium | Harper VanSteenhouse, Yumi Kasai, David Gray, Carol Clugston, Anna Dominiczak and Alex Alderton, Roberto Amato, Sonia Goncalves, Ewan Harrison, David K. Jackson, Ian Johnston, Dominic Kwiatkowski, Cordelia Langford, John Sillitoe on behalf of the Wellcome Sanger Institute COVID-19 Surveillance Team |
| EPI_ISL_777611 | Lighthouse Lab in Cambridge | Wellcome Sanger Institute for the COVID-19 Genomics UK (COG-UK) Consortium | Rob Howes, The Lighthouse Lab in Cambridge and Alex Alderton, Roberto Amato, Sonia Goncalves, Ewan Harrison, David K. Jackson, Ian Johnston, Dominic Kwiatkowski, Cordelia Langford, John Sillitoe on behalf of the Wellcome Sanger Institute COVID-19 Surveillance Team |
| EPI_ISL_777612 | Lighthouse Lab in Glasgow | Wellcome Sanger Institute for the COVID-19 Genomics UK (COG-UK) Consortium | Harper VanSteenhouse, Yumi Kasai, David Gray, Carol Clugston, Anna Dominiczak and Alex Alderton, Roberto Amato, Sonia Goncalves, Ewan Harrison, David K. Jackson, Ian Johnston, Dominic Kwiatkowski, Cordelia Langford, John Sillitoe on behalf of the Wellcome Sanger Institute COVID-19 Surveillance Team |
| EPI_ISL_777613, EPI_ISL_777614, EPI_ISL_777615, EPI_ISL_777616, EPI_ISL_777617, EPI_ISL_777618, EPI_ISL_777619, EPI_ISL_777620 | Lighthouse Lab in Cambridge | Wellcome Sanger Institute for the COVID-19 Genomics UK (COG-UK) Consortium | Rob Howes, The Lighthouse Lab in Cambridge and Alex Alderton, Roberto Amato, Sonia Goncalves, Ewan Harrison, David K. Jackson, Ian Johnston, Dominic Kwiatkowski, Cordelia Langford, John Sillitoe on behalf of the Wellcome Sanger Institute COVID-19 Surveillance Team |
| EPI_ISL_777622, EPI_ISL_777623 | Lighthouse Lab in Glasgow | Wellcome Sanger Institute for the COVID-19 Genomics UK (COG-UK) Consortium | Harper VanSteenhouse, Yumi Kasai, David Gray, Carol Clugston, Anna Dominiczak and Alex Alderton, Roberto Amato, Sonia Goncalves, Ewan Harrison, David K. Jackson, Ian Johnston, Dominic Kwiatkowski, Cordelia Langford, John Sillitoe on behalf of the Wellcome Sanger Institute COVID-19 Surveillance Team |
| EPI_ISL_777624, EPI_ISL_777625, EPI_ISL_777626 | Lighthouse Lab in Cambridge | Wellcome Sanger Institute for the COVID-19 Genomics UK (COG-UK) Consortium | Rob Howes, The Lighthouse Lab in Cambridge and Alex Alderton, Roberto Amato, Sonia Goncalves, Ewan Harrison, David K. Jackson, Ian Johnston, Dominic Kwiatkowski, Cordelia Langford, John Sillitoe on behalf of the Wellcome Sanger Institute COVID-19 Surveillance Team |
| EPI_ISL_777628, EPI_ISL_777630 | Lighthouse Lab in Glasgow | Wellcome Sanger Institute for the COVID-19 Genomics UK (COG-UK) Consortium | Harper VanSteenhouse, Yumi Kasai, David Gray, Carol Clugston, Anna Dominiczak and Alex Alderton, Roberto Amato, Sonia Goncalves, Ewan Harrison, David K. Jackson, Ian Johnston, Dominic Kwiatkowski, Cordelia Langford, John Sillitoe on behalf of the Wellcome Sanger Institute COVID-19 Surveillance Team |
| EPI_ISL_777631, EPI_ISL_777632, EPI_ISL_777635, EPI_ISL_777636, EPI_ISL_777637, EPI_ISL_777640 | Lighthouse Lab in Cambridge | Wellcome Sanger Institute for the COVID-19 Genomics UK (COG-UK) Consortium | Rob Howes, The Lighthouse Lab in Cambridge and Alex Alderton, Roberto Amato, Sonia Goncalves, Ewan Harrison, David K. Jackson, Ian Johnston, Dominic Kwiatkowski, Cordelia Langford, John Sillitoe on behalf of the Wellcome Sanger Institute COVID-19 Surveillance Team |
| EPI_ISL_777641 | Lighthouse Lab in Glasgow | Wellcome Sanger Institute for the COVID-19 Genomics UK (COG-UK) Consortium | Harper VanSteenhouse, Yumi Kasai, David Gray, Carol Clugston, Anna Dominiczak and Alex Alderton, Roberto Amato, Sonia Goncalves, Ewan Harrison, David K. Jackson, Ian Johnston, Dominic Kwiatkowski, Cordelia Langford, John Sillitoe on behalf of the Wellcome Sanger Institute COVID-19 Surveillance Team |
| EPI_ISL_777642, EPI_ISL_777643, EPI_ISL_777645 | Lighthouse Lab in Cambridge | Wellcome Sanger Institute for the COVID-19 Genomics UK (COG-UK) Consortium | Rob Howes, The Lighthouse Lab in Cambridge and Alex Alderton, Roberto Amato, Sonia Goncalves, Ewan Harrison, David K. Jackson, Ian Johnston, Dominic Kwiatkowski, Cordelia Langford, John Sillitoe on behalf of the Wellcome Sanger Institute COVID-19 Surveillance Team |
| EPI_ISL_777646 | Lighthouse Lab in Glasgow | Wellcome Sanger Institute for the COVID-19 Genomics UK (COG-UK) Consortium | Harper VanSteenhouse, Yumi Kasai, David Gray, Carol Clugston, Anna Dominiczak and Alex Alderton, Roberto Amato, Sonia Goncalves, Ewan Harrison, David K. Jackson, Ian Johnston, Dominic Kwiatkowski, Cordelia Langford, John Sillitoe on behalf of the Wellcome Sanger Institute COVID-19 Surveillance Team |
| EPI_ISL_777647, EPI_ISL_777648, EPI_ISL_777652 | Lighthouse Lab in Cambridge | Wellcome Sanger Institute for the COVID-19 Genomics UK (COG-UK) Consortium | Rob Howes, The Lighthouse Lab in Cambridge and Alex Alderton, Roberto Amato, Sonia Goncalves, Ewan Harrison, David K. Jackson, Ian Johnston, Dominic Kwiatkowski, Cordelia Langford, John Sillitoe on behalf of the Wellcome Sanger Institute COVID-19 Surveillance Team |
| EPI_ISL_777653, EPI_ISL_777654 | Lighthouse Lab in Glasgow | Wellcome Sanger Institute for the COVID-19 Genomics UK (COG-UK) Consortium | Harper VanSteenhouse, Yumi Kasai, David Gray, Carol Clugston, Anna Dominiczak and Alex Alderton, Roberto Amato, Sonia Goncalves, Ewan Harrison, David K. Jackson, Ian Johnston, Dominic Kwiatkowski, Cordelia Langford, John Sillitoe on behalf of the Wellcome Sanger Institute COVID-19 Surveillance Team |
| EPI_ISL_777656 | Lighthouse Lab in Cambridge | Wellcome Sanger Institute for the COVID-19 Genomics UK (COG-UK) Consortium | Rob Howes, The Lighthouse Lab in Cambridge and Alex Alderton, Roberto Amato, Sonia Goncalves, Ewan Harrison, David K. Jackson, Ian Johnston, Dominic Kwiatkowski, Cordelia Langford, John Sillitoe on behalf of the Wellcome Sanger Institute COVID-19 Surveillance Team |
| EPI_ISL_777657, EPI_ISL_777658, EPI_ISL_777660 | Lighthouse Lab in Glasgow | Wellcome Sanger Institute for the COVID-19 Genomics UK (COG-UK) Consortium | Harper VanSteenhouse, Yumi Kasai, David Gray, Carol Clugston, Anna Dominiczak and Alex Alderton, Roberto Amato, Sonia Goncalves, Ewan Harrison, David K. Jackson, Ian Johnston, Dominic Kwiatkowski, Cordelia Langford, John Sillitoe on behalf of the Wellcome Sanger Institute COVID-19 Surveillance Team |
| EPI_ISL_777661 | Lighthouse Lab in Cambridge | Wellcome Sanger Institute for the COVID-19 Genomics UK (COG-UK) Consortium | Rob Howes, The Lighthouse Lab in Cambridge and Alex Alderton, Roberto Amato, Sonia Goncalves, Ewan Harrison, David K. Jackson, Ian Johnston, Dominic Kwiatkowski, Cordelia Langford, John Sillitoe on behalf of the Wellcome Sanger Institute COVID-19 Surveillance Team |
| EPI_ISL_777662, EPI_ISL_777663 | Lighthouse Lab in Glasgow | Wellcome Sanger Institute for the COVID-19 Genomics UK (COG-UK) Consortium | Harper VanSteenhouse, Yumi Kasai, David Gray, Carol Clugston, Anna Dominiczak and Alex Alderton, Roberto Amato, Sonia Goncalves, Ewan Harrison, David K. Jackson, Ian Johnston, Dominic Kwiatkowski, Cordelia Langford, John Sillitoe on behalf of the Wellcome Sanger Institute COVID-19 Surveillance Team |
| EPI_ISL_777664, EPI_ISL_777665, EPI_ISL_777666 | Lighthouse Lab in Cambridge | Wellcome Sanger Institute for the COVID-19 Genomics UK (COG-UK) Consortium | Rob Howes, The Lighthouse Lab in Cambridge and Alex Alderton, Roberto Amato, Sonia Goncalves, Ewan Harrison, David K. Jackson, Ian Johnston, Dominic Kwiatkowski, Cordelia Langford, John Sillitoe on behalf of the Wellcome Sanger Institute COVID-19 Surveillance Team |
| EPI_ISL_777667 | Lighthouse Lab in Glasgow | Wellcome Sanger Institute for the COVID-19 Genomics UK (COG-UK) Consortium | Harper VanSteenhouse, Yumi Kasai, David Gray, Carol Clugston, Anna Dominiczak and Alex Alderton, Roberto Amato, Sonia Goncalves, Ewan Harrison, David K. Jackson, Ian Johnston, Dominic Kwiatkowski, Cordelia Langford, John Sillitoe on behalf of the Wellcome Sanger Institute COVID-19 Surveillance Team |
| EPI_ISL_777668 | Lighthouse Lab in Cambridge | Wellcome Sanger Institute for the COVID-19 Genomics UK (COG-UK) Consortium | Rob Howes, The Lighthouse Lab in Cambridge and Alex Alderton, Roberto Amato, Sonia Goncalves, Ewan Harrison, David K. Jackson, Ian Johnston, Dominic Kwiatkowski, Cordelia Langford, John Sillitoe on behalf of the Wellcome Sanger Institute COVID-19 Surveillance Team |
| EPI_ISL_777670 | Lighthouse Lab in Glasgow | Wellcome Sanger Institute for the COVID-19 Genomics UK (COG-UK) Consortium | Harper VanSteenhouse, Yumi Kasai, David Gray, Carol Clugston, Anna Dominiczak and Alex Alderton, Roberto Amato, Sonia Goncalves, Ewan Harrison, David K. Jackson, Ian Johnston, Dominic Kwiatkowski, Cordelia Langford, John Sillitoe on behalf of the Wellcome Sanger Institute COVID-19 Surveillance Team |

| | | | |
|---|---|---|---|
| EPI_ISL_777671, EPI_ISL_777673, EPI_ISL_777674, EPI_ISL_777677, EPI_ISL_777678, EPI_ISL_777680, EPI_ISL_777682, EPI_ISL_777683, EPI_ISL_777685, EPI_ISL_777687, EPI_ISL_777688, EPI_ISL_777689, EPI_ISL_777690, EPI_ISL_777693, EPI_ISL_777698, EPI_ISL_777699, EPI_ISL_777701, EPI_ISL_777702, EPI_ISL_777704, EPI_ISL_777705, EPI_ISL_777707, EPI_ISL_777708, EPI_ISL_777710, EPI_ISL_777711, EPI_ISL_777714, EPI_ISL_777715, EPI_ISL_777716, EPI_ISL_777717, EPI_ISL_777718, EPI_ISL_777719, EPI_ISL_777720, EPI_ISL_777721, EPI_ISL_777722, EPI_ISL_777724, EPI_ISL_777727, EPI_ISL_777729, EPI_ISL_777730, EPI_ISL_777732, EPI_ISL_777735, EPI_ISL_777738, EPI_ISL_777739, EPI_ISL_777740, EPI_ISL_777741, EPI_ISL_777743, EPI_ISL_777744, EPI_ISL_777746, EPI_ISL_777747, EPI_ISL_777749, EPI_ISL_777750, EPI_ISL_777751, EPI_ISL_777752, EPI_ISL_777753, EPI_ISL_777755, EPI_ISL_777756, EPI_ISL_777757, EPI_ISL_777758, EPI_ISL_777759, EPI_ISL_777761, EPI_ISL_777763, EPI_ISL_777764, EPI_ISL_777765, EPI_ISL_777766, EPI_ISL_777770, EPI_ISL_777771, EPI_ISL_777772, EPI_ISL_777774, EPI_ISL_777775, EPI_ISL_777777, EPI_ISL_777778, EPI_ISL_777779, EPI_ISL_777780, EPI_ISL_777781, EPI_ISL_777782, EPI_ISL_777783, EPI_ISL_777784, EPI_ISL_777785, EPI_ISL_777787, EPI_ISL_777788, EPI_ISL_777789, EPI_ISL_777790, EPI_ISL_777791, EPI_ISL_777793, EPI_ISL_777794, EPI_ISL_777797, EPI_ISL_777798, EPI_ISL_777800, EPI_ISL_777803, EPI_ISL_777804, EPI_ISL_777805, EPI_ISL_777806, EPI_ISL_777809, EPI_ISL_777810, EPI_ISL_777811, EPI_ISL_777813, EPI_ISL_777814, EPI_ISL_777815, EPI_ISL_777817, EPI_ISL_777818, EPI_ISL_777819, EPI_ISL_777820, EPI_ISL_777822, EPI_ISL_777823, EPI_ISL_777825, EPI_ISL_777827, EPI_ISL_777828, EPI_ISL_777829, EPI_ISL_777831, EPI_ISL_777832, EPI_ISL_777834, EPI_ISL_777835, EPI_ISL_777837, EPI_ISL_777838, EPI_ISL_777839, EPI_ISL_777842, EPI_ISL_777844, EPI_ISL_777848, EPI_ISL_777850, EPI_ISL_777851, EPI_ISL_777852, EPI_ISL_777853, EPI_ISL_777854, EPI_ISL_777855, EPI_ISL_777856, EPI_ISL_777857, EPI_ISL_777858, EPI_ISL_777859, EPI_ISL_777860, EPI_ISL_777861, EPI_ISL_777862, EPI_ISL_777863, EPI_ISL_777866, EPI_ISL_777867, EPI_ISL_777868, EPI_ISL_777869, EPI_ISL_777870, EPI_ISL_777871, EPI_ISL_777872, EPI_ISL_777873, EPI_ISL_777874, EPI_ISL_777875, EPI_ISL_777878, EPI_ISL_777879, EPI_ISL_777880, EPI_ISL_777884, EPI_ISL_777885, EPI_ISL_777888, EPI_ISL_777889, EPI_ISL_777890, EPI_ISL_777894, EPI_ISL_777895, EPI_ISL_777896, EPI_ISL_777897, EPI_ISL_777899, EPI_ISL_777900, EPI_ISL_777901, EPI_ISL_777903, EPI_ISL_777904, EPI_ISL_777906, EPI_ISL_777907, EPI_ISL_777908, EPI_ISL_777909, EPI_ISL_777911, EPI_ISL_777912, EPI_ISL_777914, EPI_ISL_777916, EPI_ISL_777917, EPI_ISL_777918, EPI_ISL_777919, EPI_ISL_777920, EPI_ISL_777921, EPI_ISL_777922, EPI_ISL_777926, EPI_ISL_777927, EPI_ISL_777929, EPI_ISL_777930, EPI_ISL_777931, EPI_ISL_777932, EPI_ISL_777933, EPI_ISL_777935, EPI_ISL_777936, EPI_ISL_777937, EPI_ISL_777938, EPI_ISL_777939, EPI_ISL_777941, EPI_ISL_777943, EPI_ISL_777944, EPI_ISL_777945, EPI_ISL_777946, EPI_ISL_777947, EPI_ISL_777948, EPI_ISL_777949, EPI_ISL_777950, EPI_ISL_777951, EPI_ISL_777952, EPI_ISL_777954, EPI_ISL_777956, EPI_ISL_777958, EPI_ISL_777959, EPI_ISL_777961, EPI_ISL_777964, EPI_ISL_777965, EPI_ISL_777966, EPI_ISL_777968, EPI_ISL_777970, EPI_ISL_777971, EPI_ISL_777973, EPI_ISL_777974, EPI_ISL_777975, EPI_ISL_777977, EPI_ISL_777978, EPI_ISL_777981, EPI_ISL_777983, EPI_ISL_777984, EPI_ISL_777987, EPI_ISL_777988, EPI_ISL_777990, EPI_ISL_777992, EPI_ISL_777996, EPI_ISL_777999, EPI_ISL_778000, EPI_ISL_778001, EPI_ISL_778002, EPI_ISL_778010, EPI_ISL_778011, EPI_ISL_778012, EPI_ISL_778013, EPI_ISL_778015, EPI_ISL_778017, EPI_ISL_778019, EPI_ISL_778020, EPI_ISL_778023, EPI_ISL_778027, EPI_ISL_778028, EPI_ISL_778030, EPI_ISL_778037, EPI_ISL_778038, EPI_ISL_778039, EPI_ISL_778041 | | | |
| see above | Lighthouse Lab in Cambridge | Wellcome Sanger Institute for the COVID-19 Genomics UK (COG-UK) Consortium | Rob Howes, The Lighthouse Lab in Cambridge and Alex Alderton, Roberto Amato, Sonia Goncalves, Ewan Harrison, David K. Jackson, Ian Johnston, Dominic Kwiatkowski, Cordelia Langford, John Sillitoe on behalf of the Wellcome Sanger Institute COVID-19 Surveillance Team |
| EPI_ISL_778051 | Lighthouse Lab in Alderley Park | Wellcome Sanger Institute for the COVID-19 Genomics UK (COG-UK) Consortium | Jacquelyn Wynn, Mairead Hyland, The Lighthouse Lab in Alderley Park and Alex Alderton, Roberto Amato, Sonia Goncalves, Ewan Harrison, David K. Jackson, Ian Johnston, Dominic Kwiatkowski, Cordelia Langford, John Sillitoe on behalf of the Wellcome Sanger Institute COVID-19 Surveillance Team |
| EPI_ISL_778056, EPI_ISL_778057, EPI_ISL_778061, EPI_ISL_778066, EPI_ISL_778067, EPI_ISL_778071, EPI_ISL_778072, EPI_ISL_778075, EPI_ISL_778081, EPI_ISL_778087, EPI_ISL_778091, EPI_ISL_778094, EPI_ISL_778099, EPI_ISL_778103, EPI_ISL_778105, EPI_ISL_778110, EPI_ISL_778112 | | | |
| see above | Lighthouse Lab in Cambridge | Wellcome Sanger Institute for the COVID-19 Genomics UK (COG-UK) Consortium | Rob Howes, The Lighthouse Lab in Cambridge and Alex Alderton, Roberto Amato, Sonia Goncalves, Ewan Harrison, David K. Jackson, Ian Johnston, Dominic Kwiatkowski, Cordelia Langford, John Sillitoe on behalf of the Wellcome Sanger Institute COVID-19 Surveillance Team |
| EPI_ISL_778113 | Lighthouse Lab in Alderley Park | Wellcome Sanger Institute for the COVID-19 Genomics UK (COG-UK) Consortium | Jacquelyn Wynn, Mairead Hyland, The Lighthouse Lab in Alderley Park and Alex Alderton, Roberto Amato, Sonia Goncalves, Ewan Harrison, David K. Jackson, Ian Johnston, Dominic Kwiatkowski, Cordelia Langford, John Sillitoe on behalf of the Wellcome Sanger Institute COVID-19 Surveillance Team |
| EPI_ISL_778120, EPI_ISL_778125, EPI_ISL_778126, EPI_ISL_778130, EPI_ISL_778135, EPI_ISL_778137, EPI_ISL_778139, EPI_ISL_778143, EPI_ISL_778150, EPI_ISL_778152, EPI_ISL_778169, EPI_ISL_778172, EPI_ISL_778173, EPI_ISL_778174, EPI_ISL_778175, EPI_ISL_778177, EPI_ISL_778178, EPI_ISL_778182, EPI_ISL_778185, EPI_ISL_778188, EPI_ISL_778194, EPI_ISL_778197, EPI_ISL_778199, EPI_ISL_778200, EPI_ISL_778206, EPI_ISL_778210, EPI_ISL_778213, EPI_ISL_778214, EPI_ISL_778221, EPI_ISL_778222, EPI_ISL_778232, EPI_ISL_778235, EPI_ISL_778241, EPI_ISL_778243 | | | |
| see above | Lighthouse Lab in Cambridge | Wellcome Sanger Institute for the COVID-19 Genomics UK (COG-UK) Consortium | Rob Howes, The Lighthouse Lab in Cambridge and Alex Alderton, Roberto Amato, Sonia Goncalves, Ewan Harrison, David K. Jackson, Ian Johnston, Dominic Kwiatkowski, Cordelia Langford, John Sillitoe on behalf of the Wellcome Sanger Institute COVID-19 Surveillance Team |
| EPI_ISL_778246, EPI_ISL_778256 | Lighthouse Lab in Alderley Park | Wellcome Sanger Institute for the COVID-19 Genomics UK (COG-UK) Consortium | Jacquelyn Wynn, Mairead Hyland, The Lighthouse Lab in Alderley Park and Alex Alderton, Roberto Amato, Sonia Goncalves, Ewan Harrison, David K. Jackson, Ian Johnston, Dominic Kwiatkowski, Cordelia Langford, John Sillitoe on behalf of the Wellcome Sanger Institute COVID-19 Surveillance Team |
| EPI_ISL_778257, EPI_ISL_778262, EPI_ISL_778265, EPI_ISL_778267, EPI_ISL_778271, EPI_ISL_778273 | Lighthouse Lab in Cambridge | Wellcome Sanger Institute for the COVID-19 Genomics UK (COG-UK) Consortium | Rob Howes, The Lighthouse Lab in Cambridge and Alex Alderton, Roberto Amato, Sonia Goncalves, Ewan Harrison, David K. Jackson, Ian Johnston, Dominic Kwiatkowski, Cordelia Langford, John Sillitoe on behalf of the Wellcome Sanger Institute COVID-19 Surveillance Team |
| EPI_ISL_778282 | Lighthouse Lab in Alderley Park | Wellcome Sanger Institute for the COVID-19 Genomics UK (COG-UK) Consortium | Jacquelyn Wynn, Mairead Hyland, The Lighthouse Lab in Alderley Park and Alex Alderton, Roberto Amato, Sonia Goncalves, Ewan Harrison, David K. Jackson, Ian Johnston, Dominic Kwiatkowski, Cordelia Langford, John Sillitoe on behalf of the Wellcome Sanger Institute COVID-19 Surveillance Team |
| EPI_ISL_778284, EPI_ISL_778285, EPI_ISL_778291 | Lighthouse Lab in Cambridge | Wellcome Sanger Institute for the COVID-19 Genomics UK (COG-UK) Consortium | Rob Howes, The Lighthouse Lab in Cambridge and Alex Alderton, Roberto Amato, Sonia Goncalves, Ewan Harrison, David K. Jackson, Ian Johnston, Dominic Kwiatkowski, Cordelia Langford, John Sillitoe on behalf of the Wellcome Sanger Institute COVID-19 Surveillance Team |
| EPI_ISL_778295 | Lighthouse Lab in Alderley Park | Wellcome Sanger Institute for the COVID-19 Genomics UK (COG-UK) Consortium | Jacquelyn Wynn, Mairead Hyland, The Lighthouse Lab in Alderley Park and Alex Alderton, Roberto Amato, Sonia Goncalves, Ewan Harrison, David K. Jackson, Ian Johnston, Dominic Kwiatkowski, Cordelia Langford, John Sillitoe on behalf of the Wellcome Sanger Institute COVID-19 Surveillance Team |
| EPI_ISL_778296 | Lighthouse Lab in Cambridge | Wellcome Sanger Institute for the COVID-19 Genomics UK (COG-UK) Consortium | Rob Howes, The Lighthouse Lab in Cambridge and Alex Alderton, Roberto Amato, Sonia Goncalves, Ewan Harrison, David K. Jackson, Ian Johnston, Dominic Kwiatkowski, Cordelia Langford, John Sillitoe on behalf of the Wellcome Sanger Institute COVID-19 Surveillance Team |
| EPI_ISL_778830, EPI_ISL_778832, EPI_ISL_778833, EPI_ISL_778839 | AIID | Irish Coronavirus Sequencing Consortium-Teagasc Grange | Matthew McCabe, Aljandro Abner Garcia Leon, Fiona Crispie, Calum Walsh, Michael Carr, John Kenny, Paul Cotter, Patrick Mallon, Gabriel Gonzalez |
| EPI_ISL_778868 | Virology Lab, Ospedali Riuniti di Ancona | Dipartimento di Scienze Biomediche e Cliniche, L.Sacco, Università di Milano | Alessia Lai, Annalisa Bergna, Gianni Zehender, Claudia Balotta, Sara Caucci, Laura Di Sante, Roberta Longo, Sofia Maria Luigia Tiano, Massimo Galli, Patrizia Bagnarelli, Stefano Menzo |
| EPI_ISL_778869 | Virology Lab, Ospedali Riuniti, Ancona | Dipartimento di Scenze Biomediche e Cliniche, L. Sacco, Università di Milano | Alessia Lai, Annalisa Bergna, Gianni Zehender, Claudia Balotta, Sara Caucci, Laura Di Sante, Roberta Longo, Sofia Maria Luigia Tiano, Massimo Galli, Patrizia Bagnarelli, Stefano Menzo |
| EPI_ISL_779109, EPI_ISL_779110, EPI_ISL_779111, EPI_ISL_779112, EPI_ISL_779113, EPI_ISL_779114, EPI_ISL_779115, EPI_ISL_779116 | LSUHS Emerging Viral Threat Laboratory | Microbial Genome Sequencing Center | Jennifer L. Carroll, Jeremy P. Kamil, Camille F. Abshire, Maarten Van Diest, Andrew D. Yurochko, Martin J. Sapp, Rona S. Scott, Christopher G. Kevil, Daniel J. Snyder, Vaughn S. Cooper, John A. Vanchiere |
| EPI_ISL_779117, EPI_ISL_779118, EPI_ISL_779119, EPI_ISL_779120, EPI_ISL_779121, EPI_ISL_779122, EPI_ISL_779123, EPI_ISL_779124, EPI_ISL_779125, EPI_ISL_779126, EPI_ISL_779127, EPI_ISL_779128 | | | |
| see above | LSUHS Emerging Viral Threat Laboratory | Microbial Genome Sequencing Center | Maarten Van Diest, Jeremy P. Kamil, Jennifer L. Carroll, Camille F. Abshire, Andrew D. Yurochko, Martin J. Sapp, Rona S. Scott, Christopher G. Kevil, Daniel J. Snyder, Vaughn S. Cooper, John A. Vanchiere |
| EPI_ISL_779396 | Pathology North - Royal North Shore Hospital - NSW Health Pathology | NSW Health Pathology - Institute of Clinical Pathology and Medical Research; Westmead Hospital; University of Sydney | CIDM-PH et al. |
| EPI_ISL_779408 | Royal Darwin Hospital Pathology | MDU-PHL | Meumann, E., Caly L., Seemann T., Sait, M.L., Druce J., Sherry, N.L. |
| EPI_ISL_779615 | Victorian Infectious Diseases Reference Laboratory (VIDRL) | VIDRL and MDU-PHL | Caly L., Seemann T., Sait, M.L., Druce J., Sherry, N.L. |
| EPI_ISL_779616 | Microbiological Diagnostic Unit - Public Health Laboratory (MDU-PHL) | MDU-PHL | Seemann T., Sait, M.L., Sherry, N.L. |
| EPI_ISL_779617 | Victorian Infectious Diseases Reference Laboratory (VIDRL) | VIDRL and MDU-PHL | Caly L., Seemann T., Sait, M.L., Druce J., Sherry, N.L. |
| EPI_ISL_779627 | Microbiological Diagnostic Unit - Public Health Laboratory (MDU-PHL) | MDU-PHL | Seemann T., Sait, M.L., Sherry, N.L. |
| EPI_ISL_779785 | CHU Poitiers | CNR Virus des Infections Respiratoires - France SUD | Antonin Bal, Gregory Destras, Gwendolyne Burfin, Hadrien Règue, Quentin Semanas, Martine Valette, Bruno Lina, Agnès Beby-Defaux, Magali Garcia, Clément Jousselin, Nicolas Lévêque, Laurence Josset |
| EPI_ISL_779786 | CHU Bordeaux | CNR Virus des Infections Respiratoires - France SUD | Antonin Bal, Gregory Destras, Gwendolyne Burfin, Hadrien Règue, Quentin Semanas, Martine Valette, Bruno Lina, Pantxika Bellecave, Camille Ciccone, Isabelle Garrigue, Marie-Edith Lafon, Pascale Trimoulet, Laurence Josset |
| EPI_ISL_779787, EPI_ISL_779788, EPI_ISL_779789, EPI_ISL_779790, EPI_ISL_779791, EPI_ISL_779792, EPI_ISL_779793 | CNR Virus des Infections Respiratoires - France SUD | CNR Virus des Infections Respiratoires - France SUD | Antonin Bal, Gregory Destras, Gwendolyne Burfin, Hadrien Règue, Quentin Semanas, Martine Valette, Bruno Lina, Laurence Josset |
| EPI_ISL_779839, EPI_ISL_779840, EPI_ISL_779841, EPI_ISL_779842, EPI_ISL_779843, EPI_ISL_779844 | CHU Bordeaux | CNR Virus des Infections Respiratoires - France SUD | Antonin Bal, Gregory Destras, Gwendolyne Burfin, Hadrien Règue, Quentin Semanas, Martine Valette, Bruno Lina, Pantxika Bellecave, Camille Ciccone, Isabelle Garrigue, Marie-Edith Lafon, Pascale Trimoulet, Laurence Josset |

| | | | |
|---|---|---|---|
| EPI_ISL_779866, EPI_ISL_779891, EPI_ISL_779908, EPI_ISL_779909, EPI_ISL_779921 | Servicio de Microbiología, Hospital Universitario Son Espases | SeqCOVID-SPAIN consortium/IBV(CSIC) | Carla López-Causapé, Jordi Reina, Antonio Oliver and SeqCOVID-SPAIN consortium |
| EPI_ISL_779931, EPI_ISL_779952, EPI_ISL_779953 | Center of Medical Microbiology, Virology, and Hospital Hygiene, University of Duesseldorf | Center of Medical Microbiology, Virology, and Hospital Hygiene, University of Duesseldorf | Maximilian Damagnez, Alexander Dilthey, Ashley-Jane Duplessis, Torsten Houwaart, Lisanna Hülse, Malte Kohns Vasconcelos, Nadine Lübke, Jessica Nicolai, Klaus Pfeffer, Daniel Strelow, Teresa Tamayo, Jörg Timm, Andreas Walker, Tobias Wienemann |
| EPI_ISL_781185, EPI_ISL_781188, EPI_ISL_781199, EPI_ISL_781221, EPI_ISL_781223, EPI_ISL_781260, EPI_ISL_781263, EPI_ISL_781279, EPI_ISL_781281, EPI_ISL_781287, EPI_ISL_781299, EPI_ISL_781304, EPI_ISL_781306, EPI_ISL_781308, EPI_ISL_781327, EPI_ISL_781332 | | | |
| see above | Lighthouse Lab in Cambridge | Wellcome Sanger Institute for the COVID-19 Genomics UK (COG-UK) Consortium | Rob Howes, The Lighthouse Lab in Cambridge and Alex Alderton, Roberto Amato, Sonia Goncalves, Ewan Harrison, David K. Jackson, Ian Johnston, Dominic Kwiatkowski, Cordelia Langford, John Sillitoe on behalf of the Wellcome Sanger Institute COVID-19 Surveillance Team |
| EPI_ISL_781334 | Lighthouse Lab in Milton Keynes | Wellcome Sanger Institute for the COVID-19 Genomics UK (COG-UK) Consortium | The Lighthouse Lab in Milton Keynes and Alex Alderton, Roberto Amato, Sonia Goncalves, Ewan Harrison, David K. Jackson, Ian Johnston, Dominic Kwiatkowski, Cordelia Langford, John Sillitoe on behalf of the Wellcome Sanger Institute COVID-19 Surveillance Team |
| EPI_ISL_781340, EPI_ISL_781342, EPI_ISL_781358, EPI_ISL_781364, EPI_ISL_781368 | Lighthouse Lab in Cambridge | Wellcome Sanger Institute for the COVID-19 Genomics UK (COG-UK) Consortium | Rob Howes, The Lighthouse Lab in Cambridge and Alex Alderton, Roberto Amato, Sonia Goncalves, Ewan Harrison, David K. Jackson, Ian Johnston, Dominic Kwiatkowski, Cordelia Langford, John Sillitoe on behalf of the Wellcome Sanger Institute COVID-19 Surveillance Team |
| EPI_ISL_781373 | Lighthouse Lab in Milton Keynes | Wellcome Sanger Institute for the COVID-19 Genomics UK (COG-UK) Consortium | The Lighthouse Lab in Milton Keynes and Alex Alderton, Roberto Amato, Sonia Goncalves, Ewan Harrison, David K. Jackson, Ian Johnston, Dominic Kwiatkowski, Cordelia Langford, John Sillitoe on behalf of the Wellcome Sanger Institute COVID-19 Surveillance Team |
| EPI_ISL_781376, EPI_ISL_781385, EPI_ISL_781396, EPI_ISL_781402, EPI_ISL_781407, EPI_ISL_781409, EPI_ISL_781413 | Lighthouse Lab in Cambridge | Wellcome Sanger Institute for the COVID-19 Genomics UK (COG-UK) Consortium | Rob Howes, The Lighthouse Lab in Cambridge and Alex Alderton, Roberto Amato, Sonia Goncalves, Ewan Harrison, David K. Jackson, Ian Johnston, Dominic Kwiatkowski, Cordelia Langford, John Sillitoe on behalf of the Wellcome Sanger Institute COVID-19 Surveillance Team |
| EPI_ISL_781421 | Lighthouse Lab in Milton Keynes | Wellcome Sanger Institute for the COVID-19 Genomics UK (COG-UK) Consortium | The Lighthouse Lab in Milton Keynes and Alex Alderton, Roberto Amato, Sonia Goncalves, Ewan Harrison, David K. Jackson, Ian Johnston, Dominic Kwiatkowski, Cordelia Langford, John Sillitoe on behalf of the Wellcome Sanger Institute COVID-19 Surveillance Team |
| EPI_ISL_781426, EPI_ISL_781432, EPI_ISL_781436, EPI_ISL_781438, EPI_ISL_781440, EPI_ISL_781441, EPI_ISL_781442, EPI_ISL_781466, EPI_ISL_781471, EPI_ISL_781472, EPI_ISL_781490, EPI_ISL_781494 | | | |
| see above | Lighthouse Lab in Cambridge | Wellcome Sanger Institute for the COVID-19 Genomics UK (COG-UK) Consortium | Rob Howes, The Lighthouse Lab in Cambridge and Alex Alderton, Roberto Amato, Sonia Goncalves, Ewan Harrison, David K. Jackson, Ian Johnston, Dominic Kwiatkowski, Cordelia Langford, John Sillitoe on behalf of the Wellcome Sanger Institute COVID-19 Surveillance Team |
| EPI_ISL_781496 | Lighthouse Lab in Milton Keynes | Wellcome Sanger Institute for the COVID-19 Genomics UK (COG-UK) Consortium | The Lighthouse Lab in Milton Keynes and Alex Alderton, Roberto Amato, Sonia Goncalves, Ewan Harrison, David K. Jackson, Ian Johnston, Dominic Kwiatkowski, Cordelia Langford, John Sillitoe on behalf of the Wellcome Sanger Institute COVID-19 Surveillance Team |
| EPI_ISL_781498, EPI_ISL_781512, EPI_ISL_781514, EPI_ISL_781519, EPI_ISL_781523, EPI_ISL_781524, EPI_ISL_781528, EPI_ISL_781538, EPI_ISL_781542, EPI_ISL_781543, EPI_ISL_781545, EPI_ISL_781546, EPI_ISL_781552, EPI_ISL_781553, EPI_ISL_781557, EPI_ISL_781565, EPI_ISL_781566, EPI_ISL_781581, EPI_ISL_781587, EPI_ISL_781590, EPI_ISL_781591, EPI_ISL_781593, EPI_ISL_781595, EPI_ISL_781603, EPI_ISL_781606, EPI_ISL_781610, EPI_ISL_781620, EPI_ISL_781622, EPI_ISL_781637, EPI_ISL_781638, EPI_ISL_781642, EPI_ISL_781645, EPI_ISL_781656 | | | |
| see above | Lighthouse Lab in Cambridge | Wellcome Sanger Institute for the COVID-19 Genomics UK (COG-UK) Consortium | Rob Howes, The Lighthouse Lab in Cambridge and Alex Alderton, Roberto Amato, Sonia Goncalves, Ewan Harrison, David K. Jackson, Ian Johnston, Dominic Kwiatkowski, Cordelia Langford, John Sillitoe on behalf of the Wellcome Sanger Institute COVID-19 Surveillance Team |
| EPI_ISL_781665 | Lighthouse Lab in Milton Keynes | Wellcome Sanger Institute for the COVID-19 Genomics UK (COG-UK) Consortium | The Lighthouse Lab in Milton Keynes and Alex Alderton, Roberto Amato, Sonia Goncalves, Ewan Harrison, David K. Jackson, Ian Johnston, Dominic Kwiatkowski, Cordelia Langford, John Sillitoe on behalf of the Wellcome Sanger Institute COVID-19 Surveillance Team |
| EPI_ISL_781666 | Lighthouse Lab in Cambridge | Wellcome Sanger Institute for the COVID-19 Genomics UK (COG-UK) Consortium | Rob Howes, The Lighthouse Lab in Cambridge and Alex Alderton, Roberto Amato, Sonia Goncalves, Ewan Harrison, David K. Jackson, Ian Johnston, Dominic Kwiatkowski, Cordelia Langford, John Sillitoe on behalf of the Wellcome Sanger Institute COVID-19 Surveillance Team |
| EPI_ISL_781669 | Lighthouse Lab in Milton Keynes | Wellcome Sanger Institute for the COVID-19 Genomics UK (COG-UK) Consortium | The Lighthouse Lab in Milton Keynes and Alex Alderton, Roberto Amato, Sonia Goncalves, Ewan Harrison, David K. Jackson, Ian Johnston, Dominic Kwiatkowski, Cordelia Langford, John Sillitoe on behalf of the Wellcome Sanger Institute COVID-19 Surveillance Team |
| EPI_ISL_781678, EPI_ISL_781685, EPI_ISL_781691, EPI_ISL_781698, EPI_ISL_781703, EPI_ISL_781706, EPI_ISL_781707, EPI_ISL_781711, EPI_ISL_781712, EPI_ISL_781713, EPI_ISL_781715, EPI_ISL_781716, EPI_ISL_781728, EPI_ISL_781729, EPI_ISL_781734, EPI_ISL_781736, EPI_ISL_781741, EPI_ISL_781745, EPI_ISL_781759, EPI_ISL_781760, EPI_ISL_781762, EPI_ISL_781764, EPI_ISL_781766, EPI_ISL_781771, EPI_ISL_781772, EPI_ISL_781780, EPI_ISL_781794, EPI_ISL_781797, EPI_ISL_781805, EPI_ISL_781808, EPI_ISL_781809, EPI_ISL_781815, EPI_ISL_781822, EPI_ISL_781838, EPI_ISL_781842, EPI_ISL_781843, EPI_ISL_781844, EPI_ISL_781848, EPI_ISL_781867 | | | |
| see above | Lighthouse Lab in Cambridge | Wellcome Sanger Institute for the COVID-19 Genomics UK (COG-UK) Consortium | Rob Howes, The Lighthouse Lab in Cambridge and Alex Alderton, Roberto Amato, Sonia Goncalves, Ewan Harrison, David K. Jackson, Ian Johnston, Dominic Kwiatkowski, Cordelia Langford, John Sillitoe on behalf of the Wellcome Sanger Institute COVID-19 Surveillance Team |
| EPI_ISL_781872, EPI_ISL_781874, EPI_ISL_781877, EPI_ISL_781879, EPI_ISL_781881, EPI_ISL_781883, EPI_ISL_781884, EPI_ISL_781887, EPI_ISL_781889, EPI_ISL_781892, EPI_ISL_781893, EPI_ISL_781894, EPI_ISL_781896, EPI_ISL_781899, EPI_ISL_781900, EPI_ISL_781901, EPI_ISL_781902, EPI_ISL_781904, EPI_ISL_781905, EPI_ISL_781906, EPI_ISL_781911, EPI_ISL_781912, EPI_ISL_781917, EPI_ISL_781923, EPI_ISL_781926, EPI_ISL_781927, EPI_ISL_781929, EPI_ISL_781931, EPI_ISL_781932, EPI_ISL_781934, EPI_ISL_781935, EPI_ISL_781936, EPI_ISL_781937, EPI_ISL_781939, EPI_ISL_781943, EPI_ISL_781945, EPI_ISL_781949, EPI_ISL_781951, EPI_ISL_781954, EPI_ISL_781958, EPI_ISL_781959, EPI_ISL_781961, EPI_ISL_781962, EPI_ISL_781963, EPI_ISL_781964, EPI_ISL_781965, EPI_ISL_781966, EPI_ISL_781969, EPI_ISL_781970, EPI_ISL_781972, EPI_ISL_781974, EPI_ISL_781975, EPI_ISL_781976, EPI_ISL_781981, EPI_ISL_781983, EPI_ISL_781985, EPI_ISL_781986, EPI_ISL_781989, EPI_ISL_781991, EPI_ISL_781993, EPI_ISL_781995, EPI_ISL_781996, EPI_ISL_782002, EPI_ISL_782005, EPI_ISL_782006, EPI_ISL_782007, EPI_ISL_782010, EPI_ISL_782011, EPI_ISL_782012, EPI_ISL_782013, EPI_ISL_782016, EPI_ISL_782019, EPI_ISL_782020, EPI_ISL_782022, EPI_ISL_782023, EPI_ISL_782024, EPI_ISL_782025, EPI_ISL_782029, EPI_ISL_782033, EPI_ISL_782034, EPI_ISL_782041, EPI_ISL_782043, EPI_ISL_782045, EPI_ISL_782047, EPI_ISL_782048, EPI_ISL_782050, EPI_ISL_782053, EPI_ISL_782057, EPI_ISL_782060 | | | |
| see above | Lighthouse Lab in Alderley Park | Wellcome Sanger Institute for the COVID-19 Genomics UK (COG-UK) Consortium | Jacquelyn Wynn, Mairead Hyland, The Lighthouse Lab in Alderley Park and Alex Alderton, Roberto Amato, Sonia Goncalves, Ewan Harrison, David K. Jackson, Ian Johnston, Dominic Kwiatkowski, Cordelia Langford, John Sillitoe on behalf of the Wellcome Sanger Institute COVID-19 Surveillance Team |
| EPI_ISL_782061 | Lighthouse Lab in Milton Keynes | Wellcome Sanger Institute for the COVID-19 Genomics UK (COG-UK) Consortium | The Lighthouse Lab in Milton Keynes and Alex Alderton, Roberto Amato, Sonia Goncalves, Ewan Harrison, David K. Jackson, Ian Johnston, Dominic Kwiatkowski, Cordelia Langford, John Sillitoe on behalf of the Wellcome Sanger Institute COVID-19 Surveillance Team |
| EPI_ISL_782063, EPI_ISL_782068, EPI_ISL_782070, EPI_ISL_782071, EPI_ISL_782072, EPI_ISL_782076, EPI_ISL_782078, EPI_ISL_782082, EPI_ISL_782083, EPI_ISL_782085, EPI_ISL_782086, EPI_ISL_782089, EPI_ISL_782090, EPI_ISL_782091, EPI_ISL_782095, EPI_ISL_782096, EPI_ISL_782097, EPI_ISL_782099, EPI_ISL_782100, EPI_ISL_782102, EPI_ISL_782104, EPI_ISL_782108, EPI_ISL_782109, EPI_ISL_782110, EPI_ISL_782111, EPI_ISL_782117, EPI_ISL_782119, EPI_ISL_782120, EPI_ISL_782122, EPI_ISL_782124, EPI_ISL_782126, EPI_ISL_782132, EPI_ISL_782135, EPI_ISL_782139, EPI_ISL_782140, EPI_ISL_782141, EPI_ISL_782142, EPI_ISL_782148, EPI_ISL_782149, EPI_ISL_782150 | | | |
| see above | Lighthouse Lab in Alderley Park | Wellcome Sanger Institute for the COVID-19 Genomics UK (COG-UK) Consortium | Jacquelyn Wynn, Mairead Hyland, The Lighthouse Lab in Alderley Park and Alex Alderton, Roberto Amato, Sonia Goncalves, Ewan Harrison, David K. Jackson, Ian Johnston, Dominic Kwiatkowski, Cordelia Langford, John Sillitoe on behalf of the Wellcome Sanger Institute COVID-19 Surveillance Team |
| EPI_ISL_782327 | Lighthouse Lab in Glasgow | Wellcome Sanger Institute for the COVID-19 Genomics UK (COG-UK) Consortium | Harper VanSteenhouse, Yumi Kasai, David Gray, Carol Clugston, Anna Dominiczak and Alex Alderton, Roberto Amato, Sonia Goncalves, Ewan Harrison, David K. Jackson, Ian Johnston, Dominic Kwiatkowski, Cordelia Langford, John Sillitoe on behalf of the Wellcome Sanger Institute COVID-19 Surveillance Team |
| EPI_ISL_789012, EPI_ISL_789013 | Klinisk mikrobiologi, Laboratoriemedicin | The Public Health Agency of Sweden | Department of Microbiology, The Public Health Agency of Sweden |
| EPI_ISL_789017 | Klinisk Mikrobiologi | The Public Health Agency of Sweden | Department of Microbiology, The Public Health Agency of Sweden |
| EPI_ISL_789029, EPI_ISL_789030, EPI_ISL_789031, EPI_ISL_789032, EPI_ISL_789033, EPI_ISL_789034, EPI_ISL_789035 | Klinisk mikrobiologi, Laboratoriemedicin | The Public Health Agency of Sweden | Department of Microbiology, The Public Health Agency of Sweden |
| EPI_ISL_789036 | Klinisk mikrobiologi | The Public Health Agency of Sweden | Department of Microbiology, The Public Health Agency of Sweden |
| EPI_ISL_789048, EPI_ISL_789049, EPI_ISL_789050, EPI_ISL_789051, EPI_ISL_789052, EPI_ISL_789053, EPI_ISL_789055, EPI_ISL_789056, EPI_ISL_789057 | Klinisk mikrobiologi, Viruslab | The Public Health Agency of Sweden | Department of Microbiology, The Public Health Agency of Sweden |
| EPI_ISL_790570, EPI_ISL_790572, EPI_ISL_790576, EPI_ISL_790577, EPI_ISL_790578, EPI_ISL_790589, EPI_ISL_790594, EPI_ISL_790632, EPI_ISL_790656, EPI_ISL_790680, EPI_ISL_790681, EPI_ISL_790682, EPI_ISL_790683, EPI_ISL_790684, EPI_ISL_790685, EPI_ISL_790696, EPI_ISL_790701, EPI_ISL_790702, EPI_ISL_790706, EPI_ISL_790707, EPI_ISL_790708, EPI_ISL_790722, EPI_ISL_790723, EPI_ISL_790724, EPI_ISL_790730, EPI_ISL_790731, EPI_ISL_790732, EPI_ISL_790733, EPI_ISL_790738, EPI_ISL_790739, EPI_ISL_790744, EPI_ISL_790763, EPI_ISL_790764, EPI_ISL_790766, EPI_ISL_790785, EPI_ISL_790786, EPI_ISL_790789, EPI_ISL_790790, EPI_ISL_790798, EPI_ISL_790799, EPI_ISL_790815, EPI_ISL_790833, EPI_ISL_790834, EPI_ISL_790835, EPI_ISL_790836, EPI_ISL_790838, EPI_ISL_790851, EPI_ISL_790855, EPI_ISL_790874, EPI_ISL_790875, EPI_ISL_790893, EPI_ISL_790894, EPI_ISL_790895, EPI_ISL_790896, | | | |

| | | | |
|---|---|---|---|
| EPI_ISL_790897, EPI_ISL_790898, EPI_ISL_790899, EPI_ISL_790902, EPI_ISL_790913, EPI_ISL_790915, EPI_ISL_790916, EPI_ISL_790917, EPI_ISL_790919, EPI_ISL_790921, EPI_ISL_790922, EPI_ISL_790926, EPI_ISL_790933, EPI_ISL_790934, EPI_ISL_790938, EPI_ISL_790939, EPI_ISL_790954, EPI_ISL_790968, EPI_ISL_791044, EPI_ISL_791071, EPI_ISL_791072 | | | |
| see above | Dutch COVID-19 response team | National Institute for Public Health and the Environment (RIVM) | Adam Meijer, Harry Vennema, Jeroen Cremer, Sharon van den Brink, Bas van der Veer, AnneMarie van den Brandt, Florian Zwagemaker, Dennis Schmitz, Chantal Reusken, on behalf of the national COVID-19 response team |
| EPI_ISL_791183, EPI_ISL_791187, EPI_ISL_791188, EPI_ISL_791189, EPI_ISL_791204, EPI_ISL_791205, EPI_ISL_791206, EPI_ISL_791207, EPI_ISL_791210, EPI_ISL_791212, EPI_ISL_791213, EPI_ISL_791215, EPI_ISL_791217, EPI_ISL_791218, EPI_ISL_791219, EPI_ISL_791221 | | | |
| see above | Respiratory Virus Unit, National Infection Service, Public Health England | COVID-19 Genomics UK (COG-UK) Consortium | PHE Covid Sequencing Team |
| EPI_ISL_791280, EPI_ISL_791282, EPI_ISL_791283, EPI_ISL_791285, EPI_ISL_791290, EPI_ISL_791291, EPI_ISL_791292, EPI_ISL_791295, EPI_ISL_791296, EPI_ISL_791302, EPI_ISL_791309, EPI_ISL_791310, EPI_ISL_791311, EPI_ISL_791312, EPI_ISL_791313 | | | |
| see above | National Virus Reference Laboratory | Irish Coronavirus Sequencing Consortium - Teagasc Moorepark | Alejandro Abner Garcia Leon, Paul Cotter, Fiona Crispie, John Kenny, Paddy Mallon, Calum Walsh |
| EPI_ISL_791416 | Johns Hopkins Hospital Department of Pathology | Johns Hopkins Hospital Department of Pathology | C. Paul Morris, Chun Huai Luo, Adannaya Amadi, Nicholas Gallagher, Heba H. Mostafa |
| EPI_ISL_791999 | CHU - Hôpital Cavale Blanche | National Reference Center for Viruses of Respiratory Infections, Institut Pasteur, Paris | Marion Barbet, Sylvie Behillil, Méline Bizard, Angela Brisebarre, Camille Capel, Etienne Simon-Lorière, Vincent Enouf, Maud Vanpeene, Sylvie van der Werf, Léa Pilorgé |
| EPI_ISL_792000, EPI_ISL_792001, EPI_ISL_792002, EPI_ISL_792003, EPI_ISL_792004, EPI_ISL_792005, EPI_ISL_792006, EPI_ISL_792007 | Laboratoire de Virologie Hôpital Robert Debré | National Reference Center for Viruses of Respiratory Infections, Institut Pasteur, Paris | Marion Barbet, Sylvie Behillil, Méline Bizard, Angela Brisebarre, Camille Capel, Etienne Simon-Lorière, Vincent Enouf, Maud Vanpeene, Sylvie van der Werf, Andreoletti Laurent |
| EPI_ISL_792008, EPI_ISL_792009, EPI_ISL_792010, EPI_ISL_792011, EPI_ISL_792012, EPI_ISL_792013, EPI_ISL_792014, EPI_ISL_792015 | CH de Bethune - Laboratoire de Biologie Médicale | National Reference Center for Viruses of Respiratory Infections, Institut Pasteur, Paris | Marion Barbet, Sylvie Behillil, Méline Bizard, Angela Brisebarre, Camille Capel, Etienne Simon-Lorière, Vincent Enouf, Maud Vanpeene, Sylvie van der Werf, Léa Pilorgé |
| EPI_ISL_792024, EPI_ISL_792025, EPI_ISL_792027 | Pathology North - Royal North Shore Hospital - NSW Health Pathology | NSW Health Pathology - Institute of Clinical Pathology and Medical Research; Westmead Hospital; University of Sydney | CIDM-PH et al. |
| EPI_ISL_792028 | Laverty Pathology | NSW Health Pathology - Institute of Clinical Pathology and Medical Research; Westmead Hospital; University of Sydney | CIDM-PH et al. |
| EPI_ISL_792035 | South Eastern Area Laboratory Services (SEALS) | NSW Health Pathology - Institute of Clinical Pathology and Medical Research; Westmead Hospital; University of Sydney | CIDM-PH et al. |
| EPI_ISL_792040 | hospital | National Reference Center for Viruses of Respiratory Infections, Institut Pasteur, Paris | Marion Barbet, Sylvie Behillil, Méline Bizard, Angela Brisebarre, Camille Capel, Etienne Simon-Lorière, Vincent Enouf, Maud Vanpeene, Sylvie van der Werf,Laurent Roudière |
| EPI_ISL_792523, EPI_ISL_792524 | Laboratorio de Virología del Hospital de Niños Dr. Ricardo Gutierrez | Grupo de Genómica y Bioinformática del Instituto de Investigación de la Cadena Láctea CONICET-INTA on behalf of 'Proyecto Argentino Interinstitucional de genomica de SARS-CoV-2' (PAIS Consortium) | Amadio, AF, Eberhardt, MF; Irazoqui, M; Torres, C; Aulicino, P; König, G; Acevedo, ME; Alvarez Lopez, C; Alexay, S; Jacques, O; Mistchenko, AS, Goya, S; Nabaes Jodar, MS; Viegas, M. |
| EPI_ISL_794283, EPI_ISL_794284, EPI_ISL_794285, EPI_ISL_794286, EPI_ISL_794287, EPI_ISL_794288, EPI_ISL_794289, EPI_ISL_794290, EPI_ISL_794291, EPI_ISL_794292, EPI_ISL_794293, EPI_ISL_794294, EPI_ISL_794295 | | | |
| see above | Columbia University Irving Medical Center | Wadsworth Center, New York State Department.of Health | Kirsten St. George, Daryl M. Lamson, Alexis Russel, Matthew Shudt, Melissa A Leisner, Jonathan Plitnick, Navjot Singh, John Kelly, Sara Griesemer, Erasmus Schneider, Erica Lasek-Nesselquist |
| EPI_ISL_794724, EPI_ISL_794727, EPI_ISL_794729, EPI_ISL_794731, EPI_ISL_794732, EPI_ISL_794733 | PathWest Laboratory Medicine WA | PathWest Laboratory Medicine WA Microbial Surveillance Unit | PathWest Laboratory Medicine WA Microbial Surveillance Unit |
| EPI_ISL_794754, EPI_ISL_794774, EPI_ISL_794775, EPI_ISL_794776, EPI_ISL_794777, EPI_ISL_794778, EPI_ISL_794784, EPI_ISL_794794, EPI_ISL_794795, EPI_ISL_794796, EPI_ISL_794797, EPI_ISL_794798, EPI_ISL_794799, EPI_ISL_794800, EPI_ISL_794801, EPI_ISL_794802, EPI_ISL_794809, EPI_ISL_794810, EPI_ISL_794811, EPI_ISL_794812, EPI_ISL_794813, EPI_ISL_794814, EPI_ISL_794815, EPI_ISL_794816, EPI_ISL_794817 | | | |
| see above | Istituto Zooprofilattico Sperimentale della Puglia e della Basilicata | Istituto Zooprofilattico Sperimentale della Puglia e della Basilicata | Parisi A., Bianco A., Capozzi L., Del Sambro L., Manzulli V, Rondinone V., Pace L., Cipolletta D., Galante D. |
| EPI_ISL_796653 | Department of Medical Microbiology - section Molde, Molde Hospital | Norwegian Institute of Public Health, Department of Virology | Kathrine Stene-Johansen, Kamilla Heddeland Instefjord, Hilde Elshaug, Atiya R Ali,Marie Paulsen Madsen, Rasmus Riis Kopperud, Hilde Vollan, Karoline Bragstad, Olav Hungnes |
| EPI_ISL_796655, EPI_ISL_796656, EPI_ISL_796657 | Norwegian Institute of Public Health, Department of Virology | Norwegian Institute of Public Health, Department of Virology | Kathrine Stene-Johansen, Kamilla Heddeland Instefjord, Hilde Elshaug, Atiya R Ali,Marie Paulsen Madsen, Rasmus Riis Kopperud, Hilde Vollan, Karoline Bragstad, Olav Hungnes |
| EPI_ISL_796671 | Furst Medical Laboratory | Norwegian Institute of Public Health, Department of Virology | Kathrine Stene-Johansen, Kamilla Heddeland Instefjord, Hilde Elshaug, Atiya R Ali,Marie Paulsen Madsen, Rasmus Riis Kopperud, Hilde Vollan, Karoline Bragstad, Olav Hungnes |
| EPI_ISL_796676 | Norwegian Institute of Public Health, Department of Virology | Norwegian Institute of Public Health, Department of Virology | Kathrine Stene-Johansen, Kamilla Heddeland Instefjord, Hilde Elshaug, Atiya R Ali,Marie Paulsen Madsen, Rasmus Riis Kopperud, Hilde Vollan, Karoline Bragstad, Olav Hungnes |
| EPI_ISL_796681, EPI_ISL_796682 | Department of Medical Microbiology - section Molde, Molde Hospital | Norwegian Institute of Public Health, Department of Virology | Kathrine Stene-Johansen, Kamilla Heddeland Instefjord, Hilde Elshaug, Atiya R Ali,Marie Paulsen Madsen, Rasmus Riis Kopperud, Hilde Vollan, Karoline Bragstad, Olav Hungnes |
| EPI_ISL_796687 | Department of Medical Microbiology, St. Olavs hospital | Norwegian Institute of Public Health, Department of Virology | Kathrine Stene-Johansen, Kamilla Heddeland Instefjord, Hilde Elshaug, Atiya R Ali,Marie Paulsen Madsen, Rasmus Riis Kopperud, Hilde Vollan, Karoline Bragstad, Olav Hungnes |
| EPI_ISL_796714, EPI_ISL_796716 | Hospital of Southern Norway - Kristiansand, Department of Medical Microbiology | Norwegian Institute of Public Health, Department of Virology | Kathrine Stene-Johansen, Kamilla Heddeland Instefjord, Hilde Elshaug, Atiya R Ali,Marie Paulsen Madsen, Rasmus Riis Kopperud, Hilde Vollan, Karoline Bragstad, Olav Hungnes |
| EPI_ISL_796719, EPI_ISL_796720, EPI_ISL_796721 | Department of Medical Microbiology, St. Olavs hospital | Norwegian Institute of Public Health, Department of Virology | Kathrine Stene-Johansen, Kamilla Heddeland Instefjord, Hilde Elshaug, Atiya R Ali,Marie Paulsen Madsen, Rasmus Riis Kopperud, Hilde Vollan, Karoline Bragstad, Olav Hungnes |
| EPI_ISL_796724, EPI_ISL_796726 | Furst Medical Laboratory | Norwegian Institute of Public Health, Department of Virology | Kathrine Stene-Johansen, Kamilla Heddeland Instefjord, Hilde Elshaug, Atiya R Ali,Marie Paulsen Madsen, Rasmus Riis Kopperud, Hilde Vollan, Karoline Bragstad, Olav Hungnes |
| EPI_ISL_796776 | Instituto Nacional de Saude (INSA) | Instituto Nacional de Saude (INSA) | Borges et al |
| EPI_ISL_799637 | Lighthouse Lab in Alderley Park | Wellcome Sanger Institute for the COVID-19 Genomics UK (COG-UK) Consortium | Jacquelyn Wynn, Mairead Hyland, The Lighthouse Lab in Alderley Park and Alex Alderton, Roberto Amato, Sonia Goncalves, Ewan Harrison, David K. Jackson, Ian Johnston, Dominic Kwiatkowski, Cordelia Langford, John Sillitoe on behalf of the Wellcome Sanger Institute COVID-19 Surveillance Team |
| EPI_ISL_799740 | Lighthouse Lab in Milton Keynes | Wellcome Sanger Institute for the COVID-19 Genomics UK (COG-UK) Consortium | The Lighthouse Lab in Milton Keynes and Alex Alderton, Roberto Amato, Sonia Goncalves, Ewan Harrison, David K. Jackson, Ian Johnston, Dominic Kwiatkowski, Cordelia Langford, John Sillitoe on behalf of the Wellcome Sanger Institute COVID-19 Surveillance Team |
| EPI_ISL_799877, EPI_ISL_799898, EPI_ISL_799908, EPI_ISL_799925, EPI_ISL_800575, EPI_ISL_800577, EPI_ISL_800578, EPI_ISL_800579 | Lighthouse Lab in Alderley Park | Wellcome Sanger Institute for the COVID-19 Genomics UK (COG-UK) Consortium | Jacquelyn Wynn, Mairead Hyland, The Lighthouse Lab in Alderley Park and Alex Alderton, Roberto Amato, Sonia Goncalves, Ewan Harrison, David K. Jackson, Ian Johnston, Dominic Kwiatkowski, Cordelia Langford, John Sillitoe on behalf of the Wellcome Sanger Institute COVID-19 Surveillance Team |
| EPI_ISL_801444, EPI_ISL_801445, | Dutch COVID-19 response team | Erasmus Medical Center | Bas Oude Munnink, Reina Sikkema, David Nieuwenhuijse, Irina Chestakova, Anne van der Linden, Marjan Boter, Emmanuelle Munger, Corine |

| | | | |
|---|---|---|---|
| EPI_ISL_801447, EPI_ISL_801449 | | | GeurtsvanKessel, Annemiek van der Eijk, Richard Molenkamp, Marion Koopmans, on behalf of the Dutch national COVID-19 response team. |
| EPI_ISL_801523, EPI_ISL_801536 | Sonic Reference Laboratory | Pathogen Discovery, Respiratory Viruses Branch, Division of Viral Diseases, Centers for Disease Control and Prevention | Ying Tao, Yan Li, Jing Zhang, Krista Queen, Anna Uehara, Peter Cook, Clinton R. Paden, Haibin Wang, Suxiang Tong |
| EPI_ISL_802520, EPI_ISL_802524, EPI_ISL_802525, EPI_ISL_802533 | Dutch COVID-19 response team | Erasmus Medical Center | Bas Oude Munnink, Reina Sikkema, David Nieuwenhuijse, Irina Chestakova, Anne van der Linden, Marjan Boter, Emmanuelle Munger, Corine GeurtsvanKessel, Annemiek van der Eijk, Richard Molenkamp, Marion Koopmans, on behalf of the Dutch national COVID-19 response team. |
| EPI_ISL_802573 | M Health Fairview | Minnesota Department of Health, Public Health Laboratory | Alexandra Lorentz, Jacob Garfin, Matt Plumb, and Xiong Wang |
| EPI_ISL_803902 | BBMP Urban PHC | Department of Neurovirology, National Institute of Mental Health and Neurosciences (NIMHANS) | Chitra Pattabiraman, Pramada Prasad, Anita S Desai, V Ravi |
| EPI_ISL_804038, EPI_ISL_804041, EPI_ISL_804042, EPI_ISL_804044, EPI_ISL_804048, EPI_ISL_804050, EPI_ISL_804051, EPI_ISL_804052 | SC (UCO) Igiene e Sanità Pubblica (funzione integrata con SC Microbiologia e Virologia) e Laboratory of Molecular Virology of the International Centre for Genetic Engineering and Biotechnology (ICGEB) | ARGO Laboratorio Genomica ed Epigenomica | Licastro D, Dal Monego S, Degasperi M, Marcello A, D'Agaro P |
| EPI_ISL_804227, EPI_ISL_804228, EPI_ISL_804229, EPI_ISL_804230, EPI_ISL_804231, EPI_ISL_804266, EPI_ISL_804267, EPI_ISL_804268, EPI_ISL_804269, EPI_ISL_804270 | Respiratory Virus Unit, National Infection Service, Public Health England | COVID-19 Genomics UK (COG-UK) Consortium | PHE Covid Sequencing Team |
| EPI_ISL_804370 | CHU Purpan - Laboratoire de Virologie - Institut Fédératif de Biologie | CHU Purpan - Laboratoire de Virologie - Institut Fédératif de Biologie | Latour J., Ranger N., Dubois M., Carcenac R., Harter A., Boyer P., Tremeaux P., Izopet J. |
| EPI_ISL_804610, EPI_ISL_804622, EPI_ISL_804626, EPI_ISL_804649, EPI_ISL_804709 | Michigan Department of Health and Human Services, Bureau of Laboratories | Michigan Department of Health and Human Services, Bureau of Laboratories | Blankenship HM, Riner D, Soehnlen MK |
| EPI_ISL_804815, EPI_ISL_804817, EPI_ISL_804818, EPI_ISL_804819, EPI_ISL_804822, EPI_ISL_804823, EPI_ISL_804824, EPI_ISL_804826, EPI_ISL_804828, EPI_ISL_804829, EPI_ISL_804832, EPI_ISL_804835, EPI_ISL_804838, EPI_ISL_804839, EPI_ISL_804842 | | | |
| see above | DB Diagnosticos do Brasil | Laboratório de Parasitologia Médica - Instituto de Medicina Tropical - Universidade de São Paulo | Nuno Faria, Ingra Morales Claro, Darlan Candido, Lucas A. Moyses Franco, Pamela dos Santos Andrade, Thais de Moura Coletti, Camila A. Maia da Silva, Flavia Cristina Sales, Erika Regina Manuli, Renato A. Santana, Nelson Gaburo, Cecília da Cunha Camilo, Nelson Abrahim Fraiji, Myuki Alfaia Esashika Crispim, Maria do Perpétuo Socorro Sampaio Carvalho, Andrew Rambaut, Nick Loman, Oliver G. Pybus, Ester C. Sabino; DB; HEMOAM; CDL; CADDE Genomic Network. |
| EPI_ISL_804856, EPI_ISL_804857, EPI_ISL_804858, EPI_ISL_804895, EPI_ISL_804942, EPI_ISL_804943, EPI_ISL_804944 | DC Public Health Lab/ Dept. of Forensic Sciences | DC Public Health Lab/ Dept. of Forensic Sciences | Scott Nguyen, Elizabeth Zelaya, Connie Maza, Monica Mann, Brittany Hamilton, David Payne, Jocelyn Hauser |
| EPI_ISL_806730 | Presidio Ospedaliero S.Liberatore Atri | Istituto Zooprofilattico Sperimentale dell'Abruzzo e Molise "G. Caporale" | Lorusso A, Marcacci M, Di Domenico M, Ancora M, Curini V, Mangone I, Rinaldi A, Di Pasquale A, Cammà C, Puglia I, Calistri P, Savini G |
| EPI_ISL_806731, EPI_ISL_806732 | SIESP DIPARTIMENTO DI PREVENZIONE CHIETI | Istituto Zooprofilattico Sperimentale dell'Abruzzo e Molise "G. Caporale" | Lorusso A, Marcacci M, Di Domenico M, Ancora M, Curini V, Mangone I, Rinaldi A, Di Pasquale A, Cammà C, Puglia I, Calistri P, Savini G |
| EPI_ISL_806733 | SIESP CHIETI - DRIVE IN CHIETI | Istituto Zooprofilattico Sperimentale dell'Abruzzo e Molise "G. Caporale" | Lorusso A, Marcacci M, Di Domenico M, Ancora M, Curini V, Mangone I, Rinaldi A, Di Pasquale A, Cammà C, Puglia I, Calistri P, Savini G |
| EPI_ISL_806734 | SIESP CHIETI - DRIVE IN LANCIANO | Istituto Zooprofilattico Sperimentale dell'Abruzzo e Molise "G. Caporale" | Lorusso A, Marcacci M, Di Domenico M, Ancora M, Curini V, Mangone I, Rinaldi A, Di Pasquale A, Cammà C, Puglia I, Calistri P, Savini G |
| EPI_ISL_806735, EPI_ISL_806736, EPI_ISL_806737, EPI_ISL_806738, EPI_ISL_806739, EPI_ISL_806740, EPI_ISL_806741, EPI_ISL_806742, EPI_ISL_806743, EPI_ISL_806744 | SIESP DIPARTIMENTO DI PREVENZIONE TERAMO | Istituto Zooprofilattico Sperimentale dell'Abruzzo e Molise "G. Caporale" | Lorusso A, Marcacci M, Di Domenico M, Ancora M, Curini V, Mangone I, Rinaldi A, Di Pasquale A, Cammà C, Puglia I, Calistri P, Savini G |
| EPI_ISL_806745 | RSA Giulianova | Istituto Zooprofilattico Sperimentale dell'Abruzzo e Molise "G. Caporale" | Lorusso A, Marcacci M, Di Domenico M, Ancora M, Curini V, Mangone I, Rinaldi A, Di Pasquale A, Cammà C, Puglia I, Calistri P, Savini G |
| EPI_ISL_806746, EPI_ISL_806747, EPI_ISL_806748 | SIESP DIPARTIMENTO DI PREVENZIONE TERAMO | Istituto Zooprofilattico Sperimentale dell'Abruzzo e Molise "G. Caporale" | Lorusso A, Marcacci M, Di Domenico M, Ancora M, Curini V, Mangone I, Rinaldi A, Di Pasquale A, Cammà C, Puglia I, Calistri P, Savini G |
| EPI_ISL_806749, EPI_ISL_806750, EPI_ISL_806751, EPI_ISL_806752, EPI_ISL_806753, EPI_ISL_806754, EPI_ISL_806755, EPI_ISL_806756, EPI_ISL_806757 | SIESP DIPARTIMENTO DI PREVENZIONE SULMONA | Istituto Zooprofilattico Sperimentale dell'Abruzzo e Molise "G. Caporale" | Lorusso A, Marcacci M, Di Domenico M, Ancora M, Curini V, Mangone I, Rinaldi A, Di Pasquale A, Cammà C, Puglia I, Calistri P, Savini G |
| EPI_ISL_806758, EPI_ISL_806759, EPI_ISL_806760, EPI_ISL_806761 | DIPARTIMENTO PREVENZIONE AVEZZANO-SERVIZIO DI IGIENE EPIDEMIOLOGIA E SANITA' PUBBLICA | Istituto Zooprofilattico Sperimentale dell'Abruzzo e Molise "G. Caporale" | Lorusso A, Marcacci M, Di Domenico M, Ancora M, Curini V, Mangone I, Rinaldi A, Di Pasquale A, Cammà C, Puglia I, Calistri P, Savini G |
| EPI_ISL_806762 | SIESP CHIETI - DRIVE IN ORTONA | Istituto Zooprofilattico Sperimentale dell'Abruzzo e Molise "G. Caporale" | Lorusso A, Marcacci M, Di Domenico M, Ancora M, Curini V, Mangone I, Rinaldi A, Di Pasquale A, Cammà C, Puglia I, Calistri P, Savini G |
| EPI_ISL_806763, EPI_ISL_806764, EPI_ISL_806765 | DIPARTIMENTO PREVENZIONE AVEZZANO-SERVIZIO DI IGIENE EPIDEMIOLOGIA E SANITA' PUBBLICA | Istituto Zooprofilattico Sperimentale dell'Abruzzo e Molise "G. Caporale" | Lorusso A, Marcacci M, Di Domenico M, Ancora M, Curini V, Mangone I, Rinaldi A, Di Pasquale A, Cammà C, Puglia I, Calistri P, Savini G |
| EPI_ISL_806766, EPI_ISL_806767, EPI_ISL_806768, EPI_ISL_806769, EPI_ISL_806770, EPI_ISL_806771, EPI_ISL_806772 | SIESP DIPARTIMENTO DI PREVENZIONE TERAMO | Istituto Zooprofilattico Sperimentale dell'Abruzzo e Molise "G. Caporale" | Lorusso A, Marcacci M, Di Domenico M, Ancora M, Curini V, Mangone I, Rinaldi A, Di Pasquale A, Cammà C, Puglia I, Calistri P, Savini G |
| EPI_ISL_806773, EPI_ISL_806774 | Ospedale S.Salvatore-Medicina Interna L'Aquila | Istituto Zooprofilattico Sperimentale dell'Abruzzo e Molise "G. Caporale" | Lorusso A, Marcacci M, Di Domenico M, Ancora M, Curini V, Mangone I, Rinaldi A, Di Pasquale A, Cammà C, Puglia I, Calistri P, Savini G |
| EPI_ISL_806775, EPI_ISL_806776 | SIESP CHIETI - DRIVE IN LANCIANO | Istituto Zooprofilattico Sperimentale dell'Abruzzo e Molise "G. Caporale" | Lorusso A, Marcacci M, Di Domenico M, Ancora M, Curini V, Mangone I, Rinaldi A, Di Pasquale A, Cammà C, Puglia I, Calistri P, Savini G |
| EPI_ISL_806777, EPI_ISL_806778, EPI_ISL_806779, EPI_ISL_806780 | SIESP DIPARTIMENTO DI PREVENZIONE TERAMO | Istituto Zooprofilattico Sperimentale dell'Abruzzo e Molise "G. Caporale" | Lorusso A, Marcacci M, Di Domenico M, Ancora M, Curini V, Mangone I, Rinaldi A, Di Pasquale A, Cammà C, Puglia I, Calistri P, Savini G |
| EPI_ISL_806781, EPI_ISL_806782 | DIPARTIMENTO PREVENZIONE AVEZZANO-SERVIZIO DI IGIENE EPIDEMIOLOGIA E SANITA' PUBBLICA | Istituto Zooprofilattico Sperimentale dell'Abruzzo e Molise "G. Caporale" | Lorusso A, Marcacci M, Di Domenico M, Ancora M, Curini V, Mangone I, Rinaldi A, Di Pasquale A, Cammà C, Puglia I, Calistri P, Savini G |
| EPI_ISL_806783 | Casa di cura Di Lorenzo- Avezzano | Istituto Zooprofilattico Sperimentale dell'Abruzzo e Molise "G. Caporale" | Lorusso A, Marcacci M, Di Domenico M, Ancora M, Curini V, Mangone I, Rinaldi A, Di Pasquale A, Cammà C, Puglia I, Calistri P, Savini G |

| | | | |
|---|---|---|---|
| EPI_ISL_806784, EPI_ISL_806785 | Ospedale Civile G.Mazzini-Teramo | Istituto Zooprofilattico Sperimentale dell'Abruzzo e Molise "G. Caporale" | Lorusso A, Marcacci M, Di Domenico M, Ancora M, Curini V, Mangone I, Rinaldi A, Di Pasquale A, Cammà C, Puglia I, Calistri P, Savini G |
| EPI_ISL_806786, EPI_ISL_806787, EPI_ISL_806788 | SIESP DIPARTIMENTO DI PREVENZIONE TERAMO | Istituto Zooprofilattico Sperimentale dell'Abruzzo e Molise "G. Caporale" | Lorusso A, Marcacci M, Di Domenico M, Ancora M, Curini V, Mangone I, Rinaldi A, Di Pasquale A, Cammà C, Puglia I, Calistri P, Savini G |
| EPI_ISL_806790 | SIESP DIPARTIMENTO DI PREVENZIONE DELL'AQUILA | Istituto Zooprofilattico Sperimentale dell'Abruzzo e Molise "G. Caporale" | Lorusso A, Marcacci M, Di Domenico M, Ancora M, Curini V, Mangone I, Rinaldi A, Di Pasquale A, Cammà C, Puglia I, Calistri P, Savini G |
| EPI_ISL_806791, EPI_ISL_806792, EPI_ISL_806793, EPI_ISL_806794, EPI_ISL_806795, EPI_ISL_806796, EPI_ISL_806797, EPI_ISL_806798 | SIESP CHIETI - DRIVE IN ORTONA | Istituto Zooprofilattico Sperimentale dell'Abruzzo e Molise "G. Caporale" | Lorusso A, Marcacci M, Di Domenico M, Ancora M, Curini V, Mangone I, Rinaldi A, Di Pasquale A, Cammà C, Puglia I, Calistri P, Savini G |
| EPI_ISL_806799 | SIESP CHIETI - DRIVE IN LANCIANO | Istituto Zooprofilattico Sperimentale dell'Abruzzo e Molise "G. Caporale" | Lorusso A, Marcacci M, Di Domenico M, Ancora M, Curini V, Mangone I, Rinaldi A, Di Pasquale A, Cammà C, Puglia I, Calistri P, Savini G |
| EPI_ISL_806800, EPI_ISL_806801, EPI_ISL_806802, EPI_ISL_806803 | SIESP CHIETI - DRIVE IN GISSI | Istituto Zooprofilattico Sperimentale dell'Abruzzo e Molise "G. Caporale" | Lorusso A, Marcacci M, Di Domenico M, Ancora M, Curini V, Mangone I, Rinaldi A, Di Pasquale A, Cammà C, Puglia I, Calistri P, Savini G |
| EPI_ISL_806804, EPI_ISL_806805 | SIESP CHIETI - DISTRETTO SANITARIO CHIETI | Istituto Zooprofilattico Sperimentale dell'Abruzzo e Molise "G. Caporale" | Lorusso A, Marcacci M, Di Domenico M, Ancora M, Curini V, Mangone I, Rinaldi A, Di Pasquale A, Cammà C, Puglia I, Calistri P, Savini G |
| EPI_ISL_806806 | SIESP DIPARTIMENTO DI PREVENZIONE CHIETI | Istituto Zooprofilattico Sperimentale dell'Abruzzo e Molise "G. Caporale" | Lorusso A, Marcacci M, Di Domenico M, Ancora M, Curini V, Mangone I, Rinaldi A, Di Pasquale A, Cammà C, Puglia I, Calistri P, Savini G |
| EPI_ISL_806846, EPI_ISL_806852 | Alaska State Virology Laboratory | Alaska State Virology Laboratory | Stephanie DeRonde, Lisa Smith, Ph.D., Devin M. Drown, Ph.D., Jack Chen, Ph.D. |
| EPI_ISL_811129 | Viollier AG | Department of Biosystems Science and Engineering, ETH Zürich | Chaoran Chen, Sarah Nadeau, Catharine Aquino, Ivan Topolsky, Philipp Jablonski, Lara Fuhrmann, David Dreifuss, Katharina Jahn, Andreia Cabral de Gouvea, Maria Domenica Moccia, Simon Grüter, Timothy Sykes, Lennart Opitz, Griffin White, Laura Neff, Doris Popovic, Andrea Patrignani, Jay Tracy, Ralph Schlapbach, Christiane Beckmann, Maurice Redondo, Olivier Kobel, Christoph Noppen, Sophie Seidel, Noemie Santamaria de Souza, Niko Beerenwinkel, Tanja Stadler |
| EPI_ISL_812382, EPI_ISL_812383, EPI_ISL_812396 | Utah Public Health Laboratory | Utah Public Health Laboratory | Erin L. Young, Kelly F. Oakeson, Tara Gallagher |
| EPI_ISL_812518 | Sydney South West Pathology Service (SSWPS) - Concord Repatriation General Hospital - NSW Health Pathology | NSW Health Pathology - Institute of Clinical Pathology and Medical Research; Westmead Hospital; University of Sydney | CIDM-PH et al. |
| EPI_ISL_812745 | DOHMH Riverside | New York City Public Health Laboratory | Jade Wang, et al. |
| EPI_ISL_812746, EPI_ISL_812747, EPI_ISL_812748 | DOHMH Corona | New York City Public Health Laboratory | Jade Wang, et al. |
| EPI_ISL_812749 | DOHMH Central Harlem | New York City Public Health Laboratory | Jade Wang, et al. |
| EPI_ISL_812750, EPI_ISL_812751 | DOHMH Corona | New York City Public Health Laboratory | Jade Wang, et al. |
| EPI_ISL_812752 | DOHMH Crown Heights | New York City Public Health Laboratory | Jade Wang, et al. |
| EPI_ISL_812753, EPI_ISL_812754, EPI_ISL_812755, EPI_ISL_812756, EPI_ISL_812757 | DOHMH Morrisania | New York City Public Health Laboratory | Jade Wang, et al. |
| EPI_ISL_812758 | DOHMH Crown Heights | New York City Public Health Laboratory | Jade Wang, et al. |
| EPI_ISL_812759 | DOHMH Riverside | New York City Public Health Laboratory | Jade Wang, et al. |
| EPI_ISL_812760 | DOHMH Corona | New York City Public Health Laboratory | Jade Wang, et al. |
| EPI_ISL_813062, EPI_ISL_813087, EPI_ISL_813144, EPI_ISL_813145, EPI_ISL_813146, EPI_ISL_813147, EPI_ISL_813148, EPI_ISL_813149, EPI_ISL_813150, EPI_ISL_813151, EPI_ISL_813152 | | | |
| see above | University of Birmingham | COVID-19 Genomics UK (COG-UK) Consortium | Institute of Microbiology, University of Birmingham: Claire McMurray, Joanne Stockton, Samuel Nicholls, Radoslaw Poplawski, Will Rowe, Josh Quick, Nicholas Loman. University of Birmingham Testing Laboratory: Celina M Whalley, Andrew Bosworth, Charlotte Poxon, Kasun Wanigasooriya, Oliver Pickles, Mike Kidd, Alex Richter, Andrew D Beggs PHE Heartlands Lab: Husam Osman, Andrew Bosworth. Queen Elizabeth Hospital: Anna Casey |
| EPI_ISL_813172, EPI_ISL_813196, EPI_ISL_813232, EPI_ISL_813235, EPI_ISL_813242, EPI_ISL_813297, EPI_ISL_813300, EPI_ISL_813301, EPI_ISL_813302, EPI_ISL_813303, EPI_ISL_813304, EPI_ISL_813321, EPI_ISL_813333, EPI_ISL_813334, EPI_ISL_813336, EPI_ISL_813338, EPI_ISL_813340, EPI_ISL_813342, EPI_ISL_813345, EPI_ISL_813347, EPI_ISL_813348, EPI_ISL_813349, EPI_ISL_813350, EPI_ISL_813351, EPI_ISL_813352, EPI_ISL_813353, EPI_ISL_813354, EPI_ISL_813355 | | | |
| see above | Department of Pathology, University of Cambridge | COVID-19 Genomics UK (COG-UK) Consortium | Aminu S. Jahun, Yasmin Chaudhry, Grant Hall, Iliana Georgana, Myra Hosmillo, Martin D. Curran, Malte Pinckert, Surendra Parmar, Ian Goodfellow |
| EPI_ISL_813357, EPI_ISL_813358, EPI_ISL_813359, EPI_ISL_813360, EPI_ISL_813361, EPI_ISL_813362, EPI_ISL_813363, EPI_ISL_813364, EPI_ISL_813365, EPI_ISL_813366, EPI_ISL_813367, EPI_ISL_813368, EPI_ISL_813369, EPI_ISL_813370, EPI_ISL_813371, EPI_ISL_813372, EPI_ISL_813373, EPI_ISL_813374, EPI_ISL_813375, EPI_ISL_813376, EPI_ISL_813377, EPI_ISL_813378, EPI_ISL_813379, EPI_ISL_813380, EPI_ISL_813381, EPI_ISL_813382, EPI_ISL_813383, EPI_ISL_813384, EPI_ISL_813385, EPI_ISL_813386, EPI_ISL_813387, EPI_ISL_813388, EPI_ISL_813389, EPI_ISL_813390, EPI_ISL_813391, EPI_ISL_813392, EPI_ISL_813393, EPI_ISL_813394, EPI_ISL_813395, EPI_ISL_813396, EPI_ISL_813397, EPI_ISL_813398, EPI_ISL_813399, EPI_ISL_813400, EPI_ISL_813401, EPI_ISL_813402, EPI_ISL_813403, EPI_ISL_813404, EPI_ISL_813405, EPI_ISL_813406, EPI_ISL_813407, EPI_ISL_813408, EPI_ISL_813409, EPI_ISL_813410, EPI_ISL_813411, EPI_ISL_813412, EPI_ISL_813413, EPI_ISL_813414, EPI_ISL_813415, EPI_ISL_813416, EPI_ISL_813427, EPI_ISL_813430, EPI_ISL_813431 | | | |
| see above | University of Exeter | COVID-19 Genomics UK (COG-UK) Consortium | Ben Temperton,Aaron Jeffries,Michelle Michelsen,Joanna Warwick-Dugdale,Audrey Farbos,Robyn Manley,Stephen Michell,Jane Masoli |
| EPI_ISL_813656 | Kettering General Hospital | COVID-19 Genomics UK (COG-UK) Consortium | Patrick McClure, Joseph Chappell, Theocharis Tsoleridis, Jonathan Ball, Nadine Holmes, Matthew Carlisle, Christopher Moore, Fei Sang, Johnny Debebe, Victoria Wright, Matthew Loose |
| EPI_ISL_813803, EPI_ISL_813805, EPI_ISL_813807, EPI_ISL_813808 | Liverpool Clinical Laboratories | COVID-19 Genomics UK (COG-UK) Consortium | Sam Haldenby, Anita Lucaci, Steve Paterson, Julian Hiscox, Alistair Darby, M Almsaud, A Alrezaihi, Muhannad Alruwaili, Stuart D Armstrong, Jones Benjamin, Eleanor G Bentley, Anu Chawla, Jordan J Clark, Angela Cowell, Richard Eccles, Isabel García-Dorival, Matthew Gemmell, Alessandro Gerada, PKF Gilmore, Richard Gregory, Ximeng Han, Catherine Hartley, Margaret Hughes, Miren Iturriza-Gomara, James Johnson, L Luu, Jenifer Manson, Charlotte Nelson, Elaine O'Toole, Cassie Olateju, Rebekah Penrice-Randal , Lucille Rainbow, N.P Randle, Trevor Ian Robinson, Parul Sharma, Ghada T Shawli, James P Stewart, Neil Swainston, Ecaterina Vamos, Joanne Watts, Mark Whitehead |
| EPI_ISL_813832, EPI_ISL_813833, EPI_ISL_813834, EPI_ISL_813835, EPI_ISL_813836, EPI_ISL_813837, EPI_ISL_813838, EPI_ISL_813839, EPI_ISL_813840, EPI_ISL_813907, EPI_ISL_813908, EPI_ISL_813911, EPI_ISL_813919 | | | |
| see above | University College London, Great Ormond Street Hospital for Children NHS Foundation Trust, Imperial College Healthcare NHS Trust | COVID-19 Genomics UK (COG-UK) Consortium | Sergi Castellano, Rachel Williams, Mark Kristiansen, Paola Resende Silva, Sunando Roy, Tony Brooks, Helena Tutill, Paola Niola, Patricia Dyal, Charlotte Williams, Leysa Forrest, Yasmin Panchbhaya, Jacqueline Findlay, Samuel Weeks, Julianne Brown, Kathryn Harris, Paul Randell, James Price, Alison Holmes, Judith Breuer |
| EPI_ISL_813976 | Norwegian Institute of Public Health, Department of Virology | Norwegian Institute of Public Health, Department of Virology | Kathrine Stene-Johansen, Kamilla Heddeland Instefjord, Hilde Elshaug, Atiya R Ali,Marie Paulsen Madsen, Rasmus Riis Kopperud, Hilde Vollan, Karoline Bragstad, Olav Hungnes |
| EPI_ISL_814288, EPI_ISL_814289, EPI_ISL_814290 | Bioinformatics and Biostatistics Lab, Advanced Sequencing Facility | COVID-19 Genomics UK (COG-UK) Consortium | Aengus Stewart,Jerome Nicod,Chelsea Sawyer,Laura Cubitt,Harshil Patel,Margaret Crawford |
| EPI_ISL_814303, EPI_ISL_814310, EPI_ISL_814317, EPI_ISL_814329, EPI_ISL_814334 | Wales Specialist Virology Centre Sequencing lab: Pathogen Genomics Unit | COVID-19 Genomics UK (COG-UK) Consortium | Catherine Moore, Johnathan Evans, Laura Gifford, Malorie Perry, Simon Cottrell, Angela Marchbank, Alec Birchley, Alexander Adams, Amy Gaskin, Bree Gatica-Wilcox, Jason Coombes, Joel Southgate, Lauren Gilbert, Lee Graham, Nicole Pacchiarini, Sara Kumziene-Summerhayes, Sarah Taylor, Sophie Jones, Sara Rey, Matthew Bull, Joanne Watkins, Sally Corden, Tom Connor |
| EPI_ISL_814339, EPI_ISL_814345, EPI_ISL_814348 | West of Scotland Specialist Virology Centre, NHSGGC / MRC-University of Glasgow Centre for Virus Research | COVID-19 Genomics UK (COG-UK) Consortium | Ana da Silva Filipe, Natasha Johnson, Kathy Smollett, Daniel Mair, Stephen Carmichael, Alice Broos, Lily Tong, Jenna Nichols, Kyriaki Nomikou; Sarah McDonald; Richard Orton, Joseph Hughes, Sreenu Vattipally, David L Robertson; Alasdair MacLean, Rory Gunson; Sharif Shaaban, Matthew Holden; |

| | | | |
|---|---|---|---|
| | | | Rachel Blacow, Guy Mollett, Kathy Li, James Shepherd, Antonia Ho, Emma Thomson |
| EPI_ISL_814351 | Bioinformatics and Biostatistics Lab, Advanced Sequencing Facility | COVID-19 Genomics UK (COG-UK) Consortium | Aengus Stewart,Jerome Nicod,Chelsea Sawyer,Laura Cubitt,Harshil Patel,Margaret Crawford |
| EPI_ISL_814353 | Wales Specialist Virology Centre Sequencing lab: Pathogen Genomics Unit | COVID-19 Genomics UK (COG-UK) Consortium | Catherine Moore, Johnathan Evans, Laura Gifford, Malorie Perry, Simon Cottrell, Angela Marchbank, Alec Birchley, Alexander Adams, Amy Gaskin, Bree Gatica-Wilcox, Jason Coombes, Joel Southgate, Lauren Gilbert, Lee Graham, Nicole Pacchiarini, Sara Kumziene-Summerhayes, Sarah Taylor, Sophie Jones, Sara Rey, Matthew Bull, Joanne Watkins, Sally Corden, Tom Connor |
| EPI_ISL_814355 | West of Scotland Specialist Virology Centre, NHSGGC / MRC-University of Glasgow Centre for Virus Research | COVID-19 Genomics UK (COG-UK) Consortium | Ana da Silva Filipe, Natasha Johnson, Kathy Smollett, Daniel Mair, Stephen Carmichael, Alice Broos, Lily Tong, Jenna Nichols, Kyriaki Nomikou; Sarah McDonald; Richard Orton, Joseph Hughes, Sreenu Vattipally, David L Robertson; Alasdair MacLean, Rory Gunson; Sharif Shaaban, Matthew Holden; Rachel Blacow, Guy Mollett, Kathy Li, James Shepherd, Antonia Ho, Emma Thomson |
| EPI_ISL_814359, EPI_ISL_814360, EPI_ISL_814361 | Bioinformatics and Biostatistics Lab, Advanced Sequencing Facility | COVID-19 Genomics UK (COG-UK) Consortium | Aengus Stewart,Jerome Nicod,Chelsea Sawyer,Laura Cubitt,Harshil Patel,Margaret Crawford |
| EPI_ISL_814370, EPI_ISL_814380, EPI_ISL_814396, EPI_ISL_814397, EPI_ISL_814403, EPI_ISL_814406, EPI_ISL_814410, EPI_ISL_814414 | Wales Specialist Virology Centre Sequencing lab: Pathogen Genomics Unit | COVID-19 Genomics UK (COG-UK) Consortium | Catherine Moore, Johnathan Evans, Laura Gifford, Malorie Perry, Simon Cottrell, Angela Marchbank, Alec Birchley, Alexander Adams, Amy Gaskin, Bree Gatica-Wilcox, Jason Coombes, Joel Southgate, Lauren Gilbert, Lee Graham, Nicole Pacchiarini, Sara Kumziene-Summerhayes, Sarah Taylor, Sophie Jones, Sara Rey, Matthew Bull, Joanne Watkins, Sally Corden, Tom Connor |
| EPI_ISL_814417, EPI_ISL_814419, EPI_ISL_814424, EPI_ISL_814425, EPI_ISL_814426, EPI_ISL_814427, EPI_ISL_814428, EPI_ISL_814435 | Bioinformatics and Biostatistics Lab, Advanced Sequencing Facility | COVID-19 Genomics UK (COG-UK) Consortium | Aengus Stewart,Jerome Nicod,Chelsea Sawyer,Laura Cubitt,Harshil Patel,Margaret Crawford |
| EPI_ISL_814444, EPI_ISL_814445, EPI_ISL_814446, EPI_ISL_814447, EPI_ISL_814448, EPI_ISL_814450, EPI_ISL_814451, EPI_ISL_814452, EPI_ISL_814453, EPI_ISL_814455, EPI_ISL_814458 | | | |
| see above | West of Scotland Specialist Virology Centre, NHSGGC / MRC-University of Glasgow Centre for Virus Research | COVID-19 Genomics UK (COG-UK) Consortium | Ana da Silva Filipe, Natasha Johnson, Kathy Smollett, Daniel Mair, Stephen Carmichael, Alice Broos, Lily Tong, Jenna Nichols, Kyriaki Nomikou; Sarah McDonald; Richard Orton, Joseph Hughes, Sreenu Vattipally, David L Robertson; Alasdair MacLean, Rory Gunson; Sharif Shaaban, Matthew Holden; Rachel Blacow, Guy Mollett, Kathy Li, James Shepherd, Antonia Ho, Emma Thomson |
| EPI_ISL_814467 | Oxford Viromics, NDM, University of Oxford; Oxford University Hospitals; Basingstoke and North Hampshire Hospital | COVID-19 Genomics UK (COG-UK) Consortium | Tanya Golubchik, David Bonsall, George Macintyre, Amy Trebes, Mariateresa de Cesare, Catrin Moore, Alex Mobbs, Anita Justice, Robert Shaw, Monique Andersson, Timothy Peto, Emma Wise, Nathan Moore, Jessica Lynch, Nick Cortes, Matilde Mori, Stephen Kidd, David Buck, John Todd, Christophe Fraser |
| EPI_ISL_814517, EPI_ISL_814518, EPI_ISL_814520, EPI_ISL_814521, EPI_ISL_814524, EPI_ISL_814525, EPI_ISL_814539 | Bioinformatics and Biostatistics Lab, Advanced Sequencing Facility | COVID-19 Genomics UK (COG-UK) Consortium | Aengus Stewart,Jerome Nicod,Chelsea Sawyer,Laura Cubitt,Harshil Patel,Margaret Crawford |
| EPI_ISL_814542 | Wales Specialist Virology Centre Sequencing lab: Pathogen Genomics Unit | COVID-19 Genomics UK (COG-UK) Consortium | Catherine Moore, Johnathan Evans, Laura Gifford, Malorie Perry, Simon Cottrell, Angela Marchbank, Alec Birchley, Alexander Adams, Amy Gaskin, Bree Gatica-Wilcox, Jason Coombes, Joel Southgate, Lauren Gilbert, Lee Graham, Nicole Pacchiarini, Sara Kumziene-Summerhayes, Sarah Taylor, Sophie Jones, Sara Rey, Matthew Bull, Joanne Watkins, Sally Corden, Tom Connor |
| EPI_ISL_814544, EPI_ISL_814548 | Bioinformatics and Biostatistics Lab, Advanced Sequencing Facility | COVID-19 Genomics UK (COG-UK) Consortium | Aengus Stewart,Jerome Nicod,Chelsea Sawyer,Laura Cubitt,Harshil Patel,Margaret Crawford |
| EPI_ISL_814549 | West of Scotland Specialist Virology Centre, NHSGGC / MRC-University of Glasgow Centre for Virus Research | COVID-19 Genomics UK (COG-UK) Consortium | Ana da Silva Filipe, Natasha Johnson, Kathy Smollett, Daniel Mair, Stephen Carmichael, Alice Broos, Lily Tong, Jenna Nichols, Kyriaki Nomikou; Sarah McDonald; Richard Orton, Joseph Hughes, Sreenu Vattipally, David L Robertson; Alasdair MacLean, Rory Gunson; Sharif Shaaban, Matthew Holden; Rachel Blacow, Guy Mollett, Kathy Li, James Shepherd, Antonia Ho, Emma Thomson |
| EPI_ISL_814550, EPI_ISL_814553 | Wales Specialist Virology Centre Sequencing lab: Pathogen Genomics Unit | COVID-19 Genomics UK (COG-UK) Consortium | Catherine Moore, Johnathan Evans, Laura Gifford, Malorie Perry, Simon Cottrell, Angela Marchbank, Alec Birchley, Alexander Adams, Amy Gaskin, Bree Gatica-Wilcox, Jason Coombes, Joel Southgate, Lauren Gilbert, Lee Graham, Nicole Pacchiarini, Sara Kumziene-Summerhayes, Sarah Taylor, Sophie Jones, Sara Rey, Matthew Bull, Joanne Watkins, Sally Corden, Tom Connor |
| EPI_ISL_814568 | West of Scotland Specialist Virology Centre, NHSGGC / MRC-University of Glasgow Centre for Virus Research | COVID-19 Genomics UK (COG-UK) Consortium | Ana da Silva Filipe, Natasha Johnson, Kathy Smollett, Daniel Mair, Stephen Carmichael, Alice Broos, Lily Tong, Jenna Nichols, Kyriaki Nomikou; Sarah McDonald; Richard Orton, Joseph Hughes, Sreenu Vattipally, David L Robertson; Alasdair MacLean, Rory Gunson; Sharif Shaaban, Matthew Holden; Rachel Blacow, Guy Mollett, Kathy Li, James Shepherd, Antonia Ho, Emma Thomson |
| EPI_ISL_814572, EPI_ISL_814594 | Oxford Viromics, NDM, University of Oxford; Oxford University Hospitals; Basingstoke and North Hampshire Hospital | COVID-19 Genomics UK (COG-UK) Consortium | Tanya Golubchik, David Bonsall, George Macintyre, Amy Trebes, Mariateresa de Cesare, Catrin Moore, Alex Mobbs, Anita Justice, Robert Shaw, Monique Andersson, Timothy Peto, Emma Wise, Nathan Moore, Jessica Lynch, Nick Cortes, Matilde Mori, Stephen Kidd, David Buck, John Todd, Christophe Fraser |
| EPI_ISL_814597 | Wales Specialist Virology Centre Sequencing lab: Pathogen Genomics Unit | COVID-19 Genomics UK (COG-UK) Consortium | Catherine Moore, Johnathan Evans, Laura Gifford, Malorie Perry, Simon Cottrell, Angela Marchbank, Alec Birchley, Alexander Adams, Amy Gaskin, Bree Gatica-Wilcox, Jason Coombes, Joel Southgate, Lauren Gilbert, Lee Graham, Nicole Pacchiarini, Sara Kumziene-Summerhayes, Sarah Taylor, Sophie Jones, Sara Rey, Matthew Bull, Joanne Watkins, Sally Corden, Tom Connor |
| EPI_ISL_814604, EPI_ISL_814605 | Oxford Viromics, NDM, University of Oxford; Oxford University Hospitals; Basingstoke and North Hampshire Hospital | COVID-19 Genomics UK (COG-UK) Consortium | Tanya Golubchik, David Bonsall, George Macintyre, Amy Trebes, Mariateresa de Cesare, Catrin Moore, Alex Mobbs, Anita Justice, Robert Shaw, Monique Andersson, Timothy Peto, Emma Wise, Nathan Moore, Jessica Lynch, Nick Cortes, Matilde Mori, Stephen Kidd, David Buck, John Todd, Christophe Fraser |
| EPI_ISL_814609 | West of Scotland Specialist Virology Centre, NHSGGC / MRC-University of Glasgow Centre for Virus Research | COVID-19 Genomics UK (COG-UK) Consortium | Ana da Silva Filipe, Natasha Johnson, Kathy Smollett, Daniel Mair, Stephen Carmichael, Alice Broos, Lily Tong, Jenna Nichols, Kyriaki Nomikou; Sarah McDonald; Richard Orton, Joseph Hughes, Sreenu Vattipally, David L Robertson; Alasdair MacLean, Rory Gunson; Sharif Shaaban, Matthew Holden; Rachel Blacow, Guy Mollett, Kathy Li, James Shepherd, Antonia Ho, Emma Thomson |
| EPI_ISL_814610, EPI_ISL_814612, EPI_ISL_814614, EPI_ISL_814616, EPI_ISL_814618 | Bioinformatics and Biostatistics Lab, Advanced Sequencing Facility | COVID-19 Genomics UK (COG-UK) Consortium | Aengus Stewart,Jerome Nicod,Chelsea Sawyer,Laura Cubitt,Harshil Patel,Margaret Crawford |
| EPI_ISL_814620, EPI_ISL_814621, EPI_ISL_814622, EPI_ISL_814624, EPI_ISL_814627, EPI_ISL_814628, EPI_ISL_814631, EPI_ISL_814633, EPI_ISL_814634, EPI_ISL_814636, EPI_ISL_814638, EPI_ISL_814639, EPI_ISL_814642, EPI_ISL_814644, EPI_ISL_814645, EPI_ISL_814650, EPI_ISL_814651, EPI_ISL_814653, EPI_ISL_814654, EPI_ISL_814655, EPI_ISL_814656, EPI_ISL_814657, EPI_ISL_814658, EPI_ISL_814661, EPI_ISL_814664 | | | |
| see above | West of Scotland Specialist Virology Centre, NHSGGC / MRC-University of Glasgow Centre for Virus Research | COVID-19 Genomics UK (COG-UK) Consortium | Ana da Silva Filipe, Natasha Johnson, Kathy Smollett, Daniel Mair, Stephen Carmichael, Alice Broos, Lily Tong, Jenna Nichols, Kyriaki Nomikou; Sarah McDonald; Richard Orton, Joseph Hughes, Sreenu Vattipally, David L Robertson; Alasdair MacLean, Rory Gunson; Sharif Shaaban, Matthew Holden; Rachel Blacow, Guy Mollett, Kathy Li, James Shepherd, Antonia Ho, Emma Thomson |
| EPI_ISL_814688, EPI_ISL_814690, EPI_ISL_814692, EPI_ISL_814693, EPI_ISL_814694 | Oxford Viromics, NDM, University of Oxford; Oxford University Hospitals; Basingstoke and North Hampshire Hospital | COVID-19 Genomics UK (COG-UK) Consortium | Tanya Golubchik, David Bonsall, George Macintyre, Amy Trebes, Mariateresa de Cesare, Catrin Moore, Alex Mobbs, Anita Justice, Robert Shaw, Monique Andersson, Timothy Peto, Emma Wise, Nathan Moore, Jessica Lynch, Nick Cortes, Matilde Mori, Stephen Kidd, David Buck, John Todd, Christophe Fraser |
| EPI_ISL_814941, EPI_ISL_814954, EPI_ISL_814955, EPI_ISL_814956, EPI_ISL_814957, EPI_ISL_814958, EPI_ISL_814959, EPI_ISL_814960, EPI_ISL_814961, EPI_ISL_814962, EPI_ISL_814963, EPI_ISL_814964, EPI_ISL_814965, EPI_ISL_814968, EPI_ISL_815039, EPI_ISL_815040, EPI_ISL_815041, EPI_ISL_815042, EPI_ISL_815090, EPI_ISL_815138, EPI_ISL_815139, EPI_ISL_815140, EPI_ISL_815141, EPI_ISL_815142, EPI_ISL_815143, EPI_ISL_815145, EPI_ISL_815147, EPI_ISL_815148, EPI_ISL_815149, EPI_ISL_815151, EPI_ISL_815244, EPI_ISL_815245, EPI_ISL_815246 | | | |
| see above | Wales Specialist Virology Centre Sequencing lab: Pathogen Genomics Unit | COVID-19 Genomics UK (COG-UK) Consortium | Catherine Moore, Johnathan Evans, Laura Gifford, Malorie Perry, Simon Cottrell, Angela Marchbank, Alec Birchley, Alexander Adams, Amy Gaskin, Bree Gatica-Wilcox, Jason Coombes, Joel Southgate, Lauren Gilbert, Lee Graham, Nicole Pacchiarini, Sara Kumziene-Summerhayes, Sarah Taylor, Sophie Jones, Sara Rey, Matthew Bull, Joanne Watkins, Sally Corden, Tom Connor |
| EPI_ISL_816272, EPI_ISL_816446, EPI_ISL_816481, EPI_ISL_816488, EPI_ISL_816511, EPI_ISL_816567, EPI_ISL_816580 | Virology Department, Sheffield Teaching Hospitals NHS Foundation Trust/Department of Infection, Immunity and Cardiovascular Disease, The Medical School, University of Sheffield | COVID-19 Genomics UK (COG-UK) Consortium | Thushan de Silva, Matthew Parker, Nikki Smith, Adri Angyal, Rebecca Brown, Luke Green, Rachel Tucker, Paul Parsons, Danielle Groves, Katie Johnson, Laura Carrilero, Alex Keeley, Dave Partridge, Matthew Wyles, Benjamin Lindsey, Mehmet Yavuz, Mohammad Raza, Cariad Evans |
| EPI_ISL_817050, EPI_ISL_817051, EPI_ISL_817053, EPI_ISL_817054, EPI_ISL_817057, EPI_ISL_817062, EPI_ISL_817063, EPI_ISL_817066, EPI_ISL_817067, EPI_ISL_817069, EPI_ISL_817070, EPI_ISL_817071, EPI_ISL_817072, EPI_ISL_817073, EPI_ISL_817074, EPI_ISL_817075, EPI_ISL_817076, EPI_ISL_817078, | | | |

| | | | |
|---|---|---|---|
| EPI_ISL_817079, EPI_ISL_817080, EPI_ISL_817081, EPI_ISL_817082, EPI_ISL_817083, EPI_ISL_817084, EPI_ISL_817085, EPI_ISL_817086, EPI_ISL_817087, EPI_ISL_817088, EPI_ISL_817089, EPI_ISL_817090, EPI_ISL_817091, EPI_ISL_817092, EPI_ISL_817093, EPI_ISL_817094, EPI_ISL_817095, EPI_ISL_817096, EPI_ISL_817097, EPI_ISL_817098, EPI_ISL_817099, EPI_ISL_817100, EPI_ISL_817101, EPI_ISL_817102, EPI_ISL_817103, EPI_ISL_817104, EPI_ISL_817105, EPI_ISL_817106, EPI_ISL_817107, EPI_ISL_817108, EPI_ISL_817109, EPI_ISL_817110, EPI_ISL_817111, EPI_ISL_817112, EPI_ISL_817113, EPI_ISL_817114, EPI_ISL_817115, EPI_ISL_817116, EPI_ISL_817117, EPI_ISL_817118, EPI_ISL_817119, EPI_ISL_817120, EPI_ISL_817121 | | | |
| see above | Bioinformatics and Biostatistics Lab, Advanced Sequencing Facility | COVID-19 Genomics UK (COG-UK) Consortium | Aengus Stewart,Jerome Nicod,Chelsea Sawyer,Laura Cubitt,Harshil Patel,Margaret Crawford |
| EPI_ISL_819138, EPI_ISL_819190, EPI_ISL_819197 | Servicio de Microbiología, Hospital Universitario Son Espases | SeqCOVID-SPAIN consortium/IBV(CSIC) | Carla López-Causapé, Jordi Reina, Antonio Oliver and SeqCOVID-SPAIN consortium |
| EPI_ISL_819609, EPI_ISL_819610, EPI_ISL_819611 | Queens Medical Centre, Clinical Microbiology Department / DeepSeq Nottingham | COVID-19 Genomics UK (COG-UK) Consortium | Gemma Clark, Wendy Smith, Manjinder Khakh, Vicki M Fleming, Michelle M Lister, Hannah Howson-Wells, Jonathan Ball, Patrick McClure, Joseph Chappell, Theocharis Tsoleridis, Nadine Holmes, Matthew Carlisle, Christopher Moore, Fei Sang, Johnny Debebe, Victoria Wright, Matthew Loose |
| EPI_ISL_819612, EPI_ISL_819613, EPI_ISL_819614, EPI_ISL_819615, EPI_ISL_819616, EPI_ISL_819617, EPI_ISL_819618, EPI_ISL_819619, EPI_ISL_819620, EPI_ISL_819621, EPI_ISL_819622, EPI_ISL_819623, EPI_ISL_819624, EPI_ISL_819625, EPI_ISL_819626, EPI_ISL_819627, EPI_ISL_819628, EPI_ISL_819629, EPI_ISL_819630, EPI_ISL_819631, EPI_ISL_819632, EPI_ISL_819633, EPI_ISL_819634, EPI_ISL_819635, EPI_ISL_819636, EPI_ISL_819637, EPI_ISL_819638, EPI_ISL_819639, EPI_ISL_819640, EPI_ISL_819641, EPI_ISL_819642, EPI_ISL_819643, EPI_ISL_819644, EPI_ISL_819645, EPI_ISL_819646 | | | |
| see above | Oxford Viromics, NDM, University of Oxford; Oxford University Hospitals; Basingstoke and North Hampshire Hospital | COVID-19 Genomics UK (COG-UK) Consortium | Tanya Golubchik, David Bonsall, George Macintyre, Amy Trebes, Mariateresa de Cesare, Catrin Moore, Alex Mobbs, Anita Justice, Robert Shaw, Monique Andersson, Timothy Peto, Emma Wise, Nathan Moore, Jessica Lynch, Nick Cortes, Matilde Mori, Stephen Kidd, David Buck, John Todd, Christophe Fraser |
| EPI_ISL_819647, EPI_ISL_819648, EPI_ISL_819649, EPI_ISL_819650, EPI_ISL_819651, EPI_ISL_819652 | Queens Medical Centre, Clinical Microbiology Department / DeepSeq Nottingham | COVID-19 Genomics UK (COG-UK) Consortium | Gemma Clark, Wendy Smith, Manjinder Khakh, Vicki M Fleming, Michelle M Lister, Hannah Howson-Wells, Jonathan Ball, Patrick McClure, Joseph Chappell, Theocharis Tsoleridis, Nadine Holmes, Matthew Carlisle, Christopher Moore, Fei Sang, Johnny Debebe, Victoria Wright, Matthew Loose |
| EPI_ISL_819653, EPI_ISL_819654, EPI_ISL_819655, EPI_ISL_819656, EPI_ISL_819657, EPI_ISL_819658, EPI_ISL_819659, EPI_ISL_819660, EPI_ISL_819661, EPI_ISL_819662, EPI_ISL_819663, EPI_ISL_819664, EPI_ISL_819665, EPI_ISL_819666, EPI_ISL_819667, EPI_ISL_819668, EPI_ISL_819669, EPI_ISL_819670 | | | |
| see above | Oxford Viromics, NDM, University of Oxford; Oxford University Hospitals; Basingstoke and North Hampshire Hospital | COVID-19 Genomics UK (COG-UK) Consortium | Tanya Golubchik, David Bonsall, George Macintyre, Amy Trebes, Mariateresa de Cesare, Catrin Moore, Alex Mobbs, Anita Justice, Robert Shaw, Monique Andersson, Timothy Peto, Emma Wise, Nathan Moore, Jessica Lynch, Nick Cortes, Matilde Mori, Stephen Kidd, David Buck, John Todd, Christophe Fraser |
| EPI_ISL_819671, EPI_ISL_819672, EPI_ISL_819673, EPI_ISL_819674 | Queens Medical Centre, Clinical Microbiology Department / DeepSeq Nottingham | COVID-19 Genomics UK (COG-UK) Consortium | Gemma Clark, Wendy Smith, Manjinder Khakh, Vicki M Fleming, Michelle M Lister, Hannah Howson-Wells, Jonathan Ball, Patrick McClure, Joseph Chappell, Theocharis Tsoleridis, Nadine Holmes, Matthew Carlisle, Christopher Moore, Fei Sang, Johnny Debebe, Victoria Wright, Matthew Loose |
| EPI_ISL_819675, EPI_ISL_819676, EPI_ISL_819677, EPI_ISL_819678, EPI_ISL_819679, EPI_ISL_819680, EPI_ISL_819681, EPI_ISL_819682, EPI_ISL_819683, EPI_ISL_819684, EPI_ISL_819685 | | | |
| see above | Oxford Viromics, NDM, University of Oxford; Oxford University Hospitals; Basingstoke and North Hampshire Hospital | COVID-19 Genomics UK (COG-UK) Consortium | Tanya Golubchik, David Bonsall, George Macintyre, Amy Trebes, Mariateresa de Cesare, Catrin Moore, Alex Mobbs, Anita Justice, Robert Shaw, Monique Andersson, Timothy Peto, Emma Wise, Nathan Moore, Jessica Lynch, Nick Cortes, Matilde Mori, Stephen Kidd, David Buck, John Todd, Christophe Fraser |
| EPI_ISL_819686, EPI_ISL_819687, EPI_ISL_819688 | Queens Medical Centre, Clinical Microbiology Department / DeepSeq Nottingham | COVID-19 Genomics UK (COG-UK) Consortium | Gemma Clark, Wendy Smith, Manjinder Khakh, Vicki M Fleming, Michelle M Lister, Hannah Howson-Wells, Jonathan Ball, Patrick McClure, Joseph Chappell, Theocharis Tsoleridis, Nadine Holmes, Matthew Carlisle, Christopher Moore, Fei Sang, Johnny Debebe, Victoria Wright, Matthew Loose |
| EPI_ISL_819689, EPI_ISL_819690, EPI_ISL_819691, EPI_ISL_819692, EPI_ISL_819693, EPI_ISL_819694, EPI_ISL_819695, EPI_ISL_819696, EPI_ISL_819697, EPI_ISL_819698, EPI_ISL_819699, EPI_ISL_819700, EPI_ISL_819701, EPI_ISL_819702, EPI_ISL_819703, EPI_ISL_819704, EPI_ISL_819705, EPI_ISL_819706, EPI_ISL_819707, EPI_ISL_819708, EPI_ISL_819709, EPI_ISL_819710, EPI_ISL_819711, EPI_ISL_819712, EPI_ISL_819713, EPI_ISL_819714, EPI_ISL_819715, EPI_ISL_819716, EPI_ISL_819717, EPI_ISL_819718, EPI_ISL_819719, EPI_ISL_819720, EPI_ISL_819721, EPI_ISL_819722, EPI_ISL_819723 | | | |
| see above | Oxford Viromics, NDM, University of Oxford; Oxford University Hospitals; Basingstoke and North Hampshire Hospital | COVID-19 Genomics UK (COG-UK) Consortium | Tanya Golubchik, David Bonsall, George Macintyre, Amy Trebes, Mariateresa de Cesare, Catrin Moore, Alex Mobbs, Anita Justice, Robert Shaw, Monique Andersson, Timothy Peto, Emma Wise, Nathan Moore, Jessica Lynch, Nick Cortes, Matilde Mori, Stephen Kidd, David Buck, John Todd, Christophe Fraser |
| EPI_ISL_819724, EPI_ISL_819725, EPI_ISL_819726, EPI_ISL_819727, EPI_ISL_819728, EPI_ISL_819729, EPI_ISL_819730, EPI_ISL_819731 | Queens Medical Centre, Clinical Microbiology Department / DeepSeq Nottingham | COVID-19 Genomics UK (COG-UK) Consortium | Gemma Clark, Wendy Smith, Manjinder Khakh, Vicki M Fleming, Michelle M Lister, Hannah Howson-Wells, Jonathan Ball, Patrick McClure, Joseph Chappell, Theocharis Tsoleridis, Nadine Holmes, Matthew Carlisle, Christopher Moore, Fei Sang, Johnny Debebe, Victoria Wright, Matthew Loose |
| EPI_ISL_819732, EPI_ISL_819733, EPI_ISL_819734, EPI_ISL_819735, EPI_ISL_819736, EPI_ISL_819737, EPI_ISL_819738, EPI_ISL_819739, EPI_ISL_819740, EPI_ISL_819741, EPI_ISL_819742, EPI_ISL_819743, EPI_ISL_819744, EPI_ISL_819745, EPI_ISL_819746, EPI_ISL_819747, EPI_ISL_819748, EPI_ISL_819749, EPI_ISL_819750, EPI_ISL_819751, EPI_ISL_819752, EPI_ISL_819753, EPI_ISL_819754, EPI_ISL_819755, EPI_ISL_819756 | | | |
| see above | Oxford Viromics, NDM, University of Oxford; Oxford University Hospitals; Basingstoke and North Hampshire Hospital | COVID-19 Genomics UK (COG-UK) Consortium | Tanya Golubchik, David Bonsall, George Macintyre, Amy Trebes, Mariateresa de Cesare, Catrin Moore, Alex Mobbs, Anita Justice, Robert Shaw, Monique Andersson, Timothy Peto, Emma Wise, Nathan Moore, Jessica Lynch, Nick Cortes, Matilde Mori, Stephen Kidd, David Buck, John Todd, Christophe Fraser |
| EPI_ISL_819757, EPI_ISL_819758 | Queens Medical Centre, Clinical Microbiology Department / DeepSeq Nottingham | COVID-19 Genomics UK (COG-UK) Consortium | Gemma Clark, Wendy Smith, Manjinder Khakh, Vicki M Fleming, Michelle M Lister, Hannah Howson-Wells, Jonathan Ball, Patrick McClure, Joseph Chappell, Theocharis Tsoleridis, Nadine Holmes, Matthew Carlisle, Christopher Moore, Fei Sang, Johnny Debebe, Victoria Wright, Matthew Loose |
| EPI_ISL_819759, EPI_ISL_819760, EPI_ISL_819761, EPI_ISL_819762 | Oxford Viromics, NDM, University of Oxford; Oxford University Hospitals; Basingstoke and North Hampshire Hospital | COVID-19 Genomics UK (COG-UK) Consortium | Tanya Golubchik, David Bonsall, George Macintyre, Amy Trebes, Mariateresa de Cesare, Catrin Moore, Alex Mobbs, Anita Justice, Robert Shaw, Monique Andersson, Timothy Peto, Emma Wise, Nathan Moore, Jessica Lynch, Nick Cortes, Matilde Mori, Stephen Kidd, David Buck, John Todd, Christophe Fraser |
| EPI_ISL_819763 | Queens Medical Centre, Clinical Microbiology Department / DeepSeq Nottingham | COVID-19 Genomics UK (COG-UK) Consortium | Gemma Clark, Wendy Smith, Manjinder Khakh, Vicki M Fleming, Michelle M Lister, Hannah Howson-Wells, Jonathan Ball, Patrick McClure, Joseph Chappell, Theocharis Tsoleridis, Nadine Holmes, Matthew Carlisle, Christopher Moore, Fei Sang, Johnny Debebe, Victoria Wright, Matthew Loose |
| EPI_ISL_819764, EPI_ISL_819765, EPI_ISL_819766, EPI_ISL_819767, EPI_ISL_819768, EPI_ISL_819769, EPI_ISL_819770, EPI_ISL_819771, EPI_ISL_819772, EPI_ISL_819773, EPI_ISL_819774, EPI_ISL_819775, EPI_ISL_819776, EPI_ISL_819777, EPI_ISL_819778, EPI_ISL_819779, EPI_ISL_819780, EPI_ISL_819781, EPI_ISL_819782, EPI_ISL_819783, EPI_ISL_819784, EPI_ISL_819785, EPI_ISL_819786, EPI_ISL_819787, EPI_ISL_819788, EPI_ISL_819789, EPI_ISL_819790, EPI_ISL_819791, EPI_ISL_819792, EPI_ISL_819793, EPI_ISL_819794 | | | |
| see above | Oxford Viromics, NDM, University of Oxford; Oxford University Hospitals; Basingstoke and North Hampshire Hospital | COVID-19 Genomics UK (COG-UK) Consortium | Tanya Golubchik, David Bonsall, George Macintyre, Amy Trebes, Mariateresa de Cesare, Catrin Moore, Alex Mobbs, Anita Justice, Robert Shaw, Monique Andersson, Timothy Peto, Emma Wise, Nathan Moore, Jessica Lynch, Nick Cortes, Matilde Mori, Stephen Kidd, David Buck, John Todd, Christophe Fraser |
| EPI_ISL_819795 | Queens Medical Centre, Clinical Microbiology Department / DeepSeq Nottingham | COVID-19 Genomics UK (COG-UK) Consortium | Gemma Clark, Wendy Smith, Manjinder Khakh, Vicki M Fleming, Michelle M Lister, Hannah Howson-Wells, Jonathan Ball, Patrick McClure, Joseph Chappell, Theocharis Tsoleridis, Nadine Holmes, Matthew Carlisle, Christopher Moore, Fei Sang, Johnny Debebe, Victoria Wright, Matthew Loose |
| EPI_ISL_819796, EPI_ISL_819797, EPI_ISL_819798 | Oxford Viromics, NDM, University of Oxford; Oxford University Hospitals; Basingstoke and North Hampshire Hospital | COVID-19 Genomics UK (COG-UK) Consortium | Tanya Golubchik, David Bonsall, George Macintyre, Amy Trebes, Mariateresa de Cesare, Catrin Moore, Alex Mobbs, Anita Justice, Robert Shaw, Monique Andersson, Timothy Peto, Emma Wise, Nathan Moore, Jessica Lynch, Nick Cortes, Matilde Mori, Stephen Kidd, David Buck, John Todd, Christophe Fraser |
| EPI_ISL_819799, EPI_ISL_819800, EPI_ISL_819801, EPI_ISL_819802 | Queens Medical Centre, Clinical Microbiology Department / DeepSeq Nottingham | COVID-19 Genomics UK (COG-UK) Consortium | Gemma Clark, Wendy Smith, Manjinder Khakh, Vicki M Fleming, Michelle M Lister, Hannah Howson-Wells, Jonathan Ball, Patrick McClure, Joseph Chappell, Theocharis Tsoleridis, Nadine Holmes, Matthew Carlisle, Christopher Moore, Fei Sang, Johnny Debebe, Victoria Wright, Matthew Loose |
| EPI_ISL_819803, EPI_ISL_819804, EPI_ISL_819805, EPI_ISL_819806, EPI_ISL_819807, EPI_ISL_819808, EPI_ISL_819809, EPI_ISL_819810, EPI_ISL_819811, EPI_ISL_819812, EPI_ISL_819813, EPI_ISL_819814, EPI_ISL_819815, EPI_ISL_819816, EPI_ISL_819817, EPI_ISL_819818, EPI_ISL_819819, EPI_ISL_819820 | | | |
| see above | Oxford Viromics, NDM, University of Oxford; Oxford University Hospitals; Basingstoke and North Hampshire Hospital | COVID-19 Genomics UK (COG-UK) Consortium | Tanya Golubchik, David Bonsall, George Macintyre, Amy Trebes, Mariateresa de Cesare, Catrin Moore, Alex Mobbs, Anita Justice, Robert Shaw, Monique Andersson, Timothy Peto, Emma Wise, Nathan Moore, Jessica Lynch, Nick Cortes, Matilde Mori, Stephen Kidd, David Buck, John Todd, Christophe Fraser |
| EPI_ISL_822347, EPI_ISL_822348, EPI_ISL_822349, EPI_ISL_822350, EPI_ISL_822351, EPI_ISL_822352, EPI_ISL_822353, EPI_ISL_822354, EPI_ISL_822355, EPI_ISL_822356, EPI_ISL_822357, EPI_ISL_822358, EPI_ISL_822359, EPI_ISL_822360, EPI_ISL_822361, EPI_ISL_822362, EPI_ISL_822363, EPI_ISL_822364, EPI_ISL_822365, EPI_ISL_822840, EPI_ISL_822846, EPI_ISL_822847, EPI_ISL_822851, EPI_ISL_822852, EPI_ISL_822854, EPI_ISL_822855, EPI_ISL_822856, EPI_ISL_822857, EPI_ISL_822858, EPI_ISL_822861, EPI_ISL_822862, EPI_ISL_822870, EPI_ISL_822871, EPI_ISL_822873, EPI_ISL_822876, EPI_ISL_822890, EPI_ISL_822893, EPI_ISL_822904, EPI_ISL_822906, EPI_ISL_822910, EPI_ISL_822915, EPI_ISL_822919, EPI_ISL_822929, EPI_ISL_822935, EPI_ISL_822945, EPI_ISL_822956, EPI_ISL_822960, EPI_ISL_822965, EPI_ISL_822966, EPI_ISL_822969, EPI_ISL_822976, EPI_ISL_822981, EPI_ISL_822982, EPI_ISL_822984, EPI_ISL_822986, EPI_ISL_822987, EPI_ISL_822989, EPI_ISL_822990, EPI_ISL_822991, EPI_ISL_822992, EPI_ISL_822993, EPI_ISL_822994, EPI_ISL_822995, EPI_ISL_822996, EPI_ISL_822999, EPI_ISL_823006, EPI_ISL_823009, EPI_ISL_823010, EPI_ISL_823011, EPI_ISL_823020, EPI_ISL_823025, EPI_ISL_823026, EPI_ISL_823027, EPI_ISL_823028, EPI_ISL_823031, EPI_ISL_823032, EPI_ISL_823033, EPI_ISL_823034, EPI_ISL_823111, EPI_ISL_823114, EPI_ISL_823120, EPI_ISL_823122, EPI_ISL_823123, EPI_ISL_823125, EPI_ISL_823126, EPI_ISL_823128, EPI_ISL_823130, EPI_ISL_823134, EPI_ISL_823144, EPI_ISL_823157, EPI_ISL_823163, EPI_ISL_823174, EPI_ISL_823178, EPI_ISL_823179, EPI_ISL_823180, EPI_ISL_823181, EPI_ISL_823182, EPI_ISL_823184, EPI_ISL_823185, EPI_ISL_823186, EPI_ISL_823187, EPI_ISL_823188, EPI_ISL_823197, EPI_ISL_823198, EPI_ISL_823199, EPI_ISL_823234, EPI_ISL_823246, EPI_ISL_823247, EPI_ISL_823248, EPI_ISL_823249, EPI_ISL_823250, EPI_ISL_823251, EPI_ISL_823252, EPI_ISL_823253, EPI_ISL_823254, EPI_ISL_823255, EPI_ISL_823256, EPI_ISL_823257, EPI_ISL_823258, EPI_ISL_823259, EPI_ISL_823260, EPI_ISL_823261, EPI_ISL_823262, EPI_ISL_823263, EPI_ISL_823264, EPI_ISL_823265, EPI_ISL_823266, EPI_ISL_823267, EPI_ISL_823268, EPI_ISL_823269, EPI_ISL_823270, EPI_ISL_823271, EPI_ISL_823272, EPI_ISL_823273, EPI_ISL_823274, EPI_ISL_823275, EPI_ISL_823276, EPI_ISL_823277, EPI_ISL_823278, EPI_ISL_823279, EPI_ISL_823280, EPI_ISL_823281, EPI_ISL_823282, EPI_ISL_823283, EPI_ISL_823284, EPI_ISL_823285, EPI_ISL_823286, EPI_ISL_823287, EPI_ISL_823288, EPI_ISL_823289, EPI_ISL_823290, EPI_ISL_823291, EPI_ISL_823292, EPI_ISL_823293, EPI_ISL_823294, EPI_ISL_823295, EPI_ISL_823296, EPI_ISL_823297, EPI_ISL_823298, EPI_ISL_823299, EPI_ISL_823300, EPI_ISL_823301, EPI_ISL_823302, EPI_ISL_823303, EPI_ISL_823304, EPI_ISL_823305, EPI_ISL_823306, EPI_ISL_823307, EPI_ISL_823308, EPI_ISL_823309, EPI_ISL_823310, EPI_ISL_823311, EPI_ISL_823312, EPI_ISL_823313, EPI_ISL_823314, EPI_ISL_823315, EPI_ISL_823316, EPI_ISL_823318, EPI_ISL_823319, EPI_ISL_823321, EPI_ISL_823322, EPI_ISL_823325, EPI_ISL_823337, EPI_ISL_823340, EPI_ISL_823342, EPI_ISL_823366, EPI_ISL_823370, EPI_ISL_823384, EPI_ISL_823385, EPI_ISL_823389, EPI_ISL_823404, EPI_ISL_823405, EPI_ISL_823406, EPI_ISL_823407, EPI_ISL_823408, EPI_ISL_823409, EPI_ISL_823410, EPI_ISL_823411, EPI_ISL_823412, EPI_ISL_823413, EPI_ISL_823414, EPI_ISL_823415, EPI_ISL_823416, EPI_ISL_823417, EPI_ISL_823418, EPI_ISL_823419, EPI_ISL_823420, EPI_ISL_823421, EPI_ISL_823422, EPI_ISL_823423, EPI_ISL_823424, EPI_ISL_823425, EPI_ISL_823426, EPI_ISL_823427, EPI_ISL_823428, EPI_ISL_823429, EPI_ISL_823430, EPI_ISL_823431, EPI_ISL_823432, EPI_ISL_823433, EPI_ISL_823434, EPI_ISL_823436, EPI_ISL_823437 | | | |
| see above | Wales Specialist Virology Centre Sequencing lab: Pathogen Genomics Unit | COVID-19 Genomics UK (COG-UK) Consortium | Catherine Moore, Johnathan Evans, Laura Gifford, Malorie Perry, Simon Cottrell, Angela Marchbank, Alec Birchley, Alexander Adams, Amy Gaskin, Bree Gatica-Wilcox, Jason Coombes, Joel Southgate, Lauren Gilbert, Lee Graham, Nicole Pacchiarini, Sara Kumziene-Summerhayes, Sarah Taylor, Sophie Jones, Sara Rey, Matthew Bull, Joanne Watkins, Sally Corden, Tom Connor |

| | | | |
|---|---|---|---|
| EPI_ISL_824101, EPI_ISL_824183, EPI_ISL_824184, EPI_ISL_824218, EPI_ISL_824219, EPI_ISL_824220, EPI_ISL_824241, EPI_ISL_824258 | Dutch COVID-19 response team | National Institute for Public Health and the Environment (RIVM) | Adam Meijer, Harry Vennema, Jeroen Cremer, Sharon van den Brink, Bas van der Veer, AnneMarie van den Brandt, Florian Zwagemaker, Dennis Schmitz, Chantal Reusken, on behalf of the national COVID-19 response team |
| EPI_ISL_824337, EPI_ISL_824340, EPI_ISL_824349, EPI_ISL_824381 | Michigan Department of Health and Human Services, Bureau of Laboratories | Michigan Department of Health and Human Services, Bureau of Laboratories | Blankenship HM, Riner D, Soehnlen MK |
| EPI_ISL_824404, EPI_ISL_824405, EPI_ISL_824406 | California Department of Public Health | California Department of Public Health | CDPH IDLB COVIDNet |
| EPI_ISL_824922, EPI_ISL_824923, EPI_ISL_824924, EPI_ISL_824925, EPI_ISL_824930 | Arizona State Public Health Laboratory | Arizona State Public Health Laboratory | Trung Huynh, Jessica Escobar, Katherine Fullerton, Nobuko Fukushima, Stacy White, Linda Getsinger, Victor Waddell |
| EPI_ISL_824981 | Maryland Public Health Laboratory | Maryland Public Health Laboratory | Maryland Department of Health Laboratories Administration |
| EPI_ISL_825008, EPI_ISL_825009, EPI_ISL_825010 | Arizona State Public Health Laboratory | Arizona State Public Health Laboratory | Trung Huynh, Jessica Escobar, Katherine Fullerton, Nobuko Fukushima, Stacy White, Linda Getsinger, Victor Waddell |
| EPI_ISL_825132, EPI_ISL_825133, EPI_ISL_825134, EPI_ISL_825135, EPI_ISL_825137, EPI_ISL_825138, EPI_ISL_825139 | NHLS-IALCH | KRISP, KZN Research Innovation and Sequencing Platform | Giandhari J, Pillay S, Lessells R, Mdlalose K, York D, Khan S, Tegally H, Wilkinson E, de Oliveira T |
| EPI_ISL_825328, EPI_ISL_825379, EPI_ISL_825380, EPI_ISL_825381 | Hospital Universitari Vall d'Hebron - Vall d'Hebron Institut de Rercerca | Hospital Universitari Vall d'Hebron | Cristina Andrés, Maria Piñana, Josep F Abril, Damir Garcia-Cehic, Ariadna Rando, Juliana Esperalba, Maria Gema Codina, Carla Castillo, Maria Carmen Martin, Tomàs Pumarola, Josep Quer, Andrés Antón |
| EPI_ISL_825438, EPI_ISL_825439, EPI_ISL_825440, EPI_ISL_825441, EPI_ISL_825442, EPI_ISL_825443, EPI_ISL_825444, EPI_ISL_825445, EPI_ISL_825461, EPI_ISL_825462, EPI_ISL_825463, EPI_ISL_825464 | | | |
| see above | NHLS-IALCH | KRISP, KZN Research Innovation and Sequencing Platform | Giandhari J, Pillay S, Lessells R, Mdlalose K, York D, Khan S, Tegally H, Wilkinson E, de Oliveira T |
| EPI_ISL_825807 | Nigeria Centre For Disease Control | National reference Laboratory, NCDC, Gaduwa, Abuja | Dr Ndodo Nnaemeka, Olusola Anuoluwapo Akanbi, Chimaobi Chukwu, Dr Omoare Adesuyi, Esebanmen Grace, Anthony Ahumibe, Naidoo Dhamari, Nwando Mba, Dr Chikwe Ihekweazu |
| EPI_ISL_826286, EPI_ISL_826291, EPI_ISL_826293, EPI_ISL_826294, EPI_ISL_826295 | The Jackson Laboratory | The Jackson Laboratory | Lloyd M, Maurya R, Renzette N, Omerza G, Kelly K, Li L, Wei C L, Adams M |
| EPI_ISL_826852, EPI_ISL_826864, EPI_ISL_826867, EPI_ISL_826889, EPI_ISL_826890, EPI_ISL_826891 | deCODE genetics | deCODE genetics | Daniel F Gudbjartsson; Agnar Helgason; Hakon Jonsson; Olafur T Magnusson; Pall Melsted; Gudmundur L Norddahl; Jona Saemundsdottir; Asgeir Sigurdsson; Patrick Sulem; Arna B Agustsdottir; Hannes Eggertsson; Berglind Eiriksdottir; Run Fridriksdottir; Elisabet E Gardarsdottir; Gudmundur Georgsson; Olafia S Gretarsdottir; Kjartan R Gudmundsson; Thora R Gunnarsdottir; Arnaldur Gylfason; Hilma Holm; Brynjar O Jensson; Aslaug Jonasdottir; Kamilla S Josefsdottir; Thordur Kristjansson; Droplaug N Magnusdottir; Solvi Rognvaldsson; Louise le Roux; Gudrun Sigmundsdottir; Gardar Sveinbjornsson; Kristin E Sveinsdottir; Maney Sveinsdottir; Emil A Thorarensen; Bjarni Thorbjornsson; Gisli Masson; Ingileif Jonsdottir; Alma Moller; Thorolfur Gudnason; Karl G Kristinsson; Unnur Thorsteinsdottir; Kari Stefansson |
| EPI_ISL_826893 | The National University Hospital of Iceland | deCODE genetics | Daniel F Gudbjartsson; Agnar Helgason; Hakon Jonsson; Olafur T Magnusson; Pall Melsted; Gudmundur L Norddahl; Jona Saemundsdottir; Asgeir Sigurdsson; Patrick Sulem; Arna B Agustsdottir; Hannes Eggertsson; Berglind Eiriksdottir; Run Fridriksdottir; Elisabet E Gardarsdottir; Gudmundur Georgsson; Olafia S Gretarsdottir; Kjartan R Gudmundsson; Thora R Gunnarsdottir; Arnaldur Gylfason; Hilma Holm; Brynjar O Jensson; Aslaug Jonasdottir; Kamilla S Josefsdottir; Thordur Kristjansson; Droplaug N Magnusdottir; Solvi Rognvaldsson; Louise le Roux; Gudrun Sigmundsdottir; Gardar Sveinbjornsson; Kristin E Sveinsdottir; Maney Sveinsdottir; Emil A Thorarensen; Bjarni Thorbjornsson; Gisli Masson; Ingileif Jonsdottir; Alma Moller; Thorolfur Gudnason; Karl G Kristinsson; Unnur Thorsteinsdottir; Kari Stefansson |
| EPI_ISL_827486, EPI_ISL_827487, EPI_ISL_827488, EPI_ISL_827489, EPI_ISL_827490, EPI_ISL_827491, EPI_ISL_827507, EPI_ISL_827511, EPI_ISL_827524, EPI_ISL_827998, EPI_ISL_828380, EPI_ISL_828385, EPI_ISL_828558, EPI_ISL_828797, EPI_ISL_828806, EPI_ISL_829135, EPI_ISL_829141, EPI_ISL_829427, EPI_ISL_829431, EPI_ISL_829447, EPI_ISL_829664, EPI_ISL_829693, EPI_ISL_829694, EPI_ISL_829695, EPI_ISL_829696, EPI_ISL_829716, EPI_ISL_829884, EPI_ISL_829918 | | | |
| see above | deCODE genetics | deCODE genetics | Daniel F Gudbjartsson; Agnar Helgason; Hakon Jonsson; Olafur T Magnusson; Pall Melsted; Gudmundur L Norddahl; Jona Saemundsdottir; Asgeir Sigurdsson; Patrick Sulem; Arna B Agustsdottir; Hannes Eggertsson; Berglind Eiriksdottir; Run Fridriksdottir; Elisabet E Gardarsdottir; Gudmundur Georgsson; Olafia S Gretarsdottir; Kjartan R Gudmundsson; Thora R Gunnarsdottir; Arnaldur Gylfason; Hilma Holm; Brynjar O Jensson; Aslaug Jonasdottir; Kamilla S Josefsdottir; Thordur Kristjansson; Droplaug N Magnusdottir; Solvi Rognvaldsson; Louise le Roux; Gudrun Sigmundsdottir; Gardar Sveinbjornsson; Kristin E Sveinsdottir; Maney Sveinsdottir; Emil A Thorarensen; Bjarni Thorbjornsson; Gisli Masson; Ingileif Jonsdottir; Alma Moller; Thorolfur Gudnason; Karl G Kristinsson; Unnur Thorsteinsdottir; Kari Stefansson |
| EPI_ISL_829989 | The National University Hospital of Iceland | deCODE genetics | Daniel F Gudbjartsson; Agnar Helgason; Hakon Jonsson; Olafur T Magnusson; Pall Melsted; Gudmundur L Norddahl; Jona Saemundsdottir; Asgeir Sigurdsson; Patrick Sulem; Arna B Agustsdottir; Hannes Eggertsson; Berglind Eiriksdottir; Run Fridriksdottir; Elisabet E Gardarsdottir; Gudmundur Georgsson; Olafia S Gretarsdottir; Kjartan R Gudmundsson; Thora R Gunnarsdottir; Arnaldur Gylfason; Hilma Holm; Brynjar O Jensson; Aslaug Jonasdottir; Kamilla S Josefsdottir; Thordur Kristjansson; Droplaug N Magnusdottir; Solvi Rognvaldsson; Louise le Roux; Gudrun Sigmundsdottir; Gardar Sveinbjornsson; Kristin E Sveinsdottir; Maney Sveinsdottir; Emil A Thorarensen; Bjarni Thorbjornsson; Gisli Masson; Ingileif Jonsdottir; Alma Moller; Thorolfur Gudnason; Karl G Kristinsson; Unnur Thorsteinsdottir; Kari Stefansson |
| EPI_ISL_830187, EPI_ISL_830188, EPI_ISL_830409 | deCODE genetics | deCODE genetics | Daniel F Gudbjartsson; Agnar Helgason; Hakon Jonsson; Olafur T Magnusson; Pall Melsted; Gudmundur L Norddahl; Jona Saemundsdottir; Asgeir Sigurdsson; Patrick Sulem; Arna B Agustsdottir; Hannes Eggertsson; Berglind Eiriksdottir; Run Fridriksdottir; Elisabet E Gardarsdottir; Gudmundur Georgsson; Olafia S Gretarsdottir; Kjartan R Gudmundsson; Thora R Gunnarsdottir; Arnaldur Gylfason; Hilma Holm; Brynjar O Jensson; Aslaug Jonasdottir; Kamilla S Josefsdottir; Thordur Kristjansson; Droplaug N Magnusdottir; Solvi Rognvaldsson; Louise le Roux; Gudrun Sigmundsdottir; Gardar Sveinbjornsson; Kristin E Sveinsdottir; Maney Sveinsdottir; Emil A Thorarensen; Bjarni Thorbjornsson; Gisli Masson; Ingileif Jonsdottir; Alma Moller; Thorolfur Gudnason; Karl G Kristinsson; Unnur Thorsteinsdottir; Kari Stefansson |
| EPI_ISL_831028, EPI_ISL_831037, EPI_ISL_831038, EPI_ISL_831264, EPI_ISL_831267, EPI_ISL_831268, EPI_ISL_831269, EPI_ISL_831270, EPI_ISL_831271, EPI_ISL_831272, EPI_ISL_831273, EPI_ISL_831274, EPI_ISL_831281, EPI_ISL_831283, EPI_ISL_831284 | | | |
| see above | Hospital Universitario La Paz (Madrid) | SeqCOVID-SPAIN consortium/IBV(CSIC) | Fernando Lázaro-Perona, María Rodríguez-Tejedor, Elias Dahdouh, Jesús Mingorance and SeqCOVID-SPAIN consortium |
| EPI_ISL_831375, EPI_ISL_831376, EPI_ISL_831377, EPI_ISL_831378 | Limbach - MVZ Labor Dr. Limbach & Kollegen | Robert Koch Institute, Influenza and respiratory viruses FG17 & Bioinformatics MF1, Berlin, Germany | Dr. Konrad Bode, Stephan Fuchs, Stefan Kroeger, Marianne Wedde, Oliver Drechsel, Aleksandar Radonic, Rene Kmiecinski, Ralf Duerrwald, Thorsten Wolff |
| EPI_ISL_831652, EPI_ISL_831653, EPI_ISL_831657, EPI_ISL_831658 | Institute for Infectious Diseases, University of Bern, Switzerland | Institute for Infectious Diseases, University of Bern, Switzerland | Michel C Koch, Christian Baumann, Miguel A Terrazos Miani, Cora Sägesser, Pascal Bittel, Stephen L Leib, Peter Keller, Franziska Suter-Riniker, Alban Ramette |
| EPI_ISL_831734, EPI_ISL_831747, EPI_ISL_831751, EPI_ISL_831874, EPI_ISL_831875, EPI_ISL_831876, EPI_ISL_831882, EPI_ISL_831883, EPI_ISL_831884, EPI_ISL_831885, EPI_ISL_831886 | | | |
| see above | United States Air Force School of Aerospace Medicine | United States Air Force School of Aerospace Medicine | Anthony Fries, Jennifer Meyer, William Gruner, Amanda Javorina, Sarah Purves, Clarise Starr, Elizabeth Macias |
| EPI_ISL_831958 | Klinisk mikrobiologi | The Public Health Agency of Sweden | Department of Microbiology, The Public Health Agency of Sweden |
| EPI_ISL_831993, EPI_ISL_831995 | TATAA Biocenter | The Public Health Agency of Sweden | Department of Microbiology, The Public Health Agency of Sweden |
| EPI_ISL_832001, EPI_ISL_832002 | Klinisk mikrobiologi | The Public Health Agency of Sweden | Department of Microbiology, The Public Health Agency of Sweden |
| EPI_ISL_832105, EPI_ISL_832110, EPI_ISL_832111, EPI_ISL_832112, EPI_ISL_832113, EPI_ISL_832114, EPI_ISL_832115, EPI_ISL_832117, EPI_ISL_832118, EPI_ISL_832119, EPI_ISL_832120 | | | |
| see above | MD Laboratories | Los Angeles County PHL | P. Hemarajata et al. |
| EPI_ISL_832130, EPI_ISL_832131, EPI_ISL_832132, EPI_ISL_832133, EPI_ISL_832134, EPI_ISL_832135, EPI_ISL_832136, EPI_ISL_832137, EPI_ISL_832138, EPI_ISL_832139, EPI_ISL_832140, EPI_ISL_832141, EPI_ISL_832142, EPI_ISL_832143 | | | |

| | | | |
|---|---|---|---|
| see above | Hospital | National Reference Center for Viruses of Respiratory Infections, Institut Pasteur, Paris | Marion Barbet, Sylvie Behillil, Méline Bizard, Angela Brisebarre, Camille Capel, Etienne Simon-Lorière, Vincent Enouf, Maud Vanpeene, Sylvie van der Werf, Laurent Andreoletti |
| EPI_ISL_832167 | Sentinelles Province | National Reference Center for Viruses of Respiratory Infections, Institut Pasteur, Paris | Marion Barbet, Sylvie Behillil, Méline Bizard, Angela Brisebarre, Camille Capel, Etienne Simon-Lorière, Vincent Enouf, Maud Vanpeene, Sylvie van der Werf, Jean-Michel Monnier |
| EPI_ISL_832179 | Hospital | National Reference Center for Viruses of Respiratory Infections, Institut Pasteur, Paris | Marion Barbet, Sylvie Behillil, Méline Bizard, Angela Brisebarre, Camille Capel, Etienne Simon-Lorière, Vincent Enouf, Maud Vanpeene, Sylvie van der Werf, Laurent Roudiere |
| EPI_ISL_832244 | DOHMH Crown Heights | New York City Public Health Laboratory | Jade Wang, et al. |
| EPI_ISL_832250 | DOHMH Jamaica | New York City Public Health Laboratory | Jade Wang, et al. |
| EPI_ISL_832255 | DOHMH Morrisania | New York City Public Health Laboratory | Jade Wang, et al. |
| EPI_ISL_832258, EPI_ISL_832259 | DOHMH PHL | New York City Public Health Laboratory | Jade Wang, et al. |
| EPI_ISL_832260, EPI_ISL_832261, EPI_ISL_832262 | DOHMH Corona | New York City Public Health Laboratory | Jade Wang, et al. |
| EPI_ISL_832272, EPI_ISL_832273, EPI_ISL_832274 | DOHMH Jamaica | New York City Public Health Laboratory | Jade Wang, et al. |
| EPI_ISL_832275 | DOHMH Crown Heights | New York City Public Health Laboratory | Jade Wang, et al. |
| EPI_ISL_832276 | DOHMH Jamaica | New York City Public Health Laboratory | Jade Wang, et al. |
| EPI_ISL_832277, EPI_ISL_832278, EPI_ISL_832279 | DOHMH Corona | New York City Public Health Laboratory | Jade Wang, et al. |
| EPI_ISL_832280, EPI_ISL_832281 | DOHMH Morrisania | New York City Public Health Laboratory | Jade Wang, et al. |
| EPI_ISL_832282 | DOHMH Chelsea | New York City Public Health Laboratory | Jade Wang, et al. |
| EPI_ISL_832283 | DOHMH Jamaica | New York City Public Health Laboratory | Jade Wang, et al. |
| EPI_ISL_832284 | DOHMH Central Harlem | New York City Public Health Laboratory | Jade Wang, et al. |
| EPI_ISL_832285 | DOHMH Morrisania | New York City Public Health Laboratory | Jade Wang, et al. |
| EPI_ISL_832286 | DOHMH Fort Greene | New York City Public Health Laboratory | Jade Wang, et al. |
| EPI_ISL_832287, EPI_ISL_832288 | DOHMH Crown Heights | New York City Public Health Laboratory | Jade Wang, et al. |
| EPI_ISL_832289 | DOHMH Morrisania | New York City Public Health Laboratory | Jade Wang, et al. |
| EPI_ISL_832290, EPI_ISL_832291 | DOHMH Corona | New York City Public Health Laboratory | Jade Wang, et al. |
| EPI_ISL_832292, EPI_ISL_832293 | DOHMH Crown Heights | New York City Public Health Laboratory | Jade Wang, et al. |
| EPI_ISL_832294 | DOHMH Jamaica | New York City Public Health Laboratory | Jade Wang, et al. |
| EPI_ISL_832295 | DOHMH Crown Heights | New York City Public Health Laboratory | Jade Wang, et al. |
| EPI_ISL_832296 | DOHMH Riverside | New York City Public Health Laboratory | Jade Wang, et al. |
| EPI_ISL_832297, EPI_ISL_832298 | DOHMH Jamaica | New York City Public Health Laboratory | Jade Wang, et al. |
| EPI_ISL_832387, EPI_ISL_832388 | Santa Clara County Public Health Laboratory | Santa Clara County Public Health Laboratory | Santa Clara County Public Health Department |
| EPI_ISL_832815, EPI_ISL_832816 | SIESP CHIETI - DRIVE IN LANCIANO | Istitituto Zooprofilattico Sperimentale dell'Abruzzo e Molise "G.Caporale" | Lorusso A, Marcacci M, Di Domenico M, Curini V, Ancora M, Cammà C, Rinaldi A, Mangone I, Di Pasquale A, Puglia I, Calistri P, Savini G. |
| EPI_ISL_832818 | Ospedale Civile Atri | Istitituto Zooprofilattico Sperimentale dell'Abruzzo e Molise "G.Caporale" | Lorusso A, Marcacci M, Di Domenico M, Curini V, Ancora M, Cammà C, Rinaldi A, Mangone I, Di Pasquale A, Puglia I, Calistri P, Savini G. |
| EPI_ISL_832819 | SIESP DIPARTIMENTO DI PREVENZIONE SULMONA | Istitituto Zooprofilattico Sperimentale dell'Abruzzo e Molise "G.Caporale" | Lorusso A, Marcacci M, Di Domenico M, Curini V, Ancora M, Cammà C, Rinaldi A, Mangone I, Di Pasquale A, Puglia I, Calistri P, Savini G. |
| EPI_ISL_833133, EPI_ISL_833135 | Laboratorio de Ecologia de Doencas Transmissiveis na Amazonia, Instituto Leonidas e Maria Deane - Fiocruz Amazonia | Laboratorio de Ecologia de Doencas Transmissiveis na Amazonia, Instituto Leonidas e Maria Deane - Fiocruz Amazonia | Valdinete Nascimento, Victor Souza, André Corado, Fernanda Nascimento, George Silva, Ágatha Costa, Debora Duarte, Karina Pessoa, Matilde Mejía, Luciana Gonçalves, Maria Júlia Brandão, Michele Jesus, Felipe Naveca |
| EPI_ISL_833167, EPI_ISL_833171, EPI_ISL_833172, EPI_ISL_833173, EPI_ISL_833176 | DB Diagnosticos do Brasil | Instituto Adolfo Lutz, Interdiciplinary Procedures Center, Strategic Laboratory | Claudio Tavares Sacchi, Claudia Regina Gonçalves, Erica Valessa Ramos Gomes, Karoline Rodrigues Campos |
| EPI_ISL_833250 | SIESP CHIETI - DRIVE IN ORTONA | Istituto Zooprofilattico Sperimentale dell'Abruzzo e Molise "G. Caporale" | Lorusso A, Marcacci M, Di Domenico M, Ancora M, Curini V, Mangone I, Rinaldi A, Di Pasquale A, Cammà C, Puglia I, Calistri P, Savini G |
| EPI_ISL_836981, EPI_ISL_836982, EPI_ISL_836983, EPI_ISL_836984, EPI_ISL_836985, EPI_ISL_836986, EPI_ISL_836987, EPI_ISL_837066, EPI_ISL_837068, EPI_ISL_837082, EPI_ISL_837084, EPI_ISL_837085, EPI_ISL_837097, EPI_ISL_837098, EPI_ISL_837099, EPI_ISL_837101, EPI_ISL_837102, EPI_ISL_837116, EPI_ISL_837117, EPI_ISL_837118, EPI_ISL_837119, EPI_ISL_837203, EPI_ISL_837204, EPI_ISL_837205, EPI_ISL_837206 | | | |
| see above | Respiratory Virus Unit, National Infection Service, Public Health England | COVID-19 Genomics UK (COG-UK) Consortium | PHE Covid Sequencing Team |
| EPI_ISL_837318, EPI_ISL_837449 | Istituto Zooprofilattico Sperimentale del Mezzogiorno | TIGEM | Patrizia Annunziata, Andrea Ballabio, Valentina Bouche, Davide Cacchiarelli (CorrespAuthor), Pellegrino Cerino, Chiara Colantuono, Lucio Di Filippo, Antonio Grimaldi, Antonio Limone, Gabriella Loconte, Anna Manfredi, Francesco Panariello, Biancamaria Pierri, Marcello Salvi, Lucia Vassallo |
| EPI_ISL_837837, EPI_ISL_837838, EPI_ISL_837839, EPI_ISL_837840, EPI_ISL_837841, EPI_ISL_837842 | Department of Pathology, University of Cambridge | COVID-19 Genomics UK (COG-UK) Consortium | Aminu S. Jahun, Yasmin Chaudhry, Grant Hall, Iliana Georgana, Myra Hosmillo, Martin D. Curran, Malte Pinckert, Surendra Parmar, Ian Goodfellow |
| EPI_ISL_838116, EPI_ISL_838118, EPI_ISL_838125, EPI_ISL_838128, EPI_ISL_838133 | West of Scotland Specialist Virology Centre, NHSGGC / MRC-University of Glasgow Centre for Virus Research | COVID-19 Genomics UK (COG-UK) Consortium | Ana da Silva Filipe, Natasha Johnson, Kathy Smollett, Daniel Mair, Stephen Carmichael, Alice Broos, Lily Tong, Jenna Nichols, Kyriaki Nomikou; Sarah McDonald; Richard Orton, Joseph Hughes, Sreenu Vattipally, David L Robertson; Alasdair MacLean, Rory Gunson; Sharif Shaaban, Matthew Holden; Rachel Blacow, Guy Mollett, Kathy Li, James Shepherd, Antonia Ho, Emma Thomson |
| EPI_ISL_838227, EPI_ISL_838228, EPI_ISL_838229, EPI_ISL_838230, EPI_ISL_838231, EPI_ISL_838232, EPI_ISL_838235, EPI_ISL_838236 | Virology Department, Royal Infirmary of Edinburgh, NHS Lothian / School of Biological Sciences, University of Edinburgh / Institute of Genetics and Molecular Medicine, University of Edinburgh | COVID-19 Genomics UK (COG-UK) Consortium | McHugh M, Dewar R, Rooke S, Gallagher M, Balcaza C, O'Toole Á, Scher E, Hill V, McCrone JT, Colquhoun R, Yu X, Jackson B, Rambaut A, Williams TC, Templeton K |
| EPI_ISL_838529, EPI_ISL_838531, EPI_ISL_838532, EPI_ISL_838533, EPI_ISL_838534, EPI_ISL_838535, EPI_ISL_838536, EPI_ISL_838537, EPI_ISL_838542, EPI_ISL_838545, EPI_ISL_838547, EPI_ISL_838549, EPI_ISL_838551, EPI_ISL_838552, EPI_ISL_838553, EPI_ISL_838554, EPI_ISL_838555, EPI_ISL_838556, EPI_ISL_838558, EPI_ISL_838559, EPI_ISL_838564, EPI_ISL_838568, EPI_ISL_838569, EPI_ISL_838570, EPI_ISL_838578, EPI_ISL_838579, EPI_ISL_838646, EPI_ISL_838666 | | | |
| see above | Liverpool Clinical Laboratories | COVID-19 Genomics UK (COG-UK) Consortium | Sam Haldenby, Anita Lucaci, Steve Paterson, Julian Hiscox, Alistair Darby, M Almsaud, A Alrezaihi, Muhannad Alruwaili, Stuart D Armstrong, Jones Benjamin, Eleanor G Bentley, Anu Chawla, Jordan J Clark, Angela Cowell, Richard Eccles, Isabel García-Dorival, Matthew Gemmell, Alessandro Gerada, PKF Gilmore, Richard Gregory, Ximeng Han, Catherine Hartley, Margaret Hughes, Miren Iturriza-Gomara, James Johnson, L Luu, Jenifer Manson, |

| | | | |
|---|---|---|---|
| | | | Charlotte Nelson, Elaine O'Toole, Cassie Olateju, Rebekah Penrice-Randal , Lucille Rainbow, N.P Randle, Trevor Ian Robinson, Parul Sharma, Ghada T Shawli, James P Stewart, Neil Swainston, Ecaterina Vamos, Joanne Watts, Mark Whitehead |
| EPI_ISL_838705, EPI_ISL_838708, EPI_ISL_838709, EPI_ISL_838711, EPI_ISL_838712, EPI_ISL_838713, EPI_ISL_838715, EPI_ISL_838718, EPI_ISL_838721, EPI_ISL_838723, EPI_ISL_838724, EPI_ISL_838725, EPI_ISL_838726, EPI_ISL_838727, EPI_ISL_838730, EPI_ISL_838731, EPI_ISL_838732, EPI_ISL_838733, EPI_ISL_839342, EPI_ISL_839343, EPI_ISL_839344, EPI_ISL_839347, EPI_ISL_839354, EPI_ISL_839355, EPI_ISL_839356, EPI_ISL_839357 | | | |
| see above | University College London, Great Ormond Street Hospital for Children NHS Foundation Trust, Imperial College Healthcare NHS Trust | COVID-19 Genomics UK (COG-UK) Consortium | Sergi Castellano, Rachel Williams, Mark Kristiansen, Paola Resende Silva, Sunando Roy, Tony Brooks, Helena Tutill, Paola Niola, Patricia Dyal, Charlotte Williams, Leysa Forrest, Yasmin Panchbhaya, Jacqueline Findlay, Samuel Weeks, Julianne Brown, Kathryn Harris, Paul Randell, James Price, Alison Holmes, Judith Breuer |
| EPI_ISL_839484, EPI_ISL_839508, EPI_ISL_839509, EPI_ISL_839510, EPI_ISL_839516, EPI_ISL_839517, EPI_ISL_839518, EPI_ISL_839582, EPI_ISL_839583, EPI_ISL_839584, EPI_ISL_839585, EPI_ISL_839586, EPI_ISL_839587, EPI_ISL_839588, EPI_ISL_839589, EPI_ISL_839591, EPI_ISL_839592, EPI_ISL_839600, EPI_ISL_839603, EPI_ISL_839636, EPI_ISL_839638, EPI_ISL_839640, EPI_ISL_839644, EPI_ISL_839645, EPI_ISL_839646, EPI_ISL_839648, EPI_ISL_839666, EPI_ISL_839731 | | | |
| see above | Northumbria University / South Tees Hospitals NHS Foundation Trust / North Cumbria Integrated Care NHS Foundation Trust / North Tees and Hartlepool NHS Foundation Trust / Newcastle Hospitals NHS Foundation Trust | COVID-19 Genomics UK (COG-UK) Consortium | Darren L Smith,Andrew Nelson,Matthew Bashton,Greg R Young,Joshua Loh,John Allan,Mohammad A Tariq,Giles S Holt,Gary Black,Wen C Yew,Lynn Dover,Paul Baker,Steve Liggett,Sarah Essex,Jane Greenaway,Debra Padgett,Clive Graham,Garren Scott,Edward Barton,Emma Swindells,Brendan Payne,Jennifer Collins,Yusri Taha,Gary Eltringham |
| EPI_ISL_840062, EPI_ISL_840063, EPI_ISL_840064, EPI_ISL_840065, EPI_ISL_840066, EPI_ISL_840067, EPI_ISL_840068, EPI_ISL_840069, EPI_ISL_840180, EPI_ISL_840181, EPI_ISL_840182, EPI_ISL_840183, EPI_ISL_840184, EPI_ISL_840185 | | | |
| see above | Lincolnshire Hospitals and DeepSeq Nottingham | COVID-19 Genomics UK (COG-UK) Consortium | Nichola Duckworth, Tim Sloan, Sarah Walsh, Jonathan Ball, Patrick McClure, Joeseph Chappell, Nadine Holmes, Matthew Carlisle, Christopher Moore, Fei Sang, Johnny Debebe, Victoria Wright, Matthew Loose |
| EPI_ISL_840256, EPI_ISL_840257, EPI_ISL_840260, EPI_ISL_840261, EPI_ISL_840263, EPI_ISL_840264, EPI_ISL_840267, EPI_ISL_840269, EPI_ISL_840270, EPI_ISL_840272, EPI_ISL_840274, EPI_ISL_840276, EPI_ISL_840278, EPI_ISL_840279, EPI_ISL_840280, EPI_ISL_840349, EPI_ISL_840350, EPI_ISL_840353, EPI_ISL_840357, EPI_ISL_840359, EPI_ISL_840360, EPI_ISL_840361 | | | |
| see above | Oxford Viromics, NDM, University of Oxford; Oxford University Hospitals; Basingstoke and North Hampshire Hospital | COVID-19 Genomics UK (COG-UK) Consortium | Tanya Golubchik, David Bonsall, George Macintyre, Amy Trebes, Mariateresa de Cesare, Catrin Moore, Alex Mobbs, Anita Justice, Robert Shaw, Monique Andersson, Timothy Peto, Emma Wise, Nathan Moore, Jessica Lynch, Nick Cortes, Matilde Mori, Stephen Kidd, David Buck, John Todd, Christophe Fraser |
| EPI_ISL_840712, EPI_ISL_840713, EPI_ISL_840714, EPI_ISL_840716 | Originating lab: Wales Specialist Virology Centre Sequencing lab: Pathogen Genomics Unit | Public Health Wales Microbiology Cardiff Wales Specialist Virology Centre | Catherine Moore, Johnathan Evans, Laura Gifford, Malorie Perry, Simon Cottrell, Angela Marchbank, Alec Birchley, Alexander Adams, Amy Gaskin, Bree Gatica-Wilcox, Jason Coombes, Joel Southgate, Lauren Gilbert, Lee Graham, Nicole Pacchiarini, Sara Kumziene-Summerhayes, Sarah Taylor, Sophie Jones, Sara Rey, Matthew Bull, Joanne Watkins, Sally Corden, Tom Connor |
| EPI_ISL_840836, EPI_ISL_840887 | Wales Specialist Virology Centre Sequencing lab: Pathogen Genomics Unit | Public Health Wales Microbiology Cardiff Wales Specialist Virology Centre | Catherine Moore, Johnathan Evans, Laura Gifford, Malorie Perry, Simon Cottrell, Angela Marchbank, Alec Birchley, Alexander Adams, Amy Gaskin, Bree Gatica-Wilcox, Jason Coombes, Joel Southgate, Lauren Gilbert, Lee Graham, Nicole Pacchiarini, Sara Kumziene-Summerhayes, Sarah Taylor, Sophie Jones, Sara Rey, Matthew Bull, Joanne Watkins, Sally Corden, Tom Connor |
| EPI_ISL_841505 | Originating lab: Wales Specialist Virology Centre Sequencing lab: Pathogen Genomics Unit | Public Health Wales Microbiology Cardiff Wales Specialist Virology Centre | Catherine Moore, Johnathan Evans, Laura Gifford, Malorie Perry, Simon Cottrell, Angela Marchbank, Alec Birchley, Alexander Adams, Amy Gaskin, Bree Gatica-Wilcox, Jason Coombes, Joel Southgate, Lauren Gilbert, Lee Graham, Nicole Pacchiarini, Sara Kumziene-Summerhayes, Sarah Taylor, Sophie Jones, Sara Rey, Matthew Bull, Joanne Watkins, Sally Corden, Tom Connor |
| EPI_ISL_841759, EPI_ISL_841770, EPI_ISL_841771, EPI_ISL_841772, EPI_ISL_841773, EPI_ISL_841775, EPI_ISL_841777, EPI_ISL_841778, EPI_ISL_841781, EPI_ISL_841783, EPI_ISL_841784, EPI_ISL_841785, EPI_ISL_841787, EPI_ISL_841789, EPI_ISL_841790, EPI_ISL_841791, EPI_ISL_841792, EPI_ISL_841793, EPI_ISL_841795, EPI_ISL_841797, EPI_ISL_841798, EPI_ISL_841799, EPI_ISL_841800, EPI_ISL_841801, EPI_ISL_841802, EPI_ISL_841803, EPI_ISL_841804, EPI_ISL_841805, EPI_ISL_841806, EPI_ISL_841808, EPI_ISL_841809, EPI_ISL_841810, EPI_ISL_841811, EPI_ISL_841813, EPI_ISL_841830, EPI_ISL_841831, EPI_ISL_841832, EPI_ISL_841833, EPI_ISL_841834, EPI_ISL_841835, EPI_ISL_841836, EPI_ISL_841837, EPI_ISL_841838, EPI_ISL_841839, EPI_ISL_841840, EPI_ISL_841841, EPI_ISL_841842, EPI_ISL_841843, EPI_ISL_841844, EPI_ISL_841845, EPI_ISL_841855, EPI_ISL_841856, EPI_ISL_841857, EPI_ISL_841858, EPI_ISL_841859, EPI_ISL_841860, EPI_ISL_841862, EPI_ISL_841863, EPI_ISL_841864, EPI_ISL_841866, EPI_ISL_841867, EPI_ISL_841868, EPI_ISL_841869, EPI_ISL_841870 | | | |
| see above | Centre for Enzyme Innovation, University of Portsmouth / Translational Research Laboratory, Portsmouth Hospitals NHS Trust | COVID-19 Genomics UK (COG-UK) Consortium | Angela Beckett,Yann Bourgeois,Garry Scarlett,Sharon Glaysher,Scott Elliott,Kelly Bicknell,Robert Impey,Allyson Lloyd,Sarah Wyllie,Ethan Butcher,Anoop Chauhan,Samuel Robson |
| EPI_ISL_842313, EPI_ISL_842317 | Virology Department, Sheffield Teaching Hospitals NHS Foundation Trust/Department of Infection, Immunity and Cardiovascular Disease, The Medical School, University of Sheffield | COVID-19 Genomics UK (COG-UK) Consortium | Thushan de Silva, Matthew Parker, Nikki Smith, Adri Angyal, Rebecca Brown, Luke Green, Rachel Tucker, Paul Parsons, Danielle Groves, Katie Johnson, Laura Carrilero, Alex Keeley, Dave Partridge, Matthew Wyles, Benjamin Lindsey, Mehmet Yavuz, Mohammad Raza, Cariad Evans |
| EPI_ISL_842633 | Norwegian Institute of Public Health, Department of Virology | Norwegian Institute of Public Health, Department of Virology | Kathrine Stene-Johansen, Kamilla Heddeland Instefjord, Hilde Elshaug, Atiya R Ali,Marie Paulsen Madsen, Rasmus Riis Kopperud, Hilde Vollan, Karoline Bragstad, Olav Hungnes |
| EPI_ISL_842686, EPI_ISL_842687, EPI_ISL_842689, EPI_ISL_842690, EPI_ISL_842698, EPI_ISL_842699, EPI_ISL_842701, EPI_ISL_842702, EPI_ISL_842703, EPI_ISL_842719, EPI_ISL_842721, EPI_ISL_842722, EPI_ISL_842723, EPI_ISL_842725 | | | |
| see above | University College London Hospital | COVID-19 Genomics UK (COG-UK) Consortium | Judith Heaney, Matthew Byott, Catherine Houlihan, Dan Frampton, Stuart Kirk, Moira Spyer and Eleni Nastouli |
| EPI_ISL_843091, EPI_ISL_843092, EPI_ISL_843105, EPI_ISL_843106, EPI_ISL_843107, EPI_ISL_843108, EPI_ISL_843109, EPI_ISL_843110, EPI_ISL_843116, EPI_ISL_843117, EPI_ISL_843118 | | | |
| see above | Barts Health NHS Trust | COVID-19 Genomics UK (COG-UK) Consortium | CUTINO-MOGUEL, Maria-Teresa; HARRINGTON, David; OWOYEMI, Dola; SHYLINI, Raghavendran; BROAD, Claire; KELE, Beatrix |
| EPI_ISL_845565 | National Public Health Laboratory, Cameroon | African Centre of Excellence for Genomics of Infectious Diseases (ACEGID), Redeemer's University | Oluniyi P.E. et al |
| EPI_ISL_845621, EPI_ISL_845624, EPI_ISL_845625 | Dirección de Sanidad Ejército | Instituto Nacional de Salud - Dirección de Investigación en Salud Pública | Katherine Laiton-Donato, Diego A. Álvarez-Díaz, Carlos Franco-Muñoz, Mauricio Pacheco-Montealegre, Maria T. Herrera-Sepúlveda, Jonathan Reales, Sheryll Corchuelo, Julian Naizaque, Gerardo Santamaría, Paola Muñoz-Laiton, Diego Andrés Prada, Magdalena Wiesner, Martha Lucia Ospina Martinez, Marcela Mercado-Reyes |
| EPI_ISL_845628, EPI_ISL_845629 | Laboratorio Médico Echavarria | Instituto Nacional de Salud - Dirección de Investigación en Salud Pública | Katherine Laiton-Donato, Diego A. Álvarez-Díaz, Carlos Franco-Muñoz, Mauricio Pacheco-Montealegre, Maria T. Herrera-Sepúlveda, Jonathan Reales, Sheryll Corchuelo, Julian Naizaque, Gerardo Santamaría, Paola Muñoz-Laiton, Diego Andrés Prada, Magdalena Wiesner, Martha Lucia Ospina Martinez, Marcela Mercado-Reyes |
| EPI_ISL_845656 | UNIDAD HEMATOLÓGICA ESPECIALIZADA | Instituto Nacional de Salud - Dirección de Investigación en Salud Pública | Katherine Laiton-Donato, Diego A. Álvarez-Díaz, Carlos Franco-Muñoz, Mauricio Pacheco-Montealegre, Maria T. Herrera-Sepúlveda, Jonathan Reales, Sheryll Corchuelo, Julian Naizaque, Gerardo Santamaría, Paola Muñoz-Laiton, Diego Andrés Prada, Magdalena Wiesner, Martha Lucia Ospina Martinez, Marcela Mercado-Reyes |
| EPI_ISL_845796 | Histopath | NSW Health Pathology - Institute of Clinical Pathology and Medical Research; Westmead Hospital; University of Sydney | CIDM-PH et al. |
| EPI_ISL_845809 | TGen North | TGen North | Jolene Bowers, Megan Folkerts, Chris French, Hayley Yaglom, Ashlyn Pfeiffer, Darrin Lemmer, Dave Engelthaler, The Arizona COVID Genomics Union (ACGU) |
| EPI_ISL_846587 | National Institute of Public Health - National Institute of Hygiene | National Institute of Public Health - National Institute of Hygiene | Wokowicz Tomasz, Zacharczuk Katarzyna, Gawor Jan |
| EPI_ISL_846589, EPI_ISL_846593 | Respiratory Virus Unit, National Infection Service, Public Health England | COVID-19 Genomics UK (COG-UK) Consortium | PHE Covid Sequencing Team |
| EPI_ISL_847527, EPI_ISL_847529, EPI_ISL_847531, EPI_ISL_847534, EPI_ISL_847548, EPI_ISL_847550, EPI_ISL_847557, EPI_ISL_847570, EPI_ISL_847575, EPI_ISL_847577, EPI_ISL_847623, EPI_ISL_847624, EPI_ISL_847625, EPI_ISL_847626, EPI_ISL_847627, EPI_ISL_847628, EPI_ISL_847629, EPI_ISL_847630, EPI_ISL_847631, EPI_ISL_847632, EPI_ISL_847633, EPI_ISL_847634, EPI_ISL_847635, EPI_ISL_847636, EPI_ISL_847637, EPI_ISL_847710, EPI_ISL_847711, EPI_ISL_847712, EPI_ISL_847717, EPI_ISL_847721, EPI_ISL_847722, EPI_ISL_847726, EPI_ISL_847734, EPI_ISL_847737, EPI_ISL_847750, EPI_ISL_847777, EPI_ISL_847778, EPI_ISL_847779, EPI_ISL_847780 | | | |
| see above | California Department of Public Health | Chiu Laboratory, University of California, San Francisco | Charles Chiu, Xianding (Wayne) Deng, Candace Wang, Brian Bushnell, Scot Federman, Jill Hacker, Debra Wadford |
| EPI_ISL_848048 | Michigan Department of Health and Human Services, Bureau of Laboratories | Michigan Department of Health and Human Services, Bureau of Laboratories | Blankenship HM, Riner D, Soehnlen MK |
| EPI_ISL_848206, EPI_ISL_848216, EPI_ISL_848218, EPI_ISL_848237, EPI_ISL_848265, EPI_ISL_848431, EPI_ISL_848432, EPI_ISL_848433, EPI_ISL_848434, EPI_ISL_848435, EPI_ISL_848436, EPI_ISL_848437, EPI_ISL_848438, EPI_ISL_848445, EPI_ISL_848446, EPI_ISL_848448, EPI_ISL_848459, EPI_ISL_848538, | | | |

| | | | |
|---|---|---|---|
| EPI_ISL_848539, EPI_ISL_848540, EPI_ISL_848541, EPI_ISL_848542, EPI_ISL_848543, EPI_ISL_848544, EPI_ISL_848545 | | | |
| see above | Illinois Department of Public Health | Gagnon Lab, Southern Illinois University | Keith Gagnon |
| EPI_ISL_849210, EPI_ISL_849211, EPI_ISL_849240, EPI_ISL_849246, EPI_ISL_849268 | Utah Public Health Laboratory | Utah Public Health Laboratory | Erin L. Young, Kelly F. Oakeson, Tara Gallagher |
| EPI_ISL_849732, EPI_ISL_849733, EPI_ISL_849734 | unknown | PHV-FSS | Son Nguyen et al. |
| EPI_ISL_850666, EPI_ISL_850669 | The National Institute of Public Health | State Veterinary Institute Prague | Nagy,A;Jirincova,H;Trnka,D;Vecerova,J;Trinklova,M |
| EPI_ISL_852990, EPI_ISL_852991 | Hospital General Universitario Gregorio Marañón | SeqCOVID-SPAIN consortium/IBV(CSIC) | Darío García de Viedma, Laura Pérez-Lago, Pedro J Sola-Campoy, Sergio Buenestado-Serrano, Marta Herranz, Victor Manuel de la Cueva, Julia Suárez, Pilar Catalán, Patricia Muñoz and SeqCOVID-SPAIN consortium |
| EPI_ISL_853363, EPI_ISL_853372 | UPMC Clinical Microbiology Laboratory | Microbial Genome Sequencing Center; Microbial Genomic Epidemiology Laboratory | Mustapha M. Mustapha, Jane W. Marsh, Dan Snyder, Marissa P. Griffith, Stephanie L. Mitchell, Vatsala R. Srinivasa, Kady D. Waggle, Chinelo Ezeonwuku, Vaughn S. Cooper, Lee H. Harrison |
| EPI_ISL_853750, EPI_ISL_853756, EPI_ISL_853763, EPI_ISL_853786 | Department of Microbiology, University Innsbruck | Bergthaler laboratory, CeMM Research Center for Molecular Medicine of the Austrian Academy of Sciences | Lukas Endler, Alexandra Popa, Benedikt Agerer, Jakob-Wendelin Genger, Alexander Lercher, Anna Schedl, Thomas Penz, Michael Schuster, Jan Laine, Martin Senekowitsch, Christoph Bock, Andreas Bergthaler |
| EPI_ISL_854749 | Victorian Infectious Diseases Reference Laboratory (VIDRL) | VIDRL and MDU-PHL | Caly L., Seemann T., Sait, M.L., Druce J., Sherry, N.L. |
| EPI_ISL_855395, EPI_ISL_855396, EPI_ISL_855397 | California Department of Public Health | Chiu Laboratory, University of California, San Francisco | Charles Chiu, Xianding (Wayne) Deng, Candace Wang, Brian Bushnell, Scot Federman, Jill Hacker, Debra Wadford |
| EPI_ISL_855430, EPI_ISL_855431, EPI_ISL_855432, EPI_ISL_855433, EPI_ISL_855434, EPI_ISL_855435, EPI_ISL_855436, EPI_ISL_855437, EPI_ISL_855438, EPI_ISL_855439, EPI_ISL_855440, EPI_ISL_855441, EPI_ISL_855442, EPI_ISL_855443, EPI_ISL_855444, EPI_ISL_855445, EPI_ISL_855446, EPI_ISL_855447, EPI_ISL_855448, EPI_ISL_855449, EPI_ISL_855450, EPI_ISL_855451, EPI_ISL_855452, EPI_ISL_855453, EPI_ISL_855454, EPI_ISL_855455, EPI_ISL_855456, EPI_ISL_855457, EPI_ISL_855458, EPI_ISL_855459, EPI_ISL_855460, EPI_ISL_855461, EPI_ISL_855462, EPI_ISL_855463, EPI_ISL_855464, EPI_ISL_855465, EPI_ISL_855466, EPI_ISL_855467, EPI_ISL_855468, EPI_ISL_855469, EPI_ISL_855470, EPI_ISL_855471, EPI_ISL_855472, EPI_ISL_855473, EPI_ISL_855474, EPI_ISL_855475, EPI_ISL_855476, EPI_ISL_855477, EPI_ISL_855478 | | | |
| see above | Servicio de Microbiología, Laboratori Clínic Metropolitana Nord. Hospital Universitari Germans Trias i Pujol. Institut d'Investigació en Ciències de la Salut Germans Trias i Pujol (IGTP) | SeqCOVID-SPAIN consortium/IBV(CSIC) | Elisa Martró, Antoni E. Bordoy, Anna Not, Adrián Antuori, Anabel Fernández, Nona Romaní, Verónica Saludes, Cristina Casañ and SeqCOVID-SPAIN consortium |
| EPI_ISL_855503, EPI_ISL_855504, EPI_ISL_855514, EPI_ISL_855546 | KEMRI-Wellcome Trust Research Programme/KEMRI-CGMR-C Kilifi | KEMRI-Wellcome Trust Research Programme/KEMRI-CGMR-C Kilifi | Githinji et al |
| EPI_ISL_855914, EPI_ISL_855916, EPI_ISL_855919 | Lab voor klinische biologie | Onderzoeksgroep Virologie | Laurens Lambrechts, Nick Vereecke, Marthe Pauwels, Bruno Verhasselt, Linos Vandekerckhove, Hans Nauwynck, Sebastiaan Theuns |
| EPI_ISL_856702, EPI_ISL_856755 | Ohio Department of Health Laboratory | Ohio Department of Health Laboratory | Holmes, Jennifer; Eric Brandt, Keoni Omura, Glen McGillivary, Caitlin McDonnell, Kirtana Ramadugu, Erica Leasure, Kelsey Florek, Heather Blankenship, Quanta Brown, and Tammy Bannerman |
| EPI_ISL_856788 | Servicio Virosis Respiratorias-Departamento Virología-INEI | Instituto Nacional Enfermedades Infecciosas C.G.Malbran | Baumeister E., Avaro M., Benedetti E., Russo M., Dattero ME, Pontoriero A., Cisterna D., Molina V., Perandones C., Tuduri E., Lorenzo F., Poklepovich T., Campos J. |
| EPI_ISL_856802, EPI_ISL_856803, EPI_ISL_856804, EPI_ISL_856805, EPI_ISL_856806, EPI_ISL_856807, EPI_ISL_856808, EPI_ISL_856809, EPI_ISL_856812, EPI_ISL_856813, EPI_ISL_856815, EPI_ISL_856816, EPI_ISL_856817, EPI_ISL_856818, EPI_ISL_856819, EPI_ISL_856820, EPI_ISL_856822, EPI_ISL_856824, EPI_ISL_856825, EPI_ISL_856826, EPI_ISL_856828, EPI_ISL_856829, EPI_ISL_856830, EPI_ISL_856833, EPI_ISL_856834, EPI_ISL_856837, EPI_ISL_856838, EPI_ISL_856839, EPI_ISL_856841, EPI_ISL_856842, EPI_ISL_856844, EPI_ISL_856845, EPI_ISL_856848, EPI_ISL_856849, EPI_ISL_856850, EPI_ISL_856852, EPI_ISL_856853, EPI_ISL_856855, EPI_ISL_856856, EPI_ISL_856858, EPI_ISL_856859, EPI_ISL_856860, EPI_ISL_856861, EPI_ISL_856862, EPI_ISL_856863, EPI_ISL_856864, EPI_ISL_856865, EPI_ISL_856866, EPI_ISL_856867, EPI_ISL_856868 | | | |
| see above | Utah Public Health Laboratory | Utah Public Health Laboratory | Erin L. Young, Kelly F. Oakeson, Tara Gallagher |
| EPI_ISL_856910, EPI_ISL_856911 | National Institute of Public Health - National Institute of Hygiene | National Institute of Public Health - National Institute of Hygiene | Wokowicz Tomasz, Zacharczuk Katarzyna, Gawor Jan |
| EPI_ISL_857307 | OCME Office Of Chief Medical Examiner | New York City Public Health Laboratory | Jade Wang, et al. |
| EPI_ISL_857541 | Swiss National Reference Centre for Influenza | Swiss National Reference Centre for Influenza | Tim Roloff, Ana Rita Gonçalves, Madlen Stange, Helena MB Seth-Smith, Alfredo Mari, Karoline Leuzinger, Julia Bielicki, Manuel Battegay, Hans Hirsch, Laurent Kaiser, Adrian Egli |
| EPI_ISL_857891, EPI_ISL_858902, EPI_ISL_858905, EPI_ISL_858907, EPI_ISL_858909, EPI_ISL_858913, EPI_ISL_858914, EPI_ISL_858915, EPI_ISL_858917, EPI_ISL_858918, EPI_ISL_858923, EPI_ISL_858924, EPI_ISL_858925, EPI_ISL_858926, EPI_ISL_858931, EPI_ISL_858932, EPI_ISL_858933, EPI_ISL_858934, EPI_ISL_858938, EPI_ISL_858939, EPI_ISL_858940, EPI_ISL_858941, EPI_ISL_858942, EPI_ISL_858944, EPI_ISL_858946, EPI_ISL_858947, EPI_ISL_858948, EPI_ISL_858949, EPI_ISL_858950, EPI_ISL_858951, EPI_ISL_858952, EPI_ISL_858954, EPI_ISL_858955, EPI_ISL_858957, EPI_ISL_858959, EPI_ISL_858961, EPI_ISL_858962, EPI_ISL_858963, EPI_ISL_858964, EPI_ISL_858965, EPI_ISL_858966, EPI_ISL_858967, EPI_ISL_858968, EPI_ISL_858971, EPI_ISL_858972, EPI_ISL_858976, EPI_ISL_858977, EPI_ISL_858978 | | | |
| see above | Lighthouse Lab in Alderley Park | Wellcome Sanger Institute for the COVID-19 Genomics UK (COG-UK) Consortium | Jacquelyn Wynn, Mairead Hyland, The Lighthouse Lab in Alderley Park and Alex Alderton, Roberto Amato, Sonia Goncalves, Ewan Harrison, David K. Jackson, Ian Johnston, Dominic Kwiatkowski, Cordelia Langford, John Sillitoe on behalf of the Wellcome Sanger Institute COVID-19 Surveillance Team |
| EPI_ISL_858997, EPI_ISL_859003, EPI_ISL_859012 | Lighthouse Lab in Glasgow | Wellcome Sanger Institute for the COVID-19 Genomics UK (COG-UK) Consortium | Harper VanSteenhouse, Yumi Kasai, David Gray, Carol Clugston, Anna Dominiczak and Alex Alderton, Roberto Amato, Sonia Goncalves, Ewan Harrison, David K. Jackson, Ian Johnston, Dominic Kwiatkowski, Cordelia Langford, John Sillitoe on behalf of the Wellcome Sanger Institute COVID-19 Surveillance Team |
| EPI_ISL_859035, EPI_ISL_859043, EPI_ISL_859049, EPI_ISL_859051, EPI_ISL_859061, EPI_ISL_859063, EPI_ISL_859066, EPI_ISL_859067, EPI_ISL_859070, EPI_ISL_859073, EPI_ISL_859075, EPI_ISL_859076, EPI_ISL_859080, EPI_ISL_859082, EPI_ISL_859085, EPI_ISL_859086, EPI_ISL_859089, EPI_ISL_859090, EPI_ISL_859091, EPI_ISL_859092, EPI_ISL_859096, EPI_ISL_859097, EPI_ISL_859099, EPI_ISL_859101, EPI_ISL_859103, EPI_ISL_859104, EPI_ISL_859107, EPI_ISL_859109, EPI_ISL_859110, EPI_ISL_859111, EPI_ISL_859113, EPI_ISL_859115, EPI_ISL_859117, EPI_ISL_859121, EPI_ISL_859122, EPI_ISL_859124, EPI_ISL_859126, EPI_ISL_859128, EPI_ISL_859129, EPI_ISL_859132, EPI_ISL_859134, EPI_ISL_859135, EPI_ISL_859136, EPI_ISL_859138, EPI_ISL_859141, EPI_ISL_859142, EPI_ISL_859143, EPI_ISL_859144, EPI_ISL_859145, EPI_ISL_859147, EPI_ISL_859179, EPI_ISL_859207, EPI_ISL_859212, EPI_ISL_859217, EPI_ISL_859226, EPI_ISL_859275 | | | |
| see above | Lighthouse Lab in Alderley Park | Wellcome Sanger Institute for the COVID-19 Genomics UK (COG-UK) Consortium | Jacquelyn Wynn, Mairead Hyland, The Lighthouse Lab in Alderley Park and Alex Alderton, Roberto Amato, Sonia Goncalves, Ewan Harrison, David K. Jackson, Ian Johnston, Dominic Kwiatkowski, Cordelia Langford, John Sillitoe on behalf of the Wellcome Sanger Institute COVID-19 Surveillance Team |
| EPI_ISL_859320, EPI_ISL_859321, EPI_ISL_859322, EPI_ISL_859323, EPI_ISL_859324, EPI_ISL_859325 | Lighthouse Lab in Glasgow | Wellcome Sanger Institute for the COVID-19 Genomics UK (COG-UK) Consortium | Harper VanSteenhouse, Yumi Kasai, David Gray, Carol Clugston, Anna Dominiczak and Alex Alderton, Roberto Amato, Sonia Goncalves, Ewan Harrison, David K. Jackson, Ian Johnston, Dominic Kwiatkowski, Cordelia Langford, John Sillitoe on behalf of the Wellcome Sanger Institute COVID-19 Surveillance Team |
| EPI_ISL_859327 | Lighthouse Lab in Milton Keynes | Wellcome Sanger Institute for the COVID-19 Genomics UK (COG-UK) Consortium | The Lighthouse Lab in Milton Keynes and Alex Alderton, Roberto Amato, Sonia Goncalves, Ewan Harrison, David K. Jackson, Ian Johnston, Dominic Kwiatkowski, Cordelia Langford, John Sillitoe on behalf of the Wellcome Sanger Institute COVID-19 Surveillance Team |
| EPI_ISL_859328, EPI_ISL_859329, EPI_ISL_859330, EPI_ISL_859331, EPI_ISL_859332, EPI_ISL_859333, EPI_ISL_859335, EPI_ISL_859336, EPI_ISL_859337, EPI_ISL_859338, EPI_ISL_859339, EPI_ISL_859340, EPI_ISL_859341, EPI_ISL_859342, EPI_ISL_859343, EPI_ISL_859345, EPI_ISL_859347, EPI_ISL_859348, EPI_ISL_859350, EPI_ISL_859351, EPI_ISL_859354, EPI_ISL_859355, EPI_ISL_859356, EPI_ISL_859358, EPI_ISL_859359, EPI_ISL_859360, EPI_ISL_859361, EPI_ISL_859362, EPI_ISL_859363, EPI_ISL_859364, EPI_ISL_859367, EPI_ISL_859368, EPI_ISL_859370, EPI_ISL_859371, EPI_ISL_859372, EPI_ISL_859373, EPI_ISL_859374, EPI_ISL_859375, EPI_ISL_859378, EPI_ISL_859379, EPI_ISL_859381, EPI_ISL_859382, EPI_ISL_859386, EPI_ISL_859388, EPI_ISL_859389, EPI_ISL_859390, EPI_ISL_859391, EPI_ISL_859392, EPI_ISL_859393, EPI_ISL_859394, EPI_ISL_859395, EPI_ISL_859396, EPI_ISL_859397, EPI_ISL_859398, EPI_ISL_859399, EPI_ISL_859400, EPI_ISL_859402, EPI_ISL_859403, EPI_ISL_859404, EPI_ISL_859405, EPI_ISL_859406, EPI_ISL_859407, EPI_ISL_859408, EPI_ISL_859409, EPI_ISL_859410, EPI_ISL_859411, EPI_ISL_859412, EPI_ISL_859413, EPI_ISL_859414, EPI_ISL_859415, EPI_ISL_859416, EPI_ISL_859417, EPI_ISL_859418, EPI_ISL_859419, EPI_ISL_859420, EPI_ISL_859421 | | | |
| see above | Lighthouse Lab in Alderley Park | Wellcome Sanger Institute for the COVID-19 Genomics UK (COG-UK) Consortium | Jacquelyn Wynn, Mairead Hyland, The Lighthouse Lab in Alderley Park and Alex Alderton, Roberto Amato, Sonia Goncalves, Ewan Harrison, David K. Jackson, Ian Johnston, Dominic Kwiatkowski, Cordelia Langford, John Sillitoe on behalf of the Wellcome Sanger Institute COVID-19 Surveillance Team |
| EPI_ISL_859422 | Lighthouse Lab in Glasgow | Wellcome Sanger Institute for the COVID-19 Genomics UK (COG-UK) Consortium | Harper VanSteenhouse, Yumi Kasai, David Gray, Carol Clugston, Anna Dominiczak and Alex Alderton, Roberto Amato, Sonia Goncalves, Ewan Harrison, David K. Jackson, Ian Johnston, Dominic Kwiatkowski, Cordelia Langford, John Sillitoe on behalf of the Wellcome Sanger Institute COVID-19 Surveillance Team |
| EPI_ISL_859423, EPI_ISL_859424, EPI_ISL_859425, EPI_ISL_859426, | Lighthouse Lab in Alderley Park | Wellcome Sanger Institute for the COVID-19 Genomics UK (COG-UK) Consortium | Jacquelyn Wynn, Mairead Hyland, The Lighthouse Lab in Alderley Park and Alex Alderton, Roberto Amato, Sonia Goncalves, Ewan Harrison, David K. Jackson, Ian Johnston, Dominic Kwiatkowski, Cordelia Langford, John Sillitoe on behalf of the Wellcome Sanger Institute COVID-19 Surveillance Team |

| | | | |
|---|---|---|---|
| EPI_ISL_859428, EPI_ISL_859429, EPI_ISL_859430, EPI_ISL_859431, EPI_ISL_859434 | | | |
| EPI_ISL_859435 | Lighthouse Lab in Glasgow | Wellcome Sanger Institute for the COVID-19 Genomics UK (COG-UK) Consortium | Harper VanSteenhouse, Yumi Kasai, David Gray, Carol Clugston, Anna Dominiczak and Alex Alderton, Roberto Amato, Sonia Goncalves, Ewan Harrison, David K. Jackson, Ian Johnston, Dominic Kwiatkowski, Cordelia Langford, John Sillitoe on behalf of the Wellcome Sanger Institute COVID-19 Surveillance Team |
| EPI_ISL_859436, EPI_ISL_859437, EPI_ISL_859438, EPI_ISL_859439, EPI_ISL_859440, EPI_ISL_859441, EPI_ISL_859442, EPI_ISL_859443, EPI_ISL_859444, EPI_ISL_859445, EPI_ISL_859446, EPI_ISL_859447, EPI_ISL_859448, EPI_ISL_859449, EPI_ISL_859450, EPI_ISL_859451, EPI_ISL_859452, EPI_ISL_859453, EPI_ISL_859454, EPI_ISL_859455 | | | |
| see above | Lighthouse Lab in Alderley Park | Wellcome Sanger Institute for the COVID-19 Genomics UK (COG-UK) Consortium | Jacquelyn Wynn, Mairead Hyland, The Lighthouse Lab in Alderley Park and Alex Alderton, Roberto Amato, Sonia Goncalves, Ewan Harrison, David K. Jackson, Ian Johnston, Dominic Kwiatkowski, Cordelia Langford, John Sillitoe on behalf of the Wellcome Sanger Institute COVID-19 Surveillance Team |
| EPI_ISL_860116 | Medical Microbiology Unit, Department for Laboratory Medicine, Drammen Hospital, Vestre Viken Health Trust, | Norwegian Institute of Public Health, Department of Virology | Kathrine Stene-Johansen, Kamilla Heddeland Instefjord, Hilde Elshaug, Atiya R Ali,Marie Paulsen Madsen, Rasmus Riis Kopperud, Hilde Vollan, Karoline Bragstad, Olav Hungnes |
| EPI_ISL_860206, EPI_ISL_860207 | Ostfold Hospital Trust - Kalnes, Centre for Laboratory Medicine, Section for gene technology and infection serology | Norwegian Institute of Public Health, Department of Virology | Kathrine Stene-Johansen, Kamilla Heddeland Instefjord, Hilde Elshaug, Atiya R Ali,Marie Paulsen Madsen, Rasmus Riis Kopperud, Hilde Vollan, Karoline Bragstad, Olav Hungnes |
| EPI_ISL_860209 | Norwegian Institute of Public Health, Department of Virology | Norwegian Institute of Public Health, Department of Virology | Kathrine Stene-Johansen, Kamilla Heddeland Instefjord, Hilde Elshaug, Atiya R Ali,Marie Paulsen Madsen, Rasmus Riis Kopperud, Hilde Vollan, Karoline Bragstad, Olav Hungnes |
| EPI_ISL_860291 | Innlandet Hospital Trust, Division Lillehammer, Department for Medical Microbiology | Norwegian Institute of Public Health, Department of Virology | Kathrine Stene-Johansen, Kamilla Heddeland Instefjord, Hilde Elshaug, Atiya R Ali,Marie Paulsen Madsen, Rasmus Riis Kopperud, Hilde Vollan, Karoline Bragstad, Olav Hungnes |
| EPI_ISL_860600, EPI_ISL_860611, EPI_ISL_860612, EPI_ISL_860626, EPI_ISL_860627, EPI_ISL_860628 | NHLS-IALCH | KRISP, KZn Research Innovation and Sequencing Platform | Giandhari J, Pillay S, Lessells R, Mdlalose K, York D, Khan S, Tegally H, Wilkinson E, de Oliveira T |
| EPI_ISL_860791, EPI_ISL_860792, EPI_ISL_860793 | Ohio Department of Health Laboratory | Ohio Department of Health Laboratory | Holmes, Jennifer; Eric Brandt, Keoni Omura, Glen McGillivary, Caitlin McDonnell, Kirtana Ramadugu, Erica Leasure, Kelsey Florek, Heather Blankenship, Quanta Brown, and Tammy Bannerman |
| EPI_ISL_860935 | Johns Hopkins Hospital Department of Pathology | Johns Hopkins Hospital Department of Pathology | C. Paul Morris, Chun Huai Luo, Adannaya Amadi, Nicholas Gallagher, Heba H. Mostafa |
| EPI_ISL_861509, EPI_ISL_861510, EPI_ISL_861511, EPI_ISL_861512, EPI_ISL_861513, EPI_ISL_861514, EPI_ISL_861515, EPI_ISL_861516, EPI_ISL_861517, EPI_ISL_861518, EPI_ISL_861519, EPI_ISL_861520, EPI_ISL_861521, EPI_ISL_861522, EPI_ISL_861523, EPI_ISL_861524, EPI_ISL_861525, EPI_ISL_861526, EPI_ISL_861527 | | | |
| see above | ZOTZ KLIMAS MVZ Düsseldorf-Centrum GbR ÜBAG für Labormedizin, Genetik, Zytologie, Pathologie | Center of Medical Microbiology, Virology, and Hospital Hygiene, University of Duesseldorf | Maximilian Damagnez, Alexander Dilthey, Ashley-Jane Duplessis, Patrick Finzer, Katrin Hoffmann, Torsten Houwaart, Lisanna Hülse, Malte Kohns Vasconcelos, Marek Korencak, Nadine Lübke, Jessica Nicolai, Klaus Pfeffer, Daniel Strelow, Jörg Timm, Andreas Walker, Tobias Wienemann, Rainer Zotz |
| EPI_ISL_861674, EPI_ISL_861675 | UPA Central de Caraguatatuba | Instituto Adolfo Lutz, Interdiciplinary Procedures Center, Strategic Laboratory | Claudio Tavares Sacchi, Claudia Regina Gonçalves, Erica Valessa Ramos Gomes, Karoline Rodrigues Campos |
| EPI_ISL_861778, EPI_ISL_861779, EPI_ISL_861788, EPI_ISL_861793, EPI_ISL_861811, EPI_ISL_861824 | Hospital General Universitario Gregorio Marañón | SeqCOVID-SPAIN consortium/IBV(CSIC) | Darío García de Viedma, Laura Pérez-Lago, Pedro J Sola-Campoy, Sergio Buenestado-Serrano, Marta Herranz, Victor Manuel de la Cueva, Julia Suárez, Pilar Catalán, Patricia Muñoz and SeqCOVID-SPAIN consortium |
| EPI_ISL_862127, EPI_ISL_862137 | Charité Universitätsmedizin Berlin, Institut fur Virologie/Labor Berlin | Charité Universitätsmedizin Berlin, Institut für Virologie | Victor M Corman, Barbara Mühlemann, Jörn Beheim-Schwarzbach, Tobias Bleicker, Julia Tesch, Talitha Veith, Julia Schneider, Terry Jones, Christian Drosten |
| EPI_ISL_862781 | Utah Public Health Laboratory, Utah Public Health Laboratory Infectious Disease submission group | Utah Public Health Laboratory, Utah Public Health Laboratory Infectious Disease submission group | Young,E.L., Oakeson,K.F., Gallagher,T. |
| EPI_ISL_864547, EPI_ISL_864548, EPI_ISL_864554, EPI_ISL_864555, EPI_ISL_864556, EPI_ISL_864558, EPI_ISL_864560, EPI_ISL_864561, EPI_ISL_864562 | Knappschaftskrankenhaus Bochum | Bundeswehr Institute of Microbiology | Markus Antwerpen, Mustafa Özçürümez, Antonios Katsounas, Alexandra Rehn, Mathias Walter, Malena Bestehorn-Willmann, Sabine Zange, Enrico Georgi, Roman Wölfel |
| EPI_ISL_864833, EPI_ISL_864837, EPI_ISL_864839, EPI_ISL_864854 | Department of Pathology, University of Cambridge | COVID-19 Genomics UK (COG-UK) Consortium | Aminu S. Jahun, Yasmin Chaudhry, Grant Hall, Iliana Georgana, Myra Hosmillo, Martin D. Curran, Malte Pinckert, Surendra Parmar, Ian Goodfellow |
| EPI_ISL_864960 | West of Scotland Specialist Virology Centre, NHSGGC / MRC-University of Glasgow Centre for Virus Research | COVID-19 Genomics UK (COG-UK) Consortium | Ana da Silva Filipe, Natasha Johnson, Kathy Smollett, Daniel Mair, Stephen Carmichael, Alice Broos, Lily Tong, Jenna Nichols, Kyriaki Nomikou; Sarah McDonald; Richard Orton, Joseph Hughes, Sreenu Vattipally, David L Robertson; Alasdair MacLean, Rory Gunson; Sharif Shaaban, Matthew Holden; Rachel Blacow, Guy Mollett, Kathy Li, James Shepherd, Antonia Ho, Emma Thomson |
| EPI_ISL_865072 | Virology Department, Royal Infirmary of Edinburgh, NHS Lothian / School of Biological Sciences, University of Edinburgh / Institute of Genetics and Molecular Medicine, University of Edinburgh | COVID-19 Genomics UK (COG-UK) Consortium | McHugh M, Dewar R, Rooke S, Gallagher M, Balcaza C, O'Toole Á, Scher E, Hill V, McCrone JT, Colquhoun R, Yu X, Jackson B, Rambaut A, Williams TC, Templeton K |
| EPI_ISL_865242, EPI_ISL_865243, EPI_ISL_865244, EPI_ISL_865245, EPI_ISL_865247, EPI_ISL_865251, EPI_ISL_865252, EPI_ISL_865254, EPI_ISL_865278, EPI_ISL_865448, EPI_ISL_865459, EPI_ISL_865460, EPI_ISL_865461, EPI_ISL_865462, EPI_ISL_865463, EPI_ISL_865464, EPI_ISL_865465, EPI_ISL_865466 | | | |
| see above | Liverpool Clinical Laboratories | COVID-19 Genomics UK (COG-UK) Consortium | Sam Haldenby, Anita Lucaci, Steve Paterson, Julian Hiscox, Alistair Darby, M Almsaud, A Alrezaihi, Muhannad Alruwaili, Stuart D Armstrong, Jones Benjamin, Eleanor G Bentley, Anu Chawla, Jordan J Clark, Angela Cowell, Richard Eccles, Isabel García-Dorival, Matthew Gemmell, Alessandro Gerada, PKF Gilmore, Richard Gregory, Ximeng Han, Catherine Hartley, Margaret Hughes, Miren Iturriza-Gomara, James Johnson, L Luu, Jenifer Manson, Charlotte Nelson, Elaine O'Toole, Cassie Olateju, Rebekah Penrice-Randal , Lucille Rainbow, N.P Randle, Trevor Ian Robinson, Parul Sharma, Ghada T Shawli, James P Stewart, Neil Swainston, Ecaterina Vamos, Joanne Watts, Mark Whitehead |
| EPI_ISL_865690, EPI_ISL_865721, EPI_ISL_865722, EPI_ISL_865723, EPI_ISL_865724, EPI_ISL_865725, EPI_ISL_865726, EPI_ISL_865727, EPI_ISL_865728, EPI_ISL_865729, EPI_ISL_865730, EPI_ISL_865731, EPI_ISL_865746 | | | |
| see above | University College London, Great Ormond Street Hospital for Children NHS Foundation Trust, Imperial College Healthcare NHS Trust | COVID-19 Genomics UK (COG-UK) Consortium | Sergi Castellano, Rachel Williams, Mark Kristiansen, Paola Resende Silva, Sunando Roy, Tony Brooks, Helena Tutill, Paola Niola, Patricia Dyal, Charlotte Williams, Leysa Forrest, Yasmin Panchbhaya, Jacqueline Findlay, Samuel Weeks, Julianne Brown, Kathryn Harris, Paul Randell, James Price, Alison Holmes, Judith Breuer |
| EPI_ISL_866050, EPI_ISL_866051, EPI_ISL_866068, EPI_ISL_866069, EPI_ISL_866118, EPI_ISL_866119, EPI_ISL_866132, EPI_ISL_866133, EPI_ISL_866149, EPI_ISL_866151, EPI_ISL_866164 | | | |
| see above | University College London Hospital | COVID-19 Genomics UK (COG-UK) Consortium | Judith Heaney, Matthew Byott, Catherine Houlihan, Dan Frampton, Stuart Kirk, Moira Spyer and Eleni Nastouli |
| EPI_ISL_866577, EPI_ISL_866578, EPI_ISL_866586, EPI_ISL_866802, EPI_ISL_866807, EPI_ISL_866808, EPI_ISL_866812, EPI_ISL_866814 | Quadram Institute Bioscience | COVID-19 Genomics UK (COG-UK) Consortium | Dave J. Baker, Gemma L. Kay, Alp Aydin, Thanh Le-Viet, Steven Rudder, Ana P. Tedim, Anastasia Kolyva, Maria Diaz, Leonardo de Oliveira Martins, Nabil-Fareed Alikhan, Lizzie Meadows, Rachael Stanley, Ngozi Elumogo, Muhammed Yasir, Nicholas M. Thomson, Alexander J Trotter, Rachel Gilroy, Samuel Bloomfield, Claire Stuart, Andrew Bell, Reenesh Prakash, Samir Dervisevic, Alison E. Mather, John Wain, Mark Webber, Andrew J. Page, Justin O'Grady |
| EPI_ISL_866891, EPI_ISL_866892, EPI_ISL_866893, EPI_ISL_866894, EPI_ISL_866895, EPI_ISL_866904 | Queens Medical Centre, Clinical Microbiology Department / DeepSeq Nottingham | COVID-19 Genomics UK (COG-UK) Consortium | Gemma Clark, Wendy Smith, Manjinder Khakh, Vicki M Fleming, Michelle M Lister, Hannah Howson-Wells, Jonathan Ball, Patrick McClure, Joseph Chappell, Theocharis Tsoleridis, Nadine Holmes, Matthew Carlisle, Christopher Moore, Fei Sang, Johnny Debebe, Victoria Wright, Matthew Loose |
| EPI_ISL_867035 | Oxford Viromics, NDM, University of Oxford; Oxford University Hospitals; Basingstoke and North Hampshire Hospital | COVID-19 Genomics UK (COG-UK) Consortium | Tanya Golubchik, David Bonsall, George Macintyre, Amy Trebes, Mariateresa de Cesare, Catrin Moore, Alex Mobbs, Anita Justice, Robert Shaw, Monique Andersson, Timothy Peto, Emma Wise, Nathan Moore, Jessica Lynch, Nick Cortes, Matilde Mori, Stephen Kidd, David Buck, John Todd, Christophe Fraser |
| EPI_ISL_867500, EPI_ISL_867501 | Originating lab: Wales Specialist Virology Centre Sequencing | Public Health Wales Microbiology Cardiff Wales Specialist | Catherine Moore, Johnathan Evans, Laura Gifford, Malorie Perry, Simon Cottrell, Angela Marchbank, Alec Birchley, Alexander Adams, Amy Gaskin, Bree |

| | | | |
|---|---|---|---|
| | lab: Pathogen Genomics Unit | Virology Centre | Gatica-Wilcox, Jason Coombes, Joel Southgate, Lauren Gilbert, Lee Graham, Nicole Pacchiarini, Sara Kumziene-Summerhayes, Sarah Taylor, Sophie Jones, Sara Rey, Matthew Bull, Joanne Watkins, Sally Corden, Tom Connor |
| EPI_ISL_871891, EPI_ISL_871914, EPI_ISL_871918, EPI_ISL_871922, EPI_ISL_871927, EPI_ISL_871928, EPI_ISL_871930, EPI_ISL_871933, EPI_ISL_871937 | Servicio de Microbiología, Laboratori Clínic Metropolitana Nord. Hospital Universitari Germans Trias i Pujol. Institut d'Investigació en Ciències de la Salut Germans Trias i Pujol (IGTP) | SeqCOVID-SPAIN consortium/IBV(CSIC) | Elisa Martró, Antoni E. Bordoy, Anna Not, Adrián Antuori, Anabel Fernández, Nona Romaní, Verónica Saludes, Cristina Casañ and SeqCOVID-SPAIN consortium |
| EPI_ISL_872101 | Chu Tivoli | GIGA Medical Genomics | Keith Durkin, Maria Artesi, Sébastien Bontems, Raphaël Boreux, Bouchra Boujemla, Cécile Meex, Pierrette Melin, Marie-Pierre Hayette, Vincent Bours |
| EPI_ISL_872196 | Caribbean Public Health Agency | Carrington Lab, Department of PreClinical Sciences, Faculty of Medical Sciences, The University of the West Indies | Nikita S. D. Sahadeo, Arianne Brown-Jordan, Vernie Ramkissoon, Sarah Hill, Naresh Nandram, Avery Hinds, Jerome Foster, Stanley Giddings, Karla Georges, Marsha Ivey, Rahul Naidu, Risha Singh, SueMin Nathaniel, Rajini Haraksingh, Jaya Jayaraman, Chinna Chinnadurai, Adesh Ramsubhag, Nuno Faria, Oliver Pybus, Christopher Oura, Gabriel Escobar, Christine V. F. Carrington |
| EPI_ISL_872369 | Texas Department of State Health Services (TXDSHS) | Texas Department of State Health Services (TXDSHS) | Bonnie Oh, Anita Pokharel, James Daniel Bonser, Myong Koag, Chung Wang, Rachel Lee, Grace Kubin, Rashmi Tuladhar, Mayela Pedrueza, Maliha Rahman, Jenny Zhang |
| EPI_ISL_872585 | Australian Clinical Labs | NSW Health Pathology - Institute of Clinical Pathology and Medical Research; Westmead Hospital; University of Sydney | CIDM-PH et al. |
| EPI_ISL_872937, EPI_ISL_872943, EPI_ISL_872946, EPI_ISL_872949, EPI_ISL_872950, EPI_ISL_872952, EPI_ISL_872953, EPI_ISL_872954, EPI_ISL_872955, EPI_ISL_872956, EPI_ISL_872957, EPI_ISL_872958 | | | |
| see above | WHO National Influenza Centre Russian Federation | WHO National Influenza Centre Russian Federation | Andrey Komissarov, Artem Fadeev, Anna Ivanova, Kseniya Komissarova, Dmitry Bazhenov, Mikhail Bakaev, Daria Danilenko, Ksenia Safina, Elena Nabieva, Georgii Bazykin, Dmitry Lioznov |
| EPI_ISL_873155 | University of Michigan Clinical Microbiology Laboratory | Lauring Lab, University of Michigan, Department of Microbiology and Immunology | Valesano |
| EPI_ISL_873221 | M Health Fairview | Minnesota Department of Health, Public Health Laboratory | Alexandra Lorentz, Jacob Garfin, Matt Plumb, and Xiong Wang |
| EPI_ISL_876045, EPI_ISL_876047, EPI_ISL_876066, EPI_ISL_876068, EPI_ISL_876071, EPI_ISL_876090, EPI_ISL_876110, EPI_ISL_876148, EPI_ISL_876165, EPI_ISL_876175, EPI_ISL_876181, EPI_ISL_876288, EPI_ISL_876289, EPI_ISL_876290, EPI_ISL_876291, EPI_ISL_876292, EPI_ISL_876293, EPI_ISL_876294, EPI_ISL_876295, EPI_ISL_876296 | | | |
| see above | Massachusetts State Public Health Laboratory | Massachusetts State Public Health Laboratory | Andrew Lang, Timelia Fink, Glen Gallagher, Sandra Smole |
| EPI_ISL_876533 | Florida Bureau of Public Health Laboratories | Florida Bureau of Public Health Laboratories | Sarah Schmedes, Jason Blanton |
| EPI_ISL_876592 | TN Division of Laboratory Services | Pathogen Discovery, Respiratory Viruses Branch, Division of Viral Diseases, Centers for Disease Control and Prevention | Ying Tao, Yan Li, Jing Zhang, Krista Queen, Anna Uehara, Peter Cook, Clinton R. Paden, Haibin Wang, Suxiang Tong |
| EPI_ISL_876776, EPI_ISL_876777, EPI_ISL_876778, EPI_ISL_876779, EPI_ISL_876780, EPI_ISL_876781, EPI_ISL_876782, EPI_ISL_876783 | Istituto Zooprofilattico Sperimentale della Puglia e della Basilicata | Istituto Zooprofilattico Sperimentale della Puglia e della Basilicata | Parisi A., Bianco A., Capozzi L., Del Sambro L., Manzulli V, Rondinone V., Pace L., Cipolletta D., Galante D. |
| EPI_ISL_877423 | Primasatya Husada Citra Hospital | Institute of Tropical Disease, Universitas Airlangga | Maria I Lusida, Krisnoadi Rahardjo, Aldise M Nastri, Jezzy R Dewantari, Rima R Prasetya, Pudji Djanuartono, Gatot Soegiarto, Laksmi Wulandari, Resti Yudhawati, Soetjipto, Yasuko Mori, Kazufumi Shimizu |
| EPI_ISL_877456 | Primasatya Husada Citra Hospital | Institute of Tropical Disease, Universitas Airlangga | Krisnoadi Rahardjo, Aldise M Nastri, Jezzy R Dewantari, Rima R Prasetya, Pudji Djanuartono, Gatot Soegiarto, Laksmi Wulandari, Resti Yudhawati, Soetjipto, Yasuko Mori, Maria I Lusida, Kazufumi Shimizu |
| EPI_ISL_877457 | Primasatya Husada Citra Hospital | Institute of Tropical Disease, Universitas Airlangga | Aldise M Nastri, Jezzy R Dewantari, Rima R Prasetya, Krisnoadi Rahardjo, Pudji Djanuartono, Gatot Soegiarto, Laksmi Wulandari, Resti Yudhawati, Soetjipto, Yasuko Mori, Maria I Lusida, Kazufumi Shimizu |
| EPI_ISL_877458 | Anwar Medika General Hospital | Institute of Tropical Disease, Universitas Airlangga | Jezzy R Dewantari, Rima R Prasetya, Krisnoadi Rahardjo, Aldise M Nastri, Nungky Taniasari, Gatot Soegiarto, Laksmi Wulandari, Resti Yudhawati, Soetjipto, Yasuko Mori, Maria I Lusida, Kazufumi Shimizu |
| EPI_ISL_877459 | Darmo Hospital | Institute of Tropical Disease, Universitas Airlangga | Rima R Prasetya, Krisnoadi Rahardjo, Aldise M Nastri, Jezzy R Dewantari, Sulung Budianto, Gatot Soegiarto, Laksmi Wulandari, Resti Yudhawati, Soetjipto, Yasuko Mori, Maria I Lusida, Kazufumi Shimizu |
| EPI_ISL_878589, EPI_ISL_878592, EPI_ISL_878595, EPI_ISL_878597, EPI_ISL_878603, EPI_ISL_878605, EPI_ISL_878608, EPI_ISL_878619, EPI_ISL_878624, EPI_ISL_878629, EPI_ISL_878632 | | | |
| see above | San Diego County Public Health Laboratory | Andersen lab at Scripps Research | SEARCH Alliance San Diego with Tracy Basler, Jovan Shephard, Brett Austin |
| EPI_ISL_878747, EPI_ISL_878791, EPI_ISL_880185 | Rady's Childrens Hospital | Andersen lab at Scripps Research | SEARCH Alliance San Diego with Nanda Radamchar, David Dimmock, Linda Luo, Christina Clarke, Kathryn Bouic, Teresa Mueller, Denise Malicki |
| EPI_ISL_882769 | Institute of Tropical Disease | Institute of Tropical Disease, Universitas Airlangga | Kazufumi Shimizu, Krisnoadi Rahardjo, Aldise M Nastri, Jezzy R Dewantari, Rima R Prasetya, Gatot Soegiarto, Laksmi Wulandari, Resti Yudhawati, Yasuko Mori, Soetjipto, Maria I Lusida |
| EPI_ISL_882962 | National Institute of Public Health - National Institute of Hygiene | National Institute of Public Health - National Institute of Hygiene | Wokowicz Tomasz, Zacharczuk Katarzyna, Gawor Jan |
| EPI_ISL_884266, EPI_ISL_884278 | Institute of Medical Microbiology and Hospital Hygiene | Institute of Medical Microbiology and Hospital Hygiene | Prof. Dr. Achim Kaasch, Aljoscha Tersteegen |
| EPI_ISL_884920, EPI_ISL_884923, EPI_ISL_884924, EPI_ISL_884925, EPI_ISL_884926, EPI_ISL_884927, EPI_ISL_884928, EPI_ISL_884929, EPI_ISL_884930, EPI_ISL_884931, EPI_ISL_884932, EPI_ISL_884933, EPI_ISL_884934, EPI_ISL_884935, EPI_ISL_884936, EPI_ISL_884940, EPI_ISL_884942, EPI_ISL_884972, EPI_ISL_884978, EPI_ISL_885051 | | | |
| see above | Santa Clara County Public Health Laboratory | Chan-Zuckerberg Biohub | CZB Cliahub Consortium |
| EPI_ISL_887437, EPI_ISL_887444, EPI_ISL_887472, EPI_ISL_887493 | Instituto Nacional de Saude (INS), Mozambique | KRISP, KZN Research Innovation and Sequencing Platform | Nalia Ismael, Nadia Sitoe, Paulo Arnaldo, Nedio Mabunda, Giandhari J, Pillay S, Tegally H, Wilkinson E, de Oliveira T |
| EPI_ISL_888739, EPI_ISL_888742, EPI_ISL_888744, EPI_ISL_888745, EPI_ISL_888746, EPI_ISL_888747, EPI_ISL_888749, EPI_ISL_888750, EPI_ISL_888771, EPI_ISL_888776 | KU Leuven, Rega Institute, Clinical and Epidemiological Virology | KU Leuven, Rega Institute, Clinical and Epidemiological Virology | Tony Wawina-Bokalanga, Bert Vanmechelen, Joan Marti-Carerras, Piet Maes |
| EPI_ISL_888858, EPI_ISL_888859, EPI_ISL_888874 | Michigan Department of Health and Human Services, Bureau of Laboratories | Michigan Department of Health and Human Services, Bureau of Laboratories | Blankenship HM, Riner D, Soehnlen MK |
| EPI_ISL_888981 | RS Omni Pulomas | Eijkman Institute for Molecular Biology, Ministry of Research and Technology/National Agency for Research and Innovation | Iskandar Adnan, Lydia V. Panggalo, Sukma Oktavianthi, Willy Agustine, Edison Johar, Hidayat Trimarsanto, Frilasita A Yudhaputri, Safarina G Malik, Khin Saw Myint, Amin Soebandrio |
| EPI_ISL_889014 | RSU Medika Dramaga | Eijkman Institute for Molecular Biology, Ministry of Research and Technology/National Agency for Research and Innovation | Iskandar Adnan, Lydia V. Panggalo, Sukma Oktavianthi, Willy Agustine, Edison Johar, Hidayat Trimarsanto, Frilasita A Yudhaputri, Safarina G Malik, Khin Saw Myint, Amin Soebandrio |
| EPI_ISL_889015 | RS Anna | Eijkman Institute for Molecular Biology, Ministry of Research and Technology/National Agency for Research and Innovation | Iskandar Adnan, Lydia V. Panggalo, Sukma Oktavianthi, Willy Agustine, Edison Johar, Hidayat Trimarsanto, Frilasita A Yudhaputri, Safarina G Malik, Khin Saw Myint, Amin Soebandrio |
| EPI_ISL_889016 | RS Mitra Husada | Eijkman Institute for Molecular Biology, Ministry of Research and Technology/National Agency for Research and Innovation | Iskandar Adnan, Lydia V. Panggalo, Sukma Oktavianthi, Willy Agustine, Edison Johar, Hidayat Trimarsanto, Frilasita A Yudhaputri, Safarina G Malik, Khin Saw Myint, Amin Soebandrio |

| | | | |
|---|---|---|---|
| EPI_ISL_890239, EPI_ISL_890240, EPI_ISL_890249, EPI_ISL_890254, EPI_ISL_890255, EPI_ISL_890259, EPI_ISL_890260, EPI_ISL_890264, EPI_ISL_890265, EPI_ISL_890266, EPI_ISL_890270, EPI_ISL_890271, EPI_ISL_890274, EPI_ISL_890275, EPI_ISL_890276, EPI_ISL_890292 | | | |
| see above | KU Leuven, Rega Institute, Clinical and Epidemiological Virology | KU Leuven, Rega Institute, Clinical and Epidemiological Virology | Tony Wawina-Bokalanga, Bert Vanmechelen, Joan Marti-Carerras, Piet Maes |
| EPI_ISL_890889 | Hospital | National Reference Center for Viruses of Respiratory Infections, Institut Pasteur, Paris | Marion Barbet, Sylvie Behillil, Méline Bizard, Angela Brisebarre, Camille Capel, Etienne Simon-Lorière, Vincent Enouf, Maud Vanpeene, Sylvie van der Werf, Laurent Andreoletti |
| EPI_ISL_891199 | DPH, Massachusetts State Public Health Lab | DPH, Massachusetts State Public Health Lab | Lang,A.S., Fink,T., Gallagher,G.R., Smole,S.C. |
| EPI_ISL_894162 | Institute of Medical Microbiology and Hospital Hygiene | Institute of Medical Microbiology and Hospital Hygiene | Prof. Dr. Achim Kaasch, Aljoscha Tersteegen |
| EPI_ISL_896140, EPI_ISL_896154, EPI_ISL_896161, EPI_ISL_896189, EPI_ISL_896190, EPI_ISL_896194, EPI_ISL_896195, EPI_ISL_896196 | MEPHI, Aix Marseille University | MEPHI, Aix Marseille University | Anthony LEVASSEUR |
| EPI_ISL_896295, EPI_ISL_896326, EPI_ISL_896327, EPI_ISL_896328, EPI_ISL_896329, EPI_ISL_896330, EPI_ISL_896331, EPI_ISL_896332, EPI_ISL_896333, EPI_ISL_896334, EPI_ISL_896335, EPI_ISL_896336, EPI_ISL_896337, EPI_ISL_896343, EPI_ISL_896344, EPI_ISL_896345, EPI_ISL_896346, EPI_ISL_896349, EPI_ISL_896498, EPI_ISL_896499, EPI_ISL_896521, EPI_ISL_896522, EPI_ISL_896523, EPI_ISL_896524, EPI_ISL_896562, EPI_ISL_896570 | | | |
| see above | MEMORIAL SLOAN KETTERING CANCER CENTER | Wadsworth Center, New York State Department of Health | Kirsten St. George, Daryl M. Lamson, Alexis Russel, Matthew Shudt, Melissa A Leisner, Jonathan Plitnick, Navjot Singh, John Kelly, Erasmus Schneider, Erica Lasek-Nesselquist |
| EPI_ISL_900064, EPI_ISL_900116, EPI_ISL_900159, EPI_ISL_900313, EPI_ISL_900381, EPI_ISL_900427, EPI_ISL_900445 | MEPHI, Aix Marseille University | MEPHI, Aix Marseille University | Anthony LEVASSEUR |
| EPI_ISL_900577, EPI_ISL_900578, EPI_ISL_900581, EPI_ISL_900582, EPI_ISL_900584, EPI_ISL_900592, EPI_ISL_900594, EPI_ISL_900595, EPI_ISL_900596, EPI_ISL_900597, EPI_ISL_900598, EPI_ISL_900599, EPI_ISL_900600, EPI_ISL_900601, EPI_ISL_900602, EPI_ISL_900603, EPI_ISL_900604, EPI_ISL_900617, EPI_ISL_900618, EPI_ISL_900619, EPI_ISL_900620, EPI_ISL_900621, EPI_ISL_900622, EPI_ISL_900623, EPI_ISL_900624, EPI_ISL_900625, EPI_ISL_900626, EPI_ISL_900627, EPI_ISL_900628, EPI_ISL_900629, EPI_ISL_900630, EPI_ISL_900632, EPI_ISL_900653, EPI_ISL_900654, EPI_ISL_900655, EPI_ISL_900656, EPI_ISL_900657, EPI_ISL_900658, EPI_ISL_900659, EPI_ISL_900660, EPI_ISL_900671, EPI_ISL_900672, EPI_ISL_900673, EPI_ISL_900674, EPI_ISL_900675, EPI_ISL_900676, EPI_ISL_900677, EPI_ISL_900678, EPI_ISL_900679, EPI_ISL_900680, EPI_ISL_900681, EPI_ISL_900682, EPI_ISL_900683, EPI_ISL_900684 | | | |
| see above | IZSM | TIGEM | Patrizia Annunziata, Andrea Ballabio, Valentina Bouche, Davide Cacchiarelli (CorrespAuthor), Pellegrino Cerino, Chiara Colantuono, Maria Concetta Cuomo, Denise Di Concilio, Lucio Di Filippo, Antonio Grimaldi, Antonio Limone, Anna Manfredi, Francesco Panariello, Biancamaria Pierri, Marcello Salvi |
| EPI_ISL_902758 | Department of Virology and Immunology, University of Helsinki and Helsinki University Hospital, Huslab Finland | Department of Virology, Faculty of Medicine, University of Helsinki, Helsinki, Finland | Teemu Smura, Ravi Kant, Phuoc Truong, Hussein Alburkat, Hannimari Kallio-Kokko, Jenni Virtanen, Maija Suvanto, Essi Korhonen, Sari Hannula, Harri Kangas, Hanna Liimatainen, Satu Kurkela, Hanna Jarva, Maija Lappalainen, Pekka Ellonen, Olli Vapalahti |
| EPI_ISL_903254, EPI_ISL_903255 | M Health Fairview | Minnesota Department of Health, Public Health Laboratory | Alexandra Lorentz, Jacob Garfin, Matt Plumb, and Xiong Wang |
| EPI_ISL_903575 | MI - Michigan Department of Health and Human Services - Bureau of Laboratories | Genomics and Discovery, Respiratory Viruses Branch, Division of Viral Diseases, Centers for Disease Control and Prevention | Krista Queen, Yan Li, Ying Tao, Jing Zhang, Anna Uehara, Anna Montmayeur, Clinton R. Paden, Peter W. Cook, Rachel Marine, Mili Sheth, Jasmine Padilla, Sarah Nobles, Mark Burroughs, Lori Rowe, Haibin Wang, Ben L. Rambo-Martin, Dhwani Batra, Justin Lee, Suxiang Tong |
| EPI_ISL_903581 | CO Dept. of Public Health and Environment, Lab Services Division | Genomics and Discovery, Respiratory Viruses Branch, Division of Viral Diseases, Centers for Disease Control and Prevention | Krista Queen, Yan Li, Ying Tao, Jing Zhang, Anna Uehara, Anna Montmayeur, Clinton R. Paden, Peter W. Cook, Rachel Marine, Mili Sheth, Jasmine Padilla, Sarah Nobles, Mark Burroughs, Lori Rowe, Haibin Wang, Ben L. Rambo-Martin, Dhwani Batra, Justin Lee, Suxiang Tong |
| EPI_ISL_903589, EPI_ISL_903605 | MI - Michigan Department of Health and Human Services - Bureau of Laboratories | Genomics and Discovery, Respiratory Viruses Branch, Division of Viral Diseases, Centers for Disease Control and Prevention | Krista Queen, Yan Li, Ying Tao, Jing Zhang, Anna Uehara, Anna Montmayeur, Clinton R. Paden, Peter W. Cook, Rachel Marine, Mili Sheth, Jasmine Padilla, Sarah Nobles, Mark Burroughs, Lori Rowe, Haibin Wang, Ben L. Rambo-Martin, Dhwani Batra, Justin Lee, Suxiang Tong |
| EPI_ISL_903606 | CO Dept. of Public Health and Environment, Lab Services Division | Genomics and Discovery, Respiratory Viruses Branch, Division of Viral Diseases, Centers for Disease Control and Prevention | Krista Queen, Yan Li, Ying Tao, Jing Zhang, Anna Uehara, Anna Montmayeur, Clinton R. Paden, Peter W. Cook, Rachel Marine, Mili Sheth, Jasmine Padilla, Sarah Nobles, Mark Burroughs, Lori Rowe, Haibin Wang, Ben L. Rambo-Martin, Dhwani Batra, Justin Lee, Suxiang Tong |
| EPI_ISL_903611 | OR State PHL-Virology/Immunology Section | Genomics and Discovery, Respiratory Viruses Branch, Division of Viral Diseases, Centers for Disease Control and Prevention | Krista Queen, Yan Li, Ying Tao, Jing Zhang, Anna Uehara, Anna Montmayeur, Clinton R. Paden, Peter W. Cook, Rachel Marine, Mili Sheth, Jasmine Padilla, Sarah Nobles, Mark Burroughs, Lori Rowe, Haibin Wang, Ben L. Rambo-Martin, Dhwani Batra, Justin Lee, Suxiang Tong |
| EPI_ISL_903629 | IA State Hygienic Laboratory | Genomics and Discovery, Respiratory Viruses Branch, Division of Viral Diseases, Centers for Disease Control and Prevention | Krista Queen, Yan Li, Ying Tao, Jing Zhang, Anna Uehara, Anna Montmayeur, Clinton R. Paden, Peter W. Cook, Rachel Marine, Mili Sheth, Jasmine Padilla, Sarah Nobles, Mark Burroughs, Lori Rowe, Haibin Wang, Ben L. Rambo-Martin, Dhwani Batra, Justin Lee, Suxiang Tong |
| EPI_ISL_903648 | CO Dept. of Public Health and Environment, Lab Services Division | Genomics and Discovery, Respiratory Viruses Branch, Division of Viral Diseases, Centers for Disease Control and Prevention | Krista Queen, Yan Li, Ying Tao, Jing Zhang, Anna Uehara, Anna Montmayeur, Clinton R. Paden, Peter W. Cook, Rachel Marine, Mili Sheth, Jasmine Padilla, Sarah Nobles, Mark Burroughs, Lori Rowe, Haibin Wang, Ben L. Rambo-Martin, Dhwani Batra, Justin Lee, Suxiang Tong |
| EPI_ISL_903649 | OR State PHL-Virology/Immunology Section | Genomics and Discovery, Respiratory Viruses Branch, Division of Viral Diseases, Centers for Disease Control and Prevention | Krista Queen, Yan Li, Ying Tao, Jing Zhang, Anna Uehara, Anna Montmayeur, Clinton R. Paden, Peter W. Cook, Rachel Marine, Mili Sheth, Jasmine Padilla, Sarah Nobles, Mark Burroughs, Lori Rowe, Haibin Wang, Ben L. Rambo-Martin, Dhwani Batra, Justin Lee, Suxiang Tong |
| EPI_ISL_903660 | MN PHL Division, Minnesota Department of Health | Genomics and Discovery, Respiratory Viruses Branch, Division of Viral Diseases, Centers for Disease Control and Prevention | Krista Queen, Yan Li, Ying Tao, Jing Zhang, Anna Uehara, Anna Montmayeur, Clinton R. Paden, Peter W. Cook, Rachel Marine, Mili Sheth, Jasmine Padilla, Sarah Nobles, Mark Burroughs, Lori Rowe, Haibin Wang, Ben L. Rambo-Martin, Dhwani Batra, Justin Lee, Suxiang Tong |
| EPI_ISL_903675 | MI - Michigan Department of Health and Human Services - Bureau of Laboratories | Genomics and Discovery, Respiratory Viruses Branch, Division of Viral Diseases, Centers for Disease Control and Prevention | Krista Queen, Yan Li, Ying Tao, Jing Zhang, Anna Uehara, Anna Montmayeur, Clinton R. Paden, Peter W. Cook, Rachel Marine, Mili Sheth, Jasmine Padilla, Sarah Nobles, Mark Burroughs, Lori Rowe, Haibin Wang, Ben L. Rambo-Martin, Dhwani Batra, Justin Lee, Suxiang Tong |
| EPI_ISL_903687 | OR State PHL-Virology/Immunology Section | Genomics and Discovery, Respiratory Viruses Branch, Division of Viral Diseases, Centers for Disease Control and Prevention | Krista Queen, Yan Li, Ying Tao, Jing Zhang, Anna Uehara, Anna Montmayeur, Clinton R. Paden, Peter W. Cook, Rachel Marine, Mili Sheth, Jasmine Padilla, Sarah Nobles, Mark Burroughs, Lori Rowe, Haibin Wang, Ben L. Rambo-Martin, Dhwani Batra, Justin Lee, Suxiang Tong |
| EPI_ISL_903691 | MN PHL Division, Minnesota Department of Health | Genomics and Discovery, Respiratory Viruses Branch, Division of Viral Diseases, Centers for Disease Control and Prevention | Krista Queen, Yan Li, Ying Tao, Jing Zhang, Anna Uehara, Anna Montmayeur, Clinton R. Paden, Peter W. Cook, Rachel Marine, Mili Sheth, Jasmine Padilla, Sarah Nobles, Mark Burroughs, Lori Rowe, Haibin Wang, Ben L. Rambo-Martin, Dhwani Batra, Justin Lee, Suxiang Tong |
| EPI_ISL_903703 | NV State Public Health Laboratory | Genomics and Discovery, Respiratory Viruses Branch, Division of Viral Diseases, Centers for Disease Control and Prevention | Krista Queen, Yan Li, Ying Tao, Jing Zhang, Anna Uehara, Anna Montmayeur, Clinton R. Paden, Peter W. Cook, Rachel Marine, Mili Sheth, Jasmine Padilla, Sarah Nobles, Mark Burroughs, Lori Rowe, Haibin Wang, Ben L. Rambo-Martin, Dhwani Batra, Justin Lee, Suxiang Tong |
| EPI_ISL_903705, EPI_ISL_903708, EPI_ISL_903710, EPI_ISL_903712 | MI - Michigan Department of Health and Human Services - Bureau of Laboratories | Genomics and Discovery, Respiratory Viruses Branch, Division of Viral Diseases, Centers for Disease Control and Prevention | Krista Queen, Yan Li, Ying Tao, Jing Zhang, Anna Uehara, Anna Montmayeur, Clinton R. Paden, Peter W. Cook, Rachel Marine, Mili Sheth, Jasmine Padilla, Sarah Nobles, Mark Burroughs, Lori Rowe, Haibin Wang, Ben L. Rambo-Martin, Dhwani Batra, Justin Lee, Suxiang Tong |
| EPI_ISL_903719 | NV State Public Health Laboratory | Genomics and Discovery, Respiratory Viruses Branch, Division of Viral Diseases, Centers for Disease Control and Prevention | Krista Queen, Yan Li, Ying Tao, Jing Zhang, Anna Uehara, Anna Montmayeur, Clinton R. Paden, Peter W. Cook, Rachel Marine, Mili Sheth, Jasmine Padilla, Sarah Nobles, Mark Burroughs, Lori Rowe, Haibin Wang, Ben L. Rambo-Martin, Dhwani Batra, Justin Lee, Suxiang Tong |
| EPI_ISL_903724 | MI - Michigan Department of Health and Human Services - | Genomics and Discovery, Respiratory Viruses Branch, | Krista Queen, Yan Li, Ying Tao, Jing Zhang, Anna Uehara, Anna Montmayeur, Clinton R. Paden, Peter W. Cook, Rachel Marine, Mili Sheth, Jasmine |

| | | | |
|---|---|---|---|
| | Bureau of Laboratories | Division of Viral Diseases, Centers for Disease Control and Prevention | Padilla, Sarah Nobles, Mark Burroughs, Lori Rowe, Haibin Wang, Ben L. Rambo-Martin, Dhwani Batra, Justin Lee, Suxiang Tong |
| EPI_ISL_903745 | MS Public Health Laboratory | Genomics and Discovery, Respiratory Viruses Branch, Division of Viral Diseases, Centers for Disease Control and Prevention | Krista Queen, Yan Li, Ying Tao, Jing Zhang, Anna Uehara, Anna Montmayeur, Clinton R. Paden, Peter W. Cook, Rachel Marine, Mili Sheth, Jasmine Padilla, Sarah Nobles, Mark Burroughs, Lori Rowe, Haibin Wang, Ben L. Rambo-Martin, Dhwani Batra, Justin Lee, Suxiang Tong |
| EPI_ISL_903753 | ID Bureau of Laboratories | Genomics and Discovery, Respiratory Viruses Branch, Division of Viral Diseases, Centers for Disease Control and Prevention | Krista Queen, Yan Li, Ying Tao, Jing Zhang, Anna Uehara, Anna Montmayeur, Clinton R. Paden, Peter W. Cook, Rachel Marine, Mili Sheth, Jasmine Padilla, Sarah Nobles, Mark Burroughs, Lori Rowe, Haibin Wang, Ben L. Rambo-Martin, Dhwani Batra, Justin Lee, Suxiang Tong |
| EPI_ISL_903768 | TX DSHS, Lab Services Section MC 1947 | Genomics and Discovery, Respiratory Viruses Branch, Division of Viral Diseases, Centers for Disease Control and Prevention | Krista Queen, Yan Li, Ying Tao, Jing Zhang, Anna Uehara, Anna Montmayeur, Clinton R. Paden, Peter W. Cook, Rachel Marine, Mili Sheth, Jasmine Padilla, Sarah Nobles, Mark Burroughs, Lori Rowe, Haibin Wang, Ben L. Rambo-Martin, Dhwani Batra, Justin Lee, Suxiang Tong |
| EPI_ISL_903780, EPI_ISL_903781, EPI_ISL_903798, EPI_ISL_903799, EPI_ISL_903818 | CO Dept. of Public Health and Environment, Lab Services Division | Genomics and Discovery, Respiratory Viruses Branch, Division of Viral Diseases, Centers for Disease Control and Prevention | Krista Queen, Yan Li, Ying Tao, Jing Zhang, Anna Uehara, Anna Montmayeur, Clinton R. Paden, Peter W. Cook, Rachel Marine, Mili Sheth, Jasmine Padilla, Sarah Nobles, Mark Burroughs, Lori Rowe, Haibin Wang, Ben L. Rambo-Martin, Dhwani Batra, Justin Lee, Suxiang Tong |
| EPI_ISL_903845 | NE Public Health Laboratory | Genomics and Discovery, Respiratory Viruses Branch, Division of Viral Diseases, Centers for Disease Control and Prevention | Krista Queen, Yan Li, Ying Tao, Jing Zhang, Anna Uehara, Anna Montmayeur, Clinton R. Paden, Peter W. Cook, Rachel Marine, Mili Sheth, Jasmine Padilla, Sarah Nobles, Mark Burroughs, Lori Rowe, Haibin Wang, Ben L. Rambo-Martin, Dhwani Batra, Justin Lee, Suxiang Tong |
| EPI_ISL_903850 | MS Public Health Laboratory | Genomics and Discovery, Respiratory Viruses Branch, Division of Viral Diseases, Centers for Disease Control and Prevention | Krista Queen, Yan Li, Ying Tao, Jing Zhang, Anna Uehara, Anna Montmayeur, Clinton R. Paden, Peter W. Cook, Rachel Marine, Mili Sheth, Jasmine Padilla, Sarah Nobles, Mark Burroughs, Lori Rowe, Haibin Wang, Ben L. Rambo-Martin, Dhwani Batra, Justin Lee, Suxiang Tong |
| EPI_ISL_903887, EPI_ISL_903908, EPI_ISL_903923 | ID Bureau of Laboratories | Genomics and Discovery, Respiratory Viruses Branch, Division of Viral Diseases, Centers for Disease Control and Prevention | Krista Queen, Yan Li, Ying Tao, Jing Zhang, Anna Uehara, Anna Montmayeur, Clinton R. Paden, Peter W. Cook, Rachel Marine, Mili Sheth, Jasmine Padilla, Sarah Nobles, Mark Burroughs, Lori Rowe, Haibin Wang, Ben L. Rambo-Martin, Dhwani Batra, Justin Lee, Suxiang Tong |
| EPI_ISL_903936 | CO Dept. of Public Health and Environment, Lab Services Division | Genomics and Discovery, Respiratory Viruses Branch, Division of Viral Diseases, Centers for Disease Control and Prevention | Krista Queen, Yan Li, Ying Tao, Jing Zhang, Anna Uehara, Anna Montmayeur, Clinton R. Paden, Peter W. Cook, Rachel Marine, Mili Sheth, Jasmine Padilla, Sarah Nobles, Mark Burroughs, Lori Rowe, Haibin Wang, Ben L. Rambo-Martin, Dhwani Batra, Justin Lee, Suxiang Tong |
| EPI_ISL_904243, EPI_ISL_904283, EPI_ISL_904284, EPI_ISL_904285, EPI_ISL_904286, EPI_ISL_904287, EPI_ISL_904361, EPI_ISL_904495 | Dutch COVID-19 response team | Erasmus Medical Center | Bas Oude Munnink, Reina Sikkema, David Nieuwenhuijse, Irina Chestakova, Anne van der Linden, Marjan Boter, Emmanuelle Munger, Corine GeurtsvanKessel, Annemiek van der Eijk, Richard Molenkamp, Marion Koopmans, on behalf of the Dutch national COVID-19 response team. |
| EPI_ISL_904765, EPI_ISL_904885, EPI_ISL_904890, EPI_ISL_905097, EPI_ISL_905124, EPI_ISL_905125, EPI_ISL_905129, EPI_ISL_905130, EPI_ISL_905132, EPI_ISL_905133, EPI_ISL_905137, EPI_ISL_905138, EPI_ISL_905139, EPI_ISL_905150, EPI_ISL_905160, EPI_ISL_905161, EPI_ISL_905605, EPI_ISL_905647, EPI_ISL_905648, EPI_ISL_905649, EPI_ISL_905657, EPI_ISL_905685 | | | |
| see above | Dutch COVID-19 response team | National Institute for Public Health and the Environment (RIVM) | Adam Meijer, Harry Vennema, Dirk Eggink, Jeroen Cremer, Sharon van den Brink, Bas van der Veer, AnneMarie van den Brandt, Florian Zwagemaker, Dennis Schmitz, Chantal Reusken, on behalf of the national COVID-19 response team |
| EPI_ISL_905821, EPI_ISL_905822, EPI_ISL_905823, EPI_ISL_905824, EPI_ISL_905825, EPI_ISL_905826, EPI_ISL_905827, EPI_ISL_905828, EPI_ISL_905829, EPI_ISL_905830, EPI_ISL_905831, EPI_ISL_905832, EPI_ISL_905833, EPI_ISL_905834, EPI_ISL_905835, EPI_ISL_905836, EPI_ISL_905837, EPI_ISL_905838, EPI_ISL_905839, EPI_ISL_905840, EPI_ISL_905841, EPI_ISL_905842, EPI_ISL_905843, EPI_ISL_905844, EPI_ISL_905845, EPI_ISL_905846 | | | |
| see above | OHSU Lab Services Molecular Microbiology Lab | Oregon SARS-CoV-2 Genome Sequencing Center | Brendan L. O'Connell, Sally Grindstaff, Kayla Carter, Ruth V. Nichols, Alec J. Hirsch, Donna Hansel, Guang Fan, Xuan, Qin, Daniel N. Streblow, William B. Messer, Andrew C. Adey, Benjamin N. Bimber, Brian J. O'Roak |
| EPI_ISL_906295, EPI_ISL_906296, EPI_ISL_906297, EPI_ISL_906298, EPI_ISL_906299, EPI_ISL_906300, EPI_ISL_906301 | Nigeria Centre for Disease Control (NCDC) | African Centre of Excellence for Genomics of Infectious Diseases (ACEGID), Redeemer's University | Oluniyi P.E. et al |
| EPI_ISL_906565, EPI_ISL_906566, EPI_ISL_906567, EPI_ISL_906568 | Maine Health and Environmental Testing Laboratory (Maine HETL) | Tewhey Lab, The Jackson Laboratory | Matluk,N., Dewey,H., Iosue,F., Barter,M., Lynch,R., Munger,H. and Tewhey,R. |
| EPI_ISL_906850 | Respiratory Viruses Branch, Centers for Disease Control and Prevention | Respiratory Viruses Branch, Centers for Disease Control and Prevention | Tao,Y., Li,Y., Zhang,J., Queen,K., Uehara,A., Cook,P., Paden,C.R., Wang,H., Tong,S. |
| EPI_ISL_909948 | CUSL/UCLouvain COVID testing federal platform | UCLouvain/IREC/MBLG | Jean Ruelle, Lysa Pinsmaye, Benoit Kabamba Mukadi |
| EPI_ISL_909970, EPI_ISL_909976, EPI_ISL_909983, EPI_ISL_909984, EPI_ISL_909985, EPI_ISL_909986, EPI_ISL_909987, EPI_ISL_909988, EPI_ISL_909989, EPI_ISL_909990, EPI_ISL_910002 | | | |
| see above | A. Krumbholz, Labor Dr. Krause und Kollegen MVZ GmbH, Kiel | Charité Universitätsmedizin Berlin, Institut für Virologie | Victor M Corman, Tobias Bleicker, Julia Tesch, Barbara Mühlemann, Jörn Beheim-Schwarzbach, Talitha Veith, Julia Schneider, Cornelia Schlee, Tomasz Zemojtel, Terry Jones, Christian Drosten |
| EPI_ISL_910549, EPI_ISL_910570, EPI_ISL_910578, EPI_ISL_910579, EPI_ISL_911067, EPI_ISL_911068, EPI_ISL_911069, EPI_ISL_911070, EPI_ISL_911071, EPI_ISL_911074, EPI_ISL_911075, EPI_ISL_911076, EPI_ISL_911077, EPI_ISL_911078, EPI_ISL_911079, EPI_ISL_911080, EPI_ISL_911081, EPI_ISL_911085, EPI_ISL_911086, EPI_ISL_911087, EPI_ISL_911088, EPI_ISL_911089, EPI_ISL_911090, EPI_ISL_911091, EPI_ISL_911092, EPI_ISL_911093, EPI_ISL_911094, EPI_ISL_911095, EPI_ISL_911096, EPI_ISL_911097, EPI_ISL_911098, EPI_ISL_911099, EPI_ISL_911100, EPI_ISL_911101, EPI_ISL_911102, EPI_ISL_911103 | | | |
| see above | Laboratoire national de sante, Microbiology, Virology | Laboratoire national de sante, Microbiology, Microbial Genomics Platform | Anke Wienecke-Baldacchino, Catherine Ragimbeau,Jessica Tapp, Fatu Djabi, Lise Pignon, Raoul Salmon, Tamir Abdelrahman |
| EPI_ISL_912270, EPI_ISL_912283, EPI_ISL_912331 | Hospital General Universitario Gregorio Marañón | SeqCOVID-SPAIN consortium / IBV (CSIC) | Darío García de Viedma, Laura Pérez-Lago, Pedro J Sola-Campoy, Sergio Buenestado-Serrano, Marta Herranz, Victor Manuel de la Cueva, Julia Suárez, Pilar Catalán, Patricia Muñoz and SeqCOVID-SPAIN consortium |
| EPI_ISL_912361, EPI_ISL_912362, EPI_ISL_912363, EPI_ISL_912394 | Fondation Congolaise pour la recherche medicale (FCRM), Francine Ntoumi | NGS Competence Center Tuebingen, Institut für Medizinische Mikrobiologie und Hygiene, Universitaetsklinikum Tübingen | Angel Angelov |
| EPI_ISL_912477, EPI_ISL_912490 | NHLS Universitas Academic | UFS Virology | PA Bester, MM Nyaga, P Nthiga, MT Mogotsi, D Goedhals, T de Oliveira |
| EPI_ISL_913103, EPI_ISL_913108, EPI_ISL_913109, EPI_ISL_913110, EPI_ISL_913112 | CHU Purpan - Laboratoire de Virologie - Institut Fédératif de Biologie | CHU Purpan - Laboratoire de Virologie - Institut Fédératif de Biologie | Latour J., Ranger N., Dubois M., Carcenac R., Harter A., Boyer P., Tremeaux P., Izopet J. |
| EPI_ISL_913295 | Klinisk mikrobiologi | The Public Health Agency of Sweden | Anna-Malin Linde, Maria Lind Karlberg, Carlo Berg, Oskar Karlsson Lindsjo, Sofia Stamouli, Reza Advani, Mattias Haukland, Petra Holmstrom, Noura Walai, Petra Edquist, Mia Brytting, Anna Risberg, Karin Tegmark-Wisell |
| EPI_ISL_913315 | Klinisk Mikrobiologi | The Public Health Agency of Sweden | Anna-Malin Linde, Maria Lind Karlberg, Carlo Berg, Oskar Karlsson Lindsjo, Sofia Stamouli, Reza Advani, Mattias Haukland, Petra Holmstrom, Noura Walai, Petra Edquist, Mia Brytting, Anna Risberg, Karin Tegmark-Wisell |
| EPI_ISL_913320, EPI_ISL_913321, EPI_ISL_913322 | Klinisk mikrobiologi | The Public Health Agency of Sweden | Anna-Malin Linde, Maria Lind Karlberg, Carlo Berg, Oskar Karlsson Lindsjo, Sofia Stamouli, Reza Advani, Mattias Haukland, Petra Holmstrom, Noura Walai, Petra Edquist, Mia Brytting, Anna Risberg, Karin Tegmark-Wisell |
| EPI_ISL_913621, EPI_ISL_913622, EPI_ISL_913623, EPI_ISL_913624 | Michigan Department of Health and Human Services, Bureau of Laboratories | Michigan Department of Health and Human Services, Bureau of Laboratories | Blankenship HM, Riner D, Soehnlen MK |
| EPI_ISL_913781, EPI_ISL_913801 | KU Leuven, Rega Institute, Clinical and Epidemiological Virology | KU Leuven, Rega Institute, Clinical and Epidemiological Virology | Tony Wawina-Bokalanga, Bert Vanmechelen, Joan Marti-Carerras, Piet Maes |

| | | | |
|---|---|---|---|
| EPI_ISL_913953, EPI_ISL_913954, EPI_ISL_913955, EPI_ISL_913968, EPI_ISL_913983 | Instituto de Diagnostico y Referencia Epidemiologicos INDRE_RNLSP | Instituto de Diagnostico y Referencia Epidemiologicos (INDRE) | Claudia Wong-Arambula, Abril Rodriguez-Maldonado, Fabiola Garces-Ayala, Adnan Araiza-Rodriguez, David Fragoso-Fonseca, Sergio Rangel-Guerrero, Mayra Jimenez-Morales, Nancy Munoz-Hernandez, Natividad Cruz-Ortiz, Tatiana Nunez-Garcia, Gisela Barrera-Badillo, Lucia Hernandez-Rivas, Irma Lopez-Martinez, Ernesto Ramirez-Gonzalez. |
| EPI_ISL_914802 | AREA DE SALUD CATEDRAL NORESTE | Inciensa, Instituto Costarricense de Investigación y Enseñanza en Nutrición y Salud | Francisco Duarte, Hebleen Porras, Claudio Soto-Garita, Estela Cordero, Adriana Godínez, Melany Calderón & Mariel López |
| EPI_ISL_914803 | AREA DE SALUD LA UNION | Inciensa, Instituto Costarricense de Investigación y Enseñanza en Nutrición y Salud | Francisco Duarte, Hebleen Porras, Claudio Soto-Garita, Estela Cordero, Adriana Godínez, Melany Calderón & Mónica Charpentier-Artavia |
| EPI_ISL_914805 | AREA DE SALUD CARTAGO | Inciensa, Instituto Costarricense de Investigación y Enseñanza en Nutrición y Salud | Francisco Duarte, Hebleen Porras, Claudio Soto-Garita, Estela Cordero, Adriana Godínez, Melany Calderón & Mónica Charpentier-Artavia |
| EPI_ISL_914806 | AREA DE SALUD CORONADO | Inciensa, Instituto Costarricense de Investigación y Enseñanza en Nutrición y Salud | Francisco Duarte, Hebleen Porras, Claudio Soto-Garita, Estela Cordero, Adriana Godínez & Melany Calderón |
| EPI_ISL_914809 | AREA DE SALUD OREAMUNO-PACAYAS-TIERRA BLANCA | Inciensa, Instituto Costarricense de Investigación y Enseñanza en Nutrición y Salud | Francisco Duarte, Hebleen Porras, Claudio Soto-Garita, Estela Cordero, Adriana Godínez, Melany Calderón & Mariel López |
| EPI_ISL_918170 | Department of Infectious Diseases and Immunology, National Hospital Organization Nagoya Medical Center | Clinical Research Center, National Hospital Organization Nagoya Medical Center | Yoshihiro Nakata, Hirotaka Ode, Mai Kubota, Masakazu Matsuda, Kazuhiro Matsuoka, Miho Nakasuji, Mikiko Mori, Mayumi Imahashi, Yoshiyuki Yokomaku, Yasumasa Iwatani |
| EPI_ISL_918443 | AIID | Irish Coronavirus Sequencing Consortium-Teagasc Grange | Matthew McCabe, Aljandro Abner Garcia Leon, Fiona Crispie, Calum Walsh, Michael Carr, John Kenny, Paul Cotter, Patrick Mallon, Gabriel Gonzalez |
| EPI_ISL_918554 | LACEN - Laboratório Central de Saúde Pública do Amapa | Evandro Chagas Institute | Santos, M.C.; Silva, A.M.; Junior, W.D.C.; Barbagelata, L.S.; Ferreira, J.A.; Sousa, E.M.A.; da Silva, P.S.; Pinheiro, K.C.; L.C.; Sousa Junior, E.C. |
| EPI_ISL_918828, EPI_ISL_918829, EPI_ISL_918830, EPI_ISL_918833, EPI_ISL_918834, EPI_ISL_918835, EPI_ISL_918836, EPI_ISL_918837, EPI_ISL_918838, EPI_ISL_918839, EPI_ISL_918840, EPI_ISL_918969, EPI_ISL_918970, EPI_ISL_918971, EPI_ISL_918972 | | | |
| see above | University of Birmingham | COVID-19 Genomics UK (COG-UK) Consortium | Institute of Microbiology, University of Birmingham: Claire McMurray, Joanne Stockton, Samuel Nicholls, Radoslaw Poplawski, Will Rowe, Josh Quick, Nicholas Loman. University of Birmingham Testing Laboratory: Celina M Whalley, Andrew Bosworth, Charlotte Poxon, Kasun Wanigasooriya, Oliver Pickles, Mike Kidd, Alex Richter, Andrew D Beggs PHE Heartlands Lab: Husam Osman, Andrew Bosworth. Queen Elizabeth Hospital: Anna Casey |
| EPI_ISL_919241, EPI_ISL_919242, EPI_ISL_919246, EPI_ISL_919251, EPI_ISL_919262, EPI_ISL_919264, EPI_ISL_919265 | West of Scotland Specialist Virology Centre, NHSGGC / MRC-University of Glasgow Centre for Virus Research | COVID-19 Genomics UK (COG-UK) Consortium | Ana da Silva Filipe, Natasha Johnson, Kathy Smollett, Daniel Mair, Stephen Carmichael, Alice Broos, Lily Tong, Jenna Nichols, Kyriaki Nomikou; Sarah McDonald; Richard Orton, Joseph Hughes, Sreenu Vattipally, David L Robertson; Alasdair MacLean, Rory Gunson; Sharif Shaaban, Matthew Holden; Rachel Blacow, Guy Mollett, Kathy Li, James Shepherd, Antonia Ho, Emma Thomson |
| EPI_ISL_919314, EPI_ISL_919315, EPI_ISL_919317, EPI_ISL_919371, EPI_ISL_919372, EPI_ISL_919373, EPI_ISL_919374, EPI_ISL_919375, EPI_ISL_919399 | Virology Department, Royal Infirmary of Edinburgh, NHS Lothian / School of Biological Sciences, University of Edinburgh / Institute of Genetics and Molecular Medicine, University of Edinburgh | COVID-19 Genomics UK (COG-UK) Consortium | McHugh M, Dewar R, Rooke S, Gallagher M, Balcaza C, O'Toole Á, Scher E, Hill V, McCrone JT, Colquhoun R, Yu X, Jackson B, Rambaut A, Williams TC, Templeton K |
| EPI_ISL_919446, EPI_ISL_919447 | Liverpool Clinical Laboratories | COVID-19 Genomics UK (COG-UK) Consortium | Sam Haldenby, Anita Lucaci, Steve Paterson, Julian Hiscox, Alistair Darby, M Almsaud, A Alrezaihi, Muhannad Alruwaili, Stuart D Armstrong, Jones Benjamin, Eleanor G Bentley, Anu Chawla, Jordan J Clark, Angela Cowell, Richard Eccles, Isabel García-Dorival, Matthew Gemmell, Alessandro Gerada, PKF Gilmore, Richard Gregory, Ximeng Han, Catherine Hartley, Margaret Hughes, Miren Iturriza-Gomara, James Johnson, L Luu, Jenifer Manson, Charlotte Nelson, Elaine O'Toole, Cassie Olateju, Rebekah Penrice-Randal , Lucille Rainbow, N.P Randle, Trevor Ian Robinson, Parul Sharma, Ghada T Shawli, James P Stewart, Neil Swainston, Ecaterina Vamos, Joanne Watts, Mark Whitehead |
| EPI_ISL_919725, EPI_ISL_919726 | University College London, Great Ormond Street Hospital for Children NHS Foundation Trust, Imperial College Healthcare NHS Trust | COVID-19 Genomics UK (COG-UK) Consortium | Sergi Castellano, Rachel Williams, Mark Kristiansen, Paola Resende Silva, Sunando Roy, Tony Brooks, Helena Tutill, Paola Niola, Patricia Dyal, Charlotte Williams, Leysa Forrest, Yasmin Panchbhaya, Jacqueline Findlay, Samuel Weeks, Julianne Brown, Kathryn Harris, Paul Randell, James Price, Alison Holmes, Judith Breuer |
| EPI_ISL_920176, EPI_ISL_920221, EPI_ISL_920240, EPI_ISL_920243, EPI_ISL_920248, EPI_ISL_920249, EPI_ISL_920251, EPI_ISL_920259, EPI_ISL_920260, EPI_ISL_920269, EPI_ISL_920270, EPI_ISL_920281, EPI_ISL_920291, EPI_ISL_920301, EPI_ISL_920321, EPI_ISL_920461, EPI_ISL_920468, EPI_ISL_920476, EPI_ISL_920477, EPI_ISL_920478, EPI_ISL_920485, EPI_ISL_920486, EPI_ISL_920495, EPI_ISL_920502, EPI_ISL_920505, EPI_ISL_920508, EPI_ISL_920513, EPI_ISL_920520, EPI_ISL_920521, EPI_ISL_920526, EPI_ISL_920540, EPI_ISL_920547, EPI_ISL_920548, EPI_ISL_920549, EPI_ISL_920550, EPI_ISL_920551, EPI_ISL_920558, EPI_ISL_920560, EPI_ISL_920582, EPI_ISL_920584, EPI_ISL_920585, EPI_ISL_920596, EPI_ISL_920647, EPI_ISL_920666, EPI_ISL_920674, EPI_ISL_920688, EPI_ISL_920696, EPI_ISL_920709, EPI_ISL_920710, EPI_ISL_920717, EPI_ISL_920721, EPI_ISL_920746 | | | |
| see above | University College London Hospital | COVID-19 Genomics UK (COG-UK) Consortium | Judith Heaney, Matthew Byott, Catherine Houlihan, Dan Frampton, Stuart Kirk, Moira Spyer and Eleni Nastouli |
| EPI_ISL_921137, EPI_ISL_921138, EPI_ISL_921140, EPI_ISL_921141, EPI_ISL_921142, EPI_ISL_921143, EPI_ISL_921144, EPI_ISL_921145, EPI_ISL_921146, EPI_ISL_921147, EPI_ISL_921148, EPI_ISL_921150, EPI_ISL_921158, EPI_ISL_921159, EPI_ISL_921160, EPI_ISL_921162, EPI_ISL_921163, EPI_ISL_921164, EPI_ISL_921165, EPI_ISL_921167, EPI_ISL_921170, EPI_ISL_921171, EPI_ISL_921173, EPI_ISL_921178, EPI_ISL_921179, EPI_ISL_921180, EPI_ISL_921182, EPI_ISL_921183, EPI_ISL_921184, EPI_ISL_921699, EPI_ISL_921700, EPI_ISL_921701, EPI_ISL_921702, EPI_ISL_921704, EPI_ISL_921705, EPI_ISL_921708, EPI_ISL_921709, EPI_ISL_921710, EPI_ISL_921711, EPI_ISL_921713, EPI_ISL_921714, EPI_ISL_921715, EPI_ISL_921716, EPI_ISL_921717, EPI_ISL_921720, EPI_ISL_921721, EPI_ISL_921722, EPI_ISL_921723, EPI_ISL_921725, EPI_ISL_921753, EPI_ISL_921756, EPI_ISL_921758 | | | |
| see above | Northumbria University / South Tees Hospitals NHS Foundation Trust / North Cumbria Integrated Care NHS Foundation Trust / North Tees and Hartlepool NHS Foundation Trust / Newcastle Hospitals NHS Foundation Trust | COVID-19 Genomics UK (COG-UK) Consortium | Darren L Smith,Andrew Nelson,Matthew Bashton,Greg R Young,Joshua Loh,John Allan,Mohammad A Tariq,Giles S Holt,Gary Black,Wen C Yew,Lynn Dover,Paul Baker,Steve Liggett,Sarah Essex,Jane Greenaway,Debra Padgett,Clive Graham,Garren Scott,Edward Barton,Emma Swindells,Brendan Payne,Jennifer Collins,Yusri Taha,Gary Eltringham |
| EPI_ISL_925082, EPI_ISL_925087, EPI_ISL_925088, EPI_ISL_925092 | Wyoming Public Health Laboratory | Wyoming Public Health Laboratory | Noah Hull, Taylor Fearing, Lynette Gumbleton, Channing Weber, Ashley Norberg, Bailey Bowcutt, and Wanda Manley |
| EPI_ISL_925390, EPI_ISL_925391, EPI_ISL_925392, EPI_ISL_925393, EPI_ISL_925394, EPI_ISL_925467, EPI_ISL_925468 | Department of Clinical Microbiology | GIGA Medical Genomics | Keith Durkin, Maria Artesi, Sébastien Bontems, Raphaël Boreux, Bouchra Boujemla, Cécile Meex, Pierrette Melin, Marie-Pierre Hayette, Vincent Bours |
| EPI_ISL_925498 | 1.AO Universitaria 'S. Giovanni di Dio e Ruggi D'Aragona, Scuola Medica Salernitana' Hospital / 2.UOC di Virologia e Microbiologia, Università della Campania 'L. Vanvitelli' / 3.AO Universitaria 'Federico II' Napoli Hospital / 4.AORN 'San Giuseppe Moscati' Avellino Hospital / 5.AO 'San Pio - presidio G. Rummo' Benevento Hospital / 6.AO 'Sant'Anna e San Sebastiano' Caserta Hospital / 7.PO 'Maria Santissima Addolorata' Eboli Hospital / 8.Biogem Istituto di Ricerche Genetiche | 1. Genome Research Center for Health (CRGS) / 2. Laboratory of Molecular Medicine and Genomics(LMMGe) / 3. Center for Research in Pure and Applied Mathematics (CRMPA) | Giorgio Giurato, Francesca Rizzo, Alessandro Weisz, Gianluigi Franci, Giovanni Nassa, Pasquale Pagliano, Roberta Tarallo, Elena Alexandrova, Ylenia D'Agostino, Carlo Ferravante, Jessica Lamberti, Viola Melone, Domenico Memoli, Valeria Mirici Cappa, Domenico Palumbo, Giovanni Pecoraro, Assunta Sellitto, Oriana Strianese, Ilaria Terenzi, Giuseppe Fenza, Aniello Gentile, Antonello Saccomanno, Sonia Amabile, Teresa Rocco, Annamaria Salvati, Emilia Vaccaro, Massimiliano Galdiero, Michele Cennamo, Giuseppe Portella, Maria Grazia Foti, Mariarosaria Ingino, Maria Landi, Maurizio Fumi, Vincenzo Rocco, Rita Greco, Vittoria Letizia, Arnolfo Petruzziello, Maddalena Schioppa, Gregorio Goffredi, Francesca Marciano, Michele Caraglia, Alessia Cossu, Marianna Scrima |
| EPI_ISL_925847, EPI_ISL_925849, EPI_ISL_925850, EPI_ISL_925855, EPI_ISL_925865, EPI_ISL_925866, EPI_ISL_925867, EPI_ISL_925868, EPI_ISL_925869, EPI_ISL_925870, EPI_ISL_925871, EPI_ISL_925872, EPI_ISL_925873, EPI_ISL_925874, EPI_ISL_925875, EPI_ISL_925876, EPI_ISL_925877, EPI_ISL_925878, EPI_ISL_925879, EPI_ISL_925880, EPI_ISL_925881, EPI_ISL_925893, EPI_ISL_925897, EPI_ISL_925898, EPI_ISL_925899, EPI_ISL_930634, EPI_ISL_930853 | | | |
| see above | Nucleic Acid Testing, National Reference Laboratory | GIGA Medical Genomics | Yvan Butera, Keith Durkin, Maria Artesi, Bouchra Boujemla, Robert Rutayisire, Patrick Tuyisenge, Esperence Umumararungu, Sébastien Bontems, Marie-Pierre Hayette, Nathalie Renotte, Swaibu Gatare, Jacob Souopgui, Sabin Nsanzimana, Vincent Bours, Léon Mutesa |
| EPI_ISL_931394 | University Hospital Basel, Clinical Virology | University Hospital Basel, Clinical Bacteriology | Tim Roloff, Madlen Stange, Helena MB Seth-Smith, Alfredo Mari, Karoline Leuzinger, Julia Bielicki, Manuel Battegay, Hans Hirsch, Adrian Egli |
| EPI_ISL_933481 | Lighthouse Lab in Milton Keynes | Wellcome Sanger Institute for the COVID-19 Genomics UK (COG-UK) Consortium | The Lighthouse Lab in Milton Keynes and Alex Alderton, Roberto Amato, Sonia Goncalves, Ewan Harrison, David K. Jackson, Ian Johnston, Dominic Kwiatkowski, Cordelia Langford, John Sillitoe on behalf of the Wellcome Sanger Institute COVID-19 Surveillance Team |
| EPI_ISL_933670, EPI_ISL_933671, EPI_ISL_933672, EPI_ISL_933673, EPI_ISL_933678, EPI_ISL_933699, EPI_ISL_933700, EPI_ISL_933708, EPI_ISL_933709, EPI_ISL_933710, EPI_ISL_933712 | | | |
| see above | Instituto de Diagnostico y Referencia Epidemiologicos INDRE_RNLSP | Instituto de Diagnostico y Referencia Epidemiologicos (INDRE) | Claudia Wong-Arambula, Abril Rodriguez-Maldonado, Fabiola Garces-Ayala, Adnan Araiza-Rodriguez, David Fragoso-Fonseca, Sergio Rangel-Guerrero, Mayra Jimenez-Morales, Nancy Munoz-Hernandez, Natividad Cruz-Ortiz, Tatiana Nunez-Garcia, Gisela Barrera-Badillo, Lucia Hernandez-Rivas, Irma |

| | | | |
|---|---|---|---|
| | | | Lopez-Martinez, Ernesto Ramirez-Gonzalez. |
| EPI_ISL_934171, EPI_ISL_934172, EPI_ISL_934173 | Vilnius university hospital Santaros Klinikos, Center of Laboratory Medicine | Vilnius university hospital Santaros Klinikos, Center of Laboratory Medicine | Ingrida Olendraite, Daniel Naumovas, Rimvydas Norvilas, Dovile Ezerskyte, Justinas Slikas, Gytis Dudas |
| EPI_ISL_934385, EPI_ISL_934386 | Klinisk mikrobiologi | The Public Health Agency of Sweden | Anna-Malin Linde, Maria Lind Karlberg, Carlo Berg, Oskar Karlsson Lindsjo, Sofia Stamouli, Reza Advani, Mattias Haukland, Petra Holmstrom, Noura Walai, Petra Edquist, Mia Brytting, Anna Risberg, Karin Tegmark-Wisell |
| EPI_ISL_934389 | Klinisk Mikrobiologi | The Public Health Agency of Sweden | Anna-Malin Linde, Maria Lind Karlberg, Carlo Berg, Oskar Karlsson Lindsjo, Sofia Stamouli, Reza Advani, Mattias Haukland, Petra Holmstrom, Noura Walai, Petra Edquist, Mia Brytting, Anna Risberg, Karin Tegmark-Wisell |
| EPI_ISL_934572, EPI_ISL_934573, EPI_ISL_934574, EPI_ISL_934575 | Department of Laboratory Medicine, Division of Clinical Virology, University of Medicine, Vienna | Bergthaler laboratory, CeMM Research Center for Molecular Medicine of the Austrian Academy of Sciences | Lukas Endler, Anna Schedl, Thomas Penz, Benedikt Agerer, Maelle Le Moing, Michael Schuster, Bekir Erguner, Jan Laine, Martin Senekowitsch, Christoph Bock, Andreas Bergthaler |
| EPI_ISL_934638 | Department of Microbiology, University Innsbruck | Bergthaler laboratory, CeMM Research Center for Molecular Medicine of the Austrian Academy of Sciences | Lukas Endler, Anna Schedl, Thomas Penz, Benedikt Agerer, Maelle Le Moing, Michael Schuster, Bekir Erguner, Jan Laine, Martin Senekowitsch, Christoph Bock, Andreas Bergthaler |
| EPI_ISL_935207 | KU Leuven, Rega Institute, Clinical and Epidemiological Virology | KU Leuven, Rega Institute, Clinical and Epidemiological Virology | Tony Wawina-Bokalanga, Bert Vanmechelen, Joan Marti-Carerras, Piet Maes |
| EPI_ISL_936131, EPI_ISL_936132, EPI_ISL_936134, EPI_ISL_936135, EPI_ISL_936136, EPI_ISL_936137 | New York Presbyterian Hospital | Wadsworth Center, New York State Department of Health | Kirsten St. George, Daryl M. Lamson, Alexis Russel, Matthew Shudt, Melissa A Leisner, Jonathan Plitnick, Navjot Singh, John Kelly, Erasmus Schneider, Erica Lasek-Nesselquist |
| EPI_ISL_936646, EPI_ISL_936647, EPI_ISL_936648, EPI_ISL_936649, EPI_ISL_936650, EPI_ISL_936651, EPI_ISL_936652, EPI_ISL_936685, EPI_ISL_936686, EPI_ISL_936689, EPI_ISL_936693, EPI_ISL_936694, EPI_ISL_936695 | | | |
| see above | Northwestern Memorial Hospital | Ozer Lab | Ramon Lorenzo-Redondo, Lacy M. Simons, Chad J. Achenbach, Lawrence J. Jennings, Michael G. Ison, Judd F. Hultquist, Egon A. Ozer |
| EPI_ISL_937353 | Utah Public Health Laboratory | Utah Public Health Laboratory | Erin L. Young, Kelly F. Oakeson, Tara Gallagher |
| EPI_ISL_940081, EPI_ISL_940082, EPI_ISL_940085, EPI_ISL_940088, EPI_ISL_940089, EPI_ISL_940090, EPI_ISL_940091, EPI_ISL_940092, EPI_ISL_940096, EPI_ISL_940100, EPI_ISL_940103, EPI_ISL_940110, EPI_ISL_940111, EPI_ISL_940124, EPI_ISL_940126, EPI_ISL_940131, EPI_ISL_940137 | | | |
| see above | Charlotte Maxeke Johannesburg Academic Hospital, National Health Laboratory Services, Gauteng, South Africa | National Institute for Communicable Diseases of the National Health Laboratory Service | Amoako DG, Mohale T, Ntuli N, Mahlangu B, Allam M, Ismail A, Bhiman JN |
| EPI_ISL_940827, EPI_ISL_940828, EPI_ISL_940829, EPI_ISL_940833, EPI_ISL_940834, EPI_ISL_940835, EPI_ISL_940836, EPI_ISL_940837, EPI_ISL_940838 | Virginia DCLS | Virginia DCLS | Virginia DCLS |
| EPI_ISL_940880, EPI_ISL_940881 | Vaccines and Infectious Diseases Analytics Research Unit (VIDA) | KRISP, KZN Research Innovation and Sequencing Platform | Baillie Vicky, du Plessis Jeanine, Giandhari Jennifer, Pillay Sureshnee, Naidoo Yeshnee, Tegally Houriiyah, de Oliveira Tulio, Madhi Shabir |
| EPI_ISL_941282 | Nigeria Centre for Disease Control (NCDC) | African Centre of Excellence for Genomics of Infectious Diseases (ACEGID), Redeemer's University | Oluniyi P.E. et al |
| EPI_ISL_942262, EPI_ISL_942263, EPI_ISL_942266, EPI_ISL_942267, EPI_ISL_942268, EPI_ISL_942269, EPI_ISL_942270, EPI_ISL_942271, EPI_ISL_942272, EPI_ISL_942273, EPI_ISL_942274, EPI_ISL_942275, EPI_ISL_942276, EPI_ISL_942277, EPI_ISL_942278, EPI_ISL_942279, EPI_ISL_942280, EPI_ISL_942281, EPI_ISL_942282, EPI_ISL_942283, EPI_ISL_942284, EPI_ISL_942285, EPI_ISL_942286, EPI_ISL_942308, EPI_ISL_942309, EPI_ISL_942310, EPI_ISL_942311, EPI_ISL_942312, EPI_ISL_942314, EPI_ISL_942315, EPI_ISL_942317, EPI_ISL_942318, EPI_ISL_942319, EPI_ISL_942320, EPI_ISL_942321, EPI_ISL_942322, EPI_ISL_942323, EPI_ISL_942324, EPI_ISL_942325 | | | |
| see above | Wisconsin State Laboratory of Hygiene Communicable Disease Division | Wisconsin State Laboratory of Hygiene Communicable Disease Division | Kelsey R. Florek, Abigail C. Shockey |
| EPI_ISL_942707, EPI_ISL_942708, EPI_ISL_942709, EPI_ISL_942710, EPI_ISL_942711, EPI_ISL_942712, EPI_ISL_942713, EPI_ISL_942714, EPI_ISL_942715, EPI_ISL_942716, EPI_ISL_942717, EPI_ISL_942718, EPI_ISL_942719 | | | |
| see above | Gundersen Molecular Diagnostics Laboratory | Kabara Cancer Research Institute | Craig S. Richmond, Paraic A. Kenny |
| EPI_ISL_943015, EPI_ISL_943359 | Dutch COVID-19 response team | National Institute for Public Health and the Environment (RIVM) | Adam Meijer, Harry Vennema, Dirk Eggink, Jeroen Cremer, Sharon van den Brink, Bas van der Veer, AnneMarie van den Brandt, Florian Zwagemaker, Dennis Schmitz, Chantal Reusken, on behalf of the national COVID-19 response team |
| EPI_ISL_943989 | LACEN do Estado de Goias | Instituto Adolfo Lutz, Interdiciplinary Procedures Center, Strategic Laboratory | Claudio Tavares Sacchi, Claudia Regina Gonçalves, Erica Valessa Ramos Gomes, Karoline Rodrigues Campos |
| EPI_ISL_943995 | General Hospital - Veles | Research Center for Genetic Engineering and Biotechnology "Georgi D. Efremov" , Macedonian Academy of Sciences and Arts | Aleksandar J. Dimovski, Dijana Plasheska-Karanfilska, Predrag Noveski, Gjorgji Bozinovski, Milena Jakimovska |
| EPI_ISL_943996, EPI_ISL_943997 | General Hospital - Tetovo | Research Center for Genetic Engineering and Biotechnology "Georgi D. Efremov" , Macedonian Academy of Sciences and Arts | Aleksandar J. Dimovski, Dijana Plasheska-Karanfilska, Predrag Noveski, Gjorgji Bozinovski, Milena Jakimovska |
| EPI_ISL_943998 | General Hospital - Strumica | Research Center for Genetic Engineering and Biotechnology "Georgi D. Efremov" , Macedonian Academy of Sciences and Arts | Aleksandar J. Dimovski, Dijana Plasheska-Karanfilska, Predrag Noveski, Gjorgji Bozinovski, Milena Jakimovska |
| EPI_ISL_944102, EPI_ISL_944103, EPI_ISL_944104 | Institute for Medical Research, Infectious Disease Research Centre, National Institutes of Health, Ministry of Health Malaysia | Institute for Medical Research, Infectious Disease Research Centre, National Institutes of Health, Ministry of Health Malaysia | Suppiah J, Kamel K, Azizan MA, Thayan R |
| EPI_ISL_944105, EPI_ISL_944106 | General Hospital - Strumica | Research Center for Genetic Engineering and Biotechnology "Georgi D. Efremov" , Macedonian Academy of Sciences and Arts | Aleksandar J. Dimovski, Dijana Plasheska-Karanfilska, Predrag Noveski, Gjorgji Bozinovski, Milena Jakimovska |
| EPI_ISL_945054, EPI_ISL_945085 | Lighthouse Lab in Cambridge | Wellcome Sanger Institute for the COVID-19 Genomics UK (COG-UK) Consortium | Rob Howes, The Lighthouse Lab in Cambridge and Alex Alderton, Roberto Amato, Sonia Goncalves, Ewan Harrison, David K. Jackson, Ian Johnston, Dominic Kwiatkowski, Cordelia Langford, John Sillitoe on behalf of the Wellcome Sanger Institute COVID-19 Surveillance Team |
| EPI_ISL_945088 | Lighthouse Lab in Alderley Park | Wellcome Sanger Institute for the COVID-19 Genomics UK (COG-UK) Consortium | Jacquelyn Wynn, Mairead Hyland, The Lighthouse Lab in Alderley Park and Alex Alderton, Roberto Amato, Sonia Goncalves, Ewan Harrison, David K. Jackson, Ian Johnston, Dominic Kwiatkowski, Cordelia Langford, John Sillitoe on behalf of the Wellcome Sanger Institute COVID-19 Surveillance Team |
| EPI_ISL_945089 | Lighthouse Lab in Cambridge | Wellcome Sanger Institute for the COVID-19 Genomics UK (COG-UK) Consortium | Rob Howes, The Lighthouse Lab in Cambridge and Alex Alderton, Roberto Amato, Sonia Goncalves, Ewan Harrison, David K. Jackson, Ian Johnston, Dominic Kwiatkowski, Cordelia Langford, John Sillitoe on behalf of the Wellcome Sanger Institute COVID-19 Surveillance Team |
| EPI_ISL_945099 | Lighthouse Lab in Alderley Park | Wellcome Sanger Institute for the COVID-19 Genomics UK (COG-UK) Consortium | Jacquelyn Wynn, Mairead Hyland, The Lighthouse Lab in Alderley Park and Alex Alderton, Roberto Amato, Sonia Goncalves, Ewan Harrison, David K. Jackson, Ian Johnston, Dominic Kwiatkowski, Cordelia Langford, John Sillitoe on behalf of the Wellcome Sanger Institute COVID-19 Surveillance Team |
| EPI_ISL_945111 | Lighthouse Lab in Milton Keynes | Wellcome Sanger Institute for the COVID-19 Genomics UK (COG-UK) Consortium | The Lighthouse Lab in Milton Keynes and Alex Alderton, Roberto Amato, Sonia Goncalves, Ewan Harrison, David K. Jackson, Ian Johnston, Dominic Kwiatkowski, Cordelia Langford, John Sillitoe on behalf of the Wellcome Sanger Institute COVID-19 Surveillance Team |
| EPI_ISL_945113, EPI_ISL_945117 | Lighthouse Lab in Alderley Park | Wellcome Sanger Institute for the COVID-19 Genomics UK (COG-UK) Consortium | Jacquelyn Wynn, Mairead Hyland, The Lighthouse Lab in Alderley Park and Alex Alderton, Roberto Amato, Sonia Goncalves, Ewan Harrison, David K. Jackson, Ian Johnston, Dominic Kwiatkowski, Cordelia Langford, John Sillitoe on behalf of the Wellcome Sanger Institute COVID-19 Surveillance Team |
| EPI_ISL_945119 | Lighthouse Lab in Milton Keynes | Wellcome Sanger Institute for the COVID-19 Genomics UK (COG-UK) Consortium | The Lighthouse Lab in Milton Keynes and Alex Alderton, Roberto Amato, Sonia Goncalves, Ewan Harrison, David K. Jackson, Ian Johnston, Dominic Kwiatkowski, Cordelia Langford, John Sillitoe on behalf of the Wellcome Sanger Institute COVID-19 Surveillance Team |

| | | | |
|---|---|---|---|
| EPI_ISL_945160 | Lighthouse Lab in Alderley Park | Wellcome Sanger Institute for the COVID-19 Genomics UK (COG-UK) Consortium | Jacquelyn Wynn, Mairead Hyland, The Lighthouse Lab in Alderley Park and Alex Alderton, Roberto Amato, Sonia Goncalves, Ewan Harrison, David K. Jackson, Ian Johnston, Dominic Kwiatkowski, Cordelia Langford, John Sillitoe on behalf of the Wellcome Sanger Institute COVID-19 Surveillance Team |
| EPI_ISL_945170 | Lighthouse Lab in Cambridge | Wellcome Sanger Institute for the COVID-19 Genomics UK (COG-UK) Consortium | Rob Howes, The Lighthouse Lab in Cambridge and Alex Alderton, Roberto Amato, Sonia Goncalves, Ewan Harrison, David K. Jackson, Ian Johnston, Dominic Kwiatkowski, Cordelia Langford, John Sillitoe on behalf of the Wellcome Sanger Institute COVID-19 Surveillance Team |
| EPI_ISL_945171 | Lighthouse Lab in Milton Keynes | Wellcome Sanger Institute for the COVID-19 Genomics UK (COG-UK) Consortium | The Lighthouse Lab in Milton Keynes and Alex Alderton, Roberto Amato, Sonia Goncalves, Ewan Harrison, David K. Jackson, Ian Johnston, Dominic Kwiatkowski, Cordelia Langford, John Sillitoe on behalf of the Wellcome Sanger Institute COVID-19 Surveillance Team |
| EPI_ISL_945188 | Lighthouse Lab in Cambridge | Wellcome Sanger Institute for the COVID-19 Genomics UK (COG-UK) Consortium | Rob Howes, The Lighthouse Lab in Cambridge and Alex Alderton, Roberto Amato, Sonia Goncalves, Ewan Harrison, David K. Jackson, Ian Johnston, Dominic Kwiatkowski, Cordelia Langford, John Sillitoe on behalf of the Wellcome Sanger Institute COVID-19 Surveillance Team |
| EPI_ISL_945201 | Lighthouse Lab in Alderley Park | Wellcome Sanger Institute for the COVID-19 Genomics UK (COG-UK) Consortium | Jacquelyn Wynn, Mairead Hyland, The Lighthouse Lab in Alderley Park and Alex Alderton, Roberto Amato, Sonia Goncalves, Ewan Harrison, David K. Jackson, Ian Johnston, Dominic Kwiatkowski, Cordelia Langford, John Sillitoe on behalf of the Wellcome Sanger Institute COVID-19 Surveillance Team |
| EPI_ISL_945202 | Lighthouse Lab in Cambridge | Wellcome Sanger Institute for the COVID-19 Genomics UK (COG-UK) Consortium | Rob Howes, The Lighthouse Lab in Cambridge and Alex Alderton, Roberto Amato, Sonia Goncalves, Ewan Harrison, David K. Jackson, Ian Johnston, Dominic Kwiatkowski, Cordelia Langford, John Sillitoe on behalf of the Wellcome Sanger Institute COVID-19 Surveillance Team |
| EPI_ISL_945204 | Lighthouse Lab in Alderley Park | Wellcome Sanger Institute for the COVID-19 Genomics UK (COG-UK) Consortium | Jacquelyn Wynn, Mairead Hyland, The Lighthouse Lab in Alderley Park and Alex Alderton, Roberto Amato, Sonia Goncalves, Ewan Harrison, David K. Jackson, Ian Johnston, Dominic Kwiatkowski, Cordelia Langford, John Sillitoe on behalf of the Wellcome Sanger Institute COVID-19 Surveillance Team |
| EPI_ISL_945209 | Lighthouse Lab in Glasgow | Wellcome Sanger Institute for the COVID-19 Genomics UK (COG-UK) Consortium | Harper VanSteenhouse, Yumi Kasai, David Gray, Carol Clugston, Anna Dominiczak and Alex Alderton, Roberto Amato, Sonia Goncalves, Ewan Harrison, David K. Jackson, Ian Johnston, Dominic Kwiatkowski, Cordelia Langford, John Sillitoe on behalf of the Wellcome Sanger Institute COVID-19 Surveillance Team |
| EPI_ISL_945223 | Lighthouse Lab in Milton Keynes | Wellcome Sanger Institute for the COVID-19 Genomics UK (COG-UK) Consortium | The Lighthouse Lab in Milton Keynes and Alex Alderton, Roberto Amato, Sonia Goncalves, Ewan Harrison, David K. Jackson, Ian Johnston, Dominic Kwiatkowski, Cordelia Langford, John Sillitoe on behalf of the Wellcome Sanger Institute COVID-19 Surveillance Team |
| EPI_ISL_945247, EPI_ISL_945249 | Lighthouse Lab in Cambridge | Wellcome Sanger Institute for the COVID-19 Genomics UK (COG-UK) Consortium | Rob Howes, The Lighthouse Lab in Cambridge and Alex Alderton, Roberto Amato, Sonia Goncalves, Ewan Harrison, David K. Jackson, Ian Johnston, Dominic Kwiatkowski, Cordelia Langford, John Sillitoe on behalf of the Wellcome Sanger Institute COVID-19 Surveillance Team |
| EPI_ISL_945270 | Lighthouse Lab in Milton Keynes | Wellcome Sanger Institute for the COVID-19 Genomics UK (COG-UK) Consortium | The Lighthouse Lab in Milton Keynes and Alex Alderton, Roberto Amato, Sonia Goncalves, Ewan Harrison, David K. Jackson, Ian Johnston, Dominic Kwiatkowski, Cordelia Langford, John Sillitoe on behalf of the Wellcome Sanger Institute COVID-19 Surveillance Team |
| EPI_ISL_945282 | Lighthouse Lab in Alderley Park | Wellcome Sanger Institute for the COVID-19 Genomics UK (COG-UK) Consortium | Jacquelyn Wynn, Mairead Hyland, The Lighthouse Lab in Alderley Park and Alex Alderton, Roberto Amato, Sonia Goncalves, Ewan Harrison, David K. Jackson, Ian Johnston, Dominic Kwiatkowski, Cordelia Langford, John Sillitoe on behalf of the Wellcome Sanger Institute COVID-19 Surveillance Team |
| EPI_ISL_945299 | Lighthouse Lab in Milton Keynes | Wellcome Sanger Institute for the COVID-19 Genomics UK (COG-UK) Consortium | The Lighthouse Lab in Milton Keynes and Alex Alderton, Roberto Amato, Sonia Goncalves, Ewan Harrison, David K. Jackson, Ian Johnston, Dominic Kwiatkowski, Cordelia Langford, John Sillitoe on behalf of the Wellcome Sanger Institute COVID-19 Surveillance Team |
| EPI_ISL_945305 | Lighthouse Lab in Cambridge | Wellcome Sanger Institute for the COVID-19 Genomics UK (COG-UK) Consortium | Rob Howes, The Lighthouse Lab in Cambridge and Alex Alderton, Roberto Amato, Sonia Goncalves, Ewan Harrison, David K. Jackson, Ian Johnston, Dominic Kwiatkowski, Cordelia Langford, John Sillitoe on behalf of the Wellcome Sanger Institute COVID-19 Surveillance Team |
| EPI_ISL_945306, EPI_ISL_945310 | Lighthouse Lab in Alderley Park | Wellcome Sanger Institute for the COVID-19 Genomics UK (COG-UK) Consortium | Jacquelyn Wynn, Mairead Hyland, The Lighthouse Lab in Alderley Park and Alex Alderton, Roberto Amato, Sonia Goncalves, Ewan Harrison, David K. Jackson, Ian Johnston, Dominic Kwiatkowski, Cordelia Langford, John Sillitoe on behalf of the Wellcome Sanger Institute COVID-19 Surveillance Team |
| EPI_ISL_945320 | Lighthouse Lab in Cambridge | Wellcome Sanger Institute for the COVID-19 Genomics UK (COG-UK) Consortium | Rob Howes, The Lighthouse Lab in Cambridge and Alex Alderton, Roberto Amato, Sonia Goncalves, Ewan Harrison, David K. Jackson, Ian Johnston, Dominic Kwiatkowski, Cordelia Langford, John Sillitoe on behalf of the Wellcome Sanger Institute COVID-19 Surveillance Team |
| EPI_ISL_945323 | Lighthouse Lab in Alderley Park | Wellcome Sanger Institute for the COVID-19 Genomics UK (COG-UK) Consortium | Jacquelyn Wynn, Mairead Hyland, The Lighthouse Lab in Alderley Park and Alex Alderton, Roberto Amato, Sonia Goncalves, Ewan Harrison, David K. Jackson, Ian Johnston, Dominic Kwiatkowski, Cordelia Langford, John Sillitoe on behalf of the Wellcome Sanger Institute COVID-19 Surveillance Team |
| EPI_ISL_945340 | Lighthouse Lab in Milton Keynes | Wellcome Sanger Institute for the COVID-19 Genomics UK (COG-UK) Consortium | The Lighthouse Lab in Milton Keynes and Alex Alderton, Roberto Amato, Sonia Goncalves, Ewan Harrison, David K. Jackson, Ian Johnston, Dominic Kwiatkowski, Cordelia Langford, John Sillitoe on behalf of the Wellcome Sanger Institute COVID-19 Surveillance Team |
| EPI_ISL_945350 | Lighthouse Lab in Cambridge | Wellcome Sanger Institute for the COVID-19 Genomics UK (COG-UK) Consortium | Rob Howes, The Lighthouse Lab in Cambridge and Alex Alderton, Roberto Amato, Sonia Goncalves, Ewan Harrison, David K. Jackson, Ian Johnston, Dominic Kwiatkowski, Cordelia Langford, John Sillitoe on behalf of the Wellcome Sanger Institute COVID-19 Surveillance Team |
| EPI_ISL_945361 | Lighthouse Lab in Alderley Park | Wellcome Sanger Institute for the COVID-19 Genomics UK (COG-UK) Consortium | Jacquelyn Wynn, Mairead Hyland, The Lighthouse Lab in Alderley Park and Alex Alderton, Roberto Amato, Sonia Goncalves, Ewan Harrison, David K. Jackson, Ian Johnston, Dominic Kwiatkowski, Cordelia Langford, John Sillitoe on behalf of the Wellcome Sanger Institute COVID-19 Surveillance Team |
| EPI_ISL_945367 | Lighthouse Lab in Cambridge | Wellcome Sanger Institute for the COVID-19 Genomics UK (COG-UK) Consortium | Rob Howes, The Lighthouse Lab in Cambridge and Alex Alderton, Roberto Amato, Sonia Goncalves, Ewan Harrison, David K. Jackson, Ian Johnston, Dominic Kwiatkowski, Cordelia Langford, John Sillitoe on behalf of the Wellcome Sanger Institute COVID-19 Surveillance Team |
| EPI_ISL_947312 | RSU Bunda Mulia | Eijkman Institute for Molecular Biology, Ministry of Research and Technology/National Agency for Research and Innovation | Lydia V. Panggalo, Sukma Oktavianthi, Willy Agustine, Edison Johar, Hidayat Trimarsanto, Iskandar Adnan, Frilasita A Yudhaputri, Safarina G Malik, Khin Saw Myint, Amin Soebandrio |
| EPI_ISL_947315 | RSU Harapan Bunda | Eijkman Institute for Molecular Biology, Ministry of Research and Technology/National Agency for Research and Innovation | Lydia V. Panggalo, Sukma Oktavianthi, Willy Agustine, Edison Johar, Hidayat Trimarsanto, Iskandar Adnan, Frilasita A Yudhaputri, Safarina G Malik, Khin Saw Myint, Amin Soebandrio |
| EPI_ISL_949785 | University College London, Great Ormond Street Hospital for Children NHS Foundation Trust, Imperial College Healthcare NHS Trust | COVID-19 Genomics UK (COG-UK) Consortium | Sergi Castellano, Rachel Williams, Mark Kristiansen, Paola Resende Silva, Sunando Roy, Tony Brooks, Helena Tutill, Paola Niola, Patricia Dyal, Charlotte Williams, Leysa Forrest, Yasmin Panchbhaya, Jacqueline Findlay, Samuel Weeks, Julianne Brown, Kathryn Harris, Paul Randell, James Price, Alison Holmes, Judith Breuer |
| EPI_ISL_950592 | Quadram Institute Bioscience | COVID-19 Genomics UK (COG-UK) Consortium | Dave J. Baker, Gemma L. Kay, Alp Aydin, Thanh Le-Viet, Steven Rudder, Ana P. Tedim, Anastasia Kolyva, Maria Diaz, Leonardo de Oliveira Martins, Nabil-Fareed Alikhan, Lizzie Meadows, Rachael Stanley, Ngozi Elumogo, Muhammed Yasir, Nicholas M. Thomson, Alexander J Trotter, Rachel Gilroy, Samuel Bloomfield, Claire Stuart, Andrew Bell, Reenesh Prakash, Samir Dervisevic, Alison E. Mather, John Wain, Mark Webber, Andrew J. Page, Justin O'Grady |
| EPI_ISL_950705 | Lincolnshire Hospitals and DeepSeq Nottingham | COVID-19 Genomics UK (COG-UK) Consortium | Nichola Duckworth, Tim Sloan, Sarah Walsh, Jonathan Ball, Patrick McClure, Joeseph Chappell, Nadine Holmes, Matthew Carlisle, Christopher Moore, Fei Sang, Johnny Debebe, Victoria Wright, Matthew Loose |
| EPI_ISL_950900, EPI_ISL_950901, EPI_ISL_950905, EPI_ISL_950908, EPI_ISL_950909, EPI_ISL_950910, EPI_ISL_950912, EPI_ISL_950913, EPI_ISL_950917, EPI_ISL_950918 | Oxford Viromics, NDM, University of Oxford; Oxford University Hospitals; Basingstoke and North Hampshire Hospital | COVID-19 Genomics UK (COG-UK) Consortium | Tanya Golubchik, David Bonsall, George Macintyre, Amy Trebes, Mariateresa de Cesare, Catrin Moore, Alex Mobbs, Anita Justice, Robert Shaw, Monique Andersson, Timothy Peto, Emma Wise, Nathan Moore, Jessica Lynch, Nick Cortes, Matilde Mori, Stephen Kidd, David Buck, John Todd, Christophe Fraser |
| EPI_ISL_951751 | Originating lab: Wales Specialist Virology Centre Sequencing lab: Pathogen Genomics Unit | Public Health Wales Microbiology Cardiff Wales Specialist Virology Centre | Catherine Moore, Johnathan Evans, Laura Gifford, Malorie Perry, Simon Cottrell, Angela Marchbank, Alec Birchley, Alexander Adams, Amy Gaskin, Bree Gatica-Wilcox, Jason Coombes, Joel Southgate, Lauren Gilbert, Lee Graham, Nicole Pacchiarini, Sara Kumziene-Summerhayes, Sarah Taylor, Sophie Jones, Sara Rey, Matthew Bull, Joanne Watkins, Sally Corden, Tom Connor |
| EPI_ISL_954198 | 1.AO Universitaria 'S. Giovanni di Dio e Ruggi D'Aragona, Scuola Medica Salernitana' Hospital / 2.UOC di Virologia e Microbiologia, Università della Campania 'L. Vanvitelli' / 3.AO Universitaria 'Federico II' Napoli Hospital / 4.AORN 'San Giuseppe Moscati' Avellino Hospital / 5.AO 'San Pio - presidio G. Rummo' Benevento Hospital / 6.AO 'Sant'Anna e San Sebastiano' Caserta Hospital / 7.PO 'Maria Santissima Addolorata' Eboli Hospital / 8.Biogem Istituto di Ricerche Genetiche | 1. Genome Research Center for Health (CRGS) / 2. Laboratory of Molecular Medicine and Genomics(LMMGe) / 3. Center for Research in Pure and Applied Mathematics (CRMPA) | Giorgio Giurato, Francesca Rizzo, Alessandro Weisz, Gianluigi Franci, Giovanni Nassa, Pasquale Pagliano, Roberta Tarallo, Elena Alexandrova, Ylenia D'Agostino, Carlo Ferravante, Jessica Lamberti, Viola Melone, Domenico Memoli, Valeria Mirici Cappa, Domenico Palumbo, Giovanni Pecoraro, Assunta Sellitto, Oriana Strianese, Ilaria Terenzi, Giuseppe Fenza, Aniello Gentile, Antonello Saccomanno, Sonia Amabile, Teresa Rocco, Annamaria Salvati, Emilia Vaccaro, Massimiliano Galdiero, Michele Cennamo, Giuseppe Portella, Maria Grazia Foti, Mariarosaria Ingino, Maria Landi, Maurizio Fumi, Vincenzo Rocco, Rita Greco, Vittoria Letizia, Arnolfo Petruzziello, Maddalena Schioppa, Gregorio Goffredi, Francesca Marciano, Michele Caraglia, Alessia Cossu, Marianna Scrima, Edmondo Adorisio, Morena D'Avenia, Michela Iacobellis, Rosanna Piluscio, Giorgio Dirani, Vittorio Sambri, Simona Semprini, Silvia Zanoli, Francesco Curcio, Stefania Marzinotto, Andreina Baj, Fausto Sessa. |

| | | | |
|---|---|---|---|
| EPI_ISL_955171 | University of Sarajevo, Veterinary Faculty, Laboratory for Molecular Diagnostic and Research Laboratory | University of Sarajevo, Veterinary Faculty, Laboratory for Molecular Diagnostic and Research Laboratory | Goleti Š., Goleti T., Ali-Šeho A., Softi A., Nicevi M., Šabi E., Jaži A., Hodži A., Terzi I. |
| EPI_ISL_955316, EPI_ISL_955320 | GA Department of Public Health Laboratory | Pathogen Discovery, Respiratory Viruses Branch, Division of Viral Diseases, Centers for Disease Control and Prevention | Ying Tao, Jing Zhang, Yan Li, Krista Queen, Anna Uehara, Peter Cook, Clinton R. Paden, Haibin Wang, Suxiang Tong |
| EPI_ISL_955572, EPI_ISL_955573, EPI_ISL_955574, EPI_ISL_955575, EPI_ISL_955576, EPI_ISL_955577, EPI_ISL_955578, EPI_ISL_955579, EPI_ISL_955580, EPI_ISL_955581, EPI_ISL_955582, EPI_ISL_955583, EPI_ISL_955584, EPI_ISL_955585, EPI_ISL_955586, EPI_ISL_955587, EPI_ISL_955588, EPI_ISL_955589, EPI_ISL_955590, EPI_ISL_955591, EPI_ISL_955592, EPI_ISL_955593, EPI_ISL_955594, EPI_ISL_955595, EPI_ISL_955596, EPI_ISL_955597, EPI_ISL_955598, EPI_ISL_955599, EPI_ISL_955600, EPI_ISL_955601, EPI_ISL_955602, EPI_ISL_955603, EPI_ISL_955604, EPI_ISL_955605, EPI_ISL_955606, EPI_ISL_955607, EPI_ISL_955608, EPI_ISL_955609, EPI_ISL_955610, EPI_ISL_955611, EPI_ISL_955612, EPI_ISL_955613, EPI_ISL_955614, EPI_ISL_955615, EPI_ISL_955616, EPI_ISL_955617, EPI_ISL_955618, EPI_ISL_955619, EPI_ISL_955620, EPI_ISL_955621, EPI_ISL_955622, EPI_ISL_955623, EPI_ISL_955624, EPI_ISL_955625, EPI_ISL_955626, EPI_ISL_955627 | | | |
| see above | Santa Clara County Public Health Laboratory | Chan-Zuckerberg Biohub | CZB Cliahub Consortium |
| EPI_ISL_959849 | National Virus Reference Laboratory | National Virus Reference Laboratory | Michael Carr, Gabriel Gonzalez, Jonathan Dean, Cillian F De Gascun |
| EPI_ISL_960188, EPI_ISL_960190, EPI_ISL_960191, EPI_ISL_960201, EPI_ISL_960202, EPI_ISL_960203, EPI_ISL_960204, EPI_ISL_960205, EPI_ISL_960206, EPI_ISL_960207, EPI_ISL_960208, EPI_ISL_960209, EPI_ISL_960210, EPI_ISL_960211, EPI_ISL_960212, EPI_ISL_960213, EPI_ISL_960214, EPI_ISL_960215, EPI_ISL_960216, EPI_ISL_960217, EPI_ISL_960218, EPI_ISL_960219, EPI_ISL_960220, EPI_ISL_960221, EPI_ISL_960222 | | | |
| see above | Virginia Division of Consolidated Laboratory Services | Virginia Division of Consolidated Laboratory Services | Virginia DCLS |
| EPI_ISL_960443, EPI_ISL_960444, EPI_ISL_960457, EPI_ISL_960473, EPI_ISL_960531, EPI_ISL_960532, EPI_ISL_960533, EPI_ISL_960534, EPI_ISL_960535, EPI_ISL_960536, EPI_ISL_960537, EPI_ISL_960538, EPI_ISL_960539, EPI_ISL_960540, EPI_ISL_960541, EPI_ISL_960542, EPI_ISL_960544, EPI_ISL_960545, EPI_ISL_960546, EPI_ISL_960558, EPI_ISL_960559, EPI_ISL_960576, EPI_ISL_960577, EPI_ISL_960619, EPI_ISL_960620, EPI_ISL_960621, EPI_ISL_960622, EPI_ISL_960623, EPI_ISL_960624, EPI_ISL_960625, EPI_ISL_960626, EPI_ISL_960627, EPI_ISL_960628, EPI_ISL_960629, EPI_ISL_960630, EPI_ISL_960631, EPI_ISL_960632, EPI_ISL_960633, EPI_ISL_960634, EPI_ISL_960635, EPI_ISL_960636, EPI_ISL_960637, EPI_ISL_960638, EPI_ISL_960639, EPI_ISL_960648 | | | |
| see above | Istituto Zooprofilattico Sperimentale del Mezzogiorno | TIGEM | Patrizia Annunziata, Andrea Ballabio, Valentina Bouche, Davide Cacchiarelli, Pellegrino Cerino, Chiara Colantuono, Maria Concetta Cuomo, Denise Di Concilio, Lucio Di Filippo, Antonio Grimaldi, Antonio Limone, Anna Manfredi, Francesco Panariello, Biancamaria Pierri, Marcello Salvi |
| EPI_ISL_962457 | Seattle Flu Study | Seattle Flu Study | Deborah A. Nickerson, Chris D. Frazar, Jover Lee, Benjamin Pelle, Erica Ryke, Matthew Richardson, Amanda Adler, Elisabeth Brandstetter, Peter D. Han, Kairsten Fay, Misja Ilcisin, Kirsten Lacombe, Thomas R. Sibley, Melissa Truong, Caitlin R. Wolf, Michael Boeckh, Janet A. Englund, Michael Famulare, Barry R. Lutz, Mark J. Rieder, Lea M. Starita, Matthew Thompson, Jay Shendure, Trevor Bedford, Helen Y. Chu |
| EPI_ISL_964990 | Department of Medical Microbiology, St. Olavs hospital | Norwegian Institute of Public Health, Department of Virology | Kathrine Stene-Johansen, Kamilla Heddeland Instefjord, Hilde Elshaug, Ignacio Garcia Llorente, Serina B Engebretsen, Atiya R Ali,Marie Paulsen Madsen, Rasmus Riis Kopperud, Hilde Vollan, Karoline Bragstad, Olav Hungnes |
| EPI_ISL_965059, EPI_ISL_965060, EPI_ISL_965061, EPI_ISL_965062, EPI_ISL_965063, EPI_ISL_965064, EPI_ISL_965065, EPI_ISL_965066, EPI_ISL_965067, EPI_ISL_965068, EPI_ISL_965069, EPI_ISL_965070 | | | |
| see above | Wyoming Public Health Laboratory | Wyoming Public Health Laboratory | Noah Hull, Taylor Fearing, Lynette Gumbleton, Channing Weber, Ashley Norberg, Bailey Bowcutt, and Wanda Manley |
| EPI_ISL_965387 | University of Liège COVID-19 testing center | GIGA Medical Genomics | Keith Durkin, Maria Artesi, Bouchra Boujemla, Emmanuel André, Marc Van Ranst, Fabrice Bureau, Laurent Gillet, Wouter Coppieters, Vincent Bours |
| EPI_ISL_965558, EPI_ISL_965638, EPI_ISL_965675, EPI_ISL_965680 | Dutch COVID-19 response team | Medical Microbiology, Maastricht University Medical Centre | Jozef Dingemans*, Brian van der Veer*, Erik Beuken, Carmen Reumkens, Lieke van Alphen, Christian Hoebe, Paul Savelkoul |
| EPI_ISL_966761, EPI_ISL_966762, EPI_ISL_966763, EPI_ISL_966764 | Maine HETL | Tewhey Lab, The Jackson Laboratory | Matluk,N., Dewey,H., Iosue,F., Barter,M., Lynch,R., Munger,H. and Tewhey,R. |
| EPI_ISL_967527, EPI_ISL_967528, EPI_ISL_967529, EPI_ISL_967530, EPI_ISL_967532, EPI_ISL_967533, EPI_ISL_967534, EPI_ISL_967535, EPI_ISL_967536, EPI_ISL_967539, EPI_ISL_967540, EPI_ISL_967541, EPI_ISL_967542, EPI_ISL_967543, EPI_ISL_967544, EPI_ISL_967545, EPI_ISL_967546, EPI_ISL_967548, EPI_ISL_967549, EPI_ISL_967550, EPI_ISL_967557, EPI_ISL_967558, EPI_ISL_967559, EPI_ISL_967560, EPI_ISL_967561, EPI_ISL_967562, EPI_ISL_967571, EPI_ISL_967573, EPI_ISL_967574, EPI_ISL_967575, EPI_ISL_967577, EPI_ISL_967578, EPI_ISL_967579, EPI_ISL_967580, EPI_ISL_967581, EPI_ISL_967582, EPI_ISL_967584, EPI_ISL_967585, EPI_ISL_967600, EPI_ISL_967601, EPI_ISL_967602, EPI_ISL_967603, EPI_ISL_967604, EPI_ISL_967605, EPI_ISL_967606, EPI_ISL_967607, EPI_ISL_967608, EPI_ISL_967610, EPI_ISL_967611 | | | |
| see above | State Laboratories Division, Hawaii State Department of Health | State Laboratories Division, Hawaii State Department of Health | Pamela O'Brien, Drew Kuwazaki, Ayana Garnet, Razvan Sultana, Edward Desmond |
| EPI_ISL_967884 | Arizona State Public Health Laboratory | Arizona State Public Health Laboratory | Trung Huynh, Jessica Escobar, Katherine Fullerton, Nobuko Fukushima, Stacy White, Linda Getsinger, Victor Waddell |
| EPI_ISL_968834 | KEMRI-Wellcome Trust Research Programme/KEMRI-CGMR-C Kilifi | KEMRI-Wellcome Trust Research Programme/KEMRI-CGMR-C Kilifi | Githinji et al |
| EPI_ISL_976521, EPI_ISL_976522, EPI_ISL_976523, EPI_ISL_976524, EPI_ISL_976525, EPI_ISL_976526, EPI_ISL_976527, EPI_ISL_976528, EPI_ISL_976529, EPI_ISL_976530, EPI_ISL_976531, EPI_ISL_976532, EPI_ISL_976533, EPI_ISL_976534, EPI_ISL_976535, EPI_ISL_976536, EPI_ISL_976537, EPI_ISL_976538, EPI_ISL_976539, EPI_ISL_976540, EPI_ISL_976541, EPI_ISL_976542, EPI_ISL_976543, EPI_ISL_976544, EPI_ISL_976545, EPI_ISL_976546, EPI_ISL_976547, EPI_ISL_976548, EPI_ISL_976549, EPI_ISL_976550, EPI_ISL_976551, EPI_ISL_976552, EPI_ISL_976553, EPI_ISL_976554, EPI_ISL_976555, EPI_ISL_976556, EPI_ISL_976557, EPI_ISL_976558, EPI_ISL_976559, EPI_ISL_976560, EPI_ISL_976561, EPI_ISL_976562, EPI_ISL_976563, EPI_ISL_976564, EPI_ISL_976565, EPI_ISL_976566, EPI_ISL_976567, EPI_ISL_976568, EPI_ISL_976569, EPI_ISL_976570, EPI_ISL_976571, EPI_ISL_976572, EPI_ISL_976573, EPI_ISL_976574, EPI_ISL_976575, EPI_ISL_976576, EPI_ISL_976577, EPI_ISL_976578, EPI_ISL_976579, EPI_ISL_976580, EPI_ISL_976581, EPI_ISL_976582, EPI_ISL_976583, EPI_ISL_976584, EPI_ISL_976585, EPI_ISL_976586, EPI_ISL_976587, EPI_ISL_976588, EPI_ISL_976589, EPI_ISL_976590, EPI_ISL_976591, EPI_ISL_976592 | | | |
| see above | BCCDC Public Health Laboratory | BCCDC Public Health Laboratory | Prystajecky Natalie, Linda Hoang, Dan Fornika, John Tyson, Shannon Russell, Kim Macdonald, Kimia Kamelian, Ana Pacagnella, Corrinne Ng, Loretta Janz, Robert Azana Terry Snutch, Mel Krajden |
| EPI_ISL_977161 | ULSS 03 Venezia | Istituto Zooprofilattico Sperimentale delle Venezie | Adelaide Milani, Alessia Schivo, Annalisa Salviato, Erika Giorgia Quaranta, Ambra Pastori, Bianca Zecchin, Alice Fusaro, Isabella Monne, Calogero Terregino, Antonia Ricci |
| EPI_ISL_977163 | ULSS 2 Treviso | Istituto Zooprofilattico Sperimentale delle Venezie | Adelaide Milani, Alessia Schivo, Annalisa Salviato, Erika Giorgia Quaranta, Ambra Pastori, Bianca Zecchin, Alice Fusaro, Isabella Monne, Calogero Terregino, Antonia Ricci |
| EPI_ISL_977165, EPI_ISL_977166, EPI_ISL_977180, EPI_ISL_977181, EPI_ISL_977182, EPI_ISL_977184, EPI_ISL_977185 | ULSS 6 Euganea | Istituto Zooprofilattico Sperimentale delle Venezie | Adelaide Milani, Alessia Schivo, Annalisa Salviato, Erika Giorgia Quaranta, Ambra Pastori, Bianca Zecchin, Alice Fusaro, Isabella Monne, Calogero Terregino, Antonia Ricci |
| EPI_ISL_977195, EPI_ISL_977196, EPI_ISL_977197, EPI_ISL_977198, EPI_ISL_977199 | ULSS 2 Treviso | Istituto Zooprofilattico Sperimentale delle Venezie | Adelaide Milani, Alessia Schivo, Annalisa Salviato, Erika Giorgia Quaranta, Ambra Pastori, Bianca Zecchin, Alice Fusaro, Isabella Monne, Calogero Terregino, Antonia Ricci |
| EPI_ISL_977205, EPI_ISL_977206 | ULSS 7 Pedemontana - Distretto 1 | Istituto Zooprofilattico Sperimentale delle Venezie | Adelaide Milani, Alessia Schivo, Annalisa Salviato, Erika Giorgia Quaranta, Ambra Pastori, Bianca Zecchin, Alice Fusaro, Isabella Monne, Calogero Terregino, Antonia Ricci |
| EPI_ISL_977242, EPI_ISL_977243, EPI_ISL_977244, EPI_ISL_977245 | ULSS 03 Venezia | Istituto Zooprofilattico Sperimentale delle Venezie | Adelaide Milani, Alessia Schivo, Annalisa Salviato, Erika Giorgia Quaranta, Ambra Pastori, Bianca Zecchin, Alice Fusaro, Isabella Monne, Calogero Terregino, Antonia Ricci |
| EPI_ISL_977259, EPI_ISL_977286, EPI_ISL_977337, EPI_ISL_977338, EPI_ISL_977339, EPI_ISL_977340 | University of Zambia, School of Veterinary Medicine | UNZAVET and PATH | Mulenga Mwenda-Chimfwembe, Ngonda Saasa, Daniel Bridges |
| EPI_ISL_977696, EPI_ISL_977702, EPI_ISL_977732, EPI_ISL_977737, EPI_ISL_977738, EPI_ISL_977743, EPI_ISL_977744, EPI_ISL_977745, EPI_ISL_977746, EPI_ISL_977747, EPI_ISL_977748, EPI_ISL_977751, EPI_ISL_977752, EPI_ISL_977753, EPI_ISL_977754, EPI_ISL_977755, EPI_ISL_977756, EPI_ISL_977757, EPI_ISL_977758, EPI_ISL_977759, EPI_ISL_977760, EPI_ISL_977765, EPI_ISL_977766, EPI_ISL_977767, EPI_ISL_977768, EPI_ISL_977769, EPI_ISL_977770, EPI_ISL_977771, EPI_ISL_977772, EPI_ISL_977773, EPI_ISL_977774, EPI_ISL_977775, EPI_ISL_977776, EPI_ISL_977779, EPI_ISL_977780, EPI_ISL_977781, EPI_ISL_977782, EPI_ISL_977783, EPI_ISL_977785, EPI_ISL_977786, EPI_ISL_977787, EPI_ISL_977788, EPI_ISL_977789, EPI_ISL_977790, EPI_ISL_977792, EPI_ISL_977794, EPI_ISL_977795, EPI_ISL_977796, EPI_ISL_977797, EPI_ISL_977798, EPI_ISL_977800, EPI_ISL_977802, EPI_ISL_977804, EPI_ISL_977806, EPI_ISL_977807, EPI_ISL_977808, EPI_ISL_977809, EPI_ISL_977810, EPI_ISL_977811, EPI_ISL_977812, EPI_ISL_977813, EPI_ISL_977814, EPI_ISL_977815, EPI_ISL_977816, EPI_ISL_977817, EPI_ISL_977818, EPI_ISL_977820, EPI_ISL_977821, EPI_ISL_977822, EPI_ISL_977823, EPI_ISL_977825, EPI_ISL_977826, EPI_ISL_977827, EPI_ISL_977828, EPI_ISL_977829, EPI_ISL_977830, EPI_ISL_977831, EPI_ISL_977832, EPI_ISL_977833, EPI_ISL_977834, EPI_ISL_977835, EPI_ISL_977836, EPI_ISL_977837, EPI_ISL_977838, EPI_ISL_977839, EPI_ISL_977840, EPI_ISL_977841, EPI_ISL_977842, EPI_ISL_977843, EPI_ISL_977844, EPI_ISL_977845, EPI_ISL_977846, EPI_ISL_977847, EPI_ISL_977848, EPI_ISL_977849, EPI_ISL_977850, EPI_ISL_977851, EPI_ISL_977852, EPI_ISL_977853, EPI_ISL_977854, EPI_ISL_977855, EPI_ISL_977856, EPI_ISL_977857, EPI_ISL_977858, EPI_ISL_977859, EPI_ISL_977860, EPI_ISL_977861, EPI_ISL_977862, EPI_ISL_977863, EPI_ISL_977864, EPI_ISL_977865, EPI_ISL_977866, EPI_ISL_977867, EPI_ISL_977868, EPI_ISL_977869, EPI_ISL_977870, EPI_ISL_977872, EPI_ISL_977873, EPI_ISL_977874, EPI_ISL_977876, EPI_ISL_977877, EPI_ISL_977878, EPI_ISL_977879, EPI_ISL_977880, EPI_ISL_977882, EPI_ISL_977883, EPI_ISL_977884 | | | |
| see above | California Department of Public Health | Chiu Laboratory, University of California, San Francisco | Charles Chiu, Xianding (Wayne) Deng, Candace Wang, Venice Servellita, Jill Hacker, Debra Wadford |
| EPI_ISL_977985, EPI_ISL_977986, | Chiu Laboratory, University of California, San Francisco | Chiu Laboratory, University of California, San Francisco | Charles Chiu, Xianding (Wayne) Deng, Candace Wang, Venice Servellita, Jill Hacker, Debra Wadford |

| | | | |
|---|---|---|---|
| EPI_ISL_977987, EPI_ISL_977989, EPI_ISL_977990, EPI_ISL_977992, EPI_ISL_977995, EPI_ISL_977997, EPI_ISL_977998 | | | |
| EPI_ISL_978201, EPI_ISL_978202, EPI_ISL_978203, EPI_ISL_978204, EPI_ISL_978205, EPI_ISL_978206, EPI_ISL_978207, EPI_ISL_978208, EPI_ISL_978209, EPI_ISL_978210, EPI_ISL_978211, EPI_ISL_978212 | | | |
| see above | Virginia Division of Consolidated Laboratory Services | Virginia Division of Consolidated Laboratory Services | Virginia DCLS |
| EPI_ISL_978977, EPI_ISL_978978, EPI_ISL_978979, EPI_ISL_978980, EPI_ISL_978981, EPI_ISL_978982, EPI_ISL_978983 | California Department of Public Health | Chiu Laboratory, University of California, San Francisco | Charles Chiu, Xianding (Wayne) Deng, Candace Wang, Venice Servellita, Jill Hacker, Debra Wadford |
| EPI_ISL_979183, EPI_ISL_979184, EPI_ISL_979187, EPI_ISL_979188, EPI_ISL_979189, EPI_ISL_979190, EPI_ISL_979191, EPI_ISL_979195, EPI_ISL_979196, EPI_ISL_979197, EPI_ISL_979198, EPI_ISL_979199, EPI_ISL_979200, EPI_ISL_979201, EPI_ISL_979202, EPI_ISL_979203, EPI_ISL_979204, EPI_ISL_979205, EPI_ISL_979206, EPI_ISL_979207, EPI_ISL_979208, EPI_ISL_979209, EPI_ISL_979210, EPI_ISL_979211, EPI_ISL_979212, EPI_ISL_979213, EPI_ISL_979214, EPI_ISL_979215 | | | |
| see above | Humboldt County Public Health Laboratory | Chan-Zuckerberg Biohub | CZB Cliahub Consortium |
| EPI_ISL_979294, EPI_ISL_979300, EPI_ISL_979304, EPI_ISL_979308, EPI_ISL_979309 | Cadham Provincial laboratory | National Microbiology Laboratory (NML) | Anna Majer, Shari Tyson, Grace Seo, Philip Mabon, Elsie Grudeski, Rhiannon Huzarewich, Russell Mandes, Anneliese Landgraff, Jennifer Tanner, Natalie Knox, Morag Graham, Gary Van Domselaar, Paul Van Caeseele, Jared Bullard, David Alexander, Kerry Dust, Nathalie Bastien, Yan Li, Timothy Booth, Darian Hole, Madison Chapel, Kirsten Biggar, CanCOGeN's metadata curation team, Public Health Agency of Canada CanCOGeN team |
| EPI_ISL_981920, EPI_ISL_981921, EPI_ISL_981922, EPI_ISL_981923, EPI_ISL_981924, EPI_ISL_981925, EPI_ISL_981926, EPI_ISL_981927, EPI_ISL_981928, EPI_ISL_981929, EPI_ISL_981930, EPI_ISL_981931, EPI_ISL_981932, EPI_ISL_981933, EPI_ISL_981934, EPI_ISL_981935, EPI_ISL_981936, EPI_ISL_981937, EPI_ISL_981938, EPI_ISL_981939, EPI_ISL_981940 | | | |
| see above | Microbiology Service, Hospital Universitario Clinico San Cecilio, Granada | Microbiology Service, Hospital Universitario Clinico San Cecilio, Granada | Adolfo de Salazar, Natalia Chueca, Laura Viñuela, Ana Fuentes, Federico García |
| EPI_ISL_982302, EPI_ISL_982304, EPI_ISL_982311, EPI_ISL_982324, EPI_ISL_982325, EPI_ISL_982326, EPI_ISL_982327, EPI_ISL_982328, EPI_ISL_982329, EPI_ISL_982330, EPI_ISL_982331, EPI_ISL_982332, EPI_ISL_982333, EPI_ISL_982334, EPI_ISL_982335, EPI_ISL_982336, EPI_ISL_982337, EPI_ISL_982338, EPI_ISL_982339, EPI_ISL_982340, EPI_ISL_982341, EPI_ISL_982342, EPI_ISL_982343 | | | |
| see above | Hôpital Henri Mondor | Department of Virology, Henri Mondor University Hospital, Assistance Publique Hôpitaux de Paris, Université Paris-Est Créteil, INSERM U955 | Christophe Rodriguez, Slim Fourati, Vanessa Demontant, Guillaume Gricourt, Melissa N'Debi, Alexandre Soulier, Elisabeth Trawinski, Jean-Michel Pawlotsky |
| EPI_ISL_982851 | UK Healthcare Clinical Microbiology | Kentucky State Public Health Lab | Stephanie Lunn, Karim George, Joshua Tobias, William Grooms, Vaneet Arora, Matthew Johnson, Rachel Zinner, Rhonda Lucas |
