## Supplementary material for "Detection and characterization of the SARS-CoV-2 lineage B.1.526 in New York": Supp. Table 3 GISAID Acknowledgment part 3

We gratefully acknowledge the following Authors from the Originating laboratories responsible for obtaining the specimens, as well as the Submitting laboratories where the genome data were generated and shared via GISAID, on which this research is based.

All Submitters of data may be contacted directly via www.gisaid.org

Authors are sorted alphabetically.

| Accession ID | Originating Laboratory | Submitting Laboratory | Authors |
|---|---|---|---|
| EPI_ISL_1001460 | Clinical Microbiology Laboratory, NewYork Presbyterian Hospital/Columbia University Irving Medical Center | Uhlemann Laboratory, Columbia University Irving Medical Center | Medini K. Annavajhala, Anne-Catrin Uhlemann, Hiroshi Mori, David Ho |
| EPI_ISL_1009188, EPI_ISL_1009189, EPI_ISL_1009190, EPI_ISL_1009192, EPI_ISL_1009193 | American Esoteric Laboratories | Colleen B. Jonsson | Mariah K. Taylor, Walter Reichard, Jyothi Parvathareddy, Colleen B. Jonsson |
| EPI_ISL_1009289, EPI_ISL_1009290, EPI_ISL_1009291, EPI_ISL_1009292, EPI_ISL_1009293, EPI_ISL_1009294, EPI_ISL_1009295, EPI_ISL_1009296, EPI_ISL_1009297, EPI_ISL_1009298, EPI_ISL_1009299, EPI_ISL_1009300, EPI_ISL_1009301, EPI_ISL_1009302, EPI_ISL_1009303, EPI_ISL_1009304, EPI_ISL_1009305, EPI_ISL_1009306, EPI_ISL_1009307, EPI_ISL_1009308, EPI_ISL_1009309, EPI_ISL_1009310, EPI_ISL_1009311, EPI_ISL_1009312, EPI_ISL_1009313, EPI_ISL_1009314, EPI_ISL_1009315, EPI_ISL_1009316, EPI_ISL_1009317, EPI_ISL_1009318, EPI_ISL_1009319, EPI_ISL_1009320, EPI_ISL_1009321, EPI_ISL_1009322, EPI_ISL_1009323, EPI_ISL_1009324, EPI_ISL_1009325, EPI_ISL_1009326, EPI_ISL_1009327, EPI_ISL_1009328, EPI_ISL_1009329, EPI_ISL_1009330, EPI_ISL_1009331, EPI_ISL_1009332, EPI_ISL_1009333, EPI_ISL_1009334, EPI_ISL_1009335, EPI_ISL_1009336, EPI_ISL_1009337, EPI_ISL_1009338, EPI_ISL_1009339, EPI_ISL_1009340, EPI_ISL_1009341, EPI_ISL_1009342, EPI_ISL_1009343, EPI_ISL_1009344, EPI_ISL_1009345, EPI_ISL_1009346, EPI_ISL_1009347, EPI_ISL_1009348, EPI_ISL_1009349, EPI_ISL_1009350, EPI_ISL_1009351, EPI_ISL_1009352, EPI_ISL_1009353, EPI_ISL_1009354, EPI_ISL_1009355, EPI_ISL_1009356, EPI_ISL_1009357, EPI_ISL_1009358, EPI_ISL_1009359, EPI_ISL_1009360, EPI_ISL_1009361, EPI_ISL_1009362, EPI_ISL_1009363, EPI_ISL_1009364, EPI_ISL_1009365, EPI_ISL_1009366, EPI_ISL_1009367, EPI_ISL_1009368, EPI_ISL_1009369, EPI_ISL_1009370, EPI_ISL_1009371, EPI_ISL_1009372, EPI_ISL_1009373, EPI_ISL_1009374, EPI_ISL_1009375, EPI_ISL_1009376, EPI_ISL_1009377, EPI_ISL_1009378, EPI_ISL_1009379, EPI_ISL_1009380, EPI_ISL_1009381, EPI_ISL_1009382, EPI_ISL_1009383, EPI_ISL_1009384, EPI_ISL_1009385, EPI_ISL_1009386, EPI_ISL_1009387, EPI_ISL_1009388, EPI_ISL_1009389, EPI_ISL_1009390, EPI_ISL_1009391, EPI_ISL_1009392, EPI_ISL_1009393, EPI_ISL_1009394, EPI_ISL_1009395, EPI_ISL_1009396, EPI_ISL_1009397, EPI_ISL_1009398, EPI_ISL_1009399, EPI_ISL_1009400, EPI_ISL_1009401, EPI_ISL_1009402, EPI_ISL_1009403, EPI_ISL_1009404, EPI_ISL_1009405, EPI_ISL_1009406, EPI_ISL_1009407, EPI_ISL_1009408, EPI_ISL_1009409, EPI_ISL_1009410, EPI_ISL_1009411, EPI_ISL_1009412, EPI_ISL_1009413, EPI_ISL_1009414, EPI_ISL_1009415, EPI_ISL_1009416, EPI_ISL_1009417, EPI_ISL_1009418, EPI_ISL_1009419, EPI_ISL_1009420, EPI_ISL_1009421, EPI_ISL_1009422, EPI_ISL_1009423, EPI_ISL_1009424, EPI_ISL_1009425, EPI_ISL_1009426, EPI_ISL_1009427, EPI_ISL_1009428, EPI_ISL_1009429, EPI_ISL_1009430, EPI_ISL_1009431, EPI_ISL_1009432, EPI_ISL_1009433, EPI_ISL_1009434, EPI_ISL_1009435, EPI_ISL_1009436, EPI_ISL_1009437, EPI_ISL_1009438, EPI_ISL_1009439, EPI_ISL_1009440, EPI_ISL_1009441, EPI_ISL_1009442, EPI_ISL_1009443, EPI_ISL_1009444, EPI_ISL_1009445, EPI_ISL_1009446, EPI_ISL_1009447, EPI_ISL_1009448, EPI_ISL_1009449, EPI_ISL_1009450, EPI_ISL_1009451, EPI_ISL_1009452, EPI_ISL_1009453, EPI_ISL_1009454, EPI_ISL_1009455, EPI_ISL_1009456, EPI_ISL_1009457, EPI_ISL_1009458, EPI_ISL_1009459, EPI_ISL_1009460, EPI_ISL_1009461, EPI_ISL_1009462, EPI_ISL_1009463, EPI_ISL_1009464, EPI_ISL_1009465, EPI_ISL_1009466, EPI_ISL_1009467, EPI_ISL_1009468, EPI_ISL_1009469, EPI_ISL_1009470, EPI_ISL_1009471, EPI_ISL_1009472, EPI_ISL_1009473, EPI_ISL_1009474, EPI_ISL_1009475, EPI_ISL_1009476, EPI_ISL_1009477, EPI_ISL_1009478, EPI_ISL_1009479, EPI_ISL_1009480, EPI_ISL_1009481, EPI_ISL_1009482, EPI_ISL_1009483, EPI_ISL_1009484, EPI_ISL_1009485, EPI_ISL_1009486, EPI_ISL_1009487, EPI_ISL_1009488, EPI_ISL_1009489, EPI_ISL_1009490, EPI_ISL_1009491, EPI_ISL_1009492, EPI_ISL_1009493, EPI_ISL_1009494, EPI_ISL_1009495, EPI_ISL_1009496, EPI_ISL_1009497, EPI_ISL_1009498, EPI_ISL_1009499, EPI_ISL_1009500, EPI_ISL_1009501, EPI_ISL_1009502, EPI_ISL_1009503, EPI_ISL_1009504, EPI_ISL_1009505, EPI_ISL_1009506, EPI_ISL_1009507, EPI_ISL_1009508, EPI_ISL_1009509, EPI_ISL_1009510, EPI_ISL_1009511, EPI_ISL_1009512, EPI_ISL_1009513, EPI_ISL_1009514, EPI_ISL_1009515, EPI_ISL_1009516, EPI_ISL_1009517, EPI_ISL_1009518, EPI_ISL_1009519, EPI_ISL_1009520, EPI_ISL_1009521, EPI_ISL_1009522, EPI_ISL_1009523, EPI_ISL_1009524, EPI_ISL_1009525, EPI_ISL_1009526, EPI_ISL_1009527, EPI_ISL_1009528, EPI_ISL_1009529, EPI_ISL_1009530, EPI_ISL_1009531, EPI_ISL_1009532, EPI_ISL_1009533, EPI_ISL_1009534, EPI_ISL_1009535, EPI_ISL_1009536, EPI_ISL_1009537, EPI_ISL_1009538, EPI_ISL_1009539, EPI_ISL_1009540, EPI_ISL_1009541, EPI_ISL_1009542, EPI_ISL_1009543, EPI_ISL_1009544, EPI_ISL_1009545, EPI_ISL_1009546, EPI_ISL_1009547, EPI_ISL_1009548, EPI_ISL_1009549, EPI_ISL_1009550, EPI_ISL_1009551, EPI_ISL_1009552, EPI_ISL_1009553, EPI_ISL_1009554, EPI_ISL_1009555, EPI_ISL_1009556, EPI_ISL_1009557, EPI_ISL_1009558, EPI_ISL_1009559, EPI_ISL_1009560, EPI_ISL_1009561, EPI_ISL_1009562, EPI_ISL_1009563, EPI_ISL_1009564, EPI_ISL_1009565, EPI_ISL_1009566, EPI_ISL_1009567, EPI_ISL_1009568, EPI_ISL_1009569, EPI_ISL_1009570, EPI_ISL_1009571, EPI_ISL_1009572, EPI_ISL_1009573, EPI_ISL_1009574, EPI_ISL_1009575, EPI_ISL_1009576, EPI_ISL_1009577, EPI_ISL_1009578, EPI_ISL_1009579, EPI_ISL_1009580, EPI_ISL_1009581, EPI_ISL_1009582, EPI_ISL_1009583, EPI_ISL_1009584, EPI_ISL_1009585, EPI_ISL_1009586, EPI_ISL_1009587, EPI_ISL_1009588, EPI_ISL_1009589, EPI_ISL_1009590, EPI_ISL_1009591, EPI_ISL_1009592, EPI_ISL_1009593, EPI_ISL_1009594, EPI_ISL_1009595, EPI_ISL_1009596, EPI_ISL_1009597, EPI_ISL_1009598, EPI_ISL_1009599, EPI_ISL_1009600, EPI_ISL_1009601, EPI_ISL_1009602, EPI_ISL_1009603, EPI_ISL_1009604, EPI_ISL_1009605, EPI_ISL_1009606, EPI_ISL_1009607, EPI_ISL_1009608, EPI_ISL_1009609, EPI_ISL_1009610, EPI_ISL_1009611, EPI_ISL_1009612, EPI_ISL_1009613, EPI_ISL_1009614, EPI_ISL_1009615, EPI_ISL_1009616 | | | |
| see above | Erie County Public Health (ECPHL) | University at Buffalo Genomics and Bioinformatics Core | Jonathan Bard, Natalie Lamb, Alyssa Pohlman, Brandon Marzullo, Amanda Boccolucci, Norma Nowak, Donald Yergeau, Jennifer Surtees |
| EPI_ISL_1015653, EPI_ISL_1015654, EPI_ISL_1015655, EPI_ISL_1015656, EPI_ISL_1015657, EPI_ISL_1015658, EPI_ISL_1015659, EPI_ISL_1015660, EPI_ISL_1015661, EPI_ISL_1015662, EPI_ISL_1015663, EPI_ISL_1015664, EPI_ISL_1015665, EPI_ISL_1015666, EPI_ISL_1015667, EPI_ISL_1015668, EPI_ISL_1015669, EPI_ISL_1015670, EPI_ISL_1015671, EPI_ISL_1015672, EPI_ISL_1015673, EPI_ISL_1015674, EPI_ISL_1015675, EPI_ISL_1015676, EPI_ISL_1015677, EPI_ISL_1015678, EPI_ISL_1015679, EPI_ISL_1015680, EPI_ISL_1015681, EPI_ISL_1015682, EPI_ISL_1015683, EPI_ISL_1015684, EPI_ISL_1015685, EPI_ISL_1015686, EPI_ISL_1015687, EPI_ISL_1015688, EPI_ISL_1015689, EPI_ISL_1015690, EPI_ISL_1015691, EPI_ISL_1015692, EPI_ISL_1015693, EPI_ISL_1015694, EPI_ISL_1015695, EPI_ISL_1015696, EPI_ISL_1015697, EPI_ISL_1015698, EPI_ISL_1015699, EPI_ISL_1015700, EPI_ISL_1015701, EPI_ISL_1015702, EPI_ISL_1015703, EPI_ISL_1015704, EPI_ISL_1015705, EPI_ISL_1015706, EPI_ISL_1015707, EPI_ISL_1015708, EPI_ISL_1015709, EPI_ISL_1015710, EPI_ISL_1015711, EPI_ISL_1015712 | | | |
| see above | Columbia University Irving Medical Center | Wadsworth Center, New York State Department of Health | Kirsten St. George, Daryl M. Lamson, Alexis Russel, Matthew Shudt, Melissa A Leisner, Jonathan Plitnick, Navjot Singh, John Kelly, Erasmus Schneider, Erica Lasek-Nesselquist |
| EPI_ISL_1016013, EPI_ISL_1016014, EPI_ISL_1016015, EPI_ISL_1016016, EPI_ISL_1016017, EPI_ISL_1016018, EPI_ISL_1016075, EPI_ISL_1016100 | NYU Langone Health | Departments of Pathology and Medicine, New York University School of Medicine | Adriana Heguy, Dacia Dimartino, Emily Guzman, Christian Marier, Peter Meyn, Sitharam Ramaswami, Gael Westby, Paul Zappile, Yutong Zhang, Paolo Cotzia, Guiqing Wang |
| EPI_ISL_1016101, EPI_ISL_1016102, EPI_ISL_1016103, EPI_ISL_1016104, EPI_ISL_1016105, EPI_ISL_1016106, EPI_ISL_1016107, EPI_ISL_1016108, EPI_ISL_1016109, EPI_ISL_1016110 | URMC LABS | Wadsworth Center, New York State Department of Health | Kirsten St. George, Daryl M. Lamson, Alexis Russel, Matthew Shudt, Melissa A Leisner, Jonathan Plitnick, Navjot Singh, John Kelly, Erasmus Schneider, Erica Lasek-Nesselquist |
| EPI_ISL_1016111, EPI_ISL_1016112, EPI_ISL_1016113, EPI_ISL_1016114, EPI_ISL_1016115, EPI_ISL_1016116, EPI_ISL_1016117 | Columbia University Irving Medical Center | Wadsworth Center, New York State Department of Health | Kirsten St. George, Daryl M. Lamson, Alexis Russel, Matthew Shudt, Melissa A Leisner, Jonathan Plitnick, Navjot Singh, John Kelly, Erasmus Schneider, Erica Lasek-Nesselquist |
| EPI_ISL_1016118, EPI_ISL_1016119, EPI_ISL_1016120, EPI_ISL_1016121, EPI_ISL_1016122, EPI_ISL_1016123, EPI_ISL_1016124, EPI_ISL_1016125, EPI_ISL_1016126 | URMC LABS | Wadsworth Center, New York State Department of Health | Kirsten St. George, Daryl M. Lamson, Alexis Russel, Matthew Shudt, Melissa A Leisner, Jonathan Plitnick, Navjot Singh, John Kelly, Erasmus Schneider, Erica Lasek-Nesselquist |
| EPI_ISL_1016127, EPI_ISL_1016128, EPI_ISL_1016129, EPI_ISL_1016130, EPI_ISL_1016131, EPI_ISL_1016132, EPI_ISL_1016133, EPI_ISL_1016134, EPI_ISL_1016135 | THE MARY IMOGENE BASSETT HOSPITAL | Wadsworth Center, New York State Department of Health | Kirsten St. George, Daryl M. Lamson, Alexis Russel, Matthew Shudt, Melissa A Leisner, Jonathan Plitnick, Navjot Singh, John Kelly, Erasmus Schneider, Erica Lasek-Nesselquist |
| EPI_ISL_1016136 | NORTHWELL HEALTH LABORATORIES | Wadsworth Center, New York State Department of Health | Kirsten St. George, Daryl M. Lamson, Alexis Russel, Matthew Shudt, Melissa A Leisner, Jonathan Plitnick, Navjot Singh, John Kelly, Erasmus Schneider, Erica Lasek-Nesselquist |
| EPI_ISL_1016137 | THE MARY IMOGENE BASSETT HOSPITAL | Wadsworth Center, New York State Department of Health | Kirsten St. George, Daryl M. Lamson, Alexis Russel, Matthew Shudt, Melissa A Leisner, Jonathan Plitnick, Navjot Singh, John Kelly, Erasmus Schneider, Erica Lasek-Nesselquist |
| EPI_ISL_1016138 | NORTHWELL HEALTH LABORATORIES | Wadsworth Center, New York State Department of Health | Kirsten St. George, Daryl M. Lamson, Alexis Russel, Matthew Shudt, Melissa A Leisner, Jonathan Plitnick, Navjot Singh, John Kelly, Erasmus Schneider, Erica Lasek-Nesselquist |
| EPI_ISL_1016139 | THE MARY IMOGENE BASSETT HOSPITAL | Wadsworth Center, New York State Department of Health | Kirsten St. George, Daryl M. Lamson, Alexis Russel, Matthew Shudt, Melissa A Leisner, Jonathan Plitnick, Navjot Singh, John Kelly, Erasmus Schneider, Erica Lasek-Nesselquist |
| EPI_ISL_1016140 | NORTHWELL HEALTH LABORATORIES | Wadsworth Center, New York State Department of Health | Kirsten St. George, Daryl M. Lamson, Alexis Russel, Matthew Shudt, Melissa A Leisner, Jonathan Plitnick, Navjot Singh, John Kelly, Erasmus Schneider, Erica Lasek-Nesselquist |
| EPI_ISL_1016141, EPI_ISL_1016142 | THE MARY IMOGENE BASSETT HOSPITAL | Wadsworth Center, New York State Department of Health | Kirsten St. George, Daryl M. Lamson, Alexis Russel, Matthew Shudt, Melissa A Leisner, Jonathan Plitnick, Navjot Singh, John Kelly, Erasmus Schneider, Erica Lasek-Nesselquist |

| | | | |
|---|---|---|---|
| EPI_ISL_1016143, EPI_ISL_1016144, EPI_ISL_1016145, EPI_ISL_1016146, EPI_ISL_1016147, EPI_ISL_1016148, EPI_ISL_1016149, EPI_ISL_1016150, EPI_ISL_1016151, EPI_ISL_1016152, EPI_ISL_1016153 | | | |
| see above | NORTHWELL HEALTH LABORATORIES | Wadsworth Center, New York State Department of Health | Kirsten St. George, Daryl M. Lamson, Alexis Russel, Matthew Shudt, Melissa A Leisner, Jonathan Plitnick, Navjot Singh, John Kelly, Erasmus Schneider, Erica Lasek-Nesselquist |
| EPI_ISL_1016154, EPI_ISL_1016155 | THE MARY IMOGENE BASSETT HOSPITAL | Wadsworth Center, New York State Department of Health | Kirsten St. George, Daryl M. Lamson, Alexis Russel, Matthew Shudt, Melissa A Leisner, Jonathan Plitnick, Navjot Singh, John Kelly, Erasmus Schneider, Erica Lasek-Nesselquist |
| EPI_ISL_1016156, EPI_ISL_1016157, EPI_ISL_1016158, EPI_ISL_1016159, EPI_ISL_1016160, EPI_ISL_1016161, EPI_ISL_1016162, EPI_ISL_1016163, EPI_ISL_1016164, EPI_ISL_1016165, EPI_ISL_1016166, EPI_ISL_1016167, EPI_ISL_1016168 | | | |
| see above | NORTHWELL HEALTH LABORATORIES | Wadsworth Center, New York State Department of Health | Kirsten St. George, Daryl M. Lamson, Alexis Russel, Matthew Shudt, Melissa A Leisner, Jonathan Plitnick, Navjot Singh, John Kelly, Erasmus Schneider, Erica Lasek-Nesselquist |
| EPI_ISL_1016169 | TEMPUS LABS INC | Wadsworth Center, New York State Department of Health | Kirsten St. George, Daryl M. Lamson, Alexis Russel, Matthew Shudt, Melissa A Leisner, Jonathan Plitnick, Navjot Singh, John Kelly, Erasmus Schneider, Erica Lasek-Nesselquist |
| EPI_ISL_1016170 | NYU Langone Health | Departments of Pathology and Medicine, New York University School of Medicine | Adriana Heguy, Dacia Dimartino, Emily Guzman, Christian Marier, Peter Meyn, Sitharam Ramaswami, Gael Westby, Paul Zappile, Yutong Zhang, Paolo Cotzia, Guiqing Wang |
| EPI_ISL_1016171, EPI_ISL_1016172, EPI_ISL_1016173, EPI_ISL_1016174, EPI_ISL_1016175, EPI_ISL_1016176, EPI_ISL_1016177, EPI_ISL_1016178, EPI_ISL_1016179, EPI_ISL_1016180, EPI_ISL_1016181, EPI_ISL_1016182, EPI_ISL_1016183, EPI_ISL_1016184, EPI_ISL_1016185, EPI_ISL_1016186, EPI_ISL_1016187, EPI_ISL_1016188, EPI_ISL_1016189, EPI_ISL_1016190, EPI_ISL_1016191, EPI_ISL_1016192, EPI_ISL_1016193, EPI_ISL_1016194, EPI_ISL_1016195, EPI_ISL_1016196, EPI_ISL_1016197, EPI_ISL_1016198, EPI_ISL_1016199, EPI_ISL_1016200, EPI_ISL_1016201, EPI_ISL_1016202, EPI_ISL_1016203, EPI_ISL_1016204, EPI_ISL_1016205, EPI_ISL_1016206, EPI_ISL_1016207, EPI_ISL_1016208, EPI_ISL_1016209, EPI_ISL_1016210, EPI_ISL_1016211, EPI_ISL_1016212, EPI_ISL_1016213, EPI_ISL_1016214, EPI_ISL_1016215, EPI_ISL_1016216, EPI_ISL_1016217, EPI_ISL_1016218, EPI_ISL_1016219 | | | |
| see above | MONTEFIORE MEDICAL CENTER LABORATORIES | Wadsworth Center, New York State Department of Health | Kirsten St. George, Daryl M. Lamson, Alexis Russel, Matthew Shudt, Melissa A Leisner, Jonathan Plitnick, Navjot Singh, John Kelly, Erasmus Schneider, Erica Lasek-Nesselquist |
| EPI_ISL_1016220, EPI_ISL_1016221, EPI_ISL_1016222, EPI_ISL_1016223, EPI_ISL_1016224, EPI_ISL_1016225, EPI_ISL_1016226, EPI_ISL_1016227, EPI_ISL_1016228, EPI_ISL_1016229, EPI_ISL_1016230, EPI_ISL_1016231, EPI_ISL_1016232, EPI_ISL_1016233, EPI_ISL_1016234, EPI_ISL_1016235, EPI_ISL_1016236, EPI_ISL_1016237, EPI_ISL_1016238, EPI_ISL_1016239, EPI_ISL_1016240, EPI_ISL_1016241, EPI_ISL_1016242, EPI_ISL_1016243, EPI_ISL_1016244, EPI_ISL_1016245, EPI_ISL_1016246, EPI_ISL_1016247, EPI_ISL_1016248 | | | |
| see above | New York Presbyterian Hospital | Wadsworth Center, New York State Department of Health | Kirsten St. George, Daryl M. Lamson, Alexis Russel, Matthew Shudt, Melissa A Leisner, Jonathan Plitnick, Navjot Singh, John Kelly, Erasmus Schneider, Erica Lasek-Nesselquist |
| EPI_ISL_1016249, EPI_ISL_1016250, EPI_ISL_1016251, EPI_ISL_1016252, EPI_ISL_1016253, EPI_ISL_1016254, EPI_ISL_1016255, EPI_ISL_1016256 | TEMPUS LABS INC | Wadsworth Center, New York State Department of Health | Kirsten St. George, Daryl M. Lamson, Alexis Russel, Matthew Shudt, Melissa A Leisner, Jonathan Plitnick, Navjot Singh, John Kelly, Erasmus Schneider, Erica Lasek-Nesselquist |
| EPI_ISL_1016257 | THE MARY IMOGENE BASSETT HOSPITAL | Wadsworth Center, New York State Department of Health | Kirsten St. George, Daryl M. Lamson, Alexis Russel, Matthew Shudt, Melissa A Leisner, Jonathan Plitnick, Navjot Singh, John Kelly, Erasmus Schneider, Erica Lasek-Nesselquist |
| EPI_ISL_1016258, EPI_ISL_1016259 | New York Presbyterian Hospital | Wadsworth Center, New York State Department of Health | Kirsten St. George, Daryl M. Lamson, Alexis Russel, Matthew Shudt, Melissa A Leisner, Jonathan Plitnick, Navjot Singh, John Kelly, Erasmus Schneider, Erica Lasek-Nesselquist |
| EPI_ISL_1016260, EPI_ISL_1016261 | Columbia University Irving Medical Center | Wadsworth Center, New York State Department of Health | Kirsten St. George, Daryl M. Lamson, Alexis Russel, Matthew Shudt, Melissa A Leisner, Jonathan Plitnick, Navjot Singh, John Kelly, Erasmus Schneider, Erica Lasek-Nesselquist |
| EPI_ISL_1016262, EPI_ISL_1016263, EPI_ISL_1016264, EPI_ISL_1016265, EPI_ISL_1016266 | New York Presbyterian Hospital | Wadsworth Center, New York State Department of Health | Kirsten St. George, Daryl M. Lamson, Alexis Russel, Matthew Shudt, Melissa A Leisner, Jonathan Plitnick, Navjot Singh, John Kelly, Erasmus Schneider, Erica Lasek-Nesselquist |
| EPI_ISL_1016267, EPI_ISL_1016268 | Columbia University Irving Medical Center | Wadsworth Center, New York State Department of Health | Kirsten St. George, Daryl M. Lamson, Alexis Russel, Matthew Shudt, Melissa A Leisner, Jonathan Plitnick, Navjot Singh, John Kelly, Erasmus Schneider, Erica Lasek-Nesselquist |
| EPI_ISL_1016269 | New York Presbyterian Hospital | Wadsworth Center, New York State Department of Health | Kirsten St. George, Daryl M. Lamson, Alexis Russel, Matthew Shudt, Melissa A Leisner, Jonathan Plitnick, Navjot Singh, John Kelly, Erasmus Schneider, Erica Lasek-Nesselquist |
| EPI_ISL_1016270 | Columbia University Irving Medical Center | Wadsworth Center, New York State Department of Health | Kirsten St. George, Daryl M. Lamson, Alexis Russel, Matthew Shudt, Melissa A Leisner, Jonathan Plitnick, Navjot Singh, John Kelly, Erasmus Schneider, Erica Lasek-Nesselquist |
| EPI_ISL_1016271, EPI_ISL_1016272, EPI_ISL_1016273, EPI_ISL_1016274, EPI_ISL_1016275, EPI_ISL_1016276, EPI_ISL_1016277, EPI_ISL_1016278, EPI_ISL_1016279, EPI_ISL_1016280 | New York Presbyterian Hospital | Wadsworth Center, New York State Department of Health | Kirsten St. George, Daryl M. Lamson, Alexis Russel, Matthew Shudt, Melissa A Leisner, Jonathan Plitnick, Navjot Singh, John Kelly, Erasmus Schneider, Erica Lasek-Nesselquist |
| EPI_ISL_1016281, EPI_ISL_1016282, EPI_ISL_1016283, EPI_ISL_1016284 | Columbia University Irving Medical Center | Wadsworth Center, New York State Department of Health | Kirsten St. George, Daryl M. Lamson, Alexis Russel, Matthew Shudt, Melissa A Leisner, Jonathan Plitnick, Navjot Singh, John Kelly, Erasmus Schneider, Erica Lasek-Nesselquist |
| EPI_ISL_1016285, EPI_ISL_1016286 | SUNY UPSTATE MEDICAL UNIVERSITY | Wadsworth Center, New York State Department of Health | Kirsten St. George, Daryl M. Lamson, Alexis Russel, Matthew Shudt, Melissa A Leisner, Jonathan Plitnick, Navjot Singh, John Kelly, Erasmus Schneider, Erica Lasek-Nesselquist |
| EPI_ISL_1016287, EPI_ISL_1016288, EPI_ISL_1016289, EPI_ISL_1016290, EPI_ISL_1016291, EPI_ISL_1016292, EPI_ISL_1016293, EPI_ISL_1016294, EPI_ISL_1016295, EPI_ISL_1016296, EPI_ISL_1016297 | | | |
| see above | Columbia University Irving Medical Center | Wadsworth Center, New York State Department of Health | Kirsten St. George, Daryl M. Lamson, Alexis Russel, Matthew Shudt, Melissa A Leisner, Jonathan Plitnick, Navjot Singh, John Kelly, Erasmus Schneider, Erica Lasek-Nesselquist |
| EPI_ISL_1016298, EPI_ISL_1016299, EPI_ISL_1016300, EPI_ISL_1016301, EPI_ISL_1016302, EPI_ISL_1016303 | SUNY UPSTATE MEDICAL UNIVERSITY | Wadsworth Center, New York State Department of Health | Kirsten St. George, Daryl M. Lamson, Alexis Russel, Matthew Shudt, Melissa A Leisner, Jonathan Plitnick, Navjot Singh, John Kelly, Erasmus Schneider, Erica Lasek-Nesselquist |
| EPI_ISL_1016304, EPI_ISL_1016305, EPI_ISL_1016306, EPI_ISL_1016307, EPI_ISL_1016308, EPI_ISL_1016309, EPI_ISL_1016310 | New York Presbyterian Hospital | Wadsworth Center, New York State Department of Health | Kirsten St. George, Daryl M. Lamson, Alexis Russel, Matthew Shudt, Melissa A Leisner, Jonathan Plitnick, Navjot Singh, John Kelly, Erasmus Schneider, Erica Lasek-Nesselquist |
| EPI_ISL_1016311, EPI_ISL_1016312, EPI_ISL_1016313, EPI_ISL_1016314, EPI_ISL_1016315, EPI_ISL_1016316, EPI_ISL_1016317, EPI_ISL_1016318, EPI_ISL_1016319, EPI_ISL_1016320, EPI_ISL_1016321, EPI_ISL_1016322 | | | |
| see above | SUNY UPSTATE MEDICAL UNIVERSITY | Wadsworth Center, New York State Department of Health | Kirsten St. George, Daryl M. Lamson, Alexis Russel, Matthew Shudt, Melissa A Leisner, Jonathan Plitnick, Navjot Singh, John Kelly, Erasmus Schneider, Erica Lasek-Nesselquist |
| EPI_ISL_1016323, EPI_ISL_1016324, EPI_ISL_1016325, EPI_ISL_1016326 | Columbia University Irving Medical Center | Wadsworth Center, New York State Department of Health | Kirsten St. George, Daryl M. Lamson, Alexis Russel, Matthew Shudt, Melissa A Leisner, Jonathan Plitnick, Navjot Singh, John Kelly, Erasmus Schneider, Erica Lasek-Nesselquist |
| EPI_ISL_1016327 | SUNY UPSTATE MEDICAL UNIVERSITY | Wadsworth Center, New York State Department of Health | Kirsten St. George, Daryl M. Lamson, Alexis Russel, Matthew Shudt, Melissa A Leisner, Jonathan Plitnick, Navjot Singh, John Kelly, Erasmus Schneider, Erica Lasek-Nesselquist |
| EPI_ISL_1016328, EPI_ISL_1016329, EPI_ISL_1016330, EPI_ISL_1016331 | Columbia University Irving Medical Center | Wadsworth Center, New York State Department of Health | Kirsten St. George, Daryl M. Lamson, Alexis Russel, Matthew Shudt, Melissa A Leisner, Jonathan Plitnick, Navjot Singh, John Kelly, Erasmus Schneider, Erica Lasek-Nesselquist |
| EPI_ISL_1016332 | WESTCHESTER MEDICAL CENTER | Wadsworth Center, New York State Department of Health | Kirsten St. George, Daryl M. Lamson, Alexis Russel, Matthew Shudt, Melissa A Leisner, Jonathan Plitnick, Navjot Singh, John Kelly, Erasmus Schneider, Erica Lasek-Nesselquist |
| EPI_ISL_1016333, EPI_ISL_1016334 | URMC LABS | Wadsworth Center, New York State Department of Health | Kirsten St. George, Daryl M. Lamson, Alexis Russel, Matthew Shudt, Melissa A Leisner, Jonathan Plitnick, Navjot Singh, John Kelly, Erasmus Schneider, Erica Lasek-Nesselquist |

| | | | |
|---|---|---|---|
| EPI_ISL_1016335, EPI_ISL_1016336 | New York Presbyterian Hospital | Wadsworth Center, New York State Department of Health | Kirsten St. George, Daryl M. Lamson, Alexis Russel, Matthew Shudt, Melissa A Leisner, Jonathan Plitnick, Navjot Singh, John Kelly, Erasmus Schneider, Erica Lasek-Nesselquist |
| EPI_ISL_1016337, EPI_ISL_1016338 | MEMORIAL SLOAN KETTERING CANCER CENTER | Wadsworth Center, New York State Department of Health | Kirsten St. George, Daryl M. Lamson, Alexis Russel, Matthew Shudt, Melissa A Leisner, Jonathan Plitnick, Navjot Singh, John Kelly, Erasmus Schneider, Erica Lasek-Nesselquist |
| EPI_ISL_1016339 | URMC LABS | Wadsworth Center, New York State Department of Health | Kirsten St. George, Daryl M. Lamson, Alexis Russel, Matthew Shudt, Melissa A Leisner, Jonathan Plitnick, Navjot Singh, John Kelly, Erasmus Schneider, Erica Lasek-Nesselquist |
| EPI_ISL_1016340 | WESTCHESTER MEDICAL CENTER | Wadsworth Center, New York State Department of Health | Kirsten St. George, Daryl M. Lamson, Alexis Russel, Matthew Shudt, Melissa A Leisner, Jonathan Plitnick, Navjot Singh, John Kelly, Erasmus Schneider, Erica Lasek-Nesselquist |
| EPI_ISL_1016341 | New York Presbyterian Hospital | Wadsworth Center, New York State Department of Health | Kirsten St. George, Daryl M. Lamson, Alexis Russel, Matthew Shudt, Melissa A Leisner, Jonathan Plitnick, Navjot Singh, John Kelly, Erasmus Schneider, Erica Lasek-Nesselquist |
| EPI_ISL_1016342, EPI_ISL_1016343 | MEMORIAL SLOAN KETTERING CANCER CENTER | Wadsworth Center, New York State Department of Health | Kirsten St. George, Daryl M. Lamson, Alexis Russel, Matthew Shudt, Melissa A Leisner, Jonathan Plitnick, Navjot Singh, John Kelly, Erasmus Schneider, Erica Lasek-Nesselquist |
| EPI_ISL_1016344, EPI_ISL_1016345, EPI_ISL_1016346 | WESTCHESTER MEDICAL CENTER | Wadsworth Center, New York State Department of Health | Kirsten St. George, Daryl M. Lamson, Alexis Russel, Matthew Shudt, Melissa A Leisner, Jonathan Plitnick, Navjot Singh, John Kelly, Erasmus Schneider, Erica Lasek-Nesselquist |
| EPI_ISL_1016347, EPI_ISL_1016348, EPI_ISL_1016349, EPI_ISL_1016350, EPI_ISL_1016351, EPI_ISL_1016352, EPI_ISL_1016353, EPI_ISL_1016354, EPI_ISL_1016355, EPI_ISL_1016356, EPI_ISL_1016357, EPI_ISL_1016358, EPI_ISL_1016359, EPI_ISL_1016360, EPI_ISL_1016361, EPI_ISL_1016362, EPI_ISL_1016363, EPI_ISL_1016364 | | | |
| see above | URMC LABS | Wadsworth Center, New York State Department of Health | Kirsten St. George, Daryl M. Lamson, Alexis Russel, Matthew Shudt, Melissa A Leisner, Jonathan Plitnick, Navjot Singh, John Kelly, Erasmus Schneider, Erica Lasek-Nesselquist |
| EPI_ISL_1016365 | WESTCHESTER MEDICAL CENTER | Wadsworth Center, New York State Department of Health | Kirsten St. George, Daryl M. Lamson, Alexis Russel, Matthew Shudt, Melissa A Leisner, Jonathan Plitnick, Navjot Singh, John Kelly, Erasmus Schneider, Erica Lasek-Nesselquist |
| EPI_ISL_1016366, EPI_ISL_1016367, EPI_ISL_1016368, EPI_ISL_1016369, EPI_ISL_1016370, EPI_ISL_1016371, EPI_ISL_1016372 | New York Presbyterian Hospital | Wadsworth Center, New York State Department of Health | Kirsten St. George, Daryl M. Lamson, Alexis Russel, Matthew Shudt, Melissa A Leisner, Jonathan Plitnick, Navjot Singh, John Kelly, Erasmus Schneider, Erica Lasek-Nesselquist |
| EPI_ISL_1016373, EPI_ISL_1016374, EPI_ISL_1016375, EPI_ISL_1016376, EPI_ISL_1016377, EPI_ISL_1016378, EPI_ISL_1016379, EPI_ISL_1016380 | WESTCHESTER MEDICAL CENTER | Wadsworth Center, New York State Department of Health | Kirsten St. George, Daryl M. Lamson, Alexis Russel, Matthew Shudt, Melissa A Leisner, Jonathan Plitnick, Navjot Singh, John Kelly, Erasmus Schneider, Erica Lasek-Nesselquist |
| EPI_ISL_1016381, EPI_ISL_1016382, EPI_ISL_1016383, EPI_ISL_1016384, EPI_ISL_1016385, EPI_ISL_1016386, EPI_ISL_1016387, EPI_ISL_1016388, EPI_ISL_1016389, EPI_ISL_1016390, EPI_ISL_1016391, EPI_ISL_1016392, EPI_ISL_1016393, EPI_ISL_1016394, EPI_ISL_1016395, EPI_ISL_1016396, EPI_ISL_1016397, EPI_ISL_1016398, EPI_ISL_1016399, EPI_ISL_1016400, EPI_ISL_1016401 | | | |
| see above | New York Presbyterian Hospital | Wadsworth Center, New York State Department of Health | Kirsten St. George, Daryl M. Lamson, Alexis Russel, Matthew Shudt, Melissa A Leisner, Jonathan Plitnick, Navjot Singh, John Kelly, Erasmus Schneider, Erica Lasek-Nesselquist |
| EPI_ISL_1016402, EPI_ISL_1016403, EPI_ISL_1016404, EPI_ISL_1016405, EPI_ISL_1016406, EPI_ISL_1016407, EPI_ISL_1016408, EPI_ISL_1016409, EPI_ISL_1016410, EPI_ISL_1016411 | MEMORIAL SLOAN KETTERING CANCER CENTER | Wadsworth Center, New York State Department of Health | Kirsten St. George, Daryl M. Lamson, Alexis Russel, Matthew Shudt, Melissa A Leisner, Jonathan Plitnick, Navjot Singh, John Kelly, Erasmus Schneider, Erica Lasek-Nesselquist |
| EPI_ISL_1016412 | STONY BROOK UNIVERSITY HOSPITAL | Wadsworth Center, New York State Department of Health | Kirsten St. George, Daryl M. Lamson, Alexis Russel, Matthew Shudt, Melissa A Leisner, Jonathan Plitnick, Navjot Singh, John Kelly, Erasmus Schneider, Erica Lasek-Nesselquist |
| EPI_ISL_1016413 | NORTHWELL HEALTH LABORATORIES | Wadsworth Center, New York State Department of Health | Kirsten St. George, Daryl M. Lamson, Alexis Russel, Matthew Shudt, Melissa A Leisner, Jonathan Plitnick, Navjot Singh, John Kelly, Erasmus Schneider, Erica Lasek-Nesselquist |
| EPI_ISL_1016414, EPI_ISL_1016415 | URMC LABS | Wadsworth Center, New York State Department of Health | Kirsten St. George, Daryl M. Lamson, Alexis Russel, Matthew Shudt, Melissa A Leisner, Jonathan Plitnick, Navjot Singh, John Kelly, Erasmus Schneider, Erica Lasek-Nesselquist |
| EPI_ISL_1016416 | STONY BROOK UNIVERSITY HOSPITAL | Wadsworth Center, New York State Department of Health | Kirsten St. George, Daryl M. Lamson, Alexis Russel, Matthew Shudt, Melissa A Leisner, Jonathan Plitnick, Navjot Singh, John Kelly, Erasmus Schneider, Erica Lasek-Nesselquist |
| EPI_ISL_1016417 | ALBANY MEDICAL CENTER HOSPITAL CLINICAL LABORATORIES | Wadsworth Center, New York State Department of Health | Kirsten St. George, Daryl M. Lamson, Alexis Russel, Matthew Shudt, Melissa A Leisner, Jonathan Plitnick, Navjot Singh, John Kelly, Erasmus Schneider, Erica Lasek-Nesselquist |
| EPI_ISL_1016418, EPI_ISL_1016419, EPI_ISL_1016420, EPI_ISL_1016421, EPI_ISL_1016422, EPI_ISL_1016423, EPI_ISL_1016424, EPI_ISL_1016425, EPI_ISL_1016426, EPI_ISL_1016427, EPI_ISL_1016428, EPI_ISL_1016429, EPI_ISL_1016430, EPI_ISL_1016431, EPI_ISL_1016432, EPI_ISL_1016433, EPI_ISL_1016434, EPI_ISL_1016435, EPI_ISL_1016436, EPI_ISL_1016437, EPI_ISL_1016438, EPI_ISL_1016439, EPI_ISL_1016440, EPI_ISL_1016441, EPI_ISL_1016442, EPI_ISL_1016443, EPI_ISL_1016444, EPI_ISL_1016445, EPI_ISL_1016446, EPI_ISL_1016447, EPI_ISL_1016448, EPI_ISL_1016449, EPI_ISL_1016450, EPI_ISL_1016451, EPI_ISL_1016452, EPI_ISL_1016453, EPI_ISL_1016454, EPI_ISL_1016455, EPI_ISL_1016456, EPI_ISL_1016457 | | | |
| see above | URMC LABS | Wadsworth Center, New York State Department of Health | Kirsten St. George, Daryl M. Lamson, Alexis Russel, Matthew Shudt, Melissa A Leisner, Jonathan Plitnick, Navjot Singh, John Kelly, Erasmus Schneider, Erica Lasek-Nesselquist |
| EPI_ISL_1016458 | NORTHWELL HEALTH LABORATORIES | Wadsworth Center, New York State Department of Health | Kirsten St. George, Daryl M. Lamson, Alexis Russel, Matthew Shudt, Melissa A Leisner, Jonathan Plitnick, Navjot Singh, John Kelly, Erasmus Schneider, Erica Lasek-Nesselquist |
| EPI_ISL_1016459, EPI_ISL_1016460, EPI_ISL_1016461, EPI_ISL_1016462, EPI_ISL_1016463, EPI_ISL_1016464, EPI_ISL_1016465, EPI_ISL_1016466, EPI_ISL_1016467 | STONY BROOK UNIVERSITY HOSPITAL | Wadsworth Center, New York State Department of Health | Kirsten St. George, Daryl M. Lamson, Alexis Russel, Matthew Shudt, Melissa A Leisner, Jonathan Plitnick, Navjot Singh, John Kelly, Erasmus Schneider, Erica Lasek-Nesselquist |
| EPI_ISL_1016468, EPI_ISL_1016469, EPI_ISL_1016470, EPI_ISL_1016471, EPI_ISL_1016472, EPI_ISL_1016473, EPI_ISL_1016474, EPI_ISL_1016475 | ALBANY MEDICAL CENTER HOSPITAL CLINICAL LABORATORIES | Wadsworth Center, New York State Department of Health | Kirsten St. George, Daryl M. Lamson, Alexis Russel, Matthew Shudt, Melissa A Leisner, Jonathan Plitnick, Navjot Singh, John Kelly, Erasmus Schneider, Erica Lasek-Nesselquist |
| EPI_ISL_1016476, EPI_ISL_1016477, EPI_ISL_1016478 | STONY BROOK UNIVERSITY HOSPITAL | Wadsworth Center, New York State Department of Health | Kirsten St. George, Daryl M. Lamson, Alexis Russel, Matthew Shudt, Melissa A Leisner, Jonathan Plitnick, Navjot Singh, John Kelly, Erasmus Schneider, Erica Lasek-Nesselquist |
| EPI_ISL_1016565, EPI_ISL_1016602, EPI_ISL_1016643 | Helix/Illumina | Respiratory Viruses Branch, Division of Viral Diseases, Centers for Disease Control and Prevention | Peter W. Cook, Dakota Howard, Dhwani Batra, Ben L. Rambo-Martin, Eileen de Feo, Jan Antico, Christine Tran, Matthew Tolentino, Shannon Wickline, Kim Gietzen, Brad Sickler, Jingtao Liu, Eric Allen, Phil Febbo, Summer Galloway, Nicole L. Washington, Simon White, Geraint Levan, Kelly Schiabor Barrett, Elizabeth Cirulli, Alexandre Bolze, Ary Ascencio, Chantal Rivera-Garcia, Ryan Cho, Jason Nguyen, Sherry Wang, Jimmy Ramirez, Tyler Cassens, Efren Sandoval, Magnus Isaksson, William Lee, David Becker, Marc Laurent, James Lu, Clinton R. Paden, Suxiang Tong, Duncan MacCannell |
| EPI_ISL_1017489, EPI_ISL_1017491, EPI_ISL_1017500, EPI_ISL_1017502, EPI_ISL_1017514, EPI_ISL_1017525, EPI_ISL_1017526, EPI_ISL_1017536, EPI_ISL_1017541, EPI_ISL_1017547, EPI_ISL_1017566, EPI_ISL_1017568, EPI_ISL_1017571, EPI_ISL_1017575, EPI_ISL_1017576, EPI_ISL_1017577, EPI_ISL_1017582, EPI_ISL_1017583, EPI_ISL_1017584 | | | |
| see above | Murphy Medical Associates | Grubaugh Lab - Yale School of Public Health | Mary Petrone, Joseph Fauver, Caleb Neal, Steven Murphy, Chantal Vogels, Mallery Breban, Annie Watkins, Tara Alpert, Nathan Grubaugh |
| EPI_ISL_1017969, EPI_ISL_1017970 | DOHMH PHL | New York City Public Health Laboratory | Jade Wang, et al. |

| | | | |
|---|---|---|---|
| EPI_ISL_1017971, EPI_ISL_1017972, EPI_ISL_1017973 | DOHMH Riverside | New York City Public Health Laboratory | Jade Wang, et al. |
| EPI_ISL_1017974 | DOHMH Morrisania | New York City Public Health Laboratory | Jade Wang, et al. |
| EPI_ISL_1017975 | DOHMH Corona | New York City Public Health Laboratory | Jade Wang, et al. |
| EPI_ISL_1017976, EPI_ISL_1017977 | DOHMH Jamaica | New York City Public Health Laboratory | Jade Wang, et al. |
| EPI_ISL_1017978 | DOHMH Chelsea | New York City Public Health Laboratory | Jade Wang, et al. |
| EPI_ISL_1017979 | DOHMH PHL | New York City Public Health Laboratory | Jade Wang, et al. |
| EPI_ISL_1017980, EPI_ISL_1017981 | DOHMH Morrisania | New York City Public Health Laboratory | Jade Wang, et al. |
| EPI_ISL_1017982, EPI_ISL_1017983, EPI_ISL_1017984, EPI_ISL_1017985 | DOHMH Central Harlem | New York City Public Health Laboratory | Jade Wang, et al. |
| EPI_ISL_1017986, EPI_ISL_1017987, EPI_ISL_1017988 | DOHMH Corona | New York City Public Health Laboratory | Jade Wang, et al. |
| EPI_ISL_1017989, EPI_ISL_1017990 | Department of Homeless Services | New York City Public Health Laboratory | Jade Wang, et al. |
| EPI_ISL_1017991 | DOHMH PHL | New York City Public Health Laboratory | Jade Wang, et al. |
| EPI_ISL_1017992, EPI_ISL_1017993, EPI_ISL_1017994 | DOHMH Jamaica | New York City Public Health Laboratory | Jade Wang, et al. |
| EPI_ISL_1017995, EPI_ISL_1017996, EPI_ISL_1017997, EPI_ISL_1017998, EPI_ISL_1017999 | DOHMH Corona | New York City Public Health Laboratory | Jade Wang, et al. |
| EPI_ISL_1018000 | DOHMH Morrisania | New York City Public Health Laboratory | Jade Wang, et al. |
| EPI_ISL_1018001 | DOHMH Central Harlem | New York City Public Health Laboratory | Jade Wang, et al. |
| EPI_ISL_1018002, EPI_ISL_1018003, EPI_ISL_1018004 | DOHMH PHL | New York City Public Health Laboratory | Jade Wang, et al. |
| EPI_ISL_1018005 | DOHMH Corona | New York City Public Health Laboratory | Jade Wang, et al. |
| EPI_ISL_1018006, EPI_ISL_1018007, EPI_ISL_1018008, EPI_ISL_1018009, EPI_ISL_1018010 | DOHMH Morrisania | New York City Public Health Laboratory | Jade Wang, et al. |
| EPI_ISL_1018011 | DOHMH Chelsea | New York City Public Health Laboratory | Jade Wang, et al. |
| EPI_ISL_1018012, EPI_ISL_1018013, EPI_ISL_1018014, EPI_ISL_1018015, EPI_ISL_1018016, EPI_ISL_1018017, EPI_ISL_1018018, EPI_ISL_1018019, EPI_ISL_1018020 | DOHMH Jamaica | New York City Public Health Laboratory | Jade Wang, et al. |
| EPI_ISL_1018021 | DOHMH Crown Heights | New York City Public Health Laboratory | Jade Wang, et al. |
| EPI_ISL_1018022, EPI_ISL_1018023 | DOHMH Jamaica | New York City Public Health Laboratory | Jade Wang, et al. |
| EPI_ISL_1018024 | DOHMH Crown Heights | New York City Public Health Laboratory | Jade Wang, et al. |
| EPI_ISL_1018025, EPI_ISL_1018026 | DOHMH Morrisania | New York City Public Health Laboratory | Jade Wang, et al. |
| EPI_ISL_1018027, EPI_ISL_1018028, EPI_ISL_1018029, EPI_ISL_1018030, EPI_ISL_1018031 | DOHMH Corona | New York City Public Health Laboratory | Jade Wang, et al. |
| EPI_ISL_1018032, EPI_ISL_1018033, EPI_ISL_1018034, EPI_ISL_1018035, EPI_ISL_1018036, EPI_ISL_1018037, EPI_ISL_1018038, EPI_ISL_1018039, EPI_ISL_1018040, EPI_ISL_1018041, EPI_ISL_1018042, EPI_ISL_1018043, EPI_ISL_1018044, EPI_ISL_1018045, EPI_ISL_1018046, EPI_ISL_1018047, EPI_ISL_1018048, EPI_ISL_1018049 | | | |
| see above | OCME Office Of Chief Medical Examiner | New York City Public Health Laboratory | Jade Wang, et al. |
| EPI_ISL_1020487, EPI_ISL_1020488, EPI_ISL_1020489, EPI_ISL_1020490, EPI_ISL_1020491, EPI_ISL_1020492, EPI_ISL_1020493, EPI_ISL_1020494, EPI_ISL_1020495, EPI_ISL_1020496, EPI_ISL_1020497, EPI_ISL_1020498, EPI_ISL_1020499, EPI_ISL_1020500, EPI_ISL_1020501, EPI_ISL_1020502, EPI_ISL_1020503, EPI_ISL_1020504, EPI_ISL_1020505, EPI_ISL_1020506, EPI_ISL_1020507, EPI_ISL_1020508, EPI_ISL_1020509, EPI_ISL_1020510, EPI_ISL_1020511, EPI_ISL_1020512, EPI_ISL_1020513, EPI_ISL_1020514, EPI_ISL_1020515, EPI_ISL_1020516, EPI_ISL_1020517, EPI_ISL_1020518, EPI_ISL_1020519, EPI_ISL_1020520, EPI_ISL_1020521, EPI_ISL_1020522, EPI_ISL_1020523, EPI_ISL_1020524, EPI_ISL_1020525, EPI_ISL_1020526, EPI_ISL_1020527, EPI_ISL_1020528, EPI_ISL_1020529, EPI_ISL_1020530, EPI_ISL_1020531, EPI_ISL_1020532, EPI_ISL_1020533, EPI_ISL_1020534, EPI_ISL_1020535, EPI_ISL_1020536, EPI_ISL_1020537, EPI_ISL_1020538, EPI_ISL_1020539, EPI_ISL_1020540, EPI_ISL_1020541, EPI_ISL_1020542, EPI_ISL_1020543, EPI_ISL_1020544, EPI_ISL_1020545, EPI_ISL_1020546, EPI_ISL_1020547, EPI_ISL_1020548, EPI_ISL_1020549, EPI_ISL_1020550 | | | |
| see above | Columbia University Irving Medical Center | Wadsworth Center, New York State Department of Health | Kirsten St. George, Daryl M. Lamson, Alexis Russel, Matthew Shudt, Melissa A Leisner, Jonathan Plitnick, Navjot Singh, John Kelly, Erasmus Schneider, Erica Lasek-Nesselquist |
| EPI_ISL_1020593, EPI_ISL_1020774, EPI_ISL_1020778, EPI_ISL_1020810, EPI_ISL_1020839, EPI_ISL_1020872, EPI_ISL_1020911, EPI_ISL_1020934, EPI_ISL_1020989, EPI_ISL_1020994, EPI_ISL_1021030, EPI_ISL_1021033, EPI_ISL_1021041, EPI_ISL_1021054, EPI_ISL_1021057, EPI_ISL_1021066, EPI_ISL_1021081, EPI_ISL_1021094, EPI_ISL_1021098, EPI_ISL_1021121, EPI_ISL_1021123, EPI_ISL_1021125, EPI_ISL_1021128, EPI_ISL_1021135, EPI_ISL_1021136, EPI_ISL_1021168, EPI_ISL_1021181, EPI_ISL_1021210, EPI_ISL_1021229, EPI_ISL_1021257, EPI_ISL_1021265, EPI_ISL_1021359, EPI_ISL_1021360, EPI_ISL_1021361, EPI_ISL_1021368, EPI_ISL_1021383, EPI_ISL_1021384, EPI_ISL_1021394, EPI_ISL_1021413, EPI_ISL_1021428, EPI_ISL_1021437, EPI_ISL_1021501, EPI_ISL_1021507, EPI_ISL_1021508, EPI_ISL_1021540, EPI_ISL_1021542, EPI_ISL_1021561, EPI_ISL_1021562, EPI_ISL_1021592, EPI_ISL_1021630, EPI_ISL_1021655, EPI_ISL_1021669, EPI_ISL_1021722, EPI_ISL_1021735, EPI_ISL_1021736, EPI_ISL_1021750, EPI_ISL_1021751, EPI_ISL_1021761, EPI_ISL_1021803, EPI_ISL_1021819, EPI_ISL_1021824, EPI_ISL_1021832, EPI_ISL_1021857, EPI_ISL_1021865, EPI_ISL_1021868, EPI_ISL_1021928, EPI_ISL_1021963, EPI_ISL_1021969, EPI_ISL_1021970, EPI_ISL_1021981, EPI_ISL_1022042, EPI_ISL_1022048, EPI_ISL_1022085, EPI_ISL_1022107, EPI_ISL_1022108, EPI_ISL_1022130, EPI_ISL_1022131, EPI_ISL_1022132, EPI_ISL_1022133, EPI_ISL_1022167, EPI_ISL_1022176, EPI_ISL_1022205, EPI_ISL_1022209, EPI_ISL_1022214, EPI_ISL_1022227, EPI_ISL_1022261, EPI_ISL_1022269, EPI_ISL_1022288, EPI_ISL_1022311, EPI_ISL_1022325, EPI_ISL_1022336, EPI_ISL_1022371, EPI_ISL_1022372, EPI_ISL_1022373, EPI_ISL_1022374, EPI_ISL_1022395, EPI_ISL_1022396, EPI_ISL_1026002, EPI_ISL_1026006, EPI_ISL_1026023, EPI_ISL_1026103, EPI_ISL_1026107, EPI_ISL_1026212, EPI_ISL_1026215, EPI_ISL_1026241, EPI_ISL_1026300, EPI_ISL_1026306, EPI_ISL_1026346, EPI_ISL_1026443, EPI_ISL_1026456, EPI_ISL_1026490, EPI_ISL_1026532, EPI_ISL_1026540, EPI_ISL_1026613, EPI_ISL_1026630, EPI_ISL_1026672, EPI_ISL_1026695, EPI_ISL_1026726, EPI_ISL_1026732, EPI_ISL_1026854, EPI_ISL_1027684, EPI_ISL_1027727, EPI_ISL_1027736, EPI_ISL_1027771, EPI_ISL_1027772, EPI_ISL_1027774, EPI_ISL_1027777, EPI_ISL_1027793, EPI_ISL_1027812, EPI_ISL_1027837, EPI_ISL_1027866, EPI_ISL_1027919, EPI_ISL_1027979, EPI_ISL_1027990, EPI_ISL_1028017, EPI_ISL_1028026, EPI_ISL_1028037, EPI_ISL_1028113, EPI_ISL_1028119, EPI_ISL_1028181, EPI_ISL_1028310, EPI_ISL_1028315, EPI_ISL_1028330, EPI_ISL_1028384, EPI_ISL_1028401, EPI_ISL_1028439, EPI_ISL_1028460, EPI_ISL_1028469, EPI_ISL_1028524, EPI_ISL_1028539, EPI_ISL_1028544, EPI_ISL_1028547, EPI_ISL_1028558, EPI_ISL_1028572, EPI_ISL_1028615, EPI_ISL_1028632, EPI_ISL_1028640, EPI_ISL_1028645, EPI_ISL_1028656, EPI_ISL_1028687, EPI_ISL_1028690, EPI_ISL_1028716, EPI_ISL_1028743, EPI_ISL_1028773, EPI_ISL_1028812, EPI_ISL_1028847, EPI_ISL_1028870, EPI_ISL_1028936, EPI_ISL_1028951, EPI_ISL_1028954, EPI_ISL_1029021, EPI_ISL_1029031, EPI_ISL_1029056, EPI_ISL_1029070, EPI_ISL_1029074, EPI_ISL_1029084, EPI_ISL_1029086, EPI_ISL_1029087, EPI_ISL_1029088, EPI_ISL_1029244, EPI_ISL_1029245, EPI_ISL_1029287, EPI_ISL_1029288, EPI_ISL_1029289, EPI_ISL_1029328, EPI_ISL_1029401, EPI_ISL_1029411, EPI_ISL_1029440, EPI_ISL_1029441, EPI_ISL_1029442, EPI_ISL_1029485, EPI_ISL_1029486, EPI_ISL_1029487, EPI_ISL_1029488, EPI_ISL_1029501, EPI_ISL_1029502, EPI_ISL_1029917, EPI_ISL_1029939, EPI_ISL_1029941, EPI_ISL_1029943, EPI_ISL_1030530, EPI_ISL_1030622, EPI_ISL_1030672, EPI_ISL_1030682, EPI_ISL_1030748, EPI_ISL_1030785, EPI_ISL_1030833, EPI_ISL_1030880, EPI_ISL_1030908, EPI_ISL_1030946, EPI_ISL_1030998, EPI_ISL_1031026, EPI_ISL_1031031, EPI_ISL_1031049, EPI_ISL_1031122, EPI_ISL_1031224, EPI_ISL_1031246, EPI_ISL_1031247, EPI_ISL_1031251, EPI_ISL_1031276, EPI_ISL_1031295, EPI_ISL_1031374, EPI_ISL_1031503, EPI_ISL_1031511, EPI_ISL_1031544, EPI_ISL_1031563, EPI_ISL_1031660, EPI_ISL_1031678, EPI_ISL_1031687, EPI_ISL_1031765, EPI_ISL_1031766, EPI_ISL_1031797, EPI_ISL_1031873, EPI_ISL_1032050, EPI_ISL_1032060, EPI_ISL_1032067, EPI_ISL_1032163, EPI_ISL_1032196, EPI_ISL_1032197, EPI_ISL_1032201, EPI_ISL_1032205, EPI_ISL_1032344, EPI_ISL_1032393, EPI_ISL_1032394, EPI_ISL_1032395, EPI_ISL_1032399, EPI_ISL_1032401, EPI_ISL_1032405, EPI_ISL_1032567, EPI_ISL_1032580, EPI_ISL_1032581 | | | |
| see above | Laboratory Corporation of America | Respiratory Viruses Branch, Division of Viral Diseases, Centers for Disease Control and Prevention | Peter W. Cook, Dakota Howard, Dhwani Batra, Ben L. Rambo-Martin, Clinton R. Paden, Suxiang Tong, Duncan MacCannell |
| EPI_ISL_1034279 | Northwest Pathology | UW Virology Lab | Pavitra Roychoudhury, Lasata Shrestha, Shah A Mohamed Bakhash, Michelle Lin, Hong Xie, Meei-Li Huang, Keith R Jerome, Alexander Greninger |
| EPI_ISL_1036954, EPI_ISL_1037009, EPI_ISL_1037086, EPI_ISL_1037167, EPI_ISL_1037195, EPI_ISL_1037232, EPI_ISL_1037250, EPI_ISL_1037255, EPI_ISL_1037306, EPI_ISL_1037475, EPI_ISL_1037497, EPI_ISL_1037510, EPI_ISL_1037640, EPI_ISL_1037650, EPI_ISL_1037655, EPI_ISL_1037790, EPI_ISL_1037859, EPI_ISL_1037868, EPI_ISL_1037980, EPI_ISL_1037988, EPI_ISL_1038042, EPI_ISL_1038074, EPI_ISL_1038121, EPI_ISL_1038179, EPI_ISL_1038215, EPI_ISL_1038223, EPI_ISL_1038235, EPI_ISL_1038302, EPI_ISL_1038344, EPI_ISL_1038361, EPI_ISL_1038367, EPI_ISL_1038368, EPI_ISL_1038406, EPI_ISL_1038485, EPI_ISL_1038490, EPI_ISL_1038491, EPI_ISL_1038492, EPI_ISL_1038493, EPI_ISL_1038784, EPI_ISL_1038785 | | | |
| see above | Laboratory Corporation of America | Respiratory Viruses Branch, Division of Viral Diseases, | Peter W. Cook, Dakota Howard, Dhwani Batra, Ben L. Rambo-Martin, Clinton R. Paden, Suxiang Tong, Duncan MacCannell |

Centers for Disease Control and Prevention

EPI_ISL_1038971, EPI_ISL_1038972, EPI_ISL_1038973, EPI_ISL_1038974, EPI_ISL_1038975, EPI_ISL_1038976, EPI_ISL_1038977, EPI_ISL_1038978, EPI_ISL_1038979, EPI_ISL_1038980, EPI_ISL_1038981

| | | | |
|---|---|---|---|
| see above | Tempus | Grubaugh Lab - Yale School of Public Health | Joseph Fauver, Tara Alpert, Anderson Brito, Mallery Breban, Anne Wyllie, Chantal Vogels, Mary Petrone, Annie Watkins, Chaney Kalinich, Isabel Ott, Nathan Grubaugh |
| EPI_ISL_1038996 | Murphy Medical Associates | Grubaugh Lab - Yale School of Public Health | Joseph Fauver, Tara Alpert, Anderson Brito, Mallery Breban, Anne Wyllie, Chantal Vogels, Mary Petrone, Annie Watkins, Chaney Kalinich, Isabel Ott, Nathan Grubaugh |
| EPI_ISL_1040480 | Pandemic Response Lab - NYC | Pandemic Response Lab, R&D | Henry Lee, Michael Hammerling, Melissa Hopkins, Cybill del Castillo, William Ward, Pradeep Bugga, Haiping Hao, Jon Laurent |

EPI_ISL_1040853, EPI_ISL_1040854, EPI_ISL_1040855, EPI_ISL_1040856, EPI_ISL_1040857, EPI_ISL_1040858, EPI_ISL_1040859, EPI_ISL_1040861, EPI_ISL_1040862, EPI_ISL_1040863, EPI_ISL_1040864, EPI_ISL_1040865, EPI_ISL_1040866, EPI_ISL_1040867, EPI_ISL_1040869, EPI_ISL_1040870, EPI_ISL_1040871, EPI_ISL_1040872, EPI_ISL_1040873, EPI_ISL_1040874, EPI_ISL_1040875, EPI_ISL_1040876, EPI_ISL_1040877, EPI_ISL_1040878, EPI_ISL_1040879, EPI_ISL_1040880, EPI_ISL_1040881, EPI_ISL_1040882, EPI_ISL_1040883, EPI_ISL_1040884, EPI_ISL_1040885, EPI_ISL_1040886, EPI_ISL_1040887, EPI_ISL_1040888, EPI_ISL_1040889, EPI_ISL_1040890, EPI_ISL_1040891, EPI_ISL_1040892, EPI_ISL_1040893, EPI_ISL_1040894, EPI_ISL_1040895, EPI_ISL_1040896, EPI_ISL_1040897, EPI_ISL_1040898, EPI_ISL_1040899, EPI_ISL_1040900, EPI_ISL_1040901, EPI_ISL_1040902, EPI_ISL_1040903, EPI_ISL_1040904, EPI_ISL_1040905, EPI_ISL_1040906, EPI_ISL_1040908, EPI_ISL_1040909, EPI_ISL_1040910, EPI_ISL_1040911, EPI_ISL_1040913, EPI_ISL_1040914

| | | | |
|---|---|---|---|
| see above | NYU Langone Health | Departments of Pathology and Medicine, New York University School of Medicine | Adriana Heguy, Dacia Dimartino, Emily Guzman, Christian Marier, Peter Meyn, Sitharam Ramaswami, Gael Westby, Paul Zappile, Yutong Zhang, Paolo Cotzia, Guiqing Wang |

EPI_ISL_1041213, EPI_ISL_1041214, EPI_ISL_1041215, EPI_ISL_1041216, EPI_ISL_1041217, EPI_ISL_1041218, EPI_ISL_1041219, EPI_ISL_1041220, EPI_ISL_1041221, EPI_ISL_1041222, EPI_ISL_1041223, EPI_ISL_1041224, EPI_ISL_1041225, EPI_ISL_1041226, EPI_ISL_1041227, EPI_ISL_1041228, EPI_ISL_1041229, EPI_ISL_1041230, EPI_ISL_1041231, EPI_ISL_1041232, EPI_ISL_1041233, EPI_ISL_1041234, EPI_ISL_1041235, EPI_ISL_1041236, EPI_ISL_1041237, EPI_ISL_1041238, EPI_ISL_1041239, EPI_ISL_1041240, EPI_ISL_1041241, EPI_ISL_1041242, EPI_ISL_1041243, EPI_ISL_1041244, EPI_ISL_1041245, EPI_ISL_1041246, EPI_ISL_1041247, EPI_ISL_1041248, EPI_ISL_1041249, EPI_ISL_1041250, EPI_ISL_1041251, EPI_ISL_1041252, EPI_ISL_1041253, EPI_ISL_1041254, EPI_ISL_1041255, EPI_ISL_1041256, EPI_ISL_1041257, EPI_ISL_1041258, EPI_ISL_1041259, EPI_ISL_1041260, EPI_ISL_1041261, EPI_ISL_1041262, EPI_ISL_1041263, EPI_ISL_1041264, EPI_ISL_1041265, EPI_ISL_1041266, EPI_ISL_1041267, EPI_ISL_1041268, EPI_ISL_1041269, EPI_ISL_1041270, EPI_ISL_1041271, EPI_ISL_1041272, EPI_ISL_1041273, EPI_ISL_1041274, EPI_ISL_1041275, EPI_ISL_1041276, EPI_ISL_1041277, EPI_ISL_1041278, EPI_ISL_1041279, EPI_ISL_1041280, EPI_ISL_1041281, EPI_ISL_1041282, EPI_ISL_1041283, EPI_ISL_1041284, EPI_ISL_1041285, EPI_ISL_1041286, EPI_ISL_1041287, EPI_ISL_1041288, EPI_ISL_1041289, EPI_ISL_1041290, EPI_ISL_1041291, EPI_ISL_1041292, EPI_ISL_1041293, EPI_ISL_1041294, EPI_ISL_1041295, EPI_ISL_1041296, EPI_ISL_1041297, EPI_ISL_1041298, EPI_ISL_1041299, EPI_ISL_1041300, EPI_ISL_1041301, EPI_ISL_1041302, EPI_ISL_1041303, EPI_ISL_1041304, EPI_ISL_1041305, EPI_ISL_1041306, EPI_ISL_1041307, EPI_ISL_1041308, EPI_ISL_1041309, EPI_ISL_1041310, EPI_ISL_1041311, EPI_ISL_1041312, EPI_ISL_1041313, EPI_ISL_1041314, EPI_ISL_1041315, EPI_ISL_1041316, EPI_ISL_1041317, EPI_ISL_1041318, EPI_ISL_1041319, EPI_ISL_1041320, EPI_ISL_1041321, EPI_ISL_1041322, EPI_ISL_1041323, EPI_ISL_1041324, EPI_ISL_1041325, EPI_ISL_1041326, EPI_ISL_1041327, EPI_ISL_1041328, EPI_ISL_1041329, EPI_ISL_1041330, EPI_ISL_1041331, EPI_ISL_1041332, EPI_ISL_1041333, EPI_ISL_1041334, EPI_ISL_1041335, EPI_ISL_1041336, EPI_ISL_1041337, EPI_ISL_1041338, EPI_ISL_1041339, EPI_ISL_1041340, EPI_ISL_1041341, EPI_ISL_1041342, EPI_ISL_1041343, EPI_ISL_1041344, EPI_ISL_1041345, EPI_ISL_1041346, EPI_ISL_1041347, EPI_ISL_1041348, EPI_ISL_1041349, EPI_ISL_1041350, EPI_ISL_1041351, EPI_ISL_1041352, EPI_ISL_1041353, EPI_ISL_1041354, EPI_ISL_1041355, EPI_ISL_1041356, EPI_ISL_1041357, EPI_ISL_1041358, EPI_ISL_1041359, EPI_ISL_1041360, EPI_ISL_1041361, EPI_ISL_1041362, EPI_ISL_1041363, EPI_ISL_1041364, EPI_ISL_1041365, EPI_ISL_1041366, EPI_ISL_1041367, EPI_ISL_1041368, EPI_ISL_1041369, EPI_ISL_1041370, EPI_ISL_1041371, EPI_ISL_1041372, EPI_ISL_1041373, EPI_ISL_1041374, EPI_ISL_1041375, EPI_ISL_1041376, EPI_ISL_1041377, EPI_ISL_1041378, EPI_ISL_1041379, EPI_ISL_1041380, EPI_ISL_1041381, EPI_ISL_1041382, EPI_ISL_1041383, EPI_ISL_1041384, EPI_ISL_1041385, EPI_ISL_1041386, EPI_ISL_1041387, EPI_ISL_1041388, EPI_ISL_1041389, EPI_ISL_1041390, EPI_ISL_1041391, EPI_ISL_1041392, EPI_ISL_1041393, EPI_ISL_1041394, EPI_ISL_1041395, EPI_ISL_1041396, EPI_ISL_1041397, EPI_ISL_1041398, EPI_ISL_1041399, EPI_ISL_1041400, EPI_ISL_1041401, EPI_ISL_1041402, EPI_ISL_1041403, EPI_ISL_1041404, EPI_ISL_1041405, EPI_ISL_1041406, EPI_ISL_1041407, EPI_ISL_1041408, EPI_ISL_1041409, EPI_ISL_1041410, EPI_ISL_1041411, EPI_ISL_1041412, EPI_ISL_1041413, EPI_ISL_1041414, EPI_ISL_1041415, EPI_ISL_1041416, EPI_ISL_1041417, EPI_ISL_1041418, EPI_ISL_1041419, EPI_ISL_1041420, EPI_ISL_1041421, EPI_ISL_1041422, EPI_ISL_1041423, EPI_ISL_1041424, EPI_ISL_1041425, EPI_ISL_1041426, EPI_ISL_1041427, EPI_ISL_1041428, EPI_ISL_1041429, EPI_ISL_1041430, EPI_ISL_1041431, EPI_ISL_1041432, EPI_ISL_1041433, EPI_ISL_1041434, EPI_ISL_1041435, EPI_ISL_1041436, EPI_ISL_1041437, EPI_ISL_1041438, EPI_ISL_1041439, EPI_ISL_1041440, EPI_ISL_1041441, EPI_ISL_1041442, EPI_ISL_1041443, EPI_ISL_1041444, EPI_ISL_1041445, EPI_ISL_1041446, EPI_ISL_1041447, EPI_ISL_1041448, EPI_ISL_1041449, EPI_ISL_1041450, EPI_ISL_1041451, EPI_ISL_1041452, EPI_ISL_1041453, EPI_ISL_1041454, EPI_ISL_1041455, EPI_ISL_1041456, EPI_ISL_1041457, EPI_ISL_1041458, EPI_ISL_1041459, EPI_ISL_1041460, EPI_ISL_1041461, EPI_ISL_1041462, EPI_ISL_1041463, EPI_ISL_1041464, EPI_ISL_1041465, EPI_ISL_1041466, EPI_ISL_1041467, EPI_ISL_1041468, EPI_ISL_1041469, EPI_ISL_1041470, EPI_ISL_1041471, EPI_ISL_1041472, EPI_ISL_1041473, EPI_ISL_1041474, EPI_ISL_1041475, EPI_ISL_1041476, EPI_ISL_1041477, EPI_ISL_1041478, EPI_ISL_1041479, EPI_ISL_1041480, EPI_ISL_1041481, EPI_ISL_1041482, EPI_ISL_1041483, EPI_ISL_1041484, EPI_ISL_1041485, EPI_ISL_1041486, EPI_ISL_1041487, EPI_ISL_1041488, EPI_ISL_1041489, EPI_ISL_1041490, EPI_ISL_1041491, EPI_ISL_1041492, EPI_ISL_1041493, EPI_ISL_1041494, EPI_ISL_1041495, EPI_ISL_1041496, EPI_ISL_1041497, EPI_ISL_1041498, EPI_ISL_1041499, EPI_ISL_1041500, EPI_ISL_1041501, EPI_ISL_1041502, EPI_ISL_1041503, EPI_ISL_1041504, EPI_ISL_1041505, EPI_ISL_1041506, EPI_ISL_1041507, EPI_ISL_1041508, EPI_ISL_1041510, EPI_ISL_1041511, EPI_ISL_1041512, EPI_ISL_1041513, EPI_ISL_1041514, EPI_ISL_1041515, EPI_ISL_1041516, EPI_ISL_1041517, EPI_ISL_1041518, EPI_ISL_1041519, EPI_ISL_1041520, EPI_ISL_1041521, EPI_ISL_1041522, EPI_ISL_1041523, EPI_ISL_1041524, EPI_ISL_1041525, EPI_ISL_1041526, EPI_ISL_1041527, EPI_ISL_1041528, EPI_ISL_1041529, EPI_ISL_1041530, EPI_ISL_1041531, EPI_ISL_1041532, EPI_ISL_1041533, EPI_ISL_1041534, EPI_ISL_1041535, EPI_ISL_1041536, EPI_ISL_1041537, EPI_ISL_1041538, EPI_ISL_1041539, EPI_ISL_1041540, EPI_ISL_1041541, EPI_ISL_1041542, EPI_ISL_1041543, EPI_ISL_1041544, EPI_ISL_1041545, EPI_ISL_1041546, EPI_ISL_1041547, EPI_ISL_1041548, EPI_ISL_1041549, EPI_ISL_1041550, EPI_ISL_1041551, EPI_ISL_1041552, EPI_ISL_1041553, EPI_ISL_1041554, EPI_ISL_1041555, EPI_ISL_1041556, EPI_ISL_1041557, EPI_ISL_1041558, EPI_ISL_1041559, EPI_ISL_1041560, EPI_ISL_1041561, EPI_ISL_1041562, EPI_ISL_1041563, EPI_ISL_1041564, EPI_ISL_1041565, EPI_ISL_1041566, EPI_ISL_1041567, EPI_ISL_1041568, EPI_ISL_1041569, EPI_ISL_1041570, EPI_ISL_1041571, EPI_ISL_1041572, EPI_ISL_1041573, EPI_ISL_1041574, EPI_ISL_1041575, EPI_ISL_1041576, EPI_ISL_1041577, EPI_ISL_1041578, EPI_ISL_1041579, EPI_ISL_1041580, EPI_ISL_1041581, EPI_ISL_1041582, EPI_ISL_1041583, EPI_ISL_1041584, EPI_ISL_1041585, EPI_ISL_1041586, EPI_ISL_1041587, EPI_ISL_1041588, EPI_ISL_1041589, EPI_ISL_1041590, EPI_ISL_1041591, EPI_ISL_1041592, EPI_ISL_1041593, EPI_ISL_1041594, EPI_ISL_1041595, EPI_ISL_1041596, EPI_ISL_1041597, EPI_ISL_1041598, EPI_ISL_1041599, EPI_ISL_1041600, EPI_ISL_1041601, EPI_ISL_1041602, EPI_ISL_1041603, EPI_ISL_1041604, EPI_ISL_1041605, EPI_ISL_1041606, EPI_ISL_1041607, EPI_ISL_1041608, EPI_ISL_1041609, EPI_ISL_1041610, EPI_ISL_1041611, EPI_ISL_1041612, EPI_ISL_1041613, EPI_ISL_1041614, EPI_ISL_1041615, EPI_ISL_1041616, EPI_ISL_1041617, EPI_ISL_1041618, EPI_ISL_1041619, EPI_ISL_1041620, EPI_ISL_1041621, EPI_ISL_1041622, EPI_ISL_1041623, EPI_ISL_1041624, EPI_ISL_1041625, EPI_ISL_1041626, EPI_ISL_1041627, EPI_ISL_1041628, EPI_ISL_1041629, EPI_ISL_1041630, EPI_ISL_1041631, EPI_ISL_1041632, EPI_ISL_1041633, EPI_ISL_1041634, EPI_ISL_1041635, EPI_ISL_1041636, EPI_ISL_1041637, EPI_ISL_1041638, EPI_ISL_1041639, EPI_ISL_1041640, EPI_ISL_1041641, EPI_ISL_1041642, EPI_ISL_1041643, EPI_ISL_1041644, EPI_ISL_1041645, EPI_ISL_1041646, EPI_ISL_1041647, EPI_ISL_1041648, EPI_ISL_1041649, EPI_ISL_1041650, EPI_ISL_1041651, EPI_ISL_1041652, EPI_ISL_1041653, EPI_ISL_1041654, EPI_ISL_1041655, EPI_ISL_1041656, EPI_ISL_1041657, EPI_ISL_1041658, EPI_ISL_1041659, EPI_ISL_1041660, EPI_ISL_1041661, EPI_ISL_1041662, EPI_ISL_1041663, EPI_ISL_1041664, EPI_ISL_1041665, EPI_ISL_1041666, EPI_ISL_1041667, EPI_ISL_1041668, EPI_ISL_1041669, EPI_ISL_1041670, EPI_ISL_1041671, EPI_ISL_1041672, EPI_ISL_1041673, EPI_ISL_1041674, EPI_ISL_1041675, EPI_ISL_1041676, EPI_ISL_1041677, EPI_ISL_1041678, EPI_ISL_1041679, EPI_ISL_1041680, EPI_ISL_1041681, EPI_ISL_1041682, EPI_ISL_1041683, EPI_ISL_1041684, EPI_ISL_1041685, EPI_ISL_1041686, EPI_ISL_1041687, EPI_ISL_1041688, EPI_ISL_1041689, EPI_ISL_1041690, EPI_ISL_1041691, EPI_ISL_1041692, EPI_ISL_1041693, EPI_ISL_1041694, EPI_ISL_1041695, EPI_ISL_1041696, EPI_ISL_1041697, EPI_ISL_1041698, EPI_ISL_1041699, EPI_ISL_1041700, EPI_ISL_1041701, EPI_ISL_1041702, EPI_ISL_1041703, EPI_ISL_1041704, EPI_ISL_1041705, EPI_ISL_1041706, EPI_ISL_1041707, EPI_ISL_1041708, EPI_ISL_1041709, EPI_ISL_1041710, EPI_ISL_1041711, EPI_ISL_1041712, EPI_ISL_1041713, EPI_ISL_1041714, EPI_ISL_1041715, EPI_ISL_1041716, EPI_ISL_1041717, EPI_ISL_1041718, EPI_ISL_1041719, EPI_ISL_1041720, EPI_ISL_1041721, EPI_ISL_1041722, EPI_ISL_1041723, EPI_ISL_1041724, EPI_ISL_1041725, EPI_ISL_1041726, EPI_ISL_1041727, EPI_ISL_1041728, EPI_ISL_1041729, EPI_ISL_1041730, EPI_ISL_1041731, EPI_ISL_1041732, EPI_ISL_1041733, EPI_ISL_1041734, EPI_ISL_1041735, EPI_ISL_1041736, EPI_ISL_1041737, EPI_ISL_1041738, EPI_ISL_1041739, EPI_ISL_1041740, EPI_ISL_1041741, EPI_ISL_1041742, EPI_ISL_1041743, EPI_ISL_1041744, EPI_ISL_1041745, EPI_ISL_1041746, EPI_ISL_1041747, EPI_ISL_1041748, EPI_ISL_1041749, EPI_ISL_1041750, EPI_ISL_1041751, EPI_ISL_1041752, EPI_ISL_1041753, EPI_ISL_1041754, EPI_ISL_1041755, EPI_ISL_1041756, EPI_ISL_1041757, EPI_ISL_1041758, EPI_ISL_1041759, EPI_ISL_1041760, EPI_ISL_1041761, EPI_ISL_1041762, EPI_ISL_1041763, EPI_ISL_1041764, EPI_ISL_1041765, EPI_ISL_1041766, EPI_ISL_1041767, EPI_ISL_1041768, EPI_ISL_1041769, EPI_ISL_1041770, EPI_ISL_1041771, EPI_ISL_1041772, EPI_ISL_1041773, EPI_ISL_1041774, EPI_ISL_1041775, EPI_ISL_1041776, EPI_ISL_1041777, EPI_ISL_1041778, EPI_ISL_1041779, EPI_ISL_1041780, EPI_ISL_1041781, EPI_ISL_1041782, EPI_ISL_1041783, EPI_ISL_1041784, EPI_ISL_1041785, EPI_ISL_1041786, EPI_ISL_1041787, EPI_ISL_1041788, EPI_ISL_1041789, EPI_ISL_1041790, EPI_ISL_1041791, EPI_ISL_1041792, EPI_ISL_1041793, EPI_ISL_1041794, EPI_ISL_1041795, EPI_ISL_1041796, EPI_ISL_1041797, EPI_ISL_1041798, EPI_ISL_1041799, EPI_ISL_1041800, EPI_ISL_1041801, EPI_ISL_1041802, EPI_ISL_1041803, EPI_ISL_1041804, EPI_ISL_1041805, EPI_ISL_1041806, EPI_ISL_1041807, EPI_ISL_1041808, EPI_ISL_1041809, EPI_ISL_1041810, EPI_ISL_1041811, EPI_ISL_1041812, EPI_ISL_1041813, EPI_ISL_1041814, EPI_ISL_1041815, EPI_ISL_1041816, EPI_ISL_1041817, EPI_ISL_1041818, EPI_ISL_1041819, EPI_ISL_1041820, EPI_ISL_1041821, EPI_ISL_1041822, EPI_ISL_1041823, EPI_ISL_1041824, EPI_ISL_1041825, EPI_ISL_1041826, EPI_ISL_1041827, EPI_ISL_1041828, EPI_ISL_1041829, EPI_ISL_1041830, EPI_ISL_1041831, EPI_ISL_1041832, EPI_ISL_1041833, EPI_ISL_1041834, EPI_ISL_1041835, EPI_ISL_1041836, EPI_ISL_1041837, EPI_ISL_1041838, EPI_ISL_1041839, EPI_ISL_1041840, EPI_ISL_1041841, EPI_ISL_1041842, EPI_ISL_1041843, EPI_ISL_1041844, EPI_ISL_1041845, EPI_ISL_1041846, EPI_ISL_1041847, EPI_ISL_1041848, EPI_ISL_1041849, EPI_ISL_1041850, EPI_ISL_1041851, EPI_ISL_1041852, EPI_ISL_1041853, EPI_ISL_1041854, EPI_ISL_1041855, EPI_ISL_1041856, EPI_ISL_1041857, EPI_ISL_1041858, EPI_ISL_1041859, EPI_ISL_1041860, EPI_ISL_1041861, EPI_ISL_1041862, EPI_ISL_1041863, EPI_ISL_1041864, EPI_ISL_1041865, EPI_ISL_1041866, EPI_ISL_1041867, EPI_ISL_1041868, EPI_ISL_1041869, EPI_ISL_1041870, EPI_ISL_1041871, EPI_ISL_1041872, EPI_ISL_1041873, EPI_ISL_1041874, EPI_ISL_1041875, EPI_ISL_1041876, EPI_ISL_1041877, EPI_ISL_1041878, EPI_ISL_1041879, EPI_ISL_1041880, EPI_ISL_1041881, EPI_ISL_1041882, EPI_ISL_1041883, EPI_ISL_1041884, EPI_ISL_1041885, EPI_ISL_1041886, EPI_ISL_1041887, EPI_ISL_1041888, EPI_ISL_1041889, EPI_ISL_1041890, EPI_ISL_1041891, EPI_ISL_1041892, EPI_ISL_1041893, EPI_ISL_1041894, EPI_ISL_1041895, EPI_ISL_1041896, EPI_ISL_1041897, EPI_ISL_1041898, EPI_ISL_1041899, EPI_ISL_1041900, EPI_ISL_1041901, EPI_ISL_1041902, EPI_ISL_1041903, EPI_ISL_1041904, EPI_ISL_1041905, EPI_ISL_1041906, EPI_ISL_1041907, EPI_ISL_1041908, EPI_ISL_1041909, EPI_ISL_1041910, EPI_ISL_1041911, EPI_ISL_1041912, EPI_ISL_1041913, EPI_ISL_1041914, EPI_ISL_1041915, EPI_ISL_1041916, EPI_ISL_1041917, EPI_ISL_1041918, EPI_ISL_1041919, EPI_ISL_1041920, EPI_ISL_1041921, EPI_ISL_1041922, EPI_ISL_1041923, EPI_ISL_1041924, EPI_ISL_1041925, EPI_ISL_1041926, EPI_ISL_1041927, EPI_ISL_1041928, EPI_ISL_1041929, EPI_ISL_1041930

| | | | |
|---|---|---|---|
| see above | Pandemic Response Lab - NYC | Pandemic Response Lab, R&D | Henry Lee, Michael Hammerling, Melissa Hopkins, Cybill del Castillo, William Ward, Pradeep Bugga, Haiping Hao, Jon Laurent |
| EPI_ISL_1049338, EPI_ISL_1049339 | Wadsworth Center, New York State Department.of Health | Wadsworth Center, New York State Department.of Health | Kirsten St. George, Daryl M. Lamson, Alexis Russel, Matthew Shudt, Melissa A Leisner, Jonathan Plitnick, Navjot Singh, John Kelly, Erasmus Schneider, Erica Lasek-Nesselquist |
| EPI_ISL_1049482 | MSHS Clinical Microbiology Laboratories | MSHS Pathogen Surveillance Program | Ana S. Gonzalez-Reiche, Hala Alshammary, Mitchell J. Sullivan, Brianne Ciferri, Ajay Obla, Angela Amoako, Mahmoud Awawda, Daniel Floda, Julia Matthews, Ashley Salimbangon, Levy Sominsky, Katherine Beach, Kayla Russo, Charles Gleason, Shelcie Fabre, Giulio Kleiner, Zenab Khan, Bremy Alburquerque, Adriana van de Guchte, Komal Srivastava, Matthew M. Hernandez, Jayeeta Dutta, Denise Jurczyszak, Nancy Francoeur, Betsaida Salom Melo, Irina Oussenko, Gintaras Deikus, Juan Soto, Shwetha Hara Sridhar, Ying-Chih Wang, Kathryn Twyman, Deena R. Altman, Robert Sebra, Adolfo García-Sastre, Marta Luksza, Gopi Patel, Sarah Schaefer, Melissa Gitman, Michael D. Nowak, Alberto Paniz-Mondolfi, Emilia Mia Sordillo, Viviana Simon, Harm van Bakel |

EPI_ISL_1050008, EPI_ISL_1050009, EPI_ISL_1050010, EPI_ISL_1050011, EPI_ISL_1050012, EPI_ISL_1050013, EPI_ISL_1050014, EPI_ISL_1050015, EPI_ISL_1050016, EPI_ISL_1050017, EPI_ISL_1050018, EPI_ISL_1050019, EPI_ISL_1050020, EPI_ISL_1050021, EPI_ISL_1050022, EPI_ISL_1050023, EPI_ISL_1050024, EPI_ISL_1050025, EPI_ISL_1050026, EPI_ISL_1050027, EPI_ISL_1050028, EPI_ISL_1050029, EPI_ISL_1050030, EPI_ISL_1050031, EPI_ISL_1050032, EPI_ISL_1050033, EPI_ISL_1050034, EPI_ISL_1050035, EPI_ISL_1050036, EPI_ISL_1050037, EPI_ISL_1050038, EPI_ISL_1050039, EPI_ISL_1050040, EPI_ISL_1050041, EPI_ISL_1050043, EPI_ISL_1050044, EPI_ISL_1050045, EPI_ISL_1050046, EPI_ISL_1050048, EPI_ISL_1050049, EPI_ISL_1050050, EPI_ISL_1050051, EPI_ISL_1050052, EPI_ISL_1050053, EPI_ISL_1050054, EPI_ISL_1050055, EPI_ISL_1050056, EPI_ISL_1050057, EPI_ISL_1050058, EPI_ISL_1050059, EPI_ISL_1050060, EPI_ISL_1050061, EPI_ISL_1050062, EPI_ISL_1050063, EPI_ISL_1050064

EPI_ISL_861369, EPI_ISL_861370, EPI_ISL_861371, EPI_ISL_861372, EPI_ISL_861373, EPI_ISL_861374, EPI_ISL_861375, EPI_ISL_861376, EPI_ISL_861377, EPI_ISL_861378, EPI_ISL_861379, EPI_ISL_861380, EPI_ISL_861381, EPI_ISL_861382, EPI_ISL_861383, EPI_ISL_861384, EPI_ISL_861385, EPI_ISL_861386, EPI_ISL_861387, EPI_ISL_861388, EPI_ISL_861389, EPI_ISL_861390, EPI_ISL_861391, EPI_ISL_861392, EPI_ISL_861393, EPI_ISL_861394, EPI_ISL_861395, EPI_ISL_861396, EPI_ISL_861397, EPI_ISL_861398, EPI_ISL_861399, EPI_ISL_861400, EPI_ISL_861401, EPI_ISL_861402, EPI_ISL_861403, EPI_ISL_861404, EPI_ISL_861405, EPI_ISL_861406, EPI_ISL_861407, EPI_ISL_861408, EPI_ISL_861409, EPI_ISL_861410, EPI_ISL_861411, EPI_ISL_861412

| | | | |
|---|---|---|---|
| | | | Erica Lasek-Nesselquist |
| EPI_ISL_896534 | New York Presbyterian Hospital | Wadsworth Center, New York State Department of Health | Kirsten St. George, Daryl M. Lamson, Alexis Russel, Matthew Shudt, Melissa A Leisner, Jonathan Plitnick, Navjot Singh, John Kelly, Erasmus Schneider, Erica Lasek-Nesselquist |
| EPI_ISL_896535, EPI_ISL_896536 | MEMORIAL SLOAN KETTERING CANCER CENTER | Wadsworth Center, New York State Department of Health | Kirsten St. George, Daryl M. Lamson, Alexis Russel, Matthew Shudt, Melissa A Leisner, Jonathan Plitnick, Navjot Singh, John Kelly, Erasmus Schneider, Erica Lasek-Nesselquist |
| EPI_ISL_896537, EPI_ISL_896538, EPI_ISL_896539, EPI_ISL_896540 | New York Presbyterian Hospital | Wadsworth Center, New York State Department of Health | Kirsten St. George, Daryl M. Lamson, Alexis Russel, Matthew Shudt, Melissa A Leisner, Jonathan Plitnick, Navjot Singh, John Kelly, Erasmus Schneider, Erica Lasek-Nesselquist |
| EPI_ISL_896541, EPI_ISL_896542, EPI_ISL_896543, EPI_ISL_896544, EPI_ISL_896545, EPI_ISL_896546, EPI_ISL_896547, EPI_ISL_896548 | MEMORIAL SLOAN KETTERING CANCER CENTER | Wadsworth Center, New York State Department of Health | Kirsten St. George, Daryl M. Lamson, Alexis Russel, Matthew Shudt, Melissa A Leisner, Jonathan Plitnick, Navjot Singh, John Kelly, Erasmus Schneider, Erica Lasek-Nesselquist |
| EPI_ISL_896549, EPI_ISL_896550, EPI_ISL_896551 | New York Presbyterian Hospital | Wadsworth Center, New York State Department of Health | Kirsten St. George, Daryl M. Lamson, Alexis Russel, Matthew Shudt, Melissa A Leisner, Jonathan Plitnick, Navjot Singh, John Kelly, Erasmus Schneider, Erica Lasek-Nesselquist |
| EPI_ISL_896552 | MEMORIAL SLOAN KETTERING CANCER CENTER | Wadsworth Center, New York State Department of Health | Kirsten St. George, Daryl M. Lamson, Alexis Russel, Matthew Shudt, Melissa A Leisner, Jonathan Plitnick, Navjot Singh, John Kelly, Erasmus Schneider, Erica Lasek-Nesselquist |
| EPI_ISL_896553, EPI_ISL_896554 | New York Presbyterian Hospital | Wadsworth Center, New York State Department of Health | Kirsten St. George, Daryl M. Lamson, Alexis Russel, Matthew Shudt, Melissa A Leisner, Jonathan Plitnick, Navjot Singh, John Kelly, Erasmus Schneider, Erica Lasek-Nesselquist |
| EPI_ISL_896555 | MEMORIAL SLOAN KETTERING CANCER CENTER | Wadsworth Center, New York State Department of Health | Kirsten St. George, Daryl M. Lamson, Alexis Russel, Matthew Shudt, Melissa A Leisner, Jonathan Plitnick, Navjot Singh, John Kelly, Erasmus Schneider, Erica Lasek-Nesselquist |
| EPI_ISL_896556, EPI_ISL_896557, EPI_ISL_896558, EPI_ISL_896559, EPI_ISL_896560, EPI_ISL_896561 | New York Presbyterian Hospital | Wadsworth Center, New York State Department of Health | Kirsten St. George, Daryl M. Lamson, Alexis Russel, Matthew Shudt, Melissa A Leisner, Jonathan Plitnick, Navjot Singh, John Kelly, Erasmus Schneider, Erica Lasek-Nesselquist |
| EPI_ISL_896562 | MEMORIAL SLOAN KETTERING CANCER CENTER | Wadsworth Center, New York State Department of Health | Kirsten St. George, Daryl M. Lamson, Alexis Russel, Matthew Shudt, Melissa A Leisner, Jonathan Plitnick, Navjot Singh, John Kelly, Erasmus Schneider, Erica Lasek-Nesselquist |
| EPI_ISL_896563, EPI_ISL_896564, EPI_ISL_896565, EPI_ISL_896566, EPI_ISL_896567, EPI_ISL_896568, EPI_ISL_896569 | URMC LABS | Wadsworth Center, New York State Department of Health | Kirsten St. George, Daryl M. Lamson, Alexis Russel, Matthew Shudt, Melissa A Leisner, Jonathan Plitnick, Navjot Singh, John Kelly, Erasmus Schneider, Erica Lasek-Nesselquist |
| EPI_ISL_896570, EPI_ISL_896571 | MEMORIAL SLOAN KETTERING CANCER CENTER | Wadsworth Center, New York State Department of Health | Kirsten St. George, Daryl M. Lamson, Alexis Russel, Matthew Shudt, Melissa A Leisner, Jonathan Plitnick, Navjot Singh, John Kelly, Erasmus Schneider, Erica Lasek-Nesselquist |
| EPI_ISL_903950, EPI_ISL_903977 | NYSDOH Wadsworth Center, Virology Lab | Genomics and Discovery, Respiratory Viruses Branch, Division of Viral Diseases, Centers for Disease Control and Prevention | Krista Queen, Yan Li, Ying Tao, Jing Zhang, Anna Uehara, Anna Montmayeur, Clinton R. Paden, Peter W. Cook, Rachel Marine, Mili Sheth, Jasmine Padilla, Sarah Nobles, Mark Burroughs, Lori Rowe, Haibin Wang, Ben L. Rambo-Martin, Dhwani Batra, Justin Lee, Suxiang Tong |
| EPI_ISL_912169 | East Gene | Grubaugh Lab - Yale School of Public Health | Tara Alpert, Joseph Fauver, Anderson Brito, Mallery Breban, Anne Wyllie, Chantal Vogels, Mary Petrone, Annie Watkins, Chaney Kalinich, Isabel Ott, Nathan Grubaugh |
| EPI_ISL_915340, EPI_ISL_915341, EPI_ISL_915342, EPI_ISL_915343, EPI_ISL_915344, EPI_ISL_915345 | Quest Diagnostics | Quest Diagnostics | Rosenthal,S.H., Gerasimova,A., Kagan,R.M., Anderson, B., Hua, M., Liu Y., Bernstein, L.E., Livingston, K.E., Perez, A., Shalhout, D.F., Shlyakhter, I.A., Owen, R., Tanpaiboon, P., Lacbawan, F. |
| EPI_ISL_935973, EPI_ISL_935974, EPI_ISL_935975, EPI_ISL_935976, EPI_ISL_935977, EPI_ISL_935978, EPI_ISL_935979, EPI_ISL_935980, EPI_ISL_935981, EPI_ISL_935982, EPI_ISL_935983, EPI_ISL_935984, EPI_ISL_935985, EPI_ISL_935986, EPI_ISL_935987, EPI_ISL_935988, EPI_ISL_935989, EPI_ISL_935990, EPI_ISL_935991, EPI_ISL_935992 | | | |
| see above | GLENS FALLS HOSPITAL LABORATORY | Wadsworth Center, New York State Department of Health | Kirsten St. George, Daryl M. Lamson, Alexis Russel, Matthew Shudt, Melissa A Leisner, Jonathan Plitnick, Navjot Singh, John Kelly, Erasmus Schneider, Erica Lasek-Nesselquist |
| EPI_ISL_935993, EPI_ISL_935994, EPI_ISL_935995, EPI_ISL_935996, EPI_ISL_935997, EPI_ISL_935998, EPI_ISL_935999, EPI_ISL_936000, EPI_ISL_936001, EPI_ISL_936002, EPI_ISL_936003, EPI_ISL_936004, EPI_ISL_936005, EPI_ISL_936006, EPI_ISL_936007, EPI_ISL_936008, EPI_ISL_936009, EPI_ISL_936010, EPI_ISL_936011, EPI_ISL_936012, EPI_ISL_936013, EPI_ISL_936014, EPI_ISL_936015, EPI_ISL_936016, EPI_ISL_936017, EPI_ISL_936018, EPI_ISL_936019, EPI_ISL_936020, EPI_ISL_936021, EPI_ISL_936022 | | | |
| see above | SUNY UPSTATE MEDICAL UNIVERSITY | Wadsworth Center, New York State Department of Health | Kirsten St. George, Daryl M. Lamson, Alexis Russel, Matthew Shudt, Melissa A Leisner, Jonathan Plitnick, Navjot Singh, John Kelly, Erasmus Schneider, Erica Lasek-Nesselquist |
| EPI_ISL_936023, EPI_ISL_936024, EPI_ISL_936025, EPI_ISL_936026, EPI_ISL_936027, EPI_ISL_936028, EPI_ISL_936029, EPI_ISL_936030, EPI_ISL_936031, EPI_ISL_936032, EPI_ISL_936033, EPI_ISL_936034, EPI_ISL_936035 | | | |
| see above | THE MARY IMOGENE BASSETT HOSPITAL | Wadsworth Center, New York State Department of Health | Kirsten St. George, Daryl M. Lamson, Alexis Russel, Matthew Shudt, Melissa A Leisner, Jonathan Plitnick, Navjot Singh, John Kelly, Erasmus Schneider, Erica Lasek-Nesselquist |
| EPI_ISL_936036, EPI_ISL_936037, EPI_ISL_936038, EPI_ISL_936039, EPI_ISL_936040, EPI_ISL_936041, EPI_ISL_936042, EPI_ISL_936043, EPI_ISL_936044, EPI_ISL_936045 | Columbia University Irving Medical Center | Wadsworth Center, New York State Department of Health | Kirsten St. George, Daryl M. Lamson, Alexis Russel, Matthew Shudt, Melissa A Leisner, Jonathan Plitnick, Navjot Singh, John Kelly, Erasmus Schneider, Erica Lasek-Nesselquist |
| EPI_ISL_936046, EPI_ISL_936047, EPI_ISL_936048, EPI_ISL_936049, EPI_ISL_936050, EPI_ISL_936051, EPI_ISL_936052, EPI_ISL_936053, EPI_ISL_936054, EPI_ISL_936055, EPI_ISL_936056, EPI_ISL_936057, EPI_ISL_936058, EPI_ISL_936059 | | | |
| see above | ADIRONDACK MEDICAL CENTER | Wadsworth Center, New York State Department of Health | Kirsten St. George, Daryl M. Lamson, Alexis Russel, Matthew Shudt, Melissa A Leisner, Jonathan Plitnick, Navjot Singh, John Kelly, Erasmus Schneider, Erica Lasek-Nesselquist |
| EPI_ISL_936060, EPI_ISL_936061, EPI_ISL_936062 | KALEIDA CENTER FOR LABORATORY MEDICINE | Wadsworth Center, New York State Department of Health | Kirsten St. George, Daryl M. Lamson, Alexis Russel, Matthew Shudt, Melissa A Leisner, Jonathan Plitnick, Navjot Singh, John Kelly, Erasmus Schneider, Erica Lasek-Nesselquist |
| EPI_ISL_936063, EPI_ISL_936064, EPI_ISL_936065, EPI_ISL_936066 | Columbia University Irving Medical Center | Wadsworth Center, New York State Department of Health | Kirsten St. George, Daryl M. Lamson, Alexis Russel, Matthew Shudt, Melissa A Leisner, Jonathan Plitnick, Navjot Singh, John Kelly, Erasmus Schneider, Erica Lasek-Nesselquist |
| EPI_ISL_936067, EPI_ISL_936068 | KALEIDA CENTER FOR LABORATORY MEDICINE | Wadsworth Center, New York State Department of Health | Kirsten St. George, Daryl M. Lamson, Alexis Russel, Matthew Shudt, Melissa A Leisner, Jonathan Plitnick, Navjot Singh, John Kelly, Erasmus Schneider, Erica Lasek-Nesselquist |
| EPI_ISL_936069 | Columbia University Irving Medical Center | Wadsworth Center, New York State Department of Health | Kirsten St. George, Daryl M. Lamson, Alexis Russel, Matthew Shudt, Melissa A Leisner, Jonathan Plitnick, Navjot Singh, John Kelly, Erasmus Schneider, Erica Lasek-Nesselquist |
| EPI_ISL_936070, EPI_ISL_936071, EPI_ISL_936072, EPI_ISL_936073, EPI_ISL_936074, EPI_ISL_936075, EPI_ISL_936076, EPI_ISL_936077, EPI_ISL_936078, EPI_ISL_936079, EPI_ISL_936080, EPI_ISL_936081 | | | |
| see above | KALEIDA CENTER FOR LABORATORY MEDICINE | Wadsworth Center, New York State Department of Health | Kirsten St. George, Daryl M. Lamson, Alexis Russel, Matthew Shudt, Melissa A Leisner, Jonathan Plitnick, Navjot Singh, John Kelly, Erasmus Schneider, Erica Lasek-Nesselquist |
| EPI_ISL_936082 | Columbia University Irving Medical Center | Wadsworth Center, New York State Department of Health | Kirsten St. George, Daryl M. Lamson, Alexis Russel, Matthew Shudt, Melissa A Leisner, Jonathan Plitnick, Navjot Singh, John Kelly, Erasmus Schneider, Erica Lasek-Nesselquist |
| EPI_ISL_936083, EPI_ISL_936084, EPI_ISL_936085 | KALEIDA CENTER FOR LABORATORY MEDICINE | Wadsworth Center, New York State Department of Health | Kirsten St. George, Daryl M. Lamson, Alexis Russel, Matthew Shudt, Melissa A Leisner, Jonathan Plitnick, Navjot Singh, John Kelly, Erasmus Schneider, Erica Lasek-Nesselquist |

EPI_ISL_937256, EPI_ISL_937257, EPI_ISL_937258, EPI_ISL_937259, EPI_ISL_937260, EPI_ISL_937261, EPI_ISL_937262, EPI_ISL_937263, EPI_ISL_937264, EPI_ISL_937265, EPI_ISL_937266, EPI_ISL_937267, EPI_ISL_937268, EPI_ISL_937269, EPI_ISL_937270, EPI_ISL_937271, EPI_ISL_937272

| | | | |
|---|---|---|---|
| see above | NYC HH Elmhurst Hospital Medical Center | New York City Public Health Laboratory | Jade Wang, et al. |

| | | | |
|---|---|---|---|
| EPI_ISL_955265 | Sunrise Medical Laboratory | Pathogen Discovery, Respiratory Viruses Branch, Division of Viral Diseases, Centers for Disease Control and Prevention | Ying Tao, Jing Zhang, Yan Li, Krista Queen, Anna Uehara, Peter Cook, Clinton R. Paden, Haibin Wang, Suxiang Tong |
| EPI_ISL_955271 | American Esoteric Laboratory | Pathogen Discovery, Respiratory Viruses Branch, Division of Viral Diseases, Centers for Disease Control and Prevention | Ying Tao, Jing Zhang, Yan Li, Krista Queen, Anna Uehara, Peter Cook, Clinton R. Paden, Haibin Wang, Suxiang Tong |
| EPI_ISL_955300 | Sunrise Medical Laboratory | Pathogen Discovery, Respiratory Viruses Branch, Division of Viral Diseases, Centers for Disease Control and Prevention | Ying Tao, Jing Zhang, Yan Li, Krista Queen, Anna Uehara, Peter Cook, Clinton R. Paden, Haibin Wang, Suxiang Tong |
| EPI_ISL_961289, EPI_ISL_961290, EPI_ISL_961291, EPI_ISL_961292, EPI_ISL_961293, EPI_ISL_961294, EPI_ISL_961295, EPI_ISL_961296, EPI_ISL_961297, EPI_ISL_961298, EPI_ISL_961299, EPI_ISL_961300, EPI_ISL_961301, EPI_ISL_961302, EPI_ISL_961303, EPI_ISL_961304, EPI_ISL_961305, EPI_ISL_961306, EPI_ISL_961307, EPI_ISL_961308, EPI_ISL_961309, EPI_ISL_961310, EPI_ISL_961311, EPI_ISL_961312, EPI_ISL_961313, EPI_ISL_961314, EPI_ISL_961315, EPI_ISL_961328, EPI_ISL_961329, EPI_ISL_961330, EPI_ISL_961331, EPI_ISL_961332, EPI_ISL_961333, EPI_ISL_961334, EPI_ISL_961335, EPI_ISL_961336, EPI_ISL_961337, EPI_ISL_961338, EPI_ISL_961339, EPI_ISL_961340, EPI_ISL_961341, EPI_ISL_961342, EPI_ISL_961343, EPI_ISL_961344, EPI_ISL_961345, EPI_ISL_961346, EPI_ISL_961347, EPI_ISL_961348, EPI_ISL_961349, EPI_ISL_961350, EPI_ISL_961351 | | | |
| see above | Biotia | Biotia | Dorottya Nagy-Szakal, Mara Couto-Rodriguez, Heather Wells, Joseph Barrows, Marilyne Debieu, Kristin Butcher, Siyuan Chen, Agnes Berki, Courteny Hager, Robert Boorstein, Mariah Taylor, Colleen Jonsson, Christopher Mason, Niamh O'Hara |
| EPI_ISL_965303, EPI_ISL_965528, EPI_ISL_965770, EPI_ISL_965812, EPI_ISL_965814, EPI_ISL_965826, EPI_ISL_965837, EPI_ISL_965838, EPI_ISL_965839, EPI_ISL_965910, EPI_ISL_965948, EPI_ISL_965962, EPI_ISL_965963, EPI_ISL_965964, EPI_ISL_965965, EPI_ISL_965967, EPI_ISL_965968, EPI_ISL_965969, EPI_ISL_965970, EPI_ISL_965971, EPI_ISL_965972, EPI_ISL_965973, EPI_ISL_965977, EPI_ISL_965980, EPI_ISL_965989, EPI_ISL_966020, EPI_ISL_966044, EPI_ISL_966062, EPI_ISL_966084, EPI_ISL_966108, EPI_ISL_966160, EPI_ISL_966184, EPI_ISL_966204, EPI_ISL_966227, EPI_ISL_966245, EPI_ISL_966264, EPI_ISL_966289, EPI_ISL_966308, EPI_ISL_966311, EPI_ISL_966333, EPI_ISL_966357, EPI_ISL_966375, EPI_ISL_966376, EPI_ISL_966377, EPI_ISL_966378 | | | |
| see above | NYU Langone Health | Departments of Pathology and Medicine, New York University School of Medicine | Adriana Heguy, Dacia Dimartino, Emily Guzman, Christian Marier, Peter Meyn, Sitharam Ramaswami, Gael Westby, Paul Zappile, Yutong Zhang, Paolo Cotzia, Guiqing Wang |
| EPI_ISL_966379 | OCME Office Of Chief Medical Examiner | New York City Public Health Laboratory | Jade Wang, et al. |
| EPI_ISL_966380 | NYU Langone Health | Departments of Pathology and Medicine, New York University School of Medicine | Adriana Heguy, Dacia Dimartino, Emily Guzman, Christian Marier, Peter Meyn, Sitharam Ramaswami, Gael Westby, Paul Zappile, Yutong Zhang, Paolo Cotzia, Guiqing Wang |
| EPI_ISL_966381 | OCME Office Of Chief Medical Examiner | New York City Public Health Laboratory | Jade Wang, et al. |
| EPI_ISL_966382 | DOHMH Jamaica | New York City Public Health Laboratory | Jade Wang, et al. |
| EPI_ISL_966383 | OCME Office Of Chief Medical Examiner | New York City Public Health Laboratory | Jade Wang, et al. |
| EPI_ISL_966384 | DOHMH Jamaica | New York City Public Health Laboratory | Jade Wang, et al. |
| EPI_ISL_966385 | Department of Homeless Services | New York City Public Health Laboratory | Jade Wang, et al. |
| EPI_ISL_966386 | DOHMH Corona | New York City Public Health Laboratory | Jade Wang, et al. |
| EPI_ISL_966387, EPI_ISL_966388 | DOHMH Jamaica | New York City Public Health Laboratory | Jade Wang, et al. |
| EPI_ISL_966389 | DOHMH Central Harlem | New York City Public Health Laboratory | Jade Wang, et al. |
| EPI_ISL_966390, EPI_ISL_966391 | OCME Office Of Chief Medical Examiner | New York City Public Health Laboratory | Jade Wang, et al. |
| EPI_ISL_966392 | DOHMH Corona | New York City Public Health Laboratory | Jade Wang, et al. |
| EPI_ISL_966393 | OCME Office Of Chief Medical Examiner | New York City Public Health Laboratory | Jade Wang, et al. |
| EPI_ISL_966394 | DOHMH Corona | New York City Public Health Laboratory | Jade Wang, et al. |
| EPI_ISL_966395 | DOHMH Jamaica | New York City Public Health Laboratory | Jade Wang, et al. |
| EPI_ISL_966396, EPI_ISL_966397 | OCME Office Of Chief Medical Examiner | New York City Public Health Laboratory | Jade Wang, et al. |
| EPI_ISL_966398 | DOHMH Jamaica | New York City Public Health Laboratory | Jade Wang, et al. |
| EPI_ISL_966399 | DOHMH Corona | New York City Public Health Laboratory | Jade Wang, et al. |
| EPI_ISL_966400 | OCME Office Of Chief Medical Examiner | New York City Public Health Laboratory | Jade Wang, et al. |
| EPI_ISL_966401 | Department of Homeless Services | New York City Public Health Laboratory | Jade Wang, et al. |
| EPI_ISL_966402, EPI_ISL_966403 | OCME Office Of Chief Medical Examiner | New York City Public Health Laboratory | Jade Wang, et al. |
| EPI_ISL_966404 | DOHMH PHL | New York City Public Health Laboratory | Jade Wang, et al. |
| EPI_ISL_966405 | DOHMH Jamaica | New York City Public Health Laboratory | Jade Wang, et al. |
| EPI_ISL_966406 | NYU Langone Health | Departments of Pathology and Medicine, New York University School of Medicine | Adriana Heguy, Dacia Dimartino, Emily Guzman, Christian Marier, Peter Meyn, Sitharam Ramaswami, Gael Westby, Paul Zappile, Yutong Zhang, Paolo Cotzia, Guiqing Wang |
| EPI_ISL_966407, EPI_ISL_966408 | DOHMH Morrisania | New York City Public Health Laboratory | Jade Wang, et al. |
| EPI_ISL_966409, EPI_ISL_966410 | DOHMH Corona | New York City Public Health Laboratory | Jade Wang, et al. |
| EPI_ISL_966411 | OCME Office Of Chief Medical Examiner | New York City Public Health Laboratory | Jade Wang, et al. |
| EPI_ISL_966412 | DOHMH Jamaica | New York City Public Health Laboratory | Jade Wang, et al. |
| EPI_ISL_966413 | DOHMH Central Harlem | New York City Public Health Laboratory | Jade Wang, et al. |
| EPI_ISL_966414 | DOHMH Morrisania | New York City Public Health Laboratory | Jade Wang, et al. |
| EPI_ISL_966415 | DOHMH Corona | New York City Public Health Laboratory | Jade Wang, et al. |
| EPI_ISL_966416 | DOHMH Central Harlem | New York City Public Health Laboratory | Jade Wang, et al. |
| EPI_ISL_966417, EPI_ISL_966418, EPI_ISL_966419 | OCME Office Of Chief Medical Examiner | New York City Public Health Laboratory | Jade Wang, et al. |
| EPI_ISL_966420 | DOHMH Morrisania | New York City Public Health Laboratory | Jade Wang, et al. |
| EPI_ISL_966421, EPI_ISL_966422 | DOHMH Jamaica | New York City Public Health Laboratory | Jade Wang, et al. |
| EPI_ISL_966423 | DOHMH Morrisania | New York City Public Health Laboratory | Jade Wang, et al. |
| EPI_ISL_966424 | DOHMH Jamaica | New York City Public Health Laboratory | Jade Wang, et al. |
| EPI_ISL_966425 | DOHMH Corona | New York City Public Health Laboratory | Jade Wang, et al. |
| EPI_ISL_966426 | DOHMH PHL | New York City Public Health Laboratory | Jade Wang, et al. |
| EPI_ISL_966427 | OCME Office Of Chief Medical Examiner | New York City Public Health Laboratory | Jade Wang, et al. |
| EPI_ISL_966428 | NYU Langone Health | Departments of Pathology and Medicine, New York University School of Medicine | Adriana Heguy, Dacia Dimartino, Emily Guzman, Christian Marier, Peter Meyn, Sitharam Ramaswami, Gael Westby, Paul Zappile, Yutong Zhang, Paolo Cotzia, Guiqing Wang |
| EPI_ISL_966429, EPI_ISL_966430, EPI_ISL_966431 | OCME Office Of Chief Medical Examiner | New York City Public Health Laboratory | Jade Wang, et al. |

EPI_ISL_966952, EPI_ISL_967055, EPI_ISL_967085, EPI_ISL_967137, EPI_ISL_967207, EPI_ISL_967230, EPI_ISL_967296, EPI_ISL_967332, EPI_ISL_967358, EPI_ISL_967678, EPI_ISL_967789, EPI_ISL_967880

| | | | |
|---|---|---|---|
| see above | Helix/Illumina | Respiratory Viruses Branch, Division of Viral Diseases, Centers for Disease Control and Prevention | Peter W. Cook,Dakota Howard,Dhwani Batra,Ben L. Rambo-Martin,Eileen de Feo,Jan Antico,Christine Tran,Matthew Tolentino,Shannon Wickline,Kim Gietzen,Brad Sickler,Jingtao Liu,Eric Allen,Phil Febbo,Summer Galloway,Nicole L. Washington,Simon White,Geraint Levan,Kelly Schiabor Barrett,Elizabeth Cirulli,Alexandre Bolze,Ary Ascencio,Charlotte Rivera-Garcia,Ryan Cho,Jason Nguyen,Sherry Wang,Jimmy Ramirez,Tyler Cassens,Efren Sandoval,Magnus Isaksson,William Lee,David Becker,Marc Laurent,James Lu,Clinton R. Paden,Suxiang Tong,Duncan MacCannell, |
| EPI_ISL_968080 | Pandemic Response Lab - NYC | Pandemic Response Lab, R&D | Henry Lee, Michael Hammerling, Melissa Hopkins, Cybill del Castillo, William Ward, Pradeep Bugga, Haiping Hao, Jon Laurent |
| EPI_ISL_977666 | NYU Langone Health | Departments of Pathology and Medicine, New York University School of Medicine | Adriana Heguy, Dacia Dimartino, Emily Guzman, Christian Marier, Peter Meyn, Sitharam Ramaswami, Gael Westby, Paul Zappile, Yutong Zhang, Paolo Cotzia, Guiqing Wang |
| EPI_ISL_978603, EPI_ISL_978726, EPI_ISL_978736, EPI_ISL_978741 | Helix/Illumina | Respiratory Viruses Branch, Division of Viral Diseases, Centers for Disease Control and Prevention | Peter W. Cook,Dakota Howard,Dhwani Batra,Ben L. Rambo-Martin,Eileen de Feo,Jan Antico,Christine Tran,Matthew Tolentino,Shannon Wickline,Kim Gietzen,Brad Sickler,Jingtao Liu,Eric Allen,Phil Febbo,Summer Galloway,Nicole L. Washington,Simon White,Geraint Levan,Kelly Schiabor Barrett,Elizabeth Cirulli,Alexandre Bolze,Ary Ascencio,Charlotte Rivera-Garcia,Ryan Cho,Jason Nguyen,Sherry Wang,Jimmy Ramirez,Tyler Cassens,Efren Sandoval,Magnus Isaksson,William Lee,David Becker,Marc Laurent,James Lu,Clinton R. Paden,Suxiang Tong,Duncan MacCannell, |
| EPI_ISL_982425, EPI_ISL_982426, EPI_ISL_982427, EPI_ISL_982428, EPI_ISL_982429, EPI_ISL_982430, EPI_ISL_982431, EPI_ISL_982432, EPI_ISL_982433, EPI_ISL_982434, EPI_ISL_982435, EPI_ISL_982436, EPI_ISL_982437, EPI_ISL_982438, EPI_ISL_982439, EPI_ISL_982440, EPI_ISL_982441, EPI_ISL_982442, EPI_ISL_982443, EPI_ISL_982444, EPI_ISL_982445, EPI_ISL_982446, EPI_ISL_982447, EPI_ISL_982448, EPI_ISL_982449, EPI_ISL_982450, EPI_ISL_982451, EPI_ISL_982452, EPI_ISL_982453, EPI_ISL_982454, EPI_ISL_982455, EPI_ISL_982456, EPI_ISL_982457, EPI_ISL_982458, EPI_ISL_982459, EPI_ISL_982460, EPI_ISL_982461, EPI_ISL_982462, EPI_ISL_982463, EPI_ISL_982464, EPI_ISL_982465, EPI_ISL_982466, EPI_ISL_982467, EPI_ISL_982468, EPI_ISL_982469, EPI_ISL_982470, EPI_ISL_982471, EPI_ISL_982472, EPI_ISL_982473, EPI_ISL_982474, EPI_ISL_982475, EPI_ISL_982476, EPI_ISL_982477, EPI_ISL_982478, EPI_ISL_982479, EPI_ISL_982480, EPI_ISL_982481, EPI_ISL_982482, EPI_ISL_982483, EPI_ISL_982484, EPI_ISL_982485, EPI_ISL_982486, EPI_ISL_982487, EPI_ISL_982488, EPI_ISL_982489, EPI_ISL_982490, EPI_ISL_982491, EPI_ISL_982492, EPI_ISL_982493, EPI_ISL_982494, EPI_ISL_982495, EPI_ISL_982496, EPI_ISL_982497, EPI_ISL_982498 | | | |
| see above | MONTEFIORE MEDICAL CENTER LABORATORIES | Wadsworth Center, New York State Department of Health | Kirsten St. George, Daryl M. Lamson, Alexis Russel, Matthew Shudt, Melissa A Leisner, Jonathan Plitnick, Navjot Singh, John Kelly, Erasmus Schneider, Erica Lasek-Nesselquist |
| EPI_ISL_982499, EPI_ISL_982500, EPI_ISL_982501, EPI_ISL_982502, EPI_ISL_982503, EPI_ISL_982504 | ADIRONDACK MEDICAL CENTER | Wadsworth Center, New York State Department of Health | Kirsten St. George, Daryl M. Lamson, Alexis Russel, Matthew Shudt, Melissa A Leisner, Jonathan Plitnick, Navjot Singh, John Kelly, Erasmus Schneider, Erica Lasek-Nesselquist |
| EPI_ISL_983099, EPI_ISL_983100, EPI_ISL_983101, EPI_ISL_983102, EPI_ISL_983103, EPI_ISL_983104, EPI_ISL_983105, EPI_ISL_983106, EPI_ISL_983107, EPI_ISL_983108, EPI_ISL_983109, EPI_ISL_983110, EPI_ISL_983111, EPI_ISL_983112, EPI_ISL_983113, EPI_ISL_983114, EPI_ISL_983115 | | | |
| see above | MONTEFIORE MEDICAL CENTER LABORATORIES | Wadsworth Center, New York State Department of Health | Kirsten St. George, Daryl M. Lamson, Alexis Russel, Matthew Shudt, Melissa A Leisner, Jonathan Plitnick, Navjot Singh, John Kelly, Erasmus Schneider, Erica Lasek-Nesselquist |
| EPI_ISL_983116, EPI_ISL_983117, EPI_ISL_983118 | SUNY UPSTATE MEDICAL UNIVERSITY | Wadsworth Center, New York State Department of Health | Kirsten St. George, Daryl M. Lamson, Alexis Russel, Matthew Shudt, Melissa A Leisner, Jonathan Plitnick, Navjot Singh, John Kelly, Erasmus Schneider, Erica Lasek-Nesselquist |
| EPI_ISL_983119, EPI_ISL_983120, EPI_ISL_983121, EPI_ISL_983122, EPI_ISL_983123, EPI_ISL_983124, EPI_ISL_983125, EPI_ISL_983126, EPI_ISL_983127, EPI_ISL_983128, EPI_ISL_983129, EPI_ISL_983130, EPI_ISL_983131 | | | |
| see above | THE MARY IMOGENE BASSETT HOSPITAL | Wadsworth Center, New York State Department of Health | Kirsten St. George, Daryl M. Lamson, Alexis Russel, Matthew Shudt, Melissa A Leisner, Jonathan Plitnick, Navjot Singh, John Kelly, Erasmus Schneider, Erica Lasek-Nesselquist |
| EPI_ISL_983132, EPI_ISL_983133, EPI_ISL_983134, EPI_ISL_983135, EPI_ISL_983136, EPI_ISL_983137, EPI_ISL_983138, EPI_ISL_983139, EPI_ISL_983140, EPI_ISL_983141, EPI_ISL_983142, EPI_ISL_983143, EPI_ISL_983144, EPI_ISL_983145, EPI_ISL_983146, EPI_ISL_983147, EPI_ISL_983148, EPI_ISL_983149, EPI_ISL_983150, EPI_ISL_983151, EPI_ISL_983152, EPI_ISL_983153, EPI_ISL_983154, EPI_ISL_983155, EPI_ISL_983156, EPI_ISL_983157, EPI_ISL_983158, EPI_ISL_983159, EPI_ISL_983160, EPI_ISL_983161, EPI_ISL_983162, EPI_ISL_983163, EPI_ISL_983164, EPI_ISL_983165, EPI_ISL_983166, EPI_ISL_983167, EPI_ISL_983168, EPI_ISL_983169, EPI_ISL_983170 | | | |
| see above | MONTEFIORE MEDICAL CENTER LABORATORIES | Wadsworth Center, New York State Department of Health | Kirsten St. George, Daryl M. Lamson, Alexis Russel, Matthew Shudt, Melissa A Leisner, Jonathan Plitnick, Navjot Singh, John Kelly, Erasmus Schneider, Erica Lasek-Nesselquist |
| EPI_ISL_983240, EPI_ISL_983241, EPI_ISL_983242 | KALEIDA CENTER FOR LABORATORY MEDICINE | Wadsworth Center, New York State Department of Health | Kirsten St. George, Daryl M. Lamson, Alexis Russel, Matthew Shudt, Melissa A Leisner, Jonathan Plitnick, Navjot Singh, John Kelly, Erasmus Schneider, Erica Lasek-Nesselquist |
| EPI_ISL_983243 | SUNY UPSTATE MEDICAL UNIVERSITY | Wadsworth Center, New York State Department of Health | Kirsten St. George, Daryl M. Lamson, Alexis Russel, Matthew Shudt, Melissa A Leisner, Jonathan Plitnick, Navjot Singh, John Kelly, Erasmus Schneider, Erica Lasek-Nesselquist |
| EPI_ISL_983244 | KALEIDA CENTER FOR LABORATORY MEDICINE | Wadsworth Center, New York State Department of Health | Kirsten St. George, Daryl M. Lamson, Alexis Russel, Matthew Shudt, Melissa A Leisner, Jonathan Plitnick, Navjot Singh, John Kelly, Erasmus Schneider, Erica Lasek-Nesselquist |
| EPI_ISL_983245 | SUNY UPSTATE MEDICAL UNIVERSITY | Wadsworth Center, New York State Department of Health | Kirsten St. George, Daryl M. Lamson, Alexis Russel, Matthew Shudt, Melissa A Leisner, Jonathan Plitnick, Navjot Singh, John Kelly, Erasmus Schneider, Erica Lasek-Nesselquist |
| EPI_ISL_983246, EPI_ISL_983247 | KALEIDA CENTER FOR LABORATORY MEDICINE | Wadsworth Center, New York State Department of Health | Kirsten St. George, Daryl M. Lamson, Alexis Russel, Matthew Shudt, Melissa A Leisner, Jonathan Plitnick, Navjot Singh, John Kelly, Erasmus Schneider, Erica Lasek-Nesselquist |
| EPI_ISL_983248, EPI_ISL_983249 | SUNY UPSTATE MEDICAL UNIVERSITY | Wadsworth Center, New York State Department of Health | Kirsten St. George, Daryl M. Lamson, Alexis Russel, Matthew Shudt, Melissa A Leisner, Jonathan Plitnick, Navjot Singh, John Kelly, Erasmus Schneider, Erica Lasek-Nesselquist |
| EPI_ISL_983250, EPI_ISL_983251, EPI_ISL_983252, EPI_ISL_983253, EPI_ISL_983254, EPI_ISL_983255, EPI_ISL_983256, EPI_ISL_983257, EPI_ISL_983258, EPI_ISL_983259, EPI_ISL_983260, EPI_ISL_983261, EPI_ISL_983262, EPI_ISL_983263, EPI_ISL_983264, EPI_ISL_983265, EPI_ISL_983266, EPI_ISL_983267, EPI_ISL_983268, EPI_ISL_983269, EPI_ISL_983270, EPI_ISL_983271, EPI_ISL_983272, EPI_ISL_983273, EPI_ISL_983274, EPI_ISL_983275, EPI_ISL_983276, EPI_ISL_983277, EPI_ISL_983278, EPI_ISL_983279, EPI_ISL_983280, EPI_ISL_983281, EPI_ISL_983282 | | | |
| see above | KALEIDA CENTER FOR LABORATORY MEDICINE | Wadsworth Center, New York State Department of Health | Kirsten St. George, Daryl M. Lamson, Alexis Russel, Matthew Shudt, Melissa A Leisner, Jonathan Plitnick, Navjot Singh, John Kelly, Erasmus Schneider, Erica Lasek-Nesselquist |
| EPI_ISL_983283 | SUNY UPSTATE MEDICAL UNIVERSITY | Wadsworth Center, New York State Department of Health | Kirsten St. George, Daryl M. Lamson, Alexis Russel, Matthew Shudt, Melissa A Leisner, Jonathan Plitnick, Navjot Singh, John Kelly, Erasmus Schneider, Erica Lasek-Nesselquist |
| EPI_ISL_983284, EPI_ISL_983285, EPI_ISL_983286, EPI_ISL_983287, EPI_ISL_983288 | KALEIDA CENTER FOR LABORATORY MEDICINE | Wadsworth Center, New York State Department of Health | Kirsten St. George, Daryl M. Lamson, Alexis Russel, Matthew Shudt, Melissa A Leisner, Jonathan Plitnick, Navjot Singh, John Kelly, Erasmus Schneider, Erica Lasek-Nesselquist |
| EPI_ISL_983289, EPI_ISL_983290, EPI_ISL_983291, EPI_ISL_983292, EPI_ISL_983293, EPI_ISL_983294, EPI_ISL_983295 | SUNY UPSTATE MEDICAL UNIVERSITY | Wadsworth Center, New York State Department of Health | Kirsten St. George, Daryl M. Lamson, Alexis Russel, Matthew Shudt, Melissa A Leisner, Jonathan Plitnick, Navjot Singh, John Kelly, Erasmus Schneider, Erica Lasek-Nesselquist |
| EPI_ISL_983296, EPI_ISL_983297, EPI_ISL_983298, EPI_ISL_983299 | KALEIDA CENTER FOR LABORATORY MEDICINE | Wadsworth Center, New York State Department of Health | Kirsten St. George, Daryl M. Lamson, Alexis Russel, Matthew Shudt, Melissa A Leisner, Jonathan Plitnick, Navjot Singh, John Kelly, Erasmus Schneider, Erica Lasek-Nesselquist |
| EPI_ISL_983300, EPI_ISL_983301, EPI_ISL_983302, EPI_ISL_983303, EPI_ISL_983304, EPI_ISL_983305, EPI_ISL_983306, EPI_ISL_983307, EPI_ISL_983308, EPI_ISL_983309, EPI_ISL_983310, EPI_ISL_983311, EPI_ISL_983312, EPI_ISL_983313, EPI_ISL_983314, EPI_ISL_983315, EPI_ISL_983316, EPI_ISL_983317, EPI_ISL_983318, EPI_ISL_983319 | | | |
| see above | SUNY UPSTATE MEDICAL UNIVERSITY | Wadsworth Center, New York State Department of Health | Kirsten St. George, Daryl M. Lamson, Alexis Russel, Matthew Shudt, Melissa A Leisner, Jonathan Plitnick, Navjot Singh, John Kelly, Erasmus Schneider, Erica Lasek-Nesselquist |
| EPI_ISL_983438, EPI_ISL_983439, EPI_ISL_983440, EPI_ISL_983441, EPI_ISL_983442, EPI_ISL_983443, EPI_ISL_983444 | URMC LABS | Wadsworth Center, New York State Department of Health | Kirsten St. George, Daryl M. Lamson, Alexis Russel, Matthew Shudt, Melissa A Leisner, Jonathan Plitnick, Navjot Singh, John Kelly, Erasmus Schneider, Erica Lasek-Nesselquist |
| EPI_ISL_983445, EPI_ISL_983446 | THE MARY IMOGENE BASSETT HOSPITAL | Wadsworth Center, New York State Department of Health | Kirsten St. George, Daryl M. Lamson, Alexis Russel, Matthew Shudt, Melissa A Leisner, Jonathan Plitnick, Navjot Singh, John Kelly, Erasmus Schneider, Erica Lasek-Nesselquist |
| EPI_ISL_983447, EPI_ISL_983448, EPI_ISL_983449, EPI_ISL_983450, EPI_ISL_983451, EPI_ISL_983452, EPI_ISL_983453, EPI_ISL_983454, EPI_ISL_983455, EPI_ISL_983456, EPI_ISL_983457, EPI_ISL_983458, EPI_ISL_983459, EPI_ISL_983460, EPI_ISL_983461, EPI_ISL_983462, EPI_ISL_983463, EPI_ISL_983464, EPI_ISL_983465, EPI_ISL_983466, EPI_ISL_983467, EPI_ISL_983468, EPI_ISL_983469, EPI_ISL_983470, EPI_ISL_983471, EPI_ISL_983472, EPI_ISL_983473, EPI_ISL_983474, EPI_ISL_983475 | | | |

| | | | |
|---|---|---|---|
| see above | URMC LABS | Wadsworth Center, New York State Department of Health | Kirsten St. George, Daryl M. Lamson, Alexis Russel, Matthew Shudt, Melissa A Leisner, Jonathan Plitnick, Navjot Singh, John Kelly, Erasmus Schneider, Erica Lasek-Nesselquist |
| EPI_ISL_983476 | Wadsworth Center, New York State Department of Health | Wadsworth Center, New York State Department of Health | Kirsten St. George, Daryl M. Lamson, Alexis Russel, Matthew Shudt, Melissa A Leisner, Jonathan Plitnick, Navjot Singh, John Kelly, Erasmus Schneider, Erica Lasek-Nesselquist |
| EPI_ISL_983477 | THE MARY IMOGENE BASSETT HOSPITAL | Wadsworth Center, New York State Department of Health | Kirsten St. George, Daryl M. Lamson, Alexis Russel, Matthew Shudt, Melissa A Leisner, Jonathan Plitnick, Navjot Singh, John Kelly, Erasmus Schneider, Erica Lasek-Nesselquist |
| EPI_ISL_983478, EPI_ISL_983479, EPI_ISL_983480, EPI_ISL_983481, EPI_ISL_983482, EPI_ISL_983483, EPI_ISL_983484 | URMC LABS | Wadsworth Center, New York State Department of Health | Kirsten St. George, Daryl M. Lamson, Alexis Russel, Matthew Shudt, Melissa A Leisner, Jonathan Plitnick, Navjot Singh, John Kelly, Erasmus Schneider, Erica Lasek-Nesselquist |
| EPI_ISL_983485, EPI_ISL_983486, EPI_ISL_983487, EPI_ISL_983488, EPI_ISL_983489, EPI_ISL_983490, EPI_ISL_983491, EPI_ISL_983492 | THE MARY IMOGENE BASSETT HOSPITAL | Wadsworth Center, New York State Department of Health | Kirsten St. George, Daryl M. Lamson, Alexis Russel, Matthew Shudt, Melissa A Leisner, Jonathan Plitnick, Navjot Singh, John Kelly, Erasmus Schneider, Erica Lasek-Nesselquist |

EPI_ISL_984756, EPI_ISL_984757, EPI_ISL_984758, EPI_ISL_984759, EPI_ISL_984760, EPI_ISL_984761, EPI_ISL_984762, EPI_ISL_984763, EPI_ISL_984764, EPI_ISL_984765, EPI_ISL_984766, EPI_ISL_984767, EPI_ISL_984768, EPI_ISL_984769, EPI_ISL_984770, EPI_ISL_984771, EPI_ISL_984772, EPI_ISL_984773, EPI_ISL_984774, EPI_ISL_984775, EPI_ISL_984776, EPI_ISL_984777, EPI_ISL_984778, EPI_ISL_984779, EPI_ISL_984780, EPI_ISL_984781, EPI_ISL_984782, EPI_ISL_984783, EPI_ISL_984784, EPI_ISL_984785, EPI_ISL_984786, EPI_ISL_984787, EPI_ISL_984788, EPI_ISL_984789, EPI_ISL_984790, EPI_ISL_984791, EPI_ISL_984792, EPI_ISL_984793, EPI_ISL_984794, EPI_ISL_984795, EPI_ISL_984796, EPI_ISL_984797, EPI_ISL_984798, EPI_ISL_984799, EPI_ISL_984800, EPI_ISL_984801, EPI_ISL_984802, EPI_ISL_984803, EPI_ISL_984804, EPI_ISL_984805, EPI_ISL_984806, EPI_ISL_984807, EPI_ISL_984808, EPI_ISL_984809, EPI_ISL_984810, EPI_ISL_984811, EPI_ISL_984812, EPI_ISL_984813, EPI_ISL_984814, EPI_ISL_984815, EPI_ISL_984816, EPI_ISL_984817, EPI_ISL_984818, EPI_ISL_984819, EPI_ISL_984820, EPI_ISL_984821, EPI_ISL_984822, EPI_ISL_984823, EPI_ISL_984824, EPI_ISL_984825, EPI_ISL_984826, EPI_ISL_984827, EPI_ISL_984828, EPI_ISL_984829, EPI_ISL_984830, EPI_ISL_984831, EPI_ISL_984832, EPI_ISL_984833, EPI_ISL_984834, EPI_ISL_984835, EPI_ISL_984836, EPI_ISL_984837, EPI_ISL_984838, EPI_ISL_984839, EPI_ISL_984840, EPI_ISL_984841, EPI_ISL_984842, EPI_ISL_984843, EPI_ISL_984844, EPI_ISL_984845, EPI_ISL_984846, EPI_ISL_984847, EPI_ISL_984848, EPI_ISL_984849, EPI_ISL_984850, EPI_ISL_984851, EPI_ISL_984852, EPI_ISL_984853, EPI_ISL_984854, EPI_ISL_984855, EPI_ISL_984856, EPI_ISL_984857, EPI_ISL_984858, EPI_ISL_984859, EPI_ISL_984860, EPI_ISL_984861, EPI_ISL_984862, EPI_ISL_984863, EPI_ISL_984864, EPI_ISL_984865, EPI_ISL_984866, EPI_ISL_984867, EPI_ISL_984868, EPI_ISL_984869, EPI_ISL_984870, EPI_ISL_984871, EPI_ISL_984872, EPI_ISL_984873, EPI_ISL_984874, EPI_ISL_984875, EPI_ISL_984876, EPI_ISL_984877, EPI_ISL_984878, EPI_ISL_984879, EPI_ISL_984880, EPI_ISL_984881, EPI_ISL_984882, EPI_ISL_984883, EPI_ISL_984884, EPI_ISL_984885, EPI_ISL_984886, EPI_ISL_984887, EPI_ISL_984888, EPI_ISL_984889, EPI_ISL_984890, EPI_ISL_984891, EPI_ISL_984892, EPI_ISL_984893, EPI_ISL_984894, EPI_ISL_984895, EPI_ISL_984896, EPI_ISL_984897, EPI_ISL_984898, EPI_ISL_984899, EPI_ISL_984900, EPI_ISL_984901, EPI_ISL_984902, EPI_ISL_984903, EPI_ISL_984904, EPI_ISL_984905, EPI_ISL_984906, EPI_ISL_984907, EPI_ISL_984908, EPI_ISL_984909, EPI_ISL_984910, EPI_ISL_984911, EPI_ISL_984912, EPI_ISL_984913, EPI_ISL_984914, EPI_ISL_984915, EPI_ISL_984916, EPI_ISL_984917, EPI_ISL_984918, EPI_ISL_984919, EPI_ISL_984920, EPI_ISL_984921, EPI_ISL_984922, EPI_ISL_984923, EPI_ISL_984924, EPI_ISL_984925, EPI_ISL_984926, EPI_ISL_984927, EPI_ISL_984928, EPI_ISL_984929, EPI_ISL_984930, EPI_ISL_984931, EPI_ISL_984932, EPI_ISL_984933, EPI_ISL_984934, EPI_ISL_984935, EPI_ISL_984936, EPI_ISL_984937, EPI_ISL_984938, EPI_ISL_984939, EPI_ISL_984940, EPI_ISL_984941, EPI_ISL_984942, EPI_ISL_984943, EPI_ISL_984944, EPI_ISL_984945, EPI_ISL_984946, EPI_ISL_984947, EPI_ISL_994802, EPI_ISL_994803, EPI_ISL_994804, EPI_ISL_994805, EPI_ISL_994806, EPI_ISL_994807, EPI_ISL_994808, EPI_ISL_994809, EPI_ISL_994810, EPI_ISL_994811, EPI_ISL_994812, EPI_ISL_994813, EPI_ISL_994814, EPI_ISL_994815, EPI_ISL_994816, EPI_ISL_994817, EPI_ISL_994818, EPI_ISL_994819, EPI_ISL_994820, EPI_ISL_994821, EPI_ISL_994822, EPI_ISL_994823, EPI_ISL_994824, EPI_ISL_994825, EPI_ISL_994826, EPI_ISL_994827, EPI_ISL_994828, EPI_ISL_994829, EPI_ISL_994830, EPI_ISL_994831, EPI_ISL_994832, EPI_ISL_994833, EPI_ISL_994834, EPI_ISL_994835, EPI_ISL_994836, EPI_ISL_994837, EPI_ISL_994838, EPI_ISL_994839, EPI_ISL_994840, EPI_ISL_994841, EPI_ISL_994842, EPI_ISL_994843, EPI_ISL_994844, EPI_ISL_994845, EPI_ISL_994846, EPI_ISL_994847, EPI_ISL_994848, EPI_ISL_994849, EPI_ISL_994850, EPI_ISL_994851, EPI_ISL_994852, EPI_ISL_994853, EPI_ISL_994854, EPI_ISL_994855, EPI_ISL_994856, EPI_ISL_994857, EPI_ISL_994858, EPI_ISL_994859, EPI_ISL_994860, EPI_ISL_994861, EPI_ISL_994862, EPI_ISL_994863, EPI_ISL_994864, EPI_ISL_994865, EPI_ISL_994866, EPI_ISL_994867, EPI_ISL_994868, EPI_ISL_994869, EPI_ISL_994870, EPI_ISL_994871, EPI_ISL_994872, EPI_ISL_994873, EPI_ISL_994874, EPI_ISL_994875, EPI_ISL_994876, EPI_ISL_994877, EPI_ISL_994878, EPI_ISL_994879, EPI_ISL_994880, EPI_ISL_994881, EPI_ISL_994882, EPI_ISL_994883, EPI_ISL_994884, EPI_ISL_994885, EPI_ISL_994886, EPI_ISL_994887, EPI_ISL_994888, EPI_ISL_994889, EPI_ISL_994890, EPI_ISL_994891, EPI_ISL_994892, EPI_ISL_994893, EPI_ISL_994894, EPI_ISL_994895, EPI_ISL_994896, EPI_ISL_994897, EPI_ISL_994898, EPI_ISL_994899, EPI_ISL_994900, EPI_ISL_994901, EPI_ISL_994902, EPI_ISL_994903, EPI_ISL_994904, EPI_ISL_994905, EPI_ISL_994906, EPI_ISL_994907, EPI_ISL_994908, EPI_ISL_994909, EPI_ISL_994910, EPI_ISL_994911, EPI_ISL_994912, EPI_ISL_994913, EPI_ISL_994914, EPI_ISL_994915, EPI_ISL_994916, EPI_ISL_994917, EPI_ISL_994918, EPI_ISL_994919, EPI_ISL_994920, EPI_ISL_994921, EPI_ISL_994922, EPI_ISL_994923, EPI_ISL_994924, EPI_ISL_994925, EPI_ISL_994926, EPI_ISL_994927, EPI_ISL_994928, EPI_ISL_994929, EPI_ISL_994930, EPI_ISL_994931, EPI_ISL_994932, EPI_ISL_994933, EPI_ISL_994934, EPI_ISL_994935, EPI_ISL_994936, EPI_ISL_994937, EPI_ISL_994938, EPI_ISL_994939, EPI_ISL_994940, EPI_ISL_994941, EPI_ISL_994942, EPI_ISL_994943, EPI_ISL_994944, EPI_ISL_994945, EPI_ISL_994946, EPI_ISL_994947, EPI_ISL_994948, EPI_ISL_994949, EPI_ISL_994950, EPI_ISL_994951, EPI_ISL_994952, EPI_ISL_994953, EPI_ISL_994954, EPI_ISL_994955, EPI_ISL_994956, EPI_ISL_994957, EPI_ISL_994958, EPI_ISL_994959, EPI_ISL_994960, EPI_ISL_994961, EPI_ISL_994962, EPI_ISL_994963, EPI_ISL_994964, EPI_ISL_994965, EPI_ISL_994966, EPI_ISL_994967, EPI_ISL_994968, EPI_ISL_994969, EPI_ISL_994970, EPI_ISL_994971, EPI_ISL_994972, EPI_ISL_994973, EPI_ISL_994974, EPI_ISL_994975, EPI_ISL_994976, EPI_ISL_994977, EPI_ISL_994978, EPI_ISL_994979, EPI_ISL_994980, EPI_ISL_994981, EPI_ISL_994982, EPI_ISL_994983, EPI_ISL_994984, EPI_ISL_994985, EPI_ISL_994986, EPI_ISL_994987, EPI_ISL_994988, EPI_ISL_994989, EPI_ISL_994990, EPI_ISL_994991, EPI_ISL_994992, EPI_ISL_994993, EPI_ISL_994994, EPI_ISL_994995, EPI_ISL_994996, EPI_ISL_994997, EPI_ISL_994998, EPI_ISL_994999, EPI_ISL_995000, EPI_ISL_995001, EPI_ISL_995002, EPI_ISL_995003, EPI_ISL_995004, EPI_ISL_995005, EPI_ISL_995006, EPI_ISL_995007, EPI_ISL_995008, EPI_ISL_995009, EPI_ISL_995010, EPI_ISL_995011, EPI_ISL_995012, EPI_ISL_995013, EPI_ISL_995014, EPI_ISL_995015, EPI_ISL_995016, EPI_ISL_995017, EPI_ISL_995018, EPI_ISL_995019, EPI_ISL_995020, EPI_ISL_995021, EPI_ISL_995022, EPI_ISL_995023, EPI_ISL_995024, EPI_ISL_995025, EPI_ISL_995026, EPI_ISL_995027, EPI_ISL_995028, EPI_ISL_995029, EPI_ISL_995030, EPI_ISL_995031, EPI_ISL_995032, EPI_ISL_995033, EPI_ISL_995034, EPI_ISL_995035, EPI_ISL_995036, EPI_ISL_995037, EPI_ISL_995038, EPI_ISL_995039, EPI_ISL_995040, EPI_ISL_995041, EPI_ISL_995042, EPI_ISL_995043, EPI_ISL_995044, EPI_ISL_995045, EPI_ISL_995046, EPI_ISL_995047, EPI_ISL_995048, EPI_ISL_995049, EPI_ISL_995050, EPI_ISL_995051, EPI_ISL_995052, EPI_ISL_995053, EPI_ISL_995054, EPI_ISL_995055, EPI_ISL_995056, EPI_ISL_995057, EPI_ISL_995058, EPI_ISL_995059, EPI_ISL_995060, EPI_ISL_995061, EPI_ISL_995062, EPI_ISL_995063, EPI_ISL_995064, EPI_ISL_995065, EPI_ISL_995066, EPI_ISL_995067, EPI_ISL_995068, EPI_ISL_995069, EPI_ISL_995070, EPI_ISL_995071, EPI_ISL_995072, EPI_ISL_995073, EPI_ISL_995074, EPI_ISL_995075, EPI_ISL_995076, EPI_ISL_995077, EPI_ISL_995078, EPI_ISL_995079, EPI_ISL_995080, EPI_ISL_995081, EPI_ISL_995082, EPI_ISL_995083, EPI_ISL_995084, EPI_ISL_995085, EPI_ISL_995086, EPI_ISL_995087, EPI_ISL_995088, EPI_ISL_995089, EPI_ISL_995090, EPI_ISL_995091, EPI_ISL_995092, EPI_ISL_995093, EPI_ISL_995094, EPI_ISL_995095, EPI_ISL_995096, EPI_ISL_995097, EPI_ISL_995098, EPI_ISL_995099, EPI_ISL_995100, EPI_ISL_995101, EPI_ISL_995102, EPI_ISL_995103, EPI_ISL_995104, EPI_ISL_995105, EPI_ISL_995106, EPI_ISL_995107, EPI_ISL_995108, EPI_ISL_995109, EPI_ISL_995110, EPI_ISL_995111, EPI_ISL_995112, EPI_ISL_995113, EPI_ISL_995114, EPI_ISL_995115, EPI_ISL_995116, EPI_ISL_995117, EPI_ISL_995118, EPI_ISL_995119, EPI_ISL_995120, EPI_ISL_995121, EPI_ISL_995122, EPI_ISL_995123, EPI_ISL_995124, EPI_ISL_995125, EPI_ISL_995126, EPI_ISL_995127, EPI_ISL_995128, EPI_ISL_995129, EPI_ISL_995130, EPI_ISL_995131, EPI_ISL_995132, EPI_ISL_995133, EPI_ISL_995134, EPI_ISL_995135, EPI_ISL_995136, EPI_ISL_995137, EPI_ISL_995138, EPI_ISL_995139, EPI_ISL_995140, EPI_ISL_995141, EPI_ISL_995142, EPI_ISL_995143, EPI_ISL_995144, EPI_ISL_995145, EPI_ISL_995146, EPI_ISL_995147, EPI_ISL_995148, EPI_ISL_995149, EPI_ISL_995150, EPI_ISL_995151, EPI_ISL_995152, EPI_ISL_995153, EPI_ISL_995154, EPI_ISL_995155, EPI_ISL_995156, EPI_ISL_995157, EPI_ISL_995158, EPI_ISL_995159, EPI_ISL_995160, EPI_ISL_995161, EPI_ISL_995162, EPI_ISL_995163, EPI_ISL_995164, EPI_ISL_995165, EPI_ISL_995166, EPI_ISL_995167, EPI_ISL_995168, EPI_ISL_995169, EPI_ISL_995170, EPI_ISL_995171

| | | | |
|---|---|---|---|
| see above | Pandemic Response Lab - NYC | Pandemic Response Lab, R&D | Henry Lee, Michael Hammerling, Melissa Hopkins, Cybill del Castillo, William Ward, Pradeep Bugga, Haiping Hao, Jon Laurent |
