## Supplementary material for "Detection and characterization of the SARS-CoV-2 lineage B.1.526 in New York": Supp. Table 2

**Supplementary Table 2a. Individual vaccinee characteristics and plasma titers**

| | Demographic characteristics | | | | Vaccination details | | | | | Neutralization titer (reciprocal dilution) NT50 | | | |
|---|---|---|---|---|---|---|---|---|---|---|---|---|---|
| | | | | | | | Time between (days) | | | Experiment 1 | | Experiment 2 | |
| Participant ID | Age (years) | Sex | Race § | Ethnicity | Vaccine platform | # of doses received | 1st & 2nd dose | 1st dose & blood draw | 2nd dose & blood draw | B.1.526 v.1† | D614G | B.1.526 v.2‡ | D614G |
| MOD1 | 51 | Male | W | Hispanic | Moderna | 2 | 28 | 74 | 46 | 364 | 2176 | 497 | 917 |
| MOD2 | 35 | Male | W | Non-Hispanic | Moderna | 2 | 34 | 62 | 28 | 265 | 975 | 205 | 573 |
| MOD3 | 32 | Male | W | Non-Hispanic | Moderna | 2 | 31 | 78 | 47 | 269 | 1412 | 460 | 1762 |
| MOD6 | 41 | Female | W | Non-Hispanic | Moderna | 2 | 28 | 89 | 61 | 223 | 679 | 201 | 983 |
| MOD7 | 62 | Male | A | Non-Hispanic | Moderna | 2 | 35 | 84 | 49 | 563 | 2412 | 746 | 2981 |
| MOD8 | 69 | Female | W | Non-Hispanic | Moderna | 2 | 35 | 96 | 63 | 214 | 1713 | 381 | 1660 |
| PFZ12 | 53 | Female | W | Non-Hispanic | Pfizer/BioNTech | 2 | 21 | 83 | 62 | 345 | 1516 | 733 | 1278 |
| PFZ16 | 67 | Female | W | Non-Hispanic | Pfizer/BioNTech | 2 | 22 | 119 | 97 | 109 | 566 | 188 | 718 |
| C001 | 38 | Female | W | Non-Hispanic | Pfizer/BioNTech | 2 | 22 | 57 | 35 | 884 | 2652 | 1035 | 1738 |
| C004 | 37 | Male | W | Non-Hispanic | Pfizer/BioNTech | 2 | 21 | 91 | 70 | 254 | 1603 | 860 | 1991 |

§ = White (W); Asian (A)
† = B.1.516 v.1 spike mutations L5F, T95I, D253G, E484K, D614G, and A701V
‡ = B.1.526 v.2 spike mutations: L5F, T95I, D253G, S477N, D614G, and Q957R
Characteristics from Supplementary Table 1 of Wang, et al., "mRNA vaccine-elicited antibodies to SARS-CoV-2 and circulating variants", Nature (2021)

**Supplementary Table 2b. Individual convalescent participant characteristics and plasma titers**

| | | | | | Temporal dynamics (days) | | | | | | | Neutralization titer (reciprocal dilution) NT50 | | | | | | | |
|---|---|---|---|---|---|---|---|---|---|---|---|---|---|---|---|---|---|---|---|
| | | | | | | | | | | | | Timepoint T1 - 1.3 months | | | | Timepoint T2 - 6.2 months | | | |
| | | | | | | | | | | | | Experiment 1 | | Experiment 2 | | Experiment 1 | | Experiment 2 | |
| ID | Age (years) | Sex | Race | Ethnicity | Sx duration during acute disease | Sx onset to initial visit (T1) | Sx onset to follow-up visit (T2) | Time between visits | # of solicited comorbidities § | Acute disease severity by WHO (0-8) ¶ | Post-acute Sx persistence # | B.1.526 v.1† | D614G | B.1.526 v.2‡ | D614G | B.1.526 v.1† | D614G | B.1.526 v.2‡ | D614G |
| **38** | 57 | F | White | Non-Hispanic | 10 | 38 | 211 | 173 | 0 | 2 | N | 27 | 148 | 255 | 385 | 154 | 891 | 775 | 2597 |
| **40** | 44 | M | White | Non-Hispanic | 7 | 23 | 195 | 172 | 0 | 2 | N | 13 | 37 | 72 | 90 | N.N. | 10 | N.N. | N.N. |
| **76** | 49 | F | White | Non-Hispanic | 28 | 34 | 204 | 170 | 0 | 1 | Y | 33 | 105 | 105 | 184 | 10 | 23 | 13 | 24 |
| **99** | 36 | F | White | Non-Hispanic | 13 | 29 | 204 | 175 | 0 | 2 | N | 180 | 973 | 794 | 1055 | 22 | 358 | 176 | 240 |
| **233** | 55 | M | White | Non-Hispanic | 20 | 41 | 206 | 165 | 0 | 2 | N | 55 | 194 | 108 | 165 | 19 | 50 | 17 | 46 |
| **328** | 54 | F | White | Non-Hispanic | 22 | 62 | 203 | 141 | 0 | 2 | N | 24 | 347 | 50 | 131 | 22 | 135 | 29 | 77 |
| **352** | 44 | M | White | Non-Hispanic | 16 | 43 | 197 | 154 | 0 | 2 | N | 129 | 1684 | 587 | 744 | 27 | 148 | 40 | 107 |
| **403*** | 52 | M | Asian | Non-Hispanic | 18 | 39 | 174 | 135 | 1 | 4 | Y | 555 | 3223 | 1067 | 2433 | 89 | 463 | 45 | 183 |
| **437** | 43 | F | Asian | Non-Hispanic | 14 | 34 | 192 | 158 | 1 | 2 | N | 122 | 525 | 276 | 996 | 56 | 132 | 65 | 270 |
| **537** | 52 | M | White | Non-Hispanic | 15 | 45 | 178 | 133 | 2 | 2 | Y | 650 | 3528 | 1610 | 4291 | 240 | 853 | 334 | 957 |

Sx = symptoms, N.N = non-neutralizing

* = hospitalized

§ = Arterial hypertension (HTN), obesity (OB), diabetes mellitus (DM), asthma (A), chronic obstructive pulmonary disease (COPD), coronary artery disease (CAD), cancer (CX)

¶ = WHO Ordinal Scale for Clinical Improvement, COVID-19 Trial Design Synopsis

### = Persistent fatigue, dyspnea, athletic deficit, or ≥ 3 other solicited symptoms beyond 6 weeks from Sx onset

† = B.1.516 v.1 spike mutations L5F, T95I, D253G, E484K, D614G, and A701V

‡ = B.1.526 v.2 spike mutations: L5F, T95I, D253G, S477N, D614G, and Q957R

Characteristics from Supplementary Table 2 of Gaebler, et al., "Evolution of antibody immunity to SARS-CoV-2" Nature 591, 639–644 (2021)
